# CircRNA may not be “circular”

**DOI:** 10.1101/2020.09.27.315275

**Authors:** Handong Sun, Zijuan Wu, Ming Liu, Liang Yu, Jianyong Li, Xiangming Ding, Hui Jin

## Abstract

Circular RNA (circRNA) is a novel regulatory non-coding RNA and participates in diverse physiological and pathological processes. However, the structures and molecular mechanisms of circRNAs remain unclear. In this study, we took advantage of circRNA array and bioinformatics analysis and found lots of internal complementary base-pairing sequences (ICBPS) existed in plenty of circRNAs, especially in extremely long circRNAs (el-circRNAs, > 5,000 nt). The result indicated that circRNA may not be a simple circular structure. In addition, we put forward the hypothesis of “open-close effect “ in the transition for specific circRNA from normal state to morbid state. Taken together, our results not only expand the knowledge of circRNAs, but also highlight the potential molecular mechanism of circRNAs.

## 1. Introduction

CircRNA has long been characterized as a single strand covalently closed continuous loop without 5’ to 3’ polarity and a polyadenylated tail^[1]^. The expression of circRNAs has spatio-temporal specificity, and they may exhibit distinct expression patterns in different diseases or at different stages^[2]^. Although accumulating evidence reveals that circRNAs could exert vital biological functions and serve as novel biomarkers as well as providing promising therapeutic approaches for various human diseases^[3]^, much has not yet to be elucidated about their molecular mechanisms. Here, we propose a brand new hypothesis that circRNAs may not be a single strand continuous loop, they probably have double-strand structure, which may be dynamically reversible and have impact on their mechanism and functions.

## 2. Results

### 2.1 The ICBPS existed in circRNAs

In our study, we firstly performed circRNA microarray analysis using CapitalBio Technology Human CircRNA Array V2 to detect the circRNAs expression of human plasma. Surprisingly, we discovered that there are lots of internal complementary base-pairing sequences (ICBPS) existed in plenty of circRNAs, especially in extremely long circRNAs (el-circRNAs, > 5,000 nt). Based on the bioinformatics analysis of human circRNA database (http://www.circbank.cn/)^[4]^ that containing about 140,790 circRNAs, we got similar results. The maximum length (maxLen) of the ICBPS is diverse. For most circRNAs, the maxLen is under 15 or even 10 nt, while we were surprised to find that there are near 6 thousand circRNAs with ICBPS ≥10 nt and 206 circRNAs which contain continuous ICBPS over 100 nt (Figure 1a). Next in this study, we emphasized on the analysis of 6,155 circRNAs (4.37% of total circRNAs) with ICBPS ≥20 nt, most of which contain more than 1 pair of ICBPS. The number of circRNAs with different amounts of ICBPS was analyzed statistically, and there are over 2,000 circRNAs containing more than 20 pairs of ICBPS (Figure 1b). Through bioinformatics analysis we found that the 6,155 circRNAs contain more than 14 million ICBPS in total, and for 90% circRNAs, the median length (medianLen) of the ICBPS is between 20 and 31 nt (Appendix Figure S1a). Also, the number and maxLen of ICBPS were closely correlated with the total length of circRNAs (circLen) (Figure 1c, 1d).

**Figure 1.**
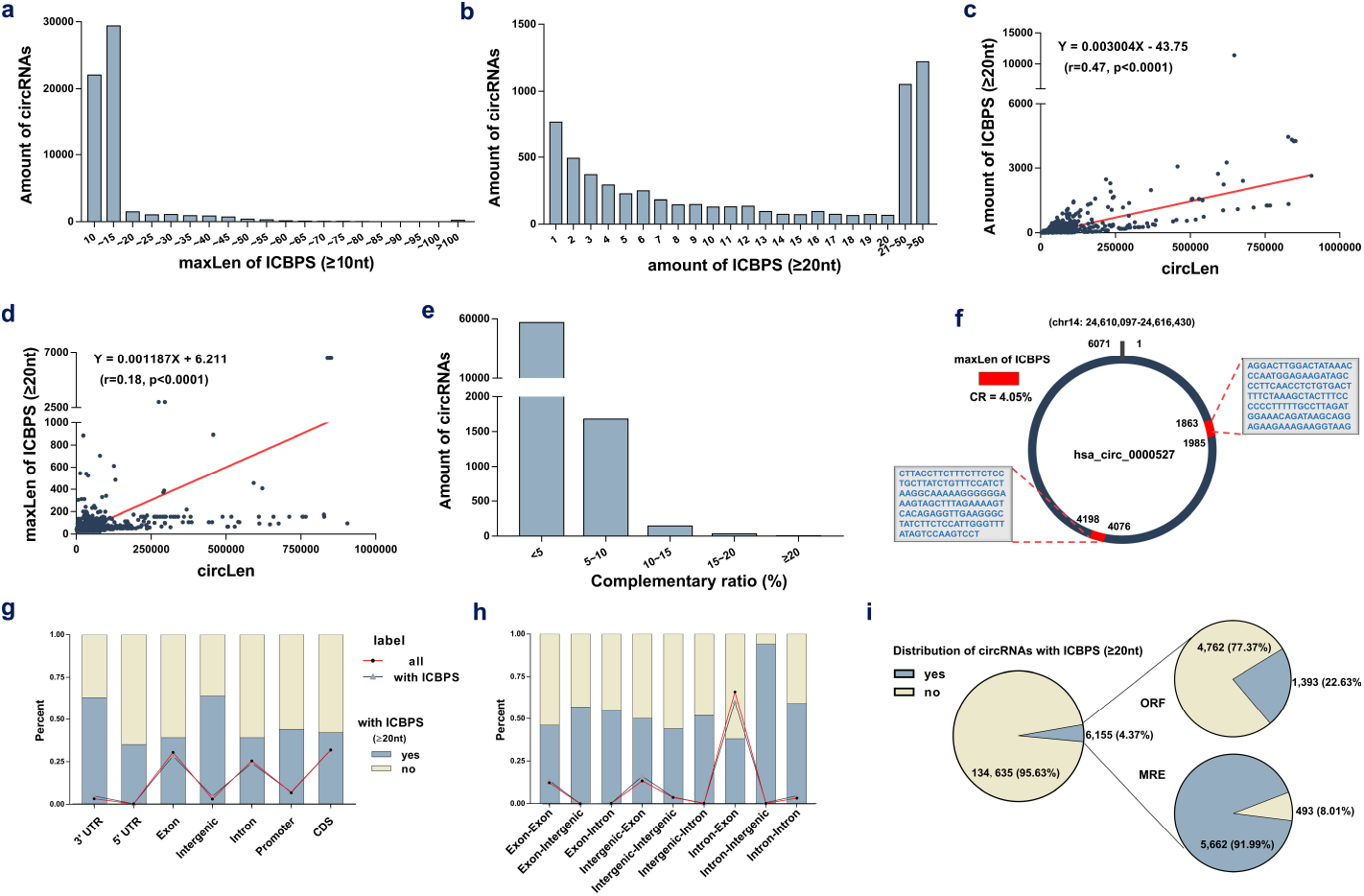
Characteristics of circRNAs. (a) The amount of circRNAs with different maxLen of ICBPS (≥10 nt). (b) The numbers of circRNAs with different amount of ICBPS. (c, d) The correlation analysis between the amount/ maxLen of ICBPS (≥20 nt) and the total length of circRNAs. (e) The amount of circRNAs with different complementary ratio. (f) Hsa_circ_0000527 was taken as an example. (g) The percentage of circRNAs (with ICBPS ≥ 20nt or not) that transcribed from different regions of their parental gene. The polyline graph shows the number distribution of circRNAs in different types. (h) The percentage of circRNAs (with ICBPS ≥ 20 nt or not) with different components originated from the parental genes. The polyline graph shows the number distribution of circRNAs in different types. (i) The distribution of circRNAs with ICBPS ≥ 20nt, and the distribution of these circRNAs with ORF/MRE at the same time.

### 2.2 CR indicates the probability of internal base pairing in circRNA

In addition, we counted the amounts of circRNAs with different complementary ratio (CR) in the database. Here in this study, we defined CR as [(maxLen of ICBPS × 2) /circLen] × 100%. A higher CR would indicate a greater probability of internal base pairing in a circRNA. We found that CR of most circRNAs is under 10% or even 5%. However, to our surprise, for circRNAs with CR ≥ 20%, their circLens are all under 200 nt (Figure 1e, Appendix Table S1). To make an example, hsa_circ_0000527, an el-circRNA originated from exon 24 of chromodomain 14, contains 3 pairs of ICBPS (≥ 20nt) and the maxLen is 123 nt, circLen is 6071 nt, thus its CR is 4.05% (Figure 1f, Appendix Data S1).

### 2.3 Characteristics of ICBPS in circRNA

Also, we calculated the percentage of circRNAs (with ICBPS ≥ 20nt or not) that transcribed from different regions of their parental gene. For circRNAs that derived from 3’-UTR of parental genes, about 60% of them contain ICBPS ≥ 20nt (Figure 1g). Similar statistics analysis was conducted according to different chromosome origins as well as different components originated from the parental gene (Appendix Figure S1b, Figure 1h).

Studies have confirmed that, functioning as competitive endogenous RNA (ceRNA), circRNAs can competitively sponge miRNAs through miRNA recognition elements (MREs)^[5]^. According to the microarray analysis data, there are 206 circRNAs whose maxLen of ICBPS >100nt, and about 64% of them have overlap between ICBPS and MREs (Appendix Figure S1c). Meanwhile, circRNA contains internal ribozyme entry site (IRES) and has the potential to translate proteins, which can be predicted through bioinformatics analysis of its open reading frame (ORF)^[6]^. Based on the analysis of circBank database, there are 6,155 circRNAs contain ICBPS ≥ 20 nt, and about 23%/ 92% of them have overlap with ORF/MRE at the same time (Figure 1i).

### 2.4 Possible structures and molecular mechanisms of circRNAs

Synthesizing the above analytical results, we proposed a brave and reasonable conjecture that circRNA may not be a simple circular structure. It probably contains double-strand structure internally because of the presence of ICBPS (shown as A and A’ in Figure 2a). Moreover, some special cases may also exist. For example, there may be one segment of ICBPS that can be complementary paired with multiple ICBPS (shown as B and B’/B” in Figure 2a). Or for one continuous sequence, it may have different complementary sequences that set close together or overlap on the same RNA chain (shown as C, D and C’, D’ in Figure 2a), but in fact, these complementary pairings can hardly exist at the same time. Thus we inferred that the “open” or “close” state of the double-strand structures in circRNAs would be a sophisticated dynamic process which might be reversible and regulated by micro-environment or other internal factors such as the length of ICBPS, the binding free energy, the distance between pairing fragments, the secondary structure of RNA, or relevant RNA modification like N6-methyladenosine (m6A), etc. And this process might be an important way of regulating the circRNAs degradation, translation and adsorption of miRNAs and so on. This dynamic process is demonstrated in Video 1. RNA-binding proteins (RBPs) act as trans-factors involved in circRNAs biogenesis^[7]^. The formation of double-stranded structure makes circRNAs compressed in space, which may help to make the bond with RBPs more firmly and thus facilitate circRNAs being exported into the cytoplasma from nucleus. In addition, circRNAs as miRNA sponge and its translation behaviors would be prevented when relevant sites on circRNAs are “closed” due to the occurrence of base-pairing. Furthermore, circRNAs are highly stable and the degradation mechanism of circRNAs has not been clarified yet. The double-strand structure in circRNA may make them easier to be degraded by relevant enzymes, and this will probably explain how cells eliminate circRNAs (Figure 2b).

**Figure 2.**
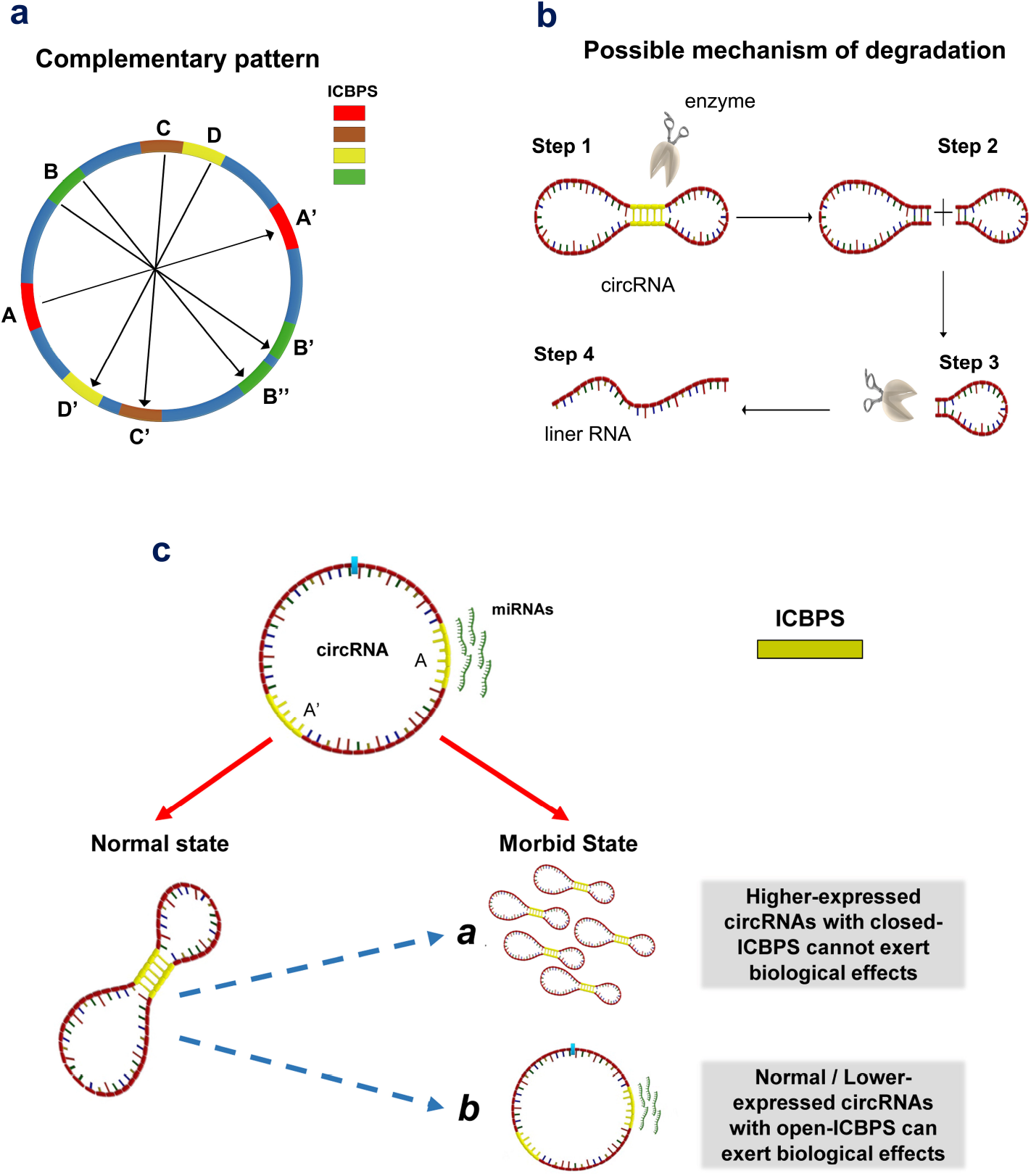
Possible structures and molecular mechanisms of circRNAs. (a) The complementary patterns of ICBPS in circRNA. (b) Possible mechanism of circRNA degradation due to the presence of double-strand structure. (c) The “open-close effect” in the transition for specific circRNA from normal state to morbid state.

Actually, several researches have questioned about the mechanism of ceRNA^[8]^: some circRNAs were predicted to be able to bind a certain miRNA and thus regulate the downstream target genes, while this cannot be well proved through related experiments. On the other hand, the abundance changes of some circRNAs cannot effectively regulate the targeted genes. Based on this phenomenon, we put forward the hypothesis of “open-close effect “ in the transition for specific circRNA from normal state to morbid state: for those circRNAs with closed ICBPS, even if they are highly expressed, they may not exert corresponding biological functions; for those circRNAs with similar or even lower expression, they may also play important roles through “opening” relevant ICBPS (Figure 2c), and vice versa.

## 3. Discussion

In conclusion, we speculated that circRNA may not be a simple circular structure, which probably contains double-strand structure internally, and the “open” or “close” of the double-strand structures would be a sophisticated dynamic process. If this hypothesis is correct, it indicates that circRNA may play roles in the occurrence and development of disease, not necessarily through its aberrant expression change, but also through the “open-close effect” of related sites on the sequence. If a certain circRNA has both oncogenic and tumor suppressor miRNA binding sites, it may selectively “open” or “close” specific miRNA binding sites, consequently leads to different effects. In addition, this hypothesis may help to provide new ideas and clues for the questions as follows: 1) certain circRNAs cannot be amplified and validated using primers designed according to the design principle; 2) why some circRNAs have no effects on their predicted target miRNAs; 3) how can circRNAs be exported from nucleus into cytoplasma; 4) how can circRNAs be degraded; 5) how does m6A regulate circRNA encoding proteins and 6) this may help to fill the research gap of el-circRNAs, which may have more complex structures. This theory may change the existing circRNA research mode, and more importantly, extend the underlying logic of selecting indicators based on circRNA-sequencing or circRNA array.

## 4. Materials and Methods

### 4.1 CircRNA microarray

The circRNA microarray was conducted using CapitalBio Technology Human CircRNA Array V2 (CapitalBio Technology, Beijing, China) containing probes interrogating about 170,340 humuan circRNAs. And those circRNA target sequences were all from Circbase, Deepbase and Rybak-Wolf 2015.

### 4.2 Database

Database used in our research containing circBank (http://www.circbank.cn/) and circBase (http://www.circbase.org/).

### 4.3 Statistical analysis

Data analysis was performed using GraphPad Prism7 software (GraphPad Software Inc., La Jolla, CA) and SPSS 20.0 software (SPSS Inc. Chicago, IL, USA). Correlations were analysed by Pearson’s correlation test. P values of < 0.05 were considered significant.

## Acknowledgments

We thank the circBank Database (http://www.circbank.cn) for providing the bioinformation of circRNAs in this study.

## Funding

This work was supported by National Natural Science Foundation of China (Grant no. 81700155, 81720108002), National Science and Technology Major Project (2018ZX09734-007), Scientific research support project of Jiangsu Commission of Health (H2018085).

## Conflict of interest

The authors declare no competing interests.

## Notes

### Competing Interest Statement

The authors have declared no competing interest.

