## Supplementary for "CircRNA may not be “circular”"

**Supplementary information, Data S1**

**Information of the instanced circRNA**

circBase ID: hsa_circ_0000527

circBank ID: hsa_circFAM158A_001

Position: chr14: 24610097-24616430

Length: 6071

RNA sequence:

CTGGACCAGTCCGGTACGCTCCGGGCCTCACCGAGGAAGTCCCACCCTGC
AGGGGAACCGACTCTGGCCGGCCCTAACCCCTCTCGAGCGAGTCCCGCCC
CATCTCCGGTCTGTCCTGAAGGCAGAGGGAAGACCCTCACCCAACCCCTG
CCCGCTCTCTTCTAAGAGCTGCTAGTCCTTCTCCACCCCAGGCGCTCCCTCT
GGCTGCCCCTCCCACTCGCTCTGATCTCTCTCCGTCCGCTACTTGCGGTCGA
GCCTTCTTGGGGGAGCCCCGCCACTGAATACCCCCTTTGCCATACTCCCAC
CTCTCTTACCCCAACCCTGGACTTCCAGTTGGCCCTAGCTATAAGCCCCGCC
CCTCTCCGAGGGACAGCCCCCGCTCCAGTCTCCGCGGTCCCACTTACCCTA
GCCCCTCCCTCGGGCAGCCCCTGGACTGTGACCCAGCCAGCACCGCCTCT
CCTAGTGGCAACTGGCTGGGAATGGCTGAGGGGCTACCTACTGCTTGCGGC
CTGCCAGCGGGAGGAGGGGGGCGGAAAGAAGAAAGGGGGCGGGGTTGA
GGGGGGCGGGCCTGGACCTGGGGAGTGAAAGCGAAAGCCCGGGCGACTA
GCCGGGAGACCAGAGATCTAGCGACTGAAGCAGCATGGCCAAGCCGTGTG
GGGTGCGCCTGAGCGGGGAAGCCCGCAAACAGGTAATGGGGAATCGGAG
GCCAGGAAGGGGTCATAGAACCCTCTTGGGTGGAGGCCAAGCTGGGTGGT
TCAGGGGACCAAGCAGTGCCAGAGAACTCCCCCAATCTCGCTAGTGCCTT
CACTTCTCGGGGATTCCCAGATTCCTGCCTTCTGACTGCAGTGAGCAGGAG
CTTCCAGGGAAAGCTAGTTTGTGGTGAGAAAGGCCTGAGCCACAGTTCGA
GGGCACCCTGTTGTGGGTGGAAGCTTTTGACAACCCAGACCCTTAACCAA
CATCCGGGTACTGCTCTCTGTGTAGGTGGAGGTCTTCAGACAGAATCTTTT
CCAGGAGGTAAGTCTCTGGATTTGAGCGATTTGACCCAGAATCTTGGACCA
ATACTTGAGGATAACAGCCTCTCAGAATGACCCCCCCAAGTCACCCTAGAC
TTAACCCTACTTCCCTTACCAAGCCTTCACCCAAAGTAACCCCGGAATTATT
GATAGCTACCTTTAATCTGACCTTTCTTATGACTCTCCCGACTTCTGGCACTT
CTCCCGTCTTTTGATGTGGAGGAAAAAAACCATGGCTTCCAAAGCAGGAAT
ATCACAGTCCTAACCTTCGTTCTGCCATTTACCAGCTTGGGCAAGTCCTTTC
ATTTTCAGGTCTTCAGTTTCCTCATTTGCAAGTCTTGGACCCAGTGATTTCC
AAAGGCCCTTCCAGTTCCATTTGTCAGACATCTCATTGAGCATATTCTGTGA

TTCCCCTCCCTTGCAATGTAACCCACAGGCTGAGGAATTCCTCTACAGATTC
TTGCCACAGAAAATCATATACCTGAATCAGCTCTTGCAAGTGAGTAGTGTG
GGCCCCTTCTGCCCCATCCTATGTCCTCTATCCCCTTTGCCCACCCTCTCTAG
CTAACAACTGGATTCAGCATGCAGAGAATAGTTTCAAATAGTGGATTAGGA
ACATTAGGATGGCGTGAGGCATGAGCCTAGGTTGGGAGGCAGAGTGCAAG
TGGGATGTATGGAGGGAAGGCTGTGACCTAGGATGAAGGGAAGAGGTTGG
GCAGCCCTTGACAGTGAGCTCTGTGCAGGAGGACTCCCTCAATGTGGCTG
ACTTGACTTCCCTCCGGGCCCCACTGGACATCCCCATCCCAGACCCTCCAC
CCAAGGATGATGAGGTGAGG[CACTCAGGGTGC**AGGACTTGGACTATAAA**
**CCCAATGGAGAAGATAGCCCTTCAACCTCTGTGACTTTTCTAAAGCTA**
**CTTTCCCCCCTTTTTGCCTTAGATGGAAACAGATAAGCAGGAGAAGAA**
**AGAAGGTAAG**AGAGATAAGAGGTXXXXXXXXXXXXXXXXXXXXXXXXX
XXXXXGGATTCAGGCCCTTTCTCATCCAGTAGTCAATGTGCCATCTCCCCTT
CCCTAGTCACCTCTTATCTCA**CTTACCTTCTTTCTTCTCCTGCTTATCTGT**
**TTCCATCTAAGGCAAAAAGGGGGGAAAGTAGCTTTAGAAAAGTCACAG**
**AGGTTGAAGGGCTATCTTCTCCATTGGGTTTATAGTCCAAGTCCT**TCAC
CCTGAGTGCCTAGGGTG]GGGATTGTTACATGCCATGGCAGCCATGCAGATA
ACCTGTATTTTGGAATGTAACAGTATTATTTTCATCCAGAGGTCTCAGGAAT
TATTGGAAGAGATGGCATTTGCACAGAGCCCTTATGGGAATAATCAGGCTTT
GGATACACAAGGGGAAAGTTCTGTGGTCAACATCAGGAATGAAGAAAGAA
AGAATCCTCATCAGTTCAGGAAAAGCCTAGAAGGAGAATAAGGTTTCACT
GGGAGAACTATGGAGAACTACGGATGAATGATTGGACTCCCCCCACCCTTT
TTTTTTTTTTTTGAGATGGAGTCTTGCTCTGTTGCCCAGGCTGGAGTGCAA
TGGCACGATCTCAGCTTACTGCTACCTCCGTCTCCCGGGTTCAAGCAATTCT
CCTGCCTCAGCCTCCCGAGTAGCTGGAATTACAGGCATGTGCCACCACACC
TGGCTAATTTTTTTGTATTTTTAATAGAGACGGGGTTTCGCCATGTTGGCCA
GGCTGGTCTCGAACTCCTGACCTCAGGCGATCCACCCACCTCGGCCTCCCA
AAGTGCTGGGATTACAAGGCGTGAGCCACCGCGCCGGGCCTGGACTCCCT
CCTTAAGGGCTAGGAGGGGAAAAAATAATGCTTCCAAGGCACTGTGGTGA
AGGAAATACATCGCAGGACATTATGGGAAGGATGAATCTGTAAGTTTTTCTT
ACTGTCCTCCCCCATATCCCCTTCAGCCAATCACTGTCTTTATCTCTGCTTTT
CAAGATTCAGCCCTGGGCTGGTGATAGAGACAAGGGTTCCTTTGCTGTGAT
CTTGCTGTAAACTCCATGGGATTGTTCTTTAATTCAGGGCCTTATCCTGTCAT
AAAATATGAGATAAGAATGTGCTGCGAAGCACTAGACTGGCTGGAAGCCTG
CCCCCATACATATCTCATTAAATCCTTAAAGGAACACGGGCAAAAGATAAAC

CAAGGGCAAGAAAACCTGCAGTCGCGAAGGCAACACGGCCCATTAGTAAG
TGACAGAGCCGATCTAGAGTACAGAGCCACTCAACCCATAATAGGCGCAA
GGCGGTCTGCTTTGCCCGCAGCGCTCCGGAACACCCGGCTACGGGATAGC
GATCCATCCTTGGGGCTCGGCGGAGCGAGTGGTAGTGGGCAACACAGGTA
ACAGGCAAAGCCAACCACCAACCGAGAGCGGAAGTCGGGGAACCTAGCC
AATAAGAGTGCGGGCAATTTTCTGTGCTTCACAATGGTGGGTTCTTTCCAG
TCCGCTGGAGACAAAGGCGGGGCAACGTTGGGGCGCGGCCGCTGATTGGA
CAAATTGAAAGCCTTTAGCCCTATCAAGGAGCACCACGTGGGCGGGACTC
CAGGAGCCAGTTCTGCAGGGAGCTGCAGTTTGGTCTGTCTCCCAGGCAGG
CGCAGAGTCGCGGCCGCCAGCTAGGGGCGCGGGAAGGCGGGGCTCGGAT
GCAATCGGGACCTCCTCCTGGACTGGGCCGGGGGCGGACTCCGGGACCCA
GGGCGCCGGGAGCCGGCGGGCTACCTGCGAGTCGAGTTAGCGTTGTCGCC
GAACCGAAGCCTCGCTCGCCATGGGGGAGGTGGAGATCTCGGCCCTGGCC
TACGTGAAGATGTGCCTGCATGCTGCCCGGTACCCACACGCCGCAGTCAAC
GGGCTGTTTTTGGCGCCAGCGCCGCGGTCTGGAGAATGCCTGTGCCTCACC
GACTGTGTGCCCCTCTTCCACAGCCACCTGGCCCTGTCCGTCATGTTGGAG
GTCGCCCTCAACCAGGTGGATGTGTGGGGAGCACAGGCCGGTCTGGTGGT
GGCTGGTTACTACCATGCCAATGCAGCTGTGAACGATCAGAG

***ICBPS is noted with red color and XXXXXX indicate an omission in the sequence***

**Supplementary information, Data S2**

### Materials and Methods

#### CircRNA microarray

The circRNA microarray was conducted using CapitalBio Technology Human CircRNA Array V2 (CapitalBio Technology, Beijing, China) containing probes interrogating about 170,340 humuan circRNAs. And those circRNA target sequences were all from Circbase, Deepbase and Rybak-Wolf 2015.

#### Database

Database used in our research containing circBank (http://www.circbank.cn/) and circBase (http://www.circbase.org/).

#### Statistical analysis

Data analysis was performed using GraphPad Prism7 software (GraphPad Software Inc., La Jolla, CA) and SPSS 20.0 software (SPSS Inc. Chicago, IL, USA). Correlations were analysed by Pearson's correlation test. P values of $< 0.05$ were considered significant.

**Supplementary information, Figure S1**

Figure S1. (a) The median length of 6,155 circRNAs' ICBPS. (b) The distribution of circRNAs (with ICBPS ≥100 nt) with or without MRE. (c) The percentage of circRNAs (with ICBPS ≥ 20 nt or not) that transcribed from different chromosome origins. The polyline graph shows the number distribution of circRNAs in different types.

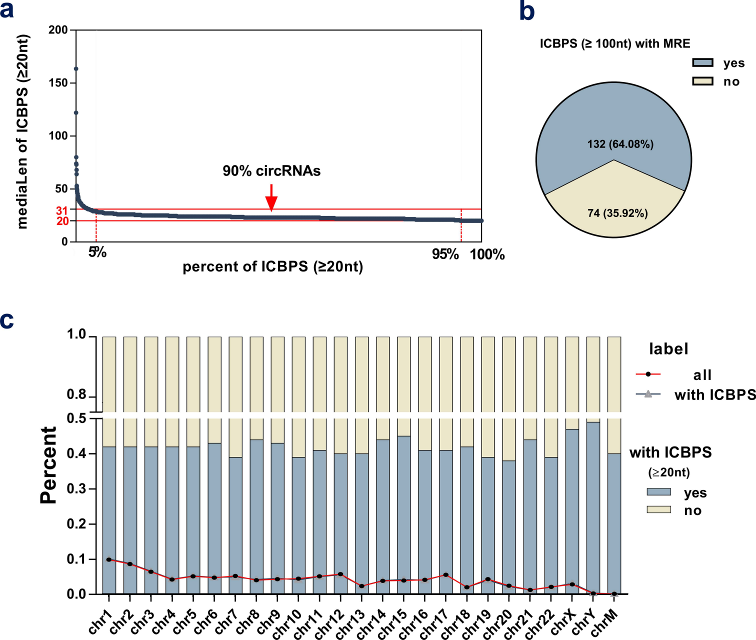
a
200
150
100
50
31
20
0
mediaLen of ICBPS (≥20nt)
90% circRNAs
5%
95%
100%
percent of ICBPS (≥20nt)
b
ICBPS (≥ 100nt) with MRE
yes
no
132 (64.08%)
74 (35.92%)
c
1.0
0.8
0.5
0.4
0.3
0.2
0.1
0.0
Percent
label
all
with ICBPS
with ICBPS
(≥20nt)
yes
no
chr1
chr2
chr3
chr4
chr5
chr6
chr7
chr8
chr9
chr10
chr11
chr12
chr13
chr14
chr15
chr16
chr17
chr18
chr19
chr20
chr21
chr22
chrX
chrY
chrM

| circBase ID | circLen | maxLen of ICBPS | complementary ratio (CR) |
|---|---|---|---|
| **hsa_circ_0028307** | **103** | **11** | **21.36%** |
| **hsa_circ_0139448** | **105** | **11** | **20.95%** |
| **hsa_circ_0136574** | **182** | **19** | **20.88%** |
| **hsa_circ_0119061** | **116** | **12** | **20.69%** |
| **hsa_circ_0001288** | **100** | **10** | **20.00%** |
| hsa_circ_0095976 | 121 | 12 | 19.83% |
| hsa_circ_0124016 | 111 | 11 | 19.82% |
| hsa_circ_0120351 | 102 | 10 | 19.61% |
| hsa_circ_0105594 | 104 | 10 | 19.23% |
| hsa_circ_0096742 | 115 | 11 | 19.13% |
| hsa_circ_0067581 | 105 | 10 | 19.05% |
| hsa_circ_0108431 | 107 | 10 | 18.69% |
| hsa_circ_0061141 | 140 | 13 | 18.57% |
| hsa_circ_0119650 | 110 | 10 | 18.18% |
| hsa_circ_0037908 | 112 | 10 | 17.86% |
| hsa_circ_0090089 | 113 | 10 | 17.70% |
| hsa_circ_0003355 | 114 | 10 | 17.54% |
| hsa_circ_0050263 | 115 | 10 | 17.39% |
| hsa_circ_0059479 | 115 | 10 | 17.39% |
| hsa_circ_0100405 | 115 | 10 | 17.39% |
| hsa_circ_0123756 | 140 | 12 | 17.14% |
| hsa_circ_0114661 | 141 | 12 | 17.02% |
| hsa_circ_0023669 | 118 | 10 | 16.95% |
| hsa_circ_0050921 | 118 | 10 | 16.95% |
| hsa_circ_0078939 | 130 | 11 | 16.92% |
| hsa_circ_0084954 | 119 | 10 | 16.81% |
| hsa_circ_0087815 | 119 | 10 | 16.81% |
| hsa_circ_0103084 | 120 | 10 | 16.67% |
| hsa_circ_0047307 | 134 | 11 | 16.42% |
| hsa_circ_0036320 | 122 | 10 | 16.39% |
| hsa_circ_0106687 | 122 | 10 | 16.39% |
| hsa_circ_0124647 | 125 | 10 | 16.00% |
| hsa_circ_0128158 | 127 | 10 | 15.75% |
| hsa_circ_0124978 | 128 | 10 | 15.63% |
| hsa_circ_0001881 | 129 | 10 | 15.50% |
| hsa_circ_0135955 | 130 | 10 | 15.38% |
| hsa_circ_0022467 | 157 | 12 | 15.29% |
| hsa_circ_0012800 | 132 | 10 | 15.15% |
| hsa_circ_0073493 | 146 | 11 | 15.07% |
| hsa_circ_0109128 | 148 | 11 | 14.86% |
| hsa_circ_0139218 | 365 | 27 | 14.79% |
| hsa_circ_0096081 | 136 | 10 | 14.71% |
| hsa_circ_0120270 | 177 | 13 | 14.69% |
| hsa_circ_0015375 | 137 | 10 | 14.60% |
| hsa_circ_0109147 | 137 | 10 | 14.60% |
| hsa_circ_0132954 | 137 | 10 | 14.60% |
| hsa_circ_0009247 | 139 | 10 | 14.39% |

| | | | |
|---|---|---|---|
| hsa_circ_0104901 | 167 | 12 | 14.37% |
| hsa_circ_0133149 | 196 | 14 | 14.29% |
| hsa_circ_0096583 | 140 | 10 | 14.29% |
| hsa_circ_0021440 | 169 | 12 | 14.20% |
| hsa_circ_0131254 | 155 | 11 | 14.19% |
| hsa_circ_0033540 | 142 | 10 | 14.08% |
| hsa_circ_0106029 | 142 | 10 | 14.08% |
| hsa_circ_0109883 | 142 | 10 | 14.08% |
| hsa_circ_0042763 | 143 | 10 | 13.99% |
| hsa_circ_0011062 | 187 | 13 | 13.90% |
| hsa_circ_0016311 | 144 | 10 | 13.89% |
| hsa_circ_0072102 | 145 | 10 | 13.79% |
| hsa_circ_0100386 | 146 | 10 | 13.70% |
| hsa_circ_0090088 | 148 | 10 | 13.51% |
| hsa_circ_0112666 | 148 | 10 | 13.51% |
| hsa_circ_0079205 | 149 | 10 | 13.42% |
| hsa_circ_0133030 | 149 | 10 | 13.42% |
| hsa_circ_0075727 | 180 | 12 | 13.33% |
| hsa_circ_0020189 | 166 | 11 | 13.25% |
| hsa_circ_0103325 | 154 | 10 | 12.99% |
| hsa_circ_0107188 | 170 | 11 | 12.94% |
| hsa_circ_0138818 | 155 | 10 | 12.90% |
| hsa_circ_0097559 | 156 | 10 | 12.82% |
| hsa_circ_0092753 | 157 | 10 | 12.74% |
| hsa_circ_0121677 | 315 | 20 | 12.70% |
| hsa_circ_0044815 | 159 | 10 | 12.58% |
| hsa_circ_0083303 | 162 | 10 | 12.35% |
| hsa_circ_0127164 | 162 | 10 | 12.35% |
| hsa_circ_0080000 | 244 | 15 | 12.30% |
| hsa_circ_0067850 | 163 | 10 | 12.27% |
| hsa_circ_0069478 | 163 | 10 | 12.27% |
| hsa_circ_0092017 | 163 | 10 | 12.27% |
| hsa_circ_0050641 | 295 | 18 | 12.20% |
| hsa_circ_0087658 | 164 | 10 | 12.20% |
| hsa_circ_0129462 | 164 | 10 | 12.20% |
| hsa_circ_0012599 | 165 | 10 | 12.12% |
| hsa_circ_0025781 | 165 | 10 | 12.12% |
| hsa_circ_0036373 | 165 | 10 | 12.12% |
| hsa_circ_0124317 | 165 | 10 | 12.12% |
| hsa_circ_0112881 | 166 | 10 | 12.05% |
| hsa_circ_0135447 | 166 | 10 | 12.05% |
| hsa_circ_0008883 | 167 | 10 | 11.98% |
| hsa_circ_0042058 | 184 | 11 | 11.96% |
| hsa_circ_0133522 | 201 | 12 | 11.94% |
| hsa_circ_0072550 | 168 | 10 | 11.90% |
| hsa_circ_0096370 | 168 | 10 | 11.90% |
| hsa_circ_0106172 | 168 | 10 | 11.90% |
| hsa_circ_0109877 | 168 | 10 | 11.90% |
| hsa_circ_0127073 | 169 | 10 | 11.83% |
| hsa_circ_0099074 | 186 | 11 | 11.83% |

| | | | |
|---|---|---|---|
| hsa_circ_0138009 | 220 | 13 | 11.82% |
| hsa_circ_0102750 | 237 | 14 | 11.81% |
| hsa_circ_0010241 | 187 | 11 | 11.76% |
| hsa_circ_0037473 | 170 | 10 | 11.76% |
| hsa_circ_0052999 | 170 | 10 | 11.76% |
| hsa_circ_0055690 | 170 | 10 | 11.76% |
| hsa_circ_0097557 | 171 | 10 | 11.70% |
| hsa_circ_0118818 | 171 | 10 | 11.70% |
| hsa_circ_0129084 | 171 | 10 | 11.70% |
| hsa_circ_0015098 | 189 | 11 | 11.64% |
| hsa_circ_0042275 | 189 | 11 | 11.64% |
| hsa_circ_0081871 | 189 | 11 | 11.64% |
| hsa_circ_0024221 | 207 | 12 | 11.59% |
| hsa_circ_0049155 | 190 | 11 | 11.58% |
| hsa_circ_0057819 | 173 | 10 | 11.56% |
| hsa_circ_0060601 | 191 | 11 | 11.52% |
| hsa_circ_0092093 | 191 | 11 | 11.52% |
| hsa_circ_0056361 | 174 | 10 | 11.49% |
| hsa_circ_0096891 | 174 | 10 | 11.49% |
| hsa_circ_0004341 | 175 | 10 | 11.43% |
| hsa_circ_0077574 | 246 | 14 | 11.38% |
| hsa_circ_0038204 | 176 | 10 | 11.36% |
| hsa_circ_0055543 | 176 | 10 | 11.36% |
| hsa_circ_0056013 | 176 | 10 | 11.36% |
| hsa_circ_0056022 | 176 | 10 | 11.36% |
| hsa_circ_0115683 | 176 | 10 | 11.36% |
| hsa_circ_0123718 | 176 | 10 | 11.36% |
| hsa_circ_0023670 | 177 | 10 | 11.30% |
| hsa_circ_0039530 | 177 | 10 | 11.30% |
| hsa_circ_0101281 | 177 | 10 | 11.30% |
| hsa_circ_0046498 | 179 | 10 | 11.17% |
| hsa_circ_0080502 | 179 | 10 | 11.17% |
| hsa_circ_0045850 | 198 | 11 | 11.11% |
| hsa_circ_0010651 | 180 | 10 | 11.11% |
| hsa_circ_0088159 | 180 | 10 | 11.11% |
| hsa_circ_0101454 | 218 | 12 | 11.01% |
| hsa_circ_0064564 | 182 | 10 | 10.99% |
| hsa_circ_0084802 | 182 | 10 | 10.99% |
| hsa_circ_0096719 | 183 | 10 | 10.93% |
| hsa_circ_0109338 | 348 | 19 | 10.92% |
| hsa_circ_0095755 | 239 | 13 | 10.88% |
| hsa_circ_0127251 | 184 | 10 | 10.87% |
| hsa_circ_0127362 | 184 | 10 | 10.87% |
| hsa_circ_0064319 | 204 | 11 | 10.78% |
| hsa_circ_0095447 | 204 | 11 | 10.78% |
| hsa_circ_0051770 | 186 | 10 | 10.75% |
| hsa_circ_0115045 | 205 | 11 | 10.73% |
| hsa_circ_0066923 | 187 | 10 | 10.70% |
| hsa_circ_0128807 | 187 | 10 | 10.70% |
| hsa_circ_0094453 | 225 | 12 | 10.67% |

| | | | |
|---|---|---|---|
| hsa_circ_0012003 | 188 | 10 | 10.64% |
| hsa_circ_0014989 | 188 | 10 | 10.64% |
| hsa_circ_0026069 | 188 | 10 | 10.64% |
| hsa_circ_0096070 | 188 | 10 | 10.64% |
| hsa_circ_0113680 | 188 | 10 | 10.64% |
| hsa_circ_0018754 | 207 | 11 | 10.63% |
| hsa_circ_0062808 | 207 | 11 | 10.63% |
| hsa_circ_0009323 | 189 | 10 | 10.58% |
| hsa_circ_0106348 | 189 | 10 | 10.58% |
| hsa_circ_0043520 | 208 | 11 | 10.58% |
| hsa_circ_0089184 | 228 | 12 | 10.53% |
| hsa_circ_0082784 | 209 | 11 | 10.53% |
| hsa_circ_0059182 | 248 | 13 | 10.48% |
| hsa_circ_0121667 | 229 | 12 | 10.48% |
| hsa_circ_0014826 | 191 | 10 | 10.47% |
| hsa_circ_0102267 | 191 | 10 | 10.47% |
| hsa_circ_0015932 | 211 | 11 | 10.43% |
| hsa_circ_0094600 | 211 | 11 | 10.43% |
| hsa_circ_0050719 | 232 | 12 | 10.34% |
| hsa_circ_0014655 | 194 | 10 | 10.31% |
| hsa_circ_0085115 | 214 | 11 | 10.28% |
| hsa_circ_0115478 | 214 | 11 | 10.28% |
| hsa_circ_0116157 | 312 | 16 | 10.26% |
| hsa_circ_0003254 | 195 | 10 | 10.26% |
| hsa_circ_0015373 | 195 | 10 | 10.26% |
| hsa_circ_0064699 | 195 | 10 | 10.26% |
| hsa_circ_0106393 | 195 | 10 | 10.26% |
| hsa_circ_0041903 | 196 | 10 | 10.20% |
| hsa_circ_0082087 | 197 | 10 | 10.15% |
| hsa_circ_0000819 | 217 | 11 | 10.14% |
| hsa_circ_0022537 | 198 | 10 | 10.10% |
| hsa_circ_0136259 | 198 | 10 | 10.10% |
| hsa_circ_0140471 | 198 | 10 | 10.10% |
| hsa_circ_0066058 | 259 | 13 | 10.04% |
| hsa_circ_0002870 | 220 | 11 | 10.00% |
| hsa_circ_0003215 | 220 | 11 | 10.00% |
| hsa_circ_0044623 | 220 | 11 | 10.00% |
| hsa_circ_0139450 | 220 | 11 | 10.00% |
| hsa_circ_0055687 | 201 | 10 | 9.95% |
| hsa_circ_0069334 | 201 | 10 | 9.95% |
| hsa_circ_0101834 | 201 | 10 | 9.95% |
| hsa_circ_0108495 | 201 | 10 | 9.95% |
| hsa_circ_0102665 | 744 | 37 | 9.95% |
| hsa_circ_0123978 | 242 | 12 | 9.92% |
| hsa_circ_0047669 | 222 | 11 | 9.91% |
| hsa_circ_0044433 | 202 | 10 | 9.90% |
| hsa_circ_0068238 | 202 | 10 | 9.90% |
| hsa_circ_0103940 | 202 | 10 | 9.90% |
| hsa_circ_0109764 | 202 | 10 | 9.90% |
| hsa_circ_0120179 | 202 | 10 | 9.90% |

| | | | |
|---|---|---|---|
| hsa_circ_0010910 | 243 | 12 | 9.88% |
| hsa_circ_0089974 | 287 | 14 | 9.76% |
| hsa_circ_0088526 | 246 | 12 | 9.76% |
| hsa_circ_0089860 | 205 | 10 | 9.76% |
| hsa_circ_0113148 | 205 | 10 | 9.76% |
| hsa_circ_0114312 | 205 | 10 | 9.76% |
| hsa_circ_0051772 | 288 | 14 | 9.72% |
| hsa_circ_0005872 | 206 | 10 | 9.71% |
| hsa_circ_0014875 | 206 | 10 | 9.71% |
| hsa_circ_0015935 | 206 | 10 | 9.71% |
| hsa_circ_0094250 | 206 | 10 | 9.71% |
| hsa_circ_0124591 | 206 | 10 | 9.71% |
| hsa_circ_0128267 | 227 | 11 | 9.69% |
| hsa_circ_0036371 | 207 | 10 | 9.66% |
| hsa_circ_0096447 | 207 | 10 | 9.66% |
| hsa_circ_0124011 | 207 | 10 | 9.66% |
| hsa_circ_0017590 | 270 | 13 | 9.63% |
| hsa_circ_0001277 | 229 | 11 | 9.61% |
| hsa_circ_0106946 | 250 | 12 | 9.60% |
| hsa_circ_0133402 | 480 | 23 | 9.58% |
| hsa_circ_0046660 | 209 | 10 | 9.57% |
| hsa_circ_0076426 | 209 | 10 | 9.57% |
| hsa_circ_0086653 | 209 | 10 | 9.57% |
| hsa_circ_0031539 | 251 | 12 | 9.56% |
| hsa_circ_0019543 | 210 | 10 | 9.52% |
| hsa_circ_0128207 | 210 | 10 | 9.52% |
| hsa_circ_0132160 | 210 | 10 | 9.52% |
| hsa_circ_0055875 | 274 | 13 | 9.49% |
| hsa_circ_0075249 | 232 | 11 | 9.48% |
| hsa_circ_0079175 | 211 | 10 | 9.48% |
| hsa_circ_0131133 | 254 | 12 | 9.45% |
| hsa_circ_0060558 | 233 | 11 | 9.44% |
| hsa_circ_0083163 | 233 | 11 | 9.44% |
| hsa_circ_0009108 | 212 | 10 | 9.43% |
| hsa_circ_0106948 | 212 | 10 | 9.43% |
| hsa_circ_0109403 | 212 | 10 | 9.43% |
| hsa_circ_0113882 | 212 | 10 | 9.43% |
| hsa_circ_0006836 | 234 | 11 | 9.40% |
| hsa_circ_0009870 | 234 | 11 | 9.40% |
| hsa_circ_0139451 | 234 | 11 | 9.40% |
| hsa_circ_0002910 | 213 | 10 | 9.39% |
| hsa_circ_0089818 | 213 | 10 | 9.39% |
| hsa_circ_0120825 | 213 | 10 | 9.39% |
| hsa_circ_0134541 | 213 | 10 | 9.39% |
| hsa_circ_0001594 | 6513 | 305 | 9.37% |
| hsa_circ_0092743 | 235 | 11 | 9.36% |
| hsa_circ_0004946 | 214 | 10 | 9.35% |
| hsa_circ_0128326 | 257 | 12 | 9.34% |
| hsa_circ_0045148 | 258 | 12 | 9.30% |
| hsa_circ_0054835 | 215 | 10 | 9.30% |

| | | | |
|---|---|---|---|
| hsa_circ_0066677 | 215 | 10 | 9.30% |
| hsa_circ_0082157 | 215 | 10 | 9.30% |
| hsa_circ_0097465 | 215 | 10 | 9.30% |
| hsa_circ_0110306 | 215 | 10 | 9.30% |
| hsa_circ_0113687 | 215 | 10 | 9.30% |
| hsa_circ_0067771 | 216 | 10 | 9.26% |
| hsa_circ_0083421 | 216 | 10 | 9.26% |
| hsa_circ_0095910 | 216 | 10 | 9.26% |
| hsa_circ_0138912 | 216 | 10 | 9.26% |
| hsa_circ_0005994 | 239 | 11 | 9.21% |
| hsa_circ_0114778 | 218 | 10 | 9.17% |
| hsa_circ_0063748 | 240 | 11 | 9.17% |
| hsa_circ_0119558 | 262 | 12 | 9.16% |
| hsa_circ_0098583 | 219 | 10 | 9.13% |
| hsa_circ_0061578 | 241 | 11 | 9.13% |
| hsa_circ_0000880 | 263 | 12 | 9.13% |
| hsa_circ_0000597 | 242 | 11 | 9.09% |
| hsa_circ_0080100 | 220 | 10 | 9.09% |
| hsa_circ_0098565 | 220 | 10 | 9.09% |
| hsa_circ_0105601 | 220 | 10 | 9.09% |
| hsa_circ_0128564 | 287 | 13 | 9.06% |
| hsa_circ_0092143 | 265 | 12 | 9.06% |
| hsa_circ_0037992 | 243 | 11 | 9.05% |
| hsa_circ_0077577 | 243 | 11 | 9.05% |
| hsa_circ_0088231 | 221 | 10 | 9.05% |
| hsa_circ_0096193 | 221 | 10 | 9.05% |
| hsa_circ_0116473 | 221 | 10 | 9.05% |
| hsa_circ_0130384 | 244 | 11 | 9.02% |
| hsa_circ_0068636 | 222 | 10 | 9.01% |
| hsa_circ_0093400 | 222 | 10 | 9.01% |
| hsa_circ_0095382 | 222 | 10 | 9.01% |
| hsa_circ_0112842 | 222 | 10 | 9.01% |
| hsa_circ_0118328 | 222 | 10 | 9.01% |
| hsa_circ_0127379 | 222 | 10 | 9.01% |
| hsa_circ_0008292 | 223 | 10 | 8.97% |
| hsa_circ_0065957 | 223 | 10 | 8.97% |
| hsa_circ_0082950 | 223 | 10 | 8.97% |
| hsa_circ_0098692 | 268 | 12 | 8.96% |
| hsa_circ_0076709 | 246 | 11 | 8.94% |
| hsa_circ_0051845 | 224 | 10 | 8.93% |
| hsa_circ_0085003 | 224 | 10 | 8.93% |
| hsa_circ_0050583 | 247 | 11 | 8.91% |
| hsa_circ_0087335 | 247 | 11 | 8.91% |
| hsa_circ_0011817 | 225 | 10 | 8.89% |
| hsa_circ_0040972 | 225 | 10 | 8.89% |
| hsa_circ_0050377 | 225 | 10 | 8.89% |
| hsa_circ_0071272 | 225 | 10 | 8.89% |
| hsa_circ_0119377 | 225 | 10 | 8.89% |
| hsa_circ_0074895 | 248 | 11 | 8.87% |
| hsa_circ_0135197 | 248 | 11 | 8.87% |

| | | | |
|---|---|---|---|
| hsa_circ_0027002 | 226 | 10 | 8.85% |
| hsa_circ_0079940 | 249 | 11 | 8.84% |
| hsa_circ_0121466 | 249 | 11 | 8.84% |
| hsa_circ_0002666 | 227 | 10 | 8.81% |
| hsa_circ_0008263 | 227 | 10 | 8.81% |
| hsa_circ_0011843 | 227 | 10 | 8.81% |
| hsa_circ_0012992 | 227 | 10 | 8.81% |
| hsa_circ_0038718 | 227 | 10 | 8.81% |
| hsa_circ_0069316 | 227 | 10 | 8.81% |
| hsa_circ_0091418 | 227 | 10 | 8.81% |
| hsa_circ_0095972 | 227 | 10 | 8.81% |
| hsa_circ_0101006 | 227 | 10 | 8.81% |
| hsa_circ_0113313 | 227 | 10 | 8.81% |
| hsa_circ_0076796 | 250 | 11 | 8.80% |
| hsa_circ_0020934 | 228 | 10 | 8.77% |
| hsa_circ_0022931 | 228 | 10 | 8.77% |
| hsa_circ_0052130 | 228 | 10 | 8.77% |
| hsa_circ_0056081 | 228 | 10 | 8.77% |
| hsa_circ_0070807 | 228 | 10 | 8.77% |
| hsa_circ_0124125 | 228 | 10 | 8.77% |
| hsa_circ_0126298 | 228 | 10 | 8.77% |
| hsa_circ_0139449 | 251 | 11 | 8.76% |
| hsa_circ_0000217 | 229 | 10 | 8.73% |
| hsa_circ_0023410 | 229 | 10 | 8.73% |
| hsa_circ_0050683 | 229 | 10 | 8.73% |
| hsa_circ_0072468 | 229 | 10 | 8.73% |
| hsa_circ_0129116 | 229 | 10 | 8.73% |
| hsa_circ_0139333 | 229 | 10 | 8.73% |
| hsa_circ_0057451 | 252 | 11 | 8.73% |
| hsa_circ_0023667 | 230 | 10 | 8.70% |
| hsa_circ_0118981 | 230 | 10 | 8.70% |
| hsa_circ_0129951 | 230 | 10 | 8.70% |
| hsa_circ_0140174 | 230 | 10 | 8.70% |
| hsa_circ_0140513 | 230 | 10 | 8.70% |
| hsa_circ_0001202 | 231 | 10 | 8.66% |
| hsa_circ_0081668 | 231 | 10 | 8.66% |
| hsa_circ_0004468 | 255 | 11 | 8.63% |
| hsa_circ_0041892 | 255 | 11 | 8.63% |
| hsa_circ_0081426 | 255 | 11 | 8.63% |
| hsa_circ_0135948 | 255 | 11 | 8.63% |
| hsa_circ_0101548 | 232 | 10 | 8.62% |
| hsa_circ_0129273 | 232 | 10 | 8.62% |
| hsa_circ_0009109 | 279 | 12 | 8.60% |
| hsa_circ_0009435 | 279 | 12 | 8.60% |
| hsa_circ_0048841 | 535 | 23 | 8.60% |
| hsa_circ_0024516 | 256 | 11 | 8.59% |
| hsa_circ_0039562 | 256 | 11 | 8.59% |
| hsa_circ_0011113 | 233 | 10 | 8.58% |
| hsa_circ_0137448 | 233 | 10 | 8.58% |
| hsa_circ_0055611 | 257 | 11 | 8.56% |

| | | | |
|---|---|---|---|
| hsa_circ_0089104 | 234 | 10 | 8.55% |
| hsa_circ_0109438 | 234 | 10 | 8.55% |
| hsa_circ_0129203 | 234 | 10 | 8.55% |
| hsa_circ_0137093 | 234 | 10 | 8.55% |
| hsa_circ_0032963 | 235 | 10 | 8.51% |
| hsa_circ_0079029 | 235 | 10 | 8.51% |
| hsa_circ_0072103 | 283 | 12 | 8.48% |
| hsa_circ_0112527 | 236 | 10 | 8.47% |
| hsa_circ_0112818 | 236 | 10 | 8.47% |
| hsa_circ_0018637 | 260 | 11 | 8.46% |
| hsa_circ_0046945 | 260 | 11 | 8.46% |
| hsa_circ_0049856 | 260 | 11 | 8.46% |
| hsa_circ_0100649 | 260 | 11 | 8.46% |
| hsa_circ_0139976 | 260 | 11 | 8.46% |
| hsa_circ_0119669 | 331 | 14 | 8.46% |
| hsa_circ_0096363 | 284 | 12 | 8.45% |
| hsa_circ_0000446 | 237 | 10 | 8.44% |
| hsa_circ_0089486 | 237 | 10 | 8.44% |
| hsa_circ_0102771 | 237 | 10 | 8.44% |
| hsa_circ_0103236 | 237 | 10 | 8.44% |
| hsa_circ_0092938 | 261 | 11 | 8.43% |
| hsa_circ_0011092 | 238 | 10 | 8.40% |
| hsa_circ_0078517 | 238 | 10 | 8.40% |
| hsa_circ_0114631 | 238 | 10 | 8.40% |
| hsa_circ_0086035 | 262 | 11 | 8.40% |
| hsa_circ_0124409 | 262 | 11 | 8.40% |
| hsa_circ_0138008 | 310 | 13 | 8.39% |
| hsa_circ_0005725 | 239 | 10 | 8.37% |
| hsa_circ_0061199 | 239 | 10 | 8.37% |
| hsa_circ_0116715 | 239 | 10 | 8.37% |
| hsa_circ_0125738 | 239 | 10 | 8.37% |
| hsa_circ_0139510 | 239 | 10 | 8.37% |
| hsa_circ_0096068 | 1028 | 43 | 8.37% |
| hsa_circ_0112442 | 336 | 14 | 8.33% |
| hsa_circ_0075077 | 264 | 11 | 8.33% |
| hsa_circ_0122229 | 264 | 11 | 8.33% |
| hsa_circ_0000057 | 240 | 10 | 8.33% |
| hsa_circ_0000383 | 240 | 10 | 8.33% |
| hsa_circ_0027475 | 240 | 10 | 8.33% |
| hsa_circ_0090973 | 240 | 10 | 8.33% |
| hsa_circ_0105732 | 240 | 10 | 8.33% |
| hsa_circ_0006608 | 241 | 10 | 8.30% |
| hsa_circ_0073208 | 241 | 10 | 8.30% |
| hsa_circ_0092018 | 241 | 10 | 8.30% |
| hsa_circ_0096011 | 241 | 10 | 8.30% |
| hsa_circ_0106018 | 241 | 10 | 8.30% |
| hsa_circ_0114677 | 241 | 10 | 8.30% |
| hsa_circ_0119064 | 458 | 19 | 8.30% |
| hsa_circ_0108382 | 13182 | 546 | 8.28% |
| hsa_circ_0012338 | 290 | 12 | 8.28% |

| | | | |
|---|---|---|---|
| hsa_circ_0040321 | 290 | 12 | 8.28% |
| hsa_circ_0008063 | 242 | 10 | 8.26% |
| hsa_circ_0098700 | 242 | 10 | 8.26% |
| hsa_circ_0101549 | 242 | 10 | 8.26% |
| hsa_circ_0034893 | 243 | 10 | 8.23% |
| hsa_circ_0039612 | 243 | 10 | 8.23% |
| hsa_circ_0040765 | 243 | 10 | 8.23% |
| hsa_circ_0095913 | 243 | 10 | 8.23% |
| hsa_circ_0004700 | 292 | 12 | 8.22% |
| hsa_circ_0082713 | 268 | 11 | 8.21% |
| hsa_circ_0001576 | 244 | 10 | 8.20% |
| hsa_circ_0023441 | 244 | 10 | 8.20% |
| hsa_circ_0034324 | 244 | 10 | 8.20% |
| hsa_circ_0067427 | 244 | 10 | 8.20% |
| hsa_circ_0098883 | 244 | 10 | 8.20% |
| hsa_circ_0105088 | 244 | 10 | 8.20% |
| hsa_circ_0124117 | 269 | 11 | 8.18% |
| hsa_circ_0005967 | 245 | 10 | 8.16% |
| hsa_circ_0007840 | 245 | 10 | 8.16% |
| hsa_circ_0058910 | 245 | 10 | 8.16% |
| hsa_circ_0074868 | 245 | 10 | 8.16% |
| hsa_circ_0113429 | 245 | 10 | 8.16% |
| hsa_circ_0121211 | 245 | 10 | 8.16% |
| hsa_circ_0072640 | 319 | 13 | 8.15% |
| hsa_circ_0022619 | 246 | 10 | 8.13% |
| hsa_circ_0051816 | 246 | 10 | 8.13% |
| hsa_circ_0124001 | 246 | 10 | 8.13% |
| hsa_circ_0128075 | 320 | 13 | 8.13% |
| hsa_circ_0008348 | 271 | 11 | 8.12% |
| hsa_circ_0008753 | 271 | 11 | 8.12% |
| hsa_circ_0073351 | 296 | 12 | 8.11% |
| hsa_circ_0018218 | 247 | 10 | 8.10% |
| hsa_circ_0019207 | 247 | 10 | 8.10% |
| hsa_circ_0035848 | 247 | 10 | 8.10% |
| hsa_circ_0073109 | 247 | 10 | 8.10% |
| hsa_circ_0074264 | 247 | 10 | 8.10% |
| hsa_circ_0032028 | 272 | 11 | 8.09% |
| hsa_circ_0064143 | 248 | 10 | 8.06% |
| hsa_circ_0064821 | 248 | 10 | 8.06% |
| hsa_circ_0095179 | 248 | 10 | 8.06% |
| hsa_circ_0109169 | 248 | 10 | 8.06% |
| hsa_circ_0116771 | 273 | 11 | 8.06% |
| hsa_circ_0128335 | 273 | 11 | 8.06% |
| hsa_circ_0004586 | 249 | 10 | 8.03% |
| hsa_circ_0033429 | 249 | 10 | 8.03% |
| hsa_circ_0076866 | 249 | 10 | 8.03% |
| hsa_circ_0090466 | 249 | 10 | 8.03% |
| hsa_circ_0093693 | 249 | 10 | 8.03% |
| hsa_circ_0006142 | 299 | 12 | 8.03% |
| hsa_circ_0075030 | 250 | 10 | 8.00% |

| | | | |
|---|---|---|---|
| hsa_circ_0097899 | 250 | 10 | 8.00% |
| hsa_circ_0118815 | 250 | 10 | 8.00% |
| hsa_circ_0130771 | 250 | 10 | 8.00% |
| hsa_circ_0099435 | 326 | 13 | 7.98% |
| hsa_circ_0003224 | 276 | 11 | 7.97% |
| hsa_circ_0044721 | 276 | 11 | 7.97% |
| hsa_circ_0001340 | 251 | 10 | 7.97% |
| hsa_circ_0110566 | 251 | 10 | 7.97% |
| hsa_circ_0131630 | 251 | 10 | 7.97% |
| hsa_circ_0136819 | 251 | 10 | 7.97% |
| hsa_circ_0087331 | 277 | 11 | 7.94% |
| hsa_circ_0055120 | 252 | 10 | 7.94% |
| hsa_circ_0058408 | 252 | 10 | 7.94% |
| hsa_circ_0077627 | 252 | 10 | 7.94% |
| hsa_circ_0109565 | 252 | 10 | 7.94% |
| hsa_circ_0119335 | 252 | 10 | 7.94% |
| hsa_circ_0096046 | 278 | 11 | 7.91% |
| hsa_circ_0035482 | 253 | 10 | 7.91% |
| hsa_circ_0036258 | 253 | 10 | 7.91% |
| hsa_circ_0116918 | 279 | 11 | 7.89% |
| hsa_circ_0131701 | 330 | 13 | 7.88% |
| hsa_circ_0077457 | 254 | 10 | 7.87% |
| hsa_circ_0115869 | 254 | 10 | 7.87% |
| hsa_circ_0121864 | 254 | 10 | 7.87% |
| hsa_circ_0135867 | 254 | 10 | 7.87% |
| hsa_circ_0119065 | 305 | 12 | 7.87% |
| hsa_circ_0030734 | 255 | 10 | 7.84% |
| hsa_circ_0037708 | 255 | 10 | 7.84% |
| hsa_circ_0058840 | 255 | 10 | 7.84% |
| hsa_circ_0079362 | 255 | 10 | 7.84% |
| hsa_circ_0105128 | 255 | 10 | 7.84% |
| hsa_circ_0119891 | 255 | 10 | 7.84% |
| hsa_circ_0089958 | 281 | 11 | 7.83% |
| hsa_circ_0000688 | 256 | 10 | 7.81% |
| hsa_circ_0006277 | 256 | 10 | 7.81% |
| hsa_circ_0021184 | 256 | 10 | 7.81% |
| hsa_circ_0070888 | 256 | 10 | 7.81% |
| hsa_circ_0100948 | 256 | 10 | 7.81% |
| hsa_circ_0103929 | 256 | 10 | 7.81% |
| hsa_circ_0138242 | 256 | 10 | 7.81% |
| hsa_circ_0052856 | 282 | 11 | 7.80% |
| hsa_circ_0107809 | 282 | 11 | 7.80% |
| hsa_circ_0128427 | 282 | 11 | 7.80% |
| hsa_circ_0078145 | 257 | 10 | 7.78% |
| hsa_circ_0089335 | 257 | 10 | 7.78% |
| hsa_circ_0115113 | 257 | 10 | 7.78% |
| hsa_circ_0133245 | 773 | 30 | 7.76% |
| hsa_circ_0107566 | 258 | 10 | 7.75% |
| hsa_circ_0124318 | 258 | 10 | 7.75% |
| hsa_circ_0137028 | 258 | 10 | 7.75% |

| | | | |
|---|---|---|---|
| hsa_circ_0127723 | 284 | 11 | 7.75% |
| hsa_circ_0027377 | 259 | 10 | 7.72% |
| hsa_circ_0032482 | 259 | 10 | 7.72% |
| hsa_circ_0134168 | 259 | 10 | 7.72% |
| hsa_circ_0139630 | 259 | 10 | 7.72% |
| hsa_circ_0121170 | 285 | 11 | 7.72% |
| hsa_circ_0032661 | 442 | 17 | 7.69% |
| hsa_circ_0028979 | 260 | 10 | 7.69% |
| hsa_circ_0049659 | 260 | 10 | 7.69% |
| hsa_circ_0072052 | 260 | 10 | 7.69% |
| hsa_circ_0072155 | 260 | 10 | 7.69% |
| hsa_circ_0078134 | 260 | 10 | 7.69% |
| hsa_circ_0112403 | 260 | 10 | 7.69% |
| hsa_circ_0123071 | 260 | 10 | 7.69% |
| hsa_circ_0139878 | 260 | 10 | 7.69% |
| hsa_circ_0105056 | 287 | 11 | 7.67% |
| hsa_circ_0069900 | 288 | 11 | 7.64% |
| hsa_circ_0097349 | 288 | 11 | 7.64% |
| hsa_circ_0130302 | 288 | 11 | 7.64% |
| hsa_circ_0045283 | 262 | 10 | 7.63% |
| hsa_circ_0099541 | 262 | 10 | 7.63% |
| hsa_circ_0019146 | 315 | 12 | 7.62% |
| hsa_circ_0004400 | 289 | 11 | 7.61% |
| hsa_circ_0088804 | 289 | 11 | 7.61% |
| hsa_circ_0117015 | 263 | 10 | 7.60% |
| hsa_circ_0116420 | 290 | 11 | 7.59% |
| hsa_circ_0015374 | 264 | 10 | 7.58% |
| hsa_circ_0025728 | 264 | 10 | 7.58% |
| hsa_circ_0041704 | 264 | 10 | 7.58% |
| hsa_circ_0088756 | 264 | 10 | 7.58% |
| hsa_circ_0097059 | 264 | 10 | 7.58% |
| hsa_circ_0123760 | 264 | 10 | 7.58% |
| hsa_circ_0087713 | 317 | 12 | 7.57% |
| hsa_circ_0011748 | 291 | 11 | 7.56% |
| hsa_circ_0008659 | 318 | 12 | 7.55% |
| hsa_circ_0037041 | 265 | 10 | 7.55% |
| hsa_circ_0065915 | 265 | 10 | 7.55% |
| hsa_circ_0070929 | 265 | 10 | 7.55% |
| hsa_circ_0077532 | 265 | 10 | 7.55% |
| hsa_circ_0117534 | 265 | 10 | 7.55% |
| hsa_circ_0058172 | 292 | 11 | 7.53% |
| hsa_circ_0089565 | 292 | 11 | 7.53% |
| hsa_circ_0049740 | 372 | 14 | 7.53% |
| hsa_circ_0001065 | 266 | 10 | 7.52% |
| hsa_circ_0057205 | 266 | 10 | 7.52% |
| hsa_circ_0060537 | 266 | 10 | 7.52% |
| hsa_circ_0005594 | 320 | 12 | 7.50% |
| hsa_circ_0047091 | 347 | 13 | 7.49% |
| hsa_circ_0033432 | 267 | 10 | 7.49% |
| hsa_circ_0035128 | 267 | 10 | 7.49% |

| | | | |
|---|---|---|---|
| hsa_circ_0064014 | 267 | 10 | 7.49% |
| hsa_circ_0135169 | 267 | 10 | 7.49% |
| hsa_circ_0137511 | 267 | 10 | 7.49% |
| hsa_circ_0006303 | 294 | 11 | 7.48% |
| hsa_circ_0051348 | 294 | 11 | 7.48% |
| hsa_circ_0101039 | 348 | 13 | 7.47% |
| hsa_circ_0122361 | 23704 | 885 | 7.47% |
| hsa_circ_0045887 | 268 | 10 | 7.46% |
| hsa_circ_0046661 | 268 | 10 | 7.46% |
| hsa_circ_0091156 | 295 | 11 | 7.46% |
| hsa_circ_0000491 | 269 | 10 | 7.43% |
| hsa_circ_0000732 | 269 | 10 | 7.43% |
| hsa_circ_0001003 | 269 | 10 | 7.43% |
| hsa_circ_0008698 | 269 | 10 | 7.43% |
| hsa_circ_0049355 | 269 | 10 | 7.43% |
| hsa_circ_0090101 | 269 | 10 | 7.43% |
| hsa_circ_0109194 | 269 | 10 | 7.43% |
| hsa_circ_0127074 | 269 | 10 | 7.43% |
| hsa_circ_0133729 | 269 | 10 | 7.43% |
| hsa_circ_0105743 | 296 | 11 | 7.43% |
| hsa_circ_0105356 | 324 | 12 | 7.41% |
| hsa_circ_0076792 | 297 | 11 | 7.41% |
| hsa_circ_0119987 | 297 | 11 | 7.41% |
| hsa_circ_0119992 | 297 | 11 | 7.41% |
| hsa_circ_0056715 | 270 | 10 | 7.41% |
| hsa_circ_0073203 | 270 | 10 | 7.41% |
| hsa_circ_0081336 | 270 | 10 | 7.41% |
| hsa_circ_0090324 | 270 | 10 | 7.41% |
| hsa_circ_0016942 | 298 | 11 | 7.38% |
| hsa_circ_0038849 | 271 | 10 | 7.38% |
| hsa_circ_0064890 | 271 | 10 | 7.38% |
| hsa_circ_0136155 | 271 | 10 | 7.38% |
| hsa_circ_0087584 | 299 | 11 | 7.36% |
| hsa_circ_0104245 | 299 | 11 | 7.36% |
| hsa_circ_0004614 | 272 | 10 | 7.35% |
| hsa_circ_0024773 | 272 | 10 | 7.35% |
| hsa_circ_0037812 | 272 | 10 | 7.35% |
| hsa_circ_0065127 | 272 | 10 | 7.35% |
| hsa_circ_0072838 | 272 | 10 | 7.35% |
| hsa_circ_0114146 | 272 | 10 | 7.35% |
| hsa_circ_0075728 | 327 | 12 | 7.34% |
| hsa_circ_0041760 | 300 | 11 | 7.33% |
| hsa_circ_0096979 | 300 | 11 | 7.33% |
| hsa_circ_0002761 | 273 | 10 | 7.33% |
| hsa_circ_0007350 | 273 | 10 | 7.33% |
| hsa_circ_0053404 | 273 | 10 | 7.33% |
| hsa_circ_0072023 | 273 | 10 | 7.33% |
| hsa_circ_0072799 | 273 | 10 | 7.33% |
| hsa_circ_0072880 | 273 | 10 | 7.33% |
| hsa_circ_0076537 | 273 | 10 | 7.33% |

| | | | |
|---|---|---|---|
| hsa_circ_0076916 | 273 | 10 | 7.33% |
| hsa_circ_0083051 | 273 | 10 | 7.33% |
| hsa_circ_0130481 | 273 | 10 | 7.33% |
| hsa_circ_0076100 | 328 | 12 | 7.32% |
| hsa_circ_0018281 | 301 | 11 | 7.31% |
| hsa_circ_0045094 | 274 | 10 | 7.30% |
| hsa_circ_0074502 | 274 | 10 | 7.30% |
| hsa_circ_0076037 | 274 | 10 | 7.30% |
| hsa_circ_0086153 | 274 | 10 | 7.30% |
| hsa_circ_0097448 | 274 | 10 | 7.30% |
| hsa_circ_0106859 | 274 | 10 | 7.30% |
| hsa_circ_0124833 | 274 | 10 | 7.30% |
| hsa_circ_0109007 | 329 | 12 | 7.29% |
| hsa_circ_0098553 | 330 | 12 | 7.27% |
| hsa_circ_0067163 | 275 | 10 | 7.27% |
| hsa_circ_0111061 | 275 | 10 | 7.27% |
| hsa_circ_0114145 | 275 | 10 | 7.27% |
| hsa_circ_0136734 | 275 | 10 | 7.27% |
| hsa_circ_0005588 | 276 | 10 | 7.25% |
| hsa_circ_0026592 | 276 | 10 | 7.25% |
| hsa_circ_0028491 | 276 | 10 | 7.25% |
| hsa_circ_0040971 | 276 | 10 | 7.25% |
| hsa_circ_0051123 | 276 | 10 | 7.25% |
| hsa_circ_0069926 | 276 | 10 | 7.25% |
| hsa_circ_0116561 | 276 | 10 | 7.25% |
| hsa_circ_0138291 | 276 | 10 | 7.25% |
| hsa_circ_0112087 | 304 | 11 | 7.24% |
| hsa_circ_0136124 | 304 | 11 | 7.24% |
| hsa_circ_0075804 | 720 | 26 | 7.22% |
| hsa_circ_0030889 | 277 | 10 | 7.22% |
| hsa_circ_0031150 | 277 | 10 | 7.22% |
| hsa_circ_0035647 | 277 | 10 | 7.22% |
| hsa_circ_0038508 | 277 | 10 | 7.22% |
| hsa_circ_0055721 | 277 | 10 | 7.22% |
| hsa_circ_0080866 | 277 | 10 | 7.22% |
| hsa_circ_0106860 | 277 | 10 | 7.22% |
| hsa_circ_0115047 | 277 | 10 | 7.22% |
| hsa_circ_0115809 | 277 | 10 | 7.22% |
| hsa_circ_0119125 | 277 | 10 | 7.22% |
| hsa_circ_0122113 | 277 | 10 | 7.22% |
| hsa_circ_0133590 | 277 | 10 | 7.22% |
| hsa_circ_0084746 | 305 | 11 | 7.21% |
| hsa_circ_0047657 | 278 | 10 | 7.19% |
| hsa_circ_0073054 | 278 | 10 | 7.19% |
| hsa_circ_0057452 | 306 | 11 | 7.19% |
| hsa_circ_0130885 | 334 | 12 | 7.19% |
| hsa_circ_0002637 | 279 | 10 | 7.17% |
| hsa_circ_0045904 | 279 | 10 | 7.17% |
| hsa_circ_0048762 | 279 | 10 | 7.17% |
| hsa_circ_0079759 | 279 | 10 | 7.17% |

| | | | |
|---|---|---|---|
| hsa_circ_0090087 | 279 | 10 | 7.17% |
| hsa_circ_0104285 | 279 | 10 | 7.17% |
| hsa_circ_0111036 | 279 | 10 | 7.17% |
| hsa_circ_0113032 | 279 | 10 | 7.17% |
| hsa_circ_0117582 | 279 | 10 | 7.17% |
| hsa_circ_0119601 | 279 | 10 | 7.17% |
| hsa_circ_0131804 | 279 | 10 | 7.17% |
| hsa_circ_0003147 | 307 | 11 | 7.17% |
| hsa_circ_0014850 | 335 | 12 | 7.16% |
| hsa_circ_0124871 | 391 | 14 | 7.16% |
| hsa_circ_0078053 | 280 | 10 | 7.14% |
| hsa_circ_0082948 | 280 | 10 | 7.14% |
| hsa_circ_0092300 | 280 | 10 | 7.14% |
| hsa_circ_0126025 | 280 | 10 | 7.14% |
| hsa_circ_0051506 | 393 | 14 | 7.12% |
| hsa_circ_0013223 | 337 | 12 | 7.12% |
| hsa_circ_0006917 | 281 | 10 | 7.12% |
| hsa_circ_0012612 | 281 | 10 | 7.12% |
| hsa_circ_0017827 | 281 | 10 | 7.12% |
| hsa_circ_0053118 | 281 | 10 | 7.12% |
| hsa_circ_0106812 | 281 | 10 | 7.12% |
| hsa_circ_0115756 | 281 | 10 | 7.12% |
| hsa_circ_0030009 | 282 | 10 | 7.09% |
| hsa_circ_0094763 | 282 | 10 | 7.09% |
| hsa_circ_0101577 | 282 | 10 | 7.09% |
| hsa_circ_0111669 | 282 | 10 | 7.09% |
| hsa_circ_0121616 | 282 | 10 | 7.09% |
| hsa_circ_0135993 | 282 | 10 | 7.09% |
| hsa_circ_0097672 | 311 | 11 | 7.07% |
| hsa_circ_0003320 | 283 | 10 | 7.07% |
| hsa_circ_0005135 | 283 | 10 | 7.07% |
| hsa_circ_0008137 | 283 | 10 | 7.07% |
| hsa_circ_0009248 | 283 | 10 | 7.07% |
| hsa_circ_0024756 | 283 | 10 | 7.07% |
| hsa_circ_0048519 | 283 | 10 | 7.07% |
| hsa_circ_0134578 | 283 | 10 | 7.07% |
| hsa_circ_0042790 | 312 | 11 | 7.05% |
| hsa_circ_0070673 | 312 | 11 | 7.05% |
| hsa_circ_0092937 | 312 | 11 | 7.05% |
| hsa_circ_0027827 | 284 | 10 | 7.04% |
| hsa_circ_0081492 | 284 | 10 | 7.04% |
| hsa_circ_0092481 | 284 | 10 | 7.04% |
| hsa_circ_0106373 | 284 | 10 | 7.04% |
| hsa_circ_0000154 | 341 | 12 | 7.04% |
| hsa_circ_0137920 | 342 | 12 | 7.02% |
| hsa_circ_0021342 | 285 | 10 | 7.02% |
| hsa_circ_0070703 | 285 | 10 | 7.02% |
| hsa_circ_0079030 | 285 | 10 | 7.02% |
| hsa_circ_0031971 | 343 | 12 | 7.00% |
| hsa_circ_0046193 | 286 | 10 | 6.99% |

| | | | |
|---|---|---|---|
| hsa_circ_0061746 | 286 | 10 | 6.99% |
| hsa_circ_0101402 | 286 | 10 | 6.99% |
| hsa_circ_0104724 | 286 | 10 | 6.99% |
| hsa_circ_0131858 | 286 | 10 | 6.99% |
| hsa_circ_0104246 | 315 | 11 | 6.98% |
| hsa_circ_0109785 | 344 | 12 | 6.98% |
| hsa_circ_0004146 | 287 | 10 | 6.97% |
| hsa_circ_0005844 | 287 | 10 | 6.97% |
| hsa_circ_0044470 | 287 | 10 | 6.97% |
| hsa_circ_0065958 | 287 | 10 | 6.97% |
| hsa_circ_0066594 | 287 | 10 | 6.97% |
| hsa_circ_0082698 | 287 | 10 | 6.97% |
| hsa_circ_0100400 | 287 | 10 | 6.97% |
| hsa_circ_0111715 | 287 | 10 | 6.97% |
| hsa_circ_0135940 | 316 | 11 | 6.96% |
| hsa_circ_0025322 | 288 | 10 | 6.94% |
| hsa_circ_0025959 | 288 | 10 | 6.94% |
| hsa_circ_0060569 | 288 | 10 | 6.94% |
| hsa_circ_0083069 | 288 | 10 | 6.94% |
| hsa_circ_0050640 | 519 | 18 | 6.94% |
| hsa_circ_0013221 | 318 | 11 | 6.92% |
| hsa_circ_0018289 | 348 | 12 | 6.90% |
| hsa_circ_0097616 | 348 | 12 | 6.90% |
| hsa_circ_0037367 | 319 | 11 | 6.90% |
| hsa_circ_0039565 | 290 | 10 | 6.90% |
| hsa_circ_0102495 | 290 | 10 | 6.90% |
| hsa_circ_0113343 | 290 | 10 | 6.90% |
| hsa_circ_0119174 | 290 | 10 | 6.90% |
| hsa_circ_0132161 | 290 | 10 | 6.90% |
| hsa_circ_0122144 | 349 | 12 | 6.88% |
| hsa_circ_0112229 | 320 | 11 | 6.88% |
| hsa_circ_0008592 | 291 | 10 | 6.87% |
| hsa_circ_0047398 | 291 | 10 | 6.87% |
| hsa_circ_0056713 | 291 | 10 | 6.87% |
| hsa_circ_0086723 | 291 | 10 | 6.87% |
| hsa_circ_0105132 | 291 | 10 | 6.87% |
| hsa_circ_0131110 | 291 | 10 | 6.87% |
| hsa_circ_0133367 | 291 | 10 | 6.87% |
| hsa_circ_0060748 | 350 | 12 | 6.86% |
| hsa_circ_0105838 | 321 | 11 | 6.85% |
| hsa_circ_0039515 | 292 | 10 | 6.85% |
| hsa_circ_0040687 | 292 | 10 | 6.85% |
| hsa_circ_0044111 | 292 | 10 | 6.85% |
| hsa_circ_0119519 | 292 | 10 | 6.85% |
| hsa_circ_0138025 | 292 | 10 | 6.85% |
| hsa_circ_0055548 | 322 | 11 | 6.83% |
| hsa_circ_0078940 | 322 | 11 | 6.83% |
| hsa_circ_0081427 | 322 | 11 | 6.83% |
| hsa_circ_0082015 | 322 | 11 | 6.83% |
| hsa_circ_0084667 | 322 | 11 | 6.83% |

| | | | |
|---|---|---|---|
| hsa_circ_0097495 | 322 | 11 | 6.83% |
| hsa_circ_0119693 | 322 | 11 | 6.83% |
| hsa_circ_0120704 | 322 | 11 | 6.83% |
| hsa_circ_0000334 | 293 | 10 | 6.83% |
| hsa_circ_0027117 | 293 | 10 | 6.83% |
| hsa_circ_0039158 | 293 | 10 | 6.83% |
| hsa_circ_0079563 | 293 | 10 | 6.83% |
| hsa_circ_0007042 | 323 | 11 | 6.81% |
| hsa_circ_0005158 | 294 | 10 | 6.80% |
| hsa_circ_0005285 | 294 | 10 | 6.80% |
| hsa_circ_0023606 | 294 | 10 | 6.80% |
| hsa_circ_0050618 | 294 | 10 | 6.80% |
| hsa_circ_0066679 | 294 | 10 | 6.80% |
| hsa_circ_0096102 | 294 | 10 | 6.80% |
| hsa_circ_0106019 | 294 | 10 | 6.80% |
| hsa_circ_0108522 | 294 | 10 | 6.80% |
| hsa_circ_0111973 | 294 | 10 | 6.80% |
| hsa_circ_0132460 | 294 | 10 | 6.80% |
| hsa_circ_0095444 | 324 | 11 | 6.79% |
| hsa_circ_0098880 | 354 | 12 | 6.78% |
| hsa_circ_0115372 | 354 | 12 | 6.78% |
| hsa_circ_0006087 | 295 | 10 | 6.78% |
| hsa_circ_0034842 | 295 | 10 | 6.78% |
| hsa_circ_0090998 | 295 | 10 | 6.78% |
| hsa_circ_0115284 | 295 | 10 | 6.78% |
| hsa_circ_0126589 | 295 | 10 | 6.78% |
| hsa_circ_0140408 | 295 | 10 | 6.78% |
| hsa_circ_0014799 | 296 | 10 | 6.76% |
| hsa_circ_0030822 | 296 | 10 | 6.76% |
| hsa_circ_0036732 | 296 | 10 | 6.76% |
| hsa_circ_0043817 | 296 | 10 | 6.76% |
| hsa_circ_0066401 | 296 | 10 | 6.76% |
| hsa_circ_0101298 | 296 | 10 | 6.76% |
| hsa_circ_0130715 | 296 | 10 | 6.76% |
| hsa_circ_0131756 | 296 | 10 | 6.76% |
| hsa_circ_0135954 | 296 | 10 | 6.76% |
| hsa_circ_0138246 | 296 | 10 | 6.76% |
| hsa_circ_0000392 | 326 | 11 | 6.75% |
| hsa_circ_0048973 | 326 | 11 | 6.75% |
| hsa_circ_0041391 | 297 | 10 | 6.73% |
| hsa_circ_0049242 | 297 | 10 | 6.73% |
| hsa_circ_0066119 | 297 | 10 | 6.73% |
| hsa_circ_0110693 | 297 | 10 | 6.73% |
| hsa_circ_0130441 | 297 | 10 | 6.73% |
| hsa_circ_0066459 | 327 | 11 | 6.73% |
| hsa_circ_0095445 | 327 | 11 | 6.73% |
| hsa_circ_0107040 | 327 | 11 | 6.73% |
| hsa_circ_0018989 | 298 | 10 | 6.71% |
| hsa_circ_0022965 | 298 | 10 | 6.71% |
| hsa_circ_0043107 | 298 | 10 | 6.71% |

| | | | |
|---|---|---|---|
| hsa_circ_0048268 | 298 | 10 | 6.71% |
| hsa_circ_0089939 | 298 | 10 | 6.71% |
| hsa_circ_0104112 | 298 | 10 | 6.71% |
| hsa_circ_0107628 | 298 | 10 | 6.71% |
| hsa_circ_0110538 | 298 | 10 | 6.71% |
| hsa_circ_0114641 | 298 | 10 | 6.71% |
| hsa_circ_0022889 | 299 | 10 | 6.69% |
| hsa_circ_0039465 | 299 | 10 | 6.69% |
| hsa_circ_0052142 | 299 | 10 | 6.69% |
| hsa_circ_0089911 | 299 | 10 | 6.69% |
| hsa_circ_0116474 | 299 | 10 | 6.69% |
| hsa_circ_0130863 | 299 | 10 | 6.69% |
| hsa_circ_0097531 | 329 | 11 | 6.69% |
| hsa_circ_0011509 | 330 | 11 | 6.67% |
| hsa_circ_0065422 | 330 | 11 | 6.67% |
| hsa_circ_0085711 | 330 | 11 | 6.67% |
| hsa_circ_0089686 | 330 | 11 | 6.67% |
| hsa_circ_0120240 | 330 | 11 | 6.67% |
| hsa_circ_0004532 | 300 | 10 | 6.67% |
| hsa_circ_0007787 | 300 | 10 | 6.67% |
| hsa_circ_0034891 | 300 | 10 | 6.67% |
| hsa_circ_0064613 | 300 | 10 | 6.67% |
| hsa_circ_0070145 | 300 | 10 | 6.67% |
| hsa_circ_0081863 | 300 | 10 | 6.67% |
| hsa_circ_0083123 | 300 | 10 | 6.67% |
| hsa_circ_0089643 | 300 | 10 | 6.67% |
| hsa_circ_0092283 | 300 | 10 | 6.67% |
| hsa_circ_0092289 | 300 | 10 | 6.67% |
| hsa_circ_0030847 | 361 | 12 | 6.65% |
| hsa_circ_0132293 | 361 | 12 | 6.65% |
| hsa_circ_0000250 | 331 | 11 | 6.65% |
| hsa_circ_0007171 | 331 | 11 | 6.65% |
| hsa_circ_0051525 | 331 | 11 | 6.65% |
| hsa_circ_0000849 | 301 | 10 | 6.64% |
| hsa_circ_0007039 | 301 | 10 | 6.64% |
| hsa_circ_0063983 | 301 | 10 | 6.64% |
| hsa_circ_0090959 | 301 | 10 | 6.64% |
| hsa_circ_0115555 | 301 | 10 | 6.64% |
| hsa_circ_0117802 | 301 | 10 | 6.64% |
| hsa_circ_0122351 | 301 | 10 | 6.64% |
| hsa_circ_0139715 | 392 | 13 | 6.63% |
| hsa_circ_0021441 | 362 | 12 | 6.63% |
| hsa_circ_0114961 | 362 | 12 | 6.63% |
| hsa_circ_0026026 | 332 | 11 | 6.63% |
| hsa_circ_0096672 | 332 | 11 | 6.63% |
| hsa_circ_0024067 | 302 | 10 | 6.62% |
| hsa_circ_0082040 | 302 | 10 | 6.62% |
| hsa_circ_0097126 | 302 | 10 | 6.62% |
| hsa_circ_0110424 | 302 | 10 | 6.62% |
| hsa_circ_0125933 | 302 | 10 | 6.62% |

| | | | |
|---|---|---|---|
| hsa_circ_0137730 | 333 | 11 | 6.61% |
| hsa_circ_0002904 | 424 | 14 | 6.60% |
| hsa_circ_0044709 | 424 | 14 | 6.60% |
| hsa_circ_0010181 | 303 | 10 | 6.60% |
| hsa_circ_0031151 | 303 | 10 | 6.60% |
| hsa_circ_0036505 | 303 | 10 | 6.60% |
| hsa_circ_0038073 | 303 | 10 | 6.60% |
| hsa_circ_0059044 | 303 | 10 | 6.60% |
| hsa_circ_0061363 | 303 | 10 | 6.60% |
| hsa_circ_0094958 | 303 | 10 | 6.60% |
| hsa_circ_0107481 | 303 | 10 | 6.60% |
| hsa_circ_0138006 | 394 | 13 | 6.60% |
| hsa_circ_0048750 | 364 | 12 | 6.59% |
| hsa_circ_0013745 | 334 | 11 | 6.59% |
| hsa_circ_0135808 | 334 | 11 | 6.59% |
| hsa_circ_0081490 | 395 | 13 | 6.58% |
| hsa_circ_0008933 | 304 | 10 | 6.58% |
| hsa_circ_0023723 | 304 | 10 | 6.58% |
| hsa_circ_0054401 | 304 | 10 | 6.58% |
| hsa_circ_0066078 | 304 | 10 | 6.58% |
| hsa_circ_0066131 | 304 | 10 | 6.58% |
| hsa_circ_0067326 | 304 | 10 | 6.58% |
| hsa_circ_0110325 | 304 | 10 | 6.58% |
| hsa_circ_0140543 | 304 | 10 | 6.58% |
| hsa_circ_0095167 | 365 | 12 | 6.58% |
| hsa_circ_0117886 | 335 | 11 | 6.57% |
| hsa_circ_0006344 | 305 | 10 | 6.56% |
| hsa_circ_0013509 | 305 | 10 | 6.56% |
| hsa_circ_0072991 | 305 | 10 | 6.56% |
| hsa_circ_0103795 | 305 | 10 | 6.56% |
| hsa_circ_0109401 | 305 | 10 | 6.56% |
| hsa_circ_0111068 | 305 | 10 | 6.56% |
| hsa_circ_0115033 | 305 | 10 | 6.56% |
| hsa_circ_0134748 | 305 | 10 | 6.56% |
| hsa_circ_0136755 | 305 | 10 | 6.56% |
| hsa_circ_0037481 | 336 | 11 | 6.55% |
| hsa_circ_0077259 | 336 | 11 | 6.55% |
| hsa_circ_0115855 | 336 | 11 | 6.55% |
| hsa_circ_0015210 | 367 | 12 | 6.54% |
| hsa_circ_0065315 | 367 | 12 | 6.54% |
| hsa_circ_0008325 | 306 | 10 | 6.54% |
| hsa_circ_0011447 | 306 | 10 | 6.54% |
| hsa_circ_0104568 | 306 | 10 | 6.54% |
| hsa_circ_0113093 | 306 | 10 | 6.54% |
| hsa_circ_0114884 | 306 | 10 | 6.54% |
| hsa_circ_0135173 | 306 | 10 | 6.54% |
| hsa_circ_0049924 | 337 | 11 | 6.53% |
| hsa_circ_0021998 | 429 | 14 | 6.53% |
| hsa_circ_0063390 | 368 | 12 | 6.52% |
| hsa_circ_0023715 | 307 | 10 | 6.51% |

| | | | |
|---|---|---|---|
| hsa_circ_0109765 | 307 | 10 | 6.51% |
| hsa_circ_0127601 | 307 | 10 | 6.51% |
| hsa_circ_0060843 | 369 | 12 | 6.50% |
| hsa_circ_0116542 | 400 | 13 | 6.50% |
| hsa_circ_0053028 | 462 | 15 | 6.49% |
| hsa_circ_0005672 | 308 | 10 | 6.49% |
| hsa_circ_0039368 | 308 | 10 | 6.49% |
| hsa_circ_0060728 | 308 | 10 | 6.49% |
| hsa_circ_0065574 | 308 | 10 | 6.49% |
| hsa_circ_0102223 | 308 | 10 | 6.49% |
| hsa_circ_0109350 | 308 | 10 | 6.49% |
| hsa_circ_0112896 | 308 | 10 | 6.49% |
| hsa_circ_0128532 | 308 | 10 | 6.49% |
| hsa_circ_0129261 | 308 | 10 | 6.49% |
| hsa_circ_0133542 | 308 | 10 | 6.49% |
| hsa_circ_0134511 | 339 | 11 | 6.49% |
| hsa_circ_0090309 | 370 | 12 | 6.49% |
| hsa_circ_0014988 | 309 | 10 | 6.47% |
| hsa_circ_0027486 | 309 | 10 | 6.47% |
| hsa_circ_0102980 | 309 | 10 | 6.47% |
| hsa_circ_0126031 | 309 | 10 | 6.47% |
| hsa_circ_0075074 | 340 | 11 | 6.47% |
| hsa_circ_0106042 | 340 | 11 | 6.47% |
| hsa_circ_0019404 | 372 | 12 | 6.45% |
| hsa_circ_0030286 | 372 | 12 | 6.45% |
| hsa_circ_0031301 | 341 | 11 | 6.45% |
| hsa_circ_0051427 | 341 | 11 | 6.45% |
| hsa_circ_0134063 | 341 | 11 | 6.45% |
| hsa_circ_0004414 | 310 | 10 | 6.45% |
| hsa_circ_0004967 | 310 | 10 | 6.45% |
| hsa_circ_0005261 | 310 | 10 | 6.45% |
| hsa_circ_0090506 | 310 | 10 | 6.45% |
| hsa_circ_0097062 | 310 | 10 | 6.45% |
| hsa_circ_0111753 | 310 | 10 | 6.45% |
| hsa_circ_0113723 | 310 | 10 | 6.45% |
| hsa_circ_0130322 | 310 | 10 | 6.45% |
| hsa_circ_0106433 | 342 | 11 | 6.43% |
| hsa_circ_0006849 | 311 | 10 | 6.43% |
| hsa_circ_0028488 | 311 | 10 | 6.43% |
| hsa_circ_0057987 | 311 | 10 | 6.43% |
| hsa_circ_0082343 | 311 | 10 | 6.43% |
| hsa_circ_0090328 | 311 | 10 | 6.43% |
| hsa_circ_0103360 | 311 | 10 | 6.43% |
| hsa_circ_0105637 | 311 | 10 | 6.43% |
| hsa_circ_0018264 | 312 | 10 | 6.41% |
| hsa_circ_0029944 | 312 | 10 | 6.41% |
| hsa_circ_0046496 | 312 | 10 | 6.41% |
| hsa_circ_0106822 | 312 | 10 | 6.41% |
| hsa_circ_0108093 | 312 | 10 | 6.41% |
| hsa_circ_0113009 | 312 | 10 | 6.41% |

| | | | |
|---|---|---|---|
| hsa_circ_0049438 | 344 | 11 | 6.40% |
| hsa_circ_0091435 | 344 | 11 | 6.40% |
| hsa_circ_0118664 | 344 | 11 | 6.40% |
| hsa_circ_0029076 | 313 | 10 | 6.39% |
| hsa_circ_0039053 | 313 | 10 | 6.39% |
| hsa_circ_0046229 | 313 | 10 | 6.39% |
| hsa_circ_0059906 | 313 | 10 | 6.39% |
| hsa_circ_0071125 | 313 | 10 | 6.39% |
| hsa_circ_0090445 | 313 | 10 | 6.39% |
| hsa_circ_0127830 | 313 | 10 | 6.39% |
| hsa_circ_0055197 | 470 | 15 | 6.38% |
| hsa_circ_0071923 | 345 | 11 | 6.38% |
| hsa_circ_0001555 | 314 | 10 | 6.37% |
| hsa_circ_0004682 | 314 | 10 | 6.37% |
| hsa_circ_0022451 | 314 | 10 | 6.37% |
| hsa_circ_0037474 | 314 | 10 | 6.37% |
| hsa_circ_0080101 | 314 | 10 | 6.37% |
| hsa_circ_0129570 | 314 | 10 | 6.37% |
| hsa_circ_0077205 | 346 | 11 | 6.36% |
| hsa_circ_0119571 | 378 | 12 | 6.35% |
| hsa_circ_0008646 | 315 | 10 | 6.35% |
| hsa_circ_0022536 | 315 | 10 | 6.35% |
| hsa_circ_0077868 | 315 | 10 | 6.35% |
| hsa_circ_0102480 | 315 | 10 | 6.35% |
| hsa_circ_0137416 | 315 | 10 | 6.35% |
| hsa_circ_0062804 | 347 | 11 | 6.34% |
| hsa_circ_0134066 | 347 | 11 | 6.34% |
| hsa_circ_0011061 | 316 | 10 | 6.33% |
| hsa_circ_0038329 | 316 | 10 | 6.33% |
| hsa_circ_0072984 | 316 | 10 | 6.33% |
| hsa_circ_0102727 | 411 | 13 | 6.33% |
| hsa_circ_0131880 | 348 | 11 | 6.32% |
| hsa_circ_0042758 | 317 | 10 | 6.31% |
| hsa_circ_0050172 | 317 | 10 | 6.31% |
| hsa_circ_0112383 | 317 | 10 | 6.31% |
| hsa_circ_0116744 | 317 | 10 | 6.31% |
| hsa_circ_0119961 | 317 | 10 | 6.31% |
| hsa_circ_0003424 | 349 | 11 | 6.30% |
| hsa_circ_0118678 | 349 | 11 | 6.30% |
| hsa_circ_0120912 | 349 | 11 | 6.30% |
| hsa_circ_0131386 | 381 | 12 | 6.30% |
| hsa_circ_0046776 | 318 | 10 | 6.29% |
| hsa_circ_0061567 | 318 | 10 | 6.29% |
| hsa_circ_0068001 | 318 | 10 | 6.29% |
| hsa_circ_0126543 | 318 | 10 | 6.29% |
| hsa_circ_0011693 | 350 | 11 | 6.29% |
| hsa_circ_0109556 | 350 | 11 | 6.29% |
| hsa_circ_0103972 | 382 | 12 | 6.28% |
| hsa_circ_0105196 | 382 | 12 | 6.28% |
| hsa_circ_0061528 | 414 | 13 | 6.28% |

| | | | |
|---|---|---|---|
| hsa_circ_0012844 | 319 | 10 | 6.27% |
| hsa_circ_0087161 | 319 | 10 | 6.27% |
| hsa_circ_0096176 | 319 | 10 | 6.27% |
| hsa_circ_0136536 | 319 | 10 | 6.27% |
| hsa_circ_0138040 | 319 | 10 | 6.27% |
| hsa_circ_0002199 | 351 | 11 | 6.27% |
| hsa_circ_0115299 | 351 | 11 | 6.27% |
| hsa_circ_0127724 | 351 | 11 | 6.27% |
| hsa_circ_0072104 | 383 | 12 | 6.27% |
| hsa_circ_0069812 | 384 | 12 | 6.25% |
| hsa_circ_0039381 | 352 | 11 | 6.25% |
| hsa_circ_0004271 | 320 | 10 | 6.25% |
| hsa_circ_0012127 | 320 | 10 | 6.25% |
| hsa_circ_0040766 | 320 | 10 | 6.25% |
| hsa_circ_0054404 | 320 | 10 | 6.25% |
| hsa_circ_0101546 | 320 | 10 | 6.25% |
| hsa_circ_0139361 | 320 | 10 | 6.25% |
| hsa_circ_0015957 | 353 | 11 | 6.23% |
| hsa_circ_0026160 | 353 | 11 | 6.23% |
| hsa_circ_0074211 | 353 | 11 | 6.23% |
| hsa_circ_0011311 | 321 | 10 | 6.23% |
| hsa_circ_0032334 | 321 | 10 | 6.23% |
| hsa_circ_0049231 | 321 | 10 | 6.23% |
| hsa_circ_0135994 | 321 | 10 | 6.23% |
| hsa_circ_0087162 | 386 | 12 | 6.22% |
| hsa_circ_0099232 | 386 | 12 | 6.22% |
| hsa_circ_0004738 | 354 | 11 | 6.21% |
| hsa_circ_0072583 | 676 | 21 | 6.21% |
| hsa_circ_0024023 | 323 | 10 | 6.19% |
| hsa_circ_0065338 | 323 | 10 | 6.19% |
| hsa_circ_0108767 | 323 | 10 | 6.19% |
| hsa_circ_0023430 | 356 | 11 | 6.18% |
| hsa_circ_0091304 | 356 | 11 | 6.18% |
| hsa_circ_0135341 | 356 | 11 | 6.18% |
| hsa_circ_0000560 | 324 | 10 | 6.17% |
| hsa_circ_0026120 | 324 | 10 | 6.17% |
| hsa_circ_0029422 | 324 | 10 | 6.17% |
| hsa_circ_0064072 | 324 | 10 | 6.17% |
| hsa_circ_0079093 | 324 | 10 | 6.17% |
| hsa_circ_0124057 | 324 | 10 | 6.17% |
| hsa_circ_0127373 | 324 | 10 | 6.17% |
| hsa_circ_0052214 | 357 | 11 | 6.16% |
| hsa_circ_0098497 | 357 | 11 | 6.16% |
| hsa_circ_0016997 | 325 | 10 | 6.15% |
| hsa_circ_0040037 | 325 | 10 | 6.15% |
| hsa_circ_0113381 | 325 | 10 | 6.15% |
| hsa_circ_0114367 | 325 | 10 | 6.15% |
| hsa_circ_0131403 | 325 | 10 | 6.15% |
| hsa_circ_0010412 | 358 | 11 | 6.15% |
| hsa_circ_0115430 | 391 | 12 | 6.14% |

| | | | |
|---|---|---|---|
| hsa_circ_0012598 | 326 | 10 | 6.13% |
| hsa_circ_0026174 | 326 | 10 | 6.13% |
| hsa_circ_0058244 | 326 | 10 | 6.13% |
| hsa_circ_0080322 | 326 | 10 | 6.13% |
| hsa_circ_0116333 | 326 | 10 | 6.13% |
| hsa_circ_0055613 | 359 | 11 | 6.13% |
| hsa_circ_0023490 | 392 | 12 | 6.12% |
| hsa_circ_0005845 | 327 | 10 | 6.12% |
| hsa_circ_0052045 | 327 | 10 | 6.12% |
| hsa_circ_0052166 | 327 | 10 | 6.12% |
| hsa_circ_0080514 | 327 | 10 | 6.12% |
| hsa_circ_0110040 | 327 | 10 | 6.12% |
| hsa_circ_0110198 | 327 | 10 | 6.12% |
| hsa_circ_0119332 | 327 | 10 | 6.12% |
| hsa_circ_0127228 | 327 | 10 | 6.12% |
| hsa_circ_0016733 | 360 | 11 | 6.11% |
| hsa_circ_0037477 | 360 | 11 | 6.11% |
| hsa_circ_0057395 | 360 | 11 | 6.11% |
| hsa_circ_0092019 | 360 | 11 | 6.11% |
| hsa_circ_0108716 | 360 | 11 | 6.11% |
| hsa_circ_0124133 | 360 | 11 | 6.11% |
| hsa_circ_0077659 | 426 | 13 | 6.10% |
| hsa_circ_0002160 | 328 | 10 | 6.10% |
| hsa_circ_0004819 | 328 | 10 | 6.10% |
| hsa_circ_0038507 | 328 | 10 | 6.10% |
| hsa_circ_0058490 | 328 | 10 | 6.10% |
| hsa_circ_0082947 | 328 | 10 | 6.10% |
| hsa_circ_0124116 | 328 | 10 | 6.10% |
| hsa_circ_0131216 | 328 | 10 | 6.10% |
| hsa_circ_0133562 | 361 | 11 | 6.09% |
| hsa_circ_0045905 | 329 | 10 | 6.08% |
| hsa_circ_0105998 | 329 | 10 | 6.08% |
| hsa_circ_0109337 | 329 | 10 | 6.08% |
| hsa_circ_0001870 | 362 | 11 | 6.08% |
| hsa_circ_0007080 | 362 | 11 | 6.08% |
| hsa_circ_0126613 | 429 | 13 | 6.06% |
| hsa_circ_0092142 | 396 | 12 | 6.06% |
| hsa_circ_0004477 | 330 | 10 | 6.06% |
| hsa_circ_0010778 | 330 | 10 | 6.06% |
| hsa_circ_0011848 | 330 | 10 | 6.06% |
| hsa_circ_0051687 | 330 | 10 | 6.06% |
| hsa_circ_0067251 | 330 | 10 | 6.06% |
| hsa_circ_0092196 | 330 | 10 | 6.06% |
| hsa_circ_0105877 | 330 | 10 | 6.06% |
| hsa_circ_0057941 | 397 | 12 | 6.05% |
| hsa_circ_0018635 | 364 | 11 | 6.04% |
| hsa_circ_0060778 | 364 | 11 | 6.04% |
| hsa_circ_0002431 | 331 | 10 | 6.04% |
| hsa_circ_0004457 | 331 | 10 | 6.04% |
| hsa_circ_0007591 | 331 | 10 | 6.04% |

| | | | |
|---|---|---|---|
| hsa_circ_0009249 | 331 | 10 | 6.04% |
| hsa_circ_0013353 | 331 | 10 | 6.04% |
| hsa_circ_0042947 | 331 | 10 | 6.04% |
| hsa_circ_0057795 | 331 | 10 | 6.04% |
| hsa_circ_0068233 | 331 | 10 | 6.04% |
| hsa_circ_0079031 | 331 | 10 | 6.04% |
| hsa_circ_0091419 | 331 | 10 | 6.04% |
| hsa_circ_0095333 | 331 | 10 | 6.04% |
| hsa_circ_0109161 | 431 | 13 | 6.03% |
| hsa_circ_0092752 | 398 | 12 | 6.03% |
| hsa_circ_0040832 | 365 | 11 | 6.03% |
| hsa_circ_0102655 | 365 | 11 | 6.03% |
| hsa_circ_0031424 | 332 | 10 | 6.02% |
| hsa_circ_0045362 | 332 | 10 | 6.02% |
| hsa_circ_0095035 | 332 | 10 | 6.02% |
| hsa_circ_0108458 | 332 | 10 | 6.02% |
| hsa_circ_0114345 | 332 | 10 | 6.02% |
| hsa_circ_0137446 | 332 | 10 | 6.02% |
| hsa_circ_0140512 | 332 | 10 | 6.02% |
| hsa_circ_0100899 | 399 | 12 | 6.02% |
| hsa_circ_0015982 | 499 | 15 | 6.01% |
| hsa_circ_0003506 | 366 | 11 | 6.01% |
| hsa_circ_0004811 | 366 | 11 | 6.01% |
| hsa_circ_0076467 | 366 | 11 | 6.01% |
| hsa_circ_0051685 | 333 | 10 | 6.01% |
| hsa_circ_0076769 | 333 | 10 | 6.01% |
| hsa_circ_0087580 | 333 | 10 | 6.01% |
| hsa_circ_0072641 | 433 | 13 | 6.00% |
| hsa_circ_0062812 | 367 | 11 | 5.99% |
| hsa_circ_0107659 | 367 | 11 | 5.99% |
| hsa_circ_0128268 | 367 | 11 | 5.99% |
| hsa_circ_0137152 | 367 | 11 | 5.99% |
| hsa_circ_0036810 | 334 | 10 | 5.99% |
| hsa_circ_0058736 | 334 | 10 | 5.99% |
| hsa_circ_0124768 | 334 | 10 | 5.99% |
| hsa_circ_0064625 | 368 | 11 | 5.98% |
| hsa_circ_0136885 | 402 | 12 | 5.97% |
| hsa_circ_0010930 | 335 | 10 | 5.97% |
| hsa_circ_0044825 | 335 | 10 | 5.97% |
| hsa_circ_0111971 | 335 | 10 | 5.97% |
| hsa_circ_0108023 | 369 | 11 | 5.96% |
| hsa_circ_0040852 | 336 | 10 | 5.95% |
| hsa_circ_0047089 | 336 | 10 | 5.95% |
| hsa_circ_0077605 | 336 | 10 | 5.95% |
| hsa_circ_0109881 | 336 | 10 | 5.95% |
| hsa_circ_0113091 | 336 | 10 | 5.95% |
| hsa_circ_0126099 | 336 | 10 | 5.95% |
| hsa_circ_0116998 | 370 | 11 | 5.95% |
| hsa_circ_0105424 | 707 | 21 | 5.94% |
| hsa_circ_0045330 | 404 | 12 | 5.94% |

| | | | |
|---|---|---|---|
| hsa_circ_0066705 | 404 | 12 | 5.94% |
| hsa_circ_0098832 | 404 | 12 | 5.94% |
| hsa_circ_0102571 | 1550 | 46 | 5.94% |
| hsa_circ_0009850 | 337 | 10 | 5.93% |
| hsa_circ_0029170 | 337 | 10 | 5.93% |
| hsa_circ_0048279 | 337 | 10 | 5.93% |
| hsa_circ_0096023 | 337 | 10 | 5.93% |
| hsa_circ_0109542 | 337 | 10 | 5.93% |
| hsa_circ_0114925 | 337 | 10 | 5.93% |
| hsa_circ_0122300 | 337 | 10 | 5.93% |
| hsa_circ_0138011 | 337 | 10 | 5.93% |
| hsa_circ_0109153 | 573 | 17 | 5.93% |
| hsa_circ_0014982 | 371 | 11 | 5.93% |
| hsa_circ_0120447 | 371 | 11 | 5.93% |
| hsa_circ_0128085 | 371 | 11 | 5.93% |
| hsa_circ_0058585 | 405 | 12 | 5.93% |
| hsa_circ_0026053 | 338 | 10 | 5.92% |
| hsa_circ_0032547 | 338 | 10 | 5.92% |
| hsa_circ_0037282 | 338 | 10 | 5.92% |
| hsa_circ_0057768 | 338 | 10 | 5.92% |
| hsa_circ_0084952 | 338 | 10 | 5.92% |
| hsa_circ_0089371 | 338 | 10 | 5.92% |
| hsa_circ_0096369 | 338 | 10 | 5.92% |
| hsa_circ_0108565 | 338 | 10 | 5.92% |
| hsa_circ_0113003 | 338 | 10 | 5.92% |
| hsa_circ_0113430 | 338 | 10 | 5.92% |
| hsa_circ_0016110 | 372 | 11 | 5.91% |
| hsa_circ_0117764 | 372 | 11 | 5.91% |
| hsa_circ_0120859 | 372 | 11 | 5.91% |
| hsa_circ_0085818 | 575 | 17 | 5.91% |
| hsa_circ_0129577 | 406 | 12 | 5.91% |
| hsa_circ_0077846 | 339 | 10 | 5.90% |
| hsa_circ_0107236 | 339 | 10 | 5.90% |
| hsa_circ_0129571 | 339 | 10 | 5.90% |
| hsa_circ_0134504 | 339 | 10 | 5.90% |
| hsa_circ_0066780 | 373 | 11 | 5.90% |
| hsa_circ_0104769 | 373 | 11 | 5.90% |
| hsa_circ_0028503 | 441 | 13 | 5.90% |
| hsa_circ_0077591 | 475 | 14 | 5.89% |
| hsa_circ_0055820 | 374 | 11 | 5.88% |
| hsa_circ_0037909 | 340 | 10 | 5.88% |
| hsa_circ_0065617 | 340 | 10 | 5.88% |
| hsa_circ_0067278 | 340 | 10 | 5.88% |
| hsa_circ_0081301 | 340 | 10 | 5.88% |
| hsa_circ_0086656 | 340 | 10 | 5.88% |
| hsa_circ_0131712 | 340 | 10 | 5.88% |
| hsa_circ_0027977 | 443 | 13 | 5.87% |
| hsa_circ_0081512 | 443 | 13 | 5.87% |
| hsa_circ_0072453 | 375 | 11 | 5.87% |
| hsa_circ_0076793 | 375 | 11 | 5.87% |

| | | | |
|---|---|---|---|
| hsa_circ_0123024 | 375 | 11 | 5.87% |
| hsa_circ_0131375 | 375 | 11 | 5.87% |
| hsa_circ_0015094 | 341 | 10 | 5.87% |
| hsa_circ_0085782 | 341 | 10 | 5.87% |
| hsa_circ_0140532 | 341 | 10 | 5.87% |
| hsa_circ_0004786 | 478 | 14 | 5.86% |
| hsa_circ_0023052 | 444 | 13 | 5.86% |
| hsa_circ_0043860 | 444 | 13 | 5.86% |
| hsa_circ_0008847 | 376 | 11 | 5.85% |
| hsa_circ_0024480 | 376 | 11 | 5.85% |
| hsa_circ_0085116 | 376 | 11 | 5.85% |
| hsa_circ_0100035 | 376 | 11 | 5.85% |
| hsa_circ_0003218 | 342 | 10 | 5.85% |
| hsa_circ_0016112 | 342 | 10 | 5.85% |
| hsa_circ_0026683 | 342 | 10 | 5.85% |
| hsa_circ_0050645 | 342 | 10 | 5.85% |
| hsa_circ_0059480 | 342 | 10 | 5.85% |
| hsa_circ_0073978 | 342 | 10 | 5.85% |
| hsa_circ_0092075 | 342 | 10 | 5.85% |
| hsa_circ_0109015 | 342 | 10 | 5.85% |
| hsa_circ_0118903 | 342 | 10 | 5.85% |
| hsa_circ_0130292 | 342 | 10 | 5.85% |
| hsa_circ_0140197 | 342 | 10 | 5.85% |
| hsa_circ_0138004 | 445 | 13 | 5.84% |
| hsa_circ_0053235 | 377 | 11 | 5.84% |
| hsa_circ_0116418 | 377 | 11 | 5.84% |
| hsa_circ_0005609 | 343 | 10 | 5.83% |
| hsa_circ_0005774 | 343 | 10 | 5.83% |
| hsa_circ_0020933 | 343 | 10 | 5.83% |
| hsa_circ_0023467 | 343 | 10 | 5.83% |
| hsa_circ_0093127 | 343 | 10 | 5.83% |
| hsa_circ_0095918 | 481 | 14 | 5.82% |
| hsa_circ_0002980 | 378 | 11 | 5.82% |
| hsa_circ_0013222 | 378 | 11 | 5.82% |
| hsa_circ_0046906 | 378 | 11 | 5.82% |
| hsa_circ_0087332 | 378 | 11 | 5.82% |
| hsa_circ_0000455 | 344 | 10 | 5.81% |
| hsa_circ_0072154 | 344 | 10 | 5.81% |
| hsa_circ_0102157 | 344 | 10 | 5.81% |
| hsa_circ_0120987 | 344 | 10 | 5.81% |
| hsa_circ_0133285 | 344 | 10 | 5.81% |
| hsa_circ_0060528 | 448 | 13 | 5.80% |
| hsa_circ_0026050 | 345 | 10 | 5.80% |
| hsa_circ_0030543 | 345 | 10 | 5.80% |
| hsa_circ_0098585 | 345 | 10 | 5.80% |
| hsa_circ_0103631 | 380 | 11 | 5.79% |
| hsa_circ_0073810 | 415 | 12 | 5.78% |
| hsa_circ_0001307 | 346 | 10 | 5.78% |
| hsa_circ_0001848 | 346 | 10 | 5.78% |
| hsa_circ_0005763 | 346 | 10 | 5.78% |

| | | | |
|---|---|---|---|
| hsa_circ_0006988 | 346 | 10 | 5.78% |
| hsa_circ_0035633 | 346 | 10 | 5.78% |
| hsa_circ_0039017 | 346 | 10 | 5.78% |
| hsa_circ_0053131 | 346 | 10 | 5.78% |
| hsa_circ_0077893 | 346 | 10 | 5.78% |
| hsa_circ_0111707 | 346 | 10 | 5.78% |
| hsa_circ_0050111 | 381 | 11 | 5.77% |
| hsa_circ_0093889 | 381 | 11 | 5.77% |
| hsa_circ_0075034 | 416 | 12 | 5.77% |
| hsa_circ_0115373 | 416 | 12 | 5.77% |
| hsa_circ_0012109 | 347 | 10 | 5.76% |
| hsa_circ_0053038 | 347 | 10 | 5.76% |
| hsa_circ_0082304 | 347 | 10 | 5.76% |
| hsa_circ_0085282 | 347 | 10 | 5.76% |
| hsa_circ_0125816 | 347 | 10 | 5.76% |
| hsa_circ_0138241 | 347 | 10 | 5.76% |
| hsa_circ_0081198 | 382 | 11 | 5.76% |
| hsa_circ_0053953 | 417 | 12 | 5.76% |
| hsa_circ_0009152 | 348 | 10 | 5.75% |
| hsa_circ_0055892 | 348 | 10 | 5.75% |
| hsa_circ_0065916 | 383 | 11 | 5.74% |
| hsa_circ_0074995 | 383 | 11 | 5.74% |
| hsa_circ_0122838 | 383 | 11 | 5.74% |
| hsa_circ_0113244 | 453 | 13 | 5.74% |
| hsa_circ_0006174 | 349 | 10 | 5.73% |
| hsa_circ_0010100 | 349 | 10 | 5.73% |
| hsa_circ_0018068 | 349 | 10 | 5.73% |
| hsa_circ_0064967 | 349 | 10 | 5.73% |
| hsa_circ_0137946 | 349 | 10 | 5.73% |
| hsa_circ_0078477 | 384 | 11 | 5.73% |
| hsa_circ_0039555 | 385 | 11 | 5.71% |
| hsa_circ_0068379 | 385 | 11 | 5.71% |
| hsa_circ_0001292 | 350 | 10 | 5.71% |
| hsa_circ_0004716 | 350 | 10 | 5.71% |
| hsa_circ_0009861 | 350 | 10 | 5.71% |
| hsa_circ_0012963 | 350 | 10 | 5.71% |
| hsa_circ_0050668 | 350 | 10 | 5.71% |
| hsa_circ_0067580 | 350 | 10 | 5.71% |
| hsa_circ_0068863 | 350 | 10 | 5.71% |
| hsa_circ_0105164 | 350 | 10 | 5.71% |
| hsa_circ_0110966 | 350 | 10 | 5.71% |
| hsa_circ_0003644 | 421 | 12 | 5.70% |
| hsa_circ_0130826 | 421 | 12 | 5.70% |
| hsa_circ_0006185 | 386 | 11 | 5.70% |
| hsa_circ_0114220 | 386 | 11 | 5.70% |
| hsa_circ_0120756 | 386 | 11 | 5.70% |
| hsa_circ_0003584 | 351 | 10 | 5.70% |
| hsa_circ_0003864 | 351 | 10 | 5.70% |
| hsa_circ_0018455 | 351 | 10 | 5.70% |
| hsa_circ_0049356 | 351 | 10 | 5.70% |

| | | | |
|---|---|---|---|
| hsa_circ_0057446 | 351 | 10 | 5.70% |
| hsa_circ_0060352 | 351 | 10 | 5.70% |
| hsa_circ_0114709 | 351 | 10 | 5.70% |
| hsa_circ_0117296 | 351 | 10 | 5.70% |
| hsa_circ_0125496 | 351 | 10 | 5.70% |
| hsa_circ_0126723 | 351 | 10 | 5.70% |
| hsa_circ_0129631 | 351 | 10 | 5.70% |
| hsa_circ_0136920 | 351 | 10 | 5.70% |
| hsa_circ_0138524 | 351 | 10 | 5.70% |
| hsa_circ_0140195 | 351 | 10 | 5.70% |
| hsa_circ_0032662 | 598 | 17 | 5.69% |
| hsa_circ_0020846 | 387 | 11 | 5.68% |
| hsa_circ_0082783 | 387 | 11 | 5.68% |
| hsa_circ_0104778 | 387 | 11 | 5.68% |
| hsa_circ_0105496 | 387 | 11 | 5.68% |
| hsa_circ_0016806 | 352 | 10 | 5.68% |
| hsa_circ_0031935 | 352 | 10 | 5.68% |
| hsa_circ_0062875 | 352 | 10 | 5.68% |
| hsa_circ_0068423 | 352 | 10 | 5.68% |
| hsa_circ_0101085 | 352 | 10 | 5.68% |
| hsa_circ_0126299 | 352 | 10 | 5.68% |
| hsa_circ_0130436 | 352 | 10 | 5.68% |
| hsa_circ_0116158 | 564 | 16 | 5.67% |
| hsa_circ_0098879 | 423 | 12 | 5.67% |
| hsa_circ_0027672 | 388 | 11 | 5.67% |
| hsa_circ_0009189 | 353 | 10 | 5.67% |
| hsa_circ_0026145 | 353 | 10 | 5.67% |
| hsa_circ_0040764 | 353 | 10 | 5.67% |
| hsa_circ_0041726 | 353 | 10 | 5.67% |
| hsa_circ_0062637 | 389 | 11 | 5.66% |
| hsa_circ_0006515 | 354 | 10 | 5.65% |
| hsa_circ_0047655 | 354 | 10 | 5.65% |
| hsa_circ_0051844 | 354 | 10 | 5.65% |
| hsa_circ_0059982 | 354 | 10 | 5.65% |
| hsa_circ_0087504 | 354 | 10 | 5.65% |
| hsa_circ_0096470 | 354 | 10 | 5.65% |
| hsa_circ_0098172 | 354 | 10 | 5.65% |
| hsa_circ_0099088 | 354 | 10 | 5.65% |
| hsa_circ_0125623 | 354 | 10 | 5.65% |
| hsa_circ_0134875 | 354 | 10 | 5.65% |
| hsa_circ_0081849 | 390 | 11 | 5.64% |
| hsa_circ_0023615 | 461 | 13 | 5.64% |
| hsa_circ_0005443 | 355 | 10 | 5.63% |
| hsa_circ_0036733 | 355 | 10 | 5.63% |
| hsa_circ_0105653 | 355 | 10 | 5.63% |
| hsa_circ_0127564 | 355 | 10 | 5.63% |
| hsa_circ_0134372 | 355 | 10 | 5.63% |
| hsa_circ_0010426 | 391 | 11 | 5.63% |
| hsa_circ_0083161 | 391 | 11 | 5.63% |
| hsa_circ_0110687 | 391 | 11 | 5.63% |

| | | | |
|---|---|---|---|
| hsa_circ_0114893 | 391 | 11 | 5.63% |
| hsa_circ_0120165 | 391 | 11 | 5.63% |
| hsa_circ_0122049 | 391 | 11 | 5.63% |
| hsa_circ_0043427 | 356 | 10 | 5.62% |
| hsa_circ_0052419 | 356 | 10 | 5.62% |
| hsa_circ_0064514 | 356 | 10 | 5.62% |
| hsa_circ_0107404 | 356 | 10 | 5.62% |
| hsa_circ_0121000 | 356 | 10 | 5.62% |
| hsa_circ_0137719 | 356 | 10 | 5.62% |
| hsa_circ_0093726 | 392 | 11 | 5.61% |
| hsa_circ_0106833 | 392 | 11 | 5.61% |
| hsa_circ_0124550 | 499 | 14 | 5.61% |
| hsa_circ_0075065 | 464 | 13 | 5.60% |
| hsa_circ_0001398 | 357 | 10 | 5.60% |
| hsa_circ_0011758 | 357 | 10 | 5.60% |
| hsa_circ_0080107 | 357 | 10 | 5.60% |
| hsa_circ_0080805 | 357 | 10 | 5.60% |
| hsa_circ_0096131 | 357 | 10 | 5.60% |
| hsa_circ_0102900 | 357 | 10 | 5.60% |
| hsa_circ_0134179 | 357 | 10 | 5.60% |
| hsa_circ_0002989 | 358 | 10 | 5.59% |
| hsa_circ_0017680 | 358 | 10 | 5.59% |
| hsa_circ_0053268 | 358 | 10 | 5.59% |
| hsa_circ_0073209 | 358 | 10 | 5.59% |
| hsa_circ_0020285 | 394 | 11 | 5.58% |
| hsa_circ_0046944 | 394 | 11 | 5.58% |
| hsa_circ_0063812 | 394 | 11 | 5.58% |
| hsa_circ_0074652 | 394 | 11 | 5.58% |
| hsa_circ_0085849 | 394 | 11 | 5.58% |
| hsa_circ_0109203 | 394 | 11 | 5.58% |
| hsa_circ_0020532 | 466 | 13 | 5.58% |
| hsa_circ_0005559 | 359 | 10 | 5.57% |
| hsa_circ_0008378 | 359 | 10 | 5.57% |
| hsa_circ_0013373 | 359 | 10 | 5.57% |
| hsa_circ_0013694 | 359 | 10 | 5.57% |
| hsa_circ_0019259 | 359 | 10 | 5.57% |
| hsa_circ_0042761 | 359 | 10 | 5.57% |
| hsa_circ_0059726 | 359 | 10 | 5.57% |
| hsa_circ_0093883 | 359 | 10 | 5.57% |
| hsa_circ_0104110 | 359 | 10 | 5.57% |
| hsa_circ_0119599 | 359 | 10 | 5.57% |
| hsa_circ_0126106 | 359 | 10 | 5.57% |
| hsa_circ_0127077 | 359 | 10 | 5.57% |
| hsa_circ_0107011 | 395 | 11 | 5.57% |
| hsa_circ_0116546 | 395 | 11 | 5.57% |
| hsa_circ_0114713 | 432 | 12 | 5.56% |
| hsa_circ_0005740 | 396 | 11 | 5.56% |
| hsa_circ_0066912 | 396 | 11 | 5.56% |
| hsa_circ_0090451 | 396 | 11 | 5.56% |
| hsa_circ_0122228 | 396 | 11 | 5.56% |

| | | | |
|---|---|---|---|
| hsa_circ_0000061 | 360 | 10 | 5.56% |
| hsa_circ_0038716 | 360 | 10 | 5.56% |
| hsa_circ_0048049 | 360 | 10 | 5.56% |
| hsa_circ_0058324 | 360 | 10 | 5.56% |
| hsa_circ_0059105 | 360 | 10 | 5.56% |
| hsa_circ_0070479 | 360 | 10 | 5.56% |
| hsa_circ_0083533 | 360 | 10 | 5.56% |
| hsa_circ_0091188 | 360 | 10 | 5.56% |
| hsa_circ_0105215 | 360 | 10 | 5.56% |
| hsa_circ_0119357 | 360 | 10 | 5.56% |
| hsa_circ_0123686 | 360 | 10 | 5.56% |
| hsa_circ_0135934 | 360 | 10 | 5.56% |
| hsa_circ_0113017 | 433 | 12 | 5.54% |
| hsa_circ_0099177 | 1191 | 33 | 5.54% |
| hsa_circ_0042738 | 397 | 11 | 5.54% |
| hsa_circ_0077112 | 397 | 11 | 5.54% |
| hsa_circ_0005990 | 361 | 10 | 5.54% |
| hsa_circ_0120053 | 361 | 10 | 5.54% |
| hsa_circ_0135221 | 361 | 10 | 5.54% |
| hsa_circ_0044777 | 434 | 12 | 5.53% |
| hsa_circ_0094451 | 434 | 12 | 5.53% |
| hsa_circ_0082014 | 398 | 11 | 5.53% |
| hsa_circ_0094843 | 398 | 11 | 5.53% |
| hsa_circ_0101882 | 398 | 11 | 5.53% |
| hsa_circ_0006810 | 362 | 10 | 5.52% |
| hsa_circ_0022432 | 362 | 10 | 5.52% |
| hsa_circ_0076764 | 362 | 10 | 5.52% |
| hsa_circ_0094381 | 362 | 10 | 5.52% |
| hsa_circ_0110578 | 362 | 10 | 5.52% |
| hsa_circ_0112077 | 362 | 10 | 5.52% |
| hsa_circ_0063750 | 399 | 11 | 5.51% |
| hsa_circ_0064318 | 399 | 11 | 5.51% |
| hsa_circ_0121164 | 399 | 11 | 5.51% |
| hsa_circ_0010151 | 363 | 10 | 5.51% |
| hsa_circ_0032642 | 363 | 10 | 5.51% |
| hsa_circ_0046136 | 363 | 10 | 5.51% |
| hsa_circ_0103895 | 363 | 10 | 5.51% |
| hsa_circ_0109097 | 363 | 10 | 5.51% |
| hsa_circ_0126102 | 363 | 10 | 5.51% |
| hsa_circ_0134749 | 363 | 10 | 5.51% |
| hsa_circ_0115847 | 472 | 13 | 5.51% |
| hsa_circ_0060491 | 436 | 12 | 5.50% |
| hsa_circ_0118158 | 436 | 12 | 5.50% |
| hsa_circ_0089956 | 400 | 11 | 5.50% |
| hsa_circ_0023830 | 364 | 10 | 5.49% |
| hsa_circ_0030382 | 364 | 10 | 5.49% |
| hsa_circ_0050355 | 364 | 10 | 5.49% |
| hsa_circ_0056188 | 364 | 10 | 5.49% |
| hsa_circ_0072099 | 364 | 10 | 5.49% |
| hsa_circ_0118953 | 364 | 10 | 5.49% |

| | | | |
|---|---|---|---|
| hsa_circ_0137469 | 364 | 10 | 5.49% |
| hsa_circ_0001203 | 437 | 12 | 5.49% |
| hsa_circ_0076099 | 437 | 12 | 5.49% |
| hsa_circ_0050954 | 401 | 11 | 5.49% |
| hsa_circ_0084376 | 401 | 11 | 5.49% |
| hsa_circ_0124899 | 401 | 11 | 5.49% |
| hsa_circ_0014804 | 438 | 12 | 5.48% |
| hsa_circ_0102192 | 438 | 12 | 5.48% |
| hsa_circ_0125857 | 438 | 12 | 5.48% |
| hsa_circ_0010615 | 365 | 10 | 5.48% |
| hsa_circ_0090467 | 365 | 10 | 5.48% |
| hsa_circ_0122441 | 365 | 10 | 5.48% |
| hsa_circ_0140753 | 365 | 10 | 5.48% |
| hsa_circ_0000832 | 402 | 11 | 5.47% |
| hsa_circ_0028305 | 402 | 11 | 5.47% |
| hsa_circ_0038822 | 402 | 11 | 5.47% |
| hsa_circ_0007328 | 366 | 10 | 5.46% |
| hsa_circ_0028655 | 366 | 10 | 5.46% |
| hsa_circ_0048125 | 366 | 10 | 5.46% |
| hsa_circ_0080947 | 366 | 10 | 5.46% |
| hsa_circ_0098764 | 366 | 10 | 5.46% |
| hsa_circ_0119894 | 366 | 10 | 5.46% |
| hsa_circ_0126187 | 366 | 10 | 5.46% |
| hsa_circ_0127378 | 366 | 10 | 5.46% |
| hsa_circ_0070478 | 403 | 11 | 5.46% |
| hsa_circ_0093681 | 403 | 11 | 5.46% |
| hsa_circ_0015723 | 440 | 12 | 5.45% |
| hsa_circ_0136415 | 917 | 25 | 5.45% |
| hsa_circ_0048832 | 367 | 10 | 5.45% |
| hsa_circ_0106368 | 367 | 10 | 5.45% |
| hsa_circ_0110619 | 367 | 10 | 5.45% |
| hsa_circ_0119201 | 367 | 10 | 5.45% |
| hsa_circ_0135493 | 367 | 10 | 5.45% |
| hsa_circ_0025156 | 404 | 11 | 5.45% |
| hsa_circ_0085705 | 404 | 11 | 5.45% |
| hsa_circ_0027660 | 441 | 12 | 5.44% |
| hsa_circ_0004894 | 478 | 13 | 5.44% |
| hsa_circ_0006170 | 368 | 10 | 5.43% |
| hsa_circ_0088360 | 368 | 10 | 5.43% |
| hsa_circ_0107717 | 368 | 10 | 5.43% |
| hsa_circ_0114112 | 368 | 10 | 5.43% |
| hsa_circ_0118964 | 368 | 10 | 5.43% |
| hsa_circ_0122916 | 368 | 10 | 5.43% |
| hsa_circ_0040078 | 405 | 11 | 5.43% |
| hsa_circ_0058170 | 405 | 11 | 5.43% |
| hsa_circ_0060313 | 405 | 11 | 5.43% |
| hsa_circ_0029286 | 369 | 10 | 5.42% |
| hsa_circ_0129827 | 369 | 10 | 5.42% |
| hsa_circ_0137512 | 369 | 10 | 5.42% |
| hsa_circ_0046505 | 406 | 11 | 5.42% |

| | | | |
|---|---|---|---|
| hsa_circ_0033074 | 443 | 12 | 5.42% |
| hsa_circ_0050639 | 665 | 18 | 5.41% |
| hsa_circ_0041807 | 407 | 11 | 5.41% |
| hsa_circ_0042242 | 370 | 10 | 5.41% |
| hsa_circ_0061704 | 370 | 10 | 5.41% |
| hsa_circ_0087900 | 370 | 10 | 5.41% |
| hsa_circ_0106020 | 370 | 10 | 5.41% |
| hsa_circ_0126094 | 370 | 10 | 5.41% |
| hsa_circ_0137609 | 370 | 10 | 5.41% |
| hsa_circ_0139614 | 370 | 10 | 5.41% |
| hsa_circ_0020190 | 408 | 11 | 5.39% |
| hsa_circ_0042268 | 408 | 11 | 5.39% |
| hsa_circ_0052857 | 408 | 11 | 5.39% |
| hsa_circ_0096115 | 408 | 11 | 5.39% |
| hsa_circ_0013508 | 371 | 10 | 5.39% |
| hsa_circ_0076683 | 371 | 10 | 5.39% |
| hsa_circ_0116177 | 371 | 10 | 5.39% |
| hsa_circ_0116560 | 371 | 10 | 5.39% |
| hsa_circ_0101476 | 520 | 14 | 5.38% |
| hsa_circ_0030287 | 446 | 12 | 5.38% |
| hsa_circ_0137030 | 409 | 11 | 5.38% |
| hsa_circ_0014766 | 372 | 10 | 5.38% |
| hsa_circ_0022535 | 372 | 10 | 5.38% |
| hsa_circ_0024217 | 372 | 10 | 5.38% |
| hsa_circ_0028657 | 372 | 10 | 5.38% |
| hsa_circ_0045574 | 372 | 10 | 5.38% |
| hsa_circ_0072192 | 372 | 10 | 5.38% |
| hsa_circ_0081663 | 372 | 10 | 5.38% |
| hsa_circ_0086129 | 372 | 10 | 5.38% |
| hsa_circ_0096261 | 372 | 10 | 5.38% |
| hsa_circ_0096975 | 372 | 10 | 5.38% |
| hsa_circ_0100401 | 372 | 10 | 5.38% |
| hsa_circ_0101887 | 447 | 12 | 5.37% |
| hsa_circ_0001935 | 410 | 11 | 5.37% |
| hsa_circ_0005758 | 373 | 10 | 5.36% |
| hsa_circ_0027093 | 373 | 10 | 5.36% |
| hsa_circ_0035481 | 373 | 10 | 5.36% |
| hsa_circ_0043815 | 373 | 10 | 5.36% |
| hsa_circ_0053115 | 373 | 10 | 5.36% |
| hsa_circ_0054630 | 373 | 10 | 5.36% |
| hsa_circ_0066204 | 373 | 10 | 5.36% |
| hsa_circ_0072985 | 373 | 10 | 5.36% |
| hsa_circ_0079990 | 373 | 10 | 5.36% |
| hsa_circ_0080503 | 373 | 10 | 5.36% |
| hsa_circ_0081716 | 373 | 10 | 5.36% |
| hsa_circ_0091074 | 373 | 10 | 5.36% |
| hsa_circ_0106215 | 373 | 10 | 5.36% |
| hsa_circ_0113504 | 373 | 10 | 5.36% |
| hsa_circ_0114304 | 373 | 10 | 5.36% |
| hsa_circ_0137026 | 373 | 10 | 5.36% |

| | | | |
|---|---|---|---|
| hsa_circ_0019147 | 448 | 12 | 5.36% |
| hsa_circ_0058582 | 448 | 12 | 5.36% |
| hsa_circ_0002993 | 411 | 11 | 5.35% |
| hsa_circ_0035073 | 411 | 11 | 5.35% |
| hsa_circ_0083676 | 411 | 11 | 5.35% |
| hsa_circ_0006116 | 561 | 15 | 5.35% |
| hsa_circ_0000045 | 374 | 10 | 5.35% |
| hsa_circ_0004164 | 374 | 10 | 5.35% |
| hsa_circ_0026051 | 374 | 10 | 5.35% |
| hsa_circ_0068866 | 374 | 10 | 5.35% |
| hsa_circ_0104448 | 374 | 10 | 5.35% |
| hsa_circ_0137449 | 374 | 10 | 5.35% |
| hsa_circ_0005795 | 524 | 14 | 5.34% |
| hsa_circ_0016387 | 412 | 11 | 5.34% |
| hsa_circ_0122131 | 412 | 11 | 5.34% |
| hsa_circ_0106802 | 450 | 12 | 5.33% |
| hsa_circ_0005986 | 375 | 10 | 5.33% |
| hsa_circ_0065785 | 375 | 10 | 5.33% |
| hsa_circ_0070281 | 375 | 10 | 5.33% |
| hsa_circ_0074262 | 375 | 10 | 5.33% |
| hsa_circ_0078340 | 375 | 10 | 5.33% |
| hsa_circ_0023428 | 413 | 11 | 5.33% |
| hsa_circ_0091250 | 413 | 11 | 5.33% |
| hsa_circ_0033073 | 451 | 12 | 5.32% |
| hsa_circ_0050718 | 451 | 12 | 5.32% |
| hsa_circ_0101888 | 451 | 12 | 5.32% |
| hsa_circ_0002194 | 376 | 10 | 5.32% |
| hsa_circ_0006634 | 376 | 10 | 5.32% |
| hsa_circ_0023442 | 376 | 10 | 5.32% |
| hsa_circ_0035445 | 376 | 10 | 5.32% |
| hsa_circ_0052412 | 376 | 10 | 5.32% |
| hsa_circ_0054212 | 376 | 10 | 5.32% |
| hsa_circ_0069688 | 376 | 10 | 5.32% |
| hsa_circ_0106858 | 376 | 10 | 5.32% |
| hsa_circ_0115684 | 376 | 10 | 5.32% |
| hsa_circ_0125922 | 376 | 10 | 5.32% |
| hsa_circ_0132157 | 376 | 10 | 5.32% |
| hsa_circ_0013357 | 414 | 11 | 5.31% |
| hsa_circ_0051146 | 414 | 11 | 5.31% |
| hsa_circ_0025336 | 377 | 10 | 5.31% |
| hsa_circ_0038866 | 377 | 10 | 5.31% |
| hsa_circ_0047658 | 377 | 10 | 5.31% |
| hsa_circ_0098063 | 377 | 10 | 5.31% |
| hsa_circ_0133547 | 377 | 10 | 5.31% |
| hsa_circ_0025386 | 453 | 12 | 5.30% |
| hsa_circ_0102077 | 453 | 12 | 5.30% |
| hsa_circ_0112805 | 453 | 12 | 5.30% |
| hsa_circ_0008465 | 378 | 10 | 5.29% |
| hsa_circ_0011816 | 378 | 10 | 5.29% |
| hsa_circ_0022839 | 378 | 10 | 5.29% |

| | | | |
|---|---|---|---|
| hsa_circ_0034885 | 378 | 10 | 5.29% |
| hsa_circ_0046215 | 378 | 10 | 5.29% |
| hsa_circ_0066970 | 378 | 10 | 5.29% |
| hsa_circ_0079363 | 378 | 10 | 5.29% |
| hsa_circ_0080313 | 378 | 10 | 5.29% |
| hsa_circ_0080964 | 378 | 10 | 5.29% |
| hsa_circ_0106200 | 378 | 10 | 5.29% |
| hsa_circ_0111785 | 378 | 10 | 5.29% |
| hsa_circ_0123778 | 378 | 10 | 5.29% |
| hsa_circ_0043320 | 416 | 11 | 5.29% |
| hsa_circ_0084030 | 492 | 13 | 5.28% |
| hsa_circ_0051505 | 530 | 14 | 5.28% |
| hsa_circ_0014027 | 379 | 10 | 5.28% |
| hsa_circ_0015371 | 379 | 10 | 5.28% |
| hsa_circ_0029301 | 379 | 10 | 5.28% |
| hsa_circ_0029618 | 379 | 10 | 5.28% |
| hsa_circ_0043519 | 379 | 10 | 5.28% |
| hsa_circ_0057569 | 379 | 10 | 5.28% |
| hsa_circ_0070295 | 379 | 10 | 5.28% |
| hsa_circ_0087594 | 379 | 10 | 5.28% |
| hsa_circ_0089487 | 379 | 10 | 5.28% |
| hsa_circ_0106813 | 379 | 10 | 5.28% |
| hsa_circ_0015933 | 417 | 11 | 5.28% |
| hsa_circ_0016944 | 417 | 11 | 5.28% |
| hsa_circ_0026027 | 417 | 11 | 5.28% |
| hsa_circ_0076651 | 417 | 11 | 5.28% |
| hsa_circ_0095966 | 417 | 11 | 5.28% |
| hsa_circ_0102611 | 645 | 17 | 5.27% |
| hsa_circ_0118932 | 570 | 15 | 5.26% |
| hsa_circ_0133941 | 456 | 12 | 5.26% |
| hsa_circ_0045200 | 418 | 11 | 5.26% |
| hsa_circ_0089644 | 418 | 11 | 5.26% |
| hsa_circ_0004377 | 380 | 10 | 5.26% |
| hsa_circ_0010927 | 380 | 10 | 5.26% |
| hsa_circ_0019667 | 380 | 10 | 5.26% |
| hsa_circ_0019785 | 380 | 10 | 5.26% |
| hsa_circ_0023031 | 380 | 10 | 5.26% |
| hsa_circ_0054299 | 380 | 10 | 5.26% |
| hsa_circ_0066098 | 380 | 10 | 5.26% |
| hsa_circ_0077456 | 380 | 10 | 5.26% |
| hsa_circ_0098118 | 380 | 10 | 5.26% |
| hsa_circ_0103160 | 380 | 10 | 5.26% |
| hsa_circ_0122309 | 380 | 10 | 5.26% |
| hsa_circ_0009740 | 495 | 13 | 5.25% |
| hsa_circ_0026149 | 457 | 12 | 5.25% |
| hsa_circ_0005468 | 419 | 11 | 5.25% |
| hsa_circ_0028109 | 381 | 10 | 5.25% |
| hsa_circ_0042192 | 381 | 10 | 5.25% |
| hsa_circ_0043595 | 381 | 10 | 5.25% |
| hsa_circ_0104113 | 381 | 10 | 5.25% |

| | | | |
|---|---|---|---|
| hsa_circ_0106355 | 381 | 10 | 5.25% |
| hsa_circ_0092259 | 496 | 13 | 5.24% |
| hsa_circ_0140302 | 496 | 13 | 5.24% |
| hsa_circ_0004047 | 382 | 10 | 5.24% |
| hsa_circ_0022769 | 382 | 10 | 5.24% |
| hsa_circ_0066624 | 382 | 10 | 5.24% |
| hsa_circ_0104412 | 382 | 10 | 5.24% |
| hsa_circ_0120891 | 382 | 10 | 5.24% |
| hsa_circ_0129828 | 382 | 10 | 5.24% |
| hsa_circ_0137740 | 382 | 10 | 5.24% |
| hsa_circ_0140085 | 382 | 10 | 5.24% |
| hsa_circ_0016765 | 421 | 11 | 5.23% |
| hsa_circ_0109205 | 421 | 11 | 5.23% |
| hsa_circ_0088245 | 383 | 10 | 5.22% |
| hsa_circ_0099700 | 383 | 10 | 5.22% |
| hsa_circ_0128713 | 383 | 10 | 5.22% |
| hsa_circ_0108763 | 422 | 11 | 5.21% |
| hsa_circ_0120798 | 422 | 11 | 5.21% |
| hsa_circ_0122129 | 422 | 11 | 5.21% |
| hsa_circ_0133144 | 422 | 11 | 5.21% |
| hsa_circ_0003449 | 384 | 10 | 5.21% |
| hsa_circ_0008519 | 384 | 10 | 5.21% |
| hsa_circ_0023409 | 384 | 10 | 5.21% |
| hsa_circ_0039214 | 384 | 10 | 5.21% |
| hsa_circ_0067282 | 384 | 10 | 5.21% |
| hsa_circ_0070702 | 384 | 10 | 5.21% |
| hsa_circ_0073490 | 384 | 10 | 5.21% |
| hsa_circ_0096200 | 384 | 10 | 5.21% |
| hsa_circ_0098338 | 384 | 10 | 5.21% |
| hsa_circ_0117601 | 384 | 10 | 5.21% |
| hsa_circ_0037211 | 423 | 11 | 5.20% |
| hsa_circ_0114788 | 423 | 11 | 5.20% |
| hsa_circ_0115523 | 423 | 11 | 5.20% |
| hsa_circ_0119993 | 423 | 11 | 5.20% |
| hsa_circ_0003931 | 385 | 10 | 5.19% |
| hsa_circ_0008466 | 385 | 10 | 5.19% |
| hsa_circ_0038509 | 385 | 10 | 5.19% |
| hsa_circ_0040854 | 385 | 10 | 5.19% |
| hsa_circ_0044471 | 385 | 10 | 5.19% |
| hsa_circ_0056652 | 385 | 10 | 5.19% |
| hsa_circ_0082340 | 385 | 10 | 5.19% |
| hsa_circ_0082620 | 385 | 10 | 5.19% |
| hsa_circ_0096443 | 385 | 10 | 5.19% |
| hsa_circ_0111374 | 385 | 10 | 5.19% |
| hsa_circ_0111823 | 385 | 10 | 5.19% |
| hsa_circ_0134233 | 385 | 10 | 5.19% |
| hsa_circ_0122258 | 424 | 11 | 5.19% |
| hsa_circ_0128627 | 424 | 11 | 5.19% |
| hsa_circ_0128735 | 463 | 12 | 5.18% |
| hsa_circ_0001802 | 386 | 10 | 5.18% |

| | | | |
|---|---|---|---|
| hsa_circ_0011866 | 386 | 10 | 5.18% |
| hsa_circ_0011960 | 386 | 10 | 5.18% |
| hsa_circ_0026115 | 386 | 10 | 5.18% |
| hsa_circ_0070269 | 386 | 10 | 5.18% |
| hsa_circ_0077626 | 386 | 10 | 5.18% |
| hsa_circ_0099887 | 386 | 10 | 5.18% |
| hsa_circ_0103736 | 386 | 10 | 5.18% |
| hsa_circ_0138950 | 386 | 10 | 5.18% |
| hsa_circ_0008657 | 387 | 10 | 5.17% |
| hsa_circ_0014019 | 387 | 10 | 5.17% |
| hsa_circ_0025026 | 387 | 10 | 5.17% |
| hsa_circ_0034088 | 387 | 10 | 5.17% |
| hsa_circ_0055333 | 387 | 10 | 5.17% |
| hsa_circ_0078711 | 387 | 10 | 5.17% |
| hsa_circ_0100295 | 387 | 10 | 5.17% |
| hsa_circ_0116348 | 387 | 10 | 5.17% |
| hsa_circ_0029374 | 426 | 11 | 5.16% |
| hsa_circ_0117391 | 426 | 11 | 5.16% |
| hsa_circ_0016340 | 388 | 10 | 5.15% |
| hsa_circ_0039150 | 388 | 10 | 5.15% |
| hsa_circ_0046656 | 388 | 10 | 5.15% |
| hsa_circ_0068749 | 388 | 10 | 5.15% |
| hsa_circ_0075235 | 388 | 10 | 5.15% |
| hsa_circ_0075835 | 388 | 10 | 5.15% |
| hsa_circ_0103358 | 388 | 10 | 5.15% |
| hsa_circ_0105261 | 388 | 10 | 5.15% |
| hsa_circ_0116076 | 388 | 10 | 5.15% |
| hsa_circ_0095933 | 427 | 11 | 5.15% |
| hsa_circ_0116116 | 427 | 11 | 5.15% |
| hsa_circ_0118159 | 466 | 12 | 5.15% |
| hsa_circ_0004000 | 389 | 10 | 5.14% |
| hsa_circ_0034063 | 389 | 10 | 5.14% |
| hsa_circ_0072727 | 389 | 10 | 5.14% |
| hsa_circ_0079192 | 389 | 10 | 5.14% |
| hsa_circ_0116440 | 389 | 10 | 5.14% |
| hsa_circ_0122659 | 389 | 10 | 5.14% |
| hsa_circ_0130670 | 389 | 10 | 5.14% |
| hsa_circ_0057868 | 428 | 11 | 5.14% |
| hsa_circ_0008198 | 507 | 13 | 5.13% |
| hsa_circ_0088527 | 468 | 12 | 5.13% |
| hsa_circ_0126480 | 468 | 12 | 5.13% |
| hsa_circ_0057458 | 429 | 11 | 5.13% |
| hsa_circ_0079938 | 429 | 11 | 5.13% |
| hsa_circ_0000570 | 390 | 10 | 5.13% |
| hsa_circ_0012613 | 390 | 10 | 5.13% |
| hsa_circ_0018620 | 390 | 10 | 5.13% |
| hsa_circ_0037751 | 390 | 10 | 5.13% |
| hsa_circ_0051441 | 390 | 10 | 5.13% |
| hsa_circ_0068092 | 390 | 10 | 5.13% |
| hsa_circ_0096458 | 390 | 10 | 5.13% |

| | | | |
|---|---|---|---|
| hsa_circ_0122249 | 390 | 10 | 5.13% |
| hsa_circ_0136645 | 390 | 10 | 5.13% |
| hsa_circ_0140366 | 390 | 10 | 5.13% |
| hsa_circ_0103570 | 430 | 11 | 5.12% |
| hsa_circ_0045773 | 391 | 10 | 5.12% |
| hsa_circ_0050119 | 391 | 10 | 5.12% |
| hsa_circ_0079032 | 391 | 10 | 5.12% |
| hsa_circ_0083767 | 391 | 10 | 5.12% |
| hsa_circ_0107017 | 391 | 10 | 5.12% |
| hsa_circ_0119892 | 391 | 10 | 5.12% |
| hsa_circ_0120894 | 391 | 10 | 5.12% |
| hsa_circ_0128770 | 391 | 10 | 5.12% |
| hsa_circ_0135772 | 391 | 10 | 5.12% |
| hsa_circ_0011112 | 509 | 13 | 5.11% |
| hsa_circ_0116765 | 666 | 17 | 5.11% |
| hsa_circ_0005079 | 431 | 11 | 5.10% |
| hsa_circ_0055194 | 588 | 15 | 5.10% |
| hsa_circ_0003281 | 392 | 10 | 5.10% |
| hsa_circ_0042228 | 392 | 10 | 5.10% |
| hsa_circ_0062448 | 392 | 10 | 5.10% |
| hsa_circ_0094482 | 392 | 10 | 5.10% |
| hsa_circ_0106091 | 392 | 10 | 5.10% |
| hsa_circ_0107907 | 392 | 10 | 5.10% |
| hsa_circ_0116457 | 392 | 10 | 5.10% |
| hsa_circ_0138646 | 392 | 10 | 5.10% |
| hsa_circ_0138702 | 392 | 10 | 5.10% |
| hsa_circ_0116294 | 471 | 12 | 5.10% |
| hsa_circ_0068378 | 432 | 11 | 5.09% |
| hsa_circ_0093356 | 432 | 11 | 5.09% |
| hsa_circ_0021818 | 393 | 10 | 5.09% |
| hsa_circ_0029945 | 393 | 10 | 5.09% |
| hsa_circ_0041095 | 393 | 10 | 5.09% |
| hsa_circ_0065121 | 393 | 10 | 5.09% |
| hsa_circ_0068746 | 393 | 10 | 5.09% |
| hsa_circ_0094478 | 393 | 10 | 5.09% |
| hsa_circ_0128348 | 393 | 10 | 5.09% |
| hsa_circ_0027228 | 433 | 11 | 5.08% |
| hsa_circ_0099012 | 433 | 11 | 5.08% |
| hsa_circ_0108602 | 433 | 11 | 5.08% |
| hsa_circ_0000914 | 394 | 10 | 5.08% |
| hsa_circ_0006646 | 394 | 10 | 5.08% |
| hsa_circ_0023665 | 394 | 10 | 5.08% |
| hsa_circ_0038194 | 394 | 10 | 5.08% |
| hsa_circ_0050365 | 394 | 10 | 5.08% |
| hsa_circ_0068643 | 394 | 10 | 5.08% |
| hsa_circ_0074683 | 394 | 10 | 5.08% |
| hsa_circ_0085404 | 394 | 10 | 5.08% |
| hsa_circ_0087192 | 394 | 10 | 5.08% |
| hsa_circ_0133192 | 394 | 10 | 5.08% |
| hsa_circ_0067434 | 473 | 12 | 5.07% |

| | | | |
|---|---|---|---|
| hsa_circ_0136170 | 552 | 14 | 5.07% |
| hsa_circ_0001158 | 395 | 10 | 5.06% |
| hsa_circ_0006121 | 395 | 10 | 5.06% |
| hsa_circ_0031942 | 395 | 10 | 5.06% |
| hsa_circ_0031948 | 395 | 10 | 5.06% |
| hsa_circ_0043816 | 395 | 10 | 5.06% |
| hsa_circ_0073559 | 395 | 10 | 5.06% |
| hsa_circ_0080291 | 395 | 10 | 5.06% |
| hsa_circ_0081979 | 395 | 10 | 5.06% |
| hsa_circ_0082178 | 395 | 10 | 5.06% |
| hsa_circ_0093888 | 395 | 10 | 5.06% |
| hsa_circ_0108859 | 395 | 10 | 5.06% |
| hsa_circ_0116856 | 395 | 10 | 5.06% |
| hsa_circ_0015102 | 435 | 11 | 5.06% |
| hsa_circ_0098265 | 435 | 11 | 5.06% |
| hsa_circ_0129760 | 435 | 11 | 5.06% |
| hsa_circ_0000266 | 396 | 10 | 5.05% |
| hsa_circ_0056596 | 396 | 10 | 5.05% |
| hsa_circ_0092411 | 396 | 10 | 5.05% |
| hsa_circ_0093634 | 396 | 10 | 5.05% |
| hsa_circ_0094520 | 396 | 10 | 5.05% |
| hsa_circ_0095690 | 396 | 10 | 5.05% |
| hsa_circ_0096095 | 396 | 10 | 5.05% |
| hsa_circ_0117237 | 396 | 10 | 5.05% |
| hsa_circ_0121713 | 396 | 10 | 5.05% |
| hsa_circ_0123362 | 396 | 10 | 5.05% |
| hsa_circ_0127597 | 396 | 10 | 5.05% |
| hsa_circ_0135987 | 396 | 10 | 5.05% |
| hsa_circ_0097630 | 436 | 11 | 5.05% |
| hsa_circ_0110567 | 436 | 11 | 5.05% |
| hsa_circ_0085819 | 674 | 17 | 5.04% |
| hsa_circ_0098849 | 476 | 12 | 5.04% |
| hsa_circ_0055632 | 1468 | 37 | 5.04% |
| hsa_circ_0006701 | 397 | 10 | 5.04% |
| hsa_circ_0038939 | 397 | 10 | 5.04% |
| hsa_circ_0099586 | 397 | 10 | 5.04% |
| hsa_circ_0066428 | 437 | 11 | 5.03% |
| hsa_circ_0093505 | 437 | 11 | 5.03% |
| hsa_circ_0133113 | 437 | 11 | 5.03% |
| hsa_circ_0016452 | 477 | 12 | 5.03% |
| hsa_circ_0020078 | 477 | 12 | 5.03% |
| hsa_circ_0012052 | 398 | 10 | 5.03% |
| hsa_circ_0025797 | 398 | 10 | 5.03% |
| hsa_circ_0048199 | 398 | 10 | 5.03% |
| hsa_circ_0113979 | 398 | 10 | 5.03% |
| hsa_circ_0126554 | 398 | 10 | 5.03% |
| hsa_circ_0139056 | 398 | 10 | 5.03% |
| hsa_circ_0015410 | 438 | 11 | 5.02% |
| hsa_circ_0020964 | 438 | 11 | 5.02% |
| hsa_circ_0013011 | 478 | 12 | 5.02% |

| | | | |
|---|---|---|---|
| hsa_circ_0003191 | 558 | 14 | 5.02% |
| hsa_circ_0003925 | 399 | 10 | 5.01% |
| hsa_circ_0004940 | 399 | 10 | 5.01% |
| hsa_circ_0024147 | 399 | 10 | 5.01% |
| hsa_circ_0039157 | 399 | 10 | 5.01% |
| hsa_circ_0066966 | 399 | 10 | 5.01% |
| hsa_circ_0071756 | 399 | 10 | 5.01% |
| hsa_circ_0104881 | 399 | 10 | 5.01% |
| hsa_circ_0122350 | 399 | 10 | 5.01% |
| hsa_circ_0130728 | 399 | 10 | 5.01% |
| hsa_circ_0036240 | 439 | 11 | 5.01% |
| hsa_circ_0075073 | 439 | 11 | 5.01% |
| hsa_circ_0001608 | 440 | 11 | 5.00% |
| hsa_circ_0006848 | 400 | 10 | 5.00% |
| hsa_circ_0018145 | 400 | 10 | 5.00% |
| hsa_circ_0119246 | 400 | 10 | 5.00% |
| hsa_circ_0138758 | 400 | 10 | 5.00% |
| hsa_circ_0010634 | 441 | 11 | 4.99% |
| hsa_circ_0039467 | 441 | 11 | 4.99% |
| hsa_circ_0099137 | 441 | 11 | 4.99% |
| hsa_circ_0009190 | 401 | 10 | 4.99% |
| hsa_circ_0012331 | 401 | 10 | 4.99% |
| hsa_circ_0012904 | 401 | 10 | 4.99% |
| hsa_circ_0015686 | 401 | 10 | 4.99% |
| hsa_circ_0055691 | 401 | 10 | 4.99% |
| hsa_circ_0056362 | 401 | 10 | 4.99% |
| hsa_circ_0065735 | 401 | 10 | 4.99% |
| hsa_circ_0091420 | 401 | 10 | 4.99% |
| hsa_circ_0095059 | 401 | 10 | 4.99% |
| hsa_circ_0095296 | 401 | 10 | 4.99% |
| hsa_circ_0118784 | 401 | 10 | 4.99% |
| hsa_circ_0117541 | 522 | 13 | 4.98% |
| hsa_circ_0001801 | 402 | 10 | 4.98% |
| hsa_circ_0001917 | 402 | 10 | 4.98% |
| hsa_circ_0031655 | 402 | 10 | 4.98% |
| hsa_circ_0038505 | 402 | 10 | 4.98% |
| hsa_circ_0042511 | 402 | 10 | 4.98% |
| hsa_circ_0083246 | 402 | 10 | 4.98% |
| hsa_circ_0090086 | 402 | 10 | 4.98% |
| hsa_circ_0092448 | 402 | 10 | 4.98% |
| hsa_circ_0093650 | 402 | 10 | 4.98% |
| hsa_circ_0101507 | 402 | 10 | 4.98% |
| hsa_circ_0109256 | 402 | 10 | 4.98% |
| hsa_circ_0112608 | 402 | 10 | 4.98% |
| hsa_circ_0125382 | 402 | 10 | 4.98% |
| hsa_circ_0131942 | 402 | 10 | 4.98% |
| hsa_circ_0137005 | 402 | 10 | 4.98% |
| hsa_circ_0106418 | 1329 | 33 | 4.97% |
| hsa_circ_0021347 | 443 | 11 | 4.97% |
| hsa_circ_0006310 | 403 | 10 | 4.96% |

| | | | |
|---|---|---|---|
| hsa_circ_0008566 | 403 | 10 | 4.96% |
| hsa_circ_0011506 | 403 | 10 | 4.96% |
| hsa_circ_0023384 | 403 | 10 | 4.96% |
| hsa_circ_0089916 | 403 | 10 | 4.96% |
| hsa_circ_0107245 | 403 | 10 | 4.96% |
| hsa_circ_0123792 | 403 | 10 | 4.96% |
| hsa_circ_0119572 | 484 | 12 | 4.96% |
| hsa_circ_0012960 | 444 | 11 | 4.95% |
| hsa_circ_0066460 | 444 | 11 | 4.95% |
| hsa_circ_0043455 | 525 | 13 | 4.95% |
| hsa_circ_0001316 | 404 | 10 | 4.95% |
| hsa_circ_0003522 | 404 | 10 | 4.95% |
| hsa_circ_0058652 | 404 | 10 | 4.95% |
| hsa_circ_0059122 | 404 | 10 | 4.95% |
| hsa_circ_0072581 | 404 | 10 | 4.95% |
| hsa_circ_0094724 | 404 | 10 | 4.95% |
| hsa_circ_0101602 | 404 | 10 | 4.95% |
| hsa_circ_0113260 | 404 | 10 | 4.95% |
| hsa_circ_0134215 | 404 | 10 | 4.95% |
| hsa_circ_0134473 | 404 | 10 | 4.95% |
| hsa_circ_0070666 | 485 | 12 | 4.95% |
| hsa_circ_0095789 | 486 | 12 | 4.94% |
| hsa_circ_0109923 | 486 | 12 | 4.94% |
| hsa_circ_0001371 | 405 | 10 | 4.94% |
| hsa_circ_0006242 | 405 | 10 | 4.94% |
| hsa_circ_0065409 | 405 | 10 | 4.94% |
| hsa_circ_0087581 | 405 | 10 | 4.94% |
| hsa_circ_0087857 | 405 | 10 | 4.94% |
| hsa_circ_0101864 | 405 | 10 | 4.94% |
| hsa_circ_0109605 | 405 | 10 | 4.94% |
| hsa_circ_0120576 | 405 | 10 | 4.94% |
| hsa_circ_0123687 | 405 | 10 | 4.94% |
| hsa_circ_0007412 | 446 | 11 | 4.93% |
| hsa_circ_0031299 | 446 | 11 | 4.93% |
| hsa_circ_0071018 | 446 | 11 | 4.93% |
| hsa_circ_0047127 | 487 | 12 | 4.93% |
| hsa_circ_0020935 | 406 | 10 | 4.93% |
| hsa_circ_0026946 | 406 | 10 | 4.93% |
| hsa_circ_0027674 | 406 | 10 | 4.93% |
| hsa_circ_0044049 | 406 | 10 | 4.93% |
| hsa_circ_0050127 | 406 | 10 | 4.93% |
| hsa_circ_0081636 | 406 | 10 | 4.93% |
| hsa_circ_0086780 | 406 | 10 | 4.93% |
| hsa_circ_0116745 | 406 | 10 | 4.93% |
| hsa_circ_0135562 | 406 | 10 | 4.93% |
| hsa_circ_0002499 | 528 | 13 | 4.92% |
| hsa_circ_0084762 | 447 | 11 | 4.92% |
| hsa_circ_0129630 | 447 | 11 | 4.92% |
| hsa_circ_0009381 | 407 | 10 | 4.91% |
| hsa_circ_0015026 | 407 | 10 | 4.91% |

| | | | |
|---|---|---|---|
| hsa_circ_0018764 | 407 | 10 | 4.91% |
| hsa_circ_0084509 | 407 | 10 | 4.91% |
| hsa_circ_0121859 | 407 | 10 | 4.91% |
| hsa_circ_0006138 | 448 | 11 | 4.91% |
| hsa_circ_0122465 | 448 | 11 | 4.91% |
| hsa_circ_0080358 | 530 | 13 | 4.91% |
| hsa_circ_0113956 | 530 | 13 | 4.91% |
| hsa_circ_0011108 | 408 | 10 | 4.90% |
| hsa_circ_0036541 | 408 | 10 | 4.90% |
| hsa_circ_0055739 | 408 | 10 | 4.90% |
| hsa_circ_0056498 | 408 | 10 | 4.90% |
| hsa_circ_0100902 | 408 | 10 | 4.90% |
| hsa_circ_0102601 | 408 | 10 | 4.90% |
| hsa_circ_0104564 | 408 | 10 | 4.90% |
| hsa_circ_0112332 | 408 | 10 | 4.90% |
| hsa_circ_0130865 | 408 | 10 | 4.90% |
| hsa_circ_0040833 | 449 | 11 | 4.90% |
| hsa_circ_0095931 | 449 | 11 | 4.90% |
| hsa_circ_0138225 | 449 | 11 | 4.90% |
| hsa_circ_0102841 | 531 | 13 | 4.90% |
| hsa_circ_0132405 | 572 | 14 | 4.90% |
| hsa_circ_0005389 | 409 | 10 | 4.89% |
| hsa_circ_0045025 | 409 | 10 | 4.89% |
| hsa_circ_0100985 | 409 | 10 | 4.89% |
| hsa_circ_0106219 | 409 | 10 | 4.89% |
| hsa_circ_0117604 | 409 | 10 | 4.89% |
| hsa_circ_0038048 | 450 | 11 | 4.89% |
| hsa_circ_0051088 | 450 | 11 | 4.89% |
| hsa_circ_0096436 | 450 | 11 | 4.89% |
| hsa_circ_0104057 | 491 | 12 | 4.89% |
| hsa_circ_0130247 | 573 | 14 | 4.89% |
| hsa_circ_0088710 | 696 | 17 | 4.89% |
| hsa_circ_0060048 | 820 | 20 | 4.88% |
| hsa_circ_0102683 | 492 | 12 | 4.88% |
| hsa_circ_0055377 | 451 | 11 | 4.88% |
| hsa_circ_0008095 | 410 | 10 | 4.88% |
| hsa_circ_0059372 | 410 | 10 | 4.88% |
| hsa_circ_0079470 | 410 | 10 | 4.88% |
| hsa_circ_0083666 | 410 | 10 | 4.88% |
| hsa_circ_0094722 | 410 | 10 | 4.88% |
| hsa_circ_0103928 | 410 | 10 | 4.88% |
| hsa_circ_0105264 | 410 | 10 | 4.88% |
| hsa_circ_0117468 | 410 | 10 | 4.88% |
| hsa_circ_0121860 | 410 | 10 | 4.88% |
| hsa_circ_0128429 | 410 | 10 | 4.88% |
| hsa_circ_0137417 | 410 | 10 | 4.88% |
| hsa_circ_0111244 | 534 | 13 | 4.87% |
| hsa_circ_0034343 | 452 | 11 | 4.87% |
| hsa_circ_0055612 | 452 | 11 | 4.87% |
| hsa_circ_0074212 | 452 | 11 | 4.87% |

| | | | |
|---|---|---|---|
| hsa_circ_0077040 | 411 | 10 | 4.87% |
| hsa_circ_0084902 | 411 | 10 | 4.87% |
| hsa_circ_0097125 | 411 | 10 | 4.87% |
| hsa_circ_0108094 | 411 | 10 | 4.87% |
| hsa_circ_0113707 | 411 | 10 | 4.87% |
| hsa_circ_0041990 | 535 | 13 | 4.86% |
| hsa_circ_0039104 | 494 | 12 | 4.86% |
| hsa_circ_0028308 | 453 | 11 | 4.86% |
| hsa_circ_0051801 | 453 | 11 | 4.86% |
| hsa_circ_0105069 | 453 | 11 | 4.86% |
| hsa_circ_0125383 | 453 | 11 | 4.86% |
| hsa_circ_0136460 | 453 | 11 | 4.86% |
| hsa_circ_0139925 | 453 | 11 | 4.86% |
| hsa_circ_0016093 | 412 | 10 | 4.85% |
| hsa_circ_0039308 | 412 | 10 | 4.85% |
| hsa_circ_0106585 | 412 | 10 | 4.85% |
| hsa_circ_0010411 | 454 | 11 | 4.85% |
| hsa_circ_0023426 | 413 | 10 | 4.84% |
| hsa_circ_0034745 | 413 | 10 | 4.84% |
| hsa_circ_0049910 | 413 | 10 | 4.84% |
| hsa_circ_0050890 | 413 | 10 | 4.84% |
| hsa_circ_0060302 | 413 | 10 | 4.84% |
| hsa_circ_0062762 | 413 | 10 | 4.84% |
| hsa_circ_0080501 | 413 | 10 | 4.84% |
| hsa_circ_0127218 | 413 | 10 | 4.84% |
| hsa_circ_0036992 | 579 | 14 | 4.84% |
| hsa_circ_0093747 | 455 | 11 | 4.84% |
| hsa_circ_0109979 | 455 | 11 | 4.84% |
| hsa_circ_0125165 | 455 | 11 | 4.84% |
| hsa_circ_0130173 | 455 | 11 | 4.84% |
| hsa_circ_0018347 | 414 | 10 | 4.83% |
| hsa_circ_0023387 | 414 | 10 | 4.83% |
| hsa_circ_0024287 | 414 | 10 | 4.83% |
| hsa_circ_0032853 | 414 | 10 | 4.83% |
| hsa_circ_0036723 | 414 | 10 | 4.83% |
| hsa_circ_0057592 | 414 | 10 | 4.83% |
| hsa_circ_0059912 | 414 | 10 | 4.83% |
| hsa_circ_0061660 | 414 | 10 | 4.83% |
| hsa_circ_0100380 | 414 | 10 | 4.83% |
| hsa_circ_0101013 | 414 | 10 | 4.83% |
| hsa_circ_0127340 | 414 | 10 | 4.83% |
| hsa_circ_0051581 | 456 | 11 | 4.82% |
| hsa_circ_0059284 | 456 | 11 | 4.82% |
| hsa_circ_0087835 | 456 | 11 | 4.82% |
| hsa_circ_0092489 | 456 | 11 | 4.82% |
| hsa_circ_0090575 | 539 | 13 | 4.82% |
| hsa_circ_0002791 | 1120 | 27 | 4.82% |
| hsa_circ_0066112 | 498 | 12 | 4.82% |
| hsa_circ_0134419 | 498 | 12 | 4.82% |
| hsa_circ_0000255 | 415 | 10 | 4.82% |

| | | | |
|---|---|---|---|
| hsa_circ_0033516 | 415 | 10 | 4.82% |
| hsa_circ_0056240 | 415 | 10 | 4.82% |
| hsa_circ_0079118 | 415 | 10 | 4.82% |
| hsa_circ_0080915 | 415 | 10 | 4.82% |
| hsa_circ_0098946 | 415 | 10 | 4.82% |
| hsa_circ_0106838 | 415 | 10 | 4.82% |
| hsa_circ_0107722 | 415 | 10 | 4.82% |
| hsa_circ_0115283 | 415 | 10 | 4.82% |
| hsa_circ_0130672 | 415 | 10 | 4.82% |
| hsa_circ_0131755 | 415 | 10 | 4.82% |
| hsa_circ_0139348 | 415 | 10 | 4.82% |
| hsa_circ_0002954 | 457 | 11 | 4.81% |
| hsa_circ_0002795 | 416 | 10 | 4.81% |
| hsa_circ_0030008 | 416 | 10 | 4.81% |
| hsa_circ_0051576 | 416 | 10 | 4.81% |
| hsa_circ_0065470 | 416 | 10 | 4.81% |
| hsa_circ_0079492 | 416 | 10 | 4.81% |
| hsa_circ_0080457 | 416 | 10 | 4.81% |
| hsa_circ_0082460 | 416 | 10 | 4.81% |
| hsa_circ_0133560 | 416 | 10 | 4.81% |
| hsa_circ_0134169 | 416 | 10 | 4.81% |
| hsa_circ_0004787 | 458 | 11 | 4.80% |
| hsa_circ_0036693 | 458 | 11 | 4.80% |
| hsa_circ_0066212 | 458 | 11 | 4.80% |
| hsa_circ_0086032 | 458 | 11 | 4.80% |
| hsa_circ_0133595 | 458 | 11 | 4.80% |
| hsa_circ_0119611 | 625 | 15 | 4.80% |
| hsa_circ_0072105 | 500 | 12 | 4.80% |
| hsa_circ_0011259 | 417 | 10 | 4.80% |
| hsa_circ_0012155 | 417 | 10 | 4.80% |
| hsa_circ_0023233 | 417 | 10 | 4.80% |
| hsa_circ_0034449 | 417 | 10 | 4.80% |
| hsa_circ_0059185 | 417 | 10 | 4.80% |
| hsa_circ_0079356 | 417 | 10 | 4.80% |
| hsa_circ_0095152 | 417 | 10 | 4.80% |
| hsa_circ_0096055 | 417 | 10 | 4.80% |
| hsa_circ_0097560 | 417 | 10 | 4.80% |
| hsa_circ_0117434 | 417 | 10 | 4.80% |
| hsa_circ_0131752 | 417 | 10 | 4.80% |
| hsa_circ_0132458 | 417 | 10 | 4.80% |
| hsa_circ_0123060 | 459 | 11 | 4.79% |
| hsa_circ_0000352 | 418 | 10 | 4.78% |
| hsa_circ_0000411 | 418 | 10 | 4.78% |
| hsa_circ_0001701 | 418 | 10 | 4.78% |
| hsa_circ_0005231 | 418 | 10 | 4.78% |
| hsa_circ_0042848 | 418 | 10 | 4.78% |
| hsa_circ_0079239 | 418 | 10 | 4.78% |
| hsa_circ_0106890 | 418 | 10 | 4.78% |
| hsa_circ_0107468 | 418 | 10 | 4.78% |
| hsa_circ_0109965 | 418 | 10 | 4.78% |

| | | | |
|---|---|---|---|
| hsa_circ_0010368 | 419 | 10 | 4.77% |
| hsa_circ_0069148 | 419 | 10 | 4.77% |
| hsa_circ_0094224 | 419 | 10 | 4.77% |
| hsa_circ_0096849 | 419 | 10 | 4.77% |
| hsa_circ_0120382 | 419 | 10 | 4.77% |
| hsa_circ_0130738 | 419 | 10 | 4.77% |
| hsa_circ_0006733 | 461 | 11 | 4.77% |
| hsa_circ_0061415 | 461 | 11 | 4.77% |
| hsa_circ_0079673 | 461 | 11 | 4.77% |
| hsa_circ_0105741 | 461 | 11 | 4.77% |
| hsa_circ_0132980 | 461 | 11 | 4.77% |
| hsa_circ_0134441 | 461 | 11 | 4.77% |
| hsa_circ_0015980 | 630 | 15 | 4.76% |
| hsa_circ_0033148 | 504 | 12 | 4.76% |
| hsa_circ_0138256 | 504 | 12 | 4.76% |
| hsa_circ_0002928 | 462 | 11 | 4.76% |
| hsa_circ_0007168 | 462 | 11 | 4.76% |
| hsa_circ_0011076 | 462 | 11 | 4.76% |
| hsa_circ_0085481 | 462 | 11 | 4.76% |
| hsa_circ_0120067 | 462 | 11 | 4.76% |
| hsa_circ_0126283 | 462 | 11 | 4.76% |
| hsa_circ_0135118 | 462 | 11 | 4.76% |
| hsa_circ_0000923 | 420 | 10 | 4.76% |
| hsa_circ_0034304 | 420 | 10 | 4.76% |
| hsa_circ_0042098 | 420 | 10 | 4.76% |
| hsa_circ_0074756 | 420 | 10 | 4.76% |
| hsa_circ_0080428 | 420 | 10 | 4.76% |
| hsa_circ_0092303 | 420 | 10 | 4.76% |
| hsa_circ_0092305 | 420 | 10 | 4.76% |
| hsa_circ_0092479 | 420 | 10 | 4.76% |
| hsa_circ_0101547 | 420 | 10 | 4.76% |
| hsa_circ_0104725 | 420 | 10 | 4.76% |
| hsa_circ_0109300 | 420 | 10 | 4.76% |
| hsa_circ_0111247 | 420 | 10 | 4.76% |
| hsa_circ_0118892 | 420 | 10 | 4.76% |
| hsa_circ_0121559 | 420 | 10 | 4.76% |
| hsa_circ_0129443 | 505 | 12 | 4.75% |
| hsa_circ_0003513 | 463 | 11 | 4.75% |
| hsa_circ_0008514 | 463 | 11 | 4.75% |
| hsa_circ_0013506 | 463 | 11 | 4.75% |
| hsa_circ_0116368 | 463 | 11 | 4.75% |
| hsa_circ_0071337 | 421 | 10 | 4.75% |
| hsa_circ_0078146 | 421 | 10 | 4.75% |
| hsa_circ_0096596 | 421 | 10 | 4.75% |
| hsa_circ_0099747 | 421 | 10 | 4.75% |
| hsa_circ_0112737 | 421 | 10 | 4.75% |
| hsa_circ_0125286 | 421 | 10 | 4.75% |
| hsa_circ_0132514 | 421 | 10 | 4.75% |
| hsa_circ_0135577 | 421 | 10 | 4.75% |
| hsa_circ_0055786 | 548 | 13 | 4.74% |

| | | | |
|---|---|---|---|
| hsa_circ_0131808 | 548 | 13 | 4.74% |
| hsa_circ_0043276 | 506 | 12 | 4.74% |
| hsa_circ_0020701 | 464 | 11 | 4.74% |
| hsa_circ_0054906 | 464 | 11 | 4.74% |
| hsa_circ_0063200 | 464 | 11 | 4.74% |
| hsa_circ_0114258 | 464 | 11 | 4.74% |
| hsa_circ_0007567 | 422 | 10 | 4.74% |
| hsa_circ_0013540 | 422 | 10 | 4.74% |
| hsa_circ_0067451 | 422 | 10 | 4.74% |
| hsa_circ_0103809 | 422 | 10 | 4.74% |
| hsa_circ_0115032 | 422 | 10 | 4.74% |
| hsa_circ_0048069 | 591 | 14 | 4.74% |
| hsa_circ_0026150 | 507 | 12 | 4.73% |
| hsa_circ_0010985 | 465 | 11 | 4.73% |
| hsa_circ_0040310 | 465 | 11 | 4.73% |
| hsa_circ_0074648 | 465 | 11 | 4.73% |
| hsa_circ_0138043 | 465 | 11 | 4.73% |
| hsa_circ_0006288 | 423 | 10 | 4.73% |
| hsa_circ_0007298 | 423 | 10 | 4.73% |
| hsa_circ_0007660 | 423 | 10 | 4.73% |
| hsa_circ_0008370 | 423 | 10 | 4.73% |
| hsa_circ_0021071 | 423 | 10 | 4.73% |
| hsa_circ_0034779 | 423 | 10 | 4.73% |
| hsa_circ_0036259 | 423 | 10 | 4.73% |
| hsa_circ_0046979 | 423 | 10 | 4.73% |
| hsa_circ_0051807 | 423 | 10 | 4.73% |
| hsa_circ_0076060 | 423 | 10 | 4.73% |
| hsa_circ_0081752 | 423 | 10 | 4.73% |
| hsa_circ_0005656 | 466 | 11 | 4.72% |
| hsa_circ_0063471 | 466 | 11 | 4.72% |
| hsa_circ_0075248 | 466 | 11 | 4.72% |
| hsa_circ_0023053 | 551 | 13 | 4.72% |
| hsa_circ_0007193 | 424 | 10 | 4.72% |
| hsa_circ_0008786 | 424 | 10 | 4.72% |
| hsa_circ_0021240 | 424 | 10 | 4.72% |
| hsa_circ_0044570 | 424 | 10 | 4.72% |
| hsa_circ_0045108 | 424 | 10 | 4.72% |
| hsa_circ_0045506 | 424 | 10 | 4.72% |
| hsa_circ_0068641 | 424 | 10 | 4.72% |
| hsa_circ_0078937 | 424 | 10 | 4.72% |
| hsa_circ_0054797 | 509 | 12 | 4.72% |
| hsa_circ_0037839 | 467 | 11 | 4.71% |
| hsa_circ_0051753 | 467 | 11 | 4.71% |
| hsa_circ_0063172 | 467 | 11 | 4.71% |
| hsa_circ_0067291 | 467 | 11 | 4.71% |
| hsa_circ_0117022 | 510 | 12 | 4.71% |
| hsa_circ_0129578 | 510 | 12 | 4.71% |
| hsa_circ_0000957 | 425 | 10 | 4.71% |
| hsa_circ_0017835 | 425 | 10 | 4.71% |
| hsa_circ_0021764 | 425 | 10 | 4.71% |

| | | | |
|---|---|---|---|
| hsa_circ_0023719 | 425 | 10 | 4.71% |
| hsa_circ_0037509 | 425 | 10 | 4.71% |
| hsa_circ_0062304 | 425 | 10 | 4.71% |
| hsa_circ_0070655 | 425 | 10 | 4.71% |
| hsa_circ_0082799 | 425 | 10 | 4.71% |
| hsa_circ_0127706 | 425 | 10 | 4.71% |
| hsa_circ_0134644 | 425 | 10 | 4.71% |
| hsa_circ_0135578 | 425 | 10 | 4.71% |
| hsa_circ_0092691 | 553 | 13 | 4.70% |
| hsa_circ_0036781 | 468 | 11 | 4.70% |
| hsa_circ_0096785 | 468 | 11 | 4.70% |
| hsa_circ_0128897 | 468 | 11 | 4.70% |
| hsa_circ_0008052 | 426 | 10 | 4.69% |
| hsa_circ_0013510 | 426 | 10 | 4.69% |
| hsa_circ_0014643 | 426 | 10 | 4.69% |
| hsa_circ_0017078 | 426 | 10 | 4.69% |
| hsa_circ_0034031 | 426 | 10 | 4.69% |
| hsa_circ_0035034 | 426 | 10 | 4.69% |
| hsa_circ_0078261 | 426 | 10 | 4.69% |
| hsa_circ_0082288 | 426 | 10 | 4.69% |
| hsa_circ_0099519 | 426 | 10 | 4.69% |
| hsa_circ_0111175 | 426 | 10 | 4.69% |
| hsa_circ_0020533 | 554 | 13 | 4.69% |
| hsa_circ_0050869 | 469 | 11 | 4.69% |
| hsa_circ_0107395 | 469 | 11 | 4.69% |
| hsa_circ_0118819 | 469 | 11 | 4.69% |
| hsa_circ_0021816 | 427 | 10 | 4.68% |
| hsa_circ_0044800 | 427 | 10 | 4.68% |
| hsa_circ_0050395 | 427 | 10 | 4.68% |
| hsa_circ_0052288 | 427 | 10 | 4.68% |
| hsa_circ_0054646 | 427 | 10 | 4.68% |
| hsa_circ_0082151 | 427 | 10 | 4.68% |
| hsa_circ_0093079 | 427 | 10 | 4.68% |
| hsa_circ_0099163 | 427 | 10 | 4.68% |
| hsa_circ_0105197 | 427 | 10 | 4.68% |
| hsa_circ_0105363 | 427 | 10 | 4.68% |
| hsa_circ_0108515 | 427 | 10 | 4.68% |
| hsa_circ_0112545 | 427 | 10 | 4.68% |
| hsa_circ_0126432 | 427 | 10 | 4.68% |
| hsa_circ_0032873 | 470 | 11 | 4.68% |
| hsa_circ_0077816 | 470 | 11 | 4.68% |
| hsa_circ_0133899 | 470 | 11 | 4.68% |
| hsa_circ_0044482 | 513 | 12 | 4.68% |
| hsa_circ_0023614 | 556 | 13 | 4.68% |
| hsa_circ_0086132 | 599 | 14 | 4.67% |
| hsa_circ_0053036 | 428 | 10 | 4.67% |
| hsa_circ_0057465 | 428 | 10 | 4.67% |
| hsa_circ_0064362 | 428 | 10 | 4.67% |
| hsa_circ_0079524 | 428 | 10 | 4.67% |
| hsa_circ_0097898 | 428 | 10 | 4.67% |

| | | | |
|---|---|---|---|
| hsa_circ_0111142 | 428 | 10 | 4.67% |
| hsa_circ_0118194 | 428 | 10 | 4.67% |
| hsa_circ_0012177 | 471 | 11 | 4.67% |
| hsa_circ_0105326 | 471 | 11 | 4.67% |
| hsa_circ_0048489 | 514 | 12 | 4.67% |
| hsa_circ_0093882 | 514 | 12 | 4.67% |
| hsa_circ_0113473 | 514 | 12 | 4.67% |
| hsa_circ_0027831 | 429 | 10 | 4.66% |
| hsa_circ_0051124 | 429 | 10 | 4.66% |
| hsa_circ_0064882 | 429 | 10 | 4.66% |
| hsa_circ_0065117 | 429 | 10 | 4.66% |
| hsa_circ_0069612 | 429 | 10 | 4.66% |
| hsa_circ_0074428 | 429 | 10 | 4.66% |
| hsa_circ_0081568 | 429 | 10 | 4.66% |
| hsa_circ_0093275 | 429 | 10 | 4.66% |
| hsa_circ_0097974 | 429 | 10 | 4.66% |
| hsa_circ_0098668 | 429 | 10 | 4.66% |
| hsa_circ_0124321 | 429 | 10 | 4.66% |
| hsa_circ_0127021 | 429 | 10 | 4.66% |
| hsa_circ_0130442 | 429 | 10 | 4.66% |
| hsa_circ_0006590 | 472 | 11 | 4.66% |
| hsa_circ_0027683 | 472 | 11 | 4.66% |
| hsa_circ_0074896 | 472 | 11 | 4.66% |
| hsa_circ_0120565 | 472 | 11 | 4.66% |
| hsa_circ_0122130 | 472 | 11 | 4.66% |
| hsa_circ_0060640 | 515 | 12 | 4.66% |
| hsa_circ_0000705 | 559 | 13 | 4.65% |
| hsa_circ_0059446 | 430 | 10 | 4.65% |
| hsa_circ_0104108 | 430 | 10 | 4.65% |
| hsa_circ_0107061 | 430 | 10 | 4.65% |
| hsa_circ_0115295 | 430 | 10 | 4.65% |
| hsa_circ_0122164 | 430 | 10 | 4.65% |
| hsa_circ_0136746 | 430 | 10 | 4.65% |
| hsa_circ_0136830 | 517 | 12 | 4.64% |
| hsa_circ_0098448 | 474 | 11 | 4.64% |
| hsa_circ_0042759 | 431 | 10 | 4.64% |
| hsa_circ_0046216 | 431 | 10 | 4.64% |
| hsa_circ_0077711 | 431 | 10 | 4.64% |
| hsa_circ_0087855 | 431 | 10 | 4.64% |
| hsa_circ_0101119 | 431 | 10 | 4.64% |
| hsa_circ_0118624 | 431 | 10 | 4.64% |
| hsa_circ_0129952 | 431 | 10 | 4.64% |
| hsa_circ_0030613 | 561 | 13 | 4.63% |
| hsa_circ_0139339 | 561 | 13 | 4.63% |
| hsa_circ_0017194 | 518 | 12 | 4.63% |
| hsa_circ_0087081 | 518 | 12 | 4.63% |
| hsa_circ_0068144 | 475 | 11 | 4.63% |
| hsa_circ_0069901 | 475 | 11 | 4.63% |
| hsa_circ_0138202 | 1166 | 27 | 4.63% |
| hsa_circ_0001806 | 432 | 10 | 4.63% |

| | | | |
|---|---|---|---|
| hsa_circ_0027020 | 432 | 10 | 4.63% |
| hsa_circ_0034700 | 432 | 10 | 4.63% |
| hsa_circ_0075764 | 432 | 10 | 4.63% |
| hsa_circ_0098199 | 432 | 10 | 4.63% |
| hsa_circ_0103361 | 432 | 10 | 4.63% |
| hsa_circ_0127388 | 432 | 10 | 4.63% |
| hsa_circ_0047789 | 519 | 12 | 4.62% |
| hsa_circ_0120440 | 519 | 12 | 4.62% |
| hsa_circ_0010756 | 476 | 11 | 4.62% |
| hsa_circ_0059589 | 476 | 11 | 4.62% |
| hsa_circ_0113104 | 476 | 11 | 4.62% |
| hsa_circ_0005921 | 433 | 10 | 4.62% |
| hsa_circ_0010126 | 433 | 10 | 4.62% |
| hsa_circ_0040224 | 433 | 10 | 4.62% |
| hsa_circ_0050394 | 433 | 10 | 4.62% |
| hsa_circ_0061011 | 433 | 10 | 4.62% |
| hsa_circ_0062228 | 433 | 10 | 4.62% |
| hsa_circ_0072492 | 433 | 10 | 4.62% |
| hsa_circ_0083716 | 433 | 10 | 4.62% |
| hsa_circ_0097459 | 433 | 10 | 4.62% |
| hsa_circ_0100193 | 433 | 10 | 4.62% |
| hsa_circ_0113939 | 433 | 10 | 4.62% |
| hsa_circ_0118963 | 433 | 10 | 4.62% |
| hsa_circ_0137692 | 433 | 10 | 4.62% |
| hsa_circ_0023532 | 520 | 12 | 4.62% |
| hsa_circ_0103173 | 520 | 12 | 4.62% |
| hsa_circ_0117021 | 520 | 12 | 4.62% |
| hsa_circ_0039056 | 477 | 11 | 4.61% |
| hsa_circ_0115126 | 477 | 11 | 4.61% |
| hsa_circ_0001338 | 434 | 10 | 4.61% |
| hsa_circ_0001400 | 434 | 10 | 4.61% |
| hsa_circ_0006311 | 434 | 10 | 4.61% |
| hsa_circ_0008297 | 434 | 10 | 4.61% |
| hsa_circ_0054213 | 434 | 10 | 4.61% |
| hsa_circ_0067711 | 434 | 10 | 4.61% |
| hsa_circ_0087219 | 434 | 10 | 4.61% |
| hsa_circ_0100397 | 434 | 10 | 4.61% |
| hsa_circ_0100448 | 434 | 10 | 4.61% |
| hsa_circ_0103844 | 434 | 10 | 4.61% |
| hsa_circ_0107814 | 434 | 10 | 4.61% |
| hsa_circ_0126427 | 434 | 10 | 4.61% |
| hsa_circ_0139372 | 434 | 10 | 4.61% |
| hsa_circ_0052226 | 478 | 11 | 4.60% |
| hsa_circ_0083159 | 478 | 11 | 4.60% |
| hsa_circ_0112085 | 478 | 11 | 4.60% |
| hsa_circ_0102303 | 522 | 12 | 4.60% |
| hsa_circ_0001262 | 435 | 10 | 4.60% |
| hsa_circ_0011978 | 435 | 10 | 4.60% |
| hsa_circ_0130937 | 435 | 10 | 4.60% |
| hsa_circ_0092936 | 479 | 11 | 4.59% |

| | | | |
|---|---|---|---|
| hsa_circ_0097532 | 479 | 11 | 4.59% |
| hsa_circ_0092140 | 523 | 12 | 4.59% |
| hsa_circ_0005530 | 436 | 10 | 4.59% |
| hsa_circ_0027090 | 436 | 10 | 4.59% |
| hsa_circ_0072493 | 436 | 10 | 4.59% |
| hsa_circ_0080513 | 436 | 10 | 4.59% |
| hsa_circ_0082154 | 436 | 10 | 4.59% |
| hsa_circ_0089757 | 436 | 10 | 4.59% |
| hsa_circ_0099713 | 436 | 10 | 4.59% |
| hsa_circ_0131308 | 436 | 10 | 4.59% |
| hsa_circ_0131402 | 436 | 10 | 4.59% |
| hsa_circ_0103536 | 480 | 11 | 4.58% |
| hsa_circ_0121436 | 480 | 11 | 4.58% |
| hsa_circ_0134100 | 480 | 11 | 4.58% |
| hsa_circ_0114069 | 1048 | 24 | 4.58% |
| hsa_circ_0100562 | 524 | 12 | 4.58% |
| hsa_circ_0002014 | 437 | 10 | 4.58% |
| hsa_circ_0006274 | 437 | 10 | 4.58% |
| hsa_circ_0006846 | 437 | 10 | 4.58% |
| hsa_circ_0020295 | 437 | 10 | 4.58% |
| hsa_circ_0048243 | 437 | 10 | 4.58% |
| hsa_circ_0068141 | 437 | 10 | 4.58% |
| hsa_circ_0068677 | 437 | 10 | 4.58% |
| hsa_circ_0068750 | 437 | 10 | 4.58% |
| hsa_circ_0077161 | 437 | 10 | 4.58% |
| hsa_circ_0080279 | 437 | 10 | 4.58% |
| hsa_circ_0094111 | 437 | 10 | 4.58% |
| hsa_circ_0128852 | 612 | 14 | 4.58% |
| hsa_circ_0044908 | 481 | 11 | 4.57% |
| hsa_circ_0137072 | 481 | 11 | 4.57% |
| hsa_circ_0073802 | 525 | 12 | 4.57% |
| hsa_circ_0090168 | 525 | 12 | 4.57% |
| hsa_circ_0023051 | 569 | 13 | 4.57% |
| hsa_circ_0030526 | 569 | 13 | 4.57% |
| hsa_circ_0006995 | 438 | 10 | 4.57% |
| hsa_circ_0018860 | 438 | 10 | 4.57% |
| hsa_circ_0035444 | 438 | 10 | 4.57% |
| hsa_circ_0059970 | 438 | 10 | 4.57% |
| hsa_circ_0068413 | 438 | 10 | 4.57% |
| hsa_circ_0086442 | 438 | 10 | 4.57% |
| hsa_circ_0092688 | 438 | 10 | 4.57% |
| hsa_circ_0108493 | 438 | 10 | 4.57% |
| hsa_circ_0127602 | 438 | 10 | 4.57% |
| hsa_circ_0133780 | 438 | 10 | 4.57% |
| hsa_circ_0138645 | 438 | 10 | 4.57% |
| hsa_circ_0026876 | 482 | 11 | 4.56% |
| hsa_circ_0030839 | 482 | 11 | 4.56% |
| hsa_circ_0089592 | 482 | 11 | 4.56% |
| hsa_circ_0096716 | 482 | 11 | 4.56% |
| hsa_circ_0020960 | 439 | 10 | 4.56% |

| | | | |
|---|---|---|---|
| hsa_circ_0043782 | 439 | 10 | 4.56% |
| hsa_circ_0058179 | 439 | 10 | 4.56% |
| hsa_circ_0060727 | 439 | 10 | 4.56% |
| hsa_circ_0065725 | 439 | 10 | 4.56% |
| hsa_circ_0089940 | 439 | 10 | 4.56% |
| hsa_circ_0104283 | 439 | 10 | 4.56% |
| hsa_circ_0107629 | 439 | 10 | 4.56% |
| hsa_circ_0114049 | 439 | 10 | 4.56% |
| hsa_circ_0127341 | 439 | 10 | 4.56% |
| hsa_circ_0026564 | 483 | 11 | 4.55% |
| hsa_circ_0057457 | 483 | 11 | 4.55% |
| hsa_circ_0067243 | 483 | 11 | 4.55% |
| hsa_circ_0081428 | 483 | 11 | 4.55% |
| hsa_circ_0083496 | 483 | 11 | 4.55% |
| hsa_circ_0118816 | 483 | 11 | 4.55% |
| hsa_circ_0007320 | 572 | 13 | 4.55% |
| hsa_circ_0076101 | 528 | 12 | 4.55% |
| hsa_circ_0109204 | 484 | 11 | 4.55% |
| hsa_circ_0126717 | 484 | 11 | 4.55% |
| hsa_circ_0001649 | 440 | 10 | 4.55% |
| hsa_circ_0003643 | 440 | 10 | 4.55% |
| hsa_circ_0006364 | 440 | 10 | 4.55% |
| hsa_circ_0036503 | 440 | 10 | 4.55% |
| hsa_circ_0059071 | 440 | 10 | 4.55% |
| hsa_circ_0098892 | 440 | 10 | 4.55% |
| hsa_circ_0115891 | 440 | 10 | 4.55% |
| hsa_circ_0032243 | 485 | 11 | 4.54% |
| hsa_circ_0048806 | 485 | 11 | 4.54% |
| hsa_circ_0140498 | 485 | 11 | 4.54% |
| hsa_circ_0011985 | 441 | 10 | 4.54% |
| hsa_circ_0014768 | 441 | 10 | 4.54% |
| hsa_circ_0016122 | 441 | 10 | 4.54% |
| hsa_circ_0021873 | 441 | 10 | 4.54% |
| hsa_circ_0028141 | 441 | 10 | 4.54% |
| hsa_circ_0038074 | 441 | 10 | 4.54% |
| hsa_circ_0044265 | 441 | 10 | 4.54% |
| hsa_circ_0055432 | 441 | 10 | 4.54% |
| hsa_circ_0094220 | 441 | 10 | 4.54% |
| hsa_circ_0098702 | 441 | 10 | 4.54% |
| hsa_circ_0107616 | 441 | 10 | 4.54% |
| hsa_circ_0110571 | 441 | 10 | 4.54% |
| hsa_circ_0112900 | 441 | 10 | 4.54% |
| hsa_circ_0064457 | 574 | 13 | 4.53% |
| hsa_circ_0109165 | 530 | 12 | 4.53% |
| hsa_circ_0051524 | 486 | 11 | 4.53% |
| hsa_circ_0060312 | 486 | 11 | 4.53% |
| hsa_circ_0090578 | 486 | 11 | 4.53% |
| hsa_circ_0125227 | 486 | 11 | 4.53% |
| hsa_circ_0135703 | 486 | 11 | 4.53% |
| hsa_circ_0004840 | 442 | 10 | 4.52% |

| | | | |
|---|---|---|---|
| hsa_circ_0008674 | 442 | 10 | 4.52% |
| hsa_circ_0008724 | 442 | 10 | 4.52% |
| hsa_circ_0012875 | 442 | 10 | 4.52% |
| hsa_circ_0017737 | 442 | 10 | 4.52% |
| hsa_circ_0031947 | 442 | 10 | 4.52% |
| hsa_circ_0032916 | 442 | 10 | 4.52% |
| hsa_circ_0035648 | 442 | 10 | 4.52% |
| hsa_circ_0037379 | 442 | 10 | 4.52% |
| hsa_circ_0039998 | 442 | 10 | 4.52% |
| hsa_circ_0044463 | 442 | 10 | 4.52% |
| hsa_circ_0063655 | 442 | 10 | 4.52% |
| hsa_circ_0083422 | 442 | 10 | 4.52% |
| hsa_circ_0099085 | 442 | 10 | 4.52% |
| hsa_circ_0118812 | 442 | 10 | 4.52% |
| hsa_circ_0040319 | 531 | 12 | 4.52% |
| hsa_circ_0100898 | 531 | 12 | 4.52% |
| hsa_circ_0122143 | 531 | 12 | 4.52% |
| hsa_circ_0001061 | 487 | 11 | 4.52% |
| hsa_circ_0088921 | 487 | 11 | 4.52% |
| hsa_circ_0130895 | 487 | 11 | 4.52% |
| hsa_circ_0131405 | 487 | 11 | 4.52% |
| hsa_circ_0004204 | 443 | 10 | 4.51% |
| hsa_circ_0006965 | 443 | 10 | 4.51% |
| hsa_circ_0080876 | 443 | 10 | 4.51% |
| hsa_circ_0105786 | 443 | 10 | 4.51% |
| hsa_circ_0113002 | 443 | 10 | 4.51% |
| hsa_circ_0113029 | 443 | 10 | 4.51% |
| hsa_circ_0129193 | 443 | 10 | 4.51% |
| hsa_circ_0089794 | 4123 | 93 | 4.51% |
| hsa_circ_0125452 | 532 | 12 | 4.51% |
| hsa_circ_0139518 | 532 | 12 | 4.51% |
| hsa_circ_0043855 | 488 | 11 | 4.51% |
| hsa_circ_0123943 | 488 | 11 | 4.51% |
| hsa_circ_0060523 | 577 | 13 | 4.51% |
| hsa_circ_0005335 | 444 | 10 | 4.50% |
| hsa_circ_0005469 | 444 | 10 | 4.50% |
| hsa_circ_0013686 | 444 | 10 | 4.50% |
| hsa_circ_0039226 | 444 | 10 | 4.50% |
| hsa_circ_0045363 | 444 | 10 | 4.50% |
| hsa_circ_0075035 | 444 | 10 | 4.50% |
| hsa_circ_0076802 | 444 | 10 | 4.50% |
| hsa_circ_0097568 | 444 | 10 | 4.50% |
| hsa_circ_0126305 | 444 | 10 | 4.50% |
| hsa_circ_0126479 | 444 | 10 | 4.50% |
| hsa_circ_0005457 | 533 | 12 | 4.50% |
| hsa_circ_0049915 | 489 | 11 | 4.50% |
| hsa_circ_0067627 | 489 | 11 | 4.50% |
| hsa_circ_0075246 | 578 | 13 | 4.50% |
| hsa_circ_0006430 | 445 | 10 | 4.49% |
| hsa_circ_0014974 | 445 | 10 | 4.49% |

| | | | |
|---|---|---|---|
| hsa_circ_0058647 | 445 | 10 | 4.49% |
| hsa_circ_0067185 | 445 | 10 | 4.49% |
| hsa_circ_0083047 | 445 | 10 | 4.49% |
| hsa_circ_0091812 | 445 | 10 | 4.49% |
| hsa_circ_0118873 | 445 | 10 | 4.49% |
| hsa_circ_0119142 | 445 | 10 | 4.49% |
| hsa_circ_0133290 | 445 | 10 | 4.49% |
| hsa_circ_0005380 | 490 | 11 | 4.49% |
| hsa_circ_0059728 | 490 | 11 | 4.49% |
| hsa_circ_0023152 | 535 | 12 | 4.49% |
| hsa_circ_0027115 | 535 | 12 | 4.49% |
| hsa_circ_0047618 | 535 | 12 | 4.49% |
| hsa_circ_0003572 | 446 | 10 | 4.48% |
| hsa_circ_0043428 | 446 | 10 | 4.48% |
| hsa_circ_0056311 | 446 | 10 | 4.48% |
| hsa_circ_0079925 | 446 | 10 | 4.48% |
| hsa_circ_0081474 | 446 | 10 | 4.48% |
| hsa_circ_0086769 | 446 | 10 | 4.48% |
| hsa_circ_0101313 | 446 | 10 | 4.48% |
| hsa_circ_0110481 | 446 | 10 | 4.48% |
| hsa_circ_0112092 | 446 | 10 | 4.48% |
| hsa_circ_0118951 | 446 | 10 | 4.48% |
| hsa_circ_0121414 | 446 | 10 | 4.48% |
| hsa_circ_0126088 | 446 | 10 | 4.48% |
| hsa_circ_0113245 | 580 | 13 | 4.48% |
| hsa_circ_0014680 | 491 | 11 | 4.48% |
| hsa_circ_0066779 | 491 | 11 | 4.48% |
| hsa_circ_0005560 | 447 | 10 | 4.47% |
| hsa_circ_0021126 | 447 | 10 | 4.47% |
| hsa_circ_0045413 | 447 | 10 | 4.47% |
| hsa_circ_0045560 | 447 | 10 | 4.47% |
| hsa_circ_0046657 | 447 | 10 | 4.47% |
| hsa_circ_0057705 | 447 | 10 | 4.47% |
| hsa_circ_0064126 | 447 | 10 | 4.47% |
| hsa_circ_0070146 | 447 | 10 | 4.47% |
| hsa_circ_0092454 | 447 | 10 | 4.47% |
| hsa_circ_0106954 | 447 | 10 | 4.47% |
| hsa_circ_0113135 | 447 | 10 | 4.47% |
| hsa_circ_0121464 | 447 | 10 | 4.47% |
| hsa_circ_0134815 | 447 | 10 | 4.47% |
| hsa_circ_0095142 | 626 | 14 | 4.47% |
| hsa_circ_0063283 | 492 | 11 | 4.47% |
| hsa_circ_0117888 | 492 | 11 | 4.47% |
| hsa_circ_0055489 | 537 | 12 | 4.47% |
| hsa_circ_0100700 | 582 | 13 | 4.47% |
| hsa_circ_0105707 | 1120 | 25 | 4.46% |
| hsa_circ_0000303 | 448 | 10 | 4.46% |
| hsa_circ_0051731 | 448 | 10 | 4.46% |
| hsa_circ_0080906 | 448 | 10 | 4.46% |
| hsa_circ_0082875 | 448 | 10 | 4.46% |

| | | | |
|---|---|---|---|
| hsa_circ_0087684 | 448 | 10 | 4.46% |
| hsa_circ_0114595 | 448 | 10 | 4.46% |
| hsa_circ_0115242 | 448 | 10 | 4.46% |
| hsa_circ_0121958 | 448 | 10 | 4.46% |
| hsa_circ_0128780 | 448 | 10 | 4.46% |
| hsa_circ_0137947 | 448 | 10 | 4.46% |
| hsa_circ_0107750 | 493 | 11 | 4.46% |
| hsa_circ_0002346 | 538 | 12 | 4.46% |
| hsa_circ_0042889 | 583 | 13 | 4.46% |
| hsa_circ_0007275 | 449 | 10 | 4.45% |
| hsa_circ_0010028 | 449 | 10 | 4.45% |
| hsa_circ_0019053 | 449 | 10 | 4.45% |
| hsa_circ_0032158 | 449 | 10 | 4.45% |
| hsa_circ_0037442 | 449 | 10 | 4.45% |
| hsa_circ_0068420 | 449 | 10 | 4.45% |
| hsa_circ_0072989 | 449 | 10 | 4.45% |
| hsa_circ_0099595 | 449 | 10 | 4.45% |
| hsa_circ_0133954 | 449 | 10 | 4.45% |
| hsa_circ_0094107 | 494 | 11 | 4.45% |
| hsa_circ_0113999 | 494 | 11 | 4.45% |
| hsa_circ_0041215 | 539 | 12 | 4.45% |
| hsa_circ_0043273 | 539 | 12 | 4.45% |
| hsa_circ_0124950 | 584 | 13 | 4.45% |
| hsa_circ_0130017 | 810 | 18 | 4.44% |
| hsa_circ_0052499 | 495 | 11 | 4.44% |
| hsa_circ_0100036 | 495 | 11 | 4.44% |
| hsa_circ_0110116 | 495 | 11 | 4.44% |
| hsa_circ_0122732 | 495 | 11 | 4.44% |
| hsa_circ_0015251 | 450 | 10 | 4.44% |
| hsa_circ_0021182 | 450 | 10 | 4.44% |
| hsa_circ_0031642 | 450 | 10 | 4.44% |
| hsa_circ_0039978 | 450 | 10 | 4.44% |
| hsa_circ_0047675 | 450 | 10 | 4.44% |
| hsa_circ_0075105 | 450 | 10 | 4.44% |
| hsa_circ_0081714 | 450 | 10 | 4.44% |
| hsa_circ_0085163 | 450 | 10 | 4.44% |
| hsa_circ_0094039 | 450 | 10 | 4.44% |
| hsa_circ_0101628 | 450 | 10 | 4.44% |
| hsa_circ_0105802 | 450 | 10 | 4.44% |
| hsa_circ_0113762 | 450 | 10 | 4.44% |
| hsa_circ_0116426 | 450 | 10 | 4.44% |
| hsa_circ_0121662 | 450 | 10 | 4.44% |
| hsa_circ_0129452 | 450 | 10 | 4.44% |
| hsa_circ_0133548 | 450 | 10 | 4.44% |
| hsa_circ_0134367 | 450 | 10 | 4.44% |
| hsa_circ_0001432 | 496 | 11 | 4.44% |
| hsa_circ_0018634 | 496 | 11 | 4.44% |
| hsa_circ_0085410 | 496 | 11 | 4.44% |
| hsa_circ_0110378 | 496 | 11 | 4.44% |
| hsa_circ_0133801 | 496 | 11 | 4.44% |

| | | | |
|---|---|---|---|
| hsa_circ_0006486 | 451 | 10 | 4.43% |
| hsa_circ_0007392 | 451 | 10 | 4.43% |
| hsa_circ_0014099 | 451 | 10 | 4.43% |
| hsa_circ_0020576 | 451 | 10 | 4.43% |
| hsa_circ_0028873 | 451 | 10 | 4.43% |
| hsa_circ_0040892 | 451 | 10 | 4.43% |
| hsa_circ_0042762 | 451 | 10 | 4.43% |
| hsa_circ_0047185 | 451 | 10 | 4.43% |
| hsa_circ_0051590 | 451 | 10 | 4.43% |
| hsa_circ_0053320 | 451 | 10 | 4.43% |
| hsa_circ_0056594 | 451 | 10 | 4.43% |
| hsa_circ_0060532 | 451 | 10 | 4.43% |
| hsa_circ_0063758 | 451 | 10 | 4.43% |
| hsa_circ_0081605 | 451 | 10 | 4.43% |
| hsa_circ_0086933 | 451 | 10 | 4.43% |
| hsa_circ_0097478 | 451 | 10 | 4.43% |
| hsa_circ_0102897 | 451 | 10 | 4.43% |
| hsa_circ_0123882 | 451 | 10 | 4.43% |
| hsa_circ_0127049 | 451 | 10 | 4.43% |
| hsa_circ_0128597 | 451 | 10 | 4.43% |
| hsa_circ_0135138 | 451 | 10 | 4.43% |
| hsa_circ_0018855 | 542 | 12 | 4.43% |
| hsa_circ_0061539 | 497 | 11 | 4.43% |
| hsa_circ_0096495 | 497 | 11 | 4.43% |
| hsa_circ_0110432 | 497 | 11 | 4.43% |
| hsa_circ_0002550 | 452 | 10 | 4.42% |
| hsa_circ_0010098 | 452 | 10 | 4.42% |
| hsa_circ_0048486 | 452 | 10 | 4.42% |
| hsa_circ_0057018 | 452 | 10 | 4.42% |
| hsa_circ_0099983 | 452 | 10 | 4.42% |
| hsa_circ_0104028 | 452 | 10 | 4.42% |
| hsa_circ_0107246 | 452 | 10 | 4.42% |
| hsa_circ_0123467 | 452 | 10 | 4.42% |
| hsa_circ_0045325 | 588 | 13 | 4.42% |
| hsa_circ_0114714 | 588 | 13 | 4.42% |
| hsa_circ_0069470 | 543 | 12 | 4.42% |
| hsa_circ_0075729 | 543 | 12 | 4.42% |
| hsa_circ_0002727 | 498 | 11 | 4.42% |
| hsa_circ_0036092 | 498 | 11 | 4.42% |
| hsa_circ_0055976 | 498 | 11 | 4.42% |
| hsa_circ_0068056 | 498 | 11 | 4.42% |
| hsa_circ_0077618 | 498 | 11 | 4.42% |
| hsa_circ_0111706 | 498 | 11 | 4.42% |
| hsa_circ_0107171 | 634 | 14 | 4.42% |
| hsa_circ_0011448 | 453 | 10 | 4.42% |
| hsa_circ_0019619 | 453 | 10 | 4.42% |
| hsa_circ_0024774 | 453 | 10 | 4.42% |
| hsa_circ_0026687 | 453 | 10 | 4.42% |
| hsa_circ_0027935 | 453 | 10 | 4.42% |
| hsa_circ_0039879 | 453 | 10 | 4.42% |

| | | | |
|---|---|---|---|
| hsa_circ_0046226 | 453 | 10 | 4.42% |
| hsa_circ_0047683 | 453 | 10 | 4.42% |
| hsa_circ_0077586 | 453 | 10 | 4.42% |
| hsa_circ_0082060 | 453 | 10 | 4.42% |
| hsa_circ_0095896 | 453 | 10 | 4.42% |
| hsa_circ_0096882 | 453 | 10 | 4.42% |
| hsa_circ_0098956 | 453 | 10 | 4.42% |
| hsa_circ_0105876 | 453 | 10 | 4.42% |
| hsa_circ_0114113 | 453 | 10 | 4.42% |
| hsa_circ_0134073 | 453 | 10 | 4.42% |
| hsa_circ_0134117 | 453 | 10 | 4.42% |
| hsa_circ_0002957 | 544 | 12 | 4.41% |
| hsa_circ_0105862 | 544 | 12 | 4.41% |
| hsa_circ_0043673 | 499 | 11 | 4.41% |
| hsa_circ_0080040 | 499 | 11 | 4.41% |
| hsa_circ_0084029 | 590 | 13 | 4.41% |
| hsa_circ_0022966 | 454 | 10 | 4.41% |
| hsa_circ_0044814 | 454 | 10 | 4.41% |
| hsa_circ_0064664 | 454 | 10 | 4.41% |
| hsa_circ_0077898 | 454 | 10 | 4.41% |
| hsa_circ_0099587 | 454 | 10 | 4.41% |
| hsa_circ_0132315 | 454 | 10 | 4.41% |
| hsa_circ_0016453 | 545 | 12 | 4.40% |
| hsa_circ_0107266 | 545 | 12 | 4.40% |
| hsa_circ_0017939 | 500 | 11 | 4.40% |
| hsa_circ_0022809 | 500 | 11 | 4.40% |
| hsa_circ_0130110 | 591 | 13 | 4.40% |
| hsa_circ_0041214 | 455 | 10 | 4.40% |
| hsa_circ_0064073 | 455 | 10 | 4.40% |
| hsa_circ_0067745 | 455 | 10 | 4.40% |
| hsa_circ_0098168 | 455 | 10 | 4.40% |
| hsa_circ_0102857 | 455 | 10 | 4.40% |
| hsa_circ_0106356 | 455 | 10 | 4.40% |
| hsa_circ_0114123 | 455 | 10 | 4.40% |
| hsa_circ_0114816 | 455 | 10 | 4.40% |
| hsa_circ_0126844 | 455 | 10 | 4.40% |
| hsa_circ_0130301 | 455 | 10 | 4.40% |
| hsa_circ_0019755 | 501 | 11 | 4.39% |
| hsa_circ_0106292 | 501 | 11 | 4.39% |
| hsa_circ_0135829 | 501 | 11 | 4.39% |
| hsa_circ_0138065 | 501 | 11 | 4.39% |
| hsa_circ_0140500 | 501 | 11 | 4.39% |
| hsa_circ_0012002 | 456 | 10 | 4.39% |
| hsa_circ_0013491 | 456 | 10 | 4.39% |
| hsa_circ_0081976 | 456 | 10 | 4.39% |
| hsa_circ_0096024 | 456 | 10 | 4.39% |
| hsa_circ_0099679 | 456 | 10 | 4.39% |
| hsa_circ_0100944 | 456 | 10 | 4.39% |
| hsa_circ_0101919 | 456 | 10 | 4.39% |
| hsa_circ_0029273 | 502 | 11 | 4.38% |

| | | | |
|---|---|---|---|
| hsa_circ_0044397 | 502 | 11 | 4.38% |
| hsa_circ_0123851 | 502 | 11 | 4.38% |
| hsa_circ_0129758 | 502 | 11 | 4.38% |
| hsa_circ_0027912 | 594 | 13 | 4.38% |
| hsa_circ_0037215 | 457 | 10 | 4.38% |
| hsa_circ_0048763 | 457 | 10 | 4.38% |
| hsa_circ_0049536 | 457 | 10 | 4.38% |
| hsa_circ_0057256 | 457 | 10 | 4.38% |
| hsa_circ_0076967 | 457 | 10 | 4.38% |
| hsa_circ_0079171 | 457 | 10 | 4.38% |
| hsa_circ_0135421 | 457 | 10 | 4.38% |
| hsa_circ_0048807 | 503 | 11 | 4.37% |
| hsa_circ_0074996 | 503 | 11 | 4.37% |
| hsa_circ_0094105 | 503 | 11 | 4.37% |
| hsa_circ_0104765 | 503 | 11 | 4.37% |
| hsa_circ_0136837 | 503 | 11 | 4.37% |
| hsa_circ_0019144 | 549 | 12 | 4.37% |
| hsa_circ_0007779 | 458 | 10 | 4.37% |
| hsa_circ_0013351 | 458 | 10 | 4.37% |
| hsa_circ_0044441 | 458 | 10 | 4.37% |
| hsa_circ_0045647 | 458 | 10 | 4.37% |
| hsa_circ_0092513 | 458 | 10 | 4.37% |
| hsa_circ_0112943 | 458 | 10 | 4.37% |
| hsa_circ_0126098 | 458 | 10 | 4.37% |
| hsa_circ_0129896 | 458 | 10 | 4.37% |
| hsa_circ_0137972 | 458 | 10 | 4.37% |
| hsa_circ_0057455 | 504 | 11 | 4.37% |
| hsa_circ_0092896 | 504 | 11 | 4.37% |
| hsa_circ_0134512 | 504 | 11 | 4.37% |
| hsa_circ_0002418 | 459 | 10 | 4.36% |
| hsa_circ_0022071 | 459 | 10 | 4.36% |
| hsa_circ_0029554 | 459 | 10 | 4.36% |
| hsa_circ_0030524 | 459 | 10 | 4.36% |
| hsa_circ_0038989 | 459 | 10 | 4.36% |
| hsa_circ_0062236 | 459 | 10 | 4.36% |
| hsa_circ_0087442 | 459 | 10 | 4.36% |
| hsa_circ_0096847 | 459 | 10 | 4.36% |
| hsa_circ_0113312 | 459 | 10 | 4.36% |
| hsa_circ_0116077 | 459 | 10 | 4.36% |
| hsa_circ_0124242 | 459 | 10 | 4.36% |
| hsa_circ_0136832 | 459 | 10 | 4.36% |
| hsa_circ_0139362 | 459 | 10 | 4.36% |
| hsa_circ_0100824 | 551 | 12 | 4.36% |
| hsa_circ_0069919 | 597 | 13 | 4.36% |
| hsa_circ_0099924 | 597 | 13 | 4.36% |
| hsa_circ_0030288 | 552 | 12 | 4.35% |
| hsa_circ_0061697 | 506 | 11 | 4.35% |
| hsa_circ_0081199 | 506 | 11 | 4.35% |
| hsa_circ_0021281 | 460 | 10 | 4.35% |
| hsa_circ_0029782 | 460 | 10 | 4.35% |

| | | | |
|---|---|---|---|
| hsa_circ_0056237 | 460 | 10 | 4.35% |
| hsa_circ_0060479 | 460 | 10 | 4.35% |
| hsa_circ_0079811 | 460 | 10 | 4.35% |
| hsa_circ_0084523 | 460 | 10 | 4.35% |
| hsa_circ_0089489 | 460 | 10 | 4.35% |
| hsa_circ_0109471 | 460 | 10 | 4.35% |
| hsa_circ_0116416 | 460 | 10 | 4.35% |
| hsa_circ_0127595 | 460 | 10 | 4.35% |
| hsa_circ_0140196 | 460 | 10 | 4.35% |
| hsa_circ_0004908 | 645 | 14 | 4.34% |
| hsa_circ_0030124 | 553 | 12 | 4.34% |
| hsa_circ_0049859 | 507 | 11 | 4.34% |
| hsa_circ_0050955 | 507 | 11 | 4.34% |
| hsa_circ_0138171 | 507 | 11 | 4.34% |
| hsa_circ_0004416 | 461 | 10 | 4.34% |
| hsa_circ_0011133 | 461 | 10 | 4.34% |
| hsa_circ_0014228 | 461 | 10 | 4.34% |
| hsa_circ_0019133 | 461 | 10 | 4.34% |
| hsa_circ_0027088 | 461 | 10 | 4.34% |
| hsa_circ_0032507 | 461 | 10 | 4.34% |
| hsa_circ_0043966 | 461 | 10 | 4.34% |
| hsa_circ_0054227 | 461 | 10 | 4.34% |
| hsa_circ_0057769 | 461 | 10 | 4.34% |
| hsa_circ_0078339 | 461 | 10 | 4.34% |
| hsa_circ_0083234 | 461 | 10 | 4.34% |
| hsa_circ_0084553 | 461 | 10 | 4.34% |
| hsa_circ_0085779 | 461 | 10 | 4.34% |
| hsa_circ_0111264 | 461 | 10 | 4.34% |
| hsa_circ_0116227 | 461 | 10 | 4.34% |
| hsa_circ_0005064 | 554 | 12 | 4.33% |
| hsa_circ_0041629 | 508 | 11 | 4.33% |
| hsa_circ_0089210 | 508 | 11 | 4.33% |
| hsa_circ_0092020 | 508 | 11 | 4.33% |
| hsa_circ_0107203 | 508 | 11 | 4.33% |
| hsa_circ_0015277 | 462 | 10 | 4.33% |
| hsa_circ_0031610 | 462 | 10 | 4.33% |
| hsa_circ_0049353 | 462 | 10 | 4.33% |
| hsa_circ_0049404 | 462 | 10 | 4.33% |
| hsa_circ_0083300 | 462 | 10 | 4.33% |
| hsa_circ_0097512 | 462 | 10 | 4.33% |
| hsa_circ_0097659 | 462 | 10 | 4.33% |
| hsa_circ_0129565 | 462 | 10 | 4.33% |
| hsa_circ_0020325 | 601 | 13 | 4.33% |
| hsa_circ_0058716 | 555 | 12 | 4.32% |
| hsa_circ_0049661 | 509 | 11 | 4.32% |
| hsa_circ_0068771 | 509 | 11 | 4.32% |
| hsa_circ_0097415 | 509 | 11 | 4.32% |
| hsa_circ_0004025 | 463 | 10 | 4.32% |
| hsa_circ_0015452 | 463 | 10 | 4.32% |
| hsa_circ_0015665 | 463 | 10 | 4.32% |

| | | | |
|---|---|---|---|
| hsa_circ_0020640 | 463 | 10 | 4.32% |
| hsa_circ_0022838 | 463 | 10 | 4.32% |
| hsa_circ_0029703 | 463 | 10 | 4.32% |
| hsa_circ_0043572 | 463 | 10 | 4.32% |
| hsa_circ_0056874 | 463 | 10 | 4.32% |
| hsa_circ_0081391 | 463 | 10 | 4.32% |
| hsa_circ_0091365 | 463 | 10 | 4.32% |
| hsa_circ_0109177 | 463 | 10 | 4.32% |
| hsa_circ_0112889 | 463 | 10 | 4.32% |
| hsa_circ_0115727 | 463 | 10 | 4.32% |
| hsa_circ_0118622 | 463 | 10 | 4.32% |
| hsa_circ_0125409 | 463 | 10 | 4.32% |
| hsa_circ_0128848 | 463 | 10 | 4.32% |
| hsa_circ_0138915 | 556 | 12 | 4.32% |
| hsa_circ_0022324 | 464 | 10 | 4.31% |
| hsa_circ_0066711 | 464 | 10 | 4.31% |
| hsa_circ_0074793 | 464 | 10 | 4.31% |
| hsa_circ_0114368 | 464 | 10 | 4.31% |
| hsa_circ_0125805 | 464 | 10 | 4.31% |
| hsa_circ_0125855 | 464 | 10 | 4.31% |
| hsa_circ_0129133 | 464 | 10 | 4.31% |
| hsa_circ_0003040 | 511 | 11 | 4.31% |
| hsa_circ_0026758 | 511 | 11 | 4.31% |
| hsa_circ_0106240 | 511 | 11 | 4.31% |
| hsa_circ_0139938 | 511 | 11 | 4.31% |
| hsa_circ_0105397 | 558 | 12 | 4.30% |
| hsa_circ_0129442 | 558 | 12 | 4.30% |
| hsa_circ_0138363 | 558 | 12 | 4.30% |
| hsa_circ_0003916 | 465 | 10 | 4.30% |
| hsa_circ_0003936 | 465 | 10 | 4.30% |
| hsa_circ_0007595 | 465 | 10 | 4.30% |
| hsa_circ_0023411 | 465 | 10 | 4.30% |
| hsa_circ_0029397 | 465 | 10 | 4.30% |
| hsa_circ_0035527 | 465 | 10 | 4.30% |
| hsa_circ_0057358 | 465 | 10 | 4.30% |
| hsa_circ_0076302 | 465 | 10 | 4.30% |
| hsa_circ_0122662 | 465 | 10 | 4.30% |
| hsa_circ_0131146 | 465 | 10 | 4.30% |
| hsa_circ_0136144 | 465 | 10 | 4.30% |
| hsa_circ_0026477 | 512 | 11 | 4.30% |
| hsa_circ_0037840 | 512 | 11 | 4.30% |
| hsa_circ_0050870 | 512 | 11 | 4.30% |
| hsa_circ_0132528 | 559 | 12 | 4.29% |
| hsa_circ_0005960 | 466 | 10 | 4.29% |
| hsa_circ_0006273 | 466 | 10 | 4.29% |
| hsa_circ_0031148 | 466 | 10 | 4.29% |
| hsa_circ_0062876 | 466 | 10 | 4.29% |
| hsa_circ_0067467 | 466 | 10 | 4.29% |
| hsa_circ_0068623 | 466 | 10 | 4.29% |
| hsa_circ_0079094 | 466 | 10 | 4.29% |

| | | | |
|---|---|---|---|
| hsa_circ_0079203 | 466 | 10 | 4.29% |
| hsa_circ_0090468 | 466 | 10 | 4.29% |
| hsa_circ_0092407 | 466 | 10 | 4.29% |
| hsa_circ_0105973 | 466 | 10 | 4.29% |
| hsa_circ_0117230 | 466 | 10 | 4.29% |
| hsa_circ_0039325 | 513 | 11 | 4.29% |
| hsa_circ_0074213 | 513 | 11 | 4.29% |
| hsa_circ_0088057 | 513 | 11 | 4.29% |
| hsa_circ_0134870 | 513 | 11 | 4.29% |
| hsa_circ_0137182 | 513 | 11 | 4.29% |
| hsa_circ_0138635 | 513 | 11 | 4.29% |
| hsa_circ_0004661 | 560 | 12 | 4.29% |
| hsa_circ_0112471 | 560 | 12 | 4.29% |
| hsa_circ_0003854 | 467 | 10 | 4.28% |
| hsa_circ_0020799 | 467 | 10 | 4.28% |
| hsa_circ_0026945 | 467 | 10 | 4.28% |
| hsa_circ_0096764 | 467 | 10 | 4.28% |
| hsa_circ_0115034 | 467 | 10 | 4.28% |
| hsa_circ_0083138 | 654 | 14 | 4.28% |
| hsa_circ_0016938 | 514 | 11 | 4.28% |
| hsa_circ_0001506 | 608 | 13 | 4.28% |
| hsa_circ_0131700 | 608 | 13 | 4.28% |
| hsa_circ_0003179 | 468 | 10 | 4.27% |
| hsa_circ_0003445 | 468 | 10 | 4.27% |
| hsa_circ_0049127 | 468 | 10 | 4.27% |
| hsa_circ_0063963 | 468 | 10 | 4.27% |
| hsa_circ_0069925 | 468 | 10 | 4.27% |
| hsa_circ_0072050 | 468 | 10 | 4.27% |
| hsa_circ_0093642 | 468 | 10 | 4.27% |
| hsa_circ_0098302 | 468 | 10 | 4.27% |
| hsa_circ_0112476 | 468 | 10 | 4.27% |
| hsa_circ_0135988 | 468 | 10 | 4.27% |
| hsa_circ_0003703 | 515 | 11 | 4.27% |
| hsa_circ_0103571 | 515 | 11 | 4.27% |
| hsa_circ_0002056 | 562 | 12 | 4.27% |
| hsa_circ_0124994 | 562 | 12 | 4.27% |
| hsa_circ_0110893 | 1078 | 23 | 4.27% |
| hsa_circ_0112680 | 1453 | 31 | 4.27% |
| hsa_circ_0129074 | 1172 | 25 | 4.27% |
| hsa_circ_0005772 | 469 | 10 | 4.26% |
| hsa_circ_0014750 | 469 | 10 | 4.26% |
| hsa_circ_0045095 | 469 | 10 | 4.26% |
| hsa_circ_0073976 | 469 | 10 | 4.26% |
| hsa_circ_0077813 | 469 | 10 | 4.26% |
| hsa_circ_0078240 | 469 | 10 | 4.26% |
| hsa_circ_0138412 | 469 | 10 | 4.26% |
| hsa_circ_0036241 | 516 | 11 | 4.26% |
| hsa_circ_0098567 | 516 | 11 | 4.26% |
| hsa_circ_0113685 | 516 | 11 | 4.26% |
| hsa_circ_0109668 | 563 | 12 | 4.26% |

| | | | |
|---|---|---|---|
| hsa_circ_0117676 | 563 | 12 | 4.26% |
| hsa_circ_0124243 | 1927 | 41 | 4.26% |
| hsa_circ_0037490 | 611 | 13 | 4.26% |
| hsa_circ_0043773 | 564 | 12 | 4.26% |
| hsa_circ_0075724 | 564 | 12 | 4.26% |
| hsa_circ_0088525 | 564 | 12 | 4.26% |
| hsa_circ_0020205 | 517 | 11 | 4.26% |
| hsa_circ_0039871 | 517 | 11 | 4.26% |
| hsa_circ_0046295 | 517 | 11 | 4.26% |
| hsa_circ_0079130 | 517 | 11 | 4.26% |
| hsa_circ_0022242 | 470 | 10 | 4.26% |
| hsa_circ_0022768 | 470 | 10 | 4.26% |
| hsa_circ_0053116 | 470 | 10 | 4.26% |
| hsa_circ_0063872 | 470 | 10 | 4.26% |
| hsa_circ_0073241 | 470 | 10 | 4.26% |
| hsa_circ_0084783 | 470 | 10 | 4.26% |
| hsa_circ_0097807 | 470 | 10 | 4.26% |
| hsa_circ_0107405 | 470 | 10 | 4.26% |
| hsa_circ_0127377 | 470 | 10 | 4.26% |
| hsa_circ_0002707 | 612 | 13 | 4.25% |
| hsa_circ_0043469 | 565 | 12 | 4.25% |
| hsa_circ_0095291 | 565 | 12 | 4.25% |
| hsa_circ_0127850 | 565 | 12 | 4.25% |
| hsa_circ_0077113 | 518 | 11 | 4.25% |
| hsa_circ_0087582 | 518 | 11 | 4.25% |
| hsa_circ_0120757 | 518 | 11 | 4.25% |
| hsa_circ_0128266 | 518 | 11 | 4.25% |
| hsa_circ_0005553 | 471 | 10 | 4.25% |
| hsa_circ_0008408 | 471 | 10 | 4.25% |
| hsa_circ_0017686 | 471 | 10 | 4.25% |
| hsa_circ_0038207 | 471 | 10 | 4.25% |
| hsa_circ_0056758 | 471 | 10 | 4.25% |
| hsa_circ_0080707 | 471 | 10 | 4.25% |
| hsa_circ_0096086 | 471 | 10 | 4.25% |
| hsa_circ_0122041 | 471 | 10 | 4.25% |
| hsa_circ_0137067 | 471 | 10 | 4.25% |
| hsa_circ_0041961 | 707 | 15 | 4.24% |
| hsa_circ_0085820 | 802 | 17 | 4.24% |
| hsa_circ_0008689 | 519 | 11 | 4.24% |
| hsa_circ_0015926 | 519 | 11 | 4.24% |
| hsa_circ_0116772 | 519 | 11 | 4.24% |
| hsa_circ_0006791 | 472 | 10 | 4.24% |
| hsa_circ_0038726 | 472 | 10 | 4.24% |
| hsa_circ_0085771 | 472 | 10 | 4.24% |
| hsa_circ_0100875 | 472 | 10 | 4.24% |
| hsa_circ_0105258 | 472 | 10 | 4.24% |
| hsa_circ_0114919 | 472 | 10 | 4.24% |
| hsa_circ_0115466 | 472 | 10 | 4.24% |
| hsa_circ_0119338 | 756 | 16 | 4.23% |
| hsa_circ_0094452 | 567 | 12 | 4.23% |

| | | | |
|---|---|---|---|
| hsa_circ_0002981 | 520 | 11 | 4.23% |
| hsa_circ_0038503 | 520 | 11 | 4.23% |
| hsa_circ_0132902 | 520 | 11 | 4.23% |
| hsa_circ_0088711 | 804 | 17 | 4.23% |
| hsa_circ_0048831 | 473 | 10 | 4.23% |
| hsa_circ_0065784 | 473 | 10 | 4.23% |
| hsa_circ_0079564 | 473 | 10 | 4.23% |
| hsa_circ_0081303 | 473 | 10 | 4.23% |
| hsa_circ_0115437 | 473 | 10 | 4.23% |
| hsa_circ_0116493 | 473 | 10 | 4.23% |
| hsa_circ_0134772 | 473 | 10 | 4.23% |
| hsa_circ_0050178 | 521 | 11 | 4.22% |
| hsa_circ_0077260 | 521 | 11 | 4.22% |
| hsa_circ_0110526 | 521 | 11 | 4.22% |
| hsa_circ_0001075 | 474 | 10 | 4.22% |
| hsa_circ_0008074 | 474 | 10 | 4.22% |
| hsa_circ_0020929 | 474 | 10 | 4.22% |
| hsa_circ_0024246 | 474 | 10 | 4.22% |
| hsa_circ_0051843 | 474 | 10 | 4.22% |
| hsa_circ_0061481 | 474 | 10 | 4.22% |
| hsa_circ_0068642 | 474 | 10 | 4.22% |
| hsa_circ_0073766 | 474 | 10 | 4.22% |
| hsa_circ_0082305 | 474 | 10 | 4.22% |
| hsa_circ_0096220 | 474 | 10 | 4.22% |
| hsa_circ_0100450 | 474 | 10 | 4.22% |
| hsa_circ_0104852 | 474 | 10 | 4.22% |
| hsa_circ_0108556 | 474 | 10 | 4.22% |
| hsa_circ_0111072 | 474 | 10 | 4.22% |
| hsa_circ_0113231 | 474 | 10 | 4.22% |
| hsa_circ_0116569 | 474 | 10 | 4.22% |
| hsa_circ_0126100 | 474 | 10 | 4.22% |
| hsa_circ_0127051 | 474 | 10 | 4.22% |
| hsa_circ_0138948 | 474 | 10 | 4.22% |
| hsa_circ_0140229 | 474 | 10 | 4.22% |
| hsa_circ_0034764 | 569 | 12 | 4.22% |
| hsa_circ_0068376 | 522 | 11 | 4.21% |
| hsa_circ_0081872 | 522 | 11 | 4.21% |
| hsa_circ_0089110 | 522 | 11 | 4.21% |
| hsa_circ_0131236 | 522 | 11 | 4.21% |
| hsa_circ_0071569 | 570 | 12 | 4.21% |
| hsa_circ_0006234 | 475 | 10 | 4.21% |
| hsa_circ_0010613 | 475 | 10 | 4.21% |
| hsa_circ_0016807 | 475 | 10 | 4.21% |
| hsa_circ_0019670 | 475 | 10 | 4.21% |
| hsa_circ_0024812 | 475 | 10 | 4.21% |
| hsa_circ_0026947 | 475 | 10 | 4.21% |
| hsa_circ_0041978 | 475 | 10 | 4.21% |
| hsa_circ_0105379 | 475 | 10 | 4.21% |
| hsa_circ_0108006 | 475 | 10 | 4.21% |
| hsa_circ_0121661 | 475 | 10 | 4.21% |

| | | | |
|---|---|---|---|
| hsa_circ_0021985 | 618 | 13 | 4.21% |
| hsa_circ_0029322 | 523 | 11 | 4.21% |
| hsa_circ_0062710 | 523 | 11 | 4.21% |
| hsa_circ_0090450 | 523 | 11 | 4.21% |
| hsa_circ_0103804 | 523 | 11 | 4.21% |
| hsa_circ_0027592 | 571 | 12 | 4.20% |
| hsa_circ_0029476 | 476 | 10 | 4.20% |
| hsa_circ_0036593 | 476 | 10 | 4.20% |
| hsa_circ_0042241 | 476 | 10 | 4.20% |
| hsa_circ_0047294 | 476 | 10 | 4.20% |
| hsa_circ_0047837 | 476 | 10 | 4.20% |
| hsa_circ_0054610 | 476 | 10 | 4.20% |
| hsa_circ_0055485 | 476 | 10 | 4.20% |
| hsa_circ_0060478 | 476 | 10 | 4.20% |
| hsa_circ_0087014 | 476 | 10 | 4.20% |
| hsa_circ_0096459 | 476 | 10 | 4.20% |
| hsa_circ_0128670 | 476 | 10 | 4.20% |
| hsa_circ_0011745 | 524 | 11 | 4.20% |
| hsa_circ_0108277 | 524 | 11 | 4.20% |
| hsa_circ_0114259 | 524 | 11 | 4.20% |
| hsa_circ_0123645 | 524 | 11 | 4.20% |
| hsa_circ_0095917 | 667 | 14 | 4.20% |
| hsa_circ_0107621 | 572 | 12 | 4.20% |
| hsa_circ_0121049 | 572 | 12 | 4.20% |
| hsa_circ_0006676 | 477 | 10 | 4.19% |
| hsa_circ_0047066 | 477 | 10 | 4.19% |
| hsa_circ_0048126 | 477 | 10 | 4.19% |
| hsa_circ_0056880 | 477 | 10 | 4.19% |
| hsa_circ_0076809 | 477 | 10 | 4.19% |
| hsa_circ_0079760 | 477 | 10 | 4.19% |
| hsa_circ_0099384 | 477 | 10 | 4.19% |
| hsa_circ_0099699 | 477 | 10 | 4.19% |
| hsa_circ_0102367 | 477 | 10 | 4.19% |
| hsa_circ_0106453 | 477 | 10 | 4.19% |
| hsa_circ_0133871 | 477 | 10 | 4.19% |
| hsa_circ_0000457 | 668 | 14 | 4.19% |
| hsa_circ_0002410 | 525 | 11 | 4.19% |
| hsa_circ_0009257 | 525 | 11 | 4.19% |
| hsa_circ_0031267 | 525 | 11 | 4.19% |
| hsa_circ_0049857 | 525 | 11 | 4.19% |
| hsa_circ_0101714 | 525 | 11 | 4.19% |
| hsa_circ_0010747 | 478 | 10 | 4.18% |
| hsa_circ_0032759 | 478 | 10 | 4.18% |
| hsa_circ_0046422 | 478 | 10 | 4.18% |
| hsa_circ_0081753 | 478 | 10 | 4.18% |
| hsa_circ_0114921 | 478 | 10 | 4.18% |
| hsa_circ_0121442 | 478 | 10 | 4.18% |
| hsa_circ_0057383 | 765 | 16 | 4.18% |
| hsa_circ_0102071 | 526 | 11 | 4.18% |
| hsa_circ_0104546 | 526 | 11 | 4.18% |

| | | | |
|---|---|---|---|
| hsa_circ_0015272 | 574 | 12 | 4.18% |
| hsa_circ_0115262 | 574 | 12 | 4.18% |
| hsa_circ_0005793 | 479 | 10 | 4.18% |
| hsa_circ_0036321 | 479 | 10 | 4.18% |
| hsa_circ_0058970 | 479 | 10 | 4.18% |
| hsa_circ_0062169 | 479 | 10 | 4.18% |
| hsa_circ_0080323 | 479 | 10 | 4.18% |
| hsa_circ_0080639 | 479 | 10 | 4.18% |
| hsa_circ_0099842 | 479 | 10 | 4.18% |
| hsa_circ_0102926 | 479 | 10 | 4.18% |
| hsa_circ_0107046 | 479 | 10 | 4.18% |
| hsa_circ_0125314 | 479 | 10 | 4.18% |
| hsa_circ_0130227 | 479 | 10 | 4.18% |
| hsa_circ_0028146 | 527 | 11 | 4.17% |
| hsa_circ_0069073 | 527 | 11 | 4.17% |
| hsa_circ_0074893 | 527 | 11 | 4.17% |
| hsa_circ_0089572 | 527 | 11 | 4.17% |
| hsa_circ_0054513 | 624 | 13 | 4.17% |
| hsa_circ_0019805 | 528 | 11 | 4.17% |
| hsa_circ_0038754 | 528 | 11 | 4.17% |
| hsa_circ_0044258 | 528 | 11 | 4.17% |
| hsa_circ_0046375 | 528 | 11 | 4.17% |
| hsa_circ_0074649 | 528 | 11 | 4.17% |
| hsa_circ_0079204 | 528 | 11 | 4.17% |
| hsa_circ_0122230 | 528 | 11 | 4.17% |
| hsa_circ_0122551 | 528 | 11 | 4.17% |
| hsa_circ_0003742 | 480 | 10 | 4.17% |
| hsa_circ_0050121 | 480 | 10 | 4.17% |
| hsa_circ_0076599 | 480 | 10 | 4.17% |
| hsa_circ_0078832 | 480 | 10 | 4.17% |
| hsa_circ_0102734 | 480 | 10 | 4.17% |
| hsa_circ_0139628 | 480 | 10 | 4.17% |
| hsa_circ_0102220 | 1009 | 21 | 4.16% |
| hsa_circ_0004495 | 529 | 11 | 4.16% |
| hsa_circ_0066458 | 529 | 11 | 4.16% |
| hsa_circ_0129761 | 529 | 11 | 4.16% |
| hsa_circ_0008089 | 481 | 10 | 4.16% |
| hsa_circ_0021753 | 481 | 10 | 4.16% |
| hsa_circ_0022850 | 481 | 10 | 4.16% |
| hsa_circ_0024235 | 481 | 10 | 4.16% |
| hsa_circ_0072193 | 481 | 10 | 4.16% |
| hsa_circ_0086139 | 481 | 10 | 4.16% |
| hsa_circ_0086717 | 481 | 10 | 4.16% |
| hsa_circ_0115937 | 481 | 10 | 4.16% |
| hsa_circ_0126628 | 481 | 10 | 4.16% |
| hsa_circ_0061230 | 530 | 11 | 4.15% |
| hsa_circ_0127725 | 530 | 11 | 4.15% |
| hsa_circ_0002483 | 482 | 10 | 4.15% |
| hsa_circ_0009023 | 482 | 10 | 4.15% |
| hsa_circ_0018762 | 482 | 10 | 4.15% |

| | | | |
|---|---|---|---|
| hsa_circ_0040591 | 482 | 10 | 4.15% |
| hsa_circ_0109255 | 482 | 10 | 4.15% |
| hsa_circ_0113338 | 482 | 10 | 4.15% |
| hsa_circ_0115282 | 482 | 10 | 4.15% |
| hsa_circ_0067474 | 627 | 13 | 4.15% |
| hsa_circ_0131699 | 627 | 13 | 4.15% |
| hsa_circ_0043277 | 579 | 12 | 4.15% |
| hsa_circ_0003008 | 531 | 11 | 4.14% |
| hsa_circ_0048248 | 531 | 11 | 4.14% |
| hsa_circ_0052392 | 531 | 11 | 4.14% |
| hsa_circ_0076808 | 531 | 11 | 4.14% |
| hsa_circ_0084668 | 531 | 11 | 4.14% |
| hsa_circ_0104070 | 531 | 11 | 4.14% |
| hsa_circ_0105343 | 531 | 11 | 4.14% |
| hsa_circ_0113724 | 531 | 11 | 4.14% |
| hsa_circ_0139659 | 531 | 11 | 4.14% |
| hsa_circ_0089766 | 966 | 20 | 4.14% |
| hsa_circ_0004657 | 483 | 10 | 4.14% |
| hsa_circ_0006421 | 483 | 10 | 4.14% |
| hsa_circ_0026645 | 483 | 10 | 4.14% |
| hsa_circ_0028969 | 483 | 10 | 4.14% |
| hsa_circ_0030065 | 483 | 10 | 4.14% |
| hsa_circ_0047888 | 483 | 10 | 4.14% |
| hsa_circ_0070701 | 483 | 10 | 4.14% |
| hsa_circ_0081234 | 483 | 10 | 4.14% |
| hsa_circ_0101605 | 483 | 10 | 4.14% |
| hsa_circ_0117255 | 483 | 10 | 4.14% |
| hsa_circ_0055198 | 725 | 15 | 4.14% |
| hsa_circ_0023651 | 532 | 11 | 4.14% |
| hsa_circ_0037128 | 532 | 11 | 4.14% |
| hsa_circ_0133343 | 532 | 11 | 4.14% |
| hsa_circ_0011957 | 484 | 10 | 4.13% |
| hsa_circ_0038563 | 484 | 10 | 4.13% |
| hsa_circ_0050229 | 484 | 10 | 4.13% |
| hsa_circ_0096538 | 484 | 10 | 4.13% |
| hsa_circ_0110905 | 484 | 10 | 4.13% |
| hsa_circ_0116531 | 484 | 10 | 4.13% |
| hsa_circ_0123407 | 484 | 10 | 4.13% |
| hsa_circ_0110633 | 1599 | 33 | 4.13% |
| hsa_circ_0006441 | 533 | 11 | 4.13% |
| hsa_circ_0049447 | 533 | 11 | 4.13% |
| hsa_circ_0062063 | 533 | 11 | 4.13% |
| hsa_circ_0002020 | 485 | 10 | 4.12% |
| hsa_circ_0010020 | 485 | 10 | 4.12% |
| hsa_circ_0062344 | 485 | 10 | 4.12% |
| hsa_circ_0064604 | 485 | 10 | 4.12% |
| hsa_circ_0069259 | 485 | 10 | 4.12% |
| hsa_circ_0118644 | 485 | 10 | 4.12% |
| hsa_circ_0126089 | 485 | 10 | 4.12% |
| hsa_circ_0126166 | 485 | 10 | 4.12% |

| | | | |
|---|---|---|---|
| hsa_circ_0132814 | 485 | 10 | 4.12% |
| hsa_circ_0112948 | 631 | 13 | 4.12% |
| hsa_circ_0020374 | 534 | 11 | 4.12% |
| hsa_circ_0026007 | 534 | 11 | 4.12% |
| hsa_circ_0089593 | 534 | 11 | 4.12% |
| hsa_circ_0096191 | 534 | 11 | 4.12% |
| hsa_circ_0112667 | 534 | 11 | 4.12% |
| hsa_circ_0048588 | 583 | 12 | 4.12% |
| hsa_circ_0012720 | 486 | 10 | 4.12% |
| hsa_circ_0020767 | 486 | 10 | 4.12% |
| hsa_circ_0027300 | 486 | 10 | 4.12% |
| hsa_circ_0046524 | 486 | 10 | 4.12% |
| hsa_circ_0050000 | 486 | 10 | 4.12% |
| hsa_circ_0051859 | 486 | 10 | 4.12% |
| hsa_circ_0060990 | 486 | 10 | 4.12% |
| hsa_circ_0068900 | 486 | 10 | 4.12% |
| hsa_circ_0088735 | 486 | 10 | 4.12% |
| hsa_circ_0090306 | 486 | 10 | 4.12% |
| hsa_circ_0094782 | 486 | 10 | 4.12% |
| hsa_circ_0098314 | 486 | 10 | 4.12% |
| hsa_circ_0112329 | 486 | 10 | 4.12% |
| hsa_circ_0116518 | 486 | 10 | 4.12% |
| hsa_circ_0130652 | 486 | 10 | 4.12% |
| hsa_circ_0134366 | 486 | 10 | 4.12% |
| hsa_circ_0058973 | 535 | 11 | 4.11% |
| hsa_circ_0081414 | 535 | 11 | 4.11% |
| hsa_circ_0106772 | 535 | 11 | 4.11% |
| hsa_circ_0007948 | 487 | 10 | 4.11% |
| hsa_circ_0008812 | 487 | 10 | 4.11% |
| hsa_circ_0015279 | 487 | 10 | 4.11% |
| hsa_circ_0025235 | 487 | 10 | 4.11% |
| hsa_circ_0041506 | 487 | 10 | 4.11% |
| hsa_circ_0043167 | 487 | 10 | 4.11% |
| hsa_circ_0051814 | 487 | 10 | 4.11% |
| hsa_circ_0064354 | 487 | 10 | 4.11% |
| hsa_circ_0102272 | 487 | 10 | 4.11% |
| hsa_circ_0134542 | 1315 | 27 | 4.11% |
| hsa_circ_0133942 | 585 | 12 | 4.10% |
| hsa_circ_0109633 | 1268 | 26 | 4.10% |
| hsa_circ_0009382 | 488 | 10 | 4.10% |
| hsa_circ_0048581 | 488 | 10 | 4.10% |
| hsa_circ_0062181 | 488 | 10 | 4.10% |
| hsa_circ_0072996 | 488 | 10 | 4.10% |
| hsa_circ_0103930 | 488 | 10 | 4.10% |
| hsa_circ_0122053 | 488 | 10 | 4.10% |
| hsa_circ_0123838 | 488 | 10 | 4.10% |
| hsa_circ_0130330 | 488 | 10 | 4.10% |
| hsa_circ_0114511 | 586 | 12 | 4.10% |
| hsa_circ_0004449 | 489 | 10 | 4.09% |
| hsa_circ_0006412 | 489 | 10 | 4.09% |

| | | | |
|---|---|---|---|
| hsa_circ_0009163 | 489 | 10 | 4.09% |
| hsa_circ_0009599 | 489 | 10 | 4.09% |
| hsa_circ_0021075 | 489 | 10 | 4.09% |
| hsa_circ_0039141 | 489 | 10 | 4.09% |
| hsa_circ_0087690 | 489 | 10 | 4.09% |
| hsa_circ_0114593 | 489 | 10 | 4.09% |
| hsa_circ_0115129 | 489 | 10 | 4.09% |
| hsa_circ_0120762 | 489 | 10 | 4.09% |
| hsa_circ_0139678 | 489 | 10 | 4.09% |
| hsa_circ_0059660 | 538 | 11 | 4.09% |
| hsa_circ_0063809 | 538 | 11 | 4.09% |
| hsa_circ_0094106 | 538 | 11 | 4.09% |
| hsa_circ_0130948 | 538 | 11 | 4.09% |
| hsa_circ_0103856 | 587 | 12 | 4.09% |
| hsa_circ_0043670 | 588 | 12 | 4.08% |
| hsa_circ_0117677 | 588 | 12 | 4.08% |
| hsa_circ_0133943 | 588 | 12 | 4.08% |
| hsa_circ_0138834 | 588 | 12 | 4.08% |
| hsa_circ_0011406 | 490 | 10 | 4.08% |
| hsa_circ_0047891 | 490 | 10 | 4.08% |
| hsa_circ_0065738 | 490 | 10 | 4.08% |
| hsa_circ_0077138 | 490 | 10 | 4.08% |
| hsa_circ_0094546 | 3921 | 80 | 4.08% |
| hsa_circ_0128743 | 589 | 12 | 4.07% |
| hsa_circ_0080362 | 1080 | 22 | 4.07% |
| hsa_circ_0013466 | 540 | 11 | 4.07% |
| hsa_circ_0043528 | 540 | 11 | 4.07% |
| hsa_circ_0089914 | 540 | 11 | 4.07% |
| hsa_circ_0129825 | 540 | 11 | 4.07% |
| hsa_circ_0008912 | 491 | 10 | 4.07% |
| hsa_circ_0011635 | 491 | 10 | 4.07% |
| hsa_circ_0068637 | 491 | 10 | 4.07% |
| hsa_circ_0070446 | 491 | 10 | 4.07% |
| hsa_circ_0072490 | 491 | 10 | 4.07% |
| hsa_circ_0081980 | 491 | 10 | 4.07% |
| hsa_circ_0095691 | 491 | 10 | 4.07% |
| hsa_circ_0119768 | 491 | 10 | 4.07% |
| hsa_circ_0122825 | 491 | 10 | 4.07% |
| hsa_circ_0126556 | 491 | 10 | 4.07% |
| hsa_circ_0140321 | 491 | 10 | 4.07% |
| hsa_circ_0084023 | 639 | 13 | 4.07% |
| hsa_circ_0009512 | 541 | 11 | 4.07% |
| hsa_circ_0045425 | 541 | 11 | 4.07% |
| hsa_circ_0055502 | 541 | 11 | 4.07% |
| hsa_circ_0060938 | 541 | 11 | 4.07% |
| hsa_circ_0081054 | 541 | 11 | 4.07% |
| hsa_circ_0095771 | 541 | 11 | 4.07% |
| hsa_circ_0101135 | 541 | 11 | 4.07% |
| hsa_circ_0137640 | 541 | 11 | 4.07% |
| hsa_circ_0001999 | 492 | 10 | 4.07% |

| | | | |
|---|---|---|---|
| hsa_circ_0005030 | 492 | 10 | 4.07% |
| hsa_circ_0006724 | 492 | 10 | 4.07% |
| hsa_circ_0007543 | 492 | 10 | 4.07% |
| hsa_circ_0015129 | 492 | 10 | 4.07% |
| hsa_circ_0033139 | 492 | 10 | 4.07% |
| hsa_circ_0045138 | 492 | 10 | 4.07% |
| hsa_circ_0077831 | 492 | 10 | 4.07% |
| hsa_circ_0089101 | 492 | 10 | 4.07% |
| hsa_circ_0099098 | 492 | 10 | 4.07% |
| hsa_circ_0099681 | 492 | 10 | 4.07% |
| hsa_circ_0105078 | 492 | 10 | 4.07% |
| hsa_circ_0111756 | 492 | 10 | 4.07% |
| hsa_circ_0118886 | 492 | 10 | 4.07% |
| hsa_circ_0134730 | 492 | 10 | 4.07% |
| hsa_circ_0092908 | 542 | 11 | 4.06% |
| hsa_circ_0140301 | 542 | 11 | 4.06% |
| hsa_circ_0029257 | 493 | 10 | 4.06% |
| hsa_circ_0068753 | 493 | 10 | 4.06% |
| hsa_circ_0095029 | 493 | 10 | 4.06% |
| hsa_circ_0100296 | 493 | 10 | 4.06% |
| hsa_circ_0112815 | 493 | 10 | 4.06% |
| hsa_circ_0113524 | 493 | 10 | 4.06% |
| hsa_circ_0116210 | 493 | 10 | 4.06% |
| hsa_circ_0126103 | 493 | 10 | 4.06% |
| hsa_circ_0140113 | 493 | 10 | 4.06% |
| hsa_circ_0064083 | 641 | 13 | 4.06% |
| hsa_circ_0000527 | 6071 | 123 | 4.05% |
| hsa_circ_0011149 | 543 | 11 | 4.05% |
| hsa_circ_0121165 | 543 | 11 | 4.05% |
| hsa_circ_0002775 | 494 | 10 | 4.05% |
| hsa_circ_0018348 | 494 | 10 | 4.05% |
| hsa_circ_0022630 | 494 | 10 | 4.05% |
| hsa_circ_0025945 | 494 | 10 | 4.05% |
| hsa_circ_0027462 | 494 | 10 | 4.05% |
| hsa_circ_0065867 | 494 | 10 | 4.05% |
| hsa_circ_0072491 | 494 | 10 | 4.05% |
| hsa_circ_0095909 | 494 | 10 | 4.05% |
| hsa_circ_0106242 | 494 | 10 | 4.05% |
| hsa_circ_0124616 | 494 | 10 | 4.05% |
| hsa_circ_0034722 | 544 | 11 | 4.04% |
| hsa_circ_0042739 | 544 | 11 | 4.04% |
| hsa_circ_0097296 | 544 | 11 | 4.04% |
| hsa_circ_0116419 | 544 | 11 | 4.04% |
| hsa_circ_0123641 | 643 | 13 | 4.04% |
| hsa_circ_0018493 | 594 | 12 | 4.04% |
| hsa_circ_0006405 | 495 | 10 | 4.04% |
| hsa_circ_0022650 | 495 | 10 | 4.04% |
| hsa_circ_0064030 | 495 | 10 | 4.04% |
| hsa_circ_0089667 | 495 | 10 | 4.04% |
| hsa_circ_0118327 | 495 | 10 | 4.04% |

| | | | |
|---|---|---|---|
| hsa_circ_0132094 | 495 | 10 | 4.04% |
| hsa_circ_0133157 | 495 | 10 | 4.04% |
| hsa_circ_0000029 | 496 | 10 | 4.03% |
| hsa_circ_0019029 | 496 | 10 | 4.03% |
| hsa_circ_0023087 | 496 | 10 | 4.03% |
| hsa_circ_0088427 | 496 | 10 | 4.03% |
| hsa_circ_0132534 | 496 | 10 | 4.03% |
| hsa_circ_0023054 | 645 | 13 | 4.03% |
| hsa_circ_0012355 | 546 | 11 | 4.03% |
| hsa_circ_0107396 | 546 | 11 | 4.03% |
| hsa_circ_0135214 | 546 | 11 | 4.03% |
| hsa_circ_0095381 | 596 | 12 | 4.03% |
| hsa_circ_0059418 | 646 | 13 | 4.02% |
| hsa_circ_0000415 | 497 | 10 | 4.02% |
| hsa_circ_0002186 | 497 | 10 | 4.02% |
| hsa_circ_0008820 | 497 | 10 | 4.02% |
| hsa_circ_0026302 | 497 | 10 | 4.02% |
| hsa_circ_0051128 | 497 | 10 | 4.02% |
| hsa_circ_0056360 | 497 | 10 | 4.02% |
| hsa_circ_0072549 | 497 | 10 | 4.02% |
| hsa_circ_0081990 | 497 | 10 | 4.02% |
| hsa_circ_0085129 | 497 | 10 | 4.02% |
| hsa_circ_0091194 | 497 | 10 | 4.02% |
| hsa_circ_0117991 | 497 | 10 | 4.02% |
| hsa_circ_0119328 | 497 | 10 | 4.02% |
| hsa_circ_0124732 | 497 | 10 | 4.02% |
| hsa_circ_0135935 | 497 | 10 | 4.02% |
| hsa_circ_0019987 | 547 | 11 | 4.02% |
| hsa_circ_0046499 | 547 | 11 | 4.02% |
| hsa_circ_0093680 | 547 | 11 | 4.02% |
| hsa_circ_0122050 | 547 | 11 | 4.02% |
| hsa_circ_0138649 | 547 | 11 | 4.02% |
| hsa_circ_0007354 | 597 | 12 | 4.02% |
| hsa_circ_0079225 | 1045 | 21 | 4.02% |
| hsa_circ_0009739 | 647 | 13 | 4.02% |
| hsa_circ_0099923 | 647 | 13 | 4.02% |
| hsa_circ_0093227 | 498 | 10 | 4.02% |
| hsa_circ_0105245 | 498 | 10 | 4.02% |
| hsa_circ_0117809 | 498 | 10 | 4.02% |
| hsa_circ_0121161 | 498 | 10 | 4.02% |
| hsa_circ_0123684 | 498 | 10 | 4.02% |
| hsa_circ_0131943 | 498 | 10 | 4.02% |
| hsa_circ_0131946 | 498 | 10 | 4.02% |
| hsa_circ_0133368 | 498 | 10 | 4.02% |
| hsa_circ_0000180 | 548 | 11 | 4.01% |
| hsa_circ_0005490 | 548 | 11 | 4.01% |
| hsa_circ_0006222 | 548 | 11 | 4.01% |
| hsa_circ_0042785 | 548 | 11 | 4.01% |
| hsa_circ_0097761 | 548 | 11 | 4.01% |
| hsa_circ_0118817 | 548 | 11 | 4.01% |

| | | | |
|---|---|---|---|
| hsa_circ_0009383 | 598 | 12 | 4.01% |
| hsa_circ_0024788 | 598 | 12 | 4.01% |
| hsa_circ_0010366 | 499 | 10 | 4.01% |
| hsa_circ_0013512 | 499 | 10 | 4.01% |
| hsa_circ_0015027 | 499 | 10 | 4.01% |
| hsa_circ_0048904 | 499 | 10 | 4.01% |
| hsa_circ_0069613 | 499 | 10 | 4.01% |
| hsa_circ_0082461 | 499 | 10 | 4.01% |
| hsa_circ_0088227 | 499 | 10 | 4.01% |
| hsa_circ_0093399 | 499 | 10 | 4.01% |
| hsa_circ_0102679 | 499 | 10 | 4.01% |
| hsa_circ_0132535 | 499 | 10 | 4.01% |
| hsa_circ_0000778 | 549 | 11 | 4.01% |
| hsa_circ_0035074 | 549 | 11 | 4.01% |
| hsa_circ_0038819 | 549 | 11 | 4.01% |
| hsa_circ_0057447 | 549 | 11 | 4.01% |
| hsa_circ_0087842 | 549 | 11 | 4.01% |
| hsa_circ_0089957 | 549 | 11 | 4.01% |
| hsa_circ_0095494 | 549 | 11 | 4.01% |
| hsa_circ_0073626 | 599 | 12 | 4.01% |
| hsa_circ_0125925 | 599 | 12 | 4.01% |
| hsa_circ_0137530 | 599 | 12 | 4.01% |
| hsa_circ_0116385 | 3694 | 74 | 4.01% |
| hsa_circ_0047092 | 649 | 13 | 4.01% |
| hsa_circ_0105458 | 649 | 13 | 4.01% |
| hsa_circ_0023612 | 650 | 13 | 4.00% |
| hsa_circ_0029957 | 550 | 11 | 4.00% |
| hsa_circ_0068118 | 550 | 11 | 4.00% |
| hsa_circ_0098133 | 550 | 11 | 4.00% |
| hsa_circ_0123026 | 550 | 11 | 4.00% |
| hsa_circ_0002060 | 500 | 10 | 4.00% |
| hsa_circ_0012058 | 500 | 10 | 4.00% |
| hsa_circ_0014986 | 500 | 10 | 4.00% |
| hsa_circ_0081734 | 500 | 10 | 4.00% |
| hsa_circ_0092296 | 500 | 10 | 4.00% |
| hsa_circ_0096647 | 500 | 10 | 4.00% |
| hsa_circ_0109697 | 500 | 10 | 4.00% |
| hsa_circ_0119514 | 500 | 10 | 4.00% |
| hsa_circ_0005054 | 551 | 11 | 3.99% |
| hsa_circ_0018636 | 551 | 11 | 3.99% |
| hsa_circ_0029375 | 551 | 11 | 3.99% |
| hsa_circ_0130812 | 551 | 11 | 3.99% |
| hsa_circ_0003557 | 501 | 10 | 3.99% |
| hsa_circ_0003979 | 501 | 10 | 3.99% |
| hsa_circ_0034554 | 501 | 10 | 3.99% |
| hsa_circ_0054402 | 501 | 10 | 3.99% |
| hsa_circ_0059792 | 501 | 10 | 3.99% |
| hsa_circ_0063493 | 501 | 10 | 3.99% |
| hsa_circ_0077995 | 501 | 10 | 3.99% |
| hsa_circ_0095210 | 501 | 10 | 3.99% |

| | | | |
|---|---|---|---|
| hsa_circ_0102493 | 501 | 10 | 3.99% |
| hsa_circ_0104284 | 501 | 10 | 3.99% |
| hsa_circ_0111266 | 501 | 10 | 3.99% |
| hsa_circ_0112097 | 501 | 10 | 3.99% |
| hsa_circ_0116701 | 501 | 10 | 3.99% |
| hsa_circ_0101626 | 602 | 12 | 3.99% |
| hsa_circ_0109163 | 602 | 12 | 3.99% |
| hsa_circ_0056412 | 552 | 11 | 3.99% |
| hsa_circ_0076288 | 552 | 11 | 3.99% |
| hsa_circ_0081055 | 552 | 11 | 3.99% |
| hsa_circ_0095934 | 552 | 11 | 3.99% |
| hsa_circ_0004861 | 502 | 10 | 3.98% |
| hsa_circ_0006245 | 502 | 10 | 3.98% |
| hsa_circ_0028506 | 502 | 10 | 3.98% |
| hsa_circ_0034339 | 502 | 10 | 3.98% |
| hsa_circ_0036929 | 502 | 10 | 3.98% |
| hsa_circ_0037363 | 502 | 10 | 3.98% |
| hsa_circ_0047087 | 502 | 10 | 3.98% |
| hsa_circ_0062877 | 502 | 10 | 3.98% |
| hsa_circ_0068794 | 502 | 10 | 3.98% |
| hsa_circ_0071361 | 502 | 10 | 3.98% |
| hsa_circ_0094222 | 502 | 10 | 3.98% |
| hsa_circ_0098898 | 502 | 10 | 3.98% |
| hsa_circ_0103359 | 502 | 10 | 3.98% |
| hsa_circ_0106672 | 502 | 10 | 3.98% |
| hsa_circ_0106908 | 502 | 10 | 3.98% |
| hsa_circ_0117602 | 502 | 10 | 3.98% |
| hsa_circ_0127000 | 502 | 10 | 3.98% |
| hsa_circ_0128062 | 502 | 10 | 3.98% |
| hsa_circ_0128879 | 502 | 10 | 3.98% |
| hsa_circ_0138214 | 502 | 10 | 3.98% |
| hsa_circ_0140692 | 502 | 10 | 3.98% |
| hsa_circ_0052824 | 653 | 13 | 3.98% |
| hsa_circ_0115324 | 3165 | 63 | 3.98% |
| hsa_circ_0034080 | 603 | 12 | 3.98% |
| hsa_circ_0043270 | 603 | 12 | 3.98% |
| hsa_circ_0050402 | 603 | 12 | 3.98% |
| hsa_circ_0096364 | 603 | 12 | 3.98% |
| hsa_circ_0014544 | 553 | 11 | 3.98% |
| hsa_circ_0047884 | 553 | 11 | 3.98% |
| hsa_circ_0061955 | 553 | 11 | 3.98% |
| hsa_circ_0062712 | 553 | 11 | 3.98% |
| hsa_circ_0063173 | 553 | 11 | 3.98% |
| hsa_circ_0099606 | 553 | 11 | 3.98% |
| hsa_circ_0021221 | 503 | 10 | 3.98% |
| hsa_circ_0021482 | 503 | 10 | 3.98% |
| hsa_circ_0028209 | 503 | 10 | 3.98% |
| hsa_circ_0076762 | 503 | 10 | 3.98% |
| hsa_circ_0090984 | 503 | 10 | 3.98% |
| hsa_circ_0107582 | 503 | 10 | 3.98% |

| | | | |
|---|---|---|---|
| hsa_circ_0109415 | 503 | 10 | 3.98% |
| hsa_circ_0137008 | 503 | 10 | 3.98% |
| hsa_circ_0048226 | 654 | 13 | 3.98% |
| hsa_circ_0009055 | 554 | 11 | 3.97% |
| hsa_circ_0057055 | 554 | 11 | 3.97% |
| hsa_circ_0099371 | 554 | 11 | 3.97% |
| hsa_circ_0111011 | 554 | 11 | 3.97% |
| hsa_circ_0121974 | 554 | 11 | 3.97% |
| hsa_circ_0002066 | 504 | 10 | 3.97% |
| hsa_circ_0007578 | 504 | 10 | 3.97% |
| hsa_circ_0016471 | 504 | 10 | 3.97% |
| hsa_circ_0030567 | 504 | 10 | 3.97% |
| hsa_circ_0031391 | 504 | 10 | 3.97% |
| hsa_circ_0054026 | 504 | 10 | 3.97% |
| hsa_circ_0062081 | 504 | 10 | 3.97% |
| hsa_circ_0075229 | 504 | 10 | 3.97% |
| hsa_circ_0080108 | 504 | 10 | 3.97% |
| hsa_circ_0100520 | 504 | 10 | 3.97% |
| hsa_circ_0109113 | 504 | 10 | 3.97% |
| hsa_circ_0109627 | 504 | 10 | 3.97% |
| hsa_circ_0115595 | 504 | 10 | 3.97% |
| hsa_circ_0131618 | 504 | 10 | 3.97% |
| hsa_circ_0137510 | 504 | 10 | 3.97% |
| hsa_circ_0061350 | 555 | 11 | 3.96% |
| hsa_circ_0109755 | 555 | 11 | 3.96% |
| hsa_circ_0128456 | 555 | 11 | 3.96% |
| hsa_circ_0020323 | 656 | 13 | 3.96% |
| hsa_circ_0022841 | 808 | 16 | 3.96% |
| hsa_circ_0001302 | 505 | 10 | 3.96% |
| hsa_circ_0004644 | 505 | 10 | 3.96% |
| hsa_circ_0008007 | 505 | 10 | 3.96% |
| hsa_circ_0010988 | 505 | 10 | 3.96% |
| hsa_circ_0016359 | 505 | 10 | 3.96% |
| hsa_circ_0019786 | 505 | 10 | 3.96% |
| hsa_circ_0024566 | 505 | 10 | 3.96% |
| hsa_circ_0041561 | 505 | 10 | 3.96% |
| hsa_circ_0043463 | 505 | 10 | 3.96% |
| hsa_circ_0044266 | 505 | 10 | 3.96% |
| hsa_circ_0077241 | 505 | 10 | 3.96% |
| hsa_circ_0085403 | 505 | 10 | 3.96% |
| hsa_circ_0100772 | 505 | 10 | 3.96% |
| hsa_circ_0107231 | 505 | 10 | 3.96% |
| hsa_circ_0116817 | 505 | 10 | 3.96% |
| hsa_circ_0120892 | 505 | 10 | 3.96% |
| hsa_circ_0136721 | 505 | 10 | 3.96% |
| hsa_circ_0139221 | 505 | 10 | 3.96% |
| hsa_circ_0117542 | 657 | 13 | 3.96% |
| hsa_circ_0002382 | 556 | 11 | 3.96% |
| hsa_circ_0019213 | 556 | 11 | 3.96% |
| hsa_circ_0023496 | 556 | 11 | 3.96% |

| | | | |
|---|---|---|---|
| hsa_circ_0132905 | 556 | 11 | 3.96% |
| hsa_circ_0140341 | 556 | 11 | 3.96% |
| hsa_circ_0063905 | 607 | 12 | 3.95% |
| hsa_circ_0002231 | 506 | 10 | 3.95% |
| hsa_circ_0051249 | 506 | 10 | 3.95% |
| hsa_circ_0062167 | 506 | 10 | 3.95% |
| hsa_circ_0066203 | 506 | 10 | 3.95% |
| hsa_circ_0095974 | 506 | 10 | 3.95% |
| hsa_circ_0112427 | 506 | 10 | 3.95% |
| hsa_circ_0123496 | 506 | 10 | 3.95% |
| hsa_circ_0115266 | 658 | 13 | 3.95% |
| hsa_circ_0009011 | 557 | 11 | 3.95% |
| hsa_circ_0124088 | 557 | 11 | 3.95% |
| hsa_circ_0128593 | 557 | 11 | 3.95% |
| hsa_circ_0135108 | 557 | 11 | 3.95% |
| hsa_circ_0060164 | 608 | 12 | 3.95% |
| hsa_circ_0112563 | 608 | 12 | 3.95% |
| hsa_circ_0003361 | 507 | 10 | 3.94% |
| hsa_circ_0005663 | 507 | 10 | 3.94% |
| hsa_circ_0020142 | 507 | 10 | 3.94% |
| hsa_circ_0030010 | 507 | 10 | 3.94% |
| hsa_circ_0040763 | 507 | 10 | 3.94% |
| hsa_circ_0042551 | 507 | 10 | 3.94% |
| hsa_circ_0043596 | 507 | 10 | 3.94% |
| hsa_circ_0066737 | 507 | 10 | 3.94% |
| hsa_circ_0075031 | 507 | 10 | 3.94% |
| hsa_circ_0102687 | 507 | 10 | 3.94% |
| hsa_circ_0106034 | 507 | 10 | 3.94% |
| hsa_circ_0119765 | 507 | 10 | 3.94% |
| hsa_circ_0125050 | 507 | 10 | 3.94% |
| hsa_circ_0130714 | 507 | 10 | 3.94% |
| hsa_circ_0133556 | 507 | 10 | 3.94% |
| hsa_circ_0022409 | 558 | 11 | 3.94% |
| hsa_circ_0064315 | 558 | 11 | 3.94% |
| hsa_circ_0096246 | 558 | 11 | 3.94% |
| hsa_circ_0139661 | 558 | 11 | 3.94% |
| hsa_circ_0026017 | 609 | 12 | 3.94% |
| hsa_circ_0027542 | 609 | 12 | 3.94% |
| hsa_circ_0047615 | 609 | 12 | 3.94% |
| hsa_circ_0102728 | 660 | 13 | 3.94% |
| hsa_circ_0011490 | 508 | 10 | 3.94% |
| hsa_circ_0011633 | 508 | 10 | 3.94% |
| hsa_circ_0016251 | 508 | 10 | 3.94% |
| hsa_circ_0038637 | 508 | 10 | 3.94% |
| hsa_circ_0065224 | 508 | 10 | 3.94% |
| hsa_circ_0076444 | 508 | 10 | 3.94% |
| hsa_circ_0106147 | 508 | 10 | 3.94% |
| hsa_circ_0106349 | 508 | 10 | 3.94% |
| hsa_circ_0107117 | 508 | 10 | 3.94% |
| hsa_circ_0115251 | 508 | 10 | 3.94% |

| | | | |
|---|---|---|---|
| hsa_circ_0117650 | 508 | 10 | 3.94% |
| hsa_circ_0123574 | 508 | 10 | 3.94% |
| hsa_circ_0127576 | 508 | 10 | 3.94% |
| hsa_circ_0127715 | 508 | 10 | 3.94% |
| hsa_circ_0129371 | 508 | 10 | 3.94% |
| hsa_circ_0062107 | 813 | 16 | 3.94% |
| hsa_circ_0026010 | 559 | 11 | 3.94% |
| hsa_circ_0100512 | 559 | 11 | 3.94% |
| hsa_circ_0112411 | 559 | 11 | 3.94% |
| hsa_circ_0064155 | 610 | 12 | 3.93% |
| hsa_circ_0102681 | 610 | 12 | 3.93% |
| hsa_circ_0129903 | 661 | 13 | 3.93% |
| hsa_circ_0024771 | 509 | 10 | 3.93% |
| hsa_circ_0031313 | 509 | 10 | 3.93% |
| hsa_circ_0038961 | 509 | 10 | 3.93% |
| hsa_circ_0075128 | 509 | 10 | 3.93% |
| hsa_circ_0116398 | 509 | 10 | 3.93% |
| hsa_circ_0012813 | 560 | 11 | 3.93% |
| hsa_circ_0014889 | 560 | 11 | 3.93% |
| hsa_circ_0078603 | 560 | 11 | 3.93% |
| hsa_circ_0132644 | 560 | 11 | 3.93% |
| hsa_circ_0020112 | 867 | 17 | 3.92% |
| hsa_circ_0083249 | 816 | 16 | 3.92% |
| hsa_circ_0019403 | 612 | 12 | 3.92% |
| hsa_circ_0108537 | 612 | 12 | 3.92% |
| hsa_circ_0023000 | 561 | 11 | 3.92% |
| hsa_circ_0062064 | 561 | 11 | 3.92% |
| hsa_circ_0088775 | 561 | 11 | 3.92% |
| hsa_circ_0090680 | 561 | 11 | 3.92% |
| hsa_circ_0008362 | 510 | 10 | 3.92% |
| hsa_circ_0008399 | 510 | 10 | 3.92% |
| hsa_circ_0012594 | 510 | 10 | 3.92% |
| hsa_circ_0014229 | 510 | 10 | 3.92% |
| hsa_circ_0023034 | 510 | 10 | 3.92% |
| hsa_circ_0033617 | 510 | 10 | 3.92% |
| hsa_circ_0041389 | 510 | 10 | 3.92% |
| hsa_circ_0069857 | 510 | 10 | 3.92% |
| hsa_circ_0080512 | 510 | 10 | 3.92% |
| hsa_circ_0086164 | 510 | 10 | 3.92% |
| hsa_circ_0086461 | 510 | 10 | 3.92% |
| hsa_circ_0090401 | 510 | 10 | 3.92% |
| hsa_circ_0090958 | 510 | 10 | 3.92% |
| hsa_circ_0119763 | 510 | 10 | 3.92% |
| hsa_circ_0123043 | 510 | 10 | 3.92% |
| hsa_circ_0137808 | 510 | 10 | 3.92% |
| hsa_circ_0048047 | 664 | 13 | 3.92% |
| hsa_circ_0079530 | 664 | 13 | 3.92% |
| hsa_circ_0126645 | 613 | 12 | 3.92% |
| hsa_circ_0011749 | 562 | 11 | 3.91% |
| hsa_circ_0067157 | 562 | 11 | 3.91% |

| | | | |
|---|---|---|---|
| hsa_circ_0140637 | 562 | 11 | 3.91% |
| hsa_circ_0003559 | 511 | 10 | 3.91% |
| hsa_circ_0010779 | 511 | 10 | 3.91% |
| hsa_circ_0012126 | 511 | 10 | 3.91% |
| hsa_circ_0012750 | 511 | 10 | 3.91% |
| hsa_circ_0016329 | 511 | 10 | 3.91% |
| hsa_circ_0055685 | 511 | 10 | 3.91% |
| hsa_circ_0059631 | 511 | 10 | 3.91% |
| hsa_circ_0065446 | 511 | 10 | 3.91% |
| hsa_circ_0068620 | 511 | 10 | 3.91% |
| hsa_circ_0096339 | 511 | 10 | 3.91% |
| hsa_circ_0102258 | 511 | 10 | 3.91% |
| hsa_circ_0112895 | 511 | 10 | 3.91% |
| hsa_circ_0135418 | 511 | 10 | 3.91% |
| hsa_circ_0000747 | 614 | 12 | 3.91% |
| hsa_circ_0071734 | 614 | 12 | 3.91% |
| hsa_circ_0037841 | 563 | 11 | 3.91% |
| hsa_circ_0063176 | 563 | 11 | 3.91% |
| hsa_circ_0098059 | 563 | 11 | 3.91% |
| hsa_circ_0118813 | 563 | 11 | 3.91% |
| hsa_circ_0002411 | 512 | 10 | 3.91% |
| hsa_circ_0013654 | 512 | 10 | 3.91% |
| hsa_circ_0015484 | 512 | 10 | 3.91% |
| hsa_circ_0016220 | 512 | 10 | 3.91% |
| hsa_circ_0022967 | 512 | 10 | 3.91% |
| hsa_circ_0023093 | 512 | 10 | 3.91% |
| hsa_circ_0035462 | 512 | 10 | 3.91% |
| hsa_circ_0036088 | 512 | 10 | 3.91% |
| hsa_circ_0067943 | 512 | 10 | 3.91% |
| hsa_circ_0086259 | 512 | 10 | 3.91% |
| hsa_circ_0087347 | 512 | 10 | 3.91% |
| hsa_circ_0093884 | 512 | 10 | 3.91% |
| hsa_circ_0103090 | 512 | 10 | 3.91% |
| hsa_circ_0106369 | 512 | 10 | 3.91% |
| hsa_circ_0110304 | 512 | 10 | 3.91% |
| hsa_circ_0113379 | 512 | 10 | 3.91% |
| hsa_circ_0119759 | 512 | 10 | 3.91% |
| hsa_circ_0125557 | 512 | 10 | 3.91% |
| hsa_circ_0140026 | 512 | 10 | 3.91% |
| hsa_circ_0123532 | 666 | 13 | 3.90% |
| hsa_circ_0043669 | 615 | 12 | 3.90% |
| hsa_circ_0102301 | 615 | 12 | 3.90% |
| hsa_circ_0121924 | 615 | 12 | 3.90% |
| hsa_circ_0055978 | 564 | 11 | 3.90% |
| hsa_circ_0058530 | 564 | 11 | 3.90% |
| hsa_circ_0061541 | 564 | 11 | 3.90% |
| hsa_circ_0074214 | 564 | 11 | 3.90% |
| hsa_circ_0095442 | 564 | 11 | 3.90% |
| hsa_circ_0139810 | 564 | 11 | 3.90% |
| hsa_circ_0016358 | 513 | 10 | 3.90% |

| | | | |
|---|---|---|---|
| hsa_circ_0025773 | 513 | 10 | 3.90% |
| hsa_circ_0032984 | 513 | 10 | 3.90% |
| hsa_circ_0043818 | 513 | 10 | 3.90% |
| hsa_circ_0047915 | 513 | 10 | 3.90% |
| hsa_circ_0054687 | 513 | 10 | 3.90% |
| hsa_circ_0060570 | 513 | 10 | 3.90% |
| hsa_circ_0064435 | 513 | 10 | 3.90% |
| hsa_circ_0069342 | 513 | 10 | 3.90% |
| hsa_circ_0093018 | 513 | 10 | 3.90% |
| hsa_circ_0093540 | 513 | 10 | 3.90% |
| hsa_circ_0094221 | 513 | 10 | 3.90% |
| hsa_circ_0104871 | 513 | 10 | 3.90% |
| hsa_circ_0122660 | 513 | 10 | 3.90% |
| hsa_circ_0123497 | 513 | 10 | 3.90% |
| hsa_circ_0126300 | 513 | 10 | 3.90% |
| hsa_circ_0126923 | 513 | 10 | 3.90% |
| hsa_circ_0055785 | 667 | 13 | 3.90% |
| hsa_circ_0006893 | 565 | 11 | 3.89% |
| hsa_circ_0008055 | 565 | 11 | 3.89% |
| hsa_circ_0130673 | 565 | 11 | 3.89% |
| hsa_circ_0140782 | 565 | 11 | 3.89% |
| hsa_circ_0005572 | 514 | 10 | 3.89% |
| hsa_circ_0014098 | 514 | 10 | 3.89% |
| hsa_circ_0021177 | 514 | 10 | 3.89% |
| hsa_circ_0023597 | 514 | 10 | 3.89% |
| hsa_circ_0045775 | 514 | 10 | 3.89% |
| hsa_circ_0058180 | 514 | 10 | 3.89% |
| hsa_circ_0068747 | 514 | 10 | 3.89% |
| hsa_circ_0079810 | 514 | 10 | 3.89% |
| hsa_circ_0089027 | 514 | 10 | 3.89% |
| hsa_circ_0089988 | 514 | 10 | 3.89% |
| hsa_circ_0096012 | 514 | 10 | 3.89% |
| hsa_circ_0117423 | 514 | 10 | 3.89% |
| hsa_circ_0119621 | 514 | 10 | 3.89% |
| hsa_circ_0003399 | 566 | 11 | 3.89% |
| hsa_circ_0023574 | 566 | 11 | 3.89% |
| hsa_circ_0036370 | 566 | 11 | 3.89% |
| hsa_circ_0095446 | 566 | 11 | 3.89% |
| hsa_circ_0090165 | 618 | 12 | 3.88% |
| hsa_circ_0099446 | 618 | 12 | 3.88% |
| hsa_circ_0005058 | 515 | 10 | 3.88% |
| hsa_circ_0008085 | 515 | 10 | 3.88% |
| hsa_circ_0015366 | 515 | 10 | 3.88% |
| hsa_circ_0034705 | 515 | 10 | 3.88% |
| hsa_circ_0063729 | 515 | 10 | 3.88% |
| hsa_circ_0075610 | 515 | 10 | 3.88% |
| hsa_circ_0076506 | 515 | 10 | 3.88% |
| hsa_circ_0107237 | 515 | 10 | 3.88% |
| hsa_circ_0129632 | 515 | 10 | 3.88% |
| hsa_circ_0121450 | 567 | 11 | 3.88% |

| | | | |
|---|---|---|---|
| hsa_circ_0134547 | 567 | 11 | 3.88% |
| hsa_circ_0094707 | 619 | 12 | 3.88% |
| hsa_circ_0001681 | 516 | 10 | 3.88% |
| hsa_circ_0012596 | 516 | 10 | 3.88% |
| hsa_circ_0012755 | 516 | 10 | 3.88% |
| hsa_circ_0014717 | 516 | 10 | 3.88% |
| hsa_circ_0035824 | 516 | 10 | 3.88% |
| hsa_circ_0045548 | 516 | 10 | 3.88% |
| hsa_circ_0050378 | 516 | 10 | 3.88% |
| hsa_circ_0063359 | 516 | 10 | 3.88% |
| hsa_circ_0079761 | 516 | 10 | 3.88% |
| hsa_circ_0099808 | 516 | 10 | 3.88% |
| hsa_circ_0103925 | 516 | 10 | 3.88% |
| hsa_circ_0106366 | 516 | 10 | 3.88% |
| hsa_circ_0126036 | 516 | 10 | 3.88% |
| hsa_circ_0127447 | 516 | 10 | 3.88% |
| hsa_circ_0129585 | 516 | 10 | 3.88% |
| hsa_circ_0134909 | 516 | 10 | 3.88% |
| hsa_circ_0003779 | 568 | 11 | 3.87% |
| hsa_circ_0028889 | 568 | 11 | 3.87% |
| hsa_circ_0093890 | 568 | 11 | 3.87% |
| hsa_circ_0014963 | 517 | 10 | 3.87% |
| hsa_circ_0025798 | 517 | 10 | 3.87% |
| hsa_circ_0031011 | 517 | 10 | 3.87% |
| hsa_circ_0044244 | 517 | 10 | 3.87% |
| hsa_circ_0066661 | 517 | 10 | 3.87% |
| hsa_circ_0081554 | 517 | 10 | 3.87% |
| hsa_circ_0082142 | 517 | 10 | 3.87% |
| hsa_circ_0083420 | 517 | 10 | 3.87% |
| hsa_circ_0090441 | 517 | 10 | 3.87% |
| hsa_circ_0092074 | 517 | 10 | 3.87% |
| hsa_circ_0107342 | 517 | 10 | 3.87% |
| hsa_circ_0112385 | 517 | 10 | 3.87% |
| hsa_circ_0024178 | 569 | 11 | 3.87% |
| hsa_circ_0055215 | 569 | 11 | 3.87% |
| hsa_circ_0082013 | 569 | 11 | 3.87% |
| hsa_circ_0118296 | 569 | 11 | 3.87% |
| hsa_circ_0004953 | 518 | 10 | 3.86% |
| hsa_circ_0032338 | 518 | 10 | 3.86% |
| hsa_circ_0032546 | 518 | 10 | 3.86% |
| hsa_circ_0057878 | 518 | 10 | 3.86% |
| hsa_circ_0066632 | 518 | 10 | 3.86% |
| hsa_circ_0078336 | 518 | 10 | 3.86% |
| hsa_circ_0079757 | 518 | 10 | 3.86% |
| hsa_circ_0126254 | 518 | 10 | 3.86% |
| hsa_circ_0130443 | 518 | 10 | 3.86% |
| hsa_circ_0139360 | 518 | 10 | 3.86% |
| hsa_circ_0139641 | 518 | 10 | 3.86% |
| hsa_circ_0057350 | 570 | 11 | 3.86% |
| hsa_circ_0085990 | 570 | 11 | 3.86% |

| | | | |
|---|---|---|---|
| hsa_circ_0089645 | 570 | 11 | 3.86% |
| hsa_circ_0127268 | 570 | 11 | 3.86% |
| hsa_circ_0086046 | 674 | 13 | 3.86% |
| hsa_circ_0001350 | 519 | 10 | 3.85% |
| hsa_circ_0002265 | 519 | 10 | 3.85% |
| hsa_circ_0011466 | 519 | 10 | 3.85% |
| hsa_circ_0030549 | 519 | 10 | 3.85% |
| hsa_circ_0055536 | 519 | 10 | 3.85% |
| hsa_circ_0085843 | 519 | 10 | 3.85% |
| hsa_circ_0090085 | 519 | 10 | 3.85% |
| hsa_circ_0091895 | 519 | 10 | 3.85% |
| hsa_circ_0093318 | 519 | 10 | 3.85% |
| hsa_circ_0100398 | 519 | 10 | 3.85% |
| hsa_circ_0104282 | 519 | 10 | 3.85% |
| hsa_circ_0113094 | 519 | 10 | 3.85% |
| hsa_circ_0135503 | 519 | 10 | 3.85% |
| hsa_circ_0012174 | 571 | 11 | 3.85% |
| hsa_circ_0048933 | 571 | 11 | 3.85% |
| hsa_circ_0070127 | 571 | 11 | 3.85% |
| hsa_circ_0103317 | 571 | 11 | 3.85% |
| hsa_circ_0119180 | 571 | 11 | 3.85% |
| hsa_circ_0137074 | 623 | 12 | 3.85% |
| hsa_circ_0057194 | 624 | 12 | 3.85% |
| hsa_circ_0116293 | 624 | 12 | 3.85% |
| hsa_circ_0091157 | 572 | 11 | 3.85% |
| hsa_circ_0092898 | 572 | 11 | 3.85% |
| hsa_circ_0096002 | 572 | 11 | 3.85% |
| hsa_circ_0096003 | 572 | 11 | 3.85% |
| hsa_circ_0125670 | 572 | 11 | 3.85% |
| hsa_circ_0035402 | 520 | 10 | 3.85% |
| hsa_circ_0047659 | 520 | 10 | 3.85% |
| hsa_circ_0075018 | 520 | 10 | 3.85% |
| hsa_circ_0079594 | 520 | 10 | 3.85% |
| hsa_circ_0090847 | 520 | 10 | 3.85% |
| hsa_circ_0100495 | 520 | 10 | 3.85% |
| hsa_circ_0118165 | 520 | 10 | 3.85% |
| hsa_circ_0022185 | 729 | 14 | 3.84% |
| hsa_circ_0042387 | 729 | 14 | 3.84% |
| hsa_circ_0127848 | 625 | 12 | 3.84% |
| hsa_circ_0039886 | 573 | 11 | 3.84% |
| hsa_circ_0059421 | 573 | 11 | 3.84% |
| hsa_circ_0063572 | 573 | 11 | 3.84% |
| hsa_circ_0089751 | 573 | 11 | 3.84% |
| hsa_circ_0130478 | 573 | 11 | 3.84% |
| hsa_circ_0017771 | 521 | 10 | 3.84% |
| hsa_circ_0022930 | 521 | 10 | 3.84% |
| hsa_circ_0023901 | 521 | 10 | 3.84% |
| hsa_circ_0038330 | 521 | 10 | 3.84% |
| hsa_circ_0038927 | 521 | 10 | 3.84% |
| hsa_circ_0044158 | 521 | 10 | 3.84% |

| | | | |
|---|---|---|---|
| hsa_circ_0057921 | 521 | 10 | 3.84% |
| hsa_circ_0064605 | 521 | 10 | 3.84% |
| hsa_circ_0084507 | 521 | 10 | 3.84% |
| hsa_circ_0140482 | 521 | 10 | 3.84% |
| hsa_circ_0134640 | 834 | 16 | 3.84% |
| hsa_circ_0062316 | 782 | 15 | 3.84% |
| hsa_circ_0035346 | 626 | 12 | 3.83% |
| hsa_circ_0050797 | 626 | 12 | 3.83% |
| hsa_circ_0009929 | 574 | 11 | 3.83% |
| hsa_circ_0028405 | 574 | 11 | 3.83% |
| hsa_circ_0108079 | 574 | 11 | 3.83% |
| hsa_circ_0116510 | 574 | 11 | 3.83% |
| hsa_circ_0062315 | 783 | 15 | 3.83% |
| hsa_circ_0025302 | 522 | 10 | 3.83% |
| hsa_circ_0029028 | 522 | 10 | 3.83% |
| hsa_circ_0043609 | 522 | 10 | 3.83% |
| hsa_circ_0052321 | 522 | 10 | 3.83% |
| hsa_circ_0055073 | 522 | 10 | 3.83% |
| hsa_circ_0055481 | 522 | 10 | 3.83% |
| hsa_circ_0096677 | 522 | 10 | 3.83% |
| hsa_circ_0096684 | 522 | 10 | 3.83% |
| hsa_circ_0126095 | 522 | 10 | 3.83% |
| hsa_circ_0135499 | 522 | 10 | 3.83% |
| hsa_circ_0079462 | 627 | 12 | 3.83% |
| hsa_circ_0092525 | 575 | 11 | 3.83% |
| hsa_circ_0100788 | 575 | 11 | 3.83% |
| hsa_circ_0123061 | 575 | 11 | 3.83% |
| hsa_circ_0136858 | 575 | 11 | 3.83% |
| hsa_circ_0048081 | 732 | 14 | 3.83% |
| hsa_circ_0055365 | 523 | 10 | 3.82% |
| hsa_circ_0064345 | 523 | 10 | 3.82% |
| hsa_circ_0087665 | 523 | 10 | 3.82% |
| hsa_circ_0093180 | 523 | 10 | 3.82% |
| hsa_circ_0106474 | 523 | 10 | 3.82% |
| hsa_circ_0118649 | 523 | 10 | 3.82% |
| hsa_circ_0125287 | 523 | 10 | 3.82% |
| hsa_circ_0047927 | 576 | 11 | 3.82% |
| hsa_circ_0052096 | 576 | 11 | 3.82% |
| hsa_circ_0057776 | 576 | 11 | 3.82% |
| hsa_circ_0127345 | 576 | 11 | 3.82% |
| hsa_circ_0139681 | 576 | 11 | 3.82% |
| hsa_circ_0068412 | 524 | 10 | 3.82% |
| hsa_circ_0090557 | 524 | 10 | 3.82% |
| hsa_circ_0100675 | 524 | 10 | 3.82% |
| hsa_circ_0116740 | 524 | 10 | 3.82% |
| hsa_circ_0128090 | 524 | 10 | 3.82% |
| hsa_circ_0042683 | 629 | 12 | 3.82% |
| hsa_circ_0132330 | 629 | 12 | 3.82% |
| hsa_circ_0061017 | 577 | 11 | 3.81% |
| hsa_circ_0076373 | 577 | 11 | 3.81% |

| | | | |
|---|---|---|---|
| hsa_circ_0120562 | 577 | 11 | 3.81% |
| hsa_circ_0061477 | 630 | 12 | 3.81% |
| hsa_circ_0027301 | 525 | 10 | 3.81% |
| hsa_circ_0069725 | 525 | 10 | 3.81% |
| hsa_circ_0077892 | 525 | 10 | 3.81% |
| hsa_circ_0085973 | 525 | 10 | 3.81% |
| hsa_circ_0094091 | 525 | 10 | 3.81% |
| hsa_circ_0100770 | 525 | 10 | 3.81% |
| hsa_circ_0123679 | 525 | 10 | 3.81% |
| hsa_circ_0130618 | 525 | 10 | 3.81% |
| hsa_circ_0132381 | 525 | 10 | 3.81% |
| hsa_circ_0132621 | 525 | 10 | 3.81% |
| hsa_circ_0134437 | 525 | 10 | 3.81% |
| hsa_circ_0138762 | 525 | 10 | 3.81% |
| hsa_circ_0002181 | 578 | 11 | 3.81% |
| hsa_circ_0033544 | 578 | 11 | 3.81% |
| hsa_circ_0037478 | 578 | 11 | 3.81% |
| hsa_circ_0006156 | 526 | 10 | 3.80% |
| hsa_circ_0015989 | 526 | 10 | 3.80% |
| hsa_circ_0038070 | 526 | 10 | 3.80% |
| hsa_circ_0057537 | 526 | 10 | 3.80% |
| hsa_circ_0092829 | 526 | 10 | 3.80% |
| hsa_circ_0126104 | 526 | 10 | 3.80% |
| hsa_circ_0130195 | 526 | 10 | 3.80% |
| hsa_circ_0135422 | 526 | 10 | 3.80% |
| hsa_circ_0099436 | 684 | 13 | 3.80% |
| hsa_circ_0003027 | 579 | 11 | 3.80% |
| hsa_circ_0004528 | 579 | 11 | 3.80% |
| hsa_circ_0035075 | 579 | 11 | 3.80% |
| hsa_circ_0038133 | 579 | 11 | 3.80% |
| hsa_circ_0049916 | 579 | 11 | 3.80% |
| hsa_circ_0063866 | 632 | 12 | 3.80% |
| hsa_circ_0029868 | 527 | 10 | 3.80% |
| hsa_circ_0059913 | 527 | 10 | 3.80% |
| hsa_circ_0061747 | 527 | 10 | 3.80% |
| hsa_circ_0062874 | 527 | 10 | 3.80% |
| hsa_circ_0084998 | 527 | 10 | 3.80% |
| hsa_circ_0127598 | 527 | 10 | 3.80% |
| hsa_circ_0065054 | 580 | 11 | 3.79% |
| hsa_circ_0048802 | 633 | 12 | 3.79% |
| hsa_circ_0114715 | 686 | 13 | 3.79% |
| hsa_circ_0004090 | 528 | 10 | 3.79% |
| hsa_circ_0022618 | 528 | 10 | 3.79% |
| hsa_circ_0023676 | 528 | 10 | 3.79% |
| hsa_circ_0045916 | 528 | 10 | 3.79% |
| hsa_circ_0049150 | 528 | 10 | 3.79% |
| hsa_circ_0054034 | 528 | 10 | 3.79% |
| hsa_circ_0072021 | 528 | 10 | 3.79% |
| hsa_circ_0081733 | 528 | 10 | 3.79% |
| hsa_circ_0116492 | 528 | 10 | 3.79% |

| | | | |
|---|---|---|---|
| hsa_circ_0124446 | 528 | 10 | 3.79% |
| hsa_circ_0023452 | 581 | 11 | 3.79% |
| hsa_circ_0027456 | 581 | 11 | 3.79% |
| hsa_circ_0059586 | 581 | 11 | 3.79% |
| hsa_circ_0100627 | 581 | 11 | 3.79% |
| hsa_circ_0011865 | 529 | 10 | 3.78% |
| hsa_circ_0021340 | 529 | 10 | 3.78% |
| hsa_circ_0025619 | 529 | 10 | 3.78% |
| hsa_circ_0027987 | 529 | 10 | 3.78% |
| hsa_circ_0036317 | 529 | 10 | 3.78% |
| hsa_circ_0058755 | 529 | 10 | 3.78% |
| hsa_circ_0100177 | 529 | 10 | 3.78% |
| hsa_circ_0101043 | 529 | 10 | 3.78% |
| hsa_circ_0109756 | 529 | 10 | 3.78% |
| hsa_circ_0112407 | 529 | 10 | 3.78% |
| hsa_circ_0122114 | 529 | 10 | 3.78% |
| hsa_circ_0125405 | 529 | 10 | 3.78% |
| hsa_circ_0125680 | 529 | 10 | 3.78% |
| hsa_circ_0128644 | 529 | 10 | 3.78% |
| hsa_circ_0132231 | 529 | 10 | 3.78% |
| hsa_circ_0008266 | 582 | 11 | 3.78% |
| hsa_circ_0027800 | 582 | 11 | 3.78% |
| hsa_circ_0132903 | 582 | 11 | 3.78% |
| hsa_circ_0060873 | 688 | 13 | 3.78% |
| hsa_circ_0001448 | 636 | 12 | 3.77% |
| hsa_circ_0067437 | 636 | 12 | 3.77% |
| hsa_circ_0051400 | 583 | 11 | 3.77% |
| hsa_circ_0092264 | 583 | 11 | 3.77% |
| hsa_circ_0116221 | 583 | 11 | 3.77% |
| hsa_circ_0126354 | 583 | 11 | 3.77% |
| hsa_circ_0133798 | 583 | 11 | 3.77% |
| hsa_circ_0008861 | 530 | 10 | 3.77% |
| hsa_circ_0037156 | 530 | 10 | 3.77% |
| hsa_circ_0080084 | 530 | 10 | 3.77% |
| hsa_circ_0135774 | 530 | 10 | 3.77% |
| hsa_circ_0102074 | 637 | 12 | 3.77% |
| hsa_circ_0057456 | 584 | 11 | 3.77% |
| hsa_circ_0083162 | 584 | 11 | 3.77% |
| hsa_circ_0113154 | 584 | 11 | 3.77% |
| hsa_circ_0007926 | 531 | 10 | 3.77% |
| hsa_circ_0023030 | 531 | 10 | 3.77% |
| hsa_circ_0036268 | 531 | 10 | 3.77% |
| hsa_circ_0039631 | 531 | 10 | 3.77% |
| hsa_circ_0040211 | 531 | 10 | 3.77% |
| hsa_circ_0062997 | 531 | 10 | 3.77% |
| hsa_circ_0074144 | 531 | 10 | 3.77% |
| hsa_circ_0083463 | 531 | 10 | 3.77% |
| hsa_circ_0108491 | 531 | 10 | 3.77% |
| hsa_circ_0110442 | 531 | 10 | 3.77% |
| hsa_circ_0110773 | 531 | 10 | 3.77% |

| | | | |
|---|---|---|---|
| hsa_circ_0117215 | 531 | 10 | 3.77% |
| hsa_circ_0135196 | 531 | 10 | 3.77% |
| hsa_circ_0136233 | 531 | 10 | 3.77% |
| hsa_circ_0140710 | 531 | 10 | 3.77% |
| hsa_circ_0021999 | 744 | 14 | 3.76% |
| hsa_circ_0074081 | 691 | 13 | 3.76% |
| hsa_circ_0097857 | 638 | 12 | 3.76% |
| hsa_circ_0111130 | 638 | 12 | 3.76% |
| hsa_circ_0033368 | 585 | 11 | 3.76% |
| hsa_circ_0131879 | 585 | 11 | 3.76% |
| hsa_circ_0138836 | 585 | 11 | 3.76% |
| hsa_circ_0012297 | 532 | 10 | 3.76% |
| hsa_circ_0035451 | 532 | 10 | 3.76% |
| hsa_circ_0037380 | 532 | 10 | 3.76% |
| hsa_circ_0038854 | 532 | 10 | 3.76% |
| hsa_circ_0048466 | 532 | 10 | 3.76% |
| hsa_circ_0051018 | 532 | 10 | 3.76% |
| hsa_circ_0072839 | 532 | 10 | 3.76% |
| hsa_circ_0085289 | 532 | 10 | 3.76% |
| hsa_circ_0113178 | 532 | 10 | 3.76% |
| hsa_circ_0127619 | 532 | 10 | 3.76% |
| hsa_circ_0137900 | 532 | 10 | 3.76% |
| hsa_circ_0006714 | 586 | 11 | 3.75% |
| hsa_circ_0014632 | 586 | 11 | 3.75% |
| hsa_circ_0037212 | 586 | 11 | 3.75% |
| hsa_circ_0051656 | 586 | 11 | 3.75% |
| hsa_circ_0116629 | 586 | 11 | 3.75% |
| hsa_circ_0011936 | 533 | 10 | 3.75% |
| hsa_circ_0020639 | 533 | 10 | 3.75% |
| hsa_circ_0034251 | 533 | 10 | 3.75% |
| hsa_circ_0046389 | 533 | 10 | 3.75% |
| hsa_circ_0051074 | 533 | 10 | 3.75% |
| hsa_circ_0079600 | 533 | 10 | 3.75% |
| hsa_circ_0087858 | 533 | 10 | 3.75% |
| hsa_circ_0100983 | 533 | 10 | 3.75% |
| hsa_circ_0108764 | 533 | 10 | 3.75% |
| hsa_circ_0121617 | 533 | 10 | 3.75% |
| hsa_circ_0133618 | 693 | 13 | 3.75% |
| hsa_circ_0008047 | 640 | 12 | 3.75% |
| hsa_circ_0089332 | 640 | 12 | 3.75% |
| hsa_circ_0052188 | 747 | 14 | 3.75% |
| hsa_circ_0034094 | 587 | 11 | 3.75% |
| hsa_circ_0040303 | 587 | 11 | 3.75% |
| hsa_circ_0099369 | 587 | 11 | 3.75% |
| hsa_circ_0133572 | 587 | 11 | 3.75% |
| hsa_circ_0135613 | 587 | 11 | 3.75% |
| hsa_circ_0003983 | 534 | 10 | 3.75% |
| hsa_circ_0009150 | 534 | 10 | 3.75% |
| hsa_circ_0037770 | 534 | 10 | 3.75% |
| hsa_circ_0043992 | 534 | 10 | 3.75% |

| | | | |
|---|---|---|---|
| hsa_circ_0050889 | 534 | 10 | 3.75% |
| hsa_circ_0082344 | 534 | 10 | 3.75% |
| hsa_circ_0082457 | 534 | 10 | 3.75% |
| hsa_circ_0107334 | 534 | 10 | 3.75% |
| hsa_circ_0134738 | 534 | 10 | 3.75% |
| hsa_circ_0073359 | 641 | 12 | 3.74% |
| hsa_circ_0111729 | 641 | 12 | 3.74% |
| hsa_circ_0107172 | 748 | 14 | 3.74% |
| hsa_circ_0000744 | 588 | 11 | 3.74% |
| hsa_circ_0008306 | 588 | 11 | 3.74% |
| hsa_circ_0038840 | 588 | 11 | 3.74% |
| hsa_circ_0128474 | 588 | 11 | 3.74% |
| hsa_circ_0040949 | 856 | 16 | 3.74% |
| hsa_circ_0130246 | 749 | 14 | 3.74% |
| hsa_circ_0001680 | 642 | 12 | 3.74% |
| hsa_circ_0024754 | 642 | 12 | 3.74% |
| hsa_circ_0032049 | 642 | 12 | 3.74% |
| hsa_circ_0082614 | 642 | 12 | 3.74% |
| hsa_circ_0091297 | 642 | 12 | 3.74% |
| hsa_circ_0002708 | 535 | 10 | 3.74% |
| hsa_circ_0014485 | 535 | 10 | 3.74% |
| hsa_circ_0048689 | 535 | 10 | 3.74% |
| hsa_circ_0077366 | 535 | 10 | 3.74% |
| hsa_circ_0088542 | 535 | 10 | 3.74% |
| hsa_circ_0112032 | 535 | 10 | 3.74% |
| hsa_circ_0118785 | 535 | 10 | 3.74% |
| hsa_circ_0123294 | 535 | 10 | 3.74% |
| hsa_circ_0084020 | 696 | 13 | 3.74% |
| hsa_circ_0018523 | 589 | 11 | 3.74% |
| hsa_circ_0029579 | 589 | 11 | 3.74% |
| hsa_circ_0051800 | 589 | 11 | 3.74% |
| hsa_circ_0068261 | 589 | 11 | 3.74% |
| hsa_circ_0128961 | 750 | 14 | 3.73% |
| hsa_circ_0016454 | 643 | 12 | 3.73% |
| hsa_circ_0003141 | 536 | 10 | 3.73% |
| hsa_circ_0014973 | 536 | 10 | 3.73% |
| hsa_circ_0032235 | 536 | 10 | 3.73% |
| hsa_circ_0033541 | 536 | 10 | 3.73% |
| hsa_circ_0046795 | 536 | 10 | 3.73% |
| hsa_circ_0124568 | 536 | 10 | 3.73% |
| hsa_circ_0125497 | 536 | 10 | 3.73% |
| hsa_circ_0137282 | 536 | 10 | 3.73% |
| hsa_circ_0139224 | 536 | 10 | 3.73% |
| hsa_circ_0121357 | 2037 | 38 | 3.73% |
| hsa_circ_0029553 | 590 | 11 | 3.73% |
| hsa_circ_0034654 | 590 | 11 | 3.73% |
| hsa_circ_0103167 | 590 | 11 | 3.73% |
| hsa_circ_0007458 | 537 | 10 | 3.72% |
| hsa_circ_0017091 | 537 | 10 | 3.72% |
| hsa_circ_0018079 | 537 | 10 | 3.72% |

| | | | |
|---|---|---|---|
| hsa_circ_0021772 | 537 | 10 | 3.72% |
| hsa_circ_0023848 | 537 | 10 | 3.72% |
| hsa_circ_0032904 | 537 | 10 | 3.72% |
| hsa_circ_0051993 | 537 | 10 | 3.72% |
| hsa_circ_0057217 | 537 | 10 | 3.72% |
| hsa_circ_0065733 | 537 | 10 | 3.72% |
| hsa_circ_0082152 | 537 | 10 | 3.72% |
| hsa_circ_0102660 | 537 | 10 | 3.72% |
| hsa_circ_0109004 | 537 | 10 | 3.72% |
| hsa_circ_0114108 | 537 | 10 | 3.72% |
| hsa_circ_0116349 | 537 | 10 | 3.72% |
| hsa_circ_0123789 | 537 | 10 | 3.72% |
| hsa_circ_0129030 | 537 | 10 | 3.72% |
| hsa_circ_0136735 | 537 | 10 | 3.72% |
| hsa_circ_0007975 | 591 | 11 | 3.72% |
| hsa_circ_0010709 | 591 | 11 | 3.72% |
| hsa_circ_0057454 | 591 | 11 | 3.72% |
| hsa_circ_0060597 | 591 | 11 | 3.72% |
| hsa_circ_0063813 | 591 | 11 | 3.72% |
| hsa_circ_0089739 | 591 | 11 | 3.72% |
| hsa_circ_0100376 | 591 | 11 | 3.72% |
| hsa_circ_0130309 | 591 | 11 | 3.72% |
| hsa_circ_0004302 | 538 | 10 | 3.72% |
| hsa_circ_0007769 | 538 | 10 | 3.72% |
| hsa_circ_0089571 | 538 | 10 | 3.72% |
| hsa_circ_0006832 | 592 | 11 | 3.72% |
| hsa_circ_0109762 | 1561 | 29 | 3.72% |
| hsa_circ_0074672 | 646 | 12 | 3.72% |
| hsa_circ_0004337 | 539 | 10 | 3.71% |
| hsa_circ_0036044 | 539 | 10 | 3.71% |
| hsa_circ_0039335 | 539 | 10 | 3.71% |
| hsa_circ_0067072 | 539 | 10 | 3.71% |
| hsa_circ_0084807 | 539 | 10 | 3.71% |
| hsa_circ_0111824 | 539 | 10 | 3.71% |
| hsa_circ_0118884 | 539 | 10 | 3.71% |
| hsa_circ_0126909 | 539 | 10 | 3.71% |
| hsa_circ_0133895 | 539 | 10 | 3.71% |
| hsa_circ_0140713 | 539 | 10 | 3.71% |
| hsa_circ_0024816 | 593 | 11 | 3.71% |
| hsa_circ_0055975 | 593 | 11 | 3.71% |
| hsa_circ_0098263 | 593 | 11 | 3.71% |
| hsa_circ_0115300 | 593 | 11 | 3.71% |
| hsa_circ_0120753 | 593 | 11 | 3.71% |
| hsa_circ_0140501 | 593 | 11 | 3.71% |
| hsa_circ_0005625 | 647 | 12 | 3.71% |
| hsa_circ_0072100 | 647 | 12 | 3.71% |
| hsa_circ_0024778 | 594 | 11 | 3.70% |
| hsa_circ_0040834 | 594 | 11 | 3.70% |
| hsa_circ_0044252 | 594 | 11 | 3.70% |
| hsa_circ_0049925 | 594 | 11 | 3.70% |

| | | | |
|---|---|---|---|
| hsa_circ_0060776 | 594 | 11 | 3.70% |
| hsa_circ_0072447 | 594 | 11 | 3.70% |
| hsa_circ_0107041 | 594 | 11 | 3.70% |
| hsa_circ_0002238 | 540 | 10 | 3.70% |
| hsa_circ_0004190 | 540 | 10 | 3.70% |
| hsa_circ_0021872 | 540 | 10 | 3.70% |
| hsa_circ_0025448 | 540 | 10 | 3.70% |
| hsa_circ_0037732 | 540 | 10 | 3.70% |
| hsa_circ_0048338 | 540 | 10 | 3.70% |
| hsa_circ_0048979 | 540 | 10 | 3.70% |
| hsa_circ_0055692 | 540 | 10 | 3.70% |
| hsa_circ_0063984 | 540 | 10 | 3.70% |
| hsa_circ_0076063 | 540 | 10 | 3.70% |
| hsa_circ_0079357 | 540 | 10 | 3.70% |
| hsa_circ_0079599 | 540 | 10 | 3.70% |
| hsa_circ_0097922 | 540 | 10 | 3.70% |
| hsa_circ_0101736 | 540 | 10 | 3.70% |
| hsa_circ_0103067 | 540 | 10 | 3.70% |
| hsa_circ_0103733 | 540 | 10 | 3.70% |
| hsa_circ_0106606 | 540 | 10 | 3.70% |
| hsa_circ_0138280 | 540 | 10 | 3.70% |
| hsa_circ_0060844 | 649 | 12 | 3.70% |
| hsa_circ_0050643 | 595 | 11 | 3.70% |
| hsa_circ_0080159 | 595 | 11 | 3.70% |
| hsa_circ_0101009 | 595 | 11 | 3.70% |
| hsa_circ_0103890 | 595 | 11 | 3.70% |
| hsa_circ_0002236 | 541 | 10 | 3.70% |
| hsa_circ_0026681 | 541 | 10 | 3.70% |
| hsa_circ_0028041 | 541 | 10 | 3.70% |
| hsa_circ_0050617 | 541 | 10 | 3.70% |
| hsa_circ_0075470 | 541 | 10 | 3.70% |
| hsa_circ_0076303 | 541 | 10 | 3.70% |
| hsa_circ_0089551 | 541 | 10 | 3.70% |
| hsa_circ_0092455 | 541 | 10 | 3.70% |
| hsa_circ_0099513 | 541 | 10 | 3.70% |
| hsa_circ_0109250 | 541 | 10 | 3.70% |
| hsa_circ_0113998 | 541 | 10 | 3.70% |
| hsa_circ_0128205 | 541 | 10 | 3.70% |
| hsa_circ_0131890 | 541 | 10 | 3.70% |
| hsa_circ_0124300 | 1300 | 24 | 3.69% |
| hsa_circ_0100825 | 650 | 12 | 3.69% |
| hsa_circ_0040304 | 596 | 11 | 3.69% |
| hsa_circ_0096980 | 596 | 11 | 3.69% |
| hsa_circ_0000374 | 542 | 10 | 3.69% |
| hsa_circ_0005649 | 542 | 10 | 3.69% |
| hsa_circ_0024693 | 542 | 10 | 3.69% |
| hsa_circ_0032006 | 542 | 10 | 3.69% |
| hsa_circ_0080099 | 542 | 10 | 3.69% |
| hsa_circ_0100424 | 542 | 10 | 3.69% |
| hsa_circ_0112486 | 542 | 10 | 3.69% |

| | | | |
|---|---|---|---|
| hsa_circ_0123283 | 542 | 10 | 3.69% |
| hsa_circ_0128505 | 542 | 10 | 3.69% |
| hsa_circ_0009191 | 759 | 14 | 3.69% |
| hsa_circ_0041339 | 651 | 12 | 3.69% |
| hsa_circ_0115890 | 651 | 12 | 3.69% |
| hsa_circ_0022770 | 597 | 11 | 3.69% |
| hsa_circ_0032612 | 597 | 11 | 3.69% |
| hsa_circ_0072199 | 597 | 11 | 3.69% |
| hsa_circ_0123999 | 597 | 11 | 3.69% |
| hsa_circ_0130133 | 597 | 11 | 3.69% |
| hsa_circ_0136458 | 597 | 11 | 3.69% |
| hsa_circ_0000070 | 543 | 10 | 3.68% |
| hsa_circ_0009250 | 543 | 10 | 3.68% |
| hsa_circ_0014022 | 543 | 10 | 3.68% |
| hsa_circ_0042750 | 543 | 10 | 3.68% |
| hsa_circ_0044687 | 543 | 10 | 3.68% |
| hsa_circ_0074142 | 543 | 10 | 3.68% |
| hsa_circ_0097956 | 543 | 10 | 3.68% |
| hsa_circ_0101545 | 543 | 10 | 3.68% |
| hsa_circ_0113183 | 543 | 10 | 3.68% |
| hsa_circ_0114521 | 543 | 10 | 3.68% |
| hsa_circ_0120395 | 543 | 10 | 3.68% |
| hsa_circ_0127603 | 543 | 10 | 3.68% |
| hsa_circ_0015560 | 706 | 13 | 3.68% |
| hsa_circ_0075721 | 652 | 12 | 3.68% |
| hsa_circ_0045424 | 598 | 11 | 3.68% |
| hsa_circ_0076663 | 707 | 13 | 3.68% |
| hsa_circ_0053029 | 816 | 15 | 3.68% |
| hsa_circ_0003304 | 544 | 10 | 3.68% |
| hsa_circ_0009860 | 544 | 10 | 3.68% |
| hsa_circ_0011093 | 544 | 10 | 3.68% |
| hsa_circ_0038667 | 544 | 10 | 3.68% |
| hsa_circ_0051072 | 544 | 10 | 3.68% |
| hsa_circ_0079727 | 544 | 10 | 3.68% |
| hsa_circ_0089545 | 544 | 10 | 3.68% |
| hsa_circ_0113145 | 544 | 10 | 3.68% |
| hsa_circ_0116075 | 544 | 10 | 3.68% |
| hsa_circ_0122898 | 544 | 10 | 3.68% |
| hsa_circ_0128432 | 544 | 10 | 3.68% |
| hsa_circ_0137027 | 544 | 10 | 3.68% |
| hsa_circ_0137955 | 544 | 10 | 3.68% |
| hsa_circ_0050120 | 653 | 12 | 3.68% |
| hsa_circ_0036105 | 599 | 11 | 3.67% |
| hsa_circ_0105807 | 599 | 11 | 3.67% |
| hsa_circ_0106299 | 599 | 11 | 3.67% |
| hsa_circ_0128149 | 817 | 15 | 3.67% |
| hsa_circ_0088524 | 654 | 12 | 3.67% |
| hsa_circ_0005563 | 545 | 10 | 3.67% |
| hsa_circ_0006971 | 545 | 10 | 3.67% |
| hsa_circ_0011958 | 545 | 10 | 3.67% |

| | | | |
|---|---|---|---|
| hsa_circ_0030725 | 545 | 10 | 3.67% |
| hsa_circ_0047676 | 545 | 10 | 3.67% |
| hsa_circ_0047894 | 545 | 10 | 3.67% |
| hsa_circ_0068751 | 545 | 10 | 3.67% |
| hsa_circ_0110255 | 545 | 10 | 3.67% |
| hsa_circ_0118679 | 545 | 10 | 3.67% |
| hsa_circ_0122474 | 545 | 10 | 3.67% |
| hsa_circ_0069969 | 709 | 13 | 3.67% |
| hsa_circ_0069055 | 600 | 11 | 3.67% |
| hsa_circ_0005298 | 655 | 12 | 3.66% |
| hsa_circ_0005260 | 546 | 10 | 3.66% |
| hsa_circ_0007056 | 546 | 10 | 3.66% |
| hsa_circ_0013348 | 546 | 10 | 3.66% |
| hsa_circ_0018725 | 546 | 10 | 3.66% |
| hsa_circ_0032564 | 546 | 10 | 3.66% |
| hsa_circ_0041431 | 546 | 10 | 3.66% |
| hsa_circ_0042296 | 546 | 10 | 3.66% |
| hsa_circ_0047498 | 546 | 10 | 3.66% |
| hsa_circ_0050361 | 546 | 10 | 3.66% |
| hsa_circ_0090469 | 546 | 10 | 3.66% |
| hsa_circ_0091790 | 546 | 10 | 3.66% |
| hsa_circ_0097123 | 546 | 10 | 3.66% |
| hsa_circ_0099380 | 546 | 10 | 3.66% |
| hsa_circ_0102412 | 546 | 10 | 3.66% |
| hsa_circ_0111201 | 546 | 10 | 3.66% |
| hsa_circ_0123683 | 546 | 10 | 3.66% |
| hsa_circ_0128206 | 546 | 10 | 3.66% |
| hsa_circ_0134231 | 546 | 10 | 3.66% |
| hsa_circ_0034624 | 601 | 11 | 3.66% |
| hsa_circ_0053344 | 601 | 11 | 3.66% |
| hsa_circ_0076445 | 601 | 11 | 3.66% |
| hsa_circ_0027128 | 656 | 12 | 3.66% |
| hsa_circ_0034424 | 656 | 12 | 3.66% |
| hsa_circ_0067312 | 656 | 12 | 3.66% |
| hsa_circ_0014860 | 547 | 10 | 3.66% |
| hsa_circ_0057995 | 547 | 10 | 3.66% |
| hsa_circ_0080069 | 547 | 10 | 3.66% |
| hsa_circ_0097122 | 547 | 10 | 3.66% |
| hsa_circ_0100792 | 547 | 10 | 3.66% |
| hsa_circ_0106891 | 547 | 10 | 3.66% |
| hsa_circ_0114863 | 547 | 10 | 3.66% |
| hsa_circ_0036374 | 602 | 11 | 3.65% |
| hsa_circ_0044241 | 602 | 11 | 3.65% |
| hsa_circ_0125214 | 602 | 11 | 3.65% |
| hsa_circ_0091305 | 821 | 15 | 3.65% |
| hsa_circ_0015095 | 657 | 12 | 3.65% |
| hsa_circ_0030845 | 657 | 12 | 3.65% |
| hsa_circ_0075110 | 657 | 12 | 3.65% |
| hsa_circ_0128073 | 657 | 12 | 3.65% |
| hsa_circ_0076668 | 712 | 13 | 3.65% |

| | | | |
|---|---|---|---|
| hsa_circ_0004175 | 548 | 10 | 3.65% |
| hsa_circ_0005416 | 548 | 10 | 3.65% |
| hsa_circ_0015445 | 548 | 10 | 3.65% |
| hsa_circ_0041904 | 548 | 10 | 3.65% |
| hsa_circ_0041987 | 548 | 10 | 3.65% |
| hsa_circ_0047507 | 548 | 10 | 3.65% |
| hsa_circ_0049354 | 548 | 10 | 3.65% |
| hsa_circ_0054948 | 548 | 10 | 3.65% |
| hsa_circ_0069149 | 548 | 10 | 3.65% |
| hsa_circ_0093017 | 548 | 10 | 3.65% |
| hsa_circ_0101709 | 548 | 10 | 3.65% |
| hsa_circ_0138064 | 548 | 10 | 3.65% |
| hsa_circ_0015164 | 603 | 11 | 3.65% |
| hsa_circ_0055281 | 603 | 11 | 3.65% |
| hsa_circ_0055378 | 603 | 11 | 3.65% |
| hsa_circ_0057448 | 603 | 11 | 3.65% |
| hsa_circ_0088058 | 603 | 11 | 3.65% |
| hsa_circ_0089441 | 603 | 11 | 3.65% |
| hsa_circ_0090449 | 603 | 11 | 3.65% |
| hsa_circ_0005228 | 658 | 12 | 3.65% |
| hsa_circ_0071567 | 658 | 12 | 3.65% |
| hsa_circ_0006874 | 549 | 10 | 3.64% |
| hsa_circ_0033241 | 549 | 10 | 3.64% |
| hsa_circ_0034883 | 549 | 10 | 3.64% |
| hsa_circ_0047055 | 549 | 10 | 3.64% |
| hsa_circ_0053130 | 549 | 10 | 3.64% |
| hsa_circ_0058413 | 549 | 10 | 3.64% |
| hsa_circ_0067164 | 549 | 10 | 3.64% |
| hsa_circ_0079212 | 549 | 10 | 3.64% |
| hsa_circ_0082830 | 549 | 10 | 3.64% |
| hsa_circ_0102031 | 549 | 10 | 3.64% |
| hsa_circ_0124766 | 549 | 10 | 3.64% |
| hsa_circ_0000298 | 604 | 11 | 3.64% |
| hsa_circ_0106773 | 604 | 11 | 3.64% |
| hsa_circ_0090307 | 659 | 12 | 3.64% |
| hsa_circ_0132406 | 769 | 14 | 3.64% |
| hsa_circ_0060524 | 715 | 13 | 3.64% |
| hsa_circ_0005579 | 660 | 12 | 3.64% |
| hsa_circ_0014149 | 605 | 11 | 3.64% |
| hsa_circ_0057134 | 605 | 11 | 3.64% |
| hsa_circ_0068058 | 605 | 11 | 3.64% |
| hsa_circ_0081850 | 605 | 11 | 3.64% |
| hsa_circ_0102033 | 605 | 11 | 3.64% |
| hsa_circ_0014824 | 550 | 10 | 3.64% |
| hsa_circ_0017411 | 550 | 10 | 3.64% |
| hsa_circ_0019012 | 550 | 10 | 3.64% |
| hsa_circ_0020795 | 550 | 10 | 3.64% |
| hsa_circ_0038199 | 550 | 10 | 3.64% |
| hsa_circ_0049954 | 550 | 10 | 3.64% |
| hsa_circ_0050684 | 550 | 10 | 3.64% |

| | | | |
|---|---|---|---|
| hsa_circ_0061674 | 550 | 10 | 3.64% |
| hsa_circ_0103537 | 550 | 10 | 3.64% |
| hsa_circ_0115276 | 550 | 10 | 3.64% |
| hsa_circ_0122966 | 550 | 10 | 3.64% |
| hsa_circ_0132894 | 550 | 10 | 3.64% |
| hsa_circ_0023491 | 661 | 12 | 3.63% |
| hsa_circ_0060022 | 661 | 12 | 3.63% |
| hsa_circ_0118229 | 661 | 12 | 3.63% |
| hsa_circ_0015654 | 606 | 11 | 3.63% |
| hsa_circ_0046296 | 606 | 11 | 3.63% |
| hsa_circ_0110897 | 606 | 11 | 3.63% |
| hsa_circ_0012953 | 551 | 10 | 3.63% |
| hsa_circ_0014497 | 551 | 10 | 3.63% |
| hsa_circ_0023370 | 551 | 10 | 3.63% |
| hsa_circ_0050413 | 551 | 10 | 3.63% |
| hsa_circ_0064767 | 551 | 10 | 3.63% |
| hsa_circ_0108324 | 551 | 10 | 3.63% |
| hsa_circ_0076665 | 717 | 13 | 3.63% |
| hsa_circ_0084024 | 717 | 13 | 3.63% |
| hsa_circ_0012855 | 662 | 12 | 3.63% |
| hsa_circ_0052236 | 662 | 12 | 3.63% |
| hsa_circ_0036738 | 607 | 11 | 3.62% |
| hsa_circ_0031352 | 552 | 10 | 3.62% |
| hsa_circ_0066721 | 552 | 10 | 3.62% |
| hsa_circ_0080934 | 552 | 10 | 3.62% |
| hsa_circ_0114050 | 552 | 10 | 3.62% |
| hsa_circ_0115133 | 552 | 10 | 3.62% |
| hsa_circ_0135624 | 552 | 10 | 3.62% |
| hsa_circ_0073497 | 718 | 13 | 3.62% |
| hsa_circ_0103973 | 663 | 12 | 3.62% |
| hsa_circ_0026478 | 608 | 11 | 3.62% |
| hsa_circ_0040282 | 608 | 11 | 3.62% |
| hsa_circ_0107982 | 608 | 11 | 3.62% |
| hsa_circ_0113489 | 608 | 11 | 3.62% |
| hsa_circ_0121148 | 608 | 11 | 3.62% |
| hsa_circ_0126284 | 608 | 11 | 3.62% |
| hsa_circ_0107516 | 774 | 14 | 3.62% |
| hsa_circ_0002631 | 553 | 10 | 3.62% |
| hsa_circ_0004346 | 553 | 10 | 3.62% |
| hsa_circ_0019386 | 553 | 10 | 3.62% |
| hsa_circ_0024454 | 553 | 10 | 3.62% |
| hsa_circ_0026320 | 553 | 10 | 3.62% |
| hsa_circ_0059608 | 553 | 10 | 3.62% |
| hsa_circ_0096475 | 553 | 10 | 3.62% |
| hsa_circ_0097553 | 553 | 10 | 3.62% |
| hsa_circ_0110560 | 553 | 10 | 3.62% |
| hsa_circ_0112937 | 553 | 10 | 3.62% |
| hsa_circ_0115334 | 553 | 10 | 3.62% |
| hsa_circ_0118150 | 553 | 10 | 3.62% |
| hsa_circ_0119619 | 553 | 10 | 3.62% |

| | | | |
|---|---|---|---|
| hsa_circ_0124537 | 553 | 10 | 3.62% |
| hsa_circ_0089904 | 719 | 13 | 3.62% |
| hsa_circ_0071534 | 664 | 12 | 3.61% |
| hsa_circ_0096321 | 664 | 12 | 3.61% |
| hsa_circ_0024775 | 609 | 11 | 3.61% |
| hsa_circ_0074215 | 609 | 11 | 3.61% |
| hsa_circ_0107840 | 609 | 11 | 3.61% |
| hsa_circ_0121872 | 609 | 11 | 3.61% |
| hsa_circ_0005023 | 720 | 13 | 3.61% |
| hsa_circ_0004557 | 554 | 10 | 3.61% |
| hsa_circ_0008558 | 554 | 10 | 3.61% |
| hsa_circ_0013603 | 554 | 10 | 3.61% |
| hsa_circ_0026042 | 554 | 10 | 3.61% |
| hsa_circ_0060434 | 554 | 10 | 3.61% |
| hsa_circ_0060531 | 554 | 10 | 3.61% |
| hsa_circ_0110739 | 554 | 10 | 3.61% |
| hsa_circ_0117424 | 554 | 10 | 3.61% |
| hsa_circ_0130739 | 554 | 10 | 3.61% |
| hsa_circ_0050807 | 665 | 12 | 3.61% |
| hsa_circ_0072962 | 665 | 12 | 3.61% |
| hsa_circ_0080912 | 665 | 12 | 3.61% |
| hsa_circ_0082739 | 610 | 11 | 3.61% |
| hsa_circ_0108698 | 610 | 11 | 3.61% |
| hsa_circ_0138208 | 666 | 12 | 3.60% |
| hsa_circ_0026941 | 555 | 10 | 3.60% |
| hsa_circ_0037362 | 555 | 10 | 3.60% |
| hsa_circ_0046130 | 555 | 10 | 3.60% |
| hsa_circ_0079798 | 555 | 10 | 3.60% |
| hsa_circ_0095101 | 555 | 10 | 3.60% |
| hsa_circ_0098659 | 555 | 10 | 3.60% |
| hsa_circ_0099674 | 555 | 10 | 3.60% |
| hsa_circ_0102206 | 555 | 10 | 3.60% |
| hsa_circ_0109156 | 555 | 10 | 3.60% |
| hsa_circ_0118166 | 555 | 10 | 3.60% |
| hsa_circ_0123465 | 555 | 10 | 3.60% |
| hsa_circ_0125902 | 555 | 10 | 3.60% |
| hsa_circ_0126785 | 555 | 10 | 3.60% |
| hsa_circ_0072374 | 722 | 13 | 3.60% |
| hsa_circ_0071003 | 611 | 11 | 3.60% |
| hsa_circ_0001875 | 556 | 10 | 3.60% |
| hsa_circ_0009211 | 556 | 10 | 3.60% |
| hsa_circ_0010496 | 556 | 10 | 3.60% |
| hsa_circ_0021604 | 556 | 10 | 3.60% |
| hsa_circ_0021874 | 556 | 10 | 3.60% |
| hsa_circ_0032082 | 556 | 10 | 3.60% |
| hsa_circ_0035649 | 556 | 10 | 3.60% |
| hsa_circ_0038019 | 556 | 10 | 3.60% |
| hsa_circ_0052664 | 556 | 10 | 3.60% |
| hsa_circ_0056205 | 556 | 10 | 3.60% |
| hsa_circ_0058715 | 556 | 10 | 3.60% |

| | | | |
|---|---|---|---|
| hsa_circ_0061163 | 556 | 10 | 3.60% |
| hsa_circ_0068860 | 556 | 10 | 3.60% |
| hsa_circ_0071434 | 556 | 10 | 3.60% |
| hsa_circ_0071757 | 556 | 10 | 3.60% |
| hsa_circ_0080504 | 556 | 10 | 3.60% |
| hsa_circ_0084882 | 556 | 10 | 3.60% |
| hsa_circ_0099705 | 556 | 10 | 3.60% |
| hsa_circ_0110157 | 556 | 10 | 3.60% |
| hsa_circ_0122962 | 556 | 10 | 3.60% |
| hsa_circ_0131041 | 556 | 10 | 3.60% |
| hsa_circ_0051523 | 612 | 11 | 3.59% |
| hsa_circ_0076421 | 612 | 11 | 3.59% |
| hsa_circ_0129762 | 612 | 11 | 3.59% |
| hsa_circ_0132900 | 612 | 11 | 3.59% |
| hsa_circ_0068269 | 7624 | 137 | 3.59% |
| hsa_circ_0022135 | 668 | 12 | 3.59% |
| hsa_circ_0001717 | 557 | 10 | 3.59% |
| hsa_circ_0038665 | 557 | 10 | 3.59% |
| hsa_circ_0039613 | 557 | 10 | 3.59% |
| hsa_circ_0065471 | 557 | 10 | 3.59% |
| hsa_circ_0083822 | 557 | 10 | 3.59% |
| hsa_circ_0085065 | 557 | 10 | 3.59% |
| hsa_circ_0094683 | 557 | 10 | 3.59% |
| hsa_circ_0116455 | 557 | 10 | 3.59% |
| hsa_circ_0128899 | 557 | 10 | 3.59% |
| hsa_circ_0138227 | 557 | 10 | 3.59% |
| hsa_circ_0031639 | 613 | 11 | 3.59% |
| hsa_circ_0055106 | 613 | 11 | 3.59% |
| hsa_circ_0074225 | 613 | 11 | 3.59% |
| hsa_circ_0024222 | 669 | 12 | 3.59% |
| hsa_circ_0054442 | 669 | 12 | 3.59% |
| hsa_circ_0069968 | 725 | 13 | 3.59% |
| hsa_circ_0128076 | 725 | 13 | 3.59% |
| hsa_circ_0012125 | 558 | 10 | 3.58% |
| hsa_circ_0015249 | 558 | 10 | 3.58% |
| hsa_circ_0022243 | 558 | 10 | 3.58% |
| hsa_circ_0043424 | 558 | 10 | 3.58% |
| hsa_circ_0045888 | 558 | 10 | 3.58% |
| hsa_circ_0058911 | 558 | 10 | 3.58% |
| hsa_circ_0062802 | 558 | 10 | 3.58% |
| hsa_circ_0066040 | 558 | 10 | 3.58% |
| hsa_circ_0071755 | 558 | 10 | 3.58% |
| hsa_circ_0072780 | 558 | 10 | 3.58% |
| hsa_circ_0073257 | 558 | 10 | 3.58% |
| hsa_circ_0076427 | 558 | 10 | 3.58% |
| hsa_circ_0080865 | 558 | 10 | 3.58% |
| hsa_circ_0092883 | 558 | 10 | 3.58% |
| hsa_circ_0116475 | 558 | 10 | 3.58% |
| hsa_circ_0100513 | 614 | 11 | 3.58% |
| hsa_circ_0125548 | 614 | 11 | 3.58% |

| | | | |
|---|---|---|---|
| hsa_circ_0139290 | 614 | 11 | 3.58% |
| hsa_circ_0139356 | 670 | 12 | 3.58% |
| hsa_circ_0003299 | 559 | 10 | 3.58% |
| hsa_circ_0012926 | 559 | 10 | 3.58% |
| hsa_circ_0015246 | 559 | 10 | 3.58% |
| hsa_circ_0025238 | 559 | 10 | 3.58% |
| hsa_circ_0032213 | 559 | 10 | 3.58% |
| hsa_circ_0032917 | 559 | 10 | 3.58% |
| hsa_circ_0041238 | 559 | 10 | 3.58% |
| hsa_circ_0055478 | 559 | 10 | 3.58% |
| hsa_circ_0082683 | 559 | 10 | 3.58% |
| hsa_circ_0086779 | 559 | 10 | 3.58% |
| hsa_circ_0111265 | 559 | 10 | 3.58% |
| hsa_circ_0112191 | 559 | 10 | 3.58% |
| hsa_circ_0114412 | 559 | 10 | 3.58% |
| hsa_circ_0119770 | 559 | 10 | 3.58% |
| hsa_circ_0005068 | 615 | 11 | 3.58% |
| hsa_circ_0052508 | 615 | 11 | 3.58% |
| hsa_circ_0068120 | 615 | 11 | 3.58% |
| hsa_circ_0071017 | 615 | 11 | 3.58% |
| hsa_circ_0104640 | 615 | 11 | 3.58% |
| hsa_circ_0124484 | 615 | 11 | 3.58% |
| hsa_circ_0130636 | 615 | 11 | 3.58% |
| hsa_circ_0131472 | 2408 | 43 | 3.57% |
| hsa_circ_0008236 | 784 | 14 | 3.57% |
| hsa_circ_0133762 | 728 | 13 | 3.57% |
| hsa_circ_0009371 | 672 | 12 | 3.57% |
| hsa_circ_0002582 | 616 | 11 | 3.57% |
| hsa_circ_0069240 | 616 | 11 | 3.57% |
| hsa_circ_0091573 | 616 | 11 | 3.57% |
| hsa_circ_0115776 | 616 | 11 | 3.57% |
| hsa_circ_0138571 | 616 | 11 | 3.57% |
| hsa_circ_0139482 | 616 | 11 | 3.57% |
| hsa_circ_0002292 | 560 | 10 | 3.57% |
| hsa_circ_0006673 | 560 | 10 | 3.57% |
| hsa_circ_0026798 | 560 | 10 | 3.57% |
| hsa_circ_0046224 | 560 | 10 | 3.57% |
| hsa_circ_0066194 | 560 | 10 | 3.57% |
| hsa_circ_0066733 | 560 | 10 | 3.57% |
| hsa_circ_0084981 | 560 | 10 | 3.57% |
| hsa_circ_0094165 | 560 | 10 | 3.57% |
| hsa_circ_0096817 | 560 | 10 | 3.57% |
| hsa_circ_0105371 | 560 | 10 | 3.57% |
| hsa_circ_0112319 | 560 | 10 | 3.57% |
| hsa_circ_0113611 | 560 | 10 | 3.57% |
| hsa_circ_0118965 | 560 | 10 | 3.57% |
| hsa_circ_0121657 | 560 | 10 | 3.57% |
| hsa_circ_0026378 | 17473 | 312 | 3.57% |
| hsa_circ_0011290 | 617 | 11 | 3.57% |
| hsa_circ_0051398 | 617 | 11 | 3.57% |

| | | | |
|---|---|---|---|
| hsa_circ_0053141 | 617 | 11 | 3.57% |
| hsa_circ_0097639 | 617 | 11 | 3.57% |
| hsa_circ_0113871 | 617 | 11 | 3.57% |
| hsa_circ_0134823 | 617 | 11 | 3.57% |
| hsa_circ_0136123 | 617 | 11 | 3.57% |
| hsa_circ_0009212 | 561 | 10 | 3.57% |
| hsa_circ_0020809 | 561 | 10 | 3.57% |
| hsa_circ_0024990 | 561 | 10 | 3.57% |
| hsa_circ_0052447 | 561 | 10 | 3.57% |
| hsa_circ_0063551 | 561 | 10 | 3.57% |
| hsa_circ_0070283 | 561 | 10 | 3.57% |
| hsa_circ_0072545 | 561 | 10 | 3.57% |
| hsa_circ_0079922 | 561 | 10 | 3.57% |
| hsa_circ_0093658 | 561 | 10 | 3.57% |
| hsa_circ_0096305 | 561 | 10 | 3.57% |
| hsa_circ_0111267 | 561 | 10 | 3.57% |
| hsa_circ_0117183 | 561 | 10 | 3.57% |
| hsa_circ_0120647 | 561 | 10 | 3.57% |
| hsa_circ_0120918 | 561 | 10 | 3.57% |
| hsa_circ_0122042 | 561 | 10 | 3.57% |
| hsa_circ_0130020 | 561 | 10 | 3.57% |
| hsa_circ_0052825 | 730 | 13 | 3.56% |
| hsa_circ_0063222 | 1685 | 30 | 3.56% |
| hsa_circ_0106293 | 618 | 11 | 3.56% |
| hsa_circ_0111705 | 618 | 11 | 3.56% |
| hsa_circ_0120944 | 843 | 15 | 3.56% |
| hsa_circ_0006299 | 562 | 10 | 3.56% |
| hsa_circ_0013318 | 562 | 10 | 3.56% |
| hsa_circ_0021745 | 562 | 10 | 3.56% |
| hsa_circ_0029390 | 562 | 10 | 3.56% |
| hsa_circ_0034145 | 562 | 10 | 3.56% |
| hsa_circ_0035288 | 562 | 10 | 3.56% |
| hsa_circ_0051578 | 562 | 10 | 3.56% |
| hsa_circ_0056049 | 562 | 10 | 3.56% |
| hsa_circ_0061607 | 562 | 10 | 3.56% |
| hsa_circ_0064003 | 562 | 10 | 3.56% |
| hsa_circ_0080635 | 562 | 10 | 3.56% |
| hsa_circ_0080674 | 562 | 10 | 3.56% |
| hsa_circ_0095552 | 562 | 10 | 3.56% |
| hsa_circ_0115496 | 562 | 10 | 3.56% |
| hsa_circ_0140112 | 562 | 10 | 3.56% |
| hsa_circ_0122070 | 731 | 13 | 3.56% |
| hsa_circ_0035269 | 675 | 12 | 3.56% |
| hsa_circ_0052190 | 675 | 12 | 3.56% |
| hsa_circ_0130260 | 675 | 12 | 3.56% |
| hsa_circ_0010424 | 619 | 11 | 3.55% |
| hsa_circ_0060314 | 619 | 11 | 3.55% |
| hsa_circ_0085847 | 619 | 11 | 3.55% |
| hsa_circ_0100305 | 619 | 11 | 3.55% |
| hsa_circ_0108124 | 619 | 11 | 3.55% |

| | | | |
|---|---|---|---|
| hsa_circ_0000251 | 563 | 10 | 3.55% |
| hsa_circ_0003338 | 563 | 10 | 3.55% |
| hsa_circ_0008336 | 563 | 10 | 3.55% |
| hsa_circ_0024442 | 563 | 10 | 3.55% |
| hsa_circ_0030542 | 563 | 10 | 3.55% |
| hsa_circ_0049537 | 563 | 10 | 3.55% |
| hsa_circ_0050198 | 563 | 10 | 3.55% |
| hsa_circ_0051873 | 563 | 10 | 3.55% |
| hsa_circ_0053413 | 563 | 10 | 3.55% |
| hsa_circ_0066950 | 563 | 10 | 3.55% |
| hsa_circ_0068032 | 563 | 10 | 3.55% |
| hsa_circ_0110199 | 563 | 10 | 3.55% |
| hsa_circ_0112354 | 563 | 10 | 3.55% |
| hsa_circ_0116659 | 563 | 10 | 3.55% |
| hsa_circ_0129109 | 563 | 10 | 3.55% |
| hsa_circ_0130437 | 563 | 10 | 3.55% |
| hsa_circ_0046276 | 676 | 12 | 3.55% |
| hsa_circ_0068369 | 620 | 11 | 3.55% |
| hsa_circ_0131748 | 620 | 11 | 3.55% |
| hsa_circ_0117543 | 733 | 13 | 3.55% |
| hsa_circ_0000329 | 564 | 10 | 3.55% |
| hsa_circ_0041941 | 564 | 10 | 3.55% |
| hsa_circ_0084708 | 564 | 10 | 3.55% |
| hsa_circ_0090414 | 564 | 10 | 3.55% |
| hsa_circ_0094223 | 564 | 10 | 3.55% |
| hsa_circ_0098430 | 564 | 10 | 3.55% |
| hsa_circ_0099680 | 564 | 10 | 3.55% |
| hsa_circ_0102887 | 564 | 10 | 3.55% |
| hsa_circ_0116425 | 564 | 10 | 3.55% |
| hsa_circ_0116729 | 564 | 10 | 3.55% |
| hsa_circ_0117207 | 564 | 10 | 3.55% |
| hsa_circ_0117557 | 564 | 10 | 3.55% |
| hsa_circ_0120096 | 564 | 10 | 3.55% |
| hsa_circ_0000489 | 677 | 12 | 3.55% |
| hsa_circ_0057305 | 677 | 12 | 3.55% |
| hsa_circ_0073486 | 677 | 12 | 3.55% |
| hsa_circ_0030220 | 790 | 14 | 3.54% |
| hsa_circ_0009217 | 621 | 11 | 3.54% |
| hsa_circ_0049004 | 621 | 11 | 3.54% |
| hsa_circ_0061969 | 621 | 11 | 3.54% |
| hsa_circ_0104501 | 621 | 11 | 3.54% |
| hsa_circ_0108019 | 621 | 11 | 3.54% |
| hsa_circ_0138267 | 621 | 11 | 3.54% |
| hsa_circ_0133763 | 734 | 13 | 3.54% |
| hsa_circ_0049977 | 678 | 12 | 3.54% |
| hsa_circ_0017455 | 565 | 10 | 3.54% |
| hsa_circ_0021900 | 565 | 10 | 3.54% |
| hsa_circ_0023930 | 565 | 10 | 3.54% |
| hsa_circ_0052802 | 565 | 10 | 3.54% |
| hsa_circ_0066167 | 565 | 10 | 3.54% |

| | | | |
|---|---|---|---|
| hsa_circ_0078658 | 565 | 10 | 3.54% |
| hsa_circ_0079557 | 565 | 10 | 3.54% |
| hsa_circ_0081566 | 565 | 10 | 3.54% |
| hsa_circ_0082378 | 565 | 10 | 3.54% |
| hsa_circ_0082770 | 565 | 10 | 3.54% |
| hsa_circ_0091492 | 565 | 10 | 3.54% |
| hsa_circ_0100566 | 565 | 10 | 3.54% |
| hsa_circ_0118088 | 565 | 10 | 3.54% |
| hsa_circ_0126083 | 565 | 10 | 3.54% |
| hsa_circ_0132176 | 565 | 10 | 3.54% |
| hsa_circ_0099102 | 2092 | 37 | 3.54% |
| hsa_circ_0016386 | 622 | 11 | 3.54% |
| hsa_circ_0140111 | 622 | 11 | 3.54% |
| hsa_circ_0021576 | 792 | 14 | 3.54% |
| hsa_circ_0019720 | 679 | 12 | 3.53% |
| hsa_circ_0037636 | 679 | 12 | 3.53% |
| hsa_circ_0057602 | 679 | 12 | 3.53% |
| hsa_circ_0011907 | 566 | 10 | 3.53% |
| hsa_circ_0013271 | 566 | 10 | 3.53% |
| hsa_circ_0026552 | 566 | 10 | 3.53% |
| hsa_circ_0031340 | 566 | 10 | 3.53% |
| hsa_circ_0040017 | 566 | 10 | 3.53% |
| hsa_circ_0050551 | 566 | 10 | 3.53% |
| hsa_circ_0064700 | 566 | 10 | 3.53% |
| hsa_circ_0098451 | 566 | 10 | 3.53% |
| hsa_circ_0099537 | 566 | 10 | 3.53% |
| hsa_circ_0103697 | 566 | 10 | 3.53% |
| hsa_circ_0105073 | 566 | 10 | 3.53% |
| hsa_circ_0112430 | 566 | 10 | 3.53% |
| hsa_circ_0118893 | 566 | 10 | 3.53% |
| hsa_circ_0124315 | 566 | 10 | 3.53% |
| hsa_circ_0127148 | 566 | 10 | 3.53% |
| hsa_circ_0131799 | 566 | 10 | 3.53% |
| hsa_circ_0137847 | 566 | 10 | 3.53% |
| hsa_circ_0018471 | 623 | 11 | 3.53% |
| hsa_circ_0023420 | 623 | 11 | 3.53% |
| hsa_circ_0065365 | 623 | 11 | 3.53% |
| hsa_circ_0109607 | 623 | 11 | 3.53% |
| hsa_circ_0015981 | 850 | 15 | 3.53% |
| hsa_circ_0133944 | 680 | 12 | 3.53% |
| hsa_circ_0000676 | 567 | 10 | 3.53% |
| hsa_circ_0032935 | 567 | 10 | 3.53% |
| hsa_circ_0050257 | 567 | 10 | 3.53% |
| hsa_circ_0057652 | 567 | 10 | 3.53% |
| hsa_circ_0070883 | 567 | 10 | 3.53% |
| hsa_circ_0074324 | 567 | 10 | 3.53% |
| hsa_circ_0083247 | 567 | 10 | 3.53% |
| hsa_circ_0103898 | 567 | 10 | 3.53% |
| hsa_circ_0104517 | 567 | 10 | 3.53% |
| hsa_circ_0106668 | 567 | 10 | 3.53% |

| | | | |
|---|---|---|---|
| hsa_circ_0118651 | 567 | 10 | 3.53% |
| hsa_circ_0124103 | 567 | 10 | 3.53% |
| hsa_circ_0124720 | 567 | 10 | 3.53% |
| hsa_circ_0129566 | 567 | 10 | 3.53% |
| hsa_circ_0067212 | 624 | 11 | 3.53% |
| hsa_circ_0102390 | 624 | 11 | 3.53% |
| hsa_circ_0118666 | 624 | 11 | 3.53% |
| hsa_circ_0000063 | 568 | 10 | 3.52% |
| hsa_circ_0006719 | 568 | 10 | 3.52% |
| hsa_circ_0007123 | 568 | 10 | 3.52% |
| hsa_circ_0014202 | 568 | 10 | 3.52% |
| hsa_circ_0045050 | 568 | 10 | 3.52% |
| hsa_circ_0051234 | 568 | 10 | 3.52% |
| hsa_circ_0051346 | 568 | 10 | 3.52% |
| hsa_circ_0060353 | 568 | 10 | 3.52% |
| hsa_circ_0065618 | 568 | 10 | 3.52% |
| hsa_circ_0066760 | 568 | 10 | 3.52% |
| hsa_circ_0081229 | 568 | 10 | 3.52% |
| hsa_circ_0122886 | 568 | 10 | 3.52% |
| hsa_circ_0125285 | 568 | 10 | 3.52% |
| hsa_circ_0129259 | 568 | 10 | 3.52% |
| hsa_circ_0019804 | 625 | 11 | 3.52% |
| hsa_circ_0023974 | 625 | 11 | 3.52% |
| hsa_circ_0076056 | 625 | 11 | 3.52% |
| hsa_circ_0110604 | 625 | 11 | 3.52% |
| hsa_circ_0129518 | 625 | 11 | 3.52% |
| hsa_circ_0133890 | 625 | 11 | 3.52% |
| hsa_circ_0137073 | 625 | 11 | 3.52% |
| hsa_circ_0065713 | 682 | 12 | 3.52% |
| hsa_circ_0007529 | 569 | 10 | 3.51% |
| hsa_circ_0010238 | 569 | 10 | 3.51% |
| hsa_circ_0015645 | 569 | 10 | 3.51% |
| hsa_circ_0018658 | 569 | 10 | 3.51% |
| hsa_circ_0040806 | 569 | 10 | 3.51% |
| hsa_circ_0051805 | 569 | 10 | 3.51% |
| hsa_circ_0064676 | 569 | 10 | 3.51% |
| hsa_circ_0095320 | 569 | 10 | 3.51% |
| hsa_circ_0103923 | 569 | 10 | 3.51% |
| hsa_circ_0104427 | 569 | 10 | 3.51% |
| hsa_circ_0126388 | 569 | 10 | 3.51% |
| hsa_circ_0116997 | 626 | 11 | 3.51% |
| hsa_circ_0121440 | 626 | 11 | 3.51% |
| hsa_circ_0132461 | 626 | 11 | 3.51% |
| hsa_circ_0002777 | 1139 | 20 | 3.51% |
| hsa_circ_0062268 | 855 | 15 | 3.51% |
| hsa_circ_0065317 | 684 | 12 | 3.51% |
| hsa_circ_0121086 | 684 | 12 | 3.51% |
| hsa_circ_0139768 | 684 | 12 | 3.51% |
| hsa_circ_0048916 | 627 | 11 | 3.51% |
| hsa_circ_0001831 | 570 | 10 | 3.51% |

| | | | |
|---|---|---|---|
| hsa_circ_0002038 | 570 | 10 | 3.51% |
| hsa_circ_0002501 | 570 | 10 | 3.51% |
| hsa_circ_0017089 | 570 | 10 | 3.51% |
| hsa_circ_0028654 | 570 | 10 | 3.51% |
| hsa_circ_0035040 | 570 | 10 | 3.51% |
| hsa_circ_0043717 | 570 | 10 | 3.51% |
| hsa_circ_0051806 | 570 | 10 | 3.51% |
| hsa_circ_0052020 | 570 | 10 | 3.51% |
| hsa_circ_0052198 | 570 | 10 | 3.51% |
| hsa_circ_0064031 | 570 | 10 | 3.51% |
| hsa_circ_0095715 | 570 | 10 | 3.51% |
| hsa_circ_0104019 | 570 | 10 | 3.51% |
| hsa_circ_0107728 | 570 | 10 | 3.51% |
| hsa_circ_0123378 | 570 | 10 | 3.51% |
| hsa_circ_0129161 | 570 | 10 | 3.51% |
| hsa_circ_0132229 | 570 | 10 | 3.51% |
| hsa_circ_0138798 | 570 | 10 | 3.51% |
| hsa_circ_0095655 | 742 | 13 | 3.50% |
| hsa_circ_0067914 | 685 | 12 | 3.50% |
| hsa_circ_0005802 | 628 | 11 | 3.50% |
| hsa_circ_0038504 | 628 | 11 | 3.50% |
| hsa_circ_0039560 | 628 | 11 | 3.50% |
| hsa_circ_0061574 | 628 | 11 | 3.50% |
| hsa_circ_0091658 | 628 | 11 | 3.50% |
| hsa_circ_0099481 | 628 | 11 | 3.50% |
| hsa_circ_0121733 | 628 | 11 | 3.50% |
| hsa_circ_0140053 | 628 | 11 | 3.50% |
| hsa_circ_0008778 | 571 | 10 | 3.50% |
| hsa_circ_0010173 | 571 | 10 | 3.50% |
| hsa_circ_0014511 | 571 | 10 | 3.50% |
| hsa_circ_0032835 | 571 | 10 | 3.50% |
| hsa_circ_0059121 | 571 | 10 | 3.50% |
| hsa_circ_0072049 | 571 | 10 | 3.50% |
| hsa_circ_0078034 | 571 | 10 | 3.50% |
| hsa_circ_0081683 | 571 | 10 | 3.50% |
| hsa_circ_0081711 | 571 | 10 | 3.50% |
| hsa_circ_0086980 | 571 | 10 | 3.50% |
| hsa_circ_0101739 | 571 | 10 | 3.50% |
| hsa_circ_0104720 | 571 | 10 | 3.50% |
| hsa_circ_0105593 | 571 | 10 | 3.50% |
| hsa_circ_0115028 | 571 | 10 | 3.50% |
| hsa_circ_0123502 | 571 | 10 | 3.50% |
| hsa_circ_0126072 | 571 | 10 | 3.50% |
| hsa_circ_0140786 | 571 | 10 | 3.50% |
| hsa_circ_0122002 | 686 | 12 | 3.50% |
| hsa_circ_0013739 | 629 | 11 | 3.50% |
| hsa_circ_0023573 | 629 | 11 | 3.50% |
| hsa_circ_0040562 | 572 | 10 | 3.50% |
| hsa_circ_0041786 | 572 | 10 | 3.50% |
| hsa_circ_0069347 | 572 | 10 | 3.50% |

| | | | |
|---|---|---|---|
| hsa_circ_0088801 | 572 | 10 | 3.50% |
| hsa_circ_0129222 | 572 | 10 | 3.50% |
| hsa_circ_0105671 | 744 | 13 | 3.49% |
| hsa_circ_0039101 | 687 | 12 | 3.49% |
| hsa_circ_0106014 | 687 | 12 | 3.49% |
| hsa_circ_0136178 | 687 | 12 | 3.49% |
| hsa_circ_0012521 | 630 | 11 | 3.49% |
| hsa_circ_0110950 | 630 | 11 | 3.49% |
| hsa_circ_0130251 | 802 | 14 | 3.49% |
| hsa_circ_0007907 | 573 | 10 | 3.49% |
| hsa_circ_0074259 | 573 | 10 | 3.49% |
| hsa_circ_0075125 | 573 | 10 | 3.49% |
| hsa_circ_0079206 | 573 | 10 | 3.49% |
| hsa_circ_0085777 | 573 | 10 | 3.49% |
| hsa_circ_0092685 | 573 | 10 | 3.49% |
| hsa_circ_0100828 | 573 | 10 | 3.49% |
| hsa_circ_0111927 | 573 | 10 | 3.49% |
| hsa_circ_0124516 | 573 | 10 | 3.49% |
| hsa_circ_0126439 | 573 | 10 | 3.49% |
| hsa_circ_0129572 | 573 | 10 | 3.49% |
| hsa_circ_0131794 | 1204 | 21 | 3.49% |
| hsa_circ_0071536 | 688 | 12 | 3.49% |
| hsa_circ_0129441 | 688 | 12 | 3.49% |
| hsa_circ_0121535 | 975 | 17 | 3.49% |
| hsa_circ_0064438 | 631 | 11 | 3.49% |
| hsa_circ_0115339 | 631 | 11 | 3.49% |
| hsa_circ_0126677 | 631 | 11 | 3.49% |
| hsa_circ_0057373 | 918 | 16 | 3.49% |
| hsa_circ_0010673 | 746 | 13 | 3.49% |
| hsa_circ_0115831 | 746 | 13 | 3.49% |
| hsa_circ_0003292 | 574 | 10 | 3.48% |
| hsa_circ_0005406 | 574 | 10 | 3.48% |
| hsa_circ_0007244 | 574 | 10 | 3.48% |
| hsa_circ_0012955 | 574 | 10 | 3.48% |
| hsa_circ_0012976 | 574 | 10 | 3.48% |
| hsa_circ_0015597 | 574 | 10 | 3.48% |
| hsa_circ_0029263 | 574 | 10 | 3.48% |
| hsa_circ_0042714 | 574 | 10 | 3.48% |
| hsa_circ_0043308 | 574 | 10 | 3.48% |
| hsa_circ_0049250 | 574 | 10 | 3.48% |
| hsa_circ_0096552 | 574 | 10 | 3.48% |
| hsa_circ_0101983 | 574 | 10 | 3.48% |
| hsa_circ_0104780 | 574 | 10 | 3.48% |
| hsa_circ_0117105 | 574 | 10 | 3.48% |
| hsa_circ_0054626 | 689 | 12 | 3.48% |
| hsa_circ_0085709 | 632 | 11 | 3.48% |
| hsa_circ_0120758 | 632 | 11 | 3.48% |
| hsa_circ_0135339 | 632 | 11 | 3.48% |
| hsa_circ_0140032 | 632 | 11 | 3.48% |
| hsa_circ_0001971 | 575 | 10 | 3.48% |

| | | | |
|---|---|---|---|
| hsa_circ_0002106 | 575 | 10 | 3.48% |
| hsa_circ_0019393 | 575 | 10 | 3.48% |
| hsa_circ_0020181 | 575 | 10 | 3.48% |
| hsa_circ_0022583 | 575 | 10 | 3.48% |
| hsa_circ_0034065 | 575 | 10 | 3.48% |
| hsa_circ_0061815 | 575 | 10 | 3.48% |
| hsa_circ_0073111 | 575 | 10 | 3.48% |
| hsa_circ_0126164 | 575 | 10 | 3.48% |
| hsa_circ_0126640 | 575 | 10 | 3.48% |
| hsa_circ_0128950 | 575 | 10 | 3.48% |
| hsa_circ_0132955 | 575 | 10 | 3.48% |
| hsa_circ_0136547 | 575 | 10 | 3.48% |
| hsa_circ_0005806 | 748 | 13 | 3.48% |
| hsa_circ_0010984 | 633 | 11 | 3.48% |
| hsa_circ_0050619 | 633 | 11 | 3.48% |
| hsa_circ_0051005 | 633 | 11 | 3.48% |
| hsa_circ_0115521 | 633 | 11 | 3.48% |
| hsa_circ_0119991 | 633 | 11 | 3.48% |
| hsa_circ_0087159 | 691 | 12 | 3.47% |
| hsa_circ_0132529 | 691 | 12 | 3.47% |
| hsa_circ_0003400 | 576 | 10 | 3.47% |
| hsa_circ_0005741 | 576 | 10 | 3.47% |
| hsa_circ_0007970 | 576 | 10 | 3.47% |
| hsa_circ_0014100 | 576 | 10 | 3.47% |
| hsa_circ_0016302 | 576 | 10 | 3.47% |
| hsa_circ_0029564 | 576 | 10 | 3.47% |
| hsa_circ_0036363 | 576 | 10 | 3.47% |
| hsa_circ_0060005 | 576 | 10 | 3.47% |
| hsa_circ_0070073 | 576 | 10 | 3.47% |
| hsa_circ_0080109 | 576 | 10 | 3.47% |
| hsa_circ_0089756 | 576 | 10 | 3.47% |
| hsa_circ_0102098 | 576 | 10 | 3.47% |
| hsa_circ_0110660 | 576 | 10 | 3.47% |
| hsa_circ_0111176 | 576 | 10 | 3.47% |
| hsa_circ_0134739 | 576 | 10 | 3.47% |
| hsa_circ_0137399 | 576 | 10 | 3.47% |
| hsa_circ_0113881 | 692 | 12 | 3.47% |
| hsa_circ_0005066 | 577 | 10 | 3.47% |
| hsa_circ_0008301 | 577 | 10 | 3.47% |
| hsa_circ_0008331 | 577 | 10 | 3.47% |
| hsa_circ_0020994 | 577 | 10 | 3.47% |
| hsa_circ_0022120 | 577 | 10 | 3.47% |
| hsa_circ_0078909 | 577 | 10 | 3.47% |
| hsa_circ_0084518 | 577 | 10 | 3.47% |
| hsa_circ_0096594 | 577 | 10 | 3.47% |
| hsa_circ_0102856 | 577 | 10 | 3.47% |
| hsa_circ_0113257 | 577 | 10 | 3.47% |
| hsa_circ_0123550 | 577 | 10 | 3.47% |
| hsa_circ_0124061 | 577 | 10 | 3.47% |
| hsa_circ_0133969 | 577 | 10 | 3.47% |

| | | | |
|---|---|---|---|
| hsa_circ_0134892 | 577 | 10 | 3.47% |
| hsa_circ_0049804 | 635 | 11 | 3.46% |
| hsa_circ_0079939 | 635 | 11 | 3.46% |
| hsa_circ_0133072 | 635 | 11 | 3.46% |
| hsa_circ_0105822 | 1097 | 19 | 3.46% |
| hsa_circ_0089185 | 693 | 12 | 3.46% |
| hsa_circ_0002055 | 578 | 10 | 3.46% |
| hsa_circ_0002183 | 578 | 10 | 3.46% |
| hsa_circ_0003511 | 578 | 10 | 3.46% |
| hsa_circ_0012600 | 578 | 10 | 3.46% |
| hsa_circ_0019556 | 578 | 10 | 3.46% |
| hsa_circ_0042659 | 578 | 10 | 3.46% |
| hsa_circ_0081521 | 578 | 10 | 3.46% |
| hsa_circ_0081637 | 578 | 10 | 3.46% |
| hsa_circ_0103566 | 578 | 10 | 3.46% |
| hsa_circ_0106674 | 578 | 10 | 3.46% |
| hsa_circ_0048339 | 636 | 11 | 3.46% |
| hsa_circ_0089115 | 636 | 11 | 3.46% |
| hsa_circ_0106783 | 636 | 11 | 3.46% |
| hsa_circ_0112094 | 636 | 11 | 3.46% |
| hsa_circ_0018135 | 694 | 12 | 3.46% |
| hsa_circ_0029195 | 694 | 12 | 3.46% |
| hsa_circ_0006050 | 579 | 10 | 3.45% |
| hsa_circ_0008700 | 579 | 10 | 3.45% |
| hsa_circ_0010686 | 579 | 10 | 3.45% |
| hsa_circ_0012730 | 579 | 10 | 3.45% |
| hsa_circ_0013772 | 579 | 10 | 3.45% |
| hsa_circ_0026786 | 579 | 10 | 3.45% |
| hsa_circ_0044819 | 579 | 10 | 3.45% |
| hsa_circ_0063685 | 579 | 10 | 3.45% |
| hsa_circ_0069894 | 579 | 10 | 3.45% |
| hsa_circ_0088361 | 579 | 10 | 3.45% |
| hsa_circ_0104726 | 579 | 10 | 3.45% |
| hsa_circ_0112063 | 579 | 10 | 3.45% |
| hsa_circ_0116978 | 579 | 10 | 3.45% |
| hsa_circ_0119318 | 579 | 10 | 3.45% |
| hsa_circ_0120789 | 579 | 10 | 3.45% |
| hsa_circ_0010635 | 637 | 11 | 3.45% |
| hsa_circ_0014233 | 637 | 11 | 3.45% |
| hsa_circ_0114741 | 637 | 11 | 3.45% |
| hsa_circ_0031031 | 695 | 12 | 3.45% |
| hsa_circ_0035750 | 695 | 12 | 3.45% |
| hsa_circ_0133884 | 695 | 12 | 3.45% |
| hsa_circ_0007346 | 2087 | 36 | 3.45% |
| hsa_circ_0036394 | 696 | 12 | 3.45% |
| hsa_circ_0055362 | 696 | 12 | 3.45% |
| hsa_circ_0015927 | 638 | 11 | 3.45% |
| hsa_circ_0050179 | 638 | 11 | 3.45% |
| hsa_circ_0059137 | 638 | 11 | 3.45% |
| hsa_circ_0001956 | 580 | 10 | 3.45% |

| | | | |
|---|---|---|---|
| hsa_circ_0005894 | 580 | 10 | 3.45% |
| hsa_circ_0017491 | 580 | 10 | 3.45% |
| hsa_circ_0025686 | 580 | 10 | 3.45% |
| hsa_circ_0027467 | 580 | 10 | 3.45% |
| hsa_circ_0032999 | 580 | 10 | 3.45% |
| hsa_circ_0036750 | 580 | 10 | 3.45% |
| hsa_circ_0073878 | 580 | 10 | 3.45% |
| hsa_circ_0091733 | 580 | 10 | 3.45% |
| hsa_circ_0099588 | 580 | 10 | 3.45% |
| hsa_circ_0114114 | 580 | 10 | 3.45% |
| hsa_circ_0115292 | 580 | 10 | 3.45% |
| hsa_circ_0119333 | 580 | 10 | 3.45% |
| hsa_circ_0067432 | 697 | 12 | 3.44% |
| hsa_circ_0140620 | 697 | 12 | 3.44% |
| hsa_circ_0007374 | 639 | 11 | 3.44% |
| hsa_circ_0076572 | 639 | 11 | 3.44% |
| hsa_circ_0114154 | 639 | 11 | 3.44% |
| hsa_circ_0126846 | 639 | 11 | 3.44% |
| hsa_circ_0127008 | 639 | 11 | 3.44% |
| hsa_circ_0000737 | 581 | 10 | 3.44% |
| hsa_circ_0022693 | 581 | 10 | 3.44% |
| hsa_circ_0046518 | 581 | 10 | 3.44% |
| hsa_circ_0059971 | 581 | 10 | 3.44% |
| hsa_circ_0088726 | 581 | 10 | 3.44% |
| hsa_circ_0099520 | 581 | 10 | 3.44% |
| hsa_circ_0105194 | 581 | 10 | 3.44% |
| hsa_circ_0129884 | 581 | 10 | 3.44% |
| hsa_circ_0038820 | 640 | 11 | 3.44% |
| hsa_circ_0055979 | 640 | 11 | 3.44% |
| hsa_circ_0085669 | 640 | 11 | 3.44% |
| hsa_circ_0089594 | 640 | 11 | 3.44% |
| hsa_circ_0106481 | 640 | 11 | 3.44% |
| hsa_circ_0018454 | 582 | 10 | 3.44% |
| hsa_circ_0020798 | 582 | 10 | 3.44% |
| hsa_circ_0032851 | 582 | 10 | 3.44% |
| hsa_circ_0042251 | 582 | 10 | 3.44% |
| hsa_circ_0060374 | 582 | 10 | 3.44% |
| hsa_circ_0081600 | 582 | 10 | 3.44% |
| hsa_circ_0093362 | 582 | 10 | 3.44% |
| hsa_circ_0098301 | 582 | 10 | 3.44% |
| hsa_circ_0100381 | 582 | 10 | 3.44% |
| hsa_circ_0100755 | 582 | 10 | 3.44% |
| hsa_circ_0102735 | 582 | 10 | 3.44% |
| hsa_circ_0115304 | 582 | 10 | 3.44% |
| hsa_circ_0117155 | 582 | 10 | 3.44% |
| hsa_circ_0117432 | 582 | 10 | 3.44% |
| hsa_circ_0133919 | 582 | 10 | 3.44% |
| hsa_circ_0072755 | 699 | 12 | 3.43% |
| hsa_circ_0132329 | 699 | 12 | 3.43% |
| hsa_circ_0002655 | 641 | 11 | 3.43% |

| | | | |
|---|---|---|---|
| hsa_circ_0009370 | 641 | 11 | 3.43% |
| hsa_circ_0058290 | 641 | 11 | 3.43% |
| hsa_circ_0109647 | 641 | 11 | 3.43% |
| hsa_circ_0120854 | 641 | 11 | 3.43% |
| hsa_circ_0127369 | 641 | 11 | 3.43% |
| hsa_circ_0026380 | 18185 | 312 | 3.43% |
| hsa_circ_0102612 | 991 | 17 | 3.43% |
| hsa_circ_0007590 | 583 | 10 | 3.43% |
| hsa_circ_0012407 | 583 | 10 | 3.43% |
| hsa_circ_0047736 | 583 | 10 | 3.43% |
| hsa_circ_0048334 | 583 | 10 | 3.43% |
| hsa_circ_0062303 | 583 | 10 | 3.43% |
| hsa_circ_0073491 | 583 | 10 | 3.43% |
| hsa_circ_0082462 | 583 | 10 | 3.43% |
| hsa_circ_0094336 | 583 | 10 | 3.43% |
| hsa_circ_0095991 | 583 | 10 | 3.43% |
| hsa_circ_0107238 | 583 | 10 | 3.43% |
| hsa_circ_0125928 | 583 | 10 | 3.43% |
| hsa_circ_0129502 | 583 | 10 | 3.43% |
| hsa_circ_0092101 | 700 | 12 | 3.43% |
| hsa_circ_0039997 | 642 | 11 | 3.43% |
| hsa_circ_0067137 | 642 | 11 | 3.43% |
| hsa_circ_0075746 | 642 | 11 | 3.43% |
| hsa_circ_0091397 | 642 | 11 | 3.43% |
| hsa_circ_0130671 | 642 | 11 | 3.43% |
| hsa_circ_0007181 | 584 | 10 | 3.42% |
| hsa_circ_0009721 | 584 | 10 | 3.42% |
| hsa_circ_0020936 | 584 | 10 | 3.42% |
| hsa_circ_0039995 | 584 | 10 | 3.42% |
| hsa_circ_0043102 | 584 | 10 | 3.42% |
| hsa_circ_0043742 | 584 | 10 | 3.42% |
| hsa_circ_0055884 | 584 | 10 | 3.42% |
| hsa_circ_0064923 | 584 | 10 | 3.42% |
| hsa_circ_0065426 | 584 | 10 | 3.42% |
| hsa_circ_0072877 | 584 | 10 | 3.42% |
| hsa_circ_0076507 | 584 | 10 | 3.42% |
| hsa_circ_0076598 | 584 | 10 | 3.42% |
| hsa_circ_0087371 | 584 | 10 | 3.42% |
| hsa_circ_0097966 | 584 | 10 | 3.42% |
| hsa_circ_0131003 | 584 | 10 | 3.42% |
| hsa_circ_0132013 | 584 | 10 | 3.42% |
| hsa_circ_0140704 | 584 | 10 | 3.42% |
| hsa_circ_0043274 | 701 | 12 | 3.42% |
| hsa_circ_0015954 | 643 | 11 | 3.42% |
| hsa_circ_0040984 | 643 | 11 | 3.42% |
| hsa_circ_0073220 | 643 | 11 | 3.42% |
| hsa_circ_0122902 | 643 | 11 | 3.42% |
| hsa_circ_0047740 | 760 | 13 | 3.42% |
| hsa_circ_0061064 | 702 | 12 | 3.42% |
| hsa_circ_0109014 | 702 | 12 | 3.42% |

| | | | |
|---|---|---|---|
| hsa_circ_0118157 | 702 | 12 | 3.42% |
| hsa_circ_0000140 | 585 | 10 | 3.42% |
| hsa_circ_0001358 | 585 | 10 | 3.42% |
| hsa_circ_0001362 | 585 | 10 | 3.42% |
| hsa_circ_0011815 | 585 | 10 | 3.42% |
| hsa_circ_0027997 | 585 | 10 | 3.42% |
| hsa_circ_0040893 | 585 | 10 | 3.42% |
| hsa_circ_0048241 | 585 | 10 | 3.42% |
| hsa_circ_0073204 | 585 | 10 | 3.42% |
| hsa_circ_0078033 | 585 | 10 | 3.42% |
| hsa_circ_0091760 | 585 | 10 | 3.42% |
| hsa_circ_0118808 | 585 | 10 | 3.42% |
| hsa_circ_0120794 | 585 | 10 | 3.42% |
| hsa_circ_0127824 | 585 | 10 | 3.42% |
| hsa_circ_0062269 | 878 | 15 | 3.42% |
| hsa_circ_0020306 | 644 | 11 | 3.42% |
| hsa_circ_0024795 | 644 | 11 | 3.42% |
| hsa_circ_0076794 | 644 | 11 | 3.42% |
| hsa_circ_0080452 | 644 | 11 | 3.42% |
| hsa_circ_0048843 | 2401 | 41 | 3.42% |
| hsa_circ_0107173 | 820 | 14 | 3.41% |
| hsa_circ_0057721 | 703 | 12 | 3.41% |
| hsa_circ_0001916 | 586 | 10 | 3.41% |
| hsa_circ_0030550 | 586 | 10 | 3.41% |
| hsa_circ_0043624 | 586 | 10 | 3.41% |
| hsa_circ_0072794 | 586 | 10 | 3.41% |
| hsa_circ_0072864 | 586 | 10 | 3.41% |
| hsa_circ_0080115 | 586 | 10 | 3.41% |
| hsa_circ_0095391 | 586 | 10 | 3.41% |
| hsa_circ_0105972 | 586 | 10 | 3.41% |
| hsa_circ_0109637 | 586 | 10 | 3.41% |
| hsa_circ_0122116 | 586 | 10 | 3.41% |
| hsa_circ_0022210 | 645 | 11 | 3.41% |
| hsa_circ_0034723 | 645 | 11 | 3.41% |
| hsa_circ_0077815 | 645 | 11 | 3.41% |
| hsa_circ_0085902 | 645 | 11 | 3.41% |
| hsa_circ_0104565 | 645 | 11 | 3.41% |
| hsa_circ_0105141 | 645 | 11 | 3.41% |
| hsa_circ_0118086 | 645 | 11 | 3.41% |
| hsa_circ_0120754 | 645 | 11 | 3.41% |
| hsa_circ_0007333 | 587 | 10 | 3.41% |
| hsa_circ_0007653 | 587 | 10 | 3.41% |
| hsa_circ_0013547 | 587 | 10 | 3.41% |
| hsa_circ_0022229 | 587 | 10 | 3.41% |
| hsa_circ_0071358 | 587 | 10 | 3.41% |
| hsa_circ_0082697 | 587 | 10 | 3.41% |
| hsa_circ_0094314 | 587 | 10 | 3.41% |
| hsa_circ_0107630 | 587 | 10 | 3.41% |
| hsa_circ_0119760 | 587 | 10 | 3.41% |
| hsa_circ_0121787 | 587 | 10 | 3.41% |

| | | | |
|---|---|---|---|
| hsa_circ_0132224 | 587 | 10 | 3.41% |
| hsa_circ_0017824 | 646 | 11 | 3.41% |
| hsa_circ_0016764 | 705 | 12 | 3.40% |
| hsa_circ_0000532 | 588 | 10 | 3.40% |
| hsa_circ_0003369 | 588 | 10 | 3.40% |
| hsa_circ_0004924 | 588 | 10 | 3.40% |
| hsa_circ_0023388 | 588 | 10 | 3.40% |
| hsa_circ_0037765 | 588 | 10 | 3.40% |
| hsa_circ_0038942 | 588 | 10 | 3.40% |
| hsa_circ_0039278 | 588 | 10 | 3.40% |
| hsa_circ_0042726 | 588 | 10 | 3.40% |
| hsa_circ_0054399 | 588 | 10 | 3.40% |
| hsa_circ_0065868 | 588 | 10 | 3.40% |
| hsa_circ_0081664 | 588 | 10 | 3.40% |
| hsa_circ_0083717 | 588 | 10 | 3.40% |
| hsa_circ_0089186 | 588 | 10 | 3.40% |
| hsa_circ_0102578 | 588 | 10 | 3.40% |
| hsa_circ_0128763 | 588 | 10 | 3.40% |
| hsa_circ_0104601 | 647 | 11 | 3.40% |
| hsa_circ_0067915 | 706 | 12 | 3.40% |
| hsa_circ_0049079 | 765 | 13 | 3.40% |
| hsa_circ_0003160 | 589 | 10 | 3.40% |
| hsa_circ_0006626 | 589 | 10 | 3.40% |
| hsa_circ_0037904 | 589 | 10 | 3.40% |
| hsa_circ_0066625 | 589 | 10 | 3.40% |
| hsa_circ_0075256 | 589 | 10 | 3.40% |
| hsa_circ_0075919 | 589 | 10 | 3.40% |
| hsa_circ_0095610 | 589 | 10 | 3.40% |
| hsa_circ_0102916 | 589 | 10 | 3.40% |
| hsa_circ_0103696 | 589 | 10 | 3.40% |
| hsa_circ_0106893 | 589 | 10 | 3.40% |
| hsa_circ_0109416 | 589 | 10 | 3.40% |
| hsa_circ_0132973 | 589 | 10 | 3.40% |
| hsa_circ_0140388 | 589 | 10 | 3.40% |
| hsa_circ_0000323 | 648 | 11 | 3.40% |
| hsa_circ_0017838 | 648 | 11 | 3.40% |
| hsa_circ_0017947 | 648 | 11 | 3.40% |
| hsa_circ_0052209 | 648 | 11 | 3.40% |
| hsa_circ_0100804 | 648 | 11 | 3.40% |
| hsa_circ_0123018 | 648 | 11 | 3.40% |
| hsa_circ_0066570 | 707 | 12 | 3.39% |
| hsa_circ_0045399 | 826 | 14 | 3.39% |
| hsa_circ_0045328 | 708 | 12 | 3.39% |
| hsa_circ_0058714 | 708 | 12 | 3.39% |
| hsa_circ_0086944 | 708 | 12 | 3.39% |
| hsa_circ_0042382 | 649 | 11 | 3.39% |
| hsa_circ_0089573 | 649 | 11 | 3.39% |
| hsa_circ_0001493 | 590 | 10 | 3.39% |
| hsa_circ_0005070 | 590 | 10 | 3.39% |
| hsa_circ_0005139 | 590 | 10 | 3.39% |

| | | | |
|---|---|---|---|
| hsa_circ_0015967 | 590 | 10 | 3.39% |
| hsa_circ_0024627 | 590 | 10 | 3.39% |
| hsa_circ_0025960 | 590 | 10 | 3.39% |
| hsa_circ_0031233 | 590 | 10 | 3.39% |
| hsa_circ_0037733 | 590 | 10 | 3.39% |
| hsa_circ_0043954 | 590 | 10 | 3.39% |
| hsa_circ_0046194 | 590 | 10 | 3.39% |
| hsa_circ_0046697 | 590 | 10 | 3.39% |
| hsa_circ_0062329 | 590 | 10 | 3.39% |
| hsa_circ_0093491 | 590 | 10 | 3.39% |
| hsa_circ_0095295 | 590 | 10 | 3.39% |
| hsa_circ_0105203 | 590 | 10 | 3.39% |
| hsa_circ_0131107 | 590 | 10 | 3.39% |
| hsa_circ_0136792 | 590 | 10 | 3.39% |
| hsa_circ_0009943 | 709 | 12 | 3.39% |
| hsa_circ_0094689 | 709 | 12 | 3.39% |
| hsa_circ_0027516 | 650 | 11 | 3.38% |
| hsa_circ_0055908 | 650 | 11 | 3.38% |
| hsa_circ_0059328 | 650 | 11 | 3.38% |
| hsa_circ_0093377 | 650 | 11 | 3.38% |
| hsa_circ_0100032 | 650 | 11 | 3.38% |
| hsa_circ_0130794 | 650 | 11 | 3.38% |
| hsa_circ_0022968 | 591 | 10 | 3.38% |
| hsa_circ_0024097 | 591 | 10 | 3.38% |
| hsa_circ_0034450 | 591 | 10 | 3.38% |
| hsa_circ_0067165 | 591 | 10 | 3.38% |
| hsa_circ_0070186 | 591 | 10 | 3.38% |
| hsa_circ_0079528 | 591 | 10 | 3.38% |
| hsa_circ_0081335 | 591 | 10 | 3.38% |
| hsa_circ_0100764 | 591 | 10 | 3.38% |
| hsa_circ_0102840 | 591 | 10 | 3.38% |
| hsa_circ_0104449 | 591 | 10 | 3.38% |
| hsa_circ_0122905 | 591 | 10 | 3.38% |
| hsa_circ_0132834 | 591 | 10 | 3.38% |
| hsa_circ_0137598 | 591 | 10 | 3.38% |
| hsa_circ_0012380 | 3487 | 59 | 3.38% |
| hsa_circ_0043460 | 828 | 14 | 3.38% |
| hsa_circ_0139665 | 769 | 13 | 3.38% |
| hsa_circ_0009757 | 710 | 12 | 3.38% |
| hsa_circ_0006525 | 651 | 11 | 3.38% |
| hsa_circ_0007884 | 651 | 11 | 3.38% |
| hsa_circ_0024723 | 651 | 11 | 3.38% |
| hsa_circ_0031302 | 651 | 11 | 3.38% |
| hsa_circ_0034572 | 651 | 11 | 3.38% |
| hsa_circ_0063751 | 651 | 11 | 3.38% |
| hsa_circ_0105490 | 651 | 11 | 3.38% |
| hsa_circ_0119990 | 651 | 11 | 3.38% |
| hsa_circ_0138169 | 651 | 11 | 3.38% |
| hsa_circ_0000470 | 592 | 10 | 3.38% |
| hsa_circ_0012727 | 592 | 10 | 3.38% |

| | | | |
|---|---|---|---|
| hsa_circ_0019139 | 592 | 10 | 3.38% |
| hsa_circ_0022767 | 592 | 10 | 3.38% |
| hsa_circ_0025865 | 592 | 10 | 3.38% |
| hsa_circ_0046137 | 592 | 10 | 3.38% |
| hsa_circ_0057783 | 592 | 10 | 3.38% |
| hsa_circ_0078237 | 592 | 10 | 3.38% |
| hsa_circ_0091791 | 592 | 10 | 3.38% |
| hsa_circ_0102981 | 592 | 10 | 3.38% |
| hsa_circ_0110944 | 592 | 10 | 3.38% |
| hsa_circ_0042684 | 711 | 12 | 3.38% |
| hsa_circ_0057444 | 711 | 12 | 3.38% |
| hsa_circ_0050016 | 652 | 11 | 3.37% |
| hsa_circ_0107204 | 652 | 11 | 3.37% |
| hsa_circ_0135675 | 652 | 11 | 3.37% |
| hsa_circ_0040581 | 593 | 10 | 3.37% |
| hsa_circ_0047488 | 593 | 10 | 3.37% |
| hsa_circ_0051842 | 593 | 10 | 3.37% |
| hsa_circ_0070912 | 593 | 10 | 3.37% |
| hsa_circ_0080903 | 593 | 10 | 3.37% |
| hsa_circ_0094010 | 593 | 10 | 3.37% |
| hsa_circ_0101776 | 593 | 10 | 3.37% |
| hsa_circ_0104975 | 593 | 10 | 3.37% |
| hsa_circ_0135959 | 593 | 10 | 3.37% |
| hsa_circ_0099617 | 890 | 15 | 3.37% |
| hsa_circ_0108923 | 890 | 15 | 3.37% |
| hsa_circ_0113955 | 890 | 15 | 3.37% |
| hsa_circ_0025112 | 653 | 11 | 3.37% |
| hsa_circ_0122147 | 653 | 11 | 3.37% |
| hsa_circ_0015446 | 594 | 10 | 3.37% |
| hsa_circ_0018539 | 594 | 10 | 3.37% |
| hsa_circ_0030283 | 594 | 10 | 3.37% |
| hsa_circ_0043518 | 594 | 10 | 3.37% |
| hsa_circ_0062312 | 594 | 10 | 3.37% |
| hsa_circ_0077495 | 594 | 10 | 3.37% |
| hsa_circ_0088309 | 594 | 10 | 3.37% |
| hsa_circ_0109170 | 594 | 10 | 3.37% |
| hsa_circ_0118002 | 594 | 10 | 3.37% |
| hsa_circ_0118931 | 594 | 10 | 3.37% |
| hsa_circ_0125498 | 594 | 10 | 3.37% |
| hsa_circ_0126433 | 594 | 10 | 3.37% |
| hsa_circ_0129737 | 594 | 10 | 3.37% |
| hsa_circ_0138058 | 594 | 10 | 3.37% |
| hsa_circ_0007637 | 713 | 12 | 3.37% |
| hsa_circ_0017034 | 713 | 12 | 3.37% |
| hsa_circ_0091815 | 832 | 14 | 3.37% |
| hsa_circ_0135538 | 832 | 14 | 3.37% |
| hsa_circ_0115991 | 654 | 11 | 3.36% |
| hsa_circ_0115366 | 714 | 12 | 3.36% |
| hsa_circ_0007456 | 595 | 10 | 3.36% |
| hsa_circ_0015658 | 595 | 10 | 3.36% |

| | | | |
|---|---|---|---|
| hsa_circ_0035326 | 595 | 10 | 3.36% |
| hsa_circ_0043429 | 595 | 10 | 3.36% |
| hsa_circ_0074431 | 595 | 10 | 3.36% |
| hsa_circ_0081304 | 595 | 10 | 3.36% |
| hsa_circ_0094931 | 595 | 10 | 3.36% |
| hsa_circ_0108729 | 595 | 10 | 3.36% |
| hsa_circ_0117288 | 595 | 10 | 3.36% |
| hsa_circ_0057389 | 774 | 13 | 3.36% |
| hsa_circ_0085802 | 655 | 11 | 3.36% |
| hsa_circ_0102645 | 655 | 11 | 3.36% |
| hsa_circ_0051773 | 834 | 14 | 3.36% |
| hsa_circ_0069030 | 715 | 12 | 3.36% |
| hsa_circ_0102682 | 715 | 12 | 3.36% |
| hsa_circ_0006056 | 596 | 10 | 3.36% |
| hsa_circ_0019783 | 596 | 10 | 3.36% |
| hsa_circ_0026558 | 596 | 10 | 3.36% |
| hsa_circ_0027986 | 596 | 10 | 3.36% |
| hsa_circ_0030523 | 596 | 10 | 3.36% |
| hsa_circ_0065626 | 596 | 10 | 3.36% |
| hsa_circ_0069165 | 596 | 10 | 3.36% |
| hsa_circ_0070330 | 596 | 10 | 3.36% |
| hsa_circ_0082050 | 596 | 10 | 3.36% |
| hsa_circ_0085101 | 596 | 10 | 3.36% |
| hsa_circ_0094397 | 596 | 10 | 3.36% |
| hsa_circ_0098574 | 596 | 10 | 3.36% |
| hsa_circ_0099711 | 596 | 10 | 3.36% |
| hsa_circ_0113378 | 596 | 10 | 3.36% |
| hsa_circ_0114305 | 596 | 10 | 3.36% |
| hsa_circ_0119758 | 596 | 10 | 3.36% |
| hsa_circ_0121827 | 596 | 10 | 3.36% |
| hsa_circ_0125731 | 596 | 10 | 3.36% |
| hsa_circ_0134954 | 596 | 10 | 3.36% |
| hsa_circ_0136756 | 596 | 10 | 3.36% |
| hsa_circ_0080817 | 656 | 11 | 3.35% |
| hsa_circ_0096000 | 656 | 11 | 3.35% |
| hsa_circ_0101998 | 656 | 11 | 3.35% |
| hsa_circ_0127720 | 656 | 11 | 3.35% |
| hsa_circ_0095924 | 895 | 15 | 3.35% |
| hsa_circ_0102977 | 716 | 12 | 3.35% |
| hsa_circ_0014026 | 597 | 10 | 3.35% |
| hsa_circ_0032977 | 597 | 10 | 3.35% |
| hsa_circ_0035528 | 597 | 10 | 3.35% |
| hsa_circ_0044120 | 597 | 10 | 3.35% |
| hsa_circ_0053125 | 597 | 10 | 3.35% |
| hsa_circ_0060212 | 597 | 10 | 3.35% |
| hsa_circ_0067087 | 597 | 10 | 3.35% |
| hsa_circ_0079830 | 597 | 10 | 3.35% |
| hsa_circ_0080276 | 597 | 10 | 3.35% |
| hsa_circ_0100493 | 597 | 10 | 3.35% |
| hsa_circ_0110579 | 597 | 10 | 3.35% |

| | | | |
|---|---|---|---|
| hsa_circ_0115728 | 597 | 10 | 3.35% |
| hsa_circ_0130172 | 597 | 10 | 3.35% |
| hsa_circ_0134426 | 597 | 10 | 3.35% |
| hsa_circ_0036989 | 836 | 14 | 3.35% |
| hsa_circ_0006060 | 657 | 11 | 3.35% |
| hsa_circ_0038622 | 657 | 11 | 3.35% |
| hsa_circ_0042751 | 657 | 11 | 3.35% |
| hsa_circ_0057445 | 657 | 11 | 3.35% |
| hsa_circ_0058233 | 657 | 11 | 3.35% |
| hsa_circ_0104766 | 657 | 11 | 3.35% |
| hsa_circ_0052170 | 896 | 15 | 3.35% |
| hsa_circ_0009600 | 598 | 10 | 3.34% |
| hsa_circ_0020565 | 598 | 10 | 3.34% |
| hsa_circ_0039858 | 598 | 10 | 3.34% |
| hsa_circ_0047923 | 598 | 10 | 3.34% |
| hsa_circ_0050569 | 598 | 10 | 3.34% |
| hsa_circ_0051868 | 598 | 10 | 3.34% |
| hsa_circ_0060536 | 598 | 10 | 3.34% |
| hsa_circ_0069381 | 598 | 10 | 3.34% |
| hsa_circ_0073114 | 598 | 10 | 3.34% |
| hsa_circ_0086942 | 598 | 10 | 3.34% |
| hsa_circ_0090042 | 598 | 10 | 3.34% |
| hsa_circ_0090462 | 598 | 10 | 3.34% |
| hsa_circ_0091984 | 598 | 10 | 3.34% |
| hsa_circ_0109171 | 598 | 10 | 3.34% |
| hsa_circ_0138980 | 598 | 10 | 3.34% |
| hsa_circ_0046506 | 658 | 11 | 3.34% |
| hsa_circ_0081002 | 658 | 11 | 3.34% |
| hsa_circ_0121169 | 658 | 11 | 3.34% |
| hsa_circ_0097562 | 718 | 12 | 3.34% |
| hsa_circ_0117678 | 718 | 12 | 3.34% |
| hsa_circ_0095656 | 778 | 13 | 3.34% |
| hsa_circ_0097262 | 778 | 13 | 3.34% |
| hsa_circ_0113243 | 778 | 13 | 3.34% |
| hsa_circ_0006823 | 599 | 10 | 3.34% |
| hsa_circ_0039143 | 599 | 10 | 3.34% |
| hsa_circ_0045492 | 599 | 10 | 3.34% |
| hsa_circ_0069145 | 599 | 10 | 3.34% |
| hsa_circ_0069333 | 599 | 10 | 3.34% |
| hsa_circ_0093641 | 599 | 10 | 3.34% |
| hsa_circ_0094721 | 599 | 10 | 3.34% |
| hsa_circ_0112393 | 599 | 10 | 3.34% |
| hsa_circ_0115400 | 599 | 10 | 3.34% |
| hsa_circ_0129736 | 599 | 10 | 3.34% |
| hsa_circ_0132106 | 599 | 10 | 3.34% |
| hsa_circ_0133784 | 599 | 10 | 3.34% |
| hsa_circ_0134414 | 599 | 10 | 3.34% |
| hsa_circ_0020154 | 659 | 11 | 3.34% |
| hsa_circ_0028632 | 659 | 11 | 3.34% |
| hsa_circ_0094103 | 659 | 11 | 3.34% |

| | | | |
|---|---|---|---|
| hsa_circ_0003111 | 719 | 12 | 3.34% |
| hsa_circ_0109082 | 719 | 12 | 3.34% |
| hsa_circ_0064702 | 779 | 13 | 3.34% |
| hsa_circ_0062861 | 780 | 13 | 3.33% |
| hsa_circ_0074046 | 780 | 13 | 3.33% |
| hsa_circ_0122140 | 720 | 12 | 3.33% |
| hsa_circ_0072643 | 660 | 11 | 3.33% |
| hsa_circ_0092134 | 660 | 11 | 3.33% |
| hsa_circ_0092722 | 660 | 11 | 3.33% |
| hsa_circ_0104537 | 660 | 11 | 3.33% |
| hsa_circ_0002894 | 600 | 10 | 3.33% |
| hsa_circ_0021715 | 600 | 10 | 3.33% |
| hsa_circ_0045172 | 600 | 10 | 3.33% |
| hsa_circ_0048207 | 600 | 10 | 3.33% |
| hsa_circ_0079838 | 600 | 10 | 3.33% |
| hsa_circ_0082827 | 600 | 10 | 3.33% |
| hsa_circ_0094040 | 600 | 10 | 3.33% |
| hsa_circ_0101907 | 600 | 10 | 3.33% |
| hsa_circ_0117238 | 600 | 10 | 3.33% |
| hsa_circ_0134096 | 600 | 10 | 3.33% |
| hsa_circ_0138525 | 600 | 10 | 3.33% |
| hsa_circ_0023610 | 781 | 13 | 3.33% |
| hsa_circ_0102704 | 781 | 13 | 3.33% |
| hsa_circ_0073621 | 721 | 12 | 3.33% |
| hsa_circ_0060182 | 661 | 11 | 3.33% |
| hsa_circ_0111976 | 661 | 11 | 3.33% |
| hsa_circ_0117573 | 661 | 11 | 3.33% |
| hsa_circ_0134079 | 661 | 11 | 3.33% |
| hsa_circ_0023663 | 601 | 10 | 3.33% |
| hsa_circ_0034992 | 601 | 10 | 3.33% |
| hsa_circ_0042334 | 601 | 10 | 3.33% |
| hsa_circ_0048694 | 601 | 10 | 3.33% |
| hsa_circ_0048926 | 601 | 10 | 3.33% |
| hsa_circ_0064346 | 601 | 10 | 3.33% |
| hsa_circ_0064683 | 601 | 10 | 3.33% |
| hsa_circ_0075228 | 601 | 10 | 3.33% |
| hsa_circ_0075964 | 601 | 10 | 3.33% |
| hsa_circ_0095547 | 601 | 10 | 3.33% |
| hsa_circ_0095562 | 601 | 10 | 3.33% |
| hsa_circ_0096104 | 601 | 10 | 3.33% |
| hsa_circ_0099940 | 601 | 10 | 3.33% |
| hsa_circ_0110586 | 601 | 10 | 3.33% |
| hsa_circ_0129958 | 601 | 10 | 3.33% |
| hsa_circ_0130399 | 601 | 10 | 3.33% |
| hsa_circ_0139225 | 601 | 10 | 3.33% |
| hsa_circ_0026379 | 18766 | 312 | 3.33% |
| hsa_circ_0109763 | 782 | 13 | 3.32% |
| hsa_circ_0016450 | 722 | 12 | 3.32% |
| hsa_circ_0019281 | 722 | 12 | 3.32% |
| hsa_circ_0005925 | 662 | 11 | 3.32% |

| | | | |
|---|---|---|---|
| hsa_circ_0048258 | 662 | 11 | 3.32% |
| hsa_circ_0063558 | 662 | 11 | 3.32% |
| hsa_circ_0116881 | 662 | 11 | 3.32% |
| hsa_circ_0019384 | 602 | 10 | 3.32% |
| hsa_circ_0028916 | 602 | 10 | 3.32% |
| hsa_circ_0036319 | 602 | 10 | 3.32% |
| hsa_circ_0069608 | 602 | 10 | 3.32% |
| hsa_circ_0089097 | 602 | 10 | 3.32% |
| hsa_circ_0094723 | 602 | 10 | 3.32% |
| hsa_circ_0096317 | 602 | 10 | 3.32% |
| hsa_circ_0105357 | 602 | 10 | 3.32% |
| hsa_circ_0120522 | 602 | 10 | 3.32% |
| hsa_circ_0121900 | 602 | 10 | 3.32% |
| hsa_circ_0123206 | 602 | 10 | 3.32% |
| hsa_circ_0126435 | 602 | 10 | 3.32% |
| hsa_circ_0126588 | 602 | 10 | 3.32% |
| hsa_circ_0131868 | 602 | 10 | 3.32% |
| hsa_circ_0132456 | 602 | 10 | 3.32% |
| hsa_circ_0137165 | 602 | 10 | 3.32% |
| hsa_circ_0138464 | 602 | 10 | 3.32% |
| hsa_circ_0040226 | 783 | 13 | 3.32% |
| hsa_circ_0092256 | 783 | 13 | 3.32% |
| hsa_circ_0005978 | 663 | 11 | 3.32% |
| hsa_circ_0029159 | 663 | 11 | 3.32% |
| hsa_circ_0030159 | 663 | 11 | 3.32% |
| hsa_circ_0034653 | 663 | 11 | 3.32% |
| hsa_circ_0038611 | 663 | 11 | 3.32% |
| hsa_circ_0040307 | 663 | 11 | 3.32% |
| hsa_circ_0043492 | 663 | 11 | 3.32% |
| hsa_circ_0058527 | 663 | 11 | 3.32% |
| hsa_circ_0094844 | 663 | 11 | 3.32% |
| hsa_circ_0130119 | 663 | 11 | 3.32% |
| hsa_circ_0007304 | 603 | 10 | 3.32% |
| hsa_circ_0015170 | 603 | 10 | 3.32% |
| hsa_circ_0018721 | 603 | 10 | 3.32% |
| hsa_circ_0025620 | 603 | 10 | 3.32% |
| hsa_circ_0035401 | 603 | 10 | 3.32% |
| hsa_circ_0043570 | 603 | 10 | 3.32% |
| hsa_circ_0056879 | 603 | 10 | 3.32% |
| hsa_circ_0067985 | 603 | 10 | 3.32% |
| hsa_circ_0070132 | 603 | 10 | 3.32% |
| hsa_circ_0081992 | 603 | 10 | 3.32% |
| hsa_circ_0085031 | 603 | 10 | 3.32% |
| hsa_circ_0088178 | 603 | 10 | 3.32% |
| hsa_circ_0097638 | 603 | 10 | 3.32% |
| hsa_circ_0115076 | 603 | 10 | 3.32% |
| hsa_circ_0124193 | 603 | 10 | 3.32% |
| hsa_circ_0135549 | 603 | 10 | 3.32% |
| hsa_circ_0035806 | 724 | 12 | 3.31% |
| hsa_circ_0008229 | 664 | 11 | 3.31% |

| | | | |
|---|---|---|---|
| hsa_circ_0050323 | 664 | 11 | 3.31% |
| hsa_circ_0091150 | 664 | 11 | 3.31% |
| hsa_circ_0116630 | 664 | 11 | 3.31% |
| hsa_circ_0129764 | 664 | 11 | 3.31% |
| hsa_circ_0025681 | 785 | 13 | 3.31% |
| hsa_circ_0087953 | 785 | 13 | 3.31% |
| hsa_circ_0094122 | 785 | 13 | 3.31% |
| hsa_circ_0014245 | 604 | 10 | 3.31% |
| hsa_circ_0019668 | 604 | 10 | 3.31% |
| hsa_circ_0040564 | 604 | 10 | 3.31% |
| hsa_circ_0043933 | 604 | 10 | 3.31% |
| hsa_circ_0046525 | 604 | 10 | 3.31% |
| hsa_circ_0049089 | 604 | 10 | 3.31% |
| hsa_circ_0063030 | 604 | 10 | 3.31% |
| hsa_circ_0069311 | 604 | 10 | 3.31% |
| hsa_circ_0093395 | 604 | 10 | 3.31% |
| hsa_circ_0095208 | 604 | 10 | 3.31% |
| hsa_circ_0108516 | 604 | 10 | 3.31% |
| hsa_circ_0126101 | 604 | 10 | 3.31% |
| hsa_circ_0138114 | 604 | 10 | 3.31% |
| hsa_circ_0095443 | 725 | 12 | 3.31% |
| hsa_circ_0024423 | 967 | 16 | 3.31% |
| hsa_circ_0022266 | 665 | 11 | 3.31% |
| hsa_circ_0025502 | 665 | 11 | 3.31% |
| hsa_circ_0067663 | 665 | 11 | 3.31% |
| hsa_circ_0098030 | 665 | 11 | 3.31% |
| hsa_circ_0105329 | 665 | 11 | 3.31% |
| hsa_circ_0089587 | 786 | 13 | 3.31% |
| hsa_circ_0087158 | 726 | 12 | 3.31% |
| hsa_circ_0099444 | 726 | 12 | 3.31% |
| hsa_circ_0000159 | 605 | 10 | 3.31% |
| hsa_circ_0013176 | 605 | 10 | 3.31% |
| hsa_circ_0015111 | 605 | 10 | 3.31% |
| hsa_circ_0037510 | 605 | 10 | 3.31% |
| hsa_circ_0037907 | 605 | 10 | 3.31% |
| hsa_circ_0041819 | 605 | 10 | 3.31% |
| hsa_circ_0045046 | 605 | 10 | 3.31% |
| hsa_circ_0067276 | 605 | 10 | 3.31% |
| hsa_circ_0082952 | 605 | 10 | 3.31% |
| hsa_circ_0121957 | 605 | 10 | 3.31% |
| hsa_circ_0133975 | 605 | 10 | 3.31% |
| hsa_circ_0135825 | 605 | 10 | 3.31% |
| hsa_circ_0136368 | 605 | 10 | 3.31% |
| hsa_circ_0042955 | 1332 | 22 | 3.30% |
| hsa_circ_0029955 | 666 | 11 | 3.30% |
| hsa_circ_0044661 | 666 | 11 | 3.30% |
| hsa_circ_0057064 | 666 | 11 | 3.30% |
| hsa_circ_0065707 | 666 | 11 | 3.30% |
| hsa_circ_0124949 | 666 | 11 | 3.30% |
| hsa_circ_0133797 | 666 | 11 | 3.30% |

| | | | |
|---|---|---|---|
| hsa_circ_0140300 | 666 | 11 | 3.30% |
| hsa_circ_0004679 | 606 | 10 | 3.30% |
| hsa_circ_0015970 | 606 | 10 | 3.30% |
| hsa_circ_0035803 | 606 | 10 | 3.30% |
| hsa_circ_0039977 | 606 | 10 | 3.30% |
| hsa_circ_0066799 | 606 | 10 | 3.30% |
| hsa_circ_0082345 | 606 | 10 | 3.30% |
| hsa_circ_0084781 | 606 | 10 | 3.30% |
| hsa_circ_0086831 | 606 | 10 | 3.30% |
| hsa_circ_0095307 | 606 | 10 | 3.30% |
| hsa_circ_0107169 | 606 | 10 | 3.30% |
| hsa_circ_0109191 | 606 | 10 | 3.30% |
| hsa_circ_0125051 | 606 | 10 | 3.30% |
| hsa_circ_0130401 | 606 | 10 | 3.30% |
| hsa_circ_0132510 | 606 | 10 | 3.30% |
| hsa_circ_0023973 | 667 | 11 | 3.30% |
| hsa_circ_0031251 | 667 | 11 | 3.30% |
| hsa_circ_0039553 | 667 | 11 | 3.30% |
| hsa_circ_0079086 | 667 | 11 | 3.30% |
| hsa_circ_0119447 | 667 | 11 | 3.30% |
| hsa_circ_0118320 | 910 | 15 | 3.30% |
| hsa_circ_0015203 | 728 | 12 | 3.30% |
| hsa_circ_0135128 | 728 | 12 | 3.30% |
| hsa_circ_0002827 | 607 | 10 | 3.29% |
| hsa_circ_0009089 | 607 | 10 | 3.29% |
| hsa_circ_0020971 | 607 | 10 | 3.29% |
| hsa_circ_0087800 | 607 | 10 | 3.29% |
| hsa_circ_0093645 | 607 | 10 | 3.29% |
| hsa_circ_0096809 | 607 | 10 | 3.29% |
| hsa_circ_0109809 | 607 | 10 | 3.29% |
| hsa_circ_0105932 | 1093 | 18 | 3.29% |
| hsa_circ_0061347 | 668 | 11 | 3.29% |
| hsa_circ_0116421 | 668 | 11 | 3.29% |
| hsa_circ_0132906 | 668 | 11 | 3.29% |
| hsa_circ_0004257 | 729 | 12 | 3.29% |
| hsa_circ_0057387 | 729 | 12 | 3.29% |
| hsa_circ_0058926 | 729 | 12 | 3.29% |
| hsa_circ_0113899 | 729 | 12 | 3.29% |
| hsa_circ_0022453 | 608 | 10 | 3.29% |
| hsa_circ_0023926 | 608 | 10 | 3.29% |
| hsa_circ_0028846 | 608 | 10 | 3.29% |
| hsa_circ_0031418 | 608 | 10 | 3.29% |
| hsa_circ_0036498 | 608 | 10 | 3.29% |
| hsa_circ_0052623 | 608 | 10 | 3.29% |
| hsa_circ_0067058 | 608 | 10 | 3.29% |
| hsa_circ_0080498 | 608 | 10 | 3.29% |
| hsa_circ_0082341 | 608 | 10 | 3.29% |
| hsa_circ_0087017 | 608 | 10 | 3.29% |
| hsa_circ_0093396 | 608 | 10 | 3.29% |
| hsa_circ_0109577 | 608 | 10 | 3.29% |

| | | | |
|---|---|---|---|
| hsa_circ_0127281 | 608 | 10 | 3.29% |
| hsa_circ_0026479 | 669 | 11 | 3.29% |
| hsa_circ_0058059 | 669 | 11 | 3.29% |
| hsa_circ_0115388 | 669 | 11 | 3.29% |
| hsa_circ_0057038 | 730 | 12 | 3.29% |
| hsa_circ_0134658 | 913 | 15 | 3.29% |
| hsa_circ_0001322 | 609 | 10 | 3.28% |
| hsa_circ_0004672 | 609 | 10 | 3.28% |
| hsa_circ_0017780 | 609 | 10 | 3.28% |
| hsa_circ_0020654 | 609 | 10 | 3.28% |
| hsa_circ_0024146 | 609 | 10 | 3.28% |
| hsa_circ_0035653 | 609 | 10 | 3.28% |
| hsa_circ_0059907 | 609 | 10 | 3.28% |
| hsa_circ_0073092 | 609 | 10 | 3.28% |
| hsa_circ_0119378 | 609 | 10 | 3.28% |
| hsa_circ_0127446 | 609 | 10 | 3.28% |
| hsa_circ_0131660 | 609 | 10 | 3.28% |
| hsa_circ_0133207 | 609 | 10 | 3.28% |
| hsa_circ_0006895 | 670 | 11 | 3.28% |
| hsa_circ_0095353 | 670 | 11 | 3.28% |
| hsa_circ_0095875 | 670 | 11 | 3.28% |
| hsa_circ_0114260 | 670 | 11 | 3.28% |
| hsa_circ_0035345 | 731 | 12 | 3.28% |
| hsa_circ_0050117 | 731 | 12 | 3.28% |
| hsa_circ_0094449 | 731 | 12 | 3.28% |
| hsa_circ_0115826 | 792 | 13 | 3.28% |
| hsa_circ_0004017 | 2988 | 49 | 3.28% |
| hsa_circ_0087976 | 671 | 11 | 3.28% |
| hsa_circ_0126285 | 671 | 11 | 3.28% |
| hsa_circ_0003246 | 610 | 10 | 3.28% |
| hsa_circ_0023385 | 610 | 10 | 3.28% |
| hsa_circ_0027678 | 610 | 10 | 3.28% |
| hsa_circ_0035825 | 610 | 10 | 3.28% |
| hsa_circ_0039472 | 610 | 10 | 3.28% |
| hsa_circ_0048120 | 610 | 10 | 3.28% |
| hsa_circ_0052609 | 610 | 10 | 3.28% |
| hsa_circ_0068293 | 610 | 10 | 3.28% |
| hsa_circ_0075124 | 610 | 10 | 3.28% |
| hsa_circ_0081211 | 610 | 10 | 3.28% |
| hsa_circ_0084121 | 610 | 10 | 3.28% |
| hsa_circ_0101987 | 610 | 10 | 3.28% |
| hsa_circ_0107094 | 610 | 10 | 3.28% |
| hsa_circ_0118162 | 610 | 10 | 3.28% |
| hsa_circ_0121411 | 610 | 10 | 3.28% |
| hsa_circ_0128772 | 610 | 10 | 3.28% |
| hsa_circ_0137913 | 610 | 10 | 3.28% |
| hsa_circ_0053129 | 855 | 14 | 3.27% |
| hsa_circ_0012360 | 733 | 12 | 3.27% |
| hsa_circ_0047756 | 733 | 12 | 3.27% |
| hsa_circ_0008927 | 672 | 11 | 3.27% |

| | | | |
|---|---|---|---|
| hsa_circ_0009111 | 672 | 11 | 3.27% |
| hsa_circ_0024582 | 672 | 11 | 3.27% |
| hsa_circ_0037470 | 672 | 11 | 3.27% |
| hsa_circ_0051318 | 672 | 11 | 3.27% |
| hsa_circ_0081958 | 672 | 11 | 3.27% |
| hsa_circ_0085676 | 672 | 11 | 3.27% |
| hsa_circ_0096001 | 672 | 11 | 3.27% |
| hsa_circ_0009162 | 611 | 10 | 3.27% |
| hsa_circ_0011104 | 611 | 10 | 3.27% |
| hsa_circ_0019094 | 611 | 10 | 3.27% |
| hsa_circ_0071356 | 611 | 10 | 3.27% |
| hsa_circ_0072952 | 611 | 10 | 3.27% |
| hsa_circ_0084798 | 611 | 10 | 3.27% |
| hsa_circ_0094892 | 611 | 10 | 3.27% |
| hsa_circ_0128954 | 611 | 10 | 3.27% |
| hsa_circ_0140299 | 611 | 10 | 3.27% |
| hsa_circ_0021979 | 795 | 13 | 3.27% |
| hsa_circ_0048589 | 734 | 12 | 3.27% |
| hsa_circ_0068707 | 673 | 11 | 3.27% |
| hsa_circ_0109206 | 673 | 11 | 3.27% |
| hsa_circ_0122550 | 673 | 11 | 3.27% |
| hsa_circ_0004564 | 612 | 10 | 3.27% |
| hsa_circ_0012363 | 612 | 10 | 3.27% |
| hsa_circ_0051668 | 612 | 10 | 3.27% |
| hsa_circ_0052164 | 612 | 10 | 3.27% |
| hsa_circ_0052473 | 612 | 10 | 3.27% |
| hsa_circ_0055222 | 612 | 10 | 3.27% |
| hsa_circ_0055475 | 612 | 10 | 3.27% |
| hsa_circ_0073055 | 612 | 10 | 3.27% |
| hsa_circ_0079058 | 612 | 10 | 3.27% |
| hsa_circ_0080637 | 612 | 10 | 3.27% |
| hsa_circ_0080670 | 612 | 10 | 3.27% |
| hsa_circ_0106839 | 612 | 10 | 3.27% |
| hsa_circ_0122052 | 612 | 10 | 3.27% |
| hsa_circ_0133286 | 612 | 10 | 3.27% |
| hsa_circ_0133447 | 612 | 10 | 3.27% |
| hsa_circ_0017033 | 735 | 12 | 3.27% |
| hsa_circ_0090166 | 735 | 12 | 3.27% |
| hsa_circ_0060469 | 674 | 11 | 3.26% |
| hsa_circ_0081429 | 674 | 11 | 3.26% |
| hsa_circ_0048070 | 858 | 14 | 3.26% |
| hsa_circ_0128709 | 1839 | 30 | 3.26% |
| hsa_circ_0003771 | 613 | 10 | 3.26% |
| hsa_circ_0008948 | 613 | 10 | 3.26% |
| hsa_circ_0009934 | 613 | 10 | 3.26% |
| hsa_circ_0028042 | 613 | 10 | 3.26% |
| hsa_circ_0046707 | 613 | 10 | 3.26% |
| hsa_circ_0050681 | 613 | 10 | 3.26% |
| hsa_circ_0068644 | 613 | 10 | 3.26% |
| hsa_circ_0082331 | 613 | 10 | 3.26% |

| | | | |
|---|---|---|---|
| hsa_circ_0087441 | 613 | 10 | 3.26% |
| hsa_circ_0105680 | 613 | 10 | 3.26% |
| hsa_circ_0108007 | 613 | 10 | 3.26% |
| hsa_circ_0109504 | 613 | 10 | 3.26% |
| hsa_circ_0110662 | 613 | 10 | 3.26% |
| hsa_circ_0126162 | 613 | 10 | 3.26% |
| hsa_circ_0006503 | 736 | 12 | 3.26% |
| hsa_circ_0054799 | 736 | 12 | 3.26% |
| hsa_circ_0073205 | 675 | 11 | 3.26% |
| hsa_circ_0081424 | 675 | 11 | 3.26% |
| hsa_circ_0097077 | 675 | 11 | 3.26% |
| hsa_circ_0109732 | 675 | 11 | 3.26% |
| hsa_circ_0131061 | 675 | 11 | 3.26% |
| hsa_circ_0003791 | 614 | 10 | 3.26% |
| hsa_circ_0008148 | 614 | 10 | 3.26% |
| hsa_circ_0011090 | 614 | 10 | 3.26% |
| hsa_circ_0022890 | 614 | 10 | 3.26% |
| hsa_circ_0042397 | 614 | 10 | 3.26% |
| hsa_circ_0043791 | 614 | 10 | 3.26% |
| hsa_circ_0048336 | 614 | 10 | 3.26% |
| hsa_circ_0055540 | 614 | 10 | 3.26% |
| hsa_circ_0064322 | 614 | 10 | 3.26% |
| hsa_circ_0086718 | 614 | 10 | 3.26% |
| hsa_circ_0100399 | 614 | 10 | 3.26% |
| hsa_circ_0115467 | 614 | 10 | 3.26% |
| hsa_circ_0117639 | 614 | 10 | 3.26% |
| hsa_circ_0122658 | 614 | 10 | 3.26% |
| hsa_circ_0120699 | 737 | 12 | 3.26% |
| hsa_circ_0134458 | 2642 | 43 | 3.26% |
| hsa_circ_0076357 | 676 | 11 | 3.25% |
| hsa_circ_0134291 | 676 | 11 | 3.25% |
| hsa_circ_0034082 | 738 | 12 | 3.25% |
| hsa_circ_0006964 | 615 | 10 | 3.25% |
| hsa_circ_0018173 | 615 | 10 | 3.25% |
| hsa_circ_0021139 | 615 | 10 | 3.25% |
| hsa_circ_0021486 | 615 | 10 | 3.25% |
| hsa_circ_0058649 | 615 | 10 | 3.25% |
| hsa_circ_0073242 | 615 | 10 | 3.25% |
| hsa_circ_0079167 | 615 | 10 | 3.25% |
| hsa_circ_0082306 | 615 | 10 | 3.25% |
| hsa_circ_0085892 | 615 | 10 | 3.25% |
| hsa_circ_0114850 | 615 | 10 | 3.25% |
| hsa_circ_0125917 | 615 | 10 | 3.25% |
| hsa_circ_0137652 | 615 | 10 | 3.25% |
| hsa_circ_0139137 | 615 | 10 | 3.25% |
| hsa_circ_0051670 | 677 | 11 | 3.25% |
| hsa_circ_0125209 | 677 | 11 | 3.25% |
| hsa_circ_0127403 | 677 | 11 | 3.25% |
| hsa_circ_0011551 | 616 | 10 | 3.25% |
| hsa_circ_0022037 | 616 | 10 | 3.25% |

| | | | |
|---|---|---|---|
| hsa_circ_0036455 | 616 | 10 | 3.25% |
| hsa_circ_0042227 | 616 | 10 | 3.25% |
| hsa_circ_0064957 | 616 | 10 | 3.25% |
| hsa_circ_0071394 | 616 | 10 | 3.25% |
| hsa_circ_0096822 | 616 | 10 | 3.25% |
| hsa_circ_0134322 | 616 | 10 | 3.25% |
| hsa_circ_0008575 | 678 | 11 | 3.24% |
| hsa_circ_0012173 | 678 | 11 | 3.24% |
| hsa_circ_0027620 | 678 | 11 | 3.24% |
| hsa_circ_0061673 | 678 | 11 | 3.24% |
| hsa_circ_0082780 | 678 | 11 | 3.24% |
| hsa_circ_0091038 | 678 | 11 | 3.24% |
| hsa_circ_0097217 | 678 | 11 | 3.24% |
| hsa_circ_0109224 | 678 | 11 | 3.24% |
| hsa_circ_0093495 | 33358 | 541 | 3.24% |
| hsa_circ_0115799 | 740 | 12 | 3.24% |
| hsa_circ_0006776 | 4564 | 74 | 3.24% |
| hsa_circ_0001496 | 617 | 10 | 3.24% |
| hsa_circ_0019637 | 617 | 10 | 3.24% |
| hsa_circ_0029704 | 617 | 10 | 3.24% |
| hsa_circ_0038847 | 617 | 10 | 3.24% |
| hsa_circ_0056208 | 617 | 10 | 3.24% |
| hsa_circ_0063795 | 617 | 10 | 3.24% |
| hsa_circ_0067656 | 617 | 10 | 3.24% |
| hsa_circ_0087889 | 617 | 10 | 3.24% |
| hsa_circ_0123498 | 617 | 10 | 3.24% |
| hsa_circ_0002474 | 679 | 11 | 3.24% |
| hsa_circ_0042786 | 679 | 11 | 3.24% |
| hsa_circ_0052455 | 679 | 11 | 3.24% |
| hsa_circ_0061575 | 679 | 11 | 3.24% |
| hsa_circ_0089705 | 679 | 11 | 3.24% |
| hsa_circ_0098608 | 2963 | 48 | 3.24% |
| hsa_circ_0012675 | 741 | 12 | 3.24% |
| hsa_circ_0031542 | 741 | 12 | 3.24% |
| hsa_circ_0000084 | 618 | 10 | 3.24% |
| hsa_circ_0008078 | 618 | 10 | 3.24% |
| hsa_circ_0013216 | 618 | 10 | 3.24% |
| hsa_circ_0050387 | 618 | 10 | 3.24% |
| hsa_circ_0055839 | 618 | 10 | 3.24% |
| hsa_circ_0061482 | 618 | 10 | 3.24% |
| hsa_circ_0065950 | 618 | 10 | 3.24% |
| hsa_circ_0075195 | 618 | 10 | 3.24% |
| hsa_circ_0077713 | 618 | 10 | 3.24% |
| hsa_circ_0088538 | 618 | 10 | 3.24% |
| hsa_circ_0091392 | 618 | 10 | 3.24% |
| hsa_circ_0096304 | 618 | 10 | 3.24% |
| hsa_circ_0107709 | 618 | 10 | 3.24% |
| hsa_circ_0116746 | 618 | 10 | 3.24% |
| hsa_circ_0124667 | 618 | 10 | 3.24% |
| hsa_circ_0138110 | 618 | 10 | 3.24% |

| | | | |
|---|---|---|---|
| hsa_circ_0032970 | 680 | 11 | 3.24% |
| hsa_circ_0046497 | 680 | 11 | 3.24% |
| hsa_circ_0099201 | 680 | 11 | 3.24% |
| hsa_circ_0116304 | 680 | 11 | 3.24% |
| hsa_circ_0122671 | 680 | 11 | 3.24% |
| hsa_circ_0039013 | 742 | 12 | 3.23% |
| hsa_circ_0009455 | 619 | 10 | 3.23% |
| hsa_circ_0033592 | 619 | 10 | 3.23% |
| hsa_circ_0038964 | 619 | 10 | 3.23% |
| hsa_circ_0046410 | 619 | 10 | 3.23% |
| hsa_circ_0055886 | 619 | 10 | 3.23% |
| hsa_circ_0064831 | 619 | 10 | 3.23% |
| hsa_circ_0078751 | 619 | 10 | 3.23% |
| hsa_circ_0112738 | 619 | 10 | 3.23% |
| hsa_circ_0116565 | 619 | 10 | 3.23% |
| hsa_circ_0118693 | 619 | 10 | 3.23% |
| hsa_circ_0119249 | 619 | 10 | 3.23% |
| hsa_circ_0120763 | 619 | 10 | 3.23% |
| hsa_circ_0123044 | 619 | 10 | 3.23% |
| hsa_circ_0125815 | 619 | 10 | 3.23% |
| hsa_circ_0133187 | 619 | 10 | 3.23% |
| hsa_circ_0138462 | 619 | 10 | 3.23% |
| hsa_circ_0139038 | 619 | 10 | 3.23% |
| hsa_circ_0068059 | 681 | 11 | 3.23% |
| hsa_circ_0085552 | 681 | 11 | 3.23% |
| hsa_circ_0124181 | 681 | 11 | 3.23% |
| hsa_circ_0128563 | 681 | 11 | 3.23% |
| hsa_circ_0027913 | 805 | 13 | 3.23% |
| hsa_circ_0097283 | 805 | 13 | 3.23% |
| hsa_circ_0029244 | 1549 | 25 | 3.23% |
| hsa_circ_0128708 | 1859 | 30 | 3.23% |
| hsa_circ_0138922 | 868 | 14 | 3.23% |
| hsa_circ_0042891 | 806 | 13 | 3.23% |
| hsa_circ_0107781 | 806 | 13 | 3.23% |
| hsa_circ_0002615 | 744 | 12 | 3.23% |
| hsa_circ_0071353 | 744 | 12 | 3.23% |
| hsa_circ_0017274 | 682 | 11 | 3.23% |
| hsa_circ_0055500 | 682 | 11 | 3.23% |
| hsa_circ_0066144 | 682 | 11 | 3.23% |
| hsa_circ_0003909 | 620 | 10 | 3.23% |
| hsa_circ_0008152 | 620 | 10 | 3.23% |
| hsa_circ_0014673 | 620 | 10 | 3.23% |
| hsa_circ_0016080 | 620 | 10 | 3.23% |
| hsa_circ_0022469 | 620 | 10 | 3.23% |
| hsa_circ_0036776 | 620 | 10 | 3.23% |
| hsa_circ_0046654 | 620 | 10 | 3.23% |
| hsa_circ_0088093 | 620 | 10 | 3.23% |
| hsa_circ_0092641 | 620 | 10 | 3.23% |
| hsa_circ_0099404 | 620 | 10 | 3.23% |
| hsa_circ_0105989 | 620 | 10 | 3.23% |

| | | | |
|---|---|---|---|
| hsa_circ_0123499 | 620 | 10 | 3.23% |
| hsa_circ_0006852 | 683 | 11 | 3.22% |
| hsa_circ_0019099 | 683 | 11 | 3.22% |
| hsa_circ_0025205 | 683 | 11 | 3.22% |
| hsa_circ_0041628 | 683 | 11 | 3.22% |
| hsa_circ_0107803 | 683 | 11 | 3.22% |
| hsa_circ_0115408 | 683 | 11 | 3.22% |
| hsa_circ_0032590 | 621 | 10 | 3.22% |
| hsa_circ_0042712 | 621 | 10 | 3.22% |
| hsa_circ_0075501 | 621 | 10 | 3.22% |
| hsa_circ_0075599 | 621 | 10 | 3.22% |
| hsa_circ_0079353 | 621 | 10 | 3.22% |
| hsa_circ_0088334 | 621 | 10 | 3.22% |
| hsa_circ_0094232 | 621 | 10 | 3.22% |
| hsa_circ_0111727 | 621 | 10 | 3.22% |
| hsa_circ_0112744 | 621 | 10 | 3.22% |
| hsa_circ_0114267 | 621 | 10 | 3.22% |
| hsa_circ_0030501 | 932 | 15 | 3.22% |
| hsa_circ_0078756 | 870 | 14 | 3.22% |
| hsa_circ_0050264 | 808 | 13 | 3.22% |
| hsa_circ_0033146 | 746 | 12 | 3.22% |
| hsa_circ_0048101 | 684 | 11 | 3.22% |
| hsa_circ_0052681 | 684 | 11 | 3.22% |
| hsa_circ_0065410 | 684 | 11 | 3.22% |
| hsa_circ_0105292 | 684 | 11 | 3.22% |
| hsa_circ_0106635 | 684 | 11 | 3.22% |
| hsa_circ_0000127 | 622 | 10 | 3.22% |
| hsa_circ_0025969 | 622 | 10 | 3.22% |
| hsa_circ_0047601 | 622 | 10 | 3.22% |
| hsa_circ_0059965 | 622 | 10 | 3.22% |
| hsa_circ_0065734 | 622 | 10 | 3.22% |
| hsa_circ_0068748 | 622 | 10 | 3.22% |
| hsa_circ_0071419 | 622 | 10 | 3.22% |
| hsa_circ_0087853 | 622 | 10 | 3.22% |
| hsa_circ_0089490 | 622 | 10 | 3.22% |
| hsa_circ_0089521 | 622 | 10 | 3.22% |
| hsa_circ_0092681 | 622 | 10 | 3.22% |
| hsa_circ_0140778 | 622 | 10 | 3.22% |
| hsa_circ_0068787 | 1742 | 28 | 3.21% |
| hsa_circ_0123437 | 809 | 13 | 3.21% |
| hsa_circ_0128069 | 809 | 13 | 3.21% |
| hsa_circ_0086120 | 747 | 12 | 3.21% |
| hsa_circ_0121319 | 747 | 12 | 3.21% |
| hsa_circ_0001186 | 685 | 11 | 3.21% |
| hsa_circ_0022539 | 685 | 11 | 3.21% |
| hsa_circ_0123993 | 685 | 11 | 3.21% |
| hsa_circ_0000523 | 623 | 10 | 3.21% |
| hsa_circ_0007026 | 623 | 10 | 3.21% |
| hsa_circ_0007876 | 623 | 10 | 3.21% |
| hsa_circ_0008800 | 623 | 10 | 3.21% |

| | | | |
|---|---|---|---|
| hsa_circ_0010590 | 623 | 10 | 3.21% |
| hsa_circ_0018074 | 623 | 10 | 3.21% |
| hsa_circ_0018146 | 623 | 10 | 3.21% |
| hsa_circ_0024628 | 623 | 10 | 3.21% |
| hsa_circ_0026121 | 623 | 10 | 3.21% |
| hsa_circ_0036439 | 623 | 10 | 3.21% |
| hsa_circ_0041597 | 623 | 10 | 3.21% |
| hsa_circ_0047652 | 623 | 10 | 3.21% |
| hsa_circ_0049618 | 623 | 10 | 3.21% |
| hsa_circ_0059979 | 623 | 10 | 3.21% |
| hsa_circ_0069456 | 623 | 10 | 3.21% |
| hsa_circ_0085795 | 623 | 10 | 3.21% |
| hsa_circ_0096720 | 623 | 10 | 3.21% |
| hsa_circ_0108752 | 623 | 10 | 3.21% |
| hsa_circ_0110337 | 623 | 10 | 3.21% |
| hsa_circ_0116491 | 623 | 10 | 3.21% |
| hsa_circ_0125017 | 623 | 10 | 3.21% |
| hsa_circ_0126167 | 623 | 10 | 3.21% |
| hsa_circ_0029828 | 810 | 13 | 3.21% |
| hsa_circ_0031838 | 1371 | 22 | 3.21% |
| hsa_circ_0016050 | 748 | 12 | 3.21% |
| hsa_circ_0074673 | 748 | 12 | 3.21% |
| hsa_circ_0057246 | 686 | 11 | 3.21% |
| hsa_circ_0096491 | 686 | 11 | 3.21% |
| hsa_circ_0114298 | 686 | 11 | 3.21% |
| hsa_circ_0007555 | 624 | 10 | 3.21% |
| hsa_circ_0009246 | 624 | 10 | 3.21% |
| hsa_circ_0019634 | 624 | 10 | 3.21% |
| hsa_circ_0031146 | 624 | 10 | 3.21% |
| hsa_circ_0050396 | 624 | 10 | 3.21% |
| hsa_circ_0069343 | 624 | 10 | 3.21% |
| hsa_circ_0091544 | 624 | 10 | 3.21% |
| hsa_circ_0110441 | 624 | 10 | 3.21% |
| hsa_circ_0111670 | 624 | 10 | 3.21% |
| hsa_circ_0126230 | 624 | 10 | 3.21% |
| hsa_circ_0110074 | 1373 | 22 | 3.20% |
| hsa_circ_0038623 | 687 | 11 | 3.20% |
| hsa_circ_0068377 | 687 | 11 | 3.20% |
| hsa_circ_0073272 | 687 | 11 | 3.20% |
| hsa_circ_0077186 | 875 | 14 | 3.20% |
| hsa_circ_0002884 | 625 | 10 | 3.20% |
| hsa_circ_0003221 | 625 | 10 | 3.20% |
| hsa_circ_0039671 | 625 | 10 | 3.20% |
| hsa_circ_0064734 | 625 | 10 | 3.20% |
| hsa_circ_0078403 | 625 | 10 | 3.20% |
| hsa_circ_0079176 | 625 | 10 | 3.20% |
| hsa_circ_0091085 | 625 | 10 | 3.20% |
| hsa_circ_0122927 | 625 | 10 | 3.20% |
| hsa_circ_0013923 | 688 | 11 | 3.20% |
| hsa_circ_0070534 | 688 | 11 | 3.20% |

| | | | |
|---|---|---|---|
| hsa_circ_0081047 | 688 | 11 | 3.20% |
| hsa_circ_0133561 | 688 | 11 | 3.20% |
| hsa_circ_0135236 | 688 | 11 | 3.20% |
| hsa_circ_0004388 | 626 | 10 | 3.19% |
| hsa_circ_0007768 | 626 | 10 | 3.19% |
| hsa_circ_0012797 | 626 | 10 | 3.19% |
| hsa_circ_0025303 | 626 | 10 | 3.19% |
| hsa_circ_0037791 | 626 | 10 | 3.19% |
| hsa_circ_0062800 | 626 | 10 | 3.19% |
| hsa_circ_0064434 | 626 | 10 | 3.19% |
| hsa_circ_0067619 | 626 | 10 | 3.19% |
| hsa_circ_0103126 | 626 | 10 | 3.19% |
| hsa_circ_0109638 | 626 | 10 | 3.19% |
| hsa_circ_0116575 | 626 | 10 | 3.19% |
| hsa_circ_0127919 | 626 | 10 | 3.19% |
| hsa_circ_0138413 | 626 | 10 | 3.19% |
| hsa_circ_0008208 | 689 | 11 | 3.19% |
| hsa_circ_0015958 | 689 | 11 | 3.19% |
| hsa_circ_0031876 | 689 | 11 | 3.19% |
| hsa_circ_0058933 | 689 | 11 | 3.19% |
| hsa_circ_0131758 | 689 | 11 | 3.19% |
| hsa_circ_0021624 | 752 | 12 | 3.19% |
| hsa_circ_0081802 | 752 | 12 | 3.19% |
| hsa_circ_0031849 | 627 | 10 | 3.19% |
| hsa_circ_0045865 | 627 | 10 | 3.19% |
| hsa_circ_0046186 | 627 | 10 | 3.19% |
| hsa_circ_0050550 | 627 | 10 | 3.19% |
| hsa_circ_0082143 | 627 | 10 | 3.19% |
| hsa_circ_0098688 | 627 | 10 | 3.19% |
| hsa_circ_0107207 | 627 | 10 | 3.19% |
| hsa_circ_0110065 | 627 | 10 | 3.19% |
| hsa_circ_0113467 | 627 | 10 | 3.19% |
| hsa_circ_0125527 | 627 | 10 | 3.19% |
| hsa_circ_0127897 | 627 | 10 | 3.19% |
| hsa_circ_0132831 | 627 | 10 | 3.19% |
| hsa_circ_0077592 | 878 | 14 | 3.19% |
| hsa_circ_0110627 | 2070 | 33 | 3.19% |
| hsa_circ_0037853 | 690 | 11 | 3.19% |
| hsa_circ_0074650 | 690 | 11 | 3.19% |
| hsa_circ_0104599 | 690 | 11 | 3.19% |
| hsa_circ_0044845 | 753 | 12 | 3.19% |
| hsa_circ_0108553 | 753 | 12 | 3.19% |
| hsa_circ_0002658 | 628 | 10 | 3.18% |
| hsa_circ_0012124 | 628 | 10 | 3.18% |
| hsa_circ_0062790 | 628 | 10 | 3.18% |
| hsa_circ_0092939 | 628 | 10 | 3.18% |
| hsa_circ_0097787 | 628 | 10 | 3.18% |
| hsa_circ_0123434 | 628 | 10 | 3.18% |
| hsa_circ_0133894 | 628 | 10 | 3.18% |
| hsa_circ_0139130 | 628 | 10 | 3.18% |

| | | | |
|---|---|---|---|
| hsa_circ_0038158 | 691 | 11 | 3.18% |
| hsa_circ_0069561 | 691 | 11 | 3.18% |
| hsa_circ_0106040 | 691 | 11 | 3.18% |
| hsa_circ_0007712 | 629 | 10 | 3.18% |
| hsa_circ_0010021 | 629 | 10 | 3.18% |
| hsa_circ_0011872 | 629 | 10 | 3.18% |
| hsa_circ_0012589 | 629 | 10 | 3.18% |
| hsa_circ_0026636 | 629 | 10 | 3.18% |
| hsa_circ_0034086 | 629 | 10 | 3.18% |
| hsa_circ_0038075 | 629 | 10 | 3.18% |
| hsa_circ_0060654 | 629 | 10 | 3.18% |
| hsa_circ_0094037 | 629 | 10 | 3.18% |
| hsa_circ_0137708 | 629 | 10 | 3.18% |
| hsa_circ_0137849 | 629 | 10 | 3.18% |
| hsa_circ_0002086 | 692 | 11 | 3.18% |
| hsa_circ_0016939 | 692 | 11 | 3.18% |
| hsa_circ_0055232 | 692 | 11 | 3.18% |
| hsa_circ_0088280 | 692 | 11 | 3.18% |
| hsa_circ_0047754 | 755 | 12 | 3.18% |
| hsa_circ_0058918 | 755 | 12 | 3.18% |
| hsa_circ_0004648 | 693 | 11 | 3.17% |
| hsa_circ_0023975 | 693 | 11 | 3.17% |
| hsa_circ_0037586 | 693 | 11 | 3.17% |
| hsa_circ_0042938 | 693 | 11 | 3.17% |
| hsa_circ_0077578 | 693 | 11 | 3.17% |
| hsa_circ_0086628 | 693 | 11 | 3.17% |
| hsa_circ_0106988 | 693 | 11 | 3.17% |
| hsa_circ_0109844 | 693 | 11 | 3.17% |
| hsa_circ_0119586 | 693 | 11 | 3.17% |
| hsa_circ_0028507 | 630 | 10 | 3.17% |
| hsa_circ_0036960 | 630 | 10 | 3.17% |
| hsa_circ_0046355 | 630 | 10 | 3.17% |
| hsa_circ_0057440 | 630 | 10 | 3.17% |
| hsa_circ_0062447 | 630 | 10 | 3.17% |
| hsa_circ_0066971 | 630 | 10 | 3.17% |
| hsa_circ_0072272 | 630 | 10 | 3.17% |
| hsa_circ_0094781 | 630 | 10 | 3.17% |
| hsa_circ_0103105 | 630 | 10 | 3.17% |
| hsa_circ_0107232 | 630 | 10 | 3.17% |
| hsa_circ_0127593 | 630 | 10 | 3.17% |
| hsa_circ_0133242 | 630 | 10 | 3.17% |
| hsa_circ_0137447 | 630 | 10 | 3.17% |
| hsa_circ_0137872 | 630 | 10 | 3.17% |
| hsa_circ_0139222 | 630 | 10 | 3.17% |
| hsa_circ_0005616 | 883 | 14 | 3.17% |
| hsa_circ_0050796 | 757 | 12 | 3.17% |
| hsa_circ_0091095 | 757 | 12 | 3.17% |
| hsa_circ_0120140 | 757 | 12 | 3.17% |
| hsa_circ_0011073 | 694 | 11 | 3.17% |
| hsa_circ_0043343 | 694 | 11 | 3.17% |

| | | | |
|---|---|---|---|
| hsa_circ_0059041 | 694 | 11 | 3.17% |
| hsa_circ_0069562 | 694 | 11 | 3.17% |
| hsa_circ_0078523 | 694 | 11 | 3.17% |
| hsa_circ_0080471 | 694 | 11 | 3.17% |
| hsa_circ_0081679 | 694 | 11 | 3.17% |
| hsa_circ_0131912 | 694 | 11 | 3.17% |
| hsa_circ_0138104 | 694 | 11 | 3.17% |
| hsa_circ_0140502 | 694 | 11 | 3.17% |
| hsa_circ_0000024 | 631 | 10 | 3.17% |
| hsa_circ_0001487 | 631 | 10 | 3.17% |
| hsa_circ_0028212 | 631 | 10 | 3.17% |
| hsa_circ_0037804 | 631 | 10 | 3.17% |
| hsa_circ_0043101 | 631 | 10 | 3.17% |
| hsa_circ_0065020 | 631 | 10 | 3.17% |
| hsa_circ_0077367 | 631 | 10 | 3.17% |
| hsa_circ_0088543 | 631 | 10 | 3.17% |
| hsa_circ_0123198 | 631 | 10 | 3.17% |
| hsa_circ_0006775 | 695 | 11 | 3.17% |
| hsa_circ_0007744 | 695 | 11 | 3.17% |
| hsa_circ_0026489 | 695 | 11 | 3.17% |
| hsa_circ_0039023 | 695 | 11 | 3.17% |
| hsa_circ_0062108 | 1011 | 16 | 3.17% |
| hsa_circ_0000542 | 632 | 10 | 3.16% |
| hsa_circ_0013104 | 632 | 10 | 3.16% |
| hsa_circ_0043897 | 632 | 10 | 3.16% |
| hsa_circ_0044685 | 632 | 10 | 3.16% |
| hsa_circ_0045778 | 632 | 10 | 3.16% |
| hsa_circ_0050379 | 632 | 10 | 3.16% |
| hsa_circ_0052174 | 632 | 10 | 3.16% |
| hsa_circ_0056313 | 632 | 10 | 3.16% |
| hsa_circ_0064221 | 632 | 10 | 3.16% |
| hsa_circ_0081392 | 632 | 10 | 3.16% |
| hsa_circ_0087854 | 632 | 10 | 3.16% |
| hsa_circ_0109400 | 632 | 10 | 3.16% |
| hsa_circ_0127113 | 632 | 10 | 3.16% |
| hsa_circ_0127599 | 632 | 10 | 3.16% |
| hsa_circ_0074269 | 759 | 12 | 3.16% |
| hsa_circ_0077234 | 759 | 12 | 3.16% |
| hsa_circ_0103053 | 759 | 12 | 3.16% |
| hsa_circ_0121925 | 759 | 12 | 3.16% |
| hsa_circ_0006190 | 696 | 11 | 3.16% |
| hsa_circ_0121162 | 696 | 11 | 3.16% |
| hsa_circ_0139658 | 696 | 11 | 3.16% |
| hsa_circ_0039628 | 633 | 10 | 3.16% |
| hsa_circ_0039991 | 633 | 10 | 3.16% |
| hsa_circ_0042509 | 633 | 10 | 3.16% |
| hsa_circ_0048330 | 633 | 10 | 3.16% |
| hsa_circ_0050884 | 633 | 10 | 3.16% |
| hsa_circ_0058177 | 633 | 10 | 3.16% |
| hsa_circ_0061757 | 633 | 10 | 3.16% |

| | | | |
|---|---|---|---|
| hsa_circ_0081279 | 633 | 10 | 3.16% |
| hsa_circ_0088240 | 633 | 10 | 3.16% |
| hsa_circ_0107167 | 633 | 10 | 3.16% |
| hsa_circ_0122529 | 633 | 10 | 3.16% |
| hsa_circ_0134175 | 633 | 10 | 3.16% |
| hsa_circ_0136962 | 633 | 10 | 3.16% |
| hsa_circ_0103127 | 760 | 12 | 3.16% |
| hsa_circ_0133882 | 760 | 12 | 3.16% |
| hsa_circ_0037366 | 697 | 11 | 3.16% |
| hsa_circ_0037476 | 697 | 11 | 3.16% |
| hsa_circ_0099885 | 697 | 11 | 3.16% |
| hsa_circ_0106882 | 697 | 11 | 3.16% |
| hsa_circ_0110947 | 697 | 11 | 3.16% |
| hsa_circ_0114371 | 697 | 11 | 3.16% |
| hsa_circ_0002339 | 634 | 10 | 3.15% |
| hsa_circ_0004791 | 634 | 10 | 3.15% |
| hsa_circ_0006431 | 634 | 10 | 3.15% |
| hsa_circ_0015629 | 634 | 10 | 3.15% |
| hsa_circ_0015939 | 634 | 10 | 3.15% |
| hsa_circ_0024920 | 634 | 10 | 3.15% |
| hsa_circ_0035089 | 634 | 10 | 3.15% |
| hsa_circ_0046313 | 634 | 10 | 3.15% |
| hsa_circ_0048240 | 634 | 10 | 3.15% |
| hsa_circ_0054935 | 634 | 10 | 3.15% |
| hsa_circ_0063185 | 634 | 10 | 3.15% |
| hsa_circ_0071071 | 634 | 10 | 3.15% |
| hsa_circ_0101542 | 634 | 10 | 3.15% |
| hsa_circ_0109236 | 634 | 10 | 3.15% |
| hsa_circ_0110075 | 634 | 10 | 3.15% |
| hsa_circ_0112683 | 634 | 10 | 3.15% |
| hsa_circ_0128148 | 634 | 10 | 3.15% |
| hsa_circ_0133263 | 634 | 10 | 3.15% |
| hsa_circ_0087025 | 761 | 12 | 3.15% |
| hsa_circ_0002663 | 698 | 11 | 3.15% |
| hsa_circ_0081200 | 698 | 11 | 3.15% |
| hsa_circ_0125540 | 698 | 11 | 3.15% |
| hsa_circ_0055146 | 825 | 13 | 3.15% |
| hsa_circ_0126644 | 762 | 12 | 3.15% |
| hsa_circ_0137603 | 762 | 12 | 3.15% |
| hsa_circ_0022898 | 635 | 10 | 3.15% |
| hsa_circ_0033361 | 635 | 10 | 3.15% |
| hsa_circ_0047968 | 635 | 10 | 3.15% |
| hsa_circ_0048836 | 635 | 10 | 3.15% |
| hsa_circ_0060444 | 635 | 10 | 3.15% |
| hsa_circ_0070296 | 635 | 10 | 3.15% |
| hsa_circ_0081369 | 635 | 10 | 3.15% |
| hsa_circ_0081685 | 635 | 10 | 3.15% |
| hsa_circ_0099086 | 635 | 10 | 3.15% |
| hsa_circ_0124526 | 635 | 10 | 3.15% |
| hsa_circ_0076666 | 826 | 13 | 3.15% |

| | | | |
|---|---|---|---|
| hsa_circ_0023952 | 699 | 11 | 3.15% |
| hsa_circ_0040306 | 699 | 11 | 3.15% |
| hsa_circ_0117885 | 699 | 11 | 3.15% |
| hsa_circ_0135109 | 699 | 11 | 3.15% |
| hsa_circ_0139616 | 890 | 14 | 3.15% |
| hsa_circ_0042685 | 763 | 12 | 3.15% |
| hsa_circ_0117755 | 763 | 12 | 3.15% |
| hsa_circ_0006281 | 636 | 10 | 3.14% |
| hsa_circ_0013032 | 636 | 10 | 3.14% |
| hsa_circ_0042558 | 636 | 10 | 3.14% |
| hsa_circ_0048714 | 636 | 10 | 3.14% |
| hsa_circ_0050496 | 636 | 10 | 3.14% |
| hsa_circ_0058945 | 636 | 10 | 3.14% |
| hsa_circ_0093228 | 636 | 10 | 3.14% |
| hsa_circ_0096678 | 636 | 10 | 3.14% |
| hsa_circ_0097884 | 636 | 10 | 3.14% |
| hsa_circ_0098955 | 636 | 10 | 3.14% |
| hsa_circ_0101692 | 636 | 10 | 3.14% |
| hsa_circ_0118001 | 636 | 10 | 3.14% |
| hsa_circ_0127416 | 636 | 10 | 3.14% |
| hsa_circ_0128362 | 636 | 10 | 3.14% |
| hsa_circ_0130474 | 636 | 10 | 3.14% |
| hsa_circ_0136936 | 636 | 10 | 3.14% |
| hsa_circ_0138311 | 636 | 10 | 3.14% |
| hsa_circ_0029173 | 827 | 13 | 3.14% |
| hsa_circ_0119445 | 827 | 13 | 3.14% |
| hsa_circ_0021014 | 700 | 11 | 3.14% |
| hsa_circ_0034623 | 700 | 11 | 3.14% |
| hsa_circ_0069774 | 700 | 11 | 3.14% |
| hsa_circ_0092474 | 700 | 11 | 3.14% |
| hsa_circ_0125215 | 700 | 11 | 3.14% |
| hsa_circ_0013585 | 891 | 14 | 3.14% |
| hsa_circ_0014594 | 828 | 13 | 3.14% |
| hsa_circ_0042114 | 828 | 13 | 3.14% |
| hsa_circ_0043865 | 828 | 13 | 3.14% |
| hsa_circ_0005601 | 637 | 10 | 3.14% |
| hsa_circ_0017732 | 637 | 10 | 3.14% |
| hsa_circ_0025050 | 637 | 10 | 3.14% |
| hsa_circ_0032669 | 637 | 10 | 3.14% |
| hsa_circ_0045490 | 637 | 10 | 3.14% |
| hsa_circ_0046777 | 637 | 10 | 3.14% |
| hsa_circ_0047892 | 637 | 10 | 3.14% |
| hsa_circ_0047999 | 637 | 10 | 3.14% |
| hsa_circ_0055042 | 637 | 10 | 3.14% |
| hsa_circ_0090084 | 637 | 10 | 3.14% |
| hsa_circ_0100459 | 637 | 10 | 3.14% |
| hsa_circ_0133549 | 637 | 10 | 3.14% |
| hsa_circ_0024777 | 701 | 11 | 3.14% |
| hsa_circ_0042953 | 701 | 11 | 3.14% |
| hsa_circ_0058972 | 701 | 11 | 3.14% |

| | | | |
|---|---|---|---|
| hsa_circ_0118510 | 701 | 11 | 3.14% |
| hsa_circ_0051970 | 765 | 12 | 3.14% |
| hsa_circ_0000085 | 638 | 10 | 3.13% |
| hsa_circ_0000115 | 638 | 10 | 3.13% |
| hsa_circ_0005269 | 638 | 10 | 3.13% |
| hsa_circ_0007393 | 638 | 10 | 3.13% |
| hsa_circ_0021547 | 638 | 10 | 3.13% |
| hsa_circ_0037734 | 638 | 10 | 3.13% |
| hsa_circ_0054776 | 638 | 10 | 3.13% |
| hsa_circ_0064344 | 638 | 10 | 3.13% |
| hsa_circ_0064737 | 638 | 10 | 3.13% |
| hsa_circ_0066522 | 638 | 10 | 3.13% |
| hsa_circ_0083016 | 638 | 10 | 3.13% |
| hsa_circ_0114909 | 638 | 10 | 3.13% |
| hsa_circ_0058377 | 702 | 11 | 3.13% |
| hsa_circ_0068871 | 702 | 11 | 3.13% |
| hsa_circ_0090924 | 702 | 11 | 3.13% |
| hsa_circ_0103282 | 702 | 11 | 3.13% |
| hsa_circ_0118665 | 702 | 11 | 3.13% |
| hsa_circ_0129654 | 702 | 11 | 3.13% |
| hsa_circ_0043668 | 766 | 12 | 3.13% |
| hsa_circ_0135308 | 766 | 12 | 3.13% |
| hsa_circ_0032652 | 1086 | 17 | 3.13% |
| hsa_circ_0003249 | 639 | 10 | 3.13% |
| hsa_circ_0004119 | 639 | 10 | 3.13% |
| hsa_circ_0004728 | 639 | 10 | 3.13% |
| hsa_circ_0007295 | 639 | 10 | 3.13% |
| hsa_circ_0011945 | 639 | 10 | 3.13% |
| hsa_circ_0020577 | 639 | 10 | 3.13% |
| hsa_circ_0032339 | 639 | 10 | 3.13% |
| hsa_circ_0048690 | 639 | 10 | 3.13% |
| hsa_circ_0058906 | 639 | 10 | 3.13% |
| hsa_circ_0063459 | 639 | 10 | 3.13% |
| hsa_circ_0065724 | 639 | 10 | 3.13% |
| hsa_circ_0087572 | 639 | 10 | 3.13% |
| hsa_circ_0103202 | 639 | 10 | 3.13% |
| hsa_circ_0103762 | 639 | 10 | 3.13% |
| hsa_circ_0106425 | 639 | 10 | 3.13% |
| hsa_circ_0108311 | 639 | 10 | 3.13% |
| hsa_circ_0110305 | 639 | 10 | 3.13% |
| hsa_circ_0119620 | 639 | 10 | 3.13% |
| hsa_circ_0132662 | 639 | 10 | 3.13% |
| hsa_circ_0133128 | 639 | 10 | 3.13% |
| hsa_circ_0135497 | 639 | 10 | 3.13% |
| hsa_circ_0064312 | 703 | 11 | 3.13% |
| hsa_circ_0080161 | 703 | 11 | 3.13% |
| hsa_circ_0107472 | 960 | 15 | 3.13% |
| hsa_circ_0061475 | 768 | 12 | 3.13% |
| hsa_circ_0071251 | 768 | 12 | 3.13% |
| hsa_circ_0036372 | 704 | 11 | 3.13% |

| | | | |
|---|---|---|---|
| hsa_circ_0037167 | 704 | 11 | 3.13% |
| hsa_circ_0046500 | 704 | 11 | 3.13% |
| hsa_circ_0048582 | 704 | 11 | 3.13% |
| hsa_circ_0059282 | 704 | 11 | 3.13% |
| hsa_circ_0074997 | 704 | 11 | 3.13% |
| hsa_circ_0092016 | 704 | 11 | 3.13% |
| hsa_circ_0122839 | 704 | 11 | 3.13% |
| hsa_circ_0010096 | 640 | 10 | 3.13% |
| hsa_circ_0012723 | 640 | 10 | 3.13% |
| hsa_circ_0032592 | 640 | 10 | 3.13% |
| hsa_circ_0043164 | 640 | 10 | 3.13% |
| hsa_circ_0049908 | 640 | 10 | 3.13% |
| hsa_circ_0066797 | 640 | 10 | 3.13% |
| hsa_circ_0072899 | 640 | 10 | 3.13% |
| hsa_circ_0112078 | 640 | 10 | 3.13% |
| hsa_circ_0118732 | 640 | 10 | 3.13% |
| hsa_circ_0123857 | 640 | 10 | 3.13% |
| hsa_circ_0132372 | 640 | 10 | 3.13% |
| hsa_circ_0078852 | 3330 | 52 | 3.12% |
| hsa_circ_0069917 | 897 | 14 | 3.12% |
| hsa_circ_0119564 | 769 | 12 | 3.12% |
| hsa_circ_0008401 | 705 | 11 | 3.12% |
| hsa_circ_0009285 | 705 | 11 | 3.12% |
| hsa_circ_0012172 | 705 | 11 | 3.12% |
| hsa_circ_0059846 | 705 | 11 | 3.12% |
| hsa_circ_0063740 | 705 | 11 | 3.12% |
| hsa_circ_0086551 | 705 | 11 | 3.12% |
| hsa_circ_0094849 | 705 | 11 | 3.12% |
| hsa_circ_0126627 | 705 | 11 | 3.12% |
| hsa_circ_0006972 | 641 | 10 | 3.12% |
| hsa_circ_0007027 | 641 | 10 | 3.12% |
| hsa_circ_0011864 | 641 | 10 | 3.12% |
| hsa_circ_0016713 | 641 | 10 | 3.12% |
| hsa_circ_0051294 | 641 | 10 | 3.12% |
| hsa_circ_0061323 | 641 | 10 | 3.12% |
| hsa_circ_0066626 | 641 | 10 | 3.12% |
| hsa_circ_0071942 | 641 | 10 | 3.12% |
| hsa_circ_0075479 | 641 | 10 | 3.12% |
| hsa_circ_0077083 | 641 | 10 | 3.12% |
| hsa_circ_0090910 | 641 | 10 | 3.12% |
| hsa_circ_0091375 | 641 | 10 | 3.12% |
| hsa_circ_0094112 | 641 | 10 | 3.12% |
| hsa_circ_0107969 | 641 | 10 | 3.12% |
| hsa_circ_0113420 | 641 | 10 | 3.12% |
| hsa_circ_0120166 | 641 | 10 | 3.12% |
| hsa_circ_0077246 | 770 | 12 | 3.12% |
| hsa_circ_0075951 | 706 | 11 | 3.12% |
| hsa_circ_0118809 | 706 | 11 | 3.12% |
| hsa_circ_0000149 | 642 | 10 | 3.12% |
| hsa_circ_0008974 | 642 | 10 | 3.12% |

| | | | |
|---|---|---|---|
| hsa_circ_0014964 | 642 | 10 | 3.12% |
| hsa_circ_0023371 | 642 | 10 | 3.12% |
| hsa_circ_0032548 | 642 | 10 | 3.12% |
| hsa_circ_0048051 | 642 | 10 | 3.12% |
| hsa_circ_0055397 | 642 | 10 | 3.12% |
| hsa_circ_0060906 | 642 | 10 | 3.12% |
| hsa_circ_0064343 | 642 | 10 | 3.12% |
| hsa_circ_0065862 | 642 | 10 | 3.12% |
| hsa_circ_0066788 | 642 | 10 | 3.12% |
| hsa_circ_0072046 | 642 | 10 | 3.12% |
| hsa_circ_0076800 | 642 | 10 | 3.12% |
| hsa_circ_0084977 | 642 | 10 | 3.12% |
| hsa_circ_0114184 | 642 | 10 | 3.12% |
| hsa_circ_0133369 | 642 | 10 | 3.12% |
| hsa_circ_0134833 | 642 | 10 | 3.12% |
| hsa_circ_0116381 | 4755 | 74 | 3.11% |
| hsa_circ_0004698 | 707 | 11 | 3.11% |
| hsa_circ_0034573 | 707 | 11 | 3.11% |
| hsa_circ_0039861 | 707 | 11 | 3.11% |
| hsa_circ_0060315 | 707 | 11 | 3.11% |
| hsa_circ_0092907 | 707 | 11 | 3.11% |
| hsa_circ_0093974 | 707 | 11 | 3.11% |
| hsa_circ_0111278 | 707 | 11 | 3.11% |
| hsa_circ_0111701 | 707 | 11 | 3.11% |
| hsa_circ_0001995 | 643 | 10 | 3.11% |
| hsa_circ_0011114 | 643 | 10 | 3.11% |
| hsa_circ_0018373 | 643 | 10 | 3.11% |
| hsa_circ_0022668 | 643 | 10 | 3.11% |
| hsa_circ_0043909 | 643 | 10 | 3.11% |
| hsa_circ_0059118 | 643 | 10 | 3.11% |
| hsa_circ_0094519 | 643 | 10 | 3.11% |
| hsa_circ_0097124 | 643 | 10 | 3.11% |
| hsa_circ_0098763 | 643 | 10 | 3.11% |
| hsa_circ_0100277 | 643 | 10 | 3.11% |
| hsa_circ_0104111 | 643 | 10 | 3.11% |
| hsa_circ_0118762 | 643 | 10 | 3.11% |
| hsa_circ_0126696 | 643 | 10 | 3.11% |
| hsa_circ_0133325 | 643 | 10 | 3.11% |
| hsa_circ_0136454 | 643 | 10 | 3.11% |
| hsa_circ_0009117 | 708 | 11 | 3.11% |
| hsa_circ_0038796 | 708 | 11 | 3.11% |
| hsa_circ_0058338 | 708 | 11 | 3.11% |
| hsa_circ_0060936 | 708 | 11 | 3.11% |
| hsa_circ_0072466 | 708 | 11 | 3.11% |
| hsa_circ_0113488 | 708 | 11 | 3.11% |
| hsa_circ_0057394 | 837 | 13 | 3.11% |
| hsa_circ_0080359 | 837 | 13 | 3.11% |
| hsa_circ_0005943 | 644 | 10 | 3.11% |
| hsa_circ_0008794 | 644 | 10 | 3.11% |
| hsa_circ_0022239 | 644 | 10 | 3.11% |

| | | | |
|---|---|---|---|
| hsa_circ_0044112 | 644 | 10 | 3.11% |
| hsa_circ_0048200 | 644 | 10 | 3.11% |
| hsa_circ_0056123 | 644 | 10 | 3.11% |
| hsa_circ_0066757 | 644 | 10 | 3.11% |
| hsa_circ_0124897 | 644 | 10 | 3.11% |
| hsa_circ_0065350 | 773 | 12 | 3.10% |
| hsa_circ_0088559 | 773 | 12 | 3.10% |
| hsa_circ_0022000 | 902 | 14 | 3.10% |
| hsa_circ_0009930 | 709 | 11 | 3.10% |
| hsa_circ_0033369 | 709 | 11 | 3.10% |
| hsa_circ_0037699 | 709 | 11 | 3.10% |
| hsa_circ_0043200 | 709 | 11 | 3.10% |
| hsa_circ_0072870 | 709 | 11 | 3.10% |
| hsa_circ_0137653 | 709 | 11 | 3.10% |
| hsa_circ_0032799 | 774 | 12 | 3.10% |
| hsa_circ_0041004 | 774 | 12 | 3.10% |
| hsa_circ_0005320 | 645 | 10 | 3.10% |
| hsa_circ_0006438 | 645 | 10 | 3.10% |
| hsa_circ_0028523 | 645 | 10 | 3.10% |
| hsa_circ_0041486 | 645 | 10 | 3.10% |
| hsa_circ_0044823 | 645 | 10 | 3.10% |
| hsa_circ_0047588 | 645 | 10 | 3.10% |
| hsa_circ_0051172 | 645 | 10 | 3.10% |
| hsa_circ_0052249 | 645 | 10 | 3.10% |
| hsa_circ_0058710 | 645 | 10 | 3.10% |
| hsa_circ_0069729 | 645 | 10 | 3.10% |
| hsa_circ_0112328 | 645 | 10 | 3.10% |
| hsa_circ_0112596 | 645 | 10 | 3.10% |
| hsa_circ_0117643 | 645 | 10 | 3.10% |
| hsa_circ_0122531 | 645 | 10 | 3.10% |
| hsa_circ_0126477 | 645 | 10 | 3.10% |
| hsa_circ_0132125 | 645 | 10 | 3.10% |
| hsa_circ_0134951 | 645 | 10 | 3.10% |
| hsa_circ_0016945 | 710 | 11 | 3.10% |
| hsa_circ_0017983 | 710 | 11 | 3.10% |
| hsa_circ_0048337 | 710 | 11 | 3.10% |
| hsa_circ_0054629 | 710 | 11 | 3.10% |
| hsa_circ_0076795 | 710 | 11 | 3.10% |
| hsa_circ_0082851 | 710 | 11 | 3.10% |
| hsa_circ_0097832 | 710 | 11 | 3.10% |
| hsa_circ_0046230 | 904 | 14 | 3.10% |
| hsa_circ_0019140 | 775 | 12 | 3.10% |
| hsa_circ_0002005 | 646 | 10 | 3.10% |
| hsa_circ_0015125 | 646 | 10 | 3.10% |
| hsa_circ_0016721 | 646 | 10 | 3.10% |
| hsa_circ_0044005 | 646 | 10 | 3.10% |
| hsa_circ_0045491 | 646 | 10 | 3.10% |
| hsa_circ_0049251 | 646 | 10 | 3.10% |
| hsa_circ_0073081 | 646 | 10 | 3.10% |
| hsa_circ_0093073 | 646 | 10 | 3.10% |

| | | | |
|---|---|---|---|
| hsa_circ_0102361 | 646 | 10 | 3.10% |
| hsa_circ_0103068 | 646 | 10 | 3.10% |
| hsa_circ_0110793 | 646 | 10 | 3.10% |
| hsa_circ_0117952 | 646 | 10 | 3.10% |
| hsa_circ_0118650 | 646 | 10 | 3.10% |
| hsa_circ_0133824 | 646 | 10 | 3.10% |
| hsa_circ_0004099 | 840 | 13 | 3.10% |
| hsa_circ_0041731 | 840 | 13 | 3.10% |
| hsa_circ_0010425 | 711 | 11 | 3.09% |
| hsa_circ_0011075 | 711 | 11 | 3.09% |
| hsa_circ_0036692 | 711 | 11 | 3.09% |
| hsa_circ_0091570 | 711 | 11 | 3.09% |
| hsa_circ_0100377 | 711 | 11 | 3.09% |
| hsa_circ_0005278 | 776 | 12 | 3.09% |
| hsa_circ_0028504 | 776 | 12 | 3.09% |
| hsa_circ_0023609 | 841 | 13 | 3.09% |
| hsa_circ_0058386 | 841 | 13 | 3.09% |
| hsa_circ_0019324 | 647 | 10 | 3.09% |
| hsa_circ_0020835 | 647 | 10 | 3.09% |
| hsa_circ_0042099 | 647 | 10 | 3.09% |
| hsa_circ_0081404 | 647 | 10 | 3.09% |
| hsa_circ_0108499 | 647 | 10 | 3.09% |
| hsa_circ_0117232 | 647 | 10 | 3.09% |
| hsa_circ_0123884 | 647 | 10 | 3.09% |
| hsa_circ_0009683 | 712 | 11 | 3.09% |
| hsa_circ_0010758 | 712 | 11 | 3.09% |
| hsa_circ_0081182 | 712 | 11 | 3.09% |
| hsa_circ_0083874 | 712 | 11 | 3.09% |
| hsa_circ_0107983 | 712 | 11 | 3.09% |
| hsa_circ_0114758 | 712 | 11 | 3.09% |
| hsa_circ_0072332 | 777 | 12 | 3.09% |
| hsa_circ_0101961 | 842 | 13 | 3.09% |
| hsa_circ_0003128 | 648 | 10 | 3.09% |
| hsa_circ_0019826 | 648 | 10 | 3.09% |
| hsa_circ_0023598 | 648 | 10 | 3.09% |
| hsa_circ_0043610 | 648 | 10 | 3.09% |
| hsa_circ_0051143 | 648 | 10 | 3.09% |
| hsa_circ_0055994 | 648 | 10 | 3.09% |
| hsa_circ_0057413 | 648 | 10 | 3.09% |
| hsa_circ_0059320 | 648 | 10 | 3.09% |
| hsa_circ_0071943 | 648 | 10 | 3.09% |
| hsa_circ_0075129 | 648 | 10 | 3.09% |
| hsa_circ_0078187 | 648 | 10 | 3.09% |
| hsa_circ_0087576 | 648 | 10 | 3.09% |
| hsa_circ_0094328 | 648 | 10 | 3.09% |
| hsa_circ_0106785 | 648 | 10 | 3.09% |
| hsa_circ_0037592 | 713 | 11 | 3.09% |
| hsa_circ_0047164 | 713 | 11 | 3.09% |
| hsa_circ_0048660 | 713 | 11 | 3.09% |
| hsa_circ_0088734 | 713 | 11 | 3.09% |

| | | | |
|---|---|---|---|
| hsa_circ_0023879 | 778 | 12 | 3.08% |
| hsa_circ_0122145 | 778 | 12 | 3.08% |
| hsa_circ_0127545 | 2919 | 45 | 3.08% |
| hsa_circ_0004937 | 649 | 10 | 3.08% |
| hsa_circ_0018425 | 649 | 10 | 3.08% |
| hsa_circ_0046131 | 649 | 10 | 3.08% |
| hsa_circ_0056359 | 649 | 10 | 3.08% |
| hsa_circ_0065109 | 649 | 10 | 3.08% |
| hsa_circ_0068780 | 649 | 10 | 3.08% |
| hsa_circ_0071509 | 649 | 10 | 3.08% |
| hsa_circ_0075855 | 649 | 10 | 3.08% |
| hsa_circ_0076485 | 649 | 10 | 3.08% |
| hsa_circ_0081461 | 649 | 10 | 3.08% |
| hsa_circ_0082705 | 649 | 10 | 3.08% |
| hsa_circ_0094057 | 649 | 10 | 3.08% |
| hsa_circ_0095070 | 649 | 10 | 3.08% |
| hsa_circ_0112317 | 649 | 10 | 3.08% |
| hsa_circ_0117970 | 649 | 10 | 3.08% |
| hsa_circ_0131634 | 649 | 10 | 3.08% |
| hsa_circ_0089740 | 714 | 11 | 3.08% |
| hsa_circ_0088889 | 779 | 12 | 3.08% |
| hsa_circ_0134703 | 779 | 12 | 3.08% |
| hsa_circ_0050344 | 1948 | 30 | 3.08% |
| hsa_circ_0128182 | 974 | 15 | 3.08% |
| hsa_circ_0067534 | 780 | 12 | 3.08% |
| hsa_circ_0075725 | 780 | 12 | 3.08% |
| hsa_circ_0017823 | 715 | 11 | 3.08% |
| hsa_circ_0038466 | 715 | 11 | 3.08% |
| hsa_circ_0043674 | 715 | 11 | 3.08% |
| hsa_circ_0023454 | 650 | 10 | 3.08% |
| hsa_circ_0062168 | 650 | 10 | 3.08% |
| hsa_circ_0066407 | 650 | 10 | 3.08% |
| hsa_circ_0069808 | 650 | 10 | 3.08% |
| hsa_circ_0079558 | 650 | 10 | 3.08% |
| hsa_circ_0082491 | 650 | 10 | 3.08% |
| hsa_circ_0115406 | 650 | 10 | 3.08% |
| hsa_circ_0116453 | 650 | 10 | 3.08% |
| hsa_circ_0120365 | 650 | 10 | 3.08% |
| hsa_circ_0121615 | 650 | 10 | 3.08% |
| hsa_circ_0134675 | 650 | 10 | 3.08% |
| hsa_circ_0081441 | 781 | 12 | 3.07% |
| hsa_circ_0029157 | 716 | 11 | 3.07% |
| hsa_circ_0044659 | 716 | 11 | 3.07% |
| hsa_circ_0008546 | 651 | 10 | 3.07% |
| hsa_circ_0010150 | 651 | 10 | 3.07% |
| hsa_circ_0013789 | 651 | 10 | 3.07% |
| hsa_circ_0017490 | 651 | 10 | 3.07% |
| hsa_circ_0027131 | 651 | 10 | 3.07% |
| hsa_circ_0028489 | 651 | 10 | 3.07% |
| hsa_circ_0028917 | 651 | 10 | 3.07% |

| | | | |
|---|---|---|---|
| hsa_circ_0046367 | 651 | 10 | 3.07% |
| hsa_circ_0048671 | 651 | 10 | 3.07% |
| hsa_circ_0051607 | 651 | 10 | 3.07% |
| hsa_circ_0058079 | 651 | 10 | 3.07% |
| hsa_circ_0070282 | 651 | 10 | 3.07% |
| hsa_circ_0071541 | 651 | 10 | 3.07% |
| hsa_circ_0078950 | 651 | 10 | 3.07% |
| hsa_circ_0087005 | 651 | 10 | 3.07% |
| hsa_circ_0087061 | 651 | 10 | 3.07% |
| hsa_circ_0087065 | 651 | 10 | 3.07% |
| hsa_circ_0090912 | 651 | 10 | 3.07% |
| hsa_circ_0091792 | 651 | 10 | 3.07% |
| hsa_circ_0108797 | 651 | 10 | 3.07% |
| hsa_circ_0135551 | 651 | 10 | 3.07% |
| hsa_circ_0135800 | 651 | 10 | 3.07% |
| hsa_circ_0033410 | 977 | 15 | 3.07% |
| hsa_circ_0050456 | 977 | 15 | 3.07% |
| hsa_circ_0131795 | 1368 | 21 | 3.07% |
| hsa_circ_0037968 | 717 | 11 | 3.07% |
| hsa_circ_0058027 | 717 | 11 | 3.07% |
| hsa_circ_0065966 | 717 | 11 | 3.07% |
| hsa_circ_0081043 | 717 | 11 | 3.07% |
| hsa_circ_0093759 | 717 | 11 | 3.07% |
| hsa_circ_0095965 | 717 | 11 | 3.07% |
| hsa_circ_0028572 | 652 | 10 | 3.07% |
| hsa_circ_0031525 | 652 | 10 | 3.07% |
| hsa_circ_0043310 | 652 | 10 | 3.07% |
| hsa_circ_0070921 | 652 | 10 | 3.07% |
| hsa_circ_0079522 | 652 | 10 | 3.07% |
| hsa_circ_0098455 | 652 | 10 | 3.07% |
| hsa_circ_0110588 | 652 | 10 | 3.07% |
| hsa_circ_0130438 | 652 | 10 | 3.07% |
| hsa_circ_0134927 | 652 | 10 | 3.07% |
| hsa_circ_0073496 | 848 | 13 | 3.07% |
| hsa_circ_0043972 | 718 | 11 | 3.06% |
| hsa_circ_0064624 | 718 | 11 | 3.06% |
| hsa_circ_0067459 | 718 | 11 | 3.06% |
| hsa_circ_0073534 | 718 | 11 | 3.06% |
| hsa_circ_0087193 | 718 | 11 | 3.06% |
| hsa_circ_0095608 | 718 | 11 | 3.06% |
| hsa_circ_0127368 | 718 | 11 | 3.06% |
| hsa_circ_0140503 | 718 | 11 | 3.06% |
| hsa_circ_0003120 | 653 | 10 | 3.06% |
| hsa_circ_0038206 | 653 | 10 | 3.06% |
| hsa_circ_0042849 | 653 | 10 | 3.06% |
| hsa_circ_0047916 | 653 | 10 | 3.06% |
| hsa_circ_0062211 | 653 | 10 | 3.06% |
| hsa_circ_0062625 | 653 | 10 | 3.06% |
| hsa_circ_0070454 | 653 | 10 | 3.06% |
| hsa_circ_0125558 | 653 | 10 | 3.06% |

| | | | |
|---|---|---|---|
| hsa_circ_0137471 | 653 | 10 | 3.06% |
| hsa_circ_0138442 | 849 | 13 | 3.06% |
| hsa_circ_0041216 | 784 | 12 | 3.06% |
| hsa_circ_0061062 | 784 | 12 | 3.06% |
| hsa_circ_0057347 | 719 | 11 | 3.06% |
| hsa_circ_0076012 | 719 | 11 | 3.06% |
| hsa_circ_0119584 | 719 | 11 | 3.06% |
| hsa_circ_0137060 | 719 | 11 | 3.06% |
| hsa_circ_0110289 | 1177 | 18 | 3.06% |
| hsa_circ_0005251 | 654 | 10 | 3.06% |
| hsa_circ_0006552 | 654 | 10 | 3.06% |
| hsa_circ_0022837 | 654 | 10 | 3.06% |
| hsa_circ_0043421 | 654 | 10 | 3.06% |
| hsa_circ_0049405 | 654 | 10 | 3.06% |
| hsa_circ_0071456 | 654 | 10 | 3.06% |
| hsa_circ_0076470 | 654 | 10 | 3.06% |
| hsa_circ_0085972 | 654 | 10 | 3.06% |
| hsa_circ_0096848 | 654 | 10 | 3.06% |
| hsa_circ_0100297 | 654 | 10 | 3.06% |
| hsa_circ_0103475 | 654 | 10 | 3.06% |
| hsa_circ_0119766 | 654 | 10 | 3.06% |
| hsa_circ_0125771 | 654 | 10 | 3.06% |
| hsa_circ_0127419 | 654 | 10 | 3.06% |
| hsa_circ_0134098 | 654 | 10 | 3.06% |
| hsa_circ_0137613 | 654 | 10 | 3.06% |
| hsa_circ_0012525 | 785 | 12 | 3.06% |
| hsa_circ_0073922 | 785 | 12 | 3.06% |
| hsa_circ_0013173 | 720 | 11 | 3.06% |
| hsa_circ_0044260 | 720 | 11 | 3.06% |
| hsa_circ_0068368 | 720 | 11 | 3.06% |
| hsa_circ_0110896 | 720 | 11 | 3.06% |
| hsa_circ_0127726 | 720 | 11 | 3.06% |
| hsa_circ_0128699 | 720 | 11 | 3.06% |
| hsa_circ_0138837 | 720 | 11 | 3.06% |
| hsa_circ_0052826 | 851 | 13 | 3.06% |
| hsa_circ_0130365 | 851 | 13 | 3.06% |
| hsa_circ_0138751 | 786 | 12 | 3.05% |
| hsa_circ_0006499 | 655 | 10 | 3.05% |
| hsa_circ_0008205 | 655 | 10 | 3.05% |
| hsa_circ_0027019 | 655 | 10 | 3.05% |
| hsa_circ_0070806 | 655 | 10 | 3.05% |
| hsa_circ_0072194 | 655 | 10 | 3.05% |
| hsa_circ_0087152 | 655 | 10 | 3.05% |
| hsa_circ_0089657 | 655 | 10 | 3.05% |
| hsa_circ_0131087 | 655 | 10 | 3.05% |
| hsa_circ_0097674 | 1704 | 26 | 3.05% |
| hsa_circ_0012207 | 721 | 11 | 3.05% |
| hsa_circ_0048260 | 721 | 11 | 3.05% |
| hsa_circ_0065545 | 721 | 11 | 3.05% |
| hsa_circ_0130635 | 721 | 11 | 3.05% |

| | | | |
|---|---|---|---|
| hsa_circ_0057368 | 918 | 14 | 3.05% |
| hsa_circ_0096252 | 1246 | 19 | 3.05% |
| hsa_circ_0116289 | 787 | 12 | 3.05% |
| hsa_circ_0000041 | 656 | 10 | 3.05% |
| hsa_circ_0009720 | 656 | 10 | 3.05% |
| hsa_circ_0015442 | 656 | 10 | 3.05% |
| hsa_circ_0016079 | 656 | 10 | 3.05% |
| hsa_circ_0032934 | 656 | 10 | 3.05% |
| hsa_circ_0039587 | 656 | 10 | 3.05% |
| hsa_circ_0046515 | 656 | 10 | 3.05% |
| hsa_circ_0060349 | 656 | 10 | 3.05% |
| hsa_circ_0081475 | 656 | 10 | 3.05% |
| hsa_circ_0089638 | 656 | 10 | 3.05% |
| hsa_circ_0092072 | 656 | 10 | 3.05% |
| hsa_circ_0108095 | 656 | 10 | 3.05% |
| hsa_circ_0119202 | 656 | 10 | 3.05% |
| hsa_circ_0119396 | 656 | 10 | 3.05% |
| hsa_circ_0125466 | 656 | 10 | 3.05% |
| hsa_circ_0130291 | 656 | 10 | 3.05% |
| hsa_circ_0130320 | 656 | 10 | 3.05% |
| hsa_circ_0006260 | 722 | 11 | 3.05% |
| hsa_circ_0019548 | 722 | 11 | 3.05% |
| hsa_circ_0023798 | 722 | 11 | 3.05% |
| hsa_circ_0120591 | 788 | 12 | 3.05% |
| hsa_circ_0006287 | 657 | 10 | 3.04% |
| hsa_circ_0007783 | 657 | 10 | 3.04% |
| hsa_circ_0011396 | 657 | 10 | 3.04% |
| hsa_circ_0018910 | 657 | 10 | 3.04% |
| hsa_circ_0023231 | 657 | 10 | 3.04% |
| hsa_circ_0026641 | 657 | 10 | 3.04% |
| hsa_circ_0057367 | 657 | 10 | 3.04% |
| hsa_circ_0066791 | 657 | 10 | 3.04% |
| hsa_circ_0072712 | 657 | 10 | 3.04% |
| hsa_circ_0092514 | 657 | 10 | 3.04% |
| hsa_circ_0102218 | 657 | 10 | 3.04% |
| hsa_circ_0105092 | 657 | 10 | 3.04% |
| hsa_circ_0107335 | 657 | 10 | 3.04% |
| hsa_circ_0118331 | 657 | 10 | 3.04% |
| hsa_circ_0119764 | 657 | 10 | 3.04% |
| hsa_circ_0122560 | 657 | 10 | 3.04% |
| hsa_circ_0124040 | 657 | 10 | 3.04% |
| hsa_circ_0129456 | 657 | 10 | 3.04% |
| hsa_circ_0132093 | 657 | 10 | 3.04% |
| hsa_circ_0019916 | 723 | 11 | 3.04% |
| hsa_circ_0053340 | 723 | 11 | 3.04% |
| hsa_circ_0100511 | 723 | 11 | 3.04% |
| hsa_circ_0105142 | 723 | 11 | 3.04% |
| hsa_circ_0012856 | 789 | 12 | 3.04% |
| hsa_circ_0027009 | 789 | 12 | 3.04% |
| hsa_circ_0057304 | 789 | 12 | 3.04% |

| | | | |
|---|---|---|---|
| hsa_circ_0004963 | 658 | 10 | 3.04% |
| hsa_circ_0006491 | 658 | 10 | 3.04% |
| hsa_circ_0035801 | 658 | 10 | 3.04% |
| hsa_circ_0038499 | 658 | 10 | 3.04% |
| hsa_circ_0038553 | 658 | 10 | 3.04% |
| hsa_circ_0051994 | 658 | 10 | 3.04% |
| hsa_circ_0067180 | 658 | 10 | 3.04% |
| hsa_circ_0068594 | 658 | 10 | 3.04% |
| hsa_circ_0081712 | 658 | 10 | 3.04% |
| hsa_circ_0085170 | 658 | 10 | 3.04% |
| hsa_circ_0092845 | 658 | 10 | 3.04% |
| hsa_circ_0101701 | 658 | 10 | 3.04% |
| hsa_circ_0112513 | 658 | 10 | 3.04% |
| hsa_circ_0132787 | 658 | 10 | 3.04% |
| hsa_circ_0133958 | 658 | 10 | 3.04% |
| hsa_circ_0022086 | 724 | 11 | 3.04% |
| hsa_circ_0053277 | 724 | 11 | 3.04% |
| hsa_circ_0085939 | 724 | 11 | 3.04% |
| hsa_circ_0089002 | 724 | 11 | 3.04% |
| hsa_circ_0128594 | 724 | 11 | 3.04% |
| hsa_circ_0138636 | 724 | 11 | 3.04% |
| hsa_circ_0074666 | 790 | 12 | 3.04% |
| hsa_circ_0103807 | 790 | 12 | 3.04% |
| hsa_circ_0096957 | 856 | 13 | 3.04% |
| hsa_circ_0007589 | 659 | 10 | 3.03% |
| hsa_circ_0012467 | 659 | 10 | 3.03% |
| hsa_circ_0024357 | 659 | 10 | 3.03% |
| hsa_circ_0025936 | 659 | 10 | 3.03% |
| hsa_circ_0029577 | 659 | 10 | 3.03% |
| hsa_circ_0051212 | 659 | 10 | 3.03% |
| hsa_circ_0053410 | 659 | 10 | 3.03% |
| hsa_circ_0057734 | 659 | 10 | 3.03% |
| hsa_circ_0068096 | 659 | 10 | 3.03% |
| hsa_circ_0108714 | 659 | 10 | 3.03% |
| hsa_circ_0120013 | 659 | 10 | 3.03% |
| hsa_circ_0001724 | 725 | 11 | 3.03% |
| hsa_circ_0008935 | 725 | 11 | 3.03% |
| hsa_circ_0021988 | 725 | 11 | 3.03% |
| hsa_circ_0029790 | 725 | 11 | 3.03% |
| hsa_circ_0031951 | 725 | 11 | 3.03% |
| hsa_circ_0044253 | 725 | 11 | 3.03% |
| hsa_circ_0056325 | 725 | 11 | 3.03% |
| hsa_circ_0076284 | 725 | 11 | 3.03% |
| hsa_circ_0113840 | 725 | 11 | 3.03% |
| hsa_circ_0082613 | 791 | 12 | 3.03% |
| hsa_circ_0112474 | 791 | 12 | 3.03% |
| hsa_circ_0115588 | 791 | 12 | 3.03% |
| hsa_circ_0106547 | 1122 | 17 | 3.03% |
| hsa_circ_0000247 | 792 | 12 | 3.03% |
| hsa_circ_0070371 | 792 | 12 | 3.03% |

| | | | |
|---|---|---|---|
| hsa_circ_0072335 | 792 | 12 | 3.03% |
| hsa_circ_0094321 | 726 | 11 | 3.03% |
| hsa_circ_0110070 | 726 | 11 | 3.03% |
| hsa_circ_0000098 | 660 | 10 | 3.03% |
| hsa_circ_0008463 | 660 | 10 | 3.03% |
| hsa_circ_0011027 | 660 | 10 | 3.03% |
| hsa_circ_0019557 | 660 | 10 | 3.03% |
| hsa_circ_0022929 | 660 | 10 | 3.03% |
| hsa_circ_0023176 | 660 | 10 | 3.03% |
| hsa_circ_0025392 | 660 | 10 | 3.03% |
| hsa_circ_0027279 | 660 | 10 | 3.03% |
| hsa_circ_0027608 | 660 | 10 | 3.03% |
| hsa_circ_0032593 | 660 | 10 | 3.03% |
| hsa_circ_0042151 | 660 | 10 | 3.03% |
| hsa_circ_0049476 | 660 | 10 | 3.03% |
| hsa_circ_0050434 | 660 | 10 | 3.03% |
| hsa_circ_0051442 | 660 | 10 | 3.03% |
| hsa_circ_0084325 | 660 | 10 | 3.03% |
| hsa_circ_0093885 | 660 | 10 | 3.03% |
| hsa_circ_0104134 | 660 | 10 | 3.03% |
| hsa_circ_0105990 | 660 | 10 | 3.03% |
| hsa_circ_0120887 | 660 | 10 | 3.03% |
| hsa_circ_0126096 | 660 | 10 | 3.03% |
| hsa_circ_0140277 | 660 | 10 | 3.03% |
| hsa_circ_0130244 | 925 | 14 | 3.03% |
| hsa_circ_0094954 | 859 | 13 | 3.03% |
| hsa_circ_0136336 | 859 | 13 | 3.03% |
| hsa_circ_0050808 | 793 | 12 | 3.03% |
| hsa_circ_0001742 | 727 | 11 | 3.03% |
| hsa_circ_0021807 | 727 | 11 | 3.03% |
| hsa_circ_0098657 | 727 | 11 | 3.03% |
| hsa_circ_0121219 | 727 | 11 | 3.03% |
| hsa_circ_0016121 | 661 | 10 | 3.03% |
| hsa_circ_0020752 | 661 | 10 | 3.03% |
| hsa_circ_0031284 | 661 | 10 | 3.03% |
| hsa_circ_0032277 | 661 | 10 | 3.03% |
| hsa_circ_0039586 | 661 | 10 | 3.03% |
| hsa_circ_0048830 | 661 | 10 | 3.03% |
| hsa_circ_0058712 | 661 | 10 | 3.03% |
| hsa_circ_0062406 | 661 | 10 | 3.03% |
| hsa_circ_0066754 | 661 | 10 | 3.03% |
| hsa_circ_0069723 | 661 | 10 | 3.03% |
| hsa_circ_0118445 | 661 | 10 | 3.03% |
| hsa_circ_0135984 | 661 | 10 | 3.03% |
| hsa_circ_0084017 | 860 | 13 | 3.02% |
| hsa_circ_0092307 | 860 | 13 | 3.02% |
| hsa_circ_0019666 | 728 | 11 | 3.02% |
| hsa_circ_0036177 | 728 | 11 | 3.02% |
| hsa_circ_0076446 | 728 | 11 | 3.02% |
| hsa_circ_0001953 | 662 | 10 | 3.02% |

| | | | |
|---|---|---|---|
| hsa_circ_0004874 | 662 | 10 | 3.02% |
| hsa_circ_0011762 | 662 | 10 | 3.02% |
| hsa_circ_0013055 | 662 | 10 | 3.02% |
| hsa_circ_0035245 | 662 | 10 | 3.02% |
| hsa_circ_0071758 | 662 | 10 | 3.02% |
| hsa_circ_0071817 | 662 | 10 | 3.02% |
| hsa_circ_0088132 | 662 | 10 | 3.02% |
| hsa_circ_0049741 | 927 | 14 | 3.02% |
| hsa_circ_0115271 | 2253 | 34 | 3.02% |
| hsa_circ_0036777 | 729 | 11 | 3.02% |
| hsa_circ_0066826 | 729 | 11 | 3.02% |
| hsa_circ_0073492 | 729 | 11 | 3.02% |
| hsa_circ_0130630 | 729 | 11 | 3.02% |
| hsa_circ_0140636 | 729 | 11 | 3.02% |
| hsa_circ_0005353 | 663 | 10 | 3.02% |
| hsa_circ_0017553 | 663 | 10 | 3.02% |
| hsa_circ_0026676 | 663 | 10 | 3.02% |
| hsa_circ_0028978 | 663 | 10 | 3.02% |
| hsa_circ_0042944 | 663 | 10 | 3.02% |
| hsa_circ_0052163 | 663 | 10 | 3.02% |
| hsa_circ_0057359 | 663 | 10 | 3.02% |
| hsa_circ_0064397 | 663 | 10 | 3.02% |
| hsa_circ_0066660 | 663 | 10 | 3.02% |
| hsa_circ_0068743 | 663 | 10 | 3.02% |
| hsa_circ_0080325 | 663 | 10 | 3.02% |
| hsa_circ_0085032 | 663 | 10 | 3.02% |
| hsa_circ_0099675 | 663 | 10 | 3.02% |
| hsa_circ_0103159 | 663 | 10 | 3.02% |
| hsa_circ_0134150 | 663 | 10 | 3.02% |
| hsa_circ_0003633 | 796 | 12 | 3.02% |
| hsa_circ_0014137 | 796 | 12 | 3.02% |
| hsa_circ_0037272 | 796 | 12 | 3.02% |
| hsa_circ_0090009 | 796 | 12 | 3.02% |
| hsa_circ_0139769 | 796 | 12 | 3.02% |
| hsa_circ_0087207 | 929 | 14 | 3.01% |
| hsa_circ_0046297 | 730 | 11 | 3.01% |
| hsa_circ_0068662 | 730 | 11 | 3.01% |
| hsa_circ_0094365 | 730 | 11 | 3.01% |
| hsa_circ_0108031 | 730 | 11 | 3.01% |
| hsa_circ_0016728 | 664 | 10 | 3.01% |
| hsa_circ_0017077 | 664 | 10 | 3.01% |
| hsa_circ_0034688 | 664 | 10 | 3.01% |
| hsa_circ_0038717 | 664 | 10 | 3.01% |
| hsa_circ_0040739 | 664 | 10 | 3.01% |
| hsa_circ_0049357 | 664 | 10 | 3.01% |
| hsa_circ_0100240 | 664 | 10 | 3.01% |
| hsa_circ_0105582 | 664 | 10 | 3.01% |
| hsa_circ_0119889 | 664 | 10 | 3.01% |
| hsa_circ_0131108 | 664 | 10 | 3.01% |
| hsa_circ_0140491 | 797 | 12 | 3.01% |

| | | | |
|---|---|---|---|
| hsa_circ_0001404 | 731 | 11 | 3.01% |
| hsa_circ_0011746 | 731 | 11 | 3.01% |
| hsa_circ_0029391 | 731 | 11 | 3.01% |
| hsa_circ_0037844 | 731 | 11 | 3.01% |
| hsa_circ_0074727 | 731 | 11 | 3.01% |
| hsa_circ_0096202 | 731 | 11 | 3.01% |
| hsa_circ_0099437 | 864 | 13 | 3.01% |
| hsa_circ_0007120 | 798 | 12 | 3.01% |
| hsa_circ_0122141 | 798 | 12 | 3.01% |
| hsa_circ_0003485 | 665 | 10 | 3.01% |
| hsa_circ_0005966 | 665 | 10 | 3.01% |
| hsa_circ_0029055 | 665 | 10 | 3.01% |
| hsa_circ_0041381 | 665 | 10 | 3.01% |
| hsa_circ_0041612 | 665 | 10 | 3.01% |
| hsa_circ_0046526 | 665 | 10 | 3.01% |
| hsa_circ_0070227 | 665 | 10 | 3.01% |
| hsa_circ_0082463 | 665 | 10 | 3.01% |
| hsa_circ_0089652 | 665 | 10 | 3.01% |
| hsa_circ_0099972 | 665 | 10 | 3.01% |
| hsa_circ_0110181 | 665 | 10 | 3.01% |
| hsa_circ_0123788 | 665 | 10 | 3.01% |
| hsa_circ_0124445 | 665 | 10 | 3.01% |
| hsa_circ_0136135 | 665 | 10 | 3.01% |
| hsa_circ_0003855 | 732 | 11 | 3.01% |
| hsa_circ_0009938 | 732 | 11 | 3.01% |
| hsa_circ_0084669 | 732 | 11 | 3.01% |
| hsa_circ_0090448 | 732 | 11 | 3.01% |
| hsa_circ_0099998 | 732 | 11 | 3.01% |
| hsa_circ_0049739 | 932 | 14 | 3.00% |
| hsa_circ_0128766 | 799 | 12 | 3.00% |
| hsa_circ_0014715 | 666 | 10 | 3.00% |
| hsa_circ_0017020 | 666 | 10 | 3.00% |
| hsa_circ_0018332 | 666 | 10 | 3.00% |
| hsa_circ_0023675 | 666 | 10 | 3.00% |
| hsa_circ_0066505 | 666 | 10 | 3.00% |
| hsa_circ_0066521 | 666 | 10 | 3.00% |
| hsa_circ_0074675 | 666 | 10 | 3.00% |
| hsa_circ_0090402 | 666 | 10 | 3.00% |
| hsa_circ_0096341 | 666 | 10 | 3.00% |
| hsa_circ_0100771 | 666 | 10 | 3.00% |
| hsa_circ_0104087 | 666 | 10 | 3.00% |
| hsa_circ_0114926 | 666 | 10 | 3.00% |
| hsa_circ_0122349 | 666 | 10 | 3.00% |
| hsa_circ_0126306 | 666 | 10 | 3.00% |
| hsa_circ_0129567 | 666 | 10 | 3.00% |
| hsa_circ_0084545 | 866 | 13 | 3.00% |
| hsa_circ_0004926 | 733 | 11 | 3.00% |
| hsa_circ_0041321 | 733 | 11 | 3.00% |
| hsa_circ_0041454 | 733 | 11 | 3.00% |
| hsa_circ_0045410 | 733 | 11 | 3.00% |

| | | | |
|---|---|---|---|
| hsa_circ_0050781 | 733 | 11 | 3.00% |
| hsa_circ_0061775 | 733 | 11 | 3.00% |
| hsa_circ_0063355 | 733 | 11 | 3.00% |
| hsa_circ_0072781 | 733 | 11 | 3.00% |
| hsa_circ_0099372 | 733 | 11 | 3.00% |
| hsa_circ_0108358 | 733 | 11 | 3.00% |
| hsa_circ_0115967 | 733 | 11 | 3.00% |
| hsa_circ_0140775 | 733 | 11 | 3.00% |
| hsa_circ_0132404 | 933 | 14 | 3.00% |
| hsa_circ_0122262 | 867 | 13 | 3.00% |
| hsa_circ_0005489 | 667 | 10 | 3.00% |
| hsa_circ_0020857 | 667 | 10 | 3.00% |
| hsa_circ_0023971 | 667 | 10 | 3.00% |
| hsa_circ_0030610 | 667 | 10 | 3.00% |
| hsa_circ_0034439 | 667 | 10 | 3.00% |
| hsa_circ_0045061 | 667 | 10 | 3.00% |
| hsa_circ_0069238 | 667 | 10 | 3.00% |
| hsa_circ_0074836 | 667 | 10 | 3.00% |
| hsa_circ_0080491 | 667 | 10 | 3.00% |
| hsa_circ_0089618 | 667 | 10 | 3.00% |
| hsa_circ_0091062 | 667 | 10 | 3.00% |
| hsa_circ_0098945 | 667 | 10 | 3.00% |
| hsa_circ_0106265 | 667 | 10 | 3.00% |
| hsa_circ_0107373 | 667 | 10 | 3.00% |
| hsa_circ_0112033 | 667 | 10 | 3.00% |
| hsa_circ_0118728 | 667 | 10 | 3.00% |
| hsa_circ_0122250 | 667 | 10 | 3.00% |
| hsa_circ_0123069 | 667 | 10 | 3.00% |
| hsa_circ_0125832 | 667 | 10 | 3.00% |
| hsa_circ_0130639 | 667 | 10 | 3.00% |
| hsa_circ_0134509 | 667 | 10 | 3.00% |
| hsa_circ_0084063 | 934 | 14 | 3.00% |
| hsa_circ_0048570 | 734 | 11 | 3.00% |
| hsa_circ_0056822 | 734 | 11 | 3.00% |
| hsa_circ_0067835 | 734 | 11 | 3.00% |
| hsa_circ_0020039 | 801 | 12 | 3.00% |
| hsa_circ_0116303 | 1736 | 26 | 3.00% |
| hsa_circ_0077575 | 935 | 14 | 2.99% |
| hsa_circ_0004043 | 668 | 10 | 2.99% |
| hsa_circ_0007415 | 668 | 10 | 2.99% |
| hsa_circ_0013228 | 668 | 10 | 2.99% |
| hsa_circ_0015855 | 668 | 10 | 2.99% |
| hsa_circ_0022649 | 668 | 10 | 2.99% |
| hsa_circ_0023931 | 668 | 10 | 2.99% |
| hsa_circ_0102127 | 668 | 10 | 2.99% |
| hsa_circ_0114109 | 668 | 10 | 2.99% |
| hsa_circ_0128714 | 668 | 10 | 2.99% |
| hsa_circ_0135995 | 668 | 10 | 2.99% |
| hsa_circ_0098256 | 1470 | 22 | 2.99% |
| hsa_circ_0004073 | 735 | 11 | 2.99% |

| | | | |
|---|---|---|---|
| hsa_circ_0016385 | 735 | 11 | 2.99% |
| hsa_circ_0038469 | 735 | 11 | 2.99% |
| hsa_circ_0050801 | 735 | 11 | 2.99% |
| hsa_circ_0074216 | 735 | 11 | 2.99% |
| hsa_circ_0077442 | 735 | 11 | 2.99% |
| hsa_circ_0114856 | 735 | 11 | 2.99% |
| hsa_circ_0006246 | 802 | 12 | 2.99% |
| hsa_circ_0057036 | 802 | 12 | 2.99% |
| hsa_circ_0008667 | 669 | 10 | 2.99% |
| hsa_circ_0014956 | 669 | 10 | 2.99% |
| hsa_circ_0025931 | 669 | 10 | 2.99% |
| hsa_circ_0027936 | 669 | 10 | 2.99% |
| hsa_circ_0042948 | 669 | 10 | 2.99% |
| hsa_circ_0043597 | 669 | 10 | 2.99% |
| hsa_circ_0043623 | 669 | 10 | 2.99% |
| hsa_circ_0055834 | 669 | 10 | 2.99% |
| hsa_circ_0082034 | 669 | 10 | 2.99% |
| hsa_circ_0082768 | 669 | 10 | 2.99% |
| hsa_circ_0082804 | 669 | 10 | 2.99% |
| hsa_circ_0084054 | 669 | 10 | 2.99% |
| hsa_circ_0093393 | 669 | 10 | 2.99% |
| hsa_circ_0095290 | 669 | 10 | 2.99% |
| hsa_circ_0096271 | 669 | 10 | 2.99% |
| hsa_circ_0109172 | 669 | 10 | 2.99% |
| hsa_circ_0122963 | 669 | 10 | 2.99% |
| hsa_circ_0140714 | 669 | 10 | 2.99% |
| hsa_circ_0103980 | 1405 | 21 | 2.99% |
| hsa_circ_0005134 | 736 | 11 | 2.99% |
| hsa_circ_0046493 | 736 | 11 | 2.99% |
| hsa_circ_0051799 | 736 | 11 | 2.99% |
| hsa_circ_0081384 | 736 | 11 | 2.99% |
| hsa_circ_0092265 | 736 | 11 | 2.99% |
| hsa_circ_0097416 | 736 | 11 | 2.99% |
| hsa_circ_0120563 | 736 | 11 | 2.99% |
| hsa_circ_0122378 | 736 | 11 | 2.99% |
| hsa_circ_0048648 | 803 | 12 | 2.99% |
| hsa_circ_0090586 | 804 | 12 | 2.99% |
| hsa_circ_0096859 | 804 | 12 | 2.99% |
| hsa_circ_0129210 | 804 | 12 | 2.99% |
| hsa_circ_0055216 | 737 | 11 | 2.99% |
| hsa_circ_0002138 | 670 | 10 | 2.99% |
| hsa_circ_0027094 | 670 | 10 | 2.99% |
| hsa_circ_0042060 | 670 | 10 | 2.99% |
| hsa_circ_0044813 | 670 | 10 | 2.99% |
| hsa_circ_0088822 | 670 | 10 | 2.99% |
| hsa_circ_0094059 | 670 | 10 | 2.99% |
| hsa_circ_0110156 | 670 | 10 | 2.99% |
| hsa_circ_0125265 | 670 | 10 | 2.99% |
| hsa_circ_0035014 | 805 | 12 | 2.98% |
| hsa_circ_0051424 | 805 | 12 | 2.98% |

| | | | |
|---|---|---|---|
| hsa_circ_0015322 | 738 | 11 | 2.98% |
| hsa_circ_0017273 | 738 | 11 | 2.98% |
| hsa_circ_0081465 | 738 | 11 | 2.98% |
| hsa_circ_0094591 | 738 | 11 | 2.98% |
| hsa_circ_0120855 | 738 | 11 | 2.98% |
| hsa_circ_0015278 | 671 | 10 | 2.98% |
| hsa_circ_0021867 | 671 | 10 | 2.98% |
| hsa_circ_0025048 | 671 | 10 | 2.98% |
| hsa_circ_0029251 | 671 | 10 | 2.98% |
| hsa_circ_0038564 | 671 | 10 | 2.98% |
| hsa_circ_0065472 | 671 | 10 | 2.98% |
| hsa_circ_0073648 | 671 | 10 | 2.98% |
| hsa_circ_0077076 | 671 | 10 | 2.98% |
| hsa_circ_0086158 | 671 | 10 | 2.98% |
| hsa_circ_0087738 | 671 | 10 | 2.98% |
| hsa_circ_0129738 | 671 | 10 | 2.98% |
| hsa_circ_0138887 | 671 | 10 | 2.98% |
| hsa_circ_0139118 | 671 | 10 | 2.98% |
| hsa_circ_0138281 | 806 | 12 | 2.98% |
| hsa_circ_0056913 | 739 | 11 | 2.98% |
| hsa_circ_0074795 | 739 | 11 | 2.98% |
| hsa_circ_0089114 | 739 | 11 | 2.98% |
| hsa_circ_0111114 | 739 | 11 | 2.98% |
| hsa_circ_0114185 | 739 | 11 | 2.98% |
| hsa_circ_0133592 | 739 | 11 | 2.98% |
| hsa_circ_0140033 | 739 | 11 | 2.98% |
| hsa_circ_0008280 | 672 | 10 | 2.98% |
| hsa_circ_0016046 | 672 | 10 | 2.98% |
| hsa_circ_0021937 | 672 | 10 | 2.98% |
| hsa_circ_0029026 | 672 | 10 | 2.98% |
| hsa_circ_0030886 | 672 | 10 | 2.98% |
| hsa_circ_0043686 | 672 | 10 | 2.98% |
| hsa_circ_0079837 | 672 | 10 | 2.98% |
| hsa_circ_0094483 | 672 | 10 | 2.98% |
| hsa_circ_0098286 | 672 | 10 | 2.98% |
| hsa_circ_0099322 | 672 | 10 | 2.98% |
| hsa_circ_0110001 | 672 | 10 | 2.98% |
| hsa_circ_0122928 | 672 | 10 | 2.98% |
| hsa_circ_0126418 | 672 | 10 | 2.98% |
| hsa_circ_0132661 | 672 | 10 | 2.98% |
| hsa_circ_0133529 | 672 | 10 | 2.98% |
| hsa_circ_0135556 | 672 | 10 | 2.98% |
| hsa_circ_0132493 | 807 | 12 | 2.97% |
| hsa_circ_0056589 | 740 | 11 | 2.97% |
| hsa_circ_0061692 | 740 | 11 | 2.97% |
| hsa_circ_0093886 | 740 | 11 | 2.97% |
| hsa_circ_0136174 | 740 | 11 | 2.97% |
| hsa_circ_0001234 | 673 | 10 | 2.97% |
| hsa_circ_0014767 | 673 | 10 | 2.97% |
| hsa_circ_0022323 | 673 | 10 | 2.97% |

| | | | |
|---|---|---|---|
| hsa_circ_0026939 | 673 | 10 | 2.97% |
| hsa_circ_0036296 | 673 | 10 | 2.97% |
| hsa_circ_0066031 | 673 | 10 | 2.97% |
| hsa_circ_0069382 | 673 | 10 | 2.97% |
| hsa_circ_0086676 | 673 | 10 | 2.97% |
| hsa_circ_0093394 | 673 | 10 | 2.97% |
| hsa_circ_0093767 | 673 | 10 | 2.97% |
| hsa_circ_0117219 | 673 | 10 | 2.97% |
| hsa_circ_0117228 | 673 | 10 | 2.97% |
| hsa_circ_0120238 | 673 | 10 | 2.97% |
| hsa_circ_0129395 | 673 | 10 | 2.97% |
| hsa_circ_0139052 | 673 | 10 | 2.97% |
| hsa_circ_0072632 | 875 | 13 | 2.97% |
| hsa_circ_0004121 | 741 | 11 | 2.97% |
| hsa_circ_0019801 | 741 | 11 | 2.97% |
| hsa_circ_0028025 | 741 | 11 | 2.97% |
| hsa_circ_0029271 | 741 | 11 | 2.97% |
| hsa_circ_0087554 | 741 | 11 | 2.97% |
| hsa_circ_0009405 | 876 | 13 | 2.97% |
| hsa_circ_0003268 | 674 | 10 | 2.97% |
| hsa_circ_0007159 | 674 | 10 | 2.97% |
| hsa_circ_0025441 | 674 | 10 | 2.97% |
| hsa_circ_0038196 | 674 | 10 | 2.97% |
| hsa_circ_0055790 | 674 | 10 | 2.97% |
| hsa_circ_0058368 | 674 | 10 | 2.97% |
| hsa_circ_0067786 | 674 | 10 | 2.97% |
| hsa_circ_0073752 | 674 | 10 | 2.97% |
| hsa_circ_0082476 | 674 | 10 | 2.97% |
| hsa_circ_0116099 | 674 | 10 | 2.97% |
| hsa_circ_0123680 | 674 | 10 | 2.97% |
| hsa_circ_0049753 | 809 | 12 | 2.97% |
| hsa_circ_0104831 | 809 | 12 | 2.97% |
| hsa_circ_0098606 | 3237 | 48 | 2.97% |
| hsa_circ_0072446 | 742 | 11 | 2.96% |
| hsa_circ_0084747 | 742 | 11 | 2.96% |
| hsa_circ_0091723 | 742 | 11 | 2.96% |
| hsa_circ_0096496 | 742 | 11 | 2.96% |
| hsa_circ_0103802 | 742 | 11 | 2.96% |
| hsa_circ_0108275 | 742 | 11 | 2.96% |
| hsa_circ_0118307 | 742 | 11 | 2.96% |
| hsa_circ_0138170 | 742 | 11 | 2.96% |
| hsa_circ_0061497 | 877 | 13 | 2.96% |
| hsa_circ_0031540 | 810 | 12 | 2.96% |
| hsa_circ_0088206 | 810 | 12 | 2.96% |
| hsa_circ_0126033 | 810 | 12 | 2.96% |
| hsa_circ_0007698 | 675 | 10 | 2.96% |
| hsa_circ_0010369 | 675 | 10 | 2.96% |
| hsa_circ_0022072 | 675 | 10 | 2.96% |
| hsa_circ_0022496 | 675 | 10 | 2.96% |
| hsa_circ_0029187 | 675 | 10 | 2.96% |

| | | | |
|---|---|---|---|
| hsa_circ_0038344 | 675 | 10 | 2.96% |
| hsa_circ_0038984 | 675 | 10 | 2.96% |
| hsa_circ_0039827 | 675 | 10 | 2.96% |
| hsa_circ_0051972 | 675 | 10 | 2.96% |
| hsa_circ_0054100 | 675 | 10 | 2.96% |
| hsa_circ_0066821 | 675 | 10 | 2.96% |
| hsa_circ_0070889 | 675 | 10 | 2.96% |
| hsa_circ_0075612 | 675 | 10 | 2.96% |
| hsa_circ_0091192 | 675 | 10 | 2.96% |
| hsa_circ_0095692 | 675 | 10 | 2.96% |
| hsa_circ_0106892 | 675 | 10 | 2.96% |
| hsa_circ_0108514 | 675 | 10 | 2.96% |
| hsa_circ_0132339 | 675 | 10 | 2.96% |
| hsa_circ_0081769 | 878 | 13 | 2.96% |
| hsa_circ_0002596 | 743 | 11 | 2.96% |
| hsa_circ_0086185 | 743 | 11 | 2.96% |
| hsa_circ_0091090 | 743 | 11 | 2.96% |
| hsa_circ_0043272 | 811 | 12 | 2.96% |
| hsa_circ_0064156 | 811 | 12 | 2.96% |
| hsa_circ_0090202 | 811 | 12 | 2.96% |
| hsa_circ_0130083 | 811 | 12 | 2.96% |
| hsa_circ_0021494 | 676 | 10 | 2.96% |
| hsa_circ_0021603 | 676 | 10 | 2.96% |
| hsa_circ_0030377 | 676 | 10 | 2.96% |
| hsa_circ_0039829 | 676 | 10 | 2.96% |
| hsa_circ_0042332 | 676 | 10 | 2.96% |
| hsa_circ_0051821 | 676 | 10 | 2.96% |
| hsa_circ_0051960 | 676 | 10 | 2.96% |
| hsa_circ_0059648 | 676 | 10 | 2.96% |
| hsa_circ_0078514 | 676 | 10 | 2.96% |
| hsa_circ_0079598 | 676 | 10 | 2.96% |
| hsa_circ_0092229 | 676 | 10 | 2.96% |
| hsa_circ_0092408 | 676 | 10 | 2.96% |
| hsa_circ_0096287 | 676 | 10 | 2.96% |
| hsa_circ_0121658 | 676 | 10 | 2.96% |
| hsa_circ_0051519 | 744 | 11 | 2.96% |
| hsa_circ_0055404 | 744 | 11 | 2.96% |
| hsa_circ_0060698 | 880 | 13 | 2.95% |
| hsa_circ_0012744 | 677 | 10 | 2.95% |
| hsa_circ_0014972 | 677 | 10 | 2.95% |
| hsa_circ_0045047 | 677 | 10 | 2.95% |
| hsa_circ_0062471 | 677 | 10 | 2.95% |
| hsa_circ_0072875 | 677 | 10 | 2.95% |
| hsa_circ_0075330 | 677 | 10 | 2.95% |
| hsa_circ_0076118 | 677 | 10 | 2.95% |
| hsa_circ_0079257 | 677 | 10 | 2.95% |
| hsa_circ_0081878 | 677 | 10 | 2.95% |
| hsa_circ_0094625 | 677 | 10 | 2.95% |
| hsa_circ_0094660 | 677 | 10 | 2.95% |
| hsa_circ_0122491 | 677 | 10 | 2.95% |

| | | | |
|---|---|---|---|
| hsa_circ_0128951 | 677 | 10 | 2.95% |
| hsa_circ_0134170 | 677 | 10 | 2.95% |
| hsa_circ_0026159 | 745 | 11 | 2.95% |
| hsa_circ_0125400 | 745 | 11 | 2.95% |
| hsa_circ_0139660 | 745 | 11 | 2.95% |
| hsa_circ_0064020 | 1016 | 15 | 2.95% |
| hsa_circ_0001325 | 678 | 10 | 2.95% |
| hsa_circ_0005311 | 678 | 10 | 2.95% |
| hsa_circ_0014577 | 678 | 10 | 2.95% |
| hsa_circ_0017561 | 678 | 10 | 2.95% |
| hsa_circ_0036757 | 678 | 10 | 2.95% |
| hsa_circ_0037170 | 678 | 10 | 2.95% |
| hsa_circ_0038394 | 678 | 10 | 2.95% |
| hsa_circ_0040155 | 678 | 10 | 2.95% |
| hsa_circ_0050384 | 678 | 10 | 2.95% |
| hsa_circ_0057760 | 678 | 10 | 2.95% |
| hsa_circ_0059016 | 678 | 10 | 2.95% |
| hsa_circ_0059834 | 678 | 10 | 2.95% |
| hsa_circ_0065436 | 678 | 10 | 2.95% |
| hsa_circ_0066148 | 678 | 10 | 2.95% |
| hsa_circ_0075426 | 678 | 10 | 2.95% |
| hsa_circ_0075879 | 678 | 10 | 2.95% |
| hsa_circ_0083604 | 678 | 10 | 2.95% |
| hsa_circ_0089720 | 678 | 10 | 2.95% |
| hsa_circ_0096249 | 678 | 10 | 2.95% |
| hsa_circ_0116072 | 678 | 10 | 2.95% |
| hsa_circ_0123649 | 678 | 10 | 2.95% |
| hsa_circ_0131950 | 678 | 10 | 2.95% |
| hsa_circ_0134935 | 678 | 10 | 2.95% |
| hsa_circ_0052389 | 746 | 11 | 2.95% |
| hsa_circ_0069906 | 746 | 11 | 2.95% |
| hsa_circ_0107205 | 746 | 11 | 2.95% |
| hsa_circ_0053217 | 882 | 13 | 2.95% |
| hsa_circ_0057392 | 882 | 13 | 2.95% |
| hsa_circ_0007148 | 679 | 10 | 2.95% |
| hsa_circ_0011718 | 679 | 10 | 2.95% |
| hsa_circ_0048733 | 679 | 10 | 2.95% |
| hsa_circ_0053075 | 679 | 10 | 2.95% |
| hsa_circ_0085281 | 679 | 10 | 2.95% |
| hsa_circ_0116125 | 679 | 10 | 2.95% |
| hsa_circ_0118203 | 679 | 10 | 2.95% |
| hsa_circ_0119425 | 679 | 10 | 2.95% |
| hsa_circ_0122794 | 679 | 10 | 2.95% |
| hsa_circ_0022810 | 747 | 11 | 2.95% |
| hsa_circ_0031970 | 747 | 11 | 2.95% |
| hsa_circ_0068057 | 747 | 11 | 2.95% |
| hsa_circ_0074644 | 747 | 11 | 2.95% |
| hsa_circ_0087432 | 747 | 11 | 2.95% |
| hsa_circ_0122771 | 747 | 11 | 2.95% |
| hsa_circ_0134824 | 747 | 11 | 2.95% |

| | | | |
|---|---|---|---|
| hsa_circ_0058923 | 815 | 12 | 2.94% |
| hsa_circ_0073465 | 815 | 12 | 2.94% |
| hsa_circ_0074082 | 883 | 13 | 2.94% |
| hsa_circ_0082311 | 883 | 13 | 2.94% |
| hsa_circ_0049252 | 952 | 14 | 2.94% |
| hsa_circ_0051120 | 952 | 14 | 2.94% |
| hsa_circ_0132916 | 952 | 14 | 2.94% |
| hsa_circ_0098232 | 884 | 13 | 2.94% |
| hsa_circ_0033346 | 816 | 12 | 2.94% |
| hsa_circ_0116562 | 816 | 12 | 2.94% |
| hsa_circ_0031637 | 748 | 11 | 2.94% |
| hsa_circ_0037738 | 748 | 11 | 2.94% |
| hsa_circ_0071873 | 748 | 11 | 2.94% |
| hsa_circ_0092675 | 748 | 11 | 2.94% |
| hsa_circ_0115397 | 748 | 11 | 2.94% |
| hsa_circ_0002072 | 680 | 10 | 2.94% |
| hsa_circ_0004870 | 680 | 10 | 2.94% |
| hsa_circ_0005128 | 680 | 10 | 2.94% |
| hsa_circ_0007786 | 680 | 10 | 2.94% |
| hsa_circ_0018726 | 680 | 10 | 2.94% |
| hsa_circ_0031910 | 680 | 10 | 2.94% |
| hsa_circ_0037721 | 680 | 10 | 2.94% |
| hsa_circ_0048441 | 680 | 10 | 2.94% |
| hsa_circ_0058987 | 680 | 10 | 2.94% |
| hsa_circ_0068634 | 680 | 10 | 2.94% |
| hsa_circ_0087270 | 680 | 10 | 2.94% |
| hsa_circ_0101861 | 680 | 10 | 2.94% |
| hsa_circ_0103389 | 680 | 10 | 2.94% |
| hsa_circ_0116741 | 680 | 10 | 2.94% |
| hsa_circ_0119756 | 680 | 10 | 2.94% |
| hsa_circ_0138981 | 680 | 10 | 2.94% |
| hsa_circ_0000675 | 953 | 14 | 2.94% |
| hsa_circ_0123768 | 885 | 13 | 2.94% |
| hsa_circ_0058369 | 817 | 12 | 2.94% |
| hsa_circ_0038214 | 749 | 11 | 2.94% |
| hsa_circ_0106469 | 749 | 11 | 2.94% |
| hsa_circ_0113843 | 749 | 11 | 2.94% |
| hsa_circ_0129423 | 749 | 11 | 2.94% |
| hsa_circ_0132459 | 749 | 11 | 2.94% |
| hsa_circ_0003667 | 681 | 10 | 2.94% |
| hsa_circ_0017456 | 681 | 10 | 2.94% |
| hsa_circ_0024893 | 681 | 10 | 2.94% |
| hsa_circ_0027125 | 681 | 10 | 2.94% |
| hsa_circ_0029066 | 681 | 10 | 2.94% |
| hsa_circ_0031254 | 681 | 10 | 2.94% |
| hsa_circ_0035422 | 681 | 10 | 2.94% |
| hsa_circ_0035443 | 681 | 10 | 2.94% |
| hsa_circ_0089668 | 681 | 10 | 2.94% |
| hsa_circ_0102598 | 681 | 10 | 2.94% |
| hsa_circ_0106227 | 681 | 10 | 2.94% |

| | | | |
|---|---|---|---|
| hsa_circ_0108661 | 681 | 10 | 2.94% |
| hsa_circ_0112906 | 681 | 10 | 2.94% |
| hsa_circ_0120074 | 681 | 10 | 2.94% |
| hsa_circ_0134216 | 681 | 10 | 2.94% |
| hsa_circ_0135625 | 681 | 10 | 2.94% |
| hsa_circ_0005634 | 750 | 11 | 2.93% |
| hsa_circ_0023421 | 750 | 11 | 2.93% |
| hsa_circ_0044156 | 750 | 11 | 2.93% |
| hsa_circ_0064316 | 750 | 11 | 2.93% |
| hsa_circ_0081056 | 750 | 11 | 2.93% |
| hsa_circ_0131749 | 750 | 11 | 2.93% |
| hsa_circ_0131750 | 750 | 11 | 2.93% |
| hsa_circ_0003371 | 682 | 10 | 2.93% |
| hsa_circ_0006924 | 682 | 10 | 2.93% |
| hsa_circ_0029262 | 682 | 10 | 2.93% |
| hsa_circ_0044472 | 682 | 10 | 2.93% |
| hsa_circ_0045774 | 682 | 10 | 2.93% |
| hsa_circ_0049230 | 682 | 10 | 2.93% |
| hsa_circ_0050734 | 682 | 10 | 2.93% |
| hsa_circ_0056621 | 682 | 10 | 2.93% |
| hsa_circ_0067445 | 682 | 10 | 2.93% |
| hsa_circ_0072547 | 682 | 10 | 2.93% |
| hsa_circ_0090187 | 682 | 10 | 2.93% |
| hsa_circ_0101041 | 682 | 10 | 2.93% |
| hsa_circ_0114294 | 682 | 10 | 2.93% |
| hsa_circ_0126443 | 682 | 10 | 2.93% |
| hsa_circ_0036575 | 1502 | 22 | 2.93% |
| hsa_circ_0044239 | 751 | 11 | 2.93% |
| hsa_circ_0064392 | 751 | 11 | 2.93% |
| hsa_circ_0074894 | 751 | 11 | 2.93% |
| hsa_circ_0076652 | 751 | 11 | 2.93% |
| hsa_circ_0114187 | 751 | 11 | 2.93% |
| hsa_circ_0106347 | 956 | 14 | 2.93% |
| hsa_circ_0006748 | 683 | 10 | 2.93% |
| hsa_circ_0008218 | 683 | 10 | 2.93% |
| hsa_circ_0023607 | 683 | 10 | 2.93% |
| hsa_circ_0053408 | 683 | 10 | 2.93% |
| hsa_circ_0064682 | 683 | 10 | 2.93% |
| hsa_circ_0079813 | 683 | 10 | 2.93% |
| hsa_circ_0086459 | 683 | 10 | 2.93% |
| hsa_circ_0095206 | 683 | 10 | 2.93% |
| hsa_circ_0096940 | 683 | 10 | 2.93% |
| hsa_circ_0116334 | 683 | 10 | 2.93% |
| hsa_circ_0116799 | 683 | 10 | 2.93% |
| hsa_circ_0123494 | 683 | 10 | 2.93% |
| hsa_circ_0132312 | 683 | 10 | 2.93% |
| hsa_circ_0132537 | 683 | 10 | 2.93% |
| hsa_circ_0092736 | 820 | 12 | 2.93% |
| hsa_circ_0102300 | 820 | 12 | 2.93% |
| hsa_circ_0139855 | 820 | 12 | 2.93% |

| | | | |
|---|---|---|---|
| hsa_circ_0031382 | 752 | 11 | 2.93% |
| hsa_circ_0093832 | 752 | 11 | 2.93% |
| hsa_circ_0067271 | 889 | 13 | 2.92% |
| hsa_circ_0014749 | 684 | 10 | 2.92% |
| hsa_circ_0021310 | 684 | 10 | 2.92% |
| hsa_circ_0027230 | 684 | 10 | 2.92% |
| hsa_circ_0027249 | 684 | 10 | 2.92% |
| hsa_circ_0031755 | 684 | 10 | 2.92% |
| hsa_circ_0044689 | 684 | 10 | 2.92% |
| hsa_circ_0057029 | 684 | 10 | 2.92% |
| hsa_circ_0059072 | 684 | 10 | 2.92% |
| hsa_circ_0070290 | 684 | 10 | 2.92% |
| hsa_circ_0077956 | 684 | 10 | 2.92% |
| hsa_circ_0079923 | 684 | 10 | 2.92% |
| hsa_circ_0080288 | 684 | 10 | 2.92% |
| hsa_circ_0080904 | 684 | 10 | 2.92% |
| hsa_circ_0087324 | 684 | 10 | 2.92% |
| hsa_circ_0090470 | 684 | 10 | 2.92% |
| hsa_circ_0092406 | 684 | 10 | 2.92% |
| hsa_circ_0095389 | 684 | 10 | 2.92% |
| hsa_circ_0099701 | 684 | 10 | 2.92% |
| hsa_circ_0103500 | 684 | 10 | 2.92% |
| hsa_circ_0109402 | 684 | 10 | 2.92% |
| hsa_circ_0126949 | 684 | 10 | 2.92% |
| hsa_circ_0127856 | 684 | 10 | 2.92% |
| hsa_circ_0076607 | 821 | 12 | 2.92% |
| hsa_circ_0081796 | 821 | 12 | 2.92% |
| hsa_circ_0096101 | 821 | 12 | 2.92% |
| hsa_circ_0048132 | 753 | 11 | 2.92% |
| hsa_circ_0065753 | 753 | 11 | 2.92% |
| hsa_circ_0125211 | 753 | 11 | 2.92% |
| hsa_circ_0140488 | 753 | 11 | 2.92% |
| hsa_circ_0000203 | 685 | 10 | 2.92% |
| hsa_circ_0005221 | 685 | 10 | 2.92% |
| hsa_circ_0015888 | 685 | 10 | 2.92% |
| hsa_circ_0021687 | 685 | 10 | 2.92% |
| hsa_circ_0034886 | 685 | 10 | 2.92% |
| hsa_circ_0039905 | 685 | 10 | 2.92% |
| hsa_circ_0045026 | 685 | 10 | 2.92% |
| hsa_circ_0059963 | 685 | 10 | 2.92% |
| hsa_circ_0062180 | 685 | 10 | 2.92% |
| hsa_circ_0063402 | 685 | 10 | 2.92% |
| hsa_circ_0087838 | 685 | 10 | 2.92% |
| hsa_circ_0090186 | 685 | 10 | 2.92% |
| hsa_circ_0095028 | 685 | 10 | 2.92% |
| hsa_circ_0114047 | 685 | 10 | 2.92% |
| hsa_circ_0120452 | 685 | 10 | 2.92% |
| hsa_circ_0111243 | 891 | 13 | 2.92% |
| hsa_circ_0001702 | 754 | 11 | 2.92% |
| hsa_circ_0031121 | 754 | 11 | 2.92% |

| | | | |
|---|---|---|---|
| hsa_circ_0079131 | 754 | 11 | 2.92% |
| hsa_circ_0107934 | 823 | 12 | 2.92% |
| hsa_circ_0005449 | 686 | 10 | 2.92% |
| hsa_circ_0013317 | 686 | 10 | 2.92% |
| hsa_circ_0014244 | 686 | 10 | 2.92% |
| hsa_circ_0023886 | 686 | 10 | 2.92% |
| hsa_circ_0026175 | 686 | 10 | 2.92% |
| hsa_circ_0026384 | 686 | 10 | 2.92% |
| hsa_circ_0031976 | 686 | 10 | 2.92% |
| hsa_circ_0036477 | 686 | 10 | 2.92% |
| hsa_circ_0044824 | 686 | 10 | 2.92% |
| hsa_circ_0047546 | 686 | 10 | 2.92% |
| hsa_circ_0047600 | 686 | 10 | 2.92% |
| hsa_circ_0064761 | 686 | 10 | 2.92% |
| hsa_circ_0066377 | 686 | 10 | 2.92% |
| hsa_circ_0081756 | 686 | 10 | 2.92% |
| hsa_circ_0095970 | 686 | 10 | 2.92% |
| hsa_circ_0112746 | 686 | 10 | 2.92% |
| hsa_circ_0125903 | 686 | 10 | 2.92% |
| hsa_circ_0011536 | 755 | 11 | 2.91% |
| hsa_circ_0049048 | 755 | 11 | 2.91% |
| hsa_circ_0065706 | 755 | 11 | 2.91% |
| hsa_circ_0105555 | 755 | 11 | 2.91% |
| hsa_circ_0117781 | 755 | 11 | 2.91% |
| hsa_circ_0077837 | 824 | 12 | 2.91% |
| hsa_circ_0105533 | 893 | 13 | 2.91% |
| hsa_circ_0017410 | 687 | 10 | 2.91% |
| hsa_circ_0020748 | 687 | 10 | 2.91% |
| hsa_circ_0058999 | 687 | 10 | 2.91% |
| hsa_circ_0072045 | 687 | 10 | 2.91% |
| hsa_circ_0084966 | 687 | 10 | 2.91% |
| hsa_circ_0087685 | 687 | 10 | 2.91% |
| hsa_circ_0094299 | 687 | 10 | 2.91% |
| hsa_circ_0106282 | 687 | 10 | 2.91% |
| hsa_circ_0110661 | 687 | 10 | 2.91% |
| hsa_circ_0131944 | 687 | 10 | 2.91% |
| hsa_circ_0134173 | 687 | 10 | 2.91% |
| hsa_circ_0135419 | 687 | 10 | 2.91% |
| hsa_circ_0001254 | 2268 | 33 | 2.91% |
| hsa_circ_0019662 | 756 | 11 | 2.91% |
| hsa_circ_0022985 | 756 | 11 | 2.91% |
| hsa_circ_0055376 | 756 | 11 | 2.91% |
| hsa_circ_0058528 | 756 | 11 | 2.91% |
| hsa_circ_0062295 | 756 | 11 | 2.91% |
| hsa_circ_0095999 | 756 | 11 | 2.91% |
| hsa_circ_0125700 | 756 | 11 | 2.91% |
| hsa_circ_0133329 | 756 | 11 | 2.91% |
| hsa_circ_0067279 | 1032 | 15 | 2.91% |
| hsa_circ_0007017 | 688 | 10 | 2.91% |
| hsa_circ_0024573 | 688 | 10 | 2.91% |

| | | | |
|---|---|---|---|
| hsa_circ_0038580 | 688 | 10 | 2.91% |
| hsa_circ_0038719 | 688 | 10 | 2.91% |
| hsa_circ_0039365 | 688 | 10 | 2.91% |
| hsa_circ_0051841 | 688 | 10 | 2.91% |
| hsa_circ_0070736 | 688 | 10 | 2.91% |
| hsa_circ_0084202 | 688 | 10 | 2.91% |
| hsa_circ_0111104 | 688 | 10 | 2.91% |
| hsa_circ_0126620 | 688 | 10 | 2.91% |
| hsa_circ_0137132 | 688 | 10 | 2.91% |
| hsa_circ_0062297 | 757 | 11 | 2.91% |
| hsa_circ_0105138 | 757 | 11 | 2.91% |
| hsa_circ_0111708 | 757 | 11 | 2.91% |
| hsa_circ_0019027 | 826 | 12 | 2.91% |
| hsa_circ_0072376 | 826 | 12 | 2.91% |
| hsa_circ_0131610 | 2272 | 33 | 2.90% |
| hsa_circ_0005112 | 689 | 10 | 2.90% |
| hsa_circ_0010949 | 689 | 10 | 2.90% |
| hsa_circ_0046698 | 689 | 10 | 2.90% |
| hsa_circ_0051710 | 689 | 10 | 2.90% |
| hsa_circ_0099706 | 689 | 10 | 2.90% |
| hsa_circ_0101395 | 689 | 10 | 2.90% |
| hsa_circ_0104450 | 689 | 10 | 2.90% |
| hsa_circ_0104856 | 689 | 10 | 2.90% |
| hsa_circ_0114115 | 689 | 10 | 2.90% |
| hsa_circ_0134218 | 689 | 10 | 2.90% |
| hsa_circ_0003220 | 758 | 11 | 2.90% |
| hsa_circ_0089003 | 758 | 11 | 2.90% |
| hsa_circ_0003788 | 827 | 12 | 2.90% |
| hsa_circ_0062513 | 2275 | 33 | 2.90% |
| hsa_circ_0019988 | 897 | 13 | 2.90% |
| hsa_circ_0042686 | 828 | 12 | 2.90% |
| hsa_circ_0111348 | 828 | 12 | 2.90% |
| hsa_circ_0121074 | 828 | 12 | 2.90% |
| hsa_circ_0087583 | 759 | 11 | 2.90% |
| hsa_circ_0120755 | 759 | 11 | 2.90% |
| hsa_circ_0133327 | 759 | 11 | 2.90% |
| hsa_circ_0018080 | 690 | 10 | 2.90% |
| hsa_circ_0023147 | 690 | 10 | 2.90% |
| hsa_circ_0027420 | 690 | 10 | 2.90% |
| hsa_circ_0031094 | 690 | 10 | 2.90% |
| hsa_circ_0037431 | 690 | 10 | 2.90% |
| hsa_circ_0088306 | 690 | 10 | 2.90% |
| hsa_circ_0096765 | 690 | 10 | 2.90% |
| hsa_circ_0110200 | 690 | 10 | 2.90% |
| hsa_circ_0135208 | 690 | 10 | 2.90% |
| hsa_circ_0135378 | 690 | 10 | 2.90% |
| hsa_circ_0135996 | 690 | 10 | 2.90% |
| hsa_circ_0015983 | 1036 | 15 | 2.90% |
| hsa_circ_0113952 | 1036 | 15 | 2.90% |
| hsa_circ_0014853 | 829 | 12 | 2.90% |

| | | | |
|---|---|---|---|
| hsa_circ_0008116 | 760 | 11 | 2.89% |
| hsa_circ_0048335 | 760 | 11 | 2.89% |
| hsa_circ_0002583 | 691 | 10 | 2.89% |
| hsa_circ_0005244 | 691 | 10 | 2.89% |
| hsa_circ_0046324 | 691 | 10 | 2.89% |
| hsa_circ_0062878 | 691 | 10 | 2.89% |
| hsa_circ_0066241 | 691 | 10 | 2.89% |
| hsa_circ_0067839 | 691 | 10 | 2.89% |
| hsa_circ_0080314 | 691 | 10 | 2.89% |
| hsa_circ_0083121 | 691 | 10 | 2.89% |
| hsa_circ_0088426 | 691 | 10 | 2.89% |
| hsa_circ_0104127 | 691 | 10 | 2.89% |
| hsa_circ_0105487 | 691 | 10 | 2.89% |
| hsa_circ_0134724 | 691 | 10 | 2.89% |
| hsa_circ_0001982 | 899 | 13 | 2.89% |
| hsa_circ_0115344 | 899 | 13 | 2.89% |
| hsa_circ_0039499 | 830 | 12 | 2.89% |
| hsa_circ_0039753 | 761 | 11 | 2.89% |
| hsa_circ_0058789 | 761 | 11 | 2.89% |
| hsa_circ_0067455 | 761 | 11 | 2.89% |
| hsa_circ_0078602 | 761 | 11 | 2.89% |
| hsa_circ_0087577 | 761 | 11 | 2.89% |
| hsa_circ_0104069 | 761 | 11 | 2.89% |
| hsa_circ_0115492 | 761 | 11 | 2.89% |
| hsa_circ_0129786 | 761 | 11 | 2.89% |
| hsa_circ_0006540 | 692 | 10 | 2.89% |
| hsa_circ_0030551 | 692 | 10 | 2.89% |
| hsa_circ_0040894 | 692 | 10 | 2.89% |
| hsa_circ_0048531 | 692 | 10 | 2.89% |
| hsa_circ_0049137 | 692 | 10 | 2.89% |
| hsa_circ_0056709 | 692 | 10 | 2.89% |
| hsa_circ_0056744 | 692 | 10 | 2.89% |
| hsa_circ_0063346 | 692 | 10 | 2.89% |
| hsa_circ_0066771 | 692 | 10 | 2.89% |
| hsa_circ_0067767 | 692 | 10 | 2.89% |
| hsa_circ_0074239 | 692 | 10 | 2.89% |
| hsa_circ_0081497 | 692 | 10 | 2.89% |
| hsa_circ_0085790 | 692 | 10 | 2.89% |
| hsa_circ_0087901 | 692 | 10 | 2.89% |
| hsa_circ_0091187 | 692 | 10 | 2.89% |
| hsa_circ_0101703 | 692 | 10 | 2.89% |
| hsa_circ_0106226 | 692 | 10 | 2.89% |
| hsa_circ_0112758 | 900 | 13 | 2.89% |
| hsa_circ_0007973 | 762 | 11 | 2.89% |
| hsa_circ_0092889 | 762 | 11 | 2.89% |
| hsa_circ_0100808 | 762 | 11 | 2.89% |
| hsa_circ_0121537 | 762 | 11 | 2.89% |
| hsa_circ_0004093 | 693 | 10 | 2.89% |
| hsa_circ_0004530 | 693 | 10 | 2.89% |
| hsa_circ_0018220 | 693 | 10 | 2.89% |

| | | | |
|---|---|---|---|
| hsa_circ_0020056 | 693 | 10 | 2.89% |
| hsa_circ_0030999 | 693 | 10 | 2.89% |
| hsa_circ_0069863 | 693 | 10 | 2.89% |
| hsa_circ_0079735 | 693 | 10 | 2.89% |
| hsa_circ_0084327 | 693 | 10 | 2.89% |
| hsa_circ_0085021 | 693 | 10 | 2.89% |
| hsa_circ_0087248 | 693 | 10 | 2.89% |
| hsa_circ_0099670 | 693 | 10 | 2.89% |
| hsa_circ_0118847 | 693 | 10 | 2.89% |
| hsa_circ_0127048 | 693 | 10 | 2.89% |
| hsa_circ_0113039 | 2288 | 33 | 2.88% |
| hsa_circ_0090167 | 832 | 12 | 2.88% |
| hsa_circ_0113478 | 832 | 12 | 2.88% |
| hsa_circ_0130033 | 971 | 14 | 2.88% |
| hsa_circ_0022270 | 763 | 11 | 2.88% |
| hsa_circ_0065544 | 763 | 11 | 2.88% |
| hsa_circ_0092014 | 763 | 11 | 2.88% |
| hsa_circ_0102069 | 763 | 11 | 2.88% |
| hsa_circ_0103569 | 763 | 11 | 2.88% |
| hsa_circ_0123168 | 763 | 11 | 2.88% |
| hsa_circ_0125879 | 1249 | 18 | 2.88% |
| hsa_circ_0000796 | 694 | 10 | 2.88% |
| hsa_circ_0002929 | 694 | 10 | 2.88% |
| hsa_circ_0004409 | 694 | 10 | 2.88% |
| hsa_circ_0005841 | 694 | 10 | 2.88% |
| hsa_circ_0007182 | 694 | 10 | 2.88% |
| hsa_circ_0024570 | 694 | 10 | 2.88% |
| hsa_circ_0028911 | 694 | 10 | 2.88% |
| hsa_circ_0052778 | 694 | 10 | 2.88% |
| hsa_circ_0059708 | 694 | 10 | 2.88% |
| hsa_circ_0067880 | 694 | 10 | 2.88% |
| hsa_circ_0100626 | 694 | 10 | 2.88% |
| hsa_circ_0103040 | 694 | 10 | 2.88% |
| hsa_circ_0111088 | 694 | 10 | 2.88% |
| hsa_circ_0111820 | 694 | 10 | 2.88% |
| hsa_circ_0123040 | 694 | 10 | 2.88% |
| hsa_circ_0136571 | 694 | 10 | 2.88% |
| hsa_circ_0138647 | 694 | 10 | 2.88% |
| hsa_circ_0016389 | 833 | 12 | 2.88% |
| hsa_circ_0000776 | 764 | 11 | 2.88% |
| hsa_circ_0036778 | 764 | 11 | 2.88% |
| hsa_circ_0095441 | 834 | 12 | 2.88% |
| hsa_circ_0009748 | 695 | 10 | 2.88% |
| hsa_circ_0014232 | 695 | 10 | 2.88% |
| hsa_circ_0038020 | 695 | 10 | 2.88% |
| hsa_circ_0043082 | 695 | 10 | 2.88% |
| hsa_circ_0046132 | 695 | 10 | 2.88% |
| hsa_circ_0049658 | 695 | 10 | 2.88% |
| hsa_circ_0066165 | 695 | 10 | 2.88% |
| hsa_circ_0068795 | 695 | 10 | 2.88% |

| | | | |
|---|---|---|---|
| hsa_circ_0087391 | 695 | 10 | 2.88% |
| hsa_circ_0092417 | 695 | 10 | 2.88% |
| hsa_circ_0094307 | 695 | 10 | 2.88% |
| hsa_circ_0097724 | 695 | 10 | 2.88% |
| hsa_circ_0131645 | 695 | 10 | 2.88% |
| hsa_circ_0138393 | 695 | 10 | 2.88% |
| hsa_circ_0138935 | 695 | 10 | 2.88% |
| hsa_circ_0025203 | 765 | 11 | 2.88% |
| hsa_circ_0033543 | 765 | 11 | 2.88% |
| hsa_circ_0076573 | 765 | 11 | 2.88% |
| hsa_circ_0094904 | 765 | 11 | 2.88% |
| hsa_circ_0012568 | 696 | 10 | 2.87% |
| hsa_circ_0022969 | 696 | 10 | 2.87% |
| hsa_circ_0034553 | 696 | 10 | 2.87% |
| hsa_circ_0037922 | 696 | 10 | 2.87% |
| hsa_circ_0049999 | 696 | 10 | 2.87% |
| hsa_circ_0061085 | 696 | 10 | 2.87% |
| hsa_circ_0066995 | 696 | 10 | 2.87% |
| hsa_circ_0067851 | 696 | 10 | 2.87% |
| hsa_circ_0071860 | 696 | 10 | 2.87% |
| hsa_circ_0075332 | 696 | 10 | 2.87% |
| hsa_circ_0079799 | 696 | 10 | 2.87% |
| hsa_circ_0079973 | 696 | 10 | 2.87% |
| hsa_circ_0080898 | 696 | 10 | 2.87% |
| hsa_circ_0093976 | 696 | 10 | 2.87% |
| hsa_circ_0101278 | 696 | 10 | 2.87% |
| hsa_circ_0107243 | 696 | 10 | 2.87% |
| hsa_circ_0112028 | 696 | 10 | 2.87% |
| hsa_circ_0115863 | 696 | 10 | 2.87% |
| hsa_circ_0118079 | 696 | 10 | 2.87% |
| hsa_circ_0121158 | 696 | 10 | 2.87% |
| hsa_circ_0122561 | 696 | 10 | 2.87% |
| hsa_circ_0127221 | 696 | 10 | 2.87% |
| hsa_circ_0127596 | 696 | 10 | 2.87% |
| hsa_circ_0128091 | 696 | 10 | 2.87% |
| hsa_circ_0134893 | 696 | 10 | 2.87% |
| hsa_circ_0018765 | 905 | 13 | 2.87% |
| hsa_circ_0090960 | 766 | 11 | 2.87% |
| hsa_circ_0116295 | 836 | 12 | 2.87% |
| hsa_circ_0004765 | 697 | 10 | 2.87% |
| hsa_circ_0009713 | 697 | 10 | 2.87% |
| hsa_circ_0013639 | 697 | 10 | 2.87% |
| hsa_circ_0014005 | 697 | 10 | 2.87% |
| hsa_circ_0039306 | 697 | 10 | 2.87% |
| hsa_circ_0041718 | 697 | 10 | 2.87% |
| hsa_circ_0051925 | 697 | 10 | 2.87% |
| hsa_circ_0081487 | 697 | 10 | 2.87% |
| hsa_circ_0094152 | 697 | 10 | 2.87% |
| hsa_circ_0114678 | 697 | 10 | 2.87% |
| hsa_circ_0138885 | 697 | 10 | 2.87% |

| | | | |
|---|---|---|---|
| hsa_circ_0005302 | 767 | 11 | 2.87% |
| hsa_circ_0038156 | 767 | 11 | 2.87% |
| hsa_circ_0050956 | 767 | 11 | 2.87% |
| hsa_circ_0063559 | 767 | 11 | 2.87% |
| hsa_circ_0094837 | 767 | 11 | 2.87% |
| hsa_circ_0005603 | 698 | 10 | 2.87% |
| hsa_circ_0015648 | 698 | 10 | 2.87% |
| hsa_circ_0021127 | 698 | 10 | 2.87% |
| hsa_circ_0051160 | 698 | 10 | 2.87% |
| hsa_circ_0061657 | 698 | 10 | 2.87% |
| hsa_circ_0064352 | 698 | 10 | 2.87% |
| hsa_circ_0080908 | 698 | 10 | 2.87% |
| hsa_circ_0083840 | 698 | 10 | 2.87% |
| hsa_circ_0084510 | 698 | 10 | 2.87% |
| hsa_circ_0105445 | 698 | 10 | 2.87% |
| hsa_circ_0107129 | 698 | 10 | 2.87% |
| hsa_circ_0107233 | 698 | 10 | 2.87% |
| hsa_circ_0004026 | 768 | 11 | 2.86% |
| hsa_circ_0060773 | 768 | 11 | 2.86% |
| hsa_circ_0076773 | 768 | 11 | 2.86% |
| hsa_circ_0099482 | 768 | 11 | 2.86% |
| hsa_circ_0124046 | 768 | 11 | 2.86% |
| hsa_circ_0016579 | 838 | 12 | 2.86% |
| hsa_circ_0017193 | 838 | 12 | 2.86% |
| hsa_circ_0005697 | 699 | 10 | 2.86% |
| hsa_circ_0011875 | 699 | 10 | 2.86% |
| hsa_circ_0024234 | 699 | 10 | 2.86% |
| hsa_circ_0025255 | 699 | 10 | 2.86% |
| hsa_circ_0032274 | 699 | 10 | 2.86% |
| hsa_circ_0037247 | 699 | 10 | 2.86% |
| hsa_circ_0042744 | 699 | 10 | 2.86% |
| hsa_circ_0042760 | 699 | 10 | 2.86% |
| hsa_circ_0049266 | 699 | 10 | 2.86% |
| hsa_circ_0053109 | 699 | 10 | 2.86% |
| hsa_circ_0062418 | 699 | 10 | 2.86% |
| hsa_circ_0063358 | 699 | 10 | 2.86% |
| hsa_circ_0067812 | 699 | 10 | 2.86% |
| hsa_circ_0084521 | 699 | 10 | 2.86% |
| hsa_circ_0085402 | 699 | 10 | 2.86% |
| hsa_circ_0098953 | 699 | 10 | 2.86% |
| hsa_circ_0106357 | 699 | 10 | 2.86% |
| hsa_circ_0109734 | 699 | 10 | 2.86% |
| hsa_circ_0110180 | 699 | 10 | 2.86% |
| hsa_circ_0116057 | 699 | 10 | 2.86% |
| hsa_circ_0125672 | 699 | 10 | 2.86% |
| hsa_circ_0133078 | 699 | 10 | 2.86% |
| hsa_circ_0133227 | 699 | 10 | 2.86% |
| hsa_circ_0133713 | 699 | 10 | 2.86% |
| hsa_circ_0140230 | 699 | 10 | 2.86% |
| hsa_circ_0009088 | 769 | 11 | 2.86% |

| | | | |
|---|---|---|---|
| hsa_circ_0025245 | 769 | 11 | 2.86% |
| hsa_circ_0026025 | 769 | 11 | 2.86% |
| hsa_circ_0032833 | 769 | 11 | 2.86% |
| hsa_circ_0063352 | 769 | 11 | 2.86% |
| hsa_circ_0074998 | 769 | 11 | 2.86% |
| hsa_circ_0117235 | 769 | 11 | 2.86% |
| hsa_circ_0020821 | 909 | 13 | 2.86% |
| hsa_circ_0010980 | 840 | 12 | 2.86% |
| hsa_circ_0132520 | 840 | 12 | 2.86% |
| hsa_circ_0013062 | 770 | 11 | 2.86% |
| hsa_circ_0027686 | 770 | 11 | 2.86% |
| hsa_circ_0050012 | 770 | 11 | 2.86% |
| hsa_circ_0085421 | 770 | 11 | 2.86% |
| hsa_circ_0114962 | 770 | 11 | 2.86% |
| hsa_circ_0118084 | 770 | 11 | 2.86% |
| hsa_circ_0007088 | 700 | 10 | 2.86% |
| hsa_circ_0028038 | 700 | 10 | 2.86% |
| hsa_circ_0042294 | 700 | 10 | 2.86% |
| hsa_circ_0062394 | 700 | 10 | 2.86% |
| hsa_circ_0089473 | 700 | 10 | 2.86% |
| hsa_circ_0114513 | 700 | 10 | 2.86% |
| hsa_circ_0131976 | 700 | 10 | 2.86% |
| hsa_circ_0023097 | 841 | 12 | 2.85% |
| hsa_circ_0058655 | 771 | 11 | 2.85% |
| hsa_circ_0083497 | 771 | 11 | 2.85% |
| hsa_circ_0083669 | 771 | 11 | 2.85% |
| hsa_circ_0099926 | 771 | 11 | 2.85% |
| hsa_circ_0108014 | 771 | 11 | 2.85% |
| hsa_circ_0114489 | 771 | 11 | 2.85% |
| hsa_circ_0128258 | 771 | 11 | 2.85% |
| hsa_circ_0004947 | 701 | 10 | 2.85% |
| hsa_circ_0017280 | 701 | 10 | 2.85% |
| hsa_circ_0055593 | 701 | 10 | 2.85% |
| hsa_circ_0067751 | 701 | 10 | 2.85% |
| hsa_circ_0069167 | 701 | 10 | 2.85% |
| hsa_circ_0072273 | 701 | 10 | 2.85% |
| hsa_circ_0076443 | 701 | 10 | 2.85% |
| hsa_circ_0081565 | 701 | 10 | 2.85% |
| hsa_circ_0081921 | 701 | 10 | 2.85% |
| hsa_circ_0084919 | 701 | 10 | 2.85% |
| hsa_circ_0094431 | 701 | 10 | 2.85% |
| hsa_circ_0101040 | 701 | 10 | 2.85% |
| hsa_circ_0115130 | 701 | 10 | 2.85% |
| hsa_circ_0126440 | 701 | 10 | 2.85% |
| hsa_circ_0021545 | 1052 | 15 | 2.85% |
| hsa_circ_0106398 | 912 | 13 | 2.85% |
| hsa_circ_0116393 | 2386 | 34 | 2.85% |
| hsa_circ_0022203 | 772 | 11 | 2.85% |
| hsa_circ_0029578 | 772 | 11 | 2.85% |
| hsa_circ_0065089 | 772 | 11 | 2.85% |

| | | | |
|---|---|---|---|
| hsa_circ_0093714 | 772 | 11 | 2.85% |
| hsa_circ_0107344 | 772 | 11 | 2.85% |
| hsa_circ_0000106 | 702 | 10 | 2.85% |
| hsa_circ_0012049 | 702 | 10 | 2.85% |
| hsa_circ_0012918 | 702 | 10 | 2.85% |
| hsa_circ_0023205 | 702 | 10 | 2.85% |
| hsa_circ_0023927 | 702 | 10 | 2.85% |
| hsa_circ_0025196 | 702 | 10 | 2.85% |
| hsa_circ_0038146 | 702 | 10 | 2.85% |
| hsa_circ_0045721 | 702 | 10 | 2.85% |
| hsa_circ_0051904 | 702 | 10 | 2.85% |
| hsa_circ_0054191 | 702 | 10 | 2.85% |
| hsa_circ_0059380 | 702 | 10 | 2.85% |
| hsa_circ_0116067 | 702 | 10 | 2.85% |
| hsa_circ_0119108 | 702 | 10 | 2.85% |
| hsa_circ_0132107 | 702 | 10 | 2.85% |
| hsa_circ_0134387 | 702 | 10 | 2.85% |
| hsa_circ_0106555 | 913 | 13 | 2.85% |
| hsa_circ_0036395 | 843 | 12 | 2.85% |
| hsa_circ_0051510 | 843 | 12 | 2.85% |
| hsa_circ_0053954 | 843 | 12 | 2.85% |
| hsa_circ_0070535 | 773 | 11 | 2.85% |
| hsa_circ_0074781 | 773 | 11 | 2.85% |
| hsa_circ_0020479 | 703 | 10 | 2.84% |
| hsa_circ_0050552 | 703 | 10 | 2.84% |
| hsa_circ_0062963 | 703 | 10 | 2.84% |
| hsa_circ_0064504 | 703 | 10 | 2.84% |
| hsa_circ_0064593 | 703 | 10 | 2.84% |
| hsa_circ_0068736 | 703 | 10 | 2.84% |
| hsa_circ_0086834 | 703 | 10 | 2.84% |
| hsa_circ_0101629 | 703 | 10 | 2.84% |
| hsa_circ_0117512 | 703 | 10 | 2.84% |
| hsa_circ_0120485 | 703 | 10 | 2.84% |
| hsa_circ_0121452 | 703 | 10 | 2.84% |
| hsa_circ_0130171 | 703 | 10 | 2.84% |
| hsa_circ_0140484 | 703 | 10 | 2.84% |
| hsa_circ_0100107 | 844 | 12 | 2.84% |
| hsa_circ_0014702 | 774 | 11 | 2.84% |
| hsa_circ_0051601 | 774 | 11 | 2.84% |
| hsa_circ_0055609 | 774 | 11 | 2.84% |
| hsa_circ_0073246 | 774 | 11 | 2.84% |
| hsa_circ_0076613 | 774 | 11 | 2.84% |
| hsa_circ_0124698 | 774 | 11 | 2.84% |
| hsa_circ_0123325 | 4856 | 69 | 2.84% |
| hsa_circ_0119779 | 915 | 13 | 2.84% |
| hsa_circ_0009862 | 704 | 10 | 2.84% |
| hsa_circ_0056908 | 704 | 10 | 2.84% |
| hsa_circ_0070598 | 704 | 10 | 2.84% |
| hsa_circ_0076938 | 704 | 10 | 2.84% |
| hsa_circ_0083400 | 704 | 10 | 2.84% |

| | | | |
|---|---|---|---|
| hsa_circ_0084997 | 704 | 10 | 2.84% |
| hsa_circ_0087549 | 704 | 10 | 2.84% |
| hsa_circ_0105338 | 704 | 10 | 2.84% |
| hsa_circ_0116197 | 704 | 10 | 2.84% |
| hsa_circ_0120607 | 704 | 10 | 2.84% |
| hsa_circ_0121768 | 704 | 10 | 2.84% |
| hsa_circ_0123836 | 704 | 10 | 2.84% |
| hsa_circ_0011459 | 986 | 14 | 2.84% |
| hsa_circ_0021969 | 986 | 14 | 2.84% |
| hsa_circ_0026391 | 775 | 11 | 2.84% |
| hsa_circ_0042766 | 775 | 11 | 2.84% |
| hsa_circ_0063752 | 775 | 11 | 2.84% |
| hsa_circ_0095438 | 775 | 11 | 2.84% |
| hsa_circ_0106981 | 775 | 11 | 2.84% |
| hsa_circ_0135304 | 775 | 11 | 2.84% |
| hsa_circ_0101962 | 916 | 13 | 2.84% |
| hsa_circ_0015539 | 705 | 10 | 2.84% |
| hsa_circ_0016204 | 705 | 10 | 2.84% |
| hsa_circ_0028483 | 705 | 10 | 2.84% |
| hsa_circ_0089278 | 705 | 10 | 2.84% |
| hsa_circ_0092661 | 705 | 10 | 2.84% |
| hsa_circ_0118962 | 705 | 10 | 2.84% |
| hsa_circ_0123559 | 705 | 10 | 2.84% |
| hsa_circ_0135309 | 705 | 10 | 2.84% |
| hsa_circ_0006555 | 776 | 11 | 2.84% |
| hsa_circ_0025214 | 776 | 11 | 2.84% |
| hsa_circ_0029470 | 776 | 11 | 2.84% |
| hsa_circ_0035553 | 776 | 11 | 2.84% |
| hsa_circ_0051156 | 776 | 11 | 2.84% |
| hsa_circ_0083070 | 776 | 11 | 2.84% |
| hsa_circ_0121833 | 776 | 11 | 2.84% |
| hsa_circ_0123058 | 776 | 11 | 2.84% |
| hsa_circ_0100981 | 847 | 12 | 2.83% |
| hsa_circ_0015126 | 706 | 10 | 2.83% |
| hsa_circ_0024549 | 706 | 10 | 2.83% |
| hsa_circ_0031312 | 706 | 10 | 2.83% |
| hsa_circ_0082302 | 706 | 10 | 2.83% |
| hsa_circ_0104027 | 706 | 10 | 2.83% |
| hsa_circ_0109681 | 706 | 10 | 2.83% |
| hsa_circ_0115868 | 706 | 10 | 2.83% |
| hsa_circ_0123466 | 706 | 10 | 2.83% |
| hsa_circ_0123548 | 706 | 10 | 2.83% |
| hsa_circ_0129974 | 706 | 10 | 2.83% |
| hsa_circ_0021532 | 777 | 11 | 2.83% |
| hsa_circ_0040274 | 777 | 11 | 2.83% |
| hsa_circ_0054178 | 777 | 11 | 2.83% |
| hsa_circ_0003263 | 848 | 12 | 2.83% |
| hsa_circ_0008807 | 848 | 12 | 2.83% |
| hsa_circ_0017292 | 848 | 12 | 2.83% |
| hsa_circ_0026157 | 848 | 12 | 2.83% |

| | | | |
|---|---|---|---|
| hsa_circ_0036538 | 848 | 12 | 2.83% |
| hsa_circ_0105281 | 848 | 12 | 2.83% |
| hsa_circ_0038400 | 1979 | 28 | 2.83% |
| hsa_circ_0004460 | 707 | 10 | 2.83% |
| hsa_circ_0007796 | 707 | 10 | 2.83% |
| hsa_circ_0008967 | 707 | 10 | 2.83% |
| hsa_circ_0015532 | 707 | 10 | 2.83% |
| hsa_circ_0027236 | 707 | 10 | 2.83% |
| hsa_circ_0037735 | 707 | 10 | 2.83% |
| hsa_circ_0042552 | 707 | 10 | 2.83% |
| hsa_circ_0047397 | 707 | 10 | 2.83% |
| hsa_circ_0053888 | 707 | 10 | 2.83% |
| hsa_circ_0065722 | 707 | 10 | 2.83% |
| hsa_circ_0067772 | 707 | 10 | 2.83% |
| hsa_circ_0072836 | 707 | 10 | 2.83% |
| hsa_circ_0080652 | 707 | 10 | 2.83% |
| hsa_circ_0092731 | 707 | 10 | 2.83% |
| hsa_circ_0102370 | 707 | 10 | 2.83% |
| hsa_circ_0102786 | 707 | 10 | 2.83% |
| hsa_circ_0104212 | 707 | 10 | 2.83% |
| hsa_circ_0110541 | 707 | 10 | 2.83% |
| hsa_circ_0128645 | 707 | 10 | 2.83% |
| hsa_circ_0131538 | 707 | 10 | 2.83% |
| hsa_circ_0139626 | 707 | 10 | 2.83% |
| hsa_circ_0005010 | 778 | 11 | 2.83% |
| hsa_circ_0015928 | 778 | 11 | 2.83% |
| hsa_circ_0030730 | 778 | 11 | 2.83% |
| hsa_circ_0062711 | 778 | 11 | 2.83% |
| hsa_circ_0079133 | 778 | 11 | 2.83% |
| hsa_circ_0100298 | 778 | 11 | 2.83% |
| hsa_circ_0137031 | 778 | 11 | 2.83% |
| hsa_circ_0117528 | 1981 | 28 | 2.83% |
| hsa_circ_0126035 | 849 | 12 | 2.83% |
| hsa_circ_0023158 | 708 | 10 | 2.82% |
| hsa_circ_0032557 | 708 | 10 | 2.82% |
| hsa_circ_0035109 | 708 | 10 | 2.82% |
| hsa_circ_0054403 | 708 | 10 | 2.82% |
| hsa_circ_0058152 | 708 | 10 | 2.82% |
| hsa_circ_0068518 | 708 | 10 | 2.82% |
| hsa_circ_0071247 | 708 | 10 | 2.82% |
| hsa_circ_0081493 | 708 | 10 | 2.82% |
| hsa_circ_0105156 | 708 | 10 | 2.82% |
| hsa_circ_0125768 | 708 | 10 | 2.82% |
| hsa_circ_0131884 | 708 | 10 | 2.82% |
| hsa_circ_0132095 | 708 | 10 | 2.82% |
| hsa_circ_0139238 | 708 | 10 | 2.82% |
| hsa_circ_0009582 | 779 | 11 | 2.82% |
| hsa_circ_0041077 | 779 | 11 | 2.82% |
| hsa_circ_0044748 | 779 | 11 | 2.82% |
| hsa_circ_0060468 | 779 | 11 | 2.82% |

| | | | |
|---|---|---|---|
| hsa_circ_0061763 | 779 | 11 | 2.82% |
| hsa_circ_0087908 | 779 | 11 | 2.82% |
| hsa_circ_0140589 | 1204 | 17 | 2.82% |
| hsa_circ_0039393 | 850 | 12 | 2.82% |
| hsa_circ_0111241 | 921 | 13 | 2.82% |
| hsa_circ_0071041 | 992 | 14 | 2.82% |
| hsa_circ_0022743 | 709 | 10 | 2.82% |
| hsa_circ_0023633 | 709 | 10 | 2.82% |
| hsa_circ_0036521 | 709 | 10 | 2.82% |
| hsa_circ_0040007 | 709 | 10 | 2.82% |
| hsa_circ_0047402 | 709 | 10 | 2.82% |
| hsa_circ_0058909 | 709 | 10 | 2.82% |
| hsa_circ_0082088 | 709 | 10 | 2.82% |
| hsa_circ_0086838 | 709 | 10 | 2.82% |
| hsa_circ_0091761 | 709 | 10 | 2.82% |
| hsa_circ_0096224 | 709 | 10 | 2.82% |
| hsa_circ_0098762 | 709 | 10 | 2.82% |
| hsa_circ_0101737 | 709 | 10 | 2.82% |
| hsa_circ_0122649 | 709 | 10 | 2.82% |
| hsa_circ_0098857 | 780 | 11 | 2.82% |
| hsa_circ_0126544 | 780 | 11 | 2.82% |
| hsa_circ_0076957 | 993 | 14 | 2.82% |
| hsa_circ_0115827 | 923 | 13 | 2.82% |
| hsa_circ_0043470 | 852 | 12 | 2.82% |
| hsa_circ_0085874 | 852 | 12 | 2.82% |
| hsa_circ_0120362 | 852 | 12 | 2.82% |
| hsa_circ_0019428 | 781 | 11 | 2.82% |
| hsa_circ_0047075 | 781 | 11 | 2.82% |
| hsa_circ_0091574 | 781 | 11 | 2.82% |
| hsa_circ_0104774 | 781 | 11 | 2.82% |
| hsa_circ_0129756 | 781 | 11 | 2.82% |
| hsa_circ_0132138 | 781 | 11 | 2.82% |
| hsa_circ_0011097 | 710 | 10 | 2.82% |
| hsa_circ_0056347 | 710 | 10 | 2.82% |
| hsa_circ_0060949 | 710 | 10 | 2.82% |
| hsa_circ_0102882 | 710 | 10 | 2.82% |
| hsa_circ_0108982 | 710 | 10 | 2.82% |
| hsa_circ_0116708 | 710 | 10 | 2.82% |
| hsa_circ_0118626 | 710 | 10 | 2.82% |
| hsa_circ_0131518 | 710 | 10 | 2.82% |
| hsa_circ_0131856 | 710 | 10 | 2.82% |
| hsa_circ_0136331 | 710 | 10 | 2.82% |
| hsa_circ_0137445 | 710 | 10 | 2.82% |
| hsa_circ_0021986 | 924 | 13 | 2.81% |
| hsa_circ_0054841 | 853 | 12 | 2.81% |
| hsa_circ_0060743 | 853 | 12 | 2.81% |
| hsa_circ_0123991 | 853 | 12 | 2.81% |
| hsa_circ_0072612 | 782 | 11 | 2.81% |
| hsa_circ_0073430 | 782 | 11 | 2.81% |
| hsa_circ_0094362 | 782 | 11 | 2.81% |

| | | | |
|---|---|---|---|
| hsa_circ_0004523 | 711 | 10 | 2.81% |
| hsa_circ_0018760 | 711 | 10 | 2.81% |
| hsa_circ_0028847 | 711 | 10 | 2.81% |
| hsa_circ_0039521 | 711 | 10 | 2.81% |
| hsa_circ_0058907 | 711 | 10 | 2.81% |
| hsa_circ_0087991 | 711 | 10 | 2.81% |
| hsa_circ_0088992 | 711 | 10 | 2.81% |
| hsa_circ_0091283 | 711 | 10 | 2.81% |
| hsa_circ_0123837 | 711 | 10 | 2.81% |
| hsa_circ_0132830 | 711 | 10 | 2.81% |
| hsa_circ_0123438 | 925 | 13 | 2.81% |
| hsa_circ_0000087 | 783 | 11 | 2.81% |
| hsa_circ_0001430 | 783 | 11 | 2.81% |
| hsa_circ_0004286 | 783 | 11 | 2.81% |
| hsa_circ_0022805 | 783 | 11 | 2.81% |
| hsa_circ_0006689 | 712 | 10 | 2.81% |
| hsa_circ_0007095 | 712 | 10 | 2.81% |
| hsa_circ_0015367 | 712 | 10 | 2.81% |
| hsa_circ_0022899 | 712 | 10 | 2.81% |
| hsa_circ_0025752 | 712 | 10 | 2.81% |
| hsa_circ_0026468 | 712 | 10 | 2.81% |
| hsa_circ_0032858 | 712 | 10 | 2.81% |
| hsa_circ_0037381 | 712 | 10 | 2.81% |
| hsa_circ_0045282 | 712 | 10 | 2.81% |
| hsa_circ_0047913 | 712 | 10 | 2.81% |
| hsa_circ_0067281 | 712 | 10 | 2.81% |
| hsa_circ_0068174 | 712 | 10 | 2.81% |
| hsa_circ_0072443 | 712 | 10 | 2.81% |
| hsa_circ_0075126 | 712 | 10 | 2.81% |
| hsa_circ_0095682 | 712 | 10 | 2.81% |
| hsa_circ_0101976 | 712 | 10 | 2.81% |
| hsa_circ_0104438 | 712 | 10 | 2.81% |
| hsa_circ_0107170 | 712 | 10 | 2.81% |
| hsa_circ_0111522 | 712 | 10 | 2.81% |
| hsa_circ_0117080 | 712 | 10 | 2.81% |
| hsa_circ_0118829 | 712 | 10 | 2.81% |
| hsa_circ_0133893 | 712 | 10 | 2.81% |
| hsa_circ_0015552 | 926 | 13 | 2.81% |
| hsa_circ_0008640 | 855 | 12 | 2.81% |
| hsa_circ_0071522 | 855 | 12 | 2.81% |
| hsa_circ_0027682 | 784 | 11 | 2.81% |
| hsa_circ_0042272 | 784 | 11 | 2.81% |
| hsa_circ_0085804 | 784 | 11 | 2.81% |
| hsa_circ_0101999 | 784 | 11 | 2.81% |
| hsa_circ_0009094 | 713 | 10 | 2.81% |
| hsa_circ_0010510 | 713 | 10 | 2.81% |
| hsa_circ_0043481 | 713 | 10 | 2.81% |
| hsa_circ_0045885 | 713 | 10 | 2.81% |
| hsa_circ_0066319 | 713 | 10 | 2.81% |
| hsa_circ_0079986 | 713 | 10 | 2.81% |

| | | | |
|---|---|---|---|
| hsa_circ_0089520 | 713 | 10 | 2.81% |
| hsa_circ_0093229 | 713 | 10 | 2.81% |
| hsa_circ_0098968 | 713 | 10 | 2.81% |
| hsa_circ_0102331 | 713 | 10 | 2.81% |
| hsa_circ_0113636 | 713 | 10 | 2.81% |
| hsa_circ_0126165 | 713 | 10 | 2.81% |
| hsa_circ_0113900 | 856 | 12 | 2.80% |
| hsa_circ_0004824 | 785 | 11 | 2.80% |
| hsa_circ_0026569 | 785 | 11 | 2.80% |
| hsa_circ_0037969 | 785 | 11 | 2.80% |
| hsa_circ_0046423 | 785 | 11 | 2.80% |
| hsa_circ_0019709 | 714 | 10 | 2.80% |
| hsa_circ_0025024 | 714 | 10 | 2.80% |
| hsa_circ_0046372 | 714 | 10 | 2.80% |
| hsa_circ_0055342 | 714 | 10 | 2.80% |
| hsa_circ_0076763 | 714 | 10 | 2.80% |
| hsa_circ_0082634 | 714 | 10 | 2.80% |
| hsa_circ_0084187 | 714 | 10 | 2.80% |
| hsa_circ_0115046 | 714 | 10 | 2.80% |
| hsa_circ_0117182 | 714 | 10 | 2.80% |
| hsa_circ_0125410 | 714 | 10 | 2.80% |
| hsa_circ_0130236 | 714 | 10 | 2.80% |
| hsa_circ_0130900 | 714 | 10 | 2.80% |
| hsa_circ_0135254 | 714 | 10 | 2.80% |
| hsa_circ_0135500 | 714 | 10 | 2.80% |
| hsa_circ_0135927 | 714 | 10 | 2.80% |
| hsa_circ_0136340 | 714 | 10 | 2.80% |
| hsa_circ_0127851 | 857 | 12 | 2.80% |
| hsa_circ_0048271 | 1000 | 14 | 2.80% |
| hsa_circ_0004843 | 786 | 11 | 2.80% |
| hsa_circ_0030507 | 786 | 11 | 2.80% |
| hsa_circ_0062638 | 786 | 11 | 2.80% |
| hsa_circ_0081430 | 786 | 11 | 2.80% |
| hsa_circ_0131558 | 2430 | 34 | 2.80% |
| hsa_circ_0000073 | 715 | 10 | 2.80% |
| hsa_circ_0010701 | 715 | 10 | 2.80% |
| hsa_circ_0014969 | 715 | 10 | 2.80% |
| hsa_circ_0016553 | 715 | 10 | 2.80% |
| hsa_circ_0033219 | 715 | 10 | 2.80% |
| hsa_circ_0035055 | 715 | 10 | 2.80% |
| hsa_circ_0037124 | 715 | 10 | 2.80% |
| hsa_circ_0037213 | 715 | 10 | 2.80% |
| hsa_circ_0046464 | 715 | 10 | 2.80% |
| hsa_circ_0072428 | 715 | 10 | 2.80% |
| hsa_circ_0076624 | 715 | 10 | 2.80% |
| hsa_circ_0096476 | 715 | 10 | 2.80% |
| hsa_circ_0098602 | 715 | 10 | 2.80% |
| hsa_circ_0104434 | 715 | 10 | 2.80% |
| hsa_circ_0110191 | 715 | 10 | 2.80% |
| hsa_circ_0112210 | 715 | 10 | 2.80% |

| | | | |
|---|---|---|---|
| hsa_circ_0113380 | 715 | 10 | 2.80% |
| hsa_circ_0113632 | 715 | 10 | 2.80% |
| hsa_circ_0116914 | 715 | 10 | 2.80% |
| hsa_circ_0117208 | 715 | 10 | 2.80% |
| hsa_circ_0122479 | 715 | 10 | 2.80% |
| hsa_circ_0140693 | 715 | 10 | 2.80% |
| hsa_circ_0039176 | 787 | 11 | 2.80% |
| hsa_circ_0095959 | 787 | 11 | 2.80% |
| hsa_circ_0115522 | 787 | 11 | 2.80% |
| hsa_circ_0042258 | 859 | 12 | 2.79% |
| hsa_circ_0088712 | 1217 | 17 | 2.79% |
| hsa_circ_0004612 | 716 | 10 | 2.79% |
| hsa_circ_0007700 | 716 | 10 | 2.79% |
| hsa_circ_0032483 | 716 | 10 | 2.79% |
| hsa_circ_0053767 | 716 | 10 | 2.79% |
| hsa_circ_0064515 | 716 | 10 | 2.79% |
| hsa_circ_0069853 | 716 | 10 | 2.79% |
| hsa_circ_0077624 | 716 | 10 | 2.79% |
| hsa_circ_0079844 | 716 | 10 | 2.79% |
| hsa_circ_0081393 | 716 | 10 | 2.79% |
| hsa_circ_0105174 | 716 | 10 | 2.79% |
| hsa_circ_0107752 | 716 | 10 | 2.79% |
| hsa_circ_0108051 | 716 | 10 | 2.79% |
| hsa_circ_0125278 | 716 | 10 | 2.79% |
| hsa_circ_0129699 | 716 | 10 | 2.79% |
| hsa_circ_0029102 | 788 | 11 | 2.79% |
| hsa_circ_0049434 | 788 | 11 | 2.79% |
| hsa_circ_0051798 | 788 | 11 | 2.79% |
| hsa_circ_0066778 | 788 | 11 | 2.79% |
| hsa_circ_0079605 | 788 | 11 | 2.79% |
| hsa_circ_0093679 | 788 | 11 | 2.79% |
| hsa_circ_0043838 | 860 | 12 | 2.79% |
| hsa_circ_0094899 | 860 | 12 | 2.79% |
| hsa_circ_0139605 | 860 | 12 | 2.79% |
| hsa_circ_0003212 | 932 | 13 | 2.79% |
| hsa_circ_0020819 | 932 | 13 | 2.79% |
| hsa_circ_0132296 | 932 | 13 | 2.79% |
| hsa_circ_0003045 | 717 | 10 | 2.79% |
| hsa_circ_0019583 | 717 | 10 | 2.79% |
| hsa_circ_0042031 | 717 | 10 | 2.79% |
| hsa_circ_0046512 | 717 | 10 | 2.79% |
| hsa_circ_0048137 | 717 | 10 | 2.79% |
| hsa_circ_0063241 | 717 | 10 | 2.79% |
| hsa_circ_0073056 | 717 | 10 | 2.79% |
| hsa_circ_0085930 | 717 | 10 | 2.79% |
| hsa_circ_0092425 | 717 | 10 | 2.79% |
| hsa_circ_0095297 | 717 | 10 | 2.79% |
| hsa_circ_0107844 | 717 | 10 | 2.79% |
| hsa_circ_0131539 | 717 | 10 | 2.79% |
| hsa_circ_0012386 | 789 | 11 | 2.79% |

| | | | |
|---|---|---|---|
| hsa_circ_0041625 | 789 | 11 | 2.79% |
| hsa_circ_0051426 | 789 | 11 | 2.79% |
| hsa_circ_0057453 | 789 | 11 | 2.79% |
| hsa_circ_0095431 | 789 | 11 | 2.79% |
| hsa_circ_0110951 | 789 | 11 | 2.79% |
| hsa_circ_0117441 | 789 | 11 | 2.79% |
| hsa_circ_0050403 | 861 | 12 | 2.79% |
| hsa_circ_0021980 | 933 | 13 | 2.79% |
| hsa_circ_0081922 | 933 | 13 | 2.79% |
| hsa_circ_0122261 | 933 | 13 | 2.79% |
| hsa_circ_0001616 | 718 | 10 | 2.79% |
| hsa_circ_0006155 | 718 | 10 | 2.79% |
| hsa_circ_0007652 | 718 | 10 | 2.79% |
| hsa_circ_0008892 | 718 | 10 | 2.79% |
| hsa_circ_0013548 | 718 | 10 | 2.79% |
| hsa_circ_0013786 | 718 | 10 | 2.79% |
| hsa_circ_0071641 | 718 | 10 | 2.79% |
| hsa_circ_0103669 | 718 | 10 | 2.79% |
| hsa_circ_0119203 | 718 | 10 | 2.79% |
| hsa_circ_0015180 | 790 | 11 | 2.78% |
| hsa_circ_0026689 | 790 | 11 | 2.78% |
| hsa_circ_0050970 | 790 | 11 | 2.78% |
| hsa_circ_0102187 | 790 | 11 | 2.78% |
| hsa_circ_0102763 | 790 | 11 | 2.78% |
| hsa_circ_0104764 | 790 | 11 | 2.78% |
| hsa_circ_0121457 | 790 | 11 | 2.78% |
| hsa_circ_0039011 | 862 | 12 | 2.78% |
| hsa_circ_0025765 | 1006 | 14 | 2.78% |
| hsa_circ_0002256 | 719 | 10 | 2.78% |
| hsa_circ_0022584 | 719 | 10 | 2.78% |
| hsa_circ_0025035 | 719 | 10 | 2.78% |
| hsa_circ_0028888 | 719 | 10 | 2.78% |
| hsa_circ_0031022 | 719 | 10 | 2.78% |
| hsa_circ_0043261 | 719 | 10 | 2.78% |
| hsa_circ_0043908 | 719 | 10 | 2.78% |
| hsa_circ_0066351 | 719 | 10 | 2.78% |
| hsa_circ_0071760 | 719 | 10 | 2.78% |
| hsa_circ_0114043 | 719 | 10 | 2.78% |
| hsa_circ_0114306 | 719 | 10 | 2.78% |
| hsa_circ_0127045 | 719 | 10 | 2.78% |
| hsa_circ_0138113 | 719 | 10 | 2.78% |
| hsa_circ_0083022 | 791 | 11 | 2.78% |
| hsa_circ_0084377 | 791 | 11 | 2.78% |
| hsa_circ_0129763 | 791 | 11 | 2.78% |
| hsa_circ_0116286 | 863 | 12 | 2.78% |
| hsa_circ_0132327 | 863 | 12 | 2.78% |
| hsa_circ_0046077 | 936 | 13 | 2.78% |
| hsa_circ_0019143 | 864 | 12 | 2.78% |
| hsa_circ_0072101 | 864 | 12 | 2.78% |
| hsa_circ_0004320 | 792 | 11 | 2.78% |

| | | | |
|---|---|---|---|
| hsa_circ_0005690 | 792 | 11 | 2.78% |
| hsa_circ_0007867 | 792 | 11 | 2.78% |
| hsa_circ_0039236 | 792 | 11 | 2.78% |
| hsa_circ_0043583 | 792 | 11 | 2.78% |
| hsa_circ_0005014 | 720 | 10 | 2.78% |
| hsa_circ_0023550 | 720 | 10 | 2.78% |
| hsa_circ_0046527 | 720 | 10 | 2.78% |
| hsa_circ_0069084 | 720 | 10 | 2.78% |
| hsa_circ_0070020 | 720 | 10 | 2.78% |
| hsa_circ_0072840 | 720 | 10 | 2.78% |
| hsa_circ_0083891 | 720 | 10 | 2.78% |
| hsa_circ_0088228 | 720 | 10 | 2.78% |
| hsa_circ_0103737 | 720 | 10 | 2.78% |
| hsa_circ_0112057 | 720 | 10 | 2.78% |
| hsa_circ_0114369 | 720 | 10 | 2.78% |
| hsa_circ_0123301 | 720 | 10 | 2.78% |
| hsa_circ_0135494 | 720 | 10 | 2.78% |
| hsa_circ_0072584 | 1513 | 21 | 2.78% |
| hsa_circ_0036993 | 1009 | 14 | 2.78% |
| hsa_circ_0074491 | 1009 | 14 | 2.78% |
| hsa_circ_0076129 | 865 | 12 | 2.77% |
| hsa_circ_0026385 | 721 | 10 | 2.77% |
| hsa_circ_0031526 | 721 | 10 | 2.77% |
| hsa_circ_0078019 | 721 | 10 | 2.77% |
| hsa_circ_0112309 | 721 | 10 | 2.77% |
| hsa_circ_0118891 | 721 | 10 | 2.77% |
| hsa_circ_0069138 | 866 | 12 | 2.77% |
| hsa_circ_0086662 | 866 | 12 | 2.77% |
| hsa_circ_0135424 | 866 | 12 | 2.77% |
| hsa_circ_0015408 | 794 | 11 | 2.77% |
| hsa_circ_0024583 | 794 | 11 | 2.77% |
| hsa_circ_0057660 | 794 | 11 | 2.77% |
| hsa_circ_0121819 | 794 | 11 | 2.77% |
| hsa_circ_0134307 | 794 | 11 | 2.77% |
| hsa_circ_0006362 | 722 | 10 | 2.77% |
| hsa_circ_0010866 | 722 | 10 | 2.77% |
| hsa_circ_0031773 | 722 | 10 | 2.77% |
| hsa_circ_0050215 | 722 | 10 | 2.77% |
| hsa_circ_0059908 | 722 | 10 | 2.77% |
| hsa_circ_0068292 | 722 | 10 | 2.77% |
| hsa_circ_0069609 | 722 | 10 | 2.77% |
| hsa_circ_0079833 | 722 | 10 | 2.77% |
| hsa_circ_0083692 | 722 | 10 | 2.77% |
| hsa_circ_0106335 | 722 | 10 | 2.77% |
| hsa_circ_0113095 | 722 | 10 | 2.77% |
| hsa_circ_0116517 | 722 | 10 | 2.77% |
| hsa_circ_0128913 | 722 | 10 | 2.77% |
| hsa_circ_0008210 | 795 | 11 | 2.77% |
| hsa_circ_0050180 | 795 | 11 | 2.77% |
| hsa_circ_0130666 | 795 | 11 | 2.77% |

| | | | |
|---|---|---|---|
| hsa_circ_0007324 | 723 | 10 | 2.77% |
| hsa_circ_0010174 | 723 | 10 | 2.77% |
| hsa_circ_0048439 | 723 | 10 | 2.77% |
| hsa_circ_0052866 | 723 | 10 | 2.77% |
| hsa_circ_0054327 | 723 | 10 | 2.77% |
| hsa_circ_0056763 | 723 | 10 | 2.77% |
| hsa_circ_0058635 | 723 | 10 | 2.77% |
| hsa_circ_0061209 | 723 | 10 | 2.77% |
| hsa_circ_0063785 | 723 | 10 | 2.77% |
| hsa_circ_0066239 | 723 | 10 | 2.77% |
| hsa_circ_0069733 | 723 | 10 | 2.77% |
| hsa_circ_0087305 | 723 | 10 | 2.77% |
| hsa_circ_0101540 | 723 | 10 | 2.77% |
| hsa_circ_0112058 | 723 | 10 | 2.77% |
| hsa_circ_0114902 | 723 | 10 | 2.77% |
| hsa_circ_0121112 | 723 | 10 | 2.77% |
| hsa_circ_0133370 | 723 | 10 | 2.77% |
| hsa_circ_0134436 | 723 | 10 | 2.77% |
| hsa_circ_0061501 | 940 | 13 | 2.77% |
| hsa_circ_0075722 | 868 | 12 | 2.76% |
| hsa_circ_0013224 | 796 | 11 | 2.76% |
| hsa_circ_0042237 | 796 | 11 | 2.76% |
| hsa_circ_0067517 | 796 | 11 | 2.76% |
| hsa_circ_0076057 | 796 | 11 | 2.76% |
| hsa_circ_0096492 | 796 | 11 | 2.76% |
| hsa_circ_0117289 | 796 | 11 | 2.76% |
| hsa_circ_0127129 | 796 | 11 | 2.76% |
| hsa_circ_0012876 | 724 | 10 | 2.76% |
| hsa_circ_0025442 | 724 | 10 | 2.76% |
| hsa_circ_0046778 | 724 | 10 | 2.76% |
| hsa_circ_0066270 | 724 | 10 | 2.76% |
| hsa_circ_0085337 | 724 | 10 | 2.76% |
| hsa_circ_0098081 | 724 | 10 | 2.76% |
| hsa_circ_0104731 | 724 | 10 | 2.76% |
| hsa_circ_0111343 | 724 | 10 | 2.76% |
| hsa_circ_0111554 | 724 | 10 | 2.76% |
| hsa_circ_0114149 | 724 | 10 | 2.76% |
| hsa_circ_0131900 | 724 | 10 | 2.76% |
| hsa_circ_0132833 | 724 | 10 | 2.76% |
| hsa_circ_0140719 | 724 | 10 | 2.76% |
| hsa_circ_0014136 | 869 | 12 | 2.76% |
| hsa_circ_0038536 | 797 | 11 | 2.76% |
| hsa_circ_0039022 | 797 | 11 | 2.76% |
| hsa_circ_0054075 | 797 | 11 | 2.76% |
| hsa_circ_0073263 | 797 | 11 | 2.76% |
| hsa_circ_0005481 | 870 | 12 | 2.76% |
| hsa_circ_0070624 | 870 | 12 | 2.76% |
| hsa_circ_0092102 | 870 | 12 | 2.76% |
| hsa_circ_0105391 | 870 | 12 | 2.76% |
| hsa_circ_0012670 | 725 | 10 | 2.76% |

| | | | |
|---|---|---|---|
| hsa_circ_0030244 | 725 | 10 | 2.76% |
| hsa_circ_0038947 | 725 | 10 | 2.76% |
| hsa_circ_0052810 | 725 | 10 | 2.76% |
| hsa_circ_0066901 | 725 | 10 | 2.76% |
| hsa_circ_0073762 | 725 | 10 | 2.76% |
| hsa_circ_0085743 | 725 | 10 | 2.76% |
| hsa_circ_0096810 | 725 | 10 | 2.76% |
| hsa_circ_0097112 | 725 | 10 | 2.76% |
| hsa_circ_0106755 | 725 | 10 | 2.76% |
| hsa_circ_0061954 | 798 | 11 | 2.76% |
| hsa_circ_0086552 | 798 | 11 | 2.76% |
| hsa_circ_0101472 | 798 | 11 | 2.76% |
| hsa_circ_0126685 | 798 | 11 | 2.76% |
| hsa_circ_0133636 | 798 | 11 | 2.76% |
| hsa_circ_0137729 | 798 | 11 | 2.76% |
| hsa_circ_0138752 | 798 | 11 | 2.76% |
| hsa_circ_0139015 | 798 | 11 | 2.76% |
| hsa_circ_0005463 | 2613 | 36 | 2.76% |
| hsa_circ_0025444 | 871 | 12 | 2.76% |
| hsa_circ_0007419 | 726 | 10 | 2.75% |
| hsa_circ_0017448 | 726 | 10 | 2.75% |
| hsa_circ_0017828 | 726 | 10 | 2.75% |
| hsa_circ_0045173 | 726 | 10 | 2.75% |
| hsa_circ_0055273 | 726 | 10 | 2.75% |
| hsa_circ_0067059 | 726 | 10 | 2.75% |
| hsa_circ_0077597 | 726 | 10 | 2.75% |
| hsa_circ_0080199 | 726 | 10 | 2.75% |
| hsa_circ_0090305 | 726 | 10 | 2.75% |
| hsa_circ_0113468 | 726 | 10 | 2.75% |
| hsa_circ_0124136 | 726 | 10 | 2.75% |
| hsa_circ_0136080 | 726 | 10 | 2.75% |
| hsa_circ_0051345 | 944 | 13 | 2.75% |
| hsa_circ_0006000 | 799 | 11 | 2.75% |
| hsa_circ_0070932 | 799 | 11 | 2.75% |
| hsa_circ_0081464 | 799 | 11 | 2.75% |
| hsa_circ_0085117 | 799 | 11 | 2.75% |
| hsa_circ_0091407 | 799 | 11 | 2.75% |
| hsa_circ_0129782 | 799 | 11 | 2.75% |
| hsa_circ_0071733 | 1017 | 14 | 2.75% |
| hsa_circ_0032663 | 1235 | 17 | 2.75% |
| hsa_circ_0032340 | 727 | 10 | 2.75% |
| hsa_circ_0051767 | 727 | 10 | 2.75% |
| hsa_circ_0069790 | 727 | 10 | 2.75% |
| hsa_circ_0073461 | 727 | 10 | 2.75% |
| hsa_circ_0075583 | 727 | 10 | 2.75% |
| hsa_circ_0079033 | 727 | 10 | 2.75% |
| hsa_circ_0080070 | 727 | 10 | 2.75% |
| hsa_circ_0081370 | 727 | 10 | 2.75% |
| hsa_circ_0082448 | 727 | 10 | 2.75% |
| hsa_circ_0101418 | 727 | 10 | 2.75% |

| | | | |
|---|---|---|---|
| hsa_circ_0109470 | 727 | 10 | 2.75% |
| hsa_circ_0113398 | 727 | 10 | 2.75% |
| hsa_circ_0114086 | 727 | 10 | 2.75% |
| hsa_circ_0125018 | 727 | 10 | 2.75% |
| hsa_circ_0126810 | 727 | 10 | 2.75% |
| hsa_circ_0127365 | 727 | 10 | 2.75% |
| hsa_circ_0139275 | 727 | 10 | 2.75% |
| hsa_circ_0036990 | 1018 | 14 | 2.75% |
| hsa_circ_0009601 | 800 | 11 | 2.75% |
| hsa_circ_0014701 | 800 | 11 | 2.75% |
| hsa_circ_0043998 | 800 | 11 | 2.75% |
| hsa_circ_0092351 | 800 | 11 | 2.75% |
| hsa_circ_0115436 | 800 | 11 | 2.75% |
| hsa_circ_0135198 | 800 | 11 | 2.75% |
| hsa_circ_0138534 | 800 | 11 | 2.75% |
| hsa_circ_0005410 | 873 | 12 | 2.75% |
| hsa_circ_0083401 | 873 | 12 | 2.75% |
| hsa_circ_0113463 | 873 | 12 | 2.75% |
| hsa_circ_0052487 | 728 | 10 | 2.75% |
| hsa_circ_0069146 | 728 | 10 | 2.75% |
| hsa_circ_0079991 | 728 | 10 | 2.75% |
| hsa_circ_0096311 | 728 | 10 | 2.75% |
| hsa_circ_0098449 | 728 | 10 | 2.75% |
| hsa_circ_0099381 | 728 | 10 | 2.75% |
| hsa_circ_0105814 | 728 | 10 | 2.75% |
| hsa_circ_0105991 | 728 | 10 | 2.75% |
| hsa_circ_0113873 | 728 | 10 | 2.75% |
| hsa_circ_0114250 | 728 | 10 | 2.75% |
| hsa_circ_0119622 | 728 | 10 | 2.75% |
| hsa_circ_0120627 | 728 | 10 | 2.75% |
| hsa_circ_0128177 | 728 | 10 | 2.75% |
| hsa_circ_0010520 | 801 | 11 | 2.75% |
| hsa_circ_0042064 | 801 | 11 | 2.75% |
| hsa_circ_0063362 | 801 | 11 | 2.75% |
| hsa_circ_0074145 | 801 | 11 | 2.75% |
| hsa_circ_0081476 | 801 | 11 | 2.75% |
| hsa_circ_0082553 | 801 | 11 | 2.75% |
| hsa_circ_0088848 | 801 | 11 | 2.75% |
| hsa_circ_0096283 | 801 | 11 | 2.75% |
| hsa_circ_0113152 | 801 | 11 | 2.75% |
| hsa_circ_0123015 | 801 | 11 | 2.75% |
| hsa_circ_0124699 | 801 | 11 | 2.75% |
| hsa_circ_0093154 | 1238 | 17 | 2.75% |
| hsa_circ_0079820 | 874 | 12 | 2.75% |
| hsa_circ_0113920 | 2040 | 28 | 2.75% |
| hsa_circ_0136577 | 1239 | 17 | 2.74% |
| hsa_circ_0005385 | 729 | 10 | 2.74% |
| hsa_circ_0013656 | 729 | 10 | 2.74% |
| hsa_circ_0025256 | 729 | 10 | 2.74% |
| hsa_circ_0052469 | 729 | 10 | 2.74% |

| | | | |
|---|---|---|---|
| hsa_circ_0058245 | 729 | 10 | 2.74% |
| hsa_circ_0059373 | 729 | 10 | 2.74% |
| hsa_circ_0060989 | 729 | 10 | 2.74% |
| hsa_circ_0061194 | 729 | 10 | 2.74% |
| hsa_circ_0065222 | 729 | 10 | 2.74% |
| hsa_circ_0074337 | 729 | 10 | 2.74% |
| hsa_circ_0086984 | 729 | 10 | 2.74% |
| hsa_circ_0087774 | 729 | 10 | 2.74% |
| hsa_circ_0099544 | 729 | 10 | 2.74% |
| hsa_circ_0103598 | 729 | 10 | 2.74% |
| hsa_circ_0118205 | 729 | 10 | 2.74% |
| hsa_circ_0121214 | 729 | 10 | 2.74% |
| hsa_circ_0123045 | 729 | 10 | 2.74% |
| hsa_circ_0126642 | 729 | 10 | 2.74% |
| hsa_circ_0130844 | 729 | 10 | 2.74% |
| hsa_circ_0028461 | 802 | 11 | 2.74% |
| hsa_circ_0043199 | 802 | 11 | 2.74% |
| hsa_circ_0047039 | 802 | 11 | 2.74% |
| hsa_circ_0089709 | 802 | 11 | 2.74% |
| hsa_circ_0135375 | 802 | 11 | 2.74% |
| hsa_circ_0067205 | 875 | 12 | 2.74% |
| hsa_circ_0067225 | 875 | 12 | 2.74% |
| hsa_circ_0119565 | 875 | 12 | 2.74% |
| hsa_circ_0129056 | 875 | 12 | 2.74% |
| hsa_circ_0140037 | 875 | 12 | 2.74% |
| hsa_circ_0077853 | 948 | 13 | 2.74% |
| hsa_circ_0100466 | 948 | 13 | 2.74% |
| hsa_circ_0117674 | 1167 | 16 | 2.74% |
| hsa_circ_0077990 | 949 | 13 | 2.74% |
| hsa_circ_0084472 | 876 | 12 | 2.74% |
| hsa_circ_0006803 | 803 | 11 | 2.74% |
| hsa_circ_0045414 | 803 | 11 | 2.74% |
| hsa_circ_0063844 | 803 | 11 | 2.74% |
| hsa_circ_0067792 | 803 | 11 | 2.74% |
| hsa_circ_0070482 | 803 | 11 | 2.74% |
| hsa_circ_0087909 | 803 | 11 | 2.74% |
| hsa_circ_0009244 | 730 | 10 | 2.74% |
| hsa_circ_0014332 | 730 | 10 | 2.74% |
| hsa_circ_0042945 | 730 | 10 | 2.74% |
| hsa_circ_0047910 | 730 | 10 | 2.74% |
| hsa_circ_0080927 | 730 | 10 | 2.74% |
| hsa_circ_0096238 | 730 | 10 | 2.74% |
| hsa_circ_0099138 | 730 | 10 | 2.74% |
| hsa_circ_0100793 | 730 | 10 | 2.74% |
| hsa_circ_0104721 | 730 | 10 | 2.74% |
| hsa_circ_0105498 | 730 | 10 | 2.74% |
| hsa_circ_0105690 | 730 | 10 | 2.74% |
| hsa_circ_0112023 | 730 | 10 | 2.74% |
| hsa_circ_0126629 | 730 | 10 | 2.74% |
| hsa_circ_0128711 | 730 | 10 | 2.74% |

| | | | |
|---|---|---|---|
| hsa_circ_0132342 | 730 | 10 | 2.74% |
| hsa_circ_0134166 | 730 | 10 | 2.74% |
| hsa_circ_0038730 | 950 | 13 | 2.74% |
| hsa_circ_0099793 | 950 | 13 | 2.74% |
| hsa_circ_0027621 | 804 | 11 | 2.74% |
| hsa_circ_0040918 | 804 | 11 | 2.74% |
| hsa_circ_0070404 | 804 | 11 | 2.74% |
| hsa_circ_0120617 | 804 | 11 | 2.74% |
| hsa_circ_0009003 | 731 | 10 | 2.74% |
| hsa_circ_0016909 | 731 | 10 | 2.74% |
| hsa_circ_0027468 | 731 | 10 | 2.74% |
| hsa_circ_0033626 | 731 | 10 | 2.74% |
| hsa_circ_0043567 | 731 | 10 | 2.74% |
| hsa_circ_0052448 | 731 | 10 | 2.74% |
| hsa_circ_0056036 | 731 | 10 | 2.74% |
| hsa_circ_0056597 | 731 | 10 | 2.74% |
| hsa_circ_0060905 | 731 | 10 | 2.74% |
| hsa_circ_0069402 | 731 | 10 | 2.74% |
| hsa_circ_0076914 | 731 | 10 | 2.74% |
| hsa_circ_0084806 | 731 | 10 | 2.74% |
| hsa_circ_0087302 | 731 | 10 | 2.74% |
| hsa_circ_0093646 | 731 | 10 | 2.74% |
| hsa_circ_0097028 | 731 | 10 | 2.74% |
| hsa_circ_0106198 | 731 | 10 | 2.74% |
| hsa_circ_0096215 | 2340 | 32 | 2.74% |
| hsa_circ_0094240 | 878 | 12 | 2.73% |
| hsa_circ_0004423 | 805 | 11 | 2.73% |
| hsa_circ_0048808 | 805 | 11 | 2.73% |
| hsa_circ_0053107 | 805 | 11 | 2.73% |
| hsa_circ_0106770 | 805 | 11 | 2.73% |
| hsa_circ_0113869 | 805 | 11 | 2.73% |
| hsa_circ_0005613 | 732 | 10 | 2.73% |
| hsa_circ_0009251 | 732 | 10 | 2.73% |
| hsa_circ_0009617 | 732 | 10 | 2.73% |
| hsa_circ_0020558 | 732 | 10 | 2.73% |
| hsa_circ_0027280 | 732 | 10 | 2.73% |
| hsa_circ_0031028 | 732 | 10 | 2.73% |
| hsa_circ_0032332 | 732 | 10 | 2.73% |
| hsa_circ_0037382 | 732 | 10 | 2.73% |
| hsa_circ_0039125 | 732 | 10 | 2.73% |
| hsa_circ_0062393 | 732 | 10 | 2.73% |
| hsa_circ_0071962 | 732 | 10 | 2.73% |
| hsa_circ_0071964 | 732 | 10 | 2.73% |
| hsa_circ_0072889 | 732 | 10 | 2.73% |
| hsa_circ_0079172 | 732 | 10 | 2.73% |
| hsa_circ_0082640 | 732 | 10 | 2.73% |
| hsa_circ_0084974 | 732 | 10 | 2.73% |
| hsa_circ_0095831 | 732 | 10 | 2.73% |
| hsa_circ_0098122 | 732 | 10 | 2.73% |
| hsa_circ_0100765 | 732 | 10 | 2.73% |

| | | | |
|---|---|---|---|
| hsa_circ_0116730 | 732 | 10 | 2.73% |
| hsa_circ_0123877 | 732 | 10 | 2.73% |
| hsa_circ_0128004 | 732 | 10 | 2.73% |
| hsa_circ_0130400 | 732 | 10 | 2.73% |
| hsa_circ_0098848 | 879 | 12 | 2.73% |
| hsa_circ_0003204 | 806 | 11 | 2.73% |
| hsa_circ_0097182 | 806 | 11 | 2.73% |
| hsa_circ_0127270 | 806 | 11 | 2.73% |
| hsa_circ_0078755 | 1026 | 14 | 2.73% |
| hsa_circ_0003237 | 733 | 10 | 2.73% |
| hsa_circ_0003373 | 733 | 10 | 2.73% |
| hsa_circ_0008524 | 733 | 10 | 2.73% |
| hsa_circ_0008813 | 733 | 10 | 2.73% |
| hsa_circ_0010333 | 733 | 10 | 2.73% |
| hsa_circ_0021483 | 733 | 10 | 2.73% |
| hsa_circ_0039992 | 733 | 10 | 2.73% |
| hsa_circ_0041510 | 733 | 10 | 2.73% |
| hsa_circ_0097023 | 733 | 10 | 2.73% |
| hsa_circ_0104446 | 733 | 10 | 2.73% |
| hsa_circ_0115029 | 733 | 10 | 2.73% |
| hsa_circ_0117684 | 733 | 10 | 2.73% |
| hsa_circ_0118763 | 733 | 10 | 2.73% |
| hsa_circ_0123520 | 733 | 10 | 2.73% |
| hsa_circ_0130737 | 733 | 10 | 2.73% |
| hsa_circ_0009577 | 807 | 11 | 2.73% |
| hsa_circ_0014690 | 807 | 11 | 2.73% |
| hsa_circ_0040895 | 807 | 11 | 2.73% |
| hsa_circ_0043628 | 807 | 11 | 2.73% |
| hsa_circ_0054881 | 807 | 11 | 2.73% |
| hsa_circ_0085814 | 807 | 11 | 2.73% |
| hsa_circ_0091575 | 807 | 11 | 2.73% |
| hsa_circ_0001481 | 954 | 13 | 2.73% |
| hsa_circ_0025916 | 954 | 13 | 2.73% |
| hsa_circ_0007102 | 734 | 10 | 2.72% |
| hsa_circ_0018738 | 734 | 10 | 2.72% |
| hsa_circ_0034252 | 734 | 10 | 2.72% |
| hsa_circ_0048198 | 734 | 10 | 2.72% |
| hsa_circ_0050969 | 734 | 10 | 2.72% |
| hsa_circ_0061756 | 734 | 10 | 2.72% |
| hsa_circ_0072384 | 734 | 10 | 2.72% |
| hsa_circ_0082029 | 734 | 10 | 2.72% |
| hsa_circ_0088626 | 734 | 10 | 2.72% |
| hsa_circ_0102104 | 734 | 10 | 2.72% |
| hsa_circ_0112777 | 734 | 10 | 2.72% |
| hsa_circ_0132098 | 734 | 10 | 2.72% |
| hsa_circ_0040316 | 881 | 12 | 2.72% |
| hsa_circ_0050398 | 881 | 12 | 2.72% |
| hsa_circ_0088882 | 881 | 12 | 2.72% |
| hsa_circ_0025879 | 808 | 11 | 2.72% |
| hsa_circ_0052506 | 808 | 11 | 2.72% |

| | | | |
|---|---|---|---|
| hsa_circ_0117857 | 808 | 11 | 2.72% |
| hsa_circ_0123752 | 808 | 11 | 2.72% |
| hsa_circ_0138838 | 808 | 11 | 2.72% |
| hsa_circ_0036624 | 882 | 12 | 2.72% |
| hsa_circ_0050655 | 882 | 12 | 2.72% |
| hsa_circ_0076469 | 882 | 12 | 2.72% |
| hsa_circ_0085634 | 882 | 12 | 2.72% |
| hsa_circ_0127037 | 882 | 12 | 2.72% |
| hsa_circ_0140492 | 882 | 12 | 2.72% |
| hsa_circ_0007771 | 735 | 10 | 2.72% |
| hsa_circ_0012991 | 735 | 10 | 2.72% |
| hsa_circ_0014375 | 735 | 10 | 2.72% |
| hsa_circ_0034534 | 735 | 10 | 2.72% |
| hsa_circ_0035160 | 735 | 10 | 2.72% |
| hsa_circ_0038495 | 735 | 10 | 2.72% |
| hsa_circ_0070941 | 735 | 10 | 2.72% |
| hsa_circ_0072388 | 735 | 10 | 2.72% |
| hsa_circ_0079728 | 735 | 10 | 2.72% |
| hsa_circ_0084930 | 735 | 10 | 2.72% |
| hsa_circ_0093230 | 735 | 10 | 2.72% |
| hsa_circ_0108312 | 735 | 10 | 2.72% |
| hsa_circ_0109718 | 735 | 10 | 2.72% |
| hsa_circ_0125531 | 735 | 10 | 2.72% |
| hsa_circ_0126301 | 735 | 10 | 2.72% |
| hsa_circ_0135416 | 735 | 10 | 2.72% |
| hsa_circ_0011410 | 809 | 11 | 2.72% |
| hsa_circ_0037831 | 809 | 11 | 2.72% |
| hsa_circ_0043986 | 809 | 11 | 2.72% |
| hsa_circ_0077539 | 809 | 11 | 2.72% |
| hsa_circ_0117009 | 809 | 11 | 2.72% |
| hsa_circ_0118186 | 809 | 11 | 2.72% |
| hsa_circ_0120752 | 809 | 11 | 2.72% |
| hsa_circ_0122907 | 809 | 11 | 2.72% |
| hsa_circ_0018254 | 1030 | 14 | 2.72% |
| hsa_circ_0044598 | 883 | 12 | 2.72% |
| hsa_circ_0019598 | 736 | 10 | 2.72% |
| hsa_circ_0022069 | 736 | 10 | 2.72% |
| hsa_circ_0058424 | 736 | 10 | 2.72% |
| hsa_circ_0077037 | 736 | 10 | 2.72% |
| hsa_circ_0077214 | 736 | 10 | 2.72% |
| hsa_circ_0079882 | 736 | 10 | 2.72% |
| hsa_circ_0095013 | 736 | 10 | 2.72% |
| hsa_circ_0097647 | 736 | 10 | 2.72% |
| hsa_circ_0097660 | 736 | 10 | 2.72% |
| hsa_circ_0098957 | 736 | 10 | 2.72% |
| hsa_circ_0104829 | 736 | 10 | 2.72% |
| hsa_circ_0111432 | 736 | 10 | 2.72% |
| hsa_circ_0118163 | 736 | 10 | 2.72% |
| hsa_circ_0126090 | 736 | 10 | 2.72% |
| hsa_circ_0130168 | 736 | 10 | 2.72% |

| | | | |
|---|---|---|---|
| hsa_circ_0077656 | 957 | 13 | 2.72% |
| hsa_circ_0006645 | 810 | 11 | 2.72% |
| hsa_circ_0007572 | 810 | 11 | 2.72% |
| hsa_circ_0026392 | 810 | 11 | 2.72% |
| hsa_circ_0048331 | 810 | 11 | 2.72% |
| hsa_circ_0057249 | 810 | 11 | 2.72% |
| hsa_circ_0057430 | 810 | 11 | 2.72% |
| hsa_circ_0059585 | 810 | 11 | 2.72% |
| hsa_circ_0060765 | 810 | 11 | 2.72% |
| hsa_circ_0075250 | 810 | 11 | 2.72% |
| hsa_circ_0090325 | 810 | 11 | 2.72% |
| hsa_circ_0099613 | 810 | 11 | 2.72% |
| hsa_circ_0103821 | 810 | 11 | 2.72% |
| hsa_circ_0123013 | 810 | 11 | 2.72% |
| hsa_circ_0131754 | 810 | 11 | 2.72% |
| hsa_circ_0107517 | 1031 | 14 | 2.72% |
| hsa_circ_0096148 | 1105 | 15 | 2.71% |
| hsa_circ_0057369 | 1179 | 16 | 2.71% |
| hsa_circ_0018722 | 737 | 10 | 2.71% |
| hsa_circ_0038565 | 737 | 10 | 2.71% |
| hsa_circ_0043312 | 737 | 10 | 2.71% |
| hsa_circ_0045889 | 737 | 10 | 2.71% |
| hsa_circ_0054463 | 737 | 10 | 2.71% |
| hsa_circ_0062799 | 737 | 10 | 2.71% |
| hsa_circ_0074357 | 737 | 10 | 2.71% |
| hsa_circ_0098575 | 737 | 10 | 2.71% |
| hsa_circ_0111057 | 737 | 10 | 2.71% |
| hsa_circ_0120050 | 737 | 10 | 2.71% |
| hsa_circ_0122925 | 737 | 10 | 2.71% |
| hsa_circ_0066077 | 811 | 11 | 2.71% |
| hsa_circ_0091415 | 811 | 11 | 2.71% |
| hsa_circ_0105789 | 1475 | 20 | 2.71% |
| hsa_circ_0014775 | 885 | 12 | 2.71% |
| hsa_circ_0032289 | 885 | 12 | 2.71% |
| hsa_circ_0140420 | 1033 | 14 | 2.71% |
| hsa_circ_0001052 | 738 | 10 | 2.71% |
| hsa_circ_0001893 | 738 | 10 | 2.71% |
| hsa_circ_0007714 | 738 | 10 | 2.71% |
| hsa_circ_0024294 | 738 | 10 | 2.71% |
| hsa_circ_0032581 | 738 | 10 | 2.71% |
| hsa_circ_0082068 | 738 | 10 | 2.71% |
| hsa_circ_0104847 | 738 | 10 | 2.71% |
| hsa_circ_0125647 | 738 | 10 | 2.71% |
| hsa_circ_0129823 | 738 | 10 | 2.71% |
| hsa_circ_0137597 | 738 | 10 | 2.71% |
| hsa_circ_0009928 | 812 | 11 | 2.71% |
| hsa_circ_0082012 | 812 | 11 | 2.71% |
| hsa_circ_0032869 | 886 | 12 | 2.71% |
| hsa_circ_0049546 | 886 | 12 | 2.71% |
| hsa_circ_0109798 | 960 | 13 | 2.71% |

| | | | |
|---|---|---|---|
| hsa_circ_0001394 | 739 | 10 | 2.71% |
| hsa_circ_0021817 | 739 | 10 | 2.71% |
| hsa_circ_0033284 | 739 | 10 | 2.71% |
| hsa_circ_0040743 | 739 | 10 | 2.71% |
| hsa_circ_0042229 | 739 | 10 | 2.71% |
| hsa_circ_0046019 | 739 | 10 | 2.71% |
| hsa_circ_0072352 | 739 | 10 | 2.71% |
| hsa_circ_0099378 | 739 | 10 | 2.71% |
| hsa_circ_0112511 | 739 | 10 | 2.71% |
| hsa_circ_0134424 | 739 | 10 | 2.71% |
| hsa_circ_0138954 | 739 | 10 | 2.71% |
| hsa_circ_0000672 | 813 | 11 | 2.71% |
| hsa_circ_0026756 | 813 | 11 | 2.71% |
| hsa_circ_0028026 | 813 | 11 | 2.71% |
| hsa_circ_0041779 | 813 | 11 | 2.71% |
| hsa_circ_0076625 | 813 | 11 | 2.71% |
| hsa_circ_0082740 | 813 | 11 | 2.71% |
| hsa_circ_0087336 | 813 | 11 | 2.71% |
| hsa_circ_0136349 | 813 | 11 | 2.71% |
| hsa_circ_0136389 | 1700 | 23 | 2.71% |
| hsa_circ_0082610 | 887 | 12 | 2.71% |
| hsa_circ_0025225 | 961 | 13 | 2.71% |
| hsa_circ_0045770 | 1035 | 14 | 2.71% |
| hsa_circ_0107083 | 962 | 13 | 2.70% |
| hsa_circ_0050105 | 888 | 12 | 2.70% |
| hsa_circ_0010899 | 814 | 11 | 2.70% |
| hsa_circ_0036375 | 814 | 11 | 2.70% |
| hsa_circ_0111702 | 814 | 11 | 2.70% |
| hsa_circ_0115968 | 814 | 11 | 2.70% |
| hsa_circ_0117051 | 814 | 11 | 2.70% |
| hsa_circ_0015431 | 740 | 10 | 2.70% |
| hsa_circ_0018797 | 740 | 10 | 2.70% |
| hsa_circ_0022433 | 740 | 10 | 2.70% |
| hsa_circ_0031608 | 740 | 10 | 2.70% |
| hsa_circ_0043194 | 740 | 10 | 2.70% |
| hsa_circ_0057464 | 740 | 10 | 2.70% |
| hsa_circ_0068095 | 740 | 10 | 2.70% |
| hsa_circ_0073093 | 740 | 10 | 2.70% |
| hsa_circ_0084280 | 740 | 10 | 2.70% |
| hsa_circ_0085422 | 740 | 10 | 2.70% |
| hsa_circ_0085842 | 740 | 10 | 2.70% |
| hsa_circ_0087890 | 740 | 10 | 2.70% |
| hsa_circ_0096818 | 740 | 10 | 2.70% |
| hsa_circ_0099123 | 740 | 10 | 2.70% |
| hsa_circ_0104830 | 740 | 10 | 2.70% |
| hsa_circ_0116053 | 740 | 10 | 2.70% |
| hsa_circ_0124668 | 740 | 10 | 2.70% |
| hsa_circ_0106442 | 963 | 13 | 2.70% |
| hsa_circ_0088411 | 1778 | 24 | 2.70% |
| hsa_circ_0027911 | 889 | 12 | 2.70% |

| | | | |
|---|---|---|---|
| hsa_circ_0140687 | 889 | 12 | 2.70% |
| hsa_circ_0025922 | 815 | 11 | 2.70% |
| hsa_circ_0039725 | 815 | 11 | 2.70% |
| hsa_circ_0048341 | 815 | 11 | 2.70% |
| hsa_circ_0064578 | 815 | 11 | 2.70% |
| hsa_circ_0083903 | 815 | 11 | 2.70% |
| hsa_circ_0089204 | 815 | 11 | 2.70% |
| hsa_circ_0122578 | 815 | 11 | 2.70% |
| hsa_circ_0133802 | 815 | 11 | 2.70% |
| hsa_circ_0012569 | 741 | 10 | 2.70% |
| hsa_circ_0015023 | 741 | 10 | 2.70% |
| hsa_circ_0015910 | 741 | 10 | 2.70% |
| hsa_circ_0016687 | 741 | 10 | 2.70% |
| hsa_circ_0025006 | 741 | 10 | 2.70% |
| hsa_circ_0035175 | 741 | 10 | 2.70% |
| hsa_circ_0050664 | 741 | 10 | 2.70% |
| hsa_circ_0057919 | 741 | 10 | 2.70% |
| hsa_circ_0067855 | 741 | 10 | 2.70% |
| hsa_circ_0072690 | 741 | 10 | 2.70% |
| hsa_circ_0079520 | 741 | 10 | 2.70% |
| hsa_circ_0085840 | 741 | 10 | 2.70% |
| hsa_circ_0094517 | 741 | 10 | 2.70% |
| hsa_circ_0117992 | 741 | 10 | 2.70% |
| hsa_circ_0132225 | 741 | 10 | 2.70% |
| hsa_circ_0132829 | 741 | 10 | 2.70% |
| hsa_circ_0136314 | 741 | 10 | 2.70% |
| hsa_circ_0138354 | 741 | 10 | 2.70% |
| hsa_circ_0077775 | 1260 | 17 | 2.70% |
| hsa_circ_0112964 | 1409 | 19 | 2.70% |
| hsa_circ_0021625 | 890 | 12 | 2.70% |
| hsa_circ_0037682 | 890 | 12 | 2.70% |
| hsa_circ_0133903 | 890 | 12 | 2.70% |
| hsa_circ_0004014 | 816 | 11 | 2.70% |
| hsa_circ_0039869 | 816 | 11 | 2.70% |
| hsa_circ_0074888 | 816 | 11 | 2.70% |
| hsa_circ_0128680 | 816 | 11 | 2.70% |
| hsa_circ_0132265 | 816 | 11 | 2.70% |
| hsa_circ_0132974 | 816 | 11 | 2.70% |
| hsa_circ_0011858 | 742 | 10 | 2.70% |
| hsa_circ_0012152 | 742 | 10 | 2.70% |
| hsa_circ_0025237 | 742 | 10 | 2.70% |
| hsa_circ_0028848 | 742 | 10 | 2.70% |
| hsa_circ_0034523 | 742 | 10 | 2.70% |
| hsa_circ_0035086 | 742 | 10 | 2.70% |
| hsa_circ_0037373 | 742 | 10 | 2.70% |
| hsa_circ_0046106 | 742 | 10 | 2.70% |
| hsa_circ_0049474 | 742 | 10 | 2.70% |
| hsa_circ_0049960 | 742 | 10 | 2.70% |
| hsa_circ_0066789 | 742 | 10 | 2.70% |
| hsa_circ_0077564 | 742 | 10 | 2.70% |

| | | | |
|---|---|---|---|
| hsa_circ_0078752 | 742 | 10 | 2.70% |
| hsa_circ_0088389 | 742 | 10 | 2.70% |
| hsa_circ_0121659 | 742 | 10 | 2.70% |
| hsa_circ_0122505 | 742 | 10 | 2.70% |
| hsa_circ_0127689 | 742 | 10 | 2.70% |
| hsa_circ_0023843 | 965 | 13 | 2.69% |
| hsa_circ_0009384 | 891 | 12 | 2.69% |
| hsa_circ_0049754 | 891 | 12 | 2.69% |
| hsa_circ_0021055 | 817 | 11 | 2.69% |
| hsa_circ_0054074 | 817 | 11 | 2.69% |
| hsa_circ_0058523 | 817 | 11 | 2.69% |
| hsa_circ_0079754 | 817 | 11 | 2.69% |
| hsa_circ_0007686 | 743 | 10 | 2.69% |
| hsa_circ_0010124 | 743 | 10 | 2.69% |
| hsa_circ_0019293 | 743 | 10 | 2.69% |
| hsa_circ_0021499 | 743 | 10 | 2.69% |
| hsa_circ_0049134 | 743 | 10 | 2.69% |
| hsa_circ_0055043 | 743 | 10 | 2.69% |
| hsa_circ_0064262 | 743 | 10 | 2.69% |
| hsa_circ_0079704 | 743 | 10 | 2.69% |
| hsa_circ_0084423 | 743 | 10 | 2.69% |
| hsa_circ_0091935 | 743 | 10 | 2.69% |
| hsa_circ_0094029 | 743 | 10 | 2.69% |
| hsa_circ_0108798 | 743 | 10 | 2.69% |
| hsa_circ_0114051 | 743 | 10 | 2.69% |
| hsa_circ_0122452 | 743 | 10 | 2.69% |
| hsa_circ_0074667 | 892 | 12 | 2.69% |
| hsa_circ_0105396 | 892 | 12 | 2.69% |
| hsa_circ_0133558 | 892 | 12 | 2.69% |
| hsa_circ_0041613 | 818 | 11 | 2.69% |
| hsa_circ_0118810 | 818 | 11 | 2.69% |
| hsa_circ_0076664 | 967 | 13 | 2.69% |
| hsa_circ_0016031 | 744 | 10 | 2.69% |
| hsa_circ_0018716 | 744 | 10 | 2.69% |
| hsa_circ_0034313 | 744 | 10 | 2.69% |
| hsa_circ_0035037 | 744 | 10 | 2.69% |
| hsa_circ_0036930 | 744 | 10 | 2.69% |
| hsa_circ_0039837 | 744 | 10 | 2.69% |
| hsa_circ_0041166 | 744 | 10 | 2.69% |
| hsa_circ_0050191 | 744 | 10 | 2.69% |
| hsa_circ_0050362 | 744 | 10 | 2.69% |
| hsa_circ_0056144 | 744 | 10 | 2.69% |
| hsa_circ_0064032 | 744 | 10 | 2.69% |
| hsa_circ_0070654 | 744 | 10 | 2.69% |
| hsa_circ_0075852 | 744 | 10 | 2.69% |
| hsa_circ_0079157 | 744 | 10 | 2.69% |
| hsa_circ_0083746 | 744 | 10 | 2.69% |
| hsa_circ_0095392 | 744 | 10 | 2.69% |
| hsa_circ_0101872 | 744 | 10 | 2.69% |
| hsa_circ_0102366 | 744 | 10 | 2.69% |

| | | | |
|---|---|---|---|
| hsa_circ_0117430 | 744 | 10 | 2.69% |
| hsa_circ_0118752 | 744 | 10 | 2.69% |
| hsa_circ_0121714 | 744 | 10 | 2.69% |
| hsa_circ_0129491 | 744 | 10 | 2.69% |
| hsa_circ_0016310 | 893 | 12 | 2.69% |
| hsa_circ_0105409 | 893 | 12 | 2.69% |
| hsa_circ_0016510 | 819 | 11 | 2.69% |
| hsa_circ_0083875 | 819 | 11 | 2.69% |
| hsa_circ_0118043 | 819 | 11 | 2.69% |
| hsa_circ_0122459 | 819 | 11 | 2.69% |
| hsa_circ_0131310 | 819 | 11 | 2.69% |
| hsa_circ_0031533 | 894 | 12 | 2.68% |
| hsa_circ_0067060 | 894 | 12 | 2.68% |
| hsa_circ_0117394 | 894 | 12 | 2.68% |
| hsa_circ_0006536 | 745 | 10 | 2.68% |
| hsa_circ_0015028 | 745 | 10 | 2.68% |
| hsa_circ_0029946 | 745 | 10 | 2.68% |
| hsa_circ_0030082 | 745 | 10 | 2.68% |
| hsa_circ_0049126 | 745 | 10 | 2.68% |
| hsa_circ_0057170 | 745 | 10 | 2.68% |
| hsa_circ_0057924 | 745 | 10 | 2.68% |
| hsa_circ_0061808 | 745 | 10 | 2.68% |
| hsa_circ_0063271 | 745 | 10 | 2.68% |
| hsa_circ_0074819 | 745 | 10 | 2.68% |
| hsa_circ_0077085 | 745 | 10 | 2.68% |
| hsa_circ_0080560 | 745 | 10 | 2.68% |
| hsa_circ_0094161 | 745 | 10 | 2.68% |
| hsa_circ_0121286 | 745 | 10 | 2.68% |
| hsa_circ_0121372 | 745 | 10 | 2.68% |
| hsa_circ_0135359 | 745 | 10 | 2.68% |
| hsa_circ_0012027 | 1118 | 15 | 2.68% |
| hsa_circ_0090138 | 969 | 13 | 2.68% |
| hsa_circ_0034939 | 820 | 11 | 2.68% |
| hsa_circ_0052271 | 820 | 11 | 2.68% |
| hsa_circ_0053345 | 820 | 11 | 2.68% |
| hsa_circ_0059195 | 820 | 11 | 2.68% |
| hsa_circ_0082283 | 820 | 11 | 2.68% |
| hsa_circ_0098027 | 820 | 11 | 2.68% |
| hsa_circ_0101748 | 820 | 11 | 2.68% |
| hsa_circ_0122257 | 820 | 11 | 2.68% |
| hsa_circ_0051265 | 895 | 12 | 2.68% |
| hsa_circ_0043377 | 2014 | 27 | 2.68% |
| hsa_circ_0019782 | 746 | 10 | 2.68% |
| hsa_circ_0038055 | 746 | 10 | 2.68% |
| hsa_circ_0039254 | 746 | 10 | 2.68% |
| hsa_circ_0076943 | 746 | 10 | 2.68% |
| hsa_circ_0080634 | 746 | 10 | 2.68% |
| hsa_circ_0080675 | 746 | 10 | 2.68% |
| hsa_circ_0087529 | 746 | 10 | 2.68% |
| hsa_circ_0111890 | 746 | 10 | 2.68% |

| | | | |
|---|---|---|---|
| hsa_circ_0116378 | 746 | 10 | 2.68% |
| hsa_circ_0118887 | 746 | 10 | 2.68% |
| hsa_circ_0122252 | 746 | 10 | 2.68% |
| hsa_circ_0133379 | 746 | 10 | 2.68% |
| hsa_circ_0134674 | 746 | 10 | 2.68% |
| hsa_circ_0135454 | 746 | 10 | 2.68% |
| hsa_circ_0140104 | 746 | 10 | 2.68% |
| hsa_circ_0019401 | 821 | 11 | 2.68% |
| hsa_circ_0038514 | 821 | 11 | 2.68% |
| hsa_circ_0048971 | 821 | 11 | 2.68% |
| hsa_circ_0067135 | 821 | 11 | 2.68% |
| hsa_circ_0076422 | 821 | 11 | 2.68% |
| hsa_circ_0080453 | 821 | 11 | 2.68% |
| hsa_circ_0129897 | 821 | 11 | 2.68% |
| hsa_circ_0137353 | 821 | 11 | 2.68% |
| hsa_circ_0023395 | 896 | 12 | 2.68% |
| hsa_circ_0080862 | 896 | 12 | 2.68% |
| hsa_circ_0003192 | 747 | 10 | 2.68% |
| hsa_circ_0011395 | 747 | 10 | 2.68% |
| hsa_circ_0012495 | 747 | 10 | 2.68% |
| hsa_circ_0018393 | 747 | 10 | 2.68% |
| hsa_circ_0020459 | 747 | 10 | 2.68% |
| hsa_circ_0022796 | 747 | 10 | 2.68% |
| hsa_circ_0028658 | 747 | 10 | 2.68% |
| hsa_circ_0031563 | 747 | 10 | 2.68% |
| hsa_circ_0032589 | 747 | 10 | 2.68% |
| hsa_circ_0041702 | 747 | 10 | 2.68% |
| hsa_circ_0043010 | 747 | 10 | 2.68% |
| hsa_circ_0044803 | 747 | 10 | 2.68% |
| hsa_circ_0057362 | 747 | 10 | 2.68% |
| hsa_circ_0060186 | 747 | 10 | 2.68% |
| hsa_circ_0073102 | 747 | 10 | 2.68% |
| hsa_circ_0086137 | 747 | 10 | 2.68% |
| hsa_circ_0100613 | 747 | 10 | 2.68% |
| hsa_circ_0105079 | 747 | 10 | 2.68% |
| hsa_circ_0111816 | 747 | 10 | 2.68% |
| hsa_circ_0112428 | 747 | 10 | 2.68% |
| hsa_circ_0114150 | 747 | 10 | 2.68% |
| hsa_circ_0122027 | 747 | 10 | 2.68% |
| hsa_circ_0123396 | 747 | 10 | 2.68% |
| hsa_circ_0137799 | 747 | 10 | 2.68% |
| hsa_circ_0139246 | 747 | 10 | 2.68% |
| hsa_circ_0010636 | 822 | 11 | 2.68% |
| hsa_circ_0020188 | 822 | 11 | 2.68% |
| hsa_circ_0021084 | 822 | 11 | 2.68% |
| hsa_circ_0056994 | 822 | 11 | 2.68% |
| hsa_circ_0061773 | 822 | 11 | 2.68% |
| hsa_circ_0074226 | 822 | 11 | 2.68% |
| hsa_circ_0078600 | 822 | 11 | 2.68% |
| hsa_circ_0091680 | 822 | 11 | 2.68% |

| | | | |
|---|---|---|---|
| hsa_circ_0101878 | 822 | 11 | 2.68% |
| hsa_circ_0119746 | 822 | 11 | 2.68% |
| hsa_circ_0126830 | 2317 | 31 | 2.68% |
| hsa_circ_0037779 | 2992 | 40 | 2.67% |
| hsa_circ_0011636 | 748 | 10 | 2.67% |
| hsa_circ_0018065 | 748 | 10 | 2.67% |
| hsa_circ_0018129 | 748 | 10 | 2.67% |
| hsa_circ_0021263 | 748 | 10 | 2.67% |
| hsa_circ_0076105 | 748 | 10 | 2.67% |
| hsa_circ_0078820 | 748 | 10 | 2.67% |
| hsa_circ_0078875 | 748 | 10 | 2.67% |
| hsa_circ_0078941 | 748 | 10 | 2.67% |
| hsa_circ_0096821 | 748 | 10 | 2.67% |
| hsa_circ_0097645 | 748 | 10 | 2.67% |
| hsa_circ_0099538 | 748 | 10 | 2.67% |
| hsa_circ_0103926 | 748 | 10 | 2.67% |
| hsa_circ_0105450 | 748 | 10 | 2.67% |
| hsa_circ_0117435 | 748 | 10 | 2.67% |
| hsa_circ_0007979 | 823 | 11 | 2.67% |
| hsa_circ_0044480 | 898 | 12 | 2.67% |
| hsa_circ_0096036 | 1423 | 19 | 2.67% |
| hsa_circ_0014639 | 749 | 10 | 2.67% |
| hsa_circ_0021591 | 749 | 10 | 2.67% |
| hsa_circ_0028287 | 749 | 10 | 2.67% |
| hsa_circ_0052327 | 749 | 10 | 2.67% |
| hsa_circ_0054248 | 749 | 10 | 2.67% |
| hsa_circ_0064665 | 749 | 10 | 2.67% |
| hsa_circ_0066029 | 749 | 10 | 2.67% |
| hsa_circ_0077629 | 749 | 10 | 2.67% |
| hsa_circ_0088094 | 749 | 10 | 2.67% |
| hsa_circ_0091359 | 749 | 10 | 2.67% |
| hsa_circ_0110675 | 749 | 10 | 2.67% |
| hsa_circ_0112014 | 749 | 10 | 2.67% |
| hsa_circ_0114177 | 749 | 10 | 2.67% |
| hsa_circ_0116073 | 749 | 10 | 2.67% |
| hsa_circ_0125939 | 749 | 10 | 2.67% |
| hsa_circ_0140703 | 749 | 10 | 2.67% |
| hsa_circ_0140762 | 749 | 10 | 2.67% |
| hsa_circ_0036265 | 824 | 11 | 2.67% |
| hsa_circ_0052682 | 824 | 11 | 2.67% |
| hsa_circ_0031754 | 899 | 12 | 2.67% |
| hsa_circ_0077242 | 899 | 12 | 2.67% |
| hsa_circ_0104992 | 1350 | 18 | 2.67% |
| hsa_circ_0076435 | 975 | 13 | 2.67% |
| hsa_circ_0102724 | 975 | 13 | 2.67% |
| hsa_circ_0105579 | 975 | 13 | 2.67% |
| hsa_circ_0001986 | 900 | 12 | 2.67% |
| hsa_circ_0017476 | 825 | 11 | 2.67% |
| hsa_circ_0019268 | 825 | 11 | 2.67% |
| hsa_circ_0047373 | 825 | 11 | 2.67% |

| | | | |
|---|---|---|---|
| hsa_circ_0057228 | 825 | 11 | 2.67% |
| hsa_circ_0108265 | 825 | 11 | 2.67% |
| hsa_circ_0119828 | 825 | 11 | 2.67% |
| hsa_circ_0032476 | 750 | 10 | 2.67% |
| hsa_circ_0036922 | 750 | 10 | 2.67% |
| hsa_circ_0041453 | 750 | 10 | 2.67% |
| hsa_circ_0041943 | 750 | 10 | 2.67% |
| hsa_circ_0055995 | 750 | 10 | 2.67% |
| hsa_circ_0074478 | 750 | 10 | 2.67% |
| hsa_circ_0075013 | 750 | 10 | 2.67% |
| hsa_circ_0078876 | 750 | 10 | 2.67% |
| hsa_circ_0088224 | 750 | 10 | 2.67% |
| hsa_circ_0089584 | 750 | 10 | 2.67% |
| hsa_circ_0091793 | 750 | 10 | 2.67% |
| hsa_circ_0093167 | 750 | 10 | 2.67% |
| hsa_circ_0102270 | 750 | 10 | 2.67% |
| hsa_circ_0103850 | 750 | 10 | 2.67% |
| hsa_circ_0106873 | 750 | 10 | 2.67% |
| hsa_circ_0114664 | 750 | 10 | 2.67% |
| hsa_circ_0118204 | 750 | 10 | 2.67% |
| hsa_circ_0120386 | 750 | 10 | 2.67% |
| hsa_circ_0123373 | 750 | 10 | 2.67% |
| hsa_circ_0128688 | 750 | 10 | 2.67% |
| hsa_circ_0131188 | 750 | 10 | 2.67% |
| hsa_circ_0135360 | 750 | 10 | 2.67% |
| hsa_circ_0138659 | 750 | 10 | 2.67% |
| hsa_circ_0106148 | 1501 | 20 | 2.66% |
| hsa_circ_0051509 | 901 | 12 | 2.66% |
| hsa_circ_0061476 | 901 | 12 | 2.66% |
| hsa_circ_0005624 | 826 | 11 | 2.66% |
| hsa_circ_0018282 | 826 | 11 | 2.66% |
| hsa_circ_0030527 | 826 | 11 | 2.66% |
| hsa_circ_0068704 | 826 | 11 | 2.66% |
| hsa_circ_0120560 | 826 | 11 | 2.66% |
| hsa_circ_0002241 | 751 | 10 | 2.66% |
| hsa_circ_0005961 | 751 | 10 | 2.66% |
| hsa_circ_0013341 | 751 | 10 | 2.66% |
| hsa_circ_0025022 | 751 | 10 | 2.66% |
| hsa_circ_0046187 | 751 | 10 | 2.66% |
| hsa_circ_0057908 | 751 | 10 | 2.66% |
| hsa_circ_0064432 | 751 | 10 | 2.66% |
| hsa_circ_0087856 | 751 | 10 | 2.66% |
| hsa_circ_0089989 | 751 | 10 | 2.66% |
| hsa_circ_0112606 | 751 | 10 | 2.66% |
| hsa_circ_0032655 | 1277 | 17 | 2.66% |
| hsa_circ_0110916 | 2780 | 37 | 2.66% |
| hsa_circ_0055482 | 902 | 12 | 2.66% |
| hsa_circ_0065714 | 902 | 12 | 2.66% |
| hsa_circ_0001863 | 827 | 11 | 2.66% |
| hsa_circ_0016863 | 827 | 11 | 2.66% |

| | | | |
|---|---|---|---|
| hsa_circ_0050435 | 827 | 11 | 2.66% |
| hsa_circ_0072200 | 827 | 11 | 2.66% |
| hsa_circ_0121146 | 827 | 11 | 2.66% |
| hsa_circ_0007969 | 752 | 10 | 2.66% |
| hsa_circ_0019869 | 752 | 10 | 2.66% |
| hsa_circ_0021871 | 752 | 10 | 2.66% |
| hsa_circ_0030264 | 752 | 10 | 2.66% |
| hsa_circ_0034706 | 752 | 10 | 2.66% |
| hsa_circ_0049565 | 752 | 10 | 2.66% |
| hsa_circ_0052590 | 752 | 10 | 2.66% |
| hsa_circ_0053135 | 752 | 10 | 2.66% |
| hsa_circ_0059696 | 752 | 10 | 2.66% |
| hsa_circ_0062797 | 752 | 10 | 2.66% |
| hsa_circ_0077712 | 752 | 10 | 2.66% |
| hsa_circ_0080320 | 752 | 10 | 2.66% |
| hsa_circ_0100607 | 752 | 10 | 2.66% |
| hsa_circ_0104147 | 752 | 10 | 2.66% |
| hsa_circ_0112311 | 752 | 10 | 2.66% |
| hsa_circ_0121899 | 752 | 10 | 2.66% |
| hsa_circ_0127587 | 752 | 10 | 2.66% |
| hsa_circ_0139776 | 752 | 10 | 2.66% |
| hsa_circ_0001906 | 828 | 11 | 2.66% |
| hsa_circ_0023422 | 828 | 11 | 2.66% |
| hsa_circ_0071289 | 828 | 11 | 2.66% |
| hsa_circ_0076465 | 828 | 11 | 2.66% |
| hsa_circ_0089459 | 828 | 11 | 2.66% |
| hsa_circ_0006075 | 753 | 10 | 2.66% |
| hsa_circ_0011938 | 753 | 10 | 2.66% |
| hsa_circ_0032609 | 753 | 10 | 2.66% |
| hsa_circ_0037360 | 753 | 10 | 2.66% |
| hsa_circ_0040287 | 753 | 10 | 2.66% |
| hsa_circ_0046733 | 753 | 10 | 2.66% |
| hsa_circ_0050662 | 753 | 10 | 2.66% |
| hsa_circ_0056260 | 753 | 10 | 2.66% |
| hsa_circ_0059018 | 753 | 10 | 2.66% |
| hsa_circ_0059914 | 753 | 10 | 2.66% |
| hsa_circ_0063987 | 753 | 10 | 2.66% |
| hsa_circ_0065469 | 753 | 10 | 2.66% |
| hsa_circ_0094716 | 753 | 10 | 2.66% |
| hsa_circ_0107835 | 753 | 10 | 2.66% |
| hsa_circ_0107968 | 753 | 10 | 2.66% |
| hsa_circ_0118124 | 753 | 10 | 2.66% |
| hsa_circ_0123462 | 753 | 10 | 2.66% |
| hsa_circ_0126571 | 753 | 10 | 2.66% |
| hsa_circ_0134726 | 753 | 10 | 2.66% |
| hsa_circ_0137329 | 753 | 10 | 2.66% |
| hsa_circ_0138394 | 753 | 10 | 2.66% |
| hsa_circ_0058573 | 904 | 12 | 2.65% |
| hsa_circ_0088428 | 904 | 12 | 2.65% |
| hsa_circ_0130090 | 904 | 12 | 2.65% |

| | | | |
|---|---|---|---|
| hsa_circ_0004065 | 829 | 11 | 2.65% |
| hsa_circ_0011508 | 829 | 11 | 2.65% |
| hsa_circ_0020968 | 829 | 11 | 2.65% |
| hsa_circ_0041324 | 829 | 11 | 2.65% |
| hsa_circ_0043408 | 829 | 11 | 2.65% |
| hsa_circ_0044660 | 829 | 11 | 2.65% |
| hsa_circ_0051579 | 829 | 11 | 2.65% |
| hsa_circ_0103235 | 829 | 11 | 2.65% |
| hsa_circ_0140515 | 829 | 11 | 2.65% |
| hsa_circ_0058680 | 980 | 13 | 2.65% |
| hsa_circ_0099614 | 1131 | 15 | 2.65% |
| hsa_circ_0005403 | 754 | 10 | 2.65% |
| hsa_circ_0011863 | 754 | 10 | 2.65% |
| hsa_circ_0013511 | 754 | 10 | 2.65% |
| hsa_circ_0027062 | 754 | 10 | 2.65% |
| hsa_circ_0037108 | 754 | 10 | 2.65% |
| hsa_circ_0037722 | 754 | 10 | 2.65% |
| hsa_circ_0043099 | 754 | 10 | 2.65% |
| hsa_circ_0048206 | 754 | 10 | 2.65% |
| hsa_circ_0056695 | 754 | 10 | 2.65% |
| hsa_circ_0103674 | 754 | 10 | 2.65% |
| hsa_circ_0104861 | 754 | 10 | 2.65% |
| hsa_circ_0106290 | 754 | 10 | 2.65% |
| hsa_circ_0129919 | 754 | 10 | 2.65% |
| hsa_circ_0135771 | 754 | 10 | 2.65% |
| hsa_circ_0136272 | 754 | 10 | 2.65% |
| hsa_circ_0002622 | 905 | 12 | 2.65% |
| hsa_circ_0098674 | 2263 | 30 | 2.65% |
| hsa_circ_0003012 | 830 | 11 | 2.65% |
| hsa_circ_0013316 | 830 | 11 | 2.65% |
| hsa_circ_0027361 | 830 | 11 | 2.65% |
| hsa_circ_0059281 | 830 | 11 | 2.65% |
| hsa_circ_0091314 | 830 | 11 | 2.65% |
| hsa_circ_0094104 | 830 | 11 | 2.65% |
| hsa_circ_0115389 | 1811 | 24 | 2.65% |
| hsa_circ_0108479 | 1283 | 17 | 2.65% |
| hsa_circ_0031839 | 1661 | 22 | 2.65% |
| hsa_circ_0081511 | 1057 | 14 | 2.65% |
| hsa_circ_0039372 | 906 | 12 | 2.65% |
| hsa_circ_0033669 | 755 | 10 | 2.65% |
| hsa_circ_0037615 | 755 | 10 | 2.65% |
| hsa_circ_0050374 | 755 | 10 | 2.65% |
| hsa_circ_0051851 | 755 | 10 | 2.65% |
| hsa_circ_0070890 | 755 | 10 | 2.65% |
| hsa_circ_0077162 | 755 | 10 | 2.65% |
| hsa_circ_0096315 | 755 | 10 | 2.65% |
| hsa_circ_0096718 | 755 | 10 | 2.65% |
| hsa_circ_0104970 | 755 | 10 | 2.65% |
| hsa_circ_0114492 | 755 | 10 | 2.65% |
| hsa_circ_0116656 | 755 | 10 | 2.65% |

| | | | |
|---|---|---|---|
| hsa_circ_0117156 | 755 | 10 | 2.65% |
| hsa_circ_0118593 | 755 | 10 | 2.65% |
| hsa_circ_0118924 | 755 | 10 | 2.65% |
| hsa_circ_0121374 | 755 | 10 | 2.65% |
| hsa_circ_0127924 | 755 | 10 | 2.65% |
| hsa_circ_0137437 | 755 | 10 | 2.65% |
| hsa_circ_0114059 | 1284 | 17 | 2.65% |
| hsa_circ_0132770 | 1133 | 15 | 2.65% |
| hsa_circ_0002788 | 831 | 11 | 2.65% |
| hsa_circ_0019917 | 831 | 11 | 2.65% |
| hsa_circ_0042273 | 831 | 11 | 2.65% |
| hsa_circ_0049917 | 831 | 11 | 2.65% |
| hsa_circ_0064317 | 831 | 11 | 2.65% |
| hsa_circ_0083723 | 831 | 11 | 2.65% |
| hsa_circ_0087939 | 831 | 11 | 2.65% |
| hsa_circ_0114155 | 831 | 11 | 2.65% |
| hsa_circ_0003095 | 907 | 12 | 2.65% |
| hsa_circ_0055486 | 907 | 12 | 2.65% |
| hsa_circ_0019083 | 756 | 10 | 2.65% |
| hsa_circ_0019306 | 756 | 10 | 2.65% |
| hsa_circ_0019660 | 756 | 10 | 2.65% |
| hsa_circ_0025304 | 756 | 10 | 2.65% |
| hsa_circ_0045507 | 756 | 10 | 2.65% |
| hsa_circ_0062970 | 756 | 10 | 2.65% |
| hsa_circ_0064075 | 756 | 10 | 2.65% |
| hsa_circ_0068314 | 756 | 10 | 2.65% |
| hsa_circ_0080506 | 756 | 10 | 2.65% |
| hsa_circ_0081434 | 756 | 10 | 2.65% |
| hsa_circ_0086650 | 756 | 10 | 2.65% |
| hsa_circ_0090957 | 756 | 10 | 2.65% |
| hsa_circ_0100640 | 756 | 10 | 2.65% |
| hsa_circ_0101868 | 756 | 10 | 2.65% |
| hsa_circ_0120648 | 756 | 10 | 2.65% |
| hsa_circ_0122117 | 756 | 10 | 2.65% |
| hsa_circ_0130123 | 756 | 10 | 2.65% |
| hsa_circ_0134149 | 756 | 10 | 2.65% |
| hsa_circ_0121267 | 983 | 13 | 2.64% |
| hsa_circ_0019663 | 832 | 11 | 2.64% |
| hsa_circ_0080042 | 832 | 11 | 2.64% |
| hsa_circ_0123057 | 832 | 11 | 2.64% |
| hsa_circ_0107712 | 1816 | 24 | 2.64% |
| hsa_circ_0067311 | 908 | 12 | 2.64% |
| hsa_circ_0072963 | 908 | 12 | 2.64% |
| hsa_circ_0103805 | 908 | 12 | 2.64% |
| hsa_circ_0109797 | 984 | 13 | 2.64% |
| hsa_circ_0004639 | 757 | 10 | 2.64% |
| hsa_circ_0015987 | 757 | 10 | 2.64% |
| hsa_circ_0030615 | 757 | 10 | 2.64% |
| hsa_circ_0034712 | 757 | 10 | 2.64% |
| hsa_circ_0044464 | 757 | 10 | 2.64% |

| | | | |
|---|---|---|---|
| hsa_circ_0050182 | 757 | 10 | 2.64% |
| hsa_circ_0055225 | 757 | 10 | 2.64% |
| hsa_circ_0077709 | 757 | 10 | 2.64% |
| hsa_circ_0100460 | 757 | 10 | 2.64% |
| hsa_circ_0124569 | 757 | 10 | 2.64% |
| hsa_circ_0133129 | 757 | 10 | 2.64% |
| hsa_circ_0137009 | 757 | 10 | 2.64% |
| hsa_circ_0107174 | 1060 | 14 | 2.64% |
| hsa_circ_0014150 | 833 | 11 | 2.64% |
| hsa_circ_0097012 | 833 | 11 | 2.64% |
| hsa_circ_0121975 | 833 | 11 | 2.64% |
| hsa_circ_0096687 | 1364 | 18 | 2.64% |
| hsa_circ_0053380 | 39787 | 525 | 2.64% |
| hsa_circ_0007288 | 758 | 10 | 2.64% |
| hsa_circ_0008143 | 758 | 10 | 2.64% |
| hsa_circ_0010097 | 758 | 10 | 2.64% |
| hsa_circ_0041418 | 758 | 10 | 2.64% |
| hsa_circ_0049705 | 758 | 10 | 2.64% |
| hsa_circ_0050070 | 758 | 10 | 2.64% |
| hsa_circ_0055265 | 758 | 10 | 2.64% |
| hsa_circ_0066166 | 758 | 10 | 2.64% |
| hsa_circ_0066169 | 758 | 10 | 2.64% |
| hsa_circ_0067201 | 758 | 10 | 2.64% |
| hsa_circ_0083252 | 758 | 10 | 2.64% |
| hsa_circ_0120875 | 758 | 10 | 2.64% |
| hsa_circ_0121276 | 758 | 10 | 2.64% |
| hsa_circ_0125464 | 758 | 10 | 2.64% |
| hsa_circ_0030231 | 834 | 11 | 2.64% |
| hsa_circ_0065419 | 834 | 11 | 2.64% |
| hsa_circ_0068717 | 834 | 11 | 2.64% |
| hsa_circ_0112409 | 834 | 11 | 2.64% |
| hsa_circ_0131142 | 834 | 11 | 2.64% |
| hsa_circ_0021606 | 910 | 12 | 2.64% |
| hsa_circ_0077588 | 1062 | 14 | 2.64% |
| hsa_circ_0003663 | 759 | 10 | 2.64% |
| hsa_circ_0006741 | 759 | 10 | 2.64% |
| hsa_circ_0029428 | 759 | 10 | 2.64% |
| hsa_circ_0039001 | 759 | 10 | 2.64% |
| hsa_circ_0039339 | 759 | 10 | 2.64% |
| hsa_circ_0070319 | 759 | 10 | 2.64% |
| hsa_circ_0075254 | 759 | 10 | 2.64% |
| hsa_circ_0079055 | 759 | 10 | 2.64% |
| hsa_circ_0095548 | 759 | 10 | 2.64% |
| hsa_circ_0101988 | 759 | 10 | 2.64% |
| hsa_circ_0103278 | 759 | 10 | 2.64% |
| hsa_circ_0110795 | 759 | 10 | 2.64% |
| hsa_circ_0111786 | 759 | 10 | 2.64% |
| hsa_circ_0120764 | 759 | 10 | 2.64% |
| hsa_circ_0123960 | 759 | 10 | 2.64% |
| hsa_circ_0127898 | 759 | 10 | 2.64% |

| | | | |
|---|---|---|---|
| hsa_circ_0136682 | 759 | 10 | 2.64% |
| hsa_circ_0012119 | 835 | 11 | 2.63% |
| hsa_circ_0034622 | 835 | 11 | 2.63% |
| hsa_circ_0037125 | 835 | 11 | 2.63% |
| hsa_circ_0042337 | 835 | 11 | 2.63% |
| hsa_circ_0099572 | 835 | 11 | 2.63% |
| hsa_circ_0045254 | 911 | 12 | 2.63% |
| hsa_circ_0083847 | 911 | 12 | 2.63% |
| hsa_circ_0124830 | 911 | 12 | 2.63% |
| hsa_circ_0096561 | 2356 | 31 | 2.63% |
| hsa_circ_0017293 | 912 | 12 | 2.63% |
| hsa_circ_0085466 | 912 | 12 | 2.63% |
| hsa_circ_0011289 | 836 | 11 | 2.63% |
| hsa_circ_0022407 | 836 | 11 | 2.63% |
| hsa_circ_0078627 | 836 | 11 | 2.63% |
| hsa_circ_0080935 | 836 | 11 | 2.63% |
| hsa_circ_0085279 | 836 | 11 | 2.63% |
| hsa_circ_0122549 | 836 | 11 | 2.63% |
| hsa_circ_0037906 | 760 | 10 | 2.63% |
| hsa_circ_0057097 | 760 | 10 | 2.63% |
| hsa_circ_0126643 | 913 | 12 | 2.63% |
| hsa_circ_0042787 | 837 | 11 | 2.63% |
| hsa_circ_0085009 | 837 | 11 | 2.63% |
| hsa_circ_0118856 | 837 | 11 | 2.63% |
| hsa_circ_0132904 | 837 | 11 | 2.63% |
| hsa_circ_0006401 | 761 | 10 | 2.63% |
| hsa_circ_0007611 | 761 | 10 | 2.63% |
| hsa_circ_0008487 | 761 | 10 | 2.63% |
| hsa_circ_0012741 | 761 | 10 | 2.63% |
| hsa_circ_0022316 | 761 | 10 | 2.63% |
| hsa_circ_0027626 | 761 | 10 | 2.63% |
| hsa_circ_0038076 | 761 | 10 | 2.63% |
| hsa_circ_0038245 | 761 | 10 | 2.63% |
| hsa_circ_0038345 | 761 | 10 | 2.63% |
| hsa_circ_0039668 | 761 | 10 | 2.63% |
| hsa_circ_0041756 | 761 | 10 | 2.63% |
| hsa_circ_0045866 | 761 | 10 | 2.63% |
| hsa_circ_0048662 | 761 | 10 | 2.63% |
| hsa_circ_0059879 | 761 | 10 | 2.63% |
| hsa_circ_0074326 | 761 | 10 | 2.63% |
| hsa_circ_0079816 | 761 | 10 | 2.63% |
| hsa_circ_0083112 | 761 | 10 | 2.63% |
| hsa_circ_0137064 | 761 | 10 | 2.63% |
| hsa_circ_0133911 | 1066 | 14 | 2.63% |
| hsa_circ_0022694 | 838 | 11 | 2.63% |
| hsa_circ_0038182 | 838 | 11 | 2.63% |
| hsa_circ_0038358 | 838 | 11 | 2.63% |
| hsa_circ_0051150 | 838 | 11 | 2.63% |
| hsa_circ_0086770 | 838 | 11 | 2.63% |
| hsa_circ_0090597 | 838 | 11 | 2.63% |

| | | | |
|---|---|---|---|
| hsa_circ_0096107 | 838 | 11 | 2.63% |
| hsa_circ_0004879 | 762 | 10 | 2.62% |
| hsa_circ_0012595 | 762 | 10 | 2.62% |
| hsa_circ_0013336 | 762 | 10 | 2.62% |
| hsa_circ_0015439 | 762 | 10 | 2.62% |
| hsa_circ_0016252 | 762 | 10 | 2.62% |
| hsa_circ_0022373 | 762 | 10 | 2.62% |
| hsa_circ_0023492 | 762 | 10 | 2.62% |
| hsa_circ_0026799 | 762 | 10 | 2.62% |
| hsa_circ_0035611 | 762 | 10 | 2.62% |
| hsa_circ_0051902 | 762 | 10 | 2.62% |
| hsa_circ_0057561 | 762 | 10 | 2.62% |
| hsa_circ_0080102 | 762 | 10 | 2.62% |
| hsa_circ_0083052 | 762 | 10 | 2.62% |
| hsa_circ_0084522 | 762 | 10 | 2.62% |
| hsa_circ_0085730 | 762 | 10 | 2.62% |
| hsa_circ_0087686 | 762 | 10 | 2.62% |
| hsa_circ_0096758 | 762 | 10 | 2.62% |
| hsa_circ_0099881 | 762 | 10 | 2.62% |
| hsa_circ_0102474 | 762 | 10 | 2.62% |
| hsa_circ_0105513 | 762 | 10 | 2.62% |
| hsa_circ_0106726 | 762 | 10 | 2.62% |
| hsa_circ_0108794 | 762 | 10 | 2.62% |
| hsa_circ_0114638 | 762 | 10 | 2.62% |
| hsa_circ_0121988 | 762 | 10 | 2.62% |
| hsa_circ_0122348 | 762 | 10 | 2.62% |
| hsa_circ_0125279 | 762 | 10 | 2.62% |
| hsa_circ_0140409 | 762 | 10 | 2.62% |
| hsa_circ_0057342 | 915 | 12 | 2.62% |
| hsa_circ_0011148 | 839 | 11 | 2.62% |
| hsa_circ_0018626 | 839 | 11 | 2.62% |
| hsa_circ_0049835 | 839 | 11 | 2.62% |
| hsa_circ_0060638 | 839 | 11 | 2.62% |
| hsa_circ_0088288 | 839 | 11 | 2.62% |
| hsa_circ_0089878 | 839 | 11 | 2.62% |
| hsa_circ_0095495 | 839 | 11 | 2.62% |
| hsa_circ_0109584 | 839 | 11 | 2.62% |
| hsa_circ_0049253 | 1068 | 14 | 2.62% |
| hsa_circ_0019541 | 763 | 10 | 2.62% |
| hsa_circ_0038990 | 763 | 10 | 2.62% |
| hsa_circ_0088544 | 763 | 10 | 2.62% |
| hsa_circ_0101775 | 763 | 10 | 2.62% |
| hsa_circ_0124527 | 992 | 13 | 2.62% |
| hsa_circ_0125923 | 916 | 12 | 2.62% |
| hsa_circ_0090418 | 1069 | 14 | 2.62% |
| hsa_circ_0003768 | 840 | 11 | 2.62% |
| hsa_circ_0009462 | 840 | 11 | 2.62% |
| hsa_circ_0020598 | 840 | 11 | 2.62% |
| hsa_circ_0034796 | 840 | 11 | 2.62% |
| hsa_circ_0069054 | 840 | 11 | 2.62% |

| | | | |
|---|---|---|---|
| hsa_circ_0090681 | 840 | 11 | 2.62% |
| hsa_circ_0092348 | 840 | 11 | 2.62% |
| hsa_circ_0098334 | 840 | 11 | 2.62% |
| hsa_circ_0101091 | 840 | 11 | 2.62% |
| hsa_circ_0113156 | 840 | 11 | 2.62% |
| hsa_circ_0120672 | 840 | 11 | 2.62% |
| hsa_circ_0121163 | 840 | 11 | 2.62% |
| hsa_circ_0126282 | 840 | 11 | 2.62% |
| hsa_circ_0135110 | 840 | 11 | 2.62% |
| hsa_circ_0048044 | 993 | 13 | 2.62% |
| hsa_circ_0016108 | 764 | 10 | 2.62% |
| hsa_circ_0037898 | 764 | 10 | 2.62% |
| hsa_circ_0038545 | 764 | 10 | 2.62% |
| hsa_circ_0068621 | 764 | 10 | 2.62% |
| hsa_circ_0116747 | 764 | 10 | 2.62% |
| hsa_circ_0126621 | 764 | 10 | 2.62% |
| hsa_circ_0127223 | 764 | 10 | 2.62% |
| hsa_circ_0136175 | 764 | 10 | 2.62% |
| hsa_circ_0139397 | 764 | 10 | 2.62% |
| hsa_circ_0031268 | 917 | 12 | 2.62% |
| hsa_circ_0060488 | 917 | 12 | 2.62% |
| hsa_circ_0023088 | 841 | 11 | 2.62% |
| hsa_circ_0025142 | 841 | 11 | 2.62% |
| hsa_circ_0044030 | 841 | 11 | 2.62% |
| hsa_circ_0057202 | 841 | 11 | 2.62% |
| hsa_circ_0070809 | 841 | 11 | 2.62% |
| hsa_circ_0075950 | 841 | 11 | 2.62% |
| hsa_circ_0076255 | 841 | 11 | 2.62% |
| hsa_circ_0105805 | 841 | 11 | 2.62% |
| hsa_circ_0118775 | 841 | 11 | 2.62% |
| hsa_circ_0033477 | 918 | 12 | 2.61% |
| hsa_circ_0037639 | 918 | 12 | 2.61% |
| hsa_circ_0088523 | 918 | 12 | 2.61% |
| hsa_circ_0013028 | 765 | 10 | 2.61% |
| hsa_circ_0033668 | 765 | 10 | 2.61% |
| hsa_circ_0036098 | 765 | 10 | 2.61% |
| hsa_circ_0041899 | 765 | 10 | 2.61% |
| hsa_circ_0060480 | 765 | 10 | 2.61% |
| hsa_circ_0063979 | 765 | 10 | 2.61% |
| hsa_circ_0073618 | 765 | 10 | 2.61% |
| hsa_circ_0084875 | 765 | 10 | 2.61% |
| hsa_circ_0120043 | 765 | 10 | 2.61% |
| hsa_circ_0132788 | 765 | 10 | 2.61% |
| hsa_circ_0019756 | 842 | 11 | 2.61% |
| hsa_circ_0020155 | 842 | 11 | 2.61% |
| hsa_circ_0044259 | 842 | 11 | 2.61% |
| hsa_circ_0068241 | 842 | 11 | 2.61% |
| hsa_circ_0091151 | 842 | 11 | 2.61% |
| hsa_circ_0077187 | 1072 | 14 | 2.61% |
| hsa_circ_0037435 | 919 | 12 | 2.61% |

| | | | |
|---|---|---|---|
| hsa_circ_0133885 | 919 | 12 | 2.61% |
| hsa_circ_0002029 | 766 | 10 | 2.61% |
| hsa_circ_0007357 | 766 | 10 | 2.61% |
| hsa_circ_0012745 | 766 | 10 | 2.61% |
| hsa_circ_0014683 | 766 | 10 | 2.61% |
| hsa_circ_0029689 | 766 | 10 | 2.61% |
| hsa_circ_0037040 | 766 | 10 | 2.61% |
| hsa_circ_0039110 | 766 | 10 | 2.61% |
| hsa_circ_0046449 | 766 | 10 | 2.61% |
| hsa_circ_0069866 | 766 | 10 | 2.61% |
| hsa_circ_0080874 | 766 | 10 | 2.61% |
| hsa_circ_0103413 | 766 | 10 | 2.61% |
| hsa_circ_0128133 | 766 | 10 | 2.61% |
| hsa_circ_0132040 | 766 | 10 | 2.61% |
| hsa_circ_0133477 | 766 | 10 | 2.61% |
| hsa_circ_0019950 | 996 | 13 | 2.61% |
| hsa_circ_0021981 | 996 | 13 | 2.61% |
| hsa_circ_0067927 | 996 | 13 | 2.61% |
| hsa_circ_0002059 | 843 | 11 | 2.61% |
| hsa_circ_0037129 | 843 | 11 | 2.61% |
| hsa_circ_0042752 | 843 | 11 | 2.61% |
| hsa_circ_0062414 | 843 | 11 | 2.61% |
| hsa_circ_0067456 | 843 | 11 | 2.61% |
| hsa_circ_0081241 | 843 | 11 | 2.61% |
| hsa_circ_0087333 | 843 | 11 | 2.61% |
| hsa_circ_0104514 | 843 | 11 | 2.61% |
| hsa_circ_0047786 | 2683 | 35 | 2.61% |
| hsa_circ_0103552 | 920 | 12 | 2.61% |
| hsa_circ_0106710 | 920 | 12 | 2.61% |
| hsa_circ_0003421 | 767 | 10 | 2.61% |
| hsa_circ_0014713 | 767 | 10 | 2.61% |
| hsa_circ_0041818 | 767 | 10 | 2.61% |
| hsa_circ_0074345 | 767 | 10 | 2.61% |
| hsa_circ_0091664 | 767 | 10 | 2.61% |
| hsa_circ_0105486 | 767 | 10 | 2.61% |
| hsa_circ_0106837 | 767 | 10 | 2.61% |
| hsa_circ_0121869 | 767 | 10 | 2.61% |
| hsa_circ_0125257 | 767 | 10 | 2.61% |
| hsa_circ_0130048 | 767 | 10 | 2.61% |
| hsa_circ_0134952 | 767 | 10 | 2.61% |
| hsa_circ_0084884 | 1074 | 14 | 2.61% |
| hsa_circ_0045056 | 844 | 11 | 2.61% |
| hsa_circ_0076354 | 844 | 11 | 2.61% |
| hsa_circ_0102764 | 844 | 11 | 2.61% |
| hsa_circ_0120449 | 844 | 11 | 2.61% |
| hsa_circ_0093961 | 2073 | 27 | 2.60% |
| hsa_circ_0002249 | 768 | 10 | 2.60% |
| hsa_circ_0006883 | 768 | 10 | 2.60% |
| hsa_circ_0058375 | 768 | 10 | 2.60% |
| hsa_circ_0061948 | 768 | 10 | 2.60% |

| | | | |
|---|---|---|---|
| hsa_circ_0064822 | 768 | 10 | 2.60% |
| hsa_circ_0069619 | 768 | 10 | 2.60% |
| hsa_circ_0076931 | 768 | 10 | 2.60% |
| hsa_circ_0086719 | 768 | 10 | 2.60% |
| hsa_circ_0087395 | 768 | 10 | 2.60% |
| hsa_circ_0094612 | 768 | 10 | 2.60% |
| hsa_circ_0098425 | 768 | 10 | 2.60% |
| hsa_circ_0100294 | 768 | 10 | 2.60% |
| hsa_circ_0107073 | 768 | 10 | 2.60% |
| hsa_circ_0108566 | 768 | 10 | 2.60% |
| hsa_circ_0113721 | 768 | 10 | 2.60% |
| hsa_circ_0117216 | 768 | 10 | 2.60% |
| hsa_circ_0118000 | 768 | 10 | 2.60% |
| hsa_circ_0133651 | 768 | 10 | 2.60% |
| hsa_circ_0133708 | 768 | 10 | 2.60% |
| hsa_circ_0052065 | 1690 | 22 | 2.60% |
| hsa_circ_0072128 | 845 | 11 | 2.60% |
| hsa_circ_0078030 | 845 | 11 | 2.60% |
| hsa_circ_0088349 | 845 | 11 | 2.60% |
| hsa_circ_0029482 | 922 | 12 | 2.60% |
| hsa_circ_0053264 | 922 | 12 | 2.60% |
| hsa_circ_0089092 | 922 | 12 | 2.60% |
| hsa_circ_0103336 | 922 | 12 | 2.60% |
| hsa_circ_0104475 | 922 | 12 | 2.60% |
| hsa_circ_0015596 | 769 | 10 | 2.60% |
| hsa_circ_0028760 | 769 | 10 | 2.60% |
| hsa_circ_0030806 | 769 | 10 | 2.60% |
| hsa_circ_0036318 | 769 | 10 | 2.60% |
| hsa_circ_0054287 | 769 | 10 | 2.60% |
| hsa_circ_0075965 | 769 | 10 | 2.60% |
| hsa_circ_0079059 | 769 | 10 | 2.60% |
| hsa_circ_0083248 | 769 | 10 | 2.60% |
| hsa_circ_0089225 | 769 | 10 | 2.60% |
| hsa_circ_0101732 | 769 | 10 | 2.60% |
| hsa_circ_0103763 | 769 | 10 | 2.60% |
| hsa_circ_0109006 | 769 | 10 | 2.60% |
| hsa_circ_0110279 | 769 | 10 | 2.60% |
| hsa_circ_0134944 | 769 | 10 | 2.60% |
| hsa_circ_0135926 | 769 | 10 | 2.60% |
| hsa_circ_0139539 | 769 | 10 | 2.60% |
| hsa_circ_0009216 | 846 | 11 | 2.60% |
| hsa_circ_0030726 | 846 | 11 | 2.60% |
| hsa_circ_0081180 | 846 | 11 | 2.60% |
| hsa_circ_0094380 | 846 | 11 | 2.60% |
| hsa_circ_0094762 | 846 | 11 | 2.60% |
| hsa_circ_0110059 | 846 | 11 | 2.60% |
| hsa_circ_0033417 | 923 | 12 | 2.60% |
| hsa_circ_0031771 | 1078 | 14 | 2.60% |
| hsa_circ_0060433 | 1001 | 13 | 2.60% |
| hsa_circ_0006212 | 924 | 12 | 2.60% |

| | | | |
|---|---|---|---|
| hsa_circ_0036391 | 924 | 12 | 2.60% |
| hsa_circ_0058919 | 924 | 12 | 2.60% |
| hsa_circ_0072191 | 924 | 12 | 2.60% |
| hsa_circ_0100108 | 924 | 12 | 2.60% |
| hsa_circ_0104476 | 924 | 12 | 2.60% |
| hsa_circ_0023701 | 847 | 11 | 2.60% |
| hsa_circ_0074992 | 847 | 11 | 2.60% |
| hsa_circ_0091513 | 847 | 11 | 2.60% |
| hsa_circ_0134288 | 847 | 11 | 2.60% |
| hsa_circ_0135828 | 847 | 11 | 2.60% |
| hsa_circ_0014484 | 770 | 10 | 2.60% |
| hsa_circ_0014506 | 770 | 10 | 2.60% |
| hsa_circ_0028476 | 770 | 10 | 2.60% |
| hsa_circ_0044327 | 770 | 10 | 2.60% |
| hsa_circ_0056638 | 770 | 10 | 2.60% |
| hsa_circ_0075437 | 770 | 10 | 2.60% |
| hsa_circ_0093777 | 770 | 10 | 2.60% |
| hsa_circ_0111402 | 770 | 10 | 2.60% |
| hsa_circ_0115173 | 770 | 10 | 2.60% |
| hsa_circ_0119204 | 770 | 10 | 2.60% |
| hsa_circ_0130290 | 770 | 10 | 2.60% |
| hsa_circ_0131815 | 770 | 10 | 2.60% |
| hsa_circ_0048751 | 925 | 12 | 2.59% |
| hsa_circ_0116757 | 925 | 12 | 2.59% |
| hsa_circ_0009467 | 848 | 11 | 2.59% |
| hsa_circ_0013463 | 848 | 11 | 2.59% |
| hsa_circ_0029272 | 848 | 11 | 2.59% |
| hsa_circ_0031636 | 848 | 11 | 2.59% |
| hsa_circ_0088031 | 848 | 11 | 2.59% |
| hsa_circ_0002649 | 771 | 10 | 2.59% |
| hsa_circ_0016682 | 771 | 10 | 2.59% |
| hsa_circ_0040092 | 771 | 10 | 2.59% |
| hsa_circ_0054511 | 771 | 10 | 2.59% |
| hsa_circ_0055434 | 771 | 10 | 2.59% |
| hsa_circ_0059321 | 771 | 10 | 2.59% |
| hsa_circ_0066202 | 771 | 10 | 2.59% |
| hsa_circ_0068744 | 771 | 10 | 2.59% |
| hsa_circ_0082030 | 771 | 10 | 2.59% |
| hsa_circ_0084572 | 771 | 10 | 2.59% |
| hsa_circ_0093225 | 771 | 10 | 2.59% |
| hsa_circ_0097878 | 771 | 10 | 2.59% |
| hsa_circ_0115468 | 771 | 10 | 2.59% |
| hsa_circ_0126441 | 771 | 10 | 2.59% |
| hsa_circ_0133959 | 771 | 10 | 2.59% |
| hsa_circ_0135131 | 771 | 10 | 2.59% |
| hsa_circ_0043015 | 3779 | 49 | 2.59% |
| hsa_circ_0101963 | 1003 | 13 | 2.59% |
| hsa_circ_0003520 | 849 | 11 | 2.59% |
| hsa_circ_0036866 | 849 | 11 | 2.59% |
| hsa_circ_0064418 | 849 | 11 | 2.59% |

| | | | |
|---|---|---|---|
| hsa_circ_0104566 | 849 | 11 | 2.59% |
| hsa_circ_0107861 | 849 | 11 | 2.59% |
| hsa_circ_0110069 | 849 | 11 | 2.59% |
| hsa_circ_0017453 | 772 | 10 | 2.59% |
| hsa_circ_0067125 | 772 | 10 | 2.59% |
| hsa_circ_0070919 | 772 | 10 | 2.59% |
| hsa_circ_0082464 | 772 | 10 | 2.59% |
| hsa_circ_0110458 | 772 | 10 | 2.59% |
| hsa_circ_0124185 | 927 | 12 | 2.59% |
| hsa_circ_0085548 | 850 | 11 | 2.59% |
| hsa_circ_0119211 | 850 | 11 | 2.59% |
| hsa_circ_0131271 | 850 | 11 | 2.59% |
| hsa_circ_0032965 | 773 | 10 | 2.59% |
| hsa_circ_0043004 | 773 | 10 | 2.59% |
| hsa_circ_0047329 | 773 | 10 | 2.59% |
| hsa_circ_0060135 | 773 | 10 | 2.59% |
| hsa_circ_0076726 | 773 | 10 | 2.59% |
| hsa_circ_0076779 | 773 | 10 | 2.59% |
| hsa_circ_0092221 | 773 | 10 | 2.59% |
| hsa_circ_0112095 | 773 | 10 | 2.59% |
| hsa_circ_0115497 | 773 | 10 | 2.59% |
| hsa_circ_0134171 | 773 | 10 | 2.59% |
| hsa_circ_0136548 | 773 | 10 | 2.59% |
| hsa_circ_0132548 | 2474 | 32 | 2.59% |
| hsa_circ_0084749 | 928 | 12 | 2.59% |
| hsa_circ_0018786 | 851 | 11 | 2.59% |
| hsa_circ_0042982 | 851 | 11 | 2.59% |
| hsa_circ_0096493 | 851 | 11 | 2.59% |
| hsa_circ_0119508 | 851 | 11 | 2.59% |
| hsa_circ_0125212 | 851 | 11 | 2.59% |
| hsa_circ_0138951 | 1006 | 13 | 2.58% |
| hsa_circ_0045744 | 1161 | 15 | 2.58% |
| hsa_circ_0007659 | 774 | 10 | 2.58% |
| hsa_circ_0018646 | 774 | 10 | 2.58% |
| hsa_circ_0020765 | 774 | 10 | 2.58% |
| hsa_circ_0030738 | 774 | 10 | 2.58% |
| hsa_circ_0038958 | 774 | 10 | 2.58% |
| hsa_circ_0054036 | 774 | 10 | 2.58% |
| hsa_circ_0057328 | 774 | 10 | 2.58% |
| hsa_circ_0061327 | 774 | 10 | 2.58% |
| hsa_circ_0074005 | 774 | 10 | 2.58% |
| hsa_circ_0091312 | 774 | 10 | 2.58% |
| hsa_circ_0095250 | 774 | 10 | 2.58% |
| hsa_circ_0112064 | 774 | 10 | 2.58% |
| hsa_circ_0112242 | 774 | 10 | 2.58% |
| hsa_circ_0122305 | 774 | 10 | 2.58% |
| hsa_circ_0125671 | 774 | 10 | 2.58% |
| hsa_circ_0026316 | 852 | 11 | 2.58% |
| hsa_circ_0026488 | 852 | 11 | 2.58% |
| hsa_circ_0030875 | 852 | 11 | 2.58% |

| | | | |
|---|---|---|---|
| hsa_circ_0044605 | 852 | 11 | 2.58% |
| hsa_circ_0081183 | 852 | 11 | 2.58% |
| hsa_circ_0088289 | 852 | 11 | 2.58% |
| hsa_circ_0089294 | 852 | 11 | 2.58% |
| hsa_circ_0037176 | 1007 | 13 | 2.58% |
| hsa_circ_0042687 | 930 | 12 | 2.58% |
| hsa_circ_0096603 | 930 | 12 | 2.58% |
| hsa_circ_0112472 | 930 | 12 | 2.58% |
| hsa_circ_0021935 | 775 | 10 | 2.58% |
| hsa_circ_0035849 | 775 | 10 | 2.58% |
| hsa_circ_0047782 | 775 | 10 | 2.58% |
| hsa_circ_0055921 | 775 | 10 | 2.58% |
| hsa_circ_0063500 | 775 | 10 | 2.58% |
| hsa_circ_0066504 | 775 | 10 | 2.58% |
| hsa_circ_0077806 | 775 | 10 | 2.58% |
| hsa_circ_0087902 | 775 | 10 | 2.58% |
| hsa_circ_0094148 | 775 | 10 | 2.58% |
| hsa_circ_0094714 | 775 | 10 | 2.58% |
| hsa_circ_0106374 | 775 | 10 | 2.58% |
| hsa_circ_0119604 | 775 | 10 | 2.58% |
| hsa_circ_0137472 | 775 | 10 | 2.58% |
| hsa_circ_0140042 | 775 | 10 | 2.58% |
| hsa_circ_0137237 | 1938 | 25 | 2.58% |
| hsa_circ_0009513 | 853 | 11 | 2.58% |
| hsa_circ_0031120 | 853 | 11 | 2.58% |
| hsa_circ_0041320 | 853 | 11 | 2.58% |
| hsa_circ_0113807 | 853 | 11 | 2.58% |
| hsa_circ_0135602 | 853 | 11 | 2.58% |
| hsa_circ_0136120 | 853 | 11 | 2.58% |
| hsa_circ_0003600 | 776 | 10 | 2.58% |
| hsa_circ_0005678 | 776 | 10 | 2.58% |
| hsa_circ_0014004 | 776 | 10 | 2.58% |
| hsa_circ_0020230 | 776 | 10 | 2.58% |
| hsa_circ_0023466 | 776 | 10 | 2.58% |
| hsa_circ_0042093 | 776 | 10 | 2.58% |
| hsa_circ_0060607 | 776 | 10 | 2.58% |
| hsa_circ_0101616 | 776 | 10 | 2.58% |
| hsa_circ_0104148 | 776 | 10 | 2.58% |
| hsa_circ_0128246 | 776 | 10 | 2.58% |
| hsa_circ_0005209 | 932 | 12 | 2.58% |
| hsa_circ_0014854 | 932 | 12 | 2.58% |
| hsa_circ_0034425 | 932 | 12 | 2.58% |
| hsa_circ_0118183 | 932 | 12 | 2.58% |
| hsa_circ_0006795 | 777 | 10 | 2.57% |
| hsa_circ_0006815 | 777 | 10 | 2.57% |
| hsa_circ_0009798 | 777 | 10 | 2.57% |
| hsa_circ_0014299 | 777 | 10 | 2.57% |
| hsa_circ_0034747 | 777 | 10 | 2.57% |
| hsa_circ_0035923 | 777 | 10 | 2.57% |
| hsa_circ_0039672 | 777 | 10 | 2.57% |

| | | | |
|---|---|---|---|
| hsa_circ_0041739 | 777 | 10 | 2.57% |
| hsa_circ_0044242 | 777 | 10 | 2.57% |
| hsa_circ_0046222 | 777 | 10 | 2.57% |
| hsa_circ_0048691 | 777 | 10 | 2.57% |
| hsa_circ_0081473 | 777 | 10 | 2.57% |
| hsa_circ_0090456 | 777 | 10 | 2.57% |
| hsa_circ_0099020 | 777 | 10 | 2.57% |
| hsa_circ_0099510 | 777 | 10 | 2.57% |
| hsa_circ_0111854 | 777 | 10 | 2.57% |
| hsa_circ_0112257 | 777 | 10 | 2.57% |
| hsa_circ_0117408 | 777 | 10 | 2.57% |
| hsa_circ_0124462 | 777 | 10 | 2.57% |
| hsa_circ_0127560 | 777 | 10 | 2.57% |
| hsa_circ_0132532 | 777 | 10 | 2.57% |
| hsa_circ_0078821 | 855 | 11 | 2.57% |
| hsa_circ_0092021 | 855 | 11 | 2.57% |
| hsa_circ_0103463 | 855 | 11 | 2.57% |
| hsa_circ_0121437 | 855 | 11 | 2.57% |
| hsa_circ_0127015 | 855 | 11 | 2.57% |
| hsa_circ_0121506 | 1166 | 15 | 2.57% |
| hsa_circ_0029402 | 2566 | 33 | 2.57% |
| hsa_circ_0111242 | 1011 | 13 | 2.57% |
| hsa_circ_0018845 | 778 | 10 | 2.57% |
| hsa_circ_0026984 | 778 | 10 | 2.57% |
| hsa_circ_0029505 | 778 | 10 | 2.57% |
| hsa_circ_0048818 | 778 | 10 | 2.57% |
| hsa_circ_0087367 | 778 | 10 | 2.57% |
| hsa_circ_0089533 | 778 | 10 | 2.57% |
| hsa_circ_0095785 | 778 | 10 | 2.57% |
| hsa_circ_0103837 | 778 | 10 | 2.57% |
| hsa_circ_0104832 | 778 | 10 | 2.57% |
| hsa_circ_0106745 | 778 | 10 | 2.57% |
| hsa_circ_0132454 | 778 | 10 | 2.57% |
| hsa_circ_0140212 | 778 | 10 | 2.57% |
| hsa_circ_0005536 | 856 | 11 | 2.57% |
| hsa_circ_0009118 | 856 | 11 | 2.57% |
| hsa_circ_0035944 | 856 | 11 | 2.57% |
| hsa_circ_0051089 | 856 | 11 | 2.57% |
| hsa_circ_0054067 | 856 | 11 | 2.57% |
| hsa_circ_0087555 | 856 | 11 | 2.57% |
| hsa_circ_0117297 | 856 | 11 | 2.57% |
| hsa_circ_0103374 | 934 | 12 | 2.57% |
| hsa_circ_0000342 | 779 | 10 | 2.57% |
| hsa_circ_0009698 | 779 | 10 | 2.57% |
| hsa_circ_0014338 | 779 | 10 | 2.57% |
| hsa_circ_0025252 | 779 | 10 | 2.57% |
| hsa_circ_0041589 | 779 | 10 | 2.57% |
| hsa_circ_0053432 | 779 | 10 | 2.57% |
| hsa_circ_0059688 | 779 | 10 | 2.57% |
| hsa_circ_0067999 | 779 | 10 | 2.57% |

| | | | |
|---|---|---|---|
| hsa_circ_0073115 | 779 | 10 | 2.57% |
| hsa_circ_0085162 | 779 | 10 | 2.57% |
| hsa_circ_0087669 | 779 | 10 | 2.57% |
| hsa_circ_0093979 | 779 | 10 | 2.57% |
| hsa_circ_0094038 | 779 | 10 | 2.57% |
| hsa_circ_0096704 | 779 | 10 | 2.57% |
| hsa_circ_0100088 | 779 | 10 | 2.57% |
| hsa_circ_0114918 | 779 | 10 | 2.57% |
| hsa_circ_0116345 | 779 | 10 | 2.57% |
| hsa_circ_0127487 | 779 | 10 | 2.57% |
| hsa_circ_0133061 | 779 | 10 | 2.57% |
| hsa_circ_0029624 | 857 | 11 | 2.57% |
| hsa_circ_0051157 | 857 | 11 | 2.57% |
| hsa_circ_0075934 | 857 | 11 | 2.57% |
| hsa_circ_0053948 | 935 | 12 | 2.57% |
| hsa_circ_0018429 | 936 | 12 | 2.56% |
| hsa_circ_0024218 | 936 | 12 | 2.56% |
| hsa_circ_0049117 | 936 | 12 | 2.56% |
| hsa_circ_0124620 | 936 | 12 | 2.56% |
| hsa_circ_0013269 | 858 | 11 | 2.56% |
| hsa_circ_0016521 | 858 | 11 | 2.56% |
| hsa_circ_0019918 | 858 | 11 | 2.56% |
| hsa_circ_0037748 | 858 | 11 | 2.56% |
| hsa_circ_0049921 | 858 | 11 | 2.56% |
| hsa_circ_0089293 | 858 | 11 | 2.56% |
| hsa_circ_0089561 | 858 | 11 | 2.56% |
| hsa_circ_0101836 | 858 | 11 | 2.56% |
| hsa_circ_0122460 | 858 | 11 | 2.56% |
| hsa_circ_0003024 | 780 | 10 | 2.56% |
| hsa_circ_0003200 | 780 | 10 | 2.56% |
| hsa_circ_0003606 | 780 | 10 | 2.56% |
| hsa_circ_0006384 | 780 | 10 | 2.56% |
| hsa_circ_0008747 | 780 | 10 | 2.56% |
| hsa_circ_0025261 | 780 | 10 | 2.56% |
| hsa_circ_0031521 | 780 | 10 | 2.56% |
| hsa_circ_0032278 | 780 | 10 | 2.56% |
| hsa_circ_0034930 | 780 | 10 | 2.56% |
| hsa_circ_0035845 | 780 | 10 | 2.56% |
| hsa_circ_0036163 | 780 | 10 | 2.56% |
| hsa_circ_0040669 | 780 | 10 | 2.56% |
| hsa_circ_0058425 | 780 | 10 | 2.56% |
| hsa_circ_0064004 | 780 | 10 | 2.56% |
| hsa_circ_0068586 | 780 | 10 | 2.56% |
| hsa_circ_0068593 | 780 | 10 | 2.56% |
| hsa_circ_0079137 | 780 | 10 | 2.56% |
| hsa_circ_0080914 | 780 | 10 | 2.56% |
| hsa_circ_0087002 | 780 | 10 | 2.56% |
| hsa_circ_0092852 | 780 | 10 | 2.56% |
| hsa_circ_0108101 | 780 | 10 | 2.56% |
| hsa_circ_0125900 | 780 | 10 | 2.56% |

| | | | |
|---|---|---|---|
| hsa_circ_0127562 | 780 | 10 | 2.56% |
| hsa_circ_0133679 | 780 | 10 | 2.56% |
| hsa_circ_0138526 | 780 | 10 | 2.56% |
| hsa_circ_0138811 | 780 | 10 | 2.56% |
| hsa_circ_0094545 | 6241 | 80 | 2.56% |
| hsa_circ_0067280 | 1171 | 15 | 2.56% |
| hsa_circ_0002494 | 937 | 12 | 2.56% |
| hsa_circ_0089042 | 937 | 12 | 2.56% |
| hsa_circ_0044151 | 859 | 11 | 2.56% |
| hsa_circ_0059847 | 859 | 11 | 2.56% |
| hsa_circ_0065542 | 859 | 11 | 2.56% |
| hsa_circ_0087578 | 859 | 11 | 2.56% |
| hsa_circ_0016794 | 781 | 10 | 2.56% |
| hsa_circ_0023050 | 781 | 10 | 2.56% |
| hsa_circ_0057238 | 781 | 10 | 2.56% |
| hsa_circ_0069728 | 781 | 10 | 2.56% |
| hsa_circ_0072472 | 781 | 10 | 2.56% |
| hsa_circ_0072868 | 781 | 10 | 2.56% |
| hsa_circ_0076677 | 781 | 10 | 2.56% |
| hsa_circ_0087359 | 781 | 10 | 2.56% |
| hsa_circ_0102927 | 781 | 10 | 2.56% |
| hsa_circ_0110132 | 781 | 10 | 2.56% |
| hsa_circ_0111257 | 781 | 10 | 2.56% |
| hsa_circ_0113179 | 781 | 10 | 2.56% |
| hsa_circ_0117265 | 781 | 10 | 2.56% |
| hsa_circ_0121393 | 781 | 10 | 2.56% |
| hsa_circ_0125960 | 781 | 10 | 2.56% |
| hsa_circ_0133264 | 781 | 10 | 2.56% |
| hsa_circ_0111830 | 1328 | 17 | 2.56% |
| hsa_circ_0138282 | 1016 | 13 | 2.56% |
| hsa_circ_0106581 | 938 | 12 | 2.56% |
| hsa_circ_0089706 | 860 | 11 | 2.56% |
| hsa_circ_0011946 | 782 | 10 | 2.56% |
| hsa_circ_0016078 | 782 | 10 | 2.56% |
| hsa_circ_0020619 | 782 | 10 | 2.56% |
| hsa_circ_0025018 | 782 | 10 | 2.56% |
| hsa_circ_0026419 | 782 | 10 | 2.56% |
| hsa_circ_0029968 | 782 | 10 | 2.56% |
| hsa_circ_0033136 | 782 | 10 | 2.56% |
| hsa_circ_0045041 | 782 | 10 | 2.56% |
| hsa_circ_0049907 | 782 | 10 | 2.56% |
| hsa_circ_0064681 | 782 | 10 | 2.56% |
| hsa_circ_0076061 | 782 | 10 | 2.56% |
| hsa_circ_0086370 | 782 | 10 | 2.56% |
| hsa_circ_0087062 | 782 | 10 | 2.56% |
| hsa_circ_0108500 | 782 | 10 | 2.56% |
| hsa_circ_0111928 | 782 | 10 | 2.56% |
| hsa_circ_0132009 | 782 | 10 | 2.56% |
| hsa_circ_0133778 | 782 | 10 | 2.56% |
| hsa_circ_0140211 | 782 | 10 | 2.56% |

| | | | |
|---|---|---|---|
| hsa_circ_0100885 | 1017 | 13 | 2.56% |
| hsa_circ_0136384 | 1017 | 13 | 2.56% |
| hsa_circ_0087104 | 939 | 12 | 2.56% |
| hsa_circ_0022312 | 861 | 11 | 2.56% |
| hsa_circ_0053335 | 861 | 11 | 2.56% |
| hsa_circ_0057051 | 861 | 11 | 2.56% |
| hsa_circ_0074890 | 861 | 11 | 2.56% |
| hsa_circ_0084664 | 861 | 11 | 2.56% |
| hsa_circ_0087799 | 861 | 11 | 2.56% |
| hsa_circ_0105146 | 2270 | 29 | 2.56% |
| hsa_circ_0072097 | 1096 | 14 | 2.55% |
| hsa_circ_0130252 | 1096 | 14 | 2.55% |
| hsa_circ_0015659 | 783 | 10 | 2.55% |
| hsa_circ_0016033 | 783 | 10 | 2.55% |
| hsa_circ_0032591 | 783 | 10 | 2.55% |
| hsa_circ_0037576 | 783 | 10 | 2.55% |
| hsa_circ_0055831 | 783 | 10 | 2.55% |
| hsa_circ_0067541 | 783 | 10 | 2.55% |
| hsa_circ_0083236 | 783 | 10 | 2.55% |
| hsa_circ_0097990 | 783 | 10 | 2.55% |
| hsa_circ_0102483 | 783 | 10 | 2.55% |
| hsa_circ_0139233 | 783 | 10 | 2.55% |
| hsa_circ_0038453 | 1488 | 19 | 2.55% |
| hsa_circ_0021324 | 940 | 12 | 2.55% |
| hsa_circ_0069474 | 940 | 12 | 2.55% |
| hsa_circ_0097247 | 940 | 12 | 2.55% |
| hsa_circ_0058760 | 1097 | 14 | 2.55% |
| hsa_circ_0001805 | 862 | 11 | 2.55% |
| hsa_circ_0003722 | 862 | 11 | 2.55% |
| hsa_circ_0006974 | 862 | 11 | 2.55% |
| hsa_circ_0035909 | 862 | 11 | 2.55% |
| hsa_circ_0098777 | 862 | 11 | 2.55% |
| hsa_circ_0025334 | 784 | 10 | 2.55% |
| hsa_circ_0033670 | 784 | 10 | 2.55% |
| hsa_circ_0048684 | 784 | 10 | 2.55% |
| hsa_circ_0066725 | 784 | 10 | 2.55% |
| hsa_circ_0087323 | 784 | 10 | 2.55% |
| hsa_circ_0096154 | 784 | 10 | 2.55% |
| hsa_circ_0140393 | 784 | 10 | 2.55% |
| hsa_circ_0031269 | 941 | 12 | 2.55% |
| hsa_circ_0033624 | 941 | 12 | 2.55% |
| hsa_circ_0088660 | 941 | 12 | 2.55% |
| hsa_circ_0122003 | 941 | 12 | 2.55% |
| hsa_circ_0006070 | 863 | 11 | 2.55% |
| hsa_circ_0022046 | 863 | 11 | 2.55% |
| hsa_circ_0031856 | 863 | 11 | 2.55% |
| hsa_circ_0040550 | 863 | 11 | 2.55% |
| hsa_circ_0046140 | 863 | 11 | 2.55% |
| hsa_circ_0074808 | 863 | 11 | 2.55% |
| hsa_circ_0108228 | 863 | 11 | 2.55% |

| | | | |
|---|---|---|---|
| hsa_circ_0135676 | 863 | 11 | 2.55% |
| hsa_circ_0081358 | 942 | 12 | 2.55% |
| hsa_circ_0139354 | 942 | 12 | 2.55% |
| hsa_circ_0004493 | 785 | 10 | 2.55% |
| hsa_circ_0007291 | 785 | 10 | 2.55% |
| hsa_circ_0009458 | 785 | 10 | 2.55% |
| hsa_circ_0010898 | 785 | 10 | 2.55% |
| hsa_circ_0012859 | 785 | 10 | 2.55% |
| hsa_circ_0012874 | 785 | 10 | 2.55% |
| hsa_circ_0025799 | 785 | 10 | 2.55% |
| hsa_circ_0033662 | 785 | 10 | 2.55% |
| hsa_circ_0047972 | 785 | 10 | 2.55% |
| hsa_circ_0051044 | 785 | 10 | 2.55% |
| hsa_circ_0051926 | 785 | 10 | 2.55% |
| hsa_circ_0055876 | 785 | 10 | 2.55% |
| hsa_circ_0072548 | 785 | 10 | 2.55% |
| hsa_circ_0075032 | 785 | 10 | 2.55% |
| hsa_circ_0086666 | 785 | 10 | 2.55% |
| hsa_circ_0088151 | 785 | 10 | 2.55% |
| hsa_circ_0092447 | 785 | 10 | 2.55% |
| hsa_circ_0103041 | 785 | 10 | 2.55% |
| hsa_circ_0103125 | 785 | 10 | 2.55% |
| hsa_circ_0118180 | 785 | 10 | 2.55% |
| hsa_circ_0121385 | 785 | 10 | 2.55% |
| hsa_circ_0122043 | 785 | 10 | 2.55% |
| hsa_circ_0124997 | 785 | 10 | 2.55% |
| hsa_circ_0010713 | 864 | 11 | 2.55% |
| hsa_circ_0022849 | 864 | 11 | 2.55% |
| hsa_circ_0023782 | 864 | 11 | 2.55% |
| hsa_circ_0026609 | 864 | 11 | 2.55% |
| hsa_circ_0057431 | 864 | 11 | 2.55% |
| hsa_circ_0060180 | 864 | 11 | 2.55% |
| hsa_circ_0076358 | 864 | 11 | 2.55% |
| hsa_circ_0084748 | 864 | 11 | 2.55% |
| hsa_circ_0094734 | 864 | 11 | 2.55% |
| hsa_circ_0100378 | 864 | 11 | 2.55% |
| hsa_circ_0021626 | 943 | 12 | 2.55% |
| hsa_circ_0039317 | 943 | 12 | 2.55% |
| hsa_circ_0042837 | 943 | 12 | 2.55% |
| hsa_circ_0012937 | 786 | 10 | 2.54% |
| hsa_circ_0033665 | 786 | 10 | 2.54% |
| hsa_circ_0046368 | 786 | 10 | 2.54% |
| hsa_circ_0056878 | 786 | 10 | 2.54% |
| hsa_circ_0057545 | 786 | 10 | 2.54% |
| hsa_circ_0059494 | 786 | 10 | 2.54% |
| hsa_circ_0062234 | 786 | 10 | 2.54% |
| hsa_circ_0079812 | 786 | 10 | 2.54% |
| hsa_circ_0087642 | 786 | 10 | 2.54% |
| hsa_circ_0127044 | 786 | 10 | 2.54% |
| hsa_circ_0132513 | 786 | 10 | 2.54% |

| | | | |
|---|---|---|---|
| hsa_circ_0133104 | 786 | 10 | 2.54% |
| hsa_circ_0128070 | 1022 | 13 | 2.54% |
| hsa_circ_0060459 | 865 | 11 | 2.54% |
| hsa_circ_0078926 | 865 | 11 | 2.54% |
| hsa_circ_0007655 | 944 | 12 | 2.54% |
| hsa_circ_0020346 | 944 | 12 | 2.54% |
| hsa_circ_0063494 | 944 | 12 | 2.54% |
| hsa_circ_0074052 | 944 | 12 | 2.54% |
| hsa_circ_0013497 | 787 | 10 | 2.54% |
| hsa_circ_0058216 | 787 | 10 | 2.54% |
| hsa_circ_0099310 | 787 | 10 | 2.54% |
| hsa_circ_0099521 | 787 | 10 | 2.54% |
| hsa_circ_0109232 | 787 | 10 | 2.54% |
| hsa_circ_0137610 | 787 | 10 | 2.54% |
| hsa_circ_0137656 | 787 | 10 | 2.54% |
| hsa_circ_0095323 | 2125 | 27 | 2.54% |
| hsa_circ_0113140 | 2204 | 28 | 2.54% |
| hsa_circ_0006812 | 866 | 11 | 2.54% |
| hsa_circ_0018391 | 866 | 11 | 2.54% |
| hsa_circ_0021202 | 866 | 11 | 2.54% |
| hsa_circ_0059338 | 866 | 11 | 2.54% |
| hsa_circ_0063684 | 866 | 11 | 2.54% |
| hsa_circ_0071514 | 866 | 11 | 2.54% |
| hsa_circ_0095003 | 866 | 11 | 2.54% |
| hsa_circ_0108229 | 866 | 11 | 2.54% |
| hsa_circ_0007473 | 945 | 12 | 2.54% |
| hsa_circ_0081959 | 945 | 12 | 2.54% |
| hsa_circ_0002783 | 788 | 10 | 2.54% |
| hsa_circ_0012591 | 788 | 10 | 2.54% |
| hsa_circ_0023382 | 788 | 10 | 2.54% |
| hsa_circ_0062589 | 788 | 10 | 2.54% |
| hsa_circ_0066782 | 788 | 10 | 2.54% |
| hsa_circ_0071550 | 788 | 10 | 2.54% |
| hsa_circ_0071576 | 788 | 10 | 2.54% |
| hsa_circ_0116127 | 788 | 10 | 2.54% |
| hsa_circ_0124232 | 788 | 10 | 2.54% |
| hsa_circ_0129503 | 788 | 10 | 2.54% |
| hsa_circ_0011074 | 867 | 11 | 2.54% |
| hsa_circ_0093809 | 867 | 11 | 2.54% |
| hsa_circ_0127346 | 867 | 11 | 2.54% |
| hsa_circ_0026151 | 946 | 12 | 2.54% |
| hsa_circ_0041221 | 946 | 12 | 2.54% |
| hsa_circ_0080618 | 946 | 12 | 2.54% |
| hsa_circ_0065531 | 1025 | 13 | 2.54% |
| hsa_circ_0067632 | 1025 | 13 | 2.54% |
| hsa_circ_0048071 | 1104 | 14 | 2.54% |
| hsa_circ_0104931 | 1104 | 14 | 2.54% |
| hsa_circ_0008460 | 789 | 10 | 2.53% |
| hsa_circ_0008664 | 789 | 10 | 2.53% |
| hsa_circ_0024995 | 789 | 10 | 2.53% |

| | | | |
|---|---|---|---|
| hsa_circ_0068268 | 789 | 10 | 2.53% |
| hsa_circ_0073110 | 789 | 10 | 2.53% |
| hsa_circ_0075657 | 789 | 10 | 2.53% |
| hsa_circ_0090510 | 789 | 10 | 2.53% |
| hsa_circ_0094324 | 789 | 10 | 2.53% |
| hsa_circ_0107721 | 789 | 10 | 2.53% |
| hsa_circ_0113342 | 789 | 10 | 2.53% |
| hsa_circ_0122873 | 789 | 10 | 2.53% |
| hsa_circ_0125317 | 789 | 10 | 2.53% |
| hsa_circ_0128504 | 789 | 10 | 2.53% |
| hsa_circ_0137443 | 789 | 10 | 2.53% |
| hsa_circ_0137673 | 789 | 10 | 2.53% |
| hsa_circ_0138965 | 789 | 10 | 2.53% |
| hsa_circ_0007684 | 868 | 11 | 2.53% |
| hsa_circ_0059275 | 868 | 11 | 2.53% |
| hsa_circ_0080936 | 868 | 11 | 2.53% |
| hsa_circ_0085938 | 868 | 11 | 2.53% |
| hsa_circ_0117392 | 868 | 11 | 2.53% |
| hsa_circ_0121003 | 868 | 11 | 2.53% |
| hsa_circ_0122292 | 868 | 11 | 2.53% |
| hsa_circ_0025680 | 1026 | 13 | 2.53% |
| hsa_circ_0022978 | 869 | 11 | 2.53% |
| hsa_circ_0063174 | 869 | 11 | 2.53% |
| hsa_circ_0083023 | 869 | 11 | 2.53% |
| hsa_circ_0106043 | 869 | 11 | 2.53% |
| hsa_circ_0114261 | 869 | 11 | 2.53% |
| hsa_circ_0012123 | 790 | 10 | 2.53% |
| hsa_circ_0015247 | 790 | 10 | 2.53% |
| hsa_circ_0017376 | 790 | 10 | 2.53% |
| hsa_circ_0019381 | 790 | 10 | 2.53% |
| hsa_circ_0023887 | 790 | 10 | 2.53% |
| hsa_circ_0038071 | 790 | 10 | 2.53% |
| hsa_circ_0040727 | 790 | 10 | 2.53% |
| hsa_circ_0056712 | 790 | 10 | 2.53% |
| hsa_circ_0057877 | 790 | 10 | 2.53% |
| hsa_circ_0070595 | 790 | 10 | 2.53% |
| hsa_circ_0090508 | 790 | 10 | 2.53% |
| hsa_circ_0093024 | 790 | 10 | 2.53% |
| hsa_circ_0096089 | 790 | 10 | 2.53% |
| hsa_circ_0111589 | 790 | 10 | 2.53% |
| hsa_circ_0112519 | 790 | 10 | 2.53% |
| hsa_circ_0115908 | 790 | 10 | 2.53% |
| hsa_circ_0126097 | 790 | 10 | 2.53% |
| hsa_circ_0127114 | 790 | 10 | 2.53% |
| hsa_circ_0133976 | 790 | 10 | 2.53% |
| hsa_circ_0140442 | 790 | 10 | 2.53% |
| hsa_circ_0063581 | 3163 | 40 | 2.53% |
| hsa_circ_0003546 | 1028 | 13 | 2.53% |
| hsa_circ_0050199 | 949 | 12 | 2.53% |
| hsa_circ_0092735 | 949 | 12 | 2.53% |

| | | | |
|---|---|---|---|
| hsa_circ_0067213 | 870 | 11 | 2.53% |
| hsa_circ_0092390 | 870 | 11 | 2.53% |
| hsa_circ_0095166 | 870 | 11 | 2.53% |
| hsa_circ_0097494 | 870 | 11 | 2.53% |
| hsa_circ_0000023 | 791 | 10 | 2.53% |
| hsa_circ_0009613 | 791 | 10 | 2.53% |
| hsa_circ_0026336 | 791 | 10 | 2.53% |
| hsa_circ_0026948 | 791 | 10 | 2.53% |
| hsa_circ_0036460 | 791 | 10 | 2.53% |
| hsa_circ_0040740 | 791 | 10 | 2.53% |
| hsa_circ_0064347 | 791 | 10 | 2.53% |
| hsa_circ_0066352 | 791 | 10 | 2.53% |
| hsa_circ_0067109 | 791 | 10 | 2.53% |
| hsa_circ_0092884 | 791 | 10 | 2.53% |
| hsa_circ_0095421 | 791 | 10 | 2.53% |
| hsa_circ_0106201 | 791 | 10 | 2.53% |
| hsa_circ_0124551 | 791 | 10 | 2.53% |
| hsa_circ_0059868 | 1108 | 14 | 2.53% |
| hsa_circ_0010665 | 1029 | 13 | 2.53% |
| hsa_circ_0013380 | 1029 | 13 | 2.53% |
| hsa_circ_0080024 | 950 | 12 | 2.53% |
| hsa_circ_0012832 | 871 | 11 | 2.53% |
| hsa_circ_0015320 | 871 | 11 | 2.53% |
| hsa_circ_0091724 | 871 | 11 | 2.53% |
| hsa_circ_0117133 | 871 | 11 | 2.53% |
| hsa_circ_0023181 | 792 | 10 | 2.53% |
| hsa_circ_0057399 | 792 | 10 | 2.53% |
| hsa_circ_0065674 | 792 | 10 | 2.53% |
| hsa_circ_0066758 | 792 | 10 | 2.53% |
| hsa_circ_0071035 | 792 | 10 | 2.53% |
| hsa_circ_0073512 | 792 | 10 | 2.53% |
| hsa_circ_0080085 | 792 | 10 | 2.53% |
| hsa_circ_0083521 | 792 | 10 | 2.53% |
| hsa_circ_0088590 | 792 | 10 | 2.53% |
| hsa_circ_0095992 | 792 | 10 | 2.53% |
| hsa_circ_0097033 | 792 | 10 | 2.53% |
| hsa_circ_0112055 | 792 | 10 | 2.53% |
| hsa_circ_0112298 | 792 | 10 | 2.53% |
| hsa_circ_0126084 | 792 | 10 | 2.53% |
| hsa_circ_0127894 | 792 | 10 | 2.53% |
| hsa_circ_0138376 | 792 | 10 | 2.53% |
| hsa_circ_0011331 | 2615 | 33 | 2.52% |
| hsa_circ_0066976 | 951 | 12 | 2.52% |
| hsa_circ_0090161 | 951 | 12 | 2.52% |
| hsa_circ_0137344 | 951 | 12 | 2.52% |
| hsa_circ_0010036 | 872 | 11 | 2.52% |
| hsa_circ_0028298 | 872 | 11 | 2.52% |
| hsa_circ_0055707 | 872 | 11 | 2.52% |
| hsa_circ_0062998 | 872 | 11 | 2.52% |
| hsa_circ_0086254 | 872 | 11 | 2.52% |

| | | | |
|---|---|---|---|
| hsa_circ_0115973 | 872 | 11 | 2.52% |
| hsa_circ_0135337 | 872 | 11 | 2.52% |
| hsa_circ_0140106 | 872 | 11 | 2.52% |
| hsa_circ_0024828 | 793 | 10 | 2.52% |
| hsa_circ_0030887 | 793 | 10 | 2.52% |
| hsa_circ_0037511 | 793 | 10 | 2.52% |
| hsa_circ_0046997 | 793 | 10 | 2.52% |
| hsa_circ_0061192 | 793 | 10 | 2.52% |
| hsa_circ_0078238 | 793 | 10 | 2.52% |
| hsa_circ_0099524 | 793 | 10 | 2.52% |
| hsa_circ_0104727 | 793 | 10 | 2.52% |
| hsa_circ_0105443 | 793 | 10 | 2.52% |
| hsa_circ_0118925 | 793 | 10 | 2.52% |
| hsa_circ_0126233 | 793 | 10 | 2.52% |
| hsa_circ_0131086 | 793 | 10 | 2.52% |
| hsa_circ_0135277 | 793 | 10 | 2.52% |
| hsa_circ_0060525 | 1031 | 13 | 2.52% |
| hsa_circ_0010707 | 873 | 11 | 2.52% |
| hsa_circ_0020731 | 873 | 11 | 2.52% |
| hsa_circ_0071281 | 873 | 11 | 2.52% |
| hsa_circ_0075363 | 873 | 11 | 2.52% |
| hsa_circ_0103411 | 873 | 11 | 2.52% |
| hsa_circ_0123016 | 873 | 11 | 2.52% |
| hsa_circ_0127013 | 873 | 11 | 2.52% |
| hsa_circ_0004201 | 794 | 10 | 2.52% |
| hsa_circ_0006632 | 794 | 10 | 2.52% |
| hsa_circ_0019137 | 794 | 10 | 2.52% |
| hsa_circ_0031419 | 794 | 10 | 2.52% |
| hsa_circ_0031783 | 794 | 10 | 2.52% |
| hsa_circ_0033664 | 794 | 10 | 2.52% |
| hsa_circ_0035032 | 794 | 10 | 2.52% |
| hsa_circ_0051768 | 794 | 10 | 2.52% |
| hsa_circ_0072234 | 794 | 10 | 2.52% |
| hsa_circ_0087220 | 794 | 10 | 2.52% |
| hsa_circ_0091545 | 794 | 10 | 2.52% |
| hsa_circ_0098754 | 794 | 10 | 2.52% |
| hsa_circ_0111822 | 794 | 10 | 2.52% |
| hsa_circ_0126557 | 794 | 10 | 2.52% |
| hsa_circ_0129697 | 794 | 10 | 2.52% |
| hsa_circ_0130842 | 794 | 10 | 2.52% |
| hsa_circ_0048752 | 953 | 12 | 2.52% |
| hsa_circ_0067910 | 953 | 12 | 2.52% |
| hsa_circ_0089093 | 953 | 12 | 2.52% |
| hsa_circ_0062422 | 874 | 11 | 2.52% |
| hsa_circ_0080156 | 874 | 11 | 2.52% |
| hsa_circ_0120759 | 874 | 11 | 2.52% |
| hsa_circ_0122672 | 874 | 11 | 2.52% |
| hsa_circ_0127503 | 874 | 11 | 2.52% |
| hsa_circ_0103347 | 954 | 12 | 2.52% |
| hsa_circ_0008704 | 795 | 10 | 2.52% |

| | | | |
|---|---|---|---|
| hsa_circ_0026310 | 795 | 10 | 2.52% |
| hsa_circ_0040592 | 795 | 10 | 2.52% |
| hsa_circ_0042736 | 795 | 10 | 2.52% |
| hsa_circ_0043953 | 795 | 10 | 2.52% |
| hsa_circ_0064744 | 795 | 10 | 2.52% |
| hsa_circ_0067413 | 795 | 10 | 2.52% |
| hsa_circ_0083048 | 795 | 10 | 2.52% |
| hsa_circ_0102892 | 795 | 10 | 2.52% |
| hsa_circ_0103764 | 795 | 10 | 2.52% |
| hsa_circ_0113268 | 795 | 10 | 2.52% |
| hsa_circ_0137398 | 795 | 10 | 2.52% |
| hsa_circ_0139164 | 795 | 10 | 2.52% |
| hsa_circ_0026246 | 1034 | 13 | 2.51% |
| hsa_circ_0027193 | 875 | 11 | 2.51% |
| hsa_circ_0074146 | 875 | 11 | 2.51% |
| hsa_circ_0080162 | 875 | 11 | 2.51% |
| hsa_circ_0093315 | 875 | 11 | 2.51% |
| hsa_circ_0025582 | 955 | 12 | 2.51% |
| hsa_circ_0014345 | 796 | 10 | 2.51% |
| hsa_circ_0018739 | 796 | 10 | 2.51% |
| hsa_circ_0029407 | 796 | 10 | 2.51% |
| hsa_circ_0058711 | 796 | 10 | 2.51% |
| hsa_circ_0065931 | 796 | 10 | 2.51% |
| hsa_circ_0071459 | 796 | 10 | 2.51% |
| hsa_circ_0073929 | 796 | 10 | 2.51% |
| hsa_circ_0102234 | 796 | 10 | 2.51% |
| hsa_circ_0108873 | 796 | 10 | 2.51% |
| hsa_circ_0115048 | 796 | 10 | 2.51% |
| hsa_circ_0119722 | 796 | 10 | 2.51% |
| hsa_circ_0122379 | 796 | 10 | 2.51% |
| hsa_circ_0123068 | 796 | 10 | 2.51% |
| hsa_circ_0122064 | 1035 | 13 | 2.51% |
| hsa_circ_0006922 | 2946 | 37 | 2.51% |
| hsa_circ_0025054 | 876 | 11 | 2.51% |
| hsa_circ_0063663 | 876 | 11 | 2.51% |
| hsa_circ_0091571 | 876 | 11 | 2.51% |
| hsa_circ_0093535 | 876 | 11 | 2.51% |
| hsa_circ_0108021 | 876 | 11 | 2.51% |
| hsa_circ_0117918 | 876 | 11 | 2.51% |
| hsa_circ_0001408 | 797 | 10 | 2.51% |
| hsa_circ_0007343 | 797 | 10 | 2.51% |
| hsa_circ_0009133 | 797 | 10 | 2.51% |
| hsa_circ_0024741 | 797 | 10 | 2.51% |
| hsa_circ_0039108 | 797 | 10 | 2.51% |
| hsa_circ_0055044 | 797 | 10 | 2.51% |
| hsa_circ_0063883 | 797 | 10 | 2.51% |
| hsa_circ_0064603 | 797 | 10 | 2.51% |
| hsa_circ_0076260 | 797 | 10 | 2.51% |
| hsa_circ_0076508 | 797 | 10 | 2.51% |
| hsa_circ_0085834 | 797 | 10 | 2.51% |

| | | | |
|---|---|---|---|
| hsa_circ_0095563 | 797 | 10 | 2.51% |
| hsa_circ_0101984 | 797 | 10 | 2.51% |
| hsa_circ_0122727 | 797 | 10 | 2.51% |
| hsa_circ_0125224 | 797 | 10 | 2.51% |
| hsa_circ_0133853 | 797 | 10 | 2.51% |
| hsa_circ_0042980 | 877 | 11 | 2.51% |
| hsa_circ_0046703 | 877 | 11 | 2.51% |
| hsa_circ_0081620 | 877 | 11 | 2.51% |
| hsa_circ_0094457 | 877 | 11 | 2.51% |
| hsa_circ_0125935 | 957 | 12 | 2.51% |
| hsa_circ_0000600 | 798 | 10 | 2.51% |
| hsa_circ_0009404 | 798 | 10 | 2.51% |
| hsa_circ_0010364 | 798 | 10 | 2.51% |
| hsa_circ_0026781 | 798 | 10 | 2.51% |
| hsa_circ_0071814 | 798 | 10 | 2.51% |
| hsa_circ_0088785 | 798 | 10 | 2.51% |
| hsa_circ_0101702 | 798 | 10 | 2.51% |
| hsa_circ_0116069 | 798 | 10 | 2.51% |
| hsa_circ_0134371 | 798 | 10 | 2.51% |
| hsa_circ_0140114 | 798 | 10 | 2.51% |
| hsa_circ_0030683 | 878 | 11 | 2.51% |
| hsa_circ_0105836 | 878 | 11 | 2.51% |
| hsa_circ_0059242 | 958 | 12 | 2.51% |
| hsa_circ_0129104 | 1038 | 13 | 2.50% |
| hsa_circ_0057374 | 1278 | 16 | 2.50% |
| hsa_circ_0011552 | 799 | 10 | 2.50% |
| hsa_circ_0011959 | 799 | 10 | 2.50% |
| hsa_circ_0020206 | 799 | 10 | 2.50% |
| hsa_circ_0041567 | 799 | 10 | 2.50% |
| hsa_circ_0044109 | 799 | 10 | 2.50% |
| hsa_circ_0046219 | 799 | 10 | 2.50% |
| hsa_circ_0067465 | 799 | 10 | 2.50% |
| hsa_circ_0067829 | 799 | 10 | 2.50% |
| hsa_circ_0079358 | 799 | 10 | 2.50% |
| hsa_circ_0093398 | 799 | 10 | 2.50% |
| hsa_circ_0094474 | 799 | 10 | 2.50% |
| hsa_circ_0094964 | 799 | 10 | 2.50% |
| hsa_circ_0095790 | 799 | 10 | 2.50% |
| hsa_circ_0098456 | 799 | 10 | 2.50% |
| hsa_circ_0100142 | 799 | 10 | 2.50% |
| hsa_circ_0110131 | 799 | 10 | 2.50% |
| hsa_circ_0119178 | 799 | 10 | 2.50% |
| hsa_circ_0127457 | 799 | 10 | 2.50% |
| hsa_circ_0135997 | 799 | 10 | 2.50% |
| hsa_circ_0126700 | 1678 | 21 | 2.50% |
| hsa_circ_0013103 | 879 | 11 | 2.50% |
| hsa_circ_0021201 | 879 | 11 | 2.50% |
| hsa_circ_0027534 | 879 | 11 | 2.50% |
| hsa_circ_0029317 | 879 | 11 | 2.50% |
| hsa_circ_0034562 | 879 | 11 | 2.50% |

| | | | |
|---|---|---|---|
| hsa_circ_0046773 | 879 | 11 | 2.50% |
| hsa_circ_0084870 | 879 | 11 | 2.50% |
| hsa_circ_0120212 | 879 | 11 | 2.50% |
| hsa_circ_0008641 | 959 | 12 | 2.50% |
| hsa_circ_0034431 | 959 | 12 | 2.50% |
| hsa_circ_0036625 | 959 | 12 | 2.50% |
| hsa_circ_0111578 | 1599 | 20 | 2.50% |
| hsa_circ_0116382 | 960 | 12 | 2.50% |
| hsa_circ_0012201 | 880 | 11 | 2.50% |
| hsa_circ_0059590 | 880 | 11 | 2.50% |
| hsa_circ_0065719 | 880 | 11 | 2.50% |
| hsa_circ_0128560 | 880 | 11 | 2.50% |
| hsa_circ_0009438 | 800 | 10 | 2.50% |
| hsa_circ_0016599 | 800 | 10 | 2.50% |
| hsa_circ_0031060 | 800 | 10 | 2.50% |
| hsa_circ_0033625 | 800 | 10 | 2.50% |
| hsa_circ_0041096 | 800 | 10 | 2.50% |
| hsa_circ_0041446 | 800 | 10 | 2.50% |
| hsa_circ_0049348 | 800 | 10 | 2.50% |
| hsa_circ_0053401 | 800 | 10 | 2.50% |
| hsa_circ_0064503 | 800 | 10 | 2.50% |
| hsa_circ_0073144 | 800 | 10 | 2.50% |
| hsa_circ_0073262 | 800 | 10 | 2.50% |
| hsa_circ_0073629 | 800 | 10 | 2.50% |
| hsa_circ_0080110 | 800 | 10 | 2.50% |
| hsa_circ_0095716 | 800 | 10 | 2.50% |
| hsa_circ_0098761 | 800 | 10 | 2.50% |
| hsa_circ_0101604 | 800 | 10 | 2.50% |
| hsa_circ_0108215 | 800 | 10 | 2.50% |
| hsa_circ_0117925 | 800 | 10 | 2.50% |
| hsa_circ_0135423 | 800 | 10 | 2.50% |
| hsa_circ_0024105 | 1121 | 14 | 2.50% |
| hsa_circ_0120589 | 961 | 12 | 2.50% |
| hsa_circ_0025200 | 881 | 11 | 2.50% |
| hsa_circ_0070715 | 881 | 11 | 2.50% |
| hsa_circ_0088055 | 881 | 11 | 2.50% |
| hsa_circ_0116907 | 881 | 11 | 2.50% |
| hsa_circ_0118686 | 881 | 11 | 2.50% |
| hsa_circ_0138850 | 881 | 11 | 2.50% |
| hsa_circ_0008144 | 801 | 10 | 2.50% |
| hsa_circ_0026638 | 801 | 10 | 2.50% |
| hsa_circ_0032341 | 801 | 10 | 2.50% |
| hsa_circ_0041562 | 801 | 10 | 2.50% |
| hsa_circ_0053103 | 801 | 10 | 2.50% |
| hsa_circ_0057255 | 801 | 10 | 2.50% |
| hsa_circ_0069100 | 801 | 10 | 2.50% |
| hsa_circ_0099671 | 801 | 10 | 2.50% |
| hsa_circ_0100023 | 801 | 10 | 2.50% |
| hsa_circ_0106353 | 801 | 10 | 2.50% |
| hsa_circ_0128850 | 801 | 10 | 2.50% |

| | | | |
|---|---|---|---|
| hsa_circ_0136720 | 801 | 10 | 2.50% |
| hsa_circ_0130034 | 1122 | 14 | 2.50% |
| hsa_circ_0023055 | 1042 | 13 | 2.50% |
| hsa_circ_0027914 | 1042 | 13 | 2.50% |
| hsa_circ_0046078 | 1042 | 13 | 2.50% |
| hsa_circ_0061168 | 962 | 12 | 2.49% |
| hsa_circ_0010040 | 882 | 11 | 2.49% |
| hsa_circ_0023972 | 882 | 11 | 2.49% |
| hsa_circ_0042238 | 882 | 11 | 2.49% |
| hsa_circ_0058971 | 882 | 11 | 2.49% |
| hsa_circ_0059923 | 882 | 11 | 2.49% |
| hsa_circ_0079935 | 882 | 11 | 2.49% |
| hsa_circ_0102147 | 882 | 11 | 2.49% |
| hsa_circ_0114963 | 882 | 11 | 2.49% |
| hsa_circ_0027452 | 802 | 10 | 2.49% |
| hsa_circ_0042460 | 802 | 10 | 2.49% |
| hsa_circ_0045504 | 802 | 10 | 2.49% |
| hsa_circ_0047784 | 802 | 10 | 2.49% |
| hsa_circ_0057600 | 802 | 10 | 2.49% |
| hsa_circ_0058298 | 802 | 10 | 2.49% |
| hsa_circ_0059236 | 802 | 10 | 2.49% |
| hsa_circ_0081684 | 802 | 10 | 2.49% |
| hsa_circ_0081715 | 802 | 10 | 2.49% |
| hsa_circ_0100652 | 802 | 10 | 2.49% |
| hsa_circ_0110875 | 802 | 10 | 2.49% |
| hsa_circ_0133970 | 802 | 10 | 2.49% |
| hsa_circ_0137006 | 802 | 10 | 2.49% |
| hsa_circ_0138135 | 802 | 10 | 2.49% |
| hsa_circ_0077794 | 1043 | 13 | 2.49% |
| hsa_circ_0003946 | 883 | 11 | 2.49% |
| hsa_circ_0017212 | 883 | 11 | 2.49% |
| hsa_circ_0045622 | 883 | 11 | 2.49% |
| hsa_circ_0055556 | 883 | 11 | 2.49% |
| hsa_circ_0092719 | 883 | 11 | 2.49% |
| hsa_circ_0102292 | 883 | 11 | 2.49% |
| hsa_circ_0118811 | 883 | 11 | 2.49% |
| hsa_circ_0000459 | 803 | 10 | 2.49% |
| hsa_circ_0011585 | 803 | 10 | 2.49% |
| hsa_circ_0014838 | 803 | 10 | 2.49% |
| hsa_circ_0015432 | 803 | 10 | 2.49% |
| hsa_circ_0019010 | 803 | 10 | 2.49% |
| hsa_circ_0052449 | 803 | 10 | 2.49% |
| hsa_circ_0059833 | 803 | 10 | 2.49% |
| hsa_circ_0062327 | 803 | 10 | 2.49% |
| hsa_circ_0063209 | 803 | 10 | 2.49% |
| hsa_circ_0068389 | 803 | 10 | 2.49% |
| hsa_circ_0082641 | 803 | 10 | 2.49% |
| hsa_circ_0101098 | 803 | 10 | 2.49% |
| hsa_circ_0102088 | 803 | 10 | 2.49% |
| hsa_circ_0106462 | 803 | 10 | 2.49% |

| | | | |
|---|---|---|---|
| hsa_circ_0107341 | 803 | 10 | 2.49% |
| hsa_circ_0128111 | 803 | 10 | 2.49% |
| hsa_circ_0138272 | 803 | 10 | 2.49% |
| hsa_circ_0010340 | 964 | 12 | 2.49% |
| hsa_circ_0113796 | 1848 | 23 | 2.49% |
| hsa_circ_0136392 | 1848 | 23 | 2.49% |
| hsa_circ_0006872 | 884 | 11 | 2.49% |
| hsa_circ_0019477 | 884 | 11 | 2.49% |
| hsa_circ_0029373 | 884 | 11 | 2.49% |
| hsa_circ_0029576 | 884 | 11 | 2.49% |
| hsa_circ_0081621 | 884 | 11 | 2.49% |
| hsa_circ_0083835 | 884 | 11 | 2.49% |
| hsa_circ_0100216 | 884 | 11 | 2.49% |
| hsa_circ_0115984 | 884 | 11 | 2.49% |
| hsa_circ_0012949 | 1045 | 13 | 2.49% |
| hsa_circ_0105583 | 1045 | 13 | 2.49% |
| hsa_circ_0001471 | 804 | 10 | 2.49% |
| hsa_circ_0023138 | 804 | 10 | 2.49% |
| hsa_circ_0066452 | 804 | 10 | 2.49% |
| hsa_circ_0098201 | 804 | 10 | 2.49% |
| hsa_circ_0113629 | 804 | 10 | 2.49% |
| hsa_circ_0124972 | 804 | 10 | 2.49% |
| hsa_circ_0126161 | 804 | 10 | 2.49% |
| hsa_circ_0126478 | 804 | 10 | 2.49% |
| hsa_circ_0127674 | 804 | 10 | 2.49% |
| hsa_circ_0133158 | 804 | 10 | 2.49% |
| hsa_circ_0134773 | 804 | 10 | 2.49% |
| hsa_circ_0136745 | 804 | 10 | 2.49% |
| hsa_circ_0137199 | 804 | 10 | 2.49% |
| hsa_circ_0096782 | 965 | 12 | 2.49% |
| hsa_circ_0102072 | 965 | 12 | 2.49% |
| hsa_circ_0001139 | 885 | 11 | 2.49% |
| hsa_circ_0028005 | 885 | 11 | 2.49% |
| hsa_circ_0037829 | 885 | 11 | 2.49% |
| hsa_circ_0040325 | 885 | 11 | 2.49% |
| hsa_circ_0064332 | 885 | 11 | 2.49% |
| hsa_circ_0123937 | 885 | 11 | 2.49% |
| hsa_circ_0133197 | 885 | 11 | 2.49% |
| hsa_circ_0010940 | 805 | 10 | 2.48% |
| hsa_circ_0011098 | 805 | 10 | 2.48% |
| hsa_circ_0025167 | 805 | 10 | 2.48% |
| hsa_circ_0027235 | 805 | 10 | 2.48% |
| hsa_circ_0038577 | 805 | 10 | 2.48% |
| hsa_circ_0042407 | 805 | 10 | 2.48% |
| hsa_circ_0061755 | 805 | 10 | 2.48% |
| hsa_circ_0071794 | 805 | 10 | 2.48% |
| hsa_circ_0075175 | 805 | 10 | 2.48% |
| hsa_circ_0079989 | 805 | 10 | 2.48% |
| hsa_circ_0086511 | 805 | 10 | 2.48% |
| hsa_circ_0124918 | 805 | 10 | 2.48% |

| | | | |
|---|---|---|---|
| hsa_circ_0009692 | 886 | 11 | 2.48% |
| hsa_circ_0055705 | 886 | 11 | 2.48% |
| hsa_circ_0069253 | 886 | 11 | 2.48% |
| hsa_circ_0090800 | 886 | 11 | 2.48% |
| hsa_circ_0030407 | 967 | 12 | 2.48% |
| hsa_circ_0114523 | 1209 | 15 | 2.48% |
| hsa_circ_0012614 | 806 | 10 | 2.48% |
| hsa_circ_0015609 | 806 | 10 | 2.48% |
| hsa_circ_0020507 | 806 | 10 | 2.48% |
| hsa_circ_0022894 | 806 | 10 | 2.48% |
| hsa_circ_0023921 | 806 | 10 | 2.48% |
| hsa_circ_0051436 | 806 | 10 | 2.48% |
| hsa_circ_0051951 | 806 | 10 | 2.48% |
| hsa_circ_0069255 | 806 | 10 | 2.48% |
| hsa_circ_0076442 | 806 | 10 | 2.48% |
| hsa_circ_0079258 | 806 | 10 | 2.48% |
| hsa_circ_0081460 | 806 | 10 | 2.48% |
| hsa_circ_0082490 | 806 | 10 | 2.48% |
| hsa_circ_0084797 | 806 | 10 | 2.48% |
| hsa_circ_0094439 | 806 | 10 | 2.48% |
| hsa_circ_0094489 | 806 | 10 | 2.48% |
| hsa_circ_0110485 | 806 | 10 | 2.48% |
| hsa_circ_0121456 | 806 | 10 | 2.48% |
| hsa_circ_0125769 | 806 | 10 | 2.48% |
| hsa_circ_0086047 | 1048 | 13 | 2.48% |
| hsa_circ_0043490 | 887 | 11 | 2.48% |
| hsa_circ_0068055 | 887 | 11 | 2.48% |
| hsa_circ_0096753 | 887 | 11 | 2.48% |
| hsa_circ_0020079 | 968 | 12 | 2.48% |
| hsa_circ_0091594 | 968 | 12 | 2.48% |
| hsa_circ_0132530 | 968 | 12 | 2.48% |
| hsa_circ_0001888 | 807 | 10 | 2.48% |
| hsa_circ_0013790 | 807 | 10 | 2.48% |
| hsa_circ_0016720 | 807 | 10 | 2.48% |
| hsa_circ_0019008 | 807 | 10 | 2.48% |
| hsa_circ_0019818 | 807 | 10 | 2.48% |
| hsa_circ_0033663 | 807 | 10 | 2.48% |
| hsa_circ_0037647 | 807 | 10 | 2.48% |
| hsa_circ_0039623 | 807 | 10 | 2.48% |
| hsa_circ_0053076 | 807 | 10 | 2.48% |
| hsa_circ_0065442 | 807 | 10 | 2.48% |
| hsa_circ_0068508 | 807 | 10 | 2.48% |
| hsa_circ_0086669 | 807 | 10 | 2.48% |
| hsa_circ_0107227 | 807 | 10 | 2.48% |
| hsa_circ_0108062 | 807 | 10 | 2.48% |
| hsa_circ_0114246 | 807 | 10 | 2.48% |
| hsa_circ_0114922 | 807 | 10 | 2.48% |
| hsa_circ_0135256 | 807 | 10 | 2.48% |
| hsa_circ_0140105 | 807 | 10 | 2.48% |
| hsa_circ_0082294 | 1130 | 14 | 2.48% |

| | | | |
|---|---|---|---|
| hsa_circ_0018966 | 888 | 11 | 2.48% |
| hsa_circ_0025721 | 888 | 11 | 2.48% |
| hsa_circ_0030400 | 888 | 11 | 2.48% |
| hsa_circ_0055228 | 888 | 11 | 2.48% |
| hsa_circ_0103990 | 888 | 11 | 2.48% |
| hsa_circ_0108829 | 888 | 11 | 2.48% |
| hsa_circ_0126545 | 888 | 11 | 2.48% |
| hsa_circ_0127292 | 888 | 11 | 2.48% |
| hsa_circ_0082612 | 969 | 12 | 2.48% |
| hsa_circ_0118426 | 969 | 12 | 2.48% |
| hsa_circ_0119021 | 969 | 12 | 2.48% |
| hsa_circ_0007235 | 1050 | 13 | 2.48% |
| hsa_circ_0018452 | 808 | 10 | 2.48% |
| hsa_circ_0031355 | 808 | 10 | 2.48% |
| hsa_circ_0036309 | 808 | 10 | 2.48% |
| hsa_circ_0043011 | 808 | 10 | 2.48% |
| hsa_circ_0043907 | 808 | 10 | 2.48% |
| hsa_circ_0050040 | 808 | 10 | 2.48% |
| hsa_circ_0062535 | 808 | 10 | 2.48% |
| hsa_circ_0072594 | 808 | 10 | 2.48% |
| hsa_circ_0079273 | 808 | 10 | 2.48% |
| hsa_circ_0096766 | 808 | 10 | 2.48% |
| hsa_circ_0104278 | 808 | 10 | 2.48% |
| hsa_circ_0111004 | 808 | 10 | 2.48% |
| hsa_circ_0131536 | 808 | 10 | 2.48% |
| hsa_circ_0132102 | 808 | 10 | 2.48% |
| hsa_circ_0140069 | 808 | 10 | 2.48% |
| hsa_circ_0038470 | 889 | 11 | 2.47% |
| hsa_circ_0055375 | 889 | 11 | 2.47% |
| hsa_circ_0098580 | 889 | 11 | 2.47% |
| hsa_circ_0104703 | 889 | 11 | 2.47% |
| hsa_circ_0111012 | 889 | 11 | 2.47% |
| hsa_circ_0127727 | 889 | 11 | 2.47% |
| hsa_circ_0028707 | 970 | 12 | 2.47% |
| hsa_circ_0055399 | 970 | 12 | 2.47% |
| hsa_circ_0123149 | 970 | 12 | 2.47% |
| hsa_circ_0095026 | 1294 | 16 | 2.47% |
| hsa_circ_0015242 | 809 | 10 | 2.47% |
| hsa_circ_0020904 | 809 | 10 | 2.47% |
| hsa_circ_0022647 | 809 | 10 | 2.47% |
| hsa_circ_0023688 | 809 | 10 | 2.47% |
| hsa_circ_0025191 | 809 | 10 | 2.47% |
| hsa_circ_0031242 | 809 | 10 | 2.47% |
| hsa_circ_0031566 | 809 | 10 | 2.47% |
| hsa_circ_0043563 | 809 | 10 | 2.47% |
| hsa_circ_0053037 | 809 | 10 | 2.47% |
| hsa_circ_0054394 | 809 | 10 | 2.47% |
| hsa_circ_0055093 | 809 | 10 | 2.47% |
| hsa_circ_0060136 | 809 | 10 | 2.47% |
| hsa_circ_0079364 | 809 | 10 | 2.47% |

| | | | |
|---|---|---|---|
| hsa_circ_0081305 | 809 | 10 | 2.47% |
| hsa_circ_0101769 | 809 | 10 | 2.47% |
| hsa_circ_0116660 | 809 | 10 | 2.47% |
| hsa_circ_0134439 | 809 | 10 | 2.47% |
| hsa_circ_0039561 | 890 | 11 | 2.47% |
| hsa_circ_0082921 | 890 | 11 | 2.47% |
| hsa_circ_0099913 | 890 | 11 | 2.47% |
| hsa_circ_0106979 | 890 | 11 | 2.47% |
| hsa_circ_0121054 | 890 | 11 | 2.47% |
| hsa_circ_0016875 | 971 | 12 | 2.47% |
| hsa_circ_0004335 | 1134 | 14 | 2.47% |
| hsa_circ_0076434 | 1053 | 13 | 2.47% |
| hsa_circ_0118427 | 972 | 12 | 2.47% |
| hsa_circ_0003209 | 891 | 11 | 2.47% |
| hsa_circ_0051006 | 891 | 11 | 2.47% |
| hsa_circ_0051797 | 891 | 11 | 2.47% |
| hsa_circ_0061666 | 891 | 11 | 2.47% |
| hsa_circ_0085411 | 891 | 11 | 2.47% |
| hsa_circ_0107401 | 891 | 11 | 2.47% |
| hsa_circ_0112735 | 891 | 11 | 2.47% |
| hsa_circ_0132617 | 891 | 11 | 2.47% |
| hsa_circ_0135486 | 891 | 11 | 2.47% |
| hsa_circ_0015368 | 810 | 10 | 2.47% |
| hsa_circ_0027281 | 810 | 10 | 2.47% |
| hsa_circ_0032635 | 810 | 10 | 2.47% |
| hsa_circ_0034555 | 810 | 10 | 2.47% |
| hsa_circ_0041420 | 810 | 10 | 2.47% |
| hsa_circ_0043611 | 810 | 10 | 2.47% |
| hsa_circ_0048275 | 810 | 10 | 2.47% |
| hsa_circ_0051711 | 810 | 10 | 2.47% |
| hsa_circ_0080324 | 810 | 10 | 2.47% |
| hsa_circ_0088817 | 810 | 10 | 2.47% |
| hsa_circ_0098595 | 810 | 10 | 2.47% |
| hsa_circ_0101918 | 810 | 10 | 2.47% |
| hsa_circ_0102368 | 810 | 10 | 2.47% |
| hsa_circ_0111924 | 810 | 10 | 2.47% |
| hsa_circ_0113078 | 810 | 10 | 2.47% |
| hsa_circ_0114092 | 810 | 10 | 2.47% |
| hsa_circ_0131067 | 810 | 10 | 2.47% |
| hsa_circ_0139337 | 1054 | 13 | 2.47% |
| hsa_circ_0010088 | 973 | 12 | 2.47% |
| hsa_circ_0006876 | 892 | 11 | 2.47% |
| hsa_circ_0017956 | 892 | 11 | 2.47% |
| hsa_circ_0019546 | 892 | 11 | 2.47% |
| hsa_circ_0023399 | 892 | 11 | 2.47% |
| hsa_circ_0107754 | 892 | 11 | 2.47% |
| hsa_circ_0003210 | 811 | 10 | 2.47% |
| hsa_circ_0004285 | 811 | 10 | 2.47% |
| hsa_circ_0004544 | 811 | 10 | 2.47% |
| hsa_circ_0007329 | 811 | 10 | 2.47% |

| | | | |
|---|---|---|---|
| hsa_circ_0018641 | 811 | 10 | 2.47% |
| hsa_circ_0022765 | 811 | 10 | 2.47% |
| hsa_circ_0023094 | 811 | 10 | 2.47% |
| hsa_circ_0034840 | 811 | 10 | 2.47% |
| hsa_circ_0035646 | 811 | 10 | 2.47% |
| hsa_circ_0037312 | 811 | 10 | 2.47% |
| hsa_circ_0039131 | 811 | 10 | 2.47% |
| hsa_circ_0063025 | 811 | 10 | 2.47% |
| hsa_circ_0113232 | 811 | 10 | 2.47% |
| hsa_circ_0119769 | 811 | 10 | 2.47% |
| hsa_circ_0126424 | 811 | 10 | 2.47% |
| hsa_circ_0126907 | 811 | 10 | 2.47% |
| hsa_circ_0134816 | 811 | 10 | 2.47% |
| hsa_circ_0106558 | 1055 | 13 | 2.46% |
| hsa_circ_0040085 | 974 | 12 | 2.46% |
| hsa_circ_0024055 | 893 | 11 | 2.46% |
| hsa_circ_0052953 | 893 | 11 | 2.46% |
| hsa_circ_0006913 | 812 | 10 | 2.46% |
| hsa_circ_0024768 | 812 | 10 | 2.46% |
| hsa_circ_0024798 | 812 | 10 | 2.46% |
| hsa_circ_0043165 | 812 | 10 | 2.46% |
| hsa_circ_0049467 | 812 | 10 | 2.46% |
| hsa_circ_0051839 | 812 | 10 | 2.46% |
| hsa_circ_0055355 | 812 | 10 | 2.46% |
| hsa_circ_0056356 | 812 | 10 | 2.46% |
| hsa_circ_0067127 | 812 | 10 | 2.46% |
| hsa_circ_0072494 | 812 | 10 | 2.46% |
| hsa_circ_0089617 | 812 | 10 | 2.46% |
| hsa_circ_0123112 | 812 | 10 | 2.46% |
| hsa_circ_0124893 | 812 | 10 | 2.46% |
| hsa_circ_0130456 | 812 | 10 | 2.46% |
| hsa_circ_0131062 | 812 | 10 | 2.46% |
| hsa_circ_0133314 | 812 | 10 | 2.46% |
| hsa_circ_0136750 | 812 | 10 | 2.46% |
| hsa_circ_0137614 | 812 | 10 | 2.46% |
| hsa_circ_0025919 | 1056 | 13 | 2.46% |
| hsa_circ_0040227 | 1056 | 13 | 2.46% |
| hsa_circ_0045149 | 975 | 12 | 2.46% |
| hsa_circ_0056680 | 975 | 12 | 2.46% |
| hsa_circ_0086099 | 975 | 12 | 2.46% |
| hsa_circ_0106949 | 975 | 12 | 2.46% |
| hsa_circ_0118424 | 975 | 12 | 2.46% |
| hsa_circ_0047706 | 3982 | 49 | 2.46% |
| hsa_circ_0067277 | 1219 | 15 | 2.46% |
| hsa_circ_0000974 | 894 | 11 | 2.46% |
| hsa_circ_0002085 | 894 | 11 | 2.46% |
| hsa_circ_0055773 | 894 | 11 | 2.46% |
| hsa_circ_0086927 | 894 | 11 | 2.46% |
| hsa_circ_0105003 | 894 | 11 | 2.46% |
| hsa_circ_0106883 | 894 | 11 | 2.46% |

| | | | |
|---|---|---|---|
| hsa_circ_0115520 | 894 | 11 | 2.46% |
| hsa_circ_0117842 | 894 | 11 | 2.46% |
| hsa_circ_0004070 | 1138 | 14 | 2.46% |
| hsa_circ_0005612 | 813 | 10 | 2.46% |
| hsa_circ_0012131 | 813 | 10 | 2.46% |
| hsa_circ_0032670 | 813 | 10 | 2.46% |
| hsa_circ_0039305 | 813 | 10 | 2.46% |
| hsa_circ_0040602 | 813 | 10 | 2.46% |
| hsa_circ_0045096 | 813 | 10 | 2.46% |
| hsa_circ_0046133 | 813 | 10 | 2.46% |
| hsa_circ_0057882 | 813 | 10 | 2.46% |
| hsa_circ_0065855 | 813 | 10 | 2.46% |
| hsa_circ_0066164 | 813 | 10 | 2.46% |
| hsa_circ_0071359 | 813 | 10 | 2.46% |
| hsa_circ_0090617 | 813 | 10 | 2.46% |
| hsa_circ_0104247 | 813 | 10 | 2.46% |
| hsa_circ_0113654 | 813 | 10 | 2.46% |
| hsa_circ_0117469 | 813 | 10 | 2.46% |
| hsa_circ_0129093 | 813 | 10 | 2.46% |
| hsa_circ_0131977 | 813 | 10 | 2.46% |
| hsa_circ_0132226 | 813 | 10 | 2.46% |
| hsa_circ_0120315 | 976 | 12 | 2.46% |
| hsa_circ_0031933 | 895 | 11 | 2.46% |
| hsa_circ_0043362 | 895 | 11 | 2.46% |
| hsa_circ_0073206 | 895 | 11 | 2.46% |
| hsa_circ_0113838 | 895 | 11 | 2.46% |
| hsa_circ_0120446 | 895 | 11 | 2.46% |
| hsa_circ_0134603 | 895 | 11 | 2.46% |
| hsa_circ_0003076 | 814 | 10 | 2.46% |
| hsa_circ_0004959 | 814 | 10 | 2.46% |
| hsa_circ_0023220 | 814 | 10 | 2.46% |
| hsa_circ_0027324 | 814 | 10 | 2.46% |
| hsa_circ_0034320 | 814 | 10 | 2.46% |
| hsa_circ_0041390 | 814 | 10 | 2.46% |
| hsa_circ_0047969 | 814 | 10 | 2.46% |
| hsa_circ_0056402 | 814 | 10 | 2.46% |
| hsa_circ_0081333 | 814 | 10 | 2.46% |
| hsa_circ_0083304 | 814 | 10 | 2.46% |
| hsa_circ_0096312 | 814 | 10 | 2.46% |
| hsa_circ_0102602 | 814 | 10 | 2.46% |
| hsa_circ_0106540 | 814 | 10 | 2.46% |
| hsa_circ_0110792 | 814 | 10 | 2.46% |
| hsa_circ_0140272 | 814 | 10 | 2.46% |
| hsa_circ_0049717 | 1140 | 14 | 2.46% |
| hsa_circ_0025498 | 896 | 11 | 2.46% |
| hsa_circ_0064177 | 896 | 11 | 2.46% |
| hsa_circ_0065745 | 896 | 11 | 2.46% |
| hsa_circ_0067202 | 978 | 12 | 2.45% |
| hsa_circ_0099931 | 978 | 12 | 2.45% |
| hsa_circ_0118425 | 978 | 12 | 2.45% |

| | | | |
|---|---|---|---|
| hsa_circ_0001955 | 815 | 10 | 2.45% |
| hsa_circ_0009102 | 815 | 10 | 2.45% |
| hsa_circ_0022240 | 815 | 10 | 2.45% |
| hsa_circ_0030616 | 815 | 10 | 2.45% |
| hsa_circ_0030995 | 815 | 10 | 2.45% |
| hsa_circ_0033666 | 815 | 10 | 2.45% |
| hsa_circ_0046519 | 815 | 10 | 2.45% |
| hsa_circ_0048876 | 815 | 10 | 2.45% |
| hsa_circ_0059630 | 815 | 10 | 2.45% |
| hsa_circ_0075333 | 815 | 10 | 2.45% |
| hsa_circ_0080289 | 815 | 10 | 2.45% |
| hsa_circ_0089480 | 815 | 10 | 2.45% |
| hsa_circ_0100114 | 815 | 10 | 2.45% |
| hsa_circ_0101865 | 815 | 10 | 2.45% |
| hsa_circ_0101989 | 815 | 10 | 2.45% |
| hsa_circ_0104987 | 815 | 10 | 2.45% |
| hsa_circ_0138858 | 815 | 10 | 2.45% |
| hsa_circ_0092295 | 1060 | 13 | 2.45% |
| hsa_circ_0030946 | 897 | 11 | 2.45% |
| hsa_circ_0080785 | 897 | 11 | 2.45% |
| hsa_circ_0041724 | 1142 | 14 | 2.45% |
| hsa_circ_0072957 | 979 | 12 | 2.45% |
| hsa_circ_0002714 | 816 | 10 | 2.45% |
| hsa_circ_0011079 | 816 | 10 | 2.45% |
| hsa_circ_0012601 | 816 | 10 | 2.45% |
| hsa_circ_0021259 | 816 | 10 | 2.45% |
| hsa_circ_0024358 | 816 | 10 | 2.45% |
| hsa_circ_0030598 | 816 | 10 | 2.45% |
| hsa_circ_0032958 | 816 | 10 | 2.45% |
| hsa_circ_0051971 | 816 | 10 | 2.45% |
| hsa_circ_0066772 | 816 | 10 | 2.45% |
| hsa_circ_0071859 | 816 | 10 | 2.45% |
| hsa_circ_0074835 | 816 | 10 | 2.45% |
| hsa_circ_0081569 | 816 | 10 | 2.45% |
| hsa_circ_0081898 | 816 | 10 | 2.45% |
| hsa_circ_0082144 | 816 | 10 | 2.45% |
| hsa_circ_0084118 | 816 | 10 | 2.45% |
| hsa_circ_0104154 | 816 | 10 | 2.45% |
| hsa_circ_0108555 | 816 | 10 | 2.45% |
| hsa_circ_0110281 | 816 | 10 | 2.45% |
| hsa_circ_0118894 | 816 | 10 | 2.45% |
| hsa_circ_0137393 | 816 | 10 | 2.45% |
| hsa_circ_0107258 | 2775 | 34 | 2.45% |
| hsa_circ_0052279 | 898 | 11 | 2.45% |
| hsa_circ_0111365 | 898 | 11 | 2.45% |
| hsa_circ_0122633 | 898 | 11 | 2.45% |
| hsa_circ_0102302 | 980 | 12 | 2.45% |
| hsa_circ_0040296 | 1062 | 13 | 2.45% |
| hsa_circ_0011028 | 817 | 10 | 2.45% |
| hsa_circ_0011237 | 817 | 10 | 2.45% |

| | | | |
|---|---|---|---|
| hsa_circ_0012939 | 817 | 10 | 2.45% |
| hsa_circ_0015599 | 817 | 10 | 2.45% |
| hsa_circ_0016931 | 817 | 10 | 2.45% |
| hsa_circ_0023307 | 817 | 10 | 2.45% |
| hsa_circ_0026420 | 817 | 10 | 2.45% |
| hsa_circ_0050071 | 817 | 10 | 2.45% |
| hsa_circ_0060611 | 817 | 10 | 2.45% |
| hsa_circ_0068453 | 817 | 10 | 2.45% |
| hsa_circ_0072981 | 817 | 10 | 2.45% |
| hsa_circ_0082233 | 817 | 10 | 2.45% |
| hsa_circ_0096272 | 817 | 10 | 2.45% |
| hsa_circ_0098833 | 817 | 10 | 2.45% |
| hsa_circ_0099702 | 817 | 10 | 2.45% |
| hsa_circ_0100761 | 817 | 10 | 2.45% |
| hsa_circ_0103237 | 817 | 10 | 2.45% |
| hsa_circ_0105765 | 817 | 10 | 2.45% |
| hsa_circ_0111739 | 817 | 10 | 2.45% |
| hsa_circ_0117076 | 817 | 10 | 2.45% |
| hsa_circ_0120326 | 817 | 10 | 2.45% |
| hsa_circ_0121928 | 817 | 10 | 2.45% |
| hsa_circ_0129054 | 817 | 10 | 2.45% |
| hsa_circ_0033634 | 899 | 11 | 2.45% |
| hsa_circ_0037971 | 899 | 11 | 2.45% |
| hsa_circ_0050957 | 899 | 11 | 2.45% |
| hsa_circ_0055379 | 899 | 11 | 2.45% |
| hsa_circ_0063996 | 899 | 11 | 2.45% |
| hsa_circ_0065633 | 899 | 11 | 2.45% |
| hsa_circ_0107819 | 899 | 11 | 2.45% |
| hsa_circ_0113745 | 899 | 11 | 2.45% |
| hsa_circ_0129131 | 899 | 11 | 2.45% |
| hsa_circ_0129653 | 899 | 11 | 2.45% |
| hsa_circ_0130206 | 981 | 12 | 2.45% |
| hsa_circ_0138952 | 1063 | 13 | 2.45% |
| hsa_circ_0008690 | 818 | 10 | 2.44% |
| hsa_circ_0020296 | 818 | 10 | 2.44% |
| hsa_circ_0029835 | 818 | 10 | 2.44% |
| hsa_circ_0048092 | 818 | 10 | 2.44% |
| hsa_circ_0053402 | 818 | 10 | 2.44% |
| hsa_circ_0055134 | 818 | 10 | 2.44% |
| hsa_circ_0067973 | 818 | 10 | 2.44% |
| hsa_circ_0080067 | 818 | 10 | 2.44% |
| hsa_circ_0081327 | 818 | 10 | 2.44% |
| hsa_circ_0087000 | 818 | 10 | 2.44% |
| hsa_circ_0087772 | 818 | 10 | 2.44% |
| hsa_circ_0087822 | 818 | 10 | 2.44% |
| hsa_circ_0088988 | 818 | 10 | 2.44% |
| hsa_circ_0106511 | 818 | 10 | 2.44% |
| hsa_circ_0108339 | 818 | 10 | 2.44% |
| hsa_circ_0111134 | 818 | 10 | 2.44% |
| hsa_circ_0111184 | 818 | 10 | 2.44% |

| | | | |
|---|---|---|---|
| hsa_circ_0113667 | 818 | 10 | 2.44% |
| hsa_circ_0114242 | 818 | 10 | 2.44% |
| hsa_circ_0116942 | 818 | 10 | 2.44% |
| hsa_circ_0125620 | 818 | 10 | 2.44% |
| hsa_circ_0126395 | 818 | 10 | 2.44% |
| hsa_circ_0135093 | 818 | 10 | 2.44% |
| hsa_circ_0137944 | 818 | 10 | 2.44% |
| hsa_circ_0023160 | 900 | 11 | 2.44% |
| hsa_circ_0061542 | 900 | 11 | 2.44% |
| hsa_circ_0073290 | 900 | 11 | 2.44% |
| hsa_circ_0105806 | 900 | 11 | 2.44% |
| hsa_circ_0048057 | 982 | 12 | 2.44% |
| hsa_circ_0004359 | 819 | 10 | 2.44% |
| hsa_circ_0011132 | 819 | 10 | 2.44% |
| hsa_circ_0022848 | 819 | 10 | 2.44% |
| hsa_circ_0028477 | 819 | 10 | 2.44% |
| hsa_circ_0037393 | 819 | 10 | 2.44% |
| hsa_circ_0043159 | 819 | 10 | 2.44% |
| hsa_circ_0044817 | 819 | 10 | 2.44% |
| hsa_circ_0072117 | 819 | 10 | 2.44% |
| hsa_circ_0091756 | 819 | 10 | 2.44% |
| hsa_circ_0103766 | 819 | 10 | 2.44% |
| hsa_circ_0108092 | 819 | 10 | 2.44% |
| hsa_circ_0124767 | 819 | 10 | 2.44% |
| hsa_circ_0126438 | 819 | 10 | 2.44% |
| hsa_circ_0129196 | 819 | 10 | 2.44% |
| hsa_circ_0004902 | 901 | 11 | 2.44% |
| hsa_circ_0034375 | 901 | 11 | 2.44% |
| hsa_circ_0051514 | 901 | 11 | 2.44% |
| hsa_circ_0059561 | 901 | 11 | 2.44% |
| hsa_circ_0069213 | 901 | 11 | 2.44% |
| hsa_circ_0076561 | 901 | 11 | 2.44% |
| hsa_circ_0093222 | 901 | 11 | 2.44% |
| hsa_circ_0104379 | 901 | 11 | 2.44% |
| hsa_circ_0051927 | 983 | 12 | 2.44% |
| hsa_circ_0074049 | 983 | 12 | 2.44% |
| hsa_circ_0078266 | 1065 | 13 | 2.44% |
| hsa_circ_0069134 | 984 | 12 | 2.44% |
| hsa_circ_0058841 | 902 | 11 | 2.44% |
| hsa_circ_0061776 | 902 | 11 | 2.44% |
| hsa_circ_0080050 | 902 | 11 | 2.44% |
| hsa_circ_0091572 | 902 | 11 | 2.44% |
| hsa_circ_0097105 | 902 | 11 | 2.44% |
| hsa_circ_0133799 | 902 | 11 | 2.44% |
| hsa_circ_0022742 | 820 | 10 | 2.44% |
| hsa_circ_0065096 | 820 | 10 | 2.44% |
| hsa_circ_0068290 | 820 | 10 | 2.44% |
| hsa_circ_0068618 | 820 | 10 | 2.44% |
| hsa_circ_0069375 | 820 | 10 | 2.44% |
| hsa_circ_0086159 | 820 | 10 | 2.44% |

| | | | |
|---|---|---|---|
| hsa_circ_0086297 | 820 | 10 | 2.44% |
| hsa_circ_0102613 | 820 | 10 | 2.44% |
| hsa_circ_0115307 | 820 | 10 | 2.44% |
| hsa_circ_0130072 | 820 | 10 | 2.44% |
| hsa_circ_0140751 | 820 | 10 | 2.44% |
| hsa_circ_0118804 | 1149 | 14 | 2.44% |
| hsa_circ_0017573 | 985 | 12 | 2.44% |
| hsa_circ_0088481 | 985 | 12 | 2.44% |
| hsa_circ_0108338 | 985 | 12 | 2.44% |
| hsa_circ_0027008 | 903 | 11 | 2.44% |
| hsa_circ_0051397 | 903 | 11 | 2.44% |
| hsa_circ_0081057 | 903 | 11 | 2.44% |
| hsa_circ_0087134 | 903 | 11 | 2.44% |
| hsa_circ_0096120 | 903 | 11 | 2.44% |
| hsa_circ_0103724 | 903 | 11 | 2.44% |
| hsa_circ_0112265 | 903 | 11 | 2.44% |
| hsa_circ_0120713 | 903 | 11 | 2.44% |
| hsa_circ_0126678 | 903 | 11 | 2.44% |
| hsa_circ_0137001 | 903 | 11 | 2.44% |
| hsa_circ_0007573 | 821 | 10 | 2.44% |
| hsa_circ_0025236 | 821 | 10 | 2.44% |
| hsa_circ_0031946 | 821 | 10 | 2.44% |
| hsa_circ_0050677 | 821 | 10 | 2.44% |
| hsa_circ_0054700 | 821 | 10 | 2.44% |
| hsa_circ_0066753 | 821 | 10 | 2.44% |
| hsa_circ_0081589 | 821 | 10 | 2.44% |
| hsa_circ_0083837 | 821 | 10 | 2.44% |
| hsa_circ_0100391 | 821 | 10 | 2.44% |
| hsa_circ_0107075 | 821 | 10 | 2.44% |
| hsa_circ_0113393 | 821 | 10 | 2.44% |
| hsa_circ_0120999 | 821 | 10 | 2.44% |
| hsa_circ_0072629 | 1068 | 13 | 2.43% |
| hsa_circ_0067530 | 986 | 12 | 2.43% |
| hsa_circ_0001284 | 904 | 11 | 2.43% |
| hsa_circ_0029622 | 904 | 11 | 2.43% |
| hsa_circ_0030508 | 904 | 11 | 2.43% |
| hsa_circ_0122333 | 904 | 11 | 2.43% |
| hsa_circ_0008900 | 822 | 10 | 2.43% |
| hsa_circ_0016768 | 822 | 10 | 2.43% |
| hsa_circ_0022216 | 822 | 10 | 2.43% |
| hsa_circ_0041065 | 822 | 10 | 2.43% |
| hsa_circ_0054639 | 822 | 10 | 2.43% |
| hsa_circ_0057986 | 822 | 10 | 2.43% |
| hsa_circ_0061339 | 822 | 10 | 2.43% |
| hsa_circ_0062373 | 822 | 10 | 2.43% |
| hsa_circ_0087515 | 822 | 10 | 2.43% |
| hsa_circ_0091896 | 822 | 10 | 2.43% |
| hsa_circ_0102392 | 822 | 10 | 2.43% |
| hsa_circ_0103767 | 822 | 10 | 2.43% |
| hsa_circ_0105252 | 822 | 10 | 2.43% |

| | | | |
|---|---|---|---|
| hsa_circ_0109920 | 822 | 10 | 2.43% |
| hsa_circ_0112774 | 822 | 10 | 2.43% |
| hsa_circ_0115021 | 822 | 10 | 2.43% |
| hsa_circ_0045979 | 1151 | 14 | 2.43% |
| hsa_circ_0014141 | 987 | 12 | 2.43% |
| hsa_circ_0069475 | 987 | 12 | 2.43% |
| hsa_circ_0104190 | 987 | 12 | 2.43% |
| hsa_circ_0124134 | 2468 | 30 | 2.43% |
| hsa_circ_0034940 | 905 | 11 | 2.43% |
| hsa_circ_0049743 | 905 | 11 | 2.43% |
| hsa_circ_0065710 | 905 | 11 | 2.43% |
| hsa_circ_0082775 | 905 | 11 | 2.43% |
| hsa_circ_0090650 | 905 | 11 | 2.43% |
| hsa_circ_0095769 | 905 | 11 | 2.43% |
| hsa_circ_0096243 | 905 | 11 | 2.43% |
| hsa_circ_0022237 | 823 | 10 | 2.43% |
| hsa_circ_0023963 | 823 | 10 | 2.43% |
| hsa_circ_0024658 | 823 | 10 | 2.43% |
| hsa_circ_0068816 | 823 | 10 | 2.43% |
| hsa_circ_0069385 | 823 | 10 | 2.43% |
| hsa_circ_0070331 | 823 | 10 | 2.43% |
| hsa_circ_0073365 | 823 | 10 | 2.43% |
| hsa_circ_0075223 | 823 | 10 | 2.43% |
| hsa_circ_0088788 | 823 | 10 | 2.43% |
| hsa_circ_0111523 | 823 | 10 | 2.43% |
| hsa_circ_0118618 | 823 | 10 | 2.43% |
| hsa_circ_0122944 | 823 | 10 | 2.43% |
| hsa_circ_0132811 | 823 | 10 | 2.43% |
| hsa_circ_0059778 | 1070 | 13 | 2.43% |
| hsa_circ_0043499 | 1235 | 15 | 2.43% |
| hsa_circ_0037637 | 988 | 12 | 2.43% |
| hsa_circ_0131647 | 988 | 12 | 2.43% |
| hsa_circ_0019092 | 906 | 11 | 2.43% |
| hsa_circ_0024578 | 906 | 11 | 2.43% |
| hsa_circ_0028027 | 906 | 11 | 2.43% |
| hsa_circ_0038823 | 906 | 11 | 2.43% |
| hsa_circ_0039233 | 906 | 11 | 2.43% |
| hsa_circ_0052211 | 906 | 11 | 2.43% |
| hsa_circ_0072197 | 906 | 11 | 2.43% |
| hsa_circ_0077146 | 906 | 11 | 2.43% |
| hsa_circ_0120750 | 906 | 11 | 2.43% |
| hsa_circ_0101964 | 1071 | 13 | 2.43% |
| hsa_circ_0007574 | 824 | 10 | 2.43% |
| hsa_circ_0011422 | 824 | 10 | 2.43% |
| hsa_circ_0017820 | 824 | 10 | 2.43% |
| hsa_circ_0018727 | 824 | 10 | 2.43% |
| hsa_circ_0029995 | 824 | 10 | 2.43% |
| hsa_circ_0032790 | 824 | 10 | 2.43% |
| hsa_circ_0041699 | 824 | 10 | 2.43% |
| hsa_circ_0067416 | 824 | 10 | 2.43% |

| | | | |
|---|---|---|---|
| hsa_circ_0069143 | 824 | 10 | 2.43% |
| hsa_circ_0087422 | 824 | 10 | 2.43% |
| hsa_circ_0088193 | 824 | 10 | 2.43% |
| hsa_circ_0089721 | 824 | 10 | 2.43% |
| hsa_circ_0099569 | 824 | 10 | 2.43% |
| hsa_circ_0104096 | 824 | 10 | 2.43% |
| hsa_circ_0135907 | 824 | 10 | 2.43% |
| hsa_circ_0098482 | 989 | 12 | 2.43% |
| hsa_circ_0031300 | 907 | 11 | 2.43% |
| hsa_circ_0056834 | 907 | 11 | 2.43% |
| hsa_circ_0065227 | 907 | 11 | 2.43% |
| hsa_circ_0099798 | 907 | 11 | 2.43% |
| hsa_circ_0028500 | 1072 | 13 | 2.43% |
| hsa_circ_0092689 | 1072 | 13 | 2.43% |
| hsa_circ_0090832 | 1155 | 14 | 2.42% |
| hsa_circ_0004671 | 825 | 10 | 2.42% |
| hsa_circ_0013626 | 825 | 10 | 2.42% |
| hsa_circ_0027688 | 825 | 10 | 2.42% |
| hsa_circ_0047589 | 825 | 10 | 2.42% |
| hsa_circ_0047998 | 825 | 10 | 2.42% |
| hsa_circ_0048532 | 825 | 10 | 2.42% |
| hsa_circ_0050258 | 825 | 10 | 2.42% |
| hsa_circ_0052223 | 825 | 10 | 2.42% |
| hsa_circ_0076622 | 825 | 10 | 2.42% |
| hsa_circ_0083566 | 825 | 10 | 2.42% |
| hsa_circ_0095259 | 825 | 10 | 2.42% |
| hsa_circ_0096126 | 825 | 10 | 2.42% |
| hsa_circ_0101120 | 825 | 10 | 2.42% |
| hsa_circ_0107438 | 825 | 10 | 2.42% |
| hsa_circ_0111335 | 825 | 10 | 2.42% |
| hsa_circ_0128208 | 825 | 10 | 2.42% |
| hsa_circ_0128312 | 825 | 10 | 2.42% |
| hsa_circ_0042200 | 908 | 11 | 2.42% |
| hsa_circ_0044125 | 908 | 11 | 2.42% |
| hsa_circ_0049799 | 908 | 11 | 2.42% |
| hsa_circ_0059326 | 908 | 11 | 2.42% |
| hsa_circ_0062289 | 908 | 11 | 2.42% |
| hsa_circ_0079941 | 908 | 11 | 2.42% |
| hsa_circ_0096240 | 908 | 11 | 2.42% |
| hsa_circ_0114381 | 908 | 11 | 2.42% |
| hsa_circ_0048487 | 991 | 12 | 2.42% |
| hsa_circ_0088883 | 991 | 12 | 2.42% |
| hsa_circ_0128949 | 991 | 12 | 2.42% |
| hsa_circ_0019870 | 826 | 10 | 2.42% |
| hsa_circ_0019998 | 826 | 10 | 2.42% |
| hsa_circ_0041595 | 826 | 10 | 2.42% |
| hsa_circ_0042394 | 826 | 10 | 2.42% |
| hsa_circ_0043598 | 826 | 10 | 2.42% |
| hsa_circ_0043654 | 826 | 10 | 2.42% |
| hsa_circ_0051684 | 826 | 10 | 2.42% |

| | | | |
|---|---|---|---|
| hsa_circ_0063833 | 826 | 10 | 2.42% |
| hsa_circ_0070904 | 826 | 10 | 2.42% |
| hsa_circ_0082706 | 826 | 10 | 2.42% |
| hsa_circ_0085788 | 826 | 10 | 2.42% |
| hsa_circ_0090471 | 826 | 10 | 2.42% |
| hsa_circ_0091732 | 826 | 10 | 2.42% |
| hsa_circ_0094530 | 826 | 10 | 2.42% |
| hsa_circ_0096241 | 826 | 10 | 2.42% |
| hsa_circ_0116074 | 826 | 10 | 2.42% |
| hsa_circ_0125681 | 826 | 10 | 2.42% |
| hsa_circ_0129223 | 826 | 10 | 2.42% |
| hsa_circ_0137205 | 826 | 10 | 2.42% |
| hsa_circ_0072624 | 1074 | 13 | 2.42% |
| hsa_circ_0002083 | 909 | 11 | 2.42% |
| hsa_circ_0011044 | 909 | 11 | 2.42% |
| hsa_circ_0034672 | 909 | 11 | 2.42% |
| hsa_circ_0066776 | 909 | 11 | 2.42% |
| hsa_circ_0102768 | 909 | 11 | 2.42% |
| hsa_circ_0034821 | 992 | 12 | 2.42% |
| hsa_circ_0042293 | 992 | 12 | 2.42% |
| hsa_circ_0010950 | 827 | 10 | 2.42% |
| hsa_circ_0015171 | 827 | 10 | 2.42% |
| hsa_circ_0019730 | 827 | 10 | 2.42% |
| hsa_circ_0040843 | 827 | 10 | 2.42% |
| hsa_circ_0057690 | 827 | 10 | 2.42% |
| hsa_circ_0070699 | 827 | 10 | 2.42% |
| hsa_circ_0073578 | 827 | 10 | 2.42% |
| hsa_circ_0081743 | 827 | 10 | 2.42% |
| hsa_circ_0087719 | 827 | 10 | 2.42% |
| hsa_circ_0088236 | 827 | 10 | 2.42% |
| hsa_circ_0103476 | 827 | 10 | 2.42% |
| hsa_circ_0110911 | 827 | 10 | 2.42% |
| hsa_circ_0112845 | 827 | 10 | 2.42% |
| hsa_circ_0119150 | 827 | 10 | 2.42% |
| hsa_circ_0119757 | 827 | 10 | 2.42% |
| hsa_circ_0017113 | 910 | 11 | 2.42% |
| hsa_circ_0036178 | 910 | 11 | 2.42% |
| hsa_circ_0058524 | 910 | 11 | 2.42% |
| hsa_circ_0108558 | 910 | 11 | 2.42% |
| hsa_circ_0114183 | 910 | 11 | 2.42% |
| hsa_circ_0102615 | 1241 | 15 | 2.42% |
| hsa_circ_0099447 | 993 | 12 | 2.42% |
| hsa_circ_0047687 | 2070 | 25 | 2.42% |
| hsa_circ_0010591 | 828 | 10 | 2.42% |
| hsa_circ_0012508 | 828 | 10 | 2.42% |
| hsa_circ_0021590 | 828 | 10 | 2.42% |
| hsa_circ_0021690 | 828 | 10 | 2.42% |
| hsa_circ_0044468 | 828 | 10 | 2.42% |
| hsa_circ_0051142 | 828 | 10 | 2.42% |
| hsa_circ_0060581 | 828 | 10 | 2.42% |

| | | | |
|---|---|---|---|
| hsa_circ_0071579 | 828 | 10 | 2.42% |
| hsa_circ_0086142 | 828 | 10 | 2.42% |
| hsa_circ_0088173 | 828 | 10 | 2.42% |
| hsa_circ_0089128 | 828 | 10 | 2.42% |
| hsa_circ_0092388 | 828 | 10 | 2.42% |
| hsa_circ_0100541 | 828 | 10 | 2.42% |
| hsa_circ_0101097 | 828 | 10 | 2.42% |
| hsa_circ_0112029 | 828 | 10 | 2.42% |
| hsa_circ_0113823 | 828 | 10 | 2.42% |
| hsa_circ_0117723 | 828 | 10 | 2.42% |
| hsa_circ_0128712 | 828 | 10 | 2.42% |
| hsa_circ_0134178 | 828 | 10 | 2.42% |
| hsa_circ_0120851 | 911 | 11 | 2.41% |
| hsa_circ_0107946 | 1574 | 19 | 2.41% |
| hsa_circ_0081923 | 1077 | 13 | 2.41% |
| hsa_circ_0022886 | 1243 | 15 | 2.41% |
| hsa_circ_0012108 | 829 | 10 | 2.41% |
| hsa_circ_0021135 | 829 | 10 | 2.41% |
| hsa_circ_0021337 | 829 | 10 | 2.41% |
| hsa_circ_0026517 | 829 | 10 | 2.41% |
| hsa_circ_0028547 | 829 | 10 | 2.41% |
| hsa_circ_0028659 | 829 | 10 | 2.41% |
| hsa_circ_0061330 | 829 | 10 | 2.41% |
| hsa_circ_0068712 | 829 | 10 | 2.41% |
| hsa_circ_0080047 | 829 | 10 | 2.41% |
| hsa_circ_0082816 | 829 | 10 | 2.41% |
| hsa_circ_0085682 | 829 | 10 | 2.41% |
| hsa_circ_0087322 | 829 | 10 | 2.41% |
| hsa_circ_0096239 | 829 | 10 | 2.41% |
| hsa_circ_0096806 | 829 | 10 | 2.41% |
| hsa_circ_0099366 | 829 | 10 | 2.41% |
| hsa_circ_0100918 | 829 | 10 | 2.41% |
| hsa_circ_0114928 | 829 | 10 | 2.41% |
| hsa_circ_0118040 | 829 | 10 | 2.41% |
| hsa_circ_0124314 | 829 | 10 | 2.41% |
| hsa_circ_0127927 | 829 | 10 | 2.41% |
| hsa_circ_0129392 | 829 | 10 | 2.41% |
| hsa_circ_0135628 | 829 | 10 | 2.41% |
| hsa_circ_0052505 | 912 | 11 | 2.41% |
| hsa_circ_0057351 | 912 | 11 | 2.41% |
| hsa_circ_0060316 | 912 | 11 | 2.41% |
| hsa_circ_0093484 | 912 | 11 | 2.41% |
| hsa_circ_0095518 | 912 | 11 | 2.41% |
| hsa_circ_0002027 | 995 | 12 | 2.41% |
| hsa_circ_0028697 | 1161 | 14 | 2.41% |
| hsa_circ_0096688 | 1576 | 19 | 2.41% |
| hsa_circ_0115951 | 2572 | 31 | 2.41% |
| hsa_circ_0015984 | 1245 | 15 | 2.41% |
| hsa_circ_0027167 | 996 | 12 | 2.41% |
| hsa_circ_0042688 | 996 | 12 | 2.41% |

| | | | |
|---|---|---|---|
| hsa_circ_0044479 | 996 | 12 | 2.41% |
| hsa_circ_0063339 | 996 | 12 | 2.41% |
| hsa_circ_0067415 | 996 | 12 | 2.41% |
| hsa_circ_0118423 | 996 | 12 | 2.41% |
| hsa_circ_0133004 | 996 | 12 | 2.41% |
| hsa_circ_0134727 | 996 | 12 | 2.41% |
| hsa_circ_0003974 | 913 | 11 | 2.41% |
| hsa_circ_0043854 | 913 | 11 | 2.41% |
| hsa_circ_0107535 | 913 | 11 | 2.41% |
| hsa_circ_0120061 | 913 | 11 | 2.41% |
| hsa_circ_0006875 | 830 | 10 | 2.41% |
| hsa_circ_0020930 | 830 | 10 | 2.41% |
| hsa_circ_0028211 | 830 | 10 | 2.41% |
| hsa_circ_0036471 | 830 | 10 | 2.41% |
| hsa_circ_0048828 | 830 | 10 | 2.41% |
| hsa_circ_0065739 | 830 | 10 | 2.41% |
| hsa_circ_0103765 | 830 | 10 | 2.41% |
| hsa_circ_0107239 | 830 | 10 | 2.41% |
| hsa_circ_0119753 | 830 | 10 | 2.41% |
| hsa_circ_0123562 | 830 | 10 | 2.41% |
| hsa_circ_0137299 | 830 | 10 | 2.41% |
| hsa_circ_0061907 | 1163 | 14 | 2.41% |
| hsa_circ_0081463 | 1080 | 13 | 2.41% |
| hsa_circ_0090010 | 997 | 12 | 2.41% |
| hsa_circ_0015959 | 914 | 11 | 2.41% |
| hsa_circ_0030713 | 914 | 11 | 2.41% |
| hsa_circ_0043494 | 914 | 11 | 2.41% |
| hsa_circ_0063353 | 914 | 11 | 2.41% |
| hsa_circ_0111472 | 914 | 11 | 2.41% |
| hsa_circ_0113841 | 914 | 11 | 2.41% |
| hsa_circ_0124852 | 914 | 11 | 2.41% |
| hsa_circ_0008740 | 831 | 10 | 2.41% |
| hsa_circ_0015718 | 831 | 10 | 2.41% |
| hsa_circ_0028039 | 831 | 10 | 2.41% |
| hsa_circ_0036045 | 831 | 10 | 2.41% |
| hsa_circ_0050978 | 831 | 10 | 2.41% |
| hsa_circ_0059737 | 831 | 10 | 2.41% |
| hsa_circ_0066740 | 831 | 10 | 2.41% |
| hsa_circ_0067162 | 831 | 10 | 2.41% |
| hsa_circ_0068173 | 831 | 10 | 2.41% |
| hsa_circ_0070651 | 831 | 10 | 2.41% |
| hsa_circ_0071861 | 831 | 10 | 2.41% |
| hsa_circ_0091373 | 831 | 10 | 2.41% |
| hsa_circ_0104447 | 831 | 10 | 2.41% |
| hsa_circ_0123973 | 831 | 10 | 2.41% |
| hsa_circ_0125445 | 831 | 10 | 2.41% |
| hsa_circ_0132797 | 831 | 10 | 2.41% |
| hsa_circ_0135498 | 831 | 10 | 2.41% |
| hsa_circ_0137941 | 831 | 10 | 2.41% |
| hsa_circ_0074492 | 1164 | 14 | 2.41% |

| | | | |
|---|---|---|---|
| hsa_circ_0035809 | 998 | 12 | 2.40% |
| hsa_circ_0120697 | 998 | 12 | 2.40% |
| hsa_circ_0021719 | 915 | 11 | 2.40% |
| hsa_circ_0058529 | 915 | 11 | 2.40% |
| hsa_circ_0060310 | 915 | 11 | 2.40% |
| hsa_circ_0060346 | 915 | 11 | 2.40% |
| hsa_circ_0085813 | 915 | 11 | 2.40% |
| hsa_circ_0001976 | 832 | 10 | 2.40% |
| hsa_circ_0011812 | 832 | 10 | 2.40% |
| hsa_circ_0063998 | 832 | 10 | 2.40% |
| hsa_circ_0079448 | 832 | 10 | 2.40% |
| hsa_circ_0082126 | 832 | 10 | 2.40% |
| hsa_circ_0087249 | 832 | 10 | 2.40% |
| hsa_circ_0106714 | 832 | 10 | 2.40% |
| hsa_circ_0119890 | 832 | 10 | 2.40% |
| hsa_circ_0137948 | 832 | 10 | 2.40% |
| hsa_circ_0097443 | 2414 | 29 | 2.40% |
| hsa_circ_0057370 | 1332 | 16 | 2.40% |
| hsa_circ_0057379 | 1332 | 16 | 2.40% |
| hsa_circ_0047649 | 999 | 12 | 2.40% |
| hsa_circ_0134729 | 999 | 12 | 2.40% |
| hsa_circ_0021061 | 916 | 11 | 2.40% |
| hsa_circ_0083724 | 916 | 11 | 2.40% |
| hsa_circ_0090446 | 916 | 11 | 2.40% |
| hsa_circ_0093485 | 916 | 11 | 2.40% |
| hsa_circ_0001220 | 833 | 10 | 2.40% |
| hsa_circ_0015024 | 833 | 10 | 2.40% |
| hsa_circ_0020897 | 833 | 10 | 2.40% |
| hsa_circ_0034535 | 833 | 10 | 2.40% |
| hsa_circ_0038879 | 833 | 10 | 2.40% |
| hsa_circ_0041725 | 833 | 10 | 2.40% |
| hsa_circ_0045133 | 833 | 10 | 2.40% |
| hsa_circ_0049468 | 833 | 10 | 2.40% |
| hsa_circ_0050546 | 833 | 10 | 2.40% |
| hsa_circ_0112310 | 833 | 10 | 2.40% |
| hsa_circ_0126833 | 833 | 10 | 2.40% |
| hsa_circ_0129927 | 833 | 10 | 2.40% |
| hsa_circ_0137811 | 833 | 10 | 2.40% |
| hsa_circ_0138049 | 833 | 10 | 2.40% |
| hsa_circ_0069444 | 917 | 11 | 2.40% |
| hsa_circ_0096065 | 917 | 11 | 2.40% |
| hsa_circ_0107860 | 917 | 11 | 2.40% |
| hsa_circ_0111321 | 917 | 11 | 2.40% |
| hsa_circ_0004641 | 834 | 10 | 2.40% |
| hsa_circ_0007835 | 834 | 10 | 2.40% |
| hsa_circ_0012616 | 834 | 10 | 2.40% |
| hsa_circ_0016799 | 834 | 10 | 2.40% |
| hsa_circ_0026373 | 834 | 10 | 2.40% |
| hsa_circ_0030764 | 834 | 10 | 2.40% |
| hsa_circ_0033355 | 834 | 10 | 2.40% |

| | | | |
|---|---|---|---|
| hsa_circ_0048732 | 834 | 10 | 2.40% |
| hsa_circ_0071269 | 834 | 10 | 2.40% |
| hsa_circ_0071437 | 834 | 10 | 2.40% |
| hsa_circ_0078337 | 834 | 10 | 2.40% |
| hsa_circ_0081486 | 834 | 10 | 2.40% |
| hsa_circ_0084519 | 834 | 10 | 2.40% |
| hsa_circ_0088408 | 834 | 10 | 2.40% |
| hsa_circ_0102599 | 834 | 10 | 2.40% |
| hsa_circ_0108008 | 834 | 10 | 2.40% |
| hsa_circ_0113993 | 834 | 10 | 2.40% |
| hsa_circ_0133442 | 834 | 10 | 2.40% |
| hsa_circ_0104827 | 1001 | 12 | 2.40% |
| hsa_circ_0024776 | 918 | 11 | 2.40% |
| hsa_circ_0041614 | 918 | 11 | 2.40% |
| hsa_circ_0048288 | 918 | 11 | 2.40% |
| hsa_circ_0089203 | 918 | 11 | 2.40% |
| hsa_circ_0123903 | 918 | 11 | 2.40% |
| hsa_circ_0139900 | 918 | 11 | 2.40% |
| hsa_circ_0039571 | 1002 | 12 | 2.40% |
| hsa_circ_0050717 | 1002 | 12 | 2.40% |
| hsa_circ_0012300 | 835 | 10 | 2.40% |
| hsa_circ_0014348 | 835 | 10 | 2.40% |
| hsa_circ_0021325 | 835 | 10 | 2.40% |
| hsa_circ_0046513 | 835 | 10 | 2.40% |
| hsa_circ_0052199 | 835 | 10 | 2.40% |
| hsa_circ_0065218 | 835 | 10 | 2.40% |
| hsa_circ_0086258 | 835 | 10 | 2.40% |
| hsa_circ_0091134 | 835 | 10 | 2.40% |
| hsa_circ_0093489 | 835 | 10 | 2.40% |
| hsa_circ_0106500 | 835 | 10 | 2.40% |
| hsa_circ_0110201 | 835 | 10 | 2.40% |
| hsa_circ_0111825 | 835 | 10 | 2.40% |
| hsa_circ_0117990 | 835 | 10 | 2.40% |
| hsa_circ_0121945 | 835 | 10 | 2.40% |
| hsa_circ_0135599 | 835 | 10 | 2.40% |
| hsa_circ_0020147 | 1086 | 13 | 2.39% |
| hsa_circ_0060699 | 1086 | 13 | 2.39% |
| hsa_circ_0095832 | 919 | 11 | 2.39% |
| hsa_circ_0102811 | 919 | 11 | 2.39% |
| hsa_circ_0004301 | 836 | 10 | 2.39% |
| hsa_circ_0004742 | 836 | 10 | 2.39% |
| hsa_circ_0017405 | 836 | 10 | 2.39% |
| hsa_circ_0017733 | 836 | 10 | 2.39% |
| hsa_circ_0031626 | 836 | 10 | 2.39% |
| hsa_circ_0033425 | 836 | 10 | 2.39% |
| hsa_circ_0035855 | 836 | 10 | 2.39% |
| hsa_circ_0039976 | 836 | 10 | 2.39% |
| hsa_circ_0041523 | 836 | 10 | 2.39% |
| hsa_circ_0060094 | 836 | 10 | 2.39% |
| hsa_circ_0062522 | 836 | 10 | 2.39% |

| | | | |
|---|---|---|---|
| hsa_circ_0066242 | 836 | 10 | 2.39% |
| hsa_circ_0084056 | 836 | 10 | 2.39% |
| hsa_circ_0098610 | 836 | 10 | 2.39% |
| hsa_circ_0102674 | 836 | 10 | 2.39% |
| hsa_circ_0123113 | 836 | 10 | 2.39% |
| hsa_circ_0134007 | 836 | 10 | 2.39% |
| hsa_circ_0137392 | 836 | 10 | 2.39% |
| hsa_circ_0007989 | 1087 | 13 | 2.39% |
| hsa_circ_0131662 | 1756 | 21 | 2.39% |
| hsa_circ_0065704 | 920 | 11 | 2.39% |
| hsa_circ_0118019 | 920 | 11 | 2.39% |
| hsa_circ_0033418 | 1004 | 12 | 2.39% |
| hsa_circ_0049755 | 1004 | 12 | 2.39% |
| hsa_circ_0074668 | 1004 | 12 | 2.39% |
| hsa_circ_0080617 | 1004 | 12 | 2.39% |
| hsa_circ_0126034 | 1004 | 12 | 2.39% |
| hsa_circ_0013493 | 837 | 10 | 2.39% |
| hsa_circ_0020458 | 837 | 10 | 2.39% |
| hsa_circ_0021752 | 837 | 10 | 2.39% |
| hsa_circ_0045131 | 837 | 10 | 2.39% |
| hsa_circ_0050370 | 837 | 10 | 2.39% |
| hsa_circ_0052195 | 837 | 10 | 2.39% |
| hsa_circ_0056963 | 837 | 10 | 2.39% |
| hsa_circ_0095137 | 837 | 10 | 2.39% |
| hsa_circ_0100886 | 837 | 10 | 2.39% |
| hsa_circ_0101601 | 837 | 10 | 2.39% |
| hsa_circ_0102888 | 837 | 10 | 2.39% |
| hsa_circ_0104421 | 837 | 10 | 2.39% |
| hsa_circ_0124440 | 837 | 10 | 2.39% |
| hsa_circ_0129955 | 837 | 10 | 2.39% |
| hsa_circ_0017685 | 921 | 11 | 2.39% |
| hsa_circ_0122665 | 921 | 11 | 2.39% |
| hsa_circ_0138103 | 921 | 11 | 2.39% |
| hsa_circ_0079194 | 1340 | 16 | 2.39% |
| hsa_circ_0123143 | 1005 | 12 | 2.39% |
| hsa_circ_0013584 | 1173 | 14 | 2.39% |
| hsa_circ_0000131 | 838 | 10 | 2.39% |
| hsa_circ_0015765 | 838 | 10 | 2.39% |
| hsa_circ_0036295 | 838 | 10 | 2.39% |
| hsa_circ_0038332 | 838 | 10 | 2.39% |
| hsa_circ_0041777 | 838 | 10 | 2.39% |
| hsa_circ_0042190 | 838 | 10 | 2.39% |
| hsa_circ_0058636 | 838 | 10 | 2.39% |
| hsa_circ_0060735 | 838 | 10 | 2.39% |
| hsa_circ_0069018 | 838 | 10 | 2.39% |
| hsa_circ_0072114 | 838 | 10 | 2.39% |
| hsa_circ_0079561 | 838 | 10 | 2.39% |
| hsa_circ_0080230 | 838 | 10 | 2.39% |
| hsa_circ_0102360 | 838 | 10 | 2.39% |
| hsa_circ_0118400 | 838 | 10 | 2.39% |

| | | | |
|---|---|---|---|
| hsa_circ_0122594 | 838 | 10 | 2.39% |
| hsa_circ_0124765 | 838 | 10 | 2.39% |
| hsa_circ_0131682 | 838 | 10 | 2.39% |
| hsa_circ_0137023 | 838 | 10 | 2.39% |
| hsa_circ_0036312 | 922 | 11 | 2.39% |
| hsa_circ_0056996 | 922 | 11 | 2.39% |
| hsa_circ_0087573 | 922 | 11 | 2.39% |
| hsa_circ_0102765 | 922 | 11 | 2.39% |
| hsa_circ_0108845 | 922 | 11 | 2.39% |
| hsa_circ_0138610 | 922 | 11 | 2.39% |
| hsa_circ_0127656 | 1006 | 12 | 2.39% |
| hsa_circ_0087468 | 1090 | 13 | 2.39% |
| hsa_circ_0013549 | 839 | 10 | 2.38% |
| hsa_circ_0016465 | 839 | 10 | 2.38% |
| hsa_circ_0023046 | 839 | 10 | 2.38% |
| hsa_circ_0023458 | 839 | 10 | 2.38% |
| hsa_circ_0042074 | 839 | 10 | 2.38% |
| hsa_circ_0042191 | 839 | 10 | 2.38% |
| hsa_circ_0054167 | 839 | 10 | 2.38% |
| hsa_circ_0057704 | 839 | 10 | 2.38% |
| hsa_circ_0058322 | 839 | 10 | 2.38% |
| hsa_circ_0064127 | 839 | 10 | 2.38% |
| hsa_circ_0099539 | 839 | 10 | 2.38% |
| hsa_circ_0114151 | 839 | 10 | 2.38% |
| hsa_circ_0117229 | 839 | 10 | 2.38% |
| hsa_circ_0118262 | 839 | 10 | 2.38% |
| hsa_circ_0137819 | 839 | 10 | 2.38% |
| hsa_circ_0008237 | 923 | 11 | 2.38% |
| hsa_circ_0019117 | 923 | 11 | 2.38% |
| hsa_circ_0043292 | 923 | 11 | 2.38% |
| hsa_circ_0074524 | 923 | 11 | 2.38% |
| hsa_circ_0092720 | 923 | 11 | 2.38% |
| hsa_circ_0063865 | 1007 | 12 | 2.38% |
| hsa_circ_0011307 | 1092 | 13 | 2.38% |
| hsa_circ_0092257 | 1092 | 13 | 2.38% |
| hsa_circ_0110190 | 1092 | 13 | 2.38% |
| hsa_circ_0064834 | 1008 | 12 | 2.38% |
| hsa_circ_0025574 | 924 | 11 | 2.38% |
| hsa_circ_0027713 | 924 | 11 | 2.38% |
| hsa_circ_0031375 | 924 | 11 | 2.38% |
| hsa_circ_0060839 | 924 | 11 | 2.38% |
| hsa_circ_0077142 | 924 | 11 | 2.38% |
| hsa_circ_0078190 | 924 | 11 | 2.38% |
| hsa_circ_0126286 | 924 | 11 | 2.38% |
| hsa_circ_0010474 | 840 | 10 | 2.38% |
| hsa_circ_0013026 | 840 | 10 | 2.38% |
| hsa_circ_0022626 | 840 | 10 | 2.38% |
| hsa_circ_0028134 | 840 | 10 | 2.38% |
| hsa_circ_0034699 | 840 | 10 | 2.38% |
| hsa_circ_0039529 | 840 | 10 | 2.38% |

| | | | |
|---|---|---|---|
| hsa_circ_0057070 | 840 | 10 | 2.38% |
| hsa_circ_0072874 | 840 | 10 | 2.38% |
| hsa_circ_0095775 | 840 | 10 | 2.38% |
| hsa_circ_0097212 | 840 | 10 | 2.38% |
| hsa_circ_0106264 | 840 | 10 | 2.38% |
| hsa_circ_0134174 | 840 | 10 | 2.38% |
| hsa_circ_0138898 | 840 | 10 | 2.38% |
| hsa_circ_0025271 | 1177 | 14 | 2.38% |
| hsa_circ_0029178 | 925 | 11 | 2.38% |
| hsa_circ_0029323 | 925 | 11 | 2.38% |
| hsa_circ_0042753 | 925 | 11 | 2.38% |
| hsa_circ_0108032 | 925 | 11 | 2.38% |
| hsa_circ_0105192 | 2607 | 31 | 2.38% |
| hsa_circ_0016421 | 841 | 10 | 2.38% |
| hsa_circ_0024245 | 841 | 10 | 2.38% |
| hsa_circ_0026783 | 841 | 10 | 2.38% |
| hsa_circ_0041559 | 841 | 10 | 2.38% |
| hsa_circ_0055410 | 841 | 10 | 2.38% |
| hsa_circ_0069171 | 841 | 10 | 2.38% |
| hsa_circ_0074422 | 841 | 10 | 2.38% |
| hsa_circ_0079454 | 841 | 10 | 2.38% |
| hsa_circ_0094787 | 841 | 10 | 2.38% |
| hsa_circ_0095153 | 841 | 10 | 2.38% |
| hsa_circ_0108029 | 841 | 10 | 2.38% |
| hsa_circ_0109407 | 841 | 10 | 2.38% |
| hsa_circ_0111129 | 841 | 10 | 2.38% |
| hsa_circ_0116731 | 841 | 10 | 2.38% |
| hsa_circ_0131246 | 841 | 10 | 2.38% |
| hsa_circ_0140334 | 841 | 10 | 2.38% |
| hsa_circ_0080360 | 1094 | 13 | 2.38% |
| hsa_circ_0110085 | 3114 | 37 | 2.38% |
| hsa_circ_0048911 | 1010 | 12 | 2.38% |
| hsa_circ_0007035 | 926 | 11 | 2.38% |
| hsa_circ_0018521 | 926 | 11 | 2.38% |
| hsa_circ_0027515 | 926 | 11 | 2.38% |
| hsa_circ_0104321 | 926 | 11 | 2.38% |
| hsa_circ_0017449 | 842 | 10 | 2.38% |
| hsa_circ_0017457 | 842 | 10 | 2.38% |
| hsa_circ_0030458 | 842 | 10 | 2.38% |
| hsa_circ_0038346 | 842 | 10 | 2.38% |
| hsa_circ_0060577 | 842 | 10 | 2.38% |
| hsa_circ_0065657 | 842 | 10 | 2.38% |
| hsa_circ_0070942 | 842 | 10 | 2.38% |
| hsa_circ_0071835 | 842 | 10 | 2.38% |
| hsa_circ_0075658 | 842 | 10 | 2.38% |
| hsa_circ_0076756 | 842 | 10 | 2.38% |
| hsa_circ_0077163 | 842 | 10 | 2.38% |
| hsa_circ_0104165 | 842 | 10 | 2.38% |
| hsa_circ_0116666 | 842 | 10 | 2.38% |
| hsa_circ_0123492 | 842 | 10 | 2.38% |

| | | | |
|---|---|---|---|
| hsa_circ_0127150 | 842 | 10 | 2.38% |
| hsa_circ_0035910 | 927 | 11 | 2.37% |
| hsa_circ_0093887 | 927 | 11 | 2.37% |
| hsa_circ_0115969 | 927 | 11 | 2.37% |
| hsa_circ_0130657 | 927 | 11 | 2.37% |
| hsa_circ_0133112 | 927 | 11 | 2.37% |
| hsa_circ_0139867 | 927 | 11 | 2.37% |
| hsa_circ_0135328 | 1180 | 14 | 2.37% |
| hsa_circ_0007553 | 843 | 10 | 2.37% |
| hsa_circ_0018075 | 843 | 10 | 2.37% |
| hsa_circ_0026785 | 843 | 10 | 2.37% |
| hsa_circ_0034725 | 843 | 10 | 2.37% |
| hsa_circ_0040011 | 843 | 10 | 2.37% |
| hsa_circ_0045644 | 843 | 10 | 2.37% |
| hsa_circ_0054209 | 843 | 10 | 2.37% |
| hsa_circ_0064601 | 843 | 10 | 2.37% |
| hsa_circ_0066408 | 843 | 10 | 2.37% |
| hsa_circ_0085725 | 843 | 10 | 2.37% |
| hsa_circ_0089098 | 843 | 10 | 2.37% |
| hsa_circ_0099760 | 843 | 10 | 2.37% |
| hsa_circ_0118125 | 843 | 10 | 2.37% |
| hsa_circ_0123941 | 843 | 10 | 2.37% |
| hsa_circ_0127276 | 843 | 10 | 2.37% |
| hsa_circ_0134708 | 843 | 10 | 2.37% |
| hsa_circ_0021577 | 1181 | 14 | 2.37% |
| hsa_circ_0049084 | 1181 | 14 | 2.37% |
| hsa_circ_0009500 | 928 | 11 | 2.37% |
| hsa_circ_0018385 | 928 | 11 | 2.37% |
| hsa_circ_0038610 | 928 | 11 | 2.37% |
| hsa_circ_0053300 | 928 | 11 | 2.37% |
| hsa_circ_0055546 | 928 | 11 | 2.37% |
| hsa_circ_0073463 | 928 | 11 | 2.37% |
| hsa_circ_0099804 | 928 | 11 | 2.37% |
| hsa_circ_0020578 | 844 | 10 | 2.37% |
| hsa_circ_0028208 | 844 | 10 | 2.37% |
| hsa_circ_0034440 | 844 | 10 | 2.37% |
| hsa_circ_0042439 | 844 | 10 | 2.37% |
| hsa_circ_0045845 | 844 | 10 | 2.37% |
| hsa_circ_0057169 | 844 | 10 | 2.37% |
| hsa_circ_0075010 | 844 | 10 | 2.37% |
| hsa_circ_0094337 | 844 | 10 | 2.37% |
| hsa_circ_0095176 | 844 | 10 | 2.37% |
| hsa_circ_0103191 | 844 | 10 | 2.37% |
| hsa_circ_0106871 | 844 | 10 | 2.37% |
| hsa_circ_0118454 | 844 | 10 | 2.37% |
| hsa_circ_0121706 | 844 | 10 | 2.37% |
| hsa_circ_0101397 | 1013 | 12 | 2.37% |
| hsa_circ_0042336 | 929 | 11 | 2.37% |
| hsa_circ_0051931 | 929 | 11 | 2.37% |
| hsa_circ_0058516 | 929 | 11 | 2.37% |

| | | | |
|---|---|---|---|
| hsa_circ_0058653 | 929 | 11 | 2.37% |
| hsa_circ_0125439 | 929 | 11 | 2.37% |
| hsa_circ_0125733 | 929 | 11 | 2.37% |
| hsa_circ_0039496 | 1014 | 12 | 2.37% |
| hsa_circ_0133945 | 1014 | 12 | 2.37% |
| hsa_circ_0015029 | 845 | 10 | 2.37% |
| hsa_circ_0024440 | 845 | 10 | 2.37% |
| hsa_circ_0035770 | 845 | 10 | 2.37% |
| hsa_circ_0042245 | 845 | 10 | 2.37% |
| hsa_circ_0054245 | 845 | 10 | 2.37% |
| hsa_circ_0074718 | 845 | 10 | 2.37% |
| hsa_circ_0083846 | 845 | 10 | 2.37% |
| hsa_circ_0091762 | 845 | 10 | 2.37% |
| hsa_circ_0106137 | 845 | 10 | 2.37% |
| hsa_circ_0118164 | 845 | 10 | 2.37% |
| hsa_circ_0127577 | 845 | 10 | 2.37% |
| hsa_circ_0130287 | 845 | 10 | 2.37% |
| hsa_circ_0132682 | 845 | 10 | 2.37% |
| hsa_circ_0021865 | 3635 | 43 | 2.37% |
| hsa_circ_0054512 | 1099 | 13 | 2.37% |
| hsa_circ_0064455 | 1099 | 13 | 2.37% |
| hsa_circ_0066832 | 930 | 11 | 2.37% |
| hsa_circ_0084739 | 930 | 11 | 2.37% |
| hsa_circ_0100048 | 930 | 11 | 2.37% |
| hsa_circ_0119888 | 930 | 11 | 2.37% |
| hsa_circ_0127678 | 930 | 11 | 2.37% |
| hsa_circ_0086768 | 1015 | 12 | 2.36% |
| hsa_circ_0000558 | 1269 | 15 | 2.36% |
| hsa_circ_0014494 | 846 | 10 | 2.36% |
| hsa_circ_0038636 | 846 | 10 | 2.36% |
| hsa_circ_0046188 | 846 | 10 | 2.36% |
| hsa_circ_0049824 | 846 | 10 | 2.36% |
| hsa_circ_0058153 | 846 | 10 | 2.36% |
| hsa_circ_0058306 | 846 | 10 | 2.36% |
| hsa_circ_0062385 | 846 | 10 | 2.36% |
| hsa_circ_0083665 | 846 | 10 | 2.36% |
| hsa_circ_0085474 | 846 | 10 | 2.36% |
| hsa_circ_0092445 | 846 | 10 | 2.36% |
| hsa_circ_0095862 | 846 | 10 | 2.36% |
| hsa_circ_0096302 | 846 | 10 | 2.36% |
| hsa_circ_0110017 | 846 | 10 | 2.36% |
| hsa_circ_0120614 | 846 | 10 | 2.36% |
| hsa_circ_0063738 | 931 | 11 | 2.36% |
| hsa_circ_0089610 | 931 | 11 | 2.36% |
| hsa_circ_0105328 | 931 | 11 | 2.36% |
| hsa_circ_0114842 | 1016 | 12 | 2.36% |
| hsa_circ_0115260 | 1016 | 12 | 2.36% |
| hsa_circ_0021982 | 1101 | 13 | 2.36% |
| hsa_circ_0005883 | 847 | 10 | 2.36% |
| hsa_circ_0011790 | 847 | 10 | 2.36% |

| | | | |
|---|---|---|---|
| hsa_circ_0012150 | 847 | 10 | 2.36% |
| hsa_circ_0012494 | 847 | 10 | 2.36% |
| hsa_circ_0015159 | 847 | 10 | 2.36% |
| hsa_circ_0022416 | 847 | 10 | 2.36% |
| hsa_circ_0036946 | 847 | 10 | 2.36% |
| hsa_circ_0048945 | 847 | 10 | 2.36% |
| hsa_circ_0059452 | 847 | 10 | 2.36% |
| hsa_circ_0062438 | 847 | 10 | 2.36% |
| hsa_circ_0081367 | 847 | 10 | 2.36% |
| hsa_circ_0099166 | 847 | 10 | 2.36% |
| hsa_circ_0099748 | 847 | 10 | 2.36% |
| hsa_circ_0109398 | 847 | 10 | 2.36% |
| hsa_circ_0111918 | 847 | 10 | 2.36% |
| hsa_circ_0124309 | 847 | 10 | 2.36% |
| hsa_circ_0132006 | 847 | 10 | 2.36% |
| hsa_circ_0120296 | 1186 | 14 | 2.36% |
| hsa_circ_0010397 | 1525 | 18 | 2.36% |
| hsa_circ_0005750 | 932 | 11 | 2.36% |
| hsa_circ_0009283 | 932 | 11 | 2.36% |
| hsa_circ_0037832 | 932 | 11 | 2.36% |
| hsa_circ_0077579 | 932 | 11 | 2.36% |
| hsa_circ_0085803 | 932 | 11 | 2.36% |
| hsa_circ_0095330 | 932 | 11 | 2.36% |
| hsa_circ_0105340 | 932 | 11 | 2.36% |
| hsa_circ_0127402 | 932 | 11 | 2.36% |
| hsa_circ_0130536 | 932 | 11 | 2.36% |
| hsa_circ_0130822 | 932 | 11 | 2.36% |
| hsa_circ_0096216 | 2712 | 32 | 2.36% |
| hsa_circ_0016675 | 848 | 10 | 2.36% |
| hsa_circ_0029984 | 848 | 10 | 2.36% |
| hsa_circ_0051901 | 848 | 10 | 2.36% |
| hsa_circ_0061324 | 848 | 10 | 2.36% |
| hsa_circ_0064341 | 848 | 10 | 2.36% |
| hsa_circ_0081563 | 848 | 10 | 2.36% |
| hsa_circ_0107965 | 848 | 10 | 2.36% |
| hsa_circ_0108148 | 848 | 10 | 2.36% |
| hsa_circ_0119500 | 848 | 10 | 2.36% |
| hsa_circ_0121788 | 848 | 10 | 2.36% |
| hsa_circ_0131654 | 848 | 10 | 2.36% |
| hsa_circ_0102701 | 2290 | 27 | 2.36% |
| hsa_circ_0019829 | 933 | 11 | 2.36% |
| hsa_circ_0039427 | 933 | 11 | 2.36% |
| hsa_circ_0039567 | 933 | 11 | 2.36% |
| hsa_circ_0042949 | 933 | 11 | 2.36% |
| hsa_circ_0046424 | 933 | 11 | 2.36% |
| hsa_circ_0070386 | 933 | 11 | 2.36% |
| hsa_circ_0097177 | 933 | 11 | 2.36% |
| hsa_circ_0133271 | 933 | 11 | 2.36% |
| hsa_circ_0063760 | 1103 | 13 | 2.36% |
| hsa_circ_0063582 | 3396 | 40 | 2.36% |

| | | | |
|---|---|---|---|
| hsa_circ_0005854 | 849 | 10 | 2.36% |
| hsa_circ_0006912 | 849 | 10 | 2.36% |
| hsa_circ_0011443 | 849 | 10 | 2.36% |
| hsa_circ_0017772 | 849 | 10 | 2.36% |
| hsa_circ_0027282 | 849 | 10 | 2.36% |
| hsa_circ_0027740 | 849 | 10 | 2.36% |
| hsa_circ_0031522 | 849 | 10 | 2.36% |
| hsa_circ_0036924 | 849 | 10 | 2.36% |
| hsa_circ_0060328 | 849 | 10 | 2.36% |
| hsa_circ_0062123 | 849 | 10 | 2.36% |
| hsa_circ_0063057 | 849 | 10 | 2.36% |
| hsa_circ_0065619 | 849 | 10 | 2.36% |
| hsa_circ_0066588 | 849 | 10 | 2.36% |
| hsa_circ_0077341 | 849 | 10 | 2.36% |
| hsa_circ_0100362 | 849 | 10 | 2.36% |
| hsa_circ_0116088 | 849 | 10 | 2.36% |
| hsa_circ_0116238 | 849 | 10 | 2.36% |
| hsa_circ_0133474 | 849 | 10 | 2.36% |
| hsa_circ_0136332 | 849 | 10 | 2.36% |
| hsa_circ_0137709 | 849 | 10 | 2.36% |
| hsa_circ_0001140 | 934 | 11 | 2.36% |
| hsa_circ_0054116 | 934 | 11 | 2.36% |
| hsa_circ_0125206 | 934 | 11 | 2.36% |
| hsa_circ_0062839 | 1019 | 12 | 2.36% |
| hsa_circ_0042887 | 1104 | 13 | 2.36% |
| hsa_circ_0036305 | 1105 | 13 | 2.35% |
| hsa_circ_0076662 | 1105 | 13 | 2.35% |
| hsa_circ_0136483 | 1105 | 13 | 2.35% |
| hsa_circ_0040140 | 1020 | 12 | 2.35% |
| hsa_circ_0065712 | 1020 | 12 | 2.35% |
| hsa_circ_0092137 | 1020 | 12 | 2.35% |
| hsa_circ_0099599 | 1020 | 12 | 2.35% |
| hsa_circ_0003344 | 935 | 11 | 2.35% |
| hsa_circ_0026486 | 935 | 11 | 2.35% |
| hsa_circ_0044153 | 935 | 11 | 2.35% |
| hsa_circ_0059741 | 935 | 11 | 2.35% |
| hsa_circ_0114188 | 935 | 11 | 2.35% |
| hsa_circ_0017898 | 850 | 10 | 2.35% |
| hsa_circ_0021140 | 850 | 10 | 2.35% |
| hsa_circ_0027901 | 850 | 10 | 2.35% |
| hsa_circ_0041545 | 850 | 10 | 2.35% |
| hsa_circ_0043417 | 850 | 10 | 2.35% |
| hsa_circ_0058997 | 850 | 10 | 2.35% |
| hsa_circ_0082158 | 850 | 10 | 2.35% |
| hsa_circ_0085390 | 850 | 10 | 2.35% |
| hsa_circ_0085604 | 850 | 10 | 2.35% |
| hsa_circ_0093586 | 850 | 10 | 2.35% |
| hsa_circ_0101296 | 850 | 10 | 2.35% |
| hsa_circ_0116415 | 850 | 10 | 2.35% |
| hsa_circ_0121765 | 850 | 10 | 2.35% |

| | | | |
|---|---|---|---|
| hsa_circ_0127417 | 850 | 10 | 2.35% |
| hsa_circ_0096556 | 1106 | 13 | 2.35% |
| hsa_circ_0006115 | 936 | 11 | 2.35% |
| hsa_circ_0093677 | 936 | 11 | 2.35% |
| hsa_circ_0096981 | 936 | 11 | 2.35% |
| hsa_circ_0120075 | 936 | 11 | 2.35% |
| hsa_circ_0124482 | 936 | 11 | 2.35% |
| hsa_circ_0134630 | 936 | 11 | 2.35% |
| hsa_circ_0136121 | 936 | 11 | 2.35% |
| hsa_circ_0032771 | 851 | 10 | 2.35% |
| hsa_circ_0038195 | 851 | 10 | 2.35% |
| hsa_circ_0045720 | 851 | 10 | 2.35% |
| hsa_circ_0077843 | 851 | 10 | 2.35% |
| hsa_circ_0086371 | 851 | 10 | 2.35% |
| hsa_circ_0092220 | 851 | 10 | 2.35% |
| hsa_circ_0096300 | 851 | 10 | 2.35% |
| hsa_circ_0103639 | 851 | 10 | 2.35% |
| hsa_circ_0108067 | 851 | 10 | 2.35% |
| hsa_circ_0113814 | 851 | 10 | 2.35% |
| hsa_circ_0128298 | 851 | 10 | 2.35% |
| hsa_circ_0132310 | 851 | 10 | 2.35% |
| hsa_circ_0045745 | 1277 | 15 | 2.35% |
| hsa_circ_0035400 | 1107 | 13 | 2.35% |
| hsa_circ_0101957 | 1107 | 13 | 2.35% |
| hsa_circ_0121503 | 1022 | 12 | 2.35% |
| hsa_circ_0029953 | 937 | 11 | 2.35% |
| hsa_circ_0081622 | 937 | 11 | 2.35% |
| hsa_circ_0100382 | 937 | 11 | 2.35% |
| hsa_circ_0021661 | 1363 | 16 | 2.35% |
| hsa_circ_0007362 | 852 | 10 | 2.35% |
| hsa_circ_0011890 | 852 | 10 | 2.35% |
| hsa_circ_0026882 | 852 | 10 | 2.35% |
| hsa_circ_0032880 | 852 | 10 | 2.35% |
| hsa_circ_0046465 | 852 | 10 | 2.35% |
| hsa_circ_0055009 | 852 | 10 | 2.35% |
| hsa_circ_0060115 | 852 | 10 | 2.35% |
| hsa_circ_0069400 | 852 | 10 | 2.35% |
| hsa_circ_0076263 | 852 | 10 | 2.35% |
| hsa_circ_0091200 | 852 | 10 | 2.35% |
| hsa_circ_0103038 | 852 | 10 | 2.35% |
| hsa_circ_0105451 | 852 | 10 | 2.35% |
| hsa_circ_0116086 | 852 | 10 | 2.35% |
| hsa_circ_0117950 | 852 | 10 | 2.35% |
| hsa_circ_0131258 | 852 | 10 | 2.35% |
| hsa_circ_0136256 | 852 | 10 | 2.35% |
| hsa_circ_0136665 | 852 | 10 | 2.35% |
| hsa_circ_0083481 | 1108 | 13 | 2.35% |
| hsa_circ_0043296 | 2131 | 25 | 2.35% |
| hsa_circ_0019758 | 1023 | 12 | 2.35% |
| hsa_circ_0043637 | 938 | 11 | 2.35% |

| | | | |
|---|---|---|---|
| hsa_circ_0073046 | 938 | 11 | 2.35% |
| hsa_circ_0098028 | 938 | 11 | 2.35% |
| hsa_circ_0110601 | 938 | 11 | 2.35% |
| hsa_circ_0014765 | 853 | 10 | 2.34% |
| hsa_circ_0024352 | 853 | 10 | 2.34% |
| hsa_circ_0026337 | 853 | 10 | 2.34% |
| hsa_circ_0027611 | 853 | 10 | 2.34% |
| hsa_circ_0030066 | 853 | 10 | 2.34% |
| hsa_circ_0031258 | 853 | 10 | 2.34% |
| hsa_circ_0033656 | 853 | 10 | 2.34% |
| hsa_circ_0035961 | 853 | 10 | 2.34% |
| hsa_circ_0037870 | 853 | 10 | 2.34% |
| hsa_circ_0100578 | 853 | 10 | 2.34% |
| hsa_circ_0102160 | 853 | 10 | 2.34% |
| hsa_circ_0105091 | 853 | 10 | 2.34% |
| hsa_circ_0109655 | 853 | 10 | 2.34% |
| hsa_circ_0121583 | 853 | 10 | 2.34% |
| hsa_circ_0128005 | 853 | 10 | 2.34% |
| hsa_circ_0132932 | 853 | 10 | 2.34% |
| hsa_circ_0087339 | 1024 | 12 | 2.34% |
| hsa_circ_0004234 | 939 | 11 | 2.34% |
| hsa_circ_0024045 | 939 | 11 | 2.34% |
| hsa_circ_0027248 | 939 | 11 | 2.34% |
| hsa_circ_0047666 | 939 | 11 | 2.34% |
| hsa_circ_0051549 | 939 | 11 | 2.34% |
| hsa_circ_0077814 | 939 | 11 | 2.34% |
| hsa_circ_0139648 | 939 | 11 | 2.34% |
| hsa_circ_0041429 | 1110 | 13 | 2.34% |
| hsa_circ_0036596 | 854 | 10 | 2.34% |
| hsa_circ_0038566 | 854 | 10 | 2.34% |
| hsa_circ_0053414 | 854 | 10 | 2.34% |
| hsa_circ_0087523 | 854 | 10 | 2.34% |
| hsa_circ_0093705 | 854 | 10 | 2.34% |
| hsa_circ_0108795 | 854 | 10 | 2.34% |
| hsa_circ_0120014 | 854 | 10 | 2.34% |
| hsa_circ_0120167 | 854 | 10 | 2.34% |
| hsa_circ_0125526 | 854 | 10 | 2.34% |
| hsa_circ_0135989 | 854 | 10 | 2.34% |
| hsa_circ_0135429 | 1025 | 12 | 2.34% |
| hsa_circ_0010005 | 940 | 11 | 2.34% |
| hsa_circ_0011409 | 940 | 11 | 2.34% |
| hsa_circ_0031237 | 940 | 11 | 2.34% |
| hsa_circ_0034922 | 940 | 11 | 2.34% |
| hsa_circ_0104612 | 940 | 11 | 2.34% |
| hsa_circ_0132772 | 940 | 11 | 2.34% |
| hsa_circ_0140145 | 1111 | 13 | 2.34% |
| hsa_circ_0126224 | 11114 | 130 | 2.34% |
| hsa_circ_0011904 | 1026 | 12 | 2.34% |
| hsa_circ_0057443 | 1026 | 12 | 2.34% |
| hsa_circ_0097748 | 1026 | 12 | 2.34% |

| | | | |
|---|---|---|---|
| hsa_circ_0119257 | 1026 | 12 | 2.34% |
| hsa_circ_0000791 | 855 | 10 | 2.34% |
| hsa_circ_0004569 | 855 | 10 | 2.34% |
| hsa_circ_0041088 | 855 | 10 | 2.34% |
| hsa_circ_0044816 | 855 | 10 | 2.34% |
| hsa_circ_0057047 | 855 | 10 | 2.34% |
| hsa_circ_0068791 | 855 | 10 | 2.34% |
| hsa_circ_0069727 | 855 | 10 | 2.34% |
| hsa_circ_0086200 | 855 | 10 | 2.34% |
| hsa_circ_0094036 | 855 | 10 | 2.34% |
| hsa_circ_0112240 | 855 | 10 | 2.34% |
| hsa_circ_0125775 | 855 | 10 | 2.34% |
| hsa_circ_0126085 | 855 | 10 | 2.34% |
| hsa_circ_0128516 | 855 | 10 | 2.34% |
| hsa_circ_0132783 | 855 | 10 | 2.34% |
| hsa_circ_0135330 | 855 | 10 | 2.34% |
| hsa_circ_0067928 | 1112 | 13 | 2.34% |
| hsa_circ_0069967 | 1112 | 13 | 2.34% |
| hsa_circ_0021710 | 941 | 11 | 2.34% |
| hsa_circ_0106209 | 941 | 11 | 2.34% |
| hsa_circ_0126370 | 941 | 11 | 2.34% |
| hsa_circ_0073622 | 1027 | 12 | 2.34% |
| hsa_circ_0121052 | 1027 | 12 | 2.34% |
| hsa_circ_0004684 | 856 | 10 | 2.34% |
| hsa_circ_0015143 | 856 | 10 | 2.34% |
| hsa_circ_0025778 | 856 | 10 | 2.34% |
| hsa_circ_0035864 | 856 | 10 | 2.34% |
| hsa_circ_0039363 | 856 | 10 | 2.34% |
| hsa_circ_0051093 | 856 | 10 | 2.34% |
| hsa_circ_0097677 | 856 | 10 | 2.34% |
| hsa_circ_0100055 | 856 | 10 | 2.34% |
| hsa_circ_0115959 | 856 | 10 | 2.34% |
| hsa_circ_0129453 | 856 | 10 | 2.34% |
| hsa_circ_0005120 | 942 | 11 | 2.34% |
| hsa_circ_0119716 | 942 | 11 | 2.34% |
| hsa_circ_0047753 | 1028 | 12 | 2.33% |
| hsa_circ_0080905 | 1028 | 12 | 2.33% |
| hsa_circ_0133723 | 10453 | 122 | 2.33% |
| hsa_circ_0010539 | 857 | 10 | 2.33% |
| hsa_circ_0016205 | 857 | 10 | 2.33% |
| hsa_circ_0017018 | 857 | 10 | 2.33% |
| hsa_circ_0032764 | 857 | 10 | 2.33% |
| hsa_circ_0061560 | 857 | 10 | 2.33% |
| hsa_circ_0075480 | 857 | 10 | 2.33% |
| hsa_circ_0076117 | 857 | 10 | 2.33% |
| hsa_circ_0087038 | 857 | 10 | 2.33% |
| hsa_circ_0121804 | 857 | 10 | 2.33% |
| hsa_circ_0133077 | 857 | 10 | 2.33% |
| hsa_circ_0012220 | 943 | 11 | 2.33% |
| hsa_circ_0040266 | 943 | 11 | 2.33% |

| | | | |
|---|---|---|---|
| hsa_circ_0042747 | 943 | 11 | 2.33% |
| hsa_circ_0122908 | 943 | 11 | 2.33% |
| hsa_circ_0136115 | 943 | 11 | 2.33% |
| hsa_circ_0017294 | 1029 | 12 | 2.33% |
| hsa_circ_0054899 | 1029 | 12 | 2.33% |
| hsa_circ_0083852 | 2144 | 25 | 2.33% |
| hsa_circ_0008961 | 1201 | 14 | 2.33% |
| hsa_circ_0112965 | 1630 | 19 | 2.33% |
| hsa_circ_0011393 | 858 | 10 | 2.33% |
| hsa_circ_0014578 | 858 | 10 | 2.33% |
| hsa_circ_0017387 | 858 | 10 | 2.33% |
| hsa_circ_0032668 | 858 | 10 | 2.33% |
| hsa_circ_0036143 | 858 | 10 | 2.33% |
| hsa_circ_0043262 | 858 | 10 | 2.33% |
| hsa_circ_0044811 | 858 | 10 | 2.33% |
| hsa_circ_0064658 | 858 | 10 | 2.33% |
| hsa_circ_0096231 | 858 | 10 | 2.33% |
| hsa_circ_0098799 | 858 | 10 | 2.33% |
| hsa_circ_0102365 | 858 | 10 | 2.33% |
| hsa_circ_0105167 | 858 | 10 | 2.33% |
| hsa_circ_0105716 | 858 | 10 | 2.33% |
| hsa_circ_0106370 | 858 | 10 | 2.33% |
| hsa_circ_0115644 | 858 | 10 | 2.33% |
| hsa_circ_0116021 | 858 | 10 | 2.33% |
| hsa_circ_0130106 | 858 | 10 | 2.33% |
| hsa_circ_0132335 | 858 | 10 | 2.33% |
| hsa_circ_0135296 | 858 | 10 | 2.33% |
| hsa_circ_0059270 | 944 | 11 | 2.33% |
| hsa_circ_0060011 | 944 | 11 | 2.33% |
| hsa_circ_0060602 | 944 | 11 | 2.33% |
| hsa_circ_0075744 | 944 | 11 | 2.33% |
| hsa_circ_0094832 | 944 | 11 | 2.33% |
| hsa_circ_0131762 | 944 | 11 | 2.33% |
| hsa_circ_0132139 | 944 | 11 | 2.33% |
| hsa_circ_0136670 | 944 | 11 | 2.33% |
| hsa_circ_0009431 | 1030 | 12 | 2.33% |
| hsa_circ_0042109 | 1030 | 12 | 2.33% |
| hsa_circ_0054900 | 1030 | 12 | 2.33% |
| hsa_circ_0102191 | 1030 | 12 | 2.33% |
| hsa_circ_0112470 | 1030 | 12 | 2.33% |
| hsa_circ_0046245 | 1116 | 13 | 2.33% |
| hsa_circ_0013136 | 2319 | 27 | 2.33% |
| hsa_circ_0003680 | 859 | 10 | 2.33% |
| hsa_circ_0019581 | 859 | 10 | 2.33% |
| hsa_circ_0025874 | 859 | 10 | 2.33% |
| hsa_circ_0072865 | 859 | 10 | 2.33% |
| hsa_circ_0083948 | 859 | 10 | 2.33% |
| hsa_circ_0086460 | 859 | 10 | 2.33% |
| hsa_circ_0089519 | 859 | 10 | 2.33% |
| hsa_circ_0090463 | 859 | 10 | 2.33% |

| | | | |
|---|---|---|---|
| hsa_circ_0096759 | 859 | 10 | 2.33% |
| hsa_circ_0116597 | 859 | 10 | 2.33% |
| hsa_circ_0117745 | 859 | 10 | 2.33% |
| hsa_circ_0122484 | 859 | 10 | 2.33% |
| hsa_circ_0126502 | 859 | 10 | 2.33% |
| hsa_circ_0138270 | 859 | 10 | 2.33% |
| hsa_circ_0002255 | 945 | 11 | 2.33% |
| hsa_circ_0003726 | 945 | 11 | 2.33% |
| hsa_circ_0007450 | 945 | 11 | 2.33% |
| hsa_circ_0044606 | 945 | 11 | 2.33% |
| hsa_circ_0063307 | 945 | 11 | 2.33% |
| hsa_circ_0070393 | 945 | 11 | 2.33% |
| hsa_circ_0071464 | 945 | 11 | 2.33% |
| hsa_circ_0114632 | 945 | 11 | 2.33% |
| hsa_circ_0131293 | 945 | 11 | 2.33% |
| hsa_circ_0033342 | 1031 | 12 | 2.33% |
| hsa_circ_0036196 | 1117 | 13 | 2.33% |
| hsa_circ_0067558 | 1117 | 13 | 2.33% |
| hsa_circ_0133878 | 1032 | 12 | 2.33% |
| hsa_circ_0011967 | 946 | 11 | 2.33% |
| hsa_circ_0055384 | 946 | 11 | 2.33% |
| hsa_circ_0067756 | 946 | 11 | 2.33% |
| hsa_circ_0133635 | 946 | 11 | 2.33% |
| hsa_circ_0011905 | 860 | 10 | 2.33% |
| hsa_circ_0012742 | 860 | 10 | 2.33% |
| hsa_circ_0030882 | 860 | 10 | 2.33% |
| hsa_circ_0032330 | 860 | 10 | 2.33% |
| hsa_circ_0035062 | 860 | 10 | 2.33% |
| hsa_circ_0052219 | 860 | 10 | 2.33% |
| hsa_circ_0056696 | 860 | 10 | 2.33% |
| hsa_circ_0064764 | 860 | 10 | 2.33% |
| hsa_circ_0067452 | 860 | 10 | 2.33% |
| hsa_circ_0069101 | 860 | 10 | 2.33% |
| hsa_circ_0073888 | 860 | 10 | 2.33% |
| hsa_circ_0094879 | 860 | 10 | 2.33% |
| hsa_circ_0120124 | 860 | 10 | 2.33% |
| hsa_circ_0011332 | 2839 | 33 | 2.32% |
| hsa_circ_0060137 | 1635 | 19 | 2.32% |
| hsa_circ_0057375 | 1377 | 16 | 2.32% |
| hsa_circ_0045980 | 1205 | 14 | 2.32% |
| hsa_circ_0103381 | 1033 | 12 | 2.32% |
| hsa_circ_0016379 | 947 | 11 | 2.32% |
| hsa_circ_0041590 | 947 | 11 | 2.32% |
| hsa_circ_0012756 | 861 | 10 | 2.32% |
| hsa_circ_0025005 | 861 | 10 | 2.32% |
| hsa_circ_0026171 | 861 | 10 | 2.32% |
| hsa_circ_0031260 | 861 | 10 | 2.32% |
| hsa_circ_0035027 | 861 | 10 | 2.32% |
| hsa_circ_0041236 | 861 | 10 | 2.32% |
| hsa_circ_0049538 | 861 | 10 | 2.32% |

| | | | |
|---|---|---|---|
| hsa_circ_0051417 | 861 | 10 | 2.32% |
| hsa_circ_0052519 | 861 | 10 | 2.32% |
| hsa_circ_0055343 | 861 | 10 | 2.32% |
| hsa_circ_0076304 | 861 | 10 | 2.32% |
| hsa_circ_0099664 | 861 | 10 | 2.32% |
| hsa_circ_0117605 | 861 | 10 | 2.32% |
| hsa_circ_0119895 | 861 | 10 | 2.32% |
| hsa_circ_0122469 | 861 | 10 | 2.32% |
| hsa_circ_0130444 | 861 | 10 | 2.32% |
| hsa_circ_0135130 | 861 | 10 | 2.32% |
| hsa_circ_0117055 | 1636 | 19 | 2.32% |
| hsa_circ_0035232 | 1034 | 12 | 2.32% |
| hsa_circ_0040027 | 1034 | 12 | 2.32% |
| hsa_circ_0022983 | 948 | 11 | 2.32% |
| hsa_circ_0040326 | 948 | 11 | 2.32% |
| hsa_circ_0044658 | 948 | 11 | 2.32% |
| hsa_circ_0052849 | 948 | 11 | 2.32% |
| hsa_circ_0055217 | 948 | 11 | 2.32% |
| hsa_circ_0088849 | 948 | 11 | 2.32% |
| hsa_circ_0009889 | 862 | 10 | 2.32% |
| hsa_circ_0017196 | 862 | 10 | 2.32% |
| hsa_circ_0026635 | 862 | 10 | 2.32% |
| hsa_circ_0043906 | 862 | 10 | 2.32% |
| hsa_circ_0045933 | 862 | 10 | 2.32% |
| hsa_circ_0047956 | 862 | 10 | 2.32% |
| hsa_circ_0065944 | 862 | 10 | 2.32% |
| hsa_circ_0070075 | 862 | 10 | 2.32% |
| hsa_circ_0077024 | 862 | 10 | 2.32% |
| hsa_circ_0086316 | 862 | 10 | 2.32% |
| hsa_circ_0089045 | 862 | 10 | 2.32% |
| hsa_circ_0105262 | 862 | 10 | 2.32% |
| hsa_circ_0106350 | 862 | 10 | 2.32% |
| hsa_circ_0107596 | 862 | 10 | 2.32% |
| hsa_circ_0110540 | 862 | 10 | 2.32% |
| hsa_circ_0116557 | 862 | 10 | 2.32% |
| hsa_circ_0130917 | 862 | 10 | 2.32% |
| hsa_circ_0136365 | 862 | 10 | 2.32% |
| hsa_circ_0081856 | 1207 | 14 | 2.32% |
| hsa_circ_0034734 | 949 | 11 | 2.32% |
| hsa_circ_0040571 | 949 | 11 | 2.32% |
| hsa_circ_0051727 | 949 | 11 | 2.32% |
| hsa_circ_0067664 | 949 | 11 | 2.32% |
| hsa_circ_0117290 | 949 | 11 | 2.32% |
| hsa_circ_0136400 | 949 | 11 | 2.32% |
| hsa_circ_0008108 | 863 | 10 | 2.32% |
| hsa_circ_0016600 | 863 | 10 | 2.32% |
| hsa_circ_0017596 | 863 | 10 | 2.32% |
| hsa_circ_0025617 | 863 | 10 | 2.32% |
| hsa_circ_0038331 | 863 | 10 | 2.32% |
| hsa_circ_0044473 | 863 | 10 | 2.32% |

| | | | |
|---|---|---|---|
| hsa_circ_0057746 | 863 | 10 | 2.32% |
| hsa_circ_0071079 | 863 | 10 | 2.32% |
| hsa_circ_0075262 | 863 | 10 | 2.32% |
| hsa_circ_0113481 | 863 | 10 | 2.32% |
| hsa_circ_0129648 | 863 | 10 | 2.32% |
| hsa_circ_0134040 | 863 | 10 | 2.32% |
| hsa_circ_0101953 | 1122 | 13 | 2.32% |
| hsa_circ_0078483 | 3885 | 45 | 2.32% |
| hsa_circ_0101345 | 1036 | 12 | 2.32% |
| hsa_circ_0130234 | 1036 | 12 | 2.32% |
| hsa_circ_0038355 | 950 | 11 | 2.32% |
| hsa_circ_0052807 | 950 | 11 | 2.32% |
| hsa_circ_0069806 | 950 | 11 | 2.32% |
| hsa_circ_0082741 | 950 | 11 | 2.32% |
| hsa_circ_0093309 | 950 | 11 | 2.32% |
| hsa_circ_0102309 | 950 | 11 | 2.32% |
| hsa_circ_0021568 | 5010 | 58 | 2.32% |
| hsa_circ_0002983 | 864 | 10 | 2.31% |
| hsa_circ_0010660 | 864 | 10 | 2.31% |
| hsa_circ_0012658 | 864 | 10 | 2.31% |
| hsa_circ_0019258 | 864 | 10 | 2.31% |
| hsa_circ_0023749 | 864 | 10 | 2.31% |
| hsa_circ_0027601 | 864 | 10 | 2.31% |
| hsa_circ_0039519 | 864 | 10 | 2.31% |
| hsa_circ_0054637 | 864 | 10 | 2.31% |
| hsa_circ_0057408 | 864 | 10 | 2.31% |
| hsa_circ_0061771 | 864 | 10 | 2.31% |
| hsa_circ_0062429 | 864 | 10 | 2.31% |
| hsa_circ_0068624 | 864 | 10 | 2.31% |
| hsa_circ_0082828 | 864 | 10 | 2.31% |
| hsa_circ_0096127 | 864 | 10 | 2.31% |
| hsa_circ_0099199 | 864 | 10 | 2.31% |
| hsa_circ_0123368 | 864 | 10 | 2.31% |
| hsa_circ_0135259 | 864 | 10 | 2.31% |
| hsa_circ_0065842 | 1037 | 12 | 2.31% |
| hsa_circ_0020681 | 1210 | 14 | 2.31% |
| hsa_circ_0107518 | 1210 | 14 | 2.31% |
| hsa_circ_0002648 | 951 | 11 | 2.31% |
| hsa_circ_0020998 | 951 | 11 | 2.31% |
| hsa_circ_0021394 | 951 | 11 | 2.31% |
| hsa_circ_0021395 | 951 | 11 | 2.31% |
| hsa_circ_0038246 | 951 | 11 | 2.31% |
| hsa_circ_0051796 | 951 | 11 | 2.31% |
| hsa_circ_0065439 | 951 | 11 | 2.31% |
| hsa_circ_0071452 | 1038 | 12 | 2.31% |
| hsa_circ_0107801 | 1038 | 12 | 2.31% |
| hsa_circ_0000072 | 865 | 10 | 2.31% |
| hsa_circ_0003234 | 865 | 10 | 2.31% |
| hsa_circ_0003758 | 865 | 10 | 2.31% |
| hsa_circ_0004275 | 865 | 10 | 2.31% |

| | | | |
|---|---|---|---|
| hsa_circ_0007832 | 865 | 10 | 2.31% |
| hsa_circ_0016541 | 865 | 10 | 2.31% |
| hsa_circ_0018526 | 865 | 10 | 2.31% |
| hsa_circ_0023372 | 865 | 10 | 2.31% |
| hsa_circ_0025775 | 865 | 10 | 2.31% |
| hsa_circ_0038938 | 865 | 10 | 2.31% |
| hsa_circ_0045825 | 865 | 10 | 2.31% |
| hsa_circ_0050604 | 865 | 10 | 2.31% |
| hsa_circ_0050862 | 865 | 10 | 2.31% |
| hsa_circ_0055791 | 865 | 10 | 2.31% |
| hsa_circ_0059877 | 865 | 10 | 2.31% |
| hsa_circ_0069835 | 865 | 10 | 2.31% |
| hsa_circ_0075025 | 865 | 10 | 2.31% |
| hsa_circ_0086338 | 865 | 10 | 2.31% |
| hsa_circ_0094230 | 865 | 10 | 2.31% |
| hsa_circ_0100663 | 865 | 10 | 2.31% |
| hsa_circ_0101543 | 865 | 10 | 2.31% |
| hsa_circ_0110455 | 865 | 10 | 2.31% |
| hsa_circ_0122255 | 865 | 10 | 2.31% |
| hsa_circ_0041635 | 1125 | 13 | 2.31% |
| hsa_circ_0044155 | 952 | 11 | 2.31% |
| hsa_circ_0064008 | 952 | 11 | 2.31% |
| hsa_circ_0080043 | 952 | 11 | 2.31% |
| hsa_circ_0105112 | 952 | 11 | 2.31% |
| hsa_circ_0114253 | 952 | 11 | 2.31% |
| hsa_circ_0120711 | 952 | 11 | 2.31% |
| hsa_circ_0051210 | 1039 | 12 | 2.31% |
| hsa_circ_0109090 | 1039 | 12 | 2.31% |
| hsa_circ_0002972 | 866 | 10 | 2.31% |
| hsa_circ_0030177 | 866 | 10 | 2.31% |
| hsa_circ_0036061 | 866 | 10 | 2.31% |
| hsa_circ_0069383 | 866 | 10 | 2.31% |
| hsa_circ_0079595 | 866 | 10 | 2.31% |
| hsa_circ_0079814 | 866 | 10 | 2.31% |
| hsa_circ_0088164 | 866 | 10 | 2.31% |
| hsa_circ_0096267 | 866 | 10 | 2.31% |
| hsa_circ_0109353 | 866 | 10 | 2.31% |
| hsa_circ_0019269 | 953 | 11 | 2.31% |
| hsa_circ_0038560 | 953 | 11 | 2.31% |
| hsa_circ_0049918 | 953 | 11 | 2.31% |
| hsa_circ_0104600 | 953 | 11 | 2.31% |
| hsa_circ_0114759 | 953 | 11 | 2.31% |
| hsa_circ_0117551 | 953 | 11 | 2.31% |
| hsa_circ_0015211 | 1040 | 12 | 2.31% |
| hsa_circ_0000522 | 867 | 10 | 2.31% |
| hsa_circ_0010696 | 867 | 10 | 2.31% |
| hsa_circ_0011759 | 867 | 10 | 2.31% |
| hsa_circ_0023599 | 867 | 10 | 2.31% |
| hsa_circ_0031607 | 867 | 10 | 2.31% |
| hsa_circ_0038720 | 867 | 10 | 2.31% |

| | | | |
|---|---|---|---|
| hsa_circ_0050490 | 867 | 10 | 2.31% |
| hsa_circ_0065909 | 867 | 10 | 2.31% |
| hsa_circ_0070850 | 867 | 10 | 2.31% |
| hsa_circ_0072769 | 867 | 10 | 2.31% |
| hsa_circ_0076538 | 867 | 10 | 2.31% |
| hsa_circ_0077565 | 867 | 10 | 2.31% |
| hsa_circ_0082695 | 867 | 10 | 2.31% |
| hsa_circ_0084720 | 867 | 10 | 2.31% |
| hsa_circ_0085001 | 867 | 10 | 2.31% |
| hsa_circ_0094962 | 867 | 10 | 2.31% |
| hsa_circ_0105689 | 867 | 10 | 2.31% |
| hsa_circ_0111087 | 867 | 10 | 2.31% |
| hsa_circ_0125686 | 867 | 10 | 2.31% |
| hsa_circ_0137658 | 867 | 10 | 2.31% |
| hsa_circ_0009042 | 954 | 11 | 2.31% |
| hsa_circ_0020995 | 954 | 11 | 2.31% |
| hsa_circ_0025232 | 954 | 11 | 2.31% |
| hsa_circ_0038956 | 954 | 11 | 2.31% |
| hsa_circ_0088053 | 954 | 11 | 2.31% |
| hsa_circ_0111703 | 954 | 11 | 2.31% |
| hsa_circ_0129625 | 954 | 11 | 2.31% |
| hsa_circ_0131348 | 954 | 11 | 2.31% |
| hsa_circ_0140524 | 954 | 11 | 2.31% |
| hsa_circ_0002348 | 1041 | 12 | 2.31% |
| hsa_circ_0066113 | 1041 | 12 | 2.31% |
| hsa_circ_0070500 | 1041 | 12 | 2.31% |
| hsa_circ_0072964 | 1041 | 12 | 2.31% |
| hsa_circ_0049080 | 1128 | 13 | 2.30% |
| hsa_circ_0017450 | 868 | 10 | 2.30% |
| hsa_circ_0020376 | 868 | 10 | 2.30% |
| hsa_circ_0021128 | 868 | 10 | 2.30% |
| hsa_circ_0029394 | 868 | 10 | 2.30% |
| hsa_circ_0036303 | 868 | 10 | 2.30% |
| hsa_circ_0046124 | 868 | 10 | 2.30% |
| hsa_circ_0047547 | 868 | 10 | 2.30% |
| hsa_circ_0048111 | 868 | 10 | 2.30% |
| hsa_circ_0077756 | 868 | 10 | 2.30% |
| hsa_circ_0101673 | 868 | 10 | 2.30% |
| hsa_circ_0108002 | 868 | 10 | 2.30% |
| hsa_circ_0111340 | 868 | 10 | 2.30% |
| hsa_circ_0000468 | 955 | 11 | 2.30% |
| hsa_circ_0006034 | 955 | 11 | 2.30% |
| hsa_circ_0008170 | 955 | 11 | 2.30% |
| hsa_circ_0075169 | 955 | 11 | 2.30% |
| hsa_circ_0045772 | 1042 | 12 | 2.30% |
| hsa_circ_0050623 | 1042 | 12 | 2.30% |
| hsa_circ_0091490 | 1042 | 12 | 2.30% |
| hsa_circ_0138283 | 1129 | 13 | 2.30% |
| hsa_circ_0051463 | 1216 | 14 | 2.30% |
| hsa_circ_0017839 | 869 | 10 | 2.30% |

| | | | |
|---|---|---|---|
| hsa_circ_0033659 | 869 | 10 | 2.30% |
| hsa_circ_0041564 | 869 | 10 | 2.30% |
| hsa_circ_0043425 | 869 | 10 | 2.30% |
| hsa_circ_0055720 | 869 | 10 | 2.30% |
| hsa_circ_0061833 | 869 | 10 | 2.30% |
| hsa_circ_0081665 | 869 | 10 | 2.30% |
| hsa_circ_0084116 | 869 | 10 | 2.30% |
| hsa_circ_0103633 | 869 | 10 | 2.30% |
| hsa_circ_0106874 | 869 | 10 | 2.30% |
| hsa_circ_0117734 | 869 | 10 | 2.30% |
| hsa_circ_0057757 | 956 | 11 | 2.30% |
| hsa_circ_0090621 | 956 | 11 | 2.30% |
| hsa_circ_0105398 | 956 | 11 | 2.30% |
| hsa_circ_0106917 | 956 | 11 | 2.30% |
| hsa_circ_0110603 | 956 | 11 | 2.30% |
| hsa_circ_0131723 | 1043 | 12 | 2.30% |
| hsa_circ_0087952 | 1130 | 13 | 2.30% |
| hsa_circ_0026370 | 27122 | 312 | 2.30% |
| hsa_circ_0125423 | 1218 | 14 | 2.30% |
| hsa_circ_0127949 | 1131 | 13 | 2.30% |
| hsa_circ_0004158 | 957 | 11 | 2.30% |
| hsa_circ_0012120 | 957 | 11 | 2.30% |
| hsa_circ_0017570 | 957 | 11 | 2.30% |
| hsa_circ_0035939 | 957 | 11 | 2.30% |
| hsa_circ_0129437 | 957 | 11 | 2.30% |
| hsa_circ_0005631 | 870 | 10 | 2.30% |
| hsa_circ_0006186 | 870 | 10 | 2.30% |
| hsa_circ_0008245 | 870 | 10 | 2.30% |
| hsa_circ_0013500 | 870 | 10 | 2.30% |
| hsa_circ_0015369 | 870 | 10 | 2.30% |
| hsa_circ_0022434 | 870 | 10 | 2.30% |
| hsa_circ_0024182 | 870 | 10 | 2.30% |
| hsa_circ_0029304 | 870 | 10 | 2.30% |
| hsa_circ_0033106 | 870 | 10 | 2.30% |
| hsa_circ_0034929 | 870 | 10 | 2.30% |
| hsa_circ_0035529 | 870 | 10 | 2.30% |
| hsa_circ_0047423 | 870 | 10 | 2.30% |
| hsa_circ_0051803 | 870 | 10 | 2.30% |
| hsa_circ_0051824 | 870 | 10 | 2.30% |
| hsa_circ_0060511 | 870 | 10 | 2.30% |
| hsa_circ_0070069 | 870 | 10 | 2.30% |
| hsa_circ_0073630 | 870 | 10 | 2.30% |
| hsa_circ_0080561 | 870 | 10 | 2.30% |
| hsa_circ_0080872 | 870 | 10 | 2.30% |
| hsa_circ_0081941 | 870 | 10 | 2.30% |
| hsa_circ_0089858 | 870 | 10 | 2.30% |
| hsa_circ_0090579 | 870 | 10 | 2.30% |
| hsa_circ_0095062 | 870 | 10 | 2.30% |
| hsa_circ_0106678 | 870 | 10 | 2.30% |
| hsa_circ_0120044 | 870 | 10 | 2.30% |

| | | | |
|---|---|---|---|
| hsa_circ_0124194 | 870 | 10 | 2.30% |
| hsa_circ_0134311 | 870 | 10 | 2.30% |
| hsa_circ_0119742 | 2002 | 23 | 2.30% |
| hsa_circ_0063966 | 1045 | 12 | 2.30% |
| hsa_circ_0069476 | 1045 | 12 | 2.30% |
| hsa_circ_0140036 | 1045 | 12 | 2.30% |
| hsa_circ_0000078 | 958 | 11 | 2.30% |
| hsa_circ_0048099 | 958 | 11 | 2.30% |
| hsa_circ_0105700 | 958 | 11 | 2.30% |
| hsa_circ_0108022 | 958 | 11 | 2.30% |
| hsa_circ_0013886 | 871 | 10 | 2.30% |
| hsa_circ_0020957 | 871 | 10 | 2.30% |
| hsa_circ_0037252 | 871 | 10 | 2.30% |
| hsa_circ_0051804 | 871 | 10 | 2.30% |
| hsa_circ_0067992 | 871 | 10 | 2.30% |
| hsa_circ_0072053 | 871 | 10 | 2.30% |
| hsa_circ_0076558 | 871 | 10 | 2.30% |
| hsa_circ_0080262 | 871 | 10 | 2.30% |
| hsa_circ_0082496 | 871 | 10 | 2.30% |
| hsa_circ_0087503 | 871 | 10 | 2.30% |
| hsa_circ_0096025 | 871 | 10 | 2.30% |
| hsa_circ_0097024 | 871 | 10 | 2.30% |
| hsa_circ_0103673 | 871 | 10 | 2.30% |
| hsa_circ_0108003 | 871 | 10 | 2.30% |
| hsa_circ_0127306 | 871 | 10 | 2.30% |
| hsa_circ_0060107 | 1220 | 14 | 2.30% |
| hsa_circ_0074432 | 1046 | 12 | 2.29% |
| hsa_circ_0069460 | 959 | 11 | 2.29% |
| hsa_circ_0094366 | 959 | 11 | 2.29% |
| hsa_circ_0014636 | 872 | 10 | 2.29% |
| hsa_circ_0018999 | 872 | 10 | 2.29% |
| hsa_circ_0034780 | 872 | 10 | 2.29% |
| hsa_circ_0039015 | 872 | 10 | 2.29% |
| hsa_circ_0048284 | 872 | 10 | 2.29% |
| hsa_circ_0059909 | 872 | 10 | 2.29% |
| hsa_circ_0063154 | 872 | 10 | 2.29% |
| hsa_circ_0072955 | 872 | 10 | 2.29% |
| hsa_circ_0086793 | 872 | 10 | 2.29% |
| hsa_circ_0087891 | 872 | 10 | 2.29% |
| hsa_circ_0088942 | 872 | 10 | 2.29% |
| hsa_circ_0092123 | 872 | 10 | 2.29% |
| hsa_circ_0119106 | 872 | 10 | 2.29% |
| hsa_circ_0122186 | 872 | 10 | 2.29% |
| hsa_circ_0019949 | 960 | 11 | 2.29% |
| hsa_circ_0059966 | 960 | 11 | 2.29% |
| hsa_circ_0077204 | 960 | 11 | 2.29% |
| hsa_circ_0089736 | 960 | 11 | 2.29% |
| hsa_circ_0010140 | 873 | 10 | 2.29% |
| hsa_circ_0011573 | 873 | 10 | 2.29% |
| hsa_circ_0015643 | 873 | 10 | 2.29% |

| | | | |
|---|---|---|---|
| hsa_circ_0024763 | 873 | 10 | 2.29% |
| hsa_circ_0025014 | 873 | 10 | 2.29% |
| hsa_circ_0027680 | 873 | 10 | 2.29% |
| hsa_circ_0029687 | 873 | 10 | 2.29% |
| hsa_circ_0032584 | 873 | 10 | 2.29% |
| hsa_circ_0045489 | 873 | 10 | 2.29% |
| hsa_circ_0060613 | 873 | 10 | 2.29% |
| hsa_circ_0076441 | 873 | 10 | 2.29% |
| hsa_circ_0078224 | 873 | 10 | 2.29% |
| hsa_circ_0084139 | 873 | 10 | 2.29% |
| hsa_circ_0097029 | 873 | 10 | 2.29% |
| hsa_circ_0112834 | 873 | 10 | 2.29% |
| hsa_circ_0138780 | 873 | 10 | 2.29% |
| hsa_circ_0009421 | 1310 | 15 | 2.29% |
| hsa_circ_0026750 | 961 | 11 | 2.29% |
| hsa_circ_0048420 | 961 | 11 | 2.29% |
| hsa_circ_0097411 | 961 | 11 | 2.29% |
| hsa_circ_0106973 | 961 | 11 | 2.29% |
| hsa_circ_0121217 | 961 | 11 | 2.29% |
| hsa_circ_0121390 | 961 | 11 | 2.29% |
| hsa_circ_0081514 | 1136 | 13 | 2.29% |
| hsa_circ_0000615 | 874 | 10 | 2.29% |
| hsa_circ_0009622 | 874 | 10 | 2.29% |
| hsa_circ_0026639 | 874 | 10 | 2.29% |
| hsa_circ_0035033 | 874 | 10 | 2.29% |
| hsa_circ_0054032 | 874 | 10 | 2.29% |
| hsa_circ_0067986 | 874 | 10 | 2.29% |
| hsa_circ_0100716 | 874 | 10 | 2.29% |
| hsa_circ_0126847 | 874 | 10 | 2.29% |
| hsa_circ_0127286 | 874 | 10 | 2.29% |
| hsa_circ_0135895 | 874 | 10 | 2.29% |
| hsa_circ_0019033 | 1049 | 12 | 2.29% |
| hsa_circ_0060749 | 1049 | 12 | 2.29% |
| hsa_circ_0086743 | 1049 | 12 | 2.29% |
| hsa_circ_0014234 | 962 | 11 | 2.29% |
| hsa_circ_0021082 | 962 | 11 | 2.29% |
| hsa_circ_0040076 | 962 | 11 | 2.29% |
| hsa_circ_0076147 | 962 | 11 | 2.29% |
| hsa_circ_0135022 | 962 | 11 | 2.29% |
| hsa_circ_0140642 | 962 | 11 | 2.29% |
| hsa_circ_0103181 | 1137 | 13 | 2.29% |
| hsa_circ_0040995 | 1050 | 12 | 2.29% |
| hsa_circ_0132521 | 1050 | 12 | 2.29% |
| hsa_circ_0008486 | 875 | 10 | 2.29% |
| hsa_circ_0013604 | 875 | 10 | 2.29% |
| hsa_circ_0021162 | 875 | 10 | 2.29% |
| hsa_circ_0033657 | 875 | 10 | 2.29% |
| hsa_circ_0039854 | 875 | 10 | 2.29% |
| hsa_circ_0052450 | 875 | 10 | 2.29% |
| hsa_circ_0055859 | 875 | 10 | 2.29% |

| | | | |
|---|---|---|---|
| hsa_circ_0060421 | 875 | 10 | 2.29% |
| hsa_circ_0071683 | 875 | 10 | 2.29% |
| hsa_circ_0079924 | 875 | 10 | 2.29% |
| hsa_circ_0081686 | 875 | 10 | 2.29% |
| hsa_circ_0084480 | 875 | 10 | 2.29% |
| hsa_circ_0086161 | 875 | 10 | 2.29% |
| hsa_circ_0112747 | 875 | 10 | 2.29% |
| hsa_circ_0116337 | 875 | 10 | 2.29% |
| hsa_circ_0123714 | 875 | 10 | 2.29% |
| hsa_circ_0134177 | 875 | 10 | 2.29% |
| hsa_circ_0136643 | 875 | 10 | 2.29% |
| hsa_circ_0138528 | 875 | 10 | 2.29% |
| hsa_circ_0139239 | 875 | 10 | 2.29% |
| hsa_circ_0024177 | 963 | 11 | 2.28% |
| hsa_circ_0037414 | 963 | 11 | 2.28% |
| hsa_circ_0039234 | 963 | 11 | 2.28% |
| hsa_circ_0058297 | 963 | 11 | 2.28% |
| hsa_circ_0070099 | 963 | 11 | 2.28% |
| hsa_circ_0086542 | 963 | 11 | 2.28% |
| hsa_circ_0090651 | 963 | 11 | 2.28% |
| hsa_circ_0090674 | 963 | 11 | 2.28% |
| hsa_circ_0091396 | 963 | 11 | 2.28% |
| hsa_circ_0026633 | 1051 | 12 | 2.28% |
| hsa_circ_0108052 | 1051 | 12 | 2.28% |
| hsa_circ_0016219 | 876 | 10 | 2.28% |
| hsa_circ_0016932 | 876 | 10 | 2.28% |
| hsa_circ_0035452 | 876 | 10 | 2.28% |
| hsa_circ_0051166 | 876 | 10 | 2.28% |
| hsa_circ_0058299 | 876 | 10 | 2.28% |
| hsa_circ_0064165 | 876 | 10 | 2.28% |
| hsa_circ_0072878 | 876 | 10 | 2.28% |
| hsa_circ_0091561 | 876 | 10 | 2.28% |
| hsa_circ_0107330 | 876 | 10 | 2.28% |
| hsa_circ_0111173 | 876 | 10 | 2.28% |
| hsa_circ_0119259 | 876 | 10 | 2.28% |
| hsa_circ_0123798 | 876 | 10 | 2.28% |
| hsa_circ_0128276 | 876 | 10 | 2.28% |
| hsa_circ_0135770 | 876 | 10 | 2.28% |
| hsa_circ_0135990 | 876 | 10 | 2.28% |
| hsa_circ_0138097 | 876 | 10 | 2.28% |
| hsa_circ_0020816 | 1139 | 13 | 2.28% |
| hsa_circ_0072627 | 1139 | 13 | 2.28% |
| hsa_circ_0038131 | 964 | 11 | 2.28% |
| hsa_circ_0043629 | 964 | 11 | 2.28% |
| hsa_circ_0083671 | 964 | 11 | 2.28% |
| hsa_circ_0099927 | 964 | 11 | 2.28% |
| hsa_circ_0058859 | 1052 | 12 | 2.28% |
| hsa_circ_0009161 | 877 | 10 | 2.28% |
| hsa_circ_0031412 | 877 | 10 | 2.28% |
| hsa_circ_0033448 | 877 | 10 | 2.28% |

| | | | |
|---|---|---|---|
| hsa_circ_0040291 | 877 | 10 | 2.28% |
| hsa_circ_0049676 | 877 | 10 | 2.28% |
| hsa_circ_0055045 | 877 | 10 | 2.28% |
| hsa_circ_0062419 | 877 | 10 | 2.28% |
| hsa_circ_0073566 | 877 | 10 | 2.28% |
| hsa_circ_0080671 | 877 | 10 | 2.28% |
| hsa_circ_0081269 | 877 | 10 | 2.28% |
| hsa_circ_0093063 | 877 | 10 | 2.28% |
| hsa_circ_0108793 | 877 | 10 | 2.28% |
| hsa_circ_0073368 | 1228 | 14 | 2.28% |
| hsa_circ_0036242 | 965 | 11 | 2.28% |
| hsa_circ_0037842 | 965 | 11 | 2.28% |
| hsa_circ_0063560 | 965 | 11 | 2.28% |
| hsa_circ_0076567 | 965 | 11 | 2.28% |
| hsa_circ_0093327 | 965 | 11 | 2.28% |
| hsa_circ_0101996 | 965 | 11 | 2.28% |
| hsa_circ_0018717 | 878 | 10 | 2.28% |
| hsa_circ_0020317 | 878 | 10 | 2.28% |
| hsa_circ_0062962 | 878 | 10 | 2.28% |
| hsa_circ_0063880 | 878 | 10 | 2.28% |
| hsa_circ_0068737 | 878 | 10 | 2.28% |
| hsa_circ_0094030 | 878 | 10 | 2.28% |
| hsa_circ_0098342 | 878 | 10 | 2.28% |
| hsa_circ_0100651 | 878 | 10 | 2.28% |
| hsa_circ_0124919 | 878 | 10 | 2.28% |
| hsa_circ_0129158 | 878 | 10 | 2.28% |
| hsa_circ_0132822 | 878 | 10 | 2.28% |
| hsa_circ_0134454 | 878 | 10 | 2.28% |
| hsa_circ_0010519 | 966 | 11 | 2.28% |
| hsa_circ_0060281 | 966 | 11 | 2.28% |
| hsa_circ_0068373 | 966 | 11 | 2.28% |
| hsa_circ_0078211 | 966 | 11 | 2.28% |
| hsa_circ_0092648 | 966 | 11 | 2.28% |
| hsa_circ_0067406 | 1054 | 12 | 2.28% |
| hsa_circ_0090928 | 1054 | 12 | 2.28% |
| hsa_circ_0139357 | 1054 | 12 | 2.28% |
| hsa_circ_0001272 | 879 | 10 | 2.28% |
| hsa_circ_0005235 | 879 | 10 | 2.28% |
| hsa_circ_0024359 | 879 | 10 | 2.28% |
| hsa_circ_0024543 | 879 | 10 | 2.28% |
| hsa_circ_0039993 | 879 | 10 | 2.28% |
| hsa_circ_0050880 | 879 | 10 | 2.28% |
| hsa_circ_0061881 | 879 | 10 | 2.28% |
| hsa_circ_0069809 | 879 | 10 | 2.28% |
| hsa_circ_0077237 | 879 | 10 | 2.28% |
| hsa_circ_0085751 | 879 | 10 | 2.28% |
| hsa_circ_0086720 | 879 | 10 | 2.28% |
| hsa_circ_0099589 | 879 | 10 | 2.28% |
| hsa_circ_0108173 | 879 | 10 | 2.28% |
| hsa_circ_0113301 | 879 | 10 | 2.28% |

| | | | |
|---|---|---|---|
| hsa_circ_0115622 | 879 | 10 | 2.28% |
| hsa_circ_0126661 | 879 | 10 | 2.28% |
| hsa_circ_0129132 | 879 | 10 | 2.28% |
| hsa_circ_0133717 | 879 | 10 | 2.28% |
| hsa_circ_0020965 | 967 | 11 | 2.28% |
| hsa_circ_0080013 | 967 | 11 | 2.28% |
| hsa_circ_0080163 | 967 | 11 | 2.28% |
| hsa_circ_0123899 | 967 | 11 | 2.28% |
| hsa_circ_0019759 | 1055 | 12 | 2.27% |
| hsa_circ_0030921 | 1055 | 12 | 2.27% |
| hsa_circ_0092462 | 2638 | 30 | 2.27% |
| hsa_circ_0112673 | 1407 | 16 | 2.27% |
| hsa_circ_0043779 | 1144 | 13 | 2.27% |
| hsa_circ_0011184 | 1056 | 12 | 2.27% |
| hsa_circ_0033560 | 1056 | 12 | 2.27% |
| hsa_circ_0064804 | 1056 | 12 | 2.27% |
| hsa_circ_0040676 | 968 | 11 | 2.27% |
| hsa_circ_0041624 | 968 | 11 | 2.27% |
| hsa_circ_0042065 | 968 | 11 | 2.27% |
| hsa_circ_0050866 | 968 | 11 | 2.27% |
| hsa_circ_0094387 | 968 | 11 | 2.27% |
| hsa_circ_0140646 | 968 | 11 | 2.27% |
| hsa_circ_0013180 | 880 | 10 | 2.27% |
| hsa_circ_0035970 | 880 | 10 | 2.27% |
| hsa_circ_0040985 | 880 | 10 | 2.27% |
| hsa_circ_0080071 | 880 | 10 | 2.27% |
| hsa_circ_0088818 | 880 | 10 | 2.27% |
| hsa_circ_0095740 | 880 | 10 | 2.27% |
| hsa_circ_0101031 | 880 | 10 | 2.27% |
| hsa_circ_0108564 | 880 | 10 | 2.27% |
| hsa_circ_0124098 | 880 | 10 | 2.27% |
| hsa_circ_0136833 | 880 | 10 | 2.27% |
| hsa_circ_0139835 | 880 | 10 | 2.27% |
| hsa_circ_0140155 | 880 | 10 | 2.27% |
| hsa_circ_0026247 | 1145 | 13 | 2.27% |
| hsa_circ_0060007 | 1145 | 13 | 2.27% |
| hsa_circ_0067262 | 1145 | 13 | 2.27% |
| hsa_circ_0100487 | 1057 | 12 | 2.27% |
| hsa_circ_0023259 | 969 | 11 | 2.27% |
| hsa_circ_0057661 | 969 | 11 | 2.27% |
| hsa_circ_0077528 | 969 | 11 | 2.27% |
| hsa_circ_0077869 | 969 | 11 | 2.27% |
| hsa_circ_0088764 | 969 | 11 | 2.27% |
| hsa_circ_0121076 | 969 | 11 | 2.27% |
| hsa_circ_0129375 | 969 | 11 | 2.27% |
| hsa_circ_0001078 | 881 | 10 | 2.27% |
| hsa_circ_0001165 | 881 | 10 | 2.27% |
| hsa_circ_0012607 | 881 | 10 | 2.27% |
| hsa_circ_0014839 | 881 | 10 | 2.27% |
| hsa_circ_0018374 | 881 | 10 | 2.27% |

| | | | |
|---|---|---|---|
| hsa_circ_0018723 | 881 | 10 | 2.27% |
| hsa_circ_0020796 | 881 | 10 | 2.27% |
| hsa_circ_0055528 | 881 | 10 | 2.27% |
| hsa_circ_0058215 | 881 | 10 | 2.27% |
| hsa_circ_0060032 | 881 | 10 | 2.27% |
| hsa_circ_0065088 | 881 | 10 | 2.27% |
| hsa_circ_0077135 | 881 | 10 | 2.27% |
| hsa_circ_0095739 | 881 | 10 | 2.27% |
| hsa_circ_0103308 | 881 | 10 | 2.27% |
| hsa_circ_0110921 | 881 | 10 | 2.27% |
| hsa_circ_0113136 | 881 | 10 | 2.27% |
| hsa_circ_0118830 | 881 | 10 | 2.27% |
| hsa_circ_0127429 | 1146 | 13 | 2.27% |
| hsa_circ_0005902 | 1058 | 12 | 2.27% |
| hsa_circ_0003628 | 970 | 11 | 2.27% |
| hsa_circ_0013314 | 970 | 11 | 2.27% |
| hsa_circ_0034646 | 970 | 11 | 2.27% |
| hsa_circ_0055708 | 970 | 11 | 2.27% |
| hsa_circ_0068054 | 970 | 11 | 2.27% |
| hsa_circ_0077288 | 970 | 11 | 2.27% |
| hsa_circ_0094848 | 970 | 11 | 2.27% |
| hsa_circ_0126845 | 970 | 11 | 2.27% |
| hsa_circ_0030996 | 882 | 10 | 2.27% |
| hsa_circ_0048351 | 882 | 10 | 2.27% |
| hsa_circ_0061745 | 882 | 10 | 2.27% |
| hsa_circ_0072623 | 882 | 10 | 2.27% |
| hsa_circ_0076041 | 882 | 10 | 2.27% |
| hsa_circ_0077486 | 882 | 10 | 2.27% |
| hsa_circ_0078636 | 882 | 10 | 2.27% |
| hsa_circ_0084089 | 882 | 10 | 2.27% |
| hsa_circ_0087320 | 882 | 10 | 2.27% |
| hsa_circ_0089415 | 882 | 10 | 2.27% |
| hsa_circ_0091794 | 882 | 10 | 2.27% |
| hsa_circ_0092126 | 882 | 10 | 2.27% |
| hsa_circ_0100308 | 882 | 10 | 2.27% |
| hsa_circ_0102099 | 882 | 10 | 2.27% |
| hsa_circ_0115704 | 882 | 10 | 2.27% |
| hsa_circ_0120282 | 882 | 10 | 2.27% |
| hsa_circ_0137331 | 882 | 10 | 2.27% |
| hsa_circ_0130144 | 1147 | 13 | 2.27% |
| hsa_circ_0135882 | 2206 | 25 | 2.27% |
| hsa_circ_0054428 | 1059 | 12 | 2.27% |
| hsa_circ_0132516 | 1059 | 12 | 2.27% |
| hsa_circ_0022271 | 971 | 11 | 2.27% |
| hsa_circ_0040678 | 971 | 11 | 2.27% |
| hsa_circ_0056590 | 971 | 11 | 2.27% |
| hsa_circ_0114573 | 971 | 11 | 2.27% |
| hsa_circ_0138971 | 971 | 11 | 2.27% |
| hsa_circ_0028478 | 883 | 10 | 2.27% |
| hsa_circ_0075252 | 883 | 10 | 2.27% |

| | | | |
|---|---|---|---|
| hsa_circ_0078007 | 883 | 10 | 2.27% |
| hsa_circ_0097055 | 883 | 10 | 2.27% |
| hsa_circ_0103845 | 883 | 10 | 2.27% |
| hsa_circ_0105725 | 883 | 10 | 2.27% |
| hsa_circ_0107322 | 883 | 10 | 2.27% |
| hsa_circ_0111245 | 883 | 10 | 2.27% |
| hsa_circ_0126811 | 883 | 10 | 2.27% |
| hsa_circ_0127588 | 883 | 10 | 2.27% |
| hsa_circ_0039885 | 1060 | 12 | 2.26% |
| hsa_circ_0007031 | 972 | 11 | 2.26% |
| hsa_circ_0025197 | 972 | 11 | 2.26% |
| hsa_circ_0004083 | 884 | 10 | 2.26% |
| hsa_circ_0050430 | 884 | 10 | 2.26% |
| hsa_circ_0051995 | 884 | 10 | 2.26% |
| hsa_circ_0062302 | 884 | 10 | 2.26% |
| hsa_circ_0072268 | 884 | 10 | 2.26% |
| hsa_circ_0074427 | 884 | 10 | 2.26% |
| hsa_circ_0077861 | 884 | 10 | 2.26% |
| hsa_circ_0101765 | 884 | 10 | 2.26% |
| hsa_circ_0118642 | 884 | 10 | 2.26% |
| hsa_circ_0120486 | 884 | 10 | 2.26% |
| hsa_circ_0128404 | 884 | 10 | 2.26% |
| hsa_circ_0133117 | 884 | 10 | 2.26% |
| hsa_circ_0003576 | 1061 | 12 | 2.26% |
| hsa_circ_0058370 | 1061 | 12 | 2.26% |
| hsa_circ_0011727 | 973 | 11 | 2.26% |
| hsa_circ_0051158 | 973 | 11 | 2.26% |
| hsa_circ_0087910 | 973 | 11 | 2.26% |
| hsa_circ_0090838 | 973 | 11 | 2.26% |
| hsa_circ_0113752 | 973 | 11 | 2.26% |
| hsa_circ_0132255 | 973 | 11 | 2.26% |
| hsa_circ_0110179 | 1239 | 14 | 2.26% |
| hsa_circ_0014712 | 1062 | 12 | 2.26% |
| hsa_circ_0004338 | 885 | 10 | 2.26% |
| hsa_circ_0010444 | 885 | 10 | 2.26% |
| hsa_circ_0013021 | 885 | 10 | 2.26% |
| hsa_circ_0028044 | 885 | 10 | 2.26% |
| hsa_circ_0031188 | 885 | 10 | 2.26% |
| hsa_circ_0041419 | 885 | 10 | 2.26% |
| hsa_circ_0064272 | 885 | 10 | 2.26% |
| hsa_circ_0069376 | 885 | 10 | 2.26% |
| hsa_circ_0089990 | 885 | 10 | 2.26% |
| hsa_circ_0092732 | 885 | 10 | 2.26% |
| hsa_circ_0097965 | 885 | 10 | 2.26% |
| hsa_circ_0101076 | 885 | 10 | 2.26% |
| hsa_circ_0101123 | 885 | 10 | 2.26% |
| hsa_circ_0112571 | 885 | 10 | 2.26% |
| hsa_circ_0116229 | 885 | 10 | 2.26% |
| hsa_circ_0119188 | 885 | 10 | 2.26% |
| hsa_circ_0121889 | 885 | 10 | 2.26% |

| | | | |
|---|---|---|---|
| hsa_circ_0122894 | 885 | 10 | 2.26% |
| hsa_circ_0128006 | 885 | 10 | 2.26% |
| hsa_circ_0130761 | 885 | 10 | 2.26% |
| hsa_circ_0131378 | 885 | 10 | 2.26% |
| hsa_circ_0028541 | 1151 | 13 | 2.26% |
| hsa_circ_0054041 | 1151 | 13 | 2.26% |
| hsa_circ_0005599 | 974 | 11 | 2.26% |
| hsa_circ_0018158 | 974 | 11 | 2.26% |
| hsa_circ_0023423 | 974 | 11 | 2.26% |
| hsa_circ_0042523 | 974 | 11 | 2.26% |
| hsa_circ_0058167 | 974 | 11 | 2.26% |
| hsa_circ_0092909 | 974 | 11 | 2.26% |
| hsa_circ_0115640 | 974 | 11 | 2.26% |
| hsa_circ_0117054 | 974 | 11 | 2.26% |
| hsa_circ_0127269 | 974 | 11 | 2.26% |
| hsa_circ_0030840 | 1240 | 14 | 2.26% |
| hsa_circ_0033362 | 886 | 10 | 2.26% |
| hsa_circ_0038925 | 886 | 10 | 2.26% |
| hsa_circ_0047194 | 886 | 10 | 2.26% |
| hsa_circ_0059830 | 886 | 10 | 2.26% |
| hsa_circ_0065669 | 886 | 10 | 2.26% |
| hsa_circ_0066980 | 886 | 10 | 2.26% |
| hsa_circ_0087271 | 886 | 10 | 2.26% |
| hsa_circ_0108517 | 886 | 10 | 2.26% |
| hsa_circ_0123538 | 886 | 10 | 2.26% |
| hsa_circ_0133059 | 886 | 10 | 2.26% |
| hsa_circ_0015178 | 975 | 11 | 2.26% |
| hsa_circ_0024006 | 975 | 11 | 2.26% |
| hsa_circ_0071279 | 975 | 11 | 2.26% |
| hsa_circ_0120341 | 975 | 11 | 2.26% |
| hsa_circ_0011517 | 1064 | 12 | 2.26% |
| hsa_circ_0041686 | 1064 | 12 | 2.26% |
| hsa_circ_0005801 | 887 | 10 | 2.25% |
| hsa_circ_0023932 | 887 | 10 | 2.25% |
| hsa_circ_0028759 | 887 | 10 | 2.25% |
| hsa_circ_0031064 | 887 | 10 | 2.25% |
| hsa_circ_0033661 | 887 | 10 | 2.25% |
| hsa_circ_0041776 | 887 | 10 | 2.25% |
| hsa_circ_0052218 | 887 | 10 | 2.25% |
| hsa_circ_0070007 | 887 | 10 | 2.25% |
| hsa_circ_0074272 | 887 | 10 | 2.25% |
| hsa_circ_0074436 | 887 | 10 | 2.25% |
| hsa_circ_0088535 | 887 | 10 | 2.25% |
| hsa_circ_0125180 | 887 | 10 | 2.25% |
| hsa_circ_0132511 | 887 | 10 | 2.25% |
| hsa_circ_0034704 | 976 | 11 | 2.25% |
| hsa_circ_0062256 | 976 | 11 | 2.25% |
| hsa_circ_0064313 | 976 | 11 | 2.25% |
| hsa_circ_0111279 | 976 | 11 | 2.25% |
| hsa_circ_0043861 | 1154 | 13 | 2.25% |

| | | | |
|---|---|---|---|
| hsa_circ_0052950 | 1154 | 13 | 2.25% |
| hsa_circ_0010352 | 888 | 10 | 2.25% |
| hsa_circ_0012611 | 888 | 10 | 2.25% |
| hsa_circ_0045058 | 888 | 10 | 2.25% |
| hsa_circ_0054242 | 888 | 10 | 2.25% |
| hsa_circ_0070133 | 888 | 10 | 2.25% |
| hsa_circ_0084072 | 888 | 10 | 2.25% |
| hsa_circ_0100309 | 888 | 10 | 2.25% |
| hsa_circ_0123386 | 888 | 10 | 2.25% |
| hsa_circ_0126009 | 888 | 10 | 2.25% |
| hsa_circ_0128310 | 888 | 10 | 2.25% |
| hsa_circ_0013526 | 977 | 11 | 2.25% |
| hsa_circ_0030978 | 977 | 11 | 2.25% |
| hsa_circ_0036779 | 977 | 11 | 2.25% |
| hsa_circ_0052387 | 977 | 11 | 2.25% |
| hsa_circ_0055008 | 977 | 11 | 2.25% |
| hsa_circ_0076944 | 977 | 11 | 2.25% |
| hsa_circ_0093159 | 977 | 11 | 2.25% |
| hsa_circ_0097907 | 977 | 11 | 2.25% |
| hsa_circ_0117852 | 977 | 11 | 2.25% |
| hsa_circ_0065954 | 1066 | 12 | 2.25% |
| hsa_circ_0125918 | 1066 | 12 | 2.25% |
| hsa_circ_0021264 | 889 | 10 | 2.25% |
| hsa_circ_0022322 | 889 | 10 | 2.25% |
| hsa_circ_0024852 | 889 | 10 | 2.25% |
| hsa_circ_0032104 | 889 | 10 | 2.25% |
| hsa_circ_0034838 | 889 | 10 | 2.25% |
| hsa_circ_0044113 | 889 | 10 | 2.25% |
| hsa_circ_0067970 | 889 | 10 | 2.25% |
| hsa_circ_0070591 | 889 | 10 | 2.25% |
| hsa_circ_0074429 | 889 | 10 | 2.25% |
| hsa_circ_0088383 | 889 | 10 | 2.25% |
| hsa_circ_0097861 | 889 | 10 | 2.25% |
| hsa_circ_0097876 | 889 | 10 | 2.25% |
| hsa_circ_0105655 | 889 | 10 | 2.25% |
| hsa_circ_0116529 | 889 | 10 | 2.25% |
| hsa_circ_0123885 | 889 | 10 | 2.25% |
| hsa_circ_0011744 | 978 | 11 | 2.25% |
| hsa_circ_0023047 | 978 | 11 | 2.25% |
| hsa_circ_0044050 | 978 | 11 | 2.25% |
| hsa_circ_0060933 | 978 | 11 | 2.25% |
| hsa_circ_0104364 | 978 | 11 | 2.25% |
| hsa_circ_0116539 | 978 | 11 | 2.25% |
| hsa_circ_0120705 | 978 | 11 | 2.25% |
| hsa_circ_0039373 | 1067 | 12 | 2.25% |
| hsa_circ_0059241 | 1067 | 12 | 2.25% |
| hsa_circ_0092056 | 1067 | 12 | 2.25% |
| hsa_circ_0007863 | 1068 | 12 | 2.25% |
| hsa_circ_0039501 | 1068 | 12 | 2.25% |
| hsa_circ_0015955 | 979 | 11 | 2.25% |

| | | | |
|---|---|---|---|
| hsa_circ_0040597 | 979 | 11 | 2.25% |
| hsa_circ_0082483 | 979 | 11 | 2.25% |
| hsa_circ_0108231 | 979 | 11 | 2.25% |
| hsa_circ_0116582 | 979 | 11 | 2.25% |
| hsa_circ_0124335 | 979 | 11 | 2.25% |
| hsa_circ_0006113 | 890 | 10 | 2.25% |
| hsa_circ_0014003 | 890 | 10 | 2.25% |
| hsa_circ_0018615 | 890 | 10 | 2.25% |
| hsa_circ_0021262 | 890 | 10 | 2.25% |
| hsa_circ_0021477 | 890 | 10 | 2.25% |
| hsa_circ_0032021 | 890 | 10 | 2.25% |
| hsa_circ_0045716 | 890 | 10 | 2.25% |
| hsa_circ_0050472 | 890 | 10 | 2.25% |
| hsa_circ_0114588 | 890 | 10 | 2.25% |
| hsa_circ_0119186 | 890 | 10 | 2.25% |
| hsa_circ_0123988 | 890 | 10 | 2.25% |
| hsa_circ_0126784 | 890 | 10 | 2.25% |
| hsa_circ_0137065 | 890 | 10 | 2.25% |
| hsa_circ_0125800 | 2761 | 31 | 2.25% |
| hsa_circ_0048056 | 1069 | 12 | 2.25% |
| hsa_circ_0019664 | 980 | 11 | 2.24% |
| hsa_circ_0045672 | 980 | 11 | 2.24% |
| hsa_circ_0064879 | 980 | 11 | 2.24% |
| hsa_circ_0066910 | 980 | 11 | 2.24% |
| hsa_circ_0101567 | 980 | 11 | 2.24% |
| hsa_circ_0004193 | 891 | 10 | 2.24% |
| hsa_circ_0005788 | 891 | 10 | 2.24% |
| hsa_circ_0008573 | 891 | 10 | 2.24% |
| hsa_circ_0009714 | 891 | 10 | 2.24% |
| hsa_circ_0032061 | 891 | 10 | 2.24% |
| hsa_circ_0032829 | 891 | 10 | 2.24% |
| hsa_circ_0033653 | 891 | 10 | 2.24% |
| hsa_circ_0041097 | 891 | 10 | 2.24% |
| hsa_circ_0053181 | 891 | 10 | 2.24% |
| hsa_circ_0057799 | 891 | 10 | 2.24% |
| hsa_circ_0059068 | 891 | 10 | 2.24% |
| hsa_circ_0073644 | 891 | 10 | 2.24% |
| hsa_circ_0082155 | 891 | 10 | 2.24% |
| hsa_circ_0087851 | 891 | 10 | 2.24% |
| hsa_circ_0095535 | 891 | 10 | 2.24% |
| hsa_circ_0100281 | 891 | 10 | 2.24% |
| hsa_circ_0112495 | 891 | 10 | 2.24% |
| hsa_circ_0118733 | 891 | 10 | 2.24% |
| hsa_circ_0137440 | 891 | 10 | 2.24% |
| hsa_circ_0138063 | 891 | 10 | 2.24% |
| hsa_circ_0013275 | 1070 | 12 | 2.24% |
| hsa_circ_0076708 | 1070 | 12 | 2.24% |
| hsa_circ_0097824 | 1070 | 12 | 2.24% |
| hsa_circ_0100822 | 1070 | 12 | 2.24% |
| hsa_circ_0023225 | 981 | 11 | 2.24% |

| | | | |
|---|---|---|---|
| hsa_circ_0046793 | 981 | 11 | 2.24% |
| hsa_circ_0091657 | 981 | 11 | 2.24% |
| hsa_circ_0096187 | 981 | 11 | 2.24% |
| hsa_circ_0019338 | 892 | 10 | 2.24% |
| hsa_circ_0029261 | 892 | 10 | 2.24% |
| hsa_circ_0050213 | 892 | 10 | 2.24% |
| hsa_circ_0065328 | 892 | 10 | 2.24% |
| hsa_circ_0074358 | 892 | 10 | 2.24% |
| hsa_circ_0086839 | 892 | 10 | 2.24% |
| hsa_circ_0089483 | 892 | 10 | 2.24% |
| hsa_circ_0097846 | 892 | 10 | 2.24% |
| hsa_circ_0108102 | 892 | 10 | 2.24% |
| hsa_circ_0111524 | 892 | 10 | 2.24% |
| hsa_circ_0116742 | 892 | 10 | 2.24% |
| hsa_circ_0129795 | 892 | 10 | 2.24% |
| hsa_circ_0138979 | 892 | 10 | 2.24% |
| hsa_circ_0021567 | 5174 | 58 | 2.24% |
| hsa_circ_0124636 | 2587 | 29 | 2.24% |
| hsa_circ_0025358 | 1071 | 12 | 2.24% |
| hsa_circ_0122001 | 1071 | 12 | 2.24% |
| hsa_circ_0080665 | 2946 | 33 | 2.24% |
| hsa_circ_0013732 | 982 | 11 | 2.24% |
| hsa_circ_0016235 | 982 | 11 | 2.24% |
| hsa_circ_0043203 | 982 | 11 | 2.24% |
| hsa_circ_0051518 | 982 | 11 | 2.24% |
| hsa_circ_0055229 | 982 | 11 | 2.24% |
| hsa_circ_0058436 | 982 | 11 | 2.24% |
| hsa_circ_0086025 | 982 | 11 | 2.24% |
| hsa_circ_0129784 | 982 | 11 | 2.24% |
| hsa_circ_0042877 | 1250 | 14 | 2.24% |
| hsa_circ_0098196 | 1250 | 14 | 2.24% |
| hsa_circ_0089793 | 2679 | 30 | 2.24% |
| hsa_circ_0012975 | 893 | 10 | 2.24% |
| hsa_circ_0026938 | 893 | 10 | 2.24% |
| hsa_circ_0045288 | 893 | 10 | 2.24% |
| hsa_circ_0092840 | 893 | 10 | 2.24% |
| hsa_circ_0099878 | 893 | 10 | 2.24% |
| hsa_circ_0107224 | 893 | 10 | 2.24% |
| hsa_circ_0116029 | 893 | 10 | 2.24% |
| hsa_circ_0128912 | 893 | 10 | 2.24% |
| hsa_circ_0134720 | 893 | 10 | 2.24% |
| hsa_circ_0059394 | 1072 | 12 | 2.24% |
| hsa_circ_0113664 | 1072 | 12 | 2.24% |
| hsa_circ_0050953 | 983 | 11 | 2.24% |
| hsa_circ_0052567 | 983 | 11 | 2.24% |
| hsa_circ_0083686 | 983 | 11 | 2.24% |
| hsa_circ_0089748 | 983 | 11 | 2.24% |
| hsa_circ_0094732 | 983 | 11 | 2.24% |
| hsa_circ_0105066 | 983 | 11 | 2.24% |
| hsa_circ_0105585 | 983 | 11 | 2.24% |

| | | | |
|---|---|---|---|
| hsa_circ_0112377 | 983 | 11 | 2.24% |
| hsa_circ_0118154 | 983 | 11 | 2.24% |
| hsa_circ_0106556 | 1162 | 13 | 2.24% |
| hsa_circ_0002045 | 894 | 10 | 2.24% |
| hsa_circ_0015275 | 894 | 10 | 2.24% |
| hsa_circ_0021836 | 894 | 10 | 2.24% |
| hsa_circ_0037216 | 894 | 10 | 2.24% |
| hsa_circ_0042473 | 894 | 10 | 2.24% |
| hsa_circ_0050864 | 894 | 10 | 2.24% |
| hsa_circ_0066353 | 894 | 10 | 2.24% |
| hsa_circ_0068132 | 894 | 10 | 2.24% |
| hsa_circ_0078978 | 894 | 10 | 2.24% |
| hsa_circ_0081496 | 894 | 10 | 2.24% |
| hsa_circ_0081741 | 894 | 10 | 2.24% |
| hsa_circ_0094731 | 894 | 10 | 2.24% |
| hsa_circ_0108770 | 894 | 10 | 2.24% |
| hsa_circ_0124953 | 894 | 10 | 2.24% |
| hsa_circ_0131867 | 894 | 10 | 2.24% |
| hsa_circ_0137360 | 894 | 10 | 2.24% |
| hsa_circ_0140376 | 894 | 10 | 2.24% |
| hsa_circ_0035408 | 1073 | 12 | 2.24% |
| hsa_circ_0035865 | 1073 | 12 | 2.24% |
| hsa_circ_0102800 | 1073 | 12 | 2.24% |
| hsa_circ_0139314 | 984 | 11 | 2.24% |
| hsa_circ_0033145 | 1074 | 12 | 2.23% |
| hsa_circ_0105595 | 1074 | 12 | 2.23% |
| hsa_circ_0004432 | 895 | 10 | 2.23% |
| hsa_circ_0009657 | 895 | 10 | 2.23% |
| hsa_circ_0030263 | 895 | 10 | 2.23% |
| hsa_circ_0035480 | 895 | 10 | 2.23% |
| hsa_circ_0041700 | 895 | 10 | 2.23% |
| hsa_circ_0046189 | 895 | 10 | 2.23% |
| hsa_circ_0046243 | 895 | 10 | 2.23% |
| hsa_circ_0048216 | 895 | 10 | 2.23% |
| hsa_circ_0082465 | 895 | 10 | 2.23% |
| hsa_circ_0102565 | 895 | 10 | 2.23% |
| hsa_circ_0106533 | 895 | 10 | 2.23% |
| hsa_circ_0064749 | 1164 | 13 | 2.23% |
| hsa_circ_0122885 | 985 | 11 | 2.23% |
| hsa_circ_0012697 | 1075 | 12 | 2.23% |
| hsa_circ_0014652 | 1075 | 12 | 2.23% |
| hsa_circ_0007494 | 896 | 10 | 2.23% |
| hsa_circ_0014114 | 896 | 10 | 2.23% |
| hsa_circ_0019491 | 896 | 10 | 2.23% |
| hsa_circ_0023017 | 896 | 10 | 2.23% |
| hsa_circ_0026537 | 896 | 10 | 2.23% |
| hsa_circ_0031014 | 896 | 10 | 2.23% |
| hsa_circ_0050061 | 896 | 10 | 2.23% |
| hsa_circ_0062311 | 896 | 10 | 2.23% |
| hsa_circ_0085971 | 896 | 10 | 2.23% |

| | | | |
|---|---|---|---|
| hsa_circ_0094217 | 896 | 10 | 2.23% |
| hsa_circ_0099083 | 896 | 10 | 2.23% |
| hsa_circ_0112745 | 896 | 10 | 2.23% |
| hsa_circ_0117217 | 896 | 10 | 2.23% |
| hsa_circ_0132011 | 896 | 10 | 2.23% |
| hsa_circ_0029154 | 1165 | 13 | 2.23% |
| hsa_circ_0079952 | 1165 | 13 | 2.23% |
| hsa_circ_0007663 | 986 | 11 | 2.23% |
| hsa_circ_0024227 | 986 | 11 | 2.23% |
| hsa_circ_0037471 | 986 | 11 | 2.23% |
| hsa_circ_0037687 | 986 | 11 | 2.23% |
| hsa_circ_0050210 | 986 | 11 | 2.23% |
| hsa_circ_0051830 | 986 | 11 | 2.23% |
| hsa_circ_0054230 | 986 | 11 | 2.23% |
| hsa_circ_0081181 | 986 | 11 | 2.23% |
| hsa_circ_0085517 | 986 | 11 | 2.23% |
| hsa_circ_0089877 | 986 | 11 | 2.23% |
| hsa_circ_0101726 | 986 | 11 | 2.23% |
| hsa_circ_0105067 | 986 | 11 | 2.23% |
| hsa_circ_0022174 | 1255 | 14 | 2.23% |
| hsa_circ_0060131 | 1076 | 12 | 2.23% |
| hsa_circ_0013483 | 897 | 10 | 2.23% |
| hsa_circ_0031604 | 897 | 10 | 2.23% |
| hsa_circ_0032475 | 897 | 10 | 2.23% |
| hsa_circ_0039574 | 897 | 10 | 2.23% |
| hsa_circ_0044357 | 897 | 10 | 2.23% |
| hsa_circ_0049030 | 897 | 10 | 2.23% |
| hsa_circ_0065467 | 897 | 10 | 2.23% |
| hsa_circ_0079880 | 897 | 10 | 2.23% |
| hsa_circ_0092496 | 897 | 10 | 2.23% |
| hsa_circ_0104948 | 897 | 10 | 2.23% |
| hsa_circ_0122442 | 897 | 10 | 2.23% |
| hsa_circ_0127819 | 897 | 10 | 2.23% |
| hsa_circ_0092331 | 3140 | 35 | 2.23% |
| hsa_circ_0017948 | 987 | 11 | 2.23% |
| hsa_circ_0041802 | 987 | 11 | 2.23% |
| hsa_circ_0046507 | 987 | 11 | 2.23% |
| hsa_circ_0059339 | 987 | 11 | 2.23% |
| hsa_circ_0086924 | 987 | 11 | 2.23% |
| hsa_circ_0104642 | 987 | 11 | 2.23% |
| hsa_circ_0119989 | 987 | 11 | 2.23% |
| hsa_circ_0134286 | 987 | 11 | 2.23% |
| hsa_circ_0003113 | 1077 | 12 | 2.23% |
| hsa_circ_0041611 | 1077 | 12 | 2.23% |
| hsa_circ_0077986 | 1167 | 13 | 2.23% |
| hsa_circ_0001281 | 898 | 10 | 2.23% |
| hsa_circ_0008941 | 898 | 10 | 2.23% |
| hsa_circ_0028737 | 898 | 10 | 2.23% |
| hsa_circ_0032960 | 898 | 10 | 2.23% |
| hsa_circ_0048616 | 898 | 10 | 2.23% |

| | | | |
|---|---|---|---|
| hsa_circ_0057895 | 898 | 10 | 2.23% |
| hsa_circ_0075187 | 898 | 10 | 2.23% |
| hsa_circ_0078256 | 898 | 10 | 2.23% |
| hsa_circ_0085783 | 898 | 10 | 2.23% |
| hsa_circ_0087218 | 898 | 10 | 2.23% |
| hsa_circ_0115558 | 898 | 10 | 2.23% |
| hsa_circ_0123925 | 898 | 10 | 2.23% |
| hsa_circ_0129051 | 898 | 10 | 2.23% |
| hsa_circ_0006604 | 2784 | 31 | 2.23% |
| hsa_circ_0012222 | 2425 | 27 | 2.23% |
| hsa_circ_0009138 | 988 | 11 | 2.23% |
| hsa_circ_0019398 | 988 | 11 | 2.23% |
| hsa_circ_0029372 | 988 | 11 | 2.23% |
| hsa_circ_0086253 | 988 | 11 | 2.23% |
| hsa_circ_0096087 | 988 | 11 | 2.23% |
| hsa_circ_0106231 | 988 | 11 | 2.23% |
| hsa_circ_0108462 | 1078 | 12 | 2.23% |
| hsa_circ_0025753 | 1168 | 13 | 2.23% |
| hsa_circ_0005982 | 899 | 10 | 2.22% |
| hsa_circ_0008621 | 899 | 10 | 2.22% |
| hsa_circ_0014025 | 899 | 10 | 2.22% |
| hsa_circ_0016336 | 899 | 10 | 2.22% |
| hsa_circ_0031021 | 899 | 10 | 2.22% |
| hsa_circ_0031905 | 899 | 10 | 2.22% |
| hsa_circ_0069027 | 899 | 10 | 2.22% |
| hsa_circ_0069384 | 899 | 10 | 2.22% |
| hsa_circ_0072725 | 899 | 10 | 2.22% |
| hsa_circ_0076939 | 899 | 10 | 2.22% |
| hsa_circ_0082843 | 899 | 10 | 2.22% |
| hsa_circ_0085983 | 899 | 10 | 2.22% |
| hsa_circ_0088422 | 899 | 10 | 2.22% |
| hsa_circ_0108562 | 899 | 10 | 2.22% |
| hsa_circ_0114293 | 899 | 10 | 2.22% |
| hsa_circ_0122238 | 899 | 10 | 2.22% |
| hsa_circ_0125595 | 899 | 10 | 2.22% |
| hsa_circ_0076058 | 989 | 11 | 2.22% |
| hsa_circ_0095429 | 989 | 11 | 2.22% |
| hsa_circ_0117442 | 989 | 11 | 2.22% |
| hsa_circ_0102748 | 1169 | 13 | 2.22% |
| hsa_circ_0009192 | 1259 | 14 | 2.22% |
| hsa_circ_0057393 | 1530 | 17 | 2.22% |
| hsa_circ_0101955 | 1170 | 13 | 2.22% |
| hsa_circ_0057442 | 1080 | 12 | 2.22% |
| hsa_circ_0070623 | 1080 | 12 | 2.22% |
| hsa_circ_0134412 | 1080 | 12 | 2.22% |
| hsa_circ_0021083 | 990 | 11 | 2.22% |
| hsa_circ_0067214 | 990 | 11 | 2.22% |
| hsa_circ_0070994 | 990 | 11 | 2.22% |
| hsa_circ_0014142 | 900 | 10 | 2.22% |
| hsa_circ_0022875 | 900 | 10 | 2.22% |

| | | | |
|---|---|---|---|
| hsa_circ_0025982 | 900 | 10 | 2.22% |
| hsa_circ_0042205 | 900 | 10 | 2.22% |
| hsa_circ_0048238 | 900 | 10 | 2.22% |
| hsa_circ_0056733 | 900 | 10 | 2.22% |
| hsa_circ_0068033 | 900 | 10 | 2.22% |
| hsa_circ_0070379 | 900 | 10 | 2.22% |
| hsa_circ_0081570 | 900 | 10 | 2.22% |
| hsa_circ_0090410 | 900 | 10 | 2.22% |
| hsa_circ_0092332 | 900 | 10 | 2.22% |
| hsa_circ_0093936 | 900 | 10 | 2.22% |
| hsa_circ_0097509 | 900 | 10 | 2.22% |
| hsa_circ_0099698 | 900 | 10 | 2.22% |
| hsa_circ_0112322 | 900 | 10 | 2.22% |
| hsa_circ_0131534 | 900 | 10 | 2.22% |
| hsa_circ_0008571 | 1081 | 12 | 2.22% |
| hsa_circ_0027512 | 1081 | 12 | 2.22% |
| hsa_circ_0113464 | 1081 | 12 | 2.22% |
| hsa_circ_0037850 | 991 | 11 | 2.22% |
| hsa_circ_0049805 | 991 | 11 | 2.22% |
| hsa_circ_0061774 | 991 | 11 | 2.22% |
| hsa_circ_0063467 | 991 | 11 | 2.22% |
| hsa_circ_0074791 | 991 | 11 | 2.22% |
| hsa_circ_0098655 | 991 | 11 | 2.22% |
| hsa_circ_0121715 | 991 | 11 | 2.22% |
| hsa_circ_0122840 | 991 | 11 | 2.22% |
| hsa_circ_0013342 | 901 | 10 | 2.22% |
| hsa_circ_0052867 | 901 | 10 | 2.22% |
| hsa_circ_0095118 | 901 | 10 | 2.22% |
| hsa_circ_0096313 | 901 | 10 | 2.22% |
| hsa_circ_0101985 | 901 | 10 | 2.22% |
| hsa_circ_0109781 | 901 | 10 | 2.22% |
| hsa_circ_0111286 | 901 | 10 | 2.22% |
| hsa_circ_0116748 | 901 | 10 | 2.22% |
| hsa_circ_0123651 | 901 | 10 | 2.22% |
| hsa_circ_0132174 | 901 | 10 | 2.22% |
| hsa_circ_0132455 | 901 | 10 | 2.22% |
| hsa_circ_0139559 | 901 | 10 | 2.22% |
| hsa_circ_0007593 | 1082 | 12 | 2.22% |
| hsa_circ_0034418 | 1082 | 12 | 2.22% |
| hsa_circ_0098693 | 1082 | 12 | 2.22% |
| hsa_circ_0086609 | 992 | 11 | 2.22% |
| hsa_circ_0124178 | 2345 | 26 | 2.22% |
| hsa_circ_0014708 | 902 | 10 | 2.22% |
| hsa_circ_0017409 | 902 | 10 | 2.22% |
| hsa_circ_0017678 | 902 | 10 | 2.22% |
| hsa_circ_0060337 | 902 | 10 | 2.22% |
| hsa_circ_0067801 | 902 | 10 | 2.22% |
| hsa_circ_0069332 | 902 | 10 | 2.22% |
| hsa_circ_0080335 | 902 | 10 | 2.22% |
| hsa_circ_0102235 | 902 | 10 | 2.22% |

| | | | |
|---|---|---|---|
| hsa_circ_0105082 | 902 | 10 | 2.22% |
| hsa_circ_0117644 | 902 | 10 | 2.22% |
| hsa_circ_0120073 | 902 | 10 | 2.22% |
| hsa_circ_0125288 | 902 | 10 | 2.22% |
| hsa_circ_0140386 | 902 | 10 | 2.22% |
| hsa_circ_0140694 | 902 | 10 | 2.22% |
| hsa_circ_0124268 | 1263 | 14 | 2.22% |
| hsa_circ_0010674 | 1173 | 13 | 2.22% |
| hsa_circ_0137239 | 2707 | 30 | 2.22% |
| hsa_circ_0055218 | 1083 | 12 | 2.22% |
| hsa_circ_0090162 | 1083 | 12 | 2.22% |
| hsa_circ_0117209 | 1083 | 12 | 2.22% |
| hsa_circ_0032029 | 993 | 11 | 2.22% |
| hsa_circ_0048975 | 993 | 11 | 2.22% |
| hsa_circ_0051863 | 993 | 11 | 2.22% |
| hsa_circ_0057905 | 993 | 11 | 2.22% |
| hsa_circ_0060467 | 993 | 11 | 2.22% |
| hsa_circ_0070909 | 993 | 11 | 2.22% |
| hsa_circ_0136116 | 993 | 11 | 2.22% |
| hsa_circ_0010439 | 903 | 10 | 2.21% |
| hsa_circ_0016196 | 903 | 10 | 2.21% |
| hsa_circ_0016398 | 903 | 10 | 2.21% |
| hsa_circ_0043313 | 903 | 10 | 2.21% |
| hsa_circ_0044613 | 903 | 10 | 2.21% |
| hsa_circ_0046450 | 903 | 10 | 2.21% |
| hsa_circ_0070225 | 903 | 10 | 2.21% |
| hsa_circ_0070821 | 903 | 10 | 2.21% |
| hsa_circ_0071401 | 903 | 10 | 2.21% |
| hsa_circ_0075147 | 903 | 10 | 2.21% |
| hsa_circ_0079377 | 903 | 10 | 2.21% |
| hsa_circ_0081717 | 903 | 10 | 2.21% |
| hsa_circ_0092452 | 903 | 10 | 2.21% |
| hsa_circ_0094788 | 903 | 10 | 2.21% |
| hsa_circ_0102733 | 903 | 10 | 2.21% |
| hsa_circ_0111925 | 903 | 10 | 2.21% |
| hsa_circ_0121961 | 903 | 10 | 2.21% |
| hsa_circ_0122480 | 903 | 10 | 2.21% |
| hsa_circ_0127899 | 903 | 10 | 2.21% |
| hsa_circ_0132462 | 903 | 10 | 2.21% |
| hsa_circ_0108621 | 1174 | 13 | 2.21% |
| hsa_circ_0126851 | 1084 | 12 | 2.21% |
| hsa_circ_0033370 | 994 | 11 | 2.21% |
| hsa_circ_0034644 | 994 | 11 | 2.21% |
| hsa_circ_0041618 | 994 | 11 | 2.21% |
| hsa_circ_0080160 | 994 | 11 | 2.21% |
| hsa_circ_0080576 | 994 | 11 | 2.21% |
| hsa_circ_0012667 | 904 | 10 | 2.21% |
| hsa_circ_0021845 | 904 | 10 | 2.21% |
| hsa_circ_0025581 | 904 | 10 | 2.21% |
| hsa_circ_0030710 | 904 | 10 | 2.21% |

| | | | |
|---|---|---|---|
| hsa_circ_0032503 | 904 | 10 | 2.21% |
| hsa_circ_0094785 | 904 | 10 | 2.21% |
| hsa_circ_0107289 | 904 | 10 | 2.21% |
| hsa_circ_0109480 | 904 | 10 | 2.21% |
| hsa_circ_0121711 | 904 | 10 | 2.21% |
| hsa_circ_0123463 | 904 | 10 | 2.21% |
| hsa_circ_0124970 | 904 | 10 | 2.21% |
| hsa_circ_0133265 | 904 | 10 | 2.21% |
| hsa_circ_0138763 | 904 | 10 | 2.21% |
| hsa_circ_0109185 | 1085 | 12 | 2.21% |
| hsa_circ_0047860 | 995 | 11 | 2.21% |
| hsa_circ_0053156 | 995 | 11 | 2.21% |
| hsa_circ_0074404 | 995 | 11 | 2.21% |
| hsa_circ_0082922 | 995 | 11 | 2.21% |
| hsa_circ_0104229 | 995 | 11 | 2.21% |
| hsa_circ_0116685 | 995 | 11 | 2.21% |
| hsa_circ_0130633 | 995 | 11 | 2.21% |
| hsa_circ_0106850 | 2714 | 30 | 2.21% |
| hsa_circ_0073471 | 1086 | 12 | 2.21% |
| hsa_circ_0104700 | 1086 | 12 | 2.21% |
| hsa_circ_0001754 | 905 | 10 | 2.21% |
| hsa_circ_0002966 | 905 | 10 | 2.21% |
| hsa_circ_0008171 | 905 | 10 | 2.21% |
| hsa_circ_0016688 | 905 | 10 | 2.21% |
| hsa_circ_0021341 | 905 | 10 | 2.21% |
| hsa_circ_0021944 | 905 | 10 | 2.21% |
| hsa_circ_0037487 | 905 | 10 | 2.21% |
| hsa_circ_0049566 | 905 | 10 | 2.21% |
| hsa_circ_0050046 | 905 | 10 | 2.21% |
| hsa_circ_0055413 | 905 | 10 | 2.21% |
| hsa_circ_0063639 | 905 | 10 | 2.21% |
| hsa_circ_0065782 | 905 | 10 | 2.21% |
| hsa_circ_0080894 | 905 | 10 | 2.21% |
| hsa_circ_0090472 | 905 | 10 | 2.21% |
| hsa_circ_0095388 | 905 | 10 | 2.21% |
| hsa_circ_0097604 | 905 | 10 | 2.21% |
| hsa_circ_0103638 | 905 | 10 | 2.21% |
| hsa_circ_0115294 | 905 | 10 | 2.21% |
| hsa_circ_0116111 | 905 | 10 | 2.21% |
| hsa_circ_0128794 | 905 | 10 | 2.21% |
| hsa_circ_0139986 | 905 | 10 | 2.21% |
| hsa_circ_0037482 | 996 | 11 | 2.21% |
| hsa_circ_0052954 | 996 | 11 | 2.21% |
| hsa_circ_0089502 | 996 | 11 | 2.21% |
| hsa_circ_0095332 | 996 | 11 | 2.21% |
| hsa_circ_0067429 | 1268 | 14 | 2.21% |
| hsa_circ_0011421 | 906 | 10 | 2.21% |
| hsa_circ_0014333 | 906 | 10 | 2.21% |
| hsa_circ_0022238 | 906 | 10 | 2.21% |
| hsa_circ_0027770 | 906 | 10 | 2.21% |

| | | | |
|---|---|---|---|
| hsa_circ_0051837 | 906 | 10 | 2.21% |
| hsa_circ_0062558 | 906 | 10 | 2.21% |
| hsa_circ_0067108 | 906 | 10 | 2.21% |
| hsa_circ_0073971 | 906 | 10 | 2.21% |
| hsa_circ_0076673 | 906 | 10 | 2.21% |
| hsa_circ_0077623 | 906 | 10 | 2.21% |
| hsa_circ_0087348 | 906 | 10 | 2.21% |
| hsa_circ_0089653 | 906 | 10 | 2.21% |
| hsa_circ_0103124 | 906 | 10 | 2.21% |
| hsa_circ_0060133 | 997 | 11 | 2.21% |
| hsa_circ_0080049 | 997 | 11 | 2.21% |
| hsa_circ_0085937 | 997 | 11 | 2.21% |
| hsa_circ_0122634 | 997 | 11 | 2.21% |
| hsa_circ_0120897 | 1632 | 18 | 2.21% |
| hsa_circ_0042689 | 1088 | 12 | 2.21% |
| hsa_circ_0087087 | 3174 | 35 | 2.21% |
| hsa_circ_0004365 | 907 | 10 | 2.21% |
| hsa_circ_0012147 | 907 | 10 | 2.21% |
| hsa_circ_0019185 | 907 | 10 | 2.21% |
| hsa_circ_0029695 | 907 | 10 | 2.21% |
| hsa_circ_0037359 | 907 | 10 | 2.21% |
| hsa_circ_0079056 | 907 | 10 | 2.21% |
| hsa_circ_0084951 | 907 | 10 | 2.21% |
| hsa_circ_0090358 | 907 | 10 | 2.21% |
| hsa_circ_0117537 | 907 | 10 | 2.21% |
| hsa_circ_0118828 | 907 | 10 | 2.21% |
| hsa_circ_0119761 | 907 | 10 | 2.21% |
| hsa_circ_0140706 | 907 | 10 | 2.21% |
| hsa_circ_0009602 | 998 | 11 | 2.20% |
| hsa_circ_0022871 | 998 | 11 | 2.20% |
| hsa_circ_0047926 | 998 | 11 | 2.20% |
| hsa_circ_0007758 | 1089 | 12 | 2.20% |
| hsa_circ_0069780 | 1089 | 12 | 2.20% |
| hsa_circ_0120441 | 1089 | 12 | 2.20% |
| hsa_circ_0016076 | 908 | 10 | 2.20% |
| hsa_circ_0022970 | 908 | 10 | 2.20% |
| hsa_circ_0023775 | 908 | 10 | 2.20% |
| hsa_circ_0025443 | 908 | 10 | 2.20% |
| hsa_circ_0029863 | 908 | 10 | 2.20% |
| hsa_circ_0033132 | 908 | 10 | 2.20% |
| hsa_circ_0033660 | 908 | 10 | 2.20% |
| hsa_circ_0037901 | 908 | 10 | 2.20% |
| hsa_circ_0039303 | 908 | 10 | 2.20% |
| hsa_circ_0045042 | 908 | 10 | 2.20% |
| hsa_circ_0052401 | 908 | 10 | 2.20% |
| hsa_circ_0057794 | 908 | 10 | 2.20% |
| hsa_circ_0058541 | 908 | 10 | 2.20% |
| hsa_circ_0065377 | 908 | 10 | 2.20% |
| hsa_circ_0074898 | 908 | 10 | 2.20% |
| hsa_circ_0075230 | 908 | 10 | 2.20% |

| | | | |
|---|---|---|---|
| hsa_circ_0083823 | 908 | 10 | 2.20% |
| hsa_circ_0098164 | 908 | 10 | 2.20% |
| hsa_circ_0103156 | 908 | 10 | 2.20% |
| hsa_circ_0103935 | 908 | 10 | 2.20% |
| hsa_circ_0105166 | 908 | 10 | 2.20% |
| hsa_circ_0110537 | 908 | 10 | 2.20% |
| hsa_circ_0110768 | 908 | 10 | 2.20% |
| hsa_circ_0119501 | 908 | 10 | 2.20% |
| hsa_circ_0122496 | 908 | 10 | 2.20% |
| hsa_circ_0125306 | 908 | 10 | 2.20% |
| hsa_circ_0130068 | 908 | 10 | 2.20% |
| hsa_circ_0138760 | 908 | 10 | 2.20% |
| hsa_circ_0138856 | 908 | 10 | 2.20% |
| hsa_circ_0007117 | 999 | 11 | 2.20% |
| hsa_circ_0054876 | 999 | 11 | 2.20% |
| hsa_circ_0057469 | 999 | 11 | 2.20% |
| hsa_circ_0067244 | 999 | 11 | 2.20% |
| hsa_circ_0089004 | 999 | 11 | 2.20% |
| hsa_circ_0089464 | 999 | 11 | 2.20% |
| hsa_circ_0097831 | 999 | 11 | 2.20% |
| hsa_circ_0135376 | 999 | 11 | 2.20% |
| hsa_circ_0021586 | 1090 | 12 | 2.20% |
| hsa_circ_0037633 | 1090 | 12 | 2.20% |
| hsa_circ_0080900 | 1090 | 12 | 2.20% |
| hsa_circ_0086742 | 1090 | 12 | 2.20% |
| hsa_circ_0093503 | 1090 | 12 | 2.20% |
| hsa_circ_0097249 | 1090 | 12 | 2.20% |
| hsa_circ_0132531 | 1090 | 12 | 2.20% |
| hsa_circ_0134704 | 1090 | 12 | 2.20% |
| hsa_circ_0108916 | 1454 | 16 | 2.20% |
| hsa_circ_0009896 | 909 | 10 | 2.20% |
| hsa_circ_0017485 | 909 | 10 | 2.20% |
| hsa_circ_0019635 | 909 | 10 | 2.20% |
| hsa_circ_0023922 | 909 | 10 | 2.20% |
| hsa_circ_0025185 | 909 | 10 | 2.20% |
| hsa_circ_0043599 | 909 | 10 | 2.20% |
| hsa_circ_0045291 | 909 | 10 | 2.20% |
| hsa_circ_0052868 | 909 | 10 | 2.20% |
| hsa_circ_0060185 | 909 | 10 | 2.20% |
| hsa_circ_0077708 | 909 | 10 | 2.20% |
| hsa_circ_0085131 | 909 | 10 | 2.20% |
| hsa_circ_0089634 | 909 | 10 | 2.20% |
| hsa_circ_0093552 | 909 | 10 | 2.20% |
| hsa_circ_0095788 | 909 | 10 | 2.20% |
| hsa_circ_0099987 | 909 | 10 | 2.20% |
| hsa_circ_0100392 | 909 | 10 | 2.20% |
| hsa_circ_0104451 | 909 | 10 | 2.20% |
| hsa_circ_0127430 | 909 | 10 | 2.20% |
| hsa_circ_0128132 | 909 | 10 | 2.20% |
| hsa_circ_0133820 | 909 | 10 | 2.20% |

| | | | |
|---|---|---|---|
| hsa_circ_0011566 | 1000 | 11 | 2.20% |
| hsa_circ_0047169 | 1000 | 11 | 2.20% |
| hsa_circ_0055957 | 1000 | 11 | 2.20% |
| hsa_circ_0100033 | 1000 | 11 | 2.20% |
| hsa_circ_0100239 | 1000 | 11 | 2.20% |
| hsa_circ_0117032 | 1000 | 11 | 2.20% |
| hsa_circ_0117121 | 1000 | 11 | 2.20% |
| hsa_circ_0078853 | 4728 | 52 | 2.20% |
| hsa_circ_0036213 | 1273 | 14 | 2.20% |
| hsa_circ_0015141 | 2274 | 25 | 2.20% |
| hsa_circ_0133308 | 2548 | 28 | 2.20% |
| hsa_circ_0107515 | 1274 | 14 | 2.20% |
| hsa_circ_0106397 | 1183 | 13 | 2.20% |
| hsa_circ_0074552 | 1092 | 12 | 2.20% |
| hsa_circ_0053787 | 1001 | 11 | 2.20% |
| hsa_circ_0098408 | 1001 | 11 | 2.20% |
| hsa_circ_0135601 | 1001 | 11 | 2.20% |
| hsa_circ_0006784 | 910 | 10 | 2.20% |
| hsa_circ_0010093 | 910 | 10 | 2.20% |
| hsa_circ_0015580 | 910 | 10 | 2.20% |
| hsa_circ_0016182 | 910 | 10 | 2.20% |
| hsa_circ_0029207 | 910 | 10 | 2.20% |
| hsa_circ_0036709 | 910 | 10 | 2.20% |
| hsa_circ_0045500 | 910 | 10 | 2.20% |
| hsa_circ_0049593 | 910 | 10 | 2.20% |
| hsa_circ_0056917 | 910 | 10 | 2.20% |
| hsa_circ_0066885 | 910 | 10 | 2.20% |
| hsa_circ_0068002 | 910 | 10 | 2.20% |
| hsa_circ_0084126 | 910 | 10 | 2.20% |
| hsa_circ_0085767 | 910 | 10 | 2.20% |
| hsa_circ_0090628 | 910 | 10 | 2.20% |
| hsa_circ_0094607 | 910 | 10 | 2.20% |
| hsa_circ_0096318 | 910 | 10 | 2.20% |
| hsa_circ_0104425 | 910 | 10 | 2.20% |
| hsa_circ_0111611 | 910 | 10 | 2.20% |
| hsa_circ_0133472 | 910 | 10 | 2.20% |
| hsa_circ_0137477 | 910 | 10 | 2.20% |
| hsa_circ_0140231 | 910 | 10 | 2.20% |
| hsa_circ_0081462 | 1184 | 13 | 2.20% |
| hsa_circ_0021893 | 1093 | 12 | 2.20% |
| hsa_circ_0079649 | 1093 | 12 | 2.20% |
| hsa_circ_0016933 | 1002 | 11 | 2.20% |
| hsa_circ_0022403 | 1002 | 11 | 2.20% |
| hsa_circ_0043489 | 1002 | 11 | 2.20% |
| hsa_circ_0045422 | 1002 | 11 | 2.20% |
| hsa_circ_0064977 | 1002 | 11 | 2.20% |
| hsa_circ_0084455 | 1002 | 11 | 2.20% |
| hsa_circ_0087579 | 1002 | 11 | 2.20% |
| hsa_circ_0094441 | 1002 | 11 | 2.20% |
| hsa_circ_0131814 | 1002 | 11 | 2.20% |

| | | | |
|---|---|---|---|
| hsa_circ_0061384 | 1822 | 20 | 2.20% |
| hsa_circ_0037311 | 911 | 10 | 2.20% |
| hsa_circ_0044914 | 911 | 10 | 2.20% |
| hsa_circ_0054035 | 911 | 10 | 2.20% |
| hsa_circ_0055996 | 911 | 10 | 2.20% |
| hsa_circ_0061658 | 911 | 10 | 2.20% |
| hsa_circ_0069610 | 911 | 10 | 2.20% |
| hsa_circ_0076942 | 911 | 10 | 2.20% |
| hsa_circ_0096247 | 911 | 10 | 2.20% |
| hsa_circ_0100501 | 911 | 10 | 2.20% |
| hsa_circ_0102899 | 911 | 10 | 2.20% |
| hsa_circ_0130819 | 911 | 10 | 2.20% |
| hsa_circ_0132367 | 911 | 10 | 2.20% |
| hsa_circ_0136963 | 911 | 10 | 2.20% |
| hsa_circ_0033622 | 1094 | 12 | 2.19% |
| hsa_circ_0117185 | 1094 | 12 | 2.19% |
| hsa_circ_0132522 | 1094 | 12 | 2.19% |
| hsa_circ_0015101 | 1003 | 11 | 2.19% |
| hsa_circ_0026469 | 1003 | 11 | 2.19% |
| hsa_circ_0028004 | 1003 | 11 | 2.19% |
| hsa_circ_0035423 | 1003 | 11 | 2.19% |
| hsa_circ_0046068 | 1003 | 11 | 2.19% |
| hsa_circ_0001175 | 912 | 10 | 2.19% |
| hsa_circ_0025930 | 912 | 10 | 2.19% |
| hsa_circ_0038324 | 912 | 10 | 2.19% |
| hsa_circ_0038845 | 912 | 10 | 2.19% |
| hsa_circ_0039669 | 912 | 10 | 2.19% |
| hsa_circ_0050363 | 912 | 10 | 2.19% |
| hsa_circ_0053136 | 912 | 10 | 2.19% |
| hsa_circ_0062759 | 912 | 10 | 2.19% |
| hsa_circ_0091133 | 912 | 10 | 2.19% |
| hsa_circ_0092835 | 912 | 10 | 2.19% |
| hsa_circ_0095501 | 912 | 10 | 2.19% |
| hsa_circ_0095883 | 912 | 10 | 2.19% |
| hsa_circ_0116737 | 912 | 10 | 2.19% |
| hsa_circ_0128479 | 912 | 10 | 2.19% |
| hsa_circ_0129204 | 912 | 10 | 2.19% |
| hsa_circ_0058107 | 1095 | 12 | 2.19% |
| hsa_circ_0068543 | 1095 | 12 | 2.19% |
| hsa_circ_0007205 | 1004 | 11 | 2.19% |
| hsa_circ_0023777 | 1004 | 11 | 2.19% |
| hsa_circ_0039192 | 1004 | 11 | 2.19% |
| hsa_circ_0055706 | 1004 | 11 | 2.19% |
| hsa_circ_0055765 | 1004 | 11 | 2.19% |
| hsa_circ_0068115 | 1004 | 11 | 2.19% |
| hsa_circ_0103567 | 1004 | 11 | 2.19% |
| hsa_circ_0003245 | 913 | 10 | 2.19% |
| hsa_circ_0011099 | 913 | 10 | 2.19% |
| hsa_circ_0011932 | 913 | 10 | 2.19% |
| hsa_circ_0015243 | 913 | 10 | 2.19% |

| | | | |
|---|---|---|---|
| hsa_circ_0024501 | 913 | 10 | 2.19% |
| hsa_circ_0031980 | 913 | 10 | 2.19% |
| hsa_circ_0042399 | 913 | 10 | 2.19% |
| hsa_circ_0044311 | 913 | 10 | 2.19% |
| hsa_circ_0056082 | 913 | 10 | 2.19% |
| hsa_circ_0056147 | 913 | 10 | 2.19% |
| hsa_circ_0077086 | 913 | 10 | 2.19% |
| hsa_circ_0098489 | 913 | 10 | 2.19% |
| hsa_circ_0104541 | 913 | 10 | 2.19% |
| hsa_circ_0117433 | 913 | 10 | 2.19% |
| hsa_circ_0119295 | 913 | 10 | 2.19% |
| hsa_circ_0123041 | 913 | 10 | 2.19% |
| hsa_circ_0133613 | 913 | 10 | 2.19% |
| hsa_circ_0137938 | 913 | 10 | 2.19% |
| hsa_circ_0126506 | 1735 | 19 | 2.19% |
| hsa_circ_0090011 | 1096 | 12 | 2.19% |
| hsa_circ_0128947 | 1096 | 12 | 2.19% |
| hsa_circ_0130207 | 1096 | 12 | 2.19% |
| hsa_circ_0031195 | 2284 | 25 | 2.19% |
| hsa_circ_0006797 | 1005 | 11 | 2.19% |
| hsa_circ_0030944 | 1005 | 11 | 2.19% |
| hsa_circ_0037479 | 1005 | 11 | 2.19% |
| hsa_circ_0044254 | 1005 | 11 | 2.19% |
| hsa_circ_0048328 | 1005 | 11 | 2.19% |
| hsa_circ_0053142 | 1005 | 11 | 2.19% |
| hsa_circ_0078893 | 1005 | 11 | 2.19% |
| hsa_circ_0080164 | 1005 | 11 | 2.19% |
| hsa_circ_0095471 | 1005 | 11 | 2.19% |
| hsa_circ_0097762 | 1005 | 11 | 2.19% |
| hsa_circ_0110233 | 1005 | 11 | 2.19% |
| hsa_circ_0138166 | 1005 | 11 | 2.19% |
| hsa_circ_0090521 | 1188 | 13 | 2.19% |
| hsa_circ_0009101 | 914 | 10 | 2.19% |
| hsa_circ_0014241 | 914 | 10 | 2.19% |
| hsa_circ_0014987 | 914 | 10 | 2.19% |
| hsa_circ_0019394 | 914 | 10 | 2.19% |
| hsa_circ_0033658 | 914 | 10 | 2.19% |
| hsa_circ_0043453 | 914 | 10 | 2.19% |
| hsa_circ_0045212 | 914 | 10 | 2.19% |
| hsa_circ_0047123 | 914 | 10 | 2.19% |
| hsa_circ_0062328 | 914 | 10 | 2.19% |
| hsa_circ_0064500 | 914 | 10 | 2.19% |
| hsa_circ_0068097 | 914 | 10 | 2.19% |
| hsa_circ_0068234 | 914 | 10 | 2.19% |
| hsa_circ_0075293 | 914 | 10 | 2.19% |
| hsa_circ_0078480 | 914 | 10 | 2.19% |
| hsa_circ_0081742 | 914 | 10 | 2.19% |
| hsa_circ_0087723 | 914 | 10 | 2.19% |
| hsa_circ_0101979 | 914 | 10 | 2.19% |
| hsa_circ_0111341 | 914 | 10 | 2.19% |

| | | | |
|---|---|---|---|
| hsa_circ_0114035 | 914 | 10 | 2.19% |
| hsa_circ_0116706 | 914 | 10 | 2.19% |
| hsa_circ_0126018 | 914 | 10 | 2.19% |
| hsa_circ_0127561 | 914 | 10 | 2.19% |
| hsa_circ_0137637 | 914 | 10 | 2.19% |
| hsa_circ_0115693 | 3108 | 34 | 2.19% |
| hsa_circ_0051663 | 1097 | 12 | 2.19% |
| hsa_circ_0002678 | 1006 | 11 | 2.19% |
| hsa_circ_0018968 | 1006 | 11 | 2.19% |
| hsa_circ_0095767 | 1006 | 11 | 2.19% |
| hsa_circ_0100299 | 1006 | 11 | 2.19% |
| hsa_circ_0060526 | 1189 | 13 | 2.19% |
| hsa_circ_0043768 | 1098 | 12 | 2.19% |
| hsa_circ_0072190 | 1098 | 12 | 2.19% |
| hsa_circ_0014287 | 915 | 10 | 2.19% |
| hsa_circ_0027283 | 915 | 10 | 2.19% |
| hsa_circ_0027481 | 915 | 10 | 2.19% |
| hsa_circ_0037486 | 915 | 10 | 2.19% |
| hsa_circ_0051073 | 915 | 10 | 2.19% |
| hsa_circ_0056950 | 915 | 10 | 2.19% |
| hsa_circ_0062392 | 915 | 10 | 2.19% |
| hsa_circ_0065214 | 915 | 10 | 2.19% |
| hsa_circ_0066735 | 915 | 10 | 2.19% |
| hsa_circ_0071552 | 915 | 10 | 2.19% |
| hsa_circ_0079918 | 915 | 10 | 2.19% |
| hsa_circ_0081276 | 915 | 10 | 2.19% |
| hsa_circ_0085053 | 915 | 10 | 2.19% |
| hsa_circ_0111673 | 915 | 10 | 2.19% |
| hsa_circ_0114639 | 915 | 10 | 2.19% |
| hsa_circ_0121890 | 915 | 10 | 2.19% |
| hsa_circ_0126105 | 915 | 10 | 2.19% |
| hsa_circ_0130303 | 915 | 10 | 2.19% |
| hsa_circ_0136343 | 915 | 10 | 2.19% |
| hsa_circ_0036576 | 2014 | 22 | 2.18% |
| hsa_circ_0014976 | 1007 | 11 | 2.18% |
| hsa_circ_0014983 | 1007 | 11 | 2.18% |
| hsa_circ_0038460 | 1007 | 11 | 2.18% |
| hsa_circ_0059531 | 1007 | 11 | 2.18% |
| hsa_circ_0130134 | 1007 | 11 | 2.18% |
| hsa_circ_0132901 | 1007 | 11 | 2.18% |
| hsa_circ_0074674 | 1099 | 12 | 2.18% |
| hsa_circ_0128031 | 1099 | 12 | 2.18% |
| hsa_circ_0015030 | 916 | 10 | 2.18% |
| hsa_circ_0031763 | 916 | 10 | 2.18% |
| hsa_circ_0042021 | 916 | 10 | 2.18% |
| hsa_circ_0098239 | 916 | 10 | 2.18% |
| hsa_circ_0099093 | 916 | 10 | 2.18% |
| hsa_circ_0100492 | 916 | 10 | 2.18% |
| hsa_circ_0110430 | 916 | 10 | 2.18% |
| hsa_circ_0125772 | 916 | 10 | 2.18% |

| | | | |
|---|---|---|---|
| hsa_circ_0008434 | 1924 | 21 | 2.18% |
| hsa_circ_0131938 | 2016 | 22 | 2.18% |
| hsa_circ_0013661 | 1008 | 11 | 2.18% |
| hsa_circ_0052393 | 1008 | 11 | 2.18% |
| hsa_circ_0055118 | 1008 | 11 | 2.18% |
| hsa_circ_0067661 | 1008 | 11 | 2.18% |
| hsa_circ_0108367 | 1008 | 11 | 2.18% |
| hsa_circ_0050975 | 1283 | 14 | 2.18% |
| hsa_circ_0102820 | 1375 | 15 | 2.18% |
| hsa_circ_0104791 | 1100 | 12 | 2.18% |
| hsa_circ_0082084 | 1192 | 13 | 2.18% |
| hsa_circ_0004471 | 917 | 10 | 2.18% |
| hsa_circ_0023689 | 917 | 10 | 2.18% |
| hsa_circ_0033469 | 917 | 10 | 2.18% |
| hsa_circ_0034701 | 917 | 10 | 2.18% |
| hsa_circ_0036462 | 917 | 10 | 2.18% |
| hsa_circ_0040207 | 917 | 10 | 2.18% |
| hsa_circ_0063345 | 917 | 10 | 2.18% |
| hsa_circ_0064915 | 917 | 10 | 2.18% |
| hsa_circ_0065075 | 917 | 10 | 2.18% |
| hsa_circ_0075834 | 917 | 10 | 2.18% |
| hsa_circ_0081779 | 917 | 10 | 2.18% |
| hsa_circ_0085445 | 917 | 10 | 2.18% |
| hsa_circ_0096853 | 917 | 10 | 2.18% |
| hsa_circ_0100496 | 917 | 10 | 2.18% |
| hsa_circ_0107362 | 917 | 10 | 2.18% |
| hsa_circ_0133371 | 917 | 10 | 2.18% |
| hsa_circ_0137827 | 917 | 10 | 2.18% |
| hsa_circ_0094905 | 1009 | 11 | 2.18% |
| hsa_circ_0102759 | 1009 | 11 | 2.18% |
| hsa_circ_0123996 | 1009 | 11 | 2.18% |
| hsa_circ_0135826 | 1009 | 11 | 2.18% |
| hsa_circ_0048560 | 1101 | 12 | 2.18% |
| hsa_circ_0088048 | 1101 | 12 | 2.18% |
| hsa_circ_0099445 | 1101 | 12 | 2.18% |
| hsa_circ_0112473 | 1101 | 12 | 2.18% |
| hsa_circ_0012938 | 918 | 10 | 2.18% |
| hsa_circ_0024291 | 918 | 10 | 2.18% |
| hsa_circ_0027574 | 918 | 10 | 2.18% |
| hsa_circ_0029238 | 918 | 10 | 2.18% |
| hsa_circ_0040020 | 918 | 10 | 2.18% |
| hsa_circ_0049829 | 918 | 10 | 2.18% |
| hsa_circ_0050335 | 918 | 10 | 2.18% |
| hsa_circ_0061765 | 918 | 10 | 2.18% |
| hsa_circ_0064740 | 918 | 10 | 2.18% |
| hsa_circ_0069862 | 918 | 10 | 2.18% |
| hsa_circ_0078066 | 918 | 10 | 2.18% |
| hsa_circ_0079596 | 918 | 10 | 2.18% |
| hsa_circ_0098222 | 918 | 10 | 2.18% |
| hsa_circ_0099506 | 918 | 10 | 2.18% |

| | | | |
|---|---|---|---|
| hsa_circ_0114110 | 918 | 10 | 2.18% |
| hsa_circ_0114124 | 918 | 10 | 2.18% |
| hsa_circ_0122812 | 918 | 10 | 2.18% |
| hsa_circ_0125730 | 918 | 10 | 2.18% |
| hsa_circ_0137942 | 918 | 10 | 2.18% |
| hsa_circ_0016946 | 1010 | 11 | 2.18% |
| hsa_circ_0044237 | 1010 | 11 | 2.18% |
| hsa_circ_0049734 | 1010 | 11 | 2.18% |
| hsa_circ_0081422 | 1010 | 11 | 2.18% |
| hsa_circ_0084670 | 1010 | 11 | 2.18% |
| hsa_circ_0100217 | 1010 | 11 | 2.18% |
| hsa_circ_0003026 | 1102 | 12 | 2.18% |
| hsa_circ_0014138 | 1102 | 12 | 2.18% |
| hsa_circ_0126977 | 1102 | 12 | 2.18% |
| hsa_circ_0016651 | 919 | 10 | 2.18% |
| hsa_circ_0020769 | 919 | 10 | 2.18% |
| hsa_circ_0029226 | 919 | 10 | 2.18% |
| hsa_circ_0031243 | 919 | 10 | 2.18% |
| hsa_circ_0049853 | 919 | 10 | 2.18% |
| hsa_circ_0066920 | 919 | 10 | 2.18% |
| hsa_circ_0068451 | 919 | 10 | 2.18% |
| hsa_circ_0072747 | 919 | 10 | 2.18% |
| hsa_circ_0079591 | 919 | 10 | 2.18% |
| hsa_circ_0122900 | 919 | 10 | 2.18% |
| hsa_circ_0125833 | 919 | 10 | 2.18% |
| hsa_circ_0127220 | 919 | 10 | 2.18% |
| hsa_circ_0133650 | 919 | 10 | 2.18% |
| hsa_circ_0139811 | 919 | 10 | 2.18% |
| hsa_circ_0140627 | 919 | 10 | 2.18% |
| hsa_circ_0015165 | 1011 | 11 | 2.18% |
| hsa_circ_0023949 | 1011 | 11 | 2.18% |
| hsa_circ_0052277 | 1011 | 11 | 2.18% |
| hsa_circ_0052365 | 1011 | 11 | 2.18% |
| hsa_circ_0062097 | 1011 | 11 | 2.18% |
| hsa_circ_0071833 | 1011 | 11 | 2.18% |
| hsa_circ_0078474 | 1011 | 11 | 2.18% |
| hsa_circ_0093223 | 1011 | 11 | 2.18% |
| hsa_circ_0098416 | 1011 | 11 | 2.18% |
| hsa_circ_0124122 | 1011 | 11 | 2.18% |
| hsa_circ_0003690 | 1103 | 12 | 2.18% |
| hsa_circ_0060304 | 1103 | 12 | 2.18% |
| hsa_circ_0038731 | 1195 | 13 | 2.18% |
| hsa_circ_0117332 | 1287 | 14 | 2.18% |
| hsa_circ_0046135 | 1104 | 12 | 2.17% |
| hsa_circ_0046603 | 1012 | 11 | 2.17% |
| hsa_circ_0060926 | 1012 | 11 | 2.17% |
| hsa_circ_0099799 | 1012 | 11 | 2.17% |
| hsa_circ_0106482 | 1012 | 11 | 2.17% |
| hsa_circ_0003627 | 920 | 10 | 2.17% |
| hsa_circ_0003683 | 920 | 10 | 2.17% |

| | | | |
|---|---|---|---|
| hsa_circ_0006387 | 920 | 10 | 2.17% |
| hsa_circ_0006981 | 920 | 10 | 2.17% |
| hsa_circ_0007501 | 920 | 10 | 2.17% |
| hsa_circ_0033654 | 920 | 10 | 2.17% |
| hsa_circ_0046694 | 920 | 10 | 2.17% |
| hsa_circ_0057762 | 920 | 10 | 2.17% |
| hsa_circ_0059739 | 920 | 10 | 2.17% |
| hsa_circ_0068506 | 920 | 10 | 2.17% |
| hsa_circ_0068720 | 920 | 10 | 2.17% |
| hsa_circ_0075176 | 920 | 10 | 2.17% |
| hsa_circ_0086160 | 920 | 10 | 2.17% |
| hsa_circ_0090005 | 920 | 10 | 2.17% |
| hsa_circ_0100292 | 920 | 10 | 2.17% |
| hsa_circ_0109233 | 920 | 10 | 2.17% |
| hsa_circ_0112034 | 920 | 10 | 2.17% |
| hsa_circ_0000319 | 1197 | 13 | 2.17% |
| hsa_circ_0047088 | 1197 | 13 | 2.17% |
| hsa_circ_0010179 | 1013 | 11 | 2.17% |
| hsa_circ_0025553 | 1013 | 11 | 2.17% |
| hsa_circ_0032244 | 1013 | 11 | 2.17% |
| hsa_circ_0069701 | 1013 | 11 | 2.17% |
| hsa_circ_0087542 | 1013 | 11 | 2.17% |
| hsa_circ_0094319 | 1013 | 11 | 2.17% |
| hsa_circ_0100214 | 1013 | 11 | 2.17% |
| hsa_circ_0123477 | 1013 | 11 | 2.17% |
| hsa_circ_0124925 | 1013 | 11 | 2.17% |
| hsa_circ_0129009 | 1013 | 11 | 2.17% |
| hsa_circ_0132256 | 1013 | 11 | 2.17% |
| hsa_circ_0113791 | 4605 | 50 | 2.17% |
| hsa_circ_0015447 | 921 | 10 | 2.17% |
| hsa_circ_0027685 | 921 | 10 | 2.17% |
| hsa_circ_0034084 | 921 | 10 | 2.17% |
| hsa_circ_0034352 | 921 | 10 | 2.17% |
| hsa_circ_0039058 | 921 | 10 | 2.17% |
| hsa_circ_0047486 | 921 | 10 | 2.17% |
| hsa_circ_0048257 | 921 | 10 | 2.17% |
| hsa_circ_0051390 | 921 | 10 | 2.17% |
| hsa_circ_0067787 | 921 | 10 | 2.17% |
| hsa_circ_0072110 | 921 | 10 | 2.17% |
| hsa_circ_0088406 | 921 | 10 | 2.17% |
| hsa_circ_0090082 | 921 | 10 | 2.17% |
| hsa_circ_0105604 | 921 | 10 | 2.17% |
| hsa_circ_0106983 | 921 | 10 | 2.17% |
| hsa_circ_0116191 | 921 | 10 | 2.17% |
| hsa_circ_0120677 | 921 | 10 | 2.17% |
| hsa_circ_0131078 | 921 | 10 | 2.17% |
| hsa_circ_0059869 | 1290 | 14 | 2.17% |
| hsa_circ_0058183 | 1106 | 12 | 2.17% |
| hsa_circ_0092103 | 1106 | 12 | 2.17% |
| hsa_circ_0019919 | 1014 | 11 | 2.17% |

| | | | |
|---|---|---|---|
| hsa_circ_0022124 | 1014 | 11 | 2.17% |
| hsa_circ_0035106 | 1014 | 11 | 2.17% |
| hsa_circ_0038456 | 1014 | 11 | 2.17% |
| hsa_circ_0076772 | 1014 | 11 | 2.17% |
| hsa_circ_0086545 | 1014 | 11 | 2.17% |
| hsa_circ_0013494 | 922 | 10 | 2.17% |
| hsa_circ_0017090 | 922 | 10 | 2.17% |
| hsa_circ_0020231 | 922 | 10 | 2.17% |
| hsa_circ_0023508 | 922 | 10 | 2.17% |
| hsa_circ_0026439 | 922 | 10 | 2.17% |
| hsa_circ_0027311 | 922 | 10 | 2.17% |
| hsa_circ_0035827 | 922 | 10 | 2.17% |
| hsa_circ_0037374 | 922 | 10 | 2.17% |
| hsa_circ_0039952 | 922 | 10 | 2.17% |
| hsa_circ_0050582 | 922 | 10 | 2.17% |
| hsa_circ_0053558 | 922 | 10 | 2.17% |
| hsa_circ_0071022 | 922 | 10 | 2.17% |
| hsa_circ_0079669 | 922 | 10 | 2.17% |
| hsa_circ_0081520 | 922 | 10 | 2.17% |
| hsa_circ_0081829 | 922 | 10 | 2.17% |
| hsa_circ_0088509 | 922 | 10 | 2.17% |
| hsa_circ_0110790 | 922 | 10 | 2.17% |
| hsa_circ_0112536 | 922 | 10 | 2.17% |
| hsa_circ_0114048 | 922 | 10 | 2.17% |
| hsa_circ_0071623 | 1199 | 13 | 2.17% |
| hsa_circ_0014855 | 1107 | 12 | 2.17% |
| hsa_circ_0086366 | 1107 | 12 | 2.17% |
| hsa_circ_0031635 | 1015 | 11 | 2.17% |
| hsa_circ_0045407 | 1015 | 11 | 2.17% |
| hsa_circ_0051361 | 1015 | 11 | 2.17% |
| hsa_circ_0058321 | 1015 | 11 | 2.17% |
| hsa_circ_0073653 | 1015 | 11 | 2.17% |
| hsa_circ_0076350 | 1015 | 11 | 2.17% |
| hsa_circ_0081231 | 1015 | 11 | 2.17% |
| hsa_circ_0114474 | 1015 | 11 | 2.17% |
| hsa_circ_0117285 | 1015 | 11 | 2.17% |
| hsa_circ_0030423 | 923 | 10 | 2.17% |
| hsa_circ_0036257 | 923 | 10 | 2.17% |
| hsa_circ_0041855 | 923 | 10 | 2.17% |
| hsa_circ_0057825 | 923 | 10 | 2.17% |
| hsa_circ_0074757 | 923 | 10 | 2.17% |
| hsa_circ_0081713 | 923 | 10 | 2.17% |
| hsa_circ_0085787 | 923 | 10 | 2.17% |
| hsa_circ_0106801 | 923 | 10 | 2.17% |
| hsa_circ_0120527 | 923 | 10 | 2.17% |
| hsa_circ_0121826 | 1385 | 15 | 2.17% |
| hsa_circ_0018390 | 1108 | 12 | 2.17% |
| hsa_circ_0065963 | 1108 | 12 | 2.17% |
| hsa_circ_0068539 | 1108 | 12 | 2.17% |
| hsa_circ_0054177 | 1016 | 11 | 2.17% |

| | | | |
|---|---|---|---|
| hsa_circ_0058668 | 1016 | 11 | 2.17% |
| hsa_circ_0061173 | 1016 | 11 | 2.17% |
| hsa_circ_0099607 | 1016 | 11 | 2.17% |
| hsa_circ_0133361 | 1016 | 11 | 2.17% |
| hsa_circ_0009456 | 924 | 10 | 2.16% |
| hsa_circ_0009649 | 924 | 10 | 2.16% |
| hsa_circ_0023182 | 924 | 10 | 2.16% |
| hsa_circ_0023380 | 924 | 10 | 2.16% |
| hsa_circ_0025003 | 924 | 10 | 2.16% |
| hsa_circ_0038988 | 924 | 10 | 2.16% |
| hsa_circ_0040691 | 924 | 10 | 2.16% |
| hsa_circ_0048352 | 924 | 10 | 2.16% |
| hsa_circ_0048498 | 924 | 10 | 2.16% |
| hsa_circ_0074174 | 924 | 10 | 2.16% |
| hsa_circ_0074500 | 924 | 10 | 2.16% |
| hsa_circ_0075290 | 924 | 10 | 2.16% |
| hsa_circ_0081352 | 924 | 10 | 2.16% |
| hsa_circ_0118068 | 924 | 10 | 2.16% |
| hsa_circ_0137413 | 924 | 10 | 2.16% |
| hsa_circ_0139057 | 924 | 10 | 2.16% |
| hsa_circ_0107560 | 1109 | 12 | 2.16% |
| hsa_circ_0024344 | 1017 | 11 | 2.16% |
| hsa_circ_0041591 | 1017 | 11 | 2.16% |
| hsa_circ_0057404 | 1017 | 11 | 2.16% |
| hsa_circ_0059895 | 1017 | 11 | 2.16% |
| hsa_circ_0069409 | 1017 | 11 | 2.16% |
| hsa_circ_0085665 | 1017 | 11 | 2.16% |
| hsa_circ_0105044 | 1017 | 11 | 2.16% |
| hsa_circ_0121256 | 1017 | 11 | 2.16% |
| hsa_circ_0060432 | 1202 | 13 | 2.16% |
| hsa_circ_0034407 | 1110 | 12 | 2.16% |
| hsa_circ_0127912 | 1110 | 12 | 2.16% |
| hsa_circ_0009614 | 925 | 10 | 2.16% |
| hsa_circ_0012513 | 925 | 10 | 2.16% |
| hsa_circ_0026116 | 925 | 10 | 2.16% |
| hsa_circ_0034061 | 925 | 10 | 2.16% |
| hsa_circ_0042094 | 925 | 10 | 2.16% |
| hsa_circ_0051068 | 925 | 10 | 2.16% |
| hsa_circ_0051823 | 925 | 10 | 2.16% |
| hsa_circ_0062793 | 925 | 10 | 2.16% |
| hsa_circ_0096850 | 925 | 10 | 2.16% |
| hsa_circ_0098422 | 925 | 10 | 2.16% |
| hsa_circ_0099938 | 925 | 10 | 2.16% |
| hsa_circ_0103201 | 925 | 10 | 2.16% |
| hsa_circ_0115533 | 925 | 10 | 2.16% |
| hsa_circ_0127050 | 925 | 10 | 2.16% |
| hsa_circ_0011300 | 1018 | 11 | 2.16% |
| hsa_circ_0045617 | 1018 | 11 | 2.16% |
| hsa_circ_0087569 | 1018 | 11 | 2.16% |
| hsa_circ_0094182 | 1018 | 11 | 2.16% |

| | | | |
|---|---|---|---|
| hsa_circ_0117333 | 1296 | 14 | 2.16% |
| hsa_circ_0097617 | 1111 | 12 | 2.16% |
| hsa_circ_0055353 | 926 | 10 | 2.16% |
| hsa_circ_0061744 | 926 | 10 | 2.16% |
| hsa_circ_0064342 | 926 | 10 | 2.16% |
| hsa_circ_0091539 | 926 | 10 | 2.16% |
| hsa_circ_0099072 | 926 | 10 | 2.16% |
| hsa_circ_0103705 | 926 | 10 | 2.16% |
| hsa_circ_0103998 | 926 | 10 | 2.16% |
| hsa_circ_0112537 | 926 | 10 | 2.16% |
| hsa_circ_0114116 | 926 | 10 | 2.16% |
| hsa_circ_0131074 | 926 | 10 | 2.16% |
| hsa_circ_0135409 | 926 | 10 | 2.16% |
| hsa_circ_0031099 | 1019 | 11 | 2.16% |
| hsa_circ_0090786 | 1019 | 11 | 2.16% |
| hsa_circ_0108242 | 1019 | 11 | 2.16% |
| hsa_circ_0117109 | 1019 | 11 | 2.16% |
| hsa_circ_0128108 | 1019 | 11 | 2.16% |
| hsa_circ_0140597 | 1019 | 11 | 2.16% |
| hsa_circ_0061056 | 1112 | 12 | 2.16% |
| hsa_circ_0008164 | 927 | 10 | 2.16% |
| hsa_circ_0012615 | 927 | 10 | 2.16% |
| hsa_circ_0014781 | 927 | 10 | 2.16% |
| hsa_circ_0015244 | 927 | 10 | 2.16% |
| hsa_circ_0033652 | 927 | 10 | 2.16% |
| hsa_circ_0057737 | 927 | 10 | 2.16% |
| hsa_circ_0059491 | 927 | 10 | 2.16% |
| hsa_circ_0060494 | 927 | 10 | 2.16% |
| hsa_circ_0063396 | 927 | 10 | 2.16% |
| hsa_circ_0069017 | 927 | 10 | 2.16% |
| hsa_circ_0073580 | 927 | 10 | 2.16% |
| hsa_circ_0073904 | 927 | 10 | 2.16% |
| hsa_circ_0091280 | 927 | 10 | 2.16% |
| hsa_circ_0095222 | 927 | 10 | 2.16% |
| hsa_circ_0101738 | 927 | 10 | 2.16% |
| hsa_circ_0104164 | 927 | 10 | 2.16% |
| hsa_circ_0106868 | 927 | 10 | 2.16% |
| hsa_circ_0111920 | 927 | 10 | 2.16% |
| hsa_circ_0127972 | 927 | 10 | 2.16% |
| hsa_circ_0133516 | 927 | 10 | 2.16% |
| hsa_circ_0133319 | 2318 | 25 | 2.16% |
| hsa_circ_0009501 | 1020 | 11 | 2.16% |
| hsa_circ_0012113 | 1020 | 11 | 2.16% |
| hsa_circ_0022791 | 1020 | 11 | 2.16% |
| hsa_circ_0028443 | 1020 | 11 | 2.16% |
| hsa_circ_0043000 | 1020 | 11 | 2.16% |
| hsa_circ_0049926 | 1020 | 11 | 2.16% |
| hsa_circ_0052683 | 1020 | 11 | 2.16% |
| hsa_circ_0055545 | 1020 | 11 | 2.16% |
| hsa_circ_0056069 | 1020 | 11 | 2.16% |

| | | | |
|---|---|---|---|
| hsa_circ_0058409 | 1020 | 11 | 2.16% |
| hsa_circ_0066876 | 1020 | 11 | 2.16% |
| hsa_circ_0071924 | 1020 | 11 | 2.16% |
| hsa_circ_0136159 | 1020 | 11 | 2.16% |
| hsa_circ_0109282 | 2504 | 27 | 2.16% |
| hsa_circ_0032562 | 1113 | 12 | 2.16% |
| hsa_circ_0118416 | 1113 | 12 | 2.16% |
| hsa_circ_0134733 | 1299 | 14 | 2.16% |
| hsa_circ_0011942 | 928 | 10 | 2.16% |
| hsa_circ_0014186 | 928 | 10 | 2.16% |
| hsa_circ_0031519 | 928 | 10 | 2.16% |
| hsa_circ_0036437 | 928 | 10 | 2.16% |
| hsa_circ_0042654 | 928 | 10 | 2.16% |
| hsa_circ_0087989 | 928 | 10 | 2.16% |
| hsa_circ_0088423 | 928 | 10 | 2.16% |
| hsa_circ_0103759 | 928 | 10 | 2.16% |
| hsa_circ_0119681 | 928 | 10 | 2.16% |
| hsa_circ_0134997 | 928 | 10 | 2.16% |
| hsa_circ_0038834 | 1021 | 11 | 2.15% |
| hsa_circ_0043638 | 1021 | 11 | 2.15% |
| hsa_circ_0058974 | 1021 | 11 | 2.15% |
| hsa_circ_0064424 | 1021 | 11 | 2.15% |
| hsa_circ_0069691 | 1021 | 11 | 2.15% |
| hsa_circ_0122733 | 1021 | 11 | 2.15% |
| hsa_circ_0003316 | 1114 | 12 | 2.15% |
| hsa_circ_0058679 | 1207 | 13 | 2.15% |
| hsa_circ_0120214 | 1207 | 13 | 2.15% |
| hsa_circ_0011015 | 929 | 10 | 2.15% |
| hsa_circ_0017016 | 929 | 10 | 2.15% |
| hsa_circ_0018665 | 929 | 10 | 2.15% |
| hsa_circ_0019218 | 929 | 10 | 2.15% |
| hsa_circ_0025506 | 929 | 10 | 2.15% |
| hsa_circ_0026356 | 929 | 10 | 2.15% |
| hsa_circ_0037375 | 929 | 10 | 2.15% |
| hsa_circ_0038817 | 929 | 10 | 2.15% |
| hsa_circ_0045132 | 929 | 10 | 2.15% |
| hsa_circ_0064731 | 929 | 10 | 2.15% |
| hsa_circ_0068354 | 929 | 10 | 2.15% |
| hsa_circ_0075836 | 929 | 10 | 2.15% |
| hsa_circ_0083974 | 929 | 10 | 2.15% |
| hsa_circ_0084055 | 929 | 10 | 2.15% |
| hsa_circ_0085365 | 929 | 10 | 2.15% |
| hsa_circ_0085830 | 929 | 10 | 2.15% |
| hsa_circ_0096306 | 929 | 10 | 2.15% |
| hsa_circ_0098784 | 929 | 10 | 2.15% |
| hsa_circ_0102775 | 929 | 10 | 2.15% |
| hsa_circ_0104439 | 929 | 10 | 2.15% |
| hsa_circ_0111887 | 929 | 10 | 2.15% |
| hsa_circ_0112190 | 929 | 10 | 2.15% |
| hsa_circ_0112330 | 929 | 10 | 2.15% |

| | | | |
|---|---|---|---|
| hsa_circ_0125463 | 929 | 10 | 2.15% |
| hsa_circ_0131379 | 929 | 10 | 2.15% |
| hsa_circ_0138395 | 929 | 10 | 2.15% |
| hsa_circ_0005670 | 1022 | 11 | 2.15% |
| hsa_circ_0021708 | 1022 | 11 | 2.15% |
| hsa_circ_0039559 | 1022 | 11 | 2.15% |
| hsa_circ_0060746 | 1115 | 12 | 2.15% |
| hsa_circ_0092054 | 1115 | 12 | 2.15% |
| hsa_circ_0125924 | 1115 | 12 | 2.15% |
| hsa_circ_0140530 | 1301 | 14 | 2.15% |
| hsa_circ_0033172 | 1209 | 13 | 2.15% |
| hsa_circ_0091504 | 1116 | 12 | 2.15% |
| hsa_circ_0007314 | 1023 | 11 | 2.15% |
| hsa_circ_0071904 | 1023 | 11 | 2.15% |
| hsa_circ_0082801 | 1023 | 11 | 2.15% |
| hsa_circ_0104762 | 1023 | 11 | 2.15% |
| hsa_circ_0111322 | 1023 | 11 | 2.15% |
| hsa_circ_0136907 | 1023 | 11 | 2.15% |
| hsa_circ_0002575 | 930 | 10 | 2.15% |
| hsa_circ_0031259 | 930 | 10 | 2.15% |
| hsa_circ_0033655 | 930 | 10 | 2.15% |
| hsa_circ_0039713 | 930 | 10 | 2.15% |
| hsa_circ_0057557 | 930 | 10 | 2.15% |
| hsa_circ_0060092 | 930 | 10 | 2.15% |
| hsa_circ_0082134 | 930 | 10 | 2.15% |
| hsa_circ_0088782 | 930 | 10 | 2.15% |
| hsa_circ_0097880 | 930 | 10 | 2.15% |
| hsa_circ_0118871 | 930 | 10 | 2.15% |
| hsa_circ_0129398 | 930 | 10 | 2.15% |
| hsa_circ_0136311 | 930 | 10 | 2.15% |
| hsa_circ_0139215 | 930 | 10 | 2.15% |
| hsa_circ_0008459 | 1117 | 12 | 2.15% |
| hsa_circ_0065824 | 1117 | 12 | 2.15% |
| hsa_circ_0116807 | 1117 | 12 | 2.15% |
| hsa_circ_0125448 | 1117 | 12 | 2.15% |
| hsa_circ_0036787 | 1024 | 11 | 2.15% |
| hsa_circ_0113808 | 1024 | 11 | 2.15% |
| hsa_circ_0010121 | 931 | 10 | 2.15% |
| hsa_circ_0014821 | 931 | 10 | 2.15% |
| hsa_circ_0057767 | 931 | 10 | 2.15% |
| hsa_circ_0067414 | 931 | 10 | 2.15% |
| hsa_circ_0072768 | 931 | 10 | 2.15% |
| hsa_circ_0073197 | 931 | 10 | 2.15% |
| hsa_circ_0075714 | 931 | 10 | 2.15% |
| hsa_circ_0089648 | 931 | 10 | 2.15% |
| hsa_circ_0106262 | 931 | 10 | 2.15% |
| hsa_circ_0128975 | 931 | 10 | 2.15% |
| hsa_circ_0138111 | 931 | 10 | 2.15% |
| hsa_circ_0013642 | 1304 | 14 | 2.15% |
| hsa_circ_0010396 | 1677 | 18 | 2.15% |

| | | | |
|---|---|---|---|
| hsa_circ_0022611 | 1118 | 12 | 2.15% |
| hsa_circ_0024784 | 1118 | 12 | 2.15% |
| hsa_circ_0044846 | 1118 | 12 | 2.15% |
| hsa_circ_0111864 | 1118 | 12 | 2.15% |
| hsa_circ_0122654 | 1584 | 17 | 2.15% |
| hsa_circ_0050181 | 1025 | 11 | 2.15% |
| hsa_circ_0057062 | 1025 | 11 | 2.15% |
| hsa_circ_0086714 | 1025 | 11 | 2.15% |
| hsa_circ_0010938 | 932 | 10 | 2.15% |
| hsa_circ_0035960 | 932 | 10 | 2.15% |
| hsa_circ_0036665 | 932 | 10 | 2.15% |
| hsa_circ_0060582 | 932 | 10 | 2.15% |
| hsa_circ_0070884 | 932 | 10 | 2.15% |
| hsa_circ_0074447 | 932 | 10 | 2.15% |
| hsa_circ_0095407 | 932 | 10 | 2.15% |
| hsa_circ_0113144 | 932 | 10 | 2.15% |
| hsa_circ_0113485 | 932 | 10 | 2.15% |
| hsa_circ_0120611 | 932 | 10 | 2.15% |
| hsa_circ_0121986 | 932 | 10 | 2.15% |
| hsa_circ_0122346 | 932 | 10 | 2.15% |
| hsa_circ_0130845 | 932 | 10 | 2.15% |
| hsa_circ_0140634 | 932 | 10 | 2.15% |
| hsa_circ_0072827 | 275244 | 2953 | 2.15% |
| hsa_circ_0049993 | 2144 | 23 | 2.15% |
| hsa_circ_0030849 | 1212 | 13 | 2.15% |
| hsa_circ_0033035 | 1212 | 13 | 2.15% |
| hsa_circ_0051438 | 1212 | 13 | 2.15% |
| hsa_circ_0071352 | 1119 | 12 | 2.14% |
| hsa_circ_0116651 | 1119 | 12 | 2.14% |
| hsa_circ_0014980 | 1026 | 11 | 2.14% |
| hsa_circ_0056997 | 1026 | 11 | 2.14% |
| hsa_circ_0065228 | 1026 | 11 | 2.14% |
| hsa_circ_0082281 | 1026 | 11 | 2.14% |
| hsa_circ_0089640 | 1026 | 11 | 2.14% |
| hsa_circ_0101136 | 1026 | 11 | 2.14% |
| hsa_circ_0118513 | 1026 | 11 | 2.14% |
| hsa_circ_0130806 | 1026 | 11 | 2.14% |
| hsa_circ_0011637 | 933 | 10 | 2.14% |
| hsa_circ_0014316 | 933 | 10 | 2.14% |
| hsa_circ_0026461 | 933 | 10 | 2.14% |
| hsa_circ_0030015 | 933 | 10 | 2.14% |
| hsa_circ_0032005 | 933 | 10 | 2.14% |
| hsa_circ_0056454 | 933 | 10 | 2.14% |
| hsa_circ_0058510 | 933 | 10 | 2.14% |
| hsa_circ_0061931 | 933 | 10 | 2.14% |
| hsa_circ_0066236 | 933 | 10 | 2.14% |
| hsa_circ_0073434 | 933 | 10 | 2.14% |
| hsa_circ_0083348 | 933 | 10 | 2.14% |
| hsa_circ_0104368 | 933 | 10 | 2.14% |
| hsa_circ_0109502 | 933 | 10 | 2.14% |

| | | | |
|---|---|---|---|
| hsa_circ_0113358 | 933 | 10 | 2.14% |
| hsa_circ_0114477 | 933 | 10 | 2.14% |
| hsa_circ_0129019 | 933 | 10 | 2.14% |
| hsa_circ_0036744 | 1120 | 12 | 2.14% |
| hsa_circ_0062825 | 1120 | 12 | 2.14% |
| hsa_circ_0067206 | 1120 | 12 | 2.14% |
| hsa_circ_0004782 | 1027 | 11 | 2.14% |
| hsa_circ_0019426 | 1027 | 11 | 2.14% |
| hsa_circ_0025222 | 1027 | 11 | 2.14% |
| hsa_circ_0029392 | 1027 | 11 | 2.14% |
| hsa_circ_0030194 | 1027 | 11 | 2.14% |
| hsa_circ_0035940 | 1027 | 11 | 2.14% |
| hsa_circ_0052874 | 1027 | 11 | 2.14% |
| hsa_circ_0081845 | 1027 | 11 | 2.14% |
| hsa_circ_0113421 | 1027 | 11 | 2.14% |
| hsa_circ_0113872 | 1027 | 11 | 2.14% |
| hsa_circ_0128071 | 1214 | 13 | 2.14% |
| hsa_circ_0022646 | 934 | 10 | 2.14% |
| hsa_circ_0053415 | 934 | 10 | 2.14% |
| hsa_circ_0087364 | 934 | 10 | 2.14% |
| hsa_circ_0089759 | 934 | 10 | 2.14% |
| hsa_circ_0090357 | 934 | 10 | 2.14% |
| hsa_circ_0095508 | 934 | 10 | 2.14% |
| hsa_circ_0102928 | 934 | 10 | 2.14% |
| hsa_circ_0105466 | 934 | 10 | 2.14% |
| hsa_circ_0107062 | 934 | 10 | 2.14% |
| hsa_circ_0114441 | 934 | 10 | 2.14% |
| hsa_circ_0122306 | 934 | 10 | 2.14% |
| hsa_circ_0068793 | 1121 | 12 | 2.14% |
| hsa_circ_0085903 | 1121 | 12 | 2.14% |
| hsa_circ_0007962 | 1028 | 11 | 2.14% |
| hsa_circ_0024005 | 1028 | 11 | 2.14% |
| hsa_circ_0074809 | 1028 | 11 | 2.14% |
| hsa_circ_0127370 | 1028 | 11 | 2.14% |
| hsa_circ_0047413 | 1215 | 13 | 2.14% |
| hsa_circ_0086781 | 1215 | 13 | 2.14% |
| hsa_circ_0115315 | 1589 | 17 | 2.14% |
| hsa_circ_0071056 | 1776 | 19 | 2.14% |
| hsa_circ_0047758 | 1122 | 12 | 2.14% |
| hsa_circ_0088522 | 1122 | 12 | 2.14% |
| hsa_circ_0003665 | 935 | 10 | 2.14% |
| hsa_circ_0005827 | 935 | 10 | 2.14% |
| hsa_circ_0021137 | 935 | 10 | 2.14% |
| hsa_circ_0022651 | 935 | 10 | 2.14% |
| hsa_circ_0073139 | 935 | 10 | 2.14% |
| hsa_circ_0079286 | 935 | 10 | 2.14% |
| hsa_circ_0080046 | 935 | 10 | 2.14% |
| hsa_circ_0080739 | 935 | 10 | 2.14% |
| hsa_circ_0091979 | 935 | 10 | 2.14% |
| hsa_circ_0112030 | 935 | 10 | 2.14% |

| | | | |
|---|---|---|---|
| hsa_circ_0131404 | 935 | 10 | 2.14% |
| hsa_circ_0131901 | 935 | 10 | 2.14% |
| hsa_circ_0025157 | 1029 | 11 | 2.14% |
| hsa_circ_0026751 | 1029 | 11 | 2.14% |
| hsa_circ_0031770 | 1029 | 11 | 2.14% |
| hsa_circ_0052561 | 1029 | 11 | 2.14% |
| hsa_circ_0057803 | 1029 | 11 | 2.14% |
| hsa_circ_0092144 | 1029 | 11 | 2.14% |
| hsa_circ_0125698 | 1029 | 11 | 2.14% |
| hsa_circ_0131913 | 1029 | 11 | 2.14% |
| hsa_circ_0138529 | 1029 | 11 | 2.14% |
| hsa_circ_0001632 | 936 | 10 | 2.14% |
| hsa_circ_0005305 | 936 | 10 | 2.14% |
| hsa_circ_0009618 | 936 | 10 | 2.14% |
| hsa_circ_0012570 | 936 | 10 | 2.14% |
| hsa_circ_0061800 | 936 | 10 | 2.14% |
| hsa_circ_0064696 | 936 | 10 | 2.14% |
| hsa_circ_0065550 | 936 | 10 | 2.14% |
| hsa_circ_0077807 | 936 | 10 | 2.14% |
| hsa_circ_0081368 | 936 | 10 | 2.14% |
| hsa_circ_0089589 | 936 | 10 | 2.14% |
| hsa_circ_0096178 | 936 | 10 | 2.14% |
| hsa_circ_0101279 | 936 | 10 | 2.14% |
| hsa_circ_0107755 | 936 | 10 | 2.14% |
| hsa_circ_0109378 | 936 | 10 | 2.14% |
| hsa_circ_0120408 | 936 | 10 | 2.14% |
| hsa_circ_0124894 | 936 | 10 | 2.14% |
| hsa_circ_0126071 | 936 | 10 | 2.14% |
| hsa_circ_0129719 | 936 | 10 | 2.14% |
| hsa_circ_0066955 | 1217 | 13 | 2.14% |
| hsa_circ_0008293 | 1030 | 11 | 2.14% |
| hsa_circ_0009465 | 1030 | 11 | 2.14% |
| hsa_circ_0046380 | 1030 | 11 | 2.14% |
| hsa_circ_0054958 | 1030 | 11 | 2.14% |
| hsa_circ_0091725 | 1030 | 11 | 2.14% |
| hsa_circ_0124198 | 1030 | 11 | 2.14% |
| hsa_circ_0125213 | 1030 | 11 | 2.14% |
| hsa_circ_0052706 | 1405 | 15 | 2.14% |
| hsa_circ_0034913 | 1124 | 12 | 2.14% |
| hsa_circ_0000261 | 937 | 10 | 2.13% |
| hsa_circ_0033648 | 937 | 10 | 2.13% |
| hsa_circ_0041341 | 937 | 10 | 2.13% |
| hsa_circ_0062224 | 937 | 10 | 2.13% |
| hsa_circ_0075422 | 937 | 10 | 2.13% |
| hsa_circ_0089961 | 937 | 10 | 2.13% |
| hsa_circ_0092784 | 937 | 10 | 2.13% |
| hsa_circ_0101486 | 937 | 10 | 2.13% |
| hsa_circ_0126307 | 937 | 10 | 2.13% |
| hsa_circ_0132483 | 937 | 10 | 2.13% |
| hsa_circ_0133776 | 937 | 10 | 2.13% |

| | | | |
|---|---|---|---|
| hsa_circ_0026393 | 1031 | 11 | 2.13% |
| hsa_circ_0051317 | 1031 | 11 | 2.13% |
| hsa_circ_0057061 | 1031 | 11 | 2.13% |
| hsa_circ_0062386 | 1031 | 11 | 2.13% |
| hsa_circ_0064394 | 1031 | 11 | 2.13% |
| hsa_circ_0072398 | 1031 | 11 | 2.13% |
| hsa_circ_0111362 | 1031 | 11 | 2.13% |
| hsa_circ_0097439 | 2156 | 23 | 2.13% |
| hsa_circ_0037043 | 1125 | 12 | 2.13% |
| hsa_circ_0050712 | 1125 | 12 | 2.13% |
| hsa_circ_0052175 | 1125 | 12 | 2.13% |
| hsa_circ_0104141 | 1125 | 12 | 2.13% |
| hsa_circ_0133880 | 1125 | 12 | 2.13% |
| hsa_circ_0121584 | 1688 | 18 | 2.13% |
| hsa_circ_0016159 | 938 | 10 | 2.13% |
| hsa_circ_0018909 | 938 | 10 | 2.13% |
| hsa_circ_0039301 | 938 | 10 | 2.13% |
| hsa_circ_0039366 | 938 | 10 | 2.13% |
| hsa_circ_0044372 | 938 | 10 | 2.13% |
| hsa_circ_0050665 | 938 | 10 | 2.13% |
| hsa_circ_0067842 | 938 | 10 | 2.13% |
| hsa_circ_0075502 | 938 | 10 | 2.13% |
| hsa_circ_0084114 | 938 | 10 | 2.13% |
| hsa_circ_0097585 | 938 | 10 | 2.13% |
| hsa_circ_0097885 | 938 | 10 | 2.13% |
| hsa_circ_0098298 | 938 | 10 | 2.13% |
| hsa_circ_0100388 | 938 | 10 | 2.13% |
| hsa_circ_0130482 | 938 | 10 | 2.13% |
| hsa_circ_0137578 | 938 | 10 | 2.13% |
| hsa_circ_0137720 | 938 | 10 | 2.13% |
| hsa_circ_0138458 | 938 | 10 | 2.13% |
| hsa_circ_0024175 | 1032 | 11 | 2.13% |
| hsa_circ_0055547 | 1032 | 11 | 2.13% |
| hsa_circ_0061508 | 1032 | 11 | 2.13% |
| hsa_circ_0068242 | 1032 | 11 | 2.13% |
| hsa_circ_0089197 | 1032 | 11 | 2.13% |
| hsa_circ_0130631 | 1032 | 11 | 2.13% |
| hsa_circ_0136405 | 1032 | 11 | 2.13% |
| hsa_circ_0029783 | 1126 | 12 | 2.13% |
| hsa_circ_0135125 | 1126 | 12 | 2.13% |
| hsa_circ_0139623 | 1126 | 12 | 2.13% |
| hsa_circ_0003251 | 939 | 10 | 2.13% |
| hsa_circ_0006340 | 939 | 10 | 2.13% |
| hsa_circ_0010061 | 939 | 10 | 2.13% |
| hsa_circ_0014002 | 939 | 10 | 2.13% |
| hsa_circ_0016025 | 939 | 10 | 2.13% |
| hsa_circ_0034391 | 939 | 10 | 2.13% |
| hsa_circ_0047970 | 939 | 10 | 2.13% |
| hsa_circ_0055046 | 939 | 10 | 2.13% |
| hsa_circ_0058155 | 939 | 10 | 2.13% |

| | | | |
|---|---|---|---|
| hsa_circ_0058417 | 939 | 10 | 2.13% |
| hsa_circ_0060077 | 939 | 10 | 2.13% |
| hsa_circ_0072941 | 939 | 10 | 2.13% |
| hsa_circ_0078186 | 939 | 10 | 2.13% |
| hsa_circ_0083514 | 939 | 10 | 2.13% |
| hsa_circ_0087110 | 939 | 10 | 2.13% |
| hsa_circ_0093631 | 939 | 10 | 2.13% |
| hsa_circ_0111696 | 939 | 10 | 2.13% |
| hsa_circ_0121927 | 939 | 10 | 2.13% |
| hsa_circ_0126225 | 939 | 10 | 2.13% |
| hsa_circ_0131548 | 939 | 10 | 2.13% |
| hsa_circ_0131593 | 939 | 10 | 2.13% |
| hsa_circ_0014340 | 1033 | 11 | 2.13% |
| hsa_circ_0046501 | 1033 | 11 | 2.13% |
| hsa_circ_0110721 | 1033 | 11 | 2.13% |
| hsa_circ_0117485 | 1033 | 11 | 2.13% |
| hsa_circ_0027285 | 1221 | 13 | 2.13% |
| hsa_circ_0072312 | 1221 | 13 | 2.13% |
| hsa_circ_0034686 | 1786 | 19 | 2.13% |
| hsa_circ_0009235 | 1128 | 12 | 2.13% |
| hsa_circ_0018627 | 1128 | 12 | 2.13% |
| hsa_circ_0071085 | 1128 | 12 | 2.13% |
| hsa_circ_0082637 | 1128 | 12 | 2.13% |
| hsa_circ_0109103 | 1128 | 12 | 2.13% |
| hsa_circ_0138914 | 1128 | 12 | 2.13% |
| hsa_circ_0006386 | 1034 | 11 | 2.13% |
| hsa_circ_0020952 | 1034 | 11 | 2.13% |
| hsa_circ_0035621 | 1034 | 11 | 2.13% |
| hsa_circ_0095507 | 1034 | 11 | 2.13% |
| hsa_circ_0109460 | 1034 | 11 | 2.13% |
| hsa_circ_0006307 | 940 | 10 | 2.13% |
| hsa_circ_0027063 | 940 | 10 | 2.13% |
| hsa_circ_0048742 | 940 | 10 | 2.13% |
| hsa_circ_0052794 | 940 | 10 | 2.13% |
| hsa_circ_0081657 | 940 | 10 | 2.13% |
| hsa_circ_0086345 | 940 | 10 | 2.13% |
| hsa_circ_0086445 | 940 | 10 | 2.13% |
| hsa_circ_0116709 | 940 | 10 | 2.13% |
| hsa_circ_0122254 | 940 | 10 | 2.13% |
| hsa_circ_0124218 | 940 | 10 | 2.13% |
| hsa_circ_0134814 | 940 | 10 | 2.13% |
| hsa_circ_0017275 | 1035 | 11 | 2.13% |
| hsa_circ_0034193 | 1035 | 11 | 2.13% |
| hsa_circ_0045612 | 1035 | 11 | 2.13% |
| hsa_circ_0057862 | 1035 | 11 | 2.13% |
| hsa_circ_0081045 | 1035 | 11 | 2.13% |
| hsa_circ_0083958 | 1035 | 11 | 2.13% |
| hsa_circ_0094846 | 1035 | 11 | 2.13% |
| hsa_circ_0100908 | 1035 | 11 | 2.13% |
| hsa_circ_0103233 | 1035 | 11 | 2.13% |

| | | | |
|---|---|---|---|
| hsa_circ_0112859 | 1035 | 11 | 2.13% |
| hsa_circ_0113137 | 1035 | 11 | 2.13% |
| hsa_circ_0009895 | 941 | 10 | 2.13% |
| hsa_circ_0017719 | 941 | 10 | 2.13% |
| hsa_circ_0017897 | 941 | 10 | 2.13% |
| hsa_circ_0018560 | 941 | 10 | 2.13% |
| hsa_circ_0020363 | 941 | 10 | 2.13% |
| hsa_circ_0022108 | 941 | 10 | 2.13% |
| hsa_circ_0025305 | 941 | 10 | 2.13% |
| hsa_circ_0029146 | 941 | 10 | 2.13% |
| hsa_circ_0053067 | 941 | 10 | 2.13% |
| hsa_circ_0065910 | 941 | 10 | 2.13% |
| hsa_circ_0076239 | 941 | 10 | 2.13% |
| hsa_circ_0114303 | 941 | 10 | 2.13% |
| hsa_circ_0127810 | 941 | 10 | 2.13% |
| hsa_circ_0137024 | 941 | 10 | 2.13% |
| hsa_circ_0031196 | 2353 | 25 | 2.12% |
| hsa_circ_0102725 | 1224 | 13 | 2.12% |
| hsa_circ_0023700 | 1036 | 11 | 2.12% |
| hsa_circ_0044029 | 1036 | 11 | 2.12% |
| hsa_circ_0063137 | 1036 | 11 | 2.12% |
| hsa_circ_0089418 | 1036 | 11 | 2.12% |
| hsa_circ_0089876 | 1036 | 11 | 2.12% |
| hsa_circ_0096233 | 1036 | 11 | 2.12% |
| hsa_circ_0110967 | 1036 | 11 | 2.12% |
| hsa_circ_0124598 | 1036 | 11 | 2.12% |
| hsa_circ_0005586 | 942 | 10 | 2.12% |
| hsa_circ_0012609 | 942 | 10 | 2.12% |
| hsa_circ_0014319 | 942 | 10 | 2.12% |
| hsa_circ_0016435 | 942 | 10 | 2.12% |
| hsa_circ_0020860 | 942 | 10 | 2.12% |
| hsa_circ_0026386 | 942 | 10 | 2.12% |
| hsa_circ_0030599 | 942 | 10 | 2.12% |
| hsa_circ_0031723 | 942 | 10 | 2.12% |
| hsa_circ_0034451 | 942 | 10 | 2.12% |
| hsa_circ_0037376 | 942 | 10 | 2.12% |
| hsa_circ_0044289 | 942 | 10 | 2.12% |
| hsa_circ_0064656 | 942 | 10 | 2.12% |
| hsa_circ_0077614 | 942 | 10 | 2.12% |
| hsa_circ_0096152 | 942 | 10 | 2.12% |
| hsa_circ_0096225 | 942 | 10 | 2.12% |
| hsa_circ_0103877 | 942 | 10 | 2.12% |
| hsa_circ_0107209 | 942 | 10 | 2.12% |
| hsa_circ_0109168 | 942 | 10 | 2.12% |
| hsa_circ_0117159 | 942 | 10 | 2.12% |
| hsa_circ_0119327 | 942 | 10 | 2.12% |
| hsa_circ_0122818 | 942 | 10 | 2.12% |
| hsa_circ_0123878 | 942 | 10 | 2.12% |
| hsa_circ_0138347 | 942 | 10 | 2.12% |
| hsa_circ_0012128 | 1225 | 13 | 2.12% |

| | | | |
|---|---|---|---|
| hsa_circ_0025226 | 1225 | 13 | 2.12% |
| hsa_circ_0037680 | 1131 | 12 | 2.12% |
| hsa_circ_0020992 | 1037 | 11 | 2.12% |
| hsa_circ_0037843 | 1037 | 11 | 2.12% |
| hsa_circ_0044654 | 1037 | 11 | 2.12% |
| hsa_circ_0124662 | 1037 | 11 | 2.12% |
| hsa_circ_0014714 | 943 | 10 | 2.12% |
| hsa_circ_0028353 | 943 | 10 | 2.12% |
| hsa_circ_0042380 | 943 | 10 | 2.12% |
| hsa_circ_0045711 | 943 | 10 | 2.12% |
| hsa_circ_0062395 | 943 | 10 | 2.12% |
| hsa_circ_0088221 | 943 | 10 | 2.12% |
| hsa_circ_0103039 | 943 | 10 | 2.12% |
| hsa_circ_0103538 | 943 | 10 | 2.12% |
| hsa_circ_0119967 | 943 | 10 | 2.12% |
| hsa_circ_0132782 | 943 | 10 | 2.12% |
| hsa_circ_0137973 | 943 | 10 | 2.12% |
| hsa_circ_0140271 | 943 | 10 | 2.12% |
| hsa_circ_0046247 | 1226 | 13 | 2.12% |
| hsa_circ_0067634 | 1226 | 13 | 2.12% |
| hsa_circ_0000295 | 1132 | 12 | 2.12% |
| hsa_circ_0062082 | 3114 | 33 | 2.12% |
| hsa_circ_0021838 | 1038 | 11 | 2.12% |
| hsa_circ_0028144 | 1038 | 11 | 2.12% |
| hsa_circ_0030033 | 1038 | 11 | 2.12% |
| hsa_circ_0063814 | 1038 | 11 | 2.12% |
| hsa_circ_0101663 | 1038 | 11 | 2.12% |
| hsa_circ_0063757 | 1227 | 13 | 2.12% |
| hsa_circ_0097498 | 1699 | 18 | 2.12% |
| hsa_circ_0029983 | 944 | 10 | 2.12% |
| hsa_circ_0033645 | 944 | 10 | 2.12% |
| hsa_circ_0054020 | 944 | 10 | 2.12% |
| hsa_circ_0062564 | 944 | 10 | 2.12% |
| hsa_circ_0071815 | 944 | 10 | 2.12% |
| hsa_circ_0075150 | 944 | 10 | 2.12% |
| hsa_circ_0076248 | 944 | 10 | 2.12% |
| hsa_circ_0087036 | 944 | 10 | 2.12% |
| hsa_circ_0087531 | 944 | 10 | 2.12% |
| hsa_circ_0099367 | 944 | 10 | 2.12% |
| hsa_circ_0104114 | 944 | 10 | 2.12% |
| hsa_circ_0104722 | 944 | 10 | 2.12% |
| hsa_circ_0110456 | 944 | 10 | 2.12% |
| hsa_circ_0119474 | 944 | 10 | 2.12% |
| hsa_circ_0121898 | 944 | 10 | 2.12% |
| hsa_circ_0127594 | 944 | 10 | 2.12% |
| hsa_circ_0130285 | 944 | 10 | 2.12% |
| hsa_circ_0134743 | 944 | 10 | 2.12% |
| hsa_circ_0057449 | 1133 | 12 | 2.12% |
| hsa_circ_0074669 | 1133 | 12 | 2.12% |
| hsa_circ_0081048 | 1133 | 12 | 2.12% |

| | | | |
|---|---|---|---|
| hsa_circ_0104193 | 1133 | 12 | 2.12% |
| hsa_circ_0118184 | 1133 | 12 | 2.12% |
| hsa_circ_0053346 | 1039 | 11 | 2.12% |
| hsa_circ_0074914 | 1039 | 11 | 2.12% |
| hsa_circ_0077870 | 1039 | 11 | 2.12% |
| hsa_circ_0083154 | 1039 | 11 | 2.12% |
| hsa_circ_0085991 | 1039 | 11 | 2.12% |
| hsa_circ_0116148 | 1039 | 11 | 2.12% |
| hsa_circ_0127267 | 1039 | 11 | 2.12% |
| hsa_circ_0110187 | 1228 | 13 | 2.12% |
| hsa_circ_0102755 | 2362 | 25 | 2.12% |
| hsa_circ_0010897 | 1134 | 12 | 2.12% |
| hsa_circ_0080546 | 1134 | 12 | 2.12% |
| hsa_circ_0089730 | 1134 | 12 | 2.12% |
| hsa_circ_0018001 | 945 | 10 | 2.12% |
| hsa_circ_0018279 | 945 | 10 | 2.12% |
| hsa_circ_0045353 | 945 | 10 | 2.12% |
| hsa_circ_0056181 | 945 | 10 | 2.12% |
| hsa_circ_0057164 | 945 | 10 | 2.12% |
| hsa_circ_0062971 | 945 | 10 | 2.12% |
| hsa_circ_0088595 | 945 | 10 | 2.12% |
| hsa_circ_0105446 | 945 | 10 | 2.12% |
| hsa_circ_0107234 | 945 | 10 | 2.12% |
| hsa_circ_0109493 | 945 | 10 | 2.12% |
| hsa_circ_0112646 | 945 | 10 | 2.12% |
| hsa_circ_0115347 | 945 | 10 | 2.12% |
| hsa_circ_0115508 | 945 | 10 | 2.12% |
| hsa_circ_0119767 | 945 | 10 | 2.12% |
| hsa_circ_0125256 | 945 | 10 | 2.12% |
| hsa_circ_0129698 | 945 | 10 | 2.12% |
| hsa_circ_0012111 | 1229 | 13 | 2.12% |
| hsa_circ_0048613 | 1040 | 11 | 2.12% |
| hsa_circ_0051024 | 1040 | 11 | 2.12% |
| hsa_circ_0068309 | 1040 | 11 | 2.12% |
| hsa_circ_0074889 | 1040 | 11 | 2.12% |
| hsa_circ_0007672 | 946 | 10 | 2.11% |
| hsa_circ_0012132 | 946 | 10 | 2.11% |
| hsa_circ_0039796 | 946 | 10 | 2.11% |
| hsa_circ_0042851 | 946 | 10 | 2.11% |
| hsa_circ_0047451 | 946 | 10 | 2.11% |
| hsa_circ_0056932 | 946 | 10 | 2.11% |
| hsa_circ_0069428 | 946 | 10 | 2.11% |
| hsa_circ_0087675 | 946 | 10 | 2.11% |
| hsa_circ_0097988 | 946 | 10 | 2.11% |
| hsa_circ_0098893 | 946 | 10 | 2.11% |
| hsa_circ_0105770 | 946 | 10 | 2.11% |
| hsa_circ_0133079 | 946 | 10 | 2.11% |
| hsa_circ_0088624 | 1230 | 13 | 2.11% |
| hsa_circ_0010041 | 1041 | 11 | 2.11% |
| hsa_circ_0057468 | 1041 | 11 | 2.11% |

| | | | |
|---|---|---|---|
| hsa_circ_0084871 | 1041 | 11 | 2.11% |
| hsa_circ_0090326 | 1041 | 11 | 2.11% |
| hsa_circ_0018183 | 1136 | 12 | 2.11% |
| hsa_circ_0083598 | 1136 | 12 | 2.11% |
| hsa_circ_0104413 | 3125 | 33 | 2.11% |
| hsa_circ_0004125 | 947 | 10 | 2.11% |
| hsa_circ_0008414 | 947 | 10 | 2.11% |
| hsa_circ_0013427 | 947 | 10 | 2.11% |
| hsa_circ_0015284 | 947 | 10 | 2.11% |
| hsa_circ_0023605 | 947 | 10 | 2.11% |
| hsa_circ_0039036 | 947 | 10 | 2.11% |
| hsa_circ_0043862 | 947 | 10 | 2.11% |
| hsa_circ_0052460 | 947 | 10 | 2.11% |
| hsa_circ_0055740 | 947 | 10 | 2.11% |
| hsa_circ_0061752 | 947 | 10 | 2.11% |
| hsa_circ_0069614 | 947 | 10 | 2.11% |
| hsa_circ_0078135 | 947 | 10 | 2.11% |
| hsa_circ_0087852 | 947 | 10 | 2.11% |
| hsa_circ_0094432 | 947 | 10 | 2.11% |
| hsa_circ_0105250 | 947 | 10 | 2.11% |
| hsa_circ_0128849 | 947 | 10 | 2.11% |
| hsa_circ_0134770 | 947 | 10 | 2.11% |
| hsa_circ_0139126 | 947 | 10 | 2.11% |
| hsa_circ_0014979 | 1042 | 11 | 2.11% |
| hsa_circ_0067273 | 1421 | 15 | 2.11% |
| hsa_circ_0107333 | 1516 | 16 | 2.11% |
| hsa_circ_0019509 | 1137 | 12 | 2.11% |
| hsa_circ_0064069 | 1137 | 12 | 2.11% |
| hsa_circ_0043866 | 1232 | 13 | 2.11% |
| hsa_circ_0134025 | 1232 | 13 | 2.11% |
| hsa_circ_0093388 | 1327 | 14 | 2.11% |
| hsa_circ_0014848 | 948 | 10 | 2.11% |
| hsa_circ_0015105 | 948 | 10 | 2.11% |
| hsa_circ_0023443 | 948 | 10 | 2.11% |
| hsa_circ_0026835 | 948 | 10 | 2.11% |
| hsa_circ_0032427 | 948 | 10 | 2.11% |
| hsa_circ_0041167 | 948 | 10 | 2.11% |
| hsa_circ_0059291 | 948 | 10 | 2.11% |
| hsa_circ_0059910 | 948 | 10 | 2.11% |
| hsa_circ_0060049 | 948 | 10 | 2.11% |
| hsa_circ_0062691 | 948 | 10 | 2.11% |
| hsa_circ_0070801 | 948 | 10 | 2.11% |
| hsa_circ_0083004 | 948 | 10 | 2.11% |
| hsa_circ_0084803 | 948 | 10 | 2.11% |
| hsa_circ_0092063 | 948 | 10 | 2.11% |
| hsa_circ_0096342 | 948 | 10 | 2.11% |
| hsa_circ_0112013 | 948 | 10 | 2.11% |
| hsa_circ_0140405 | 948 | 10 | 2.11% |
| hsa_circ_0006076 | 1043 | 11 | 2.11% |
| hsa_circ_0012388 | 1043 | 11 | 2.11% |

| | | | |
|---|---|---|---|
| hsa_circ_0021059 | 1043 | 11 | 2.11% |
| hsa_circ_0030947 | 1043 | 11 | 2.11% |
| hsa_circ_0067836 | 1043 | 11 | 2.11% |
| hsa_circ_0073161 | 1043 | 11 | 2.11% |
| hsa_circ_0075043 | 1043 | 11 | 2.11% |
| hsa_circ_0081978 | 1043 | 11 | 2.11% |
| hsa_circ_0089291 | 1043 | 11 | 2.11% |
| hsa_circ_0102766 | 1043 | 11 | 2.11% |
| hsa_circ_0129050 | 1043 | 11 | 2.11% |
| hsa_circ_0025583 | 1138 | 12 | 2.11% |
| hsa_circ_0110542 | 1138 | 12 | 2.11% |
| hsa_circ_0063583 | 3794 | 40 | 2.11% |
| hsa_circ_0015389 | 1328 | 14 | 2.11% |
| hsa_circ_0000624 | 949 | 10 | 2.11% |
| hsa_circ_0005164 | 949 | 10 | 2.11% |
| hsa_circ_0008944 | 949 | 10 | 2.11% |
| hsa_circ_0010138 | 949 | 10 | 2.11% |
| hsa_circ_0038901 | 949 | 10 | 2.11% |
| hsa_circ_0039130 | 949 | 10 | 2.11% |
| hsa_circ_0053802 | 949 | 10 | 2.11% |
| hsa_circ_0054464 | 949 | 10 | 2.11% |
| hsa_circ_0055435 | 949 | 10 | 2.11% |
| hsa_circ_0087985 | 949 | 10 | 2.11% |
| hsa_circ_0093933 | 949 | 10 | 2.11% |
| hsa_circ_0099514 | 949 | 10 | 2.11% |
| hsa_circ_0108309 | 949 | 10 | 2.11% |
| hsa_circ_0110572 | 949 | 10 | 2.11% |
| hsa_circ_0111896 | 949 | 10 | 2.11% |
| hsa_circ_0126414 | 949 | 10 | 2.11% |
| hsa_circ_0024172 | 1044 | 11 | 2.11% |
| hsa_circ_0076605 | 1044 | 11 | 2.11% |
| hsa_circ_0136431 | 1044 | 11 | 2.11% |
| hsa_circ_0036627 | 1139 | 12 | 2.11% |
| hsa_circ_0053123 | 1139 | 12 | 2.11% |
| hsa_circ_0058013 | 1139 | 12 | 2.11% |
| hsa_circ_0112258 | 1139 | 12 | 2.11% |
| hsa_circ_0135975 | 1424 | 15 | 2.11% |
| hsa_circ_0015398 | 1330 | 14 | 2.11% |
| hsa_circ_0045146 | 1140 | 12 | 2.11% |
| hsa_circ_0010714 | 1045 | 11 | 2.11% |
| hsa_circ_0025787 | 1045 | 11 | 2.11% |
| hsa_circ_0029265 | 1045 | 11 | 2.11% |
| hsa_circ_0058362 | 1045 | 11 | 2.11% |
| hsa_circ_0067886 | 1045 | 11 | 2.11% |
| hsa_circ_0123904 | 1045 | 11 | 2.11% |
| hsa_circ_0132536 | 1045 | 11 | 2.11% |
| hsa_circ_0013179 | 950 | 10 | 2.11% |
| hsa_circ_0035179 | 950 | 10 | 2.11% |
| hsa_circ_0050863 | 950 | 10 | 2.11% |
| hsa_circ_0051898 | 950 | 10 | 2.11% |

| | | | |
|---|---|---|---|
| hsa_circ_0053411 | 950 | 10 | 2.11% |
| hsa_circ_0055510 | 950 | 10 | 2.11% |
| hsa_circ_0058054 | 950 | 10 | 2.11% |
| hsa_circ_0061768 | 950 | 10 | 2.11% |
| hsa_circ_0062366 | 950 | 10 | 2.11% |
| hsa_circ_0087368 | 950 | 10 | 2.11% |
| hsa_circ_0100604 | 950 | 10 | 2.11% |
| hsa_circ_0112518 | 950 | 10 | 2.11% |
| hsa_circ_0120650 | 950 | 10 | 2.11% |
| hsa_circ_0137659 | 950 | 10 | 2.11% |
| hsa_circ_0009422 | 1426 | 15 | 2.10% |
| hsa_circ_0044459 | 1141 | 12 | 2.10% |
| hsa_circ_0055592 | 1046 | 11 | 2.10% |
| hsa_circ_0080157 | 1046 | 11 | 2.10% |
| hsa_circ_0083499 | 1046 | 11 | 2.10% |
| hsa_circ_0097636 | 1046 | 11 | 2.10% |
| hsa_circ_0122780 | 1046 | 11 | 2.10% |
| hsa_circ_0003653 | 951 | 10 | 2.10% |
| hsa_circ_0011503 | 951 | 10 | 2.10% |
| hsa_circ_0016043 | 951 | 10 | 2.10% |
| hsa_circ_0020655 | 951 | 10 | 2.10% |
| hsa_circ_0023542 | 951 | 10 | 2.10% |
| hsa_circ_0039975 | 951 | 10 | 2.10% |
| hsa_circ_0048579 | 951 | 10 | 2.10% |
| hsa_circ_0049062 | 951 | 10 | 2.10% |
| hsa_circ_0057560 | 951 | 10 | 2.10% |
| hsa_circ_0060371 | 951 | 10 | 2.10% |
| hsa_circ_0061152 | 951 | 10 | 2.10% |
| hsa_circ_0071072 | 951 | 10 | 2.10% |
| hsa_circ_0072041 | 951 | 10 | 2.10% |
| hsa_circ_0072269 | 951 | 10 | 2.10% |
| hsa_circ_0104144 | 951 | 10 | 2.10% |
| hsa_circ_0108753 | 951 | 10 | 2.10% |
| hsa_circ_0110669 | 951 | 10 | 2.10% |
| hsa_circ_0113902 | 951 | 10 | 2.10% |
| hsa_circ_0118261 | 951 | 10 | 2.10% |
| hsa_circ_0130891 | 951 | 10 | 2.10% |
| hsa_circ_0135660 | 951 | 10 | 2.10% |
| hsa_circ_0069314 | 5994 | 63 | 2.10% |
| hsa_circ_0019989 | 1237 | 13 | 2.10% |
| hsa_circ_0103184 | 1713 | 18 | 2.10% |
| hsa_circ_0001757 | 1142 | 12 | 2.10% |
| hsa_circ_0015197 | 1142 | 12 | 2.10% |
| hsa_circ_0043630 | 1047 | 11 | 2.10% |
| hsa_circ_0125183 | 1047 | 11 | 2.10% |
| hsa_circ_0129626 | 1047 | 11 | 2.10% |
| hsa_circ_0137164 | 1047 | 11 | 2.10% |
| hsa_circ_0015985 | 1428 | 15 | 2.10% |
| hsa_circ_0068551 | 1428 | 15 | 2.10% |
| hsa_circ_0014738 | 952 | 10 | 2.10% |

| | | | |
|---|---|---|---|
| hsa_circ_0034530 | 952 | 10 | 2.10% |
| hsa_circ_0052777 | 952 | 10 | 2.10% |
| hsa_circ_0053069 | 952 | 10 | 2.10% |
| hsa_circ_0064825 | 952 | 10 | 2.10% |
| hsa_circ_0068614 | 952 | 10 | 2.10% |
| hsa_circ_0071969 | 952 | 10 | 2.10% |
| hsa_circ_0079574 | 952 | 10 | 2.10% |
| hsa_circ_0081634 | 952 | 10 | 2.10% |
| hsa_circ_0085605 | 952 | 10 | 2.10% |
| hsa_circ_0087770 | 952 | 10 | 2.10% |
| hsa_circ_0104794 | 952 | 10 | 2.10% |
| hsa_circ_0111403 | 952 | 10 | 2.10% |
| hsa_circ_0116732 | 952 | 10 | 2.10% |
| hsa_circ_0139345 | 952 | 10 | 2.10% |
| hsa_circ_0031527 | 1143 | 12 | 2.10% |
| hsa_circ_0060741 | 1143 | 12 | 2.10% |
| hsa_circ_0004574 | 1048 | 11 | 2.10% |
| hsa_circ_0052454 | 1048 | 11 | 2.10% |
| hsa_circ_0080937 | 1048 | 11 | 2.10% |
| hsa_circ_0012749 | 953 | 10 | 2.10% |
| hsa_circ_0012757 | 953 | 10 | 2.10% |
| hsa_circ_0015487 | 953 | 10 | 2.10% |
| hsa_circ_0021100 | 953 | 10 | 2.10% |
| hsa_circ_0023081 | 953 | 10 | 2.10% |
| hsa_circ_0028233 | 953 | 10 | 2.10% |
| hsa_circ_0041472 | 953 | 10 | 2.10% |
| hsa_circ_0044647 | 953 | 10 | 2.10% |
| hsa_circ_0063846 | 953 | 10 | 2.10% |
| hsa_circ_0064464 | 953 | 10 | 2.10% |
| hsa_circ_0094529 | 953 | 10 | 2.10% |
| hsa_circ_0100605 | 953 | 10 | 2.10% |
| hsa_circ_0107843 | 953 | 10 | 2.10% |
| hsa_circ_0114809 | 953 | 10 | 2.10% |
| hsa_circ_0116054 | 953 | 10 | 2.10% |
| hsa_circ_0125266 | 953 | 10 | 2.10% |
| hsa_circ_0128263 | 953 | 10 | 2.10% |
| hsa_circ_0133296 | 953 | 10 | 2.10% |
| hsa_circ_0106553 | 1239 | 13 | 2.10% |
| hsa_circ_0008567 | 1049 | 11 | 2.10% |
| hsa_circ_0011408 | 1049 | 11 | 2.10% |
| hsa_circ_0029376 | 1049 | 11 | 2.10% |
| hsa_circ_0051615 | 1049 | 11 | 2.10% |
| hsa_circ_0059924 | 1049 | 11 | 2.10% |
| hsa_circ_0066395 | 1049 | 11 | 2.10% |
| hsa_circ_0096804 | 1049 | 11 | 2.10% |
| hsa_circ_0047290 | 1240 | 13 | 2.10% |
| hsa_circ_0018621 | 954 | 10 | 2.10% |
| hsa_circ_0047803 | 954 | 10 | 2.10% |
| hsa_circ_0053768 | 954 | 10 | 2.10% |
| hsa_circ_0067446 | 954 | 10 | 2.10% |

| | | | |
|---|---|---|---|
| hsa_circ_0075659 | 954 | 10 | 2.10% |
| hsa_circ_0083995 | 954 | 10 | 2.10% |
| hsa_circ_0094149 | 954 | 10 | 2.10% |
| hsa_circ_0103541 | 954 | 10 | 2.10% |
| hsa_circ_0103760 | 954 | 10 | 2.10% |
| hsa_circ_0104274 | 954 | 10 | 2.10% |
| hsa_circ_0110794 | 954 | 10 | 2.10% |
| hsa_circ_0117603 | 954 | 10 | 2.10% |
| hsa_circ_0117953 | 954 | 10 | 2.10% |
| hsa_circ_0121955 | 954 | 10 | 2.10% |
| hsa_circ_0132336 | 954 | 10 | 2.10% |
| hsa_circ_0136839 | 954 | 10 | 2.10% |
| hsa_circ_0118572 | 3435 | 36 | 2.10% |
| hsa_circ_0026147 | 1145 | 12 | 2.10% |
| hsa_circ_0097486 | 1145 | 12 | 2.10% |
| hsa_circ_0006028 | 1050 | 11 | 2.10% |
| hsa_circ_0062005 | 1050 | 11 | 2.10% |
| hsa_circ_0115386 | 1050 | 11 | 2.10% |
| hsa_circ_0134968 | 1050 | 11 | 2.10% |
| hsa_circ_0137629 | 1050 | 11 | 2.10% |
| hsa_circ_0043857 | 1241 | 13 | 2.10% |
| hsa_circ_0021566 | 5538 | 58 | 2.09% |
| hsa_circ_0010385 | 1146 | 12 | 2.09% |
| hsa_circ_0029019 | 1146 | 12 | 2.09% |
| hsa_circ_0109404 | 1146 | 12 | 2.09% |
| hsa_circ_0127456 | 1146 | 12 | 2.09% |
| hsa_circ_0025124 | 955 | 10 | 2.09% |
| hsa_circ_0027415 | 955 | 10 | 2.09% |
| hsa_circ_0028423 | 955 | 10 | 2.09% |
| hsa_circ_0031034 | 955 | 10 | 2.09% |
| hsa_circ_0032694 | 955 | 10 | 2.09% |
| hsa_circ_0044723 | 955 | 10 | 2.09% |
| hsa_circ_0046041 | 955 | 10 | 2.09% |
| hsa_circ_0046239 | 955 | 10 | 2.09% |
| hsa_circ_0050979 | 955 | 10 | 2.09% |
| hsa_circ_0057014 | 955 | 10 | 2.09% |
| hsa_circ_0057098 | 955 | 10 | 2.09% |
| hsa_circ_0082456 | 955 | 10 | 2.09% |
| hsa_circ_0086840 | 955 | 10 | 2.09% |
| hsa_circ_0098337 | 955 | 10 | 2.09% |
| hsa_circ_0109469 | 955 | 10 | 2.09% |
| hsa_circ_0113540 | 955 | 10 | 2.09% |
| hsa_circ_0135635 | 955 | 10 | 2.09% |
| hsa_circ_0009572 | 1051 | 11 | 2.09% |
| hsa_circ_0058378 | 1051 | 11 | 2.09% |
| hsa_circ_0074883 | 1051 | 11 | 2.09% |
| hsa_circ_0102807 | 1051 | 11 | 2.09% |
| hsa_circ_0104567 | 1051 | 11 | 2.09% |
| hsa_circ_0110602 | 1051 | 11 | 2.09% |
| hsa_circ_0120163 | 1051 | 11 | 2.09% |

| | | | |
|---|---|---|---|
| hsa_circ_0042292 | 1147 | 12 | 2.09% |
| hsa_circ_0085942 | 1147 | 12 | 2.09% |
| hsa_circ_0118362 | 1147 | 12 | 2.09% |
| hsa_circ_0005869 | 956 | 10 | 2.09% |
| hsa_circ_0017407 | 956 | 10 | 2.09% |
| hsa_circ_0031546 | 956 | 10 | 2.09% |
| hsa_circ_0036440 | 956 | 10 | 2.09% |
| hsa_circ_0043867 | 956 | 10 | 2.09% |
| hsa_circ_0054243 | 956 | 10 | 2.09% |
| hsa_circ_0056620 | 956 | 10 | 2.09% |
| hsa_circ_0061840 | 956 | 10 | 2.09% |
| hsa_circ_0064597 | 956 | 10 | 2.09% |
| hsa_circ_0064680 | 956 | 10 | 2.09% |
| hsa_circ_0068289 | 956 | 10 | 2.09% |
| hsa_circ_0068891 | 956 | 10 | 2.09% |
| hsa_circ_0098576 | 956 | 10 | 2.09% |
| hsa_circ_0103065 | 956 | 10 | 2.09% |
| hsa_circ_0106602 | 956 | 10 | 2.09% |
| hsa_circ_0132392 | 956 | 10 | 2.09% |
| hsa_circ_0009286 | 1243 | 13 | 2.09% |
| hsa_circ_0088850 | 1052 | 11 | 2.09% |
| hsa_circ_0092012 | 1052 | 11 | 2.09% |
| hsa_circ_0106977 | 1052 | 11 | 2.09% |
| hsa_circ_0122800 | 1052 | 11 | 2.09% |
| hsa_circ_0073520 | 1913 | 20 | 2.09% |
| hsa_circ_0046283 | 1148 | 12 | 2.09% |
| hsa_circ_0072960 | 1148 | 12 | 2.09% |
| hsa_circ_0074499 | 1148 | 12 | 2.09% |
| hsa_circ_0013273 | 957 | 10 | 2.09% |
| hsa_circ_0017349 | 957 | 10 | 2.09% |
| hsa_circ_0019518 | 957 | 10 | 2.09% |
| hsa_circ_0023457 | 957 | 10 | 2.09% |
| hsa_circ_0027126 | 957 | 10 | 2.09% |
| hsa_circ_0033399 | 957 | 10 | 2.09% |
| hsa_circ_0054836 | 957 | 10 | 2.09% |
| hsa_circ_0081738 | 957 | 10 | 2.09% |
| hsa_circ_0096232 | 957 | 10 | 2.09% |
| hsa_circ_0097590 | 957 | 10 | 2.09% |
| hsa_circ_0100321 | 957 | 10 | 2.09% |
| hsa_circ_0100457 | 957 | 10 | 2.09% |
| hsa_circ_0101021 | 957 | 10 | 2.09% |
| hsa_circ_0106371 | 957 | 10 | 2.09% |
| hsa_circ_0138662 | 957 | 10 | 2.09% |
| hsa_circ_0031578 | 2680 | 28 | 2.09% |
| hsa_circ_0027276 | 1053 | 11 | 2.09% |
| hsa_circ_0036602 | 1053 | 11 | 2.09% |
| hsa_circ_0060317 | 1053 | 11 | 2.09% |
| hsa_circ_0078637 | 1053 | 11 | 2.09% |
| hsa_circ_0078757 | 1053 | 11 | 2.09% |
| hsa_circ_0105896 | 1053 | 11 | 2.09% |

| | | | |
|---|---|---|---|
| hsa_circ_0121438 | 1053 | 11 | 2.09% |
| hsa_circ_0121483 | 1053 | 11 | 2.09% |
| hsa_circ_0125555 | 1053 | 11 | 2.09% |
| hsa_circ_0006516 | 1149 | 12 | 2.09% |
| hsa_circ_0034426 | 1149 | 12 | 2.09% |
| hsa_circ_0082231 | 1149 | 12 | 2.09% |
| hsa_circ_0088884 | 1149 | 12 | 2.09% |
| hsa_circ_0131702 | 1245 | 13 | 2.09% |
| hsa_circ_0008635 | 958 | 10 | 2.09% |
| hsa_circ_0010865 | 958 | 10 | 2.09% |
| hsa_circ_0019686 | 958 | 10 | 2.09% |
| hsa_circ_0021245 | 958 | 10 | 2.09% |
| hsa_circ_0027374 | 958 | 10 | 2.09% |
| hsa_circ_0046314 | 958 | 10 | 2.09% |
| hsa_circ_0052489 | 958 | 10 | 2.09% |
| hsa_circ_0054223 | 958 | 10 | 2.09% |
| hsa_circ_0058998 | 958 | 10 | 2.09% |
| hsa_circ_0075471 | 958 | 10 | 2.09% |
| hsa_circ_0082145 | 958 | 10 | 2.09% |
| hsa_circ_0113883 | 958 | 10 | 2.09% |
| hsa_circ_0114518 | 958 | 10 | 2.09% |
| hsa_circ_0115706 | 958 | 10 | 2.09% |
| hsa_circ_0139138 | 3354 | 35 | 2.09% |
| hsa_circ_0060869 | 1246 | 13 | 2.09% |
| hsa_circ_0071474 | 1246 | 13 | 2.09% |
| hsa_circ_0081697 | 1342 | 14 | 2.09% |
| hsa_circ_0005838 | 959 | 10 | 2.09% |
| hsa_circ_0005975 | 959 | 10 | 2.09% |
| hsa_circ_0007900 | 959 | 10 | 2.09% |
| hsa_circ_0031149 | 959 | 10 | 2.09% |
| hsa_circ_0033649 | 959 | 10 | 2.09% |
| hsa_circ_0037346 | 959 | 10 | 2.09% |
| hsa_circ_0041837 | 959 | 10 | 2.09% |
| hsa_circ_0062529 | 959 | 10 | 2.09% |
| hsa_circ_0081409 | 959 | 10 | 2.09% |
| hsa_circ_0082684 | 959 | 10 | 2.09% |
| hsa_circ_0110178 | 959 | 10 | 2.09% |
| hsa_circ_0118263 | 959 | 10 | 2.09% |
| hsa_circ_0130299 | 959 | 10 | 2.09% |
| hsa_circ_0136757 | 959 | 10 | 2.09% |
| hsa_circ_0040980 | 1055 | 11 | 2.09% |
| hsa_circ_0127265 | 1055 | 11 | 2.09% |
| hsa_circ_0128124 | 1055 | 11 | 2.09% |
| hsa_circ_0116137 | 1920 | 20 | 2.08% |
| hsa_circ_0099495 | 1152 | 12 | 2.08% |
| hsa_circ_0116290 | 1152 | 12 | 2.08% |
| hsa_circ_0125920 | 1152 | 12 | 2.08% |
| hsa_circ_0021062 | 1056 | 11 | 2.08% |
| hsa_circ_0039550 | 1056 | 11 | 2.08% |
| hsa_circ_0042075 | 1056 | 11 | 2.08% |

| | | | |
|---|---|---|---|
| hsa_circ_0060335 | 1056 | 11 | 2.08% |
| hsa_circ_0068312 | 1056 | 11 | 2.08% |
| hsa_circ_0115519 | 1056 | 11 | 2.08% |
| hsa_circ_0117841 | 1056 | 11 | 2.08% |
| hsa_circ_0011057 | 960 | 10 | 2.08% |
| hsa_circ_0025731 | 960 | 10 | 2.08% |
| hsa_circ_0026374 | 960 | 10 | 2.08% |
| hsa_circ_0030067 | 960 | 10 | 2.08% |
| hsa_circ_0039963 | 960 | 10 | 2.08% |
| hsa_circ_0044686 | 960 | 10 | 2.08% |
| hsa_circ_0045421 | 960 | 10 | 2.08% |
| hsa_circ_0045731 | 960 | 10 | 2.08% |
| hsa_circ_0047518 | 960 | 10 | 2.08% |
| hsa_circ_0047590 | 960 | 10 | 2.08% |
| hsa_circ_0049955 | 960 | 10 | 2.08% |
| hsa_circ_0057360 | 960 | 10 | 2.08% |
| hsa_circ_0070662 | 960 | 10 | 2.08% |
| hsa_circ_0071080 | 960 | 10 | 2.08% |
| hsa_circ_0072112 | 960 | 10 | 2.08% |
| hsa_circ_0076646 | 960 | 10 | 2.08% |
| hsa_circ_0077664 | 960 | 10 | 2.08% |
| hsa_circ_0078647 | 960 | 10 | 2.08% |
| hsa_circ_0086304 | 960 | 10 | 2.08% |
| hsa_circ_0090464 | 960 | 10 | 2.08% |
| hsa_circ_0094608 | 960 | 10 | 2.08% |
| hsa_circ_0094810 | 960 | 10 | 2.08% |
| hsa_circ_0098359 | 960 | 10 | 2.08% |
| hsa_circ_0100056 | 960 | 10 | 2.08% |
| hsa_circ_0108799 | 960 | 10 | 2.08% |
| hsa_circ_0114034 | 960 | 10 | 2.08% |
| hsa_circ_0124973 | 960 | 10 | 2.08% |
| hsa_circ_0089837 | 2785 | 29 | 2.08% |
| hsa_circ_0089839 | 2785 | 29 | 2.08% |
| hsa_circ_0032494 | 2113 | 22 | 2.08% |
| hsa_circ_0059102 | 1537 | 16 | 2.08% |
| hsa_circ_0102821 | 1441 | 15 | 2.08% |
| hsa_circ_0002754 | 1057 | 11 | 2.08% |
| hsa_circ_0006811 | 1057 | 11 | 2.08% |
| hsa_circ_0015960 | 1057 | 11 | 2.08% |
| hsa_circ_0082857 | 1057 | 11 | 2.08% |
| hsa_circ_0122548 | 1057 | 11 | 2.08% |
| hsa_circ_0125982 | 1057 | 11 | 2.08% |
| hsa_circ_0136364 | 1057 | 11 | 2.08% |
| hsa_circ_0024236 | 961 | 10 | 2.08% |
| hsa_circ_0025180 | 961 | 10 | 2.08% |
| hsa_circ_0025866 | 961 | 10 | 2.08% |
| hsa_circ_0060614 | 961 | 10 | 2.08% |
| hsa_circ_0063441 | 961 | 10 | 2.08% |
| hsa_circ_0081495 | 961 | 10 | 2.08% |
| hsa_circ_0081669 | 961 | 10 | 2.08% |

| | | | |
|---|---|---|---|
| hsa_circ_0084281 | 961 | 10 | 2.08% |
| hsa_circ_0135420 | 961 | 10 | 2.08% |
| hsa_circ_0009432 | 1250 | 13 | 2.08% |
| hsa_circ_0025223 | 1250 | 13 | 2.08% |
| hsa_circ_0003013 | 1154 | 12 | 2.08% |
| hsa_circ_0077496 | 1154 | 12 | 2.08% |
| hsa_circ_0097701 | 1154 | 12 | 2.08% |
| hsa_circ_0015916 | 1058 | 11 | 2.08% |
| hsa_circ_0017567 | 1058 | 11 | 2.08% |
| hsa_circ_0070811 | 1058 | 11 | 2.08% |
| hsa_circ_0093029 | 1058 | 11 | 2.08% |
| hsa_circ_0057376 | 1539 | 16 | 2.08% |
| hsa_circ_0019163 | 962 | 10 | 2.08% |
| hsa_circ_0041940 | 962 | 10 | 2.08% |
| hsa_circ_0045846 | 962 | 10 | 2.08% |
| hsa_circ_0052779 | 962 | 10 | 2.08% |
| hsa_circ_0055248 | 962 | 10 | 2.08% |
| hsa_circ_0062023 | 962 | 10 | 2.08% |
| hsa_circ_0071559 | 962 | 10 | 2.08% |
| hsa_circ_0071640 | 962 | 10 | 2.08% |
| hsa_circ_0073880 | 962 | 10 | 2.08% |
| hsa_circ_0075429 | 962 | 10 | 2.08% |
| hsa_circ_0079921 | 962 | 10 | 2.08% |
| hsa_circ_0084481 | 962 | 10 | 2.08% |
| hsa_circ_0094473 | 962 | 10 | 2.08% |
| hsa_circ_0095693 | 962 | 10 | 2.08% |
| hsa_circ_0098648 | 962 | 10 | 2.08% |
| hsa_circ_0113557 | 962 | 10 | 2.08% |
| hsa_circ_0133676 | 962 | 10 | 2.08% |
| hsa_circ_0027927 | 1155 | 12 | 2.08% |
| hsa_circ_0059393 | 1155 | 12 | 2.08% |
| hsa_circ_0125919 | 1155 | 12 | 2.08% |
| hsa_circ_0019651 | 1059 | 11 | 2.08% |
| hsa_circ_0051654 | 1059 | 11 | 2.08% |
| hsa_circ_0063660 | 1059 | 11 | 2.08% |
| hsa_circ_0065700 | 1059 | 11 | 2.08% |
| hsa_circ_0083319 | 1059 | 11 | 2.08% |
| hsa_circ_0093975 | 1059 | 11 | 2.08% |
| hsa_circ_0104388 | 1059 | 11 | 2.08% |
| hsa_circ_0138360 | 1059 | 11 | 2.08% |
| hsa_circ_0001348 | 963 | 10 | 2.08% |
| hsa_circ_0009452 | 963 | 10 | 2.08% |
| hsa_circ_0024328 | 963 | 10 | 2.08% |
| hsa_circ_0043830 | 963 | 10 | 2.08% |
| hsa_circ_0050691 | 963 | 10 | 2.08% |
| hsa_circ_0051211 | 963 | 10 | 2.08% |
| hsa_circ_0056768 | 963 | 10 | 2.08% |
| hsa_circ_0076685 | 963 | 10 | 2.08% |
| hsa_circ_0099043 | 963 | 10 | 2.08% |
| hsa_circ_0106786 | 963 | 10 | 2.08% |

| | | | |
|---|---|---|---|
| hsa_circ_0140655 | 963 | 10 | 2.08% |
| hsa_circ_0022470 | 1156 | 12 | 2.08% |
| hsa_circ_0063967 | 1156 | 12 | 2.08% |
| hsa_circ_0088555 | 1156 | 12 | 2.08% |
| hsa_circ_0135425 | 1156 | 12 | 2.08% |
| hsa_circ_0017112 | 1060 | 11 | 2.08% |
| hsa_circ_0017949 | 1060 | 11 | 2.08% |
| hsa_circ_0022975 | 1060 | 11 | 2.08% |
| hsa_circ_0046152 | 1060 | 11 | 2.08% |
| hsa_circ_0057113 | 1060 | 11 | 2.08% |
| hsa_circ_0082772 | 1060 | 11 | 2.08% |
| hsa_circ_0083018 | 1060 | 11 | 2.08% |
| hsa_circ_0119113 | 1060 | 11 | 2.08% |
| hsa_circ_0000112 | 964 | 10 | 2.07% |
| hsa_circ_0025648 | 964 | 10 | 2.07% |
| hsa_circ_0030617 | 964 | 10 | 2.07% |
| hsa_circ_0032456 | 964 | 10 | 2.07% |
| hsa_circ_0036597 | 964 | 10 | 2.07% |
| hsa_circ_0064673 | 964 | 10 | 2.07% |
| hsa_circ_0071752 | 964 | 10 | 2.07% |
| hsa_circ_0075007 | 964 | 10 | 2.07% |
| hsa_circ_0077193 | 964 | 10 | 2.07% |
| hsa_circ_0079518 | 964 | 10 | 2.07% |
| hsa_circ_0101812 | 964 | 10 | 2.07% |
| hsa_circ_0122473 | 964 | 10 | 2.07% |
| hsa_circ_0131793 | 3953 | 41 | 2.07% |
| hsa_circ_0032611 | 1157 | 12 | 2.07% |
| hsa_circ_0079513 | 1157 | 12 | 2.07% |
| hsa_circ_0083446 | 1157 | 12 | 2.07% |
| hsa_circ_0123141 | 1157 | 12 | 2.07% |
| hsa_circ_0002079 | 1061 | 11 | 2.07% |
| hsa_circ_0011898 | 1061 | 11 | 2.07% |
| hsa_circ_0049039 | 1061 | 11 | 2.07% |
| hsa_circ_0094363 | 1061 | 11 | 2.07% |
| hsa_circ_0099993 | 1061 | 11 | 2.07% |
| hsa_circ_0133087 | 1447 | 15 | 2.07% |
| hsa_circ_0006417 | 2605 | 27 | 2.07% |
| hsa_circ_0029475 | 1158 | 12 | 2.07% |
| hsa_circ_0085872 | 1158 | 12 | 2.07% |
| hsa_circ_0011175 | 965 | 10 | 2.07% |
| hsa_circ_0017444 | 965 | 10 | 2.07% |
| hsa_circ_0018790 | 965 | 10 | 2.07% |
| hsa_circ_0030280 | 965 | 10 | 2.07% |
| hsa_circ_0032705 | 965 | 10 | 2.07% |
| hsa_circ_0057236 | 965 | 10 | 2.07% |
| hsa_circ_0073734 | 965 | 10 | 2.07% |
| hsa_circ_0076913 | 965 | 10 | 2.07% |
| hsa_circ_0085744 | 965 | 10 | 2.07% |
| hsa_circ_0117030 | 965 | 10 | 2.07% |
| hsa_circ_0127668 | 965 | 10 | 2.07% |

| | | | |
|---|---|---|---|
| hsa_circ_0139560 | 965 | 10 | 2.07% |
| hsa_circ_0018285 | 1448 | 15 | 2.07% |
| hsa_circ_0064092 | 1448 | 15 | 2.07% |
| hsa_circ_0018009 | 1062 | 11 | 2.07% |
| hsa_circ_0032734 | 1062 | 11 | 2.07% |
| hsa_circ_0044000 | 1062 | 11 | 2.07% |
| hsa_circ_0055498 | 1062 | 11 | 2.07% |
| hsa_circ_0063879 | 1062 | 11 | 2.07% |
| hsa_circ_0134880 | 1062 | 11 | 2.07% |
| hsa_circ_0013683 | 1159 | 12 | 2.07% |
| hsa_circ_0026551 | 966 | 10 | 2.07% |
| hsa_circ_0031784 | 966 | 10 | 2.07% |
| hsa_circ_0033646 | 966 | 10 | 2.07% |
| hsa_circ_0041380 | 966 | 10 | 2.07% |
| hsa_circ_0041600 | 966 | 10 | 2.07% |
| hsa_circ_0050375 | 966 | 10 | 2.07% |
| hsa_circ_0054288 | 966 | 10 | 2.07% |
| hsa_circ_0083274 | 966 | 10 | 2.07% |
| hsa_circ_0088377 | 966 | 10 | 2.07% |
| hsa_circ_0093937 | 966 | 10 | 2.07% |
| hsa_circ_0094598 | 966 | 10 | 2.07% |
| hsa_circ_0097239 | 966 | 10 | 2.07% |
| hsa_circ_0122917 | 966 | 10 | 2.07% |
| hsa_circ_0130659 | 966 | 10 | 2.07% |
| hsa_circ_0137825 | 966 | 10 | 2.07% |
| hsa_circ_0095262 | 3188 | 33 | 2.07% |
| hsa_circ_0017479 | 1063 | 11 | 2.07% |
| hsa_circ_0024226 | 1063 | 11 | 2.07% |
| hsa_circ_0063610 | 1063 | 11 | 2.07% |
| hsa_circ_0086709 | 1063 | 11 | 2.07% |
| hsa_circ_0119825 | 1063 | 11 | 2.07% |
| hsa_circ_0133803 | 1063 | 11 | 2.07% |
| hsa_circ_0140643 | 1063 | 11 | 2.07% |
| hsa_circ_0026352 | 1160 | 12 | 2.07% |
| hsa_circ_0027511 | 1160 | 12 | 2.07% |
| hsa_circ_0128944 | 1160 | 12 | 2.07% |
| hsa_circ_0043612 | 967 | 10 | 2.07% |
| hsa_circ_0044408 | 967 | 10 | 2.07% |
| hsa_circ_0051811 | 967 | 10 | 2.07% |
| hsa_circ_0060659 | 967 | 10 | 2.07% |
| hsa_circ_0067687 | 967 | 10 | 2.07% |
| hsa_circ_0073107 | 967 | 10 | 2.07% |
| hsa_circ_0086655 | 967 | 10 | 2.07% |
| hsa_circ_0095922 | 967 | 10 | 2.07% |
| hsa_circ_0106314 | 967 | 10 | 2.07% |
| hsa_circ_0108103 | 967 | 10 | 2.07% |
| hsa_circ_0108417 | 967 | 10 | 2.07% |
| hsa_circ_0110459 | 967 | 10 | 2.07% |
| hsa_circ_0111298 | 967 | 10 | 2.07% |
| hsa_circ_0111520 | 967 | 10 | 2.07% |

| | | | |
|---|---|---|---|
| hsa_circ_0115509 | 967 | 10 | 2.07% |
| hsa_circ_0122650 | 967 | 10 | 2.07% |
| hsa_circ_0126416 | 967 | 10 | 2.07% |
| hsa_circ_0014748 | 1064 | 11 | 2.07% |
| hsa_circ_0028003 | 1064 | 11 | 2.07% |
| hsa_circ_0028066 | 1064 | 11 | 2.07% |
| hsa_circ_0069399 | 1064 | 11 | 2.07% |
| hsa_circ_0089690 | 1064 | 11 | 2.07% |
| hsa_circ_0123997 | 1064 | 11 | 2.07% |
| hsa_circ_0128033 | 1064 | 11 | 2.07% |
| hsa_circ_0130310 | 1064 | 11 | 2.07% |
| hsa_circ_0021824 | 1935 | 20 | 2.07% |
| hsa_circ_0018666 | 1161 | 12 | 2.07% |
| hsa_circ_0080944 | 1161 | 12 | 2.07% |
| hsa_circ_0010353 | 968 | 10 | 2.07% |
| hsa_circ_0018421 | 968 | 10 | 2.07% |
| hsa_circ_0022368 | 968 | 10 | 2.07% |
| hsa_circ_0036451 | 968 | 10 | 2.07% |
| hsa_circ_0038198 | 968 | 10 | 2.07% |
| hsa_circ_0038543 | 968 | 10 | 2.07% |
| hsa_circ_0042329 | 968 | 10 | 2.07% |
| hsa_circ_0076940 | 968 | 10 | 2.07% |
| hsa_circ_0079375 | 968 | 10 | 2.07% |
| hsa_circ_0087859 | 968 | 10 | 2.07% |
| hsa_circ_0100596 | 968 | 10 | 2.07% |
| hsa_circ_0115131 | 968 | 10 | 2.07% |
| hsa_circ_0118643 | 968 | 10 | 2.07% |
| hsa_circ_0122208 | 968 | 10 | 2.07% |
| hsa_circ_0126712 | 968 | 10 | 2.07% |
| hsa_circ_0131311 | 968 | 10 | 2.07% |
| hsa_circ_0078962 | 5421 | 56 | 2.07% |
| hsa_circ_0021940 | 1065 | 11 | 2.07% |
| hsa_circ_0057352 | 1065 | 11 | 2.07% |
| hsa_circ_0061553 | 1065 | 11 | 2.07% |
| hsa_circ_0082883 | 1065 | 11 | 2.07% |
| hsa_circ_0083323 | 1065 | 11 | 2.07% |
| hsa_circ_0084200 | 1065 | 11 | 2.07% |
| hsa_circ_0124151 | 1065 | 11 | 2.07% |
| hsa_circ_0041217 | 1162 | 12 | 2.07% |
| hsa_circ_0068082 | 5232 | 54 | 2.06% |
| hsa_circ_0000121 | 969 | 10 | 2.06% |
| hsa_circ_0010112 | 969 | 10 | 2.06% |
| hsa_circ_0015766 | 969 | 10 | 2.06% |
| hsa_circ_0036913 | 969 | 10 | 2.06% |
| hsa_circ_0046662 | 969 | 10 | 2.06% |
| hsa_circ_0060903 | 969 | 10 | 2.06% |
| hsa_circ_0064633 | 969 | 10 | 2.06% |
| hsa_circ_0079817 | 969 | 10 | 2.06% |
| hsa_circ_0086825 | 969 | 10 | 2.06% |
| hsa_circ_0099665 | 969 | 10 | 2.06% |

| | | | |
|---|---|---|---|
| hsa_circ_0103757 | 969 | 10 | 2.06% |
| hsa_circ_0112730 | 969 | 10 | 2.06% |
| hsa_circ_0113024 | 969 | 10 | 2.06% |
| hsa_circ_0116642 | 969 | 10 | 2.06% |
| hsa_circ_0129156 | 969 | 10 | 2.06% |
| hsa_circ_0029678 | 1066 | 11 | 2.06% |
| hsa_circ_0036706 | 1066 | 11 | 2.06% |
| hsa_circ_0040267 | 1066 | 11 | 2.06% |
| hsa_circ_0044149 | 1066 | 11 | 2.06% |
| hsa_circ_0072031 | 1066 | 11 | 2.06% |
| hsa_circ_0097412 | 1066 | 11 | 2.06% |
| hsa_circ_0122148 | 1066 | 11 | 2.06% |
| hsa_circ_0047277 | 1163 | 12 | 2.06% |
| hsa_circ_0050259 | 1260 | 13 | 2.06% |
| hsa_circ_0030838 | 1746 | 18 | 2.06% |
| hsa_circ_0072094 | 1358 | 14 | 2.06% |
| hsa_circ_0095047 | 1358 | 14 | 2.06% |
| hsa_circ_0128604 | 1358 | 14 | 2.06% |
| hsa_circ_0041634 | 1261 | 13 | 2.06% |
| hsa_circ_0007753 | 1067 | 11 | 2.06% |
| hsa_circ_0025037 | 1067 | 11 | 2.06% |
| hsa_circ_0041328 | 1067 | 11 | 2.06% |
| hsa_circ_0058880 | 1067 | 11 | 2.06% |
| hsa_circ_0064176 | 1067 | 11 | 2.06% |
| hsa_circ_0009802 | 970 | 10 | 2.06% |
| hsa_circ_0020315 | 970 | 10 | 2.06% |
| hsa_circ_0023915 | 970 | 10 | 2.06% |
| hsa_circ_0029169 | 970 | 10 | 2.06% |
| hsa_circ_0035028 | 970 | 10 | 2.06% |
| hsa_circ_0042188 | 970 | 10 | 2.06% |
| hsa_circ_0050547 | 970 | 10 | 2.06% |
| hsa_circ_0059132 | 970 | 10 | 2.06% |
| hsa_circ_0060608 | 970 | 10 | 2.06% |
| hsa_circ_0062246 | 970 | 10 | 2.06% |
| hsa_circ_0077751 | 970 | 10 | 2.06% |
| hsa_circ_0078239 | 970 | 10 | 2.06% |
| hsa_circ_0098279 | 970 | 10 | 2.06% |
| hsa_circ_0111612 | 970 | 10 | 2.06% |
| hsa_circ_0114815 | 970 | 10 | 2.06% |
| hsa_circ_0114923 | 970 | 10 | 2.06% |
| hsa_circ_0131261 | 970 | 10 | 2.06% |
| hsa_circ_0024794 | 1068 | 11 | 2.06% |
| hsa_circ_0073045 | 1068 | 11 | 2.06% |
| hsa_circ_0080745 | 1068 | 11 | 2.06% |
| hsa_circ_0083056 | 1068 | 11 | 2.06% |
| hsa_circ_0088054 | 1068 | 11 | 2.06% |
| hsa_circ_0105121 | 1068 | 11 | 2.06% |
| hsa_circ_0123167 | 1068 | 11 | 2.06% |
| hsa_circ_0135484 | 1068 | 11 | 2.06% |
| hsa_circ_0132803 | 1942 | 20 | 2.06% |

| | | | |
|---|---|---|---|
| hsa_circ_0006080 | 971 | 10 | 2.06% |
| hsa_circ_0016328 | 971 | 10 | 2.06% |
| hsa_circ_0019731 | 971 | 10 | 2.06% |
| hsa_circ_0022064 | 971 | 10 | 2.06% |
| hsa_circ_0030692 | 971 | 10 | 2.06% |
| hsa_circ_0041098 | 971 | 10 | 2.06% |
| hsa_circ_0042189 | 971 | 10 | 2.06% |
| hsa_circ_0046860 | 971 | 10 | 2.06% |
| hsa_circ_0061917 | 971 | 10 | 2.06% |
| hsa_circ_0063376 | 971 | 10 | 2.06% |
| hsa_circ_0079359 | 971 | 10 | 2.06% |
| hsa_circ_0087892 | 971 | 10 | 2.06% |
| hsa_circ_0091602 | 971 | 10 | 2.06% |
| hsa_circ_0111664 | 971 | 10 | 2.06% |
| hsa_circ_0112772 | 971 | 10 | 2.06% |
| hsa_circ_0090599 | 1360 | 14 | 2.06% |
| hsa_circ_0020817 | 1263 | 13 | 2.06% |
| hsa_circ_0011288 | 1069 | 11 | 2.06% |
| hsa_circ_0016943 | 1069 | 11 | 2.06% |
| hsa_circ_0037164 | 1069 | 11 | 2.06% |
| hsa_circ_0058525 | 1069 | 11 | 2.06% |
| hsa_circ_0136432 | 1069 | 11 | 2.06% |
| hsa_circ_0007157 | 972 | 10 | 2.06% |
| hsa_circ_0007690 | 972 | 10 | 2.06% |
| hsa_circ_0020579 | 972 | 10 | 2.06% |
| hsa_circ_0023902 | 972 | 10 | 2.06% |
| hsa_circ_0034062 | 972 | 10 | 2.06% |
| hsa_circ_0057510 | 972 | 10 | 2.06% |
| hsa_circ_0072488 | 972 | 10 | 2.06% |
| hsa_circ_0073077 | 972 | 10 | 2.06% |
| hsa_circ_0074713 | 972 | 10 | 2.06% |
| hsa_circ_0076595 | 972 | 10 | 2.06% |
| hsa_circ_0098989 | 972 | 10 | 2.06% |
| hsa_circ_0106336 | 972 | 10 | 2.06% |
| hsa_circ_0106537 | 972 | 10 | 2.06% |
| hsa_circ_0116083 | 972 | 10 | 2.06% |
| hsa_circ_0117638 | 972 | 10 | 2.06% |
| hsa_circ_0124669 | 972 | 10 | 2.06% |
| hsa_circ_0127046 | 972 | 10 | 2.06% |
| hsa_circ_0134717 | 972 | 10 | 2.06% |
| hsa_circ_0126173 | 2431 | 25 | 2.06% |
| hsa_circ_0028706 | 1167 | 12 | 2.06% |
| hsa_circ_0042690 | 1167 | 12 | 2.06% |
| hsa_circ_0051394 | 1070 | 11 | 2.06% |
| hsa_circ_0058566 | 1070 | 11 | 2.06% |
| hsa_circ_0120213 | 1070 | 11 | 2.06% |
| hsa_circ_0003302 | 973 | 10 | 2.06% |
| hsa_circ_0010534 | 973 | 10 | 2.06% |
| hsa_circ_0014596 | 973 | 10 | 2.06% |
| hsa_circ_0024567 | 973 | 10 | 2.06% |

| | | | |
|---|---|---|---|
| hsa_circ_0034536 | 973 | 10 | 2.06% |
| hsa_circ_0035837 | 973 | 10 | 2.06% |
| hsa_circ_0037851 | 973 | 10 | 2.06% |
| hsa_circ_0044898 | 973 | 10 | 2.06% |
| hsa_circ_0046320 | 973 | 10 | 2.06% |
| hsa_circ_0062436 | 973 | 10 | 2.06% |
| hsa_circ_0064738 | 973 | 10 | 2.06% |
| hsa_circ_0065128 | 973 | 10 | 2.06% |
| hsa_circ_0076301 | 973 | 10 | 2.06% |
| hsa_circ_0082466 | 973 | 10 | 2.06% |
| hsa_circ_0084389 | 973 | 10 | 2.06% |
| hsa_circ_0104907 | 973 | 10 | 2.06% |
| hsa_circ_0111105 | 973 | 10 | 2.06% |
| hsa_circ_0128868 | 973 | 10 | 2.06% |
| hsa_circ_0135581 | 973 | 10 | 2.06% |
| hsa_circ_0045782 | 1168 | 12 | 2.05% |
| hsa_circ_0068792 | 1168 | 12 | 2.05% |
| hsa_circ_0090562 | 1168 | 12 | 2.05% |
| hsa_circ_0078262 | 1363 | 14 | 2.05% |
| hsa_circ_0013465 | 1071 | 11 | 2.05% |
| hsa_circ_0042514 | 1071 | 11 | 2.05% |
| hsa_circ_0053108 | 1071 | 11 | 2.05% |
| hsa_circ_0110228 | 1071 | 11 | 2.05% |
| hsa_circ_0112069 | 1071 | 11 | 2.05% |
| hsa_circ_0114488 | 1071 | 11 | 2.05% |
| hsa_circ_0117566 | 1071 | 11 | 2.05% |
| hsa_circ_0123157 | 1071 | 11 | 2.05% |
| hsa_circ_0135392 | 1071 | 11 | 2.05% |
| hsa_circ_0038897 | 2629 | 27 | 2.05% |
| hsa_circ_0049081 | 1266 | 13 | 2.05% |
| hsa_circ_0119647 | 1266 | 13 | 2.05% |
| hsa_circ_0132297 | 1461 | 15 | 2.05% |
| hsa_circ_0026769 | 974 | 10 | 2.05% |
| hsa_circ_0033643 | 974 | 10 | 2.05% |
| hsa_circ_0038581 | 974 | 10 | 2.05% |
| hsa_circ_0043111 | 974 | 10 | 2.05% |
| hsa_circ_0056747 | 974 | 10 | 2.05% |
| hsa_circ_0057091 | 974 | 10 | 2.05% |
| hsa_circ_0061759 | 974 | 10 | 2.05% |
| hsa_circ_0066615 | 974 | 10 | 2.05% |
| hsa_circ_0067345 | 974 | 10 | 2.05% |
| hsa_circ_0086130 | 974 | 10 | 2.05% |
| hsa_circ_0093977 | 974 | 10 | 2.05% |
| hsa_circ_0106136 | 974 | 10 | 2.05% |
| hsa_circ_0110197 | 974 | 10 | 2.05% |
| hsa_circ_0123491 | 974 | 10 | 2.05% |
| hsa_circ_0125949 | 974 | 10 | 2.05% |
| hsa_circ_0127212 | 974 | 10 | 2.05% |
| hsa_circ_0137557 | 974 | 10 | 2.05% |
| hsa_circ_0042108 | 1169 | 12 | 2.05% |

| | | | |
|---|---|---|---|
| hsa_circ_0102078 | 1169 | 12 | 2.05% |
| hsa_circ_0015382 | 1072 | 11 | 2.05% |
| hsa_circ_0019270 | 1072 | 11 | 2.05% |
| hsa_circ_0036470 | 1072 | 11 | 2.05% |
| hsa_circ_0038839 | 1072 | 11 | 2.05% |
| hsa_circ_0087879 | 1072 | 11 | 2.05% |
| hsa_circ_0095496 | 1072 | 11 | 2.05% |
| hsa_circ_0119919 | 1072 | 11 | 2.05% |
| hsa_circ_0053955 | 1170 | 12 | 2.05% |
| hsa_circ_0006511 | 975 | 10 | 2.05% |
| hsa_circ_0018915 | 975 | 10 | 2.05% |
| hsa_circ_0033420 | 975 | 10 | 2.05% |
| hsa_circ_0041967 | 975 | 10 | 2.05% |
| hsa_circ_0048043 | 975 | 10 | 2.05% |
| hsa_circ_0060366 | 975 | 10 | 2.05% |
| hsa_circ_0061419 | 975 | 10 | 2.05% |
| hsa_circ_0096831 | 975 | 10 | 2.05% |
| hsa_circ_0098036 | 975 | 10 | 2.05% |
| hsa_circ_0098495 | 975 | 10 | 2.05% |
| hsa_circ_0104145 | 975 | 10 | 2.05% |
| hsa_circ_0111469 | 975 | 10 | 2.05% |
| hsa_circ_0017210 | 1073 | 11 | 2.05% |
| hsa_circ_0041145 | 1073 | 11 | 2.05% |
| hsa_circ_0058522 | 1073 | 11 | 2.05% |
| hsa_circ_0067129 | 1073 | 11 | 2.05% |
| hsa_circ_0024260 | 1561 | 16 | 2.05% |
| hsa_circ_0040698 | 1171 | 12 | 2.05% |
| hsa_circ_0093962 | 2635 | 27 | 2.05% |
| hsa_circ_0025239 | 976 | 10 | 2.05% |
| hsa_circ_0029724 | 976 | 10 | 2.05% |
| hsa_circ_0035448 | 976 | 10 | 2.05% |
| hsa_circ_0040292 | 976 | 10 | 2.05% |
| hsa_circ_0042353 | 976 | 10 | 2.05% |
| hsa_circ_0051736 | 976 | 10 | 2.05% |
| hsa_circ_0058365 | 976 | 10 | 2.05% |
| hsa_circ_0066163 | 976 | 10 | 2.05% |
| hsa_circ_0072687 | 976 | 10 | 2.05% |
| hsa_circ_0074423 | 976 | 10 | 2.05% |
| hsa_circ_0084624 | 976 | 10 | 2.05% |
| hsa_circ_0085127 | 976 | 10 | 2.05% |
| hsa_circ_0089910 | 976 | 10 | 2.05% |
| hsa_circ_0090987 | 976 | 10 | 2.05% |
| hsa_circ_0098156 | 976 | 10 | 2.05% |
| hsa_circ_0098159 | 976 | 10 | 2.05% |
| hsa_circ_0112544 | 976 | 10 | 2.05% |
| hsa_circ_0112846 | 976 | 10 | 2.05% |
| hsa_circ_0118734 | 976 | 10 | 2.05% |
| hsa_circ_0133865 | 976 | 10 | 2.05% |
| hsa_circ_0135597 | 976 | 10 | 2.05% |
| hsa_circ_0035911 | 1074 | 11 | 2.05% |

| | | | |
|---|---|---|---|
| hsa_circ_0065358 | 1074 | 11 | 2.05% |
| hsa_circ_0092721 | 1074 | 11 | 2.05% |
| hsa_circ_0119988 | 1074 | 11 | 2.05% |
| hsa_circ_0026158 | 1172 | 12 | 2.05% |
| hsa_circ_0119305 | 1172 | 12 | 2.05% |
| hsa_circ_0033171 | 1270 | 13 | 2.05% |
| hsa_circ_0027849 | 977 | 10 | 2.05% |
| hsa_circ_0030386 | 977 | 10 | 2.05% |
| hsa_circ_0040055 | 977 | 10 | 2.05% |
| hsa_circ_0040505 | 977 | 10 | 2.05% |
| hsa_circ_0045648 | 977 | 10 | 2.05% |
| hsa_circ_0048669 | 977 | 10 | 2.05% |
| hsa_circ_0048944 | 977 | 10 | 2.05% |
| hsa_circ_0062420 | 977 | 10 | 2.05% |
| hsa_circ_0067800 | 977 | 10 | 2.05% |
| hsa_circ_0076932 | 977 | 10 | 2.05% |
| hsa_circ_0090831 | 977 | 10 | 2.05% |
| hsa_circ_0100039 | 977 | 10 | 2.05% |
| hsa_circ_0101323 | 977 | 10 | 2.05% |
| hsa_circ_0117908 | 977 | 10 | 2.05% |
| hsa_circ_0137486 | 977 | 10 | 2.05% |
| hsa_circ_0140399 | 977 | 10 | 2.05% |
| hsa_circ_0109041 | 10552 | 108 | 2.05% |
| hsa_circ_0085035 | 1368 | 14 | 2.05% |
| hsa_circ_0009255 | 1075 | 11 | 2.05% |
| hsa_circ_0062707 | 1075 | 11 | 2.05% |
| hsa_circ_0066001 | 1075 | 11 | 2.05% |
| hsa_circ_0070115 | 1075 | 11 | 2.05% |
| hsa_circ_0104927 | 1075 | 11 | 2.05% |
| hsa_circ_0119804 | 1075 | 11 | 2.05% |
| hsa_circ_0115370 | 1173 | 12 | 2.05% |
| hsa_circ_0106790 | 2738 | 28 | 2.05% |
| hsa_circ_0025270 | 1369 | 14 | 2.05% |
| hsa_circ_0012922 | 978 | 10 | 2.04% |
| hsa_circ_0013266 | 978 | 10 | 2.04% |
| hsa_circ_0016497 | 978 | 10 | 2.04% |
| hsa_circ_0018288 | 978 | 10 | 2.04% |
| hsa_circ_0026980 | 978 | 10 | 2.04% |
| hsa_circ_0038949 | 978 | 10 | 2.04% |
| hsa_circ_0039826 | 978 | 10 | 2.04% |
| hsa_circ_0044616 | 978 | 10 | 2.04% |
| hsa_circ_0053939 | 978 | 10 | 2.04% |
| hsa_circ_0057629 | 978 | 10 | 2.04% |
| hsa_circ_0081405 | 978 | 10 | 2.04% |
| hsa_circ_0084113 | 978 | 10 | 2.04% |
| hsa_circ_0087365 | 978 | 10 | 2.04% |
| hsa_circ_0098340 | 978 | 10 | 2.04% |
| hsa_circ_0114147 | 978 | 10 | 2.04% |
| hsa_circ_0119309 | 978 | 10 | 2.04% |
| hsa_circ_0044238 | 1076 | 11 | 2.04% |

| | | | |
|---|---|---|---|
| hsa_circ_0072322 | 1076 | 11 | 2.04% |
| hsa_circ_0097168 | 1076 | 11 | 2.04% |
| hsa_circ_0117286 | 1076 | 11 | 2.04% |
| hsa_circ_0128758 | 1076 | 11 | 2.04% |
| hsa_circ_0123990 | 1174 | 12 | 2.04% |
| hsa_circ_0087473 | 1272 | 13 | 2.04% |
| hsa_circ_0095102 | 1370 | 14 | 2.04% |
| hsa_circ_0012259 | 979 | 10 | 2.04% |
| hsa_circ_0026281 | 979 | 10 | 2.04% |
| hsa_circ_0049469 | 979 | 10 | 2.04% |
| hsa_circ_0064101 | 979 | 10 | 2.04% |
| hsa_circ_0069286 | 979 | 10 | 2.04% |
| hsa_circ_0083119 | 979 | 10 | 2.04% |
| hsa_circ_0088510 | 979 | 10 | 2.04% |
| hsa_circ_0117227 | 979 | 10 | 2.04% |
| hsa_circ_0129417 | 979 | 10 | 2.04% |
| hsa_circ_0135811 | 979 | 10 | 2.04% |
| hsa_circ_0136145 | 979 | 10 | 2.04% |
| hsa_circ_0006824 | 1077 | 11 | 2.04% |
| hsa_circ_0009218 | 1077 | 11 | 2.04% |
| hsa_circ_0036179 | 1077 | 11 | 2.04% |
| hsa_circ_0072714 | 1077 | 11 | 2.04% |
| hsa_circ_0083144 | 1077 | 11 | 2.04% |
| hsa_circ_0023349 | 1175 | 12 | 2.04% |
| hsa_circ_0086587 | 2448 | 25 | 2.04% |
| hsa_circ_0121416 | 11168 | 114 | 2.04% |
| hsa_circ_0057116 | 1274 | 13 | 2.04% |
| hsa_circ_0069035 | 1274 | 13 | 2.04% |
| hsa_circ_0085822 | 1176 | 12 | 2.04% |
| hsa_circ_0107114 | 1176 | 12 | 2.04% |
| hsa_circ_0011467 | 1078 | 11 | 2.04% |
| hsa_circ_0021989 | 1078 | 11 | 2.04% |
| hsa_circ_0111460 | 1078 | 11 | 2.04% |
| hsa_circ_0136281 | 1078 | 11 | 2.04% |
| hsa_circ_0022458 | 980 | 10 | 2.04% |
| hsa_circ_0024455 | 980 | 10 | 2.04% |
| hsa_circ_0029670 | 980 | 10 | 2.04% |
| hsa_circ_0031989 | 980 | 10 | 2.04% |
| hsa_circ_0042777 | 980 | 10 | 2.04% |
| hsa_circ_0047462 | 980 | 10 | 2.04% |
| hsa_circ_0056710 | 980 | 10 | 2.04% |
| hsa_circ_0066245 | 980 | 10 | 2.04% |
| hsa_circ_0071961 | 980 | 10 | 2.04% |
| hsa_circ_0121111 | 980 | 10 | 2.04% |
| hsa_circ_0130977 | 980 | 10 | 2.04% |
| hsa_circ_0133462 | 980 | 10 | 2.04% |
| hsa_circ_0043666 | 1177 | 12 | 2.04% |
| hsa_circ_0025192 | 1079 | 11 | 2.04% |
| hsa_circ_0058060 | 1079 | 11 | 2.04% |
| hsa_circ_0063522 | 1079 | 11 | 2.04% |

| | | | |
|---|---|---|---|
| hsa_circ_0089673 | 1079 | 11 | 2.04% |
| hsa_circ_0101284 | 1079 | 11 | 2.04% |
| hsa_circ_0112374 | 1079 | 11 | 2.04% |
| hsa_circ_0113806 | 1079 | 11 | 2.04% |
| hsa_circ_0127400 | 1079 | 11 | 2.04% |
| hsa_circ_0003896 | 981 | 10 | 2.04% |
| hsa_circ_0014237 | 981 | 10 | 2.04% |
| hsa_circ_0023773 | 981 | 10 | 2.04% |
| hsa_circ_0029836 | 981 | 10 | 2.04% |
| hsa_circ_0033651 | 981 | 10 | 2.04% |
| hsa_circ_0038941 | 981 | 10 | 2.04% |
| hsa_circ_0050909 | 981 | 10 | 2.04% |
| hsa_circ_0058996 | 981 | 10 | 2.04% |
| hsa_circ_0059011 | 981 | 10 | 2.04% |
| hsa_circ_0061193 | 981 | 10 | 2.04% |
| hsa_circ_0063912 | 981 | 10 | 2.04% |
| hsa_circ_0068510 | 981 | 10 | 2.04% |
| hsa_circ_0077936 | 981 | 10 | 2.04% |
| hsa_circ_0086590 | 981 | 10 | 2.04% |
| hsa_circ_0087399 | 981 | 10 | 2.04% |
| hsa_circ_0088769 | 981 | 10 | 2.04% |
| hsa_circ_0094514 | 981 | 10 | 2.04% |
| hsa_circ_0098202 | 981 | 10 | 2.04% |
| hsa_circ_0099969 | 981 | 10 | 2.04% |
| hsa_circ_0100278 | 981 | 10 | 2.04% |
| hsa_circ_0112239 | 981 | 10 | 2.04% |
| hsa_circ_0115355 | 981 | 10 | 2.04% |
| hsa_circ_0121995 | 981 | 10 | 2.04% |
| hsa_circ_0129739 | 981 | 10 | 2.04% |
| hsa_circ_0132092 | 981 | 10 | 2.04% |
| hsa_circ_0136788 | 981 | 10 | 2.04% |
| hsa_circ_0138907 | 981 | 10 | 2.04% |
| hsa_circ_0123964 | 2846 | 29 | 2.04% |
| hsa_circ_0044154 | 1374 | 14 | 2.04% |
| hsa_circ_0047947 | 1276 | 13 | 2.04% |
| hsa_circ_0102938 | 1178 | 12 | 2.04% |
| hsa_circ_0137727 | 1178 | 12 | 2.04% |
| hsa_circ_0046425 | 1080 | 11 | 2.04% |
| hsa_circ_0048096 | 1080 | 11 | 2.04% |
| hsa_circ_0051832 | 1080 | 11 | 2.04% |
| hsa_circ_0053336 | 1080 | 11 | 2.04% |
| hsa_circ_0063564 | 1080 | 11 | 2.04% |
| hsa_circ_0064193 | 1080 | 11 | 2.04% |
| hsa_circ_0106466 | 1080 | 11 | 2.04% |
| hsa_circ_0087088 | 3437 | 35 | 2.04% |
| hsa_circ_0019041 | 982 | 10 | 2.04% |
| hsa_circ_0032895 | 982 | 10 | 2.04% |
| hsa_circ_0040728 | 982 | 10 | 2.04% |
| hsa_circ_0046973 | 982 | 10 | 2.04% |
| hsa_circ_0058591 | 982 | 10 | 2.04% |

| | | | |
|---|---|---|---|
| hsa_circ_0061834 | 982 | 10 | 2.04% |
| hsa_circ_0061853 | 982 | 10 | 2.04% |
| hsa_circ_0068410 | 982 | 10 | 2.04% |
| hsa_circ_0073174 | 982 | 10 | 2.04% |
| hsa_circ_0088989 | 982 | 10 | 2.04% |
| hsa_circ_0113821 | 982 | 10 | 2.04% |
| hsa_circ_0131304 | 982 | 10 | 2.04% |
| hsa_circ_0135915 | 982 | 10 | 2.04% |
| hsa_circ_0013472 | 1375 | 14 | 2.04% |
| hsa_circ_0123435 | 1277 | 13 | 2.04% |
| hsa_circ_0013006 | 1179 | 12 | 2.04% |
| hsa_circ_0037683 | 1179 | 12 | 2.04% |
| hsa_circ_0053267 | 1179 | 12 | 2.04% |
| hsa_circ_0104076 | 1179 | 12 | 2.04% |
| hsa_circ_0108858 | 1179 | 12 | 2.04% |
| hsa_circ_0078932 | 1474 | 15 | 2.04% |
| hsa_circ_0078971 | 1474 | 15 | 2.04% |
| hsa_circ_0022267 | 1081 | 11 | 2.04% |
| hsa_circ_0027908 | 1081 | 11 | 2.04% |
| hsa_circ_0105744 | 1081 | 11 | 2.04% |
| hsa_circ_0008118 | 983 | 10 | 2.03% |
| hsa_circ_0012496 | 983 | 10 | 2.03% |
| hsa_circ_0014169 | 983 | 10 | 2.03% |
| hsa_circ_0025934 | 983 | 10 | 2.03% |
| hsa_circ_0030110 | 983 | 10 | 2.03% |
| hsa_circ_0030783 | 983 | 10 | 2.03% |
| hsa_circ_0034916 | 983 | 10 | 2.03% |
| hsa_circ_0044812 | 983 | 10 | 2.03% |
| hsa_circ_0074271 | 983 | 10 | 2.03% |
| hsa_circ_0076116 | 983 | 10 | 2.03% |
| hsa_circ_0135317 | 983 | 10 | 2.03% |
| hsa_circ_0140410 | 983 | 10 | 2.03% |
| hsa_circ_0137042 | 1278 | 13 | 2.03% |
| hsa_circ_0004895 | 1180 | 12 | 2.03% |
| hsa_circ_0022909 | 1082 | 11 | 2.03% |
| hsa_circ_0031817 | 1082 | 11 | 2.03% |
| hsa_circ_0086252 | 1082 | 11 | 2.03% |
| hsa_circ_0003137 | 984 | 10 | 2.03% |
| hsa_circ_0012262 | 984 | 10 | 2.03% |
| hsa_circ_0022880 | 984 | 10 | 2.03% |
| hsa_circ_0067838 | 984 | 10 | 2.03% |
| hsa_circ_0073082 | 984 | 10 | 2.03% |
| hsa_circ_0081994 | 984 | 10 | 2.03% |
| hsa_circ_0082346 | 984 | 10 | 2.03% |
| hsa_circ_0103768 | 984 | 10 | 2.03% |
| hsa_circ_0108921 | 984 | 10 | 2.03% |
| hsa_circ_0123046 | 984 | 10 | 2.03% |
| hsa_circ_0126419 | 984 | 10 | 2.03% |
| hsa_circ_0127375 | 984 | 10 | 2.03% |
| hsa_circ_0128433 | 984 | 10 | 2.03% |

| | | | |
|---|---|---|---|
| hsa_circ_0129260 | 984 | 10 | 2.03% |
| hsa_circ_0140337 | 984 | 10 | 2.03% |
| hsa_circ_0002225 | 1083 | 11 | 2.03% |
| hsa_circ_0038347 | 1083 | 11 | 2.03% |
| hsa_circ_0125022 | 1083 | 11 | 2.03% |
| hsa_circ_0135305 | 1083 | 11 | 2.03% |
| hsa_circ_0119059 | 1280 | 13 | 2.03% |
| hsa_circ_0045482 | 1182 | 12 | 2.03% |
| hsa_circ_0111177 | 1182 | 12 | 2.03% |
| hsa_circ_0016087 | 985 | 10 | 2.03% |
| hsa_circ_0020351 | 985 | 10 | 2.03% |
| hsa_circ_0020801 | 985 | 10 | 2.03% |
| hsa_circ_0033193 | 985 | 10 | 2.03% |
| hsa_circ_0040761 | 985 | 10 | 2.03% |
| hsa_circ_0096658 | 985 | 10 | 2.03% |
| hsa_circ_0117651 | 985 | 10 | 2.03% |
| hsa_circ_0120487 | 985 | 10 | 2.03% |
| hsa_circ_0134214 | 985 | 10 | 2.03% |
| hsa_circ_0135991 | 985 | 10 | 2.03% |
| hsa_circ_0021272 | 1084 | 11 | 2.03% |
| hsa_circ_0047308 | 1084 | 11 | 2.03% |
| hsa_circ_0051085 | 1084 | 11 | 2.03% |
| hsa_circ_0053559 | 1084 | 11 | 2.03% |
| hsa_circ_0086015 | 1084 | 11 | 2.03% |
| hsa_circ_0095435 | 1084 | 11 | 2.03% |
| hsa_circ_0113839 | 1084 | 11 | 2.03% |
| hsa_circ_0126638 | 1084 | 11 | 2.03% |
| hsa_circ_0024782 | 1183 | 12 | 2.03% |
| hsa_circ_0025543 | 986 | 10 | 2.03% |
| hsa_circ_0033114 | 986 | 10 | 2.03% |
| hsa_circ_0055041 | 986 | 10 | 2.03% |
| hsa_circ_0063115 | 986 | 10 | 2.03% |
| hsa_circ_0068738 | 986 | 10 | 2.03% |
| hsa_circ_0085475 | 986 | 10 | 2.03% |
| hsa_circ_0089663 | 986 | 10 | 2.03% |
| hsa_circ_0101731 | 986 | 10 | 2.03% |
| hsa_circ_0102577 | 986 | 10 | 2.03% |
| hsa_circ_0103889 | 986 | 10 | 2.03% |
| hsa_circ_0119412 | 986 | 10 | 2.03% |
| hsa_circ_0046079 | 1282 | 13 | 2.03% |
| hsa_circ_0081924 | 1282 | 13 | 2.03% |
| hsa_circ_0120643 | 3649 | 37 | 2.03% |
| hsa_circ_0012637 | 1085 | 11 | 2.03% |
| hsa_circ_0014672 | 1085 | 11 | 2.03% |
| hsa_circ_0074577 | 1085 | 11 | 2.03% |
| hsa_circ_0089713 | 1085 | 11 | 2.03% |
| hsa_circ_0089847 | 1085 | 11 | 2.03% |
| hsa_circ_0096754 | 1085 | 11 | 2.03% |
| hsa_circ_0105045 | 1085 | 11 | 2.03% |
| hsa_circ_0117746 | 1085 | 11 | 2.03% |

| | | | |
|---|---|---|---|
| hsa_circ_0132357 | 1085 | 11 | 2.03% |
| hsa_circ_0086350 | 1480 | 15 | 2.03% |
| hsa_circ_0051465 | 1184 | 12 | 2.03% |
| hsa_circ_0099575 | 1184 | 12 | 2.03% |
| hsa_circ_0115020 | 1184 | 12 | 2.03% |
| hsa_circ_0049077 | 1283 | 13 | 2.03% |
| hsa_circ_0101954 | 1283 | 13 | 2.03% |
| hsa_circ_0044502 | 2270 | 23 | 2.03% |
| hsa_circ_0133268 | 2270 | 23 | 2.03% |
| hsa_circ_0010122 | 987 | 10 | 2.03% |
| hsa_circ_0022666 | 987 | 10 | 2.03% |
| hsa_circ_0028936 | 987 | 10 | 2.03% |
| hsa_circ_0030545 | 987 | 10 | 2.03% |
| hsa_circ_0033578 | 987 | 10 | 2.03% |
| hsa_circ_0053124 | 987 | 10 | 2.03% |
| hsa_circ_0060571 | 987 | 10 | 2.03% |
| hsa_circ_0062231 | 987 | 10 | 2.03% |
| hsa_circ_0065445 | 987 | 10 | 2.03% |
| hsa_circ_0065661 | 987 | 10 | 2.03% |
| hsa_circ_0065945 | 987 | 10 | 2.03% |
| hsa_circ_0070317 | 987 | 10 | 2.03% |
| hsa_circ_0091042 | 987 | 10 | 2.03% |
| hsa_circ_0099540 | 987 | 10 | 2.03% |
| hsa_circ_0101733 | 987 | 10 | 2.03% |
| hsa_circ_0102964 | 987 | 10 | 2.03% |
| hsa_circ_0108042 | 987 | 10 | 2.03% |
| hsa_circ_0111246 | 987 | 10 | 2.03% |
| hsa_circ_0138989 | 987 | 10 | 2.03% |
| hsa_circ_0139892 | 987 | 10 | 2.03% |
| hsa_circ_0039428 | 1086 | 11 | 2.03% |
| hsa_circ_0060457 | 1086 | 11 | 2.03% |
| hsa_circ_0065327 | 1086 | 11 | 2.03% |
| hsa_circ_0117571 | 1086 | 11 | 2.03% |
| hsa_circ_0049756 | 1185 | 12 | 2.03% |
| hsa_circ_0054443 | 1185 | 12 | 2.03% |
| hsa_circ_0082204 | 1185 | 12 | 2.03% |
| hsa_circ_0035971 | 988 | 10 | 2.02% |
| hsa_circ_0039836 | 988 | 10 | 2.02% |
| hsa_circ_0043620 | 988 | 10 | 2.02% |
| hsa_circ_0054608 | 988 | 10 | 2.02% |
| hsa_circ_0066723 | 988 | 10 | 2.02% |
| hsa_circ_0076264 | 988 | 10 | 2.02% |
| hsa_circ_0085766 | 988 | 10 | 2.02% |
| hsa_circ_0094168 | 988 | 10 | 2.02% |
| hsa_circ_0095150 | 988 | 10 | 2.02% |
| hsa_circ_0097782 | 988 | 10 | 2.02% |
| hsa_circ_0107966 | 988 | 10 | 2.02% |
| hsa_circ_0122964 | 988 | 10 | 2.02% |
| hsa_circ_0123278 | 988 | 10 | 2.02% |
| hsa_circ_0127589 | 988 | 10 | 2.02% |

| | | | |
|---|---|---|---|
| hsa_circ_0026317 | 1087 | 11 | 2.02% |
| hsa_circ_0060019 | 1087 | 11 | 2.02% |
| hsa_circ_0074518 | 1087 | 11 | 2.02% |
| hsa_circ_0074776 | 1087 | 11 | 2.02% |
| hsa_circ_0102812 | 1087 | 11 | 2.02% |
| hsa_circ_0128615 | 1087 | 11 | 2.02% |
| hsa_circ_0134064 | 1087 | 11 | 2.02% |
| hsa_circ_0125453 | 1186 | 12 | 2.02% |
| hsa_circ_0009193 | 1384 | 14 | 2.02% |
| hsa_circ_0031780 | 989 | 10 | 2.02% |
| hsa_circ_0032607 | 989 | 10 | 2.02% |
| hsa_circ_0054947 | 989 | 10 | 2.02% |
| hsa_circ_0066734 | 989 | 10 | 2.02% |
| hsa_circ_0068235 | 989 | 10 | 2.02% |
| hsa_circ_0072681 | 989 | 10 | 2.02% |
| hsa_circ_0080705 | 989 | 10 | 2.02% |
| hsa_circ_0110089 | 989 | 10 | 2.02% |
| hsa_circ_0114720 | 989 | 10 | 2.02% |
| hsa_circ_0119595 | 989 | 10 | 2.02% |
| hsa_circ_0139054 | 989 | 10 | 2.02% |
| hsa_circ_0038421 | 1088 | 11 | 2.02% |
| hsa_circ_0045975 | 1088 | 11 | 2.02% |
| hsa_circ_0079125 | 1088 | 11 | 2.02% |
| hsa_circ_0098149 | 1088 | 11 | 2.02% |
| hsa_circ_0100638 | 1088 | 11 | 2.02% |
| hsa_circ_0115992 | 1088 | 11 | 2.02% |
| hsa_circ_0074433 | 1187 | 12 | 2.02% |
| hsa_circ_0031101 | 1484 | 15 | 2.02% |
| hsa_circ_0003816 | 1089 | 11 | 2.02% |
| hsa_circ_0022313 | 1089 | 11 | 2.02% |
| hsa_circ_0030731 | 1089 | 11 | 2.02% |
| hsa_circ_0034936 | 1089 | 11 | 2.02% |
| hsa_circ_0055625 | 1089 | 11 | 2.02% |
| hsa_circ_0077761 | 1089 | 11 | 2.02% |
| hsa_circ_0136122 | 1089 | 11 | 2.02% |
| hsa_circ_0005700 | 990 | 10 | 2.02% |
| hsa_circ_0006753 | 990 | 10 | 2.02% |
| hsa_circ_0006772 | 990 | 10 | 2.02% |
| hsa_circ_0010445 | 990 | 10 | 2.02% |
| hsa_circ_0014949 | 990 | 10 | 2.02% |
| hsa_circ_0020786 | 990 | 10 | 2.02% |
| hsa_circ_0032877 | 990 | 10 | 2.02% |
| hsa_circ_0037580 | 990 | 10 | 2.02% |
| hsa_circ_0038056 | 990 | 10 | 2.02% |
| hsa_circ_0061580 | 990 | 10 | 2.02% |
| hsa_circ_0063881 | 990 | 10 | 2.02% |
| hsa_circ_0064431 | 990 | 10 | 2.02% |
| hsa_circ_0064692 | 990 | 10 | 2.02% |
| hsa_circ_0067766 | 990 | 10 | 2.02% |
| hsa_circ_0068313 | 990 | 10 | 2.02% |

| | | | |
|---|---|---|---|
| hsa_circ_0075656 | 990 | 10 | 2.02% |
| hsa_circ_0077449 | 990 | 10 | 2.02% |
| hsa_circ_0079270 | 990 | 10 | 2.02% |
| hsa_circ_0091940 | 990 | 10 | 2.02% |
| hsa_circ_0094261 | 990 | 10 | 2.02% |
| hsa_circ_0103015 | 990 | 10 | 2.02% |
| hsa_circ_0103966 | 990 | 10 | 2.02% |
| hsa_circ_0104728 | 990 | 10 | 2.02% |
| hsa_circ_0118648 | 990 | 10 | 2.02% |
| hsa_circ_0121942 | 990 | 10 | 2.02% |
| hsa_circ_0123196 | 990 | 10 | 2.02% |
| hsa_circ_0134162 | 990 | 10 | 2.02% |
| hsa_circ_0136505 | 990 | 10 | 2.02% |
| hsa_circ_0047686 | 2476 | 25 | 2.02% |
| hsa_circ_0138676 | 3368 | 34 | 2.02% |
| hsa_circ_0114170 | 1189 | 12 | 2.02% |
| hsa_circ_0002555 | 1090 | 11 | 2.02% |
| hsa_circ_0013016 | 1090 | 11 | 2.02% |
| hsa_circ_0044223 | 1090 | 11 | 2.02% |
| hsa_circ_0062999 | 1090 | 11 | 2.02% |
| hsa_circ_0095490 | 1090 | 11 | 2.02% |
| hsa_circ_0095833 | 1090 | 11 | 2.02% |
| hsa_circ_0097176 | 1090 | 11 | 2.02% |
| hsa_circ_0105139 | 1090 | 11 | 2.02% |
| hsa_circ_0124434 | 1090 | 11 | 2.02% |
| hsa_circ_0011947 | 991 | 10 | 2.02% |
| hsa_circ_0014651 | 991 | 10 | 2.02% |
| hsa_circ_0018349 | 991 | 10 | 2.02% |
| hsa_circ_0039667 | 991 | 10 | 2.02% |
| hsa_circ_0043263 | 991 | 10 | 2.02% |
| hsa_circ_0043413 | 991 | 10 | 2.02% |
| hsa_circ_0044104 | 991 | 10 | 2.02% |
| hsa_circ_0054925 | 991 | 10 | 2.02% |
| hsa_circ_0066921 | 991 | 10 | 2.02% |
| hsa_circ_0088815 | 991 | 10 | 2.02% |
| hsa_circ_0091797 | 991 | 10 | 2.02% |
| hsa_circ_0105512 | 991 | 10 | 2.02% |
| hsa_circ_0108834 | 991 | 10 | 2.02% |
| hsa_circ_0121763 | 991 | 10 | 2.02% |
| hsa_circ_0134476 | 991 | 10 | 2.02% |
| hsa_circ_0036660 | 1190 | 12 | 2.02% |
| hsa_circ_0126690 | 1190 | 12 | 2.02% |
| hsa_circ_0030475 | 1091 | 11 | 2.02% |
| hsa_circ_0055493 | 1091 | 11 | 2.02% |
| hsa_circ_0059196 | 1091 | 11 | 2.02% |
| hsa_circ_0107841 | 1091 | 11 | 2.02% |
| hsa_circ_0013613 | 992 | 10 | 2.02% |
| hsa_circ_0015895 | 992 | 10 | 2.02% |
| hsa_circ_0036294 | 992 | 10 | 2.02% |
| hsa_circ_0037899 | 992 | 10 | 2.02% |

| | | | |
|---|---|---|---|
| hsa_circ_0039263 | 992 | 10 | 2.02% |
| hsa_circ_0048121 | 992 | 10 | 2.02% |
| hsa_circ_0050591 | 992 | 10 | 2.02% |
| hsa_circ_0052861 | 992 | 10 | 2.02% |
| hsa_circ_0066027 | 992 | 10 | 2.02% |
| hsa_circ_0071506 | 992 | 10 | 2.02% |
| hsa_circ_0082839 | 992 | 10 | 2.02% |
| hsa_circ_0122280 | 992 | 10 | 2.02% |
| hsa_circ_0124968 | 992 | 10 | 2.02% |
| hsa_circ_0129895 | 992 | 10 | 2.02% |
| hsa_circ_0024597 | 1191 | 12 | 2.02% |
| hsa_circ_0057306 | 1191 | 12 | 2.02% |
| hsa_circ_0064805 | 1191 | 12 | 2.02% |
| hsa_circ_0021419 | 1092 | 11 | 2.01% |
| hsa_circ_0022847 | 1092 | 11 | 2.01% |
| hsa_circ_0030949 | 1092 | 11 | 2.01% |
| hsa_circ_0052383 | 1092 | 11 | 2.01% |
| hsa_circ_0053905 | 1092 | 11 | 2.01% |
| hsa_circ_0054117 | 1092 | 11 | 2.01% |
| hsa_circ_0064892 | 1092 | 11 | 2.01% |
| hsa_circ_0086928 | 1092 | 11 | 2.01% |
| hsa_circ_0103199 | 1092 | 11 | 2.01% |
| hsa_circ_0007638 | 993 | 10 | 2.01% |
| hsa_circ_0023221 | 993 | 10 | 2.01% |
| hsa_circ_0023292 | 993 | 10 | 2.01% |
| hsa_circ_0024859 | 993 | 10 | 2.01% |
| hsa_circ_0030207 | 993 | 10 | 2.01% |
| hsa_circ_0031552 | 993 | 10 | 2.01% |
| hsa_circ_0035056 | 993 | 10 | 2.01% |
| hsa_circ_0038573 | 993 | 10 | 2.01% |
| hsa_circ_0045823 | 993 | 10 | 2.01% |
| hsa_circ_0047484 | 993 | 10 | 2.01% |
| hsa_circ_0049022 | 993 | 10 | 2.01% |
| hsa_circ_0055531 | 993 | 10 | 2.01% |
| hsa_circ_0058287 | 993 | 10 | 2.01% |
| hsa_circ_0072935 | 993 | 10 | 2.01% |
| hsa_circ_0091763 | 993 | 10 | 2.01% |
| hsa_circ_0092219 | 993 | 10 | 2.01% |
| hsa_circ_0102732 | 993 | 10 | 2.01% |
| hsa_circ_0103416 | 993 | 10 | 2.01% |
| hsa_circ_0108449 | 993 | 10 | 2.01% |
| hsa_circ_0099641 | 1291 | 13 | 2.01% |
| hsa_circ_0121901 | 10528 | 106 | 2.01% |
| hsa_circ_0113571 | 1093 | 11 | 2.01% |
| hsa_circ_0122278 | 1093 | 11 | 2.01% |
| hsa_circ_0127080 | 1093 | 11 | 2.01% |
| hsa_circ_0002306 | 994 | 10 | 2.01% |
| hsa_circ_0008555 | 994 | 10 | 2.01% |
| hsa_circ_0013343 | 994 | 10 | 2.01% |
| hsa_circ_0024799 | 994 | 10 | 2.01% |

| | | | |
|---|---|---|---|
| hsa_circ_0033637 | 994 | 10 | 2.01% |
| hsa_circ_0043097 | 994 | 10 | 2.01% |
| hsa_circ_0049293 | 994 | 10 | 2.01% |
| hsa_circ_0050189 | 994 | 10 | 2.01% |
| hsa_circ_0051248 | 994 | 10 | 2.01% |
| hsa_circ_0066673 | 994 | 10 | 2.01% |
| hsa_circ_0075242 | 994 | 10 | 2.01% |
| hsa_circ_0076586 | 994 | 10 | 2.01% |
| hsa_circ_0079589 | 994 | 10 | 2.01% |
| hsa_circ_0080928 | 994 | 10 | 2.01% |
| hsa_circ_0095657 | 994 | 10 | 2.01% |
| hsa_circ_0103434 | 994 | 10 | 2.01% |
| hsa_circ_0104025 | 994 | 10 | 2.01% |
| hsa_circ_0104133 | 994 | 10 | 2.01% |
| hsa_circ_0105658 | 994 | 10 | 2.01% |
| hsa_circ_0107130 | 994 | 10 | 2.01% |
| hsa_circ_0119937 | 994 | 10 | 2.01% |
| hsa_circ_0130108 | 994 | 10 | 2.01% |
| hsa_circ_0007381 | 1193 | 12 | 2.01% |
| hsa_circ_0031629 | 1193 | 12 | 2.01% |
| hsa_circ_0048490 | 1193 | 12 | 2.01% |
| hsa_circ_0088042 | 1193 | 12 | 2.01% |
| hsa_circ_0038217 | 1094 | 11 | 2.01% |
| hsa_circ_0038835 | 1094 | 11 | 2.01% |
| hsa_circ_0041623 | 1094 | 11 | 2.01% |
| hsa_circ_0052569 | 1094 | 11 | 2.01% |
| hsa_circ_0074227 | 1094 | 11 | 2.01% |
| hsa_circ_0104596 | 1094 | 11 | 2.01% |
| hsa_circ_0123900 | 1094 | 11 | 2.01% |
| hsa_circ_0099686 | 4277 | 43 | 2.01% |
| hsa_circ_0088713 | 1691 | 17 | 2.01% |
| hsa_circ_0138735 | 1791 | 18 | 2.01% |
| hsa_circ_0039394 | 1194 | 12 | 2.01% |
| hsa_circ_0040989 | 1194 | 12 | 2.01% |
| hsa_circ_0054789 | 1194 | 12 | 2.01% |
| hsa_circ_0132494 | 1194 | 12 | 2.01% |
| hsa_circ_0011616 | 995 | 10 | 2.01% |
| hsa_circ_0015436 | 995 | 10 | 2.01% |
| hsa_circ_0025319 | 995 | 10 | 2.01% |
| hsa_circ_0031820 | 995 | 10 | 2.01% |
| hsa_circ_0048419 | 995 | 10 | 2.01% |
| hsa_circ_0075276 | 995 | 10 | 2.01% |
| hsa_circ_0079834 | 995 | 10 | 2.01% |
| hsa_circ_0084993 | 995 | 10 | 2.01% |
| hsa_circ_0085844 | 995 | 10 | 2.01% |
| hsa_circ_0095975 | 995 | 10 | 2.01% |
| hsa_circ_0105015 | 995 | 10 | 2.01% |
| hsa_circ_0113000 | 995 | 10 | 2.01% |
| hsa_circ_0115053 | 995 | 10 | 2.01% |
| hsa_circ_0120047 | 995 | 10 | 2.01% |

| | | | |
|---|---|---|---|
| hsa_circ_0125732 | 995 | 10 | 2.01% |
| hsa_circ_0130440 | 995 | 10 | 2.01% |
| hsa_circ_0133521 | 995 | 10 | 2.01% |
| hsa_circ_0137066 | 995 | 10 | 2.01% |
| hsa_circ_0010688 | 1294 | 13 | 2.01% |
| hsa_circ_0019372 | 1294 | 13 | 2.01% |
| hsa_circ_0006699 | 1095 | 11 | 2.01% |
| hsa_circ_0011481 | 1095 | 11 | 2.01% |
| hsa_circ_0066137 | 1095 | 11 | 2.01% |
| hsa_circ_0106116 | 1095 | 11 | 2.01% |
| hsa_circ_0110903 | 1095 | 11 | 2.01% |
| hsa_circ_0011522 | 3783 | 38 | 2.01% |
| hsa_circ_0014774 | 1195 | 12 | 2.01% |
| hsa_circ_0026632 | 1195 | 12 | 2.01% |
| hsa_circ_0018219 | 996 | 10 | 2.01% |
| hsa_circ_0024760 | 996 | 10 | 2.01% |
| hsa_circ_0056182 | 996 | 10 | 2.01% |
| hsa_circ_0059645 | 996 | 10 | 2.01% |
| hsa_circ_0063155 | 996 | 10 | 2.01% |
| hsa_circ_0066731 | 996 | 10 | 2.01% |
| hsa_circ_0068005 | 996 | 10 | 2.01% |
| hsa_circ_0070305 | 996 | 10 | 2.01% |
| hsa_circ_0077291 | 996 | 10 | 2.01% |
| hsa_circ_0080895 | 996 | 10 | 2.01% |
| hsa_circ_0082829 | 996 | 10 | 2.01% |
| hsa_circ_0090188 | 996 | 10 | 2.01% |
| hsa_circ_0118015 | 3288 | 33 | 2.01% |
| hsa_circ_0005708 | 1096 | 11 | 2.01% |
| hsa_circ_0011537 | 1096 | 11 | 2.01% |
| hsa_circ_0025609 | 1096 | 11 | 2.01% |
| hsa_circ_0079123 | 1096 | 11 | 2.01% |
| hsa_circ_0093758 | 1096 | 11 | 2.01% |
| hsa_circ_0097686 | 1096 | 11 | 2.01% |
| hsa_circ_0139409 | 5282 | 53 | 2.01% |
| hsa_circ_0044010 | 1196 | 12 | 2.01% |
| hsa_circ_0070366 | 1196 | 12 | 2.01% |
| hsa_circ_0103338 | 1196 | 12 | 2.01% |
| hsa_circ_0130208 | 1196 | 12 | 2.01% |
| hsa_circ_0013695 | 997 | 10 | 2.01% |
| hsa_circ_0020898 | 997 | 10 | 2.01% |
| hsa_circ_0022098 | 997 | 10 | 2.01% |
| hsa_circ_0043465 | 997 | 10 | 2.01% |
| hsa_circ_0046244 | 997 | 10 | 2.01% |
| hsa_circ_0048497 | 997 | 10 | 2.01% |
| hsa_circ_0104387 | 997 | 10 | 2.01% |
| hsa_circ_0105609 | 997 | 10 | 2.01% |
| hsa_circ_0131876 | 997 | 10 | 2.01% |
| hsa_circ_0021393 | 1097 | 11 | 2.01% |
| hsa_circ_0028068 | 1097 | 11 | 2.01% |
| hsa_circ_0043988 | 1097 | 11 | 2.01% |

| | | | |
|---|---|---|---|
| hsa_circ_0059193 | 1097 | 11 | 2.01% |
| hsa_circ_0083493 | 1097 | 11 | 2.01% |
| hsa_circ_0091149 | 1097 | 11 | 2.01% |
| hsa_circ_0114960 | 1097 | 11 | 2.01% |
| hsa_circ_0033051 | 1197 | 12 | 2.01% |
| hsa_circ_0016533 | 998 | 10 | 2.00% |
| hsa_circ_0016689 | 998 | 10 | 2.00% |
| hsa_circ_0017602 | 998 | 10 | 2.00% |
| hsa_circ_0018139 | 998 | 10 | 2.00% |
| hsa_circ_0033650 | 998 | 10 | 2.00% |
| hsa_circ_0039144 | 998 | 10 | 2.00% |
| hsa_circ_0044688 | 998 | 10 | 2.00% |
| hsa_circ_0067542 | 998 | 10 | 2.00% |
| hsa_circ_0068864 | 998 | 10 | 2.00% |
| hsa_circ_0082557 | 998 | 10 | 2.00% |
| hsa_circ_0082685 | 998 | 10 | 2.00% |
| hsa_circ_0090473 | 998 | 10 | 2.00% |
| hsa_circ_0091193 | 998 | 10 | 2.00% |
| hsa_circ_0095424 | 998 | 10 | 2.00% |
| hsa_circ_0110020 | 998 | 10 | 2.00% |
| hsa_circ_0133517 | 998 | 10 | 2.00% |
| hsa_circ_0137738 | 998 | 10 | 2.00% |
| hsa_circ_0028803 | 1098 | 11 | 2.00% |
| hsa_circ_0055956 | 1098 | 11 | 2.00% |
| hsa_circ_0065543 | 1098 | 11 | 2.00% |
| hsa_circ_0089454 | 1098 | 11 | 2.00% |
| hsa_circ_0114635 | 1098 | 11 | 2.00% |
| hsa_circ_0123799 | 1098 | 11 | 2.00% |
| hsa_circ_0095405 | 1198 | 12 | 2.00% |
| hsa_circ_0099932 | 1198 | 12 | 2.00% |
| hsa_circ_0108174 | 1298 | 13 | 2.00% |
| hsa_circ_0108620 | 1298 | 13 | 2.00% |
| hsa_circ_0137795 | 1298 | 13 | 2.00% |
| hsa_circ_0097634 | 3495 | 35 | 2.00% |
| hsa_circ_0014288 | 999 | 10 | 2.00% |
| hsa_circ_0043418 | 999 | 10 | 2.00% |
| hsa_circ_0049196 | 999 | 10 | 2.00% |
| hsa_circ_0081588 | 999 | 10 | 2.00% |
| hsa_circ_0082330 | 999 | 10 | 2.00% |
| hsa_circ_0103811 | 999 | 10 | 2.00% |
| hsa_circ_0104556 | 999 | 10 | 2.00% |
| hsa_circ_0105157 | 999 | 10 | 2.00% |
| hsa_circ_0108096 | 999 | 10 | 2.00% |
| hsa_circ_0110518 | 999 | 10 | 2.00% |
| hsa_circ_0126952 | 999 | 10 | 2.00% |
| hsa_circ_0132507 | 999 | 10 | 2.00% |
| hsa_circ_0138936 | 999 | 10 | 2.00% |
| hsa_circ_0026470 | 1099 | 11 | 2.00% |
| hsa_circ_0037881 | 1099 | 11 | 2.00% |
| hsa_circ_0043406 | 1099 | 11 | 2.00% |

| | | | |
|---|---|---|---|
| hsa_circ_0056843 | 1099 | 11 | 2.00% |
| hsa_circ_0094329 | 1199 | 12 | 2.00% |
| hsa_circ_0111422 | 1299 | 13 | 2.00% |
| hsa_circ_0029164 | 1399 | 14 | 2.00% |
| hsa_circ_0096206 | 4200 | 42 | 2.00% |
| hsa_circ_0122386 | 1300 | 13 | 2.00% |
| hsa_circ_0066710 | 1200 | 12 | 2.00% |
| hsa_circ_0114829 | 1200 | 12 | 2.00% |
| hsa_circ_0120298 | 1200 | 12 | 2.00% |
| hsa_circ_0140160 | 1200 | 12 | 2.00% |
| hsa_circ_0031169 | 1100 | 11 | 2.00% |
| hsa_circ_0035240 | 1100 | 11 | 2.00% |
| hsa_circ_0056409 | 1100 | 11 | 2.00% |
| hsa_circ_0069807 | 1100 | 11 | 2.00% |
| hsa_circ_0089005 | 1100 | 11 | 2.00% |
| hsa_circ_0091308 | 1100 | 11 | 2.00% |
| hsa_circ_0131641 | 1100 | 11 | 2.00% |
| hsa_circ_0002508 | 1000 | 10 | 2.00% |
| hsa_circ_0009068 | 1000 | 10 | 2.00% |
| hsa_circ_0010016 | 1000 | 10 | 2.00% |
| hsa_circ_0027349 | 1000 | 10 | 2.00% |
| hsa_circ_0027419 | 1000 | 10 | 2.00% |
| hsa_circ_0028996 | 1000 | 10 | 2.00% |
| hsa_circ_0029922 | 1000 | 10 | 2.00% |
| hsa_circ_0040449 | 1000 | 10 | 2.00% |
| hsa_circ_0045136 | 1000 | 10 | 2.00% |
| hsa_circ_0051619 | 1000 | 10 | 2.00% |
| hsa_circ_0073121 | 1000 | 10 | 2.00% |
| hsa_circ_0130838 | 1000 | 10 | 2.00% |
| hsa_circ_0026595 | 1601 | 16 | 2.00% |
| hsa_circ_0037905 | 1501 | 15 | 2.00% |
| hsa_circ_0139713 | 1301 | 13 | 2.00% |
| hsa_circ_0013385 | 1201 | 12 | 2.00% |
| hsa_circ_0044465 | 1201 | 12 | 2.00% |
| hsa_circ_0060015 | 1201 | 12 | 2.00% |
| hsa_circ_0057052 | 1101 | 11 | 2.00% |
| hsa_circ_0065764 | 1101 | 11 | 2.00% |
| hsa_circ_0079878 | 1101 | 11 | 2.00% |
| hsa_circ_0001973 | 1001 | 10 | 2.00% |
| hsa_circ_0017773 | 1001 | 10 | 2.00% |
| hsa_circ_0037300 | 1001 | 10 | 2.00% |
| hsa_circ_0039333 | 1001 | 10 | 2.00% |
| hsa_circ_0047560 | 1001 | 10 | 2.00% |
| hsa_circ_0051141 | 1001 | 10 | 2.00% |
| hsa_circ_0063631 | 1001 | 10 | 2.00% |
| hsa_circ_0089769 | 1001 | 10 | 2.00% |
| hsa_circ_0097788 | 1001 | 10 | 2.00% |
| hsa_circ_0101124 | 1001 | 10 | 2.00% |
| hsa_circ_0114068 | 1001 | 10 | 2.00% |
| hsa_circ_0120141 | 1001 | 10 | 2.00% |

| | | | |
|---|---|---|---|
| hsa_circ_0140424 | 1001 | 10 | 2.00% |
| hsa_circ_0042291 | 1202 | 12 | 2.00% |
| hsa_circ_0059240 | 1202 | 12 | 2.00% |
| hsa_circ_0106722 | 2304 | 23 | 2.00% |
| hsa_circ_0026588 | 1102 | 11 | 2.00% |
| hsa_circ_0036685 | 1102 | 11 | 2.00% |
| hsa_circ_0037443 | 1102 | 11 | 2.00% |
| hsa_circ_0037657 | 1102 | 11 | 2.00% |
| hsa_circ_0038530 | 1102 | 11 | 2.00% |
| hsa_circ_0063557 | 1102 | 11 | 2.00% |
| hsa_circ_0085181 | 1102 | 11 | 2.00% |
| hsa_circ_0089153 | 1102 | 11 | 2.00% |
| hsa_circ_0102881 | 1102 | 11 | 2.00% |
| hsa_circ_0115960 | 1102 | 11 | 2.00% |
| hsa_circ_0127728 | 1102 | 11 | 2.00% |
| hsa_circ_0131231 | 1102 | 11 | 2.00% |
| hsa_circ_0138079 | 1102 | 11 | 2.00% |
| hsa_circ_0001110 | 1002 | 10 | 2.00% |
| hsa_circ_0013550 | 1002 | 10 | 2.00% |
| hsa_circ_0022184 | 1002 | 10 | 2.00% |
| hsa_circ_0045598 | 1002 | 10 | 2.00% |
| hsa_circ_0062683 | 1002 | 10 | 2.00% |
| hsa_circ_0067957 | 1002 | 10 | 2.00% |
| hsa_circ_0069387 | 1002 | 10 | 2.00% |
| hsa_circ_0072365 | 1002 | 10 | 2.00% |
| hsa_circ_0073176 | 1002 | 10 | 2.00% |
| hsa_circ_0085060 | 1002 | 10 | 2.00% |
| hsa_circ_0086213 | 1002 | 10 | 2.00% |
| hsa_circ_0089044 | 1002 | 10 | 2.00% |
| hsa_circ_0096294 | 1002 | 10 | 2.00% |
| hsa_circ_0118786 | 1002 | 10 | 2.00% |
| hsa_circ_0072829 | 296025 | 2953 | 2.00% |
| hsa_circ_0039876 | 1203 | 12 | 2.00% |
| hsa_circ_0112469 | 1203 | 12 | 2.00% |
| hsa_circ_0017213 | 1103 | 11 | 1.99% |
| hsa_circ_0038105 | 1103 | 11 | 1.99% |
| hsa_circ_0042773 | 1103 | 11 | 1.99% |
| hsa_circ_0059720 | 1103 | 11 | 1.99% |
| hsa_circ_0069951 | 1103 | 11 | 1.99% |
| hsa_circ_0072444 | 1103 | 11 | 1.99% |
| hsa_circ_0073170 | 1103 | 11 | 1.99% |
| hsa_circ_0083200 | 1103 | 11 | 1.99% |
| hsa_circ_0111211 | 1103 | 11 | 1.99% |
| hsa_circ_0139645 | 1103 | 11 | 1.99% |
| hsa_circ_0089933 | 1705 | 17 | 1.99% |
| hsa_circ_0002099 | 1003 | 10 | 1.99% |
| hsa_circ_0002289 | 1003 | 10 | 1.99% |
| hsa_circ_0010692 | 1003 | 10 | 1.99% |
| hsa_circ_0026889 | 1003 | 10 | 1.99% |
| hsa_circ_0033644 | 1003 | 10 | 1.99% |

| | | | |
|---|---|---|---|
| hsa_circ_0047468 | 1003 | 10 | 1.99% |
| hsa_circ_0052021 | 1003 | 10 | 1.99% |
| hsa_circ_0059436 | 1003 | 10 | 1.99% |
| hsa_circ_0069958 | 1003 | 10 | 1.99% |
| hsa_circ_0087840 | 1003 | 10 | 1.99% |
| hsa_circ_0092062 | 1003 | 10 | 1.99% |
| hsa_circ_0122492 | 1003 | 10 | 1.99% |
| hsa_circ_0123096 | 1003 | 10 | 1.99% |
| hsa_circ_0125311 | 1003 | 10 | 1.99% |
| hsa_circ_0127860 | 1003 | 10 | 1.99% |
| hsa_circ_0137010 | 1003 | 10 | 1.99% |
| hsa_circ_0138810 | 1003 | 10 | 1.99% |
| hsa_circ_0140254 | 2809 | 28 | 1.99% |
| hsa_circ_0025281 | 1204 | 12 | 1.99% |
| hsa_circ_0042876 | 1405 | 14 | 1.99% |
| hsa_circ_0014645 | 1104 | 11 | 1.99% |
| hsa_circ_0022404 | 1104 | 11 | 1.99% |
| hsa_circ_0028033 | 1104 | 11 | 1.99% |
| hsa_circ_0039328 | 1104 | 11 | 1.99% |
| hsa_circ_0057967 | 1104 | 11 | 1.99% |
| hsa_circ_0123025 | 1104 | 11 | 1.99% |
| hsa_circ_0037488 | 1305 | 13 | 1.99% |
| hsa_circ_0008928 | 1004 | 10 | 1.99% |
| hsa_circ_0009890 | 1004 | 10 | 1.99% |
| hsa_circ_0033640 | 1004 | 10 | 1.99% |
| hsa_circ_0034746 | 1004 | 10 | 1.99% |
| hsa_circ_0040010 | 1004 | 10 | 1.99% |
| hsa_circ_0069453 | 1004 | 10 | 1.99% |
| hsa_circ_0085100 | 1004 | 10 | 1.99% |
| hsa_circ_0100393 | 1004 | 10 | 1.99% |
| hsa_circ_0108625 | 1004 | 10 | 1.99% |
| hsa_circ_0113267 | 1004 | 10 | 1.99% |
| hsa_circ_0123565 | 1004 | 10 | 1.99% |
| hsa_circ_0034419 | 1205 | 12 | 1.99% |
| hsa_circ_0048564 | 1205 | 12 | 1.99% |
| hsa_circ_0101350 | 1205 | 12 | 1.99% |
| hsa_circ_0014744 | 1406 | 14 | 1.99% |
| hsa_circ_0089503 | 1105 | 11 | 1.99% |
| hsa_circ_0095437 | 1105 | 11 | 1.99% |
| hsa_circ_0113792 | 5023 | 50 | 1.99% |
| hsa_circ_0106443 | 4421 | 44 | 1.99% |
| hsa_circ_0033620 | 1206 | 12 | 1.99% |
| hsa_circ_0086147 | 1206 | 12 | 1.99% |
| hsa_circ_0003074 | 1005 | 10 | 1.99% |
| hsa_circ_0033647 | 1005 | 10 | 1.99% |
| hsa_circ_0035650 | 1005 | 10 | 1.99% |
| hsa_circ_0043792 | 1005 | 10 | 1.99% |
| hsa_circ_0050473 | 1005 | 10 | 1.99% |
| hsa_circ_0059972 | 1005 | 10 | 1.99% |
| hsa_circ_0061207 | 1005 | 10 | 1.99% |

| | | | |
|---|---|---|---|
| hsa_circ_0067389 | 1005 | 10 | 1.99% |
| hsa_circ_0075167 | 1005 | 10 | 1.99% |
| hsa_circ_0078324 | 1005 | 10 | 1.99% |
| hsa_circ_0089715 | 1005 | 10 | 1.99% |
| hsa_circ_0091251 | 1005 | 10 | 1.99% |
| hsa_circ_0105813 | 1005 | 10 | 1.99% |
| hsa_circ_0122014 | 1005 | 10 | 1.99% |
| hsa_circ_0138355 | 1005 | 10 | 1.99% |
| hsa_circ_0058195 | 1307 | 13 | 1.99% |
| hsa_circ_0094484 | 1307 | 13 | 1.99% |
| hsa_circ_0000244 | 1106 | 11 | 1.99% |
| hsa_circ_0019364 | 1106 | 11 | 1.99% |
| hsa_circ_0045609 | 1106 | 11 | 1.99% |
| hsa_circ_0060927 | 1106 | 11 | 1.99% |
| hsa_circ_0065129 | 1106 | 11 | 1.99% |
| hsa_circ_0089148 | 1106 | 11 | 1.99% |
| hsa_circ_0097994 | 1106 | 11 | 1.99% |
| hsa_circ_0117439 | 1106 | 11 | 1.99% |
| hsa_circ_0124456 | 1106 | 11 | 1.99% |
| hsa_circ_0050622 | 1207 | 12 | 1.99% |
| hsa_circ_0087073 | 1207 | 12 | 1.99% |
| hsa_circ_0117679 | 1207 | 12 | 1.99% |
| hsa_circ_0010687 | 1006 | 10 | 1.99% |
| hsa_circ_0021773 | 1006 | 10 | 1.99% |
| hsa_circ_0031740 | 1006 | 10 | 1.99% |
| hsa_circ_0034847 | 1006 | 10 | 1.99% |
| hsa_circ_0037919 | 1006 | 10 | 1.99% |
| hsa_circ_0040741 | 1006 | 10 | 1.99% |
| hsa_circ_0042400 | 1006 | 10 | 1.99% |
| hsa_circ_0047230 | 1006 | 10 | 1.99% |
| hsa_circ_0064732 | 1006 | 10 | 1.99% |
| hsa_circ_0065658 | 1006 | 10 | 1.99% |
| hsa_circ_0076163 | 1006 | 10 | 1.99% |
| hsa_circ_0086106 | 1006 | 10 | 1.99% |
| hsa_circ_0132162 | 1006 | 10 | 1.99% |
| hsa_circ_0138389 | 1006 | 10 | 1.99% |
| hsa_circ_0026572 | 1107 | 11 | 1.99% |
| hsa_circ_0041791 | 1107 | 11 | 1.99% |
| hsa_circ_0099800 | 1107 | 11 | 1.99% |
| hsa_circ_0115357 | 1107 | 11 | 1.99% |
| hsa_circ_0120566 | 1107 | 11 | 1.99% |
| hsa_circ_0122736 | 1107 | 11 | 1.99% |
| hsa_circ_0121793 | 1409 | 14 | 1.99% |
| hsa_circ_0122437 | 1208 | 12 | 1.99% |
| hsa_circ_0041440 | 1309 | 13 | 1.99% |
| hsa_circ_0002159 | 1007 | 10 | 1.99% |
| hsa_circ_0018050 | 1007 | 10 | 1.99% |
| hsa_circ_0024133 | 1007 | 10 | 1.99% |
| hsa_circ_0066409 | 1007 | 10 | 1.99% |
| hsa_circ_0066627 | 1007 | 10 | 1.99% |

| | | | |
|---|---|---|---|
| hsa_circ_0069271 | 1007 | 10 | 1.99% |
| hsa_circ_0082718 | 1007 | 10 | 1.99% |
| hsa_circ_0084127 | 1007 | 10 | 1.99% |
| hsa_circ_0086298 | 1007 | 10 | 1.99% |
| hsa_circ_0094399 | 1007 | 10 | 1.99% |
| hsa_circ_0096414 | 1007 | 10 | 1.99% |
| hsa_circ_0110451 | 1007 | 10 | 1.99% |
| hsa_circ_0123208 | 1007 | 10 | 1.99% |
| hsa_circ_0123617 | 1007 | 10 | 1.99% |
| hsa_circ_0133825 | 1007 | 10 | 1.99% |
| hsa_circ_0139175 | 1007 | 10 | 1.99% |
| hsa_circ_0042545 | 1108 | 11 | 1.99% |
| hsa_circ_0092015 | 1108 | 11 | 1.99% |
| hsa_circ_0120946 | 1108 | 11 | 1.99% |
| hsa_circ_0130605 | 1108 | 11 | 1.99% |
| hsa_circ_0097241 | 1209 | 12 | 1.99% |
| hsa_circ_0011304 | 1008 | 10 | 1.98% |
| hsa_circ_0017384 | 1008 | 10 | 1.98% |
| hsa_circ_0026258 | 1008 | 10 | 1.98% |
| hsa_circ_0043687 | 1008 | 10 | 1.98% |
| hsa_circ_0043903 | 1008 | 10 | 1.98% |
| hsa_circ_0048533 | 1008 | 10 | 1.98% |
| hsa_circ_0056877 | 1008 | 10 | 1.98% |
| hsa_circ_0056921 | 1008 | 10 | 1.98% |
| hsa_circ_0058110 | 1008 | 10 | 1.98% |
| hsa_circ_0064009 | 1008 | 10 | 1.98% |
| hsa_circ_0064468 | 1008 | 10 | 1.98% |
| hsa_circ_0066429 | 1008 | 10 | 1.98% |
| hsa_circ_0083456 | 1008 | 10 | 1.98% |
| hsa_circ_0086640 | 1008 | 10 | 1.98% |
| hsa_circ_0102831 | 1008 | 10 | 1.98% |
| hsa_circ_0124192 | 1008 | 10 | 1.98% |
| hsa_circ_0124202 | 1008 | 10 | 1.98% |
| hsa_circ_0126436 | 1008 | 10 | 1.98% |
| hsa_circ_0060763 | 1109 | 11 | 1.98% |
| hsa_circ_0120748 | 1109 | 11 | 1.98% |
| hsa_circ_0017026 | 1210 | 12 | 1.98% |
| hsa_circ_0021983 | 1311 | 13 | 1.98% |
| hsa_circ_0097248 | 1311 | 13 | 1.98% |
| hsa_circ_0109262 | 2724 | 27 | 1.98% |
| hsa_circ_0008072 | 1009 | 10 | 1.98% |
| hsa_circ_0058709 | 1009 | 10 | 1.98% |
| hsa_circ_0068071 | 1009 | 10 | 1.98% |
| hsa_circ_0074755 | 1009 | 10 | 1.98% |
| hsa_circ_0108501 | 1009 | 10 | 1.98% |
| hsa_circ_0121280 | 1009 | 10 | 1.98% |
| hsa_circ_0127283 | 1009 | 10 | 1.98% |
| hsa_circ_0068083 | 5449 | 54 | 1.98% |
| hsa_circ_0011575 | 1110 | 11 | 1.98% |
| hsa_circ_0054467 | 1110 | 11 | 1.98% |

| | | | |
|---|---|---|---|
| hsa_circ_0058517 | 1110 | 11 | 1.98% |
| hsa_circ_0073474 | 1211 | 12 | 1.98% |
| hsa_circ_0075277 | 1211 | 12 | 1.98% |
| hsa_circ_0132860 | 1211 | 12 | 1.98% |
| hsa_circ_0104930 | 1413 | 14 | 1.98% |
| hsa_circ_0022842 | 1615 | 16 | 1.98% |
| hsa_circ_0133940 | 1212 | 12 | 1.98% |
| hsa_circ_0011983 | 1111 | 11 | 1.98% |
| hsa_circ_0031634 | 1111 | 11 | 1.98% |
| hsa_circ_0045557 | 1111 | 11 | 1.98% |
| hsa_circ_0047369 | 1111 | 11 | 1.98% |
| hsa_circ_0108978 | 1111 | 11 | 1.98% |
| hsa_circ_0136883 | 1111 | 11 | 1.98% |
| hsa_circ_0138168 | 1111 | 11 | 1.98% |
| hsa_circ_0015795 | 1010 | 10 | 1.98% |
| hsa_circ_0017587 | 1010 | 10 | 1.98% |
| hsa_circ_0026117 | 1010 | 10 | 1.98% |
| hsa_circ_0026880 | 1010 | 10 | 1.98% |
| hsa_circ_0044924 | 1010 | 10 | 1.98% |
| hsa_circ_0065079 | 1010 | 10 | 1.98% |
| hsa_circ_0067324 | 1010 | 10 | 1.98% |
| hsa_circ_0073478 | 1010 | 10 | 1.98% |
| hsa_circ_0081666 | 1010 | 10 | 1.98% |
| hsa_circ_0081729 | 1010 | 10 | 1.98% |
| hsa_circ_0090355 | 1010 | 10 | 1.98% |
| hsa_circ_0102161 | 1010 | 10 | 1.98% |
| hsa_circ_0115293 | 1010 | 10 | 1.98% |
| hsa_circ_0130612 | 1010 | 10 | 1.98% |
| hsa_circ_0134097 | 1010 | 10 | 1.98% |
| hsa_circ_0103611 | 3132 | 31 | 1.98% |
| hsa_circ_0104924 | 2829 | 28 | 1.98% |
| hsa_circ_0031534 | 1213 | 12 | 1.98% |
| hsa_circ_0049124 | 1213 | 12 | 1.98% |
| hsa_circ_0084777 | 1213 | 12 | 1.98% |
| hsa_circ_0013527 | 1112 | 11 | 1.98% |
| hsa_circ_0019134 | 1112 | 11 | 1.98% |
| hsa_circ_0025246 | 1112 | 11 | 1.98% |
| hsa_circ_0058437 | 1112 | 11 | 1.98% |
| hsa_circ_0068806 | 1112 | 11 | 1.98% |
| hsa_circ_0091450 | 1112 | 11 | 1.98% |
| hsa_circ_0091514 | 1112 | 11 | 1.98% |
| hsa_circ_0001224 | 1011 | 10 | 1.98% |
| hsa_circ_0019630 | 1011 | 10 | 1.98% |
| hsa_circ_0036154 | 1011 | 10 | 1.98% |
| hsa_circ_0043122 | 1011 | 10 | 1.98% |
| hsa_circ_0045351 | 1011 | 10 | 1.98% |
| hsa_circ_0047255 | 1011 | 10 | 1.98% |
| hsa_circ_0068613 | 1011 | 10 | 1.98% |
| hsa_circ_0070070 | 1011 | 10 | 1.98% |
| hsa_circ_0072240 | 1011 | 10 | 1.98% |

| | | | |
|---|---|---|---|
| hsa_circ_0081586 | 1011 | 10 | 1.98% |
| hsa_circ_0086661 | 1011 | 10 | 1.98% |
| hsa_circ_0087676 | 1011 | 10 | 1.98% |
| hsa_circ_0088237 | 1011 | 10 | 1.98% |
| hsa_circ_0117733 | 1011 | 10 | 1.98% |
| hsa_circ_0120327 | 1011 | 10 | 1.98% |
| hsa_circ_0137758 | 1011 | 10 | 1.98% |
| hsa_circ_0042388 | 1416 | 14 | 1.98% |
| hsa_circ_0007506 | 1214 | 12 | 1.98% |
| hsa_circ_0103967 | 1214 | 12 | 1.98% |
| hsa_circ_0023646 | 1113 | 11 | 1.98% |
| hsa_circ_0027514 | 1113 | 11 | 1.98% |
| hsa_circ_0082593 | 1113 | 11 | 1.98% |
| hsa_circ_0099912 | 1113 | 11 | 1.98% |
| hsa_circ_0107343 | 1113 | 11 | 1.98% |
| hsa_circ_0127342 | 1113 | 11 | 1.98% |
| hsa_circ_0023038 | 1012 | 10 | 1.98% |
| hsa_circ_0036046 | 1012 | 10 | 1.98% |
| hsa_circ_0039188 | 1012 | 10 | 1.98% |
| hsa_circ_0056705 | 1012 | 10 | 1.98% |
| hsa_circ_0064476 | 1012 | 10 | 1.98% |
| hsa_circ_0070877 | 1012 | 10 | 1.98% |
| hsa_circ_0072783 | 1012 | 10 | 1.98% |
| hsa_circ_0081328 | 1012 | 10 | 1.98% |
| hsa_circ_0104569 | 1012 | 10 | 1.98% |
| hsa_circ_0125626 | 1012 | 10 | 1.98% |
| hsa_circ_0138069 | 1012 | 10 | 1.98% |
| hsa_circ_0031005 | 1316 | 13 | 1.98% |
| hsa_circ_0018628 | 1215 | 12 | 1.98% |
| hsa_circ_0021488 | 1215 | 12 | 1.98% |
| hsa_circ_0026049 | 1215 | 12 | 1.98% |
| hsa_circ_0026599 | 1215 | 12 | 1.98% |
| hsa_circ_0112122 | 1215 | 12 | 1.98% |
| hsa_circ_0000975 | 1114 | 11 | 1.97% |
| hsa_circ_0016234 | 1114 | 11 | 1.97% |
| hsa_circ_0040354 | 1114 | 11 | 1.97% |
| hsa_circ_0069895 | 1114 | 11 | 1.97% |
| hsa_circ_0090192 | 1114 | 11 | 1.97% |
| hsa_circ_0139613 | 1114 | 11 | 1.97% |
| hsa_circ_0006101 | 1013 | 10 | 1.97% |
| hsa_circ_0015016 | 1013 | 10 | 1.97% |
| hsa_circ_0051835 | 1013 | 10 | 1.97% |
| hsa_circ_0071398 | 1013 | 10 | 1.97% |
| hsa_circ_0078057 | 1013 | 10 | 1.97% |
| hsa_circ_0081472 | 1013 | 10 | 1.97% |
| hsa_circ_0081627 | 1013 | 10 | 1.97% |
| hsa_circ_0084277 | 1013 | 10 | 1.97% |
| hsa_circ_0091706 | 1013 | 10 | 1.97% |
| hsa_circ_0097127 | 1013 | 10 | 1.97% |
| hsa_circ_0099382 | 1013 | 10 | 1.97% |

| | | | |
|---|---|---|---|
| hsa_circ_0121852 | 1013 | 10 | 1.97% |
| hsa_circ_0056271 | 1216 | 12 | 1.97% |
| hsa_circ_0067309 | 1216 | 12 | 1.97% |
| hsa_circ_0010983 | 1115 | 11 | 1.97% |
| hsa_circ_0054822 | 1115 | 11 | 1.97% |
| hsa_circ_0085276 | 1115 | 11 | 1.97% |
| hsa_circ_0112405 | 1115 | 11 | 1.97% |
| hsa_circ_0114355 | 1115 | 11 | 1.97% |
| hsa_circ_0000965 | 3447 | 34 | 1.97% |
| hsa_circ_0021115 | 1318 | 13 | 1.97% |
| hsa_circ_0007722 | 1014 | 10 | 1.97% |
| hsa_circ_0021143 | 1014 | 10 | 1.97% |
| hsa_circ_0022435 | 1014 | 10 | 1.97% |
| hsa_circ_0022631 | 1014 | 10 | 1.97% |
| hsa_circ_0025166 | 1014 | 10 | 1.97% |
| hsa_circ_0030456 | 1014 | 10 | 1.97% |
| hsa_circ_0031758 | 1014 | 10 | 1.97% |
| hsa_circ_0034524 | 1014 | 10 | 1.97% |
| hsa_circ_0038924 | 1014 | 10 | 1.97% |
| hsa_circ_0045934 | 1014 | 10 | 1.97% |
| hsa_circ_0052773 | 1014 | 10 | 1.97% |
| hsa_circ_0073695 | 1014 | 10 | 1.97% |
| hsa_circ_0078708 | 1014 | 10 | 1.97% |
| hsa_circ_0080458 | 1014 | 10 | 1.97% |
| hsa_circ_0081036 | 1014 | 10 | 1.97% |
| hsa_circ_0082494 | 1014 | 10 | 1.97% |
| hsa_circ_0096268 | 1014 | 10 | 1.97% |
| hsa_circ_0096309 | 1014 | 10 | 1.97% |
| hsa_circ_0096648 | 1014 | 10 | 1.97% |
| hsa_circ_0101459 | 1014 | 10 | 1.97% |
| hsa_circ_0103432 | 1014 | 10 | 1.97% |
| hsa_circ_0106431 | 1014 | 10 | 1.97% |
| hsa_circ_0114111 | 1014 | 10 | 1.97% |
| hsa_circ_0131434 | 1014 | 10 | 1.97% |
| hsa_circ_0137796 | 1014 | 10 | 1.97% |
| hsa_circ_0139710 | 1014 | 10 | 1.97% |
| hsa_circ_0075602 | 1217 | 12 | 1.97% |
| hsa_circ_0098964 | 1217 | 12 | 1.97% |
| hsa_circ_0041295 | 1116 | 11 | 1.97% |
| hsa_circ_0044603 | 1116 | 11 | 1.97% |
| hsa_circ_0081046 | 1116 | 11 | 1.97% |
| hsa_circ_0083013 | 1116 | 11 | 1.97% |
| hsa_circ_0083498 | 1116 | 11 | 1.97% |
| hsa_circ_0106413 | 1116 | 11 | 1.97% |
| hsa_circ_0137453 | 1116 | 11 | 1.97% |
| hsa_circ_0041427 | 1319 | 13 | 1.97% |
| hsa_circ_0047738 | 1319 | 13 | 1.97% |
| hsa_circ_0011520 | 3856 | 38 | 1.97% |
| hsa_circ_0010395 | 1827 | 18 | 1.97% |
| hsa_circ_0018978 | 1218 | 12 | 1.97% |

| | | | |
|---|---|---|---|
| hsa_circ_0067601 | 1218 | 12 | 1.97% |
| hsa_circ_0104136 | 1218 | 12 | 1.97% |
| hsa_circ_0024268 | 1015 | 10 | 1.97% |
| hsa_circ_0025382 | 1015 | 10 | 1.97% |
| hsa_circ_0025981 | 1015 | 10 | 1.97% |
| hsa_circ_0027473 | 1015 | 10 | 1.97% |
| hsa_circ_0030614 | 1015 | 10 | 1.97% |
| hsa_circ_0033000 | 1015 | 10 | 1.97% |
| hsa_circ_0042453 | 1015 | 10 | 1.97% |
| hsa_circ_0049650 | 1015 | 10 | 1.97% |
| hsa_circ_0053072 | 1015 | 10 | 1.97% |
| hsa_circ_0070584 | 1015 | 10 | 1.97% |
| hsa_circ_0083892 | 1015 | 10 | 1.97% |
| hsa_circ_0100766 | 1015 | 10 | 1.97% |
| hsa_circ_0118782 | 1015 | 10 | 1.97% |
| hsa_circ_0125307 | 1015 | 10 | 1.97% |
| hsa_circ_0027286 | 1320 | 13 | 1.97% |
| hsa_circ_0039378 | 1320 | 13 | 1.97% |
| hsa_circ_0032735 | 1117 | 11 | 1.97% |
| hsa_circ_0038161 | 1117 | 11 | 1.97% |
| hsa_circ_0038794 | 1117 | 11 | 1.97% |
| hsa_circ_0080165 | 1117 | 11 | 1.97% |
| hsa_circ_0124771 | 1117 | 11 | 1.97% |
| hsa_circ_0139697 | 1117 | 11 | 1.97% |
| hsa_circ_0011100 | 1219 | 12 | 1.97% |
| hsa_circ_0011291 | 1219 | 12 | 1.97% |
| hsa_circ_0015112 | 1219 | 12 | 1.97% |
| hsa_circ_0047401 | 1219 | 12 | 1.97% |
| hsa_circ_0097250 | 1219 | 12 | 1.97% |
| hsa_circ_0140035 | 1219 | 12 | 1.97% |
| hsa_circ_0019553 | 1016 | 10 | 1.97% |
| hsa_circ_0020102 | 1016 | 10 | 1.97% |
| hsa_circ_0021587 | 1016 | 10 | 1.97% |
| hsa_circ_0023506 | 1016 | 10 | 1.97% |
| hsa_circ_0031499 | 1016 | 10 | 1.97% |
| hsa_circ_0032539 | 1016 | 10 | 1.97% |
| hsa_circ_0040109 | 1016 | 10 | 1.97% |
| hsa_circ_0051893 | 1016 | 10 | 1.97% |
| hsa_circ_0073020 | 1016 | 10 | 1.97% |
| hsa_circ_0074717 | 1016 | 10 | 1.97% |
| hsa_circ_0078305 | 1016 | 10 | 1.97% |
| hsa_circ_0083860 | 1016 | 10 | 1.97% |
| hsa_circ_0087018 | 1016 | 10 | 1.97% |
| hsa_circ_0095385 | 1016 | 10 | 1.97% |
| hsa_circ_0101464 | 1016 | 10 | 1.97% |
| hsa_circ_0104275 | 1016 | 10 | 1.97% |
| hsa_circ_0108189 | 1016 | 10 | 1.97% |
| hsa_circ_0121985 | 1016 | 10 | 1.97% |
| hsa_circ_0004011 | 1118 | 11 | 1.97% |
| hsa_circ_0051520 | 1118 | 11 | 1.97% |

| | | | |
|---|---|---|---|
| hsa_circ_0063240 | 1118 | 11 | 1.97% |
| hsa_circ_0108020 | 1118 | 11 | 1.97% |
| hsa_circ_0131093 | 1118 | 11 | 1.97% |
| hsa_circ_0036254 | 1423 | 14 | 1.97% |
| hsa_circ_0130248 | 1423 | 14 | 1.97% |
| hsa_circ_0007467 | 2948 | 29 | 1.97% |
| hsa_circ_0016451 | 1220 | 12 | 1.97% |
| hsa_circ_0048753 | 1220 | 12 | 1.97% |
| hsa_circ_0064074 | 1220 | 12 | 1.97% |
| hsa_circ_0127654 | 1220 | 12 | 1.97% |
| hsa_circ_0089660 | 1322 | 13 | 1.97% |
| hsa_circ_0000174 | 1017 | 10 | 1.97% |
| hsa_circ_0008671 | 1017 | 10 | 1.97% |
| hsa_circ_0012281 | 1017 | 10 | 1.97% |
| hsa_circ_0022351 | 1017 | 10 | 1.97% |
| hsa_circ_0027489 | 1017 | 10 | 1.97% |
| hsa_circ_0038373 | 1017 | 10 | 1.97% |
| hsa_circ_0041160 | 1017 | 10 | 1.97% |
| hsa_circ_0041237 | 1017 | 10 | 1.97% |
| hsa_circ_0073760 | 1017 | 10 | 1.97% |
| hsa_circ_0090629 | 1017 | 10 | 1.97% |
| hsa_circ_0091482 | 1017 | 10 | 1.97% |
| hsa_circ_0096423 | 1017 | 10 | 1.97% |
| hsa_circ_0099253 | 1017 | 10 | 1.97% |
| hsa_circ_0101858 | 1017 | 10 | 1.97% |
| hsa_circ_0122112 | 1017 | 10 | 1.97% |
| hsa_circ_0135353 | 1017 | 10 | 1.97% |
| hsa_circ_0136652 | 1017 | 10 | 1.97% |
| hsa_circ_0138537 | 1017 | 10 | 1.97% |
| hsa_circ_0139234 | 1017 | 10 | 1.97% |
| hsa_circ_0006136 | 1119 | 11 | 1.97% |
| hsa_circ_0024661 | 1119 | 11 | 1.97% |
| hsa_circ_0050368 | 1119 | 11 | 1.97% |
| hsa_circ_0079224 | 1119 | 11 | 1.97% |
| hsa_circ_0081840 | 1119 | 11 | 1.97% |
| hsa_circ_0083672 | 1119 | 11 | 1.97% |
| hsa_circ_0089435 | 1119 | 11 | 1.97% |
| hsa_circ_0111213 | 1119 | 11 | 1.97% |
| hsa_circ_0136372 | 1119 | 11 | 1.97% |
| hsa_circ_0005862 | 1221 | 12 | 1.97% |
| hsa_circ_0006984 | 1221 | 12 | 1.97% |
| hsa_circ_0089661 | 1323 | 13 | 1.97% |
| hsa_circ_0000781 | 3461 | 34 | 1.96% |
| hsa_circ_0080469 | 2036 | 20 | 1.96% |
| hsa_circ_0018642 | 1018 | 10 | 1.96% |
| hsa_circ_0020848 | 1018 | 10 | 1.96% |
| hsa_circ_0034993 | 1018 | 10 | 1.96% |
| hsa_circ_0041066 | 1018 | 10 | 1.96% |
| hsa_circ_0041717 | 1018 | 10 | 1.96% |
| hsa_circ_0065911 | 1018 | 10 | 1.96% |

| | | | |
|---|---|---|---|
| hsa_circ_0087986 | 1018 | 10 | 1.96% |
| hsa_circ_0089683 | 1018 | 10 | 1.96% |
| hsa_circ_0101980 | 1018 | 10 | 1.96% |
| hsa_circ_0103523 | 1018 | 10 | 1.96% |
| hsa_circ_0113166 | 1018 | 10 | 1.96% |
| hsa_circ_0122449 | 1018 | 10 | 1.96% |
| hsa_circ_0128577 | 1018 | 10 | 1.96% |
| hsa_circ_0003043 | 1120 | 11 | 1.96% |
| hsa_circ_0029067 | 1120 | 11 | 1.96% |
| hsa_circ_0090675 | 1120 | 11 | 1.96% |
| hsa_circ_0096227 | 1120 | 11 | 1.96% |
| hsa_circ_0034911 | 1222 | 12 | 1.96% |
| hsa_circ_0119536 | 1222 | 12 | 1.96% |
| hsa_circ_0076661 | 1324 | 13 | 1.96% |
| hsa_circ_0087363 | 1528 | 15 | 1.96% |
| hsa_circ_0008951 | 1019 | 10 | 1.96% |
| hsa_circ_0015567 | 1019 | 10 | 1.96% |
| hsa_circ_0017594 | 1019 | 10 | 1.96% |
| hsa_circ_0022483 | 1019 | 10 | 1.96% |
| hsa_circ_0023521 | 1019 | 10 | 1.96% |
| hsa_circ_0048234 | 1019 | 10 | 1.96% |
| hsa_circ_0057550 | 1019 | 10 | 1.96% |
| hsa_circ_0059736 | 1019 | 10 | 1.96% |
| hsa_circ_0064104 | 1019 | 10 | 1.96% |
| hsa_circ_0077175 | 1019 | 10 | 1.96% |
| hsa_circ_0085397 | 1019 | 10 | 1.96% |
| hsa_circ_0086135 | 1019 | 10 | 1.96% |
| hsa_circ_0111609 | 1019 | 10 | 1.96% |
| hsa_circ_0130432 | 1019 | 10 | 1.96% |
| hsa_circ_0024765 | 1121 | 11 | 1.96% |
| hsa_circ_0058041 | 1121 | 11 | 1.96% |
| hsa_circ_0064308 | 1121 | 11 | 1.96% |
| hsa_circ_0085966 | 1121 | 11 | 1.96% |
| hsa_circ_0099571 | 1121 | 11 | 1.96% |
| hsa_circ_0131816 | 1121 | 11 | 1.96% |
| hsa_circ_0028607 | 1223 | 12 | 1.96% |
| hsa_circ_0054796 | 1223 | 12 | 1.96% |
| hsa_circ_0135124 | 1223 | 12 | 1.96% |
| hsa_circ_0138212 | 1223 | 12 | 1.96% |
| hsa_circ_0067263 | 1325 | 13 | 1.96% |
| hsa_circ_0103616 | 5199 | 51 | 1.96% |
| hsa_circ_0074553 | 1224 | 12 | 1.96% |
| hsa_circ_0093270 | 1224 | 12 | 1.96% |
| hsa_circ_0011022 | 1122 | 11 | 1.96% |
| hsa_circ_0030334 | 1122 | 11 | 1.96% |
| hsa_circ_0040551 | 1122 | 11 | 1.96% |
| hsa_circ_0072752 | 1122 | 11 | 1.96% |
| hsa_circ_0076086 | 1122 | 11 | 1.96% |
| hsa_circ_0000150 | 1020 | 10 | 1.96% |
| hsa_circ_0006291 | 1020 | 10 | 1.96% |

| | | | |
|---|---|---|---|
| hsa_circ_0021925 | 1020 | 10 | 1.96% |
| hsa_circ_0022900 | 1020 | 10 | 1.96% |
| hsa_circ_0037719 | 1020 | 10 | 1.96% |
| hsa_circ_0046695 | 1020 | 10 | 1.96% |
| hsa_circ_0050250 | 1020 | 10 | 1.96% |
| hsa_circ_0057593 | 1020 | 10 | 1.96% |
| hsa_circ_0057893 | 1020 | 10 | 1.96% |
| hsa_circ_0059738 | 1020 | 10 | 1.96% |
| hsa_circ_0087321 | 1020 | 10 | 1.96% |
| hsa_circ_0132218 | 1020 | 10 | 1.96% |
| hsa_circ_0075400 | 1939 | 19 | 1.96% |
| hsa_circ_0123986 | 2858 | 28 | 1.96% |
| hsa_circ_0014754 | 1429 | 14 | 1.96% |
| hsa_circ_0106940 | 4798 | 47 | 1.96% |
| hsa_circ_0009254 | 1123 | 11 | 1.96% |
| hsa_circ_0043994 | 1123 | 11 | 1.96% |
| hsa_circ_0054249 | 1123 | 11 | 1.96% |
| hsa_circ_0095069 | 1123 | 11 | 1.96% |
| hsa_circ_0119581 | 1123 | 11 | 1.96% |
| hsa_circ_0134614 | 3165 | 31 | 1.96% |
| hsa_circ_0001423 | 1021 | 10 | 1.96% |
| hsa_circ_0014876 | 1021 | 10 | 1.96% |
| hsa_circ_0018248 | 1021 | 10 | 1.96% |
| hsa_circ_0019592 | 1021 | 10 | 1.96% |
| hsa_circ_0022994 | 1021 | 10 | 1.96% |
| hsa_circ_0025306 | 1021 | 10 | 1.96% |
| hsa_circ_0033641 | 1021 | 10 | 1.96% |
| hsa_circ_0049852 | 1021 | 10 | 1.96% |
| hsa_circ_0052186 | 1021 | 10 | 1.96% |
| hsa_circ_0066855 | 1021 | 10 | 1.96% |
| hsa_circ_0076440 | 1021 | 10 | 1.96% |
| hsa_circ_0082427 | 1021 | 10 | 1.96% |
| hsa_circ_0085602 | 1021 | 10 | 1.96% |
| hsa_circ_0101544 | 1021 | 10 | 1.96% |
| hsa_circ_0108187 | 1021 | 10 | 1.96% |
| hsa_circ_0112869 | 1736 | 17 | 1.96% |
| hsa_circ_0052684 | 1532 | 15 | 1.96% |
| hsa_circ_0105103 | 2043 | 20 | 1.96% |
| hsa_circ_0120146 | 1226 | 12 | 1.96% |
| hsa_circ_0000592 | 1124 | 11 | 1.96% |
| hsa_circ_0048159 | 1124 | 11 | 1.96% |
| hsa_circ_0069864 | 1124 | 11 | 1.96% |
| hsa_circ_0085549 | 1124 | 11 | 1.96% |
| hsa_circ_0090372 | 1124 | 11 | 1.96% |
| hsa_circ_0095440 | 1124 | 11 | 1.96% |
| hsa_circ_0118776 | 1124 | 11 | 1.96% |
| hsa_circ_0018416 | 1022 | 10 | 1.96% |
| hsa_circ_0025040 | 1022 | 10 | 1.96% |
| hsa_circ_0038860 | 1022 | 10 | 1.96% |
| hsa_circ_0038869 | 1022 | 10 | 1.96% |

| | | | |
|---|---|---|---|
| hsa_circ_0045516 | 1022 | 10 | 1.96% |
| hsa_circ_0065986 | 1022 | 10 | 1.96% |
| hsa_circ_0069861 | 1022 | 10 | 1.96% |
| hsa_circ_0100237 | 1022 | 10 | 1.96% |
| hsa_circ_0103563 | 1022 | 10 | 1.96% |
| hsa_circ_0110460 | 1022 | 10 | 1.96% |
| hsa_circ_0114817 | 1022 | 10 | 1.96% |
| hsa_circ_0118780 | 1022 | 10 | 1.96% |
| hsa_circ_0122935 | 1022 | 10 | 1.96% |
| hsa_circ_0092885 | 3578 | 35 | 1.96% |
| hsa_circ_0026011 | 1329 | 13 | 1.96% |
| hsa_circ_0088623 | 1329 | 13 | 1.96% |
| hsa_circ_0088039 | 1227 | 12 | 1.96% |
| hsa_circ_0011301 | 1125 | 11 | 1.96% |
| hsa_circ_0033206 | 1125 | 11 | 1.96% |
| hsa_circ_0054739 | 1125 | 11 | 1.96% |
| hsa_circ_0102391 | 1125 | 11 | 1.96% |
| hsa_circ_0122284 | 1125 | 11 | 1.96% |
| hsa_circ_0139901 | 1125 | 11 | 1.96% |
| hsa_circ_0009794 | 1023 | 10 | 1.96% |
| hsa_circ_0010370 | 1023 | 10 | 1.96% |
| hsa_circ_0012830 | 1023 | 10 | 1.96% |
| hsa_circ_0017129 | 1023 | 10 | 1.96% |
| hsa_circ_0018815 | 1023 | 10 | 1.96% |
| hsa_circ_0022826 | 1023 | 10 | 1.96% |
| hsa_circ_0034269 | 1023 | 10 | 1.96% |
| hsa_circ_0037109 | 1023 | 10 | 1.96% |
| hsa_circ_0050542 | 1023 | 10 | 1.96% |
| hsa_circ_0057133 | 1023 | 10 | 1.96% |
| hsa_circ_0059297 | 1023 | 10 | 1.96% |
| hsa_circ_0069431 | 1023 | 10 | 1.96% |
| hsa_circ_0075890 | 1023 | 10 | 1.96% |
| hsa_circ_0097718 | 1023 | 10 | 1.96% |
| hsa_circ_0100389 | 1023 | 10 | 1.96% |
| hsa_circ_0114044 | 1023 | 10 | 1.96% |
| hsa_circ_0125315 | 1023 | 10 | 1.96% |
| hsa_circ_0131244 | 1023 | 10 | 1.96% |
| hsa_circ_0103185 | 1842 | 18 | 1.95% |
| hsa_circ_0068243 | 1228 | 12 | 1.95% |
| hsa_circ_0125894 | 1228 | 12 | 1.95% |
| hsa_circ_0015467 | 1126 | 11 | 1.95% |
| hsa_circ_0044624 | 1126 | 11 | 1.95% |
| hsa_circ_0049806 | 1126 | 11 | 1.95% |
| hsa_circ_0052871 | 1126 | 11 | 1.95% |
| hsa_circ_0067460 | 1126 | 11 | 1.95% |
| hsa_circ_0079858 | 1126 | 11 | 1.95% |
| hsa_circ_0083955 | 1126 | 11 | 1.95% |
| hsa_circ_0106477 | 1126 | 11 | 1.95% |
| hsa_circ_0111306 | 1126 | 11 | 1.95% |
| hsa_circ_0002024 | 1024 | 10 | 1.95% |

| | | | |
|---|---|---|---|
| hsa_circ_0023928 | 1024 | 10 | 1.95% |
| hsa_circ_0028844 | 1024 | 10 | 1.95% |
| hsa_circ_0038430 | 1024 | 10 | 1.95% |
| hsa_circ_0039215 | 1024 | 10 | 1.95% |
| hsa_circ_0042833 | 1024 | 10 | 1.95% |
| hsa_circ_0045265 | 1024 | 10 | 1.95% |
| hsa_circ_0075772 | 1024 | 10 | 1.95% |
| hsa_circ_0117428 | 1024 | 10 | 1.95% |
| hsa_circ_0133598 | 1024 | 10 | 1.95% |
| hsa_circ_0139940 | 1024 | 10 | 1.95% |
| hsa_circ_0012074 | 1434 | 14 | 1.95% |
| hsa_circ_0018242 | 1127 | 11 | 1.95% |
| hsa_circ_0019551 | 1127 | 11 | 1.95% |
| hsa_circ_0023032 | 1127 | 11 | 1.95% |
| hsa_circ_0037394 | 1127 | 11 | 1.95% |
| hsa_circ_0090610 | 1127 | 11 | 1.95% |
| hsa_circ_0122267 | 1127 | 11 | 1.95% |
| hsa_circ_0139289 | 1127 | 11 | 1.95% |
| hsa_circ_0042882 | 1332 | 13 | 1.95% |
| hsa_circ_0067435 | 1230 | 12 | 1.95% |
| hsa_circ_0020364 | 1025 | 10 | 1.95% |
| hsa_circ_0031015 | 1025 | 10 | 1.95% |
| hsa_circ_0031973 | 1025 | 10 | 1.95% |
| hsa_circ_0041582 | 1025 | 10 | 1.95% |
| hsa_circ_0041944 | 1025 | 10 | 1.95% |
| hsa_circ_0046369 | 1025 | 10 | 1.95% |
| hsa_circ_0049678 | 1025 | 10 | 1.95% |
| hsa_circ_0072385 | 1025 | 10 | 1.95% |
| hsa_circ_0077427 | 1025 | 10 | 1.95% |
| hsa_circ_0077615 | 1025 | 10 | 1.95% |
| hsa_circ_0110265 | 1025 | 10 | 1.95% |
| hsa_circ_0125618 | 1025 | 10 | 1.95% |
| hsa_circ_0140426 | 1025 | 10 | 1.95% |
| hsa_circ_0031376 | 1128 | 11 | 1.95% |
| hsa_circ_0048571 | 1128 | 11 | 1.95% |
| hsa_circ_0054877 | 1128 | 11 | 1.95% |
| hsa_circ_0131746 | 1128 | 11 | 1.95% |
| hsa_circ_0011811 | 1026 | 10 | 1.95% |
| hsa_circ_0035338 | 1026 | 10 | 1.95% |
| hsa_circ_0042005 | 1026 | 10 | 1.95% |
| hsa_circ_0042209 | 1026 | 10 | 1.95% |
| hsa_circ_0054240 | 1026 | 10 | 1.95% |
| hsa_circ_0073105 | 1026 | 10 | 1.95% |
| hsa_circ_0080807 | 1026 | 10 | 1.95% |
| hsa_circ_0102915 | 1026 | 10 | 1.95% |
| hsa_circ_0127712 | 1026 | 10 | 1.95% |
| hsa_circ_0133709 | 1026 | 10 | 1.95% |
| hsa_circ_0019705 | 1129 | 11 | 1.95% |
| hsa_circ_0026610 | 1129 | 11 | 1.95% |
| hsa_circ_0041636 | 1129 | 11 | 1.95% |

| | | | |
|---|---|---|---|
| hsa_circ_0067215 | 1129 | 11 | 1.95% |
| hsa_circ_0087550 | 1129 | 11 | 1.95% |
| hsa_circ_0093340 | 1129 | 11 | 1.95% |
| hsa_circ_0096761 | 1129 | 11 | 1.95% |
| hsa_circ_0098259 | 1129 | 11 | 1.95% |
| hsa_circ_0104638 | 1129 | 11 | 1.95% |
| hsa_circ_0104823 | 1129 | 11 | 1.95% |
| hsa_circ_0045813 | 1335 | 13 | 1.95% |
| hsa_circ_0003733 | 1027 | 10 | 1.95% |
| hsa_circ_0014018 | 1027 | 10 | 1.95% |
| hsa_circ_0022764 | 1027 | 10 | 1.95% |
| hsa_circ_0043569 | 1027 | 10 | 1.95% |
| hsa_circ_0053746 | 1027 | 10 | 1.95% |
| hsa_circ_0064763 | 1027 | 10 | 1.95% |
| hsa_circ_0069024 | 1027 | 10 | 1.95% |
| hsa_circ_0071646 | 1027 | 10 | 1.95% |
| hsa_circ_0080899 | 1027 | 10 | 1.95% |
| hsa_circ_0081459 | 1027 | 10 | 1.95% |
| hsa_circ_0103846 | 1027 | 10 | 1.95% |
| hsa_circ_0104440 | 1027 | 10 | 1.95% |
| hsa_circ_0104755 | 1027 | 10 | 1.95% |
| hsa_circ_0113708 | 1027 | 10 | 1.95% |
| hsa_circ_0119953 | 1027 | 10 | 1.95% |
| hsa_circ_0057388 | 1746 | 17 | 1.95% |
| hsa_circ_0025500 | 1130 | 11 | 1.95% |
| hsa_circ_0054848 | 1130 | 11 | 1.95% |
| hsa_circ_0058499 | 1130 | 11 | 1.95% |
| hsa_circ_0089140 | 1130 | 11 | 1.95% |
| hsa_circ_0089562 | 1130 | 11 | 1.95% |
| hsa_circ_0095468 | 1130 | 11 | 1.95% |
| hsa_circ_0104593 | 1130 | 11 | 1.95% |
| hsa_circ_0121449 | 1130 | 11 | 1.95% |
| hsa_circ_0005370 | 1233 | 12 | 1.95% |
| hsa_circ_0028501 | 1336 | 13 | 1.95% |
| hsa_circ_0052944 | 1336 | 13 | 1.95% |
| hsa_circ_0011355 | 1028 | 10 | 1.95% |
| hsa_circ_0015190 | 1028 | 10 | 1.95% |
| hsa_circ_0017052 | 1028 | 10 | 1.95% |
| hsa_circ_0028676 | 1028 | 10 | 1.95% |
| hsa_circ_0044902 | 1028 | 10 | 1.95% |
| hsa_circ_0054686 | 1028 | 10 | 1.95% |
| hsa_circ_0059857 | 1028 | 10 | 1.95% |
| hsa_circ_0065210 | 1028 | 10 | 1.95% |
| hsa_circ_0068745 | 1028 | 10 | 1.95% |
| hsa_circ_0078335 | 1028 | 10 | 1.95% |
| hsa_circ_0084392 | 1028 | 10 | 1.95% |
| hsa_circ_0084542 | 1028 | 10 | 1.95% |
| hsa_circ_0087443 | 1028 | 10 | 1.95% |
| hsa_circ_0088241 | 1028 | 10 | 1.95% |
| hsa_circ_0092922 | 1028 | 10 | 1.95% |

| | | | |
|---|---|---|---|
| hsa_circ_0107210 | 1028 | 10 | 1.95% |
| hsa_circ_0124657 | 1028 | 10 | 1.95% |
| hsa_circ_0017096 | 1131 | 11 | 1.95% |
| hsa_circ_0017946 | 1131 | 11 | 1.95% |
| hsa_circ_0020661 | 1131 | 11 | 1.95% |
| hsa_circ_0025610 | 1131 | 11 | 1.95% |
| hsa_circ_0040072 | 1131 | 11 | 1.95% |
| hsa_circ_0082919 | 1131 | 11 | 1.95% |
| hsa_circ_0086710 | 1131 | 11 | 1.95% |
| hsa_circ_0123023 | 1131 | 11 | 1.95% |
| hsa_circ_0123924 | 1131 | 11 | 1.95% |
| hsa_circ_0134331 | 1131 | 11 | 1.95% |
| hsa_circ_0137696 | 1131 | 11 | 1.95% |
| hsa_circ_0036389 | 1234 | 12 | 1.94% |
| hsa_circ_0100896 | 1543 | 15 | 1.94% |
| hsa_circ_0013334 | 1029 | 10 | 1.94% |
| hsa_circ_0013422 | 1029 | 10 | 1.94% |
| hsa_circ_0015443 | 1029 | 10 | 1.94% |
| hsa_circ_0027478 | 1029 | 10 | 1.94% |
| hsa_circ_0041168 | 1029 | 10 | 1.94% |
| hsa_circ_0043743 | 1029 | 10 | 1.94% |
| hsa_circ_0044373 | 1029 | 10 | 1.94% |
| hsa_circ_0047258 | 1029 | 10 | 1.94% |
| hsa_circ_0059478 | 1029 | 10 | 1.94% |
| hsa_circ_0072482 | 1029 | 10 | 1.94% |
| hsa_circ_0081267 | 1029 | 10 | 1.94% |
| hsa_circ_0081830 | 1029 | 10 | 1.94% |
| hsa_circ_0085981 | 1029 | 10 | 1.94% |
| hsa_circ_0090322 | 1029 | 10 | 1.94% |
| hsa_circ_0108397 | 1029 | 10 | 1.94% |
| hsa_circ_0119375 | 1029 | 10 | 1.94% |
| hsa_circ_0133775 | 1029 | 10 | 1.94% |
| hsa_circ_0135491 | 1029 | 10 | 1.94% |
| hsa_circ_0138978 | 1029 | 10 | 1.94% |
| hsa_circ_0001675 | 1132 | 11 | 1.94% |
| hsa_circ_0012683 | 1132 | 11 | 1.94% |
| hsa_circ_0013251 | 1132 | 11 | 1.94% |
| hsa_circ_0044054 | 1132 | 11 | 1.94% |
| hsa_circ_0093481 | 1132 | 11 | 1.94% |
| hsa_circ_0071560 | 1235 | 12 | 1.94% |
| hsa_circ_0081016 | 1235 | 12 | 1.94% |
| hsa_circ_0037725 | 1338 | 13 | 1.94% |
| hsa_circ_0056258 | 1339 | 13 | 1.94% |
| hsa_circ_0072754 | 1236 | 12 | 1.94% |
| hsa_circ_0085876 | 1236 | 12 | 1.94% |
| hsa_circ_0086999 | 1236 | 12 | 1.94% |
| hsa_circ_0023049 | 1133 | 11 | 1.94% |
| hsa_circ_0052212 | 1133 | 11 | 1.94% |
| hsa_circ_0060466 | 1133 | 11 | 1.94% |
| hsa_circ_0099570 | 1133 | 11 | 1.94% |

| | | | |
|---|---|---|---|
| hsa_circ_0132848 | 1133 | 11 | 1.94% |
| hsa_circ_0140600 | 1133 | 11 | 1.94% |
| hsa_circ_0009956 | 1030 | 10 | 1.94% |
| hsa_circ_0011987 | 1030 | 10 | 1.94% |
| hsa_circ_0012059 | 1030 | 10 | 1.94% |
| hsa_circ_0016603 | 1030 | 10 | 1.94% |
| hsa_circ_0026381 | 1030 | 10 | 1.94% |
| hsa_circ_0032309 | 1030 | 10 | 1.94% |
| hsa_circ_0045899 | 1030 | 10 | 1.94% |
| hsa_circ_0057619 | 1030 | 10 | 1.94% |
| hsa_circ_0071510 | 1030 | 10 | 1.94% |
| hsa_circ_0080225 | 1030 | 10 | 1.94% |
| hsa_circ_0088810 | 1030 | 10 | 1.94% |
| hsa_circ_0102860 | 1030 | 10 | 1.94% |
| hsa_circ_0104441 | 1030 | 10 | 1.94% |
| hsa_circ_0127826 | 1030 | 10 | 1.94% |
| hsa_circ_0131862 | 1030 | 10 | 1.94% |
| hsa_circ_0135015 | 1030 | 10 | 1.94% |
| hsa_circ_0113133 | 1237 | 12 | 1.94% |
| hsa_circ_0008385 | 1134 | 11 | 1.94% |
| hsa_circ_0028304 | 1134 | 11 | 1.94% |
| hsa_circ_0046451 | 1134 | 11 | 1.94% |
| hsa_circ_0050005 | 1134 | 11 | 1.94% |
| hsa_circ_0055774 | 1134 | 11 | 1.94% |
| hsa_circ_0066063 | 1134 | 11 | 1.94% |
| hsa_circ_0120669 | 1134 | 11 | 1.94% |
| hsa_circ_0133330 | 1134 | 11 | 1.94% |
| hsa_circ_0005422 | 1031 | 10 | 1.94% |
| hsa_circ_0013177 | 1031 | 10 | 1.94% |
| hsa_circ_0022645 | 1031 | 10 | 1.94% |
| hsa_circ_0024211 | 1031 | 10 | 1.94% |
| hsa_circ_0027941 | 1031 | 10 | 1.94% |
| hsa_circ_0033642 | 1031 | 10 | 1.94% |
| hsa_circ_0038381 | 1031 | 10 | 1.94% |
| hsa_circ_0055081 | 1031 | 10 | 1.94% |
| hsa_circ_0058475 | 1031 | 10 | 1.94% |
| hsa_circ_0058853 | 1031 | 10 | 1.94% |
| hsa_circ_0066453 | 1031 | 10 | 1.94% |
| hsa_circ_0071135 | 1031 | 10 | 1.94% |
| hsa_circ_0073916 | 1031 | 10 | 1.94% |
| hsa_circ_0085806 | 1031 | 10 | 1.94% |
| hsa_circ_0086794 | 1031 | 10 | 1.94% |
| hsa_circ_0129953 | 1031 | 10 | 1.94% |
| hsa_circ_0057384 | 1341 | 13 | 1.94% |
| hsa_circ_0066402 | 1238 | 12 | 1.94% |
| hsa_circ_0113457 | 1238 | 12 | 1.94% |
| hsa_circ_0036786 | 1135 | 11 | 1.94% |
| hsa_circ_0036855 | 1135 | 11 | 1.94% |
| hsa_circ_0050916 | 1135 | 11 | 1.94% |
| hsa_circ_0056220 | 1135 | 11 | 1.94% |

| | | | |
|---|---|---|---|
| hsa_circ_0091269 | 1135 | 11 | 1.94% |
| hsa_circ_0006765 | 1032 | 10 | 1.94% |
| hsa_circ_0013327 | 1032 | 10 | 1.94% |
| hsa_circ_0023895 | 1032 | 10 | 1.94% |
| hsa_circ_0037484 | 1032 | 10 | 1.94% |
| hsa_circ_0043063 | 1032 | 10 | 1.94% |
| hsa_circ_0050091 | 1032 | 10 | 1.94% |
| hsa_circ_0087532 | 1032 | 10 | 1.94% |
| hsa_circ_0115878 | 1032 | 10 | 1.94% |
| hsa_circ_0124119 | 1032 | 10 | 1.94% |
| hsa_circ_0128952 | 1032 | 10 | 1.94% |
| hsa_circ_0130156 | 1032 | 10 | 1.94% |
| hsa_circ_0137119 | 1032 | 10 | 1.94% |
| hsa_circ_0138240 | 1032 | 10 | 1.94% |
| hsa_circ_0136479 | 1342 | 13 | 1.94% |
| hsa_circ_0084750 | 1239 | 12 | 1.94% |
| hsa_circ_0138828 | 1239 | 12 | 1.94% |
| hsa_circ_0023964 | 1136 | 11 | 1.94% |
| hsa_circ_0031235 | 1136 | 11 | 1.94% |
| hsa_circ_0065912 | 1136 | 11 | 1.94% |
| hsa_circ_0071458 | 1136 | 11 | 1.94% |
| hsa_circ_0092008 | 1136 | 11 | 1.94% |
| hsa_circ_0115491 | 1136 | 11 | 1.94% |
| hsa_circ_0086131 | 1446 | 14 | 1.94% |
| hsa_circ_0016760 | 1033 | 10 | 1.94% |
| hsa_circ_0022678 | 1033 | 10 | 1.94% |
| hsa_circ_0033358 | 1033 | 10 | 1.94% |
| hsa_circ_0061315 | 1033 | 10 | 1.94% |
| hsa_circ_0063344 | 1033 | 10 | 1.94% |
| hsa_circ_0076727 | 1033 | 10 | 1.94% |
| hsa_circ_0079729 | 1033 | 10 | 1.94% |
| hsa_circ_0091621 | 1033 | 10 | 1.94% |
| hsa_circ_0096533 | 1033 | 10 | 1.94% |
| hsa_circ_0104206 | 1033 | 10 | 1.94% |
| hsa_circ_0118531 | 1033 | 10 | 1.94% |
| hsa_circ_0126010 | 1033 | 10 | 1.94% |
| hsa_circ_0137820 | 1033 | 10 | 1.94% |
| hsa_circ_0060700 | 1343 | 13 | 1.94% |
| hsa_circ_0100694 | 1343 | 13 | 1.94% |
| hsa_circ_0027066 | 1137 | 11 | 1.93% |
| hsa_circ_0066136 | 1137 | 11 | 1.93% |
| hsa_circ_0074810 | 1137 | 11 | 1.93% |
| hsa_circ_0064750 | 1344 | 13 | 1.93% |
| hsa_circ_0133235 | 1551 | 15 | 1.93% |
| hsa_circ_0008184 | 1034 | 10 | 1.93% |
| hsa_circ_0009884 | 1034 | 10 | 1.93% |
| hsa_circ_0012106 | 1034 | 10 | 1.93% |
| hsa_circ_0013267 | 1034 | 10 | 1.93% |
| hsa_circ_0022227 | 1034 | 10 | 1.93% |
| hsa_circ_0032906 | 1034 | 10 | 1.93% |

| | | | |
|---|---|---|---|
| hsa_circ_0036277 | 1034 | 10 | 1.93% |
| hsa_circ_0049995 | 1034 | 10 | 1.93% |
| hsa_circ_0059840 | 1034 | 10 | 1.93% |
| hsa_circ_0060780 | 1034 | 10 | 1.93% |
| hsa_circ_0064222 | 1034 | 10 | 1.93% |
| hsa_circ_0075039 | 1034 | 10 | 1.93% |
| hsa_circ_0117066 | 1034 | 10 | 1.93% |
| hsa_circ_0037833 | 1138 | 11 | 1.93% |
| hsa_circ_0042047 | 1138 | 11 | 1.93% |
| hsa_circ_0108233 | 1138 | 11 | 1.93% |
| hsa_circ_0130860 | 1138 | 11 | 1.93% |
| hsa_circ_0132619 | 1138 | 11 | 1.93% |
| hsa_circ_0138448 | 1138 | 11 | 1.93% |
| hsa_circ_0025641 | 1035 | 10 | 1.93% |
| hsa_circ_0029561 | 1035 | 10 | 1.93% |
| hsa_circ_0039581 | 1035 | 10 | 1.93% |
| hsa_circ_0040683 | 1035 | 10 | 1.93% |
| hsa_circ_0049135 | 1035 | 10 | 1.93% |
| hsa_circ_0052864 | 1035 | 10 | 1.93% |
| hsa_circ_0066354 | 1035 | 10 | 1.93% |
| hsa_circ_0071848 | 1035 | 10 | 1.93% |
| hsa_circ_0082682 | 1035 | 10 | 1.93% |
| hsa_circ_0088222 | 1035 | 10 | 1.93% |
| hsa_circ_0091189 | 1035 | 10 | 1.93% |
| hsa_circ_0103676 | 1035 | 10 | 1.93% |
| hsa_circ_0107240 | 1035 | 10 | 1.93% |
| hsa_circ_0107962 | 1035 | 10 | 1.93% |
| hsa_circ_0113233 | 1035 | 10 | 1.93% |
| hsa_circ_0121062 | 1035 | 10 | 1.93% |
| hsa_circ_0017492 | 1139 | 11 | 1.93% |
| hsa_circ_0026382 | 1139 | 11 | 1.93% |
| hsa_circ_0027501 | 1139 | 11 | 1.93% |
| hsa_circ_0039412 | 1139 | 11 | 1.93% |
| hsa_circ_0076433 | 1139 | 11 | 1.93% |
| hsa_circ_0121528 | 1139 | 11 | 1.93% |
| hsa_circ_0074670 | 1243 | 12 | 1.93% |
| hsa_circ_0040557 | 1036 | 10 | 1.93% |
| hsa_circ_0047785 | 1036 | 10 | 1.93% |
| hsa_circ_0063834 | 1036 | 10 | 1.93% |
| hsa_circ_0064961 | 1036 | 10 | 1.93% |
| hsa_circ_0070661 | 1036 | 10 | 1.93% |
| hsa_circ_0071046 | 1036 | 10 | 1.93% |
| hsa_circ_0088541 | 1036 | 10 | 1.93% |
| hsa_circ_0108872 | 1036 | 10 | 1.93% |
| hsa_circ_0129061 | 1036 | 10 | 1.93% |
| hsa_circ_0129499 | 1036 | 10 | 1.93% |
| hsa_circ_0011306 | 1347 | 13 | 1.93% |
| hsa_circ_0014981 | 1140 | 11 | 1.93% |
| hsa_circ_0026024 | 1140 | 11 | 1.93% |
| hsa_circ_0037165 | 1140 | 11 | 1.93% |

| | | | |
|---|---|---|---|
| hsa_circ_0043529 | 1140 | 11 | 1.93% |
| hsa_circ_0051264 | 1140 | 11 | 1.93% |
| hsa_circ_0055380 | 1140 | 11 | 1.93% |
| hsa_circ_0060766 | 1140 | 11 | 1.93% |
| hsa_circ_0061953 | 1140 | 11 | 1.93% |
| hsa_circ_0067666 | 1140 | 11 | 1.93% |
| hsa_circ_0076653 | 1140 | 11 | 1.93% |
| hsa_circ_0092317 | 1140 | 11 | 1.93% |
| hsa_circ_0094479 | 1140 | 11 | 1.93% |
| hsa_circ_0118240 | 1140 | 11 | 1.93% |
| hsa_circ_0136125 | 1140 | 11 | 1.93% |
| hsa_circ_0039121 | 1451 | 14 | 1.93% |
| hsa_circ_0130726 | 1244 | 12 | 1.93% |
| hsa_circ_0021659 | 1659 | 16 | 1.93% |
| hsa_circ_0001315 | 1037 | 10 | 1.93% |
| hsa_circ_0022719 | 1037 | 10 | 1.93% |
| hsa_circ_0037095 | 1037 | 10 | 1.93% |
| hsa_circ_0041738 | 1037 | 10 | 1.93% |
| hsa_circ_0051530 | 1037 | 10 | 1.93% |
| hsa_circ_0061753 | 1037 | 10 | 1.93% |
| hsa_circ_0066161 | 1037 | 10 | 1.93% |
| hsa_circ_0083975 | 1037 | 10 | 1.93% |
| hsa_circ_0112501 | 1037 | 10 | 1.93% |
| hsa_circ_0122502 | 1037 | 10 | 1.93% |
| hsa_circ_0122943 | 1037 | 10 | 1.93% |
| hsa_circ_0133372 | 1037 | 10 | 1.93% |
| hsa_circ_0131263 | 2800 | 27 | 1.93% |
| hsa_circ_0123932 | 4252 | 41 | 1.93% |
| hsa_circ_0003928 | 1141 | 11 | 1.93% |
| hsa_circ_0054468 | 1141 | 11 | 1.93% |
| hsa_circ_0077365 | 1141 | 11 | 1.93% |
| hsa_circ_0125280 | 1141 | 11 | 1.93% |
| hsa_circ_0031927 | 1245 | 12 | 1.93% |
| hsa_circ_0037679 | 1245 | 12 | 1.93% |
| hsa_circ_0071535 | 1245 | 12 | 1.93% |
| hsa_circ_0082187 | 1245 | 12 | 1.93% |
| hsa_circ_0088560 | 1245 | 12 | 1.93% |
| hsa_circ_0123062 | 1349 | 13 | 1.93% |
| hsa_circ_0049254 | 1453 | 14 | 1.93% |
| hsa_circ_0008467 | 1038 | 10 | 1.93% |
| hsa_circ_0026411 | 1038 | 10 | 1.93% |
| hsa_circ_0026421 | 1038 | 10 | 1.93% |
| hsa_circ_0038815 | 1038 | 10 | 1.93% |
| hsa_circ_0043884 | 1038 | 10 | 1.93% |
| hsa_circ_0070315 | 1038 | 10 | 1.93% |
| hsa_circ_0080892 | 1038 | 10 | 1.93% |
| hsa_circ_0116743 | 1038 | 10 | 1.93% |
| hsa_circ_0117181 | 1038 | 10 | 1.93% |
| hsa_circ_0130454 | 1038 | 10 | 1.93% |
| hsa_circ_0047014 | 1142 | 11 | 1.93% |

| | | | |
|---|---|---|---|
| hsa_circ_0048809 | 1142 | 11 | 1.93% |
| hsa_circ_0057257 | 1142 | 11 | 1.93% |
| hsa_circ_0085452 | 1142 | 11 | 1.93% |
| hsa_circ_0136350 | 1142 | 11 | 1.93% |
| hsa_circ_0139369 | 1142 | 11 | 1.93% |
| hsa_circ_0056039 | 1246 | 12 | 1.93% |
| hsa_circ_0080467 | 3323 | 32 | 1.93% |
| hsa_circ_0080663 | 3323 | 32 | 1.93% |
| hsa_circ_0039624 | 1039 | 10 | 1.92% |
| hsa_circ_0047360 | 1039 | 10 | 1.92% |
| hsa_circ_0050536 | 1039 | 10 | 1.92% |
| hsa_circ_0060029 | 1039 | 10 | 1.92% |
| hsa_circ_0067840 | 1039 | 10 | 1.92% |
| hsa_circ_0071577 | 1039 | 10 | 1.92% |
| hsa_circ_0077084 | 1039 | 10 | 1.92% |
| hsa_circ_0094058 | 1039 | 10 | 1.92% |
| hsa_circ_0097848 | 1039 | 10 | 1.92% |
| hsa_circ_0105803 | 2286 | 22 | 1.92% |
| hsa_circ_0002405 | 1143 | 11 | 1.92% |
| hsa_circ_0003093 | 1143 | 11 | 1.92% |
| hsa_circ_0012387 | 1143 | 11 | 1.92% |
| hsa_circ_0055766 | 1143 | 11 | 1.92% |
| hsa_circ_0087814 | 1143 | 11 | 1.92% |
| hsa_circ_0096629 | 1143 | 11 | 1.92% |
| hsa_circ_0072958 | 1247 | 12 | 1.92% |
| hsa_circ_0132523 | 1247 | 12 | 1.92% |
| hsa_circ_0074208 | 1351 | 13 | 1.92% |
| hsa_circ_0103986 | 2287 | 22 | 1.92% |
| hsa_circ_0096506 | 1352 | 13 | 1.92% |
| hsa_circ_0009234 | 1248 | 12 | 1.92% |
| hsa_circ_0046184 | 1144 | 11 | 1.92% |
| hsa_circ_0066828 | 1144 | 11 | 1.92% |
| hsa_circ_0080972 | 1144 | 11 | 1.92% |
| hsa_circ_0085949 | 1144 | 11 | 1.92% |
| hsa_circ_0103232 | 1144 | 11 | 1.92% |
| hsa_circ_0105143 | 1144 | 11 | 1.92% |
| hsa_circ_0138953 | 1144 | 11 | 1.92% |
| hsa_circ_0011045 | 1040 | 10 | 1.92% |
| hsa_circ_0025176 | 1040 | 10 | 1.92% |
| hsa_circ_0025264 | 1040 | 10 | 1.92% |
| hsa_circ_0050429 | 1040 | 10 | 1.92% |
| hsa_circ_0057689 | 1040 | 10 | 1.92% |
| hsa_circ_0083305 | 1040 | 10 | 1.92% |
| hsa_circ_0092287 | 1040 | 10 | 1.92% |
| hsa_circ_0111137 | 1040 | 10 | 1.92% |
| hsa_circ_0118869 | 1040 | 10 | 1.92% |
| hsa_circ_0122108 | 1040 | 10 | 1.92% |
| hsa_circ_0122399 | 1040 | 10 | 1.92% |
| hsa_circ_0047685 | 2601 | 25 | 1.92% |
| hsa_circ_0053218 | 1353 | 13 | 1.92% |

| | | | |
|---|---|---|---|
| hsa_circ_0106554 | 1353 | 13 | 1.92% |
| hsa_circ_0076300 | 1249 | 12 | 1.92% |
| hsa_circ_0004646 | 1145 | 11 | 1.92% |
| hsa_circ_0111363 | 1145 | 11 | 1.92% |
| hsa_circ_0132549 | 1145 | 11 | 1.92% |
| hsa_circ_0001748 | 1041 | 10 | 1.92% |
| hsa_circ_0051019 | 1041 | 10 | 1.92% |
| hsa_circ_0051036 | 1041 | 10 | 1.92% |
| hsa_circ_0096226 | 1041 | 10 | 1.92% |
| hsa_circ_0097515 | 1041 | 10 | 1.92% |
| hsa_circ_0108474 | 1041 | 10 | 1.92% |
| hsa_circ_0110280 | 1041 | 10 | 1.92% |
| hsa_circ_0127583 | 1041 | 10 | 1.92% |
| hsa_circ_0135417 | 1041 | 10 | 1.92% |
| hsa_circ_0136333 | 1041 | 10 | 1.92% |
| hsa_circ_0012847 | 1354 | 13 | 1.92% |
| hsa_circ_0090274 | 1354 | 13 | 1.92% |
| hsa_circ_0133287 | 1354 | 13 | 1.92% |
| hsa_circ_0083853 | 2604 | 25 | 1.92% |
| hsa_circ_0029435 | 1250 | 12 | 1.92% |
| hsa_circ_0132524 | 1250 | 12 | 1.92% |
| hsa_circ_0016906 | 1146 | 11 | 1.92% |
| hsa_circ_0022263 | 1146 | 11 | 1.92% |
| hsa_circ_0065418 | 1146 | 11 | 1.92% |
| hsa_circ_0071645 | 1146 | 11 | 1.92% |
| hsa_circ_0018912 | 1042 | 10 | 1.92% |
| hsa_circ_0037191 | 1042 | 10 | 1.92% |
| hsa_circ_0037621 | 1042 | 10 | 1.92% |
| hsa_circ_0051065 | 1042 | 10 | 1.92% |
| hsa_circ_0058942 | 1042 | 10 | 1.92% |
| hsa_circ_0060501 | 1042 | 10 | 1.92% |
| hsa_circ_0079828 | 1042 | 10 | 1.92% |
| hsa_circ_0110284 | 1042 | 10 | 1.92% |
| hsa_circ_0111765 | 1042 | 10 | 1.92% |
| hsa_circ_0121665 | 1042 | 10 | 1.92% |
| hsa_circ_0123429 | 1042 | 10 | 1.92% |
| hsa_circ_0126048 | 1042 | 10 | 1.92% |
| hsa_circ_0051769 | 1459 | 14 | 1.92% |
| hsa_circ_0055249 | 1355 | 13 | 1.92% |
| hsa_circ_0060673 | 1355 | 13 | 1.92% |
| hsa_circ_0003634 | 1251 | 12 | 1.92% |
| hsa_circ_0091183 | 1251 | 12 | 1.92% |
| hsa_circ_0044418 | 1147 | 11 | 1.92% |
| hsa_circ_0055709 | 1147 | 11 | 1.92% |
| hsa_circ_0019262 | 1043 | 10 | 1.92% |
| hsa_circ_0026518 | 1043 | 10 | 1.92% |
| hsa_circ_0030249 | 1043 | 10 | 1.92% |
| hsa_circ_0031764 | 1043 | 10 | 1.92% |
| hsa_circ_0033043 | 1043 | 10 | 1.92% |
| hsa_circ_0048453 | 1043 | 10 | 1.92% |

| | | | |
|---|---|---|---|
| hsa_circ_0058964 | 1043 | 10 | 1.92% |
| hsa_circ_0077613 | 1043 | 10 | 1.92% |
| hsa_circ_0078242 | 1043 | 10 | 1.92% |
| hsa_circ_0080327 | 1043 | 10 | 1.92% |
| hsa_circ_0117599 | 1043 | 10 | 1.92% |
| hsa_circ_0130519 | 1043 | 10 | 1.92% |
| hsa_circ_0055866 | 1356 | 13 | 1.92% |
| hsa_circ_0022579 | 1148 | 11 | 1.92% |
| hsa_circ_0041803 | 1148 | 11 | 1.92% |
| hsa_circ_0047858 | 1148 | 11 | 1.92% |
| hsa_circ_0065698 | 1148 | 11 | 1.92% |
| hsa_circ_0117747 | 1148 | 11 | 1.92% |
| hsa_circ_0082307 | 1357 | 13 | 1.92% |
| hsa_circ_0009070 | 1044 | 10 | 1.92% |
| hsa_circ_0015882 | 1044 | 10 | 1.92% |
| hsa_circ_0022497 | 1044 | 10 | 1.92% |
| hsa_circ_0032409 | 1044 | 10 | 1.92% |
| hsa_circ_0039338 | 1044 | 10 | 1.92% |
| hsa_circ_0041100 | 1044 | 10 | 1.92% |
| hsa_circ_0050389 | 1044 | 10 | 1.92% |
| hsa_circ_0051173 | 1044 | 10 | 1.92% |
| hsa_circ_0057307 | 1044 | 10 | 1.92% |
| hsa_circ_0061720 | 1044 | 10 | 1.92% |
| hsa_circ_0062186 | 1044 | 10 | 1.92% |
| hsa_circ_0062662 | 1044 | 10 | 1.92% |
| hsa_circ_0073334 | 1044 | 10 | 1.92% |
| hsa_circ_0075002 | 1044 | 10 | 1.92% |
| hsa_circ_0083777 | 1044 | 10 | 1.92% |
| hsa_circ_0099038 | 1044 | 10 | 1.92% |
| hsa_circ_0102373 | 1044 | 10 | 1.92% |
| hsa_circ_0109409 | 1044 | 10 | 1.92% |
| hsa_circ_0111545 | 1044 | 10 | 1.92% |
| hsa_circ_0128526 | 1044 | 10 | 1.92% |
| hsa_circ_0133986 | 1044 | 10 | 1.92% |
| hsa_circ_0135501 | 1044 | 10 | 1.92% |
| hsa_circ_0107932 | 1253 | 12 | 1.92% |
| hsa_circ_0029788 | 1149 | 11 | 1.91% |
| hsa_circ_0038471 | 1149 | 11 | 1.91% |
| hsa_circ_0044607 | 1149 | 11 | 1.91% |
| hsa_circ_0067442 | 1149 | 11 | 1.91% |
| hsa_circ_0116180 | 1149 | 11 | 1.91% |
| hsa_circ_0138268 | 1149 | 11 | 1.91% |
| hsa_circ_0028124 | 1254 | 12 | 1.91% |
| hsa_circ_0047759 | 1254 | 12 | 1.91% |
| hsa_circ_0062873 | 1254 | 12 | 1.91% |
| hsa_circ_0063882 | 1254 | 12 | 1.91% |
| hsa_circ_0106239 | 1254 | 12 | 1.91% |
| hsa_circ_0115358 | 1254 | 12 | 1.91% |
| hsa_circ_0013384 | 1045 | 10 | 1.91% |
| hsa_circ_0022954 | 1045 | 10 | 1.91% |

| | | | |
|---|---|---|---|
| hsa_circ_0029981 | 1045 | 10 | 1.91% |
| hsa_circ_0038827 | 1045 | 10 | 1.91% |
| hsa_circ_0040751 | 1045 | 10 | 1.91% |
| hsa_circ_0041784 | 1045 | 10 | 1.91% |
| hsa_circ_0051090 | 1045 | 10 | 1.91% |
| hsa_circ_0062361 | 1045 | 10 | 1.91% |
| hsa_circ_0090407 | 1045 | 10 | 1.91% |
| hsa_circ_0101157 | 1045 | 10 | 1.91% |
| hsa_circ_0108211 | 1045 | 10 | 1.91% |
| hsa_circ_0115610 | 1045 | 10 | 1.91% |
| hsa_circ_0001773 | 1150 | 11 | 1.91% |
| hsa_circ_0026058 | 1150 | 11 | 1.91% |
| hsa_circ_0051766 | 1150 | 11 | 1.91% |
| hsa_circ_0042389 | 1464 | 14 | 1.91% |
| hsa_circ_0074662 | 1255 | 12 | 1.91% |
| hsa_circ_0090585 | 1255 | 12 | 1.91% |
| hsa_circ_0004673 | 1046 | 10 | 1.91% |
| hsa_circ_0014482 | 1046 | 10 | 1.91% |
| hsa_circ_0014954 | 1046 | 10 | 1.91% |
| hsa_circ_0049550 | 1046 | 10 | 1.91% |
| hsa_circ_0063913 | 1046 | 10 | 1.91% |
| hsa_circ_0092066 | 1046 | 10 | 1.91% |
| hsa_circ_0106869 | 1046 | 10 | 1.91% |
| hsa_circ_0107214 | 1046 | 10 | 1.91% |
| hsa_circ_0112320 | 1046 | 10 | 1.91% |
| hsa_circ_0116061 | 1046 | 10 | 1.91% |
| hsa_circ_0131939 | 1046 | 10 | 1.91% |
| hsa_circ_0065519 | 1360 | 13 | 1.91% |
| hsa_circ_0068316 | 1360 | 13 | 1.91% |
| hsa_circ_0018518 | 1151 | 11 | 1.91% |
| hsa_circ_0020202 | 1151 | 11 | 1.91% |
| hsa_circ_0043913 | 1151 | 11 | 1.91% |
| hsa_circ_0046502 | 1151 | 11 | 1.91% |
| hsa_circ_0048827 | 1151 | 11 | 1.91% |
| hsa_circ_0049213 | 1151 | 11 | 1.91% |
| hsa_circ_0051064 | 1151 | 11 | 1.91% |
| hsa_circ_0110229 | 1151 | 11 | 1.91% |
| hsa_circ_0115791 | 1151 | 11 | 1.91% |
| hsa_circ_0119461 | 1151 | 11 | 1.91% |
| hsa_circ_0121712 | 1151 | 11 | 1.91% |
| hsa_circ_0068084 | 5651 | 54 | 1.91% |
| hsa_circ_0014097 | 1047 | 10 | 1.91% |
| hsa_circ_0018731 | 1047 | 10 | 1.91% |
| hsa_circ_0028751 | 1047 | 10 | 1.91% |
| hsa_circ_0030083 | 1047 | 10 | 1.91% |
| hsa_circ_0031496 | 1047 | 10 | 1.91% |
| hsa_circ_0043559 | 1047 | 10 | 1.91% |
| hsa_circ_0058994 | 1047 | 10 | 1.91% |
| hsa_circ_0065906 | 1047 | 10 | 1.91% |
| hsa_circ_0068507 | 1047 | 10 | 1.91% |

| | | | |
|---|---|---|---|
| hsa_circ_0074181 | 1047 | 10 | 1.91% |
| hsa_circ_0086954 | 1047 | 10 | 1.91% |
| hsa_circ_0088739 | 1047 | 10 | 1.91% |
| hsa_circ_0102910 | 1047 | 10 | 1.91% |
| hsa_circ_0107060 | 1047 | 10 | 1.91% |
| hsa_circ_0109186 | 1047 | 10 | 1.91% |
| hsa_circ_0026743 | 1152 | 11 | 1.91% |
| hsa_circ_0029122 | 1152 | 11 | 1.91% |
| hsa_circ_0052380 | 1152 | 11 | 1.91% |
| hsa_circ_0057059 | 1152 | 11 | 1.91% |
| hsa_circ_0067662 | 1152 | 11 | 1.91% |
| hsa_circ_0117368 | 1152 | 11 | 1.91% |
| hsa_circ_0129627 | 1152 | 11 | 1.91% |
| hsa_circ_0088049 | 1257 | 12 | 1.91% |
| hsa_circ_0026242 | 1362 | 13 | 1.91% |
| hsa_circ_0090519 | 1362 | 13 | 1.91% |
| hsa_circ_0111903 | 4401 | 42 | 1.91% |
| hsa_circ_0081440 | 1467 | 14 | 1.91% |
| hsa_circ_0022681 | 1048 | 10 | 1.91% |
| hsa_circ_0026440 | 1048 | 10 | 1.91% |
| hsa_circ_0028234 | 1048 | 10 | 1.91% |
| hsa_circ_0055099 | 1048 | 10 | 1.91% |
| hsa_circ_0057251 | 1048 | 10 | 1.91% |
| hsa_circ_0062958 | 1048 | 10 | 1.91% |
| hsa_circ_0092873 | 1048 | 10 | 1.91% |
| hsa_circ_0121950 | 1048 | 10 | 1.91% |
| hsa_circ_0130022 | 1048 | 10 | 1.91% |
| hsa_circ_0036193 | 1153 | 11 | 1.91% |
| hsa_circ_0044071 | 1153 | 11 | 1.91% |
| hsa_circ_0066188 | 1153 | 11 | 1.91% |
| hsa_circ_0066600 | 1153 | 11 | 1.91% |
| hsa_circ_0079515 | 1153 | 11 | 1.91% |
| hsa_circ_0086393 | 1153 | 11 | 1.91% |
| hsa_circ_0011358 | 1049 | 10 | 1.91% |
| hsa_circ_0018078 | 1049 | 10 | 1.91% |
| hsa_circ_0040348 | 1049 | 10 | 1.91% |
| hsa_circ_0043054 | 1049 | 10 | 1.91% |
| hsa_circ_0050059 | 1049 | 10 | 1.91% |
| hsa_circ_0104435 | 1049 | 10 | 1.91% |
| hsa_circ_0122029 | 1049 | 10 | 1.91% |
| hsa_circ_0137048 | 1049 | 10 | 1.91% |
| hsa_circ_0138778 | 1049 | 10 | 1.91% |
| hsa_circ_0020996 | 1154 | 11 | 1.91% |
| hsa_circ_0022040 | 1154 | 11 | 1.91% |
| hsa_circ_0059745 | 1154 | 11 | 1.91% |
| hsa_circ_0068571 | 1154 | 11 | 1.91% |
| hsa_circ_0073064 | 1154 | 11 | 1.91% |
| hsa_circ_0077599 | 1154 | 11 | 1.91% |
| hsa_circ_0094825 | 1154 | 11 | 1.91% |
| hsa_circ_0002822 | 1259 | 12 | 1.91% |

| | | | |
|---|---|---|---|
| hsa_circ_0024738 | 1259 | 12 | 1.91% |
| hsa_circ_0080255 | 1259 | 12 | 1.91% |
| hsa_circ_0088885 | 1259 | 12 | 1.91% |
| hsa_circ_0100273 | 1259 | 12 | 1.91% |
| hsa_circ_0072637 | 1364 | 13 | 1.91% |
| hsa_circ_0065406 | 1365 | 13 | 1.90% |
| hsa_circ_0140486 | 1260 | 12 | 1.90% |
| hsa_circ_0009045 | 1155 | 11 | 1.90% |
| hsa_circ_0014339 | 1155 | 11 | 1.90% |
| hsa_circ_0023386 | 1155 | 11 | 1.90% |
| hsa_circ_0027533 | 1155 | 11 | 1.90% |
| hsa_circ_0047183 | 1155 | 11 | 1.90% |
| hsa_circ_0052043 | 1155 | 11 | 1.90% |
| hsa_circ_0056701 | 1155 | 11 | 1.90% |
| hsa_circ_0063412 | 1155 | 11 | 1.90% |
| hsa_circ_0064302 | 1155 | 11 | 1.90% |
| hsa_circ_0084453 | 1155 | 11 | 1.90% |
| hsa_circ_0098519 | 1155 | 11 | 1.90% |
| hsa_circ_0006204 | 1050 | 10 | 1.90% |
| hsa_circ_0007271 | 1050 | 10 | 1.90% |
| hsa_circ_0014597 | 1050 | 10 | 1.90% |
| hsa_circ_0017014 | 1050 | 10 | 1.90% |
| hsa_circ_0035616 | 1050 | 10 | 1.90% |
| hsa_circ_0043613 | 1050 | 10 | 1.90% |
| hsa_circ_0050392 | 1050 | 10 | 1.90% |
| hsa_circ_0060090 | 1050 | 10 | 1.90% |
| hsa_circ_0068175 | 1050 | 10 | 1.90% |
| hsa_circ_0071676 | 1050 | 10 | 1.90% |
| hsa_circ_0079825 | 1050 | 10 | 1.90% |
| hsa_circ_0080273 | 1050 | 10 | 1.90% |
| hsa_circ_0087304 | 1050 | 10 | 1.90% |
| hsa_circ_0092181 | 1050 | 10 | 1.90% |
| hsa_circ_0110555 | 1050 | 10 | 1.90% |
| hsa_circ_0114938 | 1050 | 10 | 1.90% |
| hsa_circ_0129796 | 1050 | 10 | 1.90% |
| hsa_circ_0137870 | 1050 | 10 | 1.90% |
| hsa_circ_0140683 | 1050 | 10 | 1.90% |
| hsa_circ_0129080 | 1366 | 13 | 1.90% |
| hsa_circ_0132515 | 1261 | 12 | 1.90% |
| hsa_circ_0041751 | 1156 | 11 | 1.90% |
| hsa_circ_0098469 | 1156 | 11 | 1.90% |
| hsa_circ_0117858 | 1156 | 11 | 1.90% |
| hsa_circ_0131435 | 1156 | 11 | 1.90% |
| hsa_circ_0006046 | 1051 | 10 | 1.90% |
| hsa_circ_0007387 | 1051 | 10 | 1.90% |
| hsa_circ_0014314 | 1051 | 10 | 1.90% |
| hsa_circ_0014334 | 1051 | 10 | 1.90% |
| hsa_circ_0023888 | 1051 | 10 | 1.90% |
| hsa_circ_0031266 | 1051 | 10 | 1.90% |
| hsa_circ_0034356 | 1051 | 10 | 1.90% |

| | | | |
|---|---|---|---|
| hsa_circ_0034928 | 1051 | 10 | 1.90% |
| hsa_circ_0052465 | 1051 | 10 | 1.90% |
| hsa_circ_0085980 | 1051 | 10 | 1.90% |
| hsa_circ_0088155 | 1051 | 10 | 1.90% |
| hsa_circ_0095038 | 1051 | 10 | 1.90% |
| hsa_circ_0107228 | 1051 | 10 | 1.90% |
| hsa_circ_0109332 | 1051 | 10 | 1.90% |
| hsa_circ_0123398 | 1051 | 10 | 1.90% |
| hsa_circ_0126308 | 1051 | 10 | 1.90% |
| hsa_circ_0131737 | 1051 | 10 | 1.90% |
| hsa_circ_0137846 | 1051 | 10 | 1.90% |
| hsa_circ_0060554 | 2734 | 26 | 1.90% |
| hsa_circ_0009463 | 1157 | 11 | 1.90% |
| hsa_circ_0026394 | 1157 | 11 | 1.90% |
| hsa_circ_0029163 | 1157 | 11 | 1.90% |
| hsa_circ_0096965 | 1157 | 11 | 1.90% |
| hsa_circ_0119920 | 1157 | 11 | 1.90% |
| hsa_circ_0022136 | 1052 | 10 | 1.90% |
| hsa_circ_0028656 | 1052 | 10 | 1.90% |
| hsa_circ_0035811 | 1052 | 10 | 1.90% |
| hsa_circ_0047452 | 1052 | 10 | 1.90% |
| hsa_circ_0065781 | 1052 | 10 | 1.90% |
| hsa_circ_0108340 | 1052 | 10 | 1.90% |
| hsa_circ_0112509 | 1052 | 10 | 1.90% |
| hsa_circ_0117736 | 1052 | 10 | 1.90% |
| hsa_circ_0133551 | 1052 | 10 | 1.90% |
| hsa_circ_0134711 | 1052 | 10 | 1.90% |
| hsa_circ_0136794 | 1052 | 10 | 1.90% |
| hsa_circ_0138493 | 1052 | 10 | 1.90% |
| hsa_circ_0040584 | 3157 | 30 | 1.90% |
| hsa_circ_0031090 | 1263 | 12 | 1.90% |
| hsa_circ_0041563 | 1263 | 12 | 1.90% |
| hsa_circ_0073005 | 1263 | 12 | 1.90% |
| hsa_circ_0126641 | 1263 | 12 | 1.90% |
| hsa_circ_0009134 | 1158 | 11 | 1.90% |
| hsa_circ_0009936 | 1158 | 11 | 1.90% |
| hsa_circ_0037345 | 1158 | 11 | 1.90% |
| hsa_circ_0040058 | 1158 | 11 | 1.90% |
| hsa_circ_0056993 | 1158 | 11 | 1.90% |
| hsa_circ_0059517 | 1158 | 11 | 1.90% |
| hsa_circ_0062098 | 1158 | 11 | 1.90% |
| hsa_circ_0073961 | 1158 | 11 | 1.90% |
| hsa_circ_0086036 | 1158 | 11 | 1.90% |
| hsa_circ_0114832 | 1158 | 11 | 1.90% |
| hsa_circ_0132547 | 1158 | 11 | 1.90% |
| hsa_circ_0133747 | 1158 | 11 | 1.90% |
| hsa_circ_0135342 | 1158 | 11 | 1.90% |
| hsa_circ_0011813 | 1053 | 10 | 1.90% |
| hsa_circ_0031389 | 1053 | 10 | 1.90% |
| hsa_circ_0039273 | 1053 | 10 | 1.90% |

| | | | |
|---|---|---|---|
| hsa_circ_0056748 | 1053 | 10 | 1.90% |
| hsa_circ_0059322 | 1053 | 10 | 1.90% |
| hsa_circ_0062190 | 1053 | 10 | 1.90% |
| hsa_circ_0068456 | 1053 | 10 | 1.90% |
| hsa_circ_0070866 | 1053 | 10 | 1.90% |
| hsa_circ_0071850 | 1053 | 10 | 1.90% |
| hsa_circ_0071949 | 1053 | 10 | 1.90% |
| hsa_circ_0083811 | 1053 | 10 | 1.90% |
| hsa_circ_0096273 | 1053 | 10 | 1.90% |
| hsa_circ_0104385 | 1053 | 10 | 1.90% |
| hsa_circ_0107350 | 1053 | 10 | 1.90% |
| hsa_circ_0120678 | 1053 | 10 | 1.90% |
| hsa_circ_0137693 | 1053 | 10 | 1.90% |
| hsa_circ_0013429 | 1369 | 13 | 1.90% |
| hsa_circ_0039080 | 1264 | 12 | 1.90% |
| hsa_circ_0007299 | 1159 | 11 | 1.90% |
| hsa_circ_0038176 | 1159 | 11 | 1.90% |
| hsa_circ_0122835 | 1159 | 11 | 1.90% |
| hsa_circ_0140327 | 1159 | 11 | 1.90% |
| hsa_circ_0000831 | 1054 | 10 | 1.90% |
| hsa_circ_0011183 | 1054 | 10 | 1.90% |
| hsa_circ_0019683 | 1054 | 10 | 1.90% |
| hsa_circ_0023519 | 1054 | 10 | 1.90% |
| hsa_circ_0030367 | 1054 | 10 | 1.90% |
| hsa_circ_0041233 | 1054 | 10 | 1.90% |
| hsa_circ_0047985 | 1054 | 10 | 1.90% |
| hsa_circ_0060615 | 1054 | 10 | 1.90% |
| hsa_circ_0063699 | 1054 | 10 | 1.90% |
| hsa_circ_0071862 | 1054 | 10 | 1.90% |
| hsa_circ_0089226 | 1054 | 10 | 1.90% |
| hsa_circ_0094303 | 1054 | 10 | 1.90% |
| hsa_circ_0112880 | 1054 | 10 | 1.90% |
| hsa_circ_0115421 | 1054 | 10 | 1.90% |
| hsa_circ_0116055 | 1054 | 10 | 1.90% |
| hsa_circ_0117558 | 1054 | 10 | 1.90% |
| hsa_circ_0117647 | 1054 | 10 | 1.90% |
| hsa_circ_0123927 | 1054 | 10 | 1.90% |
| hsa_circ_0127945 | 1054 | 10 | 1.90% |
| hsa_circ_0130073 | 1054 | 10 | 1.90% |
| hsa_circ_0136987 | 1054 | 10 | 1.90% |
| hsa_circ_0070367 | 1265 | 12 | 1.90% |
| hsa_circ_0026471 | 1160 | 11 | 1.90% |
| hsa_circ_0029188 | 1160 | 11 | 1.90% |
| hsa_circ_0029556 | 1160 | 11 | 1.90% |
| hsa_circ_0054076 | 1160 | 11 | 1.90% |
| hsa_circ_0063555 | 1160 | 11 | 1.90% |
| hsa_circ_0018428 | 1266 | 12 | 1.90% |
| hsa_circ_0048114 | 1266 | 12 | 1.90% |
| hsa_circ_0086143 | 1266 | 12 | 1.90% |
| hsa_circ_0014203 | 1055 | 10 | 1.90% |

| | | | |
|---|---|---|---|
| hsa_circ_0037917 | 1055 | 10 | 1.90% |
| hsa_circ_0056593 | 1055 | 10 | 1.90% |
| hsa_circ_0057945 | 1055 | 10 | 1.90% |
| hsa_circ_0088269 | 1055 | 10 | 1.90% |
| hsa_circ_0092763 | 1055 | 10 | 1.90% |
| hsa_circ_0099234 | 1055 | 10 | 1.90% |
| hsa_circ_0129277 | 1055 | 10 | 1.90% |
| hsa_circ_0139216 | 1055 | 10 | 1.90% |
| hsa_circ_0029633 | 1161 | 11 | 1.89% |
| hsa_circ_0037786 | 1161 | 11 | 1.89% |
| hsa_circ_0047309 | 1161 | 11 | 1.89% |
| hsa_circ_0052248 | 1161 | 11 | 1.89% |
| hsa_circ_0071288 | 1161 | 11 | 1.89% |
| hsa_circ_0098654 | 1161 | 11 | 1.89% |
| hsa_circ_0098812 | 1161 | 11 | 1.89% |
| hsa_circ_0099070 | 1161 | 11 | 1.89% |
| hsa_circ_0104510 | 1161 | 11 | 1.89% |
| hsa_circ_0111102 | 1161 | 11 | 1.89% |
| hsa_circ_0047644 | 1267 | 12 | 1.89% |
| hsa_circ_0057033 | 1267 | 12 | 1.89% |
| hsa_circ_0101260 | 1267 | 12 | 1.89% |
| hsa_circ_0103770 | 1267 | 12 | 1.89% |
| hsa_circ_0020658 | 1584 | 15 | 1.89% |
| hsa_circ_0012632 | 1056 | 10 | 1.89% |
| hsa_circ_0023768 | 1056 | 10 | 1.89% |
| hsa_circ_0032319 | 1056 | 10 | 1.89% |
| hsa_circ_0042813 | 1056 | 10 | 1.89% |
| hsa_circ_0043739 | 1056 | 10 | 1.89% |
| hsa_circ_0069140 | 1056 | 10 | 1.89% |
| hsa_circ_0086207 | 1056 | 10 | 1.89% |
| hsa_circ_0090387 | 1056 | 10 | 1.89% |
| hsa_circ_0090580 | 1056 | 10 | 1.89% |
| hsa_circ_0093893 | 1056 | 10 | 1.89% |
| hsa_circ_0114434 | 1056 | 10 | 1.89% |
| hsa_circ_0127112 | 1056 | 10 | 1.89% |
| hsa_circ_0029134 | 1373 | 13 | 1.89% |
| hsa_circ_0098850 | 1162 | 11 | 1.89% |
| hsa_circ_0128757 | 1162 | 11 | 1.89% |
| hsa_circ_0024598 | 1268 | 12 | 1.89% |
| hsa_circ_0048912 | 1268 | 12 | 1.89% |
| hsa_circ_0107615 | 1268 | 12 | 1.89% |
| hsa_circ_0008199 | 1057 | 10 | 1.89% |
| hsa_circ_0010587 | 1057 | 10 | 1.89% |
| hsa_circ_0014483 | 1057 | 10 | 1.89% |
| hsa_circ_0015448 | 1057 | 10 | 1.89% |
| hsa_circ_0037665 | 1057 | 10 | 1.89% |
| hsa_circ_0040762 | 1057 | 10 | 1.89% |
| hsa_circ_0057812 | 1057 | 10 | 1.89% |
| hsa_circ_0064011 | 1057 | 10 | 1.89% |
| hsa_circ_0089379 | 1057 | 10 | 1.89% |

| | | | |
|---|---|---|---|
| hsa_circ_0098215 | 1057 | 10 | 1.89% |
| hsa_circ_0099176 | 1057 | 10 | 1.89% |
| hsa_circ_0103640 | 1057 | 10 | 1.89% |
| hsa_circ_0111815 | 1057 | 10 | 1.89% |
| hsa_circ_0130298 | 1057 | 10 | 1.89% |
| hsa_circ_0140374 | 1057 | 10 | 1.89% |
| hsa_circ_0002647 | 1163 | 11 | 1.89% |
| hsa_circ_0009903 | 1163 | 11 | 1.89% |
| hsa_circ_0063246 | 1163 | 11 | 1.89% |
| hsa_circ_0064175 | 1163 | 11 | 1.89% |
| hsa_circ_0032047 | 1269 | 12 | 1.89% |
| hsa_circ_0049278 | 1269 | 12 | 1.89% |
| hsa_circ_0057193 | 1269 | 12 | 1.89% |
| hsa_circ_0068847 | 1269 | 12 | 1.89% |
| hsa_circ_0089574 | 1269 | 12 | 1.89% |
| hsa_circ_0116248 | 1269 | 12 | 1.89% |
| hsa_circ_0046071 | 1375 | 13 | 1.89% |
| hsa_circ_0122071 | 1375 | 13 | 1.89% |
| hsa_circ_0018103 | 1058 | 10 | 1.89% |
| hsa_circ_0035922 | 1058 | 10 | 1.89% |
| hsa_circ_0036513 | 1058 | 10 | 1.89% |
| hsa_circ_0050338 | 1058 | 10 | 1.89% |
| hsa_circ_0069617 | 1058 | 10 | 1.89% |
| hsa_circ_0071612 | 1058 | 10 | 1.89% |
| hsa_circ_0092439 | 1058 | 10 | 1.89% |
| hsa_circ_0096245 | 1058 | 10 | 1.89% |
| hsa_circ_0100679 | 1058 | 10 | 1.89% |
| hsa_circ_0124692 | 1058 | 10 | 1.89% |
| hsa_circ_0124694 | 1058 | 10 | 1.89% |
| hsa_circ_0126832 | 1058 | 10 | 1.89% |
| hsa_circ_0134438 | 1058 | 10 | 1.89% |
| hsa_circ_0135971 | 1058 | 10 | 1.89% |
| hsa_circ_0136989 | 1058 | 10 | 1.89% |
| hsa_circ_0036244 | 1164 | 11 | 1.89% |
| hsa_circ_0042186 | 1164 | 11 | 1.89% |
| hsa_circ_0051316 | 1164 | 11 | 1.89% |
| hsa_circ_0069063 | 1164 | 11 | 1.89% |
| hsa_circ_0078960 | 1164 | 11 | 1.89% |
| hsa_circ_0091207 | 1164 | 11 | 1.89% |
| hsa_circ_0002616 | 2540 | 24 | 1.89% |
| hsa_circ_0007276 | 1270 | 12 | 1.89% |
| hsa_circ_0010087 | 1270 | 12 | 1.89% |
| hsa_circ_0063284 | 1270 | 12 | 1.89% |
| hsa_circ_0039142 | 2011 | 19 | 1.89% |
| hsa_circ_0072313 | 1376 | 13 | 1.89% |
| hsa_circ_0046234 | 1482 | 14 | 1.89% |
| hsa_circ_0068085 | 5718 | 54 | 1.89% |
| hsa_circ_0046318 | 1059 | 10 | 1.89% |
| hsa_circ_0060628 | 1059 | 10 | 1.89% |
| hsa_circ_0075227 | 1059 | 10 | 1.89% |

| | | | |
|---|---|---|---|
| hsa_circ_0101595 | 1059 | 10 | 1.89% |
| hsa_circ_0113185 | 1059 | 10 | 1.89% |
| hsa_circ_0114478 | 1059 | 10 | 1.89% |
| hsa_circ_0117013 | 1059 | 10 | 1.89% |
| hsa_circ_0123366 | 1059 | 10 | 1.89% |
| hsa_circ_0130778 | 1059 | 10 | 1.89% |
| hsa_circ_0131106 | 1059 | 10 | 1.89% |
| hsa_circ_0137389 | 1059 | 10 | 1.89% |
| hsa_circ_0081184 | 1165 | 11 | 1.89% |
| hsa_circ_0086016 | 1165 | 11 | 1.89% |
| hsa_circ_0094658 | 1165 | 11 | 1.89% |
| hsa_circ_0111280 | 1165 | 11 | 1.89% |
| hsa_circ_0124172 | 2754 | 26 | 1.89% |
| hsa_circ_0057353 | 1695 | 16 | 1.89% |
| hsa_circ_0005752 | 2543 | 24 | 1.89% |
| hsa_circ_0111051 | 4557 | 43 | 1.89% |
| hsa_circ_0012110 | 1378 | 13 | 1.89% |
| hsa_circ_0067254 | 1378 | 13 | 1.89% |
| hsa_circ_0015273 | 1272 | 12 | 1.89% |
| hsa_circ_0097485 | 1272 | 12 | 1.89% |
| hsa_circ_0014747 | 1166 | 11 | 1.89% |
| hsa_circ_0016732 | 1166 | 11 | 1.89% |
| hsa_circ_0033593 | 1166 | 11 | 1.89% |
| hsa_circ_0052235 | 1166 | 11 | 1.89% |
| hsa_circ_0101842 | 1166 | 11 | 1.89% |
| hsa_circ_0103991 | 1166 | 11 | 1.89% |
| hsa_circ_0129433 | 1166 | 11 | 1.89% |
| hsa_circ_0000736 | 1060 | 10 | 1.89% |
| hsa_circ_0011862 | 1060 | 10 | 1.89% |
| hsa_circ_0014242 | 1060 | 10 | 1.89% |
| hsa_circ_0021178 | 1060 | 10 | 1.89% |
| hsa_circ_0027284 | 1060 | 10 | 1.89% |
| hsa_circ_0038950 | 1060 | 10 | 1.89% |
| hsa_circ_0055149 | 1060 | 10 | 1.89% |
| hsa_circ_0061511 | 1060 | 10 | 1.89% |
| hsa_circ_0062573 | 1060 | 10 | 1.89% |
| hsa_circ_0073982 | 1060 | 10 | 1.89% |
| hsa_circ_0078124 | 1060 | 10 | 1.89% |
| hsa_circ_0086592 | 1060 | 10 | 1.89% |
| hsa_circ_0086978 | 1060 | 10 | 1.89% |
| hsa_circ_0092326 | 1060 | 10 | 1.89% |
| hsa_circ_0098341 | 1060 | 10 | 1.89% |
| hsa_circ_0121275 | 1060 | 10 | 1.89% |
| hsa_circ_0083572 | 1591 | 15 | 1.89% |
| hsa_circ_0002801 | 2758 | 26 | 1.89% |
| hsa_circ_0009756 | 1273 | 12 | 1.89% |
| hsa_circ_0046277 | 1273 | 12 | 1.89% |
| hsa_circ_0041651 | 1167 | 11 | 1.89% |
| hsa_circ_0059246 | 1167 | 11 | 1.89% |
| hsa_circ_0068052 | 1167 | 11 | 1.89% |

| | | | |
|---|---|---|---|
| hsa_circ_0076233 | 1167 | 11 | 1.89% |
| hsa_circ_0108830 | 1167 | 11 | 1.89% |
| hsa_circ_0110192 | 1167 | 11 | 1.89% |
| hsa_circ_0123995 | 1167 | 11 | 1.89% |
| hsa_circ_0131143 | 1167 | 11 | 1.89% |
| hsa_circ_0005518 | 1061 | 10 | 1.89% |
| hsa_circ_0016241 | 1061 | 10 | 1.89% |
| hsa_circ_0040510 | 1061 | 10 | 1.89% |
| hsa_circ_0047895 | 1061 | 10 | 1.89% |
| hsa_circ_0055594 | 1061 | 10 | 1.89% |
| hsa_circ_0056194 | 1061 | 10 | 1.89% |
| hsa_circ_0097827 | 1061 | 10 | 1.89% |
| hsa_circ_0111372 | 1061 | 10 | 1.89% |
| hsa_circ_0127669 | 1061 | 10 | 1.89% |
| hsa_circ_0129971 | 1061 | 10 | 1.89% |
| hsa_circ_0063389 | 1274 | 12 | 1.88% |
| hsa_circ_0004018 | 1168 | 11 | 1.88% |
| hsa_circ_0044656 | 1168 | 11 | 1.88% |
| hsa_circ_0063167 | 1168 | 11 | 1.88% |
| hsa_circ_0068053 | 1168 | 11 | 1.88% |
| hsa_circ_0072767 | 1168 | 11 | 1.88% |
| hsa_circ_0100218 | 1168 | 11 | 1.88% |
| hsa_circ_0012900 | 1062 | 10 | 1.88% |
| hsa_circ_0029048 | 1062 | 10 | 1.88% |
| hsa_circ_0035817 | 1062 | 10 | 1.88% |
| hsa_circ_0047431 | 1062 | 10 | 1.88% |
| hsa_circ_0063507 | 1062 | 10 | 1.88% |
| hsa_circ_0063980 | 1062 | 10 | 1.88% |
| hsa_circ_0069102 | 1062 | 10 | 1.88% |
| hsa_circ_0086107 | 1062 | 10 | 1.88% |
| hsa_circ_0087893 | 1062 | 10 | 1.88% |
| hsa_circ_0093488 | 1062 | 10 | 1.88% |
| hsa_circ_0097120 | 1062 | 10 | 1.88% |
| hsa_circ_0097240 | 1062 | 10 | 1.88% |
| hsa_circ_0104302 | 1062 | 10 | 1.88% |
| hsa_circ_0105499 | 1062 | 10 | 1.88% |
| hsa_circ_0127210 | 1062 | 10 | 1.88% |
| hsa_circ_0128072 | 1381 | 13 | 1.88% |
| hsa_circ_0103776 | 1275 | 12 | 1.88% |
| hsa_circ_0107115 | 1275 | 12 | 1.88% |
| hsa_circ_0010708 | 1169 | 11 | 1.88% |
| hsa_circ_0011254 | 1169 | 11 | 1.88% |
| hsa_circ_0057758 | 1169 | 11 | 1.88% |
| hsa_circ_0064027 | 1169 | 11 | 1.88% |
| hsa_circ_0083904 | 1169 | 11 | 1.88% |
| hsa_circ_0114932 | 1169 | 11 | 1.88% |
| hsa_circ_0001480 | 1063 | 10 | 1.88% |
| hsa_circ_0005456 | 1063 | 10 | 1.88% |
| hsa_circ_0014064 | 1063 | 10 | 1.88% |
| hsa_circ_0018450 | 1063 | 10 | 1.88% |

| | | | |
|---|---|---|---|
| hsa_circ_0023449 | 1063 | 10 | 1.88% |
| hsa_circ_0043437 | 1063 | 10 | 1.88% |
| hsa_circ_0066581 | 1063 | 10 | 1.88% |
| hsa_circ_0067589 | 1063 | 10 | 1.88% |
| hsa_circ_0068236 | 1063 | 10 | 1.88% |
| hsa_circ_0075611 | 1063 | 10 | 1.88% |
| hsa_circ_0086512 | 1063 | 10 | 1.88% |
| hsa_circ_0096316 | 1063 | 10 | 1.88% |
| hsa_circ_0121896 | 1063 | 10 | 1.88% |
| hsa_circ_0138026 | 1063 | 10 | 1.88% |
| hsa_circ_0027287 | 1382 | 13 | 1.88% |
| hsa_circ_0057364 | 1701 | 16 | 1.88% |
| hsa_circ_0037902 | 1595 | 15 | 1.88% |
| hsa_circ_0012064 | 1170 | 11 | 1.88% |
| hsa_circ_0015464 | 1170 | 11 | 1.88% |
| hsa_circ_0086536 | 1170 | 11 | 1.88% |
| hsa_circ_0090156 | 1170 | 11 | 1.88% |
| hsa_circ_0031004 | 1383 | 13 | 1.88% |
| hsa_circ_0015485 | 1064 | 10 | 1.88% |
| hsa_circ_0021682 | 1064 | 10 | 1.88% |
| hsa_circ_0026881 | 1064 | 10 | 1.88% |
| hsa_circ_0030568 | 1064 | 10 | 1.88% |
| hsa_circ_0037688 | 1064 | 10 | 1.88% |
| hsa_circ_0051611 | 1064 | 10 | 1.88% |
| hsa_circ_0055887 | 1064 | 10 | 1.88% |
| hsa_circ_0055915 | 1064 | 10 | 1.88% |
| hsa_circ_0071118 | 1064 | 10 | 1.88% |
| hsa_circ_0074159 | 1064 | 10 | 1.88% |
| hsa_circ_0086778 | 1064 | 10 | 1.88% |
| hsa_circ_0090006 | 1064 | 10 | 1.88% |
| hsa_circ_0090995 | 1064 | 10 | 1.88% |
| hsa_circ_0104082 | 1064 | 10 | 1.88% |
| hsa_circ_0114292 | 1064 | 10 | 1.88% |
| hsa_circ_0129418 | 1064 | 10 | 1.88% |
| hsa_circ_0133972 | 1064 | 10 | 1.88% |
| hsa_circ_0057378 | 1809 | 17 | 1.88% |
| hsa_circ_0000875 | 1277 | 12 | 1.88% |
| hsa_circ_0010042 | 1171 | 11 | 1.88% |
| hsa_circ_0017206 | 1171 | 11 | 1.88% |
| hsa_circ_0031298 | 1171 | 11 | 1.88% |
| hsa_circ_0044255 | 1171 | 11 | 1.88% |
| hsa_circ_0061668 | 1171 | 11 | 1.88% |
| hsa_circ_0068116 | 1171 | 11 | 1.88% |
| hsa_circ_0113842 | 1171 | 11 | 1.88% |
| hsa_circ_0123338 | 1171 | 11 | 1.88% |
| hsa_circ_0053128 | 1491 | 14 | 1.88% |
| hsa_circ_0083708 | 1278 | 12 | 1.88% |
| hsa_circ_0000080 | 1065 | 10 | 1.88% |
| hsa_circ_0011926 | 1065 | 10 | 1.88% |
| hsa_circ_0014216 | 1065 | 10 | 1.88% |

| | | | |
|---|---|---|---|
| hsa_circ_0021318 | 1065 | 10 | 1.88% |
| hsa_circ_0026934 | 1065 | 10 | 1.88% |
| hsa_circ_0031994 | 1065 | 10 | 1.88% |
| hsa_circ_0037706 | 1065 | 10 | 1.88% |
| hsa_circ_0039050 | 1065 | 10 | 1.88% |
| hsa_circ_0043096 | 1065 | 10 | 1.88% |
| hsa_circ_0054346 | 1065 | 10 | 1.88% |
| hsa_circ_0058886 | 1065 | 10 | 1.88% |
| hsa_circ_0060392 | 1065 | 10 | 1.88% |
| hsa_circ_0062365 | 1065 | 10 | 1.88% |
| hsa_circ_0070585 | 1065 | 10 | 1.88% |
| hsa_circ_0080318 | 1065 | 10 | 1.88% |
| hsa_circ_0085891 | 1065 | 10 | 1.88% |
| hsa_circ_0088821 | 1065 | 10 | 1.88% |
| hsa_circ_0094511 | 1065 | 10 | 1.88% |
| hsa_circ_0102196 | 1065 | 10 | 1.88% |
| hsa_circ_0102785 | 1065 | 10 | 1.88% |
| hsa_circ_0103238 | 1065 | 10 | 1.88% |
| hsa_circ_0105177 | 1065 | 10 | 1.88% |
| hsa_circ_0120387 | 1065 | 10 | 1.88% |
| hsa_circ_0124099 | 1065 | 10 | 1.88% |
| hsa_circ_0125289 | 1065 | 10 | 1.88% |
| hsa_circ_0125296 | 1065 | 10 | 1.88% |
| hsa_circ_0137558 | 1065 | 10 | 1.88% |
| hsa_circ_0035472 | 1385 | 13 | 1.88% |
| hsa_circ_0025110 | 1172 | 11 | 1.88% |
| hsa_circ_0029551 | 1172 | 11 | 1.88% |
| hsa_circ_0080577 | 1172 | 11 | 1.88% |
| hsa_circ_0081846 | 1172 | 11 | 1.88% |
| hsa_circ_0121004 | 1172 | 11 | 1.88% |
| hsa_circ_0136959 | 1172 | 11 | 1.88% |
| hsa_circ_0004395 | 1066 | 10 | 1.88% |
| hsa_circ_0013369 | 1066 | 10 | 1.88% |
| hsa_circ_0016614 | 1066 | 10 | 1.88% |
| hsa_circ_0077174 | 1066 | 10 | 1.88% |
| hsa_circ_0091897 | 1066 | 10 | 1.88% |
| hsa_circ_0098423 | 1066 | 10 | 1.88% |
| hsa_circ_0100290 | 1066 | 10 | 1.88% |
| hsa_circ_0112012 | 1066 | 10 | 1.88% |
| hsa_circ_0118704 | 1066 | 10 | 1.88% |
| hsa_circ_0119358 | 1066 | 10 | 1.88% |
| hsa_circ_0121786 | 1066 | 10 | 1.88% |
| hsa_circ_0138024 | 1066 | 10 | 1.88% |
| hsa_circ_0004744 | 1173 | 11 | 1.88% |
| hsa_circ_0008920 | 1173 | 11 | 1.88% |
| hsa_circ_0047384 | 1173 | 11 | 1.88% |
| hsa_circ_0059559 | 1173 | 11 | 1.88% |
| hsa_circ_0128379 | 1173 | 11 | 1.88% |
| hsa_circ_0024219 | 1280 | 12 | 1.88% |
| hsa_circ_0027390 | 1280 | 12 | 1.88% |

| | | | |
|---|---|---|---|
| hsa_circ_0073624 | 1280 | 12 | 1.88% |
| hsa_circ_0092105 | 1387 | 13 | 1.87% |
| hsa_circ_0064735 | 1067 | 10 | 1.87% |
| hsa_circ_0067987 | 1067 | 10 | 1.87% |
| hsa_circ_0077789 | 1067 | 10 | 1.87% |
| hsa_circ_0091017 | 1067 | 10 | 1.87% |
| hsa_circ_0097213 | 1067 | 10 | 1.87% |
| hsa_circ_0114900 | 1067 | 10 | 1.87% |
| hsa_circ_0138888 | 1067 | 10 | 1.87% |
| hsa_circ_0140080 | 1067 | 10 | 1.87% |
| hsa_circ_0022261 | 1174 | 11 | 1.87% |
| hsa_circ_0024579 | 1174 | 11 | 1.87% |
| hsa_circ_0061693 | 1174 | 11 | 1.87% |
| hsa_circ_0074342 | 1174 | 11 | 1.87% |
| hsa_circ_0106811 | 1174 | 11 | 1.87% |
| hsa_circ_0073947 | 1281 | 12 | 1.87% |
| hsa_circ_0101897 | 1281 | 12 | 1.87% |
| hsa_circ_0101943 | 1922 | 18 | 1.87% |
| hsa_circ_0012329 | 1495 | 14 | 1.87% |
| hsa_circ_0107519 | 1495 | 14 | 1.87% |
| hsa_circ_0010864 | 1068 | 10 | 1.87% |
| hsa_circ_0014239 | 1068 | 10 | 1.87% |
| hsa_circ_0016463 | 1068 | 10 | 1.87% |
| hsa_circ_0021305 | 1068 | 10 | 1.87% |
| hsa_circ_0029295 | 1068 | 10 | 1.87% |
| hsa_circ_0042861 | 1068 | 10 | 1.87% |
| hsa_circ_0054340 | 1068 | 10 | 1.87% |
| hsa_circ_0056178 | 1068 | 10 | 1.87% |
| hsa_circ_0079271 | 1068 | 10 | 1.87% |
| hsa_circ_0083113 | 1068 | 10 | 1.87% |
| hsa_circ_0090567 | 1068 | 10 | 1.87% |
| hsa_circ_0098299 | 1068 | 10 | 1.87% |
| hsa_circ_0100491 | 1068 | 10 | 1.87% |
| hsa_circ_0109274 | 1068 | 10 | 1.87% |
| hsa_circ_0115707 | 1068 | 10 | 1.87% |
| hsa_circ_0123886 | 1068 | 10 | 1.87% |
| hsa_circ_0125770 | 1068 | 10 | 1.87% |
| hsa_circ_0126539 | 1068 | 10 | 1.87% |
| hsa_circ_0019472 | 1175 | 11 | 1.87% |
| hsa_circ_0023774 | 1175 | 11 | 1.87% |
| hsa_circ_0028931 | 1175 | 11 | 1.87% |
| hsa_circ_0041622 | 1175 | 11 | 1.87% |
| hsa_circ_0073525 | 1175 | 11 | 1.87% |
| hsa_circ_0073765 | 1175 | 11 | 1.87% |
| hsa_circ_0098147 | 1175 | 11 | 1.87% |
| hsa_circ_0136459 | 1175 | 11 | 1.87% |
| hsa_circ_0040690 | 1282 | 12 | 1.87% |
| hsa_circ_0013029 | 1069 | 10 | 1.87% |
| hsa_circ_0014801 | 1069 | 10 | 1.87% |
| hsa_circ_0028203 | 1069 | 10 | 1.87% |

| | | | |
|---|---|---|---|
| hsa_circ_0039129 | 1069 | 10 | 1.87% |
| hsa_circ_0044808 | 1069 | 10 | 1.87% |
| hsa_circ_0049268 | 1069 | 10 | 1.87% |
| hsa_circ_0049296 | 1069 | 10 | 1.87% |
| hsa_circ_0049470 | 1069 | 10 | 1.87% |
| hsa_circ_0050038 | 1069 | 10 | 1.87% |
| hsa_circ_0050380 | 1069 | 10 | 1.87% |
| hsa_circ_0075715 | 1069 | 10 | 1.87% |
| hsa_circ_0084965 | 1069 | 10 | 1.87% |
| hsa_circ_0085825 | 1069 | 10 | 1.87% |
| hsa_circ_0103927 | 1069 | 10 | 1.87% |
| hsa_circ_0118453 | 1069 | 10 | 1.87% |
| hsa_circ_0119260 | 1069 | 10 | 1.87% |
| hsa_circ_0135144 | 1069 | 10 | 1.87% |
| hsa_circ_0136018 | 1069 | 10 | 1.87% |
| hsa_circ_0011333 | 3528 | 33 | 1.87% |
| hsa_circ_0015869 | 1176 | 11 | 1.87% |
| hsa_circ_0019940 | 1176 | 11 | 1.87% |
| hsa_circ_0031271 | 1176 | 11 | 1.87% |
| hsa_circ_0035945 | 1176 | 11 | 1.87% |
| hsa_circ_0041805 | 1176 | 11 | 1.87% |
| hsa_circ_0045961 | 1176 | 11 | 1.87% |
| hsa_circ_0069053 | 1176 | 11 | 1.87% |
| hsa_circ_0110679 | 1176 | 11 | 1.87% |
| hsa_circ_0135117 | 1176 | 11 | 1.87% |
| hsa_circ_0140756 | 1176 | 11 | 1.87% |
| hsa_circ_0044474 | 1283 | 12 | 1.87% |
| hsa_circ_0100063 | 4705 | 44 | 1.87% |
| hsa_circ_0116433 | 2675 | 25 | 1.87% |
| hsa_circ_0051464 | 1284 | 12 | 1.87% |
| hsa_circ_0071845 | 1284 | 12 | 1.87% |
| hsa_circ_0089307 | 1284 | 12 | 1.87% |
| hsa_circ_0135129 | 1284 | 12 | 1.87% |
| hsa_circ_0009502 | 1177 | 11 | 1.87% |
| hsa_circ_0069896 | 1177 | 11 | 1.87% |
| hsa_circ_0108806 | 1177 | 11 | 1.87% |
| hsa_circ_0117853 | 1177 | 11 | 1.87% |
| hsa_circ_0128909 | 1177 | 11 | 1.87% |
| hsa_circ_0031212 | 1070 | 10 | 1.87% |
| hsa_circ_0034452 | 1070 | 10 | 1.87% |
| hsa_circ_0058201 | 1070 | 10 | 1.87% |
| hsa_circ_0067149 | 1070 | 10 | 1.87% |
| hsa_circ_0071110 | 1070 | 10 | 1.87% |
| hsa_circ_0085982 | 1070 | 10 | 1.87% |
| hsa_circ_0095947 | 1070 | 10 | 1.87% |
| hsa_circ_0106396 | 1070 | 10 | 1.87% |
| hsa_circ_0126121 | 1070 | 10 | 1.87% |
| hsa_circ_0130764 | 1070 | 10 | 1.87% |
| hsa_circ_0048754 | 1285 | 12 | 1.87% |
| hsa_circ_0013172 | 1178 | 11 | 1.87% |

| | | | |
|---|---|---|---|
| hsa_circ_0037356 | 1178 | 11 | 1.87% |
| hsa_circ_0002552 | 1071 | 10 | 1.87% |
| hsa_circ_0011238 | 1071 | 10 | 1.87% |
| hsa_circ_0022109 | 1071 | 10 | 1.87% |
| hsa_circ_0023455 | 1071 | 10 | 1.87% |
| hsa_circ_0029425 | 1071 | 10 | 1.87% |
| hsa_circ_0051377 | 1071 | 10 | 1.87% |
| hsa_circ_0055664 | 1071 | 10 | 1.87% |
| hsa_circ_0067346 | 1071 | 10 | 1.87% |
| hsa_circ_0070270 | 1071 | 10 | 1.87% |
| hsa_circ_0075350 | 1071 | 10 | 1.87% |
| hsa_circ_0081920 | 1071 | 10 | 1.87% |
| hsa_circ_0090025 | 1071 | 10 | 1.87% |
| hsa_circ_0100622 | 1071 | 10 | 1.87% |
| hsa_circ_0103874 | 1071 | 10 | 1.87% |
| hsa_circ_0108796 | 1071 | 10 | 1.87% |
| hsa_circ_0113361 | 1071 | 10 | 1.87% |
| hsa_circ_0121839 | 1071 | 10 | 1.87% |
| hsa_circ_0048977 | 1821 | 17 | 1.87% |
| hsa_circ_0055423 | 1393 | 13 | 1.87% |
| hsa_circ_0010977 | 1286 | 12 | 1.87% |
| hsa_circ_0061214 | 1286 | 12 | 1.87% |
| hsa_circ_0097353 | 1286 | 12 | 1.87% |
| hsa_circ_0023424 | 1179 | 11 | 1.87% |
| hsa_circ_0025186 | 1179 | 11 | 1.87% |
| hsa_circ_0039554 | 1179 | 11 | 1.87% |
| hsa_circ_0062639 | 1179 | 11 | 1.87% |
| hsa_circ_0083673 | 1179 | 11 | 1.87% |
| hsa_circ_0091656 | 1179 | 11 | 1.87% |
| hsa_circ_0122334 | 1179 | 11 | 1.87% |
| hsa_circ_0131547 | 1179 | 11 | 1.87% |
| hsa_circ_0131743 | 1179 | 11 | 1.87% |
| hsa_circ_0000350 | 1072 | 10 | 1.87% |
| hsa_circ_0003548 | 1072 | 10 | 1.87% |
| hsa_circ_0018643 | 1072 | 10 | 1.87% |
| hsa_circ_0024571 | 1072 | 10 | 1.87% |
| hsa_circ_0027605 | 1072 | 10 | 1.87% |
| hsa_circ_0027609 | 1072 | 10 | 1.87% |
| hsa_circ_0031765 | 1072 | 10 | 1.87% |
| hsa_circ_0042868 | 1072 | 10 | 1.87% |
| hsa_circ_0090474 | 1072 | 10 | 1.87% |
| hsa_circ_0104771 | 1072 | 10 | 1.87% |
| hsa_circ_0123693 | 1072 | 10 | 1.87% |
| hsa_circ_0132838 | 1072 | 10 | 1.87% |
| hsa_circ_0136791 | 1072 | 10 | 1.87% |
| hsa_circ_0120148 | 3110 | 29 | 1.86% |
| hsa_circ_0064993 | 1287 | 12 | 1.86% |
| hsa_circ_0081017 | 1287 | 12 | 1.86% |
| hsa_circ_0002810 | 1180 | 11 | 1.86% |
| hsa_circ_0025497 | 1180 | 11 | 1.86% |

| | | | |
|---|---|---|---|
| hsa_circ_0037783 | 1180 | 11 | 1.86% |
| hsa_circ_0038836 | 1180 | 11 | 1.86% |
| hsa_circ_0051730 | 1180 | 11 | 1.86% |
| hsa_circ_0053073 | 1180 | 11 | 1.86% |
| hsa_circ_0083725 | 1180 | 11 | 1.86% |
| hsa_circ_0085352 | 1180 | 11 | 1.86% |
| hsa_circ_0139612 | 1180 | 11 | 1.86% |
| hsa_circ_0010393 | 1931 | 18 | 1.86% |
| hsa_circ_0000566 | 1073 | 10 | 1.86% |
| hsa_circ_0008363 | 1073 | 10 | 1.86% |
| hsa_circ_0015271 | 1073 | 10 | 1.86% |
| hsa_circ_0022513 | 1073 | 10 | 1.86% |
| hsa_circ_0024699 | 1073 | 10 | 1.86% |
| hsa_circ_0032909 | 1073 | 10 | 1.86% |
| hsa_circ_0054765 | 1073 | 10 | 1.86% |
| hsa_circ_0064698 | 1073 | 10 | 1.86% |
| hsa_circ_0073145 | 1073 | 10 | 1.86% |
| hsa_circ_0079200 | 1073 | 10 | 1.86% |
| hsa_circ_0086446 | 1073 | 10 | 1.86% |
| hsa_circ_0101167 | 1073 | 10 | 1.86% |
| hsa_circ_0107338 | 1073 | 10 | 1.86% |
| hsa_circ_0108481 | 1073 | 10 | 1.86% |
| hsa_circ_0110142 | 1073 | 10 | 1.86% |
| hsa_circ_0132096 | 1073 | 10 | 1.86% |
| hsa_circ_0017257 | 1395 | 13 | 1.86% |
| hsa_circ_0036492 | 1395 | 13 | 1.86% |
| hsa_circ_0038552 | 1395 | 13 | 1.86% |
| hsa_circ_0097510 | 1932 | 18 | 1.86% |
| hsa_circ_0021322 | 1288 | 12 | 1.86% |
| hsa_circ_0049666 | 1288 | 12 | 1.86% |
| hsa_circ_0051664 | 1288 | 12 | 1.86% |
| hsa_circ_0076254 | 1181 | 11 | 1.86% |
| hsa_circ_0040583 | 3221 | 30 | 1.86% |
| hsa_circ_0122957 | 3973 | 37 | 1.86% |
| hsa_circ_0008859 | 1074 | 10 | 1.86% |
| hsa_circ_0010935 | 1074 | 10 | 1.86% |
| hsa_circ_0011239 | 1074 | 10 | 1.86% |
| hsa_circ_0012734 | 1074 | 10 | 1.86% |
| hsa_circ_0023896 | 1074 | 10 | 1.86% |
| hsa_circ_0041515 | 1074 | 10 | 1.86% |
| hsa_circ_0042222 | 1074 | 10 | 1.86% |
| hsa_circ_0051588 | 1074 | 10 | 1.86% |
| hsa_circ_0055047 | 1074 | 10 | 1.86% |
| hsa_circ_0056788 | 1074 | 10 | 1.86% |
| hsa_circ_0072301 | 1074 | 10 | 1.86% |
| hsa_circ_0075438 | 1074 | 10 | 1.86% |
| hsa_circ_0083040 | 1074 | 10 | 1.86% |
| hsa_circ_0084896 | 1074 | 10 | 1.86% |
| hsa_circ_0085681 | 1074 | 10 | 1.86% |
| hsa_circ_0085823 | 1074 | 10 | 1.86% |

| | | | |
|---|---|---|---|
| hsa_circ_0087677 | 1074 | 10 | 1.86% |
| hsa_circ_0121875 | 1074 | 10 | 1.86% |
| hsa_circ_0135032 | 1074 | 10 | 1.86% |
| hsa_circ_0140628 | 1074 | 10 | 1.86% |
| hsa_circ_0123727 | 2900 | 27 | 1.86% |
| hsa_circ_0133437 | 1289 | 12 | 1.86% |
| hsa_circ_0014470 | 1182 | 11 | 1.86% |
| hsa_circ_0048768 | 1182 | 11 | 1.86% |
| hsa_circ_0063606 | 1182 | 11 | 1.86% |
| hsa_circ_0080116 | 1182 | 11 | 1.86% |
| hsa_circ_0087936 | 1182 | 11 | 1.86% |
| hsa_circ_0056849 | 1290 | 12 | 1.86% |
| hsa_circ_0086004 | 1290 | 12 | 1.86% |
| hsa_circ_0091411 | 1290 | 12 | 1.86% |
| hsa_circ_0125449 | 1290 | 12 | 1.86% |
| hsa_circ_0018618 | 1075 | 10 | 1.86% |
| hsa_circ_0020028 | 1075 | 10 | 1.86% |
| hsa_circ_0028842 | 1075 | 10 | 1.86% |
| hsa_circ_0065736 | 1075 | 10 | 1.86% |
| hsa_circ_0066129 | 1075 | 10 | 1.86% |
| hsa_circ_0089532 | 1075 | 10 | 1.86% |
| hsa_circ_0092785 | 1075 | 10 | 1.86% |
| hsa_circ_0096539 | 1075 | 10 | 1.86% |
| hsa_circ_0099970 | 1075 | 10 | 1.86% |
| hsa_circ_0116940 | 1075 | 10 | 1.86% |
| hsa_circ_0117431 | 1075 | 10 | 1.86% |
| hsa_circ_0118824 | 1075 | 10 | 1.86% |
| hsa_circ_0125612 | 1075 | 10 | 1.86% |
| hsa_circ_0134118 | 1075 | 10 | 1.86% |
| hsa_circ_0025575 | 1183 | 11 | 1.86% |
| hsa_circ_0044745 | 1183 | 11 | 1.86% |
| hsa_circ_0066000 | 1183 | 11 | 1.86% |
| hsa_circ_0015835 | 1291 | 12 | 1.86% |
| hsa_circ_0034855 | 1291 | 12 | 1.86% |
| hsa_circ_0126835 | 3120 | 29 | 1.86% |
| hsa_circ_0007944 | 1076 | 10 | 1.86% |
| hsa_circ_0008060 | 1076 | 10 | 1.86% |
| hsa_circ_0009621 | 1076 | 10 | 1.86% |
| hsa_circ_0016677 | 1076 | 10 | 1.86% |
| hsa_circ_0022924 | 1076 | 10 | 1.86% |
| hsa_circ_0027123 | 1076 | 10 | 1.86% |
| hsa_circ_0029941 | 1076 | 10 | 1.86% |
| hsa_circ_0033635 | 1076 | 10 | 1.86% |
| hsa_circ_0035212 | 1076 | 10 | 1.86% |
| hsa_circ_0037900 | 1076 | 10 | 1.86% |
| hsa_circ_0042271 | 1076 | 10 | 1.86% |
| hsa_circ_0052275 | 1076 | 10 | 1.86% |
| hsa_circ_0067210 | 1076 | 10 | 1.86% |
| hsa_circ_0068288 | 1076 | 10 | 1.86% |
| hsa_circ_0075917 | 1076 | 10 | 1.86% |

| | | | |
|---|---|---|---|
| hsa_circ_0078856 | 1076 | 10 | 1.86% |
| hsa_circ_0090907 | 1076 | 10 | 1.86% |
| hsa_circ_0104900 | 1076 | 10 | 1.86% |
| hsa_circ_0111296 | 1076 | 10 | 1.86% |
| hsa_circ_0113059 | 1076 | 10 | 1.86% |
| hsa_circ_0125216 | 1076 | 10 | 1.86% |
| hsa_circ_0135255 | 1076 | 10 | 1.86% |
| hsa_circ_0136204 | 1076 | 10 | 1.86% |
| hsa_circ_0021282 | 1399 | 13 | 1.86% |
| hsa_circ_0122202 | 3229 | 30 | 1.86% |
| hsa_circ_0077143 | 1184 | 11 | 1.86% |
| hsa_circ_0026937 | 1507 | 14 | 1.86% |
| hsa_circ_0029369 | 1507 | 14 | 1.86% |
| hsa_circ_0022235 | 1292 | 12 | 1.86% |
| hsa_circ_0031778 | 1292 | 12 | 1.86% |
| hsa_circ_0105280 | 1292 | 12 | 1.86% |
| hsa_circ_0021794 | 1077 | 10 | 1.86% |
| hsa_circ_0022299 | 1077 | 10 | 1.86% |
| hsa_circ_0023313 | 1077 | 10 | 1.86% |
| hsa_circ_0023416 | 1077 | 10 | 1.86% |
| hsa_circ_0037155 | 1077 | 10 | 1.86% |
| hsa_circ_0050961 | 1077 | 10 | 1.86% |
| hsa_circ_0058706 | 1077 | 10 | 1.86% |
| hsa_circ_0063181 | 1077 | 10 | 1.86% |
| hsa_circ_0065598 | 1077 | 10 | 1.86% |
| hsa_circ_0067081 | 1077 | 10 | 1.86% |
| hsa_circ_0086591 | 1077 | 10 | 1.86% |
| hsa_circ_0087116 | 1077 | 10 | 1.86% |
| hsa_circ_0110439 | 1077 | 10 | 1.86% |
| hsa_circ_0115352 | 1077 | 10 | 1.86% |
| hsa_circ_0124952 | 1077 | 10 | 1.86% |
| hsa_circ_0132097 | 1077 | 10 | 1.86% |
| hsa_circ_0013464 | 1185 | 11 | 1.86% |
| hsa_circ_0019946 | 1185 | 11 | 1.86% |
| hsa_circ_0026494 | 1185 | 11 | 1.86% |
| hsa_circ_0062103 | 1185 | 11 | 1.86% |
| hsa_circ_0074576 | 1185 | 11 | 1.86% |
| hsa_circ_0098407 | 1185 | 11 | 1.86% |
| hsa_circ_0013004 | 1293 | 12 | 1.86% |
| hsa_circ_0076495 | 1293 | 12 | 1.86% |
| hsa_circ_0092104 | 1293 | 12 | 1.86% |
| hsa_circ_0068549 | 1617 | 15 | 1.86% |
| hsa_circ_0013617 | 1078 | 10 | 1.86% |
| hsa_circ_0017267 | 1078 | 10 | 1.86% |
| hsa_circ_0019015 | 1078 | 10 | 1.86% |
| hsa_circ_0021326 | 1078 | 10 | 1.86% |
| hsa_circ_0022452 | 1078 | 10 | 1.86% |
| hsa_circ_0024446 | 1078 | 10 | 1.86% |
| hsa_circ_0028419 | 1078 | 10 | 1.86% |
| hsa_circ_0034887 | 1078 | 10 | 1.86% |

| | | | |
|---|---|---|---|
| hsa_circ_0056697 | 1078 | 10 | 1.86% |
| hsa_circ_0078834 | 1078 | 10 | 1.86% |
| hsa_circ_0093179 | 1078 | 10 | 1.86% |
| hsa_circ_0116228 | 1078 | 10 | 1.86% |
| hsa_circ_0022806 | 1186 | 11 | 1.85% |
| hsa_circ_0043158 | 1186 | 11 | 1.85% |
| hsa_circ_0053683 | 1186 | 11 | 1.85% |
| hsa_circ_0011518 | 4098 | 38 | 1.85% |
| hsa_circ_0003509 | 1510 | 14 | 1.85% |
| hsa_circ_0002520 | 1079 | 10 | 1.85% |
| hsa_circ_0013329 | 1079 | 10 | 1.85% |
| hsa_circ_0017162 | 1079 | 10 | 1.85% |
| hsa_circ_0051097 | 1079 | 10 | 1.85% |
| hsa_circ_0059309 | 1079 | 10 | 1.85% |
| hsa_circ_0068433 | 1079 | 10 | 1.85% |
| hsa_circ_0070718 | 1079 | 10 | 1.85% |
| hsa_circ_0071023 | 1079 | 10 | 1.85% |
| hsa_circ_0077841 | 1079 | 10 | 1.85% |
| hsa_circ_0085786 | 1079 | 10 | 1.85% |
| hsa_circ_0097599 | 1079 | 10 | 1.85% |
| hsa_circ_0097667 | 1079 | 10 | 1.85% |
| hsa_circ_0107359 | 1079 | 10 | 1.85% |
| hsa_circ_0109176 | 1079 | 10 | 1.85% |
| hsa_circ_0110990 | 1079 | 10 | 1.85% |
| hsa_circ_0127156 | 1079 | 10 | 1.85% |
| hsa_circ_0135661 | 1079 | 10 | 1.85% |
| hsa_circ_0012114 | 1187 | 11 | 1.85% |
| hsa_circ_0059192 | 1187 | 11 | 1.85% |
| hsa_circ_0068205 | 1187 | 11 | 1.85% |
| hsa_circ_0098529 | 1187 | 11 | 1.85% |
| hsa_circ_0104835 | 1187 | 11 | 1.85% |
| hsa_circ_0130821 | 1187 | 11 | 1.85% |
| hsa_circ_0135371 | 1187 | 11 | 1.85% |
| hsa_circ_0013425 | 1295 | 12 | 1.85% |
| hsa_circ_0016962 | 1295 | 12 | 1.85% |
| hsa_circ_0031630 | 1295 | 12 | 1.85% |
| hsa_circ_0044593 | 1295 | 12 | 1.85% |
| hsa_circ_0086098 | 1295 | 12 | 1.85% |
| hsa_circ_0108009 | 1295 | 12 | 1.85% |
| hsa_circ_0032537 | 1296 | 12 | 1.85% |
| hsa_circ_0063204 | 1296 | 12 | 1.85% |
| hsa_circ_0075464 | 1296 | 12 | 1.85% |
| hsa_circ_0136510 | 1296 | 12 | 1.85% |
| hsa_circ_0025867 | 1188 | 11 | 1.85% |
| hsa_circ_0035948 | 1188 | 11 | 1.85% |
| hsa_circ_0062102 | 1188 | 11 | 1.85% |
| hsa_circ_0064719 | 1188 | 11 | 1.85% |
| hsa_circ_0066142 | 1188 | 11 | 1.85% |
| hsa_circ_0129774 | 1188 | 11 | 1.85% |
| hsa_circ_0020478 | 1080 | 10 | 1.85% |

| | | | |
|---|---|---|---|
| hsa_circ_0020570 | 1080 | 10 | 1.85% |
| hsa_circ_0027345 | 1080 | 10 | 1.85% |
| hsa_circ_0030759 | 1080 | 10 | 1.85% |
| hsa_circ_0038635 | 1080 | 10 | 1.85% |
| hsa_circ_0039079 | 1080 | 10 | 1.85% |
| hsa_circ_0045168 | 1080 | 10 | 1.85% |
| hsa_circ_0054188 | 1080 | 10 | 1.85% |
| hsa_circ_0066560 | 1080 | 10 | 1.85% |
| hsa_circ_0069909 | 1080 | 10 | 1.85% |
| hsa_circ_0075151 | 1080 | 10 | 1.85% |
| hsa_circ_0075459 | 1080 | 10 | 1.85% |
| hsa_circ_0085454 | 1080 | 10 | 1.85% |
| hsa_circ_0128787 | 1080 | 10 | 1.85% |
| hsa_circ_0129815 | 1080 | 10 | 1.85% |
| hsa_circ_0132393 | 1080 | 10 | 1.85% |
| hsa_circ_0139168 | 1080 | 10 | 1.85% |
| hsa_circ_0110878 | 8425 | 78 | 1.85% |
| hsa_circ_0125422 | 1513 | 14 | 1.85% |
| hsa_circ_0132713 | 7241 | 67 | 1.85% |
| hsa_circ_0004859 | 1189 | 11 | 1.85% |
| hsa_circ_0043360 | 1189 | 11 | 1.85% |
| hsa_circ_0058521 | 1189 | 11 | 1.85% |
| hsa_circ_0065252 | 1189 | 11 | 1.85% |
| hsa_circ_0070401 | 1189 | 11 | 1.85% |
| hsa_circ_0101722 | 1189 | 11 | 1.85% |
| hsa_circ_0106145 | 1189 | 11 | 1.85% |
| hsa_circ_0139534 | 1189 | 11 | 1.85% |
| hsa_circ_0002437 | 1081 | 10 | 1.85% |
| hsa_circ_0019132 | 1081 | 10 | 1.85% |
| hsa_circ_0019638 | 1081 | 10 | 1.85% |
| hsa_circ_0033638 | 1081 | 10 | 1.85% |
| hsa_circ_0043645 | 1081 | 10 | 1.85% |
| hsa_circ_0077093 | 1081 | 10 | 1.85% |
| hsa_circ_0088443 | 1081 | 10 | 1.85% |
| hsa_circ_0094166 | 1081 | 10 | 1.85% |
| hsa_circ_0097808 | 1081 | 10 | 1.85% |
| hsa_circ_0112031 | 1081 | 10 | 1.85% |
| hsa_circ_0121709 | 1081 | 10 | 1.85% |
| hsa_circ_0136148 | 1081 | 10 | 1.85% |
| hsa_circ_0047414 | 1406 | 13 | 1.85% |
| hsa_circ_0086943 | 1298 | 12 | 1.85% |
| hsa_circ_0009276 | 1190 | 11 | 1.85% |
| hsa_circ_0058420 | 1190 | 11 | 1.85% |
| hsa_circ_0061507 | 1190 | 11 | 1.85% |
| hsa_circ_0064311 | 1190 | 11 | 1.85% |
| hsa_circ_0090194 | 1190 | 11 | 1.85% |
| hsa_circ_0132760 | 1190 | 11 | 1.85% |
| hsa_circ_0017370 | 1082 | 10 | 1.85% |
| hsa_circ_0019194 | 1082 | 10 | 1.85% |
| hsa_circ_0022230 | 1082 | 10 | 1.85% |

| | | | |
|---|---|---|---|
| hsa_circ_0040209 | 1082 | 10 | 1.85% |
| hsa_circ_0047709 | 1082 | 10 | 1.85% |
| hsa_circ_0047737 | 1082 | 10 | 1.85% |
| hsa_circ_0071328 | 1082 | 10 | 1.85% |
| hsa_circ_0075177 | 1082 | 10 | 1.85% |
| hsa_circ_0091604 | 1082 | 10 | 1.85% |
| hsa_circ_0097730 | 1082 | 10 | 1.85% |
| hsa_circ_0102107 | 1082 | 10 | 1.85% |
| hsa_circ_0102600 | 1082 | 10 | 1.85% |
| hsa_circ_0114806 | 1082 | 10 | 1.85% |
| hsa_circ_0120233 | 1082 | 10 | 1.85% |
| hsa_circ_0120405 | 1082 | 10 | 1.85% |
| hsa_circ_0127977 | 1082 | 10 | 1.85% |
| hsa_circ_0080941 | 1407 | 13 | 1.85% |
| hsa_circ_0001168 | 1299 | 12 | 1.85% |
| hsa_circ_0044022 | 1299 | 12 | 1.85% |
| hsa_circ_0079644 | 1299 | 12 | 1.85% |
| hsa_circ_0104191 | 1299 | 12 | 1.85% |
| hsa_circ_0113462 | 1299 | 12 | 1.85% |
| hsa_circ_0029574 | 1191 | 11 | 1.85% |
| hsa_circ_0036675 | 1191 | 11 | 1.85% |
| hsa_circ_0047180 | 1191 | 11 | 1.85% |
| hsa_circ_0080118 | 1191 | 11 | 1.85% |
| hsa_circ_0085270 | 1191 | 11 | 1.85% |
| hsa_circ_0026942 | 1516 | 14 | 1.85% |
| hsa_circ_0003814 | 1083 | 10 | 1.85% |
| hsa_circ_0013659 | 1083 | 10 | 1.85% |
| hsa_circ_0015947 | 1083 | 10 | 1.85% |
| hsa_circ_0020800 | 1083 | 10 | 1.85% |
| hsa_circ_0023720 | 1083 | 10 | 1.85% |
| hsa_circ_0042210 | 1083 | 10 | 1.85% |
| hsa_circ_0043322 | 1083 | 10 | 1.85% |
| hsa_circ_0047268 | 1083 | 10 | 1.85% |
| hsa_circ_0055261 | 1083 | 10 | 1.85% |
| hsa_circ_0069234 | 1083 | 10 | 1.85% |
| hsa_circ_0088702 | 1083 | 10 | 1.85% |
| hsa_circ_0089380 | 1083 | 10 | 1.85% |
| hsa_circ_0094647 | 1083 | 10 | 1.85% |
| hsa_circ_0024879 | 1300 | 12 | 1.85% |
| hsa_circ_0035866 | 1300 | 12 | 1.85% |
| hsa_circ_0032656 | 1842 | 17 | 1.85% |
| hsa_circ_0031197 | 2709 | 25 | 1.85% |
| hsa_circ_0098007 | 1192 | 11 | 1.85% |
| hsa_circ_0101837 | 1192 | 11 | 1.85% |
| hsa_circ_0123166 | 1192 | 11 | 1.85% |
| hsa_circ_0003751 | 1084 | 10 | 1.85% |
| hsa_circ_0009665 | 1084 | 10 | 1.85% |
| hsa_circ_0065928 | 1084 | 10 | 1.85% |
| hsa_circ_0066130 | 1084 | 10 | 1.85% |
| hsa_circ_0079592 | 1084 | 10 | 1.85% |

| | | | |
|---|---|---|---|
| hsa_circ_0082389 | 1084 | 10 | 1.85% |
| hsa_circ_0088296 | 1084 | 10 | 1.85% |
| hsa_circ_0103634 | 1084 | 10 | 1.85% |
| hsa_circ_0042691 | 1301 | 12 | 1.84% |
| hsa_circ_0076364 | 1301 | 12 | 1.84% |
| hsa_circ_0109279 | 1518 | 14 | 1.84% |
| hsa_circ_0000602 | 1193 | 11 | 1.84% |
| hsa_circ_0068632 | 1193 | 11 | 1.84% |
| hsa_circ_0079124 | 1193 | 11 | 1.84% |
| hsa_circ_0128097 | 1193 | 11 | 1.84% |
| hsa_circ_0017574 | 1302 | 12 | 1.84% |
| hsa_circ_0061473 | 1302 | 12 | 1.84% |
| hsa_circ_0119120 | 1302 | 12 | 1.84% |
| hsa_circ_0010104 | 1085 | 10 | 1.84% |
| hsa_circ_0012433 | 1085 | 10 | 1.84% |
| hsa_circ_0054853 | 1085 | 10 | 1.84% |
| hsa_circ_0060784 | 1085 | 10 | 1.84% |
| hsa_circ_0061760 | 1085 | 10 | 1.84% |
| hsa_circ_0074691 | 1085 | 10 | 1.84% |
| hsa_circ_0078427 | 1085 | 10 | 1.84% |
| hsa_circ_0084508 | 1085 | 10 | 1.84% |
| hsa_circ_0091225 | 1085 | 10 | 1.84% |
| hsa_circ_0098665 | 1085 | 10 | 1.84% |
| hsa_circ_0136146 | 1085 | 10 | 1.84% |
| hsa_circ_0012287 | 3256 | 30 | 1.84% |
| hsa_circ_0009219 | 1194 | 11 | 1.84% |
| hsa_circ_0014516 | 1194 | 11 | 1.84% |
| hsa_circ_0014703 | 1194 | 11 | 1.84% |
| hsa_circ_0017271 | 1194 | 11 | 1.84% |
| hsa_circ_0021297 | 1194 | 11 | 1.84% |
| hsa_circ_0027971 | 1194 | 11 | 1.84% |
| hsa_circ_0048913 | 1194 | 11 | 1.84% |
| hsa_circ_0082429 | 1194 | 11 | 1.84% |
| hsa_circ_0113570 | 1194 | 11 | 1.84% |
| hsa_circ_0132075 | 1194 | 11 | 1.84% |
| hsa_circ_0139686 | 1194 | 11 | 1.84% |
| hsa_circ_0048647 | 1303 | 12 | 1.84% |
| hsa_circ_0065955 | 1303 | 12 | 1.84% |
| hsa_circ_0011939 | 1086 | 10 | 1.84% |
| hsa_circ_0016337 | 1086 | 10 | 1.84% |
| hsa_circ_0030262 | 1086 | 10 | 1.84% |
| hsa_circ_0032770 | 1086 | 10 | 1.84% |
| hsa_circ_0065437 | 1086 | 10 | 1.84% |
| hsa_circ_0070071 | 1086 | 10 | 1.84% |
| hsa_circ_0071858 | 1086 | 10 | 1.84% |
| hsa_circ_0080738 | 1086 | 10 | 1.84% |
| hsa_circ_0088203 | 1086 | 10 | 1.84% |
| hsa_circ_0102105 | 1086 | 10 | 1.84% |
| hsa_circ_0111863 | 1086 | 10 | 1.84% |
| hsa_circ_0116718 | 1086 | 10 | 1.84% |

| | | | |
|---|---|---|---|
| hsa_circ_0118083 | 1086 | 10 | 1.84% |
| hsa_circ_0131958 | 1086 | 10 | 1.84% |
| hsa_circ_0105580 | 1413 | 13 | 1.84% |
| hsa_circ_0004732 | 1087 | 10 | 1.84% |
| hsa_circ_0011519 | 1087 | 10 | 1.84% |
| hsa_circ_0014393 | 1087 | 10 | 1.84% |
| hsa_circ_0016250 | 1087 | 10 | 1.84% |
| hsa_circ_0028424 | 1087 | 10 | 1.84% |
| hsa_circ_0029126 | 1087 | 10 | 1.84% |
| hsa_circ_0058797 | 1087 | 10 | 1.84% |
| hsa_circ_0060652 | 1087 | 10 | 1.84% |
| hsa_circ_0073175 | 1087 | 10 | 1.84% |
| hsa_circ_0087109 | 1087 | 10 | 1.84% |
| hsa_circ_0101753 | 1087 | 10 | 1.84% |
| hsa_circ_0106607 | 1087 | 10 | 1.84% |
| hsa_circ_0110843 | 1087 | 10 | 1.84% |
| hsa_circ_0115030 | 1087 | 10 | 1.84% |
| hsa_circ_0123207 | 1087 | 10 | 1.84% |
| hsa_circ_0124279 | 1087 | 10 | 1.84% |
| hsa_circ_0129647 | 1087 | 10 | 1.84% |
| hsa_circ_0135592 | 1087 | 10 | 1.84% |
| hsa_circ_0021425 | 1196 | 11 | 1.84% |
| hsa_circ_0043207 | 1196 | 11 | 1.84% |
| hsa_circ_0135116 | 1196 | 11 | 1.84% |
| hsa_circ_0069135 | 1305 | 12 | 1.84% |
| hsa_circ_0085904 | 1305 | 12 | 1.84% |
| hsa_circ_0140674 | 1305 | 12 | 1.84% |
| hsa_circ_0114929 | 2719 | 25 | 1.84% |
| hsa_circ_0014357 | 1088 | 10 | 1.84% |
| hsa_circ_0016496 | 1088 | 10 | 1.84% |
| hsa_circ_0016706 | 1088 | 10 | 1.84% |
| hsa_circ_0016797 | 1088 | 10 | 1.84% |
| hsa_circ_0017895 | 1088 | 10 | 1.84% |
| hsa_circ_0025019 | 1088 | 10 | 1.84% |
| hsa_circ_0030111 | 1088 | 10 | 1.84% |
| hsa_circ_0043546 | 1088 | 10 | 1.84% |
| hsa_circ_0046511 | 1088 | 10 | 1.84% |
| hsa_circ_0049714 | 1088 | 10 | 1.84% |
| hsa_circ_0051070 | 1088 | 10 | 1.84% |
| hsa_circ_0057828 | 1088 | 10 | 1.84% |
| hsa_circ_0068520 | 1088 | 10 | 1.84% |
| hsa_circ_0084079 | 1088 | 10 | 1.84% |
| hsa_circ_0087285 | 1088 | 10 | 1.84% |
| hsa_circ_0092862 | 1088 | 10 | 1.84% |
| hsa_circ_0096151 | 1088 | 10 | 1.84% |
| hsa_circ_0077503 | 3917 | 36 | 1.84% |
| hsa_circ_0052143 | 1197 | 11 | 1.84% |
| hsa_circ_0057604 | 1197 | 11 | 1.84% |
| hsa_circ_0064309 | 1197 | 11 | 1.84% |
| hsa_circ_0080044 | 1197 | 11 | 1.84% |

| | | | |
|---|---|---|---|
| hsa_circ_0088995 | 1197 | 11 | 1.84% |
| hsa_circ_0107524 | 1197 | 11 | 1.84% |
| hsa_circ_0031012 | 1306 | 12 | 1.84% |
| hsa_circ_0078048 | 1306 | 12 | 1.84% |
| hsa_circ_0117210 | 1306 | 12 | 1.84% |
| hsa_circ_0122061 | 1415 | 13 | 1.84% |
| hsa_circ_0003540 | 1089 | 10 | 1.84% |
| hsa_circ_0018594 | 1089 | 10 | 1.84% |
| hsa_circ_0039830 | 1089 | 10 | 1.84% |
| hsa_circ_0047233 | 1089 | 10 | 1.84% |
| hsa_circ_0051233 | 1089 | 10 | 1.84% |
| hsa_circ_0051891 | 1089 | 10 | 1.84% |
| hsa_circ_0056385 | 1089 | 10 | 1.84% |
| hsa_circ_0070653 | 1089 | 10 | 1.84% |
| hsa_circ_0083213 | 1089 | 10 | 1.84% |
| hsa_circ_0092005 | 1089 | 10 | 1.84% |
| hsa_circ_0097540 | 1089 | 10 | 1.84% |
| hsa_circ_0102526 | 1089 | 10 | 1.84% |
| hsa_circ_0108518 | 1089 | 10 | 1.84% |
| hsa_circ_0119297 | 1089 | 10 | 1.84% |
| hsa_circ_0124826 | 1089 | 10 | 1.84% |
| hsa_circ_0005699 | 1198 | 11 | 1.84% |
| hsa_circ_0030395 | 1198 | 11 | 1.84% |
| hsa_circ_0045739 | 1198 | 11 | 1.84% |
| hsa_circ_0059269 | 1198 | 11 | 1.84% |
| hsa_circ_0065965 | 1198 | 11 | 1.84% |
| hsa_circ_0072818 | 1198 | 11 | 1.84% |
| hsa_circ_0083495 | 1198 | 11 | 1.84% |
| hsa_circ_0085935 | 1198 | 11 | 1.84% |
| hsa_circ_0012695 | 1307 | 12 | 1.84% |
| hsa_circ_0074434 | 1307 | 12 | 1.84% |
| hsa_circ_0118547 | 1853 | 17 | 1.83% |
| hsa_circ_0137962 | 1417 | 13 | 1.83% |
| hsa_circ_0024158 | 1308 | 12 | 1.83% |
| hsa_circ_0053122 | 1308 | 12 | 1.83% |
| hsa_circ_0069658 | 1308 | 12 | 1.83% |
| hsa_circ_0100711 | 1308 | 12 | 1.83% |
| hsa_circ_0140034 | 1308 | 12 | 1.83% |
| hsa_circ_0038510 | 1199 | 11 | 1.83% |
| hsa_circ_0099408 | 1199 | 11 | 1.83% |
| hsa_circ_0112660 | 1199 | 11 | 1.83% |
| hsa_circ_0008595 | 1090 | 10 | 1.83% |
| hsa_circ_0025466 | 1090 | 10 | 1.83% |
| hsa_circ_0032504 | 1090 | 10 | 1.83% |
| hsa_circ_0078379 | 1090 | 10 | 1.83% |
| hsa_circ_0082467 | 1090 | 10 | 1.83% |
| hsa_circ_0084991 | 1090 | 10 | 1.83% |
| hsa_circ_0101481 | 1090 | 10 | 1.83% |
| hsa_circ_0108854 | 1090 | 10 | 1.83% |
| hsa_circ_0118769 | 1090 | 10 | 1.83% |

| | | | |
|---|---|---|---|
| hsa_circ_0037631 | 1309 | 12 | 1.83% |
| hsa_circ_0047693 | 1309 | 12 | 1.83% |
| hsa_circ_0066183 | 1309 | 12 | 1.83% |
| hsa_circ_0097251 | 1309 | 12 | 1.83% |
| hsa_circ_0101351 | 1309 | 12 | 1.83% |
| hsa_circ_0020564 | 1200 | 11 | 1.83% |
| hsa_circ_0028438 | 1200 | 11 | 1.83% |
| hsa_circ_0052955 | 1200 | 11 | 1.83% |
| hsa_circ_0055955 | 1200 | 11 | 1.83% |
| hsa_circ_0065696 | 1200 | 11 | 1.83% |
| hsa_circ_0069398 | 1200 | 11 | 1.83% |
| hsa_circ_0077145 | 1200 | 11 | 1.83% |
| hsa_circ_0091682 | 1200 | 11 | 1.83% |
| hsa_circ_0113346 | 1200 | 11 | 1.83% |
| hsa_circ_0118680 | 1200 | 11 | 1.83% |
| hsa_circ_0005162 | 1091 | 10 | 1.83% |
| hsa_circ_0018128 | 1091 | 10 | 1.83% |
| hsa_circ_0037357 | 1091 | 10 | 1.83% |
| hsa_circ_0041817 | 1091 | 10 | 1.83% |
| hsa_circ_0045090 | 1091 | 10 | 1.83% |
| hsa_circ_0062530 | 1091 | 10 | 1.83% |
| hsa_circ_0070022 | 1091 | 10 | 1.83% |
| hsa_circ_0086007 | 1091 | 10 | 1.83% |
| hsa_circ_0094934 | 1091 | 10 | 1.83% |
| hsa_circ_0097420 | 1091 | 10 | 1.83% |
| hsa_circ_0111998 | 1091 | 10 | 1.83% |
| hsa_circ_0115060 | 1091 | 10 | 1.83% |
| hsa_circ_0116082 | 1091 | 10 | 1.83% |
| hsa_circ_0127209 | 1091 | 10 | 1.83% |
| hsa_circ_0024556 | 1419 | 13 | 1.83% |
| hsa_circ_0074515 | 1310 | 12 | 1.83% |
| hsa_circ_0135080 | 1310 | 12 | 1.83% |
| hsa_circ_0034934 | 1201 | 11 | 1.83% |
| hsa_circ_0040565 | 1201 | 11 | 1.83% |
| hsa_circ_0041319 | 1201 | 11 | 1.83% |
| hsa_circ_0045053 | 1201 | 11 | 1.83% |
| hsa_circ_0053305 | 1201 | 11 | 1.83% |
| hsa_circ_0069298 | 1201 | 11 | 1.83% |
| hsa_circ_0099008 | 1201 | 11 | 1.83% |
| hsa_circ_0014296 | 1092 | 10 | 1.83% |
| hsa_circ_0014759 | 1092 | 10 | 1.83% |
| hsa_circ_0024443 | 1092 | 10 | 1.83% |
| hsa_circ_0032993 | 1092 | 10 | 1.83% |
| hsa_circ_0035816 | 1092 | 10 | 1.83% |
| hsa_circ_0037251 | 1092 | 10 | 1.83% |
| hsa_circ_0040448 | 1092 | 10 | 1.83% |
| hsa_circ_0050545 | 1092 | 10 | 1.83% |
| hsa_circ_0061921 | 1092 | 10 | 1.83% |
| hsa_circ_0072936 | 1092 | 10 | 1.83% |
| hsa_circ_0084007 | 1092 | 10 | 1.83% |

| | | | |
|---|---|---|---|
| hsa_circ_0097648 | 1092 | 10 | 1.83% |
| hsa_circ_0114716 | 1092 | 10 | 1.83% |
| hsa_circ_0132158 | 1092 | 10 | 1.83% |
| hsa_circ_0075773 | 2512 | 23 | 1.83% |
| hsa_circ_0082085 | 1420 | 13 | 1.83% |
| hsa_circ_0054903 | 1311 | 12 | 1.83% |
| hsa_circ_0111131 | 1311 | 12 | 1.83% |
| hsa_circ_0036892 | 1202 | 11 | 1.83% |
| hsa_circ_0045486 | 1202 | 11 | 1.83% |
| hsa_circ_0046639 | 1202 | 11 | 1.83% |
| hsa_circ_0060318 | 1202 | 11 | 1.83% |
| hsa_circ_0002134 | 1093 | 10 | 1.83% |
| hsa_circ_0008416 | 1093 | 10 | 1.83% |
| hsa_circ_0010986 | 1093 | 10 | 1.83% |
| hsa_circ_0011827 | 1093 | 10 | 1.83% |
| hsa_circ_0016800 | 1093 | 10 | 1.83% |
| hsa_circ_0039510 | 1093 | 10 | 1.83% |
| hsa_circ_0045741 | 1093 | 10 | 1.83% |
| hsa_circ_0070476 | 1093 | 10 | 1.83% |
| hsa_circ_0084128 | 1093 | 10 | 1.83% |
| hsa_circ_0086715 | 1093 | 10 | 1.83% |
| hsa_circ_0094963 | 1093 | 10 | 1.83% |
| hsa_circ_0104849 | 1093 | 10 | 1.83% |
| hsa_circ_0120998 | 1093 | 10 | 1.83% |
| hsa_circ_0087566 | 2405 | 22 | 1.83% |
| hsa_circ_0065037 | 1312 | 12 | 1.83% |
| hsa_circ_0118398 | 1312 | 12 | 1.83% |
| hsa_circ_0032736 | 1203 | 11 | 1.83% |
| hsa_circ_0075952 | 1203 | 11 | 1.83% |
| hsa_circ_0091036 | 1203 | 11 | 1.83% |
| hsa_circ_0114833 | 1203 | 11 | 1.83% |
| hsa_circ_0137694 | 1203 | 11 | 1.83% |
| hsa_circ_0049513 | 2625 | 24 | 1.83% |
| hsa_circ_0029156 | 1422 | 13 | 1.83% |
| hsa_circ_0129309 | 1422 | 13 | 1.83% |
| hsa_circ_0012739 | 1094 | 10 | 1.83% |
| hsa_circ_0027487 | 1094 | 10 | 1.83% |
| hsa_circ_0028739 | 1094 | 10 | 1.83% |
| hsa_circ_0030574 | 1094 | 10 | 1.83% |
| hsa_circ_0042071 | 1094 | 10 | 1.83% |
| hsa_circ_0067788 | 1094 | 10 | 1.83% |
| hsa_circ_0075028 | 1094 | 10 | 1.83% |
| hsa_circ_0075865 | 1094 | 10 | 1.83% |
| hsa_circ_0083690 | 1094 | 10 | 1.83% |
| hsa_circ_0091764 | 1094 | 10 | 1.83% |
| hsa_circ_0096362 | 1094 | 10 | 1.83% |
| hsa_circ_0098026 | 1094 | 10 | 1.83% |
| hsa_circ_0137848 | 1094 | 10 | 1.83% |
| hsa_circ_0014649 | 1313 | 12 | 1.83% |
| hsa_circ_0074554 | 1313 | 12 | 1.83% |

| | | | |
|---|---|---|---|
| hsa_circ_0109800 | 2845 | 26 | 1.83% |
| hsa_circ_0025339 | 1204 | 11 | 1.83% |
| hsa_circ_0111709 | 1204 | 11 | 1.83% |
| hsa_circ_0127343 | 1204 | 11 | 1.83% |
| hsa_circ_0026901 | 1314 | 12 | 1.83% |
| hsa_circ_0042692 | 1314 | 12 | 1.83% |
| hsa_circ_0045145 | 1314 | 12 | 1.83% |
| hsa_circ_0050704 | 1314 | 12 | 1.83% |
| hsa_circ_0053262 | 1314 | 12 | 1.83% |
| hsa_circ_0114203 | 1314 | 12 | 1.83% |
| hsa_circ_0012738 | 1095 | 10 | 1.83% |
| hsa_circ_0019620 | 1095 | 10 | 1.83% |
| hsa_circ_0052153 | 1095 | 10 | 1.83% |
| hsa_circ_0058317 | 1095 | 10 | 1.83% |
| hsa_circ_0070373 | 1095 | 10 | 1.83% |
| hsa_circ_0089633 | 1095 | 10 | 1.83% |
| hsa_circ_0095945 | 1095 | 10 | 1.83% |
| hsa_circ_0111485 | 1095 | 10 | 1.83% |
| hsa_circ_0122743 | 1095 | 10 | 1.83% |
| hsa_circ_0129793 | 1095 | 10 | 1.83% |
| hsa_circ_0131077 | 1095 | 10 | 1.83% |
| hsa_circ_0133543 | 1095 | 10 | 1.83% |
| hsa_circ_0136524 | 1095 | 10 | 1.83% |
| hsa_circ_0137873 | 1095 | 10 | 1.83% |
| hsa_circ_0035125 | 1205 | 11 | 1.83% |
| hsa_circ_0065359 | 1205 | 11 | 1.83% |
| hsa_circ_0097041 | 1205 | 11 | 1.83% |
| hsa_circ_0111473 | 1205 | 11 | 1.83% |
| hsa_circ_0123826 | 1205 | 11 | 1.83% |
| hsa_circ_0014856 | 1315 | 12 | 1.83% |
| hsa_circ_0015505 | 1315 | 12 | 1.83% |
| hsa_circ_0001944 | 1096 | 10 | 1.82% |
| hsa_circ_0007139 | 1096 | 10 | 1.82% |
| hsa_circ_0045913 | 1096 | 10 | 1.82% |
| hsa_circ_0047249 | 1096 | 10 | 1.82% |
| hsa_circ_0068733 | 1096 | 10 | 1.82% |
| hsa_circ_0080632 | 1096 | 10 | 1.82% |
| hsa_circ_0088699 | 1096 | 10 | 1.82% |
| hsa_circ_0091963 | 1096 | 10 | 1.82% |
| hsa_circ_0118095 | 1096 | 10 | 1.82% |
| hsa_circ_0126118 | 1096 | 10 | 1.82% |
| hsa_circ_0135644 | 1096 | 10 | 1.82% |
| hsa_circ_0137939 | 1096 | 10 | 1.82% |
| hsa_circ_0038467 | 1206 | 11 | 1.82% |
| hsa_circ_0042155 | 1206 | 11 | 1.82% |
| hsa_circ_0123080 | 1206 | 11 | 1.82% |
| hsa_circ_0123164 | 1206 | 11 | 1.82% |
| hsa_circ_0129053 | 1206 | 11 | 1.82% |
| hsa_circ_0137330 | 1206 | 11 | 1.82% |
| hsa_circ_0058240 | 1645 | 15 | 1.82% |

| | | | |
|---|---|---|---|
| hsa_circ_0080863 | 1316 | 12 | 1.82% |
| hsa_circ_0057366 | 1755 | 16 | 1.82% |
| hsa_circ_0062017 | 1426 | 13 | 1.82% |
| hsa_circ_0003724 | 1097 | 10 | 1.82% |
| hsa_circ_0005915 | 1097 | 10 | 1.82% |
| hsa_circ_0047567 | 1097 | 10 | 1.82% |
| hsa_circ_0048417 | 1097 | 10 | 1.82% |
| hsa_circ_0063491 | 1097 | 10 | 1.82% |
| hsa_circ_0072562 | 1097 | 10 | 1.82% |
| hsa_circ_0086721 | 1097 | 10 | 1.82% |
| hsa_circ_0088703 | 1097 | 10 | 1.82% |
| hsa_circ_0090780 | 1097 | 10 | 1.82% |
| hsa_circ_0109408 | 1097 | 10 | 1.82% |
| hsa_circ_0113631 | 1097 | 10 | 1.82% |
| hsa_circ_0122109 | 1097 | 10 | 1.82% |
| hsa_circ_0022696 | 1207 | 11 | 1.82% |
| hsa_circ_0087911 | 1207 | 11 | 1.82% |
| hsa_circ_0104389 | 1207 | 11 | 1.82% |
| hsa_circ_0132211 | 1207 | 11 | 1.82% |
| hsa_circ_0032695 | 1317 | 12 | 1.82% |
| hsa_circ_0128945 | 1317 | 12 | 1.82% |
| hsa_circ_0004500 | 1098 | 10 | 1.82% |
| hsa_circ_0016026 | 1098 | 10 | 1.82% |
| hsa_circ_0016652 | 1098 | 10 | 1.82% |
| hsa_circ_0033636 | 1098 | 10 | 1.82% |
| hsa_circ_0034892 | 1098 | 10 | 1.82% |
| hsa_circ_0036144 | 1098 | 10 | 1.82% |
| hsa_circ_0037120 | 1098 | 10 | 1.82% |
| hsa_circ_0042327 | 1098 | 10 | 1.82% |
| hsa_circ_0047445 | 1098 | 10 | 1.82% |
| hsa_circ_0064695 | 1098 | 10 | 1.82% |
| hsa_circ_0076305 | 1098 | 10 | 1.82% |
| hsa_circ_0082119 | 1098 | 10 | 1.82% |
| hsa_circ_0095309 | 1098 | 10 | 1.82% |
| hsa_circ_0111912 | 1098 | 10 | 1.82% |
| hsa_circ_0112540 | 1098 | 10 | 1.82% |
| hsa_circ_0122645 | 1098 | 10 | 1.82% |
| hsa_circ_0125249 | 1098 | 10 | 1.82% |
| hsa_circ_0021378 | 1208 | 11 | 1.82% |
| hsa_circ_0034340 | 1208 | 11 | 1.82% |
| hsa_circ_0037235 | 1208 | 11 | 1.82% |
| hsa_circ_0058061 | 1208 | 11 | 1.82% |
| hsa_circ_0073767 | 1208 | 11 | 1.82% |
| hsa_circ_0086030 | 1208 | 11 | 1.82% |
| hsa_circ_0089504 | 1208 | 11 | 1.82% |
| hsa_circ_0099771 | 1208 | 11 | 1.82% |
| hsa_circ_0122845 | 1208 | 11 | 1.82% |
| hsa_circ_0129214 | 1208 | 11 | 1.82% |
| hsa_circ_0136975 | 4395 | 40 | 1.82% |
| hsa_circ_0001099 | 1099 | 10 | 1.82% |

| | | | |
|---|---|---|---|
| hsa_circ_0002826 | 1099 | 10 | 1.82% |
| hsa_circ_0010152 | 1099 | 10 | 1.82% |
| hsa_circ_0011602 | 1099 | 10 | 1.82% |
| hsa_circ_0029662 | 1099 | 10 | 1.82% |
| hsa_circ_0048023 | 1099 | 10 | 1.82% |
| hsa_circ_0076040 | 1099 | 10 | 1.82% |
| hsa_circ_0087792 | 1099 | 10 | 1.82% |
| hsa_circ_0094544 | 1099 | 10 | 1.82% |
| hsa_circ_0107655 | 1099 | 10 | 1.82% |
| hsa_circ_0123097 | 1099 | 10 | 1.82% |
| hsa_circ_0134172 | 1099 | 10 | 1.82% |
| hsa_circ_0136729 | 1099 | 10 | 1.82% |
| hsa_circ_0029175 | 1209 | 11 | 1.82% |
| hsa_circ_0072764 | 1209 | 11 | 1.82% |
| hsa_circ_0083831 | 1209 | 11 | 1.82% |
| hsa_circ_0094833 | 1209 | 11 | 1.82% |
| hsa_circ_0112483 | 1209 | 11 | 1.82% |
| hsa_circ_0124301 | 2638 | 24 | 1.82% |
| hsa_circ_0037623 | 1319 | 12 | 1.82% |
| hsa_circ_0110562 | 1319 | 12 | 1.82% |
| hsa_circ_0013137 | 1320 | 12 | 1.82% |
| hsa_circ_0047645 | 1320 | 12 | 1.82% |
| hsa_circ_0052979 | 1320 | 12 | 1.82% |
| hsa_circ_0055889 | 1320 | 12 | 1.82% |
| hsa_circ_0071553 | 1320 | 12 | 1.82% |
| hsa_circ_0135265 | 1320 | 12 | 1.82% |
| hsa_circ_0015263 | 1210 | 11 | 1.82% |
| hsa_circ_0027003 | 1210 | 11 | 1.82% |
| hsa_circ_0028271 | 1210 | 11 | 1.82% |
| hsa_circ_0051831 | 1210 | 11 | 1.82% |
| hsa_circ_0002591 | 1100 | 10 | 1.82% |
| hsa_circ_0007535 | 1100 | 10 | 1.82% |
| hsa_circ_0019642 | 1100 | 10 | 1.82% |
| hsa_circ_0035029 | 1100 | 10 | 1.82% |
| hsa_circ_0037039 | 1100 | 10 | 1.82% |
| hsa_circ_0071016 | 1100 | 10 | 1.82% |
| hsa_circ_0073057 | 1100 | 10 | 1.82% |
| hsa_circ_0099565 | 1100 | 10 | 1.82% |
| hsa_circ_0100762 | 1100 | 10 | 1.82% |
| hsa_circ_0128129 | 1100 | 10 | 1.82% |
| hsa_circ_0112959 | 5503 | 50 | 1.82% |
| hsa_circ_0106199 | 1541 | 14 | 1.82% |
| hsa_circ_0076667 | 1431 | 13 | 1.82% |
| hsa_circ_0062838 | 2642 | 24 | 1.82% |
| hsa_circ_0097162 | 2642 | 24 | 1.82% |
| hsa_circ_0036658 | 1321 | 12 | 1.82% |
| hsa_circ_0048894 | 1321 | 12 | 1.82% |
| hsa_circ_0056267 | 1321 | 12 | 1.82% |
| hsa_circ_0127238 | 1321 | 12 | 1.82% |
| hsa_circ_0089735 | 1211 | 11 | 1.82% |

| | | | |
|---|---|---|---|
| hsa_circ_0129757 | 1211 | 11 | 1.82% |
| hsa_circ_0134012 | 1211 | 11 | 1.82% |
| hsa_circ_0068086 | 5945 | 54 | 1.82% |
| hsa_circ_0017164 | 1101 | 10 | 1.82% |
| hsa_circ_0018814 | 1101 | 10 | 1.82% |
| hsa_circ_0028150 | 1101 | 10 | 1.82% |
| hsa_circ_0043902 | 1101 | 10 | 1.82% |
| hsa_circ_0045124 | 1101 | 10 | 1.82% |
| hsa_circ_0053290 | 1101 | 10 | 1.82% |
| hsa_circ_0053412 | 1101 | 10 | 1.82% |
| hsa_circ_0054297 | 1101 | 10 | 1.82% |
| hsa_circ_0055900 | 1101 | 10 | 1.82% |
| hsa_circ_0079984 | 1101 | 10 | 1.82% |
| hsa_circ_0087015 | 1101 | 10 | 1.82% |
| hsa_circ_0098278 | 1101 | 10 | 1.82% |
| hsa_circ_0101763 | 1101 | 10 | 1.82% |
| hsa_circ_0120460 | 1101 | 10 | 1.82% |
| hsa_circ_0122244 | 1101 | 10 | 1.82% |
| hsa_circ_0125052 | 1101 | 10 | 1.82% |
| hsa_circ_0128239 | 1101 | 10 | 1.82% |
| hsa_circ_0138906 | 1101 | 10 | 1.82% |
| hsa_circ_0083973 | 1432 | 13 | 1.82% |
| hsa_circ_0089659 | 1432 | 13 | 1.82% |
| hsa_circ_0099624 | 1322 | 12 | 1.82% |
| hsa_circ_0010742 | 1212 | 11 | 1.82% |
| hsa_circ_0025596 | 1212 | 11 | 1.82% |
| hsa_circ_0032511 | 1212 | 11 | 1.82% |
| hsa_circ_0033371 | 1212 | 11 | 1.82% |
| hsa_circ_0053337 | 1212 | 11 | 1.82% |
| hsa_circ_0064777 | 1212 | 11 | 1.82% |
| hsa_circ_0091270 | 1212 | 11 | 1.82% |
| hsa_circ_0120564 | 1212 | 11 | 1.82% |
| hsa_circ_0131818 | 1212 | 11 | 1.82% |
| hsa_circ_0030046 | 3416 | 31 | 1.81% |
| hsa_circ_0011990 | 1102 | 10 | 1.81% |
| hsa_circ_0025515 | 1102 | 10 | 1.81% |
| hsa_circ_0025687 | 1102 | 10 | 1.81% |
| hsa_circ_0060118 | 1102 | 10 | 1.81% |
| hsa_circ_0064704 | 1102 | 10 | 1.81% |
| hsa_circ_0080315 | 1102 | 10 | 1.81% |
| hsa_circ_0090415 | 1102 | 10 | 1.81% |
| hsa_circ_0091601 | 1102 | 10 | 1.81% |
| hsa_circ_0091730 | 1102 | 10 | 1.81% |
| hsa_circ_0092389 | 1102 | 10 | 1.81% |
| hsa_circ_0096682 | 1102 | 10 | 1.81% |
| hsa_circ_0104423 | 1102 | 10 | 1.81% |
| hsa_circ_0106287 | 1102 | 10 | 1.81% |
| hsa_circ_0033170 | 1433 | 13 | 1.81% |
| hsa_circ_0018629 | 1323 | 12 | 1.81% |
| hsa_circ_0031528 | 1323 | 12 | 1.81% |

| | | | |
|---|---|---|---|
| hsa_circ_0037634 | 1323 | 12 | 1.81% |
| hsa_circ_0040137 | 1323 | 12 | 1.81% |
| hsa_circ_0052803 | 1323 | 12 | 1.81% |
| hsa_circ_0064390 | 1323 | 12 | 1.81% |
| hsa_circ_0005141 | 3639 | 33 | 1.81% |
| hsa_circ_0016947 | 1213 | 11 | 1.81% |
| hsa_circ_0021973 | 1213 | 11 | 1.81% |
| hsa_circ_0038609 | 1213 | 11 | 1.81% |
| hsa_circ_0053144 | 1213 | 11 | 1.81% |
| hsa_circ_0056467 | 1213 | 11 | 1.81% |
| hsa_circ_0063747 | 1213 | 11 | 1.81% |
| hsa_circ_0117437 | 1213 | 11 | 1.81% |
| hsa_circ_0128614 | 1213 | 11 | 1.81% |
| hsa_circ_0009681 | 3198 | 29 | 1.81% |
| hsa_circ_0101944 | 1985 | 18 | 1.81% |
| hsa_circ_0006866 | 1103 | 10 | 1.81% |
| hsa_circ_0013033 | 1103 | 10 | 1.81% |
| hsa_circ_0025605 | 1103 | 10 | 1.81% |
| hsa_circ_0028754 | 1103 | 10 | 1.81% |
| hsa_circ_0046628 | 1103 | 10 | 1.81% |
| hsa_circ_0047424 | 1103 | 10 | 1.81% |
| hsa_circ_0059973 | 1103 | 10 | 1.81% |
| hsa_circ_0060757 | 1103 | 10 | 1.81% |
| hsa_circ_0073501 | 1103 | 10 | 1.81% |
| hsa_circ_0122922 | 1103 | 10 | 1.81% |
| hsa_circ_0123005 | 1103 | 10 | 1.81% |
| hsa_circ_0133974 | 1103 | 10 | 1.81% |
| hsa_circ_0140232 | 1103 | 10 | 1.81% |
| hsa_circ_0029356 | 1324 | 12 | 1.81% |
| hsa_circ_0037638 | 1324 | 12 | 1.81% |
| hsa_circ_0040996 | 1324 | 12 | 1.81% |
| hsa_circ_0128355 | 1324 | 12 | 1.81% |
| hsa_circ_0007919 | 1545 | 14 | 1.81% |
| hsa_circ_0016718 | 1214 | 11 | 1.81% |
| hsa_circ_0025208 | 1214 | 11 | 1.81% |
| hsa_circ_0025643 | 1214 | 11 | 1.81% |
| hsa_circ_0025949 | 1214 | 11 | 1.81% |
| hsa_circ_0051682 | 1214 | 11 | 1.81% |
| hsa_circ_0082002 | 1214 | 11 | 1.81% |
| hsa_circ_0103449 | 1214 | 11 | 1.81% |
| hsa_circ_0117847 | 1214 | 11 | 1.81% |
| hsa_circ_0124492 | 1214 | 11 | 1.81% |
| hsa_circ_0114892 | 3312 | 30 | 1.81% |
| hsa_circ_0006207 | 1104 | 10 | 1.81% |
| hsa_circ_0029205 | 1104 | 10 | 1.81% |
| hsa_circ_0030390 | 1104 | 10 | 1.81% |
| hsa_circ_0052196 | 1104 | 10 | 1.81% |
| hsa_circ_0062021 | 1104 | 10 | 1.81% |
| hsa_circ_0071740 | 1104 | 10 | 1.81% |
| hsa_circ_0078649 | 1104 | 10 | 1.81% |

| | | | |
|---|---|---|---|
| hsa_circ_0101335 | 1104 | 10 | 1.81% |
| hsa_circ_0109399 | 1104 | 10 | 1.81% |
| hsa_circ_0110283 | 1104 | 10 | 1.81% |
| hsa_circ_0117324 | 1104 | 10 | 1.81% |
| hsa_circ_0121232 | 1104 | 10 | 1.81% |
| hsa_circ_0124964 | 1104 | 10 | 1.81% |
| hsa_circ_0128089 | 1104 | 10 | 1.81% |
| hsa_circ_0129691 | 1104 | 10 | 1.81% |
| hsa_circ_0039096 | 1325 | 12 | 1.81% |
| hsa_circ_0121075 | 1325 | 12 | 1.81% |
| hsa_circ_0008446 | 1215 | 11 | 1.81% |
| hsa_circ_0020644 | 1215 | 11 | 1.81% |
| hsa_circ_0040183 | 1215 | 11 | 1.81% |
| hsa_circ_0057662 | 1215 | 11 | 1.81% |
| hsa_circ_0088851 | 1215 | 11 | 1.81% |
| hsa_circ_0031606 | 1326 | 12 | 1.81% |
| hsa_circ_0006444 | 1105 | 10 | 1.81% |
| hsa_circ_0009801 | 1105 | 10 | 1.81% |
| hsa_circ_0016550 | 1105 | 10 | 1.81% |
| hsa_circ_0033494 | 1105 | 10 | 1.81% |
| hsa_circ_0035854 | 1105 | 10 | 1.81% |
| hsa_circ_0046195 | 1105 | 10 | 1.81% |
| hsa_circ_0063182 | 1105 | 10 | 1.81% |
| hsa_circ_0068739 | 1105 | 10 | 1.81% |
| hsa_circ_0075472 | 1105 | 10 | 1.81% |
| hsa_circ_0077160 | 1105 | 10 | 1.81% |
| hsa_circ_0077935 | 1105 | 10 | 1.81% |
| hsa_circ_0095422 | 1105 | 10 | 1.81% |
| hsa_circ_0099806 | 1105 | 10 | 1.81% |
| hsa_circ_0102475 | 1105 | 10 | 1.81% |
| hsa_circ_0126946 | 1105 | 10 | 1.81% |
| hsa_circ_0134769 | 1105 | 10 | 1.81% |
| hsa_circ_0023841 | 1437 | 13 | 1.81% |
| hsa_circ_0039329 | 1216 | 11 | 1.81% |
| hsa_circ_0091122 | 1216 | 11 | 1.81% |
| hsa_circ_0096801 | 1216 | 11 | 1.81% |
| hsa_circ_0103445 | 1216 | 11 | 1.81% |
| hsa_circ_0010391 | 1990 | 18 | 1.81% |
| hsa_circ_0013156 | 1548 | 14 | 1.81% |
| hsa_circ_0010896 | 1327 | 12 | 1.81% |
| hsa_circ_0053956 | 1327 | 12 | 1.81% |
| hsa_circ_0000702 | 1106 | 10 | 1.81% |
| hsa_circ_0025307 | 1106 | 10 | 1.81% |
| hsa_circ_0028484 | 1106 | 10 | 1.81% |
| hsa_circ_0028843 | 1106 | 10 | 1.81% |
| hsa_circ_0030600 | 1106 | 10 | 1.81% |
| hsa_circ_0031825 | 1106 | 10 | 1.81% |
| hsa_circ_0035525 | 1106 | 10 | 1.81% |
| hsa_circ_0044619 | 1106 | 10 | 1.81% |
| hsa_circ_0089279 | 1106 | 10 | 1.81% |

| | | | |
|---|---|---|---|
| hsa_circ_0095676 | 1106 | 10 | 1.81% |
| hsa_circ_0098447 | 1106 | 10 | 1.81% |
| hsa_circ_0107614 | 1106 | 10 | 1.81% |
| hsa_circ_0117731 | 1106 | 10 | 1.81% |
| hsa_circ_0118181 | 1106 | 10 | 1.81% |
| hsa_circ_0121959 | 1106 | 10 | 1.81% |
| hsa_circ_0140406 | 1106 | 10 | 1.81% |
| hsa_circ_0100470 | 1438 | 13 | 1.81% |
| hsa_circ_0042246 | 1770 | 16 | 1.81% |
| hsa_circ_0013857 | 1217 | 11 | 1.81% |
| hsa_circ_0042715 | 1217 | 11 | 1.81% |
| hsa_circ_0052811 | 1217 | 11 | 1.81% |
| hsa_circ_0092899 | 1217 | 11 | 1.81% |
| hsa_circ_0122959 | 5644 | 51 | 1.81% |
| hsa_circ_0035807 | 1328 | 12 | 1.81% |
| hsa_circ_0100109 | 1328 | 12 | 1.81% |
| hsa_circ_0114079 | 1328 | 12 | 1.81% |
| hsa_circ_0119015 | 1328 | 12 | 1.81% |
| hsa_circ_0123197 | 1439 | 13 | 1.81% |
| hsa_circ_0010360 | 1107 | 10 | 1.81% |
| hsa_circ_0021619 | 1107 | 10 | 1.81% |
| hsa_circ_0057472 | 1107 | 10 | 1.81% |
| hsa_circ_0058733 | 1107 | 10 | 1.81% |
| hsa_circ_0058802 | 1107 | 10 | 1.81% |
| hsa_circ_0065267 | 1107 | 10 | 1.81% |
| hsa_circ_0086722 | 1107 | 10 | 1.81% |
| hsa_circ_0115746 | 1107 | 10 | 1.81% |
| hsa_circ_0119355 | 1107 | 10 | 1.81% |
| hsa_circ_0119593 | 1107 | 10 | 1.81% |
| hsa_circ_0122040 | 1107 | 10 | 1.81% |
| hsa_circ_0138977 | 1107 | 10 | 1.81% |
| hsa_circ_0051169 | 1218 | 11 | 1.81% |
| hsa_circ_0058310 | 1218 | 11 | 1.81% |
| hsa_circ_0094204 | 1329 | 12 | 1.81% |
| hsa_circ_0117277 | 1329 | 12 | 1.81% |
| hsa_circ_0020110 | 1883 | 17 | 1.81% |
| hsa_circ_0004690 | 1108 | 10 | 1.81% |
| hsa_circ_0007749 | 1108 | 10 | 1.81% |
| hsa_circ_0011370 | 1108 | 10 | 1.81% |
| hsa_circ_0022345 | 1108 | 10 | 1.81% |
| hsa_circ_0029534 | 1108 | 10 | 1.81% |
| hsa_circ_0036725 | 1108 | 10 | 1.81% |
| hsa_circ_0038187 | 1108 | 10 | 1.81% |
| hsa_circ_0040512 | 1108 | 10 | 1.81% |
| hsa_circ_0041045 | 1108 | 10 | 1.81% |
| hsa_circ_0055860 | 1108 | 10 | 1.81% |
| hsa_circ_0063026 | 1108 | 10 | 1.81% |
| hsa_circ_0070228 | 1108 | 10 | 1.81% |
| hsa_circ_0073140 | 1108 | 10 | 1.81% |
| hsa_circ_0075783 | 1108 | 10 | 1.81% |

| | | | |
|---|---|---|---|
| hsa_circ_0088218 | 1108 | 10 | 1.81% |
| hsa_circ_0088928 | 1108 | 10 | 1.81% |
| hsa_circ_0114389 | 1108 | 10 | 1.81% |
| hsa_circ_0115291 | 1108 | 10 | 1.81% |
| hsa_circ_0121664 | 1108 | 10 | 1.81% |
| hsa_circ_0122918 | 1108 | 10 | 1.81% |
| hsa_circ_0123361 | 1108 | 10 | 1.81% |
| hsa_circ_0123464 | 1108 | 10 | 1.81% |
| hsa_circ_0131147 | 1108 | 10 | 1.81% |
| hsa_circ_0054105 | 2881 | 26 | 1.80% |
| hsa_circ_0081775 | 1219 | 11 | 1.80% |
| hsa_circ_0090769 | 1219 | 11 | 1.80% |
| hsa_circ_0111324 | 1219 | 11 | 1.80% |
| hsa_circ_0113226 | 3214 | 29 | 1.80% |
| hsa_circ_0015167 | 1330 | 12 | 1.80% |
| hsa_circ_0101624 | 1330 | 12 | 1.80% |
| hsa_circ_0130736 | 1330 | 12 | 1.80% |
| hsa_circ_0067265 | 1663 | 15 | 1.80% |
| hsa_circ_0022547 | 1885 | 17 | 1.80% |
| hsa_circ_0013692 | 1109 | 10 | 1.80% |
| hsa_circ_0021797 | 1109 | 10 | 1.80% |
| hsa_circ_0053034 | 1109 | 10 | 1.80% |
| hsa_circ_0066424 | 1109 | 10 | 1.80% |
| hsa_circ_0067543 | 1109 | 10 | 1.80% |
| hsa_circ_0069278 | 1109 | 10 | 1.80% |
| hsa_circ_0091376 | 1109 | 10 | 1.80% |
| hsa_circ_0091466 | 1109 | 10 | 1.80% |
| hsa_circ_0094918 | 1109 | 10 | 1.80% |
| hsa_circ_0100019 | 1109 | 10 | 1.80% |
| hsa_circ_0100185 | 1109 | 10 | 1.80% |
| hsa_circ_0111959 | 1109 | 10 | 1.80% |
| hsa_circ_0123618 | 1109 | 10 | 1.80% |
| hsa_circ_0126805 | 1109 | 10 | 1.80% |
| hsa_circ_0028956 | 1220 | 11 | 1.80% |
| hsa_circ_0040595 | 1220 | 11 | 1.80% |
| hsa_circ_0078570 | 1220 | 11 | 1.80% |
| hsa_circ_0085288 | 1220 | 11 | 1.80% |
| hsa_circ_0096762 | 1220 | 11 | 1.80% |
| hsa_circ_0097763 | 1220 | 11 | 1.80% |
| hsa_circ_0110776 | 1220 | 11 | 1.80% |
| hsa_circ_0132773 | 1220 | 11 | 1.80% |
| hsa_circ_0009044 | 1331 | 12 | 1.80% |
| hsa_circ_0048833 | 1331 | 12 | 1.80% |
| hsa_circ_0071354 | 1331 | 12 | 1.80% |
| hsa_circ_0091367 | 1331 | 12 | 1.80% |
| hsa_circ_0132738 | 1331 | 12 | 1.80% |
| hsa_circ_0131834 | 1664 | 15 | 1.80% |
| hsa_circ_0057432 | 1332 | 12 | 1.80% |
| hsa_circ_0009603 | 1221 | 11 | 1.80% |
| hsa_circ_0020557 | 1221 | 11 | 1.80% |

| | | | |
|---|---|---|---|
| hsa_circ_0065161 | 1221 | 11 | 1.80% |
| hsa_circ_0136402 | 1221 | 11 | 1.80% |
| hsa_circ_0001328 | 1110 | 10 | 1.80% |
| hsa_circ_0028462 | 1110 | 10 | 1.80% |
| hsa_circ_0039580 | 1110 | 10 | 1.80% |
| hsa_circ_0057630 | 1110 | 10 | 1.80% |
| hsa_circ_0060401 | 1110 | 10 | 1.80% |
| hsa_circ_0061661 | 1110 | 10 | 1.80% |
| hsa_circ_0072093 | 1110 | 10 | 1.80% |
| hsa_circ_0076631 | 1110 | 10 | 1.80% |
| hsa_circ_0079590 | 1110 | 10 | 1.80% |
| hsa_circ_0081394 | 1110 | 10 | 1.80% |
| hsa_circ_0085677 | 1110 | 10 | 1.80% |
| hsa_circ_0093837 | 1110 | 10 | 1.80% |
| hsa_circ_0105216 | 1110 | 10 | 1.80% |
| hsa_circ_0111724 | 1110 | 10 | 1.80% |
| hsa_circ_0124312 | 1110 | 10 | 1.80% |
| hsa_circ_0126510 | 1110 | 10 | 1.80% |
| hsa_circ_0067270 | 1666 | 15 | 1.80% |
| hsa_circ_0005593 | 1444 | 13 | 1.80% |
| hsa_circ_0117680 | 1333 | 12 | 1.80% |
| hsa_circ_0090664 | 1222 | 11 | 1.80% |
| hsa_circ_0135393 | 1222 | 11 | 1.80% |
| hsa_circ_0010772 | 1111 | 10 | 1.80% |
| hsa_circ_0022459 | 1111 | 10 | 1.80% |
| hsa_circ_0024237 | 1111 | 10 | 1.80% |
| hsa_circ_0059915 | 1111 | 10 | 1.80% |
| hsa_circ_0064585 | 1111 | 10 | 1.80% |
| hsa_circ_0070770 | 1111 | 10 | 1.80% |
| hsa_circ_0074578 | 1111 | 10 | 1.80% |
| hsa_circ_0079457 | 1111 | 10 | 1.80% |
| hsa_circ_0084067 | 3334 | 30 | 1.80% |
| hsa_circ_0014764 | 1445 | 13 | 1.80% |
| hsa_circ_0012183 | 1334 | 12 | 1.80% |
| hsa_circ_0103337 | 1334 | 12 | 1.80% |
| hsa_circ_0051208 | 1223 | 11 | 1.80% |
| hsa_circ_0076318 | 1223 | 11 | 1.80% |
| hsa_circ_0076431 | 1223 | 11 | 1.80% |
| hsa_circ_0000795 | 1112 | 10 | 1.80% |
| hsa_circ_0019020 | 1112 | 10 | 1.80% |
| hsa_circ_0029864 | 1112 | 10 | 1.80% |
| hsa_circ_0048942 | 1112 | 10 | 1.80% |
| hsa_circ_0064502 | 1112 | 10 | 1.80% |
| hsa_circ_0069008 | 1112 | 10 | 1.80% |
| hsa_circ_0075654 | 1112 | 10 | 1.80% |
| hsa_circ_0077616 | 1112 | 10 | 1.80% |
| hsa_circ_0079465 | 1112 | 10 | 1.80% |
| hsa_circ_0088632 | 1112 | 10 | 1.80% |
| hsa_circ_0119923 | 1112 | 10 | 1.80% |
| hsa_circ_0089724 | 1335 | 12 | 1.80% |

| | | | |
|---|---|---|---|
| hsa_circ_0104184 | 1335 | 12 | 1.80% |
| hsa_circ_0132517 | 1335 | 12 | 1.80% |
| hsa_circ_0022043 | 1224 | 11 | 1.80% |
| hsa_circ_0027307 | 1224 | 11 | 1.80% |
| hsa_circ_0049598 | 1224 | 11 | 1.80% |
| hsa_circ_0054118 | 1224 | 11 | 1.80% |
| hsa_circ_0056991 | 1224 | 11 | 1.80% |
| hsa_circ_0090545 | 1224 | 11 | 1.80% |
| hsa_circ_0106003 | 1224 | 11 | 1.80% |
| hsa_circ_0025944 | 1113 | 10 | 1.80% |
| hsa_circ_0037189 | 1113 | 10 | 1.80% |
| hsa_circ_0051979 | 1113 | 10 | 1.80% |
| hsa_circ_0078016 | 1113 | 10 | 1.80% |
| hsa_circ_0080060 | 1113 | 10 | 1.80% |
| hsa_circ_0081458 | 1113 | 10 | 1.80% |
| hsa_circ_0086998 | 1113 | 10 | 1.80% |
| hsa_circ_0090584 | 1113 | 10 | 1.80% |
| hsa_circ_0127205 | 1113 | 10 | 1.80% |
| hsa_circ_0136663 | 1113 | 10 | 1.80% |
| hsa_circ_0139629 | 1113 | 10 | 1.80% |
| hsa_circ_0066111 | 1336 | 12 | 1.80% |
| hsa_circ_0085467 | 1336 | 12 | 1.80% |
| hsa_circ_0101372 | 3675 | 33 | 1.80% |
| hsa_circ_0048870 | 1225 | 11 | 1.80% |
| hsa_circ_0053702 | 1225 | 11 | 1.80% |
| hsa_circ_0099815 | 1225 | 11 | 1.80% |
| hsa_circ_0082303 | 1448 | 13 | 1.80% |
| hsa_circ_0006370 | 1114 | 10 | 1.80% |
| hsa_circ_0020896 | 1114 | 10 | 1.80% |
| hsa_circ_0025266 | 1114 | 10 | 1.80% |
| hsa_circ_0029306 | 1114 | 10 | 1.80% |
| hsa_circ_0031490 | 1114 | 10 | 1.80% |
| hsa_circ_0034311 | 1114 | 10 | 1.80% |
| hsa_circ_0035213 | 1114 | 10 | 1.80% |
| hsa_circ_0049349 | 1114 | 10 | 1.80% |
| hsa_circ_0049726 | 1114 | 10 | 1.80% |
| hsa_circ_0065946 | 1114 | 10 | 1.80% |
| hsa_circ_0078503 | 1114 | 10 | 1.80% |
| hsa_circ_0099035 | 1114 | 10 | 1.80% |
| hsa_circ_0100359 | 1114 | 10 | 1.80% |
| hsa_circ_0127418 | 1114 | 10 | 1.80% |
| hsa_circ_0132485 | 1114 | 10 | 1.80% |
| hsa_circ_0136786 | 1114 | 10 | 1.80% |
| hsa_circ_0130235 | 1337 | 12 | 1.80% |
| hsa_circ_0076958 | 1560 | 14 | 1.79% |
| hsa_circ_0000079 | 1226 | 11 | 1.79% |
| hsa_circ_0016362 | 1226 | 11 | 1.79% |
| hsa_circ_0023206 | 1226 | 11 | 1.79% |
| hsa_circ_0029384 | 1226 | 11 | 1.79% |
| hsa_circ_0042261 | 1226 | 11 | 1.79% |

| | | | |
|---|---|---|---|
| hsa_circ_0059556 | 1226 | 11 | 1.79% |
| hsa_circ_0072713 | 1226 | 11 | 1.79% |
| hsa_circ_0088805 | 1226 | 11 | 1.79% |
| hsa_circ_0095436 | 1226 | 11 | 1.79% |
| hsa_circ_0086703 | 1449 | 13 | 1.79% |
| hsa_circ_0024150 | 1338 | 12 | 1.79% |
| hsa_circ_0066114 | 1338 | 12 | 1.79% |
| hsa_circ_0006146 | 1115 | 10 | 1.79% |
| hsa_circ_0014009 | 1115 | 10 | 1.79% |
| hsa_circ_0014861 | 1115 | 10 | 1.79% |
| hsa_circ_0035090 | 1115 | 10 | 1.79% |
| hsa_circ_0049969 | 1115 | 10 | 1.79% |
| hsa_circ_0067876 | 1115 | 10 | 1.79% |
| hsa_circ_0075225 | 1115 | 10 | 1.79% |
| hsa_circ_0077933 | 1115 | 10 | 1.79% |
| hsa_circ_0079162 | 1115 | 10 | 1.79% |
| hsa_circ_0091043 | 1115 | 10 | 1.79% |
| hsa_circ_0055787 | 1450 | 13 | 1.79% |
| hsa_circ_0136480 | 1450 | 13 | 1.79% |
| hsa_circ_0017945 | 1227 | 11 | 1.79% |
| hsa_circ_0031390 | 1227 | 11 | 1.79% |
| hsa_circ_0083615 | 1227 | 11 | 1.79% |
| hsa_circ_0102667 | 1227 | 11 | 1.79% |
| hsa_circ_0136284 | 1562 | 14 | 1.79% |
| hsa_circ_0022105 | 1339 | 12 | 1.79% |
| hsa_circ_0030925 | 1339 | 12 | 1.79% |
| hsa_circ_0031877 | 1339 | 12 | 1.79% |
| hsa_circ_0039901 | 1339 | 12 | 1.79% |
| hsa_circ_0056848 | 1339 | 12 | 1.79% |
| hsa_circ_0019590 | 1116 | 10 | 1.79% |
| hsa_circ_0020350 | 1116 | 10 | 1.79% |
| hsa_circ_0025774 | 1116 | 10 | 1.79% |
| hsa_circ_0033421 | 1116 | 10 | 1.79% |
| hsa_circ_0064124 | 1116 | 10 | 1.79% |
| hsa_circ_0064505 | 1116 | 10 | 1.79% |
| hsa_circ_0076291 | 1116 | 10 | 1.79% |
| hsa_circ_0105977 | 1116 | 10 | 1.79% |
| hsa_circ_0116222 | 1116 | 10 | 1.79% |
| hsa_circ_0131826 | 1116 | 10 | 1.79% |
| hsa_circ_0083830 | 1228 | 11 | 1.79% |
| hsa_circ_0127721 | 1228 | 11 | 1.79% |
| hsa_circ_0138542 | 1228 | 11 | 1.79% |
| hsa_circ_0016539 | 1340 | 12 | 1.79% |
| hsa_circ_0022414 | 1340 | 12 | 1.79% |
| hsa_circ_0057942 | 1340 | 12 | 1.79% |
| hsa_circ_0039454 | 1452 | 13 | 1.79% |
| hsa_circ_0014510 | 1117 | 10 | 1.79% |
| hsa_circ_0016338 | 1117 | 10 | 1.79% |
| hsa_circ_0024247 | 1117 | 10 | 1.79% |
| hsa_circ_0032286 | 1117 | 10 | 1.79% |

| | | | |
|---|---|---|---|
| hsa_circ_0048483 | 1117 | 10 | 1.79% |
| hsa_circ_0056191 | 1117 | 10 | 1.79% |
| hsa_circ_0058630 | 1117 | 10 | 1.79% |
| hsa_circ_0097469 | 1117 | 10 | 1.79% |
| hsa_circ_0118080 | 1117 | 10 | 1.79% |
| hsa_circ_0132891 | 1117 | 10 | 1.79% |
| hsa_circ_0137049 | 1117 | 10 | 1.79% |
| hsa_circ_0140684 | 1117 | 10 | 1.79% |
| hsa_circ_0012328 | 1564 | 14 | 1.79% |
| hsa_circ_0072091 | 1564 | 14 | 1.79% |
| hsa_circ_0083650 | 1564 | 14 | 1.79% |
| hsa_circ_0010006 | 1229 | 11 | 1.79% |
| hsa_circ_0013903 | 1229 | 11 | 1.79% |
| hsa_circ_0020923 | 1229 | 11 | 1.79% |
| hsa_circ_0021829 | 1229 | 11 | 1.79% |
| hsa_circ_0064561 | 1229 | 11 | 1.79% |
| hsa_circ_0067216 | 1229 | 11 | 1.79% |
| hsa_circ_0067863 | 1229 | 11 | 1.79% |
| hsa_circ_0115993 | 1229 | 11 | 1.79% |
| hsa_circ_0062104 | 1676 | 15 | 1.79% |
| hsa_circ_0030935 | 1341 | 12 | 1.79% |
| hsa_circ_0063615 | 1341 | 12 | 1.79% |
| hsa_circ_0093978 | 1341 | 12 | 1.79% |
| hsa_circ_0086581 | 1677 | 15 | 1.79% |
| hsa_circ_0016022 | 1118 | 10 | 1.79% |
| hsa_circ_0017451 | 1118 | 10 | 1.79% |
| hsa_circ_0020753 | 1118 | 10 | 1.79% |
| hsa_circ_0025004 | 1118 | 10 | 1.79% |
| hsa_circ_0029454 | 1118 | 10 | 1.79% |
| hsa_circ_0031016 | 1118 | 10 | 1.79% |
| hsa_circ_0035671 | 1118 | 10 | 1.79% |
| hsa_circ_0071578 | 1118 | 10 | 1.79% |
| hsa_circ_0079165 | 1118 | 10 | 1.79% |
| hsa_circ_0115054 | 1118 | 10 | 1.79% |
| hsa_circ_0010178 | 1230 | 11 | 1.79% |
| hsa_circ_0030934 | 1230 | 11 | 1.79% |
| hsa_circ_0036180 | 1230 | 11 | 1.79% |
| hsa_circ_0039568 | 1230 | 11 | 1.79% |
| hsa_circ_0048917 | 1230 | 11 | 1.79% |
| hsa_circ_0051129 | 1230 | 11 | 1.79% |
| hsa_circ_0093173 | 1230 | 11 | 1.79% |
| hsa_circ_0121149 | 1230 | 11 | 1.79% |
| hsa_circ_0136640 | 1230 | 11 | 1.79% |
| hsa_circ_0075620 | 1342 | 12 | 1.79% |
| hsa_circ_0097755 | 1342 | 12 | 1.79% |
| hsa_circ_0001832 | 1119 | 10 | 1.79% |
| hsa_circ_0014832 | 1119 | 10 | 1.79% |
| hsa_circ_0020454 | 1119 | 10 | 1.79% |
| hsa_circ_0036501 | 1119 | 10 | 1.79% |
| hsa_circ_0045599 | 1119 | 10 | 1.79% |

| | | | |
|---|---|---|---|
| hsa_circ_0057613 | 1119 | 10 | 1.79% |
| hsa_circ_0060184 | 1119 | 10 | 1.79% |
| hsa_circ_0073693 | 1119 | 10 | 1.79% |
| hsa_circ_0093226 | 1119 | 10 | 1.79% |
| hsa_circ_0106024 | 1119 | 10 | 1.79% |
| hsa_circ_0109942 | 1119 | 10 | 1.79% |
| hsa_circ_0112429 | 1119 | 10 | 1.79% |
| hsa_circ_0113309 | 1119 | 10 | 1.79% |
| hsa_circ_0135550 | 1119 | 10 | 1.79% |
| hsa_circ_0013428 | 1231 | 11 | 1.79% |
| hsa_circ_0025181 | 1231 | 11 | 1.79% |
| hsa_circ_0063356 | 1231 | 11 | 1.79% |
| hsa_circ_0078550 | 1231 | 11 | 1.79% |
| hsa_circ_0118024 | 1231 | 11 | 1.79% |
| hsa_circ_0006682 | 1343 | 12 | 1.79% |
| hsa_circ_0031753 | 1343 | 12 | 1.79% |
| hsa_circ_0083291 | 1567 | 14 | 1.79% |
| hsa_circ_0084065 | 1567 | 14 | 1.79% |
| hsa_circ_0042875 | 1568 | 14 | 1.79% |
| hsa_circ_0012294 | 1456 | 13 | 1.79% |
| hsa_circ_0084544 | 1456 | 13 | 1.79% |
| hsa_circ_0059640 | 1344 | 12 | 1.79% |
| hsa_circ_0120857 | 1344 | 12 | 1.79% |
| hsa_circ_0022508 | 1232 | 11 | 1.79% |
| hsa_circ_0022580 | 1232 | 11 | 1.79% |
| hsa_circ_0049697 | 1232 | 11 | 1.79% |
| hsa_circ_0051515 | 1232 | 11 | 1.79% |
| hsa_circ_0062308 | 1232 | 11 | 1.79% |
| hsa_circ_0113155 | 1232 | 11 | 1.79% |
| hsa_circ_0119267 | 1232 | 11 | 1.79% |
| hsa_circ_0014378 | 1120 | 10 | 1.79% |
| hsa_circ_0017309 | 1120 | 10 | 1.79% |
| hsa_circ_0024132 | 1120 | 10 | 1.79% |
| hsa_circ_0033639 | 1120 | 10 | 1.79% |
| hsa_circ_0039037 | 1120 | 10 | 1.79% |
| hsa_circ_0051416 | 1120 | 10 | 1.79% |
| hsa_circ_0057146 | 1120 | 10 | 1.79% |
| hsa_circ_0064067 | 1120 | 10 | 1.79% |
| hsa_circ_0068978 | 1120 | 10 | 1.79% |
| hsa_circ_0072480 | 1120 | 10 | 1.79% |
| hsa_circ_0074184 | 1120 | 10 | 1.79% |
| hsa_circ_0077650 | 1120 | 10 | 1.79% |
| hsa_circ_0095897 | 1120 | 10 | 1.79% |
| hsa_circ_0107058 | 1120 | 10 | 1.79% |
| hsa_circ_0068087 | 6052 | 54 | 1.78% |
| hsa_circ_0100159 | 2914 | 26 | 1.78% |
| hsa_circ_0056265 | 1345 | 12 | 1.78% |
| hsa_circ_0013313 | 1233 | 11 | 1.78% |
| hsa_circ_0060307 | 1233 | 11 | 1.78% |
| hsa_circ_0064093 | 1233 | 11 | 1.78% |

| | | | |
|---|---|---|---|
| hsa_circ_0071457 | 1233 | 11 | 1.78% |
| hsa_circ_0124782 | 1233 | 11 | 1.78% |
| hsa_circ_0128561 | 1233 | 11 | 1.78% |
| hsa_circ_0130044 | 1233 | 11 | 1.78% |
| hsa_circ_0011128 | 1121 | 10 | 1.78% |
| hsa_circ_0016755 | 1121 | 10 | 1.78% |
| hsa_circ_0022971 | 1121 | 10 | 1.78% |
| hsa_circ_0055274 | 1121 | 10 | 1.78% |
| hsa_circ_0057558 | 1121 | 10 | 1.78% |
| hsa_circ_0062972 | 1121 | 10 | 1.78% |
| hsa_circ_0073917 | 1121 | 10 | 1.78% |
| hsa_circ_0096649 | 1121 | 10 | 1.78% |
| hsa_circ_0117620 | 1121 | 10 | 1.78% |
| hsa_circ_0105572 | 1458 | 13 | 1.78% |
| hsa_circ_0017282 | 1346 | 12 | 1.78% |
| hsa_circ_0046821 | 1234 | 11 | 1.78% |
| hsa_circ_0101137 | 1234 | 11 | 1.78% |
| hsa_circ_0118020 | 1234 | 11 | 1.78% |
| hsa_circ_0004327 | 1122 | 10 | 1.78% |
| hsa_circ_0007065 | 1122 | 10 | 1.78% |
| hsa_circ_0011659 | 1122 | 10 | 1.78% |
| hsa_circ_0014971 | 1122 | 10 | 1.78% |
| hsa_circ_0020656 | 1122 | 10 | 1.78% |
| hsa_circ_0023677 | 1122 | 10 | 1.78% |
| hsa_circ_0034519 | 1122 | 10 | 1.78% |
| hsa_circ_0039062 | 1122 | 10 | 1.78% |
| hsa_circ_0053241 | 1122 | 10 | 1.78% |
| hsa_circ_0068503 | 1122 | 10 | 1.78% |
| hsa_circ_0079762 | 1122 | 10 | 1.78% |
| hsa_circ_0110769 | 1122 | 10 | 1.78% |
| hsa_circ_0111106 | 1122 | 10 | 1.78% |
| hsa_circ_0114603 | 1122 | 10 | 1.78% |
| hsa_circ_0130091 | 1122 | 10 | 1.78% |
| hsa_circ_0133842 | 1122 | 10 | 1.78% |
| hsa_circ_0138555 | 1122 | 10 | 1.78% |
| hsa_circ_0095919 | 1571 | 14 | 1.78% |
| hsa_circ_0034638 | 1347 | 12 | 1.78% |
| hsa_circ_0034961 | 1347 | 12 | 1.78% |
| hsa_circ_0070796 | 1347 | 12 | 1.78% |
| hsa_circ_0133886 | 1347 | 12 | 1.78% |
| hsa_circ_0138129 | 1347 | 12 | 1.78% |
| hsa_circ_0100794 | 2133 | 19 | 1.78% |
| hsa_circ_0024727 | 1235 | 11 | 1.78% |
| hsa_circ_0043582 | 1235 | 11 | 1.78% |
| hsa_circ_0064739 | 1235 | 11 | 1.78% |
| hsa_circ_0085832 | 1235 | 11 | 1.78% |
| hsa_circ_0103777 | 1235 | 11 | 1.78% |
| hsa_circ_0029822 | 1123 | 10 | 1.78% |
| hsa_circ_0035921 | 1123 | 10 | 1.78% |
| hsa_circ_0038582 | 1123 | 10 | 1.78% |

| | | | |
|---|---|---|---|
| hsa_circ_0039572 | 1123 | 10 | 1.78% |
| hsa_circ_0054546 | 1123 | 10 | 1.78% |
| hsa_circ_0065271 | 1123 | 10 | 1.78% |
| hsa_circ_0065863 | 1123 | 10 | 1.78% |
| hsa_circ_0086305 | 1123 | 10 | 1.78% |
| hsa_circ_0103636 | 1123 | 10 | 1.78% |
| hsa_circ_0112245 | 1123 | 10 | 1.78% |
| hsa_circ_0131300 | 1123 | 10 | 1.78% |
| hsa_circ_0020077 | 1348 | 12 | 1.78% |
| hsa_circ_0117422 | 79210 | 705 | 1.78% |
| hsa_circ_0022052 | 1236 | 11 | 1.78% |
| hsa_circ_0024485 | 1236 | 11 | 1.78% |
| hsa_circ_0027642 | 1236 | 11 | 1.78% |
| hsa_circ_0065630 | 1236 | 11 | 1.78% |
| hsa_circ_0071290 | 1236 | 11 | 1.78% |
| hsa_circ_0076555 | 1236 | 11 | 1.78% |
| hsa_circ_0087419 | 1236 | 11 | 1.78% |
| hsa_circ_0091681 | 1236 | 11 | 1.78% |
| hsa_circ_0123576 | 1236 | 11 | 1.78% |
| hsa_circ_0063958 | 1124 | 10 | 1.78% |
| hsa_circ_0069571 | 1124 | 10 | 1.78% |
| hsa_circ_0081494 | 1124 | 10 | 1.78% |
| hsa_circ_0083886 | 1124 | 10 | 1.78% |
| hsa_circ_0125901 | 1124 | 10 | 1.78% |
| hsa_circ_0126639 | 1124 | 10 | 1.78% |
| hsa_circ_0065639 | 1237 | 11 | 1.78% |
| hsa_circ_0096288 | 1237 | 11 | 1.78% |
| hsa_circ_0096717 | 1237 | 11 | 1.78% |
| hsa_circ_0103438 | 1237 | 11 | 1.78% |
| hsa_circ_0097214 | 1687 | 15 | 1.78% |
| hsa_circ_0119942 | 1350 | 12 | 1.78% |
| hsa_circ_0004120 | 1125 | 10 | 1.78% |
| hsa_circ_0007521 | 1125 | 10 | 1.78% |
| hsa_circ_0014023 | 1125 | 10 | 1.78% |
| hsa_circ_0016034 | 1125 | 10 | 1.78% |
| hsa_circ_0026375 | 1125 | 10 | 1.78% |
| hsa_circ_0031194 | 1125 | 10 | 1.78% |
| hsa_circ_0037059 | 1125 | 10 | 1.78% |
| hsa_circ_0044282 | 1125 | 10 | 1.78% |
| hsa_circ_0058403 | 1125 | 10 | 1.78% |
| hsa_circ_0070685 | 1125 | 10 | 1.78% |
| hsa_circ_0089681 | 1125 | 10 | 1.78% |
| hsa_circ_0094619 | 1125 | 10 | 1.78% |
| hsa_circ_0108402 | 1125 | 10 | 1.78% |
| hsa_circ_0110672 | 1125 | 10 | 1.78% |
| hsa_circ_0036856 | 1238 | 11 | 1.78% |
| hsa_circ_0085943 | 1238 | 11 | 1.78% |
| hsa_circ_0089732 | 1238 | 11 | 1.78% |
| hsa_circ_0100538 | 1238 | 11 | 1.78% |
| hsa_circ_0114835 | 1238 | 11 | 1.78% |

| | | | |
|---|---|---|---|
| hsa_circ_0012070 | 1576 | 14 | 1.78% |
| hsa_circ_0047271 | 1351 | 12 | 1.78% |
| hsa_circ_0069129 | 1351 | 12 | 1.78% |
| hsa_circ_0107371 | 1351 | 12 | 1.78% |
| hsa_circ_0000780 | 1126 | 10 | 1.78% |
| hsa_circ_0001884 | 1126 | 10 | 1.78% |
| hsa_circ_0051887 | 1126 | 10 | 1.78% |
| hsa_circ_0059837 | 1126 | 10 | 1.78% |
| hsa_circ_0073584 | 1126 | 10 | 1.78% |
| hsa_circ_0077368 | 1126 | 10 | 1.78% |
| hsa_circ_0114519 | 1126 | 10 | 1.78% |
| hsa_circ_0117837 | 1126 | 10 | 1.78% |
| hsa_circ_0114186 | 1464 | 13 | 1.78% |
| hsa_circ_0011286 | 1239 | 11 | 1.78% |
| hsa_circ_0025154 | 1239 | 11 | 1.78% |
| hsa_circ_0028401 | 1239 | 11 | 1.78% |
| hsa_circ_0083887 | 1239 | 11 | 1.78% |
| hsa_circ_0097924 | 1239 | 11 | 1.78% |
| hsa_circ_0114499 | 1239 | 11 | 1.78% |
| hsa_circ_0057506 | 1577 | 14 | 1.78% |
| hsa_circ_0072095 | 1577 | 14 | 1.78% |
| hsa_circ_0087070 | 1352 | 12 | 1.78% |
| hsa_circ_0089228 | 1352 | 12 | 1.78% |
| hsa_circ_0122004 | 1352 | 12 | 1.78% |
| hsa_circ_0100416 | 2254 | 20 | 1.77% |
| hsa_circ_0001425 | 1127 | 10 | 1.77% |
| hsa_circ_0003541 | 1127 | 10 | 1.77% |
| hsa_circ_0020314 | 1127 | 10 | 1.77% |
| hsa_circ_0027577 | 1127 | 10 | 1.77% |
| hsa_circ_0031158 | 1127 | 10 | 1.77% |
| hsa_circ_0036722 | 1127 | 10 | 1.77% |
| hsa_circ_0037863 | 1127 | 10 | 1.77% |
| hsa_circ_0039814 | 1127 | 10 | 1.77% |
| hsa_circ_0042553 | 1127 | 10 | 1.77% |
| hsa_circ_0063207 | 1127 | 10 | 1.77% |
| hsa_circ_0081064 | 1127 | 10 | 1.77% |
| hsa_circ_0085061 | 1127 | 10 | 1.77% |
| hsa_circ_0029451 | 1916 | 17 | 1.77% |
| hsa_circ_0046180 | 1240 | 11 | 1.77% |
| hsa_circ_0046466 | 1240 | 11 | 1.77% |
| hsa_circ_0089744 | 1240 | 11 | 1.77% |
| hsa_circ_0099801 | 1240 | 11 | 1.77% |
| hsa_circ_0007420 | 1353 | 12 | 1.77% |
| hsa_circ_0012522 | 1353 | 12 | 1.77% |
| hsa_circ_0099598 | 1353 | 12 | 1.77% |
| hsa_circ_0044503 | 2594 | 23 | 1.77% |
| hsa_circ_0023923 | 1128 | 10 | 1.77% |
| hsa_circ_0025439 | 1128 | 10 | 1.77% |
| hsa_circ_0046340 | 1128 | 10 | 1.77% |
| hsa_circ_0047653 | 1128 | 10 | 1.77% |

| | | | |
|---|---|---|---|
| hsa_circ_0058076 | 1128 | 10 | 1.77% |
| hsa_circ_0058150 | 1128 | 10 | 1.77% |
| hsa_circ_0061888 | 1128 | 10 | 1.77% |
| hsa_circ_0063183 | 1128 | 10 | 1.77% |
| hsa_circ_0082122 | 1128 | 10 | 1.77% |
| hsa_circ_0084907 | 1128 | 10 | 1.77% |
| hsa_circ_0112346 | 1128 | 10 | 1.77% |
| hsa_circ_0118846 | 1128 | 10 | 1.77% |
| hsa_circ_0120604 | 1128 | 10 | 1.77% |
| hsa_circ_0132382 | 1128 | 10 | 1.77% |
| hsa_circ_0140682 | 1128 | 10 | 1.77% |
| hsa_circ_0004308 | 1241 | 11 | 1.77% |
| hsa_circ_0055710 | 1241 | 11 | 1.77% |
| hsa_circ_0080253 | 1354 | 12 | 1.77% |
| hsa_circ_0092053 | 1354 | 12 | 1.77% |
| hsa_circ_0103334 | 1354 | 12 | 1.77% |
| hsa_circ_0067631 | 1467 | 13 | 1.77% |
| hsa_circ_0130460 | 1467 | 13 | 1.77% |
| hsa_circ_0052068 | 1580 | 14 | 1.77% |
| hsa_circ_0004194 | 1129 | 10 | 1.77% |
| hsa_circ_0006317 | 1129 | 10 | 1.77% |
| hsa_circ_0012872 | 1129 | 10 | 1.77% |
| hsa_circ_0016795 | 1129 | 10 | 1.77% |
| hsa_circ_0023618 | 1129 | 10 | 1.77% |
| hsa_circ_0024473 | 1129 | 10 | 1.77% |
| hsa_circ_0049648 | 1129 | 10 | 1.77% |
| hsa_circ_0061946 | 1129 | 10 | 1.77% |
| hsa_circ_0073138 | 1129 | 10 | 1.77% |
| hsa_circ_0083557 | 1129 | 10 | 1.77% |
| hsa_circ_0097717 | 1129 | 10 | 1.77% |
| hsa_circ_0113053 | 1129 | 10 | 1.77% |
| hsa_circ_0009279 | 1242 | 11 | 1.77% |
| hsa_circ_0040312 | 1242 | 11 | 1.77% |
| hsa_circ_0044055 | 1242 | 11 | 1.77% |
| hsa_circ_0046150 | 1242 | 11 | 1.77% |
| hsa_circ_0064436 | 1242 | 11 | 1.77% |
| hsa_circ_0086699 | 1242 | 11 | 1.77% |
| hsa_circ_0114382 | 1242 | 11 | 1.77% |
| hsa_circ_0123507 | 1242 | 11 | 1.77% |
| hsa_circ_0125207 | 1242 | 11 | 1.77% |
| hsa_circ_0133795 | 1242 | 11 | 1.77% |
| hsa_circ_0136061 | 1242 | 11 | 1.77% |
| hsa_circ_0123476 | 1355 | 12 | 1.77% |
| hsa_circ_0064446 | 1468 | 13 | 1.77% |
| hsa_circ_0067602 | 1356 | 12 | 1.77% |
| hsa_circ_0088204 | 1356 | 12 | 1.77% |
| hsa_circ_0108990 | 1356 | 12 | 1.77% |
| hsa_circ_0000355 | 1243 | 11 | 1.77% |
| hsa_circ_0010208 | 1243 | 11 | 1.77% |
| hsa_circ_0023201 | 1243 | 11 | 1.77% |

| | | | |
|---|---|---|---|
| hsa_circ_0042737 | 1243 | 11 | 1.77% |
| hsa_circ_0044419 | 1243 | 11 | 1.77% |
| hsa_circ_0052366 | 1243 | 11 | 1.77% |
| hsa_circ_0075944 | 1243 | 11 | 1.77% |
| hsa_circ_0089669 | 1243 | 11 | 1.77% |
| hsa_circ_0090720 | 1243 | 11 | 1.77% |
| hsa_circ_0093620 | 1243 | 11 | 1.77% |
| hsa_circ_0099805 | 1243 | 11 | 1.77% |
| hsa_circ_0003577 | 1130 | 10 | 1.77% |
| hsa_circ_0012507 | 1130 | 10 | 1.77% |
| hsa_circ_0041518 | 1130 | 10 | 1.77% |
| hsa_circ_0060332 | 1130 | 10 | 1.77% |
| hsa_circ_0060942 | 1130 | 10 | 1.77% |
| hsa_circ_0074034 | 1130 | 10 | 1.77% |
| hsa_circ_0077705 | 1130 | 10 | 1.77% |
| hsa_circ_0080558 | 1130 | 10 | 1.77% |
| hsa_circ_0112847 | 1130 | 10 | 1.77% |
| hsa_circ_0127145 | 1130 | 10 | 1.77% |
| hsa_circ_0011583 | 4974 | 44 | 1.77% |
| hsa_circ_0057365 | 1809 | 16 | 1.77% |
| hsa_circ_0052827 | 1470 | 13 | 1.77% |
| hsa_circ_0076133 | 1470 | 13 | 1.77% |
| hsa_circ_0071490 | 1357 | 12 | 1.77% |
| hsa_circ_0115371 | 1357 | 12 | 1.77% |
| hsa_circ_0047374 | 1244 | 11 | 1.77% |
| hsa_circ_0091241 | 1244 | 11 | 1.77% |
| hsa_circ_0096866 | 1244 | 11 | 1.77% |
| hsa_circ_0105323 | 1244 | 11 | 1.77% |
| hsa_circ_0118232 | 1244 | 11 | 1.77% |
| hsa_circ_0106033 | 2375 | 21 | 1.77% |
| hsa_circ_0019927 | 1131 | 10 | 1.77% |
| hsa_circ_0023229 | 1131 | 10 | 1.77% |
| hsa_circ_0024107 | 1131 | 10 | 1.77% |
| hsa_circ_0032080 | 1131 | 10 | 1.77% |
| hsa_circ_0040944 | 1131 | 10 | 1.77% |
| hsa_circ_0043651 | 1131 | 10 | 1.77% |
| hsa_circ_0048814 | 1131 | 10 | 1.77% |
| hsa_circ_0048816 | 1131 | 10 | 1.77% |
| hsa_circ_0051809 | 1131 | 10 | 1.77% |
| hsa_circ_0052775 | 1131 | 10 | 1.77% |
| hsa_circ_0059735 | 1131 | 10 | 1.77% |
| hsa_circ_0063114 | 1131 | 10 | 1.77% |
| hsa_circ_0063988 | 1131 | 10 | 1.77% |
| hsa_circ_0086299 | 1131 | 10 | 1.77% |
| hsa_circ_0095132 | 1131 | 10 | 1.77% |
| hsa_circ_0126442 | 1131 | 10 | 1.77% |
| hsa_circ_0132890 | 1131 | 10 | 1.77% |
| hsa_circ_0021656 | 1810 | 16 | 1.77% |
| hsa_circ_0002875 | 1471 | 13 | 1.77% |
| hsa_circ_0052193 | 1471 | 13 | 1.77% |

| | | | |
|---|---|---|---|
| hsa_circ_0040692 | 1358 | 12 | 1.77% |
| hsa_circ_0101346 | 1358 | 12 | 1.77% |
| hsa_circ_0102799 | 1358 | 12 | 1.77% |
| hsa_circ_0005849 | 1245 | 11 | 1.77% |
| hsa_circ_0034673 | 1245 | 11 | 1.77% |
| hsa_circ_0066621 | 1245 | 11 | 1.77% |
| hsa_circ_0101796 | 1245 | 11 | 1.77% |
| hsa_circ_0116519 | 1245 | 11 | 1.77% |
| hsa_circ_0116993 | 1245 | 11 | 1.77% |
| hsa_circ_0138643 | 1245 | 11 | 1.77% |
| hsa_circ_0004570 | 1132 | 10 | 1.77% |
| hsa_circ_0020456 | 1132 | 10 | 1.77% |
| hsa_circ_0027479 | 1132 | 10 | 1.77% |
| hsa_circ_0030879 | 1132 | 10 | 1.77% |
| hsa_circ_0052259 | 1132 | 10 | 1.77% |
| hsa_circ_0085439 | 1132 | 10 | 1.77% |
| hsa_circ_0094826 | 1132 | 10 | 1.77% |
| hsa_circ_0096383 | 1132 | 10 | 1.77% |
| hsa_circ_0097586 | 1132 | 10 | 1.77% |
| hsa_circ_0116859 | 1132 | 10 | 1.77% |
| hsa_circ_0123856 | 1132 | 10 | 1.77% |
| hsa_circ_0000176 | 1359 | 12 | 1.77% |
| hsa_circ_0002779 | 1359 | 12 | 1.77% |
| hsa_circ_0012274 | 1359 | 12 | 1.77% |
| hsa_circ_0105742 | 1246 | 11 | 1.77% |
| hsa_circ_0114354 | 1246 | 11 | 1.77% |
| hsa_circ_0138474 | 1246 | 11 | 1.77% |
| hsa_circ_0140480 | 1246 | 11 | 1.77% |
| hsa_circ_0023907 | 1133 | 10 | 1.77% |
| hsa_circ_0029358 | 1133 | 10 | 1.77% |
| hsa_circ_0047973 | 1133 | 10 | 1.77% |
| hsa_circ_0050592 | 1133 | 10 | 1.77% |
| hsa_circ_0057252 | 1133 | 10 | 1.77% |
| hsa_circ_0061361 | 1133 | 10 | 1.77% |
| hsa_circ_0063118 | 1133 | 10 | 1.77% |
| hsa_circ_0064423 | 1133 | 10 | 1.77% |
| hsa_circ_0064655 | 1133 | 10 | 1.77% |
| hsa_circ_0085740 | 1133 | 10 | 1.77% |
| hsa_circ_0137959 | 1473 | 13 | 1.77% |
| hsa_circ_0014857 | 1360 | 12 | 1.76% |
| hsa_circ_0091184 | 1360 | 12 | 1.76% |
| hsa_circ_0037867 | 1247 | 11 | 1.76% |
| hsa_circ_0052568 | 1247 | 11 | 1.76% |
| hsa_circ_0069892 | 1247 | 11 | 1.76% |
| hsa_circ_0085719 | 1247 | 11 | 1.76% |
| hsa_circ_0102813 | 1247 | 11 | 1.76% |
| hsa_circ_0109776 | 1247 | 11 | 1.76% |
| hsa_circ_0134065 | 1247 | 11 | 1.76% |
| hsa_circ_0075401 | 2154 | 19 | 1.76% |
| hsa_circ_0108843 | 1474 | 13 | 1.76% |

| | | | |
|---|---|---|---|
| hsa_circ_0039115 | 1134 | 10 | 1.76% |
| hsa_circ_0048388 | 1134 | 10 | 1.76% |
| hsa_circ_0050169 | 1134 | 10 | 1.76% |
| hsa_circ_0056749 | 1134 | 10 | 1.76% |
| hsa_circ_0056944 | 1134 | 10 | 1.76% |
| hsa_circ_0068223 | 1134 | 10 | 1.76% |
| hsa_circ_0078766 | 1134 | 10 | 1.76% |
| hsa_circ_0090032 | 1134 | 10 | 1.76% |
| hsa_circ_0133373 | 1134 | 10 | 1.76% |
| hsa_circ_0009583 | 1248 | 11 | 1.76% |
| hsa_circ_0040355 | 1248 | 11 | 1.76% |
| hsa_circ_0052079 | 1248 | 11 | 1.76% |
| hsa_circ_0060785 | 1248 | 11 | 1.76% |
| hsa_circ_0064405 | 1248 | 11 | 1.76% |
| hsa_circ_0089442 | 1248 | 11 | 1.76% |
| hsa_circ_0090131 | 1248 | 11 | 1.76% |
| hsa_circ_0097611 | 1248 | 11 | 1.76% |
| hsa_circ_0116084 | 1248 | 11 | 1.76% |
| hsa_circ_0120184 | 1248 | 11 | 1.76% |
| hsa_circ_0052693 | 1702 | 15 | 1.76% |
| hsa_circ_0020347 | 1362 | 12 | 1.76% |
| hsa_circ_0067409 | 1362 | 12 | 1.76% |
| hsa_circ_0002487 | 1135 | 10 | 1.76% |
| hsa_circ_0005160 | 1135 | 10 | 1.76% |
| hsa_circ_0005615 | 1135 | 10 | 1.76% |
| hsa_circ_0019017 | 1135 | 10 | 1.76% |
| hsa_circ_0028082 | 1135 | 10 | 1.76% |
| hsa_circ_0044446 | 1135 | 10 | 1.76% |
| hsa_circ_0049883 | 1135 | 10 | 1.76% |
| hsa_circ_0065215 | 1135 | 10 | 1.76% |
| hsa_circ_0096656 | 1135 | 10 | 1.76% |
| hsa_circ_0112318 | 1135 | 10 | 1.76% |
| hsa_circ_0030732 | 1249 | 11 | 1.76% |
| hsa_circ_0035372 | 1249 | 11 | 1.76% |
| hsa_circ_0059136 | 1249 | 11 | 1.76% |
| hsa_circ_0076028 | 1249 | 11 | 1.76% |
| hsa_circ_0091317 | 1249 | 11 | 1.76% |
| hsa_circ_0093224 | 1249 | 11 | 1.76% |
| hsa_circ_0136351 | 1249 | 11 | 1.76% |
| hsa_circ_0068533 | 1363 | 12 | 1.76% |
| hsa_circ_0138125 | 1363 | 12 | 1.76% |
| hsa_circ_0011389 | 1136 | 10 | 1.76% |
| hsa_circ_0013390 | 1136 | 10 | 1.76% |
| hsa_circ_0016719 | 1136 | 10 | 1.76% |
| hsa_circ_0037404 | 1136 | 10 | 1.76% |
| hsa_circ_0065427 | 1136 | 10 | 1.76% |
| hsa_circ_0075331 | 1136 | 10 | 1.76% |
| hsa_circ_0081321 | 1136 | 10 | 1.76% |
| hsa_circ_0091503 | 1136 | 10 | 1.76% |
| hsa_circ_0093386 | 1136 | 10 | 1.76% |

| | | | |
|---|---|---|---|
| hsa_circ_0112174 | 1136 | 10 | 1.76% |
| hsa_circ_0116915 | 1136 | 10 | 1.76% |
| hsa_circ_0121803 | 1136 | 10 | 1.76% |
| hsa_circ_0125139 | 1136 | 10 | 1.76% |
| hsa_circ_0136601 | 1136 | 10 | 1.76% |
| hsa_circ_0137043 | 1477 | 13 | 1.76% |
| hsa_circ_0031633 | 1250 | 11 | 1.76% |
| hsa_circ_0074189 | 1250 | 11 | 1.76% |
| hsa_circ_0086711 | 1250 | 11 | 1.76% |
| hsa_circ_0115211 | 1250 | 11 | 1.76% |
| hsa_circ_0116989 | 1250 | 11 | 1.76% |
| hsa_circ_0135827 | 1250 | 11 | 1.76% |
| hsa_circ_0089680 | 1364 | 12 | 1.76% |
| hsa_circ_0139766 | 1364 | 12 | 1.76% |
| hsa_circ_0002998 | 1137 | 10 | 1.76% |
| hsa_circ_0007890 | 1137 | 10 | 1.76% |
| hsa_circ_0015909 | 1137 | 10 | 1.76% |
| hsa_circ_0017793 | 1137 | 10 | 1.76% |
| hsa_circ_0021841 | 1137 | 10 | 1.76% |
| hsa_circ_0026974 | 1137 | 10 | 1.76% |
| hsa_circ_0028810 | 1137 | 10 | 1.76% |
| hsa_circ_0029674 | 1137 | 10 | 1.76% |
| hsa_circ_0038567 | 1137 | 10 | 1.76% |
| hsa_circ_0041560 | 1137 | 10 | 1.76% |
| hsa_circ_0048032 | 1137 | 10 | 1.76% |
| hsa_circ_0061360 | 1137 | 10 | 1.76% |
| hsa_circ_0069386 | 1137 | 10 | 1.76% |
| hsa_circ_0076266 | 1137 | 10 | 1.76% |
| hsa_circ_0091160 | 1137 | 10 | 1.76% |
| hsa_circ_0102965 | 1137 | 10 | 1.76% |
| hsa_circ_0107325 | 1137 | 10 | 1.76% |
| hsa_circ_0114435 | 1137 | 10 | 1.76% |
| hsa_circ_0132128 | 1137 | 10 | 1.76% |
| hsa_circ_0040429 | 1251 | 11 | 1.76% |
| hsa_circ_0044070 | 1251 | 11 | 1.76% |
| hsa_circ_0089582 | 1251 | 11 | 1.76% |
| hsa_circ_0108127 | 1251 | 11 | 1.76% |
| hsa_circ_0110221 | 1251 | 11 | 1.76% |
| hsa_circ_0140601 | 1251 | 11 | 1.76% |
| hsa_circ_0023528 | 1365 | 12 | 1.76% |
| hsa_circ_0067426 | 1365 | 12 | 1.76% |
| hsa_circ_0069953 | 1365 | 12 | 1.76% |
| hsa_circ_0049699 | 1479 | 13 | 1.76% |
| hsa_circ_0000423 | 1138 | 10 | 1.76% |
| hsa_circ_0022158 | 1138 | 10 | 1.76% |
| hsa_circ_0023015 | 1138 | 10 | 1.76% |
| hsa_circ_0053747 | 1138 | 10 | 1.76% |
| hsa_circ_0060089 | 1138 | 10 | 1.76% |
| hsa_circ_0095039 | 1138 | 10 | 1.76% |
| hsa_circ_0097113 | 1138 | 10 | 1.76% |

| | | | |
|---|---|---|---|
| hsa_circ_0107374 | 1138 | 10 | 1.76% |
| hsa_circ_0123692 | 1138 | 10 | 1.76% |
| hsa_circ_0008685 | 1252 | 11 | 1.76% |
| hsa_circ_0030866 | 1252 | 11 | 1.76% |
| hsa_circ_0050871 | 1252 | 11 | 1.76% |
| hsa_circ_0111212 | 1252 | 11 | 1.76% |
| hsa_circ_0124769 | 1252 | 11 | 1.76% |
| hsa_circ_0133328 | 1252 | 11 | 1.76% |
| hsa_circ_0021442 | 1366 | 12 | 1.76% |
| hsa_circ_0071351 | 1366 | 12 | 1.76% |
| hsa_circ_0080519 | 1366 | 12 | 1.76% |
| hsa_circ_0089662 | 1366 | 12 | 1.76% |
| hsa_circ_0122960 | 5806 | 51 | 1.76% |
| hsa_circ_0007927 | 2960 | 26 | 1.76% |
| hsa_circ_0001902 | 1139 | 10 | 1.76% |
| hsa_circ_0009816 | 1139 | 10 | 1.76% |
| hsa_circ_0017818 | 1139 | 10 | 1.76% |
| hsa_circ_0026640 | 1139 | 10 | 1.76% |
| hsa_circ_0031720 | 1139 | 10 | 1.76% |
| hsa_circ_0036908 | 1139 | 10 | 1.76% |
| hsa_circ_0039048 | 1139 | 10 | 1.76% |
| hsa_circ_0041798 | 1139 | 10 | 1.76% |
| hsa_circ_0043699 | 1139 | 10 | 1.76% |
| hsa_circ_0064474 | 1139 | 10 | 1.76% |
| hsa_circ_0081939 | 1139 | 10 | 1.76% |
| hsa_circ_0096125 | 1139 | 10 | 1.76% |
| hsa_circ_0099466 | 1139 | 10 | 1.76% |
| hsa_circ_0135209 | 1139 | 10 | 1.76% |
| hsa_circ_0074252 | 1253 | 11 | 1.76% |
| hsa_circ_0110631 | 1253 | 11 | 1.76% |
| hsa_circ_0117119 | 1253 | 11 | 1.76% |
| hsa_circ_0058381 | 1481 | 13 | 1.76% |
| hsa_circ_0131549 | 4444 | 39 | 1.76% |
| hsa_circ_0133821 | 2052 | 18 | 1.75% |
| hsa_circ_0010854 | 1368 | 12 | 1.75% |
| hsa_circ_0029890 | 1368 | 12 | 1.75% |
| hsa_circ_0098305 | 1368 | 12 | 1.75% |
| hsa_circ_0114204 | 1368 | 12 | 1.75% |
| hsa_circ_0011299 | 1254 | 11 | 1.75% |
| hsa_circ_0049983 | 1254 | 11 | 1.75% |
| hsa_circ_0050324 | 1254 | 11 | 1.75% |
| hsa_circ_0064774 | 1254 | 11 | 1.75% |
| hsa_circ_0071910 | 1254 | 11 | 1.75% |
| hsa_circ_0076423 | 1254 | 11 | 1.75% |
| hsa_circ_0083677 | 1254 | 11 | 1.75% |
| hsa_circ_0103694 | 1254 | 11 | 1.75% |
| hsa_circ_0108403 | 1254 | 11 | 1.75% |
| hsa_circ_0002142 | 1140 | 10 | 1.75% |
| hsa_circ_0016601 | 1140 | 10 | 1.75% |
| hsa_circ_0028641 | 1140 | 10 | 1.75% |

| | | | |
|---|---|---|---|
| hsa_circ_0047519 | 1140 | 10 | 1.75% |
| hsa_circ_0048688 | 1140 | 10 | 1.75% |
| hsa_circ_0066964 | 1140 | 10 | 1.75% |
| hsa_circ_0078908 | 1140 | 10 | 1.75% |
| hsa_circ_0098183 | 1140 | 10 | 1.75% |
| hsa_circ_0106987 | 1140 | 10 | 1.75% |
| hsa_circ_0120893 | 1140 | 10 | 1.75% |
| hsa_circ_0131070 | 1140 | 10 | 1.75% |
| hsa_circ_0135998 | 1140 | 10 | 1.75% |
| hsa_circ_0140163 | 1140 | 10 | 1.75% |
| hsa_circ_0054104 | 2966 | 26 | 1.75% |
| hsa_circ_0047942 | 1483 | 13 | 1.75% |
| hsa_circ_0022735 | 1369 | 12 | 1.75% |
| hsa_circ_0061331 | 1369 | 12 | 1.75% |
| hsa_circ_0100821 | 1369 | 12 | 1.75% |
| hsa_circ_0115962 | 1369 | 12 | 1.75% |
| hsa_circ_0133883 | 1369 | 12 | 1.75% |
| hsa_circ_0038122 | 1255 | 11 | 1.75% |
| hsa_circ_0052029 | 1255 | 11 | 1.75% |
| hsa_circ_0052224 | 1255 | 11 | 1.75% |
| hsa_circ_0056990 | 1255 | 11 | 1.75% |
| hsa_circ_0061460 | 1255 | 11 | 1.75% |
| hsa_circ_0106478 | 1255 | 11 | 1.75% |
| hsa_circ_0126401 | 1255 | 11 | 1.75% |
| hsa_circ_0131270 | 1255 | 11 | 1.75% |
| hsa_circ_0024456 | 1141 | 10 | 1.75% |
| hsa_circ_0033522 | 1141 | 10 | 1.75% |
| hsa_circ_0047463 | 1141 | 10 | 1.75% |
| hsa_circ_0056658 | 1141 | 10 | 1.75% |
| hsa_circ_0059496 | 1141 | 10 | 1.75% |
| hsa_circ_0062524 | 1141 | 10 | 1.75% |
| hsa_circ_0071807 | 1141 | 10 | 1.75% |
| hsa_circ_0086651 | 1141 | 10 | 1.75% |
| hsa_circ_0093367 | 1141 | 10 | 1.75% |
| hsa_circ_0116598 | 1141 | 10 | 1.75% |
| hsa_circ_0121350 | 1141 | 10 | 1.75% |
| hsa_circ_0122958 | 5821 | 51 | 1.75% |
| hsa_circ_0045477 | 1370 | 12 | 1.75% |
| hsa_circ_0050804 | 1370 | 12 | 1.75% |
| hsa_circ_0093090 | 1256 | 11 | 1.75% |
| hsa_circ_0093319 | 1256 | 11 | 1.75% |
| hsa_circ_0109898 | 1256 | 11 | 1.75% |
| hsa_circ_0118066 | 1256 | 11 | 1.75% |
| hsa_circ_0014693 | 1142 | 10 | 1.75% |
| hsa_circ_0026826 | 1142 | 10 | 1.75% |
| hsa_circ_0026879 | 1142 | 10 | 1.75% |
| hsa_circ_0038951 | 1142 | 10 | 1.75% |
| hsa_circ_0045174 | 1142 | 10 | 1.75% |
| hsa_circ_0077087 | 1142 | 10 | 1.75% |
| hsa_circ_0079826 | 1142 | 10 | 1.75% |

| | | | |
|---|---|---|---|
| hsa_circ_0102477 | 1142 | 10 | 1.75% |
| hsa_circ_0119762 | 1142 | 10 | 1.75% |
| hsa_circ_0123647 | 1142 | 10 | 1.75% |
| hsa_circ_0124231 | 1142 | 10 | 1.75% |
| hsa_circ_0130905 | 1142 | 10 | 1.75% |
| hsa_circ_0029450 | 1485 | 13 | 1.75% |
| hsa_circ_0094777 | 1485 | 13 | 1.75% |
| hsa_circ_0072328 | 1371 | 12 | 1.75% |
| hsa_circ_0088050 | 1371 | 12 | 1.75% |
| hsa_circ_0120587 | 1371 | 12 | 1.75% |
| hsa_circ_0132525 | 1371 | 12 | 1.75% |
| hsa_circ_0037015 | 2628 | 23 | 1.75% |
| hsa_circ_0136596 | 3771 | 33 | 1.75% |
| hsa_circ_0011021 | 1257 | 11 | 1.75% |
| hsa_circ_0017097 | 1257 | 11 | 1.75% |
| hsa_circ_0059285 | 1257 | 11 | 1.75% |
| hsa_circ_0064736 | 1257 | 11 | 1.75% |
| hsa_circ_0080151 | 1257 | 11 | 1.75% |
| hsa_circ_0080413 | 1257 | 11 | 1.75% |
| hsa_circ_0112266 | 1257 | 11 | 1.75% |
| hsa_circ_0131747 | 1257 | 11 | 1.75% |
| hsa_circ_0136347 | 1257 | 11 | 1.75% |
| hsa_circ_0016183 | 1143 | 10 | 1.75% |
| hsa_circ_0017441 | 1143 | 10 | 1.75% |
| hsa_circ_0023293 | 1143 | 10 | 1.75% |
| hsa_circ_0024919 | 1143 | 10 | 1.75% |
| hsa_circ_0025320 | 1143 | 10 | 1.75% |
| hsa_circ_0035796 | 1143 | 10 | 1.75% |
| hsa_circ_0044267 | 1143 | 10 | 1.75% |
| hsa_circ_0055223 | 1143 | 10 | 1.75% |
| hsa_circ_0069593 | 1143 | 10 | 1.75% |
| hsa_circ_0083775 | 1143 | 10 | 1.75% |
| hsa_circ_0085159 | 1143 | 10 | 1.75% |
| hsa_circ_0088811 | 1143 | 10 | 1.75% |
| hsa_circ_0092253 | 1143 | 10 | 1.75% |
| hsa_circ_0103761 | 1143 | 10 | 1.75% |
| hsa_circ_0132474 | 1143 | 10 | 1.75% |
| hsa_circ_0075942 | 3316 | 29 | 1.75% |
| hsa_circ_0032653 | 1944 | 17 | 1.75% |
| hsa_circ_0106276 | 1258 | 11 | 1.75% |
| hsa_circ_0122669 | 1258 | 11 | 1.75% |
| hsa_circ_0062109 | 1830 | 16 | 1.75% |
| hsa_circ_0047415 | 1487 | 13 | 1.75% |
| hsa_circ_0040890 | 1144 | 10 | 1.75% |
| hsa_circ_0049471 | 1144 | 10 | 1.75% |
| hsa_circ_0068888 | 1144 | 10 | 1.75% |
| hsa_circ_0072476 | 1144 | 10 | 1.75% |
| hsa_circ_0074150 | 1144 | 10 | 1.75% |
| hsa_circ_0082037 | 1144 | 10 | 1.75% |
| hsa_circ_0097591 | 1144 | 10 | 1.75% |

| | | | |
|---|---|---|---|
| hsa_circ_0122317 | 1144 | 10 | 1.75% |
| hsa_circ_0137974 | 1144 | 10 | 1.75% |
| hsa_circ_0138889 | 1144 | 10 | 1.75% |
| hsa_circ_0040086 | 1373 | 12 | 1.75% |
| hsa_circ_0048755 | 1373 | 12 | 1.75% |
| hsa_circ_0122142 | 1373 | 12 | 1.75% |
| hsa_circ_0011505 | 1259 | 11 | 1.75% |
| hsa_circ_0030945 | 1259 | 11 | 1.75% |
| hsa_circ_0071693 | 1259 | 11 | 1.75% |
| hsa_circ_0099821 | 1259 | 11 | 1.75% |
| hsa_circ_0111364 | 1259 | 11 | 1.75% |
| hsa_circ_0135111 | 1259 | 11 | 1.75% |
| hsa_circ_0056253 | 1488 | 13 | 1.75% |
| hsa_circ_0067907 | 1488 | 13 | 1.75% |
| hsa_circ_0016460 | 1603 | 14 | 1.75% |
| hsa_circ_0065614 | 1374 | 12 | 1.75% |
| hsa_circ_0114174 | 1374 | 12 | 1.75% |
| hsa_circ_0006041 | 1145 | 10 | 1.75% |
| hsa_circ_0023026 | 1145 | 10 | 1.75% |
| hsa_circ_0023134 | 1145 | 10 | 1.75% |
| hsa_circ_0036361 | 1145 | 10 | 1.75% |
| hsa_circ_0036800 | 1145 | 10 | 1.75% |
| hsa_circ_0040666 | 1145 | 10 | 1.75% |
| hsa_circ_0049019 | 1145 | 10 | 1.75% |
| hsa_circ_0050537 | 1145 | 10 | 1.75% |
| hsa_circ_0057216 | 1145 | 10 | 1.75% |
| hsa_circ_0059224 | 1145 | 10 | 1.75% |
| hsa_circ_0065456 | 1145 | 10 | 1.75% |
| hsa_circ_0068814 | 1145 | 10 | 1.75% |
| hsa_circ_0072489 | 1145 | 10 | 1.75% |
| hsa_circ_0081949 | 1145 | 10 | 1.75% |
| hsa_circ_0091579 | 1145 | 10 | 1.75% |
| hsa_circ_0106072 | 1145 | 10 | 1.75% |
| hsa_circ_0107970 | 1145 | 10 | 1.75% |
| hsa_circ_0128814 | 1145 | 10 | 1.75% |
| hsa_circ_0137657 | 1145 | 10 | 1.75% |
| hsa_circ_0139208 | 1145 | 10 | 1.75% |
| hsa_circ_0035079 | 1489 | 13 | 1.75% |
| hsa_circ_0064703 | 1489 | 13 | 1.75% |
| hsa_circ_0014948 | 1260 | 11 | 1.75% |
| hsa_circ_0016927 | 1260 | 11 | 1.75% |
| hsa_circ_0020226 | 1260 | 11 | 1.75% |
| hsa_circ_0022081 | 1260 | 11 | 1.75% |
| hsa_circ_0072122 | 1260 | 11 | 1.75% |
| hsa_circ_0087533 | 1260 | 11 | 1.75% |
| hsa_circ_0087844 | 1260 | 11 | 1.75% |
| hsa_circ_0068661 | 1375 | 12 | 1.75% |
| hsa_circ_0031704 | 1719 | 15 | 1.75% |
| hsa_circ_0008176 | 1146 | 10 | 1.75% |
| hsa_circ_0015533 | 1146 | 10 | 1.75% |

| | | | |
|---|---|---|---|
| hsa_circ_0034441 | 1146 | 10 | 1.75% |
| hsa_circ_0048128 | 1146 | 10 | 1.75% |
| hsa_circ_0049169 | 1146 | 10 | 1.75% |
| hsa_circ_0061252 | 1146 | 10 | 1.75% |
| hsa_circ_0062404 | 1146 | 10 | 1.75% |
| hsa_circ_0074475 | 1146 | 10 | 1.75% |
| hsa_circ_0084112 | 1146 | 10 | 1.75% |
| hsa_circ_0092059 | 1146 | 10 | 1.75% |
| hsa_circ_0116081 | 1146 | 10 | 1.75% |
| hsa_circ_0131542 | 1146 | 10 | 1.75% |
| hsa_circ_0135411 | 1146 | 10 | 1.75% |
| hsa_circ_0068088 | 6190 | 54 | 1.74% |
| hsa_circ_0010764 | 1261 | 11 | 1.74% |
| hsa_circ_0052391 | 1261 | 11 | 1.74% |
| hsa_circ_0098417 | 1261 | 11 | 1.74% |
| hsa_circ_0103458 | 1261 | 11 | 1.74% |
| hsa_circ_0017867 | 1376 | 12 | 1.74% |
| hsa_circ_0081356 | 1376 | 12 | 1.74% |
| hsa_circ_0106784 | 3899 | 34 | 1.74% |
| hsa_circ_0068617 | 1491 | 13 | 1.74% |
| hsa_circ_0122259 | 1491 | 13 | 1.74% |
| hsa_circ_0001610 | 1147 | 10 | 1.74% |
| hsa_circ_0011754 | 1147 | 10 | 1.74% |
| hsa_circ_0012182 | 1147 | 10 | 1.74% |
| hsa_circ_0036167 | 1147 | 10 | 1.74% |
| hsa_circ_0037643 | 1147 | 10 | 1.74% |
| hsa_circ_0040135 | 1147 | 10 | 1.74% |
| hsa_circ_0057909 | 1147 | 10 | 1.74% |
| hsa_circ_0101077 | 1147 | 10 | 1.74% |
| hsa_circ_0111891 | 1147 | 10 | 1.74% |
| hsa_circ_0112205 | 1147 | 10 | 1.74% |
| hsa_circ_0113012 | 1147 | 10 | 1.74% |
| hsa_circ_0117214 | 1147 | 10 | 1.74% |
| hsa_circ_0124100 | 1147 | 10 | 1.74% |
| hsa_circ_0030031 | 1262 | 11 | 1.74% |
| hsa_circ_0038301 | 1262 | 11 | 1.74% |
| hsa_circ_0048805 | 1262 | 11 | 1.74% |
| hsa_circ_0052502 | 1262 | 11 | 1.74% |
| hsa_circ_0062125 | 1262 | 11 | 1.74% |
| hsa_circ_0063281 | 1262 | 11 | 1.74% |
| hsa_circ_0067049 | 1262 | 11 | 1.74% |
| hsa_circ_0097696 | 1262 | 11 | 1.74% |
| hsa_circ_0102760 | 1262 | 11 | 1.74% |
| hsa_circ_0059534 | 1377 | 12 | 1.74% |
| hsa_circ_0011356 | 1148 | 10 | 1.74% |
| hsa_circ_0022331 | 1148 | 10 | 1.74% |
| hsa_circ_0050381 | 1148 | 10 | 1.74% |
| hsa_circ_0057976 | 1148 | 10 | 1.74% |
| hsa_circ_0075215 | 1148 | 10 | 1.74% |
| hsa_circ_0081658 | 1148 | 10 | 1.74% |

| | | | |
|---|---|---|---|
| hsa_circ_0085456 | 1148 | 10 | 1.74% |
| hsa_circ_0089991 | 1148 | 10 | 1.74% |
| hsa_circ_0111292 | 1148 | 10 | 1.74% |
| hsa_circ_0118635 | 1148 | 10 | 1.74% |
| hsa_circ_0123563 | 1148 | 10 | 1.74% |
| hsa_circ_0125312 | 1148 | 10 | 1.74% |
| hsa_circ_0135780 | 1148 | 10 | 1.74% |
| hsa_circ_0139046 | 1148 | 10 | 1.74% |
| hsa_circ_0014827 | 1263 | 11 | 1.74% |
| hsa_circ_0019476 | 1263 | 11 | 1.74% |
| hsa_circ_0021200 | 1263 | 11 | 1.74% |
| hsa_circ_0031760 | 1263 | 11 | 1.74% |
| hsa_circ_0035493 | 1263 | 11 | 1.74% |
| hsa_circ_0049210 | 1263 | 11 | 1.74% |
| hsa_circ_0054861 | 1263 | 11 | 1.74% |
| hsa_circ_0054870 | 1263 | 11 | 1.74% |
| hsa_circ_0069064 | 1263 | 11 | 1.74% |
| hsa_circ_0076775 | 1263 | 11 | 1.74% |
| hsa_circ_0105144 | 1263 | 11 | 1.74% |
| hsa_circ_0118292 | 1263 | 11 | 1.74% |
| hsa_circ_0138472 | 1263 | 11 | 1.74% |
| hsa_circ_0023165 | 1378 | 12 | 1.74% |
| hsa_circ_0102188 | 1378 | 12 | 1.74% |
| hsa_circ_0025075 | 22967 | 200 | 1.74% |
| hsa_circ_0028542 | 1493 | 13 | 1.74% |
| hsa_circ_0076658 | 1493 | 13 | 1.74% |
| hsa_circ_0086782 | 1493 | 13 | 1.74% |
| hsa_circ_0048976 | 1953 | 17 | 1.74% |
| hsa_circ_0013627 | 1149 | 10 | 1.74% |
| hsa_circ_0014598 | 1149 | 10 | 1.74% |
| hsa_circ_0024970 | 1149 | 10 | 1.74% |
| hsa_circ_0025397 | 1149 | 10 | 1.74% |
| hsa_circ_0032267 | 1149 | 10 | 1.74% |
| hsa_circ_0032757 | 1149 | 10 | 1.74% |
| hsa_circ_0037343 | 1149 | 10 | 1.74% |
| hsa_circ_0044407 | 1149 | 10 | 1.74% |
| hsa_circ_0048001 | 1149 | 10 | 1.74% |
| hsa_circ_0065276 | 1149 | 10 | 1.74% |
| hsa_circ_0076473 | 1149 | 10 | 1.74% |
| hsa_circ_0116371 | 1149 | 10 | 1.74% |
| hsa_circ_0133055 | 1149 | 10 | 1.74% |
| hsa_circ_0012180 | 1264 | 11 | 1.74% |
| hsa_circ_0014059 | 1264 | 11 | 1.74% |
| hsa_circ_0015673 | 1264 | 11 | 1.74% |
| hsa_circ_0019107 | 1264 | 11 | 1.74% |
| hsa_circ_0031247 | 1264 | 11 | 1.74% |
| hsa_circ_0049085 | 1264 | 11 | 1.74% |
| hsa_circ_0067646 | 1264 | 11 | 1.74% |
| hsa_circ_0089563 | 1264 | 11 | 1.74% |
| hsa_circ_0130358 | 1264 | 11 | 1.74% |

| | | | |
|---|---|---|---|
| hsa_circ_0046286 | 1379 | 12 | 1.74% |
| hsa_circ_0049793 | 1379 | 12 | 1.74% |
| hsa_circ_0052284 | 1379 | 12 | 1.74% |
| hsa_circ_0082424 | 1379 | 12 | 1.74% |
| hsa_circ_0032664 | 1954 | 17 | 1.74% |
| hsa_circ_0060701 | 1495 | 13 | 1.74% |
| hsa_circ_0074077 | 1495 | 13 | 1.74% |
| hsa_circ_0088040 | 1380 | 12 | 1.74% |
| hsa_circ_0028795 | 1265 | 11 | 1.74% |
| hsa_circ_0035576 | 1265 | 11 | 1.74% |
| hsa_circ_0083145 | 1265 | 11 | 1.74% |
| hsa_circ_0016077 | 1150 | 10 | 1.74% |
| hsa_circ_0023600 | 1150 | 10 | 1.74% |
| hsa_circ_0025403 | 1150 | 10 | 1.74% |
| hsa_circ_0047561 | 1150 | 10 | 1.74% |
| hsa_circ_0063021 | 1150 | 10 | 1.74% |
| hsa_circ_0064105 | 1150 | 10 | 1.74% |
| hsa_circ_0067783 | 1150 | 10 | 1.74% |
| hsa_circ_0086790 | 1150 | 10 | 1.74% |
| hsa_circ_0089190 | 1150 | 10 | 1.74% |
| hsa_circ_0095709 | 1150 | 10 | 1.74% |
| hsa_circ_0102745 | 1150 | 10 | 1.74% |
| hsa_circ_0104436 | 1150 | 10 | 1.74% |
| hsa_circ_0107235 | 1150 | 10 | 1.74% |
| hsa_circ_0112543 | 1150 | 10 | 1.74% |
| hsa_circ_0132709 | 1150 | 10 | 1.74% |
| hsa_circ_0105662 | 1496 | 13 | 1.74% |
| hsa_circ_0012088 | 1381 | 12 | 1.74% |
| hsa_circ_0115374 | 1381 | 12 | 1.74% |
| hsa_circ_0130096 | 1381 | 12 | 1.74% |
| hsa_circ_0001896 | 1266 | 11 | 1.74% |
| hsa_circ_0061428 | 1266 | 11 | 1.74% |
| hsa_circ_0063027 | 1266 | 11 | 1.74% |
| hsa_circ_0112447 | 1266 | 11 | 1.74% |
| hsa_circ_0120688 | 1266 | 11 | 1.74% |
| hsa_circ_0122772 | 1266 | 11 | 1.74% |
| hsa_circ_0053237 | 1151 | 10 | 1.74% |
| hsa_circ_0064141 | 1151 | 10 | 1.74% |
| hsa_circ_0078058 | 1151 | 10 | 1.74% |
| hsa_circ_0092616 | 1151 | 10 | 1.74% |
| hsa_circ_0110119 | 1151 | 10 | 1.74% |
| hsa_circ_0125666 | 1151 | 10 | 1.74% |
| hsa_circ_0127558 | 1151 | 10 | 1.74% |
| hsa_circ_0044475 | 1382 | 12 | 1.74% |
| hsa_circ_0080893 | 1382 | 12 | 1.74% |
| hsa_circ_0127696 | 1382 | 12 | 1.74% |
| hsa_circ_0129579 | 1382 | 12 | 1.74% |
| hsa_circ_0019131 | 1267 | 11 | 1.74% |
| hsa_circ_0037834 | 1267 | 11 | 1.74% |
| hsa_circ_0040327 | 1267 | 11 | 1.74% |

| | | | |
|---|---|---|---|
| hsa_circ_0100300 | 1267 | 11 | 1.74% |
| hsa_circ_0106257 | 1267 | 11 | 1.74% |
| hsa_circ_0117799 | 1267 | 11 | 1.74% |
| hsa_circ_0126747 | 1267 | 11 | 1.74% |
| hsa_circ_0004654 | 1728 | 15 | 1.74% |
| hsa_circ_0022643 | 1152 | 10 | 1.74% |
| hsa_circ_0022928 | 1152 | 10 | 1.74% |
| hsa_circ_0033426 | 1152 | 10 | 1.74% |
| hsa_circ_0042211 | 1152 | 10 | 1.74% |
| hsa_circ_0046052 | 1152 | 10 | 1.74% |
| hsa_circ_0049823 | 1152 | 10 | 1.74% |
| hsa_circ_0051096 | 1152 | 10 | 1.74% |
| hsa_circ_0061259 | 1152 | 10 | 1.74% |
| hsa_circ_0077092 | 1152 | 10 | 1.74% |
| hsa_circ_0081946 | 1152 | 10 | 1.74% |
| hsa_circ_0096222 | 1152 | 10 | 1.74% |
| hsa_circ_0111127 | 1152 | 10 | 1.74% |
| hsa_circ_0111375 | 1152 | 10 | 1.74% |
| hsa_circ_0111515 | 1152 | 10 | 1.74% |
| hsa_circ_0119177 | 1152 | 10 | 1.74% |
| hsa_circ_0126288 | 1152 | 10 | 1.74% |
| hsa_circ_0126925 | 1152 | 10 | 1.74% |
| hsa_circ_0134036 | 1152 | 10 | 1.74% |
| hsa_circ_0137401 | 1152 | 10 | 1.74% |
| hsa_circ_0056256 | 1498 | 13 | 1.74% |
| hsa_circ_0021822 | 2305 | 20 | 1.74% |
| hsa_circ_0009901 | 1268 | 11 | 1.74% |
| hsa_circ_0023159 | 1268 | 11 | 1.74% |
| hsa_circ_0033158 | 1268 | 11 | 1.74% |
| hsa_circ_0101084 | 1268 | 11 | 1.74% |
| hsa_circ_0119507 | 1268 | 11 | 1.74% |
| hsa_circ_0138041 | 1268 | 11 | 1.74% |
| hsa_circ_0039271 | 1153 | 10 | 1.73% |
| hsa_circ_0046114 | 1153 | 10 | 1.73% |
| hsa_circ_0046528 | 1153 | 10 | 1.73% |
| hsa_circ_0061962 | 1153 | 10 | 1.73% |
| hsa_circ_0080272 | 1153 | 10 | 1.73% |
| hsa_circ_0093727 | 1153 | 10 | 1.73% |
| hsa_circ_0113824 | 1153 | 10 | 1.73% |
| hsa_circ_0129445 | 1153 | 10 | 1.73% |
| hsa_circ_0129740 | 1153 | 10 | 1.73% |
| hsa_circ_0132479 | 1153 | 10 | 1.73% |
| hsa_circ_0139321 | 1153 | 10 | 1.73% |
| hsa_circ_0092179 | 5304 | 46 | 1.73% |
| hsa_circ_0029938 | 1384 | 12 | 1.73% |
| hsa_circ_0003149 | 1269 | 11 | 1.73% |
| hsa_circ_0037845 | 1269 | 11 | 1.73% |
| hsa_circ_0039856 | 1269 | 11 | 1.73% |
| hsa_circ_0062382 | 1269 | 11 | 1.73% |
| hsa_circ_0093968 | 1269 | 11 | 1.73% |

| | | | |
|---|---|---|---|
| hsa_circ_0068615 | 1500 | 13 | 1.73% |
| hsa_circ_0026307 | 1154 | 10 | 1.73% |
| hsa_circ_0037994 | 1154 | 10 | 1.73% |
| hsa_circ_0038326 | 1154 | 10 | 1.73% |
| hsa_circ_0047252 | 1154 | 10 | 1.73% |
| hsa_circ_0054341 | 1154 | 10 | 1.73% |
| hsa_circ_0058185 | 1154 | 10 | 1.73% |
| hsa_circ_0062051 | 1154 | 10 | 1.73% |
| hsa_circ_0071660 | 1154 | 10 | 1.73% |
| hsa_circ_0075966 | 1154 | 10 | 1.73% |
| hsa_circ_0076910 | 1154 | 10 | 1.73% |
| hsa_circ_0100318 | 1154 | 10 | 1.73% |
| hsa_circ_0120858 | 1154 | 10 | 1.73% |
| hsa_circ_0122233 | 1154 | 10 | 1.73% |
| hsa_circ_0133463 | 1154 | 10 | 1.73% |
| hsa_circ_0079283 | 1385 | 12 | 1.73% |
| hsa_circ_0039938 | 1270 | 11 | 1.73% |
| hsa_circ_0045051 | 1270 | 11 | 1.73% |
| hsa_circ_0090533 | 1270 | 11 | 1.73% |
| hsa_circ_0128381 | 1270 | 11 | 1.73% |
| hsa_circ_0137695 | 1270 | 11 | 1.73% |
| hsa_circ_0024104 | 1617 | 14 | 1.73% |
| hsa_circ_0129438 | 1386 | 12 | 1.73% |
| hsa_circ_0000204 | 1155 | 10 | 1.73% |
| hsa_circ_0010690 | 1155 | 10 | 1.73% |
| hsa_circ_0013295 | 1155 | 10 | 1.73% |
| hsa_circ_0015675 | 1155 | 10 | 1.73% |
| hsa_circ_0026123 | 1155 | 10 | 1.73% |
| hsa_circ_0032410 | 1155 | 10 | 1.73% |
| hsa_circ_0036047 | 1155 | 10 | 1.73% |
| hsa_circ_0036162 | 1155 | 10 | 1.73% |
| hsa_circ_0061854 | 1155 | 10 | 1.73% |
| hsa_circ_0068645 | 1155 | 10 | 1.73% |
| hsa_circ_0079168 | 1155 | 10 | 1.73% |
| hsa_circ_0081073 | 1155 | 10 | 1.73% |
| hsa_circ_0086876 | 1155 | 10 | 1.73% |
| hsa_circ_0089484 | 1155 | 10 | 1.73% |
| hsa_circ_0093674 | 1155 | 10 | 1.73% |
| hsa_circ_0096828 | 1155 | 10 | 1.73% |
| hsa_circ_0127151 | 1155 | 10 | 1.73% |
| hsa_circ_0132081 | 1155 | 10 | 1.73% |
| hsa_circ_0133062 | 1155 | 10 | 1.73% |
| hsa_circ_0135520 | 1155 | 10 | 1.73% |
| hsa_circ_0040657 | 1271 | 11 | 1.73% |
| hsa_circ_0049481 | 1271 | 11 | 1.73% |
| hsa_circ_0064303 | 1271 | 11 | 1.73% |
| hsa_circ_0114178 | 1271 | 11 | 1.73% |
| hsa_circ_0136766 | 1271 | 11 | 1.73% |
| hsa_circ_0048565 | 1387 | 12 | 1.73% |
| hsa_circ_0128941 | 1387 | 12 | 1.73% |

| | | | |
|---|---|---|---|
| hsa_circ_0010773 | 1156 | 10 | 1.73% |
| hsa_circ_0011948 | 1156 | 10 | 1.73% |
| hsa_circ_0012724 | 1156 | 10 | 1.73% |
| hsa_circ_0020937 | 1156 | 10 | 1.73% |
| hsa_circ_0023690 | 1156 | 10 | 1.73% |
| hsa_circ_0024304 | 1156 | 10 | 1.73% |
| hsa_circ_0044840 | 1156 | 10 | 1.73% |
| hsa_circ_0046034 | 1156 | 10 | 1.73% |
| hsa_circ_0057894 | 1156 | 10 | 1.73% |
| hsa_circ_0058055 | 1156 | 10 | 1.73% |
| hsa_circ_0060085 | 1156 | 10 | 1.73% |
| hsa_circ_0065125 | 1156 | 10 | 1.73% |
| hsa_circ_0065947 | 1156 | 10 | 1.73% |
| hsa_circ_0071822 | 1156 | 10 | 1.73% |
| hsa_circ_0101312 | 1156 | 10 | 1.73% |
| hsa_circ_0127827 | 1156 | 10 | 1.73% |
| hsa_circ_0135919 | 1156 | 10 | 1.73% |
| hsa_circ_0001672 | 4971 | 43 | 1.73% |
| hsa_circ_0029155 | 1503 | 13 | 1.73% |
| hsa_circ_0031561 | 1272 | 11 | 1.73% |
| hsa_circ_0056600 | 1272 | 11 | 1.73% |
| hsa_circ_0064301 | 1272 | 11 | 1.73% |
| hsa_circ_0068872 | 1272 | 11 | 1.73% |
| hsa_circ_0103230 | 1272 | 11 | 1.73% |
| hsa_circ_0131092 | 1272 | 11 | 1.73% |
| hsa_circ_0030917 | 1388 | 12 | 1.73% |
| hsa_circ_0043858 | 1504 | 13 | 1.73% |
| hsa_circ_0136098 | 1504 | 13 | 1.73% |
| hsa_circ_0022436 | 1157 | 10 | 1.73% |
| hsa_circ_0031553 | 1157 | 10 | 1.73% |
| hsa_circ_0083890 | 1157 | 10 | 1.73% |
| hsa_circ_0092440 | 1157 | 10 | 1.73% |
| hsa_circ_0098470 | 1157 | 10 | 1.73% |
| hsa_circ_0104756 | 1157 | 10 | 1.73% |
| hsa_circ_0106535 | 1157 | 10 | 1.73% |
| hsa_circ_0118387 | 1157 | 10 | 1.73% |
| hsa_circ_0130105 | 1157 | 10 | 1.73% |
| hsa_circ_0115970 | 1273 | 11 | 1.73% |
| hsa_circ_0106081 | 3125 | 27 | 1.73% |
| hsa_circ_0040990 | 1389 | 12 | 1.73% |
| hsa_circ_0119602 | 2431 | 21 | 1.73% |
| hsa_circ_0106343 | 1621 | 14 | 1.73% |
| hsa_circ_0001196 | 1158 | 10 | 1.73% |
| hsa_circ_0014820 | 1158 | 10 | 1.73% |
| hsa_circ_0016761 | 1158 | 10 | 1.73% |
| hsa_circ_0028652 | 1158 | 10 | 1.73% |
| hsa_circ_0032582 | 1158 | 10 | 1.73% |
| hsa_circ_0035030 | 1158 | 10 | 1.73% |
| hsa_circ_0046309 | 1158 | 10 | 1.73% |
| hsa_circ_0056772 | 1158 | 10 | 1.73% |

| | | | |
|---|---|---|---|
| hsa_circ_0058646 | 1158 | 10 | 1.73% |
| hsa_circ_0064733 | 1158 | 10 | 1.73% |
| hsa_circ_0069880 | 1158 | 10 | 1.73% |
| hsa_circ_0097789 | 1158 | 10 | 1.73% |
| hsa_circ_0100661 | 1158 | 10 | 1.73% |
| hsa_circ_0104109 | 1158 | 10 | 1.73% |
| hsa_circ_0117635 | 1158 | 10 | 1.73% |
| hsa_circ_0122413 | 1158 | 10 | 1.73% |
| hsa_circ_0001226 | 1274 | 11 | 1.73% |
| hsa_circ_0010716 | 1274 | 11 | 1.73% |
| hsa_circ_0021426 | 1274 | 11 | 1.73% |
| hsa_circ_0025029 | 1274 | 11 | 1.73% |
| hsa_circ_0015113 | 1390 | 12 | 1.73% |
| hsa_circ_0056037 | 1390 | 12 | 1.73% |
| hsa_circ_0110216 | 1738 | 15 | 1.73% |
| hsa_circ_0035483 | 1159 | 10 | 1.73% |
| hsa_circ_0060292 | 1159 | 10 | 1.73% |
| hsa_circ_0061623 | 1159 | 10 | 1.73% |
| hsa_circ_0070931 | 1159 | 10 | 1.73% |
| hsa_circ_0072042 | 1159 | 10 | 1.73% |
| hsa_circ_0076439 | 1159 | 10 | 1.73% |
| hsa_circ_0078380 | 1159 | 10 | 1.73% |
| hsa_circ_0089227 | 1159 | 10 | 1.73% |
| hsa_circ_0103387 | 1159 | 10 | 1.73% |
| hsa_circ_0109815 | 1159 | 10 | 1.73% |
| hsa_circ_0136790 | 1159 | 10 | 1.73% |
| hsa_circ_0019138 | 1275 | 11 | 1.73% |
| hsa_circ_0051393 | 1275 | 11 | 1.73% |
| hsa_circ_0054914 | 1275 | 11 | 1.73% |
| hsa_circ_0064174 | 1275 | 11 | 1.73% |
| hsa_circ_0074884 | 1275 | 11 | 1.73% |
| hsa_circ_0074907 | 1275 | 11 | 1.73% |
| hsa_circ_0080593 | 1275 | 11 | 1.73% |
| hsa_circ_0091016 | 1275 | 11 | 1.73% |
| hsa_circ_0132561 | 1275 | 11 | 1.73% |
| hsa_circ_0135489 | 1275 | 11 | 1.73% |
| hsa_circ_0131385 | 1391 | 12 | 1.73% |
| hsa_circ_0080205 | 1508 | 13 | 1.72% |
| hsa_circ_0005776 | 1392 | 12 | 1.72% |
| hsa_circ_0028940 | 1392 | 12 | 1.72% |
| hsa_circ_0080901 | 1392 | 12 | 1.72% |
| hsa_circ_0026355 | 1276 | 11 | 1.72% |
| hsa_circ_0048555 | 1276 | 11 | 1.72% |
| hsa_circ_0084671 | 1276 | 11 | 1.72% |
| hsa_circ_0133804 | 1276 | 11 | 1.72% |
| hsa_circ_0010118 | 1160 | 10 | 1.72% |
| hsa_circ_0015019 | 1160 | 10 | 1.72% |
| hsa_circ_0016457 | 1160 | 10 | 1.72% |
| hsa_circ_0027602 | 1160 | 10 | 1.72% |
| hsa_circ_0031560 | 1160 | 10 | 1.72% |

| | | | |
|---|---|---|---|
| hsa_circ_0034901 | 1160 | 10 | 1.72% |
| hsa_circ_0050543 | 1160 | 10 | 1.72% |
| hsa_circ_0057601 | 1160 | 10 | 1.72% |
| hsa_circ_0058644 | 1160 | 10 | 1.72% |
| hsa_circ_0068471 | 1160 | 10 | 1.72% |
| hsa_circ_0084990 | 1160 | 10 | 1.72% |
| hsa_circ_0088540 | 1160 | 10 | 1.72% |
| hsa_circ_0091374 | 1160 | 10 | 1.72% |
| hsa_circ_0106444 | 1160 | 10 | 1.72% |
| hsa_circ_0107620 | 1160 | 10 | 1.72% |
| hsa_circ_0110361 | 1160 | 10 | 1.72% |
| hsa_circ_0113566 | 1160 | 10 | 1.72% |
| hsa_circ_0031003 | 1509 | 13 | 1.72% |
| hsa_circ_0044356 | 1509 | 13 | 1.72% |
| hsa_circ_0010895 | 1393 | 12 | 1.72% |
| hsa_circ_0024769 | 1277 | 11 | 1.72% |
| hsa_circ_0046708 | 1277 | 11 | 1.72% |
| hsa_circ_0049200 | 1277 | 11 | 1.72% |
| hsa_circ_0066145 | 1277 | 11 | 1.72% |
| hsa_circ_0086796 | 1277 | 11 | 1.72% |
| hsa_circ_0104320 | 1277 | 11 | 1.72% |
| hsa_circ_0008068 | 1161 | 10 | 1.72% |
| hsa_circ_0009907 | 1161 | 10 | 1.72% |
| hsa_circ_0040752 | 1161 | 10 | 1.72% |
| hsa_circ_0071857 | 1161 | 10 | 1.72% |
| hsa_circ_0080027 | 1161 | 10 | 1.72% |
| hsa_circ_0097030 | 1161 | 10 | 1.72% |
| hsa_circ_0100289 | 1161 | 10 | 1.72% |
| hsa_circ_0107230 | 1161 | 10 | 1.72% |
| hsa_circ_0134673 | 1161 | 10 | 1.72% |
| hsa_circ_0128857 | 2671 | 23 | 1.72% |
| hsa_circ_0050100 | 1510 | 13 | 1.72% |
| hsa_circ_0067310 | 1394 | 12 | 1.72% |
| hsa_circ_0004626 | 1278 | 11 | 1.72% |
| hsa_circ_0019735 | 1278 | 11 | 1.72% |
| hsa_circ_0052273 | 1278 | 11 | 1.72% |
| hsa_circ_0096421 | 1278 | 11 | 1.72% |
| hsa_circ_0116632 | 1278 | 11 | 1.72% |
| hsa_circ_0123897 | 1278 | 11 | 1.72% |
| hsa_circ_0017486 | 1162 | 10 | 1.72% |
| hsa_circ_0051296 | 1162 | 10 | 1.72% |
| hsa_circ_0056075 | 1162 | 10 | 1.72% |
| hsa_circ_0056750 | 1162 | 10 | 1.72% |
| hsa_circ_0059529 | 1162 | 10 | 1.72% |
| hsa_circ_0061328 | 1162 | 10 | 1.72% |
| hsa_circ_0066719 | 1162 | 10 | 1.72% |
| hsa_circ_0075178 | 1162 | 10 | 1.72% |
| hsa_circ_0078143 | 1162 | 10 | 1.72% |
| hsa_circ_0079035 | 1162 | 10 | 1.72% |
| hsa_circ_0090391 | 1162 | 10 | 1.72% |

| | | | |
|---|---|---|---|
| hsa_circ_0091898 | 1162 | 10 | 1.72% |
| hsa_circ_0122239 | 1162 | 10 | 1.72% |
| hsa_circ_0123397 | 1162 | 10 | 1.72% |
| hsa_circ_0126249 | 1162 | 10 | 1.72% |
| hsa_circ_0129908 | 1162 | 10 | 1.72% |
| hsa_circ_0132163 | 1162 | 10 | 1.72% |
| hsa_circ_0039453 | 1511 | 13 | 1.72% |
| hsa_circ_0116436 | 1511 | 13 | 1.72% |
| hsa_circ_0074540 | 1395 | 12 | 1.72% |
| hsa_circ_0136390 | 2674 | 23 | 1.72% |
| hsa_circ_0124876 | 2558 | 22 | 1.72% |
| hsa_circ_0049061 | 1279 | 11 | 1.72% |
| hsa_circ_0053288 | 1279 | 11 | 1.72% |
| hsa_circ_0078191 | 1279 | 11 | 1.72% |
| hsa_circ_0088391 | 1279 | 11 | 1.72% |
| hsa_circ_0104760 | 1279 | 11 | 1.72% |
| hsa_circ_0130202 | 2093 | 18 | 1.72% |
| hsa_circ_0007647 | 1163 | 10 | 1.72% |
| hsa_circ_0010234 | 1163 | 10 | 1.72% |
| hsa_circ_0020770 | 1163 | 10 | 1.72% |
| hsa_circ_0037622 | 1163 | 10 | 1.72% |
| hsa_circ_0039705 | 1163 | 10 | 1.72% |
| hsa_circ_0044131 | 1163 | 10 | 1.72% |
| hsa_circ_0062674 | 1163 | 10 | 1.72% |
| hsa_circ_0066759 | 1163 | 10 | 1.72% |
| hsa_circ_0081581 | 1163 | 10 | 1.72% |
| hsa_circ_0091360 | 1163 | 10 | 1.72% |
| hsa_circ_0098860 | 1163 | 10 | 1.72% |
| hsa_circ_0109001 | 1163 | 10 | 1.72% |
| hsa_circ_0067692 | 1512 | 13 | 1.72% |
| hsa_circ_0115061 | 1512 | 13 | 1.72% |
| hsa_circ_0035772 | 1396 | 12 | 1.72% |
| hsa_circ_0075716 | 1396 | 12 | 1.72% |
| hsa_circ_0097284 | 3607 | 31 | 1.72% |
| hsa_circ_0012993 | 1280 | 11 | 1.72% |
| hsa_circ_0036216 | 1280 | 11 | 1.72% |
| hsa_circ_0083201 | 1280 | 11 | 1.72% |
| hsa_circ_0119635 | 1280 | 11 | 1.72% |
| hsa_circ_0137627 | 1280 | 11 | 1.72% |
| hsa_circ_0001172 | 1513 | 13 | 1.72% |
| hsa_circ_0092106 | 1513 | 13 | 1.72% |
| hsa_circ_0015761 | 1164 | 10 | 1.72% |
| hsa_circ_0026412 | 1164 | 10 | 1.72% |
| hsa_circ_0026422 | 1164 | 10 | 1.72% |
| hsa_circ_0039857 | 1164 | 10 | 1.72% |
| hsa_circ_0068171 | 1164 | 10 | 1.72% |
| hsa_circ_0073327 | 1164 | 10 | 1.72% |
| hsa_circ_0077509 | 1164 | 10 | 1.72% |
| hsa_circ_0091941 | 1164 | 10 | 1.72% |
| hsa_circ_0094308 | 1164 | 10 | 1.72% |

| | | | |
|---|---|---|---|
| hsa_circ_0103036 | 1164 | 10 | 1.72% |
| hsa_circ_0126587 | 1164 | 10 | 1.72% |
| hsa_circ_0040018 | 1397 | 12 | 1.72% |
| hsa_circ_0043833 | 1397 | 12 | 1.72% |
| hsa_circ_0115356 | 1397 | 12 | 1.72% |
| hsa_circ_0125929 | 1397 | 12 | 1.72% |
| hsa_circ_0057377 | 1863 | 16 | 1.72% |
| hsa_circ_0009868 | 1281 | 11 | 1.72% |
| hsa_circ_0012821 | 1281 | 11 | 1.72% |
| hsa_circ_0021941 | 1281 | 11 | 1.72% |
| hsa_circ_0039966 | 1281 | 11 | 1.72% |
| hsa_circ_0052247 | 1281 | 11 | 1.72% |
| hsa_circ_0091311 | 1281 | 11 | 1.72% |
| hsa_circ_0106154 | 1281 | 11 | 1.72% |
| hsa_circ_0113120 | 1281 | 11 | 1.72% |
| hsa_circ_0023344 | 1398 | 12 | 1.72% |
| hsa_circ_0099147 | 1398 | 12 | 1.72% |
| hsa_circ_0005711 | 1165 | 10 | 1.72% |
| hsa_circ_0010025 | 1165 | 10 | 1.72% |
| hsa_circ_0011240 | 1165 | 10 | 1.72% |
| hsa_circ_0011528 | 1165 | 10 | 1.72% |
| hsa_circ_0012610 | 1165 | 10 | 1.72% |
| hsa_circ_0035022 | 1165 | 10 | 1.72% |
| hsa_circ_0051037 | 1165 | 10 | 1.72% |
| hsa_circ_0051913 | 1165 | 10 | 1.72% |
| hsa_circ_0066752 | 1165 | 10 | 1.72% |
| hsa_circ_0093268 | 1165 | 10 | 1.72% |
| hsa_circ_0098297 | 1165 | 10 | 1.72% |
| hsa_circ_0108351 | 1165 | 10 | 1.72% |
| hsa_circ_0124222 | 1165 | 10 | 1.72% |
| hsa_circ_0139752 | 1165 | 10 | 1.72% |
| hsa_circ_0026012 | 1282 | 11 | 1.72% |
| hsa_circ_0055494 | 1282 | 11 | 1.72% |
| hsa_circ_0063258 | 1282 | 11 | 1.72% |
| hsa_circ_0084091 | 1282 | 11 | 1.72% |
| hsa_circ_0073948 | 1399 | 12 | 1.72% |
| hsa_circ_0017406 | 1166 | 10 | 1.72% |
| hsa_circ_0033616 | 1166 | 10 | 1.72% |
| hsa_circ_0058558 | 1166 | 10 | 1.72% |
| hsa_circ_0065572 | 1166 | 10 | 1.72% |
| hsa_circ_0065670 | 1166 | 10 | 1.72% |
| hsa_circ_0078257 | 1166 | 10 | 1.72% |
| hsa_circ_0084701 | 1166 | 10 | 1.72% |
| hsa_circ_0091534 | 1166 | 10 | 1.72% |
| hsa_circ_0104085 | 1166 | 10 | 1.72% |
| hsa_circ_0118765 | 1166 | 10 | 1.72% |
| hsa_circ_0002750 | 1283 | 11 | 1.71% |
| hsa_circ_0027648 | 1283 | 11 | 1.71% |
| hsa_circ_0028306 | 1283 | 11 | 1.71% |
| hsa_circ_0038971 | 1283 | 11 | 1.71% |

| | | | |
|---|---|---|---|
| hsa_circ_0058518 | 1283 | 11 | 1.71% |
| hsa_circ_0104817 | 1283 | 11 | 1.71% |
| hsa_circ_0108867 | 1283 | 11 | 1.71% |
| hsa_circ_0126761 | 1283 | 11 | 1.71% |
| hsa_circ_0071561 | 1400 | 12 | 1.71% |
| hsa_circ_0101583 | 1400 | 12 | 1.71% |
| hsa_circ_0013256 | 1167 | 10 | 1.71% |
| hsa_circ_0014047 | 1167 | 10 | 1.71% |
| hsa_circ_0015283 | 1167 | 10 | 1.71% |
| hsa_circ_0023010 | 1167 | 10 | 1.71% |
| hsa_circ_0027738 | 1167 | 10 | 1.71% |
| hsa_circ_0029350 | 1167 | 10 | 1.71% |
| hsa_circ_0030232 | 1167 | 10 | 1.71% |
| hsa_circ_0032474 | 1167 | 10 | 1.71% |
| hsa_circ_0038276 | 1167 | 10 | 1.71% |
| hsa_circ_0054755 | 1167 | 10 | 1.71% |
| hsa_circ_0062367 | 1167 | 10 | 1.71% |
| hsa_circ_0070128 | 1167 | 10 | 1.71% |
| hsa_circ_0086895 | 1167 | 10 | 1.71% |
| hsa_circ_0095154 | 1167 | 10 | 1.71% |
| hsa_circ_0100178 | 1167 | 10 | 1.71% |
| hsa_circ_0100892 | 1167 | 10 | 1.71% |
| hsa_circ_0103088 | 1167 | 10 | 1.71% |
| hsa_circ_0119729 | 1167 | 10 | 1.71% |
| hsa_circ_0126926 | 1167 | 10 | 1.71% |
| hsa_circ_0132388 | 1167 | 10 | 1.71% |
| hsa_circ_0135891 | 1167 | 10 | 1.71% |
| hsa_circ_0012078 | 1634 | 14 | 1.71% |
| hsa_circ_0077600 | 1284 | 11 | 1.71% |
| hsa_circ_0079193 | 1284 | 11 | 1.71% |
| hsa_circ_0101697 | 5137 | 44 | 1.71% |
| hsa_circ_0033515 | 1401 | 12 | 1.71% |
| hsa_circ_0094442 | 1401 | 12 | 1.71% |
| hsa_circ_0096590 | 1401 | 12 | 1.71% |
| hsa_circ_0128930 | 1401 | 12 | 1.71% |
| hsa_circ_0093911 | 1518 | 13 | 1.71% |
| hsa_circ_0108175 | 1518 | 13 | 1.71% |
| hsa_circ_0001164 | 1168 | 10 | 1.71% |
| hsa_circ_0005202 | 1168 | 10 | 1.71% |
| hsa_circ_0018375 | 1168 | 10 | 1.71% |
| hsa_circ_0020781 | 1168 | 10 | 1.71% |
| hsa_circ_0026114 | 1168 | 10 | 1.71% |
| hsa_circ_0029626 | 1168 | 10 | 1.71% |
| hsa_circ_0031756 | 1168 | 10 | 1.71% |
| hsa_circ_0048515 | 1168 | 10 | 1.71% |
| hsa_circ_0057594 | 1168 | 10 | 1.71% |
| hsa_circ_0068450 | 1168 | 10 | 1.71% |
| hsa_circ_0079705 | 1168 | 10 | 1.71% |
| hsa_circ_0079898 | 1168 | 10 | 1.71% |
| hsa_circ_0086317 | 1168 | 10 | 1.71% |

| | | | |
|---|---|---|---|
| hsa_circ_0086513 | 1168 | 10 | 1.71% |
| hsa_circ_0092833 | 1168 | 10 | 1.71% |
| hsa_circ_0094068 | 1168 | 10 | 1.71% |
| hsa_circ_0098221 | 1168 | 10 | 1.71% |
| hsa_circ_0113001 | 1168 | 10 | 1.71% |
| hsa_circ_0118690 | 1168 | 10 | 1.71% |
| hsa_circ_0123042 | 1168 | 10 | 1.71% |
| hsa_circ_0038036 | 2570 | 22 | 1.71% |
| hsa_circ_0014630 | 1285 | 11 | 1.71% |
| hsa_circ_0016940 | 1285 | 11 | 1.71% |
| hsa_circ_0045511 | 1285 | 11 | 1.71% |
| hsa_circ_0054462 | 1285 | 11 | 1.71% |
| hsa_circ_0055919 | 1285 | 11 | 1.71% |
| hsa_circ_0071925 | 1285 | 11 | 1.71% |
| hsa_circ_0096219 | 1285 | 11 | 1.71% |
| hsa_circ_0098592 | 1285 | 11 | 1.71% |
| hsa_circ_0100475 | 1285 | 11 | 1.71% |
| hsa_circ_0129376 | 1285 | 11 | 1.71% |
| hsa_circ_0133989 | 1285 | 11 | 1.71% |
| hsa_circ_0136296 | 1285 | 11 | 1.71% |
| hsa_circ_0059927 | 1402 | 12 | 1.71% |
| hsa_circ_0085799 | 1402 | 12 | 1.71% |
| hsa_circ_0108005 | 1402 | 12 | 1.71% |
| hsa_circ_0108336 | 1402 | 12 | 1.71% |
| hsa_circ_0014477 | 1169 | 10 | 1.71% |
| hsa_circ_0053314 | 1169 | 10 | 1.71% |
| hsa_circ_0055411 | 1169 | 10 | 1.71% |
| hsa_circ_0064353 | 1169 | 10 | 1.71% |
| hsa_circ_0070799 | 1169 | 10 | 1.71% |
| hsa_circ_0083174 | 1169 | 10 | 1.71% |
| hsa_circ_0086795 | 1169 | 10 | 1.71% |
| hsa_circ_0088442 | 1169 | 10 | 1.71% |
| hsa_circ_0096819 | 1169 | 10 | 1.71% |
| hsa_circ_0108742 | 1169 | 10 | 1.71% |
| hsa_circ_0126247 | 1169 | 10 | 1.71% |
| hsa_circ_0137553 | 1169 | 10 | 1.71% |
| hsa_circ_0024676 | 1286 | 11 | 1.71% |
| hsa_circ_0029275 | 1286 | 11 | 1.71% |
| hsa_circ_0037781 | 1286 | 11 | 1.71% |
| hsa_circ_0052390 | 1286 | 11 | 1.71% |
| hsa_circ_0102767 | 1286 | 11 | 1.71% |
| hsa_circ_0105745 | 1286 | 11 | 1.71% |
| hsa_circ_0132849 | 1286 | 11 | 1.71% |
| hsa_circ_0133138 | 1286 | 11 | 1.71% |
| hsa_circ_0005270 | 1403 | 12 | 1.71% |
| hsa_circ_0024391 | 1403 | 12 | 1.71% |
| hsa_circ_0062908 | 1403 | 12 | 1.71% |
| hsa_circ_0086144 | 1403 | 12 | 1.71% |
| hsa_circ_0017258 | 1520 | 13 | 1.71% |
| hsa_circ_0067264 | 1520 | 13 | 1.71% |

| | | | |
|---|---|---|---|
| hsa_circ_0101951 | 1521 | 13 | 1.71% |
| hsa_circ_0034432 | 1404 | 12 | 1.71% |
| hsa_circ_0049118 | 1404 | 12 | 1.71% |
| hsa_circ_0113347 | 1404 | 12 | 1.71% |
| hsa_circ_0117914 | 1404 | 12 | 1.71% |
| hsa_circ_0124394 | 1404 | 12 | 1.71% |
| hsa_circ_0099409 | 1287 | 11 | 1.71% |
| hsa_circ_0130667 | 1287 | 11 | 1.71% |
| hsa_circ_0020580 | 1170 | 10 | 1.71% |
| hsa_circ_0021246 | 1170 | 10 | 1.71% |
| hsa_circ_0022254 | 1170 | 10 | 1.71% |
| hsa_circ_0044932 | 1170 | 10 | 1.71% |
| hsa_circ_0052862 | 1170 | 10 | 1.71% |
| hsa_circ_0066035 | 1170 | 10 | 1.71% |
| hsa_circ_0075582 | 1170 | 10 | 1.71% |
| hsa_circ_0078497 | 1170 | 10 | 1.71% |
| hsa_circ_0081300 | 1170 | 10 | 1.71% |
| hsa_circ_0117001 | 1170 | 10 | 1.71% |
| hsa_circ_0124711 | 1170 | 10 | 1.71% |
| hsa_circ_0128998 | 1170 | 10 | 1.71% |
| hsa_circ_0131396 | 1170 | 10 | 1.71% |
| hsa_circ_0130249 | 1639 | 14 | 1.71% |
| hsa_circ_0053121 | 1405 | 12 | 1.71% |
| hsa_circ_0061055 | 1405 | 12 | 1.71% |
| hsa_circ_0079510 | 1405 | 12 | 1.71% |
| hsa_circ_0006180 | 1288 | 11 | 1.71% |
| hsa_circ_0080014 | 1288 | 11 | 1.71% |
| hsa_circ_0007407 | 1171 | 10 | 1.71% |
| hsa_circ_0010316 | 1171 | 10 | 1.71% |
| hsa_circ_0013105 | 1171 | 10 | 1.71% |
| hsa_circ_0014312 | 1171 | 10 | 1.71% |
| hsa_circ_0017190 | 1171 | 10 | 1.71% |
| hsa_circ_0030302 | 1171 | 10 | 1.71% |
| hsa_circ_0032787 | 1171 | 10 | 1.71% |
| hsa_circ_0036931 | 1171 | 10 | 1.71% |
| hsa_circ_0064949 | 1171 | 10 | 1.71% |
| hsa_circ_0069907 | 1171 | 10 | 1.71% |
| hsa_circ_0094069 | 1171 | 10 | 1.71% |
| hsa_circ_0137150 | 1171 | 10 | 1.71% |
| hsa_circ_0138232 | 1171 | 10 | 1.71% |
| hsa_circ_0138660 | 1171 | 10 | 1.71% |
| hsa_circ_0103186 | 2108 | 18 | 1.71% |
| hsa_circ_0063618 | 1406 | 12 | 1.71% |
| hsa_circ_0018194 | 1289 | 11 | 1.71% |
| hsa_circ_0094361 | 1289 | 11 | 1.71% |
| hsa_circ_0045292 | 1172 | 10 | 1.71% |
| hsa_circ_0065465 | 1172 | 10 | 1.71% |
| hsa_circ_0068778 | 1172 | 10 | 1.71% |
| hsa_circ_0072473 | 1172 | 10 | 1.71% |
| hsa_circ_0076933 | 1172 | 10 | 1.71% |

| | | | |
|---|---|---|---|
| hsa_circ_0088780 | 1172 | 10 | 1.71% |
| hsa_circ_0046162 | 1524 | 13 | 1.71% |
| hsa_circ_0052407 | 1524 | 13 | 1.71% |
| hsa_circ_0028182 | 1407 | 12 | 1.71% |
| hsa_circ_0047275 | 1407 | 12 | 1.71% |
| hsa_circ_0123147 | 1407 | 12 | 1.71% |
| hsa_circ_0019358 | 1290 | 11 | 1.71% |
| hsa_circ_0052027 | 1290 | 11 | 1.71% |
| hsa_circ_0082431 | 1290 | 11 | 1.71% |
| hsa_circ_0086702 | 1290 | 11 | 1.71% |
| hsa_circ_0086805 | 1290 | 11 | 1.71% |
| hsa_circ_0087688 | 1290 | 11 | 1.71% |
| hsa_circ_0094364 | 1290 | 11 | 1.71% |
| hsa_circ_0130261 | 1290 | 11 | 1.71% |
| hsa_circ_0012735 | 1173 | 10 | 1.71% |
| hsa_circ_0014217 | 1173 | 10 | 1.71% |
| hsa_circ_0014320 | 1173 | 10 | 1.71% |
| hsa_circ_0023381 | 1173 | 10 | 1.71% |
| hsa_circ_0058634 | 1173 | 10 | 1.71% |
| hsa_circ_0062718 | 1173 | 10 | 1.71% |
| hsa_circ_0064598 | 1173 | 10 | 1.71% |
| hsa_circ_0071073 | 1173 | 10 | 1.71% |
| hsa_circ_0090978 | 1173 | 10 | 1.71% |
| hsa_circ_0095058 | 1173 | 10 | 1.71% |
| hsa_circ_0095502 | 1173 | 10 | 1.71% |
| hsa_circ_0117466 | 1173 | 10 | 1.71% |
| hsa_circ_0131019 | 1173 | 10 | 1.71% |
| hsa_circ_0132129 | 1173 | 10 | 1.71% |
| hsa_circ_0135564 | 1173 | 10 | 1.71% |
| hsa_circ_0124175 | 3050 | 26 | 1.70% |
| hsa_circ_0140605 | 1408 | 12 | 1.70% |
| hsa_circ_0083139 | 1643 | 14 | 1.70% |
| hsa_circ_0031932 | 1291 | 11 | 1.70% |
| hsa_circ_0046452 | 1291 | 11 | 1.70% |
| hsa_circ_0108225 | 1291 | 11 | 1.70% |
| hsa_circ_0046005 | 1526 | 13 | 1.70% |
| hsa_circ_0080860 | 1526 | 13 | 1.70% |
| hsa_circ_0101034 | 1526 | 13 | 1.70% |
| hsa_circ_0009259 | 1174 | 10 | 1.70% |
| hsa_circ_0066475 | 1174 | 10 | 1.70% |
| hsa_circ_0071414 | 1174 | 10 | 1.70% |
| hsa_circ_0076728 | 1174 | 10 | 1.70% |
| hsa_circ_0095386 | 1174 | 10 | 1.70% |
| hsa_circ_0097155 | 1174 | 10 | 1.70% |
| hsa_circ_0123715 | 1174 | 10 | 1.70% |
| hsa_circ_0012087 | 1409 | 12 | 1.70% |
| hsa_circ_0061212 | 1409 | 12 | 1.70% |
| hsa_circ_0038337 | 1292 | 11 | 1.70% |
| hsa_circ_0086540 | 1292 | 11 | 1.70% |
| hsa_circ_0087717 | 1292 | 11 | 1.70% |

| | | | |
|---|---|---|---|
| hsa_circ_0135306 | 1292 | 11 | 1.70% |
| hsa_circ_0074041 | 1527 | 13 | 1.70% |
| hsa_circ_0014785 | 1762 | 15 | 1.70% |
| hsa_circ_0140531 | 1645 | 14 | 1.70% |
| hsa_circ_0037472 | 1410 | 12 | 1.70% |
| hsa_circ_0057343 | 1410 | 12 | 1.70% |
| hsa_circ_0074265 | 1410 | 12 | 1.70% |
| hsa_circ_0078091 | 1410 | 12 | 1.70% |
| hsa_circ_0013115 | 1175 | 10 | 1.70% |
| hsa_circ_0022533 | 1175 | 10 | 1.70% |
| hsa_circ_0040349 | 1175 | 10 | 1.70% |
| hsa_circ_0076909 | 1175 | 10 | 1.70% |
| hsa_circ_0136752 | 1175 | 10 | 1.70% |
| hsa_circ_0013539 | 1763 | 15 | 1.70% |
| hsa_circ_0020657 | 1293 | 11 | 1.70% |
| hsa_circ_0035101 | 1293 | 11 | 1.70% |
| hsa_circ_0048421 | 1293 | 11 | 1.70% |
| hsa_circ_0091755 | 1293 | 11 | 1.70% |
| hsa_circ_0113850 | 1293 | 11 | 1.70% |
| hsa_circ_0119829 | 1293 | 11 | 1.70% |
| hsa_circ_0014043 | 1411 | 12 | 1.70% |
| hsa_circ_0036666 | 1411 | 12 | 1.70% |
| hsa_circ_0106451 | 3998 | 34 | 1.70% |
| hsa_circ_0015814 | 1176 | 10 | 1.70% |
| hsa_circ_0020849 | 1176 | 10 | 1.70% |
| hsa_circ_0034888 | 1176 | 10 | 1.70% |
| hsa_circ_0038542 | 1176 | 10 | 1.70% |
| hsa_circ_0054238 | 1176 | 10 | 1.70% |
| hsa_circ_0064263 | 1176 | 10 | 1.70% |
| hsa_circ_0066312 | 1176 | 10 | 1.70% |
| hsa_circ_0075226 | 1176 | 10 | 1.70% |
| hsa_circ_0076113 | 1176 | 10 | 1.70% |
| hsa_circ_0083634 | 1176 | 10 | 1.70% |
| hsa_circ_0083718 | 1176 | 10 | 1.70% |
| hsa_circ_0121922 | 1176 | 10 | 1.70% |
| hsa_circ_0123430 | 1176 | 10 | 1.70% |
| hsa_circ_0125308 | 1176 | 10 | 1.70% |
| hsa_circ_0126412 | 1176 | 10 | 1.70% |
| hsa_circ_0127893 | 1176 | 10 | 1.70% |
| hsa_circ_0130617 | 1176 | 10 | 1.70% |
| hsa_circ_0131277 | 1176 | 10 | 1.70% |
| hsa_circ_0132121 | 1176 | 10 | 1.70% |
| hsa_circ_0134013 | 1176 | 10 | 1.70% |
| hsa_circ_0140322 | 1176 | 10 | 1.70% |
| hsa_circ_0117330 | 5293 | 45 | 1.70% |
| hsa_circ_0017941 | 1294 | 11 | 1.70% |
| hsa_circ_0025499 | 1294 | 11 | 1.70% |
| hsa_circ_0051673 | 1294 | 11 | 1.70% |
| hsa_circ_0077598 | 1294 | 11 | 1.70% |
| hsa_circ_0114386 | 1294 | 11 | 1.70% |

| | | | |
|---|---|---|---|
| hsa_circ_0117443 | 1294 | 11 | 1.70% |
| hsa_circ_0138875 | 1294 | 11 | 1.70% |
| hsa_circ_0043772 | 1412 | 12 | 1.70% |
| hsa_circ_0101340 | 1412 | 12 | 1.70% |
| hsa_circ_0135766 | 1412 | 12 | 1.70% |
| hsa_circ_0014335 | 1177 | 10 | 1.70% |
| hsa_circ_0078311 | 1177 | 10 | 1.70% |
| hsa_circ_0087982 | 1177 | 10 | 1.70% |
| hsa_circ_0093669 | 1177 | 10 | 1.70% |
| hsa_circ_0100674 | 1177 | 10 | 1.70% |
| hsa_circ_0123439 | 1177 | 10 | 1.70% |
| hsa_circ_0135898 | 1177 | 10 | 1.70% |
| hsa_circ_0136106 | 1177 | 10 | 1.70% |
| hsa_circ_0044655 | 1295 | 11 | 1.70% |
| hsa_circ_0079554 | 1295 | 11 | 1.70% |
| hsa_circ_0116646 | 1295 | 11 | 1.70% |
| hsa_circ_0125553 | 1295 | 11 | 1.70% |
| hsa_circ_0007185 | 3179 | 27 | 1.70% |
| hsa_circ_0077233 | 1413 | 12 | 1.70% |
| hsa_circ_0135960 | 1531 | 13 | 1.70% |
| hsa_circ_0008924 | 1178 | 10 | 1.70% |
| hsa_circ_0042070 | 1178 | 10 | 1.70% |
| hsa_circ_0060207 | 1178 | 10 | 1.70% |
| hsa_circ_0071931 | 1178 | 10 | 1.70% |
| hsa_circ_0109417 | 1178 | 10 | 1.70% |
| hsa_circ_0115178 | 1178 | 10 | 1.70% |
| hsa_circ_0116030 | 1178 | 10 | 1.70% |
| hsa_circ_0038952 | 1296 | 11 | 1.70% |
| hsa_circ_0060578 | 1296 | 11 | 1.70% |
| hsa_circ_0074790 | 1296 | 11 | 1.70% |
| hsa_circ_0129434 | 1296 | 11 | 1.70% |
| hsa_circ_0135963 | 1296 | 11 | 1.70% |
| hsa_circ_0069139 | 1414 | 12 | 1.70% |
| hsa_circ_0084865 | 1532 | 13 | 1.70% |
| hsa_circ_0007833 | 1179 | 10 | 1.70% |
| hsa_circ_0015767 | 1179 | 10 | 1.70% |
| hsa_circ_0017350 | 1179 | 10 | 1.70% |
| hsa_circ_0018689 | 1179 | 10 | 1.70% |
| hsa_circ_0023889 | 1179 | 10 | 1.70% |
| hsa_circ_0067802 | 1179 | 10 | 1.70% |
| hsa_circ_0081083 | 1179 | 10 | 1.70% |
| hsa_circ_0082200 | 1179 | 10 | 1.70% |
| hsa_circ_0090963 | 1179 | 10 | 1.70% |
| hsa_circ_0092877 | 1179 | 10 | 1.70% |
| hsa_circ_0094921 | 1179 | 10 | 1.70% |
| hsa_circ_0102673 | 1179 | 10 | 1.70% |
| hsa_circ_0107328 | 1179 | 10 | 1.70% |
| hsa_circ_0120598 | 1179 | 10 | 1.70% |
| hsa_circ_0121960 | 1179 | 10 | 1.70% |
| hsa_circ_0129348 | 1179 | 10 | 1.70% |

| | | | |
|---|---|---|---|
| hsa_circ_0136782 | 1179 | 10 | 1.70% |
| hsa_circ_0138401 | 1179 | 10 | 1.70% |
| hsa_circ_0009315 | 1297 | 11 | 1.70% |
| hsa_circ_0029897 | 1297 | 11 | 1.70% |
| hsa_circ_0038096 | 1297 | 11 | 1.70% |
| hsa_circ_0042754 | 1297 | 11 | 1.70% |
| hsa_circ_0060929 | 1297 | 11 | 1.70% |
| hsa_circ_0079073 | 1297 | 11 | 1.70% |
| hsa_circ_0117762 | 1297 | 11 | 1.70% |
| hsa_circ_0087366 | 1415 | 12 | 1.70% |
| hsa_circ_0102341 | 1415 | 12 | 1.70% |
| hsa_circ_0051023 | 1533 | 13 | 1.70% |
| hsa_circ_0000322 | 1887 | 16 | 1.70% |
| hsa_circ_0056200 | 1534 | 13 | 1.69% |
| hsa_circ_0108939 | 1534 | 13 | 1.69% |
| hsa_circ_0013630 | 1416 | 12 | 1.69% |
| hsa_circ_0019808 | 1416 | 12 | 1.69% |
| hsa_circ_0090185 | 1416 | 12 | 1.69% |
| hsa_circ_0112259 | 1416 | 12 | 1.69% |
| hsa_circ_0032737 | 1298 | 11 | 1.69% |
| hsa_circ_0035905 | 1298 | 11 | 1.69% |
| hsa_circ_0064880 | 1298 | 11 | 1.69% |
| hsa_circ_0065695 | 1298 | 11 | 1.69% |
| hsa_circ_0079560 | 1298 | 11 | 1.69% |
| hsa_circ_0104897 | 1298 | 11 | 1.69% |
| hsa_circ_0105503 | 1298 | 11 | 1.69% |
| hsa_circ_0025440 | 1180 | 10 | 1.69% |
| hsa_circ_0025646 | 1180 | 10 | 1.69% |
| hsa_circ_0034253 | 1180 | 10 | 1.69% |
| hsa_circ_0036673 | 1180 | 10 | 1.69% |
| hsa_circ_0040742 | 1180 | 10 | 1.69% |
| hsa_circ_0058053 | 1180 | 10 | 1.69% |
| hsa_circ_0062616 | 1180 | 10 | 1.69% |
| hsa_circ_0063977 | 1180 | 10 | 1.69% |
| hsa_circ_0064010 | 1180 | 10 | 1.69% |
| hsa_circ_0069103 | 1180 | 10 | 1.69% |
| hsa_circ_0078985 | 1180 | 10 | 1.69% |
| hsa_circ_0111859 | 1180 | 10 | 1.69% |
| hsa_circ_0124622 | 1180 | 10 | 1.69% |
| hsa_circ_0127973 | 1180 | 10 | 1.69% |
| hsa_circ_0060672 | 1535 | 13 | 1.69% |
| hsa_circ_0075278 | 1417 | 12 | 1.69% |
| hsa_circ_0119427 | 1417 | 12 | 1.69% |
| hsa_circ_0011407 | 1299 | 11 | 1.69% |
| hsa_circ_0013017 | 1299 | 11 | 1.69% |
| hsa_circ_0013098 | 1299 | 11 | 1.69% |
| hsa_circ_0050988 | 1299 | 11 | 1.69% |
| hsa_circ_0083622 | 1299 | 11 | 1.69% |
| hsa_circ_0085992 | 1299 | 11 | 1.69% |
| hsa_circ_0111474 | 1299 | 11 | 1.69% |

| | | | |
|---|---|---|---|
| hsa_circ_0112392 | 1299 | 11 | 1.69% |
| hsa_circ_0114784 | 1299 | 11 | 1.69% |
| hsa_circ_0006850 | 1181 | 10 | 1.69% |
| hsa_circ_0031420 | 1181 | 10 | 1.69% |
| hsa_circ_0047217 | 1181 | 10 | 1.69% |
| hsa_circ_0075555 | 1181 | 10 | 1.69% |
| hsa_circ_0078504 | 1181 | 10 | 1.69% |
| hsa_circ_0088146 | 1181 | 10 | 1.69% |
| hsa_circ_0088378 | 1181 | 10 | 1.69% |
| hsa_circ_0001911 | 1536 | 13 | 1.69% |
| hsa_circ_0017679 | 1536 | 13 | 1.69% |
| hsa_circ_0062862 | 1536 | 13 | 1.69% |
| hsa_circ_0102073 | 1418 | 12 | 1.69% |
| hsa_circ_0016527 | 1300 | 11 | 1.69% |
| hsa_circ_0017748 | 1300 | 11 | 1.69% |
| hsa_circ_0021389 | 1300 | 11 | 1.69% |
| hsa_circ_0032874 | 1300 | 11 | 1.69% |
| hsa_circ_0061999 | 1300 | 11 | 1.69% |
| hsa_circ_0098280 | 1891 | 16 | 1.69% |
| hsa_circ_0008525 | 1182 | 10 | 1.69% |
| hsa_circ_0009647 | 1182 | 10 | 1.69% |
| hsa_circ_0019586 | 1182 | 10 | 1.69% |
| hsa_circ_0027412 | 1182 | 10 | 1.69% |
| hsa_circ_0028879 | 1182 | 10 | 1.69% |
| hsa_circ_0043064 | 1182 | 10 | 1.69% |
| hsa_circ_0049715 | 1182 | 10 | 1.69% |
| hsa_circ_0054851 | 1182 | 10 | 1.69% |
| hsa_circ_0066231 | 1182 | 10 | 1.69% |
| hsa_circ_0084726 | 1182 | 10 | 1.69% |
| hsa_circ_0127928 | 1182 | 10 | 1.69% |
| hsa_circ_0066960 | 1655 | 14 | 1.69% |
| hsa_circ_0103490 | 1655 | 14 | 1.69% |
| hsa_circ_0098930 | 4493 | 38 | 1.69% |
| hsa_circ_0026902 | 1419 | 12 | 1.69% |
| hsa_circ_0074236 | 1419 | 12 | 1.69% |
| hsa_circ_0089656 | 1419 | 12 | 1.69% |
| hsa_circ_0136511 | 1419 | 12 | 1.69% |
| hsa_circ_0051025 | 1301 | 11 | 1.69% |
| hsa_circ_0058062 | 1301 | 11 | 1.69% |
| hsa_circ_0086561 | 1301 | 11 | 1.69% |
| hsa_circ_0098131 | 1301 | 11 | 1.69% |
| hsa_circ_0104635 | 1301 | 11 | 1.69% |
| hsa_circ_0139284 | 1301 | 11 | 1.69% |
| hsa_circ_0014315 | 1183 | 10 | 1.69% |
| hsa_circ_0019780 | 1183 | 10 | 1.69% |
| hsa_circ_0023014 | 1183 | 10 | 1.69% |
| hsa_circ_0030757 | 1183 | 10 | 1.69% |
| hsa_circ_0035722 | 1183 | 10 | 1.69% |
| hsa_circ_0042484 | 1183 | 10 | 1.69% |
| hsa_circ_0042508 | 1183 | 10 | 1.69% |

| | | | |
|---|---|---|---|
| hsa_circ_0043718 | 1183 | 10 | 1.69% |
| hsa_circ_0054029 | 1183 | 10 | 1.69% |
| hsa_circ_0065638 | 1183 | 10 | 1.69% |
| hsa_circ_0065856 | 1183 | 10 | 1.69% |
| hsa_circ_0074017 | 1183 | 10 | 1.69% |
| hsa_circ_0087641 | 1183 | 10 | 1.69% |
| hsa_circ_0090917 | 1183 | 10 | 1.69% |
| hsa_circ_0111124 | 1183 | 10 | 1.69% |
| hsa_circ_0111371 | 1183 | 10 | 1.69% |
| hsa_circ_0116720 | 1183 | 10 | 1.69% |
| hsa_circ_0128055 | 1183 | 10 | 1.69% |
| hsa_circ_0132397 | 1183 | 10 | 1.69% |
| hsa_circ_0023450 | 1302 | 11 | 1.69% |
| hsa_circ_0034659 | 1302 | 11 | 1.69% |
| hsa_circ_0038422 | 1302 | 11 | 1.69% |
| hsa_circ_0057229 | 1302 | 11 | 1.69% |
| hsa_circ_0066061 | 1302 | 11 | 1.69% |
| hsa_circ_0073426 | 1302 | 11 | 1.69% |
| hsa_circ_0119917 | 1302 | 11 | 1.69% |
| hsa_circ_0006399 | 1184 | 10 | 1.69% |
| hsa_circ_0011638 | 1184 | 10 | 1.69% |
| hsa_circ_0023474 | 1184 | 10 | 1.69% |
| hsa_circ_0023800 | 1184 | 10 | 1.69% |
| hsa_circ_0026779 | 1184 | 10 | 1.69% |
| hsa_circ_0033202 | 1184 | 10 | 1.69% |
| hsa_circ_0044278 | 1184 | 10 | 1.69% |
| hsa_circ_0048935 | 1184 | 10 | 1.69% |
| hsa_circ_0057235 | 1184 | 10 | 1.69% |
| hsa_circ_0063508 | 1184 | 10 | 1.69% |
| hsa_circ_0063640 | 1184 | 10 | 1.69% |
| hsa_circ_0064897 | 1184 | 10 | 1.69% |
| hsa_circ_0068211 | 1184 | 10 | 1.69% |
| hsa_circ_0071933 | 1184 | 10 | 1.69% |
| hsa_circ_0079695 | 1184 | 10 | 1.69% |
| hsa_circ_0105200 | 1184 | 10 | 1.69% |
| hsa_circ_0116124 | 1184 | 10 | 1.69% |
| hsa_circ_0122411 | 1184 | 10 | 1.69% |
| hsa_circ_0124621 | 1184 | 10 | 1.69% |
| hsa_circ_0107783 | 1421 | 12 | 1.69% |
| hsa_circ_0115259 | 1421 | 12 | 1.69% |
| hsa_circ_0015677 | 1303 | 11 | 1.69% |
| hsa_circ_0039211 | 1303 | 11 | 1.69% |
| hsa_circ_0066827 | 1303 | 11 | 1.69% |
| hsa_circ_0093757 | 1303 | 11 | 1.69% |
| hsa_circ_0127253 | 1303 | 11 | 1.69% |
| hsa_circ_0136478 | 1540 | 13 | 1.69% |
| hsa_circ_0032658 | 2014 | 17 | 1.69% |
| hsa_circ_0069418 | 4265 | 36 | 1.69% |
| hsa_circ_0106749 | 2488 | 21 | 1.69% |
| hsa_circ_0134009 | 5095 | 43 | 1.69% |

| | | | |
|---|---|---|---|
| hsa_circ_0021996 | 1659 | 14 | 1.69% |
| hsa_circ_0010384 | 1422 | 12 | 1.69% |
| hsa_circ_0055293 | 1422 | 12 | 1.69% |
| hsa_circ_0005093 | 1185 | 10 | 1.69% |
| hsa_circ_0016801 | 1185 | 10 | 1.69% |
| hsa_circ_0024447 | 1185 | 10 | 1.69% |
| hsa_circ_0034262 | 1185 | 10 | 1.69% |
| hsa_circ_0038197 | 1185 | 10 | 1.69% |
| hsa_circ_0046190 | 1185 | 10 | 1.69% |
| hsa_circ_0065987 | 1185 | 10 | 1.69% |
| hsa_circ_0077080 | 1185 | 10 | 1.69% |
| hsa_circ_0085446 | 1185 | 10 | 1.69% |
| hsa_circ_0093549 | 1185 | 10 | 1.69% |
| hsa_circ_0114436 | 1185 | 10 | 1.69% |
| hsa_circ_0122485 | 1185 | 10 | 1.69% |
| hsa_circ_0128723 | 1185 | 10 | 1.69% |
| hsa_circ_0140072 | 1185 | 10 | 1.69% |
| hsa_circ_0038335 | 1304 | 11 | 1.69% |
| hsa_circ_0047385 | 1304 | 11 | 1.69% |
| hsa_circ_0048379 | 1304 | 11 | 1.69% |
| hsa_circ_0125281 | 1304 | 11 | 1.69% |
| hsa_circ_0036435 | 1423 | 12 | 1.69% |
| hsa_circ_0074053 | 1423 | 12 | 1.69% |
| hsa_circ_0018813 | 1186 | 10 | 1.69% |
| hsa_circ_0029962 | 1186 | 10 | 1.69% |
| hsa_circ_0035302 | 1186 | 10 | 1.69% |
| hsa_circ_0048819 | 1186 | 10 | 1.69% |
| hsa_circ_0051531 | 1186 | 10 | 1.69% |
| hsa_circ_0051538 | 1186 | 10 | 1.69% |
| hsa_circ_0060502 | 1186 | 10 | 1.69% |
| hsa_circ_0064580 | 1186 | 10 | 1.69% |
| hsa_circ_0087687 | 1186 | 10 | 1.69% |
| hsa_circ_0112404 | 1186 | 10 | 1.69% |
| hsa_circ_0112879 | 1186 | 10 | 1.69% |
| hsa_circ_0119936 | 1186 | 10 | 1.69% |
| hsa_circ_0078656 | 1542 | 13 | 1.69% |
| hsa_circ_0091474 | 1542 | 13 | 1.69% |
| hsa_circ_0015162 | 1305 | 11 | 1.69% |
| hsa_circ_0019009 | 1305 | 11 | 1.69% |
| hsa_circ_0030397 | 1305 | 11 | 1.69% |
| hsa_circ_0049037 | 1305 | 11 | 1.69% |
| hsa_circ_0074801 | 1305 | 11 | 1.69% |
| hsa_circ_0094137 | 1305 | 11 | 1.69% |
| hsa_circ_0104698 | 1305 | 11 | 1.69% |
| hsa_circ_0111700 | 1305 | 11 | 1.69% |
| hsa_circ_0034101 | 4627 | 39 | 1.69% |
| hsa_circ_0107514 | 1661 | 14 | 1.69% |
| hsa_circ_0035808 | 1424 | 12 | 1.69% |
| hsa_circ_0069091 | 1424 | 12 | 1.69% |
| hsa_circ_0083178 | 1424 | 12 | 1.69% |

| | | | |
|---|---|---|---|
| hsa_circ_0100467 | 1543 | 13 | 1.69% |
| hsa_circ_0028149 | 1187 | 10 | 1.68% |
| hsa_circ_0032957 | 1187 | 10 | 1.68% |
| hsa_circ_0033356 | 1187 | 10 | 1.68% |
| hsa_circ_0043112 | 1187 | 10 | 1.68% |
| hsa_circ_0060183 | 1187 | 10 | 1.68% |
| hsa_circ_0105090 | 1187 | 10 | 1.68% |
| hsa_circ_0107126 | 1187 | 10 | 1.68% |
| hsa_circ_0124644 | 1187 | 10 | 1.68% |
| hsa_circ_0129639 | 1187 | 10 | 1.68% |
| hsa_circ_0001014 | 2493 | 21 | 1.68% |
| hsa_circ_0003913 | 1306 | 11 | 1.68% |
| hsa_circ_0038106 | 1306 | 11 | 1.68% |
| hsa_circ_0049204 | 1306 | 11 | 1.68% |
| hsa_circ_0056591 | 1306 | 11 | 1.68% |
| hsa_circ_0068307 | 1306 | 11 | 1.68% |
| hsa_circ_0095488 | 1306 | 11 | 1.68% |
| hsa_circ_0113722 | 1306 | 11 | 1.68% |
| hsa_circ_0127177 | 1306 | 11 | 1.68% |
| hsa_circ_0128159 | 1306 | 11 | 1.68% |
| hsa_circ_0136472 | 1306 | 11 | 1.68% |
| hsa_circ_0010892 | 1425 | 12 | 1.68% |
| hsa_circ_0085875 | 1425 | 12 | 1.68% |
| hsa_circ_0104187 | 1425 | 12 | 1.68% |
| hsa_circ_0120694 | 1425 | 12 | 1.68% |
| hsa_circ_0122387 | 1544 | 13 | 1.68% |
| hsa_circ_0070168 | 1663 | 14 | 1.68% |
| hsa_circ_0010264 | 1188 | 10 | 1.68% |
| hsa_circ_0065627 | 1188 | 10 | 1.68% |
| hsa_circ_0074684 | 1188 | 10 | 1.68% |
| hsa_circ_0075423 | 1188 | 10 | 1.68% |
| hsa_circ_0107831 | 1188 | 10 | 1.68% |
| hsa_circ_0113815 | 1188 | 10 | 1.68% |
| hsa_circ_0120577 | 1188 | 10 | 1.68% |
| hsa_circ_0130577 | 1188 | 10 | 1.68% |
| hsa_circ_0134461 | 1188 | 10 | 1.68% |
| hsa_circ_0138807 | 1188 | 10 | 1.68% |
| hsa_circ_0139916 | 1188 | 10 | 1.68% |
| hsa_circ_0019061 | 1307 | 11 | 1.68% |
| hsa_circ_0021432 | 1307 | 11 | 1.68% |
| hsa_circ_0024830 | 1307 | 11 | 1.68% |
| hsa_circ_0025587 | 1307 | 11 | 1.68% |
| hsa_circ_0031295 | 1307 | 11 | 1.68% |
| hsa_circ_0064775 | 1307 | 11 | 1.68% |
| hsa_circ_0067769 | 1307 | 11 | 1.68% |
| hsa_circ_0136513 | 1307 | 11 | 1.68% |
| hsa_circ_0107691 | 3446 | 29 | 1.68% |
| hsa_circ_0014786 | 1783 | 15 | 1.68% |
| hsa_circ_0011435 | 1189 | 10 | 1.68% |
| hsa_circ_0013689 | 1189 | 10 | 1.68% |

| | | | |
|---|---|---|---|
| hsa_circ_0020477 | 1189 | 10 | 1.68% |
| hsa_circ_0034267 | 1189 | 10 | 1.68% |
| hsa_circ_0039888 | 1189 | 10 | 1.68% |
| hsa_circ_0045554 | 1189 | 10 | 1.68% |
| hsa_circ_0047932 | 1189 | 10 | 1.68% |
| hsa_circ_0049014 | 1189 | 10 | 1.68% |
| hsa_circ_0058949 | 1189 | 10 | 1.68% |
| hsa_circ_0083233 | 1189 | 10 | 1.68% |
| hsa_circ_0092456 | 1189 | 10 | 1.68% |
| hsa_circ_0100420 | 1189 | 10 | 1.68% |
| hsa_circ_0110530 | 1189 | 10 | 1.68% |
| hsa_circ_0112607 | 1189 | 10 | 1.68% |
| hsa_circ_0128017 | 1189 | 10 | 1.68% |
| hsa_circ_0140397 | 1189 | 10 | 1.68% |
| hsa_circ_0010825 | 1308 | 11 | 1.68% |
| hsa_circ_0041181 | 1308 | 11 | 1.68% |
| hsa_circ_0048158 | 1308 | 11 | 1.68% |
| hsa_circ_0069554 | 1308 | 11 | 1.68% |
| hsa_circ_0087903 | 1308 | 11 | 1.68% |
| hsa_circ_0124895 | 1308 | 11 | 1.68% |
| hsa_circ_0009318 | 1427 | 12 | 1.68% |
| hsa_circ_0076171 | 1427 | 12 | 1.68% |
| hsa_circ_0135849 | 1665 | 14 | 1.68% |
| hsa_circ_0058159 | 3213 | 27 | 1.68% |
| hsa_circ_0009667 | 1428 | 12 | 1.68% |
| hsa_circ_0011101 | 1428 | 12 | 1.68% |
| hsa_circ_0018426 | 1428 | 12 | 1.68% |
| hsa_circ_0053051 | 1428 | 12 | 1.68% |
| hsa_circ_0036165 | 1309 | 11 | 1.68% |
| hsa_circ_0045043 | 1309 | 11 | 1.68% |
| hsa_circ_0052858 | 1309 | 11 | 1.68% |
| hsa_circ_0089333 | 1309 | 11 | 1.68% |
| hsa_circ_0099980 | 1309 | 11 | 1.68% |
| hsa_circ_0014061 | 1190 | 10 | 1.68% |
| hsa_circ_0017776 | 1190 | 10 | 1.68% |
| hsa_circ_0043257 | 1190 | 10 | 1.68% |
| hsa_circ_0052795 | 1190 | 10 | 1.68% |
| hsa_circ_0079158 | 1190 | 10 | 1.68% |
| hsa_circ_0092114 | 1190 | 10 | 1.68% |
| hsa_circ_0095177 | 1190 | 10 | 1.68% |
| hsa_circ_0104088 | 1190 | 10 | 1.68% |
| hsa_circ_0050506 | 2143 | 18 | 1.68% |
| hsa_circ_0011334 | 3929 | 33 | 1.68% |
| hsa_circ_0090012 | 1429 | 12 | 1.68% |
| hsa_circ_0108581 | 1429 | 12 | 1.68% |
| hsa_circ_0025177 | 1310 | 11 | 1.68% |
| hsa_circ_0027563 | 1310 | 11 | 1.68% |
| hsa_circ_0028064 | 1310 | 11 | 1.68% |
| hsa_circ_0061943 | 1310 | 11 | 1.68% |
| hsa_circ_0063877 | 1310 | 11 | 1.68% |

| | | | |
|---|---|---|---|
| hsa_circ_0093486 | 1310 | 11 | 1.68% |
| hsa_circ_0111024 | 1310 | 11 | 1.68% |
| hsa_circ_0000107 | 1191 | 10 | 1.68% |
| hsa_circ_0011550 | 1191 | 10 | 1.68% |
| hsa_circ_0012472 | 1191 | 10 | 1.68% |
| hsa_circ_0022137 | 1191 | 10 | 1.68% |
| hsa_circ_0026129 | 1191 | 10 | 1.68% |
| hsa_circ_0034894 | 1191 | 10 | 1.68% |
| hsa_circ_0038414 | 1191 | 10 | 1.68% |
| hsa_circ_0051069 | 1191 | 10 | 1.68% |
| hsa_circ_0056684 | 1191 | 10 | 1.68% |
| hsa_circ_0058610 | 1191 | 10 | 1.68% |
| hsa_circ_0066742 | 1191 | 10 | 1.68% |
| hsa_circ_0069754 | 1191 | 10 | 1.68% |
| hsa_circ_0078679 | 1191 | 10 | 1.68% |
| hsa_circ_0081268 | 1191 | 10 | 1.68% |
| hsa_circ_0085283 | 1191 | 10 | 1.68% |
| hsa_circ_0085969 | 1191 | 10 | 1.68% |
| hsa_circ_0089129 | 1191 | 10 | 1.68% |
| hsa_circ_0091230 | 1191 | 10 | 1.68% |
| hsa_circ_0093284 | 1191 | 10 | 1.68% |
| hsa_circ_0126008 | 1191 | 10 | 1.68% |
| hsa_circ_0140754 | 1191 | 10 | 1.68% |
| hsa_circ_0011369 | 3932 | 33 | 1.68% |
| hsa_circ_0027290 | 1549 | 13 | 1.68% |
| hsa_circ_0010086 | 1430 | 12 | 1.68% |
| hsa_circ_0076608 | 1430 | 12 | 1.68% |
| hsa_circ_0096541 | 1430 | 12 | 1.68% |
| hsa_circ_0008289 | 1311 | 11 | 1.68% |
| hsa_circ_0022509 | 1311 | 11 | 1.68% |
| hsa_circ_0084735 | 1311 | 11 | 1.68% |
| hsa_circ_0085934 | 1311 | 11 | 1.68% |
| hsa_circ_0125034 | 1311 | 11 | 1.68% |
| hsa_circ_0138080 | 1311 | 11 | 1.68% |
| hsa_circ_0139682 | 1311 | 11 | 1.68% |
| hsa_circ_0103123 | 4768 | 40 | 1.68% |
| hsa_circ_0011348 | 1192 | 10 | 1.68% |
| hsa_circ_0013435 | 1192 | 10 | 1.68% |
| hsa_circ_0017371 | 1192 | 10 | 1.68% |
| hsa_circ_0021260 | 1192 | 10 | 1.68% |
| hsa_circ_0024958 | 1192 | 10 | 1.68% |
| hsa_circ_0052765 | 1192 | 10 | 1.68% |
| hsa_circ_0065991 | 1192 | 10 | 1.68% |
| hsa_circ_0081607 | 1192 | 10 | 1.68% |
| hsa_circ_0090398 | 1192 | 10 | 1.68% |
| hsa_circ_0095072 | 1192 | 10 | 1.68% |
| hsa_circ_0095533 | 1192 | 10 | 1.68% |
| hsa_circ_0127528 | 1192 | 10 | 1.68% |
| hsa_circ_0137574 | 1192 | 10 | 1.68% |
| hsa_circ_0053127 | 1669 | 14 | 1.68% |

| | | | |
|---|---|---|---|
| hsa_circ_0068404 | 1550 | 13 | 1.68% |
| hsa_circ_0082086 | 1550 | 13 | 1.68% |
| hsa_circ_0035268 | 1431 | 12 | 1.68% |
| hsa_circ_0057433 | 1431 | 12 | 1.68% |
| hsa_circ_0057441 | 1431 | 12 | 1.68% |
| hsa_circ_0034180 | 1312 | 11 | 1.68% |
| hsa_circ_0066427 | 1312 | 11 | 1.68% |
| hsa_circ_0100903 | 1312 | 11 | 1.68% |
| hsa_circ_0083651 | 1670 | 14 | 1.68% |
| hsa_circ_0009734 | 1193 | 10 | 1.68% |
| hsa_circ_0016047 | 1193 | 10 | 1.68% |
| hsa_circ_0036736 | 1193 | 10 | 1.68% |
| hsa_circ_0065272 | 1193 | 10 | 1.68% |
| hsa_circ_0068360 | 1193 | 10 | 1.68% |
| hsa_circ_0070885 | 1193 | 10 | 1.68% |
| hsa_circ_0072389 | 1193 | 10 | 1.68% |
| hsa_circ_0073977 | 1193 | 10 | 1.68% |
| hsa_circ_0075029 | 1193 | 10 | 1.68% |
| hsa_circ_0083821 | 1193 | 10 | 1.68% |
| hsa_circ_0084254 | 1193 | 10 | 1.68% |
| hsa_circ_0090603 | 1193 | 10 | 1.68% |
| hsa_circ_0093215 | 1193 | 10 | 1.68% |
| hsa_circ_0100188 | 1193 | 10 | 1.68% |
| hsa_circ_0104083 | 1193 | 10 | 1.68% |
| hsa_circ_0112578 | 1193 | 10 | 1.68% |
| hsa_circ_0123094 | 1193 | 10 | 1.68% |
| hsa_circ_0123516 | 1193 | 10 | 1.68% |
| hsa_circ_0131578 | 1193 | 10 | 1.68% |
| hsa_circ_0101369 | 3937 | 33 | 1.68% |
| hsa_circ_0036082 | 4057 | 34 | 1.68% |
| hsa_circ_0087972 | 1432 | 12 | 1.68% |
| hsa_circ_0097202 | 1432 | 12 | 1.68% |
| hsa_circ_0102726 | 1432 | 12 | 1.68% |
| hsa_circ_0125915 | 1432 | 12 | 1.68% |
| hsa_circ_0049255 | 1671 | 14 | 1.68% |
| hsa_circ_0066930 | 1671 | 14 | 1.68% |
| hsa_circ_0004485 | 1313 | 11 | 1.68% |
| hsa_circ_0031436 | 1313 | 11 | 1.68% |
| hsa_circ_0065913 | 1313 | 11 | 1.68% |
| hsa_circ_0074202 | 1313 | 11 | 1.68% |
| hsa_circ_0074042 | 1552 | 13 | 1.68% |
| hsa_circ_0001022 | 1194 | 10 | 1.68% |
| hsa_circ_0031311 | 1194 | 10 | 1.68% |
| hsa_circ_0036923 | 1194 | 10 | 1.68% |
| hsa_circ_0043831 | 1194 | 10 | 1.68% |
| hsa_circ_0047783 | 1194 | 10 | 1.68% |
| hsa_circ_0050337 | 1194 | 10 | 1.68% |
| hsa_circ_0079829 | 1194 | 10 | 1.68% |
| hsa_circ_0081395 | 1194 | 10 | 1.68% |
| hsa_circ_0093283 | 1194 | 10 | 1.68% |

| | | | |
|---|---|---|---|
| hsa_circ_0095734 | 1194 | 10 | 1.68% |
| hsa_circ_0105311 | 1194 | 10 | 1.68% |
| hsa_circ_0133879 | 1433 | 12 | 1.67% |
| hsa_circ_0062809 | 1314 | 11 | 1.67% |
| hsa_circ_0063409 | 1314 | 11 | 1.67% |
| hsa_circ_0063989 | 1314 | 11 | 1.67% |
| hsa_circ_0089514 | 1314 | 11 | 1.67% |
| hsa_circ_0101178 | 1314 | 11 | 1.67% |
| hsa_circ_0093745 | 2868 | 24 | 1.67% |
| hsa_circ_0019399 | 1434 | 12 | 1.67% |
| hsa_circ_0058026 | 1434 | 12 | 1.67% |
| hsa_circ_0068848 | 1434 | 12 | 1.67% |
| hsa_circ_0002925 | 1195 | 10 | 1.67% |
| hsa_circ_0034392 | 1195 | 10 | 1.67% |
| hsa_circ_0039300 | 1195 | 10 | 1.67% |
| hsa_circ_0051620 | 1195 | 10 | 1.67% |
| hsa_circ_0059237 | 1195 | 10 | 1.67% |
| hsa_circ_0078081 | 1195 | 10 | 1.67% |
| hsa_circ_0090200 | 1195 | 10 | 1.67% |
| hsa_circ_0091088 | 1195 | 10 | 1.67% |
| hsa_circ_0091362 | 1195 | 10 | 1.67% |
| hsa_circ_0103300 | 1195 | 10 | 1.67% |
| hsa_circ_0115052 | 1195 | 10 | 1.67% |
| hsa_circ_0119946 | 1195 | 10 | 1.67% |
| hsa_circ_0107095 | 3227 | 27 | 1.67% |
| hsa_circ_0019176 | 1315 | 11 | 1.67% |
| hsa_circ_0042148 | 1315 | 11 | 1.67% |
| hsa_circ_0058028 | 1315 | 11 | 1.67% |
| hsa_circ_0061694 | 1315 | 11 | 1.67% |
| hsa_circ_0068835 | 1315 | 11 | 1.67% |
| hsa_circ_0073763 | 1315 | 11 | 1.67% |
| hsa_circ_0089601 | 1315 | 11 | 1.67% |
| hsa_circ_0091451 | 1315 | 11 | 1.67% |
| hsa_circ_0103459 | 1315 | 11 | 1.67% |
| hsa_circ_0107390 | 1315 | 11 | 1.67% |
| hsa_circ_0029225 | 1435 | 12 | 1.67% |
| hsa_circ_0134725 | 1435 | 12 | 1.67% |
| hsa_circ_0001428 | 1196 | 10 | 1.67% |
| hsa_circ_0023913 | 1196 | 10 | 1.67% |
| hsa_circ_0024700 | 1196 | 10 | 1.67% |
| hsa_circ_0057687 | 1196 | 10 | 1.67% |
| hsa_circ_0064511 | 1196 | 10 | 1.67% |
| hsa_circ_0080889 | 1196 | 10 | 1.67% |
| hsa_circ_0087811 | 1196 | 10 | 1.67% |
| hsa_circ_0094933 | 1196 | 10 | 1.67% |
| hsa_circ_0112833 | 1196 | 10 | 1.67% |
| hsa_circ_0135158 | 1196 | 10 | 1.67% |
| hsa_circ_0140681 | 1196 | 10 | 1.67% |
| hsa_circ_0018151 | 1316 | 11 | 1.67% |
| hsa_circ_0031377 | 1316 | 11 | 1.67% |

| | | | |
|---|---|---|---|
| hsa_circ_0033372 | 1316 | 11 | 1.67% |
| hsa_circ_0039416 | 1316 | 11 | 1.67% |
| hsa_circ_0039551 | 1316 | 11 | 1.67% |
| hsa_circ_0063191 | 1316 | 11 | 1.67% |
| hsa_circ_0064577 | 1316 | 11 | 1.67% |
| hsa_circ_0076420 | 1316 | 11 | 1.67% |
| hsa_circ_0091446 | 1316 | 11 | 1.67% |
| hsa_circ_0097873 | 1316 | 11 | 1.67% |
| hsa_circ_0100304 | 1316 | 11 | 1.67% |
| hsa_circ_0101994 | 1316 | 11 | 1.67% |
| hsa_circ_0114636 | 1316 | 11 | 1.67% |
| hsa_circ_0050572 | 1436 | 12 | 1.67% |
| hsa_circ_0001543 | 1197 | 10 | 1.67% |
| hsa_circ_0012973 | 1197 | 10 | 1.67% |
| hsa_circ_0019179 | 1197 | 10 | 1.67% |
| hsa_circ_0023959 | 1197 | 10 | 1.67% |
| hsa_circ_0037161 | 1197 | 10 | 1.67% |
| hsa_circ_0040054 | 1197 | 10 | 1.67% |
| hsa_circ_0040670 | 1197 | 10 | 1.67% |
| hsa_circ_0068357 | 1197 | 10 | 1.67% |
| hsa_circ_0071625 | 1197 | 10 | 1.67% |
| hsa_circ_0080590 | 1197 | 10 | 1.67% |
| hsa_circ_0097121 | 1197 | 10 | 1.67% |
| hsa_circ_0098736 | 1197 | 10 | 1.67% |
| hsa_circ_0110629 | 1197 | 10 | 1.67% |
| hsa_circ_0116017 | 1197 | 10 | 1.67% |
| hsa_circ_0121943 | 1197 | 10 | 1.67% |
| hsa_circ_0001926 | 2873 | 24 | 1.67% |
| hsa_circ_0023001 | 1317 | 11 | 1.67% |
| hsa_circ_0025598 | 1317 | 11 | 1.67% |
| hsa_circ_0026465 | 1317 | 11 | 1.67% |
| hsa_circ_0028176 | 1317 | 11 | 1.67% |
| hsa_circ_0052859 | 1317 | 11 | 1.67% |
| hsa_circ_0053899 | 1317 | 11 | 1.67% |
| hsa_circ_0112425 | 1317 | 11 | 1.67% |
| hsa_circ_0036537 | 1437 | 12 | 1.67% |
| hsa_circ_0085462 | 1437 | 12 | 1.67% |
| hsa_circ_0087277 | 1437 | 12 | 1.67% |
| hsa_circ_0112260 | 1437 | 12 | 1.67% |
| hsa_circ_0116903 | 1437 | 12 | 1.67% |
| hsa_circ_0019371 | 1557 | 13 | 1.67% |
| hsa_circ_0084514 | 1557 | 13 | 1.67% |
| hsa_circ_0128859 | 2755 | 23 | 1.67% |
| hsa_circ_0048272 | 1677 | 14 | 1.67% |
| hsa_circ_0024800 | 1198 | 10 | 1.67% |
| hsa_circ_0055899 | 1198 | 10 | 1.67% |
| hsa_circ_0060269 | 1198 | 10 | 1.67% |
| hsa_circ_0060482 | 1198 | 10 | 1.67% |
| hsa_circ_0069019 | 1198 | 10 | 1.67% |
| hsa_circ_0080563 | 1198 | 10 | 1.67% |

| | | | |
|---|---|---|---|
| hsa_circ_0089099 | 1198 | 10 | 1.67% |
| hsa_circ_0098934 | 1198 | 10 | 1.67% |
| hsa_circ_0103268 | 1198 | 10 | 1.67% |
| hsa_circ_0137007 | 1198 | 10 | 1.67% |
| hsa_circ_0011717 | 1318 | 11 | 1.67% |
| hsa_circ_0036703 | 1318 | 11 | 1.67% |
| hsa_circ_0039964 | 1318 | 11 | 1.67% |
| hsa_circ_0104371 | 1318 | 11 | 1.67% |
| hsa_circ_0113804 | 1318 | 11 | 1.67% |
| hsa_circ_0039081 | 1438 | 12 | 1.67% |
| hsa_circ_0044466 | 1438 | 12 | 1.67% |
| hsa_circ_0131658 | 1438 | 12 | 1.67% |
| hsa_circ_0072560 | 1558 | 13 | 1.67% |
| hsa_circ_0076659 | 1558 | 13 | 1.67% |
| hsa_circ_0013051 | 1199 | 10 | 1.67% |
| hsa_circ_0017021 | 1199 | 10 | 1.67% |
| hsa_circ_0017063 | 1199 | 10 | 1.67% |
| hsa_circ_0024259 | 1199 | 10 | 1.67% |
| hsa_circ_0038969 | 1199 | 10 | 1.67% |
| hsa_circ_0090839 | 1199 | 10 | 1.67% |
| hsa_circ_0105500 | 1199 | 10 | 1.67% |
| hsa_circ_0124971 | 1199 | 10 | 1.67% |
| hsa_circ_0132316 | 1199 | 10 | 1.67% |
| hsa_circ_0136602 | 1199 | 10 | 1.67% |
| hsa_circ_0020924 | 1319 | 11 | 1.67% |
| hsa_circ_0052030 | 1319 | 11 | 1.67% |
| hsa_circ_0095566 | 1319 | 11 | 1.67% |
| hsa_circ_0123165 | 1319 | 11 | 1.67% |
| hsa_circ_0071491 | 1439 | 12 | 1.67% |
| hsa_circ_0111090 | 1439 | 12 | 1.67% |
| hsa_circ_0057231 | 1559 | 13 | 1.67% |
| hsa_circ_0068616 | 1559 | 13 | 1.67% |
| hsa_circ_0081410 | 1559 | 13 | 1.67% |
| hsa_circ_0078808 | 3240 | 27 | 1.67% |
| hsa_circ_0089039 | 1560 | 13 | 1.67% |
| hsa_circ_0013386 | 1440 | 12 | 1.67% |
| hsa_circ_0014710 | 1440 | 12 | 1.67% |
| hsa_circ_0077342 | 1440 | 12 | 1.67% |
| hsa_circ_0110002 | 1440 | 12 | 1.67% |
| hsa_circ_0013676 | 1320 | 11 | 1.67% |
| hsa_circ_0036192 | 1320 | 11 | 1.67% |
| hsa_circ_0042316 | 1320 | 11 | 1.67% |
| hsa_circ_0063401 | 1320 | 11 | 1.67% |
| hsa_circ_0091033 | 1320 | 11 | 1.67% |
| hsa_circ_0093158 | 1320 | 11 | 1.67% |
| hsa_circ_0101843 | 1320 | 11 | 1.67% |
| hsa_circ_0129783 | 1320 | 11 | 1.67% |
| hsa_circ_0135603 | 1320 | 11 | 1.67% |
| hsa_circ_0003562 | 1200 | 10 | 1.67% |
| hsa_circ_0023764 | 1200 | 10 | 1.67% |

| | | | |
|---|---|---|---|
| hsa_circ_0024077 | 1200 | 10 | 1.67% |
| hsa_circ_0044268 | 1200 | 10 | 1.67% |
| hsa_circ_0056307 | 1200 | 10 | 1.67% |
| hsa_circ_0056876 | 1200 | 10 | 1.67% |
| hsa_circ_0061932 | 1200 | 10 | 1.67% |
| hsa_circ_0062024 | 1200 | 10 | 1.67% |
| hsa_circ_0063765 | 1200 | 10 | 1.67% |
| hsa_circ_0069998 | 1200 | 10 | 1.67% |
| hsa_circ_0096939 | 1200 | 10 | 1.67% |
| hsa_circ_0102773 | 1200 | 10 | 1.67% |
| hsa_circ_0138804 | 1200 | 10 | 1.67% |
| hsa_circ_0078230 | 3001 | 25 | 1.67% |
| hsa_circ_0064144 | 2761 | 23 | 1.67% |
| hsa_circ_0134922 | 1561 | 13 | 1.67% |
| hsa_circ_0079103 | 1441 | 12 | 1.67% |
| hsa_circ_0021837 | 1321 | 11 | 1.67% |
| hsa_circ_0118021 | 1321 | 11 | 1.67% |
| hsa_circ_0119115 | 1321 | 11 | 1.67% |
| hsa_circ_0136070 | 1321 | 11 | 1.67% |
| hsa_circ_0018169 | 1201 | 10 | 1.67% |
| hsa_circ_0038303 | 1201 | 10 | 1.67% |
| hsa_circ_0047747 | 1201 | 10 | 1.67% |
| hsa_circ_0058590 | 1201 | 10 | 1.67% |
| hsa_circ_0066128 | 1201 | 10 | 1.67% |
| hsa_circ_0070192 | 1201 | 10 | 1.67% |
| hsa_circ_0085294 | 1201 | 10 | 1.67% |
| hsa_circ_0085784 | 1201 | 10 | 1.67% |
| hsa_circ_0088910 | 1201 | 10 | 1.67% |
| hsa_circ_0099416 | 1201 | 10 | 1.67% |
| hsa_circ_0116193 | 1201 | 10 | 1.67% |
| hsa_circ_0136787 | 1201 | 10 | 1.67% |
| hsa_circ_0137367 | 1201 | 10 | 1.67% |
| hsa_circ_0044828 | 1802 | 15 | 1.66% |
| hsa_circ_0071505 | 1442 | 12 | 1.66% |
| hsa_circ_0111178 | 1442 | 12 | 1.66% |
| hsa_circ_0026395 | 1322 | 11 | 1.66% |
| hsa_circ_0028175 | 1322 | 11 | 1.66% |
| hsa_circ_0035937 | 1322 | 11 | 1.66% |
| hsa_circ_0046364 | 1322 | 11 | 1.66% |
| hsa_circ_0063002 | 1322 | 11 | 1.66% |
| hsa_circ_0085592 | 1322 | 11 | 1.66% |
| hsa_circ_0089419 | 1322 | 11 | 1.66% |
| hsa_circ_0018265 | 1202 | 10 | 1.66% |
| hsa_circ_0021144 | 1202 | 10 | 1.66% |
| hsa_circ_0024506 | 1202 | 10 | 1.66% |
| hsa_circ_0060849 | 1202 | 10 | 1.66% |
| hsa_circ_0072483 | 1202 | 10 | 1.66% |
| hsa_circ_0080459 | 1202 | 10 | 1.66% |
| hsa_circ_0103001 | 1202 | 10 | 1.66% |
| hsa_circ_0110533 | 1202 | 10 | 1.66% |

| | | | |
|---|---|---|---|
| hsa_circ_0129052 | 1202 | 10 | 1.66% |
| hsa_circ_0024399 | 1563 | 13 | 1.66% |
| hsa_circ_0078229 | 3006 | 25 | 1.66% |
| hsa_circ_0029565 | 1323 | 11 | 1.66% |
| hsa_circ_0003448 | 1203 | 10 | 1.66% |
| hsa_circ_0008830 | 1203 | 10 | 1.66% |
| hsa_circ_0016240 | 1203 | 10 | 1.66% |
| hsa_circ_0018147 | 1203 | 10 | 1.66% |
| hsa_circ_0021679 | 1203 | 10 | 1.66% |
| hsa_circ_0025640 | 1203 | 10 | 1.66% |
| hsa_circ_0039726 | 1203 | 10 | 1.66% |
| hsa_circ_0042829 | 1203 | 10 | 1.66% |
| hsa_circ_0044447 | 1203 | 10 | 1.66% |
| hsa_circ_0048665 | 1203 | 10 | 1.66% |
| hsa_circ_0072835 | 1203 | 10 | 1.66% |
| hsa_circ_0078147 | 1203 | 10 | 1.66% |
| hsa_circ_0079876 | 1203 | 10 | 1.66% |
| hsa_circ_0088993 | 1203 | 10 | 1.66% |
| hsa_circ_0099345 | 1203 | 10 | 1.66% |
| hsa_circ_0101071 | 1203 | 10 | 1.66% |
| hsa_circ_0103847 | 1203 | 10 | 1.66% |
| hsa_circ_0126309 | 1203 | 10 | 1.66% |
| hsa_circ_0138123 | 1203 | 10 | 1.66% |
| hsa_circ_0095403 | 1444 | 12 | 1.66% |
| hsa_circ_0029826 | 1324 | 11 | 1.66% |
| hsa_circ_0038506 | 1324 | 11 | 1.66% |
| hsa_circ_0040184 | 1324 | 11 | 1.66% |
| hsa_circ_0061364 | 1324 | 11 | 1.66% |
| hsa_circ_0065249 | 1324 | 11 | 1.66% |
| hsa_circ_0071402 | 1324 | 11 | 1.66% |
| hsa_circ_0081227 | 1324 | 11 | 1.66% |
| hsa_circ_0113947 | 1324 | 11 | 1.66% |
| hsa_circ_0117999 | 1324 | 11 | 1.66% |
| hsa_circ_0137662 | 1324 | 11 | 1.66% |
| hsa_circ_0001098 | 1204 | 10 | 1.66% |
| hsa_circ_0004145 | 1204 | 10 | 1.66% |
| hsa_circ_0004630 | 1204 | 10 | 1.66% |
| hsa_circ_0051091 | 1204 | 10 | 1.66% |
| hsa_circ_0072518 | 1204 | 10 | 1.66% |
| hsa_circ_0089217 | 1204 | 10 | 1.66% |
| hsa_circ_0090781 | 1204 | 10 | 1.66% |
| hsa_circ_0092567 | 1204 | 10 | 1.66% |
| hsa_circ_0097826 | 1204 | 10 | 1.66% |
| hsa_circ_0109769 | 1204 | 10 | 1.66% |
| hsa_circ_0119751 | 1204 | 10 | 1.66% |
| hsa_circ_0122520 | 1204 | 10 | 1.66% |
| hsa_circ_0132127 | 1204 | 10 | 1.66% |
| hsa_circ_0048055 | 1445 | 12 | 1.66% |
| hsa_circ_0079526 | 1445 | 12 | 1.66% |
| hsa_circ_0006967 | 1325 | 11 | 1.66% |

| | | | |
|---|---|---|---|
| hsa_circ_0045057 | 1325 | 11 | 1.66% |
| hsa_circ_0063168 | 1325 | 11 | 1.66% |
| hsa_circ_0078436 | 1325 | 11 | 1.66% |
| hsa_circ_0091098 | 1325 | 11 | 1.66% |
| hsa_circ_0094120 | 1566 | 13 | 1.66% |
| hsa_circ_0050621 | 1446 | 12 | 1.66% |
| hsa_circ_0091096 | 1446 | 12 | 1.66% |
| hsa_circ_0011512 | 1205 | 10 | 1.66% |
| hsa_circ_0060848 | 1205 | 10 | 1.66% |
| hsa_circ_0075458 | 1205 | 10 | 1.66% |
| hsa_circ_0084721 | 1205 | 10 | 1.66% |
| hsa_circ_0089072 | 1205 | 10 | 1.66% |
| hsa_circ_0093768 | 1205 | 10 | 1.66% |
| hsa_circ_0100015 | 1205 | 10 | 1.66% |
| hsa_circ_0101524 | 1205 | 10 | 1.66% |
| hsa_circ_0103013 | 1205 | 10 | 1.66% |
| hsa_circ_0121389 | 1205 | 10 | 1.66% |
| hsa_circ_0135448 | 1205 | 10 | 1.66% |
| hsa_circ_0042713 | 1326 | 11 | 1.66% |
| hsa_circ_0044406 | 1326 | 11 | 1.66% |
| hsa_circ_0054234 | 1326 | 11 | 1.66% |
| hsa_circ_0059584 | 1326 | 11 | 1.66% |
| hsa_circ_0077810 | 1326 | 11 | 1.66% |
| hsa_circ_0100521 | 1326 | 11 | 1.66% |
| hsa_circ_0139676 | 1326 | 11 | 1.66% |
| hsa_circ_0098110 | 4341 | 36 | 1.66% |
| hsa_circ_0115982 | 1447 | 12 | 1.66% |
| hsa_circ_0021723 | 1206 | 10 | 1.66% |
| hsa_circ_0022150 | 1206 | 10 | 1.66% |
| hsa_circ_0028651 | 1206 | 10 | 1.66% |
| hsa_circ_0036907 | 1206 | 10 | 1.66% |
| hsa_circ_0047234 | 1206 | 10 | 1.66% |
| hsa_circ_0048298 | 1206 | 10 | 1.66% |
| hsa_circ_0051223 | 1206 | 10 | 1.66% |
| hsa_circ_0054523 | 1206 | 10 | 1.66% |
| hsa_circ_0059573 | 1206 | 10 | 1.66% |
| hsa_circ_0072689 | 1206 | 10 | 1.66% |
| hsa_circ_0073019 | 1206 | 10 | 1.66% |
| hsa_circ_0074279 | 1206 | 10 | 1.66% |
| hsa_circ_0076097 | 1206 | 10 | 1.66% |
| hsa_circ_0077225 | 1206 | 10 | 1.66% |
| hsa_circ_0085449 | 1206 | 10 | 1.66% |
| hsa_circ_0088196 | 1206 | 10 | 1.66% |
| hsa_circ_0090313 | 1206 | 10 | 1.66% |
| hsa_circ_0092117 | 1206 | 10 | 1.66% |
| hsa_circ_0092672 | 1206 | 10 | 1.66% |
| hsa_circ_0096218 | 1206 | 10 | 1.66% |
| hsa_circ_0100324 | 1206 | 10 | 1.66% |
| hsa_circ_0101280 | 1206 | 10 | 1.66% |
| hsa_circ_0106337 | 1206 | 10 | 1.66% |

| | | | |
|---|---|---|---|
| hsa_circ_0114995 | 1206 | 10 | 1.66% |
| hsa_circ_0126155 | 1206 | 10 | 1.66% |
| hsa_circ_0040005 | 1327 | 11 | 1.66% |
| hsa_circ_0068568 | 1327 | 11 | 1.66% |
| hsa_circ_0078470 | 1327 | 11 | 1.66% |
| hsa_circ_0099802 | 1327 | 11 | 1.66% |
| hsa_circ_0112357 | 1327 | 11 | 1.66% |
| hsa_circ_0066417 | 1448 | 12 | 1.66% |
| hsa_circ_0083791 | 1448 | 12 | 1.66% |
| hsa_circ_0118905 | 1448 | 12 | 1.66% |
| hsa_circ_0137496 | 1448 | 12 | 1.66% |
| hsa_circ_0087469 | 1569 | 13 | 1.66% |
| hsa_circ_0011010 | 1207 | 10 | 1.66% |
| hsa_circ_0014187 | 1207 | 10 | 1.66% |
| hsa_circ_0024339 | 1207 | 10 | 1.66% |
| hsa_circ_0036811 | 1207 | 10 | 1.66% |
| hsa_circ_0041352 | 1207 | 10 | 1.66% |
| hsa_circ_0047293 | 1207 | 10 | 1.66% |
| hsa_circ_0094512 | 1207 | 10 | 1.66% |
| hsa_circ_0094741 | 1207 | 10 | 1.66% |
| hsa_circ_0110329 | 1207 | 10 | 1.66% |
| hsa_circ_0115303 | 1207 | 10 | 1.66% |
| hsa_circ_0123619 | 1207 | 10 | 1.66% |
| hsa_circ_0130498 | 1207 | 10 | 1.66% |
| hsa_circ_0131824 | 1207 | 10 | 1.66% |
| hsa_circ_0119706 | 3259 | 27 | 1.66% |
| hsa_circ_0050483 | 1328 | 11 | 1.66% |
| hsa_circ_0083296 | 1328 | 11 | 1.66% |
| hsa_circ_0099988 | 1328 | 11 | 1.66% |
| hsa_circ_0122294 | 1328 | 11 | 1.66% |
| hsa_circ_0000077 | 3622 | 30 | 1.66% |
| hsa_circ_0010697 | 1449 | 12 | 1.66% |
| hsa_circ_0013138 | 1449 | 12 | 1.66% |
| hsa_circ_0137426 | 1449 | 12 | 1.66% |
| hsa_circ_0056286 | 1570 | 13 | 1.66% |
| hsa_circ_0005947 | 1208 | 10 | 1.66% |
| hsa_circ_0006039 | 1208 | 10 | 1.66% |
| hsa_circ_0073631 | 1208 | 10 | 1.66% |
| hsa_circ_0083114 | 1208 | 10 | 1.66% |
| hsa_circ_0099710 | 1208 | 10 | 1.66% |
| hsa_circ_0110328 | 1208 | 10 | 1.66% |
| hsa_circ_0129712 | 1208 | 10 | 1.66% |
| hsa_circ_0010419 | 1329 | 11 | 1.66% |
| hsa_circ_0023769 | 1329 | 11 | 1.66% |
| hsa_circ_0033307 | 1329 | 11 | 1.66% |
| hsa_circ_0036194 | 1329 | 11 | 1.66% |
| hsa_circ_0065709 | 1329 | 11 | 1.66% |
| hsa_circ_0129130 | 1329 | 11 | 1.66% |
| hsa_circ_0050393 | 1450 | 12 | 1.66% |
| hsa_circ_0089229 | 1450 | 12 | 1.66% |

| | | | |
|---|---|---|---|
| hsa_circ_0024545 | 1571 | 13 | 1.65% |
| hsa_circ_0009071 | 1209 | 10 | 1.65% |
| hsa_circ_0017434 | 1209 | 10 | 1.65% |
| hsa_circ_0019174 | 1209 | 10 | 1.65% |
| hsa_circ_0031187 | 1209 | 10 | 1.65% |
| hsa_circ_0044305 | 1209 | 10 | 1.65% |
| hsa_circ_0045696 | 1209 | 10 | 1.65% |
| hsa_circ_0046658 | 1209 | 10 | 1.65% |
| hsa_circ_0054544 | 1209 | 10 | 1.65% |
| hsa_circ_0056380 | 1209 | 10 | 1.65% |
| hsa_circ_0060088 | 1209 | 10 | 1.65% |
| hsa_circ_0060660 | 1209 | 10 | 1.65% |
| hsa_circ_0063156 | 1209 | 10 | 1.65% |
| hsa_circ_0066038 | 1209 | 10 | 1.65% |
| hsa_circ_0068094 | 1209 | 10 | 1.65% |
| hsa_circ_0068980 | 1209 | 10 | 1.65% |
| hsa_circ_0085603 | 1209 | 10 | 1.65% |
| hsa_circ_0113994 | 1209 | 10 | 1.65% |
| hsa_circ_0140629 | 1209 | 10 | 1.65% |
| hsa_circ_0010710 | 1330 | 11 | 1.65% |
| hsa_circ_0010751 | 1330 | 11 | 1.65% |
| hsa_circ_0040596 | 1330 | 11 | 1.65% |
| hsa_circ_0076483 | 1330 | 11 | 1.65% |
| hsa_circ_0121981 | 1330 | 11 | 1.65% |
| hsa_circ_0058571 | 1451 | 12 | 1.65% |
| hsa_circ_0098306 | 1451 | 12 | 1.65% |
| hsa_circ_0015561 | 1572 | 13 | 1.65% |
| hsa_circ_0024261 | 1936 | 16 | 1.65% |
| hsa_circ_0050705 | 1452 | 12 | 1.65% |
| hsa_circ_0079514 | 1452 | 12 | 1.65% |
| hsa_circ_0053338 | 1331 | 11 | 1.65% |
| hsa_circ_0069011 | 1331 | 11 | 1.65% |
| hsa_circ_0110222 | 1331 | 11 | 1.65% |
| hsa_circ_0115792 | 1331 | 11 | 1.65% |
| hsa_circ_0018280 | 1210 | 10 | 1.65% |
| hsa_circ_0036293 | 1210 | 10 | 1.65% |
| hsa_circ_0038921 | 1210 | 10 | 1.65% |
| hsa_circ_0045302 | 1210 | 10 | 1.65% |
| hsa_circ_0064359 | 1210 | 10 | 1.65% |
| hsa_circ_0080197 | 1210 | 10 | 1.65% |
| hsa_circ_0085953 | 1210 | 10 | 1.65% |
| hsa_circ_0086236 | 1210 | 10 | 1.65% |
| hsa_circ_0093766 | 1210 | 10 | 1.65% |
| hsa_circ_0103544 | 1210 | 10 | 1.65% |
| hsa_circ_0133527 | 1210 | 10 | 1.65% |
| hsa_circ_0138857 | 1210 | 10 | 1.65% |
| hsa_circ_0140081 | 1210 | 10 | 1.65% |
| hsa_circ_0040876 | 1695 | 14 | 1.65% |
| hsa_circ_0052069 | 1695 | 14 | 1.65% |
| hsa_circ_0047948 | 1574 | 13 | 1.65% |

| | | | |
|---|---|---|---|
| hsa_circ_0009284 | 1332 | 11 | 1.65% |
| hsa_circ_0032250 | 1332 | 11 | 1.65% |
| hsa_circ_0049327 | 1332 | 11 | 1.65% |
| hsa_circ_0026932 | 1211 | 10 | 1.65% |
| hsa_circ_0037204 | 1211 | 10 | 1.65% |
| hsa_circ_0063489 | 1211 | 10 | 1.65% |
| hsa_circ_0072351 | 1211 | 10 | 1.65% |
| hsa_circ_0089381 | 1211 | 10 | 1.65% |
| hsa_circ_0106285 | 1211 | 10 | 1.65% |
| hsa_circ_0112331 | 1211 | 10 | 1.65% |
| hsa_circ_0129674 | 1211 | 10 | 1.65% |
| hsa_circ_0130775 | 1211 | 10 | 1.65% |
| hsa_circ_0135664 | 1211 | 10 | 1.65% |
| hsa_circ_0019948 | 1575 | 13 | 1.65% |
| hsa_circ_0067635 | 1575 | 13 | 1.65% |
| hsa_circ_0024381 | 1454 | 12 | 1.65% |
| hsa_circ_0037116 | 1454 | 12 | 1.65% |
| hsa_circ_0122135 | 1454 | 12 | 1.65% |
| hsa_circ_0116403 | 4968 | 41 | 1.65% |
| hsa_circ_0030098 | 1333 | 11 | 1.65% |
| hsa_circ_0037332 | 1333 | 11 | 1.65% |
| hsa_circ_0083571 | 1818 | 15 | 1.65% |
| hsa_circ_0003696 | 1212 | 10 | 1.65% |
| hsa_circ_0009763 | 1212 | 10 | 1.65% |
| hsa_circ_0012184 | 1212 | 10 | 1.65% |
| hsa_circ_0013851 | 1212 | 10 | 1.65% |
| hsa_circ_0015482 | 1212 | 10 | 1.65% |
| hsa_circ_0023619 | 1212 | 10 | 1.65% |
| hsa_circ_0035851 | 1212 | 10 | 1.65% |
| hsa_circ_0037769 | 1212 | 10 | 1.65% |
| hsa_circ_0040566 | 1212 | 10 | 1.65% |
| hsa_circ_0041157 | 1212 | 10 | 1.65% |
| hsa_circ_0047489 | 1212 | 10 | 1.65% |
| hsa_circ_0048902 | 1212 | 10 | 1.65% |
| hsa_circ_0049311 | 1212 | 10 | 1.65% |
| hsa_circ_0065438 | 1212 | 10 | 1.65% |
| hsa_circ_0076425 | 1212 | 10 | 1.65% |
| hsa_circ_0085241 | 1212 | 10 | 1.65% |
| hsa_circ_0087502 | 1212 | 10 | 1.65% |
| hsa_circ_0088700 | 1212 | 10 | 1.65% |
| hsa_circ_0091923 | 1212 | 10 | 1.65% |
| hsa_circ_0094609 | 1212 | 10 | 1.65% |
| hsa_circ_0131020 | 1212 | 10 | 1.65% |
| hsa_circ_0131148 | 1212 | 10 | 1.65% |
| hsa_circ_0134810 | 1212 | 10 | 1.65% |
| hsa_circ_0132728 | 2061 | 17 | 1.65% |
| hsa_circ_0015138 | 3031 | 25 | 1.65% |
| hsa_circ_0048556 | 1455 | 12 | 1.65% |
| hsa_circ_0058162 | 1455 | 12 | 1.65% |
| hsa_circ_0088181 | 1455 | 12 | 1.65% |

| | | | |
|---|---|---|---|
| hsa_circ_0127219 | 1455 | 12 | 1.65% |
| hsa_circ_0019069 | 1334 | 11 | 1.65% |
| hsa_circ_0019265 | 1334 | 11 | 1.65% |
| hsa_circ_0021972 | 1334 | 11 | 1.65% |
| hsa_circ_0049315 | 1334 | 11 | 1.65% |
| hsa_circ_0052270 | 1334 | 11 | 1.65% |
| hsa_circ_0099995 | 1334 | 11 | 1.65% |
| hsa_circ_0026441 | 1213 | 10 | 1.65% |
| hsa_circ_0026797 | 1213 | 10 | 1.65% |
| hsa_circ_0030261 | 1213 | 10 | 1.65% |
| hsa_circ_0032010 | 1213 | 10 | 1.65% |
| hsa_circ_0044722 | 1213 | 10 | 1.65% |
| hsa_circ_0048246 | 1213 | 10 | 1.65% |
| hsa_circ_0076550 | 1213 | 10 | 1.65% |
| hsa_circ_0086955 | 1213 | 10 | 1.65% |
| hsa_circ_0088812 | 1213 | 10 | 1.65% |
| hsa_circ_0118130 | 1213 | 10 | 1.65% |
| hsa_circ_0047272 | 1456 | 12 | 1.65% |
| hsa_circ_0043617 | 1335 | 11 | 1.65% |
| hsa_circ_0069448 | 1335 | 11 | 1.65% |
| hsa_circ_0077144 | 1335 | 11 | 1.65% |
| hsa_circ_0106039 | 1335 | 11 | 1.65% |
| hsa_circ_0102537 | 4249 | 35 | 1.65% |
| hsa_circ_0009769 | 1214 | 10 | 1.65% |
| hsa_circ_0045169 | 1214 | 10 | 1.65% |
| hsa_circ_0055080 | 1214 | 10 | 1.65% |
| hsa_circ_0055828 | 1214 | 10 | 1.65% |
| hsa_circ_0068224 | 1214 | 10 | 1.65% |
| hsa_circ_0071010 | 1214 | 10 | 1.65% |
| hsa_circ_0073094 | 1214 | 10 | 1.65% |
| hsa_circ_0083124 | 1214 | 10 | 1.65% |
| hsa_circ_0088819 | 1214 | 10 | 1.65% |
| hsa_circ_0116446 | 1214 | 10 | 1.65% |
| hsa_circ_0132659 | 1214 | 10 | 1.65% |
| hsa_circ_0137351 | 1214 | 10 | 1.65% |
| hsa_circ_0002617 | 1457 | 12 | 1.65% |
| hsa_circ_0053043 | 1457 | 12 | 1.65% |
| hsa_circ_0074575 | 1457 | 12 | 1.65% |
| hsa_circ_0043206 | 1336 | 11 | 1.65% |
| hsa_circ_0052215 | 1336 | 11 | 1.65% |
| hsa_circ_0053703 | 1336 | 11 | 1.65% |
| hsa_circ_0067461 | 1336 | 11 | 1.65% |
| hsa_circ_0106970 | 1336 | 11 | 1.65% |
| hsa_circ_0069810 | 1458 | 12 | 1.65% |
| hsa_circ_0028677 | 1215 | 10 | 1.65% |
| hsa_circ_0036145 | 1215 | 10 | 1.65% |
| hsa_circ_0039302 | 1215 | 10 | 1.65% |
| hsa_circ_0058811 | 1215 | 10 | 1.65% |
| hsa_circ_0064422 | 1215 | 10 | 1.65% |
| hsa_circ_0079057 | 1215 | 10 | 1.65% |

| | | | |
|---|---|---|---|
| hsa_circ_0080929 | 1215 | 10 | 1.65% |
| hsa_circ_0082388 | 1215 | 10 | 1.65% |
| hsa_circ_0084949 | 1215 | 10 | 1.65% |
| hsa_circ_0085255 | 1215 | 10 | 1.65% |
| hsa_circ_0086950 | 1215 | 10 | 1.65% |
| hsa_circ_0087768 | 1215 | 10 | 1.65% |
| hsa_circ_0095946 | 1215 | 10 | 1.65% |
| hsa_circ_0118011 | 1215 | 10 | 1.65% |
| hsa_circ_0123213 | 1215 | 10 | 1.65% |
| hsa_circ_0127431 | 1215 | 10 | 1.65% |
| hsa_circ_0129389 | 1215 | 10 | 1.65% |
| hsa_circ_0131306 | 1215 | 10 | 1.65% |
| hsa_circ_0136841 | 1215 | 10 | 1.65% |
| hsa_circ_0021984 | 1580 | 13 | 1.65% |
| hsa_circ_0038967 | 1337 | 11 | 1.65% |
| hsa_circ_0046351 | 1337 | 11 | 1.65% |
| hsa_circ_0057800 | 1337 | 11 | 1.65% |
| hsa_circ_0121247 | 1337 | 11 | 1.65% |
| hsa_circ_0052127 | 1702 | 14 | 1.65% |
| hsa_circ_0010855 | 1459 | 12 | 1.64% |
| hsa_circ_0012698 | 1459 | 12 | 1.64% |
| hsa_circ_0003650 | 1216 | 10 | 1.64% |
| hsa_circ_0018841 | 1216 | 10 | 1.64% |
| hsa_circ_0029215 | 1216 | 10 | 1.64% |
| hsa_circ_0039845 | 1216 | 10 | 1.64% |
| hsa_circ_0056021 | 1216 | 10 | 1.64% |
| hsa_circ_0074505 | 1216 | 10 | 1.64% |
| hsa_circ_0111405 | 1216 | 10 | 1.64% |
| hsa_circ_0132377 | 1216 | 10 | 1.64% |
| hsa_circ_0138956 | 1216 | 10 | 1.64% |
| hsa_circ_0042883 | 1581 | 13 | 1.64% |
| hsa_circ_0120215 | 1581 | 13 | 1.64% |
| hsa_circ_0007846 | 1338 | 11 | 1.64% |
| hsa_circ_0009397 | 1338 | 11 | 1.64% |
| hsa_circ_0013501 | 1338 | 11 | 1.64% |
| hsa_circ_0024764 | 1338 | 11 | 1.64% |
| hsa_circ_0038348 | 1338 | 11 | 1.64% |
| hsa_circ_0077827 | 1338 | 11 | 1.64% |
| hsa_circ_0083158 | 1338 | 11 | 1.64% |
| hsa_circ_0083646 | 1338 | 11 | 1.64% |
| hsa_circ_0107527 | 1338 | 11 | 1.64% |
| hsa_circ_0128203 | 1338 | 11 | 1.64% |
| hsa_circ_0135862 | 1338 | 11 | 1.64% |
| hsa_circ_0136117 | 1338 | 11 | 1.64% |
| hsa_circ_0131015 | 1825 | 15 | 1.64% |
| hsa_circ_0024599 | 1460 | 12 | 1.64% |
| hsa_circ_0088740 | 1460 | 12 | 1.64% |
| hsa_circ_0052210 | 1582 | 13 | 1.64% |
| hsa_circ_0018486 | 1217 | 10 | 1.64% |
| hsa_circ_0021476 | 1217 | 10 | 1.64% |

| | | | |
|---|---|---|---|
| hsa_circ_0026259 | 1217 | 10 | 1.64% |
| hsa_circ_0031273 | 1217 | 10 | 1.64% |
| hsa_circ_0038861 | 1217 | 10 | 1.64% |
| hsa_circ_0049131 | 1217 | 10 | 1.64% |
| hsa_circ_0060350 | 1217 | 10 | 1.64% |
| hsa_circ_0065528 | 1217 | 10 | 1.64% |
| hsa_circ_0067378 | 1217 | 10 | 1.64% |
| hsa_circ_0079567 | 1217 | 10 | 1.64% |
| hsa_circ_0085084 | 1217 | 10 | 1.64% |
| hsa_circ_0090351 | 1217 | 10 | 1.64% |
| hsa_circ_0095042 | 1217 | 10 | 1.64% |
| hsa_circ_0107375 | 1217 | 10 | 1.64% |
| hsa_circ_0116194 | 1217 | 10 | 1.64% |
| hsa_circ_0122936 | 1217 | 10 | 1.64% |
| hsa_circ_0103122 | 4869 | 40 | 1.64% |
| hsa_circ_0046820 | 1339 | 11 | 1.64% |
| hsa_circ_0049205 | 1339 | 11 | 1.64% |
| hsa_circ_0083950 | 1339 | 11 | 1.64% |
| hsa_circ_0013057 | 1461 | 12 | 1.64% |
| hsa_circ_0030401 | 1461 | 12 | 1.64% |
| hsa_circ_0039374 | 1461 | 12 | 1.64% |
| hsa_circ_0121580 | 2557 | 21 | 1.64% |
| hsa_circ_0057390 | 2070 | 17 | 1.64% |
| hsa_circ_0003051 | 1218 | 10 | 1.64% |
| hsa_circ_0016668 | 1218 | 10 | 1.64% |
| hsa_circ_0020385 | 1218 | 10 | 1.64% |
| hsa_circ_0039367 | 1218 | 10 | 1.64% |
| hsa_circ_0068287 | 1218 | 10 | 1.64% |
| hsa_circ_0081485 | 1218 | 10 | 1.64% |
| hsa_circ_0082174 | 1218 | 10 | 1.64% |
| hsa_circ_0102861 | 1218 | 10 | 1.64% |
| hsa_circ_0112580 | 1218 | 10 | 1.64% |
| hsa_circ_0123292 | 1218 | 10 | 1.64% |
| hsa_circ_0126800 | 1218 | 10 | 1.64% |
| hsa_circ_0124173 | 3167 | 26 | 1.64% |
| hsa_circ_0021385 | 1340 | 11 | 1.64% |
| hsa_circ_0040043 | 1340 | 11 | 1.64% |
| hsa_circ_0069805 | 1340 | 11 | 1.64% |
| hsa_circ_0108246 | 1340 | 11 | 1.64% |
| hsa_circ_0119460 | 1340 | 11 | 1.64% |
| hsa_circ_0124599 | 1340 | 11 | 1.64% |
| hsa_circ_0035867 | 1462 | 12 | 1.64% |
| hsa_circ_0101623 | 1462 | 12 | 1.64% |
| hsa_circ_0103547 | 1462 | 12 | 1.64% |
| hsa_circ_0044432 | 1584 | 13 | 1.64% |
| hsa_circ_0072620 | 1584 | 13 | 1.64% |
| hsa_circ_0013890 | 1219 | 10 | 1.64% |
| hsa_circ_0018919 | 1219 | 10 | 1.64% |
| hsa_circ_0024928 | 1219 | 10 | 1.64% |
| hsa_circ_0032053 | 1219 | 10 | 1.64% |

| | | | |
|---|---|---|---|
| hsa_circ_0043172 | 1219 | 10 | 1.64% |
| hsa_circ_0055287 | 1219 | 10 | 1.64% |
| hsa_circ_0056446 | 1219 | 10 | 1.64% |
| hsa_circ_0068396 | 1219 | 10 | 1.64% |
| hsa_circ_0130824 | 1219 | 10 | 1.64% |
| hsa_circ_0019216 | 1341 | 11 | 1.64% |
| hsa_circ_0045632 | 1341 | 11 | 1.64% |
| hsa_circ_0048422 | 1341 | 11 | 1.64% |
| hsa_circ_0085266 | 1341 | 11 | 1.64% |
| hsa_circ_0099817 | 1341 | 11 | 1.64% |
| hsa_circ_0088043 | 1463 | 12 | 1.64% |
| hsa_circ_0099622 | 1463 | 12 | 1.64% |
| hsa_circ_0024779 | 1464 | 12 | 1.64% |
| hsa_circ_0033090 | 1464 | 12 | 1.64% |
| hsa_circ_0048786 | 1464 | 12 | 1.64% |
| hsa_circ_0054839 | 1464 | 12 | 1.64% |
| hsa_circ_0089731 | 1464 | 12 | 1.64% |
| hsa_circ_0027970 | 1342 | 11 | 1.64% |
| hsa_circ_0041434 | 1342 | 11 | 1.64% |
| hsa_circ_0053303 | 1342 | 11 | 1.64% |
| hsa_circ_0087137 | 1342 | 11 | 1.64% |
| hsa_circ_0108144 | 1342 | 11 | 1.64% |
| hsa_circ_0130651 | 1342 | 11 | 1.64% |
| hsa_circ_0136669 | 1342 | 11 | 1.64% |
| hsa_circ_0015270 | 1220 | 10 | 1.64% |
| hsa_circ_0016690 | 1220 | 10 | 1.64% |
| hsa_circ_0028479 | 1220 | 10 | 1.64% |
| hsa_circ_0029416 | 1220 | 10 | 1.64% |
| hsa_circ_0034520 | 1220 | 10 | 1.64% |
| hsa_circ_0043152 | 1220 | 10 | 1.64% |
| hsa_circ_0058418 | 1220 | 10 | 1.64% |
| hsa_circ_0065779 | 1220 | 10 | 1.64% |
| hsa_circ_0074417 | 1220 | 10 | 1.64% |
| hsa_circ_0075042 | 1220 | 10 | 1.64% |
| hsa_circ_0076630 | 1220 | 10 | 1.64% |
| hsa_circ_0080869 | 1220 | 10 | 1.64% |
| hsa_circ_0080994 | 1220 | 10 | 1.64% |
| hsa_circ_0086493 | 1220 | 10 | 1.64% |
| hsa_circ_0120122 | 1220 | 10 | 1.64% |
| hsa_circ_0136176 | 1220 | 10 | 1.64% |
| hsa_circ_0137798 | 1220 | 10 | 1.64% |
| hsa_circ_0139240 | 1220 | 10 | 1.64% |
| hsa_circ_0058239 | 1831 | 15 | 1.64% |
| hsa_circ_0102790 | 1465 | 12 | 1.64% |
| hsa_circ_0003152 | 1343 | 11 | 1.64% |
| hsa_circ_0037236 | 1343 | 11 | 1.64% |
| hsa_circ_0089733 | 1343 | 11 | 1.64% |
| hsa_circ_0106915 | 1343 | 11 | 1.64% |
| hsa_circ_0021306 | 1221 | 10 | 1.64% |
| hsa_circ_0022437 | 1221 | 10 | 1.64% |

| | | | |
|---|---|---|---|
| hsa_circ_0026376 | 1221 | 10 | 1.64% |
| hsa_circ_0033404 | 1221 | 10 | 1.64% |
| hsa_circ_0047135 | 1221 | 10 | 1.64% |
| hsa_circ_0058016 | 1221 | 10 | 1.64% |
| hsa_circ_0059444 | 1221 | 10 | 1.64% |
| hsa_circ_0078669 | 1221 | 10 | 1.64% |
| hsa_circ_0079226 | 1221 | 10 | 1.64% |
| hsa_circ_0086826 | 1221 | 10 | 1.64% |
| hsa_circ_0089517 | 1221 | 10 | 1.64% |
| hsa_circ_0122795 | 1221 | 10 | 1.64% |
| hsa_circ_0123744 | 1221 | 10 | 1.64% |
| hsa_circ_0125958 | 1221 | 10 | 1.64% |
| hsa_circ_0127753 | 1221 | 10 | 1.64% |
| hsa_circ_0135523 | 1221 | 10 | 1.64% |
| hsa_circ_0140017 | 1221 | 10 | 1.64% |
| hsa_circ_0114454 | 2809 | 23 | 1.64% |
| hsa_circ_0118417 | 1466 | 12 | 1.64% |
| hsa_circ_0028302 | 1344 | 11 | 1.64% |
| hsa_circ_0039429 | 1344 | 11 | 1.64% |
| hsa_circ_0087865 | 1344 | 11 | 1.64% |
| hsa_circ_0092022 | 1344 | 11 | 1.64% |
| hsa_circ_0101008 | 1344 | 11 | 1.64% |
| hsa_circ_0106004 | 1344 | 11 | 1.64% |
| hsa_circ_0139724 | 1344 | 11 | 1.64% |
| hsa_circ_0106984 | 1833 | 15 | 1.64% |
| hsa_circ_0015880 | 1222 | 10 | 1.64% |
| hsa_circ_0034506 | 1222 | 10 | 1.64% |
| hsa_circ_0036488 | 1222 | 10 | 1.64% |
| hsa_circ_0054091 | 1222 | 10 | 1.64% |
| hsa_circ_0071469 | 1222 | 10 | 1.64% |
| hsa_circ_0100356 | 1222 | 10 | 1.64% |
| hsa_circ_0102671 | 1222 | 10 | 1.64% |
| hsa_circ_0105665 | 1222 | 10 | 1.64% |
| hsa_circ_0108656 | 1222 | 10 | 1.64% |
| hsa_circ_0114861 | 1222 | 10 | 1.64% |
| hsa_circ_0137718 | 1222 | 10 | 1.64% |
| hsa_circ_0038060 | 1711 | 14 | 1.64% |
| hsa_circ_0019990 | 1589 | 13 | 1.64% |
| hsa_circ_0020144 | 1589 | 13 | 1.64% |
| hsa_circ_0071453 | 1467 | 12 | 1.64% |
| hsa_circ_0102801 | 1467 | 12 | 1.64% |
| hsa_circ_0133504 | 1467 | 12 | 1.64% |
| hsa_circ_0035104 | 1345 | 11 | 1.64% |
| hsa_circ_0058795 | 1345 | 11 | 1.64% |
| hsa_circ_0063004 | 1345 | 11 | 1.64% |
| hsa_circ_0070005 | 1345 | 11 | 1.64% |
| hsa_circ_0081531 | 1345 | 11 | 1.64% |
| hsa_circ_0090797 | 1345 | 11 | 1.64% |
| hsa_circ_0092682 | 1345 | 11 | 1.64% |
| hsa_circ_0096259 | 1345 | 11 | 1.64% |

| | | | |
|---|---|---|---|
| hsa_circ_0106483 | 1345 | 11 | 1.64% |
| hsa_circ_0107820 | 1345 | 11 | 1.64% |
| hsa_circ_0123905 | 1345 | 11 | 1.64% |
| hsa_circ_0042874 | 1712 | 14 | 1.64% |
| hsa_circ_0020452 | 1223 | 10 | 1.64% |
| hsa_circ_0024087 | 1223 | 10 | 1.64% |
| hsa_circ_0034453 | 1223 | 10 | 1.64% |
| hsa_circ_0035087 | 1223 | 10 | 1.64% |
| hsa_circ_0035601 | 1223 | 10 | 1.64% |
| hsa_circ_0047446 | 1223 | 10 | 1.64% |
| hsa_circ_0050845 | 1223 | 10 | 1.64% |
| hsa_circ_0052159 | 1223 | 10 | 1.64% |
| hsa_circ_0055847 | 1223 | 10 | 1.64% |
| hsa_circ_0056698 | 1223 | 10 | 1.64% |
| hsa_circ_0061809 | 1223 | 10 | 1.64% |
| hsa_circ_0086619 | 1223 | 10 | 1.64% |
| hsa_circ_0101981 | 1223 | 10 | 1.64% |
| hsa_circ_0117073 | 1223 | 10 | 1.64% |
| hsa_circ_0123790 | 1223 | 10 | 1.64% |
| hsa_circ_0140209 | 1223 | 10 | 1.64% |
| hsa_circ_0089658 | 1590 | 13 | 1.64% |
| hsa_circ_0040889 | 1346 | 11 | 1.63% |
| hsa_circ_0068206 | 1346 | 11 | 1.63% |
| hsa_circ_0087120 | 1346 | 11 | 1.63% |
| hsa_circ_0098409 | 1346 | 11 | 1.63% |
| hsa_circ_0100533 | 1346 | 11 | 1.63% |
| hsa_circ_0137770 | 1346 | 11 | 1.63% |
| hsa_circ_0007399 | 1224 | 10 | 1.63% |
| hsa_circ_0032161 | 1224 | 10 | 1.63% |
| hsa_circ_0052780 | 1224 | 10 | 1.63% |
| hsa_circ_0064159 | 1224 | 10 | 1.63% |
| hsa_circ_0069897 | 1224 | 10 | 1.63% |
| hsa_circ_0082262 | 1224 | 10 | 1.63% |
| hsa_circ_0083893 | 1224 | 10 | 1.63% |
| hsa_circ_0085862 | 1224 | 10 | 1.63% |
| hsa_circ_0088437 | 1224 | 10 | 1.63% |
| hsa_circ_0089095 | 1224 | 10 | 1.63% |
| hsa_circ_0091199 | 1224 | 10 | 1.63% |
| hsa_circ_0117645 | 1224 | 10 | 1.63% |
| hsa_circ_0120406 | 1224 | 10 | 1.63% |
| hsa_circ_0124951 | 1224 | 10 | 1.63% |
| hsa_circ_0125134 | 1224 | 10 | 1.63% |
| hsa_circ_0125140 | 1224 | 10 | 1.63% |
| hsa_circ_0132952 | 1224 | 10 | 1.63% |
| hsa_circ_0139711 | 1224 | 10 | 1.63% |
| hsa_circ_0022415 | 1469 | 12 | 1.63% |
| hsa_circ_0045209 | 1469 | 12 | 1.63% |
| hsa_circ_0128788 | 2571 | 21 | 1.63% |
| hsa_circ_0015956 | 1347 | 11 | 1.63% |
| hsa_circ_0019819 | 1347 | 11 | 1.63% |

| | | | |
|---|---|---|---|
| hsa_circ_0032215 | 1347 | 11 | 1.63% |
| hsa_circ_0050223 | 1347 | 11 | 1.63% |
| hsa_circ_0052388 | 1347 | 11 | 1.63% |
| hsa_circ_0064417 | 1347 | 11 | 1.63% |
| hsa_circ_0105065 | 1347 | 11 | 1.63% |
| hsa_circ_0121496 | 1347 | 11 | 1.63% |
| hsa_circ_0122770 | 1347 | 11 | 1.63% |
| hsa_circ_0127309 | 1347 | 11 | 1.63% |
| hsa_circ_0134189 | 1470 | 12 | 1.63% |
| hsa_circ_0001352 | 1225 | 10 | 1.63% |
| hsa_circ_0015440 | 1225 | 10 | 1.63% |
| hsa_circ_0016094 | 1225 | 10 | 1.63% |
| hsa_circ_0022878 | 1225 | 10 | 1.63% |
| hsa_circ_0041012 | 1225 | 10 | 1.63% |
| hsa_circ_0050855 | 1225 | 10 | 1.63% |
| hsa_circ_0052176 | 1225 | 10 | 1.63% |
| hsa_circ_0058000 | 1225 | 10 | 1.63% |
| hsa_circ_0060625 | 1225 | 10 | 1.63% |
| hsa_circ_0060898 | 1225 | 10 | 1.63% |
| hsa_circ_0065763 | 1225 | 10 | 1.63% |
| hsa_circ_0073258 | 1225 | 10 | 1.63% |
| hsa_circ_0088147 | 1225 | 10 | 1.63% |
| hsa_circ_0095795 | 1225 | 10 | 1.63% |
| hsa_circ_0107609 | 1225 | 10 | 1.63% |
| hsa_circ_0120402 | 1225 | 10 | 1.63% |
| hsa_circ_0021856 | 5268 | 43 | 1.63% |
| hsa_circ_0100251 | 1593 | 13 | 1.63% |
| hsa_circ_0132140 | 1348 | 11 | 1.63% |
| hsa_circ_0026949 | 1226 | 10 | 1.63% |
| hsa_circ_0048115 | 1226 | 10 | 1.63% |
| hsa_circ_0052260 | 1226 | 10 | 1.63% |
| hsa_circ_0063782 | 1226 | 10 | 1.63% |
| hsa_circ_0065610 | 1226 | 10 | 1.63% |
| hsa_circ_0074361 | 1226 | 10 | 1.63% |
| hsa_circ_0101789 | 1226 | 10 | 1.63% |
| hsa_circ_0114536 | 1226 | 10 | 1.63% |
| hsa_circ_0119382 | 1226 | 10 | 1.63% |
| hsa_circ_0124285 | 1226 | 10 | 1.63% |
| hsa_circ_0135721 | 1226 | 10 | 1.63% |
| hsa_circ_0010681 | 1594 | 13 | 1.63% |
| hsa_circ_0083350 | 2698 | 22 | 1.63% |
| hsa_circ_0010135 | 1349 | 11 | 1.63% |
| hsa_circ_0037784 | 1349 | 11 | 1.63% |
| hsa_circ_0065229 | 1349 | 11 | 1.63% |
| hsa_circ_0075361 | 1349 | 11 | 1.63% |
| hsa_circ_0096550 | 1349 | 11 | 1.63% |
| hsa_circ_0126758 | 1349 | 11 | 1.63% |
| hsa_circ_0039490 | 1472 | 12 | 1.63% |
| hsa_circ_0048895 | 1472 | 12 | 1.63% |
| hsa_circ_0091405 | 1472 | 12 | 1.63% |

| | | | |
|---|---|---|---|
| hsa_circ_0001451 | 1227 | 10 | 1.63% |
| hsa_circ_0014833 | 1227 | 10 | 1.63% |
| hsa_circ_0014874 | 1227 | 10 | 1.63% |
| hsa_circ_0047988 | 1227 | 10 | 1.63% |
| hsa_circ_0057551 | 1227 | 10 | 1.63% |
| hsa_circ_0075243 | 1227 | 10 | 1.63% |
| hsa_circ_0099417 | 1227 | 10 | 1.63% |
| hsa_circ_0106313 | 1227 | 10 | 1.63% |
| hsa_circ_0107348 | 1227 | 10 | 1.63% |
| hsa_circ_0120742 | 1227 | 10 | 1.63% |
| hsa_circ_0127143 | 1227 | 10 | 1.63% |
| hsa_circ_0129321 | 1227 | 10 | 1.63% |
| hsa_circ_0131075 | 1227 | 10 | 1.63% |
| hsa_circ_0140253 | 1227 | 10 | 1.63% |
| hsa_circ_0051086 | 1350 | 11 | 1.63% |
| hsa_circ_0052246 | 1350 | 11 | 1.63% |
| hsa_circ_0064299 | 1350 | 11 | 1.63% |
| hsa_circ_0065538 | 1350 | 11 | 1.63% |
| hsa_circ_0134408 | 1350 | 11 | 1.63% |
| hsa_circ_0020349 | 1473 | 12 | 1.63% |
| hsa_circ_0034420 | 1473 | 12 | 1.63% |
| hsa_circ_0097709 | 2824 | 23 | 1.63% |
| hsa_circ_0014640 | 1228 | 10 | 1.63% |
| hsa_circ_0031855 | 1228 | 10 | 1.63% |
| hsa_circ_0044761 | 1228 | 10 | 1.63% |
| hsa_circ_0056952 | 1228 | 10 | 1.63% |
| hsa_circ_0057735 | 1228 | 10 | 1.63% |
| hsa_circ_0059820 | 1228 | 10 | 1.63% |
| hsa_circ_0078059 | 1228 | 10 | 1.63% |
| hsa_circ_0085697 | 1228 | 10 | 1.63% |
| hsa_circ_0085952 | 1228 | 10 | 1.63% |
| hsa_circ_0089948 | 1228 | 10 | 1.63% |
| hsa_circ_0107610 | 1228 | 10 | 1.63% |
| hsa_circ_0125304 | 1228 | 10 | 1.63% |
| hsa_circ_0125406 | 1228 | 10 | 1.63% |
| hsa_circ_0130023 | 1228 | 10 | 1.63% |
| hsa_circ_0047859 | 1351 | 11 | 1.63% |
| hsa_circ_0058030 | 1351 | 11 | 1.63% |
| hsa_circ_0062065 | 1351 | 11 | 1.63% |
| hsa_circ_0072613 | 1351 | 11 | 1.63% |
| hsa_circ_0039902 | 1474 | 12 | 1.63% |
| hsa_circ_0097615 | 1474 | 12 | 1.63% |
| hsa_circ_0045395 | 1720 | 14 | 1.63% |
| hsa_circ_0001503 | 1229 | 10 | 1.63% |
| hsa_circ_0014391 | 1229 | 10 | 1.63% |
| hsa_circ_0026792 | 1229 | 10 | 1.63% |
| hsa_circ_0037746 | 1229 | 10 | 1.63% |
| hsa_circ_0077487 | 1229 | 10 | 1.63% |
| hsa_circ_0077906 | 1229 | 10 | 1.63% |
| hsa_circ_0079919 | 1229 | 10 | 1.63% |

| | | | |
|---|---|---|---|
| hsa_circ_0083258 | 1229 | 10 | 1.63% |
| hsa_circ_0097668 | 1229 | 10 | 1.63% |
| hsa_circ_0120385 | 1229 | 10 | 1.63% |
| hsa_circ_0121894 | 1229 | 10 | 1.63% |
| hsa_circ_0138459 | 1229 | 10 | 1.63% |
| hsa_circ_0005363 | 1352 | 11 | 1.63% |
| hsa_circ_0020497 | 1352 | 11 | 1.63% |
| hsa_circ_0029721 | 1352 | 11 | 1.63% |
| hsa_circ_0042746 | 1352 | 11 | 1.63% |
| hsa_circ_0063876 | 1352 | 11 | 1.63% |
| hsa_circ_0070782 | 1352 | 11 | 1.63% |
| hsa_circ_0102805 | 1352 | 11 | 1.63% |
| hsa_circ_0119448 | 1352 | 11 | 1.63% |
| hsa_circ_0131540 | 1352 | 11 | 1.63% |
| hsa_circ_0133787 | 1352 | 11 | 1.63% |
| hsa_circ_0071350 | 1475 | 12 | 1.63% |
| hsa_circ_0018284 | 1845 | 15 | 1.63% |
| hsa_circ_0072096 | 1722 | 14 | 1.63% |
| hsa_circ_0015553 | 1599 | 13 | 1.63% |
| hsa_circ_0107042 | 1476 | 12 | 1.63% |
| hsa_circ_0012837 | 1353 | 11 | 1.63% |
| hsa_circ_0016363 | 1353 | 11 | 1.63% |
| hsa_circ_0026484 | 1353 | 11 | 1.63% |
| hsa_circ_0046181 | 1353 | 11 | 1.63% |
| hsa_circ_0062412 | 1353 | 11 | 1.63% |
| hsa_circ_0074678 | 1353 | 11 | 1.63% |
| hsa_circ_0096517 | 1353 | 11 | 1.63% |
| hsa_circ_0009039 | 1230 | 10 | 1.63% |
| hsa_circ_0029657 | 1230 | 10 | 1.63% |
| hsa_circ_0031663 | 1230 | 10 | 1.63% |
| hsa_circ_0048127 | 1230 | 10 | 1.63% |
| hsa_circ_0051387 | 1230 | 10 | 1.63% |
| hsa_circ_0060339 | 1230 | 10 | 1.63% |
| hsa_circ_0071855 | 1230 | 10 | 1.63% |
| hsa_circ_0073103 | 1230 | 10 | 1.63% |
| hsa_circ_0075998 | 1230 | 10 | 1.63% |
| hsa_circ_0091765 | 1230 | 10 | 1.63% |
| hsa_circ_0096183 | 1230 | 10 | 1.63% |
| hsa_circ_0102588 | 1230 | 10 | 1.63% |
| hsa_circ_0110167 | 1230 | 10 | 1.63% |
| hsa_circ_0110791 | 1230 | 10 | 1.63% |
| hsa_circ_0115075 | 1230 | 10 | 1.63% |
| hsa_circ_0122393 | 1230 | 10 | 1.63% |
| hsa_circ_0125985 | 1230 | 10 | 1.63% |
| hsa_circ_0135645 | 1230 | 10 | 1.63% |
| hsa_circ_0013680 | 1477 | 12 | 1.62% |
| hsa_circ_0060178 | 1354 | 11 | 1.62% |
| hsa_circ_0074190 | 1354 | 11 | 1.62% |
| hsa_circ_0083923 | 1354 | 11 | 1.62% |
| hsa_circ_0087970 | 1354 | 11 | 1.62% |

| | | | |
|---|---|---|---|
| hsa_circ_0107823 | 1354 | 11 | 1.62% |
| hsa_circ_0115031 | 1354 | 11 | 1.62% |
| hsa_circ_0117371 | 1354 | 11 | 1.62% |
| hsa_circ_0122547 | 1354 | 11 | 1.62% |
| hsa_circ_0122666 | 1354 | 11 | 1.62% |
| hsa_circ_0012721 | 1231 | 10 | 1.62% |
| hsa_circ_0042454 | 1231 | 10 | 1.62% |
| hsa_circ_0046107 | 1231 | 10 | 1.62% |
| hsa_circ_0085344 | 1231 | 10 | 1.62% |
| hsa_circ_0094127 | 1231 | 10 | 1.62% |
| hsa_circ_0107145 | 1231 | 10 | 1.62% |
| hsa_circ_0121293 | 1231 | 10 | 1.62% |
| hsa_circ_0067475 | 1601 | 13 | 1.62% |
| hsa_circ_0072622 | 1601 | 13 | 1.62% |
| hsa_circ_0114321 | 1478 | 12 | 1.62% |
| hsa_circ_0009604 | 1355 | 11 | 1.62% |
| hsa_circ_0014797 | 1355 | 11 | 1.62% |
| hsa_circ_0041025 | 1355 | 11 | 1.62% |
| hsa_circ_0046343 | 1355 | 11 | 1.62% |
| hsa_circ_0101164 | 1355 | 11 | 1.62% |
| hsa_circ_0011344 | 1232 | 10 | 1.62% |
| hsa_circ_0057989 | 1232 | 10 | 1.62% |
| hsa_circ_0058992 | 1232 | 10 | 1.62% |
| hsa_circ_0059374 | 1232 | 10 | 1.62% |
| hsa_circ_0068266 | 1232 | 10 | 1.62% |
| hsa_circ_0083316 | 1232 | 10 | 1.62% |
| hsa_circ_0120070 | 1232 | 10 | 1.62% |
| hsa_circ_0089655 | 1725 | 14 | 1.62% |
| hsa_circ_0017716 | 1479 | 12 | 1.62% |
| hsa_circ_0025742 | 1479 | 12 | 1.62% |
| hsa_circ_0131384 | 1479 | 12 | 1.62% |
| hsa_circ_0014947 | 1356 | 11 | 1.62% |
| hsa_circ_0044775 | 1356 | 11 | 1.62% |
| hsa_circ_0064772 | 1356 | 11 | 1.62% |
| hsa_circ_0090528 | 1356 | 11 | 1.62% |
| hsa_circ_0110945 | 1356 | 11 | 1.62% |
| hsa_circ_0129429 | 1356 | 11 | 1.62% |
| hsa_circ_0014065 | 1233 | 10 | 1.62% |
| hsa_circ_0024967 | 1233 | 10 | 1.62% |
| hsa_circ_0025871 | 1233 | 10 | 1.62% |
| hsa_circ_0026387 | 1233 | 10 | 1.62% |
| hsa_circ_0043419 | 1233 | 10 | 1.62% |
| hsa_circ_0068731 | 1233 | 10 | 1.62% |
| hsa_circ_0075641 | 1233 | 10 | 1.62% |
| hsa_circ_0078022 | 1233 | 10 | 1.62% |
| hsa_circ_0084383 | 1233 | 10 | 1.62% |
| hsa_circ_0102923 | 1233 | 10 | 1.62% |
| hsa_circ_0110242 | 1233 | 10 | 1.62% |
| hsa_circ_0118706 | 1233 | 10 | 1.62% |
| hsa_circ_0122945 | 1233 | 10 | 1.62% |

| | | | |
|---|---|---|---|
| hsa_circ_0132998 | 1233 | 10 | 1.62% |
| hsa_circ_0097706 | 5302 | 43 | 1.62% |
| hsa_circ_0036608 | 2836 | 23 | 1.62% |
| hsa_circ_0121261 | 3083 | 25 | 1.62% |
| hsa_circ_0052707 | 1850 | 15 | 1.62% |
| hsa_circ_0009199 | 1480 | 12 | 1.62% |
| hsa_circ_0127652 | 1480 | 12 | 1.62% |
| hsa_circ_0024581 | 1357 | 11 | 1.62% |
| hsa_circ_0039382 | 1357 | 11 | 1.62% |
| hsa_circ_0058067 | 1357 | 11 | 1.62% |
| hsa_circ_0062388 | 1357 | 11 | 1.62% |
| hsa_circ_0116352 | 1357 | 11 | 1.62% |
| hsa_circ_0125953 | 1357 | 11 | 1.62% |
| hsa_circ_0139286 | 1357 | 11 | 1.62% |
| hsa_circ_0073814 | 4688 | 38 | 1.62% |
| hsa_circ_0098931 | 4688 | 38 | 1.62% |
| hsa_circ_0001795 | 1234 | 10 | 1.62% |
| hsa_circ_0020660 | 1234 | 10 | 1.62% |
| hsa_circ_0024757 | 1234 | 10 | 1.62% |
| hsa_circ_0031283 | 1234 | 10 | 1.62% |
| hsa_circ_0094943 | 1234 | 10 | 1.62% |
| hsa_circ_0101892 | 1234 | 10 | 1.62% |
| hsa_circ_0114268 | 1234 | 10 | 1.62% |
| hsa_circ_0117625 | 1234 | 10 | 1.62% |
| hsa_circ_0140115 | 1234 | 10 | 1.62% |
| hsa_circ_0072807 | 2098 | 17 | 1.62% |
| hsa_circ_0072826 | 2098 | 17 | 1.62% |
| hsa_circ_0090157 | 1481 | 12 | 1.62% |
| hsa_circ_0092052 | 1481 | 12 | 1.62% |
| hsa_circ_0015674 | 1358 | 11 | 1.62% |
| hsa_circ_0028528 | 1358 | 11 | 1.62% |
| hsa_circ_0038423 | 1358 | 11 | 1.62% |
| hsa_circ_0040070 | 1358 | 11 | 1.62% |
| hsa_circ_0056232 | 1358 | 11 | 1.62% |
| hsa_circ_0068050 | 1358 | 11 | 1.62% |
| hsa_circ_0071692 | 1358 | 11 | 1.62% |
| hsa_circ_0104060 | 1358 | 11 | 1.62% |
| hsa_circ_0026388 | 1605 | 13 | 1.62% |
| hsa_circ_0072849 | 2099 | 17 | 1.62% |
| hsa_circ_0051728 | 1482 | 12 | 1.62% |
| hsa_circ_0036525 | 1235 | 10 | 1.62% |
| hsa_circ_0061488 | 1235 | 10 | 1.62% |
| hsa_circ_0064412 | 1235 | 10 | 1.62% |
| hsa_circ_0070226 | 1235 | 10 | 1.62% |
| hsa_circ_0070764 | 1235 | 10 | 1.62% |
| hsa_circ_0075439 | 1235 | 10 | 1.62% |
| hsa_circ_0077947 | 1235 | 10 | 1.62% |
| hsa_circ_0084476 | 1235 | 10 | 1.62% |
| hsa_circ_0087058 | 1235 | 10 | 1.62% |
| hsa_circ_0094162 | 1235 | 10 | 1.62% |

| | | | |
|---|---|---|---|
| hsa_circ_0122647 | 1235 | 10 | 1.62% |
| hsa_circ_0123521 | 1235 | 10 | 1.62% |
| hsa_circ_0136109 | 1235 | 10 | 1.62% |
| hsa_circ_0064447 | 1606 | 13 | 1.62% |
| hsa_circ_0068401 | 1606 | 13 | 1.62% |
| hsa_circ_0102842 | 1606 | 13 | 1.62% |
| hsa_circ_0020561 | 1359 | 11 | 1.62% |
| hsa_circ_0038134 | 1359 | 11 | 1.62% |
| hsa_circ_0065694 | 1359 | 11 | 1.62% |
| hsa_circ_0081847 | 1359 | 11 | 1.62% |
| hsa_circ_0128907 | 1359 | 11 | 1.62% |
| hsa_circ_0107839 | 1730 | 14 | 1.62% |
| hsa_circ_0125176 | 1483 | 12 | 1.62% |
| hsa_circ_0010133 | 1236 | 10 | 1.62% |
| hsa_circ_0010153 | 1236 | 10 | 1.62% |
| hsa_circ_0022632 | 1236 | 10 | 1.62% |
| hsa_circ_0023373 | 1236 | 10 | 1.62% |
| hsa_circ_0026778 | 1236 | 10 | 1.62% |
| hsa_circ_0030801 | 1236 | 10 | 1.62% |
| hsa_circ_0033449 | 1236 | 10 | 1.62% |
| hsa_circ_0034689 | 1236 | 10 | 1.62% |
| hsa_circ_0065430 | 1236 | 10 | 1.62% |
| hsa_circ_0067780 | 1236 | 10 | 1.62% |
| hsa_circ_0082169 | 1236 | 10 | 1.62% |
| hsa_circ_0092200 | 1236 | 10 | 1.62% |
| hsa_circ_0101127 | 1236 | 10 | 1.62% |
| hsa_circ_0122447 | 1236 | 10 | 1.62% |
| hsa_circ_0123354 | 5439 | 44 | 1.62% |
| hsa_circ_0034643 | 1360 | 11 | 1.62% |
| hsa_circ_0044420 | 1360 | 11 | 1.62% |
| hsa_circ_0052031 | 1360 | 11 | 1.62% |
| hsa_circ_0054738 | 1360 | 11 | 1.62% |
| hsa_circ_0083614 | 1360 | 11 | 1.62% |
| hsa_circ_0139902 | 1360 | 11 | 1.62% |
| hsa_circ_0078202 | 3957 | 32 | 1.62% |
| hsa_circ_0134705 | 1484 | 12 | 1.62% |
| hsa_circ_0039380 | 1608 | 13 | 1.62% |
| hsa_circ_0013616 | 1237 | 10 | 1.62% |
| hsa_circ_0018944 | 1237 | 10 | 1.62% |
| hsa_circ_0023679 | 1237 | 10 | 1.62% |
| hsa_circ_0037037 | 1237 | 10 | 1.62% |
| hsa_circ_0047223 | 1237 | 10 | 1.62% |
| hsa_circ_0063973 | 1237 | 10 | 1.62% |
| hsa_circ_0064267 | 1237 | 10 | 1.62% |
| hsa_circ_0066159 | 1237 | 10 | 1.62% |
| hsa_circ_0067752 | 1237 | 10 | 1.62% |
| hsa_circ_0068729 | 1237 | 10 | 1.62% |
| hsa_circ_0079104 | 1237 | 10 | 1.62% |
| hsa_circ_0092478 | 1237 | 10 | 1.62% |
| hsa_circ_0104795 | 1237 | 10 | 1.62% |

| | | | |
|---|---|---|---|
| hsa_circ_0105497 | 1237 | 10 | 1.62% |
| hsa_circ_0110636 | 1237 | 10 | 1.62% |
| hsa_circ_0117928 | 1237 | 10 | 1.62% |
| hsa_circ_0140353 | 1237 | 10 | 1.62% |
| hsa_circ_0009921 | 1361 | 11 | 1.62% |
| hsa_circ_0029677 | 1361 | 11 | 1.62% |
| hsa_circ_0034735 | 1361 | 11 | 1.62% |
| hsa_circ_0038808 | 1361 | 11 | 1.62% |
| hsa_circ_0049807 | 1361 | 11 | 1.62% |
| hsa_circ_0097609 | 1361 | 11 | 1.62% |
| hsa_circ_0108125 | 1361 | 11 | 1.62% |
| hsa_circ_0127371 | 1361 | 11 | 1.62% |
| hsa_circ_0130295 | 1361 | 11 | 1.62% |
| hsa_circ_0136295 | 1361 | 11 | 1.62% |
| hsa_circ_0136442 | 1361 | 11 | 1.62% |
| hsa_circ_0101250 | 2475 | 20 | 1.62% |
| hsa_circ_0054781 | 1485 | 12 | 1.62% |
| hsa_circ_0065879 | 1485 | 12 | 1.62% |
| hsa_circ_0071554 | 1485 | 12 | 1.62% |
| hsa_circ_0085798 | 1485 | 12 | 1.62% |
| hsa_circ_0128503 | 1485 | 12 | 1.62% |
| hsa_circ_0077959 | 1609 | 13 | 1.62% |
| hsa_circ_0002742 | 1238 | 10 | 1.62% |
| hsa_circ_0006679 | 1238 | 10 | 1.62% |
| hsa_circ_0020905 | 1238 | 10 | 1.62% |
| hsa_circ_0032422 | 1238 | 10 | 1.62% |
| hsa_circ_0038830 | 1238 | 10 | 1.62% |
| hsa_circ_0042872 | 1238 | 10 | 1.62% |
| hsa_circ_0043005 | 1238 | 10 | 1.62% |
| hsa_circ_0072542 | 1238 | 10 | 1.62% |
| hsa_circ_0091130 | 1238 | 10 | 1.62% |
| hsa_circ_0095124 | 1238 | 10 | 1.62% |
| hsa_circ_0107339 | 1238 | 10 | 1.62% |
| hsa_circ_0124139 | 1238 | 10 | 1.62% |
| hsa_circ_0125297 | 1238 | 10 | 1.62% |
| hsa_circ_0130520 | 1238 | 10 | 1.62% |
| hsa_circ_0133211 | 1238 | 10 | 1.62% |
| hsa_circ_0005107 | 1362 | 11 | 1.62% |
| hsa_circ_0012678 | 1362 | 11 | 1.62% |
| hsa_circ_0024007 | 1362 | 11 | 1.62% |
| hsa_circ_0053892 | 1362 | 11 | 1.62% |
| hsa_circ_0061966 | 1362 | 11 | 1.62% |
| hsa_circ_0077845 | 1362 | 11 | 1.62% |
| hsa_circ_0079134 | 1362 | 11 | 1.62% |
| hsa_circ_0088852 | 1362 | 11 | 1.62% |
| hsa_circ_0096832 | 1362 | 11 | 1.62% |
| hsa_circ_0118013 | 1362 | 11 | 1.62% |
| hsa_circ_0130625 | 1362 | 11 | 1.62% |
| hsa_circ_0079195 | 1486 | 12 | 1.62% |
| hsa_circ_0010871 | 1239 | 10 | 1.61% |

| | | | |
|---|---|---|---|
| hsa_circ_0017597 | 1239 | 10 | 1.61% |
| hsa_circ_0021897 | 1239 | 10 | 1.61% |
| hsa_circ_0032594 | 1239 | 10 | 1.61% |
| hsa_circ_0036548 | 1239 | 10 | 1.61% |
| hsa_circ_0046116 | 1239 | 10 | 1.61% |
| hsa_circ_0065379 | 1239 | 10 | 1.61% |
| hsa_circ_0070812 | 1239 | 10 | 1.61% |
| hsa_circ_0088006 | 1239 | 10 | 1.61% |
| hsa_circ_0092765 | 1239 | 10 | 1.61% |
| hsa_circ_0097131 | 1239 | 10 | 1.61% |
| hsa_circ_0099939 | 1239 | 10 | 1.61% |
| hsa_circ_0100282 | 1239 | 10 | 1.61% |
| hsa_circ_0129646 | 1239 | 10 | 1.61% |
| hsa_circ_0135413 | 1239 | 10 | 1.61% |
| hsa_circ_0040656 | 1363 | 11 | 1.61% |
| hsa_circ_0080916 | 1363 | 11 | 1.61% |
| hsa_circ_0133800 | 1363 | 11 | 1.61% |
| hsa_circ_0129620 | 1487 | 12 | 1.61% |
| hsa_circ_0039526 | 1611 | 13 | 1.61% |
| hsa_circ_0135141 | 1611 | 13 | 1.61% |
| hsa_circ_0100897 | 1859 | 15 | 1.61% |
| hsa_circ_0005983 | 4958 | 40 | 1.61% |
| hsa_circ_0105136 | 4587 | 37 | 1.61% |
| hsa_circ_0036208 | 1488 | 12 | 1.61% |
| hsa_circ_0114205 | 1488 | 12 | 1.61% |
| hsa_circ_0010045 | 1364 | 11 | 1.61% |
| hsa_circ_0010410 | 1364 | 11 | 1.61% |
| hsa_circ_0041296 | 1364 | 11 | 1.61% |
| hsa_circ_0095830 | 1364 | 11 | 1.61% |
| hsa_circ_0118179 | 1364 | 11 | 1.61% |
| hsa_circ_0136348 | 1364 | 11 | 1.61% |
| hsa_circ_0023063 | 1240 | 10 | 1.61% |
| hsa_circ_0032665 | 1240 | 10 | 1.61% |
| hsa_circ_0035771 | 1240 | 10 | 1.61% |
| hsa_circ_0039255 | 1240 | 10 | 1.61% |
| hsa_circ_0067377 | 1240 | 10 | 1.61% |
| hsa_circ_0072241 | 1240 | 10 | 1.61% |
| hsa_circ_0079585 | 1240 | 10 | 1.61% |
| hsa_circ_0089481 | 1240 | 10 | 1.61% |
| hsa_circ_0090824 | 1240 | 10 | 1.61% |
| hsa_circ_0123006 | 1240 | 10 | 1.61% |
| hsa_circ_0135809 | 1240 | 10 | 1.61% |
| hsa_circ_0085036 | 1737 | 14 | 1.61% |
| hsa_circ_0085624 | 1489 | 12 | 1.61% |
| hsa_circ_0089306 | 1489 | 12 | 1.61% |
| hsa_circ_0119943 | 1489 | 12 | 1.61% |
| hsa_circ_0128942 | 1489 | 12 | 1.61% |
| hsa_circ_0010074 | 1365 | 11 | 1.61% |
| hsa_circ_0011250 | 1365 | 11 | 1.61% |
| hsa_circ_0089295 | 1365 | 11 | 1.61% |

| | | | |
|---|---|---|---|
| hsa_circ_0090408 | 1365 | 11 | 1.61% |
| hsa_circ_0096694 | 1365 | 11 | 1.61% |
| hsa_circ_0114537 | 1365 | 11 | 1.61% |
| hsa_circ_0116984 | 1365 | 11 | 1.61% |
| hsa_circ_0130030 | 1365 | 11 | 1.61% |
| hsa_circ_0018101 | 1241 | 10 | 1.61% |
| hsa_circ_0020632 | 1241 | 10 | 1.61% |
| hsa_circ_0056328 | 1241 | 10 | 1.61% |
| hsa_circ_0060033 | 1241 | 10 | 1.61% |
| hsa_circ_0071584 | 1241 | 10 | 1.61% |
| hsa_circ_0079332 | 1241 | 10 | 1.61% |
| hsa_circ_0111725 | 1241 | 10 | 1.61% |
| hsa_circ_0117771 | 1241 | 10 | 1.61% |
| hsa_circ_0100468 | 1614 | 13 | 1.61% |
| hsa_circ_0121387 | 1490 | 12 | 1.61% |
| hsa_circ_0007585 | 1366 | 11 | 1.61% |
| hsa_circ_0028002 | 1366 | 11 | 1.61% |
| hsa_circ_0051729 | 1366 | 11 | 1.61% |
| hsa_circ_0053552 | 1366 | 11 | 1.61% |
| hsa_circ_0013436 | 1242 | 10 | 1.61% |
| hsa_circ_0015131 | 1242 | 10 | 1.61% |
| hsa_circ_0015945 | 1242 | 10 | 1.61% |
| hsa_circ_0022512 | 1242 | 10 | 1.61% |
| hsa_circ_0042212 | 1242 | 10 | 1.61% |
| hsa_circ_0043048 | 1242 | 10 | 1.61% |
| hsa_circ_0045701 | 1242 | 10 | 1.61% |
| hsa_circ_0061961 | 1242 | 10 | 1.61% |
| hsa_circ_0066800 | 1242 | 10 | 1.61% |
| hsa_circ_0071816 | 1242 | 10 | 1.61% |
| hsa_circ_0075634 | 1242 | 10 | 1.61% |
| hsa_circ_0079587 | 1242 | 10 | 1.61% |
| hsa_circ_0094822 | 1242 | 10 | 1.61% |
| hsa_circ_0095419 | 1242 | 10 | 1.61% |
| hsa_circ_0097450 | 1242 | 10 | 1.61% |
| hsa_circ_0109546 | 1242 | 10 | 1.61% |
| hsa_circ_0116456 | 1242 | 10 | 1.61% |
| hsa_circ_0131071 | 1242 | 10 | 1.61% |
| hsa_circ_0134904 | 1242 | 10 | 1.61% |
| hsa_circ_0012010 | 1615 | 13 | 1.61% |
| hsa_circ_0069777 | 1491 | 12 | 1.61% |
| hsa_circ_0047181 | 1367 | 11 | 1.61% |
| hsa_circ_0071806 | 1367 | 11 | 1.61% |
| hsa_circ_0110204 | 1367 | 11 | 1.61% |
| hsa_circ_0117135 | 1367 | 11 | 1.61% |
| hsa_circ_0139864 | 1367 | 11 | 1.61% |
| hsa_circ_0019291 | 1243 | 10 | 1.61% |
| hsa_circ_0019684 | 1243 | 10 | 1.61% |
| hsa_circ_0044612 | 1243 | 10 | 1.61% |
| hsa_circ_0053840 | 1243 | 10 | 1.61% |
| hsa_circ_0055633 | 1243 | 10 | 1.61% |

| | | | |
|---|---|---|---|
| hsa_circ_0056241 | 1243 | 10 | 1.61% |
| hsa_circ_0068740 | 1243 | 10 | 1.61% |
| hsa_circ_0080492 | 1243 | 10 | 1.61% |
| hsa_circ_0090392 | 1243 | 10 | 1.61% |
| hsa_circ_0110573 | 1243 | 10 | 1.61% |
| hsa_circ_0136822 | 1243 | 10 | 1.61% |
| hsa_circ_0075774 | 2859 | 23 | 1.61% |
| hsa_circ_0093242 | 2984 | 24 | 1.61% |
| hsa_circ_0022887 | 1865 | 15 | 1.61% |
| hsa_circ_0094897 | 1492 | 12 | 1.61% |
| hsa_circ_0022546 | 2114 | 17 | 1.61% |
| hsa_circ_0019424 | 1368 | 11 | 1.61% |
| hsa_circ_0019910 | 1368 | 11 | 1.61% |
| hsa_circ_0059194 | 1368 | 11 | 1.61% |
| hsa_circ_0060276 | 1368 | 11 | 1.61% |
| hsa_circ_0065640 | 1368 | 11 | 1.61% |
| hsa_circ_0094845 | 1368 | 11 | 1.61% |
| hsa_circ_0139370 | 1368 | 11 | 1.61% |
| hsa_circ_0021283 | 1617 | 13 | 1.61% |
| hsa_circ_0124678 | 1617 | 13 | 1.61% |
| hsa_circ_0024389 | 1244 | 10 | 1.61% |
| hsa_circ_0024572 | 1244 | 10 | 1.61% |
| hsa_circ_0027344 | 1244 | 10 | 1.61% |
| hsa_circ_0031319 | 1244 | 10 | 1.61% |
| hsa_circ_0034564 | 1244 | 10 | 1.61% |
| hsa_circ_0043264 | 1244 | 10 | 1.61% |
| hsa_circ_0050062 | 1244 | 10 | 1.61% |
| hsa_circ_0065073 | 1244 | 10 | 1.61% |
| hsa_circ_0065737 | 1244 | 10 | 1.61% |
| hsa_circ_0078006 | 1244 | 10 | 1.61% |
| hsa_circ_0082825 | 1244 | 10 | 1.61% |
| hsa_circ_0087981 | 1244 | 10 | 1.61% |
| hsa_circ_0107996 | 1244 | 10 | 1.61% |
| hsa_circ_0113048 | 1244 | 10 | 1.61% |
| hsa_circ_0128668 | 1244 | 10 | 1.61% |
| hsa_circ_0137390 | 1244 | 10 | 1.61% |
| hsa_circ_0019545 | 1369 | 11 | 1.61% |
| hsa_circ_0026472 | 1369 | 11 | 1.61% |
| hsa_circ_0032872 | 1369 | 11 | 1.61% |
| hsa_circ_0041127 | 1369 | 11 | 1.61% |
| hsa_circ_0053347 | 1369 | 11 | 1.61% |
| hsa_circ_0083416 | 1618 | 13 | 1.61% |
| hsa_circ_0010540 | 1245 | 10 | 1.61% |
| hsa_circ_0017483 | 1245 | 10 | 1.61% |
| hsa_circ_0021253 | 1245 | 10 | 1.61% |
| hsa_circ_0027225 | 1245 | 10 | 1.61% |
| hsa_circ_0028492 | 1245 | 10 | 1.61% |
| hsa_circ_0039243 | 1245 | 10 | 1.61% |
| hsa_circ_0044003 | 1245 | 10 | 1.61% |
| hsa_circ_0044047 | 1245 | 10 | 1.61% |

| | | | |
|---|---|---|---|
| hsa_circ_0055259 | 1245 | 10 | 1.61% |
| hsa_circ_0058154 | 1245 | 10 | 1.61% |
| hsa_circ_0060119 | 1245 | 10 | 1.61% |
| hsa_circ_0060566 | 1245 | 10 | 1.61% |
| hsa_circ_0070377 | 1245 | 10 | 1.61% |
| hsa_circ_0090189 | 1245 | 10 | 1.61% |
| hsa_circ_0095941 | 1245 | 10 | 1.61% |
| hsa_circ_0096409 | 1245 | 10 | 1.61% |
| hsa_circ_0122340 | 1245 | 10 | 1.61% |
| hsa_circ_0129735 | 1245 | 10 | 1.61% |
| hsa_circ_0129993 | 1245 | 10 | 1.61% |
| hsa_circ_0131544 | 1245 | 10 | 1.61% |
| hsa_circ_0014675 | 1370 | 11 | 1.61% |
| hsa_circ_0016120 | 1370 | 11 | 1.61% |
| hsa_circ_0035967 | 1370 | 11 | 1.61% |
| hsa_circ_0052208 | 1370 | 11 | 1.61% |
| hsa_circ_0056960 | 1370 | 11 | 1.61% |
| hsa_circ_0107406 | 1370 | 11 | 1.61% |
| hsa_circ_0022761 | 1495 | 12 | 1.61% |
| hsa_circ_0048561 | 1495 | 12 | 1.61% |
| hsa_circ_0075717 | 1495 | 12 | 1.61% |
| hsa_circ_0103971 | 1495 | 12 | 1.61% |
| hsa_circ_0024937 | 1246 | 10 | 1.61% |
| hsa_circ_0025638 | 1246 | 10 | 1.61% |
| hsa_circ_0031450 | 1246 | 10 | 1.61% |
| hsa_circ_0045163 | 1246 | 10 | 1.61% |
| hsa_circ_0049842 | 1246 | 10 | 1.61% |
| hsa_circ_0051874 | 1246 | 10 | 1.61% |
| hsa_circ_0056304 | 1246 | 10 | 1.61% |
| hsa_circ_0079835 | 1246 | 10 | 1.61% |
| hsa_circ_0090986 | 1246 | 10 | 1.61% |
| hsa_circ_0109468 | 1246 | 10 | 1.61% |
| hsa_circ_0119146 | 1246 | 10 | 1.61% |
| hsa_circ_0050318 | 3365 | 27 | 1.60% |
| hsa_circ_0026760 | 1371 | 11 | 1.60% |
| hsa_circ_0044056 | 1371 | 11 | 1.60% |
| hsa_circ_0053341 | 1371 | 11 | 1.60% |
| hsa_circ_0066619 | 1371 | 11 | 1.60% |
| hsa_circ_0089001 | 1371 | 11 | 1.60% |
| hsa_circ_0095935 | 1371 | 11 | 1.60% |
| hsa_circ_0130306 | 1371 | 11 | 1.60% |
| hsa_circ_0077493 | 1496 | 12 | 1.60% |
| hsa_circ_0086145 | 1496 | 12 | 1.60% |
| hsa_circ_0135074 | 1496 | 12 | 1.60% |
| hsa_circ_0099457 | 1621 | 13 | 1.60% |
| hsa_circ_0005384 | 1247 | 10 | 1.60% |
| hsa_circ_0012509 | 1247 | 10 | 1.60% |
| hsa_circ_0021129 | 1247 | 10 | 1.60% |
| hsa_circ_0023553 | 1247 | 10 | 1.60% |
| hsa_circ_0031498 | 1247 | 10 | 1.60% |

| | | | |
|---|---|---|---|
| hsa_circ_0035709 | 1247 | 10 | 1.60% |
| hsa_circ_0035810 | 1247 | 10 | 1.60% |
| hsa_circ_0040851 | 1247 | 10 | 1.60% |
| hsa_circ_0042814 | 1247 | 10 | 1.60% |
| hsa_circ_0047317 | 1247 | 10 | 1.60% |
| hsa_circ_0047358 | 1247 | 10 | 1.60% |
| hsa_circ_0064261 | 1247 | 10 | 1.60% |
| hsa_circ_0103213 | 1247 | 10 | 1.60% |
| hsa_circ_0116112 | 1247 | 10 | 1.60% |
| hsa_circ_0128242 | 1247 | 10 | 1.60% |
| hsa_circ_0136528 | 1247 | 10 | 1.60% |
| hsa_circ_0037700 | 1372 | 11 | 1.60% |
| hsa_circ_0038201 | 1372 | 11 | 1.60% |
| hsa_circ_0056605 | 1372 | 11 | 1.60% |
| hsa_circ_0074147 | 1372 | 11 | 1.60% |
| hsa_circ_0090931 | 1372 | 11 | 1.60% |
| hsa_circ_0027543 | 1497 | 12 | 1.60% |
| hsa_circ_0076166 | 1497 | 12 | 1.60% |
| hsa_circ_0107372 | 1497 | 12 | 1.60% |
| hsa_circ_0127661 | 1497 | 12 | 1.60% |
| hsa_circ_0046080 | 1622 | 13 | 1.60% |
| hsa_circ_0016653 | 1248 | 10 | 1.60% |
| hsa_circ_0031423 | 1248 | 10 | 1.60% |
| hsa_circ_0063055 | 1248 | 10 | 1.60% |
| hsa_circ_0065211 | 1248 | 10 | 1.60% |
| hsa_circ_0065527 | 1248 | 10 | 1.60% |
| hsa_circ_0072372 | 1248 | 10 | 1.60% |
| hsa_circ_0073700 | 1248 | 10 | 1.60% |
| hsa_circ_0073901 | 1248 | 10 | 1.60% |
| hsa_circ_0086675 | 1248 | 10 | 1.60% |
| hsa_circ_0096157 | 1248 | 10 | 1.60% |
| hsa_circ_0110655 | 1248 | 10 | 1.60% |
| hsa_circ_0120091 | 1248 | 10 | 1.60% |
| hsa_circ_0121874 | 1248 | 10 | 1.60% |
| hsa_circ_0132159 | 1248 | 10 | 1.60% |
| hsa_circ_0133374 | 1248 | 10 | 1.60% |
| hsa_circ_0136783 | 1248 | 10 | 1.60% |
| hsa_circ_0138377 | 1248 | 10 | 1.60% |
| hsa_circ_0037658 | 1373 | 11 | 1.60% |
| hsa_circ_0059742 | 1373 | 11 | 1.60% |
| hsa_circ_0129410 | 1373 | 11 | 1.60% |
| hsa_circ_0129435 | 1498 | 12 | 1.60% |
| hsa_circ_0135848 | 1748 | 14 | 1.60% |
| hsa_circ_0037999 | 1249 | 10 | 1.60% |
| hsa_circ_0055321 | 1249 | 10 | 1.60% |
| hsa_circ_0067650 | 1249 | 10 | 1.60% |
| hsa_circ_0120633 | 1249 | 10 | 1.60% |
| hsa_circ_0122188 | 1249 | 10 | 1.60% |
| hsa_circ_0138396 | 1249 | 10 | 1.60% |
| hsa_circ_0030989 | 1374 | 11 | 1.60% |

| | | | |
|---|---|---|---|
| hsa_circ_0053100 | 1374 | 11 | 1.60% |
| hsa_circ_0053654 | 1374 | 11 | 1.60% |
| hsa_circ_0057283 | 1374 | 11 | 1.60% |
| hsa_circ_0065983 | 1374 | 11 | 1.60% |
| hsa_circ_0068074 | 1374 | 11 | 1.60% |
| hsa_circ_0116142 | 1374 | 11 | 1.60% |
| hsa_circ_0016734 | 1499 | 12 | 1.60% |
| hsa_circ_0039094 | 1499 | 12 | 1.60% |
| hsa_circ_0084012 | 1499 | 12 | 1.60% |
| hsa_circ_0119304 | 1499 | 12 | 1.60% |
| hsa_circ_0056254 | 1624 | 13 | 1.60% |
| hsa_circ_0012093 | 1874 | 15 | 1.60% |
| hsa_circ_0051451 | 1750 | 14 | 1.60% |
| hsa_circ_0139160 | 1625 | 13 | 1.60% |
| hsa_circ_0016737 | 1500 | 12 | 1.60% |
| hsa_circ_0034427 | 1500 | 12 | 1.60% |
| hsa_circ_0047760 | 1500 | 12 | 1.60% |
| hsa_circ_0061516 | 1500 | 12 | 1.60% |
| hsa_circ_0078915 | 1500 | 12 | 1.60% |
| hsa_circ_0089723 | 1500 | 12 | 1.60% |
| hsa_circ_0013733 | 1375 | 11 | 1.60% |
| hsa_circ_0024103 | 1375 | 11 | 1.60% |
| hsa_circ_0043361 | 1375 | 11 | 1.60% |
| hsa_circ_0055954 | 1375 | 11 | 1.60% |
| hsa_circ_0067457 | 1375 | 11 | 1.60% |
| hsa_circ_0093802 | 1375 | 11 | 1.60% |
| hsa_circ_0015025 | 1250 | 10 | 1.60% |
| hsa_circ_0032936 | 1250 | 10 | 1.60% |
| hsa_circ_0043166 | 1250 | 10 | 1.60% |
| hsa_circ_0064759 | 1250 | 10 | 1.60% |
| hsa_circ_0065628 | 1250 | 10 | 1.60% |
| hsa_circ_0066470 | 1250 | 10 | 1.60% |
| hsa_circ_0079877 | 1250 | 10 | 1.60% |
| hsa_circ_0088690 | 1250 | 10 | 1.60% |
| hsa_circ_0096269 | 1250 | 10 | 1.60% |
| hsa_circ_0101850 | 1250 | 10 | 1.60% |
| hsa_circ_0105883 | 1250 | 10 | 1.60% |
| hsa_circ_0113746 | 1250 | 10 | 1.60% |
| hsa_circ_0115864 | 1250 | 10 | 1.60% |
| hsa_circ_0121465 | 1250 | 10 | 1.60% |
| hsa_circ_0125220 | 1250 | 10 | 1.60% |
| hsa_circ_0125764 | 1250 | 10 | 1.60% |
| hsa_circ_0121551 | 1626 | 13 | 1.60% |
| hsa_circ_0042288 | 1501 | 12 | 1.60% |
| hsa_circ_0069732 | 1501 | 12 | 1.60% |
| hsa_circ_0019109 | 1376 | 11 | 1.60% |
| hsa_circ_0043889 | 1376 | 11 | 1.60% |
| hsa_circ_0047370 | 1376 | 11 | 1.60% |
| hsa_circ_0052000 | 1376 | 11 | 1.60% |
| hsa_circ_0076112 | 1376 | 11 | 1.60% |

| | | | |
|---|---|---|---|
| hsa_circ_0077871 | 1376 | 11 | 1.60% |
| hsa_circ_0103140 | 3878 | 31 | 1.60% |
| hsa_circ_0019356 | 1251 | 10 | 1.60% |
| hsa_circ_0026359 | 1251 | 10 | 1.60% |
| hsa_circ_0037264 | 1251 | 10 | 1.60% |
| hsa_circ_0039169 | 1251 | 10 | 1.60% |
| hsa_circ_0045137 | 1251 | 10 | 1.60% |
| hsa_circ_0051232 | 1251 | 10 | 1.60% |
| hsa_circ_0087016 | 1251 | 10 | 1.60% |
| hsa_circ_0092248 | 1251 | 10 | 1.60% |
| hsa_circ_0106673 | 1251 | 10 | 1.60% |
| hsa_circ_0120674 | 1251 | 10 | 1.60% |
| hsa_circ_0127321 | 1251 | 10 | 1.60% |
| hsa_circ_0138724 | 1251 | 10 | 1.60% |
| hsa_circ_0027737 | 2628 | 21 | 1.60% |
| hsa_circ_0081027 | 1627 | 13 | 1.60% |
| hsa_circ_0000658 | 1502 | 12 | 1.60% |
| hsa_circ_0097751 | 1502 | 12 | 1.60% |
| hsa_circ_0135326 | 2128 | 17 | 1.60% |
| hsa_circ_0011285 | 1377 | 11 | 1.60% |
| hsa_circ_0015870 | 1377 | 11 | 1.60% |
| hsa_circ_0039479 | 1377 | 11 | 1.60% |
| hsa_circ_0059027 | 1377 | 11 | 1.60% |
| hsa_circ_0059587 | 1377 | 11 | 1.60% |
| hsa_circ_0084454 | 1377 | 11 | 1.60% |
| hsa_circ_0096619 | 1377 | 11 | 1.60% |
| hsa_circ_0006118 | 1252 | 10 | 1.60% |
| hsa_circ_0034525 | 1252 | 10 | 1.60% |
| hsa_circ_0039855 | 1252 | 10 | 1.60% |
| hsa_circ_0061743 | 1252 | 10 | 1.60% |
| hsa_circ_0072561 | 1252 | 10 | 1.60% |
| hsa_circ_0089213 | 1252 | 10 | 1.60% |
| hsa_circ_0098661 | 1252 | 10 | 1.60% |
| hsa_circ_0110102 | 1252 | 10 | 1.60% |
| hsa_circ_0114332 | 1252 | 10 | 1.60% |
| hsa_circ_0122189 | 1252 | 10 | 1.60% |
| hsa_circ_0049584 | 1628 | 13 | 1.60% |
| hsa_circ_0037624 | 1503 | 12 | 1.60% |
| hsa_circ_0058991 | 1503 | 12 | 1.60% |
| hsa_circ_0080616 | 1503 | 12 | 1.60% |
| hsa_circ_0137427 | 1503 | 12 | 1.60% |
| hsa_circ_0023109 | 1378 | 11 | 1.60% |
| hsa_circ_0044069 | 1378 | 11 | 1.60% |
| hsa_circ_0049927 | 1378 | 11 | 1.60% |
| hsa_circ_0104809 | 1378 | 11 | 1.60% |
| hsa_circ_0011135 | 1253 | 10 | 1.60% |
| hsa_circ_0057892 | 1253 | 10 | 1.60% |
| hsa_circ_0076974 | 1253 | 10 | 1.60% |
| hsa_circ_0077611 | 1253 | 10 | 1.60% |
| hsa_circ_0094793 | 1253 | 10 | 1.60% |

| | | | |
|---|---|---|---|
| hsa_circ_0113160 | 1253 | 10 | 1.60% |
| hsa_circ_0117939 | 1253 | 10 | 1.60% |
| hsa_circ_0126245 | 1253 | 10 | 1.60% |
| hsa_circ_0110189 | 1629 | 13 | 1.60% |
| hsa_circ_0054115 | 1880 | 15 | 1.60% |
| hsa_circ_0040087 | 1504 | 12 | 1.60% |
| hsa_circ_0051136 | 1504 | 12 | 1.60% |
| hsa_circ_0058583 | 1504 | 12 | 1.60% |
| hsa_circ_0099597 | 1504 | 12 | 1.60% |
| hsa_circ_0049919 | 1379 | 11 | 1.60% |
| hsa_circ_0065489 | 1379 | 11 | 1.60% |
| hsa_circ_0125663 | 1379 | 11 | 1.60% |
| hsa_circ_0002052 | 1254 | 10 | 1.59% |
| hsa_circ_0005118 | 1254 | 10 | 1.59% |
| hsa_circ_0013375 | 1254 | 10 | 1.59% |
| hsa_circ_0036937 | 1254 | 10 | 1.59% |
| hsa_circ_0038325 | 1254 | 10 | 1.59% |
| hsa_circ_0053937 | 1254 | 10 | 1.59% |
| hsa_circ_0056902 | 1254 | 10 | 1.59% |
| hsa_circ_0066659 | 1254 | 10 | 1.59% |
| hsa_circ_0067782 | 1254 | 10 | 1.59% |
| hsa_circ_0068887 | 1254 | 10 | 1.59% |
| hsa_circ_0073565 | 1254 | 10 | 1.59% |
| hsa_circ_0074600 | 1254 | 10 | 1.59% |
| hsa_circ_0092529 | 1254 | 10 | 1.59% |
| hsa_circ_0104267 | 1254 | 10 | 1.59% |
| hsa_circ_0135029 | 1254 | 10 | 1.59% |
| hsa_circ_0137871 | 1254 | 10 | 1.59% |
| hsa_circ_0124286 | 3386 | 27 | 1.59% |
| hsa_circ_0039149 | 1505 | 12 | 1.59% |
| hsa_circ_0073619 | 1505 | 12 | 1.59% |
| hsa_circ_0133518 | 1505 | 12 | 1.59% |
| hsa_circ_0042386 | 1756 | 14 | 1.59% |
| hsa_circ_0014410 | 1380 | 11 | 1.59% |
| hsa_circ_0020645 | 1380 | 11 | 1.59% |
| hsa_circ_0029149 | 1380 | 11 | 1.59% |
| hsa_circ_0076134 | 1380 | 11 | 1.59% |
| hsa_circ_0089436 | 1380 | 11 | 1.59% |
| hsa_circ_0066648 | 2259 | 18 | 1.59% |
| hsa_circ_0017868 | 1506 | 12 | 1.59% |
| hsa_circ_0000051 | 1255 | 10 | 1.59% |
| hsa_circ_0022642 | 1255 | 10 | 1.59% |
| hsa_circ_0029279 | 1255 | 10 | 1.59% |
| hsa_circ_0044925 | 1255 | 10 | 1.59% |
| hsa_circ_0047464 | 1255 | 10 | 1.59% |
| hsa_circ_0052009 | 1255 | 10 | 1.59% |
| hsa_circ_0056083 | 1255 | 10 | 1.59% |
| hsa_circ_0067953 | 1255 | 10 | 1.59% |
| hsa_circ_0109503 | 1255 | 10 | 1.59% |
| hsa_circ_0113447 | 1255 | 10 | 1.59% |

| | | | |
|---|---|---|---|
| hsa_circ_0114745 | 1255 | 10 | 1.59% |
| hsa_circ_0121277 | 1255 | 10 | 1.59% |
| hsa_circ_0025137 | 1381 | 11 | 1.59% |
| hsa_circ_0036181 | 1381 | 11 | 1.59% |
| hsa_circ_0085717 | 1381 | 11 | 1.59% |
| hsa_circ_0114447 | 1381 | 11 | 1.59% |
| hsa_circ_0131715 | 1381 | 11 | 1.59% |
| hsa_circ_0118305 | 17455 | 139 | 1.59% |
| hsa_circ_0114401 | 1507 | 12 | 1.59% |
| hsa_circ_0008083 | 1256 | 10 | 1.59% |
| hsa_circ_0022110 | 1256 | 10 | 1.59% |
| hsa_circ_0041507 | 1256 | 10 | 1.59% |
| hsa_circ_0043676 | 1256 | 10 | 1.59% |
| hsa_circ_0074939 | 1256 | 10 | 1.59% |
| hsa_circ_0075460 | 1256 | 10 | 1.59% |
| hsa_circ_0076415 | 1256 | 10 | 1.59% |
| hsa_circ_0082188 | 1256 | 10 | 1.59% |
| hsa_circ_0103881 | 1256 | 10 | 1.59% |
| hsa_circ_0104276 | 1256 | 10 | 1.59% |
| hsa_circ_0123098 | 1256 | 10 | 1.59% |
| hsa_circ_0128549 | 1256 | 10 | 1.59% |
| hsa_circ_0133085 | 1256 | 10 | 1.59% |
| hsa_circ_0138505 | 1256 | 10 | 1.59% |
| hsa_circ_0101939 | 2261 | 18 | 1.59% |
| hsa_circ_0014761 | 1633 | 13 | 1.59% |
| hsa_circ_0049690 | 2387 | 19 | 1.59% |
| hsa_circ_0138678 | 2387 | 19 | 1.59% |
| hsa_circ_0022888 | 1382 | 11 | 1.59% |
| hsa_circ_0040412 | 1382 | 11 | 1.59% |
| hsa_circ_0043615 | 1382 | 11 | 1.59% |
| hsa_circ_0075585 | 1382 | 11 | 1.59% |
| hsa_circ_0090908 | 1382 | 11 | 1.59% |
| hsa_circ_0108649 | 1382 | 11 | 1.59% |
| hsa_circ_0011533 | 1508 | 12 | 1.59% |
| hsa_circ_0056272 | 1508 | 12 | 1.59% |
| hsa_circ_0088886 | 1508 | 12 | 1.59% |
| hsa_circ_0105887 | 1634 | 13 | 1.59% |
| hsa_circ_0044106 | 1257 | 10 | 1.59% |
| hsa_circ_0096807 | 1257 | 10 | 1.59% |
| hsa_circ_0104892 | 1257 | 10 | 1.59% |
| hsa_circ_0107816 | 1257 | 10 | 1.59% |
| hsa_circ_0113328 | 1257 | 10 | 1.59% |
| hsa_circ_0116863 | 1257 | 10 | 1.59% |
| hsa_circ_0049435 | 1383 | 11 | 1.59% |
| hsa_circ_0050364 | 1383 | 11 | 1.59% |
| hsa_circ_0089449 | 1383 | 11 | 1.59% |
| hsa_circ_0096726 | 1383 | 11 | 1.59% |
| hsa_circ_0099160 | 1383 | 11 | 1.59% |
| hsa_circ_0121375 | 1383 | 11 | 1.59% |
| hsa_circ_0121707 | 1383 | 11 | 1.59% |

| | | | |
|---|---|---|---|
| hsa_circ_0123944 | 1383 | 11 | 1.59% |
| hsa_circ_0085080 | 1886 | 15 | 1.59% |
| hsa_circ_0030880 | 1509 | 12 | 1.59% |
| hsa_circ_0044666 | 3270 | 26 | 1.59% |
| hsa_circ_0017720 | 1258 | 10 | 1.59% |
| hsa_circ_0026110 | 1258 | 10 | 1.59% |
| hsa_circ_0041090 | 1258 | 10 | 1.59% |
| hsa_circ_0081667 | 1258 | 10 | 1.59% |
| hsa_circ_0082468 | 1258 | 10 | 1.59% |
| hsa_circ_0108771 | 1258 | 10 | 1.59% |
| hsa_circ_0137425 | 1258 | 10 | 1.59% |
| hsa_circ_0025584 | 1384 | 11 | 1.59% |
| hsa_circ_0047667 | 1384 | 11 | 1.59% |
| hsa_circ_0049328 | 1384 | 11 | 1.59% |
| hsa_circ_0051392 | 1384 | 11 | 1.59% |
| hsa_circ_0052026 | 1384 | 11 | 1.59% |
| hsa_circ_0080098 | 1384 | 11 | 1.59% |
| hsa_circ_0120615 | 1384 | 11 | 1.59% |
| hsa_circ_0017027 | 1510 | 12 | 1.59% |
| hsa_circ_0004498 | 1259 | 10 | 1.59% |
| hsa_circ_0008886 | 1259 | 10 | 1.59% |
| hsa_circ_0032908 | 1259 | 10 | 1.59% |
| hsa_circ_0055048 | 1259 | 10 | 1.59% |
| hsa_circ_0091998 | 1259 | 10 | 1.59% |
| hsa_circ_0110210 | 1259 | 10 | 1.59% |
| hsa_circ_0122940 | 1259 | 10 | 1.59% |
| hsa_circ_0123654 | 1259 | 10 | 1.59% |
| hsa_circ_0124208 | 1259 | 10 | 1.59% |
| hsa_circ_0126804 | 1259 | 10 | 1.59% |
| hsa_circ_0127779 | 1259 | 10 | 1.59% |
| hsa_circ_0134624 | 1259 | 10 | 1.59% |
| hsa_circ_0019135 | 1385 | 11 | 1.59% |
| hsa_circ_0024420 | 1385 | 11 | 1.59% |
| hsa_circ_0027513 | 1385 | 11 | 1.59% |
| hsa_circ_0038257 | 1385 | 11 | 1.59% |
| hsa_circ_0060590 | 1385 | 11 | 1.59% |
| hsa_circ_0066073 | 1385 | 11 | 1.59% |
| hsa_circ_0072614 | 1385 | 11 | 1.59% |
| hsa_circ_0080097 | 1385 | 11 | 1.59% |
| hsa_circ_0089670 | 1385 | 11 | 1.59% |
| hsa_circ_0095524 | 1385 | 11 | 1.59% |
| hsa_circ_0114243 | 1385 | 11 | 1.59% |
| hsa_circ_0131181 | 1385 | 11 | 1.59% |
| hsa_circ_0094440 | 1511 | 12 | 1.59% |
| hsa_circ_0110326 | 1511 | 12 | 1.59% |
| hsa_circ_0118140 | 1763 | 14 | 1.59% |
| hsa_circ_0113224 | 1764 | 14 | 1.59% |
| hsa_circ_0067384 | 1386 | 11 | 1.59% |
| hsa_circ_0071905 | 1386 | 11 | 1.59% |
| hsa_circ_0073047 | 1386 | 11 | 1.59% |

| | | | |
|---|---|---|---|
| hsa_circ_0076317 | 1386 | 11 | 1.59% |
| hsa_circ_0098148 | 1386 | 11 | 1.59% |
| hsa_circ_0112048 | 1386 | 11 | 1.59% |
| hsa_circ_0013628 | 1260 | 10 | 1.59% |
| hsa_circ_0030385 | 1260 | 10 | 1.59% |
| hsa_circ_0032601 | 1260 | 10 | 1.59% |
| hsa_circ_0053854 | 1260 | 10 | 1.59% |
| hsa_circ_0062515 | 1260 | 10 | 1.59% |
| hsa_circ_0077787 | 1260 | 10 | 1.59% |
| hsa_circ_0099030 | 1260 | 10 | 1.59% |
| hsa_circ_0105879 | 1260 | 10 | 1.59% |
| hsa_circ_0124747 | 1260 | 10 | 1.59% |
| hsa_circ_0136044 | 1260 | 10 | 1.59% |
| hsa_circ_0134955 | 1639 | 13 | 1.59% |
| hsa_circ_0009464 | 1387 | 11 | 1.59% |
| hsa_circ_0021812 | 1387 | 11 | 1.59% |
| hsa_circ_0079132 | 1387 | 11 | 1.59% |
| hsa_circ_0114619 | 1387 | 11 | 1.59% |
| hsa_circ_0000790 | 1261 | 10 | 1.59% |
| hsa_circ_0027606 | 1261 | 10 | 1.59% |
| hsa_circ_0036438 | 1261 | 10 | 1.59% |
| hsa_circ_0046724 | 1261 | 10 | 1.59% |
| hsa_circ_0056683 | 1261 | 10 | 1.59% |
| hsa_circ_0062531 | 1261 | 10 | 1.59% |
| hsa_circ_0066149 | 1261 | 10 | 1.59% |
| hsa_circ_0066562 | 1261 | 10 | 1.59% |
| hsa_circ_0066884 | 1261 | 10 | 1.59% |
| hsa_circ_0091729 | 1261 | 10 | 1.59% |
| hsa_circ_0092806 | 1261 | 10 | 1.59% |
| hsa_circ_0099974 | 1261 | 10 | 1.59% |
| hsa_circ_0101631 | 1261 | 10 | 1.59% |
| hsa_circ_0103246 | 1261 | 10 | 1.59% |
| hsa_circ_0116451 | 1261 | 10 | 1.59% |
| hsa_circ_0127978 | 1261 | 10 | 1.59% |
| hsa_circ_0131565 | 1261 | 10 | 1.59% |
| hsa_circ_0132394 | 1261 | 10 | 1.59% |
| hsa_circ_0078231 | 3153 | 25 | 1.59% |
| hsa_circ_0052070 | 1766 | 14 | 1.59% |
| hsa_circ_0069036 | 1640 | 13 | 1.59% |
| hsa_circ_0114927 | 3154 | 25 | 1.59% |
| hsa_circ_0009923 | 1388 | 11 | 1.59% |
| hsa_circ_0027658 | 1388 | 11 | 1.59% |
| hsa_circ_0037835 | 1388 | 11 | 1.59% |
| hsa_circ_0038789 | 1388 | 11 | 1.59% |
| hsa_circ_0060939 | 1388 | 11 | 1.59% |
| hsa_circ_0069804 | 1388 | 11 | 1.59% |
| hsa_circ_0074897 | 1388 | 11 | 1.59% |
| hsa_circ_0084379 | 1388 | 11 | 1.59% |
| hsa_circ_0099410 | 1388 | 11 | 1.59% |
| hsa_circ_0118214 | 1388 | 11 | 1.59% |

| | | | |
|---|---|---|---|
| hsa_circ_0012732 | 1262 | 10 | 1.58% |
| hsa_circ_0013778 | 1262 | 10 | 1.58% |
| hsa_circ_0015361 | 1262 | 10 | 1.58% |
| hsa_circ_0017620 | 1262 | 10 | 1.58% |
| hsa_circ_0035522 | 1262 | 10 | 1.58% |
| hsa_circ_0038638 | 1262 | 10 | 1.58% |
| hsa_circ_0043156 | 1262 | 10 | 1.58% |
| hsa_circ_0047896 | 1262 | 10 | 1.58% |
| hsa_circ_0055110 | 1262 | 10 | 1.58% |
| hsa_circ_0079708 | 1262 | 10 | 1.58% |
| hsa_circ_0081277 | 1262 | 10 | 1.58% |
| hsa_circ_0127635 | 1262 | 10 | 1.58% |
| hsa_circ_0131021 | 1262 | 10 | 1.58% |
| hsa_circ_0046163 | 1641 | 13 | 1.58% |
| hsa_circ_0086712 | 1641 | 13 | 1.58% |
| hsa_circ_0082911 | 1515 | 12 | 1.58% |
| hsa_circ_0129439 | 1515 | 12 | 1.58% |
| hsa_circ_0043378 | 1389 | 11 | 1.58% |
| hsa_circ_0060928 | 1389 | 11 | 1.58% |
| hsa_circ_0072766 | 1389 | 11 | 1.58% |
| hsa_circ_0108404 | 1389 | 11 | 1.58% |
| hsa_circ_0110976 | 1389 | 11 | 1.58% |
| hsa_circ_0122456 | 1389 | 11 | 1.58% |
| hsa_circ_0140157 | 1389 | 11 | 1.58% |
| hsa_circ_0010390 | 2273 | 18 | 1.58% |
| hsa_circ_0027270 | 1263 | 10 | 1.58% |
| hsa_circ_0031767 | 1263 | 10 | 1.58% |
| hsa_circ_0104369 | 1263 | 10 | 1.58% |
| hsa_circ_0110975 | 1263 | 10 | 1.58% |
| hsa_circ_0115100 | 1263 | 10 | 1.58% |
| hsa_circ_0118422 | 1263 | 10 | 1.58% |
| hsa_circ_0129256 | 1263 | 10 | 1.58% |
| hsa_circ_0129386 | 1263 | 10 | 1.58% |
| hsa_circ_0129468 | 1263 | 10 | 1.58% |
| hsa_circ_0133103 | 1263 | 10 | 1.58% |
| hsa_circ_0131114 | 3790 | 30 | 1.58% |
| hsa_circ_0010792 | 1516 | 12 | 1.58% |
| hsa_circ_0040693 | 1516 | 12 | 1.58% |
| hsa_circ_0073472 | 1516 | 12 | 1.58% |
| hsa_circ_0039937 | 1769 | 14 | 1.58% |
| hsa_circ_0049060 | 1390 | 11 | 1.58% |
| hsa_circ_0068802 | 1390 | 11 | 1.58% |
| hsa_circ_0085315 | 1390 | 11 | 1.58% |
| hsa_circ_0089505 | 1390 | 11 | 1.58% |
| hsa_circ_0121141 | 1390 | 11 | 1.58% |
| hsa_circ_0059656 | 1643 | 13 | 1.58% |
| hsa_circ_0001719 | 3539 | 28 | 1.58% |
| hsa_circ_0003491 | 1264 | 10 | 1.58% |
| hsa_circ_0015396 | 1264 | 10 | 1.58% |
| hsa_circ_0032964 | 1264 | 10 | 1.58% |

| | | | |
|---|---|---|---|
| hsa_circ_0039482 | 1264 | 10 | 1.58% |
| hsa_circ_0072937 | 1264 | 10 | 1.58% |
| hsa_circ_0075650 | 1264 | 10 | 1.58% |
| hsa_circ_0080887 | 1264 | 10 | 1.58% |
| hsa_circ_0082102 | 1264 | 10 | 1.58% |
| hsa_circ_0093235 | 1264 | 10 | 1.58% |
| hsa_circ_0100602 | 1264 | 10 | 1.58% |
| hsa_circ_0111297 | 1264 | 10 | 1.58% |
| hsa_circ_0131381 | 1264 | 10 | 1.58% |
| hsa_circ_0139016 | 1264 | 10 | 1.58% |
| hsa_circ_0065852 | 1517 | 12 | 1.58% |
| hsa_circ_0091091 | 1517 | 12 | 1.58% |
| hsa_circ_0066773 | 1391 | 11 | 1.58% |
| hsa_circ_0067495 | 1391 | 11 | 1.58% |
| hsa_circ_0068694 | 1391 | 11 | 1.58% |
| hsa_circ_0079072 | 1391 | 11 | 1.58% |
| hsa_circ_0061457 | 1644 | 13 | 1.58% |
| hsa_circ_0071463 | 4048 | 32 | 1.58% |
| hsa_circ_0027313 | 1518 | 12 | 1.58% |
| hsa_circ_0034955 | 1518 | 12 | 1.58% |
| hsa_circ_0017159 | 1265 | 10 | 1.58% |
| hsa_circ_0020834 | 1265 | 10 | 1.58% |
| hsa_circ_0032300 | 1265 | 10 | 1.58% |
| hsa_circ_0033128 | 1265 | 10 | 1.58% |
| hsa_circ_0044024 | 1265 | 10 | 1.58% |
| hsa_circ_0044290 | 1265 | 10 | 1.58% |
| hsa_circ_0046241 | 1265 | 10 | 1.58% |
| hsa_circ_0048366 | 1265 | 10 | 1.58% |
| hsa_circ_0051378 | 1265 | 10 | 1.58% |
| hsa_circ_0054239 | 1265 | 10 | 1.58% |
| hsa_circ_0062384 | 1265 | 10 | 1.58% |
| hsa_circ_0086231 | 1265 | 10 | 1.58% |
| hsa_circ_0108633 | 1265 | 10 | 1.58% |
| hsa_circ_0121948 | 1265 | 10 | 1.58% |
| hsa_circ_0124955 | 1265 | 10 | 1.58% |
| hsa_circ_0126209 | 1265 | 10 | 1.58% |
| hsa_circ_0132236 | 1265 | 10 | 1.58% |
| hsa_circ_0139000 | 1265 | 10 | 1.58% |
| hsa_circ_0004997 | 1392 | 11 | 1.58% |
| hsa_circ_0011404 | 1392 | 11 | 1.58% |
| hsa_circ_0019920 | 1392 | 11 | 1.58% |
| hsa_circ_0027079 | 1392 | 11 | 1.58% |
| hsa_circ_0043508 | 1392 | 11 | 1.58% |
| hsa_circ_0060764 | 1392 | 11 | 1.58% |
| hsa_circ_0060952 | 1392 | 11 | 1.58% |
| hsa_circ_0088390 | 1392 | 11 | 1.58% |
| hsa_circ_0089463 | 1392 | 11 | 1.58% |
| hsa_circ_0097517 | 1392 | 11 | 1.58% |
| hsa_circ_0104315 | 1392 | 11 | 1.58% |
| hsa_circ_0001135 | 1266 | 10 | 1.58% |

| | | | |
|---|---|---|---|
| hsa_circ_0020807 | 1266 | 10 | 1.58% |
| hsa_circ_0021180 | 1266 | 10 | 1.58% |
| hsa_circ_0046191 | 1266 | 10 | 1.58% |
| hsa_circ_0057916 | 1266 | 10 | 1.58% |
| hsa_circ_0079378 | 1266 | 10 | 1.58% |
| hsa_circ_0082135 | 1266 | 10 | 1.58% |
| hsa_circ_0084537 | 1266 | 10 | 1.58% |
| hsa_circ_0090149 | 1266 | 10 | 1.58% |
| hsa_circ_0091600 | 1266 | 10 | 1.58% |
| hsa_circ_0096534 | 1266 | 10 | 1.58% |
| hsa_circ_0099021 | 1266 | 10 | 1.58% |
| hsa_circ_0108886 | 1266 | 10 | 1.58% |
| hsa_circ_0111855 | 1266 | 10 | 1.58% |
| hsa_circ_0137845 | 1266 | 10 | 1.58% |
| hsa_circ_0010765 | 1393 | 11 | 1.58% |
| hsa_circ_0014668 | 1393 | 11 | 1.58% |
| hsa_circ_0109308 | 4053 | 32 | 1.58% |
| hsa_circ_0073004 | 1520 | 12 | 1.58% |
| hsa_circ_0079696 | 1520 | 12 | 1.58% |
| hsa_circ_0110238 | 1520 | 12 | 1.58% |
| hsa_circ_0081406 | 1647 | 13 | 1.58% |
| hsa_circ_0014188 | 1267 | 10 | 1.58% |
| hsa_circ_0014937 | 1267 | 10 | 1.58% |
| hsa_circ_0022876 | 1267 | 10 | 1.58% |
| hsa_circ_0023629 | 1267 | 10 | 1.58% |
| hsa_circ_0030013 | 1267 | 10 | 1.58% |
| hsa_circ_0037171 | 1267 | 10 | 1.58% |
| hsa_circ_0037310 | 1267 | 10 | 1.58% |
| hsa_circ_0050971 | 1267 | 10 | 1.58% |
| hsa_circ_0063373 | 1267 | 10 | 1.58% |
| hsa_circ_0063847 | 1267 | 10 | 1.58% |
| hsa_circ_0087019 | 1267 | 10 | 1.58% |
| hsa_circ_0072823 | 2154 | 17 | 1.58% |
| hsa_circ_0012994 | 1394 | 11 | 1.58% |
| hsa_circ_0029573 | 1394 | 11 | 1.58% |
| hsa_circ_0049427 | 1394 | 11 | 1.58% |
| hsa_circ_0060381 | 1394 | 11 | 1.58% |
| hsa_circ_0138289 | 1394 | 11 | 1.58% |
| hsa_circ_0014787 | 1901 | 15 | 1.58% |
| hsa_circ_0039206 | 1521 | 12 | 1.58% |
| hsa_circ_0112252 | 1521 | 12 | 1.58% |
| hsa_circ_0116291 | 1521 | 12 | 1.58% |
| hsa_circ_0120580 | 1521 | 12 | 1.58% |
| hsa_circ_0000563 | 1268 | 10 | 1.58% |
| hsa_circ_0005914 | 1268 | 10 | 1.58% |
| hsa_circ_0025267 | 1268 | 10 | 1.58% |
| hsa_circ_0051111 | 1268 | 10 | 1.58% |
| hsa_circ_0070878 | 1268 | 10 | 1.58% |
| hsa_circ_0071263 | 1268 | 10 | 1.58% |
| hsa_circ_0079824 | 1268 | 10 | 1.58% |

| | | | |
|---|---|---|---|
| hsa_circ_0088370 | 1268 | 10 | 1.58% |
| hsa_circ_0088438 | 1268 | 10 | 1.58% |
| hsa_circ_0089787 | 1268 | 10 | 1.58% |
| hsa_circ_0127421 | 1268 | 10 | 1.58% |
| hsa_circ_0039589 | 1395 | 11 | 1.58% |
| hsa_circ_0043970 | 1395 | 11 | 1.58% |
| hsa_circ_0053145 | 1395 | 11 | 1.58% |
| hsa_circ_0066706 | 1395 | 11 | 1.58% |
| hsa_circ_0112489 | 1395 | 11 | 1.58% |
| hsa_circ_0127825 | 1395 | 11 | 1.58% |
| hsa_circ_0023529 | 1522 | 12 | 1.58% |
| hsa_circ_0054682 | 1522 | 12 | 1.58% |
| hsa_circ_0112907 | 1522 | 12 | 1.58% |
| hsa_circ_0044498 | 1649 | 13 | 1.58% |
| hsa_circ_0005976 | 1269 | 10 | 1.58% |
| hsa_circ_0030107 | 1269 | 10 | 1.58% |
| hsa_circ_0046022 | 1269 | 10 | 1.58% |
| hsa_circ_0059632 | 1269 | 10 | 1.58% |
| hsa_circ_0061967 | 1269 | 10 | 1.58% |
| hsa_circ_0068257 | 1269 | 10 | 1.58% |
| hsa_circ_0081942 | 1269 | 10 | 1.58% |
| hsa_circ_0085851 | 1269 | 10 | 1.58% |
| hsa_circ_0088583 | 1269 | 10 | 1.58% |
| hsa_circ_0090914 | 1269 | 10 | 1.58% |
| hsa_circ_0122988 | 1269 | 10 | 1.58% |
| hsa_circ_0124670 | 1269 | 10 | 1.58% |
| hsa_circ_0126336 | 1269 | 10 | 1.58% |
| hsa_circ_0133706 | 1269 | 10 | 1.58% |
| hsa_circ_0040268 | 1396 | 11 | 1.58% |
| hsa_circ_0041042 | 1396 | 11 | 1.58% |
| hsa_circ_0044163 | 1396 | 11 | 1.58% |
| hsa_circ_0044167 | 1396 | 11 | 1.58% |
| hsa_circ_0052278 | 1396 | 11 | 1.58% |
| hsa_circ_0063247 | 1396 | 11 | 1.58% |
| hsa_circ_0099109 | 1396 | 11 | 1.58% |
| hsa_circ_0029784 | 1523 | 12 | 1.58% |
| hsa_circ_0053120 | 1523 | 12 | 1.58% |
| hsa_circ_0077172 | 1523 | 12 | 1.58% |
| hsa_circ_0081960 | 1523 | 12 | 1.58% |
| hsa_circ_0024555 | 1650 | 13 | 1.58% |
| hsa_circ_0045815 | 1650 | 13 | 1.58% |
| hsa_circ_0067806 | 2666 | 21 | 1.58% |
| hsa_circ_0015664 | 1524 | 12 | 1.57% |
| hsa_circ_0067436 | 1524 | 12 | 1.57% |
| hsa_circ_0088382 | 1524 | 12 | 1.57% |
| hsa_circ_0031632 | 1397 | 11 | 1.57% |
| hsa_circ_0057968 | 1397 | 11 | 1.57% |
| hsa_circ_0097693 | 1397 | 11 | 1.57% |
| hsa_circ_0106270 | 1397 | 11 | 1.57% |
| hsa_circ_0008779 | 1270 | 10 | 1.57% |

| | | | |
|---|---|---|---|
| hsa_circ_0026720 | 1270 | 10 | 1.57% |
| hsa_circ_0048353 | 1270 | 10 | 1.57% |
| hsa_circ_0055421 | 1270 | 10 | 1.57% |
| hsa_circ_0072728 | 1270 | 10 | 1.57% |
| hsa_circ_0074503 | 1270 | 10 | 1.57% |
| hsa_circ_0090658 | 1270 | 10 | 1.57% |
| hsa_circ_0091361 | 1270 | 10 | 1.57% |
| hsa_circ_0121388 | 1270 | 10 | 1.57% |
| hsa_circ_0135972 | 1270 | 10 | 1.57% |
| hsa_circ_0138556 | 1270 | 10 | 1.57% |
| hsa_circ_0139209 | 1270 | 10 | 1.57% |
| hsa_circ_0140331 | 1652 | 13 | 1.57% |
| hsa_circ_0027164 | 1525 | 12 | 1.57% |
| hsa_circ_0031689 | 1525 | 12 | 1.57% |
| hsa_circ_0043770 | 1525 | 12 | 1.57% |
| hsa_circ_0059532 | 1525 | 12 | 1.57% |
| hsa_circ_0079419 | 1525 | 12 | 1.57% |
| hsa_circ_0010235 | 1398 | 11 | 1.57% |
| hsa_circ_0015632 | 1398 | 11 | 1.57% |
| hsa_circ_0071675 | 1398 | 11 | 1.57% |
| hsa_circ_0100436 | 1398 | 11 | 1.57% |
| hsa_circ_0029290 | 1271 | 10 | 1.57% |
| hsa_circ_0040446 | 1271 | 10 | 1.57% |
| hsa_circ_0041968 | 1271 | 10 | 1.57% |
| hsa_circ_0052470 | 1271 | 10 | 1.57% |
| hsa_circ_0056523 | 1271 | 10 | 1.57% |
| hsa_circ_0056760 | 1271 | 10 | 1.57% |
| hsa_circ_0061836 | 1271 | 10 | 1.57% |
| hsa_circ_0079311 | 1271 | 10 | 1.57% |
| hsa_circ_0083088 | 1271 | 10 | 1.57% |
| hsa_circ_0083603 | 1271 | 10 | 1.57% |
| hsa_circ_0106655 | 1271 | 10 | 1.57% |
| hsa_circ_0112187 | 1271 | 10 | 1.57% |
| hsa_circ_0125262 | 1271 | 10 | 1.57% |
| hsa_circ_0133348 | 1271 | 10 | 1.57% |
| hsa_circ_0134712 | 1271 | 10 | 1.57% |
| hsa_circ_0122969 | 6228 | 49 | 1.57% |
| hsa_circ_0105327 | 1526 | 12 | 1.57% |
| hsa_circ_0133195 | 1526 | 12 | 1.57% |
| hsa_circ_0060069 | 2798 | 22 | 1.57% |
| hsa_circ_0023953 | 1399 | 11 | 1.57% |
| hsa_circ_0066744 | 1399 | 11 | 1.57% |
| hsa_circ_0069260 | 1399 | 11 | 1.57% |
| hsa_circ_0082284 | 1399 | 11 | 1.57% |
| hsa_circ_0095953 | 1399 | 11 | 1.57% |
| hsa_circ_0110755 | 3180 | 25 | 1.57% |
| hsa_circ_0006617 | 1272 | 10 | 1.57% |
| hsa_circ_0016870 | 1272 | 10 | 1.57% |
| hsa_circ_0044667 | 1272 | 10 | 1.57% |
| hsa_circ_0053748 | 1272 | 10 | 1.57% |

| | | | |
|---|---|---|---|
| hsa_circ_0067993 | 1272 | 10 | 1.57% |
| hsa_circ_0085749 | 1272 | 10 | 1.57% |
| hsa_circ_0095100 | 1272 | 10 | 1.57% |
| hsa_circ_0100911 | 1272 | 10 | 1.57% |
| hsa_circ_0123540 | 1272 | 10 | 1.57% |
| hsa_circ_0126248 | 1272 | 10 | 1.57% |
| hsa_circ_0126302 | 1272 | 10 | 1.57% |
| hsa_circ_0064801 | 1527 | 12 | 1.57% |
| hsa_circ_0118380 | 1527 | 12 | 1.57% |
| hsa_circ_0000752 | 1400 | 11 | 1.57% |
| hsa_circ_0001283 | 1400 | 11 | 1.57% |
| hsa_circ_0057060 | 1400 | 11 | 1.57% |
| hsa_circ_0080595 | 1400 | 11 | 1.57% |
| hsa_circ_0042390 | 1782 | 14 | 1.57% |
| hsa_circ_0012705 | 1273 | 10 | 1.57% |
| hsa_circ_0017404 | 1273 | 10 | 1.57% |
| hsa_circ_0020218 | 1273 | 10 | 1.57% |
| hsa_circ_0024623 | 1273 | 10 | 1.57% |
| hsa_circ_0050336 | 1273 | 10 | 1.57% |
| hsa_circ_0054337 | 1273 | 10 | 1.57% |
| hsa_circ_0062275 | 1273 | 10 | 1.57% |
| hsa_circ_0085109 | 1273 | 10 | 1.57% |
| hsa_circ_0070390 | 1528 | 12 | 1.57% |
| hsa_circ_0009679 | 3693 | 29 | 1.57% |
| hsa_circ_0109910 | 1783 | 14 | 1.57% |
| hsa_circ_0001381 | 1401 | 11 | 1.57% |
| hsa_circ_0003025 | 1401 | 11 | 1.57% |
| hsa_circ_0071926 | 1401 | 11 | 1.57% |
| hsa_circ_0091023 | 1401 | 11 | 1.57% |
| hsa_circ_0101879 | 1401 | 11 | 1.57% |
| hsa_circ_0103464 | 1401 | 11 | 1.57% |
| hsa_circ_0133836 | 1401 | 11 | 1.57% |
| hsa_circ_0001377 | 1274 | 10 | 1.57% |
| hsa_circ_0035672 | 1274 | 10 | 1.57% |
| hsa_circ_0042223 | 1274 | 10 | 1.57% |
| hsa_circ_0050065 | 1274 | 10 | 1.57% |
| hsa_circ_0078488 | 1274 | 10 | 1.57% |
| hsa_circ_0095245 | 1274 | 10 | 1.57% |
| hsa_circ_0100673 | 1274 | 10 | 1.57% |
| hsa_circ_0108214 | 1274 | 10 | 1.57% |
| hsa_circ_0118384 | 1274 | 10 | 1.57% |
| hsa_circ_0133349 | 1274 | 10 | 1.57% |
| hsa_circ_0037117 | 1529 | 12 | 1.57% |
| hsa_circ_0045331 | 1529 | 12 | 1.57% |
| hsa_circ_0081355 | 1529 | 12 | 1.57% |
| hsa_circ_0037838 | 1402 | 11 | 1.57% |
| hsa_circ_0038119 | 1402 | 11 | 1.57% |
| hsa_circ_0063184 | 1402 | 11 | 1.57% |
| hsa_circ_0074006 | 1402 | 11 | 1.57% |
| hsa_circ_0098418 | 1402 | 11 | 1.57% |

| | | | |
|---|---|---|---|
| hsa_circ_0117801 | 1402 | 11 | 1.57% |
| hsa_circ_0072633 | 1657 | 13 | 1.57% |
| hsa_circ_0009666 | 1275 | 10 | 1.57% |
| hsa_circ_0012102 | 1275 | 10 | 1.57% |
| hsa_circ_0013289 | 1275 | 10 | 1.57% |
| hsa_circ_0026038 | 1275 | 10 | 1.57% |
| hsa_circ_0034387 | 1275 | 10 | 1.57% |
| hsa_circ_0065457 | 1275 | 10 | 1.57% |
| hsa_circ_0072486 | 1275 | 10 | 1.57% |
| hsa_circ_0078580 | 1275 | 10 | 1.57% |
| hsa_circ_0116089 | 1275 | 10 | 1.57% |
| hsa_circ_0139753 | 1275 | 10 | 1.57% |
| hsa_circ_0012950 | 2168 | 17 | 1.57% |
| hsa_circ_0002301 | 1403 | 11 | 1.57% |
| hsa_circ_0041349 | 1403 | 11 | 1.57% |
| hsa_circ_0051163 | 1403 | 11 | 1.57% |
| hsa_circ_0083049 | 1403 | 11 | 1.57% |
| hsa_circ_0100530 | 1403 | 11 | 1.57% |
| hsa_circ_0123863 | 1403 | 11 | 1.57% |
| hsa_circ_0119960 | 2424 | 19 | 1.57% |
| hsa_circ_0096212 | 3700 | 29 | 1.57% |
| hsa_circ_0012736 | 1276 | 10 | 1.57% |
| hsa_circ_0036932 | 1276 | 10 | 1.57% |
| hsa_circ_0044294 | 1276 | 10 | 1.57% |
| hsa_circ_0054013 | 1276 | 10 | 1.57% |
| hsa_circ_0072439 | 1276 | 10 | 1.57% |
| hsa_circ_0080072 | 1404 | 11 | 1.57% |
| hsa_circ_0096522 | 1404 | 11 | 1.57% |
| hsa_circ_0118797 | 1404 | 11 | 1.57% |
| hsa_circ_0122903 | 1404 | 11 | 1.57% |
| hsa_circ_0130795 | 1404 | 11 | 1.57% |
| hsa_circ_0140389 | 1404 | 11 | 1.57% |
| hsa_circ_0000240 | 1532 | 12 | 1.57% |
| hsa_circ_0045255 | 1532 | 12 | 1.57% |
| hsa_circ_0064166 | 2554 | 20 | 1.57% |
| hsa_circ_0004157 | 1277 | 10 | 1.57% |
| hsa_circ_0016912 | 1277 | 10 | 1.57% |
| hsa_circ_0019414 | 1277 | 10 | 1.57% |
| hsa_circ_0029663 | 1277 | 10 | 1.57% |
| hsa_circ_0029976 | 1277 | 10 | 1.57% |
| hsa_circ_0032261 | 1277 | 10 | 1.57% |
| hsa_circ_0064730 | 1277 | 10 | 1.57% |
| hsa_circ_0064941 | 1277 | 10 | 1.57% |
| hsa_circ_0076075 | 1277 | 10 | 1.57% |
| hsa_circ_0086237 | 1277 | 10 | 1.57% |
| hsa_circ_0087727 | 1277 | 10 | 1.57% |
| hsa_circ_0095114 | 1277 | 10 | 1.57% |
| hsa_circ_0095677 | 1277 | 10 | 1.57% |
| hsa_circ_0134650 | 3193 | 25 | 1.57% |
| hsa_circ_0011297 | 1405 | 11 | 1.57% |

| | | | |
|---|---|---|---|
| hsa_circ_0013795 | 1405 | 11 | 1.57% |
| hsa_circ_0020293 | 1405 | 11 | 1.57% |
| hsa_circ_0039735 | 1405 | 11 | 1.57% |
| hsa_circ_0069893 | 1405 | 11 | 1.57% |
| hsa_circ_0070596 | 1405 | 11 | 1.57% |
| hsa_circ_0079553 | 1405 | 11 | 1.57% |
| hsa_circ_0101825 | 1405 | 11 | 1.57% |
| hsa_circ_0102751 | 1405 | 11 | 1.57% |
| hsa_circ_0129785 | 1405 | 11 | 1.57% |
| hsa_circ_0009844 | 1533 | 12 | 1.57% |
| hsa_circ_0047993 | 1533 | 12 | 1.57% |
| hsa_circ_0054627 | 1533 | 12 | 1.57% |
| hsa_circ_0097278 | 3450 | 27 | 1.57% |
| hsa_circ_0023072 | 1278 | 10 | 1.56% |
| hsa_circ_0023918 | 1278 | 10 | 1.56% |
| hsa_circ_0041712 | 1278 | 10 | 1.56% |
| hsa_circ_0059734 | 1278 | 10 | 1.56% |
| hsa_circ_0061465 | 1278 | 10 | 1.56% |
| hsa_circ_0102628 | 1278 | 10 | 1.56% |
| hsa_circ_0109569 | 1278 | 10 | 1.56% |
| hsa_circ_0110457 | 1278 | 10 | 1.56% |
| hsa_circ_0010518 | 3451 | 27 | 1.56% |
| hsa_circ_0019357 | 1406 | 11 | 1.56% |
| hsa_circ_0035596 | 1406 | 11 | 1.56% |
| hsa_circ_0053684 | 1406 | 11 | 1.56% |
| hsa_circ_0062380 | 1406 | 11 | 1.56% |
| hsa_circ_0070398 | 1406 | 11 | 1.56% |
| hsa_circ_0086598 | 1406 | 11 | 1.56% |
| hsa_circ_0089289 | 1406 | 11 | 1.56% |
| hsa_circ_0089639 | 1406 | 11 | 1.56% |
| hsa_circ_0092092 | 1406 | 11 | 1.56% |
| hsa_circ_0122264 | 1406 | 11 | 1.56% |
| hsa_circ_0133105 | 1406 | 11 | 1.56% |
| hsa_circ_0017575 | 1534 | 12 | 1.56% |
| hsa_circ_0100847 | 1534 | 12 | 1.56% |
| hsa_circ_0077025 | 1662 | 13 | 1.56% |
| hsa_circ_0062131 | 2046 | 16 | 1.56% |
| hsa_circ_0013389 | 1279 | 10 | 1.56% |
| hsa_circ_0014095 | 1279 | 10 | 1.56% |
| hsa_circ_0015683 | 1279 | 10 | 1.56% |
| hsa_circ_0018249 | 1279 | 10 | 1.56% |
| hsa_circ_0025508 | 1279 | 10 | 1.56% |
| hsa_circ_0040053 | 1279 | 10 | 1.56% |
| hsa_circ_0040540 | 1279 | 10 | 1.56% |
| hsa_circ_0047465 | 1279 | 10 | 1.56% |
| hsa_circ_0047491 | 1279 | 10 | 1.56% |
| hsa_circ_0048617 | 1279 | 10 | 1.56% |
| hsa_circ_0056425 | 1279 | 10 | 1.56% |
| hsa_circ_0086647 | 1279 | 10 | 1.56% |
| hsa_circ_0102966 | 1279 | 10 | 1.56% |

| | | | |
|---|---|---|---|
| hsa_circ_0113215 | 1279 | 10 | 1.56% |
| hsa_circ_0121406 | 1279 | 10 | 1.56% |
| hsa_circ_0126417 | 1279 | 10 | 1.56% |
| hsa_circ_0133826 | 1279 | 10 | 1.56% |
| hsa_circ_0139556 | 1279 | 10 | 1.56% |
| hsa_circ_0010563 | 1407 | 11 | 1.56% |
| hsa_circ_0027799 | 1407 | 11 | 1.56% |
| hsa_circ_0065413 | 1407 | 11 | 1.56% |
| hsa_circ_0108754 | 1407 | 11 | 1.56% |
| hsa_circ_0127765 | 1407 | 11 | 1.56% |
| hsa_circ_0013631 | 1535 | 12 | 1.56% |
| hsa_circ_0036209 | 1535 | 12 | 1.56% |
| hsa_circ_0051721 | 1535 | 12 | 1.56% |
| hsa_circ_0055888 | 1535 | 12 | 1.56% |
| hsa_circ_0063199 | 1535 | 12 | 1.56% |
| hsa_circ_0112908 | 1535 | 12 | 1.56% |
| hsa_circ_0047946 | 1663 | 13 | 1.56% |
| hsa_circ_0080475 | 1919 | 15 | 1.56% |
| hsa_circ_0022171 | 1792 | 14 | 1.56% |
| hsa_circ_0085873 | 1536 | 12 | 1.56% |
| hsa_circ_0044744 | 1408 | 11 | 1.56% |
| hsa_circ_0045243 | 1408 | 11 | 1.56% |
| hsa_circ_0051825 | 1408 | 11 | 1.56% |
| hsa_circ_0094989 | 1408 | 11 | 1.56% |
| hsa_circ_0135126 | 1408 | 11 | 1.56% |
| hsa_circ_0018102 | 1280 | 10 | 1.56% |
| hsa_circ_0022255 | 1280 | 10 | 1.56% |
| hsa_circ_0038285 | 1280 | 10 | 1.56% |
| hsa_circ_0039853 | 1280 | 10 | 1.56% |
| hsa_circ_0075179 | 1280 | 10 | 1.56% |
| hsa_circ_0075474 | 1280 | 10 | 1.56% |
| hsa_circ_0076865 | 1280 | 10 | 1.56% |
| hsa_circ_0087722 | 1280 | 10 | 1.56% |
| hsa_circ_0089646 | 1280 | 10 | 1.56% |
| hsa_circ_0099649 | 1280 | 10 | 1.56% |
| hsa_circ_0126019 | 1280 | 10 | 1.56% |
| hsa_circ_0136834 | 1280 | 10 | 1.56% |
| hsa_circ_0019572 | 1409 | 11 | 1.56% |
| hsa_circ_0027502 | 1409 | 11 | 1.56% |
| hsa_circ_0030977 | 1409 | 11 | 1.56% |
| hsa_circ_0069910 | 1409 | 11 | 1.56% |
| hsa_circ_0099566 | 1409 | 11 | 1.56% |
| hsa_circ_0103412 | 1409 | 11 | 1.56% |
| hsa_circ_0122285 | 1409 | 11 | 1.56% |
| hsa_circ_0127261 | 1409 | 11 | 1.56% |
| hsa_circ_0140572 | 1409 | 11 | 1.56% |
| hsa_circ_0011511 | 1281 | 10 | 1.56% |
| hsa_circ_0014006 | 1281 | 10 | 1.56% |
| hsa_circ_0049307 | 1281 | 10 | 1.56% |
| hsa_circ_0062973 | 1281 | 10 | 1.56% |

| | | | |
|---|---|---|---|
| hsa_circ_0067784 | 1281 | 10 | 1.56% |
| hsa_circ_0075026 | 1281 | 10 | 1.56% |
| hsa_circ_0077238 | 1281 | 10 | 1.56% |
| hsa_circ_0089054 | 1281 | 10 | 1.56% |
| hsa_circ_0092754 | 1281 | 10 | 1.56% |
| hsa_circ_0119426 | 1281 | 10 | 1.56% |
| hsa_circ_0127226 | 1281 | 10 | 1.56% |
| hsa_circ_0135452 | 1281 | 10 | 1.56% |
| hsa_circ_0135589 | 1281 | 10 | 1.56% |
| hsa_circ_0109036 | 2947 | 23 | 1.56% |
| hsa_circ_0005548 | 1922 | 15 | 1.56% |
| hsa_circ_0013538 | 1922 | 15 | 1.56% |
| hsa_circ_0046412 | 1538 | 12 | 1.56% |
| hsa_circ_0134876 | 1538 | 12 | 1.56% |
| hsa_circ_0122481 | 2948 | 23 | 1.56% |
| hsa_circ_0039863 | 1410 | 11 | 1.56% |
| hsa_circ_0048676 | 1410 | 11 | 1.56% |
| hsa_circ_0068117 | 1410 | 11 | 1.56% |
| hsa_circ_0074250 | 1410 | 11 | 1.56% |
| hsa_circ_0078569 | 1410 | 11 | 1.56% |
| hsa_circ_0010693 | 1282 | 10 | 1.56% |
| hsa_circ_0024953 | 1282 | 10 | 1.56% |
| hsa_circ_0033539 | 1282 | 10 | 1.56% |
| hsa_circ_0040953 | 1282 | 10 | 1.56% |
| hsa_circ_0043553 | 1282 | 10 | 1.56% |
| hsa_circ_0051533 | 1282 | 10 | 1.56% |
| hsa_circ_0066716 | 1282 | 10 | 1.56% |
| hsa_circ_0090710 | 1282 | 10 | 1.56% |
| hsa_circ_0093963 | 1282 | 10 | 1.56% |
| hsa_circ_0102159 | 1282 | 10 | 1.56% |
| hsa_circ_0137874 | 1282 | 10 | 1.56% |
| hsa_circ_0077682 | 1539 | 12 | 1.56% |
| hsa_circ_0058238 | 1924 | 15 | 1.56% |
| hsa_circ_0124166 | 3335 | 26 | 1.56% |
| hsa_circ_0027562 | 1411 | 11 | 1.56% |
| hsa_circ_0037797 | 1411 | 11 | 1.56% |
| hsa_circ_0038249 | 1411 | 11 | 1.56% |
| hsa_circ_0042717 | 1411 | 11 | 1.56% |
| hsa_circ_0059554 | 1411 | 11 | 1.56% |
| hsa_circ_0063990 | 1411 | 11 | 1.56% |
| hsa_circ_0076482 | 1411 | 11 | 1.56% |
| hsa_circ_0113946 | 1411 | 11 | 1.56% |
| hsa_circ_0121259 | 3207 | 25 | 1.56% |
| hsa_circ_0028755 | 1283 | 10 | 1.56% |
| hsa_circ_0034087 | 1283 | 10 | 1.56% |
| hsa_circ_0046398 | 1283 | 10 | 1.56% |
| hsa_circ_0060208 | 1283 | 10 | 1.56% |
| hsa_circ_0062225 | 1283 | 10 | 1.56% |
| hsa_circ_0079568 | 1283 | 10 | 1.56% |
| hsa_circ_0080304 | 1283 | 10 | 1.56% |

| | | | |
|---|---|---|---|
| hsa_circ_0106312 | 1283 | 10 | 1.56% |
| hsa_circ_0108097 | 1283 | 10 | 1.56% |
| hsa_circ_0122472 | 1283 | 10 | 1.56% |
| hsa_circ_0129603 | 1283 | 10 | 1.56% |
| hsa_circ_0139532 | 1283 | 10 | 1.56% |
| hsa_circ_0015736 | 1668 | 13 | 1.56% |
| hsa_circ_0064748 | 1668 | 13 | 1.56% |
| hsa_circ_0051714 | 1540 | 12 | 1.56% |
| hsa_circ_0012520 | 1412 | 11 | 1.56% |
| hsa_circ_0024885 | 1412 | 11 | 1.56% |
| hsa_circ_0043342 | 1412 | 11 | 1.56% |
| hsa_circ_0075679 | 1412 | 11 | 1.56% |
| hsa_circ_0097923 | 1412 | 11 | 1.56% |
| hsa_circ_0139588 | 1412 | 11 | 1.56% |
| hsa_circ_0064082 | 1669 | 13 | 1.56% |
| hsa_circ_0016088 | 1284 | 10 | 1.56% |
| hsa_circ_0038886 | 1284 | 10 | 1.56% |
| hsa_circ_0056920 | 1284 | 10 | 1.56% |
| hsa_circ_0085125 | 1284 | 10 | 1.56% |
| hsa_circ_0086214 | 1284 | 10 | 1.56% |
| hsa_circ_0092165 | 1284 | 10 | 1.56% |
| hsa_circ_0126151 | 1284 | 10 | 1.56% |
| hsa_circ_0127122 | 1284 | 10 | 1.56% |
| hsa_circ_0129883 | 1284 | 10 | 1.56% |
| hsa_circ_0058327 | 1541 | 12 | 1.56% |
| hsa_circ_0110920 | 1541 | 12 | 1.56% |
| hsa_circ_0127648 | 1541 | 12 | 1.56% |
| hsa_circ_0008140 | 1413 | 11 | 1.56% |
| hsa_circ_0020891 | 1413 | 11 | 1.56% |
| hsa_circ_0038298 | 1413 | 11 | 1.56% |
| hsa_circ_0079614 | 1413 | 11 | 1.56% |
| hsa_circ_0099220 | 1413 | 11 | 1.56% |
| hsa_circ_0089725 | 1542 | 12 | 1.56% |
| hsa_circ_0007729 | 1285 | 10 | 1.56% |
| hsa_circ_0036275 | 1285 | 10 | 1.56% |
| hsa_circ_0040012 | 1285 | 10 | 1.56% |
| hsa_circ_0040514 | 1285 | 10 | 1.56% |
| hsa_circ_0061161 | 1285 | 10 | 1.56% |
| hsa_circ_0064142 | 1285 | 10 | 1.56% |
| hsa_circ_0065577 | 1285 | 10 | 1.56% |
| hsa_circ_0075216 | 1285 | 10 | 1.56% |
| hsa_circ_0088385 | 1285 | 10 | 1.56% |
| hsa_circ_0128744 | 7197 | 56 | 1.56% |
| hsa_circ_0025155 | 1414 | 11 | 1.56% |
| hsa_circ_0030948 | 1414 | 11 | 1.56% |
| hsa_circ_0046467 | 1414 | 11 | 1.56% |
| hsa_circ_0051314 | 1414 | 11 | 1.56% |
| hsa_circ_0105005 | 1414 | 11 | 1.56% |
| hsa_circ_0055682 | 2057 | 16 | 1.56% |
| hsa_circ_0035131 | 1543 | 12 | 1.56% |

| | | | |
|---|---|---|---|
| hsa_circ_0020771 | 1286 | 10 | 1.56% |
| hsa_circ_0020899 | 1286 | 10 | 1.56% |
| hsa_circ_0038668 | 1286 | 10 | 1.56% |
| hsa_circ_0045167 | 1286 | 10 | 1.56% |
| hsa_circ_0048050 | 1286 | 10 | 1.56% |
| hsa_circ_0057744 | 1286 | 10 | 1.56% |
| hsa_circ_0060246 | 1286 | 10 | 1.56% |
| hsa_circ_0113327 | 1286 | 10 | 1.56% |
| hsa_circ_0124875 | 1286 | 10 | 1.56% |
| hsa_circ_0009569 | 1415 | 11 | 1.55% |
| hsa_circ_0015047 | 1415 | 11 | 1.55% |
| hsa_circ_0016126 | 1415 | 11 | 1.55% |
| hsa_circ_0020652 | 1415 | 11 | 1.55% |
| hsa_circ_0056074 | 1415 | 11 | 1.55% |
| hsa_circ_0068711 | 1415 | 11 | 1.55% |
| hsa_circ_0077905 | 1415 | 11 | 1.55% |
| hsa_circ_0111486 | 1415 | 11 | 1.55% |
| hsa_circ_0063272 | 1544 | 12 | 1.55% |
| hsa_circ_0095402 | 1544 | 12 | 1.55% |
| hsa_circ_0006652 | 1673 | 13 | 1.55% |
| hsa_circ_0028543 | 1673 | 13 | 1.55% |
| hsa_circ_0060674 | 1673 | 13 | 1.55% |
| hsa_circ_0088573 | 1673 | 13 | 1.55% |
| hsa_circ_0039264 | 1287 | 10 | 1.55% |
| hsa_circ_0050810 | 1287 | 10 | 1.55% |
| hsa_circ_0054946 | 1287 | 10 | 1.55% |
| hsa_circ_0064218 | 1287 | 10 | 1.55% |
| hsa_circ_0065662 | 1287 | 10 | 1.55% |
| hsa_circ_0069104 | 1287 | 10 | 1.55% |
| hsa_circ_0073579 | 1287 | 10 | 1.55% |
| hsa_circ_0080271 | 1287 | 10 | 1.55% |
| hsa_circ_0090555 | 1287 | 10 | 1.55% |
| hsa_circ_0113449 | 1287 | 10 | 1.55% |
| hsa_circ_0126838 | 1287 | 10 | 1.55% |
| hsa_circ_0139639 | 1287 | 10 | 1.55% |
| hsa_circ_0140333 | 1287 | 10 | 1.55% |
| hsa_circ_0072813 | 838926 | 6518 | 1.55% |
| hsa_circ_0090600 | 1802 | 14 | 1.55% |
| hsa_circ_0020613 | 1416 | 11 | 1.55% |
| hsa_circ_0037945 | 1416 | 11 | 1.55% |
| hsa_circ_0047688 | 1416 | 11 | 1.55% |
| hsa_circ_0054114 | 1416 | 11 | 1.55% |
| hsa_circ_0080713 | 1416 | 11 | 1.55% |
| hsa_circ_0082742 | 1416 | 11 | 1.55% |
| hsa_circ_0089714 | 1416 | 11 | 1.55% |
| hsa_circ_0124783 | 1416 | 11 | 1.55% |
| hsa_circ_0135176 | 1416 | 11 | 1.55% |
| hsa_circ_0032558 | 1545 | 12 | 1.55% |
| hsa_circ_0069659 | 1545 | 12 | 1.55% |
| hsa_circ_0082229 | 1545 | 12 | 1.55% |

| | | | |
|---|---|---|---|
| hsa_circ_0084396 | 1674 | 13 | 1.55% |
| hsa_circ_0040578 | 3864 | 30 | 1.55% |
| hsa_circ_0014599 | 1288 | 10 | 1.55% |
| hsa_circ_0030795 | 1288 | 10 | 1.55% |
| hsa_circ_0044126 | 1288 | 10 | 1.55% |
| hsa_circ_0055039 | 1288 | 10 | 1.55% |
| hsa_circ_0056238 | 1288 | 10 | 1.55% |
| hsa_circ_0061968 | 1288 | 10 | 1.55% |
| hsa_circ_0066175 | 1288 | 10 | 1.55% |
| hsa_circ_0070922 | 1288 | 10 | 1.55% |
| hsa_circ_0073196 | 1288 | 10 | 1.55% |
| hsa_circ_0073974 | 1288 | 10 | 1.55% |
| hsa_circ_0081555 | 1288 | 10 | 1.55% |
| hsa_circ_0090356 | 1288 | 10 | 1.55% |
| hsa_circ_0100283 | 1288 | 10 | 1.55% |
| hsa_circ_0112686 | 1288 | 10 | 1.55% |
| hsa_circ_0117719 | 1288 | 10 | 1.55% |
| hsa_circ_0127559 | 1288 | 10 | 1.55% |
| hsa_circ_0010752 | 1417 | 11 | 1.55% |
| hsa_circ_0025089 | 1417 | 11 | 1.55% |
| hsa_circ_0028625 | 1417 | 11 | 1.55% |
| hsa_circ_0031134 | 1417 | 11 | 1.55% |
| hsa_circ_0037882 | 1417 | 11 | 1.55% |
| hsa_circ_0038607 | 1417 | 11 | 1.55% |
| hsa_circ_0109458 | 1417 | 11 | 1.55% |
| hsa_circ_0135698 | 1417 | 11 | 1.55% |
| hsa_circ_0044452 | 1546 | 12 | 1.55% |
| hsa_circ_0072732 | 1546 | 12 | 1.55% |
| hsa_circ_0076172 | 1546 | 12 | 1.55% |
| hsa_circ_0059870 | 1804 | 14 | 1.55% |
| hsa_circ_0067266 | 1933 | 15 | 1.55% |
| hsa_circ_0022498 | 1289 | 10 | 1.55% |
| hsa_circ_0024701 | 1289 | 10 | 1.55% |
| hsa_circ_0039033 | 1289 | 10 | 1.55% |
| hsa_circ_0046757 | 1289 | 10 | 1.55% |
| hsa_circ_0052037 | 1289 | 10 | 1.55% |
| hsa_circ_0062918 | 1289 | 10 | 1.55% |
| hsa_circ_0064117 | 1289 | 10 | 1.55% |
| hsa_circ_0081957 | 1289 | 10 | 1.55% |
| hsa_circ_0098929 | 1289 | 10 | 1.55% |
| hsa_circ_0113613 | 1289 | 10 | 1.55% |
| hsa_circ_0133851 | 1289 | 10 | 1.55% |
| hsa_circ_0026396 | 1418 | 11 | 1.55% |
| hsa_circ_0034396 | 1418 | 11 | 1.55% |
| hsa_circ_0076424 | 1418 | 11 | 1.55% |
| hsa_circ_0094673 | 1418 | 11 | 1.55% |
| hsa_circ_0120751 | 1418 | 11 | 1.55% |
| hsa_circ_0126731 | 1418 | 11 | 1.55% |
| hsa_circ_0116395 | 3094 | 24 | 1.55% |
| hsa_circ_0034963 | 1547 | 12 | 1.55% |

| | | | |
|---|---|---|---|
| hsa_circ_0036256 | 1547 | 12 | 1.55% |
| hsa_circ_0037635 | 1547 | 12 | 1.55% |
| hsa_circ_0028695 | 1805 | 14 | 1.55% |
| hsa_circ_0011147 | 2063 | 16 | 1.55% |
| hsa_circ_0034259 | 1548 | 12 | 1.55% |
| hsa_circ_0036616 | 1548 | 12 | 1.55% |
| hsa_circ_0072189 | 1548 | 12 | 1.55% |
| hsa_circ_0090760 | 1548 | 12 | 1.55% |
| hsa_circ_0121577 | 1548 | 12 | 1.55% |
| hsa_circ_0026166 | 1419 | 11 | 1.55% |
| hsa_circ_0027745 | 1419 | 11 | 1.55% |
| hsa_circ_0046266 | 1419 | 11 | 1.55% |
| hsa_circ_0057067 | 1419 | 11 | 1.55% |
| hsa_circ_0086946 | 1419 | 11 | 1.55% |
| hsa_circ_0002923 | 1290 | 10 | 1.55% |
| hsa_circ_0008603 | 1290 | 10 | 1.55% |
| hsa_circ_0011382 | 1290 | 10 | 1.55% |
| hsa_circ_0012924 | 1290 | 10 | 1.55% |
| hsa_circ_0014641 | 1290 | 10 | 1.55% |
| hsa_circ_0022350 | 1290 | 10 | 1.55% |
| hsa_circ_0029024 | 1290 | 10 | 1.55% |
| hsa_circ_0077717 | 1290 | 10 | 1.55% |
| hsa_circ_0079868 | 1290 | 10 | 1.55% |
| hsa_circ_0079993 | 1290 | 10 | 1.55% |
| hsa_circ_0080967 | 1290 | 10 | 1.55% |
| hsa_circ_0088332 | 1290 | 10 | 1.55% |
| hsa_circ_0119928 | 1290 | 10 | 1.55% |
| hsa_circ_0128002 | 1290 | 10 | 1.55% |
| hsa_circ_0130763 | 1290 | 10 | 1.55% |
| hsa_circ_0134160 | 1290 | 10 | 1.55% |
| hsa_circ_0139367 | 4774 | 37 | 1.55% |
| hsa_circ_0123139 | 1807 | 14 | 1.55% |
| hsa_circ_0086744 | 1549 | 12 | 1.55% |
| hsa_circ_0133273 | 1549 | 12 | 1.55% |
| hsa_circ_0027306 | 1420 | 11 | 1.55% |
| hsa_circ_0070236 | 1420 | 11 | 1.55% |
| hsa_circ_0120068 | 1420 | 11 | 1.55% |
| hsa_circ_0014321 | 1291 | 10 | 1.55% |
| hsa_circ_0021684 | 1291 | 10 | 1.55% |
| hsa_circ_0030157 | 1291 | 10 | 1.55% |
| hsa_circ_0033422 | 1291 | 10 | 1.55% |
| hsa_circ_0034174 | 1291 | 10 | 1.55% |
| hsa_circ_0036308 | 1291 | 10 | 1.55% |
| hsa_circ_0074635 | 1291 | 10 | 1.55% |
| hsa_circ_0089271 | 1291 | 10 | 1.55% |
| hsa_circ_0092453 | 1291 | 10 | 1.55% |
| hsa_circ_0095416 | 1291 | 10 | 1.55% |
| hsa_circ_0125489 | 1291 | 10 | 1.55% |
| hsa_circ_0052359 | 2583 | 20 | 1.55% |
| hsa_circ_0029937 | 1550 | 12 | 1.55% |

| | | | |
|---|---|---|---|
| hsa_circ_0054793 | 1550 | 12 | 1.55% |
| hsa_circ_0013621 | 1421 | 11 | 1.55% |
| hsa_circ_0017566 | 1421 | 11 | 1.55% |
| hsa_circ_0022578 | 1421 | 11 | 1.55% |
| hsa_circ_0026482 | 1421 | 11 | 1.55% |
| hsa_circ_0031289 | 1421 | 11 | 1.55% |
| hsa_circ_0035266 | 1421 | 11 | 1.55% |
| hsa_circ_0038150 | 1421 | 11 | 1.55% |
| hsa_circ_0038461 | 1421 | 11 | 1.55% |
| hsa_circ_0043843 | 1421 | 11 | 1.55% |
| hsa_circ_0045408 | 1421 | 11 | 1.55% |
| hsa_circ_0047656 | 1421 | 11 | 1.55% |
| hsa_circ_0071511 | 1421 | 11 | 1.55% |
| hsa_circ_0073963 | 1421 | 11 | 1.55% |
| hsa_circ_0091293 | 1421 | 11 | 1.55% |
| hsa_circ_0129780 | 1421 | 11 | 1.55% |
| hsa_circ_0049973 | 4134 | 32 | 1.55% |
| hsa_circ_0013525 | 1292 | 10 | 1.55% |
| hsa_circ_0016197 | 1292 | 10 | 1.55% |
| hsa_circ_0017420 | 1292 | 10 | 1.55% |
| hsa_circ_0019263 | 1292 | 10 | 1.55% |
| hsa_circ_0022534 | 1292 | 10 | 1.55% |
| hsa_circ_0031189 | 1292 | 10 | 1.55% |
| hsa_circ_0037933 | 1292 | 10 | 1.55% |
| hsa_circ_0039340 | 1292 | 10 | 1.55% |
| hsa_circ_0052087 | 1292 | 10 | 1.55% |
| hsa_circ_0062245 | 1292 | 10 | 1.55% |
| hsa_circ_0064886 | 1292 | 10 | 1.55% |
| hsa_circ_0065216 | 1292 | 10 | 1.55% |
| hsa_circ_0067653 | 1292 | 10 | 1.55% |
| hsa_circ_0083794 | 1292 | 10 | 1.55% |
| hsa_circ_0101754 | 1292 | 10 | 1.55% |
| hsa_circ_0109570 | 1292 | 10 | 1.55% |
| hsa_circ_0022129 | 1551 | 12 | 1.55% |
| hsa_circ_0014336 | 1422 | 11 | 1.55% |
| hsa_circ_0022272 | 1422 | 11 | 1.55% |
| hsa_circ_0027969 | 1422 | 11 | 1.55% |
| hsa_circ_0042314 | 1422 | 11 | 1.55% |
| hsa_circ_0051826 | 1422 | 11 | 1.55% |
| hsa_circ_0059848 | 1422 | 11 | 1.55% |
| hsa_circ_0065097 | 1422 | 11 | 1.55% |
| hsa_circ_0066968 | 1422 | 11 | 1.55% |
| hsa_circ_0071243 | 1422 | 11 | 1.55% |
| hsa_circ_0089452 | 1422 | 11 | 1.55% |
| hsa_circ_0089455 | 1422 | 11 | 1.55% |
| hsa_circ_0129377 | 1422 | 11 | 1.55% |
| hsa_circ_0024233 | 1293 | 10 | 1.55% |
| hsa_circ_0026499 | 1293 | 10 | 1.55% |
| hsa_circ_0028808 | 1293 | 10 | 1.55% |
| hsa_circ_0031714 | 1293 | 10 | 1.55% |

| | | | |
|---|---|---|---|
| hsa_circ_0034411 | 1293 | 10 | 1.55% |
| hsa_circ_0041980 | 1293 | 10 | 1.55% |
| hsa_circ_0058990 | 1293 | 10 | 1.55% |
| hsa_circ_0060400 | 1293 | 10 | 1.55% |
| hsa_circ_0080061 | 1293 | 10 | 1.55% |
| hsa_circ_0080379 | 1293 | 10 | 1.55% |
| hsa_circ_0105465 | 1293 | 10 | 1.55% |
| hsa_circ_0108563 | 1293 | 10 | 1.55% |
| hsa_circ_0137875 | 1293 | 10 | 1.55% |
| hsa_circ_0086704 | 1681 | 13 | 1.55% |
| hsa_circ_0109875 | 3233 | 25 | 1.55% |
| hsa_circ_0080811 | 1552 | 12 | 1.55% |
| hsa_circ_0090874 | 1552 | 12 | 1.55% |
| hsa_circ_0031981 | 1423 | 11 | 1.55% |
| hsa_circ_0052066 | 1423 | 11 | 1.55% |
| hsa_circ_0082791 | 1423 | 11 | 1.55% |
| hsa_circ_0091123 | 1423 | 11 | 1.55% |
| hsa_circ_0100201 | 1423 | 11 | 1.55% |
| hsa_circ_0123316 | 1423 | 11 | 1.55% |
| hsa_circ_0108592 | 1682 | 13 | 1.55% |
| hsa_circ_0016193 | 1294 | 10 | 1.55% |
| hsa_circ_0031653 | 1294 | 10 | 1.55% |
| hsa_circ_0067343 | 1294 | 10 | 1.55% |
| hsa_circ_0088273 | 1294 | 10 | 1.55% |
| hsa_circ_0123841 | 1294 | 10 | 1.55% |
| hsa_circ_0132308 | 1294 | 10 | 1.55% |
| hsa_circ_0074266 | 1553 | 12 | 1.55% |
| hsa_circ_0108004 | 1553 | 12 | 1.55% |
| hsa_circ_0021654 | 2071 | 16 | 1.55% |
| hsa_circ_0113800 | 4919 | 38 | 1.55% |
| hsa_circ_0040283 | 1424 | 11 | 1.54% |
| hsa_circ_0050468 | 1424 | 11 | 1.54% |
| hsa_circ_0058902 | 1424 | 11 | 1.54% |
| hsa_circ_0062676 | 1424 | 11 | 1.54% |
| hsa_circ_0065138 | 1424 | 11 | 1.54% |
| hsa_circ_0093501 | 1424 | 11 | 1.54% |
| hsa_circ_0099803 | 1424 | 11 | 1.54% |
| hsa_circ_0138362 | 1424 | 11 | 1.54% |
| hsa_circ_0034358 | 1683 | 13 | 1.54% |
| hsa_circ_0122999 | 4402 | 34 | 1.54% |
| hsa_circ_0052141 | 1554 | 12 | 1.54% |
| hsa_circ_0098809 | 1554 | 12 | 1.54% |
| hsa_circ_0013552 | 1295 | 10 | 1.54% |
| hsa_circ_0092119 | 1295 | 10 | 1.54% |
| hsa_circ_0104655 | 1295 | 10 | 1.54% |
| hsa_circ_0112599 | 1295 | 10 | 1.54% |
| hsa_circ_0117065 | 1295 | 10 | 1.54% |
| hsa_circ_0125231 | 1295 | 10 | 1.54% |
| hsa_circ_0128315 | 1295 | 10 | 1.54% |
| hsa_circ_0129153 | 1295 | 10 | 1.54% |

| | | | |
|---|---|---|---|
| hsa_circ_0009229 | 1425 | 11 | 1.54% |
| hsa_circ_0009280 | 1425 | 11 | 1.54% |
| hsa_circ_0029176 | 1425 | 11 | 1.54% |
| hsa_circ_0113138 | 1425 | 11 | 1.54% |
| hsa_circ_0117567 | 1425 | 11 | 1.54% |
| hsa_circ_0122846 | 1425 | 11 | 1.54% |
| hsa_circ_0139355 | 1425 | 11 | 1.54% |
| hsa_circ_0139770 | 1555 | 12 | 1.54% |
| hsa_circ_0008226 | 1296 | 10 | 1.54% |
| hsa_circ_0016035 | 1296 | 10 | 1.54% |
| hsa_circ_0020451 | 1296 | 10 | 1.54% |
| hsa_circ_0024293 | 1296 | 10 | 1.54% |
| hsa_circ_0036359 | 1296 | 10 | 1.54% |
| hsa_circ_0040014 | 1296 | 10 | 1.54% |
| hsa_circ_0040916 | 1296 | 10 | 1.54% |
| hsa_circ_0045964 | 1296 | 10 | 1.54% |
| hsa_circ_0049064 | 1296 | 10 | 1.54% |
| hsa_circ_0065750 | 1296 | 10 | 1.54% |
| hsa_circ_0066784 | 1296 | 10 | 1.54% |
| hsa_circ_0094860 | 1296 | 10 | 1.54% |
| hsa_circ_0100020 | 1296 | 10 | 1.54% |
| hsa_circ_0047949 | 1685 | 13 | 1.54% |
| hsa_circ_0040063 | 1426 | 11 | 1.54% |
| hsa_circ_0056736 | 1426 | 11 | 1.54% |
| hsa_circ_0070100 | 1426 | 11 | 1.54% |
| hsa_circ_0094401 | 1426 | 11 | 1.54% |
| hsa_circ_0029449 | 1556 | 12 | 1.54% |
| hsa_circ_0043771 | 1556 | 12 | 1.54% |
| hsa_circ_0093251 | 1686 | 13 | 1.54% |
| hsa_circ_0009761 | 1297 | 10 | 1.54% |
| hsa_circ_0024497 | 1297 | 10 | 1.54% |
| hsa_circ_0028288 | 1297 | 10 | 1.54% |
| hsa_circ_0035339 | 1297 | 10 | 1.54% |
| hsa_circ_0047069 | 1297 | 10 | 1.54% |
| hsa_circ_0047490 | 1297 | 10 | 1.54% |
| hsa_circ_0056076 | 1297 | 10 | 1.54% |
| hsa_circ_0061187 | 1297 | 10 | 1.54% |
| hsa_circ_0103848 | 1297 | 10 | 1.54% |
| hsa_circ_0115981 | 1297 | 10 | 1.54% |
| hsa_circ_0129467 | 1297 | 10 | 1.54% |
| hsa_circ_0010736 | 1427 | 11 | 1.54% |
| hsa_circ_0056001 | 1427 | 11 | 1.54% |
| hsa_circ_0066250 | 1427 | 11 | 1.54% |
| hsa_circ_0073764 | 1427 | 11 | 1.54% |
| hsa_circ_0124460 | 1427 | 11 | 1.54% |
| hsa_circ_0025220 | 1557 | 12 | 1.54% |
| hsa_circ_0110618 | 1557 | 12 | 1.54% |
| hsa_circ_0005502 | 1687 | 13 | 1.54% |
| hsa_circ_0000711 | 1298 | 10 | 1.54% |
| hsa_circ_0014899 | 1298 | 10 | 1.54% |

| | | | |
|---|---|---|---|
| hsa_circ_0052010 | 1298 | 10 | 1.54% |
| hsa_circ_0055595 | 1298 | 10 | 1.54% |
| hsa_circ_0060120 | 1298 | 10 | 1.54% |
| hsa_circ_0062363 | 1298 | 10 | 1.54% |
| hsa_circ_0063526 | 1298 | 10 | 1.54% |
| hsa_circ_0064246 | 1298 | 10 | 1.54% |
| hsa_circ_0071406 | 1298 | 10 | 1.54% |
| hsa_circ_0085758 | 1298 | 10 | 1.54% |
| hsa_circ_0104119 | 1298 | 10 | 1.54% |
| hsa_circ_0112056 | 1298 | 10 | 1.54% |
| hsa_circ_0118781 | 1298 | 10 | 1.54% |
| hsa_circ_0134737 | 1298 | 10 | 1.54% |
| hsa_circ_0036474 | 1428 | 11 | 1.54% |
| hsa_circ_0100212 | 1428 | 11 | 1.54% |
| hsa_circ_0130118 | 1428 | 11 | 1.54% |
| hsa_circ_0138469 | 1428 | 11 | 1.54% |
| hsa_circ_0032608 | 1558 | 12 | 1.54% |
| hsa_circ_0052144 | 1558 | 12 | 1.54% |
| hsa_circ_0119013 | 1558 | 12 | 1.54% |
| hsa_circ_0127222 | 1558 | 12 | 1.54% |
| hsa_circ_0021498 | 1688 | 13 | 1.54% |
| hsa_circ_0072810 | 846530 | 6518 | 1.54% |
| hsa_circ_0009153 | 1299 | 10 | 1.54% |
| hsa_circ_0024882 | 1299 | 10 | 1.54% |
| hsa_circ_0034442 | 1299 | 10 | 1.54% |
| hsa_circ_0038998 | 1299 | 10 | 1.54% |
| hsa_circ_0040290 | 1299 | 10 | 1.54% |
| hsa_circ_0046696 | 1299 | 10 | 1.54% |
| hsa_circ_0057852 | 1299 | 10 | 1.54% |
| hsa_circ_0064599 | 1299 | 10 | 1.54% |
| hsa_circ_0065951 | 1299 | 10 | 1.54% |
| hsa_circ_0087246 | 1299 | 10 | 1.54% |
| hsa_circ_0088335 | 1299 | 10 | 1.54% |
| hsa_circ_0091746 | 1299 | 10 | 1.54% |
| hsa_circ_0098300 | 1299 | 10 | 1.54% |
| hsa_circ_0103417 | 1299 | 10 | 1.54% |
| hsa_circ_0107975 | 1299 | 10 | 1.54% |
| hsa_circ_0026464 | 1429 | 11 | 1.54% |
| hsa_circ_0029921 | 1429 | 11 | 1.54% |
| hsa_circ_0038837 | 1429 | 11 | 1.54% |
| hsa_circ_0048572 | 1429 | 11 | 1.54% |
| hsa_circ_0052503 | 1429 | 11 | 1.54% |
| hsa_circ_0070783 | 1429 | 11 | 1.54% |
| hsa_circ_0015556 | 1689 | 13 | 1.54% |
| hsa_circ_0119528 | 3638 | 28 | 1.54% |
| hsa_circ_0013826 | 2470 | 19 | 1.54% |
| hsa_circ_0068155 | 1560 | 12 | 1.54% |
| hsa_circ_0009503 | 1430 | 11 | 1.54% |
| hsa_circ_0014629 | 1430 | 11 | 1.54% |
| hsa_circ_0025842 | 1430 | 11 | 1.54% |

| | | | |
|---|---|---|---|
| hsa_circ_0048380 | 1430 | 11 | 1.54% |
| hsa_circ_0058315 | 1430 | 11 | 1.54% |
| hsa_circ_0062635 | 1430 | 11 | 1.54% |
| hsa_circ_0140647 | 1430 | 11 | 1.54% |
| hsa_circ_0008560 | 1300 | 10 | 1.54% |
| hsa_circ_0009201 | 1300 | 10 | 1.54% |
| hsa_circ_0027863 | 1300 | 10 | 1.54% |
| hsa_circ_0029136 | 1300 | 10 | 1.54% |
| hsa_circ_0075779 | 1300 | 10 | 1.54% |
| hsa_circ_0087636 | 1300 | 10 | 1.54% |
| hsa_circ_0105955 | 1300 | 10 | 1.54% |
| hsa_circ_0120071 | 1300 | 10 | 1.54% |
| hsa_circ_0012327 | 1821 | 14 | 1.54% |
| hsa_circ_0012693 | 1561 | 12 | 1.54% |
| hsa_circ_0047681 | 1561 | 12 | 1.54% |
| hsa_circ_0005308 | 1431 | 11 | 1.54% |
| hsa_circ_0022865 | 1431 | 11 | 1.54% |
| hsa_circ_0030229 | 1431 | 11 | 1.54% |
| hsa_circ_0038953 | 1431 | 11 | 1.54% |
| hsa_circ_0080282 | 1431 | 11 | 1.54% |
| hsa_circ_0096693 | 1431 | 11 | 1.54% |
| hsa_circ_0110948 | 1431 | 11 | 1.54% |
| hsa_circ_0136066 | 1431 | 11 | 1.54% |
| hsa_circ_0027940 | 1301 | 10 | 1.54% |
| hsa_circ_0056179 | 1301 | 10 | 1.54% |
| hsa_circ_0098766 | 1301 | 10 | 1.54% |
| hsa_circ_0104155 | 1301 | 10 | 1.54% |
| hsa_circ_0117927 | 1301 | 10 | 1.54% |
| hsa_circ_0126119 | 1301 | 10 | 1.54% |
| hsa_circ_0127432 | 1301 | 10 | 1.54% |
| hsa_circ_0071462 | 4164 | 32 | 1.54% |
| hsa_circ_0076698 | 1692 | 13 | 1.54% |
| hsa_circ_0024600 | 1562 | 12 | 1.54% |
| hsa_circ_0106832 | 1562 | 12 | 1.54% |
| hsa_circ_0042755 | 1432 | 11 | 1.54% |
| hsa_circ_0054871 | 1432 | 11 | 1.54% |
| hsa_circ_0063561 | 1432 | 11 | 1.54% |
| hsa_circ_0042121 | 1302 | 10 | 1.54% |
| hsa_circ_0049622 | 1302 | 10 | 1.54% |
| hsa_circ_0050332 | 1302 | 10 | 1.54% |
| hsa_circ_0052804 | 1302 | 10 | 1.54% |
| hsa_circ_0064088 | 1302 | 10 | 1.54% |
| hsa_circ_0064654 | 1302 | 10 | 1.54% |
| hsa_circ_0071027 | 1302 | 10 | 1.54% |
| hsa_circ_0075092 | 1302 | 10 | 1.54% |
| hsa_circ_0087666 | 1302 | 10 | 1.54% |
| hsa_circ_0137570 | 1302 | 10 | 1.54% |
| hsa_circ_0098399 | 4558 | 35 | 1.54% |
| hsa_circ_0056249 | 1693 | 13 | 1.54% |
| hsa_circ_0029830 | 1563 | 12 | 1.54% |

| | | | |
|---|---|---|---|
| hsa_circ_0056847 | 1563 | 12 | 1.54% |
| hsa_circ_0093761 | 1563 | 12 | 1.54% |
| hsa_circ_0002559 | 1433 | 11 | 1.54% |
| hsa_circ_0050209 | 1433 | 11 | 1.54% |
| hsa_circ_0131734 | 1433 | 11 | 1.54% |
| hsa_circ_0135135 | 1433 | 11 | 1.54% |
| hsa_circ_0009297 | 1303 | 10 | 1.53% |
| hsa_circ_0014819 | 1303 | 10 | 1.53% |
| hsa_circ_0038918 | 1303 | 10 | 1.53% |
| hsa_circ_0064122 | 1303 | 10 | 1.53% |
| hsa_circ_0068003 | 1303 | 10 | 1.53% |
| hsa_circ_0069360 | 1303 | 10 | 1.53% |
| hsa_circ_0069415 | 1303 | 10 | 1.53% |
| hsa_circ_0070266 | 1303 | 10 | 1.53% |
| hsa_circ_0070905 | 1303 | 10 | 1.53% |
| hsa_circ_0073100 | 1303 | 10 | 1.53% |
| hsa_circ_0074579 | 1303 | 10 | 1.53% |
| hsa_circ_0103649 | 1303 | 10 | 1.53% |
| hsa_circ_0128576 | 1303 | 10 | 1.53% |
| hsa_circ_0134460 | 1303 | 10 | 1.53% |
| hsa_circ_0122788 | 5082 | 39 | 1.53% |
| hsa_circ_0020011 | 1564 | 12 | 1.53% |
| hsa_circ_0113131 | 1564 | 12 | 1.53% |
| hsa_circ_0062126 | 1434 | 11 | 1.53% |
| hsa_circ_0066141 | 1434 | 11 | 1.53% |
| hsa_circ_0081868 | 1434 | 11 | 1.53% |
| hsa_circ_0117930 | 1434 | 11 | 1.53% |
| hsa_circ_0122674 | 1434 | 11 | 1.53% |
| hsa_circ_0135340 | 1434 | 11 | 1.53% |
| hsa_circ_0136352 | 1434 | 11 | 1.53% |
| hsa_circ_0013520 | 1304 | 10 | 1.53% |
| hsa_circ_0026124 | 1304 | 10 | 1.53% |
| hsa_circ_0069235 | 1304 | 10 | 1.53% |
| hsa_circ_0084478 | 1304 | 10 | 1.53% |
| hsa_circ_0092601 | 1304 | 10 | 1.53% |
| hsa_circ_0116025 | 1304 | 10 | 1.53% |
| hsa_circ_0118530 | 1304 | 10 | 1.53% |
| hsa_circ_0019067 | 1565 | 12 | 1.53% |
| hsa_circ_0085464 | 1565 | 12 | 1.53% |
| hsa_circ_0085625 | 1565 | 12 | 1.53% |
| hsa_circ_0119429 | 1565 | 12 | 1.53% |
| hsa_circ_0006404 | 1435 | 11 | 1.53% |
| hsa_circ_0078435 | 1435 | 11 | 1.53% |
| hsa_circ_0091315 | 1435 | 11 | 1.53% |
| hsa_circ_0003199 | 1566 | 12 | 1.53% |
| hsa_circ_0020791 | 1566 | 12 | 1.53% |
| hsa_circ_0065239 | 1566 | 12 | 1.53% |
| hsa_circ_0002807 | 1305 | 10 | 1.53% |
| hsa_circ_0022217 | 1305 | 10 | 1.53% |
| hsa_circ_0026172 | 1305 | 10 | 1.53% |

| | | | |
|---|---|---|---|
| hsa_circ_0040511 | 1305 | 10 | 1.53% |
| hsa_circ_0066582 | 1305 | 10 | 1.53% |
| hsa_circ_0089966 | 1305 | 10 | 1.53% |
| hsa_circ_0100016 | 1305 | 10 | 1.53% |
| hsa_circ_0113626 | 1305 | 10 | 1.53% |
| hsa_circ_0132156 | 1305 | 10 | 1.53% |
| hsa_circ_0031002 | 1697 | 13 | 1.53% |
| hsa_circ_0000053 | 1436 | 11 | 1.53% |
| hsa_circ_0003982 | 1436 | 11 | 1.53% |
| hsa_circ_0015260 | 1436 | 11 | 1.53% |
| hsa_circ_0037339 | 1436 | 11 | 1.53% |
| hsa_circ_0040275 | 1436 | 11 | 1.53% |
| hsa_circ_0074424 | 1436 | 11 | 1.53% |
| hsa_circ_0072811 | 851140 | 6518 | 1.53% |
| hsa_circ_0053949 | 1567 | 12 | 1.53% |
| hsa_circ_0026113 | 1306 | 10 | 1.53% |
| hsa_circ_0032011 | 1306 | 10 | 1.53% |
| hsa_circ_0035651 | 1306 | 10 | 1.53% |
| hsa_circ_0038026 | 1306 | 10 | 1.53% |
| hsa_circ_0043151 | 1306 | 10 | 1.53% |
| hsa_circ_0047457 | 1306 | 10 | 1.53% |
| hsa_circ_0047537 | 1306 | 10 | 1.53% |
| hsa_circ_0064899 | 1306 | 10 | 1.53% |
| hsa_circ_0071154 | 1306 | 10 | 1.53% |
| hsa_circ_0083759 | 1306 | 10 | 1.53% |
| hsa_circ_0023039 | 1437 | 11 | 1.53% |
| hsa_circ_0030179 | 1437 | 11 | 1.53% |
| hsa_circ_0038336 | 1437 | 11 | 1.53% |
| hsa_circ_0040074 | 1437 | 11 | 1.53% |
| hsa_circ_0059333 | 1437 | 11 | 1.53% |
| hsa_circ_0080223 | 1437 | 11 | 1.53% |
| hsa_circ_0080875 | 1437 | 11 | 1.53% |
| hsa_circ_0089091 | 1437 | 11 | 1.53% |
| hsa_circ_0104631 | 1437 | 11 | 1.53% |
| hsa_circ_0117800 | 1437 | 11 | 1.53% |
| hsa_circ_0052801 | 1568 | 12 | 1.53% |
| hsa_circ_0058023 | 1568 | 12 | 1.53% |
| hsa_circ_0092435 | 1568 | 12 | 1.53% |
| hsa_circ_0121621 | 1568 | 12 | 1.53% |
| hsa_circ_0000099 | 1307 | 10 | 1.53% |
| hsa_circ_0016083 | 1307 | 10 | 1.53% |
| hsa_circ_0022499 | 1307 | 10 | 1.53% |
| hsa_circ_0034288 | 1307 | 10 | 1.53% |
| hsa_circ_0036463 | 1307 | 10 | 1.53% |
| hsa_circ_0037760 | 1307 | 10 | 1.53% |
| hsa_circ_0068093 | 1307 | 10 | 1.53% |
| hsa_circ_0074572 | 1307 | 10 | 1.53% |
| hsa_circ_0081456 | 1307 | 10 | 1.53% |
| hsa_circ_0082264 | 1307 | 10 | 1.53% |
| hsa_circ_0096314 | 1307 | 10 | 1.53% |

| | | | |
|---|---|---|---|
| hsa_circ_0097725 | 1307 | 10 | 1.53% |
| hsa_circ_0105752 | 1307 | 10 | 1.53% |
| hsa_circ_0069043 | 1830 | 14 | 1.53% |
| hsa_circ_0020997 | 1438 | 11 | 1.53% |
| hsa_circ_0040180 | 1438 | 11 | 1.53% |
| hsa_circ_0043832 | 1438 | 11 | 1.53% |
| hsa_circ_0055284 | 1438 | 11 | 1.53% |
| hsa_circ_0074779 | 1438 | 11 | 1.53% |
| hsa_circ_0084931 | 1438 | 11 | 1.53% |
| hsa_circ_0087543 | 1438 | 11 | 1.53% |
| hsa_circ_0032657 | 2223 | 17 | 1.53% |
| hsa_circ_0072621 | 1700 | 13 | 1.53% |
| hsa_circ_0119334 | 1700 | 13 | 1.53% |
| hsa_circ_0080092 | 1831 | 14 | 1.53% |
| hsa_circ_0016482 | 1308 | 10 | 1.53% |
| hsa_circ_0022333 | 1308 | 10 | 1.53% |
| hsa_circ_0025529 | 1308 | 10 | 1.53% |
| hsa_circ_0026500 | 1308 | 10 | 1.53% |
| hsa_circ_0079736 | 1308 | 10 | 1.53% |
| hsa_circ_0102274 | 1308 | 10 | 1.53% |
| hsa_circ_0109356 | 1308 | 10 | 1.53% |
| hsa_circ_0117606 | 1308 | 10 | 1.53% |
| hsa_circ_0118209 | 1308 | 10 | 1.53% |
| hsa_circ_0120232 | 1308 | 10 | 1.53% |
| hsa_circ_0140068 | 1308 | 10 | 1.53% |
| hsa_circ_0010458 | 1439 | 11 | 1.53% |
| hsa_circ_0024112 | 1439 | 11 | 1.53% |
| hsa_circ_0026109 | 1439 | 11 | 1.53% |
| hsa_circ_0028929 | 1439 | 11 | 1.53% |
| hsa_circ_0031316 | 1439 | 11 | 1.53% |
| hsa_circ_0035376 | 1439 | 11 | 1.53% |
| hsa_circ_0052850 | 1439 | 11 | 1.53% |
| hsa_circ_0075217 | 1439 | 11 | 1.53% |
| hsa_circ_0091447 | 1439 | 11 | 1.53% |
| hsa_circ_0093339 | 1439 | 11 | 1.53% |
| hsa_circ_0108232 | 1439 | 11 | 1.53% |
| hsa_circ_0123021 | 1439 | 11 | 1.53% |
| hsa_circ_0139008 | 1439 | 11 | 1.53% |
| hsa_circ_0033478 | 1570 | 12 | 1.53% |
| hsa_circ_0071566 | 1570 | 12 | 1.53% |
| hsa_circ_0104613 | 31664 | 242 | 1.53% |
| hsa_circ_0026442 | 1309 | 10 | 1.53% |
| hsa_circ_0046588 | 1309 | 10 | 1.53% |
| hsa_circ_0046803 | 1309 | 10 | 1.53% |
| hsa_circ_0049961 | 1309 | 10 | 1.53% |
| hsa_circ_0060503 | 1309 | 10 | 1.53% |
| hsa_circ_0069374 | 1309 | 10 | 1.53% |
| hsa_circ_0082954 | 1309 | 10 | 1.53% |
| hsa_circ_0104870 | 1309 | 10 | 1.53% |
| hsa_circ_0117646 | 1309 | 10 | 1.53% |

| | | | |
|---|---|---|---|
| hsa_circ_0135257 | 1309 | 10 | 1.53% |
| hsa_circ_0138890 | 1309 | 10 | 1.53% |
| hsa_circ_0083531 | 1440 | 11 | 1.53% |
| hsa_circ_0065477 | 1571 | 12 | 1.53% |
| hsa_circ_0138451 | 1571 | 12 | 1.53% |
| hsa_circ_0011146 | 2096 | 16 | 1.53% |
| hsa_circ_0022104 | 1703 | 13 | 1.53% |
| hsa_circ_0061455 | 1703 | 13 | 1.53% |
| hsa_circ_0014709 | 1572 | 12 | 1.53% |
| hsa_circ_0018524 | 1572 | 12 | 1.53% |
| hsa_circ_0043483 | 1572 | 12 | 1.53% |
| hsa_circ_0064070 | 1572 | 12 | 1.53% |
| hsa_circ_0077823 | 1572 | 12 | 1.53% |
| hsa_circ_0011327 | 1441 | 11 | 1.53% |
| hsa_circ_0033536 | 1441 | 11 | 1.53% |
| hsa_circ_0037864 | 1441 | 11 | 1.53% |
| hsa_circ_0055711 | 1441 | 11 | 1.53% |
| hsa_circ_0070121 | 1441 | 11 | 1.53% |
| hsa_circ_0081541 | 1441 | 11 | 1.53% |
| hsa_circ_0136784 | 1441 | 11 | 1.53% |
| hsa_circ_0010684 | 1310 | 10 | 1.53% |
| hsa_circ_0018777 | 1310 | 10 | 1.53% |
| hsa_circ_0021961 | 1310 | 10 | 1.53% |
| hsa_circ_0022943 | 1310 | 10 | 1.53% |
| hsa_circ_0023543 | 1310 | 10 | 1.53% |
| hsa_circ_0023765 | 1310 | 10 | 1.53% |
| hsa_circ_0062789 | 1310 | 10 | 1.53% |
| hsa_circ_0074415 | 1310 | 10 | 1.53% |
| hsa_circ_0088990 | 1310 | 10 | 1.53% |
| hsa_circ_0095203 | 1310 | 10 | 1.53% |
| hsa_circ_0112463 | 1310 | 10 | 1.53% |
| hsa_circ_0115177 | 1310 | 10 | 1.53% |
| hsa_circ_0112224 | 4848 | 37 | 1.53% |
| hsa_circ_0009316 | 1573 | 12 | 1.53% |
| hsa_circ_0025740 | 1573 | 12 | 1.53% |
| hsa_circ_0076192 | 1573 | 12 | 1.53% |
| hsa_circ_0106047 | 1573 | 12 | 1.53% |
| hsa_circ_0106955 | 1573 | 12 | 1.53% |
| hsa_circ_0016106 | 1442 | 11 | 1.53% |
| hsa_circ_0020185 | 1442 | 11 | 1.53% |
| hsa_circ_0047950 | 1442 | 11 | 1.53% |
| hsa_circ_0058519 | 1442 | 11 | 1.53% |
| hsa_circ_0065930 | 1442 | 11 | 1.53% |
| hsa_circ_0082594 | 1442 | 11 | 1.53% |
| hsa_circ_0083494 | 1442 | 11 | 1.53% |
| hsa_circ_0094603 | 1442 | 11 | 1.53% |
| hsa_circ_0117291 | 1442 | 11 | 1.53% |
| hsa_circ_0120185 | 1442 | 11 | 1.53% |
| hsa_circ_0011305 | 1311 | 10 | 1.53% |
| hsa_circ_0031245 | 1311 | 10 | 1.53% |

| | | | |
|---|---|---|---|
| hsa_circ_0031854 | 1311 | 10 | 1.53% |
| hsa_circ_0033137 | 1311 | 10 | 1.53% |
| hsa_circ_0040131 | 1311 | 10 | 1.53% |
| hsa_circ_0040588 | 1311 | 10 | 1.53% |
| hsa_circ_0071853 | 1311 | 10 | 1.53% |
| hsa_circ_0072656 | 1311 | 10 | 1.53% |
| hsa_circ_0112455 | 1311 | 10 | 1.53% |
| hsa_circ_0118128 | 1311 | 10 | 1.53% |
| hsa_circ_0124140 | 1311 | 10 | 1.53% |
| hsa_circ_0035473 | 1705 | 13 | 1.52% |
| hsa_circ_0012857 | 1574 | 12 | 1.52% |
| hsa_circ_0071647 | 1574 | 12 | 1.52% |
| hsa_circ_0096783 | 1574 | 12 | 1.52% |
| hsa_circ_0045822 | 2230 | 17 | 1.52% |
| hsa_circ_0015929 | 1443 | 11 | 1.52% |
| hsa_circ_0023202 | 1443 | 11 | 1.52% |
| hsa_circ_0032753 | 1443 | 11 | 1.52% |
| hsa_circ_0073184 | 1443 | 11 | 1.52% |
| hsa_circ_0129723 | 1443 | 11 | 1.52% |
| hsa_circ_0037916 | 1312 | 10 | 1.52% |
| hsa_circ_0038934 | 1312 | 10 | 1.52% |
| hsa_circ_0039067 | 1312 | 10 | 1.52% |
| hsa_circ_0042815 | 1312 | 10 | 1.52% |
| hsa_circ_0047251 | 1312 | 10 | 1.52% |
| hsa_circ_0062321 | 1312 | 10 | 1.52% |
| hsa_circ_0079455 | 1312 | 10 | 1.52% |
| hsa_circ_0084417 | 1312 | 10 | 1.52% |
| hsa_circ_0126697 | 1312 | 10 | 1.52% |
| hsa_circ_0139312 | 1312 | 10 | 1.52% |
| hsa_circ_0139346 | 1312 | 10 | 1.52% |
| hsa_circ_0084406 | 1706 | 13 | 1.52% |
| hsa_circ_0119242 | 1706 | 13 | 1.52% |
| hsa_circ_0009290 | 1444 | 11 | 1.52% |
| hsa_circ_0017211 | 1444 | 11 | 1.52% |
| hsa_circ_0063248 | 1444 | 11 | 1.52% |
| hsa_circ_0087894 | 1444 | 11 | 1.52% |
| hsa_circ_0105313 | 1444 | 11 | 1.52% |
| hsa_circ_0013225 | 1313 | 10 | 1.52% |
| hsa_circ_0014990 | 1313 | 10 | 1.52% |
| hsa_circ_0027795 | 1313 | 10 | 1.52% |
| hsa_circ_0038943 | 1313 | 10 | 1.52% |
| hsa_circ_0059477 | 1313 | 10 | 1.52% |
| hsa_circ_0071836 | 1313 | 10 | 1.52% |
| hsa_circ_0080266 | 1313 | 10 | 1.52% |
| hsa_circ_0099357 | 1313 | 10 | 1.52% |
| hsa_circ_0115117 | 1313 | 10 | 1.52% |
| hsa_circ_0119090 | 1313 | 10 | 1.52% |
| hsa_circ_0128130 | 1313 | 10 | 1.52% |
| hsa_circ_0039102 | 1576 | 12 | 1.52% |
| hsa_circ_0067733 | 1576 | 12 | 1.52% |

| | | | |
|---|---|---|---|
| hsa_circ_0135368 | 1839 | 14 | 1.52% |
| hsa_circ_0004807 | 1445 | 11 | 1.52% |
| hsa_circ_0016478 | 1445 | 11 | 1.52% |
| hsa_circ_0025108 | 1445 | 11 | 1.52% |
| hsa_circ_0027001 | 1445 | 11 | 1.52% |
| hsa_circ_0029312 | 1445 | 11 | 1.52% |
| hsa_circ_0060767 | 1445 | 11 | 1.52% |
| hsa_circ_0070374 | 1445 | 11 | 1.52% |
| hsa_circ_0108226 | 1445 | 11 | 1.52% |
| hsa_circ_0134223 | 1445 | 11 | 1.52% |
| hsa_circ_0139899 | 1445 | 11 | 1.52% |
| hsa_circ_0026950 | 1314 | 10 | 1.52% |
| hsa_circ_0027000 | 1314 | 10 | 1.52% |
| hsa_circ_0038928 | 1314 | 10 | 1.52% |
| hsa_circ_0039706 | 1314 | 10 | 1.52% |
| hsa_circ_0056337 | 1314 | 10 | 1.52% |
| hsa_circ_0057937 | 1314 | 10 | 1.52% |
| hsa_circ_0058903 | 1314 | 10 | 1.52% |
| hsa_circ_0059974 | 1314 | 10 | 1.52% |
| hsa_circ_0064506 | 1314 | 10 | 1.52% |
| hsa_circ_0066787 | 1314 | 10 | 1.52% |
| hsa_circ_0076546 | 1314 | 10 | 1.52% |
| hsa_circ_0081831 | 1314 | 10 | 1.52% |
| hsa_circ_0081950 | 1314 | 10 | 1.52% |
| hsa_circ_0088233 | 1314 | 10 | 1.52% |
| hsa_circ_0130989 | 1314 | 10 | 1.52% |
| hsa_circ_0051665 | 1577 | 12 | 1.52% |
| hsa_circ_0080193 | 1577 | 12 | 1.52% |
| hsa_circ_0118428 | 1577 | 12 | 1.52% |
| hsa_circ_0032545 | 1446 | 11 | 1.52% |
| hsa_circ_0097370 | 1446 | 11 | 1.52% |
| hsa_circ_0108126 | 1446 | 11 | 1.52% |
| hsa_circ_0121371 | 1446 | 11 | 1.52% |
| hsa_circ_0046081 | 1709 | 13 | 1.52% |
| hsa_circ_0090512 | 1709 | 13 | 1.52% |
| hsa_circ_0076124 | 3944 | 30 | 1.52% |
| hsa_circ_0041826 | 1578 | 12 | 1.52% |
| hsa_circ_0043667 | 1578 | 12 | 1.52% |
| hsa_circ_0051726 | 1578 | 12 | 1.52% |
| hsa_circ_0089230 | 1578 | 12 | 1.52% |
| hsa_circ_0097242 | 1578 | 12 | 1.52% |
| hsa_circ_0036149 | 1315 | 10 | 1.52% |
| hsa_circ_0036961 | 1315 | 10 | 1.52% |
| hsa_circ_0056502 | 1315 | 10 | 1.52% |
| hsa_circ_0063906 | 1315 | 10 | 1.52% |
| hsa_circ_0069087 | 1315 | 10 | 1.52% |
| hsa_circ_0073173 | 1315 | 10 | 1.52% |
| hsa_circ_0074735 | 1315 | 10 | 1.52% |
| hsa_circ_0091158 | 1315 | 10 | 1.52% |
| hsa_circ_0108680 | 1315 | 10 | 1.52% |

| | | | |
|---|---|---|---|
| hsa_circ_0122744 | 1315 | 10 | 1.52% |
| hsa_circ_0129879 | 1315 | 10 | 1.52% |
| hsa_circ_0078388 | 3025 | 23 | 1.52% |
| hsa_circ_0090988 | 1973 | 15 | 1.52% |
| hsa_circ_0059655 | 1710 | 13 | 1.52% |
| hsa_circ_0022095 | 1447 | 11 | 1.52% |
| hsa_circ_0028441 | 1447 | 11 | 1.52% |
| hsa_circ_0114722 | 1447 | 11 | 1.52% |
| hsa_circ_0121934 | 1447 | 11 | 1.52% |
| hsa_circ_0078232 | 3289 | 25 | 1.52% |
| hsa_circ_0105933 | 2105 | 16 | 1.52% |
| hsa_circ_0058010 | 1579 | 12 | 1.52% |
| hsa_circ_0111449 | 1579 | 12 | 1.52% |
| hsa_circ_0001067 | 1316 | 10 | 1.52% |
| hsa_circ_0011357 | 1316 | 10 | 1.52% |
| hsa_circ_0037149 | 1316 | 10 | 1.52% |
| hsa_circ_0039511 | 1316 | 10 | 1.52% |
| hsa_circ_0043963 | 1316 | 10 | 1.52% |
| hsa_circ_0052177 | 1316 | 10 | 1.52% |
| hsa_circ_0053257 | 1316 | 10 | 1.52% |
| hsa_circ_0057640 | 1316 | 10 | 1.52% |
| hsa_circ_0057793 | 1316 | 10 | 1.52% |
| hsa_circ_0091406 | 1316 | 10 | 1.52% |
| hsa_circ_0123522 | 1316 | 10 | 1.52% |
| hsa_circ_0126293 | 1316 | 10 | 1.52% |
| hsa_circ_0133342 | 1316 | 10 | 1.52% |
| hsa_circ_0105231 | 5923 | 45 | 1.52% |
| hsa_circ_0009899 | 1448 | 11 | 1.52% |
| hsa_circ_0084934 | 1448 | 11 | 1.52% |
| hsa_circ_0104207 | 1448 | 11 | 1.52% |
| hsa_circ_0120612 | 1448 | 11 | 1.52% |
| hsa_circ_0089678 | 1580 | 12 | 1.52% |
| hsa_circ_0099621 | 1580 | 12 | 1.52% |
| hsa_circ_0117487 | 1580 | 12 | 1.52% |
| hsa_circ_0136594 | 1580 | 12 | 1.52% |
| hsa_circ_0021066 | 1712 | 13 | 1.52% |
| hsa_circ_0064448 | 1712 | 13 | 1.52% |
| hsa_circ_0004987 | 1317 | 10 | 1.52% |
| hsa_circ_0014792 | 1317 | 10 | 1.52% |
| hsa_circ_0028682 | 1317 | 10 | 1.52% |
| hsa_circ_0044100 | 1317 | 10 | 1.52% |
| hsa_circ_0044306 | 1317 | 10 | 1.52% |
| hsa_circ_0045626 | 1317 | 10 | 1.52% |
| hsa_circ_0046659 | 1317 | 10 | 1.52% |
| hsa_circ_0047562 | 1317 | 10 | 1.52% |
| hsa_circ_0056150 | 1317 | 10 | 1.52% |
| hsa_circ_0069881 | 1317 | 10 | 1.52% |
| hsa_circ_0085049 | 1317 | 10 | 1.52% |
| hsa_circ_0087381 | 1317 | 10 | 1.52% |
| hsa_circ_0088421 | 1317 | 10 | 1.52% |

| | | | |
|---|---|---|---|
| hsa_circ_0096413 | 1317 | 10 | 1.52% |
| hsa_circ_0131112 | 1317 | 10 | 1.52% |
| hsa_circ_0137206 | 1317 | 10 | 1.52% |
| hsa_circ_0110023 | 1844 | 14 | 1.52% |
| hsa_circ_0023254 | 1449 | 11 | 1.52% |
| hsa_circ_0045048 | 1449 | 11 | 1.52% |
| hsa_circ_0063237 | 1449 | 11 | 1.52% |
| hsa_circ_0126505 | 2503 | 19 | 1.52% |
| hsa_circ_0027705 | 1581 | 12 | 1.52% |
| hsa_circ_0035163 | 1581 | 12 | 1.52% |
| hsa_circ_0040320 | 1581 | 12 | 1.52% |
| hsa_circ_0040829 | 1581 | 12 | 1.52% |
| hsa_circ_0073560 | 1581 | 12 | 1.52% |
| hsa_circ_0090154 | 1581 | 12 | 1.52% |
| hsa_circ_0052951 | 1713 | 13 | 1.52% |
| hsa_circ_0074039 | 1713 | 13 | 1.52% |
| hsa_circ_0123671 | 5799 | 44 | 1.52% |
| hsa_circ_0106985 | 1977 | 15 | 1.52% |
| hsa_circ_0014253 | 1318 | 10 | 1.52% |
| hsa_circ_0035246 | 1318 | 10 | 1.52% |
| hsa_circ_0052047 | 1318 | 10 | 1.52% |
| hsa_circ_0058553 | 1318 | 10 | 1.52% |
| hsa_circ_0077426 | 1318 | 10 | 1.52% |
| hsa_circ_0078481 | 1318 | 10 | 1.52% |
| hsa_circ_0085968 | 1318 | 10 | 1.52% |
| hsa_circ_0089992 | 1318 | 10 | 1.52% |
| hsa_circ_0090475 | 1318 | 10 | 1.52% |
| hsa_circ_0114289 | 1318 | 10 | 1.52% |
| hsa_circ_0121156 | 1318 | 10 | 1.52% |
| hsa_circ_0133458 | 1318 | 10 | 1.52% |
| hsa_circ_0140233 | 1318 | 10 | 1.52% |
| hsa_circ_0046426 | 1450 | 11 | 1.52% |
| hsa_circ_0048149 | 1450 | 11 | 1.52% |
| hsa_circ_0076127 | 1450 | 11 | 1.52% |
| hsa_circ_0100200 | 1450 | 11 | 1.52% |
| hsa_circ_0112366 | 1450 | 11 | 1.52% |
| hsa_circ_0137625 | 1450 | 11 | 1.52% |
| hsa_circ_0137639 | 1450 | 11 | 1.52% |
| hsa_circ_0010588 | 1582 | 12 | 1.52% |
| hsa_circ_0115088 | 1582 | 12 | 1.52% |
| hsa_circ_0121209 | 1582 | 12 | 1.52% |
| hsa_circ_0036490 | 1714 | 13 | 1.52% |
| hsa_circ_0014788 | 1978 | 15 | 1.52% |
| hsa_circ_0023213 | 1319 | 10 | 1.52% |
| hsa_circ_0029852 | 1319 | 10 | 1.52% |
| hsa_circ_0128777 | 1319 | 10 | 1.52% |
| hsa_circ_0129087 | 1319 | 10 | 1.52% |
| hsa_circ_0129985 | 1319 | 10 | 1.52% |
| hsa_circ_0002472 | 1451 | 11 | 1.52% |
| hsa_circ_0026023 | 1451 | 11 | 1.52% |

| | | | |
|---|---|---|---|
| hsa_circ_0043655 | 1451 | 11 | 1.52% |
| hsa_circ_0065998 | 1451 | 11 | 1.52% |
| hsa_circ_0067824 | 1451 | 11 | 1.52% |
| hsa_circ_0081841 | 1451 | 11 | 1.52% |
| hsa_circ_0087552 | 1451 | 11 | 1.52% |
| hsa_circ_0094733 | 1451 | 11 | 1.52% |
| hsa_circ_0083569 | 1979 | 15 | 1.52% |
| hsa_circ_0002688 | 1584 | 12 | 1.52% |
| hsa_circ_0051722 | 1584 | 12 | 1.52% |
| hsa_circ_0055483 | 1584 | 12 | 1.52% |
| hsa_circ_0075660 | 1584 | 12 | 1.52% |
| hsa_circ_0135426 | 1584 | 12 | 1.52% |
| hsa_circ_0010136 | 1452 | 11 | 1.52% |
| hsa_circ_0014704 | 1452 | 11 | 1.52% |
| hsa_circ_0031781 | 1320 | 10 | 1.52% |
| hsa_circ_0045894 | 1320 | 10 | 1.52% |
| hsa_circ_0056378 | 1320 | 10 | 1.52% |
| hsa_circ_0064128 | 1320 | 10 | 1.52% |
| hsa_circ_0071157 | 1320 | 10 | 1.52% |
| hsa_circ_0076535 | 1320 | 10 | 1.52% |
| hsa_circ_0090465 | 1320 | 10 | 1.52% |
| hsa_circ_0092061 | 1320 | 10 | 1.52% |
| hsa_circ_0104149 | 1320 | 10 | 1.52% |
| hsa_circ_0124890 | 1320 | 10 | 1.52% |
| hsa_circ_0129145 | 1320 | 10 | 1.52% |
| hsa_circ_0027291 | 1717 | 13 | 1.51% |
| hsa_circ_0102432 | 1585 | 12 | 1.51% |
| hsa_circ_0118394 | 1585 | 12 | 1.51% |
| hsa_circ_0010209 | 1453 | 11 | 1.51% |
| hsa_circ_0018236 | 1453 | 11 | 1.51% |
| hsa_circ_0052855 | 1453 | 11 | 1.51% |
| hsa_circ_0095228 | 1453 | 11 | 1.51% |
| hsa_circ_0134199 | 1453 | 11 | 1.51% |
| hsa_circ_0011634 | 1321 | 10 | 1.51% |
| hsa_circ_0012434 | 1321 | 10 | 1.51% |
| hsa_circ_0015255 | 1321 | 10 | 1.51% |
| hsa_circ_0024924 | 1321 | 10 | 1.51% |
| hsa_circ_0041484 | 1321 | 10 | 1.51% |
| hsa_circ_0099208 | 1321 | 10 | 1.51% |
| hsa_circ_0101862 | 1321 | 10 | 1.51% |
| hsa_circ_0114862 | 1321 | 10 | 1.51% |
| hsa_circ_0117069 | 1321 | 10 | 1.51% |
| hsa_circ_0119727 | 1321 | 10 | 1.51% |
| hsa_circ_0043851 | 1586 | 12 | 1.51% |
| hsa_circ_0065012 | 1586 | 12 | 1.51% |
| hsa_circ_0083695 | 1586 | 12 | 1.51% |
| hsa_circ_0059579 | 1454 | 11 | 1.51% |
| hsa_circ_0076108 | 1454 | 11 | 1.51% |
| hsa_circ_0095834 | 1454 | 11 | 1.51% |
| hsa_circ_0124674 | 1454 | 11 | 1.51% |

| | | | |
|---|---|---|---|
| hsa_circ_0007368 | 1322 | 10 | 1.51% |
| hsa_circ_0011755 | 1322 | 10 | 1.51% |
| hsa_circ_0012740 | 1322 | 10 | 1.51% |
| hsa_circ_0018062 | 1322 | 10 | 1.51% |
| hsa_circ_0026890 | 1322 | 10 | 1.51% |
| hsa_circ_0031821 | 1322 | 10 | 1.51% |
| hsa_circ_0034257 | 1322 | 10 | 1.51% |
| hsa_circ_0078448 | 1322 | 10 | 1.51% |
| hsa_circ_0096319 | 1322 | 10 | 1.51% |
| hsa_circ_0108413 | 1322 | 10 | 1.51% |
| hsa_circ_0117212 | 1322 | 10 | 1.51% |
| hsa_circ_0098225 | 2380 | 18 | 1.51% |
| hsa_circ_0036436 | 1587 | 12 | 1.51% |
| hsa_circ_0049980 | 1587 | 12 | 1.51% |
| hsa_circ_0089851 | 1587 | 12 | 1.51% |
| hsa_circ_0125910 | 1587 | 12 | 1.51% |
| hsa_circ_0126852 | 1587 | 12 | 1.51% |
| hsa_circ_0019921 | 1455 | 11 | 1.51% |
| hsa_circ_0027791 | 1455 | 11 | 1.51% |
| hsa_circ_0054233 | 1455 | 11 | 1.51% |
| hsa_circ_0083940 | 1455 | 11 | 1.51% |
| hsa_circ_0100012 | 1455 | 11 | 1.51% |
| hsa_circ_0104390 | 1455 | 11 | 1.51% |
| hsa_circ_0070129 | 1323 | 10 | 1.51% |
| hsa_circ_0073058 | 1323 | 10 | 1.51% |
| hsa_circ_0075649 | 1323 | 10 | 1.51% |
| hsa_circ_0084947 | 1323 | 10 | 1.51% |
| hsa_circ_0087011 | 1323 | 10 | 1.51% |
| hsa_circ_0088405 | 1323 | 10 | 1.51% |
| hsa_circ_0097669 | 1323 | 10 | 1.51% |
| hsa_circ_0099393 | 1323 | 10 | 1.51% |
| hsa_circ_0110963 | 1323 | 10 | 1.51% |
| hsa_circ_0113060 | 1323 | 10 | 1.51% |
| hsa_circ_0133544 | 1323 | 10 | 1.51% |
| hsa_circ_0053025 | 1720 | 13 | 1.51% |
| hsa_circ_0066649 | 2382 | 18 | 1.51% |
| hsa_circ_0025772 | 1588 | 12 | 1.51% |
| hsa_circ_0037396 | 1588 | 12 | 1.51% |
| hsa_circ_0042257 | 1588 | 12 | 1.51% |
| hsa_circ_0083592 | 1588 | 12 | 1.51% |
| hsa_circ_0001282 | 1853 | 14 | 1.51% |
| hsa_circ_0040877 | 1853 | 14 | 1.51% |
| hsa_circ_0010075 | 1456 | 11 | 1.51% |
| hsa_circ_0012057 | 1324 | 10 | 1.51% |
| hsa_circ_0014115 | 1324 | 10 | 1.51% |
| hsa_circ_0035458 | 1324 | 10 | 1.51% |
| hsa_circ_0038981 | 1324 | 10 | 1.51% |
| hsa_circ_0041581 | 1324 | 10 | 1.51% |
| hsa_circ_0042729 | 1324 | 10 | 1.51% |
| hsa_circ_0050068 | 1324 | 10 | 1.51% |

| | | | |
|---|---|---|---|
| hsa_circ_0064357 | 1324 | 10 | 1.51% |
| hsa_circ_0126420 | 1324 | 10 | 1.51% |
| hsa_circ_0049494 | 5032 | 38 | 1.51% |
| hsa_circ_0131471 | 5297 | 40 | 1.51% |
| hsa_circ_0011284 | 1457 | 11 | 1.51% |
| hsa_circ_0012515 | 1457 | 11 | 1.51% |
| hsa_circ_0116600 | 1457 | 11 | 1.51% |
| hsa_circ_0079105 | 1722 | 13 | 1.51% |
| hsa_circ_0113630 | 4107 | 31 | 1.51% |
| hsa_circ_0031523 | 1590 | 12 | 1.51% |
| hsa_circ_0011313 | 1325 | 10 | 1.51% |
| hsa_circ_0046090 | 1325 | 10 | 1.51% |
| hsa_circ_0050935 | 1325 | 10 | 1.51% |
| hsa_circ_0076729 | 1325 | 10 | 1.51% |
| hsa_circ_0084482 | 1325 | 10 | 1.51% |
| hsa_circ_0126752 | 1325 | 10 | 1.51% |
| hsa_circ_0138402 | 1325 | 10 | 1.51% |
| hsa_circ_0140656 | 1325 | 10 | 1.51% |
| hsa_circ_0042147 | 1458 | 11 | 1.51% |
| hsa_circ_0073627 | 1458 | 11 | 1.51% |
| hsa_circ_0083639 | 1458 | 11 | 1.51% |
| hsa_circ_0091421 | 1458 | 11 | 1.51% |
| hsa_circ_0020348 | 1591 | 12 | 1.51% |
| hsa_circ_0041211 | 1591 | 12 | 1.51% |
| hsa_circ_0121386 | 1591 | 12 | 1.51% |
| hsa_circ_0008084 | 1326 | 10 | 1.51% |
| hsa_circ_0013034 | 1326 | 10 | 1.51% |
| hsa_circ_0027124 | 1326 | 10 | 1.51% |
| hsa_circ_0038086 | 1326 | 10 | 1.51% |
| hsa_circ_0043650 | 1326 | 10 | 1.51% |
| hsa_circ_0049904 | 1326 | 10 | 1.51% |
| hsa_circ_0077124 | 1326 | 10 | 1.51% |
| hsa_circ_0092237 | 1326 | 10 | 1.51% |
| hsa_circ_0127284 | 1326 | 10 | 1.51% |
| hsa_circ_0130751 | 1326 | 10 | 1.51% |
| hsa_circ_0135663 | 1326 | 10 | 1.51% |
| hsa_circ_0135992 | 1326 | 10 | 1.51% |
| hsa_circ_0016236 | 1459 | 11 | 1.51% |
| hsa_circ_0053301 | 1459 | 11 | 1.51% |
| hsa_circ_0064170 | 1459 | 11 | 1.51% |
| hsa_circ_0081177 | 1459 | 11 | 1.51% |
| hsa_circ_0093360 | 1459 | 11 | 1.51% |
| hsa_circ_0106479 | 1459 | 11 | 1.51% |
| hsa_circ_0138165 | 1459 | 11 | 1.51% |
| hsa_circ_0052703 | 1990 | 15 | 1.51% |
| hsa_circ_0085797 | 1592 | 12 | 1.51% |
| hsa_circ_0116203 | 3981 | 30 | 1.51% |
| hsa_circ_0012255 | 1327 | 10 | 1.51% |
| hsa_circ_0019040 | 1327 | 10 | 1.51% |
| hsa_circ_0019288 | 1327 | 10 | 1.51% |

| | | | |
|---|---|---|---|
| hsa_circ_0024457 | 1327 | 10 | 1.51% |
| hsa_circ_0034917 | 1327 | 10 | 1.51% |
| hsa_circ_0068983 | 1327 | 10 | 1.51% |
| hsa_circ_0076989 | 1327 | 10 | 1.51% |
| hsa_circ_0081484 | 1327 | 10 | 1.51% |
| hsa_circ_0099989 | 1327 | 10 | 1.51% |
| hsa_circ_0105462 | 1327 | 10 | 1.51% |
| hsa_circ_0110243 | 1327 | 10 | 1.51% |
| hsa_circ_0123539 | 1327 | 10 | 1.51% |
| hsa_circ_0067793 | 3716 | 28 | 1.51% |
| hsa_circ_0028151 | 1460 | 11 | 1.51% |
| hsa_circ_0032875 | 1460 | 11 | 1.51% |
| hsa_circ_0056841 | 1460 | 11 | 1.51% |
| hsa_circ_0059717 | 1460 | 11 | 1.51% |
| hsa_circ_0068804 | 1460 | 11 | 1.51% |
| hsa_circ_0092402 | 1460 | 11 | 1.51% |
| hsa_circ_0103345 | 1460 | 11 | 1.51% |
| hsa_circ_0084688 | 1593 | 12 | 1.51% |
| hsa_circ_0009733 | 1328 | 10 | 1.51% |
| hsa_circ_0018602 | 1328 | 10 | 1.51% |
| hsa_circ_0024321 | 1328 | 10 | 1.51% |
| hsa_circ_0025932 | 1328 | 10 | 1.51% |
| hsa_circ_0026256 | 1328 | 10 | 1.51% |
| hsa_circ_0027677 | 1328 | 10 | 1.51% |
| hsa_circ_0029969 | 1328 | 10 | 1.51% |
| hsa_circ_0041158 | 1328 | 10 | 1.51% |
| hsa_circ_0061256 | 1328 | 10 | 1.51% |
| hsa_circ_0085175 | 1328 | 10 | 1.51% |
| hsa_circ_0089402 | 1328 | 10 | 1.51% |
| hsa_circ_0123270 | 1328 | 10 | 1.51% |
| hsa_circ_0137831 | 1328 | 10 | 1.51% |
| hsa_circ_0138384 | 1328 | 10 | 1.51% |
| hsa_circ_0140276 | 1328 | 10 | 1.51% |
| hsa_circ_0055926 | 2922 | 22 | 1.51% |
| hsa_circ_0018507 | 1461 | 11 | 1.51% |
| hsa_circ_0024761 | 1461 | 11 | 1.51% |
| hsa_circ_0025090 | 1461 | 11 | 1.51% |
| hsa_circ_0029097 | 1461 | 11 | 1.51% |
| hsa_circ_0120561 | 1461 | 11 | 1.51% |
| hsa_circ_0135598 | 1461 | 11 | 1.51% |
| hsa_circ_0022011 | 1594 | 12 | 1.51% |
| hsa_circ_0037469 | 1594 | 12 | 1.51% |
| hsa_circ_0068387 | 1594 | 12 | 1.51% |
| hsa_circ_0007624 | 1329 | 10 | 1.50% |
| hsa_circ_0009628 | 1329 | 10 | 1.50% |
| hsa_circ_0017388 | 1329 | 10 | 1.50% |
| hsa_circ_0019612 | 1329 | 10 | 1.50% |
| hsa_circ_0020581 | 1329 | 10 | 1.50% |
| hsa_circ_0022111 | 1329 | 10 | 1.50% |
| hsa_circ_0026423 | 1329 | 10 | 1.50% |

| | | | |
|---|---|---|---|
| hsa_circ_0028241 | 1329 | 10 | 1.50% |
| hsa_circ_0031992 | 1329 | 10 | 1.50% |
| hsa_circ_0047447 | 1329 | 10 | 1.50% |
| hsa_circ_0048059 | 1329 | 10 | 1.50% |
| hsa_circ_0048666 | 1329 | 10 | 1.50% |
| hsa_circ_0048851 | 1329 | 10 | 1.50% |
| hsa_circ_0049802 | 1329 | 10 | 1.50% |
| hsa_circ_0071810 | 1329 | 10 | 1.50% |
| hsa_circ_0084989 | 1329 | 10 | 1.50% |
| hsa_circ_0086127 | 1329 | 10 | 1.50% |
| hsa_circ_0094922 | 1329 | 10 | 1.50% |
| hsa_circ_0097022 | 1329 | 10 | 1.50% |
| hsa_circ_0100658 | 1329 | 10 | 1.50% |
| hsa_circ_0106989 | 1329 | 10 | 1.50% |
| hsa_circ_0108142 | 1329 | 10 | 1.50% |
| hsa_circ_0114269 | 1329 | 10 | 1.50% |
| hsa_circ_0035119 | 5582 | 42 | 1.50% |
| hsa_circ_0026167 | 1462 | 11 | 1.50% |
| hsa_circ_0030509 | 1462 | 11 | 1.50% |
| hsa_circ_0102890 | 1462 | 11 | 1.50% |
| hsa_circ_0108463 | 1462 | 11 | 1.50% |
| hsa_circ_0013218 | 1595 | 12 | 1.50% |
| hsa_circ_0102265 | 1994 | 15 | 1.50% |
| hsa_circ_0109093 | 2926 | 22 | 1.50% |
| hsa_circ_0073797 | 1596 | 12 | 1.50% |
| hsa_circ_0019538 | 1463 | 11 | 1.50% |
| hsa_circ_0021811 | 1463 | 11 | 1.50% |
| hsa_circ_0046934 | 1463 | 11 | 1.50% |
| hsa_circ_0075358 | 1463 | 11 | 1.50% |
| hsa_circ_0076554 | 1463 | 11 | 1.50% |
| hsa_circ_0077119 | 1463 | 11 | 1.50% |
| hsa_circ_0118250 | 1463 | 11 | 1.50% |
| hsa_circ_0023123 | 1330 | 10 | 1.50% |
| hsa_circ_0044926 | 1330 | 10 | 1.50% |
| hsa_circ_0061848 | 1330 | 10 | 1.50% |
| hsa_circ_0101114 | 1330 | 10 | 1.50% |
| hsa_circ_0118467 | 1330 | 10 | 1.50% |
| hsa_circ_0127554 | 1330 | 10 | 1.50% |
| hsa_circ_0135262 | 1330 | 10 | 1.50% |
| hsa_circ_0138654 | 4923 | 37 | 1.50% |
| hsa_circ_0117173 | 1730 | 13 | 1.50% |
| hsa_circ_0016767 | 1597 | 12 | 1.50% |
| hsa_circ_0074498 | 1597 | 12 | 1.50% |
| hsa_circ_0075930 | 1597 | 12 | 1.50% |
| hsa_circ_0115367 | 1597 | 12 | 1.50% |
| hsa_circ_0129589 | 1597 | 12 | 1.50% |
| hsa_circ_0049619 | 1464 | 11 | 1.50% |
| hsa_circ_0116680 | 1464 | 11 | 1.50% |
| hsa_circ_0119206 | 1464 | 11 | 1.50% |
| hsa_circ_0125660 | 1464 | 11 | 1.50% |

| | | | |
|---|---|---|---|
| hsa_circ_0126728 | 1464 | 11 | 1.50% |
| hsa_circ_0021847 | 5723 | 43 | 1.50% |
| hsa_circ_0055257 | 1331 | 10 | 1.50% |
| hsa_circ_0065114 | 1331 | 10 | 1.50% |
| hsa_circ_0065671 | 1331 | 10 | 1.50% |
| hsa_circ_0071673 | 1331 | 10 | 1.50% |
| hsa_circ_0072354 | 1331 | 10 | 1.50% |
| hsa_circ_0079376 | 1331 | 10 | 1.50% |
| hsa_circ_0124984 | 1331 | 10 | 1.50% |
| hsa_circ_0129078 | 1331 | 10 | 1.50% |
| hsa_circ_0135683 | 1331 | 10 | 1.50% |
| hsa_circ_0069832 | 1997 | 15 | 1.50% |
| hsa_circ_0022608 | 1598 | 12 | 1.50% |
| hsa_circ_0073350 | 1598 | 12 | 1.50% |
| hsa_circ_0084620 | 1598 | 12 | 1.50% |
| hsa_circ_0090158 | 1598 | 12 | 1.50% |
| hsa_circ_0101478 | 1598 | 12 | 1.50% |
| hsa_circ_0018625 | 1465 | 11 | 1.50% |
| hsa_circ_0021535 | 1465 | 11 | 1.50% |
| hsa_circ_0046453 | 1465 | 11 | 1.50% |
| hsa_circ_0054733 | 1465 | 11 | 1.50% |
| hsa_circ_0059086 | 1465 | 11 | 1.50% |
| hsa_circ_0062968 | 1465 | 11 | 1.50% |
| hsa_circ_0066616 | 1465 | 11 | 1.50% |
| hsa_circ_0088825 | 1465 | 11 | 1.50% |
| hsa_circ_0134247 | 1465 | 11 | 1.50% |
| hsa_circ_0000712 | 1332 | 10 | 1.50% |
| hsa_circ_0005226 | 1332 | 10 | 1.50% |
| hsa_circ_0026260 | 1332 | 10 | 1.50% |
| hsa_circ_0028675 | 1332 | 10 | 1.50% |
| hsa_circ_0041161 | 1332 | 10 | 1.50% |
| hsa_circ_0062413 | 1332 | 10 | 1.50% |
| hsa_circ_0063131 | 1332 | 10 | 1.50% |
| hsa_circ_0071108 | 1332 | 10 | 1.50% |
| hsa_circ_0074918 | 1332 | 10 | 1.50% |
| hsa_circ_0075632 | 1332 | 10 | 1.50% |
| hsa_circ_0094179 | 1332 | 10 | 1.50% |
| hsa_circ_0094470 | 1332 | 10 | 1.50% |
| hsa_circ_0098018 | 1332 | 10 | 1.50% |
| hsa_circ_0102589 | 1332 | 10 | 1.50% |
| hsa_circ_0040694 | 1599 | 12 | 1.50% |
| hsa_circ_0082638 | 1599 | 12 | 1.50% |
| hsa_circ_0108582 | 1599 | 12 | 1.50% |
| hsa_circ_0016658 | 1466 | 11 | 1.50% |
| hsa_circ_0044228 | 1466 | 11 | 1.50% |
| hsa_circ_0045411 | 1466 | 11 | 1.50% |
| hsa_circ_0016322 | 1333 | 10 | 1.50% |
| hsa_circ_0020780 | 1333 | 10 | 1.50% |
| hsa_circ_0057910 | 1333 | 10 | 1.50% |
| hsa_circ_0066837 | 1333 | 10 | 1.50% |

| | | | |
|---|---|---|---|
| hsa_circ_0074541 | 1333 | 10 | 1.50% |
| hsa_circ_0112148 | 1333 | 10 | 1.50% |
| hsa_circ_0017272 | 1733 | 13 | 1.50% |
| hsa_circ_0118654 | 1733 | 13 | 1.50% |
| hsa_circ_0021653 | 2133 | 16 | 1.50% |
| hsa_circ_0075928 | 2000 | 15 | 1.50% |
| hsa_circ_0078931 | 2000 | 15 | 1.50% |
| hsa_circ_0001605 | 1600 | 12 | 1.50% |
| hsa_circ_0024213 | 1600 | 12 | 1.50% |
| hsa_circ_0036661 | 1600 | 12 | 1.50% |
| hsa_circ_0058580 | 1600 | 12 | 1.50% |
| hsa_circ_0070415 | 1600 | 12 | 1.50% |
| hsa_circ_0097252 | 1600 | 12 | 1.50% |
| hsa_circ_0131898 | 1600 | 12 | 1.50% |
| hsa_circ_0026483 | 1467 | 11 | 1.50% |
| hsa_circ_0062633 | 1467 | 11 | 1.50% |
| hsa_circ_0073650 | 1467 | 11 | 1.50% |
| hsa_circ_0076351 | 1467 | 11 | 1.50% |
| hsa_circ_0076518 | 1467 | 11 | 1.50% |
| hsa_circ_0089741 | 1467 | 11 | 1.50% |
| hsa_circ_0015480 | 1334 | 10 | 1.50% |
| hsa_circ_0016042 | 1334 | 10 | 1.50% |
| hsa_circ_0021142 | 1334 | 10 | 1.50% |
| hsa_circ_0065948 | 1334 | 10 | 1.50% |
| hsa_circ_0079053 | 1334 | 10 | 1.50% |
| hsa_circ_0087807 | 1334 | 10 | 1.50% |
| hsa_circ_0136664 | 1334 | 10 | 1.50% |
| hsa_circ_0018742 | 2935 | 22 | 1.50% |
| hsa_circ_0051521 | 1468 | 11 | 1.50% |
| hsa_circ_0087797 | 1468 | 11 | 1.50% |
| hsa_circ_0096864 | 1468 | 11 | 1.50% |
| hsa_circ_0105756 | 1468 | 11 | 1.50% |
| hsa_circ_0113528 | 1468 | 11 | 1.50% |
| hsa_circ_0113809 | 1468 | 11 | 1.50% |
| hsa_circ_0117369 | 1468 | 11 | 1.50% |
| hsa_circ_0120450 | 1468 | 11 | 1.50% |
| hsa_circ_0022048 | 1869 | 14 | 1.50% |
| hsa_circ_0003617 | 1335 | 10 | 1.50% |
| hsa_circ_0006026 | 1335 | 10 | 1.50% |
| hsa_circ_0013860 | 1335 | 10 | 1.50% |
| hsa_circ_0014760 | 1335 | 10 | 1.50% |
| hsa_circ_0017055 | 1335 | 10 | 1.50% |
| hsa_circ_0020972 | 1335 | 10 | 1.50% |
| hsa_circ_0030569 | 1335 | 10 | 1.50% |
| hsa_circ_0038853 | 1335 | 10 | 1.50% |
| hsa_circ_0045266 | 1335 | 10 | 1.50% |
| hsa_circ_0047043 | 1335 | 10 | 1.50% |
| hsa_circ_0048062 | 1335 | 10 | 1.50% |
| hsa_circ_0052516 | 1335 | 10 | 1.50% |
| hsa_circ_0055344 | 1335 | 10 | 1.50% |

| | | | |
|---|---|---|---|
| hsa_circ_0057614 | 1335 | 10 | 1.50% |
| hsa_circ_0077934 | 1335 | 10 | 1.50% |
| hsa_circ_0089946 | 1335 | 10 | 1.50% |
| hsa_circ_0105958 | 1335 | 10 | 1.50% |
| hsa_circ_0129733 | 1335 | 10 | 1.50% |
| hsa_circ_0139210 | 1335 | 10 | 1.50% |
| hsa_circ_0093109 | 1736 | 13 | 1.50% |
| hsa_circ_0026567 | 1469 | 11 | 1.50% |
| hsa_circ_0030727 | 1469 | 11 | 1.50% |
| hsa_circ_0034674 | 1469 | 11 | 1.50% |
| hsa_circ_0038157 | 1469 | 11 | 1.50% |
| hsa_circ_0048423 | 1469 | 11 | 1.50% |
| hsa_circ_0084456 | 1469 | 11 | 1.50% |
| hsa_circ_0103182 | 2805 | 21 | 1.50% |
| hsa_circ_0002900 | 1603 | 12 | 1.50% |
| hsa_circ_0041570 | 1603 | 12 | 1.50% |
| hsa_circ_0091707 | 1603 | 12 | 1.50% |
| hsa_circ_0107386 | 1603 | 12 | 1.50% |
| hsa_circ_0138126 | 1603 | 12 | 1.50% |
| hsa_circ_0010347 | 1336 | 10 | 1.50% |
| hsa_circ_0038568 | 1336 | 10 | 1.50% |
| hsa_circ_0063157 | 1336 | 10 | 1.50% |
| hsa_circ_0092655 | 1336 | 10 | 1.50% |
| hsa_circ_0114514 | 1336 | 10 | 1.50% |
| hsa_circ_0116012 | 1336 | 10 | 1.50% |
| hsa_circ_0116047 | 1336 | 10 | 1.50% |
| hsa_circ_0122231 | 1336 | 10 | 1.50% |
| hsa_circ_0131823 | 1336 | 10 | 1.50% |
| hsa_circ_0046164 | 1737 | 13 | 1.50% |
| hsa_circ_0073076 | 1737 | 13 | 1.50% |
| hsa_circ_0011255 | 1470 | 11 | 1.50% |
| hsa_circ_0018410 | 1470 | 11 | 1.50% |
| hsa_circ_0019006 | 1470 | 11 | 1.50% |
| hsa_circ_0020830 | 1470 | 11 | 1.50% |
| hsa_circ_0059017 | 1470 | 11 | 1.50% |
| hsa_circ_0087547 | 1470 | 11 | 1.50% |
| hsa_circ_0009423 | 2005 | 15 | 1.50% |
| hsa_circ_0053027 | 2005 | 15 | 1.50% |
| hsa_circ_0080742 | 1604 | 12 | 1.50% |
| hsa_circ_0124384 | 1604 | 12 | 1.50% |
| hsa_circ_0015049 | 1337 | 10 | 1.50% |
| hsa_circ_0058631 | 1337 | 10 | 1.50% |
| hsa_circ_0073141 | 1337 | 10 | 1.50% |
| hsa_circ_0078390 | 1337 | 10 | 1.50% |
| hsa_circ_0089816 | 1337 | 10 | 1.50% |
| hsa_circ_0091007 | 1337 | 10 | 1.50% |
| hsa_circ_0108879 | 1337 | 10 | 1.50% |
| hsa_circ_0123214 | 1337 | 10 | 1.50% |
| hsa_circ_0087787 | 1872 | 14 | 1.50% |
| hsa_circ_0028962 | 1471 | 11 | 1.50% |

| | | | |
|---|---|---|---|
| hsa_circ_0044057 | 1471 | 11 | 1.50% |
| hsa_circ_0072123 | 1471 | 11 | 1.50% |
| hsa_circ_0089290 | 1471 | 11 | 1.50% |
| hsa_circ_0101990 | 1471 | 11 | 1.50% |
| hsa_circ_0005332 | 1605 | 12 | 1.50% |
| hsa_circ_0075136 | 1605 | 12 | 1.50% |
| hsa_circ_0112870 | 2274 | 17 | 1.50% |
| hsa_circ_0073367 | 1873 | 14 | 1.49% |
| hsa_circ_0010505 | 1338 | 10 | 1.49% |
| hsa_circ_0042000 | 1338 | 10 | 1.49% |
| hsa_circ_0059225 | 1338 | 10 | 1.49% |
| hsa_circ_0062421 | 1338 | 10 | 1.49% |
| hsa_circ_0066355 | 1338 | 10 | 1.49% |
| hsa_circ_0067470 | 1338 | 10 | 1.49% |
| hsa_circ_0079285 | 1338 | 10 | 1.49% |
| hsa_circ_0084381 | 1338 | 10 | 1.49% |
| hsa_circ_0085511 | 1338 | 10 | 1.49% |
| hsa_circ_0086766 | 1338 | 10 | 1.49% |
| hsa_circ_0013322 | 1472 | 11 | 1.49% |
| hsa_circ_0100301 | 1472 | 11 | 1.49% |
| hsa_circ_0047197 | 1606 | 12 | 1.49% |
| hsa_circ_0020233 | 1339 | 10 | 1.49% |
| hsa_circ_0031228 | 1339 | 10 | 1.49% |
| hsa_circ_0042873 | 1339 | 10 | 1.49% |
| hsa_circ_0049646 | 1339 | 10 | 1.49% |
| hsa_circ_0062705 | 1339 | 10 | 1.49% |
| hsa_circ_0064962 | 1339 | 10 | 1.49% |
| hsa_circ_0084499 | 1339 | 10 | 1.49% |
| hsa_circ_0088439 | 1339 | 10 | 1.49% |
| hsa_circ_0092031 | 1339 | 10 | 1.49% |
| hsa_circ_0099515 | 1339 | 10 | 1.49% |
| hsa_circ_0121351 | 1339 | 10 | 1.49% |
| hsa_circ_0126501 | 1339 | 10 | 1.49% |
| hsa_circ_0136134 | 1339 | 10 | 1.49% |
| hsa_circ_0137197 | 1339 | 10 | 1.49% |
| hsa_circ_0140046 | 1339 | 10 | 1.49% |
| hsa_circ_0002071 | 1473 | 11 | 1.49% |
| hsa_circ_0023258 | 1473 | 11 | 1.49% |
| hsa_circ_0035373 | 1473 | 11 | 1.49% |
| hsa_circ_0038087 | 1473 | 11 | 1.49% |
| hsa_circ_0039027 | 1473 | 11 | 1.49% |
| hsa_circ_0045280 | 1473 | 11 | 1.49% |
| hsa_circ_0052268 | 1473 | 11 | 1.49% |
| hsa_circ_0066774 | 1473 | 11 | 1.49% |
| hsa_circ_0092077 | 1473 | 11 | 1.49% |
| hsa_circ_0128925 | 1473 | 11 | 1.49% |
| hsa_circ_0079506 | 1607 | 12 | 1.49% |
| hsa_circ_0091309 | 1607 | 12 | 1.49% |
| hsa_circ_0005430 | 2009 | 15 | 1.49% |
| hsa_circ_0137776 | 2277 | 17 | 1.49% |

| | | | |
|---|---|---|---|
| hsa_circ_0009487 | 4288 | 32 | 1.49% |
| hsa_circ_0029973 | 1474 | 11 | 1.49% |
| hsa_circ_0059036 | 1474 | 11 | 1.49% |
| hsa_circ_0063590 | 1474 | 11 | 1.49% |
| hsa_circ_0075091 | 1474 | 11 | 1.49% |
| hsa_circ_0078546 | 1474 | 11 | 1.49% |
| hsa_circ_0099226 | 1474 | 11 | 1.49% |
| hsa_circ_0101844 | 1474 | 11 | 1.49% |
| hsa_circ_0108405 | 1474 | 11 | 1.49% |
| hsa_circ_0001907 | 1340 | 10 | 1.49% |
| hsa_circ_0026130 | 1340 | 10 | 1.49% |
| hsa_circ_0036682 | 1340 | 10 | 1.49% |
| hsa_circ_0057026 | 1340 | 10 | 1.49% |
| hsa_circ_0077948 | 1340 | 10 | 1.49% |
| hsa_circ_0138619 | 1340 | 10 | 1.49% |
| hsa_circ_0139568 | 1340 | 10 | 1.49% |
| hsa_circ_0108171 | 1743 | 13 | 1.49% |
| hsa_circ_0029084 | 1475 | 11 | 1.49% |
| hsa_circ_0058956 | 1475 | 11 | 1.49% |
| hsa_circ_0077812 | 1475 | 11 | 1.49% |
| hsa_circ_0095493 | 1475 | 11 | 1.49% |
| hsa_circ_0111482 | 1475 | 11 | 1.49% |
| hsa_circ_0035485 | 1341 | 10 | 1.49% |
| hsa_circ_0035891 | 1341 | 10 | 1.49% |
| hsa_circ_0041359 | 1341 | 10 | 1.49% |
| hsa_circ_0049190 | 1341 | 10 | 1.49% |
| hsa_circ_0049421 | 1341 | 10 | 1.49% |
| hsa_circ_0072543 | 1341 | 10 | 1.49% |
| hsa_circ_0074009 | 1341 | 10 | 1.49% |
| hsa_circ_0086956 | 1341 | 10 | 1.49% |
| hsa_circ_0089871 | 1341 | 10 | 1.49% |
| hsa_circ_0096768 | 1341 | 10 | 1.49% |
| hsa_circ_0097501 | 1341 | 10 | 1.49% |
| hsa_circ_0116240 | 1341 | 10 | 1.49% |
| hsa_circ_0125096 | 1341 | 10 | 1.49% |
| hsa_circ_0136188 | 1341 | 10 | 1.49% |
| hsa_circ_0119263 | 4024 | 30 | 1.49% |
| hsa_circ_0032283 | 1878 | 14 | 1.49% |
| hsa_circ_0062860 | 1744 | 13 | 1.49% |
| hsa_circ_0029970 | 1476 | 11 | 1.49% |
| hsa_circ_0074247 | 1476 | 11 | 1.49% |
| hsa_circ_0107394 | 1476 | 11 | 1.49% |
| hsa_circ_0011549 | 1342 | 10 | 1.49% |
| hsa_circ_0021338 | 1342 | 10 | 1.49% |
| hsa_circ_0023018 | 1342 | 10 | 1.49% |
| hsa_circ_0051127 | 1342 | 10 | 1.49% |
| hsa_circ_0062022 | 1342 | 10 | 1.49% |
| hsa_circ_0064938 | 1342 | 10 | 1.49% |
| hsa_circ_0087791 | 1342 | 10 | 1.49% |
| hsa_circ_0094747 | 1342 | 10 | 1.49% |

| | | | |
|---|---|---|---|
| hsa_circ_0094909 | 1342 | 10 | 1.49% |
| hsa_circ_0104442 | 1342 | 10 | 1.49% |
| hsa_circ_0118825 | 1342 | 10 | 1.49% |
| hsa_circ_0130843 | 1342 | 10 | 1.49% |
| hsa_circ_0137840 | 1342 | 10 | 1.49% |
| hsa_circ_0139276 | 1342 | 10 | 1.49% |
| hsa_circ_0070906 | 1745 | 13 | 1.49% |
| hsa_circ_0023389 | 1611 | 12 | 1.49% |
| hsa_circ_0035902 | 1477 | 11 | 1.49% |
| hsa_circ_0080930 | 1477 | 11 | 1.49% |
| hsa_circ_0092023 | 1477 | 11 | 1.49% |
| hsa_circ_0105897 | 1477 | 11 | 1.49% |
| hsa_circ_0125552 | 1477 | 11 | 1.49% |
| hsa_circ_0016224 | 1343 | 10 | 1.49% |
| hsa_circ_0016802 | 1343 | 10 | 1.49% |
| hsa_circ_0020386 | 1343 | 10 | 1.49% |
| hsa_circ_0024362 | 1343 | 10 | 1.49% |
| hsa_circ_0044586 | 1343 | 10 | 1.49% |
| hsa_circ_0057631 | 1343 | 10 | 1.49% |
| hsa_circ_0060030 | 1343 | 10 | 1.49% |
| hsa_circ_0070652 | 1343 | 10 | 1.49% |
| hsa_circ_0071688 | 1343 | 10 | 1.49% |
| hsa_circ_0092724 | 1343 | 10 | 1.49% |
| hsa_circ_0110958 | 1343 | 10 | 1.49% |
| hsa_circ_0119983 | 1343 | 10 | 1.49% |
| hsa_circ_0127943 | 1343 | 10 | 1.49% |
| hsa_circ_0128636 | 1343 | 10 | 1.49% |
| hsa_circ_0133054 | 1343 | 10 | 1.49% |
| hsa_circ_0133464 | 1343 | 10 | 1.49% |
| hsa_circ_0135892 | 1343 | 10 | 1.49% |
| hsa_circ_0131609 | 2821 | 21 | 1.49% |
| hsa_circ_0075718 | 1612 | 12 | 1.49% |
| hsa_circ_0131470 | 5374 | 40 | 1.49% |
| hsa_circ_0038127 | 1478 | 11 | 1.49% |
| hsa_circ_0043319 | 1478 | 11 | 1.49% |
| hsa_circ_0050735 | 1478 | 11 | 1.49% |
| hsa_circ_0051655 | 1478 | 11 | 1.49% |
| hsa_circ_0066251 | 1478 | 11 | 1.49% |
| hsa_circ_0077872 | 1478 | 11 | 1.49% |
| hsa_circ_0103662 | 1478 | 11 | 1.49% |
| hsa_circ_0121902 | 14245 | 106 | 1.49% |
| hsa_circ_0011809 | 1344 | 10 | 1.49% |
| hsa_circ_0063924 | 1344 | 10 | 1.49% |
| hsa_circ_0073914 | 1344 | 10 | 1.49% |
| hsa_circ_0086972 | 1344 | 10 | 1.49% |
| hsa_circ_0088244 | 1344 | 10 | 1.49% |
| hsa_circ_0089130 | 1344 | 10 | 1.49% |
| hsa_circ_0098545 | 1344 | 10 | 1.49% |
| hsa_circ_0107562 | 1344 | 10 | 1.49% |
| hsa_circ_0112036 | 1344 | 10 | 1.49% |

| | | | |
|---|---|---|---|
| hsa_circ_0120218 | 1344 | 10 | 1.49% |
| hsa_circ_0122621 | 1344 | 10 | 1.49% |
| hsa_circ_0124738 | 1344 | 10 | 1.49% |
| hsa_circ_0009265 | 1613 | 12 | 1.49% |
| hsa_circ_0054429 | 1613 | 12 | 1.49% |
| hsa_circ_0103968 | 1613 | 12 | 1.49% |
| hsa_circ_0130607 | 1613 | 12 | 1.49% |
| hsa_circ_0009457 | 1479 | 11 | 1.49% |
| hsa_circ_0015871 | 1479 | 11 | 1.49% |
| hsa_circ_0026397 | 1479 | 11 | 1.49% |
| hsa_circ_0030942 | 1479 | 11 | 1.49% |
| hsa_circ_0032962 | 1479 | 11 | 1.49% |
| hsa_circ_0044742 | 1479 | 11 | 1.49% |
| hsa_circ_0054878 | 1479 | 11 | 1.49% |
| hsa_circ_0071468 | 1479 | 11 | 1.49% |
| hsa_circ_0048920 | 1748 | 13 | 1.49% |
| hsa_circ_0045150 | 1614 | 12 | 1.49% |
| hsa_circ_0060744 | 1614 | 12 | 1.49% |
| hsa_circ_0078220 | 1614 | 12 | 1.49% |
| hsa_circ_0090242 | 1614 | 12 | 1.49% |
| hsa_circ_0128940 | 1614 | 12 | 1.49% |
| hsa_circ_0011343 | 1345 | 10 | 1.49% |
| hsa_circ_0012372 | 1345 | 10 | 1.49% |
| hsa_circ_0015315 | 1345 | 10 | 1.49% |
| hsa_circ_0017892 | 1345 | 10 | 1.49% |
| hsa_circ_0018002 | 1345 | 10 | 1.49% |
| hsa_circ_0018798 | 1345 | 10 | 1.49% |
| hsa_circ_0018859 | 1345 | 10 | 1.49% |
| hsa_circ_0023571 | 1345 | 10 | 1.49% |
| hsa_circ_0040871 | 1345 | 10 | 1.49% |
| hsa_circ_0055286 | 1345 | 10 | 1.49% |
| hsa_circ_0059500 | 1345 | 10 | 1.49% |
| hsa_circ_0063054 | 1345 | 10 | 1.49% |
| hsa_circ_0067841 | 1345 | 10 | 1.49% |
| hsa_circ_0068172 | 1345 | 10 | 1.49% |
| hsa_circ_0090198 | 1345 | 10 | 1.49% |
| hsa_circ_0096671 | 1345 | 10 | 1.49% |
| hsa_circ_0101771 | 1345 | 10 | 1.49% |
| hsa_circ_0127890 | 1345 | 10 | 1.49% |
| hsa_circ_0130157 | 1345 | 10 | 1.49% |
| hsa_circ_0138801 | 1345 | 10 | 1.49% |
| hsa_circ_0103401 | 1749 | 13 | 1.49% |
| hsa_circ_0003969 | 1480 | 11 | 1.49% |
| hsa_circ_0026752 | 1480 | 11 | 1.49% |
| hsa_circ_0039965 | 1480 | 11 | 1.49% |
| hsa_circ_0041125 | 1480 | 11 | 1.49% |
| hsa_circ_0070805 | 1480 | 11 | 1.49% |
| hsa_circ_0118353 | 1480 | 11 | 1.49% |
| hsa_circ_0121250 | 1480 | 11 | 1.49% |
| hsa_circ_0016958 | 1615 | 12 | 1.49% |

| | | | |
|---|---|---|---|
| hsa_circ_0045332 | 1615 | 12 | 1.49% |
| hsa_circ_0077243 | 1615 | 12 | 1.49% |
| hsa_circ_0123148 | 1615 | 12 | 1.49% |
| hsa_circ_0102011 | 10633 | 79 | 1.49% |
| hsa_circ_0030537 | 1346 | 10 | 1.49% |
| hsa_circ_0030807 | 1346 | 10 | 1.49% |
| hsa_circ_0035880 | 1346 | 10 | 1.49% |
| hsa_circ_0049360 | 1346 | 10 | 1.49% |
| hsa_circ_0073979 | 1346 | 10 | 1.49% |
| hsa_circ_0080821 | 1346 | 10 | 1.49% |
| hsa_circ_0084904 | 1346 | 10 | 1.49% |
| hsa_circ_0088102 | 1346 | 10 | 1.49% |
| hsa_circ_0090199 | 1346 | 10 | 1.49% |
| hsa_circ_0135941 | 1346 | 10 | 1.49% |
| hsa_circ_0012820 | 1481 | 11 | 1.49% |
| hsa_circ_0017431 | 1481 | 11 | 1.49% |
| hsa_circ_0021199 | 1481 | 11 | 1.49% |
| hsa_circ_0049913 | 1481 | 11 | 1.49% |
| hsa_circ_0065692 | 1481 | 11 | 1.49% |
| hsa_circ_0136143 | 1481 | 11 | 1.49% |
| hsa_circ_0079396 | 3905 | 29 | 1.49% |
| hsa_circ_0015504 | 1616 | 12 | 1.49% |
| hsa_circ_0023835 | 1616 | 12 | 1.49% |
| hsa_circ_0047640 | 1616 | 12 | 1.49% |
| hsa_circ_0089119 | 1616 | 12 | 1.49% |
| hsa_circ_0101347 | 1616 | 12 | 1.49% |
| hsa_circ_0010441 | 1347 | 10 | 1.48% |
| hsa_circ_0015350 | 1347 | 10 | 1.48% |
| hsa_circ_0026144 | 1347 | 10 | 1.48% |
| hsa_circ_0031396 | 1347 | 10 | 1.48% |
| hsa_circ_0034884 | 1347 | 10 | 1.48% |
| hsa_circ_0035842 | 1347 | 10 | 1.48% |
| hsa_circ_0046053 | 1347 | 10 | 1.48% |
| hsa_circ_0073095 | 1347 | 10 | 1.48% |
| hsa_circ_0083596 | 1347 | 10 | 1.48% |
| hsa_circ_0117160 | 1347 | 10 | 1.48% |
| hsa_circ_0122077 | 1347 | 10 | 1.48% |
| hsa_circ_0130839 | 1347 | 10 | 1.48% |
| hsa_circ_0006721 | 1482 | 11 | 1.48% |
| hsa_circ_0018773 | 1482 | 11 | 1.48% |
| hsa_circ_0065754 | 1482 | 11 | 1.48% |
| hsa_circ_0080610 | 1482 | 11 | 1.48% |
| hsa_circ_0117722 | 1482 | 11 | 1.48% |
| hsa_circ_0071555 | 1617 | 12 | 1.48% |
| hsa_circ_0128946 | 1617 | 12 | 1.48% |
| hsa_circ_0134474 | 1617 | 12 | 1.48% |
| hsa_circ_0123669 | 5931 | 44 | 1.48% |
| hsa_circ_0008816 | 1348 | 10 | 1.48% |
| hsa_circ_0014581 | 1348 | 10 | 1.48% |
| hsa_circ_0021172 | 1348 | 10 | 1.48% |

| | | | |
|---|---|---|---|
| hsa_circ_0022148 | 1348 | 10 | 1.48% |
| hsa_circ_0028838 | 1348 | 10 | 1.48% |
| hsa_circ_0043967 | 1348 | 10 | 1.48% |
| hsa_circ_0047930 | 1348 | 10 | 1.48% |
| hsa_circ_0056299 | 1348 | 10 | 1.48% |
| hsa_circ_0057215 | 1348 | 10 | 1.48% |
| hsa_circ_0079418 | 1348 | 10 | 1.48% |
| hsa_circ_0083229 | 1348 | 10 | 1.48% |
| hsa_circ_0090353 | 1348 | 10 | 1.48% |
| hsa_circ_0113988 | 1348 | 10 | 1.48% |
| hsa_circ_0118122 | 1348 | 10 | 1.48% |
| hsa_circ_0125765 | 1348 | 10 | 1.48% |
| hsa_circ_0132389 | 1348 | 10 | 1.48% |
| hsa_circ_0035614 | 1483 | 11 | 1.48% |
| hsa_circ_0087898 | 1483 | 11 | 1.48% |
| hsa_circ_0091239 | 1483 | 11 | 1.48% |
| hsa_circ_0129034 | 4854 | 36 | 1.48% |
| hsa_circ_0035868 | 1618 | 12 | 1.48% |
| hsa_circ_0095292 | 1618 | 12 | 1.48% |
| hsa_circ_0025247 | 1888 | 14 | 1.48% |
| hsa_circ_0090419 | 1888 | 14 | 1.48% |
| hsa_circ_0014784 | 1349 | 10 | 1.48% |
| hsa_circ_0026303 | 1349 | 10 | 1.48% |
| hsa_circ_0037184 | 1349 | 10 | 1.48% |
| hsa_circ_0041101 | 1349 | 10 | 1.48% |
| hsa_circ_0045412 | 1349 | 10 | 1.48% |
| hsa_circ_0064427 | 1349 | 10 | 1.48% |
| hsa_circ_0101060 | 1349 | 10 | 1.48% |
| hsa_circ_0118657 | 1349 | 10 | 1.48% |
| hsa_circ_0123644 | 1349 | 10 | 1.48% |
| hsa_circ_0005959 | 1484 | 11 | 1.48% |
| hsa_circ_0009605 | 1484 | 11 | 1.48% |
| hsa_circ_0113157 | 1484 | 11 | 1.48% |
| hsa_circ_0115247 | 1484 | 11 | 1.48% |
| hsa_circ_0072630 | 1754 | 13 | 1.48% |
| hsa_circ_0084866 | 1754 | 13 | 1.48% |
| hsa_circ_0049689 | 2564 | 19 | 1.48% |
| hsa_circ_0037415 | 1485 | 11 | 1.48% |
| hsa_circ_0043999 | 1485 | 11 | 1.48% |
| hsa_circ_0058286 | 1485 | 11 | 1.48% |
| hsa_circ_0071513 | 1485 | 11 | 1.48% |
| hsa_circ_0083612 | 1485 | 11 | 1.48% |
| hsa_circ_0083880 | 1485 | 11 | 1.48% |
| hsa_circ_0089462 | 1485 | 11 | 1.48% |
| hsa_circ_0124580 | 1485 | 11 | 1.48% |
| hsa_circ_0031557 | 1350 | 10 | 1.48% |
| hsa_circ_0055040 | 1350 | 10 | 1.48% |
| hsa_circ_0055665 | 1350 | 10 | 1.48% |
| hsa_circ_0076934 | 1350 | 10 | 1.48% |
| hsa_circ_0077488 | 1350 | 10 | 1.48% |

| | | | |
|---|---|---|---|
| hsa_circ_0083642 | 1350 | 10 | 1.48% |
| hsa_circ_0086589 | 1350 | 10 | 1.48% |
| hsa_circ_0092782 | 1350 | 10 | 1.48% |
| hsa_circ_0134220 | 1350 | 10 | 1.48% |
| hsa_circ_0140759 | 1350 | 10 | 1.48% |
| hsa_circ_0121676 | 14585 | 108 | 1.48% |
| hsa_circ_0048273 | 1891 | 14 | 1.48% |
| hsa_circ_0091274 | 1756 | 13 | 1.48% |
| hsa_circ_0036544 | 1486 | 11 | 1.48% |
| hsa_circ_0050964 | 1486 | 11 | 1.48% |
| hsa_circ_0087408 | 1486 | 11 | 1.48% |
| hsa_circ_0127064 | 1486 | 11 | 1.48% |
| hsa_circ_0129366 | 1486 | 11 | 1.48% |
| hsa_circ_0129430 | 1486 | 11 | 1.48% |
| hsa_circ_0132212 | 1486 | 11 | 1.48% |
| hsa_circ_0000363 | 1351 | 10 | 1.48% |
| hsa_circ_0016666 | 1351 | 10 | 1.48% |
| hsa_circ_0052774 | 1351 | 10 | 1.48% |
| hsa_circ_0064677 | 1351 | 10 | 1.48% |
| hsa_circ_0086332 | 1351 | 10 | 1.48% |
| hsa_circ_0090497 | 1351 | 10 | 1.48% |
| hsa_circ_0115109 | 1351 | 10 | 1.48% |
| hsa_circ_0120324 | 1351 | 10 | 1.48% |
| hsa_circ_0035107 | 1487 | 11 | 1.48% |
| hsa_circ_0046488 | 1487 | 11 | 1.48% |
| hsa_circ_0048597 | 1487 | 11 | 1.48% |
| hsa_circ_0051613 | 1487 | 11 | 1.48% |
| hsa_circ_0086553 | 1487 | 11 | 1.48% |
| hsa_circ_0127820 | 1487 | 11 | 1.48% |
| hsa_circ_0004474 | 1352 | 10 | 1.48% |
| hsa_circ_0024691 | 1352 | 10 | 1.48% |
| hsa_circ_0031204 | 1352 | 10 | 1.48% |
| hsa_circ_0042213 | 1352 | 10 | 1.48% |
| hsa_circ_0072688 | 1352 | 10 | 1.48% |
| hsa_circ_0087240 | 1352 | 10 | 1.48% |
| hsa_circ_0130820 | 1352 | 10 | 1.48% |
| hsa_circ_0048072 | 1893 | 14 | 1.48% |
| hsa_circ_0079512 | 1623 | 12 | 1.48% |
| hsa_circ_0124161 | 3517 | 26 | 1.48% |
| hsa_circ_0016941 | 1488 | 11 | 1.48% |
| hsa_circ_0044068 | 1488 | 11 | 1.48% |
| hsa_circ_0046065 | 1488 | 11 | 1.48% |
| hsa_circ_0051822 | 1488 | 11 | 1.48% |
| hsa_circ_0063400 | 1488 | 11 | 1.48% |
| hsa_circ_0070018 | 1488 | 11 | 1.48% |
| hsa_circ_0089376 | 1488 | 11 | 1.48% |
| hsa_circ_0106706 | 1488 | 11 | 1.48% |
| hsa_circ_0095423 | 1894 | 14 | 1.48% |
| hsa_circ_0001478 | 1353 | 10 | 1.48% |
| hsa_circ_0009715 | 1353 | 10 | 1.48% |

| | | | |
|---|---|---|---|
| hsa_circ_0018605 | 1353 | 10 | 1.48% |
| hsa_circ_0030373 | 1353 | 10 | 1.48% |
| hsa_circ_0037806 | 1353 | 10 | 1.48% |
| hsa_circ_0044771 | 1353 | 10 | 1.48% |
| hsa_circ_0053935 | 1353 | 10 | 1.48% |
| hsa_circ_0058050 | 1353 | 10 | 1.48% |
| hsa_circ_0095242 | 1353 | 10 | 1.48% |
| hsa_circ_0097783 | 1353 | 10 | 1.48% |
| hsa_circ_0103580 | 1353 | 10 | 1.48% |
| hsa_circ_0119853 | 1353 | 10 | 1.48% |
| hsa_circ_0125725 | 1353 | 10 | 1.48% |
| hsa_circ_0125761 | 1353 | 10 | 1.48% |
| hsa_circ_0128999 | 1353 | 10 | 1.48% |
| hsa_circ_0130024 | 1353 | 10 | 1.48% |
| hsa_circ_0134866 | 1353 | 10 | 1.48% |
| hsa_circ_0138976 | 1353 | 10 | 1.48% |
| hsa_circ_0100669 | 1759 | 13 | 1.48% |
| hsa_circ_0080466 | 3924 | 29 | 1.48% |
| hsa_circ_0079509 | 1624 | 12 | 1.48% |
| hsa_circ_0120322 | 1624 | 12 | 1.48% |
| hsa_circ_0061070 | 1489 | 11 | 1.48% |
| hsa_circ_0065641 | 1489 | 11 | 1.48% |
| hsa_circ_0084732 | 1489 | 11 | 1.48% |
| hsa_circ_0094604 | 1489 | 11 | 1.48% |
| hsa_circ_0100081 | 1489 | 11 | 1.48% |
| hsa_circ_0066762 | 1760 | 13 | 1.48% |
| hsa_circ_0114448 | 1760 | 13 | 1.48% |
| hsa_circ_0018389 | 2031 | 15 | 1.48% |
| hsa_circ_0015890 | 1354 | 10 | 1.48% |
| hsa_circ_0020365 | 1354 | 10 | 1.48% |
| hsa_circ_0025568 | 1354 | 10 | 1.48% |
| hsa_circ_0039889 | 1354 | 10 | 1.48% |
| hsa_circ_0048245 | 1354 | 10 | 1.48% |
| hsa_circ_0071929 | 1354 | 10 | 1.48% |
| hsa_circ_0072235 | 1354 | 10 | 1.48% |
| hsa_circ_0102858 | 1354 | 10 | 1.48% |
| hsa_circ_0109229 | 1354 | 10 | 1.48% |
| hsa_circ_0110103 | 1354 | 10 | 1.48% |
| hsa_circ_0110529 | 1354 | 10 | 1.48% |
| hsa_circ_0118914 | 1354 | 10 | 1.48% |
| hsa_circ_0131417 | 1354 | 10 | 1.48% |
| hsa_circ_0131572 | 1354 | 10 | 1.48% |
| hsa_circ_0006584 | 1490 | 11 | 1.48% |
| hsa_circ_0043971 | 1490 | 11 | 1.48% |
| hsa_circ_0059746 | 1490 | 11 | 1.48% |
| hsa_circ_0081999 | 1490 | 11 | 1.48% |
| hsa_circ_0092910 | 1490 | 11 | 1.48% |
| hsa_circ_0100474 | 1490 | 11 | 1.48% |
| hsa_circ_0025861 | 2439 | 18 | 1.48% |
| hsa_circ_0037195 | 1897 | 14 | 1.48% |

| | | | |
|---|---|---|---|
| hsa_circ_0041898 | 1626 | 12 | 1.48% |
| hsa_circ_0045329 | 1626 | 12 | 1.48% |
| hsa_circ_0078086 | 1626 | 12 | 1.48% |
| hsa_circ_0012514 | 1355 | 10 | 1.48% |
| hsa_circ_0036994 | 1355 | 10 | 1.48% |
| hsa_circ_0050902 | 1355 | 10 | 1.48% |
| hsa_circ_0055792 | 1355 | 10 | 1.48% |
| hsa_circ_0063914 | 1355 | 10 | 1.48% |
| hsa_circ_0070380 | 1355 | 10 | 1.48% |
| hsa_circ_0072555 | 1355 | 10 | 1.48% |
| hsa_circ_0088929 | 1355 | 10 | 1.48% |
| hsa_circ_0093050 | 1355 | 10 | 1.48% |
| hsa_circ_0117429 | 1355 | 10 | 1.48% |
| hsa_circ_0121348 | 1355 | 10 | 1.48% |
| hsa_circ_0121770 | 1355 | 10 | 1.48% |
| hsa_circ_0012431 | 1491 | 11 | 1.48% |
| hsa_circ_0014493 | 1491 | 11 | 1.48% |
| hsa_circ_0033359 | 1491 | 11 | 1.48% |
| hsa_circ_0034059 | 1491 | 11 | 1.48% |
| hsa_circ_0050597 | 1491 | 11 | 1.48% |
| hsa_circ_0070828 | 1491 | 11 | 1.48% |
| hsa_circ_0031541 | 1627 | 12 | 1.48% |
| hsa_circ_0068538 | 1627 | 12 | 1.48% |
| hsa_circ_0077694 | 1627 | 12 | 1.48% |
| hsa_circ_0003475 | 1356 | 10 | 1.47% |
| hsa_circ_0018295 | 1356 | 10 | 1.47% |
| hsa_circ_0032671 | 1356 | 10 | 1.47% |
| hsa_circ_0033238 | 1356 | 10 | 1.47% |
| hsa_circ_0039065 | 1356 | 10 | 1.47% |
| hsa_circ_0051110 | 1356 | 10 | 1.47% |
| hsa_circ_0067183 | 1356 | 10 | 1.47% |
| hsa_circ_0084709 | 1356 | 10 | 1.47% |
| hsa_circ_0088813 | 1356 | 10 | 1.47% |
| hsa_circ_0110554 | 1356 | 10 | 1.47% |
| hsa_circ_0126246 | 1356 | 10 | 1.47% |
| hsa_circ_0126669 | 1356 | 10 | 1.47% |
| hsa_circ_0052952 | 1763 | 13 | 1.47% |
| hsa_circ_0037097 | 2170 | 16 | 1.47% |
| hsa_circ_0004171 | 1492 | 11 | 1.47% |
| hsa_circ_0082587 | 1492 | 11 | 1.47% |
| hsa_circ_0061210 | 1628 | 12 | 1.47% |
| hsa_circ_0063616 | 1628 | 12 | 1.47% |
| hsa_circ_0076707 | 1628 | 12 | 1.47% |
| hsa_circ_0086146 | 1628 | 12 | 1.47% |
| hsa_circ_0039168 | 1357 | 10 | 1.47% |
| hsa_circ_0051115 | 1357 | 10 | 1.47% |
| hsa_circ_0067611 | 1357 | 10 | 1.47% |
| hsa_circ_0072696 | 1357 | 10 | 1.47% |
| hsa_circ_0082127 | 1357 | 10 | 1.47% |
| hsa_circ_0013643 | 1900 | 14 | 1.47% |

| | | | |
|---|---|---|---|
| hsa_circ_0021298 | 1493 | 11 | 1.47% |
| hsa_circ_0027641 | 1493 | 11 | 1.47% |
| hsa_circ_0063257 | 1493 | 11 | 1.47% |
| hsa_circ_0079209 | 1493 | 11 | 1.47% |
| hsa_circ_0091655 | 1493 | 11 | 1.47% |
| hsa_circ_0112484 | 1493 | 11 | 1.47% |
| hsa_circ_0126546 | 1493 | 11 | 1.47% |
| hsa_circ_0128905 | 1493 | 11 | 1.47% |
| hsa_circ_0135229 | 1493 | 11 | 1.47% |
| hsa_circ_0070537 | 1629 | 12 | 1.47% |
| hsa_circ_0100620 | 1629 | 12 | 1.47% |
| hsa_circ_0119299 | 1629 | 12 | 1.47% |
| hsa_circ_0120588 | 1629 | 12 | 1.47% |
| hsa_circ_0016246 | 1358 | 10 | 1.47% |
| hsa_circ_0019487 | 1358 | 10 | 1.47% |
| hsa_circ_0019993 | 1358 | 10 | 1.47% |
| hsa_circ_0028220 | 1358 | 10 | 1.47% |
| hsa_circ_0046966 | 1358 | 10 | 1.47% |
| hsa_circ_0055465 | 1358 | 10 | 1.47% |
| hsa_circ_0062582 | 1358 | 10 | 1.47% |
| hsa_circ_0070886 | 1358 | 10 | 1.47% |
| hsa_circ_0091190 | 1358 | 10 | 1.47% |
| hsa_circ_0095730 | 1358 | 10 | 1.47% |
| hsa_circ_0116495 | 1358 | 10 | 1.47% |
| hsa_circ_0129635 | 1358 | 10 | 1.47% |
| hsa_circ_0019360 | 1494 | 11 | 1.47% |
| hsa_circ_0044405 | 1494 | 11 | 1.47% |
| hsa_circ_0049830 | 1494 | 11 | 1.47% |
| hsa_circ_0089671 | 1494 | 11 | 1.47% |
| hsa_circ_0119830 | 1494 | 11 | 1.47% |
| hsa_circ_0124578 | 1494 | 11 | 1.47% |
| hsa_circ_0103542 | 1630 | 12 | 1.47% |
| hsa_circ_0102791 | 1766 | 13 | 1.47% |
| hsa_circ_0137984 | 2038 | 15 | 1.47% |
| hsa_circ_0000177 | 1359 | 10 | 1.47% |
| hsa_circ_0043160 | 1359 | 10 | 1.47% |
| hsa_circ_0049486 | 1359 | 10 | 1.47% |
| hsa_circ_0050266 | 1359 | 10 | 1.47% |
| hsa_circ_0053913 | 1359 | 10 | 1.47% |
| hsa_circ_0060520 | 1359 | 10 | 1.47% |
| hsa_circ_0065555 | 1359 | 10 | 1.47% |
| hsa_circ_0066812 | 1359 | 10 | 1.47% |
| hsa_circ_0082426 | 1359 | 10 | 1.47% |
| hsa_circ_0085440 | 1359 | 10 | 1.47% |
| hsa_circ_0091964 | 1359 | 10 | 1.47% |
| hsa_circ_0098210 | 1359 | 10 | 1.47% |
| hsa_circ_0098781 | 1359 | 10 | 1.47% |
| hsa_circ_0102891 | 1359 | 10 | 1.47% |
| hsa_circ_0127975 | 1359 | 10 | 1.47% |
| hsa_circ_0129640 | 1359 | 10 | 1.47% |

| | | | |
|---|---|---|---|
| hsa_circ_0133838 | 1359 | 10 | 1.47% |
| hsa_circ_0133936 | 1359 | 10 | 1.47% |
| hsa_circ_0025209 | 1495 | 11 | 1.47% |
| hsa_circ_0025608 | 1495 | 11 | 1.47% |
| hsa_circ_0030872 | 1495 | 11 | 1.47% |
| hsa_circ_0038424 | 1495 | 11 | 1.47% |
| hsa_circ_0056413 | 1495 | 11 | 1.47% |
| hsa_circ_0093385 | 1495 | 11 | 1.47% |
| hsa_circ_0100480 | 1495 | 11 | 1.47% |
| hsa_circ_0140058 | 1495 | 11 | 1.47% |
| hsa_circ_0086624 | 6116 | 45 | 1.47% |
| hsa_circ_0054842 | 1631 | 12 | 1.47% |
| hsa_circ_0083177 | 1631 | 12 | 1.47% |
| hsa_circ_0102075 | 1631 | 12 | 1.47% |
| hsa_circ_0040555 | 1903 | 14 | 1.47% |
| hsa_circ_0103981 | 2991 | 22 | 1.47% |
| hsa_circ_0014946 | 1632 | 12 | 1.47% |
| hsa_circ_0031529 | 1632 | 12 | 1.47% |
| hsa_circ_0042057 | 1632 | 12 | 1.47% |
| hsa_circ_0122245 | 1632 | 12 | 1.47% |
| hsa_circ_0009922 | 1496 | 11 | 1.47% |
| hsa_circ_0028966 | 1496 | 11 | 1.47% |
| hsa_circ_0031759 | 1496 | 11 | 1.47% |
| hsa_circ_0034977 | 1496 | 11 | 1.47% |
| hsa_circ_0043507 | 1496 | 11 | 1.47% |
| hsa_circ_0076013 | 1496 | 11 | 1.47% |
| hsa_circ_0089856 | 1496 | 11 | 1.47% |
| hsa_circ_0104318 | 1496 | 11 | 1.47% |
| hsa_circ_0120109 | 1496 | 11 | 1.47% |
| hsa_circ_0002642 | 1360 | 10 | 1.47% |
| hsa_circ_0034380 | 1360 | 10 | 1.47% |
| hsa_circ_0048850 | 1360 | 10 | 1.47% |
| hsa_circ_0051229 | 1360 | 10 | 1.47% |
| hsa_circ_0089632 | 1360 | 10 | 1.47% |
| hsa_circ_0120724 | 1360 | 10 | 1.47% |
| hsa_circ_0024503 | 4897 | 36 | 1.47% |
| hsa_circ_0045129 | 1633 | 12 | 1.47% |
| hsa_circ_0119537 | 1633 | 12 | 1.47% |
| hsa_circ_0134518 | 7757 | 57 | 1.47% |
| hsa_circ_0031791 | 1497 | 11 | 1.47% |
| hsa_circ_0088994 | 1497 | 11 | 1.47% |
| hsa_circ_0006463 | 1361 | 10 | 1.47% |
| hsa_circ_0012310 | 1361 | 10 | 1.47% |
| hsa_circ_0017921 | 1361 | 10 | 1.47% |
| hsa_circ_0018170 | 1361 | 10 | 1.47% |
| hsa_circ_0047235 | 1361 | 10 | 1.47% |
| hsa_circ_0064888 | 1361 | 10 | 1.47% |
| hsa_circ_0075435 | 1361 | 10 | 1.47% |
| hsa_circ_0084962 | 1361 | 10 | 1.47% |
| hsa_circ_0096477 | 1361 | 10 | 1.47% |

| | | | |
|---|---|---|---|
| hsa_circ_0107125 | 1361 | 10 | 1.47% |
| hsa_circ_0110112 | 1361 | 10 | 1.47% |
| hsa_circ_0010399 | 4900 | 36 | 1.47% |
| hsa_circ_0049716 | 1906 | 14 | 1.47% |
| hsa_circ_0062018 | 1770 | 13 | 1.47% |
| hsa_circ_0121425 | 1770 | 13 | 1.47% |
| hsa_circ_0021247 | 1634 | 12 | 1.47% |
| hsa_circ_0045256 | 1634 | 12 | 1.47% |
| hsa_circ_0051723 | 1634 | 12 | 1.47% |
| hsa_circ_0070868 | 1634 | 12 | 1.47% |
| hsa_circ_0090302 | 1634 | 12 | 1.47% |
| hsa_circ_0036717 | 1498 | 11 | 1.47% |
| hsa_circ_0037410 | 1498 | 11 | 1.47% |
| hsa_circ_0093499 | 1498 | 11 | 1.47% |
| hsa_circ_0094592 | 1498 | 11 | 1.47% |
| hsa_circ_0095951 | 1498 | 11 | 1.47% |
| hsa_circ_0000857 | 1362 | 10 | 1.47% |
| hsa_circ_0017706 | 1362 | 10 | 1.47% |
| hsa_circ_0030536 | 1362 | 10 | 1.47% |
| hsa_circ_0037720 | 1362 | 10 | 1.47% |
| hsa_circ_0048534 | 1362 | 10 | 1.47% |
| hsa_circ_0050385 | 1362 | 10 | 1.47% |
| hsa_circ_0065458 | 1362 | 10 | 1.47% |
| hsa_circ_0080057 | 1362 | 10 | 1.47% |
| hsa_circ_0081659 | 1362 | 10 | 1.47% |
| hsa_circ_0083772 | 1362 | 10 | 1.47% |
| hsa_circ_0129600 | 1362 | 10 | 1.47% |
| hsa_circ_0129972 | 1362 | 10 | 1.47% |
| hsa_circ_0091310 | 1907 | 14 | 1.47% |
| hsa_circ_0134034 | 1771 | 13 | 1.47% |
| hsa_circ_0008826 | 1499 | 11 | 1.47% |
| hsa_circ_0038972 | 1499 | 11 | 1.47% |
| hsa_circ_0066283 | 1499 | 11 | 1.47% |
| hsa_circ_0117997 | 1499 | 11 | 1.47% |
| hsa_circ_0010078 | 1363 | 10 | 1.47% |
| hsa_circ_0017242 | 1363 | 10 | 1.47% |
| hsa_circ_0034837 | 1363 | 10 | 1.47% |
| hsa_circ_0036364 | 1363 | 10 | 1.47% |
| hsa_circ_0039665 | 1363 | 10 | 1.47% |
| hsa_circ_0039812 | 1363 | 10 | 1.47% |
| hsa_circ_0051386 | 1363 | 10 | 1.47% |
| hsa_circ_0051930 | 1363 | 10 | 1.47% |
| hsa_circ_0063848 | 1363 | 10 | 1.47% |
| hsa_circ_0071146 | 1363 | 10 | 1.47% |
| hsa_circ_0123037 | 1363 | 10 | 1.47% |
| hsa_circ_0129387 | 1363 | 10 | 1.47% |
| hsa_circ_0094829 | 4498 | 33 | 1.47% |
| hsa_circ_0093480 | 2045 | 15 | 1.47% |
| hsa_circ_0108337 | 1636 | 12 | 1.47% |
| hsa_circ_0088853 | 1500 | 11 | 1.47% |

| | | | |
|---|---|---|---|
| hsa_circ_0136705 | 1500 | 11 | 1.47% |
| hsa_circ_0001192 | 1364 | 10 | 1.47% |
| hsa_circ_0016071 | 1364 | 10 | 1.47% |
| hsa_circ_0020594 | 1364 | 10 | 1.47% |
| hsa_circ_0029415 | 1364 | 10 | 1.47% |
| hsa_circ_0032788 | 1364 | 10 | 1.47% |
| hsa_circ_0036766 | 1364 | 10 | 1.47% |
| hsa_circ_0040444 | 1364 | 10 | 1.47% |
| hsa_circ_0042455 | 1364 | 10 | 1.47% |
| hsa_circ_0070881 | 1364 | 10 | 1.47% |
| hsa_circ_0085755 | 1364 | 10 | 1.47% |
| hsa_circ_0095425 | 1364 | 10 | 1.47% |
| hsa_circ_0120723 | 1364 | 10 | 1.47% |
| hsa_circ_0138825 | 1364 | 10 | 1.47% |
| hsa_circ_0126304 | 3683 | 27 | 1.47% |
| hsa_circ_0053258 | 1637 | 12 | 1.47% |
| hsa_circ_0130741 | 1637 | 12 | 1.47% |
| hsa_circ_0037822 | 1501 | 11 | 1.47% |
| hsa_circ_0065539 | 1501 | 11 | 1.47% |
| hsa_circ_0069299 | 1501 | 11 | 1.47% |
| hsa_circ_0077967 | 1501 | 11 | 1.47% |
| hsa_circ_0094187 | 1501 | 11 | 1.47% |
| hsa_circ_0036207 | 1638 | 12 | 1.47% |
| hsa_circ_0138831 | 1638 | 12 | 1.47% |
| hsa_circ_0008355 | 1365 | 10 | 1.47% |
| hsa_circ_0014791 | 1365 | 10 | 1.47% |
| hsa_circ_0032616 | 1365 | 10 | 1.47% |
| hsa_circ_0034097 | 1365 | 10 | 1.47% |
| hsa_circ_0038994 | 1365 | 10 | 1.47% |
| hsa_circ_0043740 | 1365 | 10 | 1.47% |
| hsa_circ_0054834 | 1365 | 10 | 1.47% |
| hsa_circ_0074122 | 1365 | 10 | 1.47% |
| hsa_circ_0082257 | 1365 | 10 | 1.47% |
| hsa_circ_0099556 | 1365 | 10 | 1.47% |
| hsa_circ_0111811 | 1365 | 10 | 1.47% |
| hsa_circ_0123670 | 6008 | 44 | 1.46% |
| hsa_circ_0028995 | 1502 | 11 | 1.46% |
| hsa_circ_0106939 | 3551 | 26 | 1.46% |
| hsa_circ_0000756 | 1366 | 10 | 1.46% |
| hsa_circ_0016660 | 1366 | 10 | 1.46% |
| hsa_circ_0047428 | 1366 | 10 | 1.46% |
| hsa_circ_0060086 | 1366 | 10 | 1.46% |
| hsa_circ_0067954 | 1366 | 10 | 1.46% |
| hsa_circ_0072559 | 1366 | 10 | 1.46% |
| hsa_circ_0081296 | 1366 | 10 | 1.46% |
| hsa_circ_0105757 | 1366 | 10 | 1.46% |
| hsa_circ_0109418 | 1366 | 10 | 1.46% |
| hsa_circ_0122303 | 1366 | 10 | 1.46% |
| hsa_circ_0123734 | 1366 | 10 | 1.46% |
| hsa_circ_0042348 | 6011 | 44 | 1.46% |

| | | | |
|---|---|---|---|
| hsa_circ_0032076 | 2186 | 16 | 1.46% |
| hsa_circ_0036050 | 1503 | 11 | 1.46% |
| hsa_circ_0046029 | 1503 | 11 | 1.46% |
| hsa_circ_0071131 | 1503 | 11 | 1.46% |
| hsa_circ_0087714 | 1503 | 11 | 1.46% |
| hsa_circ_0114473 | 1503 | 11 | 1.46% |
| hsa_circ_0115176 | 1503 | 11 | 1.46% |
| hsa_circ_0040128 | 1640 | 12 | 1.46% |
| hsa_circ_0015269 | 1367 | 10 | 1.46% |
| hsa_circ_0019631 | 1367 | 10 | 1.46% |
| hsa_circ_0027939 | 1367 | 10 | 1.46% |
| hsa_circ_0044338 | 1367 | 10 | 1.46% |
| hsa_circ_0065491 | 1367 | 10 | 1.46% |
| hsa_circ_0065507 | 1367 | 10 | 1.46% |
| hsa_circ_0077901 | 1367 | 10 | 1.46% |
| hsa_circ_0092192 | 1367 | 10 | 1.46% |
| hsa_circ_0093584 | 1367 | 10 | 1.46% |
| hsa_circ_0118987 | 1367 | 10 | 1.46% |
| hsa_circ_0140067 | 1367 | 10 | 1.46% |
| hsa_circ_0116224 | 1914 | 14 | 1.46% |
| hsa_circ_0016509 | 1504 | 11 | 1.46% |
| hsa_circ_0021038 | 1504 | 11 | 1.46% |
| hsa_circ_0023766 | 1504 | 11 | 1.46% |
| hsa_circ_0038349 | 1504 | 11 | 1.46% |
| hsa_circ_0049198 | 1504 | 11 | 1.46% |
| hsa_circ_0070123 | 1504 | 11 | 1.46% |
| hsa_circ_0080447 | 1504 | 11 | 1.46% |
| hsa_circ_0089007 | 1504 | 11 | 1.46% |
| hsa_circ_0089588 | 1504 | 11 | 1.46% |
| hsa_circ_0119205 | 1504 | 11 | 1.46% |
| hsa_circ_0128550 | 1504 | 11 | 1.46% |
| hsa_circ_0004841 | 2735 | 20 | 1.46% |
| hsa_circ_0109307 | 2735 | 20 | 1.46% |
| hsa_circ_0013700 | 1641 | 12 | 1.46% |
| hsa_circ_0037118 | 1641 | 12 | 1.46% |
| hsa_circ_0053263 | 1641 | 12 | 1.46% |
| hsa_circ_0125909 | 1778 | 13 | 1.46% |
| hsa_circ_0003836 | 1368 | 10 | 1.46% |
| hsa_circ_0018690 | 1368 | 10 | 1.46% |
| hsa_circ_0036938 | 1368 | 10 | 1.46% |
| hsa_circ_0056967 | 1368 | 10 | 1.46% |
| hsa_circ_0069105 | 1368 | 10 | 1.46% |
| hsa_circ_0095090 | 1368 | 10 | 1.46% |
| hsa_circ_0135035 | 1368 | 10 | 1.46% |
| hsa_circ_0038168 | 1505 | 11 | 1.46% |
| hsa_circ_0038457 | 1505 | 11 | 1.46% |
| hsa_circ_0048323 | 1505 | 11 | 1.46% |
| hsa_circ_0058985 | 1505 | 11 | 1.46% |
| hsa_circ_0061505 | 1505 | 11 | 1.46% |
| hsa_circ_0068807 | 1505 | 11 | 1.46% |

| | | | |
|---|---|---|---|
| hsa_circ_0017865 | 1642 | 12 | 1.46% |
| hsa_circ_0033337 | 1642 | 12 | 1.46% |
| hsa_circ_0013751 | 1369 | 10 | 1.46% |
| hsa_circ_0018819 | 1369 | 10 | 1.46% |
| hsa_circ_0026509 | 1369 | 10 | 1.46% |
| hsa_circ_0034748 | 1369 | 10 | 1.46% |
| hsa_circ_0047084 | 1369 | 10 | 1.46% |
| hsa_circ_0048496 | 1369 | 10 | 1.46% |
| hsa_circ_0122246 | 1369 | 10 | 1.46% |
| hsa_circ_0124064 | 1369 | 10 | 1.46% |
| hsa_circ_0005683 | 1506 | 11 | 1.46% |
| hsa_circ_0009568 | 1506 | 11 | 1.46% |
| hsa_circ_0013099 | 1506 | 11 | 1.46% |
| hsa_circ_0071906 | 1506 | 11 | 1.46% |
| hsa_circ_0081058 | 1506 | 11 | 1.46% |
| hsa_circ_0086537 | 1506 | 11 | 1.46% |
| hsa_circ_0092457 | 1506 | 11 | 1.46% |
| hsa_circ_0108865 | 1506 | 11 | 1.46% |
| hsa_circ_0127126 | 1506 | 11 | 1.46% |
| hsa_circ_0127229 | 1506 | 11 | 1.46% |
| hsa_circ_0016056 | 1643 | 12 | 1.46% |
| hsa_circ_0033529 | 1643 | 12 | 1.46% |
| hsa_circ_0046284 | 1643 | 12 | 1.46% |
| hsa_circ_0094893 | 1643 | 12 | 1.46% |
| hsa_circ_0113465 | 1643 | 12 | 1.46% |
| hsa_circ_0136875 | 1643 | 12 | 1.46% |
| hsa_circ_0048634 | 1780 | 13 | 1.46% |
| hsa_circ_0005642 | 2739 | 20 | 1.46% |
| hsa_circ_0025227 | 1781 | 13 | 1.46% |
| hsa_circ_0066115 | 1644 | 12 | 1.46% |
| hsa_circ_0120745 | 1507 | 11 | 1.46% |
| hsa_circ_0122667 | 1507 | 11 | 1.46% |
| hsa_circ_0026443 | 1370 | 10 | 1.46% |
| hsa_circ_0032008 | 1370 | 10 | 1.46% |
| hsa_circ_0039618 | 1370 | 10 | 1.46% |
| hsa_circ_0055788 | 1370 | 10 | 1.46% |
| hsa_circ_0062776 | 1370 | 10 | 1.46% |
| hsa_circ_0071851 | 1370 | 10 | 1.46% |
| hsa_circ_0073551 | 1370 | 10 | 1.46% |
| hsa_circ_0088536 | 1370 | 10 | 1.46% |
| hsa_circ_0092934 | 1370 | 10 | 1.46% |
| hsa_circ_0103957 | 1370 | 10 | 1.46% |
| hsa_circ_0124882 | 1370 | 10 | 1.46% |
| hsa_circ_0137688 | 3700 | 27 | 1.46% |
| hsa_circ_0048748 | 1645 | 12 | 1.46% |
| hsa_circ_0075710 | 1645 | 12 | 1.46% |
| hsa_circ_0103436 | 6032 | 44 | 1.46% |
| hsa_circ_0038838 | 1508 | 11 | 1.46% |
| hsa_circ_0054447 | 1508 | 11 | 1.46% |
| hsa_circ_0058955 | 1508 | 11 | 1.46% |

| | | | |
|---|---|---|---|
| hsa_circ_0085993 | 1508 | 11 | 1.46% |
| hsa_circ_0126759 | 1508 | 11 | 1.46% |
| hsa_circ_0033405 | 1371 | 10 | 1.46% |
| hsa_circ_0036933 | 1371 | 10 | 1.46% |
| hsa_circ_0051270 | 1371 | 10 | 1.46% |
| hsa_circ_0052657 | 1371 | 10 | 1.46% |
| hsa_circ_0060504 | 1371 | 10 | 1.46% |
| hsa_circ_0061715 | 1371 | 10 | 1.46% |
| hsa_circ_0066017 | 1371 | 10 | 1.46% |
| hsa_circ_0066158 | 1371 | 10 | 1.46% |
| hsa_circ_0070314 | 1371 | 10 | 1.46% |
| hsa_circ_0077212 | 1371 | 10 | 1.46% |
| hsa_circ_0079312 | 1371 | 10 | 1.46% |
| hsa_circ_0082716 | 1371 | 10 | 1.46% |
| hsa_circ_0082840 | 1371 | 10 | 1.46% |
| hsa_circ_0083044 | 1371 | 10 | 1.46% |
| hsa_circ_0057385 | 2331 | 17 | 1.46% |
| hsa_circ_0070167 | 1920 | 14 | 1.46% |
| hsa_circ_0012995 | 1783 | 13 | 1.46% |
| hsa_circ_0023530 | 1646 | 12 | 1.46% |
| hsa_circ_0035946 | 1509 | 11 | 1.46% |
| hsa_circ_0062121 | 1509 | 11 | 1.46% |
| hsa_circ_0099816 | 1509 | 11 | 1.46% |
| hsa_circ_0117568 | 1509 | 11 | 1.46% |
| hsa_circ_0139908 | 1509 | 11 | 1.46% |
| hsa_circ_0020683 | 1372 | 10 | 1.46% |
| hsa_circ_0020861 | 1372 | 10 | 1.46% |
| hsa_circ_0079310 | 1372 | 10 | 1.46% |
| hsa_circ_0085160 | 1372 | 10 | 1.46% |
| hsa_circ_0094078 | 1372 | 10 | 1.46% |
| hsa_circ_0103778 | 1372 | 10 | 1.46% |
| hsa_circ_0113164 | 1372 | 10 | 1.46% |
| hsa_circ_0138209 | 1372 | 10 | 1.46% |
| hsa_circ_0074899 | 1784 | 13 | 1.46% |
| hsa_circ_0087729 | 1647 | 12 | 1.46% |
| hsa_circ_0104185 | 1647 | 12 | 1.46% |
| hsa_circ_0137918 | 1647 | 12 | 1.46% |
| hsa_circ_0018372 | 1510 | 11 | 1.46% |
| hsa_circ_0019734 | 1510 | 11 | 1.46% |
| hsa_circ_0043392 | 1510 | 11 | 1.46% |
| hsa_circ_0051013 | 1510 | 11 | 1.46% |
| hsa_circ_0053018 | 1510 | 11 | 1.46% |
| hsa_circ_0085593 | 1510 | 11 | 1.46% |
| hsa_circ_0087803 | 1510 | 11 | 1.46% |
| hsa_circ_0090661 | 1510 | 11 | 1.46% |
| hsa_circ_0111755 | 1510 | 11 | 1.46% |
| hsa_circ_0118681 | 1510 | 11 | 1.46% |
| hsa_circ_0008265 | 1373 | 10 | 1.46% |
| hsa_circ_0011988 | 1373 | 10 | 1.46% |
| hsa_circ_0022551 | 1373 | 10 | 1.46% |

| | | | |
|---|---|---|---|
| hsa_circ_0023691 | 1373 | 10 | 1.46% |
| hsa_circ_0064623 | 1373 | 10 | 1.46% |
| hsa_circ_0105509 | 1373 | 10 | 1.46% |
| hsa_circ_0128284 | 1785 | 13 | 1.46% |
| hsa_circ_0010447 | 1648 | 12 | 1.46% |
| hsa_circ_0016995 | 1648 | 12 | 1.46% |
| hsa_circ_0058790 | 1648 | 12 | 1.46% |
| hsa_circ_0062840 | 1648 | 12 | 1.46% |
| hsa_circ_0007634 | 1511 | 11 | 1.46% |
| hsa_circ_0028527 | 1511 | 11 | 1.46% |
| hsa_circ_0041026 | 1511 | 11 | 1.46% |
| hsa_circ_0071823 | 1511 | 11 | 1.46% |
| hsa_circ_0085793 | 1511 | 11 | 1.46% |
| hsa_circ_0089506 | 1511 | 11 | 1.46% |
| hsa_circ_0010500 | 1374 | 10 | 1.46% |
| hsa_circ_0022138 | 1374 | 10 | 1.46% |
| hsa_circ_0026730 | 1374 | 10 | 1.46% |
| hsa_circ_0035959 | 1374 | 10 | 1.46% |
| hsa_circ_0043173 | 1374 | 10 | 1.46% |
| hsa_circ_0045175 | 1374 | 10 | 1.46% |
| hsa_circ_0060256 | 1374 | 10 | 1.46% |
| hsa_circ_0071879 | 1374 | 10 | 1.46% |
| hsa_circ_0072474 | 1374 | 10 | 1.46% |
| hsa_circ_0076034 | 1374 | 10 | 1.46% |
| hsa_circ_0086212 | 1374 | 10 | 1.46% |
| hsa_circ_0090964 | 1374 | 10 | 1.46% |
| hsa_circ_0092475 | 1374 | 10 | 1.46% |
| hsa_circ_0125567 | 1374 | 10 | 1.46% |
| hsa_circ_0131590 | 1374 | 10 | 1.46% |
| hsa_circ_0133937 | 1374 | 10 | 1.46% |
| hsa_circ_0029433 | 1649 | 12 | 1.46% |
| hsa_circ_0042564 | 1649 | 12 | 1.46% |
| hsa_circ_0059642 | 1649 | 12 | 1.46% |
| hsa_circ_0070867 | 1649 | 12 | 1.46% |
| hsa_circ_0119116 | 1649 | 12 | 1.46% |
| hsa_circ_0012202 | 1512 | 11 | 1.46% |
| hsa_circ_0028787 | 1512 | 11 | 1.46% |
| hsa_circ_0051304 | 1512 | 11 | 1.46% |
| hsa_circ_0054068 | 1512 | 11 | 1.46% |
| hsa_circ_0076564 | 1512 | 11 | 1.46% |
| hsa_circ_0079222 | 1512 | 11 | 1.46% |
| hsa_circ_0090546 | 1512 | 11 | 1.46% |
| hsa_circ_0100197 | 1512 | 11 | 1.46% |
| hsa_circ_0022544 | 2337 | 17 | 1.45% |
| hsa_circ_0019272 | 1650 | 12 | 1.45% |
| hsa_circ_0065444 | 1650 | 12 | 1.45% |
| hsa_circ_0119484 | 1650 | 12 | 1.45% |
| hsa_circ_0137724 | 1650 | 12 | 1.45% |
| hsa_circ_0040965 | 1375 | 10 | 1.45% |
| hsa_circ_0048416 | 1375 | 10 | 1.45% |

| | | | |
|---|---|---|---|
| hsa_circ_0076306 | 1375 | 10 | 1.45% |
| hsa_circ_0085091 | 1375 | 10 | 1.45% |
| hsa_circ_0086986 | 1375 | 10 | 1.45% |
| hsa_circ_0087261 | 1375 | 10 | 1.45% |
| hsa_circ_0095895 | 1375 | 10 | 1.45% |
| hsa_circ_0132219 | 1375 | 10 | 1.45% |
| hsa_circ_0014763 | 1788 | 13 | 1.45% |
| hsa_circ_0072608 | 1788 | 13 | 1.45% |
| hsa_circ_0021400 | 1513 | 11 | 1.45% |
| hsa_circ_0035241 | 1513 | 11 | 1.45% |
| hsa_circ_0046788 | 1513 | 11 | 1.45% |
| hsa_circ_0090798 | 1513 | 11 | 1.45% |
| hsa_circ_0097687 | 1513 | 11 | 1.45% |
| hsa_circ_0105556 | 1513 | 11 | 1.45% |
| hsa_circ_0114042 | 1513 | 11 | 1.45% |
| hsa_circ_0119097 | 1513 | 11 | 1.45% |
| hsa_circ_0126912 | 1513 | 11 | 1.45% |
| hsa_circ_0131228 | 1513 | 11 | 1.45% |
| hsa_circ_0113031 | 4265 | 31 | 1.45% |
| hsa_circ_0039566 | 1651 | 12 | 1.45% |
| hsa_circ_0075336 | 1651 | 12 | 1.45% |
| hsa_circ_0009157 | 1376 | 10 | 1.45% |
| hsa_circ_0016195 | 1376 | 10 | 1.45% |
| hsa_circ_0020084 | 1376 | 10 | 1.45% |
| hsa_circ_0044495 | 1376 | 10 | 1.45% |
| hsa_circ_0044893 | 1376 | 10 | 1.45% |
| hsa_circ_0066343 | 1376 | 10 | 1.45% |
| hsa_circ_0068216 | 1376 | 10 | 1.45% |
| hsa_circ_0075651 | 1376 | 10 | 1.45% |
| hsa_circ_0076502 | 1376 | 10 | 1.45% |
| hsa_circ_0084540 | 1376 | 10 | 1.45% |
| hsa_circ_0089716 | 1376 | 10 | 1.45% |
| hsa_circ_0093071 | 1376 | 10 | 1.45% |
| hsa_circ_0119817 | 1376 | 10 | 1.45% |
| hsa_circ_0125630 | 1376 | 10 | 1.45% |
| hsa_circ_0031625 | 1789 | 13 | 1.45% |
| hsa_circ_0052625 | 1789 | 13 | 1.45% |
| hsa_circ_0024421 | 1514 | 11 | 1.45% |
| hsa_circ_0043344 | 1514 | 11 | 1.45% |
| hsa_circ_0065849 | 1514 | 11 | 1.45% |
| hsa_circ_0070383 | 1514 | 11 | 1.45% |
| hsa_circ_0077455 | 1514 | 11 | 1.45% |
| hsa_circ_0086403 | 1514 | 11 | 1.45% |
| hsa_circ_0081603 | 1790 | 13 | 1.45% |
| hsa_circ_0026931 | 1377 | 10 | 1.45% |
| hsa_circ_0027768 | 1377 | 10 | 1.45% |
| hsa_circ_0040935 | 1377 | 10 | 1.45% |
| hsa_circ_0046629 | 1377 | 10 | 1.45% |
| hsa_circ_0048618 | 1377 | 10 | 1.45% |
| hsa_circ_0065665 | 1377 | 10 | 1.45% |

| | | | |
|---|---|---|---|
| hsa_circ_0071582 | 1377 | 10 | 1.45% |
| hsa_circ_0073791 | 1377 | 10 | 1.45% |
| hsa_circ_0084424 | 1377 | 10 | 1.45% |
| hsa_circ_0094923 | 1377 | 10 | 1.45% |
| hsa_circ_0101093 | 1377 | 10 | 1.45% |
| hsa_circ_0115097 | 1377 | 10 | 1.45% |
| hsa_circ_0072092 | 1928 | 14 | 1.45% |
| hsa_circ_0004050 | 1515 | 11 | 1.45% |
| hsa_circ_0016724 | 1515 | 11 | 1.45% |
| hsa_circ_0031234 | 1515 | 11 | 1.45% |
| hsa_circ_0035465 | 1515 | 11 | 1.45% |
| hsa_circ_0041347 | 1515 | 11 | 1.45% |
| hsa_circ_0060402 | 1515 | 11 | 1.45% |
| hsa_circ_0091991 | 1515 | 11 | 1.45% |
| hsa_circ_0099358 | 1515 | 11 | 1.45% |
| hsa_circ_0131377 | 1515 | 11 | 1.45% |
| hsa_circ_0138839 | 1515 | 11 | 1.45% |
| hsa_circ_0140523 | 1515 | 11 | 1.45% |
| hsa_circ_0018388 | 2066 | 15 | 1.45% |
| hsa_circ_0011145 | 2204 | 16 | 1.45% |
| hsa_circ_0102458 | 2342 | 17 | 1.45% |
| hsa_circ_0066650 | 2480 | 18 | 1.45% |
| hsa_circ_0005455 | 1378 | 10 | 1.45% |
| hsa_circ_0036549 | 1378 | 10 | 1.45% |
| hsa_circ_0041981 | 1378 | 10 | 1.45% |
| hsa_circ_0046622 | 1378 | 10 | 1.45% |
| hsa_circ_0060956 | 1378 | 10 | 1.45% |
| hsa_circ_0062243 | 1378 | 10 | 1.45% |
| hsa_circ_0063225 | 1378 | 10 | 1.45% |
| hsa_circ_0068146 | 1378 | 10 | 1.45% |
| hsa_circ_0088229 | 1378 | 10 | 1.45% |
| hsa_circ_0092224 | 1378 | 10 | 1.45% |
| hsa_circ_0018311 | 1516 | 11 | 1.45% |
| hsa_circ_0035238 | 1516 | 11 | 1.45% |
| hsa_circ_0089377 | 1516 | 11 | 1.45% |
| hsa_circ_0015986 | 2068 | 15 | 1.45% |
| hsa_circ_0110722 | 3585 | 26 | 1.45% |
| hsa_circ_0011445 | 1379 | 10 | 1.45% |
| hsa_circ_0019395 | 1379 | 10 | 1.45% |
| hsa_circ_0020168 | 1379 | 10 | 1.45% |
| hsa_circ_0032091 | 1379 | 10 | 1.45% |
| hsa_circ_0038922 | 1379 | 10 | 1.45% |
| hsa_circ_0066322 | 1379 | 10 | 1.45% |
| hsa_circ_0072212 | 1379 | 10 | 1.45% |
| hsa_circ_0104159 | 1379 | 10 | 1.45% |
| hsa_circ_0111588 | 1379 | 10 | 1.45% |
| hsa_circ_0125141 | 1379 | 10 | 1.45% |
| hsa_circ_0008913 | 1517 | 11 | 1.45% |
| hsa_circ_0015344 | 1517 | 11 | 1.45% |
| hsa_circ_0063991 | 1517 | 11 | 1.45% |

| | | | |
|---|---|---|---|
| hsa_circ_0110820 | 1517 | 11 | 1.45% |
| hsa_circ_0138411 | 1517 | 11 | 1.45% |
| hsa_circ_0045821 | 2345 | 17 | 1.45% |
| hsa_circ_0083942 | 1656 | 12 | 1.45% |
| hsa_circ_0069445 | 1518 | 11 | 1.45% |
| hsa_circ_0074880 | 1518 | 11 | 1.45% |
| hsa_circ_0090614 | 1518 | 11 | 1.45% |
| hsa_circ_0123940 | 1518 | 11 | 1.45% |
| hsa_circ_0022438 | 1380 | 10 | 1.45% |
| hsa_circ_0031715 | 1380 | 10 | 1.45% |
| hsa_circ_0069504 | 1380 | 10 | 1.45% |
| hsa_circ_0089280 | 1380 | 10 | 1.45% |
| hsa_circ_0111197 | 1380 | 10 | 1.45% |
| hsa_circ_0138718 | 1380 | 10 | 1.45% |
| hsa_circ_0062464 | 27463 | 199 | 1.45% |
| hsa_circ_0115327 | 4142 | 30 | 1.45% |
| hsa_circ_0062342 | 1795 | 13 | 1.45% |
| hsa_circ_0077653 | 1795 | 13 | 1.45% |
| hsa_circ_0099411 | 1795 | 13 | 1.45% |
| hsa_circ_0049194 | 1657 | 12 | 1.45% |
| hsa_circ_0043506 | 1519 | 11 | 1.45% |
| hsa_circ_0074832 | 1519 | 11 | 1.45% |
| hsa_circ_0080931 | 1519 | 11 | 1.45% |
| hsa_circ_0091215 | 1519 | 11 | 1.45% |
| hsa_circ_0098410 | 1519 | 11 | 1.45% |
| hsa_circ_0134760 | 1519 | 11 | 1.45% |
| hsa_circ_0002316 | 1381 | 10 | 1.45% |
| hsa_circ_0012435 | 1381 | 10 | 1.45% |
| hsa_circ_0026453 | 1381 | 10 | 1.45% |
| hsa_circ_0033134 | 1381 | 10 | 1.45% |
| hsa_circ_0069510 | 1381 | 10 | 1.45% |
| hsa_circ_0082488 | 1381 | 10 | 1.45% |
| hsa_circ_0091552 | 1381 | 10 | 1.45% |
| hsa_circ_0096349 | 1381 | 10 | 1.45% |
| hsa_circ_0100599 | 1381 | 10 | 1.45% |
| hsa_circ_0113704 | 1381 | 10 | 1.45% |
| hsa_circ_0121564 | 1381 | 10 | 1.45% |
| hsa_circ_0137019 | 1381 | 10 | 1.45% |
| hsa_circ_0139822 | 1381 | 10 | 1.45% |
| hsa_circ_0027298 | 1658 | 12 | 1.45% |
| hsa_circ_0112253 | 1658 | 12 | 1.45% |
| hsa_circ_0019468 | 1520 | 11 | 1.45% |
| hsa_circ_0027275 | 1520 | 11 | 1.45% |
| hsa_circ_0045537 | 1520 | 11 | 1.45% |
| hsa_circ_0051362 | 1520 | 11 | 1.45% |
| hsa_circ_0052374 | 1520 | 11 | 1.45% |
| hsa_circ_0068370 | 1520 | 11 | 1.45% |
| hsa_circ_0074191 | 1520 | 11 | 1.45% |
| hsa_circ_0106590 | 1520 | 11 | 1.45% |
| hsa_circ_0002820 | 1382 | 10 | 1.45% |

| | | | |
|---|---|---|---|
| hsa_circ_0012101 | 1382 | 10 | 1.45% |
| hsa_circ_0017790 | 1382 | 10 | 1.45% |
| hsa_circ_0019415 | 1382 | 10 | 1.45% |
| hsa_circ_0022641 | 1382 | 10 | 1.45% |
| hsa_circ_0050518 | 1382 | 10 | 1.45% |
| hsa_circ_0062319 | 1382 | 10 | 1.45% |
| hsa_circ_0068725 | 1382 | 10 | 1.45% |
| hsa_circ_0076928 | 1382 | 10 | 1.45% |
| hsa_circ_0082848 | 1382 | 10 | 1.45% |
| hsa_circ_0092750 | 1382 | 10 | 1.45% |
| hsa_circ_0106344 | 1935 | 14 | 1.45% |
| hsa_circ_0021148 | 1797 | 13 | 1.45% |
| hsa_circ_0037480 | 1797 | 13 | 1.45% |
| hsa_circ_0053957 | 1659 | 12 | 1.45% |
| hsa_circ_0125358 | 3180 | 23 | 1.45% |
| hsa_circ_0079447 | 2074 | 15 | 1.45% |
| hsa_circ_0071029 | 1521 | 11 | 1.45% |
| hsa_circ_0002710 | 1383 | 10 | 1.45% |
| hsa_circ_0026213 | 1383 | 10 | 1.45% |
| hsa_circ_0046577 | 1383 | 10 | 1.45% |
| hsa_circ_0050528 | 1383 | 10 | 1.45% |
| hsa_circ_0081859 | 1383 | 10 | 1.45% |
| hsa_circ_0091784 | 1383 | 10 | 1.45% |
| hsa_circ_0116102 | 1383 | 10 | 1.45% |
| hsa_circ_0120077 | 1383 | 10 | 1.45% |
| hsa_circ_0013518 | 1660 | 12 | 1.45% |
| hsa_circ_0022866 | 1522 | 11 | 1.45% |
| hsa_circ_0029625 | 1522 | 11 | 1.45% |
| hsa_circ_0047070 | 1522 | 11 | 1.45% |
| hsa_circ_0065690 | 1522 | 11 | 1.45% |
| hsa_circ_0067458 | 1522 | 11 | 1.45% |
| hsa_circ_0069699 | 1522 | 11 | 1.45% |
| hsa_circ_0098020 | 1522 | 11 | 1.45% |
| hsa_circ_0114571 | 1522 | 11 | 1.45% |
| hsa_circ_0122298 | 1522 | 11 | 1.45% |
| hsa_circ_0017809 | 1799 | 13 | 1.45% |
| hsa_circ_0012866 | 1384 | 10 | 1.45% |
| hsa_circ_0025579 | 1384 | 10 | 1.45% |
| hsa_circ_0028485 | 1384 | 10 | 1.45% |
| hsa_circ_0029853 | 1384 | 10 | 1.45% |
| hsa_circ_0037309 | 1384 | 10 | 1.45% |
| hsa_circ_0047466 | 1384 | 10 | 1.45% |
| hsa_circ_0048244 | 1384 | 10 | 1.45% |
| hsa_circ_0050072 | 1384 | 10 | 1.45% |
| hsa_circ_0055845 | 1384 | 10 | 1.45% |
| hsa_circ_0071759 | 1384 | 10 | 1.45% |
| hsa_circ_0094613 | 1384 | 10 | 1.45% |
| hsa_circ_0099785 | 1384 | 10 | 1.45% |
| hsa_circ_0107225 | 1384 | 10 | 1.45% |
| hsa_circ_0119954 | 1384 | 10 | 1.45% |

| | | | |
|---|---|---|---|
| hsa_circ_0073864 | 1661 | 12 | 1.44% |
| hsa_circ_0037827 | 1523 | 11 | 1.44% |
| hsa_circ_0112375 | 1523 | 11 | 1.44% |
| hsa_circ_0055300 | 1662 | 12 | 1.44% |
| hsa_circ_0014376 | 1385 | 10 | 1.44% |
| hsa_circ_0021261 | 1385 | 10 | 1.44% |
| hsa_circ_0048982 | 1385 | 10 | 1.44% |
| hsa_circ_0060472 | 1385 | 10 | 1.44% |
| hsa_circ_0062619 | 1385 | 10 | 1.44% |
| hsa_circ_0090252 | 1385 | 10 | 1.44% |
| hsa_circ_0092530 | 1385 | 10 | 1.44% |
| hsa_circ_0092958 | 1385 | 10 | 1.44% |
| hsa_circ_0117720 | 1385 | 10 | 1.44% |
| hsa_circ_0050102 | 7064 | 51 | 1.44% |
| hsa_circ_0082821 | 2078 | 15 | 1.44% |
| hsa_circ_0010081 | 1524 | 11 | 1.44% |
| hsa_circ_0049158 | 1524 | 11 | 1.44% |
| hsa_circ_0084487 | 1524 | 11 | 1.44% |
| hsa_circ_0051715 | 1663 | 12 | 1.44% |
| hsa_circ_0015705 | 1386 | 10 | 1.44% |
| hsa_circ_0017130 | 1386 | 10 | 1.44% |
| hsa_circ_0019733 | 1386 | 10 | 1.44% |
| hsa_circ_0031697 | 1386 | 10 | 1.44% |
| hsa_circ_0035096 | 1386 | 10 | 1.44% |
| hsa_circ_0039385 | 1386 | 10 | 1.44% |
| hsa_circ_0065083 | 1386 | 10 | 1.44% |
| hsa_circ_0067740 | 1386 | 10 | 1.44% |
| hsa_circ_0072511 | 1386 | 10 | 1.44% |
| hsa_circ_0074834 | 1386 | 10 | 1.44% |
| hsa_circ_0080111 | 1386 | 10 | 1.44% |
| hsa_circ_0096655 | 1386 | 10 | 1.44% |
| hsa_circ_0100315 | 1386 | 10 | 1.44% |
| hsa_circ_0100774 | 1386 | 10 | 1.44% |
| hsa_circ_0101059 | 1386 | 10 | 1.44% |
| hsa_circ_0107229 | 1386 | 10 | 1.44% |
| hsa_circ_0110962 | 1386 | 10 | 1.44% |
| hsa_circ_0125313 | 1386 | 10 | 1.44% |
| hsa_circ_0017259 | 1802 | 13 | 1.44% |
| hsa_circ_0025228 | 1802 | 13 | 1.44% |
| hsa_circ_0034376 | 1525 | 11 | 1.44% |
| hsa_circ_0034548 | 1525 | 11 | 1.44% |
| hsa_circ_0041498 | 1525 | 11 | 1.44% |
| hsa_circ_0067385 | 1525 | 11 | 1.44% |
| hsa_circ_0074253 | 1525 | 11 | 1.44% |
| hsa_circ_0131523 | 1525 | 11 | 1.44% |
| hsa_circ_0052968 | 2496 | 18 | 1.44% |
| hsa_circ_0037625 | 1664 | 12 | 1.44% |
| hsa_circ_0091752 | 1803 | 13 | 1.44% |
| hsa_circ_0025013 | 1387 | 10 | 1.44% |
| hsa_circ_0044134 | 1387 | 10 | 1.44% |

| | | | |
|---|---|---|---|
| hsa_circ_0026753 | 1526 | 11 | 1.44% |
| hsa_circ_0046360 | 1526 | 11 | 1.44% |
| hsa_circ_0083433 | 1526 | 11 | 1.44% |
| hsa_circ_0102809 | 1526 | 11 | 1.44% |
| hsa_circ_0107805 | 3747 | 27 | 1.44% |
| hsa_circ_0016200 | 1388 | 10 | 1.44% |
| hsa_circ_0041270 | 1388 | 10 | 1.44% |
| hsa_circ_0059815 | 1388 | 10 | 1.44% |
| hsa_circ_0064527 | 1388 | 10 | 1.44% |
| hsa_circ_0089471 | 1388 | 10 | 1.44% |
| hsa_circ_0112325 | 1388 | 10 | 1.44% |
| hsa_circ_0139031 | 1388 | 10 | 1.44% |
| hsa_circ_0124157 | 3609 | 26 | 1.44% |
| hsa_circ_0001276 | 1527 | 11 | 1.44% |
| hsa_circ_0019922 | 1527 | 11 | 1.44% |
| hsa_circ_0020889 | 1527 | 11 | 1.44% |
| hsa_circ_0026125 | 1527 | 11 | 1.44% |
| hsa_circ_0045004 | 1527 | 11 | 1.44% |
| hsa_circ_0069090 | 1527 | 11 | 1.44% |
| hsa_circ_0039873 | 1666 | 12 | 1.44% |
| hsa_circ_0086141 | 1666 | 12 | 1.44% |
| hsa_circ_0018290 | 1805 | 13 | 1.44% |
| hsa_circ_0073494 | 1805 | 13 | 1.44% |
| hsa_circ_0071040 | 1944 | 14 | 1.44% |
| hsa_circ_0011336 | 1389 | 10 | 1.44% |
| hsa_circ_0046779 | 1389 | 10 | 1.44% |
| hsa_circ_0052044 | 1389 | 10 | 1.44% |
| hsa_circ_0059405 | 1389 | 10 | 1.44% |
| hsa_circ_0061257 | 1389 | 10 | 1.44% |
| hsa_circ_0112344 | 1389 | 10 | 1.44% |
| hsa_circ_0125547 | 1389 | 10 | 1.44% |
| hsa_circ_0022790 | 1528 | 11 | 1.44% |
| hsa_circ_0033373 | 1528 | 11 | 1.44% |
| hsa_circ_0042044 | 1528 | 11 | 1.44% |
| hsa_circ_0087260 | 1528 | 11 | 1.44% |
| hsa_circ_0136465 | 1528 | 11 | 1.44% |
| hsa_circ_0048557 | 1667 | 12 | 1.44% |
| hsa_circ_0059931 | 1667 | 12 | 1.44% |
| hsa_circ_0069954 | 1667 | 12 | 1.44% |
| hsa_circ_0029382 | 1806 | 13 | 1.44% |
| hsa_circ_0071528 | 1806 | 13 | 1.44% |
| hsa_circ_0083137 | 4029 | 29 | 1.44% |
| hsa_circ_0026001 | 2502 | 18 | 1.44% |
| hsa_circ_0054616 | 1668 | 12 | 1.44% |
| hsa_circ_0056035 | 1668 | 12 | 1.44% |
| hsa_circ_0119014 | 1668 | 12 | 1.44% |
| hsa_circ_0055555 | 1529 | 11 | 1.44% |
| hsa_circ_0061420 | 1529 | 11 | 1.44% |
| hsa_circ_0091030 | 1529 | 11 | 1.44% |
| hsa_circ_0004761 | 1390 | 10 | 1.44% |

| | | | |
|---|---|---|---|
| hsa_circ_0023570 | 1390 | 10 | 1.44% |
| hsa_circ_0030591 | 1390 | 10 | 1.44% |
| hsa_circ_0036769 | 1390 | 10 | 1.44% |
| hsa_circ_0063126 | 1390 | 10 | 1.44% |
| hsa_circ_0092968 | 1390 | 10 | 1.44% |
| hsa_circ_0093165 | 2920 | 21 | 1.44% |
| hsa_circ_0122859 | 1947 | 14 | 1.44% |
| hsa_circ_0048027 | 1808 | 13 | 1.44% |
| hsa_circ_0128995 | 1808 | 13 | 1.44% |
| hsa_circ_0000055 | 1669 | 12 | 1.44% |
| hsa_circ_0072733 | 1669 | 12 | 1.44% |
| hsa_circ_0085468 | 1669 | 12 | 1.44% |
| hsa_circ_0085841 | 1669 | 12 | 1.44% |
| hsa_circ_0107999 | 1669 | 12 | 1.44% |
| hsa_circ_0024484 | 1530 | 11 | 1.44% |
| hsa_circ_0049827 | 1530 | 11 | 1.44% |
| hsa_circ_0055819 | 1530 | 11 | 1.44% |
| hsa_circ_0065751 | 1530 | 11 | 1.44% |
| hsa_circ_0077844 | 1530 | 11 | 1.44% |
| hsa_circ_0094331 | 1530 | 11 | 1.44% |
| hsa_circ_0104757 | 1530 | 11 | 1.44% |
| hsa_circ_0122295 | 1530 | 11 | 1.44% |
| hsa_circ_0129372 | 1530 | 11 | 1.44% |
| hsa_circ_0013360 | 1391 | 10 | 1.44% |
| hsa_circ_0022945 | 1391 | 10 | 1.44% |
| hsa_circ_0024692 | 1391 | 10 | 1.44% |
| hsa_circ_0043076 | 1391 | 10 | 1.44% |
| hsa_circ_0053433 | 1391 | 10 | 1.44% |
| hsa_circ_0058554 | 1391 | 10 | 1.44% |
| hsa_circ_0062209 | 1391 | 10 | 1.44% |
| hsa_circ_0066965 | 1391 | 10 | 1.44% |
| hsa_circ_0072433 | 1391 | 10 | 1.44% |
| hsa_circ_0079831 | 1391 | 10 | 1.44% |
| hsa_circ_0104084 | 1391 | 10 | 1.44% |
| hsa_circ_0107191 | 1391 | 10 | 1.44% |
| hsa_circ_0112170 | 1391 | 10 | 1.44% |
| hsa_circ_0122471 | 1391 | 10 | 1.44% |
| hsa_circ_0130101 | 1391 | 10 | 1.44% |
| hsa_circ_0133614 | 1391 | 10 | 1.44% |
| hsa_circ_0044205 | 2922 | 21 | 1.44% |
| hsa_circ_0138391 | 1948 | 14 | 1.44% |
| hsa_circ_0098105 | 5705 | 41 | 1.44% |
| hsa_circ_0052796 | 1670 | 12 | 1.44% |
| hsa_circ_0101341 | 1670 | 12 | 1.44% |
| hsa_circ_0129038 | 1670 | 12 | 1.44% |
| hsa_circ_0006800 | 1531 | 11 | 1.44% |
| hsa_circ_0021416 | 1531 | 11 | 1.44% |
| hsa_circ_0037237 | 1531 | 11 | 1.44% |
| hsa_circ_0101603 | 1531 | 11 | 1.44% |
| hsa_circ_0115383 | 1531 | 11 | 1.44% |

| | | | |
|---|---|---|---|
| hsa_circ_0125208 | 1531 | 11 | 1.44% |
| hsa_circ_0010449 | 1392 | 10 | 1.44% |
| hsa_circ_0020746 | 1392 | 10 | 1.44% |
| hsa_circ_0023028 | 1392 | 10 | 1.44% |
| hsa_circ_0030315 | 1392 | 10 | 1.44% |
| hsa_circ_0043153 | 1392 | 10 | 1.44% |
| hsa_circ_0091377 | 1392 | 10 | 1.44% |
| hsa_circ_0079106 | 1810 | 13 | 1.44% |
| hsa_circ_0059047 | 1671 | 12 | 1.44% |
| hsa_circ_0111133 | 1671 | 12 | 1.44% |
| hsa_circ_0013901 | 1532 | 11 | 1.44% |
| hsa_circ_0020553 | 1532 | 11 | 1.44% |
| hsa_circ_0029571 | 1532 | 11 | 1.44% |
| hsa_circ_0042711 | 1532 | 11 | 1.44% |
| hsa_circ_0060435 | 1532 | 11 | 1.44% |
| hsa_circ_0070965 | 1532 | 11 | 1.44% |
| hsa_circ_0082912 | 1532 | 11 | 1.44% |
| hsa_circ_0089981 | 1532 | 11 | 1.44% |
| hsa_circ_0101188 | 1532 | 11 | 1.44% |
| hsa_circ_0121720 | 1532 | 11 | 1.44% |
| hsa_circ_0027271 | 1393 | 10 | 1.44% |
| hsa_circ_0054545 | 1393 | 10 | 1.44% |
| hsa_circ_0066018 | 1393 | 10 | 1.44% |
| hsa_circ_0069463 | 1393 | 10 | 1.44% |
| hsa_circ_0070019 | 1393 | 10 | 1.44% |
| hsa_circ_0071065 | 1393 | 10 | 1.44% |
| hsa_circ_0075238 | 1393 | 10 | 1.44% |
| hsa_circ_0136544 | 1393 | 10 | 1.44% |
| hsa_circ_0009488 | 4458 | 32 | 1.44% |
| hsa_circ_0032798 | 1672 | 12 | 1.44% |
| hsa_circ_0069130 | 1672 | 12 | 1.44% |
| hsa_circ_0071837 | 1672 | 12 | 1.44% |
| hsa_circ_0113396 | 1951 | 14 | 1.44% |
| hsa_circ_0011701 | 1533 | 11 | 1.44% |
| hsa_circ_0068308 | 1533 | 11 | 1.44% |
| hsa_circ_0072573 | 1533 | 11 | 1.44% |
| hsa_circ_0089737 | 1533 | 11 | 1.44% |
| hsa_circ_0100293 | 1533 | 11 | 1.44% |
| hsa_circ_0130376 | 1533 | 11 | 1.44% |
| hsa_circ_0006011 | 1394 | 10 | 1.43% |
| hsa_circ_0009881 | 1394 | 10 | 1.43% |
| hsa_circ_0016155 | 1394 | 10 | 1.43% |
| hsa_circ_0024317 | 1394 | 10 | 1.43% |
| hsa_circ_0038634 | 1394 | 10 | 1.43% |
| hsa_circ_0045677 | 1394 | 10 | 1.43% |
| hsa_circ_0064758 | 1394 | 10 | 1.43% |
| hsa_circ_0033013 | 1673 | 12 | 1.43% |
| hsa_circ_0106634 | 1673 | 12 | 1.43% |
| hsa_circ_0020651 | 1534 | 11 | 1.43% |
| hsa_circ_0098653 | 1534 | 11 | 1.43% |

| | | | |
|---|---|---|---|
| hsa_circ_0104208 | 1534 | 11 | 1.43% |
| hsa_circ_0139903 | 1534 | 11 | 1.43% |
| hsa_circ_0030848 | 1813 | 13 | 1.43% |
| hsa_circ_0055476 | 1674 | 12 | 1.43% |
| hsa_circ_0008701 | 1395 | 10 | 1.43% |
| hsa_circ_0020449 | 1395 | 10 | 1.43% |
| hsa_circ_0029963 | 1395 | 10 | 1.43% |
| hsa_circ_0034860 | 1395 | 10 | 1.43% |
| hsa_circ_0035608 | 1395 | 10 | 1.43% |
| hsa_circ_0048817 | 1395 | 10 | 1.43% |
| hsa_circ_0049350 | 1395 | 10 | 1.43% |
| hsa_circ_0051349 | 1395 | 10 | 1.43% |
| hsa_circ_0058404 | 1395 | 10 | 1.43% |
| hsa_circ_0059831 | 1395 | 10 | 1.43% |
| hsa_circ_0073435 | 1395 | 10 | 1.43% |
| hsa_circ_0080885 | 1395 | 10 | 1.43% |
| hsa_circ_0097780 | 1395 | 10 | 1.43% |
| hsa_circ_0119310 | 1395 | 10 | 1.43% |
| hsa_circ_0133380 | 1395 | 10 | 1.43% |
| hsa_circ_0058178 | 1814 | 13 | 1.43% |
| hsa_circ_0037865 | 1535 | 11 | 1.43% |
| hsa_circ_0044257 | 1535 | 11 | 1.43% |
| hsa_circ_0046185 | 1535 | 11 | 1.43% |
| hsa_circ_0057068 | 1535 | 11 | 1.43% |
| hsa_circ_0072257 | 1535 | 11 | 1.43% |
| hsa_circ_0076103 | 1535 | 11 | 1.43% |
| hsa_circ_0126753 | 1535 | 11 | 1.43% |
| hsa_circ_0131535 | 1535 | 11 | 1.43% |
| hsa_circ_0079281 | 1675 | 12 | 1.43% |
| hsa_circ_0009913 | 1396 | 10 | 1.43% |
| hsa_circ_0011063 | 1396 | 10 | 1.43% |
| hsa_circ_0017233 | 1396 | 10 | 1.43% |
| hsa_circ_0028901 | 1396 | 10 | 1.43% |
| hsa_circ_0031205 | 1396 | 10 | 1.43% |
| hsa_circ_0080344 | 1396 | 10 | 1.43% |
| hsa_circ_0104976 | 1396 | 10 | 1.43% |
| hsa_circ_0122503 | 1396 | 10 | 1.43% |
| hsa_circ_0135414 | 1396 | 10 | 1.43% |
| hsa_circ_0019912 | 1536 | 11 | 1.43% |
| hsa_circ_0056933 | 1536 | 11 | 1.43% |
| hsa_circ_0092010 | 1536 | 11 | 1.43% |
| hsa_circ_0102719 | 1536 | 11 | 1.43% |
| hsa_circ_0049003 | 1676 | 12 | 1.43% |
| hsa_circ_0087013 | 1676 | 12 | 1.43% |
| hsa_circ_0091488 | 1676 | 12 | 1.43% |
| hsa_circ_0045533 | 1816 | 13 | 1.43% |
| hsa_circ_0011198 | 1397 | 10 | 1.43% |
| hsa_circ_0013553 | 1397 | 10 | 1.43% |
| hsa_circ_0025234 | 1397 | 10 | 1.43% |
| hsa_circ_0028529 | 1397 | 10 | 1.43% |

| | | | |
|---|---|---|---|
| hsa_circ_0033168 | 1397 | 10 | 1.43% |
| hsa_circ_0042391 | 1397 | 10 | 1.43% |
| hsa_circ_0058611 | 1397 | 10 | 1.43% |
| hsa_circ_0079373 | 1397 | 10 | 1.43% |
| hsa_circ_0089993 | 1397 | 10 | 1.43% |
| hsa_circ_0100115 | 1397 | 10 | 1.43% |
| hsa_circ_0107218 | 1397 | 10 | 1.43% |
| hsa_circ_0135292 | 1397 | 10 | 1.43% |
| hsa_circ_0137391 | 1397 | 10 | 1.43% |
| hsa_circ_0006545 | 2934 | 21 | 1.43% |
| hsa_circ_0103183 | 2934 | 21 | 1.43% |
| hsa_circ_0013888 | 1537 | 11 | 1.43% |
| hsa_circ_0017114 | 1537 | 11 | 1.43% |
| hsa_circ_0026877 | 1537 | 11 | 1.43% |
| hsa_circ_0050561 | 1537 | 11 | 1.43% |
| hsa_circ_0066785 | 1537 | 11 | 1.43% |
| hsa_circ_0068051 | 1537 | 11 | 1.43% |
| hsa_circ_0083339 | 1537 | 11 | 1.43% |
| hsa_circ_0101594 | 1537 | 11 | 1.43% |
| hsa_circ_0031509 | 1677 | 12 | 1.43% |
| hsa_circ_0048890 | 1677 | 12 | 1.43% |
| hsa_circ_0132847 | 1817 | 13 | 1.43% |
| hsa_circ_0008217 | 1398 | 10 | 1.43% |
| hsa_circ_0009731 | 1398 | 10 | 1.43% |
| hsa_circ_0011716 | 1398 | 10 | 1.43% |
| hsa_circ_0017010 | 1398 | 10 | 1.43% |
| hsa_circ_0039336 | 1398 | 10 | 1.43% |
| hsa_circ_0042797 | 1398 | 10 | 1.43% |
| hsa_circ_0049837 | 1398 | 10 | 1.43% |
| hsa_circ_0054301 | 1398 | 10 | 1.43% |
| hsa_circ_0078118 | 1398 | 10 | 1.43% |
| hsa_circ_0082469 | 1398 | 10 | 1.43% |
| hsa_circ_0083833 | 1398 | 10 | 1.43% |
| hsa_circ_0113394 | 1398 | 10 | 1.43% |
| hsa_circ_0012818 | 1538 | 11 | 1.43% |
| hsa_circ_0031135 | 1538 | 11 | 1.43% |
| hsa_circ_0049423 | 1538 | 11 | 1.43% |
| hsa_circ_0051391 | 1538 | 11 | 1.43% |
| hsa_circ_0072445 | 1538 | 11 | 1.43% |
| hsa_circ_0077975 | 1538 | 11 | 1.43% |
| hsa_circ_0089707 | 1538 | 11 | 1.43% |
| hsa_circ_0065499 | 1678 | 12 | 1.43% |
| hsa_circ_0089305 | 1678 | 12 | 1.43% |
| hsa_circ_0047226 | 1399 | 10 | 1.43% |
| hsa_circ_0075180 | 1399 | 10 | 1.43% |
| hsa_circ_0083747 | 1399 | 10 | 1.43% |
| hsa_circ_0085123 | 1399 | 10 | 1.43% |
| hsa_circ_0090347 | 1399 | 10 | 1.43% |
| hsa_circ_0122453 | 1399 | 10 | 1.43% |
| hsa_circ_0123560 | 1399 | 10 | 1.43% |

| | | | |
|---|---|---|---|
| hsa_circ_0139307 | 1399 | 10 | 1.43% |
| hsa_circ_0017043 | 1539 | 11 | 1.43% |
| hsa_circ_0020182 | 1539 | 11 | 1.43% |
| hsa_circ_0025597 | 1539 | 11 | 1.43% |
| hsa_circ_0043294 | 1539 | 11 | 1.43% |
| hsa_circ_0094400 | 1539 | 11 | 1.43% |
| hsa_circ_0139868 | 1539 | 11 | 1.43% |
| hsa_circ_0061862 | 1679 | 12 | 1.43% |
| hsa_circ_0115368 | 1679 | 12 | 1.43% |
| hsa_circ_0105265 | 1819 | 13 | 1.43% |
| hsa_circ_0113893 | 1819 | 13 | 1.43% |
| hsa_circ_0058236 | 2099 | 15 | 1.43% |
| hsa_circ_0132866 | 4480 | 32 | 1.43% |
| hsa_circ_0059724 | 3220 | 23 | 1.43% |
| hsa_circ_0074516 | 1680 | 12 | 1.43% |
| hsa_circ_0113901 | 1680 | 12 | 1.43% |
| hsa_circ_0021018 | 1540 | 11 | 1.43% |
| hsa_circ_0043938 | 1540 | 11 | 1.43% |
| hsa_circ_0053685 | 1540 | 11 | 1.43% |
| hsa_circ_0062378 | 1540 | 11 | 1.43% |
| hsa_circ_0085780 | 1540 | 11 | 1.43% |
| hsa_circ_0003288 | 1400 | 10 | 1.43% |
| hsa_circ_0027143 | 1400 | 10 | 1.43% |
| hsa_circ_0027681 | 1400 | 10 | 1.43% |
| hsa_circ_0031058 | 1400 | 10 | 1.43% |
| hsa_circ_0057912 | 1400 | 10 | 1.43% |
| hsa_circ_0066300 | 1400 | 10 | 1.43% |
| hsa_circ_0072913 | 1400 | 10 | 1.43% |
| hsa_circ_0088727 | 1400 | 10 | 1.43% |
| hsa_circ_0110025 | 1400 | 10 | 1.43% |
| hsa_circ_0123655 | 1400 | 10 | 1.43% |
| hsa_circ_0129159 | 1400 | 10 | 1.43% |
| hsa_circ_0130574 | 1400 | 10 | 1.43% |
| hsa_circ_0068386 | 1681 | 12 | 1.43% |
| hsa_circ_0069654 | 1681 | 12 | 1.43% |
| hsa_circ_0031494 | 1541 | 11 | 1.43% |
| hsa_circ_0087149 | 1541 | 11 | 1.43% |
| hsa_circ_0049761 | 1401 | 10 | 1.43% |
| hsa_circ_0055898 | 1401 | 10 | 1.43% |
| hsa_circ_0062532 | 1401 | 10 | 1.43% |
| hsa_circ_0070592 | 1401 | 10 | 1.43% |
| hsa_circ_0075294 | 1401 | 10 | 1.43% |
| hsa_circ_0079904 | 1401 | 10 | 1.43% |
| hsa_circ_0094889 | 1401 | 10 | 1.43% |
| hsa_circ_0126413 | 1401 | 10 | 1.43% |
| hsa_circ_0127926 | 1401 | 10 | 1.43% |
| hsa_circ_0084407 | 1822 | 13 | 1.43% |
| hsa_circ_0012089 | 1682 | 12 | 1.43% |
| hsa_circ_0058901 | 1682 | 12 | 1.43% |
| hsa_circ_0128752 | 1682 | 12 | 1.43% |

| | | | |
|---|---|---|---|
| hsa_circ_0017207 | 1542 | 11 | 1.43% |
| hsa_circ_0023251 | 1542 | 11 | 1.43% |
| hsa_circ_0061395 | 1542 | 11 | 1.43% |
| hsa_circ_0064406 | 1542 | 11 | 1.43% |
| hsa_circ_0096829 | 1542 | 11 | 1.43% |
| hsa_circ_0099221 | 1542 | 11 | 1.43% |
| hsa_circ_0116985 | 1542 | 11 | 1.43% |
| hsa_circ_0009214 | 1402 | 10 | 1.43% |
| hsa_circ_0019298 | 1402 | 10 | 1.43% |
| hsa_circ_0039990 | 1402 | 10 | 1.43% |
| hsa_circ_0043478 | 1402 | 10 | 1.43% |
| hsa_circ_0043677 | 1402 | 10 | 1.43% |
| hsa_circ_0063481 | 1402 | 10 | 1.43% |
| hsa_circ_0069372 | 1402 | 10 | 1.43% |
| hsa_circ_0075198 | 1402 | 10 | 1.43% |
| hsa_circ_0119284 | 1402 | 10 | 1.43% |
| hsa_circ_0067430 | 1963 | 14 | 1.43% |
| hsa_circ_0082891 | 1963 | 14 | 1.43% |
| hsa_circ_0039092 | 1683 | 12 | 1.43% |
| hsa_circ_0054786 | 1683 | 12 | 1.43% |
| hsa_circ_0015670 | 1543 | 11 | 1.43% |
| hsa_circ_0016883 | 1543 | 11 | 1.43% |
| hsa_circ_0024728 | 1543 | 11 | 1.43% |
| hsa_circ_0060591 | 1543 | 11 | 1.43% |
| hsa_circ_0075970 | 1543 | 11 | 1.43% |
| hsa_circ_0012056 | 1403 | 10 | 1.43% |
| hsa_circ_0020823 | 1403 | 10 | 1.43% |
| hsa_circ_0023472 | 1403 | 10 | 1.43% |
| hsa_circ_0024209 | 1403 | 10 | 1.43% |
| hsa_circ_0088386 | 1403 | 10 | 1.43% |
| hsa_circ_0125322 | 1403 | 10 | 1.43% |
| hsa_circ_0129278 | 1403 | 10 | 1.43% |
| hsa_circ_0129816 | 1403 | 10 | 1.43% |
| hsa_circ_0066956 | 1824 | 13 | 1.43% |
| hsa_circ_0059238 | 1684 | 12 | 1.43% |
| hsa_circ_0072785 | 1684 | 12 | 1.43% |
| hsa_circ_0136874 | 1684 | 12 | 1.43% |
| hsa_circ_0036991 | 1965 | 14 | 1.42% |
| hsa_circ_0051284 | 1544 | 11 | 1.42% |
| hsa_circ_0052668 | 1544 | 11 | 1.42% |
| hsa_circ_0076656 | 1544 | 11 | 1.42% |
| hsa_circ_0082278 | 1544 | 11 | 1.42% |
| hsa_circ_0087824 | 1544 | 11 | 1.42% |
| hsa_circ_0087912 | 1544 | 11 | 1.42% |
| hsa_circ_0107432 | 1544 | 11 | 1.42% |
| hsa_circ_0029132 | 1404 | 10 | 1.42% |
| hsa_circ_0039506 | 1404 | 10 | 1.42% |
| hsa_circ_0044860 | 1404 | 10 | 1.42% |
| hsa_circ_0049962 | 1404 | 10 | 1.42% |
| hsa_circ_0060267 | 1404 | 10 | 1.42% |

| | | | |
|---|---|---|---|
| hsa_circ_0067364 | 1404 | 10 | 1.42% |
| hsa_circ_0071404 | 1404 | 10 | 1.42% |
| hsa_circ_0083313 | 1404 | 10 | 1.42% |
| hsa_circ_0090979 | 1404 | 10 | 1.42% |
| hsa_circ_0095773 | 1404 | 10 | 1.42% |
| hsa_circ_0109383 | 1404 | 10 | 1.42% |
| hsa_circ_0111333 | 1404 | 10 | 1.42% |
| hsa_circ_0113161 | 1404 | 10 | 1.42% |
| hsa_circ_0131729 | 1404 | 10 | 1.42% |
| hsa_circ_0032659 | 2387 | 17 | 1.42% |
| hsa_circ_0013424 | 1685 | 12 | 1.42% |
| hsa_circ_0031154 | 1685 | 12 | 1.42% |
| hsa_circ_0101889 | 1685 | 12 | 1.42% |
| hsa_circ_0136942 | 3090 | 22 | 1.42% |
| hsa_circ_0020308 | 1545 | 11 | 1.42% |
| hsa_circ_0021422 | 1545 | 11 | 1.42% |
| hsa_circ_0070458 | 1545 | 11 | 1.42% |
| hsa_circ_0101611 | 1545 | 11 | 1.42% |
| hsa_circ_0136149 | 1545 | 11 | 1.42% |
| hsa_circ_0111879 | 6321 | 45 | 1.42% |
| hsa_circ_0001021 | 1967 | 14 | 1.42% |
| hsa_circ_0018246 | 1686 | 12 | 1.42% |
| hsa_circ_0034628 | 1686 | 12 | 1.42% |
| hsa_circ_0018842 | 1405 | 10 | 1.42% |
| hsa_circ_0024860 | 1405 | 10 | 1.42% |
| hsa_circ_0032285 | 1405 | 10 | 1.42% |
| hsa_circ_0076637 | 1405 | 10 | 1.42% |
| hsa_circ_0084304 | 1405 | 10 | 1.42% |
| hsa_circ_0086431 | 1405 | 10 | 1.42% |
| hsa_circ_0096242 | 1405 | 10 | 1.42% |
| hsa_circ_0122253 | 1405 | 10 | 1.42% |
| hsa_circ_0106932 | 3654 | 26 | 1.42% |
| hsa_circ_0046072 | 1827 | 13 | 1.42% |
| hsa_circ_0011979 | 1546 | 11 | 1.42% |
| hsa_circ_0027640 | 1546 | 11 | 1.42% |
| hsa_circ_0055283 | 1546 | 11 | 1.42% |
| hsa_circ_0056909 | 1546 | 11 | 1.42% |
| hsa_circ_0134596 | 2249 | 16 | 1.42% |
| hsa_circ_0030012 | 1687 | 12 | 1.42% |
| hsa_circ_0031029 | 1687 | 12 | 1.42% |
| hsa_circ_0011542 | 1406 | 10 | 1.42% |
| hsa_circ_0023908 | 1406 | 10 | 1.42% |
| hsa_circ_0025629 | 1406 | 10 | 1.42% |
| hsa_circ_0031547 | 1406 | 10 | 1.42% |
| hsa_circ_0033003 | 1406 | 10 | 1.42% |
| hsa_circ_0112610 | 1406 | 10 | 1.42% |
| hsa_circ_0112753 | 1406 | 10 | 1.42% |
| hsa_circ_0121729 | 1406 | 10 | 1.42% |
| hsa_circ_0122183 | 1406 | 10 | 1.42% |
| hsa_circ_0044147 | 1547 | 11 | 1.42% |

| | | | |
|---|---|---|---|
| hsa_circ_0058687 | 1547 | 11 | 1.42% |
| hsa_circ_0060465 | 1547 | 11 | 1.42% |
| hsa_circ_0105691 | 1547 | 11 | 1.42% |
| hsa_circ_0121005 | 1547 | 11 | 1.42% |
| hsa_circ_0076329 | 3657 | 26 | 1.42% |
| hsa_circ_0048566 | 1688 | 12 | 1.42% |
| hsa_circ_0058707 | 1688 | 12 | 1.42% |
| hsa_circ_0070389 | 1688 | 12 | 1.42% |
| hsa_circ_0079697 | 1688 | 12 | 1.42% |
| hsa_circ_0110327 | 1688 | 12 | 1.42% |
| hsa_circ_0129431 | 1688 | 12 | 1.42% |
| hsa_circ_0016067 | 1407 | 10 | 1.42% |
| hsa_circ_0032575 | 1407 | 10 | 1.42% |
| hsa_circ_0042670 | 1407 | 10 | 1.42% |
| hsa_circ_0056009 | 1407 | 10 | 1.42% |
| hsa_circ_0074833 | 1407 | 10 | 1.42% |
| hsa_circ_0086909 | 1407 | 10 | 1.42% |
| hsa_circ_0091244 | 1407 | 10 | 1.42% |
| hsa_circ_0110104 | 1407 | 10 | 1.42% |
| hsa_circ_0117556 | 1407 | 10 | 1.42% |
| hsa_circ_0014543 | 1548 | 11 | 1.42% |
| hsa_circ_0083522 | 1548 | 11 | 1.42% |
| hsa_circ_0086538 | 1548 | 11 | 1.42% |
| hsa_circ_0089564 | 1548 | 11 | 1.42% |
| hsa_circ_0089664 | 1548 | 11 | 1.42% |
| hsa_circ_0013294 | 1689 | 12 | 1.42% |
| hsa_circ_0049757 | 1689 | 12 | 1.42% |
| hsa_circ_0056903 | 1689 | 12 | 1.42% |
| hsa_circ_0083591 | 1689 | 12 | 1.42% |
| hsa_circ_0013461 | 1408 | 10 | 1.42% |
| hsa_circ_0016710 | 1408 | 10 | 1.42% |
| hsa_circ_0025453 | 1408 | 10 | 1.42% |
| hsa_circ_0031421 | 1408 | 10 | 1.42% |
| hsa_circ_0034040 | 1408 | 10 | 1.42% |
| hsa_circ_0045890 | 1408 | 10 | 1.42% |
| hsa_circ_0020698 | 1549 | 11 | 1.42% |
| hsa_circ_0041350 | 1549 | 11 | 1.42% |
| hsa_circ_0064779 | 1549 | 11 | 1.42% |
| hsa_circ_0065642 | 1549 | 11 | 1.42% |
| hsa_circ_0067894 | 1549 | 11 | 1.42% |
| hsa_circ_0071050 | 1549 | 11 | 1.42% |
| hsa_circ_0136147 | 1549 | 11 | 1.42% |
| hsa_circ_0086138 | 1690 | 12 | 1.42% |
| hsa_circ_0121421 | 1831 | 13 | 1.42% |
| hsa_circ_0012165 | 1409 | 10 | 1.42% |
| hsa_circ_0016910 | 1409 | 10 | 1.42% |
| hsa_circ_0017385 | 1409 | 10 | 1.42% |
| hsa_circ_0020605 | 1409 | 10 | 1.42% |
| hsa_circ_0026052 | 1409 | 10 | 1.42% |
| hsa_circ_0028133 | 1409 | 10 | 1.42% |

| | | | |
|---|---|---|---|
| hsa_circ_0034388 | 1409 | 10 | 1.42% |
| hsa_circ_0036721 | 1409 | 10 | 1.42% |
| hsa_circ_0044039 | 1409 | 10 | 1.42% |
| hsa_circ_0058399 | 1409 | 10 | 1.42% |
| hsa_circ_0072737 | 1409 | 10 | 1.42% |
| hsa_circ_0085028 | 1409 | 10 | 1.42% |
| hsa_circ_0085150 | 1409 | 10 | 1.42% |
| hsa_circ_0089055 | 1409 | 10 | 1.42% |
| hsa_circ_0101851 | 1409 | 10 | 1.42% |
| hsa_circ_0114945 | 1409 | 10 | 1.42% |
| hsa_circ_0129017 | 1409 | 10 | 1.42% |
| hsa_circ_0038211 | 1550 | 11 | 1.42% |
| hsa_circ_0053146 | 1550 | 11 | 1.42% |
| hsa_circ_0127310 | 1550 | 11 | 1.42% |
| hsa_circ_0040130 | 1691 | 12 | 1.42% |
| hsa_circ_0105477 | 1691 | 12 | 1.42% |
| hsa_circ_0063213 | 1833 | 13 | 1.42% |
| hsa_circ_0069551 | 1833 | 13 | 1.42% |
| hsa_circ_0024772 | 1692 | 12 | 1.42% |
| hsa_circ_0026351 | 1692 | 12 | 1.42% |
| hsa_circ_0040655 | 1692 | 12 | 1.42% |
| hsa_circ_0065604 | 1692 | 12 | 1.42% |
| hsa_circ_0069126 | 1692 | 12 | 1.42% |
| hsa_circ_0023226 | 1551 | 11 | 1.42% |
| hsa_circ_0046179 | 1551 | 11 | 1.42% |
| hsa_circ_0052562 | 1551 | 11 | 1.42% |
| hsa_circ_0064304 | 1551 | 11 | 1.42% |
| hsa_circ_0111373 | 1551 | 11 | 1.42% |
| hsa_circ_0038290 | 1410 | 10 | 1.42% |
| hsa_circ_0048523 | 1410 | 10 | 1.42% |
| hsa_circ_0058051 | 1410 | 10 | 1.42% |
| hsa_circ_0066454 | 1410 | 10 | 1.42% |
| hsa_circ_0091369 | 1410 | 10 | 1.42% |
| hsa_circ_0091899 | 1410 | 10 | 1.42% |
| hsa_circ_0093389 | 1410 | 10 | 1.42% |
| hsa_circ_0099874 | 1410 | 10 | 1.42% |
| hsa_circ_0105237 | 1410 | 10 | 1.42% |
| hsa_circ_0106608 | 1410 | 10 | 1.42% |
| hsa_circ_0113968 | 1410 | 10 | 1.42% |
| hsa_circ_0127645 | 1410 | 10 | 1.42% |
| hsa_circ_0129692 | 1410 | 10 | 1.42% |
| hsa_circ_0029524 | 1834 | 13 | 1.42% |
| hsa_circ_0034853 | 1693 | 12 | 1.42% |
| hsa_circ_0067584 | 1693 | 12 | 1.42% |
| hsa_circ_0080249 | 1693 | 12 | 1.42% |
| hsa_circ_0111179 | 1693 | 12 | 1.42% |
| hsa_circ_0134187 | 1693 | 12 | 1.42% |
| hsa_circ_0031378 | 1552 | 11 | 1.42% |
| hsa_circ_0035242 | 1552 | 11 | 1.42% |
| hsa_circ_0062478 | 1552 | 11 | 1.42% |

| | | | |
|---|---|---|---|
| hsa_circ_0074140 | 1552 | 11 | 1.42% |
| hsa_circ_0016604 | 1411 | 10 | 1.42% |
| hsa_circ_0042020 | 1411 | 10 | 1.42% |
| hsa_circ_0062193 | 1411 | 10 | 1.42% |
| hsa_circ_0076551 | 1411 | 10 | 1.42% |
| hsa_circ_0091226 | 1411 | 10 | 1.42% |
| hsa_circ_0118956 | 1411 | 10 | 1.42% |
| hsa_circ_0009514 | 4657 | 33 | 1.42% |
| hsa_circ_0038034 | 3105 | 22 | 1.42% |
| hsa_circ_0050659 | 1835 | 13 | 1.42% |
| hsa_circ_0071390 | 1694 | 12 | 1.42% |
| hsa_circ_0132495 | 1694 | 12 | 1.42% |
| hsa_circ_0086625 | 6353 | 45 | 1.42% |
| hsa_circ_0004246 | 1553 | 11 | 1.42% |
| hsa_circ_0010065 | 1553 | 11 | 1.42% |
| hsa_circ_0034309 | 1553 | 11 | 1.42% |
| hsa_circ_0038275 | 1553 | 11 | 1.42% |
| hsa_circ_0046370 | 1553 | 11 | 1.42% |
| hsa_circ_0066082 | 1553 | 11 | 1.42% |
| hsa_circ_0089844 | 1553 | 11 | 1.42% |
| hsa_circ_0110904 | 1553 | 11 | 1.42% |
| hsa_circ_0042831 | 1412 | 10 | 1.42% |
| hsa_circ_0068353 | 1412 | 10 | 1.42% |
| hsa_circ_0082173 | 1412 | 10 | 1.42% |
| hsa_circ_0090352 | 1412 | 10 | 1.42% |
| hsa_circ_0098629 | 1412 | 10 | 1.42% |
| hsa_circ_0100486 | 1412 | 10 | 1.42% |
| hsa_circ_0103198 | 1977 | 14 | 1.42% |
| hsa_circ_0093660 | 2401 | 17 | 1.42% |
| hsa_circ_0090159 | 1695 | 12 | 1.42% |
| hsa_circ_0022901 | 1554 | 11 | 1.42% |
| hsa_circ_0027968 | 1554 | 11 | 1.42% |
| hsa_circ_0064206 | 1554 | 11 | 1.42% |
| hsa_circ_0074601 | 1554 | 11 | 1.42% |
| hsa_circ_0077811 | 1554 | 11 | 1.42% |
| hsa_circ_0080716 | 1554 | 11 | 1.42% |
| hsa_circ_0084323 | 1554 | 11 | 1.42% |
| hsa_circ_0116087 | 1554 | 11 | 1.42% |
| hsa_circ_0025456 | 1978 | 14 | 1.42% |
| hsa_circ_0005897 | 1413 | 10 | 1.42% |
| hsa_circ_0026729 | 1413 | 10 | 1.42% |
| hsa_circ_0027932 | 1413 | 10 | 1.42% |
| hsa_circ_0043049 | 1413 | 10 | 1.42% |
| hsa_circ_0046089 | 1413 | 10 | 1.42% |
| hsa_circ_0049851 | 1413 | 10 | 1.42% |
| hsa_circ_0064659 | 1413 | 10 | 1.42% |
| hsa_circ_0074035 | 1413 | 10 | 1.42% |
| hsa_circ_0079809 | 1413 | 10 | 1.42% |
| hsa_circ_0091766 | 1413 | 10 | 1.42% |
| hsa_circ_0133984 | 1413 | 10 | 1.42% |

| | | | |
|---|---|---|---|
| hsa_circ_0135659 | 1413 | 10 | 1.42% |
| hsa_circ_0060876 | 1837 | 13 | 1.42% |
| hsa_circ_0080861 | 1837 | 13 | 1.42% |
| hsa_circ_0065878 | 1696 | 12 | 1.42% |
| hsa_circ_0088877 | 1696 | 12 | 1.42% |
| hsa_circ_0135369 | 1979 | 14 | 1.41% |
| hsa_circ_0002833 | 1555 | 11 | 1.41% |
| hsa_circ_0012516 | 1555 | 11 | 1.41% |
| hsa_circ_0025722 | 1555 | 11 | 1.41% |
| hsa_circ_0069898 | 1555 | 11 | 1.41% |
| hsa_circ_0097095 | 1555 | 11 | 1.41% |
| hsa_circ_0135115 | 1555 | 11 | 1.41% |
| hsa_circ_0123436 | 1838 | 13 | 1.41% |
| hsa_circ_0119228 | 6221 | 44 | 1.41% |
| hsa_circ_0020637 | 1414 | 10 | 1.41% |
| hsa_circ_0064856 | 1414 | 10 | 1.41% |
| hsa_circ_0086349 | 1414 | 10 | 1.41% |
| hsa_circ_0089768 | 1414 | 10 | 1.41% |
| hsa_circ_0092147 | 1414 | 10 | 1.41% |
| hsa_circ_0115996 | 1414 | 10 | 1.41% |
| hsa_circ_0116864 | 1414 | 10 | 1.41% |
| hsa_circ_0121944 | 1414 | 10 | 1.41% |
| hsa_circ_0089118 | 1697 | 12 | 1.41% |
| hsa_circ_0012987 | 1556 | 11 | 1.41% |
| hsa_circ_0016238 | 1556 | 11 | 1.41% |
| hsa_circ_0034060 | 1556 | 11 | 1.41% |
| hsa_circ_0037883 | 1556 | 11 | 1.41% |
| hsa_circ_0065250 | 1556 | 11 | 1.41% |
| hsa_circ_0122901 | 1556 | 11 | 1.41% |
| hsa_circ_0101035 | 1839 | 13 | 1.41% |
| hsa_circ_0086341 | 2122 | 15 | 1.41% |
| hsa_circ_0127246 | 3679 | 26 | 1.41% |
| hsa_circ_0011620 | 1698 | 12 | 1.41% |
| hsa_circ_0018367 | 1698 | 12 | 1.41% |
| hsa_circ_0052283 | 1698 | 12 | 1.41% |
| hsa_circ_0013030 | 1415 | 10 | 1.41% |
| hsa_circ_0023289 | 1415 | 10 | 1.41% |
| hsa_circ_0050075 | 1415 | 10 | 1.41% |
| hsa_circ_0057771 | 1415 | 10 | 1.41% |
| hsa_circ_0058599 | 1415 | 10 | 1.41% |
| hsa_circ_0083464 | 1415 | 10 | 1.41% |
| hsa_circ_0088064 | 1415 | 10 | 1.41% |
| hsa_circ_0129399 | 1415 | 10 | 1.41% |
| hsa_circ_0032141 | 2123 | 15 | 1.41% |
| hsa_circ_0008634 | 1557 | 11 | 1.41% |
| hsa_circ_0027778 | 1557 | 11 | 1.41% |
| hsa_circ_0031988 | 1557 | 11 | 1.41% |
| hsa_circ_0032518 | 1557 | 11 | 1.41% |
| hsa_circ_0072600 | 1557 | 11 | 1.41% |
| hsa_circ_0083678 | 1557 | 11 | 1.41% |

| | | | |
|---|---|---|---|
| hsa_circ_0009197 | 1699 | 12 | 1.41% |
| hsa_circ_0104660 | 1699 | 12 | 1.41% |
| hsa_circ_0132883 | 1699 | 12 | 1.41% |
| hsa_circ_0006576 | 1416 | 10 | 1.41% |
| hsa_circ_0019016 | 1416 | 10 | 1.41% |
| hsa_circ_0027783 | 1416 | 10 | 1.41% |
| hsa_circ_0036302 | 1416 | 10 | 1.41% |
| hsa_circ_0044648 | 1416 | 10 | 1.41% |
| hsa_circ_0045162 | 1416 | 10 | 1.41% |
| hsa_circ_0047745 | 1416 | 10 | 1.41% |
| hsa_circ_0062910 | 1416 | 10 | 1.41% |
| hsa_circ_0074420 | 1416 | 10 | 1.41% |
| hsa_circ_0075190 | 1416 | 10 | 1.41% |
| hsa_circ_0075306 | 1416 | 10 | 1.41% |
| hsa_circ_0122162 | 1416 | 10 | 1.41% |
| hsa_circ_0114445 | 1841 | 13 | 1.41% |
| hsa_circ_0101716 | 5240 | 37 | 1.41% |
| hsa_circ_0012425 | 1558 | 11 | 1.41% |
| hsa_circ_0022925 | 1558 | 11 | 1.41% |
| hsa_circ_0035494 | 1558 | 11 | 1.41% |
| hsa_circ_0040178 | 1558 | 11 | 1.41% |
| hsa_circ_0050366 | 1558 | 11 | 1.41% |
| hsa_circ_0091326 | 1558 | 11 | 1.41% |
| hsa_circ_0120182 | 1558 | 11 | 1.41% |
| hsa_circ_0052313 | 1983 | 14 | 1.41% |
| hsa_circ_0031274 | 1417 | 10 | 1.41% |
| hsa_circ_0039593 | 1417 | 10 | 1.41% |
| hsa_circ_0043552 | 1417 | 10 | 1.41% |
| hsa_circ_0088909 | 1417 | 10 | 1.41% |
| hsa_circ_0092664 | 1417 | 10 | 1.41% |
| hsa_circ_0104893 | 1417 | 10 | 1.41% |
| hsa_circ_0109419 | 1417 | 10 | 1.41% |
| hsa_circ_0118945 | 1417 | 10 | 1.41% |
| hsa_circ_0114335 | 1984 | 14 | 1.41% |
| hsa_circ_0021313 | 1559 | 11 | 1.41% |
| hsa_circ_0054953 | 1559 | 11 | 1.41% |
| hsa_circ_0069258 | 1559 | 11 | 1.41% |
| hsa_circ_0117790 | 1559 | 11 | 1.41% |
| hsa_circ_0120710 | 1559 | 11 | 1.41% |
| hsa_circ_0100208 | 3685 | 26 | 1.41% |
| hsa_circ_0011398 | 1701 | 12 | 1.41% |
| hsa_circ_0078688 | 1701 | 12 | 1.41% |
| hsa_circ_0124461 | 1701 | 12 | 1.41% |
| hsa_circ_0036276 | 1418 | 10 | 1.41% |
| hsa_circ_0041905 | 1418 | 10 | 1.41% |
| hsa_circ_0045814 | 1418 | 10 | 1.41% |
| hsa_circ_0069657 | 1418 | 10 | 1.41% |
| hsa_circ_0100288 | 1418 | 10 | 1.41% |
| hsa_circ_0113196 | 1418 | 10 | 1.41% |
| hsa_circ_0120888 | 1418 | 10 | 1.41% |

| | | | |
|---|---|---|---|
| hsa_circ_0127828 | 1418 | 10 | 1.41% |
| hsa_circ_0131727 | 1418 | 10 | 1.41% |
| hsa_circ_0138871 | 1418 | 10 | 1.41% |
| hsa_circ_0061446 | 2269 | 16 | 1.41% |
| hsa_circ_0025229 | 1560 | 11 | 1.41% |
| hsa_circ_0029949 | 1560 | 11 | 1.41% |
| hsa_circ_0034645 | 1560 | 11 | 1.41% |
| hsa_circ_0046670 | 1560 | 11 | 1.41% |
| hsa_circ_0048740 | 1560 | 11 | 1.41% |
| hsa_circ_0058430 | 1560 | 11 | 1.41% |
| hsa_circ_0059845 | 1560 | 11 | 1.41% |
| hsa_circ_0076574 | 1560 | 11 | 1.41% |
| hsa_circ_0107034 | 1560 | 11 | 1.41% |
| hsa_circ_0018856 | 1702 | 12 | 1.41% |
| hsa_circ_0016036 | 1419 | 10 | 1.41% |
| hsa_circ_0026591 | 1419 | 10 | 1.41% |
| hsa_circ_0027482 | 1419 | 10 | 1.41% |
| hsa_circ_0031722 | 1419 | 10 | 1.41% |
| hsa_circ_0057653 | 1419 | 10 | 1.41% |
| hsa_circ_0068504 | 1419 | 10 | 1.41% |
| hsa_circ_0086228 | 1419 | 10 | 1.41% |
| hsa_circ_0090388 | 1419 | 10 | 1.41% |
| hsa_circ_0099703 | 1419 | 10 | 1.41% |
| hsa_circ_0112748 | 1419 | 10 | 1.41% |
| hsa_circ_0117467 | 1419 | 10 | 1.41% |
| hsa_circ_0013794 | 1561 | 11 | 1.41% |
| hsa_circ_0038537 | 1561 | 11 | 1.41% |
| hsa_circ_0074421 | 1561 | 11 | 1.41% |
| hsa_circ_0087057 | 1561 | 11 | 1.41% |
| hsa_circ_0111349 | 1561 | 11 | 1.41% |
| hsa_circ_0129590 | 1561 | 11 | 1.41% |
| hsa_circ_0001387 | 1703 | 12 | 1.41% |
| hsa_circ_0078233 | 3548 | 25 | 1.41% |
| hsa_circ_0059654 | 1845 | 13 | 1.41% |
| hsa_circ_0013535 | 2129 | 15 | 1.41% |
| hsa_circ_0076102 | 1704 | 12 | 1.41% |
| hsa_circ_0088430 | 1704 | 12 | 1.41% |
| hsa_circ_0088758 | 1704 | 12 | 1.41% |
| hsa_circ_0017214 | 1562 | 11 | 1.41% |
| hsa_circ_0138269 | 1562 | 11 | 1.41% |
| hsa_circ_0007070 | 1420 | 10 | 1.41% |
| hsa_circ_0029611 | 1420 | 10 | 1.41% |
| hsa_circ_0042083 | 1420 | 10 | 1.41% |
| hsa_circ_0064728 | 1420 | 10 | 1.41% |
| hsa_circ_0084124 | 1420 | 10 | 1.41% |
| hsa_circ_0096310 | 1420 | 10 | 1.41% |
| hsa_circ_0097115 | 1420 | 10 | 1.41% |
| hsa_circ_0099018 | 1420 | 10 | 1.41% |
| hsa_circ_0107363 | 1420 | 10 | 1.41% |
| hsa_circ_0135412 | 1420 | 10 | 1.41% |

| | | | |
|---|---|---|---|
| hsa_circ_0135936 | 1420 | 10 | 1.41% |
| hsa_circ_0057473 | 1705 | 12 | 1.41% |
| hsa_circ_0081608 | 1705 | 12 | 1.41% |
| hsa_circ_0052038 | 1563 | 11 | 1.41% |
| hsa_circ_0113781 | 1563 | 11 | 1.41% |
| hsa_circ_0117017 | 1563 | 11 | 1.41% |
| hsa_circ_0117363 | 1563 | 11 | 1.41% |
| hsa_circ_0122778 | 1563 | 11 | 1.41% |
| hsa_circ_0009489 | 4547 | 32 | 1.41% |
| hsa_circ_0013889 | 1421 | 10 | 1.41% |
| hsa_circ_0020355 | 1421 | 10 | 1.41% |
| hsa_circ_0031362 | 1421 | 10 | 1.41% |
| hsa_circ_0038142 | 1421 | 10 | 1.41% |
| hsa_circ_0047538 | 1421 | 10 | 1.41% |
| hsa_circ_0052444 | 1421 | 10 | 1.41% |
| hsa_circ_0052466 | 1421 | 10 | 1.41% |
| hsa_circ_0103003 | 1421 | 10 | 1.41% |
| hsa_circ_0126004 | 1421 | 10 | 1.41% |
| hsa_circ_0133852 | 1421 | 10 | 1.41% |
| hsa_circ_0024546 | 1848 | 13 | 1.41% |
| hsa_circ_0033584 | 1706 | 12 | 1.41% |
| hsa_circ_0034433 | 1706 | 12 | 1.41% |
| hsa_circ_0064798 | 1706 | 12 | 1.41% |
| hsa_circ_0014985 | 1564 | 11 | 1.41% |
| hsa_circ_0021891 | 1564 | 11 | 1.41% |
| hsa_circ_0038247 | 1564 | 11 | 1.41% |
| hsa_circ_0056914 | 1564 | 11 | 1.41% |
| hsa_circ_0011919 | 1422 | 10 | 1.41% |
| hsa_circ_0023024 | 1422 | 10 | 1.41% |
| hsa_circ_0025943 | 1422 | 10 | 1.41% |
| hsa_circ_0054992 | 1422 | 10 | 1.41% |
| hsa_circ_0059319 | 1422 | 10 | 1.41% |
| hsa_circ_0068263 | 1422 | 10 | 1.41% |
| hsa_circ_0071190 | 1422 | 10 | 1.41% |
| hsa_circ_0071849 | 1422 | 10 | 1.41% |
| hsa_circ_0086221 | 1422 | 10 | 1.41% |
| hsa_circ_0098601 | 1422 | 10 | 1.41% |
| hsa_circ_0106104 | 1422 | 10 | 1.41% |
| hsa_circ_0012319 | 1849 | 13 | 1.41% |
| hsa_circ_0088282 | 1849 | 13 | 1.41% |
| hsa_circ_0018752 | 1565 | 11 | 1.41% |
| hsa_circ_0068938 | 1565 | 11 | 1.41% |
| hsa_circ_0088723 | 3842 | 27 | 1.41% |
| hsa_circ_0008594 | 1423 | 10 | 1.41% |
| hsa_circ_0011436 | 1423 | 10 | 1.41% |
| hsa_circ_0014254 | 1423 | 10 | 1.41% |
| hsa_circ_0027864 | 1423 | 10 | 1.41% |
| hsa_circ_0031017 | 1423 | 10 | 1.41% |
| hsa_circ_0039527 | 1423 | 10 | 1.41% |
| hsa_circ_0089303 | 1423 | 10 | 1.41% |

| | | | |
|---|---|---|---|
| hsa_circ_0108743 | 1423 | 10 | 1.41% |
| hsa_circ_0137522 | 1423 | 10 | 1.41% |
| hsa_circ_0137852 | 1423 | 10 | 1.41% |
| hsa_circ_0071246 | 1708 | 12 | 1.41% |
| hsa_circ_0029103 | 1566 | 11 | 1.40% |
| hsa_circ_0031200 | 1566 | 11 | 1.40% |
| hsa_circ_0038970 | 1566 | 11 | 1.40% |
| hsa_circ_0052025 | 1566 | 11 | 1.40% |
| hsa_circ_0085715 | 1566 | 11 | 1.40% |
| hsa_circ_0109752 | 1566 | 11 | 1.40% |
| hsa_circ_0109937 | 1566 | 11 | 1.40% |
| hsa_circ_0116990 | 1566 | 11 | 1.40% |
| hsa_circ_0124663 | 1566 | 11 | 1.40% |
| hsa_circ_0118595 | 1851 | 13 | 1.40% |
| hsa_circ_0016793 | 1424 | 10 | 1.40% |
| hsa_circ_0051411 | 1424 | 10 | 1.40% |
| hsa_circ_0057891 | 1424 | 10 | 1.40% |
| hsa_circ_0061974 | 1424 | 10 | 1.40% |
| hsa_circ_0067379 | 1424 | 10 | 1.40% |
| hsa_circ_0092430 | 1424 | 10 | 1.40% |
| hsa_circ_0103758 | 1424 | 10 | 1.40% |
| hsa_circ_0133716 | 1424 | 10 | 1.40% |
| hsa_circ_0036619 | 1709 | 12 | 1.40% |
| hsa_circ_0074677 | 1709 | 12 | 1.40% |
| hsa_circ_0128241 | 1709 | 12 | 1.40% |
| hsa_circ_0133904 | 1709 | 12 | 1.40% |
| hsa_circ_0041197 | 1567 | 11 | 1.40% |
| hsa_circ_0120656 | 1567 | 11 | 1.40% |
| hsa_circ_0087476 | 1852 | 13 | 1.40% |
| hsa_circ_0000388 | 5415 | 38 | 1.40% |
| hsa_circ_0058825 | 1710 | 12 | 1.40% |
| hsa_circ_0135177 | 1710 | 12 | 1.40% |
| hsa_circ_0012808 | 1425 | 10 | 1.40% |
| hsa_circ_0025265 | 1425 | 10 | 1.40% |
| hsa_circ_0026413 | 1425 | 10 | 1.40% |
| hsa_circ_0026424 | 1425 | 10 | 1.40% |
| hsa_circ_0026510 | 1425 | 10 | 1.40% |
| hsa_circ_0041580 | 1425 | 10 | 1.40% |
| hsa_circ_0045652 | 1425 | 10 | 1.40% |
| hsa_circ_0051481 | 1425 | 10 | 1.40% |
| hsa_circ_0063670 | 1425 | 10 | 1.40% |
| hsa_circ_0093202 | 1425 | 10 | 1.40% |
| hsa_circ_0098493 | 1425 | 10 | 1.40% |
| hsa_circ_0102603 | 1425 | 10 | 1.40% |
| hsa_circ_0105717 | 1425 | 10 | 1.40% |
| hsa_circ_0126660 | 1425 | 10 | 1.40% |
| hsa_circ_0132126 | 1425 | 10 | 1.40% |
| hsa_circ_0132867 | 2423 | 17 | 1.40% |
| hsa_circ_0049833 | 1568 | 11 | 1.40% |
| hsa_circ_0051341 | 1568 | 11 | 1.40% |

| | | | |
|---|---|---|---|
| hsa_circ_0066252 | 1568 | 11 | 1.40% |
| hsa_circ_0066457 | 1568 | 11 | 1.40% |
| hsa_circ_0090787 | 1568 | 11 | 1.40% |
| hsa_circ_0115971 | 1568 | 11 | 1.40% |
| hsa_circ_0127822 | 1568 | 11 | 1.40% |
| hsa_circ_0016904 | 1996 | 14 | 1.40% |
| hsa_circ_0020900 | 1426 | 10 | 1.40% |
| hsa_circ_0026190 | 1426 | 10 | 1.40% |
| hsa_circ_0031487 | 1426 | 10 | 1.40% |
| hsa_circ_0033398 | 1426 | 10 | 1.40% |
| hsa_circ_0073252 | 1426 | 10 | 1.40% |
| hsa_circ_0079869 | 1426 | 10 | 1.40% |
| hsa_circ_0083232 | 1426 | 10 | 1.40% |
| hsa_circ_0085581 | 1426 | 10 | 1.40% |
| hsa_circ_0087763 | 1426 | 10 | 1.40% |
| hsa_circ_0118910 | 1426 | 10 | 1.40% |
| hsa_circ_0038152 | 1854 | 13 | 1.40% |
| hsa_circ_0040651 | 1569 | 11 | 1.40% |
| hsa_circ_0046468 | 1569 | 11 | 1.40% |
| hsa_circ_0105137 | 1569 | 11 | 1.40% |
| hsa_circ_0138077 | 1569 | 11 | 1.40% |
| hsa_circ_0023725 | 1427 | 10 | 1.40% |
| hsa_circ_0042230 | 1427 | 10 | 1.40% |
| hsa_circ_0045844 | 1427 | 10 | 1.40% |
| hsa_circ_0089549 | 1427 | 10 | 1.40% |
| hsa_circ_0096381 | 1427 | 10 | 1.40% |
| hsa_circ_0120092 | 1427 | 10 | 1.40% |
| hsa_circ_0004152 | 1570 | 11 | 1.40% |
| hsa_circ_0010918 | 1570 | 11 | 1.40% |
| hsa_circ_0037923 | 1570 | 11 | 1.40% |
| hsa_circ_0041183 | 1570 | 11 | 1.40% |
| hsa_circ_0075586 | 1570 | 11 | 1.40% |
| hsa_circ_0138685 | 1570 | 11 | 1.40% |
| hsa_circ_0052800 | 1713 | 12 | 1.40% |
| hsa_circ_0091708 | 1713 | 12 | 1.40% |
| hsa_circ_0131875 | 1713 | 12 | 1.40% |
| hsa_circ_0138121 | 1713 | 12 | 1.40% |
| hsa_circ_0051434 | 1856 | 13 | 1.40% |
| hsa_circ_0030590 | 1428 | 10 | 1.40% |
| hsa_circ_0030736 | 1428 | 10 | 1.40% |
| hsa_circ_0040052 | 1428 | 10 | 1.40% |
| hsa_circ_0042381 | 1428 | 10 | 1.40% |
| hsa_circ_0042570 | 1428 | 10 | 1.40% |
| hsa_circ_0045021 | 1428 | 10 | 1.40% |
| hsa_circ_0090636 | 1428 | 10 | 1.40% |
| hsa_circ_0127837 | 1428 | 10 | 1.40% |
| hsa_circ_0132313 | 1428 | 10 | 1.40% |
| hsa_circ_0137394 | 1428 | 10 | 1.40% |
| hsa_circ_0011276 | 1571 | 11 | 1.40% |
| hsa_circ_0013620 | 1571 | 11 | 1.40% |

| | | | |
|---|---|---|---|
| hsa_circ_0036304 | 1571 | 11 | 1.40% |
| hsa_circ_0046489 | 1571 | 11 | 1.40% |
| hsa_circ_0092897 | 1571 | 11 | 1.40% |
| hsa_circ_0097413 | 1571 | 11 | 1.40% |
| hsa_circ_0122731 | 1571 | 11 | 1.40% |
| hsa_circ_0010791 | 1714 | 12 | 1.40% |
| hsa_circ_0038365 | 1714 | 12 | 1.40% |
| hsa_circ_0040879 | 1714 | 12 | 1.40% |
| hsa_circ_0090531 | 1714 | 12 | 1.40% |
| hsa_circ_0025164 | 1429 | 10 | 1.40% |
| hsa_circ_0054098 | 1429 | 10 | 1.40% |
| hsa_circ_0072431 | 1429 | 10 | 1.40% |
| hsa_circ_0081553 | 1429 | 10 | 1.40% |
| hsa_circ_0084905 | 1429 | 10 | 1.40% |
| hsa_circ_0131083 | 1429 | 10 | 1.40% |
| hsa_circ_0000922 | 1572 | 11 | 1.40% |
| hsa_circ_0014517 | 1572 | 11 | 1.40% |
| hsa_circ_0015581 | 1572 | 11 | 1.40% |
| hsa_circ_0020610 | 1572 | 11 | 1.40% |
| hsa_circ_0022309 | 1572 | 11 | 1.40% |
| hsa_circ_0025504 | 1572 | 11 | 1.40% |
| hsa_circ_0054883 | 1572 | 11 | 1.40% |
| hsa_circ_0076330 | 1572 | 11 | 1.40% |
| hsa_circ_0104636 | 1572 | 11 | 1.40% |
| hsa_circ_0106088 | 1572 | 11 | 1.40% |
| hsa_circ_0135593 | 1572 | 11 | 1.40% |
| hsa_circ_0008736 | 1715 | 12 | 1.40% |
| hsa_circ_0015557 | 1858 | 13 | 1.40% |
| hsa_circ_0118048 | 1858 | 13 | 1.40% |
| hsa_circ_0085037 | 2001 | 14 | 1.40% |
| hsa_circ_0099103 | 2144 | 15 | 1.40% |
| hsa_circ_0100670 | 1859 | 13 | 1.40% |
| hsa_circ_0022397 | 1716 | 12 | 1.40% |
| hsa_circ_0040088 | 1716 | 12 | 1.40% |
| hsa_circ_0084011 | 1716 | 12 | 1.40% |
| hsa_circ_0086939 | 1716 | 12 | 1.40% |
| hsa_circ_0019763 | 1573 | 11 | 1.40% |
| hsa_circ_0039996 | 1573 | 11 | 1.40% |
| hsa_circ_0049808 | 1573 | 11 | 1.40% |
| hsa_circ_0010467 | 1430 | 10 | 1.40% |
| hsa_circ_0015879 | 1430 | 10 | 1.40% |
| hsa_circ_0022708 | 1430 | 10 | 1.40% |
| hsa_circ_0028747 | 1430 | 10 | 1.40% |
| hsa_circ_0032706 | 1430 | 10 | 1.40% |
| hsa_circ_0047291 | 1430 | 10 | 1.40% |
| hsa_circ_0048438 | 1430 | 10 | 1.40% |
| hsa_circ_0052517 | 1430 | 10 | 1.40% |
| hsa_circ_0071074 | 1430 | 10 | 1.40% |
| hsa_circ_0082112 | 1430 | 10 | 1.40% |
| hsa_circ_0096650 | 1430 | 10 | 1.40% |

| | | | |
|---|---|---|---|
| hsa_circ_0117910 | 1430 | 10 | 1.40% |
| hsa_circ_0129930 | 1430 | 10 | 1.40% |
| hsa_circ_0123661 | 5865 | 41 | 1.40% |
| hsa_circ_0029266 | 1717 | 12 | 1.40% |
| hsa_circ_0026861 | 1574 | 11 | 1.40% |
| hsa_circ_0035388 | 1574 | 11 | 1.40% |
| hsa_circ_0043639 | 1574 | 11 | 1.40% |
| hsa_circ_0015426 | 1431 | 10 | 1.40% |
| hsa_circ_0016142 | 1431 | 10 | 1.40% |
| hsa_circ_0046693 | 1431 | 10 | 1.40% |
| hsa_circ_0057834 | 1431 | 10 | 1.40% |
| hsa_circ_0108352 | 1431 | 10 | 1.40% |
| hsa_circ_0109974 | 1431 | 10 | 1.40% |
| hsa_circ_0110673 | 1431 | 10 | 1.40% |
| hsa_circ_0135356 | 1431 | 10 | 1.40% |
| hsa_circ_0107723 | 3006 | 21 | 1.40% |
| hsa_circ_0080091 | 2004 | 14 | 1.40% |
| hsa_circ_0045655 | 1861 | 13 | 1.40% |
| hsa_circ_0128898 | 1861 | 13 | 1.40% |
| hsa_circ_0031631 | 1718 | 12 | 1.40% |
| hsa_circ_0113130 | 1718 | 12 | 1.40% |
| hsa_circ_0119940 | 1718 | 12 | 1.40% |
| hsa_circ_0124189 | 1718 | 12 | 1.40% |
| hsa_circ_0015633 | 1575 | 11 | 1.40% |
| hsa_circ_0022087 | 1575 | 11 | 1.40% |
| hsa_circ_0024056 | 1575 | 11 | 1.40% |
| hsa_circ_0045734 | 1575 | 11 | 1.40% |
| hsa_circ_0056851 | 1575 | 11 | 1.40% |
| hsa_circ_0060345 | 1575 | 11 | 1.40% |
| hsa_circ_0086582 | 2148 | 15 | 1.40% |
| hsa_circ_0016537 | 1432 | 10 | 1.40% |
| hsa_circ_0019971 | 1432 | 10 | 1.40% |
| hsa_circ_0034289 | 1432 | 10 | 1.40% |
| hsa_circ_0058145 | 1432 | 10 | 1.40% |
| hsa_circ_0089943 | 1432 | 10 | 1.40% |
| hsa_circ_0106375 | 1432 | 10 | 1.40% |
| hsa_circ_0116499 | 4583 | 32 | 1.40% |
| hsa_circ_0061600 | 1862 | 13 | 1.40% |
| hsa_circ_0051466 | 1719 | 12 | 1.40% |
| hsa_circ_0083107 | 1719 | 12 | 1.40% |
| hsa_circ_0018794 | 2149 | 15 | 1.40% |
| hsa_circ_0014480 | 1576 | 11 | 1.40% |
| hsa_circ_0022800 | 1576 | 11 | 1.40% |
| hsa_circ_0040260 | 1576 | 11 | 1.40% |
| hsa_circ_0049442 | 1576 | 11 | 1.40% |
| hsa_circ_0069194 | 1576 | 11 | 1.40% |
| hsa_circ_0074591 | 1576 | 11 | 1.40% |
| hsa_circ_0076122 | 1576 | 11 | 1.40% |
| hsa_circ_0078041 | 1576 | 11 | 1.40% |
| hsa_circ_0022049 | 2006 | 14 | 1.40% |

| | | | |
|---|---|---|---|
| hsa_circ_0016877 | 1433 | 10 | 1.40% |
| hsa_circ_0028593 | 1433 | 10 | 1.40% |
| hsa_circ_0056349 | 1433 | 10 | 1.40% |
| hsa_circ_0061253 | 1433 | 10 | 1.40% |
| hsa_circ_0064106 | 1433 | 10 | 1.40% |
| hsa_circ_0079413 | 1433 | 10 | 1.40% |
| hsa_circ_0110269 | 1433 | 10 | 1.40% |
| hsa_circ_0137289 | 1433 | 10 | 1.40% |
| hsa_circ_0011102 | 1720 | 12 | 1.40% |
| hsa_circ_0034501 | 1720 | 12 | 1.40% |
| hsa_circ_0067599 | 1720 | 12 | 1.40% |
| hsa_circ_0015538 | 1577 | 11 | 1.40% |
| hsa_circ_0017474 | 1577 | 11 | 1.40% |
| hsa_circ_0018934 | 1577 | 11 | 1.40% |
| hsa_circ_0065962 | 1577 | 11 | 1.40% |
| hsa_circ_0066750 | 1577 | 11 | 1.40% |
| hsa_circ_0071467 | 1577 | 11 | 1.40% |
| hsa_circ_0117854 | 1577 | 11 | 1.40% |
| hsa_circ_0022615 | 1434 | 10 | 1.39% |
| hsa_circ_0034256 | 1434 | 10 | 1.39% |
| hsa_circ_0049645 | 1434 | 10 | 1.39% |
| hsa_circ_0090381 | 1434 | 10 | 1.39% |
| hsa_circ_0108329 | 1434 | 10 | 1.39% |
| hsa_circ_0116727 | 1434 | 10 | 1.39% |
| hsa_circ_0089043 | 1721 | 12 | 1.39% |
| hsa_circ_0094715 | 1721 | 12 | 1.39% |
| hsa_circ_0113373 | 1721 | 12 | 1.39% |
| hsa_circ_0139904 | 1721 | 12 | 1.39% |
| hsa_circ_0000837 | 1578 | 11 | 1.39% |
| hsa_circ_0024327 | 1578 | 11 | 1.39% |
| hsa_circ_0044608 | 1578 | 11 | 1.39% |
| hsa_circ_0048355 | 1578 | 11 | 1.39% |
| hsa_circ_0073018 | 1578 | 11 | 1.39% |
| hsa_circ_0087691 | 1578 | 11 | 1.39% |
| hsa_circ_0136161 | 1578 | 11 | 1.39% |
| hsa_circ_0137790 | 1578 | 11 | 1.39% |
| hsa_circ_0139924 | 1578 | 11 | 1.39% |
| hsa_circ_0047416 | 1865 | 13 | 1.39% |
| hsa_circ_0086034 | 1865 | 13 | 1.39% |
| hsa_circ_0034744 | 2009 | 14 | 1.39% |
| hsa_circ_0034493 | 1722 | 12 | 1.39% |
| hsa_circ_0072956 | 1722 | 12 | 1.39% |
| hsa_circ_0040872 | 1435 | 10 | 1.39% |
| hsa_circ_0046630 | 1435 | 10 | 1.39% |
| hsa_circ_0053848 | 1435 | 10 | 1.39% |
| hsa_circ_0061837 | 1435 | 10 | 1.39% |
| hsa_circ_0069541 | 1435 | 10 | 1.39% |
| hsa_circ_0108590 | 1435 | 10 | 1.39% |
| hsa_circ_0114350 | 1435 | 10 | 1.39% |
| hsa_circ_0118455 | 1435 | 10 | 1.39% |

| | | | |
|---|---|---|---|
| hsa_circ_0132337 | 1435 | 10 | 1.39% |
| hsa_circ_0030499 | 2153 | 15 | 1.39% |
| hsa_circ_0117174 | 1866 | 13 | 1.39% |
| hsa_circ_0010561 | 1579 | 11 | 1.39% |
| hsa_circ_0039752 | 1579 | 11 | 1.39% |
| hsa_circ_0060266 | 1579 | 11 | 1.39% |
| hsa_circ_0077415 | 1579 | 11 | 1.39% |
| hsa_circ_0086654 | 1579 | 11 | 1.39% |
| hsa_circ_0025321 | 1723 | 12 | 1.39% |
| hsa_circ_0110124 | 1723 | 12 | 1.39% |
| hsa_circ_0032720 | 1436 | 10 | 1.39% |
| hsa_circ_0034835 | 1436 | 10 | 1.39% |
| hsa_circ_0035012 | 1436 | 10 | 1.39% |
| hsa_circ_0043799 | 1436 | 10 | 1.39% |
| hsa_circ_0058257 | 1436 | 10 | 1.39% |
| hsa_circ_0069817 | 1436 | 10 | 1.39% |
| hsa_circ_0087980 | 1436 | 10 | 1.39% |
| hsa_circ_0071461 | 4596 | 32 | 1.39% |
| hsa_circ_0009606 | 1580 | 11 | 1.39% |
| hsa_circ_0041448 | 1580 | 11 | 1.39% |
| hsa_circ_0046267 | 1580 | 11 | 1.39% |
| hsa_circ_0052005 | 1580 | 11 | 1.39% |
| hsa_circ_0052497 | 1580 | 11 | 1.39% |
| hsa_circ_0091721 | 1580 | 11 | 1.39% |
| hsa_circ_0036954 | 2011 | 14 | 1.39% |
| hsa_circ_0059558 | 2011 | 14 | 1.39% |
| hsa_circ_0124009 | 4166 | 29 | 1.39% |
| hsa_circ_0016951 | 1724 | 12 | 1.39% |
| hsa_circ_0035267 | 1724 | 12 | 1.39% |
| hsa_circ_0047024 | 1868 | 13 | 1.39% |
| hsa_circ_0010524 | 1437 | 10 | 1.39% |
| hsa_circ_0025370 | 1437 | 10 | 1.39% |
| hsa_circ_0025404 | 1437 | 10 | 1.39% |
| hsa_circ_0038327 | 1437 | 10 | 1.39% |
| hsa_circ_0042602 | 1437 | 10 | 1.39% |
| hsa_circ_0056699 | 1437 | 10 | 1.39% |
| hsa_circ_0058137 | 1437 | 10 | 1.39% |
| hsa_circ_0075475 | 1437 | 10 | 1.39% |
| hsa_circ_0086372 | 1437 | 10 | 1.39% |
| hsa_circ_0089026 | 1437 | 10 | 1.39% |
| hsa_circ_0123620 | 1437 | 10 | 1.39% |
| hsa_circ_0138722 | 1437 | 10 | 1.39% |
| hsa_circ_0121407 | 6611 | 46 | 1.39% |
| hsa_circ_0016669 | 1581 | 11 | 1.39% |
| hsa_circ_0019923 | 1581 | 11 | 1.39% |
| hsa_circ_0029557 | 1581 | 11 | 1.39% |
| hsa_circ_0029716 | 1581 | 11 | 1.39% |
| hsa_circ_0031751 | 1581 | 11 | 1.39% |
| hsa_circ_0045052 | 1581 | 11 | 1.39% |
| hsa_circ_0072750 | 1725 | 12 | 1.39% |

| | | | |
|---|---|---|---|
| hsa_circ_0045819 | 2444 | 17 | 1.39% |
| hsa_circ_0100722 | 1869 | 13 | 1.39% |
| hsa_circ_0086182 | 2013 | 14 | 1.39% |
| hsa_circ_0048408 | 2157 | 15 | 1.39% |
| hsa_circ_0016588 | 1438 | 10 | 1.39% |
| hsa_circ_0039139 | 1438 | 10 | 1.39% |
| hsa_circ_0061922 | 1438 | 10 | 1.39% |
| hsa_circ_0069623 | 1438 | 10 | 1.39% |
| hsa_circ_0090026 | 1438 | 10 | 1.39% |
| hsa_circ_0090411 | 1438 | 10 | 1.39% |
| hsa_circ_0091745 | 1438 | 10 | 1.39% |
| hsa_circ_0101787 | 1438 | 10 | 1.39% |
| hsa_circ_0102101 | 1438 | 10 | 1.39% |
| hsa_circ_0111510 | 1438 | 10 | 1.39% |
| hsa_circ_0057279 | 2301 | 16 | 1.39% |
| hsa_circ_0067493 | 1582 | 11 | 1.39% |
| hsa_circ_0070238 | 1582 | 11 | 1.39% |
| hsa_circ_0073158 | 1582 | 11 | 1.39% |
| hsa_circ_0076005 | 1582 | 11 | 1.39% |
| hsa_circ_0080280 | 1582 | 11 | 1.39% |
| hsa_circ_0117859 | 1582 | 11 | 1.39% |
| hsa_circ_0107309 | 2445 | 17 | 1.39% |
| hsa_circ_0018820 | 1726 | 12 | 1.39% |
| hsa_circ_0048031 | 3596 | 25 | 1.39% |
| hsa_circ_0069972 | 2014 | 14 | 1.39% |
| hsa_circ_0004156 | 1439 | 10 | 1.39% |
| hsa_circ_0019292 | 1439 | 10 | 1.39% |
| hsa_circ_0034514 | 1439 | 10 | 1.39% |
| hsa_circ_0036150 | 1439 | 10 | 1.39% |
| hsa_circ_0041713 | 1439 | 10 | 1.39% |
| hsa_circ_0060567 | 1439 | 10 | 1.39% |
| hsa_circ_0065603 | 1439 | 10 | 1.39% |
| hsa_circ_0065769 | 1439 | 10 | 1.39% |
| hsa_circ_0072938 | 1439 | 10 | 1.39% |
| hsa_circ_0082041 | 1439 | 10 | 1.39% |
| hsa_circ_0082894 | 1439 | 10 | 1.39% |
| hsa_circ_0085750 | 1439 | 10 | 1.39% |
| hsa_circ_0101891 | 1439 | 10 | 1.39% |
| hsa_circ_0104415 | 1439 | 10 | 1.39% |
| hsa_circ_0014032 | 1583 | 11 | 1.39% |
| hsa_circ_0017015 | 1583 | 11 | 1.39% |
| hsa_circ_0028526 | 1583 | 11 | 1.39% |
| hsa_circ_0052075 | 1583 | 11 | 1.39% |
| hsa_circ_0053078 | 1583 | 11 | 1.39% |
| hsa_circ_0068633 | 1583 | 11 | 1.39% |
| hsa_circ_0088841 | 1583 | 11 | 1.39% |
| hsa_circ_0016373 | 1727 | 12 | 1.39% |
| hsa_circ_0109980 | 4319 | 30 | 1.39% |
| hsa_circ_0031961 | 3456 | 24 | 1.39% |
| hsa_circ_0064013 | 3456 | 24 | 1.39% |

| | | | |
|---|---|---|---|
| hsa_circ_0075418 | 1872 | 13 | 1.39% |
| hsa_circ_0035270 | 1728 | 12 | 1.39% |
| hsa_circ_0085296 | 1728 | 12 | 1.39% |
| hsa_circ_0091464 | 1728 | 12 | 1.39% |
| hsa_circ_0034178 | 1584 | 11 | 1.39% |
| hsa_circ_0050325 | 1584 | 11 | 1.39% |
| hsa_circ_0060177 | 1584 | 11 | 1.39% |
| hsa_circ_0060483 | 1584 | 11 | 1.39% |
| hsa_circ_0064294 | 1584 | 11 | 1.39% |
| hsa_circ_0069888 | 1584 | 11 | 1.39% |
| hsa_circ_0071286 | 1584 | 11 | 1.39% |
| hsa_circ_0122753 | 1584 | 11 | 1.39% |
| hsa_circ_0016028 | 1440 | 10 | 1.39% |
| hsa_circ_0017389 | 1440 | 10 | 1.39% |
| hsa_circ_0035447 | 1440 | 10 | 1.39% |
| hsa_circ_0071573 | 1440 | 10 | 1.39% |
| hsa_circ_0078739 | 1440 | 10 | 1.39% |
| hsa_circ_0074221 | 1873 | 13 | 1.39% |
| hsa_circ_0020663 | 1729 | 12 | 1.39% |
| hsa_circ_0049472 | 1729 | 12 | 1.39% |
| hsa_circ_0049522 | 1729 | 12 | 1.39% |
| hsa_circ_0063589 | 1729 | 12 | 1.39% |
| hsa_circ_0119428 | 1729 | 12 | 1.39% |
| hsa_circ_0012959 | 1585 | 11 | 1.39% |
| hsa_circ_0058312 | 1585 | 11 | 1.39% |
| hsa_circ_0099562 | 1585 | 11 | 1.39% |
| hsa_circ_0106484 | 1585 | 11 | 1.39% |
| hsa_circ_0107821 | 1585 | 11 | 1.39% |
| hsa_circ_0112836 | 1585 | 11 | 1.39% |
| hsa_circ_0122287 | 1585 | 11 | 1.39% |
| hsa_circ_0135166 | 2882 | 20 | 1.39% |
| hsa_circ_0016744 | 1441 | 10 | 1.39% |
| hsa_circ_0037696 | 1441 | 10 | 1.39% |
| hsa_circ_0047532 | 1441 | 10 | 1.39% |
| hsa_circ_0048148 | 1441 | 10 | 1.39% |
| hsa_circ_0065206 | 1441 | 10 | 1.39% |
| hsa_circ_0081396 | 1441 | 10 | 1.39% |
| hsa_circ_0089378 | 1441 | 10 | 1.39% |
| hsa_circ_0089835 | 1441 | 10 | 1.39% |
| hsa_circ_0090237 | 1441 | 10 | 1.39% |
| hsa_circ_0107054 | 1441 | 10 | 1.39% |
| hsa_circ_0133014 | 1441 | 10 | 1.39% |
| hsa_circ_0073742 | 2883 | 20 | 1.39% |
| hsa_circ_0081915 | 1874 | 13 | 1.39% |
| hsa_circ_0083417 | 1874 | 13 | 1.39% |
| hsa_circ_0052280 | 1730 | 12 | 1.39% |
| hsa_circ_0057450 | 1730 | 12 | 1.39% |
| hsa_circ_0084471 | 1730 | 12 | 1.39% |
| hsa_circ_0129404 | 1730 | 12 | 1.39% |
| hsa_circ_0014442 | 1586 | 11 | 1.39% |

| | | | |
|---|---|---|---|
| hsa_circ_0015715 | 1586 | 11 | 1.39% |
| hsa_circ_0054879 | 1586 | 11 | 1.39% |
| hsa_circ_0065244 | 1586 | 11 | 1.39% |
| hsa_circ_0068199 | 1586 | 11 | 1.39% |
| hsa_circ_0111796 | 1586 | 11 | 1.39% |
| hsa_circ_0113025 | 1586 | 11 | 1.39% |
| hsa_circ_0014255 | 1442 | 10 | 1.39% |
| hsa_circ_0020387 | 1442 | 10 | 1.39% |
| hsa_circ_0033551 | 1442 | 10 | 1.39% |
| hsa_circ_0034454 | 1442 | 10 | 1.39% |
| hsa_circ_0071689 | 1442 | 10 | 1.39% |
| hsa_circ_0095564 | 1442 | 10 | 1.39% |
| hsa_circ_0013773 | 1587 | 11 | 1.39% |
| hsa_circ_0066070 | 1587 | 11 | 1.39% |
| hsa_circ_0078434 | 1587 | 11 | 1.39% |
| hsa_circ_0080746 | 1587 | 11 | 1.39% |
| hsa_circ_0091124 | 1587 | 11 | 1.39% |
| hsa_circ_0106929 | 1587 | 11 | 1.39% |
| hsa_circ_0066651 | 2597 | 18 | 1.39% |
| hsa_circ_0002553 | 1443 | 10 | 1.39% |
| hsa_circ_0019206 | 1443 | 10 | 1.39% |
| hsa_circ_0025308 | 1443 | 10 | 1.39% |
| hsa_circ_0047393 | 1443 | 10 | 1.39% |
| hsa_circ_0077399 | 1443 | 10 | 1.39% |
| hsa_circ_0117463 | 1443 | 10 | 1.39% |
| hsa_circ_0058943 | 1876 | 13 | 1.39% |
| hsa_circ_0024785 | 1732 | 12 | 1.39% |
| hsa_circ_0112053 | 1732 | 12 | 1.39% |
| hsa_circ_0140487 | 1732 | 12 | 1.39% |
| hsa_circ_0021771 | 1588 | 11 | 1.39% |
| hsa_circ_0035947 | 1588 | 11 | 1.39% |
| hsa_circ_0039005 | 1588 | 11 | 1.39% |
| hsa_circ_0048147 | 1588 | 11 | 1.39% |
| hsa_circ_0074026 | 1588 | 11 | 1.39% |
| hsa_circ_0084630 | 1588 | 11 | 1.39% |
| hsa_circ_0140483 | 1588 | 11 | 1.39% |
| hsa_circ_0045531 | 1877 | 13 | 1.39% |
| hsa_circ_0011943 | 1444 | 10 | 1.39% |
| hsa_circ_0014755 | 1444 | 10 | 1.39% |
| hsa_circ_0022939 | 1444 | 10 | 1.39% |
| hsa_circ_0050175 | 1444 | 10 | 1.39% |
| hsa_circ_0051638 | 1444 | 10 | 1.39% |
| hsa_circ_0068285 | 1444 | 10 | 1.39% |
| hsa_circ_0104178 | 1444 | 10 | 1.39% |
| hsa_circ_0115618 | 1444 | 10 | 1.39% |
| hsa_circ_0121560 | 1444 | 10 | 1.39% |
| hsa_circ_0017576 | 1733 | 12 | 1.38% |
| hsa_circ_0017654 | 1733 | 12 | 1.38% |
| hsa_circ_0056845 | 1733 | 12 | 1.38% |
| hsa_circ_0028928 | 1589 | 11 | 1.38% |

| | | | |
|---|---|---|---|
| hsa_circ_0041124 | 1589 | 11 | 1.38% |
| hsa_circ_0098526 | 1589 | 11 | 1.38% |
| hsa_circ_0122975 | 1589 | 11 | 1.38% |
| hsa_circ_0121222 | 4046 | 28 | 1.38% |
| hsa_circ_0135853 | 2023 | 14 | 1.38% |
| hsa_circ_0077240 | 1734 | 12 | 1.38% |
| hsa_circ_0011646 | 1445 | 10 | 1.38% |
| hsa_circ_0015020 | 1445 | 10 | 1.38% |
| hsa_circ_0018902 | 1445 | 10 | 1.38% |
| hsa_circ_0027938 | 1445 | 10 | 1.38% |
| hsa_circ_0039166 | 1445 | 10 | 1.38% |
| hsa_circ_0041853 | 1445 | 10 | 1.38% |
| hsa_circ_0064238 | 1445 | 10 | 1.38% |
| hsa_circ_0085978 | 1445 | 10 | 1.38% |
| hsa_circ_0099168 | 1445 | 10 | 1.38% |
| hsa_circ_0106395 | 1445 | 10 | 1.38% |
| hsa_circ_0127356 | 1445 | 10 | 1.38% |
| hsa_circ_0128056 | 1445 | 10 | 1.38% |
| hsa_circ_0016729 | 1590 | 11 | 1.38% |
| hsa_circ_0021583 | 1590 | 11 | 1.38% |
| hsa_circ_0051155 | 1590 | 11 | 1.38% |
| hsa_circ_0052266 | 1590 | 11 | 1.38% |
| hsa_circ_0076861 | 1590 | 11 | 1.38% |
| hsa_circ_0083647 | 1590 | 11 | 1.38% |
| hsa_circ_0094405 | 1590 | 11 | 1.38% |
| hsa_circ_0102460 | 1590 | 11 | 1.38% |
| hsa_circ_0103961 | 1590 | 11 | 1.38% |
| hsa_circ_0108060 | 1590 | 11 | 1.38% |
| hsa_circ_0140369 | 1590 | 11 | 1.38% |
| hsa_circ_0124053 | 4915 | 34 | 1.38% |
| hsa_circ_0070538 | 1735 | 12 | 1.38% |
| hsa_circ_0011009 | 1446 | 10 | 1.38% |
| hsa_circ_0011478 | 1446 | 10 | 1.38% |
| hsa_circ_0015239 | 1446 | 10 | 1.38% |
| hsa_circ_0021173 | 1446 | 10 | 1.38% |
| hsa_circ_0024568 | 1446 | 10 | 1.38% |
| hsa_circ_0030004 | 1446 | 10 | 1.38% |
| hsa_circ_0032298 | 1446 | 10 | 1.38% |
| hsa_circ_0038706 | 1446 | 10 | 1.38% |
| hsa_circ_0044026 | 1446 | 10 | 1.38% |
| hsa_circ_0045170 | 1446 | 10 | 1.38% |
| hsa_circ_0056761 | 1446 | 10 | 1.38% |
| hsa_circ_0060655 | 1446 | 10 | 1.38% |
| hsa_circ_0064669 | 1446 | 10 | 1.38% |
| hsa_circ_0070240 | 1446 | 10 | 1.38% |
| hsa_circ_0070610 | 1446 | 10 | 1.38% |
| hsa_circ_0072506 | 1446 | 10 | 1.38% |
| hsa_circ_0088407 | 1446 | 10 | 1.38% |
| hsa_circ_0094383 | 1446 | 10 | 1.38% |
| hsa_circ_0108598 | 1446 | 10 | 1.38% |

| | | | |
|---|---|---|---|
| hsa_circ_0130264 | 1446 | 10 | 1.38% |
| hsa_circ_0135252 | 1446 | 10 | 1.38% |
| hsa_circ_0010407 | 1591 | 11 | 1.38% |
| hsa_circ_0011110 | 1591 | 11 | 1.38% |
| hsa_circ_0047170 | 1591 | 11 | 1.38% |
| hsa_circ_0083541 | 1591 | 11 | 1.38% |
| hsa_circ_0037627 | 1736 | 12 | 1.38% |
| hsa_circ_0060731 | 1736 | 12 | 1.38% |
| hsa_circ_0074497 | 1736 | 12 | 1.38% |
| hsa_circ_0064621 | 1881 | 13 | 1.38% |
| hsa_circ_0019465 | 1447 | 10 | 1.38% |
| hsa_circ_0021912 | 1447 | 10 | 1.38% |
| hsa_circ_0071812 | 1447 | 10 | 1.38% |
| hsa_circ_0075694 | 1447 | 10 | 1.38% |
| hsa_circ_0107349 | 1447 | 10 | 1.38% |
| hsa_circ_0140270 | 1447 | 10 | 1.38% |
| hsa_circ_0140715 | 3039 | 21 | 1.38% |
| hsa_circ_0026935 | 2026 | 14 | 1.38% |
| hsa_circ_0079849 | 2026 | 14 | 1.38% |
| hsa_circ_0014337 | 1592 | 11 | 1.38% |
| hsa_circ_0121408 | 1592 | 11 | 1.38% |
| hsa_circ_0067267 | 2171 | 15 | 1.38% |
| hsa_circ_0014840 | 1737 | 12 | 1.38% |
| hsa_circ_0080655 | 1737 | 12 | 1.38% |
| hsa_circ_0082425 | 1737 | 12 | 1.38% |
| hsa_circ_0129775 | 1737 | 12 | 1.38% |
| hsa_circ_0084064 | 2027 | 14 | 1.38% |
| hsa_circ_0107685 | 4199 | 29 | 1.38% |
| hsa_circ_0044283 | 1448 | 10 | 1.38% |
| hsa_circ_0050523 | 1448 | 10 | 1.38% |
| hsa_circ_0055846 | 1448 | 10 | 1.38% |
| hsa_circ_0062720 | 1448 | 10 | 1.38% |
| hsa_circ_0067544 | 1448 | 10 | 1.38% |
| hsa_circ_0123819 | 1448 | 10 | 1.38% |
| hsa_circ_0077114 | 1593 | 11 | 1.38% |
| hsa_circ_0092730 | 1593 | 11 | 1.38% |
| hsa_circ_0095315 | 1593 | 11 | 1.38% |
| hsa_circ_0065956 | 1738 | 12 | 1.38% |
| hsa_circ_0017313 | 1449 | 10 | 1.38% |
| hsa_circ_0040163 | 1449 | 10 | 1.38% |
| hsa_circ_0053855 | 1449 | 10 | 1.38% |
| hsa_circ_0065992 | 1449 | 10 | 1.38% |
| hsa_circ_0073456 | 1449 | 10 | 1.38% |
| hsa_circ_0085752 | 1449 | 10 | 1.38% |
| hsa_circ_0092078 | 1449 | 10 | 1.38% |
| hsa_circ_0094008 | 1449 | 10 | 1.38% |
| hsa_circ_0111135 | 1449 | 10 | 1.38% |
| hsa_circ_0117772 | 1449 | 10 | 1.38% |
| hsa_circ_0128003 | 1449 | 10 | 1.38% |
| hsa_circ_0021187 | 1594 | 11 | 1.38% |

| | | | |
|---|---|---|---|
| hsa_circ_0023068 | 1594 | 11 | 1.38% |
| hsa_circ_0092880 | 1594 | 11 | 1.38% |
| hsa_circ_0112462 | 1594 | 11 | 1.38% |
| hsa_circ_0124966 | 1594 | 11 | 1.38% |
| hsa_circ_0023166 | 1739 | 12 | 1.38% |
| hsa_circ_0056161 | 1739 | 12 | 1.38% |
| hsa_circ_0078450 | 1739 | 12 | 1.38% |
| hsa_circ_0080886 | 1739 | 12 | 1.38% |
| hsa_circ_0094690 | 1739 | 12 | 1.38% |
| hsa_circ_0017260 | 1884 | 13 | 1.38% |
| hsa_circ_0053212 | 1884 | 13 | 1.38% |
| hsa_circ_0126614 | 2465 | 17 | 1.38% |
| hsa_circ_0011897 | 1740 | 12 | 1.38% |
| hsa_circ_0040755 | 1595 | 11 | 1.38% |
| hsa_circ_0042718 | 1595 | 11 | 1.38% |
| hsa_circ_0045301 | 1595 | 11 | 1.38% |
| hsa_circ_0063915 | 1595 | 11 | 1.38% |
| hsa_circ_0125586 | 1595 | 11 | 1.38% |
| hsa_circ_0004527 | 1450 | 10 | 1.38% |
| hsa_circ_0007725 | 1450 | 10 | 1.38% |
| hsa_circ_0011202 | 1450 | 10 | 1.38% |
| hsa_circ_0055733 | 1450 | 10 | 1.38% |
| hsa_circ_0058824 | 1450 | 10 | 1.38% |
| hsa_circ_0062893 | 1450 | 10 | 1.38% |
| hsa_circ_0070229 | 1450 | 10 | 1.38% |
| hsa_circ_0084644 | 1450 | 10 | 1.38% |
| hsa_circ_0089139 | 1450 | 10 | 1.38% |
| hsa_circ_0121795 | 1450 | 10 | 1.38% |
| hsa_circ_0129880 | 1450 | 10 | 1.38% |
| hsa_circ_0086116 | 1741 | 12 | 1.38% |
| hsa_circ_0010236 | 1596 | 11 | 1.38% |
| hsa_circ_0032598 | 1596 | 11 | 1.38% |
| hsa_circ_0053704 | 1596 | 11 | 1.38% |
| hsa_circ_0081317 | 1596 | 11 | 1.38% |
| hsa_circ_0015017 | 1451 | 10 | 1.38% |
| hsa_circ_0015564 | 1451 | 10 | 1.38% |
| hsa_circ_0032437 | 1451 | 10 | 1.38% |
| hsa_circ_0036174 | 1451 | 10 | 1.38% |
| hsa_circ_0040917 | 1451 | 10 | 1.38% |
| hsa_circ_0041610 | 1451 | 10 | 1.38% |
| hsa_circ_0081698 | 1451 | 10 | 1.38% |
| hsa_circ_0087297 | 1451 | 10 | 1.38% |
| hsa_circ_0090608 | 1451 | 10 | 1.38% |
| hsa_circ_0096729 | 1451 | 10 | 1.38% |
| hsa_circ_0130561 | 1451 | 10 | 1.38% |
| hsa_circ_0131422 | 1451 | 10 | 1.38% |
| hsa_circ_0134621 | 1451 | 10 | 1.38% |
| hsa_circ_0012996 | 1887 | 13 | 1.38% |
| hsa_circ_0055335 | 1887 | 13 | 1.38% |
| hsa_circ_0040313 | 1742 | 12 | 1.38% |

| | | | |
|---|---|---|---|
| hsa_circ_0040701 | 1742 | 12 | 1.38% |
| hsa_circ_0049748 | 1742 | 12 | 1.38% |
| hsa_circ_0071568 | 1742 | 12 | 1.38% |
| hsa_circ_0023171 | 1597 | 11 | 1.38% |
| hsa_circ_0026578 | 1597 | 11 | 1.38% |
| hsa_circ_0048367 | 1597 | 11 | 1.38% |
| hsa_circ_0059901 | 1597 | 11 | 1.38% |
| hsa_circ_0114455 | 5372 | 37 | 1.38% |
| hsa_circ_0020768 | 1452 | 10 | 1.38% |
| hsa_circ_0054647 | 1452 | 10 | 1.38% |
| hsa_circ_0057020 | 1452 | 10 | 1.38% |
| hsa_circ_0057253 | 1452 | 10 | 1.38% |
| hsa_circ_0076082 | 1452 | 10 | 1.38% |
| hsa_circ_0076730 | 1452 | 10 | 1.38% |
| hsa_circ_0093074 | 1452 | 10 | 1.38% |
| hsa_circ_0103641 | 1452 | 10 | 1.38% |
| hsa_circ_0113013 | 1452 | 10 | 1.38% |
| hsa_circ_0122967 | 1452 | 10 | 1.38% |
| hsa_circ_0124671 | 1452 | 10 | 1.38% |
| hsa_circ_0136804 | 1452 | 10 | 1.38% |
| hsa_circ_0108170 | 1888 | 13 | 1.38% |
| hsa_circ_0027176 | 1743 | 12 | 1.38% |
| hsa_circ_0087210 | 1743 | 12 | 1.38% |
| hsa_circ_0036051 | 1598 | 11 | 1.38% |
| hsa_circ_0038256 | 1598 | 11 | 1.38% |
| hsa_circ_0043197 | 1598 | 11 | 1.38% |
| hsa_circ_0072124 | 1598 | 11 | 1.38% |
| hsa_circ_0091999 | 1598 | 11 | 1.38% |
| hsa_circ_0093060 | 1598 | 11 | 1.38% |
| hsa_circ_0108580 | 1598 | 11 | 1.38% |
| hsa_circ_0114378 | 1598 | 11 | 1.38% |
| hsa_circ_0114568 | 1598 | 11 | 1.38% |
| hsa_circ_0139575 | 1598 | 11 | 1.38% |
| hsa_circ_0009886 | 1453 | 10 | 1.38% |
| hsa_circ_0012719 | 1453 | 10 | 1.38% |
| hsa_circ_0036767 | 1453 | 10 | 1.38% |
| hsa_circ_0072885 | 1453 | 10 | 1.38% |
| hsa_circ_0073910 | 1453 | 10 | 1.38% |
| hsa_circ_0082497 | 1453 | 10 | 1.38% |
| hsa_circ_0086011 | 1453 | 10 | 1.38% |
| hsa_circ_0098019 | 1453 | 10 | 1.38% |
| hsa_circ_0099990 | 1453 | 10 | 1.38% |
| hsa_circ_0106501 | 1453 | 10 | 1.38% |
| hsa_circ_0124962 | 1453 | 10 | 1.38% |
| hsa_circ_0135013 | 1453 | 10 | 1.38% |
| hsa_circ_0114905 | 4214 | 29 | 1.38% |
| hsa_circ_0062314 | 3633 | 25 | 1.38% |
| hsa_circ_0000387 | 1744 | 12 | 1.38% |
| hsa_circ_0000417 | 1599 | 11 | 1.38% |
| hsa_circ_0013502 | 1599 | 11 | 1.38% |

| | | | |
|---|---|---|---|
| hsa_circ_0025338 | 1599 | 11 | 1.38% |
| hsa_circ_0048450 | 1599 | 11 | 1.38% |
| hsa_circ_0072324 | 1599 | 11 | 1.38% |
| hsa_circ_0095193 | 1599 | 11 | 1.38% |
| hsa_circ_0096833 | 1599 | 11 | 1.38% |
| hsa_circ_0112051 | 1599 | 11 | 1.38% |
| hsa_circ_0123442 | 1599 | 11 | 1.38% |
| hsa_circ_0131888 | 1599 | 11 | 1.38% |
| hsa_circ_0088569 | 1890 | 13 | 1.38% |
| hsa_circ_0092107 | 1890 | 13 | 1.38% |
| hsa_circ_0119168 | 2908 | 20 | 1.38% |
| hsa_circ_0014897 | 1454 | 10 | 1.38% |
| hsa_circ_0015936 | 1454 | 10 | 1.38% |
| hsa_circ_0020973 | 1454 | 10 | 1.38% |
| hsa_circ_0023500 | 1454 | 10 | 1.38% |
| hsa_circ_0023801 | 1454 | 10 | 1.38% |
| hsa_circ_0052178 | 1454 | 10 | 1.38% |
| hsa_circ_0052325 | 1454 | 10 | 1.38% |
| hsa_circ_0052766 | 1454 | 10 | 1.38% |
| hsa_circ_0056077 | 1454 | 10 | 1.38% |
| hsa_circ_0081660 | 1454 | 10 | 1.38% |
| hsa_circ_0103290 | 1454 | 10 | 1.38% |
| hsa_circ_0128578 | 1454 | 10 | 1.38% |
| hsa_circ_0128813 | 1454 | 10 | 1.38% |
| hsa_circ_0134116 | 1454 | 10 | 1.38% |
| hsa_circ_0059235 | 1745 | 12 | 1.38% |
| hsa_circ_0063588 | 1745 | 12 | 1.38% |
| hsa_circ_0028407 | 1600 | 11 | 1.38% |
| hsa_circ_0037238 | 1600 | 11 | 1.38% |
| hsa_circ_0079853 | 1600 | 11 | 1.38% |
| hsa_circ_0087899 | 1600 | 11 | 1.38% |
| hsa_circ_0012951 | 2473 | 17 | 1.37% |
| hsa_circ_0064439 | 1746 | 12 | 1.37% |
| hsa_circ_0009866 | 1455 | 10 | 1.37% |
| hsa_circ_0013031 | 1455 | 10 | 1.37% |
| hsa_circ_0029147 | 1455 | 10 | 1.37% |
| hsa_circ_0050844 | 1455 | 10 | 1.37% |
| hsa_circ_0059733 | 1455 | 10 | 1.37% |
| hsa_circ_0072113 | 1455 | 10 | 1.37% |
| hsa_circ_0077670 | 1455 | 10 | 1.37% |
| hsa_circ_0098403 | 1455 | 10 | 1.37% |
| hsa_circ_0107264 | 1455 | 10 | 1.37% |
| hsa_circ_0111759 | 1455 | 10 | 1.37% |
| hsa_circ_0116625 | 1455 | 10 | 1.37% |
| hsa_circ_0129636 | 1455 | 10 | 1.37% |
| hsa_circ_0104906 | 3638 | 25 | 1.37% |
| hsa_circ_0101384 | 2183 | 15 | 1.37% |
| hsa_circ_0081407 | 1892 | 13 | 1.37% |
| hsa_circ_0080477 | 1601 | 11 | 1.37% |
| hsa_circ_0083383 | 1601 | 11 | 1.37% |

| | | | |
|---|---|---|---|
| hsa_circ_0109213 | 1601 | 11 | 1.37% |
| hsa_circ_0018422 | 1747 | 12 | 1.37% |
| hsa_circ_0056681 | 1747 | 12 | 1.37% |
| hsa_circ_0088035 | 1747 | 12 | 1.37% |
| hsa_circ_0021617 | 1456 | 10 | 1.37% |
| hsa_circ_0027824 | 1456 | 10 | 1.37% |
| hsa_circ_0036334 | 1456 | 10 | 1.37% |
| hsa_circ_0040442 | 1456 | 10 | 1.37% |
| hsa_circ_0049906 | 1456 | 10 | 1.37% |
| hsa_circ_0054606 | 1456 | 10 | 1.37% |
| hsa_circ_0070698 | 1456 | 10 | 1.37% |
| hsa_circ_0072306 | 1456 | 10 | 1.37% |
| hsa_circ_0074123 | 1456 | 10 | 1.37% |
| hsa_circ_0086320 | 1456 | 10 | 1.37% |
| hsa_circ_0091881 | 1456 | 10 | 1.37% |
| hsa_circ_0104122 | 1456 | 10 | 1.37% |
| hsa_circ_0124306 | 1456 | 10 | 1.37% |
| hsa_circ_0083480 | 1893 | 13 | 1.37% |
| hsa_circ_0088622 | 1893 | 13 | 1.37% |
| hsa_circ_0018325 | 1602 | 11 | 1.37% |
| hsa_circ_0034377 | 1602 | 11 | 1.37% |
| hsa_circ_0043656 | 1602 | 11 | 1.37% |
| hsa_circ_0067753 | 1602 | 11 | 1.37% |
| hsa_circ_0026943 | 2039 | 14 | 1.37% |
| hsa_circ_0059871 | 2039 | 14 | 1.37% |
| hsa_circ_0044504 | 3350 | 23 | 1.37% |
| hsa_circ_0035236 | 1748 | 12 | 1.37% |
| hsa_circ_0048898 | 1748 | 12 | 1.37% |
| hsa_circ_0106582 | 1748 | 12 | 1.37% |
| hsa_circ_0015486 | 1457 | 10 | 1.37% |
| hsa_circ_0017721 | 1457 | 10 | 1.37% |
| hsa_circ_0017977 | 1457 | 10 | 1.37% |
| hsa_circ_0026575 | 1457 | 10 | 1.37% |
| hsa_circ_0028620 | 1457 | 10 | 1.37% |
| hsa_circ_0050649 | 1457 | 10 | 1.37% |
| hsa_circ_0117810 | 1457 | 10 | 1.37% |
| hsa_circ_0133345 | 1457 | 10 | 1.37% |
| hsa_circ_0107027 | 5683 | 39 | 1.37% |
| hsa_circ_0011241 | 1603 | 11 | 1.37% |
| hsa_circ_0034551 | 1603 | 11 | 1.37% |
| hsa_circ_0046490 | 1603 | 11 | 1.37% |
| hsa_circ_0122804 | 1603 | 11 | 1.37% |
| hsa_circ_0128107 | 1603 | 11 | 1.37% |
| hsa_circ_0032145 | 1749 | 12 | 1.37% |
| hsa_circ_0031711 | 1458 | 10 | 1.37% |
| hsa_circ_0056379 | 1458 | 10 | 1.37% |
| hsa_circ_0056714 | 1458 | 10 | 1.37% |
| hsa_circ_0063120 | 1458 | 10 | 1.37% |
| hsa_circ_0066729 | 1458 | 10 | 1.37% |
| hsa_circ_0071231 | 1458 | 10 | 1.37% |

| | | | |
|---|---|---|---|
| hsa_circ_0090389 | 1458 | 10 | 1.37% |
| hsa_circ_0108816 | 1458 | 10 | 1.37% |
| hsa_circ_0109298 | 1458 | 10 | 1.37% |
| hsa_circ_0123795 | 1458 | 10 | 1.37% |
| hsa_circ_0058320 | 1604 | 11 | 1.37% |
| hsa_circ_0003023 | 1750 | 12 | 1.37% |
| hsa_circ_0056796 | 1750 | 12 | 1.37% |
| hsa_circ_0075138 | 1750 | 12 | 1.37% |
| hsa_circ_0092051 | 1750 | 12 | 1.37% |
| hsa_circ_0009712 | 1459 | 10 | 1.37% |
| hsa_circ_0010356 | 1459 | 10 | 1.37% |
| hsa_circ_0016714 | 1459 | 10 | 1.37% |
| hsa_circ_0038569 | 1459 | 10 | 1.37% |
| hsa_circ_0046050 | 1459 | 10 | 1.37% |
| hsa_circ_0085683 | 1459 | 10 | 1.37% |
| hsa_circ_0091525 | 1459 | 10 | 1.37% |
| hsa_circ_0099735 | 1459 | 10 | 1.37% |
| hsa_circ_0110674 | 1459 | 10 | 1.37% |
| hsa_circ_0019938 | 1605 | 11 | 1.37% |
| hsa_circ_0030178 | 1605 | 11 | 1.37% |
| hsa_circ_0038531 | 1605 | 11 | 1.37% |
| hsa_circ_0039918 | 1605 | 11 | 1.37% |
| hsa_circ_0041348 | 1605 | 11 | 1.37% |
| hsa_circ_0080500 | 1605 | 11 | 1.37% |
| hsa_circ_0088854 | 1605 | 11 | 1.37% |
| hsa_circ_0091348 | 1605 | 11 | 1.37% |
| hsa_circ_0100100 | 1605 | 11 | 1.37% |
| hsa_circ_0120252 | 1605 | 11 | 1.37% |
| hsa_circ_0110126 | 1751 | 12 | 1.37% |
| hsa_circ_0061451 | 1897 | 13 | 1.37% |
| hsa_circ_0066403 | 1752 | 12 | 1.37% |
| hsa_circ_0067603 | 1752 | 12 | 1.37% |
| hsa_circ_0028967 | 1606 | 11 | 1.37% |
| hsa_circ_0080073 | 1606 | 11 | 1.37% |
| hsa_circ_0098021 | 1606 | 11 | 1.37% |
| hsa_circ_0022616 | 1460 | 10 | 1.37% |
| hsa_circ_0027689 | 1460 | 10 | 1.37% |
| hsa_circ_0039579 | 1460 | 10 | 1.37% |
| hsa_circ_0042085 | 1460 | 10 | 1.37% |
| hsa_circ_0051980 | 1460 | 10 | 1.37% |
| hsa_circ_0067832 | 1460 | 10 | 1.37% |
| hsa_circ_0092441 | 1460 | 10 | 1.37% |
| hsa_circ_0064566 | 3505 | 24 | 1.37% |
| hsa_circ_0128066 | 1899 | 13 | 1.37% |
| hsa_circ_0094342 | 1753 | 12 | 1.37% |
| hsa_circ_0013089 | 1607 | 11 | 1.37% |
| hsa_circ_0049211 | 1607 | 11 | 1.37% |
| hsa_circ_0085817 | 1607 | 11 | 1.37% |
| hsa_circ_0091873 | 1607 | 11 | 1.37% |
| hsa_circ_0009646 | 1461 | 10 | 1.37% |

| | | | |
|---|---|---|---|
| hsa_circ_0034861 | 1461 | 10 | 1.37% |
| hsa_circ_0044307 | 1461 | 10 | 1.37% |
| hsa_circ_0051501 | 1461 | 10 | 1.37% |
| hsa_circ_0072047 | 1461 | 10 | 1.37% |
| hsa_circ_0104143 | 1461 | 10 | 1.37% |
| hsa_circ_0135206 | 1461 | 10 | 1.37% |
| hsa_circ_0005419 | 2777 | 19 | 1.37% |
| hsa_circ_0114206 | 1754 | 12 | 1.37% |
| hsa_circ_0133440 | 1754 | 12 | 1.37% |
| hsa_circ_0012394 | 1608 | 11 | 1.37% |
| hsa_circ_0046969 | 1608 | 11 | 1.37% |
| hsa_circ_0056411 | 1608 | 11 | 1.37% |
| hsa_circ_0067905 | 1608 | 11 | 1.37% |
| hsa_circ_0114244 | 1608 | 11 | 1.37% |
| hsa_circ_0125659 | 1608 | 11 | 1.37% |
| hsa_circ_0009652 | 6725 | 46 | 1.37% |
| hsa_circ_0133293 | 2924 | 20 | 1.37% |
| hsa_circ_0013534 | 2193 | 15 | 1.37% |
| hsa_circ_0011678 | 1462 | 10 | 1.37% |
| hsa_circ_0028425 | 1462 | 10 | 1.37% |
| hsa_circ_0056607 | 1462 | 10 | 1.37% |
| hsa_circ_0058048 | 1462 | 10 | 1.37% |
| hsa_circ_0068785 | 1462 | 10 | 1.37% |
| hsa_circ_0074590 | 1462 | 10 | 1.37% |
| hsa_circ_0090238 | 1462 | 10 | 1.37% |
| hsa_circ_0118452 | 1462 | 10 | 1.37% |
| hsa_circ_0008877 | 1609 | 11 | 1.37% |
| hsa_circ_0082893 | 1609 | 11 | 1.37% |
| hsa_circ_0111475 | 1609 | 11 | 1.37% |
| hsa_circ_0001511 | 1463 | 10 | 1.37% |
| hsa_circ_0036739 | 1463 | 10 | 1.37% |
| hsa_circ_0056524 | 1463 | 10 | 1.37% |
| hsa_circ_0099707 | 1463 | 10 | 1.37% |
| hsa_circ_0130817 | 1463 | 10 | 1.37% |
| hsa_circ_0134706 | 1463 | 10 | 1.37% |
| hsa_circ_0072609 | 1902 | 13 | 1.37% |
| hsa_circ_0026903 | 1756 | 12 | 1.37% |
| hsa_circ_0048984 | 1756 | 12 | 1.37% |
| hsa_circ_0004367 | 1610 | 11 | 1.37% |
| hsa_circ_0008071 | 1610 | 11 | 1.37% |
| hsa_circ_0044058 | 1610 | 11 | 1.37% |
| hsa_circ_0089420 | 1610 | 11 | 1.37% |
| hsa_circ_0107523 | 1610 | 11 | 1.37% |
| hsa_circ_0000054 | 1464 | 10 | 1.37% |
| hsa_circ_0038021 | 1464 | 10 | 1.37% |
| hsa_circ_0038828 | 1464 | 10 | 1.37% |
| hsa_circ_0043123 | 1464 | 10 | 1.37% |
| hsa_circ_0046568 | 1464 | 10 | 1.37% |
| hsa_circ_0054926 | 1464 | 10 | 1.37% |
| hsa_circ_0058480 | 1464 | 10 | 1.37% |

| | | | |
|---|---|---|---|
| hsa_circ_0064260 | 1464 | 10 | 1.37% |
| hsa_circ_0077567 | 1464 | 10 | 1.37% |
| hsa_circ_0087721 | 1464 | 10 | 1.37% |
| hsa_circ_0116890 | 1464 | 10 | 1.37% |
| hsa_circ_0123554 | 1464 | 10 | 1.37% |
| hsa_circ_0130025 | 1464 | 10 | 1.37% |
| hsa_circ_0138180 | 1464 | 10 | 1.37% |
| hsa_circ_0022732 | 1757 | 12 | 1.37% |
| hsa_circ_0050571 | 1757 | 12 | 1.37% |
| hsa_circ_0010043 | 1611 | 11 | 1.37% |
| hsa_circ_0051259 | 1611 | 11 | 1.37% |
| hsa_circ_0113407 | 1611 | 11 | 1.37% |
| hsa_circ_0009099 | 5860 | 40 | 1.37% |
| hsa_circ_0025860 | 2637 | 18 | 1.37% |
| hsa_circ_0020080 | 1758 | 12 | 1.37% |
| hsa_circ_0039371 | 1758 | 12 | 1.37% |
| hsa_circ_0066747 | 1758 | 12 | 1.37% |
| hsa_circ_0071556 | 1758 | 12 | 1.37% |
| hsa_circ_0074047 | 1758 | 12 | 1.37% |
| hsa_circ_0020219 | 1465 | 10 | 1.37% |
| hsa_circ_0057086 | 1465 | 10 | 1.37% |
| hsa_circ_0069519 | 1465 | 10 | 1.37% |
| hsa_circ_0101072 | 1465 | 10 | 1.37% |
| hsa_circ_0111897 | 1465 | 10 | 1.37% |
| hsa_circ_0086269 | 1905 | 13 | 1.36% |
| hsa_circ_0011715 | 1612 | 11 | 1.36% |
| hsa_circ_0017432 | 1612 | 11 | 1.36% |
| hsa_circ_0038977 | 1612 | 11 | 1.36% |
| hsa_circ_0040420 | 1612 | 11 | 1.36% |
| hsa_circ_0120456 | 2931 | 20 | 1.36% |
| hsa_circ_0066861 | 4837 | 33 | 1.36% |
| hsa_circ_0128971 | 1759 | 12 | 1.36% |
| hsa_circ_0014256 | 1466 | 10 | 1.36% |
| hsa_circ_0029486 | 1466 | 10 | 1.36% |
| hsa_circ_0033460 | 1466 | 10 | 1.36% |
| hsa_circ_0050511 | 1466 | 10 | 1.36% |
| hsa_circ_0053405 | 1466 | 10 | 1.36% |
| hsa_circ_0054315 | 1466 | 10 | 1.36% |
| hsa_circ_0066174 | 1466 | 10 | 1.36% |
| hsa_circ_0110357 | 1466 | 10 | 1.36% |
| hsa_circ_0119594 | 1466 | 10 | 1.36% |
| hsa_circ_0130840 | 1466 | 10 | 1.36% |
| hsa_circ_0132390 | 1466 | 10 | 1.36% |
| hsa_circ_0139823 | 1906 | 13 | 1.36% |
| hsa_circ_0031962 | 3519 | 24 | 1.36% |
| hsa_circ_0011144 | 2346 | 16 | 1.36% |
| hsa_circ_0021390 | 1613 | 11 | 1.36% |
| hsa_circ_0035317 | 1613 | 11 | 1.36% |
| hsa_circ_0020828 | 1760 | 12 | 1.36% |
| hsa_circ_0022823 | 1760 | 12 | 1.36% |

| | | | |
|---|---|---|---|
| hsa_circ_0044214 | 1760 | 12 | 1.36% |
| hsa_circ_0060747 | 1760 | 12 | 1.36% |
| hsa_circ_0027895 | 1467 | 10 | 1.36% |
| hsa_circ_0029448 | 1467 | 10 | 1.36% |
| hsa_circ_0032617 | 1467 | 10 | 1.36% |
| hsa_circ_0041816 | 1467 | 10 | 1.36% |
| hsa_circ_0043075 | 1467 | 10 | 1.36% |
| hsa_circ_0045417 | 1467 | 10 | 1.36% |
| hsa_circ_0048815 | 1467 | 10 | 1.36% |
| hsa_circ_0066116 | 1467 | 10 | 1.36% |
| hsa_circ_0072662 | 1467 | 10 | 1.36% |
| hsa_circ_0074138 | 1467 | 10 | 1.36% |
| hsa_circ_0127555 | 1467 | 10 | 1.36% |
| hsa_circ_0129731 | 1467 | 10 | 1.36% |
| hsa_circ_0133906 | 1467 | 10 | 1.36% |
| hsa_circ_0049203 | 1614 | 11 | 1.36% |
| hsa_circ_0054872 | 1614 | 11 | 1.36% |
| hsa_circ_0065687 | 1614 | 11 | 1.36% |
| hsa_circ_0070375 | 1614 | 11 | 1.36% |
| hsa_circ_0085275 | 1614 | 11 | 1.36% |
| hsa_circ_0086546 | 1614 | 11 | 1.36% |
| hsa_circ_0104270 | 1614 | 11 | 1.36% |
| hsa_circ_0134657 | 2201 | 15 | 1.36% |
| hsa_circ_0051724 | 1761 | 12 | 1.36% |
| hsa_circ_0074574 | 1761 | 12 | 1.36% |
| hsa_circ_0020103 | 1468 | 10 | 1.36% |
| hsa_circ_0021835 | 1468 | 10 | 1.36% |
| hsa_circ_0034211 | 1468 | 10 | 1.36% |
| hsa_circ_0043544 | 1468 | 10 | 1.36% |
| hsa_circ_0047448 | 1468 | 10 | 1.36% |
| hsa_circ_0055793 | 1468 | 10 | 1.36% |
| hsa_circ_0061492 | 1468 | 10 | 1.36% |
| hsa_circ_0062274 | 1468 | 10 | 1.36% |
| hsa_circ_0084382 | 1468 | 10 | 1.36% |
| hsa_circ_0087790 | 1468 | 10 | 1.36% |
| hsa_circ_0088982 | 1468 | 10 | 1.36% |
| hsa_circ_0101042 | 1468 | 10 | 1.36% |
| hsa_circ_0136986 | 1468 | 10 | 1.36% |
| hsa_circ_0031422 | 1615 | 11 | 1.36% |
| hsa_circ_0045197 | 1615 | 11 | 1.36% |
| hsa_circ_0063249 | 1615 | 11 | 1.36% |
| hsa_circ_0068111 | 1615 | 11 | 1.36% |
| hsa_circ_0122796 | 1615 | 11 | 1.36% |
| hsa_circ_0039103 | 1762 | 12 | 1.36% |
| hsa_circ_0094022 | 1762 | 12 | 1.36% |
| hsa_circ_0078234 | 3671 | 25 | 1.36% |
| hsa_circ_0094548 | 2644 | 18 | 1.36% |
| hsa_circ_0004226 | 1469 | 10 | 1.36% |
| hsa_circ_0016081 | 1469 | 10 | 1.36% |
| hsa_circ_0033088 | 1469 | 10 | 1.36% |

| | | | |
|---|---|---|---|
| hsa_circ_0037344 | 1469 | 10 | 1.36% |
| hsa_circ_0038683 | 1469 | 10 | 1.36% |
| hsa_circ_0045116 | 1469 | 10 | 1.36% |
| hsa_circ_0049552 | 1469 | 10 | 1.36% |
| hsa_circ_0060650 | 1469 | 10 | 1.36% |
| hsa_circ_0061887 | 1469 | 10 | 1.36% |
| hsa_circ_0069876 | 1469 | 10 | 1.36% |
| hsa_circ_0077888 | 1469 | 10 | 1.36% |
| hsa_circ_0079597 | 1469 | 10 | 1.36% |
| hsa_circ_0105646 | 1469 | 10 | 1.36% |
| hsa_circ_0125722 | 1469 | 10 | 1.36% |
| hsa_circ_0136200 | 1469 | 10 | 1.36% |
| hsa_circ_0038143 | 1616 | 11 | 1.36% |
| hsa_circ_0044342 | 1616 | 11 | 1.36% |
| hsa_circ_0081540 | 1616 | 11 | 1.36% |
| hsa_circ_0090660 | 1616 | 11 | 1.36% |
| hsa_circ_0096295 | 1616 | 11 | 1.36% |
| hsa_circ_0096473 | 1616 | 11 | 1.36% |
| hsa_circ_0125518 | 1616 | 11 | 1.36% |
| hsa_circ_0132423 | 1616 | 11 | 1.36% |
| hsa_circ_0134570 | 1616 | 11 | 1.36% |
| hsa_circ_0042107 | 1763 | 12 | 1.36% |
| hsa_circ_0078687 | 1763 | 12 | 1.36% |
| hsa_circ_0137339 | 1763 | 12 | 1.36% |
| hsa_circ_0021037 | 2204 | 15 | 1.36% |
| hsa_circ_0019991 | 1911 | 13 | 1.36% |
| hsa_circ_0086402 | 1911 | 13 | 1.36% |
| hsa_circ_0031596 | 1764 | 12 | 1.36% |
| hsa_circ_0075137 | 1764 | 12 | 1.36% |
| hsa_circ_0112254 | 1764 | 12 | 1.36% |
| hsa_circ_0019911 | 1617 | 11 | 1.36% |
| hsa_circ_0019924 | 1617 | 11 | 1.36% |
| hsa_circ_0024111 | 1617 | 11 | 1.36% |
| hsa_circ_0061086 | 1617 | 11 | 1.36% |
| hsa_circ_0079617 | 1617 | 11 | 1.36% |
| hsa_circ_0080039 | 1617 | 11 | 1.36% |
| hsa_circ_0108222 | 1617 | 11 | 1.36% |
| hsa_circ_0112449 | 1617 | 11 | 1.36% |
| hsa_circ_0021714 | 1470 | 10 | 1.36% |
| hsa_circ_0022699 | 1470 | 10 | 1.36% |
| hsa_circ_0024903 | 1470 | 10 | 1.36% |
| hsa_circ_0043883 | 1470 | 10 | 1.36% |
| hsa_circ_0048535 | 1470 | 10 | 1.36% |
| hsa_circ_0057138 | 1470 | 10 | 1.36% |
| hsa_circ_0058732 | 1470 | 10 | 1.36% |
| hsa_circ_0064528 | 1470 | 10 | 1.36% |
| hsa_circ_0087176 | 1470 | 10 | 1.36% |
| hsa_circ_0092776 | 1470 | 10 | 1.36% |
| hsa_circ_0101125 | 1470 | 10 | 1.36% |
| hsa_circ_0105463 | 1470 | 10 | 1.36% |

| | | | |
|---|---|---|---|
| hsa_circ_0107329 | 1470 | 10 | 1.36% |
| hsa_circ_0123737 | 1470 | 10 | 1.36% |
| hsa_circ_0139096 | 1470 | 10 | 1.36% |
| hsa_circ_0140657 | 1470 | 10 | 1.36% |
| hsa_circ_0047773 | 1765 | 12 | 1.36% |
| hsa_circ_0010248 | 1618 | 11 | 1.36% |
| hsa_circ_0017488 | 1618 | 11 | 1.36% |
| hsa_circ_0022863 | 1618 | 11 | 1.36% |
| hsa_circ_0078462 | 1618 | 11 | 1.36% |
| hsa_circ_0080571 | 1618 | 11 | 1.36% |
| hsa_circ_0086151 | 1618 | 11 | 1.36% |
| hsa_circ_0099751 | 1618 | 11 | 1.36% |
| hsa_circ_0106041 | 1618 | 11 | 1.36% |
| hsa_circ_0097712 | 6325 | 43 | 1.36% |
| hsa_circ_0016691 | 1471 | 10 | 1.36% |
| hsa_circ_0020672 | 1471 | 10 | 1.36% |
| hsa_circ_0041740 | 1471 | 10 | 1.36% |
| hsa_circ_0043560 | 1471 | 10 | 1.36% |
| hsa_circ_0043600 | 1471 | 10 | 1.36% |
| hsa_circ_0048718 | 1471 | 10 | 1.36% |
| hsa_circ_0088436 | 1471 | 10 | 1.36% |
| hsa_circ_0105255 | 1471 | 10 | 1.36% |
| hsa_circ_0126262 | 1471 | 10 | 1.36% |
| hsa_circ_0129146 | 1471 | 10 | 1.36% |
| hsa_circ_0139537 | 2501 | 17 | 1.36% |
| hsa_circ_0126014 | 3090 | 21 | 1.36% |
| hsa_circ_0015720 | 1766 | 12 | 1.36% |
| hsa_circ_0086266 | 1766 | 12 | 1.36% |
| hsa_circ_0134721 | 1766 | 12 | 1.36% |
| hsa_circ_0015924 | 1619 | 11 | 1.36% |
| hsa_circ_0035466 | 1619 | 11 | 1.36% |
| hsa_circ_0074879 | 1619 | 11 | 1.36% |
| hsa_circ_0132983 | 1619 | 11 | 1.36% |
| hsa_circ_0011703 | 1472 | 10 | 1.36% |
| hsa_circ_0012204 | 1472 | 10 | 1.36% |
| hsa_circ_0026417 | 1472 | 10 | 1.36% |
| hsa_circ_0038919 | 1472 | 10 | 1.36% |
| hsa_circ_0040237 | 1472 | 10 | 1.36% |
| hsa_circ_0041942 | 1472 | 10 | 1.36% |
| hsa_circ_0044809 | 1472 | 10 | 1.36% |
| hsa_circ_0052756 | 1472 | 10 | 1.36% |
| hsa_circ_0055082 | 1472 | 10 | 1.36% |
| hsa_circ_0059445 | 1472 | 10 | 1.36% |
| hsa_circ_0064727 | 1472 | 10 | 1.36% |
| hsa_circ_0084418 | 1472 | 10 | 1.36% |
| hsa_circ_0087174 | 1472 | 10 | 1.36% |
| hsa_circ_0098257 | 1472 | 10 | 1.36% |
| hsa_circ_0119298 | 1472 | 10 | 1.36% |
| hsa_circ_0121947 | 1472 | 10 | 1.36% |
| hsa_circ_0128531 | 1472 | 10 | 1.36% |

| | | | |
|---|---|---|---|
| hsa_circ_0139405 | 1472 | 10 | 1.36% |
| hsa_circ_0066107 | 1767 | 12 | 1.36% |
| hsa_circ_0046454 | 1620 | 11 | 1.36% |
| hsa_circ_0070239 | 1620 | 11 | 1.36% |
| hsa_circ_0052600 | 2062 | 14 | 1.36% |
| hsa_circ_0020300 | 1473 | 10 | 1.36% |
| hsa_circ_0021945 | 1473 | 10 | 1.36% |
| hsa_circ_0060955 | 1473 | 10 | 1.36% |
| hsa_circ_0075185 | 1473 | 10 | 1.36% |
| hsa_circ_0083979 | 1473 | 10 | 1.36% |
| hsa_circ_0087012 | 1473 | 10 | 1.36% |
| hsa_circ_0094556 | 1473 | 10 | 1.36% |
| hsa_circ_0112456 | 1473 | 10 | 1.36% |
| hsa_circ_0127231 | 1473 | 10 | 1.36% |
| hsa_circ_0129794 | 1473 | 10 | 1.36% |
| hsa_circ_0009325 | 1768 | 12 | 1.36% |
| hsa_circ_0020299 | 1768 | 12 | 1.36% |
| hsa_circ_0094477 | 4126 | 28 | 1.36% |
| hsa_circ_0010907 | 1621 | 11 | 1.36% |
| hsa_circ_0018519 | 1621 | 11 | 1.36% |
| hsa_circ_0127766 | 1621 | 11 | 1.36% |
| hsa_circ_0009822 | 1474 | 10 | 1.36% |
| hsa_circ_0010526 | 1474 | 10 | 1.36% |
| hsa_circ_0022843 | 1474 | 10 | 1.36% |
| hsa_circ_0035585 | 1474 | 10 | 1.36% |
| hsa_circ_0043420 | 1474 | 10 | 1.36% |
| hsa_circ_0048699 | 1474 | 10 | 1.36% |
| hsa_circ_0068168 | 1474 | 10 | 1.36% |
| hsa_circ_0092442 | 1474 | 10 | 1.36% |
| hsa_circ_0021627 | 1769 | 12 | 1.36% |
| hsa_circ_0030149 | 2064 | 14 | 1.36% |
| hsa_circ_0109278 | 2064 | 14 | 1.36% |
| hsa_circ_0041139 | 1622 | 11 | 1.36% |
| hsa_circ_0093384 | 1622 | 11 | 1.36% |
| hsa_circ_0101612 | 1622 | 11 | 1.36% |
| hsa_circ_0131497 | 1622 | 11 | 1.36% |
| hsa_circ_0051496 | 2065 | 14 | 1.36% |
| hsa_circ_0048646 | 1770 | 12 | 1.36% |
| hsa_circ_0053166 | 1770 | 12 | 1.36% |
| hsa_circ_0053259 | 1770 | 12 | 1.36% |
| hsa_circ_0008792 | 1475 | 10 | 1.36% |
| hsa_circ_0018561 | 1475 | 10 | 1.36% |
| hsa_circ_0023232 | 1475 | 10 | 1.36% |
| hsa_circ_0036906 | 1475 | 10 | 1.36% |
| hsa_circ_0084995 | 1475 | 10 | 1.36% |
| hsa_circ_0085472 | 1475 | 10 | 1.36% |
| hsa_circ_0086125 | 1475 | 10 | 1.36% |
| hsa_circ_0088533 | 1475 | 10 | 1.36% |
| hsa_circ_0098759 | 1475 | 10 | 1.36% |
| hsa_circ_0102556 | 1475 | 10 | 1.36% |

| | | | |
|---|---|---|---|
| hsa_circ_0120813 | 1475 | 10 | 1.36% |
| hsa_circ_0005106 | 1623 | 11 | 1.36% |
| hsa_circ_0021123 | 1623 | 11 | 1.36% |
| hsa_circ_0023647 | 1623 | 11 | 1.36% |
| hsa_circ_0033308 | 1623 | 11 | 1.36% |
| hsa_circ_0038350 | 1623 | 11 | 1.36% |
| hsa_circ_0051516 | 1623 | 11 | 1.36% |
| hsa_circ_0058029 | 1623 | 11 | 1.36% |
| hsa_circ_0118191 | 1623 | 11 | 1.36% |
| hsa_circ_0118549 | 1623 | 11 | 1.36% |
| hsa_circ_0135485 | 1623 | 11 | 1.36% |
| hsa_circ_0125255 | 2066 | 14 | 1.36% |
| hsa_circ_0048756 | 1771 | 12 | 1.36% |
| hsa_circ_0073945 | 1771 | 12 | 1.36% |
| hsa_circ_0077777 | 1771 | 12 | 1.36% |
| hsa_circ_0101256 | 2509 | 17 | 1.36% |
| hsa_circ_0020180 | 1476 | 10 | 1.36% |
| hsa_circ_0029982 | 1476 | 10 | 1.36% |
| hsa_circ_0048354 | 1476 | 10 | 1.36% |
| hsa_circ_0074473 | 1476 | 10 | 1.36% |
| hsa_circ_0088795 | 1476 | 10 | 1.36% |
| hsa_circ_0096026 | 1476 | 10 | 1.36% |
| hsa_circ_0099917 | 1476 | 10 | 1.36% |
| hsa_circ_0116734 | 1476 | 10 | 1.36% |
| hsa_circ_0025678 | 1919 | 13 | 1.35% |
| hsa_circ_0087470 | 1919 | 13 | 1.35% |
| hsa_circ_0128092 | 3248 | 22 | 1.35% |
| hsa_circ_0044067 | 1624 | 11 | 1.35% |
| hsa_circ_0071694 | 1624 | 11 | 1.35% |
| hsa_circ_0086286 | 1624 | 11 | 1.35% |
| hsa_circ_0043459 | 2067 | 14 | 1.35% |
| hsa_circ_0097702 | 6349 | 43 | 1.35% |
| hsa_circ_0074233 | 1772 | 12 | 1.35% |
| hsa_circ_0088342 | 1772 | 12 | 1.35% |
| hsa_circ_0027739 | 1477 | 10 | 1.35% |
| hsa_circ_0030422 | 1477 | 10 | 1.35% |
| hsa_circ_0033041 | 1477 | 10 | 1.35% |
| hsa_circ_0033546 | 1477 | 10 | 1.35% |
| hsa_circ_0042778 | 1477 | 10 | 1.35% |
| hsa_circ_0059299 | 1477 | 10 | 1.35% |
| hsa_circ_0065554 | 1477 | 10 | 1.35% |
| hsa_circ_0066336 | 1477 | 10 | 1.35% |
| hsa_circ_0085649 | 1477 | 10 | 1.35% |
| hsa_circ_0091965 | 1477 | 10 | 1.35% |
| hsa_circ_0121857 | 1477 | 10 | 1.35% |
| hsa_circ_0009504 | 1625 | 11 | 1.35% |
| hsa_circ_0026126 | 1625 | 11 | 1.35% |
| hsa_circ_0027532 | 1625 | 11 | 1.35% |
| hsa_circ_0030684 | 1625 | 11 | 1.35% |
| hsa_circ_0034191 | 1625 | 11 | 1.35% |

| | | | |
|---|---|---|---|
| hsa_circ_0051313 | 1625 | 11 | 1.35% |
| hsa_circ_0109184 | 1625 | 11 | 1.35% |
| hsa_circ_0126729 | 1625 | 11 | 1.35% |
| hsa_circ_0017788 | 1773 | 12 | 1.35% |
| hsa_circ_0065838 | 1773 | 12 | 1.35% |
| hsa_circ_0067518 | 1773 | 12 | 1.35% |
| hsa_circ_0011420 | 1478 | 10 | 1.35% |
| hsa_circ_0013075 | 1478 | 10 | 1.35% |
| hsa_circ_0023601 | 1478 | 10 | 1.35% |
| hsa_circ_0024916 | 1478 | 10 | 1.35% |
| hsa_circ_0038959 | 1478 | 10 | 1.35% |
| hsa_circ_0048495 | 1478 | 10 | 1.35% |
| hsa_circ_0093583 | 1478 | 10 | 1.35% |
| hsa_circ_0112526 | 1478 | 10 | 1.35% |
| hsa_circ_0115865 | 1478 | 10 | 1.35% |
| hsa_circ_0134930 | 1478 | 10 | 1.35% |
| hsa_circ_0019935 | 1626 | 11 | 1.35% |
| hsa_circ_0028788 | 1626 | 11 | 1.35% |
| hsa_circ_0038092 | 1626 | 11 | 1.35% |
| hsa_circ_0043997 | 1626 | 11 | 1.35% |
| hsa_circ_0049159 | 1626 | 11 | 1.35% |
| hsa_circ_0049727 | 1626 | 11 | 1.35% |
| hsa_circ_0073955 | 1626 | 11 | 1.35% |
| hsa_circ_0077890 | 1626 | 11 | 1.35% |
| hsa_circ_0086404 | 1626 | 11 | 1.35% |
| hsa_circ_0089710 | 1626 | 11 | 1.35% |
| hsa_circ_0117374 | 1626 | 11 | 1.35% |
| hsa_circ_0134303 | 1626 | 11 | 1.35% |
| hsa_circ_0014495 | 1774 | 12 | 1.35% |
| hsa_circ_0067906 | 1774 | 12 | 1.35% |
| hsa_circ_0021851 | 6062 | 41 | 1.35% |
| hsa_circ_0115058 | 4436 | 30 | 1.35% |
| hsa_circ_0045278 | 2366 | 16 | 1.35% |
| hsa_circ_0007936 | 1479 | 10 | 1.35% |
| hsa_circ_0023552 | 1479 | 10 | 1.35% |
| hsa_circ_0025935 | 1479 | 10 | 1.35% |
| hsa_circ_0047493 | 1479 | 10 | 1.35% |
| hsa_circ_0065492 | 1479 | 10 | 1.35% |
| hsa_circ_0066344 | 1479 | 10 | 1.35% |
| hsa_circ_0067861 | 1479 | 10 | 1.35% |
| hsa_circ_0088197 | 1479 | 10 | 1.35% |
| hsa_circ_0096153 | 1479 | 10 | 1.35% |
| hsa_circ_0126947 | 1479 | 10 | 1.35% |
| hsa_circ_0137357 | 1479 | 10 | 1.35% |
| hsa_circ_0010210 | 1627 | 11 | 1.35% |
| hsa_circ_0036166 | 1627 | 11 | 1.35% |
| hsa_circ_0041032 | 1627 | 11 | 1.35% |
| hsa_circ_0080278 | 1627 | 11 | 1.35% |
| hsa_circ_0117793 | 1627 | 11 | 1.35% |
| hsa_circ_0125182 | 1627 | 11 | 1.35% |

| | | | |
|---|---|---|---|
| hsa_circ_0024315 | 1775 | 12 | 1.35% |
| hsa_circ_0037106 | 1775 | 12 | 1.35% |
| hsa_circ_0052315 | 1775 | 12 | 1.35% |
| hsa_circ_0062611 | 1775 | 12 | 1.35% |
| hsa_circ_0063613 | 1775 | 12 | 1.35% |
| hsa_circ_0077678 | 1775 | 12 | 1.35% |
| hsa_circ_0089870 | 1775 | 12 | 1.35% |
| hsa_circ_0112338 | 1775 | 12 | 1.35% |
| hsa_circ_0132119 | 2663 | 18 | 1.35% |
| hsa_circ_0014546 | 3404 | 23 | 1.35% |
| hsa_circ_0016735 | 1776 | 12 | 1.35% |
| hsa_circ_0079552 | 1776 | 12 | 1.35% |
| hsa_circ_0100838 | 1776 | 12 | 1.35% |
| hsa_circ_0009220 | 1628 | 11 | 1.35% |
| hsa_circ_0039028 | 1628 | 11 | 1.35% |
| hsa_circ_0072258 | 1628 | 11 | 1.35% |
| hsa_circ_0074925 | 1628 | 11 | 1.35% |
| hsa_circ_0078545 | 1628 | 11 | 1.35% |
| hsa_circ_0086285 | 1628 | 11 | 1.35% |
| hsa_circ_0008963 | 1480 | 10 | 1.35% |
| hsa_circ_0023287 | 1480 | 10 | 1.35% |
| hsa_circ_0039707 | 1480 | 10 | 1.35% |
| hsa_circ_0041092 | 1480 | 10 | 1.35% |
| hsa_circ_0047136 | 1480 | 10 | 1.35% |
| hsa_circ_0023842 | 1925 | 13 | 1.35% |
| hsa_circ_0046819 | 1629 | 11 | 1.35% |
| hsa_circ_0051139 | 1629 | 11 | 1.35% |
| hsa_circ_0055799 | 1629 | 11 | 1.35% |
| hsa_circ_0060217 | 1629 | 11 | 1.35% |
| hsa_circ_0061394 | 1629 | 11 | 1.35% |
| hsa_circ_0032905 | 1481 | 10 | 1.35% |
| hsa_circ_0034581 | 1481 | 10 | 1.35% |
| hsa_circ_0060710 | 1481 | 10 | 1.35% |
| hsa_circ_0062786 | 1481 | 10 | 1.35% |
| hsa_circ_0067875 | 1481 | 10 | 1.35% |
| hsa_circ_0088947 | 1481 | 10 | 1.35% |
| hsa_circ_0105559 | 1481 | 10 | 1.35% |
| hsa_circ_0112600 | 1481 | 10 | 1.35% |
| hsa_circ_0128398 | 1481 | 10 | 1.35% |
| hsa_circ_0097713 | 6369 | 43 | 1.35% |
| hsa_circ_0057437 | 1926 | 13 | 1.35% |
| hsa_circ_0053117 | 1778 | 12 | 1.35% |
| hsa_circ_0058832 | 1778 | 12 | 1.35% |
| hsa_circ_0083106 | 1778 | 12 | 1.35% |
| hsa_circ_0006002 | 1630 | 11 | 1.35% |
| hsa_circ_0043531 | 1630 | 11 | 1.35% |
| hsa_circ_0117797 | 1630 | 11 | 1.35% |
| hsa_circ_0001232 | 3705 | 25 | 1.35% |
| hsa_circ_0033411 | 2223 | 15 | 1.35% |
| hsa_circ_0020469 | 1482 | 10 | 1.35% |

| | | | |
|---|---|---|---|
| hsa_circ_0024448 | 1482 | 10 | 1.35% |
| hsa_circ_0067651 | 1482 | 10 | 1.35% |
| hsa_circ_0069354 | 1482 | 10 | 1.35% |
| hsa_circ_0070957 | 1482 | 10 | 1.35% |
| hsa_circ_0073282 | 1482 | 10 | 1.35% |
| hsa_circ_0082249 | 1482 | 10 | 1.35% |
| hsa_circ_0084297 | 1482 | 10 | 1.35% |
| hsa_circ_0092233 | 1482 | 10 | 1.35% |
| hsa_circ_0112059 | 1482 | 10 | 1.35% |
| hsa_circ_0113448 | 1482 | 10 | 1.35% |
| hsa_circ_0052071 | 2075 | 14 | 1.35% |
| hsa_circ_0071529 | 1927 | 13 | 1.35% |
| hsa_circ_0013633 | 1779 | 12 | 1.35% |
| hsa_circ_0022405 | 1779 | 12 | 1.35% |
| hsa_circ_0041827 | 1779 | 12 | 1.35% |
| hsa_circ_0045147 | 1779 | 12 | 1.35% |
| hsa_circ_0120581 | 1779 | 12 | 1.35% |
| hsa_circ_0028459 | 1631 | 11 | 1.35% |
| hsa_circ_0048661 | 1631 | 11 | 1.35% |
| hsa_circ_0061996 | 1631 | 11 | 1.35% |
| hsa_circ_0079398 | 1631 | 11 | 1.35% |
| hsa_circ_0116898 | 1631 | 11 | 1.35% |
| hsa_circ_0128384 | 1631 | 11 | 1.35% |
| hsa_circ_0128755 | 1631 | 11 | 1.35% |
| hsa_circ_0009241 | 1483 | 10 | 1.35% |
| hsa_circ_0012640 | 1483 | 10 | 1.35% |
| hsa_circ_0030168 | 1483 | 10 | 1.35% |
| hsa_circ_0039606 | 1483 | 10 | 1.35% |
| hsa_circ_0057572 | 1483 | 10 | 1.35% |
| hsa_circ_0064973 | 1483 | 10 | 1.35% |
| hsa_circ_0067781 | 1483 | 10 | 1.35% |
| hsa_circ_0072386 | 1483 | 10 | 1.35% |
| hsa_circ_0078391 | 1483 | 10 | 1.35% |
| hsa_circ_0086588 | 1483 | 10 | 1.35% |
| hsa_circ_0088371 | 1483 | 10 | 1.35% |
| hsa_circ_0115534 | 1483 | 10 | 1.35% |
| hsa_circ_0126353 | 1483 | 10 | 1.35% |
| hsa_circ_0129197 | 1483 | 10 | 1.35% |
| hsa_circ_0001660 | 1928 | 13 | 1.35% |
| hsa_circ_0010656 | 1928 | 13 | 1.35% |
| hsa_circ_0106525 | 4301 | 29 | 1.35% |
| hsa_circ_0025364 | 1780 | 12 | 1.35% |
| hsa_circ_0040452 | 1780 | 12 | 1.35% |
| hsa_circ_0023194 | 1632 | 11 | 1.35% |
| hsa_circ_0062550 | 1632 | 11 | 1.35% |
| hsa_circ_0067217 | 1632 | 11 | 1.35% |
| hsa_circ_0079446 | 1632 | 11 | 1.35% |
| hsa_circ_0127864 | 3858 | 26 | 1.35% |
| hsa_circ_0029380 | 1929 | 13 | 1.35% |
| hsa_circ_0027272 | 1484 | 10 | 1.35% |

| | | | |
|---|---|---|---|
| hsa_circ_0060389 | 1484 | 10 | 1.35% |
| hsa_circ_0086957 | 1484 | 10 | 1.35% |
| hsa_circ_0091249 | 1484 | 10 | 1.35% |
| hsa_circ_0092201 | 1484 | 10 | 1.35% |
| hsa_circ_0093344 | 1484 | 10 | 1.35% |
| hsa_circ_0111425 | 1484 | 10 | 1.35% |
| hsa_circ_0114793 | 1484 | 10 | 1.35% |
| hsa_circ_0121154 | 1484 | 10 | 1.35% |
| hsa_circ_0125649 | 1484 | 10 | 1.35% |
| hsa_circ_0012084 | 1633 | 11 | 1.35% |
| hsa_circ_0019661 | 1633 | 11 | 1.35% |
| hsa_circ_0035105 | 1633 | 11 | 1.35% |
| hsa_circ_0037325 | 1633 | 11 | 1.35% |
| hsa_circ_0088705 | 1633 | 11 | 1.35% |
| hsa_circ_0089008 | 1633 | 11 | 1.35% |
| hsa_circ_0109972 | 1633 | 11 | 1.35% |
| hsa_circ_0139865 | 1633 | 11 | 1.35% |
| hsa_circ_0009427 | 1930 | 13 | 1.35% |
| hsa_circ_0060675 | 1930 | 13 | 1.35% |
| hsa_circ_0072615 | 1930 | 13 | 1.35% |
| hsa_circ_0117170 | 1930 | 13 | 1.35% |
| hsa_circ_0132298 | 2227 | 15 | 1.35% |
| hsa_circ_0072089 | 2079 | 14 | 1.35% |
| hsa_circ_0016766 | 1782 | 12 | 1.35% |
| hsa_circ_0058022 | 1782 | 12 | 1.35% |
| hsa_circ_0130696 | 1782 | 12 | 1.35% |
| hsa_circ_0020833 | 1485 | 10 | 1.35% |
| hsa_circ_0025783 | 1485 | 10 | 1.35% |
| hsa_circ_0036934 | 1485 | 10 | 1.35% |
| hsa_circ_0066703 | 1485 | 10 | 1.35% |
| hsa_circ_0077484 | 1485 | 10 | 1.35% |
| hsa_circ_0091770 | 1485 | 10 | 1.35% |
| hsa_circ_0101978 | 1485 | 10 | 1.35% |
| hsa_circ_0120323 | 1485 | 10 | 1.35% |
| hsa_circ_0132466 | 1485 | 10 | 1.35% |
| hsa_circ_0133766 | 1485 | 10 | 1.35% |
| hsa_circ_0060702 | 1931 | 13 | 1.35% |
| hsa_circ_0083284 | 1634 | 11 | 1.35% |
| hsa_circ_0098260 | 1634 | 11 | 1.35% |
| hsa_circ_0111342 | 1634 | 11 | 1.35% |
| hsa_circ_0053944 | 1783 | 12 | 1.35% |
| hsa_circ_0088741 | 1783 | 12 | 1.35% |
| hsa_circ_0125895 | 1783 | 12 | 1.35% |
| hsa_circ_0109084 | 4458 | 30 | 1.35% |
| hsa_circ_0004867 | 1486 | 10 | 1.35% |
| hsa_circ_0006827 | 1486 | 10 | 1.35% |
| hsa_circ_0010451 | 1486 | 10 | 1.35% |
| hsa_circ_0021922 | 1486 | 10 | 1.35% |
| hsa_circ_0052458 | 1486 | 10 | 1.35% |
| hsa_circ_0059441 | 1486 | 10 | 1.35% |

| | | | |
|---|---|---|---|
| hsa_circ_0074298 | 1486 | 10 | 1.35% |
| hsa_circ_0085111 | 1486 | 10 | 1.35% |
| hsa_circ_0121133 | 1486 | 10 | 1.35% |
| hsa_circ_0133545 | 1486 | 10 | 1.35% |
| hsa_circ_0135662 | 1486 | 10 | 1.35% |
| hsa_circ_0137821 | 1486 | 10 | 1.35% |
| hsa_circ_0014647 | 1932 | 13 | 1.35% |
| hsa_circ_0077289 | 1932 | 13 | 1.35% |
| hsa_circ_0013887 | 1635 | 11 | 1.35% |
| hsa_circ_0021072 | 1635 | 11 | 1.35% |
| hsa_circ_0043341 | 1635 | 11 | 1.35% |
| hsa_circ_0064770 | 1635 | 11 | 1.35% |
| hsa_circ_0084364 | 1635 | 11 | 1.35% |
| hsa_circ_0105401 | 3122 | 21 | 1.35% |
| hsa_circ_0022114 | 1784 | 12 | 1.35% |
| hsa_circ_0057192 | 1784 | 12 | 1.35% |
| hsa_circ_0067131 | 1784 | 12 | 1.35% |
| hsa_circ_0069131 | 1784 | 12 | 1.35% |
| hsa_circ_0073680 | 1784 | 12 | 1.35% |
| hsa_circ_0119009 | 1784 | 12 | 1.35% |
| hsa_circ_0008495 | 1487 | 10 | 1.34% |
| hsa_circ_0028897 | 1487 | 10 | 1.34% |
| hsa_circ_0037150 | 1487 | 10 | 1.34% |
| hsa_circ_0063125 | 1487 | 10 | 1.34% |
| hsa_circ_0071821 | 1487 | 10 | 1.34% |
| hsa_circ_0076805 | 1487 | 10 | 1.34% |
| hsa_circ_0137680 | 1487 | 10 | 1.34% |
| hsa_circ_0138460 | 1487 | 10 | 1.34% |
| hsa_circ_0057962 | 3123 | 21 | 1.34% |
| hsa_circ_0043614 | 1636 | 11 | 1.34% |
| hsa_circ_0050390 | 1785 | 12 | 1.34% |
| hsa_circ_0067501 | 1785 | 12 | 1.34% |
| hsa_circ_0103543 | 1785 | 12 | 1.34% |
| hsa_circ_0125969 | 1785 | 12 | 1.34% |
| hsa_circ_0040556 | 2083 | 14 | 1.34% |
| hsa_circ_0013214 | 1488 | 10 | 1.34% |
| hsa_circ_0028638 | 1488 | 10 | 1.34% |
| hsa_circ_0030133 | 1488 | 10 | 1.34% |
| hsa_circ_0039337 | 1488 | 10 | 1.34% |
| hsa_circ_0064266 | 1488 | 10 | 1.34% |
| hsa_circ_0071626 | 1488 | 10 | 1.34% |
| hsa_circ_0089942 | 1488 | 10 | 1.34% |
| hsa_circ_0092783 | 1488 | 10 | 1.34% |
| hsa_circ_0114270 | 1488 | 10 | 1.34% |
| hsa_circ_0121838 | 1488 | 10 | 1.34% |
| hsa_circ_0129991 | 1488 | 10 | 1.34% |
| hsa_circ_0130615 | 1488 | 10 | 1.34% |
| hsa_circ_0139170 | 1488 | 10 | 1.34% |
| hsa_circ_0025899 | 1637 | 11 | 1.34% |
| hsa_circ_0074849 | 1637 | 11 | 1.34% |

| | | | |
|---|---|---|---|
| hsa_circ_0076484 | 1637 | 11 | 1.34% |
| hsa_circ_0083611 | 1637 | 11 | 1.34% |
| hsa_circ_0106641 | 1637 | 11 | 1.34% |
| hsa_circ_0110989 | 1637 | 11 | 1.34% |
| hsa_circ_0112144 | 1637 | 11 | 1.34% |
| hsa_circ_0112364 | 1637 | 11 | 1.34% |
| hsa_circ_0118352 | 1637 | 11 | 1.34% |
| hsa_circ_0087069 | 1786 | 12 | 1.34% |
| hsa_circ_0044772 | 1489 | 10 | 1.34% |
| hsa_circ_0045927 | 1489 | 10 | 1.34% |
| hsa_circ_0075889 | 1489 | 10 | 1.34% |
| hsa_circ_0096464 | 1489 | 10 | 1.34% |
| hsa_circ_0101447 | 1489 | 10 | 1.34% |
| hsa_circ_0127891 | 1489 | 10 | 1.34% |
| hsa_circ_0020773 | 1638 | 11 | 1.34% |
| hsa_circ_0098214 | 1638 | 11 | 1.34% |
| hsa_circ_0099684 | 1638 | 11 | 1.34% |
| hsa_circ_0061137 | 1787 | 12 | 1.34% |
| hsa_circ_0071349 | 1787 | 12 | 1.34% |
| hsa_circ_0042289 | 1788 | 12 | 1.34% |
| hsa_circ_0103157 | 1788 | 12 | 1.34% |
| hsa_circ_0027015 | 1639 | 11 | 1.34% |
| hsa_circ_0030095 | 1639 | 11 | 1.34% |
| hsa_circ_0048612 | 1639 | 11 | 1.34% |
| hsa_circ_0059330 | 1639 | 11 | 1.34% |
| hsa_circ_0068045 | 1639 | 11 | 1.34% |
| hsa_circ_0019325 | 1490 | 10 | 1.34% |
| hsa_circ_0021316 | 1490 | 10 | 1.34% |
| hsa_circ_0025011 | 1490 | 10 | 1.34% |
| hsa_circ_0056568 | 1490 | 10 | 1.34% |
| hsa_circ_0059232 | 1490 | 10 | 1.34% |
| hsa_circ_0075581 | 1490 | 10 | 1.34% |
| hsa_circ_0093464 | 1490 | 10 | 1.34% |
| hsa_circ_0095034 | 1490 | 10 | 1.34% |
| hsa_circ_0096121 | 1490 | 10 | 1.34% |
| hsa_circ_0099383 | 1490 | 10 | 1.34% |
| hsa_circ_0117479 | 1490 | 10 | 1.34% |
| hsa_circ_0121367 | 1490 | 10 | 1.34% |
| hsa_circ_0126421 | 1490 | 10 | 1.34% |
| hsa_circ_0128970 | 1490 | 10 | 1.34% |
| hsa_circ_0101707 | 1938 | 13 | 1.34% |
| hsa_circ_0058063 | 1640 | 11 | 1.34% |
| hsa_circ_0072869 | 1640 | 11 | 1.34% |
| hsa_circ_0083674 | 1640 | 11 | 1.34% |
| hsa_circ_0093964 | 1640 | 11 | 1.34% |
| hsa_circ_0140075 | 1640 | 11 | 1.34% |
| hsa_circ_0022218 | 1491 | 10 | 1.34% |
| hsa_circ_0023029 | 1491 | 10 | 1.34% |
| hsa_circ_0026377 | 1491 | 10 | 1.34% |
| hsa_circ_0028696 | 1491 | 10 | 1.34% |

| | | | |
|---|---|---|---|
| hsa_circ_0037405 | 1491 | 10 | 1.34% |
| hsa_circ_0040515 | 1491 | 10 | 1.34% |
| hsa_circ_0047563 | 1491 | 10 | 1.34% |
| hsa_circ_0055461 | 1491 | 10 | 1.34% |
| hsa_circ_0060751 | 1491 | 10 | 1.34% |
| hsa_circ_0077938 | 1491 | 10 | 1.34% |
| hsa_circ_0095417 | 1491 | 10 | 1.34% |
| hsa_circ_0103779 | 1491 | 10 | 1.34% |
| hsa_circ_0117874 | 1491 | 10 | 1.34% |
| hsa_circ_0132477 | 1491 | 10 | 1.34% |
| hsa_circ_0135588 | 1491 | 10 | 1.34% |
| hsa_circ_0058757 | 1790 | 12 | 1.34% |
| hsa_circ_0044618 | 1641 | 11 | 1.34% |
| hsa_circ_0054744 | 1641 | 11 | 1.34% |
| hsa_circ_0058115 | 1641 | 11 | 1.34% |
| hsa_circ_0081238 | 1641 | 11 | 1.34% |
| hsa_circ_0120512 | 1641 | 11 | 1.34% |
| hsa_circ_0026214 | 1492 | 10 | 1.34% |
| hsa_circ_0030629 | 1492 | 10 | 1.34% |
| hsa_circ_0031743 | 1492 | 10 | 1.34% |
| hsa_circ_0038851 | 1492 | 10 | 1.34% |
| hsa_circ_0041336 | 1492 | 10 | 1.34% |
| hsa_circ_0049063 | 1492 | 10 | 1.34% |
| hsa_circ_0060719 | 1492 | 10 | 1.34% |
| hsa_circ_0064160 | 1492 | 10 | 1.34% |
| hsa_circ_0064693 | 1492 | 10 | 1.34% |
| hsa_circ_0068926 | 1492 | 10 | 1.34% |
| hsa_circ_0073436 | 1492 | 10 | 1.34% |
| hsa_circ_0090375 | 1492 | 10 | 1.34% |
| hsa_circ_0096644 | 1492 | 10 | 1.34% |
| hsa_circ_0118895 | 1492 | 10 | 1.34% |
| hsa_circ_0129501 | 1492 | 10 | 1.34% |
| hsa_circ_0138717 | 1492 | 10 | 1.34% |
| hsa_circ_0082293 | 2089 | 14 | 1.34% |
| hsa_circ_0090126 | 2089 | 14 | 1.34% |
| hsa_circ_0098212 | 2686 | 18 | 1.34% |
| hsa_circ_0042285 | 1791 | 12 | 1.34% |
| hsa_circ_0062075 | 1791 | 12 | 1.34% |
| hsa_circ_0016127 | 1642 | 11 | 1.34% |
| hsa_circ_0069300 | 1642 | 11 | 1.34% |
| hsa_circ_0076702 | 1642 | 11 | 1.34% |
| hsa_circ_0086017 | 1642 | 11 | 1.34% |
| hsa_circ_0098022 | 1642 | 11 | 1.34% |
| hsa_circ_0104316 | 1642 | 11 | 1.34% |
| hsa_circ_0113569 | 1642 | 11 | 1.34% |
| hsa_circ_0045818 | 2538 | 17 | 1.34% |
| hsa_circ_0014170 | 1493 | 10 | 1.34% |
| hsa_circ_0028221 | 1493 | 10 | 1.34% |
| hsa_circ_0035385 | 1493 | 10 | 1.34% |
| hsa_circ_0055923 | 1493 | 10 | 1.34% |

| | | | |
|---|---|---|---|
| hsa_circ_0060718 | 1493 | 10 | 1.34% |
| hsa_circ_0065988 | 1493 | 10 | 1.34% |
| hsa_circ_0084425 | 1493 | 10 | 1.34% |
| hsa_circ_0085441 | 1493 | 10 | 1.34% |
| hsa_circ_0088063 | 1493 | 10 | 1.34% |
| hsa_circ_0100600 | 1493 | 10 | 1.34% |
| hsa_circ_0114991 | 1493 | 10 | 1.34% |
| hsa_circ_0120632 | 1493 | 10 | 1.34% |
| hsa_circ_0126143 | 1493 | 10 | 1.34% |
| hsa_circ_0127208 | 1493 | 10 | 1.34% |
| hsa_circ_0022704 | 1792 | 12 | 1.34% |
| hsa_circ_0072749 | 1792 | 12 | 1.34% |
| hsa_circ_0128962 | 2091 | 14 | 1.34% |
| hsa_circ_0031364 | 1643 | 11 | 1.34% |
| hsa_circ_0052286 | 1643 | 11 | 1.34% |
| hsa_circ_0053638 | 1643 | 11 | 1.34% |
| hsa_circ_0091767 | 1643 | 11 | 1.34% |
| hsa_circ_0094188 | 1643 | 11 | 1.34% |
| hsa_circ_0020140 | 1942 | 13 | 1.34% |
| hsa_circ_0034507 | 1494 | 10 | 1.34% |
| hsa_circ_0046012 | 1494 | 10 | 1.34% |
| hsa_circ_0052761 | 1494 | 10 | 1.34% |
| hsa_circ_0057125 | 1494 | 10 | 1.34% |
| hsa_circ_0068844 | 1494 | 10 | 1.34% |
| hsa_circ_0085745 | 1494 | 10 | 1.34% |
| hsa_circ_0097801 | 1494 | 10 | 1.34% |
| hsa_circ_0108140 | 1494 | 10 | 1.34% |
| hsa_circ_0138615 | 1494 | 10 | 1.34% |
| hsa_circ_0010429 | 1793 | 12 | 1.34% |
| hsa_circ_0098669 | 5230 | 35 | 1.34% |
| hsa_circ_0010066 | 1644 | 11 | 1.34% |
| hsa_circ_0011294 | 1644 | 11 | 1.34% |
| hsa_circ_0012819 | 1644 | 11 | 1.34% |
| hsa_circ_0015759 | 1644 | 11 | 1.34% |
| hsa_circ_0023569 | 1644 | 11 | 1.34% |
| hsa_circ_0025573 | 1644 | 11 | 1.34% |
| hsa_circ_0038273 | 1644 | 11 | 1.34% |
| hsa_circ_0071512 | 1644 | 11 | 1.34% |
| hsa_circ_0087401 | 1644 | 11 | 1.34% |
| hsa_circ_0127311 | 1644 | 11 | 1.34% |
| hsa_circ_0112695 | 1943 | 13 | 1.34% |
| hsa_circ_0036985 | 2093 | 14 | 1.34% |
| hsa_circ_0080089 | 2093 | 14 | 1.34% |
| hsa_circ_0054430 | 1794 | 12 | 1.34% |
| hsa_circ_0013729 | 1495 | 10 | 1.34% |
| hsa_circ_0064472 | 1495 | 10 | 1.34% |
| hsa_circ_0107026 | 5831 | 39 | 1.34% |
| hsa_circ_0036072 | 5084 | 34 | 1.34% |
| hsa_circ_0040176 | 1645 | 11 | 1.34% |
| hsa_circ_0087904 | 1645 | 11 | 1.34% |

| | | | |
|---|---|---|---|
| hsa_circ_0087979 | 1645 | 11 | 1.34% |
| hsa_circ_0023754 | 1795 | 12 | 1.34% |
| hsa_circ_0038362 | 1795 | 12 | 1.34% |
| hsa_circ_0028032 | 1496 | 10 | 1.34% |
| hsa_circ_0041366 | 1496 | 10 | 1.34% |
| hsa_circ_0061838 | 1496 | 10 | 1.34% |
| hsa_circ_0061975 | 1496 | 10 | 1.34% |
| hsa_circ_0078015 | 1496 | 10 | 1.34% |
| hsa_circ_0088936 | 1496 | 10 | 1.34% |
| hsa_circ_0138741 | 1496 | 10 | 1.34% |
| hsa_circ_0052937 | 1945 | 13 | 1.34% |
| hsa_circ_0025230 | 1646 | 11 | 1.34% |
| hsa_circ_0069397 | 1646 | 11 | 1.34% |
| hsa_circ_0074192 | 1646 | 11 | 1.34% |
| hsa_circ_0091722 | 1646 | 11 | 1.34% |
| hsa_circ_0105975 | 1646 | 11 | 1.34% |
| hsa_circ_0106190 | 1646 | 11 | 1.34% |
| hsa_circ_0008161 | 7782 | 52 | 1.34% |
| hsa_circ_0034487 | 1796 | 12 | 1.34% |
| hsa_circ_0048562 | 1796 | 12 | 1.34% |
| hsa_circ_0013197 | 1497 | 10 | 1.34% |
| hsa_circ_0020797 | 1497 | 10 | 1.34% |
| hsa_circ_0028902 | 1497 | 10 | 1.34% |
| hsa_circ_0031782 | 1497 | 10 | 1.34% |
| hsa_circ_0073118 | 1497 | 10 | 1.34% |
| hsa_circ_0089631 | 1497 | 10 | 1.34% |
| hsa_circ_0106196 | 1497 | 10 | 1.34% |
| hsa_circ_0046235 | 2096 | 14 | 1.34% |
| hsa_circ_0010454 | 1647 | 11 | 1.34% |
| hsa_circ_0026721 | 1647 | 11 | 1.34% |
| hsa_circ_0029387 | 1647 | 11 | 1.34% |
| hsa_circ_0030084 | 1647 | 11 | 1.34% |
| hsa_circ_0033535 | 1647 | 11 | 1.34% |
| hsa_circ_0065643 | 1647 | 11 | 1.34% |
| hsa_circ_0069992 | 1647 | 11 | 1.34% |
| hsa_circ_0075523 | 1647 | 11 | 1.34% |
| hsa_circ_0091868 | 1647 | 11 | 1.34% |
| hsa_circ_0115190 | 1647 | 11 | 1.34% |
| hsa_circ_0122559 | 1647 | 11 | 1.34% |
| hsa_circ_0025406 | 1797 | 12 | 1.34% |
| hsa_circ_0126911 | 1797 | 12 | 1.34% |
| hsa_circ_0128098 | 1797 | 12 | 1.34% |
| hsa_circ_0087136 | 1947 | 13 | 1.34% |
| hsa_circ_0011756 | 1498 | 10 | 1.34% |
| hsa_circ_0016089 | 1498 | 10 | 1.34% |
| hsa_circ_0024564 | 1498 | 10 | 1.34% |
| hsa_circ_0024971 | 1498 | 10 | 1.34% |
| hsa_circ_0028480 | 1498 | 10 | 1.34% |
| hsa_circ_0039927 | 1498 | 10 | 1.34% |
| hsa_circ_0049712 | 1498 | 10 | 1.34% |

| | | | |
|---|---|---|---|
| hsa_circ_0057964 | 1498 | 10 | 1.34% |
| hsa_circ_0072699 | 1498 | 10 | 1.34% |
| hsa_circ_0081914 | 1498 | 10 | 1.34% |
| hsa_circ_0134654 | 1498 | 10 | 1.34% |
| hsa_circ_0138869 | 1498 | 10 | 1.34% |
| hsa_circ_0010446 | 1648 | 11 | 1.33% |
| hsa_circ_0011282 | 1648 | 11 | 1.33% |
| hsa_circ_0015634 | 1648 | 11 | 1.33% |
| hsa_circ_0016237 | 1648 | 11 | 1.33% |
| hsa_circ_0062383 | 1648 | 11 | 1.33% |
| hsa_circ_0087689 | 1648 | 11 | 1.33% |
| hsa_circ_0105131 | 1648 | 11 | 1.33% |
| hsa_circ_0118041 | 1648 | 11 | 1.33% |
| hsa_circ_0121870 | 1648 | 11 | 1.33% |
| hsa_circ_0035869 | 1798 | 12 | 1.33% |
| hsa_circ_0106711 | 1798 | 12 | 1.33% |
| hsa_circ_0115363 | 1798 | 12 | 1.33% |
| hsa_circ_0136767 | 1798 | 12 | 1.33% |
| hsa_circ_0058176 | 1948 | 13 | 1.33% |
| hsa_circ_0021648 | 2398 | 16 | 1.33% |
| hsa_circ_0013106 | 1499 | 10 | 1.33% |
| hsa_circ_0016084 | 1499 | 10 | 1.33% |
| hsa_circ_0026825 | 1499 | 10 | 1.33% |
| hsa_circ_0081886 | 1499 | 10 | 1.33% |
| hsa_circ_0117029 | 1499 | 10 | 1.33% |
| hsa_circ_0126708 | 1499 | 10 | 1.33% |
| hsa_circ_0136313 | 1499 | 10 | 1.33% |
| hsa_circ_0034857 | 1649 | 11 | 1.33% |
| hsa_circ_0040001 | 1649 | 11 | 1.33% |
| hsa_circ_0047842 | 1649 | 11 | 1.33% |
| hsa_circ_0048859 | 1649 | 11 | 1.33% |
| hsa_circ_0075218 | 1649 | 11 | 1.33% |
| hsa_circ_0102832 | 1649 | 11 | 1.33% |
| hsa_circ_0127262 | 1649 | 11 | 1.33% |
| hsa_circ_0018522 | 1799 | 12 | 1.33% |
| hsa_circ_0034909 | 1799 | 12 | 1.33% |
| hsa_circ_0084745 | 1949 | 13 | 1.33% |
| hsa_circ_0034258 | 1800 | 12 | 1.33% |
| hsa_circ_0091182 | 1800 | 12 | 1.33% |
| hsa_circ_0099738 | 1800 | 12 | 1.33% |
| hsa_circ_0019359 | 1650 | 11 | 1.33% |
| hsa_circ_0035329 | 1650 | 11 | 1.33% |
| hsa_circ_0080126 | 1650 | 11 | 1.33% |
| hsa_circ_0125657 | 1650 | 11 | 1.33% |
| hsa_circ_0126216 | 1650 | 11 | 1.33% |
| hsa_circ_0013791 | 1500 | 10 | 1.33% |
| hsa_circ_0027899 | 1500 | 10 | 1.33% |
| hsa_circ_0036804 | 1500 | 10 | 1.33% |
| hsa_circ_0045151 | 1500 | 10 | 1.33% |
| hsa_circ_0063598 | 1500 | 10 | 1.33% |

| | | | |
|---|---|---|---|
| hsa_circ_0073053 | 1500 | 10 | 1.33% |
| hsa_circ_0085741 | 1500 | 10 | 1.33% |
| hsa_circ_0090820 | 1500 | 10 | 1.33% |
| hsa_circ_0095961 | 1500 | 10 | 1.33% |
| hsa_circ_0078811 | 3001 | 20 | 1.33% |
| hsa_circ_0015563 | 1951 | 13 | 1.33% |
| hsa_circ_0045816 | 1951 | 13 | 1.33% |
| hsa_circ_0072610 | 1951 | 13 | 1.33% |
| hsa_circ_0123605 | 1801 | 12 | 1.33% |
| hsa_circ_0138829 | 1801 | 12 | 1.33% |
| hsa_circ_0005842 | 1651 | 11 | 1.33% |
| hsa_circ_0031246 | 1651 | 11 | 1.33% |
| hsa_circ_0036523 | 1651 | 11 | 1.33% |
| hsa_circ_0042544 | 1651 | 11 | 1.33% |
| hsa_circ_0042555 | 1651 | 11 | 1.33% |
| hsa_circ_0072253 | 1651 | 11 | 1.33% |
| hsa_circ_0090799 | 1651 | 11 | 1.33% |
| hsa_circ_0093500 | 1651 | 11 | 1.33% |
| hsa_circ_0127176 | 1651 | 11 | 1.33% |
| hsa_circ_0026688 | 1501 | 10 | 1.33% |
| hsa_circ_0029964 | 1501 | 10 | 1.33% |
| hsa_circ_0034416 | 1501 | 10 | 1.33% |
| hsa_circ_0049241 | 1501 | 10 | 1.33% |
| hsa_circ_0075947 | 1501 | 10 | 1.33% |
| hsa_circ_0086443 | 1501 | 10 | 1.33% |
| hsa_circ_0089070 | 1501 | 10 | 1.33% |
| hsa_circ_0122719 | 1501 | 10 | 1.33% |
| hsa_circ_0123551 | 1501 | 10 | 1.33% |
| hsa_circ_0139012 | 1501 | 10 | 1.33% |
| hsa_circ_0038551 | 1952 | 13 | 1.33% |
| hsa_circ_0083479 | 1952 | 13 | 1.33% |
| hsa_circ_0034169 | 1802 | 12 | 1.33% |
| hsa_circ_0009826 | 1652 | 11 | 1.33% |
| hsa_circ_0036182 | 1652 | 11 | 1.33% |
| hsa_circ_0046250 | 1652 | 11 | 1.33% |
| hsa_circ_0051674 | 1652 | 11 | 1.33% |
| hsa_circ_0079352 | 1652 | 11 | 1.33% |
| hsa_circ_0111049 | 1652 | 11 | 1.33% |
| hsa_circ_0129779 | 1652 | 11 | 1.33% |
| hsa_circ_0052708 | 2253 | 15 | 1.33% |
| hsa_circ_0024078 | 1502 | 10 | 1.33% |
| hsa_circ_0027991 | 1502 | 10 | 1.33% |
| hsa_circ_0059975 | 1502 | 10 | 1.33% |
| hsa_circ_0072685 | 1502 | 10 | 1.33% |
| hsa_circ_0076906 | 1502 | 10 | 1.33% |
| hsa_circ_0098613 | 1502 | 10 | 1.33% |
| hsa_circ_0100179 | 1502 | 10 | 1.33% |
| hsa_circ_0112541 | 1502 | 10 | 1.33% |
| hsa_circ_0118745 | 1502 | 10 | 1.33% |
| hsa_circ_0138390 | 1502 | 10 | 1.33% |

| | | | |
|---|---|---|---|
| hsa_circ_0138854 | 1502 | 10 | 1.33% |
| hsa_circ_0093046 | 4657 | 31 | 1.33% |
| hsa_circ_0100671 | 1953 | 13 | 1.33% |
| hsa_circ_0102792 | 1953 | 13 | 1.33% |
| hsa_circ_0031279 | 1803 | 12 | 1.33% |
| hsa_circ_0066972 | 1803 | 12 | 1.33% |
| hsa_circ_0129205 | 1803 | 12 | 1.33% |
| hsa_circ_0018992 | 1653 | 11 | 1.33% |
| hsa_circ_0063253 | 1653 | 11 | 1.33% |
| hsa_circ_0094866 | 1653 | 11 | 1.33% |
| hsa_circ_0120363 | 1653 | 11 | 1.33% |
| hsa_circ_0137595 | 1653 | 11 | 1.33% |
| hsa_circ_0023855 | 1503 | 10 | 1.33% |
| hsa_circ_0029148 | 1503 | 10 | 1.33% |
| hsa_circ_0032571 | 1503 | 10 | 1.33% |
| hsa_circ_0047236 | 1503 | 10 | 1.33% |
| hsa_circ_0065459 | 1503 | 10 | 1.33% |
| hsa_circ_0071994 | 1503 | 10 | 1.33% |
| hsa_circ_0074338 | 1503 | 10 | 1.33% |
| hsa_circ_0100616 | 1503 | 10 | 1.33% |
| hsa_circ_0119818 | 1503 | 10 | 1.33% |
| hsa_circ_0138886 | 1503 | 10 | 1.33% |
| hsa_circ_0140695 | 1503 | 10 | 1.33% |
| hsa_circ_0136673 | 1954 | 13 | 1.33% |
| hsa_circ_0005414 | 4059 | 27 | 1.33% |
| hsa_circ_0113653 | 1804 | 12 | 1.33% |
| hsa_circ_0125606 | 1804 | 12 | 1.33% |
| hsa_circ_0048002 | 1654 | 11 | 1.33% |
| hsa_circ_0061069 | 1654 | 11 | 1.33% |
| hsa_circ_0072871 | 1654 | 11 | 1.33% |
| hsa_circ_0096289 | 1654 | 11 | 1.33% |
| hsa_circ_0107773 | 1654 | 11 | 1.33% |
| hsa_circ_0010308 | 1504 | 10 | 1.33% |
| hsa_circ_0012867 | 1504 | 10 | 1.33% |
| hsa_circ_0019608 | 1504 | 10 | 1.33% |
| hsa_circ_0025426 | 1504 | 10 | 1.33% |
| hsa_circ_0028454 | 1504 | 10 | 1.33% |
| hsa_circ_0028689 | 1504 | 10 | 1.33% |
| hsa_circ_0029359 | 1504 | 10 | 1.33% |
| hsa_circ_0034720 | 1504 | 10 | 1.33% |
| hsa_circ_0054640 | 1504 | 10 | 1.33% |
| hsa_circ_0057708 | 1504 | 10 | 1.33% |
| hsa_circ_0072302 | 1504 | 10 | 1.33% |
| hsa_circ_0123531 | 1504 | 10 | 1.33% |
| hsa_circ_0138385 | 1504 | 10 | 1.33% |
| hsa_circ_0008282 | 6317 | 42 | 1.33% |
| hsa_circ_0062072 | 1805 | 12 | 1.33% |
| hsa_circ_0122899 | 1805 | 12 | 1.33% |
| hsa_circ_0054605 | 3761 | 25 | 1.33% |
| hsa_circ_0089812 | 6018 | 40 | 1.33% |

| | | | |
|---|---|---|---|
| hsa_circ_0010818 | 1655 | 11 | 1.33% |
| hsa_circ_0025526 | 1655 | 11 | 1.33% |
| hsa_circ_0051983 | 1655 | 11 | 1.33% |
| hsa_circ_0059147 | 1655 | 11 | 1.33% |
| hsa_circ_0071811 | 1655 | 11 | 1.33% |
| hsa_circ_0046082 | 1956 | 13 | 1.33% |
| hsa_circ_0079299 | 1956 | 13 | 1.33% |
| hsa_circ_0021678 | 1505 | 10 | 1.33% |
| hsa_circ_0062374 | 1505 | 10 | 1.33% |
| hsa_circ_0069427 | 1505 | 10 | 1.33% |
| hsa_circ_0074014 | 1505 | 10 | 1.33% |
| hsa_circ_0084123 | 1505 | 10 | 1.33% |
| hsa_circ_0089470 | 1505 | 10 | 1.33% |
| hsa_circ_0092778 | 1505 | 10 | 1.33% |
| hsa_circ_0096615 | 1505 | 10 | 1.33% |
| hsa_circ_0096936 | 1505 | 10 | 1.33% |
| hsa_circ_0097726 | 1505 | 10 | 1.33% |
| hsa_circ_0105783 | 1505 | 10 | 1.33% |
| hsa_circ_0113391 | 1505 | 10 | 1.33% |
| hsa_circ_0062852 | 1957 | 13 | 1.33% |
| hsa_circ_0015254 | 1656 | 11 | 1.33% |
| hsa_circ_0018098 | 1656 | 11 | 1.33% |
| hsa_circ_0037296 | 1656 | 11 | 1.33% |
| hsa_circ_0046310 | 1656 | 11 | 1.33% |
| hsa_circ_0049571 | 1656 | 11 | 1.33% |
| hsa_circ_0065325 | 1656 | 11 | 1.33% |
| hsa_circ_0072803 | 1656 | 11 | 1.33% |
| hsa_circ_0072845 | 1656 | 11 | 1.33% |
| hsa_circ_0130485 | 1656 | 11 | 1.33% |
| hsa_circ_0009515 | 4969 | 33 | 1.33% |
| hsa_circ_0031278 | 1807 | 12 | 1.33% |
| hsa_circ_0075711 | 1807 | 12 | 1.33% |
| hsa_circ_0126492 | 4518 | 30 | 1.33% |
| hsa_circ_0005346 | 1506 | 10 | 1.33% |
| hsa_circ_0010724 | 1506 | 10 | 1.33% |
| hsa_circ_0010924 | 1506 | 10 | 1.33% |
| hsa_circ_0019555 | 1506 | 10 | 1.33% |
| hsa_circ_0025578 | 1506 | 10 | 1.33% |
| hsa_circ_0033597 | 1506 | 10 | 1.33% |
| hsa_circ_0042033 | 1506 | 10 | 1.33% |
| hsa_circ_0048619 | 1506 | 10 | 1.33% |
| hsa_circ_0057546 | 1506 | 10 | 1.33% |
| hsa_circ_0071950 | 1506 | 10 | 1.33% |
| hsa_circ_0073898 | 1506 | 10 | 1.33% |
| hsa_circ_0093187 | 1506 | 10 | 1.33% |
| hsa_circ_0096651 | 1506 | 10 | 1.33% |
| hsa_circ_0125619 | 1506 | 10 | 1.33% |
| hsa_circ_0101945 | 1958 | 13 | 1.33% |
| hsa_circ_0015635 | 1657 | 11 | 1.33% |
| hsa_circ_0047599 | 1657 | 11 | 1.33% |

| | | | |
|---|---|---|---|
| hsa_circ_0057943 | 1808 | 12 | 1.33% |
| hsa_circ_0116132 | 1959 | 13 | 1.33% |
| hsa_circ_0018064 | 1507 | 10 | 1.33% |
| hsa_circ_0021795 | 1507 | 10 | 1.33% |
| hsa_circ_0026501 | 1507 | 10 | 1.33% |
| hsa_circ_0029712 | 1507 | 10 | 1.33% |
| hsa_circ_0058632 | 1507 | 10 | 1.33% |
| hsa_circ_0063028 | 1507 | 10 | 1.33% |
| hsa_circ_0074007 | 1507 | 10 | 1.33% |
| hsa_circ_0077048 | 1507 | 10 | 1.33% |
| hsa_circ_0093654 | 1507 | 10 | 1.33% |
| hsa_circ_0099216 | 1507 | 10 | 1.33% |
| hsa_circ_0115179 | 1507 | 10 | 1.33% |
| hsa_circ_0121884 | 1507 | 10 | 1.33% |
| hsa_circ_0135981 | 1507 | 10 | 1.33% |
| hsa_circ_0135325 | 2562 | 17 | 1.33% |
| hsa_circ_0022867 | 1658 | 11 | 1.33% |
| hsa_circ_0038647 | 1658 | 11 | 1.33% |
| hsa_circ_0056929 | 1658 | 11 | 1.33% |
| hsa_circ_0118477 | 1658 | 11 | 1.33% |
| hsa_circ_0118675 | 2261 | 15 | 1.33% |
| hsa_circ_0008675 | 1809 | 12 | 1.33% |
| hsa_circ_0085995 | 1809 | 12 | 1.33% |
| hsa_circ_0042667 | 1508 | 10 | 1.33% |
| hsa_circ_0047508 | 1508 | 10 | 1.33% |
| hsa_circ_0059083 | 1508 | 10 | 1.33% |
| hsa_circ_0061835 | 1508 | 10 | 1.33% |
| hsa_circ_0084987 | 1508 | 10 | 1.33% |
| hsa_circ_0088381 | 1508 | 10 | 1.33% |
| hsa_circ_0088930 | 1508 | 10 | 1.33% |
| hsa_circ_0094557 | 1508 | 10 | 1.33% |
| hsa_circ_0135807 | 1508 | 10 | 1.33% |
| hsa_circ_0059142 | 1659 | 11 | 1.33% |
| hsa_circ_0085994 | 1659 | 11 | 1.33% |
| hsa_circ_0010331 | 1810 | 12 | 1.33% |
| hsa_circ_0078192 | 1810 | 12 | 1.33% |
| hsa_circ_0103553 | 1810 | 12 | 1.33% |
| hsa_circ_0119560 | 1810 | 12 | 1.33% |
| hsa_circ_0099581 | 2565 | 17 | 1.33% |
| hsa_circ_0037895 | 1509 | 10 | 1.33% |
| hsa_circ_0041262 | 1509 | 10 | 1.33% |
| hsa_circ_0049132 | 1509 | 10 | 1.33% |
| hsa_circ_0049882 | 1509 | 10 | 1.33% |
| hsa_circ_0061377 | 1509 | 10 | 1.33% |
| hsa_circ_0063066 | 1509 | 10 | 1.33% |
| hsa_circ_0082258 | 1509 | 10 | 1.33% |
| hsa_circ_0102604 | 1509 | 10 | 1.33% |
| hsa_circ_0106640 | 1509 | 10 | 1.33% |
| hsa_circ_0125264 | 1509 | 10 | 1.33% |
| hsa_circ_0064295 | 1660 | 11 | 1.33% |

| | | | |
|---|---|---|---|
| hsa_circ_0065395 | 1660 | 11 | 1.33% |
| hsa_circ_0136160 | 1660 | 11 | 1.33% |
| hsa_circ_0114452 | 1811 | 12 | 1.33% |
| hsa_circ_0129910 | 2113 | 14 | 1.33% |
| hsa_circ_0012050 | 1661 | 11 | 1.32% |
| hsa_circ_0056403 | 1661 | 11 | 1.32% |
| hsa_circ_0061698 | 1661 | 11 | 1.32% |
| hsa_circ_0124898 | 1661 | 11 | 1.32% |
| hsa_circ_0011029 | 1510 | 10 | 1.32% |
| hsa_circ_0015429 | 1510 | 10 | 1.32% |
| hsa_circ_0025776 | 1510 | 10 | 1.32% |
| hsa_circ_0069653 | 1510 | 10 | 1.32% |
| hsa_circ_0077507 | 4682 | 31 | 1.32% |
| hsa_circ_0050260 | 1964 | 13 | 1.32% |
| hsa_circ_0070539 | 1813 | 12 | 1.32% |
| hsa_circ_0076079 | 1813 | 12 | 1.32% |
| hsa_circ_0119117 | 1813 | 12 | 1.32% |
| hsa_circ_0016758 | 1662 | 11 | 1.32% |
| hsa_circ_0038272 | 1662 | 11 | 1.32% |
| hsa_circ_0045910 | 1662 | 11 | 1.32% |
| hsa_circ_0076331 | 1662 | 11 | 1.32% |
| hsa_circ_0124443 | 1662 | 11 | 1.32% |
| hsa_circ_0018127 | 1511 | 10 | 1.32% |
| hsa_circ_0044559 | 1511 | 10 | 1.32% |
| hsa_circ_0067941 | 1511 | 10 | 1.32% |
| hsa_circ_0073645 | 1511 | 10 | 1.32% |
| hsa_circ_0080584 | 1511 | 10 | 1.32% |
| hsa_circ_0085231 | 1511 | 10 | 1.32% |
| hsa_circ_0099327 | 1511 | 10 | 1.32% |
| hsa_circ_0108695 | 1511 | 10 | 1.32% |
| hsa_circ_0111623 | 1511 | 10 | 1.32% |
| hsa_circ_0116133 | 1511 | 10 | 1.32% |
| hsa_circ_0123374 | 1511 | 10 | 1.32% |
| hsa_circ_0125634 | 1511 | 10 | 1.32% |
| hsa_circ_0127282 | 1511 | 10 | 1.32% |
| hsa_circ_0137545 | 1511 | 10 | 1.32% |
| hsa_circ_0062463 | 30069 | 199 | 1.32% |
| hsa_circ_0052945 | 1965 | 13 | 1.32% |
| hsa_circ_0098230 | 1965 | 13 | 1.32% |
| hsa_circ_0116190 | 1965 | 13 | 1.32% |
| hsa_circ_0024770 | 1814 | 12 | 1.32% |
| hsa_circ_0073043 | 1814 | 12 | 1.32% |
| hsa_circ_0122223 | 1814 | 12 | 1.32% |
| hsa_circ_0061503 | 1663 | 11 | 1.32% |
| hsa_circ_0062548 | 1663 | 11 | 1.32% |
| hsa_circ_0068569 | 1663 | 11 | 1.32% |
| hsa_circ_0099560 | 1663 | 11 | 1.32% |
| hsa_circ_0112206 | 1663 | 11 | 1.32% |
| hsa_circ_0140066 | 1663 | 11 | 1.32% |
| hsa_circ_0016020 | 1512 | 10 | 1.32% |

| | | | |
|---|---|---|---|
| hsa_circ_0016673 | 1512 | 10 | 1.32% |
| hsa_circ_0020169 | 1512 | 10 | 1.32% |
| hsa_circ_0037758 | 1512 | 10 | 1.32% |
| hsa_circ_0041281 | 1512 | 10 | 1.32% |
| hsa_circ_0060386 | 1512 | 10 | 1.32% |
| hsa_circ_0063482 | 1512 | 10 | 1.32% |
| hsa_circ_0094419 | 1512 | 10 | 1.32% |
| hsa_circ_0125495 | 1512 | 10 | 1.32% |
| hsa_circ_0135023 | 1512 | 10 | 1.32% |
| hsa_circ_0137844 | 1512 | 10 | 1.32% |
| hsa_circ_0074222 | 1966 | 13 | 1.32% |
| hsa_circ_0030505 | 1815 | 12 | 1.32% |
| hsa_circ_0058357 | 1815 | 12 | 1.32% |
| hsa_circ_0067598 | 1815 | 12 | 1.32% |
| hsa_circ_0128695 | 1815 | 12 | 1.32% |
| hsa_circ_0005450 | 1664 | 11 | 1.32% |
| hsa_circ_0022073 | 1664 | 11 | 1.32% |
| hsa_circ_0046087 | 1664 | 11 | 1.32% |
| hsa_circ_0056781 | 1664 | 11 | 1.32% |
| hsa_circ_0062634 | 1664 | 11 | 1.32% |
| hsa_circ_0069195 | 1664 | 11 | 1.32% |
| hsa_circ_0048073 | 2118 | 14 | 1.32% |
| hsa_circ_0044635 | 1513 | 10 | 1.32% |
| hsa_circ_0057474 | 1513 | 10 | 1.32% |
| hsa_circ_0077045 | 1513 | 10 | 1.32% |
| hsa_circ_0110609 | 1513 | 10 | 1.32% |
| hsa_circ_0124065 | 1513 | 10 | 1.32% |
| hsa_circ_0127092 | 1513 | 10 | 1.32% |
| hsa_circ_0135415 | 1513 | 10 | 1.32% |
| hsa_circ_0035474 | 1967 | 13 | 1.32% |
| hsa_circ_0123431 | 1967 | 13 | 1.32% |
| hsa_circ_0066684 | 2421 | 16 | 1.32% |
| hsa_circ_0025741 | 1816 | 12 | 1.32% |
| hsa_circ_0085962 | 1816 | 12 | 1.32% |
| hsa_circ_0113870 | 1816 | 12 | 1.32% |
| hsa_circ_0044421 | 1665 | 11 | 1.32% |
| hsa_circ_0047908 | 1665 | 11 | 1.32% |
| hsa_circ_0050856 | 1665 | 11 | 1.32% |
| hsa_circ_0053362 | 1665 | 11 | 1.32% |
| hsa_circ_0069257 | 1665 | 11 | 1.32% |
| hsa_circ_0079535 | 1665 | 11 | 1.32% |
| hsa_circ_0083384 | 1665 | 11 | 1.32% |
| hsa_circ_0083538 | 1665 | 11 | 1.32% |
| hsa_circ_0085194 | 1665 | 11 | 1.32% |
| hsa_circ_0094339 | 1665 | 11 | 1.32% |
| hsa_circ_0140525 | 1665 | 11 | 1.32% |
| hsa_circ_0136724 | 1968 | 13 | 1.32% |
| hsa_circ_0062846 | 2271 | 15 | 1.32% |
| hsa_circ_0010537 | 1514 | 10 | 1.32% |
| hsa_circ_0011639 | 1514 | 10 | 1.32% |

| | | | |
|---|---|---|---|
| hsa_circ_0027882 | 1514 | 10 | 1.32% |
| hsa_circ_0042393 | 1514 | 10 | 1.32% |
| hsa_circ_0046387 | 1514 | 10 | 1.32% |
| hsa_circ_0052659 | 1514 | 10 | 1.32% |
| hsa_circ_0054485 | 1514 | 10 | 1.32% |
| hsa_circ_0060209 | 1514 | 10 | 1.32% |
| hsa_circ_0063439 | 1514 | 10 | 1.32% |
| hsa_circ_0084282 | 1514 | 10 | 1.32% |
| hsa_circ_0058778 | 1817 | 12 | 1.32% |
| hsa_circ_0006578 | 1666 | 11 | 1.32% |
| hsa_circ_0011251 | 1666 | 11 | 1.32% |
| hsa_circ_0020887 | 1666 | 11 | 1.32% |
| hsa_circ_0021268 | 1666 | 11 | 1.32% |
| hsa_circ_0076455 | 1666 | 11 | 1.32% |
| hsa_circ_0077974 | 1666 | 11 | 1.32% |
| hsa_circ_0098144 | 1666 | 11 | 1.32% |
| hsa_circ_0139003 | 1666 | 11 | 1.32% |
| hsa_circ_0139101 | 1666 | 11 | 1.32% |
| hsa_circ_0125013 | 3636 | 24 | 1.32% |
| hsa_circ_0064508 | 1818 | 12 | 1.32% |
| hsa_circ_0088887 | 1818 | 12 | 1.32% |
| hsa_circ_0019517 | 1515 | 10 | 1.32% |
| hsa_circ_0020388 | 1515 | 10 | 1.32% |
| hsa_circ_0046327 | 1515 | 10 | 1.32% |
| hsa_circ_0064247 | 1515 | 10 | 1.32% |
| hsa_circ_0064617 | 1515 | 10 | 1.32% |
| hsa_circ_0082146 | 1515 | 10 | 1.32% |
| hsa_circ_0083986 | 1515 | 10 | 1.32% |
| hsa_circ_0099364 | 1515 | 10 | 1.32% |
| hsa_circ_0103883 | 1515 | 10 | 1.32% |
| hsa_circ_0105921 | 1515 | 10 | 1.32% |
| hsa_circ_0106372 | 1515 | 10 | 1.32% |
| hsa_circ_0046165 | 1970 | 13 | 1.32% |
| hsa_circ_0044891 | 1667 | 11 | 1.32% |
| hsa_circ_0117848 | 1667 | 11 | 1.32% |
| hsa_circ_0028180 | 1819 | 12 | 1.32% |
| hsa_circ_0052281 | 1819 | 12 | 1.32% |
| hsa_circ_0082701 | 1819 | 12 | 1.32% |
| hsa_circ_0089231 | 1819 | 12 | 1.32% |
| hsa_circ_0091306 | 2274 | 15 | 1.32% |
| hsa_circ_0012373 | 1516 | 10 | 1.32% |
| hsa_circ_0015889 | 1516 | 10 | 1.32% |
| hsa_circ_0027676 | 1516 | 10 | 1.32% |
| hsa_circ_0048040 | 1516 | 10 | 1.32% |
| hsa_circ_0049617 | 1516 | 10 | 1.32% |
| hsa_circ_0050993 | 1516 | 10 | 1.32% |
| hsa_circ_0054645 | 1516 | 10 | 1.32% |
| hsa_circ_0061799 | 1516 | 10 | 1.32% |
| hsa_circ_0073193 | 1516 | 10 | 1.32% |
| hsa_circ_0077847 | 1516 | 10 | 1.32% |

| | | | |
|---|---|---|---|
| hsa_circ_0103867 | 1516 | 10 | 1.32% |
| hsa_circ_0132651 | 1516 | 10 | 1.32% |
| hsa_circ_0010993 | 3487 | 23 | 1.32% |
| hsa_circ_0014421 | 1668 | 11 | 1.32% |
| hsa_circ_0026626 | 1668 | 11 | 1.32% |
| hsa_circ_0028533 | 1668 | 11 | 1.32% |
| hsa_circ_0080592 | 1668 | 11 | 1.32% |
| hsa_circ_0004287 | 2275 | 15 | 1.32% |
| hsa_circ_0010085 | 1820 | 12 | 1.32% |
| hsa_circ_0039564 | 1820 | 12 | 1.32% |
| hsa_circ_0067239 | 1820 | 12 | 1.32% |
| hsa_circ_0008780 | 6371 | 42 | 1.32% |
| hsa_circ_0046097 | 1972 | 13 | 1.32% |
| hsa_circ_0009730 | 1517 | 10 | 1.32% |
| hsa_circ_0017726 | 1517 | 10 | 1.32% |
| hsa_circ_0021896 | 1517 | 10 | 1.32% |
| hsa_circ_0045159 | 1517 | 10 | 1.32% |
| hsa_circ_0074956 | 1517 | 10 | 1.32% |
| hsa_circ_0113668 | 1517 | 10 | 1.32% |
| hsa_circ_0132314 | 1517 | 10 | 1.32% |
| hsa_circ_0080087 | 2124 | 14 | 1.32% |
| hsa_circ_0019830 | 1669 | 11 | 1.32% |
| hsa_circ_0047855 | 1669 | 11 | 1.32% |
| hsa_circ_0059276 | 1669 | 11 | 1.32% |
| hsa_circ_0076017 | 1669 | 11 | 1.32% |
| hsa_circ_0106470 | 1669 | 11 | 1.32% |
| hsa_circ_0115607 | 2428 | 16 | 1.32% |
| hsa_circ_0038191 | 1821 | 12 | 1.32% |
| hsa_circ_0119535 | 1821 | 12 | 1.32% |
| hsa_circ_0019002 | 1518 | 10 | 1.32% |
| hsa_circ_0029236 | 1518 | 10 | 1.32% |
| hsa_circ_0034443 | 1518 | 10 | 1.32% |
| hsa_circ_0041360 | 1518 | 10 | 1.32% |
| hsa_circ_0042554 | 1518 | 10 | 1.32% |
| hsa_circ_0079979 | 1518 | 10 | 1.32% |
| hsa_circ_0085247 | 1518 | 10 | 1.32% |
| hsa_circ_0122504 | 1518 | 10 | 1.32% |
| hsa_circ_0027746 | 1670 | 11 | 1.32% |
| hsa_circ_0048791 | 1670 | 11 | 1.32% |
| hsa_circ_0039878 | 1822 | 12 | 1.32% |
| hsa_circ_0105476 | 1822 | 12 | 1.32% |
| hsa_circ_0002999 | 1519 | 10 | 1.32% |
| hsa_circ_0006211 | 1519 | 10 | 1.32% |
| hsa_circ_0017360 | 1519 | 10 | 1.32% |
| hsa_circ_0020772 | 1519 | 10 | 1.32% |
| hsa_circ_0022099 | 1519 | 10 | 1.32% |
| hsa_circ_0055148 | 1519 | 10 | 1.32% |
| hsa_circ_0075880 | 1519 | 10 | 1.32% |
| hsa_circ_0082297 | 1519 | 10 | 1.32% |
| hsa_circ_0101908 | 1519 | 10 | 1.32% |

| | | | |
|---|---|---|---|
| hsa_circ_0111812 | 1519 | 10 | 1.32% |
| hsa_circ_0131335 | 1519 | 10 | 1.32% |
| hsa_circ_0048424 | 1671 | 11 | 1.32% |
| hsa_circ_0053287 | 1671 | 11 | 1.32% |
| hsa_circ_0034367 | 1823 | 12 | 1.32% |
| hsa_circ_0102868 | 1823 | 12 | 1.32% |
| hsa_circ_0114387 | 3950 | 26 | 1.32% |
| hsa_circ_0090756 | 1975 | 13 | 1.32% |
| hsa_circ_0108918 | 1975 | 13 | 1.32% |
| hsa_circ_0027172 | 1824 | 12 | 1.32% |
| hsa_circ_0039089 | 1824 | 12 | 1.32% |
| hsa_circ_0094021 | 1824 | 12 | 1.32% |
| hsa_circ_0023211 | 1672 | 11 | 1.32% |
| hsa_circ_0027064 | 1672 | 11 | 1.32% |
| hsa_circ_0093671 | 1672 | 11 | 1.32% |
| hsa_circ_0094493 | 1672 | 11 | 1.32% |
| hsa_circ_0112668 | 1672 | 11 | 1.32% |
| hsa_circ_0002767 | 1520 | 10 | 1.32% |
| hsa_circ_0012602 | 1520 | 10 | 1.32% |
| hsa_circ_0018343 | 1520 | 10 | 1.32% |
| hsa_circ_0020446 | 1520 | 10 | 1.32% |
| hsa_circ_0022325 | 1520 | 10 | 1.32% |
| hsa_circ_0031554 | 1520 | 10 | 1.32% |
| hsa_circ_0046623 | 1520 | 10 | 1.32% |
| hsa_circ_0047568 | 1520 | 10 | 1.32% |
| hsa_circ_0055997 | 1520 | 10 | 1.32% |
| hsa_circ_0077049 | 1520 | 10 | 1.32% |
| hsa_circ_0089401 | 1520 | 10 | 1.32% |
| hsa_circ_0075538 | 2281 | 15 | 1.32% |
| hsa_circ_0106298 | 4106 | 27 | 1.32% |
| hsa_circ_0077960 | 1977 | 13 | 1.32% |
| hsa_circ_0011846 | 1825 | 12 | 1.32% |
| hsa_circ_0040695 | 1825 | 12 | 1.32% |
| hsa_circ_0050676 | 1825 | 12 | 1.32% |
| hsa_circ_0011403 | 1673 | 11 | 1.32% |
| hsa_circ_0025210 | 1673 | 11 | 1.32% |
| hsa_circ_0054954 | 1673 | 11 | 1.32% |
| hsa_circ_0068328 | 1673 | 11 | 1.32% |
| hsa_circ_0004858 | 1521 | 10 | 1.31% |
| hsa_circ_0025840 | 1521 | 10 | 1.31% |
| hsa_circ_0037036 | 1521 | 10 | 1.31% |
| hsa_circ_0045354 | 1521 | 10 | 1.31% |
| hsa_circ_0050030 | 1521 | 10 | 1.31% |
| hsa_circ_0063524 | 1521 | 10 | 1.31% |
| hsa_circ_0074585 | 1521 | 10 | 1.31% |
| hsa_circ_0076293 | 1521 | 10 | 1.31% |
| hsa_circ_0094861 | 1521 | 10 | 1.31% |
| hsa_circ_0124654 | 1521 | 10 | 1.31% |
| hsa_circ_0048778 | 1826 | 12 | 1.31% |
| hsa_circ_0101392 | 3348 | 22 | 1.31% |

| | | | |
|---|---|---|---|
| hsa_circ_0009607 | 1674 | 11 | 1.31% |
| hsa_circ_0022747 | 1674 | 11 | 1.31% |
| hsa_circ_0029116 | 1674 | 11 | 1.31% |
| hsa_circ_0038454 | 1674 | 11 | 1.31% |
| hsa_circ_0057663 | 1674 | 11 | 1.31% |
| hsa_circ_0065261 | 1674 | 11 | 1.31% |
| hsa_circ_0083881 | 1674 | 11 | 1.31% |
| hsa_circ_0105181 | 1674 | 11 | 1.31% |
| hsa_circ_0116085 | 1674 | 11 | 1.31% |
| hsa_circ_0120072 | 1674 | 11 | 1.31% |
| hsa_circ_0121982 | 1674 | 11 | 1.31% |
| hsa_circ_0122266 | 1674 | 11 | 1.31% |
| hsa_circ_0012729 | 1522 | 10 | 1.31% |
| hsa_circ_0015018 | 1522 | 10 | 1.31% |
| hsa_circ_0034055 | 1522 | 10 | 1.31% |
| hsa_circ_0042326 | 1522 | 10 | 1.31% |
| hsa_circ_0065777 | 1522 | 10 | 1.31% |
| hsa_circ_0066080 | 1522 | 10 | 1.31% |
| hsa_circ_0090962 | 1522 | 10 | 1.31% |
| hsa_circ_0115498 | 1522 | 10 | 1.31% |
| hsa_circ_0110525 | 5175 | 34 | 1.31% |
| hsa_circ_0104040 | 2131 | 14 | 1.31% |
| hsa_circ_0074452 | 1827 | 12 | 1.31% |
| hsa_circ_0052076 | 1675 | 11 | 1.31% |
| hsa_circ_0089522 | 1675 | 11 | 1.31% |
| hsa_circ_0136118 | 1675 | 11 | 1.31% |
| hsa_circ_0120856 | 4112 | 27 | 1.31% |
| hsa_circ_0111390 | 3046 | 20 | 1.31% |
| hsa_circ_0002593 | 1523 | 10 | 1.31% |
| hsa_circ_0019289 | 1523 | 10 | 1.31% |
| hsa_circ_0028740 | 1523 | 10 | 1.31% |
| hsa_circ_0040940 | 1523 | 10 | 1.31% |
| hsa_circ_0047392 | 1523 | 10 | 1.31% |
| hsa_circ_0063123 | 1523 | 10 | 1.31% |
| hsa_circ_0068154 | 1523 | 10 | 1.31% |
| hsa_circ_0077750 | 1523 | 10 | 1.31% |
| hsa_circ_0087290 | 1523 | 10 | 1.31% |
| hsa_circ_0092431 | 1523 | 10 | 1.31% |
| hsa_circ_0111351 | 1523 | 10 | 1.31% |
| hsa_circ_0034074 | 1828 | 12 | 1.31% |
| hsa_circ_0035773 | 1828 | 12 | 1.31% |
| hsa_circ_0077601 | 1828 | 12 | 1.31% |
| hsa_circ_0010759 | 1676 | 11 | 1.31% |
| hsa_circ_0055772 | 1676 | 11 | 1.31% |
| hsa_circ_0091039 | 1676 | 11 | 1.31% |
| hsa_circ_0060290 | 2743 | 18 | 1.31% |
| hsa_circ_0015268 | 1524 | 10 | 1.31% |
| hsa_circ_0029886 | 1524 | 10 | 1.31% |
| hsa_circ_0035662 | 1524 | 10 | 1.31% |
| hsa_circ_0042597 | 1524 | 10 | 1.31% |

| | | | |
|---|---|---|---|
| hsa_circ_0043477 | 1524 | 10 | 1.31% |
| hsa_circ_0059427 | 1524 | 10 | 1.31% |
| hsa_circ_0065273 | 1524 | 10 | 1.31% |
| hsa_circ_0088737 | 1524 | 10 | 1.31% |
| hsa_circ_0089890 | 1524 | 10 | 1.31% |
| hsa_circ_0102271 | 1524 | 10 | 1.31% |
| hsa_circ_0117222 | 1524 | 10 | 1.31% |
| hsa_circ_0125990 | 1524 | 10 | 1.31% |
| hsa_circ_0135457 | 1524 | 10 | 1.31% |
| hsa_circ_0003821 | 1829 | 12 | 1.31% |
| hsa_circ_0032602 | 1829 | 12 | 1.31% |
| hsa_circ_0034167 | 1677 | 11 | 1.31% |
| hsa_circ_0087678 | 1677 | 11 | 1.31% |
| hsa_circ_0089155 | 1677 | 11 | 1.31% |
| hsa_circ_0104836 | 1677 | 11 | 1.31% |
| hsa_circ_0022102 | 1982 | 13 | 1.31% |
| hsa_circ_0063511 | 1982 | 13 | 1.31% |
| hsa_circ_0095517 | 1982 | 13 | 1.31% |
| hsa_circ_0128990 | 1982 | 13 | 1.31% |
| hsa_circ_0124147 | 3965 | 26 | 1.31% |
| hsa_circ_0091800 | 1830 | 12 | 1.31% |
| hsa_circ_0128478 | 1830 | 12 | 1.31% |
| hsa_circ_0011918 | 1525 | 10 | 1.31% |
| hsa_circ_0017403 | 1525 | 10 | 1.31% |
| hsa_circ_0024883 | 1525 | 10 | 1.31% |
| hsa_circ_0047634 | 1525 | 10 | 1.31% |
| hsa_circ_0057736 | 1525 | 10 | 1.31% |
| hsa_circ_0067122 | 1525 | 10 | 1.31% |
| hsa_circ_0073191 | 1525 | 10 | 1.31% |
| hsa_circ_0078004 | 1525 | 10 | 1.31% |
| hsa_circ_0081940 | 1525 | 10 | 1.31% |
| hsa_circ_0107045 | 1525 | 10 | 1.31% |
| hsa_circ_0120464 | 1525 | 10 | 1.31% |
| hsa_circ_0129137 | 1525 | 10 | 1.31% |
| hsa_circ_0132306 | 1525 | 10 | 1.31% |
| hsa_circ_0094114 | 6253 | 41 | 1.31% |
| hsa_circ_0016361 | 1678 | 11 | 1.31% |
| hsa_circ_0019004 | 1678 | 11 | 1.31% |
| hsa_circ_0028127 | 1678 | 11 | 1.31% |
| hsa_circ_0038128 | 1678 | 11 | 1.31% |
| hsa_circ_0038243 | 1678 | 11 | 1.31% |
| hsa_circ_0043842 | 1678 | 11 | 1.31% |
| hsa_circ_0061662 | 1678 | 11 | 1.31% |
| hsa_circ_0062282 | 1678 | 11 | 1.31% |
| hsa_circ_0087895 | 1678 | 11 | 1.31% |
| hsa_circ_0136947 | 1678 | 11 | 1.31% |
| hsa_circ_0016577 | 1831 | 12 | 1.31% |
| hsa_circ_0018691 | 1526 | 10 | 1.31% |
| hsa_circ_0021951 | 1526 | 10 | 1.31% |
| hsa_circ_0036245 | 1526 | 10 | 1.31% |

| | | | |
|---|---|---|---|
| hsa_circ_0041709 | 1526 | 10 | 1.31% |
| hsa_circ_0061094 | 1526 | 10 | 1.31% |
| hsa_circ_0064960 | 1526 | 10 | 1.31% |
| hsa_circ_0085436 | 1526 | 10 | 1.31% |
| hsa_circ_0116798 | 1526 | 10 | 1.31% |
| hsa_circ_0130479 | 1526 | 10 | 1.31% |
| hsa_circ_0132997 | 1526 | 10 | 1.31% |
| hsa_circ_0028525 | 1679 | 11 | 1.31% |
| hsa_circ_0051066 | 1679 | 11 | 1.31% |
| hsa_circ_0054147 | 1679 | 11 | 1.31% |
| hsa_circ_0066829 | 1679 | 11 | 1.31% |
| hsa_circ_0116946 | 1679 | 11 | 1.31% |
| hsa_circ_0118036 | 1679 | 11 | 1.31% |
| hsa_circ_0121952 | 1679 | 11 | 1.31% |
| hsa_circ_0127266 | 1679 | 11 | 1.31% |
| hsa_circ_0129368 | 1679 | 11 | 1.31% |
| hsa_circ_0139261 | 1679 | 11 | 1.31% |
| hsa_circ_0014040 | 3053 | 20 | 1.31% |
| hsa_circ_0075181 | 1832 | 12 | 1.31% |
| hsa_circ_0079278 | 1832 | 12 | 1.31% |
| hsa_circ_0010915 | 1527 | 10 | 1.31% |
| hsa_circ_0023137 | 1527 | 10 | 1.31% |
| hsa_circ_0031822 | 1527 | 10 | 1.31% |
| hsa_circ_0039846 | 1527 | 10 | 1.31% |
| hsa_circ_0040127 | 1527 | 10 | 1.31% |
| hsa_circ_0061512 | 1527 | 10 | 1.31% |
| hsa_circ_0065578 | 1527 | 10 | 1.31% |
| hsa_circ_0109501 | 1527 | 10 | 1.31% |
| hsa_circ_0113634 | 1527 | 10 | 1.31% |
| hsa_circ_0134008 | 1527 | 10 | 1.31% |
| hsa_circ_0022542 | 2596 | 17 | 1.31% |
| hsa_circ_0062189 | 3207 | 21 | 1.31% |
| hsa_circ_0097707 | 6567 | 43 | 1.31% |
| hsa_circ_0031112 | 1680 | 11 | 1.31% |
| hsa_circ_0035281 | 1680 | 11 | 1.31% |
| hsa_circ_0044227 | 1680 | 11 | 1.31% |
| hsa_circ_0063875 | 1680 | 11 | 1.31% |
| hsa_circ_0076359 | 1680 | 11 | 1.31% |
| hsa_circ_0092299 | 1680 | 11 | 1.31% |
| hsa_circ_0029287 | 1833 | 12 | 1.31% |
| hsa_circ_0030014 | 1528 | 10 | 1.31% |
| hsa_circ_0030129 | 1528 | 10 | 1.31% |
| hsa_circ_0033353 | 1528 | 10 | 1.31% |
| hsa_circ_0053283 | 1528 | 10 | 1.31% |
| hsa_circ_0086514 | 1528 | 10 | 1.31% |
| hsa_circ_0091524 | 1528 | 10 | 1.31% |
| hsa_circ_0094547 | 1528 | 10 | 1.31% |
| hsa_circ_0098978 | 1528 | 10 | 1.31% |
| hsa_circ_0019539 | 1681 | 11 | 1.31% |
| hsa_circ_0035843 | 1681 | 11 | 1.31% |

| | | | |
|---|---|---|---|
| hsa_circ_0067639 | 1681 | 11 | 1.31% |
| hsa_circ_0067896 | 1681 | 11 | 1.31% |
| hsa_circ_0097456 | 1987 | 13 | 1.31% |
| hsa_circ_0017386 | 1529 | 10 | 1.31% |
| hsa_circ_0036338 | 1529 | 10 | 1.31% |
| hsa_circ_0072043 | 1529 | 10 | 1.31% |
| hsa_circ_0085975 | 1529 | 10 | 1.31% |
| hsa_circ_0110118 | 1529 | 10 | 1.31% |
| hsa_circ_0137700 | 1529 | 10 | 1.31% |
| hsa_circ_0137999 | 1529 | 10 | 1.31% |
| hsa_circ_0022047 | 1682 | 11 | 1.31% |
| hsa_circ_0041621 | 1682 | 11 | 1.31% |
| hsa_circ_0082596 | 1682 | 11 | 1.31% |
| hsa_circ_0120709 | 1682 | 11 | 1.31% |
| hsa_circ_0127632 | 5199 | 34 | 1.31% |
| hsa_circ_0131708 | 2600 | 17 | 1.31% |
| hsa_circ_0067308 | 1836 | 12 | 1.31% |
| hsa_circ_0059902 | 1683 | 11 | 1.31% |
| hsa_circ_0068374 | 1683 | 11 | 1.31% |
| hsa_circ_0086216 | 1683 | 11 | 1.31% |
| hsa_circ_0117349 | 1683 | 11 | 1.31% |
| hsa_circ_0125055 | 1683 | 11 | 1.31% |
| hsa_circ_0014300 | 1530 | 10 | 1.31% |
| hsa_circ_0037768 | 1530 | 10 | 1.31% |
| hsa_circ_0043232 | 1530 | 10 | 1.31% |
| hsa_circ_0047746 | 1530 | 10 | 1.31% |
| hsa_circ_0052520 | 1530 | 10 | 1.31% |
| hsa_circ_0063978 | 1530 | 10 | 1.31% |
| hsa_circ_0074180 | 1530 | 10 | 1.31% |
| hsa_circ_0081681 | 1530 | 10 | 1.31% |
| hsa_circ_0090231 | 1530 | 10 | 1.31% |
| hsa_circ_0091959 | 1530 | 10 | 1.31% |
| hsa_circ_0107971 | 1530 | 10 | 1.31% |
| hsa_circ_0113216 | 1530 | 10 | 1.31% |
| hsa_circ_0059557 | 2143 | 14 | 1.31% |
| hsa_circ_0002817 | 1990 | 13 | 1.31% |
| hsa_circ_0082096 | 1837 | 12 | 1.31% |
| hsa_circ_0021185 | 1684 | 11 | 1.31% |
| hsa_circ_0039057 | 1684 | 11 | 1.31% |
| hsa_circ_0041601 | 1684 | 11 | 1.31% |
| hsa_circ_0065493 | 1684 | 11 | 1.31% |
| hsa_circ_0071325 | 1684 | 11 | 1.31% |
| hsa_circ_0098134 | 1684 | 11 | 1.31% |
| hsa_circ_0136379 | 1684 | 11 | 1.31% |
| hsa_circ_0023674 | 1531 | 10 | 1.31% |
| hsa_circ_0026511 | 1531 | 10 | 1.31% |
| hsa_circ_0043793 | 1531 | 10 | 1.31% |
| hsa_circ_0044911 | 1531 | 10 | 1.31% |
| hsa_circ_0076039 | 1531 | 10 | 1.31% |
| hsa_circ_0078482 | 1531 | 10 | 1.31% |

| | | | |
|---|---|---|---|
| hsa_circ_0139751 | 1531 | 10 | 1.31% |
| hsa_circ_0120454 | 3063 | 20 | 1.31% |
| hsa_circ_0034852 | 1838 | 12 | 1.31% |
| hsa_circ_0072784 | 1838 | 12 | 1.31% |
| hsa_circ_0121513 | 1838 | 12 | 1.31% |
| hsa_circ_0011325 | 1685 | 11 | 1.31% |
| hsa_circ_0029483 | 1685 | 11 | 1.31% |
| hsa_circ_0030733 | 1685 | 11 | 1.31% |
| hsa_circ_0087534 | 1685 | 11 | 1.31% |
| hsa_circ_0090177 | 1685 | 11 | 1.31% |
| hsa_circ_0117559 | 1685 | 11 | 1.31% |
| hsa_circ_0041196 | 2298 | 15 | 1.31% |
| hsa_circ_0039883 | 1532 | 10 | 1.31% |
| hsa_circ_0040841 | 1532 | 10 | 1.31% |
| hsa_circ_0049706 | 1532 | 10 | 1.31% |
| hsa_circ_0060505 | 1532 | 10 | 1.31% |
| hsa_circ_0062851 | 1532 | 10 | 1.31% |
| hsa_circ_0080269 | 1532 | 10 | 1.31% |
| hsa_circ_0084897 | 1532 | 10 | 1.31% |
| hsa_circ_0086495 | 1532 | 10 | 1.31% |
| hsa_circ_0091744 | 1532 | 10 | 1.31% |
| hsa_circ_0110419 | 1532 | 10 | 1.31% |
| hsa_circ_0120403 | 1532 | 10 | 1.31% |
| hsa_circ_0124302 | 1532 | 10 | 1.31% |
| hsa_circ_0124748 | 1532 | 10 | 1.31% |
| hsa_circ_0017261 | 1992 | 13 | 1.31% |
| hsa_circ_0066059 | 1992 | 13 | 1.31% |
| hsa_circ_0026075 | 1839 | 12 | 1.31% |
| hsa_circ_0030871 | 1839 | 12 | 1.31% |
| hsa_circ_0048152 | 1839 | 12 | 1.31% |
| hsa_circ_0028968 | 1686 | 11 | 1.30% |
| hsa_circ_0044801 | 1686 | 11 | 1.30% |
| hsa_circ_0116129 | 1686 | 11 | 1.30% |
| hsa_circ_0024154 | 1533 | 10 | 1.30% |
| hsa_circ_0035152 | 1533 | 10 | 1.30% |
| hsa_circ_0044937 | 1533 | 10 | 1.30% |
| hsa_circ_0076076 | 1533 | 10 | 1.30% |
| hsa_circ_0081079 | 1533 | 10 | 1.30% |
| hsa_circ_0082031 | 1533 | 10 | 1.30% |
| hsa_circ_0086985 | 1533 | 10 | 1.30% |
| hsa_circ_0091889 | 1533 | 10 | 1.30% |
| hsa_circ_0099094 | 1533 | 10 | 1.30% |
| hsa_circ_0108709 | 1533 | 10 | 1.30% |
| hsa_circ_0122486 | 1533 | 10 | 1.30% |
| hsa_circ_0126204 | 1533 | 10 | 1.30% |
| hsa_circ_0052213 | 1993 | 13 | 1.30% |
| hsa_circ_0119109 | 1993 | 13 | 1.30% |
| hsa_circ_0137044 | 1993 | 13 | 1.30% |
| hsa_circ_0002922 | 1840 | 12 | 1.30% |
| hsa_circ_0014104 | 1840 | 12 | 1.30% |

| | | | |
|---|---|---|---|
| hsa_circ_0083269 | 1840 | 12 | 1.30% |
| hsa_circ_0118382 | 1840 | 12 | 1.30% |
| hsa_circ_0007023 | 2147 | 14 | 1.30% |
| hsa_circ_0015046 | 1687 | 11 | 1.30% |
| hsa_circ_0016901 | 1687 | 11 | 1.30% |
| hsa_circ_0038083 | 1687 | 11 | 1.30% |
| hsa_circ_0046371 | 1687 | 11 | 1.30% |
| hsa_circ_0046469 | 1687 | 11 | 1.30% |
| hsa_circ_0085836 | 1687 | 11 | 1.30% |
| hsa_circ_0091684 | 1687 | 11 | 1.30% |
| hsa_circ_0105113 | 1687 | 11 | 1.30% |
| hsa_circ_0138081 | 1687 | 11 | 1.30% |
| hsa_circ_0052873 | 1994 | 13 | 1.30% |
| hsa_circ_0048162 | 2301 | 15 | 1.30% |
| hsa_circ_0026261 | 1534 | 10 | 1.30% |
| hsa_circ_0041735 | 1534 | 10 | 1.30% |
| hsa_circ_0046756 | 1534 | 10 | 1.30% |
| hsa_circ_0052467 | 1534 | 10 | 1.30% |
| hsa_circ_0055417 | 1534 | 10 | 1.30% |
| hsa_circ_0060717 | 1534 | 10 | 1.30% |
| hsa_circ_0061947 | 1534 | 10 | 1.30% |
| hsa_circ_0081374 | 1534 | 10 | 1.30% |
| hsa_circ_0086077 | 1534 | 10 | 1.30% |
| hsa_circ_0100352 | 1534 | 10 | 1.30% |
| hsa_circ_0102374 | 1534 | 10 | 1.30% |
| hsa_circ_0123564 | 1534 | 10 | 1.30% |
| hsa_circ_0127670 | 1534 | 10 | 1.30% |
| hsa_circ_0006154 | 1841 | 12 | 1.30% |
| hsa_circ_0016992 | 1841 | 12 | 1.30% |
| hsa_circ_0111514 | 1841 | 12 | 1.30% |
| hsa_circ_0040888 | 1688 | 11 | 1.30% |
| hsa_circ_0137058 | 1688 | 11 | 1.30% |
| hsa_circ_0022396 | 1842 | 12 | 1.30% |
| hsa_circ_0036656 | 1842 | 12 | 1.30% |
| hsa_circ_0076167 | 1842 | 12 | 1.30% |
| hsa_circ_0104186 | 1842 | 12 | 1.30% |
| hsa_circ_0013243 | 1535 | 10 | 1.30% |
| hsa_circ_0013779 | 1535 | 10 | 1.30% |
| hsa_circ_0020665 | 1535 | 10 | 1.30% |
| hsa_circ_0036935 | 1535 | 10 | 1.30% |
| hsa_circ_0063959 | 1535 | 10 | 1.30% |
| hsa_circ_0064107 | 1535 | 10 | 1.30% |
| hsa_circ_0084722 | 1535 | 10 | 1.30% |
| hsa_circ_0106236 | 1535 | 10 | 1.30% |
| hsa_circ_0115515 | 1535 | 10 | 1.30% |
| hsa_circ_0019762 | 1689 | 11 | 1.30% |
| hsa_circ_0059003 | 1689 | 11 | 1.30% |
| hsa_circ_0087977 | 1689 | 11 | 1.30% |
| hsa_circ_0102283 | 1689 | 11 | 1.30% |
| hsa_circ_0103670 | 1689 | 11 | 1.30% |

| | | | |
|---|---|---|---|
| hsa_circ_0122454 | 1689 | 11 | 1.30% |
| hsa_circ_0139315 | 1689 | 11 | 1.30% |
| hsa_circ_0051666 | 1843 | 12 | 1.30% |
| hsa_circ_0006283 | 1536 | 10 | 1.30% |
| hsa_circ_0015649 | 1536 | 10 | 1.30% |
| hsa_circ_0021913 | 1536 | 10 | 1.30% |
| hsa_circ_0055684 | 1536 | 10 | 1.30% |
| hsa_circ_0060616 | 1536 | 10 | 1.30% |
| hsa_circ_0062533 | 1536 | 10 | 1.30% |
| hsa_circ_0075633 | 1536 | 10 | 1.30% |
| hsa_circ_0079215 | 1536 | 10 | 1.30% |
| hsa_circ_0084278 | 1536 | 10 | 1.30% |
| hsa_circ_0115910 | 1536 | 10 | 1.30% |
| hsa_circ_0120265 | 1536 | 10 | 1.30% |
| hsa_circ_0137361 | 1536 | 10 | 1.30% |
| hsa_circ_0069028 | 5837 | 38 | 1.30% |
| hsa_circ_0031963 | 3687 | 24 | 1.30% |
| hsa_circ_0040650 | 1690 | 11 | 1.30% |
| hsa_circ_0064211 | 1690 | 11 | 1.30% |
| hsa_circ_0066275 | 1690 | 11 | 1.30% |
| hsa_circ_0069446 | 1690 | 11 | 1.30% |
| hsa_circ_0077139 | 1690 | 11 | 1.30% |
| hsa_circ_0126541 | 1690 | 11 | 1.30% |
| hsa_circ_0058407 | 2151 | 14 | 1.30% |
| hsa_circ_0013423 | 1844 | 12 | 1.30% |
| hsa_circ_0139721 | 1844 | 12 | 1.30% |
| hsa_circ_0086696 | 1998 | 13 | 1.30% |
| hsa_circ_0016711 | 1537 | 10 | 1.30% |
| hsa_circ_0020664 | 1537 | 10 | 1.30% |
| hsa_circ_0029233 | 1537 | 10 | 1.30% |
| hsa_circ_0064972 | 1537 | 10 | 1.30% |
| hsa_circ_0072477 | 1537 | 10 | 1.30% |
| hsa_circ_0073096 | 1537 | 10 | 1.30% |
| hsa_circ_0074322 | 1537 | 10 | 1.30% |
| hsa_circ_0090197 | 1537 | 10 | 1.30% |
| hsa_circ_0119832 | 1537 | 10 | 1.30% |
| hsa_circ_0123095 | 1537 | 10 | 1.30% |
| hsa_circ_0045049 | 1691 | 11 | 1.30% |
| hsa_circ_0094490 | 1691 | 11 | 1.30% |
| hsa_circ_0110646 | 1691 | 11 | 1.30% |
| hsa_circ_0120735 | 1691 | 11 | 1.30% |
| hsa_circ_0139642 | 1691 | 11 | 1.30% |
| hsa_circ_0035876 | 1999 | 13 | 1.30% |
| hsa_circ_0070499 | 1999 | 13 | 1.30% |
| hsa_circ_0092108 | 1999 | 13 | 1.30% |
| hsa_circ_0014568 | 1538 | 10 | 1.30% |
| hsa_circ_0028870 | 1538 | 10 | 1.30% |
| hsa_circ_0035324 | 1538 | 10 | 1.30% |
| hsa_circ_0040114 | 1538 | 10 | 1.30% |
| hsa_circ_0045891 | 1538 | 10 | 1.30% |

| | | | |
|---|---|---|---|
| hsa_circ_0056375 | 1538 | 10 | 1.30% |
| hsa_circ_0070940 | 1538 | 10 | 1.30% |
| hsa_circ_0086616 | 1538 | 10 | 1.30% |
| hsa_circ_0101073 | 1538 | 10 | 1.30% |
| hsa_circ_0107922 | 1538 | 10 | 1.30% |
| hsa_circ_0129141 | 1538 | 10 | 1.30% |
| hsa_circ_0015917 | 1692 | 11 | 1.30% |
| hsa_circ_0021451 | 1692 | 11 | 1.30% |
| hsa_circ_0041297 | 1692 | 11 | 1.30% |
| hsa_circ_0091862 | 1692 | 11 | 1.30% |
| hsa_circ_0080429 | 1846 | 12 | 1.30% |
| hsa_circ_0081700 | 1846 | 12 | 1.30% |
| hsa_circ_0115828 | 2000 | 13 | 1.30% |
| hsa_circ_0020787 | 2308 | 15 | 1.30% |
| hsa_circ_0028809 | 1539 | 10 | 1.30% |
| hsa_circ_0038633 | 1539 | 10 | 1.30% |
| hsa_circ_0058042 | 1539 | 10 | 1.30% |
| hsa_circ_0064239 | 1539 | 10 | 1.30% |
| hsa_circ_0066030 | 1539 | 10 | 1.30% |
| hsa_circ_0081084 | 1539 | 10 | 1.30% |
| hsa_circ_0082849 | 1539 | 10 | 1.30% |
| hsa_circ_0087764 | 1539 | 10 | 1.30% |
| hsa_circ_0116441 | 1539 | 10 | 1.30% |
| hsa_circ_0116494 | 1539 | 10 | 1.30% |
| hsa_circ_0125323 | 1539 | 10 | 1.30% |
| hsa_circ_0130237 | 1539 | 10 | 1.30% |
| hsa_circ_0035846 | 1693 | 11 | 1.30% |
| hsa_circ_0064286 | 1693 | 11 | 1.30% |
| hsa_circ_0083434 | 1693 | 11 | 1.30% |
| hsa_circ_0109325 | 1693 | 11 | 1.30% |
| hsa_circ_0078235 | 3848 | 25 | 1.30% |
| hsa_circ_0064614 | 4002 | 26 | 1.30% |
| hsa_circ_0013529 | 2309 | 15 | 1.30% |
| hsa_circ_0026944 | 2156 | 14 | 1.30% |
| hsa_circ_0068676 | 2002 | 13 | 1.30% |
| hsa_circ_0021280 | 1848 | 12 | 1.30% |
| hsa_circ_0112255 | 1848 | 12 | 1.30% |
| hsa_circ_0136684 | 1848 | 12 | 1.30% |
| hsa_circ_0017013 | 1694 | 11 | 1.30% |
| hsa_circ_0026398 | 1694 | 11 | 1.30% |
| hsa_circ_0029570 | 1694 | 11 | 1.30% |
| hsa_circ_0046933 | 1694 | 11 | 1.30% |
| hsa_circ_0076562 | 1694 | 11 | 1.30% |
| hsa_circ_0031746 | 1540 | 10 | 1.30% |
| hsa_circ_0066722 | 1540 | 10 | 1.30% |
| hsa_circ_0076503 | 1540 | 10 | 1.30% |
| hsa_circ_0079944 | 1540 | 10 | 1.30% |
| hsa_circ_0130124 | 1540 | 10 | 1.30% |
| hsa_circ_0138824 | 1540 | 10 | 1.30% |
| hsa_circ_0139277 | 1540 | 10 | 1.30% |

| | | | |
|---|---|---|---|
| hsa_circ_0048558 | 1849 | 12 | 1.30% |
| hsa_circ_0061466 | 1849 | 12 | 1.30% |
| hsa_circ_0013619 | 1695 | 11 | 1.30% |
| hsa_circ_0019606 | 1695 | 11 | 1.30% |
| hsa_circ_0035567 | 1695 | 11 | 1.30% |
| hsa_circ_0057535 | 1695 | 11 | 1.30% |
| hsa_circ_0084167 | 1695 | 11 | 1.30% |
| hsa_circ_0090767 | 1695 | 11 | 1.30% |
| hsa_circ_0022485 | 1541 | 10 | 1.30% |
| hsa_circ_0036172 | 1541 | 10 | 1.30% |
| hsa_circ_0045176 | 1541 | 10 | 1.30% |
| hsa_circ_0055678 | 1541 | 10 | 1.30% |
| hsa_circ_0063093 | 1541 | 10 | 1.30% |
| hsa_circ_0067447 | 1541 | 10 | 1.30% |
| hsa_circ_0095278 | 1541 | 10 | 1.30% |
| hsa_circ_0110880 | 3545 | 23 | 1.30% |
| hsa_circ_0019992 | 2004 | 13 | 1.30% |
| hsa_circ_0013678 | 1850 | 12 | 1.30% |
| hsa_circ_0021248 | 1850 | 12 | 1.30% |
| hsa_circ_0019271 | 1696 | 11 | 1.30% |
| hsa_circ_0044229 | 1696 | 11 | 1.30% |
| hsa_circ_0049206 | 1696 | 11 | 1.30% |
| hsa_circ_0061425 | 1696 | 11 | 1.30% |
| hsa_circ_0080932 | 1696 | 11 | 1.30% |
| hsa_circ_0106097 | 1696 | 11 | 1.30% |
| hsa_circ_0011055 | 1542 | 10 | 1.30% |
| hsa_circ_0019294 | 1542 | 10 | 1.30% |
| hsa_circ_0026814 | 1542 | 10 | 1.30% |
| hsa_circ_0027315 | 1542 | 10 | 1.30% |
| hsa_circ_0029127 | 1542 | 10 | 1.30% |
| hsa_circ_0083235 | 1542 | 10 | 1.30% |
| hsa_circ_0095414 | 1542 | 10 | 1.30% |
| hsa_circ_0102859 | 1542 | 10 | 1.30% |
| hsa_circ_0107364 | 1542 | 10 | 1.30% |
| hsa_circ_0119422 | 1542 | 10 | 1.30% |
| hsa_circ_0139561 | 1542 | 10 | 1.30% |
| hsa_circ_0039599 | 1851 | 12 | 1.30% |
| hsa_circ_0031170 | 1697 | 11 | 1.30% |
| hsa_circ_0128904 | 1697 | 11 | 1.30% |
| hsa_circ_0132764 | 1697 | 11 | 1.30% |
| hsa_circ_0086181 | 2160 | 14 | 1.30% |
| hsa_circ_0011203 | 1543 | 10 | 1.30% |
| hsa_circ_0019339 | 1543 | 10 | 1.30% |
| hsa_circ_0038043 | 1543 | 10 | 1.30% |
| hsa_circ_0039650 | 1543 | 10 | 1.30% |
| hsa_circ_0041047 | 1543 | 10 | 1.30% |
| hsa_circ_0046388 | 1543 | 10 | 1.30% |
| hsa_circ_0062777 | 1543 | 10 | 1.30% |
| hsa_circ_0065949 | 1543 | 10 | 1.30% |
| hsa_circ_0072736 | 1543 | 10 | 1.30% |

| | | | |
|---|---|---|---|
| hsa_circ_0075765 | 1543 | 10 | 1.30% |
| hsa_circ_0080950 | 1543 | 10 | 1.30% |
| hsa_circ_0113173 | 1543 | 10 | 1.30% |
| hsa_circ_0114349 | 1543 | 10 | 1.30% |
| hsa_circ_0129598 | 1543 | 10 | 1.30% |
| hsa_circ_0040968 | 2006 | 13 | 1.30% |
| hsa_circ_0051577 | 1852 | 12 | 1.30% |
| hsa_circ_0074555 | 1852 | 12 | 1.30% |
| hsa_circ_0037355 | 1698 | 11 | 1.30% |
| hsa_circ_0095835 | 1698 | 11 | 1.30% |
| hsa_circ_0105550 | 1698 | 11 | 1.30% |
| hsa_circ_0075684 | 2007 | 13 | 1.30% |
| hsa_circ_0100787 | 2007 | 13 | 1.30% |
| hsa_circ_0017148 | 1544 | 10 | 1.30% |
| hsa_circ_0018707 | 1544 | 10 | 1.30% |
| hsa_circ_0030587 | 1544 | 10 | 1.30% |
| hsa_circ_0043798 | 1544 | 10 | 1.30% |
| hsa_circ_0068669 | 1544 | 10 | 1.30% |
| hsa_circ_0079066 | 1544 | 10 | 1.30% |
| hsa_circ_0095104 | 1544 | 10 | 1.30% |
| hsa_circ_0120889 | 1544 | 10 | 1.30% |
| hsa_circ_0058328 | 1853 | 12 | 1.30% |
| hsa_circ_0002621 | 1699 | 11 | 1.29% |
| hsa_circ_0019706 | 1699 | 11 | 1.29% |
| hsa_circ_0055282 | 1699 | 11 | 1.29% |
| hsa_circ_0072259 | 1699 | 11 | 1.29% |
| hsa_circ_0075640 | 1699 | 11 | 1.29% |
| hsa_circ_0085796 | 1699 | 11 | 1.29% |
| hsa_circ_0061449 | 2008 | 13 | 1.29% |
| hsa_circ_0007558 | 3862 | 25 | 1.29% |
| hsa_circ_0026990 | 1854 | 12 | 1.29% |
| hsa_circ_0046413 | 1854 | 12 | 1.29% |
| hsa_circ_0107571 | 1854 | 12 | 1.29% |
| hsa_circ_0119538 | 1854 | 12 | 1.29% |
| hsa_circ_0010837 | 1545 | 10 | 1.29% |
| hsa_circ_0040661 | 1545 | 10 | 1.29% |
| hsa_circ_0058501 | 1545 | 10 | 1.29% |
| hsa_circ_0059310 | 1545 | 10 | 1.29% |
| hsa_circ_0060031 | 1545 | 10 | 1.29% |
| hsa_circ_0062721 | 1545 | 10 | 1.29% |
| hsa_circ_0064388 | 1545 | 10 | 1.29% |
| hsa_circ_0071485 | 1545 | 10 | 1.29% |
| hsa_circ_0071611 | 1545 | 10 | 1.29% |
| hsa_circ_0108475 | 1545 | 10 | 1.29% |
| hsa_circ_0113192 | 1545 | 10 | 1.29% |
| hsa_circ_0128805 | 1545 | 10 | 1.29% |
| hsa_circ_0131316 | 1545 | 10 | 1.29% |
| hsa_circ_0086740 | 5563 | 36 | 1.29% |
| hsa_circ_0022836 | 2009 | 13 | 1.29% |
| hsa_circ_0080006 | 1700 | 11 | 1.29% |

| | | | |
|---|---|---|---|
| hsa_circ_0084733 | 1700 | 11 | 1.29% |
| hsa_circ_0085794 | 1700 | 11 | 1.29% |
| hsa_circ_0127312 | 1700 | 11 | 1.29% |
| hsa_circ_0030221 | 2164 | 14 | 1.29% |
| hsa_circ_0086626 | 6957 | 45 | 1.29% |
| hsa_circ_0011413 | 1546 | 10 | 1.29% |
| hsa_circ_0034164 | 1546 | 10 | 1.29% |
| hsa_circ_0039064 | 1546 | 10 | 1.29% |
| hsa_circ_0042595 | 1546 | 10 | 1.29% |
| hsa_circ_0046620 | 1546 | 10 | 1.29% |
| hsa_circ_0046785 | 1546 | 10 | 1.29% |
| hsa_circ_0065080 | 1546 | 10 | 1.29% |
| hsa_circ_0077748 | 1546 | 10 | 1.29% |
| hsa_circ_0086124 | 1546 | 10 | 1.29% |
| hsa_circ_0086206 | 1546 | 10 | 1.29% |
| hsa_circ_0090773 | 1546 | 10 | 1.29% |
| hsa_circ_0093938 | 1546 | 10 | 1.29% |
| hsa_circ_0133375 | 1546 | 10 | 1.29% |
| hsa_circ_0086705 | 2010 | 13 | 1.29% |
| hsa_circ_0066396 | 1701 | 11 | 1.29% |
| hsa_circ_0073964 | 1701 | 11 | 1.29% |
| hsa_circ_0088387 | 1701 | 11 | 1.29% |
| hsa_circ_0089461 | 1701 | 11 | 1.29% |
| hsa_circ_0053260 | 1856 | 12 | 1.29% |
| hsa_circ_0100209 | 4022 | 26 | 1.29% |
| hsa_circ_0072616 | 2011 | 13 | 1.29% |
| hsa_circ_0000129 | 1547 | 10 | 1.29% |
| hsa_circ_0016745 | 1547 | 10 | 1.29% |
| hsa_circ_0027469 | 1547 | 10 | 1.29% |
| hsa_circ_0035900 | 1547 | 10 | 1.29% |
| hsa_circ_0041531 | 1547 | 10 | 1.29% |
| hsa_circ_0054136 | 1547 | 10 | 1.29% |
| hsa_circ_0084001 | 1547 | 10 | 1.29% |
| hsa_circ_0088230 | 1547 | 10 | 1.29% |
| hsa_circ_0032079 | 4177 | 27 | 1.29% |
| hsa_circ_0000264 | 1702 | 11 | 1.29% |
| hsa_circ_0020892 | 1702 | 11 | 1.29% |
| hsa_circ_0069865 | 1702 | 11 | 1.29% |
| hsa_circ_0096296 | 1702 | 11 | 1.29% |
| hsa_circ_0023348 | 1857 | 12 | 1.29% |
| hsa_circ_0091743 | 1857 | 12 | 1.29% |
| hsa_circ_0135690 | 1857 | 12 | 1.29% |
| hsa_circ_0024397 | 2012 | 13 | 1.29% |
| hsa_circ_0008584 | 1548 | 10 | 1.29% |
| hsa_circ_0017614 | 1548 | 10 | 1.29% |
| hsa_circ_0027449 | 1548 | 10 | 1.29% |
| hsa_circ_0031190 | 1548 | 10 | 1.29% |
| hsa_circ_0039135 | 1548 | 10 | 1.29% |
| hsa_circ_0043700 | 1548 | 10 | 1.29% |
| hsa_circ_0060888 | 1548 | 10 | 1.29% |

| | | | |
|---|---|---|---|
| hsa_circ_0064114 | 1548 | 10 | 1.29% |
| hsa_circ_0078120 | 1548 | 10 | 1.29% |
| hsa_circ_0085690 | 1548 | 10 | 1.29% |
| hsa_circ_0099991 | 1548 | 10 | 1.29% |
| hsa_circ_0105778 | 1548 | 10 | 1.29% |
| hsa_circ_0117390 | 1548 | 10 | 1.29% |
| hsa_circ_0125234 | 1548 | 10 | 1.29% |
| hsa_circ_0128968 | 1548 | 10 | 1.29% |
| hsa_circ_0003106 | 1703 | 11 | 1.29% |
| hsa_circ_0018972 | 1703 | 11 | 1.29% |
| hsa_circ_0027966 | 1703 | 11 | 1.29% |
| hsa_circ_0034378 | 1703 | 11 | 1.29% |
| hsa_circ_0052192 | 1703 | 11 | 1.29% |
| hsa_circ_0068929 | 1703 | 11 | 1.29% |
| hsa_circ_0080575 | 1703 | 11 | 1.29% |
| hsa_circ_0083523 | 1703 | 11 | 1.29% |
| hsa_circ_0024736 | 1858 | 12 | 1.29% |
| hsa_circ_0125361 | 1858 | 12 | 1.29% |
| hsa_circ_0013579 | 2168 | 14 | 1.29% |
| hsa_circ_0023909 | 1549 | 10 | 1.29% |
| hsa_circ_0030519 | 1549 | 10 | 1.29% |
| hsa_circ_0035484 | 1549 | 10 | 1.29% |
| hsa_circ_0054927 | 1549 | 10 | 1.29% |
| hsa_circ_0137893 | 1549 | 10 | 1.29% |
| hsa_circ_0049304 | 1704 | 11 | 1.29% |
| hsa_circ_0061220 | 1704 | 11 | 1.29% |
| hsa_circ_0066102 | 1704 | 11 | 1.29% |
| hsa_circ_0093668 | 1704 | 11 | 1.29% |
| hsa_circ_0103717 | 1704 | 11 | 1.29% |
| hsa_circ_0026350 | 1859 | 12 | 1.29% |
| hsa_circ_0052797 | 1859 | 12 | 1.29% |
| hsa_circ_0009194 | 2169 | 14 | 1.29% |
| hsa_circ_0031280 | 2945 | 19 | 1.29% |
| hsa_circ_0023154 | 1860 | 12 | 1.29% |
| hsa_circ_0034502 | 1860 | 12 | 1.29% |
| hsa_circ_0041766 | 1860 | 12 | 1.29% |
| hsa_circ_0050190 | 1860 | 12 | 1.29% |
| hsa_circ_0101885 | 1860 | 12 | 1.29% |
| hsa_circ_0048145 | 1705 | 11 | 1.29% |
| hsa_circ_0061365 | 1705 | 11 | 1.29% |
| hsa_circ_0072710 | 1705 | 11 | 1.29% |
| hsa_circ_0076450 | 1705 | 11 | 1.29% |
| hsa_circ_0001068 | 1550 | 10 | 1.29% |
| hsa_circ_0011603 | 1550 | 10 | 1.29% |
| hsa_circ_0015252 | 1550 | 10 | 1.29% |
| hsa_circ_0034455 | 1550 | 10 | 1.29% |
| hsa_circ_0042436 | 1550 | 10 | 1.29% |
| hsa_circ_0047137 | 1550 | 10 | 1.29% |
| hsa_circ_0098649 | 1550 | 10 | 1.29% |
| hsa_circ_0113628 | 1550 | 10 | 1.29% |

| | | | |
|---|---|---|---|
| hsa_circ_0120608 | 1550 | 10 | 1.29% |
| hsa_circ_0126474 | 1550 | 10 | 1.29% |
| hsa_circ_0134328 | 4341 | 28 | 1.29% |
| hsa_circ_0044505 | 3566 | 23 | 1.29% |
| hsa_circ_0013821 | 2946 | 19 | 1.29% |
| hsa_circ_0128502 | 2481 | 16 | 1.29% |
| hsa_circ_0009403 | 2016 | 13 | 1.29% |
| hsa_circ_0055623 | 2016 | 13 | 1.29% |
| hsa_circ_0044455 | 1861 | 12 | 1.29% |
| hsa_circ_0057759 | 1706 | 11 | 1.29% |
| hsa_circ_0064622 | 1706 | 11 | 1.29% |
| hsa_circ_0078568 | 1706 | 11 | 1.29% |
| hsa_circ_0022112 | 1551 | 10 | 1.29% |
| hsa_circ_0022639 | 1551 | 10 | 1.29% |
| hsa_circ_0026851 | 1551 | 10 | 1.29% |
| hsa_circ_0044298 | 1551 | 10 | 1.29% |
| hsa_circ_0049090 | 1551 | 10 | 1.29% |
| hsa_circ_0052918 | 1551 | 10 | 1.29% |
| hsa_circ_0059731 | 1551 | 10 | 1.29% |
| hsa_circ_0068139 | 1551 | 10 | 1.29% |
| hsa_circ_0082657 | 1551 | 10 | 1.29% |
| hsa_circ_0099691 | 1551 | 10 | 1.29% |
| hsa_circ_0105587 | 1551 | 10 | 1.29% |
| hsa_circ_0108722 | 1551 | 10 | 1.29% |
| hsa_circ_0114848 | 1551 | 10 | 1.29% |
| hsa_circ_0121983 | 1551 | 10 | 1.29% |
| hsa_circ_0125232 | 1551 | 10 | 1.29% |
| hsa_circ_0138715 | 1551 | 10 | 1.29% |
| hsa_circ_0138909 | 1551 | 10 | 1.29% |
| hsa_circ_0117093 | 2327 | 15 | 1.29% |
| hsa_circ_0017717 | 1862 | 12 | 1.29% |
| hsa_circ_0132457 | 1862 | 12 | 1.29% |
| hsa_circ_0124340 | 4500 | 29 | 1.29% |
| hsa_circ_0002558 | 1707 | 11 | 1.29% |
| hsa_circ_0016479 | 1707 | 11 | 1.29% |
| hsa_circ_0016528 | 1707 | 11 | 1.29% |
| hsa_circ_0018159 | 1707 | 11 | 1.29% |
| hsa_circ_0037701 | 1707 | 11 | 1.29% |
| hsa_circ_0038181 | 1707 | 11 | 1.29% |
| hsa_circ_0075996 | 1707 | 11 | 1.29% |
| hsa_circ_0087741 | 1707 | 11 | 1.29% |
| hsa_circ_0091924 | 1707 | 11 | 1.29% |
| hsa_circ_0129988 | 1707 | 11 | 1.29% |
| hsa_circ_0011940 | 1552 | 10 | 1.29% |
| hsa_circ_0018510 | 1552 | 10 | 1.29% |
| hsa_circ_0035026 | 1552 | 10 | 1.29% |
| hsa_circ_0075432 | 1552 | 10 | 1.29% |
| hsa_circ_0077433 | 1552 | 10 | 1.29% |
| hsa_circ_0081270 | 1552 | 10 | 1.29% |
| hsa_circ_0121311 | 1552 | 10 | 1.29% |

| | | | |
|---|---|---|---|
| hsa_circ_0127091 | 1552 | 10 | 1.29% |
| hsa_circ_0130092 | 1552 | 10 | 1.29% |
| hsa_circ_0135516 | 1552 | 10 | 1.29% |
| hsa_circ_0123261 | 5122 | 33 | 1.29% |
| hsa_circ_0015097 | 1863 | 12 | 1.29% |
| hsa_circ_0032579 | 1863 | 12 | 1.29% |
| hsa_circ_0040024 | 1863 | 12 | 1.29% |
| hsa_circ_0134590 | 2950 | 19 | 1.29% |
| hsa_circ_0041620 | 1708 | 11 | 1.29% |
| hsa_circ_0044617 | 1708 | 11 | 1.29% |
| hsa_circ_0068570 | 1708 | 11 | 1.29% |
| hsa_circ_0114920 | 1708 | 11 | 1.29% |
| hsa_circ_0137244 | 1708 | 11 | 1.29% |
| hsa_circ_0021285 | 1553 | 10 | 1.29% |
| hsa_circ_0035797 | 1553 | 10 | 1.29% |
| hsa_circ_0037877 | 1553 | 10 | 1.29% |
| hsa_circ_0046020 | 1553 | 10 | 1.29% |
| hsa_circ_0050124 | 1553 | 10 | 1.29% |
| hsa_circ_0050291 | 1553 | 10 | 1.29% |
| hsa_circ_0056521 | 1553 | 10 | 1.29% |
| hsa_circ_0057128 | 1553 | 10 | 1.29% |
| hsa_circ_0057889 | 1553 | 10 | 1.29% |
| hsa_circ_0061024 | 1553 | 10 | 1.29% |
| hsa_circ_0071960 | 1553 | 10 | 1.29% |
| hsa_circ_0094738 | 1553 | 10 | 1.29% |
| hsa_circ_0102944 | 1553 | 10 | 1.29% |
| hsa_circ_0112193 | 1553 | 10 | 1.29% |
| hsa_circ_0117379 | 1553 | 10 | 1.29% |
| hsa_circ_0123007 | 1553 | 10 | 1.29% |
| hsa_circ_0123556 | 1553 | 10 | 1.29% |
| hsa_circ_0126391 | 1553 | 10 | 1.29% |
| hsa_circ_0039452 | 2019 | 13 | 1.29% |
| hsa_circ_0117169 | 2019 | 13 | 1.29% |
| hsa_circ_0021642 | 2485 | 16 | 1.29% |
| hsa_circ_0118764 | 1864 | 12 | 1.29% |
| hsa_circ_0027886 | 1709 | 11 | 1.29% |
| hsa_circ_0032040 | 1709 | 11 | 1.29% |
| hsa_circ_0037416 | 1709 | 11 | 1.29% |
| hsa_circ_0047843 | 1709 | 11 | 1.29% |
| hsa_circ_0055372 | 1709 | 11 | 1.29% |
| hsa_circ_0078001 | 1709 | 11 | 1.29% |
| hsa_circ_0038449 | 7147 | 46 | 1.29% |
| hsa_circ_0006216 | 1554 | 10 | 1.29% |
| hsa_circ_0013652 | 1554 | 10 | 1.29% |
| hsa_circ_0016344 | 1554 | 10 | 1.29% |
| hsa_circ_0058426 | 1554 | 10 | 1.29% |
| hsa_circ_0073459 | 1554 | 10 | 1.29% |
| hsa_circ_0073509 | 1554 | 10 | 1.29% |
| hsa_circ_0111208 | 1554 | 10 | 1.29% |
| hsa_circ_0127976 | 1554 | 10 | 1.29% |

| | | | |
|---|---|---|---|
| hsa_circ_0129396 | 1554 | 10 | 1.29% |
| hsa_circ_0058165 | 1865 | 12 | 1.29% |
| hsa_circ_0073906 | 1865 | 12 | 1.29% |
| hsa_circ_0013608 | 1710 | 11 | 1.29% |
| hsa_circ_0023568 | 1710 | 11 | 1.29% |
| hsa_circ_0029620 | 1710 | 11 | 1.29% |
| hsa_circ_0030677 | 1710 | 11 | 1.29% |
| hsa_circ_0050165 | 1710 | 11 | 1.29% |
| hsa_circ_0053893 | 1710 | 11 | 1.29% |
| hsa_circ_0091029 | 1710 | 11 | 1.29% |
| hsa_circ_0098527 | 1710 | 11 | 1.29% |
| hsa_circ_0108227 | 1710 | 11 | 1.29% |
| hsa_circ_0023247 | 3731 | 24 | 1.29% |
| hsa_circ_0058939 | 2021 | 13 | 1.29% |
| hsa_circ_0041571 | 1866 | 12 | 1.29% |
| hsa_circ_0005140 | 1555 | 10 | 1.29% |
| hsa_circ_0012045 | 1555 | 10 | 1.29% |
| hsa_circ_0024886 | 1555 | 10 | 1.29% |
| hsa_circ_0032514 | 1555 | 10 | 1.29% |
| hsa_circ_0063611 | 1555 | 10 | 1.29% |
| hsa_circ_0071224 | 1555 | 10 | 1.29% |
| hsa_circ_0074512 | 1555 | 10 | 1.29% |
| hsa_circ_0119730 | 1555 | 10 | 1.29% |
| hsa_circ_0124723 | 1555 | 10 | 1.29% |
| hsa_circ_0130916 | 1555 | 10 | 1.29% |
| hsa_circ_0113518 | 7932 | 51 | 1.29% |
| hsa_circ_0083568 | 2333 | 15 | 1.29% |
| hsa_circ_0019930 | 2022 | 13 | 1.29% |
| hsa_circ_0004785 | 1711 | 11 | 1.29% |
| hsa_circ_0061421 | 1711 | 11 | 1.29% |
| hsa_circ_0081412 | 1711 | 11 | 1.29% |
| hsa_circ_0082938 | 1711 | 11 | 1.29% |
| hsa_circ_0089211 | 1711 | 11 | 1.29% |
| hsa_circ_0092928 | 1711 | 11 | 1.29% |
| hsa_circ_0093502 | 1711 | 11 | 1.29% |
| hsa_circ_0094406 | 1711 | 11 | 1.29% |
| hsa_circ_0134472 | 1867 | 12 | 1.29% |
| hsa_circ_0066990 | 2645 | 17 | 1.29% |
| hsa_circ_0036531 | 1556 | 10 | 1.29% |
| hsa_circ_0046108 | 1556 | 10 | 1.29% |
| hsa_circ_0082754 | 1556 | 10 | 1.29% |
| hsa_circ_0083602 | 1556 | 10 | 1.29% |
| hsa_circ_0128573 | 1556 | 10 | 1.29% |
| hsa_circ_0098090 | 6381 | 41 | 1.29% |
| hsa_circ_0010731 | 1712 | 11 | 1.29% |
| hsa_circ_0041039 | 1712 | 11 | 1.29% |
| hsa_circ_0074517 | 1712 | 11 | 1.29% |
| hsa_circ_0105004 | 1712 | 11 | 1.29% |
| hsa_circ_0051617 | 2179 | 14 | 1.28% |
| hsa_circ_0101396 | 1868 | 12 | 1.28% |

| | | | |
|---|---|---|---|
| hsa_circ_0118356 | 1868 | 12 | 1.28% |
| hsa_circ_0016676 | 1557 | 10 | 1.28% |
| hsa_circ_0023318 | 1557 | 10 | 1.28% |
| hsa_circ_0025942 | 1557 | 10 | 1.28% |
| hsa_circ_0038136 | 1557 | 10 | 1.28% |
| hsa_circ_0050553 | 1557 | 10 | 1.28% |
| hsa_circ_0055844 | 1557 | 10 | 1.28% |
| hsa_circ_0057876 | 1557 | 10 | 1.28% |
| hsa_circ_0106628 | 1557 | 10 | 1.28% |
| hsa_circ_0107215 | 1557 | 10 | 1.28% |
| hsa_circ_0113107 | 1557 | 10 | 1.28% |
| hsa_circ_0113820 | 1557 | 10 | 1.28% |
| hsa_circ_0125581 | 1557 | 10 | 1.28% |
| hsa_circ_0127121 | 1557 | 10 | 1.28% |
| hsa_circ_0037218 | 1713 | 11 | 1.28% |
| hsa_circ_0041322 | 1713 | 11 | 1.28% |
| hsa_circ_0051955 | 1713 | 11 | 1.28% |
| hsa_circ_0086690 | 1713 | 11 | 1.28% |
| hsa_circ_0053024 | 2336 | 15 | 1.28% |
| hsa_circ_0097243 | 1869 | 12 | 1.28% |
| hsa_circ_0014560 | 1558 | 10 | 1.28% |
| hsa_circ_0042392 | 1558 | 10 | 1.28% |
| hsa_circ_0042636 | 1558 | 10 | 1.28% |
| hsa_circ_0064259 | 1558 | 10 | 1.28% |
| hsa_circ_0065551 | 1558 | 10 | 1.28% |
| hsa_circ_0069616 | 1558 | 10 | 1.28% |
| hsa_circ_0088794 | 1558 | 10 | 1.28% |
| hsa_circ_0088999 | 1558 | 10 | 1.28% |
| hsa_circ_0090444 | 1558 | 10 | 1.28% |
| hsa_circ_0097146 | 1558 | 10 | 1.28% |
| hsa_circ_0100864 | 1558 | 10 | 1.28% |
| hsa_circ_0019463 | 1714 | 11 | 1.28% |
| hsa_circ_0085931 | 1714 | 11 | 1.28% |
| hsa_circ_0106142 | 1714 | 11 | 1.28% |
| hsa_circ_0057354 | 2649 | 17 | 1.28% |
| hsa_circ_0017869 | 1870 | 12 | 1.28% |
| hsa_circ_0030406 | 1870 | 12 | 1.28% |
| hsa_circ_0100904 | 1870 | 12 | 1.28% |
| hsa_circ_0032686 | 3741 | 24 | 1.28% |
| hsa_circ_0040906 | 1559 | 10 | 1.28% |
| hsa_circ_0078421 | 1559 | 10 | 1.28% |
| hsa_circ_0091495 | 1559 | 10 | 1.28% |
| hsa_circ_0121880 | 1559 | 10 | 1.28% |
| hsa_circ_0124070 | 1559 | 10 | 1.28% |
| hsa_circ_0073523 | 1715 | 11 | 1.28% |
| hsa_circ_0015721 | 1871 | 12 | 1.28% |
| hsa_circ_0032302 | 1871 | 12 | 1.28% |
| hsa_circ_0045478 | 1871 | 12 | 1.28% |
| hsa_circ_0067523 | 1871 | 12 | 1.28% |
| hsa_circ_0106921 | 4055 | 26 | 1.28% |

| | | | |
|---|---|---|---|
| hsa_circ_0031516 | 1872 | 12 | 1.28% |
| hsa_circ_0043761 | 1872 | 12 | 1.28% |
| hsa_circ_0049119 | 1872 | 12 | 1.28% |
| hsa_circ_0074514 | 1872 | 12 | 1.28% |
| hsa_circ_0102525 | 1872 | 12 | 1.28% |
| hsa_circ_0017019 | 1716 | 11 | 1.28% |
| hsa_circ_0019470 | 1716 | 11 | 1.28% |
| hsa_circ_0021244 | 1716 | 11 | 1.28% |
| hsa_circ_0035495 | 1716 | 11 | 1.28% |
| hsa_circ_0048356 | 1716 | 11 | 1.28% |
| hsa_circ_0052381 | 1716 | 11 | 1.28% |
| hsa_circ_0058419 | 1716 | 11 | 1.28% |
| hsa_circ_0059023 | 1716 | 11 | 1.28% |
| hsa_circ_0061129 | 1716 | 11 | 1.28% |
| hsa_circ_0068019 | 1716 | 11 | 1.28% |
| hsa_circ_0074848 | 1716 | 11 | 1.28% |
| hsa_circ_0088629 | 1716 | 11 | 1.28% |
| hsa_circ_0032982 | 1560 | 10 | 1.28% |
| hsa_circ_0044277 | 1560 | 10 | 1.28% |
| hsa_circ_0054195 | 1560 | 10 | 1.28% |
| hsa_circ_0056853 | 1560 | 10 | 1.28% |
| hsa_circ_0078319 | 1560 | 10 | 1.28% |
| hsa_circ_0079538 | 1560 | 10 | 1.28% |
| hsa_circ_0085354 | 1560 | 10 | 1.28% |
| hsa_circ_0110152 | 1560 | 10 | 1.28% |
| hsa_circ_0130616 | 1560 | 10 | 1.28% |
| hsa_circ_0135492 | 1560 | 10 | 1.28% |
| hsa_circ_0140585 | 25590 | 164 | 1.28% |
| hsa_circ_0042283 | 1873 | 12 | 1.28% |
| hsa_circ_0038338 | 1717 | 11 | 1.28% |
| hsa_circ_0043631 | 1717 | 11 | 1.28% |
| hsa_circ_0101608 | 1717 | 11 | 1.28% |
| hsa_circ_0011683 | 1561 | 10 | 1.28% |
| hsa_circ_0021166 | 1561 | 10 | 1.28% |
| hsa_circ_0023911 | 1561 | 10 | 1.28% |
| hsa_circ_0035015 | 1561 | 10 | 1.28% |
| hsa_circ_0051148 | 1561 | 10 | 1.28% |
| hsa_circ_0058883 | 1561 | 10 | 1.28% |
| hsa_circ_0063800 | 1561 | 10 | 1.28% |
| hsa_circ_0066198 | 1561 | 10 | 1.28% |
| hsa_circ_0069323 | 1561 | 10 | 1.28% |
| hsa_circ_0082044 | 1561 | 10 | 1.28% |
| hsa_circ_0085121 | 1561 | 10 | 1.28% |
| hsa_circ_0091063 | 1561 | 10 | 1.28% |
| hsa_circ_0100213 | 1561 | 10 | 1.28% |
| hsa_circ_0110086 | 1561 | 10 | 1.28% |
| hsa_circ_0123523 | 1561 | 10 | 1.28% |
| hsa_circ_0010442 | 3435 | 22 | 1.28% |
| hsa_circ_0052588 | 2186 | 14 | 1.28% |
| hsa_circ_0044499 | 2030 | 13 | 1.28% |

| | | | |
|---|---|---|---|
| hsa_circ_0037201 | 1874 | 12 | 1.28% |
| hsa_circ_0054431 | 1874 | 12 | 1.28% |
| hsa_circ_0072731 | 1874 | 12 | 1.28% |
| hsa_circ_0005500 | 1718 | 11 | 1.28% |
| hsa_circ_0015048 | 1718 | 11 | 1.28% |
| hsa_circ_0016739 | 1718 | 11 | 1.28% |
| hsa_circ_0016948 | 1718 | 11 | 1.28% |
| hsa_circ_0020921 | 1718 | 11 | 1.28% |
| hsa_circ_0055920 | 1718 | 11 | 1.28% |
| hsa_circ_0069698 | 1718 | 11 | 1.28% |
| hsa_circ_0076245 | 1718 | 11 | 1.28% |
| hsa_circ_0005575 | 1562 | 10 | 1.28% |
| hsa_circ_0023013 | 1562 | 10 | 1.28% |
| hsa_circ_0024108 | 1562 | 10 | 1.28% |
| hsa_circ_0039298 | 1562 | 10 | 1.28% |
| hsa_circ_0041271 | 1562 | 10 | 1.28% |
| hsa_circ_0049696 | 1562 | 10 | 1.28% |
| hsa_circ_0054004 | 1562 | 10 | 1.28% |
| hsa_circ_0064934 | 1562 | 10 | 1.28% |
| hsa_circ_0084742 | 1562 | 10 | 1.28% |
| hsa_circ_0107025 | 6092 | 39 | 1.28% |
| hsa_circ_0111238 | 4375 | 28 | 1.28% |
| hsa_circ_0053958 | 1875 | 12 | 1.28% |
| hsa_circ_0124186 | 1875 | 12 | 1.28% |
| hsa_circ_0024575 | 1719 | 11 | 1.28% |
| hsa_circ_0048320 | 1719 | 11 | 1.28% |
| hsa_circ_0056836 | 1719 | 11 | 1.28% |
| hsa_circ_0095475 | 1719 | 11 | 1.28% |
| hsa_circ_0123883 | 1719 | 11 | 1.28% |
| hsa_circ_0026131 | 1563 | 10 | 1.28% |
| hsa_circ_0026502 | 1563 | 10 | 1.28% |
| hsa_circ_0040164 | 1563 | 10 | 1.28% |
| hsa_circ_0042569 | 1563 | 10 | 1.28% |
| hsa_circ_0055661 | 1563 | 10 | 1.28% |
| hsa_circ_0058300 | 1563 | 10 | 1.28% |
| hsa_circ_0115944 | 1563 | 10 | 1.28% |
| hsa_circ_0057300 | 2970 | 19 | 1.28% |
| hsa_circ_0019136 | 1876 | 12 | 1.28% |
| hsa_circ_0047761 | 1876 | 12 | 1.28% |
| hsa_circ_0039475 | 1720 | 11 | 1.28% |
| hsa_circ_0059156 | 1720 | 11 | 1.28% |
| hsa_circ_0060447 | 1720 | 11 | 1.28% |
| hsa_circ_0060603 | 1720 | 11 | 1.28% |
| hsa_circ_0069256 | 1720 | 11 | 1.28% |
| hsa_circ_0066957 | 2033 | 13 | 1.28% |
| hsa_circ_0081576 | 2033 | 13 | 1.28% |
| hsa_circ_0014388 | 1564 | 10 | 1.28% |
| hsa_circ_0038555 | 1564 | 10 | 1.28% |
| hsa_circ_0040124 | 1564 | 10 | 1.28% |
| hsa_circ_0061373 | 1564 | 10 | 1.28% |

| | | | |
|---|---|---|---|
| hsa_circ_0107972 | 1564 | 10 | 1.28% |
| hsa_circ_0126507 | 1564 | 10 | 1.28% |
| hsa_circ_0025340 | 1877 | 12 | 1.28% |
| hsa_circ_0037628 | 1877 | 12 | 1.28% |
| hsa_circ_0091485 | 1877 | 12 | 1.28% |
| hsa_circ_0092050 | 1877 | 12 | 1.28% |
| hsa_circ_0115980 | 1877 | 12 | 1.28% |
| hsa_circ_0025516 | 2190 | 14 | 1.28% |
| hsa_circ_0054873 | 1721 | 11 | 1.28% |
| hsa_circ_0065925 | 1721 | 11 | 1.28% |
| hsa_circ_0090721 | 1721 | 11 | 1.28% |
| hsa_circ_0125030 | 1721 | 11 | 1.28% |
| hsa_circ_0052201 | 2034 | 13 | 1.28% |
| hsa_circ_0118359 | 2034 | 13 | 1.28% |
| hsa_circ_0104973 | 4381 | 28 | 1.28% |
| hsa_circ_0010856 | 1878 | 12 | 1.28% |
| hsa_circ_0040830 | 1878 | 12 | 1.28% |
| hsa_circ_0041579 | 1878 | 12 | 1.28% |
| hsa_circ_0054790 | 1878 | 12 | 1.28% |
| hsa_circ_0103963 | 1878 | 12 | 1.28% |
| hsa_circ_0123919 | 1878 | 12 | 1.28% |
| hsa_circ_0011529 | 1565 | 10 | 1.28% |
| hsa_circ_0037307 | 1565 | 10 | 1.28% |
| hsa_circ_0046589 | 1565 | 10 | 1.28% |
| hsa_circ_0056503 | 1565 | 10 | 1.28% |
| hsa_circ_0068984 | 1565 | 10 | 1.28% |
| hsa_circ_0074012 | 1565 | 10 | 1.28% |
| hsa_circ_0112533 | 1565 | 10 | 1.28% |
| hsa_circ_0048028 | 2035 | 13 | 1.28% |
| hsa_circ_0018235 | 1722 | 11 | 1.28% |
| hsa_circ_0032032 | 1722 | 11 | 1.28% |
| hsa_circ_0038511 | 1722 | 11 | 1.28% |
| hsa_circ_0065797 | 1722 | 11 | 1.28% |
| hsa_circ_0071239 | 1722 | 11 | 1.28% |
| hsa_circ_0093028 | 1722 | 11 | 1.28% |
| hsa_circ_0115762 | 1722 | 11 | 1.28% |
| hsa_circ_0117220 | 1722 | 11 | 1.28% |
| hsa_circ_0117905 | 1722 | 11 | 1.28% |
| hsa_circ_0037276 | 2192 | 14 | 1.28% |
| hsa_circ_0022893 | 1879 | 12 | 1.28% |
| hsa_circ_0090609 | 1879 | 12 | 1.28% |
| hsa_circ_0139228 | 1879 | 12 | 1.28% |
| hsa_circ_0022219 | 1566 | 10 | 1.28% |
| hsa_circ_0034918 | 1566 | 10 | 1.28% |
| hsa_circ_0059437 | 1566 | 10 | 1.28% |
| hsa_circ_0069088 | 1566 | 10 | 1.28% |
| hsa_circ_0089912 | 1566 | 10 | 1.28% |
| hsa_circ_0104279 | 1566 | 10 | 1.28% |
| hsa_circ_0126005 | 1566 | 10 | 1.28% |
| hsa_circ_0012816 | 1723 | 11 | 1.28% |

| | | | |
|---|---|---|---|
| hsa_circ_0040471 | 1723 | 11 | 1.28% |
| hsa_circ_0086841 | 1723 | 11 | 1.28% |
| hsa_circ_0109296 | 1723 | 11 | 1.28% |
| hsa_circ_0030403 | 1880 | 12 | 1.28% |
| hsa_circ_0115364 | 1880 | 12 | 1.28% |
| hsa_circ_0118381 | 1880 | 12 | 1.28% |
| hsa_circ_0092109 | 2037 | 13 | 1.28% |
| hsa_circ_0004796 | 1567 | 10 | 1.28% |
| hsa_circ_0044413 | 1567 | 10 | 1.28% |
| hsa_circ_0044927 | 1567 | 10 | 1.28% |
| hsa_circ_0056682 | 1567 | 10 | 1.28% |
| hsa_circ_0056773 | 1567 | 10 | 1.28% |
| hsa_circ_0056916 | 1567 | 10 | 1.28% |
| hsa_circ_0077222 | 1567 | 10 | 1.28% |
| hsa_circ_0077461 | 1567 | 10 | 1.28% |
| hsa_circ_0087372 | 1567 | 10 | 1.28% |
| hsa_circ_0088114 | 1567 | 10 | 1.28% |
| hsa_circ_0110111 | 1567 | 10 | 1.28% |
| hsa_circ_0125726 | 1567 | 10 | 1.28% |
| hsa_circ_0019042 | 1724 | 11 | 1.28% |
| hsa_circ_0046141 | 1724 | 11 | 1.28% |
| hsa_circ_0075745 | 1724 | 11 | 1.28% |
| hsa_circ_0117508 | 1724 | 11 | 1.28% |
| hsa_circ_0137452 | 1881 | 12 | 1.28% |
| hsa_circ_0048261 | 2038 | 13 | 1.28% |
| hsa_circ_0049687 | 2979 | 19 | 1.28% |
| hsa_circ_0001190 | 1568 | 10 | 1.28% |
| hsa_circ_0018953 | 1568 | 10 | 1.28% |
| hsa_circ_0019232 | 1568 | 10 | 1.28% |
| hsa_circ_0023958 | 1568 | 10 | 1.28% |
| hsa_circ_0035284 | 1568 | 10 | 1.28% |
| hsa_circ_0035523 | 1568 | 10 | 1.28% |
| hsa_circ_0060247 | 1568 | 10 | 1.28% |
| hsa_circ_0063907 | 1568 | 10 | 1.28% |
| hsa_circ_0080003 | 1568 | 10 | 1.28% |
| hsa_circ_0094196 | 1568 | 10 | 1.28% |
| hsa_circ_0101538 | 1568 | 10 | 1.28% |
| hsa_circ_0107579 | 1568 | 10 | 1.28% |
| hsa_circ_0026262 | 1725 | 11 | 1.28% |
| hsa_circ_0078431 | 1725 | 11 | 1.28% |
| hsa_circ_0098703 | 1725 | 11 | 1.28% |
| hsa_circ_0109371 | 1725 | 11 | 1.28% |
| hsa_circ_0119559 | 1725 | 11 | 1.28% |
| hsa_circ_0122296 | 1725 | 11 | 1.28% |
| hsa_circ_0004450 | 1882 | 12 | 1.28% |
| hsa_circ_0009973 | 1882 | 12 | 1.28% |
| hsa_circ_0087974 | 1882 | 12 | 1.28% |
| hsa_circ_0117915 | 1882 | 12 | 1.28% |
| hsa_circ_0093994 | 3137 | 20 | 1.28% |
| hsa_circ_0021788 | 1569 | 10 | 1.27% |

| | | | |
|---|---|---|---|
| hsa_circ_0038546 | 1569 | 10 | 1.27% |
| hsa_circ_0041997 | 1569 | 10 | 1.27% |
| hsa_circ_0057880 | 1569 | 10 | 1.27% |
| hsa_circ_0067359 | 1569 | 10 | 1.27% |
| hsa_circ_0070320 | 1569 | 10 | 1.27% |
| hsa_circ_0092148 | 1569 | 10 | 1.27% |
| hsa_circ_0014479 | 1726 | 11 | 1.27% |
| hsa_circ_0028230 | 1726 | 11 | 1.27% |
| hsa_circ_0030938 | 1726 | 11 | 1.27% |
| hsa_circ_0084102 | 1726 | 11 | 1.27% |
| hsa_circ_0085713 | 1726 | 11 | 1.27% |
| hsa_circ_0051135 | 1883 | 12 | 1.27% |
| hsa_circ_0128934 | 1883 | 12 | 1.27% |
| hsa_circ_0010675 | 2040 | 13 | 1.27% |
| hsa_circ_0022832 | 2197 | 14 | 1.27% |
| hsa_circ_0009326 | 1884 | 12 | 1.27% |
| hsa_circ_0011896 | 1884 | 12 | 1.27% |
| hsa_circ_0064071 | 1884 | 12 | 1.27% |
| hsa_circ_0090875 | 1884 | 12 | 1.27% |
| hsa_circ_0031475 | 1727 | 11 | 1.27% |
| hsa_circ_0041415 | 1727 | 11 | 1.27% |
| hsa_circ_0072323 | 1727 | 11 | 1.27% |
| hsa_circ_0089738 | 1727 | 11 | 1.27% |
| hsa_circ_0120651 | 1727 | 11 | 1.27% |
| hsa_circ_0133176 | 1727 | 11 | 1.27% |
| hsa_circ_0019223 | 1570 | 10 | 1.27% |
| hsa_circ_0029414 | 1570 | 10 | 1.27% |
| hsa_circ_0067697 | 1570 | 10 | 1.27% |
| hsa_circ_0072355 | 1570 | 10 | 1.27% |
| hsa_circ_0075027 | 1570 | 10 | 1.27% |
| hsa_circ_0089241 | 1570 | 10 | 1.27% |
| hsa_circ_0126276 | 4397 | 28 | 1.27% |
| hsa_circ_0083478 | 2042 | 13 | 1.27% |
| hsa_circ_0123186 | 2042 | 13 | 1.27% |
| hsa_circ_0001933 | 1728 | 11 | 1.27% |
| hsa_circ_0021093 | 1728 | 11 | 1.27% |
| hsa_circ_0029717 | 1728 | 11 | 1.27% |
| hsa_circ_0048322 | 1728 | 11 | 1.27% |
| hsa_circ_0067825 | 1728 | 11 | 1.27% |
| hsa_circ_0007422 | 1571 | 10 | 1.27% |
| hsa_circ_0017402 | 1571 | 10 | 1.27% |
| hsa_circ_0073326 | 1571 | 10 | 1.27% |
| hsa_circ_0083574 | 1571 | 10 | 1.27% |
| hsa_circ_0112457 | 1571 | 10 | 1.27% |
| hsa_circ_0134144 | 1571 | 10 | 1.27% |
| hsa_circ_0057434 | 2043 | 13 | 1.27% |
| hsa_circ_0045257 | 1886 | 12 | 1.27% |
| hsa_circ_0117211 | 1886 | 12 | 1.27% |
| hsa_circ_0041836 | 1729 | 11 | 1.27% |
| hsa_circ_0081398 | 1729 | 11 | 1.27% |

| | | | |
|---|---|---|---|
| hsa_circ_0104767 | 1729 | 11 | 1.27% |
| hsa_circ_0035896 | 1572 | 10 | 1.27% |
| hsa_circ_0041009 | 1572 | 10 | 1.27% |
| hsa_circ_0059222 | 1572 | 10 | 1.27% |
| hsa_circ_0066228 | 1572 | 10 | 1.27% |
| hsa_circ_0067667 | 1572 | 10 | 1.27% |
| hsa_circ_0069326 | 1572 | 10 | 1.27% |
| hsa_circ_0071801 | 1572 | 10 | 1.27% |
| hsa_circ_0088779 | 1572 | 10 | 1.27% |
| hsa_circ_0096478 | 1572 | 10 | 1.27% |
| hsa_circ_0088814 | 2044 | 13 | 1.27% |
| hsa_circ_0074425 | 1887 | 12 | 1.27% |
| hsa_circ_0008313 | 1730 | 11 | 1.27% |
| hsa_circ_0009690 | 1730 | 11 | 1.27% |
| hsa_circ_0028439 | 1730 | 11 | 1.27% |
| hsa_circ_0091155 | 1730 | 11 | 1.27% |
| hsa_circ_0132981 | 1730 | 11 | 1.27% |
| hsa_circ_0009709 | 1573 | 10 | 1.27% |
| hsa_circ_0012436 | 1573 | 10 | 1.27% |
| hsa_circ_0033591 | 1573 | 10 | 1.27% |
| hsa_circ_0034926 | 1573 | 10 | 1.27% |
| hsa_circ_0044641 | 1573 | 10 | 1.27% |
| hsa_circ_0052227 | 1573 | 10 | 1.27% |
| hsa_circ_0056445 | 1573 | 10 | 1.27% |
| hsa_circ_0059402 | 1573 | 10 | 1.27% |
| hsa_circ_0060364 | 1573 | 10 | 1.27% |
| hsa_circ_0066364 | 1573 | 10 | 1.27% |
| hsa_circ_0068447 | 1573 | 10 | 1.27% |
| hsa_circ_0076618 | 1573 | 10 | 1.27% |
| hsa_circ_0081561 | 1573 | 10 | 1.27% |
| hsa_circ_0090260 | 1573 | 10 | 1.27% |
| hsa_circ_0090711 | 1573 | 10 | 1.27% |
| hsa_circ_0123517 | 1573 | 10 | 1.27% |
| hsa_circ_0123917 | 1573 | 10 | 1.27% |
| hsa_circ_0110703 | 5349 | 34 | 1.27% |
| hsa_circ_0017516 | 1888 | 12 | 1.27% |
| hsa_circ_0037395 | 1888 | 12 | 1.27% |
| hsa_circ_0012085 | 1731 | 11 | 1.27% |
| hsa_circ_0029085 | 1731 | 11 | 1.27% |
| hsa_circ_0053560 | 1731 | 11 | 1.27% |
| hsa_circ_0001472 | 1574 | 10 | 1.27% |
| hsa_circ_0025217 | 1574 | 10 | 1.27% |
| hsa_circ_0036138 | 1574 | 10 | 1.27% |
| hsa_circ_0057024 | 1574 | 10 | 1.27% |
| hsa_circ_0063925 | 1574 | 10 | 1.27% |
| hsa_circ_0064529 | 1574 | 10 | 1.27% |
| hsa_circ_0078964 | 1574 | 10 | 1.27% |
| hsa_circ_0088261 | 1574 | 10 | 1.27% |
| hsa_circ_0121067 | 1574 | 10 | 1.27% |
| hsa_circ_0014358 | 1889 | 12 | 1.27% |

| | | | |
|---|---|---|---|
| hsa_circ_0025780 | 1732 | 11 | 1.27% |
| hsa_circ_0058396 | 1732 | 11 | 1.27% |
| hsa_circ_0062003 | 1732 | 11 | 1.27% |
| hsa_circ_0063523 | 1732 | 11 | 1.27% |
| hsa_circ_0064780 | 1732 | 11 | 1.27% |
| hsa_circ_0065324 | 1732 | 11 | 1.27% |
| hsa_circ_0068695 | 1732 | 11 | 1.27% |
| hsa_circ_0083679 | 1732 | 11 | 1.27% |
| hsa_circ_0117484 | 1732 | 11 | 1.27% |
| hsa_circ_0119826 | 1732 | 11 | 1.27% |
| hsa_circ_0095806 | 3307 | 21 | 1.27% |
| hsa_circ_0130748 | 4095 | 26 | 1.27% |
| hsa_circ_0013757 | 1575 | 10 | 1.27% |
| hsa_circ_0016672 | 1575 | 10 | 1.27% |
| hsa_circ_0019420 | 1575 | 10 | 1.27% |
| hsa_circ_0022471 | 1575 | 10 | 1.27% |
| hsa_circ_0023924 | 1575 | 10 | 1.27% |
| hsa_circ_0027726 | 1575 | 10 | 1.27% |
| hsa_circ_0033167 | 1575 | 10 | 1.27% |
| hsa_circ_0043551 | 1575 | 10 | 1.27% |
| hsa_circ_0050002 | 1575 | 10 | 1.27% |
| hsa_circ_0058138 | 1575 | 10 | 1.27% |
| hsa_circ_0070554 | 1575 | 10 | 1.27% |
| hsa_circ_0088376 | 1575 | 10 | 1.27% |
| hsa_circ_0089581 | 1575 | 10 | 1.27% |
| hsa_circ_0097731 | 1575 | 10 | 1.27% |
| hsa_circ_0121868 | 1575 | 10 | 1.27% |
| hsa_circ_0003857 | 1733 | 11 | 1.27% |
| hsa_circ_0022751 | 1733 | 11 | 1.27% |
| hsa_circ_0027190 | 1733 | 11 | 1.27% |
| hsa_circ_0032212 | 1733 | 11 | 1.27% |
| hsa_circ_0062000 | 1733 | 11 | 1.27% |
| hsa_circ_0083872 | 1733 | 11 | 1.27% |
| hsa_circ_0092438 | 1733 | 11 | 1.27% |
| hsa_circ_0127258 | 1733 | 11 | 1.27% |
| hsa_circ_0133837 | 1733 | 11 | 1.27% |
| hsa_circ_0136062 | 1733 | 11 | 1.27% |
| hsa_circ_0086627 | 7091 | 45 | 1.27% |
| hsa_circ_0058577 | 1891 | 12 | 1.27% |
| hsa_circ_0140677 | 13553 | 86 | 1.27% |
| hsa_circ_0013688 | 1576 | 10 | 1.27% |
| hsa_circ_0013730 | 1576 | 10 | 1.27% |
| hsa_circ_0030260 | 1576 | 10 | 1.27% |
| hsa_circ_0071191 | 1576 | 10 | 1.27% |
| hsa_circ_0073975 | 1576 | 10 | 1.27% |
| hsa_circ_0076935 | 1576 | 10 | 1.27% |
| hsa_circ_0093707 | 1576 | 10 | 1.27% |
| hsa_circ_0094259 | 1576 | 10 | 1.27% |
| hsa_circ_0132837 | 1576 | 10 | 1.27% |
| hsa_circ_0137930 | 1576 | 10 | 1.27% |

| | | | |
|---|---|---|---|
| hsa_circ_0037147 | 2049 | 13 | 1.27% |
| hsa_circ_0074460 | 2049 | 13 | 1.27% |
| hsa_circ_0010813 | 1734 | 11 | 1.27% |
| hsa_circ_0024807 | 1734 | 11 | 1.27% |
| hsa_circ_0082933 | 1734 | 11 | 1.27% |
| hsa_circ_0040314 | 1892 | 12 | 1.27% |
| hsa_circ_0111799 | 5361 | 34 | 1.27% |
| hsa_circ_0006678 | 1577 | 10 | 1.27% |
| hsa_circ_0016784 | 1577 | 10 | 1.27% |
| hsa_circ_0025845 | 1577 | 10 | 1.27% |
| hsa_circ_0033350 | 1577 | 10 | 1.27% |
| hsa_circ_0055727 | 1577 | 10 | 1.27% |
| hsa_circ_0063894 | 1577 | 10 | 1.27% |
| hsa_circ_0090223 | 1577 | 10 | 1.27% |
| hsa_circ_0104280 | 1577 | 10 | 1.27% |
| hsa_circ_0112491 | 1577 | 10 | 1.27% |
| hsa_circ_0127233 | 1577 | 10 | 1.27% |
| hsa_circ_0020183 | 5362 | 34 | 1.27% |
| hsa_circ_0053361 | 1735 | 11 | 1.27% |
| hsa_circ_0062001 | 1735 | 11 | 1.27% |
| hsa_circ_0085587 | 1735 | 11 | 1.27% |
| hsa_circ_0102810 | 1735 | 11 | 1.27% |
| hsa_circ_0110857 | 1735 | 11 | 1.27% |
| hsa_circ_0115748 | 1735 | 11 | 1.27% |
| hsa_circ_0105332 | 1893 | 12 | 1.27% |
| hsa_circ_0119483 | 1893 | 12 | 1.27% |
| hsa_circ_0066652 | 2840 | 18 | 1.27% |
| hsa_circ_0014895 | 1578 | 10 | 1.27% |
| hsa_circ_0018397 | 1578 | 10 | 1.27% |
| hsa_circ_0068217 | 1578 | 10 | 1.27% |
| hsa_circ_0091540 | 1578 | 10 | 1.27% |
| hsa_circ_0113633 | 1578 | 10 | 1.27% |
| hsa_circ_0119240 | 1578 | 10 | 1.27% |
| hsa_circ_0125219 | 1578 | 10 | 1.27% |
| hsa_circ_0130521 | 1578 | 10 | 1.27% |
| hsa_circ_0045538 | 1736 | 11 | 1.27% |
| hsa_circ_0079034 | 1736 | 11 | 1.27% |
| hsa_circ_0099559 | 1736 | 11 | 1.27% |
| hsa_circ_0114262 | 1736 | 11 | 1.27% |
| hsa_circ_0034488 | 1894 | 12 | 1.27% |
| hsa_circ_0112878 | 1894 | 12 | 1.27% |
| hsa_circ_0136800 | 1894 | 12 | 1.27% |
| hsa_circ_0019939 | 2052 | 13 | 1.27% |
| hsa_circ_0076660 | 2052 | 13 | 1.27% |
| hsa_circ_0072111 | 1579 | 10 | 1.27% |
| hsa_circ_0078702 | 1579 | 10 | 1.27% |
| hsa_circ_0091599 | 1579 | 10 | 1.27% |
| hsa_circ_0102224 | 1579 | 10 | 1.27% |
| hsa_circ_0119320 | 1579 | 10 | 1.27% |
| hsa_circ_0129983 | 1579 | 10 | 1.27% |

| | | | |
|---|---|---|---|
| hsa_circ_0019738 | 1737 | 11 | 1.27% |
| hsa_circ_0044066 | 1737 | 11 | 1.27% |
| hsa_circ_0094838 | 1737 | 11 | 1.27% |
| hsa_circ_0096865 | 1737 | 11 | 1.27% |
| hsa_circ_0113258 | 1737 | 11 | 1.27% |
| hsa_circ_0121702 | 1737 | 11 | 1.27% |
| hsa_circ_0127741 | 1737 | 11 | 1.27% |
| hsa_circ_0124054 | 5369 | 34 | 1.27% |
| hsa_circ_0077508 | 4897 | 31 | 1.27% |
| hsa_circ_0136238 | 2054 | 13 | 1.27% |
| hsa_circ_0025125 | 1896 | 12 | 1.27% |
| hsa_circ_0037119 | 1896 | 12 | 1.27% |
| hsa_circ_0083993 | 1896 | 12 | 1.27% |
| hsa_circ_0089531 | 1896 | 12 | 1.27% |
| hsa_circ_0137466 | 1896 | 12 | 1.27% |
| hsa_circ_0046455 | 1738 | 11 | 1.27% |
| hsa_circ_0074695 | 1738 | 11 | 1.27% |
| hsa_circ_0084450 | 1738 | 11 | 1.27% |
| hsa_circ_0101521 | 1738 | 11 | 1.27% |
| hsa_circ_0107961 | 1738 | 11 | 1.27% |
| hsa_circ_0118857 | 1738 | 11 | 1.27% |
| hsa_circ_0009151 | 1580 | 10 | 1.27% |
| hsa_circ_0012374 | 1580 | 10 | 1.27% |
| hsa_circ_0045925 | 1580 | 10 | 1.27% |
| hsa_circ_0054321 | 1580 | 10 | 1.27% |
| hsa_circ_0057254 | 1580 | 10 | 1.27% |
| hsa_circ_0060258 | 1580 | 10 | 1.27% |
| hsa_circ_0074084 | 1580 | 10 | 1.27% |
| hsa_circ_0074877 | 1580 | 10 | 1.27% |
| hsa_circ_0096858 | 1580 | 10 | 1.27% |
| hsa_circ_0117618 | 1580 | 10 | 1.27% |
| hsa_circ_0008415 | 2371 | 15 | 1.27% |
| hsa_circ_0114179 | 2055 | 13 | 1.27% |
| hsa_circ_0010789 | 1897 | 12 | 1.27% |
| hsa_circ_0045122 | 1897 | 12 | 1.27% |
| hsa_circ_0082608 | 1897 | 12 | 1.27% |
| hsa_circ_0132851 | 1897 | 12 | 1.27% |
| hsa_circ_0005254 | 1739 | 11 | 1.27% |
| hsa_circ_0026473 | 1739 | 11 | 1.27% |
| hsa_circ_0073536 | 1739 | 11 | 1.27% |
| hsa_circ_0100263 | 1739 | 11 | 1.27% |
| hsa_circ_0117292 | 1739 | 11 | 1.27% |
| hsa_circ_0013855 | 1581 | 10 | 1.27% |
| hsa_circ_0014555 | 1581 | 10 | 1.27% |
| hsa_circ_0024748 | 1581 | 10 | 1.27% |
| hsa_circ_0029569 | 1581 | 10 | 1.27% |
| hsa_circ_0036752 | 1581 | 10 | 1.27% |
| hsa_circ_0038328 | 1581 | 10 | 1.27% |
| hsa_circ_0041159 | 1581 | 10 | 1.27% |
| hsa_circ_0055511 | 1581 | 10 | 1.27% |

| | | | |
|---|---|---|---|
| hsa_circ_0064936 | 1581 | 10 | 1.27% |
| hsa_circ_0067648 | 1581 | 10 | 1.27% |
| hsa_circ_0101315 | 2372 | 15 | 1.26% |
| hsa_circ_0050270 | 6643 | 42 | 1.26% |
| hsa_circ_0102288 | 1898 | 12 | 1.26% |
| hsa_circ_0010007 | 1740 | 11 | 1.26% |
| hsa_circ_0032063 | 1740 | 11 | 1.26% |
| hsa_circ_0051225 | 1740 | 11 | 1.26% |
| hsa_circ_0059903 | 1740 | 11 | 1.26% |
| hsa_circ_0063897 | 1740 | 11 | 1.26% |
| hsa_circ_0127821 | 1740 | 11 | 1.26% |
| hsa_circ_0132544 | 1740 | 11 | 1.26% |
| hsa_circ_0030515 | 1582 | 10 | 1.26% |
| hsa_circ_0079051 | 1582 | 10 | 1.26% |
| hsa_circ_0107360 | 1582 | 10 | 1.26% |
| hsa_circ_0114818 | 1582 | 10 | 1.26% |
| hsa_circ_0129279 | 1582 | 10 | 1.26% |
| hsa_circ_0030148 | 2215 | 14 | 1.26% |
| hsa_circ_0088856 | 2057 | 13 | 1.26% |
| hsa_circ_0136579 | 2057 | 13 | 1.26% |
| hsa_circ_0022300 | 1899 | 12 | 1.26% |
| hsa_circ_0102431 | 1899 | 12 | 1.26% |
| hsa_circ_0133106 | 2374 | 15 | 1.26% |
| hsa_circ_0041558 | 1741 | 11 | 1.26% |
| hsa_circ_0045922 | 1741 | 11 | 1.26% |
| hsa_circ_0064296 | 1741 | 11 | 1.26% |
| hsa_circ_0083011 | 1741 | 11 | 1.26% |
| hsa_circ_0131475 | 1741 | 11 | 1.26% |
| hsa_circ_0009727 | 1583 | 10 | 1.26% |
| hsa_circ_0020292 | 1583 | 10 | 1.26% |
| hsa_circ_0020832 | 1583 | 10 | 1.26% |
| hsa_circ_0025371 | 1583 | 10 | 1.26% |
| hsa_circ_0051017 | 1583 | 10 | 1.26% |
| hsa_circ_0054832 | 1583 | 10 | 1.26% |
| hsa_circ_0063671 | 1583 | 10 | 1.26% |
| hsa_circ_0101163 | 1583 | 10 | 1.26% |
| hsa_circ_0132919 | 1583 | 10 | 1.26% |
| hsa_circ_0020034 | 2058 | 13 | 1.26% |
| hsa_circ_0069265 | 4117 | 26 | 1.26% |
| hsa_circ_0112381 | 2217 | 14 | 1.26% |
| hsa_circ_0052264 | 1742 | 11 | 1.26% |
| hsa_circ_0079340 | 1742 | 11 | 1.26% |
| hsa_circ_0117994 | 1742 | 11 | 1.26% |
| hsa_circ_0046285 | 2059 | 13 | 1.26% |
| hsa_circ_0013437 | 1584 | 10 | 1.26% |
| hsa_circ_0016190 | 1584 | 10 | 1.26% |
| hsa_circ_0027568 | 1584 | 10 | 1.26% |
| hsa_circ_0037154 | 1584 | 10 | 1.26% |
| hsa_circ_0048494 | 1584 | 10 | 1.26% |
| hsa_circ_0065257 | 1584 | 10 | 1.26% |

| | | | |
|---|---|---|---|
| hsa_circ_0065268 | 1584 | 10 | 1.26% |
| hsa_circ_0070707 | 1584 | 10 | 1.26% |
| hsa_circ_0071150 | 1584 | 10 | 1.26% |
| hsa_circ_0084960 | 1584 | 10 | 1.26% |
| hsa_circ_0099323 | 1584 | 10 | 1.26% |
| hsa_circ_0104606 | 1584 | 10 | 1.26% |
| hsa_circ_0112297 | 1584 | 10 | 1.26% |
| hsa_circ_0035237 | 1901 | 12 | 1.26% |
| hsa_circ_0079196 | 1901 | 12 | 1.26% |
| hsa_circ_0010076 | 1743 | 11 | 1.26% |
| hsa_circ_0019152 | 1743 | 11 | 1.26% |
| hsa_circ_0023767 | 1743 | 11 | 1.26% |
| hsa_circ_0080568 | 1743 | 11 | 1.26% |
| hsa_circ_0083146 | 1743 | 11 | 1.26% |
| hsa_circ_0083888 | 1743 | 11 | 1.26% |
| hsa_circ_0098261 | 1743 | 11 | 1.26% |
| hsa_circ_0011419 | 3170 | 20 | 1.26% |
| hsa_circ_0020687 | 1902 | 12 | 1.26% |
| hsa_circ_0057223 | 1902 | 12 | 1.26% |
| hsa_circ_0092751 | 1902 | 12 | 1.26% |
| hsa_circ_0097747 | 1902 | 12 | 1.26% |
| hsa_circ_0101293 | 1902 | 12 | 1.26% |
| hsa_circ_0025405 | 1585 | 10 | 1.26% |
| hsa_circ_0026444 | 1585 | 10 | 1.26% |
| hsa_circ_0028688 | 1585 | 10 | 1.26% |
| hsa_circ_0035791 | 1585 | 10 | 1.26% |
| hsa_circ_0040051 | 1585 | 10 | 1.26% |
| hsa_circ_0043965 | 1585 | 10 | 1.26% |
| hsa_circ_0046923 | 1585 | 10 | 1.26% |
| hsa_circ_0061717 | 1585 | 10 | 1.26% |
| hsa_circ_0078023 | 1585 | 10 | 1.26% |
| hsa_circ_0085203 | 1585 | 10 | 1.26% |
| hsa_circ_0089941 | 1585 | 10 | 1.26% |
| hsa_circ_0086102 | 4280 | 27 | 1.26% |
| hsa_circ_0045281 | 1744 | 11 | 1.26% |
| hsa_circ_0051614 | 1744 | 11 | 1.26% |
| hsa_circ_0079896 | 1744 | 11 | 1.26% |
| hsa_circ_0139004 | 1744 | 11 | 1.26% |
| hsa_circ_0099739 | 1903 | 12 | 1.26% |
| hsa_circ_0075666 | 4599 | 29 | 1.26% |
| hsa_circ_0006226 | 1586 | 10 | 1.26% |
| hsa_circ_0016154 | 1586 | 10 | 1.26% |
| hsa_circ_0046437 | 1586 | 10 | 1.26% |
| hsa_circ_0047543 | 1586 | 10 | 1.26% |
| hsa_circ_0066346 | 1586 | 10 | 1.26% |
| hsa_circ_0070855 | 1586 | 10 | 1.26% |
| hsa_circ_0072230 | 1586 | 10 | 1.26% |
| hsa_circ_0072434 | 1586 | 10 | 1.26% |
| hsa_circ_0075580 | 1586 | 10 | 1.26% |
| hsa_circ_0080316 | 1586 | 10 | 1.26% |

| | | | |
|---|---|---|---|
| hsa_circ_0087286 | 1586 | 10 | 1.26% |
| hsa_circ_0088728 | 1586 | 10 | 1.26% |
| hsa_circ_0092874 | 1586 | 10 | 1.26% |
| hsa_circ_0118438 | 1586 | 10 | 1.26% |
| hsa_circ_0123262 | 5235 | 33 | 1.26% |
| hsa_circ_0052375 | 1745 | 11 | 1.26% |
| hsa_circ_0077965 | 1745 | 11 | 1.26% |
| hsa_circ_0079615 | 1745 | 11 | 1.26% |
| hsa_circ_0086543 | 1745 | 11 | 1.26% |
| hsa_circ_0114924 | 1745 | 11 | 1.26% |
| hsa_circ_0115778 | 1745 | 11 | 1.26% |
| hsa_circ_0083696 | 1904 | 12 | 1.26% |
| hsa_circ_0118193 | 3491 | 22 | 1.26% |
| hsa_circ_0068606 | 2063 | 13 | 1.26% |
| hsa_circ_0055964 | 1587 | 10 | 1.26% |
| hsa_circ_0085736 | 1587 | 10 | 1.26% |
| hsa_circ_0127165 | 1587 | 10 | 1.26% |
| hsa_circ_0016309 | 1746 | 11 | 1.26% |
| hsa_circ_0026267 | 1746 | 11 | 1.26% |
| hsa_circ_0036326 | 1746 | 11 | 1.26% |
| hsa_circ_0038797 | 1746 | 11 | 1.26% |
| hsa_circ_0103503 | 1746 | 11 | 1.26% |
| hsa_circ_0104305 | 1746 | 11 | 1.26% |
| hsa_circ_0111845 | 1746 | 11 | 1.26% |
| hsa_circ_0117507 | 1746 | 11 | 1.26% |
| hsa_circ_0119801 | 1746 | 11 | 1.26% |
| hsa_circ_0043561 | 1905 | 12 | 1.26% |
| hsa_circ_0069955 | 1905 | 12 | 1.26% |
| hsa_circ_0056862 | 2699 | 17 | 1.26% |
| hsa_circ_0088570 | 2064 | 13 | 1.26% |
| hsa_circ_0065893 | 2223 | 14 | 1.26% |
| hsa_circ_0010365 | 1588 | 10 | 1.26% |
| hsa_circ_0021406 | 1588 | 10 | 1.26% |
| hsa_circ_0023869 | 1588 | 10 | 1.26% |
| hsa_circ_0027410 | 1588 | 10 | 1.26% |
| hsa_circ_0028898 | 1588 | 10 | 1.26% |
| hsa_circ_0045679 | 1588 | 10 | 1.26% |
| hsa_circ_0084419 | 1588 | 10 | 1.26% |
| hsa_circ_0090551 | 1588 | 10 | 1.26% |
| hsa_circ_0097691 | 1588 | 10 | 1.26% |
| hsa_circ_0103005 | 1588 | 10 | 1.26% |
| hsa_circ_0126709 | 1588 | 10 | 1.26% |
| hsa_circ_0135327 | 1588 | 10 | 1.26% |
| hsa_circ_0138867 | 1588 | 10 | 1.26% |
| hsa_circ_0110358 | 6035 | 38 | 1.26% |
| hsa_circ_0006822 | 1747 | 11 | 1.26% |
| hsa_circ_0031437 | 1747 | 11 | 1.26% |
| hsa_circ_0046427 | 1747 | 11 | 1.26% |
| hsa_circ_0061695 | 1747 | 11 | 1.26% |
| hsa_circ_0063135 | 1747 | 11 | 1.26% |

| | | | |
|---|---|---|---|
| hsa_circ_0105971 | 3812 | 24 | 1.26% |
| hsa_circ_0044019 | 1906 | 12 | 1.26% |
| hsa_circ_0051716 | 1906 | 12 | 1.26% |
| hsa_circ_0076883 | 1906 | 12 | 1.26% |
| hsa_circ_0122554 | 1906 | 12 | 1.26% |
| hsa_circ_0043579 | 2065 | 13 | 1.26% |
| hsa_circ_0098307 | 2065 | 13 | 1.26% |
| hsa_circ_0131806 | 2065 | 13 | 1.26% |
| hsa_circ_0015974 | 1589 | 10 | 1.26% |
| hsa_circ_0026577 | 1589 | 10 | 1.26% |
| hsa_circ_0039980 | 1589 | 10 | 1.26% |
| hsa_circ_0048524 | 1589 | 10 | 1.26% |
| hsa_circ_0050812 | 1589 | 10 | 1.26% |
| hsa_circ_0054111 | 1589 | 10 | 1.26% |
| hsa_circ_0073857 | 1589 | 10 | 1.26% |
| hsa_circ_0090348 | 1589 | 10 | 1.26% |
| hsa_circ_0092158 | 1589 | 10 | 1.26% |
| hsa_circ_0093655 | 1589 | 10 | 1.26% |
| hsa_circ_0131280 | 1589 | 10 | 1.26% |
| hsa_circ_0137876 | 1589 | 10 | 1.26% |
| hsa_circ_0011402 | 1748 | 11 | 1.26% |
| hsa_circ_0019847 | 1748 | 11 | 1.26% |
| hsa_circ_0027617 | 1748 | 11 | 1.26% |
| hsa_circ_0122543 | 1748 | 11 | 1.26% |
| hsa_circ_0134385 | 1748 | 11 | 1.26% |
| hsa_circ_0063528 | 2543 | 16 | 1.26% |
| hsa_circ_0029936 | 2067 | 13 | 1.26% |
| hsa_circ_0045530 | 2067 | 13 | 1.26% |
| hsa_circ_0085947 | 2067 | 13 | 1.26% |
| hsa_circ_0026803 | 1908 | 12 | 1.26% |
| hsa_circ_0040702 | 1908 | 12 | 1.26% |
| hsa_circ_0058096 | 1908 | 12 | 1.26% |
| hsa_circ_0041745 | 1749 | 11 | 1.26% |
| hsa_circ_0042038 | 1749 | 11 | 1.26% |
| hsa_circ_0054862 | 1749 | 11 | 1.26% |
| hsa_circ_0088017 | 1749 | 11 | 1.26% |
| hsa_circ_0089742 | 1749 | 11 | 1.26% |
| hsa_circ_0109935 | 1749 | 11 | 1.26% |
| hsa_circ_0116255 | 1749 | 11 | 1.26% |
| hsa_circ_0124664 | 1749 | 11 | 1.26% |
| hsa_circ_0011136 | 1590 | 10 | 1.26% |
| hsa_circ_0018547 | 1590 | 10 | 1.26% |
| hsa_circ_0020196 | 1590 | 10 | 1.26% |
| hsa_circ_0024610 | 1590 | 10 | 1.26% |
| hsa_circ_0027897 | 1590 | 10 | 1.26% |
| hsa_circ_0053364 | 1590 | 10 | 1.26% |
| hsa_circ_0067554 | 1590 | 10 | 1.26% |
| hsa_circ_0078442 | 1590 | 10 | 1.26% |
| hsa_circ_0078524 | 1590 | 10 | 1.26% |
| hsa_circ_0094919 | 1590 | 10 | 1.26% |

| | | | |
|---|---|---|---|
| hsa_circ_0139531 | 1590 | 10 | 1.26% |
| hsa_circ_0114385 | 4136 | 26 | 1.26% |
| hsa_circ_0010084 | 1909 | 12 | 1.26% |
| hsa_circ_0020213 | 1909 | 12 | 1.26% |
| hsa_circ_0093972 | 1909 | 12 | 1.26% |
| hsa_circ_0043800 | 3500 | 22 | 1.26% |
| hsa_circ_0022262 | 1750 | 11 | 1.26% |
| hsa_circ_0036164 | 1750 | 11 | 1.26% |
| hsa_circ_0066983 | 1750 | 11 | 1.26% |
| hsa_circ_0087402 | 1750 | 11 | 1.26% |
| hsa_circ_0020170 | 1591 | 10 | 1.26% |
| hsa_circ_0020464 | 1591 | 10 | 1.26% |
| hsa_circ_0035573 | 1591 | 10 | 1.26% |
| hsa_circ_0081397 | 1591 | 10 | 1.26% |
| hsa_circ_0099704 | 1591 | 10 | 1.26% |
| hsa_circ_0118369 | 1591 | 10 | 1.26% |
| hsa_circ_0125283 | 1591 | 10 | 1.26% |
| hsa_circ_0128388 | 1591 | 10 | 1.26% |
| hsa_circ_0133567 | 1591 | 10 | 1.26% |
| hsa_circ_0139063 | 1591 | 10 | 1.26% |
| hsa_circ_0062110 | 2546 | 16 | 1.26% |
| hsa_circ_0120294 | 2069 | 13 | 1.26% |
| hsa_circ_0128996 | 2069 | 13 | 1.26% |
| hsa_circ_0005881 | 1910 | 12 | 1.26% |
| hsa_circ_0034956 | 1910 | 12 | 1.26% |
| hsa_circ_0077489 | 1910 | 12 | 1.26% |
| hsa_circ_0012517 | 1751 | 11 | 1.26% |
| hsa_circ_0026127 | 1751 | 11 | 1.26% |
| hsa_circ_0027659 | 1751 | 11 | 1.26% |
| hsa_circ_0050897 | 1751 | 11 | 1.26% |
| hsa_circ_0056992 | 1751 | 11 | 1.26% |
| hsa_circ_0001726 | 1592 | 10 | 1.26% |
| hsa_circ_0019732 | 1592 | 10 | 1.26% |
| hsa_circ_0034508 | 1592 | 10 | 1.26% |
| hsa_circ_0036720 | 1592 | 10 | 1.26% |
| hsa_circ_0043712 | 1592 | 10 | 1.26% |
| hsa_circ_0063051 | 1592 | 10 | 1.26% |
| hsa_circ_0067545 | 1592 | 10 | 1.26% |
| hsa_circ_0089527 | 1592 | 10 | 1.26% |
| hsa_circ_0022016 | 2070 | 13 | 1.26% |
| hsa_circ_0063487 | 1911 | 12 | 1.26% |
| hsa_circ_0073799 | 1911 | 12 | 1.26% |
| hsa_circ_0028804 | 1752 | 11 | 1.26% |
| hsa_circ_0038137 | 1752 | 11 | 1.26% |
| hsa_circ_0049942 | 1752 | 11 | 1.26% |
| hsa_circ_0063942 | 1752 | 11 | 1.26% |
| hsa_circ_0065679 | 1752 | 11 | 1.26% |
| hsa_circ_0071691 | 1752 | 11 | 1.26% |
| hsa_circ_0072195 | 1752 | 11 | 1.26% |
| hsa_circ_0096820 | 1752 | 11 | 1.26% |

| | | | |
|---|---|---|---|
| hsa_circ_0110230 | 1752 | 11 | 1.26% |
| hsa_circ_0026655 | 2230 | 14 | 1.26% |
| hsa_circ_0035887 | 2230 | 14 | 1.26% |
| hsa_circ_0015587 | 1593 | 10 | 1.26% |
| hsa_circ_0052767 | 1593 | 10 | 1.26% |
| hsa_circ_0078392 | 1593 | 10 | 1.26% |
| hsa_circ_0083601 | 1593 | 10 | 1.26% |
| hsa_circ_0095151 | 1593 | 10 | 1.26% |
| hsa_circ_0111738 | 1593 | 10 | 1.26% |
| hsa_circ_0113710 | 1593 | 10 | 1.26% |
| hsa_circ_0132668 | 1593 | 10 | 1.26% |
| hsa_circ_0025407 | 1912 | 12 | 1.26% |
| hsa_circ_0090029 | 1912 | 12 | 1.26% |
| hsa_circ_0069973 | 2231 | 14 | 1.26% |
| hsa_circ_0010079 | 1753 | 11 | 1.25% |
| hsa_circ_0016545 | 1753 | 11 | 1.25% |
| hsa_circ_0028927 | 1753 | 11 | 1.25% |
| hsa_circ_0030728 | 1753 | 11 | 1.25% |
| hsa_circ_0044987 | 1753 | 11 | 1.25% |
| hsa_circ_0135112 | 1753 | 11 | 1.25% |
| hsa_circ_0091167 | 2072 | 13 | 1.25% |
| hsa_circ_0003623 | 1594 | 10 | 1.25% |
| hsa_circ_0059823 | 1594 | 10 | 1.25% |
| hsa_circ_0077506 | 1594 | 10 | 1.25% |
| hsa_circ_0094558 | 1594 | 10 | 1.25% |
| hsa_circ_0102982 | 1594 | 10 | 1.25% |
| hsa_circ_0107008 | 1594 | 10 | 1.25% |
| hsa_circ_0123215 | 1594 | 10 | 1.25% |
| hsa_circ_0124121 | 1594 | 10 | 1.25% |
| hsa_circ_0126039 | 1594 | 10 | 1.25% |
| hsa_circ_0027711 | 1913 | 12 | 1.25% |
| hsa_circ_0049831 | 1754 | 11 | 1.25% |
| hsa_circ_0121448 | 1754 | 11 | 1.25% |
| hsa_circ_0091105 | 2871 | 18 | 1.25% |
| hsa_circ_0093971 | 1914 | 12 | 1.25% |
| hsa_circ_0117157 | 1914 | 12 | 1.25% |
| hsa_circ_0015719 | 1595 | 10 | 1.25% |
| hsa_circ_0023890 | 1595 | 10 | 1.25% |
| hsa_circ_0058044 | 1595 | 10 | 1.25% |
| hsa_circ_0086321 | 1595 | 10 | 1.25% |
| hsa_circ_0093934 | 1595 | 10 | 1.25% |
| hsa_circ_0096874 | 1595 | 10 | 1.25% |
| hsa_circ_0108205 | 1595 | 10 | 1.25% |
| hsa_circ_0115788 | 1595 | 10 | 1.25% |
| hsa_circ_0134126 | 1595 | 10 | 1.25% |
| hsa_circ_0052704 | 2393 | 15 | 1.25% |
| hsa_circ_0032618 | 1755 | 11 | 1.25% |
| hsa_circ_0057396 | 1755 | 11 | 1.25% |
| hsa_circ_0060257 | 1755 | 11 | 1.25% |
| hsa_circ_0083720 | 1755 | 11 | 1.25% |

| | | | |
|---|---|---|---|
| hsa_circ_0116434 | 1755 | 11 | 1.25% |
| hsa_circ_0131819 | 1755 | 11 | 1.25% |
| hsa_circ_0105969 | 3830 | 24 | 1.25% |
| hsa_circ_0063849 | 1915 | 12 | 1.25% |
| hsa_circ_0129432 | 1915 | 12 | 1.25% |
| hsa_circ_0132309 | 1915 | 12 | 1.25% |
| hsa_circ_0010862 | 1596 | 10 | 1.25% |
| hsa_circ_0012656 | 1596 | 10 | 1.25% |
| hsa_circ_0036707 | 1596 | 10 | 1.25% |
| hsa_circ_0051621 | 1596 | 10 | 1.25% |
| hsa_circ_0061254 | 1596 | 10 | 1.25% |
| hsa_circ_0061310 | 1596 | 10 | 1.25% |
| hsa_circ_0071177 | 1596 | 10 | 1.25% |
| hsa_circ_0077088 | 1596 | 10 | 1.25% |
| hsa_circ_0077863 | 1596 | 10 | 1.25% |
| hsa_circ_0088004 | 1596 | 10 | 1.25% |
| hsa_circ_0100980 | 6544 | 41 | 1.25% |
| hsa_circ_0003188 | 1756 | 11 | 1.25% |
| hsa_circ_0030939 | 1756 | 11 | 1.25% |
| hsa_circ_0097714 | 1756 | 11 | 1.25% |
| hsa_circ_0119676 | 1756 | 11 | 1.25% |
| hsa_circ_0126754 | 1756 | 11 | 1.25% |
| hsa_circ_0140391 | 1756 | 11 | 1.25% |
| hsa_circ_0010992 | 3672 | 23 | 1.25% |
| hsa_circ_0091404 | 1916 | 12 | 1.25% |
| hsa_circ_0102420 | 1916 | 12 | 1.25% |
| hsa_circ_0031000 | 2076 | 13 | 1.25% |
| hsa_circ_0086697 | 2076 | 13 | 1.25% |
| hsa_circ_0058719 | 1597 | 10 | 1.25% |
| hsa_circ_0093414 | 1597 | 10 | 1.25% |
| hsa_circ_0102721 | 1597 | 10 | 1.25% |
| hsa_circ_0118278 | 1597 | 10 | 1.25% |
| hsa_circ_0123681 | 1597 | 10 | 1.25% |
| hsa_circ_0131566 | 1597 | 10 | 1.25% |
| hsa_circ_0134354 | 1597 | 10 | 1.25% |
| hsa_circ_0137971 | 1597 | 10 | 1.25% |
| hsa_circ_0000215 | 3354 | 21 | 1.25% |
| hsa_circ_0024401 | 1757 | 11 | 1.25% |
| hsa_circ_0039133 | 1757 | 11 | 1.25% |
| hsa_circ_0040649 | 1757 | 11 | 1.25% |
| hsa_circ_0046251 | 1757 | 11 | 1.25% |
| hsa_circ_0055896 | 1757 | 11 | 1.25% |
| hsa_circ_0057664 | 1757 | 11 | 1.25% |
| hsa_circ_0064063 | 1757 | 11 | 1.25% |
| hsa_circ_0073662 | 1757 | 11 | 1.25% |
| hsa_circ_0089431 | 1757 | 11 | 1.25% |
| hsa_circ_0031510 | 1917 | 12 | 1.25% |
| hsa_circ_0050805 | 1917 | 12 | 1.25% |
| hsa_circ_0012369 | 1598 | 10 | 1.25% |
| hsa_circ_0027990 | 1598 | 10 | 1.25% |

| | | | |
|---|---|---|---|
| hsa_circ_0032320 | 1598 | 10 | 1.25% |
| hsa_circ_0035088 | 1598 | 10 | 1.25% |
| hsa_circ_0059887 | 1598 | 10 | 1.25% |
| hsa_circ_0065563 | 1598 | 10 | 1.25% |
| hsa_circ_0067338 | 1598 | 10 | 1.25% |
| hsa_circ_0074329 | 1598 | 10 | 1.25% |
| hsa_circ_0090040 | 1598 | 10 | 1.25% |
| hsa_circ_0095093 | 1598 | 10 | 1.25% |
| hsa_circ_0107588 | 1598 | 10 | 1.25% |
| hsa_circ_0129415 | 1598 | 10 | 1.25% |
| hsa_circ_0137654 | 1598 | 10 | 1.25% |
| hsa_circ_0137853 | 1598 | 10 | 1.25% |
| hsa_circ_0018213 | 1758 | 11 | 1.25% |
| hsa_circ_0066749 | 1758 | 11 | 1.25% |
| hsa_circ_0113153 | 5434 | 34 | 1.25% |
| hsa_circ_0049523 | 1918 | 12 | 1.25% |
| hsa_circ_0071367 | 1918 | 12 | 1.25% |
| hsa_circ_0087551 | 1918 | 12 | 1.25% |
| hsa_circ_0064753 | 2078 | 13 | 1.25% |
| hsa_circ_0094043 | 8792 | 55 | 1.25% |
| hsa_circ_0079120 | 2238 | 14 | 1.25% |
| hsa_circ_0042529 | 1599 | 10 | 1.25% |
| hsa_circ_0059148 | 1599 | 10 | 1.25% |
| hsa_circ_0084207 | 1599 | 10 | 1.25% |
| hsa_circ_0087298 | 1599 | 10 | 1.25% |
| hsa_circ_0093525 | 1599 | 10 | 1.25% |
| hsa_circ_0109572 | 1599 | 10 | 1.25% |
| hsa_circ_0118707 | 1599 | 10 | 1.25% |
| hsa_circ_0130688 | 1599 | 10 | 1.25% |
| hsa_circ_0010554 | 1759 | 11 | 1.25% |
| hsa_circ_0017472 | 1759 | 11 | 1.25% |
| hsa_circ_0040269 | 1759 | 11 | 1.25% |
| hsa_circ_0054232 | 1759 | 11 | 1.25% |
| hsa_circ_0054957 | 1759 | 11 | 1.25% |
| hsa_circ_0057082 | 1759 | 11 | 1.25% |
| hsa_circ_0058313 | 1759 | 11 | 1.25% |
| hsa_circ_0062845 | 1759 | 11 | 1.25% |
| hsa_circ_0098523 | 1759 | 11 | 1.25% |
| hsa_circ_0103115 | 1759 | 11 | 1.25% |
| hsa_circ_0041746 | 1919 | 12 | 1.25% |
| hsa_circ_0105475 | 1919 | 12 | 1.25% |
| hsa_circ_0037329 | 2079 | 13 | 1.25% |
| hsa_circ_0081916 | 2079 | 13 | 1.25% |
| hsa_circ_0081498 | 2559 | 16 | 1.25% |
| hsa_circ_0053114 | 1920 | 12 | 1.25% |
| hsa_circ_0061128 | 1920 | 12 | 1.25% |
| hsa_circ_0075127 | 1920 | 12 | 1.25% |
| hsa_circ_0027520 | 1760 | 11 | 1.25% |
| hsa_circ_0031405 | 1760 | 11 | 1.25% |
| hsa_circ_0052776 | 1760 | 11 | 1.25% |

| | | | |
|---|---|---|---|
| hsa_circ_0086018 | 1760 | 11 | 1.25% |
| hsa_circ_0086283 | 1760 | 11 | 1.25% |
| hsa_circ_0098130 | 1760 | 11 | 1.25% |
| hsa_circ_0123675 | 1760 | 11 | 1.25% |
| hsa_circ_0131281 | 1760 | 11 | 1.25% |
| hsa_circ_0006726 | 1600 | 10 | 1.25% |
| hsa_circ_0017890 | 1600 | 10 | 1.25% |
| hsa_circ_0043149 | 1600 | 10 | 1.25% |
| hsa_circ_0053846 | 1600 | 10 | 1.25% |
| hsa_circ_0054183 | 1600 | 10 | 1.25% |
| hsa_circ_0060893 | 1600 | 10 | 1.25% |
| hsa_circ_0077609 | 1600 | 10 | 1.25% |
| hsa_circ_0077011 | 7842 | 49 | 1.25% |
| hsa_circ_0062626 | 1921 | 12 | 1.25% |
| hsa_circ_0080019 | 1921 | 12 | 1.25% |
| hsa_circ_0095491 | 1761 | 11 | 1.25% |
| hsa_circ_0113181 | 1761 | 11 | 1.25% |
| hsa_circ_0137714 | 3202 | 20 | 1.25% |
| hsa_circ_0045600 | 1601 | 10 | 1.25% |
| hsa_circ_0047539 | 1601 | 10 | 1.25% |
| hsa_circ_0067240 | 1601 | 10 | 1.25% |
| hsa_circ_0072698 | 1601 | 10 | 1.25% |
| hsa_circ_0079153 | 1601 | 10 | 1.25% |
| hsa_circ_0113554 | 1601 | 10 | 1.25% |
| hsa_circ_0125298 | 1601 | 10 | 1.25% |
| hsa_circ_0127946 | 1601 | 10 | 1.25% |
| hsa_circ_0135585 | 1601 | 10 | 1.25% |
| hsa_circ_0060676 | 2082 | 13 | 1.25% |
| hsa_circ_0022395 | 1922 | 12 | 1.25% |
| hsa_circ_0034899 | 1922 | 12 | 1.25% |
| hsa_circ_0037261 | 1922 | 12 | 1.25% |
| hsa_circ_0063533 | 6407 | 40 | 1.25% |
| hsa_circ_0011204 | 1762 | 11 | 1.25% |
| hsa_circ_0015060 | 1762 | 11 | 1.25% |
| hsa_circ_0020774 | 1762 | 11 | 1.25% |
| hsa_circ_0031826 | 1762 | 11 | 1.25% |
| hsa_circ_0055214 | 1762 | 11 | 1.25% |
| hsa_circ_0070376 | 1762 | 11 | 1.25% |
| hsa_circ_0103081 | 1762 | 11 | 1.25% |
| hsa_circ_0113848 | 1762 | 11 | 1.25% |
| hsa_circ_0123363 | 1762 | 11 | 1.25% |
| hsa_circ_0116347 | 3364 | 21 | 1.25% |
| hsa_circ_0002650 | 1602 | 10 | 1.25% |
| hsa_circ_0017727 | 1602 | 10 | 1.25% |
| hsa_circ_0017738 | 1602 | 10 | 1.25% |
| hsa_circ_0032913 | 1602 | 10 | 1.25% |
| hsa_circ_0043826 | 1602 | 10 | 1.25% |
| hsa_circ_0069604 | 1602 | 10 | 1.25% |
| hsa_circ_0075780 | 1602 | 10 | 1.25% |
| hsa_circ_0075936 | 1602 | 10 | 1.25% |

| | | | |
|---|---|---|---|
| hsa_circ_0082655 | 1602 | 10 | 1.25% |
| hsa_circ_0118691 | 1602 | 10 | 1.25% |
| hsa_circ_0121885 | 1602 | 10 | 1.25% |
| hsa_circ_0119118 | 1923 | 12 | 1.25% |
| hsa_circ_0014661 | 1763 | 11 | 1.25% |
| hsa_circ_0021174 | 1763 | 11 | 1.25% |
| hsa_circ_0048324 | 1763 | 11 | 1.25% |
| hsa_circ_0070277 | 1763 | 11 | 1.25% |
| hsa_circ_0074379 | 1763 | 11 | 1.25% |
| hsa_circ_0117798 | 1763 | 11 | 1.25% |
| hsa_circ_0123935 | 1763 | 11 | 1.25% |
| hsa_circ_0050512 | 1603 | 10 | 1.25% |
| hsa_circ_0074501 | 1603 | 10 | 1.25% |
| hsa_circ_0094929 | 1603 | 10 | 1.25% |
| hsa_circ_0101463 | 1603 | 10 | 1.25% |
| hsa_circ_0107923 | 1603 | 10 | 1.25% |
| hsa_circ_0004865 | 1764 | 11 | 1.25% |
| hsa_circ_0043888 | 1764 | 11 | 1.25% |
| hsa_circ_0050966 | 1764 | 11 | 1.25% |
| hsa_circ_0070017 | 1764 | 11 | 1.25% |
| hsa_circ_0084378 | 1764 | 11 | 1.25% |
| hsa_circ_0053022 | 2406 | 15 | 1.25% |
| hsa_circ_0016368 | 1604 | 10 | 1.25% |
| hsa_circ_0030531 | 1604 | 10 | 1.25% |
| hsa_circ_0055897 | 1604 | 10 | 1.25% |
| hsa_circ_0066227 | 1604 | 10 | 1.25% |
| hsa_circ_0016085 | 1925 | 12 | 1.25% |
| hsa_circ_0056038 | 1925 | 12 | 1.25% |
| hsa_circ_0064799 | 1925 | 12 | 1.25% |
| hsa_circ_0038806 | 1765 | 11 | 1.25% |
| hsa_circ_0041198 | 1765 | 11 | 1.25% |
| hsa_circ_0051652 | 1765 | 11 | 1.25% |
| hsa_circ_0133508 | 1765 | 11 | 1.25% |
| hsa_circ_0083473 | 2086 | 13 | 1.25% |
| hsa_circ_0019790 | 1605 | 10 | 1.25% |
| hsa_circ_0021680 | 1605 | 10 | 1.25% |
| hsa_circ_0036918 | 1605 | 10 | 1.25% |
| hsa_circ_0053914 | 1605 | 10 | 1.25% |
| hsa_circ_0068361 | 1605 | 10 | 1.25% |
| hsa_circ_0101785 | 1605 | 10 | 1.25% |
| hsa_circ_0112767 | 1605 | 10 | 1.25% |
| hsa_circ_0015931 | 1766 | 11 | 1.25% |
| hsa_circ_0023155 | 1766 | 11 | 1.25% |
| hsa_circ_0026968 | 1766 | 11 | 1.25% |
| hsa_circ_0101838 | 1766 | 11 | 1.25% |
| hsa_circ_0102947 | 14290 | 89 | 1.25% |
| hsa_circ_0051881 | 1927 | 12 | 1.25% |
| hsa_circ_0061068 | 1927 | 12 | 1.25% |
| hsa_circ_0076224 | 1927 | 12 | 1.25% |
| hsa_circ_0079656 | 1927 | 12 | 1.25% |

| | | | |
|---|---|---|---|
| hsa_circ_0085626 | 1927 | 12 | 1.25% |
| hsa_circ_0133241 | 1927 | 12 | 1.25% |
| hsa_circ_0029628 | 1606 | 10 | 1.25% |
| hsa_circ_0069430 | 1606 | 10 | 1.25% |
| hsa_circ_0073562 | 1606 | 10 | 1.25% |
| hsa_circ_0083230 | 1606 | 10 | 1.25% |
| hsa_circ_0089630 | 1606 | 10 | 1.25% |
| hsa_circ_0115866 | 1606 | 10 | 1.25% |
| hsa_circ_0125564 | 1606 | 10 | 1.25% |
| hsa_circ_0018237 | 1767 | 11 | 1.25% |
| hsa_circ_0019003 | 1767 | 11 | 1.25% |
| hsa_circ_0035239 | 1767 | 11 | 1.25% |
| hsa_circ_0035331 | 1767 | 11 | 1.25% |
| hsa_circ_0049107 | 1767 | 11 | 1.25% |
| hsa_circ_0060305 | 1767 | 11 | 1.25% |
| hsa_circ_0082693 | 1767 | 11 | 1.25% |
| hsa_circ_0104039 | 1767 | 11 | 1.25% |
| hsa_circ_0052282 | 1928 | 12 | 1.24% |
| hsa_circ_0009744 | 1607 | 10 | 1.24% |
| hsa_circ_0015608 | 1607 | 10 | 1.24% |
| hsa_circ_0015832 | 1607 | 10 | 1.24% |
| hsa_circ_0016624 | 1607 | 10 | 1.24% |
| hsa_circ_0023012 | 1607 | 10 | 1.24% |
| hsa_circ_0048041 | 1607 | 10 | 1.24% |
| hsa_circ_0051686 | 1607 | 10 | 1.24% |
| hsa_circ_0057875 | 4982 | 31 | 1.24% |
| hsa_circ_0010766 | 1768 | 11 | 1.24% |
| hsa_circ_0048143 | 1768 | 11 | 1.24% |
| hsa_circ_0050632 | 1768 | 11 | 1.24% |
| hsa_circ_0059729 | 1768 | 11 | 1.24% |
| hsa_circ_0063509 | 1768 | 11 | 1.24% |
| hsa_circ_0026800 | 1929 | 12 | 1.24% |
| hsa_circ_0049758 | 1929 | 12 | 1.24% |
| hsa_circ_0060490 | 5948 | 37 | 1.24% |
| hsa_circ_0118404 | 2090 | 13 | 1.24% |
| hsa_circ_0052317 | 4341 | 27 | 1.24% |
| hsa_circ_0004892 | 5306 | 33 | 1.24% |
| hsa_circ_0016499 | 1608 | 10 | 1.24% |
| hsa_circ_0024522 | 1608 | 10 | 1.24% |
| hsa_circ_0043125 | 1608 | 10 | 1.24% |
| hsa_circ_0073915 | 1608 | 10 | 1.24% |
| hsa_circ_0084831 | 1608 | 10 | 1.24% |
| hsa_circ_0095420 | 1608 | 10 | 1.24% |
| hsa_circ_0108348 | 1608 | 10 | 1.24% |
| hsa_circ_0109252 | 1608 | 10 | 1.24% |
| hsa_circ_0113597 | 1608 | 10 | 1.24% |
| hsa_circ_0056516 | 1769 | 11 | 1.24% |
| hsa_circ_0078571 | 1769 | 11 | 1.24% |
| hsa_circ_0086383 | 1769 | 11 | 1.24% |
| hsa_circ_0090529 | 1769 | 11 | 1.24% |

| | | | |
|---|---|---|---|
| hsa_circ_0098411 | 1769 | 11 | 1.24% |
| hsa_circ_0024453 | 1930 | 12 | 1.24% |
| hsa_circ_0062841 | 1930 | 12 | 1.24% |
| hsa_circ_0086339 | 2413 | 15 | 1.24% |
| hsa_circ_0109474 | 3861 | 24 | 1.24% |
| hsa_circ_0022449 | 1609 | 10 | 1.24% |
| hsa_circ_0024530 | 1609 | 10 | 1.24% |
| hsa_circ_0030802 | 1609 | 10 | 1.24% |
| hsa_circ_0069563 | 1609 | 10 | 1.24% |
| hsa_circ_0074980 | 1609 | 10 | 1.24% |
| hsa_circ_0085375 | 1609 | 10 | 1.24% |
| hsa_circ_0092656 | 1609 | 10 | 1.24% |
| hsa_circ_0126801 | 1609 | 10 | 1.24% |
| hsa_circ_0031018 | 1770 | 11 | 1.24% |
| hsa_circ_0038351 | 1770 | 11 | 1.24% |
| hsa_circ_0099916 | 1770 | 11 | 1.24% |
| hsa_circ_0105083 | 1770 | 11 | 1.24% |
| hsa_circ_0118049 | 2092 | 13 | 1.24% |
| hsa_circ_0030502 | 2415 | 15 | 1.24% |
| hsa_circ_0078332 | 2093 | 13 | 1.24% |
| hsa_circ_0022822 | 1932 | 12 | 1.24% |
| hsa_circ_0089116 | 1932 | 12 | 1.24% |
| hsa_circ_0103624 | 1932 | 12 | 1.24% |
| hsa_circ_0012312 | 1771 | 11 | 1.24% |
| hsa_circ_0021391 | 1771 | 11 | 1.24% |
| hsa_circ_0031549 | 1771 | 11 | 1.24% |
| hsa_circ_0038296 | 1771 | 11 | 1.24% |
| hsa_circ_0046470 | 1771 | 11 | 1.24% |
| hsa_circ_0053536 | 1771 | 11 | 1.24% |
| hsa_circ_0119986 | 1771 | 11 | 1.24% |
| hsa_circ_0019613 | 1610 | 10 | 1.24% |
| hsa_circ_0061023 | 1610 | 10 | 1.24% |
| hsa_circ_0080468 | 1610 | 10 | 1.24% |
| hsa_circ_0081661 | 1610 | 10 | 1.24% |
| hsa_circ_0082946 | 1610 | 10 | 1.24% |
| hsa_circ_0085088 | 1610 | 10 | 1.24% |
| hsa_circ_0088750 | 1610 | 10 | 1.24% |
| hsa_circ_0116424 | 1610 | 10 | 1.24% |
| hsa_circ_0137898 | 1610 | 10 | 1.24% |
| hsa_circ_0074381 | 2577 | 16 | 1.24% |
| hsa_circ_0083567 | 2416 | 15 | 1.24% |
| hsa_circ_0016903 | 2255 | 14 | 1.24% |
| hsa_circ_0046024 | 1933 | 12 | 1.24% |
| hsa_circ_0082097 | 1933 | 12 | 1.24% |
| hsa_circ_0097913 | 1933 | 12 | 1.24% |
| hsa_circ_0125605 | 1933 | 12 | 1.24% |
| hsa_circ_0014469 | 1772 | 11 | 1.24% |
| hsa_circ_0015483 | 1772 | 11 | 1.24% |
| hsa_circ_0034379 | 1772 | 11 | 1.24% |
| hsa_circ_0074193 | 1772 | 11 | 1.24% |

| | | | |
|---|---|---|---|
| hsa_circ_0113253 | 1772 | 11 | 1.24% |
| hsa_circ_0016029 | 1611 | 10 | 1.24% |
| hsa_circ_0065672 | 1611 | 10 | 1.24% |
| hsa_circ_0075526 | 1611 | 10 | 1.24% |
| hsa_circ_0083231 | 1611 | 10 | 1.24% |
| hsa_circ_0098219 | 1611 | 10 | 1.24% |
| hsa_circ_0105460 | 1611 | 10 | 1.24% |
| hsa_circ_0123271 | 1611 | 10 | 1.24% |
| hsa_circ_0125392 | 1611 | 10 | 1.24% |
| hsa_circ_0137701 | 1611 | 10 | 1.24% |
| hsa_circ_0047943 | 2095 | 13 | 1.24% |
| hsa_circ_0139820 | 2095 | 13 | 1.24% |
| hsa_circ_0055294 | 1934 | 12 | 1.24% |
| hsa_circ_0018515 | 1773 | 11 | 1.24% |
| hsa_circ_0057278 | 1773 | 11 | 1.24% |
| hsa_circ_0080228 | 1773 | 11 | 1.24% |
| hsa_circ_0106189 | 1773 | 11 | 1.24% |
| hsa_circ_0014257 | 1612 | 10 | 1.24% |
| hsa_circ_0040951 | 1612 | 10 | 1.24% |
| hsa_circ_0047742 | 1612 | 10 | 1.24% |
| hsa_circ_0052901 | 1612 | 10 | 1.24% |
| hsa_circ_0064509 | 1612 | 10 | 1.24% |
| hsa_circ_0069538 | 1612 | 10 | 1.24% |
| hsa_circ_0078075 | 1612 | 10 | 1.24% |
| hsa_circ_0138822 | 1612 | 10 | 1.24% |
| hsa_circ_0046069 | 2096 | 13 | 1.24% |
| hsa_circ_0051224 | 1774 | 11 | 1.24% |
| hsa_circ_0076657 | 1774 | 11 | 1.24% |
| hsa_circ_0085775 | 1774 | 11 | 1.24% |
| hsa_circ_0038906 | 6290 | 39 | 1.24% |
| hsa_circ_0022541 | 2742 | 17 | 1.24% |
| hsa_circ_0009908 | 1613 | 10 | 1.24% |
| hsa_circ_0022288 | 1613 | 10 | 1.24% |
| hsa_circ_0035526 | 1613 | 10 | 1.24% |
| hsa_circ_0058612 | 1613 | 10 | 1.24% |
| hsa_circ_0064586 | 1613 | 10 | 1.24% |
| hsa_circ_0074408 | 1613 | 10 | 1.24% |
| hsa_circ_0077929 | 1613 | 10 | 1.24% |
| hsa_circ_0078116 | 1613 | 10 | 1.24% |
| hsa_circ_0083264 | 1613 | 10 | 1.24% |
| hsa_circ_0086234 | 1613 | 10 | 1.24% |
| hsa_circ_0088272 | 1613 | 10 | 1.24% |
| hsa_circ_0092649 | 1613 | 10 | 1.24% |
| hsa_circ_0127285 | 1613 | 10 | 1.24% |
| hsa_circ_0046074 | 2097 | 13 | 1.24% |
| hsa_circ_0046166 | 2097 | 13 | 1.24% |
| hsa_circ_0043852 | 1936 | 12 | 1.24% |
| hsa_circ_0028093 | 5809 | 36 | 1.24% |
| hsa_circ_0072120 | 1775 | 11 | 1.24% |
| hsa_circ_0085118 | 1775 | 11 | 1.24% |

| | | | |
|---|---|---|---|
| hsa_circ_0085967 | 1775 | 11 | 1.24% |
| hsa_circ_0089198 | 1775 | 11 | 1.24% |
| hsa_circ_0089879 | 1775 | 11 | 1.24% |
| hsa_circ_0104991 | 1775 | 11 | 1.24% |
| hsa_circ_0114379 | 1775 | 11 | 1.24% |
| hsa_circ_0058276 | 2905 | 18 | 1.24% |
| hsa_circ_0069830 | 2421 | 15 | 1.24% |
| hsa_circ_0024629 | 1614 | 10 | 1.24% |
| hsa_circ_0028481 | 1614 | 10 | 1.24% |
| hsa_circ_0031911 | 1614 | 10 | 1.24% |
| hsa_circ_0032585 | 1614 | 10 | 1.24% |
| hsa_circ_0039116 | 1614 | 10 | 1.24% |
| hsa_circ_0042018 | 1614 | 10 | 1.24% |
| hsa_circ_0069369 | 1614 | 10 | 1.24% |
| hsa_circ_0080122 | 1614 | 10 | 1.24% |
| hsa_circ_0086301 | 1614 | 10 | 1.24% |
| hsa_circ_0109518 | 1614 | 10 | 1.24% |
| hsa_circ_0125375 | 1614 | 10 | 1.24% |
| hsa_circ_0129000 | 1614 | 10 | 1.24% |
| hsa_circ_0135242 | 1614 | 10 | 1.24% |
| hsa_circ_0138530 | 1614 | 10 | 1.24% |
| hsa_circ_0140401 | 1614 | 10 | 1.24% |
| hsa_circ_0046640 | 1937 | 12 | 1.24% |
| hsa_circ_0037278 | 2260 | 14 | 1.24% |
| hsa_circ_0000677 | 1776 | 11 | 1.24% |
| hsa_circ_0033360 | 1776 | 11 | 1.24% |
| hsa_circ_0034660 | 1776 | 11 | 1.24% |
| hsa_circ_0037121 | 1776 | 11 | 1.24% |
| hsa_circ_0039967 | 1776 | 11 | 1.24% |
| hsa_circ_0079189 | 1776 | 11 | 1.24% |
| hsa_circ_0089437 | 1776 | 11 | 1.24% |
| hsa_circ_0126208 | 1776 | 11 | 1.24% |
| hsa_circ_0015114 | 2099 | 13 | 1.24% |
| hsa_circ_0114182 | 2099 | 13 | 1.24% |
| hsa_circ_0018307 | 2745 | 17 | 1.24% |
| hsa_circ_0072086 | 2261 | 14 | 1.24% |
| hsa_circ_0009426 | 1938 | 12 | 1.24% |
| hsa_circ_0094896 | 1938 | 12 | 1.24% |
| hsa_circ_0097765 | 1938 | 12 | 1.24% |
| hsa_circ_0021596 | 1615 | 10 | 1.24% |
| hsa_circ_0022479 | 1615 | 10 | 1.24% |
| hsa_circ_0042410 | 1615 | 10 | 1.24% |
| hsa_circ_0047509 | 1615 | 10 | 1.24% |
| hsa_circ_0105781 | 1615 | 10 | 1.24% |
| hsa_circ_0120743 | 1615 | 10 | 1.24% |
| hsa_circ_0128530 | 1615 | 10 | 1.24% |
| hsa_circ_0006013 | 1777 | 11 | 1.24% |
| hsa_circ_0019233 | 1777 | 11 | 1.24% |
| hsa_circ_0019827 | 1777 | 11 | 1.24% |
| hsa_circ_0033057 | 1777 | 11 | 1.24% |

| | | | |
|---|---|---|---|
| hsa_circ_0105130 | 1777 | 11 | 1.24% |
| hsa_circ_0117754 | 1777 | 11 | 1.24% |
| hsa_circ_0047304 | 2262 | 14 | 1.24% |
| hsa_circ_0090125 | 2262 | 14 | 1.24% |
| hsa_circ_0010584 | 1939 | 12 | 1.24% |
| hsa_circ_0067218 | 1939 | 12 | 1.24% |
| hsa_circ_0124188 | 1939 | 12 | 1.24% |
| hsa_circ_0018795 | 2424 | 15 | 1.24% |
| hsa_circ_0009984 | 1616 | 10 | 1.24% |
| hsa_circ_0011640 | 1616 | 10 | 1.24% |
| hsa_circ_0014936 | 1616 | 10 | 1.24% |
| hsa_circ_0039353 | 1616 | 10 | 1.24% |
| hsa_circ_0039509 | 1616 | 10 | 1.24% |
| hsa_circ_0053743 | 1616 | 10 | 1.24% |
| hsa_circ_0054641 | 1616 | 10 | 1.24% |
| hsa_circ_0056927 | 1616 | 10 | 1.24% |
| hsa_circ_0057005 | 1616 | 10 | 1.24% |
| hsa_circ_0061822 | 1616 | 10 | 1.24% |
| hsa_circ_0073549 | 1616 | 10 | 1.24% |
| hsa_circ_0088589 | 1616 | 10 | 1.24% |
| hsa_circ_0115229 | 1616 | 10 | 1.24% |
| hsa_circ_0118783 | 1616 | 10 | 1.24% |
| hsa_circ_0139839 | 1616 | 10 | 1.24% |
| hsa_circ_0030068 | 2101 | 13 | 1.24% |
| hsa_circ_0047834 | 3879 | 24 | 1.24% |
| hsa_circ_0018150 | 1778 | 11 | 1.24% |
| hsa_circ_0034549 | 1778 | 11 | 1.24% |
| hsa_circ_0035570 | 1778 | 11 | 1.24% |
| hsa_circ_0079612 | 1778 | 11 | 1.24% |
| hsa_circ_0137055 | 1778 | 11 | 1.24% |
| hsa_circ_0126277 | 4526 | 28 | 1.24% |
| hsa_circ_0040654 | 1940 | 12 | 1.24% |
| hsa_circ_0071119 | 1940 | 12 | 1.24% |
| hsa_circ_0051342 | 2102 | 13 | 1.24% |
| hsa_circ_0056255 | 2102 | 13 | 1.24% |
| hsa_circ_0086114 | 2102 | 13 | 1.24% |
| hsa_circ_0014218 | 1617 | 10 | 1.24% |
| hsa_circ_0036905 | 1617 | 10 | 1.24% |
| hsa_circ_0065460 | 1617 | 10 | 1.24% |
| hsa_circ_0093015 | 3396 | 21 | 1.24% |
| hsa_circ_0010832 | 1779 | 11 | 1.24% |
| hsa_circ_0044004 | 1779 | 11 | 1.24% |
| hsa_circ_0065412 | 1779 | 11 | 1.24% |
| hsa_circ_0074461 | 1779 | 11 | 1.24% |
| hsa_circ_0140064 | 1779 | 11 | 1.24% |
| hsa_circ_0135943 | 3720 | 23 | 1.24% |
| hsa_circ_0015387 | 1941 | 12 | 1.24% |
| hsa_circ_0045667 | 1941 | 12 | 1.24% |
| hsa_circ_0058329 | 1941 | 12 | 1.24% |
| hsa_circ_0071530 | 2103 | 13 | 1.24% |

| | | | |
|---|---|---|---|
| hsa_circ_0025547 | 1618 | 10 | 1.24% |
| hsa_circ_0032428 | 1618 | 10 | 1.24% |
| hsa_circ_0032487 | 1618 | 10 | 1.24% |
| hsa_circ_0034767 | 1618 | 10 | 1.24% |
| hsa_circ_0070958 | 1618 | 10 | 1.24% |
| hsa_circ_0100947 | 1618 | 10 | 1.24% |
| hsa_circ_0109357 | 1618 | 10 | 1.24% |
| hsa_circ_0110968 | 1618 | 10 | 1.24% |
| hsa_circ_0008509 | 1780 | 11 | 1.24% |
| hsa_circ_0029575 | 1780 | 11 | 1.24% |
| hsa_circ_0044422 | 1780 | 11 | 1.24% |
| hsa_circ_0070586 | 1780 | 11 | 1.24% |
| hsa_circ_0074402 | 1780 | 11 | 1.24% |
| hsa_circ_0076352 | 1780 | 11 | 1.24% |
| hsa_circ_0114353 | 1780 | 11 | 1.24% |
| hsa_circ_0122675 | 1780 | 11 | 1.24% |
| hsa_circ_0123938 | 1780 | 11 | 1.24% |
| hsa_circ_0061067 | 1942 | 12 | 1.24% |
| hsa_circ_0054169 | 2428 | 15 | 1.24% |
| hsa_circ_0091302 | 2428 | 15 | 1.24% |
| hsa_circ_0024708 | 1619 | 10 | 1.24% |
| hsa_circ_0029942 | 1619 | 10 | 1.24% |
| hsa_circ_0036563 | 1619 | 10 | 1.24% |
| hsa_circ_0099937 | 1619 | 10 | 1.24% |
| hsa_circ_0111813 | 1619 | 10 | 1.24% |
| hsa_circ_0138872 | 1619 | 10 | 1.24% |
| hsa_circ_0139074 | 1619 | 10 | 1.24% |
| hsa_circ_0018623 | 1781 | 11 | 1.24% |
| hsa_circ_0136686 | 1781 | 11 | 1.24% |
| hsa_circ_0023153 | 1943 | 12 | 1.24% |
| hsa_circ_0050713 | 1943 | 12 | 1.24% |
| hsa_circ_0121422 | 2105 | 13 | 1.24% |
| hsa_circ_0123549 | 2105 | 13 | 1.24% |
| hsa_circ_0136484 | 2105 | 13 | 1.24% |
| hsa_circ_0030146 | 2267 | 14 | 1.24% |
| hsa_circ_0090505 | 4211 | 26 | 1.23% |
| hsa_circ_0043871 | 2430 | 15 | 1.23% |
| hsa_circ_0013632 | 1944 | 12 | 1.23% |
| hsa_circ_0034368 | 1944 | 12 | 1.23% |
| hsa_circ_0010044 | 1782 | 11 | 1.23% |
| hsa_circ_0031492 | 1782 | 11 | 1.23% |
| hsa_circ_0043939 | 1782 | 11 | 1.23% |
| hsa_circ_0078291 | 1782 | 11 | 1.23% |
| hsa_circ_0080045 | 1782 | 11 | 1.23% |
| hsa_circ_0125056 | 1782 | 11 | 1.23% |
| hsa_circ_0014509 | 1620 | 10 | 1.23% |
| hsa_circ_0054362 | 1620 | 10 | 1.23% |
| hsa_circ_0066345 | 1620 | 10 | 1.23% |
| hsa_circ_0081483 | 1620 | 10 | 1.23% |
| hsa_circ_0082189 | 1620 | 10 | 1.23% |

| | | | |
|---|---|---|---|
| hsa_circ_0102152 | 1620 | 10 | 1.23% |
| hsa_circ_0072617 | 2107 | 13 | 1.23% |
| hsa_circ_0074330 | 1945 | 12 | 1.23% |
| hsa_circ_0014057 | 1783 | 11 | 1.23% |
| hsa_circ_0082658 | 1783 | 11 | 1.23% |
| hsa_circ_0094193 | 1783 | 11 | 1.23% |
| hsa_circ_0121338 | 1783 | 11 | 1.23% |
| hsa_circ_0006192 | 1621 | 10 | 1.23% |
| hsa_circ_0006321 | 1621 | 10 | 1.23% |
| hsa_circ_0029137 | 1621 | 10 | 1.23% |
| hsa_circ_0032667 | 1621 | 10 | 1.23% |
| hsa_circ_0035120 | 1621 | 10 | 1.23% |
| hsa_circ_0038022 | 1621 | 10 | 1.23% |
| hsa_circ_0048242 | 1621 | 10 | 1.23% |
| hsa_circ_0061741 | 1621 | 10 | 1.23% |
| hsa_circ_0064827 | 1621 | 10 | 1.23% |
| hsa_circ_0073431 | 1621 | 10 | 1.23% |
| hsa_circ_0095196 | 1621 | 10 | 1.23% |
| hsa_circ_0106819 | 1621 | 10 | 1.23% |
| hsa_circ_0108149 | 1621 | 10 | 1.23% |
| hsa_circ_0046731 | 2108 | 13 | 1.23% |
| hsa_circ_0020298 | 1946 | 12 | 1.23% |
| hsa_circ_0032144 | 1946 | 12 | 1.23% |
| hsa_circ_0021106 | 1784 | 11 | 1.23% |
| hsa_circ_0036085 | 1784 | 11 | 1.23% |
| hsa_circ_0049436 | 1784 | 11 | 1.23% |
| hsa_circ_0064778 | 1784 | 11 | 1.23% |
| hsa_circ_0067155 | 1784 | 11 | 1.23% |
| hsa_circ_0069886 | 1784 | 11 | 1.23% |
| hsa_circ_0116254 | 1784 | 11 | 1.23% |
| hsa_circ_0012017 | 1622 | 10 | 1.23% |
| hsa_circ_0012603 | 1622 | 10 | 1.23% |
| hsa_circ_0038493 | 1622 | 10 | 1.23% |
| hsa_circ_0046624 | 1622 | 10 | 1.23% |
| hsa_circ_0055842 | 1622 | 10 | 1.23% |
| hsa_circ_0078709 | 1622 | 10 | 1.23% |
| hsa_circ_0097056 | 1622 | 10 | 1.23% |
| hsa_circ_0139435 | 1622 | 10 | 1.23% |
| hsa_circ_0052539 | 4704 | 29 | 1.23% |
| hsa_circ_0030846 | 2109 | 13 | 1.23% |
| hsa_circ_0100337 | 2109 | 13 | 1.23% |
| hsa_circ_0018085 | 1947 | 12 | 1.23% |
| hsa_circ_0048465 | 1947 | 12 | 1.23% |
| hsa_circ_0067237 | 1947 | 12 | 1.23% |
| hsa_circ_0019610 | 1785 | 11 | 1.23% |
| hsa_circ_0049562 | 1785 | 11 | 1.23% |
| hsa_circ_0075219 | 1785 | 11 | 1.23% |
| hsa_circ_0003479 | 1623 | 10 | 1.23% |
| hsa_circ_0030383 | 1623 | 10 | 1.23% |
| hsa_circ_0052463 | 1623 | 10 | 1.23% |

| | | | |
|---|---|---|---|
| hsa_circ_0062230 | 1623 | 10 | 1.23% |
| hsa_circ_0067075 | 1623 | 10 | 1.23% |
| hsa_circ_0103418 | 1623 | 10 | 1.23% |
| hsa_circ_0106538 | 1623 | 10 | 1.23% |
| hsa_circ_0111960 | 1623 | 10 | 1.23% |
| hsa_circ_0123414 | 1623 | 10 | 1.23% |
| hsa_circ_0131068 | 1623 | 10 | 1.23% |
| hsa_circ_0134961 | 1623 | 10 | 1.23% |
| hsa_circ_0081399 | 2110 | 13 | 1.23% |
| hsa_circ_0021484 | 1948 | 12 | 1.23% |
| hsa_circ_0041681 | 1948 | 12 | 1.23% |
| hsa_circ_0013842 | 1786 | 11 | 1.23% |
| hsa_circ_0034770 | 1786 | 11 | 1.23% |
| hsa_circ_0069301 | 1786 | 11 | 1.23% |
| hsa_circ_0022942 | 1624 | 10 | 1.23% |
| hsa_circ_0024884 | 1624 | 10 | 1.23% |
| hsa_circ_0045704 | 1624 | 10 | 1.23% |
| hsa_circ_0082841 | 1624 | 10 | 1.23% |
| hsa_circ_0091900 | 1624 | 10 | 1.23% |
| hsa_circ_0096307 | 1624 | 10 | 1.23% |
| hsa_circ_0105860 | 1624 | 10 | 1.23% |
| hsa_circ_0115192 | 1624 | 10 | 1.23% |
| hsa_circ_0071241 | 1949 | 12 | 1.23% |
| hsa_circ_0019736 | 1787 | 11 | 1.23% |
| hsa_circ_0063745 | 1787 | 11 | 1.23% |
| hsa_circ_0103408 | 1787 | 11 | 1.23% |
| hsa_circ_0118110 | 2112 | 13 | 1.23% |
| hsa_circ_0005981 | 2762 | 17 | 1.23% |
| hsa_circ_0030841 | 2275 | 14 | 1.23% |
| hsa_circ_0012796 | 1950 | 12 | 1.23% |
| hsa_circ_0018363 | 1950 | 12 | 1.23% |
| hsa_circ_0047114 | 1950 | 12 | 1.23% |
| hsa_circ_0068156 | 1950 | 12 | 1.23% |
| hsa_circ_0016494 | 1625 | 10 | 1.23% |
| hsa_circ_0025858 | 1625 | 10 | 1.23% |
| hsa_circ_0035881 | 1625 | 10 | 1.23% |
| hsa_circ_0038192 | 1625 | 10 | 1.23% |
| hsa_circ_0061976 | 1625 | 10 | 1.23% |
| hsa_circ_0114475 | 1625 | 10 | 1.23% |
| hsa_circ_0138403 | 1625 | 10 | 1.23% |
| hsa_circ_0047939 | 2113 | 13 | 1.23% |
| hsa_circ_0010211 | 1788 | 11 | 1.23% |
| hsa_circ_0023648 | 1788 | 11 | 1.23% |
| hsa_circ_0049621 | 1788 | 11 | 1.23% |
| hsa_circ_0071319 | 1788 | 11 | 1.23% |
| hsa_circ_0079637 | 1788 | 11 | 1.23% |
| hsa_circ_0080083 | 2276 | 14 | 1.23% |
| hsa_circ_0024085 | 1626 | 10 | 1.23% |
| hsa_circ_0041361 | 1626 | 10 | 1.23% |
| hsa_circ_0076290 | 1626 | 10 | 1.23% |

| | | | |
|---|---|---|---|
| hsa_circ_0085085 | 1626 | 10 | 1.23% |
| hsa_circ_0039552 | 1789 | 11 | 1.23% |
| hsa_circ_0054849 | 1789 | 11 | 1.23% |
| hsa_circ_0077094 | 1789 | 11 | 1.23% |
| hsa_circ_0078633 | 1789 | 11 | 1.23% |
| hsa_circ_0030918 | 1952 | 12 | 1.23% |
| hsa_circ_0034421 | 1952 | 12 | 1.23% |
| hsa_circ_0067153 | 1952 | 12 | 1.23% |
| hsa_circ_0071562 | 1952 | 12 | 1.23% |
| hsa_circ_0115642 | 1952 | 12 | 1.23% |
| hsa_circ_0127322 | 1952 | 12 | 1.23% |
| hsa_circ_0041609 | 1627 | 10 | 1.23% |
| hsa_circ_0067376 | 1627 | 10 | 1.23% |
| hsa_circ_0076265 | 1627 | 10 | 1.23% |
| hsa_circ_0076731 | 1627 | 10 | 1.23% |
| hsa_circ_0104684 | 1627 | 10 | 1.23% |
| hsa_circ_0136685 | 1627 | 10 | 1.23% |
| hsa_circ_0034056 | 1790 | 11 | 1.23% |
| hsa_circ_0064407 | 1790 | 11 | 1.23% |
| hsa_circ_0074556 | 1953 | 12 | 1.23% |
| hsa_circ_0112256 | 1953 | 12 | 1.23% |
| hsa_circ_0115359 | 1953 | 12 | 1.23% |
| hsa_circ_0040969 | 2116 | 13 | 1.23% |
| hsa_circ_0078203 | 2116 | 13 | 1.23% |
| hsa_circ_0081602 | 2116 | 13 | 1.23% |
| hsa_circ_0088796 | 2279 | 14 | 1.23% |
| hsa_circ_0129424 | 3093 | 19 | 1.23% |
| hsa_circ_0020228 | 1628 | 10 | 1.23% |
| hsa_circ_0027121 | 1628 | 10 | 1.23% |
| hsa_circ_0039383 | 1628 | 10 | 1.23% |
| hsa_circ_0056522 | 1628 | 10 | 1.23% |
| hsa_circ_0064544 | 1628 | 10 | 1.23% |
| hsa_circ_0068996 | 1628 | 10 | 1.23% |
| hsa_circ_0082342 | 1628 | 10 | 1.23% |
| hsa_circ_0103939 | 1628 | 10 | 1.23% |
| hsa_circ_0106452 | 1628 | 10 | 1.23% |
| hsa_circ_0139923 | 1628 | 10 | 1.23% |
| hsa_circ_0010829 | 1791 | 11 | 1.23% |
| hsa_circ_0053900 | 1791 | 11 | 1.23% |
| hsa_circ_0060768 | 1791 | 11 | 1.23% |
| hsa_circ_0062119 | 1791 | 11 | 1.23% |
| hsa_circ_0136353 | 1791 | 11 | 1.23% |
| hsa_circ_0011513 | 1629 | 10 | 1.23% |
| hsa_circ_0012302 | 1629 | 10 | 1.23% |
| hsa_circ_0015593 | 1629 | 10 | 1.23% |
| hsa_circ_0016321 | 1629 | 10 | 1.23% |
| hsa_circ_0026191 | 1629 | 10 | 1.23% |
| hsa_circ_0041085 | 1629 | 10 | 1.23% |
| hsa_circ_0054342 | 1629 | 10 | 1.23% |
| hsa_circ_0082136 | 1629 | 10 | 1.23% |

| | | | |
|---|---|---|---|
| hsa_circ_0098490 | 1629 | 10 | 1.23% |
| hsa_circ_0106609 | 1629 | 10 | 1.23% |
| hsa_circ_0026874 | 1792 | 11 | 1.23% |
| hsa_circ_0032961 | 1792 | 11 | 1.23% |
| hsa_circ_0109370 | 1792 | 11 | 1.23% |
| hsa_circ_0113115 | 1792 | 11 | 1.23% |
| hsa_circ_0052798 | 1955 | 12 | 1.23% |
| hsa_circ_0056268 | 1955 | 12 | 1.23% |
| hsa_circ_0062114 | 1955 | 12 | 1.23% |
| hsa_circ_0069266 | 4236 | 26 | 1.23% |
| hsa_circ_0020307 | 1793 | 11 | 1.23% |
| hsa_circ_0062966 | 1793 | 11 | 1.23% |
| hsa_circ_0112211 | 1793 | 11 | 1.23% |
| hsa_circ_0021946 | 1630 | 10 | 1.23% |
| hsa_circ_0039797 | 1630 | 10 | 1.23% |
| hsa_circ_0044291 | 1630 | 10 | 1.23% |
| hsa_circ_0050414 | 1630 | 10 | 1.23% |
| hsa_circ_0085747 | 1630 | 10 | 1.23% |
| hsa_circ_0091726 | 1630 | 10 | 1.23% |
| hsa_circ_0106750 | 1630 | 10 | 1.23% |
| hsa_circ_0109492 | 1630 | 10 | 1.23% |
| hsa_circ_0127090 | 1630 | 10 | 1.23% |
| hsa_circ_0042655 | 2120 | 13 | 1.23% |
| hsa_circ_0009974 | 1957 | 12 | 1.23% |
| hsa_circ_0072346 | 1957 | 12 | 1.23% |
| hsa_circ_0083267 | 1957 | 12 | 1.23% |
| hsa_circ_0111939 | 1957 | 12 | 1.23% |
| hsa_circ_0013125 | 1794 | 11 | 1.23% |
| hsa_circ_0022066 | 1794 | 11 | 1.23% |
| hsa_circ_0022860 | 1794 | 11 | 1.23% |
| hsa_circ_0037392 | 1794 | 11 | 1.23% |
| hsa_circ_0063992 | 1794 | 11 | 1.23% |
| hsa_circ_0133791 | 1794 | 11 | 1.23% |
| hsa_circ_0020812 | 1631 | 10 | 1.23% |
| hsa_circ_0025416 | 1631 | 10 | 1.23% |
| hsa_circ_0032910 | 1631 | 10 | 1.23% |
| hsa_circ_0055749 | 1631 | 10 | 1.23% |
| hsa_circ_0073923 | 1631 | 10 | 1.23% |
| hsa_circ_0102277 | 1631 | 10 | 1.23% |
| hsa_circ_0121216 | 1631 | 10 | 1.23% |
| hsa_circ_0126716 | 1631 | 10 | 1.23% |
| hsa_circ_0134907 | 1631 | 10 | 1.23% |
| hsa_circ_0093918 | 6199 | 38 | 1.23% |
| hsa_circ_0102438 | 1958 | 12 | 1.23% |
| hsa_circ_0048318 | 1795 | 11 | 1.23% |
| hsa_circ_0087480 | 1795 | 11 | 1.23% |
| hsa_circ_0087896 | 1795 | 11 | 1.23% |
| hsa_circ_0081259 | 2448 | 15 | 1.23% |
| hsa_circ_0029478 | 1632 | 10 | 1.23% |
| hsa_circ_0035580 | 1632 | 10 | 1.23% |

| | | | |
|---|---|---|---|
| hsa_circ_0065374 | 1632 | 10 | 1.23% |
| hsa_circ_0085785 | 1632 | 10 | 1.23% |
| hsa_circ_0085914 | 1632 | 10 | 1.23% |
| hsa_circ_0096520 | 1632 | 10 | 1.23% |
| hsa_circ_0116735 | 1632 | 10 | 1.23% |
| hsa_circ_0135232 | 1632 | 10 | 1.23% |
| hsa_circ_0106421 | 2938 | 18 | 1.23% |
| hsa_circ_0076130 | 2122 | 13 | 1.23% |
| hsa_circ_0027372 | 1959 | 12 | 1.23% |
| hsa_circ_0011328 | 1796 | 11 | 1.22% |
| hsa_circ_0020754 | 1796 | 11 | 1.22% |
| hsa_circ_0026474 | 1796 | 11 | 1.22% |
| hsa_circ_0034754 | 1796 | 11 | 1.22% |
| hsa_circ_0041043 | 1796 | 11 | 1.22% |
| hsa_circ_0045044 | 1796 | 11 | 1.22% |
| hsa_circ_0069631 | 1796 | 11 | 1.22% |
| hsa_circ_0021130 | 1633 | 10 | 1.22% |
| hsa_circ_0021853 | 1633 | 10 | 1.22% |
| hsa_circ_0029765 | 1633 | 10 | 1.22% |
| hsa_circ_0040729 | 1633 | 10 | 1.22% |
| hsa_circ_0040910 | 1633 | 10 | 1.22% |
| hsa_circ_0100617 | 1633 | 10 | 1.22% |
| hsa_circ_0113661 | 1633 | 10 | 1.22% |
| hsa_circ_0121568 | 1633 | 10 | 1.22% |
| hsa_circ_0138463 | 1633 | 10 | 1.22% |
| hsa_circ_0067859 | 2123 | 13 | 1.22% |
| hsa_circ_0037629 | 1960 | 12 | 1.22% |
| hsa_circ_0050920 | 1960 | 12 | 1.22% |
| hsa_circ_0058574 | 1960 | 12 | 1.22% |
| hsa_circ_0043801 | 3594 | 22 | 1.22% |
| hsa_circ_0004133 | 1797 | 11 | 1.22% |
| hsa_circ_0009474 | 1797 | 11 | 1.22% |
| hsa_circ_0036583 | 1797 | 11 | 1.22% |
| hsa_circ_0046268 | 1797 | 11 | 1.22% |
| hsa_circ_0056838 | 1797 | 11 | 1.22% |
| hsa_circ_0037200 | 1634 | 10 | 1.22% |
| hsa_circ_0072242 | 1634 | 10 | 1.22% |
| hsa_circ_0084694 | 1634 | 10 | 1.22% |
| hsa_circ_0097900 | 1634 | 10 | 1.22% |
| hsa_circ_0010224 | 1961 | 12 | 1.22% |
| hsa_circ_0022606 | 1961 | 12 | 1.22% |
| hsa_circ_0048746 | 1961 | 12 | 1.22% |
| hsa_circ_0103823 | 1961 | 12 | 1.22% |
| hsa_circ_0010067 | 1798 | 11 | 1.22% |
| hsa_circ_0035113 | 1798 | 11 | 1.22% |
| hsa_circ_0042756 | 1798 | 11 | 1.22% |
| hsa_circ_0096234 | 1798 | 11 | 1.22% |
| hsa_circ_0100916 | 1798 | 11 | 1.22% |
| hsa_circ_0106485 | 1798 | 11 | 1.22% |
| hsa_circ_0026389 | 2125 | 13 | 1.22% |

| | | | |
|---|---|---|---|
| hsa_circ_0062859 | 2125 | 13 | 1.22% |
| hsa_circ_0110704 | 5558 | 34 | 1.22% |
| hsa_circ_0048074 | 2289 | 14 | 1.22% |
| hsa_circ_0085905 | 1962 | 12 | 1.22% |
| hsa_circ_0019609 | 1635 | 10 | 1.22% |
| hsa_circ_0066720 | 1635 | 10 | 1.22% |
| hsa_circ_0077103 | 1635 | 10 | 1.22% |
| hsa_circ_0079541 | 1635 | 10 | 1.22% |
| hsa_circ_0084474 | 1635 | 10 | 1.22% |
| hsa_circ_0101074 | 1635 | 10 | 1.22% |
| hsa_circ_0108883 | 1635 | 10 | 1.22% |
| hsa_circ_0111856 | 1635 | 10 | 1.22% |
| hsa_circ_0056863 | 2780 | 17 | 1.22% |
| hsa_circ_0000862 | 1799 | 11 | 1.22% |
| hsa_circ_0028457 | 1799 | 11 | 1.22% |
| hsa_circ_0043205 | 1799 | 11 | 1.22% |
| hsa_circ_0045404 | 1799 | 11 | 1.22% |
| hsa_circ_0049302 | 1799 | 11 | 1.22% |
| hsa_circ_0056601 | 1799 | 11 | 1.22% |
| hsa_circ_0062546 | 1799 | 11 | 1.22% |
| hsa_circ_0072183 | 1799 | 11 | 1.22% |
| hsa_circ_0102527 | 1799 | 11 | 1.22% |
| hsa_circ_0051504 | 2290 | 14 | 1.22% |
| hsa_circ_0072953 | 1963 | 12 | 1.22% |
| hsa_circ_0066072 | 2781 | 17 | 1.22% |
| hsa_circ_0000381 | 1636 | 10 | 1.22% |
| hsa_circ_0010834 | 1636 | 10 | 1.22% |
| hsa_circ_0032092 | 1636 | 10 | 1.22% |
| hsa_circ_0049881 | 1636 | 10 | 1.22% |
| hsa_circ_0061124 | 1636 | 10 | 1.22% |
| hsa_circ_0065559 | 1636 | 10 | 1.22% |
| hsa_circ_0090552 | 1636 | 10 | 1.22% |
| hsa_circ_0104682 | 1636 | 10 | 1.22% |
| hsa_circ_0113092 | 1636 | 10 | 1.22% |
| hsa_circ_0101305 | 2945 | 18 | 1.22% |
| hsa_circ_0018971 | 1800 | 11 | 1.22% |
| hsa_circ_0056213 | 1800 | 11 | 1.22% |
| hsa_circ_0063498 | 1800 | 11 | 1.22% |
| hsa_circ_0064284 | 1800 | 11 | 1.22% |
| hsa_circ_0079746 | 1800 | 11 | 1.22% |
| hsa_circ_0082595 | 1800 | 11 | 1.22% |
| hsa_circ_0083255 | 1800 | 11 | 1.22% |
| hsa_circ_0087535 | 1800 | 11 | 1.22% |
| hsa_circ_0052072 | 2291 | 14 | 1.22% |
| hsa_circ_0018323 | 1964 | 12 | 1.22% |
| hsa_circ_0117597 | 1964 | 12 | 1.22% |
| hsa_circ_0082900 | 2128 | 13 | 1.22% |
| hsa_circ_0123201 | 2128 | 13 | 1.22% |
| hsa_circ_0028512 | 1637 | 10 | 1.22% |
| hsa_circ_0063999 | 1637 | 10 | 1.22% |

| | | | |
|---|---|---|---|
| hsa_circ_0071225 | 1637 | 10 | 1.22% |
| hsa_circ_0086958 | 1637 | 10 | 1.22% |
| hsa_circ_0101975 | 1637 | 10 | 1.22% |
| hsa_circ_0009453 | 1801 | 11 | 1.22% |
| hsa_circ_0007109 | 1965 | 12 | 1.22% |
| hsa_circ_0020119 | 1965 | 12 | 1.22% |
| hsa_circ_0022762 | 1965 | 12 | 1.22% |
| hsa_circ_0024161 | 1965 | 12 | 1.22% |
| hsa_circ_0032605 | 1965 | 12 | 1.22% |
| hsa_circ_0040978 | 1965 | 12 | 1.22% |
| hsa_circ_0048262 | 2129 | 13 | 1.22% |
| hsa_circ_0001864 | 1638 | 10 | 1.22% |
| hsa_circ_0016796 | 1638 | 10 | 1.22% |
| hsa_circ_0019615 | 1638 | 10 | 1.22% |
| hsa_circ_0024269 | 1638 | 10 | 1.22% |
| hsa_circ_0049642 | 1638 | 10 | 1.22% |
| hsa_circ_0058301 | 1638 | 10 | 1.22% |
| hsa_circ_0078986 | 1638 | 10 | 1.22% |
| hsa_circ_0109061 | 1638 | 10 | 1.22% |
| hsa_circ_0135973 | 1638 | 10 | 1.22% |
| hsa_circ_0027877 | 1802 | 11 | 1.22% |
| hsa_circ_0029675 | 1802 | 11 | 1.22% |
| hsa_circ_0070778 | 1802 | 11 | 1.22% |
| hsa_circ_0072517 | 1802 | 11 | 1.22% |
| hsa_circ_0103655 | 1802 | 11 | 1.22% |
| hsa_circ_0016736 | 1966 | 12 | 1.22% |
| hsa_circ_0024396 | 2130 | 13 | 1.22% |
| hsa_circ_0038153 | 2130 | 13 | 1.22% |
| hsa_circ_0086787 | 2130 | 13 | 1.22% |
| hsa_circ_0055833 | 2458 | 15 | 1.22% |
| hsa_circ_0111751 | 2786 | 17 | 1.22% |
| hsa_circ_0006592 | 1639 | 10 | 1.22% |
| hsa_circ_0018980 | 1639 | 10 | 1.22% |
| hsa_circ_0073345 | 1639 | 10 | 1.22% |
| hsa_circ_0127279 | 1639 | 10 | 1.22% |
| hsa_circ_0139077 | 1639 | 10 | 1.22% |
| hsa_circ_0139278 | 1639 | 10 | 1.22% |
| hsa_circ_0012946 | 1803 | 11 | 1.22% |
| hsa_circ_0013707 | 1803 | 11 | 1.22% |
| hsa_circ_0020389 | 1803 | 11 | 1.22% |
| hsa_circ_0034370 | 1803 | 11 | 1.22% |
| hsa_circ_0046471 | 1803 | 11 | 1.22% |
| hsa_circ_0077973 | 1803 | 11 | 1.22% |
| hsa_circ_0128520 | 1803 | 11 | 1.22% |
| hsa_circ_0135482 | 1803 | 11 | 1.22% |
| hsa_circ_0039379 | 2131 | 13 | 1.22% |
| hsa_circ_0083133 | 4754 | 29 | 1.22% |
| hsa_circ_0136294 | 2296 | 14 | 1.22% |
| hsa_circ_0139053 | 2296 | 14 | 1.22% |
| hsa_circ_0011837 | 1968 | 12 | 1.22% |

| | | | |
|---|---|---|---|
| hsa_circ_0017577 | 1968 | 12 | 1.22% |
| hsa_circ_0036063 | 1968 | 12 | 1.22% |
| hsa_circ_0034566 | 1804 | 11 | 1.22% |
| hsa_circ_0039576 | 1804 | 11 | 1.22% |
| hsa_circ_0049963 | 1804 | 11 | 1.22% |
| hsa_circ_0135594 | 1804 | 11 | 1.22% |
| hsa_circ_0017399 | 1640 | 10 | 1.22% |
| hsa_circ_0054390 | 1640 | 10 | 1.22% |
| hsa_circ_0057789 | 1640 | 10 | 1.22% |
| hsa_circ_0073172 | 1640 | 10 | 1.22% |
| hsa_circ_0080221 | 1640 | 10 | 1.22% |
| hsa_circ_0098223 | 1640 | 10 | 1.22% |
| hsa_circ_0044506 | 3773 | 23 | 1.22% |
| hsa_circ_0022017 | 2133 | 13 | 1.22% |
| hsa_circ_0039450 | 2133 | 13 | 1.22% |
| hsa_circ_0051550 | 2133 | 13 | 1.22% |
| hsa_circ_0057436 | 2133 | 13 | 1.22% |
| hsa_circ_0094189 | 1805 | 11 | 1.22% |
| hsa_circ_0126578 | 1805 | 11 | 1.22% |
| hsa_circ_0016247 | 1641 | 10 | 1.22% |
| hsa_circ_0017153 | 1641 | 10 | 1.22% |
| hsa_circ_0039165 | 1641 | 10 | 1.22% |
| hsa_circ_0058481 | 1641 | 10 | 1.22% |
| hsa_circ_0071808 | 1641 | 10 | 1.22% |
| hsa_circ_0077041 | 1641 | 10 | 1.22% |
| hsa_circ_0094468 | 1641 | 10 | 1.22% |
| hsa_circ_0099209 | 1641 | 10 | 1.22% |
| hsa_circ_0010212 | 1970 | 12 | 1.22% |
| hsa_circ_0091742 | 1970 | 12 | 1.22% |
| hsa_circ_0094894 | 1970 | 12 | 1.22% |
| hsa_circ_0031792 | 1806 | 11 | 1.22% |
| hsa_circ_0055771 | 1806 | 11 | 1.22% |
| hsa_circ_0072184 | 1806 | 11 | 1.22% |
| hsa_circ_0076649 | 1806 | 11 | 1.22% |
| hsa_circ_0102371 | 1806 | 11 | 1.22% |
| hsa_circ_0121401 | 1806 | 11 | 1.22% |
| hsa_circ_0030697 | 1642 | 10 | 1.22% |
| hsa_circ_0065742 | 1642 | 10 | 1.22% |
| hsa_circ_0079174 | 1642 | 10 | 1.22% |
| hsa_circ_0130702 | 1642 | 10 | 1.22% |
| hsa_circ_0096511 | 2792 | 17 | 1.22% |
| hsa_circ_0040453 | 1971 | 12 | 1.22% |
| hsa_circ_0043466 | 1971 | 12 | 1.22% |
| hsa_circ_0063273 | 1971 | 12 | 1.22% |
| hsa_circ_0070363 | 1971 | 12 | 1.22% |
| hsa_circ_0073006 | 1971 | 12 | 1.22% |
| hsa_circ_0080611 | 1971 | 12 | 1.22% |
| hsa_circ_0095451 | 1971 | 12 | 1.22% |
| hsa_circ_0009550 | 1807 | 11 | 1.22% |
| hsa_circ_0013814 | 1807 | 11 | 1.22% |

| | | | |
|---|---|---|---|
| hsa_circ_0030446 | 1807 | 11 | 1.22% |
| hsa_circ_0046753 | 1807 | 11 | 1.22% |
| hsa_circ_0052587 | 2300 | 14 | 1.22% |
| hsa_circ_0014172 | 1643 | 10 | 1.22% |
| hsa_circ_0029208 | 1643 | 10 | 1.22% |
| hsa_circ_0046863 | 1643 | 10 | 1.22% |
| hsa_circ_0066318 | 1643 | 10 | 1.22% |
| hsa_circ_0081197 | 1643 | 10 | 1.22% |
| hsa_circ_0092779 | 1643 | 10 | 1.22% |
| hsa_circ_0125359 | 3779 | 23 | 1.22% |
| hsa_circ_0012384 | 2136 | 13 | 1.22% |
| hsa_circ_0061863 | 2136 | 13 | 1.22% |
| hsa_circ_0034850 | 1972 | 12 | 1.22% |
| hsa_circ_0043225 | 1972 | 12 | 1.22% |
| hsa_circ_0059530 | 1972 | 12 | 1.22% |
| hsa_circ_0071492 | 1972 | 12 | 1.22% |
| hsa_circ_0138945 | 1972 | 12 | 1.22% |
| hsa_circ_0010459 | 1808 | 11 | 1.22% |
| hsa_circ_0034663 | 1808 | 11 | 1.22% |
| hsa_circ_0041637 | 1808 | 11 | 1.22% |
| hsa_circ_0055712 | 1808 | 11 | 1.22% |
| hsa_circ_0063033 | 1808 | 11 | 1.22% |
| hsa_circ_0065000 | 1808 | 11 | 1.22% |
| hsa_circ_0105621 | 1808 | 11 | 1.22% |
| hsa_circ_0048932 | 2137 | 13 | 1.22% |
| hsa_circ_0012441 | 1644 | 10 | 1.22% |
| hsa_circ_0044494 | 1644 | 10 | 1.22% |
| hsa_circ_0048033 | 1644 | 10 | 1.22% |
| hsa_circ_0051622 | 1644 | 10 | 1.22% |
| hsa_circ_0056444 | 1644 | 10 | 1.22% |
| hsa_circ_0061374 | 1644 | 10 | 1.22% |
| hsa_circ_0103936 | 1644 | 10 | 1.22% |
| hsa_circ_0125942 | 1644 | 10 | 1.22% |
| hsa_circ_0034483 | 1973 | 12 | 1.22% |
| hsa_circ_0106856 | 1973 | 12 | 1.22% |
| hsa_circ_0000811 | 1809 | 11 | 1.22% |
| hsa_circ_0014238 | 1809 | 11 | 1.22% |
| hsa_circ_0019478 | 1809 | 11 | 1.22% |
| hsa_circ_0025564 | 1809 | 11 | 1.22% |
| hsa_circ_0044993 | 1809 | 11 | 1.22% |
| hsa_circ_0048860 | 1809 | 11 | 1.22% |
| hsa_circ_0081539 | 1809 | 11 | 1.22% |
| hsa_circ_0105551 | 1809 | 11 | 1.22% |
| hsa_circ_0010083 | 1974 | 12 | 1.22% |
| hsa_circ_0015535 | 1974 | 12 | 1.22% |
| hsa_circ_0056212 | 1974 | 12 | 1.22% |
| hsa_circ_0083590 | 1974 | 12 | 1.22% |
| hsa_circ_0089996 | 1974 | 12 | 1.22% |
| hsa_circ_0011438 | 1645 | 10 | 1.22% |
| hsa_circ_0028348 | 1645 | 10 | 1.22% |

| | | | |
|---|---|---|---|
| hsa_circ_0029232 | 1645 | 10 | 1.22% |
| hsa_circ_0049803 | 1645 | 10 | 1.22% |
| hsa_circ_0055724 | 1645 | 10 | 1.22% |
| hsa_circ_0056951 | 1645 | 10 | 1.22% |
| hsa_circ_0080943 | 1645 | 10 | 1.22% |
| hsa_circ_0082654 | 1645 | 10 | 1.22% |
| hsa_circ_0088532 | 1645 | 10 | 1.22% |
| hsa_circ_0095827 | 1645 | 10 | 1.22% |
| hsa_circ_0129492 | 1645 | 10 | 1.22% |
| hsa_circ_0074459 | 2139 | 13 | 1.22% |
| hsa_circ_0011281 | 1810 | 11 | 1.22% |
| hsa_circ_0049728 | 1810 | 11 | 1.22% |
| hsa_circ_0075994 | 1810 | 11 | 1.22% |
| hsa_circ_0103659 | 1810 | 11 | 1.22% |
| hsa_circ_0057427 | 2304 | 14 | 1.22% |
| hsa_circ_0041027 | 1975 | 12 | 1.22% |
| hsa_circ_0052145 | 1975 | 12 | 1.22% |
| hsa_circ_0078527 | 1975 | 12 | 1.22% |
| hsa_circ_0127236 | 1975 | 12 | 1.22% |
| hsa_circ_0044212 | 2469 | 15 | 1.22% |
| hsa_circ_0015566 | 1646 | 10 | 1.22% |
| hsa_circ_0020444 | 1646 | 10 | 1.22% |
| hsa_circ_0028286 | 1646 | 10 | 1.22% |
| hsa_circ_0036339 | 1646 | 10 | 1.22% |
| hsa_circ_0047487 | 1646 | 10 | 1.22% |
| hsa_circ_0090530 | 1646 | 10 | 1.22% |
| hsa_circ_0135263 | 1646 | 10 | 1.22% |
| hsa_circ_0136498 | 1646 | 10 | 1.22% |
| hsa_circ_0139282 | 1646 | 10 | 1.22% |
| hsa_circ_0136849 | 5103 | 31 | 1.21% |
| hsa_circ_0000232 | 4280 | 26 | 1.21% |
| hsa_circ_0089703 | 2140 | 13 | 1.21% |
| hsa_circ_0024110 | 1811 | 11 | 1.21% |
| hsa_circ_0083529 | 1811 | 11 | 1.21% |
| hsa_circ_0085535 | 1811 | 11 | 1.21% |
| hsa_circ_0089432 | 1811 | 11 | 1.21% |
| hsa_circ_0099084 | 1811 | 11 | 1.21% |
| hsa_circ_0117995 | 1811 | 11 | 1.21% |
| hsa_circ_0124416 | 1811 | 11 | 1.21% |
| hsa_circ_0125984 | 1811 | 11 | 1.21% |
| hsa_circ_0022540 | 2799 | 17 | 1.21% |
| hsa_circ_0057391 | 2799 | 17 | 1.21% |
| hsa_circ_0045827 | 1976 | 12 | 1.21% |
| hsa_circ_0065877 | 1976 | 12 | 1.21% |
| hsa_circ_0093744 | 1976 | 12 | 1.21% |
| hsa_circ_0015115 | 2141 | 13 | 1.21% |
| hsa_circ_0063512 | 2141 | 13 | 1.21% |
| hsa_circ_0032566 | 1647 | 10 | 1.21% |
| hsa_circ_0036914 | 1647 | 10 | 1.21% |
| hsa_circ_0052993 | 1647 | 10 | 1.21% |

| | | | |
|---|---|---|---|
| hsa_circ_0059888 | 1647 | 10 | 1.21% |
| hsa_circ_0091966 | 1647 | 10 | 1.21% |
| hsa_circ_0007225 | 1812 | 11 | 1.21% |
| hsa_circ_0012834 | 1812 | 11 | 1.21% |
| hsa_circ_0041615 | 1812 | 11 | 1.21% |
| hsa_circ_0047330 | 1812 | 11 | 1.21% |
| hsa_circ_0053705 | 1812 | 11 | 1.21% |
| hsa_circ_0063003 | 1812 | 11 | 1.21% |
| hsa_circ_0079351 | 1812 | 11 | 1.21% |
| hsa_circ_0101497 | 1812 | 11 | 1.21% |
| hsa_circ_0112464 | 1812 | 11 | 1.21% |
| hsa_circ_0115957 | 1812 | 11 | 1.21% |
| hsa_circ_0138731 | 6095 | 37 | 1.21% |
| hsa_circ_0085078 | 2471 | 15 | 1.21% |
| hsa_circ_0040699 | 1977 | 12 | 1.21% |
| hsa_circ_0093477 | 2801 | 17 | 1.21% |
| hsa_circ_0019936 | 2142 | 13 | 1.21% |
| hsa_circ_0046653 | 2472 | 15 | 1.21% |
| hsa_circ_0012731 | 1648 | 10 | 1.21% |
| hsa_circ_0042796 | 1648 | 10 | 1.21% |
| hsa_circ_0068899 | 1648 | 10 | 1.21% |
| hsa_circ_0082436 | 1648 | 10 | 1.21% |
| hsa_circ_0024751 | 1813 | 11 | 1.21% |
| hsa_circ_0106669 | 1813 | 11 | 1.21% |
| hsa_circ_0108131 | 1813 | 11 | 1.21% |
| hsa_circ_0118182 | 1813 | 11 | 1.21% |
| hsa_circ_0127263 | 1813 | 11 | 1.21% |
| hsa_circ_0132686 | 1978 | 12 | 1.21% |
| hsa_circ_0050248 | 2473 | 15 | 1.21% |
| hsa_circ_0023903 | 1649 | 10 | 1.21% |
| hsa_circ_0034085 | 1649 | 10 | 1.21% |
| hsa_circ_0034827 | 1649 | 10 | 1.21% |
| hsa_circ_0039364 | 1649 | 10 | 1.21% |
| hsa_circ_0054936 | 1649 | 10 | 1.21% |
| hsa_circ_0072505 | 1649 | 10 | 1.21% |
| hsa_circ_0084943 | 1649 | 10 | 1.21% |
| hsa_circ_0092437 | 1649 | 10 | 1.21% |
| hsa_circ_0099660 | 1649 | 10 | 1.21% |
| hsa_circ_0122487 | 1649 | 10 | 1.21% |
| hsa_circ_0123424 | 1649 | 10 | 1.21% |
| hsa_circ_0084953 | 1814 | 11 | 1.21% |
| hsa_circ_0088855 | 1814 | 11 | 1.21% |
| hsa_circ_0119326 | 1814 | 11 | 1.21% |
| hsa_circ_0022801 | 1979 | 12 | 1.21% |
| hsa_circ_0043762 | 1979 | 12 | 1.21% |
| hsa_circ_0078088 | 1979 | 12 | 1.21% |
| hsa_circ_0086140 | 1979 | 12 | 1.21% |
| hsa_circ_0003384 | 6267 | 38 | 1.21% |
| hsa_circ_0121072 | 2145 | 13 | 1.21% |
| hsa_circ_0006859 | 1815 | 11 | 1.21% |

| | | | |
|---|---|---|---|
| hsa_circ_0024647 | 1815 | 11 | 1.21% |
| hsa_circ_0038382 | 1815 | 11 | 1.21% |
| hsa_circ_0038425 | 1815 | 11 | 1.21% |
| hsa_circ_0098023 | 1815 | 11 | 1.21% |
| hsa_circ_0111926 | 1815 | 11 | 1.21% |
| hsa_circ_0024592 | 1650 | 10 | 1.21% |
| hsa_circ_0025104 | 1650 | 10 | 1.21% |
| hsa_circ_0084982 | 1650 | 10 | 1.21% |
| hsa_circ_0109395 | 1650 | 10 | 1.21% |
| hsa_circ_0116048 | 1650 | 10 | 1.21% |
| hsa_circ_0072606 | 2146 | 13 | 1.21% |
| hsa_circ_0087342 | 2146 | 13 | 1.21% |
| hsa_circ_0063586 | 1981 | 12 | 1.21% |
| hsa_circ_0122221 | 1981 | 12 | 1.21% |
| hsa_circ_0050161 | 1816 | 11 | 1.21% |
| hsa_circ_0024702 | 1651 | 10 | 1.21% |
| hsa_circ_0043065 | 1651 | 10 | 1.21% |
| hsa_circ_0079886 | 1651 | 10 | 1.21% |
| hsa_circ_0119237 | 1651 | 10 | 1.21% |
| hsa_circ_0014481 | 2147 | 13 | 1.21% |
| hsa_circ_0074033 | 2147 | 13 | 1.21% |
| hsa_circ_0002429 | 1982 | 12 | 1.21% |
| hsa_circ_0023347 | 1982 | 12 | 1.21% |
| hsa_circ_0034325 | 1982 | 12 | 1.21% |
| hsa_circ_0039370 | 1982 | 12 | 1.21% |
| hsa_circ_0061467 | 1982 | 12 | 1.21% |
| hsa_circ_0062810 | 1817 | 11 | 1.21% |
| hsa_circ_0101593 | 1817 | 11 | 1.21% |
| hsa_circ_0034287 | 1652 | 10 | 1.21% |
| hsa_circ_0040967 | 1652 | 10 | 1.21% |
| hsa_circ_0056084 | 1652 | 10 | 1.21% |
| hsa_circ_0058045 | 1652 | 10 | 1.21% |
| hsa_circ_0078412 | 1652 | 10 | 1.21% |
| hsa_circ_0089994 | 1652 | 10 | 1.21% |
| hsa_circ_0067138 | 2313 | 14 | 1.21% |
| hsa_circ_0025030 | 1983 | 12 | 1.21% |
| hsa_circ_0125604 | 1983 | 12 | 1.21% |
| hsa_circ_0039194 | 1818 | 11 | 1.21% |
| hsa_circ_0081529 | 1818 | 11 | 1.21% |
| hsa_circ_0134282 | 1818 | 11 | 1.21% |
| hsa_circ_0015891 | 1653 | 10 | 1.21% |
| hsa_circ_0016484 | 1653 | 10 | 1.21% |
| hsa_circ_0016534 | 1653 | 10 | 1.21% |
| hsa_circ_0018990 | 1653 | 10 | 1.21% |
| hsa_circ_0043119 | 1653 | 10 | 1.21% |
| hsa_circ_0069272 | 1653 | 10 | 1.21% |
| hsa_circ_0071629 | 1653 | 10 | 1.21% |
| hsa_circ_0114271 | 1653 | 10 | 1.21% |
| hsa_circ_0140100 | 4298 | 26 | 1.21% |
| hsa_circ_0005684 | 4794 | 29 | 1.21% |

| | | | |
|---|---|---|---|
| hsa_circ_0011397 | 1984 | 12 | 1.21% |
| hsa_circ_0067600 | 1984 | 12 | 1.21% |
| hsa_circ_0057355 | 2811 | 17 | 1.21% |
| hsa_circ_0065997 | 1819 | 11 | 1.21% |
| hsa_circ_0095247 | 1819 | 11 | 1.21% |
| hsa_circ_0120609 | 1819 | 11 | 1.21% |
| hsa_circ_0011052 | 1654 | 10 | 1.21% |
| hsa_circ_0013802 | 1654 | 10 | 1.21% |
| hsa_circ_0018598 | 1654 | 10 | 1.21% |
| hsa_circ_0046621 | 1654 | 10 | 1.21% |
| hsa_circ_0068845 | 1654 | 10 | 1.21% |
| hsa_circ_0108809 | 1654 | 10 | 1.21% |
| hsa_circ_0108974 | 1654 | 10 | 1.21% |
| hsa_circ_0114584 | 1654 | 10 | 1.21% |
| hsa_circ_0115017 | 2812 | 17 | 1.21% |
| hsa_circ_0124190 | 1985 | 12 | 1.21% |
| hsa_circ_0125226 | 1985 | 12 | 1.21% |
| hsa_circ_0011302 | 1820 | 11 | 1.21% |
| hsa_circ_0068327 | 1820 | 11 | 1.21% |
| hsa_circ_0087406 | 1820 | 11 | 1.21% |
| hsa_circ_0118342 | 1820 | 11 | 1.21% |
| hsa_circ_0021849 | 2482 | 15 | 1.21% |
| hsa_circ_0012523 | 1986 | 12 | 1.21% |
| hsa_circ_0032576 | 1986 | 12 | 1.21% |
| hsa_circ_0083896 | 1986 | 12 | 1.21% |
| hsa_circ_0137334 | 1986 | 12 | 1.21% |
| hsa_circ_0033444 | 1655 | 10 | 1.21% |
| hsa_circ_0046728 | 1655 | 10 | 1.21% |
| hsa_circ_0067951 | 1655 | 10 | 1.21% |
| hsa_circ_0074454 | 1655 | 10 | 1.21% |
| hsa_circ_0135733 | 1655 | 10 | 1.21% |
| hsa_circ_0013393 | 4635 | 28 | 1.21% |
| hsa_circ_0082054 | 2152 | 13 | 1.21% |
| hsa_circ_0110188 | 2152 | 13 | 1.21% |
| hsa_circ_0128058 | 2152 | 13 | 1.21% |
| hsa_circ_0009655 | 1821 | 11 | 1.21% |
| hsa_circ_0024811 | 1821 | 11 | 1.21% |
| hsa_circ_0025888 | 1821 | 11 | 1.21% |
| hsa_circ_0045657 | 1821 | 11 | 1.21% |
| hsa_circ_0054654 | 1821 | 11 | 1.21% |
| hsa_circ_0059024 | 1821 | 11 | 1.21% |
| hsa_circ_0065152 | 1821 | 11 | 1.21% |
| hsa_circ_0070166 | 2318 | 14 | 1.21% |
| hsa_circ_0048182 | 1987 | 12 | 1.21% |
| hsa_circ_0051725 | 1987 | 12 | 1.21% |
| hsa_circ_0080656 | 1987 | 12 | 1.21% |
| hsa_circ_0112422 | 1987 | 12 | 1.21% |
| hsa_circ_0128051 | 1987 | 12 | 1.21% |
| hsa_circ_0020029 | 1656 | 10 | 1.21% |
| hsa_circ_0074474 | 1656 | 10 | 1.21% |

| | | | |
|---|---|---|---|
| hsa_circ_0095125 | 1656 | 10 | 1.21% |
| hsa_circ_0126425 | 1656 | 10 | 1.21% |
| hsa_circ_0139184 | 1656 | 10 | 1.21% |
| hsa_circ_0108134 | 4637 | 28 | 1.21% |
| hsa_circ_0121368 | 4637 | 28 | 1.21% |
| hsa_circ_0044065 | 1822 | 11 | 1.21% |
| hsa_circ_0068081 | 1822 | 11 | 1.21% |
| hsa_circ_0123001 | 1822 | 11 | 1.21% |
| hsa_circ_0074230 | 1988 | 12 | 1.21% |
| hsa_circ_0029209 | 1657 | 10 | 1.21% |
| hsa_circ_0062178 | 1657 | 10 | 1.21% |
| hsa_circ_0065651 | 1657 | 10 | 1.21% |
| hsa_circ_0079722 | 1657 | 10 | 1.21% |
| hsa_circ_0017044 | 1823 | 11 | 1.21% |
| hsa_circ_0041081 | 1823 | 11 | 1.21% |
| hsa_circ_0057610 | 1823 | 11 | 1.21% |
| hsa_circ_0065644 | 1823 | 11 | 1.21% |
| hsa_circ_0095052 | 1823 | 11 | 1.21% |
| hsa_circ_0101606 | 1823 | 11 | 1.21% |
| hsa_circ_0118482 | 1823 | 11 | 1.21% |
| hsa_circ_0136991 | 1823 | 11 | 1.21% |
| hsa_circ_0060630 | 1989 | 12 | 1.21% |
| hsa_circ_0025926 | 2155 | 13 | 1.21% |
| hsa_circ_0085708 | 2155 | 13 | 1.21% |
| hsa_circ_0014433 | 1658 | 10 | 1.21% |
| hsa_circ_0015250 | 1658 | 10 | 1.21% |
| hsa_circ_0036171 | 1658 | 10 | 1.21% |
| hsa_circ_0052751 | 1658 | 10 | 1.21% |
| hsa_circ_0066637 | 1658 | 10 | 1.21% |
| hsa_circ_0087493 | 1658 | 10 | 1.21% |
| hsa_circ_0099328 | 1658 | 10 | 1.21% |
| hsa_circ_0139091 | 1658 | 10 | 1.21% |
| hsa_circ_0021890 | 1824 | 11 | 1.21% |
| hsa_circ_0036183 | 1824 | 11 | 1.21% |
| hsa_circ_0088279 | 1824 | 11 | 1.21% |
| hsa_circ_0117901 | 1824 | 11 | 1.21% |
| hsa_circ_0021116 | 2985 | 18 | 1.21% |
| hsa_circ_0086227 | 1990 | 12 | 1.21% |
| hsa_circ_0119423 | 1990 | 12 | 1.21% |
| hsa_circ_0091443 | 2156 | 13 | 1.21% |
| hsa_circ_0065889 | 2322 | 14 | 1.21% |
| hsa_circ_0016878 | 1659 | 10 | 1.21% |
| hsa_circ_0026141 | 1659 | 10 | 1.21% |
| hsa_circ_0031548 | 1659 | 10 | 1.21% |
| hsa_circ_0062047 | 1659 | 10 | 1.21% |
| hsa_circ_0090994 | 1659 | 10 | 1.21% |
| hsa_circ_0106836 | 1659 | 10 | 1.21% |
| hsa_circ_0029934 | 1825 | 11 | 1.21% |
| hsa_circ_0033374 | 1825 | 11 | 1.21% |
| hsa_circ_0039024 | 1825 | 11 | 1.21% |

| | | | |
|---|---|---|---|
| hsa_circ_0042710 | 1825 | 11 | 1.21% |
| hsa_circ_0046491 | 1825 | 11 | 1.21% |
| hsa_circ_0066810 | 1825 | 11 | 1.21% |
| hsa_circ_0090606 | 2821 | 17 | 1.21% |
| hsa_circ_0010706 | 1992 | 12 | 1.20% |
| hsa_circ_0085369 | 1992 | 12 | 1.20% |
| hsa_circ_0091487 | 1992 | 12 | 1.20% |
| hsa_circ_0019607 | 1826 | 11 | 1.20% |
| hsa_circ_0044423 | 1826 | 11 | 1.20% |
| hsa_circ_0017593 | 1660 | 10 | 1.20% |
| hsa_circ_0018034 | 1660 | 10 | 1.20% |
| hsa_circ_0029487 | 1660 | 10 | 1.20% |
| hsa_circ_0047835 | 1660 | 10 | 1.20% |
| hsa_circ_0073348 | 1660 | 10 | 1.20% |
| hsa_circ_0091284 | 1660 | 10 | 1.20% |
| hsa_circ_0117608 | 1660 | 10 | 1.20% |
| hsa_circ_0009669 | 1993 | 12 | 1.20% |
| hsa_circ_0012251 | 1827 | 11 | 1.20% |
| hsa_circ_0019722 | 1827 | 11 | 1.20% |
| hsa_circ_0022045 | 1827 | 11 | 1.20% |
| hsa_circ_0022510 | 1827 | 11 | 1.20% |
| hsa_circ_0052122 | 1827 | 11 | 1.20% |
| hsa_circ_0065996 | 1827 | 11 | 1.20% |
| hsa_circ_0106005 | 1827 | 11 | 1.20% |
| hsa_circ_0120059 | 1827 | 11 | 1.20% |
| hsa_circ_0130039 | 1827 | 11 | 1.20% |
| hsa_circ_0016082 | 1661 | 10 | 1.20% |
| hsa_circ_0024569 | 1661 | 10 | 1.20% |
| hsa_circ_0039352 | 1661 | 10 | 1.20% |
| hsa_circ_0047564 | 1661 | 10 | 1.20% |
| hsa_circ_0055748 | 1661 | 10 | 1.20% |
| hsa_circ_0065207 | 1661 | 10 | 1.20% |
| hsa_circ_0102269 | 1661 | 10 | 1.20% |
| hsa_circ_0115174 | 1661 | 10 | 1.20% |
| hsa_circ_0115594 | 1661 | 10 | 1.20% |
| hsa_circ_0092940 | 4153 | 25 | 1.20% |
| hsa_circ_0109122 | 5482 | 33 | 1.20% |
| hsa_circ_0068658 | 2160 | 13 | 1.20% |
| hsa_circ_0073800 | 1994 | 12 | 1.20% |
| hsa_circ_0081169 | 1994 | 12 | 1.20% |
| hsa_circ_0027730 | 4653 | 28 | 1.20% |
| hsa_circ_0124489 | 3656 | 22 | 1.20% |
| hsa_circ_0019484 | 1828 | 11 | 1.20% |
| hsa_circ_0043827 | 1828 | 11 | 1.20% |
| hsa_circ_0061910 | 1828 | 11 | 1.20% |
| hsa_circ_0063688 | 1828 | 11 | 1.20% |
| hsa_circ_0069786 | 1828 | 11 | 1.20% |
| hsa_circ_0077136 | 1828 | 11 | 1.20% |
| hsa_circ_0106974 | 1828 | 11 | 1.20% |
| hsa_circ_0107526 | 1828 | 11 | 1.20% |

| | | | |
|---|---|---|---|
| hsa_circ_0125603 | 1828 | 11 | 1.20% |
| hsa_circ_0050316 | 3490 | 21 | 1.20% |
| hsa_circ_0011139 | 1662 | 10 | 1.20% |
| hsa_circ_0039582 | 1662 | 10 | 1.20% |
| hsa_circ_0086083 | 1662 | 10 | 1.20% |
| hsa_circ_0094542 | 1662 | 10 | 1.20% |
| hsa_circ_0094925 | 1662 | 10 | 1.20% |
| hsa_circ_0114437 | 1662 | 10 | 1.20% |
| hsa_circ_0137207 | 1662 | 10 | 1.20% |
| hsa_circ_0139569 | 1662 | 10 | 1.20% |
| hsa_circ_0139708 | 1662 | 10 | 1.20% |
| hsa_circ_0106820 | 1995 | 12 | 1.20% |
| hsa_circ_0005232 | 1829 | 11 | 1.20% |
| hsa_circ_0043449 | 1829 | 11 | 1.20% |
| hsa_circ_0138437 | 1829 | 11 | 1.20% |
| hsa_circ_0033542 | 2328 | 14 | 1.20% |
| hsa_circ_0006667 | 1663 | 10 | 1.20% |
| hsa_circ_0011850 | 1663 | 10 | 1.20% |
| hsa_circ_0028832 | 1663 | 10 | 1.20% |
| hsa_circ_0031684 | 1663 | 10 | 1.20% |
| hsa_circ_0041969 | 1663 | 10 | 1.20% |
| hsa_circ_0046109 | 1663 | 10 | 1.20% |
| hsa_circ_0049306 | 1663 | 10 | 1.20% |
| hsa_circ_0051105 | 1663 | 10 | 1.20% |
| hsa_circ_0067652 | 1663 | 10 | 1.20% |
| hsa_circ_0088401 | 1663 | 10 | 1.20% |
| hsa_circ_0090690 | 1663 | 10 | 1.20% |
| hsa_circ_0135464 | 1663 | 10 | 1.20% |
| hsa_circ_0043226 | 1996 | 12 | 1.20% |
| hsa_circ_0047461 | 1996 | 12 | 1.20% |
| hsa_circ_0065010 | 1996 | 12 | 1.20% |
| hsa_circ_0083600 | 1996 | 12 | 1.20% |
| hsa_circ_0012925 | 1830 | 11 | 1.20% |
| hsa_circ_0014666 | 1830 | 11 | 1.20% |
| hsa_circ_0029673 | 1830 | 11 | 1.20% |
| hsa_circ_0032954 | 1830 | 11 | 1.20% |
| hsa_circ_0077828 | 1830 | 11 | 1.20% |
| hsa_circ_0080714 | 1830 | 11 | 1.20% |
| hsa_circ_0086920 | 1830 | 11 | 1.20% |
| hsa_circ_0120060 | 1830 | 11 | 1.20% |
| hsa_circ_0015172 | 1664 | 10 | 1.20% |
| hsa_circ_0060248 | 1664 | 10 | 1.20% |
| hsa_circ_0061886 | 1664 | 10 | 1.20% |
| hsa_circ_0062950 | 1664 | 10 | 1.20% |
| hsa_circ_0071785 | 1664 | 10 | 1.20% |
| hsa_circ_0098785 | 1664 | 10 | 1.20% |
| hsa_circ_0113545 | 1664 | 10 | 1.20% |
| hsa_circ_0032290 | 1997 | 12 | 1.20% |
| hsa_circ_0040880 | 1997 | 12 | 1.20% |
| hsa_circ_0079099 | 1997 | 12 | 1.20% |

| | | | |
|---|---|---|---|
| hsa_circ_0089676 | 1997 | 12 | 1.20% |
| hsa_circ_0030411 | 1831 | 11 | 1.20% |
| hsa_circ_0044146 | 1831 | 11 | 1.20% |
| hsa_circ_0052379 | 1831 | 11 | 1.20% |
| hsa_circ_0081869 | 1831 | 11 | 1.20% |
| hsa_circ_0020638 | 1665 | 10 | 1.20% |
| hsa_circ_0026132 | 1665 | 10 | 1.20% |
| hsa_circ_0026951 | 1665 | 10 | 1.20% |
| hsa_circ_0032477 | 1665 | 10 | 1.20% |
| hsa_circ_0032572 | 1665 | 10 | 1.20% |
| hsa_circ_0057316 | 1665 | 10 | 1.20% |
| hsa_circ_0091530 | 1665 | 10 | 1.20% |
| hsa_circ_0126789 | 1665 | 10 | 1.20% |
| hsa_circ_0139872 | 1665 | 10 | 1.20% |
| hsa_circ_0000994 | 1832 | 11 | 1.20% |
| hsa_circ_0039588 | 1832 | 11 | 1.20% |
| hsa_circ_0071644 | 1832 | 11 | 1.20% |
| hsa_circ_0084932 | 1832 | 11 | 1.20% |
| hsa_circ_0086691 | 1832 | 11 | 1.20% |
| hsa_circ_0092927 | 1832 | 11 | 1.20% |
| hsa_circ_0095828 | 1832 | 11 | 1.20% |
| hsa_circ_0101859 | 1832 | 11 | 1.20% |
| hsa_circ_0105107 | 1832 | 11 | 1.20% |
| hsa_circ_0107184 | 1832 | 11 | 1.20% |
| hsa_circ_0041763 | 1999 | 12 | 1.20% |
| hsa_circ_0015355 | 1666 | 10 | 1.20% |
| hsa_circ_0022065 | 1666 | 10 | 1.20% |
| hsa_circ_0036369 | 1666 | 10 | 1.20% |
| hsa_circ_0056157 | 1666 | 10 | 1.20% |
| hsa_circ_0072213 | 1666 | 10 | 1.20% |
| hsa_circ_0079827 | 1666 | 10 | 1.20% |
| hsa_circ_0082259 | 1666 | 10 | 1.20% |
| hsa_circ_0101772 | 1666 | 10 | 1.20% |
| hsa_circ_0129154 | 1666 | 10 | 1.20% |
| hsa_circ_0135220 | 1666 | 10 | 1.20% |
| hsa_circ_0137136 | 1666 | 10 | 1.20% |
| hsa_circ_0038777 | 1833 | 11 | 1.20% |
| hsa_circ_0043202 | 1833 | 11 | 1.20% |
| hsa_circ_0049832 | 1833 | 11 | 1.20% |
| hsa_circ_0055212 | 1833 | 11 | 1.20% |
| hsa_circ_0077930 | 1833 | 11 | 1.20% |
| hsa_circ_0125029 | 1833 | 11 | 1.20% |
| hsa_circ_0108724 | 2333 | 14 | 1.20% |
| hsa_circ_0072832 | 3833 | 23 | 1.20% |
| hsa_circ_0135561 | 7167 | 43 | 1.20% |
| hsa_circ_0056196 | 2167 | 13 | 1.20% |
| hsa_circ_0066781 | 2167 | 13 | 1.20% |
| hsa_circ_0016134 | 1667 | 10 | 1.20% |
| hsa_circ_0053853 | 1667 | 10 | 1.20% |
| hsa_circ_0090232 | 1667 | 10 | 1.20% |

| | | | |
|---|---|---|---|
| hsa_circ_0090349 | 1667 | 10 | 1.20% |
| hsa_circ_0091816 | 1667 | 10 | 1.20% |
| hsa_circ_0093838 | 1667 | 10 | 1.20% |
| hsa_circ_0019266 | 1834 | 11 | 1.20% |
| hsa_circ_0028409 | 1834 | 11 | 1.20% |
| hsa_circ_0053299 | 1834 | 11 | 1.20% |
| hsa_circ_0058633 | 1834 | 11 | 1.20% |
| hsa_circ_0102707 | 2668 | 16 | 1.20% |
| hsa_circ_0138613 | 2168 | 13 | 1.20% |
| hsa_circ_0134278 | 4670 | 28 | 1.20% |
| hsa_circ_0135700 | 2335 | 14 | 1.20% |
| hsa_circ_0039041 | 1668 | 10 | 1.20% |
| hsa_circ_0039063 | 1668 | 10 | 1.20% |
| hsa_circ_0053007 | 1668 | 10 | 1.20% |
| hsa_circ_0054756 | 1668 | 10 | 1.20% |
| hsa_circ_0094940 | 1668 | 10 | 1.20% |
| hsa_circ_0099324 | 1668 | 10 | 1.20% |
| hsa_circ_0105251 | 1668 | 10 | 1.20% |
| hsa_circ_0129044 | 1668 | 10 | 1.20% |
| hsa_circ_0132478 | 1668 | 10 | 1.20% |
| hsa_circ_0135318 | 1668 | 10 | 1.20% |
| hsa_circ_0030675 | 1835 | 11 | 1.20% |
| hsa_circ_0060460 | 1835 | 11 | 1.20% |
| hsa_circ_0075911 | 1835 | 11 | 1.20% |
| hsa_circ_0076243 | 1835 | 11 | 1.20% |
| hsa_circ_0111274 | 1835 | 11 | 1.20% |
| hsa_circ_0012699 | 2002 | 12 | 1.20% |
| hsa_circ_0046167 | 2169 | 13 | 1.20% |
| hsa_circ_0006479 | 1669 | 10 | 1.20% |
| hsa_circ_0010230 | 1669 | 10 | 1.20% |
| hsa_circ_0026503 | 1669 | 10 | 1.20% |
| hsa_circ_0030900 | 1669 | 10 | 1.20% |
| hsa_circ_0040858 | 1669 | 10 | 1.20% |
| hsa_circ_0077923 | 1669 | 10 | 1.20% |
| hsa_circ_0081652 | 1669 | 10 | 1.20% |
| hsa_circ_0126040 | 1669 | 10 | 1.20% |
| hsa_circ_0001181 | 1836 | 11 | 1.20% |
| hsa_circ_0012811 | 1836 | 11 | 1.20% |
| hsa_circ_0023207 | 1836 | 11 | 1.20% |
| hsa_circ_0037239 | 1836 | 11 | 1.20% |
| hsa_circ_0049430 | 1836 | 11 | 1.20% |
| hsa_circ_0068207 | 1836 | 11 | 1.20% |
| hsa_circ_0069254 | 1836 | 11 | 1.20% |
| hsa_circ_0092892 | 1836 | 11 | 1.20% |
| hsa_circ_0132421 | 1836 | 11 | 1.20% |
| hsa_circ_0066844 | 2003 | 12 | 1.20% |
| hsa_circ_0062333 | 2505 | 15 | 1.20% |
| hsa_circ_0067219 | 2004 | 12 | 1.20% |
| hsa_circ_0080241 | 2004 | 12 | 1.20% |
| hsa_circ_0084005 | 2004 | 12 | 1.20% |

| | | | |
|---|---|---|---|
| hsa_circ_0106866 | 2004 | 12 | 1.20% |
| hsa_circ_0022784 | 1837 | 11 | 1.20% |
| hsa_circ_0026611 | 1837 | 11 | 1.20% |
| hsa_circ_0053147 | 1837 | 11 | 1.20% |
| hsa_circ_0085588 | 1837 | 11 | 1.20% |
| hsa_circ_0126215 | 1837 | 11 | 1.20% |
| hsa_circ_0015266 | 1670 | 10 | 1.20% |
| hsa_circ_0015568 | 1670 | 10 | 1.20% |
| hsa_circ_0040438 | 1670 | 10 | 1.20% |
| hsa_circ_0056312 | 1670 | 10 | 1.20% |
| hsa_circ_0075274 | 1670 | 10 | 1.20% |
| hsa_circ_0092238 | 1670 | 10 | 1.20% |
| hsa_circ_0125838 | 1670 | 10 | 1.20% |
| hsa_circ_0127213 | 1670 | 10 | 1.20% |
| hsa_circ_0133180 | 1670 | 10 | 1.20% |
| hsa_circ_0015558 | 2172 | 13 | 1.20% |
| hsa_circ_0112136 | 2005 | 12 | 1.20% |
| hsa_circ_0013100 | 1838 | 11 | 1.20% |
| hsa_circ_0043258 | 1838 | 11 | 1.20% |
| hsa_circ_0052015 | 1838 | 11 | 1.20% |
| hsa_circ_0075324 | 1838 | 11 | 1.20% |
| hsa_circ_0107365 | 1838 | 11 | 1.20% |
| hsa_circ_0123688 | 1838 | 11 | 1.20% |
| hsa_circ_0113030 | 5180 | 31 | 1.20% |
| hsa_circ_0009972 | 1671 | 10 | 1.20% |
| hsa_circ_0022220 | 1671 | 10 | 1.20% |
| hsa_circ_0071947 | 1671 | 10 | 1.20% |
| hsa_circ_0084002 | 1671 | 10 | 1.20% |
| hsa_circ_0096645 | 1671 | 10 | 1.20% |
| hsa_circ_0105776 | 1671 | 10 | 1.20% |
| hsa_circ_0123621 | 1671 | 10 | 1.20% |
| hsa_circ_0136541 | 1671 | 10 | 1.20% |
| hsa_circ_0104677 | 2507 | 15 | 1.20% |
| hsa_circ_0029189 | 2006 | 12 | 1.20% |
| hsa_circ_0137721 | 2006 | 12 | 1.20% |
| hsa_circ_0006769 | 1839 | 11 | 1.20% |
| hsa_circ_0030064 | 1839 | 11 | 1.20% |
| hsa_circ_0032830 | 1839 | 11 | 1.20% |
| hsa_circ_0036184 | 1839 | 11 | 1.20% |
| hsa_circ_0053655 | 1839 | 11 | 1.20% |
| hsa_circ_0054129 | 1839 | 11 | 1.20% |
| hsa_circ_0061426 | 1839 | 11 | 1.20% |
| hsa_circ_0062006 | 1839 | 11 | 1.20% |
| hsa_circ_0071919 | 1839 | 11 | 1.20% |
| hsa_circ_0075089 | 1839 | 11 | 1.20% |
| hsa_circ_0092000 | 1839 | 11 | 1.20% |
| hsa_circ_0007834 | 1672 | 10 | 1.20% |
| hsa_circ_0032994 | 1672 | 10 | 1.20% |
| hsa_circ_0049879 | 1672 | 10 | 1.20% |
| hsa_circ_0051602 | 1672 | 10 | 1.20% |

| | | | |
|---|---|---|---|
| hsa_circ_0055676 | 1672 | 10 | 1.20% |
| hsa_circ_0078824 | 1672 | 10 | 1.20% |
| hsa_circ_0049686 | 3177 | 19 | 1.20% |
| hsa_circ_0127994 | 3846 | 23 | 1.20% |
| hsa_circ_0032595 | 2007 | 12 | 1.20% |
| hsa_circ_0062071 | 2007 | 12 | 1.20% |
| hsa_circ_0010888 | 1840 | 11 | 1.20% |
| hsa_circ_0019739 | 1840 | 11 | 1.20% |
| hsa_circ_0028524 | 1840 | 11 | 1.20% |
| hsa_circ_0036697 | 1840 | 11 | 1.20% |
| hsa_circ_0069867 | 1840 | 11 | 1.20% |
| hsa_circ_0092432 | 1840 | 11 | 1.20% |
| hsa_circ_0100080 | 1840 | 11 | 1.20% |
| hsa_circ_0136980 | 1840 | 11 | 1.20% |
| hsa_circ_0003360 | 5855 | 35 | 1.20% |
| hsa_circ_0039605 | 1673 | 10 | 1.20% |
| hsa_circ_0051405 | 1673 | 10 | 1.20% |
| hsa_circ_0080741 | 1673 | 10 | 1.20% |
| hsa_circ_0088418 | 1673 | 10 | 1.20% |
| hsa_circ_0093932 | 1673 | 10 | 1.20% |
| hsa_circ_0117863 | 1673 | 10 | 1.20% |
| hsa_circ_0128067 | 1673 | 10 | 1.20% |
| hsa_circ_0134684 | 1673 | 10 | 1.20% |
| hsa_circ_0074223 | 2175 | 13 | 1.20% |
| hsa_circ_0109667 | 2175 | 13 | 1.20% |
| hsa_circ_0126995 | 2175 | 13 | 1.20% |
| hsa_circ_0061456 | 2677 | 16 | 1.20% |
| hsa_circ_0076098 | 2510 | 15 | 1.20% |
| hsa_circ_0067234 | 2008 | 12 | 1.20% |
| hsa_circ_0132502 | 2008 | 12 | 1.20% |
| hsa_circ_0093092 | 5523 | 33 | 1.20% |
| hsa_circ_0112458 | 1841 | 11 | 1.20% |
| hsa_circ_0123008 | 1841 | 11 | 1.20% |
| hsa_circ_0138736 | 5021 | 30 | 1.19% |
| hsa_circ_0038200 | 2176 | 13 | 1.19% |
| hsa_circ_0013650 | 1674 | 10 | 1.19% |
| hsa_circ_0053384 | 1674 | 10 | 1.19% |
| hsa_circ_0091231 | 1674 | 10 | 1.19% |
| hsa_circ_0112003 | 1674 | 10 | 1.19% |
| hsa_circ_0001237 | 2009 | 12 | 1.19% |
| hsa_circ_0128967 | 2009 | 12 | 1.19% |
| hsa_circ_0096112 | 2679 | 16 | 1.19% |
| hsa_circ_0043802 | 3684 | 22 | 1.19% |
| hsa_circ_0000801 | 1842 | 11 | 1.19% |
| hsa_circ_0019534 | 1842 | 11 | 1.19% |
| hsa_circ_0020893 | 1842 | 11 | 1.19% |
| hsa_circ_0026725 | 1842 | 11 | 1.19% |
| hsa_circ_0048357 | 1842 | 11 | 1.19% |
| hsa_circ_0062975 | 1842 | 11 | 1.19% |
| hsa_circ_0129218 | 1842 | 11 | 1.19% |

| | | | |
|---|---|---|---|
| hsa_circ_0124049 | 7537 | 45 | 1.19% |
| hsa_circ_0075616 | 2010 | 12 | 1.19% |
| hsa_circ_0130579 | 2010 | 12 | 1.19% |
| hsa_circ_0046590 | 1675 | 10 | 1.19% |
| hsa_circ_0063536 | 1675 | 10 | 1.19% |
| hsa_circ_0063955 | 1675 | 10 | 1.19% |
| hsa_circ_0094017 | 1675 | 10 | 1.19% |
| hsa_circ_0094151 | 1675 | 10 | 1.19% |
| hsa_circ_0099727 | 1675 | 10 | 1.19% |
| hsa_circ_0102978 | 1675 | 10 | 1.19% |
| hsa_circ_0107353 | 1675 | 10 | 1.19% |
| hsa_circ_0110616 | 2178 | 13 | 1.19% |
| hsa_circ_0097408 | 1843 | 11 | 1.19% |
| hsa_circ_0113944 | 1843 | 11 | 1.19% |
| hsa_circ_0120616 | 1843 | 11 | 1.19% |
| hsa_circ_0074040 | 2346 | 14 | 1.19% |
| hsa_circ_0083697 | 2011 | 12 | 1.19% |
| hsa_circ_0007932 | 1676 | 10 | 1.19% |
| hsa_circ_0013022 | 1676 | 10 | 1.19% |
| hsa_circ_0037306 | 1676 | 10 | 1.19% |
| hsa_circ_0040660 | 1676 | 10 | 1.19% |
| hsa_circ_0041272 | 1676 | 10 | 1.19% |
| hsa_circ_0043161 | 1676 | 10 | 1.19% |
| hsa_circ_0104648 | 1676 | 10 | 1.19% |
| hsa_circ_0107715 | 1676 | 10 | 1.19% |
| hsa_circ_0112292 | 1676 | 10 | 1.19% |
| hsa_circ_0129106 | 1676 | 10 | 1.19% |
| hsa_circ_0129138 | 1676 | 10 | 1.19% |
| hsa_circ_0017262 | 2179 | 13 | 1.19% |
| hsa_circ_0029413 | 1844 | 11 | 1.19% |
| hsa_circ_0033015 | 1844 | 11 | 1.19% |
| hsa_circ_0042312 | 1844 | 11 | 1.19% |
| hsa_circ_0058954 | 1844 | 11 | 1.19% |
| hsa_circ_0112459 | 1844 | 11 | 1.19% |
| hsa_circ_0118865 | 1844 | 11 | 1.19% |
| hsa_circ_0024743 | 2012 | 12 | 1.19% |
| hsa_circ_0065500 | 2012 | 12 | 1.19% |
| hsa_circ_0090937 | 2683 | 16 | 1.19% |
| hsa_circ_0026953 | 1677 | 10 | 1.19% |
| hsa_circ_0036135 | 1677 | 10 | 1.19% |
| hsa_circ_0068908 | 1677 | 10 | 1.19% |
| hsa_circ_0071507 | 1677 | 10 | 1.19% |
| hsa_circ_0072342 | 1677 | 10 | 1.19% |
| hsa_circ_0085399 | 1677 | 10 | 1.19% |
| hsa_circ_0109628 | 1677 | 10 | 1.19% |
| hsa_circ_0130155 | 1677 | 10 | 1.19% |
| hsa_circ_0113910 | 1845 | 11 | 1.19% |
| hsa_circ_0019931 | 2181 | 13 | 1.19% |
| hsa_circ_0016785 | 1678 | 10 | 1.19% |
| hsa_circ_0036754 | 1678 | 10 | 1.19% |

| | | | |
|---|---|---|---|
| hsa_circ_0052179 | 1678 | 10 | 1.19% |
| hsa_circ_0078758 | 1678 | 10 | 1.19% |
| hsa_circ_0082818 | 1678 | 10 | 1.19% |
| hsa_circ_0137957 | 1678 | 10 | 1.19% |
| hsa_circ_0011479 | 1846 | 11 | 1.19% |
| hsa_circ_0013764 | 1846 | 11 | 1.19% |
| hsa_circ_0031171 | 1846 | 11 | 1.19% |
| hsa_circ_0039791 | 1846 | 11 | 1.19% |
| hsa_circ_0052263 | 1846 | 11 | 1.19% |
| hsa_circ_0089810 | 6713 | 40 | 1.19% |
| hsa_circ_0014174 | 1679 | 10 | 1.19% |
| hsa_circ_0024449 | 1679 | 10 | 1.19% |
| hsa_circ_0044762 | 1679 | 10 | 1.19% |
| hsa_circ_0088359 | 1679 | 10 | 1.19% |
| hsa_circ_0090940 | 1679 | 10 | 1.19% |
| hsa_circ_0120629 | 1679 | 10 | 1.19% |
| hsa_circ_0009516 | 5541 | 33 | 1.19% |
| hsa_circ_0011564 | 1847 | 11 | 1.19% |
| hsa_circ_0049567 | 1847 | 11 | 1.19% |
| hsa_circ_0087870 | 1847 | 11 | 1.19% |
| hsa_circ_0120712 | 1847 | 11 | 1.19% |
| hsa_circ_0022115 | 2015 | 12 | 1.19% |
| hsa_circ_0082240 | 2015 | 12 | 1.19% |
| hsa_circ_0010685 | 2183 | 13 | 1.19% |
| hsa_circ_0024554 | 2183 | 13 | 1.19% |
| hsa_circ_0037279 | 2351 | 14 | 1.19% |
| hsa_circ_0128395 | 2351 | 14 | 1.19% |
| hsa_circ_0009046 | 2687 | 16 | 1.19% |
| hsa_circ_0123966 | 5880 | 35 | 1.19% |
| hsa_circ_0019933 | 2184 | 13 | 1.19% |
| hsa_circ_0114668 | 2184 | 13 | 1.19% |
| hsa_circ_0025092 | 1848 | 11 | 1.19% |
| hsa_circ_0049424 | 1848 | 11 | 1.19% |
| hsa_circ_0061770 | 1848 | 11 | 1.19% |
| hsa_circ_0071284 | 1848 | 11 | 1.19% |
| hsa_circ_0097608 | 1848 | 11 | 1.19% |
| hsa_circ_0105809 | 1848 | 11 | 1.19% |
| hsa_circ_0112007 | 1848 | 11 | 1.19% |
| hsa_circ_0114456 | 1848 | 11 | 1.19% |
| hsa_circ_0021007 | 1680 | 10 | 1.19% |
| hsa_circ_0022113 | 1680 | 10 | 1.19% |
| hsa_circ_0025130 | 1680 | 10 | 1.19% |
| hsa_circ_0076412 | 1680 | 10 | 1.19% |
| hsa_circ_0076634 | 1680 | 10 | 1.19% |
| hsa_circ_0104120 | 1680 | 10 | 1.19% |
| hsa_circ_0117026 | 1680 | 10 | 1.19% |
| hsa_circ_0134670 | 1680 | 10 | 1.19% |
| hsa_circ_0013647 | 2017 | 12 | 1.19% |
| hsa_circ_0017653 | 2017 | 12 | 1.19% |
| hsa_circ_0028458 | 1849 | 11 | 1.19% |

| | | | |
|---|---|---|---|
| hsa_circ_0077038 | 1849 | 11 | 1.19% |
| hsa_circ_0086842 | 1849 | 11 | 1.19% |
| hsa_circ_0129836 | 1849 | 11 | 1.19% |
| hsa_circ_0012016 | 1681 | 10 | 1.19% |
| hsa_circ_0041378 | 1681 | 10 | 1.19% |
| hsa_circ_0047337 | 1681 | 10 | 1.19% |
| hsa_circ_0064248 | 1681 | 10 | 1.19% |
| hsa_circ_0067618 | 1681 | 10 | 1.19% |
| hsa_circ_0074471 | 1681 | 10 | 1.19% |
| hsa_circ_0115911 | 1681 | 10 | 1.19% |
| hsa_circ_0117058 | 1681 | 10 | 1.19% |
| hsa_circ_0117615 | 1681 | 10 | 1.19% |
| hsa_circ_0137349 | 1681 | 10 | 1.19% |
| hsa_circ_0074074 | 2186 | 13 | 1.19% |
| hsa_circ_0073291 | 2018 | 12 | 1.19% |
| hsa_circ_0024342 | 1850 | 11 | 1.19% |
| hsa_circ_0031019 | 1850 | 11 | 1.19% |
| hsa_circ_0046363 | 1850 | 11 | 1.19% |
| hsa_circ_0057596 | 1850 | 11 | 1.19% |
| hsa_circ_0126112 | 1850 | 11 | 1.19% |
| hsa_circ_0062462 | 33471 | 199 | 1.19% |
| hsa_circ_0012370 | 1682 | 10 | 1.19% |
| hsa_circ_0012865 | 1682 | 10 | 1.19% |
| hsa_circ_0063067 | 1682 | 10 | 1.19% |
| hsa_circ_0065853 | 1682 | 10 | 1.19% |
| hsa_circ_0066599 | 1682 | 10 | 1.19% |
| hsa_circ_0119388 | 1682 | 10 | 1.19% |
| hsa_circ_0121771 | 1682 | 10 | 1.19% |
| hsa_circ_0125957 | 1682 | 10 | 1.19% |
| hsa_circ_0127163 | 1682 | 10 | 1.19% |
| hsa_circ_0010991 | 3869 | 23 | 1.19% |
| hsa_circ_0022394 | 2019 | 12 | 1.19% |
| hsa_circ_0046830 | 2019 | 12 | 1.19% |
| hsa_circ_0110539 | 2019 | 12 | 1.19% |
| hsa_circ_0132499 | 2019 | 12 | 1.19% |
| hsa_circ_0083562 | 2524 | 15 | 1.19% |
| hsa_circ_0018417 | 1851 | 11 | 1.19% |
| hsa_circ_0022937 | 1851 | 11 | 1.19% |
| hsa_circ_0060255 | 1851 | 11 | 1.19% |
| hsa_circ_0071898 | 1851 | 11 | 1.19% |
| hsa_circ_0083608 | 1851 | 11 | 1.19% |
| hsa_circ_0123019 | 1851 | 11 | 1.19% |
| hsa_circ_0127506 | 1851 | 11 | 1.19% |
| hsa_circ_0025872 | 1683 | 10 | 1.19% |
| hsa_circ_0075760 | 1683 | 10 | 1.19% |
| hsa_circ_0078525 | 1683 | 10 | 1.19% |
| hsa_circ_0084085 | 1683 | 10 | 1.19% |
| hsa_circ_0095865 | 1683 | 10 | 1.19% |
| hsa_circ_0099379 | 1683 | 10 | 1.19% |
| hsa_circ_0100092 | 1683 | 10 | 1.19% |

| | | | |
|---|---|---|---|
| hsa_circ_0015549 | 2188 | 13 | 1.19% |
| hsa_circ_0053727 | 2188 | 13 | 1.19% |
| hsa_circ_0105584 | 2188 | 13 | 1.19% |
| hsa_circ_0039330 | 2020 | 12 | 1.19% |
| hsa_circ_0055679 | 1852 | 11 | 1.19% |
| hsa_circ_0094494 | 1852 | 11 | 1.19% |
| hsa_circ_0136569 | 5725 | 34 | 1.19% |
| hsa_circ_0067628 | 2189 | 13 | 1.19% |
| hsa_circ_0043195 | 1684 | 10 | 1.19% |
| hsa_circ_0054643 | 1684 | 10 | 1.19% |
| hsa_circ_0064762 | 1684 | 10 | 1.19% |
| hsa_circ_0073101 | 1684 | 10 | 1.19% |
| hsa_circ_0077348 | 1684 | 10 | 1.19% |
| hsa_circ_0080920 | 1684 | 10 | 1.19% |
| hsa_circ_0089889 | 1684 | 10 | 1.19% |
| hsa_circ_0120152 | 1684 | 10 | 1.19% |
| hsa_circ_0121345 | 1684 | 10 | 1.19% |
| hsa_circ_0113365 | 2021 | 12 | 1.19% |
| hsa_circ_0133731 | 2021 | 12 | 1.19% |
| hsa_circ_0028796 | 1853 | 11 | 1.19% |
| hsa_circ_0039770 | 1853 | 11 | 1.19% |
| hsa_circ_0046142 | 1853 | 11 | 1.19% |
| hsa_circ_0080096 | 1853 | 11 | 1.19% |
| hsa_circ_0047417 | 2190 | 13 | 1.19% |
| hsa_circ_0088857 | 2190 | 13 | 1.19% |
| hsa_circ_0071116 | 2022 | 12 | 1.19% |
| hsa_circ_0124907 | 2022 | 12 | 1.19% |
| hsa_circ_0020309 | 1685 | 10 | 1.19% |
| hsa_circ_0028694 | 1685 | 10 | 1.19% |
| hsa_circ_0044928 | 1685 | 10 | 1.19% |
| hsa_circ_0047353 | 1685 | 10 | 1.19% |
| hsa_circ_0072236 | 1685 | 10 | 1.19% |
| hsa_circ_0074089 | 1685 | 10 | 1.19% |
| hsa_circ_0078121 | 1685 | 10 | 1.19% |
| hsa_circ_0112615 | 1685 | 10 | 1.19% |
| hsa_circ_0127277 | 1685 | 10 | 1.19% |
| hsa_circ_0134114 | 1685 | 10 | 1.19% |
| hsa_circ_0135210 | 1685 | 10 | 1.19% |
| hsa_circ_0138721 | 1685 | 10 | 1.19% |
| hsa_circ_0042884 | 2191 | 13 | 1.19% |
| hsa_circ_0065405 | 2191 | 13 | 1.19% |
| hsa_circ_0117716 | 2191 | 13 | 1.19% |
| hsa_circ_0015161 | 1854 | 11 | 1.19% |
| hsa_circ_0025921 | 1854 | 11 | 1.19% |
| hsa_circ_0037879 | 1854 | 11 | 1.19% |
| hsa_circ_0046456 | 1854 | 11 | 1.19% |
| hsa_circ_0052024 | 1854 | 11 | 1.19% |
| hsa_circ_0059141 | 1854 | 11 | 1.19% |
| hsa_circ_0086910 | 1854 | 11 | 1.19% |
| hsa_circ_0106705 | 1854 | 11 | 1.19% |

| | | | |
|---|---|---|---|
| hsa_circ_0127224 | 1854 | 11 | 1.19% |
| hsa_circ_0119128 | 3371 | 20 | 1.19% |
| hsa_circ_0048891 | 2023 | 12 | 1.19% |
| hsa_circ_0009063 | 1686 | 10 | 1.19% |
| hsa_circ_0019694 | 1686 | 10 | 1.19% |
| hsa_circ_0050910 | 1686 | 10 | 1.19% |
| hsa_circ_0055843 | 1686 | 10 | 1.19% |
| hsa_circ_0059438 | 1686 | 10 | 1.19% |
| hsa_circ_0086333 | 1686 | 10 | 1.19% |
| hsa_circ_0094115 | 1686 | 10 | 1.19% |
| hsa_circ_0133376 | 1686 | 10 | 1.19% |
| hsa_circ_0029825 | 1855 | 11 | 1.19% |
| hsa_circ_0074343 | 1855 | 11 | 1.19% |
| hsa_circ_0087197 | 1855 | 11 | 1.19% |
| hsa_circ_0101569 | 1855 | 11 | 1.19% |
| hsa_circ_0118042 | 1855 | 11 | 1.19% |
| hsa_circ_0134759 | 1855 | 11 | 1.19% |
| hsa_circ_0140479 | 2361 | 14 | 1.19% |
| hsa_circ_0033325 | 2024 | 12 | 1.19% |
| hsa_circ_0087637 | 7254 | 43 | 1.19% |
| hsa_circ_0009795 | 1687 | 10 | 1.19% |
| hsa_circ_0010722 | 1687 | 10 | 1.19% |
| hsa_circ_0021926 | 1687 | 10 | 1.19% |
| hsa_circ_0031061 | 1687 | 10 | 1.19% |
| hsa_circ_0039213 | 1687 | 10 | 1.19% |
| hsa_circ_0049020 | 1687 | 10 | 1.19% |
| hsa_circ_0051623 | 1687 | 10 | 1.19% |
| hsa_circ_0072897 | 1687 | 10 | 1.19% |
| hsa_circ_0098564 | 1687 | 10 | 1.19% |
| hsa_circ_0116056 | 1687 | 10 | 1.19% |
| hsa_circ_0127185 | 1687 | 10 | 1.19% |
| hsa_circ_0024046 | 1856 | 11 | 1.19% |
| hsa_circ_0051858 | 1856 | 11 | 1.19% |
| hsa_circ_0073171 | 1856 | 11 | 1.19% |
| hsa_circ_0083240 | 1856 | 11 | 1.19% |
| hsa_circ_0112220 | 2025 | 12 | 1.19% |
| hsa_circ_0023611 | 1688 | 10 | 1.18% |
| hsa_circ_0046438 | 1688 | 10 | 1.18% |
| hsa_circ_0056377 | 1688 | 10 | 1.18% |
| hsa_circ_0058043 | 1688 | 10 | 1.18% |
| hsa_circ_0079946 | 1688 | 10 | 1.18% |
| hsa_circ_0047371 | 1857 | 11 | 1.18% |
| hsa_circ_0081842 | 1857 | 11 | 1.18% |
| hsa_circ_0100712 | 1857 | 11 | 1.18% |
| hsa_circ_0127708 | 1857 | 11 | 1.18% |
| hsa_circ_0136550 | 1857 | 11 | 1.18% |
| hsa_circ_0139380 | 1857 | 11 | 1.18% |
| hsa_circ_0006953 | 5065 | 30 | 1.18% |
| hsa_circ_0000799 | 2026 | 12 | 1.18% |
| hsa_circ_0102935 | 2026 | 12 | 1.18% |

| | | | |
|---|---|---|---|
| hsa_circ_0059653 | 2195 | 13 | 1.18% |
| hsa_circ_0020095 | 1689 | 10 | 1.18% |
| hsa_circ_0026733 | 1689 | 10 | 1.18% |
| hsa_circ_0042798 | 1689 | 10 | 1.18% |
| hsa_circ_0047971 | 1689 | 10 | 1.18% |
| hsa_circ_0065851 | 1689 | 10 | 1.18% |
| hsa_circ_0087587 | 1689 | 10 | 1.18% |
| hsa_circ_0099500 | 1689 | 10 | 1.18% |
| hsa_circ_0139241 | 1689 | 10 | 1.18% |
| hsa_circ_0026002 | 1858 | 11 | 1.18% |
| hsa_circ_0071809 | 1858 | 11 | 1.18% |
| hsa_circ_0074200 | 1858 | 11 | 1.18% |
| hsa_circ_0007209 | 2027 | 12 | 1.18% |
| hsa_circ_0105474 | 2027 | 12 | 1.18% |
| hsa_circ_0115963 | 2027 | 12 | 1.18% |
| hsa_circ_0138878 | 2027 | 12 | 1.18% |
| hsa_circ_0012659 | 2534 | 15 | 1.18% |
| hsa_circ_0105236 | 8787 | 52 | 1.18% |
| hsa_circ_0058938 | 2197 | 13 | 1.18% |
| hsa_circ_0011474 | 2028 | 12 | 1.18% |
| hsa_circ_0120582 | 2028 | 12 | 1.18% |
| hsa_circ_0009549 | 1859 | 11 | 1.18% |
| hsa_circ_0011713 | 1859 | 11 | 1.18% |
| hsa_circ_0019255 | 1859 | 11 | 1.18% |
| hsa_circ_0020304 | 1859 | 11 | 1.18% |
| hsa_circ_0039228 | 1859 | 11 | 1.18% |
| hsa_circ_0058511 | 1859 | 11 | 1.18% |
| hsa_circ_0083623 | 1859 | 11 | 1.18% |
| hsa_circ_0020480 | 1690 | 10 | 1.18% |
| hsa_circ_0052014 | 1690 | 10 | 1.18% |
| hsa_circ_0069022 | 1690 | 10 | 1.18% |
| hsa_circ_0129045 | 1690 | 10 | 1.18% |
| hsa_circ_0030997 | 2198 | 13 | 1.18% |
| hsa_circ_0103332 | 2198 | 13 | 1.18% |
| hsa_circ_0020012 | 2029 | 12 | 1.18% |
| hsa_circ_0086003 | 2029 | 12 | 1.18% |
| hsa_circ_0032515 | 1860 | 11 | 1.18% |
| hsa_circ_0046445 | 1860 | 11 | 1.18% |
| hsa_circ_0064203 | 1860 | 11 | 1.18% |
| hsa_circ_0083297 | 1860 | 11 | 1.18% |
| hsa_circ_0103116 | 1860 | 11 | 1.18% |
| hsa_circ_0122813 | 1860 | 11 | 1.18% |
| hsa_circ_0127817 | 1860 | 11 | 1.18% |
| hsa_circ_0029884 | 1691 | 10 | 1.18% |
| hsa_circ_0054559 | 1691 | 10 | 1.18% |
| hsa_circ_0073550 | 1691 | 10 | 1.18% |
| hsa_circ_0075258 | 1691 | 10 | 1.18% |
| hsa_circ_0132010 | 1691 | 10 | 1.18% |
| hsa_circ_0140208 | 1691 | 10 | 1.18% |
| hsa_circ_0023877 | 2030 | 12 | 1.18% |

| | | | |
|---|---|---|---|
| hsa_circ_0058827 | 2030 | 12 | 1.18% |
| hsa_circ_0012246 | 1861 | 11 | 1.18% |
| hsa_circ_0038109 | 1861 | 11 | 1.18% |
| hsa_circ_0050857 | 1861 | 11 | 1.18% |
| hsa_circ_0059277 | 1861 | 11 | 1.18% |
| hsa_circ_0076945 | 1861 | 11 | 1.18% |
| hsa_circ_0080709 | 1861 | 11 | 1.18% |
| hsa_circ_0087742 | 1861 | 11 | 1.18% |
| hsa_circ_0114834 | 1861 | 11 | 1.18% |
| hsa_circ_0063515 | 5414 | 32 | 1.18% |
| hsa_circ_0008766 | 1692 | 10 | 1.18% |
| hsa_circ_0011234 | 1692 | 10 | 1.18% |
| hsa_circ_0035520 | 1692 | 10 | 1.18% |
| hsa_circ_0038395 | 1692 | 10 | 1.18% |
| hsa_circ_0052768 | 1692 | 10 | 1.18% |
| hsa_circ_0058397 | 1692 | 10 | 1.18% |
| hsa_circ_0066081 | 1692 | 10 | 1.18% |
| hsa_circ_0071979 | 1692 | 10 | 1.18% |
| hsa_circ_0082376 | 1692 | 10 | 1.18% |
| hsa_circ_0117470 | 1692 | 10 | 1.18% |
| hsa_circ_0119899 | 1692 | 10 | 1.18% |
| hsa_circ_0128574 | 1692 | 10 | 1.18% |
| hsa_circ_0046228 | 2369 | 14 | 1.18% |
| hsa_circ_0101302 | 7784 | 46 | 1.18% |
| hsa_circ_0093912 | 2200 | 13 | 1.18% |
| hsa_circ_0129403 | 2031 | 12 | 1.18% |
| hsa_circ_0020552 | 1862 | 11 | 1.18% |
| hsa_circ_0021105 | 1862 | 11 | 1.18% |
| hsa_circ_0040284 | 1862 | 11 | 1.18% |
| hsa_circ_0016692 | 1693 | 10 | 1.18% |
| hsa_circ_0022637 | 1693 | 10 | 1.18% |
| hsa_circ_0054382 | 1693 | 10 | 1.18% |
| hsa_circ_0125715 | 1693 | 10 | 1.18% |
| hsa_circ_0102443 | 7280 | 43 | 1.18% |
| hsa_circ_0066653 | 3048 | 18 | 1.18% |
| hsa_circ_0088876 | 2032 | 12 | 1.18% |
| hsa_circ_0099175 | 2879 | 17 | 1.18% |
| hsa_circ_0029123 | 1863 | 11 | 1.18% |
| hsa_circ_0036358 | 1863 | 11 | 1.18% |
| hsa_circ_0040004 | 1863 | 11 | 1.18% |
| hsa_circ_0043150 | 1863 | 11 | 1.18% |
| hsa_circ_0067242 | 1863 | 11 | 1.18% |
| hsa_circ_0099789 | 1863 | 11 | 1.18% |
| hsa_circ_0130012 | 5590 | 33 | 1.18% |
| hsa_circ_0005658 | 1694 | 10 | 1.18% |
| hsa_circ_0040236 | 1694 | 10 | 1.18% |
| hsa_circ_0046873 | 1694 | 10 | 1.18% |
| hsa_circ_0082042 | 1694 | 10 | 1.18% |
| hsa_circ_0124518 | 1694 | 10 | 1.18% |
| hsa_circ_0125143 | 1694 | 10 | 1.18% |

| | | | |
|---|---|---|---|
| hsa_circ_0044362 | 2033 | 12 | 1.18% |
| hsa_circ_0131340 | 2033 | 12 | 1.18% |
| hsa_circ_0038077 | 1864 | 11 | 1.18% |
| hsa_circ_0051389 | 1864 | 11 | 1.18% |
| hsa_circ_0064297 | 1864 | 11 | 1.18% |
| hsa_circ_0091172 | 1864 | 11 | 1.18% |
| hsa_circ_0133435 | 1864 | 11 | 1.18% |
| hsa_circ_0131784 | 2203 | 13 | 1.18% |
| hsa_circ_0034489 | 2034 | 12 | 1.18% |
| hsa_circ_0066418 | 2034 | 12 | 1.18% |
| hsa_circ_0092006 | 2034 | 12 | 1.18% |
| hsa_circ_0120578 | 2034 | 12 | 1.18% |
| hsa_circ_0006232 | 1695 | 10 | 1.18% |
| hsa_circ_0036146 | 1695 | 10 | 1.18% |
| hsa_circ_0041690 | 1695 | 10 | 1.18% |
| hsa_circ_0062974 | 1695 | 10 | 1.18% |
| hsa_circ_0081280 | 1695 | 10 | 1.18% |
| hsa_circ_0092115 | 1695 | 10 | 1.18% |
| hsa_circ_0106595 | 1695 | 10 | 1.18% |
| hsa_circ_0138397 | 1695 | 10 | 1.18% |
| hsa_circ_0087474 | 2204 | 13 | 1.18% |
| hsa_circ_0016934 | 1865 | 11 | 1.18% |
| hsa_circ_0081998 | 1865 | 11 | 1.18% |
| hsa_circ_0020987 | 2035 | 12 | 1.18% |
| hsa_circ_0031943 | 2035 | 12 | 1.18% |
| hsa_circ_0041335 | 1696 | 10 | 1.18% |
| hsa_circ_0060270 | 1696 | 10 | 1.18% |
| hsa_circ_0071655 | 1696 | 10 | 1.18% |
| hsa_circ_0082653 | 1696 | 10 | 1.18% |
| hsa_circ_0102375 | 1696 | 10 | 1.18% |
| hsa_circ_0104158 | 1696 | 10 | 1.18% |
| hsa_circ_0105715 | 1696 | 10 | 1.18% |
| hsa_circ_0131567 | 1696 | 10 | 1.18% |
| hsa_circ_0062338 | 2205 | 13 | 1.18% |
| hsa_circ_0009272 | 1866 | 11 | 1.18% |
| hsa_circ_0010119 | 1866 | 11 | 1.18% |
| hsa_circ_0011400 | 1866 | 11 | 1.18% |
| hsa_circ_0013807 | 1866 | 11 | 1.18% |
| hsa_circ_0029672 | 1866 | 11 | 1.18% |
| hsa_circ_0046472 | 1866 | 11 | 1.18% |
| hsa_circ_0103042 | 1866 | 11 | 1.18% |
| hsa_circ_0116375 | 1866 | 11 | 1.18% |
| hsa_circ_0119814 | 1866 | 11 | 1.18% |
| hsa_circ_0010698 | 2036 | 12 | 1.18% |
| hsa_circ_0032606 | 2036 | 12 | 1.18% |
| hsa_circ_0090876 | 2036 | 12 | 1.18% |
| hsa_circ_0048201 | 1697 | 10 | 1.18% |
| hsa_circ_0061255 | 1697 | 10 | 1.18% |
| hsa_circ_0072519 | 1697 | 10 | 1.18% |
| hsa_circ_0081836 | 1697 | 10 | 1.18% |

| | | | |
|---|---|---|---|
| hsa_circ_0085691 | 1697 | 10 | 1.18% |
| hsa_circ_0086232 | 1697 | 10 | 1.18% |
| hsa_circ_0105034 | 1697 | 10 | 1.18% |
| hsa_circ_0105810 | 1697 | 10 | 1.18% |
| hsa_circ_0114805 | 1697 | 10 | 1.18% |
| hsa_circ_0117907 | 1697 | 10 | 1.18% |
| hsa_circ_0134483 | 1697 | 10 | 1.18% |
| hsa_circ_0124658 | 4582 | 27 | 1.18% |
| hsa_circ_0013044 | 1867 | 11 | 1.18% |
| hsa_circ_0031931 | 1867 | 11 | 1.18% |
| hsa_circ_0066683 | 1867 | 11 | 1.18% |
| hsa_circ_0089507 | 1867 | 11 | 1.18% |
| hsa_circ_0099108 | 1867 | 11 | 1.18% |
| hsa_circ_0122274 | 1867 | 11 | 1.18% |
| hsa_circ_0135702 | 1867 | 11 | 1.18% |
| hsa_circ_0063203 | 2037 | 12 | 1.18% |
| hsa_circ_0071557 | 2037 | 12 | 1.18% |
| hsa_circ_0132311 | 2037 | 12 | 1.18% |
| hsa_circ_0064751 | 2207 | 13 | 1.18% |
| hsa_circ_0139821 | 2207 | 13 | 1.18% |
| hsa_circ_0016006 | 1698 | 10 | 1.18% |
| hsa_circ_0016187 | 1698 | 10 | 1.18% |
| hsa_circ_0023011 | 1698 | 10 | 1.18% |
| hsa_circ_0035303 | 1698 | 10 | 1.18% |
| hsa_circ_0042409 | 1698 | 10 | 1.18% |
| hsa_circ_0063440 | 1698 | 10 | 1.18% |
| hsa_circ_0109079 | 1698 | 10 | 1.18% |
| hsa_circ_0139026 | 1698 | 10 | 1.18% |
| hsa_circ_0039931 | 1868 | 11 | 1.18% |
| hsa_circ_0063510 | 1868 | 11 | 1.18% |
| hsa_circ_0074902 | 1868 | 11 | 1.18% |
| hsa_circ_0099557 | 1868 | 11 | 1.18% |
| hsa_circ_0117262 | 1868 | 11 | 1.18% |
| hsa_circ_0064507 | 2038 | 12 | 1.18% |
| hsa_circ_0114180 | 2208 | 13 | 1.18% |
| hsa_circ_0022191 | 1699 | 10 | 1.18% |
| hsa_circ_0056093 | 1699 | 10 | 1.18% |
| hsa_circ_0066219 | 1699 | 10 | 1.18% |
| hsa_circ_0067546 | 1699 | 10 | 1.18% |
| hsa_circ_0075427 | 1699 | 10 | 1.18% |
| hsa_circ_0101374 | 1699 | 10 | 1.18% |
| hsa_circ_0116126 | 1699 | 10 | 1.18% |
| hsa_circ_0125373 | 1699 | 10 | 1.18% |
| hsa_circ_0138870 | 1699 | 10 | 1.18% |
| hsa_circ_0054863 | 1869 | 11 | 1.18% |
| hsa_circ_0099992 | 1869 | 11 | 1.18% |
| hsa_circ_0131632 | 1869 | 11 | 1.18% |
| hsa_circ_0003291 | 2039 | 12 | 1.18% |
| hsa_circ_0026989 | 2039 | 12 | 1.18% |
| hsa_circ_0035723 | 2039 | 12 | 1.18% |

| | | | |
|---|---|---|---|
| hsa_circ_0111626 | 2039 | 12 | 1.18% |
| hsa_circ_0081499 | 2720 | 16 | 1.18% |
| hsa_circ_0047944 | 2210 | 13 | 1.18% |
| hsa_circ_0015696 | 2040 | 12 | 1.18% |
| hsa_circ_0085996 | 2040 | 12 | 1.18% |
| hsa_circ_0055713 | 1870 | 11 | 1.18% |
| hsa_circ_0100262 | 1870 | 11 | 1.18% |
| hsa_circ_0104571 | 1870 | 11 | 1.18% |
| hsa_circ_0109754 | 1870 | 11 | 1.18% |
| hsa_circ_0027937 | 1700 | 10 | 1.18% |
| hsa_circ_0030424 | 1700 | 10 | 1.18% |
| hsa_circ_0038944 | 1700 | 10 | 1.18% |
| hsa_circ_0056774 | 1700 | 10 | 1.18% |
| hsa_circ_0066189 | 1700 | 10 | 1.18% |
| hsa_circ_0072484 | 1700 | 10 | 1.18% |
| hsa_circ_0093910 | 1700 | 10 | 1.18% |
| hsa_circ_0124452 | 1700 | 10 | 1.18% |
| hsa_circ_0136775 | 3401 | 20 | 1.18% |
| hsa_circ_0060677 | 2211 | 13 | 1.18% |
| hsa_circ_0091751 | 2211 | 13 | 1.18% |
| hsa_circ_0013134 | 2041 | 12 | 1.18% |
| hsa_circ_0065875 | 2041 | 12 | 1.18% |
| hsa_circ_0128891 | 1871 | 11 | 1.18% |
| hsa_circ_0017126 | 1701 | 10 | 1.18% |
| hsa_circ_0026414 | 1701 | 10 | 1.18% |
| hsa_circ_0026425 | 1701 | 10 | 1.18% |
| hsa_circ_0027480 | 1701 | 10 | 1.18% |
| hsa_circ_0030255 | 1701 | 10 | 1.18% |
| hsa_circ_0055176 | 1701 | 10 | 1.18% |
| hsa_circ_0066356 | 1701 | 10 | 1.18% |
| hsa_circ_0092643 | 1701 | 10 | 1.18% |
| hsa_circ_0094195 | 1701 | 10 | 1.18% |
| hsa_circ_0001388 | 2042 | 12 | 1.18% |
| hsa_circ_0036160 | 2042 | 12 | 1.18% |
| hsa_circ_0091458 | 2042 | 12 | 1.18% |
| hsa_circ_0016780 | 1872 | 11 | 1.18% |
| hsa_circ_0043940 | 1872 | 11 | 1.18% |
| hsa_circ_0056745 | 1872 | 11 | 1.18% |
| hsa_circ_0082555 | 1872 | 11 | 1.18% |
| hsa_circ_0114765 | 1872 | 11 | 1.18% |
| hsa_circ_0117370 | 1872 | 11 | 1.18% |
| hsa_circ_0074382 | 2723 | 16 | 1.18% |
| hsa_circ_0014579 | 1702 | 10 | 1.18% |
| hsa_circ_0016498 | 1702 | 10 | 1.18% |
| hsa_circ_0026357 | 1702 | 10 | 1.18% |
| hsa_circ_0056335 | 1702 | 10 | 1.18% |
| hsa_circ_0075188 | 1702 | 10 | 1.18% |
| hsa_circ_0134611 | 1702 | 10 | 1.18% |
| hsa_circ_0029698 | 4085 | 24 | 1.18% |
| hsa_circ_0047991 | 2043 | 12 | 1.17% |

| | | | |
|---|---|---|---|
| hsa_circ_0049313 | 2043 | 12 | 1.17% |
| hsa_circ_0061978 | 2043 | 12 | 1.17% |
| hsa_circ_0013782 | 1873 | 11 | 1.17% |
| hsa_circ_0076784 | 1873 | 11 | 1.17% |
| hsa_circ_0014871 | 1703 | 10 | 1.17% |
| hsa_circ_0033129 | 1703 | 10 | 1.17% |
| hsa_circ_0034032 | 1703 | 10 | 1.17% |
| hsa_circ_0035362 | 1703 | 10 | 1.17% |
| hsa_circ_0043678 | 1703 | 10 | 1.17% |
| hsa_circ_0059440 | 1703 | 10 | 1.17% |
| hsa_circ_0062884 | 1703 | 10 | 1.17% |
| hsa_circ_0083664 | 1703 | 10 | 1.17% |
| hsa_circ_0095546 | 1703 | 10 | 1.17% |
| hsa_circ_0121274 | 1703 | 10 | 1.17% |
| hsa_circ_0125235 | 1703 | 10 | 1.17% |
| hsa_circ_0047305 | 2385 | 14 | 1.17% |
| hsa_circ_0019473 | 1874 | 11 | 1.17% |
| hsa_circ_0100917 | 1874 | 11 | 1.17% |
| hsa_circ_0108058 | 1874 | 11 | 1.17% |
| hsa_circ_0002293 | 1704 | 10 | 1.17% |
| hsa_circ_0017418 | 1704 | 10 | 1.17% |
| hsa_circ_0019326 | 1704 | 10 | 1.17% |
| hsa_circ_0026815 | 1704 | 10 | 1.17% |
| hsa_circ_0045125 | 1704 | 10 | 1.17% |
| hsa_circ_0060759 | 1704 | 10 | 1.17% |
| hsa_circ_0066195 | 1704 | 10 | 1.17% |
| hsa_circ_0087175 | 1704 | 10 | 1.17% |
| hsa_circ_0130433 | 1704 | 10 | 1.17% |
| hsa_circ_0050411 | 2045 | 12 | 1.17% |
| hsa_circ_0077485 | 2045 | 12 | 1.17% |
| hsa_circ_0021392 | 1875 | 11 | 1.17% |
| hsa_circ_0045054 | 1875 | 11 | 1.17% |
| hsa_circ_0052916 | 1875 | 11 | 1.17% |
| hsa_circ_0062479 | 1875 | 11 | 1.17% |
| hsa_circ_0073965 | 1875 | 11 | 1.17% |
| hsa_circ_0079350 | 1875 | 11 | 1.17% |
| hsa_circ_0135113 | 1875 | 11 | 1.17% |
| hsa_circ_0048163 | 2557 | 15 | 1.17% |
| hsa_circ_0079550 | 2046 | 12 | 1.17% |
| hsa_circ_0011597 | 1705 | 10 | 1.17% |
| hsa_circ_0017785 | 1705 | 10 | 1.17% |
| hsa_circ_0020901 | 1705 | 10 | 1.17% |
| hsa_circ_0040050 | 1705 | 10 | 1.17% |
| hsa_circ_0066156 | 1705 | 10 | 1.17% |
| hsa_circ_0070696 | 1705 | 10 | 1.17% |
| hsa_circ_0087096 | 1705 | 10 | 1.17% |
| hsa_circ_0108212 | 1705 | 10 | 1.17% |
| hsa_circ_0137856 | 1705 | 10 | 1.17% |
| hsa_circ_0009412 | 1876 | 11 | 1.17% |
| hsa_circ_0014467 | 1876 | 11 | 1.17% |

| | | | |
|---|---|---|---|
| hsa_circ_0016109 | 1876 | 11 | 1.17% |
| hsa_circ_0019466 | 1876 | 11 | 1.17% |
| hsa_circ_0104575 | 2388 | 14 | 1.17% |
| hsa_circ_0018387 | 2559 | 15 | 1.17% |
| hsa_circ_0031912 | 1706 | 10 | 1.17% |
| hsa_circ_0048713 | 1706 | 10 | 1.17% |
| hsa_circ_0051327 | 1706 | 10 | 1.17% |
| hsa_circ_0056968 | 1706 | 10 | 1.17% |
| hsa_circ_0084917 | 1706 | 10 | 1.17% |
| hsa_circ_0098089 | 1706 | 10 | 1.17% |
| hsa_circ_0108166 | 1706 | 10 | 1.17% |
| hsa_circ_0108381 | 1706 | 10 | 1.17% |
| hsa_circ_0109064 | 1706 | 10 | 1.17% |
| hsa_circ_0114438 | 1706 | 10 | 1.17% |
| hsa_circ_0121248 | 1706 | 10 | 1.17% |
| hsa_circ_0125233 | 1706 | 10 | 1.17% |
| hsa_circ_0135289 | 1706 | 10 | 1.17% |
| hsa_circ_0052167 | 2218 | 13 | 1.17% |
| hsa_circ_0045086 | 1877 | 11 | 1.17% |
| hsa_circ_0123664 | 1877 | 11 | 1.17% |
| hsa_circ_0055832 | 2560 | 15 | 1.17% |
| hsa_circ_0065443 | 2048 | 12 | 1.17% |
| hsa_circ_0074448 | 2048 | 12 | 1.17% |
| hsa_circ_0044507 | 3926 | 23 | 1.17% |
| hsa_circ_0032411 | 1707 | 10 | 1.17% |
| hsa_circ_0034862 | 1707 | 10 | 1.17% |
| hsa_circ_0043295 | 1707 | 10 | 1.17% |
| hsa_circ_0048729 | 1707 | 10 | 1.17% |
| hsa_circ_0058505 | 1707 | 10 | 1.17% |
| hsa_circ_0088007 | 1707 | 10 | 1.17% |
| hsa_circ_0088578 | 1707 | 10 | 1.17% |
| hsa_circ_0124684 | 1707 | 10 | 1.17% |
| hsa_circ_0006910 | 2902 | 17 | 1.17% |
| hsa_circ_0052946 | 4780 | 28 | 1.17% |
| hsa_circ_0139516 | 13999 | 82 | 1.17% |
| hsa_circ_0022204 | 1878 | 11 | 1.17% |
| hsa_circ_0028440 | 1878 | 11 | 1.17% |
| hsa_circ_0029823 | 1878 | 11 | 1.17% |
| hsa_circ_0043657 | 1878 | 11 | 1.17% |
| hsa_circ_0065245 | 1878 | 11 | 1.17% |
| hsa_circ_0086554 | 1878 | 11 | 1.17% |
| hsa_circ_0094864 | 1878 | 11 | 1.17% |
| hsa_circ_0018444 | 1708 | 10 | 1.17% |
| hsa_circ_0038632 | 1708 | 10 | 1.17% |
| hsa_circ_0081701 | 1708 | 10 | 1.17% |
| hsa_circ_0090775 | 1708 | 10 | 1.17% |
| hsa_circ_0100316 | 1708 | 10 | 1.17% |
| hsa_circ_0113014 | 1708 | 10 | 1.17% |
| hsa_circ_0133012 | 1708 | 10 | 1.17% |
| hsa_circ_0010728 | 1879 | 11 | 1.17% |

| | | | |
|---|---|---|---|
| hsa_circ_0073416 | 1879 | 11 | 1.17% |
| hsa_circ_0081385 | 1879 | 11 | 1.17% |
| hsa_circ_0094867 | 1879 | 11 | 1.17% |
| hsa_circ_0101982 | 1879 | 11 | 1.17% |
| hsa_circ_0088403 | 4100 | 24 | 1.17% |
| hsa_circ_0059932 | 2050 | 12 | 1.17% |
| hsa_circ_0072618 | 2221 | 13 | 1.17% |
| hsa_circ_0082873 | 2221 | 13 | 1.17% |
| hsa_circ_0088324 | 2563 | 15 | 1.17% |
| hsa_circ_0009517 | 5639 | 33 | 1.17% |
| hsa_circ_0004702 | 2905 | 17 | 1.17% |
| hsa_circ_0023897 | 1709 | 10 | 1.17% |
| hsa_circ_0028653 | 1709 | 10 | 1.17% |
| hsa_circ_0032423 | 1709 | 10 | 1.17% |
| hsa_circ_0036121 | 1709 | 10 | 1.17% |
| hsa_circ_0036368 | 1709 | 10 | 1.17% |
| hsa_circ_0042869 | 1709 | 10 | 1.17% |
| hsa_circ_0047540 | 1709 | 10 | 1.17% |
| hsa_circ_0079173 | 1709 | 10 | 1.17% |
| hsa_circ_0090212 | 1709 | 10 | 1.17% |
| hsa_circ_0029200 | 1880 | 11 | 1.17% |
| hsa_circ_0050322 | 1880 | 11 | 1.17% |
| hsa_circ_0083607 | 1880 | 11 | 1.17% |
| hsa_circ_0098524 | 1880 | 11 | 1.17% |
| hsa_circ_0108218 | 1880 | 11 | 1.17% |
| hsa_circ_0031013 | 2051 | 12 | 1.17% |
| hsa_circ_0108240 | 5128 | 30 | 1.17% |
| hsa_circ_0138350 | 2564 | 15 | 1.17% |
| hsa_circ_0117134 | 2394 | 14 | 1.17% |
| hsa_circ_0136674 | 2223 | 13 | 1.17% |
| hsa_circ_0010785 | 2052 | 12 | 1.17% |
| hsa_circ_0082098 | 2052 | 12 | 1.17% |
| hsa_circ_0024167 | 1881 | 11 | 1.17% |
| hsa_circ_0036882 | 1881 | 11 | 1.17% |
| hsa_circ_0053553 | 1881 | 11 | 1.17% |
| hsa_circ_0068326 | 1881 | 11 | 1.17% |
| hsa_circ_0087740 | 1881 | 11 | 1.17% |
| hsa_circ_0102808 | 1881 | 11 | 1.17% |
| hsa_circ_0114802 | 1881 | 11 | 1.17% |
| hsa_circ_0129956 | 1881 | 11 | 1.17% |
| hsa_circ_0014624 | 1710 | 10 | 1.17% |
| hsa_circ_0016090 | 1710 | 10 | 1.17% |
| hsa_circ_0016803 | 1710 | 10 | 1.17% |
| hsa_circ_0023738 | 1710 | 10 | 1.17% |
| hsa_circ_0024821 | 1710 | 10 | 1.17% |
| hsa_circ_0034456 | 1710 | 10 | 1.17% |
| hsa_circ_0065202 | 1710 | 10 | 1.17% |
| hsa_circ_0073894 | 1710 | 10 | 1.17% |
| hsa_circ_0075431 | 1710 | 10 | 1.17% |
| hsa_circ_0091494 | 1710 | 10 | 1.17% |

| | | | |
|---|---|---|---|
| hsa_circ_0101066 | 1710 | 10 | 1.17% |
| hsa_circ_0114881 | 1710 | 10 | 1.17% |
| hsa_circ_0116452 | 1710 | 10 | 1.17% |
| hsa_circ_0136535 | 1710 | 10 | 1.17% |
| hsa_circ_0108241 | 5132 | 30 | 1.17% |
| hsa_circ_0042290 | 2053 | 12 | 1.17% |
| hsa_circ_0045921 | 2053 | 12 | 1.17% |
| hsa_circ_0017134 | 1882 | 11 | 1.17% |
| hsa_circ_0026872 | 1882 | 11 | 1.17% |
| hsa_circ_0073962 | 1882 | 11 | 1.17% |
| hsa_circ_0086997 | 1882 | 11 | 1.17% |
| hsa_circ_0134092 | 1882 | 11 | 1.17% |
| hsa_circ_0015351 | 1711 | 10 | 1.17% |
| hsa_circ_0015937 | 1711 | 10 | 1.17% |
| hsa_circ_0016009 | 1711 | 10 | 1.17% |
| hsa_circ_0016273 | 1711 | 10 | 1.17% |
| hsa_circ_0077283 | 1711 | 10 | 1.17% |
| hsa_circ_0077549 | 1711 | 10 | 1.17% |
| hsa_circ_0088820 | 1711 | 10 | 1.17% |
| hsa_circ_0118611 | 1711 | 10 | 1.17% |
| hsa_circ_0123961 | 1711 | 10 | 1.17% |
| hsa_circ_0125247 | 1711 | 10 | 1.17% |
| hsa_circ_0021117 | 3080 | 18 | 1.17% |
| hsa_circ_0076781 | 4107 | 24 | 1.17% |
| hsa_circ_0027061 | 1883 | 11 | 1.17% |
| hsa_circ_0051026 | 1883 | 11 | 1.17% |
| hsa_circ_0108219 | 1883 | 11 | 1.17% |
| hsa_circ_0113406 | 1883 | 11 | 1.17% |
| hsa_circ_0018137 | 1712 | 10 | 1.17% |
| hsa_circ_0022874 | 1712 | 10 | 1.17% |
| hsa_circ_0058400 | 1712 | 10 | 1.17% |
| hsa_circ_0065579 | 1712 | 10 | 1.17% |
| hsa_circ_0065650 | 1712 | 10 | 1.17% |
| hsa_circ_0066469 | 1712 | 10 | 1.17% |
| hsa_circ_0121999 | 1712 | 10 | 1.17% |
| hsa_circ_0081026 | 2226 | 13 | 1.17% |
| hsa_circ_0013274 | 2055 | 12 | 1.17% |
| hsa_circ_0047110 | 2055 | 12 | 1.17% |
| hsa_circ_0085627 | 2055 | 12 | 1.17% |
| hsa_circ_0018843 | 1884 | 11 | 1.17% |
| hsa_circ_0039029 | 1884 | 11 | 1.17% |
| hsa_circ_0049560 | 1884 | 11 | 1.17% |
| hsa_circ_0065599 | 1884 | 11 | 1.17% |
| hsa_circ_0096755 | 1884 | 11 | 1.17% |
| hsa_circ_0014561 | 1713 | 10 | 1.17% |
| hsa_circ_0041362 | 1713 | 10 | 1.17% |
| hsa_circ_0076275 | 1713 | 10 | 1.17% |
| hsa_circ_0080631 | 1713 | 10 | 1.17% |
| hsa_circ_0125762 | 1713 | 10 | 1.17% |
| hsa_circ_0026009 | 2227 | 13 | 1.17% |

| | | | |
|---|---|---|---|
| hsa_circ_0031308 | 2056 | 12 | 1.17% |
| hsa_circ_0088379 | 2056 | 12 | 1.17% |
| hsa_circ_0126984 | 1885 | 11 | 1.17% |
| hsa_circ_0088283 | 2228 | 13 | 1.17% |
| hsa_circ_0020085 | 1714 | 10 | 1.17% |
| hsa_circ_0028903 | 1714 | 10 | 1.17% |
| hsa_circ_0033039 | 1714 | 10 | 1.17% |
| hsa_circ_0059482 | 1714 | 10 | 1.17% |
| hsa_circ_0073874 | 1714 | 10 | 1.17% |
| hsa_circ_0099630 | 1714 | 10 | 1.17% |
| hsa_circ_0118487 | 1714 | 10 | 1.17% |
| hsa_circ_0130340 | 1714 | 10 | 1.17% |
| hsa_circ_0131573 | 1714 | 10 | 1.17% |
| hsa_circ_0036550 | 2057 | 12 | 1.17% |
| hsa_circ_0063579 | 2057 | 12 | 1.17% |
| hsa_circ_0084827 | 2057 | 12 | 1.17% |
| hsa_circ_0010828 | 1886 | 11 | 1.17% |
| hsa_circ_0034242 | 2229 | 13 | 1.17% |
| hsa_circ_0084397 | 2229 | 13 | 1.17% |
| hsa_circ_0053126 | 2401 | 14 | 1.17% |
| hsa_circ_0048254 | 2058 | 12 | 1.17% |
| hsa_circ_0115369 | 2058 | 12 | 1.17% |
| hsa_circ_0002480 | 1715 | 10 | 1.17% |
| hsa_circ_0013599 | 1715 | 10 | 1.17% |
| hsa_circ_0016495 | 1715 | 10 | 1.17% |
| hsa_circ_0032050 | 1715 | 10 | 1.17% |
| hsa_circ_0036873 | 1715 | 10 | 1.17% |
| hsa_circ_0042598 | 1715 | 10 | 1.17% |
| hsa_circ_0056351 | 1715 | 10 | 1.17% |
| hsa_circ_0088934 | 1715 | 10 | 1.17% |
| hsa_circ_0114221 | 1715 | 10 | 1.17% |
| hsa_circ_0121984 | 1715 | 10 | 1.17% |
| hsa_circ_0057363 | 2916 | 17 | 1.17% |
| hsa_circ_0055675 | 2230 | 13 | 1.17% |
| hsa_circ_0097840 | 3774 | 22 | 1.17% |
| hsa_circ_0025670 | 1887 | 11 | 1.17% |
| hsa_circ_0025838 | 1887 | 11 | 1.17% |
| hsa_circ_0053782 | 1887 | 11 | 1.17% |
| hsa_circ_0070430 | 1887 | 11 | 1.17% |
| hsa_circ_0093846 | 1887 | 11 | 1.17% |
| hsa_circ_0135019 | 1887 | 11 | 1.17% |
| hsa_circ_0057747 | 3431 | 20 | 1.17% |
| hsa_circ_0124393 | 3431 | 20 | 1.17% |
| hsa_circ_0044442 | 1716 | 10 | 1.17% |
| hsa_circ_0066313 | 1716 | 10 | 1.17% |
| hsa_circ_0069373 | 1716 | 10 | 1.17% |
| hsa_circ_0077023 | 1716 | 10 | 1.17% |
| hsa_circ_0103672 | 1716 | 10 | 1.17% |
| hsa_circ_0106675 | 1716 | 10 | 1.17% |
| hsa_circ_0117619 | 1716 | 10 | 1.17% |

| | | | |
|---|---|---|---|
| hsa_circ_0061422 | 1888 | 11 | 1.17% |
| hsa_circ_0091240 | 1888 | 11 | 1.17% |
| hsa_circ_0080870 | 2060 | 12 | 1.17% |
| hsa_circ_0071531 | 2232 | 13 | 1.16% |
| hsa_circ_0090275 | 2232 | 13 | 1.16% |
| hsa_circ_0028222 | 1717 | 10 | 1.16% |
| hsa_circ_0038554 | 1717 | 10 | 1.16% |
| hsa_circ_0077329 | 1717 | 10 | 1.16% |
| hsa_circ_0087412 | 1717 | 10 | 1.16% |
| hsa_circ_0099756 | 1717 | 10 | 1.16% |
| hsa_circ_0006674 | 1889 | 11 | 1.16% |
| hsa_circ_0010008 | 1889 | 11 | 1.16% |
| hsa_circ_0027630 | 1889 | 11 | 1.16% |
| hsa_circ_0059904 | 1889 | 11 | 1.16% |
| hsa_circ_0127230 | 1889 | 11 | 1.16% |
| hsa_circ_0127998 | 1889 | 11 | 1.16% |
| hsa_circ_0025131 | 2061 | 12 | 1.16% |
| hsa_circ_0035162 | 2061 | 12 | 1.16% |
| hsa_circ_0109789 | 4981 | 29 | 1.16% |
| hsa_circ_0062791 | 3092 | 18 | 1.16% |
| hsa_circ_0052586 | 2405 | 14 | 1.16% |
| hsa_circ_0108723 | 2405 | 14 | 1.16% |
| hsa_circ_0010260 | 1718 | 10 | 1.16% |
| hsa_circ_0026192 | 1718 | 10 | 1.16% |
| hsa_circ_0045017 | 1718 | 10 | 1.16% |
| hsa_circ_0053236 | 1718 | 10 | 1.16% |
| hsa_circ_0058607 | 1718 | 10 | 1.16% |
| hsa_circ_0073432 | 1718 | 10 | 1.16% |
| hsa_circ_0088415 | 1718 | 10 | 1.16% |
| hsa_circ_0102642 | 1718 | 10 | 1.16% |
| hsa_circ_0134407 | 1718 | 10 | 1.16% |
| hsa_circ_0136206 | 6357 | 37 | 1.16% |
| hsa_circ_0085351 | 1890 | 11 | 1.16% |
| hsa_circ_0098546 | 1890 | 11 | 1.16% |
| hsa_circ_0054482 | 2062 | 12 | 1.16% |
| hsa_circ_0132508 | 2062 | 12 | 1.16% |
| hsa_circ_0032166 | 2234 | 13 | 1.16% |
| hsa_circ_0075415 | 2578 | 15 | 1.16% |
| hsa_circ_0008796 | 1719 | 10 | 1.16% |
| hsa_circ_0024561 | 1719 | 10 | 1.16% |
| hsa_circ_0073189 | 1719 | 10 | 1.16% |
| hsa_circ_0085139 | 1719 | 10 | 1.16% |
| hsa_circ_0096698 | 1719 | 10 | 1.16% |
| hsa_circ_0131919 | 1719 | 10 | 1.16% |
| hsa_circ_0030410 | 1891 | 11 | 1.16% |
| hsa_circ_0040647 | 1891 | 11 | 1.16% |
| hsa_circ_0045740 | 1891 | 11 | 1.16% |
| hsa_circ_0075941 | 1891 | 11 | 1.16% |
| hsa_circ_0085242 | 1891 | 11 | 1.16% |
| hsa_circ_0136433 | 1891 | 11 | 1.16% |

| | | | |
|---|---|---|---|
| hsa_circ_0013387 | 2063 | 12 | 1.16% |
| hsa_circ_0029568 | 2063 | 12 | 1.16% |
| hsa_circ_0030506 | 2063 | 12 | 1.16% |
| hsa_circ_0062218 | 2063 | 12 | 1.16% |
| hsa_circ_0066736 | 2063 | 12 | 1.16% |
| hsa_circ_0123920 | 2063 | 12 | 1.16% |
| hsa_circ_0113141 | 5332 | 31 | 1.16% |
| hsa_circ_0083130 | 2408 | 14 | 1.16% |
| hsa_circ_0009230 | 2064 | 12 | 1.16% |
| hsa_circ_0069128 | 2064 | 12 | 1.16% |
| hsa_circ_0073349 | 2064 | 12 | 1.16% |
| hsa_circ_0034655 | 1892 | 11 | 1.16% |
| hsa_circ_0048430 | 1892 | 11 | 1.16% |
| hsa_circ_0076557 | 1892 | 11 | 1.16% |
| hsa_circ_0098024 | 1892 | 11 | 1.16% |
| hsa_circ_0132761 | 1892 | 11 | 1.16% |
| hsa_circ_0023319 | 1720 | 10 | 1.16% |
| hsa_circ_0078706 | 1720 | 10 | 1.16% |
| hsa_circ_0084283 | 1720 | 10 | 1.16% |
| hsa_circ_0094971 | 1720 | 10 | 1.16% |
| hsa_circ_0121812 | 1720 | 10 | 1.16% |
| hsa_circ_0132220 | 1720 | 10 | 1.16% |
| hsa_circ_0009385 | 2065 | 12 | 1.16% |
| hsa_circ_0021628 | 2065 | 12 | 1.16% |
| hsa_circ_0083588 | 2065 | 12 | 1.16% |
| hsa_circ_0054880 | 1893 | 11 | 1.16% |
| hsa_circ_0078544 | 1893 | 11 | 1.16% |
| hsa_circ_0084320 | 1893 | 11 | 1.16% |
| hsa_circ_0119802 | 1893 | 11 | 1.16% |
| hsa_circ_0138276 | 1893 | 11 | 1.16% |
| hsa_circ_0021947 | 1721 | 10 | 1.16% |
| hsa_circ_0075815 | 1721 | 10 | 1.16% |
| hsa_circ_0078816 | 1721 | 10 | 1.16% |
| hsa_circ_0081275 | 1721 | 10 | 1.16% |
| hsa_circ_0086900 | 1721 | 10 | 1.16% |
| hsa_circ_0093581 | 1721 | 10 | 1.16% |
| hsa_circ_0123868 | 1721 | 10 | 1.16% |
| hsa_circ_0129001 | 1721 | 10 | 1.16% |
| hsa_circ_0048211 | 2582 | 15 | 1.16% |
| hsa_circ_0022934 | 2238 | 13 | 1.16% |
| hsa_circ_0089035 | 2238 | 13 | 1.16% |
| hsa_circ_0000912 | 6542 | 38 | 1.16% |
| hsa_circ_0096250 | 3099 | 18 | 1.16% |
| hsa_circ_0011516 | 2066 | 12 | 1.16% |
| hsa_circ_0022958 | 2066 | 12 | 1.16% |
| hsa_circ_0034592 | 2066 | 12 | 1.16% |
| hsa_circ_0075259 | 2066 | 12 | 1.16% |
| hsa_circ_0022857 | 1894 | 11 | 1.16% |
| hsa_circ_0023172 | 1894 | 11 | 1.16% |
| hsa_circ_0052814 | 1894 | 11 | 1.16% |

| | | | |
|---|---|---|---|
| hsa_circ_0115736 | 1894 | 11 | 1.16% |
| hsa_circ_0119793 | 1894 | 11 | 1.16% |
| hsa_circ_0140062 | 1894 | 11 | 1.16% |
| hsa_circ_0140339 | 1894 | 11 | 1.16% |
| hsa_circ_0121174 | 3444 | 20 | 1.16% |
| hsa_circ_0001613 | 1722 | 10 | 1.16% |
| hsa_circ_0009413 | 1722 | 10 | 1.16% |
| hsa_circ_0037944 | 1722 | 10 | 1.16% |
| hsa_circ_0041229 | 1722 | 10 | 1.16% |
| hsa_circ_0042582 | 1722 | 10 | 1.16% |
| hsa_circ_0042637 | 1722 | 10 | 1.16% |
| hsa_circ_0055966 | 1722 | 10 | 1.16% |
| hsa_circ_0061436 | 1722 | 10 | 1.16% |
| hsa_circ_0065560 | 1722 | 10 | 1.16% |
| hsa_circ_0095305 | 1722 | 10 | 1.16% |
| hsa_circ_0096854 | 1722 | 10 | 1.16% |
| hsa_circ_0100322 | 1722 | 10 | 1.16% |
| hsa_circ_0102405 | 1722 | 10 | 1.16% |
| hsa_circ_0119401 | 1722 | 10 | 1.16% |
| hsa_circ_0074037 | 2411 | 14 | 1.16% |
| hsa_circ_0068157 | 2067 | 12 | 1.16% |
| hsa_circ_0089232 | 2067 | 12 | 1.16% |
| hsa_circ_0104784 | 2067 | 12 | 1.16% |
| hsa_circ_0036335 | 1895 | 11 | 1.16% |
| hsa_circ_0042048 | 1895 | 11 | 1.16% |
| hsa_circ_0061669 | 1895 | 11 | 1.16% |
| hsa_circ_0067895 | 1895 | 11 | 1.16% |
| hsa_circ_0073699 | 1895 | 11 | 1.16% |
| hsa_circ_0097414 | 1895 | 11 | 1.16% |
| hsa_circ_0125031 | 1895 | 11 | 1.16% |
| hsa_circ_0140641 | 1895 | 11 | 1.16% |
| hsa_circ_0011169 | 1723 | 10 | 1.16% |
| hsa_circ_0038286 | 1723 | 10 | 1.16% |
| hsa_circ_0043057 | 1723 | 10 | 1.16% |
| hsa_circ_0053837 | 1723 | 10 | 1.16% |
| hsa_circ_0068964 | 1723 | 10 | 1.16% |
| hsa_circ_0081662 | 1723 | 10 | 1.16% |
| hsa_circ_0093647 | 1723 | 10 | 1.16% |
| hsa_circ_0058173 | 2240 | 13 | 1.16% |
| hsa_circ_0002095 | 1896 | 11 | 1.16% |
| hsa_circ_0018458 | 1896 | 11 | 1.16% |
| hsa_circ_0115112 | 1896 | 11 | 1.16% |
| hsa_circ_0120670 | 1896 | 11 | 1.16% |
| hsa_circ_0128710 | 1896 | 11 | 1.16% |
| hsa_circ_0063637 | 2241 | 13 | 1.16% |
| hsa_circ_0134924 | 2241 | 13 | 1.16% |
| hsa_circ_0047067 | 1724 | 10 | 1.16% |
| hsa_circ_0052354 | 1724 | 10 | 1.16% |
| hsa_circ_0078620 | 1724 | 10 | 1.16% |
| hsa_circ_0088152 | 1724 | 10 | 1.16% |

| | | | |
|---|---|---|---|
| hsa_circ_0104078 | 1724 | 10 | 1.16% |
| hsa_circ_0041036 | 2069 | 12 | 1.16% |
| hsa_circ_0112424 | 2069 | 12 | 1.16% |
| hsa_circ_0037194 | 2414 | 14 | 1.16% |
| hsa_circ_0021634 | 2759 | 16 | 1.16% |
| hsa_circ_0026723 | 1897 | 11 | 1.16% |
| hsa_circ_0104038 | 1897 | 11 | 1.16% |
| hsa_circ_0111476 | 1897 | 11 | 1.16% |
| hsa_circ_0013956 | 2242 | 13 | 1.16% |
| hsa_circ_0121323 | 2242 | 13 | 1.16% |
| hsa_circ_0026179 | 2587 | 15 | 1.16% |
| hsa_circ_0105351 | 4657 | 27 | 1.16% |
| hsa_circ_0128845 | 2415 | 14 | 1.16% |
| hsa_circ_0022979 | 2070 | 12 | 1.16% |
| hsa_circ_0023151 | 2070 | 12 | 1.16% |
| hsa_circ_0045828 | 2070 | 12 | 1.16% |
| hsa_circ_0047633 | 2070 | 12 | 1.16% |
| hsa_circ_0078446 | 2070 | 12 | 1.16% |
| hsa_circ_0089717 | 2070 | 12 | 1.16% |
| hsa_circ_0089997 | 2070 | 12 | 1.16% |
| hsa_circ_0011632 | 1725 | 10 | 1.16% |
| hsa_circ_0021147 | 1725 | 10 | 1.16% |
| hsa_circ_0024994 | 1725 | 10 | 1.16% |
| hsa_circ_0050813 | 1725 | 10 | 1.16% |
| hsa_circ_0053734 | 1725 | 10 | 1.16% |
| hsa_circ_0064223 | 1725 | 10 | 1.16% |
| hsa_circ_0106467 | 1725 | 10 | 1.16% |
| hsa_circ_0118450 | 1725 | 10 | 1.16% |
| hsa_circ_0137569 | 2588 | 15 | 1.16% |
| hsa_circ_0118076 | 2243 | 13 | 1.16% |
| hsa_circ_0128991 | 2243 | 13 | 1.16% |
| hsa_circ_0017081 | 1898 | 11 | 1.16% |
| hsa_circ_0110909 | 1898 | 11 | 1.16% |
| hsa_circ_0077679 | 2071 | 12 | 1.16% |
| hsa_circ_0001368 | 1726 | 10 | 1.16% |
| hsa_circ_0014564 | 1726 | 10 | 1.16% |
| hsa_circ_0015356 | 1726 | 10 | 1.16% |
| hsa_circ_0018960 | 1726 | 10 | 1.16% |
| hsa_circ_0020440 | 1726 | 10 | 1.16% |
| hsa_circ_0061884 | 1726 | 10 | 1.16% |
| hsa_circ_0077431 | 1726 | 10 | 1.16% |
| hsa_circ_0088792 | 1726 | 10 | 1.16% |
| hsa_circ_0097114 | 1726 | 10 | 1.16% |
| hsa_circ_0098071 | 1726 | 10 | 1.16% |
| hsa_circ_0103808 | 1726 | 10 | 1.16% |
| hsa_circ_0108679 | 1726 | 10 | 1.16% |
| hsa_circ_0126359 | 2244 | 13 | 1.16% |
| hsa_circ_0024376 | 3625 | 21 | 1.16% |
| hsa_circ_0012112 | 1899 | 11 | 1.16% |
| hsa_circ_0037868 | 1899 | 11 | 1.16% |

| | | | |
|---|---|---|---|
| hsa_circ_0058007 | 1899 | 11 | 1.16% |
| hsa_circ_0094190 | 1899 | 11 | 1.16% |
| hsa_circ_0103527 | 1899 | 11 | 1.16% |
| hsa_circ_0137059 | 1899 | 11 | 1.16% |
| hsa_circ_0007571 | 2072 | 12 | 1.16% |
| hsa_circ_0016370 | 2072 | 12 | 1.16% |
| hsa_circ_0058756 | 2072 | 12 | 1.16% |
| hsa_circ_0064108 | 2072 | 12 | 1.16% |
| hsa_circ_0140785 | 2072 | 12 | 1.16% |
| hsa_circ_0093916 | 6562 | 38 | 1.16% |
| hsa_circ_0114232 | 5526 | 32 | 1.16% |
| hsa_circ_0051149 | 2245 | 13 | 1.16% |
| hsa_circ_0032738 | 1727 | 10 | 1.16% |
| hsa_circ_0039604 | 1727 | 10 | 1.16% |
| hsa_circ_0042002 | 1727 | 10 | 1.16% |
| hsa_circ_0047313 | 1727 | 10 | 1.16% |
| hsa_circ_0066049 | 1727 | 10 | 1.16% |
| hsa_circ_0071379 | 1727 | 10 | 1.16% |
| hsa_circ_0083577 | 1727 | 10 | 1.16% |
| hsa_circ_0121865 | 4663 | 27 | 1.16% |
| hsa_circ_0012252 | 1900 | 11 | 1.16% |
| hsa_circ_0116026 | 1900 | 11 | 1.16% |
| hsa_circ_0037673 | 2073 | 12 | 1.16% |
| hsa_circ_0066108 | 2073 | 12 | 1.16% |
| hsa_circ_0108445 | 6048 | 35 | 1.16% |
| hsa_circ_0027982 | 1728 | 10 | 1.16% |
| hsa_circ_0079903 | 1728 | 10 | 1.16% |
| hsa_circ_0108882 | 1728 | 10 | 1.16% |
| hsa_circ_0119085 | 1728 | 10 | 1.16% |
| hsa_circ_0009390 | 1901 | 11 | 1.16% |
| hsa_circ_0012159 | 1901 | 11 | 1.16% |
| hsa_circ_0019351 | 1901 | 11 | 1.16% |
| hsa_circ_0020390 | 1901 | 11 | 1.16% |
| hsa_circ_0024808 | 1901 | 11 | 1.16% |
| hsa_circ_0047673 | 1901 | 11 | 1.16% |
| hsa_circ_0103501 | 1901 | 11 | 1.16% |
| hsa_circ_0114263 | 1901 | 11 | 1.16% |
| hsa_circ_0058330 | 2074 | 12 | 1.16% |
| hsa_circ_0071583 | 2074 | 12 | 1.16% |
| hsa_circ_0100017 | 2074 | 12 | 1.16% |
| hsa_circ_0026215 | 1729 | 10 | 1.16% |
| hsa_circ_0043265 | 1729 | 10 | 1.16% |
| hsa_circ_0048536 | 1729 | 10 | 1.16% |
| hsa_circ_0084255 | 1729 | 10 | 1.16% |
| hsa_circ_0100601 | 1729 | 10 | 1.16% |
| hsa_circ_0121418 | 1729 | 10 | 1.16% |
| hsa_circ_0030940 | 1902 | 11 | 1.16% |
| hsa_circ_0059006 | 1902 | 11 | 1.16% |
| hsa_circ_0118144 | 1902 | 11 | 1.16% |
| hsa_circ_0102802 | 2075 | 12 | 1.16% |

| | | | |
|---|---|---|---|
| hsa_circ_0105276 | 2248 | 13 | 1.16% |
| hsa_circ_0034743 | 2421 | 14 | 1.16% |
| hsa_circ_0029485 | 2422 | 14 | 1.16% |
| hsa_circ_0036846 | 2249 | 13 | 1.16% |
| hsa_circ_0017578 | 2076 | 12 | 1.16% |
| hsa_circ_0013133 | 1903 | 11 | 1.16% |
| hsa_circ_0016808 | 1903 | 11 | 1.16% |
| hsa_circ_0017106 | 1903 | 11 | 1.16% |
| hsa_circ_0038183 | 1903 | 11 | 1.16% |
| hsa_circ_0045335 | 1903 | 11 | 1.16% |
| hsa_circ_0054519 | 1903 | 11 | 1.16% |
| hsa_circ_0088900 | 1903 | 11 | 1.16% |
| hsa_circ_0092203 | 1903 | 11 | 1.16% |
| hsa_circ_0011543 | 1730 | 10 | 1.16% |
| hsa_circ_0021257 | 1730 | 10 | 1.16% |
| hsa_circ_0041075 | 1730 | 10 | 1.16% |
| hsa_circ_0063121 | 1730 | 10 | 1.16% |
| hsa_circ_0101852 | 1730 | 10 | 1.16% |
| hsa_circ_0102129 | 1730 | 10 | 1.16% |
| hsa_circ_0113110 | 1730 | 10 | 1.16% |
| hsa_circ_0115543 | 1730 | 10 | 1.16% |
| hsa_circ_0127889 | 1730 | 10 | 1.16% |
| hsa_circ_0139963 | 1730 | 10 | 1.16% |
| hsa_circ_0044508 | 3980 | 23 | 1.16% |
| hsa_circ_0056864 | 2942 | 17 | 1.16% |
| hsa_circ_0056703 | 6577 | 38 | 1.16% |
| hsa_circ_0021962 | 1904 | 11 | 1.16% |
| hsa_circ_0071642 | 1904 | 11 | 1.16% |
| hsa_circ_0116135 | 1904 | 11 | 1.16% |
| hsa_circ_0064929 | 1731 | 10 | 1.16% |
| hsa_circ_0067938 | 1731 | 10 | 1.16% |
| hsa_circ_0086121 | 1731 | 10 | 1.16% |
| hsa_circ_0118888 | 1731 | 10 | 1.16% |
| hsa_circ_0127523 | 1731 | 10 | 1.16% |
| hsa_circ_0127954 | 1731 | 10 | 1.16% |
| hsa_circ_0091300 | 2597 | 15 | 1.16% |
| hsa_circ_0097653 | 2251 | 13 | 1.16% |
| hsa_circ_0010382 | 2078 | 12 | 1.15% |
| hsa_circ_0034434 | 2078 | 12 | 1.15% |
| hsa_circ_0049496 | 2078 | 12 | 1.15% |
| hsa_circ_0014028 | 1905 | 11 | 1.15% |
| hsa_circ_0082737 | 1905 | 11 | 1.15% |
| hsa_circ_0091034 | 1905 | 11 | 1.15% |
| hsa_circ_0120665 | 1905 | 11 | 1.15% |
| hsa_circ_0132210 | 2598 | 15 | 1.15% |
| hsa_circ_0049286 | 1732 | 10 | 1.15% |
| hsa_circ_0064569 | 1732 | 10 | 1.15% |
| hsa_circ_0109573 | 1732 | 10 | 1.15% |
| hsa_circ_0107607 | 2425 | 14 | 1.15% |
| hsa_circ_0052129 | 3118 | 18 | 1.15% |

| | | | |
|---|---|---|---|
| hsa_circ_0018932 | 2252 | 13 | 1.15% |
| hsa_circ_0067858 | 2252 | 13 | 1.15% |
| hsa_circ_0048844 | 7103 | 41 | 1.15% |
| hsa_circ_0027387 | 2079 | 12 | 1.15% |
| hsa_circ_0046829 | 2079 | 12 | 1.15% |
| hsa_circ_0078905 | 2079 | 12 | 1.15% |
| hsa_circ_0080396 | 2079 | 12 | 1.15% |
| hsa_circ_0081044 | 2079 | 12 | 1.15% |
| hsa_circ_0132488 | 2079 | 12 | 1.15% |
| hsa_circ_0137270 | 2079 | 12 | 1.15% |
| hsa_circ_0103915 | 2599 | 15 | 1.15% |
| hsa_circ_0012426 | 1906 | 11 | 1.15% |
| hsa_circ_0030057 | 1906 | 11 | 1.15% |
| hsa_circ_0046269 | 1906 | 11 | 1.15% |
| hsa_circ_0052851 | 1906 | 11 | 1.15% |
| hsa_circ_0073859 | 1906 | 11 | 1.15% |
| hsa_circ_0086093 | 1906 | 11 | 1.15% |
| hsa_circ_0054961 | 2426 | 14 | 1.15% |
| hsa_circ_0046125 | 1733 | 10 | 1.15% |
| hsa_circ_0072677 | 1733 | 10 | 1.15% |
| hsa_circ_0077102 | 1733 | 10 | 1.15% |
| hsa_circ_0086268 | 1733 | 10 | 1.15% |
| hsa_circ_0093841 | 1733 | 10 | 1.15% |
| hsa_circ_0102956 | 1733 | 10 | 1.15% |
| hsa_circ_0105275 | 2253 | 13 | 1.15% |
| hsa_circ_0113058 | 4507 | 26 | 1.15% |
| hsa_circ_0018993 | 1907 | 11 | 1.15% |
| hsa_circ_0061355 | 1907 | 11 | 1.15% |
| hsa_circ_0073370 | 1907 | 11 | 1.15% |
| hsa_circ_0091388 | 1907 | 11 | 1.15% |
| hsa_circ_0114166 | 1907 | 11 | 1.15% |
| hsa_circ_0128410 | 1907 | 11 | 1.15% |
| hsa_circ_0055407 | 9882 | 57 | 1.15% |
| hsa_circ_0050333 | 1734 | 10 | 1.15% |
| hsa_circ_0050371 | 1734 | 10 | 1.15% |
| hsa_circ_0059811 | 1734 | 10 | 1.15% |
| hsa_circ_0063952 | 1734 | 10 | 1.15% |
| hsa_circ_0070164 | 1734 | 10 | 1.15% |
| hsa_circ_0078839 | 1734 | 10 | 1.15% |
| hsa_circ_0079266 | 1734 | 10 | 1.15% |
| hsa_circ_0082246 | 1734 | 10 | 1.15% |
| hsa_circ_0084892 | 1734 | 10 | 1.15% |
| hsa_circ_0085360 | 1734 | 10 | 1.15% |
| hsa_circ_0109808 | 1734 | 10 | 1.15% |
| hsa_circ_0136560 | 1734 | 10 | 1.15% |
| hsa_circ_0006693 | 2081 | 12 | 1.15% |
| hsa_circ_0050780 | 2081 | 12 | 1.15% |
| hsa_circ_0134489 | 5550 | 32 | 1.15% |
| hsa_circ_0030712 | 1908 | 11 | 1.15% |
| hsa_circ_0065348 | 1908 | 11 | 1.15% |

| | | | |
|---|---|---|---|
| hsa_circ_0076332 | 1908 | 11 | 1.15% |
| hsa_circ_0097802 | 1908 | 11 | 1.15% |
| hsa_circ_0040317 | 2082 | 12 | 1.15% |
| hsa_circ_0043571 | 2082 | 12 | 1.15% |
| hsa_circ_0042280 | 1735 | 10 | 1.15% |
| hsa_circ_0048620 | 1735 | 10 | 1.15% |
| hsa_circ_0135517 | 1735 | 10 | 1.15% |
| hsa_circ_0026344 | 2603 | 15 | 1.15% |
| hsa_circ_0010616 | 2256 | 13 | 1.15% |
| hsa_circ_0046168 | 2256 | 13 | 1.15% |
| hsa_circ_0095852 | 2256 | 13 | 1.15% |
| hsa_circ_0005242 | 1909 | 11 | 1.15% |
| hsa_circ_0012200 | 1909 | 11 | 1.15% |
| hsa_circ_0035761 | 1909 | 11 | 1.15% |
| hsa_circ_0069196 | 1909 | 11 | 1.15% |
| hsa_circ_0078858 | 1909 | 11 | 1.15% |
| hsa_circ_0130792 | 1909 | 11 | 1.15% |
| hsa_circ_0017381 | 2083 | 12 | 1.15% |
| hsa_circ_0001142 | 1736 | 10 | 1.15% |
| hsa_circ_0011880 | 1736 | 10 | 1.15% |
| hsa_circ_0014386 | 1736 | 10 | 1.15% |
| hsa_circ_0017401 | 1736 | 10 | 1.15% |
| hsa_circ_0034719 | 1736 | 10 | 1.15% |
| hsa_circ_0040396 | 1736 | 10 | 1.15% |
| hsa_circ_0054084 | 1736 | 10 | 1.15% |
| hsa_circ_0061882 | 1736 | 10 | 1.15% |
| hsa_circ_0078676 | 1736 | 10 | 1.15% |
| hsa_circ_0088943 | 1736 | 10 | 1.15% |
| hsa_circ_0121365 | 1736 | 10 | 1.15% |
| hsa_circ_0135450 | 1736 | 10 | 1.15% |
| hsa_circ_0062858 | 2257 | 13 | 1.15% |
| hsa_circ_0000544 | 1910 | 11 | 1.15% |
| hsa_circ_0011881 | 1910 | 11 | 1.15% |
| hsa_circ_0041140 | 1910 | 11 | 1.15% |
| hsa_circ_0047178 | 1910 | 11 | 1.15% |
| hsa_circ_0048141 | 1910 | 11 | 1.15% |
| hsa_circ_0059727 | 1910 | 11 | 1.15% |
| hsa_circ_0090788 | 1910 | 11 | 1.15% |
| hsa_circ_0098412 | 1910 | 11 | 1.15% |
| hsa_circ_0049157 | 6772 | 39 | 1.15% |
| hsa_circ_0062842 | 2084 | 12 | 1.15% |
| hsa_circ_0069276 | 2084 | 12 | 1.15% |
| hsa_circ_0128781 | 2084 | 12 | 1.15% |
| hsa_circ_0004514 | 2258 | 13 | 1.15% |
| hsa_circ_0018445 | 1737 | 10 | 1.15% |
| hsa_circ_0022279 | 1737 | 10 | 1.15% |
| hsa_circ_0034556 | 1737 | 10 | 1.15% |
| hsa_circ_0037217 | 1737 | 10 | 1.15% |
| hsa_circ_0056165 | 1737 | 10 | 1.15% |
| hsa_circ_0059821 | 1737 | 10 | 1.15% |

| | | | |
|---|---|---|---|
| hsa_circ_0067952 | 1737 | 10 | 1.15% |
| hsa_circ_0078095 | 1737 | 10 | 1.15% |
| hsa_circ_0097940 | 1737 | 10 | 1.15% |
| hsa_circ_0098077 | 1737 | 10 | 1.15% |
| hsa_circ_0112199 | 1737 | 10 | 1.15% |
| hsa_circ_0139075 | 1737 | 10 | 1.15% |
| hsa_circ_0049794 | 2432 | 14 | 1.15% |
| hsa_circ_0090123 | 2432 | 14 | 1.15% |
| hsa_circ_0030150 | 1911 | 11 | 1.15% |
| hsa_circ_0043094 | 1911 | 11 | 1.15% |
| hsa_circ_0043841 | 1911 | 11 | 1.15% |
| hsa_circ_0053686 | 1911 | 11 | 1.15% |
| hsa_circ_0076449 | 1911 | 11 | 1.15% |
| hsa_circ_0085929 | 1911 | 11 | 1.15% |
| hsa_circ_0086689 | 1911 | 11 | 1.15% |
| hsa_circ_0068889 | 2085 | 12 | 1.15% |
| hsa_circ_0128697 | 5039 | 29 | 1.15% |
| hsa_circ_0011439 | 1738 | 10 | 1.15% |
| hsa_circ_0026628 | 1738 | 10 | 1.15% |
| hsa_circ_0036919 | 1738 | 10 | 1.15% |
| hsa_circ_0043126 | 1738 | 10 | 1.15% |
| hsa_circ_0057818 | 1738 | 10 | 1.15% |
| hsa_circ_0062241 | 1738 | 10 | 1.15% |
| hsa_circ_0081834 | 1738 | 10 | 1.15% |
| hsa_circ_0085086 | 1738 | 10 | 1.15% |
| hsa_circ_0107714 | 1738 | 10 | 1.15% |
| hsa_circ_0129961 | 1738 | 10 | 1.15% |
| hsa_circ_0132307 | 1738 | 10 | 1.15% |
| hsa_circ_0136172 | 1738 | 10 | 1.15% |
| hsa_circ_0107308 | 6431 | 37 | 1.15% |
| hsa_circ_0089685 | 1912 | 11 | 1.15% |
| hsa_circ_0096529 | 1912 | 11 | 1.15% |
| hsa_circ_0022454 | 2086 | 12 | 1.15% |
| hsa_circ_0025102 | 2086 | 12 | 1.15% |
| hsa_circ_0073473 | 2086 | 12 | 1.15% |
| hsa_circ_0014049 | 3303 | 19 | 1.15% |
| hsa_circ_0063792 | 1739 | 10 | 1.15% |
| hsa_circ_0066578 | 1739 | 10 | 1.15% |
| hsa_circ_0086773 | 1739 | 10 | 1.15% |
| hsa_circ_0129613 | 1739 | 10 | 1.15% |
| hsa_circ_0135034 | 1739 | 10 | 1.15% |
| hsa_circ_0053360 | 1913 | 11 | 1.15% |
| hsa_circ_0069556 | 1913 | 11 | 1.15% |
| hsa_circ_0071802 | 1913 | 11 | 1.15% |
| hsa_circ_0108436 | 1913 | 11 | 1.15% |
| hsa_circ_0131922 | 1913 | 11 | 1.15% |
| hsa_circ_0140749 | 1913 | 11 | 1.15% |
| hsa_circ_0002815 | 2087 | 12 | 1.15% |
| hsa_circ_0093594 | 4348 | 25 | 1.15% |
| hsa_circ_0021517 | 5392 | 31 | 1.15% |

| | | | |
|---|---|---|---|
| hsa_circ_0001873 | 2957 | 17 | 1.15% |
| hsa_circ_0039449 | 2262 | 13 | 1.15% |
| hsa_circ_0063483 | 2088 | 12 | 1.15% |
| hsa_circ_0119119 | 2088 | 12 | 1.15% |
| hsa_circ_0003190 | 1914 | 11 | 1.15% |
| hsa_circ_0009584 | 1914 | 11 | 1.15% |
| hsa_circ_0010546 | 1914 | 11 | 1.15% |
| hsa_circ_0011878 | 1914 | 11 | 1.15% |
| hsa_circ_0012885 | 1914 | 11 | 1.15% |
| hsa_circ_0022264 | 1914 | 11 | 1.15% |
| hsa_circ_0032089 | 1914 | 11 | 1.15% |
| hsa_circ_0062004 | 1914 | 11 | 1.15% |
| hsa_circ_0067887 | 1914 | 11 | 1.15% |
| hsa_circ_0094367 | 1914 | 11 | 1.15% |
| hsa_circ_0094834 | 1914 | 11 | 1.15% |
| hsa_circ_0116136 | 1914 | 11 | 1.15% |
| hsa_circ_0005063 | 1740 | 10 | 1.15% |
| hsa_circ_0044560 | 1740 | 10 | 1.15% |
| hsa_circ_0047565 | 1740 | 10 | 1.15% |
| hsa_circ_0049639 | 1740 | 10 | 1.15% |
| hsa_circ_0060249 | 1740 | 10 | 1.15% |
| hsa_circ_0072859 | 1740 | 10 | 1.15% |
| hsa_circ_0078729 | 1740 | 10 | 1.15% |
| hsa_circ_0099022 | 1740 | 10 | 1.15% |
| hsa_circ_0102386 | 1740 | 10 | 1.15% |
| hsa_circ_0128583 | 3133 | 18 | 1.15% |
| hsa_circ_0072319 | 2611 | 15 | 1.15% |
| hsa_circ_0058816 | 2263 | 13 | 1.15% |
| hsa_circ_0025571 | 1915 | 11 | 1.15% |
| hsa_circ_0040048 | 1915 | 11 | 1.15% |
| hsa_circ_0056381 | 1915 | 11 | 1.15% |
| hsa_circ_0137594 | 1915 | 11 | 1.15% |
| hsa_circ_0011012 | 1741 | 10 | 1.15% |
| hsa_circ_0025604 | 1741 | 10 | 1.15% |
| hsa_circ_0034775 | 1741 | 10 | 1.15% |
| hsa_circ_0035756 | 1741 | 10 | 1.15% |
| hsa_circ_0042306 | 1741 | 10 | 1.15% |
| hsa_circ_0056965 | 1741 | 10 | 1.15% |
| hsa_circ_0086983 | 1741 | 10 | 1.15% |
| hsa_circ_0117600 | 1741 | 10 | 1.15% |
| hsa_circ_0120230 | 1741 | 10 | 1.15% |
| hsa_circ_0136675 | 1741 | 10 | 1.15% |
| hsa_circ_0061979 | 2090 | 12 | 1.15% |
| hsa_circ_0074777 | 2090 | 12 | 1.15% |
| hsa_circ_0115396 | 2090 | 12 | 1.15% |
| hsa_circ_0017484 | 1916 | 11 | 1.15% |
| hsa_circ_0064285 | 1916 | 11 | 1.15% |
| hsa_circ_0065033 | 1916 | 11 | 1.15% |
| hsa_circ_0065677 | 1916 | 11 | 1.15% |
| hsa_circ_0066508 | 1916 | 11 | 1.15% |

| | | | |
|---|---|---|---|
| hsa_circ_0074926 | 1916 | 11 | 1.15% |
| hsa_circ_0112207 | 1916 | 11 | 1.15% |
| hsa_circ_0083560 | 2613 | 15 | 1.15% |
| hsa_circ_0017728 | 1742 | 10 | 1.15% |
| hsa_circ_0039276 | 1742 | 10 | 1.15% |
| hsa_circ_0040165 | 1742 | 10 | 1.15% |
| hsa_circ_0084259 | 1742 | 10 | 1.15% |
| hsa_circ_0126426 | 1742 | 10 | 1.15% |
| hsa_circ_0044443 | 2091 | 12 | 1.15% |
| hsa_circ_0028926 | 1917 | 11 | 1.15% |
| hsa_circ_0038115 | 1917 | 11 | 1.15% |
| hsa_circ_0038526 | 1917 | 11 | 1.15% |
| hsa_circ_0080572 | 1917 | 11 | 1.15% |
| hsa_circ_0137733 | 6971 | 40 | 1.15% |
| hsa_circ_0012604 | 1743 | 10 | 1.15% |
| hsa_circ_0025417 | 1743 | 10 | 1.15% |
| hsa_circ_0029664 | 1743 | 10 | 1.15% |
| hsa_circ_0044493 | 1743 | 10 | 1.15% |
| hsa_circ_0048089 | 1743 | 10 | 1.15% |
| hsa_circ_0068907 | 1743 | 10 | 1.15% |
| hsa_circ_0085171 | 1743 | 10 | 1.15% |
| hsa_circ_0135512 | 1743 | 10 | 1.15% |
| hsa_circ_0138921 | 1743 | 10 | 1.15% |
| hsa_circ_0139062 | 1743 | 10 | 1.15% |
| hsa_circ_0009275 | 2789 | 16 | 1.15% |
| hsa_circ_0001077 | 2092 | 12 | 1.15% |
| hsa_circ_0024878 | 2092 | 12 | 1.15% |
| hsa_circ_0039880 | 2092 | 12 | 1.15% |
| hsa_circ_0054444 | 2092 | 12 | 1.15% |
| hsa_circ_0067585 | 2092 | 12 | 1.15% |
| hsa_circ_0114461 | 2092 | 12 | 1.15% |
| hsa_circ_0120624 | 2092 | 12 | 1.15% |
| hsa_circ_0011330 | 2441 | 14 | 1.15% |
| hsa_circ_0018619 | 1918 | 11 | 1.15% |
| hsa_circ_0034192 | 1918 | 11 | 1.15% |
| hsa_circ_0054864 | 1918 | 11 | 1.15% |
| hsa_circ_0074787 | 1918 | 11 | 1.15% |
| hsa_circ_0076644 | 1918 | 11 | 1.15% |
| hsa_circ_0078634 | 1918 | 11 | 1.15% |
| hsa_circ_0083385 | 1918 | 11 | 1.15% |
| hsa_circ_0084525 | 1918 | 11 | 1.15% |
| hsa_circ_0048263 | 2267 | 13 | 1.15% |
| hsa_circ_0020375 | 1744 | 10 | 1.15% |
| hsa_circ_0033352 | 1744 | 10 | 1.15% |
| hsa_circ_0073574 | 1744 | 10 | 1.15% |
| hsa_circ_0088234 | 1744 | 10 | 1.15% |
| hsa_circ_0088684 | 1744 | 10 | 1.15% |
| hsa_circ_0120693 | 1744 | 10 | 1.15% |
| hsa_circ_0140288 | 1744 | 10 | 1.15% |
| hsa_circ_0079002 | 2093 | 12 | 1.15% |

| | | | |
|---|---|---|---|
| hsa_circ_0009442 | 1919 | 11 | 1.15% |
| hsa_circ_0017799 | 1919 | 11 | 1.15% |
| hsa_circ_0047171 | 1919 | 11 | 1.15% |
| hsa_circ_0072118 | 1919 | 11 | 1.15% |
| hsa_circ_0115814 | 1919 | 11 | 1.15% |
| hsa_circ_0058491 | 2617 | 15 | 1.15% |
| hsa_circ_0003963 | 1745 | 10 | 1.15% |
| hsa_circ_0011126 | 1745 | 10 | 1.15% |
| hsa_circ_0020014 | 1745 | 10 | 1.15% |
| hsa_circ_0022221 | 1745 | 10 | 1.15% |
| hsa_circ_0082386 | 1745 | 10 | 1.15% |
| hsa_circ_0122608 | 1745 | 10 | 1.15% |
| hsa_circ_0010990 | 4014 | 23 | 1.15% |
| hsa_circ_0086112 | 2269 | 13 | 1.15% |
| hsa_circ_0105851 | 6633 | 38 | 1.15% |
| hsa_circ_0029015 | 6460 | 37 | 1.15% |
| hsa_circ_0019459 | 1746 | 10 | 1.15% |
| hsa_circ_0071226 | 1746 | 10 | 1.15% |
| hsa_circ_0084887 | 1746 | 10 | 1.15% |
| hsa_circ_0113965 | 1746 | 10 | 1.15% |
| hsa_circ_0020141 | 2270 | 13 | 1.15% |
| hsa_circ_0092110 | 2270 | 13 | 1.15% |
| hsa_circ_0097649 | 2270 | 13 | 1.15% |
| hsa_circ_0017215 | 1921 | 11 | 1.15% |
| hsa_circ_0028805 | 1921 | 11 | 1.15% |
| hsa_circ_0042727 | 1921 | 11 | 1.15% |
| hsa_circ_0065645 | 1921 | 11 | 1.15% |
| hsa_circ_0087679 | 1921 | 11 | 1.15% |
| hsa_circ_0079399 | 3493 | 20 | 1.15% |
| hsa_circ_0032923 | 1747 | 10 | 1.14% |
| hsa_circ_0036962 | 1747 | 10 | 1.14% |
| hsa_circ_0038224 | 1747 | 10 | 1.14% |
| hsa_circ_0044936 | 1747 | 10 | 1.14% |
| hsa_circ_0049240 | 1747 | 10 | 1.14% |
| hsa_circ_0052919 | 1747 | 10 | 1.14% |
| hsa_circ_0075327 | 1747 | 10 | 1.14% |
| hsa_circ_0081624 | 1747 | 10 | 1.14% |
| hsa_circ_0083072 | 1747 | 10 | 1.14% |
| hsa_circ_0115424 | 1747 | 10 | 1.14% |
| hsa_circ_0117377 | 1747 | 10 | 1.14% |
| hsa_circ_0030870 | 2446 | 14 | 1.14% |
| hsa_circ_0009475 | 1922 | 11 | 1.14% |
| hsa_circ_0012986 | 1922 | 11 | 1.14% |
| hsa_circ_0020775 | 1922 | 11 | 1.14% |
| hsa_circ_0055894 | 1922 | 11 | 1.14% |
| hsa_circ_0005442 | 2097 | 12 | 1.14% |
| hsa_circ_0031511 | 2097 | 12 | 1.14% |
| hsa_circ_0031597 | 2097 | 12 | 1.14% |
| hsa_circ_0089674 | 2097 | 12 | 1.14% |
| hsa_circ_0011310 | 2272 | 13 | 1.14% |

| | | | |
|---|---|---|---|
| hsa_circ_0119434 | 2272 | 13 | 1.14% |
| hsa_circ_0001844 | 1748 | 10 | 1.14% |
| hsa_circ_0053731 | 1748 | 10 | 1.14% |
| hsa_circ_0058474 | 1748 | 10 | 1.14% |
| hsa_circ_0062770 | 1748 | 10 | 1.14% |
| hsa_circ_0072237 | 1748 | 10 | 1.14% |
| hsa_circ_0072667 | 1748 | 10 | 1.14% |
| hsa_circ_0086821 | 1748 | 10 | 1.14% |
| hsa_circ_0134349 | 1748 | 10 | 1.14% |
| hsa_circ_0030673 | 1923 | 11 | 1.14% |
| hsa_circ_0051681 | 1923 | 11 | 1.14% |
| hsa_circ_0055666 | 1923 | 11 | 1.14% |
| hsa_circ_0058695 | 1923 | 11 | 1.14% |
| hsa_circ_0100909 | 1923 | 11 | 1.14% |
| hsa_circ_0105544 | 1923 | 11 | 1.14% |
| hsa_circ_0114520 | 1923 | 11 | 1.14% |
| hsa_circ_0138430 | 2098 | 12 | 1.14% |
| hsa_circ_0096956 | 11718 | 67 | 1.14% |
| hsa_circ_0012783 | 1749 | 10 | 1.14% |
| hsa_circ_0030384 | 1749 | 10 | 1.14% |
| hsa_circ_0032567 | 1749 | 10 | 1.14% |
| hsa_circ_0054576 | 1749 | 10 | 1.14% |
| hsa_circ_0067328 | 1749 | 10 | 1.14% |
| hsa_circ_0109333 | 1749 | 10 | 1.14% |
| hsa_circ_0136287 | 1749 | 10 | 1.14% |
| hsa_circ_0047906 | 1924 | 11 | 1.14% |
| hsa_circ_0137744 | 1924 | 11 | 1.14% |
| hsa_circ_0006091 | 2099 | 12 | 1.14% |
| hsa_circ_0040256 | 2099 | 12 | 1.14% |
| hsa_circ_0092049 | 2099 | 12 | 1.14% |
| hsa_circ_0069267 | 4548 | 26 | 1.14% |
| hsa_circ_0119384 | 2274 | 13 | 1.14% |
| hsa_circ_0055902 | 7347 | 42 | 1.14% |
| hsa_circ_0085038 | 2449 | 14 | 1.14% |
| hsa_circ_0100210 | 6298 | 36 | 1.14% |
| hsa_circ_0007301 | 10497 | 60 | 1.14% |
| hsa_circ_0049516 | 2275 | 13 | 1.14% |
| hsa_circ_0081400 | 2275 | 13 | 1.14% |
| hsa_circ_0090272 | 2275 | 13 | 1.14% |
| hsa_circ_0078083 | 2100 | 12 | 1.14% |
| hsa_circ_0086148 | 2100 | 12 | 1.14% |
| hsa_circ_0013983 | 1925 | 11 | 1.14% |
| hsa_circ_0022807 | 1925 | 11 | 1.14% |
| hsa_circ_0017610 | 1750 | 10 | 1.14% |
| hsa_circ_0025168 | 1750 | 10 | 1.14% |
| hsa_circ_0030583 | 1750 | 10 | 1.14% |
| hsa_circ_0057127 | 1750 | 10 | 1.14% |
| hsa_circ_0078599 | 1750 | 10 | 1.14% |
| hsa_circ_0108472 | 1750 | 10 | 1.14% |
| hsa_circ_0120262 | 1750 | 10 | 1.14% |

| | | | |
|---|---|---|---|
| hsa_circ_0136656 | 1750 | 10 | 1.14% |
| hsa_circ_0102385 | 2101 | 12 | 1.14% |
| hsa_circ_0004658 | 1926 | 11 | 1.14% |
| hsa_circ_0017934 | 1926 | 11 | 1.14% |
| hsa_circ_0029718 | 1926 | 11 | 1.14% |
| hsa_circ_0043968 | 1926 | 11 | 1.14% |
| hsa_circ_0065900 | 1926 | 11 | 1.14% |
| hsa_circ_0072048 | 1926 | 11 | 1.14% |
| hsa_circ_0121150 | 1926 | 11 | 1.14% |
| hsa_circ_0127084 | 1926 | 11 | 1.14% |
| hsa_circ_0027584 | 1751 | 10 | 1.14% |
| hsa_circ_0108875 | 1751 | 10 | 1.14% |
| hsa_circ_0010637 | 2277 | 13 | 1.14% |
| hsa_circ_0025384 | 2102 | 12 | 1.14% |
| hsa_circ_0063735 | 2102 | 12 | 1.14% |
| hsa_circ_0085672 | 2102 | 12 | 1.14% |
| hsa_circ_0004326 | 1927 | 11 | 1.14% |
| hsa_circ_0034755 | 1927 | 11 | 1.14% |
| hsa_circ_0076241 | 1927 | 11 | 1.14% |
| hsa_circ_0095472 | 1927 | 11 | 1.14% |
| hsa_circ_0127254 | 1927 | 11 | 1.14% |
| hsa_circ_0016092 | 1752 | 10 | 1.14% |
| hsa_circ_0029925 | 1752 | 10 | 1.14% |
| hsa_circ_0045013 | 1752 | 10 | 1.14% |
| hsa_circ_0053919 | 1752 | 10 | 1.14% |
| hsa_circ_0069882 | 1752 | 10 | 1.14% |
| hsa_circ_0082622 | 1752 | 10 | 1.14% |
| hsa_circ_0136048 | 1752 | 10 | 1.14% |
| hsa_circ_0091486 | 2103 | 12 | 1.14% |
| hsa_circ_0138835 | 2103 | 12 | 1.14% |
| hsa_circ_0023040 | 1928 | 11 | 1.14% |
| hsa_circ_0023567 | 1928 | 11 | 1.14% |
| hsa_circ_0063943 | 1928 | 11 | 1.14% |
| hsa_circ_0067154 | 1928 | 11 | 1.14% |
| hsa_circ_0089205 | 1928 | 11 | 1.14% |
| hsa_circ_0140675 | 1928 | 11 | 1.14% |
| hsa_circ_0048075 | 2454 | 14 | 1.14% |
| hsa_circ_0075478 | 2454 | 14 | 1.14% |
| hsa_circ_0056785 | 1753 | 10 | 1.14% |
| hsa_circ_0075307 | 1753 | 10 | 1.14% |
| hsa_circ_0102961 | 1753 | 10 | 1.14% |
| hsa_circ_0107977 | 1753 | 10 | 1.14% |
| hsa_circ_0138616 | 1753 | 10 | 1.14% |
| hsa_circ_0084398 | 2279 | 13 | 1.14% |
| hsa_circ_0093208 | 4909 | 28 | 1.14% |
| hsa_circ_0049186 | 2104 | 12 | 1.14% |
| hsa_circ_0053110 | 2104 | 12 | 1.14% |
| hsa_circ_0102097 | 2104 | 12 | 1.14% |
| hsa_circ_0127211 | 2104 | 12 | 1.14% |
| hsa_circ_0132500 | 2104 | 12 | 1.14% |

| | | | |
|---|---|---|---|
| hsa_circ_0016786 | 1929 | 11 | 1.14% |
| hsa_circ_0026161 | 1929 | 11 | 1.14% |
| hsa_circ_0038750 | 1929 | 11 | 1.14% |
| hsa_circ_0058117 | 1929 | 11 | 1.14% |
| hsa_circ_0130483 | 1929 | 11 | 1.14% |
| hsa_circ_0015116 | 2280 | 13 | 1.14% |
| hsa_circ_0044621 | 2280 | 13 | 1.14% |
| hsa_circ_0132574 | 8419 | 48 | 1.14% |
| hsa_circ_0011782 | 1754 | 10 | 1.14% |
| hsa_circ_0020956 | 1754 | 10 | 1.14% |
| hsa_circ_0036139 | 1754 | 10 | 1.14% |
| hsa_circ_0039124 | 1754 | 10 | 1.14% |
| hsa_circ_0042084 | 1754 | 10 | 1.14% |
| hsa_circ_0055524 | 1754 | 10 | 1.14% |
| hsa_circ_0070046 | 1754 | 10 | 1.14% |
| hsa_circ_0084535 | 1754 | 10 | 1.14% |
| hsa_circ_0086359 | 1754 | 10 | 1.14% |
| hsa_circ_0116921 | 1754 | 10 | 1.14% |
| hsa_circ_0081210 | 2105 | 12 | 1.14% |
| hsa_circ_0124220 | 2456 | 14 | 1.14% |
| hsa_circ_0117794 | 1930 | 11 | 1.14% |
| hsa_circ_0014865 | 2281 | 13 | 1.14% |
| hsa_circ_0025679 | 2281 | 13 | 1.14% |
| hsa_circ_0002702 | 2106 | 12 | 1.14% |
| hsa_circ_0006506 | 2106 | 12 | 1.14% |
| hsa_circ_0034545 | 2106 | 12 | 1.14% |
| hsa_circ_0074267 | 2106 | 12 | 1.14% |
| hsa_circ_0052180 | 1755 | 10 | 1.14% |
| hsa_circ_0060399 | 1755 | 10 | 1.14% |
| hsa_circ_0082755 | 1755 | 10 | 1.14% |
| hsa_circ_0086302 | 1755 | 10 | 1.14% |
| hsa_circ_0090915 | 1755 | 10 | 1.14% |
| hsa_circ_0097567 | 1755 | 10 | 1.14% |
| hsa_circ_0134793 | 1755 | 10 | 1.14% |
| hsa_circ_0103628 | 2282 | 13 | 1.14% |
| hsa_circ_0108121 | 2282 | 13 | 1.14% |
| hsa_circ_0010068 | 1931 | 11 | 1.14% |
| hsa_circ_0017011 | 1931 | 11 | 1.14% |
| hsa_circ_0026869 | 1931 | 11 | 1.14% |
| hsa_circ_0042019 | 1931 | 11 | 1.14% |
| hsa_circ_0086336 | 1931 | 11 | 1.14% |
| hsa_circ_0126831 | 1931 | 11 | 1.14% |
| hsa_circ_0132489 | 1931 | 11 | 1.14% |
| hsa_circ_0069974 | 2458 | 14 | 1.14% |
| hsa_circ_0017487 | 1756 | 10 | 1.14% |
| hsa_circ_0024025 | 1756 | 10 | 1.14% |
| hsa_circ_0025527 | 1756 | 10 | 1.14% |
| hsa_circ_0029303 | 1756 | 10 | 1.14% |
| hsa_circ_0037305 | 1756 | 10 | 1.14% |
| hsa_circ_0045703 | 1756 | 10 | 1.14% |

| | | | |
|---|---|---|---|
| hsa_circ_0065899 | 1756 | 10 | 1.14% |
| hsa_circ_0068592 | 1756 | 10 | 1.14% |
| hsa_circ_0071938 | 1756 | 10 | 1.14% |
| hsa_circ_0076461 | 1756 | 10 | 1.14% |
| hsa_circ_0011711 | 1932 | 11 | 1.14% |
| hsa_circ_0037209 | 1932 | 11 | 1.14% |
| hsa_circ_0037294 | 1932 | 11 | 1.14% |
| hsa_circ_0040276 | 1932 | 11 | 1.14% |
| hsa_circ_0053537 | 1932 | 11 | 1.14% |
| hsa_circ_0061171 | 1932 | 11 | 1.14% |
| hsa_circ_0079349 | 1932 | 11 | 1.14% |
| hsa_circ_0086918 | 1932 | 11 | 1.14% |
| hsa_circ_0091102 | 1932 | 11 | 1.14% |
| hsa_circ_0099414 | 1932 | 11 | 1.14% |
| hsa_circ_0105415 | 1932 | 11 | 1.14% |
| hsa_circ_0110223 | 1932 | 11 | 1.14% |
| hsa_circ_0090602 | 2459 | 14 | 1.14% |
| hsa_circ_0130459 | 2284 | 13 | 1.14% |
| hsa_circ_0014392 | 1757 | 10 | 1.14% |
| hsa_circ_0025433 | 1757 | 10 | 1.14% |
| hsa_circ_0052658 | 1757 | 10 | 1.14% |
| hsa_circ_0066160 | 1757 | 10 | 1.14% |
| hsa_circ_0123415 | 1757 | 10 | 1.14% |
| hsa_circ_0019886 | 1933 | 11 | 1.14% |
| hsa_circ_0129458 | 2109 | 12 | 1.14% |
| hsa_circ_0043066 | 1758 | 10 | 1.14% |
| hsa_circ_0055527 | 1758 | 10 | 1.14% |
| hsa_circ_0073796 | 1758 | 10 | 1.14% |
| hsa_circ_0073900 | 1758 | 10 | 1.14% |
| hsa_circ_0076228 | 1758 | 10 | 1.14% |
| hsa_circ_0026991 | 1934 | 11 | 1.14% |
| hsa_circ_0038064 | 1934 | 11 | 1.14% |
| hsa_circ_0048527 | 1934 | 11 | 1.14% |
| hsa_circ_0091422 | 1934 | 11 | 1.14% |
| hsa_circ_0029831 | 2110 | 12 | 1.14% |
| hsa_circ_0076760 | 2110 | 12 | 1.14% |
| hsa_circ_0093909 | 6682 | 38 | 1.14% |
| hsa_circ_0046831 | 2286 | 13 | 1.14% |
| hsa_circ_0053707 | 2286 | 13 | 1.14% |
| hsa_circ_0038383 | 3869 | 22 | 1.14% |
| hsa_circ_0058234 | 2638 | 15 | 1.14% |
| hsa_circ_0019167 | 1759 | 10 | 1.14% |
| hsa_circ_0028660 | 1759 | 10 | 1.14% |
| hsa_circ_0045239 | 1759 | 10 | 1.14% |
| hsa_circ_0066124 | 1759 | 10 | 1.14% |
| hsa_circ_0071127 | 1759 | 10 | 1.14% |
| hsa_circ_0073858 | 1759 | 10 | 1.14% |
| hsa_circ_0078819 | 1759 | 10 | 1.14% |
| hsa_circ_0098404 | 1759 | 10 | 1.14% |
| hsa_circ_0133649 | 1759 | 10 | 1.14% |

| | | | |
|---|---|---|---|
| hsa_circ_0024343 | 1935 | 11 | 1.14% |
| hsa_circ_0040887 | 1935 | 11 | 1.14% |
| hsa_circ_0048611 | 1935 | 11 | 1.14% |
| hsa_circ_0042565 | 2111 | 12 | 1.14% |
| hsa_circ_0003990 | 4048 | 23 | 1.14% |
| hsa_circ_0018976 | 2992 | 17 | 1.14% |
| hsa_circ_0015979 | 2640 | 15 | 1.14% |
| hsa_circ_0013677 | 2112 | 12 | 1.14% |
| hsa_circ_0022760 | 2112 | 12 | 1.14% |
| hsa_circ_0111346 | 2112 | 12 | 1.14% |
| hsa_circ_0047833 | 1936 | 11 | 1.14% |
| hsa_circ_0063034 | 1936 | 11 | 1.14% |
| hsa_circ_0063810 | 1936 | 11 | 1.14% |
| hsa_circ_0080012 | 1936 | 11 | 1.14% |
| hsa_circ_0106178 | 1936 | 11 | 1.14% |
| hsa_circ_0118349 | 1936 | 11 | 1.14% |
| hsa_circ_0135864 | 1936 | 11 | 1.14% |
| hsa_circ_0005689 | 1760 | 10 | 1.14% |
| hsa_circ_0046591 | 1760 | 10 | 1.14% |
| hsa_circ_0060609 | 1760 | 10 | 1.14% |
| hsa_circ_0050932 | 2993 | 17 | 1.14% |
| hsa_circ_0139272 | 2993 | 17 | 1.14% |
| hsa_circ_0010630 | 2289 | 13 | 1.14% |
| hsa_circ_0042281 | 2113 | 12 | 1.14% |
| hsa_circ_0067597 | 2113 | 12 | 1.14% |
| hsa_circ_0022268 | 1937 | 11 | 1.14% |
| hsa_circ_0080921 | 1937 | 11 | 1.14% |
| hsa_circ_0091214 | 1937 | 11 | 1.14% |
| hsa_circ_0103703 | 1937 | 11 | 1.14% |
| hsa_circ_0019418 | 1761 | 10 | 1.14% |
| hsa_circ_0043760 | 1761 | 10 | 1.14% |
| hsa_circ_0047212 | 1761 | 10 | 1.14% |
| hsa_circ_0070936 | 1761 | 10 | 1.14% |
| hsa_circ_0097809 | 1761 | 10 | 1.14% |
| hsa_circ_0127608 | 1761 | 10 | 1.14% |
| hsa_circ_0057424 | 2466 | 14 | 1.14% |
| hsa_circ_0121887 | 2290 | 13 | 1.14% |
| hsa_circ_0024762 | 2114 | 12 | 1.14% |
| hsa_circ_0028522 | 1938 | 11 | 1.14% |
| hsa_circ_0128866 | 1938 | 11 | 1.14% |
| hsa_circ_0011791 | 1762 | 10 | 1.14% |
| hsa_circ_0043776 | 1762 | 10 | 1.14% |
| hsa_circ_0091005 | 1762 | 10 | 1.14% |
| hsa_circ_0095178 | 1762 | 10 | 1.14% |
| hsa_circ_0104150 | 1762 | 10 | 1.14% |
| hsa_circ_0120628 | 1762 | 10 | 1.14% |
| hsa_circ_0023413 | 2291 | 13 | 1.13% |
| hsa_circ_0050798 | 2115 | 12 | 1.13% |
| hsa_circ_0054772 | 2115 | 12 | 1.13% |
| hsa_circ_0037596 | 1939 | 11 | 1.13% |

| | | | |
|---|---|---|---|
| hsa_circ_0067897 | 1939 | 11 | 1.13% |
| hsa_circ_0074248 | 1939 | 11 | 1.13% |
| hsa_circ_0076453 | 1939 | 11 | 1.13% |
| hsa_circ_0129367 | 1939 | 11 | 1.13% |
| hsa_circ_0074846 | 2468 | 14 | 1.13% |
| hsa_circ_0009725 | 1763 | 10 | 1.13% |
| hsa_circ_0022636 | 1763 | 10 | 1.13% |
| hsa_circ_0035478 | 1763 | 10 | 1.13% |
| hsa_circ_0038664 | 1763 | 10 | 1.13% |
| hsa_circ_0041508 | 1763 | 10 | 1.13% |
| hsa_circ_0061359 | 1763 | 10 | 1.13% |
| hsa_circ_0066292 | 1763 | 10 | 1.13% |
| hsa_circ_0066690 | 1763 | 10 | 1.13% |
| hsa_circ_0078404 | 1763 | 10 | 1.13% |
| hsa_circ_0112304 | 1763 | 10 | 1.13% |
| hsa_circ_0114286 | 1763 | 10 | 1.13% |
| hsa_circ_0085075 | 2645 | 15 | 1.13% |
| hsa_circ_0077335 | 2116 | 12 | 1.13% |
| hsa_circ_0084468 | 2116 | 12 | 1.13% |
| hsa_circ_0115330 | 2116 | 12 | 1.13% |
| hsa_circ_0136768 | 2116 | 12 | 1.13% |
| hsa_circ_0074194 | 1940 | 11 | 1.13% |
| hsa_circ_0089053 | 1940 | 11 | 1.13% |
| hsa_circ_0106253 | 1940 | 11 | 1.13% |
| hsa_circ_0121738 | 1940 | 11 | 1.13% |
| hsa_circ_0016694 | 2822 | 16 | 1.13% |
| hsa_circ_0072746 | 4410 | 25 | 1.13% |
| hsa_circ_0012130 | 1764 | 10 | 1.13% |
| hsa_circ_0014373 | 1764 | 10 | 1.13% |
| hsa_circ_0134917 | 1764 | 10 | 1.13% |
| hsa_circ_0039972 | 6351 | 36 | 1.13% |
| hsa_circ_0070839 | 2117 | 12 | 1.13% |
| hsa_circ_0099643 | 9880 | 56 | 1.13% |
| hsa_circ_0065890 | 2470 | 14 | 1.13% |
| hsa_circ_0033401 | 1941 | 11 | 1.13% |
| hsa_circ_0092878 | 1941 | 11 | 1.13% |
| hsa_circ_0108057 | 1941 | 11 | 1.13% |
| hsa_circ_0032624 | 2118 | 12 | 1.13% |
| hsa_circ_0056737 | 2118 | 12 | 1.13% |
| hsa_circ_0070364 | 2118 | 12 | 1.13% |
| hsa_circ_0027314 | 1765 | 10 | 1.13% |
| hsa_circ_0041650 | 1765 | 10 | 1.13% |
| hsa_circ_0044805 | 1765 | 10 | 1.13% |
| hsa_circ_0052353 | 1765 | 10 | 1.13% |
| hsa_circ_0056648 | 1765 | 10 | 1.13% |
| hsa_circ_0057025 | 1765 | 10 | 1.13% |
| hsa_circ_0090953 | 1765 | 10 | 1.13% |
| hsa_circ_0116647 | 1765 | 10 | 1.13% |
| hsa_circ_0119534 | 1765 | 10 | 1.13% |
| hsa_circ_0009424 | 2648 | 15 | 1.13% |

| | | | |
|---|---|---|---|
| hsa_circ_0054510 | 2295 | 13 | 1.13% |
| hsa_circ_0017098 | 1942 | 11 | 1.13% |
| hsa_circ_0028802 | 1942 | 11 | 1.13% |
| hsa_circ_0102231 | 1942 | 11 | 1.13% |
| hsa_circ_0122297 | 1942 | 11 | 1.13% |
| hsa_circ_0131144 | 1942 | 11 | 1.13% |
| hsa_circ_0049495 | 5827 | 33 | 1.13% |
| hsa_circ_0007696 | 2119 | 12 | 1.13% |
| hsa_circ_0017651 | 2119 | 12 | 1.13% |
| hsa_circ_0060649 | 2119 | 12 | 1.13% |
| hsa_circ_0114775 | 2119 | 12 | 1.13% |
| hsa_circ_0015892 | 1766 | 10 | 1.13% |
| hsa_circ_0024109 | 1766 | 10 | 1.13% |
| hsa_circ_0061839 | 1766 | 10 | 1.13% |
| hsa_circ_0062229 | 1766 | 10 | 1.13% |
| hsa_circ_0074974 | 1766 | 10 | 1.13% |
| hsa_circ_0013844 | 1943 | 11 | 1.13% |
| hsa_circ_0046473 | 1943 | 11 | 1.13% |
| hsa_circ_0050858 | 1943 | 11 | 1.13% |
| hsa_circ_0084101 | 1943 | 11 | 1.13% |
| hsa_circ_0122282 | 1943 | 11 | 1.13% |
| hsa_circ_0020269 | 2120 | 12 | 1.13% |
| hsa_circ_0066010 | 2120 | 12 | 1.13% |
| hsa_circ_0018324 | 1767 | 10 | 1.13% |
| hsa_circ_0029844 | 1767 | 10 | 1.13% |
| hsa_circ_0040049 | 1767 | 10 | 1.13% |
| hsa_circ_0043564 | 1767 | 10 | 1.13% |
| hsa_circ_0054369 | 1767 | 10 | 1.13% |
| hsa_circ_0060707 | 1767 | 10 | 1.13% |
| hsa_circ_0069702 | 1767 | 10 | 1.13% |
| hsa_circ_0070879 | 1767 | 10 | 1.13% |
| hsa_circ_0094800 | 1767 | 10 | 1.13% |
| hsa_circ_0109065 | 1767 | 10 | 1.13% |
| hsa_circ_0109975 | 1767 | 10 | 1.13% |
| hsa_circ_0093182 | 8482 | 48 | 1.13% |
| hsa_circ_0112382 | 2474 | 14 | 1.13% |
| hsa_circ_0014478 | 1944 | 11 | 1.13% |
| hsa_circ_0022131 | 1944 | 11 | 1.13% |
| hsa_circ_0048692 | 1944 | 11 | 1.13% |
| hsa_circ_0052080 | 1944 | 11 | 1.13% |
| hsa_circ_0074141 | 1944 | 11 | 1.13% |
| hsa_circ_0106475 | 1944 | 11 | 1.13% |
| hsa_circ_0083778 | 2121 | 12 | 1.13% |
| hsa_circ_0068970 | 1768 | 10 | 1.13% |
| hsa_circ_0087373 | 1768 | 10 | 1.13% |
| hsa_circ_0004021 | 4774 | 27 | 1.13% |
| hsa_circ_0025051 | 1945 | 11 | 1.13% |
| hsa_circ_0040181 | 1945 | 11 | 1.13% |
| hsa_circ_0075220 | 1945 | 11 | 1.13% |
| hsa_circ_0109952 | 1945 | 11 | 1.13% |

| | | | |
|---|---|---|---|
| hsa_circ_0117218 | 1945 | 11 | 1.13% |
| hsa_circ_0124716 | 1945 | 11 | 1.13% |
| hsa_circ_0096432 | 4244 | 24 | 1.13% |
| hsa_circ_0058817 | 2299 | 13 | 1.13% |
| hsa_circ_0022940 | 1769 | 10 | 1.13% |
| hsa_circ_0030243 | 1769 | 10 | 1.13% |
| hsa_circ_0031906 | 1769 | 10 | 1.13% |
| hsa_circ_0047631 | 1769 | 10 | 1.13% |
| hsa_circ_0048034 | 1769 | 10 | 1.13% |
| hsa_circ_0053799 | 1769 | 10 | 1.13% |
| hsa_circ_0054080 | 1769 | 10 | 1.13% |
| hsa_circ_0067776 | 1769 | 10 | 1.13% |
| hsa_circ_0086429 | 1769 | 10 | 1.13% |
| hsa_circ_0127671 | 1769 | 10 | 1.13% |
| hsa_circ_0134281 | 1769 | 10 | 1.13% |
| hsa_circ_0135290 | 1769 | 10 | 1.13% |
| hsa_circ_0043547 | 1946 | 11 | 1.13% |
| hsa_circ_0059145 | 1946 | 11 | 1.13% |
| hsa_circ_0031628 | 2123 | 12 | 1.13% |
| hsa_circ_0051785 | 2123 | 12 | 1.13% |
| hsa_circ_0058896 | 2123 | 12 | 1.13% |
| hsa_circ_0046070 | 2300 | 13 | 1.13% |
| hsa_circ_0113142 | 5486 | 31 | 1.13% |
| hsa_circ_0028168 | 2301 | 13 | 1.13% |
| hsa_circ_0013715 | 2124 | 12 | 1.13% |
| hsa_circ_0016731 | 2124 | 12 | 1.13% |
| hsa_circ_0048484 | 2124 | 12 | 1.13% |
| hsa_circ_0013487 | 1947 | 11 | 1.13% |
| hsa_circ_0057902 | 1947 | 11 | 1.13% |
| hsa_circ_0061148 | 1947 | 11 | 1.13% |
| hsa_circ_0062549 | 1947 | 11 | 1.13% |
| hsa_circ_0068200 | 1947 | 11 | 1.13% |
| hsa_circ_0071911 | 1947 | 11 | 1.13% |
| hsa_circ_0117898 | 1947 | 11 | 1.13% |
| hsa_circ_0014368 | 1770 | 10 | 1.13% |
| hsa_circ_0072399 | 1770 | 10 | 1.13% |
| hsa_circ_0076823 | 1770 | 10 | 1.13% |
| hsa_circ_0139242 | 1770 | 10 | 1.13% |
| hsa_circ_0097703 | 7614 | 43 | 1.13% |
| hsa_circ_0060348 | 2479 | 14 | 1.13% |
| hsa_circ_0104179 | 2125 | 12 | 1.13% |
| hsa_circ_0132877 | 2125 | 12 | 1.13% |
| hsa_circ_0006437 | 1948 | 11 | 1.13% |
| hsa_circ_0016320 | 1948 | 11 | 1.13% |
| hsa_circ_0044064 | 1948 | 11 | 1.13% |
| hsa_circ_0052377 | 1948 | 11 | 1.13% |
| hsa_circ_0055680 | 1948 | 11 | 1.13% |
| hsa_circ_0062288 | 1948 | 11 | 1.13% |
| hsa_circ_0065153 | 1948 | 11 | 1.13% |
| hsa_circ_0083428 | 1948 | 11 | 1.13% |

| | | | |
|---|---|---|---|
| hsa_circ_0102522 | 1948 | 11 | 1.13% |
| hsa_circ_0108925 | 1948 | 11 | 1.13% |
| hsa_circ_0118606 | 1948 | 11 | 1.13% |
| hsa_circ_0018342 | 1771 | 10 | 1.13% |
| hsa_circ_0020813 | 1771 | 10 | 1.13% |
| hsa_circ_0039662 | 1771 | 10 | 1.13% |
| hsa_circ_0054379 | 1771 | 10 | 1.13% |
| hsa_circ_0066123 | 1771 | 10 | 1.13% |
| hsa_circ_0080204 | 1771 | 10 | 1.13% |
| hsa_circ_0090653 | 1771 | 10 | 1.13% |
| hsa_circ_0102851 | 1771 | 10 | 1.13% |
| hsa_circ_0111587 | 1771 | 10 | 1.13% |
| hsa_circ_0114601 | 1771 | 10 | 1.13% |
| hsa_circ_0137371 | 1771 | 10 | 1.13% |
| hsa_circ_0134365 | 4074 | 23 | 1.13% |
| hsa_circ_0108055 | 2126 | 12 | 1.13% |
| hsa_circ_0070447 | 1949 | 11 | 1.13% |
| hsa_circ_0071399 | 1949 | 11 | 1.13% |
| hsa_circ_0090041 | 1772 | 10 | 1.13% |
| hsa_circ_0091955 | 1772 | 10 | 1.13% |
| hsa_circ_0135611 | 1772 | 10 | 1.13% |
| hsa_circ_0035607 | 2481 | 14 | 1.13% |
| hsa_circ_0010652 | 2304 | 13 | 1.13% |
| hsa_circ_0033774 | 2127 | 12 | 1.13% |
| hsa_circ_0130184 | 4077 | 23 | 1.13% |
| hsa_circ_0009281 | 1950 | 11 | 1.13% |
| hsa_circ_0044851 | 1950 | 11 | 1.13% |
| hsa_circ_0061579 | 1950 | 11 | 1.13% |
| hsa_circ_0082046 | 1950 | 11 | 1.13% |
| hsa_circ_0129989 | 1950 | 11 | 1.13% |
| hsa_circ_0140465 | 1950 | 11 | 1.13% |
| hsa_circ_0091106 | 3191 | 18 | 1.13% |
| hsa_circ_0057348 | 3014 | 17 | 1.13% |
| hsa_circ_0022225 | 1773 | 10 | 1.13% |
| hsa_circ_0024030 | 1773 | 10 | 1.13% |
| hsa_circ_0024207 | 1773 | 10 | 1.13% |
| hsa_circ_0034923 | 1773 | 10 | 1.13% |
| hsa_circ_0039163 | 1773 | 10 | 1.13% |
| hsa_circ_0041493 | 1773 | 10 | 1.13% |
| hsa_circ_0056545 | 1773 | 10 | 1.13% |
| hsa_circ_0114813 | 1773 | 10 | 1.13% |
| hsa_circ_0024392 | 2128 | 12 | 1.13% |
| hsa_circ_0104181 | 2128 | 12 | 1.13% |
| hsa_circ_0017936 | 1951 | 11 | 1.13% |
| hsa_circ_0025668 | 1951 | 11 | 1.13% |
| hsa_circ_0030176 | 1951 | 11 | 1.13% |
| hsa_circ_0033363 | 1951 | 11 | 1.13% |
| hsa_circ_0062145 | 1951 | 11 | 1.13% |
| hsa_circ_0069696 | 1951 | 11 | 1.13% |
| hsa_circ_0097221 | 1951 | 11 | 1.13% |

| | | | |
|---|---|---|---|
| hsa_circ_0103643 | 1951 | 11 | 1.13% |
| hsa_circ_0139819 | 2306 | 13 | 1.13% |
| hsa_circ_0020030 | 1774 | 10 | 1.13% |
| hsa_circ_0023851 | 1774 | 10 | 1.13% |
| hsa_circ_0047256 | 1774 | 10 | 1.13% |
| hsa_circ_0054920 | 1774 | 10 | 1.13% |
| hsa_circ_0055017 | 1774 | 10 | 1.13% |
| hsa_circ_0069106 | 1774 | 10 | 1.13% |
| hsa_circ_0134490 | 5677 | 32 | 1.13% |
| hsa_circ_0104139 | 2129 | 12 | 1.13% |
| hsa_circ_0112465 | 2129 | 12 | 1.13% |
| hsa_circ_0070887 | 6920 | 39 | 1.13% |
| hsa_circ_0025150 | 1952 | 11 | 1.13% |
| hsa_circ_0115789 | 1952 | 11 | 1.13% |
| hsa_circ_0016138 | 2307 | 13 | 1.13% |
| hsa_circ_0075719 | 2130 | 12 | 1.13% |
| hsa_circ_0021597 | 1775 | 10 | 1.13% |
| hsa_circ_0031564 | 1775 | 10 | 1.13% |
| hsa_circ_0044922 | 1775 | 10 | 1.13% |
| hsa_circ_0050089 | 1775 | 10 | 1.13% |
| hsa_circ_0052906 | 1775 | 10 | 1.13% |
| hsa_circ_0117099 | 1775 | 10 | 1.13% |
| hsa_circ_0138642 | 1775 | 10 | 1.13% |
| hsa_circ_0060930 | 1953 | 11 | 1.13% |
| hsa_circ_0089460 | 1953 | 11 | 1.13% |
| hsa_circ_0105526 | 1953 | 11 | 1.13% |
| hsa_circ_0087638 | 7636 | 43 | 1.13% |
| hsa_circ_0074418 | 2131 | 12 | 1.13% |
| hsa_circ_0079551 | 2131 | 12 | 1.13% |
| hsa_circ_0131026 | 12076 | 68 | 1.13% |
| hsa_circ_0123299 | 5328 | 30 | 1.13% |
| hsa_circ_0020420 | 1776 | 10 | 1.13% |
| hsa_circ_0031707 | 1776 | 10 | 1.13% |
| hsa_circ_0045699 | 1776 | 10 | 1.13% |
| hsa_circ_0068499 | 1776 | 10 | 1.13% |
| hsa_circ_0074067 | 1776 | 10 | 1.13% |
| hsa_circ_0089635 | 1776 | 10 | 1.13% |
| hsa_circ_0110517 | 1776 | 10 | 1.13% |
| hsa_circ_0083024 | 2309 | 13 | 1.13% |
| hsa_circ_0128060 | 2309 | 13 | 1.13% |
| hsa_circ_0131785 | 2309 | 13 | 1.13% |
| hsa_circ_0016921 | 1954 | 11 | 1.13% |
| hsa_circ_0023252 | 1954 | 11 | 1.13% |
| hsa_circ_0082651 | 1954 | 11 | 1.13% |
| hsa_circ_0128903 | 1954 | 11 | 1.13% |
| hsa_circ_0104215 | 7996 | 45 | 1.13% |
| hsa_circ_0135096 | 2310 | 13 | 1.13% |
| hsa_circ_0030588 | 1777 | 10 | 1.13% |
| hsa_circ_0036133 | 1777 | 10 | 1.13% |
| hsa_circ_0042603 | 1777 | 10 | 1.13% |

| | | | |
|---|---|---|---|
| hsa_circ_0043721 | 1777 | 10 | 1.13% |
| hsa_circ_0055840 | 1777 | 10 | 1.13% |
| hsa_circ_0082916 | 1777 | 10 | 1.13% |
| hsa_circ_0089934 | 1777 | 10 | 1.13% |
| hsa_circ_0135646 | 1777 | 10 | 1.13% |
| hsa_circ_0045396 | 2488 | 14 | 1.13% |
| hsa_circ_0020888 | 1955 | 11 | 1.13% |
| hsa_circ_0022960 | 1955 | 11 | 1.13% |
| hsa_circ_0039030 | 1955 | 11 | 1.13% |
| hsa_circ_0074203 | 1955 | 11 | 1.13% |
| hsa_circ_0115759 | 1955 | 11 | 1.13% |
| hsa_circ_0137964 | 4799 | 27 | 1.13% |
| hsa_circ_0057302 | 2133 | 12 | 1.13% |
| hsa_circ_0063274 | 2133 | 12 | 1.13% |
| hsa_circ_0076490 | 2489 | 14 | 1.12% |
| hsa_circ_0069539 | 1778 | 10 | 1.12% |
| hsa_circ_0114222 | 1778 | 10 | 1.12% |
| hsa_circ_0071425 | 1956 | 11 | 1.12% |
| hsa_circ_0135114 | 1956 | 11 | 1.12% |
| hsa_circ_0061332 | 2134 | 12 | 1.12% |
| hsa_circ_0093969 | 2134 | 12 | 1.12% |
| hsa_circ_0062857 | 2312 | 13 | 1.12% |
| hsa_circ_0006424 | 2668 | 15 | 1.12% |
| hsa_circ_0102008 | 7649 | 43 | 1.12% |
| hsa_circ_0032273 | 3202 | 18 | 1.12% |
| hsa_circ_0024244 | 1779 | 10 | 1.12% |
| hsa_circ_0029915 | 1779 | 10 | 1.12% |
| hsa_circ_0035023 | 1779 | 10 | 1.12% |
| hsa_circ_0067898 | 1779 | 10 | 1.12% |
| hsa_circ_0119851 | 1779 | 10 | 1.12% |
| hsa_circ_0130356 | 1779 | 10 | 1.12% |
| hsa_circ_0137823 | 1779 | 10 | 1.12% |
| hsa_circ_0038769 | 1957 | 11 | 1.12% |
| hsa_circ_0080062 | 1957 | 11 | 1.12% |
| hsa_circ_0080152 | 1957 | 11 | 1.12% |
| hsa_circ_0091982 | 1957 | 11 | 1.12% |
| hsa_circ_0091992 | 1957 | 11 | 1.12% |
| hsa_circ_0105472 | 2135 | 12 | 1.12% |
| hsa_circ_0098638 | 4093 | 23 | 1.12% |
| hsa_circ_0034100 | 6942 | 39 | 1.12% |
| hsa_circ_0020033 | 2314 | 13 | 1.12% |
| hsa_circ_0100721 | 2314 | 13 | 1.12% |
| hsa_circ_0017866 | 2136 | 12 | 1.12% |
| hsa_circ_0111940 | 2136 | 12 | 1.12% |
| hsa_circ_0016882 | 1958 | 11 | 1.12% |
| hsa_circ_0020885 | 1958 | 11 | 1.12% |
| hsa_circ_0031493 | 1958 | 11 | 1.12% |
| hsa_circ_0041835 | 1958 | 11 | 1.12% |
| hsa_circ_0069675 | 1958 | 11 | 1.12% |
| hsa_circ_0092001 | 1958 | 11 | 1.12% |

| | | | |
|---|---|---|---|
| hsa_circ_0117287 | 1958 | 11 | 1.12% |
| hsa_circ_0137358 | 1958 | 11 | 1.12% |
| hsa_circ_0019516 | 1780 | 10 | 1.12% |
| hsa_circ_0025796 | 1780 | 10 | 1.12% |
| hsa_circ_0036915 | 1780 | 10 | 1.12% |
| hsa_circ_0040607 | 1780 | 10 | 1.12% |
| hsa_circ_0045683 | 1780 | 10 | 1.12% |
| hsa_circ_0052411 | 1780 | 10 | 1.12% |
| hsa_circ_0058254 | 1780 | 10 | 1.12% |
| hsa_circ_0096852 | 1780 | 10 | 1.12% |
| hsa_circ_0129393 | 1780 | 10 | 1.12% |
| hsa_circ_0123534 | 6765 | 38 | 1.12% |
| hsa_circ_0021848 | 2671 | 15 | 1.12% |
| hsa_circ_0089875 | 2315 | 13 | 1.12% |
| hsa_circ_0116189 | 2315 | 13 | 1.12% |
| hsa_circ_0031598 | 2137 | 12 | 1.12% |
| hsa_circ_0035235 | 2137 | 12 | 1.12% |
| hsa_circ_0060736 | 2137 | 12 | 1.12% |
| hsa_circ_0110628 | 2137 | 12 | 1.12% |
| hsa_circ_0019823 | 1959 | 11 | 1.12% |
| hsa_circ_0023302 | 1959 | 11 | 1.12% |
| hsa_circ_0055651 | 1959 | 11 | 1.12% |
| hsa_circ_0069051 | 1959 | 11 | 1.12% |
| hsa_circ_0085340 | 1959 | 11 | 1.12% |
| hsa_circ_0104235 | 5343 | 30 | 1.12% |
| hsa_circ_0000663 | 1781 | 10 | 1.12% |
| hsa_circ_0014694 | 1781 | 10 | 1.12% |
| hsa_circ_0034863 | 1781 | 10 | 1.12% |
| hsa_circ_0048239 | 1781 | 10 | 1.12% |
| hsa_circ_0050680 | 1781 | 10 | 1.12% |
| hsa_circ_0055725 | 1781 | 10 | 1.12% |
| hsa_circ_0066317 | 1781 | 10 | 1.12% |
| hsa_circ_0091901 | 1781 | 10 | 1.12% |
| hsa_circ_0113008 | 1781 | 10 | 1.12% |
| hsa_circ_0001013 | 3741 | 21 | 1.12% |
| hsa_circ_0035475 | 2316 | 13 | 1.12% |
| hsa_circ_0111937 | 2138 | 12 | 1.12% |
| hsa_circ_0128240 | 2138 | 12 | 1.12% |
| hsa_circ_0015872 | 1960 | 11 | 1.12% |
| hsa_circ_0054228 | 1960 | 11 | 1.12% |
| hsa_circ_0059331 | 1960 | 11 | 1.12% |
| hsa_circ_0083381 | 1960 | 11 | 1.12% |
| hsa_circ_0106636 | 1960 | 11 | 1.12% |
| hsa_circ_0015600 | 1782 | 10 | 1.12% |
| hsa_circ_0064755 | 1782 | 10 | 1.12% |
| hsa_circ_0066301 | 1782 | 10 | 1.12% |
| hsa_circ_0069464 | 1782 | 10 | 1.12% |
| hsa_circ_0069513 | 1782 | 10 | 1.12% |
| hsa_circ_0094916 | 1782 | 10 | 1.12% |
| hsa_circ_0049737 | 2495 | 14 | 1.12% |

| | | | |
|---|---|---|---|
| hsa_circ_0045263 | 2139 | 12 | 1.12% |
| hsa_circ_0052828 | 2139 | 12 | 1.12% |
| hsa_circ_0078528 | 2139 | 12 | 1.12% |
| hsa_circ_0052685 | 2674 | 15 | 1.12% |
| hsa_circ_0017480 | 1961 | 11 | 1.12% |
| hsa_circ_0033176 | 1961 | 11 | 1.12% |
| hsa_circ_0045040 | 1961 | 11 | 1.12% |
| hsa_circ_0065421 | 1961 | 11 | 1.12% |
| hsa_circ_0068558 | 1961 | 11 | 1.12% |
| hsa_circ_0071798 | 1961 | 11 | 1.12% |
| hsa_circ_0079756 | 1961 | 11 | 1.12% |
| hsa_circ_0085518 | 1961 | 11 | 1.12% |
| hsa_circ_0014108 | 1783 | 10 | 1.12% |
| hsa_circ_0031741 | 1783 | 10 | 1.12% |
| hsa_circ_0093630 | 1783 | 10 | 1.12% |
| hsa_circ_0116884 | 1783 | 10 | 1.12% |
| hsa_circ_0132061 | 1783 | 10 | 1.12% |
| hsa_circ_0120550 | 3745 | 21 | 1.12% |
| hsa_circ_0031928 | 2140 | 12 | 1.12% |
| hsa_circ_0104664 | 2140 | 12 | 1.12% |
| hsa_circ_0121920 | 2140 | 12 | 1.12% |
| hsa_circ_0005592 | 1962 | 11 | 1.12% |
| hsa_circ_0011326 | 1962 | 11 | 1.12% |
| hsa_circ_0023208 | 1962 | 11 | 1.12% |
| hsa_circ_0065629 | 1962 | 11 | 1.12% |
| hsa_circ_0107000 | 7135 | 40 | 1.12% |
| hsa_circ_0102413 | 4281 | 24 | 1.12% |
| hsa_circ_0012362 | 1784 | 10 | 1.12% |
| hsa_circ_0038040 | 1784 | 10 | 1.12% |
| hsa_circ_0044807 | 1784 | 10 | 1.12% |
| hsa_circ_0062754 | 1784 | 10 | 1.12% |
| hsa_circ_0067959 | 1784 | 10 | 1.12% |
| hsa_circ_0074114 | 1784 | 10 | 1.12% |
| hsa_circ_0084716 | 1784 | 10 | 1.12% |
| hsa_circ_0121660 | 1784 | 10 | 1.12% |
| hsa_circ_0029567 | 2141 | 12 | 1.12% |
| hsa_circ_0085370 | 2141 | 12 | 1.12% |
| hsa_circ_0133001 | 2141 | 12 | 1.12% |
| hsa_circ_0007406 | 3569 | 20 | 1.12% |
| hsa_circ_0014329 | 1963 | 11 | 1.12% |
| hsa_circ_0034306 | 1963 | 11 | 1.12% |
| hsa_circ_0100264 | 1963 | 11 | 1.12% |
| hsa_circ_0119820 | 1963 | 11 | 1.12% |
| hsa_circ_0138841 | 1963 | 11 | 1.12% |
| hsa_circ_0084408 | 2320 | 13 | 1.12% |
| hsa_circ_0064076 | 3034 | 17 | 1.12% |
| hsa_circ_0024375 | 3748 | 21 | 1.12% |
| hsa_circ_0016293 | 2142 | 12 | 1.12% |
| hsa_circ_0103086 | 2142 | 12 | 1.12% |
| hsa_circ_0127655 | 2142 | 12 | 1.12% |

| | | | |
|---|---|---|---|
| hsa_circ_0140602 | 2142 | 12 | 1.12% |
| hsa_circ_0009764 | 1785 | 10 | 1.12% |
| hsa_circ_0042006 | 1785 | 10 | 1.12% |
| hsa_circ_0067612 | 1785 | 10 | 1.12% |
| hsa_circ_0127442 | 1785 | 10 | 1.12% |
| hsa_circ_0128568 | 1785 | 10 | 1.12% |
| hsa_circ_0122428 | 7856 | 44 | 1.12% |
| hsa_circ_0011365 | 1964 | 11 | 1.12% |
| hsa_circ_0022729 | 1964 | 11 | 1.12% |
| hsa_circ_0074722 | 1964 | 11 | 1.12% |
| hsa_circ_0096228 | 5178 | 29 | 1.12% |
| hsa_circ_0041033 | 2143 | 12 | 1.12% |
| hsa_circ_0104322 | 3929 | 22 | 1.12% |
| hsa_circ_0025677 | 1786 | 10 | 1.12% |
| hsa_circ_0040126 | 1786 | 10 | 1.12% |
| hsa_circ_0040706 | 1786 | 10 | 1.12% |
| hsa_circ_0046312 | 1786 | 10 | 1.12% |
| hsa_circ_0068409 | 1786 | 10 | 1.12% |
| hsa_circ_0099331 | 1786 | 10 | 1.12% |
| hsa_circ_0119089 | 1786 | 10 | 1.12% |
| hsa_circ_0132395 | 1786 | 10 | 1.12% |
| hsa_circ_0031544 | 4823 | 27 | 1.12% |
| hsa_circ_0099247 | 1965 | 11 | 1.12% |
| hsa_circ_0101896 | 1965 | 11 | 1.12% |
| hsa_circ_0020943 | 2501 | 14 | 1.12% |
| hsa_circ_0029699 | 4288 | 24 | 1.12% |
| hsa_circ_0021875 | 2144 | 12 | 1.12% |
| hsa_circ_0010666 | 2323 | 13 | 1.12% |
| hsa_circ_0012339 | 1787 | 10 | 1.12% |
| hsa_circ_0068977 | 1787 | 10 | 1.12% |
| hsa_circ_0075186 | 1787 | 10 | 1.12% |
| hsa_circ_0079807 | 1787 | 10 | 1.12% |
| hsa_circ_0084129 | 1787 | 10 | 1.12% |
| hsa_circ_0089995 | 1787 | 10 | 1.12% |
| hsa_circ_0104156 | 1787 | 10 | 1.12% |
| hsa_circ_0080456 | 1966 | 11 | 1.12% |
| hsa_circ_0083721 | 1966 | 11 | 1.12% |
| hsa_circ_0124224 | 1966 | 11 | 1.12% |
| hsa_circ_0135310 | 4111 | 23 | 1.12% |
| hsa_circ_0113541 | 3039 | 17 | 1.12% |
| hsa_circ_0078693 | 2324 | 13 | 1.12% |
| hsa_circ_0104533 | 4112 | 23 | 1.12% |
| hsa_circ_0014556 | 1788 | 10 | 1.12% |
| hsa_circ_0028894 | 1788 | 10 | 1.12% |
| hsa_circ_0042425 | 1788 | 10 | 1.12% |
| hsa_circ_0061202 | 1788 | 10 | 1.12% |
| hsa_circ_0072920 | 1788 | 10 | 1.12% |
| hsa_circ_0079734 | 1788 | 10 | 1.12% |
| hsa_circ_0084191 | 1788 | 10 | 1.12% |
| hsa_circ_0086503 | 1788 | 10 | 1.12% |

| | | | |
|---|---|---|---|
| hsa_circ_0090023 | 1788 | 10 | 1.12% |
| hsa_circ_0093314 | 1788 | 10 | 1.12% |
| hsa_circ_0100336 | 1788 | 10 | 1.12% |
| hsa_circ_0104868 | 1788 | 10 | 1.12% |
| hsa_circ_0115883 | 1788 | 10 | 1.12% |
| hsa_circ_0131504 | 1788 | 10 | 1.12% |
| hsa_circ_0138405 | 1788 | 10 | 1.12% |
| hsa_circ_0030826 | 1967 | 11 | 1.12% |
| hsa_circ_0053148 | 1967 | 11 | 1.12% |
| hsa_circ_0097605 | 1967 | 11 | 1.12% |
| hsa_circ_0119253 | 1967 | 11 | 1.12% |
| hsa_circ_0071081 | 2146 | 12 | 1.12% |
| hsa_circ_0132711 | 2146 | 12 | 1.12% |
| hsa_circ_0061244 | 5544 | 31 | 1.12% |
| hsa_circ_0052757 | 2325 | 13 | 1.12% |
| hsa_circ_0020438 | 1789 | 10 | 1.12% |
| hsa_circ_0070583 | 1789 | 10 | 1.12% |
| hsa_circ_0080839 | 1789 | 10 | 1.12% |
| hsa_circ_0096958 | 1789 | 10 | 1.12% |
| hsa_circ_0114274 | 1789 | 10 | 1.12% |
| hsa_circ_0127047 | 1789 | 10 | 1.12% |
| hsa_circ_0128976 | 1789 | 10 | 1.12% |
| hsa_circ_0138868 | 1789 | 10 | 1.12% |
| hsa_circ_0010826 | 1968 | 11 | 1.12% |
| hsa_circ_0011205 | 1968 | 11 | 1.12% |
| hsa_circ_0011323 | 1968 | 11 | 1.12% |
| hsa_circ_0019929 | 1968 | 11 | 1.12% |
| hsa_circ_0028019 | 1968 | 11 | 1.12% |
| hsa_circ_0073697 | 1968 | 11 | 1.12% |
| hsa_circ_0075894 | 1968 | 11 | 1.12% |
| hsa_circ_0091131 | 1968 | 11 | 1.12% |
| hsa_circ_0121347 | 1968 | 11 | 1.12% |
| hsa_circ_0132345 | 1968 | 11 | 1.12% |
| hsa_circ_0020081 | 2147 | 12 | 1.12% |
| hsa_circ_0034037 | 2147 | 12 | 1.12% |
| hsa_circ_0034095 | 2147 | 12 | 1.12% |
| hsa_circ_0034478 | 2147 | 12 | 1.12% |
| hsa_circ_0050650 | 2147 | 12 | 1.12% |
| hsa_circ_0085628 | 2147 | 12 | 1.12% |
| hsa_circ_0034949 | 2685 | 15 | 1.12% |
| hsa_circ_0028891 | 2148 | 12 | 1.12% |
| hsa_circ_0048983 | 2148 | 12 | 1.12% |
| hsa_circ_0051014 | 2148 | 12 | 1.12% |
| hsa_circ_0072325 | 2148 | 12 | 1.12% |
| hsa_circ_0083698 | 2148 | 12 | 1.12% |
| hsa_circ_0103548 | 2148 | 12 | 1.12% |
| hsa_circ_0132463 | 2148 | 12 | 1.12% |
| hsa_circ_0075922 | 1969 | 11 | 1.12% |
| hsa_circ_0029764 | 1790 | 10 | 1.12% |
| hsa_circ_0041228 | 1790 | 10 | 1.12% |

| | | | |
|---|---|---|---|
| hsa_circ_0056155 | 1790 | 10 | 1.12% |
| hsa_circ_0099095 | 1790 | 10 | 1.12% |
| hsa_circ_0117714 | 1790 | 10 | 1.12% |
| hsa_circ_0121476 | 1790 | 10 | 1.12% |
| hsa_circ_0132652 | 1790 | 10 | 1.12% |
| hsa_circ_0084516 | 2328 | 13 | 1.12% |
| hsa_circ_0133108 | 2328 | 13 | 1.12% |
| hsa_circ_0045177 | 2149 | 12 | 1.12% |
| hsa_circ_0088902 | 2149 | 12 | 1.12% |
| hsa_circ_0089233 | 2149 | 12 | 1.12% |
| hsa_circ_0015918 | 1970 | 11 | 1.12% |
| hsa_circ_0071748 | 1970 | 11 | 1.12% |
| hsa_circ_0074847 | 1970 | 11 | 1.12% |
| hsa_circ_0101997 | 1970 | 11 | 1.12% |
| hsa_circ_0011796 | 1791 | 10 | 1.12% |
| hsa_circ_0056090 | 1791 | 10 | 1.12% |
| hsa_circ_0057886 | 1791 | 10 | 1.12% |
| hsa_circ_0058731 | 1791 | 10 | 1.12% |
| hsa_circ_0061647 | 1791 | 10 | 1.12% |
| hsa_circ_0072684 | 1791 | 10 | 1.12% |
| hsa_circ_0041834 | 2508 | 14 | 1.12% |
| hsa_circ_0092164 | 2329 | 13 | 1.12% |
| hsa_circ_0139116 | 2329 | 13 | 1.12% |
| hsa_circ_0053182 | 2150 | 12 | 1.12% |
| hsa_circ_0023620 | 1971 | 11 | 1.12% |
| hsa_circ_0029199 | 1971 | 11 | 1.12% |
| hsa_circ_0036185 | 1971 | 11 | 1.12% |
| hsa_circ_0038093 | 1971 | 11 | 1.12% |
| hsa_circ_0045276 | 1971 | 11 | 1.12% |
| hsa_circ_0048088 | 1971 | 11 | 1.12% |
| hsa_circ_0064168 | 1971 | 11 | 1.12% |
| hsa_circ_0082694 | 1971 | 11 | 1.12% |
| hsa_circ_0120340 | 1971 | 11 | 1.12% |
| hsa_circ_0017428 | 1792 | 10 | 1.12% |
| hsa_circ_0068225 | 1792 | 10 | 1.12% |
| hsa_circ_0074299 | 1792 | 10 | 1.12% |
| hsa_circ_0087813 | 1792 | 10 | 1.12% |
| hsa_circ_0088372 | 1792 | 10 | 1.12% |
| hsa_circ_0092149 | 1792 | 10 | 1.12% |
| hsa_circ_0113698 | 1792 | 10 | 1.12% |
| hsa_circ_0046112 | 2330 | 13 | 1.12% |
| hsa_circ_0076223 | 2151 | 12 | 1.12% |
| hsa_circ_0120945 | 5557 | 31 | 1.12% |
| hsa_circ_0020559 | 1972 | 11 | 1.12% |
| hsa_circ_0021774 | 1972 | 11 | 1.12% |
| hsa_circ_0071320 | 1972 | 11 | 1.12% |
| hsa_circ_0074036 | 2510 | 14 | 1.12% |
| hsa_circ_0003434 | 3048 | 17 | 1.12% |
| hsa_circ_0036170 | 1793 | 10 | 1.12% |
| hsa_circ_0054636 | 1793 | 10 | 1.12% |

| | | | |
|---|---|---|---|
| hsa_circ_0093894 | 1793 | 10 | 1.12% |
| hsa_circ_0110244 | 1793 | 10 | 1.12% |
| hsa_circ_0133083 | 1793 | 10 | 1.12% |
| hsa_circ_0058937 | 2331 | 13 | 1.12% |
| hsa_circ_0038403 | 3766 | 21 | 1.12% |
| hsa_circ_0093365 | 2152 | 12 | 1.12% |
| hsa_circ_0001201 | 1973 | 11 | 1.12% |
| hsa_circ_0034808 | 1973 | 11 | 1.12% |
| hsa_circ_0048896 | 1973 | 11 | 1.12% |
| hsa_circ_0063993 | 1973 | 11 | 1.12% |
| hsa_circ_0072250 | 1973 | 11 | 1.12% |
| hsa_circ_0063636 | 2332 | 13 | 1.11% |
| hsa_circ_0020082 | 1794 | 10 | 1.11% |
| hsa_circ_0057022 | 1794 | 10 | 1.11% |
| hsa_circ_0057972 | 1794 | 10 | 1.11% |
| hsa_circ_0061791 | 1794 | 10 | 1.11% |
| hsa_circ_0097790 | 1794 | 10 | 1.11% |
| hsa_circ_0126068 | 1794 | 10 | 1.11% |
| hsa_circ_0062978 | 2153 | 12 | 1.11% |
| hsa_circ_0104140 | 2153 | 12 | 1.11% |
| hsa_circ_0061452 | 2871 | 16 | 1.11% |
| hsa_circ_0019603 | 1974 | 11 | 1.11% |
| hsa_circ_0019711 | 1974 | 11 | 1.11% |
| hsa_circ_0040624 | 1974 | 11 | 1.11% |
| hsa_circ_0066069 | 1974 | 11 | 1.11% |
| hsa_circ_0108220 | 1974 | 11 | 1.11% |
| hsa_circ_0081572 | 2333 | 13 | 1.11% |
| hsa_circ_0000973 | 3590 | 20 | 1.11% |
| hsa_circ_0032287 | 2154 | 12 | 1.11% |
| hsa_circ_0004735 | 1795 | 10 | 1.11% |
| hsa_circ_0018268 | 1795 | 10 | 1.11% |
| hsa_circ_0023721 | 1795 | 10 | 1.11% |
| hsa_circ_0026734 | 1795 | 10 | 1.11% |
| hsa_circ_0030190 | 1795 | 10 | 1.11% |
| hsa_circ_0031506 | 1795 | 10 | 1.11% |
| hsa_circ_0040842 | 1795 | 10 | 1.11% |
| hsa_circ_0043074 | 1795 | 10 | 1.11% |
| hsa_circ_0046529 | 1795 | 10 | 1.11% |
| hsa_circ_0086090 | 1795 | 10 | 1.11% |
| hsa_circ_0029133 | 2334 | 13 | 1.11% |
| hsa_circ_0035938 | 1975 | 11 | 1.11% |
| hsa_circ_0052852 | 1975 | 11 | 1.11% |
| hsa_circ_0105997 | 1975 | 11 | 1.11% |
| hsa_circ_0130040 | 1975 | 11 | 1.11% |
| hsa_circ_0015397 | 1796 | 10 | 1.11% |
| hsa_circ_0036300 | 1796 | 10 | 1.11% |
| hsa_circ_0073304 | 1796 | 10 | 1.11% |
| hsa_circ_0083937 | 1796 | 10 | 1.11% |
| hsa_circ_0101065 | 1796 | 10 | 1.11% |
| hsa_circ_0114739 | 1796 | 10 | 1.11% |

| | | | |
|---|---|---|---|
| hsa_circ_0085997 | 2335 | 13 | 1.11% |
| hsa_circ_0031664 | 1976 | 11 | 1.11% |
| hsa_circ_0049178 | 1976 | 11 | 1.11% |
| hsa_circ_0083317 | 1976 | 11 | 1.11% |
| hsa_circ_0092433 | 1976 | 11 | 1.11% |
| hsa_circ_0023150 | 2156 | 12 | 1.11% |
| hsa_circ_0058356 | 2156 | 12 | 1.11% |
| hsa_circ_0100021 | 2156 | 12 | 1.11% |
| hsa_circ_0138830 | 2156 | 12 | 1.11% |
| hsa_circ_0049517 | 2336 | 13 | 1.11% |
| hsa_circ_0014896 | 1797 | 10 | 1.11% |
| hsa_circ_0017734 | 1797 | 10 | 1.11% |
| hsa_circ_0056569 | 1797 | 10 | 1.11% |
| hsa_circ_0063029 | 1797 | 10 | 1.11% |
| hsa_circ_0139607 | 1797 | 10 | 1.11% |
| hsa_circ_0117683 | 1977 | 11 | 1.11% |
| hsa_circ_0125404 | 1977 | 11 | 1.11% |
| hsa_circ_0014676 | 2337 | 13 | 1.11% |
| hsa_circ_0044038 | 1798 | 10 | 1.11% |
| hsa_circ_0065076 | 1798 | 10 | 1.11% |
| hsa_circ_0065776 | 1798 | 10 | 1.11% |
| hsa_circ_0076410 | 1798 | 10 | 1.11% |
| hsa_circ_0113728 | 1798 | 10 | 1.11% |
| hsa_circ_0122522 | 1798 | 10 | 1.11% |
| hsa_circ_0018162 | 1978 | 11 | 1.11% |
| hsa_circ_0030090 | 1978 | 11 | 1.11% |
| hsa_circ_0044350 | 1978 | 11 | 1.11% |
| hsa_circ_0126280 | 4496 | 25 | 1.11% |
| hsa_circ_0008746 | 3597 | 20 | 1.11% |
| hsa_circ_0089191 | 7016 | 39 | 1.11% |
| hsa_circ_0127908 | 3598 | 20 | 1.11% |
| hsa_circ_0019390 | 1799 | 10 | 1.11% |
| hsa_circ_0020435 | 1799 | 10 | 1.11% |
| hsa_circ_0057686 | 1799 | 10 | 1.11% |
| hsa_circ_0057733 | 1799 | 10 | 1.11% |
| hsa_circ_0068589 | 1799 | 10 | 1.11% |
| hsa_circ_0069603 | 1799 | 10 | 1.11% |
| hsa_circ_0011669 | 1979 | 11 | 1.11% |
| hsa_circ_0034756 | 1979 | 11 | 1.11% |
| hsa_circ_0054451 | 1979 | 11 | 1.11% |
| hsa_circ_0070681 | 1979 | 11 | 1.11% |
| hsa_circ_0091152 | 1979 | 11 | 1.11% |
| hsa_circ_0062417 | 2520 | 14 | 1.11% |
| hsa_circ_0060758 | 2340 | 13 | 1.11% |
| hsa_circ_0067626 | 2340 | 13 | 1.11% |
| hsa_circ_0023364 | 1980 | 11 | 1.11% |
| hsa_circ_0037866 | 1980 | 11 | 1.11% |
| hsa_circ_0050022 | 1980 | 11 | 1.11% |
| hsa_circ_0063689 | 1980 | 11 | 1.11% |
| hsa_circ_0066786 | 1980 | 11 | 1.11% |

| | | | |
|---|---|---|---|
| hsa_circ_0074772 | 1980 | 11 | 1.11% |
| hsa_circ_0077058 | 1980 | 11 | 1.11% |
| hsa_circ_0077829 | 1980 | 11 | 1.11% |
| hsa_circ_0098941 | 1980 | 11 | 1.11% |
| hsa_circ_0009784 | 1800 | 10 | 1.11% |
| hsa_circ_0012864 | 1800 | 10 | 1.11% |
| hsa_circ_0046775 | 1800 | 10 | 1.11% |
| hsa_circ_0059076 | 1800 | 10 | 1.11% |
| hsa_circ_0088931 | 1800 | 10 | 1.11% |
| hsa_circ_0091882 | 1800 | 10 | 1.11% |
| hsa_circ_0093332 | 1800 | 10 | 1.11% |
| hsa_circ_0138998 | 1800 | 10 | 1.11% |
| hsa_circ_0053729 | 2341 | 13 | 1.11% |
| hsa_circ_0078328 | 2161 | 12 | 1.11% |
| hsa_circ_0091595 | 2161 | 12 | 1.11% |
| hsa_circ_0001286 | 1981 | 11 | 1.11% |
| hsa_circ_0015672 | 1981 | 11 | 1.11% |
| hsa_circ_0021015 | 1981 | 11 | 1.11% |
| hsa_circ_0049420 | 1981 | 11 | 1.11% |
| hsa_circ_0075081 | 1981 | 11 | 1.11% |
| hsa_circ_0076333 | 1981 | 11 | 1.11% |
| hsa_circ_0106486 | 1981 | 11 | 1.11% |
| hsa_circ_0026084 | 1801 | 10 | 1.11% |
| hsa_circ_0032615 | 1801 | 10 | 1.11% |
| hsa_circ_0035091 | 1801 | 10 | 1.11% |
| hsa_circ_0084208 | 1801 | 10 | 1.11% |
| hsa_circ_0109981 | 1801 | 10 | 1.11% |
| hsa_circ_0113322 | 1801 | 10 | 1.11% |
| hsa_circ_0010699 | 2162 | 12 | 1.11% |
| hsa_circ_0061206 | 2162 | 12 | 1.11% |
| hsa_circ_0063202 | 2162 | 12 | 1.11% |
| hsa_circ_0067305 | 2162 | 12 | 1.11% |
| hsa_circ_0126733 | 2162 | 12 | 1.11% |
| hsa_circ_0015636 | 1982 | 11 | 1.11% |
| hsa_circ_0017988 | 1982 | 11 | 1.11% |
| hsa_circ_0079261 | 1982 | 11 | 1.11% |
| hsa_circ_0106272 | 1982 | 11 | 1.11% |
| hsa_circ_0110960 | 1982 | 11 | 1.11% |
| hsa_circ_0112669 | 1982 | 11 | 1.11% |
| hsa_circ_0122288 | 1982 | 11 | 1.11% |
| hsa_circ_0136953 | 1982 | 11 | 1.11% |
| hsa_circ_0056376 | 1802 | 10 | 1.11% |
| hsa_circ_0067055 | 1802 | 10 | 1.11% |
| hsa_circ_0067698 | 1802 | 10 | 1.11% |
| hsa_circ_0084978 | 1802 | 10 | 1.11% |
| hsa_circ_0085684 | 1802 | 10 | 1.11% |
| hsa_circ_0118129 | 1802 | 10 | 1.11% |
| hsa_circ_0121987 | 1802 | 10 | 1.11% |
| hsa_circ_0124223 | 1802 | 10 | 1.11% |
| hsa_circ_0127089 | 1802 | 10 | 1.11% |

| | | | |
|---|---|---|---|
| hsa_circ_0074911 | 2163 | 12 | 1.11% |
| hsa_circ_0075279 | 2163 | 12 | 1.11% |
| hsa_circ_0010959 | 1983 | 11 | 1.11% |
| hsa_circ_0011498 | 1983 | 11 | 1.11% |
| hsa_circ_0052853 | 1983 | 11 | 1.11% |
| hsa_circ_0063031 | 1983 | 11 | 1.11% |
| hsa_circ_0083524 | 1983 | 11 | 1.11% |
| hsa_circ_0089887 | 1983 | 11 | 1.11% |
| hsa_circ_0097929 | 1983 | 11 | 1.11% |
| hsa_circ_0112460 | 1983 | 11 | 1.11% |
| hsa_circ_0006407 | 1803 | 10 | 1.11% |
| hsa_circ_0011851 | 1803 | 10 | 1.11% |
| hsa_circ_0040288 | 1803 | 10 | 1.11% |
| hsa_circ_0057326 | 1803 | 10 | 1.11% |
| hsa_circ_0069321 | 1803 | 10 | 1.11% |
| hsa_circ_0078393 | 1803 | 10 | 1.11% |
| hsa_circ_0085110 | 1803 | 10 | 1.11% |
| hsa_circ_0099325 | 1803 | 10 | 1.11% |
| hsa_circ_0052626 | 2344 | 13 | 1.11% |
| hsa_circ_0024275 | 3787 | 21 | 1.11% |
| hsa_circ_0059233 | 2164 | 12 | 1.11% |
| hsa_circ_0080657 | 2164 | 12 | 1.11% |
| hsa_circ_0089009 | 1984 | 11 | 1.11% |
| hsa_circ_0095484 | 1984 | 11 | 1.11% |
| hsa_circ_0140061 | 1984 | 11 | 1.11% |
| hsa_circ_0007679 | 6133 | 34 | 1.11% |
| hsa_circ_0023317 | 2345 | 13 | 1.11% |
| hsa_circ_0097652 | 2345 | 13 | 1.11% |
| hsa_circ_0024801 | 1804 | 10 | 1.11% |
| hsa_circ_0039162 | 1804 | 10 | 1.11% |
| hsa_circ_0040108 | 1804 | 10 | 1.11% |
| hsa_circ_0063871 | 1804 | 10 | 1.11% |
| hsa_circ_0071747 | 1804 | 10 | 1.11% |
| hsa_circ_0091132 | 1804 | 10 | 1.11% |
| hsa_circ_0140738 | 1804 | 10 | 1.11% |
| hsa_circ_0032603 | 2165 | 12 | 1.11% |
| hsa_circ_0048549 | 2165 | 12 | 1.11% |
| hsa_circ_0088875 | 2165 | 12 | 1.11% |
| hsa_circ_0052073 | 2526 | 14 | 1.11% |
| hsa_circ_0010545 | 1985 | 11 | 1.11% |
| hsa_circ_0056027 | 1985 | 11 | 1.11% |
| hsa_circ_0091027 | 1985 | 11 | 1.11% |
| hsa_circ_0017539 | 2346 | 13 | 1.11% |
| hsa_circ_0016637 | 2166 | 12 | 1.11% |
| hsa_circ_0053195 | 2166 | 12 | 1.11% |
| hsa_circ_0014293 | 1805 | 10 | 1.11% |
| hsa_circ_0034707 | 1805 | 10 | 1.11% |
| hsa_circ_0056852 | 1805 | 10 | 1.11% |
| hsa_circ_0060393 | 1805 | 10 | 1.11% |
| hsa_circ_0068218 | 1805 | 10 | 1.11% |

| | | | |
|---|---|---|---|
| hsa_circ_0097587 | 1805 | 10 | 1.11% |
| hsa_circ_0036188 | 2347 | 13 | 1.11% |
| hsa_circ_0102793 | 2347 | 13 | 1.11% |
| hsa_circ_0083243 | 1986 | 11 | 1.11% |
| hsa_circ_0104768 | 1986 | 11 | 1.11% |
| hsa_circ_0080666 | 7764 | 43 | 1.11% |
| hsa_circ_0056742 | 2528 | 14 | 1.11% |
| hsa_circ_0090601 | 2528 | 14 | 1.11% |
| hsa_circ_0026801 | 2167 | 12 | 1.11% |
| hsa_circ_0100832 | 2167 | 12 | 1.11% |
| hsa_circ_0003402 | 1806 | 10 | 1.11% |
| hsa_circ_0010018 | 1806 | 10 | 1.11% |
| hsa_circ_0012364 | 1806 | 10 | 1.11% |
| hsa_circ_0012974 | 1806 | 10 | 1.11% |
| hsa_circ_0013023 | 1806 | 10 | 1.11% |
| hsa_circ_0018259 | 1806 | 10 | 1.11% |
| hsa_circ_0022674 | 1806 | 10 | 1.11% |
| hsa_circ_0032268 | 1806 | 10 | 1.11% |
| hsa_circ_0040907 | 1806 | 10 | 1.11% |
| hsa_circ_0058371 | 1806 | 10 | 1.11% |
| hsa_circ_0082035 | 1806 | 10 | 1.11% |
| hsa_circ_0083575 | 1806 | 10 | 1.11% |
| hsa_circ_0139331 | 1806 | 10 | 1.11% |
| hsa_circ_0120690 | 5960 | 33 | 1.11% |
| hsa_circ_0131027 | 12283 | 68 | 1.11% |
| hsa_circ_0026268 | 1987 | 11 | 1.11% |
| hsa_circ_0103656 | 1987 | 11 | 1.11% |
| hsa_circ_0087681 | 4697 | 26 | 1.11% |
| hsa_circ_0052250 | 2168 | 12 | 1.11% |
| hsa_circ_0055885 | 2168 | 12 | 1.11% |
| hsa_circ_0061463 | 2168 | 12 | 1.11% |
| hsa_circ_0067363 | 2168 | 12 | 1.11% |
| hsa_circ_0072740 | 2168 | 12 | 1.11% |
| hsa_circ_0089413 | 2168 | 12 | 1.11% |
| hsa_circ_0123393 | 2168 | 12 | 1.11% |
| hsa_circ_0099552 | 8131 | 45 | 1.11% |
| hsa_circ_0012437 | 1807 | 10 | 1.11% |
| hsa_circ_0050814 | 1807 | 10 | 1.11% |
| hsa_circ_0061977 | 1807 | 10 | 1.11% |
| hsa_circ_0076343 | 1807 | 10 | 1.11% |
| hsa_circ_0124033 | 1807 | 10 | 1.11% |
| hsa_circ_0135638 | 1807 | 10 | 1.11% |
| hsa_circ_0016117 | 1988 | 11 | 1.11% |
| hsa_circ_0030825 | 1988 | 11 | 1.11% |
| hsa_circ_0059091 | 1988 | 11 | 1.11% |
| hsa_circ_0100285 | 1988 | 11 | 1.11% |
| hsa_circ_0107679 | 1988 | 11 | 1.11% |
| hsa_circ_0029014 | 6687 | 37 | 1.11% |
| hsa_circ_0005277 | 5422 | 30 | 1.11% |
| hsa_circ_0003860 | 1808 | 10 | 1.11% |

| | | | |
|---|---|---|---|
| hsa_circ_0014898 | 1808 | 10 | 1.11% |
| hsa_circ_0030808 | 1808 | 10 | 1.11% |
| hsa_circ_0044763 | 1808 | 10 | 1.11% |
| hsa_circ_0056309 | 1808 | 10 | 1.11% |
| hsa_circ_0065032 | 1808 | 10 | 1.11% |
| hsa_circ_0072303 | 1808 | 10 | 1.11% |
| hsa_circ_0073489 | 1808 | 10 | 1.11% |
| hsa_circ_0076925 | 1808 | 10 | 1.11% |
| hsa_circ_0109571 | 1808 | 10 | 1.11% |
| hsa_circ_0133126 | 1808 | 10 | 1.11% |
| hsa_circ_0103749 | 5605 | 31 | 1.11% |
| hsa_circ_0029119 | 1989 | 11 | 1.11% |
| hsa_circ_0068191 | 1989 | 11 | 1.11% |
| hsa_circ_0071287 | 1989 | 11 | 1.11% |
| hsa_circ_0074988 | 1989 | 11 | 1.11% |
| hsa_circ_0102282 | 1989 | 11 | 1.11% |
| hsa_circ_0103635 | 1989 | 11 | 1.11% |
| hsa_circ_0137056 | 1989 | 11 | 1.11% |
| hsa_circ_0010857 | 2170 | 12 | 1.11% |
| hsa_circ_0074557 | 2170 | 12 | 1.11% |
| hsa_circ_0132781 | 2170 | 12 | 1.11% |
| hsa_circ_0083472 | 2351 | 13 | 1.11% |
| hsa_circ_0130799 | 3617 | 20 | 1.11% |
| hsa_circ_0117339 | 7596 | 42 | 1.11% |
| hsa_circ_0026933 | 2532 | 14 | 1.11% |
| hsa_circ_0018614 | 1809 | 10 | 1.11% |
| hsa_circ_0018692 | 1809 | 10 | 1.11% |
| hsa_circ_0020902 | 1809 | 10 | 1.11% |
| hsa_circ_0024382 | 1809 | 10 | 1.11% |
| hsa_circ_0060390 | 1809 | 10 | 1.11% |
| hsa_circ_0075766 | 1809 | 10 | 1.11% |
| hsa_circ_0116860 | 1809 | 10 | 1.11% |
| hsa_circ_0119184 | 1809 | 10 | 1.11% |
| hsa_circ_0132303 | 1809 | 10 | 1.11% |
| hsa_circ_0028957 | 1990 | 11 | 1.11% |
| hsa_circ_0039999 | 1990 | 11 | 1.11% |
| hsa_circ_0045426 | 1990 | 11 | 1.11% |
| hsa_circ_0059905 | 1990 | 11 | 1.11% |
| hsa_circ_0076316 | 1990 | 11 | 1.11% |
| hsa_circ_0103225 | 1990 | 11 | 1.11% |
| hsa_circ_0043763 | 2171 | 12 | 1.11% |
| hsa_circ_0080112 | 2171 | 12 | 1.11% |
| hsa_circ_0102376 | 2171 | 12 | 1.11% |
| hsa_circ_0087639 | 7780 | 43 | 1.11% |
| hsa_circ_0081596 | 7961 | 44 | 1.11% |
| hsa_circ_0077008 | 2895 | 16 | 1.11% |
| hsa_circ_0119243 | 6695 | 37 | 1.11% |
| hsa_circ_0069827 | 2715 | 15 | 1.10% |
| hsa_circ_0058171 | 2353 | 13 | 1.10% |
| hsa_circ_0056519 | 2172 | 12 | 1.10% |

| | | | |
|---|---|---|---|
| hsa_circ_0062719 | 2172 | 12 | 1.10% |
| hsa_circ_0109368 | 2172 | 12 | 1.10% |
| hsa_circ_0020391 | 1991 | 11 | 1.10% |
| hsa_circ_0024565 | 1991 | 11 | 1.10% |
| hsa_circ_0044981 | 1991 | 11 | 1.10% |
| hsa_circ_0053523 | 1991 | 11 | 1.10% |
| hsa_circ_0053528 | 1991 | 11 | 1.10% |
| hsa_circ_0084657 | 1991 | 11 | 1.10% |
| hsa_circ_0086287 | 1991 | 11 | 1.10% |
| hsa_circ_0095787 | 1991 | 11 | 1.10% |
| hsa_circ_0025673 | 1810 | 10 | 1.10% |
| hsa_circ_0049866 | 1810 | 10 | 1.10% |
| hsa_circ_0064007 | 1810 | 10 | 1.10% |
| hsa_circ_0098067 | 1810 | 10 | 1.10% |
| hsa_circ_0100319 | 1810 | 10 | 1.10% |
| hsa_circ_0101159 | 1810 | 10 | 1.10% |
| hsa_circ_0111498 | 1810 | 10 | 1.10% |
| hsa_circ_0040970 | 2354 | 13 | 1.10% |
| hsa_circ_0037702 | 1992 | 11 | 1.10% |
| hsa_circ_0055302 | 1992 | 11 | 1.10% |
| hsa_circ_0065675 | 1992 | 11 | 1.10% |
| hsa_circ_0073860 | 1992 | 11 | 1.10% |
| hsa_circ_0108747 | 1992 | 11 | 1.10% |
| hsa_circ_0112529 | 1992 | 11 | 1.10% |
| hsa_circ_0127329 | 1992 | 11 | 1.10% |
| hsa_circ_0133206 | 1992 | 11 | 1.10% |
| hsa_circ_0014575 | 1811 | 10 | 1.10% |
| hsa_circ_0059601 | 1811 | 10 | 1.10% |
| hsa_circ_0069220 | 1811 | 10 | 1.10% |
| hsa_circ_0078730 | 1811 | 10 | 1.10% |
| hsa_circ_0094920 | 1811 | 10 | 1.10% |
| hsa_circ_0104522 | 1811 | 10 | 1.10% |
| hsa_circ_0118279 | 1811 | 10 | 1.10% |
| hsa_circ_0121066 | 1811 | 10 | 1.10% |
| hsa_circ_0122652 | 1811 | 10 | 1.10% |
| hsa_circ_0024551 | 2355 | 13 | 1.10% |
| hsa_circ_0060689 | 2355 | 13 | 1.10% |
| hsa_circ_0081401 | 2355 | 13 | 1.10% |
| hsa_circ_0034428 | 2174 | 12 | 1.10% |
| hsa_circ_0041572 | 2174 | 12 | 1.10% |
| hsa_circ_0016030 | 1812 | 10 | 1.10% |
| hsa_circ_0021676 | 1812 | 10 | 1.10% |
| hsa_circ_0038678 | 1812 | 10 | 1.10% |
| hsa_circ_0060412 | 1812 | 10 | 1.10% |
| hsa_circ_0061883 | 1812 | 10 | 1.10% |
| hsa_circ_0079267 | 1812 | 10 | 1.10% |
| hsa_circ_0097879 | 1812 | 10 | 1.10% |
| hsa_circ_0105777 | 1812 | 10 | 1.10% |
| hsa_circ_0113713 | 1812 | 10 | 1.10% |
| hsa_circ_0124135 | 1812 | 10 | 1.10% |

| | | | |
|---|---|---|---|
| hsa_circ_0133982 | 1812 | 10 | 1.10% |
| hsa_circ_0117272 | 3806 | 21 | 1.10% |
| hsa_circ_0109381 | 2175 | 12 | 1.10% |
| hsa_circ_0139014 | 2175 | 12 | 1.10% |
| hsa_circ_0044991 | 1994 | 11 | 1.10% |
| hsa_circ_0046457 | 1994 | 11 | 1.10% |
| hsa_circ_0063112 | 1994 | 11 | 1.10% |
| hsa_circ_0016184 | 1813 | 10 | 1.10% |
| hsa_circ_0018920 | 1813 | 10 | 1.10% |
| hsa_circ_0036526 | 1813 | 10 | 1.10% |
| hsa_circ_0042629 | 1813 | 10 | 1.10% |
| hsa_circ_0042932 | 1813 | 10 | 1.10% |
| hsa_circ_0044284 | 1813 | 10 | 1.10% |
| hsa_circ_0046998 | 1813 | 10 | 1.10% |
| hsa_circ_0047227 | 1813 | 10 | 1.10% |
| hsa_circ_0048621 | 1813 | 10 | 1.10% |
| hsa_circ_0085687 | 1813 | 10 | 1.10% |
| hsa_circ_0089123 | 1813 | 10 | 1.10% |
| hsa_circ_0114896 | 1813 | 10 | 1.10% |
| hsa_circ_0133601 | 1813 | 10 | 1.10% |
| hsa_circ_0011819 | 6709 | 37 | 1.10% |
| hsa_circ_0083558 | 2720 | 15 | 1.10% |
| hsa_circ_0090989 | 2720 | 15 | 1.10% |
| hsa_circ_0052584 | 2539 | 14 | 1.10% |
| hsa_circ_0011025 | 1995 | 11 | 1.10% |
| hsa_circ_0021828 | 1995 | 11 | 1.10% |
| hsa_circ_0039928 | 1995 | 11 | 1.10% |
| hsa_circ_0072054 | 1995 | 11 | 1.10% |
| hsa_circ_0073613 | 1995 | 11 | 1.10% |
| hsa_circ_0075061 | 1995 | 11 | 1.10% |
| hsa_circ_0079116 | 1995 | 11 | 1.10% |
| hsa_circ_0083084 | 1995 | 11 | 1.10% |
| hsa_circ_0091863 | 1995 | 11 | 1.10% |
| hsa_circ_0092083 | 1995 | 11 | 1.10% |
| hsa_circ_0112208 | 1995 | 11 | 1.10% |
| hsa_circ_0036845 | 2358 | 13 | 1.10% |
| hsa_circ_0108257 | 2358 | 13 | 1.10% |
| hsa_circ_0011731 | 1814 | 10 | 1.10% |
| hsa_circ_0014305 | 1814 | 10 | 1.10% |
| hsa_circ_0023891 | 1814 | 10 | 1.10% |
| hsa_circ_0042585 | 1814 | 10 | 1.10% |
| hsa_circ_0061494 | 1814 | 10 | 1.10% |
| hsa_circ_0064510 | 1814 | 10 | 1.10% |
| hsa_circ_0099016 | 1814 | 10 | 1.10% |
| hsa_circ_0105711 | 1814 | 10 | 1.10% |
| hsa_circ_0011680 | 2177 | 12 | 1.10% |
| hsa_circ_0131999 | 2177 | 12 | 1.10% |
| hsa_circ_0020955 | 1996 | 11 | 1.10% |
| hsa_circ_0031930 | 1996 | 11 | 1.10% |
| hsa_circ_0048610 | 1996 | 11 | 1.10% |

| | | | |
|---|---|---|---|
| hsa_circ_0061030 | 1996 | 11 | 1.10% |
| hsa_circ_0086019 | 1996 | 11 | 1.10% |
| hsa_circ_0103021 | 1996 | 11 | 1.10% |
| hsa_circ_0106414 | 1996 | 11 | 1.10% |
| hsa_circ_0112662 | 1996 | 11 | 1.10% |
| hsa_circ_0047130 | 2178 | 12 | 1.10% |
| hsa_circ_0058092 | 2178 | 12 | 1.10% |
| hsa_circ_0036533 | 1815 | 10 | 1.10% |
| hsa_circ_0048667 | 1815 | 10 | 1.10% |
| hsa_circ_0060362 | 1815 | 10 | 1.10% |
| hsa_circ_0073510 | 1815 | 10 | 1.10% |
| hsa_circ_0097661 | 1815 | 10 | 1.10% |
| hsa_circ_0098011 | 1815 | 10 | 1.10% |
| hsa_circ_0104257 | 1815 | 10 | 1.10% |
| hsa_circ_0111814 | 1815 | 10 | 1.10% |
| hsa_circ_0047418 | 2360 | 13 | 1.10% |
| hsa_circ_0067476 | 2360 | 13 | 1.10% |
| hsa_circ_0012985 | 1997 | 11 | 1.10% |
| hsa_circ_0066523 | 1997 | 11 | 1.10% |
| hsa_circ_0067916 | 1997 | 11 | 1.10% |
| hsa_circ_0105304 | 2542 | 14 | 1.10% |
| hsa_circ_0106529 | 2542 | 14 | 1.10% |
| hsa_circ_0033349 | 2179 | 12 | 1.10% |
| hsa_circ_0070795 | 2179 | 12 | 1.10% |
| hsa_circ_0029472 | 4903 | 27 | 1.10% |
| hsa_circ_0015938 | 1816 | 10 | 1.10% |
| hsa_circ_0025398 | 1816 | 10 | 1.10% |
| hsa_circ_0054378 | 1816 | 10 | 1.10% |
| hsa_circ_0057658 | 1816 | 10 | 1.10% |
| hsa_circ_0075424 | 1816 | 10 | 1.10% |
| hsa_circ_0097650 | 2361 | 13 | 1.10% |
| hsa_circ_0135094 | 2361 | 13 | 1.10% |
| hsa_circ_0026969 | 1998 | 11 | 1.10% |
| hsa_circ_0038255 | 1998 | 11 | 1.10% |
| hsa_circ_0089006 | 1998 | 11 | 1.10% |
| hsa_circ_0103284 | 1998 | 11 | 1.10% |
| hsa_circ_0135590 | 1998 | 11 | 1.10% |
| hsa_circ_0054522 | 2180 | 12 | 1.10% |
| hsa_circ_0137746 | 3452 | 19 | 1.10% |
| hsa_circ_0050300 | 2362 | 13 | 1.10% |
| hsa_circ_0009707 | 1817 | 10 | 1.10% |
| hsa_circ_0029278 | 1817 | 10 | 1.10% |
| hsa_circ_0042960 | 1817 | 10 | 1.10% |
| hsa_circ_0046825 | 1817 | 10 | 1.10% |
| hsa_circ_0091470 | 1817 | 10 | 1.10% |
| hsa_circ_0104168 | 1817 | 10 | 1.10% |
| hsa_circ_0117305 | 1817 | 10 | 1.10% |
| hsa_circ_0129496 | 1817 | 10 | 1.10% |
| hsa_circ_0135036 | 1817 | 10 | 1.10% |
| hsa_circ_0011275 | 1999 | 11 | 1.10% |

| | | | |
|---|---|---|---|
| hsa_circ_0075196 | 1999 | 11 | 1.10% |
| hsa_circ_0077308 | 1999 | 11 | 1.10% |
| hsa_circ_0085119 | 1999 | 11 | 1.10% |
| hsa_circ_0114883 | 5816 | 32 | 1.10% |
| hsa_circ_0081500 | 2908 | 16 | 1.10% |
| hsa_circ_0017568 | 2181 | 12 | 1.10% |
| hsa_circ_0026897 | 2181 | 12 | 1.10% |
| hsa_circ_0037283 | 2181 | 12 | 1.10% |
| hsa_circ_0065873 | 2181 | 12 | 1.10% |
| hsa_circ_0067798 | 2181 | 12 | 1.10% |
| hsa_circ_0076884 | 2181 | 12 | 1.10% |
| hsa_circ_0047814 | 2363 | 13 | 1.10% |
| hsa_circ_0030867 | 2545 | 14 | 1.10% |
| hsa_circ_0083140 | 2545 | 14 | 1.10% |
| hsa_circ_0018125 | 1818 | 10 | 1.10% |
| hsa_circ_0021916 | 1818 | 10 | 1.10% |
| hsa_circ_0026845 | 1818 | 10 | 1.10% |
| hsa_circ_0028741 | 1818 | 10 | 1.10% |
| hsa_circ_0034676 | 1818 | 10 | 1.10% |
| hsa_circ_0035431 | 1818 | 10 | 1.10% |
| hsa_circ_0043059 | 1818 | 10 | 1.10% |
| hsa_circ_0056301 | 1818 | 10 | 1.10% |
| hsa_circ_0056608 | 1818 | 10 | 1.10% |
| hsa_circ_0080685 | 1818 | 10 | 1.10% |
| hsa_circ_0135357 | 1818 | 10 | 1.10% |
| hsa_circ_0027218 | 2000 | 11 | 1.10% |
| hsa_circ_0034158 | 2000 | 11 | 1.10% |
| hsa_circ_0037203 | 2000 | 11 | 1.10% |
| hsa_circ_0047854 | 2000 | 11 | 1.10% |
| hsa_circ_0063916 | 2000 | 11 | 1.10% |
| hsa_circ_0076543 | 2000 | 11 | 1.10% |
| hsa_circ_0079067 | 2000 | 11 | 1.10% |
| hsa_circ_0107361 | 2000 | 11 | 1.10% |
| hsa_circ_0036433 | 2182 | 12 | 1.10% |
| hsa_circ_0043493 | 2182 | 12 | 1.10% |
| hsa_circ_0046155 | 7093 | 39 | 1.10% |
| hsa_circ_0128501 | 2910 | 16 | 1.10% |
| hsa_circ_0001537 | 1819 | 10 | 1.10% |
| hsa_circ_0016431 | 1819 | 10 | 1.10% |
| hsa_circ_0030630 | 1819 | 10 | 1.10% |
| hsa_circ_0077462 | 1819 | 10 | 1.10% |
| hsa_circ_0048635 | 2001 | 11 | 1.10% |
| hsa_circ_0051677 | 2001 | 11 | 1.10% |
| hsa_circ_0055211 | 2001 | 11 | 1.10% |
| hsa_circ_0091673 | 2001 | 11 | 1.10% |
| hsa_circ_0103014 | 2001 | 11 | 1.10% |
| hsa_circ_0090782 | 2183 | 12 | 1.10% |
| hsa_circ_0039160 | 2365 | 13 | 1.10% |
| hsa_circ_0060870 | 2365 | 13 | 1.10% |
| hsa_circ_0025263 | 2547 | 14 | 1.10% |

| | | | |
|---|---|---|---|
| hsa_circ_0123921 | 5095 | 28 | 1.10% |
| hsa_circ_0024631 | 2366 | 13 | 1.10% |
| hsa_circ_0085534 | 2366 | 13 | 1.10% |
| hsa_circ_0026332 | 2184 | 12 | 1.10% |
| hsa_circ_0034819 | 2184 | 12 | 1.10% |
| hsa_circ_0040991 | 2184 | 12 | 1.10% |
| hsa_circ_0131997 | 2184 | 12 | 1.10% |
| hsa_circ_0018668 | 2002 | 11 | 1.10% |
| hsa_circ_0025149 | 2002 | 11 | 1.10% |
| hsa_circ_0031476 | 2002 | 11 | 1.10% |
| hsa_circ_0064273 | 2002 | 11 | 1.10% |
| hsa_circ_0033459 | 1820 | 10 | 1.10% |
| hsa_circ_0045552 | 1820 | 10 | 1.10% |
| hsa_circ_0049878 | 1820 | 10 | 1.10% |
| hsa_circ_0057641 | 1820 | 10 | 1.10% |
| hsa_circ_0061414 | 1820 | 10 | 1.10% |
| hsa_circ_0062722 | 1820 | 10 | 1.10% |
| hsa_circ_0067759 | 1820 | 10 | 1.10% |
| hsa_circ_0071944 | 1820 | 10 | 1.10% |
| hsa_circ_0082993 | 1820 | 10 | 1.10% |
| hsa_circ_0128965 | 1820 | 10 | 1.10% |
| hsa_circ_0134279 | 5097 | 28 | 1.10% |
| hsa_circ_0073739 | 3641 | 20 | 1.10% |
| hsa_circ_0036903 | 2185 | 12 | 1.10% |
| hsa_circ_0103965 | 2185 | 12 | 1.10% |
| hsa_circ_0138879 | 2185 | 12 | 1.10% |
| hsa_circ_0080143 | 2003 | 11 | 1.10% |
| hsa_circ_0089607 | 2003 | 11 | 1.10% |
| hsa_circ_0095829 | 2003 | 11 | 1.10% |
| hsa_circ_0100914 | 2003 | 11 | 1.10% |
| hsa_circ_0117855 | 2003 | 11 | 1.10% |
| hsa_circ_0118476 | 2003 | 11 | 1.10% |
| hsa_circ_0138840 | 2003 | 11 | 1.10% |
| hsa_circ_0024363 | 1821 | 10 | 1.10% |
| hsa_circ_0026843 | 1821 | 10 | 1.10% |
| hsa_circ_0029751 | 1821 | 10 | 1.10% |
| hsa_circ_0035333 | 1821 | 10 | 1.10% |
| hsa_circ_0036644 | 1821 | 10 | 1.10% |
| hsa_circ_0044363 | 1821 | 10 | 1.10% |
| hsa_circ_0055306 | 1821 | 10 | 1.10% |
| hsa_circ_0061604 | 1821 | 10 | 1.10% |
| hsa_circ_0063020 | 1821 | 10 | 1.10% |
| hsa_circ_0090553 | 1821 | 10 | 1.10% |
| hsa_circ_0099165 | 1821 | 10 | 1.10% |
| hsa_circ_0124063 | 1821 | 10 | 1.10% |
| hsa_circ_0139627 | 1821 | 10 | 1.10% |
| hsa_circ_0042127 | 2186 | 12 | 1.10% |
| hsa_circ_0069645 | 2186 | 12 | 1.10% |
| hsa_circ_0078861 | 4190 | 23 | 1.10% |
| hsa_circ_0014454 | 2004 | 11 | 1.10% |

| | | | |
|---|---|---|---|
| hsa_circ_0045558 | 2004 | 11 | 1.10% |
| hsa_circ_0069065 | 2004 | 11 | 1.10% |
| hsa_circ_0099908 | 2004 | 11 | 1.10% |
| hsa_circ_0105538 | 2004 | 11 | 1.10% |
| hsa_circ_0014382 | 1822 | 10 | 1.10% |
| hsa_circ_0028223 | 1822 | 10 | 1.10% |
| hsa_circ_0030112 | 1822 | 10 | 1.10% |
| hsa_circ_0060250 | 1822 | 10 | 1.10% |
| hsa_circ_0076968 | 1822 | 10 | 1.10% |
| hsa_circ_0086233 | 1822 | 10 | 1.10% |
| hsa_circ_0123375 | 1822 | 10 | 1.10% |
| hsa_circ_0087837 | 2369 | 13 | 1.10% |
| hsa_circ_0033585 | 2187 | 12 | 1.10% |
| hsa_circ_0050021 | 2187 | 12 | 1.10% |
| hsa_circ_0050907 | 2005 | 11 | 1.10% |
| hsa_circ_0068914 | 2005 | 11 | 1.10% |
| hsa_circ_0085585 | 2005 | 11 | 1.10% |
| hsa_circ_0099120 | 2005 | 11 | 1.10% |
| hsa_circ_0136304 | 2005 | 11 | 1.10% |
| hsa_circ_0136689 | 2005 | 11 | 1.10% |
| hsa_circ_0066057 | 2552 | 14 | 1.10% |
| hsa_circ_0032818 | 1823 | 10 | 1.10% |
| hsa_circ_0067882 | 1823 | 10 | 1.10% |
| hsa_circ_0088238 | 1823 | 10 | 1.10% |
| hsa_circ_0118889 | 1823 | 10 | 1.10% |
| hsa_circ_0086010 | 2370 | 13 | 1.10% |
| hsa_circ_0040285 | 2006 | 11 | 1.10% |
| hsa_circ_0053890 | 2006 | 11 | 1.10% |
| hsa_circ_0121834 | 2006 | 11 | 1.10% |
| hsa_circ_0023728 | 3648 | 20 | 1.10% |
| hsa_circ_0004305 | 1824 | 10 | 1.10% |
| hsa_circ_0017344 | 1824 | 10 | 1.10% |
| hsa_circ_0050548 | 1824 | 10 | 1.10% |
| hsa_circ_0079710 | 1824 | 10 | 1.10% |
| hsa_circ_0138386 | 1824 | 10 | 1.10% |
| hsa_circ_0029013 | 6749 | 37 | 1.10% |
| hsa_circ_0103824 | 2189 | 12 | 1.10% |
| hsa_circ_0018693 | 2007 | 11 | 1.10% |
| hsa_circ_0020303 | 2007 | 11 | 1.10% |
| hsa_circ_0029947 | 2007 | 11 | 1.10% |
| hsa_circ_0032424 | 2007 | 11 | 1.10% |
| hsa_circ_0042914 | 2007 | 11 | 1.10% |
| hsa_circ_0067166 | 2007 | 11 | 1.10% |
| hsa_circ_0076334 | 2007 | 11 | 1.10% |
| hsa_circ_0126310 | 2007 | 11 | 1.10% |
| hsa_circ_0081601 | 2372 | 13 | 1.10% |
| hsa_circ_0128418 | 2920 | 16 | 1.10% |
| hsa_circ_0048255 | 2190 | 12 | 1.10% |
| hsa_circ_0004465 | 1825 | 10 | 1.10% |
| hsa_circ_0011805 | 1825 | 10 | 1.10% |

| | | | |
|---|---|---|---|
| hsa_circ_0045653 | 1825 | 10 | 1.10% |
| hsa_circ_0066574 | 1825 | 10 | 1.10% |
| hsa_circ_0067487 | 1825 | 10 | 1.10% |
| hsa_circ_0078413 | 1825 | 10 | 1.10% |
| hsa_circ_0130901 | 1825 | 10 | 1.10% |
| hsa_circ_0126180 | 3468 | 19 | 1.10% |
| hsa_circ_0022851 | 2008 | 11 | 1.10% |
| hsa_circ_0076785 | 2008 | 11 | 1.10% |
| hsa_circ_0082931 | 2008 | 11 | 1.10% |
| hsa_circ_0083006 | 2008 | 11 | 1.10% |
| hsa_circ_0094186 | 2008 | 11 | 1.10% |
| hsa_circ_0016507 | 2191 | 12 | 1.10% |
| hsa_circ_0065316 | 2191 | 12 | 1.10% |
| hsa_circ_0119715 | 2191 | 12 | 1.10% |
| hsa_circ_0009882 | 1826 | 10 | 1.10% |
| hsa_circ_0035352 | 1826 | 10 | 1.10% |
| hsa_circ_0042328 | 1826 | 10 | 1.10% |
| hsa_circ_0044552 | 1826 | 10 | 1.10% |
| hsa_circ_0047425 | 1826 | 10 | 1.10% |
| hsa_circ_0089274 | 1826 | 10 | 1.10% |
| hsa_circ_0104117 | 1826 | 10 | 1.10% |
| hsa_circ_0106603 | 1826 | 10 | 1.10% |
| hsa_circ_0140735 | 1826 | 10 | 1.10% |
| hsa_circ_0120297 | 2922 | 16 | 1.10% |
| hsa_circ_0087922 | 2009 | 11 | 1.10% |
| hsa_circ_0007922 | 2557 | 14 | 1.10% |
| hsa_circ_0086180 | 2557 | 14 | 1.10% |
| hsa_circ_0091014 | 2192 | 12 | 1.09% |
| hsa_circ_0064896 | 2375 | 13 | 1.09% |
| hsa_circ_0127829 | 3654 | 20 | 1.09% |
| hsa_circ_0009840 | 1827 | 10 | 1.09% |
| hsa_circ_0026183 | 1827 | 10 | 1.09% |
| hsa_circ_0036893 | 1827 | 10 | 1.09% |
| hsa_circ_0039842 | 1827 | 10 | 1.09% |
| hsa_circ_0042591 | 1827 | 10 | 1.09% |
| hsa_circ_0047065 | 1827 | 10 | 1.09% |
| hsa_circ_0067958 | 1827 | 10 | 1.09% |
| hsa_circ_0068281 | 1827 | 10 | 1.09% |
| hsa_circ_0082123 | 1827 | 10 | 1.09% |
| hsa_circ_0090443 | 1827 | 10 | 1.09% |
| hsa_circ_0090950 | 1827 | 10 | 1.09% |
| hsa_circ_0137523 | 1827 | 10 | 1.09% |
| hsa_circ_0021168 | 2010 | 11 | 1.09% |
| hsa_circ_0036773 | 2010 | 11 | 1.09% |
| hsa_circ_0056214 | 2010 | 11 | 1.09% |
| hsa_circ_0057666 | 2010 | 11 | 1.09% |
| hsa_circ_0087254 | 2010 | 11 | 1.09% |
| hsa_circ_0087536 | 2010 | 11 | 1.09% |
| hsa_circ_0093844 | 2010 | 11 | 1.09% |
| hsa_circ_0132762 | 2010 | 11 | 1.09% |

| | | | |
|---|---|---|---|
| hsa_circ_0106978 | 2741 | 15 | 1.09% |
| hsa_circ_0009264 | 2193 | 12 | 1.09% |
| hsa_circ_0089998 | 2193 | 12 | 1.09% |
| hsa_circ_0063518 | 7677 | 42 | 1.09% |
| hsa_circ_0005873 | 4570 | 25 | 1.09% |
| hsa_circ_0024004 | 1828 | 10 | 1.09% |
| hsa_circ_0038570 | 1828 | 10 | 1.09% |
| hsa_circ_0063403 | 1828 | 10 | 1.09% |
| hsa_circ_0011480 | 2011 | 11 | 1.09% |
| hsa_circ_0024645 | 2011 | 11 | 1.09% |
| hsa_circ_0083944 | 2011 | 11 | 1.09% |
| hsa_circ_0107211 | 2011 | 11 | 1.09% |
| hsa_circ_0060913 | 2194 | 12 | 1.09% |
| hsa_circ_0087537 | 2194 | 12 | 1.09% |
| hsa_circ_0122895 | 2194 | 12 | 1.09% |
| hsa_circ_0036131 | 2560 | 14 | 1.09% |
| hsa_circ_0022879 | 2743 | 15 | 1.09% |
| hsa_circ_0084197 | 1829 | 10 | 1.09% |
| hsa_circ_0122684 | 1829 | 10 | 1.09% |
| hsa_circ_0125300 | 1829 | 10 | 1.09% |
| hsa_circ_0040167 | 2012 | 11 | 1.09% |
| hsa_circ_0049982 | 2012 | 11 | 1.09% |
| hsa_circ_0119827 | 2012 | 11 | 1.09% |
| hsa_circ_0120666 | 2012 | 11 | 1.09% |
| hsa_circ_0132356 | 2012 | 11 | 1.09% |
| hsa_circ_0049747 | 2195 | 12 | 1.09% |
| hsa_circ_0014758 | 2379 | 13 | 1.09% |
| hsa_circ_0063506 | 2196 | 12 | 1.09% |
| hsa_circ_0077680 | 2196 | 12 | 1.09% |
| hsa_circ_0085629 | 2196 | 12 | 1.09% |
| hsa_circ_0034941 | 2013 | 11 | 1.09% |
| hsa_circ_0042680 | 2013 | 11 | 1.09% |
| hsa_circ_0119947 | 2013 | 11 | 1.09% |
| hsa_circ_0035661 | 1830 | 10 | 1.09% |
| hsa_circ_0046569 | 1830 | 10 | 1.09% |
| hsa_circ_0047544 | 1830 | 10 | 1.09% |
| hsa_circ_0051604 | 1830 | 10 | 1.09% |
| hsa_circ_0064387 | 1830 | 10 | 1.09% |
| hsa_circ_0088150 | 1830 | 10 | 1.09% |
| hsa_circ_0105603 | 3478 | 19 | 1.09% |
| hsa_circ_0016300 | 2380 | 13 | 1.09% |
| hsa_circ_0013685 | 2197 | 12 | 1.09% |
| hsa_circ_0003408 | 2014 | 11 | 1.09% |
| hsa_circ_0045195 | 2014 | 11 | 1.09% |
| hsa_circ_0052332 | 2014 | 11 | 1.09% |
| hsa_circ_0066537 | 2014 | 11 | 1.09% |
| hsa_circ_0070777 | 2014 | 11 | 1.09% |
| hsa_circ_0094828 | 2014 | 11 | 1.09% |
| hsa_circ_0102806 | 2014 | 11 | 1.09% |
| hsa_circ_0042068 | 1831 | 10 | 1.09% |

| | | | |
|---|---|---|---|
| hsa_circ_0052762 | 1831 | 10 | 1.09% |
| hsa_circ_0063437 | 1831 | 10 | 1.09% |
| hsa_circ_0087789 | 1831 | 10 | 1.09% |
| hsa_circ_0134802 | 1831 | 10 | 1.09% |
| hsa_circ_0135582 | 1831 | 10 | 1.09% |
| hsa_circ_0139066 | 9340 | 51 | 1.09% |
| hsa_circ_0055929 | 2015 | 11 | 1.09% |
| hsa_circ_0087897 | 2015 | 11 | 1.09% |
| hsa_circ_0110261 | 6961 | 38 | 1.09% |
| hsa_circ_0010723 | 1832 | 10 | 1.09% |
| hsa_circ_0027978 | 1832 | 10 | 1.09% |
| hsa_circ_0058803 | 1832 | 10 | 1.09% |
| hsa_circ_0067178 | 1832 | 10 | 1.09% |
| hsa_circ_0098750 | 1832 | 10 | 1.09% |
| hsa_circ_0114515 | 1832 | 10 | 1.09% |
| hsa_circ_0023346 | 2199 | 12 | 1.09% |
| hsa_circ_0066845 | 2199 | 12 | 1.09% |
| hsa_circ_0081263 | 2749 | 15 | 1.09% |
| hsa_circ_0003924 | 2016 | 11 | 1.09% |
| hsa_circ_0051363 | 2016 | 11 | 1.09% |
| hsa_circ_0063394 | 2016 | 11 | 1.09% |
| hsa_circ_0111849 | 2016 | 11 | 1.09% |
| hsa_circ_0115760 | 2016 | 11 | 1.09% |
| hsa_circ_0126212 | 2016 | 11 | 1.09% |
| hsa_circ_0017224 | 1833 | 10 | 1.09% |
| hsa_circ_0028284 | 1833 | 10 | 1.09% |
| hsa_circ_0028910 | 1833 | 10 | 1.09% |
| hsa_circ_0059287 | 1833 | 10 | 1.09% |
| hsa_circ_0083980 | 1833 | 10 | 1.09% |
| hsa_circ_0113816 | 1833 | 10 | 1.09% |
| hsa_circ_0124490 | 1833 | 10 | 1.09% |
| hsa_circ_0052168 | 2383 | 13 | 1.09% |
| hsa_circ_0053210 | 2383 | 13 | 1.09% |
| hsa_circ_0074027 | 2383 | 13 | 1.09% |
| hsa_circ_0100672 | 2383 | 13 | 1.09% |
| hsa_circ_0016448 | 2200 | 12 | 1.09% |
| hsa_circ_0041136 | 2200 | 12 | 1.09% |
| hsa_circ_0005381 | 2017 | 11 | 1.09% |
| hsa_circ_0035572 | 2017 | 11 | 1.09% |
| hsa_circ_0059157 | 2017 | 11 | 1.09% |
| hsa_circ_0066425 | 2017 | 11 | 1.09% |
| hsa_circ_0081538 | 2017 | 11 | 1.09% |
| hsa_circ_0137778 | 2017 | 11 | 1.09% |
| hsa_circ_0036956 | 2384 | 13 | 1.09% |
| hsa_circ_0103326 | 1834 | 10 | 1.09% |
| hsa_circ_0103828 | 1834 | 10 | 1.09% |
| hsa_circ_0107002 | 1834 | 10 | 1.09% |
| hsa_circ_0116106 | 1834 | 10 | 1.09% |
| hsa_circ_0139065 | 9354 | 51 | 1.09% |
| hsa_circ_0037076 | 2201 | 12 | 1.09% |

| | | | |
|---|---|---|---|
| hsa_circ_0082295 | 2568 | 14 | 1.09% |
| hsa_circ_0009548 | 2018 | 11 | 1.09% |
| hsa_circ_0018293 | 2018 | 11 | 1.09% |
| hsa_circ_0037727 | 2018 | 11 | 1.09% |
| hsa_circ_0059352 | 2018 | 11 | 1.09% |
| hsa_circ_0062007 | 2018 | 11 | 1.09% |
| hsa_circ_0084440 | 2018 | 11 | 1.09% |
| hsa_circ_0093521 | 2018 | 11 | 1.09% |
| hsa_circ_0094464 | 2018 | 11 | 1.09% |
| hsa_circ_0127778 | 2018 | 11 | 1.09% |
| hsa_circ_0132070 | 2018 | 11 | 1.09% |
| hsa_circ_0043087 | 2385 | 13 | 1.09% |
| hsa_circ_0082226 | 2202 | 12 | 1.09% |
| hsa_circ_0022634 | 1835 | 10 | 1.09% |
| hsa_circ_0113543 | 1835 | 10 | 1.09% |
| hsa_circ_0035280 | 2019 | 11 | 1.09% |
| hsa_circ_0038065 | 2019 | 11 | 1.09% |
| hsa_circ_0043993 | 2019 | 11 | 1.09% |
| hsa_circ_0061391 | 2019 | 11 | 1.09% |
| hsa_circ_0071594 | 2019 | 11 | 1.09% |
| hsa_circ_0090558 | 2019 | 11 | 1.09% |
| hsa_circ_0098528 | 2019 | 11 | 1.09% |
| hsa_circ_0081687 | 2570 | 14 | 1.09% |
| hsa_circ_0120507 | 2570 | 14 | 1.09% |
| hsa_circ_0035602 | 2203 | 12 | 1.09% |
| hsa_circ_0011229 | 1836 | 10 | 1.09% |
| hsa_circ_0030878 | 1836 | 10 | 1.09% |
| hsa_circ_0043565 | 1836 | 10 | 1.09% |
| hsa_circ_0064600 | 1836 | 10 | 1.09% |
| hsa_circ_0078125 | 1836 | 10 | 1.09% |
| hsa_circ_0079870 | 1836 | 10 | 1.09% |
| hsa_circ_0084593 | 1836 | 10 | 1.09% |
| hsa_circ_0112577 | 1836 | 10 | 1.09% |
| hsa_circ_0114956 | 1836 | 10 | 1.09% |
| hsa_circ_0122721 | 1836 | 10 | 1.09% |
| hsa_circ_0134920 | 1836 | 10 | 1.09% |
| hsa_circ_0052202 | 2387 | 13 | 1.09% |
| hsa_circ_0021290 | 2020 | 11 | 1.09% |
| hsa_circ_0023212 | 2020 | 11 | 1.09% |
| hsa_circ_0126244 | 2020 | 11 | 1.09% |
| hsa_circ_0130221 | 2571 | 14 | 1.09% |
| hsa_circ_0022885 | 2755 | 15 | 1.09% |
| hsa_circ_0022820 | 2204 | 12 | 1.09% |
| hsa_circ_0056961 | 2204 | 12 | 1.09% |
| hsa_circ_0118739 | 2204 | 12 | 1.09% |
| hsa_circ_0001244 | 1837 | 10 | 1.09% |
| hsa_circ_0034165 | 1837 | 10 | 1.09% |
| hsa_circ_0039060 | 1837 | 10 | 1.09% |
| hsa_circ_0041906 | 1837 | 10 | 1.09% |
| hsa_circ_0059481 | 1837 | 10 | 1.09% |

| | | | |
|---|---|---|---|
| hsa_circ_0098361 | 1837 | 10 | 1.09% |
| hsa_circ_0123376 | 1837 | 10 | 1.09% |
| hsa_circ_0020864 | 2021 | 11 | 1.09% |
| hsa_circ_0053621 | 2021 | 11 | 1.09% |
| hsa_circ_0084375 | 2021 | 11 | 1.09% |
| hsa_circ_0116241 | 2021 | 11 | 1.09% |
| hsa_circ_0117795 | 2021 | 11 | 1.09% |
| hsa_circ_0016993 | 2205 | 12 | 1.09% |
| hsa_circ_0043227 | 2205 | 12 | 1.09% |
| hsa_circ_0051257 | 2205 | 12 | 1.09% |
| hsa_circ_0010532 | 1838 | 10 | 1.09% |
| hsa_circ_0012931 | 1838 | 10 | 1.09% |
| hsa_circ_0029230 | 1838 | 10 | 1.09% |
| hsa_circ_0074455 | 1838 | 10 | 1.09% |
| hsa_circ_0130026 | 1838 | 10 | 1.09% |
| hsa_circ_0004100 | 2022 | 11 | 1.09% |
| hsa_circ_0098145 | 2022 | 11 | 1.09% |
| hsa_circ_0101656 | 2022 | 11 | 1.09% |
| hsa_circ_0131028 | 12500 | 68 | 1.09% |
| hsa_circ_0022116 | 2206 | 12 | 1.09% |
| hsa_circ_0041200 | 2206 | 12 | 1.09% |
| hsa_circ_0118339 | 2206 | 12 | 1.09% |
| hsa_circ_0135930 | 4045 | 22 | 1.09% |
| hsa_circ_0006164 | 1839 | 10 | 1.09% |
| hsa_circ_0009724 | 1839 | 10 | 1.09% |
| hsa_circ_0021669 | 1839 | 10 | 1.09% |
| hsa_circ_0047974 | 1839 | 10 | 1.09% |
| hsa_circ_0051553 | 1839 | 10 | 1.09% |
| hsa_circ_0078114 | 1839 | 10 | 1.09% |
| hsa_circ_0084582 | 1839 | 10 | 1.09% |
| hsa_circ_0101627 | 1839 | 10 | 1.09% |
| hsa_circ_0116277 | 1839 | 10 | 1.09% |
| hsa_circ_0117461 | 1839 | 10 | 1.09% |
| hsa_circ_0125517 | 1839 | 10 | 1.09% |
| hsa_circ_0019887 | 2023 | 11 | 1.09% |
| hsa_circ_0022817 | 2023 | 11 | 1.09% |
| hsa_circ_0058422 | 2023 | 11 | 1.09% |
| hsa_circ_0066331 | 2023 | 11 | 1.09% |
| hsa_circ_0127965 | 2023 | 11 | 1.09% |
| hsa_circ_0082885 | 2207 | 12 | 1.09% |
| hsa_circ_0009171 | 3680 | 20 | 1.09% |
| hsa_circ_0027163 | 2208 | 12 | 1.09% |
| hsa_circ_0009363 | 2024 | 11 | 1.09% |
| hsa_circ_0047479 | 2024 | 11 | 1.09% |
| hsa_circ_0050314 | 2024 | 11 | 1.09% |
| hsa_circ_0075044 | 2024 | 11 | 1.09% |
| hsa_circ_0078042 | 2024 | 11 | 1.09% |
| hsa_circ_0084053 | 2024 | 11 | 1.09% |
| hsa_circ_0139406 | 2024 | 11 | 1.09% |
| hsa_circ_0013324 | 1840 | 10 | 1.09% |

| | | | |
|---|---|---|---|
| hsa_circ_0032395 | 1840 | 10 | 1.09% |
| hsa_circ_0035320 | 1840 | 10 | 1.09% |
| hsa_circ_0039592 | 1840 | 10 | 1.09% |
| hsa_circ_0041334 | 1840 | 10 | 1.09% |
| hsa_circ_0054715 | 1840 | 10 | 1.09% |
| hsa_circ_0061375 | 1840 | 10 | 1.09% |
| hsa_circ_0081827 | 1840 | 10 | 1.09% |
| hsa_circ_0082652 | 1840 | 10 | 1.09% |
| hsa_circ_0085161 | 1840 | 10 | 1.09% |
| hsa_circ_0104173 | 1840 | 10 | 1.09% |
| hsa_circ_0126515 | 1840 | 10 | 1.09% |
| hsa_circ_0024374 | 3865 | 21 | 1.09% |
| hsa_circ_0090122 | 2577 | 14 | 1.09% |
| hsa_circ_0046474 | 2025 | 11 | 1.09% |
| hsa_circ_0089158 | 2025 | 11 | 1.09% |
| hsa_circ_0129102 | 7364 | 40 | 1.09% |
| hsa_circ_0008643 | 3682 | 20 | 1.09% |
| hsa_circ_0016444 | 1841 | 10 | 1.09% |
| hsa_circ_0034915 | 1841 | 10 | 1.09% |
| hsa_circ_0042638 | 1841 | 10 | 1.09% |
| hsa_circ_0050811 | 1841 | 10 | 1.09% |
| hsa_circ_0055841 | 1841 | 10 | 1.09% |
| hsa_circ_0058595 | 1841 | 10 | 1.09% |
| hsa_circ_0066048 | 1841 | 10 | 1.09% |
| hsa_circ_0073907 | 1841 | 10 | 1.09% |
| hsa_circ_0084387 | 1841 | 10 | 1.09% |
| hsa_circ_0086820 | 1841 | 10 | 1.09% |
| hsa_circ_0110515 | 1841 | 10 | 1.09% |
| hsa_circ_0132806 | 1841 | 10 | 1.09% |
| hsa_circ_0134609 | 1841 | 10 | 1.09% |
| hsa_circ_0038209 | 6812 | 37 | 1.09% |
| hsa_circ_0128983 | 4603 | 25 | 1.09% |
| hsa_circ_0085039 | 2578 | 14 | 1.09% |
| hsa_circ_0016086 | 2210 | 12 | 1.09% |
| hsa_circ_0123184 | 2210 | 12 | 1.09% |
| hsa_circ_0038751 | 2026 | 11 | 1.09% |
| hsa_circ_0068325 | 2026 | 11 | 1.09% |
| hsa_circ_0081835 | 2026 | 11 | 1.09% |
| hsa_circ_0087804 | 2026 | 11 | 1.09% |
| hsa_circ_0091161 | 2026 | 11 | 1.09% |
| hsa_circ_0029011 | 6815 | 37 | 1.09% |
| hsa_circ_0012702 | 1842 | 10 | 1.09% |
| hsa_circ_0014434 | 1842 | 10 | 1.09% |
| hsa_circ_0045972 | 1842 | 10 | 1.09% |
| hsa_circ_0047632 | 1842 | 10 | 1.09% |
| hsa_circ_0056183 | 1842 | 10 | 1.09% |
| hsa_circ_0115180 | 1842 | 10 | 1.09% |
| hsa_circ_0016982 | 2211 | 12 | 1.09% |
| hsa_circ_0022082 | 2027 | 11 | 1.09% |
| hsa_circ_0040047 | 2027 | 11 | 1.09% |

| | | | |
|---|---|---|---|
| hsa_circ_0052262 | 2027 | 11 | 1.09% |
| hsa_circ_0065423 | 2027 | 11 | 1.09% |
| hsa_circ_0066929 | 2027 | 11 | 1.09% |
| hsa_circ_0101465 | 4976 | 27 | 1.09% |
| hsa_circ_0136462 | 3686 | 20 | 1.09% |
| hsa_circ_0009787 | 1843 | 10 | 1.09% |
| hsa_circ_0023124 | 1843 | 10 | 1.09% |
| hsa_circ_0032393 | 1843 | 10 | 1.09% |
| hsa_circ_0048728 | 1843 | 10 | 1.09% |
| hsa_circ_0125246 | 1843 | 10 | 1.09% |
| hsa_circ_0135399 | 1843 | 10 | 1.09% |
| hsa_circ_0137854 | 1843 | 10 | 1.09% |
| hsa_circ_0019945 | 2396 | 13 | 1.09% |
| hsa_circ_0124118 | 8847 | 48 | 1.09% |
| hsa_circ_0016137 | 2028 | 11 | 1.08% |
| hsa_circ_0016787 | 2028 | 11 | 1.08% |
| hsa_circ_0041199 | 2028 | 11 | 1.08% |
| hsa_circ_0050817 | 2028 | 11 | 1.08% |
| hsa_circ_0067892 | 2028 | 11 | 1.08% |
| hsa_circ_0081870 | 2028 | 11 | 1.08% |
| hsa_circ_0086555 | 2028 | 11 | 1.08% |
| hsa_circ_0120671 | 2028 | 11 | 1.08% |
| hsa_circ_0076408 | 1844 | 10 | 1.08% |
| hsa_circ_0078526 | 1844 | 10 | 1.08% |
| hsa_circ_0084003 | 1844 | 10 | 1.08% |
| hsa_circ_0107001 | 1844 | 10 | 1.08% |
| hsa_circ_0122338 | 1844 | 10 | 1.08% |
| hsa_circ_0131920 | 1844 | 10 | 1.08% |
| hsa_circ_0057356 | 3135 | 17 | 1.08% |
| hsa_circ_0018863 | 2213 | 12 | 1.08% |
| hsa_circ_0046252 | 2029 | 11 | 1.08% |
| hsa_circ_0075080 | 2029 | 11 | 1.08% |
| hsa_circ_0076556 | 2029 | 11 | 1.08% |
| hsa_circ_0114272 | 2029 | 11 | 1.08% |
| hsa_circ_0015690 | 2583 | 14 | 1.08% |
| hsa_circ_0009964 | 2214 | 12 | 1.08% |
| hsa_circ_0024027 | 2214 | 12 | 1.08% |
| hsa_circ_0032920 | 1845 | 10 | 1.08% |
| hsa_circ_0058139 | 1845 | 10 | 1.08% |
| hsa_circ_0072722 | 1845 | 10 | 1.08% |
| hsa_circ_0079253 | 1845 | 10 | 1.08% |
| hsa_circ_0083768 | 1845 | 10 | 1.08% |
| hsa_circ_0095303 | 1845 | 10 | 1.08% |
| hsa_circ_0120878 | 1845 | 10 | 1.08% |
| hsa_circ_0123656 | 1845 | 10 | 1.08% |
| hsa_circ_0011937 | 2030 | 11 | 1.08% |
| hsa_circ_0022864 | 2030 | 11 | 1.08% |
| hsa_circ_0038216 | 2030 | 11 | 1.08% |
| hsa_circ_0065621 | 2030 | 11 | 1.08% |
| hsa_circ_0130760 | 2030 | 11 | 1.08% |

| | | | |
|---|---|---|---|
| hsa_circ_0136898 | 2030 | 11 | 1.08% |
| hsa_circ_0037997 | 2215 | 12 | 1.08% |
| hsa_circ_0047541 | 1846 | 10 | 1.08% |
| hsa_circ_0057439 | 2400 | 13 | 1.08% |
| hsa_circ_0053788 | 2031 | 11 | 1.08% |
| hsa_circ_0057206 | 2031 | 11 | 1.08% |
| hsa_circ_0104304 | 2031 | 11 | 1.08% |
| hsa_circ_0114161 | 2031 | 11 | 1.08% |
| hsa_circ_0022044 | 2585 | 14 | 1.08% |
| hsa_circ_0047762 | 2216 | 12 | 1.08% |
| hsa_circ_0087971 | 2216 | 12 | 1.08% |
| hsa_circ_0018821 | 1847 | 10 | 1.08% |
| hsa_circ_0027238 | 1847 | 10 | 1.08% |
| hsa_circ_0042408 | 1847 | 10 | 1.08% |
| hsa_circ_0068176 | 1847 | 10 | 1.08% |
| hsa_circ_0075963 | 1847 | 10 | 1.08% |
| hsa_circ_0077177 | 1847 | 10 | 1.08% |
| hsa_circ_0097798 | 1847 | 10 | 1.08% |
| hsa_circ_0112171 | 1847 | 10 | 1.08% |
| hsa_circ_0116939 | 1847 | 10 | 1.08% |
| hsa_circ_0129280 | 1847 | 10 | 1.08% |
| hsa_circ_0129101 | 7389 | 40 | 1.08% |
| hsa_circ_0015602 | 2032 | 11 | 1.08% |
| hsa_circ_0042119 | 2032 | 11 | 1.08% |
| hsa_circ_0048138 | 2032 | 11 | 1.08% |
| hsa_circ_0054956 | 2032 | 11 | 1.08% |
| hsa_circ_0111478 | 2217 | 12 | 1.08% |
| hsa_circ_0088950 | 2402 | 13 | 1.08% |
| hsa_circ_0028762 | 1848 | 10 | 1.08% |
| hsa_circ_0050372 | 1848 | 10 | 1.08% |
| hsa_circ_0102967 | 1848 | 10 | 1.08% |
| hsa_circ_0110120 | 1848 | 10 | 1.08% |
| hsa_circ_0115616 | 1848 | 10 | 1.08% |
| hsa_circ_0030226 | 2033 | 11 | 1.08% |
| hsa_circ_0032296 | 2033 | 11 | 1.08% |
| hsa_circ_0053504 | 2033 | 11 | 1.08% |
| hsa_circ_0078111 | 2033 | 11 | 1.08% |
| hsa_circ_0091758 | 2033 | 11 | 1.08% |
| hsa_circ_0094618 | 2033 | 11 | 1.08% |
| hsa_circ_0018931 | 2403 | 13 | 1.08% |
| hsa_circ_0011910 | 1849 | 10 | 1.08% |
| hsa_circ_0012438 | 1849 | 10 | 1.08% |
| hsa_circ_0017007 | 1849 | 10 | 1.08% |
| hsa_circ_0018398 | 1849 | 10 | 1.08% |
| hsa_circ_0039658 | 1849 | 10 | 1.08% |
| hsa_circ_0076042 | 1849 | 10 | 1.08% |
| hsa_circ_0086576 | 1849 | 10 | 1.08% |
| hsa_circ_0090378 | 1849 | 10 | 1.08% |
| hsa_circ_0052023 | 2034 | 11 | 1.08% |
| hsa_circ_0073427 | 2034 | 11 | 1.08% |

| | | | |
|---|---|---|---|
| hsa_circ_0024208 | 2219 | 12 | 1.08% |
| hsa_circ_0091741 | 2219 | 12 | 1.08% |
| hsa_circ_0117779 | 2219 | 12 | 1.08% |
| hsa_circ_0025351 | 2404 | 13 | 1.08% |
| hsa_circ_0025447 | 2404 | 13 | 1.08% |
| hsa_circ_0092269 | 18123 | 98 | 1.08% |
| hsa_circ_0115089 | 5920 | 32 | 1.08% |
| hsa_circ_0012952 | 3145 | 17 | 1.08% |
| hsa_circ_0033775 | 2220 | 12 | 1.08% |
| hsa_circ_0010182 | 2035 | 11 | 1.08% |
| hsa_circ_0035114 | 2035 | 11 | 1.08% |
| hsa_circ_0049729 | 2035 | 11 | 1.08% |
| hsa_circ_0059324 | 2035 | 11 | 1.08% |
| hsa_circ_0082934 | 2035 | 11 | 1.08% |
| hsa_circ_0087680 | 2035 | 11 | 1.08% |
| hsa_circ_0136599 | 2035 | 11 | 1.08% |
| hsa_circ_0007287 | 1850 | 10 | 1.08% |
| hsa_circ_0014900 | 1850 | 10 | 1.08% |
| hsa_circ_0030245 | 1850 | 10 | 1.08% |
| hsa_circ_0057314 | 1850 | 10 | 1.08% |
| hsa_circ_0069932 | 1850 | 10 | 1.08% |
| hsa_circ_0134343 | 1850 | 10 | 1.08% |
| hsa_circ_0038151 | 2406 | 13 | 1.08% |
| hsa_circ_0037281 | 2221 | 12 | 1.08% |
| hsa_circ_0061513 | 2221 | 12 | 1.08% |
| hsa_circ_0121918 | 2221 | 12 | 1.08% |
| hsa_circ_0027707 | 2036 | 11 | 1.08% |
| hsa_circ_0029412 | 2036 | 11 | 1.08% |
| hsa_circ_0051512 | 2036 | 11 | 1.08% |
| hsa_circ_0108223 | 2036 | 11 | 1.08% |
| hsa_circ_0137583 | 2036 | 11 | 1.08% |
| hsa_circ_0019563 | 1851 | 10 | 1.08% |
| hsa_circ_0054638 | 1851 | 10 | 1.08% |
| hsa_circ_0064930 | 1851 | 10 | 1.08% |
| hsa_circ_0071001 | 1851 | 10 | 1.08% |
| hsa_circ_0077612 | 1851 | 10 | 1.08% |
| hsa_circ_0086229 | 1851 | 10 | 1.08% |
| hsa_circ_0089056 | 1851 | 10 | 1.08% |
| hsa_circ_0095005 | 1851 | 10 | 1.08% |
| hsa_circ_0110527 | 1851 | 10 | 1.08% |
| hsa_circ_0046407 | 2222 | 12 | 1.08% |
| hsa_circ_0026627 | 2037 | 11 | 1.08% |
| hsa_circ_0026754 | 2037 | 11 | 1.08% |
| hsa_circ_0039864 | 2037 | 11 | 1.08% |
| hsa_circ_0106023 | 2037 | 11 | 1.08% |
| hsa_circ_0127358 | 2037 | 11 | 1.08% |
| hsa_circ_0136485 | 2037 | 11 | 1.08% |
| hsa_circ_0103060 | 7593 | 41 | 1.08% |
| hsa_circ_0004297 | 1852 | 10 | 1.08% |
| hsa_circ_0011604 | 1852 | 10 | 1.08% |

| | | | |
|---|---|---|---|
| hsa_circ_0022280 | 1852 | 10 | 1.08% |
| hsa_circ_0052421 | 1852 | 10 | 1.08% |
| hsa_circ_0087660 | 1852 | 10 | 1.08% |
| hsa_circ_0105557 | 1852 | 10 | 1.08% |
| hsa_circ_0131223 | 1852 | 10 | 1.08% |
| hsa_circ_0047163 | 2593 | 14 | 1.08% |
| hsa_circ_0035924 | 2408 | 13 | 1.08% |
| hsa_circ_0016809 | 2038 | 11 | 1.08% |
| hsa_circ_0090406 | 2038 | 11 | 1.08% |
| hsa_circ_0113314 | 2038 | 11 | 1.08% |
| hsa_circ_0028826 | 1853 | 10 | 1.08% |
| hsa_circ_0036755 | 1853 | 10 | 1.08% |
| hsa_circ_0054483 | 1853 | 10 | 1.08% |
| hsa_circ_0075578 | 1853 | 10 | 1.08% |
| hsa_circ_0116182 | 1853 | 10 | 1.08% |
| hsa_circ_0121305 | 1853 | 10 | 1.08% |
| hsa_circ_0137932 | 1853 | 10 | 1.08% |
| hsa_circ_0036672 | 2224 | 12 | 1.08% |
| hsa_circ_0038316 | 2224 | 12 | 1.08% |
| hsa_circ_0071112 | 2224 | 12 | 1.08% |
| hsa_circ_0034665 | 2039 | 11 | 1.08% |
| hsa_circ_0038108 | 2039 | 11 | 1.08% |
| hsa_circ_0059495 | 2039 | 11 | 1.08% |
| hsa_circ_0076215 | 2039 | 11 | 1.08% |
| hsa_circ_0085586 | 2039 | 11 | 1.08% |
| hsa_circ_0121001 | 2039 | 11 | 1.08% |
| hsa_circ_0055782 | 2966 | 16 | 1.08% |
| hsa_circ_0023819 | 1854 | 10 | 1.08% |
| hsa_circ_0032455 | 1854 | 10 | 1.08% |
| hsa_circ_0064518 | 1854 | 10 | 1.08% |
| hsa_circ_0066357 | 1854 | 10 | 1.08% |
| hsa_circ_0074919 | 1854 | 10 | 1.08% |
| hsa_circ_0081089 | 1854 | 10 | 1.08% |
| hsa_circ_0090394 | 1854 | 10 | 1.08% |
| hsa_circ_0115879 | 1854 | 10 | 1.08% |
| hsa_circ_0117715 | 1854 | 10 | 1.08% |
| hsa_circ_0120366 | 1854 | 10 | 1.08% |
| hsa_circ_0138116 | 1854 | 10 | 1.08% |
| hsa_circ_0004094 | 2225 | 12 | 1.08% |
| hsa_circ_0070916 | 2225 | 12 | 1.08% |
| hsa_circ_0031350 | 2040 | 11 | 1.08% |
| hsa_circ_0031466 | 2040 | 11 | 1.08% |
| hsa_circ_0055909 | 2040 | 11 | 1.08% |
| hsa_circ_0103342 | 2040 | 11 | 1.08% |
| hsa_circ_0014700 | 2226 | 12 | 1.08% |
| hsa_circ_0063617 | 2226 | 12 | 1.08% |
| hsa_circ_0011058 | 1855 | 10 | 1.08% |
| hsa_circ_0023499 | 1855 | 10 | 1.08% |
| hsa_circ_0024630 | 1855 | 10 | 1.08% |
| hsa_circ_0038039 | 1855 | 10 | 1.08% |

| | | | |
|---|---|---|---|
| hsa_circ_0115584 | 1855 | 10 | 1.08% |
| hsa_circ_0118266 | 1855 | 10 | 1.08% |
| hsa_circ_0045221 | 7235 | 39 | 1.08% |
| hsa_circ_0032871 | 2041 | 11 | 1.08% |
| hsa_circ_0073655 | 2041 | 11 | 1.08% |
| hsa_circ_0082662 | 2041 | 11 | 1.08% |
| hsa_circ_0091423 | 2041 | 11 | 1.08% |
| hsa_circ_0104651 | 2041 | 11 | 1.08% |
| hsa_circ_0012243 | 2227 | 12 | 1.08% |
| hsa_circ_0023183 | 2227 | 12 | 1.08% |
| hsa_circ_0087544 | 2227 | 12 | 1.08% |
| hsa_circ_0017702 | 1856 | 10 | 1.08% |
| hsa_circ_0056094 | 1856 | 10 | 1.08% |
| hsa_circ_0067994 | 1856 | 10 | 1.08% |
| hsa_circ_0085153 | 1856 | 10 | 1.08% |
| hsa_circ_0085355 | 1856 | 10 | 1.08% |
| hsa_circ_0058818 | 2413 | 13 | 1.08% |
| hsa_circ_0010103 | 2042 | 11 | 1.08% |
| hsa_circ_0049809 | 2042 | 11 | 1.08% |
| hsa_circ_0064554 | 2042 | 11 | 1.08% |
| hsa_circ_0084538 | 2042 | 11 | 1.08% |
| hsa_circ_0098768 | 2042 | 11 | 1.08% |
| hsa_circ_0106328 | 2042 | 11 | 1.08% |
| hsa_circ_0116780 | 2042 | 11 | 1.08% |
| hsa_circ_0126392 | 2042 | 11 | 1.08% |
| hsa_circ_0140526 | 2042 | 11 | 1.08% |
| hsa_circ_0071389 | 2228 | 12 | 1.08% |
| hsa_circ_0113371 | 2228 | 12 | 1.08% |
| hsa_circ_0103244 | 2414 | 13 | 1.08% |
| hsa_circ_0028111 | 1857 | 10 | 1.08% |
| hsa_circ_0056567 | 1857 | 10 | 1.08% |
| hsa_circ_0059257 | 1857 | 10 | 1.08% |
| hsa_circ_0095418 | 1857 | 10 | 1.08% |
| hsa_circ_0099295 | 1857 | 10 | 1.08% |
| hsa_circ_0103327 | 1857 | 10 | 1.08% |
| hsa_circ_0107578 | 1857 | 10 | 1.08% |
| hsa_circ_0115499 | 1857 | 10 | 1.08% |
| hsa_circ_0022256 | 2043 | 11 | 1.08% |
| hsa_circ_0043941 | 2043 | 11 | 1.08% |
| hsa_circ_0116594 | 2972 | 16 | 1.08% |
| hsa_circ_0104409 | 2229 | 12 | 1.08% |
| hsa_circ_0138666 | 4087 | 22 | 1.08% |
| hsa_circ_0012082 | 1858 | 10 | 1.08% |
| hsa_circ_0039666 | 1858 | 10 | 1.08% |
| hsa_circ_0066012 | 1858 | 10 | 1.08% |
| hsa_circ_0082837 | 1858 | 10 | 1.08% |
| hsa_circ_0091002 | 1858 | 10 | 1.08% |
| hsa_circ_0012774 | 2044 | 11 | 1.08% |
| hsa_circ_0082492 | 2044 | 11 | 1.08% |
| hsa_circ_0120253 | 2044 | 11 | 1.08% |

| | | | |
|---|---|---|---|
| hsa_circ_0001915 | 2416 | 13 | 1.08% |
| hsa_circ_0011213 | 2416 | 13 | 1.08% |
| hsa_circ_0125806 | 2602 | 14 | 1.08% |
| hsa_circ_0107253 | 6320 | 34 | 1.08% |
| hsa_circ_0042366 | 1859 | 10 | 1.08% |
| hsa_circ_0079749 | 1859 | 10 | 1.08% |
| hsa_circ_0052078 | 2045 | 11 | 1.08% |
| hsa_circ_0078782 | 2045 | 11 | 1.08% |
| hsa_circ_0104370 | 2045 | 11 | 1.08% |
| hsa_circ_0115401 | 2045 | 11 | 1.08% |
| hsa_circ_0119912 | 2045 | 11 | 1.08% |
| hsa_circ_0002136 | 2231 | 12 | 1.08% |
| hsa_circ_0090783 | 2231 | 12 | 1.08% |
| hsa_circ_0048076 | 2604 | 14 | 1.08% |
| hsa_circ_0070163 | 2604 | 14 | 1.08% |
| hsa_circ_0067232 | 2232 | 12 | 1.08% |
| hsa_circ_0088871 | 2232 | 12 | 1.08% |
| hsa_circ_0043113 | 2046 | 11 | 1.08% |
| hsa_circ_0069766 | 2046 | 11 | 1.08% |
| hsa_circ_0076257 | 2046 | 11 | 1.08% |
| hsa_circ_0076335 | 2046 | 11 | 1.08% |
| hsa_circ_0089580 | 2046 | 11 | 1.08% |
| hsa_circ_0120772 | 2046 | 11 | 1.08% |
| hsa_circ_0009885 | 1860 | 10 | 1.08% |
| hsa_circ_0010475 | 1860 | 10 | 1.08% |
| hsa_circ_0041192 | 1860 | 10 | 1.08% |
| hsa_circ_0050976 | 1860 | 10 | 1.08% |
| hsa_circ_0071880 | 1860 | 10 | 1.08% |
| hsa_circ_0073079 | 1860 | 10 | 1.08% |
| hsa_circ_0082519 | 1860 | 10 | 1.08% |
| hsa_circ_0089629 | 1860 | 10 | 1.08% |
| hsa_circ_0134455 | 1860 | 10 | 1.08% |
| hsa_circ_0005988 | 2977 | 16 | 1.07% |
| hsa_circ_0118880 | 2791 | 15 | 1.07% |
| hsa_circ_0080661 | 2233 | 12 | 1.07% |
| hsa_circ_0045037 | 2047 | 11 | 1.07% |
| hsa_circ_0137088 | 2047 | 11 | 1.07% |
| hsa_circ_0010229 | 1861 | 10 | 1.07% |
| hsa_circ_0029837 | 1861 | 10 | 1.07% |
| hsa_circ_0035300 | 1861 | 10 | 1.07% |
| hsa_circ_0050187 | 1861 | 10 | 1.07% |
| hsa_circ_0061176 | 1861 | 10 | 1.07% |
| hsa_circ_0115599 | 1861 | 10 | 1.07% |
| hsa_circ_0130102 | 1861 | 10 | 1.07% |
| hsa_circ_0140587 | 1861 | 10 | 1.07% |
| hsa_circ_0033072 | 2606 | 14 | 1.07% |
| hsa_circ_0029900 | 2234 | 12 | 1.07% |
| hsa_circ_0055398 | 2234 | 12 | 1.07% |
| hsa_circ_0096591 | 2234 | 12 | 1.07% |
| hsa_circ_0048657 | 2048 | 11 | 1.07% |

| | | | |
|---|---|---|---|
| hsa_circ_0080023 | 2048 | 11 | 1.07% |
| hsa_circ_0030893 | 1862 | 10 | 1.07% |
| hsa_circ_0033351 | 1862 | 10 | 1.07% |
| hsa_circ_0082670 | 1862 | 10 | 1.07% |
| hsa_circ_0126914 | 1862 | 10 | 1.07% |
| hsa_circ_0101475 | 2607 | 14 | 1.07% |
| hsa_circ_0047021 | 2421 | 13 | 1.07% |
| hsa_circ_0022466 | 2235 | 12 | 1.07% |
| hsa_circ_0039106 | 2235 | 12 | 1.07% |
| hsa_circ_0053945 | 2235 | 12 | 1.07% |
| hsa_circ_0055295 | 2235 | 12 | 1.07% |
| hsa_circ_0078097 | 2235 | 12 | 1.07% |
| hsa_circ_0115581 | 2235 | 12 | 1.07% |
| hsa_circ_0022125 | 2049 | 11 | 1.07% |
| hsa_circ_0028545 | 2049 | 11 | 1.07% |
| hsa_circ_0029965 | 2049 | 11 | 1.07% |
| hsa_circ_0050678 | 2049 | 11 | 1.07% |
| hsa_circ_0078484 | 2049 | 11 | 1.07% |
| hsa_circ_0081235 | 2049 | 11 | 1.07% |
| hsa_circ_0100302 | 2049 | 11 | 1.07% |
| hsa_circ_0119244 | 6893 | 37 | 1.07% |
| hsa_circ_0015706 | 1863 | 10 | 1.07% |
| hsa_circ_0028214 | 1863 | 10 | 1.07% |
| hsa_circ_0045691 | 1863 | 10 | 1.07% |
| hsa_circ_0059780 | 1863 | 10 | 1.07% |
| hsa_circ_0064937 | 1863 | 10 | 1.07% |
| hsa_circ_0070308 | 1863 | 10 | 1.07% |
| hsa_circ_0108407 | 1863 | 10 | 1.07% |
| hsa_circ_0139750 | 1863 | 10 | 1.07% |
| hsa_circ_0105190 | 3913 | 21 | 1.07% |
| hsa_circ_0052686 | 2795 | 15 | 1.07% |
| hsa_circ_0022733 | 2236 | 12 | 1.07% |
| hsa_circ_0078529 | 2236 | 12 | 1.07% |
| hsa_circ_0101620 | 2236 | 12 | 1.07% |
| hsa_circ_0018270 | 2050 | 11 | 1.07% |
| hsa_circ_0026744 | 2050 | 11 | 1.07% |
| hsa_circ_0079347 | 2050 | 11 | 1.07% |
| hsa_circ_0086816 | 2050 | 11 | 1.07% |
| hsa_circ_0118779 | 2050 | 11 | 1.07% |
| hsa_circ_0044105 | 1864 | 10 | 1.07% |
| hsa_circ_0048035 | 1864 | 10 | 1.07% |
| hsa_circ_0079981 | 1864 | 10 | 1.07% |
| hsa_circ_0137208 | 1864 | 10 | 1.07% |
| hsa_circ_0021118 | 3356 | 18 | 1.07% |
| hsa_circ_0028734 | 2051 | 11 | 1.07% |
| hsa_circ_0037391 | 2051 | 11 | 1.07% |
| hsa_circ_0048381 | 2051 | 11 | 1.07% |
| hsa_circ_0101138 | 2051 | 11 | 1.07% |
| hsa_circ_0030252 | 2424 | 13 | 1.07% |
| hsa_circ_0000389 | 2611 | 14 | 1.07% |

| | | | |
|---|---|---|---|
| hsa_circ_0044597 | 2238 | 12 | 1.07% |
| hsa_circ_0086439 | 2238 | 12 | 1.07% |
| hsa_circ_0025749 | 1865 | 10 | 1.07% |
| hsa_circ_0055240 | 1865 | 10 | 1.07% |
| hsa_circ_0072509 | 1865 | 10 | 1.07% |
| hsa_circ_0086959 | 1865 | 10 | 1.07% |
| hsa_circ_0048991 | 2425 | 13 | 1.07% |
| hsa_circ_0050660 | 2425 | 13 | 1.07% |
| hsa_circ_0059032 | 2052 | 11 | 1.07% |
| hsa_circ_0108130 | 2052 | 11 | 1.07% |
| hsa_circ_0018663 | 2239 | 12 | 1.07% |
| hsa_circ_0049524 | 2239 | 12 | 1.07% |
| hsa_circ_0078447 | 2239 | 12 | 1.07% |
| hsa_circ_0080612 | 2239 | 12 | 1.07% |
| hsa_circ_0004079 | 1866 | 10 | 1.07% |
| hsa_circ_0044035 | 1866 | 10 | 1.07% |
| hsa_circ_0052815 | 1866 | 10 | 1.07% |
| hsa_circ_0121359 | 1866 | 10 | 1.07% |
| hsa_circ_0123220 | 1866 | 10 | 1.07% |
| hsa_circ_0093621 | 3359 | 18 | 1.07% |
| hsa_circ_0089819 | 5039 | 27 | 1.07% |
| hsa_circ_0009411 | 2053 | 11 | 1.07% |
| hsa_circ_0013132 | 2053 | 11 | 1.07% |
| hsa_circ_0015343 | 2053 | 11 | 1.07% |
| hsa_circ_0028408 | 2053 | 11 | 1.07% |
| hsa_circ_0028436 | 2053 | 11 | 1.07% |
| hsa_circ_0046350 | 2053 | 11 | 1.07% |
| hsa_circ_0048319 | 2053 | 11 | 1.07% |
| hsa_circ_0017862 | 2240 | 12 | 1.07% |
| hsa_circ_0087586 | 2240 | 12 | 1.07% |
| hsa_circ_0114724 | 2240 | 12 | 1.07% |
| hsa_circ_0020662 | 2427 | 13 | 1.07% |
| hsa_circ_0048264 | 2427 | 13 | 1.07% |
| hsa_circ_0033547 | 1867 | 10 | 1.07% |
| hsa_circ_0058626 | 1867 | 10 | 1.07% |
| hsa_circ_0084420 | 1867 | 10 | 1.07% |
| hsa_circ_0086812 | 1867 | 10 | 1.07% |
| hsa_circ_0113729 | 1867 | 10 | 1.07% |
| hsa_circ_0134495 | 1867 | 10 | 1.07% |
| hsa_circ_0015468 | 2054 | 11 | 1.07% |
| hsa_circ_0052837 | 2054 | 11 | 1.07% |
| hsa_circ_0080274 | 2054 | 11 | 1.07% |
| hsa_circ_0081433 | 2054 | 11 | 1.07% |
| hsa_circ_0091874 | 2054 | 11 | 1.07% |
| hsa_circ_0104317 | 2054 | 11 | 1.07% |
| hsa_circ_0117027 | 2054 | 11 | 1.07% |
| hsa_circ_0011613 | 2241 | 12 | 1.07% |
| hsa_circ_0087548 | 2241 | 12 | 1.07% |
| hsa_circ_0009877 | 1868 | 10 | 1.07% |
| hsa_circ_0063119 | 1868 | 10 | 1.07% |

| | | | |
|---|---|---|---|
| hsa_circ_0109574 | 1868 | 10 | 1.07% |
| hsa_circ_0131579 | 1868 | 10 | 1.07% |
| hsa_circ_0132299 | 1868 | 10 | 1.07% |
| hsa_circ_0016923 | 2055 | 11 | 1.07% |
| hsa_circ_0022759 | 2242 | 12 | 1.07% |
| hsa_circ_0038442 | 2242 | 12 | 1.07% |
| hsa_circ_0045576 | 2242 | 12 | 1.07% |
| hsa_circ_0089585 | 2429 | 13 | 1.07% |
| hsa_circ_0027224 | 1869 | 10 | 1.07% |
| hsa_circ_0054811 | 1869 | 10 | 1.07% |
| hsa_circ_0103835 | 1869 | 10 | 1.07% |
| hsa_circ_0017987 | 2056 | 11 | 1.07% |
| hsa_circ_0026568 | 2056 | 11 | 1.07% |
| hsa_circ_0039362 | 2056 | 11 | 1.07% |
| hsa_circ_0075221 | 2056 | 11 | 1.07% |
| hsa_circ_0140538 | 2243 | 12 | 1.07% |
| hsa_circ_0033055 | 2430 | 13 | 1.07% |
| hsa_circ_0053231 | 2617 | 14 | 1.07% |
| hsa_circ_0086179 | 2617 | 14 | 1.07% |
| hsa_circ_0087640 | 8040 | 43 | 1.07% |
| hsa_circ_0066056 | 2805 | 15 | 1.07% |
| hsa_circ_0032573 | 2244 | 12 | 1.07% |
| hsa_circ_0047115 | 2244 | 12 | 1.07% |
| hsa_circ_0058572 | 2244 | 12 | 1.07% |
| hsa_circ_0010175 | 2057 | 11 | 1.07% |
| hsa_circ_0043532 | 2057 | 11 | 1.07% |
| hsa_circ_0052006 | 2057 | 11 | 1.07% |
| hsa_circ_0076936 | 2057 | 11 | 1.07% |
| hsa_circ_0081193 | 2057 | 11 | 1.07% |
| hsa_circ_0136344 | 2057 | 11 | 1.07% |
| hsa_circ_0042118 | 1870 | 10 | 1.07% |
| hsa_circ_0056419 | 1870 | 10 | 1.07% |
| hsa_circ_0068299 | 1870 | 10 | 1.07% |
| hsa_circ_0030815 | 2432 | 13 | 1.07% |
| hsa_circ_0038441 | 2245 | 12 | 1.07% |
| hsa_circ_0013101 | 2058 | 11 | 1.07% |
| hsa_circ_0017863 | 2058 | 11 | 1.07% |
| hsa_circ_0024675 | 2058 | 11 | 1.07% |
| hsa_circ_0034828 | 2058 | 11 | 1.07% |
| hsa_circ_0038100 | 2058 | 11 | 1.07% |
| hsa_circ_0079187 | 2058 | 11 | 1.07% |
| hsa_circ_0056443 | 1871 | 10 | 1.07% |
| hsa_circ_0068915 | 1871 | 10 | 1.07% |
| hsa_circ_0071028 | 1871 | 10 | 1.07% |
| hsa_circ_0086776 | 1871 | 10 | 1.07% |
| hsa_circ_0089183 | 1871 | 10 | 1.07% |
| hsa_circ_0112566 | 1871 | 10 | 1.07% |
| hsa_circ_0135734 | 1871 | 10 | 1.07% |
| hsa_circ_0068762 | 4678 | 25 | 1.07% |
| hsa_circ_0031681 | 2433 | 13 | 1.07% |

| | | | |
|---|---|---|---|
| hsa_circ_0052627 | 2433 | 13 | 1.07% |
| hsa_circ_0039082 | 2246 | 12 | 1.07% |
| hsa_circ_0077384 | 2246 | 12 | 1.07% |
| hsa_circ_0043839 | 2059 | 11 | 1.07% |
| hsa_circ_0065494 | 2059 | 11 | 1.07% |
| hsa_circ_0086639 | 2059 | 11 | 1.07% |
| hsa_circ_0115224 | 2059 | 11 | 1.07% |
| hsa_circ_0135307 | 2059 | 11 | 1.07% |
| hsa_circ_0089771 | 7488 | 40 | 1.07% |
| hsa_circ_0028450 | 1872 | 10 | 1.07% |
| hsa_circ_0045014 | 1872 | 10 | 1.07% |
| hsa_circ_0095618 | 1872 | 10 | 1.07% |
| hsa_circ_0097721 | 1872 | 10 | 1.07% |
| hsa_circ_0025385 | 2247 | 12 | 1.07% |
| hsa_circ_0048552 | 2247 | 12 | 1.07% |
| hsa_circ_0080113 | 2247 | 12 | 1.07% |
| hsa_circ_0063474 | 7116 | 38 | 1.07% |
| hsa_circ_0019848 | 2060 | 11 | 1.07% |
| hsa_circ_0093531 | 2060 | 11 | 1.07% |
| hsa_circ_0130662 | 2060 | 11 | 1.07% |
| hsa_circ_0135699 | 2622 | 14 | 1.07% |
| hsa_circ_0093902 | 7117 | 38 | 1.07% |
| hsa_circ_0139530 | 5057 | 27 | 1.07% |
| hsa_circ_0009832 | 1873 | 10 | 1.07% |
| hsa_circ_0021854 | 1873 | 10 | 1.07% |
| hsa_circ_0043709 | 1873 | 10 | 1.07% |
| hsa_circ_0047591 | 1873 | 10 | 1.07% |
| hsa_circ_0062046 | 1873 | 10 | 1.07% |
| hsa_circ_0064660 | 1873 | 10 | 1.07% |
| hsa_circ_0110553 | 1873 | 10 | 1.07% |
| hsa_circ_0116052 | 1873 | 10 | 1.07% |
| hsa_circ_0123773 | 1873 | 10 | 1.07% |
| hsa_circ_0133957 | 1873 | 10 | 1.07% |
| hsa_circ_0062612 | 2997 | 16 | 1.07% |
| hsa_circ_0021876 | 2248 | 12 | 1.07% |
| hsa_circ_0070845 | 2248 | 12 | 1.07% |
| hsa_circ_0071245 | 2248 | 12 | 1.07% |
| hsa_circ_0080086 | 2623 | 14 | 1.07% |
| hsa_circ_0010568 | 2061 | 11 | 1.07% |
| hsa_circ_0017989 | 2061 | 11 | 1.07% |
| hsa_circ_0038264 | 2061 | 11 | 1.07% |
| hsa_circ_0042579 | 2061 | 11 | 1.07% |
| hsa_circ_0058629 | 2061 | 11 | 1.07% |
| hsa_circ_0070093 | 2061 | 11 | 1.07% |
| hsa_circ_0087111 | 2061 | 11 | 1.07% |
| hsa_circ_0087195 | 2061 | 11 | 1.07% |
| hsa_circ_0046169 | 2436 | 13 | 1.07% |
| hsa_circ_0027802 | 1874 | 10 | 1.07% |
| hsa_circ_0057632 | 1874 | 10 | 1.07% |
| hsa_circ_0072657 | 1874 | 10 | 1.07% |

| | | | |
|---|---|---|---|
| hsa_circ_0117478 | 1874 | 10 | 1.07% |
| hsa_circ_0010055 | 5060 | 27 | 1.07% |
| hsa_circ_0020631 | 2249 | 12 | 1.07% |
| hsa_circ_0115269 | 2249 | 12 | 1.07% |
| hsa_circ_0105763 | 2999 | 16 | 1.07% |
| hsa_circ_0080150 | 2062 | 11 | 1.07% |
| hsa_circ_0090287 | 2062 | 11 | 1.07% |
| hsa_circ_0091179 | 2062 | 11 | 1.07% |
| hsa_circ_0111704 | 2062 | 11 | 1.07% |
| hsa_circ_0124672 | 2062 | 11 | 1.07% |
| hsa_circ_0127776 | 5062 | 27 | 1.07% |
| hsa_circ_0023462 | 2250 | 12 | 1.07% |
| hsa_circ_0054791 | 2250 | 12 | 1.07% |
| hsa_circ_0079549 | 2250 | 12 | 1.07% |
| hsa_circ_0014380 | 1875 | 10 | 1.07% |
| hsa_circ_0021859 | 1875 | 10 | 1.07% |
| hsa_circ_0026383 | 1875 | 10 | 1.07% |
| hsa_circ_0029217 | 1875 | 10 | 1.07% |
| hsa_circ_0037767 | 1875 | 10 | 1.07% |
| hsa_circ_0043252 | 1875 | 10 | 1.07% |
| hsa_circ_0069664 | 1875 | 10 | 1.07% |
| hsa_circ_0079663 | 1875 | 10 | 1.07% |
| hsa_circ_0123302 | 1875 | 10 | 1.07% |
| hsa_circ_0126333 | 1875 | 10 | 1.07% |
| hsa_circ_0129482 | 1875 | 10 | 1.07% |
| hsa_circ_0122514 | 35630 | 190 | 1.07% |
| hsa_circ_0015698 | 2063 | 11 | 1.07% |
| hsa_circ_0028174 | 2063 | 11 | 1.07% |
| hsa_circ_0036186 | 2063 | 11 | 1.07% |
| hsa_circ_0052669 | 2063 | 11 | 1.07% |
| hsa_circ_0115127 | 2063 | 11 | 1.07% |
| hsa_circ_0136990 | 2063 | 11 | 1.07% |
| hsa_circ_0126989 | 3001 | 16 | 1.07% |
| hsa_circ_0095543 | 4877 | 26 | 1.07% |
| hsa_circ_0126178 | 2251 | 12 | 1.07% |
| hsa_circ_0136807 | 2251 | 12 | 1.07% |
| hsa_circ_0009914 | 1876 | 10 | 1.07% |
| hsa_circ_0064979 | 1876 | 10 | 1.07% |
| hsa_circ_0078593 | 1876 | 10 | 1.07% |
| hsa_circ_0118375 | 1876 | 10 | 1.07% |
| hsa_circ_0089063 | 2064 | 11 | 1.07% |
| hsa_circ_0120448 | 3753 | 20 | 1.07% |
| hsa_circ_0025399 | 2252 | 12 | 1.07% |
| hsa_circ_0068140 | 2252 | 12 | 1.07% |
| hsa_circ_0079578 | 2252 | 12 | 1.07% |
| hsa_circ_0064618 | 2440 | 13 | 1.07% |
| hsa_circ_0036130 | 1877 | 10 | 1.07% |
| hsa_circ_0065205 | 1877 | 10 | 1.07% |
| hsa_circ_0082038 | 1877 | 10 | 1.07% |
| hsa_circ_0119086 | 1877 | 10 | 1.07% |

| | | | |
|---|---|---|---|
| hsa_circ_0122356 | 1877 | 10 | 1.07% |
| hsa_circ_0131334 | 1877 | 10 | 1.07% |
| hsa_circ_0140153 | 1877 | 10 | 1.07% |
| hsa_circ_0113742 | 3379 | 18 | 1.07% |
| hsa_circ_0031244 | 2065 | 11 | 1.07% |
| hsa_circ_0101854 | 2065 | 11 | 1.07% |
| hsa_circ_0113829 | 2065 | 11 | 1.07% |
| hsa_circ_0135595 | 2065 | 11 | 1.07% |
| hsa_circ_0050410 | 2253 | 12 | 1.07% |
| hsa_circ_0053261 | 2253 | 12 | 1.07% |
| hsa_circ_0031627 | 2441 | 13 | 1.07% |
| hsa_circ_0115256 | 11642 | 62 | 1.07% |
| hsa_circ_0035035 | 1878 | 10 | 1.06% |
| hsa_circ_0055462 | 1878 | 10 | 1.06% |
| hsa_circ_0056173 | 1878 | 10 | 1.06% |
| hsa_circ_0068902 | 1878 | 10 | 1.06% |
| hsa_circ_0111720 | 1878 | 10 | 1.06% |
| hsa_circ_0119845 | 1878 | 10 | 1.06% |
| hsa_circ_0136310 | 1878 | 10 | 1.06% |
| hsa_circ_0047363 | 2066 | 11 | 1.06% |
| hsa_circ_0074900 | 2066 | 11 | 1.06% |
| hsa_circ_0018945 | 2254 | 12 | 1.06% |
| hsa_circ_0089711 | 2254 | 12 | 1.06% |
| hsa_circ_0083418 | 2442 | 13 | 1.06% |
| hsa_circ_0045727 | 1879 | 10 | 1.06% |
| hsa_circ_0055930 | 1879 | 10 | 1.06% |
| hsa_circ_0085572 | 1879 | 10 | 1.06% |
| hsa_circ_0114977 | 1879 | 10 | 1.06% |
| hsa_circ_0125316 | 1879 | 10 | 1.06% |
| hsa_circ_0002871 | 2067 | 11 | 1.06% |
| hsa_circ_0013165 | 2067 | 11 | 1.06% |
| hsa_circ_0040352 | 2067 | 11 | 1.06% |
| hsa_circ_0063917 | 2067 | 11 | 1.06% |
| hsa_circ_0077971 | 2067 | 11 | 1.06% |
| hsa_circ_0112045 | 2067 | 11 | 1.06% |
| hsa_circ_0117849 | 2067 | 11 | 1.06% |
| hsa_circ_0137593 | 2067 | 11 | 1.06% |
| hsa_circ_0029593 | 2443 | 13 | 1.06% |
| hsa_circ_0030859 | 2443 | 13 | 1.06% |
| hsa_circ_0036981 | 2631 | 14 | 1.06% |
| hsa_circ_0089654 | 2631 | 14 | 1.06% |
| hsa_circ_0139622 | 3008 | 16 | 1.06% |
| hsa_circ_0013219 | 2444 | 13 | 1.06% |
| hsa_circ_0034772 | 2256 | 12 | 1.06% |
| hsa_circ_0040257 | 2256 | 12 | 1.06% |
| hsa_circ_0054628 | 2256 | 12 | 1.06% |
| hsa_circ_0019175 | 2068 | 11 | 1.06% |
| hsa_circ_0031744 | 2068 | 11 | 1.06% |
| hsa_circ_0048425 | 2068 | 11 | 1.06% |
| hsa_circ_0036794 | 1880 | 10 | 1.06% |

| | | | |
|---|---|---|---|
| hsa_circ_0044551 | 1880 | 10 | 1.06% |
| hsa_circ_0054162 | 1880 | 10 | 1.06% |
| hsa_circ_0067974 | 1880 | 10 | 1.06% |
| hsa_circ_0068219 | 1880 | 10 | 1.06% |
| hsa_circ_0085248 | 1880 | 10 | 1.06% |
| hsa_circ_0087151 | 1880 | 10 | 1.06% |
| hsa_circ_0099329 | 1880 | 10 | 1.06% |
| hsa_circ_0125848 | 1880 | 10 | 1.06% |
| hsa_circ_0055830 | 2821 | 15 | 1.06% |
| hsa_circ_0096007 | 2821 | 15 | 1.06% |
| hsa_circ_0072026 | 4890 | 26 | 1.06% |
| hsa_circ_0086111 | 2445 | 13 | 1.06% |
| hsa_circ_0017517 | 2257 | 12 | 1.06% |
| hsa_circ_0102510 | 2257 | 12 | 1.06% |
| hsa_circ_0110512 | 2257 | 12 | 1.06% |
| hsa_circ_0041842 | 2069 | 11 | 1.06% |
| hsa_circ_0042024 | 2069 | 11 | 1.06% |
| hsa_circ_0063741 | 2069 | 11 | 1.06% |
| hsa_circ_0070372 | 2069 | 11 | 1.06% |
| hsa_circ_0034245 | 1881 | 10 | 1.06% |
| hsa_circ_0034991 | 1881 | 10 | 1.06% |
| hsa_circ_0071064 | 1881 | 10 | 1.06% |
| hsa_circ_0076744 | 1881 | 10 | 1.06% |
| hsa_circ_0101062 | 1881 | 10 | 1.06% |
| hsa_circ_0113265 | 1881 | 10 | 1.06% |
| hsa_circ_0121363 | 1881 | 10 | 1.06% |
| hsa_circ_0002061 | 2446 | 13 | 1.06% |
| hsa_circ_0005303 | 8279 | 44 | 1.06% |
| hsa_circ_0007215 | 2258 | 12 | 1.06% |
| hsa_circ_0041218 | 2258 | 12 | 1.06% |
| hsa_circ_0045178 | 2258 | 12 | 1.06% |
| hsa_circ_0017012 | 2070 | 11 | 1.06% |
| hsa_circ_0040107 | 2070 | 11 | 1.06% |
| hsa_circ_0058485 | 2070 | 11 | 1.06% |
| hsa_circ_0063461 | 2070 | 11 | 1.06% |
| hsa_circ_0085206 | 2070 | 11 | 1.06% |
| hsa_circ_0091860 | 2070 | 11 | 1.06% |
| hsa_circ_0110778 | 2070 | 11 | 1.06% |
| hsa_circ_0128204 | 2070 | 11 | 1.06% |
| hsa_circ_0007631 | 1882 | 10 | 1.06% |
| hsa_circ_0014814 | 1882 | 10 | 1.06% |
| hsa_circ_0039590 | 1882 | 10 | 1.06% |
| hsa_circ_0039828 | 1882 | 10 | 1.06% |
| hsa_circ_0070243 | 1882 | 10 | 1.06% |
| hsa_circ_0091833 | 1882 | 10 | 1.06% |
| hsa_circ_0140116 | 1882 | 10 | 1.06% |
| hsa_circ_0115365 | 2259 | 12 | 1.06% |
| hsa_circ_0018564 | 7342 | 39 | 1.06% |
| hsa_circ_0023365 | 2071 | 11 | 1.06% |
| hsa_circ_0024875 | 2071 | 11 | 1.06% |

| | | | |
|---|---|---|---|
| hsa_circ_0046999 | 2071 | 11 | 1.06% |
| hsa_circ_0054955 | 2071 | 11 | 1.06% |
| hsa_circ_0063399 | 2071 | 11 | 1.06% |
| hsa_circ_0012340 | 1883 | 10 | 1.06% |
| hsa_circ_0043797 | 1883 | 10 | 1.06% |
| hsa_circ_0054549 | 1883 | 10 | 1.06% |
| hsa_circ_0070505 | 1883 | 10 | 1.06% |
| hsa_circ_0084399 | 2448 | 13 | 1.06% |
| hsa_circ_0024372 | 3955 | 21 | 1.06% |
| hsa_circ_0011024 | 2260 | 12 | 1.06% |
| hsa_circ_0043665 | 2260 | 12 | 1.06% |
| hsa_circ_0133483 | 2260 | 12 | 1.06% |
| hsa_circ_0011274 | 2072 | 11 | 1.06% |
| hsa_circ_0026570 | 2072 | 11 | 1.06% |
| hsa_circ_0031966 | 2072 | 11 | 1.06% |
| hsa_circ_0047156 | 2072 | 11 | 1.06% |
| hsa_circ_0068877 | 2072 | 11 | 1.06% |
| hsa_circ_0076504 | 2072 | 11 | 1.06% |
| hsa_circ_0081938 | 2072 | 11 | 1.06% |
| hsa_circ_0130837 | 2072 | 11 | 1.06% |
| hsa_circ_0121866 | 3014 | 16 | 1.06% |
| hsa_circ_0088858 | 2449 | 13 | 1.06% |
| hsa_circ_0021846 | 2826 | 15 | 1.06% |
| hsa_circ_0010529 | 1884 | 10 | 1.06% |
| hsa_circ_0039293 | 1884 | 10 | 1.06% |
| hsa_circ_0064756 | 1884 | 10 | 1.06% |
| hsa_circ_0088908 | 1884 | 10 | 1.06% |
| hsa_circ_0093917 | 1884 | 10 | 1.06% |
| hsa_circ_0124310 | 1884 | 10 | 1.06% |
| hsa_circ_0136377 | 1884 | 10 | 1.06% |
| hsa_circ_0101121 | 2261 | 12 | 1.06% |
| hsa_circ_0121811 | 2638 | 14 | 1.06% |
| hsa_circ_0104249 | 3015 | 16 | 1.06% |
| hsa_circ_0022938 | 2073 | 11 | 1.06% |
| hsa_circ_0080604 | 2073 | 11 | 1.06% |
| hsa_circ_0118639 | 2073 | 11 | 1.06% |
| hsa_circ_0013684 | 2450 | 13 | 1.06% |
| hsa_circ_0136481 | 2450 | 13 | 1.06% |
| hsa_circ_0009518 | 6220 | 33 | 1.06% |
| hsa_circ_0046223 | 2639 | 14 | 1.06% |
| hsa_circ_0061980 | 2262 | 12 | 1.06% |
| hsa_circ_0010309 | 1885 | 10 | 1.06% |
| hsa_circ_0013363 | 1885 | 10 | 1.06% |
| hsa_circ_0044037 | 1885 | 10 | 1.06% |
| hsa_circ_0047338 | 1885 | 10 | 1.06% |
| hsa_circ_0060240 | 1885 | 10 | 1.06% |
| hsa_circ_0087287 | 1885 | 10 | 1.06% |
| hsa_circ_0124102 | 1885 | 10 | 1.06% |
| hsa_circ_0058936 | 2451 | 13 | 1.06% |
| hsa_circ_0121073 | 2451 | 13 | 1.06% |

| | | | |
|---|---|---|---|
| hsa_circ_0010946 | 2074 | 11 | 1.06% |
| hsa_circ_0088899 | 2074 | 11 | 1.06% |
| hsa_circ_0113114 | 2074 | 11 | 1.06% |
| hsa_circ_0113845 | 2074 | 11 | 1.06% |
| hsa_circ_0087165 | 2640 | 14 | 1.06% |
| hsa_circ_0080573 | 2263 | 12 | 1.06% |
| hsa_circ_0082702 | 2263 | 12 | 1.06% |
| hsa_circ_0032859 | 1886 | 10 | 1.06% |
| hsa_circ_0034558 | 1886 | 10 | 1.06% |
| hsa_circ_0039042 | 1886 | 10 | 1.06% |
| hsa_circ_0049822 | 1886 | 10 | 1.06% |
| hsa_circ_0090390 | 1886 | 10 | 1.06% |
| hsa_circ_0090700 | 1886 | 10 | 1.06% |
| hsa_circ_0129254 | 1886 | 10 | 1.06% |
| hsa_circ_0010821 | 2075 | 11 | 1.06% |
| hsa_circ_0015057 | 2075 | 11 | 1.06% |
| hsa_circ_0028806 | 2075 | 11 | 1.06% |
| hsa_circ_0030974 | 2075 | 11 | 1.06% |
| hsa_circ_0032814 | 2075 | 11 | 1.06% |
| hsa_circ_0106971 | 2075 | 11 | 1.06% |
| hsa_circ_0129812 | 2075 | 11 | 1.06% |
| hsa_circ_0076699 | 2453 | 13 | 1.06% |
| hsa_circ_0085008 | 5661 | 30 | 1.06% |
| hsa_circ_0032059 | 1887 | 10 | 1.06% |
| hsa_circ_0040166 | 1887 | 10 | 1.06% |
| hsa_circ_0097391 | 1887 | 10 | 1.06% |
| hsa_circ_0051596 | 2076 | 11 | 1.06% |
| hsa_circ_0058487 | 2076 | 11 | 1.06% |
| hsa_circ_0068201 | 2076 | 11 | 1.06% |
| hsa_circ_0126303 | 2076 | 11 | 1.06% |
| hsa_circ_0013681 | 2265 | 12 | 1.06% |
| hsa_circ_0061464 | 2265 | 12 | 1.06% |
| hsa_circ_0006126 | 2454 | 13 | 1.06% |
| hsa_circ_0023631 | 2454 | 13 | 1.06% |
| hsa_circ_0050261 | 2454 | 13 | 1.06% |
| hsa_circ_0018778 | 1888 | 10 | 1.06% |
| hsa_circ_0043192 | 1888 | 10 | 1.06% |
| hsa_circ_0056017 | 1888 | 10 | 1.06% |
| hsa_circ_0118890 | 1888 | 10 | 1.06% |
| hsa_circ_0127888 | 1888 | 10 | 1.06% |
| hsa_circ_0059158 | 2077 | 11 | 1.06% |
| hsa_circ_0016862 | 2266 | 12 | 1.06% |
| hsa_circ_0044489 | 2266 | 12 | 1.06% |
| hsa_circ_0061427 | 2266 | 12 | 1.06% |
| hsa_circ_0122791 | 2455 | 13 | 1.06% |
| hsa_circ_0022844 | 1889 | 10 | 1.06% |
| hsa_circ_0036947 | 1889 | 10 | 1.06% |
| hsa_circ_0052982 | 1889 | 10 | 1.06% |
| hsa_circ_0117304 | 1889 | 10 | 1.06% |
| hsa_circ_0026625 | 2078 | 11 | 1.06% |

| | | | |
|---|---|---|---|
| hsa_circ_0029197 | 2078 | 11 | 1.06% |
| hsa_circ_0041040 | 2078 | 11 | 1.06% |
| hsa_circ_0045277 | 2078 | 11 | 1.06% |
| hsa_circ_0073563 | 2078 | 11 | 1.06% |
| hsa_circ_0113225 | 2078 | 11 | 1.06% |
| hsa_circ_0124675 | 2078 | 11 | 1.06% |
| hsa_circ_0042282 | 2267 | 12 | 1.06% |
| hsa_circ_0063484 | 2268 | 12 | 1.06% |
| hsa_circ_0040280 | 2079 | 11 | 1.06% |
| hsa_circ_0044978 | 2079 | 11 | 1.06% |
| hsa_circ_0054723 | 2079 | 11 | 1.06% |
| hsa_circ_0075079 | 2079 | 11 | 1.06% |
| hsa_circ_0083684 | 2079 | 11 | 1.06% |
| hsa_circ_0100915 | 2079 | 11 | 1.06% |
| hsa_circ_0119502 | 2079 | 11 | 1.06% |
| hsa_circ_0015248 | 1890 | 10 | 1.06% |
| hsa_circ_0027198 | 1890 | 10 | 1.06% |
| hsa_circ_0036648 | 1890 | 10 | 1.06% |
| hsa_circ_0047566 | 1890 | 10 | 1.06% |
| hsa_circ_0076746 | 1890 | 10 | 1.06% |
| hsa_circ_0079623 | 1890 | 10 | 1.06% |
| hsa_circ_0082069 | 1890 | 10 | 1.06% |
| hsa_circ_0096004 | 1890 | 10 | 1.06% |
| hsa_circ_0140658 | 1890 | 10 | 1.06% |
| hsa_circ_0102414 | 4537 | 24 | 1.06% |
| hsa_circ_0101243 | 3781 | 20 | 1.06% |
| hsa_circ_0010661 | 2458 | 13 | 1.06% |
| hsa_circ_0099470 | 2458 | 13 | 1.06% |
| hsa_circ_0008351 | 4727 | 25 | 1.06% |
| hsa_circ_0086822 | 2269 | 12 | 1.06% |
| hsa_circ_0034731 | 2080 | 11 | 1.06% |
| hsa_circ_0069557 | 2080 | 11 | 1.06% |
| hsa_circ_0078885 | 2080 | 11 | 1.06% |
| hsa_circ_0078979 | 2080 | 11 | 1.06% |
| hsa_circ_0103102 | 2080 | 11 | 1.06% |
| hsa_circ_0115093 | 2080 | 11 | 1.06% |
| hsa_circ_0042279 | 1891 | 10 | 1.06% |
| hsa_circ_0052909 | 1891 | 10 | 1.06% |
| hsa_circ_0068258 | 1891 | 10 | 1.06% |
| hsa_circ_0136730 | 1891 | 10 | 1.06% |
| hsa_circ_0003118 | 5106 | 27 | 1.06% |
| hsa_circ_0029555 | 5107 | 27 | 1.06% |
| hsa_circ_0024316 | 2270 | 12 | 1.06% |
| hsa_circ_0030404 | 2270 | 12 | 1.06% |
| hsa_circ_0030054 | 2081 | 11 | 1.06% |
| hsa_circ_0038525 | 2081 | 11 | 1.06% |
| hsa_circ_0073561 | 2081 | 11 | 1.06% |
| hsa_circ_0049287 | 1892 | 10 | 1.06% |
| hsa_circ_0085726 | 1892 | 10 | 1.06% |
| hsa_circ_0092239 | 1892 | 10 | 1.06% |

| | | | |
|---|---|---|---|
| hsa_circ_0120119 | 1892 | 10 | 1.06% |
| hsa_circ_0066763 | 2460 | 13 | 1.06% |
| hsa_circ_0137960 | 2460 | 13 | 1.06% |
| hsa_circ_0033332 | 2271 | 12 | 1.06% |
| hsa_circ_0022853 | 2082 | 11 | 1.06% |
| hsa_circ_0056071 | 2082 | 11 | 1.06% |
| hsa_circ_0059723 | 2082 | 11 | 1.06% |
| hsa_circ_0107104 | 2082 | 11 | 1.06% |
| hsa_circ_0002171 | 5679 | 30 | 1.06% |
| hsa_circ_0011675 | 1893 | 10 | 1.06% |
| hsa_circ_0015230 | 1893 | 10 | 1.06% |
| hsa_circ_0017729 | 1893 | 10 | 1.06% |
| hsa_circ_0027215 | 1893 | 10 | 1.06% |
| hsa_circ_0081080 | 1893 | 10 | 1.06% |
| hsa_circ_0132227 | 1893 | 10 | 1.06% |
| hsa_circ_0139330 | 1893 | 10 | 1.06% |
| hsa_circ_0022226 | 2461 | 13 | 1.06% |
| hsa_circ_0046098 | 2461 | 13 | 1.06% |
| hsa_circ_0094898 | 2272 | 12 | 1.06% |
| hsa_circ_0019820 | 2083 | 11 | 1.06% |
| hsa_circ_0034574 | 2083 | 11 | 1.06% |
| hsa_circ_0128513 | 2083 | 11 | 1.06% |
| hsa_circ_0140750 | 2083 | 11 | 1.06% |
| hsa_circ_0018958 | 1894 | 10 | 1.06% |
| hsa_circ_0020013 | 1894 | 10 | 1.06% |
| hsa_circ_0032416 | 1894 | 10 | 1.06% |
| hsa_circ_0061376 | 1894 | 10 | 1.06% |
| hsa_circ_0066809 | 1894 | 10 | 1.06% |
| hsa_circ_0069020 | 1894 | 10 | 1.06% |
| hsa_circ_0075430 | 1894 | 10 | 1.06% |
| hsa_circ_0104966 | 1894 | 10 | 1.06% |
| hsa_circ_0114797 | 1894 | 10 | 1.06% |
| hsa_circ_0127317 | 1894 | 10 | 1.06% |
| hsa_circ_0025031 | 2273 | 12 | 1.06% |
| hsa_circ_0113149 | 2273 | 12 | 1.06% |
| hsa_circ_0052376 | 2084 | 11 | 1.06% |
| hsa_circ_0070745 | 2084 | 11 | 1.06% |
| hsa_circ_0083882 | 2084 | 11 | 1.06% |
| hsa_circ_0094388 | 2084 | 11 | 1.06% |
| hsa_circ_0049518 | 2463 | 13 | 1.06% |
| hsa_circ_0105573 | 2463 | 13 | 1.06% |
| hsa_circ_0109955 | 7011 | 37 | 1.06% |
| hsa_circ_0073346 | 1895 | 10 | 1.06% |
| hsa_circ_0114982 | 1895 | 10 | 1.06% |
| hsa_circ_0132624 | 1895 | 10 | 1.06% |
| hsa_circ_0030729 | 2085 | 11 | 1.06% |
| hsa_circ_0039132 | 2085 | 11 | 1.06% |
| hsa_circ_0051958 | 2085 | 11 | 1.06% |
| hsa_circ_0066559 | 2085 | 11 | 1.06% |
| hsa_circ_0069693 | 2085 | 11 | 1.06% |

| | | | |
|---|---|---|---|
| hsa_circ_0111477 | 2085 | 11 | 1.06% |
| hsa_circ_0052829 | 2275 | 12 | 1.05% |
| hsa_circ_0133323 | 5119 | 27 | 1.05% |
| hsa_circ_0024319 | 1896 | 10 | 1.05% |
| hsa_circ_0042672 | 1896 | 10 | 1.05% |
| hsa_circ_0068256 | 1896 | 10 | 1.05% |
| hsa_circ_0073305 | 1896 | 10 | 1.05% |
| hsa_circ_0078369 | 1896 | 10 | 1.05% |
| hsa_circ_0079539 | 1896 | 10 | 1.05% |
| hsa_circ_0112197 | 1896 | 10 | 1.05% |
| hsa_circ_0040225 | 2465 | 13 | 1.05% |
| hsa_circ_0046006 | 2465 | 13 | 1.05% |
| hsa_circ_0049556 | 2086 | 11 | 1.05% |
| hsa_circ_0062002 | 2086 | 11 | 1.05% |
| hsa_circ_0065673 | 2086 | 11 | 1.05% |
| hsa_circ_0135586 | 2086 | 11 | 1.05% |
| hsa_circ_0032565 | 2466 | 13 | 1.05% |
| hsa_circ_0087495 | 1897 | 10 | 1.05% |
| hsa_circ_0050503 | 2656 | 14 | 1.05% |
| hsa_circ_0016230 | 2087 | 11 | 1.05% |
| hsa_circ_0018870 | 2087 | 11 | 1.05% |
| hsa_circ_0054860 | 2087 | 11 | 1.05% |
| hsa_circ_0064855 | 2087 | 11 | 1.05% |
| hsa_circ_0004288 | 2467 | 13 | 1.05% |
| hsa_circ_0040587 | 2467 | 13 | 1.05% |
| hsa_circ_0133320 | 5124 | 27 | 1.05% |
| hsa_circ_0014362 | 1898 | 10 | 1.05% |
| hsa_circ_0023230 | 1898 | 10 | 1.05% |
| hsa_circ_0039708 | 1898 | 10 | 1.05% |
| hsa_circ_0062226 | 1898 | 10 | 1.05% |
| hsa_circ_0098732 | 1898 | 10 | 1.05% |
| hsa_circ_0106748 | 3986 | 21 | 1.05% |
| hsa_circ_0009547 | 2088 | 11 | 1.05% |
| hsa_circ_0062596 | 2088 | 11 | 1.05% |
| hsa_circ_0097602 | 2088 | 11 | 1.05% |
| hsa_circ_0127504 | 2088 | 11 | 1.05% |
| hsa_circ_0091675 | 2278 | 12 | 1.05% |
| hsa_circ_0104014 | 3038 | 16 | 1.05% |
| hsa_circ_0021598 | 1899 | 10 | 1.05% |
| hsa_circ_0022275 | 1899 | 10 | 1.05% |
| hsa_circ_0023316 | 1899 | 10 | 1.05% |
| hsa_circ_0033156 | 1899 | 10 | 1.05% |
| hsa_circ_0058260 | 1899 | 10 | 1.05% |
| hsa_circ_0067547 | 1899 | 10 | 1.05% |
| hsa_circ_0069429 | 1899 | 10 | 1.05% |
| hsa_circ_0072682 | 1899 | 10 | 1.05% |
| hsa_circ_0073972 | 1899 | 10 | 1.05% |
| hsa_circ_0077734 | 1899 | 10 | 1.05% |
| hsa_circ_0091967 | 1899 | 10 | 1.05% |
| hsa_circ_0055269 | 2089 | 11 | 1.05% |

| | | | |
|---|---|---|---|
| hsa_circ_0056295 | 2089 | 11 | 1.05% |
| hsa_circ_0069684 | 2089 | 11 | 1.05% |
| hsa_circ_0097366 | 2089 | 11 | 1.05% |
| hsa_circ_0098008 | 2089 | 11 | 1.05% |
| hsa_circ_0101525 | 2089 | 11 | 1.05% |
| hsa_circ_0099412 | 2850 | 15 | 1.05% |
| hsa_circ_0099471 | 2470 | 13 | 1.05% |
| hsa_circ_0023048 | 2280 | 12 | 1.05% |
| hsa_circ_0054617 | 2280 | 12 | 1.05% |
| hsa_circ_0076214 | 2090 | 11 | 1.05% |
| hsa_circ_0025410 | 1900 | 10 | 1.05% |
| hsa_circ_0030318 | 1900 | 10 | 1.05% |
| hsa_circ_0094967 | 1900 | 10 | 1.05% |
| hsa_circ_0133362 | 1900 | 10 | 1.05% |
| hsa_circ_0094266 | 3611 | 19 | 1.05% |
| hsa_circ_0134255 | 2661 | 14 | 1.05% |
| hsa_circ_0030234 | 2281 | 12 | 1.05% |
| hsa_circ_0120575 | 2281 | 12 | 1.05% |
| hsa_circ_0019928 | 2091 | 11 | 1.05% |
| hsa_circ_0046804 | 2091 | 11 | 1.05% |
| hsa_circ_0073425 | 2091 | 11 | 1.05% |
| hsa_circ_0119533 | 2091 | 11 | 1.05% |
| hsa_circ_0123381 | 2091 | 11 | 1.05% |
| hsa_circ_0125053 | 2091 | 11 | 1.05% |
| hsa_circ_0073135 | 1901 | 10 | 1.05% |
| hsa_circ_0032217 | 2472 | 13 | 1.05% |
| hsa_circ_0113372 | 2282 | 12 | 1.05% |
| hsa_circ_0020392 | 2092 | 11 | 1.05% |
| hsa_circ_0025012 | 2092 | 11 | 1.05% |
| hsa_circ_0029082 | 2092 | 11 | 1.05% |
| hsa_circ_0059028 | 2092 | 11 | 1.05% |
| hsa_circ_0095825 | 2092 | 11 | 1.05% |
| hsa_circ_0105586 | 2092 | 11 | 1.05% |
| hsa_circ_0123002 | 2092 | 11 | 1.05% |
| hsa_circ_0037022 | 1902 | 10 | 1.05% |
| hsa_circ_0047811 | 2473 | 13 | 1.05% |
| hsa_circ_0140052 | 8561 | 45 | 1.05% |
| hsa_circ_0058099 | 2283 | 12 | 1.05% |
| hsa_circ_0009818 | 2093 | 11 | 1.05% |
| hsa_circ_0010080 | 2093 | 11 | 1.05% |
| hsa_circ_0044063 | 2093 | 11 | 1.05% |
| hsa_circ_0079022 | 2093 | 11 | 1.05% |
| hsa_circ_0022477 | 1903 | 10 | 1.05% |
| hsa_circ_0041737 | 1903 | 10 | 1.05% |
| hsa_circ_0047237 | 1903 | 10 | 1.05% |
| hsa_circ_0119648 | 2474 | 13 | 1.05% |
| hsa_circ_0050917 | 2284 | 12 | 1.05% |
| hsa_circ_0070215 | 2284 | 12 | 1.05% |
| hsa_circ_0102377 | 2284 | 12 | 1.05% |
| hsa_circ_0111479 | 2284 | 12 | 1.05% |

| | | | |
|---|---|---|---|
| hsa_circ_0009446 | 2094 | 11 | 1.05% |
| hsa_circ_0019485 | 2094 | 11 | 1.05% |
| hsa_circ_0096327 | 2094 | 11 | 1.05% |
| hsa_circ_0112044 | 2094 | 11 | 1.05% |
| hsa_circ_0103134 | 4950 | 26 | 1.05% |
| hsa_circ_0032112 | 2475 | 13 | 1.05% |
| hsa_circ_0052256 | 1904 | 10 | 1.05% |
| hsa_circ_0072305 | 1904 | 10 | 1.05% |
| hsa_circ_0076411 | 1904 | 10 | 1.05% |
| hsa_circ_0100579 | 1904 | 10 | 1.05% |
| hsa_circ_0118325 | 1904 | 10 | 1.05% |
| hsa_circ_0010443 | 4189 | 22 | 1.05% |
| hsa_circ_0063505 | 2285 | 12 | 1.05% |
| hsa_circ_0013576 | 2666 | 14 | 1.05% |
| hsa_circ_0035888 | 2666 | 14 | 1.05% |
| hsa_circ_0031464 | 2095 | 11 | 1.05% |
| hsa_circ_0034307 | 2095 | 11 | 1.05% |
| hsa_circ_0072051 | 2095 | 11 | 1.05% |
| hsa_circ_0079369 | 2095 | 11 | 1.05% |
| hsa_circ_0131153 | 2095 | 11 | 1.05% |
| hsa_circ_0139260 | 2095 | 11 | 1.05% |
| hsa_circ_0140635 | 2095 | 11 | 1.05% |
| hsa_circ_0046253 | 2476 | 13 | 1.05% |
| hsa_circ_0092177 | 2476 | 13 | 1.05% |
| hsa_circ_0119885 | 2476 | 13 | 1.05% |
| hsa_circ_0128061 | 2476 | 13 | 1.05% |
| hsa_circ_0070416 | 2286 | 12 | 1.05% |
| hsa_circ_0024029 | 1905 | 10 | 1.05% |
| hsa_circ_0025418 | 1905 | 10 | 1.05% |
| hsa_circ_0030901 | 1905 | 10 | 1.05% |
| hsa_circ_0034444 | 1905 | 10 | 1.05% |
| hsa_circ_0037666 | 1905 | 10 | 1.05% |
| hsa_circ_0052197 | 1905 | 10 | 1.05% |
| hsa_circ_0055442 | 1905 | 10 | 1.05% |
| hsa_circ_0057266 | 1905 | 10 | 1.05% |
| hsa_circ_0095824 | 1905 | 10 | 1.05% |
| hsa_circ_0105310 | 6478 | 34 | 1.05% |
| hsa_circ_0081767 | 2477 | 13 | 1.05% |
| hsa_circ_0083415 | 2477 | 13 | 1.05% |
| hsa_circ_0129990 | 2477 | 13 | 1.05% |
| hsa_circ_0011329 | 2096 | 11 | 1.05% |
| hsa_circ_0049431 | 2096 | 11 | 1.05% |
| hsa_circ_0105438 | 2096 | 11 | 1.05% |
| hsa_circ_0123140 | 2096 | 11 | 1.05% |
| hsa_circ_0036126 | 2668 | 14 | 1.05% |
| hsa_circ_0022721 | 2287 | 12 | 1.05% |
| hsa_circ_0123144 | 2287 | 12 | 1.05% |
| hsa_circ_0024079 | 1906 | 10 | 1.05% |
| hsa_circ_0030534 | 1906 | 10 | 1.05% |
| hsa_circ_0034776 | 1906 | 10 | 1.05% |

| | | | |
|---|---|---|---|
| hsa_circ_0036920 | 1906 | 10 | 1.05% |
| hsa_circ_0052879 | 1906 | 10 | 1.05% |
| hsa_circ_0061646 | 1906 | 10 | 1.05% |
| hsa_circ_0098072 | 1906 | 10 | 1.05% |
| hsa_circ_0098581 | 1906 | 10 | 1.05% |
| hsa_circ_0101884 | 1906 | 10 | 1.05% |
| hsa_circ_0018148 | 2097 | 11 | 1.05% |
| hsa_circ_0044383 | 2097 | 11 | 1.05% |
| hsa_circ_0079872 | 2097 | 11 | 1.05% |
| hsa_circ_0016704 | 2288 | 12 | 1.05% |
| hsa_circ_0060631 | 2288 | 12 | 1.05% |
| hsa_circ_0073620 | 2288 | 12 | 1.05% |
| hsa_circ_0080427 | 2288 | 12 | 1.05% |
| hsa_circ_0040499 | 3814 | 20 | 1.05% |
| hsa_circ_0030075 | 1907 | 10 | 1.05% |
| hsa_circ_0065620 | 1907 | 10 | 1.05% |
| hsa_circ_0106828 | 1907 | 10 | 1.05% |
| hsa_circ_0059588 | 2098 | 11 | 1.05% |
| hsa_circ_0075794 | 2098 | 11 | 1.05% |
| hsa_circ_0133009 | 2098 | 11 | 1.05% |
| hsa_circ_0031155 | 2289 | 12 | 1.05% |
| hsa_circ_0066654 | 3434 | 18 | 1.05% |
| hsa_circ_0116004 | 2671 | 14 | 1.05% |
| hsa_circ_0016074 | 1908 | 10 | 1.05% |
| hsa_circ_0020880 | 1908 | 10 | 1.05% |
| hsa_circ_0021163 | 1908 | 10 | 1.05% |
| hsa_circ_0023489 | 1908 | 10 | 1.05% |
| hsa_circ_0025530 | 1908 | 10 | 1.05% |
| hsa_circ_0043825 | 1908 | 10 | 1.05% |
| hsa_circ_0044523 | 1908 | 10 | 1.05% |
| hsa_circ_0069014 | 1908 | 10 | 1.05% |
| hsa_circ_0076462 | 1908 | 10 | 1.05% |
| hsa_circ_0081625 | 1908 | 10 | 1.05% |
| hsa_circ_0127149 | 1908 | 10 | 1.05% |
| hsa_circ_0013207 | 2099 | 11 | 1.05% |
| hsa_circ_0053656 | 2099 | 11 | 1.05% |
| hsa_circ_0063250 | 2099 | 11 | 1.05% |
| hsa_circ_0080141 | 2099 | 11 | 1.05% |
| hsa_circ_0135228 | 2099 | 11 | 1.05% |
| hsa_circ_0069904 | 2290 | 12 | 1.05% |
| hsa_circ_0064895 | 2481 | 13 | 1.05% |
| hsa_circ_0129319 | 2481 | 13 | 1.05% |
| hsa_circ_0103816 | 5153 | 27 | 1.05% |
| hsa_circ_0017598 | 2672 | 14 | 1.05% |
| hsa_circ_0079964 | 2672 | 14 | 1.05% |
| hsa_circ_0090120 | 2672 | 14 | 1.05% |
| hsa_circ_0009765 | 1909 | 10 | 1.05% |
| hsa_circ_0036528 | 1909 | 10 | 1.05% |
| hsa_circ_0037314 | 1909 | 10 | 1.05% |
| hsa_circ_0045633 | 1909 | 10 | 1.05% |

| | | | |
|---|---|---|---|
| hsa_circ_0050029 | 1909 | 10 | 1.05% |
| hsa_circ_0111788 | 1909 | 10 | 1.05% |
| hsa_circ_0040419 | 2100 | 11 | 1.05% |
| hsa_circ_0069252 | 2100 | 11 | 1.05% |
| hsa_circ_0076647 | 2100 | 11 | 1.05% |
| hsa_circ_0079545 | 2100 | 11 | 1.05% |
| hsa_circ_0120661 | 2100 | 11 | 1.05% |
| hsa_circ_0128888 | 2100 | 11 | 1.05% |
| hsa_circ_0135332 | 2100 | 11 | 1.05% |
| hsa_circ_0041573 | 2291 | 12 | 1.05% |
| hsa_circ_0100338 | 2482 | 13 | 1.05% |
| hsa_circ_0048082 | 2673 | 14 | 1.05% |
| hsa_circ_0022884 | 2864 | 15 | 1.05% |
| hsa_circ_0131949 | 6302 | 33 | 1.05% |
| hsa_circ_0139948 | 2674 | 14 | 1.05% |
| hsa_circ_0121543 | 2483 | 13 | 1.05% |
| hsa_circ_0034458 | 2292 | 12 | 1.05% |
| hsa_circ_0046602 | 2292 | 12 | 1.05% |
| hsa_circ_0076818 | 2292 | 12 | 1.05% |
| hsa_circ_0138274 | 2292 | 12 | 1.05% |
| hsa_circ_0070766 | 2101 | 11 | 1.05% |
| hsa_circ_0072792 | 2101 | 11 | 1.05% |
| hsa_circ_0102889 | 2101 | 11 | 1.05% |
| hsa_circ_0122972 | 2101 | 11 | 1.05% |
| hsa_circ_0020690 | 1910 | 10 | 1.05% |
| hsa_circ_0041527 | 1910 | 10 | 1.05% |
| hsa_circ_0045064 | 1910 | 10 | 1.05% |
| hsa_circ_0056078 | 1910 | 10 | 1.05% |
| hsa_circ_0093839 | 1910 | 10 | 1.05% |
| hsa_circ_0111874 | 1910 | 10 | 1.05% |
| hsa_circ_0126513 | 1910 | 10 | 1.05% |
| hsa_circ_0006648 | 7832 | 41 | 1.05% |
| hsa_circ_0079994 | 3821 | 20 | 1.05% |
| hsa_circ_0036367 | 2484 | 13 | 1.05% |
| hsa_circ_0053106 | 2293 | 12 | 1.05% |
| hsa_circ_0021164 | 2102 | 11 | 1.05% |
| hsa_circ_0046747 | 2102 | 11 | 1.05% |
| hsa_circ_0055673 | 2102 | 11 | 1.05% |
| hsa_circ_0058231 | 2102 | 11 | 1.05% |
| hsa_circ_0074185 | 2102 | 11 | 1.05% |
| hsa_circ_0077596 | 2102 | 11 | 1.05% |
| hsa_circ_0089065 | 2102 | 11 | 1.05% |
| hsa_circ_0127146 | 2102 | 11 | 1.05% |
| hsa_circ_0021043 | 1911 | 10 | 1.05% |
| hsa_circ_0027565 | 1911 | 10 | 1.05% |
| hsa_circ_0039603 | 1911 | 10 | 1.05% |
| hsa_circ_0040606 | 1911 | 10 | 1.05% |
| hsa_circ_0040946 | 1911 | 10 | 1.05% |
| hsa_circ_0049637 | 1911 | 10 | 1.05% |
| hsa_circ_0119236 | 1911 | 10 | 1.05% |

| | | | |
|---|---|---|---|
| hsa_circ_0122187 | 1911 | 10 | 1.05% |
| hsa_circ_0135348 | 1911 | 10 | 1.05% |
| hsa_circ_0038032 | 6882 | 36 | 1.05% |
| hsa_circ_0030686 | 2103 | 11 | 1.05% |
| hsa_circ_0048386 | 2103 | 11 | 1.05% |
| hsa_circ_0053157 | 2103 | 11 | 1.05% |
| hsa_circ_0083939 | 2103 | 11 | 1.05% |
| hsa_circ_0087697 | 2103 | 11 | 1.05% |
| hsa_circ_0014685 | 1912 | 10 | 1.05% |
| hsa_circ_0024594 | 1912 | 10 | 1.05% |
| hsa_circ_0029607 | 1912 | 10 | 1.05% |
| hsa_circ_0058861 | 1912 | 10 | 1.05% |
| hsa_circ_0117943 | 1912 | 10 | 1.05% |
| hsa_circ_0118267 | 1912 | 10 | 1.05% |
| hsa_circ_0128529 | 1912 | 10 | 1.05% |
| hsa_circ_0035774 | 2486 | 13 | 1.05% |
| hsa_circ_0083737 | 2486 | 13 | 1.05% |
| hsa_circ_0061458 | 3060 | 16 | 1.05% |
| hsa_circ_0031139 | 2104 | 11 | 1.05% |
| hsa_circ_0075222 | 2104 | 11 | 1.05% |
| hsa_circ_0094865 | 2104 | 11 | 1.05% |
| hsa_circ_0008104 | 3826 | 20 | 1.05% |
| hsa_circ_0017198 | 1913 | 10 | 1.05% |
| hsa_circ_0033575 | 1913 | 10 | 1.05% |
| hsa_circ_0043726 | 1913 | 10 | 1.05% |
| hsa_circ_0054370 | 1913 | 10 | 1.05% |
| hsa_circ_0090257 | 1913 | 10 | 1.05% |
| hsa_circ_0113483 | 1913 | 10 | 1.05% |
| hsa_circ_0132925 | 1913 | 10 | 1.05% |
| hsa_circ_0037630 | 2296 | 12 | 1.05% |
| hsa_circ_0077672 | 2296 | 12 | 1.05% |
| hsa_circ_0046143 | 2105 | 11 | 1.05% |
| hsa_circ_0066680 | 2105 | 11 | 1.05% |
| hsa_circ_0081042 | 2105 | 11 | 1.05% |
| hsa_circ_0081166 | 2105 | 11 | 1.05% |
| hsa_circ_0022830 | 2488 | 13 | 1.05% |
| hsa_circ_0009834 | 1914 | 10 | 1.04% |
| hsa_circ_0013970 | 1914 | 10 | 1.04% |
| hsa_circ_0060417 | 1914 | 10 | 1.04% |
| hsa_circ_0064042 | 1914 | 10 | 1.04% |
| hsa_circ_0066432 | 1914 | 10 | 1.04% |
| hsa_circ_0069364 | 1914 | 10 | 1.04% |
| hsa_circ_0047990 | 2297 | 12 | 1.04% |
| hsa_circ_0050802 | 2297 | 12 | 1.04% |
| hsa_circ_0130089 | 2297 | 12 | 1.04% |
| hsa_circ_0072824 | 5934 | 31 | 1.04% |
| hsa_circ_0004647 | 2106 | 11 | 1.04% |
| hsa_circ_0011876 | 2106 | 11 | 1.04% |
| hsa_circ_0039031 | 2106 | 11 | 1.04% |
| hsa_circ_0053298 | 2106 | 11 | 1.04% |

| | | | |
|---|---|---|---|
| hsa_circ_0092002 | 2106 | 11 | 1.04% |
| hsa_circ_0097642 | 2106 | 11 | 1.04% |
| hsa_circ_0116374 | 2106 | 11 | 1.04% |
| hsa_circ_0121699 | 2106 | 11 | 1.04% |
| hsa_circ_0056210 | 2298 | 12 | 1.04% |
| hsa_circ_0081799 | 2298 | 12 | 1.04% |
| hsa_circ_0064950 | 1915 | 10 | 1.04% |
| hsa_circ_0069611 | 1915 | 10 | 1.04% |
| hsa_circ_0080417 | 1915 | 10 | 1.04% |
| hsa_circ_0100344 | 1915 | 10 | 1.04% |
| hsa_circ_0123231 | 1915 | 10 | 1.04% |
| hsa_circ_0139553 | 1915 | 10 | 1.04% |
| hsa_circ_0013783 | 2107 | 11 | 1.04% |
| hsa_circ_0022265 | 2107 | 11 | 1.04% |
| hsa_circ_0039140 | 2107 | 11 | 1.04% |
| hsa_circ_0039547 | 2107 | 11 | 1.04% |
| hsa_circ_0053017 | 2107 | 11 | 1.04% |
| hsa_circ_0065747 | 2107 | 11 | 1.04% |
| hsa_circ_0070774 | 2107 | 11 | 1.04% |
| hsa_circ_0092949 | 2107 | 11 | 1.04% |
| hsa_circ_0139453 | 2107 | 11 | 1.04% |
| hsa_circ_0033347 | 2299 | 12 | 1.04% |
| hsa_circ_0070784 | 2299 | 12 | 1.04% |
| hsa_circ_0109541 | 7855 | 41 | 1.04% |
| hsa_circ_0121101 | 1916 | 10 | 1.04% |
| hsa_circ_0014210 | 2108 | 11 | 1.04% |
| hsa_circ_0019740 | 2108 | 11 | 1.04% |
| hsa_circ_0031477 | 2108 | 11 | 1.04% |
| hsa_circ_0053149 | 2108 | 11 | 1.04% |
| hsa_circ_0071051 | 2108 | 11 | 1.04% |
| hsa_circ_0031453 | 2300 | 12 | 1.04% |
| hsa_circ_0034889 | 2300 | 12 | 1.04% |
| hsa_circ_0088738 | 2300 | 12 | 1.04% |
| hsa_circ_0029558 | 2492 | 13 | 1.04% |
| hsa_circ_0009838 | 1917 | 10 | 1.04% |
| hsa_circ_0028827 | 1917 | 10 | 1.04% |
| hsa_circ_0033552 | 1917 | 10 | 1.04% |
| hsa_circ_0056332 | 1917 | 10 | 1.04% |
| hsa_circ_0084337 | 1917 | 10 | 1.04% |
| hsa_circ_0086469 | 1917 | 10 | 1.04% |
| hsa_circ_0135250 | 1917 | 10 | 1.04% |
| hsa_circ_0041492 | 2684 | 14 | 1.04% |
| hsa_circ_0052433 | 2684 | 14 | 1.04% |
| hsa_circ_0038088 | 2109 | 11 | 1.04% |
| hsa_circ_0063113 | 2109 | 11 | 1.04% |
| hsa_circ_0068560 | 2109 | 11 | 1.04% |
| hsa_circ_0072678 | 2109 | 11 | 1.04% |
| hsa_circ_0120335 | 2109 | 11 | 1.04% |
| hsa_circ_0009785 | 1918 | 10 | 1.04% |
| hsa_circ_0100597 | 1918 | 10 | 1.04% |

| | | | |
|---|---|---|---|
| hsa_circ_0076894 | 2110 | 11 | 1.04% |
| hsa_circ_0084070 | 2110 | 11 | 1.04% |
| hsa_circ_0094460 | 2110 | 11 | 1.04% |
| hsa_circ_0100078 | 2110 | 11 | 1.04% |
| hsa_circ_0114796 | 2110 | 11 | 1.04% |
| hsa_circ_0137745 | 2110 | 11 | 1.04% |
| hsa_circ_0026321 | 5563 | 29 | 1.04% |
| hsa_circ_0048191 | 2302 | 12 | 1.04% |
| hsa_circ_0050283 | 2302 | 12 | 1.04% |
| hsa_circ_0091483 | 2302 | 12 | 1.04% |
| hsa_circ_0024652 | 6716 | 35 | 1.04% |
| hsa_circ_0020596 | 5565 | 29 | 1.04% |
| hsa_circ_0024205 | 1919 | 10 | 1.04% |
| hsa_circ_0042302 | 1919 | 10 | 1.04% |
| hsa_circ_0063788 | 1919 | 10 | 1.04% |
| hsa_circ_0069883 | 1919 | 10 | 1.04% |
| hsa_circ_0072068 | 1919 | 10 | 1.04% |
| hsa_circ_0100320 | 1919 | 10 | 1.04% |
| hsa_circ_0036262 | 2111 | 11 | 1.04% |
| hsa_circ_0052494 | 2111 | 11 | 1.04% |
| hsa_circ_0057665 | 2111 | 11 | 1.04% |
| hsa_circ_0063944 | 2111 | 11 | 1.04% |
| hsa_circ_0075088 | 2111 | 11 | 1.04% |
| hsa_circ_0086771 | 2111 | 11 | 1.04% |
| hsa_circ_0123567 | 2111 | 11 | 1.04% |
| hsa_circ_0136303 | 2111 | 11 | 1.04% |
| hsa_circ_0058702 | 2303 | 12 | 1.04% |
| hsa_circ_0088352 | 2303 | 12 | 1.04% |
| hsa_circ_0042788 | 2495 | 13 | 1.04% |
| hsa_circ_0121423 | 2495 | 13 | 1.04% |
| hsa_circ_0068761 | 4799 | 25 | 1.04% |
| hsa_circ_0005065 | 5375 | 28 | 1.04% |
| hsa_circ_0062821 | 2304 | 12 | 1.04% |
| hsa_circ_0071090 | 2304 | 12 | 1.04% |
| hsa_circ_0019110 | 2112 | 11 | 1.04% |
| hsa_circ_0072854 | 2112 | 11 | 1.04% |
| hsa_circ_0074829 | 2112 | 11 | 1.04% |
| hsa_circ_0084039 | 2112 | 11 | 1.04% |
| hsa_circ_0039277 | 1920 | 10 | 1.04% |
| hsa_circ_0039689 | 1920 | 10 | 1.04% |
| hsa_circ_0079548 | 2305 | 12 | 1.04% |
| hsa_circ_0085582 | 2113 | 11 | 1.04% |
| hsa_circ_0113182 | 2113 | 11 | 1.04% |
| hsa_circ_0009989 | 1921 | 10 | 1.04% |
| hsa_circ_0027723 | 1921 | 10 | 1.04% |
| hsa_circ_0051743 | 1921 | 10 | 1.04% |
| hsa_circ_0059497 | 1921 | 10 | 1.04% |
| hsa_circ_0069674 | 1921 | 10 | 1.04% |
| hsa_circ_0071315 | 1921 | 10 | 1.04% |
| hsa_circ_0100598 | 1921 | 10 | 1.04% |

| | | | |
|---|---|---|---|
| hsa_circ_0103219 | 1921 | 10 | 1.04% |
| hsa_circ_0113826 | 1921 | 10 | 1.04% |
| hsa_circ_0108301 | 4227 | 22 | 1.04% |
| hsa_circ_0018854 | 2306 | 12 | 1.04% |
| hsa_circ_0056171 | 2306 | 12 | 1.04% |
| hsa_circ_0019845 | 2114 | 11 | 1.04% |
| hsa_circ_0056028 | 2114 | 11 | 1.04% |
| hsa_circ_0062664 | 2114 | 11 | 1.04% |
| hsa_circ_0091985 | 2114 | 11 | 1.04% |
| hsa_circ_0127902 | 2114 | 11 | 1.04% |
| hsa_circ_0025857 | 1922 | 10 | 1.04% |
| hsa_circ_0064470 | 1922 | 10 | 1.04% |
| hsa_circ_0069275 | 1922 | 10 | 1.04% |
| hsa_circ_0069622 | 1922 | 10 | 1.04% |
| hsa_circ_0071155 | 1922 | 10 | 1.04% |
| hsa_circ_0078753 | 1922 | 10 | 1.04% |
| hsa_circ_0030393 | 2499 | 13 | 1.04% |
| hsa_circ_0130120 | 2499 | 13 | 1.04% |
| hsa_circ_0034364 | 2307 | 12 | 1.04% |
| hsa_circ_0058500 | 2307 | 12 | 1.04% |
| hsa_circ_0077434 | 2307 | 12 | 1.04% |
| hsa_circ_0082606 | 2307 | 12 | 1.04% |
| hsa_circ_0085906 | 2307 | 12 | 1.04% |
| hsa_circ_0026176 | 2884 | 15 | 1.04% |
| hsa_circ_0067255 | 2884 | 15 | 1.04% |
| hsa_circ_0102895 | 4999 | 26 | 1.04% |
| hsa_circ_0019562 | 2115 | 11 | 1.04% |
| hsa_circ_0080068 | 2115 | 11 | 1.04% |
| hsa_circ_0129710 | 5192 | 27 | 1.04% |
| hsa_circ_0005895 | 1923 | 10 | 1.04% |
| hsa_circ_0024427 | 1923 | 10 | 1.04% |
| hsa_circ_0036332 | 1923 | 10 | 1.04% |
| hsa_circ_0042214 | 1923 | 10 | 1.04% |
| hsa_circ_0053823 | 1923 | 10 | 1.04% |
| hsa_circ_0056587 | 1923 | 10 | 1.04% |
| hsa_circ_0058243 | 1923 | 10 | 1.04% |
| hsa_circ_0073080 | 1923 | 10 | 1.04% |
| hsa_circ_0134561 | 2308 | 12 | 1.04% |
| hsa_circ_0014041 | 3847 | 20 | 1.04% |
| hsa_circ_0025262 | 2693 | 14 | 1.04% |
| hsa_circ_0082003 | 2116 | 11 | 1.04% |
| hsa_circ_0086382 | 2116 | 11 | 1.04% |
| hsa_circ_0102083 | 2116 | 11 | 1.04% |
| hsa_circ_0087471 | 2501 | 13 | 1.04% |
| hsa_circ_0048706 | 1924 | 10 | 1.04% |
| hsa_circ_0088634 | 2309 | 12 | 1.04% |
| hsa_circ_0101484 | 2309 | 12 | 1.04% |
| hsa_circ_0091814 | 2694 | 14 | 1.04% |
| hsa_circ_0010767 | 2117 | 11 | 1.04% |
| hsa_circ_0019535 | 2117 | 11 | 1.04% |

| | | | |
|---|---|---|---|
| hsa_circ_0044145 | 2117 | 11 | 1.04% |
| hsa_circ_0071508 | 2117 | 11 | 1.04% |
| hsa_circ_0073698 | 2117 | 11 | 1.04% |
| hsa_circ_0090789 | 2117 | 11 | 1.04% |
| hsa_circ_0136671 | 2117 | 11 | 1.04% |
| hsa_circ_0017810 | 2502 | 13 | 1.04% |
| hsa_circ_0028036 | 2502 | 13 | 1.04% |
| hsa_circ_0106143 | 3465 | 18 | 1.04% |
| hsa_circ_0037626 | 2310 | 12 | 1.04% |
| hsa_circ_0041135 | 2310 | 12 | 1.04% |
| hsa_circ_0074496 | 2310 | 12 | 1.04% |
| hsa_circ_0080963 | 2310 | 12 | 1.04% |
| hsa_circ_0010348 | 1925 | 10 | 1.04% |
| hsa_circ_0048537 | 1925 | 10 | 1.04% |
| hsa_circ_0075847 | 1925 | 10 | 1.04% |
| hsa_circ_0086515 | 1925 | 10 | 1.04% |
| hsa_circ_0086577 | 1925 | 10 | 1.04% |
| hsa_circ_0100120 | 1925 | 10 | 1.04% |
| hsa_circ_0109327 | 1925 | 10 | 1.04% |
| hsa_circ_0024370 | 4043 | 21 | 1.04% |
| hsa_circ_0016139 | 2503 | 13 | 1.04% |
| hsa_circ_0014691 | 2118 | 11 | 1.04% |
| hsa_circ_0037412 | 2118 | 11 | 1.04% |
| hsa_circ_0074840 | 2118 | 11 | 1.04% |
| hsa_circ_0114264 | 2118 | 11 | 1.04% |
| hsa_circ_0118022 | 2118 | 11 | 1.04% |
| hsa_circ_0014745 | 2696 | 14 | 1.04% |
| hsa_circ_0012769 | 2311 | 12 | 1.04% |
| hsa_circ_0085457 | 2311 | 12 | 1.04% |
| hsa_circ_0125363 | 2311 | 12 | 1.04% |
| hsa_circ_0026063 | 1926 | 10 | 1.04% |
| hsa_circ_0030193 | 1926 | 10 | 1.04% |
| hsa_circ_0030381 | 1926 | 10 | 1.04% |
| hsa_circ_0052628 | 2504 | 13 | 1.04% |
| hsa_circ_0059652 | 2504 | 13 | 1.04% |
| hsa_circ_0110812 | 2119 | 11 | 1.04% |
| hsa_circ_0130703 | 2119 | 11 | 1.04% |
| hsa_circ_0120443 | 2312 | 12 | 1.04% |
| hsa_circ_0025437 | 1927 | 10 | 1.04% |
| hsa_circ_0027922 | 1927 | 10 | 1.04% |
| hsa_circ_0051624 | 1927 | 10 | 1.04% |
| hsa_circ_0073575 | 1927 | 10 | 1.04% |
| hsa_circ_0074116 | 1927 | 10 | 1.04% |
| hsa_circ_0081628 | 1927 | 10 | 1.04% |
| hsa_circ_0100497 | 1927 | 10 | 1.04% |
| hsa_circ_0109335 | 1927 | 10 | 1.04% |
| hsa_circ_0114731 | 1927 | 10 | 1.04% |
| hsa_circ_0128846 | 2698 | 14 | 1.04% |
| hsa_circ_0072381 | 2120 | 11 | 1.04% |
| hsa_circ_0082437 | 2120 | 11 | 1.04% |

| | | | |
|---|---|---|---|
| hsa_circ_0118152 | 2120 | 11 | 1.04% |
| hsa_circ_0088664 | 2313 | 12 | 1.04% |
| hsa_circ_0108444 | 1928 | 10 | 1.04% |
| hsa_circ_0111911 | 1928 | 10 | 1.04% |
| hsa_circ_0052094 | 7905 | 41 | 1.04% |
| hsa_circ_0000948 | 2121 | 11 | 1.04% |
| hsa_circ_0029572 | 2121 | 11 | 1.04% |
| hsa_circ_0057095 | 2121 | 11 | 1.04% |
| hsa_circ_0075098 | 2121 | 11 | 1.04% |
| hsa_circ_0094610 | 2121 | 11 | 1.04% |
| hsa_circ_0132076 | 2121 | 11 | 1.04% |
| hsa_circ_0009704 | 1929 | 10 | 1.04% |
| hsa_circ_0023290 | 1929 | 10 | 1.04% |
| hsa_circ_0081951 | 1929 | 10 | 1.04% |
| hsa_circ_0088575 | 1929 | 10 | 1.04% |
| hsa_circ_0088926 | 1929 | 10 | 1.04% |
| hsa_circ_0113709 | 1929 | 10 | 1.04% |
| hsa_circ_0010069 | 2122 | 11 | 1.04% |
| hsa_circ_0041976 | 2122 | 11 | 1.04% |
| hsa_circ_0042039 | 2122 | 11 | 1.04% |
| hsa_circ_0062128 | 2122 | 11 | 1.04% |
| hsa_circ_0079345 | 2122 | 11 | 1.04% |
| hsa_circ_0090954 | 2122 | 11 | 1.04% |
| hsa_circ_0010213 | 2315 | 12 | 1.04% |
| hsa_circ_0012247 | 2315 | 12 | 1.04% |
| hsa_circ_0050019 | 2315 | 12 | 1.04% |
| hsa_circ_0061444 | 2508 | 13 | 1.04% |
| hsa_circ_0091107 | 3473 | 18 | 1.04% |
| hsa_circ_0015554 | 2509 | 13 | 1.04% |
| hsa_circ_0071841 | 2316 | 12 | 1.04% |
| hsa_circ_0120583 | 2316 | 12 | 1.04% |
| hsa_circ_0011728 | 2123 | 11 | 1.04% |
| hsa_circ_0014790 | 2123 | 11 | 1.04% |
| hsa_circ_0020776 | 2123 | 11 | 1.04% |
| hsa_circ_0039059 | 2123 | 11 | 1.04% |
| hsa_circ_0049563 | 2123 | 11 | 1.04% |
| hsa_circ_0067333 | 2123 | 11 | 1.04% |
| hsa_circ_0086204 | 2123 | 11 | 1.04% |
| hsa_circ_0103642 | 2123 | 11 | 1.04% |
| hsa_circ_0014307 | 1930 | 10 | 1.04% |
| hsa_circ_0018260 | 1930 | 10 | 1.04% |
| hsa_circ_0053871 | 1930 | 10 | 1.04% |
| hsa_circ_0077842 | 1930 | 10 | 1.04% |
| hsa_circ_0129048 | 7528 | 39 | 1.04% |
| hsa_circ_0066685 | 3089 | 16 | 1.04% |
| hsa_circ_0009878 | 5406 | 28 | 1.04% |
| hsa_circ_0065115 | 2317 | 12 | 1.04% |
| hsa_circ_0068535 | 2317 | 12 | 1.04% |
| hsa_circ_0111938 | 2317 | 12 | 1.04% |
| hsa_circ_0042546 | 2124 | 11 | 1.04% |

| | | | |
|---|---|---|---|
| hsa_circ_0098767 | 2124 | 11 | 1.04% |
| hsa_circ_0111281 | 2124 | 11 | 1.04% |
| hsa_circ_0022480 | 1931 | 10 | 1.04% |
| hsa_circ_0030324 | 1931 | 10 | 1.04% |
| hsa_circ_0064047 | 1931 | 10 | 1.04% |
| hsa_circ_0052752 | 2511 | 13 | 1.04% |
| hsa_circ_0021124 | 1932 | 10 | 1.04% |
| hsa_circ_0021677 | 1932 | 10 | 1.04% |
| hsa_circ_0067881 | 1932 | 10 | 1.04% |
| hsa_circ_0112998 | 1932 | 10 | 1.04% |
| hsa_circ_0140733 | 1932 | 10 | 1.04% |
| hsa_circ_0087457 | 2512 | 13 | 1.04% |
| hsa_circ_0034777 | 2319 | 12 | 1.03% |
| hsa_circ_0071563 | 2319 | 12 | 1.03% |
| hsa_circ_0137170 | 2319 | 12 | 1.03% |
| hsa_circ_0026248 | 5798 | 30 | 1.03% |
| hsa_circ_0064932 | 2126 | 11 | 1.03% |
| hsa_circ_0107807 | 2126 | 11 | 1.03% |
| hsa_circ_0005942 | 1933 | 10 | 1.03% |
| hsa_circ_0034286 | 1933 | 10 | 1.03% |
| hsa_circ_0043733 | 1933 | 10 | 1.03% |
| hsa_circ_0054094 | 1933 | 10 | 1.03% |
| hsa_circ_0055075 | 1933 | 10 | 1.03% |
| hsa_circ_0073391 | 1933 | 10 | 1.03% |
| hsa_circ_0100778 | 1933 | 10 | 1.03% |
| hsa_circ_0108344 | 1933 | 10 | 1.03% |
| hsa_circ_0138163 | 1933 | 10 | 1.03% |
| hsa_circ_0089048 | 2513 | 13 | 1.03% |
| hsa_circ_0097442 | 5606 | 29 | 1.03% |
| hsa_circ_0017478 | 2127 | 11 | 1.03% |
| hsa_circ_0053687 | 2127 | 11 | 1.03% |
| hsa_circ_0079221 | 2127 | 11 | 1.03% |
| hsa_circ_0083683 | 2514 | 13 | 1.03% |
| hsa_circ_0085998 | 2514 | 13 | 1.03% |
| hsa_circ_0004748 | 1934 | 10 | 1.03% |
| hsa_circ_0076314 | 1934 | 10 | 1.03% |
| hsa_circ_0119087 | 1934 | 10 | 1.03% |
| hsa_circ_0034895 | 2321 | 12 | 1.03% |
| hsa_circ_0043658 | 2321 | 12 | 1.03% |
| hsa_circ_0081013 | 2321 | 12 | 1.03% |
| hsa_circ_0017009 | 2128 | 11 | 1.03% |
| hsa_circ_0070484 | 2128 | 11 | 1.03% |
| hsa_circ_0084660 | 2128 | 11 | 1.03% |
| hsa_circ_0126975 | 2128 | 11 | 1.03% |
| hsa_circ_0136354 | 2128 | 11 | 1.03% |
| hsa_circ_0094325 | 4837 | 25 | 1.03% |
| hsa_circ_0027173 | 2322 | 12 | 1.03% |
| hsa_circ_0028509 | 2322 | 12 | 1.03% |
| hsa_circ_0027898 | 1935 | 10 | 1.03% |
| hsa_circ_0032037 | 1935 | 10 | 1.03% |

| | | | |
|---|---|---|---|
| hsa_circ_0048283 | 1935 | 10 | 1.03% |
| hsa_circ_0063546 | 1935 | 10 | 1.03% |
| hsa_circ_0079980 | 1935 | 10 | 1.03% |
| hsa_circ_0105859 | 1935 | 10 | 1.03% |
| hsa_circ_0063513 | 2516 | 13 | 1.03% |
| hsa_circ_0023173 | 2129 | 11 | 1.03% |
| hsa_circ_0050340 | 2129 | 11 | 1.03% |
| hsa_circ_0061723 | 2129 | 11 | 1.03% |
| hsa_circ_0073522 | 2129 | 11 | 1.03% |
| hsa_circ_0080933 | 2129 | 11 | 1.03% |
| hsa_circ_0021205 | 2323 | 12 | 1.03% |
| hsa_circ_0030804 | 1936 | 10 | 1.03% |
| hsa_circ_0031565 | 1936 | 10 | 1.03% |
| hsa_circ_0058821 | 1936 | 10 | 1.03% |
| hsa_circ_0071000 | 1936 | 10 | 1.03% |
| hsa_circ_0072592 | 1936 | 10 | 1.03% |
| hsa_circ_0097978 | 1936 | 10 | 1.03% |
| hsa_circ_0073166 | 4453 | 23 | 1.03% |
| hsa_circ_0007161 | 2130 | 11 | 1.03% |
| hsa_circ_0021536 | 2130 | 11 | 1.03% |
| hsa_circ_0045556 | 2130 | 11 | 1.03% |
| hsa_circ_0049559 | 2130 | 11 | 1.03% |
| hsa_circ_0077132 | 2130 | 11 | 1.03% |
| hsa_circ_0093204 | 2130 | 11 | 1.03% |
| hsa_circ_0034422 | 2324 | 12 | 1.03% |
| hsa_circ_0131899 | 2324 | 12 | 1.03% |
| hsa_circ_0020814 | 2518 | 13 | 1.03% |
| hsa_circ_0050637 | 2518 | 13 | 1.03% |
| hsa_circ_0009723 | 1937 | 10 | 1.03% |
| hsa_circ_0041122 | 1937 | 10 | 1.03% |
| hsa_circ_0127925 | 1937 | 10 | 1.03% |
| hsa_circ_0066071 | 3293 | 17 | 1.03% |
| hsa_circ_0024902 | 2712 | 14 | 1.03% |
| hsa_circ_0010278 | 2131 | 11 | 1.03% |
| hsa_circ_0068152 | 2131 | 11 | 1.03% |
| hsa_circ_0092746 | 2131 | 11 | 1.03% |
| hsa_circ_0019741 | 2325 | 12 | 1.03% |
| hsa_circ_0048112 | 2325 | 12 | 1.03% |
| hsa_circ_0113101 | 2325 | 12 | 1.03% |
| hsa_circ_0113677 | 2519 | 13 | 1.03% |
| hsa_circ_0011046 | 1938 | 10 | 1.03% |
| hsa_circ_0036963 | 1938 | 10 | 1.03% |
| hsa_circ_0071088 | 1938 | 10 | 1.03% |
| hsa_circ_0099387 | 1938 | 10 | 1.03% |
| hsa_circ_0103533 | 1938 | 10 | 1.03% |
| hsa_circ_0108832 | 1938 | 10 | 1.03% |
| hsa_circ_0113582 | 1938 | 10 | 1.03% |
| hsa_circ_0121069 | 1938 | 10 | 1.03% |
| hsa_circ_0017800 | 2132 | 11 | 1.03% |
| hsa_circ_0021533 | 2132 | 11 | 1.03% |

| | | | |
|---|---|---|---|
| hsa_circ_0032429 | 2132 | 11 | 1.03% |
| hsa_circ_0099190 | 2132 | 11 | 1.03% |
| hsa_circ_0067082 | 2326 | 12 | 1.03% |
| hsa_circ_0080574 | 2326 | 12 | 1.03% |
| hsa_circ_0068578 | 2520 | 13 | 1.03% |
| hsa_circ_0043803 | 4265 | 22 | 1.03% |
| hsa_circ_0131283 | 5623 | 29 | 1.03% |
| hsa_circ_0069959 | 1939 | 10 | 1.03% |
| hsa_circ_0071658 | 1939 | 10 | 1.03% |
| hsa_circ_0077463 | 1939 | 10 | 1.03% |
| hsa_circ_0082156 | 1939 | 10 | 1.03% |
| hsa_circ_0105712 | 1939 | 10 | 1.03% |
| hsa_circ_0040886 | 2133 | 11 | 1.03% |
| hsa_circ_0062096 | 2133 | 11 | 1.03% |
| hsa_circ_0064482 | 2133 | 11 | 1.03% |
| hsa_circ_0076786 | 2133 | 11 | 1.03% |
| hsa_circ_0120744 | 2133 | 11 | 1.03% |
| hsa_circ_0129401 | 2327 | 12 | 1.03% |
| hsa_circ_0085007 | 5818 | 30 | 1.03% |
| hsa_circ_0052687 | 2909 | 15 | 1.03% |
| hsa_circ_0048407 | 2910 | 15 | 1.03% |
| hsa_circ_0047551 | 2716 | 14 | 1.03% |
| hsa_circ_0011874 | 2328 | 12 | 1.03% |
| hsa_circ_0018517 | 2328 | 12 | 1.03% |
| hsa_circ_0006251 | 2134 | 11 | 1.03% |
| hsa_circ_0013116 | 2134 | 11 | 1.03% |
| hsa_circ_0019602 | 2134 | 11 | 1.03% |
| hsa_circ_0036972 | 2134 | 11 | 1.03% |
| hsa_circ_0139193 | 2134 | 11 | 1.03% |
| hsa_circ_0011861 | 1940 | 10 | 1.03% |
| hsa_circ_0012927 | 1940 | 10 | 1.03% |
| hsa_circ_0131787 | 1940 | 10 | 1.03% |
| hsa_circ_0076722 | 2523 | 13 | 1.03% |
| hsa_circ_0091810 | 2523 | 13 | 1.03% |
| hsa_circ_0088263 | 2329 | 12 | 1.03% |
| hsa_circ_0118864 | 2329 | 12 | 1.03% |
| hsa_circ_0014209 | 2135 | 11 | 1.03% |
| hsa_circ_0132000 | 2135 | 11 | 1.03% |
| hsa_circ_0040682 | 1941 | 10 | 1.03% |
| hsa_circ_0045950 | 1941 | 10 | 1.03% |
| hsa_circ_0058402 | 1941 | 10 | 1.03% |
| hsa_circ_0068279 | 1941 | 10 | 1.03% |
| hsa_circ_0082260 | 1941 | 10 | 1.03% |
| hsa_circ_0092150 | 1941 | 10 | 1.03% |
| hsa_circ_0134908 | 1941 | 10 | 1.03% |
| hsa_circ_0094099 | 2912 | 15 | 1.03% |
| hsa_circ_0020824 | 2330 | 12 | 1.03% |
| hsa_circ_0023825 | 2330 | 12 | 1.03% |
| hsa_circ_0101625 | 2330 | 12 | 1.03% |
| hsa_circ_0017735 | 2136 | 11 | 1.03% |

| | | | |
|---|---|---|---|
| hsa_circ_0052077 | 2136 | 11 | 1.03% |
| hsa_circ_0080063 | 2136 | 11 | 1.03% |
| hsa_circ_0081059 | 2136 | 11 | 1.03% |
| hsa_circ_0091864 | 2136 | 11 | 1.03% |
| hsa_circ_0013714 | 1942 | 10 | 1.03% |
| hsa_circ_0016911 | 1942 | 10 | 1.03% |
| hsa_circ_0032060 | 1942 | 10 | 1.03% |
| hsa_circ_0047254 | 1942 | 10 | 1.03% |
| hsa_circ_0048997 | 1942 | 10 | 1.03% |
| hsa_circ_0070872 | 1942 | 10 | 1.03% |
| hsa_circ_0071407 | 1942 | 10 | 1.03% |
| hsa_circ_0035164 | 2331 | 12 | 1.03% |
| hsa_circ_0131896 | 2331 | 12 | 1.03% |
| hsa_circ_0016717 | 2137 | 11 | 1.03% |
| hsa_circ_0120786 | 2137 | 11 | 1.03% |
| hsa_circ_0026216 | 1943 | 10 | 1.03% |
| hsa_circ_0036821 | 1943 | 10 | 1.03% |
| hsa_circ_0053995 | 1943 | 10 | 1.03% |
| hsa_circ_0136049 | 1943 | 10 | 1.03% |
| hsa_circ_0015117 | 2526 | 13 | 1.03% |
| hsa_circ_0020529 | 2526 | 13 | 1.03% |
| hsa_circ_0088859 | 2526 | 13 | 1.03% |
| hsa_circ_0074852 | 2915 | 15 | 1.03% |
| hsa_circ_0081260 | 2915 | 15 | 1.03% |
| hsa_circ_0086485 | 4470 | 23 | 1.03% |
| hsa_circ_0070161 | 2721 | 14 | 1.03% |
| hsa_circ_0077026 | 2721 | 14 | 1.03% |
| hsa_circ_0023649 | 2138 | 11 | 1.03% |
| hsa_circ_0086497 | 2138 | 11 | 1.03% |
| hsa_circ_0136722 | 2138 | 11 | 1.03% |
| hsa_circ_0089032 | 2527 | 13 | 1.03% |
| hsa_circ_0008541 | 1944 | 10 | 1.03% |
| hsa_circ_0072432 | 1944 | 10 | 1.03% |
| hsa_circ_0091677 | 1944 | 10 | 1.03% |
| hsa_circ_0127014 | 2722 | 14 | 1.03% |
| hsa_circ_0000404 | 2139 | 11 | 1.03% |
| hsa_circ_0015317 | 2139 | 11 | 1.03% |
| hsa_circ_0022868 | 2139 | 11 | 1.03% |
| hsa_circ_0049730 | 2139 | 11 | 1.03% |
| hsa_circ_0072463 | 2139 | 11 | 1.03% |
| hsa_circ_0102579 | 2139 | 11 | 1.03% |
| hsa_circ_0023045 | 2528 | 13 | 1.03% |
| hsa_circ_0123416 | 2528 | 13 | 1.03% |
| hsa_circ_0123432 | 2528 | 13 | 1.03% |
| hsa_circ_0022465 | 2334 | 12 | 1.03% |
| hsa_circ_0053052 | 2334 | 12 | 1.03% |
| hsa_circ_0013836 | 1945 | 10 | 1.03% |
| hsa_circ_0018908 | 1945 | 10 | 1.03% |
| hsa_circ_0065066 | 1945 | 10 | 1.03% |
| hsa_circ_0079182 | 1945 | 10 | 1.03% |

| | | | |
|---|---|---|---|
| hsa_circ_0105302 | 1945 | 10 | 1.03% |
| hsa_circ_0112172 | 1945 | 10 | 1.03% |
| hsa_circ_0122184 | 1945 | 10 | 1.03% |
| hsa_circ_0132059 | 1945 | 10 | 1.03% |
| hsa_circ_0089141 | 2918 | 15 | 1.03% |
| hsa_circ_0029701 | 4669 | 24 | 1.03% |
| hsa_circ_0043985 | 2140 | 11 | 1.03% |
| hsa_circ_0045401 | 2140 | 11 | 1.03% |
| hsa_circ_0045794 | 2140 | 11 | 1.03% |
| hsa_circ_0046361 | 2140 | 11 | 1.03% |
| hsa_circ_0076336 | 2140 | 11 | 1.03% |
| hsa_circ_0089375 | 2140 | 11 | 1.03% |
| hsa_circ_0111547 | 2140 | 11 | 1.03% |
| hsa_circ_0140059 | 2140 | 11 | 1.03% |
| hsa_circ_0043254 | 3113 | 16 | 1.03% |
| hsa_circ_0016638 | 2335 | 12 | 1.03% |
| hsa_circ_0030447 | 2335 | 12 | 1.03% |
| hsa_circ_0063232 | 2335 | 12 | 1.03% |
| hsa_circ_0065601 | 2335 | 12 | 1.03% |
| hsa_circ_0106825 | 2335 | 12 | 1.03% |
| hsa_circ_0002327 | 5254 | 27 | 1.03% |
| hsa_circ_0051965 | 1946 | 10 | 1.03% |
| hsa_circ_0071959 | 1946 | 10 | 1.03% |
| hsa_circ_0078803 | 1946 | 10 | 1.03% |
| hsa_circ_0085748 | 1946 | 10 | 1.03% |
| hsa_circ_0123272 | 1946 | 10 | 1.03% |
| hsa_circ_0040605 | 2530 | 13 | 1.03% |
| hsa_circ_0062019 | 2530 | 13 | 1.03% |
| hsa_circ_0026623 | 2141 | 11 | 1.03% |
| hsa_circ_0029065 | 2141 | 11 | 1.03% |
| hsa_circ_0043204 | 2141 | 11 | 1.03% |
| hsa_circ_0065717 | 2141 | 11 | 1.03% |
| hsa_circ_0093862 | 2141 | 11 | 1.03% |
| hsa_circ_0125413 | 2920 | 15 | 1.03% |
| hsa_circ_0036434 | 2336 | 12 | 1.03% |
| hsa_circ_0039872 | 2336 | 12 | 1.03% |
| hsa_circ_0073625 | 2336 | 12 | 1.03% |
| hsa_circ_0114629 | 2336 | 12 | 1.03% |
| hsa_circ_0035387 | 1947 | 10 | 1.03% |
| hsa_circ_0052301 | 1947 | 10 | 1.03% |
| hsa_circ_0068031 | 1947 | 10 | 1.03% |
| hsa_circ_0074731 | 1947 | 10 | 1.03% |
| hsa_circ_0087169 | 1947 | 10 | 1.03% |
| hsa_circ_0013897 | 2142 | 11 | 1.03% |
| hsa_circ_0020215 | 2142 | 11 | 1.03% |
| hsa_circ_0038120 | 2142 | 11 | 1.03% |
| hsa_circ_0032521 | 2532 | 13 | 1.03% |
| hsa_circ_0001693 | 11493 | 59 | 1.03% |
| hsa_circ_0138865 | 31752 | 163 | 1.03% |
| hsa_circ_0131360 | 3896 | 20 | 1.03% |

| | | | |
|---|---|---|---|
| hsa_circ_0013625 | 1948 | 10 | 1.03% |
| hsa_circ_0031680 | 1948 | 10 | 1.03% |
| hsa_circ_0036753 | 1948 | 10 | 1.03% |
| hsa_circ_0041998 | 1948 | 10 | 1.03% |
| hsa_circ_0060259 | 1948 | 10 | 1.03% |
| hsa_circ_0061656 | 1948 | 10 | 1.03% |
| hsa_circ_0070864 | 1948 | 10 | 1.03% |
| hsa_circ_0096412 | 1948 | 10 | 1.03% |
| hsa_circ_0105530 | 1948 | 10 | 1.03% |
| hsa_circ_0123240 | 1948 | 10 | 1.03% |
| hsa_circ_0140071 | 1948 | 10 | 1.03% |
| hsa_circ_0030520 | 2143 | 11 | 1.03% |
| hsa_circ_0037819 | 2143 | 11 | 1.03% |
| hsa_circ_0041044 | 2143 | 11 | 1.03% |
| hsa_circ_0076559 | 2143 | 11 | 1.03% |
| hsa_circ_0090405 | 2143 | 11 | 1.03% |
| hsa_circ_0106916 | 2143 | 11 | 1.03% |
| hsa_circ_0000101 | 2338 | 12 | 1.03% |
| hsa_circ_0090152 | 2338 | 12 | 1.03% |
| hsa_circ_0024632 | 2533 | 13 | 1.03% |
| hsa_circ_0030868 | 2728 | 14 | 1.03% |
| hsa_circ_0008682 | 1949 | 10 | 1.03% |
| hsa_circ_0125849 | 1949 | 10 | 1.03% |
| hsa_circ_0027350 | 2144 | 11 | 1.03% |
| hsa_circ_0046475 | 2144 | 11 | 1.03% |
| hsa_circ_0116482 | 2144 | 11 | 1.03% |
| hsa_circ_0067399 | 2339 | 12 | 1.03% |
| hsa_circ_0105574 | 2535 | 13 | 1.03% |
| hsa_circ_0024851 | 2340 | 12 | 1.03% |
| hsa_circ_0009410 | 2145 | 11 | 1.03% |
| hsa_circ_0078384 | 2145 | 11 | 1.03% |
| hsa_circ_0092961 | 2145 | 11 | 1.03% |
| hsa_circ_0101657 | 2145 | 11 | 1.03% |
| hsa_circ_0122293 | 2145 | 11 | 1.03% |
| hsa_circ_0019456 | 1950 | 10 | 1.03% |
| hsa_circ_0022289 | 1950 | 10 | 1.03% |
| hsa_circ_0073787 | 1950 | 10 | 1.03% |
| hsa_circ_0091355 | 1950 | 10 | 1.03% |
| hsa_circ_0000931 | 4291 | 22 | 1.03% |
| hsa_circ_0049905 | 2731 | 14 | 1.03% |
| hsa_circ_0004062 | 2341 | 12 | 1.03% |
| hsa_circ_0023195 | 2341 | 12 | 1.03% |
| hsa_circ_0099835 | 4487 | 23 | 1.03% |
| hsa_circ_0041342 | 2146 | 11 | 1.03% |
| hsa_circ_0080011 | 2146 | 11 | 1.03% |
| hsa_circ_0036134 | 1951 | 10 | 1.03% |
| hsa_circ_0060714 | 1951 | 10 | 1.03% |
| hsa_circ_0084260 | 1951 | 10 | 1.03% |
| hsa_circ_0086313 | 1951 | 10 | 1.03% |
| hsa_circ_0089317 | 1951 | 10 | 1.03% |

| | | | |
|---|---|---|---|
| hsa_circ_0134496 | 1951 | 10 | 1.03% |
| hsa_circ_0013573 | 2732 | 14 | 1.02% |
| hsa_circ_0034170 | 2342 | 12 | 1.02% |
| hsa_circ_0071871 | 2342 | 12 | 1.02% |
| hsa_circ_0085712 | 2147 | 11 | 1.02% |
| hsa_circ_0094868 | 2147 | 11 | 1.02% |
| hsa_circ_0095079 | 2147 | 11 | 1.02% |
| hsa_circ_0132984 | 2147 | 11 | 1.02% |
| hsa_circ_0012375 | 1952 | 10 | 1.02% |
| hsa_circ_0086496 | 1952 | 10 | 1.02% |
| hsa_circ_0093585 | 1952 | 10 | 1.02% |
| hsa_circ_0081598 | 2538 | 13 | 1.02% |
| hsa_circ_0022701 | 2148 | 11 | 1.02% |
| hsa_circ_0083314 | 2148 | 11 | 1.02% |
| hsa_circ_0091424 | 2148 | 11 | 1.02% |
| hsa_circ_0048700 | 1953 | 10 | 1.02% |
| hsa_circ_0097960 | 1953 | 10 | 1.02% |
| hsa_circ_0129147 | 1953 | 10 | 1.02% |
| hsa_circ_0136074 | 1953 | 10 | 1.02% |
| hsa_circ_0098228 | 2539 | 13 | 1.02% |
| hsa_circ_0134916 | 2930 | 15 | 1.02% |
| hsa_circ_0034544 | 2344 | 12 | 1.02% |
| hsa_circ_0011273 | 2149 | 11 | 1.02% |
| hsa_circ_0049441 | 2149 | 11 | 1.02% |
| hsa_circ_0079477 | 4689 | 24 | 1.02% |
| hsa_circ_0030993 | 2540 | 13 | 1.02% |
| hsa_circ_0022679 | 1954 | 10 | 1.02% |
| hsa_circ_0053367 | 1954 | 10 | 1.02% |
| hsa_circ_0062352 | 1954 | 10 | 1.02% |
| hsa_circ_0070765 | 1954 | 10 | 1.02% |
| hsa_circ_0071024 | 1954 | 10 | 1.02% |
| hsa_circ_0093895 | 1954 | 10 | 1.02% |
| hsa_circ_0081501 | 3127 | 16 | 1.02% |
| hsa_circ_0048672 | 2150 | 11 | 1.02% |
| hsa_circ_0052670 | 2150 | 11 | 1.02% |
| hsa_circ_0047419 | 2541 | 13 | 1.02% |
| hsa_circ_0122263 | 2541 | 13 | 1.02% |
| hsa_circ_0026343 | 2932 | 15 | 1.02% |
| hsa_circ_0034435 | 2346 | 12 | 1.02% |
| hsa_circ_0039651 | 1955 | 10 | 1.02% |
| hsa_circ_0071586 | 1955 | 10 | 1.02% |
| hsa_circ_0085573 | 1955 | 10 | 1.02% |
| hsa_circ_0139039 | 2933 | 15 | 1.02% |
| hsa_circ_0018694 | 2151 | 11 | 1.02% |
| hsa_circ_0065349 | 2151 | 11 | 1.02% |
| hsa_circ_0066173 | 2151 | 11 | 1.02% |
| hsa_circ_0114252 | 2151 | 11 | 1.02% |
| hsa_circ_0032574 | 2347 | 12 | 1.02% |
| hsa_circ_0111936 | 2347 | 12 | 1.02% |
| hsa_circ_0015694 | 1956 | 10 | 1.02% |

| | | | |
|---|---|---|---|
| hsa_circ_0057421 | 1956 | 10 | 1.02% |
| hsa_circ_0069755 | 1956 | 10 | 1.02% |
| hsa_circ_0074320 | 1956 | 10 | 1.02% |
| hsa_circ_0077180 | 1956 | 10 | 1.02% |
| hsa_circ_0107358 | 1956 | 10 | 1.02% |
| hsa_circ_0134663 | 1956 | 10 | 1.02% |
| hsa_circ_0021119 | 3521 | 18 | 1.02% |
| hsa_circ_0011530 | 2152 | 11 | 1.02% |
| hsa_circ_0053524 | 2152 | 11 | 1.02% |
| hsa_circ_0053706 | 2152 | 11 | 1.02% |
| hsa_circ_0090559 | 2152 | 11 | 1.02% |
| hsa_circ_0112631 | 2152 | 11 | 1.02% |
| hsa_circ_0068831 | 2348 | 12 | 1.02% |
| hsa_circ_0040228 | 2544 | 13 | 1.02% |
| hsa_circ_0068425 | 2544 | 13 | 1.02% |
| hsa_circ_0139535 | 2544 | 13 | 1.02% |
| hsa_circ_0027865 | 1957 | 10 | 1.02% |
| hsa_circ_0056095 | 1957 | 10 | 1.02% |
| hsa_circ_0073854 | 1957 | 10 | 1.02% |
| hsa_circ_0100041 | 1957 | 10 | 1.02% |
| hsa_circ_0102722 | 1957 | 10 | 1.02% |
| hsa_circ_0133394 | 1957 | 10 | 1.02% |
| hsa_circ_0028521 | 2153 | 11 | 1.02% |
| hsa_circ_0044573 | 2153 | 11 | 1.02% |
| hsa_circ_0048598 | 2153 | 11 | 1.02% |
| hsa_circ_0043515 | 2349 | 12 | 1.02% |
| hsa_circ_0118601 | 2545 | 13 | 1.02% |
| hsa_circ_0014692 | 1958 | 10 | 1.02% |
| hsa_circ_0039709 | 1958 | 10 | 1.02% |
| hsa_circ_0046110 | 1958 | 10 | 1.02% |
| hsa_circ_0055512 | 1958 | 10 | 1.02% |
| hsa_circ_0089526 | 1958 | 10 | 1.02% |
| hsa_circ_0102837 | 1958 | 10 | 1.02% |
| hsa_circ_0115867 | 1958 | 10 | 1.02% |
| hsa_circ_0123403 | 1958 | 10 | 1.02% |
| hsa_circ_0131921 | 1958 | 10 | 1.02% |
| hsa_circ_0073293 | 2154 | 11 | 1.02% |
| hsa_circ_0093354 | 2154 | 11 | 1.02% |
| hsa_circ_0020630 | 2350 | 12 | 1.02% |
| hsa_circ_0052799 | 2350 | 12 | 1.02% |
| hsa_circ_0085630 | 2350 | 12 | 1.02% |
| hsa_circ_0094742 | 2350 | 12 | 1.02% |
| hsa_circ_0045532 | 2546 | 13 | 1.02% |
| hsa_circ_0086211 | 2938 | 15 | 1.02% |
| hsa_circ_0032168 | 1959 | 10 | 1.02% |
| hsa_circ_0056661 | 1959 | 10 | 1.02% |
| hsa_circ_0084498 | 1959 | 10 | 1.02% |
| hsa_circ_0107181 | 1959 | 10 | 1.02% |
| hsa_circ_0133160 | 1959 | 10 | 1.02% |
| hsa_circ_0010420 | 2351 | 12 | 1.02% |

| | | | |
|---|---|---|---|
| hsa_circ_0014646 | 2547 | 13 | 1.02% |
| hsa_circ_0089488 | 3135 | 16 | 1.02% |
| hsa_circ_0097171 | 3723 | 19 | 1.02% |
| hsa_circ_0109271 | 6860 | 35 | 1.02% |
| hsa_circ_0130601 | 4116 | 21 | 1.02% |
| hsa_circ_0030487 | 2940 | 15 | 1.02% |
| hsa_circ_0007566 | 2352 | 12 | 1.02% |
| hsa_circ_0082095 | 2352 | 12 | 1.02% |
| hsa_circ_0088350 | 2352 | 12 | 1.02% |
| hsa_circ_0086091 | 2156 | 11 | 1.02% |
| hsa_circ_0087698 | 2156 | 11 | 1.02% |
| hsa_circ_0052769 | 1960 | 10 | 1.02% |
| hsa_circ_0092234 | 1960 | 10 | 1.02% |
| hsa_circ_0098544 | 1960 | 10 | 1.02% |
| hsa_circ_0117048 | 1960 | 10 | 1.02% |
| hsa_circ_0119111 | 4902 | 25 | 1.02% |
| hsa_circ_0043764 | 2353 | 12 | 1.02% |
| hsa_circ_0132484 | 2353 | 12 | 1.02% |
| hsa_circ_0036886 | 2157 | 11 | 1.02% |
| hsa_circ_0053901 | 2157 | 11 | 1.02% |
| hsa_circ_0066678 | 2157 | 11 | 1.02% |
| hsa_circ_0068392 | 2157 | 11 | 1.02% |
| hsa_circ_0094042 | 2157 | 11 | 1.02% |
| hsa_circ_0110910 | 2157 | 11 | 1.02% |
| hsa_circ_0114836 | 2157 | 11 | 1.02% |
| hsa_circ_0014552 | 1961 | 10 | 1.02% |
| hsa_circ_0021220 | 1961 | 10 | 1.02% |
| hsa_circ_0055838 | 1961 | 10 | 1.02% |
| hsa_circ_0062755 | 1961 | 10 | 1.02% |
| hsa_circ_0072906 | 1961 | 10 | 1.02% |
| hsa_circ_0082632 | 1961 | 10 | 1.02% |
| hsa_circ_0104559 | 1961 | 10 | 1.02% |
| hsa_circ_0076600 | 6864 | 35 | 1.02% |
| hsa_circ_0052203 | 2550 | 13 | 1.02% |
| hsa_circ_0072300 | 2354 | 12 | 1.02% |
| hsa_circ_0089247 | 2354 | 12 | 1.02% |
| hsa_circ_0115988 | 2354 | 12 | 1.02% |
| hsa_circ_0011109 | 2158 | 11 | 1.02% |
| hsa_circ_0014094 | 2158 | 11 | 1.02% |
| hsa_circ_0032384 | 2158 | 11 | 1.02% |
| hsa_circ_0051480 | 2158 | 11 | 1.02% |
| hsa_circ_0064815 | 2158 | 11 | 1.02% |
| hsa_circ_0077998 | 2158 | 11 | 1.02% |
| hsa_circ_0139543 | 2158 | 11 | 1.02% |
| hsa_circ_0038033 | 5101 | 26 | 1.02% |
| hsa_circ_0116526 | 2943 | 15 | 1.02% |
| hsa_circ_0010127 | 1962 | 10 | 1.02% |
| hsa_circ_0011225 | 1962 | 10 | 1.02% |
| hsa_circ_0023733 | 1962 | 10 | 1.02% |
| hsa_circ_0039986 | 1962 | 10 | 1.02% |

| | | | |
|---|---|---|---|
| hsa_circ_0042224 | 1962 | 10 | 1.02% |
| hsa_circ_0050376 | 1962 | 10 | 1.02% |
| hsa_circ_0071270 | 1962 | 10 | 1.02% |
| hsa_circ_0132182 | 2747 | 14 | 1.02% |
| hsa_circ_0012043 | 2551 | 13 | 1.02% |
| hsa_circ_0103428 | 2551 | 13 | 1.02% |
| hsa_circ_0052975 | 2355 | 12 | 1.02% |
| hsa_circ_0023566 | 2159 | 11 | 1.02% |
| hsa_circ_0102830 | 4907 | 25 | 1.02% |
| hsa_circ_0012728 | 1963 | 10 | 1.02% |
| hsa_circ_0032451 | 1963 | 10 | 1.02% |
| hsa_circ_0039843 | 1963 | 10 | 1.02% |
| hsa_circ_0040744 | 1963 | 10 | 1.02% |
| hsa_circ_0074087 | 1963 | 10 | 1.02% |
| hsa_circ_0087452 | 1963 | 10 | 1.02% |
| hsa_circ_0115547 | 1963 | 10 | 1.02% |
| hsa_circ_0047020 | 2552 | 13 | 1.02% |
| hsa_circ_0083414 | 2552 | 13 | 1.02% |
| hsa_circ_0108881 | 7068 | 36 | 1.02% |
| hsa_circ_0016298 | 2356 | 12 | 1.02% |
| hsa_circ_0021249 | 2356 | 12 | 1.02% |
| hsa_circ_0036902 | 2356 | 12 | 1.02% |
| hsa_circ_0010102 | 2160 | 11 | 1.02% |
| hsa_circ_0017661 | 2160 | 11 | 1.02% |
| hsa_circ_0029671 | 2160 | 11 | 1.02% |
| hsa_circ_0091001 | 2160 | 11 | 1.02% |
| hsa_circ_0119530 | 2160 | 11 | 1.02% |
| hsa_circ_0129175 | 2160 | 11 | 1.02% |
| hsa_circ_0124280 | 4124 | 21 | 1.02% |
| hsa_circ_0043162 | 1964 | 10 | 1.02% |
| hsa_circ_0050419 | 1964 | 10 | 1.02% |
| hsa_circ_0061734 | 1964 | 10 | 1.02% |
| hsa_circ_0047102 | 2357 | 12 | 1.02% |
| hsa_circ_0019796 | 5697 | 29 | 1.02% |
| hsa_circ_0025260 | 2161 | 11 | 1.02% |
| hsa_circ_0042580 | 2161 | 11 | 1.02% |
| hsa_circ_0051864 | 2161 | 11 | 1.02% |
| hsa_circ_0067888 | 2161 | 11 | 1.02% |
| hsa_circ_0117486 | 3733 | 19 | 1.02% |
| hsa_circ_0040997 | 2358 | 12 | 1.02% |
| hsa_circ_0035334 | 1965 | 10 | 1.02% |
| hsa_circ_0057786 | 1965 | 10 | 1.02% |
| hsa_circ_0063817 | 1965 | 10 | 1.02% |
| hsa_circ_0067761 | 1965 | 10 | 1.02% |
| hsa_circ_0079719 | 1965 | 10 | 1.02% |
| hsa_circ_0097662 | 1965 | 10 | 1.02% |
| hsa_circ_0108160 | 1965 | 10 | 1.02% |
| hsa_circ_0080708 | 2162 | 11 | 1.02% |
| hsa_circ_0101826 | 2162 | 11 | 1.02% |
| hsa_circ_0040295 | 6683 | 34 | 1.02% |

| | | | |
|---|---|---|---|
| hsa_circ_0034224 | 2359 | 12 | 1.02% |
| hsa_circ_0093595 | 4915 | 25 | 1.02% |
| hsa_circ_0024528 | 1966 | 10 | 1.02% |
| hsa_circ_0111276 | 1966 | 10 | 1.02% |
| hsa_circ_0122965 | 1966 | 10 | 1.02% |
| hsa_circ_0125845 | 1966 | 10 | 1.02% |
| hsa_circ_0057435 | 2556 | 13 | 1.02% |
| hsa_circ_0020222 | 2163 | 11 | 1.02% |
| hsa_circ_0020551 | 2163 | 11 | 1.02% |
| hsa_circ_0040168 | 2163 | 11 | 1.02% |
| hsa_circ_0042719 | 2163 | 11 | 1.02% |
| hsa_circ_0045039 | 2163 | 11 | 1.02% |
| hsa_circ_0131887 | 2163 | 11 | 1.02% |
| hsa_circ_0023345 | 2360 | 12 | 1.02% |
| hsa_circ_0121569 | 3147 | 16 | 1.02% |
| hsa_circ_0053708 | 2557 | 13 | 1.02% |
| hsa_circ_0066764 | 2557 | 13 | 1.02% |
| hsa_circ_0139889 | 2557 | 13 | 1.02% |
| hsa_circ_0008699 | 1967 | 10 | 1.02% |
| hsa_circ_0034254 | 1967 | 10 | 1.02% |
| hsa_circ_0042731 | 1967 | 10 | 1.02% |
| hsa_circ_0069482 | 1967 | 10 | 1.02% |
| hsa_circ_0101061 | 1967 | 10 | 1.02% |
| hsa_circ_0131893 | 1967 | 10 | 1.02% |
| hsa_circ_0105560 | 2754 | 14 | 1.02% |
| hsa_circ_0070130 | 2164 | 11 | 1.02% |
| hsa_circ_0136345 | 2164 | 11 | 1.02% |
| hsa_circ_0038887 | 2361 | 12 | 1.02% |
| hsa_circ_0076520 | 2361 | 12 | 1.02% |
| hsa_circ_0131891 | 2361 | 12 | 1.02% |
| hsa_circ_0132545 | 2361 | 12 | 1.02% |
| hsa_circ_0105830 | 5707 | 29 | 1.02% |
| hsa_circ_0012098 | 1968 | 10 | 1.02% |
| hsa_circ_0022675 | 1968 | 10 | 1.02% |
| hsa_circ_0026838 | 1968 | 10 | 1.02% |
| hsa_circ_0033590 | 1968 | 10 | 1.02% |
| hsa_circ_0053852 | 1968 | 10 | 1.02% |
| hsa_circ_0088000 | 1968 | 10 | 1.02% |
| hsa_circ_0111665 | 1968 | 10 | 1.02% |
| hsa_circ_0007311 | 2165 | 11 | 1.02% |
| hsa_circ_0017731 | 2165 | 11 | 1.02% |
| hsa_circ_0026852 | 2165 | 11 | 1.02% |
| hsa_circ_0117735 | 2165 | 11 | 1.02% |
| hsa_circ_0133008 | 2165 | 11 | 1.02% |
| hsa_circ_0000245 | 2362 | 12 | 1.02% |
| hsa_circ_0027220 | 2362 | 12 | 1.02% |
| hsa_circ_0037328 | 2362 | 12 | 1.02% |
| hsa_circ_0045008 | 2362 | 12 | 1.02% |
| hsa_circ_0048265 | 2559 | 13 | 1.02% |
| hsa_circ_0062345 | 2559 | 13 | 1.02% |

| | | | |
|---|---|---|---|
| hsa_circ_0103626 | 2559 | 13 | 1.02% |
| hsa_circ_0114470 | 2559 | 13 | 1.02% |
| hsa_circ_0061489 | 1969 | 10 | 1.02% |
| hsa_circ_0085738 | 1969 | 10 | 1.02% |
| hsa_circ_0039865 | 2166 | 11 | 1.02% |
| hsa_circ_0043095 | 2166 | 11 | 1.02% |
| hsa_circ_0039875 | 2363 | 12 | 1.02% |
| hsa_circ_0048892 | 2363 | 12 | 1.02% |
| hsa_circ_0132300 | 2363 | 12 | 1.02% |
| hsa_circ_0046170 | 2560 | 13 | 1.02% |
| hsa_circ_0107217 | 2560 | 13 | 1.02% |
| hsa_circ_0017357 | 2561 | 13 | 1.02% |
| hsa_circ_0034469 | 2364 | 12 | 1.02% |
| hsa_circ_0030671 | 2167 | 11 | 1.02% |
| hsa_circ_0043554 | 2167 | 11 | 1.02% |
| hsa_circ_0053783 | 2167 | 11 | 1.02% |
| hsa_circ_0066327 | 2167 | 11 | 1.02% |
| hsa_circ_0133788 | 2167 | 11 | 1.02% |
| hsa_circ_0015878 | 1970 | 10 | 1.02% |
| hsa_circ_0016443 | 1970 | 10 | 1.02% |
| hsa_circ_0017400 | 1970 | 10 | 1.02% |
| hsa_circ_0025434 | 1970 | 10 | 1.02% |
| hsa_circ_0030191 | 1970 | 10 | 1.02% |
| hsa_circ_0120400 | 1970 | 10 | 1.02% |
| hsa_circ_0022819 | 2365 | 12 | 1.01% |
| hsa_circ_0041676 | 2365 | 12 | 1.01% |
| hsa_circ_0087100 | 2365 | 12 | 1.01% |
| hsa_circ_0101831 | 2365 | 12 | 1.01% |
| hsa_circ_0010819 | 2168 | 11 | 1.01% |
| hsa_circ_0022752 | 2168 | 11 | 1.01% |
| hsa_circ_0054943 | 2168 | 11 | 1.01% |
| hsa_circ_0091000 | 2168 | 11 | 1.01% |
| hsa_circ_0011927 | 1971 | 10 | 1.01% |
| hsa_circ_0046780 | 1971 | 10 | 1.01% |
| hsa_circ_0064224 | 1971 | 10 | 1.01% |
| hsa_circ_0071141 | 1971 | 10 | 1.01% |
| hsa_circ_0113825 | 1971 | 10 | 1.01% |
| hsa_circ_0101234 | 2563 | 13 | 1.01% |
| hsa_circ_0060123 | 8281 | 42 | 1.01% |
| hsa_circ_0041201 | 2366 | 12 | 1.01% |
| hsa_circ_0074490 | 2366 | 12 | 1.01% |
| hsa_circ_0016823 | 2169 | 11 | 1.01% |
| hsa_circ_0020334 | 2169 | 11 | 1.01% |
| hsa_circ_0040697 | 2169 | 11 | 1.01% |
| hsa_circ_0044165 | 2169 | 11 | 1.01% |
| hsa_circ_0080996 | 2169 | 11 | 1.01% |
| hsa_circ_0122444 | 2169 | 11 | 1.01% |
| hsa_circ_0010421 | 1972 | 10 | 1.01% |
| hsa_circ_0031766 | 1972 | 10 | 1.01% |
| hsa_circ_0080959 | 1972 | 10 | 1.01% |

| | | | |
|---|---|---|---|
| hsa_circ_0109309 | 1972 | 10 | 1.01% |
| hsa_circ_0127280 | 1972 | 10 | 1.01% |
| hsa_circ_0136057 | 1972 | 10 | 1.01% |
| hsa_circ_0029003 | 7297 | 37 | 1.01% |
| hsa_circ_0047394 | 2564 | 13 | 1.01% |
| hsa_circ_0121424 | 2564 | 13 | 1.01% |
| hsa_circ_0010383 | 2367 | 12 | 1.01% |
| hsa_circ_0048463 | 2367 | 12 | 1.01% |
| hsa_circ_0102383 | 2367 | 12 | 1.01% |
| hsa_circ_0120573 | 2367 | 12 | 1.01% |
| hsa_circ_0094438 | 2170 | 11 | 1.01% |
| hsa_circ_0138348 | 2170 | 11 | 1.01% |
| hsa_circ_0079478 | 4735 | 24 | 1.01% |
| hsa_circ_0016191 | 1973 | 10 | 1.01% |
| hsa_circ_0046826 | 1973 | 10 | 1.01% |
| hsa_circ_0085233 | 1973 | 10 | 1.01% |
| hsa_circ_0104022 | 1973 | 10 | 1.01% |
| hsa_circ_0127922 | 1973 | 10 | 1.01% |
| hsa_circ_0093091 | 6511 | 33 | 1.01% |
| hsa_circ_0067853 | 2565 | 13 | 1.01% |
| hsa_circ_0116793 | 4341 | 22 | 1.01% |
| hsa_circ_0024393 | 2368 | 12 | 1.01% |
| hsa_circ_0076885 | 2368 | 12 | 1.01% |
| hsa_circ_0086765 | 2368 | 12 | 1.01% |
| hsa_circ_0017046 | 2171 | 11 | 1.01% |
| hsa_circ_0082489 | 2171 | 11 | 1.01% |
| hsa_circ_0124225 | 2171 | 11 | 1.01% |
| hsa_circ_0017609 | 1974 | 10 | 1.01% |
| hsa_circ_0058140 | 1974 | 10 | 1.01% |
| hsa_circ_0065663 | 1974 | 10 | 1.01% |
| hsa_circ_0073540 | 1974 | 10 | 1.01% |
| hsa_circ_0076824 | 1974 | 10 | 1.01% |
| hsa_circ_0019039 | 2369 | 12 | 1.01% |
| hsa_circ_0054837 | 2369 | 12 | 1.01% |
| hsa_circ_0094998 | 4146 | 21 | 1.01% |
| hsa_circ_0036881 | 2172 | 11 | 1.01% |
| hsa_circ_0038110 | 2172 | 11 | 1.01% |
| hsa_circ_0046058 | 2172 | 11 | 1.01% |
| hsa_circ_0056559 | 2172 | 11 | 1.01% |
| hsa_circ_0071927 | 2172 | 11 | 1.01% |
| hsa_circ_0086911 | 2172 | 11 | 1.01% |
| hsa_circ_0116235 | 2172 | 11 | 1.01% |
| hsa_circ_0117902 | 2172 | 11 | 1.01% |
| hsa_circ_0009327 | 2370 | 12 | 1.01% |
| hsa_circ_0024586 | 2370 | 12 | 1.01% |
| hsa_circ_0076516 | 2370 | 12 | 1.01% |
| hsa_circ_0089726 | 2370 | 12 | 1.01% |
| hsa_circ_0134921 | 2370 | 12 | 1.01% |
| hsa_circ_0049890 | 1975 | 10 | 1.01% |
| hsa_circ_0078113 | 1975 | 10 | 1.01% |

| | | | |
|---|---|---|---|
| hsa_circ_0104650 | 8296 | 42 | 1.01% |
| hsa_circ_0122335 | 2568 | 13 | 1.01% |
| hsa_circ_0010543 | 2173 | 11 | 1.01% |
| hsa_circ_0059077 | 2173 | 11 | 1.01% |
| hsa_circ_0068805 | 2173 | 11 | 1.01% |
| hsa_circ_0083935 | 2173 | 11 | 1.01% |
| hsa_circ_0061981 | 2371 | 12 | 1.01% |
| hsa_circ_0136781 | 6323 | 32 | 1.01% |
| hsa_circ_0014574 | 1976 | 10 | 1.01% |
| hsa_circ_0045726 | 1976 | 10 | 1.01% |
| hsa_circ_0071780 | 1976 | 10 | 1.01% |
| hsa_circ_0086230 | 1976 | 10 | 1.01% |
| hsa_circ_0102461 | 1976 | 10 | 1.01% |
| hsa_circ_0108972 | 1976 | 10 | 1.01% |
| hsa_circ_0127437 | 1976 | 10 | 1.01% |
| hsa_circ_0063208 | 2569 | 13 | 1.01% |
| hsa_circ_0022897 | 2174 | 11 | 1.01% |
| hsa_circ_0046956 | 2174 | 11 | 1.01% |
| hsa_circ_0105696 | 2174 | 11 | 1.01% |
| hsa_circ_0077362 | 2372 | 12 | 1.01% |
| hsa_circ_0022682 | 1977 | 10 | 1.01% |
| hsa_circ_0038571 | 1977 | 10 | 1.01% |
| hsa_circ_0062179 | 1977 | 10 | 1.01% |
| hsa_circ_0066358 | 1977 | 10 | 1.01% |
| hsa_circ_0076822 | 1977 | 10 | 1.01% |
| hsa_circ_0082375 | 1977 | 10 | 1.01% |
| hsa_circ_0084541 | 1977 | 10 | 1.01% |
| hsa_circ_0003978 | 4350 | 22 | 1.01% |
| hsa_circ_0010137 | 2175 | 11 | 1.01% |
| hsa_circ_0040106 | 2175 | 11 | 1.01% |
| hsa_circ_0083938 | 2175 | 11 | 1.01% |
| hsa_circ_0094194 | 2175 | 11 | 1.01% |
| hsa_circ_0078912 | 2373 | 12 | 1.01% |
| hsa_circ_0015830 | 1978 | 10 | 1.01% |
| hsa_circ_0047204 | 1978 | 10 | 1.01% |
| hsa_circ_0058537 | 1978 | 10 | 1.01% |
| hsa_circ_0067942 | 1978 | 10 | 1.01% |
| hsa_circ_0033577 | 2176 | 11 | 1.01% |
| hsa_circ_0040044 | 2176 | 11 | 1.01% |
| hsa_circ_0136301 | 2176 | 11 | 1.01% |
| hsa_circ_0004753 | 5540 | 28 | 1.01% |
| hsa_circ_0013178 | 1979 | 10 | 1.01% |
| hsa_circ_0017613 | 1979 | 10 | 1.01% |
| hsa_circ_0033053 | 1979 | 10 | 1.01% |
| hsa_circ_0035598 | 1979 | 10 | 1.01% |
| hsa_circ_0063933 | 1979 | 10 | 1.01% |
| hsa_circ_0066314 | 1979 | 10 | 1.01% |
| hsa_circ_0088745 | 1979 | 10 | 1.01% |
| hsa_circ_0095006 | 1979 | 10 | 1.01% |
| hsa_circ_0130153 | 1979 | 10 | 1.01% |

| | | | |
|---|---|---|---|
| hsa_circ_0135788 | 1979 | 10 | 1.01% |
| hsa_circ_0031365 | 2177 | 11 | 1.01% |
| hsa_circ_0042522 | 2177 | 11 | 1.01% |
| hsa_circ_0044579 | 2177 | 11 | 1.01% |
| hsa_circ_0047331 | 2177 | 11 | 1.01% |
| hsa_circ_0080144 | 2177 | 11 | 1.01% |
| hsa_circ_0086667 | 2177 | 11 | 1.01% |
| hsa_circ_0110448 | 2177 | 11 | 1.01% |
| hsa_circ_0111009 | 2375 | 12 | 1.01% |
| hsa_circ_0022672 | 2178 | 11 | 1.01% |
| hsa_circ_0056404 | 2178 | 11 | 1.01% |
| hsa_circ_0063127 | 2178 | 11 | 1.01% |
| hsa_circ_0070746 | 2178 | 11 | 1.01% |
| hsa_circ_0074380 | 2178 | 11 | 1.01% |
| hsa_circ_0077924 | 2178 | 11 | 1.01% |
| hsa_circ_0103722 | 2178 | 11 | 1.01% |
| hsa_circ_0104020 | 2178 | 11 | 1.01% |
| hsa_circ_0011804 | 1980 | 10 | 1.01% |
| hsa_circ_0011840 | 1980 | 10 | 1.01% |
| hsa_circ_0035894 | 1980 | 10 | 1.01% |
| hsa_circ_0036790 | 1980 | 10 | 1.01% |
| hsa_circ_0044492 | 1980 | 10 | 1.01% |
| hsa_circ_0084741 | 1980 | 10 | 1.01% |
| hsa_circ_0116036 | 1980 | 10 | 1.01% |
| hsa_circ_0139895 | 1980 | 10 | 1.01% |
| hsa_circ_0092472 | 3367 | 17 | 1.01% |
| hsa_circ_0100549 | 2575 | 13 | 1.01% |
| hsa_circ_0021877 | 2377 | 12 | 1.01% |
| hsa_circ_0023149 | 2377 | 12 | 1.01% |
| hsa_circ_0017990 | 2179 | 11 | 1.01% |
| hsa_circ_0038778 | 2179 | 11 | 1.01% |
| hsa_circ_0074384 | 2179 | 11 | 1.01% |
| hsa_circ_0018593 | 1981 | 10 | 1.01% |
| hsa_circ_0053226 | 1981 | 10 | 1.01% |
| hsa_circ_0063538 | 1981 | 10 | 1.01% |
| hsa_circ_0074472 | 1981 | 10 | 1.01% |
| hsa_circ_0081179 | 1981 | 10 | 1.01% |
| hsa_circ_0096657 | 1981 | 10 | 1.01% |
| hsa_circ_0118740 | 1981 | 10 | 1.01% |
| hsa_circ_0139992 | 1981 | 10 | 1.01% |
| hsa_circ_0103781 | 4953 | 25 | 1.01% |
| hsa_circ_0118848 | 2774 | 14 | 1.01% |
| hsa_circ_0034848 | 2378 | 12 | 1.01% |
| hsa_circ_0034914 | 2378 | 12 | 1.01% |
| hsa_circ_0037670 | 2378 | 12 | 1.01% |
| hsa_circ_0041846 | 2378 | 12 | 1.01% |
| hsa_circ_0090239 | 2378 | 12 | 1.01% |
| hsa_circ_0092244 | 2378 | 12 | 1.01% |
| hsa_circ_0120630 | 2378 | 12 | 1.01% |
| hsa_circ_0007837 | 2180 | 11 | 1.01% |

| | | | |
|---|---|---|---|
| hsa_circ_0013709 | 2180 | 11 | 1.01% |
| hsa_circ_0059580 | 2180 | 11 | 1.01% |
| hsa_circ_0119689 | 2180 | 11 | 1.01% |
| hsa_circ_0125766 | 2180 | 11 | 1.01% |
| hsa_circ_0100998 | 3171 | 16 | 1.01% |
| hsa_circ_0000345 | 1982 | 10 | 1.01% |
| hsa_circ_0049762 | 1982 | 10 | 1.01% |
| hsa_circ_0055150 | 1982 | 10 | 1.01% |
| hsa_circ_0064864 | 1982 | 10 | 1.01% |
| hsa_circ_0075895 | 1982 | 10 | 1.01% |
| hsa_circ_0102034 | 1982 | 10 | 1.01% |
| hsa_circ_0104277 | 1982 | 10 | 1.01% |
| hsa_circ_0110906 | 1982 | 10 | 1.01% |
| hsa_circ_0074912 | 2379 | 12 | 1.01% |
| hsa_circ_0066712 | 2974 | 15 | 1.01% |
| hsa_circ_0112440 | 4362 | 22 | 1.01% |
| hsa_circ_0005661 | 2181 | 11 | 1.01% |
| hsa_circ_0011500 | 2181 | 11 | 1.01% |
| hsa_circ_0034050 | 2181 | 11 | 1.01% |
| hsa_circ_0054852 | 2181 | 11 | 1.01% |
| hsa_circ_0075708 | 2181 | 11 | 1.01% |
| hsa_circ_0012707 | 1983 | 10 | 1.01% |
| hsa_circ_0013361 | 1983 | 10 | 1.01% |
| hsa_circ_0055130 | 1983 | 10 | 1.01% |
| hsa_circ_0055789 | 1983 | 10 | 1.01% |
| hsa_circ_0088924 | 1983 | 10 | 1.01% |
| hsa_circ_0127703 | 1983 | 10 | 1.01% |
| hsa_circ_0046254 | 2578 | 13 | 1.01% |
| hsa_circ_0002763 | 2380 | 12 | 1.01% |
| hsa_circ_0048550 | 2380 | 12 | 1.01% |
| hsa_circ_0069277 | 2380 | 12 | 1.01% |
| hsa_circ_0056070 | 2182 | 11 | 1.01% |
| hsa_circ_0065581 | 2182 | 11 | 1.01% |
| hsa_circ_0073181 | 2182 | 11 | 1.01% |
| hsa_circ_0076064 | 2182 | 11 | 1.01% |
| hsa_circ_0076745 | 2182 | 11 | 1.01% |
| hsa_circ_0044246 | 9324 | 47 | 1.01% |
| hsa_circ_0044450 | 2381 | 12 | 1.01% |
| hsa_circ_0027219 | 2183 | 11 | 1.01% |
| hsa_circ_0028761 | 2183 | 11 | 1.01% |
| hsa_circ_0053614 | 2183 | 11 | 1.01% |
| hsa_circ_0101977 | 2183 | 11 | 1.01% |
| hsa_circ_0114254 | 2183 | 11 | 1.01% |
| hsa_circ_0029465 | 2580 | 13 | 1.01% |
| hsa_circ_0030497 | 2977 | 15 | 1.01% |
| hsa_circ_0039195 | 2382 | 12 | 1.01% |
| hsa_circ_0040258 | 2382 | 12 | 1.01% |
| hsa_circ_0090422 | 2382 | 12 | 1.01% |
| hsa_circ_0111180 | 2382 | 12 | 1.01% |
| hsa_circ_0035516 | 1985 | 10 | 1.01% |

| | | | |
|---|---|---|---|
| hsa_circ_0036122 | 1985 | 10 | 1.01% |
| hsa_circ_0102518 | 1985 | 10 | 1.01% |
| hsa_circ_0034713 | 2184 | 11 | 1.01% |
| hsa_circ_0049599 | 2184 | 11 | 1.01% |
| hsa_circ_0122276 | 2184 | 11 | 1.01% |
| hsa_circ_0125459 | 2184 | 11 | 1.01% |
| hsa_circ_0013618 | 2383 | 12 | 1.01% |
| hsa_circ_0016977 | 2383 | 12 | 1.01% |
| hsa_circ_0078530 | 2383 | 12 | 1.01% |
| hsa_circ_0052100 | 6752 | 34 | 1.01% |
| hsa_circ_0071368 | 4369 | 22 | 1.01% |
| hsa_circ_0027784 | 1986 | 10 | 1.01% |
| hsa_circ_0038391 | 1986 | 10 | 1.01% |
| hsa_circ_0061786 | 1986 | 10 | 1.01% |
| hsa_circ_0092060 | 1986 | 10 | 1.01% |
| hsa_circ_0114794 | 1986 | 10 | 1.01% |
| hsa_circ_0126839 | 1986 | 10 | 1.01% |
| hsa_circ_0058935 | 2582 | 13 | 1.01% |
| hsa_circ_0047905 | 2185 | 11 | 1.01% |
| hsa_circ_0068323 | 2185 | 11 | 1.01% |
| hsa_circ_0086504 | 2185 | 11 | 1.01% |
| hsa_circ_0051857 | 2384 | 12 | 1.01% |
| hsa_circ_0014881 | 1987 | 10 | 1.01% |
| hsa_circ_0033589 | 1987 | 10 | 1.01% |
| hsa_circ_0041171 | 1987 | 10 | 1.01% |
| hsa_circ_0045352 | 1987 | 10 | 1.01% |
| hsa_circ_0053735 | 1987 | 10 | 1.01% |
| hsa_circ_0065178 | 1987 | 10 | 1.01% |
| hsa_circ_0066795 | 1987 | 10 | 1.01% |
| hsa_circ_0071936 | 1987 | 10 | 1.01% |
| hsa_circ_0024957 | 2782 | 14 | 1.01% |
| hsa_circ_0041126 | 2186 | 11 | 1.01% |
| hsa_circ_0049555 | 2186 | 11 | 1.01% |
| hsa_circ_0089935 | 2186 | 11 | 1.01% |
| hsa_circ_0091147 | 2186 | 11 | 1.01% |
| hsa_circ_0124896 | 2186 | 11 | 1.01% |
| hsa_circ_0127120 | 2186 | 11 | 1.01% |
| hsa_circ_0047763 | 2385 | 12 | 1.01% |
| hsa_circ_0050706 | 2385 | 12 | 1.01% |
| hsa_circ_0083021 | 2584 | 13 | 1.01% |
| hsa_circ_0086612 | 2584 | 13 | 1.01% |
| hsa_circ_0138873 | 5765 | 29 | 1.01% |
| hsa_circ_0014379 | 1988 | 10 | 1.01% |
| hsa_circ_0051408 | 1988 | 10 | 1.01% |
| hsa_circ_0100768 | 1988 | 10 | 1.01% |
| hsa_circ_0009608 | 2187 | 11 | 1.01% |
| hsa_circ_0022932 | 2187 | 11 | 1.01% |
| hsa_circ_0067893 | 2187 | 11 | 1.01% |
| hsa_circ_0069050 | 2187 | 11 | 1.01% |
| hsa_circ_0120667 | 2187 | 11 | 1.01% |

| | | | |
|---|---|---|---|
| hsa_circ_0024212 | 2386 | 12 | 1.01% |
| hsa_circ_0103339 | 2386 | 12 | 1.01% |
| hsa_circ_0125859 | 2386 | 12 | 1.01% |
| hsa_circ_0015087 | 1989 | 10 | 1.01% |
| hsa_circ_0046951 | 1989 | 10 | 1.01% |
| hsa_circ_0070152 | 1989 | 10 | 1.01% |
| hsa_circ_0081245 | 1989 | 10 | 1.01% |
| hsa_circ_0081552 | 1989 | 10 | 1.01% |
| hsa_circ_0038254 | 2188 | 11 | 1.01% |
| hsa_circ_0044382 | 2188 | 11 | 1.01% |
| hsa_circ_0078281 | 2188 | 11 | 1.01% |
| hsa_circ_0088897 | 2188 | 11 | 1.01% |
| hsa_circ_0091678 | 2188 | 11 | 1.01% |
| hsa_circ_0104312 | 2188 | 11 | 1.01% |
| hsa_circ_0025792 | 2387 | 12 | 1.01% |
| hsa_circ_0068536 | 2387 | 12 | 1.01% |
| hsa_circ_0017811 | 2586 | 13 | 1.01% |
| hsa_circ_0050502 | 2785 | 14 | 1.01% |
| hsa_circ_0024367 | 4178 | 21 | 1.01% |
| hsa_circ_0030813 | 2587 | 13 | 1.01% |
| hsa_circ_0052169 | 2587 | 13 | 1.01% |
| hsa_circ_0028446 | 2189 | 11 | 1.01% |
| hsa_circ_0068841 | 2189 | 11 | 1.01% |
| hsa_circ_0098525 | 2189 | 11 | 1.01% |
| hsa_circ_0132416 | 2189 | 11 | 1.01% |
| hsa_circ_0003075 | 1990 | 10 | 1.01% |
| hsa_circ_0015866 | 1990 | 10 | 1.01% |
| hsa_circ_0031488 | 1990 | 10 | 1.01% |
| hsa_circ_0044034 | 1990 | 10 | 1.01% |
| hsa_circ_0054560 | 1990 | 10 | 1.01% |
| hsa_circ_0087661 | 1990 | 10 | 1.01% |
| hsa_circ_0100121 | 1990 | 10 | 1.01% |
| hsa_circ_0137372 | 1990 | 10 | 1.01% |
| hsa_circ_0046113 | 2588 | 13 | 1.00% |
| hsa_circ_0049515 | 2588 | 13 | 1.00% |
| hsa_circ_0090276 | 2588 | 13 | 1.00% |
| hsa_circ_0058020 | 2389 | 12 | 1.00% |
| hsa_circ_0064326 | 2389 | 12 | 1.00% |
| hsa_circ_0129352 | 2389 | 12 | 1.00% |
| hsa_circ_0029690 | 2190 | 11 | 1.00% |
| hsa_circ_0037707 | 2190 | 11 | 1.00% |
| hsa_circ_0045631 | 2190 | 11 | 1.00% |
| hsa_circ_0073855 | 2190 | 11 | 1.00% |
| hsa_circ_0081060 | 2190 | 11 | 1.00% |
| hsa_circ_0091817 | 2190 | 11 | 1.00% |
| hsa_circ_0115535 | 2190 | 11 | 1.00% |
| hsa_circ_0137829 | 2190 | 11 | 1.00% |
| hsa_circ_0027144 | 1991 | 10 | 1.00% |
| hsa_circ_0091078 | 1991 | 10 | 1.00% |
| hsa_circ_0103586 | 1991 | 10 | 1.00% |

| | | | |
|---|---|---|---|
| hsa_circ_0111586 | 1991 | 10 | 1.00% |
| hsa_circ_0129651 | 3584 | 18 | 1.00% |
| hsa_circ_0124946 | 2589 | 13 | 1.00% |
| hsa_circ_0100564 | 11552 | 58 | 1.00% |
| hsa_circ_0026867 | 2191 | 11 | 1.00% |
| hsa_circ_0043476 | 2191 | 11 | 1.00% |
| hsa_circ_0059327 | 2191 | 11 | 1.00% |
| hsa_circ_0056442 | 1992 | 10 | 1.00% |
| hsa_circ_0063117 | 1992 | 10 | 1.00% |
| hsa_circ_0067590 | 1992 | 10 | 1.00% |
| hsa_circ_0100767 | 1992 | 10 | 1.00% |
| hsa_circ_0107594 | 1992 | 10 | 1.00% |
| hsa_circ_0111207 | 1992 | 10 | 1.00% |
| hsa_circ_0140285 | 1992 | 10 | 1.00% |
| hsa_circ_0078092 | 2391 | 12 | 1.00% |
| hsa_circ_0031140 | 2192 | 11 | 1.00% |
| hsa_circ_0042504 | 2192 | 11 | 1.00% |
| hsa_circ_0049825 | 2192 | 11 | 1.00% |
| hsa_circ_0073875 | 2192 | 11 | 1.00% |
| hsa_circ_0110649 | 2192 | 11 | 1.00% |
| hsa_circ_0139735 | 2192 | 11 | 1.00% |
| hsa_circ_0134827 | 2790 | 14 | 1.00% |
| hsa_circ_0030444 | 1993 | 10 | 1.00% |
| hsa_circ_0097732 | 1993 | 10 | 1.00% |
| hsa_circ_0116274 | 1993 | 10 | 1.00% |
| hsa_circ_0123370 | 1993 | 10 | 1.00% |
| hsa_circ_0017017 | 2392 | 12 | 1.00% |
| hsa_circ_0072993 | 2392 | 12 | 1.00% |
| hsa_circ_0077673 | 2392 | 12 | 1.00% |
| hsa_circ_0048077 | 2791 | 14 | 1.00% |
| hsa_circ_0048685 | 2193 | 11 | 1.00% |
| hsa_circ_0099415 | 2193 | 11 | 1.00% |
| hsa_circ_0102884 | 2193 | 11 | 1.00% |
| hsa_circ_0033169 | 2592 | 13 | 1.00% |
| hsa_circ_0060945 | 2592 | 13 | 1.00% |
| hsa_circ_0071532 | 2592 | 13 | 1.00% |
| hsa_circ_0067877 | 1994 | 10 | 1.00% |
| hsa_circ_0079289 | 1994 | 10 | 1.00% |
| hsa_circ_0091715 | 1994 | 10 | 1.00% |
| hsa_circ_0067057 | 2393 | 12 | 1.00% |
| hsa_circ_0068158 | 2393 | 12 | 1.00% |
| hsa_circ_0119183 | 2393 | 12 | 1.00% |
| hsa_circ_0121698 | 2393 | 12 | 1.00% |
| hsa_circ_0016506 | 2194 | 11 | 1.00% |
| hsa_circ_0025172 | 2194 | 11 | 1.00% |
| hsa_circ_0060948 | 2194 | 11 | 1.00% |
| hsa_circ_0091037 | 2194 | 11 | 1.00% |
| hsa_circ_0099814 | 2194 | 11 | 1.00% |
| hsa_circ_0110211 | 2194 | 11 | 1.00% |
| hsa_circ_0047741 | 2593 | 13 | 1.00% |

| | | | |
|---|---|---|---|
| hsa_circ_0061074 | 2593 | 13 | 1.00% |
| hsa_circ_0016091 | 1995 | 10 | 1.00% |
| hsa_circ_0018566 | 1995 | 10 | 1.00% |
| hsa_circ_0040439 | 1995 | 10 | 1.00% |
| hsa_circ_0040934 | 1995 | 10 | 1.00% |
| hsa_circ_0055677 | 1995 | 10 | 1.00% |
| hsa_circ_0100933 | 1995 | 10 | 1.00% |
| hsa_circ_0102387 | 1995 | 10 | 1.00% |
| hsa_circ_0040218 | 2594 | 13 | 1.00% |
| hsa_circ_0043984 | 2195 | 11 | 1.00% |
| hsa_circ_0068876 | 2195 | 11 | 1.00% |
| hsa_circ_0137581 | 2195 | 11 | 1.00% |
| hsa_circ_0049497 | 2395 | 12 | 1.00% |
| hsa_circ_0090816 | 2395 | 12 | 1.00% |
| hsa_circ_0116858 | 2994 | 15 | 1.00% |
| hsa_circ_0020436 | 1996 | 10 | 1.00% |
| hsa_circ_0023910 | 1996 | 10 | 1.00% |
| hsa_circ_0045240 | 1996 | 10 | 1.00% |
| hsa_circ_0046864 | 1996 | 10 | 1.00% |
| hsa_circ_0052257 | 1996 | 10 | 1.00% |
| hsa_circ_0067101 | 1996 | 10 | 1.00% |
| hsa_circ_0076258 | 1996 | 10 | 1.00% |
| hsa_circ_0113589 | 1996 | 10 | 1.00% |
| hsa_circ_0067472 | 2595 | 13 | 1.00% |
| hsa_circ_0015925 | 2196 | 11 | 1.00% |
| hsa_circ_0027628 | 2196 | 11 | 1.00% |
| hsa_circ_0037920 | 2196 | 11 | 1.00% |
| hsa_circ_0056293 | 2196 | 11 | 1.00% |
| hsa_circ_0115490 | 2196 | 11 | 1.00% |
| hsa_circ_0056833 | 2396 | 12 | 1.00% |
| hsa_circ_0001269 | 1997 | 10 | 1.00% |
| hsa_circ_0019838 | 1997 | 10 | 1.00% |
| hsa_circ_0023125 | 1997 | 10 | 1.00% |
| hsa_circ_0051101 | 1997 | 10 | 1.00% |
| hsa_circ_0099330 | 1997 | 10 | 1.00% |
| hsa_circ_0105447 | 1997 | 10 | 1.00% |
| hsa_circ_0025834 | 6191 | 31 | 1.00% |
| hsa_circ_0016880 | 2197 | 11 | 1.00% |
| hsa_circ_0021104 | 2197 | 11 | 1.00% |
| hsa_circ_0040270 | 2197 | 11 | 1.00% |
| hsa_circ_0065995 | 2197 | 11 | 1.00% |
| hsa_circ_0066982 | 2197 | 11 | 1.00% |
| hsa_circ_0081910 | 2197 | 11 | 1.00% |
| hsa_circ_0138527 | 6192 | 31 | 1.00% |
| hsa_circ_0129791 | 6392 | 32 | 1.00% |
| hsa_circ_0011405 | 2397 | 12 | 1.00% |
| hsa_circ_0042427 | 2397 | 12 | 1.00% |
| hsa_circ_0067150 | 2397 | 12 | 1.00% |
| hsa_circ_0108364 | 2397 | 12 | 1.00% |
| hsa_circ_0016432 | 1998 | 10 | 1.00% |

| | | | |
|---|---|---|---|
| hsa_circ_0031400 | 1998 | 10 | 1.00% |
| hsa_circ_0057269 | 1998 | 10 | 1.00% |
| hsa_circ_0103000 | 1998 | 10 | 1.00% |
| hsa_circ_0133311 | 1998 | 10 | 1.00% |
| hsa_circ_0021179 | 2198 | 11 | 1.00% |
| hsa_circ_0026970 | 2198 | 11 | 1.00% |
| hsa_circ_0081319 | 5595 | 28 | 1.00% |
| hsa_circ_0060798 | 2398 | 12 | 1.00% |
| hsa_circ_0034038 | 2598 | 13 | 1.00% |
| hsa_circ_0019795 | 5796 | 29 | 1.00% |
| hsa_circ_0027664 | 2998 | 15 | 1.00% |
| hsa_circ_0058730 | 1999 | 10 | 1.00% |
| hsa_circ_0058946 | 1999 | 10 | 1.00% |
| hsa_circ_0079169 | 1999 | 10 | 1.00% |
| hsa_circ_0092602 | 1999 | 10 | 1.00% |
| hsa_circ_0120118 | 1999 | 10 | 1.00% |
| hsa_circ_0121361 | 1999 | 10 | 1.00% |
| hsa_circ_0136103 | 1999 | 10 | 1.00% |
| hsa_circ_0023209 | 2199 | 11 | 1.00% |
| hsa_circ_0121145 | 2199 | 11 | 1.00% |
| hsa_circ_0137359 | 2199 | 11 | 1.00% |
| hsa_circ_0047025 | 2599 | 13 | 1.00% |
| hsa_circ_0049668 | 2999 | 15 | 1.00% |
| hsa_circ_0116379 | 14799 | 74 | 1.00% |
| hsa_circ_0124737 | 3200 | 16 | 1.00% |
| hsa_circ_0001024 | 2200 | 11 | 1.00% |
| hsa_circ_0013712 | 2200 | 11 | 1.00% |
| hsa_circ_0015152 | 2200 | 11 | 1.00% |
| hsa_circ_0024106 | 2200 | 11 | 1.00% |
| hsa_circ_0031742 | 2200 | 11 | 1.00% |
| hsa_circ_0040333 | 2200 | 11 | 1.00% |
| hsa_circ_0043892 | 2200 | 11 | 1.00% |
| hsa_circ_0078376 | 2200 | 11 | 1.00% |
| hsa_circ_0086843 | 2200 | 11 | 1.00% |
| hsa_circ_0017730 | 2000 | 10 | 1.00% |
| hsa_circ_0026576 | 2000 | 10 | 1.00% |
| hsa_circ_0112678 | 2000 | 10 | 1.00% |
| hsa_circ_0076489 | 2801 | 14 | 1.00% |
| hsa_circ_0011867 | 2201 | 11 | 1.00% |
| hsa_circ_0018117 | 2201 | 11 | 1.00% |
| hsa_circ_0063023 | 2201 | 11 | 1.00% |
| hsa_circ_0084321 | 2201 | 11 | 1.00% |
| hsa_circ_0115536 | 2201 | 11 | 1.00% |
| hsa_circ_0132074 | 2201 | 11 | 1.00% |
| hsa_circ_0056816 | 2001 | 10 | 1.00% |
| hsa_circ_0071667 | 2001 | 10 | 1.00% |
| hsa_circ_0085105 | 2001 | 10 | 1.00% |
| hsa_circ_0085313 | 2001 | 10 | 1.00% |
| hsa_circ_0085645 | 2001 | 10 | 1.00% |
| hsa_circ_0010667 | 2602 | 13 | 1.00% |

| | | | |
|---|---|---|---|
| hsa_circ_0043679 | 2602 | 13 | 1.00% |
| hsa_circ_0074814 | 3603 | 18 | 1.00% |
| hsa_circ_0020903 | 2202 | 11 | 1.00% |
| hsa_circ_0044933 | 2202 | 11 | 1.00% |
| hsa_circ_0053894 | 2202 | 11 | 1.00% |
| hsa_circ_0073428 | 2202 | 11 | 1.00% |
| hsa_circ_0102170 | 2202 | 11 | 1.00% |
| hsa_circ_0103052 | 2202 | 11 | 1.00% |
| hsa_circ_0000632 | 2002 | 10 | 1.00% |
| hsa_circ_0009783 | 2002 | 10 | 1.00% |
| hsa_circ_0044929 | 2002 | 10 | 1.00% |
| hsa_circ_0049239 | 2002 | 10 | 1.00% |
| hsa_circ_0075876 | 2002 | 10 | 1.00% |
| hsa_circ_0079916 | 2002 | 10 | 1.00% |
| hsa_circ_0097979 | 2002 | 10 | 1.00% |
| hsa_circ_0108970 | 2002 | 10 | 1.00% |
| hsa_circ_0134709 | 2803 | 14 | 1.00% |
| hsa_circ_0131137 | 2603 | 13 | 1.00% |
| hsa_circ_0025400 | 2403 | 12 | 1.00% |
| hsa_circ_0031371 | 2203 | 11 | 1.00% |
| hsa_circ_0090904 | 2203 | 11 | 1.00% |
| hsa_circ_0014308 | 2003 | 10 | 1.00% |
| hsa_circ_0040753 | 2003 | 10 | 1.00% |
| hsa_circ_0055468 | 2003 | 10 | 1.00% |
| hsa_circ_0074704 | 2003 | 10 | 1.00% |
| hsa_circ_0119544 | 2003 | 10 | 1.00% |
| hsa_circ_0029516 | 2604 | 13 | 1.00% |
| hsa_circ_0076834 | 2404 | 12 | 1.00% |
| hsa_circ_0022753 | 2204 | 11 | 1.00% |
| hsa_circ_0030954 | 2204 | 11 | 1.00% |
| hsa_circ_0068443 | 2204 | 11 | 1.00% |
| hsa_circ_0070775 | 2204 | 11 | 1.00% |
| hsa_circ_0052098 | 6813 | 34 | 1.00% |
| hsa_circ_0093068 | 3607 | 18 | 1.00% |
| hsa_circ_0071444 | 3006 | 15 | 1.00% |
| hsa_circ_0016390 | 2004 | 10 | 1.00% |
| hsa_circ_0030584 | 2004 | 10 | 1.00% |
| hsa_circ_0045018 | 2004 | 10 | 1.00% |
| hsa_circ_0046541 | 2004 | 10 | 1.00% |
| hsa_circ_0052770 | 2004 | 10 | 1.00% |
| hsa_circ_0114153 | 2004 | 10 | 1.00% |
| hsa_circ_0045656 | 2405 | 12 | 1.00% |
| hsa_circ_0078133 | 2806 | 14 | 1.00% |
| hsa_circ_0129405 | 4410 | 22 | 1.00% |
| hsa_circ_0009546 | 2205 | 11 | 1.00% |
| hsa_circ_0010945 | 2205 | 11 | 1.00% |
| hsa_circ_0043991 | 2205 | 11 | 1.00% |
| hsa_circ_0048316 | 2205 | 11 | 1.00% |
| hsa_circ_0073935 | 2205 | 11 | 1.00% |
| hsa_circ_0075036 | 2205 | 11 | 1.00% |

| | | | |
|---|---|---|---|
| hsa_circ_0079873 | 2205 | 11 | 1.00% |
| hsa_circ_0048946 | 2606 | 13 | 1.00% |
| hsa_circ_0054718 | 2606 | 13 | 1.00% |
| hsa_circ_0090118 | 2807 | 14 | 1.00% |
| hsa_circ_0075183 | 2406 | 12 | 1.00% |
| hsa_circ_0037514 | 2005 | 10 | 1.00% |
| hsa_circ_0063224 | 2005 | 10 | 1.00% |
| hsa_circ_0099631 | 2005 | 10 | 1.00% |
| hsa_circ_0119588 | 2005 | 10 | 1.00% |
| hsa_circ_0052696 | 3008 | 15 | 1.00% |
| hsa_circ_0086398 | 2607 | 13 | 1.00% |
| hsa_circ_0054119 | 2206 | 11 | 1.00% |
| hsa_circ_0070724 | 2206 | 11 | 1.00% |
| hsa_circ_0086020 | 2206 | 11 | 1.00% |
| hsa_circ_0089250 | 2206 | 11 | 1.00% |
| hsa_circ_0113105 | 2206 | 11 | 1.00% |
| hsa_circ_0123524 | 2206 | 11 | 1.00% |
| hsa_circ_0135243 | 4212 | 21 | 1.00% |
| hsa_circ_0031524 | 2407 | 12 | 1.00% |
| hsa_circ_0023724 | 4012 | 20 | 1.00% |
| hsa_circ_0030156 | 3009 | 15 | 1.00% |
| hsa_circ_0072894 | 2006 | 10 | 1.00% |
| hsa_circ_0137824 | 2006 | 10 | 1.00% |
| hsa_circ_0024047 | 2207 | 11 | 1.00% |
| hsa_circ_0038790 | 2207 | 11 | 1.00% |
| hsa_circ_0054865 | 2207 | 11 | 1.00% |
| hsa_circ_0064766 | 2207 | 11 | 1.00% |
| hsa_circ_0070708 | 2207 | 11 | 1.00% |
| hsa_circ_0129837 | 2207 | 11 | 1.00% |
| hsa_circ_0043961 | 6221 | 31 | 1.00% |
| hsa_circ_0017812 | 2609 | 13 | 1.00% |
| hsa_circ_0103820 | 2609 | 13 | 1.00% |
| hsa_circ_0026062 | 2007 | 10 | 1.00% |
| hsa_circ_0039047 | 2007 | 10 | 1.00% |
| hsa_circ_0059301 | 2007 | 10 | 1.00% |
| hsa_circ_0082214 | 2007 | 10 | 1.00% |
| hsa_circ_0127119 | 2007 | 10 | 1.00% |
| hsa_circ_0055327 | 8631 | 43 | 1.00% |
| hsa_circ_0009915 | 2208 | 11 | 1.00% |
| hsa_circ_0065141 | 2208 | 11 | 1.00% |
| hsa_circ_0085400 | 2208 | 11 | 1.00% |
| hsa_circ_0071546 | 7026 | 35 | 1.00% |
| hsa_circ_0067573 | 2409 | 12 | 1.00% |
| hsa_circ_0070368 | 2409 | 12 | 1.00% |
| hsa_circ_0085349 | 2610 | 13 | 1.00% |
| hsa_circ_0090577 | 2811 | 14 | 1.00% |
| hsa_circ_0055438 | 3815 | 19 | 1.00% |
| hsa_circ_0049417 | 2008 | 10 | 1.00% |
| hsa_circ_0078394 | 2008 | 10 | 1.00% |
| hsa_circ_0079138 | 2008 | 10 | 1.00% |

| | | | |
|---|---|---|---|
| hsa_circ_0090694 | 2008 | 10 | 1.00% |
| hsa_circ_0011007 | 2209 | 11 | 1.00% |
| hsa_circ_0014060 | 2209 | 11 | 1.00% |
| hsa_circ_0021212 | 2209 | 11 | 1.00% |
| hsa_circ_0024888 | 2209 | 11 | 1.00% |
| hsa_circ_0052495 | 2209 | 11 | 1.00% |
| hsa_circ_0130021 | 2209 | 11 | 1.00% |
| hsa_circ_0051282 | 2410 | 12 | 1.00% |
| hsa_circ_0066192 | 2410 | 12 | 1.00% |
| hsa_circ_0071620 | 2410 | 12 | 1.00% |
| hsa_circ_0094023 | 2410 | 12 | 1.00% |
| hsa_circ_0104913 | 2410 | 12 | 1.00% |
| hsa_circ_0057334 | 2812 | 14 | 1.00% |
| hsa_circ_0116602 | 7633 | 38 | 1.00% |
| hsa_circ_0015392 | 2009 | 10 | 1.00% |
| hsa_circ_0015623 | 2009 | 10 | 1.00% |
| hsa_circ_0038291 | 2009 | 10 | 1.00% |
| hsa_circ_0056345 | 2009 | 10 | 1.00% |
| hsa_circ_0057208 | 2009 | 10 | 1.00% |
| hsa_circ_0059810 | 2009 | 10 | 1.00% |
| hsa_circ_0132941 | 2009 | 10 | 1.00% |
| hsa_circ_0017008 | 2210 | 11 | 1.00% |
| hsa_circ_0019849 | 2210 | 11 | 1.00% |
| hsa_circ_0025747 | 2210 | 11 | 1.00% |
| hsa_circ_0065124 | 2210 | 11 | 1.00% |
| hsa_circ_0067564 | 2210 | 11 | 1.00% |
| hsa_circ_0070461 | 2210 | 11 | 1.00% |
| hsa_circ_0136362 | 2210 | 11 | 1.00% |
| hsa_circ_0038945 | 2411 | 12 | 1.00% |
| hsa_circ_0083732 | 2612 | 13 | 1.00% |
| hsa_circ_0128987 | 2612 | 13 | 1.00% |
| hsa_circ_0042653 | 3014 | 15 | 1.00% |
| hsa_circ_0138148 | 4422 | 22 | 1.00% |
| hsa_circ_0059676 | 3015 | 15 | 1.00% |
| hsa_circ_0033343 | 2412 | 12 | 1.00% |
| hsa_circ_0057548 | 2412 | 12 | 1.00% |
| hsa_circ_0023621 | 2211 | 11 | 1.00% |
| hsa_circ_0055662 | 2211 | 11 | 1.00% |
| hsa_circ_0073085 | 2211 | 11 | 1.00% |
| hsa_circ_0076216 | 2211 | 11 | 1.00% |
| hsa_circ_0078703 | 2211 | 11 | 1.00% |
| hsa_circ_0083244 | 2211 | 11 | 1.00% |
| hsa_circ_0123544 | 2211 | 11 | 1.00% |
| hsa_circ_0125607 | 2211 | 11 | 1.00% |
| hsa_circ_0137559 | 2211 | 11 | 1.00% |
| hsa_circ_0030247 | 2010 | 10 | 1.00% |
| hsa_circ_0033084 | 2010 | 10 | 1.00% |
| hsa_circ_0055640 | 2010 | 10 | 1.00% |
| hsa_circ_0077762 | 2010 | 10 | 1.00% |
| hsa_circ_0077939 | 2010 | 10 | 1.00% |

| | | | |
|---|---|---|---|
| hsa_circ_0079871 | 2010 | 10 | 1.00% |
| hsa_circ_0086303 | 2010 | 10 | 1.00% |
| hsa_circ_0116389 | 4423 | 22 | 0.99% |
| hsa_circ_0082809 | 3820 | 19 | 0.99% |
| hsa_circ_0025501 | 2413 | 12 | 0.99% |
| hsa_circ_0063485 | 2413 | 12 | 0.99% |
| hsa_circ_0086556 | 2212 | 11 | 0.99% |
| hsa_circ_0134003 | 2212 | 11 | 0.99% |
| hsa_circ_0004566 | 4022 | 20 | 0.99% |
| hsa_circ_0102010 | 4022 | 20 | 0.99% |
| hsa_circ_0010473 | 2011 | 10 | 0.99% |
| hsa_circ_0034988 | 2011 | 10 | 0.99% |
| hsa_circ_0047260 | 2011 | 10 | 0.99% |
| hsa_circ_0054184 | 2011 | 10 | 0.99% |
| hsa_circ_0072892 | 2011 | 10 | 0.99% |
| hsa_circ_0093601 | 2011 | 10 | 0.99% |
| hsa_circ_0130373 | 4827 | 24 | 0.99% |
| hsa_circ_0039069 | 2615 | 13 | 0.99% |
| hsa_circ_0063746 | 2615 | 13 | 0.99% |
| hsa_circ_0031944 | 2414 | 12 | 0.99% |
| hsa_circ_0051599 | 2414 | 12 | 0.99% |
| hsa_circ_0063438 | 2414 | 12 | 0.99% |
| hsa_circ_0040293 | 6840 | 34 | 0.99% |
| hsa_circ_0041284 | 2213 | 11 | 0.99% |
| hsa_circ_0046791 | 2213 | 11 | 0.99% |
| hsa_circ_0065666 | 2213 | 11 | 0.99% |
| hsa_circ_0083543 | 2213 | 11 | 0.99% |
| hsa_circ_0084435 | 2213 | 11 | 0.99% |
| hsa_circ_0084526 | 2213 | 11 | 0.99% |
| hsa_circ_0091162 | 2213 | 11 | 0.99% |
| hsa_circ_0126297 | 2213 | 11 | 0.99% |
| hsa_circ_0014806 | 2012 | 10 | 0.99% |
| hsa_circ_0027132 | 2012 | 10 | 0.99% |
| hsa_circ_0073832 | 2012 | 10 | 0.99% |
| hsa_circ_0098360 | 2012 | 10 | 0.99% |
| hsa_circ_0109111 | 2012 | 10 | 0.99% |
| hsa_circ_0110559 | 2012 | 10 | 0.99% |
| hsa_circ_0041491 | 2817 | 14 | 0.99% |
| hsa_circ_0074558 | 2616 | 13 | 0.99% |
| hsa_circ_0130334 | 4025 | 20 | 0.99% |
| hsa_circ_0053959 | 2415 | 12 | 0.99% |
| hsa_circ_0139817 | 2415 | 12 | 0.99% |
| hsa_circ_0012492 | 2214 | 11 | 0.99% |
| hsa_circ_0039032 | 2214 | 11 | 0.99% |
| hsa_circ_0042034 | 2214 | 11 | 0.99% |
| hsa_circ_0051205 | 2214 | 11 | 0.99% |
| hsa_circ_0082010 | 2214 | 11 | 0.99% |
| hsa_circ_0016021 | 2013 | 10 | 0.99% |
| hsa_circ_0072418 | 2013 | 10 | 0.99% |
| hsa_circ_0085218 | 2013 | 10 | 0.99% |

| | | | |
|---|---|---|---|
| hsa_circ_0045285 | 2617 | 13 | 0.99% |
| hsa_circ_0032554 | 2416 | 12 | 0.99% |
| hsa_circ_0113665 | 2416 | 12 | 0.99% |
| hsa_circ_0136399 | 2819 | 14 | 0.99% |
| hsa_circ_0035758 | 2215 | 11 | 0.99% |
| hsa_circ_0067425 | 2215 | 11 | 0.99% |
| hsa_circ_0085583 | 2215 | 11 | 0.99% |
| hsa_circ_0096114 | 4833 | 24 | 0.99% |
| hsa_circ_0009687 | 2618 | 13 | 0.99% |
| hsa_circ_0012566 | 2014 | 10 | 0.99% |
| hsa_circ_0047213 | 2014 | 10 | 0.99% |
| hsa_circ_0057817 | 2014 | 10 | 0.99% |
| hsa_circ_0077916 | 2014 | 10 | 0.99% |
| hsa_circ_0083259 | 2014 | 10 | 0.99% |
| hsa_circ_0077235 | 2417 | 12 | 0.99% |
| hsa_circ_0020550 | 2216 | 11 | 0.99% |
| hsa_circ_0027104 | 2216 | 11 | 0.99% |
| hsa_circ_0054817 | 2216 | 11 | 0.99% |
| hsa_circ_0065428 | 2216 | 11 | 0.99% |
| hsa_circ_0067208 | 2216 | 11 | 0.99% |
| hsa_circ_0039446 | 2619 | 13 | 0.99% |
| hsa_circ_0059651 | 2619 | 13 | 0.99% |
| hsa_circ_0086688 | 2619 | 13 | 0.99% |
| hsa_circ_0091166 | 2619 | 13 | 0.99% |
| hsa_circ_0017518 | 2418 | 12 | 0.99% |
| hsa_circ_0018516 | 2418 | 12 | 0.99% |
| hsa_circ_0013024 | 2015 | 10 | 0.99% |
| hsa_circ_0029608 | 2015 | 10 | 0.99% |
| hsa_circ_0061441 | 2620 | 13 | 0.99% |
| hsa_circ_0012790 | 2217 | 11 | 0.99% |
| hsa_circ_0038066 | 2217 | 11 | 0.99% |
| hsa_circ_0052371 | 2217 | 11 | 0.99% |
| hsa_circ_0085816 | 2217 | 11 | 0.99% |
| hsa_circ_0091915 | 2217 | 11 | 0.99% |
| hsa_circ_0139005 | 2217 | 11 | 0.99% |
| hsa_circ_0019449 | 2016 | 10 | 0.99% |
| hsa_circ_0024982 | 2016 | 10 | 0.99% |
| hsa_circ_0025296 | 2016 | 10 | 0.99% |
| hsa_circ_0033440 | 2016 | 10 | 0.99% |
| hsa_circ_0044521 | 2016 | 10 | 0.99% |
| hsa_circ_0051551 | 2016 | 10 | 0.99% |
| hsa_circ_0056363 | 2016 | 10 | 0.99% |
| hsa_circ_0067739 | 2016 | 10 | 0.99% |
| hsa_circ_0076114 | 2016 | 10 | 0.99% |
| hsa_circ_0103264 | 2016 | 10 | 0.99% |
| hsa_circ_0100565 | 11693 | 58 | 0.99% |
| hsa_circ_0027569 | 2218 | 11 | 0.99% |
| hsa_circ_0087868 | 2218 | 11 | 0.99% |
| hsa_circ_0121470 | 2218 | 11 | 0.99% |
| hsa_circ_0069127 | 2420 | 12 | 0.99% |

| | | | |
|---|---|---|---|
| hsa_circ_0020431 | 2017 | 10 | 0.99% |
| hsa_circ_0029745 | 2017 | 10 | 0.99% |
| hsa_circ_0037151 | 2017 | 10 | 0.99% |
| hsa_circ_0067056 | 2017 | 10 | 0.99% |
| hsa_circ_0077504 | 2017 | 10 | 0.99% |
| hsa_circ_0087171 | 2017 | 10 | 0.99% |
| hsa_circ_0106534 | 2017 | 10 | 0.99% |
| hsa_circ_0060458 | 2219 | 11 | 0.99% |
| hsa_circ_0089069 | 2219 | 11 | 0.99% |
| hsa_circ_0110007 | 2421 | 12 | 0.99% |
| hsa_circ_0076252 | 2623 | 13 | 0.99% |
| hsa_circ_0036324 | 2825 | 14 | 0.99% |
| hsa_circ_0074456 | 2018 | 10 | 0.99% |
| hsa_circ_0113650 | 2018 | 10 | 0.99% |
| hsa_circ_0139343 | 2018 | 10 | 0.99% |
| hsa_circ_0019234 | 2220 | 11 | 0.99% |
| hsa_circ_0045628 | 2220 | 11 | 0.99% |
| hsa_circ_0063994 | 2220 | 11 | 0.99% |
| hsa_circ_0104894 | 2220 | 11 | 0.99% |
| hsa_circ_0119790 | 2220 | 11 | 0.99% |
| hsa_circ_0101719 | 7871 | 39 | 0.99% |
| hsa_circ_0019038 | 2422 | 12 | 0.99% |
| hsa_circ_0085040 | 2826 | 14 | 0.99% |
| hsa_circ_0007172 | 3230 | 16 | 0.99% |
| hsa_circ_0004585 | 2019 | 10 | 0.99% |
| hsa_circ_0015904 | 2019 | 10 | 0.99% |
| hsa_circ_0041534 | 2019 | 10 | 0.99% |
| hsa_circ_0049877 | 2019 | 10 | 0.99% |
| hsa_circ_0056085 | 2019 | 10 | 0.99% |
| hsa_circ_0021430 | 2221 | 11 | 0.99% |
| hsa_circ_0054812 | 2221 | 11 | 0.99% |
| hsa_circ_0073535 | 2221 | 11 | 0.99% |
| hsa_circ_0077543 | 2221 | 11 | 0.99% |
| hsa_circ_0079634 | 2221 | 11 | 0.99% |
| hsa_circ_0086279 | 2221 | 11 | 0.99% |
| hsa_circ_0101845 | 2221 | 11 | 0.99% |
| hsa_circ_0116377 | 2221 | 11 | 0.99% |
| hsa_circ_0024394 | 2423 | 12 | 0.99% |
| hsa_circ_0032142 | 2423 | 12 | 0.99% |
| hsa_circ_0058900 | 2423 | 12 | 0.99% |
| hsa_circ_0105761 | 2423 | 12 | 0.99% |
| hsa_circ_0116479 | 8077 | 40 | 0.99% |
| hsa_circ_0030837 | 3635 | 18 | 0.99% |
| hsa_circ_0112065 | 4645 | 23 | 0.99% |
| hsa_circ_0084033 | 4242 | 21 | 0.99% |
| hsa_circ_0017595 | 2828 | 14 | 0.99% |
| hsa_circ_0092161 | 2626 | 13 | 0.99% |
| hsa_circ_0061982 | 2424 | 12 | 0.99% |
| hsa_circ_0052771 | 2222 | 11 | 0.99% |
| hsa_circ_0070677 | 2222 | 11 | 0.99% |

| | | | |
|---|---|---|---|
| hsa_circ_0086498 | 2222 | 11 | 0.99% |
| hsa_circ_0089421 | 2222 | 11 | 0.99% |
| hsa_circ_0094458 | 2222 | 11 | 0.99% |
| hsa_circ_0120657 | 2222 | 11 | 0.99% |
| hsa_circ_0028828 | 2020 | 10 | 0.99% |
| hsa_circ_0063974 | 2020 | 10 | 0.99% |
| hsa_circ_0085063 | 2020 | 10 | 0.99% |
| hsa_circ_0039762 | 6869 | 34 | 0.99% |
| hsa_circ_0133248 | 3637 | 18 | 0.99% |
| hsa_circ_0062343 | 2627 | 13 | 0.99% |
| hsa_circ_0036671 | 2425 | 12 | 0.99% |
| hsa_circ_0045939 | 2425 | 12 | 0.99% |
| hsa_circ_0052318 | 2425 | 12 | 0.99% |
| hsa_circ_0139913 | 2425 | 12 | 0.99% |
| hsa_circ_0097676 | 6871 | 34 | 0.99% |
| hsa_circ_0035932 | 2223 | 11 | 0.99% |
| hsa_circ_0043174 | 2223 | 11 | 0.99% |
| hsa_circ_0051152 | 2223 | 11 | 0.99% |
| hsa_circ_0062061 | 2223 | 11 | 0.99% |
| hsa_circ_0065146 | 2223 | 11 | 0.99% |
| hsa_circ_0102523 | 2223 | 11 | 0.99% |
| hsa_circ_0021599 | 2021 | 10 | 0.99% |
| hsa_circ_0046871 | 2021 | 10 | 0.99% |
| hsa_circ_0057115 | 2021 | 10 | 0.99% |
| hsa_circ_0057633 | 2021 | 10 | 0.99% |
| hsa_circ_0061737 | 2021 | 10 | 0.99% |
| hsa_circ_0072658 | 2021 | 10 | 0.99% |
| hsa_circ_0088935 | 2021 | 10 | 0.99% |
| hsa_circ_0115244 | 2021 | 10 | 0.99% |
| hsa_circ_0115612 | 2021 | 10 | 0.99% |
| hsa_circ_0112905 | 3032 | 15 | 0.99% |
| hsa_circ_0127012 | 2830 | 14 | 0.99% |
| hsa_circ_0058366 | 2628 | 13 | 0.99% |
| hsa_circ_0092176 | 2628 | 13 | 0.99% |
| hsa_circ_0017320 | 2224 | 11 | 0.99% |
| hsa_circ_0045081 | 2224 | 11 | 0.99% |
| hsa_circ_0061527 | 2224 | 11 | 0.99% |
| hsa_circ_0086524 | 2224 | 11 | 0.99% |
| hsa_circ_0004138 | 2022 | 10 | 0.99% |
| hsa_circ_0010361 | 2022 | 10 | 0.99% |
| hsa_circ_0032276 | 2022 | 10 | 0.99% |
| hsa_circ_0071104 | 2022 | 10 | 0.99% |
| hsa_circ_0081445 | 2022 | 10 | 0.99% |
| hsa_circ_0085005 | 6067 | 30 | 0.99% |
| hsa_circ_0068858 | 2427 | 12 | 0.99% |
| hsa_circ_0083102 | 2427 | 12 | 0.99% |
| hsa_circ_0129400 | 2427 | 12 | 0.99% |
| hsa_circ_0022785 | 2225 | 11 | 0.99% |
| hsa_circ_0046746 | 2225 | 11 | 0.99% |
| hsa_circ_0092536 | 2225 | 11 | 0.99% |

| | | | |
|---|---|---|---|
| hsa_circ_0112209 | 2225 | 11 | 0.99% |
| hsa_circ_0016003 | 2023 | 10 | 0.99% |
| hsa_circ_0021005 | 2023 | 10 | 0.99% |
| hsa_circ_0085391 | 2023 | 10 | 0.99% |
| hsa_circ_0091968 | 2023 | 10 | 0.99% |
| hsa_circ_0102106 | 2023 | 10 | 0.99% |
| hsa_circ_0132295 | 2023 | 10 | 0.99% |
| hsa_circ_0032121 | 2428 | 12 | 0.99% |
| hsa_circ_0053105 | 2428 | 12 | 0.99% |
| hsa_circ_0043154 | 2226 | 11 | 0.99% |
| hsa_circ_0048599 | 2226 | 11 | 0.99% |
| hsa_circ_0048609 | 2226 | 11 | 0.99% |
| hsa_circ_0078269 | 2226 | 11 | 0.99% |
| hsa_circ_0135596 | 2226 | 11 | 0.99% |
| hsa_circ_0018138 | 2024 | 10 | 0.99% |
| hsa_circ_0070897 | 2024 | 10 | 0.99% |
| hsa_circ_0071958 | 2024 | 10 | 0.99% |
| hsa_circ_0078829 | 2024 | 10 | 0.99% |
| hsa_circ_0082824 | 2024 | 10 | 0.99% |
| hsa_circ_0085461 | 2429 | 12 | 0.99% |
| hsa_circ_0020393 | 2227 | 11 | 0.99% |
| hsa_circ_0034652 | 2227 | 11 | 0.99% |
| hsa_circ_0053487 | 2227 | 11 | 0.99% |
| hsa_circ_0012743 | 3037 | 15 | 0.99% |
| hsa_circ_0121570 | 3240 | 16 | 0.99% |
| hsa_circ_0132304 | 2430 | 12 | 0.99% |
| hsa_circ_0027724 | 2025 | 10 | 0.99% |
| hsa_circ_0028047 | 2025 | 10 | 0.99% |
| hsa_circ_0032396 | 2025 | 10 | 0.99% |
| hsa_circ_0063908 | 2025 | 10 | 0.99% |
| hsa_circ_0078880 | 2025 | 10 | 0.99% |
| hsa_circ_0090021 | 2025 | 10 | 0.99% |
| hsa_circ_0067737 | 2228 | 11 | 0.99% |
| hsa_circ_0122991 | 2228 | 11 | 0.99% |
| hsa_circ_0045155 | 2431 | 12 | 0.99% |
| hsa_circ_0018592 | 2026 | 10 | 0.99% |
| hsa_circ_0043058 | 2026 | 10 | 0.99% |
| hsa_circ_0071741 | 2026 | 10 | 0.99% |
| hsa_circ_0025390 | 2229 | 11 | 0.99% |
| hsa_circ_0054229 | 2229 | 11 | 0.99% |
| hsa_circ_0091321 | 2229 | 11 | 0.99% |
| hsa_circ_0098009 | 2229 | 11 | 0.99% |
| hsa_circ_0114159 | 2229 | 11 | 0.99% |
| hsa_circ_0130705 | 2229 | 11 | 0.99% |
| hsa_circ_0092985 | 4256 | 21 | 0.99% |
| hsa_circ_0079507 | 2432 | 12 | 0.99% |
| hsa_circ_0073033 | 2635 | 13 | 0.99% |
| hsa_circ_0079928 | 2635 | 13 | 0.99% |
| hsa_circ_0015245 | 2027 | 10 | 0.99% |
| hsa_circ_0026061 | 2027 | 10 | 0.99% |

| | | | |
|---|---|---|---|
| hsa_circ_0032412 | 2027 | 10 | 0.99% |
| hsa_circ_0044935 | 2027 | 10 | 0.99% |
| hsa_circ_0059781 | 2027 | 10 | 0.99% |
| hsa_circ_0064232 | 2027 | 10 | 0.99% |
| hsa_circ_0094128 | 2027 | 10 | 0.99% |
| hsa_circ_0036187 | 2230 | 11 | 0.99% |
| hsa_circ_0091222 | 5881 | 29 | 0.99% |
| hsa_circ_0022883 | 3042 | 15 | 0.99% |
| hsa_circ_0085238 | 2028 | 10 | 0.99% |
| hsa_circ_0095171 | 2028 | 10 | 0.99% |
| hsa_circ_0011134 | 2231 | 11 | 0.99% |
| hsa_circ_0015650 | 2231 | 11 | 0.99% |
| hsa_circ_0022067 | 2231 | 11 | 0.99% |
| hsa_circ_0038041 | 2231 | 11 | 0.99% |
| hsa_circ_0039358 | 2231 | 11 | 0.99% |
| hsa_circ_0109540 | 8316 | 41 | 0.99% |
| hsa_circ_0017470 | 2434 | 12 | 0.99% |
| hsa_circ_0127214 | 2434 | 12 | 0.99% |
| hsa_circ_0086113 | 2637 | 13 | 0.99% |
| hsa_circ_0019793 | 5883 | 29 | 0.99% |
| hsa_circ_0025647 | 2029 | 10 | 0.99% |
| hsa_circ_0045952 | 2029 | 10 | 0.99% |
| hsa_circ_0071089 | 2029 | 10 | 0.99% |
| hsa_circ_0017801 | 2232 | 11 | 0.99% |
| hsa_circ_0118605 | 2232 | 11 | 0.99% |
| hsa_circ_0029685 | 2435 | 12 | 0.99% |
| hsa_circ_0028998 | 7510 | 37 | 0.99% |
| hsa_circ_0129082 | 5481 | 27 | 0.99% |
| hsa_circ_0010787 | 4872 | 24 | 0.99% |
| hsa_circ_0062613 | 3248 | 16 | 0.99% |
| hsa_circ_0047103 | 2436 | 12 | 0.99% |
| hsa_circ_0013894 | 2233 | 11 | 0.99% |
| hsa_circ_0019888 | 2233 | 11 | 0.99% |
| hsa_circ_0038766 | 2233 | 11 | 0.99% |
| hsa_circ_0042513 | 2233 | 11 | 0.99% |
| hsa_circ_0065201 | 2233 | 11 | 0.99% |
| hsa_circ_0086516 | 2233 | 11 | 0.99% |
| hsa_circ_0072221 | 2030 | 10 | 0.99% |
| hsa_circ_0114075 | 2843 | 14 | 0.98% |
| hsa_circ_0039629 | 2640 | 13 | 0.98% |
| hsa_circ_0010214 | 2437 | 12 | 0.98% |
| hsa_circ_0052432 | 2437 | 12 | 0.98% |
| hsa_circ_0076886 | 2437 | 12 | 0.98% |
| hsa_circ_0014756 | 2234 | 11 | 0.98% |
| hsa_circ_0022070 | 2234 | 11 | 0.98% |
| hsa_circ_0059721 | 2234 | 11 | 0.98% |
| hsa_circ_0068359 | 2234 | 11 | 0.98% |
| hsa_circ_0090768 | 2234 | 11 | 0.98% |
| hsa_circ_0129281 | 2234 | 11 | 0.98% |
| hsa_circ_0130757 | 2234 | 11 | 0.98% |

| | | | |
|---|---|---|---|
| hsa_circ_0037196 | 2031 | 10 | 0.98% |
| hsa_circ_0057906 | 2031 | 10 | 0.98% |
| hsa_circ_0066996 | 2031 | 10 | 0.98% |
| hsa_circ_0069511 | 2031 | 10 | 0.98% |
| hsa_circ_0086774 | 2031 | 10 | 0.98% |
| hsa_circ_0112198 | 2031 | 10 | 0.98% |
| hsa_circ_0120622 | 2031 | 10 | 0.98% |
| hsa_circ_0013402 | 2844 | 14 | 0.98% |
| hsa_circ_0041219 | 2438 | 12 | 0.98% |
| hsa_circ_0048711 | 2438 | 12 | 0.98% |
| hsa_circ_0022852 | 2235 | 11 | 0.98% |
| hsa_circ_0054804 | 2235 | 11 | 0.98% |
| hsa_circ_0081061 | 2235 | 11 | 0.98% |
| hsa_circ_0083551 | 3048 | 15 | 0.98% |
| hsa_circ_0023868 | 2032 | 10 | 0.98% |
| hsa_circ_0041736 | 2032 | 10 | 0.98% |
| hsa_circ_0117609 | 2032 | 10 | 0.98% |
| hsa_circ_0135355 | 2032 | 10 | 0.98% |
| hsa_circ_0041664 | 5894 | 29 | 0.98% |
| hsa_circ_0040262 | 2439 | 12 | 0.98% |
| hsa_circ_0135830 | 2439 | 12 | 0.98% |
| hsa_circ_0115738 | 3049 | 15 | 0.98% |
| hsa_circ_0021013 | 2236 | 11 | 0.98% |
| hsa_circ_0046476 | 2236 | 11 | 0.98% |
| hsa_circ_0053749 | 2236 | 11 | 0.98% |
| hsa_circ_0131138 | 2236 | 11 | 0.98% |
| hsa_circ_0097290 | 5895 | 29 | 0.98% |
| hsa_circ_0058253 | 2033 | 10 | 0.98% |
| hsa_circ_0064543 | 2033 | 10 | 0.98% |
| hsa_circ_0025915 | 2643 | 13 | 0.98% |
| hsa_circ_0064620 | 2643 | 13 | 0.98% |
| hsa_circ_0130323 | 3253 | 16 | 0.98% |
| hsa_circ_0135680 | 4880 | 24 | 0.98% |
| hsa_circ_0125038 | 2440 | 12 | 0.98% |
| hsa_circ_0019471 | 2237 | 11 | 0.98% |
| hsa_circ_0065194 | 2237 | 11 | 0.98% |
| hsa_circ_0126260 | 2237 | 11 | 0.98% |
| hsa_circ_0063514 | 2644 | 13 | 0.98% |
| hsa_circ_0085781 | 2644 | 13 | 0.98% |
| hsa_circ_0137794 | 2644 | 13 | 0.98% |
| hsa_circ_0009519 | 6712 | 33 | 0.98% |
| hsa_circ_0018483 | 2034 | 10 | 0.98% |
| hsa_circ_0074965 | 2034 | 10 | 0.98% |
| hsa_circ_0085742 | 2034 | 10 | 0.98% |
| hsa_circ_0103177 | 2034 | 10 | 0.98% |
| hsa_circ_0135028 | 2034 | 10 | 0.98% |
| hsa_circ_0036127 | 2848 | 14 | 0.98% |
| hsa_circ_0017318 | 2238 | 11 | 0.98% |
| hsa_circ_0059771 | 2238 | 11 | 0.98% |
| hsa_circ_0085010 | 2238 | 11 | 0.98% |

| | | | |
|---|---|---|---|
| hsa_circ_0094540 | 2238 | 11 | 0.98% |
| hsa_circ_0122448 | 2238 | 11 | 0.98% |
| hsa_circ_0011309 | 2645 | 13 | 0.98% |
| hsa_circ_0017121 | 2442 | 12 | 0.98% |
| hsa_circ_0034429 | 2442 | 12 | 0.98% |
| hsa_circ_0036609 | 2442 | 12 | 0.98% |
| hsa_circ_0021673 | 2035 | 10 | 0.98% |
| hsa_circ_0026193 | 2035 | 10 | 0.98% |
| hsa_circ_0033154 | 2035 | 10 | 0.98% |
| hsa_circ_0037304 | 2035 | 10 | 0.98% |
| hsa_circ_0066575 | 2035 | 10 | 0.98% |
| hsa_circ_0091834 | 2035 | 10 | 0.98% |
| hsa_circ_0107734 | 2035 | 10 | 0.98% |
| hsa_circ_0115227 | 2035 | 10 | 0.98% |
| hsa_circ_0032820 | 2646 | 13 | 0.98% |
| hsa_circ_0001780 | 2239 | 11 | 0.98% |
| hsa_circ_0014468 | 2239 | 11 | 0.98% |
| hsa_circ_0054719 | 2239 | 11 | 0.98% |
| hsa_circ_0066676 | 2239 | 11 | 0.98% |
| hsa_circ_0071830 | 2239 | 11 | 0.98% |
| hsa_circ_0114653 | 8549 | 42 | 0.98% |
| hsa_circ_0124600 | 3664 | 18 | 0.98% |
| hsa_circ_0029832 | 2443 | 12 | 0.98% |
| hsa_circ_0043765 | 2443 | 12 | 0.98% |
| hsa_circ_0096308 | 2036 | 10 | 0.98% |
| hsa_circ_0059100 | 2647 | 13 | 0.98% |
| hsa_circ_0011206 | 2240 | 11 | 0.98% |
| hsa_circ_0014053 | 2240 | 11 | 0.98% |
| hsa_circ_0045973 | 2240 | 11 | 0.98% |
| hsa_circ_0046353 | 2240 | 11 | 0.98% |
| hsa_circ_0084629 | 2240 | 11 | 0.98% |
| hsa_circ_0109796 | 2240 | 11 | 0.98% |
| hsa_circ_0037275 | 2851 | 14 | 0.98% |
| hsa_circ_0055829 | 3055 | 15 | 0.98% |
| hsa_circ_0031236 | 2444 | 12 | 0.98% |
| hsa_circ_0035720 | 2444 | 12 | 0.98% |
| hsa_circ_0050088 | 2444 | 12 | 0.98% |
| hsa_circ_0069946 | 2444 | 12 | 0.98% |
| hsa_circ_0091740 | 2444 | 12 | 0.98% |
| hsa_circ_0126408 | 8962 | 44 | 0.98% |
| hsa_circ_0012696 | 2648 | 13 | 0.98% |
| hsa_circ_0102085 | 2648 | 13 | 0.98% |
| hsa_circ_0025548 | 2037 | 10 | 0.98% |
| hsa_circ_0052875 | 2037 | 10 | 0.98% |
| hsa_circ_0067348 | 2037 | 10 | 0.98% |
| hsa_circ_0073973 | 2037 | 10 | 0.98% |
| hsa_circ_0034737 | 2852 | 14 | 0.98% |
| hsa_circ_0042586 | 2445 | 12 | 0.98% |
| hsa_circ_0064052 | 2445 | 12 | 0.98% |
| hsa_circ_0047839 | 2853 | 14 | 0.98% |

| | | | |
|---|---|---|---|
| hsa_circ_0101473 | 2853 | 14 | 0.98% |
| hsa_circ_0032297 | 2038 | 10 | 0.98% |
| hsa_circ_0132676 | 2038 | 10 | 0.98% |
| hsa_circ_0135245 | 2038 | 10 | 0.98% |
| hsa_circ_0011565 | 2242 | 11 | 0.98% |
| hsa_circ_0080147 | 2242 | 11 | 0.98% |
| hsa_circ_0106471 | 2242 | 11 | 0.98% |
| hsa_circ_0118660 | 2242 | 11 | 0.98% |
| hsa_circ_0063492 | 2446 | 12 | 0.98% |
| hsa_circ_0125493 | 8563 | 42 | 0.98% |
| hsa_circ_0012298 | 2039 | 10 | 0.98% |
| hsa_circ_0038307 | 2039 | 10 | 0.98% |
| hsa_circ_0062787 | 2039 | 10 | 0.98% |
| hsa_circ_0076194 | 2039 | 10 | 0.98% |
| hsa_circ_0080510 | 2039 | 10 | 0.98% |
| hsa_circ_0088751 | 2039 | 10 | 0.98% |
| hsa_circ_0052846 | 2243 | 11 | 0.98% |
| hsa_circ_0055751 | 2243 | 11 | 0.98% |
| hsa_circ_0083019 | 2243 | 11 | 0.98% |
| hsa_circ_0106631 | 2243 | 11 | 0.98% |
| hsa_circ_0115008 | 2243 | 11 | 0.98% |
| hsa_circ_0018318 | 2447 | 12 | 0.98% |
| hsa_circ_0004160 | 11424 | 56 | 0.98% |
| hsa_circ_0040153 | 2856 | 14 | 0.98% |
| hsa_circ_0019807 | 2448 | 12 | 0.98% |
| hsa_circ_0020988 | 2448 | 12 | 0.98% |
| hsa_circ_0029737 | 2448 | 12 | 0.98% |
| hsa_circ_0093353 | 2448 | 12 | 0.98% |
| hsa_circ_0025883 | 2244 | 11 | 0.98% |
| hsa_circ_0029488 | 2244 | 11 | 0.98% |
| hsa_circ_0048673 | 2244 | 11 | 0.98% |
| hsa_circ_0070011 | 2244 | 11 | 0.98% |
| hsa_circ_0113088 | 2244 | 11 | 0.98% |
| hsa_circ_0122488 | 2244 | 11 | 0.98% |
| hsa_circ_0135810 | 2244 | 11 | 0.98% |
| hsa_circ_0138563 | 2244 | 11 | 0.98% |
| hsa_circ_0042215 | 2040 | 10 | 0.98% |
| hsa_circ_0056637 | 2040 | 10 | 0.98% |
| hsa_circ_0082153 | 2040 | 10 | 0.98% |
| hsa_circ_0085249 | 2040 | 10 | 0.98% |
| hsa_circ_0135639 | 2040 | 10 | 0.98% |
| hsa_circ_0139176 | 2040 | 10 | 0.98% |
| hsa_circ_0093203 | 5713 | 28 | 0.98% |
| hsa_circ_0091403 | 2857 | 14 | 0.98% |
| hsa_circ_0032085 | 2449 | 12 | 0.98% |
| hsa_circ_0040979 | 2449 | 12 | 0.98% |
| hsa_circ_0020870 | 2245 | 11 | 0.98% |
| hsa_circ_0023066 | 2245 | 11 | 0.98% |
| hsa_circ_0054714 | 2245 | 11 | 0.98% |
| hsa_circ_0057815 | 2245 | 11 | 0.98% |

| | | | |
|---|---|---|---|
| hsa_circ_0077969 | 2245 | 11 | 0.98% |
| hsa_circ_0091653 | 2041 | 10 | 0.98% |
| hsa_circ_0130646 | 2041 | 10 | 0.98% |
| hsa_circ_0131119 | 2041 | 10 | 0.98% |
| hsa_circ_0066686 | 3266 | 16 | 0.98% |
| hsa_circ_0025927 | 2654 | 13 | 0.98% |
| hsa_circ_0083471 | 2654 | 13 | 0.98% |
| hsa_circ_0118377 | 2654 | 13 | 0.98% |
| hsa_circ_0054766 | 2450 | 12 | 0.98% |
| hsa_circ_0058393 | 2450 | 12 | 0.98% |
| hsa_circ_0026536 | 2246 | 11 | 0.98% |
| hsa_circ_0029821 | 2246 | 11 | 0.98% |
| hsa_circ_0061014 | 2246 | 11 | 0.98% |
| hsa_circ_0064398 | 2246 | 11 | 0.98% |
| hsa_circ_0020925 | 2042 | 10 | 0.98% |
| hsa_circ_0037094 | 2042 | 10 | 0.98% |
| hsa_circ_0041333 | 2042 | 10 | 0.98% |
| hsa_circ_0060303 | 2042 | 10 | 0.98% |
| hsa_circ_0128637 | 2042 | 10 | 0.98% |
| hsa_circ_0138030 | 2042 | 10 | 0.98% |
| hsa_circ_0106495 | 2859 | 14 | 0.98% |
| hsa_circ_0046171 | 2655 | 13 | 0.98% |
| hsa_circ_0029992 | 2451 | 12 | 0.98% |
| hsa_circ_0031613 | 2451 | 12 | 0.98% |
| hsa_circ_0034851 | 2451 | 12 | 0.98% |
| hsa_circ_0099401 | 2451 | 12 | 0.98% |
| hsa_circ_0010814 | 2247 | 11 | 0.98% |
| hsa_circ_0018466 | 2247 | 11 | 0.98% |
| hsa_circ_0068544 | 2247 | 11 | 0.98% |
| hsa_circ_0077065 | 2247 | 11 | 0.98% |
| hsa_circ_0101165 | 2247 | 11 | 0.98% |
| hsa_circ_0100888 | 2860 | 14 | 0.98% |
| hsa_circ_0035149 | 2043 | 10 | 0.98% |
| hsa_circ_0053244 | 2043 | 10 | 0.98% |
| hsa_circ_0068942 | 2043 | 10 | 0.98% |
| hsa_circ_0081256 | 2043 | 10 | 0.98% |
| hsa_circ_0097654 | 2043 | 10 | 0.98% |
| hsa_circ_0069274 | 2452 | 12 | 0.98% |
| hsa_circ_0117985 | 2452 | 12 | 0.98% |
| hsa_circ_0037989 | 2248 | 11 | 0.98% |
| hsa_circ_0052016 | 2248 | 11 | 0.98% |
| hsa_circ_0060651 | 2248 | 11 | 0.98% |
| hsa_circ_0063648 | 2248 | 11 | 0.98% |
| hsa_circ_0065154 | 2248 | 11 | 0.98% |
| hsa_circ_0071805 | 2248 | 11 | 0.98% |
| hsa_circ_0075099 | 2248 | 11 | 0.98% |
| hsa_circ_0087388 | 2248 | 11 | 0.98% |
| hsa_circ_0092870 | 2248 | 11 | 0.98% |
| hsa_circ_0012365 | 2044 | 10 | 0.98% |
| hsa_circ_0050090 | 2044 | 10 | 0.98% |

| | | | |
|---|---|---|---|
| hsa_circ_0085251 | 2044 | 10 | 0.98% |
| hsa_circ_0085098 | 2453 | 12 | 0.98% |
| hsa_circ_0092003 | 2453 | 12 | 0.98% |
| hsa_circ_0050501 | 2862 | 14 | 0.98% |
| hsa_circ_0062356 | 2862 | 14 | 0.98% |
| hsa_circ_0050689 | 3271 | 16 | 0.98% |
| hsa_circ_0102520 | 6747 | 33 | 0.98% |
| hsa_circ_0015311 | 2249 | 11 | 0.98% |
| hsa_circ_0059066 | 2249 | 11 | 0.98% |
| hsa_circ_0063918 | 2249 | 11 | 0.98% |
| hsa_circ_0051678 | 2454 | 12 | 0.98% |
| hsa_circ_0076642 | 2454 | 12 | 0.98% |
| hsa_circ_0097689 | 2454 | 12 | 0.98% |
| hsa_circ_0010341 | 2045 | 10 | 0.98% |
| hsa_circ_0022633 | 2045 | 10 | 0.98% |
| hsa_circ_0024028 | 2045 | 10 | 0.98% |
| hsa_circ_0058625 | 2045 | 10 | 0.98% |
| hsa_circ_0076632 | 2045 | 10 | 0.98% |
| hsa_circ_0083594 | 2045 | 10 | 0.98% |
| hsa_circ_0137894 | 2045 | 10 | 0.98% |
| hsa_circ_0138484 | 2045 | 10 | 0.98% |
| hsa_circ_0060678 | 2659 | 13 | 0.98% |
| hsa_circ_0022869 | 2250 | 11 | 0.98% |
| hsa_circ_0044580 | 2250 | 11 | 0.98% |
| hsa_circ_0087558 | 2250 | 11 | 0.98% |
| hsa_circ_0106120 | 2250 | 11 | 0.98% |
| hsa_circ_0124601 | 2250 | 11 | 0.98% |
| hsa_circ_0021878 | 2455 | 12 | 0.98% |
| hsa_circ_0098188 | 2455 | 12 | 0.98% |
| hsa_circ_0100499 | 2455 | 12 | 0.98% |
| hsa_circ_0106085 | 3478 | 17 | 0.98% |
| hsa_circ_0068752 | 5115 | 25 | 0.98% |
| hsa_circ_0127909 | 4092 | 20 | 0.98% |
| hsa_circ_0011556 | 2046 | 10 | 0.98% |
| hsa_circ_0013716 | 2046 | 10 | 0.98% |
| hsa_circ_0091018 | 2046 | 10 | 0.98% |
| hsa_circ_0103946 | 2046 | 10 | 0.98% |
| hsa_circ_0122391 | 2046 | 10 | 0.98% |
| hsa_circ_0131788 | 2046 | 10 | 0.98% |
| hsa_circ_0014330 | 2251 | 11 | 0.98% |
| hsa_circ_0043942 | 2251 | 11 | 0.98% |
| hsa_circ_0050952 | 2251 | 11 | 0.98% |
| hsa_circ_0051008 | 2251 | 11 | 0.98% |
| hsa_circ_0062866 | 2251 | 11 | 0.98% |
| hsa_circ_0067373 | 2251 | 11 | 0.98% |
| hsa_circ_0100912 | 2251 | 11 | 0.98% |
| hsa_circ_0023148 | 2456 | 12 | 0.98% |
| hsa_circ_0079671 | 2456 | 12 | 0.98% |
| hsa_circ_0081616 | 2456 | 12 | 0.98% |
| hsa_circ_0118409 | 2456 | 12 | 0.98% |

| | | | |
|---|---|---|---|
| hsa_circ_0066765 | 2661 | 13 | 0.98% |
| hsa_circ_0050066 | 2047 | 10 | 0.98% |
| hsa_circ_0060884 | 2047 | 10 | 0.98% |
| hsa_circ_0090253 | 2047 | 10 | 0.98% |
| hsa_circ_0118241 | 2047 | 10 | 0.98% |
| hsa_circ_0010786 | 4913 | 24 | 0.98% |
| hsa_circ_0024887 | 2252 | 11 | 0.98% |
| hsa_circ_0033357 | 2252 | 11 | 0.98% |
| hsa_circ_0034649 | 2252 | 11 | 0.98% |
| hsa_circ_0035321 | 2252 | 11 | 0.98% |
| hsa_circ_0058555 | 2252 | 11 | 0.98% |
| hsa_circ_0063109 | 2252 | 11 | 0.98% |
| hsa_circ_0013303 | 2457 | 12 | 0.98% |
| hsa_circ_0010592 | 2662 | 13 | 0.98% |
| hsa_circ_0109917 | 6348 | 31 | 0.98% |
| hsa_circ_0061624 | 2048 | 10 | 0.98% |
| hsa_circ_0083981 | 2048 | 10 | 0.98% |
| hsa_circ_0086289 | 2048 | 10 | 0.98% |
| hsa_circ_0089621 | 2048 | 10 | 0.98% |
| hsa_circ_0088100 | 2253 | 11 | 0.98% |
| hsa_circ_0088976 | 2253 | 11 | 0.98% |
| hsa_circ_0061423 | 2458 | 12 | 0.98% |
| hsa_circ_0109535 | 5121 | 25 | 0.98% |
| hsa_circ_0020345 | 2663 | 13 | 0.98% |
| hsa_circ_0099450 | 2663 | 13 | 0.98% |
| hsa_circ_0050101 | 6966 | 34 | 0.98% |
| hsa_circ_0036110 | 2049 | 10 | 0.98% |
| hsa_circ_0056091 | 2049 | 10 | 0.98% |
| hsa_circ_0085739 | 2049 | 10 | 0.98% |
| hsa_circ_0135287 | 2049 | 10 | 0.98% |
| hsa_circ_0028520 | 2254 | 11 | 0.98% |
| hsa_circ_0030073 | 2254 | 11 | 0.98% |
| hsa_circ_0064015 | 2254 | 11 | 0.98% |
| hsa_circ_0069784 | 2254 | 11 | 0.98% |
| hsa_circ_0135181 | 2459 | 12 | 0.98% |
| hsa_circ_0049519 | 2664 | 13 | 0.98% |
| hsa_circ_0116665 | 4510 | 22 | 0.98% |
| hsa_circ_0083413 | 2665 | 13 | 0.98% |
| hsa_circ_0019843 | 2255 | 11 | 0.98% |
| hsa_circ_0041884 | 2255 | 11 | 0.98% |
| hsa_circ_0103486 | 2255 | 11 | 0.98% |
| hsa_circ_0088944 | 2050 | 10 | 0.98% |
| hsa_circ_0135008 | 2050 | 10 | 0.98% |
| hsa_circ_0016639 | 2461 | 12 | 0.98% |
| hsa_circ_0054540 | 2461 | 12 | 0.98% |
| hsa_circ_0059111 | 2256 | 11 | 0.98% |
| hsa_circ_0097688 | 2256 | 11 | 0.98% |
| hsa_circ_0028829 | 2051 | 10 | 0.98% |
| hsa_circ_0044632 | 2051 | 10 | 0.98% |
| hsa_circ_0062751 | 2051 | 10 | 0.98% |

| | | | |
|---|---|---|---|
| hsa_circ_0072062 | 2051 | 10 | 0.98% |
| hsa_circ_0089121 | 2051 | 10 | 0.98% |
| hsa_circ_0123405 | 2051 | 10 | 0.98% |
| hsa_circ_0052688 | 3077 | 15 | 0.97% |
| hsa_circ_0011594 | 2667 | 13 | 0.97% |
| hsa_circ_0014297 | 2052 | 10 | 0.97% |
| hsa_circ_0019014 | 2052 | 10 | 0.97% |
| hsa_circ_0030418 | 2052 | 10 | 0.97% |
| hsa_circ_0047836 | 2052 | 10 | 0.97% |
| hsa_circ_0056639 | 2052 | 10 | 0.97% |
| hsa_circ_0057888 | 2052 | 10 | 0.97% |
| hsa_circ_0062364 | 2052 | 10 | 0.97% |
| hsa_circ_0070875 | 2052 | 10 | 0.97% |
| hsa_circ_0101067 | 2052 | 10 | 0.97% |
| hsa_circ_0009282 | 2668 | 13 | 0.97% |
| hsa_circ_0139420 | 2668 | 13 | 0.97% |
| hsa_circ_0104011 | 3284 | 16 | 0.97% |
| hsa_circ_0048541 | 2463 | 12 | 0.97% |
| hsa_circ_0104833 | 5542 | 27 | 0.97% |
| hsa_circ_0015109 | 2258 | 11 | 0.97% |
| hsa_circ_0055105 | 2258 | 11 | 0.97% |
| hsa_circ_0069293 | 2258 | 11 | 0.97% |
| hsa_circ_0132982 | 2258 | 11 | 0.97% |
| hsa_circ_0028224 | 2053 | 10 | 0.97% |
| hsa_circ_0038521 | 2053 | 10 | 0.97% |
| hsa_circ_0079260 | 2053 | 10 | 0.97% |
| hsa_circ_0120776 | 2053 | 10 | 0.97% |
| hsa_circ_0125850 | 2053 | 10 | 0.97% |
| hsa_circ_0054976 | 3285 | 16 | 0.97% |
| hsa_circ_0053834 | 2464 | 12 | 0.97% |
| hsa_circ_0104547 | 2464 | 12 | 0.97% |
| hsa_circ_0132780 | 2464 | 12 | 0.97% |
| hsa_circ_0006519 | 4107 | 20 | 0.97% |
| hsa_circ_0018105 | 2875 | 14 | 0.97% |
| hsa_circ_0029594 | 2259 | 11 | 0.97% |
| hsa_circ_0041078 | 2259 | 11 | 0.97% |
| hsa_circ_0051592 | 2259 | 11 | 0.97% |
| hsa_circ_0061944 | 2259 | 11 | 0.97% |
| hsa_circ_0089060 | 2259 | 11 | 0.97% |
| hsa_circ_0135030 | 2259 | 11 | 0.97% |
| hsa_circ_0030102 | 2054 | 10 | 0.97% |
| hsa_circ_0060836 | 2054 | 10 | 0.97% |
| hsa_circ_0087403 | 2054 | 10 | 0.97% |
| hsa_circ_0110416 | 2054 | 10 | 0.97% |
| hsa_circ_0117378 | 2054 | 10 | 0.97% |
| hsa_circ_0039493 | 2465 | 12 | 0.97% |
| hsa_circ_0004939 | 5958 | 29 | 0.97% |
| hsa_circ_0021691 | 2260 | 11 | 0.97% |
| hsa_circ_0029650 | 2260 | 11 | 0.97% |
| hsa_circ_0071132 | 2260 | 11 | 0.97% |

| | | | |
|---|---|---|---|
| hsa_circ_0086501 | 2260 | 11 | 0.97% |
| hsa_circ_0118023 | 2260 | 11 | 0.97% |
| hsa_circ_0046734 | 2671 | 13 | 0.97% |
| hsa_circ_0018520 | 2466 | 12 | 0.97% |
| hsa_circ_0031507 | 2466 | 12 | 0.97% |
| hsa_circ_0039196 | 2466 | 12 | 0.97% |
| hsa_circ_0043659 | 2466 | 12 | 0.97% |
| hsa_circ_0044642 | 2466 | 12 | 0.97% |
| hsa_circ_0054828 | 2466 | 12 | 0.97% |
| hsa_circ_0036385 | 2055 | 10 | 0.97% |
| hsa_circ_0042639 | 2055 | 10 | 0.97% |
| hsa_circ_0090224 | 2055 | 10 | 0.97% |
| hsa_circ_0095781 | 2055 | 10 | 0.97% |
| hsa_circ_0139624 | 2055 | 10 | 0.97% |
| hsa_circ_0008038 | 2672 | 13 | 0.97% |
| hsa_circ_0014030 | 2261 | 11 | 0.97% |
| hsa_circ_0049553 | 2261 | 11 | 0.97% |
| hsa_circ_0087788 | 2878 | 14 | 0.97% |
| hsa_circ_0076641 | 2467 | 12 | 0.97% |
| hsa_circ_0078531 | 2467 | 12 | 0.97% |
| hsa_circ_0095406 | 2467 | 12 | 0.97% |
| hsa_circ_0097959 | 2467 | 12 | 0.97% |
| hsa_circ_0132301 | 2467 | 12 | 0.97% |
| hsa_circ_0102473 | 6579 | 32 | 0.97% |
| hsa_circ_0023722 | 2056 | 10 | 0.97% |
| hsa_circ_0051100 | 2056 | 10 | 0.97% |
| hsa_circ_0056079 | 2056 | 10 | 0.97% |
| hsa_circ_0075038 | 2056 | 10 | 0.97% |
| hsa_circ_0121304 | 2056 | 10 | 0.97% |
| hsa_circ_0052629 | 2673 | 13 | 0.97% |
| hsa_circ_0075328 | 2673 | 13 | 0.97% |
| hsa_circ_0041327 | 2262 | 11 | 0.97% |
| hsa_circ_0070857 | 2262 | 11 | 0.97% |
| hsa_circ_0125932 | 2262 | 11 | 0.97% |
| hsa_circ_0136988 | 2262 | 11 | 0.97% |
| hsa_circ_0136948 | 7404 | 36 | 0.97% |
| hsa_circ_0041574 | 2468 | 12 | 0.97% |
| hsa_circ_0117778 | 2468 | 12 | 0.97% |
| hsa_circ_0055332 | 2674 | 13 | 0.97% |
| hsa_circ_0021855 | 2057 | 10 | 0.97% |
| hsa_circ_0036760 | 2057 | 10 | 0.97% |
| hsa_circ_0079662 | 2057 | 10 | 0.97% |
| hsa_circ_0116050 | 2057 | 10 | 0.97% |
| hsa_circ_0032272 | 3703 | 18 | 0.97% |
| hsa_circ_0013206 | 2263 | 11 | 0.97% |
| hsa_circ_0018121 | 2263 | 11 | 0.97% |
| hsa_circ_0019816 | 2263 | 11 | 0.97% |
| hsa_circ_0024187 | 2263 | 11 | 0.97% |
| hsa_circ_0053150 | 2263 | 11 | 0.97% |
| hsa_circ_0024026 | 2469 | 12 | 0.97% |

| | | | |
|---|---|---|---|
| hsa_circ_0053946 | 2469 | 12 | 0.97% |
| hsa_circ_0086613 | 2675 | 13 | 0.97% |
| hsa_circ_0020425 | 2058 | 10 | 0.97% |
| hsa_circ_0048719 | 2058 | 10 | 0.97% |
| hsa_circ_0136568 | 2058 | 10 | 0.97% |
| hsa_circ_0010434 | 2264 | 11 | 0.97% |
| hsa_circ_0015921 | 2264 | 11 | 0.97% |
| hsa_circ_0022503 | 2264 | 11 | 0.97% |
| hsa_circ_0035318 | 2264 | 11 | 0.97% |
| hsa_circ_0070587 | 2264 | 11 | 0.97% |
| hsa_circ_0073374 | 2264 | 11 | 0.97% |
| hsa_circ_0085815 | 2264 | 11 | 0.97% |
| hsa_circ_0134700 | 2264 | 11 | 0.97% |
| hsa_circ_0041775 | 2470 | 12 | 0.97% |
| hsa_circ_0074092 | 2470 | 12 | 0.97% |
| hsa_circ_0132647 | 2882 | 14 | 0.97% |
| hsa_circ_0106632 | 3088 | 15 | 0.97% |
| hsa_circ_0013976 | 2059 | 10 | 0.97% |
| hsa_circ_0055526 | 2059 | 10 | 0.97% |
| hsa_circ_0070880 | 2059 | 10 | 0.97% |
| hsa_circ_0086073 | 2059 | 10 | 0.97% |
| hsa_circ_0071255 | 2265 | 11 | 0.97% |
| hsa_circ_0092426 | 2265 | 11 | 0.97% |
| hsa_circ_0093375 | 2265 | 11 | 0.97% |
| hsa_circ_0117444 | 2265 | 11 | 0.97% |
| hsa_circ_0123364 | 2265 | 11 | 0.97% |
| hsa_circ_0065602 | 2471 | 12 | 0.97% |
| hsa_circ_0082338 | 2471 | 12 | 0.97% |
| hsa_circ_0088736 | 2471 | 12 | 0.97% |
| hsa_circ_0060875 | 2677 | 13 | 0.97% |
| hsa_circ_0089783 | 10505 | 51 | 0.97% |
| hsa_circ_0103365 | 4326 | 21 | 0.97% |
| hsa_circ_0123409 | 2678 | 13 | 0.97% |
| hsa_circ_0139890 | 2678 | 13 | 0.97% |
| hsa_circ_0010225 | 2472 | 12 | 0.97% |
| hsa_circ_0082720 | 2472 | 12 | 0.97% |
| hsa_circ_0089443 | 2472 | 12 | 0.97% |
| hsa_circ_0135301 | 2472 | 12 | 0.97% |
| hsa_circ_0071520 | 2266 | 11 | 0.97% |
| hsa_circ_0023102 | 2060 | 10 | 0.97% |
| hsa_circ_0026033 | 2060 | 10 | 0.97% |
| hsa_circ_0032315 | 2060 | 10 | 0.97% |
| hsa_circ_0035518 | 2060 | 10 | 0.97% |
| hsa_circ_0057570 | 2060 | 10 | 0.97% |
| hsa_circ_0081254 | 2060 | 10 | 0.97% |
| hsa_circ_0130621 | 2060 | 10 | 0.97% |
| hsa_circ_0055813 | 126073 | 612 | 0.97% |
| hsa_circ_0006616 | 11127 | 54 | 0.97% |
| hsa_circ_0131945 | 6800 | 33 | 0.97% |
| hsa_circ_0035925 | 2679 | 13 | 0.97% |

| | | | |
|---|---|---|---|
| hsa_circ_0018092 | 2061 | 10 | 0.97% |
| hsa_circ_0023673 | 2061 | 10 | 0.97% |
| hsa_circ_0046111 | 2061 | 10 | 0.97% |
| hsa_circ_0081281 | 2061 | 10 | 0.97% |
| hsa_circ_0128831 | 4329 | 21 | 0.97% |
| hsa_circ_0115692 | 6391 | 31 | 0.97% |
| hsa_circ_0006162 | 4948 | 24 | 0.97% |
| hsa_circ_0044594 | 2474 | 12 | 0.97% |
| hsa_circ_0058570 | 2474 | 12 | 0.97% |
| hsa_circ_0026735 | 2268 | 11 | 0.97% |
| hsa_circ_0056067 | 2268 | 11 | 0.97% |
| hsa_circ_0112412 | 2268 | 11 | 0.97% |
| hsa_circ_0036486 | 2062 | 10 | 0.97% |
| hsa_circ_0069483 | 2062 | 10 | 0.97% |
| hsa_circ_0122685 | 2062 | 10 | 0.97% |
| hsa_circ_0138933 | 2062 | 10 | 0.97% |
| hsa_circ_0092160 | 2681 | 13 | 0.97% |
| hsa_circ_0017504 | 2269 | 11 | 0.97% |
| hsa_circ_0053785 | 2269 | 11 | 0.97% |
| hsa_circ_0063945 | 2269 | 11 | 0.97% |
| hsa_circ_0023482 | 2063 | 10 | 0.97% |
| hsa_circ_0034552 | 2063 | 10 | 0.97% |
| hsa_circ_0047677 | 2063 | 10 | 0.97% |
| hsa_circ_0069317 | 2063 | 10 | 0.97% |
| hsa_circ_0069673 | 2063 | 10 | 0.97% |
| hsa_circ_0077353 | 2063 | 10 | 0.97% |
| hsa_circ_0106406 | 2063 | 10 | 0.97% |
| hsa_circ_0129425 | 2063 | 10 | 0.97% |
| hsa_circ_0052306 | 2270 | 11 | 0.97% |
| hsa_circ_0058412 | 2270 | 11 | 0.97% |
| hsa_circ_0090560 | 2270 | 11 | 0.97% |
| hsa_circ_0027725 | 2064 | 10 | 0.97% |
| hsa_circ_0029885 | 2064 | 10 | 0.97% |
| hsa_circ_0034457 | 2064 | 10 | 0.97% |
| hsa_circ_0048538 | 2064 | 10 | 0.97% |
| hsa_circ_0050682 | 2064 | 10 | 0.97% |
| hsa_circ_0088225 | 2064 | 10 | 0.97% |
| hsa_circ_0104166 | 2064 | 10 | 0.97% |
| hsa_circ_0088278 | 2477 | 12 | 0.97% |
| hsa_circ_0097008 | 2477 | 12 | 0.97% |
| hsa_circ_0022349 | 2271 | 11 | 0.97% |
| hsa_circ_0068698 | 2271 | 11 | 0.97% |
| hsa_circ_0071902 | 2271 | 11 | 0.97% |
| hsa_circ_0077131 | 2271 | 11 | 0.97% |
| hsa_circ_0090150 | 2478 | 12 | 0.97% |
| hsa_circ_0090163 | 2478 | 12 | 0.97% |
| hsa_circ_0103118 | 2478 | 12 | 0.97% |
| hsa_circ_0110639 | 2478 | 12 | 0.97% |
| hsa_circ_0111668 | 2478 | 12 | 0.97% |
| hsa_circ_0132452 | 2478 | 12 | 0.97% |

| | | | |
|---|---|---|---|
| hsa_circ_0012376 | 2065 | 10 | 0.97% |
| hsa_circ_0033458 | 2065 | 10 | 0.97% |
| hsa_circ_0034864 | 2065 | 10 | 0.97% |
| hsa_circ_0043127 | 2065 | 10 | 0.97% |
| hsa_circ_0047467 | 2065 | 10 | 0.97% |
| hsa_circ_0071338 | 2065 | 10 | 0.97% |
| hsa_circ_0041663 | 5989 | 29 | 0.97% |
| hsa_circ_0021103 | 2272 | 11 | 0.97% |
| hsa_circ_0030668 | 2272 | 11 | 0.97% |
| hsa_circ_0040169 | 2272 | 11 | 0.97% |
| hsa_circ_0073743 | 2272 | 11 | 0.97% |
| hsa_circ_0133682 | 2272 | 11 | 0.97% |
| hsa_circ_0136260 | 6610 | 32 | 0.97% |
| hsa_circ_0030860 | 2892 | 14 | 0.97% |
| hsa_circ_0010567 | 2479 | 12 | 0.97% |
| hsa_circ_0060325 | 2479 | 12 | 0.97% |
| hsa_circ_0091433 | 2479 | 12 | 0.97% |
| hsa_circ_0034317 | 2066 | 10 | 0.97% |
| hsa_circ_0036340 | 2066 | 10 | 0.97% |
| hsa_circ_0047827 | 3306 | 16 | 0.97% |
| hsa_circ_0027631 | 2273 | 11 | 0.97% |
| hsa_circ_0054967 | 2273 | 11 | 0.97% |
| hsa_circ_0131262 | 2273 | 11 | 0.97% |
| hsa_circ_0132551 | 13640 | 66 | 0.97% |
| hsa_circ_0132496 | 2480 | 12 | 0.97% |
| hsa_circ_0034360 | 2687 | 13 | 0.97% |
| hsa_circ_0087836 | 2687 | 13 | 0.97% |
| hsa_circ_0000877 | 2067 | 10 | 0.97% |
| hsa_circ_0056641 | 2067 | 10 | 0.97% |
| hsa_circ_0106596 | 2067 | 10 | 0.97% |
| hsa_circ_0010162 | 2274 | 11 | 0.97% |
| hsa_circ_0061162 | 2274 | 11 | 0.97% |
| hsa_circ_0065376 | 2274 | 11 | 0.97% |
| hsa_circ_0134398 | 2481 | 12 | 0.97% |
| hsa_circ_0068579 | 2688 | 13 | 0.97% |
| hsa_circ_0125873 | 2688 | 13 | 0.97% |
| hsa_circ_0015071 | 2068 | 10 | 0.97% |
| hsa_circ_0042426 | 2068 | 10 | 0.97% |
| hsa_circ_0052934 | 2068 | 10 | 0.97% |
| hsa_circ_0054356 | 2068 | 10 | 0.97% |
| hsa_circ_0059255 | 2068 | 10 | 0.97% |
| hsa_circ_0067380 | 2068 | 10 | 0.97% |
| hsa_circ_0078554 | 2068 | 10 | 0.97% |
| hsa_circ_0082374 | 2068 | 10 | 0.97% |
| hsa_circ_0087114 | 2068 | 10 | 0.97% |
| hsa_circ_0031348 | 2275 | 11 | 0.97% |
| hsa_circ_0065526 | 2275 | 11 | 0.97% |
| hsa_circ_0089885 | 2275 | 11 | 0.97% |
| hsa_circ_0100087 | 2275 | 11 | 0.97% |
| hsa_circ_0095506 | 6205 | 30 | 0.97% |

| | | | |
|---|---|---|---|
| hsa_circ_0033044 | 2069 | 10 | 0.97% |
| hsa_circ_0117607 | 2069 | 10 | 0.97% |
| hsa_circ_0118268 | 2069 | 10 | 0.97% |
| hsa_circ_0076355 | 2276 | 11 | 0.97% |
| hsa_circ_0089422 | 2276 | 11 | 0.97% |
| hsa_circ_0131063 | 2276 | 11 | 0.97% |
| hsa_circ_0089177 | 2483 | 12 | 0.97% |
| hsa_circ_0079657 | 2690 | 13 | 0.97% |
| hsa_circ_0048383 | 2897 | 14 | 0.97% |
| hsa_circ_0041430 | 2691 | 13 | 0.97% |
| hsa_circ_0029646 | 2484 | 12 | 0.97% |
| hsa_circ_0067553 | 2484 | 12 | 0.97% |
| hsa_circ_0123987 | 2484 | 12 | 0.97% |
| hsa_circ_0023174 | 2277 | 11 | 0.97% |
| hsa_circ_0054589 | 2277 | 11 | 0.97% |
| hsa_circ_0057400 | 2277 | 11 | 0.97% |
| hsa_circ_0064209 | 2277 | 11 | 0.97% |
| hsa_circ_0114265 | 2277 | 11 | 0.97% |
| hsa_circ_0015622 | 2070 | 10 | 0.97% |
| hsa_circ_0020470 | 2070 | 10 | 0.97% |
| hsa_circ_0052302 | 2070 | 10 | 0.97% |
| hsa_circ_0057650 | 2070 | 10 | 0.97% |
| hsa_circ_0075122 | 2070 | 10 | 0.97% |
| hsa_circ_0094524 | 2485 | 12 | 0.97% |
| hsa_circ_0098679 | 2278 | 11 | 0.97% |
| hsa_circ_0120559 | 2278 | 11 | 0.97% |
| hsa_circ_0014573 | 2071 | 10 | 0.97% |
| hsa_circ_0032312 | 2071 | 10 | 0.97% |
| hsa_circ_0042919 | 2071 | 10 | 0.97% |
| hsa_circ_0074319 | 2071 | 10 | 0.97% |
| hsa_circ_0077864 | 2071 | 10 | 0.97% |
| hsa_circ_0095173 | 2071 | 10 | 0.97% |
| hsa_circ_0138568 | 4972 | 24 | 0.97% |
| hsa_circ_0058164 | 2486 | 12 | 0.97% |
| hsa_circ_0080426 | 2486 | 12 | 0.97% |
| hsa_circ_0096209 | 2486 | 12 | 0.97% |
| hsa_circ_0084099 | 2279 | 11 | 0.97% |
| hsa_circ_0011339 | 2072 | 10 | 0.97% |
| hsa_circ_0041503 | 2072 | 10 | 0.97% |
| hsa_circ_0114522 | 2072 | 10 | 0.97% |
| hsa_circ_0116861 | 2072 | 10 | 0.97% |
| hsa_circ_0134494 | 2072 | 10 | 0.97% |
| hsa_circ_0018420 | 2487 | 12 | 0.97% |
| hsa_circ_0018548 | 2487 | 12 | 0.97% |
| hsa_circ_0071543 | 9120 | 44 | 0.96% |
| hsa_circ_0016804 | 2280 | 11 | 0.96% |
| hsa_circ_0019352 | 2280 | 11 | 0.96% |
| hsa_circ_0033107 | 2280 | 11 | 0.96% |
| hsa_circ_0064403 | 2280 | 11 | 0.96% |
| hsa_circ_0010468 | 2073 | 10 | 0.96% |

| | | | |
|---|---|---|---|
| hsa_circ_0063545 | 2073 | 10 | 0.96% |
| hsa_circ_0067292 | 2073 | 10 | 0.96% |
| hsa_circ_0069313 | 2073 | 10 | 0.96% |
| hsa_circ_0073851 | 2073 | 10 | 0.96% |
| hsa_circ_0117464 | 2073 | 10 | 0.96% |
| hsa_circ_0044337 | 2695 | 13 | 0.96% |
| hsa_circ_0055784 | 2695 | 13 | 0.96% |
| hsa_circ_0004669 | 2488 | 12 | 0.96% |
| hsa_circ_0048707 | 2488 | 12 | 0.96% |
| hsa_circ_0057988 | 2488 | 12 | 0.96% |
| hsa_circ_0030861 | 2281 | 11 | 0.96% |
| hsa_circ_0047679 | 2281 | 11 | 0.96% |
| hsa_circ_0086606 | 2281 | 11 | 0.96% |
| hsa_circ_0091969 | 2281 | 11 | 0.96% |
| hsa_circ_0031677 | 2074 | 10 | 0.96% |
| hsa_circ_0073514 | 2074 | 10 | 0.96% |
| hsa_circ_0075123 | 2074 | 10 | 0.96% |
| hsa_circ_0104199 | 2074 | 10 | 0.96% |
| hsa_circ_0090877 | 2489 | 12 | 0.96% |
| hsa_circ_0024036 | 2282 | 11 | 0.96% |
| hsa_circ_0026399 | 2282 | 11 | 0.96% |
| hsa_circ_0049418 | 2282 | 11 | 0.96% |
| hsa_circ_0092893 | 2282 | 11 | 0.96% |
| hsa_circ_0100907 | 2282 | 11 | 0.96% |
| hsa_circ_0020635 | 2697 | 13 | 0.96% |
| hsa_circ_0060666 | 2697 | 13 | 0.96% |
| hsa_circ_0016557 | 2490 | 12 | 0.96% |
| hsa_circ_0058752 | 2490 | 12 | 0.96% |
| hsa_circ_0081773 | 2490 | 12 | 0.96% |
| hsa_circ_0041995 | 2075 | 10 | 0.96% |
| hsa_circ_0056584 | 2075 | 10 | 0.96% |
| hsa_circ_0071956 | 2075 | 10 | 0.96% |
| hsa_circ_0083745 | 2075 | 10 | 0.96% |
| hsa_circ_0121557 | 2075 | 10 | 0.96% |
| hsa_circ_0060695 | 2698 | 13 | 0.96% |
| hsa_circ_0086399 | 2698 | 13 | 0.96% |
| hsa_circ_0139340 | 2698 | 13 | 0.96% |
| hsa_circ_0046560 | 4981 | 24 | 0.96% |
| hsa_circ_0036880 | 2283 | 11 | 0.96% |
| hsa_circ_0075837 | 2283 | 11 | 0.96% |
| hsa_circ_0076887 | 2283 | 11 | 0.96% |
| hsa_circ_0085915 | 2283 | 11 | 0.96% |
| hsa_circ_0139381 | 2283 | 11 | 0.96% |
| hsa_circ_0101014 | 2906 | 14 | 0.96% |
| hsa_circ_0134254 | 2906 | 14 | 0.96% |
| hsa_circ_0051882 | 2491 | 12 | 0.96% |
| hsa_circ_0029665 | 2076 | 10 | 0.96% |
| hsa_circ_0122590 | 2076 | 10 | 0.96% |
| hsa_circ_0011308 | 2699 | 13 | 0.96% |
| hsa_circ_0102644 | 4360 | 21 | 0.96% |

| | | | |
|---|---|---|---|
| hsa_circ_0041326 | 2284 | 11 | 0.96% |
| hsa_circ_0048540 | 2284 | 11 | 0.96% |
| hsa_circ_0026057 | 2492 | 12 | 0.96% |
| hsa_circ_0082053 | 2700 | 13 | 0.96% |
| hsa_circ_0007132 | 2077 | 10 | 0.96% |
| hsa_circ_0030378 | 2077 | 10 | 0.96% |
| hsa_circ_0042530 | 2077 | 10 | 0.96% |
| hsa_circ_0045702 | 2077 | 10 | 0.96% |
| hsa_circ_0103831 | 2077 | 10 | 0.96% |
| hsa_circ_0106385 | 2077 | 10 | 0.96% |
| hsa_circ_0128843 | 2908 | 14 | 0.96% |
| hsa_circ_0137139 | 4570 | 22 | 0.96% |
| hsa_circ_0049456 | 2285 | 11 | 0.96% |
| hsa_circ_0051067 | 2285 | 11 | 0.96% |
| hsa_circ_0055279 | 2285 | 11 | 0.96% |
| hsa_circ_0083934 | 2285 | 11 | 0.96% |
| hsa_circ_0106402 | 2285 | 11 | 0.96% |
| hsa_circ_0044483 | 2493 | 12 | 0.96% |
| hsa_circ_0051854 | 2493 | 12 | 0.96% |
| hsa_circ_0089999 | 2493 | 12 | 0.96% |
| hsa_circ_0090676 | 2493 | 12 | 0.96% |
| hsa_circ_0103868 | 2701 | 13 | 0.96% |
| hsa_circ_0021798 | 2078 | 10 | 0.96% |
| hsa_circ_0041741 | 2078 | 10 | 0.96% |
| hsa_circ_0109275 | 2078 | 10 | 0.96% |
| hsa_circ_0035085 | 2286 | 11 | 0.96% |
| hsa_circ_0037860 | 2286 | 11 | 0.96% |
| hsa_circ_0083992 | 2286 | 11 | 0.96% |
| hsa_circ_0053950 | 2494 | 12 | 0.96% |
| hsa_circ_0128126 | 2494 | 12 | 0.96% |
| hsa_circ_0057344 | 3534 | 17 | 0.96% |
| hsa_circ_0020595 | 6029 | 29 | 0.96% |
| hsa_circ_0009813 | 2079 | 10 | 0.96% |
| hsa_circ_0011496 | 2079 | 10 | 0.96% |
| hsa_circ_0024563 | 2079 | 10 | 0.96% |
| hsa_circ_0041900 | 2079 | 10 | 0.96% |
| hsa_circ_0051559 | 2079 | 10 | 0.96% |
| hsa_circ_0058242 | 2079 | 10 | 0.96% |
| hsa_circ_0074316 | 2079 | 10 | 0.96% |
| hsa_circ_0115755 | 2079 | 10 | 0.96% |
| hsa_circ_0023300 | 2287 | 11 | 0.96% |
| hsa_circ_0042720 | 2287 | 11 | 0.96% |
| hsa_circ_0042830 | 2287 | 11 | 0.96% |
| hsa_circ_0076006 | 2287 | 11 | 0.96% |
| hsa_circ_0076068 | 2287 | 11 | 0.96% |
| hsa_circ_0102026 | 2287 | 11 | 0.96% |
| hsa_circ_0061204 | 2495 | 12 | 0.96% |
| hsa_circ_0123145 | 2495 | 12 | 0.96% |
| hsa_circ_0072211 | 2703 | 13 | 0.96% |
| hsa_circ_0007545 | 2911 | 14 | 0.96% |

| | | | |
|---|---|---|---|
| hsa_circ_0094827 | 6863 | 33 | 0.96% |
| hsa_circ_0062416 | 2912 | 14 | 0.96% |
| hsa_circ_0046172 | 2704 | 13 | 0.96% |
| hsa_circ_0053374 | 2704 | 13 | 0.96% |
| hsa_circ_0060737 | 2496 | 12 | 0.96% |
| hsa_circ_0027105 | 2288 | 11 | 0.96% |
| hsa_circ_0090612 | 2288 | 11 | 0.96% |
| hsa_circ_0100417 | 2288 | 11 | 0.96% |
| hsa_circ_0018033 | 2080 | 10 | 0.96% |
| hsa_circ_0053372 | 2080 | 10 | 0.96% |
| hsa_circ_0055616 | 2080 | 10 | 0.96% |
| hsa_circ_0067097 | 2080 | 10 | 0.96% |
| hsa_circ_0071227 | 2080 | 10 | 0.96% |
| hsa_circ_0072039 | 2080 | 10 | 0.96% |
| hsa_circ_0000175 | 4577 | 22 | 0.96% |
| hsa_circ_0052697 | 3121 | 15 | 0.96% |
| hsa_circ_0000495 | 2913 | 14 | 0.96% |
| hsa_circ_0025413 | 2705 | 13 | 0.96% |
| hsa_circ_0010944 | 2289 | 11 | 0.96% |
| hsa_circ_0127722 | 2289 | 11 | 0.96% |
| hsa_circ_0067320 | 2081 | 10 | 0.96% |
| hsa_circ_0097035 | 2081 | 10 | 0.96% |
| hsa_circ_0116508 | 2081 | 10 | 0.96% |
| hsa_circ_0134497 | 2081 | 10 | 0.96% |
| hsa_circ_0051751 | 2914 | 14 | 0.96% |
| hsa_circ_0016366 | 2706 | 13 | 0.96% |
| hsa_circ_0103623 | 2706 | 13 | 0.96% |
| hsa_circ_0022117 | 2498 | 12 | 0.96% |
| hsa_circ_0060646 | 2498 | 12 | 0.96% |
| hsa_circ_0079579 | 2498 | 12 | 0.96% |
| hsa_circ_0009982 | 2082 | 10 | 0.96% |
| hsa_circ_0024684 | 2082 | 10 | 0.96% |
| hsa_circ_0032388 | 2082 | 10 | 0.96% |
| hsa_circ_0040338 | 2082 | 10 | 0.96% |
| hsa_circ_0045793 | 2082 | 10 | 0.96% |
| hsa_circ_0087527 | 2082 | 10 | 0.96% |
| hsa_circ_0095308 | 2082 | 10 | 0.96% |
| hsa_circ_0126342 | 2707 | 13 | 0.96% |
| hsa_circ_0051439 | 2499 | 12 | 0.96% |
| hsa_circ_0031229 | 2291 | 11 | 0.96% |
| hsa_circ_0032385 | 2291 | 11 | 0.96% |
| hsa_circ_0034221 | 2291 | 11 | 0.96% |
| hsa_circ_0073292 | 2291 | 11 | 0.96% |
| hsa_circ_0078831 | 2291 | 11 | 0.96% |
| hsa_circ_0129813 | 2291 | 11 | 0.96% |
| hsa_circ_0139263 | 2291 | 11 | 0.96% |
| hsa_circ_0021218 | 2083 | 10 | 0.96% |
| hsa_circ_0035568 | 2292 | 11 | 0.96% |
| hsa_circ_0053774 | 2292 | 11 | 0.96% |
| hsa_circ_0088895 | 2292 | 11 | 0.96% |

| | | | |
|---|---|---|---|
| hsa_circ_0103210 | 2292 | 11 | 0.96% |
| hsa_circ_0031673 | 2709 | 13 | 0.96% |
| hsa_circ_0055329 | 2709 | 13 | 0.96% |
| hsa_circ_0109534 | 5210 | 25 | 0.96% |
| hsa_circ_0031162 | 2084 | 10 | 0.96% |
| hsa_circ_0054185 | 2084 | 10 | 0.96% |
| hsa_circ_0063909 | 2084 | 10 | 0.96% |
| hsa_circ_0065177 | 2084 | 10 | 0.96% |
| hsa_circ_0072304 | 2084 | 10 | 0.96% |
| hsa_circ_0132994 | 2084 | 10 | 0.96% |
| hsa_circ_0000420 | 3543 | 17 | 0.96% |
| hsa_circ_0030217 | 2501 | 12 | 0.96% |
| hsa_circ_0078814 | 2501 | 12 | 0.96% |
| hsa_circ_0122790 | 5836 | 28 | 0.96% |
| hsa_circ_0021175 | 2293 | 11 | 0.96% |
| hsa_circ_0046362 | 2293 | 11 | 0.96% |
| hsa_circ_0067424 | 2293 | 11 | 0.96% |
| hsa_circ_0128653 | 2293 | 11 | 0.96% |
| hsa_circ_0025352 | 2710 | 13 | 0.96% |
| hsa_circ_0058944 | 2710 | 13 | 0.96% |
| hsa_circ_0135343 | 2919 | 14 | 0.96% |
| hsa_circ_0065580 | 2085 | 10 | 0.96% |
| hsa_circ_0134933 | 2085 | 10 | 0.96% |
| hsa_circ_0120871 | 4379 | 21 | 0.96% |
| hsa_circ_0023366 | 2294 | 11 | 0.96% |
| hsa_circ_0030061 | 2294 | 11 | 0.96% |
| hsa_circ_0039811 | 2294 | 11 | 0.96% |
| hsa_circ_0042576 | 2294 | 11 | 0.96% |
| hsa_circ_0059088 | 2294 | 11 | 0.96% |
| hsa_circ_0138667 | 4589 | 22 | 0.96% |
| hsa_circ_0057616 | 2086 | 10 | 0.96% |
| hsa_circ_0064661 | 2086 | 10 | 0.96% |
| hsa_circ_0069015 | 2086 | 10 | 0.96% |
| hsa_circ_0084971 | 2086 | 10 | 0.96% |
| hsa_circ_0092971 | 2086 | 10 | 0.96% |
| hsa_circ_0098396 | 2086 | 10 | 0.96% |
| hsa_circ_0053709 | 2712 | 13 | 0.96% |
| hsa_circ_0065397 | 2712 | 13 | 0.96% |
| hsa_circ_0036261 | 2295 | 11 | 0.96% |
| hsa_circ_0063739 | 2295 | 11 | 0.96% |
| hsa_circ_0117043 | 2295 | 11 | 0.96% |
| hsa_circ_0044509 | 4799 | 23 | 0.96% |
| hsa_circ_0014808 | 2087 | 10 | 0.96% |
| hsa_circ_0039710 | 2087 | 10 | 0.96% |
| hsa_circ_0054575 | 2087 | 10 | 0.96% |
| hsa_circ_0055728 | 2087 | 10 | 0.96% |
| hsa_circ_0075948 | 2087 | 10 | 0.96% |
| hsa_circ_0077610 | 2087 | 10 | 0.96% |
| hsa_circ_0092178 | 2087 | 10 | 0.96% |
| hsa_circ_0096499 | 2087 | 10 | 0.96% |

| | | | |
|---|---|---|---|
| hsa_circ_0099648 | 2087 | 10 | 0.96% |
| hsa_circ_0102462 | 2087 | 10 | 0.96% |
| hsa_circ_0114065 | 2087 | 10 | 0.96% |
| hsa_circ_0137378 | 2087 | 10 | 0.96% |
| hsa_circ_0111536 | 5844 | 28 | 0.96% |
| hsa_circ_0048078 | 2922 | 14 | 0.96% |
| hsa_circ_0019529 | 2296 | 11 | 0.96% |
| hsa_circ_0040105 | 2296 | 11 | 0.96% |
| hsa_circ_0065254 | 2296 | 11 | 0.96% |
| hsa_circ_0068393 | 2296 | 11 | 0.96% |
| hsa_circ_0016987 | 2505 | 12 | 0.96% |
| hsa_circ_0030823 | 2505 | 12 | 0.96% |
| hsa_circ_0125037 | 2505 | 12 | 0.96% |
| hsa_circ_0074559 | 2714 | 13 | 0.96% |
| hsa_circ_0069361 | 2088 | 10 | 0.96% |
| hsa_circ_0136300 | 2088 | 10 | 0.96% |
| hsa_circ_0139857 | 6891 | 33 | 0.96% |
| hsa_circ_0012414 | 2297 | 11 | 0.96% |
| hsa_circ_0013292 | 2506 | 12 | 0.96% |
| hsa_circ_0017356 | 2715 | 13 | 0.96% |
| hsa_circ_0052182 | 2924 | 14 | 0.96% |
| hsa_circ_0007352 | 2089 | 10 | 0.96% |
| hsa_circ_0071793 | 2089 | 10 | 0.96% |
| hsa_circ_0094171 | 2089 | 10 | 0.96% |
| hsa_circ_0127168 | 2089 | 10 | 0.96% |
| hsa_circ_0138565 | 2089 | 10 | 0.96% |
| hsa_circ_0015621 | 2298 | 11 | 0.96% |
| hsa_circ_0020881 | 2298 | 11 | 0.96% |
| hsa_circ_0043823 | 2298 | 11 | 0.96% |
| hsa_circ_0058129 | 2298 | 11 | 0.96% |
| hsa_circ_0113911 | 2298 | 11 | 0.96% |
| hsa_circ_0131022 | 2298 | 11 | 0.96% |
| hsa_circ_0138071 | 2298 | 11 | 0.96% |
| hsa_circ_0017380 | 2507 | 12 | 0.96% |
| hsa_circ_0109567 | 4598 | 22 | 0.96% |
| hsa_circ_0062340 | 2717 | 13 | 0.96% |
| hsa_circ_0108176 | 2717 | 13 | 0.96% |
| hsa_circ_0037070 | 2508 | 12 | 0.96% |
| hsa_circ_0023699 | 2299 | 11 | 0.96% |
| hsa_circ_0073410 | 2299 | 11 | 0.96% |
| hsa_circ_0111386 | 2299 | 11 | 0.96% |
| hsa_circ_0001629 | 2090 | 10 | 0.96% |
| hsa_circ_0047542 | 2090 | 10 | 0.96% |
| hsa_circ_0052880 | 2090 | 10 | 0.96% |
| hsa_circ_0074957 | 2090 | 10 | 0.96% |
| hsa_circ_0108258 | 2718 | 13 | 0.96% |
| hsa_circ_0130758 | 2300 | 11 | 0.96% |
| hsa_circ_0012605 | 2091 | 10 | 0.96% |
| hsa_circ_0017958 | 2091 | 10 | 0.96% |
| hsa_circ_0042303 | 2091 | 10 | 0.96% |

| | | | |
|---|---|---|---|
| hsa_circ_0042999 | 2091 | 10 | 0.96% |
| hsa_circ_0057838 | 2091 | 10 | 0.96% |
| hsa_circ_0068298 | 2091 | 10 | 0.96% |
| hsa_circ_0096020 | 2091 | 10 | 0.96% |
| hsa_circ_0100498 | 2091 | 10 | 0.96% |
| hsa_circ_0132385 | 2091 | 10 | 0.96% |
| hsa_circ_0129998 | 5437 | 26 | 0.96% |
| hsa_circ_0043480 | 3137 | 15 | 0.96% |
| hsa_circ_0091442 | 2719 | 13 | 0.96% |
| hsa_circ_0039778 | 5020 | 24 | 0.96% |
| hsa_circ_0022758 | 2510 | 12 | 0.96% |
| hsa_circ_0031538 | 2510 | 12 | 0.96% |
| hsa_circ_0044380 | 2301 | 11 | 0.96% |
| hsa_circ_0057667 | 2301 | 11 | 0.96% |
| hsa_circ_0058109 | 2301 | 11 | 0.96% |
| hsa_circ_0063947 | 2301 | 11 | 0.96% |
| hsa_circ_0089365 | 2301 | 11 | 0.96% |
| hsa_circ_0099161 | 2301 | 11 | 0.96% |
| hsa_circ_0127910 | 4184 | 20 | 0.96% |
| hsa_circ_0014168 | 2092 | 10 | 0.96% |
| hsa_circ_0015821 | 2092 | 10 | 0.96% |
| hsa_circ_0071340 | 2092 | 10 | 0.96% |
| hsa_circ_0090176 | 2092 | 10 | 0.96% |
| hsa_circ_0118274 | 2092 | 10 | 0.96% |
| hsa_circ_0051600 | 2511 | 12 | 0.96% |
| hsa_circ_0048972 | 2302 | 11 | 0.96% |
| hsa_circ_0050524 | 2302 | 11 | 0.96% |
| hsa_circ_0068192 | 2302 | 11 | 0.96% |
| hsa_circ_0112548 | 2302 | 11 | 0.96% |
| hsa_circ_0041662 | 6069 | 29 | 0.96% |
| hsa_circ_0107219 | 2093 | 10 | 0.96% |
| hsa_circ_0110004 | 2093 | 10 | 0.96% |
| hsa_circ_0103637 | 2721 | 13 | 0.96% |
| hsa_circ_0123203 | 8582 | 41 | 0.96% |
| hsa_circ_0104626 | 3140 | 15 | 0.96% |
| hsa_circ_0013711 | 2303 | 11 | 0.96% |
| hsa_circ_0080153 | 2303 | 11 | 0.96% |
| hsa_circ_0100986 | 2303 | 11 | 0.96% |
| hsa_circ_0017266 | 5444 | 26 | 0.96% |
| hsa_circ_0047815 | 2722 | 13 | 0.96% |
| hsa_circ_0015828 | 2094 | 10 | 0.96% |
| hsa_circ_0132845 | 2094 | 10 | 0.96% |
| hsa_circ_0027296 | 2513 | 12 | 0.96% |
| hsa_circ_0035165 | 2513 | 12 | 0.96% |
| hsa_circ_0128681 | 9634 | 46 | 0.95% |
| hsa_circ_0132062 | 6912 | 33 | 0.95% |
| hsa_circ_0014884 | 2304 | 11 | 0.95% |
| hsa_circ_0019863 | 2304 | 11 | 0.95% |
| hsa_circ_0039813 | 2304 | 11 | 0.95% |
| hsa_circ_0045468 | 2304 | 11 | 0.95% |

| | | | |
|---|---|---|---|
| hsa_circ_0051138 | 2304 | 11 | 0.95% |
| hsa_circ_0112006 | 2304 | 11 | 0.95% |
| hsa_circ_0125040 | 6075 | 29 | 0.95% |
| hsa_circ_0033581 | 2514 | 12 | 0.95% |
| hsa_circ_0043505 | 2514 | 12 | 0.95% |
| hsa_circ_0007604 | 2095 | 10 | 0.95% |
| hsa_circ_0059360 | 2095 | 10 | 0.95% |
| hsa_circ_0059882 | 2095 | 10 | 0.95% |
| hsa_circ_0061923 | 2095 | 10 | 0.95% |
| hsa_circ_0065865 | 2095 | 10 | 0.95% |
| hsa_circ_0078325 | 2095 | 10 | 0.95% |
| hsa_circ_0111952 | 2095 | 10 | 0.95% |
| hsa_circ_0114795 | 2095 | 10 | 0.95% |
| hsa_circ_0087343 | 2724 | 13 | 0.95% |
| hsa_circ_0016747 | 2305 | 11 | 0.95% |
| hsa_circ_0044062 | 2305 | 11 | 0.95% |
| hsa_circ_0076787 | 2305 | 11 | 0.95% |
| hsa_circ_0135364 | 2305 | 11 | 0.95% |
| hsa_circ_0064806 | 3353 | 16 | 0.95% |
| hsa_circ_0018116 | 2096 | 10 | 0.95% |
| hsa_circ_0042195 | 2096 | 10 | 0.95% |
| hsa_circ_0046011 | 2096 | 10 | 0.95% |
| hsa_circ_0059570 | 2096 | 10 | 0.95% |
| hsa_circ_0075163 | 2096 | 10 | 0.95% |
| hsa_circ_0112160 | 2096 | 10 | 0.95% |
| hsa_circ_0118885 | 2096 | 10 | 0.95% |
| hsa_circ_0138562 | 2096 | 10 | 0.95% |
| hsa_circ_0017700 | 2725 | 13 | 0.95% |
| hsa_circ_0052763 | 2306 | 11 | 0.95% |
| hsa_circ_0009809 | 2935 | 14 | 0.95% |
| hsa_circ_0022896 | 2516 | 12 | 0.95% |
| hsa_circ_0072299 | 2516 | 12 | 0.95% |
| hsa_circ_0083103 | 2516 | 12 | 0.95% |
| hsa_circ_0085004 | 6291 | 30 | 0.95% |
| hsa_circ_0023041 | 2307 | 11 | 0.95% |
| hsa_circ_0032955 | 2307 | 11 | 0.95% |
| hsa_circ_0098775 | 2307 | 11 | 0.95% |
| hsa_circ_0025032 | 2517 | 12 | 0.95% |
| hsa_circ_0053050 | 2517 | 12 | 0.95% |
| hsa_circ_0105607 | 2517 | 12 | 0.95% |
| hsa_circ_0061440 | 2727 | 13 | 0.95% |
| hsa_circ_0028259 | 2098 | 10 | 0.95% |
| hsa_circ_0037424 | 2098 | 10 | 0.95% |
| hsa_circ_0048036 | 2098 | 10 | 0.95% |
| hsa_circ_0109897 | 2098 | 10 | 0.95% |
| hsa_circ_0132920 | 2098 | 10 | 0.95% |
| hsa_circ_0033608 | 2518 | 12 | 0.95% |
| hsa_circ_0049525 | 2518 | 12 | 0.95% |
| hsa_circ_0067852 | 2728 | 13 | 0.95% |
| hsa_circ_0008352 | 8605 | 41 | 0.95% |

| | | | |
|---|---|---|---|
| hsa_circ_0015710 | 2099 | 10 | 0.95% |
| hsa_circ_0056101 | 2099 | 10 | 0.95% |
| hsa_circ_0061531 | 2099 | 10 | 0.95% |
| hsa_circ_0067548 | 2099 | 10 | 0.95% |
| hsa_circ_0126534 | 2099 | 10 | 0.95% |
| hsa_circ_0027408 | 2309 | 11 | 0.95% |
| hsa_circ_0028015 | 2309 | 11 | 0.95% |
| hsa_circ_0038038 | 2309 | 11 | 0.95% |
| hsa_circ_0048627 | 2309 | 11 | 0.95% |
| hsa_circ_0052231 | 2309 | 11 | 0.95% |
| hsa_circ_0099413 | 2309 | 11 | 0.95% |
| hsa_circ_0091801 | 2519 | 12 | 0.95% |
| hsa_circ_0083753 | 2729 | 13 | 0.95% |
| hsa_circ_0085999 | 2729 | 13 | 0.95% |
| hsa_circ_0067937 | 3149 | 15 | 0.95% |
| hsa_circ_0097657 | 7140 | 34 | 0.95% |
| hsa_circ_0000289 | 3150 | 15 | 0.95% |
| hsa_circ_0122256 | 2730 | 13 | 0.95% |
| hsa_circ_0018631 | 2520 | 12 | 0.95% |
| hsa_circ_0025077 | 2520 | 12 | 0.95% |
| hsa_circ_0034902 | 2520 | 12 | 0.95% |
| hsa_circ_0092011 | 2520 | 12 | 0.95% |
| hsa_circ_0023210 | 2310 | 11 | 0.95% |
| hsa_circ_0045423 | 2310 | 11 | 0.95% |
| hsa_circ_0051137 | 2310 | 11 | 0.95% |
| hsa_circ_0065431 | 2310 | 11 | 0.95% |
| hsa_circ_0045065 | 2100 | 10 | 0.95% |
| hsa_circ_0064757 | 2100 | 10 | 0.95% |
| hsa_circ_0078577 | 2100 | 10 | 0.95% |
| hsa_circ_0119831 | 2100 | 10 | 0.95% |
| hsa_circ_0101314 | 3151 | 15 | 0.95% |
| hsa_circ_0120785 | 2941 | 14 | 0.95% |
| hsa_circ_0104174 | 2521 | 12 | 0.95% |
| hsa_circ_0009028 | 4622 | 22 | 0.95% |
| hsa_circ_0045455 | 2311 | 11 | 0.95% |
| hsa_circ_0053688 | 2311 | 11 | 0.95% |
| hsa_circ_0070460 | 2311 | 11 | 0.95% |
| hsa_circ_0015503 | 2101 | 10 | 0.95% |
| hsa_circ_0043124 | 2101 | 10 | 0.95% |
| hsa_circ_0070991 | 2101 | 10 | 0.95% |
| hsa_circ_0087241 | 2101 | 10 | 0.95% |
| hsa_circ_0118882 | 2101 | 10 | 0.95% |
| hsa_circ_0094001 | 3782 | 18 | 0.95% |
| hsa_circ_0057295 | 4833 | 23 | 0.95% |
| hsa_circ_0052312 | 2942 | 14 | 0.95% |
| hsa_circ_0007063 | 5254 | 25 | 0.95% |
| hsa_circ_0020297 | 2522 | 12 | 0.95% |
| hsa_circ_0057515 | 2522 | 12 | 0.95% |
| hsa_circ_0115747 | 2522 | 12 | 0.95% |
| hsa_circ_0018299 | 2312 | 11 | 0.95% |

| | | | |
|---|---|---|---|
| hsa_circ_0060454 | 2312 | 11 | 0.95% |
| hsa_circ_0078872 | 2102 | 10 | 0.95% |
| hsa_circ_0087662 | 2102 | 10 | 0.95% |
| hsa_circ_0105567 | 2102 | 10 | 0.95% |
| hsa_circ_0073355 | 2733 | 13 | 0.95% |
| hsa_circ_0123417 | 2733 | 13 | 0.95% |
| hsa_circ_0130324 | 3364 | 16 | 0.95% |
| hsa_circ_0063649 | 2523 | 12 | 0.95% |
| hsa_circ_0102382 | 2523 | 12 | 0.95% |
| hsa_circ_0120584 | 2523 | 12 | 0.95% |
| hsa_circ_0021094 | 2313 | 11 | 0.95% |
| hsa_circ_0038155 | 2313 | 11 | 0.95% |
| hsa_circ_0048382 | 2313 | 11 | 0.95% |
| hsa_circ_0077361 | 2313 | 11 | 0.95% |
| hsa_circ_0013974 | 2103 | 10 | 0.95% |
| hsa_circ_0021785 | 2103 | 10 | 0.95% |
| hsa_circ_0028589 | 2103 | 10 | 0.95% |
| hsa_circ_0033441 | 2103 | 10 | 0.95% |
| hsa_circ_0034064 | 2103 | 10 | 0.95% |
| hsa_circ_0067672 | 2103 | 10 | 0.95% |
| hsa_circ_0069814 | 2103 | 10 | 0.95% |
| hsa_circ_0086566 | 2103 | 10 | 0.95% |
| hsa_circ_0092385 | 2103 | 10 | 0.95% |
| hsa_circ_0129473 | 2103 | 10 | 0.95% |
| hsa_circ_0138107 | 2103 | 10 | 0.95% |
| hsa_circ_0010215 | 2524 | 12 | 0.95% |
| hsa_circ_0033491 | 2524 | 12 | 0.95% |
| hsa_circ_0037541 | 2314 | 11 | 0.95% |
| hsa_circ_0070101 | 2314 | 11 | 0.95% |
| hsa_circ_0046514 | 2104 | 10 | 0.95% |
| hsa_circ_0046801 | 2104 | 10 | 0.95% |
| hsa_circ_0055074 | 2104 | 10 | 0.95% |
| hsa_circ_0061735 | 2104 | 10 | 0.95% |
| hsa_circ_0081518 | 2104 | 10 | 0.95% |
| hsa_circ_0132223 | 2104 | 10 | 0.95% |
| hsa_circ_0075516 | 2946 | 14 | 0.95% |
| hsa_circ_0021219 | 2315 | 11 | 0.95% |
| hsa_circ_0039313 | 2315 | 11 | 0.95% |
| hsa_circ_0053657 | 2315 | 11 | 0.95% |
| hsa_circ_0071004 | 2315 | 11 | 0.95% |
| hsa_circ_0017001 | 2736 | 13 | 0.95% |
| hsa_circ_0094298 | 6946 | 33 | 0.95% |
| hsa_circ_0009835 | 2526 | 12 | 0.95% |
| hsa_circ_0057722 | 2526 | 12 | 0.95% |
| hsa_circ_0066419 | 2526 | 12 | 0.95% |
| hsa_circ_0073405 | 2105 | 10 | 0.95% |
| hsa_circ_0077290 | 2737 | 13 | 0.95% |
| hsa_circ_0085428 | 2737 | 13 | 0.95% |
| hsa_circ_0065901 | 2316 | 11 | 0.95% |
| hsa_circ_0071622 | 2316 | 11 | 0.95% |

| | | | |
|---|---|---|---|
| hsa_circ_0085394 | 2316 | 11 | 0.95% |
| hsa_circ_0101827 | 2316 | 11 | 0.95% |
| hsa_circ_0131531 | 5054 | 24 | 0.95% |
| hsa_circ_0089360 | 2527 | 12 | 0.95% |
| hsa_circ_0009369 | 2106 | 10 | 0.95% |
| hsa_circ_0076268 | 2106 | 10 | 0.95% |
| hsa_circ_0085151 | 2106 | 10 | 0.95% |
| hsa_circ_0089057 | 2106 | 10 | 0.95% |
| hsa_circ_0091773 | 2106 | 10 | 0.95% |
| hsa_circ_0126290 | 2106 | 10 | 0.95% |
| hsa_circ_0134237 | 2106 | 10 | 0.95% |
| hsa_circ_0042656 | 2738 | 13 | 0.95% |
| hsa_circ_0112829 | 5055 | 24 | 0.95% |
| hsa_circ_0017477 | 2317 | 11 | 0.95% |
| hsa_circ_0020220 | 2317 | 11 | 0.95% |
| hsa_circ_0044988 | 2317 | 11 | 0.95% |
| hsa_circ_0102833 | 2317 | 11 | 0.95% |
| hsa_circ_0122582 | 2317 | 11 | 0.95% |
| hsa_circ_0076718 | 3160 | 15 | 0.95% |
| hsa_circ_0085671 | 2528 | 12 | 0.95% |
| hsa_circ_0032995 | 2107 | 10 | 0.95% |
| hsa_circ_0037023 | 2107 | 10 | 0.95% |
| hsa_circ_0120959 | 2107 | 10 | 0.95% |
| hsa_circ_0013119 | 2318 | 11 | 0.95% |
| hsa_circ_0096071 | 2318 | 11 | 0.95% |
| hsa_circ_0098356 | 2318 | 11 | 0.95% |
| hsa_circ_0135587 | 2318 | 11 | 0.95% |
| hsa_circ_0138947 | 2318 | 11 | 0.95% |
| hsa_circ_0010858 | 2529 | 12 | 0.95% |
| hsa_circ_0063969 | 2529 | 12 | 0.95% |
| hsa_circ_0077464 | 2740 | 13 | 0.95% |
| hsa_circ_0139824 | 2740 | 13 | 0.95% |
| hsa_circ_0011592 | 2108 | 10 | 0.95% |
| hsa_circ_0090377 | 2108 | 10 | 0.95% |
| hsa_circ_0120283 | 2108 | 10 | 0.95% |
| hsa_circ_0003551 | 2319 | 11 | 0.95% |
| hsa_circ_0054868 | 2319 | 11 | 0.95% |
| hsa_circ_0065199 | 2319 | 11 | 0.95% |
| hsa_circ_0083310 | 2319 | 11 | 0.95% |
| hsa_circ_0137295 | 36684 | 174 | 0.95% |
| hsa_circ_0022818 | 2530 | 12 | 0.95% |
| hsa_circ_0063578 | 2530 | 12 | 0.95% |
| hsa_circ_0083584 | 2530 | 12 | 0.95% |
| hsa_circ_0053199 | 2109 | 10 | 0.95% |
| hsa_circ_0087410 | 2109 | 10 | 0.95% |
| hsa_circ_0109280 | 2109 | 10 | 0.95% |
| hsa_circ_0137373 | 2109 | 10 | 0.95% |
| hsa_circ_0051312 | 2320 | 11 | 0.95% |
| hsa_circ_0113565 | 2320 | 11 | 0.95% |
| hsa_circ_0115800 | 2320 | 11 | 0.95% |

| | | | |
|---|---|---|---|
| hsa_circ_0120725 | 2320 | 11 | 0.95% |
| hsa_circ_0031517 | 2531 | 12 | 0.95% |
| hsa_circ_0014500 | 2742 | 13 | 0.95% |
| hsa_circ_0114466 | 8437 | 40 | 0.95% |
| hsa_circ_0139232 | 4852 | 23 | 0.95% |
| hsa_circ_0138000 | 4220 | 20 | 0.95% |
| hsa_circ_0052710 | 3165 | 15 | 0.95% |
| hsa_circ_0091402 | 2954 | 14 | 0.95% |
| hsa_circ_0021879 | 2532 | 12 | 0.95% |
| hsa_circ_0127215 | 2532 | 12 | 0.95% |
| hsa_circ_0039038 | 2321 | 11 | 0.95% |
| hsa_circ_0086806 | 2321 | 11 | 0.95% |
| hsa_circ_0106446 | 2321 | 11 | 0.95% |
| hsa_circ_0117506 | 2321 | 11 | 0.95% |
| hsa_circ_0125899 | 2321 | 11 | 0.95% |
| hsa_circ_0017920 | 2110 | 10 | 0.95% |
| hsa_circ_0009328 | 2533 | 12 | 0.95% |
| hsa_circ_0030448 | 2533 | 12 | 0.95% |
| hsa_circ_0052295 | 2533 | 12 | 0.95% |
| hsa_circ_0137919 | 2533 | 12 | 0.95% |
| hsa_circ_0030663 | 2322 | 11 | 0.95% |
| hsa_circ_0032831 | 2322 | 11 | 0.95% |
| hsa_circ_0050967 | 2322 | 11 | 0.95% |
| hsa_circ_0069441 | 2322 | 11 | 0.95% |
| hsa_circ_0026449 | 2111 | 10 | 0.95% |
| hsa_circ_0027011 | 2111 | 10 | 0.95% |
| hsa_circ_0077502 | 2111 | 10 | 0.95% |
| hsa_circ_0081649 | 2111 | 10 | 0.95% |
| hsa_circ_0120812 | 2111 | 10 | 0.95% |
| hsa_circ_0061076 | 2745 | 13 | 0.95% |
| hsa_circ_0018664 | 2534 | 12 | 0.95% |
| hsa_circ_0084004 | 2534 | 12 | 0.95% |
| hsa_circ_0019301 | 2323 | 11 | 0.95% |
| hsa_circ_0047904 | 2323 | 11 | 0.95% |
| hsa_circ_0084050 | 2323 | 11 | 0.95% |
| hsa_circ_0119270 | 2323 | 11 | 0.95% |
| hsa_circ_0071825 | 4224 | 20 | 0.95% |
| hsa_circ_0020291 | 2112 | 10 | 0.95% |
| hsa_circ_0029727 | 2112 | 10 | 0.95% |
| hsa_circ_0054158 | 2112 | 10 | 0.95% |
| hsa_circ_0072400 | 2112 | 10 | 0.95% |
| hsa_circ_0073301 | 2112 | 10 | 0.95% |
| hsa_circ_0080683 | 2112 | 10 | 0.95% |
| hsa_circ_0131914 | 2112 | 10 | 0.95% |
| hsa_circ_0008276 | 4013 | 19 | 0.95% |
| hsa_circ_0060685 | 2746 | 13 | 0.95% |
| hsa_circ_0013076 | 2535 | 12 | 0.95% |
| hsa_circ_0013682 | 2535 | 12 | 0.95% |
| hsa_circ_0026269 | 2324 | 11 | 0.95% |
| hsa_circ_0041592 | 2324 | 11 | 0.95% |

| | | | |
|---|---|---|---|
| hsa_circ_0013713 | 2113 | 10 | 0.95% |
| hsa_circ_0094823 | 6973 | 33 | 0.95% |
| hsa_circ_0100776 | 2747 | 13 | 0.95% |
| hsa_circ_0115900 | 2747 | 13 | 0.95% |
| hsa_circ_0131132 | 3804 | 18 | 0.95% |
| hsa_circ_0067362 | 2536 | 12 | 0.95% |
| hsa_circ_0015930 | 2325 | 11 | 0.95% |
| hsa_circ_0053895 | 2325 | 11 | 0.95% |
| hsa_circ_0061670 | 2325 | 11 | 0.95% |
| hsa_circ_0066670 | 2325 | 11 | 0.95% |
| hsa_circ_0056544 | 2748 | 13 | 0.95% |
| hsa_circ_0058934 | 2748 | 13 | 0.95% |
| hsa_circ_0022222 | 2114 | 10 | 0.95% |
| hsa_circ_0045027 | 2537 | 12 | 0.95% |
| hsa_circ_0085099 | 2537 | 12 | 0.95% |
| hsa_circ_0042547 | 2326 | 11 | 0.95% |
| hsa_circ_0080145 | 2326 | 11 | 0.95% |
| hsa_circ_0083843 | 2326 | 11 | 0.95% |
| hsa_circ_0084740 | 2326 | 11 | 0.95% |
| hsa_circ_0113942 | 2326 | 11 | 0.95% |
| hsa_circ_0055780 | 3384 | 16 | 0.95% |
| hsa_circ_0016983 | 2538 | 12 | 0.95% |
| hsa_circ_0051283 | 2538 | 12 | 0.95% |
| hsa_circ_0080658 | 2538 | 12 | 0.95% |
| hsa_circ_0032394 | 2115 | 10 | 0.95% |
| hsa_circ_0036123 | 2115 | 10 | 0.95% |
| hsa_circ_0066348 | 2115 | 10 | 0.95% |
| hsa_circ_0076863 | 2115 | 10 | 0.95% |
| hsa_circ_0091070 | 2115 | 10 | 0.95% |
| hsa_circ_0086115 | 2750 | 13 | 0.95% |
| hsa_circ_0019840 | 2327 | 11 | 0.95% |
| hsa_circ_0093840 | 2327 | 11 | 0.95% |
| hsa_circ_0100913 | 2327 | 11 | 0.95% |
| hsa_circ_0116819 | 2327 | 11 | 0.95% |
| hsa_circ_0138999 | 2327 | 11 | 0.95% |
| hsa_circ_0022231 | 2539 | 12 | 0.95% |
| hsa_circ_0058100 | 2539 | 12 | 0.95% |
| hsa_circ_0115208 | 6771 | 32 | 0.95% |
| hsa_circ_0032397 | 2116 | 10 | 0.95% |
| hsa_circ_0044101 | 2116 | 10 | 0.95% |
| hsa_circ_0092535 | 2116 | 10 | 0.95% |
| hsa_circ_0021120 | 3809 | 18 | 0.95% |
| hsa_circ_0017813 | 2751 | 13 | 0.95% |
| hsa_circ_0040733 | 2751 | 13 | 0.95% |
| hsa_circ_0020563 | 2328 | 11 | 0.95% |
| hsa_circ_0040992 | 2540 | 12 | 0.94% |
| hsa_circ_0086438 | 2540 | 12 | 0.94% |
| hsa_circ_0094525 | 2540 | 12 | 0.94% |
| hsa_circ_0035517 | 2117 | 10 | 0.94% |
| hsa_circ_0042632 | 2117 | 10 | 0.94% |

| | | | |
|---|---|---|---|
| hsa_circ_0075505 | 2117 | 10 | 0.94% |
| hsa_circ_0039229 | 2329 | 11 | 0.94% |
| hsa_circ_0062016 | 2329 | 11 | 0.94% |
| hsa_circ_0070241 | 2329 | 11 | 0.94% |
| hsa_circ_0055827 | 3176 | 15 | 0.94% |
| hsa_circ_0095237 | 3388 | 16 | 0.94% |
| hsa_circ_0129842 | 3388 | 16 | 0.94% |
| hsa_circ_0138748 | 2541 | 12 | 0.94% |
| hsa_circ_0070847 | 2753 | 13 | 0.94% |
| hsa_circ_0021216 | 2118 | 10 | 0.94% |
| hsa_circ_0043701 | 2118 | 10 | 0.94% |
| hsa_circ_0048701 | 2118 | 10 | 0.94% |
| hsa_circ_0115754 | 2118 | 10 | 0.94% |
| hsa_circ_0118448 | 2118 | 10 | 0.94% |
| hsa_circ_0041285 | 2330 | 11 | 0.94% |
| hsa_circ_0052370 | 2330 | 11 | 0.94% |
| hsa_circ_0067704 | 2330 | 11 | 0.94% |
| hsa_circ_0069291 | 2330 | 11 | 0.94% |
| hsa_circ_0073861 | 2330 | 11 | 0.94% |
| hsa_circ_0085290 | 2330 | 11 | 0.94% |
| hsa_circ_0023461 | 2542 | 12 | 0.94% |
| hsa_circ_0057597 | 2542 | 12 | 0.94% |
| hsa_circ_0091484 | 2542 | 12 | 0.94% |
| hsa_circ_0020090 | 2966 | 14 | 0.94% |
| hsa_circ_0090115 | 2966 | 14 | 0.94% |
| hsa_circ_0109390 | 3602 | 17 | 0.94% |
| hsa_circ_0026822 | 2119 | 10 | 0.94% |
| hsa_circ_0027407 | 2119 | 10 | 0.94% |
| hsa_circ_0042620 | 2119 | 10 | 0.94% |
| hsa_circ_0056632 | 2119 | 10 | 0.94% |
| hsa_circ_0062785 | 2119 | 10 | 0.94% |
| hsa_circ_0086183 | 2119 | 10 | 0.94% |
| hsa_circ_0051305 | 2331 | 11 | 0.94% |
| hsa_circ_0053151 | 2331 | 11 | 0.94% |
| hsa_circ_0126347 | 2543 | 12 | 0.94% |
| hsa_circ_0049800 | 2755 | 13 | 0.94% |
| hsa_circ_0101154 | 7207 | 34 | 0.94% |
| hsa_circ_0078873 | 7632 | 36 | 0.94% |
| hsa_circ_0022829 | 2756 | 13 | 0.94% |
| hsa_circ_0056124 | 2544 | 12 | 0.94% |
| hsa_circ_0014887 | 2332 | 11 | 0.94% |
| hsa_circ_0015772 | 2332 | 11 | 0.94% |
| hsa_circ_0070242 | 2332 | 11 | 0.94% |
| hsa_circ_0125204 | 2332 | 11 | 0.94% |
| hsa_circ_0034824 | 2120 | 10 | 0.94% |
| hsa_circ_0061645 | 2120 | 10 | 0.94% |
| hsa_circ_0088148 | 2120 | 10 | 0.94% |
| hsa_circ_0090948 | 2120 | 10 | 0.94% |
| hsa_circ_0110415 | 2120 | 10 | 0.94% |
| hsa_circ_0066190 | 2969 | 14 | 0.94% |

| | | | |
|---|---|---|---|
| hsa_circ_0047945 | 2757 | 13 | 0.94% |
| hsa_circ_0127873 | 2545 | 12 | 0.94% |
| hsa_circ_0137333 | 2545 | 12 | 0.94% |
| hsa_circ_0017195 | 2333 | 11 | 0.94% |
| hsa_circ_0036631 | 2333 | 11 | 0.94% |
| hsa_circ_0059065 | 2333 | 11 | 0.94% |
| hsa_circ_0098220 | 2333 | 11 | 0.94% |
| hsa_circ_0023726 | 2121 | 10 | 0.94% |
| hsa_circ_0078543 | 2121 | 10 | 0.94% |
| hsa_circ_0102499 | 10394 | 49 | 0.94% |
| hsa_circ_0083412 | 2758 | 13 | 0.94% |
| hsa_circ_0054411 | 2546 | 12 | 0.94% |
| hsa_circ_0104192 | 2546 | 12 | 0.94% |
| hsa_circ_0030085 | 2334 | 11 | 0.94% |
| hsa_circ_0038258 | 2334 | 11 | 0.94% |
| hsa_circ_0067563 | 2334 | 11 | 0.94% |
| hsa_circ_0074571 | 2334 | 11 | 0.94% |
| hsa_circ_0084915 | 2334 | 11 | 0.94% |
| hsa_circ_0086882 | 2334 | 11 | 0.94% |
| hsa_circ_0088344 | 2334 | 11 | 0.94% |
| hsa_circ_0061738 | 2122 | 10 | 0.94% |
| hsa_circ_0072285 | 2122 | 10 | 0.94% |
| hsa_circ_0076732 | 2122 | 10 | 0.94% |
| hsa_circ_0081730 | 2122 | 10 | 0.94% |
| hsa_circ_0083302 | 2122 | 10 | 0.94% |
| hsa_circ_0014834 | 2547 | 12 | 0.94% |
| hsa_circ_0085469 | 2547 | 12 | 0.94% |
| hsa_circ_0130903 | 2547 | 12 | 0.94% |
| hsa_circ_0083550 | 3184 | 15 | 0.94% |
| hsa_circ_0005333 | 2335 | 11 | 0.94% |
| hsa_circ_0028797 | 2335 | 11 | 0.94% |
| hsa_circ_0034637 | 2335 | 11 | 0.94% |
| hsa_circ_0043753 | 2335 | 11 | 0.94% |
| hsa_circ_0065142 | 2335 | 11 | 0.94% |
| hsa_circ_0076552 | 2335 | 11 | 0.94% |
| hsa_circ_0078973 | 2335 | 11 | 0.94% |
| hsa_circ_0016369 | 2123 | 10 | 0.94% |
| hsa_circ_0044135 | 2123 | 10 | 0.94% |
| hsa_circ_0056224 | 2760 | 13 | 0.94% |
| hsa_circ_0047306 | 2973 | 14 | 0.94% |
| hsa_circ_0020394 | 2336 | 11 | 0.94% |
| hsa_circ_0118378 | 2761 | 13 | 0.94% |
| hsa_circ_0055965 | 2124 | 10 | 0.94% |
| hsa_circ_0059782 | 2124 | 10 | 0.94% |
| hsa_circ_0062227 | 2124 | 10 | 0.94% |
| hsa_circ_0078320 | 2124 | 10 | 0.94% |
| hsa_circ_0108343 | 2124 | 10 | 0.94% |
| hsa_circ_0016972 | 2549 | 12 | 0.94% |
| hsa_circ_0047012 | 2549 | 12 | 0.94% |
| hsa_circ_0010281 | 2337 | 11 | 0.94% |

| | | | |
|---|---|---|---|
| hsa_circ_0019536 | 2337 | 11 | 0.94% |
| hsa_circ_0057813 | 2337 | 11 | 0.94% |
| hsa_circ_0076814 | 2337 | 11 | 0.94% |
| hsa_circ_0048039 | 2762 | 13 | 0.94% |
| hsa_circ_0082621 | 5312 | 25 | 0.94% |
| hsa_circ_0111174 | 2550 | 12 | 0.94% |
| hsa_circ_0011440 | 2125 | 10 | 0.94% |
| hsa_circ_0066794 | 2125 | 10 | 0.94% |
| hsa_circ_0069770 | 2125 | 10 | 0.94% |
| hsa_circ_0133479 | 2125 | 10 | 0.94% |
| hsa_circ_0116185 | 2763 | 13 | 0.94% |
| hsa_circ_0020871 | 2338 | 11 | 0.94% |
| hsa_circ_0022846 | 2338 | 11 | 0.94% |
| hsa_circ_0086896 | 2338 | 11 | 0.94% |
| hsa_circ_0115186 | 2338 | 11 | 0.94% |
| hsa_circ_0129002 | 2338 | 11 | 0.94% |
| hsa_circ_0136785 | 2338 | 11 | 0.94% |
| hsa_circ_0140638 | 2338 | 11 | 0.94% |
| hsa_circ_0029229 | 2126 | 10 | 0.94% |
| hsa_circ_0066576 | 2126 | 10 | 0.94% |
| hsa_circ_0074112 | 2126 | 10 | 0.94% |
| hsa_circ_0009892 | 2339 | 11 | 0.94% |
| hsa_circ_0010101 | 2339 | 11 | 0.94% |
| hsa_circ_0023191 | 2339 | 11 | 0.94% |
| hsa_circ_0049561 | 2339 | 11 | 0.94% |
| hsa_circ_0054063 | 2339 | 11 | 0.94% |
| hsa_circ_0055907 | 2339 | 11 | 0.94% |
| hsa_circ_0083509 | 2339 | 11 | 0.94% |
| hsa_circ_0086547 | 2339 | 11 | 0.94% |
| hsa_circ_0096713 | 2339 | 11 | 0.94% |
| hsa_circ_0067142 | 2552 | 12 | 0.94% |
| hsa_circ_0109993 | 2552 | 12 | 0.94% |
| hsa_circ_0018482 | 2127 | 10 | 0.94% |
| hsa_circ_0019416 | 2127 | 10 | 0.94% |
| hsa_circ_0029353 | 2127 | 10 | 0.94% |
| hsa_circ_0036964 | 2127 | 10 | 0.94% |
| hsa_circ_0060539 | 2127 | 10 | 0.94% |
| hsa_circ_0123404 | 2127 | 10 | 0.94% |
| hsa_circ_0104770 | 2340 | 11 | 0.94% |
| hsa_circ_0122661 | 2340 | 11 | 0.94% |
| hsa_circ_0010338 | 2553 | 12 | 0.94% |
| hsa_circ_0124138 | 5532 | 26 | 0.94% |
| hsa_circ_0044336 | 2766 | 13 | 0.94% |
| hsa_circ_0093263 | 4894 | 23 | 0.94% |
| hsa_circ_0036936 | 2128 | 10 | 0.94% |
| hsa_circ_0083982 | 2128 | 10 | 0.94% |
| hsa_circ_0131574 | 2128 | 10 | 0.94% |
| hsa_circ_0046790 | 2341 | 11 | 0.94% |
| hsa_circ_0047887 | 2341 | 11 | 0.94% |
| hsa_circ_0051745 | 2341 | 11 | 0.94% |

| | | | |
|---|---|---|---|
| hsa_circ_0063960 | 2554 | 12 | 0.94% |
| hsa_circ_0103491 | 2554 | 12 | 0.94% |
| hsa_circ_0024633 | 2767 | 13 | 0.94% |
| hsa_circ_0034036 | 2767 | 13 | 0.94% |
| hsa_circ_0087166 | 2980 | 14 | 0.94% |
| hsa_circ_0104012 | 4896 | 23 | 0.94% |
| hsa_circ_0133693 | 7238 | 34 | 0.94% |
| hsa_circ_0017154 | 2129 | 10 | 0.94% |
| hsa_circ_0139279 | 2129 | 10 | 0.94% |
| hsa_circ_0108141 | 2342 | 11 | 0.94% |
| hsa_circ_0115237 | 2342 | 11 | 0.94% |
| hsa_circ_0015671 | 2555 | 12 | 0.94% |
| hsa_circ_0031616 | 2555 | 12 | 0.94% |
| hsa_circ_0050017 | 2555 | 12 | 0.94% |
| hsa_circ_0065831 | 2555 | 12 | 0.94% |
| hsa_circ_0109994 | 2555 | 12 | 0.94% |
| hsa_circ_0062020 | 2768 | 13 | 0.94% |
| hsa_circ_0044510 | 4898 | 23 | 0.94% |
| hsa_circ_0113931 | 2769 | 13 | 0.94% |
| hsa_circ_0046348 | 2556 | 12 | 0.94% |
| hsa_circ_0070692 | 2556 | 12 | 0.94% |
| hsa_circ_0132302 | 2556 | 12 | 0.94% |
| hsa_circ_0049128 | 2343 | 11 | 0.94% |
| hsa_circ_0073956 | 2343 | 11 | 0.94% |
| hsa_circ_0075078 | 2343 | 11 | 0.94% |
| hsa_circ_0085341 | 2343 | 11 | 0.94% |
| hsa_circ_0014377 | 2130 | 10 | 0.94% |
| hsa_circ_0082147 | 2130 | 10 | 0.94% |
| hsa_circ_0119568 | 4900 | 23 | 0.94% |
| hsa_circ_0097287 | 6179 | 29 | 0.94% |
| hsa_circ_0024422 | 2770 | 13 | 0.94% |
| hsa_circ_0048878 | 2770 | 13 | 0.94% |
| hsa_circ_0010452 | 2344 | 11 | 0.94% |
| hsa_circ_0021575 | 2344 | 11 | 0.94% |
| hsa_circ_0024858 | 2344 | 11 | 0.94% |
| hsa_circ_0121144 | 2344 | 11 | 0.94% |
| hsa_circ_0140757 | 2344 | 11 | 0.94% |
| hsa_circ_0023483 | 2131 | 10 | 0.94% |
| hsa_circ_0054830 | 2131 | 10 | 0.94% |
| hsa_circ_0063975 | 2131 | 10 | 0.94% |
| hsa_circ_0084940 | 2131 | 10 | 0.94% |
| hsa_circ_0000717 | 2558 | 12 | 0.94% |
| hsa_circ_0062073 | 2558 | 12 | 0.94% |
| hsa_circ_0114465 | 8527 | 40 | 0.94% |
| hsa_circ_0033290 | 2132 | 10 | 0.94% |
| hsa_circ_0079982 | 2132 | 10 | 0.94% |
| hsa_circ_0088402 | 2132 | 10 | 0.94% |
| hsa_circ_0131580 | 2132 | 10 | 0.94% |
| hsa_circ_0014686 | 2559 | 12 | 0.94% |
| hsa_circ_0076081 | 2559 | 12 | 0.94% |

| | | | |
|---|---|---|---|
| hsa_circ_0023727 | 2346 | 11 | 0.94% |
| hsa_circ_0065664 | 2346 | 11 | 0.94% |
| hsa_circ_0073280 | 2346 | 11 | 0.94% |
| hsa_circ_0090376 | 2346 | 11 | 0.94% |
| hsa_circ_0010120 | 2986 | 14 | 0.94% |
| hsa_circ_0006147 | 2133 | 10 | 0.94% |
| hsa_circ_0011802 | 2133 | 10 | 0.94% |
| hsa_circ_0012219 | 2133 | 10 | 0.94% |
| hsa_circ_0015905 | 2133 | 10 | 0.94% |
| hsa_circ_0099779 | 2133 | 10 | 0.94% |
| hsa_circ_0105018 | 9172 | 43 | 0.94% |
| hsa_circ_0062856 | 2773 | 13 | 0.94% |
| hsa_circ_0088574 | 2773 | 13 | 0.94% |
| hsa_circ_0102092 | 2773 | 13 | 0.94% |
| hsa_circ_0000248 | 3840 | 18 | 0.94% |
| hsa_circ_0007121 | 2560 | 12 | 0.94% |
| hsa_circ_0099973 | 6614 | 31 | 0.94% |
| hsa_circ_0052074 | 2987 | 14 | 0.94% |
| hsa_circ_0038752 | 2347 | 11 | 0.94% |
| hsa_circ_0041288 | 2347 | 11 | 0.94% |
| hsa_circ_0046311 | 2347 | 11 | 0.94% |
| hsa_circ_0086844 | 2347 | 11 | 0.94% |
| hsa_circ_0086981 | 2347 | 11 | 0.94% |
| hsa_circ_0131591 | 2347 | 11 | 0.94% |
| hsa_circ_0061201 | 2134 | 10 | 0.94% |
| hsa_circ_0107220 | 2134 | 10 | 0.94% |
| hsa_circ_0026736 | 2561 | 12 | 0.94% |
| hsa_circ_0077581 | 2561 | 12 | 0.94% |
| hsa_circ_0024611 | 2348 | 11 | 0.94% |
| hsa_circ_0059007 | 2348 | 11 | 0.94% |
| hsa_circ_0077129 | 2348 | 11 | 0.94% |
| hsa_circ_0079370 | 2348 | 11 | 0.94% |
| hsa_circ_0091163 | 2348 | 11 | 0.94% |
| hsa_circ_0092875 | 2348 | 11 | 0.94% |
| hsa_circ_0018480 | 2135 | 10 | 0.94% |
| hsa_circ_0130941 | 2135 | 10 | 0.94% |
| hsa_circ_0073936 | 2349 | 11 | 0.94% |
| hsa_circ_0007854 | 3417 | 16 | 0.94% |
| hsa_circ_0106216 | 7475 | 35 | 0.94% |
| hsa_circ_0009967 | 2563 | 12 | 0.94% |
| hsa_circ_0120442 | 4272 | 20 | 0.94% |
| hsa_circ_0012551 | 2136 | 10 | 0.94% |
| hsa_circ_0018341 | 2136 | 10 | 0.94% |
| hsa_circ_0024943 | 2136 | 10 | 0.94% |
| hsa_circ_0027834 | 2136 | 10 | 0.94% |
| hsa_circ_0027847 | 2136 | 10 | 0.94% |
| hsa_circ_0038561 | 2136 | 10 | 0.94% |
| hsa_circ_0050538 | 2136 | 10 | 0.94% |
| hsa_circ_0069905 | 5981 | 28 | 0.94% |
| hsa_circ_0010676 | 2777 | 13 | 0.94% |

| | | | |
|---|---|---|---|
| hsa_circ_0032026 | 4486 | 21 | 0.94% |
| hsa_circ_0031478 | 2350 | 11 | 0.94% |
| hsa_circ_0045831 | 2350 | 11 | 0.94% |
| hsa_circ_0113789 | 10682 | 50 | 0.94% |
| hsa_circ_0021810 | 3205 | 15 | 0.94% |
| hsa_circ_0033528 | 2564 | 12 | 0.94% |
| hsa_circ_0034459 | 2564 | 12 | 0.94% |
| hsa_circ_0070840 | 2564 | 12 | 0.94% |
| hsa_circ_0137444 | 5342 | 25 | 0.94% |
| hsa_circ_0083250 | 3419 | 16 | 0.94% |
| hsa_circ_0032568 | 2778 | 13 | 0.94% |
| hsa_circ_0055136 | 2778 | 13 | 0.94% |
| hsa_circ_0081911 | 2778 | 13 | 0.94% |
| hsa_circ_0021931 | 2137 | 10 | 0.94% |
| hsa_circ_0030627 | 2137 | 10 | 0.94% |
| hsa_circ_0060883 | 2137 | 10 | 0.94% |
| hsa_circ_0013708 | 2351 | 11 | 0.94% |
| hsa_circ_0023199 | 2351 | 11 | 0.94% |
| hsa_circ_0033153 | 2351 | 11 | 0.94% |
| hsa_circ_0051204 | 2351 | 11 | 0.94% |
| hsa_circ_0060195 | 2351 | 11 | 0.94% |
| hsa_circ_0061319 | 2351 | 11 | 0.94% |
| hsa_circ_0126730 | 2351 | 11 | 0.94% |
| hsa_circ_0027165 | 2565 | 12 | 0.94% |
| hsa_circ_0035157 | 2565 | 12 | 0.94% |
| hsa_circ_0140159 | 2565 | 12 | 0.94% |
| hsa_circ_0076806 | 2779 | 13 | 0.94% |
| hsa_circ_0081925 | 2779 | 13 | 0.94% |
| hsa_circ_0036128 | 2993 | 14 | 0.94% |
| hsa_circ_0042012 | 2138 | 10 | 0.94% |
| hsa_circ_0046869 | 2138 | 10 | 0.94% |
| hsa_circ_0049863 | 2138 | 10 | 0.94% |
| hsa_circ_0065895 | 2138 | 10 | 0.94% |
| hsa_circ_0099650 | 2138 | 10 | 0.94% |
| hsa_circ_0018695 | 2352 | 11 | 0.94% |
| hsa_circ_0055646 | 2352 | 11 | 0.94% |
| hsa_circ_0090785 | 2352 | 11 | 0.94% |
| hsa_circ_0012700 | 2566 | 12 | 0.94% |
| hsa_circ_0031172 | 2566 | 12 | 0.94% |
| hsa_circ_0089665 | 2566 | 12 | 0.94% |
| hsa_circ_0113751 | 2566 | 12 | 0.94% |
| hsa_circ_0026342 | 3208 | 15 | 0.94% |
| hsa_circ_0052050 | 3208 | 15 | 0.94% |
| hsa_circ_0010535 | 2139 | 10 | 0.94% |
| hsa_circ_0025295 | 2139 | 10 | 0.94% |
| hsa_circ_0078318 | 2139 | 10 | 0.94% |
| hsa_circ_0112037 | 2139 | 10 | 0.94% |
| hsa_circ_0117341 | 2139 | 10 | 0.94% |
| hsa_circ_0034163 | 2353 | 11 | 0.93% |
| hsa_circ_0095066 | 2353 | 11 | 0.93% |

| | | | |
|---|---|---|---|
| hsa_circ_0139995 | 2995 | 14 | 0.93% |
| hsa_circ_0102574 | 6419 | 30 | 0.93% |
| hsa_circ_0024419 | 2568 | 12 | 0.93% |
| hsa_circ_0090746 | 2568 | 12 | 0.93% |
| hsa_circ_0043752 | 2354 | 11 | 0.93% |
| hsa_circ_0025676 | 2140 | 10 | 0.93% |
| hsa_circ_0032716 | 2140 | 10 | 0.93% |
| hsa_circ_0094718 | 2140 | 10 | 0.93% |
| hsa_circ_0111369 | 2140 | 10 | 0.93% |
| hsa_circ_0118126 | 2140 | 10 | 0.93% |
| hsa_circ_0138062 | 2140 | 10 | 0.93% |
| hsa_circ_0059650 | 2783 | 13 | 0.93% |
| hsa_circ_0068580 | 2783 | 13 | 0.93% |
| hsa_circ_0138322 | 2783 | 13 | 0.93% |
| hsa_circ_0041034 | 2569 | 12 | 0.93% |
| hsa_circ_0041675 | 2569 | 12 | 0.93% |
| hsa_circ_0118363 | 2569 | 12 | 0.93% |
| hsa_circ_0118955 | 2569 | 12 | 0.93% |
| hsa_circ_0029090 | 2355 | 11 | 0.93% |
| hsa_circ_0054949 | 2355 | 11 | 0.93% |
| hsa_circ_0090547 | 2355 | 11 | 0.93% |
| hsa_circ_0078844 | 4496 | 21 | 0.93% |
| hsa_circ_0053371 | 2141 | 10 | 0.93% |
| hsa_circ_0090431 | 2141 | 10 | 0.93% |
| hsa_circ_0013255 | 2570 | 12 | 0.93% |
| hsa_circ_0010730 | 2356 | 11 | 0.93% |
| hsa_circ_0037406 | 2356 | 11 | 0.93% |
| hsa_circ_0039939 | 2356 | 11 | 0.93% |
| hsa_circ_0051027 | 2356 | 11 | 0.93% |
| hsa_circ_0059719 | 2356 | 11 | 0.93% |
| hsa_circ_0075798 | 2356 | 11 | 0.93% |
| hsa_circ_0105436 | 2356 | 11 | 0.93% |
| hsa_circ_0024540 | 2142 | 10 | 0.93% |
| hsa_circ_0066308 | 2142 | 10 | 0.93% |
| hsa_circ_0100279 | 2142 | 10 | 0.93% |
| hsa_circ_0129148 | 2142 | 10 | 0.93% |
| hsa_circ_0106463 | 3428 | 16 | 0.93% |
| hsa_circ_0087538 | 2571 | 12 | 0.93% |
| hsa_circ_0091993 | 2571 | 12 | 0.93% |
| hsa_circ_0122797 | 2571 | 12 | 0.93% |
| hsa_circ_0044075 | 2357 | 11 | 0.93% |
| hsa_circ_0047548 | 2357 | 11 | 0.93% |
| hsa_circ_0062919 | 2357 | 11 | 0.93% |
| hsa_circ_0123009 | 2357 | 11 | 0.93% |
| hsa_circ_0050308 | 3000 | 14 | 0.93% |
| hsa_circ_0109057 | 4286 | 20 | 0.93% |
| hsa_circ_0010928 | 2143 | 10 | 0.93% |
| hsa_circ_0035503 | 2143 | 10 | 0.93% |
| hsa_circ_0035515 | 2143 | 10 | 0.93% |
| hsa_circ_0041535 | 2143 | 10 | 0.93% |

| | | | |
|---|---|---|---|
| hsa_circ_0047205 | 2143 | 10 | 0.93% |
| hsa_circ_0083036 | 2143 | 10 | 0.93% |
| hsa_circ_0087172 | 2143 | 10 | 0.93% |
| hsa_circ_0114408 | 2143 | 10 | 0.93% |
| hsa_circ_0046099 | 2786 | 13 | 0.93% |
| hsa_circ_0064684 | 2786 | 13 | 0.93% |
| hsa_circ_0120300 | 3858 | 18 | 0.93% |
| hsa_circ_0131338 | 3001 | 14 | 0.93% |
| hsa_circ_0045512 | 2358 | 11 | 0.93% |
| hsa_circ_0046781 | 2358 | 11 | 0.93% |
| hsa_circ_0074273 | 2358 | 11 | 0.93% |
| hsa_circ_0131989 | 2358 | 11 | 0.93% |
| hsa_circ_0028035 | 2787 | 13 | 0.93% |
| hsa_circ_0125892 | 7718 | 36 | 0.93% |
| hsa_circ_0041332 | 2144 | 10 | 0.93% |
| hsa_circ_0086169 | 2144 | 10 | 0.93% |
| hsa_circ_0097066 | 2144 | 10 | 0.93% |
| hsa_circ_0032586 | 2573 | 12 | 0.93% |
| hsa_circ_0079580 | 2573 | 12 | 0.93% |
| hsa_circ_0060322 | 3002 | 14 | 0.93% |
| hsa_circ_0022348 | 2359 | 11 | 0.93% |
| hsa_circ_0061941 | 2359 | 11 | 0.93% |
| hsa_circ_0094034 | 2359 | 11 | 0.93% |
| hsa_circ_0124665 | 2359 | 11 | 0.93% |
| hsa_circ_0063308 | 3003 | 14 | 0.93% |
| hsa_circ_0090000 | 2574 | 12 | 0.93% |
| hsa_circ_0131091 | 2574 | 12 | 0.93% |
| hsa_circ_0009837 | 2145 | 10 | 0.93% |
| hsa_circ_0039839 | 2145 | 10 | 0.93% |
| hsa_circ_0081302 | 2145 | 10 | 0.93% |
| hsa_circ_0102744 | 3218 | 15 | 0.93% |
| hsa_circ_0029388 | 2789 | 13 | 0.93% |
| hsa_circ_0053659 | 2789 | 13 | 0.93% |
| hsa_circ_0007428 | 2360 | 11 | 0.93% |
| hsa_circ_0036522 | 2360 | 11 | 0.93% |
| hsa_circ_0089369 | 2360 | 11 | 0.93% |
| hsa_circ_0133963 | 2360 | 11 | 0.93% |
| hsa_circ_0115729 | 2575 | 12 | 0.93% |
| hsa_circ_0001642 | 2146 | 10 | 0.93% |
| hsa_circ_0051744 | 2146 | 10 | 0.93% |
| hsa_circ_0057709 | 2146 | 10 | 0.93% |
| hsa_circ_0084409 | 2790 | 13 | 0.93% |
| hsa_circ_0045525 | 2361 | 11 | 0.93% |
| hsa_circ_0079442 | 2361 | 11 | 0.93% |
| hsa_circ_0111082 | 2361 | 11 | 0.93% |
| hsa_circ_0115749 | 2361 | 11 | 0.93% |
| hsa_circ_0133792 | 2361 | 11 | 0.93% |
| hsa_circ_0019742 | 2576 | 12 | 0.93% |
| hsa_circ_0054371 | 2147 | 10 | 0.93% |
| hsa_circ_0057475 | 2147 | 10 | 0.93% |

| | | | |
|---|---|---|---|
| hsa_circ_0061797 | 2147 | 10 | 0.93% |
| hsa_circ_0078395 | 2147 | 10 | 0.93% |
| hsa_circ_0120367 | 2147 | 10 | 0.93% |
| hsa_circ_0016902 | 2362 | 11 | 0.93% |
| hsa_circ_0098010 | 2362 | 11 | 0.93% |
| hsa_circ_0109380 | 2362 | 11 | 0.93% |
| hsa_circ_0110107 | 2362 | 11 | 0.93% |
| hsa_circ_0139944 | 2362 | 11 | 0.93% |
| hsa_circ_0023146 | 2577 | 12 | 0.93% |
| hsa_circ_0030493 | 2577 | 12 | 0.93% |
| hsa_circ_0021009 | 2792 | 13 | 0.93% |
| hsa_circ_0002627 | 2363 | 11 | 0.93% |
| hsa_circ_0004054 | 2363 | 11 | 0.93% |
| hsa_circ_0006514 | 2363 | 11 | 0.93% |
| hsa_circ_0055626 | 2363 | 11 | 0.93% |
| hsa_circ_0059808 | 2363 | 11 | 0.93% |
| hsa_circ_0090397 | 2363 | 11 | 0.93% |
| hsa_circ_0110108 | 2363 | 11 | 0.93% |
| hsa_circ_0118640 | 2363 | 11 | 0.93% |
| hsa_circ_0118853 | 2363 | 11 | 0.93% |
| hsa_circ_0128997 | 2793 | 13 | 0.93% |
| hsa_circ_0133035 | 2793 | 13 | 0.93% |
| hsa_circ_0004779 | 7306 | 34 | 0.93% |
| hsa_circ_0109531 | 5587 | 26 | 0.93% |
| hsa_circ_0052902 | 2149 | 10 | 0.93% |
| hsa_circ_0077066 | 2364 | 11 | 0.93% |
| hsa_circ_0030216 | 2579 | 12 | 0.93% |
| hsa_circ_0053104 | 2579 | 12 | 0.93% |
| hsa_circ_0087306 | 2579 | 12 | 0.93% |
| hsa_circ_0140603 | 2579 | 12 | 0.93% |
| hsa_circ_0063517 | 2794 | 13 | 0.93% |
| hsa_circ_0092526 | 2794 | 13 | 0.93% |
| hsa_circ_0139621 | 5803 | 27 | 0.93% |
| hsa_circ_0132285 | 4515 | 21 | 0.93% |
| hsa_circ_0114003 | 2795 | 13 | 0.93% |
| hsa_circ_0011321 | 2580 | 12 | 0.93% |
| hsa_circ_0089444 | 2580 | 12 | 0.93% |
| hsa_circ_0134586 | 2580 | 12 | 0.93% |
| hsa_circ_0022754 | 2365 | 11 | 0.93% |
| hsa_circ_0043155 | 2365 | 11 | 0.93% |
| hsa_circ_0049455 | 2365 | 11 | 0.93% |
| hsa_circ_0053481 | 2365 | 11 | 0.93% |
| hsa_circ_0078574 | 2365 | 11 | 0.93% |
| hsa_circ_0121871 | 2365 | 11 | 0.93% |
| hsa_circ_0025481 | 2150 | 10 | 0.93% |
| hsa_circ_0027983 | 2150 | 10 | 0.93% |
| hsa_circ_0030902 | 2150 | 10 | 0.93% |
| hsa_circ_0062353 | 2150 | 10 | 0.93% |
| hsa_circ_0109334 | 2150 | 10 | 0.93% |
| hsa_circ_0109568 | 4731 | 22 | 0.93% |

| | | | |
|---|---|---|---|
| hsa_circ_0030155 | 3226 | 15 | 0.93% |
| hsa_circ_0078815 | 3226 | 15 | 0.93% |
| hsa_circ_0078929 | 3226 | 15 | 0.93% |
| hsa_circ_0078970 | 3226 | 15 | 0.93% |
| hsa_circ_0088667 | 2581 | 12 | 0.93% |
| hsa_circ_0000593 | 2366 | 11 | 0.93% |
| hsa_circ_0009760 | 2366 | 11 | 0.93% |
| hsa_circ_0019130 | 2366 | 11 | 0.93% |
| hsa_circ_0036464 | 2366 | 11 | 0.93% |
| hsa_circ_0087812 | 2151 | 10 | 0.93% |
| hsa_circ_0090913 | 2151 | 10 | 0.93% |
| hsa_circ_0116049 | 2151 | 10 | 0.93% |
| hsa_circ_0117942 | 2151 | 10 | 0.93% |
| hsa_circ_0036973 | 3012 | 14 | 0.93% |
| hsa_circ_0091456 | 7315 | 34 | 0.93% |
| hsa_circ_0007349 | 4303 | 20 | 0.93% |
| hsa_circ_0095367 | 2797 | 13 | 0.93% |
| hsa_circ_0044627 | 5164 | 24 | 0.93% |
| hsa_circ_0078690 | 2582 | 12 | 0.93% |
| hsa_circ_0120800 | 2582 | 12 | 0.93% |
| hsa_circ_0019926 | 2367 | 11 | 0.93% |
| hsa_circ_0021895 | 2367 | 11 | 0.93% |
| hsa_circ_0063110 | 2367 | 11 | 0.93% |
| hsa_circ_0083451 | 2367 | 11 | 0.93% |
| hsa_circ_0095456 | 2367 | 11 | 0.93% |
| hsa_circ_0131140 | 2367 | 11 | 0.93% |
| hsa_circ_0109056 | 4304 | 20 | 0.93% |
| hsa_circ_0030357 | 2152 | 10 | 0.93% |
| hsa_circ_0047791 | 2152 | 10 | 0.93% |
| hsa_circ_0059498 | 2152 | 10 | 0.93% |
| hsa_circ_0074115 | 2152 | 10 | 0.93% |
| hsa_circ_0074636 | 2152 | 10 | 0.93% |
| hsa_circ_0084192 | 2152 | 10 | 0.93% |
| hsa_circ_0119215 | 2152 | 10 | 0.93% |
| hsa_circ_0125843 | 2152 | 10 | 0.93% |
| hsa_circ_0083469 | 2798 | 13 | 0.93% |
| hsa_circ_0067936 | 3229 | 15 | 0.93% |
| hsa_circ_0120116 | 5597 | 26 | 0.93% |
| hsa_circ_0030933 | 2368 | 11 | 0.93% |
| hsa_circ_0041649 | 2368 | 11 | 0.93% |
| hsa_circ_0006983 | 5167 | 24 | 0.93% |
| hsa_circ_0039692 | 2153 | 10 | 0.93% |
| hsa_circ_0061412 | 2153 | 10 | 0.93% |
| hsa_circ_0133411 | 2153 | 10 | 0.93% |
| hsa_circ_0023661 | 2799 | 13 | 0.93% |
| hsa_circ_0058168 | 2799 | 13 | 0.93% |
| hsa_circ_0117993 | 2584 | 12 | 0.93% |
| hsa_circ_0030667 | 2369 | 11 | 0.93% |
| hsa_circ_0054562 | 2369 | 11 | 0.93% |
| hsa_circ_0014841 | 2154 | 10 | 0.93% |

| | | | |
|---|---|---|---|
| hsa_circ_0024195 | 2154 | 10 | 0.93% |
| hsa_circ_0025435 | 2154 | 10 | 0.93% |
| hsa_circ_0073913 | 2154 | 10 | 0.93% |
| hsa_circ_0062415 | 3016 | 14 | 0.93% |
| hsa_circ_0074383 | 3447 | 16 | 0.93% |
| hsa_circ_0084311 | 2370 | 11 | 0.93% |
| hsa_circ_0126409 | 5602 | 26 | 0.93% |
| hsa_circ_0013551 | 2155 | 10 | 0.93% |
| hsa_circ_0017703 | 2155 | 10 | 0.93% |
| hsa_circ_0019889 | 2371 | 11 | 0.93% |
| hsa_circ_0053570 | 2371 | 11 | 0.93% |
| hsa_circ_0081975 | 2371 | 11 | 0.93% |
| hsa_circ_0086982 | 2371 | 11 | 0.93% |
| hsa_circ_0107883 | 2371 | 11 | 0.93% |
| hsa_circ_0013391 | 6036 | 28 | 0.93% |
| hsa_circ_0061683 | 2587 | 12 | 0.93% |
| hsa_circ_0117149 | 2587 | 12 | 0.93% |
| hsa_circ_0087510 | 2156 | 10 | 0.93% |
| hsa_circ_0113193 | 2156 | 10 | 0.93% |
| hsa_circ_0134701 | 2156 | 10 | 0.93% |
| hsa_circ_0067624 | 2803 | 13 | 0.93% |
| hsa_circ_0016597 | 2372 | 11 | 0.93% |
| hsa_circ_0019530 | 2372 | 11 | 0.93% |
| hsa_circ_0071932 | 2372 | 11 | 0.93% |
| hsa_circ_0085197 | 2372 | 11 | 0.93% |
| hsa_circ_0063231 | 2588 | 12 | 0.93% |
| hsa_circ_0095238 | 3451 | 16 | 0.93% |
| hsa_circ_0015559 | 2804 | 13 | 0.93% |
| hsa_circ_0087477 | 2804 | 13 | 0.93% |
| hsa_circ_0022845 | 2157 | 10 | 0.93% |
| hsa_circ_0057552 | 2157 | 10 | 0.93% |
| hsa_circ_0064136 | 2157 | 10 | 0.93% |
| hsa_circ_0095616 | 2157 | 10 | 0.93% |
| hsa_circ_0096411 | 2157 | 10 | 0.93% |
| hsa_circ_0133011 | 2157 | 10 | 0.93% |
| hsa_circ_0052103 | 7981 | 37 | 0.93% |
| hsa_circ_0138738 | 4530 | 21 | 0.93% |
| hsa_circ_0018238 | 2373 | 11 | 0.93% |
| hsa_circ_0033562 | 2373 | 11 | 0.93% |
| hsa_circ_0048086 | 2373 | 11 | 0.93% |
| hsa_circ_0050519 | 2373 | 11 | 0.93% |
| hsa_circ_0086021 | 2373 | 11 | 0.93% |
| hsa_circ_0122432 | 6042 | 28 | 0.93% |
| hsa_circ_0114410 | 3021 | 14 | 0.93% |
| hsa_circ_0040875 | 3237 | 15 | 0.93% |
| hsa_circ_0035757 | 2158 | 10 | 0.93% |
| hsa_circ_0078354 | 2158 | 10 | 0.93% |
| hsa_circ_0092712 | 2158 | 10 | 0.93% |
| hsa_circ_0074692 | 3453 | 16 | 0.93% |
| hsa_circ_0019331 | 2374 | 11 | 0.93% |

| | | | |
|---|---|---|---|
| hsa_circ_0036977 | 2374 | 11 | 0.93% |
| hsa_circ_0045405 | 2374 | 11 | 0.93% |
| hsa_circ_0062703 | 2374 | 11 | 0.93% |
| hsa_circ_0079188 | 2374 | 11 | 0.93% |
| hsa_circ_0081909 | 2374 | 11 | 0.93% |
| hsa_circ_0031986 | 3669 | 17 | 0.93% |
| hsa_circ_0106423 | 2590 | 12 | 0.93% |
| hsa_circ_0070031 | 2159 | 10 | 0.93% |
| hsa_circ_0072144 | 2159 | 10 | 0.93% |
| hsa_circ_0081551 | 2159 | 10 | 0.93% |
| hsa_circ_0010748 | 2375 | 11 | 0.93% |
| hsa_circ_0017842 | 2375 | 11 | 0.93% |
| hsa_circ_0026745 | 2375 | 11 | 0.93% |
| hsa_circ_0037990 | 2375 | 11 | 0.93% |
| hsa_circ_0046391 | 2375 | 11 | 0.93% |
| hsa_circ_0053517 | 2375 | 11 | 0.93% |
| hsa_circ_0018962 | 2591 | 12 | 0.93% |
| hsa_circ_0082241 | 2591 | 12 | 0.93% |
| hsa_circ_0012079 | 3023 | 14 | 0.93% |
| hsa_circ_0131484 | 6695 | 31 | 0.93% |
| hsa_circ_0032860 | 2808 | 13 | 0.93% |
| hsa_circ_0029526 | 2376 | 11 | 0.93% |
| hsa_circ_0049130 | 2376 | 11 | 0.93% |
| hsa_circ_0051532 | 2376 | 11 | 0.93% |
| hsa_circ_0065561 | 2160 | 10 | 0.93% |
| hsa_circ_0088958 | 2160 | 10 | 0.93% |
| hsa_circ_0111252 | 5401 | 25 | 0.93% |
| hsa_circ_0018946 | 2593 | 12 | 0.93% |
| hsa_circ_0054541 | 2593 | 12 | 0.93% |
| hsa_circ_0031402 | 2377 | 11 | 0.93% |
| hsa_circ_0037818 | 2377 | 11 | 0.93% |
| hsa_circ_0045867 | 2377 | 11 | 0.93% |
| hsa_circ_0059350 | 2377 | 11 | 0.93% |
| hsa_circ_0100814 | 2377 | 11 | 0.93% |
| hsa_circ_0015988 | 2161 | 10 | 0.93% |
| hsa_circ_0046530 | 2161 | 10 | 0.93% |
| hsa_circ_0088320 | 2161 | 10 | 0.93% |
| hsa_circ_0110245 | 2161 | 10 | 0.93% |
| hsa_circ_0117833 | 2161 | 10 | 0.93% |
| hsa_circ_0017904 | 7132 | 33 | 0.93% |
| hsa_circ_0041843 | 9078 | 42 | 0.93% |
| hsa_circ_0111417 | 3891 | 18 | 0.93% |
| hsa_circ_0043533 | 2594 | 12 | 0.93% |
| hsa_circ_0010960 | 2378 | 11 | 0.93% |
| hsa_circ_0019844 | 2378 | 11 | 0.93% |
| hsa_circ_0042721 | 2378 | 11 | 0.93% |
| hsa_circ_0059351 | 2378 | 11 | 0.93% |
| hsa_circ_0093368 | 2378 | 11 | 0.93% |
| hsa_circ_0118777 | 2378 | 11 | 0.93% |
| hsa_circ_0126339 | 2378 | 11 | 0.93% |

| | | | |
|---|---|---|---|
| hsa_circ_0027293 | 3243 | 15 | 0.93% |
| hsa_circ_0034823 | 2162 | 10 | 0.93% |
| hsa_circ_0041575 | 2595 | 12 | 0.92% |
| hsa_circ_0046931 | 2379 | 11 | 0.92% |
| hsa_circ_0061729 | 2379 | 11 | 0.92% |
| hsa_circ_0104374 | 5407 | 25 | 0.92% |
| hsa_circ_0023314 | 2163 | 10 | 0.92% |
| hsa_circ_0086264 | 2596 | 12 | 0.92% |
| hsa_circ_0128213 | 2596 | 12 | 0.92% |
| hsa_circ_0117096 | 3029 | 14 | 0.92% |
| hsa_circ_0041447 | 2380 | 11 | 0.92% |
| hsa_circ_0068339 | 2380 | 11 | 0.92% |
| hsa_circ_0089608 | 2380 | 11 | 0.92% |
| hsa_circ_0093859 | 2380 | 11 | 0.92% |
| hsa_circ_0008851 | 4111 | 19 | 0.92% |
| hsa_circ_0011672 | 2164 | 10 | 0.92% |
| hsa_circ_0047101 | 2164 | 10 | 0.92% |
| hsa_circ_0074466 | 2164 | 10 | 0.92% |
| hsa_circ_0075428 | 2164 | 10 | 0.92% |
| hsa_circ_0080267 | 2164 | 10 | 0.92% |
| hsa_circ_0130964 | 2164 | 10 | 0.92% |
| hsa_circ_0118436 | 3679 | 17 | 0.92% |
| hsa_circ_0009329 | 2597 | 12 | 0.92% |
| hsa_circ_0057955 | 4329 | 20 | 0.92% |
| hsa_circ_0010810 | 2381 | 11 | 0.92% |
| hsa_circ_0082738 | 2381 | 11 | 0.92% |
| hsa_circ_0089154 | 2381 | 11 | 0.92% |
| hsa_circ_0010593 | 2814 | 13 | 0.92% |
| hsa_circ_0032951 | 2814 | 13 | 0.92% |
| hsa_circ_0067306 | 2598 | 12 | 0.92% |
| hsa_circ_0115360 | 2598 | 12 | 0.92% |
| hsa_circ_0064254 | 2165 | 10 | 0.92% |
| hsa_circ_0025353 | 2815 | 13 | 0.92% |
| hsa_circ_0017501 | 2382 | 11 | 0.92% |
| hsa_circ_0028742 | 2382 | 11 | 0.92% |
| hsa_circ_0054854 | 2382 | 11 | 0.92% |
| hsa_circ_0114162 | 2382 | 11 | 0.92% |
| hsa_circ_0120662 | 2382 | 11 | 0.92% |
| hsa_circ_0127144 | 2382 | 11 | 0.92% |
| hsa_circ_0102803 | 2599 | 12 | 0.92% |
| hsa_circ_0014243 | 2166 | 10 | 0.92% |
| hsa_circ_0020439 | 2166 | 10 | 0.92% |
| hsa_circ_0041400 | 2166 | 10 | 0.92% |
| hsa_circ_0049028 | 2166 | 10 | 0.92% |
| hsa_circ_0072069 | 2166 | 10 | 0.92% |
| hsa_circ_0079711 | 2166 | 10 | 0.92% |
| hsa_circ_0081216 | 2166 | 10 | 0.92% |
| hsa_circ_0105226 | 8881 | 41 | 0.92% |
| hsa_circ_0127846 | 3466 | 16 | 0.92% |
| hsa_circ_0053152 | 2383 | 11 | 0.92% |

| | | | |
|---|---|---|---|
| hsa_circ_0053358 | 2383 | 11 | 0.92% |
| hsa_circ_0107937 | 2383 | 11 | 0.92% |
| hsa_circ_0063449 | 2600 | 12 | 0.92% |
| hsa_circ_0075244 | 2600 | 12 | 0.92% |
| hsa_circ_0077582 | 2600 | 12 | 0.92% |
| hsa_circ_0033892 | 2167 | 10 | 0.92% |
| hsa_circ_0057634 | 2167 | 10 | 0.92% |
| hsa_circ_0062045 | 2167 | 10 | 0.92% |
| hsa_circ_0064662 | 2167 | 10 | 0.92% |
| hsa_circ_0074513 | 2167 | 10 | 0.92% |
| hsa_circ_0096498 | 2167 | 10 | 0.92% |
| hsa_circ_0096851 | 2167 | 10 | 0.92% |
| hsa_circ_0119545 | 2167 | 10 | 0.92% |
| hsa_circ_0052910 | 2384 | 11 | 0.92% |
| hsa_circ_0056468 | 2384 | 11 | 0.92% |
| hsa_circ_0056685 | 2384 | 11 | 0.92% |
| hsa_circ_0059101 | 2384 | 11 | 0.92% |
| hsa_circ_0060453 | 2384 | 11 | 0.92% |
| hsa_circ_0100552 | 2384 | 11 | 0.92% |
| hsa_circ_0075477 | 6720 | 31 | 0.92% |
| hsa_circ_0024018 | 4119 | 19 | 0.92% |
| hsa_circ_0137977 | 6504 | 30 | 0.92% |
| hsa_circ_0030803 | 2168 | 10 | 0.92% |
| hsa_circ_0050307 | 2168 | 10 | 0.92% |
| hsa_circ_0024386 | 2385 | 11 | 0.92% |
| hsa_circ_0076020 | 2385 | 11 | 0.92% |
| hsa_circ_0112485 | 2602 | 12 | 0.92% |
| hsa_circ_0012054 | 2819 | 13 | 0.92% |
| hsa_circ_0081917 | 2819 | 13 | 0.92% |
| hsa_circ_0083945 | 2819 | 13 | 0.92% |
| hsa_circ_0009494 | 3036 | 14 | 0.92% |
| hsa_circ_0130272 | 10843 | 50 | 0.92% |
| hsa_circ_0006165 | 5639 | 26 | 0.92% |
| hsa_circ_0010469 | 2169 | 10 | 0.92% |
| hsa_circ_0014984 | 2169 | 10 | 0.92% |
| hsa_circ_0035150 | 2169 | 10 | 0.92% |
| hsa_circ_0047339 | 2169 | 10 | 0.92% |
| hsa_circ_0056302 | 2169 | 10 | 0.92% |
| hsa_circ_0063787 | 2169 | 10 | 0.92% |
| hsa_circ_0116042 | 2169 | 10 | 0.92% |
| hsa_circ_0137890 | 2169 | 10 | 0.92% |
| hsa_circ_0089066 | 2386 | 11 | 0.92% |
| hsa_circ_0053960 | 2603 | 12 | 0.92% |
| hsa_circ_0048943 | 2820 | 13 | 0.92% |
| hsa_circ_0113790 | 10848 | 50 | 0.92% |
| hsa_circ_0012896 | 3038 | 14 | 0.92% |
| hsa_circ_0032823 | 2821 | 13 | 0.92% |
| hsa_circ_0081402 | 2821 | 13 | 0.92% |
| hsa_circ_0018419 | 2604 | 12 | 0.92% |
| hsa_circ_0058751 | 2604 | 12 | 0.92% |

| | | | |
|---|---|---|---|
| hsa_circ_0044982 | 2387 | 11 | 0.92% |
| hsa_circ_0071025 | 2387 | 11 | 0.92% |
| hsa_circ_0083082 | 2387 | 11 | 0.92% |
| hsa_circ_0073831 | 2170 | 10 | 0.92% |
| hsa_circ_0063516 | 8683 | 40 | 0.92% |
| hsa_circ_0074560 | 2822 | 13 | 0.92% |
| hsa_circ_0107933 | 2822 | 13 | 0.92% |
| hsa_circ_0031339 | 2388 | 11 | 0.92% |
| hsa_circ_0058141 | 2388 | 11 | 0.92% |
| hsa_circ_0067423 | 2388 | 11 | 0.92% |
| hsa_circ_0073086 | 2388 | 11 | 0.92% |
| hsa_circ_0117445 | 2388 | 11 | 0.92% |
| hsa_circ_0051552 | 2171 | 10 | 0.92% |
| hsa_circ_0055328 | 2171 | 10 | 0.92% |
| hsa_circ_0091127 | 2171 | 10 | 0.92% |
| hsa_circ_0097071 | 3474 | 16 | 0.92% |
| hsa_circ_0090990 | 3257 | 15 | 0.92% |
| hsa_circ_0019195 | 2606 | 12 | 0.92% |
| hsa_circ_0023627 | 2389 | 11 | 0.92% |
| hsa_circ_0043259 | 2389 | 11 | 0.92% |
| hsa_circ_0049551 | 2389 | 11 | 0.92% |
| hsa_circ_0051151 | 2389 | 11 | 0.92% |
| hsa_circ_0060175 | 2389 | 11 | 0.92% |
| hsa_circ_0124545 | 2389 | 11 | 0.92% |
| hsa_circ_0010125 | 2172 | 10 | 0.92% |
| hsa_circ_0012803 | 2172 | 10 | 0.92% |
| hsa_circ_0029729 | 2172 | 10 | 0.92% |
| hsa_circ_0095174 | 2172 | 10 | 0.92% |
| hsa_circ_0099326 | 2172 | 10 | 0.92% |
| hsa_circ_0108167 | 2172 | 10 | 0.92% |
| hsa_circ_0132221 | 2172 | 10 | 0.92% |
| hsa_circ_0127968 | 4562 | 21 | 0.92% |
| hsa_circ_0019220 | 2607 | 12 | 0.92% |
| hsa_circ_0032555 | 2607 | 12 | 0.92% |
| hsa_circ_0044040 | 2607 | 12 | 0.92% |
| hsa_circ_0083583 | 2607 | 12 | 0.92% |
| hsa_circ_0044511 | 4997 | 23 | 0.92% |
| hsa_circ_0010719 | 2390 | 11 | 0.92% |
| hsa_circ_0038321 | 2390 | 11 | 0.92% |
| hsa_circ_0074400 | 2390 | 11 | 0.92% |
| hsa_circ_0083832 | 2390 | 11 | 0.92% |
| hsa_circ_0022676 | 2173 | 10 | 0.92% |
| hsa_circ_0080489 | 2173 | 10 | 0.92% |
| hsa_circ_0086074 | 2173 | 10 | 0.92% |
| hsa_circ_0118463 | 2173 | 10 | 0.92% |
| hsa_circ_0140150 | 2173 | 10 | 0.92% |
| hsa_circ_0000346 | 6085 | 28 | 0.92% |
| hsa_circ_0002147 | 7389 | 34 | 0.92% |
| hsa_circ_0132501 | 2608 | 12 | 0.92% |
| hsa_circ_0022854 | 2391 | 11 | 0.92% |

| | | | |
|---|---|---|---|
| hsa_circ_0028452 | 2391 | 11 | 0.92% |
| hsa_circ_0032398 | 2174 | 10 | 0.92% |
| hsa_circ_0117708 | 2174 | 10 | 0.92% |
| hsa_circ_0063447 | 2609 | 12 | 0.92% |
| hsa_circ_0063310 | 3479 | 16 | 0.92% |
| hsa_circ_0032399 | 2392 | 11 | 0.92% |
| hsa_circ_0088959 | 2392 | 11 | 0.92% |
| hsa_circ_0103651 | 2392 | 11 | 0.92% |
| hsa_circ_0113660 | 2392 | 11 | 0.92% |
| hsa_circ_0112019 | 10003 | 46 | 0.92% |
| hsa_circ_0048183 | 2610 | 12 | 0.92% |
| hsa_circ_0020429 | 2175 | 10 | 0.92% |
| hsa_circ_0058957 | 2175 | 10 | 0.92% |
| hsa_circ_0005550 | 2393 | 11 | 0.92% |
| hsa_circ_0040885 | 2393 | 11 | 0.92% |
| hsa_circ_0051483 | 2393 | 11 | 0.92% |
| hsa_circ_0061725 | 2393 | 11 | 0.92% |
| hsa_circ_0083933 | 2393 | 11 | 0.92% |
| hsa_circ_0108221 | 2393 | 11 | 0.92% |
| hsa_circ_0123689 | 7832 | 36 | 0.92% |
| hsa_circ_0079119 | 3046 | 14 | 0.92% |
| hsa_circ_0085002 | 6528 | 30 | 0.92% |
| hsa_circ_0013839 | 2176 | 10 | 0.92% |
| hsa_circ_0049875 | 2176 | 10 | 0.92% |
| hsa_circ_0103452 | 2176 | 10 | 0.92% |
| hsa_circ_0107355 | 2176 | 10 | 0.92% |
| hsa_circ_0060679 | 2829 | 13 | 0.92% |
| hsa_circ_0101155 | 7399 | 34 | 0.92% |
| hsa_circ_0130717 | 8923 | 41 | 0.92% |
| hsa_circ_0019355 | 2394 | 11 | 0.92% |
| hsa_circ_0034228 | 2394 | 11 | 0.92% |
| hsa_circ_0068208 | 2394 | 11 | 0.92% |
| hsa_circ_0060909 | 2612 | 12 | 0.92% |
| hsa_circ_0063861 | 2612 | 12 | 0.92% |
| hsa_circ_0074958 | 2612 | 12 | 0.92% |
| hsa_circ_0114430 | 2612 | 12 | 0.92% |
| hsa_circ_0049868 | 2177 | 10 | 0.92% |
| hsa_circ_0052181 | 2177 | 10 | 0.92% |
| hsa_circ_0071585 | 2177 | 10 | 0.92% |
| hsa_circ_0076711 | 2177 | 10 | 0.92% |
| hsa_circ_0079915 | 2177 | 10 | 0.92% |
| hsa_circ_0103934 | 2177 | 10 | 0.92% |
| hsa_circ_0107580 | 2177 | 10 | 0.92% |
| hsa_circ_0016589 | 2395 | 11 | 0.92% |
| hsa_circ_0019710 | 2395 | 11 | 0.92% |
| hsa_circ_0129440 | 5879 | 27 | 0.92% |
| hsa_circ_0029738 | 2613 | 12 | 0.92% |
| hsa_circ_0088139 | 2613 | 12 | 0.92% |
| hsa_circ_0010371 | 2178 | 10 | 0.92% |
| hsa_circ_0046328 | 2178 | 10 | 0.92% |

| | | | |
|---|---|---|---|
| hsa_circ_0048539 | 2178 | 10 | 0.92% |
| hsa_circ_0081085 | 2178 | 10 | 0.92% |
| hsa_circ_0089368 | 2178 | 10 | 0.92% |
| hsa_circ_0019235 | 2396 | 11 | 0.92% |
| hsa_circ_0047812 | 2832 | 13 | 0.92% |
| hsa_circ_0091691 | 2832 | 13 | 0.92% |
| hsa_circ_0013847 | 2179 | 10 | 0.92% |
| hsa_circ_0038547 | 2179 | 10 | 0.92% |
| hsa_circ_0078672 | 2179 | 10 | 0.92% |
| hsa_circ_0082373 | 2179 | 10 | 0.92% |
| hsa_circ_0017658 | 2397 | 11 | 0.92% |
| hsa_circ_0023042 | 2397 | 11 | 0.92% |
| hsa_circ_0071690 | 2615 | 12 | 0.92% |
| hsa_circ_0098794 | 6975 | 32 | 0.92% |
| hsa_circ_0065501 | 4142 | 19 | 0.92% |
| hsa_circ_0057438 | 2834 | 13 | 0.92% |
| hsa_circ_0034542 | 2616 | 12 | 0.92% |
| hsa_circ_0023367 | 2398 | 11 | 0.92% |
| hsa_circ_0049858 | 2398 | 11 | 0.92% |
| hsa_circ_0056915 | 2398 | 11 | 0.92% |
| hsa_circ_0061393 | 2398 | 11 | 0.92% |
| hsa_circ_0073182 | 2398 | 11 | 0.92% |
| hsa_circ_0079343 | 2398 | 11 | 0.92% |
| hsa_circ_0112921 | 2398 | 11 | 0.92% |
| hsa_circ_0120336 | 2398 | 11 | 0.92% |
| hsa_circ_0013687 | 2180 | 10 | 0.92% |
| hsa_circ_0035082 | 2180 | 10 | 0.92% |
| hsa_circ_0048668 | 2180 | 10 | 0.92% |
| hsa_circ_0019407 | 3053 | 14 | 0.92% |
| hsa_circ_0084640 | 3053 | 14 | 0.92% |
| hsa_circ_0031136 | 2399 | 11 | 0.92% |
| hsa_circ_0077732 | 2181 | 10 | 0.92% |
| hsa_circ_0098346 | 2181 | 10 | 0.92% |
| hsa_circ_0139754 | 2181 | 10 | 0.92% |
| hsa_circ_0037605 | 2618 | 12 | 0.92% |
| hsa_circ_0045179 | 2618 | 12 | 0.92% |
| hsa_circ_0063502 | 2618 | 12 | 0.92% |
| hsa_circ_0090776 | 2618 | 12 | 0.92% |
| hsa_circ_0010717 | 2400 | 11 | 0.92% |
| hsa_circ_0033520 | 2400 | 11 | 0.92% |
| hsa_circ_0054597 | 2400 | 11 | 0.92% |
| hsa_circ_0060215 | 2400 | 11 | 0.92% |
| hsa_circ_0065582 | 2400 | 11 | 0.92% |
| hsa_circ_0114255 | 2400 | 11 | 0.92% |
| hsa_circ_0064990 | 3273 | 15 | 0.92% |
| hsa_circ_0055837 | 2182 | 10 | 0.92% |
| hsa_circ_0074734 | 2182 | 10 | 0.92% |
| hsa_circ_0091026 | 2182 | 10 | 0.92% |
| hsa_circ_0050425 | 3055 | 14 | 0.92% |
| hsa_circ_0030530 | 2837 | 13 | 0.92% |

| | | | |
|---|---|---|---|
| hsa_circ_0138441 | 2837 | 13 | 0.92% |
| hsa_circ_0033451 | 2619 | 12 | 0.92% |
| hsa_circ_0053049 | 2619 | 12 | 0.92% |
| hsa_circ_0055154 | 2619 | 12 | 0.92% |
| hsa_circ_0084117 | 2619 | 12 | 0.92% |
| hsa_circ_0108302 | 4802 | 22 | 0.92% |
| hsa_circ_0059107 | 2401 | 11 | 0.92% |
| hsa_circ_0068193 | 2401 | 11 | 0.92% |
| hsa_circ_0091835 | 2401 | 11 | 0.92% |
| hsa_circ_0131412 | 2401 | 11 | 0.92% |
| hsa_circ_0056373 | 2183 | 10 | 0.92% |
| hsa_circ_0116271 | 2183 | 10 | 0.92% |
| hsa_circ_0036189 | 2838 | 13 | 0.92% |
| hsa_circ_0099238 | 4148 | 19 | 0.92% |
| hsa_circ_0067734 | 2620 | 12 | 0.92% |
| hsa_circ_0070359 | 2620 | 12 | 0.92% |
| hsa_circ_0137532 | 2620 | 12 | 0.92% |
| hsa_circ_0038101 | 2402 | 11 | 0.92% |
| hsa_circ_0062594 | 2402 | 11 | 0.92% |
| hsa_circ_0122286 | 2402 | 11 | 0.92% |
| hsa_circ_0010655 | 2839 | 13 | 0.92% |
| hsa_circ_0053710 | 2839 | 13 | 0.92% |
| hsa_circ_0064269 | 2184 | 10 | 0.92% |
| hsa_circ_0078405 | 2184 | 10 | 0.92% |
| hsa_circ_0131606 | 2184 | 10 | 0.92% |
| hsa_circ_0014455 | 2403 | 11 | 0.92% |
| hsa_circ_0022700 | 2403 | 11 | 0.92% |
| hsa_circ_0047155 | 2403 | 11 | 0.92% |
| hsa_circ_0059012 | 2403 | 11 | 0.92% |
| hsa_circ_0068741 | 5462 | 25 | 0.92% |
| hsa_circ_0029484 | 3059 | 14 | 0.92% |
| hsa_circ_0074043 | 3059 | 14 | 0.92% |
| hsa_circ_0028225 | 2185 | 10 | 0.92% |
| hsa_circ_0064931 | 2185 | 10 | 0.92% |
| hsa_circ_0018928 | 2841 | 13 | 0.92% |
| hsa_circ_0090918 | 2841 | 13 | 0.92% |
| hsa_circ_0033177 | 2404 | 11 | 0.92% |
| hsa_circ_0043251 | 2404 | 11 | 0.92% |
| hsa_circ_0063690 | 2404 | 11 | 0.92% |
| hsa_circ_0070865 | 2404 | 11 | 0.92% |
| hsa_circ_0091425 | 2404 | 11 | 0.92% |
| hsa_circ_0120658 | 2404 | 11 | 0.92% |
| hsa_circ_0090823 | 3060 | 14 | 0.92% |
| hsa_circ_0018514 | 2623 | 12 | 0.91% |
| hsa_circ_0074773 | 2623 | 12 | 0.91% |
| hsa_circ_0078601 | 2623 | 12 | 0.91% |
| hsa_circ_0053978 | 2186 | 10 | 0.91% |
| hsa_circ_0077902 | 2186 | 10 | 0.91% |
| hsa_circ_0097346 | 6559 | 30 | 0.91% |
| hsa_circ_0016072 | 2405 | 11 | 0.91% |

| | | | |
|---|---|---|---|
| hsa_circ_0045958 | 2405 | 11 | 0.91% |
| hsa_circ_0085382 | 2405 | 11 | 0.91% |
| hsa_circ_0025132 | 2624 | 12 | 0.91% |
| hsa_circ_0052204 | 2843 | 13 | 0.91% |
| hsa_circ_0035613 | 2187 | 10 | 0.91% |
| hsa_circ_0076964 | 2187 | 10 | 0.91% |
| hsa_circ_0077482 | 2187 | 10 | 0.91% |
| hsa_circ_0113584 | 2187 | 10 | 0.91% |
| hsa_circ_0069927 | 3062 | 14 | 0.91% |
| hsa_circ_0106403 | 3062 | 14 | 0.91% |
| hsa_circ_0011494 | 2406 | 11 | 0.91% |
| hsa_circ_0022673 | 2406 | 11 | 0.91% |
| hsa_circ_0037319 | 2406 | 11 | 0.91% |
| hsa_circ_0071304 | 2406 | 11 | 0.91% |
| hsa_circ_0101658 | 2406 | 11 | 0.91% |
| hsa_circ_0086937 | 2625 | 12 | 0.91% |
| hsa_circ_0111168 | 2625 | 12 | 0.91% |
| hsa_circ_0017814 | 2844 | 13 | 0.91% |
| hsa_circ_0084511 | 7439 | 34 | 0.91% |
| hsa_circ_0024709 | 2188 | 10 | 0.91% |
| hsa_circ_0039123 | 2188 | 10 | 0.91% |
| hsa_circ_0046866 | 2188 | 10 | 0.91% |
| hsa_circ_0080998 | 2188 | 10 | 0.91% |
| hsa_circ_0087491 | 2188 | 10 | 0.91% |
| hsa_circ_0095881 | 2188 | 10 | 0.91% |
| hsa_circ_0097781 | 2188 | 10 | 0.91% |
| hsa_circ_0059130 | 3501 | 16 | 0.91% |
| hsa_circ_0020395 | 2407 | 11 | 0.91% |
| hsa_circ_0094373 | 2407 | 11 | 0.91% |
| hsa_circ_0034069 | 2626 | 12 | 0.91% |
| hsa_circ_0085907 | 2626 | 12 | 0.91% |
| hsa_circ_0060667 | 2845 | 13 | 0.91% |
| hsa_circ_0078806 | 5909 | 27 | 0.91% |
| hsa_circ_0029666 | 2189 | 10 | 0.91% |
| hsa_circ_0069021 | 2189 | 10 | 0.91% |
| hsa_circ_0072867 | 2408 | 11 | 0.91% |
| hsa_circ_0088098 | 2408 | 11 | 0.91% |
| hsa_circ_0047764 | 2627 | 12 | 0.91% |
| hsa_circ_0049334 | 2627 | 12 | 0.91% |
| hsa_circ_0083688 | 2627 | 12 | 0.91% |
| hsa_circ_0089030 | 2846 | 13 | 0.91% |
| hsa_circ_0119207 | 9852 | 45 | 0.91% |
| hsa_circ_0136566 | 8541 | 39 | 0.91% |
| hsa_circ_0080293 | 2847 | 13 | 0.91% |
| hsa_circ_0095514 | 2847 | 13 | 0.91% |
| hsa_circ_0048636 | 2409 | 11 | 0.91% |
| hsa_circ_0070235 | 2409 | 11 | 0.91% |
| hsa_circ_0071341 | 2409 | 11 | 0.91% |
| hsa_circ_0112535 | 2409 | 11 | 0.91% |
| hsa_circ_0060251 | 2190 | 10 | 0.91% |

| | | | |
|---|---|---|---|
| hsa_circ_0110443 | 2190 | 10 | 0.91% |
| hsa_circ_0011586 | 2848 | 13 | 0.91% |
| hsa_circ_0121979 | 2629 | 12 | 0.91% |
| hsa_circ_0031837 | 4820 | 22 | 0.91% |
| hsa_circ_0050356 | 2410 | 11 | 0.91% |
| hsa_circ_0066538 | 2410 | 11 | 0.91% |
| hsa_circ_0068322 | 2410 | 11 | 0.91% |
| hsa_circ_0073830 | 2410 | 11 | 0.91% |
| hsa_circ_0040543 | 6793 | 31 | 0.91% |
| hsa_circ_0047076 | 2630 | 12 | 0.91% |
| hsa_circ_0066595 | 2630 | 12 | 0.91% |
| hsa_circ_0047674 | 2192 | 10 | 0.91% |
| hsa_circ_0078731 | 2192 | 10 | 0.91% |
| hsa_circ_0100432 | 2192 | 10 | 0.91% |
| hsa_circ_0052311 | 3069 | 14 | 0.91% |
| hsa_circ_0025411 | 2850 | 13 | 0.91% |
| hsa_circ_0060690 | 2850 | 13 | 0.91% |
| hsa_circ_0139421 | 2850 | 13 | 0.91% |
| hsa_circ_0059673 | 3289 | 15 | 0.91% |
| hsa_circ_0011899 | 2412 | 11 | 0.91% |
| hsa_circ_0035140 | 2412 | 11 | 0.91% |
| hsa_circ_0087732 | 2412 | 11 | 0.91% |
| hsa_circ_0107121 | 2412 | 11 | 0.91% |
| hsa_circ_0106422 | 3070 | 14 | 0.91% |
| hsa_circ_0053920 | 2193 | 10 | 0.91% |
| hsa_circ_0064233 | 2193 | 10 | 0.91% |
| hsa_circ_0079808 | 2193 | 10 | 0.91% |
| hsa_circ_0089022 | 2193 | 10 | 0.91% |
| hsa_circ_0067469 | 2851 | 13 | 0.91% |
| hsa_circ_0114659 | 5922 | 27 | 0.91% |
| hsa_circ_0028016 | 2413 | 11 | 0.91% |
| hsa_circ_0028518 | 2413 | 11 | 0.91% |
| hsa_circ_0044378 | 2413 | 11 | 0.91% |
| hsa_circ_0044634 | 2413 | 11 | 0.91% |
| hsa_circ_0083245 | 2413 | 11 | 0.91% |
| hsa_circ_0050262 | 2852 | 13 | 0.91% |
| hsa_circ_0099449 | 2852 | 13 | 0.91% |
| hsa_circ_0033213 | 2194 | 10 | 0.91% |
| hsa_circ_0049238 | 2194 | 10 | 0.91% |
| hsa_circ_0093451 | 2194 | 10 | 0.91% |
| hsa_circ_0127234 | 2194 | 10 | 0.91% |
| hsa_circ_0027828 | 2633 | 12 | 0.91% |
| hsa_circ_0038850 | 2633 | 12 | 0.91% |
| hsa_circ_0010904 | 2414 | 11 | 0.91% |
| hsa_circ_0019469 | 2414 | 11 | 0.91% |
| hsa_circ_0035931 | 2414 | 11 | 0.91% |
| hsa_circ_0017080 | 2853 | 13 | 0.91% |
| hsa_circ_0052630 | 2853 | 13 | 0.91% |
| hsa_circ_0091307 | 3292 | 15 | 0.91% |
| hsa_circ_0055303 | 2634 | 12 | 0.91% |

| | | | |
|---|---|---|---|
| hsa_circ_0077681 | 2634 | 12 | 0.91% |
| hsa_circ_0036830 | 2195 | 10 | 0.91% |
| hsa_circ_0040377 | 2195 | 10 | 0.91% |
| hsa_circ_0054449 | 2195 | 10 | 0.91% |
| hsa_circ_0065203 | 2195 | 10 | 0.91% |
| hsa_circ_0039779 | 5269 | 24 | 0.91% |
| hsa_circ_0031231 | 2415 | 11 | 0.91% |
| hsa_circ_0066558 | 2415 | 11 | 0.91% |
| hsa_circ_0133789 | 2415 | 11 | 0.91% |
| hsa_circ_0025401 | 2635 | 12 | 0.91% |
| hsa_circ_0087170 | 2196 | 10 | 0.91% |
| hsa_circ_0031472 | 2416 | 11 | 0.91% |
| hsa_circ_0077917 | 2416 | 11 | 0.91% |
| hsa_circ_0088172 | 2856 | 13 | 0.91% |
| hsa_circ_0011553 | 2197 | 10 | 0.91% |
| hsa_circ_0043053 | 2197 | 10 | 0.91% |
| hsa_circ_0056308 | 2197 | 10 | 0.91% |
| hsa_circ_0063435 | 2197 | 10 | 0.91% |
| hsa_circ_0103829 | 2197 | 10 | 0.91% |
| hsa_circ_0055369 | 2417 | 11 | 0.91% |
| hsa_circ_0101263 | 2417 | 11 | 0.91% |
| hsa_circ_0108216 | 2417 | 11 | 0.91% |
| hsa_circ_0119269 | 2417 | 11 | 0.91% |
| hsa_circ_0032717 | 2637 | 12 | 0.91% |
| hsa_circ_0086932 | 2637 | 12 | 0.91% |
| hsa_circ_0094791 | 2637 | 12 | 0.91% |
| hsa_circ_0023133 | 2857 | 13 | 0.91% |
| hsa_circ_0085419 | 2857 | 13 | 0.91% |
| hsa_circ_0013610 | 2198 | 10 | 0.91% |
| hsa_circ_0056087 | 2198 | 10 | 0.91% |
| hsa_circ_0029651 | 2418 | 11 | 0.91% |
| hsa_circ_0131809 | 2418 | 11 | 0.91% |
| hsa_circ_0035804 | 2638 | 12 | 0.91% |
| hsa_circ_0052319 | 2638 | 12 | 0.91% |
| hsa_circ_0086152 | 2638 | 12 | 0.91% |
| hsa_circ_0091802 | 2638 | 12 | 0.91% |
| hsa_circ_0015345 | 2858 | 13 | 0.91% |
| hsa_circ_0078874 | 2199 | 10 | 0.91% |
| hsa_circ_0078914 | 2199 | 10 | 0.91% |
| hsa_circ_0082067 | 2199 | 10 | 0.91% |
| hsa_circ_0129477 | 2199 | 10 | 0.91% |
| hsa_circ_0092419 | 7037 | 32 | 0.91% |
| hsa_circ_0017319 | 2419 | 11 | 0.91% |
| hsa_circ_0038017 | 2419 | 11 | 0.91% |
| hsa_circ_0067889 | 2419 | 11 | 0.91% |
| hsa_circ_0071519 | 2419 | 11 | 0.91% |
| hsa_circ_0071804 | 2419 | 11 | 0.91% |
| hsa_circ_0072214 | 2419 | 11 | 0.91% |
| hsa_circ_0075075 | 2419 | 11 | 0.91% |
| hsa_circ_0138461 | 2419 | 11 | 0.91% |

| | | | |
|---|---|---|---|
| hsa_circ_0011437 | 2639 | 12 | 0.91% |
| hsa_circ_0041010 | 2639 | 12 | 0.91% |
| hsa_circ_0043335 | 2639 | 12 | 0.91% |
| hsa_circ_0057514 | 2639 | 12 | 0.91% |
| hsa_circ_0075743 | 2639 | 12 | 0.91% |
| hsa_circ_0044308 | 3079 | 14 | 0.91% |
| hsa_circ_0066191 | 3080 | 14 | 0.91% |
| hsa_circ_0032950 | 2860 | 13 | 0.91% |
| hsa_circ_0013777 | 2640 | 12 | 0.91% |
| hsa_circ_0032561 | 2640 | 12 | 0.91% |
| hsa_circ_0038440 | 2640 | 12 | 0.91% |
| hsa_circ_0125036 | 2640 | 12 | 0.91% |
| hsa_circ_0069319 | 2420 | 11 | 0.91% |
| hsa_circ_0120599 | 2420 | 11 | 0.91% |
| hsa_circ_0019025 | 2200 | 10 | 0.91% |
| hsa_circ_0036844 | 2861 | 13 | 0.91% |
| hsa_circ_0068581 | 2861 | 13 | 0.91% |
| hsa_circ_0044481 | 2641 | 12 | 0.91% |
| hsa_circ_0020636 | 2421 | 11 | 0.91% |
| hsa_circ_0023291 | 2421 | 11 | 0.91% |
| hsa_circ_0026746 | 2421 | 11 | 0.91% |
| hsa_circ_0011559 | 2201 | 10 | 0.91% |
| hsa_circ_0022629 | 2201 | 10 | 0.91% |
| hsa_circ_0065372 | 2201 | 10 | 0.91% |
| hsa_circ_0079256 | 2201 | 10 | 0.91% |
| hsa_circ_0090821 | 2201 | 10 | 0.91% |
| hsa_circ_0139171 | 2201 | 10 | 0.91% |
| hsa_circ_0013158 | 6164 | 28 | 0.91% |
| hsa_circ_0069947 | 2642 | 12 | 0.91% |
| hsa_circ_0099551 | 9908 | 45 | 0.91% |
| hsa_circ_0024129 | 2422 | 11 | 0.91% |
| hsa_circ_0080585 | 2422 | 11 | 0.91% |
| hsa_circ_0090939 | 2422 | 11 | 0.91% |
| hsa_circ_0112733 | 2422 | 11 | 0.91% |
| hsa_circ_0016185 | 2202 | 10 | 0.91% |
| hsa_circ_0018261 | 2202 | 10 | 0.91% |
| hsa_circ_0038522 | 2202 | 10 | 0.91% |
| hsa_circ_0058627 | 2202 | 10 | 0.91% |
| hsa_circ_0073097 | 2202 | 10 | 0.91% |
| hsa_circ_0114978 | 2202 | 10 | 0.91% |
| hsa_circ_0043495 | 2643 | 12 | 0.91% |
| hsa_circ_0054754 | 2643 | 12 | 0.91% |
| hsa_circ_0058093 | 2643 | 12 | 0.91% |
| hsa_circ_0050859 | 2423 | 11 | 0.91% |
| hsa_circ_0054470 | 2423 | 11 | 0.91% |
| hsa_circ_0082454 | 2423 | 11 | 0.91% |
| hsa_circ_0090790 | 2423 | 11 | 0.91% |
| hsa_circ_0104822 | 2423 | 11 | 0.91% |
| hsa_circ_0139463 | 2423 | 11 | 0.91% |
| hsa_circ_0027803 | 3084 | 14 | 0.91% |

| | | | |
|---|---|---|---|
| hsa_circ_0079458 | 2644 | 12 | 0.91% |
| hsa_circ_0101185 | 5068 | 23 | 0.91% |
| hsa_circ_0020938 | 3085 | 14 | 0.91% |
| hsa_circ_0069401 | 3085 | 14 | 0.91% |
| hsa_circ_0010435 | 2424 | 11 | 0.91% |
| hsa_circ_0035076 | 2424 | 11 | 0.91% |
| hsa_circ_0036716 | 2424 | 11 | 0.91% |
| hsa_circ_0043548 | 2424 | 11 | 0.91% |
| hsa_circ_0124714 | 2204 | 10 | 0.91% |
| hsa_circ_0137017 | 2204 | 10 | 0.91% |
| hsa_circ_0130992 | 29093 | 132 | 0.91% |
| hsa_circ_0137332 | 2645 | 12 | 0.91% |
| hsa_circ_0134046 | 7715 | 35 | 0.91% |
| hsa_circ_0043461 | 3086 | 14 | 0.91% |
| hsa_circ_0029943 | 2425 | 11 | 0.91% |
| hsa_circ_0046458 | 2425 | 11 | 0.91% |
| hsa_circ_0056120 | 2425 | 11 | 0.91% |
| hsa_circ_0080706 | 2425 | 11 | 0.91% |
| hsa_circ_0110946 | 2425 | 11 | 0.91% |
| hsa_circ_0037073 | 2866 | 13 | 0.91% |
| hsa_circ_0118574 | 5953 | 27 | 0.91% |
| hsa_circ_0128413 | 15876 | 72 | 0.91% |
| hsa_circ_0082224 | 3087 | 14 | 0.91% |
| hsa_circ_0041134 | 2646 | 12 | 0.91% |
| hsa_circ_0054760 | 2646 | 12 | 0.91% |
| hsa_circ_0070693 | 2646 | 12 | 0.91% |
| hsa_circ_0016442 | 2205 | 10 | 0.91% |
| hsa_circ_0035332 | 2205 | 10 | 0.91% |
| hsa_circ_0044526 | 2205 | 10 | 0.91% |
| hsa_circ_0045954 | 2205 | 10 | 0.91% |
| hsa_circ_0052881 | 2205 | 10 | 0.91% |
| hsa_circ_0053175 | 2205 | 10 | 0.91% |
| hsa_circ_0015265 | 2426 | 11 | 0.91% |
| hsa_circ_0019219 | 2426 | 11 | 0.91% |
| hsa_circ_0027409 | 2426 | 11 | 0.91% |
| hsa_circ_0046774 | 2426 | 11 | 0.91% |
| hsa_circ_0051372 | 2426 | 11 | 0.91% |
| hsa_circ_0084436 | 2426 | 11 | 0.91% |
| hsa_circ_0104503 | 5514 | 25 | 0.91% |
| hsa_circ_0017082 | 2206 | 10 | 0.91% |
| hsa_circ_0018501 | 2206 | 10 | 0.91% |
| hsa_circ_0021056 | 2206 | 10 | 0.91% |
| hsa_circ_0068405 | 2206 | 10 | 0.91% |
| hsa_circ_0087078 | 2206 | 10 | 0.91% |
| hsa_circ_0040229 | 2868 | 13 | 0.91% |
| hsa_circ_0009409 | 2427 | 11 | 0.91% |
| hsa_circ_0028534 | 2427 | 11 | 0.91% |
| hsa_circ_0077932 | 2427 | 11 | 0.91% |
| hsa_circ_0080997 | 2427 | 11 | 0.91% |
| hsa_circ_0088991 | 2427 | 11 | 0.91% |

| | | | |
|---|---|---|---|
| hsa_circ_0136180 | 2427 | 11 | 0.91% |
| hsa_circ_0060647 | 2648 | 12 | 0.91% |
| hsa_circ_0136738 | 2648 | 12 | 0.91% |
| hsa_circ_0037303 | 2207 | 10 | 0.91% |
| hsa_circ_0104928 | 3090 | 14 | 0.91% |
| hsa_circ_0047525 | 2428 | 11 | 0.91% |
| hsa_circ_0052912 | 2428 | 11 | 0.91% |
| hsa_circ_0070378 | 2428 | 11 | 0.91% |
| hsa_circ_0073969 | 2428 | 11 | 0.91% |
| hsa_circ_0085207 | 2428 | 11 | 0.91% |
| hsa_circ_0090903 | 2428 | 11 | 0.91% |
| hsa_circ_0102761 | 2428 | 11 | 0.91% |
| hsa_circ_0032113 | 3311 | 15 | 0.91% |
| hsa_circ_0065021 | 2649 | 12 | 0.91% |
| hsa_circ_0087148 | 2649 | 12 | 0.91% |
| hsa_circ_0133495 | 2649 | 12 | 0.91% |
| hsa_circ_0132277 | 6844 | 31 | 0.91% |
| hsa_circ_0016010 | 2208 | 10 | 0.91% |
| hsa_circ_0022481 | 2208 | 10 | 0.91% |
| hsa_circ_0025507 | 2208 | 10 | 0.91% |
| hsa_circ_0057845 | 2208 | 10 | 0.91% |
| hsa_circ_0067199 | 2208 | 10 | 0.91% |
| hsa_circ_0020216 | 2429 | 11 | 0.91% |
| hsa_circ_0028402 | 2429 | 11 | 0.91% |
| hsa_circ_0071876 | 2871 | 13 | 0.91% |
| hsa_circ_0004454 | 9276 | 42 | 0.91% |
| hsa_circ_0023867 | 2209 | 10 | 0.91% |
| hsa_circ_0039411 | 2209 | 10 | 0.91% |
| hsa_circ_0072914 | 2209 | 10 | 0.91% |
| hsa_circ_0086056 | 2209 | 10 | 0.91% |
| hsa_circ_0109878 | 2209 | 10 | 0.91% |
| hsa_circ_0115912 | 2209 | 10 | 0.91% |
| hsa_circ_0118451 | 2209 | 10 | 0.91% |
| hsa_circ_0129142 | 2209 | 10 | 0.91% |
| hsa_circ_0011842 | 2430 | 11 | 0.91% |
| hsa_circ_0053906 | 2430 | 11 | 0.91% |
| hsa_circ_0054767 | 2651 | 12 | 0.91% |
| hsa_circ_0118101 | 7071 | 32 | 0.91% |
| hsa_circ_0038968 | 3094 | 14 | 0.90% |
| hsa_circ_0044841 | 3094 | 14 | 0.90% |
| hsa_circ_0019743 | 2652 | 12 | 0.90% |
| hsa_circ_0061653 | 2652 | 12 | 0.90% |
| hsa_circ_0063197 | 2652 | 12 | 0.90% |
| hsa_circ_0053750 | 2431 | 11 | 0.90% |
| hsa_circ_0070384 | 2431 | 11 | 0.90% |
| hsa_circ_0075983 | 2431 | 11 | 0.90% |
| hsa_circ_0076937 | 2431 | 11 | 0.90% |
| hsa_circ_0105545 | 2431 | 11 | 0.90% |
| hsa_circ_0137029 | 2431 | 11 | 0.90% |
| hsa_circ_0065731 | 2210 | 10 | 0.90% |

| | | | |
|---|---|---|---|
| hsa_circ_0073908 | 2210 | 10 | 0.90% |
| hsa_circ_0115807 | 2210 | 10 | 0.90% |
| hsa_circ_0134213 | 2210 | 10 | 0.90% |
| hsa_circ_0140258 | 2210 | 10 | 0.90% |
| hsa_circ_0000701 | 3095 | 14 | 0.90% |
| hsa_circ_0026330 | 2653 | 12 | 0.90% |
| hsa_circ_0016315 | 2432 | 11 | 0.90% |
| hsa_circ_0084810 | 2432 | 11 | 0.90% |
| hsa_circ_0091170 | 2432 | 11 | 0.90% |
| hsa_circ_0017471 | 2211 | 10 | 0.90% |
| hsa_circ_0048873 | 2211 | 10 | 0.90% |
| hsa_circ_0050415 | 2211 | 10 | 0.90% |
| hsa_circ_0084893 | 2211 | 10 | 0.90% |
| hsa_circ_0119214 | 2211 | 10 | 0.90% |
| hsa_circ_0132467 | 2211 | 10 | 0.90% |
| hsa_circ_0137531 | 4201 | 19 | 0.90% |
| hsa_circ_0099448 | 2654 | 12 | 0.90% |
| hsa_circ_0036969 | 2433 | 11 | 0.90% |
| hsa_circ_0080142 | 2433 | 11 | 0.90% |
| hsa_circ_0102388 | 2433 | 11 | 0.90% |
| hsa_circ_0073911 | 2212 | 10 | 0.90% |
| hsa_circ_0128018 | 2212 | 10 | 0.90% |
| hsa_circ_0023315 | 2876 | 13 | 0.90% |
| hsa_circ_0083991 | 2655 | 12 | 0.90% |
| hsa_circ_0089178 | 2655 | 12 | 0.90% |
| hsa_circ_0024873 | 2434 | 11 | 0.90% |
| hsa_circ_0054069 | 2434 | 11 | 0.90% |
| hsa_circ_0116498 | 2434 | 11 | 0.90% |
| hsa_circ_0071952 | 2213 | 10 | 0.90% |
| hsa_circ_0074708 | 2213 | 10 | 0.90% |
| hsa_circ_0074975 | 2213 | 10 | 0.90% |
| hsa_circ_0014751 | 3099 | 14 | 0.90% |
| hsa_circ_0033108 | 2435 | 11 | 0.90% |
| hsa_circ_0050249 | 2435 | 11 | 0.90% |
| hsa_circ_0054060 | 2435 | 11 | 0.90% |
| hsa_circ_0061091 | 2435 | 11 | 0.90% |
| hsa_circ_0085913 | 2435 | 11 | 0.90% |
| hsa_circ_0088894 | 2435 | 11 | 0.90% |
| hsa_circ_0109515 | 2435 | 11 | 0.90% |
| hsa_circ_0052753 | 2878 | 13 | 0.90% |
| hsa_circ_0041074 | 2214 | 10 | 0.90% |
| hsa_circ_0076922 | 2214 | 10 | 0.90% |
| hsa_circ_0088416 | 2214 | 10 | 0.90% |
| hsa_circ_0042428 | 2657 | 12 | 0.90% |
| hsa_circ_0051015 | 2657 | 12 | 0.90% |
| hsa_circ_0089373 | 2657 | 12 | 0.90% |
| hsa_circ_0090001 | 2657 | 12 | 0.90% |
| hsa_circ_0013118 | 2436 | 11 | 0.90% |
| hsa_circ_0114000 | 2436 | 11 | 0.90% |
| hsa_circ_0107213 | 2879 | 13 | 0.90% |

| | | | |
|---|---|---|---|
| hsa_circ_0117672 | 5094 | 23 | 0.90% |
| hsa_circ_0035841 | 3101 | 14 | 0.90% |
| hsa_circ_0044857 | 2658 | 12 | 0.90% |
| hsa_circ_0089528 | 2658 | 12 | 0.90% |
| hsa_circ_0016615 | 2215 | 10 | 0.90% |
| hsa_circ_0047778 | 2215 | 10 | 0.90% |
| hsa_circ_0064146 | 2215 | 10 | 0.90% |
| hsa_circ_0083089 | 2215 | 10 | 0.90% |
| hsa_circ_0031619 | 2880 | 13 | 0.90% |
| hsa_circ_0087445 | 2437 | 11 | 0.90% |
| hsa_circ_0002651 | 3988 | 18 | 0.90% |
| hsa_circ_0100569 | 4210 | 19 | 0.90% |
| hsa_circ_0081007 | 2659 | 12 | 0.90% |
| hsa_circ_0022290 | 2216 | 10 | 0.90% |
| hsa_circ_0078457 | 2216 | 10 | 0.90% |
| hsa_circ_0082442 | 2216 | 10 | 0.90% |
| hsa_circ_0129969 | 2216 | 10 | 0.90% |
| hsa_circ_0046173 | 2881 | 13 | 0.90% |
| hsa_circ_0049812 | 2881 | 13 | 0.90% |
| hsa_circ_0055137 | 2881 | 13 | 0.90% |
| hsa_circ_0090271 | 2881 | 13 | 0.90% |
| hsa_circ_0053777 | 2438 | 11 | 0.90% |
| hsa_circ_0055735 | 2438 | 11 | 0.90% |
| hsa_circ_0060196 | 2438 | 11 | 0.90% |
| hsa_circ_0091389 | 2438 | 11 | 0.90% |
| hsa_circ_0017694 | 2882 | 13 | 0.90% |
| hsa_circ_0108259 | 2882 | 13 | 0.90% |
| hsa_circ_0047238 | 2217 | 10 | 0.90% |
| hsa_circ_0085363 | 2217 | 10 | 0.90% |
| hsa_circ_0013014 | 2439 | 11 | 0.90% |
| hsa_circ_0024049 | 2439 | 11 | 0.90% |
| hsa_circ_0024524 | 2439 | 11 | 0.90% |
| hsa_circ_0036718 | 2439 | 11 | 0.90% |
| hsa_circ_0051311 | 2439 | 11 | 0.90% |
| hsa_circ_0031600 | 2661 | 12 | 0.90% |
| hsa_circ_0063448 | 2661 | 12 | 0.90% |
| hsa_circ_0120312 | 2661 | 12 | 0.90% |
| hsa_circ_0076315 | 5766 | 26 | 0.90% |
| hsa_circ_0087344 | 2883 | 13 | 0.90% |
| hsa_circ_0037185 | 3105 | 14 | 0.90% |
| hsa_circ_0036527 | 2218 | 10 | 0.90% |
| hsa_circ_0131202 | 2218 | 10 | 0.90% |
| hsa_circ_0036475 | 2440 | 11 | 0.90% |
| hsa_circ_0074476 | 2440 | 11 | 0.90% |
| hsa_circ_0117020 | 2440 | 11 | 0.90% |
| hsa_circ_0102295 | 3771 | 17 | 0.90% |
| hsa_circ_0050085 | 9098 | 41 | 0.90% |
| hsa_circ_0040334 | 2441 | 11 | 0.90% |
| hsa_circ_0058666 | 2441 | 11 | 0.90% |
| hsa_circ_0083309 | 2441 | 11 | 0.90% |

| | | | |
|---|---|---|---|
| hsa_circ_0098025 | 2441 | 11 | 0.90% |
| hsa_circ_0085361 | 2663 | 12 | 0.90% |
| hsa_circ_0068426 | 2885 | 13 | 0.90% |
| hsa_circ_0090864 | 2886 | 13 | 0.90% |
| hsa_circ_0023021 | 2664 | 12 | 0.90% |
| hsa_circ_0014325 | 2442 | 11 | 0.90% |
| hsa_circ_0067167 | 2442 | 11 | 0.90% |
| hsa_circ_0080564 | 2442 | 11 | 0.90% |
| hsa_circ_0086877 | 2442 | 11 | 0.90% |
| hsa_circ_0089156 | 2442 | 11 | 0.90% |
| hsa_circ_0053864 | 2220 | 10 | 0.90% |
| hsa_circ_0058726 | 2220 | 10 | 0.90% |
| hsa_circ_0114735 | 2220 | 10 | 0.90% |
| hsa_circ_0021623 | 3553 | 16 | 0.90% |
| hsa_circ_0060108 | 2665 | 12 | 0.90% |
| hsa_circ_0088907 | 2665 | 12 | 0.90% |
| hsa_circ_0019350 | 2443 | 11 | 0.90% |
| hsa_circ_0049246 | 2443 | 11 | 0.90% |
| hsa_circ_0035597 | 2221 | 10 | 0.90% |
| hsa_circ_0042196 | 2221 | 10 | 0.90% |
| hsa_circ_0012789 | 3776 | 17 | 0.90% |
| hsa_circ_0119631 | 3999 | 18 | 0.90% |
| hsa_circ_0056432 | 7776 | 35 | 0.90% |
| hsa_circ_0065193 | 2444 | 11 | 0.90% |
| hsa_circ_0065337 | 2444 | 11 | 0.90% |
| hsa_circ_0085578 | 2444 | 11 | 0.90% |
| hsa_circ_0089516 | 2444 | 11 | 0.90% |
| hsa_circ_0023706 | 2222 | 10 | 0.90% |
| hsa_circ_0042169 | 2222 | 10 | 0.90% |
| hsa_circ_0069362 | 2222 | 10 | 0.90% |
| hsa_circ_0125648 | 2222 | 10 | 0.90% |
| hsa_circ_0037671 | 2667 | 12 | 0.90% |
| hsa_circ_0057259 | 2667 | 12 | 0.90% |
| hsa_circ_0072321 | 3334 | 15 | 0.90% |
| hsa_circ_0009828 | 2445 | 11 | 0.90% |
| hsa_circ_0046095 | 2445 | 11 | 0.90% |
| hsa_circ_0048314 | 2445 | 11 | 0.90% |
| hsa_circ_0056779 | 2445 | 11 | 0.90% |
| hsa_circ_0062375 | 2445 | 11 | 0.90% |
| hsa_circ_0070399 | 2445 | 11 | 0.90% |
| hsa_circ_0086409 | 2445 | 11 | 0.90% |
| hsa_circ_0135051 | 2445 | 11 | 0.90% |
| hsa_circ_0113460 | 3779 | 17 | 0.90% |
| hsa_circ_0030624 | 2223 | 10 | 0.90% |
| hsa_circ_0071232 | 2223 | 10 | 0.90% |
| hsa_circ_0084970 | 2223 | 10 | 0.90% |
| hsa_circ_0103836 | 2223 | 10 | 0.90% |
| hsa_circ_0134795 | 2223 | 10 | 0.90% |
| hsa_circ_0001918 | 4224 | 19 | 0.90% |
| hsa_circ_0025218 | 2668 | 12 | 0.90% |

| | | | |
|---|---|---|---|
| hsa_circ_0043660 | 2668 | 12 | 0.90% |
| hsa_circ_0018106 | 3113 | 14 | 0.90% |
| hsa_circ_0009790 | 2446 | 11 | 0.90% |
| hsa_circ_0041343 | 2446 | 11 | 0.90% |
| hsa_circ_0083306 | 2446 | 11 | 0.90% |
| hsa_circ_0089785 | 2446 | 11 | 0.90% |
| hsa_circ_0107617 | 2446 | 11 | 0.90% |
| hsa_circ_0069324 | 2224 | 10 | 0.90% |
| hsa_circ_0085054 | 2224 | 10 | 0.90% |
| hsa_circ_0102945 | 2224 | 10 | 0.90% |
| hsa_circ_0037726 | 2447 | 11 | 0.90% |
| hsa_circ_0057081 | 2447 | 11 | 0.90% |
| hsa_circ_0039773 | 2670 | 12 | 0.90% |
| hsa_circ_0021405 | 2225 | 10 | 0.90% |
| hsa_circ_0027780 | 2225 | 10 | 0.90% |
| hsa_circ_0030037 | 2225 | 10 | 0.90% |
| hsa_circ_0044633 | 2225 | 10 | 0.90% |
| hsa_circ_0070642 | 2225 | 10 | 0.90% |
| hsa_circ_0046100 | 2893 | 13 | 0.90% |
| hsa_circ_0130370 | 2893 | 13 | 0.90% |
| hsa_circ_0009215 | 2448 | 11 | 0.90% |
| hsa_circ_0021382 | 2448 | 11 | 0.90% |
| hsa_circ_0025427 | 2448 | 11 | 0.90% |
| hsa_circ_0049828 | 2448 | 11 | 0.90% |
| hsa_circ_0053769 | 2448 | 11 | 0.90% |
| hsa_circ_0057808 | 2448 | 11 | 0.90% |
| hsa_circ_0064956 | 2448 | 11 | 0.90% |
| hsa_circ_0076813 | 2448 | 11 | 0.90% |
| hsa_circ_0090327 | 2448 | 11 | 0.90% |
| hsa_circ_0131478 | 6899 | 31 | 0.90% |
| hsa_circ_0022118 | 2671 | 12 | 0.90% |
| hsa_circ_0021133 | 2894 | 13 | 0.90% |
| hsa_circ_0053153 | 2449 | 11 | 0.90% |
| hsa_circ_0069960 | 2449 | 11 | 0.90% |
| hsa_circ_0085710 | 2449 | 11 | 0.90% |
| hsa_circ_0095300 | 2449 | 11 | 0.90% |
| hsa_circ_0117825 | 2449 | 11 | 0.90% |
| hsa_circ_0113348 | 2672 | 12 | 0.90% |
| hsa_circ_0044335 | 2895 | 13 | 0.90% |
| hsa_circ_0116044 | 2227 | 10 | 0.90% |
| hsa_circ_0135961 | 2227 | 10 | 0.90% |
| hsa_circ_0026426 | 2450 | 11 | 0.90% |
| hsa_circ_0028789 | 2450 | 11 | 0.90% |
| hsa_circ_0061730 | 2450 | 11 | 0.90% |
| hsa_circ_0084916 | 2450 | 11 | 0.90% |
| hsa_circ_0086078 | 2450 | 11 | 0.90% |
| hsa_circ_0098874 | 2450 | 11 | 0.90% |
| hsa_circ_0050769 | 5346 | 24 | 0.90% |
| hsa_circ_0011895 | 2673 | 12 | 0.90% |
| hsa_circ_0034684 | 2673 | 12 | 0.90% |

| | | | |
|---|---|---|---|
| hsa_circ_0040474 | 2673 | 12 | 0.90% |
| hsa_circ_0071264 | 2673 | 12 | 0.90% |
| hsa_circ_0077363 | 2673 | 12 | 0.90% |
| hsa_circ_0093718 | 2673 | 12 | 0.90% |
| hsa_circ_0129119 | 2228 | 10 | 0.90% |
| hsa_circ_0011582 | 2451 | 11 | 0.90% |
| hsa_circ_0047844 | 2451 | 11 | 0.90% |
| hsa_circ_0069459 | 2451 | 11 | 0.90% |
| hsa_circ_0114440 | 2897 | 13 | 0.90% |
| hsa_circ_0024418 | 3120 | 14 | 0.90% |
| hsa_circ_0014571 | 2229 | 10 | 0.90% |
| hsa_circ_0016783 | 2229 | 10 | 0.90% |
| hsa_circ_0027720 | 2229 | 10 | 0.90% |
| hsa_circ_0069026 | 2229 | 10 | 0.90% |
| hsa_circ_0129481 | 2229 | 10 | 0.90% |
| hsa_circ_0009891 | 2452 | 11 | 0.90% |
| hsa_circ_0071590 | 2452 | 11 | 0.90% |
| hsa_circ_0086641 | 2452 | 11 | 0.90% |
| hsa_circ_0091960 | 2452 | 11 | 0.90% |
| hsa_circ_0104311 | 2452 | 11 | 0.90% |
| hsa_circ_0048542 | 2675 | 12 | 0.90% |
| hsa_circ_0098503 | 2898 | 13 | 0.90% |
| hsa_circ_0101578 | 3344 | 15 | 0.90% |
| hsa_circ_0050991 | 6689 | 30 | 0.90% |
| hsa_circ_0078967 | 5129 | 23 | 0.90% |
| hsa_circ_0000093 | 2676 | 12 | 0.90% |
| hsa_circ_0072070 | 2676 | 12 | 0.90% |
| hsa_circ_0010176 | 2453 | 11 | 0.90% |
| hsa_circ_0023565 | 2453 | 11 | 0.90% |
| hsa_circ_0056989 | 2453 | 11 | 0.90% |
| hsa_circ_0090204 | 2453 | 11 | 0.90% |
| hsa_circ_0056583 | 2230 | 10 | 0.90% |
| hsa_circ_0069877 | 2230 | 10 | 0.90% |
| hsa_circ_0122824 | 2230 | 10 | 0.90% |
| hsa_circ_0131171 | 2230 | 10 | 0.90% |
| hsa_circ_0135354 | 2230 | 10 | 0.90% |
| hsa_circ_0070360 | 2677 | 12 | 0.90% |
| hsa_circ_0088919 | 2677 | 12 | 0.90% |
| hsa_circ_0100951 | 2677 | 12 | 0.90% |
| hsa_circ_0107989 | 2677 | 12 | 0.90% |
| hsa_circ_0054823 | 2454 | 11 | 0.90% |
| hsa_circ_0054913 | 2454 | 11 | 0.90% |
| hsa_circ_0076733 | 2231 | 10 | 0.90% |
| hsa_circ_0078002 | 2231 | 10 | 0.90% |
| hsa_circ_0122324 | 2231 | 10 | 0.90% |
| hsa_circ_0010453 | 2455 | 11 | 0.90% |
| hsa_circ_0043356 | 2455 | 11 | 0.90% |
| hsa_circ_0055244 | 2455 | 11 | 0.90% |
| hsa_circ_0092972 | 2232 | 10 | 0.90% |
| hsa_circ_0091439 | 2902 | 13 | 0.90% |

| | | | |
|---|---|---|---|
| hsa_circ_0134005 | 5358 | 24 | 0.90% |
| hsa_circ_0065870 | 2679 | 12 | 0.90% |
| hsa_circ_0070362 | 2679 | 12 | 0.90% |
| hsa_circ_0063000 | 2456 | 11 | 0.90% |
| hsa_circ_0088752 | 2456 | 11 | 0.90% |
| hsa_circ_0096301 | 2456 | 11 | 0.90% |
| hsa_circ_0109379 | 2456 | 11 | 0.90% |
| hsa_circ_0109887 | 2456 | 11 | 0.90% |
| hsa_circ_0019429 | 2233 | 10 | 0.90% |
| hsa_circ_0036564 | 2233 | 10 | 0.90% |
| hsa_circ_0076539 | 2233 | 10 | 0.90% |
| hsa_circ_0092837 | 2233 | 10 | 0.90% |
| hsa_circ_0101049 | 2233 | 10 | 0.90% |
| hsa_circ_0117504 | 7146 | 32 | 0.90% |
| hsa_circ_0039197 | 2680 | 12 | 0.90% |
| hsa_circ_0051375 | 2457 | 11 | 0.90% |
| hsa_circ_0054598 | 2457 | 11 | 0.90% |
| hsa_circ_0062120 | 2457 | 11 | 0.90% |
| hsa_circ_0098737 | 2457 | 11 | 0.90% |
| hsa_circ_0118604 | 2457 | 11 | 0.90% |
| hsa_circ_0091108 | 4021 | 18 | 0.90% |
| hsa_circ_0050272 | 2234 | 10 | 0.90% |
| hsa_circ_0051554 | 2234 | 10 | 0.90% |
| hsa_circ_0124303 | 2234 | 10 | 0.90% |
| hsa_circ_0005491 | 2681 | 12 | 0.90% |
| hsa_circ_0016558 | 2681 | 12 | 0.90% |
| hsa_circ_0103117 | 2681 | 12 | 0.90% |
| hsa_circ_0048133 | 2458 | 11 | 0.90% |
| hsa_circ_0068394 | 2458 | 11 | 0.90% |
| hsa_circ_0139949 | 3129 | 14 | 0.89% |
| hsa_circ_0048703 | 2682 | 12 | 0.89% |
| hsa_circ_0136650 | 2682 | 12 | 0.89% |
| hsa_circ_0076629 | 2235 | 10 | 0.89% |
| hsa_circ_0140739 | 2235 | 10 | 0.89% |
| hsa_circ_0041298 | 2459 | 11 | 0.89% |
| hsa_circ_0057723 | 2459 | 11 | 0.89% |
| hsa_circ_0128878 | 6036 | 27 | 0.89% |
| hsa_circ_0033002 | 2683 | 12 | 0.89% |
| hsa_circ_0035511 | 2236 | 10 | 0.89% |
| hsa_circ_0043550 | 2236 | 10 | 0.89% |
| hsa_circ_0044806 | 2236 | 10 | 0.89% |
| hsa_circ_0088226 | 2236 | 10 | 0.89% |
| hsa_circ_0123487 | 2236 | 10 | 0.89% |
| hsa_circ_0022092 | 2460 | 11 | 0.89% |
| hsa_circ_0029667 | 2460 | 11 | 0.89% |
| hsa_circ_0042722 | 2460 | 11 | 0.89% |
| hsa_circ_0138853 | 2460 | 11 | 0.89% |
| hsa_circ_0023860 | 4026 | 18 | 0.89% |
| hsa_circ_0067932 | 3355 | 15 | 0.89% |
| hsa_circ_0032333 | 2684 | 12 | 0.89% |

| | | | |
|---|---|---|---|
| hsa_circ_0045083 | 2684 | 12 | 0.89% |
| hsa_circ_0014294 | 2237 | 10 | 0.89% |
| hsa_circ_0050529 | 2237 | 10 | 0.89% |
| hsa_circ_0066793 | 2237 | 10 | 0.89% |
| hsa_circ_0062817 | 3132 | 14 | 0.89% |
| hsa_circ_0070180 | 3132 | 14 | 0.89% |
| hsa_circ_0018964 | 2461 | 11 | 0.89% |
| hsa_circ_0071799 | 2461 | 11 | 0.89% |
| hsa_circ_0100291 | 2461 | 11 | 0.89% |
| hsa_circ_0106394 | 2461 | 11 | 0.89% |
| hsa_circ_0119587 | 2461 | 11 | 0.89% |
| hsa_circ_0023836 | 2685 | 12 | 0.89% |
| hsa_circ_0027055 | 2685 | 12 | 0.89% |
| hsa_circ_0064501 | 2685 | 12 | 0.89% |
| hsa_circ_0018091 | 2238 | 10 | 0.89% |
| hsa_circ_0044515 | 2238 | 10 | 0.89% |
| hsa_circ_0132230 | 2238 | 10 | 0.89% |
| hsa_circ_0051484 | 2462 | 11 | 0.89% |
| hsa_circ_0131761 | 2462 | 11 | 0.89% |
| hsa_circ_0117023 | 6939 | 31 | 0.89% |
| hsa_circ_0031103 | 4030 | 18 | 0.89% |
| hsa_circ_0055466 | 2239 | 10 | 0.89% |
| hsa_circ_0060711 | 2239 | 10 | 0.89% |
| hsa_circ_0108968 | 2239 | 10 | 0.89% |
| hsa_circ_0039256 | 2463 | 11 | 0.89% |
| hsa_circ_0048358 | 2463 | 11 | 0.89% |
| hsa_circ_0041954 | 2687 | 12 | 0.89% |
| hsa_circ_0057720 | 2687 | 12 | 0.89% |
| hsa_circ_0069956 | 2687 | 12 | 0.89% |
| hsa_circ_0089901 | 2911 | 13 | 0.89% |
| hsa_circ_0091400 | 3135 | 14 | 0.89% |
| hsa_circ_0070511 | 3359 | 15 | 0.89% |
| hsa_circ_0127468 | 3583 | 16 | 0.89% |
| hsa_circ_0072815 | 4032 | 18 | 0.89% |
| hsa_circ_0096855 | 2688 | 12 | 0.89% |
| hsa_circ_0038606 | 2464 | 11 | 0.89% |
| hsa_circ_0039269 | 2464 | 11 | 0.89% |
| hsa_circ_0091818 | 2464 | 11 | 0.89% |
| hsa_circ_0126340 | 2464 | 11 | 0.89% |
| hsa_circ_0027980 | 2240 | 10 | 0.89% |
| hsa_circ_0090753 | 2913 | 13 | 0.89% |
| hsa_circ_0033338 | 2689 | 12 | 0.89% |
| hsa_circ_0053951 | 2689 | 12 | 0.89% |
| hsa_circ_0072298 | 2689 | 12 | 0.89% |
| hsa_circ_0086103 | 2689 | 12 | 0.89% |
| hsa_circ_0040770 | 2465 | 11 | 0.89% |
| hsa_circ_0083153 | 2465 | 11 | 0.89% |
| hsa_circ_0059302 | 2241 | 10 | 0.89% |
| hsa_circ_0136921 | 5828 | 26 | 0.89% |
| hsa_circ_0047810 | 2914 | 13 | 0.89% |

| | | | |
|---|---|---|---|
| hsa_circ_0089248 | 2690 | 12 | 0.89% |
| hsa_circ_0091356 | 2690 | 12 | 0.89% |
| hsa_circ_0028763 | 2466 | 11 | 0.89% |
| hsa_circ_0032294 | 2466 | 11 | 0.89% |
| hsa_circ_0071329 | 2466 | 11 | 0.89% |
| hsa_circ_0135363 | 2466 | 11 | 0.89% |
| hsa_circ_0042676 | 2242 | 10 | 0.89% |
| hsa_circ_0047975 | 2242 | 10 | 0.89% |
| hsa_circ_0053838 | 2242 | 10 | 0.89% |
| hsa_circ_0067549 | 2242 | 10 | 0.89% |
| hsa_circ_0068004 | 2242 | 10 | 0.89% |
| hsa_circ_0088115 | 2242 | 10 | 0.89% |
| hsa_circ_0104331 | 2242 | 10 | 0.89% |
| hsa_circ_0016594 | 5606 | 25 | 0.89% |
| hsa_circ_0031591 | 2691 | 12 | 0.89% |
| hsa_circ_0046138 | 2691 | 12 | 0.89% |
| hsa_circ_0118548 | 6280 | 28 | 0.89% |
| hsa_circ_0049485 | 3140 | 14 | 0.89% |
| hsa_circ_0062667 | 3140 | 14 | 0.89% |
| hsa_circ_0116138 | 3140 | 14 | 0.89% |
| hsa_circ_0120115 | 2243 | 10 | 0.89% |
| hsa_circ_0022232 | 2692 | 12 | 0.89% |
| hsa_circ_0031500 | 2692 | 12 | 0.89% |
| hsa_circ_0056163 | 2692 | 12 | 0.89% |
| hsa_circ_0032650 | 3814 | 17 | 0.89% |
| hsa_circ_0021768 | 2468 | 11 | 0.89% |
| hsa_circ_0081502 | 3590 | 16 | 0.89% |
| hsa_circ_0093397 | 5834 | 26 | 0.89% |
| hsa_circ_0102021 | 2917 | 13 | 0.89% |
| hsa_circ_0039711 | 2244 | 10 | 0.89% |
| hsa_circ_0066359 | 2244 | 10 | 0.89% |
| hsa_circ_0082137 | 2244 | 10 | 0.89% |
| hsa_circ_0084195 | 2244 | 10 | 0.89% |
| hsa_circ_0090247 | 2244 | 10 | 0.89% |
| hsa_circ_0128595 | 2244 | 10 | 0.89% |
| hsa_circ_0036900 | 2693 | 12 | 0.89% |
| hsa_circ_0061203 | 2693 | 12 | 0.89% |
| hsa_circ_0088345 | 2469 | 11 | 0.89% |
| hsa_circ_0108038 | 2469 | 11 | 0.89% |
| hsa_circ_0044360 | 2918 | 13 | 0.89% |
| hsa_circ_0084210 | 2694 | 12 | 0.89% |
| hsa_circ_0050767 | 2245 | 10 | 0.89% |
| hsa_circ_0134212 | 9430 | 42 | 0.89% |
| hsa_circ_0068587 | 2919 | 13 | 0.89% |
| hsa_circ_0018118 | 2470 | 11 | 0.89% |
| hsa_circ_0050357 | 2470 | 11 | 0.89% |
| hsa_circ_0058603 | 2470 | 11 | 0.89% |
| hsa_circ_0059087 | 2470 | 11 | 0.89% |
| hsa_circ_0051750 | 3144 | 14 | 0.89% |
| hsa_circ_0074522 | 2695 | 12 | 0.89% |

| | | | |
|---|---|---|---|
| hsa_circ_0096535 | 2695 | 12 | 0.89% |
| hsa_circ_0029513 | 2246 | 10 | 0.89% |
| hsa_circ_0120908 | 2246 | 10 | 0.89% |
| hsa_circ_0035775 | 2920 | 13 | 0.89% |
| hsa_circ_0014548 | 2471 | 11 | 0.89% |
| hsa_circ_0058596 | 2471 | 11 | 0.89% |
| hsa_circ_0122025 | 2471 | 11 | 0.89% |
| hsa_circ_0125028 | 2471 | 11 | 0.89% |
| hsa_circ_0050500 | 3145 | 14 | 0.89% |
| hsa_circ_0104482 | 5841 | 26 | 0.89% |
| hsa_circ_0032056 | 3370 | 15 | 0.89% |
| hsa_circ_0037197 | 2696 | 12 | 0.89% |
| hsa_circ_0079670 | 2696 | 12 | 0.89% |
| hsa_circ_0138273 | 2696 | 12 | 0.89% |
| hsa_circ_0106510 | 3595 | 16 | 0.89% |
| hsa_circ_0087188 | 2921 | 13 | 0.89% |
| hsa_circ_0100339 | 2921 | 13 | 0.89% |
| hsa_circ_0083827 | 2247 | 10 | 0.89% |
| hsa_circ_0104151 | 2247 | 10 | 0.89% |
| hsa_circ_0123622 | 2247 | 10 | 0.89% |
| hsa_circ_0007098 | 3146 | 14 | 0.89% |
| hsa_circ_0012835 | 2472 | 11 | 0.89% |
| hsa_circ_0023222 | 2472 | 11 | 0.89% |
| hsa_circ_0049298 | 2472 | 11 | 0.89% |
| hsa_circ_0089770 | 2472 | 11 | 0.89% |
| hsa_circ_0130904 | 2697 | 12 | 0.89% |
| hsa_circ_0016272 | 2248 | 10 | 0.89% |
| hsa_circ_0068442 | 2248 | 10 | 0.89% |
| hsa_circ_0092095 | 2248 | 10 | 0.89% |
| hsa_circ_0099979 | 2248 | 10 | 0.89% |
| hsa_circ_0104170 | 2248 | 10 | 0.89% |
| hsa_circ_0113290 | 2248 | 10 | 0.89% |
| hsa_circ_0019474 | 2473 | 11 | 0.89% |
| hsa_circ_0030626 | 2473 | 11 | 0.89% |
| hsa_circ_0140748 | 2473 | 11 | 0.89% |
| hsa_circ_0041028 | 2698 | 12 | 0.89% |
| hsa_circ_0102109 | 2698 | 12 | 0.89% |
| hsa_circ_0023549 | 2249 | 10 | 0.89% |
| hsa_circ_0041394 | 2249 | 10 | 0.89% |
| hsa_circ_0019839 | 2474 | 11 | 0.89% |
| hsa_circ_0030076 | 2474 | 11 | 0.89% |
| hsa_circ_0082190 | 2474 | 11 | 0.89% |
| hsa_circ_0009968 | 2699 | 12 | 0.89% |
| hsa_circ_0044858 | 2699 | 12 | 0.89% |
| hsa_circ_0047739 | 2924 | 13 | 0.89% |
| hsa_circ_0045398 | 3149 | 14 | 0.89% |
| hsa_circ_0052434 | 3149 | 14 | 0.89% |
| hsa_circ_0109182 | 7875 | 35 | 0.89% |
| hsa_circ_0100332 | 2925 | 13 | 0.89% |
| hsa_circ_0138777 | 2925 | 13 | 0.89% |

| | | | |
|---|---|---|---|
| hsa_circ_0052680 | 2700 | 12 | 0.89% |
| hsa_circ_0070689 | 2700 | 12 | 0.89% |
| hsa_circ_0081577 | 2700 | 12 | 0.89% |
| hsa_circ_0094018 | 2700 | 12 | 0.89% |
| hsa_circ_0016824 | 2475 | 11 | 0.89% |
| hsa_circ_0017475 | 2475 | 11 | 0.89% |
| hsa_circ_0029091 | 2475 | 11 | 0.89% |
| hsa_circ_0044979 | 2475 | 11 | 0.89% |
| hsa_circ_0066323 | 2475 | 11 | 0.89% |
| hsa_circ_0089457 | 2475 | 11 | 0.89% |
| hsa_circ_0091970 | 2475 | 11 | 0.89% |
| hsa_circ_0094735 | 2475 | 11 | 0.89% |
| hsa_circ_0025165 | 2250 | 10 | 0.89% |
| hsa_circ_0078045 | 2250 | 10 | 0.89% |
| hsa_circ_0120113 | 2250 | 10 | 0.89% |
| hsa_circ_0018230 | 9901 | 44 | 0.89% |
| hsa_circ_0106092 | 7651 | 34 | 0.89% |
| hsa_circ_0029629 | 2926 | 13 | 0.89% |
| hsa_circ_0134060 | 5402 | 24 | 0.89% |
| hsa_circ_0039648 | 2701 | 12 | 0.89% |
| hsa_circ_0054833 | 2701 | 12 | 0.89% |
| hsa_circ_0074764 | 2701 | 12 | 0.89% |
| hsa_circ_0018090 | 2476 | 11 | 0.89% |
| hsa_circ_0028727 | 2476 | 11 | 0.89% |
| hsa_circ_0054730 | 2476 | 11 | 0.89% |
| hsa_circ_0069514 | 2476 | 11 | 0.89% |
| hsa_circ_0084448 | 2476 | 11 | 0.89% |
| hsa_circ_0116243 | 2476 | 11 | 0.89% |
| hsa_circ_0139830 | 2476 | 11 | 0.89% |
| hsa_circ_0122191 | 4502 | 20 | 0.89% |
| hsa_circ_0047202 | 2251 | 10 | 0.89% |
| hsa_circ_0086166 | 2251 | 10 | 0.89% |
| hsa_circ_0110088 | 2251 | 10 | 0.89% |
| hsa_circ_0077109 | 5178 | 23 | 0.89% |
| hsa_circ_0009644 | 2927 | 13 | 0.89% |
| hsa_circ_0023144 | 2702 | 12 | 0.89% |
| hsa_circ_0019850 | 2477 | 11 | 0.89% |
| hsa_circ_0068177 | 2477 | 11 | 0.89% |
| hsa_circ_0081934 | 2477 | 11 | 0.89% |
| hsa_circ_0125551 | 5855 | 26 | 0.89% |
| hsa_circ_0011544 | 2252 | 10 | 0.89% |
| hsa_circ_0132060 | 2252 | 10 | 0.89% |
| hsa_circ_0014845 | 2703 | 12 | 0.89% |
| hsa_circ_0065388 | 4956 | 22 | 0.89% |
| hsa_circ_0018160 | 2478 | 11 | 0.89% |
| hsa_circ_0018696 | 2478 | 11 | 0.89% |
| hsa_circ_0060451 | 2478 | 11 | 0.89% |
| hsa_circ_0083525 | 2478 | 11 | 0.89% |
| hsa_circ_0084365 | 2478 | 11 | 0.89% |
| hsa_circ_0086533 | 2478 | 11 | 0.89% |

| | | | |
|---|---|---|---|
| hsa_circ_0116386 | 2478 | 11 | 0.89% |
| hsa_circ_0017722 | 2253 | 10 | 0.89% |
| hsa_circ_0040730 | 2253 | 10 | 0.89% |
| hsa_circ_0094719 | 2253 | 10 | 0.89% |
| hsa_circ_0103587 | 2253 | 10 | 0.89% |
| hsa_circ_0104443 | 2253 | 10 | 0.89% |
| hsa_circ_0116278 | 2253 | 10 | 0.89% |
| hsa_circ_0128920 | 2253 | 10 | 0.89% |
| hsa_circ_0125549 | 5858 | 26 | 0.89% |
| hsa_circ_0042152 | 2704 | 12 | 0.89% |
| hsa_circ_0045326 | 2704 | 12 | 0.89% |
| hsa_circ_0059063 | 2479 | 11 | 0.89% |
| hsa_circ_0059718 | 2479 | 11 | 0.89% |
| hsa_circ_0078704 | 2479 | 11 | 0.89% |
| hsa_circ_0095465 | 2479 | 11 | 0.89% |
| hsa_circ_0105435 | 2479 | 11 | 0.89% |
| hsa_circ_0059649 | 2930 | 13 | 0.89% |
| hsa_circ_0063070 | 2930 | 13 | 0.89% |
| hsa_circ_0087475 | 2930 | 13 | 0.89% |
| hsa_circ_0005073 | 5635 | 25 | 0.89% |
| hsa_circ_0017347 | 2254 | 10 | 0.89% |
| hsa_circ_0051702 | 2254 | 10 | 0.89% |
| hsa_circ_0100161 | 2254 | 10 | 0.89% |
| hsa_circ_0105666 | 2254 | 10 | 0.89% |
| hsa_circ_0118216 | 2254 | 10 | 0.89% |
| hsa_circ_0034752 | 2705 | 12 | 0.89% |
| hsa_circ_0100500 | 2705 | 12 | 0.89% |
| hsa_circ_0062614 | 3607 | 16 | 0.89% |
| hsa_circ_0010760 | 2480 | 11 | 0.89% |
| hsa_circ_0024962 | 2480 | 11 | 0.89% |
| hsa_circ_0067562 | 2480 | 11 | 0.89% |
| hsa_circ_0072201 | 2480 | 11 | 0.89% |
| hsa_circ_0079019 | 2480 | 11 | 0.89% |
| hsa_circ_0139027 | 9696 | 43 | 0.89% |
| hsa_circ_0030862 | 3157 | 14 | 0.89% |
| hsa_circ_0039406 | 2706 | 12 | 0.89% |
| hsa_circ_0017146 | 2255 | 10 | 0.89% |
| hsa_circ_0130027 | 2255 | 10 | 0.89% |
| hsa_circ_0122160 | 6089 | 27 | 0.89% |
| hsa_circ_0045795 | 2481 | 11 | 0.89% |
| hsa_circ_0058119 | 2481 | 11 | 0.89% |
| hsa_circ_0063691 | 2481 | 11 | 0.89% |
| hsa_circ_0135045 | 4286 | 19 | 0.89% |
| hsa_circ_0009266 | 2707 | 12 | 0.89% |
| hsa_circ_0027229 | 2256 | 10 | 0.89% |
| hsa_circ_0030371 | 2256 | 10 | 0.89% |
| hsa_circ_0053818 | 2256 | 10 | 0.89% |
| hsa_circ_0066341 | 2256 | 10 | 0.89% |
| hsa_circ_0125844 | 2256 | 10 | 0.89% |
| hsa_circ_0127232 | 2256 | 10 | 0.89% |

| | | | |
|---|---|---|---|
| hsa_circ_0139960 | 2933 | 13 | 0.89% |
| hsa_circ_0042548 | 2482 | 11 | 0.89% |
| hsa_circ_0052367 | 2482 | 11 | 0.89% |
| hsa_circ_0069066 | 2482 | 11 | 0.89% |
| hsa_circ_0061916 | 2708 | 12 | 0.89% |
| hsa_circ_0062599 | 2257 | 10 | 0.89% |
| hsa_circ_0018868 | 2483 | 11 | 0.89% |
| hsa_circ_0069197 | 2483 | 11 | 0.89% |
| hsa_circ_0011023 | 2709 | 12 | 0.89% |
| hsa_circ_0066065 | 2709 | 12 | 0.89% |
| hsa_circ_0101774 | 2935 | 13 | 0.89% |
| hsa_circ_0114122 | 2935 | 13 | 0.89% |
| hsa_circ_0011598 | 2258 | 10 | 0.89% |
| hsa_circ_0057612 | 2258 | 10 | 0.89% |
| hsa_circ_0118883 | 2258 | 10 | 0.89% |
| hsa_circ_0019858 | 2484 | 11 | 0.89% |
| hsa_circ_0030021 | 2484 | 11 | 0.89% |
| hsa_circ_0037540 | 2484 | 11 | 0.89% |
| hsa_circ_0086912 | 2484 | 11 | 0.89% |
| hsa_circ_0023662 | 2710 | 12 | 0.89% |
| hsa_circ_0017355 | 2936 | 13 | 0.89% |
| hsa_circ_0027022 | 9261 | 41 | 0.89% |
| hsa_circ_0031813 | 6325 | 28 | 0.89% |
| hsa_circ_0011800 | 2259 | 10 | 0.89% |
| hsa_circ_0028575 | 2259 | 10 | 0.89% |
| hsa_circ_0035350 | 2259 | 10 | 0.89% |
| hsa_circ_0075420 | 2259 | 10 | 0.89% |
| hsa_circ_0101107 | 2259 | 10 | 0.89% |
| hsa_circ_0031479 | 2485 | 11 | 0.89% |
| hsa_circ_0037407 | 2485 | 11 | 0.89% |
| hsa_circ_0041202 | 2711 | 12 | 0.89% |
| hsa_circ_0071345 | 2711 | 12 | 0.89% |
| hsa_circ_0060181 | 2937 | 13 | 0.89% |
| hsa_circ_0085348 | 2937 | 13 | 0.89% |
| hsa_circ_0090270 | 2937 | 13 | 0.89% |
| hsa_circ_0075115 | 3163 | 14 | 0.89% |
| hsa_circ_0133075 | 3163 | 14 | 0.89% |
| hsa_circ_0017515 | 4293 | 19 | 0.89% |
| hsa_circ_0114464 | 9040 | 40 | 0.88% |
| hsa_circ_0022861 | 2938 | 13 | 0.88% |
| hsa_circ_0053373 | 2938 | 13 | 0.88% |
| hsa_circ_0054780 | 2712 | 12 | 0.88% |
| hsa_circ_0031461 | 2486 | 11 | 0.88% |
| hsa_circ_0043886 | 2486 | 11 | 0.88% |
| hsa_circ_0073828 | 2486 | 11 | 0.88% |
| hsa_circ_0074195 | 2486 | 11 | 0.88% |
| hsa_circ_0100906 | 2486 | 11 | 0.88% |
| hsa_circ_0018174 | 2260 | 10 | 0.88% |
| hsa_circ_0071656 | 2260 | 10 | 0.88% |
| hsa_circ_0098348 | 2260 | 10 | 0.88% |

| | | | |
|---|---|---|---|
| hsa_circ_0110925 | 10172 | 45 | 0.88% |
| hsa_circ_0024019 | 4295 | 19 | 0.88% |
| hsa_circ_0057773 | 4069 | 18 | 0.88% |
| hsa_circ_0097534 | 7234 | 32 | 0.88% |
| hsa_circ_0115387 | 8817 | 39 | 0.88% |
| hsa_circ_0060871 | 2939 | 13 | 0.88% |
| hsa_circ_0018512 | 2713 | 12 | 0.88% |
| hsa_circ_0043766 | 2713 | 12 | 0.88% |
| hsa_circ_0051295 | 2713 | 12 | 0.88% |
| hsa_circ_0074671 | 2713 | 12 | 0.88% |
| hsa_circ_0011775 | 2261 | 10 | 0.88% |
| hsa_circ_0109587 | 4975 | 22 | 0.88% |
| hsa_circ_0038126 | 9046 | 40 | 0.88% |
| hsa_circ_0136040 | 2488 | 11 | 0.88% |
| hsa_circ_0019118 | 3619 | 16 | 0.88% |
| hsa_circ_0024231 | 2262 | 10 | 0.88% |
| hsa_circ_0097031 | 2262 | 10 | 0.88% |
| hsa_circ_0132897 | 2262 | 10 | 0.88% |
| hsa_circ_0061450 | 3620 | 16 | 0.88% |
| hsa_circ_0012518 | 2715 | 12 | 0.88% |
| hsa_circ_0017378 | 2715 | 12 | 0.88% |
| hsa_circ_0131998 | 2715 | 12 | 0.88% |
| hsa_circ_0017502 | 2489 | 11 | 0.88% |
| hsa_circ_0045034 | 2489 | 11 | 0.88% |
| hsa_circ_0084913 | 2489 | 11 | 0.88% |
| hsa_circ_0113659 | 2489 | 11 | 0.88% |
| hsa_circ_0048384 | 3168 | 14 | 0.88% |
| hsa_circ_0059628 | 2263 | 10 | 0.88% |
| hsa_circ_0067762 | 2263 | 10 | 0.88% |
| hsa_circ_0083032 | 2263 | 10 | 0.88% |
| hsa_circ_0026245 | 2942 | 13 | 0.88% |
| hsa_circ_0137742 | 4300 | 19 | 0.88% |
| hsa_circ_0023826 | 2716 | 12 | 0.88% |
| hsa_circ_0048193 | 2716 | 12 | 0.88% |
| hsa_circ_0073666 | 2716 | 12 | 0.88% |
| hsa_circ_0083580 | 2716 | 12 | 0.88% |
| hsa_circ_0103549 | 2716 | 12 | 0.88% |
| hsa_circ_0104175 | 2716 | 12 | 0.88% |
| hsa_circ_0122234 | 2716 | 12 | 0.88% |
| hsa_circ_0131476 | 7017 | 31 | 0.88% |
| hsa_circ_0012519 | 3169 | 14 | 0.88% |
| hsa_circ_0044299 | 3169 | 14 | 0.88% |
| hsa_circ_0085641 | 3169 | 14 | 0.88% |
| hsa_circ_0061671 | 2490 | 11 | 0.88% |
| hsa_circ_0089628 | 2490 | 11 | 0.88% |
| hsa_circ_0025914 | 2943 | 13 | 0.88% |
| hsa_circ_0038229 | 2264 | 10 | 0.88% |
| hsa_circ_0125295 | 2264 | 10 | 0.88% |
| hsa_circ_0010326 | 2717 | 12 | 0.88% |
| hsa_circ_0010455 | 2717 | 12 | 0.88% |

| | | | |
|---|---|---|---|
| hsa_circ_0023167 | 2717 | 12 | 0.88% |
| hsa_circ_0029566 | 2717 | 12 | 0.88% |
| hsa_circ_0112359 | 2717 | 12 | 0.88% |
| hsa_circ_0116372 | 2491 | 11 | 0.88% |
| hsa_circ_0101018 | 4756 | 21 | 0.88% |
| hsa_circ_0120686 | 6342 | 28 | 0.88% |
| hsa_circ_0051929 | 3624 | 16 | 0.88% |
| hsa_circ_0030132 | 2718 | 12 | 0.88% |
| hsa_circ_0030486 | 2718 | 12 | 0.88% |
| hsa_circ_0041220 | 2718 | 12 | 0.88% |
| hsa_circ_0086941 | 2718 | 12 | 0.88% |
| hsa_circ_0034979 | 2265 | 10 | 0.88% |
| hsa_circ_0081615 | 2265 | 10 | 0.88% |
| hsa_circ_0133960 | 2265 | 10 | 0.88% |
| hsa_circ_0012290 | 2945 | 13 | 0.88% |
| hsa_circ_0015550 | 2945 | 13 | 0.88% |
| hsa_circ_0046735 | 2945 | 13 | 0.88% |
| hsa_circ_0060668 | 2945 | 13 | 0.88% |
| hsa_circ_0050073 | 2492 | 11 | 0.88% |
| hsa_circ_0084426 | 2492 | 11 | 0.88% |
| hsa_circ_0089577 | 2492 | 11 | 0.88% |
| hsa_circ_0025606 | 2719 | 12 | 0.88% |
| hsa_circ_0026737 | 2719 | 12 | 0.88% |
| hsa_circ_0028128 | 2719 | 12 | 0.88% |
| hsa_circ_0012260 | 2266 | 10 | 0.88% |
| hsa_circ_0126332 | 7025 | 31 | 0.88% |
| hsa_circ_0086632 | 2946 | 13 | 0.88% |
| hsa_circ_0011232 | 2493 | 11 | 0.88% |
| hsa_circ_0042081 | 2493 | 11 | 0.88% |
| hsa_circ_0048686 | 2493 | 11 | 0.88% |
| hsa_circ_0065387 | 2493 | 11 | 0.88% |
| hsa_circ_0086405 | 2493 | 11 | 0.88% |
| hsa_circ_0088893 | 2493 | 11 | 0.88% |
| hsa_circ_0093998 | 2493 | 11 | 0.88% |
| hsa_circ_0117850 | 2493 | 11 | 0.88% |
| hsa_circ_0118280 | 2493 | 11 | 0.88% |
| hsa_circ_0088323 | 3400 | 15 | 0.88% |
| hsa_circ_0047381 | 2720 | 12 | 0.88% |
| hsa_circ_0032389 | 2267 | 10 | 0.88% |
| hsa_circ_0064943 | 2267 | 10 | 0.88% |
| hsa_circ_0040718 | 2494 | 11 | 0.88% |
| hsa_circ_0055655 | 2494 | 11 | 0.88% |
| hsa_circ_0056073 | 2494 | 11 | 0.88% |
| hsa_circ_0068362 | 2494 | 11 | 0.88% |
| hsa_circ_0089149 | 2494 | 11 | 0.88% |
| hsa_circ_0055155 | 2721 | 12 | 0.88% |
| hsa_circ_0065432 | 2721 | 12 | 0.88% |
| hsa_circ_0082225 | 2721 | 12 | 0.88% |
| hsa_circ_0113298 | 2721 | 12 | 0.88% |
| hsa_circ_0073822 | 5896 | 26 | 0.88% |

| | | | |
|---|---|---|---|
| hsa_circ_0139744 | 52842 | 233 | 0.88% |
| hsa_circ_0137442 | 5670 | 25 | 0.88% |
| hsa_circ_0011418 | 4536 | 20 | 0.88% |
| hsa_circ_0033537 | 2268 | 10 | 0.88% |
| hsa_circ_0035323 | 2268 | 10 | 0.88% |
| hsa_circ_0093982 | 2268 | 10 | 0.88% |
| hsa_circ_0124296 | 2268 | 10 | 0.88% |
| hsa_circ_0131466 | 2268 | 10 | 0.88% |
| hsa_circ_0040059 | 2495 | 11 | 0.88% |
| hsa_circ_0045036 | 2495 | 11 | 0.88% |
| hsa_circ_0079255 | 2495 | 11 | 0.88% |
| hsa_circ_0094869 | 2495 | 11 | 0.88% |
| hsa_circ_0011214 | 2949 | 13 | 0.88% |
| hsa_circ_0036843 | 2949 | 13 | 0.88% |
| hsa_circ_0042923 | 2949 | 13 | 0.88% |
| hsa_circ_0044334 | 2949 | 13 | 0.88% |
| hsa_circ_0055138 | 2949 | 13 | 0.88% |
| hsa_circ_0084400 | 2949 | 13 | 0.88% |
| hsa_circ_0013159 | 6352 | 28 | 0.88% |
| hsa_circ_0047840 | 3176 | 14 | 0.88% |
| hsa_circ_0053594 | 2496 | 11 | 0.88% |
| hsa_circ_0065334 | 2496 | 11 | 0.88% |
| hsa_circ_0094881 | 2496 | 11 | 0.88% |
| hsa_circ_0054898 | 2723 | 12 | 0.88% |
| hsa_circ_0045643 | 2950 | 13 | 0.88% |
| hsa_circ_0060406 | 2950 | 13 | 0.88% |
| hsa_circ_0070255 | 3178 | 14 | 0.88% |
| hsa_circ_0010564 | 2724 | 12 | 0.88% |
| hsa_circ_0044015 | 2724 | 12 | 0.88% |
| hsa_circ_0019227 | 2497 | 11 | 0.88% |
| hsa_circ_0026865 | 2497 | 11 | 0.88% |
| hsa_circ_0081623 | 2497 | 11 | 0.88% |
| hsa_circ_0066717 | 2270 | 10 | 0.88% |
| hsa_circ_0136823 | 2270 | 10 | 0.88% |
| hsa_circ_0065398 | 2952 | 13 | 0.88% |
| hsa_circ_0019828 | 2498 | 11 | 0.88% |
| hsa_circ_0046592 | 2498 | 11 | 0.88% |
| hsa_circ_0047903 | 2498 | 11 | 0.88% |
| hsa_circ_0083285 | 2498 | 11 | 0.88% |
| hsa_circ_0101609 | 2498 | 11 | 0.88% |
| hsa_circ_0119254 | 2271 | 10 | 0.88% |
| hsa_circ_0047777 | 2953 | 13 | 0.88% |
| hsa_circ_0140718 | 2726 | 12 | 0.88% |
| hsa_circ_0006560 | 2499 | 11 | 0.88% |
| hsa_circ_0053658 | 2499 | 11 | 0.88% |
| hsa_circ_0127868 | 2499 | 11 | 0.88% |
| hsa_circ_0009803 | 2272 | 10 | 0.88% |
| hsa_circ_0066206 | 2272 | 10 | 0.88% |
| hsa_circ_0000449 | 3181 | 14 | 0.88% |
| hsa_circ_0009399 | 2727 | 12 | 0.88% |

| | | | |
|---|---|---|---|
| hsa_circ_0035161 | 2727 | 12 | 0.88% |
| hsa_circ_0010177 | 2500 | 11 | 0.88% |
| hsa_circ_0080586 | 2500 | 11 | 0.88% |
| hsa_circ_0085335 | 2500 | 11 | 0.88% |
| hsa_circ_0089145 | 2500 | 11 | 0.88% |
| hsa_circ_0036978 | 3182 | 14 | 0.88% |
| hsa_circ_0020288 | 2273 | 10 | 0.88% |
| hsa_circ_0048229 | 2273 | 10 | 0.88% |
| hsa_circ_0054092 | 2273 | 10 | 0.88% |
| hsa_circ_0076474 | 2273 | 10 | 0.88% |
| hsa_circ_0086385 | 2273 | 10 | 0.88% |
| hsa_circ_0090205 | 2273 | 10 | 0.88% |
| hsa_circ_0111119 | 2273 | 10 | 0.88% |
| hsa_circ_0102794 | 2955 | 13 | 0.88% |
| hsa_circ_0004068 | 2728 | 12 | 0.88% |
| hsa_circ_0008376 | 2728 | 12 | 0.88% |
| hsa_circ_0131733 | 2728 | 12 | 0.88% |
| hsa_circ_0002163 | 2501 | 11 | 0.88% |
| hsa_circ_0027616 | 2501 | 11 | 0.88% |
| hsa_circ_0049994 | 2501 | 11 | 0.88% |
| hsa_circ_0063108 | 2501 | 11 | 0.88% |
| hsa_circ_0114372 | 2501 | 11 | 0.88% |
| hsa_circ_0070717 | 10460 | 46 | 0.88% |
| hsa_circ_0076923 | 2274 | 10 | 0.88% |
| hsa_circ_0113213 | 2274 | 10 | 0.88% |
| hsa_circ_0078518 | 3866 | 17 | 0.88% |
| hsa_circ_0026589 | 2729 | 12 | 0.88% |
| hsa_circ_0011538 | 2502 | 11 | 0.88% |
| hsa_circ_0019537 | 2502 | 11 | 0.88% |
| hsa_circ_0073417 | 2502 | 11 | 0.88% |
| hsa_circ_0081881 | 2502 | 11 | 0.88% |
| hsa_circ_0052745 | 2957 | 13 | 0.88% |
| hsa_circ_0116902 | 2730 | 12 | 0.88% |
| hsa_circ_0044854 | 2275 | 10 | 0.88% |
| hsa_circ_0010711 | 2503 | 11 | 0.88% |
| hsa_circ_0016805 | 2503 | 11 | 0.88% |
| hsa_circ_0027777 | 2503 | 11 | 0.88% |
| hsa_circ_0127082 | 2503 | 11 | 0.88% |
| hsa_circ_0046714 | 2276 | 10 | 0.88% |
| hsa_circ_0052251 | 2276 | 10 | 0.88% |
| hsa_circ_0066347 | 2276 | 10 | 0.88% |
| hsa_circ_0071957 | 2276 | 10 | 0.88% |
| hsa_circ_0113817 | 2276 | 10 | 0.88% |
| hsa_circ_0130965 | 2276 | 10 | 0.88% |
| hsa_circ_0052101 | 8422 | 37 | 0.88% |
| hsa_circ_0007813 | 2504 | 11 | 0.88% |
| hsa_circ_0035673 | 2504 | 11 | 0.88% |
| hsa_circ_0063068 | 2504 | 11 | 0.88% |
| hsa_circ_0074723 | 2504 | 11 | 0.88% |
| hsa_circ_0033538 | 3187 | 14 | 0.88% |

| | | | |
|---|---|---|---|
| hsa_circ_0052748 | 2277 | 10 | 0.88% |
| hsa_circ_0084209 | 2277 | 10 | 0.88% |
| hsa_circ_0010249 | 2505 | 11 | 0.88% |
| hsa_circ_0021423 | 2505 | 11 | 0.88% |
| hsa_circ_0027961 | 2505 | 11 | 0.88% |
| hsa_circ_0080020 | 2505 | 11 | 0.88% |
| hsa_circ_0090712 | 2505 | 11 | 0.88% |
| hsa_circ_0013077 | 2733 | 12 | 0.88% |
| hsa_circ_0022952 | 2733 | 12 | 0.88% |
| hsa_circ_0031535 | 2733 | 12 | 0.88% |
| hsa_circ_0074959 | 2733 | 12 | 0.88% |
| hsa_circ_0135832 | 2733 | 12 | 0.88% |
| hsa_circ_0067071 | 3189 | 14 | 0.88% |
| hsa_circ_0043796 | 2278 | 10 | 0.88% |
| hsa_circ_0027575 | 2506 | 11 | 0.88% |
| hsa_circ_0085107 | 2506 | 11 | 0.88% |
| hsa_circ_0127083 | 2506 | 11 | 0.88% |
| hsa_circ_0121905 | 3873 | 17 | 0.88% |
| hsa_circ_0004076 | 2962 | 13 | 0.88% |
| hsa_circ_0042988 | 2962 | 13 | 0.88% |
| hsa_circ_0062853 | 2962 | 13 | 0.88% |
| hsa_circ_0113459 | 2962 | 13 | 0.88% |
| hsa_circ_0035642 | 4557 | 20 | 0.88% |
| hsa_circ_0071151 | 2279 | 10 | 0.88% |
| hsa_circ_0074706 | 2279 | 10 | 0.88% |
| hsa_circ_0095509 | 2279 | 10 | 0.88% |
| hsa_circ_0019841 | 2507 | 11 | 0.88% |
| hsa_circ_0020126 | 2507 | 11 | 0.88% |
| hsa_circ_0028516 | 2507 | 11 | 0.88% |
| hsa_circ_0052355 | 2507 | 11 | 0.88% |
| hsa_circ_0054968 | 2507 | 11 | 0.88% |
| hsa_circ_0078714 | 2507 | 11 | 0.88% |
| hsa_circ_0079371 | 2507 | 11 | 0.88% |
| hsa_circ_0099826 | 2507 | 11 | 0.88% |
| hsa_circ_0034683 | 2735 | 12 | 0.88% |
| hsa_circ_0034766 | 2735 | 12 | 0.88% |
| hsa_circ_0039877 | 2735 | 12 | 0.88% |
| hsa_circ_0054752 | 2735 | 12 | 0.88% |
| hsa_circ_0052060 | 3876 | 17 | 0.88% |
| hsa_circ_0137275 | 3192 | 14 | 0.88% |
| hsa_circ_0050037 | 2964 | 13 | 0.88% |
| hsa_circ_0004734 | 2736 | 12 | 0.88% |
| hsa_circ_0023184 | 2736 | 12 | 0.88% |
| hsa_circ_0031688 | 2736 | 12 | 0.88% |
| hsa_circ_0058718 | 2736 | 12 | 0.88% |
| hsa_circ_0009931 | 2508 | 11 | 0.88% |
| hsa_circ_0023484 | 2280 | 10 | 0.88% |
| hsa_circ_0029138 | 2280 | 10 | 0.88% |
| hsa_circ_0057710 | 2280 | 10 | 0.88% |
| hsa_circ_0062368 | 2280 | 10 | 0.88% |

| | | | |
|---|---|---|---|
| hsa_circ_0077019 | 2280 | 10 | 0.88% |
| hsa_circ_0129062 | 4105 | 18 | 0.88% |
| hsa_circ_0124715 | 3193 | 14 | 0.88% |
| hsa_circ_0126343 | 2965 | 13 | 0.88% |
| hsa_circ_0020186 | 2737 | 12 | 0.88% |
| hsa_circ_0126976 | 2737 | 12 | 0.88% |
| hsa_circ_0001041 | 2509 | 11 | 0.88% |
| hsa_circ_0002562 | 2509 | 11 | 0.88% |
| hsa_circ_0008425 | 2509 | 11 | 0.88% |
| hsa_circ_0024678 | 2509 | 11 | 0.88% |
| hsa_circ_0028798 | 2509 | 11 | 0.88% |
| hsa_circ_0052067 | 2509 | 11 | 0.88% |
| hsa_circ_0059581 | 2509 | 11 | 0.88% |
| hsa_circ_0065155 | 2509 | 11 | 0.88% |
| hsa_circ_0121147 | 2509 | 11 | 0.88% |
| hsa_circ_0029746 | 2281 | 10 | 0.88% |
| hsa_circ_0021243 | 2966 | 13 | 0.88% |
| hsa_circ_0034445 | 2738 | 12 | 0.88% |
| hsa_circ_0054768 | 2738 | 12 | 0.88% |
| hsa_circ_0045464 | 2510 | 11 | 0.88% |
| hsa_circ_0068077 | 2510 | 11 | 0.88% |
| hsa_circ_0089051 | 2510 | 11 | 0.88% |
| hsa_circ_0126341 | 2510 | 11 | 0.88% |
| hsa_circ_0009781 | 2282 | 10 | 0.88% |
| hsa_circ_0021008 | 2282 | 10 | 0.88% |
| hsa_circ_0076893 | 2282 | 10 | 0.88% |
| hsa_circ_0104268 | 2282 | 10 | 0.88% |
| hsa_circ_0126635 | 2282 | 10 | 0.88% |
| hsa_circ_0083141 | 3195 | 14 | 0.88% |
| hsa_circ_0128992 | 2967 | 13 | 0.88% |
| hsa_circ_0063311 | 3652 | 16 | 0.88% |
| hsa_circ_0023820 | 2739 | 12 | 0.88% |
| hsa_circ_0134899 | 2739 | 12 | 0.88% |
| hsa_circ_0078807 | 6163 | 27 | 0.88% |
| hsa_circ_0010768 | 2511 | 11 | 0.88% |
| hsa_circ_0048085 | 2511 | 11 | 0.88% |
| hsa_circ_0049557 | 2511 | 11 | 0.88% |
| hsa_circ_0051203 | 2511 | 11 | 0.88% |
| hsa_circ_0054857 | 2511 | 11 | 0.88% |
| hsa_circ_0065143 | 2511 | 11 | 0.88% |
| hsa_circ_0116066 | 3196 | 14 | 0.88% |
| hsa_circ_0115035 | 4566 | 20 | 0.88% |
| hsa_circ_0011803 | 2283 | 10 | 0.88% |
| hsa_circ_0054363 | 2283 | 10 | 0.88% |
| hsa_circ_0061742 | 2283 | 10 | 0.88% |
| hsa_circ_0070608 | 2283 | 10 | 0.88% |
| hsa_circ_0009233 | 2740 | 12 | 0.88% |
| hsa_circ_0036298 | 2740 | 12 | 0.88% |
| hsa_circ_0075189 | 2740 | 12 | 0.88% |
| hsa_circ_0108805 | 10732 | 47 | 0.88% |

| | | | |
|---|---|---|---|
| hsa_circ_0013813 | 3882 | 17 | 0.88% |
| hsa_circ_0044694 | 5024 | 22 | 0.88% |
| hsa_circ_0060541 | 2512 | 11 | 0.88% |
| hsa_circ_0086000 | 2969 | 13 | 0.88% |
| hsa_circ_0064991 | 3426 | 15 | 0.88% |
| hsa_circ_0013945 | 2284 | 10 | 0.88% |
| hsa_circ_0029040 | 2284 | 10 | 0.88% |
| hsa_circ_0082372 | 2284 | 10 | 0.88% |
| hsa_circ_0087173 | 2284 | 10 | 0.88% |
| hsa_circ_0100718 | 2284 | 10 | 0.88% |
| hsa_circ_0120904 | 2284 | 10 | 0.88% |
| hsa_circ_0124465 | 2284 | 10 | 0.88% |
| hsa_circ_0020629 | 2741 | 12 | 0.88% |
| hsa_circ_0063556 | 2741 | 12 | 0.88% |
| hsa_circ_0024417 | 3198 | 14 | 0.88% |
| hsa_circ_0027101 | 2513 | 11 | 0.88% |
| hsa_circ_0045241 | 2513 | 11 | 0.88% |
| hsa_circ_0067647 | 2513 | 11 | 0.88% |
| hsa_circ_0083842 | 2513 | 11 | 0.88% |
| hsa_circ_0101839 | 2513 | 11 | 0.88% |
| hsa_circ_0040873 | 3199 | 14 | 0.88% |
| hsa_circ_0078532 | 2742 | 12 | 0.88% |
| hsa_circ_0027840 | 2285 | 10 | 0.88% |
| hsa_circ_0067198 | 2285 | 10 | 0.88% |
| hsa_circ_0069605 | 2285 | 10 | 0.88% |
| hsa_circ_0078174 | 2285 | 10 | 0.88% |
| hsa_circ_0084261 | 2285 | 10 | 0.88% |
| hsa_circ_0114406 | 2285 | 10 | 0.88% |
| hsa_circ_0075921 | 6170 | 27 | 0.88% |
| hsa_circ_0013015 | 2514 | 11 | 0.88% |
| hsa_circ_0019531 | 2514 | 11 | 0.88% |
| hsa_circ_0020396 | 2514 | 11 | 0.88% |
| hsa_circ_0034199 | 2514 | 11 | 0.88% |
| hsa_circ_0098035 | 2514 | 11 | 0.88% |
| hsa_circ_0108036 | 2514 | 11 | 0.88% |
| hsa_circ_0120663 | 2514 | 11 | 0.88% |
| hsa_circ_0009330 | 2743 | 12 | 0.87% |
| hsa_circ_0044512 | 5258 | 23 | 0.87% |
| hsa_circ_0135240 | 4115 | 18 | 0.87% |
| hsa_circ_0036965 | 2515 | 11 | 0.87% |
| hsa_circ_0060301 | 3201 | 14 | 0.87% |
| hsa_circ_0090991 | 3430 | 15 | 0.87% |
| hsa_circ_0019760 | 2744 | 12 | 0.87% |
| hsa_circ_0024519 | 2744 | 12 | 0.87% |
| hsa_circ_0039600 | 2287 | 10 | 0.87% |
| hsa_circ_0067995 | 2287 | 10 | 0.87% |
| hsa_circ_0071283 | 2516 | 11 | 0.87% |
| hsa_circ_0076371 | 2516 | 11 | 0.87% |
| hsa_circ_0080686 | 2516 | 11 | 0.87% |
| hsa_circ_0040998 | 2745 | 12 | 0.87% |

| | | | |
|---|---|---|---|
| hsa_circ_0044786 | 2745 | 12 | 0.87% |
| hsa_circ_0138633 | 2745 | 12 | 0.87% |
| hsa_circ_0040768 | 2288 | 10 | 0.87% |
| hsa_circ_0063065 | 2288 | 10 | 0.87% |
| hsa_circ_0090999 | 2517 | 11 | 0.87% |
| hsa_circ_0135024 | 2517 | 11 | 0.87% |
| hsa_circ_0025394 | 2746 | 12 | 0.87% |
| hsa_circ_0041576 | 2746 | 12 | 0.87% |
| hsa_circ_0132577 | 7324 | 32 | 0.87% |
| hsa_circ_0026415 | 2289 | 10 | 0.87% |
| hsa_circ_0076712 | 2289 | 10 | 0.87% |
| hsa_circ_0081298 | 2289 | 10 | 0.87% |
| hsa_circ_0016319 | 2518 | 11 | 0.87% |
| hsa_circ_0055893 | 2518 | 11 | 0.87% |
| hsa_circ_0114439 | 2518 | 11 | 0.87% |
| hsa_circ_0109884 | 7327 | 32 | 0.87% |
| hsa_circ_0102296 | 3893 | 17 | 0.87% |
| hsa_circ_0037186 | 3206 | 14 | 0.87% |
| hsa_circ_0053207 | 2977 | 13 | 0.87% |
| hsa_circ_0084410 | 2977 | 13 | 0.87% |
| hsa_circ_0010216 | 2748 | 12 | 0.87% |
| hsa_circ_0043228 | 2748 | 12 | 0.87% |
| hsa_circ_0061681 | 2748 | 12 | 0.87% |
| hsa_circ_0086935 | 2748 | 12 | 0.87% |
| hsa_circ_0003077 | 2519 | 11 | 0.87% |
| hsa_circ_0017050 | 2519 | 11 | 0.87% |
| hsa_circ_0026971 | 2519 | 11 | 0.87% |
| hsa_circ_0034162 | 2519 | 11 | 0.87% |
| hsa_circ_0049237 | 2290 | 10 | 0.87% |
| hsa_circ_0082956 | 2749 | 12 | 0.87% |
| hsa_circ_0089708 | 2749 | 12 | 0.87% |
| hsa_circ_0123384 | 2749 | 12 | 0.87% |
| hsa_circ_0122809 | 8018 | 35 | 0.87% |
| hsa_circ_0082732 | 5040 | 22 | 0.87% |
| hsa_circ_0014328 | 2520 | 11 | 0.87% |
| hsa_circ_0054064 | 2520 | 11 | 0.87% |
| hsa_circ_0073180 | 2520 | 11 | 0.87% |
| hsa_circ_0073937 | 2520 | 11 | 0.87% |
| hsa_circ_0080135 | 2520 | 11 | 0.87% |
| hsa_circ_0086507 | 2520 | 11 | 0.87% |
| hsa_circ_0088097 | 2520 | 11 | 0.87% |
| hsa_circ_0091491 | 2520 | 11 | 0.87% |
| hsa_circ_0099907 | 2520 | 11 | 0.87% |
| hsa_circ_0100660 | 2520 | 11 | 0.87% |
| hsa_circ_0100754 | 2520 | 11 | 0.87% |
| hsa_circ_0074085 | 2291 | 10 | 0.87% |
| hsa_circ_0076261 | 2291 | 10 | 0.87% |
| hsa_circ_0129474 | 2291 | 10 | 0.87% |
| hsa_circ_0123326 | 15808 | 69 | 0.87% |
| hsa_circ_0018926 | 2979 | 13 | 0.87% |

| | | | |
|---|---|---|---|
| hsa_circ_0133295 | 2979 | 13 | 0.87% |
| hsa_circ_0118361 | 2750 | 12 | 0.87% |
| hsa_circ_0090902 | 2521 | 11 | 0.87% |
| hsa_circ_0127078 | 2521 | 11 | 0.87% |
| hsa_circ_0015911 | 7336 | 32 | 0.87% |
| hsa_circ_0103153 | 6419 | 28 | 0.87% |
| hsa_circ_0098002 | 3668 | 16 | 0.87% |
| hsa_circ_0115964 | 2751 | 12 | 0.87% |
| hsa_circ_0120571 | 2751 | 12 | 0.87% |
| hsa_circ_0077126 | 2522 | 11 | 0.87% |
| hsa_circ_0095800 | 2522 | 11 | 0.87% |
| hsa_circ_0010297 | 2293 | 10 | 0.87% |
| hsa_circ_0061439 | 2981 | 13 | 0.87% |
| hsa_circ_0140535 | 2981 | 13 | 0.87% |
| hsa_circ_0050015 | 2752 | 12 | 0.87% |
| hsa_circ_0071316 | 3211 | 14 | 0.87% |
| hsa_circ_0009447 | 2523 | 11 | 0.87% |
| hsa_circ_0054536 | 2523 | 11 | 0.87% |
| hsa_circ_0062010 | 2523 | 11 | 0.87% |
| hsa_circ_0064391 | 2523 | 11 | 0.87% |
| hsa_circ_0075676 | 2523 | 11 | 0.87% |
| hsa_circ_0091890 | 2523 | 11 | 0.87% |
| hsa_circ_0069630 | 2294 | 10 | 0.87% |
| hsa_circ_0022514 | 2753 | 12 | 0.87% |
| hsa_circ_0076519 | 2753 | 12 | 0.87% |
| hsa_circ_0104936 | 2524 | 11 | 0.87% |
| hsa_circ_0113480 | 19046 | 83 | 0.87% |
| hsa_circ_0057112 | 3672 | 16 | 0.87% |
| hsa_circ_0032577 | 2754 | 12 | 0.87% |
| hsa_circ_0052830 | 2754 | 12 | 0.87% |
| hsa_circ_0068790 | 2754 | 12 | 0.87% |
| hsa_circ_0074774 | 2754 | 12 | 0.87% |
| hsa_circ_0030576 | 2295 | 10 | 0.87% |
| hsa_circ_0091883 | 2295 | 10 | 0.87% |
| hsa_circ_0080789 | 3443 | 15 | 0.87% |
| hsa_circ_0100735 | 2525 | 11 | 0.87% |
| hsa_circ_0097085 | 6199 | 27 | 0.87% |
| hsa_circ_0020047 | 2296 | 10 | 0.87% |
| hsa_circ_0093452 | 2296 | 10 | 0.87% |
| hsa_circ_0013785 | 2526 | 11 | 0.87% |
| hsa_circ_0027632 | 2526 | 11 | 0.87% |
| hsa_circ_0053529 | 2526 | 11 | 0.87% |
| hsa_circ_0076547 | 2526 | 11 | 0.87% |
| hsa_circ_0016595 | 5743 | 25 | 0.87% |
| hsa_circ_0030824 | 2527 | 11 | 0.87% |
| hsa_circ_0031349 | 2527 | 11 | 0.87% |
| hsa_circ_0047262 | 2527 | 11 | 0.87% |
| hsa_circ_0107520 | 2527 | 11 | 0.87% |
| hsa_circ_0101572 | 3676 | 16 | 0.87% |
| hsa_circ_0040332 | 2757 | 12 | 0.87% |

| | | | |
|---|---|---|---|
| hsa_circ_0056797 | 2757 | 12 | 0.87% |
| hsa_circ_0088262 | 2757 | 12 | 0.87% |
| hsa_circ_0104172 | 2757 | 12 | 0.87% |
| hsa_circ_0132497 | 2757 | 12 | 0.87% |
| hsa_circ_0106472 | 3217 | 14 | 0.87% |
| hsa_circ_0010470 | 2298 | 10 | 0.87% |
| hsa_circ_0027666 | 2298 | 10 | 0.87% |
| hsa_circ_0044930 | 2298 | 10 | 0.87% |
| hsa_circ_0085731 | 2298 | 10 | 0.87% |
| hsa_circ_0014867 | 2528 | 11 | 0.87% |
| hsa_circ_0056743 | 2528 | 11 | 0.87% |
| hsa_circ_0088970 | 2528 | 11 | 0.87% |
| hsa_circ_0127887 | 2528 | 11 | 0.87% |
| hsa_circ_0035223 | 2758 | 12 | 0.87% |
| hsa_circ_0035734 | 2758 | 12 | 0.87% |
| hsa_circ_0130431 | 2758 | 12 | 0.87% |
| hsa_circ_0092090 | 2988 | 13 | 0.87% |
| hsa_circ_0041912 | 2299 | 10 | 0.87% |
| hsa_circ_0052194 | 2299 | 10 | 0.87% |
| hsa_circ_0126206 | 2299 | 10 | 0.87% |
| hsa_circ_0018671 | 2529 | 11 | 0.87% |
| hsa_circ_0021095 | 2529 | 11 | 0.87% |
| hsa_circ_0053902 | 2529 | 11 | 0.87% |
| hsa_circ_0107806 | 2529 | 11 | 0.87% |
| hsa_circ_0124673 | 2529 | 11 | 0.87% |
| hsa_circ_0054438 | 2759 | 12 | 0.87% |
| hsa_circ_0073821 | 5978 | 26 | 0.87% |
| hsa_circ_0060703 | 2989 | 13 | 0.87% |
| hsa_circ_0124389 | 8280 | 36 | 0.87% |
| hsa_circ_0021091 | 4370 | 19 | 0.87% |
| hsa_circ_0115267 | 3910 | 17 | 0.87% |
| hsa_circ_0088571 | 2990 | 13 | 0.87% |
| hsa_circ_0047885 | 2530 | 11 | 0.87% |
| hsa_circ_0085243 | 2530 | 11 | 0.87% |
| hsa_circ_0085579 | 2530 | 11 | 0.87% |
| hsa_circ_0108037 | 2530 | 11 | 0.87% |
| hsa_circ_0125126 | 2530 | 11 | 0.87% |
| hsa_circ_0039611 | 2300 | 10 | 0.87% |
| hsa_circ_0104994 | 7592 | 33 | 0.87% |
| hsa_circ_0033409 | 3451 | 15 | 0.87% |
| hsa_circ_0093997 | 2531 | 11 | 0.87% |
| hsa_circ_0041073 | 2301 | 10 | 0.87% |
| hsa_circ_0119132 | 11046 | 48 | 0.87% |
| hsa_circ_0138882 | 6444 | 28 | 0.87% |
| hsa_circ_0083017 | 2992 | 13 | 0.87% |
| hsa_circ_0017322 | 2762 | 12 | 0.87% |
| hsa_circ_0045474 | 2762 | 12 | 0.87% |
| hsa_circ_0071643 | 2762 | 12 | 0.87% |
| hsa_circ_0107990 | 2762 | 12 | 0.87% |
| hsa_circ_0027845 | 2532 | 11 | 0.87% |

| | | | |
|---|---|---|---|
| hsa_circ_0035707 | 2532 | 11 | 0.87% |
| hsa_circ_0068321 | 2532 | 11 | 0.87% |
| hsa_circ_0097803 | 2532 | 11 | 0.87% |
| hsa_circ_0018477 | 2302 | 10 | 0.87% |
| hsa_circ_0088215 | 2302 | 10 | 0.87% |
| hsa_circ_0124385 | 6446 | 28 | 0.87% |
| hsa_circ_0067816 | 2993 | 13 | 0.87% |
| hsa_circ_0039083 | 2763 | 12 | 0.87% |
| hsa_circ_0078072 | 2763 | 12 | 0.87% |
| hsa_circ_0055206 | 2533 | 11 | 0.87% |
| hsa_circ_0066981 | 2533 | 11 | 0.87% |
| hsa_circ_0087048 | 2533 | 11 | 0.87% |
| hsa_circ_0098547 | 2533 | 11 | 0.87% |
| hsa_circ_0109160 | 6218 | 27 | 0.87% |
| hsa_circ_0026445 | 2303 | 10 | 0.87% |
| hsa_circ_0117709 | 2303 | 10 | 0.87% |
| hsa_circ_0033619 | 2994 | 13 | 0.87% |
| hsa_circ_0046174 | 2994 | 13 | 0.87% |
| hsa_circ_0020949 | 2764 | 12 | 0.87% |
| hsa_circ_0047143 | 2764 | 12 | 0.87% |
| hsa_circ_0082644 | 2764 | 12 | 0.87% |
| hsa_circ_0084832 | 2764 | 12 | 0.87% |
| hsa_circ_0121690 | 2764 | 12 | 0.87% |
| hsa_circ_0089539 | 3225 | 14 | 0.87% |
| hsa_circ_0021379 | 2534 | 11 | 0.87% |
| hsa_circ_0027453 | 2534 | 11 | 0.87% |
| hsa_circ_0028907 | 2534 | 11 | 0.87% |
| hsa_circ_0031595 | 2534 | 11 | 0.87% |
| hsa_circ_0051485 | 2534 | 11 | 0.87% |
| hsa_circ_0105484 | 2534 | 11 | 0.87% |
| hsa_circ_0137579 | 2534 | 11 | 0.87% |
| hsa_circ_0039690 | 2304 | 10 | 0.87% |
| hsa_circ_0082991 | 2304 | 10 | 0.87% |
| hsa_circ_0094000 | 2304 | 10 | 0.87% |
| hsa_circ_0113549 | 2304 | 10 | 0.87% |
| hsa_circ_0122686 | 2304 | 10 | 0.87% |
| hsa_circ_0032234 | 2765 | 12 | 0.87% |
| hsa_circ_0044041 | 2765 | 12 | 0.87% |
| hsa_circ_0024547 | 2535 | 11 | 0.87% |
| hsa_circ_0028073 | 2535 | 11 | 0.87% |
| hsa_circ_0033576 | 2535 | 11 | 0.87% |
| hsa_circ_0034714 | 2535 | 11 | 0.87% |
| hsa_circ_0037009 | 2535 | 11 | 0.87% |
| hsa_circ_0031489 | 2766 | 12 | 0.87% |
| hsa_circ_0067143 | 2766 | 12 | 0.87% |
| hsa_circ_0092096 | 2305 | 10 | 0.87% |
| hsa_circ_0050086 | 9451 | 41 | 0.87% |
| hsa_circ_0028451 | 2536 | 11 | 0.87% |
| hsa_circ_0065880 | 3228 | 14 | 0.87% |
| hsa_circ_0128349 | 7609 | 33 | 0.87% |

| | | | |
|---|---|---|---|
| hsa_circ_0112927 | 2767 | 12 | 0.87% |
| hsa_circ_0027396 | 2306 | 10 | 0.87% |
| hsa_circ_0041815 | 2306 | 10 | 0.87% |
| hsa_circ_0051320 | 2306 | 10 | 0.87% |
| hsa_circ_0104153 | 2306 | 10 | 0.87% |
| hsa_circ_0076451 | 2998 | 13 | 0.87% |
| hsa_circ_0046048 | 4382 | 19 | 0.87% |
| hsa_circ_0071686 | 2537 | 11 | 0.87% |
| hsa_circ_0111167 | 2537 | 11 | 0.87% |
| hsa_circ_0041893 | 2768 | 12 | 0.87% |
| hsa_circ_0081503 | 3691 | 16 | 0.87% |
| hsa_circ_0052597 | 2999 | 13 | 0.87% |
| hsa_circ_0076447 | 2999 | 13 | 0.87% |
| hsa_circ_0084388 | 2999 | 13 | 0.87% |
| hsa_circ_0049900 | 2307 | 10 | 0.87% |
| hsa_circ_0081282 | 2307 | 10 | 0.87% |
| hsa_circ_0113577 | 2307 | 10 | 0.87% |
| hsa_circ_0016740 | 2538 | 11 | 0.87% |
| hsa_circ_0027329 | 2538 | 11 | 0.87% |
| hsa_circ_0035496 | 2538 | 11 | 0.87% |
| hsa_circ_0068181 | 2538 | 11 | 0.87% |
| hsa_circ_0074020 | 2538 | 11 | 0.87% |
| hsa_circ_0105244 | 2538 | 11 | 0.87% |
| hsa_circ_0010339 | 2769 | 12 | 0.87% |
| hsa_circ_0088668 | 2769 | 12 | 0.87% |
| hsa_circ_0021600 | 2308 | 10 | 0.87% |
| hsa_circ_0032819 | 2308 | 10 | 0.87% |
| hsa_circ_0077714 | 2308 | 10 | 0.87% |
| hsa_circ_0084190 | 2308 | 10 | 0.87% |
| hsa_circ_0092969 | 2308 | 10 | 0.87% |
| hsa_circ_0093996 | 2308 | 10 | 0.87% |
| hsa_circ_0103388 | 2308 | 10 | 0.87% |
| hsa_circ_0125012 | 2308 | 10 | 0.87% |
| hsa_circ_0036534 | 2539 | 11 | 0.87% |
| hsa_circ_0066691 | 2539 | 11 | 0.87% |
| hsa_circ_0071207 | 2539 | 11 | 0.87% |
| hsa_circ_0075791 | 2539 | 11 | 0.87% |
| hsa_circ_0025101 | 2770 | 12 | 0.87% |
| hsa_circ_0043231 | 2770 | 12 | 0.87% |
| hsa_circ_0082703 | 2770 | 12 | 0.87% |
| hsa_circ_0111258 | 7387 | 32 | 0.87% |
| hsa_circ_0123425 | 3001 | 13 | 0.87% |
| hsa_circ_0137439 | 5772 | 25 | 0.87% |
| hsa_circ_0089809 | 6003 | 26 | 0.87% |
| hsa_circ_0041660 | 6696 | 29 | 0.87% |
| hsa_circ_0079806 | 2309 | 10 | 0.87% |
| hsa_circ_0120101 | 2309 | 10 | 0.87% |
| hsa_circ_0021169 | 2540 | 11 | 0.87% |
| hsa_circ_0043255 | 2540 | 11 | 0.87% |
| hsa_circ_0048600 | 2540 | 11 | 0.87% |

| | | | |
|---|---|---|---|
| hsa_circ_0089882 | 2540 | 11 | 0.87% |
| hsa_circ_0098304 | 2771 | 12 | 0.87% |
| hsa_circ_0138175 | 2771 | 12 | 0.87% |
| hsa_circ_0044091 | 7390 | 32 | 0.87% |
| hsa_circ_0025354 | 3003 | 13 | 0.87% |
| hsa_circ_0039728 | 3003 | 13 | 0.87% |
| hsa_circ_0035287 | 2772 | 12 | 0.87% |
| hsa_circ_0049312 | 2772 | 12 | 0.87% |
| hsa_circ_0032413 | 2541 | 11 | 0.87% |
| hsa_circ_0112534 | 2541 | 11 | 0.87% |
| hsa_circ_0013211 | 2310 | 10 | 0.87% |
| hsa_circ_0015711 | 2310 | 10 | 0.87% |
| hsa_circ_0032751 | 2310 | 10 | 0.87% |
| hsa_circ_0037081 | 2310 | 10 | 0.87% |
| hsa_circ_0055013 | 2310 | 10 | 0.87% |
| hsa_circ_0062043 | 2310 | 10 | 0.87% |
| hsa_circ_0114732 | 2310 | 10 | 0.87% |
| hsa_circ_0011873 | 2773 | 12 | 0.87% |
| hsa_circ_0074510 | 2773 | 12 | 0.87% |
| hsa_circ_0121953 | 2773 | 12 | 0.87% |
| hsa_circ_0024977 | 2542 | 11 | 0.87% |
| hsa_circ_0046932 | 2542 | 11 | 0.87% |
| hsa_circ_0048214 | 2542 | 11 | 0.87% |
| hsa_circ_0052646 | 2542 | 11 | 0.87% |
| hsa_circ_0020428 | 2311 | 10 | 0.87% |
| hsa_circ_0033380 | 2311 | 10 | 0.87% |
| hsa_circ_0043120 | 2311 | 10 | 0.87% |
| hsa_circ_0053828 | 2311 | 10 | 0.87% |
| hsa_circ_0079426 | 2311 | 10 | 0.87% |
| hsa_circ_0085678 | 2311 | 10 | 0.87% |
| hsa_circ_0125011 | 2311 | 10 | 0.87% |
| hsa_circ_0084401 | 3005 | 13 | 0.87% |
| hsa_circ_0096689 | 4392 | 19 | 0.87% |
| hsa_circ_0018632 | 2774 | 12 | 0.87% |
| hsa_circ_0041133 | 2774 | 12 | 0.87% |
| hsa_circ_0088665 | 2774 | 12 | 0.87% |
| hsa_circ_0112461 | 2774 | 12 | 0.87% |
| hsa_circ_0114157 | 2774 | 12 | 0.87% |
| hsa_circ_0034160 | 2543 | 11 | 0.87% |
| hsa_circ_0073342 | 2543 | 11 | 0.87% |
| hsa_circ_0027341 | 2312 | 10 | 0.87% |
| hsa_circ_0043680 | 3006 | 13 | 0.86% |
| hsa_circ_0130081 | 3931 | 17 | 0.86% |
| hsa_circ_0031173 | 2775 | 12 | 0.86% |
| hsa_circ_0092918 | 2775 | 12 | 0.86% |
| hsa_circ_0034142 | 2544 | 11 | 0.86% |
| hsa_circ_0041260 | 2544 | 11 | 0.86% |
| hsa_circ_0073422 | 2544 | 11 | 0.86% |
| hsa_circ_0074477 | 2544 | 11 | 0.86% |
| hsa_circ_0136355 | 2544 | 11 | 0.86% |

| | | | |
|---|---|---|---|
| hsa_circ_0079797 | 2313 | 10 | 0.86% |
| hsa_circ_0091942 | 2313 | 10 | 0.86% |
| hsa_circ_0137826 | 2313 | 10 | 0.86% |
| hsa_circ_0046745 | 3007 | 13 | 0.86% |
| hsa_circ_0125197 | 3007 | 13 | 0.86% |
| hsa_circ_0071547 | 8096 | 35 | 0.86% |
| hsa_circ_0122567 | 4164 | 18 | 0.86% |
| hsa_circ_0046666 | 2776 | 12 | 0.86% |
| hsa_circ_0074350 | 2776 | 12 | 0.86% |
| hsa_circ_0019890 | 2545 | 11 | 0.86% |
| hsa_circ_0056072 | 2545 | 11 | 0.86% |
| hsa_circ_0058601 | 2545 | 11 | 0.86% |
| hsa_circ_0069302 | 2545 | 11 | 0.86% |
| hsa_circ_0088892 | 2545 | 11 | 0.86% |
| hsa_circ_0080038 | 3008 | 13 | 0.86% |
| hsa_circ_0032271 | 4165 | 18 | 0.86% |
| hsa_circ_0021160 | 2314 | 10 | 0.86% |
| hsa_circ_0041426 | 2314 | 10 | 0.86% |
| hsa_circ_0059524 | 2314 | 10 | 0.86% |
| hsa_circ_0033252 | 2777 | 12 | 0.86% |
| hsa_circ_0001624 | 36337 | 157 | 0.86% |
| hsa_circ_0032400 | 2546 | 11 | 0.86% |
| hsa_circ_0044848 | 2546 | 11 | 0.86% |
| hsa_circ_0118143 | 4167 | 18 | 0.86% |
| hsa_circ_0077663 | 2778 | 12 | 0.86% |
| hsa_circ_0125580 | 2778 | 12 | 0.86% |
| hsa_circ_0046857 | 2315 | 10 | 0.86% |
| hsa_circ_0072238 | 2315 | 10 | 0.86% |
| hsa_circ_0090430 | 2315 | 10 | 0.86% |
| hsa_circ_0012295 | 3010 | 13 | 0.86% |
| hsa_circ_0015919 | 2547 | 11 | 0.86% |
| hsa_circ_0018122 | 2547 | 11 | 0.86% |
| hsa_circ_0046392 | 2547 | 11 | 0.86% |
| hsa_circ_0053442 | 2547 | 11 | 0.86% |
| hsa_circ_0053889 | 2547 | 11 | 0.86% |
| hsa_circ_0073661 | 2547 | 11 | 0.86% |
| hsa_circ_0096763 | 2547 | 11 | 0.86% |
| hsa_circ_0121132 | 2547 | 11 | 0.86% |
| hsa_circ_0053101 | 2779 | 12 | 0.86% |
| hsa_circ_0079456 | 2779 | 12 | 0.86% |
| hsa_circ_0112569 | 2779 | 12 | 0.86% |
| hsa_circ_0012470 | 2316 | 10 | 0.86% |
| hsa_circ_0012687 | 2316 | 10 | 0.86% |
| hsa_circ_0013264 | 2316 | 10 | 0.86% |
| hsa_circ_0030420 | 2316 | 10 | 0.86% |
| hsa_circ_0030805 | 2316 | 10 | 0.86% |
| hsa_circ_0053921 | 2316 | 10 | 0.86% |
| hsa_circ_0061247 | 2316 | 10 | 0.86% |
| hsa_circ_0073541 | 2316 | 10 | 0.86% |
| hsa_circ_0134794 | 2316 | 10 | 0.86% |

| | | | |
|---|---|---|---|
| hsa_circ_0085395 | 2548 | 11 | 0.86% |
| hsa_circ_0117293 | 2548 | 11 | 0.86% |
| hsa_circ_0079694 | 5328 | 23 | 0.86% |
| hsa_circ_0096504 | 3012 | 13 | 0.86% |
| hsa_circ_0036801 | 2317 | 10 | 0.86% |
| hsa_circ_0088209 | 2317 | 10 | 0.86% |
| hsa_circ_0094859 | 2317 | 10 | 0.86% |
| hsa_circ_0039423 | 2549 | 11 | 0.86% |
| hsa_circ_0058788 | 2549 | 11 | 0.86% |
| hsa_circ_0091875 | 2549 | 11 | 0.86% |
| hsa_circ_0102762 | 2549 | 11 | 0.86% |
| hsa_circ_0070690 | 2781 | 12 | 0.86% |
| hsa_circ_0037801 | 3013 | 13 | 0.86% |
| hsa_circ_0039646 | 3013 | 13 | 0.86% |
| hsa_circ_0004191 | 3477 | 15 | 0.86% |
| hsa_circ_0021404 | 2318 | 10 | 0.86% |
| hsa_circ_0139172 | 2318 | 10 | 0.86% |
| hsa_circ_0057417 | 2550 | 11 | 0.86% |
| hsa_circ_0088977 | 2550 | 11 | 0.86% |
| hsa_circ_0139043 | 2550 | 11 | 0.86% |
| hsa_circ_0012493 | 2782 | 12 | 0.86% |
| hsa_circ_0024587 | 2782 | 12 | 0.86% |
| hsa_circ_0084211 | 2782 | 12 | 0.86% |
| hsa_circ_0095511 | 3014 | 13 | 0.86% |
| hsa_circ_0015902 | 2319 | 10 | 0.86% |
| hsa_circ_0042013 | 2319 | 10 | 0.86% |
| hsa_circ_0054307 | 2319 | 10 | 0.86% |
| hsa_circ_0086977 | 2319 | 10 | 0.86% |
| hsa_circ_0109281 | 2319 | 10 | 0.86% |
| hsa_circ_0114736 | 2319 | 10 | 0.86% |
| hsa_circ_0137018 | 2319 | 10 | 0.86% |
| hsa_circ_0030092 | 2551 | 11 | 0.86% |
| hsa_circ_0072675 | 2551 | 11 | 0.86% |
| hsa_circ_0076369 | 2551 | 11 | 0.86% |
| hsa_circ_0090863 | 2551 | 11 | 0.86% |
| hsa_circ_0126622 | 2551 | 11 | 0.86% |
| hsa_circ_0129493 | 2551 | 11 | 0.86% |
| hsa_circ_0063347 | 2783 | 12 | 0.86% |
| hsa_circ_0084816 | 2783 | 12 | 0.86% |
| hsa_circ_0104428 | 2783 | 12 | 0.86% |
| hsa_circ_0034361 | 3015 | 13 | 0.86% |
| hsa_circ_0085541 | 8814 | 38 | 0.86% |
| hsa_circ_0028061 | 2552 | 11 | 0.86% |
| hsa_circ_0051956 | 2552 | 11 | 0.86% |
| hsa_circ_0066328 | 2552 | 11 | 0.86% |
| hsa_circ_0114273 | 2552 | 11 | 0.86% |
| hsa_circ_0036124 | 2320 | 10 | 0.86% |
| hsa_circ_0109625 | 8585 | 37 | 0.86% |
| hsa_circ_0018697 | 2553 | 11 | 0.86% |
| hsa_circ_0076563 | 2553 | 11 | 0.86% |

| | | | |
|---|---|---|---|
| hsa_circ_0096103 | 2553 | 11 | 0.86% |
| hsa_circ_0032966 | 2321 | 10 | 0.86% |
| hsa_circ_0068732 | 5803 | 25 | 0.86% |
| hsa_circ_0050497 | 3250 | 14 | 0.86% |
| hsa_circ_0039265 | 2554 | 11 | 0.86% |
| hsa_circ_0059483 | 2322 | 10 | 0.86% |
| hsa_circ_0076014 | 3251 | 14 | 0.86% |
| hsa_circ_0058819 | 3019 | 13 | 0.86% |
| hsa_circ_0035229 | 2787 | 12 | 0.86% |
| hsa_circ_0048409 | 2787 | 12 | 0.86% |
| hsa_circ_0083546 | 3484 | 15 | 0.86% |
| hsa_circ_0031589 | 2555 | 11 | 0.86% |
| hsa_circ_0009993 | 2323 | 10 | 0.86% |
| hsa_circ_0099169 | 2323 | 10 | 0.86% |
| hsa_circ_0009940 | 2788 | 12 | 0.86% |
| hsa_circ_0016695 | 3718 | 16 | 0.86% |
| hsa_circ_0063519 | 3021 | 13 | 0.86% |
| hsa_circ_0070760 | 3021 | 13 | 0.86% |
| hsa_circ_0077458 | 3021 | 13 | 0.86% |
| hsa_circ_0018338 | 2324 | 10 | 0.86% |
| hsa_circ_0052984 | 2324 | 10 | 0.86% |
| hsa_circ_0092260 | 2324 | 10 | 0.86% |
| hsa_circ_0027145 | 2789 | 12 | 0.86% |
| hsa_circ_0034896 | 2789 | 12 | 0.86% |
| hsa_circ_0009799 | 2557 | 11 | 0.86% |
| hsa_circ_0054351 | 2557 | 11 | 0.86% |
| hsa_circ_0025984 | 3022 | 13 | 0.86% |
| hsa_circ_0032559 | 2790 | 12 | 0.86% |
| hsa_circ_0064978 | 2790 | 12 | 0.86% |
| hsa_circ_0015822 | 2325 | 10 | 0.86% |
| hsa_circ_0046439 | 2325 | 10 | 0.86% |
| hsa_circ_0070334 | 2325 | 10 | 0.86% |
| hsa_circ_0087217 | 2325 | 10 | 0.86% |
| hsa_circ_0112657 | 2325 | 10 | 0.86% |
| hsa_circ_0059647 | 3023 | 13 | 0.86% |
| hsa_circ_0021381 | 2558 | 11 | 0.86% |
| hsa_circ_0043248 | 2558 | 11 | 0.86% |
| hsa_circ_0063620 | 2558 | 11 | 0.86% |
| hsa_circ_0028925 | 3721 | 16 | 0.86% |
| hsa_circ_0120570 | 2791 | 12 | 0.86% |
| hsa_circ_0057197 | 2326 | 10 | 0.86% |
| hsa_circ_0060540 | 2326 | 10 | 0.86% |
| hsa_circ_0063995 | 2559 | 11 | 0.86% |
| hsa_circ_0114256 | 2559 | 11 | 0.86% |
| hsa_circ_0126294 | 2559 | 11 | 0.86% |
| hsa_circ_0034978 | 2560 | 11 | 0.86% |
| hsa_circ_0075161 | 2560 | 11 | 0.86% |
| hsa_circ_0112664 | 2560 | 11 | 0.86% |
| hsa_circ_0047259 | 2793 | 12 | 0.86% |
| hsa_circ_0063366 | 3026 | 13 | 0.86% |

| | | | |
|---|---|---|---|
| hsa_circ_0134538 | 3026 | 13 | 0.86% |
| hsa_circ_0034285 | 2328 | 10 | 0.86% |
| hsa_circ_0042323 | 2328 | 10 | 0.86% |
| hsa_circ_0012810 | 2561 | 11 | 0.86% |
| hsa_circ_0030516 | 2561 | 11 | 0.86% |
| hsa_circ_0025675 | 3027 | 13 | 0.86% |
| hsa_circ_0041143 | 3726 | 16 | 0.86% |
| hsa_circ_0067673 | 2329 | 10 | 0.86% |
| hsa_circ_0071934 | 2329 | 10 | 0.86% |
| hsa_circ_0101325 | 2329 | 10 | 0.86% |
| hsa_circ_0135640 | 2329 | 10 | 0.86% |
| hsa_circ_0023548 | 2562 | 11 | 0.86% |
| hsa_circ_0056646 | 2562 | 11 | 0.86% |
| hsa_circ_0063111 | 2562 | 11 | 0.86% |
| hsa_circ_0058785 | 2795 | 12 | 0.86% |
| hsa_circ_0082223 | 3261 | 14 | 0.86% |
| hsa_circ_0128871 | 9785 | 42 | 0.86% |
| hsa_circ_0057694 | 3728 | 16 | 0.86% |
| hsa_circ_0057335 | 3262 | 14 | 0.86% |
| hsa_circ_0061136 | 2796 | 12 | 0.86% |
| hsa_circ_0067303 | 2796 | 12 | 0.86% |
| hsa_circ_0022855 | 2563 | 11 | 0.86% |
| hsa_circ_0040170 | 2563 | 11 | 0.86% |
| hsa_circ_0102335 | 2563 | 11 | 0.86% |
| hsa_circ_0023390 | 2330 | 10 | 0.86% |
| hsa_circ_0077733 | 2330 | 10 | 0.86% |
| hsa_circ_0079732 | 2330 | 10 | 0.86% |
| hsa_circ_0101451 | 2330 | 10 | 0.86% |
| hsa_circ_0129255 | 2330 | 10 | 0.86% |
| hsa_circ_0006177 | 9321 | 40 | 0.86% |
| hsa_circ_0123809 | 3496 | 15 | 0.86% |
| hsa_circ_0100334 | 3030 | 13 | 0.86% |
| hsa_circ_0027595 | 2797 | 12 | 0.86% |
| hsa_circ_0069860 | 2797 | 12 | 0.86% |
| hsa_circ_0010106 | 2564 | 11 | 0.86% |
| hsa_circ_0058594 | 2564 | 11 | 0.86% |
| hsa_circ_0074340 | 2564 | 11 | 0.86% |
| hsa_circ_0120652 | 2564 | 11 | 0.86% |
| hsa_circ_0022602 | 2331 | 10 | 0.86% |
| hsa_circ_0049035 | 2331 | 10 | 0.86% |
| hsa_circ_0098347 | 2331 | 10 | 0.86% |
| hsa_circ_0101054 | 2331 | 10 | 0.86% |
| hsa_circ_0131275 | 2331 | 10 | 0.86% |
| hsa_circ_0048940 | 5595 | 24 | 0.86% |
| hsa_circ_0097481 | 3264 | 14 | 0.86% |
| hsa_circ_0022722 | 2798 | 12 | 0.86% |
| hsa_circ_0075351 | 2798 | 12 | 0.86% |
| hsa_circ_0009298 | 2565 | 11 | 0.86% |
| hsa_circ_0090399 | 2565 | 11 | 0.86% |
| hsa_circ_0087411 | 2332 | 10 | 0.86% |

| | | | |
|---|---|---|---|
| hsa_circ_0096497 | 2332 | 10 | 0.86% |
| hsa_circ_0139081 | 2332 | 10 | 0.86% |
| hsa_circ_0072487 | 3032 | 13 | 0.86% |
| hsa_circ_0125184 | 3032 | 13 | 0.86% |
| hsa_circ_0034075 | 2799 | 12 | 0.86% |
| hsa_circ_0057513 | 2799 | 12 | 0.86% |
| hsa_circ_0035310 | 2566 | 11 | 0.86% |
| hsa_circ_0052007 | 2566 | 11 | 0.86% |
| hsa_circ_0068716 | 2566 | 11 | 0.86% |
| hsa_circ_0083460 | 2566 | 11 | 0.86% |
| hsa_circ_0120618 | 2566 | 11 | 0.86% |
| hsa_circ_0044092 | 7466 | 32 | 0.86% |
| hsa_circ_0119465 | 2800 | 12 | 0.86% |
| hsa_circ_0032386 | 2567 | 11 | 0.86% |
| hsa_circ_0054965 | 3734 | 16 | 0.86% |
| hsa_circ_0131462 | 12369 | 53 | 0.86% |
| hsa_circ_0027657 | 2334 | 10 | 0.86% |
| hsa_circ_0121360 | 2334 | 10 | 0.86% |
| hsa_circ_0131196 | 2334 | 10 | 0.86% |
| hsa_circ_0009849 | 2801 | 12 | 0.86% |
| hsa_circ_0056806 | 2801 | 12 | 0.86% |
| hsa_circ_0127115 | 4902 | 21 | 0.86% |
| hsa_circ_0017088 | 2568 | 11 | 0.86% |
| hsa_circ_0085246 | 2568 | 11 | 0.86% |
| hsa_circ_0110775 | 2568 | 11 | 0.86% |
| hsa_circ_0117893 | 2568 | 11 | 0.86% |
| hsa_circ_0097975 | 3035 | 13 | 0.86% |
| hsa_circ_0115243 | 3969 | 17 | 0.86% |
| hsa_circ_0140024 | 7705 | 33 | 0.86% |
| hsa_circ_0015005 | 2802 | 12 | 0.86% |
| hsa_circ_0034460 | 2802 | 12 | 0.86% |
| hsa_circ_0074251 | 2802 | 12 | 0.86% |
| hsa_circ_0111654 | 2802 | 12 | 0.86% |
| hsa_circ_0014565 | 2335 | 10 | 0.86% |
| hsa_circ_0127460 | 2335 | 10 | 0.86% |
| hsa_circ_0139993 | 2335 | 10 | 0.86% |
| hsa_circ_0029128 | 3036 | 13 | 0.86% |
| hsa_circ_0065322 | 2569 | 11 | 0.86% |
| hsa_circ_0023833 | 2803 | 12 | 0.86% |
| hsa_circ_0032528 | 2803 | 12 | 0.86% |
| hsa_circ_0082007 | 2803 | 12 | 0.86% |
| hsa_circ_0131878 | 2803 | 12 | 0.86% |
| hsa_circ_0013880 | 2336 | 10 | 0.86% |
| hsa_circ_0027404 | 2336 | 10 | 0.86% |
| hsa_circ_0127525 | 2336 | 10 | 0.86% |
| hsa_circ_0042059 | 2570 | 11 | 0.86% |
| hsa_circ_0048608 | 2570 | 11 | 0.86% |
| hsa_circ_0053505 | 2570 | 11 | 0.86% |
| hsa_circ_0109700 | 4673 | 20 | 0.86% |
| hsa_circ_0033341 | 2804 | 12 | 0.86% |

| | | | |
|---|---|---|---|
| hsa_circ_0016616 | 2337 | 10 | 0.86% |
| hsa_circ_0044516 | 2337 | 10 | 0.86% |
| hsa_circ_0053843 | 2337 | 10 | 0.86% |
| hsa_circ_0074317 | 2337 | 10 | 0.86% |
| hsa_circ_0085727 | 2337 | 10 | 0.86% |
| hsa_circ_0024847 | 2571 | 11 | 0.86% |
| hsa_circ_0040104 | 2571 | 11 | 0.86% |
| hsa_circ_0091716 | 2571 | 11 | 0.86% |
| hsa_circ_0031020 | 2805 | 12 | 0.86% |
| hsa_circ_0077866 | 3039 | 13 | 0.86% |
| hsa_circ_0109701 | 4676 | 20 | 0.86% |
| hsa_circ_0103650 | 2338 | 10 | 0.86% |
| hsa_circ_0031460 | 2572 | 11 | 0.86% |
| hsa_circ_0078308 | 2572 | 11 | 0.86% |
| hsa_circ_0120909 | 2572 | 11 | 0.86% |
| hsa_circ_0012371 | 2339 | 10 | 0.86% |
| hsa_circ_0012469 | 2339 | 10 | 0.86% |
| hsa_circ_0041116 | 2339 | 10 | 0.86% |
| hsa_circ_0073508 | 2339 | 10 | 0.86% |
| hsa_circ_0140770 | 2339 | 10 | 0.86% |
| hsa_circ_0019103 | 2573 | 11 | 0.86% |
| hsa_circ_0030022 | 2573 | 11 | 0.86% |
| hsa_circ_0059582 | 2573 | 11 | 0.86% |
| hsa_circ_0020755 | 2807 | 12 | 0.86% |
| hsa_circ_0072465 | 3509 | 15 | 0.85% |
| hsa_circ_0110756 | 11465 | 49 | 0.85% |
| hsa_circ_0052631 | 3042 | 13 | 0.85% |
| hsa_circ_0081926 | 3042 | 13 | 0.85% |
| hsa_circ_0083094 | 2808 | 12 | 0.85% |
| hsa_circ_0123962 | 2808 | 12 | 0.85% |
| hsa_circ_0017006 | 2574 | 11 | 0.85% |
| hsa_circ_0133793 | 2574 | 11 | 0.85% |
| hsa_circ_0138910 | 2574 | 11 | 0.85% |
| hsa_circ_0019388 | 2340 | 10 | 0.85% |
| hsa_circ_0021510 | 2340 | 10 | 0.85% |
| hsa_circ_0061796 | 2340 | 10 | 0.85% |
| hsa_circ_0042812 | 3043 | 13 | 0.85% |
| hsa_circ_0020628 | 2809 | 12 | 0.85% |
| hsa_circ_0021601 | 2809 | 12 | 0.85% |
| hsa_circ_0091836 | 2575 | 11 | 0.85% |
| hsa_circ_0023857 | 4214 | 18 | 0.85% |
| hsa_circ_0022359 | 3044 | 13 | 0.85% |
| hsa_circ_0066516 | 3044 | 13 | 0.85% |
| hsa_circ_0125638 | 3981 | 17 | 0.85% |
| hsa_circ_0009825 | 2576 | 11 | 0.85% |
| hsa_circ_0033178 | 2576 | 11 | 0.85% |
| hsa_circ_0090909 | 2576 | 11 | 0.85% |
| hsa_circ_0127298 | 2576 | 11 | 0.85% |
| hsa_circ_0130614 | 2576 | 11 | 0.85% |
| hsa_circ_0023447 | 2342 | 10 | 0.85% |

| | | | |
|---|---|---|---|
| hsa_circ_0081614 | 2342 | 10 | 0.85% |
| hsa_circ_0125719 | 2342 | 10 | 0.85% |
| hsa_circ_0086685 | 3045 | 13 | 0.85% |
| hsa_circ_0094850 | 2811 | 12 | 0.85% |
| hsa_circ_0087459 | 2577 | 11 | 0.85% |
| hsa_circ_0122773 | 2577 | 11 | 0.85% |
| hsa_circ_0033099 | 2343 | 10 | 0.85% |
| hsa_circ_0038572 | 2343 | 10 | 0.85% |
| hsa_circ_0069509 | 2343 | 10 | 0.85% |
| hsa_circ_0077465 | 3046 | 13 | 0.85% |
| hsa_circ_0077961 | 3046 | 13 | 0.85% |
| hsa_circ_0016640 | 2812 | 12 | 0.85% |
| hsa_circ_0034110 | 2812 | 12 | 0.85% |
| hsa_circ_0129478 | 2812 | 12 | 0.85% |
| hsa_circ_0038236 | 2578 | 11 | 0.85% |
| hsa_circ_0072044 | 2578 | 11 | 0.85% |
| hsa_circ_0072742 | 2578 | 11 | 0.85% |
| hsa_circ_0102280 | 2578 | 11 | 0.85% |
| hsa_circ_0127905 | 2578 | 11 | 0.85% |
| hsa_circ_0130028 | 2578 | 11 | 0.85% |
| hsa_circ_0088399 | 2344 | 10 | 0.85% |
| hsa_circ_0032401 | 2579 | 11 | 0.85% |
| hsa_circ_0033263 | 2814 | 12 | 0.85% |
| hsa_circ_0034363 | 2814 | 12 | 0.85% |
| hsa_circ_0013848 | 2345 | 10 | 0.85% |
| hsa_circ_0017221 | 2345 | 10 | 0.85% |
| hsa_circ_0048710 | 2345 | 10 | 0.85% |
| hsa_circ_0074705 | 2345 | 10 | 0.85% |
| hsa_circ_0116279 | 2345 | 10 | 0.85% |
| hsa_circ_0123776 | 2345 | 10 | 0.85% |
| hsa_circ_0044333 | 3049 | 13 | 0.85% |
| hsa_circ_0054753 | 3049 | 13 | 0.85% |
| hsa_circ_0041308 | 2580 | 11 | 0.85% |
| hsa_circ_0061172 | 2580 | 11 | 0.85% |
| hsa_circ_0093413 | 2580 | 11 | 0.85% |
| hsa_circ_0114160 | 2580 | 11 | 0.85% |
| hsa_circ_0120310 | 5630 | 24 | 0.85% |
| hsa_circ_0014396 | 2815 | 12 | 0.85% |
| hsa_circ_0032485 | 2346 | 10 | 0.85% |
| hsa_circ_0035083 | 2346 | 10 | 0.85% |
| hsa_circ_0037058 | 2346 | 10 | 0.85% |
| hsa_circ_0129730 | 2346 | 10 | 0.85% |
| hsa_circ_0080378 | 3050 | 13 | 0.85% |
| hsa_circ_0024612 | 2581 | 11 | 0.85% |
| hsa_circ_0053284 | 2581 | 11 | 0.85% |
| hsa_circ_0066564 | 2581 | 11 | 0.85% |
| hsa_circ_0097793 | 2581 | 11 | 0.85% |
| hsa_circ_0111064 | 2581 | 11 | 0.85% |
| hsa_circ_0079581 | 2816 | 12 | 0.85% |
| hsa_circ_0006193 | 3051 | 13 | 0.85% |

| | | | |
|---|---|---|---|
| hsa_circ_0073356 | 3051 | 13 | 0.85% |
| hsa_circ_0011545 | 2347 | 10 | 0.85% |
| hsa_circ_0022100 | 2347 | 10 | 0.85% |
| hsa_circ_0045303 | 2347 | 10 | 0.85% |
| hsa_circ_0072977 | 3286 | 14 | 0.85% |
| hsa_circ_0043989 | 2582 | 11 | 0.85% |
| hsa_circ_0090225 | 2582 | 11 | 0.85% |
| hsa_circ_0120076 | 7512 | 32 | 0.85% |
| hsa_circ_0073000 | 2817 | 12 | 0.85% |
| hsa_circ_0080942 | 3052 | 13 | 0.85% |
| hsa_circ_0075940 | 4226 | 18 | 0.85% |
| hsa_circ_0028108 | 2348 | 10 | 0.85% |
| hsa_circ_0057553 | 2348 | 10 | 0.85% |
| hsa_circ_0104656 | 2348 | 10 | 0.85% |
| hsa_circ_0009812 | 2583 | 11 | 0.85% |
| hsa_circ_0019453 | 2583 | 11 | 0.85% |
| hsa_circ_0027184 | 2583 | 11 | 0.85% |
| hsa_circ_0039716 | 2583 | 11 | 0.85% |
| hsa_circ_0046789 | 2583 | 11 | 0.85% |
| hsa_circ_0085575 | 2583 | 11 | 0.85% |
| hsa_circ_0089456 | 2583 | 11 | 0.85% |
| hsa_circ_0091865 | 2583 | 11 | 0.85% |
| hsa_circ_0038753 | 2818 | 12 | 0.85% |
| hsa_circ_0109991 | 2818 | 12 | 0.85% |
| hsa_circ_0075924 | 8689 | 37 | 0.85% |
| hsa_circ_0102686 | 3053 | 13 | 0.85% |
| hsa_circ_0005885 | 7985 | 34 | 0.85% |
| hsa_circ_0012538 | 2349 | 10 | 0.85% |
| hsa_circ_0078587 | 2349 | 10 | 0.85% |
| hsa_circ_0086960 | 2349 | 10 | 0.85% |
| hsa_circ_0131508 | 2349 | 10 | 0.85% |
| hsa_circ_0026862 | 2584 | 11 | 0.85% |
| hsa_circ_0073009 | 2584 | 11 | 0.85% |
| hsa_circ_0091384 | 2584 | 11 | 0.85% |
| hsa_circ_0136981 | 2584 | 11 | 0.85% |
| hsa_circ_0022119 | 2819 | 12 | 0.85% |
| hsa_circ_0067891 | 2819 | 12 | 0.85% |
| hsa_circ_0126780 | 2819 | 12 | 0.85% |
| hsa_circ_0042985 | 3054 | 13 | 0.85% |
| hsa_circ_0012076 | 3289 | 14 | 0.85% |
| hsa_circ_0122566 | 4230 | 18 | 0.85% |
| hsa_circ_0085685 | 2820 | 12 | 0.85% |
| hsa_circ_0089438 | 2820 | 12 | 0.85% |
| hsa_circ_0026972 | 2585 | 11 | 0.85% |
| hsa_circ_0013767 | 2350 | 10 | 0.85% |
| hsa_circ_0057715 | 2350 | 10 | 0.85% |
| hsa_circ_0090822 | 2350 | 10 | 0.85% |
| hsa_circ_0047165 | 3291 | 14 | 0.85% |
| hsa_circ_0002036 | 3056 | 13 | 0.85% |
| hsa_circ_0135174 | 3056 | 13 | 0.85% |

| | | | |
|---|---|---|---|
| hsa_circ_0011917 | 2821 | 12 | 0.85% |
| hsa_circ_0029194 | 2821 | 12 | 0.85% |
| hsa_circ_0039647 | 2821 | 12 | 0.85% |
| hsa_circ_0085362 | 2821 | 12 | 0.85% |
| hsa_circ_0129850 | 2821 | 12 | 0.85% |
| hsa_circ_0017802 | 2586 | 11 | 0.85% |
| hsa_circ_0021057 | 2586 | 11 | 0.85% |
| hsa_circ_0024264 | 2586 | 11 | 0.85% |
| hsa_circ_0042217 | 2586 | 11 | 0.85% |
| hsa_circ_0117002 | 2586 | 11 | 0.85% |
| hsa_circ_0059571 | 2351 | 10 | 0.85% |
| hsa_circ_0114810 | 2351 | 10 | 0.85% |
| hsa_circ_0123433 | 3057 | 13 | 0.85% |
| hsa_circ_0028018 | 2587 | 11 | 0.85% |
| hsa_circ_0118275 | 2587 | 11 | 0.85% |
| hsa_circ_0061445 | 3763 | 16 | 0.85% |
| hsa_circ_0049614 | 2352 | 10 | 0.85% |
| hsa_circ_0062798 | 3058 | 13 | 0.85% |
| hsa_circ_0023022 | 2823 | 12 | 0.85% |
| hsa_circ_0033333 | 2823 | 12 | 0.85% |
| hsa_circ_0056029 | 2823 | 12 | 0.85% |
| hsa_circ_0091357 | 2823 | 12 | 0.85% |
| hsa_circ_0005027 | 2588 | 11 | 0.85% |
| hsa_circ_0042101 | 2588 | 11 | 0.85% |
| hsa_circ_0075980 | 2588 | 11 | 0.85% |
| hsa_circ_0080148 | 2588 | 11 | 0.85% |
| hsa_circ_0086084 | 2588 | 11 | 0.85% |
| hsa_circ_0087739 | 2588 | 11 | 0.85% |
| hsa_circ_0127882 | 2588 | 11 | 0.85% |
| hsa_circ_0134184 | 2588 | 11 | 0.85% |
| hsa_circ_0040125 | 2824 | 12 | 0.85% |
| hsa_circ_0060226 | 3295 | 14 | 0.85% |
| hsa_circ_0043702 | 2589 | 11 | 0.85% |
| hsa_circ_0048087 | 2589 | 11 | 0.85% |
| hsa_circ_0083839 | 2589 | 11 | 0.85% |
| hsa_circ_0112875 | 4472 | 19 | 0.85% |
| hsa_circ_0053660 | 3060 | 13 | 0.85% |
| hsa_circ_0057887 | 2354 | 10 | 0.85% |
| hsa_circ_0069878 | 2354 | 10 | 0.85% |
| hsa_circ_0112420 | 10358 | 44 | 0.85% |
| hsa_circ_0060197 | 2825 | 12 | 0.85% |
| hsa_circ_0054977 | 3296 | 14 | 0.85% |
| hsa_circ_0110484 | 9418 | 40 | 0.85% |
| hsa_circ_0041405 | 2590 | 11 | 0.85% |
| hsa_circ_0069689 | 2590 | 11 | 0.85% |
| hsa_circ_0077696 | 2590 | 11 | 0.85% |
| hsa_circ_0073820 | 6122 | 26 | 0.85% |
| hsa_circ_0041747 | 2826 | 12 | 0.85% |
| hsa_circ_0061983 | 2826 | 12 | 0.85% |
| hsa_circ_0110008 | 2826 | 12 | 0.85% |

| | | | |
|---|---|---|---|
| hsa_circ_0011798 | 2355 | 10 | 0.85% |
| hsa_circ_0021131 | 2355 | 10 | 0.85% |
| hsa_circ_0054343 | 2355 | 10 | 0.85% |
| hsa_circ_0055835 | 2355 | 10 | 0.85% |
| hsa_circ_0086314 | 2355 | 10 | 0.85% |
| hsa_circ_0120120 | 2355 | 10 | 0.85% |
| hsa_circ_0029658 | 2591 | 11 | 0.85% |
| hsa_circ_0049247 | 2591 | 11 | 0.85% |
| hsa_circ_0113924 | 2591 | 11 | 0.85% |
| hsa_circ_0025282 | 2827 | 12 | 0.85% |
| hsa_circ_0030215 | 2827 | 12 | 0.85% |
| hsa_circ_0136426 | 6832 | 29 | 0.85% |
| hsa_circ_0013849 | 2356 | 10 | 0.85% |
| hsa_circ_0020465 | 2356 | 10 | 0.85% |
| hsa_circ_0034778 | 2356 | 10 | 0.85% |
| hsa_circ_0058723 | 2356 | 10 | 0.85% |
| hsa_circ_0006106 | 2592 | 11 | 0.85% |
| hsa_circ_0022698 | 2592 | 11 | 0.85% |
| hsa_circ_0054070 | 2592 | 11 | 0.85% |
| hsa_circ_0062094 | 2592 | 11 | 0.85% |
| hsa_circ_0073780 | 2592 | 11 | 0.85% |
| hsa_circ_0077357 | 2592 | 11 | 0.85% |
| hsa_circ_0106980 | 5656 | 24 | 0.85% |
| hsa_circ_0045144 | 2828 | 12 | 0.85% |
| hsa_circ_0092816 | 2828 | 12 | 0.85% |
| hsa_circ_0062044 | 2357 | 10 | 0.85% |
| hsa_circ_0034736 | 2593 | 11 | 0.85% |
| hsa_circ_0041203 | 2829 | 12 | 0.85% |
| hsa_circ_0050806 | 2829 | 12 | 0.85% |
| hsa_circ_0075466 | 3301 | 14 | 0.85% |
| hsa_circ_0114336 | 3301 | 14 | 0.85% |
| hsa_circ_0131419 | 6838 | 29 | 0.85% |
| hsa_circ_0012804 | 2358 | 10 | 0.85% |
| hsa_circ_0023541 | 2358 | 10 | 0.85% |
| hsa_circ_0032390 | 2358 | 10 | 0.85% |
| hsa_circ_0064234 | 2358 | 10 | 0.85% |
| hsa_circ_0019333 | 2594 | 11 | 0.85% |
| hsa_circ_0092988 | 2594 | 11 | 0.85% |
| hsa_circ_0025928 | 3066 | 13 | 0.85% |
| hsa_circ_0048083 | 3302 | 14 | 0.85% |
| hsa_circ_0138523 | 3538 | 15 | 0.85% |
| hsa_circ_0099783 | 2359 | 10 | 0.85% |
| hsa_circ_0041283 | 2595 | 11 | 0.85% |
| hsa_circ_0049826 | 2595 | 11 | 0.85% |
| hsa_circ_0038438 | 2831 | 12 | 0.85% |
| hsa_circ_0048281 | 2831 | 12 | 0.85% |
| hsa_circ_0109152 | 6371 | 27 | 0.85% |
| hsa_circ_0027160 | 2832 | 12 | 0.85% |
| hsa_circ_0056844 | 2832 | 12 | 0.85% |
| hsa_circ_0038078 | 2596 | 11 | 0.85% |

| | | | |
|---|---|---|---|
| hsa_circ_0049236 | 2596 | 11 | 0.85% |
| hsa_circ_0117685 | 2596 | 11 | 0.85% |
| hsa_circ_0033293 | 2360 | 10 | 0.85% |
| hsa_circ_0043719 | 2360 | 10 | 0.85% |
| hsa_circ_0050271 | 9915 | 42 | 0.85% |
| hsa_circ_0024922 | 2833 | 12 | 0.85% |
| hsa_circ_0071883 | 2597 | 11 | 0.85% |
| hsa_circ_0019851 | 2361 | 10 | 0.85% |
| hsa_circ_0023227 | 2361 | 10 | 0.85% |
| hsa_circ_0058839 | 2361 | 10 | 0.85% |
| hsa_circ_0088927 | 2361 | 10 | 0.85% |
| hsa_circ_0091988 | 2834 | 12 | 0.85% |
| hsa_circ_0095272 | 9211 | 39 | 0.85% |
| hsa_circ_0024677 | 2598 | 11 | 0.85% |
| hsa_circ_0046503 | 2598 | 11 | 0.85% |
| hsa_circ_0109767 | 5905 | 25 | 0.85% |
| hsa_circ_0019724 | 2362 | 10 | 0.85% |
| hsa_circ_0034212 | 2362 | 10 | 0.85% |
| hsa_circ_0057481 | 2362 | 10 | 0.85% |
| hsa_circ_0086938 | 3071 | 13 | 0.85% |
| hsa_circ_0120986 | 3780 | 16 | 0.85% |
| hsa_circ_0102384 | 2835 | 12 | 0.85% |
| hsa_circ_0130082 | 2835 | 12 | 0.85% |
| hsa_circ_0024932 | 3544 | 15 | 0.85% |
| hsa_circ_0058556 | 2599 | 11 | 0.85% |
| hsa_circ_0062041 | 2599 | 11 | 0.85% |
| hsa_circ_0062056 | 2599 | 11 | 0.85% |
| hsa_circ_0080565 | 2599 | 11 | 0.85% |
| hsa_circ_0118778 | 2599 | 11 | 0.85% |
| hsa_circ_0014813 | 2363 | 10 | 0.85% |
| hsa_circ_0034136 | 2363 | 10 | 0.85% |
| hsa_circ_0088724 | 2363 | 10 | 0.85% |
| hsa_circ_0062854 | 3072 | 13 | 0.85% |
| hsa_circ_0052831 | 2836 | 12 | 0.85% |
| hsa_circ_0048360 | 3309 | 14 | 0.85% |
| hsa_circ_0026718 | 2600 | 11 | 0.85% |
| hsa_circ_0031588 | 2600 | 11 | 0.85% |
| hsa_circ_0055767 | 2600 | 11 | 0.85% |
| hsa_circ_0122679 | 8746 | 37 | 0.85% |
| hsa_circ_0074318 | 2364 | 10 | 0.85% |
| hsa_circ_0082726 | 2364 | 10 | 0.85% |
| hsa_circ_0083459 | 2364 | 10 | 0.85% |
| hsa_circ_0087726 | 2364 | 10 | 0.85% |
| hsa_circ_0044042 | 2837 | 12 | 0.85% |
| hsa_circ_0102103 | 2837 | 12 | 0.85% |
| hsa_circ_0115353 | 2837 | 12 | 0.85% |
| hsa_circ_0044376 | 2601 | 11 | 0.85% |
| hsa_circ_0082660 | 2601 | 11 | 0.85% |
| hsa_circ_0084628 | 2601 | 11 | 0.85% |
| hsa_circ_0127867 | 2601 | 11 | 0.85% |

| | | | |
|---|---|---|---|
| hsa_circ_0090111 | 3311 | 14 | 0.85% |
| hsa_circ_0019021 | 2838 | 12 | 0.85% |
| hsa_circ_0078533 | 2838 | 12 | 0.85% |
| hsa_circ_0011676 | 2365 | 10 | 0.85% |
| hsa_circ_0041970 | 2365 | 10 | 0.85% |
| hsa_circ_0046175 | 3075 | 13 | 0.85% |
| hsa_circ_0118668 | 9936 | 42 | 0.85% |
| hsa_circ_0044994 | 2839 | 12 | 0.85% |
| hsa_circ_0040731 | 2366 | 10 | 0.85% |
| hsa_circ_0045906 | 2366 | 10 | 0.85% |
| hsa_circ_0053488 | 2603 | 11 | 0.85% |
| hsa_circ_0056560 | 2603 | 11 | 0.85% |
| hsa_circ_0072915 | 2603 | 11 | 0.85% |
| hsa_circ_0080198 | 2603 | 11 | 0.85% |
| hsa_circ_0062666 | 3313 | 14 | 0.85% |
| hsa_circ_0097363 | 7336 | 31 | 0.85% |
| hsa_circ_0115157 | 2840 | 12 | 0.85% |
| hsa_circ_0099423 | 11361 | 48 | 0.84% |
| hsa_circ_0075898 | 7811 | 33 | 0.84% |
| hsa_circ_0116471 | 2367 | 10 | 0.84% |
| hsa_circ_0018811 | 2604 | 11 | 0.84% |
| hsa_circ_0030464 | 2604 | 11 | 0.84% |
| hsa_circ_0054805 | 2604 | 11 | 0.84% |
| hsa_circ_0055826 | 3551 | 15 | 0.84% |
| hsa_circ_0040754 | 2841 | 12 | 0.84% |
| hsa_circ_0069957 | 2841 | 12 | 0.84% |
| hsa_circ_0030854 | 3315 | 14 | 0.84% |
| hsa_circ_0035010 | 2368 | 10 | 0.84% |
| hsa_circ_0045077 | 2368 | 10 | 0.84% |
| hsa_circ_0084095 | 2368 | 10 | 0.84% |
| hsa_circ_0041499 | 2605 | 11 | 0.84% |
| hsa_circ_0084914 | 2605 | 11 | 0.84% |
| hsa_circ_0116996 | 9236 | 39 | 0.84% |
| hsa_circ_0129678 | 2842 | 12 | 0.84% |
| hsa_circ_0098913 | 3553 | 15 | 0.84% |
| hsa_circ_0012256 | 2369 | 10 | 0.84% |
| hsa_circ_0016016 | 2369 | 10 | 0.84% |
| hsa_circ_0028590 | 2369 | 10 | 0.84% |
| hsa_circ_0111107 | 2369 | 10 | 0.84% |
| hsa_circ_0032208 | 10424 | 44 | 0.84% |
| hsa_circ_0048368 | 2606 | 11 | 0.84% |
| hsa_circ_0070136 | 2606 | 11 | 0.84% |
| hsa_circ_0086800 | 2606 | 11 | 0.84% |
| hsa_circ_0087809 | 2606 | 11 | 0.84% |
| hsa_circ_0087867 | 2606 | 11 | 0.84% |
| hsa_circ_0099984 | 2606 | 11 | 0.84% |
| hsa_circ_0113106 | 2606 | 11 | 0.84% |
| hsa_circ_0115652 | 2843 | 12 | 0.84% |
| hsa_circ_0043221 | 3080 | 13 | 0.84% |
| hsa_circ_0083752 | 3080 | 13 | 0.84% |

| | | | |
|---|---|---|---|
| hsa_circ_0079693 | 5451 | 23 | 0.84% |
| hsa_circ_0135091 | 3081 | 13 | 0.84% |
| hsa_circ_0034539 | 2844 | 12 | 0.84% |
| hsa_circ_0051307 | 2844 | 12 | 0.84% |
| hsa_circ_0074249 | 2844 | 12 | 0.84% |
| hsa_circ_0010802 | 2607 | 11 | 0.84% |
| hsa_circ_0030570 | 2607 | 11 | 0.84% |
| hsa_circ_0140283 | 2370 | 10 | 0.84% |
| hsa_circ_0043503 | 2845 | 12 | 0.84% |
| hsa_circ_0063968 | 2845 | 12 | 0.84% |
| hsa_circ_0038532 | 2608 | 11 | 0.84% |
| hsa_circ_0065761 | 2608 | 11 | 0.84% |
| hsa_circ_0067335 | 2608 | 11 | 0.84% |
| hsa_circ_0079372 | 2608 | 11 | 0.84% |
| hsa_circ_0097683 | 2608 | 11 | 0.84% |
| hsa_circ_0131213 | 2608 | 11 | 0.84% |
| hsa_circ_0017140 | 2371 | 10 | 0.84% |
| hsa_circ_0064942 | 2371 | 10 | 0.84% |
| hsa_circ_0091597 | 2371 | 10 | 0.84% |
| hsa_circ_0009331 | 2846 | 12 | 0.84% |
| hsa_circ_0048981 | 2846 | 12 | 0.84% |
| hsa_circ_0123951 | 2846 | 12 | 0.84% |
| hsa_circ_0026059 | 2609 | 11 | 0.84% |
| hsa_circ_0037703 | 2609 | 11 | 0.84% |
| hsa_circ_0065332 | 2609 | 11 | 0.84% |
| hsa_circ_0083788 | 2609 | 11 | 0.84% |
| hsa_circ_0132471 | 2609 | 11 | 0.84% |
| hsa_circ_0015901 | 2372 | 10 | 0.84% |
| hsa_circ_0039045 | 2372 | 10 | 0.84% |
| hsa_circ_0042635 | 2372 | 10 | 0.84% |
| hsa_circ_0044626 | 2372 | 10 | 0.84% |
| hsa_circ_0063815 | 2372 | 10 | 0.84% |
| hsa_circ_0088746 | 2372 | 10 | 0.84% |
| hsa_circ_0088932 | 2372 | 10 | 0.84% |
| hsa_circ_0053711 | 3084 | 13 | 0.84% |
| hsa_circ_0090277 | 3084 | 13 | 0.84% |
| hsa_circ_0046992 | 2847 | 12 | 0.84% |
| hsa_circ_0091370 | 2847 | 12 | 0.84% |
| hsa_circ_0048359 | 2610 | 11 | 0.84% |
| hsa_circ_0058550 | 2610 | 11 | 0.84% |
| hsa_circ_0107208 | 2610 | 11 | 0.84% |
| hsa_circ_0120907 | 2610 | 11 | 0.84% |
| hsa_circ_0024983 | 2373 | 10 | 0.84% |
| hsa_circ_0105592 | 2373 | 10 | 0.84% |
| hsa_circ_0113433 | 5458 | 23 | 0.84% |
| hsa_circ_0036842 | 3085 | 13 | 0.84% |
| hsa_circ_0032030 | 2848 | 12 | 0.84% |
| hsa_circ_0039548 | 2848 | 12 | 0.84% |
| hsa_circ_0052357 | 2848 | 12 | 0.84% |
| hsa_circ_0053048 | 2848 | 12 | 0.84% |

| | | | |
|---|---|---|---|
| hsa_circ_0078689 | 2848 | 12 | 0.84% |
| hsa_circ_0071336 | 2611 | 11 | 0.84% |
| hsa_circ_0110709 | 2611 | 11 | 0.84% |
| hsa_circ_0105596 | 6172 | 26 | 0.84% |
| hsa_circ_0009401 | 3086 | 13 | 0.84% |
| hsa_circ_0101535 | 4273 | 18 | 0.84% |
| hsa_circ_0010471 | 2374 | 10 | 0.84% |
| hsa_circ_0011047 | 2374 | 10 | 0.84% |
| hsa_circ_0101272 | 2374 | 10 | 0.84% |
| hsa_circ_0115596 | 2374 | 10 | 0.84% |
| hsa_circ_0128019 | 2374 | 10 | 0.84% |
| hsa_circ_0128114 | 2374 | 10 | 0.84% |
| hsa_circ_0010099 | 2612 | 11 | 0.84% |
| hsa_circ_0010737 | 2612 | 11 | 0.84% |
| hsa_circ_0060542 | 2612 | 11 | 0.84% |
| hsa_circ_0068395 | 2612 | 11 | 0.84% |
| hsa_circ_0071282 | 2612 | 11 | 0.84% |
| hsa_circ_0030555 | 3562 | 15 | 0.84% |
| hsa_circ_0013575 | 2375 | 10 | 0.84% |
| hsa_circ_0071344 | 2375 | 10 | 0.84% |
| hsa_circ_0077046 | 2375 | 10 | 0.84% |
| hsa_circ_0019153 | 2613 | 11 | 0.84% |
| hsa_circ_0055852 | 2613 | 11 | 0.84% |
| hsa_circ_0058410 | 2613 | 11 | 0.84% |
| hsa_circ_0086913 | 2613 | 11 | 0.84% |
| hsa_circ_0138958 | 2613 | 11 | 0.84% |
| hsa_circ_0133152 | 3326 | 14 | 0.84% |
| hsa_circ_0098742 | 4514 | 19 | 0.84% |
| hsa_circ_0015261 | 2851 | 12 | 0.84% |
| hsa_circ_0026666 | 2851 | 12 | 0.84% |
| hsa_circ_0053203 | 2851 | 12 | 0.84% |
| hsa_circ_0109154 | 2851 | 12 | 0.84% |
| hsa_circ_0124772 | 2851 | 12 | 0.84% |
| hsa_circ_0056633 | 2376 | 10 | 0.84% |
| hsa_circ_0086355 | 2376 | 10 | 0.84% |
| hsa_circ_0113670 | 2376 | 10 | 0.84% |
| hsa_circ_0021278 | 2614 | 11 | 0.84% |
| hsa_circ_0024696 | 2614 | 11 | 0.84% |
| hsa_circ_0031111 | 2614 | 11 | 0.84% |
| hsa_circ_0057245 | 2614 | 11 | 0.84% |
| hsa_circ_0093302 | 2614 | 11 | 0.84% |
| hsa_circ_0067551 | 2852 | 12 | 0.84% |
| hsa_circ_0071265 | 2852 | 12 | 0.84% |
| hsa_circ_0106138 | 9508 | 40 | 0.84% |
| hsa_circ_0067339 | 2377 | 10 | 0.84% |
| hsa_circ_0080263 | 2377 | 10 | 0.84% |
| hsa_circ_0116035 | 2377 | 10 | 0.84% |
| hsa_circ_0123872 | 2377 | 10 | 0.84% |
| hsa_circ_0044628 | 5705 | 24 | 0.84% |
| hsa_circ_0016701 | 2615 | 11 | 0.84% |

| | | | |
|---|---|---|---|
| hsa_circ_0019228 | 2615 | 11 | 0.84% |
| hsa_circ_0020541 | 2615 | 11 | 0.84% |
| hsa_circ_0061090 | 2615 | 11 | 0.84% |
| hsa_circ_0081974 | 2615 | 11 | 0.84% |
| hsa_circ_0085576 | 2615 | 11 | 0.84% |
| hsa_circ_0007211 | 4517 | 19 | 0.84% |
| hsa_circ_0111766 | 8560 | 36 | 0.84% |
| hsa_circ_0046905 | 3329 | 14 | 0.84% |
| hsa_circ_0077498 | 2378 | 10 | 0.84% |
| hsa_circ_0080485 | 2378 | 10 | 0.84% |
| hsa_circ_0136984 | 2378 | 10 | 0.84% |
| hsa_circ_0108942 | 5232 | 22 | 0.84% |
| hsa_circ_0009887 | 2616 | 11 | 0.84% |
| hsa_circ_0044614 | 2616 | 11 | 0.84% |
| hsa_circ_0045454 | 2616 | 11 | 0.84% |
| hsa_circ_0035776 | 3092 | 13 | 0.84% |
| hsa_circ_0063367 | 3092 | 13 | 0.84% |
| hsa_circ_0035643 | 4757 | 20 | 0.84% |
| hsa_circ_0076486 | 3330 | 14 | 0.84% |
| hsa_circ_0074689 | 3806 | 16 | 0.84% |
| hsa_circ_0012862 | 2379 | 10 | 0.84% |
| hsa_circ_0028349 | 2379 | 10 | 0.84% |
| hsa_circ_0041374 | 2379 | 10 | 0.84% |
| hsa_circ_0089534 | 2379 | 10 | 0.84% |
| hsa_circ_0022504 | 2617 | 11 | 0.84% |
| hsa_circ_0046477 | 2617 | 11 | 0.84% |
| hsa_circ_0083526 | 2617 | 11 | 0.84% |
| hsa_circ_0099784 | 2617 | 11 | 0.84% |
| hsa_circ_0137748 | 16892 | 71 | 0.84% |
| hsa_circ_0135478 | 2855 | 12 | 0.84% |
| hsa_circ_0140730 | 2855 | 12 | 0.84% |
| hsa_circ_0017966 | 3093 | 13 | 0.84% |
| hsa_circ_0047890 | 3569 | 15 | 0.84% |
| hsa_circ_0110637 | 2856 | 12 | 0.84% |
| hsa_circ_0014058 | 2618 | 11 | 0.84% |
| hsa_circ_0038902 | 2618 | 11 | 0.84% |
| hsa_circ_0076158 | 2618 | 11 | 0.84% |
| hsa_circ_0084359 | 2618 | 11 | 0.84% |
| hsa_circ_0087126 | 2618 | 11 | 0.84% |
| hsa_circ_0091950 | 2380 | 10 | 0.84% |
| hsa_circ_0048543 | 2857 | 12 | 0.84% |
| hsa_circ_0083090 | 2857 | 12 | 0.84% |
| hsa_circ_0057804 | 2619 | 11 | 0.84% |
| hsa_circ_0038450 | 10953 | 46 | 0.84% |
| hsa_circ_0126920 | 4763 | 20 | 0.84% |
| hsa_circ_0112788 | 3096 | 13 | 0.84% |
| hsa_circ_0009999 | 2858 | 12 | 0.84% |
| hsa_circ_0053354 | 2620 | 11 | 0.84% |
| hsa_circ_0118603 | 2620 | 11 | 0.84% |
| hsa_circ_0037798 | 2382 | 10 | 0.84% |

| | | | |
|---|---|---|---|
| hsa_circ_0066360 | 2382 | 10 | 0.84% |
| hsa_circ_0072030 | 2382 | 10 | 0.84% |
| hsa_circ_0090778 | 3335 | 14 | 0.84% |
| hsa_circ_0079071 | 4288 | 18 | 0.84% |
| hsa_circ_0024588 | 2859 | 12 | 0.84% |
| hsa_circ_0075485 | 2859 | 12 | 0.84% |
| hsa_circ_0080605 | 2859 | 12 | 0.84% |
| hsa_circ_0023711 | 2621 | 11 | 0.84% |
| hsa_circ_0048606 | 2621 | 11 | 0.84% |
| hsa_circ_0051201 | 2621 | 11 | 0.84% |
| hsa_circ_0068209 | 2621 | 11 | 0.84% |
| hsa_circ_0119901 | 2621 | 11 | 0.84% |
| hsa_circ_0127297 | 2621 | 11 | 0.84% |
| hsa_circ_0122365 | 3336 | 14 | 0.84% |
| hsa_circ_0137374 | 2383 | 10 | 0.84% |
| hsa_circ_0063312 | 3813 | 16 | 0.84% |
| hsa_circ_0023198 | 2622 | 11 | 0.84% |
| hsa_circ_0023588 | 2622 | 11 | 0.84% |
| hsa_circ_0104213 | 3099 | 13 | 0.84% |
| hsa_circ_0020437 | 2384 | 10 | 0.84% |
| hsa_circ_0043809 | 2384 | 10 | 0.84% |
| hsa_circ_0106532 | 2384 | 10 | 0.84% |
| hsa_circ_0062811 | 3338 | 14 | 0.84% |
| hsa_circ_0004974 | 2623 | 11 | 0.84% |
| hsa_circ_0044279 | 2623 | 11 | 0.84% |
| hsa_circ_0071800 | 2623 | 11 | 0.84% |
| hsa_circ_0060109 | 2862 | 12 | 0.84% |
| hsa_circ_0081610 | 2862 | 12 | 0.84% |
| hsa_circ_0048871 | 2385 | 10 | 0.84% |
| hsa_circ_0054573 | 2385 | 10 | 0.84% |
| hsa_circ_0072145 | 2385 | 10 | 0.84% |
| hsa_circ_0077474 | 2385 | 10 | 0.84% |
| hsa_circ_0097810 | 2385 | 10 | 0.84% |
| hsa_circ_0055533 | 2624 | 11 | 0.84% |
| hsa_circ_0088156 | 2624 | 11 | 0.84% |
| hsa_circ_0065659 | 2386 | 10 | 0.84% |
| hsa_circ_0079914 | 2386 | 10 | 0.84% |
| hsa_circ_0139328 | 2386 | 10 | 0.84% |
| hsa_circ_0088978 | 2625 | 11 | 0.84% |
| hsa_circ_0134183 | 2625 | 11 | 0.84% |
| hsa_circ_0134967 | 5489 | 23 | 0.84% |
| hsa_circ_0006440 | 2864 | 12 | 0.84% |
| hsa_circ_0000731 | 3819 | 16 | 0.84% |
| hsa_circ_0046726 | 3103 | 13 | 0.84% |
| hsa_circ_0055139 | 3103 | 13 | 0.84% |
| hsa_circ_0078225 | 2387 | 10 | 0.84% |
| hsa_circ_0108206 | 2387 | 10 | 0.84% |
| hsa_circ_0137285 | 3342 | 14 | 0.84% |
| hsa_circ_0027934 | 2626 | 11 | 0.84% |
| hsa_circ_0037417 | 2626 | 11 | 0.84% |

| | | | |
|---|---|---|---|
| hsa_circ_0087129 | 2626 | 11 | 0.84% |
| hsa_circ_0044094 | 7640 | 32 | 0.84% |
| hsa_circ_0036512 | 2865 | 12 | 0.84% |
| hsa_circ_0052309 | 59928 | 251 | 0.84% |
| hsa_circ_0009778 | 2388 | 10 | 0.84% |
| hsa_circ_0030419 | 2388 | 10 | 0.84% |
| hsa_circ_0046952 | 2388 | 10 | 0.84% |
| hsa_circ_0083167 | 2388 | 10 | 0.84% |
| hsa_circ_0085219 | 2388 | 10 | 0.84% |
| hsa_circ_0096079 | 2388 | 10 | 0.84% |
| hsa_circ_0140539 | 2388 | 10 | 0.84% |
| hsa_circ_0022988 | 2627 | 11 | 0.84% |
| hsa_circ_0086326 | 2627 | 11 | 0.84% |
| hsa_circ_0076700 | 3105 | 13 | 0.84% |
| hsa_circ_0105285 | 7405 | 31 | 0.84% |
| hsa_circ_0044093 | 7644 | 32 | 0.84% |
| hsa_circ_0093220 | 7167 | 30 | 0.84% |
| hsa_circ_0048302 | 2389 | 10 | 0.84% |
| hsa_circ_0120905 | 2389 | 10 | 0.84% |
| hsa_circ_0022060 | 2628 | 11 | 0.84% |
| hsa_circ_0032034 | 2628 | 11 | 0.84% |
| hsa_circ_0052990 | 2628 | 11 | 0.84% |
| hsa_circ_0086919 | 2628 | 11 | 0.84% |
| hsa_circ_0077801 | 2867 | 12 | 0.84% |
| hsa_circ_0039070 | 3106 | 13 | 0.84% |
| hsa_circ_0091412 | 3106 | 13 | 0.84% |
| hsa_circ_0052126 | 3345 | 14 | 0.84% |
| hsa_circ_0073819 | 6213 | 26 | 0.84% |
| hsa_circ_0015118 | 3107 | 13 | 0.84% |
| hsa_circ_0070691 | 2868 | 12 | 0.84% |
| hsa_circ_0014456 | 2629 | 11 | 0.84% |
| hsa_circ_0024265 | 2629 | 11 | 0.84% |
| hsa_circ_0053538 | 2629 | 11 | 0.84% |
| hsa_circ_0082089 | 2629 | 11 | 0.84% |
| hsa_circ_0071987 | 2390 | 10 | 0.84% |
| hsa_circ_0022951 | 2869 | 12 | 0.84% |
| hsa_circ_0022727 | 2630 | 11 | 0.84% |
| hsa_circ_0080562 | 2630 | 11 | 0.84% |
| hsa_circ_0127081 | 2630 | 11 | 0.84% |
| hsa_circ_0073319 | 2391 | 10 | 0.84% |
| hsa_circ_0140736 | 2391 | 10 | 0.84% |
| hsa_circ_0111301 | 8847 | 37 | 0.84% |
| hsa_circ_0052049 | 3587 | 15 | 0.84% |
| hsa_circ_0028102 | 3348 | 14 | 0.84% |
| hsa_circ_0037123 | 3109 | 13 | 0.84% |
| hsa_circ_0123418 | 3109 | 13 | 0.84% |
| hsa_circ_0110127 | 2870 | 12 | 0.84% |
| hsa_circ_0120313 | 2870 | 12 | 0.84% |
| hsa_circ_0066333 | 2631 | 11 | 0.84% |
| hsa_circ_0069006 | 2631 | 11 | 0.84% |

| | | | |
|---|---|---|---|
| hsa_circ_0081991 | 2631 | 11 | 0.84% |
| hsa_circ_0051055 | 8850 | 37 | 0.84% |
| hsa_circ_0011018 | 2392 | 10 | 0.84% |
| hsa_circ_0066339 | 2392 | 10 | 0.84% |
| hsa_circ_0067975 | 2392 | 10 | 0.84% |
| hsa_circ_0078590 | 2392 | 10 | 0.84% |
| hsa_circ_0013388 | 6698 | 28 | 0.84% |
| hsa_circ_0033348 | 2871 | 12 | 0.84% |
| hsa_circ_0038779 | 2871 | 12 | 0.84% |
| hsa_circ_0025428 | 2632 | 11 | 0.84% |
| hsa_circ_0037466 | 2632 | 11 | 0.84% |
| hsa_circ_0042723 | 2632 | 11 | 0.84% |
| hsa_circ_0049199 | 2632 | 11 | 0.84% |
| hsa_circ_0065616 | 2632 | 11 | 0.84% |
| hsa_circ_0119821 | 2632 | 11 | 0.84% |
| hsa_circ_0037152 | 2393 | 10 | 0.84% |
| hsa_circ_0098163 | 3111 | 13 | 0.84% |
| hsa_circ_0076628 | 3590 | 15 | 0.84% |
| hsa_circ_0125896 | 2872 | 12 | 0.84% |
| hsa_circ_0041496 | 3351 | 14 | 0.84% |
| hsa_circ_0104916 | 3351 | 14 | 0.84% |
| hsa_circ_0047902 | 2633 | 11 | 0.84% |
| hsa_circ_0138359 | 2633 | 11 | 0.84% |
| hsa_circ_0053723 | 3112 | 13 | 0.84% |
| hsa_circ_0131988 | 3112 | 13 | 0.84% |
| hsa_circ_0061684 | 2394 | 10 | 0.84% |
| hsa_circ_0070149 | 2394 | 10 | 0.84% |
| hsa_circ_0114729 | 2394 | 10 | 0.84% |
| hsa_circ_0119835 | 2394 | 10 | 0.84% |
| hsa_circ_0019744 | 2873 | 12 | 0.84% |
| hsa_circ_0100797 | 2873 | 12 | 0.84% |
| hsa_circ_0032391 | 2634 | 11 | 0.84% |
| hsa_circ_0051486 | 2634 | 11 | 0.84% |
| hsa_circ_0061522 | 2634 | 11 | 0.84% |
| hsa_circ_0088835 | 2634 | 11 | 0.84% |
| hsa_circ_0058747 | 3113 | 13 | 0.84% |
| hsa_circ_0075514 | 3353 | 14 | 0.84% |
| hsa_circ_0038539 | 2395 | 10 | 0.84% |
| hsa_circ_0061989 | 2395 | 10 | 0.84% |
| hsa_circ_0064132 | 2395 | 10 | 0.84% |
| hsa_circ_0115605 | 2395 | 10 | 0.84% |
| hsa_circ_0109626 | 8862 | 37 | 0.84% |
| hsa_circ_0018123 | 2635 | 11 | 0.83% |
| hsa_circ_0026008 | 2635 | 11 | 0.83% |
| hsa_circ_0033104 | 2635 | 11 | 0.83% |
| hsa_circ_0098385 | 8146 | 34 | 0.83% |
| hsa_circ_0069629 | 2396 | 10 | 0.83% |
| hsa_circ_0079865 | 2396 | 10 | 0.83% |
| hsa_circ_0090587 | 2396 | 10 | 0.83% |
| hsa_circ_0090911 | 2396 | 10 | 0.83% |

| | | | |
|---|---|---|---|
| hsa_circ_0022683 | 2636 | 11 | 0.83% |
| hsa_circ_0125784 | 2636 | 11 | 0.83% |
| hsa_circ_0088564 | 3355 | 14 | 0.83% |
| hsa_circ_0132179 | 3355 | 14 | 0.83% |
| hsa_circ_0073167 | 11024 | 46 | 0.83% |
| hsa_circ_0058018 | 2876 | 12 | 0.83% |
| hsa_circ_0090777 | 2876 | 12 | 0.83% |
| hsa_circ_0024595 | 2397 | 10 | 0.83% |
| hsa_circ_0105817 | 10068 | 42 | 0.83% |
| hsa_circ_0041658 | 6952 | 29 | 0.83% |
| hsa_circ_0016924 | 2637 | 11 | 0.83% |
| hsa_circ_0021004 | 2637 | 11 | 0.83% |
| hsa_circ_0047361 | 2637 | 11 | 0.83% |
| hsa_circ_0089453 | 2637 | 11 | 0.83% |
| hsa_circ_0031601 | 2877 | 12 | 0.83% |
| hsa_circ_0063889 | 2877 | 12 | 0.83% |
| hsa_circ_0080527 | 2877 | 12 | 0.83% |
| hsa_circ_0081945 | 2877 | 12 | 0.83% |
| hsa_circ_0115739 | 2877 | 12 | 0.83% |
| hsa_circ_0113854 | 4076 | 17 | 0.83% |
| hsa_circ_0079070 | 5275 | 22 | 0.83% |
| hsa_circ_0014752 | 3357 | 14 | 0.83% |
| hsa_circ_0018262 | 2398 | 10 | 0.83% |
| hsa_circ_0078136 | 2398 | 10 | 0.83% |
| hsa_circ_0042757 | 2638 | 11 | 0.83% |
| hsa_circ_0059853 | 2638 | 11 | 0.83% |
| hsa_circ_0061410 | 2638 | 11 | 0.83% |
| hsa_circ_0085388 | 2638 | 11 | 0.83% |
| hsa_circ_0099876 | 2638 | 11 | 0.83% |
| hsa_circ_0041577 | 2878 | 12 | 0.83% |
| hsa_circ_0041828 | 2878 | 12 | 0.83% |
| hsa_circ_0088566 | 3118 | 13 | 0.83% |
| hsa_circ_0136866 | 31662 | 132 | 0.83% |
| hsa_circ_0041408 | 5277 | 22 | 0.83% |
| hsa_circ_0126049 | 2399 | 10 | 0.83% |
| hsa_circ_0031467 | 2639 | 11 | 0.83% |
| hsa_circ_0063221 | 12236 | 51 | 0.83% |
| hsa_circ_0012047 | 3119 | 13 | 0.83% |
| hsa_circ_0128475 | 4560 | 19 | 0.83% |
| hsa_circ_0027952 | 3360 | 14 | 0.83% |
| hsa_circ_0013774 | 2880 | 12 | 0.83% |
| hsa_circ_0045769 | 2880 | 12 | 0.83% |
| hsa_circ_0073687 | 2880 | 12 | 0.83% |
| hsa_circ_0108114 | 2880 | 12 | 0.83% |
| hsa_circ_0038782 | 2640 | 11 | 0.83% |
| hsa_circ_0040242 | 2640 | 11 | 0.83% |
| hsa_circ_0047082 | 2640 | 11 | 0.83% |
| hsa_circ_0059008 | 2640 | 11 | 0.83% |
| hsa_circ_0116027 | 2640 | 11 | 0.83% |
| hsa_circ_0019695 | 2400 | 10 | 0.83% |

| | | | |
|---|---|---|---|
| hsa_circ_0101773 | 2400 | 10 | 0.83% |
| hsa_circ_0129819 | 3361 | 14 | 0.83% |
| hsa_circ_0034845 | 2881 | 12 | 0.83% |
| hsa_circ_0061648 | 2881 | 12 | 0.83% |
| hsa_circ_0086092 | 2881 | 12 | 0.83% |
| hsa_circ_0134898 | 2881 | 12 | 0.83% |
| hsa_circ_0034373 | 2641 | 11 | 0.83% |
| hsa_circ_0068922 | 2641 | 11 | 0.83% |
| hsa_circ_0080149 | 2641 | 11 | 0.83% |
| hsa_circ_0086883 | 2641 | 11 | 0.83% |
| hsa_circ_0094936 | 2641 | 11 | 0.83% |
| hsa_circ_0032707 | 2401 | 10 | 0.83% |
| hsa_circ_0063970 | 2401 | 10 | 0.83% |
| hsa_circ_0072659 | 2401 | 10 | 0.83% |
| hsa_circ_0074732 | 2401 | 10 | 0.83% |
| hsa_circ_0009814 | 3362 | 14 | 0.83% |
| hsa_circ_0073667 | 2882 | 12 | 0.83% |
| hsa_circ_0088380 | 2882 | 12 | 0.83% |
| hsa_circ_0019354 | 2642 | 11 | 0.83% |
| hsa_circ_0027100 | 2642 | 11 | 0.83% |
| hsa_circ_0065246 | 2642 | 11 | 0.83% |
| hsa_circ_0083283 | 2642 | 11 | 0.83% |
| hsa_circ_0128154 | 2642 | 11 | 0.83% |
| hsa_circ_0109623 | 3603 | 15 | 0.83% |
| hsa_circ_0015687 | 2402 | 10 | 0.83% |
| hsa_circ_0040734 | 3123 | 13 | 0.83% |
| hsa_circ_0113862 | 2883 | 12 | 0.83% |
| hsa_circ_0014327 | 2643 | 11 | 0.83% |
| hsa_circ_0016810 | 2643 | 11 | 0.83% |
| hsa_circ_0076540 | 2643 | 11 | 0.83% |
| hsa_circ_0081880 | 2643 | 11 | 0.83% |
| hsa_circ_0021546 | 2403 | 10 | 0.83% |
| hsa_circ_0039392 | 2403 | 10 | 0.83% |
| hsa_circ_0042014 | 2403 | 10 | 0.83% |
| hsa_circ_0045019 | 2403 | 10 | 0.83% |
| hsa_circ_0099152 | 2403 | 10 | 0.83% |
| hsa_circ_0131995 | 2884 | 12 | 0.83% |
| hsa_circ_0028435 | 2644 | 11 | 0.83% |
| hsa_circ_0029462 | 2644 | 11 | 0.83% |
| hsa_circ_0029527 | 2644 | 11 | 0.83% |
| hsa_circ_0075767 | 2644 | 11 | 0.83% |
| hsa_circ_0117294 | 2644 | 11 | 0.83% |
| hsa_circ_0050139 | 2404 | 10 | 0.83% |
| hsa_circ_0073530 | 2885 | 12 | 0.83% |
| hsa_circ_0086087 | 2885 | 12 | 0.83% |
| hsa_circ_0073532 | 3366 | 14 | 0.83% |
| hsa_circ_0065023 | 4809 | 20 | 0.83% |
| hsa_circ_0010070 | 2645 | 11 | 0.83% |
| hsa_circ_0019460 | 2645 | 11 | 0.83% |
| hsa_circ_0034177 | 2645 | 11 | 0.83% |

| | | | |
|---|---|---|---|
| hsa_circ_0047080 | 2645 | 11 | 0.83% |
| hsa_circ_0054951 | 2645 | 11 | 0.83% |
| hsa_circ_0121031 | 2645 | 11 | 0.83% |
| hsa_circ_0085346 | 3126 | 13 | 0.83% |
| hsa_circ_0094003 | 4329 | 18 | 0.83% |
| hsa_circ_0075888 | 2886 | 12 | 0.83% |
| hsa_circ_0128519 | 2886 | 12 | 0.83% |
| hsa_circ_0055463 | 2405 | 10 | 0.83% |
| hsa_circ_0064226 | 2405 | 10 | 0.83% |
| hsa_circ_0027103 | 2646 | 11 | 0.83% |
| hsa_circ_0047078 | 2646 | 11 | 0.83% |
| hsa_circ_0063309 | 3368 | 14 | 0.83% |
| hsa_circ_0040281 | 8180 | 34 | 0.83% |
| hsa_circ_0114955 | 14195 | 59 | 0.83% |
| hsa_circ_0004030 | 2406 | 10 | 0.83% |
| hsa_circ_0012703 | 2406 | 10 | 0.83% |
| hsa_circ_0024119 | 2406 | 10 | 0.83% |
| hsa_circ_0024196 | 2406 | 10 | 0.83% |
| hsa_circ_0057647 | 2406 | 10 | 0.83% |
| hsa_circ_0060174 | 2647 | 11 | 0.83% |
| hsa_circ_0121175 | 4813 | 20 | 0.83% |
| hsa_circ_0038389 | 2888 | 12 | 0.83% |
| hsa_circ_0039602 | 2888 | 12 | 0.83% |
| hsa_circ_0034359 | 3129 | 13 | 0.83% |
| hsa_circ_0075759 | 2407 | 10 | 0.83% |
| hsa_circ_0107599 | 2407 | 10 | 0.83% |
| hsa_circ_0132880 | 5777 | 24 | 0.83% |
| hsa_circ_0034738 | 3370 | 14 | 0.83% |
| hsa_circ_0136739 | 2648 | 11 | 0.83% |
| hsa_circ_0128163 | 3130 | 13 | 0.83% |
| hsa_circ_0047893 | 2408 | 10 | 0.83% |
| hsa_circ_0069484 | 2408 | 10 | 0.83% |
| hsa_circ_0018263 | 2649 | 11 | 0.83% |
| hsa_circ_0012035 | 2890 | 12 | 0.83% |
| hsa_circ_0086818 | 2890 | 12 | 0.83% |
| hsa_circ_0017815 | 3131 | 13 | 0.83% |
| hsa_circ_0005300 | 2409 | 10 | 0.83% |
| hsa_circ_0114733 | 2409 | 10 | 0.83% |
| hsa_circ_0132228 | 2409 | 10 | 0.83% |
| hsa_circ_0052496 | 2650 | 11 | 0.83% |
| hsa_circ_0017120 | 2891 | 12 | 0.83% |
| hsa_circ_0057425 | 3374 | 14 | 0.83% |
| hsa_circ_0046246 | 3133 | 13 | 0.83% |
| hsa_circ_0057512 | 2892 | 12 | 0.83% |
| hsa_circ_0086934 | 2892 | 12 | 0.83% |
| hsa_circ_0010277 | 2651 | 11 | 0.83% |
| hsa_circ_0037697 | 2651 | 11 | 0.83% |
| hsa_circ_0042704 | 2651 | 11 | 0.83% |
| hsa_circ_0042673 | 2410 | 10 | 0.83% |
| hsa_circ_0042131 | 4821 | 20 | 0.83% |

| | | | |
|---|---|---|---|
| hsa_circ_0037987 | 7232 | 30 | 0.83% |
| hsa_circ_0090108 | 3375 | 14 | 0.83% |
| hsa_circ_0084391 | 3134 | 13 | 0.83% |
| hsa_circ_0018511 | 2893 | 12 | 0.83% |
| hsa_circ_0033344 | 2893 | 12 | 0.83% |
| hsa_circ_0115675 | 2893 | 12 | 0.83% |
| hsa_circ_0041257 | 5545 | 23 | 0.83% |
| hsa_circ_0019236 | 2652 | 11 | 0.83% |
| hsa_circ_0076788 | 2652 | 11 | 0.83% |
| hsa_circ_0115398 | 2652 | 11 | 0.83% |
| hsa_circ_0110598 | 5787 | 24 | 0.83% |
| hsa_circ_0081504 | 3858 | 16 | 0.83% |
| hsa_circ_0103712 | 12298 | 51 | 0.83% |
| hsa_circ_0010617 | 3135 | 13 | 0.83% |
| hsa_circ_0030656 | 2894 | 12 | 0.83% |
| hsa_circ_0047000 | 2894 | 12 | 0.83% |
| hsa_circ_0058897 | 2894 | 12 | 0.83% |
| hsa_circ_0102229 | 2894 | 12 | 0.83% |
| hsa_circ_0010009 | 2653 | 11 | 0.83% |
| hsa_circ_0067890 | 2653 | 11 | 0.83% |
| hsa_circ_0070108 | 2653 | 11 | 0.83% |
| hsa_circ_0073862 | 2653 | 11 | 0.83% |
| hsa_circ_0036822 | 2412 | 10 | 0.83% |
| hsa_circ_0084583 | 2412 | 10 | 0.83% |
| hsa_circ_0113271 | 2412 | 10 | 0.83% |
| hsa_circ_0037067 | 3377 | 14 | 0.83% |
| hsa_circ_0063368 | 3136 | 13 | 0.83% |
| hsa_circ_0088304 | 3136 | 13 | 0.83% |
| hsa_circ_0108835 | 3136 | 13 | 0.83% |
| hsa_circ_0009332 | 2895 | 12 | 0.83% |
| hsa_circ_0055768 | 2654 | 11 | 0.83% |
| hsa_circ_0061663 | 2654 | 11 | 0.83% |
| hsa_circ_0008706 | 2413 | 10 | 0.83% |
| hsa_circ_0109290 | 2413 | 10 | 0.83% |
| hsa_circ_0103506 | 8206 | 34 | 0.83% |
| hsa_circ_0086868 | 2655 | 11 | 0.83% |
| hsa_circ_0086878 | 2655 | 11 | 0.83% |
| hsa_circ_0104918 | 2414 | 10 | 0.83% |
| hsa_circ_0033269 | 2897 | 12 | 0.83% |
| hsa_circ_0026859 | 2656 | 11 | 0.83% |
| hsa_circ_0030018 | 2656 | 11 | 0.83% |
| hsa_circ_0061724 | 2656 | 11 | 0.83% |
| hsa_circ_0097270 | 2656 | 11 | 0.83% |
| hsa_circ_0127438 | 2656 | 11 | 0.83% |
| hsa_circ_0083410 | 3139 | 13 | 0.83% |
| hsa_circ_0091258 | 3139 | 13 | 0.83% |
| hsa_circ_0045180 | 2898 | 12 | 0.83% |
| hsa_circ_0061408 | 2898 | 12 | 0.83% |
| hsa_circ_0138271 | 2898 | 12 | 0.83% |
| hsa_circ_0016270 | 2415 | 10 | 0.83% |

| | | | |
|---|---|---|---|
| hsa_circ_0063220 | 12318 | 51 | 0.83% |
| hsa_circ_0052754 | 3140 | 13 | 0.83% |
| hsa_circ_0074277 | 2657 | 11 | 0.83% |
| hsa_circ_0102281 | 2657 | 11 | 0.83% |
| hsa_circ_0126295 | 2657 | 11 | 0.83% |
| hsa_circ_0127344 | 2657 | 11 | 0.83% |
| hsa_circ_0057853 | 2899 | 12 | 0.83% |
| hsa_circ_0121336 | 2899 | 12 | 0.83% |
| hsa_circ_0085140 | 2416 | 10 | 0.83% |
| hsa_circ_0087382 | 2416 | 10 | 0.83% |
| hsa_circ_0088116 | 2416 | 10 | 0.83% |
| hsa_circ_0022360 | 3141 | 13 | 0.83% |
| hsa_circ_0023284 | 3141 | 13 | 0.83% |
| hsa_circ_0075234 | 3141 | 13 | 0.83% |
| hsa_circ_0071303 | 2658 | 11 | 0.83% |
| hsa_circ_0081062 | 2658 | 11 | 0.83% |
| hsa_circ_0114001 | 2658 | 11 | 0.83% |
| hsa_circ_0115691 | 6525 | 27 | 0.83% |
| hsa_circ_0081378 | 5075 | 21 | 0.83% |
| hsa_circ_0010981 | 2900 | 12 | 0.83% |
| hsa_circ_0026070 | 2900 | 12 | 0.83% |
| hsa_circ_0063488 | 2900 | 12 | 0.83% |
| hsa_circ_0122896 | 2900 | 12 | 0.83% |
| hsa_circ_0020689 | 2417 | 10 | 0.83% |
| hsa_circ_0028456 | 2417 | 10 | 0.83% |
| hsa_circ_0100630 | 6768 | 28 | 0.83% |
| hsa_circ_0028799 | 2659 | 11 | 0.83% |
| hsa_circ_0080058 | 2659 | 11 | 0.83% |
| hsa_circ_0102494 | 2659 | 11 | 0.83% |
| hsa_circ_0121954 | 2659 | 11 | 0.83% |
| hsa_circ_0035715 | 2901 | 12 | 0.83% |
| hsa_circ_0057135 | 3143 | 13 | 0.83% |
| hsa_circ_0020088 | 3385 | 14 | 0.83% |
| hsa_circ_0035432 | 2418 | 10 | 0.83% |
| hsa_circ_0056891 | 2418 | 10 | 0.83% |
| hsa_circ_0060407 | 2418 | 10 | 0.83% |
| hsa_circ_0086307 | 2418 | 10 | 0.83% |
| hsa_circ_0094853 | 2418 | 10 | 0.83% |
| hsa_circ_0052252 | 2660 | 11 | 0.83% |
| hsa_circ_0071103 | 2902 | 12 | 0.83% |
| hsa_circ_0093387 | 2902 | 12 | 0.83% |
| hsa_circ_0010638 | 3144 | 13 | 0.83% |
| hsa_circ_0094086 | 5563 | 23 | 0.83% |
| hsa_circ_0051701 | 2419 | 10 | 0.83% |
| hsa_circ_0128087 | 2419 | 10 | 0.83% |
| hsa_circ_0129469 | 2419 | 10 | 0.83% |
| hsa_circ_0062381 | 2661 | 11 | 0.83% |
| hsa_circ_0091125 | 2661 | 11 | 0.83% |
| hsa_circ_0010217 | 2903 | 12 | 0.83% |
| hsa_circ_0081612 | 2903 | 12 | 0.83% |

| | | | |
|---|---|---|---|
| hsa_circ_0040874 | 3388 | 14 | 0.83% |
| hsa_circ_0058011 | 3388 | 14 | 0.83% |
| hsa_circ_0088860 | 3146 | 13 | 0.83% |
| hsa_circ_0129475 | 2904 | 12 | 0.83% |
| hsa_circ_0018670 | 2662 | 11 | 0.83% |
| hsa_circ_0024128 | 2662 | 11 | 0.83% |
| hsa_circ_0062040 | 2662 | 11 | 0.83% |
| hsa_circ_0100327 | 2662 | 11 | 0.83% |
| hsa_circ_0108213 | 2662 | 11 | 0.83% |
| hsa_circ_0070332 | 2420 | 10 | 0.83% |
| hsa_circ_0135224 | 9439 | 39 | 0.83% |
| hsa_circ_0083771 | 3631 | 15 | 0.83% |
| hsa_circ_0132082 | 3631 | 15 | 0.83% |
| hsa_circ_0031226 | 2663 | 11 | 0.83% |
| hsa_circ_0040235 | 2663 | 11 | 0.83% |
| hsa_circ_0040335 | 2663 | 11 | 0.83% |
| hsa_circ_0067674 | 2663 | 11 | 0.83% |
| hsa_circ_0031497 | 2421 | 10 | 0.83% |
| hsa_circ_0041858 | 2421 | 10 | 0.83% |
| hsa_circ_0058628 | 2421 | 10 | 0.83% |
| hsa_circ_0066792 | 2421 | 10 | 0.83% |
| hsa_circ_0104429 | 2906 | 12 | 0.83% |
| hsa_circ_0108136 | 2906 | 12 | 0.83% |
| hsa_circ_0058380 | 2664 | 11 | 0.83% |
| hsa_circ_0061709 | 2664 | 11 | 0.83% |
| hsa_circ_0076544 | 2664 | 11 | 0.83% |
| hsa_circ_0031672 | 2422 | 10 | 0.83% |
| hsa_circ_0046329 | 2422 | 10 | 0.83% |
| hsa_circ_0054484 | 2422 | 10 | 0.83% |
| hsa_circ_0135641 | 2422 | 10 | 0.83% |
| hsa_circ_0007950 | 4118 | 17 | 0.83% |
| hsa_circ_0106620 | 7994 | 33 | 0.83% |
| hsa_circ_0017473 | 2665 | 11 | 0.83% |
| hsa_circ_0031823 | 2665 | 11 | 0.83% |
| hsa_circ_0054596 | 2665 | 11 | 0.83% |
| hsa_circ_0060598 | 2665 | 11 | 0.83% |
| hsa_circ_0082925 | 2665 | 11 | 0.83% |
| hsa_circ_0140666 | 4846 | 20 | 0.83% |
| hsa_circ_0039691 | 2423 | 10 | 0.83% |
| hsa_circ_0043594 | 2423 | 10 | 0.83% |
| hsa_circ_0061546 | 2423 | 10 | 0.83% |
| hsa_circ_0071339 | 2423 | 10 | 0.83% |
| hsa_circ_0125005 | 2423 | 10 | 0.83% |
| hsa_circ_0126061 | 2423 | 10 | 0.83% |
| hsa_circ_0025698 | 3150 | 13 | 0.83% |
| hsa_circ_0071083 | 2908 | 12 | 0.83% |
| hsa_circ_0034247 | 2666 | 11 | 0.83% |
| hsa_circ_0084912 | 2666 | 11 | 0.83% |
| hsa_circ_0104303 | 3151 | 13 | 0.83% |
| hsa_circ_0011825 | 2424 | 10 | 0.83% |

| | | | |
|---|---|---|---|
| hsa_circ_0072222 | 2424 | 10 | 0.83% |
| hsa_circ_0007784 | 4121 | 17 | 0.83% |
| hsa_circ_0029193 | 2909 | 12 | 0.83% |
| hsa_circ_0046619 | 2909 | 12 | 0.83% |
| hsa_circ_0051011 | 2909 | 12 | 0.83% |
| hsa_circ_0068948 | 2909 | 12 | 0.83% |
| hsa_circ_0122443 | 2909 | 12 | 0.83% |
| hsa_circ_0128137 | 2667 | 11 | 0.82% |
| hsa_circ_0022491 | 3152 | 13 | 0.82% |
| hsa_circ_0020627 | 2910 | 12 | 0.82% |
| hsa_circ_0086360 | 2425 | 10 | 0.82% |
| hsa_circ_0099549 | 2425 | 10 | 0.82% |
| hsa_circ_0104264 | 2425 | 10 | 0.82% |
| hsa_circ_0115633 | 2425 | 10 | 0.82% |
| hsa_circ_0134361 | 2425 | 10 | 0.82% |
| hsa_circ_0008067 | 7518 | 31 | 0.82% |
| hsa_circ_0024197 | 2668 | 11 | 0.82% |
| hsa_circ_0046459 | 2668 | 11 | 0.82% |
| hsa_circ_0058600 | 2668 | 11 | 0.82% |
| hsa_circ_0133588 | 2668 | 11 | 0.82% |
| hsa_circ_0047510 | 2911 | 12 | 0.82% |
| hsa_circ_0039294 | 2426 | 10 | 0.82% |
| hsa_circ_0132655 | 2426 | 10 | 0.82% |
| hsa_circ_0053689 | 3154 | 13 | 0.82% |
| hsa_circ_0060680 | 3154 | 13 | 0.82% |
| hsa_circ_0069546 | 3154 | 13 | 0.82% |
| hsa_circ_0049297 | 2669 | 11 | 0.82% |
| hsa_circ_0059159 | 2669 | 11 | 0.82% |
| hsa_circ_0099954 | 2669 | 11 | 0.82% |
| hsa_circ_0136922 | 2669 | 11 | 0.82% |
| hsa_circ_0075584 | 3397 | 14 | 0.82% |
| hsa_circ_0058311 | 2912 | 12 | 0.82% |
| hsa_circ_0111900 | 6310 | 26 | 0.82% |
| hsa_circ_0030589 | 2427 | 10 | 0.82% |
| hsa_circ_0069730 | 2427 | 10 | 0.82% |
| hsa_circ_0074940 | 2427 | 10 | 0.82% |
| hsa_circ_0114524 | 2427 | 10 | 0.82% |
| hsa_circ_0019105 | 2670 | 11 | 0.82% |
| hsa_circ_0019713 | 2670 | 11 | 0.82% |
| hsa_circ_0020831 | 2670 | 11 | 0.82% |
| hsa_circ_0037800 | 2670 | 11 | 0.82% |
| hsa_circ_0059685 | 2670 | 11 | 0.82% |
| hsa_circ_0028122 | 2913 | 12 | 0.82% |
| hsa_circ_0102885 | 2913 | 12 | 0.82% |
| hsa_circ_0039644 | 3156 | 13 | 0.82% |
| hsa_circ_0009321 | 2428 | 10 | 0.82% |
| hsa_circ_0023343 | 2428 | 10 | 0.82% |
| hsa_circ_0031583 | 3885 | 16 | 0.82% |
| hsa_circ_0044946 | 3885 | 16 | 0.82% |
| hsa_circ_0010732 | 2671 | 11 | 0.82% |

| | | | |
|---|---|---|---|
| hsa_circ_0054855 | 2671 | 11 | 0.82% |
| hsa_circ_0060298 | 3400 | 14 | 0.82% |
| hsa_circ_0064463 | 4858 | 20 | 0.82% |
| hsa_circ_0053970 | 2429 | 10 | 0.82% |
| hsa_circ_0061054 | 2429 | 10 | 0.82% |
| hsa_circ_0076198 | 5344 | 22 | 0.82% |
| hsa_circ_0031747 | 2672 | 11 | 0.82% |
| hsa_circ_0080146 | 2672 | 11 | 0.82% |
| hsa_circ_0091440 | 3158 | 13 | 0.82% |
| hsa_circ_0042218 | 2673 | 11 | 0.82% |
| hsa_circ_0091654 | 2673 | 11 | 0.82% |
| hsa_circ_0110449 | 2673 | 11 | 0.82% |
| hsa_circ_0065562 | 2430 | 10 | 0.82% |
| hsa_circ_0105080 | 2430 | 10 | 0.82% |
| hsa_circ_0019789 | 7049 | 29 | 0.82% |
| hsa_circ_0053206 | 3160 | 13 | 0.82% |
| hsa_circ_0029597 | 2917 | 12 | 0.82% |
| hsa_circ_0034066 | 2917 | 12 | 0.82% |
| hsa_circ_0089548 | 2917 | 12 | 0.82% |
| hsa_circ_0027846 | 2674 | 11 | 0.82% |
| hsa_circ_0028764 | 2674 | 11 | 0.82% |
| hsa_circ_0044375 | 2674 | 11 | 0.82% |
| hsa_circ_0103989 | 2674 | 11 | 0.82% |
| hsa_circ_0070426 | 2431 | 10 | 0.82% |
| hsa_circ_0073542 | 2431 | 10 | 0.82% |
| hsa_circ_0124504 | 3890 | 16 | 0.82% |
| hsa_circ_0057006 | 2918 | 12 | 0.82% |
| hsa_circ_0091047 | 2918 | 12 | 0.82% |
| hsa_circ_0010957 | 2675 | 11 | 0.82% |
| hsa_circ_0040884 | 2675 | 11 | 0.82% |
| hsa_circ_0095339 | 2675 | 11 | 0.82% |
| hsa_circ_0068986 | 2432 | 10 | 0.82% |
| hsa_circ_0123426 | 3162 | 13 | 0.82% |
| hsa_circ_0003712 | 3892 | 16 | 0.82% |
| hsa_circ_0026146 | 2919 | 12 | 0.82% |
| hsa_circ_0052608 | 2919 | 12 | 0.82% |
| hsa_circ_0063855 | 2919 | 12 | 0.82% |
| hsa_circ_0125934 | 2919 | 12 | 0.82% |
| hsa_circ_0023708 | 2676 | 11 | 0.82% |
| hsa_circ_0051306 | 2676 | 11 | 0.82% |
| hsa_circ_0077808 | 2676 | 11 | 0.82% |
| hsa_circ_0120311 | 5839 | 24 | 0.82% |
| hsa_circ_0011194 | 2433 | 10 | 0.82% |
| hsa_circ_0081081 | 2433 | 10 | 0.82% |
| hsa_circ_0116272 | 2433 | 10 | 0.82% |
| hsa_circ_0124308 | 2433 | 10 | 0.82% |
| hsa_circ_0139173 | 2433 | 10 | 0.82% |
| hsa_circ_0134744 | 6813 | 28 | 0.82% |
| hsa_circ_0043309 | 2920 | 12 | 0.82% |
| hsa_circ_0088594 | 2677 | 11 | 0.82% |

| | | | |
|---|---|---|---|
| hsa_circ_0028924 | 3894 | 16 | 0.82% |
| hsa_circ_0002139 | 7058 | 29 | 0.82% |
| hsa_circ_0066655 | 4381 | 18 | 0.82% |
| hsa_circ_0017040 | 9249 | 38 | 0.82% |
| hsa_circ_0008323 | 6085 | 25 | 0.82% |
| hsa_circ_0072537 | 2434 | 10 | 0.82% |
| hsa_circ_0105796 | 2434 | 10 | 0.82% |
| hsa_circ_0040277 | 8276 | 34 | 0.82% |
| hsa_circ_0031602 | 2921 | 12 | 0.82% |
| hsa_circ_0047035 | 2921 | 12 | 0.82% |
| hsa_circ_0067394 | 2921 | 12 | 0.82% |
| hsa_circ_0009558 | 2678 | 11 | 0.82% |
| hsa_circ_0037752 | 2678 | 11 | 0.82% |
| hsa_circ_0083397 | 2678 | 11 | 0.82% |
| hsa_circ_0005438 | 4870 | 20 | 0.82% |
| hsa_circ_0016202 | 2922 | 12 | 0.82% |
| hsa_circ_0073506 | 2435 | 10 | 0.82% |
| hsa_circ_0088327 | 2435 | 10 | 0.82% |
| hsa_circ_0115230 | 2435 | 10 | 0.82% |
| hsa_circ_0122039 | 2435 | 10 | 0.82% |
| hsa_circ_0019778 | 3166 | 13 | 0.82% |
| hsa_circ_0030512 | 2679 | 11 | 0.82% |
| hsa_circ_0040234 | 2679 | 11 | 0.82% |
| hsa_circ_0054352 | 2679 | 11 | 0.82% |
| hsa_circ_0057682 | 2679 | 11 | 0.82% |
| hsa_circ_0140599 | 2679 | 11 | 0.82% |
| hsa_circ_0135380 | 5846 | 24 | 0.82% |
| hsa_circ_0043337 | 2923 | 12 | 0.82% |
| hsa_circ_0055824 | 3654 | 15 | 0.82% |
| hsa_circ_0027209 | 2436 | 10 | 0.82% |
| hsa_circ_0060340 | 2436 | 10 | 0.82% |
| hsa_circ_0062605 | 2436 | 10 | 0.82% |
| hsa_circ_0072232 | 2436 | 10 | 0.82% |
| hsa_circ_0073387 | 2436 | 10 | 0.82% |
| hsa_circ_0047816 | 3167 | 13 | 0.82% |
| hsa_circ_0059646 | 3167 | 13 | 0.82% |
| hsa_circ_0061442 | 3898 | 16 | 0.82% |
| hsa_circ_0019891 | 2680 | 11 | 0.82% |
| hsa_circ_0064526 | 2680 | 11 | 0.82% |
| hsa_circ_0087966 | 2680 | 11 | 0.82% |
| hsa_circ_0013291 | 2924 | 12 | 0.82% |
| hsa_circ_0026664 | 2924 | 12 | 0.82% |
| hsa_circ_0011673 | 2437 | 10 | 0.82% |
| hsa_circ_0063219 | 12430 | 51 | 0.82% |
| hsa_circ_0028067 | 2681 | 11 | 0.82% |
| hsa_circ_0035467 | 2681 | 11 | 0.82% |
| hsa_circ_0062704 | 2681 | 11 | 0.82% |
| hsa_circ_0094686 | 2681 | 11 | 0.82% |
| hsa_circ_0013421 | 2925 | 12 | 0.82% |
| hsa_circ_0042577 | 2925 | 12 | 0.82% |

| | | | |
|---|---|---|---|
| hsa_circ_0016897 | 3169 | 13 | 0.82% |
| hsa_circ_0008493 | 7070 | 29 | 0.82% |
| hsa_circ_0083545 | 3657 | 15 | 0.82% |
| hsa_circ_0055634 | 2438 | 10 | 0.82% |
| hsa_circ_0057635 | 2438 | 10 | 0.82% |
| hsa_circ_0091563 | 2438 | 10 | 0.82% |
| hsa_circ_0109880 | 2438 | 10 | 0.82% |
| hsa_circ_0105019 | 8535 | 35 | 0.82% |
| hsa_circ_0017141 | 2439 | 10 | 0.82% |
| hsa_circ_0050083 | 10000 | 41 | 0.82% |
| hsa_circ_0013050 | 2683 | 11 | 0.82% |
| hsa_circ_0024791 | 2683 | 11 | 0.82% |
| hsa_circ_0054586 | 2683 | 11 | 0.82% |
| hsa_circ_0085737 | 2683 | 11 | 0.82% |
| hsa_circ_0097057 | 2683 | 11 | 0.82% |
| hsa_circ_0127227 | 2683 | 11 | 0.82% |
| hsa_circ_0018653 | 2927 | 12 | 0.82% |
| hsa_circ_0022697 | 2927 | 12 | 0.82% |
| hsa_circ_0032870 | 2927 | 12 | 0.82% |
| hsa_circ_0075023 | 2927 | 12 | 0.82% |
| hsa_circ_0078989 | 3171 | 13 | 0.82% |
| hsa_circ_0064615 | 6344 | 26 | 0.82% |
| hsa_circ_0017394 | 3904 | 16 | 0.82% |
| hsa_circ_0035081 | 3172 | 13 | 0.82% |
| hsa_circ_0014769 | 2928 | 12 | 0.82% |
| hsa_circ_0058787 | 2684 | 11 | 0.82% |
| hsa_circ_0063407 | 2684 | 11 | 0.82% |
| hsa_circ_0131923 | 2684 | 11 | 0.82% |
| hsa_circ_0076981 | 2440 | 10 | 0.82% |
| hsa_circ_0099516 | 2440 | 10 | 0.82% |
| hsa_circ_0025221 | 3173 | 13 | 0.82% |
| hsa_circ_0052294 | 2929 | 12 | 0.82% |
| hsa_circ_0029652 | 2685 | 11 | 0.82% |
| hsa_circ_0055714 | 2685 | 11 | 0.82% |
| hsa_circ_0059317 | 2685 | 11 | 0.82% |
| hsa_circ_0064952 | 2685 | 11 | 0.82% |
| hsa_circ_0076065 | 2685 | 11 | 0.82% |
| hsa_circ_0085401 | 2685 | 11 | 0.82% |
| hsa_circ_0087460 | 2685 | 11 | 0.82% |
| hsa_circ_0137555 | 2685 | 11 | 0.82% |
| hsa_circ_0044332 | 3174 | 13 | 0.82% |
| hsa_circ_0058101 | 3174 | 13 | 0.82% |
| hsa_circ_0086633 | 3174 | 13 | 0.82% |
| hsa_circ_0034903 | 2930 | 12 | 0.82% |
| hsa_circ_0009946 | 2686 | 11 | 0.82% |
| hsa_circ_0051376 | 2686 | 11 | 0.82% |
| hsa_circ_0086517 | 2686 | 11 | 0.82% |
| hsa_circ_0120659 | 2686 | 11 | 0.82% |
| hsa_circ_0007563 | 6593 | 27 | 0.82% |
| hsa_circ_0021242 | 3419 | 14 | 0.82% |

| | | | |
|---|---|---|---|
| hsa_circ_0021065 | 3175 | 13 | 0.82% |
| hsa_circ_0045796 | 3175 | 13 | 0.82% |
| hsa_circ_0085912 | 2931 | 12 | 0.82% |
| hsa_circ_0107704 | 2931 | 12 | 0.82% |
| hsa_circ_0018698 | 2687 | 11 | 0.82% |
| hsa_circ_0043253 | 2687 | 11 | 0.82% |
| hsa_circ_0012714 | 2443 | 10 | 0.82% |
| hsa_circ_0040938 | 2443 | 10 | 0.82% |
| hsa_circ_0132046 | 2443 | 10 | 0.82% |
| hsa_circ_0102894 | 6352 | 26 | 0.82% |
| hsa_circ_0007500 | 6597 | 27 | 0.82% |
| hsa_circ_0134851 | 4398 | 18 | 0.82% |
| hsa_circ_0088669 | 2932 | 12 | 0.82% |
| hsa_circ_0129041 | 2932 | 12 | 0.82% |
| hsa_circ_0130362 | 3421 | 14 | 0.82% |
| hsa_circ_0100192 | 2688 | 11 | 0.82% |
| hsa_circ_0131924 | 2688 | 11 | 0.82% |
| hsa_circ_0140345 | 2688 | 11 | 0.82% |
| hsa_circ_0058721 | 2444 | 10 | 0.82% |
| hsa_circ_0103893 | 2444 | 10 | 0.82% |
| hsa_circ_0099174 | 4155 | 17 | 0.82% |
| hsa_circ_0039084 | 2933 | 12 | 0.82% |
| hsa_circ_0073011 | 2933 | 12 | 0.82% |
| hsa_circ_0012341 | 2689 | 11 | 0.82% |
| hsa_circ_0109896 | 2689 | 11 | 0.82% |
| hsa_circ_0037075 | 3178 | 13 | 0.82% |
| hsa_circ_0037221 | 2934 | 12 | 0.82% |
| hsa_circ_0139078 | 2445 | 10 | 0.82% |
| hsa_circ_0061437 | 3179 | 13 | 0.82% |
| hsa_circ_0063071 | 3179 | 13 | 0.82% |
| hsa_circ_0099567 | 3179 | 13 | 0.82% |
| hsa_circ_0019348 | 2690 | 11 | 0.82% |
| hsa_circ_0037320 | 2690 | 11 | 0.82% |
| hsa_circ_0115961 | 2690 | 11 | 0.82% |
| hsa_circ_0122115 | 2690 | 11 | 0.82% |
| hsa_circ_0130066 | 2690 | 11 | 0.82% |
| hsa_circ_0004497 | 6603 | 27 | 0.82% |
| hsa_circ_0130363 | 3424 | 14 | 0.82% |
| hsa_circ_0041204 | 2935 | 12 | 0.82% |
| hsa_circ_0134205 | 2935 | 12 | 0.82% |
| hsa_circ_0015324 | 2446 | 10 | 0.82% |
| hsa_circ_0073893 | 2446 | 10 | 0.82% |
| hsa_circ_0106656 | 2446 | 10 | 0.82% |
| hsa_circ_0001274 | 2691 | 11 | 0.82% |
| hsa_circ_0016788 | 2691 | 11 | 0.82% |
| hsa_circ_0048637 | 2691 | 11 | 0.82% |
| hsa_circ_0052110 | 2691 | 11 | 0.82% |
| hsa_circ_0063406 | 2691 | 11 | 0.82% |
| hsa_circ_0082619 | 2691 | 11 | 0.82% |
| hsa_circ_0011835 | 2936 | 12 | 0.82% |

| | | | |
|---|---|---|---|
| hsa_circ_0012062 | 2936 | 12 | 0.82% |
| hsa_circ_0023185 | 2936 | 12 | 0.82% |
| hsa_circ_0067799 | 2936 | 12 | 0.82% |
| hsa_circ_0138863 | 39881 | 163 | 0.82% |
| hsa_circ_0030699 | 2447 | 10 | 0.82% |
| hsa_circ_0039712 | 2447 | 10 | 0.82% |
| hsa_circ_0020137 | 2692 | 11 | 0.82% |
| hsa_circ_0105818 | 10279 | 42 | 0.82% |
| hsa_circ_0010332 | 2937 | 12 | 0.82% |
| hsa_circ_0015481 | 2937 | 12 | 0.82% |
| hsa_circ_0027199 | 2448 | 10 | 0.82% |
| hsa_circ_0030259 | 2448 | 10 | 0.82% |
| hsa_circ_0043216 | 2448 | 10 | 0.82% |
| hsa_circ_0043715 | 2448 | 10 | 0.82% |
| hsa_circ_0019462 | 2693 | 11 | 0.82% |
| hsa_circ_0023175 | 2693 | 11 | 0.82% |
| hsa_circ_0026203 | 2693 | 11 | 0.82% |
| hsa_circ_0080587 | 2693 | 11 | 0.82% |
| hsa_circ_0105629 | 7100 | 29 | 0.82% |
| hsa_circ_0000868 | 2938 | 12 | 0.82% |
| hsa_circ_0073098 | 2449 | 10 | 0.82% |
| hsa_circ_0014453 | 2694 | 11 | 0.82% |
| hsa_circ_0047028 | 2694 | 11 | 0.82% |
| hsa_circ_0058130 | 2694 | 11 | 0.82% |
| hsa_circ_0059019 | 2694 | 11 | 0.82% |
| hsa_circ_0060173 | 2694 | 11 | 0.82% |
| hsa_circ_0084313 | 2694 | 11 | 0.82% |
| hsa_circ_0024180 | 4409 | 18 | 0.82% |
| hsa_circ_0084384 | 3185 | 13 | 0.82% |
| hsa_circ_0021151 | 2940 | 12 | 0.82% |
| hsa_circ_0140737 | 2940 | 12 | 0.82% |
| hsa_circ_0023587 | 2695 | 11 | 0.82% |
| hsa_circ_0109340 | 2695 | 11 | 0.82% |
| hsa_circ_0037046 | 2450 | 10 | 0.82% |
| hsa_circ_0042633 | 2450 | 10 | 0.82% |
| hsa_circ_0105531 | 2450 | 10 | 0.82% |
| hsa_circ_0117946 | 2450 | 10 | 0.82% |
| hsa_circ_0081609 | 2941 | 12 | 0.82% |
| hsa_circ_0046393 | 2696 | 11 | 0.82% |
| hsa_circ_0050898 | 2696 | 11 | 0.82% |
| hsa_circ_0029455 | 2451 | 10 | 0.82% |
| hsa_circ_0008893 | 3187 | 13 | 0.82% |
| hsa_circ_0070209 | 3187 | 13 | 0.82% |
| hsa_circ_0012768 | 2942 | 12 | 0.82% |
| hsa_circ_0115393 | 2942 | 12 | 0.82% |
| hsa_circ_0130535 | 2942 | 12 | 0.82% |
| hsa_circ_0002735 | 2697 | 11 | 0.82% |
| hsa_circ_0053443 | 2697 | 11 | 0.82% |
| hsa_circ_0094033 | 2697 | 11 | 0.82% |
| hsa_circ_0135481 | 2697 | 11 | 0.82% |

| | | | |
|---|---|---|---|
| hsa_circ_0132222 | 2452 | 10 | 0.82% |
| hsa_circ_0031623 | 3188 | 13 | 0.82% |
| hsa_circ_0031926 | 3188 | 13 | 0.82% |
| hsa_circ_0036840 | 3188 | 13 | 0.82% |
| hsa_circ_0046101 | 3188 | 13 | 0.82% |
| hsa_circ_0081918 | 3188 | 13 | 0.82% |
| hsa_circ_0129479 | 2943 | 12 | 0.82% |
| hsa_circ_0018119 | 2698 | 11 | 0.82% |
| hsa_circ_0137706 | 2698 | 11 | 0.82% |
| hsa_circ_0039981 | 2453 | 10 | 0.82% |
| hsa_circ_0056234 | 3189 | 13 | 0.82% |
| hsa_circ_0077459 | 3189 | 13 | 0.82% |
| hsa_circ_0103151 | 5888 | 24 | 0.82% |
| hsa_circ_0050081 | 10059 | 41 | 0.82% |
| hsa_circ_0017003 | 2699 | 11 | 0.82% |
| hsa_circ_0117906 | 2699 | 11 | 0.82% |
| hsa_circ_0004737 | 3190 | 13 | 0.82% |
| hsa_circ_0016069 | 2454 | 10 | 0.81% |
| hsa_circ_0026060 | 2454 | 10 | 0.81% |
| hsa_circ_0034690 | 2454 | 10 | 0.81% |
| hsa_circ_0081283 | 2454 | 10 | 0.81% |
| hsa_circ_0117337 | 2454 | 10 | 0.81% |
| hsa_circ_0014634 | 2945 | 12 | 0.81% |
| hsa_circ_0135479 | 2945 | 12 | 0.81% |
| hsa_circ_0031459 | 2700 | 11 | 0.81% |
| hsa_circ_0054061 | 2700 | 11 | 0.81% |
| hsa_circ_0061945 | 2700 | 11 | 0.81% |
| hsa_circ_0063811 | 2700 | 11 | 0.81% |
| hsa_circ_0064064 | 2946 | 12 | 0.81% |
| hsa_circ_0039652 | 2455 | 10 | 0.81% |
| hsa_circ_0090321 | 2455 | 10 | 0.81% |
| hsa_circ_0011461 | 2701 | 11 | 0.81% |
| hsa_circ_0024063 | 2947 | 12 | 0.81% |
| hsa_circ_0048738 | 2947 | 12 | 0.81% |
| hsa_circ_0057349 | 4175 | 17 | 0.81% |
| hsa_circ_0059295 | 2456 | 10 | 0.81% |
| hsa_circ_0124948 | 3193 | 13 | 0.81% |
| hsa_circ_0131987 | 3193 | 13 | 0.81% |
| hsa_circ_0024669 | 2702 | 11 | 0.81% |
| hsa_circ_0033098 | 2702 | 11 | 0.81% |
| hsa_circ_0071828 | 2702 | 11 | 0.81% |
| hsa_circ_0093221 | 2702 | 11 | 0.81% |
| hsa_circ_0046904 | 3439 | 14 | 0.81% |
| hsa_circ_0138182 | 2948 | 12 | 0.81% |
| hsa_circ_0089023 | 2457 | 10 | 0.81% |
| hsa_circ_0107597 | 2457 | 10 | 0.81% |
| hsa_circ_0112173 | 2457 | 10 | 0.81% |
| hsa_circ_0012083 | 2703 | 11 | 0.81% |
| hsa_circ_0013094 | 2703 | 11 | 0.81% |
| hsa_circ_0036785 | 2703 | 11 | 0.81% |

| | | | |
|---|---|---|---|
| hsa_circ_0045465 | 2703 | 11 | 0.81% |
| hsa_circ_0081858 | 2703 | 11 | 0.81% |
| hsa_circ_0084808 | 2703 | 11 | 0.81% |
| hsa_circ_0075961 | 8847 | 36 | 0.81% |
| hsa_circ_0021518 | 3932 | 16 | 0.81% |
| hsa_circ_0018459 | 2949 | 12 | 0.81% |
| hsa_circ_0062628 | 2949 | 12 | 0.81% |
| hsa_circ_0122332 | 6144 | 25 | 0.81% |
| hsa_circ_0076452 | 3195 | 13 | 0.81% |
| hsa_circ_0010422 | 2458 | 10 | 0.81% |
| hsa_circ_0011599 | 2458 | 10 | 0.81% |
| hsa_circ_0073163 | 2458 | 10 | 0.81% |
| hsa_circ_0103259 | 2458 | 10 | 0.81% |
| hsa_circ_0097359 | 7620 | 31 | 0.81% |
| hsa_circ_0023501 | 2704 | 11 | 0.81% |
| hsa_circ_0023502 | 2704 | 11 | 0.81% |
| hsa_circ_0027454 | 2704 | 11 | 0.81% |
| hsa_circ_0047845 | 2704 | 11 | 0.81% |
| hsa_circ_0067171 | 2704 | 11 | 0.81% |
| hsa_circ_0077538 | 2704 | 11 | 0.81% |
| hsa_circ_0097804 | 2704 | 11 | 0.81% |
| hsa_circ_0123321 | 2704 | 11 | 0.81% |
| hsa_circ_0015852 | 2950 | 12 | 0.81% |
| hsa_circ_0044043 | 2950 | 12 | 0.81% |
| hsa_circ_0077232 | 2950 | 12 | 0.81% |
| hsa_circ_0089867 | 2950 | 12 | 0.81% |
| hsa_circ_0088303 | 3196 | 13 | 0.81% |
| hsa_circ_0088937 | 3196 | 13 | 0.81% |
| hsa_circ_0027342 | 2459 | 10 | 0.81% |
| hsa_circ_0061804 | 2459 | 10 | 0.81% |
| hsa_circ_0070328 | 2705 | 11 | 0.81% |
| hsa_circ_0093999 | 2705 | 11 | 0.81% |
| hsa_circ_0110718 | 2705 | 11 | 0.81% |
| hsa_circ_0046755 | 2951 | 12 | 0.81% |
| hsa_circ_0066334 | 2951 | 12 | 0.81% |
| hsa_circ_0138864 | 40086 | 163 | 0.81% |
| hsa_circ_0004192 | 9100 | 37 | 0.81% |
| hsa_circ_0017558 | 2952 | 12 | 0.81% |
| hsa_circ_0023827 | 2952 | 12 | 0.81% |
| hsa_circ_0079582 | 2952 | 12 | 0.81% |
| hsa_circ_0104426 | 2952 | 12 | 0.81% |
| hsa_circ_0019707 | 2706 | 11 | 0.81% |
| hsa_circ_0033103 | 2706 | 11 | 0.81% |
| hsa_circ_0040336 | 2706 | 11 | 0.81% |
| hsa_circ_0058114 | 2706 | 11 | 0.81% |
| hsa_circ_0011797 | 2460 | 10 | 0.81% |
| hsa_circ_0015691 | 2460 | 10 | 0.81% |
| hsa_circ_0020426 | 2460 | 10 | 0.81% |
| hsa_circ_0032867 | 2953 | 12 | 0.81% |
| hsa_circ_0131996 | 2953 | 12 | 0.81% |

| | | | |
|---|---|---|---|
| hsa_circ_0031338 | 2707 | 11 | 0.81% |
| hsa_circ_0038259 | 2707 | 11 | 0.81% |
| hsa_circ_0080233 | 2707 | 11 | 0.81% |
| hsa_circ_0086315 | 2461 | 10 | 0.81% |
| hsa_circ_0100946 | 2461 | 10 | 0.81% |
| hsa_circ_0124515 | 2461 | 10 | 0.81% |
| hsa_circ_0081693 | 5415 | 22 | 0.81% |
| hsa_circ_0022162 | 3200 | 13 | 0.81% |
| hsa_circ_0055140 | 3200 | 13 | 0.81% |
| hsa_circ_0010082 | 2954 | 12 | 0.81% |
| hsa_circ_0018633 | 2954 | 12 | 0.81% |
| hsa_circ_0058899 | 2954 | 12 | 0.81% |
| hsa_circ_0030476 | 2708 | 11 | 0.81% |
| hsa_circ_0043885 | 2708 | 11 | 0.81% |
| hsa_circ_0111458 | 2708 | 11 | 0.81% |
| hsa_circ_0058087 | 2462 | 10 | 0.81% |
| hsa_circ_0024475 | 3201 | 13 | 0.81% |
| hsa_circ_0030449 | 2955 | 12 | 0.81% |
| hsa_circ_0038212 | 2709 | 11 | 0.81% |
| hsa_circ_0038604 | 2709 | 11 | 0.81% |
| hsa_circ_0045032 | 2709 | 11 | 0.81% |
| hsa_circ_0066606 | 2709 | 11 | 0.81% |
| hsa_circ_0076213 | 2709 | 11 | 0.81% |
| hsa_circ_0085192 | 2709 | 11 | 0.81% |
| hsa_circ_0029918 | 2463 | 10 | 0.81% |
| hsa_circ_0072905 | 2463 | 10 | 0.81% |
| hsa_circ_0116670 | 5665 | 23 | 0.81% |
| hsa_circ_0095926 | 3695 | 15 | 0.81% |
| hsa_circ_0035279 | 2956 | 12 | 0.81% |
| hsa_circ_0035593 | 2710 | 11 | 0.81% |
| hsa_circ_0041511 | 2710 | 11 | 0.81% |
| hsa_circ_0014789 | 9855 | 40 | 0.81% |
| hsa_circ_0049245 | 3696 | 15 | 0.81% |
| hsa_circ_0101458 | 2464 | 10 | 0.81% |
| hsa_circ_0050013 | 2957 | 12 | 0.81% |
| hsa_circ_0061453 | 3943 | 16 | 0.81% |
| hsa_circ_0037802 | 2711 | 11 | 0.81% |
| hsa_circ_0051487 | 2711 | 11 | 0.81% |
| hsa_circ_0091703 | 2711 | 11 | 0.81% |
| hsa_circ_0022827 | 3204 | 13 | 0.81% |
| hsa_circ_0060642 | 3204 | 13 | 0.81% |
| hsa_circ_0084539 | 3204 | 13 | 0.81% |
| hsa_circ_0055129 | 3697 | 15 | 0.81% |
| hsa_circ_0011320 | 2958 | 12 | 0.81% |
| hsa_circ_0018371 | 2958 | 12 | 0.81% |
| hsa_circ_0106051 | 2958 | 12 | 0.81% |
| hsa_circ_0014553 | 2465 | 10 | 0.81% |
| hsa_circ_0068427 | 3205 | 13 | 0.81% |
| hsa_circ_0004729 | 2712 | 11 | 0.81% |
| hsa_circ_0009823 | 2712 | 11 | 0.81% |

| | | | |
|---|---|---|---|
| hsa_circ_0055659 | 2712 | 11 | 0.81% |
| hsa_circ_0116968 | 2712 | 11 | 0.81% |
| hsa_circ_0004475 | 4931 | 20 | 0.81% |
| hsa_circ_0119520 | 7150 | 29 | 0.81% |
| hsa_circ_0050082 | 10109 | 41 | 0.81% |
| hsa_circ_0138155 | 5671 | 23 | 0.81% |
| hsa_circ_0122787 | 18246 | 74 | 0.81% |
| hsa_circ_0011051 | 2959 | 12 | 0.81% |
| hsa_circ_0079848 | 2959 | 12 | 0.81% |
| hsa_circ_0041072 | 2466 | 10 | 0.81% |
| hsa_circ_0042216 | 2466 | 10 | 0.81% |
| hsa_circ_0061795 | 2466 | 10 | 0.81% |
| hsa_circ_0075812 | 2466 | 10 | 0.81% |
| hsa_circ_0112182 | 2466 | 10 | 0.81% |
| hsa_circ_0037817 | 2713 | 11 | 0.81% |
| hsa_circ_0060448 | 2713 | 11 | 0.81% |
| hsa_circ_0091380 | 2713 | 11 | 0.81% |
| hsa_circ_0116022 | 2713 | 11 | 0.81% |
| hsa_circ_0122177 | 2713 | 11 | 0.81% |
| hsa_circ_0055301 | 2960 | 12 | 0.81% |
| hsa_circ_0065098 | 2960 | 12 | 0.81% |
| hsa_circ_0120879 | 2467 | 10 | 0.81% |
| hsa_circ_0018171 | 2714 | 11 | 0.81% |
| hsa_circ_0042423 | 2961 | 12 | 0.81% |
| hsa_circ_0112166 | 2961 | 12 | 0.81% |
| hsa_circ_0137421 | 2961 | 12 | 0.81% |
| hsa_circ_0016216 | 2468 | 10 | 0.81% |
| hsa_circ_0057646 | 2468 | 10 | 0.81% |
| hsa_circ_0019837 | 2715 | 11 | 0.81% |
| hsa_circ_0067421 | 2715 | 11 | 0.81% |
| hsa_circ_0036897 | 2962 | 12 | 0.81% |
| hsa_circ_0060326 | 2962 | 12 | 0.81% |
| hsa_circ_0064327 | 2962 | 12 | 0.81% |
| hsa_circ_0110006 | 2962 | 12 | 0.81% |
| hsa_circ_0092764 | 3209 | 13 | 0.81% |
| hsa_circ_0020471 | 2469 | 10 | 0.81% |
| hsa_circ_0019700 | 2716 | 11 | 0.81% |
| hsa_circ_0030619 | 2716 | 11 | 0.81% |
| hsa_circ_0033366 | 2716 | 11 | 0.81% |
| hsa_circ_0088704 | 2716 | 11 | 0.81% |
| hsa_circ_0107552 | 2716 | 11 | 0.81% |
| hsa_circ_0041688 | 2963 | 12 | 0.81% |
| hsa_circ_0041748 | 2963 | 12 | 0.81% |
| hsa_circ_0035762 | 3210 | 13 | 0.81% |
| hsa_circ_0035644 | 4939 | 20 | 0.81% |
| hsa_circ_0088872 | 2964 | 12 | 0.81% |
| hsa_circ_0003885 | 2717 | 11 | 0.81% |
| hsa_circ_0014883 | 2717 | 11 | 0.81% |
| hsa_circ_0025083 | 2717 | 11 | 0.81% |
| hsa_circ_0033110 | 2717 | 11 | 0.81% |

| | | | |
|---|---|---|---|
| hsa_circ_0035069 | 2717 | 11 | 0.81% |
| hsa_circ_0058667 | 2717 | 11 | 0.81% |
| hsa_circ_0063069 | 2717 | 11 | 0.81% |
| hsa_circ_0044102 | 2470 | 10 | 0.81% |
| hsa_circ_0089112 | 2965 | 12 | 0.81% |
| hsa_circ_0011197 | 2718 | 11 | 0.81% |
| hsa_circ_0025967 | 2718 | 11 | 0.81% |
| hsa_circ_0035586 | 2718 | 11 | 0.81% |
| hsa_circ_0053571 | 2718 | 11 | 0.81% |
| hsa_circ_0076789 | 2718 | 11 | 0.81% |
| hsa_circ_0087670 | 2718 | 11 | 0.81% |
| hsa_circ_0129987 | 2718 | 11 | 0.81% |
| hsa_circ_0019447 | 2471 | 10 | 0.81% |
| hsa_circ_0078396 | 2471 | 10 | 0.81% |
| hsa_circ_0027900 | 3460 | 14 | 0.81% |
| hsa_circ_0108457 | 3460 | 14 | 0.81% |
| hsa_circ_0035777 | 3213 | 13 | 0.81% |
| hsa_circ_0058898 | 2966 | 12 | 0.81% |
| hsa_circ_0019703 | 2719 | 11 | 0.81% |
| hsa_circ_0071895 | 2719 | 11 | 0.81% |
| hsa_circ_0086249 | 2719 | 11 | 0.81% |
| hsa_circ_0063313 | 3955 | 16 | 0.81% |
| hsa_circ_0025421 | 2472 | 10 | 0.81% |
| hsa_circ_0042003 | 2472 | 10 | 0.81% |
| hsa_circ_0097728 | 2472 | 10 | 0.81% |
| hsa_circ_0135967 | 2472 | 10 | 0.81% |
| hsa_circ_0028062 | 2967 | 12 | 0.81% |
| hsa_circ_0100900 | 2967 | 12 | 0.81% |
| hsa_circ_0031558 | 2720 | 11 | 0.81% |
| hsa_circ_0085244 | 2720 | 11 | 0.81% |
| hsa_circ_0127883 | 2720 | 11 | 0.81% |
| hsa_circ_0140402 | 2720 | 11 | 0.81% |
| hsa_circ_0059303 | 2473 | 10 | 0.81% |
| hsa_circ_0053661 | 3215 | 13 | 0.81% |
| hsa_circ_0056030 | 2968 | 12 | 0.81% |
| hsa_circ_0100494 | 2968 | 12 | 0.81% |
| hsa_circ_0068561 | 2721 | 11 | 0.81% |
| hsa_circ_0070841 | 3216 | 13 | 0.81% |
| hsa_circ_0053849 | 2474 | 10 | 0.81% |
| hsa_circ_0084647 | 2969 | 12 | 0.81% |
| hsa_circ_0136463 | 4950 | 20 | 0.81% |
| hsa_circ_0140528 | 4455 | 18 | 0.81% |
| hsa_circ_0090105 | 3465 | 14 | 0.81% |
| hsa_circ_0046139 | 2970 | 12 | 0.81% |
| hsa_circ_0062221 | 2970 | 12 | 0.81% |
| hsa_circ_0073793 | 2970 | 12 | 0.81% |
| hsa_circ_0117912 | 2970 | 12 | 0.81% |
| hsa_circ_0014891 | 2475 | 10 | 0.81% |
| hsa_circ_0024984 | 2475 | 10 | 0.81% |
| hsa_circ_0035798 | 2475 | 10 | 0.81% |

| | | | |
|---|---|---|---|
| hsa_circ_0064228 | 2475 | 10 | 0.81% |
| hsa_circ_0071781 | 2475 | 10 | 0.81% |
| hsa_circ_0085735 | 2723 | 11 | 0.81% |
| hsa_circ_0009495 | 3466 | 14 | 0.81% |
| hsa_circ_0058314 | 2971 | 12 | 0.81% |
| hsa_circ_0065767 | 2476 | 10 | 0.81% |
| hsa_circ_0090701 | 2476 | 10 | 0.81% |
| hsa_circ_0035068 | 3219 | 13 | 0.81% |
| hsa_circ_0047775 | 3219 | 13 | 0.81% |
| hsa_circ_0020862 | 2724 | 11 | 0.81% |
| hsa_circ_0091211 | 2724 | 11 | 0.81% |
| hsa_circ_0038500 | 5696 | 23 | 0.81% |
| hsa_circ_0054985 | 2972 | 12 | 0.81% |
| hsa_circ_0125694 | 2972 | 12 | 0.81% |
| hsa_circ_0122643 | 3220 | 13 | 0.81% |
| hsa_circ_0065024 | 4954 | 20 | 0.81% |
| hsa_circ_0035519 | 2477 | 10 | 0.81% |
| hsa_circ_0055111 | 2477 | 10 | 0.81% |
| hsa_circ_0071165 | 2477 | 10 | 0.81% |
| hsa_circ_0078137 | 2477 | 10 | 0.81% |
| hsa_circ_0062039 | 2725 | 11 | 0.81% |
| hsa_circ_0084347 | 2725 | 11 | 0.81% |
| hsa_circ_0093300 | 2725 | 11 | 0.81% |
| hsa_circ_0114799 | 2725 | 11 | 0.81% |
| hsa_circ_0037996 | 2973 | 12 | 0.81% |
| hsa_circ_0067360 | 2973 | 12 | 0.81% |
| hsa_circ_0062847 | 3221 | 13 | 0.81% |
| hsa_circ_0017840 | 2478 | 10 | 0.81% |
| hsa_circ_0045515 | 2478 | 10 | 0.81% |
| hsa_circ_0090639 | 2478 | 10 | 0.81% |
| hsa_circ_0098632 | 2478 | 10 | 0.81% |
| hsa_circ_0128885 | 2478 | 10 | 0.81% |
| hsa_circ_0132654 | 2478 | 10 | 0.81% |
| hsa_circ_0016317 | 2726 | 11 | 0.81% |
| hsa_circ_0087446 | 2726 | 11 | 0.81% |
| hsa_circ_0136726 | 2726 | 11 | 0.81% |
| hsa_circ_0136806 | 2974 | 12 | 0.81% |
| hsa_circ_0083751 | 3222 | 13 | 0.81% |
| hsa_circ_0104558 | 20576 | 83 | 0.81% |
| hsa_circ_0034161 | 2727 | 11 | 0.81% |
| hsa_circ_0035497 | 2727 | 11 | 0.81% |
| hsa_circ_0037663 | 2975 | 12 | 0.81% |
| hsa_circ_0045467 | 2975 | 12 | 0.81% |
| hsa_circ_0112247 | 2975 | 12 | 0.81% |
| hsa_circ_0065095 | 4463 | 18 | 0.81% |
| hsa_circ_0115049 | 5455 | 22 | 0.81% |
| hsa_circ_0076921 | 4712 | 19 | 0.81% |
| hsa_circ_0064530 | 2728 | 11 | 0.81% |
| hsa_circ_0089576 | 2728 | 11 | 0.81% |
| hsa_circ_0111385 | 2728 | 11 | 0.81% |

| | | | |
|---|---|---|---|
| hsa_circ_0054364 | 2480 | 10 | 0.81% |
| hsa_circ_0114807 | 2480 | 10 | 0.81% |
| hsa_circ_0138478 | 2480 | 10 | 0.81% |
| hsa_circ_0140540 | 2480 | 10 | 0.81% |
| hsa_circ_0002497 | 6697 | 27 | 0.81% |
| hsa_circ_0082051 | 3225 | 13 | 0.81% |
| hsa_circ_0065433 | 2977 | 12 | 0.81% |
| hsa_circ_0083462 | 2729 | 11 | 0.81% |
| hsa_circ_0134181 | 2729 | 11 | 0.81% |
| hsa_circ_0046800 | 2481 | 10 | 0.81% |
| hsa_circ_0053841 | 2481 | 10 | 0.81% |
| hsa_circ_0092058 | 2481 | 10 | 0.81% |
| hsa_circ_0102349 | 2481 | 10 | 0.81% |
| hsa_circ_0123103 | 2481 | 10 | 0.81% |
| hsa_circ_0105819 | 10422 | 42 | 0.81% |
| hsa_circ_0032553 | 3226 | 13 | 0.81% |
| hsa_circ_0043987 | 2730 | 11 | 0.81% |
| hsa_circ_0097358 | 7694 | 31 | 0.81% |
| hsa_circ_0014371 | 2482 | 10 | 0.81% |
| hsa_circ_0045011 | 2482 | 10 | 0.81% |
| hsa_circ_0072136 | 2482 | 10 | 0.81% |
| hsa_circ_0119979 | 2482 | 10 | 0.81% |
| hsa_circ_0134672 | 2482 | 10 | 0.81% |
| hsa_circ_0036155 | 2979 | 12 | 0.81% |
| hsa_circ_0044044 | 2979 | 12 | 0.81% |
| hsa_circ_0035308 | 2731 | 11 | 0.81% |
| hsa_circ_0043703 | 2731 | 11 | 0.81% |
| hsa_circ_0080588 | 2731 | 11 | 0.81% |
| hsa_circ_0015589 | 2483 | 10 | 0.81% |
| hsa_circ_0072404 | 2483 | 10 | 0.81% |
| hsa_circ_0042277 | 2980 | 12 | 0.81% |
| hsa_circ_0139378 | 7699 | 31 | 0.81% |
| hsa_circ_0024899 | 2732 | 11 | 0.81% |
| hsa_circ_0063218 | 12667 | 51 | 0.81% |
| hsa_circ_0100993 | 3974 | 16 | 0.81% |
| hsa_circ_0020806 | 3229 | 13 | 0.81% |
| hsa_circ_0024976 | 2484 | 10 | 0.81% |
| hsa_circ_0077634 | 2484 | 10 | 0.81% |
| hsa_circ_0139706 | 12669 | 51 | 0.81% |
| hsa_circ_0053186 | 2981 | 12 | 0.81% |
| hsa_circ_0076517 | 2981 | 12 | 0.81% |
| hsa_circ_0085371 | 2981 | 12 | 0.81% |
| hsa_circ_0134513 | 3478 | 14 | 0.81% |
| hsa_circ_0061459 | 3975 | 16 | 0.81% |
| hsa_circ_0094345 | 3975 | 16 | 0.81% |
| hsa_circ_0012983 | 2733 | 11 | 0.80% |
| hsa_circ_0021003 | 2733 | 11 | 0.80% |
| hsa_circ_0065335 | 2733 | 11 | 0.80% |
| hsa_circ_0075160 | 2733 | 11 | 0.80% |
| hsa_circ_0136356 | 2733 | 11 | 0.80% |

| | | | |
|---|---|---|---|
| hsa_circ_0087255 | 2982 | 12 | 0.80% |
| hsa_circ_0026832 | 2485 | 10 | 0.80% |
| hsa_circ_0031567 | 2485 | 10 | 0.80% |
| hsa_circ_0082371 | 2485 | 10 | 0.80% |
| hsa_circ_0139377 | 7704 | 31 | 0.80% |
| hsa_circ_0011165 | 2734 | 11 | 0.80% |
| hsa_circ_0080138 | 2734 | 11 | 0.80% |
| hsa_circ_0033001 | 2983 | 12 | 0.80% |
| hsa_circ_0070579 | 2983 | 12 | 0.80% |
| hsa_circ_0074133 | 2983 | 12 | 0.80% |
| hsa_circ_0078618 | 2983 | 12 | 0.80% |
| hsa_circ_0042983 | 3232 | 13 | 0.80% |
| hsa_circ_0140439 | 8205 | 33 | 0.80% |
| hsa_circ_0036476 | 2735 | 11 | 0.80% |
| hsa_circ_0059716 | 2735 | 11 | 0.80% |
| hsa_circ_0079250 | 2735 | 11 | 0.80% |
| hsa_circ_0126616 | 2735 | 11 | 0.80% |
| hsa_circ_0086413 | 2487 | 10 | 0.80% |
| hsa_circ_0062369 | 2736 | 11 | 0.80% |
| hsa_circ_0086922 | 2736 | 11 | 0.80% |
| hsa_circ_0091971 | 2736 | 11 | 0.80% |
| hsa_circ_0129555 | 2736 | 11 | 0.80% |
| hsa_circ_0069731 | 2985 | 12 | 0.80% |
| hsa_circ_0103773 | 2985 | 12 | 0.80% |
| hsa_circ_0014374 | 2488 | 10 | 0.80% |
| hsa_circ_0019304 | 2488 | 10 | 0.80% |
| hsa_circ_0049503 | 2488 | 10 | 0.80% |
| hsa_circ_0086427 | 2488 | 10 | 0.80% |
| hsa_circ_0069665 | 2737 | 11 | 0.80% |
| hsa_circ_0071864 | 2737 | 11 | 0.80% |
| hsa_circ_0131161 | 2737 | 11 | 0.80% |
| hsa_circ_0129168 | 2986 | 12 | 0.80% |
| hsa_circ_0030500 | 3733 | 15 | 0.80% |
| hsa_circ_0045786 | 11200 | 45 | 0.80% |
| hsa_circ_0015502 | 2489 | 10 | 0.80% |
| hsa_circ_0049615 | 2489 | 10 | 0.80% |
| hsa_circ_0067550 | 2489 | 10 | 0.80% |
| hsa_circ_0020397 | 2738 | 11 | 0.80% |
| hsa_circ_0049940 | 2738 | 11 | 0.80% |
| hsa_circ_0079247 | 2738 | 11 | 0.80% |
| hsa_circ_0117353 | 2738 | 11 | 0.80% |
| hsa_circ_0068830 | 2987 | 12 | 0.80% |
| hsa_circ_0137328 | 2987 | 12 | 0.80% |
| hsa_circ_0096471 | 3236 | 13 | 0.80% |
| hsa_circ_0041409 | 5477 | 22 | 0.80% |
| hsa_circ_0101391 | 5477 | 22 | 0.80% |
| hsa_circ_0030554 | 3735 | 15 | 0.80% |
| hsa_circ_0127686 | 3237 | 13 | 0.80% |
| hsa_circ_0048188 | 2988 | 12 | 0.80% |
| hsa_circ_0056904 | 2988 | 12 | 0.80% |

| | | | |
|---|---|---|---|
| hsa_circ_0059426 | 2988 | 12 | 0.80% |
| hsa_circ_0129420 | 2988 | 12 | 0.80% |
| hsa_circ_0024228 | 2739 | 11 | 0.80% |
| hsa_circ_0039274 | 2490 | 10 | 0.80% |
| hsa_circ_0094549 | 9714 | 39 | 0.80% |
| hsa_circ_0028034 | 3238 | 13 | 0.80% |
| hsa_circ_0101685 | 2740 | 11 | 0.80% |
| hsa_circ_0113099 | 2740 | 11 | 0.80% |
| hsa_circ_0127886 | 2740 | 11 | 0.80% |
| hsa_circ_0086412 | 2491 | 10 | 0.80% |
| hsa_circ_0087262 | 2491 | 10 | 0.80% |
| hsa_circ_0120906 | 2491 | 10 | 0.80% |
| hsa_circ_0065106 | 2990 | 12 | 0.80% |
| hsa_circ_0067552 | 2990 | 12 | 0.80% |
| hsa_circ_0043007 | 2741 | 11 | 0.80% |
| hsa_circ_0053482 | 2741 | 11 | 0.80% |
| hsa_circ_0083798 | 2741 | 11 | 0.80% |
| hsa_circ_0138577 | 2741 | 11 | 0.80% |
| hsa_circ_0055012 | 2492 | 10 | 0.80% |
| hsa_circ_0078615 | 2492 | 10 | 0.80% |
| hsa_circ_0081265 | 2492 | 10 | 0.80% |
| hsa_circ_0017996 | 3489 | 14 | 0.80% |
| hsa_circ_0069358 | 3240 | 13 | 0.80% |
| hsa_circ_0019104 | 2991 | 12 | 0.80% |
| hsa_circ_0030212 | 2991 | 12 | 0.80% |
| hsa_circ_0045085 | 2991 | 12 | 0.80% |
| hsa_circ_0066836 | 2991 | 12 | 0.80% |
| hsa_circ_0125966 | 2991 | 12 | 0.80% |
| hsa_circ_0050670 | 2742 | 11 | 0.80% |
| hsa_circ_0059256 | 2742 | 11 | 0.80% |
| hsa_circ_0060961 | 2742 | 11 | 0.80% |
| hsa_circ_0065415 | 2742 | 11 | 0.80% |
| hsa_circ_0084366 | 2742 | 11 | 0.80% |
| hsa_circ_0087866 | 2742 | 11 | 0.80% |
| hsa_circ_0110096 | 2742 | 11 | 0.80% |
| hsa_circ_0036323 | 3490 | 14 | 0.80% |
| hsa_circ_0016469 | 2493 | 10 | 0.80% |
| hsa_circ_0091206 | 2493 | 10 | 0.80% |
| hsa_circ_0118376 | 2493 | 10 | 0.80% |
| hsa_circ_0124307 | 2493 | 10 | 0.80% |
| hsa_circ_0032000 | 3241 | 13 | 0.80% |
| hsa_circ_0076614 | 7729 | 31 | 0.80% |
| hsa_circ_0091983 | 2992 | 12 | 0.80% |
| hsa_circ_0114906 | 19199 | 77 | 0.80% |
| hsa_circ_0051198 | 2743 | 11 | 0.80% |
| hsa_circ_0062940 | 2743 | 11 | 0.80% |
| hsa_circ_0083407 | 3242 | 13 | 0.80% |
| hsa_circ_0027832 | 2494 | 10 | 0.80% |
| hsa_circ_0036125 | 2494 | 10 | 0.80% |
| hsa_circ_0119541 | 2494 | 10 | 0.80% |

| | | | |
|---|---|---|---|
| hsa_circ_0001146 | 5487 | 22 | 0.80% |
| hsa_circ_0099073 | 2993 | 12 | 0.80% |
| hsa_circ_0110487 | 3492 | 14 | 0.80% |
| hsa_circ_0112931 | 7982 | 32 | 0.80% |
| hsa_circ_0020601 | 2744 | 11 | 0.80% |
| hsa_circ_0062095 | 2744 | 11 | 0.80% |
| hsa_circ_0073418 | 2744 | 11 | 0.80% |
| hsa_circ_0090145 | 2744 | 11 | 0.80% |
| hsa_circ_0109971 | 7484 | 30 | 0.80% |
| hsa_circ_0119463 | 3493 | 14 | 0.80% |
| hsa_circ_0071335 | 2495 | 10 | 0.80% |
| hsa_circ_0081550 | 2495 | 10 | 0.80% |
| hsa_circ_0107551 | 2745 | 11 | 0.80% |
| hsa_circ_0035838 | 3494 | 14 | 0.80% |
| hsa_circ_0060612 | 2995 | 12 | 0.80% |
| hsa_circ_0082198 | 2995 | 12 | 0.80% |
| hsa_circ_0089181 | 2995 | 12 | 0.80% |
| hsa_circ_0039227 | 2496 | 10 | 0.80% |
| hsa_circ_0047976 | 2496 | 10 | 0.80% |
| hsa_circ_0030662 | 2746 | 11 | 0.80% |
| hsa_circ_0074315 | 2746 | 11 | 0.80% |
| hsa_circ_0083342 | 2746 | 11 | 0.80% |
| hsa_circ_0122279 | 2746 | 11 | 0.80% |
| hsa_circ_0024848 | 2996 | 12 | 0.80% |
| hsa_circ_0082128 | 2996 | 12 | 0.80% |
| hsa_circ_0063217 | 12736 | 51 | 0.80% |
| hsa_circ_0020333 | 2747 | 11 | 0.80% |
| hsa_circ_0075978 | 2747 | 11 | 0.80% |
| hsa_circ_0060629 | 2997 | 12 | 0.80% |
| hsa_circ_0068178 | 2997 | 12 | 0.80% |
| hsa_circ_0073531 | 2997 | 12 | 0.80% |
| hsa_circ_0073912 | 2997 | 12 | 0.80% |
| hsa_circ_0080790 | 3747 | 15 | 0.80% |
| hsa_circ_0054308 | 2498 | 10 | 0.80% |
| hsa_circ_0094429 | 2498 | 10 | 0.80% |
| hsa_circ_0116007 | 2498 | 10 | 0.80% |
| hsa_circ_0091813 | 3498 | 14 | 0.80% |
| hsa_circ_0000562 | 2499 | 10 | 0.80% |
| hsa_circ_0054047 | 2499 | 10 | 0.80% |
| hsa_circ_0071954 | 2499 | 10 | 0.80% |
| hsa_circ_0083362 | 2499 | 10 | 0.80% |
| hsa_circ_0079712 | 2749 | 11 | 0.80% |
| hsa_circ_0028164 | 3249 | 13 | 0.80% |
| hsa_circ_0128864 | 3499 | 14 | 0.80% |
| hsa_circ_0029401 | 20248 | 81 | 0.80% |
| hsa_circ_0044193 | 11750 | 47 | 0.80% |
| hsa_circ_0058592 | 3250 | 13 | 0.80% |
| hsa_circ_0041299 | 2750 | 11 | 0.80% |
| hsa_circ_0074021 | 2750 | 11 | 0.80% |
| hsa_circ_0080486 | 2500 | 10 | 0.80% |

| | | | |
|---|---|---|---|
| hsa_circ_0128139 | 2500 | 10 | 0.80% |
| hsa_circ_0036423 | 3251 | 13 | 0.80% |
| hsa_circ_0002235 | 3001 | 12 | 0.80% |
| hsa_circ_0019018 | 3001 | 12 | 0.80% |
| hsa_circ_0062891 | 3001 | 12 | 0.80% |
| hsa_circ_0052558 | 2751 | 11 | 0.80% |
| hsa_circ_0082924 | 2751 | 11 | 0.80% |
| hsa_circ_0110801 | 2751 | 11 | 0.80% |
| hsa_circ_0114002 | 2751 | 11 | 0.80% |
| hsa_circ_0020138 | 2501 | 10 | 0.80% |
| hsa_circ_0074303 | 2501 | 10 | 0.80% |
| hsa_circ_0047809 | 3252 | 13 | 0.80% |
| hsa_circ_0099172 | 4253 | 17 | 0.80% |
| hsa_circ_0015971 | 2502 | 10 | 0.80% |
| hsa_circ_0071219 | 2502 | 10 | 0.80% |
| hsa_circ_0030266 | 3503 | 14 | 0.80% |
| hsa_circ_0097535 | 8008 | 32 | 0.80% |
| hsa_circ_0039929 | 3003 | 12 | 0.80% |
| hsa_circ_0040793 | 3003 | 12 | 0.80% |
| hsa_circ_0092091 | 2753 | 11 | 0.80% |
| hsa_circ_0046615 | 11763 | 47 | 0.80% |
| hsa_circ_0069016 | 2503 | 10 | 0.80% |
| hsa_circ_0092832 | 2503 | 10 | 0.80% |
| hsa_circ_0062037 | 3004 | 12 | 0.80% |
| hsa_circ_0019327 | 2754 | 11 | 0.80% |
| hsa_circ_0061280 | 2754 | 11 | 0.80% |
| hsa_circ_0068317 | 2754 | 11 | 0.80% |
| hsa_circ_0129769 | 2754 | 11 | 0.80% |
| hsa_circ_0062518 | 3255 | 13 | 0.80% |
| hsa_circ_0056153 | 2504 | 10 | 0.80% |
| hsa_circ_0066448 | 2504 | 10 | 0.80% |
| hsa_circ_0066577 | 2504 | 10 | 0.80% |
| hsa_circ_0084969 | 2504 | 10 | 0.80% |
| hsa_circ_0088321 | 2504 | 10 | 0.80% |
| hsa_circ_0101326 | 2504 | 10 | 0.80% |
| hsa_circ_0131664 | 8013 | 32 | 0.80% |
| hsa_circ_0096351 | 6010 | 24 | 0.80% |
| hsa_circ_0057978 | 3005 | 12 | 0.80% |
| hsa_circ_0012590 | 2755 | 11 | 0.80% |
| hsa_circ_0057683 | 2755 | 11 | 0.80% |
| hsa_circ_0100280 | 2755 | 11 | 0.80% |
| hsa_circ_0107550 | 2755 | 11 | 0.80% |
| hsa_circ_0128881 | 2755 | 11 | 0.80% |
| hsa_circ_0036344 | 3006 | 12 | 0.80% |
| hsa_circ_0082642 | 3006 | 12 | 0.80% |
| hsa_circ_0082815 | 3006 | 12 | 0.80% |
| hsa_circ_0099214 | 3006 | 12 | 0.80% |
| hsa_circ_0037799 | 2505 | 10 | 0.80% |
| hsa_circ_0129497 | 2505 | 10 | 0.80% |
| hsa_circ_0048874 | 3257 | 13 | 0.80% |

| | | | |
|---|---|---|---|
| hsa_circ_0021964 | 2756 | 11 | 0.80% |
| hsa_circ_0026716 | 2756 | 11 | 0.80% |
| hsa_circ_0016696 | 4009 | 16 | 0.80% |
| hsa_circ_0015810 | 3007 | 12 | 0.80% |
| hsa_circ_0078534 | 3007 | 12 | 0.80% |
| hsa_circ_0088873 | 3007 | 12 | 0.80% |
| hsa_circ_0077478 | 2506 | 10 | 0.80% |
| hsa_circ_0105597 | 6516 | 26 | 0.80% |
| hsa_circ_0019544 | 3258 | 13 | 0.80% |
| hsa_circ_0031617 | 3258 | 13 | 0.80% |
| hsa_circ_0049709 | 3258 | 13 | 0.80% |
| hsa_circ_0067622 | 3258 | 13 | 0.80% |
| hsa_circ_0090362 | 2757 | 11 | 0.80% |
| hsa_circ_0062807 | 3509 | 14 | 0.80% |
| hsa_circ_0106391 | 9525 | 38 | 0.80% |
| hsa_circ_0068947 | 3008 | 12 | 0.80% |
| hsa_circ_0105394 | 4011 | 16 | 0.80% |
| hsa_circ_0050138 | 2507 | 10 | 0.80% |
| hsa_circ_0060241 | 2507 | 10 | 0.80% |
| hsa_circ_0043944 | 2758 | 11 | 0.80% |
| hsa_circ_0065660 | 2758 | 11 | 0.80% |
| hsa_circ_0069940 | 2758 | 11 | 0.80% |
| hsa_circ_0082022 | 2758 | 11 | 0.80% |
| hsa_circ_0136346 | 2758 | 11 | 0.80% |
| hsa_circ_0009313 | 6018 | 24 | 0.80% |
| hsa_circ_0016641 | 3009 | 12 | 0.80% |
| hsa_circ_0093474 | 3009 | 12 | 0.80% |
| hsa_circ_0009836 | 2508 | 10 | 0.80% |
| hsa_circ_0134610 | 2508 | 10 | 0.80% |
| hsa_circ_0099362 | 4013 | 16 | 0.80% |
| hsa_circ_0038745 | 2759 | 11 | 0.80% |
| hsa_circ_0077999 | 2759 | 11 | 0.80% |
| hsa_circ_0080566 | 2759 | 11 | 0.80% |
| hsa_circ_0107553 | 2759 | 11 | 0.80% |
| hsa_circ_0048151 | 3010 | 12 | 0.80% |
| hsa_circ_0060394 | 3010 | 12 | 0.80% |
| hsa_circ_0088171 | 3261 | 13 | 0.80% |
| hsa_circ_0046645 | 11791 | 47 | 0.80% |
| hsa_circ_0090591 | 2509 | 10 | 0.80% |
| hsa_circ_0074724 | 2760 | 11 | 0.80% |
| hsa_circ_0047074 | 3011 | 12 | 0.80% |
| hsa_circ_0073882 | 3011 | 12 | 0.80% |
| hsa_circ_0005630 | 3262 | 13 | 0.80% |
| hsa_circ_0035763 | 3262 | 13 | 0.80% |
| hsa_circ_0084393 | 3262 | 13 | 0.80% |
| hsa_circ_0087345 | 3262 | 13 | 0.80% |
| hsa_circ_0018069 | 3764 | 15 | 0.80% |
| hsa_circ_0067929 | 3764 | 15 | 0.80% |
| hsa_circ_0123194 | 5270 | 21 | 0.80% |
| hsa_circ_0045646 | 3263 | 13 | 0.80% |

| | | | |
|---|---|---|---|
| hsa_circ_0080424 | 3012 | 12 | 0.80% |
| hsa_circ_0122798 | 3012 | 12 | 0.80% |
| hsa_circ_0050358 | 2761 | 11 | 0.80% |
| hsa_circ_0064512 | 2761 | 11 | 0.80% |
| hsa_circ_0113575 | 2761 | 11 | 0.80% |
| hsa_circ_0025430 | 2510 | 10 | 0.80% |
| hsa_circ_0033391 | 2510 | 10 | 0.80% |
| hsa_circ_0013129 | 3515 | 14 | 0.80% |
| hsa_circ_0048361 | 3515 | 14 | 0.80% |
| hsa_circ_0060704 | 3264 | 13 | 0.80% |
| hsa_circ_0008387 | 3013 | 12 | 0.80% |
| hsa_circ_0045941 | 3013 | 12 | 0.80% |
| hsa_circ_0054712 | 2762 | 11 | 0.80% |
| hsa_circ_0084352 | 2762 | 11 | 0.80% |
| hsa_circ_0023135 | 2511 | 10 | 0.80% |
| hsa_circ_0027214 | 2511 | 10 | 0.80% |
| hsa_circ_0028226 | 2511 | 10 | 0.80% |
| hsa_circ_0036829 | 2511 | 10 | 0.80% |
| hsa_circ_0114907 | 19335 | 77 | 0.80% |
| hsa_circ_0103063 | 9041 | 36 | 0.80% |
| hsa_circ_0039642 | 3265 | 13 | 0.80% |
| hsa_circ_0042037 | 2763 | 11 | 0.80% |
| hsa_circ_0065718 | 2763 | 11 | 0.80% |
| hsa_circ_0128919 | 2763 | 11 | 0.80% |
| hsa_circ_0137346 | 2763 | 11 | 0.80% |
| hsa_circ_0112603 | 2512 | 10 | 0.80% |
| hsa_circ_0042130 | 5025 | 20 | 0.80% |
| hsa_circ_0045918 | 3015 | 12 | 0.80% |
| hsa_circ_0056358 | 2764 | 11 | 0.80% |
| hsa_circ_0057583 | 2764 | 11 | 0.80% |
| hsa_circ_0089146 | 2764 | 11 | 0.80% |
| hsa_circ_0051445 | 2513 | 10 | 0.80% |
| hsa_circ_0070634 | 2513 | 10 | 0.80% |
| hsa_circ_0072223 | 2513 | 10 | 0.80% |
| hsa_circ_0075145 | 2513 | 10 | 0.80% |
| hsa_circ_0063314 | 4021 | 16 | 0.80% |
| hsa_circ_0056206 | 3016 | 12 | 0.80% |
| hsa_circ_0065485 | 3016 | 12 | 0.80% |
| hsa_circ_0020224 | 2765 | 11 | 0.80% |
| hsa_circ_0060477 | 2765 | 11 | 0.80% |
| hsa_circ_0076770 | 2765 | 11 | 0.80% |
| hsa_circ_0111908 | 2765 | 11 | 0.80% |
| hsa_circ_0132468 | 2765 | 11 | 0.80% |
| hsa_circ_0030966 | 2514 | 10 | 0.80% |
| hsa_circ_0102428 | 5531 | 22 | 0.80% |
| hsa_circ_0122637 | 6537 | 26 | 0.80% |
| hsa_circ_0126267 | 7040 | 28 | 0.80% |
| hsa_circ_0025049 | 3520 | 14 | 0.80% |
| hsa_circ_0017035 | 2766 | 11 | 0.80% |
| hsa_circ_0057416 | 2766 | 11 | 0.80% |

| | | | |
|---|---|---|---|
| hsa_circ_0070506 | 2766 | 11 | 0.80% |
| hsa_circ_0075993 | 2766 | 11 | 0.80% |
| hsa_circ_0114564 | 5533 | 22 | 0.80% |
| hsa_circ_0034739 | 3521 | 14 | 0.80% |
| hsa_circ_0048410 | 3018 | 12 | 0.80% |
| hsa_circ_0058951 | 3018 | 12 | 0.80% |
| hsa_circ_0078170 | 2515 | 10 | 0.80% |
| hsa_circ_0090632 | 2515 | 10 | 0.80% |
| hsa_circ_0128138 | 2515 | 10 | 0.80% |
| hsa_circ_0015412 | 3773 | 15 | 0.80% |
| hsa_circ_0123419 | 3270 | 13 | 0.80% |
| hsa_circ_0053329 | 15847 | 63 | 0.80% |
| hsa_circ_0083762 | 2767 | 11 | 0.80% |
| hsa_circ_0122695 | 2767 | 11 | 0.80% |
| hsa_circ_0024601 | 3019 | 12 | 0.79% |
| hsa_circ_0039331 | 3019 | 12 | 0.79% |
| hsa_circ_0010123 | 2516 | 10 | 0.79% |
| hsa_circ_0013411 | 2516 | 10 | 0.79% |
| hsa_circ_0064940 | 2516 | 10 | 0.79% |
| hsa_circ_0139441 | 2516 | 10 | 0.79% |
| hsa_circ_0127241 | 9058 | 36 | 0.79% |
| hsa_circ_0045992 | 2768 | 11 | 0.79% |
| hsa_circ_0054584 | 2768 | 11 | 0.79% |
| hsa_circ_0071026 | 2768 | 11 | 0.79% |
| hsa_circ_0073957 | 2768 | 11 | 0.79% |
| hsa_circ_0118371 | 7046 | 28 | 0.79% |
| hsa_circ_0113861 | 3020 | 12 | 0.79% |
| hsa_circ_0046176 | 3272 | 13 | 0.79% |
| hsa_circ_0060691 | 3272 | 13 | 0.79% |
| hsa_circ_0010362 | 2517 | 10 | 0.79% |
| hsa_circ_0058546 | 2517 | 10 | 0.79% |
| hsa_circ_0112001 | 2517 | 10 | 0.79% |
| hsa_circ_0050292 | 11831 | 47 | 0.79% |
| hsa_circ_0089050 | 2769 | 11 | 0.79% |
| hsa_circ_0108893 | 2769 | 11 | 0.79% |
| hsa_circ_0047001 | 3021 | 12 | 0.79% |
| hsa_circ_0047516 | 3021 | 12 | 0.79% |
| hsa_circ_0090320 | 2518 | 10 | 0.79% |
| hsa_circ_0031550 | 2770 | 11 | 0.79% |
| hsa_circ_0091956 | 2770 | 11 | 0.79% |
| hsa_circ_0135279 | 2770 | 11 | 0.79% |
| hsa_circ_0040632 | 3022 | 12 | 0.79% |
| hsa_circ_0078076 | 3022 | 12 | 0.79% |
| hsa_circ_0083713 | 3022 | 12 | 0.79% |
| hsa_circ_0133283 | 3022 | 12 | 0.79% |
| hsa_circ_0139001 | 2519 | 10 | 0.79% |
| hsa_circ_0028722 | 2771 | 11 | 0.79% |
| hsa_circ_0084738 | 2771 | 11 | 0.79% |
| hsa_circ_0069625 | 3275 | 13 | 0.79% |
| hsa_circ_0019165 | 3276 | 13 | 0.79% |

| | | | |
|---|---|---|---|
| hsa_circ_0114004 | 3276 | 13 | 0.79% |
| hsa_circ_0118014 | 3276 | 13 | 0.79% |
| hsa_circ_0088277 | 3024 | 12 | 0.79% |
| hsa_circ_0049129 | 2772 | 11 | 0.79% |
| hsa_circ_0077927 | 2772 | 11 | 0.79% |
| hsa_circ_0105033 | 2772 | 11 | 0.79% |
| hsa_circ_0030877 | 2520 | 10 | 0.79% |
| hsa_circ_0036791 | 2520 | 10 | 0.79% |
| hsa_circ_0109652 | 7057 | 28 | 0.79% |
| hsa_circ_0031584 | 4033 | 16 | 0.79% |
| hsa_circ_0036982 | 3529 | 14 | 0.79% |
| hsa_circ_0018579 | 3277 | 13 | 0.79% |
| hsa_circ_0026071 | 3025 | 12 | 0.79% |
| hsa_circ_0082588 | 3025 | 12 | 0.79% |
| hsa_circ_0061731 | 2773 | 11 | 0.79% |
| hsa_circ_0064441 | 2773 | 11 | 0.79% |
| hsa_circ_0127841 | 2773 | 11 | 0.79% |
| hsa_circ_0132057 | 2521 | 10 | 0.79% |
| hsa_circ_0014730 | 4034 | 16 | 0.79% |
| hsa_circ_0099150 | 3782 | 15 | 0.79% |
| hsa_circ_0039074 | 3026 | 12 | 0.79% |
| hsa_circ_0138946 | 2774 | 11 | 0.79% |
| hsa_circ_0046007 | 3279 | 13 | 0.79% |
| hsa_circ_0081025 | 3279 | 13 | 0.79% |
| hsa_circ_0085679 | 3279 | 13 | 0.79% |
| hsa_circ_0095281 | 3027 | 12 | 0.79% |
| hsa_circ_0015920 | 2775 | 11 | 0.79% |
| hsa_circ_0032139 | 2775 | 11 | 0.79% |
| hsa_circ_0035438 | 2775 | 11 | 0.79% |
| hsa_circ_0040936 | 2775 | 11 | 0.79% |
| hsa_circ_0044615 | 2775 | 11 | 0.79% |
| hsa_circ_0056832 | 2775 | 11 | 0.79% |
| hsa_circ_0059025 | 2775 | 11 | 0.79% |
| hsa_circ_0088960 | 2775 | 11 | 0.79% |
| hsa_circ_0115009 | 2523 | 10 | 0.79% |
| hsa_circ_0010639 | 3280 | 13 | 0.79% |
| hsa_circ_0125221 | 3028 | 12 | 0.79% |
| hsa_circ_0071302 | 2776 | 11 | 0.79% |
| hsa_circ_0076234 | 2776 | 11 | 0.79% |
| hsa_circ_0137243 | 2776 | 11 | 0.79% |
| hsa_circ_0117811 | 3281 | 13 | 0.79% |
| hsa_circ_0052464 | 2524 | 10 | 0.79% |
| hsa_circ_0036670 | 3029 | 12 | 0.79% |
| hsa_circ_0106691 | 5301 | 21 | 0.79% |
| hsa_circ_0000587 | 2777 | 11 | 0.79% |
| hsa_circ_0108035 | 2777 | 11 | 0.79% |
| hsa_circ_0134595 | 6564 | 26 | 0.79% |
| hsa_circ_0014323 | 3535 | 14 | 0.79% |
| hsa_circ_0058567 | 3030 | 12 | 0.79% |
| hsa_circ_0094895 | 3030 | 12 | 0.79% |

| | | | |
|---|---|---|---|
| hsa_circ_0021751 | 3283 | 13 | 0.79% |
| hsa_circ_0007398 | 2778 | 11 | 0.79% |
| hsa_circ_0011131 | 2778 | 11 | 0.79% |
| hsa_circ_0034664 | 2778 | 11 | 0.79% |
| hsa_circ_0060604 | 2778 | 11 | 0.79% |
| hsa_circ_0069005 | 2778 | 11 | 0.79% |
| hsa_circ_0091509 | 2778 | 11 | 0.79% |
| hsa_circ_0127761 | 2778 | 11 | 0.79% |
| hsa_circ_0063215 | 12881 | 51 | 0.79% |
| hsa_circ_0046014 | 3031 | 12 | 0.79% |
| hsa_circ_0086300 | 6820 | 27 | 0.79% |
| hsa_circ_0063479 | 2526 | 10 | 0.79% |
| hsa_circ_0098053 | 2526 | 10 | 0.79% |
| hsa_circ_0126050 | 2526 | 10 | 0.79% |
| hsa_circ_0031137 | 2779 | 11 | 0.79% |
| hsa_circ_0050860 | 2779 | 11 | 0.79% |
| hsa_circ_0058202 | 2779 | 11 | 0.79% |
| hsa_circ_0062083 | 2779 | 11 | 0.79% |
| hsa_circ_0075705 | 2779 | 11 | 0.79% |
| hsa_circ_0045181 | 3032 | 12 | 0.79% |
| hsa_circ_0070920 | 3032 | 12 | 0.79% |
| hsa_circ_0022492 | 3285 | 13 | 0.79% |
| hsa_circ_0035926 | 3285 | 13 | 0.79% |
| hsa_circ_0070643 | 2527 | 10 | 0.79% |
| hsa_circ_0115637 | 2527 | 10 | 0.79% |
| hsa_circ_0027933 | 2780 | 11 | 0.79% |
| hsa_circ_0065416 | 2780 | 11 | 0.79% |
| hsa_circ_0084437 | 2780 | 11 | 0.79% |
| hsa_circ_0094119 | 9351 | 37 | 0.79% |
| hsa_circ_0119995 | 7835 | 31 | 0.79% |
| hsa_circ_0026581 | 3033 | 12 | 0.79% |
| hsa_circ_0034369 | 3033 | 12 | 0.79% |
| hsa_circ_0090002 | 3033 | 12 | 0.79% |
| hsa_circ_0066767 | 3286 | 13 | 0.79% |
| hsa_circ_0089244 | 3286 | 13 | 0.79% |
| hsa_circ_0079692 | 5814 | 23 | 0.79% |
| hsa_circ_0014753 | 3539 | 14 | 0.79% |
| hsa_circ_0037024 | 2528 | 10 | 0.79% |
| hsa_circ_0042614 | 2528 | 10 | 0.79% |
| hsa_circ_0056080 | 2528 | 10 | 0.79% |
| hsa_circ_0032230 | 2781 | 11 | 0.79% |
| hsa_circ_0034051 | 2781 | 11 | 0.79% |
| hsa_circ_0070455 | 2781 | 11 | 0.79% |
| hsa_circ_0105820 | 10619 | 42 | 0.79% |
| hsa_circ_0012690 | 3034 | 12 | 0.79% |
| hsa_circ_0091316 | 3034 | 12 | 0.79% |
| hsa_circ_0041373 | 3287 | 13 | 0.79% |
| hsa_circ_0079394 | 3287 | 13 | 0.79% |
| hsa_circ_0006940 | 8850 | 35 | 0.79% |
| hsa_circ_0044520 | 2529 | 10 | 0.79% |

| | | | |
|---|---|---|---|
| hsa_circ_0122687 | 2529 | 10 | 0.79% |
| hsa_circ_0047172 | 2782 | 11 | 0.79% |
| hsa_circ_0034248 | 3035 | 12 | 0.79% |
| hsa_circ_0071326 | 3035 | 12 | 0.79% |
| hsa_circ_0056223 | 3288 | 13 | 0.79% |
| hsa_circ_0122952 | 3794 | 15 | 0.79% |
| hsa_circ_0115331 | 7084 | 28 | 0.79% |
| hsa_circ_0114127 | 6072 | 24 | 0.79% |
| hsa_circ_0074686 | 4048 | 16 | 0.79% |
| hsa_circ_0089439 | 3036 | 12 | 0.79% |
| hsa_circ_0019457 | 2783 | 11 | 0.79% |
| hsa_circ_0060391 | 2783 | 11 | 0.79% |
| hsa_circ_0107591 | 2530 | 10 | 0.79% |
| hsa_circ_0016963 | 3037 | 12 | 0.79% |
| hsa_circ_0125863 | 3037 | 12 | 0.79% |
| hsa_circ_0003635 | 2784 | 11 | 0.79% |
| hsa_circ_0087808 | 2784 | 11 | 0.79% |
| hsa_circ_0044285 | 2531 | 10 | 0.79% |
| hsa_circ_0137348 | 2531 | 10 | 0.79% |
| hsa_circ_0046102 | 3291 | 13 | 0.79% |
| hsa_circ_0047128 | 3038 | 12 | 0.79% |
| hsa_circ_0076277 | 2785 | 11 | 0.79% |
| hsa_circ_0090762 | 2785 | 11 | 0.79% |
| hsa_circ_0015475 | 2532 | 10 | 0.79% |
| hsa_circ_0109046 | 2532 | 10 | 0.79% |
| hsa_circ_0131197 | 2532 | 10 | 0.79% |
| hsa_circ_0108390 | 3545 | 14 | 0.79% |
| hsa_circ_0060637 | 3039 | 12 | 0.79% |
| hsa_circ_0018120 | 2786 | 11 | 0.79% |
| hsa_circ_0037991 | 2786 | 11 | 0.79% |
| hsa_circ_0069303 | 2786 | 11 | 0.79% |
| hsa_circ_0078712 | 2786 | 11 | 0.79% |
| hsa_circ_0091118 | 2786 | 11 | 0.79% |
| hsa_circ_0123775 | 2786 | 11 | 0.79% |
| hsa_circ_0046903 | 3546 | 14 | 0.79% |
| hsa_circ_0117256 | 3546 | 14 | 0.79% |
| hsa_circ_0013651 | 2533 | 10 | 0.79% |
| hsa_circ_0035430 | 2533 | 10 | 0.79% |
| hsa_circ_0124124 | 3293 | 13 | 0.79% |
| hsa_circ_0022834 | 3800 | 15 | 0.79% |
| hsa_circ_0010310 | 3040 | 12 | 0.79% |
| hsa_circ_0042053 | 3040 | 12 | 0.79% |
| hsa_circ_0013893 | 2787 | 11 | 0.79% |
| hsa_circ_0020733 | 2787 | 11 | 0.79% |
| hsa_circ_0053839 | 2787 | 11 | 0.79% |
| hsa_circ_0044008 | 3294 | 13 | 0.79% |
| hsa_circ_0117812 | 3294 | 13 | 0.79% |
| hsa_circ_0060066 | 15711 | 62 | 0.79% |
| hsa_circ_0045466 | 2788 | 11 | 0.79% |
| hsa_circ_0062038 | 2788 | 11 | 0.79% |

| | | | |
|---|---|---|---|
| hsa_circ_0028162 | 3295 | 13 | 0.79% |
| hsa_circ_0000996 | 3042 | 12 | 0.79% |
| hsa_circ_0029029 | 2535 | 10 | 0.79% |
| hsa_circ_0078226 | 2535 | 10 | 0.79% |
| hsa_circ_0083260 | 2535 | 10 | 0.79% |
| hsa_circ_0085728 | 2535 | 10 | 0.79% |
| hsa_circ_0078823 | 10901 | 43 | 0.79% |
| hsa_circ_0048875 | 3296 | 13 | 0.79% |
| hsa_circ_0096916 | 2789 | 11 | 0.79% |
| hsa_circ_0083544 | 3804 | 15 | 0.79% |
| hsa_circ_0018328 | 2536 | 10 | 0.79% |
| hsa_circ_0052981 | 2536 | 10 | 0.79% |
| hsa_circ_0139327 | 2536 | 10 | 0.79% |
| hsa_circ_0001407 | 2790 | 11 | 0.79% |
| hsa_circ_0025660 | 2790 | 11 | 0.79% |
| hsa_circ_0057801 | 2790 | 11 | 0.79% |
| hsa_circ_0139371 | 2790 | 11 | 0.79% |
| hsa_circ_0048079 | 3551 | 14 | 0.79% |
| hsa_circ_0122277 | 3298 | 13 | 0.79% |
| hsa_circ_0093667 | 7611 | 30 | 0.79% |
| hsa_circ_0079913 | 2537 | 10 | 0.79% |
| hsa_circ_0104263 | 2537 | 10 | 0.79% |
| hsa_circ_0123750 | 2537 | 10 | 0.79% |
| hsa_circ_0086508 | 2791 | 11 | 0.79% |
| hsa_circ_0009559 | 3045 | 12 | 0.79% |
| hsa_circ_0011831 | 3045 | 12 | 0.79% |
| hsa_circ_0021602 | 3045 | 12 | 0.79% |
| hsa_circ_0081364 | 4314 | 17 | 0.79% |
| hsa_circ_0058932 | 3299 | 13 | 0.79% |
| hsa_circ_0028692 | 3553 | 14 | 0.79% |
| hsa_circ_0057608 | 2538 | 10 | 0.79% |
| hsa_circ_0034919 | 2792 | 11 | 0.79% |
| hsa_circ_0053819 | 2792 | 11 | 0.79% |
| hsa_circ_0077738 | 2792 | 11 | 0.79% |
| hsa_circ_0075024 | 3046 | 12 | 0.79% |
| hsa_circ_0064619 | 3300 | 13 | 0.79% |
| hsa_circ_0022475 | 2539 | 10 | 0.79% |
| hsa_circ_0041309 | 2793 | 11 | 0.79% |
| hsa_circ_0049294 | 2793 | 11 | 0.79% |
| hsa_circ_0070771 | 2793 | 11 | 0.79% |
| hsa_circ_0077063 | 2793 | 11 | 0.79% |
| hsa_circ_0067400 | 3047 | 12 | 0.79% |
| hsa_circ_0044778 | 4063 | 16 | 0.79% |
| hsa_circ_0090836 | 8128 | 32 | 0.79% |
| hsa_circ_0109692 | 7874 | 31 | 0.79% |
| hsa_circ_0000246 | 7620 | 30 | 0.79% |
| hsa_circ_0019892 | 2794 | 11 | 0.79% |
| hsa_circ_0071591 | 2794 | 11 | 0.79% |
| hsa_circ_0076353 | 2794 | 11 | 0.79% |
| hsa_circ_0016269 | 2540 | 10 | 0.79% |

| | | | |
|---|---|---|---|
| hsa_circ_0020123 | 2540 | 10 | 0.79% |
| hsa_circ_0030479 | 2540 | 10 | 0.79% |
| hsa_circ_0065176 | 2540 | 10 | 0.79% |
| hsa_circ_0130562 | 3049 | 12 | 0.79% |
| hsa_circ_0064204 | 2795 | 11 | 0.79% |
| hsa_circ_0040413 | 2541 | 10 | 0.79% |
| hsa_circ_0080482 | 2541 | 10 | 0.79% |
| hsa_circ_0082032 | 2541 | 10 | 0.79% |
| hsa_circ_0108400 | 3050 | 12 | 0.79% |
| hsa_circ_0125457 | 3050 | 12 | 0.79% |
| hsa_circ_0035840 | 2796 | 11 | 0.79% |
| hsa_circ_0117553 | 2796 | 11 | 0.79% |
| hsa_circ_0118676 | 9659 | 38 | 0.79% |
| hsa_circ_0061794 | 2542 | 10 | 0.79% |
| hsa_circ_0067976 | 2542 | 10 | 0.79% |
| hsa_circ_0140434 | 2797 | 11 | 0.79% |
| hsa_circ_0058401 | 3560 | 14 | 0.79% |
| hsa_circ_0126183 | 2543 | 10 | 0.79% |
| hsa_circ_0139076 | 2543 | 10 | 0.79% |
| hsa_circ_0085680 | 3306 | 13 | 0.79% |
| hsa_circ_0125973 | 8901 | 35 | 0.79% |
| hsa_circ_0021766 | 3052 | 12 | 0.79% |
| hsa_circ_0083703 | 3052 | 12 | 0.79% |
| hsa_circ_0121233 | 3561 | 14 | 0.79% |
| hsa_circ_0009445 | 2798 | 11 | 0.79% |
| hsa_circ_0126205 | 2798 | 11 | 0.79% |
| hsa_circ_0041657 | 7377 | 29 | 0.79% |
| hsa_circ_0057648 | 2544 | 10 | 0.79% |
| hsa_circ_0061605 | 2544 | 10 | 0.79% |
| hsa_circ_0105564 | 2544 | 10 | 0.79% |
| hsa_circ_0033334 | 3053 | 12 | 0.79% |
| hsa_circ_0055133 | 3053 | 12 | 0.79% |
| hsa_circ_0075943 | 10940 | 43 | 0.79% |
| hsa_circ_0030863 | 3562 | 14 | 0.79% |
| hsa_circ_0045393 | 3562 | 14 | 0.79% |
| hsa_circ_0062805 | 3562 | 14 | 0.79% |
| hsa_circ_0012342 | 2799 | 11 | 0.79% |
| hsa_circ_0053903 | 2799 | 11 | 0.79% |
| hsa_circ_0078948 | 10942 | 43 | 0.79% |
| hsa_circ_0025482 | 2545 | 10 | 0.79% |
| hsa_circ_0044918 | 3309 | 13 | 0.79% |
| hsa_circ_0069721 | 3309 | 13 | 0.79% |
| hsa_circ_0135092 | 3309 | 13 | 0.79% |
| hsa_circ_0078886 | 9418 | 37 | 0.79% |
| hsa_circ_0037537 | 2800 | 11 | 0.79% |
| hsa_circ_0114266 | 2800 | 11 | 0.79% |
| hsa_circ_0122355 | 2800 | 11 | 0.79% |
| hsa_circ_0078946 | 9420 | 37 | 0.79% |
| hsa_circ_0118399 | 5347 | 21 | 0.79% |
| hsa_circ_0123728 | 6875 | 27 | 0.79% |

| | | | |
|---|---|---|---|
| hsa_circ_0067657 | 2801 | 11 | 0.79% |
| hsa_circ_0096505 | 3311 | 13 | 0.79% |
| hsa_circ_0015712 | 2547 | 10 | 0.79% |
| hsa_circ_0019450 | 2547 | 10 | 0.79% |
| hsa_circ_0059143 | 8660 | 34 | 0.79% |
| hsa_circ_0093183 | 4585 | 18 | 0.79% |
| hsa_circ_0033242 | 2802 | 11 | 0.79% |
| hsa_circ_0091120 | 2802 | 11 | 0.79% |
| hsa_circ_0116043 | 2802 | 11 | 0.79% |
| hsa_circ_0073616 | 3057 | 12 | 0.79% |
| hsa_circ_0062042 | 2548 | 10 | 0.78% |
| hsa_circ_0038871 | 2803 | 11 | 0.78% |
| hsa_circ_0072179 | 2803 | 11 | 0.78% |
| hsa_circ_0122273 | 2803 | 11 | 0.78% |
| hsa_circ_0140398 | 2803 | 11 | 0.78% |
| hsa_circ_0037698 | 3058 | 12 | 0.78% |
| hsa_circ_0050631 | 3058 | 12 | 0.78% |
| hsa_circ_0111019 | 3313 | 13 | 0.78% |
| hsa_circ_0050493 | 3568 | 14 | 0.78% |
| hsa_circ_0096380 | 7137 | 28 | 0.78% |
| hsa_circ_0046762 | 2549 | 10 | 0.78% |
| hsa_circ_0051482 | 2804 | 11 | 0.78% |
| hsa_circ_0054595 | 2804 | 11 | 0.78% |
| hsa_circ_0101303 | 2804 | 11 | 0.78% |
| hsa_circ_0029562 | 3059 | 12 | 0.78% |
| hsa_circ_0046991 | 3059 | 12 | 0.78% |
| hsa_circ_0070686 | 3059 | 12 | 0.78% |
| hsa_circ_0087360 | 3059 | 12 | 0.78% |
| hsa_circ_0016879 | 2805 | 11 | 0.78% |
| hsa_circ_0015282 | 2550 | 10 | 0.78% |
| hsa_circ_0017321 | 2550 | 10 | 0.78% |
| hsa_circ_0028690 | 2550 | 10 | 0.78% |
| hsa_circ_0084584 | 2550 | 10 | 0.78% |
| hsa_circ_0087796 | 2550 | 10 | 0.78% |
| hsa_circ_0103880 | 2550 | 10 | 0.78% |
| hsa_circ_0105610 | 6121 | 24 | 0.78% |
| hsa_circ_0061713 | 3571 | 14 | 0.78% |
| hsa_circ_0022361 | 3316 | 13 | 0.78% |
| hsa_circ_0010859 | 3061 | 12 | 0.78% |
| hsa_circ_0113606 | 4592 | 18 | 0.78% |
| hsa_circ_0003812 | 6889 | 27 | 0.78% |
| hsa_circ_0050679 | 3062 | 12 | 0.78% |
| hsa_circ_0074758 | 3062 | 12 | 0.78% |
| hsa_circ_0086079 | 2807 | 11 | 0.78% |
| hsa_circ_0042017 | 2808 | 11 | 0.78% |
| hsa_circ_0075958 | 2808 | 11 | 0.78% |
| hsa_circ_0077128 | 2808 | 11 | 0.78% |
| hsa_circ_0083906 | 3574 | 14 | 0.78% |
| hsa_circ_0012537 | 2553 | 10 | 0.78% |
| hsa_circ_0030480 | 2553 | 10 | 0.78% |

| | | | |
|---|---|---|---|
| hsa_circ_0034173 | 2553 | 10 | 0.78% |
| hsa_circ_0081526 | 3319 | 13 | 0.78% |
| hsa_circ_0046665 | 3064 | 12 | 0.78% |
| hsa_circ_0067050 | 3064 | 12 | 0.78% |
| hsa_circ_0085396 | 2809 | 11 | 0.78% |
| hsa_circ_0039610 | 2554 | 10 | 0.78% |
| hsa_circ_0033246 | 3065 | 12 | 0.78% |
| hsa_circ_0128262 | 3065 | 12 | 0.78% |
| hsa_circ_0091811 | 3576 | 14 | 0.78% |
| hsa_circ_0137242 | 8430 | 33 | 0.78% |
| hsa_circ_0011277 | 2810 | 11 | 0.78% |
| hsa_circ_0054282 | 10475 | 41 | 0.78% |
| hsa_circ_0022949 | 3066 | 12 | 0.78% |
| hsa_circ_0115646 | 3066 | 12 | 0.78% |
| hsa_circ_0062085 | 2555 | 10 | 0.78% |
| hsa_circ_0095393 | 3322 | 13 | 0.78% |
| hsa_circ_0038280 | 2811 | 11 | 0.78% |
| hsa_circ_0039569 | 2811 | 11 | 0.78% |
| hsa_circ_0065929 | 2811 | 11 | 0.78% |
| hsa_circ_0131512 | 2811 | 11 | 0.78% |
| hsa_circ_0018369 | 3067 | 12 | 0.78% |
| hsa_circ_0069822 | 3067 | 12 | 0.78% |
| hsa_circ_0069437 | 2556 | 10 | 0.78% |
| hsa_circ_0128209 | 2812 | 11 | 0.78% |
| hsa_circ_0104972 | 12016 | 47 | 0.78% |
| hsa_circ_0109691 | 7926 | 31 | 0.78% |
| hsa_circ_0020423 | 2557 | 10 | 0.78% |
| hsa_circ_0097811 | 2557 | 10 | 0.78% |
| hsa_circ_0048362 | 3580 | 14 | 0.78% |
| hsa_circ_0014326 | 2813 | 11 | 0.78% |
| hsa_circ_0090900 | 2813 | 11 | 0.78% |
| hsa_circ_0100815 | 2813 | 11 | 0.78% |
| hsa_circ_0140478 | 2813 | 11 | 0.78% |
| hsa_circ_0053227 | 3581 | 14 | 0.78% |
| hsa_circ_0099626 | 10999 | 43 | 0.78% |
| hsa_circ_0077743 | 2558 | 10 | 0.78% |
| hsa_circ_0056060 | 2814 | 11 | 0.78% |
| hsa_circ_0071192 | 2814 | 11 | 0.78% |
| hsa_circ_0115051 | 2814 | 11 | 0.78% |
| hsa_circ_0024857 | 2559 | 10 | 0.78% |
| hsa_circ_0041602 | 2815 | 11 | 0.78% |
| hsa_circ_0061523 | 2815 | 11 | 0.78% |
| hsa_circ_0064875 | 3071 | 12 | 0.78% |
| hsa_circ_0089179 | 3071 | 12 | 0.78% |
| hsa_circ_0084402 | 3327 | 13 | 0.78% |
| hsa_circ_0048544 | 3072 | 12 | 0.78% |
| hsa_circ_0122530 | 3072 | 12 | 0.78% |
| hsa_circ_0088305 | 2816 | 11 | 0.78% |
| hsa_circ_0017142 | 2560 | 10 | 0.78% |
| hsa_circ_0130578 | 3329 | 13 | 0.78% |

| | | | |
|---|---|---|---|
| hsa_circ_0011531 | 3073 | 12 | 0.78% |
| hsa_circ_0052661 | 2817 | 11 | 0.78% |
| hsa_circ_0106101 | 3330 | 13 | 0.78% |
| hsa_circ_0117552 | 3074 | 12 | 0.78% |
| hsa_circ_0054858 | 2818 | 11 | 0.78% |
| hsa_circ_0077931 | 2818 | 11 | 0.78% |
| hsa_circ_0087990 | 2818 | 11 | 0.78% |
| hsa_circ_0091986 | 2818 | 11 | 0.78% |
| hsa_circ_0063100 | 2562 | 10 | 0.78% |
| hsa_circ_0074580 | 2562 | 10 | 0.78% |
| hsa_circ_0102316 | 2562 | 10 | 0.78% |
| hsa_circ_0100340 | 3331 | 13 | 0.78% |
| hsa_circ_0051009 | 3075 | 12 | 0.78% |
| hsa_circ_0065331 | 3075 | 12 | 0.78% |
| hsa_circ_0091972 | 3075 | 12 | 0.78% |
| hsa_circ_0010749 | 2819 | 11 | 0.78% |
| hsa_circ_0039266 | 2819 | 11 | 0.78% |
| hsa_circ_0088001 | 2819 | 11 | 0.78% |
| hsa_circ_0029752 | 2563 | 10 | 0.78% |
| hsa_circ_0039810 | 2563 | 10 | 0.78% |
| hsa_circ_0083754 | 2563 | 10 | 0.78% |
| hsa_circ_0085442 | 2563 | 10 | 0.78% |
| hsa_circ_0136983 | 2563 | 10 | 0.78% |
| hsa_circ_0027566 | 3076 | 12 | 0.78% |
| hsa_circ_0058765 | 3076 | 12 | 0.78% |
| hsa_circ_0063612 | 3076 | 12 | 0.78% |
| hsa_circ_0069687 | 3076 | 12 | 0.78% |
| hsa_circ_0106847 | 3076 | 12 | 0.78% |
| hsa_circ_0067084 | 2820 | 11 | 0.78% |
| hsa_circ_0103309 | 2820 | 11 | 0.78% |
| hsa_circ_0052632 | 3333 | 13 | 0.78% |
| hsa_circ_0095169 | 3333 | 13 | 0.78% |
| hsa_circ_0034076 | 3334 | 13 | 0.78% |
| hsa_circ_0058227 | 5130 | 20 | 0.78% |
| hsa_circ_0027221 | 3078 | 12 | 0.78% |
| hsa_circ_0057511 | 3078 | 12 | 0.78% |
| hsa_circ_0083699 | 3078 | 12 | 0.78% |
| hsa_circ_0084469 | 3078 | 12 | 0.78% |
| hsa_circ_0030964 | 2565 | 10 | 0.78% |
| hsa_circ_0065369 | 2565 | 10 | 0.78% |
| hsa_circ_0102313 | 2565 | 10 | 0.78% |
| hsa_circ_0001659 | 4361 | 17 | 0.78% |
| hsa_circ_0021152 | 3079 | 12 | 0.78% |
| hsa_circ_0034537 | 3079 | 12 | 0.78% |
| hsa_circ_0046156 | 3079 | 12 | 0.78% |
| hsa_circ_0083655 | 3079 | 12 | 0.78% |
| hsa_circ_0123755 | 3079 | 12 | 0.78% |
| hsa_circ_0135476 | 3079 | 12 | 0.78% |
| hsa_circ_0139669 | 3079 | 12 | 0.78% |
| hsa_circ_0053896 | 2823 | 11 | 0.78% |

| | | | |
|---|---|---|---|
| hsa_circ_0071280 | 2823 | 11 | 0.78% |
| hsa_circ_0051748 | 3593 | 14 | 0.78% |
| hsa_circ_0052660 | 3593 | 14 | 0.78% |
| hsa_circ_0031603 | 3080 | 12 | 0.78% |
| hsa_circ_0108741 | 7187 | 28 | 0.78% |
| hsa_circ_0041293 | 2824 | 11 | 0.78% |
| hsa_circ_0126787 | 8731 | 34 | 0.78% |
| hsa_circ_0017225 | 2568 | 10 | 0.78% |
| hsa_circ_0049897 | 2568 | 10 | 0.78% |
| hsa_circ_0053004 | 3082 | 12 | 0.78% |
| hsa_circ_0068829 | 3082 | 12 | 0.78% |
| hsa_circ_0085455 | 3082 | 12 | 0.78% |
| hsa_circ_0138716 | 3339 | 13 | 0.78% |
| hsa_circ_0091398 | 3596 | 14 | 0.78% |
| hsa_circ_0125041 | 7449 | 29 | 0.78% |
| hsa_circ_0116719 | 2569 | 10 | 0.78% |
| hsa_circ_0031138 | 2826 | 11 | 0.78% |
| hsa_circ_0124666 | 2826 | 11 | 0.78% |
| hsa_circ_0006581 | 6166 | 24 | 0.78% |
| hsa_circ_0031685 | 3083 | 12 | 0.78% |
| hsa_circ_0035594 | 3083 | 12 | 0.78% |
| hsa_circ_0047511 | 3083 | 12 | 0.78% |
| hsa_circ_0101642 | 4111 | 16 | 0.78% |
| hsa_circ_0013066 | 3084 | 12 | 0.78% |
| hsa_circ_0030654 | 3084 | 12 | 0.78% |
| hsa_circ_0086265 | 3084 | 12 | 0.78% |
| hsa_circ_0103211 | 2827 | 11 | 0.78% |
| hsa_circ_0027765 | 2570 | 10 | 0.78% |
| hsa_circ_0119542 | 2570 | 10 | 0.78% |
| hsa_circ_0065025 | 5141 | 20 | 0.78% |
| hsa_circ_0064987 | 3856 | 15 | 0.78% |
| hsa_circ_0034216 | 3342 | 13 | 0.78% |
| hsa_circ_0091413 | 3342 | 13 | 0.78% |
| hsa_circ_0050080 | 2828 | 11 | 0.78% |
| hsa_circ_0076982 | 2828 | 11 | 0.78% |
| hsa_circ_0086070 | 2828 | 11 | 0.78% |
| hsa_circ_0088117 | 2571 | 10 | 0.78% |
| hsa_circ_0139854 | 10542 | 41 | 0.78% |
| hsa_circ_0122950 | 3857 | 15 | 0.78% |
| hsa_circ_0079121 | 3600 | 14 | 0.78% |
| hsa_circ_0032649 | 3343 | 13 | 0.78% |
| hsa_circ_0098231 | 3343 | 13 | 0.78% |
| hsa_circ_0023622 | 2829 | 11 | 0.78% |
| hsa_circ_0062684 | 2829 | 11 | 0.78% |
| hsa_circ_0047257 | 2572 | 10 | 0.78% |
| hsa_circ_0062430 | 4116 | 16 | 0.78% |
| hsa_circ_0004540 | 3087 | 12 | 0.78% |
| hsa_circ_0071558 | 3087 | 12 | 0.78% |
| hsa_circ_0092749 | 3087 | 12 | 0.78% |
| hsa_circ_0075467 | 3602 | 14 | 0.78% |

| | | | |
|---|---|---|---|
| hsa_circ_0064874 | 3345 | 13 | 0.78% |
| hsa_circ_0126565 | 9007 | 35 | 0.78% |
| hsa_circ_0018699 | 2831 | 11 | 0.78% |
| hsa_circ_0039186 | 2831 | 11 | 0.78% |
| hsa_circ_0074196 | 2831 | 11 | 0.78% |
| hsa_circ_0117049 | 2831 | 11 | 0.78% |
| hsa_circ_0015555 | 3346 | 13 | 0.78% |
| hsa_circ_0020178 | 3346 | 13 | 0.78% |
| hsa_circ_0033408 | 3861 | 15 | 0.78% |
| hsa_circ_0028346 | 2574 | 10 | 0.78% |
| hsa_circ_0103406 | 2574 | 10 | 0.78% |
| hsa_circ_0140574 | 2574 | 10 | 0.78% |
| hsa_circ_0028513 | 2832 | 11 | 0.78% |
| hsa_circ_0051699 | 2832 | 11 | 0.78% |
| hsa_circ_0072527 | 2832 | 11 | 0.78% |
| hsa_circ_0013198 | 3090 | 12 | 0.78% |
| hsa_circ_0032003 | 2575 | 10 | 0.78% |
| hsa_circ_0048721 | 3348 | 13 | 0.78% |
| hsa_circ_0062700 | 2833 | 11 | 0.78% |
| hsa_circ_0087995 | 2833 | 11 | 0.78% |
| hsa_circ_0013775 | 3091 | 12 | 0.78% |
| hsa_circ_0113860 | 3091 | 12 | 0.78% |
| hsa_circ_0107456 | 4379 | 17 | 0.78% |
| hsa_circ_0021593 | 2576 | 10 | 0.78% |
| hsa_circ_0084094 | 2576 | 10 | 0.78% |
| hsa_circ_0128068 | 4895 | 19 | 0.78% |
| hsa_circ_0075976 | 2834 | 11 | 0.78% |
| hsa_circ_0076365 | 2834 | 11 | 0.78% |
| hsa_circ_0050011 | 3092 | 12 | 0.78% |
| hsa_circ_0081613 | 3092 | 12 | 0.78% |
| hsa_circ_0130864 | 3092 | 12 | 0.78% |
| hsa_circ_0053645 | 3350 | 13 | 0.78% |
| hsa_circ_0070210 | 3350 | 13 | 0.78% |
| hsa_circ_0071143 | 2577 | 10 | 0.78% |
| hsa_circ_0004535 | 6443 | 25 | 0.78% |
| hsa_circ_0030077 | 2835 | 11 | 0.78% |
| hsa_circ_0019037 | 3093 | 12 | 0.78% |
| hsa_circ_0086926 | 3093 | 12 | 0.78% |
| hsa_circ_0019893 | 2836 | 11 | 0.78% |
| hsa_circ_0024963 | 2836 | 11 | 0.78% |
| hsa_circ_0042219 | 2836 | 11 | 0.78% |
| hsa_circ_0051197 | 2836 | 11 | 0.78% |
| hsa_circ_0059160 | 2836 | 11 | 0.78% |
| hsa_circ_0031530 | 3094 | 12 | 0.78% |
| hsa_circ_0044469 | 3094 | 12 | 0.78% |
| hsa_circ_0101886 | 3094 | 12 | 0.78% |
| hsa_circ_0083373 | 3868 | 15 | 0.78% |
| hsa_circ_0005040 | 2579 | 10 | 0.78% |
| hsa_circ_0107488 | 11090 | 43 | 0.78% |
| hsa_circ_0123815 | 3095 | 12 | 0.78% |

| | | | |
|---|---|---|---|
| hsa_circ_0012068 | 3611 | 14 | 0.78% |
| hsa_circ_0000939 | 7997 | 31 | 0.78% |
| hsa_circ_0032167 | 3354 | 13 | 0.78% |
| hsa_circ_0096530 | 3354 | 13 | 0.78% |
| hsa_circ_0132990 | 3354 | 13 | 0.78% |
| hsa_circ_0033101 | 3096 | 12 | 0.78% |
| hsa_circ_0053201 | 3096 | 12 | 0.78% |
| hsa_circ_0018165 | 2838 | 11 | 0.78% |
| hsa_circ_0093166 | 2838 | 11 | 0.78% |
| hsa_circ_0071164 | 2580 | 10 | 0.78% |
| hsa_circ_0072850 | 6709 | 26 | 0.78% |
| hsa_circ_0072808 | 6710 | 26 | 0.77% |
| hsa_circ_0021197 | 3355 | 13 | 0.77% |
| hsa_circ_0099563 | 3355 | 13 | 0.77% |
| hsa_circ_0074511 | 3097 | 12 | 0.77% |
| hsa_circ_0022059 | 2839 | 11 | 0.77% |
| hsa_circ_0082923 | 2839 | 11 | 0.77% |
| hsa_circ_0072821 | 6711 | 26 | 0.77% |
| hsa_circ_0057136 | 3356 | 13 | 0.77% |
| hsa_circ_0047059 | 3098 | 12 | 0.77% |
| hsa_circ_0077328 | 3098 | 12 | 0.77% |
| hsa_circ_0019836 | 2840 | 11 | 0.77% |
| hsa_circ_0020398 | 2840 | 11 | 0.77% |
| hsa_circ_0070767 | 2840 | 11 | 0.77% |
| hsa_circ_0085245 | 2840 | 11 | 0.77% |
| hsa_circ_0111838 | 2840 | 11 | 0.77% |
| hsa_circ_0124475 | 2582 | 10 | 0.77% |
| hsa_circ_0062517 | 3357 | 13 | 0.77% |
| hsa_circ_0036342 | 3099 | 12 | 0.77% |
| hsa_circ_0043543 | 2841 | 11 | 0.77% |
| hsa_circ_0054724 | 2841 | 11 | 0.77% |
| hsa_circ_0067334 | 2841 | 11 | 0.77% |
| hsa_circ_0023707 | 2583 | 10 | 0.77% |
| hsa_circ_0063936 | 2583 | 10 | 0.77% |
| hsa_circ_0097305 | 3358 | 13 | 0.77% |
| hsa_circ_0129397 | 3100 | 12 | 0.77% |
| hsa_circ_0021002 | 2842 | 11 | 0.77% |
| hsa_circ_0068151 | 2842 | 11 | 0.77% |
| hsa_circ_0071228 | 3359 | 13 | 0.77% |
| hsa_circ_0120217 | 11369 | 44 | 0.77% |
| hsa_circ_0029351 | 2584 | 10 | 0.77% |
| hsa_circ_0128360 | 2584 | 10 | 0.77% |
| hsa_circ_0077580 | 3101 | 12 | 0.77% |
| hsa_circ_0046901 | 3618 | 14 | 0.77% |
| hsa_circ_0009493 | 2843 | 11 | 0.77% |
| hsa_circ_0041273 | 2843 | 11 | 0.77% |
| hsa_circ_0061550 | 2843 | 11 | 0.77% |
| hsa_circ_0128143 | 2843 | 11 | 0.77% |
| hsa_circ_0053633 | 3360 | 13 | 0.77% |
| hsa_circ_0078848 | 3877 | 15 | 0.77% |

| | | | |
|---|---|---|---|
| hsa_circ_0028250 | 2585 | 10 | 0.77% |
| hsa_circ_0029517 | 3361 | 13 | 0.77% |
| hsa_circ_0116970 | 2844 | 11 | 0.77% |
| hsa_circ_0059644 | 3362 | 13 | 0.77% |
| hsa_circ_0086634 | 3362 | 13 | 0.77% |
| hsa_circ_0045035 | 2845 | 11 | 0.77% |
| hsa_circ_0091837 | 2845 | 11 | 0.77% |
| hsa_circ_0035858 | 3104 | 12 | 0.77% |
| hsa_circ_0045344 | 3104 | 12 | 0.77% |
| hsa_circ_0058331 | 3104 | 12 | 0.77% |
| hsa_circ_0060395 | 3104 | 12 | 0.77% |
| hsa_circ_0084470 | 4915 | 19 | 0.77% |
| hsa_circ_0085379 | 3363 | 13 | 0.77% |
| hsa_circ_0041813 | 2587 | 10 | 0.77% |
| hsa_circ_0046440 | 2587 | 10 | 0.77% |
| hsa_circ_0054360 | 2587 | 10 | 0.77% |
| hsa_circ_0066226 | 2587 | 10 | 0.77% |
| hsa_circ_0098321 | 9314 | 36 | 0.77% |
| hsa_circ_0065333 | 2846 | 11 | 0.77% |
| hsa_circ_0091019 | 2846 | 11 | 0.77% |
| hsa_circ_0120653 | 2846 | 11 | 0.77% |
| hsa_circ_0040261 | 8797 | 34 | 0.77% |
| hsa_circ_0031520 | 3105 | 12 | 0.77% |
| hsa_circ_0066261 | 4658 | 18 | 0.77% |
| hsa_circ_0029668 | 2847 | 11 | 0.77% |
| hsa_circ_0063948 | 2847 | 11 | 0.77% |
| hsa_circ_0016925 | 3106 | 12 | 0.77% |
| hsa_circ_0075706 | 3106 | 12 | 0.77% |
| hsa_circ_0127681 | 3624 | 14 | 0.77% |
| hsa_circ_0039664 | 2589 | 10 | 0.77% |
| hsa_circ_0048702 | 2589 | 10 | 0.77% |
| hsa_circ_0082107 | 2589 | 10 | 0.77% |
| hsa_circ_0086364 | 2589 | 10 | 0.77% |
| hsa_circ_0020378 | 2848 | 11 | 0.77% |
| hsa_circ_0029655 | 2848 | 11 | 0.77% |
| hsa_circ_0065247 | 2848 | 11 | 0.77% |
| hsa_circ_0136357 | 2848 | 11 | 0.77% |
| hsa_circ_0082129 | 3107 | 12 | 0.77% |
| hsa_circ_0006079 | 4920 | 19 | 0.77% |
| hsa_circ_0136275 | 5179 | 20 | 0.77% |
| hsa_circ_0055141 | 3367 | 13 | 0.77% |
| hsa_circ_0010147 | 3108 | 12 | 0.77% |
| hsa_circ_0054048 | 2849 | 11 | 0.77% |
| hsa_circ_0084736 | 2849 | 11 | 0.77% |
| hsa_circ_0039785 | 2590 | 10 | 0.77% |
| hsa_circ_0043315 | 7771 | 30 | 0.77% |
| hsa_circ_0012767 | 3109 | 12 | 0.77% |
| hsa_circ_0018508 | 3109 | 12 | 0.77% |
| hsa_circ_0082428 | 2850 | 11 | 0.77% |
| hsa_circ_0135655 | 2850 | 11 | 0.77% |

| | | | |
|---|---|---|---|
| hsa_circ_0012007 | 2591 | 10 | 0.77% |
| hsa_circ_0041536 | 2591 | 10 | 0.77% |
| hsa_circ_0050695 | 3110 | 12 | 0.77% |
| hsa_circ_0124859 | 3110 | 12 | 0.77% |
| hsa_circ_0025505 | 2851 | 11 | 0.77% |
| hsa_circ_0045519 | 2851 | 11 | 0.77% |
| hsa_circ_0091932 | 2851 | 11 | 0.77% |
| hsa_circ_0135025 | 2851 | 11 | 0.77% |
| hsa_circ_0052121 | 2592 | 10 | 0.77% |
| hsa_circ_0053803 | 2592 | 10 | 0.77% |
| hsa_circ_0099222 | 3370 | 13 | 0.77% |
| hsa_circ_0059128 | 4148 | 16 | 0.77% |
| hsa_circ_0028121 | 3111 | 12 | 0.77% |
| hsa_circ_0032156 | 3111 | 12 | 0.77% |
| hsa_circ_0048769 | 3111 | 12 | 0.77% |
| hsa_circ_0102529 | 2593 | 10 | 0.77% |
| hsa_circ_0031156 | 3112 | 12 | 0.77% |
| hsa_circ_0043767 | 3112 | 12 | 0.77% |
| hsa_circ_0076758 | 3112 | 12 | 0.77% |
| hsa_circ_0082221 | 3631 | 14 | 0.77% |
| hsa_circ_0021165 | 2853 | 11 | 0.77% |
| hsa_circ_0028765 | 2853 | 11 | 0.77% |
| hsa_circ_0126251 | 2853 | 11 | 0.77% |
| hsa_circ_0087310 | 2594 | 10 | 0.77% |
| hsa_circ_0103700 | 2594 | 10 | 0.77% |
| hsa_circ_0097175 | 4410 | 17 | 0.77% |
| hsa_circ_0008476 | 6226 | 24 | 0.77% |
| hsa_circ_0080521 | 3113 | 12 | 0.77% |
| hsa_circ_0088208 | 3113 | 12 | 0.77% |
| hsa_circ_0079242 | 3892 | 15 | 0.77% |
| hsa_circ_0031174 | 3114 | 12 | 0.77% |
| hsa_circ_0035582 | 3114 | 12 | 0.77% |
| hsa_circ_0024681 | 2595 | 10 | 0.77% |
| hsa_circ_0111272 | 7007 | 27 | 0.77% |
| hsa_circ_0111787 | 5450 | 21 | 0.77% |
| hsa_circ_0130079 | 4931 | 19 | 0.77% |
| hsa_circ_0068418 | 3374 | 13 | 0.77% |
| hsa_circ_0011844 | 2855 | 11 | 0.77% |
| hsa_circ_0072382 | 2855 | 11 | 0.77% |
| hsa_circ_0098400 | 15054 | 58 | 0.77% |
| hsa_circ_0078535 | 3115 | 12 | 0.77% |
| hsa_circ_0035428 | 2596 | 10 | 0.77% |
| hsa_circ_0064951 | 2596 | 10 | 0.77% |
| hsa_circ_0051403 | 2856 | 11 | 0.77% |
| hsa_circ_0077061 | 2856 | 11 | 0.77% |
| hsa_circ_0017832 | 4674 | 18 | 0.77% |
| hsa_circ_0024589 | 3116 | 12 | 0.77% |
| hsa_circ_0046644 | 9608 | 37 | 0.77% |
| hsa_circ_0000606 | 3376 | 13 | 0.77% |
| hsa_circ_0062600 | 2597 | 10 | 0.77% |

| | | | |
|---|---|---|---|
| hsa_circ_0021963 | 2857 | 11 | 0.77% |
| hsa_circ_0054850 | 2857 | 11 | 0.77% |
| hsa_circ_0045919 | 3117 | 12 | 0.77% |
| hsa_circ_0022362 | 3377 | 13 | 0.77% |
| hsa_circ_0074561 | 3377 | 13 | 0.77% |
| hsa_circ_0039979 | 2598 | 10 | 0.77% |
| hsa_circ_0054319 | 2598 | 10 | 0.77% |
| hsa_circ_0137855 | 2598 | 10 | 0.77% |
| hsa_circ_0015183 | 2858 | 11 | 0.77% |
| hsa_circ_0059714 | 2858 | 11 | 0.77% |
| hsa_circ_0075369 | 2858 | 11 | 0.77% |
| hsa_circ_0025098 | 3118 | 12 | 0.77% |
| hsa_circ_0123394 | 3118 | 12 | 0.77% |
| hsa_circ_0125674 | 3118 | 12 | 0.77% |
| hsa_circ_0063214 | 13252 | 51 | 0.77% |
| hsa_circ_0080377 | 3378 | 13 | 0.77% |
| hsa_circ_0086936 | 3378 | 13 | 0.77% |
| hsa_circ_0091046 | 3378 | 13 | 0.77% |
| hsa_circ_0091783 | 3638 | 14 | 0.77% |
| hsa_circ_0055077 | 2599 | 10 | 0.77% |
| hsa_circ_0018903 | 3119 | 12 | 0.77% |
| hsa_circ_0067151 | 3119 | 12 | 0.77% |
| hsa_circ_0047502 | 3639 | 14 | 0.77% |
| hsa_circ_0097911 | 3380 | 13 | 0.77% |
| hsa_circ_0049235 | 2860 | 11 | 0.77% |
| hsa_circ_0090561 | 2860 | 11 | 0.77% |
| hsa_circ_0091925 | 2860 | 11 | 0.77% |
| hsa_circ_0022276 | 2600 | 10 | 0.77% |
| hsa_circ_0033089 | 2600 | 10 | 0.77% |
| hsa_circ_0039805 | 2600 | 10 | 0.77% |
| hsa_circ_0051444 | 2600 | 10 | 0.77% |
| hsa_circ_0055738 | 2600 | 10 | 0.77% |
| hsa_circ_0136254 | 2600 | 10 | 0.77% |
| hsa_circ_0094002 | 4681 | 18 | 0.77% |
| hsa_circ_0052832 | 3121 | 12 | 0.77% |
| hsa_circ_0062036 | 3121 | 12 | 0.77% |
| hsa_circ_0109992 | 3121 | 12 | 0.77% |
| hsa_circ_0077809 | 2861 | 11 | 0.77% |
| hsa_circ_0076734 | 2601 | 10 | 0.77% |
| hsa_circ_0082120 | 2601 | 10 | 0.77% |
| hsa_circ_0114811 | 2601 | 10 | 0.77% |
| hsa_circ_0122125 | 28097 | 108 | 0.77% |
| hsa_circ_0032752 | 3122 | 12 | 0.77% |
| hsa_circ_0045942 | 3122 | 12 | 0.77% |
| hsa_circ_0074134 | 3122 | 12 | 0.77% |
| hsa_circ_0070089 | 2862 | 11 | 0.77% |
| hsa_circ_0071498 | 2862 | 11 | 0.77% |
| hsa_circ_0091635 | 2862 | 11 | 0.77% |
| hsa_circ_0018144 | 2602 | 10 | 0.77% |
| hsa_circ_0075421 | 2602 | 10 | 0.77% |

| | | | |
|---|---|---|---|
| hsa_circ_0077020 | 2602 | 10 | 0.77% |
| hsa_circ_0104378 | 5985 | 23 | 0.77% |
| hsa_circ_0016530 | 3123 | 12 | 0.77% |
| hsa_circ_0019976 | 3123 | 12 | 0.77% |
| hsa_circ_0027166 | 3123 | 12 | 0.77% |
| hsa_circ_0053183 | 3123 | 12 | 0.77% |
| hsa_circ_0089524 | 3123 | 12 | 0.77% |
| hsa_circ_0024069 | 2863 | 11 | 0.77% |
| hsa_circ_0028800 | 2863 | 11 | 0.77% |
| hsa_circ_0044931 | 2863 | 11 | 0.77% |
| hsa_circ_0061930 | 2863 | 11 | 0.77% |
| hsa_circ_0036502 | 2603 | 10 | 0.77% |
| hsa_circ_0069462 | 2603 | 10 | 0.77% |
| hsa_circ_0128185 | 9893 | 38 | 0.77% |
| hsa_circ_0030277 | 7290 | 28 | 0.77% |
| hsa_circ_0111370 | 2864 | 11 | 0.77% |
| hsa_circ_0104971 | 7811 | 30 | 0.77% |
| hsa_circ_0051131 | 3385 | 13 | 0.77% |
| hsa_circ_0064893 | 3385 | 13 | 0.77% |
| hsa_circ_0018961 | 2604 | 10 | 0.77% |
| hsa_circ_0126291 | 2604 | 10 | 0.77% |
| hsa_circ_0015618 | 3125 | 12 | 0.77% |
| hsa_circ_0029192 | 3125 | 12 | 0.77% |
| hsa_circ_0077364 | 3125 | 12 | 0.77% |
| hsa_circ_0078115 | 3125 | 12 | 0.77% |
| hsa_circ_0135708 | 3125 | 12 | 0.77% |
| hsa_circ_0022041 | 3646 | 14 | 0.77% |
| hsa_circ_0034656 | 2865 | 11 | 0.77% |
| hsa_circ_0035933 | 2865 | 11 | 0.77% |
| hsa_circ_0057684 | 2865 | 11 | 0.77% |
| hsa_circ_0138358 | 2865 | 11 | 0.77% |
| hsa_circ_0047672 | 3386 | 13 | 0.77% |
| hsa_circ_0021344 | 6252 | 24 | 0.77% |
| hsa_circ_0087973 | 3126 | 12 | 0.77% |
| hsa_circ_0015867 | 2605 | 10 | 0.77% |
| hsa_circ_0050416 | 2605 | 10 | 0.77% |
| hsa_circ_0056563 | 2605 | 10 | 0.77% |
| hsa_circ_0026547 | 2866 | 11 | 0.77% |
| hsa_circ_0026858 | 2866 | 11 | 0.77% |
| hsa_circ_0077593 | 2866 | 11 | 0.77% |
| hsa_circ_0009830 | 3127 | 12 | 0.77% |
| hsa_circ_0087187 | 2606 | 10 | 0.77% |
| hsa_circ_0126657 | 2606 | 10 | 0.77% |
| hsa_circ_0034362 | 3388 | 13 | 0.77% |
| hsa_circ_0019831 | 2867 | 11 | 0.77% |
| hsa_circ_0031736 | 2867 | 11 | 0.77% |
| hsa_circ_0122126 | 28149 | 108 | 0.77% |
| hsa_circ_0099148 | 3910 | 15 | 0.77% |
| hsa_circ_0078071 | 3128 | 12 | 0.77% |
| hsa_circ_0090677 | 3128 | 12 | 0.77% |

| | | | |
|---|---|---|---|
| hsa_circ_0050079 | 10688 | 41 | 0.77% |
| hsa_circ_0077757 | 2607 | 10 | 0.77% |
| hsa_circ_0129470 | 2607 | 10 | 0.77% |
| hsa_circ_0027844 | 2868 | 11 | 0.77% |
| hsa_circ_0049507 | 2868 | 11 | 0.77% |
| hsa_circ_0056058 | 2868 | 11 | 0.77% |
| hsa_circ_0088961 | 2868 | 11 | 0.77% |
| hsa_circ_0065036 | 3129 | 12 | 0.77% |
| hsa_circ_0074908 | 3129 | 12 | 0.77% |
| hsa_circ_0126826 | 7562 | 29 | 0.77% |
| hsa_circ_0025985 | 3390 | 13 | 0.77% |
| hsa_circ_0042920 | 3390 | 13 | 0.77% |
| hsa_circ_0047813 | 3390 | 13 | 0.77% |
| hsa_circ_0017965 | 2608 | 10 | 0.77% |
| hsa_circ_0061785 | 2608 | 10 | 0.77% |
| hsa_circ_0072224 | 2608 | 10 | 0.77% |
| hsa_circ_0029691 | 2869 | 11 | 0.77% |
| hsa_circ_0032430 | 2869 | 11 | 0.77% |
| hsa_circ_0067561 | 2869 | 11 | 0.77% |
| hsa_circ_0075158 | 2869 | 11 | 0.77% |
| hsa_circ_0107489 | 11216 | 43 | 0.77% |
| hsa_circ_0137541 | 3391 | 13 | 0.77% |
| hsa_circ_0050491 | 3652 | 14 | 0.77% |
| hsa_circ_0019196 | 2870 | 11 | 0.77% |
| hsa_circ_0039309 | 2870 | 11 | 0.77% |
| hsa_circ_0022307 | 3131 | 12 | 0.77% |
| hsa_circ_0045521 | 3131 | 12 | 0.77% |
| hsa_circ_0058563 | 3131 | 12 | 0.77% |
| hsa_circ_0098690 | 3131 | 12 | 0.77% |
| hsa_circ_0120796 | 8352 | 32 | 0.77% |
| hsa_circ_0050972 | 3132 | 12 | 0.77% |
| hsa_circ_0067361 | 3132 | 12 | 0.77% |
| hsa_circ_0076830 | 3132 | 12 | 0.77% |
| hsa_circ_0030313 | 2871 | 11 | 0.77% |
| hsa_circ_0035138 | 2871 | 11 | 0.77% |
| hsa_circ_0016619 | 2610 | 10 | 0.77% |
| hsa_circ_0090835 | 8353 | 32 | 0.77% |
| hsa_circ_0102545 | 4960 | 19 | 0.77% |
| hsa_circ_0045797 | 3394 | 13 | 0.77% |
| hsa_circ_0101958 | 6527 | 25 | 0.77% |
| hsa_circ_0058761 | 3133 | 12 | 0.77% |
| hsa_circ_0098689 | 3133 | 12 | 0.77% |
| hsa_circ_0137209 | 2872 | 11 | 0.77% |
| hsa_circ_0051558 | 2611 | 10 | 0.77% |
| hsa_circ_0094915 | 2611 | 10 | 0.77% |
| hsa_circ_0113521 | 3656 | 14 | 0.77% |
| hsa_circ_0070041 | 3395 | 13 | 0.77% |
| hsa_circ_0041578 | 3134 | 12 | 0.77% |
| hsa_circ_0119189 | 3134 | 12 | 0.77% |
| hsa_circ_0125692 | 3134 | 12 | 0.77% |

| | | | |
|---|---|---|---|
| hsa_circ_0119711 | 10448 | 40 | 0.77% |
| hsa_circ_0114442 | 6531 | 25 | 0.77% |
| hsa_circ_0002691 | 4180 | 16 | 0.77% |
| hsa_circ_0140783 | 3135 | 12 | 0.77% |
| hsa_circ_0079443 | 2874 | 11 | 0.77% |
| hsa_circ_0020421 | 2613 | 10 | 0.77% |
| hsa_circ_0072660 | 2613 | 10 | 0.77% |
| hsa_circ_0090757 | 3397 | 13 | 0.77% |
| hsa_circ_0026072 | 3136 | 12 | 0.77% |
| hsa_circ_0023718 | 2875 | 11 | 0.77% |
| hsa_circ_0079052 | 2614 | 10 | 0.77% |
| hsa_circ_0015612 | 3137 | 12 | 0.77% |
| hsa_circ_0022695 | 3137 | 12 | 0.77% |
| hsa_circ_0024437 | 3137 | 12 | 0.77% |
| hsa_circ_0090784 | 3137 | 12 | 0.77% |
| hsa_circ_0035307 | 2876 | 11 | 0.76% |
| hsa_circ_0058983 | 2876 | 11 | 0.76% |
| hsa_circ_0091644 | 2876 | 11 | 0.76% |
| hsa_circ_0053690 | 3399 | 13 | 0.76% |
| hsa_circ_0042424 | 3138 | 12 | 0.76% |
| hsa_circ_0056807 | 3138 | 12 | 0.76% |
| hsa_circ_0130088 | 3138 | 12 | 0.76% |
| hsa_circ_0011546 | 2615 | 10 | 0.76% |
| hsa_circ_0027988 | 2877 | 11 | 0.76% |
| hsa_circ_0054824 | 2877 | 11 | 0.76% |
| hsa_circ_0122696 | 2877 | 11 | 0.76% |
| hsa_circ_0018947 | 3139 | 12 | 0.76% |
| hsa_circ_0035710 | 3139 | 12 | 0.76% |
| hsa_circ_0040006 | 3139 | 12 | 0.76% |
| hsa_circ_0015907 | 8371 | 32 | 0.76% |
| hsa_circ_0083542 | 3924 | 15 | 0.76% |
| hsa_circ_0016267 | 2616 | 10 | 0.76% |
| hsa_circ_0084585 | 2616 | 10 | 0.76% |
| hsa_circ_0052205 | 3401 | 13 | 0.76% |
| hsa_circ_0046478 | 2878 | 11 | 0.76% |
| hsa_circ_0052917 | 2878 | 11 | 0.76% |
| hsa_circ_0084403 | 3402 | 13 | 0.76% |
| hsa_circ_0053368 | 2617 | 10 | 0.76% |
| hsa_circ_0110782 | 2617 | 10 | 0.76% |
| hsa_circ_0011220 | 2879 | 11 | 0.76% |
| hsa_circ_0071187 | 2879 | 11 | 0.76% |
| hsa_circ_0094906 | 2879 | 11 | 0.76% |
| hsa_circ_0132696 | 6282 | 24 | 0.76% |
| hsa_circ_0012034 | 3403 | 13 | 0.76% |
| hsa_circ_0013130 | 3665 | 14 | 0.76% |
| hsa_circ_0065894 | 2618 | 10 | 0.76% |
| hsa_circ_0086573 | 2880 | 11 | 0.76% |
| hsa_circ_0090203 | 2880 | 11 | 0.76% |
| hsa_circ_0110055 | 2880 | 11 | 0.76% |
| hsa_circ_0021880 | 3142 | 12 | 0.76% |

| | | | |
|---|---|---|---|
| hsa_circ_0126449 | 3404 | 13 | 0.76% |
| hsa_circ_0048720 | 2619 | 10 | 0.76% |
| hsa_circ_0071988 | 2619 | 10 | 0.76% |
| hsa_circ_0114808 | 2619 | 10 | 0.76% |
| hsa_circ_0027917 | 2881 | 11 | 0.76% |
| hsa_circ_0013308 | 3143 | 12 | 0.76% |
| hsa_circ_0043314 | 9691 | 37 | 0.76% |
| hsa_circ_0006293 | 7334 | 28 | 0.76% |
| hsa_circ_0117360 | 6026 | 23 | 0.76% |
| hsa_circ_0085833 | 3406 | 13 | 0.76% |
| hsa_circ_0073883 | 3144 | 12 | 0.76% |
| hsa_circ_0052744 | 2620 | 10 | 0.76% |
| hsa_circ_0025355 | 3407 | 13 | 0.76% |
| hsa_circ_0060643 | 3407 | 13 | 0.76% |
| hsa_circ_0072449 | 3145 | 12 | 0.76% |
| hsa_circ_0083486 | 3145 | 12 | 0.76% |
| hsa_circ_0045832 | 2883 | 11 | 0.76% |
| hsa_circ_0058547 | 2883 | 11 | 0.76% |
| hsa_circ_0073067 | 2883 | 11 | 0.76% |
| hsa_circ_0060189 | 3408 | 13 | 0.76% |
| hsa_circ_0071221 | 3408 | 13 | 0.76% |
| hsa_circ_0033179 | 2884 | 11 | 0.76% |
| hsa_circ_0044866 | 2884 | 11 | 0.76% |
| hsa_circ_0055660 | 2884 | 11 | 0.76% |
| hsa_circ_0063096 | 2884 | 11 | 0.76% |
| hsa_circ_0078478 | 2884 | 11 | 0.76% |
| hsa_circ_0011795 | 2622 | 10 | 0.76% |
| hsa_circ_0013160 | 7342 | 28 | 0.76% |
| hsa_circ_0087185 | 3409 | 13 | 0.76% |
| hsa_circ_0065126 | 3147 | 12 | 0.76% |
| hsa_circ_0036478 | 3148 | 12 | 0.76% |
| hsa_circ_0121337 | 3148 | 12 | 0.76% |
| hsa_circ_0116140 | 86579 | 330 | 0.76% |
| hsa_circ_0061708 | 2886 | 11 | 0.76% |
| hsa_circ_0073357 | 3411 | 13 | 0.76% |
| hsa_circ_0027564 | 3936 | 15 | 0.76% |
| hsa_circ_0014809 | 2624 | 10 | 0.76% |
| hsa_circ_0053187 | 3149 | 12 | 0.76% |
| hsa_circ_0088867 | 3149 | 12 | 0.76% |
| hsa_circ_0139467 | 3674 | 14 | 0.76% |
| hsa_circ_0065198 | 2887 | 11 | 0.76% |
| hsa_circ_0121776 | 9711 | 37 | 0.76% |
| hsa_circ_0034152 | 3412 | 13 | 0.76% |
| hsa_circ_0139442 | 2625 | 10 | 0.76% |
| hsa_circ_0098294 | 7088 | 27 | 0.76% |
| hsa_circ_0035099 | 2888 | 11 | 0.76% |
| hsa_circ_0035351 | 2888 | 11 | 0.76% |
| hsa_circ_0135338 | 2888 | 11 | 0.76% |
| hsa_circ_0016596 | 6564 | 25 | 0.76% |
| hsa_circ_0033490 | 3151 | 12 | 0.76% |

| | | | |
|---|---|---|---|
| hsa_circ_0026820 | 2626 | 10 | 0.76% |
| hsa_circ_0029747 | 2626 | 10 | 0.76% |
| hsa_circ_0038445 | 2626 | 10 | 0.76% |
| hsa_circ_0086334 | 2626 | 10 | 0.76% |
| hsa_circ_0087346 | 3414 | 13 | 0.76% |
| hsa_circ_0027216 | 2889 | 11 | 0.76% |
| hsa_circ_0070237 | 2889 | 11 | 0.76% |
| hsa_circ_0094685 | 2889 | 11 | 0.76% |
| hsa_circ_0086823 | 3152 | 12 | 0.76% |
| hsa_circ_0024395 | 3415 | 13 | 0.76% |
| hsa_circ_0025699 | 3415 | 13 | 0.76% |
| hsa_circ_0009380 | 2627 | 10 | 0.76% |
| hsa_circ_0054344 | 2627 | 10 | 0.76% |
| hsa_circ_0061614 | 2627 | 10 | 0.76% |
| hsa_circ_0091884 | 2627 | 10 | 0.76% |
| hsa_circ_0019894 | 2890 | 11 | 0.76% |
| hsa_circ_0121100 | 2890 | 11 | 0.76% |
| hsa_circ_0057559 | 3941 | 15 | 0.76% |
| hsa_circ_0033312 | 3153 | 12 | 0.76% |
| hsa_circ_0069670 | 3153 | 12 | 0.76% |
| hsa_circ_0099066 | 3153 | 12 | 0.76% |
| hsa_circ_0012674 | 3416 | 13 | 0.76% |
| hsa_circ_0032552 | 3416 | 13 | 0.76% |
| hsa_circ_0046103 | 3416 | 13 | 0.76% |
| hsa_circ_0088861 | 3416 | 13 | 0.76% |
| hsa_circ_0076007 | 3679 | 14 | 0.76% |
| hsa_circ_0091785 | 2628 | 10 | 0.76% |
| hsa_circ_0029775 | 2891 | 11 | 0.76% |
| hsa_circ_0038274 | 2891 | 11 | 0.76% |
| hsa_circ_0060962 | 2891 | 11 | 0.76% |
| hsa_circ_0097784 | 2891 | 11 | 0.76% |
| hsa_circ_0129768 | 2891 | 11 | 0.76% |
| hsa_circ_0011894 | 3154 | 12 | 0.76% |
| hsa_circ_0031682 | 3417 | 13 | 0.76% |
| hsa_circ_0092623 | 3680 | 14 | 0.76% |
| hsa_circ_0068014 | 3943 | 15 | 0.76% |
| hsa_circ_0077633 | 2629 | 10 | 0.76% |
| hsa_circ_0091648 | 2629 | 10 | 0.76% |
| hsa_circ_0112026 | 2629 | 10 | 0.76% |
| hsa_circ_0065727 | 2892 | 11 | 0.76% |
| hsa_circ_0073852 | 2892 | 11 | 0.76% |
| hsa_circ_0046643 | 3155 | 12 | 0.76% |
| hsa_circ_0069081 | 3155 | 12 | 0.76% |
| hsa_circ_0084212 | 3155 | 12 | 0.76% |
| hsa_circ_0091980 | 3155 | 12 | 0.76% |
| hsa_circ_0001668 | 3418 | 13 | 0.76% |
| hsa_circ_0109866 | 8940 | 34 | 0.76% |
| hsa_circ_0127107 | 9204 | 35 | 0.76% |
| hsa_circ_0087829 | 3419 | 13 | 0.76% |
| hsa_circ_0048882 | 3156 | 12 | 0.76% |

| | | | |
|---|---|---|---|
| hsa_circ_0016825 | 2893 | 11 | 0.76% |
| hsa_circ_0048313 | 2893 | 11 | 0.76% |
| hsa_circ_0072180 | 2893 | 11 | 0.76% |
| hsa_circ_0082065 | 2630 | 10 | 0.76% |
| hsa_circ_0090319 | 2630 | 10 | 0.76% |
| hsa_circ_0025097 | 3157 | 12 | 0.76% |
| hsa_circ_0089111 | 3157 | 12 | 0.76% |
| hsa_circ_0040232 | 2894 | 11 | 0.76% |
| hsa_circ_0056096 | 2631 | 10 | 0.76% |
| hsa_circ_0060413 | 2631 | 10 | 0.76% |
| hsa_circ_0074304 | 2631 | 10 | 0.76% |
| hsa_circ_0074942 | 2631 | 10 | 0.76% |
| hsa_circ_0090702 | 2631 | 10 | 0.76% |
| hsa_circ_0112230 | 7104 | 27 | 0.76% |
| hsa_circ_0074562 | 3421 | 13 | 0.76% |
| hsa_circ_0048545 | 3158 | 12 | 0.76% |
| hsa_circ_0088830 | 3158 | 12 | 0.76% |
| hsa_circ_0067046 | 2895 | 11 | 0.76% |
| hsa_circ_0076021 | 2895 | 11 | 0.76% |
| hsa_circ_0085208 | 2895 | 11 | 0.76% |
| hsa_circ_0077755 | 2896 | 11 | 0.76% |
| hsa_circ_0090396 | 2896 | 11 | 0.76% |
| hsa_circ_0065081 | 3160 | 12 | 0.76% |
| hsa_circ_0081363 | 4477 | 17 | 0.76% |
| hsa_circ_0042015 | 2897 | 11 | 0.76% |
| hsa_circ_0063919 | 2897 | 11 | 0.76% |
| hsa_circ_0086406 | 2897 | 11 | 0.76% |
| hsa_circ_0089423 | 2897 | 11 | 0.76% |
| hsa_circ_0090919 | 3424 | 13 | 0.76% |
| hsa_circ_0036894 | 3161 | 12 | 0.76% |
| hsa_circ_0067613 | 3161 | 12 | 0.76% |
| hsa_circ_0131134 | 3161 | 12 | 0.76% |
| hsa_circ_0039021 | 2898 | 11 | 0.76% |
| hsa_circ_0062091 | 2898 | 11 | 0.76% |
| hsa_circ_0137571 | 2898 | 11 | 0.76% |
| hsa_circ_0077300 | 3689 | 14 | 0.76% |
| hsa_circ_0128110 | 3162 | 12 | 0.76% |
| hsa_circ_0015500 | 2635 | 10 | 0.76% |
| hsa_circ_0057642 | 2635 | 10 | 0.76% |
| hsa_circ_0012597 | 2899 | 11 | 0.76% |
| hsa_circ_0019723 | 2899 | 11 | 0.76% |
| hsa_circ_0011460 | 3163 | 12 | 0.76% |
| hsa_circ_0060644 | 3163 | 12 | 0.76% |
| hsa_circ_0119846 | 2636 | 10 | 0.76% |
| hsa_circ_0097060 | 2900 | 11 | 0.76% |
| hsa_circ_0076067 | 3164 | 12 | 0.76% |
| hsa_circ_0119838 | 3164 | 12 | 0.76% |
| hsa_circ_0018339 | 2637 | 10 | 0.76% |
| hsa_circ_0025254 | 2637 | 10 | 0.76% |
| hsa_circ_0040732 | 2637 | 10 | 0.76% |

| | | | |
|---|---|---|---|
| hsa_circ_0078511 | 4483 | 17 | 0.76% |
| hsa_circ_0002317 | 10549 | 40 | 0.76% |
| hsa_circ_0040171 | 2901 | 11 | 0.76% |
| hsa_circ_0041964 | 2901 | 11 | 0.76% |
| hsa_circ_0056553 | 2901 | 11 | 0.76% |
| hsa_circ_0134958 | 5275 | 20 | 0.76% |
| hsa_circ_0056031 | 3165 | 12 | 0.76% |
| hsa_circ_0013892 | 2902 | 11 | 0.76% |
| hsa_circ_0053483 | 2902 | 11 | 0.76% |
| hsa_circ_0060431 | 2902 | 11 | 0.76% |
| hsa_circ_0102706 | 8970 | 34 | 0.76% |
| hsa_circ_0010077 | 3430 | 13 | 0.76% |
| hsa_circ_0028546 | 3430 | 13 | 0.76% |
| hsa_circ_0014037 | 2639 | 10 | 0.76% |
| hsa_circ_0038746 | 2903 | 11 | 0.76% |
| hsa_circ_0042903 | 2903 | 11 | 0.76% |
| hsa_circ_0083301 | 2903 | 11 | 0.76% |
| hsa_circ_0042422 | 3167 | 12 | 0.76% |
| hsa_circ_0053369 | 3167 | 12 | 0.76% |
| hsa_circ_0074521 | 3167 | 12 | 0.76% |
| hsa_circ_0106223 | 4751 | 18 | 0.76% |
| hsa_circ_0102143 | 3696 | 14 | 0.76% |
| hsa_circ_0069098 | 3432 | 13 | 0.76% |
| hsa_circ_0091441 | 3432 | 13 | 0.76% |
| hsa_circ_0030211 | 3168 | 12 | 0.76% |
| hsa_circ_0064133 | 3168 | 12 | 0.76% |
| hsa_circ_0023203 | 2904 | 11 | 0.76% |
| hsa_circ_0068343 | 2904 | 11 | 0.76% |
| hsa_circ_0084308 | 2904 | 11 | 0.76% |
| hsa_circ_0009708 | 2640 | 10 | 0.76% |
| hsa_circ_0113305 | 2640 | 10 | 0.76% |
| hsa_circ_0070412 | 3169 | 12 | 0.76% |
| hsa_circ_0079861 | 3169 | 12 | 0.76% |
| hsa_circ_0062294 | 2905 | 11 | 0.76% |
| hsa_circ_0069671 | 2905 | 11 | 0.76% |
| hsa_circ_0077968 | 2905 | 11 | 0.76% |
| hsa_circ_0035512 | 2641 | 10 | 0.76% |
| hsa_circ_0140319 | 5018 | 19 | 0.76% |
| hsa_circ_0060020 | 3170 | 12 | 0.76% |
| hsa_circ_0025864 | 2906 | 11 | 0.76% |
| hsa_circ_0059680 | 2906 | 11 | 0.76% |
| hsa_circ_0121142 | 2906 | 11 | 0.76% |
| hsa_circ_0013331 | 2642 | 10 | 0.76% |
| hsa_circ_0050598 | 2642 | 10 | 0.76% |
| hsa_circ_0039640 | 3435 | 13 | 0.76% |
| hsa_circ_0033310 | 3171 | 12 | 0.76% |
| hsa_circ_0040397 | 3171 | 12 | 0.76% |
| hsa_circ_0083715 | 3171 | 12 | 0.76% |
| hsa_circ_0026992 | 2907 | 11 | 0.76% |
| hsa_circ_0054583 | 2907 | 11 | 0.76% |

| | | | |
|---|---|---|---|
| hsa_circ_0108750 | 2907 | 11 | 0.76% |
| hsa_circ_0111275 | 2907 | 11 | 0.76% |
| hsa_circ_0132702 | 5550 | 21 | 0.76% |
| hsa_circ_0046898 | 3700 | 14 | 0.76% |
| hsa_circ_0131259 | 3700 | 14 | 0.76% |
| hsa_circ_0139443 | 2643 | 10 | 0.76% |
| hsa_circ_0031622 | 3436 | 13 | 0.76% |
| hsa_circ_0126003 | 7930 | 30 | 0.76% |
| hsa_circ_0047521 | 3172 | 12 | 0.76% |
| hsa_circ_0022726 | 2908 | 11 | 0.76% |
| hsa_circ_0106700 | 2908 | 11 | 0.76% |
| hsa_circ_0087119 | 3437 | 13 | 0.76% |
| hsa_circ_0006233 | 6081 | 23 | 0.76% |
| hsa_circ_0013161 | 7403 | 28 | 0.76% |
| hsa_circ_0033392 | 2644 | 10 | 0.76% |
| hsa_circ_0029190 | 3173 | 12 | 0.76% |
| hsa_circ_0050087 | 6611 | 25 | 0.76% |
| hsa_circ_0016210 | 5818 | 22 | 0.76% |
| hsa_circ_0007670 | 2909 | 11 | 0.76% |
| hsa_circ_0017036 | 2909 | 11 | 0.76% |
| hsa_circ_0086802 | 2909 | 11 | 0.76% |
| hsa_circ_0096760 | 2909 | 11 | 0.76% |
| hsa_circ_0047016 | 3438 | 13 | 0.76% |
| hsa_circ_0081247 | 3967 | 15 | 0.76% |
| hsa_circ_0067830 | 3174 | 12 | 0.76% |
| hsa_circ_0091700 | 3174 | 12 | 0.76% |
| hsa_circ_0013112 | 2645 | 10 | 0.76% |
| hsa_circ_0099653 | 2645 | 10 | 0.76% |
| hsa_circ_0101853 | 7936 | 30 | 0.76% |
| hsa_circ_0029101 | 2910 | 11 | 0.76% |
| hsa_circ_0035735 | 3175 | 12 | 0.76% |
| hsa_circ_0039401 | 3175 | 12 | 0.76% |
| hsa_circ_0075688 | 2646 | 10 | 0.76% |
| hsa_circ_0100335 | 3440 | 13 | 0.76% |
| hsa_circ_0109497 | 14025 | 53 | 0.76% |
| hsa_circ_0019001 | 2911 | 11 | 0.76% |
| hsa_circ_0030086 | 2911 | 11 | 0.76% |
| hsa_circ_0126210 | 2911 | 11 | 0.76% |
| hsa_circ_0135331 | 7146 | 27 | 0.76% |
| hsa_circ_0087967 | 3176 | 12 | 0.76% |
| hsa_circ_0062615 | 4235 | 16 | 0.76% |
| hsa_circ_0050397 | 2647 | 10 | 0.76% |
| hsa_circ_0062223 | 2647 | 10 | 0.76% |
| hsa_circ_0001344 | 2912 | 11 | 0.76% |
| hsa_circ_0053751 | 2912 | 11 | 0.76% |
| hsa_circ_0116969 | 2912 | 11 | 0.76% |
| hsa_circ_0091511 | 3177 | 12 | 0.76% |
| hsa_circ_0032648 | 3442 | 13 | 0.76% |
| hsa_circ_0015457 | 2648 | 10 | 0.76% |
| hsa_circ_0055011 | 2648 | 10 | 0.76% |

| | | | |
|---|---|---|---|
| hsa_circ_0023866 | 2913 | 11 | 0.76% |
| hsa_circ_0082899 | 2913 | 11 | 0.76% |
| hsa_circ_0086525 | 2913 | 11 | 0.76% |
| hsa_circ_0000377 | 3178 | 12 | 0.76% |
| hsa_circ_0050792 | 3178 | 12 | 0.76% |
| hsa_circ_0120192 | 3178 | 12 | 0.76% |
| hsa_circ_0095024 | 3443 | 13 | 0.76% |
| hsa_circ_0056303 | 2649 | 10 | 0.76% |
| hsa_circ_0051194 | 2914 | 11 | 0.75% |
| hsa_circ_0135884 | 3179 | 12 | 0.75% |
| hsa_circ_0036422 | 3445 | 13 | 0.75% |
| hsa_circ_0066669 | 3180 | 12 | 0.75% |
| hsa_circ_0026856 | 2915 | 11 | 0.75% |
| hsa_circ_0044581 | 2915 | 11 | 0.75% |
| hsa_circ_0085392 | 2915 | 11 | 0.75% |
| hsa_circ_0078117 | 2650 | 10 | 0.75% |
| hsa_circ_0025913 | 3711 | 14 | 0.75% |
| hsa_circ_0073366 | 3711 | 14 | 0.75% |
| hsa_circ_0060681 | 3446 | 13 | 0.75% |
| hsa_circ_0040794 | 3181 | 12 | 0.75% |
| hsa_circ_0044467 | 3181 | 12 | 0.75% |
| hsa_circ_0062906 | 3181 | 12 | 0.75% |
| hsa_circ_0069162 | 2916 | 11 | 0.75% |
| hsa_circ_0056829 | 2651 | 10 | 0.75% |
| hsa_circ_0084778 | 2651 | 10 | 0.75% |
| hsa_circ_0130085 | 6628 | 25 | 0.75% |
| hsa_circ_0069304 | 2917 | 11 | 0.75% |
| hsa_circ_0114959 | 7956 | 30 | 0.75% |
| hsa_circ_0022473 | 2652 | 10 | 0.75% |
| hsa_circ_0109516 | 9814 | 37 | 0.75% |
| hsa_circ_0049010 | 3183 | 12 | 0.75% |
| hsa_circ_0081449 | 3183 | 12 | 0.75% |
| hsa_circ_0114411 | 3183 | 12 | 0.75% |
| hsa_circ_0085450 | 2918 | 11 | 0.75% |
| hsa_circ_0037694 | 3714 | 14 | 0.75% |
| hsa_circ_0048037 | 2653 | 10 | 0.75% |
| hsa_circ_0060187 | 3449 | 13 | 0.75% |
| hsa_circ_0090266 | 3449 | 13 | 0.75% |
| hsa_circ_0021153 | 3184 | 12 | 0.75% |
| hsa_circ_0029167 | 3184 | 12 | 0.75% |
| hsa_circ_0088373 | 3184 | 12 | 0.75% |
| hsa_circ_0024127 | 2919 | 11 | 0.75% |
| hsa_circ_0073087 | 2919 | 11 | 0.75% |
| hsa_circ_0032226 | 3450 | 13 | 0.75% |
| hsa_circ_0085345 | 3450 | 13 | 0.75% |
| hsa_circ_0020419 | 2654 | 10 | 0.75% |
| hsa_circ_0077903 | 2654 | 10 | 0.75% |
| hsa_circ_0129772 | 2654 | 10 | 0.75% |
| hsa_circ_0015851 | 3185 | 12 | 0.75% |
| hsa_circ_0126540 | 3185 | 12 | 0.75% |

| | | | |
|---|---|---|---|
| hsa_circ_0034246 | 2920 | 11 | 0.75% |
| hsa_circ_0129904 | 27612 | 104 | 0.75% |
| hsa_circ_0051302 | 3186 | 12 | 0.75% |
| hsa_circ_0116543 | 3186 | 12 | 0.75% |
| hsa_circ_0096741 | 2655 | 10 | 0.75% |
| hsa_circ_0129979 | 2921 | 11 | 0.75% |
| hsa_circ_0140152 | 2921 | 11 | 0.75% |
| hsa_circ_0075484 | 3187 | 12 | 0.75% |
| hsa_circ_0099011 | 3187 | 12 | 0.75% |
| hsa_circ_0078321 | 2656 | 10 | 0.75% |
| hsa_circ_0053662 | 3453 | 13 | 0.75% |
| hsa_circ_0124705 | 26828 | 101 | 0.75% |
| hsa_circ_0041528 | 2922 | 11 | 0.75% |
| hsa_circ_0042220 | 2922 | 11 | 0.75% |
| hsa_circ_0085805 | 2922 | 11 | 0.75% |
| hsa_circ_0086869 | 2922 | 11 | 0.75% |
| hsa_circ_0108983 | 3188 | 12 | 0.75% |
| hsa_circ_0090605 | 4251 | 16 | 0.75% |
| hsa_circ_0026331 | 3454 | 13 | 0.75% |
| hsa_circ_0039159 | 3454 | 13 | 0.75% |
| hsa_circ_0024613 | 2923 | 11 | 0.75% |
| hsa_circ_0053444 | 2923 | 11 | 0.75% |
| hsa_circ_0081673 | 9035 | 34 | 0.75% |
| hsa_circ_0114650 | 4252 | 16 | 0.75% |
| hsa_circ_0014033 | 3189 | 12 | 0.75% |
| hsa_circ_0102100 | 3189 | 12 | 0.75% |
| hsa_circ_0071229 | 3455 | 13 | 0.75% |
| hsa_circ_0051755 | 3721 | 14 | 0.75% |
| hsa_circ_0030961 | 2658 | 10 | 0.75% |
| hsa_circ_0106093 | 2658 | 10 | 0.75% |
| hsa_circ_0019832 | 2924 | 11 | 0.75% |
| hsa_circ_0028756 | 2924 | 11 | 0.75% |
| hsa_circ_0073421 | 2924 | 11 | 0.75% |
| hsa_circ_0020626 | 3190 | 12 | 0.75% |
| hsa_circ_0071348 | 3190 | 12 | 0.75% |
| hsa_circ_0118360 | 3456 | 13 | 0.75% |
| hsa_circ_0112962 | 14622 | 55 | 0.75% |
| hsa_circ_0133481 | 4786 | 18 | 0.75% |
| hsa_circ_0069292 | 2925 | 11 | 0.75% |
| hsa_circ_0072246 | 2925 | 11 | 0.75% |
| hsa_circ_0077203 | 2925 | 11 | 0.75% |
| hsa_circ_0022214 | 3457 | 13 | 0.75% |
| hsa_circ_0077299 | 3457 | 13 | 0.75% |
| hsa_circ_0112129 | 9575 | 36 | 0.75% |
| hsa_circ_0083406 | 3458 | 13 | 0.75% |
| hsa_circ_0063501 | 3192 | 12 | 0.75% |
| hsa_circ_0019895 | 2926 | 11 | 0.75% |
| hsa_circ_0047002 | 3193 | 12 | 0.75% |
| hsa_circ_0056464 | 3193 | 12 | 0.75% |
| hsa_circ_0062035 | 3193 | 12 | 0.75% |

| | | | |
|---|---|---|---|
| hsa_circ_0080194 | 3193 | 12 | 0.75% |
| hsa_circ_0013801 | 2927 | 11 | 0.75% |
| hsa_circ_0039185 | 2927 | 11 | 0.75% |
| hsa_circ_0039267 | 2927 | 11 | 0.75% |
| hsa_circ_0041375 | 2927 | 11 | 0.75% |
| hsa_circ_0005062 | 2661 | 10 | 0.75% |
| hsa_circ_0092379 | 2661 | 10 | 0.75% |
| hsa_circ_0072819 | 6919 | 26 | 0.75% |
| hsa_circ_0132175 | 3726 | 14 | 0.75% |
| hsa_circ_0016670 | 3460 | 13 | 0.75% |
| hsa_circ_0132694 | 6388 | 24 | 0.75% |
| hsa_circ_0028398 | 2928 | 11 | 0.75% |
| hsa_circ_0071217 | 2662 | 10 | 0.75% |
| hsa_circ_0104037 | 3461 | 13 | 0.75% |
| hsa_circ_0048161 | 2929 | 11 | 0.75% |
| hsa_circ_0056775 | 2929 | 11 | 0.75% |
| hsa_circ_0093298 | 2929 | 11 | 0.75% |
| hsa_circ_0131886 | 2929 | 11 | 0.75% |
| hsa_circ_0090946 | 2663 | 10 | 0.75% |
| hsa_circ_0097038 | 2663 | 10 | 0.75% |
| hsa_circ_0134626 | 10386 | 39 | 0.75% |
| hsa_circ_0078994 | 3995 | 15 | 0.75% |
| hsa_circ_0014451 | 3196 | 12 | 0.75% |
| hsa_circ_0025790 | 3196 | 12 | 0.75% |
| hsa_circ_0034089 | 3196 | 12 | 0.75% |
| hsa_circ_0119395 | 3196 | 12 | 0.75% |
| hsa_circ_0115625 | 8790 | 33 | 0.75% |
| hsa_circ_0051698 | 2930 | 11 | 0.75% |
| hsa_circ_0083461 | 2930 | 11 | 0.75% |
| hsa_circ_0113310 | 2930 | 11 | 0.75% |
| hsa_circ_0007013 | 12254 | 46 | 0.75% |
| hsa_circ_0059689 | 3197 | 12 | 0.75% |
| hsa_circ_0061919 | 3197 | 12 | 0.75% |
| hsa_circ_0013726 | 2931 | 11 | 0.75% |
| hsa_circ_0039257 | 2931 | 11 | 0.75% |
| hsa_circ_0081762 | 2931 | 11 | 0.75% |
| hsa_circ_0139843 | 4530 | 17 | 0.75% |
| hsa_circ_0018432 | 3198 | 12 | 0.75% |
| hsa_circ_0034531 | 3198 | 12 | 0.75% |
| hsa_circ_0016007 | 2665 | 10 | 0.75% |
| hsa_circ_0091951 | 2665 | 10 | 0.75% |
| hsa_circ_0083539 | 3998 | 15 | 0.75% |
| hsa_circ_0019164 | 3465 | 13 | 0.75% |
| hsa_circ_0044331 | 3465 | 13 | 0.75% |
| hsa_circ_0021011 | 2932 | 11 | 0.75% |
| hsa_circ_0024188 | 2932 | 11 | 0.75% |
| hsa_circ_0075893 | 2932 | 11 | 0.75% |
| hsa_circ_0039085 | 3199 | 12 | 0.75% |
| hsa_circ_0023654 | 2666 | 10 | 0.75% |
| hsa_circ_0128156 | 2666 | 10 | 0.75% |

| | | | |
|---|---|---|---|
| hsa_circ_0082049 | 3466 | 13 | 0.75% |
| hsa_circ_0089245 | 3466 | 13 | 0.75% |
| hsa_circ_0091255 | 3466 | 13 | 0.75% |
| hsa_circ_0081505 | 4266 | 16 | 0.75% |
| hsa_circ_0034769 | 2933 | 11 | 0.75% |
| hsa_circ_0077127 | 2933 | 11 | 0.75% |
| hsa_circ_0079424 | 2933 | 11 | 0.75% |
| hsa_circ_0034436 | 3200 | 12 | 0.75% |
| hsa_circ_0064065 | 3200 | 12 | 0.75% |
| hsa_circ_0068582 | 3467 | 13 | 0.75% |
| hsa_circ_0070291 | 2667 | 10 | 0.75% |
| hsa_circ_0078588 | 2667 | 10 | 0.75% |
| hsa_circ_0023019 | 4534 | 17 | 0.75% |
| hsa_circ_0052833 | 3201 | 12 | 0.75% |
| hsa_circ_0078080 | 3201 | 12 | 0.75% |
| hsa_circ_0119904 | 3201 | 12 | 0.75% |
| hsa_circ_0130407 | 3201 | 12 | 0.75% |
| hsa_circ_0133519 | 3201 | 12 | 0.75% |
| hsa_circ_0030532 | 2668 | 10 | 0.75% |
| hsa_circ_0071955 | 2668 | 10 | 0.75% |
| hsa_circ_0020399 | 2935 | 11 | 0.75% |
| hsa_circ_0088291 | 2935 | 11 | 0.75% |
| hsa_circ_0138449 | 2935 | 11 | 0.75% |
| hsa_circ_0055296 | 3202 | 12 | 0.75% |
| hsa_circ_0060331 | 3469 | 13 | 0.75% |
| hsa_circ_0040958 | 4003 | 15 | 0.75% |
| hsa_circ_0034990 | 2669 | 10 | 0.75% |
| hsa_circ_0041398 | 2669 | 10 | 0.75% |
| hsa_circ_0077403 | 2669 | 10 | 0.75% |
| hsa_circ_0034708 | 2936 | 11 | 0.75% |
| hsa_circ_0072035 | 2936 | 11 | 0.75% |
| hsa_circ_0028118 | 3203 | 12 | 0.75% |
| hsa_circ_0069010 | 3203 | 12 | 0.75% |
| hsa_circ_0110125 | 3203 | 12 | 0.75% |
| hsa_circ_0120703 | 3203 | 12 | 0.75% |
| hsa_circ_0022858 | 3470 | 13 | 0.75% |
| hsa_circ_0088862 | 3470 | 13 | 0.75% |
| hsa_circ_0066064 | 3204 | 12 | 0.75% |
| hsa_circ_0138438 | 2937 | 11 | 0.75% |
| hsa_circ_0019129 | 2670 | 10 | 0.75% |
| hsa_circ_0090631 | 2670 | 10 | 0.75% |
| hsa_circ_0048391 | 3739 | 14 | 0.75% |
| hsa_circ_0047404 | 3472 | 13 | 0.75% |
| hsa_circ_0013698 | 3205 | 12 | 0.75% |
| hsa_circ_0086081 | 3205 | 12 | 0.75% |
| hsa_circ_0019332 | 2938 | 11 | 0.75% |
| hsa_circ_0042173 | 2938 | 11 | 0.75% |
| hsa_circ_0139434 | 7747 | 29 | 0.75% |
| hsa_circ_0010744 | 3740 | 14 | 0.75% |
| hsa_circ_0004930 | 3473 | 13 | 0.75% |

| | | | |
|---|---|---|---|
| hsa_circ_0077468 | 2939 | 11 | 0.75% |
| hsa_circ_0025469 | 6947 | 26 | 0.75% |
| hsa_circ_0074941 | 2672 | 10 | 0.75% |
| hsa_circ_0129966 | 2672 | 10 | 0.75% |
| hsa_circ_0127018 | 3741 | 14 | 0.75% |
| hsa_circ_0022825 | 3474 | 13 | 0.75% |
| hsa_circ_0049099 | 3207 | 12 | 0.75% |
| hsa_circ_0010761 | 2940 | 11 | 0.75% |
| hsa_circ_0043819 | 2940 | 11 | 0.75% |
| hsa_circ_0108737 | 2940 | 11 | 0.75% |
| hsa_circ_0010525 | 2673 | 10 | 0.75% |
| hsa_circ_0063772 | 2673 | 10 | 0.75% |
| hsa_circ_0070309 | 2673 | 10 | 0.75% |
| hsa_circ_0070424 | 2673 | 10 | 0.75% |
| hsa_circ_0084711 | 2673 | 10 | 0.75% |
| hsa_circ_0079531 | 3475 | 13 | 0.75% |
| hsa_circ_0029653 | 2941 | 11 | 0.75% |
| hsa_circ_0081823 | 2941 | 11 | 0.75% |
| hsa_circ_0091119 | 2941 | 11 | 0.75% |
| hsa_circ_0010408 | 4011 | 15 | 0.75% |
| hsa_circ_0069628 | 2674 | 10 | 0.75% |
| hsa_circ_0094997 | 2674 | 10 | 0.75% |
| hsa_circ_0054057 | 3209 | 12 | 0.75% |
| hsa_circ_0079122 | 3744 | 14 | 0.75% |
| hsa_circ_0038712 | 11768 | 44 | 0.75% |
| hsa_circ_0031562 | 2942 | 11 | 0.75% |
| hsa_circ_0075197 | 2942 | 11 | 0.75% |
| hsa_circ_0050009 | 3210 | 12 | 0.75% |
| hsa_circ_0054819 | 3210 | 12 | 0.75% |
| hsa_circ_0067197 | 2675 | 10 | 0.75% |
| hsa_circ_0111914 | 9898 | 37 | 0.75% |
| hsa_circ_0064350 | 9899 | 37 | 0.75% |
| hsa_circ_0037401 | 2943 | 11 | 0.75% |
| hsa_circ_0085811 | 2943 | 11 | 0.75% |
| hsa_circ_0060319 | 3746 | 14 | 0.75% |
| hsa_circ_0120726 | 8830 | 33 | 0.75% |
| hsa_circ_0066745 | 3211 | 12 | 0.75% |
| hsa_circ_0084856 | 3211 | 12 | 0.75% |
| hsa_circ_0102795 | 3479 | 13 | 0.75% |
| hsa_circ_0113475 | 3212 | 12 | 0.75% |
| hsa_circ_0100751 | 3480 | 13 | 0.75% |
| hsa_circ_0085051 | 2677 | 10 | 0.75% |
| hsa_circ_0103830 | 2677 | 10 | 0.75% |
| hsa_circ_0059555 | 3748 | 14 | 0.75% |
| hsa_circ_0076199 | 5890 | 22 | 0.75% |
| hsa_circ_0010002 | 2945 | 11 | 0.75% |
| hsa_circ_0019188 | 2945 | 11 | 0.75% |
| hsa_circ_0019467 | 2945 | 11 | 0.75% |
| hsa_circ_0049125 | 3213 | 12 | 0.75% |
| hsa_circ_0094394 | 4017 | 15 | 0.75% |

| | | | |
|---|---|---|---|
| hsa_circ_0016617 | 2678 | 10 | 0.75% |
| hsa_circ_0106575 | 2678 | 10 | 0.75% |
| hsa_circ_0019443 | 2946 | 11 | 0.75% |
| hsa_circ_0065596 | 2946 | 11 | 0.75% |
| hsa_circ_0115475 | 2946 | 11 | 0.75% |
| hsa_circ_0068237 | 3214 | 12 | 0.75% |
| hsa_circ_0034033 | 3482 | 13 | 0.75% |
| hsa_circ_0030335 | 3750 | 14 | 0.75% |
| hsa_circ_0138471 | 3750 | 14 | 0.75% |
| hsa_circ_0016013 | 2679 | 10 | 0.75% |
| hsa_circ_0041908 | 2679 | 10 | 0.75% |
| hsa_circ_0085106 | 2947 | 11 | 0.75% |
| hsa_circ_0040700 | 3215 | 12 | 0.75% |
| hsa_circ_0042149 | 3215 | 12 | 0.75% |
| hsa_circ_0046827 | 3215 | 12 | 0.75% |
| hsa_circ_0068828 | 3215 | 12 | 0.75% |
| hsa_circ_0094783 | 3215 | 12 | 0.75% |
| hsa_circ_0122169 | 4020 | 15 | 0.75% |
| hsa_circ_0053639 | 3484 | 13 | 0.75% |
| hsa_circ_0074353 | 3216 | 12 | 0.75% |
| hsa_circ_0014868 | 2948 | 11 | 0.75% |
| hsa_circ_0038625 | 2948 | 11 | 0.75% |
| hsa_circ_0051671 | 2948 | 11 | 0.75% |
| hsa_circ_0076066 | 2948 | 11 | 0.75% |
| hsa_circ_0091838 | 2948 | 11 | 0.75% |
| hsa_circ_0134211 | 4289 | 16 | 0.75% |
| hsa_circ_0010632 | 3485 | 13 | 0.75% |
| hsa_circ_0083404 | 3485 | 13 | 0.75% |
| hsa_circ_0015678 | 3217 | 12 | 0.75% |
| hsa_circ_0014324 | 2949 | 11 | 0.75% |
| hsa_circ_0066928 | 2949 | 11 | 0.75% |
| hsa_circ_0053204 | 3486 | 13 | 0.75% |
| hsa_circ_0069049 | 3486 | 13 | 0.75% |
| hsa_circ_0117389 | 3486 | 13 | 0.75% |
| hsa_circ_0112617 | 10190 | 38 | 0.75% |
| hsa_circ_0108253 | 3218 | 12 | 0.75% |
| hsa_circ_0034661 | 2950 | 11 | 0.75% |
| hsa_circ_0077687 | 4291 | 16 | 0.75% |
| hsa_circ_0046865 | 2682 | 10 | 0.75% |
| hsa_circ_0104167 | 2682 | 10 | 0.75% |
| hsa_circ_0128155 | 2682 | 10 | 0.75% |
| hsa_circ_0109513 | 5901 | 22 | 0.75% |
| hsa_circ_0033082 | 3487 | 13 | 0.75% |
| hsa_circ_0010329 | 3219 | 12 | 0.75% |
| hsa_circ_0085908 | 3219 | 12 | 0.75% |
| hsa_circ_0016832 | 2951 | 11 | 0.75% |
| hsa_circ_0045033 | 2951 | 11 | 0.75% |
| hsa_circ_0044629 | 6439 | 24 | 0.75% |
| hsa_circ_0085058 | 2683 | 10 | 0.75% |
| hsa_circ_0034219 | 3488 | 13 | 0.75% |

| | | | |
|---|---|---|---|
| hsa_circ_0108373 | 4025 | 15 | 0.75% |
| hsa_circ_0033214 | 3220 | 12 | 0.75% |
| hsa_circ_0049191 | 3220 | 12 | 0.75% |
| hsa_circ_0062034 | 3220 | 12 | 0.75% |
| hsa_circ_0079452 | 3220 | 12 | 0.75% |
| hsa_circ_0034726 | 2952 | 11 | 0.75% |
| hsa_circ_0091819 | 2952 | 11 | 0.75% |
| hsa_circ_0123322 | 2952 | 11 | 0.75% |
| hsa_circ_0067982 | 2684 | 10 | 0.75% |
| hsa_circ_0078317 | 2684 | 10 | 0.75% |
| hsa_circ_0129822 | 3758 | 14 | 0.75% |
| hsa_circ_0001109 | 2953 | 11 | 0.75% |
| hsa_circ_0080134 | 2953 | 11 | 0.75% |
| hsa_circ_0030450 | 3222 | 12 | 0.74% |
| hsa_circ_0043339 | 3222 | 12 | 0.74% |
| hsa_circ_0035080 | 2685 | 10 | 0.74% |
| hsa_circ_0025986 | 3491 | 13 | 0.74% |
| hsa_circ_0125783 | 2954 | 11 | 0.74% |
| hsa_circ_0045943 | 3223 | 12 | 0.74% |
| hsa_circ_0125642 | 3223 | 12 | 0.74% |
| hsa_circ_0028573 | 2686 | 10 | 0.74% |
| hsa_circ_0084016 | 5641 | 21 | 0.74% |
| hsa_circ_0038175 | 2955 | 11 | 0.74% |
| hsa_circ_0087992 | 2955 | 11 | 0.74% |
| hsa_circ_0049526 | 3224 | 12 | 0.74% |
| hsa_circ_0084661 | 7523 | 28 | 0.74% |
| hsa_circ_0061114 | 2687 | 10 | 0.74% |
| hsa_circ_0072401 | 2687 | 10 | 0.74% |
| hsa_circ_0038178 | 2956 | 11 | 0.74% |
| hsa_circ_0059643 | 3494 | 13 | 0.74% |
| hsa_circ_0092714 | 3494 | 13 | 0.74% |
| hsa_circ_0124219 | 3763 | 14 | 0.74% |
| hsa_circ_0065370 | 2688 | 10 | 0.74% |
| hsa_circ_0041300 | 2957 | 11 | 0.74% |
| hsa_circ_0050843 | 2957 | 11 | 0.74% |
| hsa_circ_0111676 | 2957 | 11 | 0.74% |
| hsa_circ_0084404 | 3495 | 13 | 0.74% |
| hsa_circ_0012861 | 2689 | 10 | 0.74% |
| hsa_circ_0019600 | 2689 | 10 | 0.74% |
| hsa_circ_0062601 | 2689 | 10 | 0.74% |
| hsa_circ_0020867 | 2958 | 11 | 0.74% |
| hsa_circ_0036095 | 2958 | 11 | 0.74% |
| hsa_circ_0047173 | 2958 | 11 | 0.74% |
| hsa_circ_0082618 | 2958 | 11 | 0.74% |
| hsa_circ_0129986 | 2958 | 11 | 0.74% |
| hsa_circ_0010218 | 3227 | 12 | 0.74% |
| hsa_circ_0013067 | 3228 | 12 | 0.74% |
| hsa_circ_0070861 | 3228 | 12 | 0.74% |
| hsa_circ_0041971 | 2959 | 11 | 0.74% |
| hsa_circ_0052368 | 2959 | 11 | 0.74% |

| | | | |
|---|---|---|---|
| hsa_circ_0081207 | 2959 | 11 | 0.74% |
| hsa_circ_0099782 | 2959 | 11 | 0.74% |
| hsa_circ_0035359 | 2690 | 10 | 0.74% |
| hsa_circ_0064726 | 2690 | 10 | 0.74% |
| hsa_circ_0070669 | 2690 | 10 | 0.74% |
| hsa_circ_0099167 | 2690 | 10 | 0.74% |
| hsa_circ_0118669 | 10761 | 40 | 0.74% |
| hsa_circ_0019392 | 3229 | 12 | 0.74% |
| hsa_circ_0015603 | 2960 | 11 | 0.74% |
| hsa_circ_0041314 | 2960 | 11 | 0.74% |
| hsa_circ_0102524 | 2960 | 11 | 0.74% |
| hsa_circ_0131392 | 2691 | 10 | 0.74% |
| hsa_circ_0003329 | 16148 | 60 | 0.74% |
| hsa_circ_0029129 | 3499 | 13 | 0.74% |
| hsa_circ_0073236 | 2961 | 11 | 0.74% |
| hsa_circ_0135020 | 2961 | 11 | 0.74% |
| hsa_circ_0067195 | 2692 | 10 | 0.74% |
| hsa_circ_0137323 | 6731 | 25 | 0.74% |
| hsa_circ_0027596 | 3231 | 12 | 0.74% |
| hsa_circ_0034642 | 3231 | 12 | 0.74% |
| hsa_circ_0103859 | 7270 | 27 | 0.74% |
| hsa_circ_0082118 | 2693 | 10 | 0.74% |
| hsa_circ_0124304 | 2693 | 10 | 0.74% |
| hsa_circ_0125111 | 2693 | 10 | 0.74% |
| hsa_circ_0040999 | 3232 | 12 | 0.74% |
| hsa_circ_0090763 | 2963 | 11 | 0.74% |
| hsa_circ_0090278 | 3502 | 13 | 0.74% |
| hsa_circ_0042630 | 2694 | 10 | 0.74% |
| hsa_circ_0035748 | 3233 | 12 | 0.74% |
| hsa_circ_0065975 | 3233 | 12 | 0.74% |
| hsa_circ_0050485 | 3772 | 14 | 0.74% |
| hsa_circ_0079698 | 3772 | 14 | 0.74% |
| hsa_circ_0070456 | 2964 | 11 | 0.74% |
| hsa_circ_0091885 | 2964 | 11 | 0.74% |
| hsa_circ_0135656 | 2964 | 11 | 0.74% |
| hsa_circ_0110564 | 3503 | 13 | 0.74% |
| hsa_circ_0071986 | 2695 | 10 | 0.74% |
| hsa_circ_0075689 | 2695 | 10 | 0.74% |
| hsa_circ_0042886 | 3504 | 13 | 0.74% |
| hsa_circ_0024639 | 2965 | 11 | 0.74% |
| hsa_circ_0084963 | 2965 | 11 | 0.74% |
| hsa_circ_0117759 | 8356 | 31 | 0.74% |
| hsa_circ_0120585 | 3235 | 12 | 0.74% |
| hsa_circ_0059522 | 2696 | 10 | 0.74% |
| hsa_circ_0014038 | 2966 | 11 | 0.74% |
| hsa_circ_0073958 | 2966 | 11 | 0.74% |
| hsa_circ_0083374 | 4045 | 15 | 0.74% |
| hsa_circ_0113991 | 3236 | 12 | 0.74% |
| hsa_circ_0120128 | 4315 | 16 | 0.74% |
| hsa_circ_0022363 | 3506 | 13 | 0.74% |

| | | | |
|---|---|---|---|
| hsa_circ_0074139 | 3506 | 13 | 0.74% |
| hsa_circ_0135151 | 2697 | 10 | 0.74% |
| hsa_circ_0095022 | 7282 | 27 | 0.74% |
| hsa_circ_0035574 | 2967 | 11 | 0.74% |
| hsa_circ_0097403 | 2967 | 11 | 0.74% |
| hsa_circ_0056726 | 3237 | 12 | 0.74% |
| hsa_circ_0017530 | 131099 | 486 | 0.74% |
| hsa_circ_0045798 | 3507 | 13 | 0.74% |
| hsa_circ_0114366 | 12950 | 48 | 0.74% |
| hsa_circ_0036817 | 2698 | 10 | 0.74% |
| hsa_circ_0052102 | 7015 | 26 | 0.74% |
| hsa_circ_0078333 | 2968 | 11 | 0.74% |
| hsa_circ_0091973 | 3238 | 12 | 0.74% |
| hsa_circ_0042035 | 2969 | 11 | 0.74% |
| hsa_circ_0039332 | 3239 | 12 | 0.74% |
| hsa_circ_0062318 | 3509 | 13 | 0.74% |
| hsa_circ_0090369 | 3779 | 14 | 0.74% |
| hsa_circ_0046895 | 3780 | 14 | 0.74% |
| hsa_circ_0094709 | 3510 | 13 | 0.74% |
| hsa_circ_0027222 | 3240 | 12 | 0.74% |
| hsa_circ_0079504 | 3240 | 12 | 0.74% |
| hsa_circ_0009299 | 2970 | 11 | 0.74% |
| hsa_circ_0041873 | 2970 | 11 | 0.74% |
| hsa_circ_0011794 | 2700 | 10 | 0.74% |
| hsa_circ_0058540 | 2700 | 10 | 0.74% |
| hsa_circ_0119540 | 2700 | 10 | 0.74% |
| hsa_circ_0033489 | 3241 | 12 | 0.74% |
| hsa_circ_0061087 | 2971 | 11 | 0.74% |
| hsa_circ_0069941 | 2971 | 11 | 0.74% |
| hsa_circ_0059523 | 2701 | 10 | 0.74% |
| hsa_circ_0040735 | 3512 | 13 | 0.74% |
| hsa_circ_0021096 | 3242 | 12 | 0.74% |
| hsa_circ_0037513 | 3242 | 12 | 0.74% |
| hsa_circ_0010743 | 2972 | 11 | 0.74% |
| hsa_circ_0028801 | 2972 | 11 | 0.74% |
| hsa_circ_0073099 | 2702 | 10 | 0.74% |
| hsa_circ_0082369 | 2702 | 10 | 0.74% |
| hsa_circ_0097061 | 2702 | 10 | 0.74% |
| hsa_circ_0069159 | 3243 | 12 | 0.74% |
| hsa_circ_0069508 | 2973 | 11 | 0.74% |
| hsa_circ_0085191 | 2973 | 11 | 0.74% |
| hsa_circ_0005998 | 13785 | 51 | 0.74% |
| hsa_circ_0012032 | 3514 | 13 | 0.74% |
| hsa_circ_0044749 | 3514 | 13 | 0.74% |
| hsa_circ_0072726 | 3244 | 12 | 0.74% |
| hsa_circ_0078069 | 3244 | 12 | 0.74% |
| hsa_circ_0132122 | 3244 | 12 | 0.74% |
| hsa_circ_0041155 | 3515 | 13 | 0.74% |
| hsa_circ_0086386 | 2704 | 10 | 0.74% |
| hsa_circ_0130070 | 2704 | 10 | 0.74% |

| | | | |
|---|---|---|---|
| hsa_circ_0029489 | 3245 | 12 | 0.74% |
| hsa_circ_0041123 | 3245 | 12 | 0.74% |
| hsa_circ_0062272 | 2975 | 11 | 0.74% |
| hsa_circ_0109857 | 9196 | 34 | 0.74% |
| hsa_circ_0138259 | 3787 | 14 | 0.74% |
| hsa_circ_0011318 | 3246 | 12 | 0.74% |
| hsa_circ_0090667 | 3246 | 12 | 0.74% |
| hsa_circ_0027893 | 2705 | 10 | 0.74% |
| hsa_circ_0050134 | 2705 | 10 | 0.74% |
| hsa_circ_0057636 | 2705 | 10 | 0.74% |
| hsa_circ_0106574 | 2705 | 10 | 0.74% |
| hsa_circ_0069229 | 3517 | 13 | 0.74% |
| hsa_circ_0011130 | 2976 | 11 | 0.74% |
| hsa_circ_0068020 | 2976 | 11 | 0.74% |
| hsa_circ_0069515 | 2976 | 11 | 0.74% |
| hsa_circ_0077918 | 2976 | 11 | 0.74% |
| hsa_circ_0081857 | 2976 | 11 | 0.74% |
| hsa_circ_0017372 | 3247 | 12 | 0.74% |
| hsa_circ_0048681 | 3247 | 12 | 0.74% |
| hsa_circ_0134463 | 7577 | 28 | 0.74% |
| hsa_circ_0075955 | 3518 | 13 | 0.74% |
| hsa_circ_0031354 | 3248 | 12 | 0.74% |
| hsa_circ_0129234 | 4331 | 16 | 0.74% |
| hsa_circ_0019913 | 3519 | 13 | 0.74% |
| hsa_circ_0069669 | 2707 | 10 | 0.74% |
| hsa_circ_0015584 | 2978 | 11 | 0.74% |
| hsa_circ_0083380 | 2978 | 11 | 0.74% |
| hsa_circ_0120654 | 2978 | 11 | 0.74% |
| hsa_circ_0010190 | 3249 | 12 | 0.74% |
| hsa_circ_0067712 | 3249 | 12 | 0.74% |
| hsa_circ_0083484 | 3249 | 12 | 0.74% |
| hsa_circ_0113102 | 3249 | 12 | 0.74% |
| hsa_circ_0136740 | 3249 | 12 | 0.74% |
| hsa_circ_0054181 | 2708 | 10 | 0.74% |
| hsa_circ_0096692 | 5688 | 21 | 0.74% |
| hsa_circ_0017047 | 2980 | 11 | 0.74% |
| hsa_circ_0064587 | 2980 | 11 | 0.74% |
| hsa_circ_0077674 | 3251 | 12 | 0.74% |
| hsa_circ_0080902 | 5419 | 20 | 0.74% |
| hsa_circ_0131208 | 4065 | 15 | 0.74% |
| hsa_circ_0051298 | 3252 | 12 | 0.74% |
| hsa_circ_0037535 | 2981 | 11 | 0.74% |
| hsa_circ_0122099 | 2981 | 11 | 0.74% |
| hsa_circ_0084093 | 2710 | 10 | 0.74% |
| hsa_circ_0084270 | 2710 | 10 | 0.74% |
| hsa_circ_0125999 | 3253 | 12 | 0.74% |
| hsa_circ_0017175 | 2982 | 11 | 0.74% |
| hsa_circ_0018239 | 2982 | 11 | 0.74% |
| hsa_circ_0048312 | 2982 | 11 | 0.74% |
| hsa_circ_0053154 | 2982 | 11 | 0.74% |

| | | | |
|---|---|---|---|
| hsa_circ_0086202 | 2982 | 11 | 0.74% |
| hsa_circ_0016068 | 2711 | 10 | 0.74% |
| hsa_circ_0079864 | 2711 | 10 | 0.74% |
| hsa_circ_0095304 | 2711 | 10 | 0.74% |
| hsa_circ_0066998 | 3525 | 13 | 0.74% |
| hsa_circ_0028719 | 2983 | 11 | 0.74% |
| hsa_circ_0135674 | 2983 | 11 | 0.74% |
| hsa_circ_0074685 | 4339 | 16 | 0.74% |
| hsa_circ_0051557 | 2712 | 10 | 0.74% |
| hsa_circ_0116275 | 2712 | 10 | 0.74% |
| hsa_circ_0043338 | 3255 | 12 | 0.74% |
| hsa_circ_0055977 | 4069 | 15 | 0.74% |
| hsa_circ_0017905 | 2984 | 11 | 0.74% |
| hsa_circ_0030089 | 2984 | 11 | 0.74% |
| hsa_circ_0069690 | 2984 | 11 | 0.74% |
| hsa_circ_0072243 | 2984 | 11 | 0.74% |
| hsa_circ_0125473 | 3527 | 13 | 0.74% |
| hsa_circ_0018355 | 8954 | 33 | 0.74% |
| hsa_circ_0093262 | 7055 | 26 | 0.74% |
| hsa_circ_0015681 | 2985 | 11 | 0.74% |
| hsa_circ_0015894 | 2985 | 11 | 0.74% |
| hsa_circ_0072528 | 2985 | 11 | 0.74% |
| hsa_circ_0086075 | 2985 | 11 | 0.74% |
| hsa_circ_0032647 | 3528 | 13 | 0.74% |
| hsa_circ_0005888 | 4071 | 15 | 0.74% |
| hsa_circ_0046985 | 3257 | 12 | 0.74% |
| hsa_circ_0065715 | 3257 | 12 | 0.74% |
| hsa_circ_0122068 | 11671 | 43 | 0.74% |
| hsa_circ_0048080 | 3800 | 14 | 0.74% |
| hsa_circ_0066687 | 4343 | 16 | 0.74% |
| hsa_circ_0053489 | 2986 | 11 | 0.74% |
| hsa_circ_0060430 | 2986 | 11 | 0.74% |
| hsa_circ_0103644 | 2986 | 11 | 0.74% |
| hsa_circ_0133350 | 2986 | 11 | 0.74% |
| hsa_circ_0122805 | 7058 | 26 | 0.74% |
| hsa_circ_0080376 | 3529 | 13 | 0.74% |
| hsa_circ_0083010 | 3529 | 13 | 0.74% |
| hsa_circ_0010860 | 3258 | 12 | 0.74% |
| hsa_circ_0080882 | 3258 | 12 | 0.74% |
| hsa_circ_0082545 | 3258 | 12 | 0.74% |
| hsa_circ_0132884 | 3258 | 12 | 0.74% |
| hsa_circ_0123427 | 3530 | 13 | 0.74% |
| hsa_circ_0026714 | 2987 | 11 | 0.74% |
| hsa_circ_0040220 | 2987 | 11 | 0.74% |
| hsa_circ_0064399 | 2987 | 11 | 0.74% |
| hsa_circ_0071930 | 2987 | 11 | 0.74% |
| hsa_circ_0109463 | 9233 | 34 | 0.74% |
| hsa_circ_0079934 | 7877 | 29 | 0.74% |
| hsa_circ_0004567 | 5433 | 20 | 0.74% |
| hsa_circ_0102515 | 4075 | 15 | 0.74% |

| | | | |
|---|---|---|---|
| hsa_circ_0017697 | 3260 | 12 | 0.74% |
| hsa_circ_0017217 | 2717 | 10 | 0.74% |
| hsa_circ_0006347 | 2989 | 11 | 0.74% |
| hsa_circ_0042140 | 2989 | 11 | 0.74% |
| hsa_circ_0093405 | 5435 | 20 | 0.74% |
| hsa_circ_0029355 | 3261 | 12 | 0.74% |
| hsa_circ_0054081 | 2718 | 10 | 0.74% |
| hsa_circ_0081090 | 2718 | 10 | 0.74% |
| hsa_circ_0031585 | 4349 | 16 | 0.74% |
| hsa_circ_0083802 | 2990 | 11 | 0.74% |
| hsa_circ_0050289 | 11417 | 42 | 0.74% |
| hsa_circ_0074135 | 3262 | 12 | 0.74% |
| hsa_circ_0072225 | 2719 | 10 | 0.74% |
| hsa_circ_0084015 | 5710 | 21 | 0.74% |
| hsa_circ_0008279 | 2991 | 11 | 0.74% |
| hsa_circ_0017037 | 2991 | 11 | 0.74% |
| hsa_circ_0129127 | 2991 | 11 | 0.74% |
| hsa_circ_0070982 | 3263 | 12 | 0.74% |
| hsa_circ_0090054 | 3263 | 12 | 0.74% |
| hsa_circ_0107985 | 3263 | 12 | 0.74% |
| hsa_circ_0037664 | 3264 | 12 | 0.74% |
| hsa_circ_0083776 | 3264 | 12 | 0.74% |
| hsa_circ_0119842 | 3264 | 12 | 0.74% |
| hsa_circ_0020332 | 2992 | 11 | 0.74% |
| hsa_circ_0048861 | 2992 | 11 | 0.74% |
| hsa_circ_0086845 | 2992 | 11 | 0.74% |
| hsa_circ_0090374 | 2992 | 11 | 0.74% |
| hsa_circ_0034986 | 2720 | 10 | 0.74% |
| hsa_circ_0112180 | 2720 | 10 | 0.74% |
| hsa_circ_0135104 | 2720 | 10 | 0.74% |
| hsa_circ_0032392 | 2993 | 11 | 0.74% |
| hsa_circ_0065321 | 2993 | 11 | 0.74% |
| hsa_circ_0070762 | 2993 | 11 | 0.74% |
| hsa_circ_0088971 | 2993 | 11 | 0.74% |
| hsa_circ_0136437 | 2993 | 11 | 0.74% |
| hsa_circ_0073186 | 2721 | 10 | 0.74% |
| hsa_circ_0087383 | 2721 | 10 | 0.74% |
| hsa_circ_0112189 | 2721 | 10 | 0.74% |
| hsa_circ_0117769 | 2721 | 10 | 0.74% |
| hsa_circ_0090269 | 3538 | 13 | 0.73% |
| hsa_circ_0035792 | 2722 | 10 | 0.73% |
| hsa_circ_0056922 | 2722 | 10 | 0.73% |
| hsa_circ_0071212 | 2722 | 10 | 0.73% |
| hsa_circ_0026065 | 3267 | 12 | 0.73% |
| hsa_circ_0001627 | 2995 | 11 | 0.73% |
| hsa_circ_0066389 | 2995 | 11 | 0.73% |
| hsa_circ_0139745 | 63444 | 233 | 0.73% |
| hsa_circ_0100931 | 2723 | 10 | 0.73% |
| hsa_circ_0108725 | 2723 | 10 | 0.73% |
| hsa_circ_0123111 | 3540 | 13 | 0.73% |

| | | | |
|---|---|---|---|
| hsa_circ_0019806 | 3268 | 12 | 0.73% |
| hsa_circ_0067299 | 3268 | 12 | 0.73% |
| hsa_circ_0088753 | 2996 | 11 | 0.73% |
| hsa_circ_0069099 | 3541 | 13 | 0.73% |
| hsa_circ_0020466 | 2724 | 10 | 0.73% |
| hsa_circ_0016715 | 3269 | 12 | 0.73% |
| hsa_circ_0028063 | 3269 | 12 | 0.73% |
| hsa_circ_0083261 | 3269 | 12 | 0.73% |
| hsa_circ_0090219 | 3269 | 12 | 0.73% |
| hsa_circ_0120127 | 4904 | 18 | 0.73% |
| hsa_circ_0038884 | 2997 | 11 | 0.73% |
| hsa_circ_0062377 | 2997 | 11 | 0.73% |
| hsa_circ_0116967 | 2997 | 11 | 0.73% |
| hsa_circ_0021765 | 3270 | 12 | 0.73% |
| hsa_circ_0035158 | 3270 | 12 | 0.73% |
| hsa_circ_0072851 | 7086 | 26 | 0.73% |
| hsa_circ_0031483 | 2998 | 11 | 0.73% |
| hsa_circ_0036694 | 2998 | 11 | 0.73% |
| hsa_circ_0021519 | 4361 | 16 | 0.73% |
| hsa_circ_0072809 | 7087 | 26 | 0.73% |
| hsa_circ_0015619 | 3271 | 12 | 0.73% |
| hsa_circ_0035002 | 3271 | 12 | 0.73% |
| hsa_circ_0083682 | 3271 | 12 | 0.73% |
| hsa_circ_0011670 | 2726 | 10 | 0.73% |
| hsa_circ_0015499 | 2726 | 10 | 0.73% |
| hsa_circ_0027965 | 3544 | 13 | 0.73% |
| hsa_circ_0036543 | 2999 | 11 | 0.73% |
| hsa_circ_0084908 | 2999 | 11 | 0.73% |
| hsa_circ_0036545 | 3272 | 12 | 0.73% |
| hsa_circ_0076803 | 3272 | 12 | 0.73% |
| hsa_circ_0077475 | 2727 | 10 | 0.73% |
| hsa_circ_0103593 | 2727 | 10 | 0.73% |
| hsa_circ_0110249 | 2727 | 10 | 0.73% |
| hsa_circ_0070858 | 3000 | 11 | 0.73% |
| hsa_circ_0023444 | 6273 | 23 | 0.73% |
| hsa_circ_0038780 | 3273 | 12 | 0.73% |
| hsa_circ_0043089 | 3273 | 12 | 0.73% |
| hsa_circ_0049108 | 3273 | 12 | 0.73% |
| hsa_circ_0071831 | 3273 | 12 | 0.73% |
| hsa_circ_0073776 | 3273 | 12 | 0.73% |
| hsa_circ_0109902 | 2728 | 10 | 0.73% |
| hsa_circ_0024614 | 3001 | 11 | 0.73% |
| hsa_circ_0030093 | 3001 | 11 | 0.73% |
| hsa_circ_0054720 | 3001 | 11 | 0.73% |
| hsa_circ_0056557 | 3001 | 11 | 0.73% |
| hsa_circ_0100706 | 3001 | 11 | 0.73% |
| hsa_circ_0135070 | 10095 | 37 | 0.73% |
| hsa_circ_0004484 | 3547 | 13 | 0.73% |
| hsa_circ_0060692 | 3547 | 13 | 0.73% |
| hsa_circ_0062753 | 2729 | 10 | 0.73% |

| | | | |
|---|---|---|---|
| hsa_circ_0082515 | 2729 | 10 | 0.73% |
| hsa_circ_0058982 | 3002 | 11 | 0.73% |
| hsa_circ_0075704 | 3275 | 12 | 0.73% |
| hsa_circ_0086914 | 3275 | 12 | 0.73% |
| hsa_circ_0035159 | 3548 | 13 | 0.73% |
| hsa_circ_0030874 | 3549 | 13 | 0.73% |
| hsa_circ_0047022 | 3549 | 13 | 0.73% |
| hsa_circ_0047806 | 3549 | 13 | 0.73% |
| hsa_circ_0021215 | 3003 | 11 | 0.73% |
| hsa_circ_0058837 | 3003 | 11 | 0.73% |
| hsa_circ_0110212 | 3003 | 11 | 0.73% |
| hsa_circ_0011489 | 2730 | 10 | 0.73% |
| hsa_circ_0072405 | 2730 | 10 | 0.73% |
| hsa_circ_0063650 | 3277 | 12 | 0.73% |
| hsa_circ_0005377 | 3551 | 13 | 0.73% |
| hsa_circ_0061266 | 3551 | 13 | 0.73% |
| hsa_circ_0112814 | 3551 | 13 | 0.73% |
| hsa_circ_0017183 | 3278 | 12 | 0.73% |
| hsa_circ_0021154 | 3278 | 12 | 0.73% |
| hsa_circ_0046157 | 3278 | 12 | 0.73% |
| hsa_circ_0011230 | 3005 | 11 | 0.73% |
| hsa_circ_0038624 | 3005 | 11 | 0.73% |
| hsa_circ_0073423 | 3005 | 11 | 0.73% |
| hsa_circ_0041099 | 2732 | 10 | 0.73% |
| hsa_circ_0087782 | 2732 | 10 | 0.73% |
| hsa_circ_0089743 | 3279 | 12 | 0.73% |
| hsa_circ_0128972 | 3279 | 12 | 0.73% |
| hsa_circ_0020882 | 3006 | 11 | 0.73% |
| hsa_circ_0079862 | 3006 | 11 | 0.73% |
| hsa_circ_0107554 | 3006 | 11 | 0.73% |
| hsa_circ_0107770 | 3006 | 11 | 0.73% |
| hsa_circ_0062357 | 3826 | 14 | 0.73% |
| hsa_circ_0090752 | 3553 | 13 | 0.73% |
| hsa_circ_0067356 | 3280 | 12 | 0.73% |
| hsa_circ_0087184 | 3280 | 12 | 0.73% |
| hsa_circ_0103078 | 3280 | 12 | 0.73% |
| hsa_circ_0043704 | 3007 | 11 | 0.73% |
| hsa_circ_0053539 | 3007 | 11 | 0.73% |
| hsa_circ_0071193 | 3007 | 11 | 0.73% |
| hsa_circ_0036839 | 3554 | 13 | 0.73% |
| hsa_circ_0090695 | 3554 | 13 | 0.73% |
| hsa_circ_0035802 | 2734 | 10 | 0.73% |
| hsa_circ_0028723 | 3008 | 11 | 0.73% |
| hsa_circ_0080450 | 3008 | 11 | 0.73% |
| hsa_circ_0091437 | 3008 | 11 | 0.73% |
| hsa_circ_0024634 | 3555 | 13 | 0.73% |
| hsa_circ_0093345 | 3555 | 13 | 0.73% |
| hsa_circ_0068145 | 4102 | 15 | 0.73% |
| hsa_circ_0013078 | 3282 | 12 | 0.73% |
| hsa_circ_0069685 | 3282 | 12 | 0.73% |

| | | | |
|---|---|---|---|
| hsa_circ_0019023 | 2735 | 10 | 0.73% |
| hsa_circ_0069815 | 2735 | 10 | 0.73% |
| hsa_circ_0011527 | 4103 | 15 | 0.73% |
| hsa_circ_0070139 | 3009 | 11 | 0.73% |
| hsa_circ_0073576 | 3009 | 11 | 0.73% |
| hsa_circ_0033329 | 3283 | 12 | 0.73% |
| hsa_circ_0056164 | 2736 | 10 | 0.73% |
| hsa_circ_0073899 | 2736 | 10 | 0.73% |
| hsa_circ_0028895 | 3010 | 11 | 0.73% |
| hsa_circ_0051193 | 3010 | 11 | 0.73% |
| hsa_circ_0068202 | 3010 | 11 | 0.73% |
| hsa_circ_0083080 | 3010 | 11 | 0.73% |
| hsa_circ_0084353 | 3010 | 11 | 0.73% |
| hsa_circ_0090267 | 3558 | 13 | 0.73% |
| hsa_circ_0098560 | 2737 | 10 | 0.73% |
| hsa_circ_0129149 | 2737 | 10 | 0.73% |
| hsa_circ_0013484 | 3011 | 11 | 0.73% |
| hsa_circ_0056782 | 3011 | 11 | 0.73% |
| hsa_circ_0041607 | 3285 | 12 | 0.73% |
| hsa_circ_0072990 | 3285 | 12 | 0.73% |
| hsa_circ_0095742 | 7118 | 26 | 0.73% |
| hsa_circ_0087826 | 3559 | 13 | 0.73% |
| hsa_circ_0091262 | 3559 | 13 | 0.73% |
| hsa_circ_0127840 | 3833 | 14 | 0.73% |
| hsa_circ_0006262 | 3012 | 11 | 0.73% |
| hsa_circ_0048307 | 3012 | 11 | 0.73% |
| hsa_circ_0065133 | 3012 | 11 | 0.73% |
| hsa_circ_0105270 | 3012 | 11 | 0.73% |
| hsa_circ_0028389 | 3286 | 12 | 0.73% |
| hsa_circ_0125185 | 3286 | 12 | 0.73% |
| hsa_circ_0046104 | 3560 | 13 | 0.73% |
| hsa_circ_0094720 | 2739 | 10 | 0.73% |
| hsa_circ_0042621 | 3013 | 11 | 0.73% |
| hsa_circ_0091208 | 3013 | 11 | 0.73% |
| hsa_circ_0124371 | 3013 | 11 | 0.73% |
| hsa_circ_0024520 | 3287 | 12 | 0.73% |
| hsa_circ_0021195 | 3561 | 13 | 0.73% |
| hsa_circ_0040068 | 3835 | 14 | 0.73% |
| hsa_circ_0010618 | 3562 | 13 | 0.73% |
| hsa_circ_0020400 | 3014 | 11 | 0.73% |
| hsa_circ_0070768 | 3014 | 11 | 0.73% |
| hsa_circ_0083799 | 3014 | 11 | 0.73% |
| hsa_circ_0085062 | 2740 | 10 | 0.73% |
| hsa_circ_0019914 | 3015 | 11 | 0.73% |
| hsa_circ_0065375 | 3015 | 11 | 0.73% |
| hsa_circ_0091378 | 3015 | 11 | 0.73% |
| hsa_circ_0015236 | 2741 | 10 | 0.73% |
| hsa_circ_0020792 | 4112 | 15 | 0.73% |
| hsa_circ_0048722 | 3564 | 13 | 0.73% |
| hsa_circ_0032622 | 3290 | 12 | 0.73% |

| | | | |
|---|---|---|---|
| hsa_circ_0077125 | 3016 | 11 | 0.73% |
| hsa_circ_0095651 | 3016 | 11 | 0.73% |
| hsa_circ_0084250 | 2742 | 10 | 0.73% |
| hsa_circ_0077298 | 3565 | 13 | 0.73% |
| hsa_circ_0020556 | 3291 | 12 | 0.73% |
| hsa_circ_0029869 | 3291 | 12 | 0.73% |
| hsa_circ_0068138 | 3291 | 12 | 0.73% |
| hsa_circ_0093871 | 4114 | 15 | 0.73% |
| hsa_circ_0026531 | 3017 | 11 | 0.73% |
| hsa_circ_0053366 | 3017 | 11 | 0.73% |
| hsa_circ_0120231 | 5760 | 21 | 0.73% |
| hsa_circ_0016266 | 2743 | 10 | 0.73% |
| hsa_circ_0014200 | 5212 | 19 | 0.73% |
| hsa_circ_0103076 | 3292 | 12 | 0.73% |
| hsa_circ_0070015 | 3018 | 11 | 0.73% |
| hsa_circ_0096494 | 3018 | 11 | 0.73% |
| hsa_circ_0129524 | 3018 | 11 | 0.73% |
| hsa_circ_0048038 | 2744 | 10 | 0.73% |
| hsa_circ_0090427 | 2744 | 10 | 0.73% |
| hsa_circ_0052834 | 3293 | 12 | 0.73% |
| hsa_circ_0043780 | 3019 | 11 | 0.73% |
| hsa_circ_0071803 | 3019 | 11 | 0.73% |
| hsa_circ_0012891 | 4392 | 16 | 0.73% |
| hsa_circ_0041687 | 3294 | 12 | 0.73% |
| hsa_circ_0081771 | 2745 | 10 | 0.73% |
| hsa_circ_0023712 | 3020 | 11 | 0.73% |
| hsa_circ_0123592 | 7688 | 28 | 0.73% |
| hsa_circ_0066604 | 3570 | 13 | 0.73% |
| hsa_circ_0065583 | 3021 | 11 | 0.73% |
| hsa_circ_0069002 | 3021 | 11 | 0.73% |
| hsa_circ_0075984 | 3021 | 11 | 0.73% |
| hsa_circ_0078710 | 3021 | 11 | 0.73% |
| hsa_circ_0024880 | 3296 | 12 | 0.73% |
| hsa_circ_0034430 | 3296 | 12 | 0.73% |
| hsa_circ_0034749 | 3296 | 12 | 0.73% |
| hsa_circ_0054475 | 3296 | 12 | 0.73% |
| hsa_circ_0082047 | 3571 | 13 | 0.73% |
| hsa_circ_0055258 | 5769 | 21 | 0.73% |
| hsa_circ_0082984 | 3022 | 11 | 0.73% |
| hsa_circ_0111677 | 3022 | 11 | 0.73% |
| hsa_circ_0018368 | 3297 | 12 | 0.73% |
| hsa_circ_0137842 | 3297 | 12 | 0.73% |
| hsa_circ_0052245 | 3847 | 14 | 0.73% |
| hsa_circ_0052293 | 4122 | 15 | 0.73% |
| hsa_circ_0069507 | 2748 | 10 | 0.73% |
| hsa_circ_0073789 | 2748 | 10 | 0.73% |
| hsa_circ_0120046 | 2748 | 10 | 0.73% |
| hsa_circ_0111680 | 6046 | 22 | 0.73% |
| hsa_circ_0054071 | 3023 | 11 | 0.73% |
| hsa_circ_0140289 | 3573 | 13 | 0.73% |

| | | | |
|---|---|---|---|
| hsa_circ_0074976 | 3024 | 11 | 0.73% |
| hsa_circ_0085291 | 3024 | 11 | 0.73% |
| hsa_circ_0010514 | 3299 | 12 | 0.73% |
| hsa_circ_0033100 | 3299 | 12 | 0.73% |
| hsa_circ_0078868 | 4124 | 15 | 0.73% |
| hsa_circ_0078928 | 4124 | 15 | 0.73% |
| hsa_circ_0112155 | 6874 | 25 | 0.73% |
| hsa_circ_0025470 | 7150 | 26 | 0.73% |
| hsa_circ_0112844 | 4400 | 16 | 0.73% |
| hsa_circ_0035856 | 3575 | 13 | 0.73% |
| hsa_circ_0051675 | 3575 | 13 | 0.73% |
| hsa_circ_0009231 | 3300 | 12 | 0.73% |
| hsa_circ_0026545 | 3300 | 12 | 0.73% |
| hsa_circ_0105485 | 3300 | 12 | 0.73% |
| hsa_circ_0113986 | 3300 | 12 | 0.73% |
| hsa_circ_0128468 | 3300 | 12 | 0.73% |
| hsa_circ_0017004 | 3025 | 11 | 0.73% |
| hsa_circ_0083804 | 3025 | 11 | 0.73% |
| hsa_circ_0086389 | 3025 | 11 | 0.73% |
| hsa_circ_0128692 | 3025 | 11 | 0.73% |
| hsa_circ_0013410 | 2750 | 10 | 0.73% |
| hsa_circ_0052239 | 4401 | 16 | 0.73% |
| hsa_circ_0059612 | 3576 | 13 | 0.73% |
| hsa_circ_0060682 | 3576 | 13 | 0.73% |
| hsa_circ_0054820 | 3301 | 12 | 0.73% |
| hsa_circ_0067879 | 3301 | 12 | 0.73% |
| hsa_circ_0027223 | 3026 | 11 | 0.73% |
| hsa_circ_0017132 | 2751 | 10 | 0.73% |
| hsa_circ_0114152 | 2751 | 10 | 0.73% |
| hsa_circ_0095905 | 6603 | 24 | 0.73% |
| hsa_circ_0066482 | 4402 | 16 | 0.73% |
| hsa_circ_0138531 | 4127 | 15 | 0.73% |
| hsa_circ_0088962 | 3027 | 11 | 0.73% |
| hsa_circ_0109856 | 9357 | 34 | 0.73% |
| hsa_circ_0049509 | 12385 | 45 | 0.73% |
| hsa_circ_0030658 | 3028 | 11 | 0.73% |
| hsa_circ_0061793 | 2753 | 10 | 0.73% |
| hsa_circ_0061147 | 6332 | 23 | 0.73% |
| hsa_circ_0007738 | 3579 | 13 | 0.73% |
| hsa_circ_0097606 | 3579 | 13 | 0.73% |
| hsa_circ_0033335 | 3304 | 12 | 0.73% |
| hsa_circ_0047735 | 3304 | 12 | 0.73% |
| hsa_circ_0081876 | 3304 | 12 | 0.73% |
| hsa_circ_0068150 | 3029 | 11 | 0.73% |
| hsa_circ_0063315 | 4406 | 16 | 0.73% |
| hsa_circ_0020424 | 2754 | 10 | 0.73% |
| hsa_circ_0015386 | 3305 | 12 | 0.73% |
| hsa_circ_0025009 | 3305 | 12 | 0.73% |
| hsa_circ_0131713 | 3305 | 12 | 0.73% |
| hsa_circ_0079254 | 3856 | 14 | 0.73% |

| | | | |
|---|---|---|---|
| hsa_circ_0071899 | 3030 | 11 | 0.73% |
| hsa_circ_0077919 | 3030 | 11 | 0.73% |
| hsa_circ_0097881 | 3030 | 11 | 0.73% |
| hsa_circ_0131963 | 3581 | 13 | 0.73% |
| hsa_circ_0051301 | 3306 | 12 | 0.73% |
| hsa_circ_0058561 | 3306 | 12 | 0.73% |
| hsa_circ_0090226 | 3306 | 12 | 0.73% |
| hsa_circ_0132056 | 6337 | 23 | 0.73% |
| hsa_circ_0108374 | 4133 | 15 | 0.73% |
| hsa_circ_0032225 | 3582 | 13 | 0.73% |
| hsa_circ_0102398 | 3582 | 13 | 0.73% |
| hsa_circ_0038079 | 3031 | 11 | 0.73% |
| hsa_circ_0082859 | 3031 | 11 | 0.73% |
| hsa_circ_0090395 | 3031 | 11 | 0.73% |
| hsa_circ_0134316 | 3031 | 11 | 0.73% |
| hsa_circ_0044194 | 5787 | 21 | 0.73% |
| hsa_circ_0033565 | 3307 | 12 | 0.73% |
| hsa_circ_0046916 | 3307 | 12 | 0.73% |
| hsa_circ_0052698 | 4134 | 15 | 0.73% |
| hsa_circ_0117770 | 2756 | 10 | 0.73% |
| hsa_circ_0010631 | 3583 | 13 | 0.73% |
| hsa_circ_0084350 | 3032 | 11 | 0.73% |
| hsa_circ_0090548 | 3032 | 11 | 0.73% |
| hsa_circ_0123365 | 3032 | 11 | 0.73% |
| hsa_circ_0000998 | 3584 | 13 | 0.73% |
| hsa_circ_0010465 | 2757 | 10 | 0.73% |
| hsa_circ_0045433 | 2757 | 10 | 0.73% |
| hsa_circ_0089832 | 2757 | 10 | 0.73% |
| hsa_circ_0015688 | 3860 | 14 | 0.73% |
| hsa_circ_0066925 | 3033 | 11 | 0.73% |
| hsa_circ_0067642 | 3033 | 11 | 0.73% |
| hsa_circ_0098483 | 3033 | 11 | 0.73% |
| hsa_circ_0039780 | 6618 | 24 | 0.73% |
| hsa_circ_0090590 | 2758 | 10 | 0.73% |
| hsa_circ_0125565 | 2758 | 10 | 0.73% |
| hsa_circ_0135642 | 2758 | 10 | 0.73% |
| hsa_circ_0031824 | 3034 | 11 | 0.73% |
| hsa_circ_0059713 | 3034 | 11 | 0.73% |
| hsa_circ_0068182 | 3034 | 11 | 0.73% |
| hsa_circ_0034753 | 3310 | 12 | 0.73% |
| hsa_circ_0035286 | 3310 | 12 | 0.73% |
| hsa_circ_0136672 | 3586 | 13 | 0.73% |
| hsa_circ_0029748 | 2759 | 10 | 0.72% |
| hsa_circ_0080490 | 2759 | 10 | 0.72% |
| hsa_circ_0009945 | 3035 | 11 | 0.72% |
| hsa_circ_0043534 | 3311 | 12 | 0.72% |
| hsa_circ_0055520 | 3311 | 12 | 0.72% |
| hsa_circ_0138105 | 3311 | 12 | 0.72% |
| hsa_circ_0053663 | 3587 | 13 | 0.72% |
| hsa_circ_0052633 | 3588 | 13 | 0.72% |

| | | | |
|---|---|---|---|
| hsa_circ_0053646 | 3588 | 13 | 0.72% |
| hsa_circ_0058102 | 3588 | 13 | 0.72% |
| hsa_circ_0010720 | 3312 | 12 | 0.72% |
| hsa_circ_0099654 | 2760 | 10 | 0.72% |
| hsa_circ_0109854 | 9387 | 34 | 0.72% |
| hsa_circ_0026263 | 3037 | 11 | 0.72% |
| hsa_circ_0070725 | 3037 | 11 | 0.72% |
| hsa_circ_0108508 | 3037 | 11 | 0.72% |
| hsa_circ_0031401 | 2761 | 10 | 0.72% |
| hsa_circ_0008185 | 3314 | 12 | 0.72% |
| hsa_circ_0024198 | 3314 | 12 | 0.72% |
| hsa_circ_0097431 | 3314 | 12 | 0.72% |
| hsa_circ_0139733 | 3314 | 12 | 0.72% |
| hsa_circ_0093136 | 5524 | 20 | 0.72% |
| hsa_circ_0027830 | 2762 | 10 | 0.72% |
| hsa_circ_0078166 | 2762 | 10 | 0.72% |
| hsa_circ_0086318 | 2762 | 10 | 0.72% |
| hsa_circ_0044779 | 4420 | 16 | 0.72% |
| hsa_circ_0046044 | 3315 | 12 | 0.72% |
| hsa_circ_0064201 | 7183 | 26 | 0.72% |
| hsa_circ_0039258 | 3039 | 11 | 0.72% |
| hsa_circ_0130019 | 3039 | 11 | 0.72% |
| hsa_circ_0109967 | 11051 | 40 | 0.72% |
| hsa_circ_0129770 | 2763 | 10 | 0.72% |
| hsa_circ_0081549 | 2764 | 10 | 0.72% |
| hsa_circ_0091318 | 3317 | 12 | 0.72% |
| hsa_circ_0117535 | 3317 | 12 | 0.72% |
| hsa_circ_0016290 | 4423 | 16 | 0.72% |
| hsa_circ_0048747 | 3318 | 12 | 0.72% |
| hsa_circ_0078771 | 2765 | 10 | 0.72% |
| hsa_circ_0113162 | 2765 | 10 | 0.72% |
| hsa_circ_0010978 | 3595 | 13 | 0.72% |
| hsa_circ_0054405 | 3042 | 11 | 0.72% |
| hsa_circ_0071395 | 3042 | 11 | 0.72% |
| hsa_circ_0039399 | 3319 | 12 | 0.72% |
| hsa_circ_0123191 | 12170 | 44 | 0.72% |
| hsa_circ_0031144 | 3043 | 11 | 0.72% |
| hsa_circ_0044219 | 3043 | 11 | 0.72% |
| hsa_circ_0049938 | 3043 | 11 | 0.72% |
| hsa_circ_0057584 | 3043 | 11 | 0.72% |
| hsa_circ_0093295 | 3043 | 11 | 0.72% |
| hsa_circ_0052303 | 3320 | 12 | 0.72% |
| hsa_circ_0069158 | 3320 | 12 | 0.72% |
| hsa_circ_0111415 | 3320 | 12 | 0.72% |
| hsa_circ_0106804 | 7747 | 28 | 0.72% |
| hsa_circ_0139117 | 3597 | 13 | 0.72% |
| hsa_circ_0011547 | 2767 | 10 | 0.72% |
| hsa_circ_0096072 | 2767 | 10 | 0.72% |
| hsa_circ_0104160 | 2767 | 10 | 0.72% |
| hsa_circ_0118303 | 3874 | 14 | 0.72% |

| | | | |
|---|---|---|---|
| hsa_circ_0055532 | 3044 | 11 | 0.72% |
| hsa_circ_0091839 | 3044 | 11 | 0.72% |
| hsa_circ_0125874 | 6642 | 24 | 0.72% |
| hsa_circ_0032702 | 3321 | 12 | 0.72% |
| hsa_circ_0050006 | 3321 | 12 | 0.72% |
| hsa_circ_0059641 | 3598 | 13 | 0.72% |
| hsa_circ_0086635 | 3598 | 13 | 0.72% |
| hsa_circ_0108864 | 3598 | 13 | 0.72% |
| hsa_circ_0007963 | 2768 | 10 | 0.72% |
| hsa_circ_0083510 | 3045 | 11 | 0.72% |
| hsa_circ_0071346 | 3322 | 12 | 0.72% |
| hsa_circ_0075707 | 3322 | 12 | 0.72% |
| hsa_circ_0082130 | 3322 | 12 | 0.72% |
| hsa_circ_0128088 | 2769 | 10 | 0.72% |
| hsa_circ_0100844 | 8861 | 32 | 0.72% |
| hsa_circ_0056405 | 3046 | 11 | 0.72% |
| hsa_circ_0034641 | 3323 | 12 | 0.72% |
| hsa_circ_0076374 | 3323 | 12 | 0.72% |
| hsa_circ_0096690 | 5816 | 21 | 0.72% |
| hsa_circ_0070157 | 3878 | 14 | 0.72% |
| hsa_circ_0052734 | 3601 | 13 | 0.72% |
| hsa_circ_0070561 | 3601 | 13 | 0.72% |
| hsa_circ_0021881 | 3324 | 12 | 0.72% |
| hsa_circ_0014309 | 3047 | 11 | 0.72% |
| hsa_circ_0036386 | 3047 | 11 | 0.72% |
| hsa_circ_0069161 | 3047 | 11 | 0.72% |
| hsa_circ_0092085 | 3047 | 11 | 0.72% |
| hsa_circ_0060693 | 3602 | 13 | 0.72% |
| hsa_circ_0073089 | 3325 | 12 | 0.72% |
| hsa_circ_0033109 | 3048 | 11 | 0.72% |
| hsa_circ_0039230 | 3048 | 11 | 0.72% |
| hsa_circ_0041404 | 3048 | 11 | 0.72% |
| hsa_circ_0074637 | 3048 | 11 | 0.72% |
| hsa_circ_0101218 | 3048 | 11 | 0.72% |
| hsa_circ_0136054 | 3048 | 11 | 0.72% |
| hsa_circ_0119847 | 2771 | 10 | 0.72% |
| hsa_circ_0060079 | 3603 | 13 | 0.72% |
| hsa_circ_0099768 | 3326 | 12 | 0.72% |
| hsa_circ_0129394 | 3326 | 12 | 0.72% |
| hsa_circ_0031480 | 3049 | 11 | 0.72% |
| hsa_circ_0043043 | 3049 | 11 | 0.72% |
| hsa_circ_0053155 | 3049 | 11 | 0.72% |
| hsa_circ_0089223 | 3049 | 11 | 0.72% |
| hsa_circ_0027288 | 4158 | 15 | 0.72% |
| hsa_circ_0000065 | 3604 | 13 | 0.72% |
| hsa_circ_0075473 | 3604 | 13 | 0.72% |
| hsa_circ_0066039 | 3050 | 11 | 0.72% |
| hsa_circ_0086381 | 3050 | 11 | 0.72% |
| hsa_circ_0085089 | 2773 | 10 | 0.72% |
| hsa_circ_0043336 | 3328 | 12 | 0.72% |

| | | | |
|---|---|---|---|
| hsa_circ_0000282 | 4715 | 17 | 0.72% |
| hsa_circ_0010725 | 3051 | 11 | 0.72% |
| hsa_circ_0070338 | 3051 | 11 | 0.72% |
| hsa_circ_0119959 | 5825 | 21 | 0.72% |
| hsa_circ_0128539 | 3052 | 11 | 0.72% |
| hsa_circ_0073031 | 3607 | 13 | 0.72% |
| hsa_circ_0005891 | 4717 | 17 | 0.72% |
| hsa_circ_0024590 | 3330 | 12 | 0.72% |
| hsa_circ_0042612 | 2775 | 10 | 0.72% |
| hsa_circ_0077742 | 2775 | 10 | 0.72% |
| hsa_circ_0135510 | 4718 | 17 | 0.72% |
| hsa_circ_0122728 | 10824 | 39 | 0.72% |
| hsa_circ_0032229 | 3053 | 11 | 0.72% |
| hsa_circ_0087758 | 3053 | 11 | 0.72% |
| hsa_circ_0104911 | 3053 | 11 | 0.72% |
| hsa_circ_0054441 | 3886 | 14 | 0.72% |
| hsa_circ_0040795 | 3331 | 12 | 0.72% |
| hsa_circ_0041205 | 3331 | 12 | 0.72% |
| hsa_circ_0041894 | 3331 | 12 | 0.72% |
| hsa_circ_0083534 | 4164 | 15 | 0.72% |
| hsa_circ_0046330 | 2776 | 10 | 0.72% |
| hsa_circ_0069668 | 2776 | 10 | 0.72% |
| hsa_circ_0095274 | 2776 | 10 | 0.72% |
| hsa_circ_0039184 | 3054 | 11 | 0.72% |
| hsa_circ_0101133 | 3054 | 11 | 0.72% |
| hsa_circ_0045182 | 3332 | 12 | 0.72% |
| hsa_circ_0049103 | 3332 | 12 | 0.72% |
| hsa_circ_0067354 | 3332 | 12 | 0.72% |
| hsa_circ_0117028 | 3332 | 12 | 0.72% |
| hsa_circ_0106400 | 8886 | 32 | 0.72% |
| hsa_circ_0070968 | 2777 | 10 | 0.72% |
| hsa_circ_0071360 | 3055 | 11 | 0.72% |
| hsa_circ_0072244 | 3055 | 11 | 0.72% |
| hsa_circ_0017696 | 3333 | 12 | 0.72% |
| hsa_circ_0042566 | 3333 | 12 | 0.72% |
| hsa_circ_0062927 | 14166 | 51 | 0.72% |
| hsa_circ_0040850 | 3611 | 13 | 0.72% |
| hsa_circ_0106399 | 8889 | 32 | 0.72% |
| hsa_circ_0079113 | 3889 | 14 | 0.72% |
| hsa_circ_0025895 | 3056 | 11 | 0.72% |
| hsa_circ_0091820 | 3056 | 11 | 0.72% |
| hsa_circ_0116276 | 3056 | 11 | 0.72% |
| hsa_circ_0010861 | 3334 | 12 | 0.72% |
| hsa_circ_0020120 | 3334 | 12 | 0.72% |
| hsa_circ_0032845 | 3334 | 12 | 0.72% |
| hsa_circ_0037705 | 3334 | 12 | 0.72% |
| hsa_circ_0040249 | 3334 | 12 | 0.72% |
| hsa_circ_0063798 | 3334 | 12 | 0.72% |
| hsa_circ_0126174 | 6946 | 25 | 0.72% |
| hsa_circ_0140049 | 13338 | 48 | 0.72% |

| | | | |
|---|---|---|---|
| hsa_circ_0009288 | 14173 | 51 | 0.72% |
| hsa_circ_0100099 | 3057 | 11 | 0.72% |
| hsa_circ_0015411 | 4170 | 15 | 0.72% |
| hsa_circ_0045645 | 3614 | 13 | 0.72% |
| hsa_circ_0140697 | 3614 | 13 | 0.72% |
| hsa_circ_0031599 | 3336 | 12 | 0.72% |
| hsa_circ_0084213 | 3336 | 12 | 0.72% |
| hsa_circ_0045959 | 3058 | 11 | 0.72% |
| hsa_circ_0086080 | 3058 | 11 | 0.72% |
| hsa_circ_0017611 | 2780 | 10 | 0.72% |
| hsa_circ_0062602 | 2780 | 10 | 0.72% |
| hsa_circ_0113121 | 2780 | 10 | 0.72% |
| hsa_circ_0117527 | 3615 | 13 | 0.72% |
| hsa_circ_0020685 | 3337 | 12 | 0.72% |
| hsa_circ_0071884 | 3059 | 11 | 0.72% |
| hsa_circ_0086807 | 3059 | 11 | 0.72% |
| hsa_circ_0041387 | 2781 | 10 | 0.72% |
| hsa_circ_0092945 | 2781 | 10 | 0.72% |
| hsa_circ_0040067 | 3894 | 14 | 0.72% |
| hsa_circ_0035711 | 3338 | 12 | 0.72% |
| hsa_circ_0103832 | 3338 | 12 | 0.72% |
| hsa_circ_0126974 | 3338 | 12 | 0.72% |
| hsa_circ_0011164 | 3060 | 11 | 0.72% |
| hsa_circ_0128948 | 3060 | 11 | 0.72% |
| hsa_circ_0106327 | 5842 | 21 | 0.72% |
| hsa_circ_0137817 | 8624 | 31 | 0.72% |
| hsa_circ_0040467 | 2782 | 10 | 0.72% |
| hsa_circ_0121050 | 5286 | 19 | 0.72% |
| hsa_circ_0084662 | 7790 | 28 | 0.72% |
| hsa_circ_0079532 | 3617 | 13 | 0.72% |
| hsa_circ_0057846 | 3339 | 12 | 0.72% |
| hsa_circ_0081818 | 3339 | 12 | 0.72% |
| hsa_circ_0019680 | 3061 | 11 | 0.72% |
| hsa_circ_0088836 | 3061 | 11 | 0.72% |
| hsa_circ_0094910 | 2783 | 10 | 0.72% |
| hsa_circ_0010071 | 3618 | 13 | 0.72% |
| hsa_circ_0073954 | 4732 | 17 | 0.72% |
| hsa_circ_0023703 | 3062 | 11 | 0.72% |
| hsa_circ_0042368 | 3062 | 11 | 0.72% |
| hsa_circ_0045955 | 3062 | 11 | 0.72% |
| hsa_circ_0086901 | 3062 | 11 | 0.72% |
| hsa_circ_0133247 | 3062 | 11 | 0.72% |
| hsa_circ_0113610 | 5846 | 21 | 0.72% |
| hsa_circ_0003058 | 3619 | 13 | 0.72% |
| hsa_circ_0027057 | 3341 | 12 | 0.72% |
| hsa_circ_0058834 | 3341 | 12 | 0.72% |
| hsa_circ_0090148 | 3341 | 12 | 0.72% |
| hsa_circ_0034975 | 3063 | 11 | 0.72% |
| hsa_circ_0043820 | 3063 | 11 | 0.72% |
| hsa_circ_0041156 | 3620 | 13 | 0.72% |

| | | | |
|---|---|---|---|
| hsa_circ_0047420 | 3620 | 13 | 0.72% |
| hsa_circ_0001978 | 7797 | 28 | 0.72% |
| hsa_circ_0126111 | 4177 | 15 | 0.72% |
| hsa_circ_0082546 | 3342 | 12 | 0.72% |
| hsa_circ_0125456 | 3342 | 12 | 0.72% |
| hsa_circ_0132362 | 3342 | 12 | 0.72% |
| hsa_circ_0001879 | 13647 | 49 | 0.72% |
| hsa_circ_0012691 | 3621 | 13 | 0.72% |
| hsa_circ_0009926 | 3064 | 11 | 0.72% |
| hsa_circ_0050028 | 3064 | 11 | 0.72% |
| hsa_circ_0140787 | 3064 | 11 | 0.72% |
| hsa_circ_0090318 | 2786 | 10 | 0.72% |
| hsa_circ_0047174 | 3065 | 11 | 0.72% |
| hsa_circ_0105243 | 3065 | 11 | 0.72% |
| hsa_circ_0138532 | 3065 | 11 | 0.72% |
| hsa_circ_0010745 | 3901 | 14 | 0.72% |
| hsa_circ_0021882 | 3344 | 12 | 0.72% |
| hsa_circ_0134267 | 8639 | 31 | 0.72% |
| hsa_circ_0068428 | 3623 | 13 | 0.72% |
| hsa_circ_0132374 | 3623 | 13 | 0.72% |
| hsa_circ_0018166 | 3066 | 11 | 0.72% |
| hsa_circ_0032002 | 3066 | 11 | 0.72% |
| hsa_circ_0035703 | 3066 | 11 | 0.72% |
| hsa_circ_0060165 | 3066 | 11 | 0.72% |
| hsa_circ_0071827 | 3066 | 11 | 0.72% |
| hsa_circ_0128663 | 3066 | 11 | 0.72% |
| hsa_circ_0010461 | 3345 | 12 | 0.72% |
| hsa_circ_0057981 | 3345 | 12 | 0.72% |
| hsa_circ_0132649 | 3345 | 12 | 0.72% |
| hsa_circ_0077450 | 3624 | 13 | 0.72% |
| hsa_circ_0105314 | 2788 | 10 | 0.72% |
| hsa_circ_0065597 | 3067 | 11 | 0.72% |
| hsa_circ_0036153 | 3346 | 12 | 0.72% |
| hsa_circ_0083693 | 3346 | 12 | 0.72% |
| hsa_circ_0115965 | 3346 | 12 | 0.72% |
| hsa_circ_0128633 | 3625 | 13 | 0.72% |
| hsa_circ_0050078 | 11433 | 41 | 0.72% |
| hsa_circ_0105667 | 2789 | 10 | 0.72% |
| hsa_circ_0125781 | 2789 | 10 | 0.72% |
| hsa_circ_0048311 | 3068 | 11 | 0.72% |
| hsa_circ_0132555 | 22871 | 82 | 0.72% |
| hsa_circ_0055304 | 3347 | 12 | 0.72% |
| hsa_circ_0016974 | 7531 | 27 | 0.72% |
| hsa_circ_0005834 | 10321 | 37 | 0.72% |
| hsa_circ_0101517 | 32641 | 117 | 0.72% |
| hsa_circ_0089814 | 11997 | 43 | 0.72% |
| hsa_circ_0077301 | 3906 | 14 | 0.72% |
| hsa_circ_0036238 | 3348 | 12 | 0.72% |
| hsa_circ_0058082 | 3348 | 12 | 0.72% |
| hsa_circ_0128294 | 3348 | 12 | 0.72% |

| | | | |
|---|---|---|---|
| hsa_circ_0127301 | 3069 | 11 | 0.72% |
| hsa_circ_0030657 | 3070 | 11 | 0.72% |
| hsa_circ_0121245 | 3070 | 11 | 0.72% |
| hsa_circ_0121353 | 4187 | 15 | 0.72% |
| hsa_circ_0010219 | 3350 | 12 | 0.72% |
| hsa_circ_0076471 | 3071 | 11 | 0.72% |
| hsa_circ_0078024 | 2792 | 10 | 0.72% |
| hsa_circ_0095175 | 3630 | 13 | 0.72% |
| hsa_circ_0012888 | 4468 | 16 | 0.72% |
| hsa_circ_0113345 | 3351 | 12 | 0.72% |
| hsa_circ_0129471 | 3351 | 12 | 0.72% |
| hsa_circ_0035636 | 3072 | 11 | 0.72% |
| hsa_circ_0053632 | 3072 | 11 | 0.72% |
| hsa_circ_0097404 | 3072 | 11 | 0.72% |
| hsa_circ_0056928 | 2793 | 10 | 0.72% |
| hsa_circ_0126056 | 2793 | 10 | 0.72% |
| hsa_circ_0042890 | 3631 | 13 | 0.72% |
| hsa_circ_0131241 | 3352 | 12 | 0.72% |
| hsa_circ_0115476 | 3073 | 11 | 0.72% |
| hsa_circ_0122178 | 3073 | 11 | 0.72% |
| hsa_circ_0053634 | 3632 | 13 | 0.72% |
| hsa_circ_0047303 | 3353 | 12 | 0.72% |
| hsa_circ_0088346 | 3074 | 11 | 0.72% |
| hsa_circ_0090899 | 3074 | 11 | 0.72% |
| hsa_circ_0032645 | 3633 | 13 | 0.72% |
| hsa_circ_0040428 | 3913 | 14 | 0.72% |
| hsa_circ_0030700 | 3354 | 12 | 0.72% |
| hsa_circ_0067996 | 2795 | 10 | 0.72% |
| hsa_circ_0090206 | 2795 | 10 | 0.72% |
| hsa_circ_0078891 | 4193 | 15 | 0.72% |
| hsa_circ_0097325 | 4193 | 15 | 0.72% |
| hsa_circ_0076701 | 3634 | 13 | 0.72% |
| hsa_circ_0052104 | 6709 | 24 | 0.72% |
| hsa_circ_0023335 | 3075 | 11 | 0.72% |
| hsa_circ_0023709 | 3075 | 11 | 0.72% |
| hsa_circ_0032381 | 3075 | 11 | 0.72% |
| hsa_circ_0072106 | 3075 | 11 | 0.72% |
| hsa_circ_0073088 | 3075 | 11 | 0.72% |
| hsa_circ_0073424 | 3075 | 11 | 0.72% |
| hsa_circ_0120649 | 3075 | 11 | 0.72% |
| hsa_circ_0060320 | 3914 | 14 | 0.72% |
| hsa_circ_0038205 | 3355 | 12 | 0.72% |
| hsa_circ_0101468 | 3355 | 12 | 0.72% |
| hsa_circ_0118173 | 3355 | 12 | 0.72% |
| hsa_circ_0084092 | 2796 | 10 | 0.72% |
| hsa_circ_0084305 | 2796 | 10 | 0.72% |
| hsa_circ_0017395 | 4474 | 16 | 0.72% |
| hsa_circ_0027675 | 3076 | 11 | 0.72% |
| hsa_circ_0091821 | 3356 | 12 | 0.72% |
| hsa_circ_0026334 | 2797 | 10 | 0.72% |

| | | | |
|---|---|---|---|
| hsa_circ_0085220 | 2797 | 10 | 0.72% |
| hsa_circ_0004489 | 20143 | 72 | 0.71% |
| hsa_circ_0030963 | 2798 | 10 | 0.71% |
| hsa_circ_0140151 | 2798 | 10 | 0.71% |
| hsa_circ_0068342 | 3078 | 11 | 0.71% |
| hsa_circ_0010283 | 3358 | 12 | 0.71% |
| hsa_circ_0034526 | 3358 | 12 | 0.71% |
| hsa_circ_0049801 | 3358 | 12 | 0.71% |
| hsa_circ_0030000 | 3918 | 14 | 0.71% |
| hsa_circ_0030932 | 2799 | 10 | 0.71% |
| hsa_circ_0094428 | 2799 | 10 | 0.71% |
| hsa_circ_0019896 | 3079 | 11 | 0.71% |
| hsa_circ_0041278 | 3079 | 11 | 0.71% |
| hsa_circ_0049140 | 3079 | 11 | 0.71% |
| hsa_circ_0028360 | 3359 | 12 | 0.71% |
| hsa_circ_0101596 | 3359 | 12 | 0.71% |
| hsa_circ_0036838 | 3639 | 13 | 0.71% |
| hsa_circ_0137880 | 5319 | 19 | 0.71% |
| hsa_circ_0026710 | 3080 | 11 | 0.71% |
| hsa_circ_0035370 | 3080 | 11 | 0.71% |
| hsa_circ_0078583 | 2800 | 10 | 0.71% |
| hsa_circ_0121523 | 2800 | 10 | 0.71% |
| hsa_circ_0077448 | 3641 | 13 | 0.71% |
| hsa_circ_0030210 | 3361 | 12 | 0.71% |
| hsa_circ_0052538 | 3361 | 12 | 0.71% |
| hsa_circ_0124858 | 3361 | 12 | 0.71% |
| hsa_circ_0042622 | 3081 | 11 | 0.71% |
| hsa_circ_0076790 | 3081 | 11 | 0.71% |
| hsa_circ_0054574 | 2801 | 10 | 0.71% |
| hsa_circ_0103414 | 2801 | 10 | 0.71% |
| hsa_circ_0131324 | 2801 | 10 | 0.71% |
| hsa_circ_0022525 | 3642 | 13 | 0.71% |
| hsa_circ_0060669 | 3642 | 13 | 0.71% |
| hsa_circ_0014181 | 3362 | 12 | 0.71% |
| hsa_circ_0039354 | 3362 | 12 | 0.71% |
| hsa_circ_0114257 | 3082 | 11 | 0.71% |
| hsa_circ_0116847 | 17092 | 61 | 0.71% |
| hsa_circ_0081257 | 4203 | 15 | 0.71% |
| hsa_circ_0105964 | 6726 | 24 | 0.71% |
| hsa_circ_0053223 | 3083 | 11 | 0.71% |
| hsa_circ_0140400 | 3083 | 11 | 0.71% |
| hsa_circ_0036745 | 5045 | 18 | 0.71% |
| hsa_circ_0051556 | 2803 | 10 | 0.71% |
| hsa_circ_0139174 | 2803 | 10 | 0.71% |
| hsa_circ_0063072 | 3644 | 13 | 0.71% |
| hsa_circ_0056032 | 3364 | 12 | 0.71% |
| hsa_circ_0062033 | 3364 | 12 | 0.71% |
| hsa_circ_0027572 | 3084 | 11 | 0.71% |
| hsa_circ_0053445 | 3084 | 11 | 0.71% |
| hsa_circ_0086557 | 3084 | 11 | 0.71% |

| | | | |
|---|---|---|---|
| hsa_circ_0016671 | 3645 | 13 | 0.71% |
| hsa_circ_0037413 | 3645 | 13 | 0.71% |
| hsa_circ_0085343 | 3645 | 13 | 0.71% |
| hsa_circ_0096503 | 3645 | 13 | 0.71% |
| hsa_circ_0014284 | 3365 | 12 | 0.71% |
| hsa_circ_0089424 | 3365 | 12 | 0.71% |
| hsa_circ_0092758 | 3085 | 11 | 0.71% |
| hsa_circ_0113458 | 11781 | 42 | 0.71% |
| hsa_circ_0009650 | 3366 | 12 | 0.71% |
| hsa_circ_0086923 | 3366 | 12 | 0.71% |
| hsa_circ_0034243 | 2805 | 10 | 0.71% |
| hsa_circ_0044517 | 2805 | 10 | 0.71% |
| hsa_circ_0071218 | 2805 | 10 | 0.71% |
| hsa_circ_0135351 | 2805 | 10 | 0.71% |
| hsa_circ_0032866 | 3647 | 13 | 0.71% |
| hsa_circ_0078300 | 3086 | 11 | 0.71% |
| hsa_circ_0139257 | 3086 | 11 | 0.71% |
| hsa_circ_0024901 | 3367 | 12 | 0.71% |
| hsa_circ_0047405 | 3648 | 13 | 0.71% |
| hsa_circ_0068194 | 3087 | 11 | 0.71% |
| hsa_circ_0126979 | 5894 | 21 | 0.71% |
| hsa_circ_0087825 | 3649 | 13 | 0.71% |
| hsa_circ_0016175 | 2807 | 10 | 0.71% |
| hsa_circ_0057637 | 2807 | 10 | 0.71% |
| hsa_circ_0085103 | 3088 | 11 | 0.71% |
| hsa_circ_0007038 | 5896 | 21 | 0.71% |
| hsa_circ_0009496 | 3931 | 14 | 0.71% |
| hsa_circ_0039874 | 3931 | 14 | 0.71% |
| hsa_circ_0041605 | 2808 | 10 | 0.71% |
| hsa_circ_0050186 | 2808 | 10 | 0.71% |
| hsa_circ_0061490 | 2808 | 10 | 0.71% |
| hsa_circ_0066361 | 2808 | 10 | 0.71% |
| hsa_circ_0010436 | 3089 | 11 | 0.71% |
| hsa_circ_0077962 | 3089 | 11 | 0.71% |
| hsa_circ_0119810 | 3089 | 11 | 0.71% |
| hsa_circ_0054749 | 3370 | 12 | 0.71% |
| hsa_circ_0057586 | 3370 | 12 | 0.71% |
| hsa_circ_0086684 | 3651 | 13 | 0.71% |
| hsa_circ_0050480 | 3932 | 14 | 0.71% |
| hsa_circ_0026993 | 3090 | 11 | 0.71% |
| hsa_circ_0040172 | 3090 | 11 | 0.71% |
| hsa_circ_0056066 | 3090 | 11 | 0.71% |
| hsa_circ_0117813 | 3652 | 13 | 0.71% |
| hsa_circ_0038905 | 4496 | 16 | 0.71% |
| hsa_circ_0045594 | 3091 | 11 | 0.71% |
| hsa_circ_0020422 | 2810 | 10 | 0.71% |
| hsa_circ_0076735 | 2810 | 10 | 0.71% |
| hsa_circ_0079912 | 2810 | 10 | 0.71% |
| hsa_circ_0039019 | 4497 | 16 | 0.71% |
| hsa_circ_0046105 | 3654 | 13 | 0.71% |

| | | | |
|---|---|---|---|
| hsa_circ_0111368 | 3092 | 11 | 0.71% |
| hsa_circ_0047239 | 2811 | 10 | 0.71% |
| hsa_circ_0050131 | 2811 | 10 | 0.71% |
| hsa_circ_0011483 | 3655 | 13 | 0.71% |
| hsa_circ_0084411 | 3655 | 13 | 0.71% |
| hsa_circ_0018209 | 3374 | 12 | 0.71% |
| hsa_circ_0035330 | 3093 | 11 | 0.71% |
| hsa_circ_0044611 | 3093 | 11 | 0.71% |
| hsa_circ_0073419 | 3093 | 11 | 0.71% |
| hsa_circ_0086948 | 2812 | 10 | 0.71% |
| hsa_circ_0034715 | 3375 | 12 | 0.71% |
| hsa_circ_0119839 | 3375 | 12 | 0.71% |
| hsa_circ_0054052 | 3094 | 11 | 0.71% |
| hsa_circ_0030241 | 2813 | 10 | 0.71% |
| hsa_circ_0058108 | 3657 | 13 | 0.71% |
| hsa_circ_0084316 | 3376 | 12 | 0.71% |
| hsa_circ_0039376 | 3939 | 14 | 0.71% |
| hsa_circ_0137045 | 6190 | 22 | 0.71% |
| hsa_circ_0075371 | 3658 | 13 | 0.71% |
| hsa_circ_0054320 | 2814 | 10 | 0.71% |
| hsa_circ_0026073 | 3377 | 12 | 0.71% |
| hsa_circ_0045082 | 3377 | 12 | 0.71% |
| hsa_circ_0075907 | 3377 | 12 | 0.71% |
| hsa_circ_0046356 | 3096 | 11 | 0.71% |
| hsa_circ_0059678 | 3096 | 11 | 0.71% |
| hsa_circ_0015346 | 3659 | 13 | 0.71% |
| hsa_circ_0086255 | 3659 | 13 | 0.71% |
| hsa_circ_0046342 | 3378 | 12 | 0.71% |
| hsa_circ_0125897 | 3378 | 12 | 0.71% |
| hsa_circ_0087384 | 2815 | 10 | 0.71% |
| hsa_circ_0090317 | 3097 | 11 | 0.71% |
| hsa_circ_0052936 | 5068 | 18 | 0.71% |
| hsa_circ_0083663 | 2816 | 10 | 0.71% |
| hsa_circ_0100839 | 9012 | 32 | 0.71% |
| hsa_circ_0031110 | 3098 | 11 | 0.71% |
| hsa_circ_0059412 | 3098 | 11 | 0.71% |
| hsa_circ_0010721 | 3380 | 12 | 0.71% |
| hsa_circ_0027848 | 3380 | 12 | 0.71% |
| hsa_circ_0032864 | 3662 | 13 | 0.71% |
| hsa_circ_0045799 | 3662 | 13 | 0.71% |
| hsa_circ_0002013 | 5634 | 20 | 0.71% |
| hsa_circ_0087781 | 2817 | 10 | 0.71% |
| hsa_circ_0019495 | 3099 | 11 | 0.71% |
| hsa_circ_0053564 | 3099 | 11 | 0.71% |
| hsa_circ_0056973 | 3099 | 11 | 0.71% |
| hsa_circ_0078406 | 3099 | 11 | 0.71% |
| hsa_circ_0040250 | 9579 | 34 | 0.71% |
| hsa_circ_0022944 | 3381 | 12 | 0.71% |
| hsa_circ_0065110 | 3381 | 12 | 0.71% |
| hsa_circ_0031621 | 3663 | 13 | 0.71% |

| | | | |
|---|---|---|---|
| hsa_circ_0032431 | 3100 | 11 | 0.71% |
| hsa_circ_0067826 | 3100 | 11 | 0.71% |
| hsa_circ_0071896 | 3100 | 11 | 0.71% |
| hsa_circ_0123994 | 3100 | 11 | 0.71% |
| hsa_circ_0013420 | 3382 | 12 | 0.71% |
| hsa_circ_0011303 | 3664 | 13 | 0.71% |
| hsa_circ_0042978 | 3664 | 13 | 0.71% |
| hsa_circ_0036966 | 3946 | 14 | 0.71% |
| hsa_circ_0014562 | 2819 | 10 | 0.71% |
| hsa_circ_0027656 | 3101 | 11 | 0.71% |
| hsa_circ_0048212 | 3101 | 11 | 0.71% |
| hsa_circ_0086205 | 3101 | 11 | 0.71% |
| hsa_circ_0043661 | 3383 | 12 | 0.71% |
| hsa_circ_0055693 | 3383 | 12 | 0.71% |
| hsa_circ_0063693 | 3383 | 12 | 0.71% |
| hsa_circ_0101160 | 6203 | 22 | 0.71% |
| hsa_circ_0016872 | 3384 | 12 | 0.71% |
| hsa_circ_0034904 | 3384 | 12 | 0.71% |
| hsa_circ_0083579 | 3384 | 12 | 0.71% |
| hsa_circ_0083774 | 3384 | 12 | 0.71% |
| hsa_circ_0002428 | 3102 | 11 | 0.71% |
| hsa_circ_0031555 | 3102 | 11 | 0.71% |
| hsa_circ_0051187 | 3102 | 11 | 0.71% |
| hsa_circ_0091879 | 3102 | 11 | 0.71% |
| hsa_circ_0113880 | 3102 | 11 | 0.71% |
| hsa_circ_0133922 | 3102 | 11 | 0.71% |
| hsa_circ_0072317 | 4231 | 15 | 0.71% |
| hsa_circ_0039638 | 3667 | 13 | 0.71% |
| hsa_circ_0062032 | 3385 | 12 | 0.71% |
| hsa_circ_0027763 | 2821 | 10 | 0.71% |
| hsa_circ_0036806 | 2821 | 10 | 0.71% |
| hsa_circ_0133145 | 4232 | 15 | 0.71% |
| hsa_circ_0088322 | 3104 | 11 | 0.71% |
| hsa_circ_0091699 | 5644 | 20 | 0.71% |
| hsa_circ_0035695 | 3669 | 13 | 0.71% |
| hsa_circ_0021250 | 3387 | 12 | 0.71% |
| hsa_circ_0119264 | 3387 | 12 | 0.71% |
| hsa_circ_0037534 | 3105 | 11 | 0.71% |
| hsa_circ_0125027 | 3105 | 11 | 0.71% |
| hsa_circ_0016265 | 2823 | 10 | 0.71% |
| hsa_circ_0035078 | 2823 | 10 | 0.71% |
| hsa_circ_0019772 | 3670 | 13 | 0.71% |
| hsa_circ_0056364 | 3670 | 13 | 0.71% |
| hsa_circ_0066949 | 3670 | 13 | 0.71% |
| hsa_circ_0017064 | 3388 | 12 | 0.71% |
| hsa_circ_0063965 | 3388 | 12 | 0.71% |
| hsa_circ_0019897 | 3106 | 11 | 0.71% |
| hsa_circ_0025672 | 3106 | 11 | 0.71% |
| hsa_circ_0060686 | 3671 | 13 | 0.71% |
| hsa_circ_0074563 | 3671 | 13 | 0.71% |

| | | | |
|---|---|---|---|
| hsa_circ_0113938 | 3389 | 12 | 0.71% |
| hsa_circ_0119539 | 3389 | 12 | 0.71% |
| hsa_circ_0139945 | 22312 | 79 | 0.71% |
| hsa_circ_0005790 | 3107 | 11 | 0.71% |
| hsa_circ_0010800 | 3107 | 11 | 0.71% |
| hsa_circ_0086567 | 3107 | 11 | 0.71% |
| hsa_circ_0136887 | 3672 | 13 | 0.71% |
| hsa_circ_0031960 | 3955 | 14 | 0.71% |
| hsa_circ_0034249 | 3390 | 12 | 0.71% |
| hsa_circ_0039777 | 3390 | 12 | 0.71% |
| hsa_circ_0045165 | 3390 | 12 | 0.71% |
| hsa_circ_0106846 | 3390 | 12 | 0.71% |
| hsa_circ_0035579 | 2825 | 10 | 0.71% |
| hsa_circ_0052876 | 2825 | 10 | 0.71% |
| hsa_circ_0076924 | 2825 | 10 | 0.71% |
| hsa_circ_0037704 | 3108 | 11 | 0.71% |
| hsa_circ_0064134 | 3391 | 12 | 0.71% |
| hsa_circ_0078802 | 16674 | 59 | 0.71% |
| hsa_circ_0001725 | 3109 | 11 | 0.71% |
| hsa_circ_0129717 | 3392 | 12 | 0.71% |
| hsa_circ_0012293 | 3675 | 13 | 0.71% |
| hsa_circ_0035635 | 2827 | 10 | 0.71% |
| hsa_circ_0109539 | 2827 | 10 | 0.71% |
| hsa_circ_0005230 | 3958 | 14 | 0.71% |
| hsa_circ_0105525 | 3110 | 11 | 0.71% |
| hsa_circ_0010335 | 3393 | 12 | 0.71% |
| hsa_circ_0031512 | 3393 | 12 | 0.71% |
| hsa_circ_0045944 | 3393 | 12 | 0.71% |
| hsa_circ_0109851 | 9614 | 34 | 0.71% |
| hsa_circ_0128383 | 3676 | 13 | 0.71% |
| hsa_circ_0038001 | 2828 | 10 | 0.71% |
| hsa_circ_0062086 | 3111 | 11 | 0.71% |
| hsa_circ_0076736 | 3111 | 11 | 0.71% |
| hsa_circ_0128374 | 3111 | 11 | 0.71% |
| hsa_circ_0096667 | 11596 | 41 | 0.71% |
| hsa_circ_0060635 | 3394 | 12 | 0.71% |
| hsa_circ_0059639 | 3677 | 13 | 0.71% |
| hsa_circ_0103969 | 9900 | 35 | 0.71% |
| hsa_circ_0015497 | 2829 | 10 | 0.71% |
| hsa_circ_0015884 | 3112 | 11 | 0.71% |
| hsa_circ_0072121 | 3112 | 11 | 0.71% |
| hsa_circ_0026544 | 3395 | 12 | 0.71% |
| hsa_circ_0033122 | 3395 | 12 | 0.71% |
| hsa_circ_0077698 | 3395 | 12 | 0.71% |
| hsa_circ_0086352 | 3395 | 12 | 0.71% |
| hsa_circ_0014322 | 3961 | 14 | 0.71% |
| hsa_circ_0126837 | 3679 | 13 | 0.71% |
| hsa_circ_0041403 | 3113 | 11 | 0.71% |
| hsa_circ_0069452 | 3113 | 11 | 0.71% |
| hsa_circ_0016165 | 9624 | 34 | 0.71% |

| | | | |
|---|---|---|---|
| hsa_circ_0011539 | 3114 | 11 | 0.71% |
| hsa_circ_0016833 | 3114 | 11 | 0.71% |
| hsa_circ_0022677 | 3114 | 11 | 0.71% |
| hsa_circ_0027905 | 2831 | 10 | 0.71% |
| hsa_circ_0033085 | 2831 | 10 | 0.71% |
| hsa_circ_0018231 | 12457 | 44 | 0.71% |
| hsa_circ_0009975 | 3398 | 12 | 0.71% |
| hsa_circ_0139694 | 3398 | 12 | 0.71% |
| hsa_circ_0020401 | 3115 | 11 | 0.71% |
| hsa_circ_0078591 | 3115 | 11 | 0.71% |
| hsa_circ_0124297 | 3115 | 11 | 0.71% |
| hsa_circ_0014067 | 2832 | 10 | 0.71% |
| hsa_circ_0090633 | 2832 | 10 | 0.71% |
| hsa_circ_0064053 | 3399 | 12 | 0.71% |
| hsa_circ_0083483 | 3399 | 12 | 0.71% |
| hsa_circ_0015478 | 3116 | 11 | 0.71% |
| hsa_circ_0132422 | 3116 | 11 | 0.71% |
| hsa_circ_0010298 | 2833 | 10 | 0.71% |
| hsa_circ_0132480 | 3400 | 12 | 0.71% |
| hsa_circ_0011441 | 3117 | 11 | 0.71% |
| hsa_circ_0047366 | 3117 | 11 | 0.71% |
| hsa_circ_0132058 | 3117 | 11 | 0.71% |
| hsa_circ_0010594 | 3684 | 13 | 0.71% |
| hsa_circ_0016203 | 3401 | 12 | 0.71% |
| hsa_circ_0063807 | 3401 | 12 | 0.71% |
| hsa_circ_0048638 | 3118 | 11 | 0.71% |
| hsa_circ_0070747 | 3118 | 11 | 0.71% |
| hsa_circ_0119567 | 7938 | 28 | 0.71% |
| hsa_circ_0055114 | 3402 | 12 | 0.71% |
| hsa_circ_0066052 | 3402 | 12 | 0.71% |
| hsa_circ_0076819 | 3402 | 12 | 0.71% |
| hsa_circ_0032441 | 2835 | 10 | 0.71% |
| hsa_circ_0125780 | 2835 | 10 | 0.71% |
| hsa_circ_0058931 | 3686 | 13 | 0.71% |
| hsa_circ_0100329 | 3686 | 13 | 0.71% |
| hsa_circ_0083803 | 3119 | 11 | 0.71% |
| hsa_circ_0118682 | 13611 | 48 | 0.71% |
| hsa_circ_0025962 | 3403 | 12 | 0.71% |
| hsa_circ_0036479 | 3403 | 12 | 0.71% |
| hsa_circ_0113016 | 3403 | 12 | 0.71% |
| hsa_circ_0133081 | 4254 | 15 | 0.71% |
| hsa_circ_0059186 | 2836 | 10 | 0.71% |
| hsa_circ_0091564 | 3687 | 13 | 0.71% |
| hsa_circ_0062362 | 3120 | 11 | 0.71% |
| hsa_circ_0086509 | 3120 | 11 | 0.71% |
| hsa_circ_0025023 | 7659 | 27 | 0.71% |
| hsa_circ_0101573 | 6808 | 24 | 0.71% |
| hsa_circ_0018925 | 3688 | 13 | 0.70% |
| hsa_circ_0105697 | 7660 | 27 | 0.70% |
| hsa_circ_0092249 | 3121 | 11 | 0.70% |

| | | | |
|---|---|---|---|
| hsa_circ_0007599 | 5959 | 21 | 0.70% |
| hsa_circ_0092709 | 19296 | 68 | 0.70% |
| hsa_circ_0126020 | 3689 | 13 | 0.70% |
| hsa_circ_0137054 | 3689 | 13 | 0.70% |
| hsa_circ_0122726 | 11068 | 39 | 0.70% |
| hsa_circ_0016549 | 2838 | 10 | 0.70% |
| hsa_circ_0066201 | 3122 | 11 | 0.70% |
| hsa_circ_0105267 | 3122 | 11 | 0.70% |
| hsa_circ_0048411 | 3406 | 12 | 0.70% |
| hsa_circ_0000212 | 4258 | 15 | 0.70% |
| hsa_circ_0047210 | 2839 | 10 | 0.70% |
| hsa_circ_0062350 | 2839 | 10 | 0.70% |
| hsa_circ_0077351 | 2839 | 10 | 0.70% |
| hsa_circ_0140286 | 2839 | 10 | 0.70% |
| hsa_circ_0027157 | 3407 | 12 | 0.70% |
| hsa_circ_0128116 | 3407 | 12 | 0.70% |
| hsa_circ_0029510 | 3692 | 13 | 0.70% |
| hsa_circ_0032755 | 3408 | 12 | 0.70% |
| hsa_circ_0021380 | 3124 | 11 | 0.70% |
| hsa_circ_0036878 | 3124 | 11 | 0.70% |
| hsa_circ_0062603 | 3124 | 11 | 0.70% |
| hsa_circ_0107973 | 3124 | 11 | 0.70% |
| hsa_circ_0079463 | 2840 | 10 | 0.70% |
| hsa_circ_0120996 | 3693 | 13 | 0.70% |
| hsa_circ_0091081 | 3409 | 12 | 0.70% |
| hsa_circ_0018616 | 3125 | 11 | 0.70% |
| hsa_circ_0063920 | 3125 | 11 | 0.70% |
| hsa_circ_0076983 | 3125 | 11 | 0.70% |
| hsa_circ_0086058 | 3125 | 11 | 0.70% |
| hsa_circ_0025033 | 3410 | 12 | 0.70% |
| hsa_circ_0073884 | 3410 | 12 | 0.70% |
| hsa_circ_0113044 | 3410 | 12 | 0.70% |
| hsa_circ_0016339 | 3126 | 11 | 0.70% |
| hsa_circ_0086907 | 3126 | 11 | 0.70% |
| hsa_circ_0047228 | 2842 | 10 | 0.70% |
| hsa_circ_0060414 | 2842 | 10 | 0.70% |
| hsa_circ_0071334 | 2842 | 10 | 0.70% |
| hsa_circ_0087424 | 9948 | 35 | 0.70% |
| hsa_circ_0000555 | 3411 | 12 | 0.70% |
| hsa_circ_0079459 | 3411 | 12 | 0.70% |
| hsa_circ_0083189 | 3411 | 12 | 0.70% |
| hsa_circ_0077302 | 3980 | 14 | 0.70% |
| hsa_circ_0010342 | 2843 | 10 | 0.70% |
| hsa_circ_0071142 | 2843 | 10 | 0.70% |
| hsa_circ_0138277 | 3696 | 13 | 0.70% |
| hsa_circ_0112820 | 10804 | 38 | 0.70% |
| hsa_circ_0135931 | 5402 | 19 | 0.70% |
| hsa_circ_0009831 | 3412 | 12 | 0.70% |
| hsa_circ_0041384 | 3412 | 12 | 0.70% |
| hsa_circ_0065077 | 3412 | 12 | 0.70% |

| | | | |
|---|---|---|---|
| hsa_circ_0045030 | 3128 | 11 | 0.70% |
| hsa_circ_0071920 | 3128 | 11 | 0.70% |
| hsa_circ_0054182 | 2844 | 10 | 0.70% |
| hsa_circ_0127439 | 3413 | 12 | 0.70% |
| hsa_circ_0026202 | 3129 | 11 | 0.70% |
| hsa_circ_0083071 | 3129 | 11 | 0.70% |
| hsa_circ_0083878 | 3129 | 11 | 0.70% |
| hsa_circ_0090703 | 3129 | 11 | 0.70% |
| hsa_circ_0044750 | 3698 | 13 | 0.70% |
| hsa_circ_0126885 | 8250 | 29 | 0.70% |
| hsa_circ_0049036 | 3699 | 13 | 0.70% |
| hsa_circ_0013843 | 3130 | 11 | 0.70% |
| hsa_circ_0069505 | 3130 | 11 | 0.70% |
| hsa_circ_0077753 | 3130 | 11 | 0.70% |
| hsa_circ_0063778 | 4553 | 16 | 0.70% |
| hsa_circ_0006788 | 3415 | 12 | 0.70% |
| hsa_circ_0087975 | 3415 | 12 | 0.70% |
| hsa_circ_0083532 | 4269 | 15 | 0.70% |
| hsa_circ_0052741 | 3700 | 13 | 0.70% |
| hsa_circ_0129833 | 3131 | 11 | 0.70% |
| hsa_circ_0062716 | 3985 | 14 | 0.70% |
| hsa_circ_0036341 | 3416 | 12 | 0.70% |
| hsa_circ_0072345 | 3416 | 12 | 0.70% |
| hsa_circ_0089366 | 3416 | 12 | 0.70% |
| hsa_circ_0113476 | 3416 | 12 | 0.70% |
| hsa_circ_0077297 | 3701 | 13 | 0.70% |
| hsa_circ_0071137 | 2847 | 10 | 0.70% |
| hsa_circ_0028716 | 3132 | 11 | 0.70% |
| hsa_circ_0048303 | 3132 | 11 | 0.70% |
| hsa_circ_0013079 | 3417 | 12 | 0.70% |
| hsa_circ_0124183 | 7690 | 27 | 0.70% |
| hsa_circ_0037425 | 3133 | 11 | 0.70% |
| hsa_circ_0085570 | 3133 | 11 | 0.70% |
| hsa_circ_0134154 | 6267 | 22 | 0.70% |
| hsa_circ_0016171 | 5983 | 21 | 0.70% |
| hsa_circ_0117686 | 5983 | 21 | 0.70% |
| hsa_circ_0046480 | 3134 | 11 | 0.70% |
| hsa_circ_0058784 | 3134 | 11 | 0.70% |
| hsa_circ_0069942 | 3134 | 11 | 0.70% |
| hsa_circ_0071393 | 3134 | 11 | 0.70% |
| hsa_circ_0073498 | 3134 | 11 | 0.70% |
| hsa_circ_0078039 | 3419 | 12 | 0.70% |
| hsa_circ_0082339 | 3419 | 12 | 0.70% |
| hsa_circ_0089684 | 3989 | 14 | 0.70% |
| hsa_circ_0088843 | 3705 | 13 | 0.70% |
| hsa_circ_0049309 | 3420 | 12 | 0.70% |
| hsa_circ_0083700 | 3420 | 12 | 0.70% |
| hsa_circ_0049289 | 3135 | 11 | 0.70% |
| hsa_circ_0105916 | 9692 | 34 | 0.70% |
| hsa_circ_0032643 | 3706 | 13 | 0.70% |

| | | | |
|---|---|---|---|
| hsa_circ_0023585 | 3136 | 11 | 0.70% |
| hsa_circ_0056357 | 3422 | 12 | 0.70% |
| hsa_circ_0086356 | 3422 | 12 | 0.70% |
| hsa_circ_0084374 | 3137 | 11 | 0.70% |
| hsa_circ_0121403 | 4278 | 15 | 0.70% |
| hsa_circ_0051585 | 9414 | 33 | 0.70% |
| hsa_circ_0088002 | 3138 | 11 | 0.70% |
| hsa_circ_0046892 | 3994 | 14 | 0.70% |
| hsa_circ_0088167 | 3709 | 13 | 0.70% |
| hsa_circ_0086076 | 3424 | 12 | 0.70% |
| hsa_circ_0064723 | 8275 | 29 | 0.70% |
| hsa_circ_0061130 | 3995 | 14 | 0.70% |
| hsa_circ_0084789 | 3139 | 11 | 0.70% |
| hsa_circ_0066340 | 2854 | 10 | 0.70% |
| hsa_circ_0073605 | 2854 | 10 | 0.70% |
| hsa_circ_0082115 | 2854 | 10 | 0.70% |
| hsa_circ_0090771 | 3996 | 14 | 0.70% |
| hsa_circ_0118735 | 7707 | 27 | 0.70% |
| hsa_circ_0014052 | 3140 | 11 | 0.70% |
| hsa_circ_0028728 | 3140 | 11 | 0.70% |
| hsa_circ_0049923 | 3140 | 11 | 0.70% |
| hsa_circ_0131496 | 3140 | 11 | 0.70% |
| hsa_circ_0088939 | 3711 | 13 | 0.70% |
| hsa_circ_0125198 | 3711 | 13 | 0.70% |
| hsa_circ_0113502 | 3426 | 12 | 0.70% |
| hsa_circ_0059711 | 3141 | 11 | 0.70% |
| hsa_circ_0069823 | 3141 | 11 | 0.70% |
| hsa_circ_0070131 | 3141 | 11 | 0.70% |
| hsa_circ_0081284 | 3141 | 11 | 0.70% |
| hsa_circ_0089936 | 3141 | 11 | 0.70% |
| hsa_circ_0090292 | 3998 | 14 | 0.70% |
| hsa_circ_0102429 | 8853 | 31 | 0.70% |
| hsa_circ_0031341 | 3427 | 12 | 0.70% |
| hsa_circ_0088670 | 3427 | 12 | 0.70% |
| hsa_circ_0051555 | 2856 | 10 | 0.70% |
| hsa_circ_0064249 | 2856 | 10 | 0.70% |
| hsa_circ_0087425 | 9997 | 35 | 0.70% |
| hsa_circ_0035674 | 3142 | 11 | 0.70% |
| hsa_circ_0091103 | 3142 | 11 | 0.70% |
| hsa_circ_0014435 | 3428 | 12 | 0.70% |
| hsa_circ_0040475 | 3428 | 12 | 0.70% |
| hsa_circ_0040796 | 3428 | 12 | 0.70% |
| hsa_circ_0047298 | 3428 | 12 | 0.70% |
| hsa_circ_0010619 | 3714 | 13 | 0.70% |
| hsa_circ_0079503 | 3714 | 13 | 0.70% |
| hsa_circ_0053789 | 3143 | 11 | 0.70% |
| hsa_circ_0068318 | 3143 | 11 | 0.70% |
| hsa_circ_0068927 | 3143 | 11 | 0.70% |
| hsa_circ_0125471 | 3143 | 11 | 0.70% |
| hsa_circ_0136358 | 3143 | 11 | 0.70% |

| | | | |
|---|---|---|---|
| hsa_circ_0069773 | 9144 | 32 | 0.70% |
| hsa_circ_0059243 | 3429 | 12 | 0.70% |
| hsa_circ_0001323 | 4858 | 17 | 0.70% |
| hsa_circ_0053664 | 3715 | 13 | 0.70% |
| hsa_circ_0049935 | 3144 | 11 | 0.70% |
| hsa_circ_0063692 | 3144 | 11 | 0.70% |
| hsa_circ_0114245 | 3144 | 11 | 0.70% |
| hsa_circ_0082220 | 4002 | 14 | 0.70% |
| hsa_circ_0040247 | 9720 | 34 | 0.70% |
| hsa_circ_0061803 | 2859 | 10 | 0.70% |
| hsa_circ_0029079 | 3145 | 11 | 0.70% |
| hsa_circ_0093322 | 3145 | 11 | 0.70% |
| hsa_circ_0077953 | 3431 | 12 | 0.70% |
| hsa_circ_0108892 | 3431 | 12 | 0.70% |
| hsa_circ_0064616 | 7434 | 26 | 0.70% |
| hsa_circ_0082213 | 3432 | 12 | 0.70% |
| hsa_circ_0053622 | 3146 | 11 | 0.70% |
| hsa_circ_0069636 | 3146 | 11 | 0.70% |
| hsa_circ_0042304 | 2860 | 10 | 0.70% |
| hsa_circ_0087094 | 29748 | 104 | 0.70% |
| hsa_circ_0103376 | 3433 | 12 | 0.70% |
| hsa_circ_0095717 | 3147 | 11 | 0.70% |
| hsa_circ_0110624 | 3147 | 11 | 0.70% |
| hsa_circ_0034209 | 2861 | 10 | 0.70% |
| hsa_circ_0006298 | 8012 | 28 | 0.70% |
| hsa_circ_0126978 | 6009 | 21 | 0.70% |
| hsa_circ_0091394 | 4006 | 14 | 0.70% |
| hsa_circ_0012689 | 3720 | 13 | 0.70% |
| hsa_circ_0067614 | 3434 | 12 | 0.70% |
| hsa_circ_0072529 | 3148 | 11 | 0.70% |
| hsa_circ_0116362 | 2862 | 10 | 0.70% |
| hsa_circ_0057187 | 6011 | 21 | 0.70% |
| hsa_circ_0013776 | 3435 | 12 | 0.70% |
| hsa_circ_0055657 | 3435 | 12 | 0.70% |
| hsa_circ_0063651 | 3435 | 12 | 0.70% |
| hsa_circ_0066220 | 3435 | 12 | 0.70% |
| hsa_circ_0047962 | 3149 | 11 | 0.70% |
| hsa_circ_0036746 | 5153 | 18 | 0.70% |
| hsa_circ_0100943 | 2863 | 10 | 0.70% |
| hsa_circ_0111958 | 2863 | 10 | 0.70% |
| hsa_circ_0102268 | 6872 | 24 | 0.70% |
| hsa_circ_0030483 | 4295 | 15 | 0.70% |
| hsa_circ_0093254 | 6300 | 22 | 0.70% |
| hsa_circ_0053897 | 3150 | 11 | 0.70% |
| hsa_circ_0113720 | 5728 | 20 | 0.70% |
| hsa_circ_0090637 | 2864 | 10 | 0.70% |
| hsa_circ_0041206 | 3437 | 12 | 0.70% |
| hsa_circ_0015471 | 3151 | 11 | 0.70% |
| hsa_circ_0015707 | 3151 | 11 | 0.70% |
| hsa_circ_0035778 | 3724 | 13 | 0.70% |

| | | | |
|---|---|---|---|
| hsa_circ_0070408 | 3724 | 13 | 0.70% |
| hsa_circ_0040230 | 3438 | 12 | 0.70% |
| hsa_circ_0055983 | 3438 | 12 | 0.70% |
| hsa_circ_0064329 | 3438 | 12 | 0.70% |
| hsa_circ_0108115 | 3438 | 12 | 0.70% |
| hsa_circ_0049499 | 2865 | 10 | 0.70% |
| hsa_circ_0112179 | 2865 | 10 | 0.70% |
| hsa_circ_0083515 | 3152 | 11 | 0.70% |
| hsa_circ_0091434 | 3152 | 11 | 0.70% |
| hsa_circ_0023853 | 3153 | 11 | 0.70% |
| hsa_circ_0039609 | 3153 | 11 | 0.70% |
| hsa_circ_0056688 | 4013 | 14 | 0.70% |
| hsa_circ_0114512 | 3727 | 13 | 0.70% |
| hsa_circ_0126482 | 3727 | 13 | 0.70% |
| hsa_circ_0011013 | 2867 | 10 | 0.70% |
| hsa_circ_0071989 | 2867 | 10 | 0.70% |
| hsa_circ_0065200 | 3154 | 11 | 0.70% |
| hsa_circ_0122270 | 3154 | 11 | 0.70% |
| hsa_circ_0136277 | 5736 | 20 | 0.70% |
| hsa_circ_0020019 | 2868 | 10 | 0.70% |
| hsa_circ_0100837 | 9178 | 32 | 0.70% |
| hsa_circ_0026528 | 3155 | 11 | 0.70% |
| hsa_circ_0035498 | 3155 | 11 | 0.70% |
| hsa_circ_0056676 | 3442 | 12 | 0.70% |
| hsa_circ_0067141 | 3442 | 12 | 0.70% |
| hsa_circ_0118174 | 3442 | 12 | 0.70% |
| hsa_circ_0088866 | 3729 | 13 | 0.70% |
| hsa_circ_0088216 | 2869 | 10 | 0.70% |
| hsa_circ_0097980 | 2869 | 10 | 0.70% |
| hsa_circ_0140347 | 3156 | 11 | 0.70% |
| hsa_circ_0053205 | 3730 | 13 | 0.70% |
| hsa_circ_0111263 | 7748 | 27 | 0.70% |
| hsa_circ_0034591 | 3444 | 12 | 0.70% |
| hsa_circ_0054059 | 3444 | 12 | 0.70% |
| hsa_circ_0083262 | 3444 | 12 | 0.70% |
| hsa_circ_0086664 | 3444 | 12 | 0.70% |
| hsa_circ_0013485 | 3157 | 11 | 0.70% |
| hsa_circ_0020872 | 3157 | 11 | 0.70% |
| hsa_circ_0038107 | 3157 | 11 | 0.70% |
| hsa_circ_0095408 | 14639 | 51 | 0.70% |
| hsa_circ_0060838 | 4306 | 15 | 0.70% |
| hsa_circ_0070155 | 4019 | 14 | 0.70% |
| hsa_circ_0090037 | 4019 | 14 | 0.70% |
| hsa_circ_0023547 | 3158 | 11 | 0.70% |
| hsa_circ_0067645 | 3158 | 11 | 0.70% |
| hsa_circ_0057555 | 4307 | 15 | 0.70% |
| hsa_circ_0035645 | 5743 | 20 | 0.70% |
| hsa_circ_0032221 | 3733 | 13 | 0.70% |
| hsa_circ_0034423 | 3446 | 12 | 0.70% |
| hsa_circ_0135234 | 3446 | 12 | 0.70% |

| | | | |
|---|---|---|---|
| hsa_circ_0043925 | 3159 | 11 | 0.70% |
| hsa_circ_0042640 | 2872 | 10 | 0.70% |
| hsa_circ_0067146 | 3447 | 12 | 0.70% |
| hsa_circ_0033433 | 3160 | 11 | 0.70% |
| hsa_circ_0053775 | 3160 | 11 | 0.70% |
| hsa_circ_0140346 | 3160 | 11 | 0.70% |
| hsa_circ_0111919 | 4022 | 14 | 0.70% |
| hsa_circ_0128854 | 11492 | 40 | 0.70% |
| hsa_circ_0039035 | 2873 | 10 | 0.70% |
| hsa_circ_0051625 | 2873 | 10 | 0.70% |
| hsa_circ_0081065 | 2873 | 10 | 0.70% |
| hsa_circ_0125334 | 2873 | 10 | 0.70% |
| hsa_circ_0077098 | 3735 | 13 | 0.70% |
| hsa_circ_0073610 | 9769 | 34 | 0.70% |
| hsa_circ_0020125 | 4310 | 15 | 0.70% |
| hsa_circ_0001199 | 3448 | 12 | 0.70% |
| hsa_circ_0032088 | 3448 | 12 | 0.70% |
| hsa_circ_0058559 | 3448 | 12 | 0.70% |
| hsa_circ_0066332 | 3448 | 12 | 0.70% |
| hsa_circ_0067000 | 3448 | 12 | 0.70% |
| hsa_circ_0042100 | 3161 | 11 | 0.70% |
| hsa_circ_0050074 | 3161 | 11 | 0.70% |
| hsa_circ_0029560 | 3449 | 12 | 0.70% |
| hsa_circ_0048189 | 3449 | 12 | 0.70% |
| hsa_circ_0063107 | 3449 | 12 | 0.70% |
| hsa_circ_0025659 | 3162 | 11 | 0.70% |
| hsa_circ_0068195 | 3162 | 11 | 0.70% |
| hsa_circ_0070090 | 3162 | 11 | 0.70% |
| hsa_circ_0084672 | 3737 | 13 | 0.70% |
| hsa_circ_0045397 | 4025 | 14 | 0.70% |
| hsa_circ_0074523 | 3450 | 12 | 0.70% |
| hsa_circ_0015496 | 2875 | 10 | 0.70% |
| hsa_circ_0035513 | 2875 | 10 | 0.70% |
| hsa_circ_0057711 | 2875 | 10 | 0.70% |
| hsa_circ_0077585 | 15527 | 54 | 0.70% |
| hsa_circ_0038044 | 3163 | 11 | 0.70% |
| hsa_circ_0043882 | 3163 | 11 | 0.70% |
| hsa_circ_0136045 | 3163 | 11 | 0.70% |
| hsa_circ_0006159 | 4026 | 14 | 0.70% |
| hsa_circ_0060172 | 4026 | 14 | 0.70% |
| hsa_circ_0113666 | 3451 | 12 | 0.70% |
| hsa_circ_0060804 | 3739 | 13 | 0.70% |
| hsa_circ_0054966 | 4602 | 16 | 0.70% |
| hsa_circ_0011316 | 3164 | 11 | 0.70% |
| hsa_circ_0028713 | 3164 | 11 | 0.70% |
| hsa_circ_0032701 | 3452 | 12 | 0.70% |
| hsa_circ_0034155 | 3452 | 12 | 0.70% |
| hsa_circ_0066051 | 3452 | 12 | 0.70% |
| hsa_circ_0081041 | 3452 | 12 | 0.70% |
| hsa_circ_0061454 | 4603 | 16 | 0.70% |

| | | | |
|---|---|---|---|
| hsa_circ_0038084 | 3165 | 11 | 0.70% |
| hsa_circ_0065584 | 3165 | 11 | 0.70% |
| hsa_circ_0013997 | 3453 | 12 | 0.70% |
| hsa_circ_0021251 | 3453 | 12 | 0.70% |
| hsa_circ_0050793 | 3453 | 12 | 0.70% |
| hsa_circ_0052980 | 3453 | 12 | 0.70% |
| hsa_circ_0109070 | 10360 | 36 | 0.69% |
| hsa_circ_0003672 | 12664 | 44 | 0.69% |
| hsa_circ_0070049 | 3166 | 11 | 0.69% |
| hsa_circ_0073838 | 3166 | 11 | 0.69% |
| hsa_circ_0076366 | 3166 | 11 | 0.69% |
| hsa_circ_0111010 | 3166 | 11 | 0.69% |
| hsa_circ_0084496 | 3454 | 12 | 0.69% |
| hsa_circ_0089869 | 3742 | 13 | 0.69% |
| hsa_circ_0113159 | 2879 | 10 | 0.69% |
| hsa_circ_0039253 | 3167 | 11 | 0.69% |
| hsa_circ_0026089 | 3744 | 13 | 0.69% |
| hsa_circ_0078692 | 3744 | 13 | 0.69% |
| hsa_circ_0082547 | 3456 | 12 | 0.69% |
| hsa_circ_0118220 | 3456 | 12 | 0.69% |
| hsa_circ_0028766 | 3168 | 11 | 0.69% |
| hsa_circ_0016014 | 2880 | 10 | 0.69% |
| hsa_circ_0100827 | 12387 | 43 | 0.69% |
| hsa_circ_0121692 | 3457 | 12 | 0.69% |
| hsa_circ_0056064 | 3169 | 11 | 0.69% |
| hsa_circ_0091957 | 3169 | 11 | 0.69% |
| hsa_circ_0119822 | 3169 | 11 | 0.69% |
| hsa_circ_0010351 | 2881 | 10 | 0.69% |
| hsa_circ_0070485 | 2881 | 10 | 0.69% |
| hsa_circ_0077402 | 2881 | 10 | 0.69% |
| hsa_circ_0032054 | 4322 | 15 | 0.69% |
| hsa_circ_0075468 | 4034 | 14 | 0.69% |
| hsa_circ_0026255 | 3170 | 11 | 0.69% |
| hsa_circ_0100091 | 2882 | 10 | 0.69% |
| hsa_circ_0104424 | 3459 | 12 | 0.69% |
| hsa_circ_0014039 | 3171 | 11 | 0.69% |
| hsa_circ_0019328 | 3171 | 11 | 0.69% |
| hsa_circ_0022079 | 3171 | 11 | 0.69% |
| hsa_circ_0025160 | 3171 | 11 | 0.69% |
| hsa_circ_0137340 | 5478 | 19 | 0.69% |
| hsa_circ_0073617 | 3460 | 12 | 0.69% |
| hsa_circ_0074314 | 3172 | 11 | 0.69% |
| hsa_circ_0083003 | 3172 | 11 | 0.69% |
| hsa_circ_0052634 | 3749 | 13 | 0.69% |
| hsa_circ_0016790 | 2884 | 10 | 0.69% |
| hsa_circ_0036237 | 3461 | 12 | 0.69% |
| hsa_circ_0051300 | 3461 | 12 | 0.69% |
| hsa_circ_0026709 | 3173 | 11 | 0.69% |
| hsa_circ_0013641 | 3750 | 13 | 0.69% |
| hsa_circ_0098801 | 15001 | 52 | 0.69% |

| | | | |
|---|---|---|---|
| hsa_circ_0010718 | 3462 | 12 | 0.69% |
| hsa_circ_0033152 | 3462 | 12 | 0.69% |
| hsa_circ_0139353 | 4328 | 15 | 0.69% |
| hsa_circ_0020402 | 3174 | 11 | 0.69% |
| hsa_circ_0058549 | 3174 | 11 | 0.69% |
| hsa_circ_0128144 | 3174 | 11 | 0.69% |
| hsa_circ_0046344 | 3463 | 12 | 0.69% |
| hsa_circ_0056692 | 3463 | 12 | 0.69% |
| hsa_circ_0101880 | 3463 | 12 | 0.69% |
| hsa_circ_0026853 | 3175 | 11 | 0.69% |
| hsa_circ_0015418 | 4907 | 17 | 0.69% |
| hsa_circ_0025422 | 3464 | 12 | 0.69% |
| hsa_circ_0047345 | 3464 | 12 | 0.69% |
| hsa_circ_0104679 | 7217 | 25 | 0.69% |
| hsa_circ_0060835 | 2887 | 10 | 0.69% |
| hsa_circ_0021112 | 7795 | 27 | 0.69% |
| hsa_circ_0060588 | 3176 | 11 | 0.69% |
| hsa_circ_0086280 | 3176 | 11 | 0.69% |
| hsa_circ_0111158 | 3176 | 11 | 0.69% |
| hsa_circ_0034640 | 3465 | 12 | 0.69% |
| hsa_circ_0137296 | 3754 | 13 | 0.69% |
| hsa_circ_0016263 | 2888 | 10 | 0.69% |
| hsa_circ_0010712 | 3177 | 11 | 0.69% |
| hsa_circ_0060324 | 3466 | 12 | 0.69% |
| hsa_circ_0053804 | 2889 | 10 | 0.69% |
| hsa_circ_0024663 | 3178 | 11 | 0.69% |
| hsa_circ_0060599 | 3178 | 11 | 0.69% |
| hsa_circ_0030208 | 3467 | 12 | 0.69% |
| hsa_circ_0131612 | 6935 | 24 | 0.69% |
| hsa_circ_0072817 | 7513 | 26 | 0.69% |
| hsa_circ_0099149 | 11560 | 40 | 0.69% |
| hsa_circ_0094108 | 4335 | 15 | 0.69% |
| hsa_circ_0056837 | 3468 | 12 | 0.69% |
| hsa_circ_0021490 | 3179 | 11 | 0.69% |
| hsa_circ_0043881 | 3179 | 11 | 0.69% |
| hsa_circ_0066037 | 3179 | 11 | 0.69% |
| hsa_circ_0081879 | 3179 | 11 | 0.69% |
| hsa_circ_0032365 | 2890 | 10 | 0.69% |
| hsa_circ_0072226 | 2890 | 10 | 0.69% |
| hsa_circ_0123263 | 12428 | 43 | 0.69% |
| hsa_circ_0067318 | 5781 | 20 | 0.69% |
| hsa_circ_0041895 | 3469 | 12 | 0.69% |
| hsa_circ_0054888 | 3469 | 12 | 0.69% |
| hsa_circ_0048792 | 2891 | 10 | 0.69% |
| hsa_circ_0081506 | 4626 | 16 | 0.69% |
| hsa_circ_0130125 | 3470 | 12 | 0.69% |
| hsa_circ_0028807 | 3181 | 11 | 0.69% |
| hsa_circ_0031665 | 3181 | 11 | 0.69% |
| hsa_circ_0068999 | 3181 | 11 | 0.69% |
| hsa_circ_0080009 | 3181 | 11 | 0.69% |

| | | | |
|---|---|---|---|
| hsa_circ_0136985 | 3181 | 11 | 0.69% |
| hsa_circ_0057185 | 6073 | 21 | 0.69% |
| hsa_circ_0125701 | 7230 | 25 | 0.69% |
| hsa_circ_0016162 | 9833 | 34 | 0.69% |
| hsa_circ_0129855 | 3760 | 13 | 0.69% |
| hsa_circ_0078043 | 3471 | 12 | 0.69% |
| hsa_circ_0071797 | 3182 | 11 | 0.69% |
| hsa_circ_0078584 | 3182 | 11 | 0.69% |
| hsa_circ_0066445 | 2893 | 10 | 0.69% |
| hsa_circ_0088118 | 2893 | 10 | 0.69% |
| hsa_circ_0044724 | 3761 | 13 | 0.69% |
| hsa_circ_0045593 | 3183 | 11 | 0.69% |
| hsa_circ_0125366 | 8681 | 30 | 0.69% |
| hsa_circ_0044751 | 3762 | 13 | 0.69% |
| hsa_circ_0047787 | 2894 | 10 | 0.69% |
| hsa_circ_0061732 | 3184 | 11 | 0.69% |
| hsa_circ_0065197 | 3184 | 11 | 0.69% |
| hsa_circ_0070102 | 3184 | 11 | 0.69% |
| hsa_circ_0097596 | 3184 | 11 | 0.69% |
| hsa_circ_0020020 | 2895 | 10 | 0.69% |
| hsa_circ_0056297 | 3185 | 11 | 0.69% |
| hsa_circ_0060210 | 3185 | 11 | 0.69% |
| hsa_circ_0090500 | 3185 | 11 | 0.69% |
| hsa_circ_0012490 | 3475 | 12 | 0.69% |
| hsa_circ_0029185 | 3475 | 12 | 0.69% |
| hsa_circ_0071773 | 2896 | 10 | 0.69% |
| hsa_circ_0088863 | 3765 | 13 | 0.69% |
| hsa_circ_0094501 | 3765 | 13 | 0.69% |
| hsa_circ_0101493 | 3765 | 13 | 0.69% |
| hsa_circ_0096489 | 10427 | 36 | 0.69% |
| hsa_circ_0044906 | 4345 | 15 | 0.69% |
| hsa_circ_0135643 | 2897 | 10 | 0.69% |
| hsa_circ_0025007 | 3477 | 12 | 0.69% |
| hsa_circ_0123657 | 3188 | 11 | 0.69% |
| hsa_circ_0025099 | 3478 | 12 | 0.69% |
| hsa_circ_0123410 | 3768 | 13 | 0.69% |
| hsa_circ_0140776 | 5797 | 20 | 0.69% |
| hsa_circ_0030931 | 2899 | 10 | 0.69% |
| hsa_circ_0059518 | 2899 | 10 | 0.69% |
| hsa_circ_0082498 | 2899 | 10 | 0.69% |
| hsa_circ_0100835 | 9277 | 32 | 0.69% |
| hsa_circ_0038674 | 3189 | 11 | 0.69% |
| hsa_circ_0072665 | 3189 | 11 | 0.69% |
| hsa_circ_0126338 | 3769 | 13 | 0.69% |
| hsa_circ_0133663 | 3769 | 13 | 0.69% |
| hsa_circ_0049156 | 5220 | 18 | 0.69% |
| hsa_circ_0048863 | 3770 | 13 | 0.69% |
| hsa_circ_0034068 | 3480 | 12 | 0.69% |
| hsa_circ_0086824 | 3480 | 12 | 0.69% |
| hsa_circ_0019532 | 3190 | 11 | 0.69% |

| | | | |
|---|---|---|---|
| hsa_circ_0048309 | 3190 | 11 | 0.69% |
| hsa_circ_0077123 | 3190 | 11 | 0.69% |
| hsa_circ_0097785 | 3190 | 11 | 0.69% |
| hsa_circ_0064215 | 2900 | 10 | 0.69% |
| hsa_circ_0067357 | 3481 | 12 | 0.69% |
| hsa_circ_0080883 | 3481 | 12 | 0.69% |
| hsa_circ_0022257 | 3191 | 11 | 0.69% |
| hsa_circ_0028433 | 3191 | 11 | 0.69% |
| hsa_circ_0122000 | 4062 | 14 | 0.69% |
| hsa_circ_0046008 | 3772 | 13 | 0.69% |
| hsa_circ_0032211 | 12767 | 44 | 0.69% |
| hsa_circ_0015038 | 3482 | 12 | 0.69% |
| hsa_circ_0072739 | 3482 | 12 | 0.69% |
| hsa_circ_0035701 | 3192 | 11 | 0.69% |
| hsa_circ_0060670 | 3773 | 13 | 0.69% |
| hsa_circ_0083773 | 3483 | 12 | 0.69% |
| hsa_circ_0055974 | 4354 | 15 | 0.69% |
| hsa_circ_0092699 | 3193 | 11 | 0.69% |
| hsa_circ_0036747 | 5225 | 18 | 0.69% |
| hsa_circ_0016642 | 3484 | 12 | 0.69% |
| hsa_circ_0049171 | 3484 | 12 | 0.69% |
| hsa_circ_0073637 | 3484 | 12 | 0.69% |
| hsa_circ_0087748 | 4065 | 14 | 0.69% |
| hsa_circ_0088754 | 3194 | 11 | 0.69% |
| hsa_circ_0047406 | 3775 | 13 | 0.69% |
| hsa_circ_0053712 | 3775 | 13 | 0.69% |
| hsa_circ_0111380 | 2904 | 10 | 0.69% |
| hsa_circ_0060429 | 3195 | 11 | 0.69% |
| hsa_circ_0073343 | 3195 | 11 | 0.69% |
| hsa_circ_0113383 | 3195 | 11 | 0.69% |
| hsa_circ_0011584 | 3776 | 13 | 0.69% |
| hsa_circ_0054049 | 3486 | 12 | 0.69% |
| hsa_circ_0104115 | 3486 | 12 | 0.69% |
| hsa_circ_0118298 | 3486 | 12 | 0.69% |
| hsa_circ_0133726 | 7263 | 25 | 0.69% |
| hsa_circ_0041279 | 3196 | 11 | 0.69% |
| hsa_circ_0039086 | 3487 | 12 | 0.69% |
| hsa_circ_0104137 | 3487 | 12 | 0.69% |
| hsa_circ_0074714 | 4359 | 15 | 0.69% |
| hsa_circ_0030960 | 2906 | 10 | 0.69% |
| hsa_circ_0036807 | 2906 | 10 | 0.69% |
| hsa_circ_0032640 | 3778 | 13 | 0.69% |
| hsa_circ_0015641 | 3197 | 11 | 0.69% |
| hsa_circ_0041963 | 3197 | 11 | 0.69% |
| hsa_circ_0062092 | 3197 | 11 | 0.69% |
| hsa_circ_0041952 | 3488 | 12 | 0.69% |
| hsa_circ_0059637 | 3779 | 13 | 0.69% |
| hsa_circ_0121140 | 3779 | 13 | 0.69% |
| hsa_circ_0070342 | 3198 | 11 | 0.69% |
| hsa_circ_0110914 | 4652 | 16 | 0.69% |

| | | | |
|---|---|---|---|
| hsa_circ_0036837 | 3780 | 13 | 0.69% |
| hsa_circ_0132173 | 4071 | 14 | 0.69% |
| hsa_circ_0102347 | 2908 | 10 | 0.69% |
| hsa_circ_0007379 | 3199 | 11 | 0.69% |
| hsa_circ_0023103 | 3199 | 11 | 0.69% |
| hsa_circ_0126861 | 3781 | 13 | 0.69% |
| hsa_circ_0087798 | 14254 | 49 | 0.69% |
| hsa_circ_0061790 | 2909 | 10 | 0.69% |
| hsa_circ_0030955 | 3200 | 11 | 0.69% |
| hsa_circ_0072162 | 3200 | 11 | 0.69% |
| hsa_circ_0090393 | 3200 | 11 | 0.69% |
| hsa_circ_0026738 | 3491 | 12 | 0.69% |
| hsa_circ_0073344 | 3491 | 12 | 0.69% |
| hsa_circ_0034241 | 3782 | 13 | 0.69% |
| hsa_circ_0019624 | 3492 | 12 | 0.69% |
| hsa_circ_0035156 | 3492 | 12 | 0.69% |
| hsa_circ_0058980 | 3492 | 12 | 0.69% |
| hsa_circ_0079583 | 3492 | 12 | 0.69% |
| hsa_circ_0002895 | 3201 | 11 | 0.69% |
| hsa_circ_0092246 | 3201 | 11 | 0.69% |
| hsa_circ_0053842 | 2910 | 10 | 0.69% |
| hsa_circ_0075367 | 4366 | 15 | 0.69% |
| hsa_circ_0046928 | 3784 | 13 | 0.69% |
| hsa_circ_0018549 | 3493 | 12 | 0.69% |
| hsa_circ_0031275 | 3493 | 12 | 0.69% |
| hsa_circ_0014810 | 3202 | 11 | 0.69% |
| hsa_circ_0087231 | 2911 | 10 | 0.69% |
| hsa_circ_0112009 | 2911 | 10 | 0.69% |
| hsa_circ_0120803 | 5823 | 20 | 0.69% |
| hsa_circ_0085372 | 3494 | 12 | 0.69% |
| hsa_circ_0120900 | 3494 | 12 | 0.69% |
| hsa_circ_0023545 | 3203 | 11 | 0.69% |
| hsa_circ_0048641 | 3203 | 11 | 0.69% |
| hsa_circ_0090588 | 2912 | 10 | 0.69% |
| hsa_circ_0060687 | 3786 | 13 | 0.69% |
| hsa_circ_0054784 | 3495 | 12 | 0.69% |
| hsa_circ_0024283 | 9029 | 31 | 0.69% |
| hsa_circ_0030573 | 3204 | 11 | 0.69% |
| hsa_circ_0062349 | 4078 | 14 | 0.69% |
| hsa_circ_0134998 | 2913 | 10 | 0.69% |
| hsa_circ_0045068 | 3787 | 13 | 0.69% |
| hsa_circ_0136723 | 3787 | 13 | 0.69% |
| hsa_circ_0140441 | 8448 | 29 | 0.69% |
| hsa_circ_0051754 | 3496 | 12 | 0.69% |
| hsa_circ_0032001 | 3205 | 11 | 0.69% |
| hsa_circ_0090568 | 3205 | 11 | 0.69% |
| hsa_circ_0056222 | 3788 | 13 | 0.69% |
| hsa_circ_0071230 | 3788 | 13 | 0.69% |
| hsa_circ_0040432 | 3497 | 12 | 0.69% |
| hsa_circ_0045084 | 3497 | 12 | 0.69% |

| | | | |
|---|---|---|---|
| hsa_circ_0042623 | 3206 | 11 | 0.69% |
| hsa_circ_0063073 | 3789 | 13 | 0.69% |
| hsa_circ_0130541 | 3789 | 13 | 0.69% |
| hsa_circ_0052699 | 4372 | 15 | 0.69% |
| hsa_circ_0054818 | 3498 | 12 | 0.69% |
| hsa_circ_0004036 | 8163 | 28 | 0.69% |
| hsa_circ_0001334 | 3207 | 11 | 0.69% |
| hsa_circ_0021098 | 3207 | 11 | 0.69% |
| hsa_circ_0086879 | 3207 | 11 | 0.69% |
| hsa_circ_0134999 | 3207 | 11 | 0.69% |
| hsa_circ_0104792 | 3499 | 12 | 0.69% |
| hsa_circ_0028242 | 2916 | 10 | 0.69% |
| hsa_circ_0004263 | 5249 | 18 | 0.69% |
| hsa_circ_0061443 | 4666 | 16 | 0.69% |
| hsa_circ_0013205 | 3208 | 11 | 0.69% |
| hsa_circ_0019677 | 3208 | 11 | 0.69% |
| hsa_circ_0090334 | 4083 | 14 | 0.69% |
| hsa_circ_0120139 | 8459 | 29 | 0.69% |
| hsa_circ_0111784 | 99763 | 342 | 0.69% |
| hsa_circ_0019197 | 3209 | 11 | 0.69% |
| hsa_circ_0022317 | 3209 | 11 | 0.69% |
| hsa_circ_0043983 | 3209 | 11 | 0.69% |
| hsa_circ_0046953 | 3209 | 11 | 0.69% |
| hsa_circ_0054821 | 3501 | 12 | 0.69% |
| hsa_circ_0105036 | 3501 | 12 | 0.69% |
| hsa_circ_0113607 | 6127 | 21 | 0.69% |
| hsa_circ_0062354 | 4085 | 14 | 0.69% |
| hsa_circ_0084421 | 3210 | 11 | 0.69% |
| hsa_circ_0056033 | 3502 | 12 | 0.69% |
| hsa_circ_0090668 | 3502 | 12 | 0.69% |
| hsa_circ_0019045 | 12258 | 42 | 0.69% |
| hsa_circ_0018359 | 2919 | 10 | 0.69% |
| hsa_circ_0024804 | 2919 | 10 | 0.69% |
| hsa_circ_0033047 | 2919 | 10 | 0.69% |
| hsa_circ_0090944 | 2919 | 10 | 0.69% |
| hsa_circ_0052913 | 3211 | 11 | 0.69% |
| hsa_circ_0084911 | 3211 | 11 | 0.69% |
| hsa_circ_0043880 | 3503 | 12 | 0.69% |
| hsa_circ_0111074 | 3503 | 12 | 0.69% |
| hsa_circ_0132360 | 3503 | 12 | 0.69% |
| hsa_circ_0029989 | 3795 | 13 | 0.69% |
| hsa_circ_0117399 | 9635 | 33 | 0.69% |
| hsa_circ_0109283 | 9927 | 34 | 0.69% |
| hsa_circ_0066671 | 3504 | 12 | 0.68% |
| hsa_circ_0062370 | 3212 | 11 | 0.68% |
| hsa_circ_0108207 | 3212 | 11 | 0.68% |
| hsa_circ_0115643 | 3212 | 11 | 0.68% |
| hsa_circ_0126572 | 3212 | 11 | 0.68% |
| hsa_circ_0094341 | 3797 | 13 | 0.68% |
| hsa_circ_0040208 | 29208 | 100 | 0.68% |

| | | | |
|---|---|---|---|
| hsa_circ_0009227 | 3505 | 12 | 0.68% |
| hsa_circ_0064489 | 3505 | 12 | 0.68% |
| hsa_circ_0084087 | 3505 | 12 | 0.68% |
| hsa_circ_0025956 | 3213 | 11 | 0.68% |
| hsa_circ_0028535 | 3213 | 11 | 0.68% |
| hsa_circ_0075953 | 8763 | 30 | 0.68% |
| hsa_circ_0121063 | 2921 | 10 | 0.68% |
| hsa_circ_0133412 | 2921 | 10 | 0.68% |
| hsa_circ_0014687 | 3506 | 12 | 0.68% |
| hsa_circ_0027633 | 3506 | 12 | 0.68% |
| hsa_circ_0029490 | 3506 | 12 | 0.68% |
| hsa_circ_0078489 | 4967 | 17 | 0.68% |
| hsa_circ_0059296 | 2922 | 10 | 0.68% |
| hsa_circ_0122783 | 9643 | 33 | 0.68% |
| hsa_circ_0035084 | 3799 | 13 | 0.68% |
| hsa_circ_0086614 | 3799 | 13 | 0.68% |
| hsa_circ_0014667 | 3507 | 12 | 0.68% |
| hsa_circ_0024439 | 2923 | 10 | 0.68% |
| hsa_circ_0042774 | 2923 | 10 | 0.68% |
| hsa_circ_0044625 | 2923 | 10 | 0.68% |
| hsa_circ_0009944 | 3508 | 12 | 0.68% |
| hsa_circ_0043663 | 3508 | 12 | 0.68% |
| hsa_circ_0051315 | 3508 | 12 | 0.68% |
| hsa_circ_0057508 | 4093 | 14 | 0.68% |
| hsa_circ_0075959 | 3216 | 11 | 0.68% |
| hsa_circ_0119723 | 3216 | 11 | 0.68% |
| hsa_circ_0107070 | 11696 | 40 | 0.68% |
| hsa_circ_0099332 | 2924 | 10 | 0.68% |
| hsa_circ_0118872 | 2924 | 10 | 0.68% |
| hsa_circ_0131217 | 6434 | 22 | 0.68% |
| hsa_circ_0056148 | 3217 | 11 | 0.68% |
| hsa_circ_0094950 | 3802 | 13 | 0.68% |
| hsa_circ_0030504 | 3510 | 12 | 0.68% |
| hsa_circ_0057184 | 6143 | 21 | 0.68% |
| hsa_circ_0111511 | 4388 | 15 | 0.68% |
| hsa_circ_0095148 | 3803 | 13 | 0.68% |
| hsa_circ_0025586 | 3218 | 11 | 0.68% |
| hsa_circ_0066563 | 3218 | 11 | 0.68% |
| hsa_circ_0135004 | 3218 | 11 | 0.68% |
| hsa_circ_0058203 | 4096 | 14 | 0.68% |
| hsa_circ_0013833 | 3511 | 12 | 0.68% |
| hsa_circ_0029514 | 2926 | 10 | 0.68% |
| hsa_circ_0044009 | 3804 | 13 | 0.68% |
| hsa_circ_0066481 | 4682 | 16 | 0.68% |
| hsa_circ_0076984 | 3512 | 12 | 0.68% |
| hsa_circ_0039737 | 2927 | 10 | 0.68% |
| hsa_circ_0081066 | 2927 | 10 | 0.68% |
| hsa_circ_0117949 | 2927 | 10 | 0.68% |
| hsa_circ_0031481 | 3220 | 11 | 0.68% |
| hsa_circ_0046782 | 3220 | 11 | 0.68% |

| | | | |
|---|---|---|---|
| hsa_circ_0080136 | 3220 | 11 | 0.68% |
| hsa_circ_0074117 | 3513 | 12 | 0.68% |
| hsa_circ_0061784 | 2928 | 10 | 0.68% |
| hsa_circ_0053595 | 3221 | 11 | 0.68% |
| hsa_circ_0018941 | 3514 | 12 | 0.68% |
| hsa_circ_0055464 | 3514 | 12 | 0.68% |
| hsa_circ_0015458 | 3807 | 13 | 0.68% |
| hsa_circ_0102796 | 3807 | 13 | 0.68% |
| hsa_circ_0072402 | 2929 | 10 | 0.68% |
| hsa_circ_0028394 | 3222 | 11 | 0.68% |
| hsa_circ_0094536 | 3222 | 11 | 0.68% |
| hsa_circ_0049228 | 3515 | 12 | 0.68% |
| hsa_circ_0012894 | 4101 | 14 | 0.68% |
| hsa_circ_0010700 | 3516 | 12 | 0.68% |
| hsa_circ_0038771 | 3516 | 12 | 0.68% |
| hsa_circ_0038762 | 3223 | 11 | 0.68% |
| hsa_circ_0055736 | 3223 | 11 | 0.68% |
| hsa_circ_0019542 | 3810 | 13 | 0.68% |
| hsa_circ_0068429 | 3810 | 13 | 0.68% |
| hsa_circ_0094500 | 3517 | 12 | 0.68% |
| hsa_circ_0047240 | 2931 | 10 | 0.68% |
| hsa_circ_0020866 | 3225 | 11 | 0.68% |
| hsa_circ_0027728 | 3225 | 11 | 0.68% |
| hsa_circ_0089712 | 3519 | 12 | 0.68% |
| hsa_circ_0032214 | 7332 | 25 | 0.68% |
| hsa_circ_0063928 | 2933 | 10 | 0.68% |
| hsa_circ_0019078 | 3813 | 13 | 0.68% |
| hsa_circ_0011064 | 4107 | 14 | 0.68% |
| hsa_circ_0066948 | 3814 | 13 | 0.68% |
| hsa_circ_0026328 | 3521 | 12 | 0.68% |
| hsa_circ_0083994 | 3521 | 12 | 0.68% |
| hsa_circ_0119840 | 3521 | 12 | 0.68% |
| hsa_circ_0100559 | 8510 | 29 | 0.68% |
| hsa_circ_0023197 | 3228 | 11 | 0.68% |
| hsa_circ_0031582 | 4696 | 16 | 0.68% |
| hsa_circ_0013068 | 3522 | 12 | 0.68% |
| hsa_circ_0015610 | 3522 | 12 | 0.68% |
| hsa_circ_0025093 | 3522 | 12 | 0.68% |
| hsa_circ_0089900 | 12915 | 44 | 0.68% |
| hsa_circ_0002907 | 13209 | 45 | 0.68% |
| hsa_circ_0035595 | 3523 | 12 | 0.68% |
| hsa_circ_0086353 | 2936 | 10 | 0.68% |
| hsa_circ_0035637 | 3230 | 11 | 0.68% |
| hsa_circ_0033248 | 3524 | 12 | 0.68% |
| hsa_circ_0070365 | 3524 | 12 | 0.68% |
| hsa_circ_0100465 | 12629 | 43 | 0.68% |
| hsa_circ_0011354 | 6168 | 21 | 0.68% |
| hsa_circ_0057211 | 3231 | 11 | 0.68% |
| hsa_circ_0091786 | 3231 | 11 | 0.68% |
| hsa_circ_0131861 | 3231 | 11 | 0.68% |

| | | | |
|---|---|---|---|
| hsa_circ_0099647 | 12631 | 43 | 0.68% |
| hsa_circ_0097108 | 3819 | 13 | 0.68% |
| hsa_circ_0048385 | 4113 | 14 | 0.68% |
| hsa_circ_0021649 | 4407 | 15 | 0.68% |
| hsa_circ_0014623 | 2938 | 10 | 0.68% |
| hsa_circ_0026994 | 3232 | 11 | 0.68% |
| hsa_circ_0029299 | 3232 | 11 | 0.68% |
| hsa_circ_0082920 | 3232 | 11 | 0.68% |
| hsa_circ_0140614 | 3526 | 12 | 0.68% |
| hsa_circ_0010595 | 3820 | 13 | 0.68% |
| hsa_circ_0041240 | 2939 | 10 | 0.68% |
| hsa_circ_0010035 | 3233 | 11 | 0.68% |
| hsa_circ_0011226 | 3233 | 11 | 0.68% |
| hsa_circ_0048743 | 3233 | 11 | 0.68% |
| hsa_circ_0076278 | 3233 | 11 | 0.68% |
| hsa_circ_0047520 | 3527 | 12 | 0.68% |
| hsa_circ_0075062 | 3527 | 12 | 0.68% |
| hsa_circ_0127101 | 3527 | 12 | 0.68% |
| hsa_circ_0045264 | 4410 | 15 | 0.68% |
| hsa_circ_0032865 | 3822 | 13 | 0.68% |
| hsa_circ_0062516 | 3822 | 13 | 0.68% |
| hsa_circ_0086110 | 3822 | 13 | 0.68% |
| hsa_circ_0093629 | 3528 | 12 | 0.68% |
| hsa_circ_0040103 | 3234 | 11 | 0.68% |
| hsa_circ_0055641 | 3234 | 11 | 0.68% |
| hsa_circ_0105433 | 3234 | 11 | 0.68% |
| hsa_circ_0121099 | 3234 | 11 | 0.68% |
| hsa_circ_0039776 | 3529 | 12 | 0.68% |
| hsa_circ_0087301 | 3529 | 12 | 0.68% |
| hsa_circ_0068965 | 3235 | 11 | 0.68% |
| hsa_circ_0090742 | 3824 | 13 | 0.68% |
| hsa_circ_0054769 | 3530 | 12 | 0.68% |
| hsa_circ_0000276 | 6472 | 22 | 0.68% |
| hsa_circ_0035499 | 3236 | 11 | 0.68% |
| hsa_circ_0076713 | 3236 | 11 | 0.68% |
| hsa_circ_0086929 | 3825 | 13 | 0.68% |
| hsa_circ_0009963 | 3531 | 12 | 0.68% |
| hsa_circ_0030205 | 3531 | 12 | 0.68% |
| hsa_circ_0031518 | 3531 | 12 | 0.68% |
| hsa_circ_0010973 | 3826 | 13 | 0.68% |
| hsa_circ_0108396 | 7947 | 27 | 0.68% |
| hsa_circ_0024849 | 3532 | 12 | 0.68% |
| hsa_circ_0040797 | 3532 | 12 | 0.68% |
| hsa_circ_0027186 | 3238 | 11 | 0.68% |
| hsa_circ_0052108 | 2944 | 10 | 0.68% |
| hsa_circ_0067355 | 3533 | 12 | 0.68% |
| hsa_circ_0084059 | 3533 | 12 | 0.68% |
| hsa_circ_0076478 | 3239 | 11 | 0.68% |
| hsa_circ_0003631 | 10306 | 35 | 0.68% |
| hsa_circ_0131707 | 4418 | 15 | 0.68% |

| | | | |
|---|---|---|---|
| hsa_circ_0035724 | 3535 | 12 | 0.68% |
| hsa_circ_0041089 | 2946 | 10 | 0.68% |
| hsa_circ_0118447 | 2946 | 10 | 0.68% |
| hsa_circ_0062544 | 8840 | 30 | 0.68% |
| hsa_circ_0137036 | 6188 | 21 | 0.68% |
| hsa_circ_0089477 | 3536 | 12 | 0.68% |
| hsa_circ_0095859 | 3536 | 12 | 0.68% |
| hsa_circ_0104719 | 3536 | 12 | 0.68% |
| hsa_circ_0015395 | 3242 | 11 | 0.68% |
| hsa_circ_0054566 | 3242 | 11 | 0.68% |
| hsa_circ_0026074 | 3537 | 12 | 0.68% |
| hsa_circ_0054453 | 6190 | 21 | 0.68% |
| hsa_circ_0117751 | 12675 | 43 | 0.68% |
| hsa_circ_0019771 | 3832 | 13 | 0.68% |
| hsa_circ_0110421 | 7959 | 27 | 0.68% |
| hsa_circ_0061235 | 9139 | 31 | 0.68% |
| hsa_circ_0021058 | 3243 | 11 | 0.68% |
| hsa_circ_0049029 | 3243 | 11 | 0.68% |
| hsa_circ_0067675 | 3243 | 11 | 0.68% |
| hsa_circ_0135282 | 3243 | 11 | 0.68% |
| hsa_circ_0024070 | 3538 | 12 | 0.68% |
| hsa_circ_0049122 | 3538 | 12 | 0.68% |
| hsa_circ_0109197 | 3538 | 12 | 0.68% |
| hsa_circ_0122815 | 3538 | 12 | 0.68% |
| hsa_circ_0133724 | 3833 | 13 | 0.68% |
| hsa_circ_0014890 | 3244 | 11 | 0.68% |
| hsa_circ_0041070 | 3244 | 11 | 0.68% |
| hsa_circ_0084354 | 3244 | 11 | 0.68% |
| hsa_circ_0103645 | 3244 | 11 | 0.68% |
| hsa_circ_0009819 | 3539 | 12 | 0.68% |
| hsa_circ_0090053 | 3539 | 12 | 0.68% |
| hsa_circ_0080677 | 7373 | 25 | 0.68% |
| hsa_circ_0028536 | 3834 | 13 | 0.68% |
| hsa_circ_0085342 | 3834 | 13 | 0.68% |
| hsa_circ_0024176 | 5309 | 18 | 0.68% |
| hsa_circ_0126899 | 31561 | 107 | 0.68% |
| hsa_circ_0010063 | 3835 | 13 | 0.68% |
| hsa_circ_0045078 | 3835 | 13 | 0.68% |
| hsa_circ_0029895 | 3540 | 12 | 0.68% |
| hsa_circ_0067398 | 3540 | 12 | 0.68% |
| hsa_circ_0108116 | 3540 | 12 | 0.68% |
| hsa_circ_0010345 | 2950 | 10 | 0.68% |
| hsa_circ_0104680 | 7376 | 25 | 0.68% |
| hsa_circ_0094890 | 3541 | 12 | 0.68% |
| hsa_circ_0049234 | 3246 | 11 | 0.68% |
| hsa_circ_0059020 | 3246 | 11 | 0.68% |
| hsa_circ_0119439 | 3246 | 11 | 0.68% |
| hsa_circ_0029153 | 3837 | 13 | 0.68% |
| hsa_circ_0041975 | 3247 | 11 | 0.68% |
| hsa_circ_0057118 | 3247 | 11 | 0.68% |

| | | | |
|---|---|---|---|
| hsa_circ_0090316 | 3247 | 11 | 0.68% |
| hsa_circ_0136982 | 3247 | 11 | 0.68% |
| hsa_circ_0011453 | 2952 | 10 | 0.68% |
| hsa_circ_0016111 | 3543 | 12 | 0.68% |
| hsa_circ_0114610 | 3543 | 12 | 0.68% |
| hsa_circ_0007805 | 3248 | 11 | 0.68% |
| hsa_circ_0022074 | 3248 | 11 | 0.68% |
| hsa_circ_0128691 | 3248 | 11 | 0.68% |
| hsa_circ_0012071 | 4134 | 14 | 0.68% |
| hsa_circ_0025912 | 4134 | 14 | 0.68% |
| hsa_circ_0137743 | 5611 | 19 | 0.68% |
| hsa_circ_0078892 | 4430 | 15 | 0.68% |
| hsa_circ_0128969 | 3544 | 12 | 0.68% |
| hsa_circ_0140616 | 3249 | 11 | 0.68% |
| hsa_circ_0010266 | 4431 | 15 | 0.68% |
| hsa_circ_0084596 | 2954 | 10 | 0.68% |
| hsa_circ_0012666 | 3545 | 12 | 0.68% |
| hsa_circ_0041000 | 3545 | 12 | 0.68% |
| hsa_circ_0085237 | 3545 | 12 | 0.68% |
| hsa_circ_0064198 | 17430 | 59 | 0.68% |
| hsa_circ_0019519 | 7977 | 27 | 0.68% |
| hsa_circ_0052914 | 3250 | 11 | 0.68% |
| hsa_circ_0072668 | 3250 | 11 | 0.68% |
| hsa_circ_0089302 | 3250 | 11 | 0.68% |
| hsa_circ_0024624 | 3841 | 13 | 0.68% |
| hsa_circ_0034233 | 3841 | 13 | 0.68% |
| hsa_circ_0038203 | 3841 | 13 | 0.68% |
| hsa_circ_0091165 | 3841 | 13 | 0.68% |
| hsa_circ_0016146 | 4432 | 15 | 0.68% |
| hsa_circ_0128921 | 4137 | 14 | 0.68% |
| hsa_circ_0023186 | 3546 | 12 | 0.68% |
| hsa_circ_0084271 | 2955 | 10 | 0.68% |
| hsa_circ_0112601 | 2955 | 10 | 0.68% |
| hsa_circ_0001339 | 11525 | 39 | 0.68% |
| hsa_circ_0078995 | 4433 | 15 | 0.68% |
| hsa_circ_0085235 | 3251 | 11 | 0.68% |
| hsa_circ_0013355 | 2956 | 10 | 0.68% |
| hsa_circ_0043716 | 2956 | 10 | 0.68% |
| hsa_circ_0041376 | 3252 | 11 | 0.68% |
| hsa_circ_0061146 | 3252 | 11 | 0.68% |
| hsa_circ_0140617 | 3252 | 11 | 0.68% |
| hsa_circ_0036891 | 3548 | 12 | 0.68% |
| hsa_circ_0058786 | 3548 | 12 | 0.68% |
| hsa_circ_0114393 | 3844 | 13 | 0.68% |
| hsa_circ_0071985 | 2957 | 10 | 0.68% |
| hsa_circ_0085179 | 4140 | 14 | 0.68% |
| hsa_circ_0109968 | 11829 | 40 | 0.68% |
| hsa_circ_0139283 | 14195 | 48 | 0.68% |
| hsa_circ_0018364 | 3549 | 12 | 0.68% |
| hsa_circ_0054478 | 3549 | 12 | 0.68% |

| | | | |
|---|---|---|---|
| hsa_circ_0094200 | 3549 | 12 | 0.68% |
| hsa_circ_0009869 | 3845 | 13 | 0.68% |
| hsa_circ_0015219 | 3254 | 11 | 0.68% |
| hsa_circ_0065364 | 3254 | 11 | 0.68% |
| hsa_circ_0091209 | 3254 | 11 | 0.68% |
| hsa_circ_0024215 | 3550 | 12 | 0.68% |
| hsa_circ_0094430 | 2959 | 10 | 0.68% |
| hsa_circ_0016618 | 3255 | 11 | 0.68% |
| hsa_circ_0040173 | 3255 | 11 | 0.68% |
| hsa_circ_0049731 | 3255 | 11 | 0.68% |
| hsa_circ_0063921 | 3255 | 11 | 0.68% |
| hsa_circ_0073577 | 3255 | 11 | 0.68% |
| hsa_circ_0082985 | 3255 | 11 | 0.68% |
| hsa_circ_0090019 | 3255 | 11 | 0.68% |
| hsa_circ_0058411 | 3551 | 12 | 0.68% |
| hsa_circ_0095858 | 3551 | 12 | 0.68% |
| hsa_circ_0072467 | 3848 | 13 | 0.68% |
| hsa_circ_0137642 | 3552 | 12 | 0.68% |
| hsa_circ_0024667 | 3256 | 11 | 0.68% |
| hsa_circ_0085569 | 3256 | 11 | 0.68% |
| hsa_circ_0075469 | 4145 | 14 | 0.68% |
| hsa_circ_0050077 | 3849 | 13 | 0.68% |
| hsa_circ_0048644 | 3553 | 12 | 0.68% |
| hsa_circ_0088003 | 3257 | 11 | 0.68% |
| hsa_circ_0010620 | 3850 | 13 | 0.68% |
| hsa_circ_0010689 | 3850 | 13 | 0.68% |
| hsa_circ_0012005 | 3258 | 11 | 0.68% |
| hsa_circ_0062087 | 3258 | 11 | 0.68% |
| hsa_circ_0026973 | 7405 | 25 | 0.68% |
| hsa_circ_0014811 | 2962 | 10 | 0.68% |
| hsa_circ_0050025 | 3851 | 13 | 0.68% |
| hsa_circ_0051593 | 3851 | 13 | 0.68% |
| hsa_circ_0111262 | 7999 | 27 | 0.68% |
| hsa_circ_0044524 | 2963 | 10 | 0.67% |
| hsa_circ_0091698 | 3556 | 12 | 0.67% |
| hsa_circ_0099171 | 5038 | 17 | 0.67% |
| hsa_circ_0137607 | 14522 | 49 | 0.67% |
| hsa_circ_0080487 | 2964 | 10 | 0.67% |
| hsa_circ_0045183 | 3557 | 12 | 0.67% |
| hsa_circ_0135863 | 3557 | 12 | 0.67% |
| hsa_circ_0091649 | 3261 | 11 | 0.67% |
| hsa_circ_0042674 | 3558 | 12 | 0.67% |
| hsa_circ_0075969 | 2965 | 10 | 0.67% |
| hsa_circ_0025420 | 3262 | 11 | 0.67% |
| hsa_circ_0025968 | 3262 | 11 | 0.67% |
| hsa_circ_0051181 | 3262 | 11 | 0.67% |
| hsa_circ_0065187 | 3262 | 11 | 0.67% |
| hsa_circ_0035751 | 3559 | 12 | 0.67% |
| hsa_circ_0041896 | 3559 | 12 | 0.67% |
| hsa_circ_0009300 | 5042 | 17 | 0.67% |

| | | | |
|---|---|---|---|
| hsa_circ_0061792 | 2966 | 10 | 0.67% |
| hsa_circ_0004943 | 3263 | 11 | 0.67% |
| hsa_circ_0010092 | 3263 | 11 | 0.67% |
| hsa_circ_0015585 | 3263 | 11 | 0.67% |
| hsa_circ_0030336 | 4153 | 14 | 0.67% |
| hsa_circ_0081678 | 9197 | 31 | 0.67% |
| hsa_circ_0073277 | 3857 | 13 | 0.67% |
| hsa_circ_0071220 | 2967 | 10 | 0.67% |
| hsa_circ_0059552 | 4154 | 14 | 0.67% |
| hsa_circ_0056296 | 3264 | 11 | 0.67% |
| hsa_circ_0044768 | 3561 | 12 | 0.67% |
| hsa_circ_0088666 | 3561 | 12 | 0.67% |
| hsa_circ_0128980 | 3858 | 13 | 0.67% |
| hsa_circ_0020472 | 2968 | 10 | 0.67% |
| hsa_circ_0065195 | 3265 | 11 | 0.67% |
| hsa_circ_0138258 | 3562 | 12 | 0.67% |
| hsa_circ_0025700 | 3859 | 13 | 0.67% |
| hsa_circ_0047388 | 2969 | 10 | 0.67% |
| hsa_circ_0057210 | 2969 | 10 | 0.67% |
| hsa_circ_0039183 | 3266 | 11 | 0.67% |
| hsa_circ_0068486 | 3266 | 11 | 0.67% |
| hsa_circ_0072239 | 3266 | 11 | 0.67% |
| hsa_circ_0111909 | 3266 | 11 | 0.67% |
| hsa_circ_0124631 | 3563 | 12 | 0.67% |
| hsa_circ_0061077 | 3861 | 13 | 0.67% |
| hsa_circ_0028114 | 3564 | 12 | 0.67% |
| hsa_circ_0047459 | 3564 | 12 | 0.67% |
| hsa_circ_0047387 | 3267 | 11 | 0.67% |
| hsa_circ_0065866 | 3267 | 11 | 0.67% |
| hsa_circ_0018475 | 2970 | 10 | 0.67% |
| hsa_circ_0139438 | 2970 | 10 | 0.67% |
| hsa_circ_0010640 | 3862 | 13 | 0.67% |
| hsa_circ_0062994 | 3862 | 13 | 0.67% |
| hsa_circ_0025974 | 3565 | 12 | 0.67% |
| hsa_circ_0062590 | 3565 | 12 | 0.67% |
| hsa_circ_0057712 | 2971 | 10 | 0.67% |
| hsa_circ_0070154 | 4160 | 14 | 0.67% |
| hsa_circ_0068664 | 5349 | 18 | 0.67% |
| hsa_circ_0090993 | 3566 | 12 | 0.67% |
| hsa_circ_0035693 | 3269 | 11 | 0.67% |
| hsa_circ_0091622 | 3269 | 11 | 0.67% |
| hsa_circ_0099355 | 3269 | 11 | 0.67% |
| hsa_circ_0016524 | 2972 | 10 | 0.67% |
| hsa_circ_0065022 | 2972 | 10 | 0.67% |
| hsa_circ_0009372 | 4161 | 14 | 0.67% |
| hsa_circ_0009221 | 3567 | 12 | 0.67% |
| hsa_circ_0027643 | 3567 | 12 | 0.67% |
| hsa_circ_0015856 | 3270 | 11 | 0.67% |
| hsa_circ_0019329 | 3270 | 11 | 0.67% |
| hsa_circ_0043390 | 3270 | 11 | 0.67% |

| | | | |
|---|---|---|---|
| hsa_circ_0048308 | 3270 | 11 | 0.67% |
| hsa_circ_0125367 | 8919 | 30 | 0.67% |
| hsa_circ_0002447 | 2973 | 10 | 0.67% |
| hsa_circ_0060696 | 3865 | 13 | 0.67% |
| hsa_circ_0079114 | 3568 | 12 | 0.67% |
| hsa_circ_0041874 | 3271 | 11 | 0.67% |
| hsa_circ_0068356 | 3271 | 11 | 0.67% |
| hsa_circ_0029508 | 3866 | 13 | 0.67% |
| hsa_circ_0024416 | 4164 | 14 | 0.67% |
| hsa_circ_0047897 | 4164 | 14 | 0.67% |
| hsa_circ_0062941 | 3272 | 11 | 0.67% |
| hsa_circ_0080917 | 3272 | 11 | 0.67% |
| hsa_circ_0088838 | 3272 | 11 | 0.67% |
| hsa_circ_0033253 | 3570 | 12 | 0.67% |
| hsa_circ_0074901 | 3570 | 12 | 0.67% |
| hsa_circ_0035386 | 2975 | 10 | 0.67% |
| hsa_circ_0035764 | 3868 | 13 | 0.67% |
| hsa_circ_0051774 | 3273 | 11 | 0.67% |
| hsa_circ_0089187 | 3273 | 11 | 0.67% |
| hsa_circ_0091822 | 3571 | 12 | 0.67% |
| hsa_circ_0034206 | 2976 | 10 | 0.67% |
| hsa_circ_0029559 | 3572 | 12 | 0.67% |
| hsa_circ_0010672 | 3870 | 13 | 0.67% |
| hsa_circ_0127216 | 3870 | 13 | 0.67% |
| hsa_circ_0104995 | 13397 | 45 | 0.67% |
| hsa_circ_0036480 | 3573 | 12 | 0.67% |
| hsa_circ_0047522 | 3573 | 12 | 0.67% |
| hsa_circ_0098770 | 3573 | 12 | 0.67% |
| hsa_circ_0062341 | 4169 | 14 | 0.67% |
| hsa_circ_0027762 | 2978 | 10 | 0.67% |
| hsa_circ_0117947 | 2978 | 10 | 0.67% |
| hsa_circ_0053540 | 3276 | 11 | 0.67% |
| hsa_circ_0018433 | 3574 | 12 | 0.67% |
| hsa_circ_0049115 | 3574 | 12 | 0.67% |
| hsa_circ_0110613 | 5362 | 18 | 0.67% |
| hsa_circ_0059993 | 2979 | 10 | 0.67% |
| hsa_circ_0066223 | 2979 | 10 | 0.67% |
| hsa_circ_0023043 | 3277 | 11 | 0.67% |
| hsa_circ_0047407 | 3873 | 13 | 0.67% |
| hsa_circ_0063805 | 3576 | 12 | 0.67% |
| hsa_circ_0024793 | 3278 | 11 | 0.67% |
| hsa_circ_0024973 | 3278 | 11 | 0.67% |
| hsa_circ_0032414 | 3278 | 11 | 0.67% |
| hsa_circ_0038260 | 3278 | 11 | 0.67% |
| hsa_circ_0053752 | 3278 | 11 | 0.67% |
| hsa_circ_0091943 | 3278 | 11 | 0.67% |
| hsa_circ_0029210 | 2980 | 10 | 0.67% |
| hsa_circ_0108589 | 2980 | 10 | 0.67% |
| hsa_circ_0065571 | 4174 | 14 | 0.67% |
| hsa_circ_0109580 | 11032 | 37 | 0.67% |

| | | | |
|---|---|---|---|
| hsa_circ_0042607 | 2982 | 10 | 0.67% |
| hsa_circ_0094433 | 2982 | 10 | 0.67% |
| hsa_circ_0118446 | 2982 | 10 | 0.67% |
| hsa_circ_0051281 | 3877 | 13 | 0.67% |
| hsa_circ_0083749 | 3877 | 13 | 0.67% |
| hsa_circ_0022734 | 3579 | 12 | 0.67% |
| hsa_circ_0023854 | 3281 | 11 | 0.67% |
| hsa_circ_0076360 | 3281 | 11 | 0.67% |
| hsa_circ_0091623 | 3281 | 11 | 0.67% |
| hsa_circ_0109285 | 2983 | 10 | 0.67% |
| hsa_circ_0021194 | 3878 | 13 | 0.67% |
| hsa_circ_0100098 | 3282 | 11 | 0.67% |
| hsa_circ_0030873 | 3879 | 13 | 0.67% |
| hsa_circ_0100086 | 3879 | 13 | 0.67% |
| hsa_circ_0091739 | 5968 | 20 | 0.67% |
| hsa_circ_0056204 | 3581 | 12 | 0.67% |
| hsa_circ_0089425 | 3581 | 12 | 0.67% |
| hsa_circ_0120610 | 3581 | 12 | 0.67% |
| hsa_circ_0052064 | 8058 | 27 | 0.67% |
| hsa_circ_0001461 | 3283 | 11 | 0.67% |
| hsa_circ_0042164 | 3283 | 11 | 0.67% |
| hsa_circ_0126573 | 3283 | 11 | 0.67% |
| hsa_circ_0062991 | 4179 | 14 | 0.67% |
| hsa_circ_0090426 | 2985 | 10 | 0.67% |
| hsa_circ_0025987 | 3881 | 13 | 0.67% |
| hsa_circ_0036472 | 3881 | 13 | 0.67% |
| hsa_circ_0025965 | 3284 | 11 | 0.67% |
| hsa_circ_0032043 | 3284 | 11 | 0.67% |
| hsa_circ_0042615 | 3284 | 11 | 0.67% |
| hsa_circ_0064043 | 3583 | 12 | 0.67% |
| hsa_circ_0116273 | 2986 | 10 | 0.67% |
| hsa_circ_0023305 | 3882 | 13 | 0.67% |
| hsa_circ_0061447 | 4778 | 16 | 0.67% |
| hsa_circ_0029094 | 3285 | 11 | 0.67% |
| hsa_circ_0049229 | 3584 | 12 | 0.67% |
| hsa_circ_0067713 | 3584 | 12 | 0.67% |
| hsa_circ_0061022 | 2987 | 10 | 0.67% |
| hsa_circ_0029740 | 3286 | 11 | 0.67% |
| hsa_circ_0045697 | 3286 | 11 | 0.67% |
| hsa_circ_0048304 | 3286 | 11 | 0.67% |
| hsa_circ_0063824 | 4183 | 14 | 0.67% |
| hsa_circ_0039806 | 2988 | 10 | 0.67% |
| hsa_circ_0034709 | 3287 | 11 | 0.67% |
| hsa_circ_0050044 | 3287 | 11 | 0.67% |
| hsa_circ_0053790 | 3287 | 11 | 0.67% |
| hsa_circ_0062057 | 3287 | 11 | 0.67% |
| hsa_circ_0067746 | 3287 | 11 | 0.67% |
| hsa_circ_0034357 | 3885 | 13 | 0.67% |
| hsa_circ_0017152 | 2989 | 10 | 0.67% |
| hsa_circ_0033045 | 2989 | 10 | 0.67% |

| | | | |
|---|---|---|---|
| hsa_circ_0030956 | 3288 | 11 | 0.67% |
| hsa_circ_0078010 | 3288 | 11 | 0.67% |
| hsa_circ_0024481 | 3587 | 12 | 0.67% |
| hsa_circ_0075419 | 3886 | 13 | 0.67% |
| hsa_circ_0115429 | 10165 | 34 | 0.67% |
| hsa_circ_0069928 | 4186 | 14 | 0.67% |
| hsa_circ_0041344 | 3887 | 13 | 0.67% |
| hsa_circ_0038265 | 3289 | 11 | 0.67% |
| hsa_circ_0085443 | 2990 | 10 | 0.67% |
| hsa_circ_0135677 | 2990 | 10 | 0.67% |
| hsa_circ_0105800 | 4187 | 14 | 0.67% |
| hsa_circ_0035295 | 13459 | 45 | 0.67% |
| hsa_circ_0017083 | 3290 | 11 | 0.67% |
| hsa_circ_0020453 | 3290 | 11 | 0.67% |
| hsa_circ_0065726 | 3290 | 11 | 0.67% |
| hsa_circ_0138423 | 4786 | 16 | 0.67% |
| hsa_circ_0025414 | 3889 | 13 | 0.67% |
| hsa_circ_0136307 | 10172 | 34 | 0.67% |
| hsa_circ_0063116 | 3291 | 11 | 0.67% |
| hsa_circ_0074306 | 3291 | 11 | 0.67% |
| hsa_circ_0027896 | 4189 | 14 | 0.67% |
| hsa_circ_0032563 | 3890 | 13 | 0.67% |
| hsa_circ_0032639 | 3890 | 13 | 0.67% |
| hsa_circ_0073454 | 3890 | 13 | 0.67% |
| hsa_circ_0036600 | 3292 | 11 | 0.67% |
| hsa_circ_0045688 | 3292 | 11 | 0.67% |
| hsa_circ_0083386 | 3292 | 11 | 0.67% |
| hsa_circ_0091840 | 3292 | 11 | 0.67% |
| hsa_circ_0015231 | 2993 | 10 | 0.67% |
| hsa_circ_0066337 | 3592 | 12 | 0.67% |
| hsa_circ_0035368 | 3293 | 11 | 0.67% |
| hsa_circ_0054563 | 3293 | 11 | 0.67% |
| hsa_circ_0070339 | 3293 | 11 | 0.67% |
| hsa_circ_0016166 | 10179 | 34 | 0.67% |
| hsa_circ_0010653 | 3892 | 13 | 0.67% |
| hsa_circ_0049667 | 4491 | 15 | 0.67% |
| hsa_circ_0021903 | 2994 | 10 | 0.67% |
| hsa_circ_0090020 | 2994 | 10 | 0.67% |
| hsa_circ_0131317 | 2994 | 10 | 0.67% |
| hsa_circ_0050003 | 3593 | 12 | 0.67% |
| hsa_circ_0101758 | 6887 | 23 | 0.67% |
| hsa_circ_0092846 | 4791 | 16 | 0.67% |
| hsa_circ_0072125 | 3294 | 11 | 0.67% |
| hsa_circ_0090286 | 4193 | 14 | 0.67% |
| hsa_circ_0033607 | 3594 | 12 | 0.67% |
| hsa_circ_0039424 | 3594 | 12 | 0.67% |
| hsa_circ_0056691 | 3594 | 12 | 0.67% |
| hsa_circ_0073792 | 3594 | 12 | 0.67% |
| hsa_circ_0046717 | 2995 | 10 | 0.67% |
| hsa_circ_0070329 | 2995 | 10 | 0.67% |

| | | | |
|---|---|---|---|
| hsa_circ_0023583 | 3295 | 11 | 0.67% |
| hsa_circ_0030080 | 3295 | 11 | 0.67% |
| hsa_circ_0042165 | 3295 | 11 | 0.67% |
| hsa_circ_0043293 | 3295 | 11 | 0.67% |
| hsa_circ_0091891 | 3295 | 11 | 0.67% |
| hsa_circ_0076838 | 3595 | 12 | 0.67% |
| hsa_circ_0044529 | 2996 | 10 | 0.67% |
| hsa_circ_0016949 | 3596 | 12 | 0.67% |
| hsa_circ_0033434 | 3297 | 11 | 0.67% |
| hsa_circ_0036877 | 3297 | 11 | 0.67% |
| hsa_circ_0045451 | 3297 | 11 | 0.67% |
| hsa_circ_0054548 | 3297 | 11 | 0.67% |
| hsa_circ_0029700 | 7194 | 24 | 0.67% |
| hsa_circ_0063316 | 4796 | 16 | 0.67% |
| hsa_circ_0063446 | 3597 | 12 | 0.67% |
| hsa_circ_0080093 | 3597 | 12 | 0.67% |
| hsa_circ_0107503 | 3597 | 12 | 0.67% |
| hsa_circ_0092890 | 8993 | 30 | 0.67% |
| hsa_circ_0010569 | 3897 | 13 | 0.67% |
| hsa_circ_0081507 | 4797 | 16 | 0.67% |
| hsa_circ_0064588 | 3298 | 11 | 0.67% |
| hsa_circ_0031161 | 2999 | 10 | 0.67% |
| hsa_circ_0019528 | 3299 | 11 | 0.67% |
| hsa_circ_0026849 | 3299 | 11 | 0.67% |
| hsa_circ_0053490 | 3299 | 11 | 0.67% |
| hsa_circ_0026739 | 3599 | 12 | 0.67% |
| hsa_circ_0117911 | 3899 | 13 | 0.67% |
| hsa_circ_0108934 | 7800 | 26 | 0.67% |
| hsa_circ_0024591 | 3600 | 12 | 0.67% |
| hsa_circ_0094436 | 3600 | 12 | 0.67% |
| hsa_circ_0024185 | 3300 | 11 | 0.67% |
| hsa_circ_0085108 | 3300 | 11 | 0.67% |
| hsa_circ_0099062 | 3901 | 13 | 0.67% |
| hsa_circ_0010738 | 3301 | 11 | 0.67% |
| hsa_circ_0032170 | 3301 | 11 | 0.67% |
| hsa_circ_0062604 | 3301 | 11 | 0.67% |
| hsa_circ_0083379 | 3301 | 11 | 0.67% |
| hsa_circ_0125222 | 3301 | 11 | 0.67% |
| hsa_circ_0007710 | 9004 | 30 | 0.67% |
| hsa_circ_0083528 | 4502 | 15 | 0.67% |
| hsa_circ_0039608 | 3302 | 11 | 0.67% |
| hsa_circ_0102979 | 3002 | 10 | 0.67% |
| hsa_circ_0060973 | 4203 | 14 | 0.67% |
| hsa_circ_0009824 | 3603 | 12 | 0.67% |
| hsa_circ_0096775 | 3603 | 12 | 0.67% |
| hsa_circ_0104091 | 3603 | 12 | 0.67% |
| hsa_circ_0050470 | 3303 | 11 | 0.67% |
| hsa_circ_0071672 | 3303 | 11 | 0.67% |
| hsa_circ_0083809 | 3303 | 11 | 0.67% |
| hsa_circ_0012709 | 3003 | 10 | 0.67% |

| | | | |
|---|---|---|---|
| hsa_circ_0044456 | 3604 | 12 | 0.67% |
| hsa_circ_0087291 | 3604 | 12 | 0.67% |
| hsa_circ_0005843 | 3304 | 11 | 0.67% |
| hsa_circ_0043545 | 3304 | 11 | 0.67% |
| hsa_circ_0078975 | 10815 | 36 | 0.67% |
| hsa_circ_0055297 | 3605 | 12 | 0.67% |
| hsa_circ_0048642 | 3305 | 11 | 0.67% |
| hsa_circ_0080505 | 3305 | 11 | 0.67% |
| hsa_circ_0134855 | 6611 | 22 | 0.67% |
| hsa_circ_0066835 | 3606 | 12 | 0.67% |
| hsa_circ_0091961 | 3606 | 12 | 0.67% |
| hsa_circ_0057638 | 3005 | 10 | 0.67% |
| hsa_circ_0038915 | 7814 | 26 | 0.67% |
| hsa_circ_0041901 | 3306 | 11 | 0.67% |
| hsa_circ_0070175 | 3306 | 11 | 0.67% |
| hsa_circ_0086468 | 3306 | 11 | 0.67% |
| hsa_circ_0119797 | 3306 | 11 | 0.67% |
| hsa_circ_0064330 | 6312 | 21 | 0.67% |
| hsa_circ_0036958 | 4208 | 14 | 0.67% |
| hsa_circ_0075121 | 4208 | 14 | 0.67% |
| hsa_circ_0041461 | 3607 | 12 | 0.67% |
| hsa_circ_0071213 | 3006 | 10 | 0.67% |
| hsa_circ_0074565 | 3908 | 13 | 0.67% |
| hsa_circ_0047703 | 3307 | 11 | 0.67% |
| hsa_circ_0052645 | 3307 | 11 | 0.67% |
| hsa_circ_0063099 | 3307 | 11 | 0.67% |
| hsa_circ_0121818 | 3307 | 11 | 0.67% |
| hsa_circ_0034250 | 3608 | 12 | 0.67% |
| hsa_circ_0054813 | 3608 | 12 | 0.67% |
| hsa_circ_0079584 | 3608 | 12 | 0.67% |
| hsa_circ_0028923 | 4811 | 16 | 0.67% |
| hsa_circ_0090748 | 3909 | 13 | 0.67% |
| hsa_circ_0091771 | 3909 | 13 | 0.67% |
| hsa_circ_0024802 | 3007 | 10 | 0.67% |
| hsa_circ_0074104 | 3007 | 10 | 0.67% |
| hsa_circ_0107676 | 3007 | 10 | 0.67% |
| hsa_circ_0105833 | 12630 | 42 | 0.67% |
| hsa_circ_0062717 | 4210 | 14 | 0.67% |
| hsa_circ_0038210 | 3308 | 11 | 0.67% |
| hsa_circ_0086884 | 3308 | 11 | 0.67% |
| hsa_circ_0057258 | 3609 | 12 | 0.67% |
| hsa_circ_0093308 | 3609 | 12 | 0.67% |
| hsa_circ_0111050 | 3609 | 12 | 0.67% |
| hsa_circ_0129039 | 3609 | 12 | 0.67% |
| hsa_circ_0038616 | 3910 | 13 | 0.66% |
| hsa_circ_0134523 | 12633 | 42 | 0.66% |
| hsa_circ_0043789 | 4211 | 14 | 0.66% |
| hsa_circ_0000348 | 3309 | 11 | 0.66% |
| hsa_circ_0030473 | 3309 | 11 | 0.66% |
| hsa_circ_0043879 | 3610 | 12 | 0.66% |

| | | | |
|---|---|---|---|
| hsa_circ_0039649 | 3009 | 10 | 0.66% |
| hsa_circ_0007068 | 3310 | 11 | 0.66% |
| hsa_circ_0035225 | 3611 | 12 | 0.66% |
| hsa_circ_0066060 | 3611 | 12 | 0.66% |
| hsa_circ_0084060 | 3611 | 12 | 0.66% |
| hsa_circ_0079682 | 3912 | 13 | 0.66% |
| hsa_circ_0033322 | 3612 | 12 | 0.66% |
| hsa_circ_0127478 | 3612 | 12 | 0.66% |
| hsa_circ_0014055 | 3311 | 11 | 0.66% |
| hsa_circ_0016480 | 3311 | 11 | 0.66% |
| hsa_circ_0066451 | 3311 | 11 | 0.66% |
| hsa_circ_0080977 | 3311 | 11 | 0.66% |
| hsa_circ_0135347 | 3311 | 11 | 0.66% |
| hsa_circ_0133089 | 30704 | 102 | 0.66% |
| hsa_circ_0101597 | 8730 | 29 | 0.66% |
| hsa_circ_0128324 | 8730 | 29 | 0.66% |
| hsa_circ_0106549 | 3914 | 13 | 0.66% |
| hsa_circ_0114158 | 3312 | 11 | 0.66% |
| hsa_circ_0033393 | 3011 | 10 | 0.66% |
| hsa_circ_0034716 | 3614 | 12 | 0.66% |
| hsa_circ_0114847 | 3614 | 12 | 0.66% |
| hsa_circ_0019898 | 3313 | 11 | 0.66% |
| hsa_circ_0026995 | 3313 | 11 | 0.66% |
| hsa_circ_0029603 | 3313 | 11 | 0.66% |
| hsa_circ_0080448 | 3313 | 11 | 0.66% |
| hsa_circ_0140520 | 4518 | 15 | 0.66% |
| hsa_circ_0121775 | 3012 | 10 | 0.66% |
| hsa_circ_0035779 | 3916 | 13 | 0.66% |
| hsa_circ_0038732 | 3916 | 13 | 0.66% |
| hsa_circ_0104073 | 3615 | 12 | 0.66% |
| hsa_circ_0035504 | 3013 | 10 | 0.66% |
| hsa_circ_0045967 | 3917 | 13 | 0.66% |
| hsa_circ_0066605 | 3616 | 12 | 0.66% |
| hsa_circ_0122876 | 37670 | 125 | 0.66% |
| hsa_circ_0035148 | 3918 | 13 | 0.66% |
| hsa_circ_0091841 | 3316 | 11 | 0.66% |
| hsa_circ_0067181 | 3919 | 13 | 0.66% |
| hsa_circ_0031404 | 5125 | 17 | 0.66% |
| hsa_circ_0099121 | 8443 | 28 | 0.66% |
| hsa_circ_0048864 | 3920 | 13 | 0.66% |
| hsa_circ_0122994 | 3317 | 11 | 0.66% |
| hsa_circ_0084165 | 12666 | 42 | 0.66% |
| hsa_circ_0057182 | 6333 | 21 | 0.66% |
| hsa_circ_0062348 | 4222 | 14 | 0.66% |
| hsa_circ_0048813 | 3921 | 13 | 0.66% |
| hsa_circ_0000480 | 3318 | 11 | 0.66% |
| hsa_circ_0063405 | 3318 | 11 | 0.66% |
| hsa_circ_0086870 | 3318 | 11 | 0.66% |
| hsa_circ_0031501 | 3620 | 12 | 0.66% |
| hsa_circ_0065827 | 3922 | 13 | 0.66% |

| | | | |
|---|---|---|---|
| hsa_circ_0036796 | 3017 | 10 | 0.66% |
| hsa_circ_0025096 | 3621 | 12 | 0.66% |
| hsa_circ_0054053 | 3621 | 12 | 0.66% |
| hsa_circ_0056592 | 3621 | 12 | 0.66% |
| hsa_circ_0119747 | 3621 | 12 | 0.66% |
| hsa_circ_0135978 | 3621 | 12 | 0.66% |
| hsa_circ_0056300 | 3018 | 10 | 0.66% |
| hsa_circ_0015661 | 3320 | 11 | 0.66% |
| hsa_circ_0060427 | 3320 | 11 | 0.66% |
| hsa_circ_0102278 | 3320 | 11 | 0.66% |
| hsa_circ_0131367 | 5735 | 19 | 0.66% |
| hsa_circ_0071222 | 3924 | 13 | 0.66% |
| hsa_circ_0070969 | 8453 | 28 | 0.66% |
| hsa_circ_0011687 | 3321 | 11 | 0.66% |
| hsa_circ_0017792 | 3321 | 11 | 0.66% |
| hsa_circ_0077476 | 3321 | 11 | 0.66% |
| hsa_circ_0094435 | 3321 | 11 | 0.66% |
| hsa_circ_0056640 | 3926 | 13 | 0.66% |
| hsa_circ_0064752 | 3926 | 13 | 0.66% |
| hsa_circ_0060648 | 3624 | 12 | 0.66% |
| hsa_circ_0015997 | 3322 | 11 | 0.66% |
| hsa_circ_0082826 | 3020 | 10 | 0.66% |
| hsa_circ_0044203 | 11175 | 37 | 0.66% |
| hsa_circ_0135271 | 8155 | 27 | 0.66% |
| hsa_circ_0018929 | 3927 | 13 | 0.66% |
| hsa_circ_0060805 | 3927 | 13 | 0.66% |
| hsa_circ_0097107 | 3927 | 13 | 0.66% |
| hsa_circ_0058665 | 3625 | 12 | 0.66% |
| hsa_circ_0020403 | 3323 | 11 | 0.66% |
| hsa_circ_0066222 | 3323 | 11 | 0.66% |
| hsa_circ_0047828 | 4834 | 16 | 0.66% |
| hsa_circ_0055823 | 4532 | 15 | 0.66% |
| hsa_circ_0037995 | 3626 | 12 | 0.66% |
| hsa_circ_0054827 | 3626 | 12 | 0.66% |
| hsa_circ_0036420 | 3324 | 11 | 0.66% |
| hsa_circ_0041862 | 3324 | 11 | 0.66% |
| hsa_circ_0039659 | 3022 | 10 | 0.66% |
| hsa_circ_0140290 | 3929 | 13 | 0.66% |
| hsa_circ_0030498 | 4534 | 15 | 0.66% |
| hsa_circ_0045956 | 3325 | 11 | 0.66% |
| hsa_circ_0122179 | 3325 | 11 | 0.66% |
| hsa_circ_0030451 | 3628 | 12 | 0.66% |
| hsa_circ_0041999 | 3024 | 10 | 0.66% |
| hsa_circ_0139838 | 3327 | 11 | 0.66% |
| hsa_circ_0037280 | 4236 | 14 | 0.66% |
| hsa_circ_0052360 | 15432 | 51 | 0.66% |
| hsa_circ_0134856 | 6658 | 22 | 0.66% |
| hsa_circ_0046481 | 3329 | 11 | 0.66% |
| hsa_circ_0097678 | 10290 | 34 | 0.66% |
| hsa_circ_0127974 | 9988 | 33 | 0.66% |

| | | | |
|---|---|---|---|
| hsa_circ_0058568 | 3632 | 12 | 0.66% |
| hsa_circ_0085666 | 3632 | 12 | 0.66% |
| hsa_circ_0136492 | 3632 | 12 | 0.66% |
| hsa_circ_0019769 | 3935 | 13 | 0.66% |
| hsa_circ_0052729 | 3935 | 13 | 0.66% |
| hsa_circ_0066362 | 3027 | 10 | 0.66% |
| hsa_circ_0086057 | 3027 | 10 | 0.66% |
| hsa_circ_0102904 | 3330 | 11 | 0.66% |
| hsa_circ_0052055 | 3633 | 12 | 0.66% |
| hsa_circ_0048305 | 3331 | 11 | 0.66% |
| hsa_circ_0129115 | 5451 | 18 | 0.66% |
| hsa_circ_0009659 | 3634 | 12 | 0.66% |
| hsa_circ_0039105 | 3634 | 12 | 0.66% |
| hsa_circ_0062070 | 3635 | 12 | 0.66% |
| hsa_circ_0088192 | 3635 | 12 | 0.66% |
| hsa_circ_0036280 | 4241 | 14 | 0.66% |
| hsa_circ_0055607 | 4241 | 14 | 0.66% |
| hsa_circ_0083576 | 3636 | 12 | 0.66% |
| hsa_circ_0140615 | 3636 | 12 | 0.66% |
| hsa_circ_0028790 | 3333 | 11 | 0.66% |
| hsa_circ_0063101 | 3333 | 11 | 0.66% |
| hsa_circ_0081824 | 3333 | 11 | 0.66% |
| hsa_circ_0042928 | 3940 | 13 | 0.66% |
| hsa_circ_0057499 | 3637 | 12 | 0.66% |
| hsa_circ_0024125 | 3334 | 11 | 0.66% |
| hsa_circ_0124742 | 13033 | 43 | 0.66% |
| hsa_circ_0036748 | 5456 | 18 | 0.66% |
| hsa_circ_0105480 | 11216 | 37 | 0.66% |
| hsa_circ_0013070 | 3638 | 12 | 0.66% |
| hsa_circ_0054808 | 3638 | 12 | 0.66% |
| hsa_circ_0089218 | 3638 | 12 | 0.66% |
| hsa_circ_0013486 | 3335 | 11 | 0.66% |
| hsa_circ_0056298 | 3335 | 11 | 0.66% |
| hsa_circ_0056776 | 3335 | 11 | 0.66% |
| hsa_circ_0037066 | 4245 | 14 | 0.66% |
| hsa_circ_0138823 | 4245 | 14 | 0.66% |
| hsa_circ_0037016 | 5458 | 18 | 0.66% |
| hsa_circ_0039639 | 3942 | 13 | 0.66% |
| hsa_circ_0041001 | 3639 | 12 | 0.66% |
| hsa_circ_0074351 | 3639 | 12 | 0.66% |
| hsa_circ_0090147 | 3639 | 12 | 0.66% |
| hsa_circ_0086558 | 3336 | 11 | 0.66% |
| hsa_circ_0104858 | 6066 | 20 | 0.66% |
| hsa_circ_0085057 | 3033 | 10 | 0.66% |
| hsa_circ_0136094 | 3943 | 13 | 0.66% |
| hsa_circ_0002187 | 4550 | 15 | 0.66% |
| hsa_circ_0086430 | 3640 | 12 | 0.66% |
| hsa_circ_0059551 | 4247 | 14 | 0.66% |
| hsa_circ_0019824 | 3337 | 11 | 0.66% |
| hsa_circ_0085389 | 3337 | 11 | 0.66% |

| | | | |
|---|---|---|---|
| hsa_circ_0042931 | 3034 | 10 | 0.66% |
| hsa_circ_0010000 | 3641 | 12 | 0.66% |
| hsa_circ_0021775 | 3641 | 12 | 0.66% |
| hsa_circ_0028236 | 3641 | 12 | 0.66% |
| hsa_circ_0019821 | 3338 | 11 | 0.66% |
| hsa_circ_0061603 | 3945 | 13 | 0.66% |
| hsa_circ_0102447 | 3945 | 13 | 0.66% |
| hsa_circ_0094635 | 8802 | 29 | 0.66% |
| hsa_circ_0103497 | 4250 | 14 | 0.66% |
| hsa_circ_0058960 | 3643 | 12 | 0.66% |
| hsa_circ_0079460 | 3643 | 12 | 0.66% |
| hsa_circ_0065488 | 13662 | 45 | 0.66% |
| hsa_circ_0073543 | 3036 | 10 | 0.66% |
| hsa_circ_0130345 | 3036 | 10 | 0.66% |
| hsa_circ_0049437 | 3644 | 12 | 0.66% |
| hsa_circ_0100872 | 3644 | 12 | 0.66% |
| hsa_circ_0109696 | 3644 | 12 | 0.66% |
| hsa_circ_0131001 | 3644 | 12 | 0.66% |
| hsa_circ_0118493 | 3948 | 13 | 0.66% |
| hsa_circ_0120991 | 3948 | 13 | 0.66% |
| hsa_circ_0089792 | 9111 | 30 | 0.66% |
| hsa_circ_0037182 | 4252 | 14 | 0.66% |
| hsa_circ_0053820 | 3645 | 12 | 0.66% |
| hsa_circ_0087311 | 3645 | 12 | 0.66% |
| hsa_circ_0128634 | 3949 | 13 | 0.66% |
| hsa_circ_0096566 | 3342 | 11 | 0.66% |
| hsa_circ_0140707 | 3646 | 12 | 0.66% |
| hsa_circ_0085064 | 4558 | 15 | 0.66% |
| hsa_circ_0052329 | 13371 | 44 | 0.66% |
| hsa_circ_0021907 | 3039 | 10 | 0.66% |
| hsa_circ_0085087 | 3039 | 10 | 0.66% |
| hsa_circ_0098251 | 15499 | 51 | 0.66% |
| hsa_circ_0037082 | 3343 | 11 | 0.66% |
| hsa_circ_0077120 | 3343 | 11 | 0.66% |
| hsa_circ_0053713 | 3951 | 13 | 0.66% |
| hsa_circ_0131493 | 4255 | 14 | 0.66% |
| hsa_circ_0011364 | 3648 | 12 | 0.66% |
| hsa_circ_0015860 | 3648 | 12 | 0.66% |
| hsa_circ_0029563 | 3648 | 12 | 0.66% |
| hsa_circ_0037512 | 3648 | 12 | 0.66% |
| hsa_circ_0040174 | 3344 | 11 | 0.66% |
| hsa_circ_0056374 | 3344 | 11 | 0.66% |
| hsa_circ_0058548 | 3344 | 11 | 0.66% |
| hsa_circ_0131139 | 3344 | 11 | 0.66% |
| hsa_circ_0042486 | 3649 | 12 | 0.66% |
| hsa_circ_0101598 | 3649 | 12 | 0.66% |
| hsa_circ_0029118 | 3954 | 13 | 0.66% |
| hsa_circ_0068596 | 3346 | 11 | 0.66% |
| hsa_circ_0068595 | 3042 | 10 | 0.66% |
| hsa_circ_0082506 | 3042 | 10 | 0.66% |

| | | | |
|---|---|---|---|
| hsa_circ_0067319 | 6085 | 20 | 0.66% |
| hsa_circ_0091738 | 5173 | 17 | 0.66% |
| hsa_circ_0056504 | 4566 | 15 | 0.66% |
| hsa_circ_0039738 | 3044 | 10 | 0.66% |
| hsa_circ_0012636 | 3653 | 12 | 0.66% |
| hsa_circ_0010091 | 3349 | 11 | 0.66% |
| hsa_circ_0058457 | 3958 | 13 | 0.66% |
| hsa_circ_0114693 | 7308 | 24 | 0.66% |
| hsa_circ_0017226 | 3045 | 10 | 0.66% |
| hsa_circ_0073298 | 3045 | 10 | 0.66% |
| hsa_circ_0001883 | 4568 | 15 | 0.66% |
| hsa_circ_0022258 | 3350 | 11 | 0.66% |
| hsa_circ_0086502 | 3350 | 11 | 0.66% |
| hsa_circ_0097357 | 11878 | 39 | 0.66% |
| hsa_circ_0045946 | 3655 | 12 | 0.66% |
| hsa_circ_0009301 | 5178 | 17 | 0.66% |
| hsa_circ_0004407 | 5483 | 18 | 0.66% |
| hsa_circ_0010614 | 3960 | 13 | 0.66% |
| hsa_circ_0020217 | 3351 | 11 | 0.66% |
| hsa_circ_0091871 | 3351 | 11 | 0.66% |
| hsa_circ_0091809 | 4265 | 14 | 0.66% |
| hsa_circ_0040798 | 3656 | 12 | 0.66% |
| hsa_circ_0108924 | 3047 | 10 | 0.66% |
| hsa_circ_0115717 | 3047 | 10 | 0.66% |
| hsa_circ_0037426 | 3352 | 11 | 0.66% |
| hsa_circ_0054588 | 3352 | 11 | 0.66% |
| hsa_circ_0063930 | 3352 | 11 | 0.66% |
| hsa_circ_0078397 | 3352 | 11 | 0.66% |
| hsa_circ_0015545 | 3657 | 12 | 0.66% |
| hsa_circ_0018294 | 3657 | 12 | 0.66% |
| hsa_circ_0078098 | 3657 | 12 | 0.66% |
| hsa_circ_0140774 | 6400 | 21 | 0.66% |
| hsa_circ_0044518 | 3048 | 10 | 0.66% |
| hsa_circ_0072403 | 3048 | 10 | 0.66% |
| hsa_circ_0011560 | 3353 | 11 | 0.66% |
| hsa_circ_0048862 | 3353 | 11 | 0.66% |
| hsa_circ_0053732 | 3353 | 11 | 0.66% |
| hsa_circ_0119803 | 3353 | 11 | 0.66% |
| hsa_circ_0140667 | 3658 | 12 | 0.66% |
| hsa_circ_0040866 | 3963 | 13 | 0.66% |
| hsa_circ_0121774 | 3049 | 10 | 0.66% |
| hsa_circ_0120831 | 3354 | 11 | 0.66% |
| hsa_circ_0025657 | 3355 | 11 | 0.66% |
| hsa_circ_0026846 | 3355 | 11 | 0.66% |
| hsa_circ_0111581 | 4881 | 16 | 0.66% |
| hsa_circ_0027951 | 3966 | 13 | 0.66% |
| hsa_circ_0086755 | 3966 | 13 | 0.66% |
| hsa_circ_0124373 | 3661 | 12 | 0.66% |
| hsa_circ_0043234 | 3356 | 11 | 0.66% |
| hsa_circ_0055608 | 4272 | 14 | 0.66% |

| | | | |
|---|---|---|---|
| hsa_circ_0063106 | 3662 | 12 | 0.66% |
| hsa_circ_0089479 | 3662 | 12 | 0.66% |
| hsa_circ_0059013 | 3357 | 11 | 0.66% |
| hsa_circ_0105171 | 3357 | 11 | 0.66% |
| hsa_circ_0115848 | 3357 | 11 | 0.66% |
| hsa_circ_0081519 | 3052 | 10 | 0.66% |
| hsa_circ_0125812 | 3968 | 13 | 0.66% |
| hsa_circ_0016643 | 3663 | 12 | 0.66% |
| hsa_circ_0009776 | 3358 | 11 | 0.66% |
| hsa_circ_0065585 | 3358 | 11 | 0.66% |
| hsa_circ_0028531 | 3969 | 13 | 0.66% |
| hsa_circ_0051285 | 3969 | 13 | 0.66% |
| hsa_circ_0026586 | 3664 | 12 | 0.66% |
| hsa_circ_0091926 | 3664 | 12 | 0.66% |
| hsa_circ_0024229 | 3359 | 11 | 0.65% |
| hsa_circ_0071912 | 3359 | 11 | 0.65% |
| hsa_circ_0051922 | 5497 | 18 | 0.65% |
| hsa_circ_0035433 | 3054 | 10 | 0.65% |
| hsa_circ_0082111 | 3054 | 10 | 0.65% |
| hsa_circ_0093530 | 3665 | 12 | 0.65% |
| hsa_circ_0094199 | 3665 | 12 | 0.65% |
| hsa_circ_0051648 | 4276 | 14 | 0.65% |
| hsa_circ_0065360 | 3666 | 12 | 0.65% |
| hsa_circ_0033083 | 3055 | 10 | 0.65% |
| hsa_circ_0111953 | 3055 | 10 | 0.65% |
| hsa_circ_0010596 | 3972 | 13 | 0.65% |
| hsa_circ_0024116 | 3361 | 11 | 0.65% |
| hsa_circ_0061718 | 3361 | 11 | 0.65% |
| hsa_circ_0093291 | 3361 | 11 | 0.65% |
| hsa_circ_0105864 | 12528 | 41 | 0.65% |
| hsa_circ_0105728 | 3667 | 12 | 0.65% |
| hsa_circ_0079111 | 4279 | 14 | 0.65% |
| hsa_circ_0038714 | 3668 | 12 | 0.65% |
| hsa_circ_0067117 | 3668 | 12 | 0.65% |
| hsa_circ_0073228 | 3057 | 10 | 0.65% |
| hsa_circ_0012069 | 4280 | 14 | 0.65% |
| hsa_circ_0083748 | 4280 | 14 | 0.65% |
| hsa_circ_0033395 | 3363 | 11 | 0.65% |
| hsa_circ_0016837 | 3669 | 12 | 0.65% |
| hsa_circ_0024921 | 3669 | 12 | 0.65% |
| hsa_circ_0031536 | 3669 | 12 | 0.65% |
| hsa_circ_0046158 | 3669 | 12 | 0.65% |
| hsa_circ_0137526 | 3669 | 12 | 0.65% |
| hsa_circ_0029092 | 3364 | 11 | 0.65% |
| hsa_circ_0049928 | 3364 | 11 | 0.65% |
| hsa_circ_0111529 | 5811 | 19 | 0.65% |
| hsa_circ_0047524 | 4282 | 14 | 0.65% |
| hsa_circ_0102293 | 5200 | 17 | 0.65% |
| hsa_circ_0132576 | 6118 | 20 | 0.65% |
| hsa_circ_0038675 | 3365 | 11 | 0.65% |

| | | | |
|---|---|---|---|
| hsa_circ_0065420 | 3365 | 11 | 0.65% |
| hsa_circ_0044919 | 3977 | 13 | 0.65% |
| hsa_circ_0056221 | 4284 | 14 | 0.65% |
| hsa_circ_0038916 | 3978 | 13 | 0.65% |
| hsa_circ_0113970 | 3978 | 13 | 0.65% |
| hsa_circ_0035153 | 3672 | 12 | 0.65% |
| hsa_circ_0044195 | 3672 | 12 | 0.65% |
| hsa_circ_0044207 | 3673 | 12 | 0.65% |
| hsa_circ_0058948 | 3367 | 11 | 0.65% |
| hsa_circ_0100097 | 3367 | 11 | 0.65% |
| hsa_circ_0111582 | 4898 | 16 | 0.65% |
| hsa_circ_0126407 | 13470 | 44 | 0.65% |
| hsa_circ_0033302 | 3674 | 12 | 0.65% |
| hsa_circ_0048643 | 3368 | 11 | 0.65% |
| hsa_circ_0083438 | 3368 | 11 | 0.65% |
| hsa_circ_0019162 | 3981 | 13 | 0.65% |
| hsa_circ_0042971 | 3981 | 13 | 0.65% |
| hsa_circ_0027150 | 3675 | 12 | 0.65% |
| hsa_circ_0046648 | 3675 | 12 | 0.65% |
| hsa_circ_0049104 | 3675 | 12 | 0.65% |
| hsa_circ_0072320 | 4594 | 15 | 0.65% |
| hsa_circ_0023453 | 3676 | 12 | 0.65% |
| hsa_circ_0127244 | 3676 | 12 | 0.65% |
| hsa_circ_0133172 | 3676 | 12 | 0.65% |
| hsa_circ_0041529 | 3370 | 11 | 0.65% |
| hsa_circ_0073715 | 3370 | 11 | 0.65% |
| hsa_circ_0089552 | 3370 | 11 | 0.65% |
| hsa_circ_0117903 | 3370 | 11 | 0.65% |
| hsa_circ_0083527 | 4596 | 15 | 0.65% |
| hsa_circ_0029582 | 3064 | 10 | 0.65% |
| hsa_circ_0085221 | 3064 | 10 | 0.65% |
| hsa_circ_0036532 | 3371 | 11 | 0.65% |
| hsa_circ_0049819 | 3371 | 11 | 0.65% |
| hsa_circ_0084790 | 3371 | 11 | 0.65% |
| hsa_circ_0102905 | 3371 | 11 | 0.65% |
| hsa_circ_0062993 | 3984 | 13 | 0.65% |
| hsa_circ_0095817 | 12259 | 40 | 0.65% |
| hsa_circ_0069157 | 3678 | 12 | 0.65% |
| hsa_circ_0039670 | 3065 | 10 | 0.65% |
| hsa_circ_0047241 | 3065 | 10 | 0.65% |
| hsa_circ_0111379 | 3065 | 10 | 0.65% |
| hsa_circ_0066961 | 4292 | 14 | 0.65% |
| hsa_circ_0079109 | 4292 | 14 | 0.65% |
| hsa_circ_0060213 | 3679 | 12 | 0.65% |
| hsa_circ_0017859 | 4599 | 15 | 0.65% |
| hsa_circ_0016620 | 3066 | 10 | 0.65% |
| hsa_circ_0027761 | 3066 | 10 | 0.65% |
| hsa_circ_0041902 | 3066 | 10 | 0.65% |
| hsa_circ_0100714 | 14104 | 46 | 0.65% |
| hsa_circ_0045752 | 3986 | 13 | 0.65% |

| | | | |
|---|---|---|---|
| hsa_circ_0061267 | 3986 | 13 | 0.65% |
| hsa_circ_0060589 | 3373 | 11 | 0.65% |
| hsa_circ_0086682 | 3373 | 11 | 0.65% |
| hsa_circ_0136337 | 11041 | 36 | 0.65% |
| hsa_circ_0012540 | 3067 | 10 | 0.65% |
| hsa_circ_0100560 | 3067 | 10 | 0.65% |
| hsa_circ_0064400 | 3374 | 11 | 0.65% |
| hsa_circ_0090018 | 3374 | 11 | 0.65% |
| hsa_circ_0137234 | 13497 | 44 | 0.65% |
| hsa_circ_0015825 | 3681 | 12 | 0.65% |
| hsa_circ_0091974 | 3681 | 12 | 0.65% |
| hsa_circ_0104418 | 3681 | 12 | 0.65% |
| hsa_circ_0045394 | 4295 | 14 | 0.65% |
| hsa_circ_0001878 | 7670 | 25 | 0.65% |
| hsa_circ_0004889 | 3375 | 11 | 0.65% |
| hsa_circ_0049283 | 3375 | 11 | 0.65% |
| hsa_circ_0075756 | 3375 | 11 | 0.65% |
| hsa_circ_0126574 | 3375 | 11 | 0.65% |
| hsa_circ_0128662 | 3375 | 11 | 0.65% |
| hsa_circ_0073709 | 3989 | 13 | 0.65% |
| hsa_circ_0047503 | 4296 | 14 | 0.65% |
| hsa_circ_0124558 | 11661 | 38 | 0.65% |
| hsa_circ_0042631 | 3069 | 10 | 0.65% |
| hsa_circ_0042634 | 3069 | 10 | 0.65% |
| hsa_circ_0061050 | 3376 | 11 | 0.65% |
| hsa_circ_0088591 | 3683 | 12 | 0.65% |
| hsa_circ_0106468 | 7060 | 23 | 0.65% |
| hsa_circ_0086930 | 3991 | 13 | 0.65% |
| hsa_circ_0018008 | 3684 | 12 | 0.65% |
| hsa_circ_0071158 | 3070 | 10 | 0.65% |
| hsa_circ_0084251 | 3070 | 10 | 0.65% |
| hsa_circ_0087092 | 3070 | 10 | 0.65% |
| hsa_circ_0080371 | 3992 | 13 | 0.65% |
| hsa_circ_0034515 | 3685 | 12 | 0.65% |
| hsa_circ_0069003 | 3378 | 11 | 0.65% |
| hsa_circ_0044522 | 3071 | 10 | 0.65% |
| hsa_circ_0069929 | 4300 | 14 | 0.65% |
| hsa_circ_0087864 | 3686 | 12 | 0.65% |
| hsa_circ_0060918 | 3379 | 11 | 0.65% |
| hsa_circ_0015899 | 9830 | 32 | 0.65% |
| hsa_circ_0065770 | 3072 | 10 | 0.65% |
| hsa_circ_0139439 | 3072 | 10 | 0.65% |
| hsa_circ_0056819 | 3994 | 13 | 0.65% |
| hsa_circ_0132688 | 3687 | 12 | 0.65% |
| hsa_circ_0043240 | 3380 | 11 | 0.65% |
| hsa_circ_0046460 | 3380 | 11 | 0.65% |
| hsa_circ_0117862 | 3380 | 11 | 0.65% |
| hsa_circ_0018473 | 3073 | 10 | 0.65% |
| hsa_circ_0129963 | 3073 | 10 | 0.65% |
| hsa_circ_0064191 | 3688 | 12 | 0.65% |

| | | | |
|---|---|---|---|
| hsa_circ_0071208 | 3381 | 11 | 0.65% |
| hsa_circ_0073479 | 3074 | 10 | 0.65% |
| hsa_circ_0017256 | 3689 | 12 | 0.65% |
| hsa_circ_0063804 | 3689 | 12 | 0.65% |
| hsa_circ_0066050 | 3689 | 12 | 0.65% |
| hsa_circ_0127425 | 3689 | 12 | 0.65% |
| hsa_circ_0121932 | 3382 | 11 | 0.65% |
| hsa_circ_0129388 | 3690 | 12 | 0.65% |
| hsa_circ_0004227 | 5228 | 17 | 0.65% |
| hsa_circ_0015522 | 3998 | 13 | 0.65% |
| hsa_circ_0013297 | 3383 | 11 | 0.65% |
| hsa_circ_0020865 | 3383 | 11 | 0.65% |
| hsa_circ_0054072 | 3383 | 11 | 0.65% |
| hsa_circ_0084328 | 3383 | 11 | 0.65% |
| hsa_circ_0101466 | 3691 | 12 | 0.65% |
| hsa_circ_0097106 | 3999 | 13 | 0.65% |
| hsa_circ_0070153 | 4307 | 14 | 0.65% |
| hsa_circ_0064236 | 3077 | 10 | 0.65% |
| hsa_circ_0028636 | 4308 | 14 | 0.65% |
| hsa_circ_0017038 | 3385 | 11 | 0.65% |
| hsa_circ_0054291 | 3693 | 12 | 0.65% |
| hsa_circ_0079874 | 3693 | 12 | 0.65% |
| hsa_circ_0134138 | 10465 | 34 | 0.65% |
| hsa_circ_0088292 | 3386 | 11 | 0.65% |
| hsa_circ_0052911 | 3387 | 11 | 0.65% |
| hsa_circ_0062031 | 3695 | 12 | 0.65% |
| hsa_circ_0052484 | 5235 | 17 | 0.65% |
| hsa_circ_0127308 | 46196 | 150 | 0.65% |
| hsa_circ_0047523 | 4312 | 14 | 0.65% |
| hsa_circ_0002568 | 4004 | 13 | 0.65% |
| hsa_circ_0084586 | 3696 | 12 | 0.65% |
| hsa_circ_0087968 | 3696 | 12 | 0.65% |
| hsa_circ_0105613 | 3696 | 12 | 0.65% |
| hsa_circ_0041307 | 3388 | 11 | 0.65% |
| hsa_circ_0053737 | 3388 | 11 | 0.65% |
| hsa_circ_0036835 | 4005 | 13 | 0.65% |
| hsa_circ_0010702 | 3697 | 12 | 0.65% |
| hsa_circ_0014869 | 3697 | 12 | 0.65% |
| hsa_circ_0002118 | 3389 | 11 | 0.65% |
| hsa_circ_0047823 | 3389 | 11 | 0.65% |
| hsa_circ_0056615 | 3389 | 11 | 0.65% |
| hsa_circ_0135824 | 4622 | 15 | 0.65% |
| hsa_circ_0017045 | 3390 | 11 | 0.65% |
| hsa_circ_0073718 | 3390 | 11 | 0.65% |
| hsa_circ_0037028 | 3082 | 10 | 0.65% |
| hsa_circ_0092599 | 8323 | 27 | 0.65% |
| hsa_circ_0031208 | 4316 | 14 | 0.65% |
| hsa_circ_0033446 | 3700 | 12 | 0.65% |
| hsa_circ_0035439 | 3700 | 12 | 0.65% |
| hsa_circ_0074254 | 3700 | 12 | 0.65% |

| | | | |
|---|---|---|---|
| hsa_circ_0040437 | 3392 | 11 | 0.65% |
| hsa_circ_0037433 | 4009 | 13 | 0.65% |
| hsa_circ_0052974 | 3701 | 12 | 0.65% |
| hsa_circ_0124565 | 7094 | 23 | 0.65% |
| hsa_circ_0081299 | 3393 | 11 | 0.65% |
| hsa_circ_0135591 | 12031 | 39 | 0.65% |
| hsa_circ_0015898 | 9872 | 32 | 0.65% |
| hsa_circ_0085909 | 3702 | 12 | 0.65% |
| hsa_circ_0090146 | 3702 | 12 | 0.65% |
| hsa_circ_0009400 | 4011 | 13 | 0.65% |
| hsa_circ_0010668 | 4011 | 13 | 0.65% |
| hsa_circ_0066947 | 4011 | 13 | 0.65% |
| hsa_circ_0085940 | 4011 | 13 | 0.65% |
| hsa_circ_0052106 | 11725 | 38 | 0.65% |
| hsa_circ_0106561 | 3086 | 10 | 0.65% |
| hsa_circ_0061164 | 4012 | 13 | 0.65% |
| hsa_circ_0083402 | 4012 | 13 | 0.65% |
| hsa_circ_0133692 | 10494 | 34 | 0.65% |
| hsa_circ_0045184 | 3704 | 12 | 0.65% |
| hsa_circ_0108920 | 4013 | 13 | 0.65% |
| hsa_circ_0077031 | 4322 | 14 | 0.65% |
| hsa_circ_0068971 | 3396 | 11 | 0.65% |
| hsa_circ_0073403 | 3396 | 11 | 0.65% |
| hsa_circ_0075989 | 3396 | 11 | 0.65% |
| hsa_circ_0057797 | 4014 | 13 | 0.65% |
| hsa_circ_0066020 | 3397 | 11 | 0.65% |
| hsa_circ_0116971 | 3397 | 11 | 0.65% |
| hsa_circ_0115773 | 7103 | 23 | 0.65% |
| hsa_circ_0018948 | 3706 | 12 | 0.65% |
| hsa_circ_0067714 | 3706 | 12 | 0.65% |
| hsa_circ_0043032 | 4015 | 13 | 0.65% |
| hsa_circ_0058743 | 4015 | 13 | 0.65% |
| hsa_circ_0079699 | 4324 | 14 | 0.65% |
| hsa_circ_0096840 | 13281 | 43 | 0.65% |
| hsa_circ_0089138 | 3707 | 12 | 0.65% |
| hsa_circ_0023668 | 4016 | 13 | 0.65% |
| hsa_circ_0132407 | 6797 | 22 | 0.65% |
| hsa_circ_0117404 | 4017 | 13 | 0.65% |
| hsa_circ_0033354 | 3399 | 11 | 0.65% |
| hsa_circ_0094862 | 3400 | 11 | 0.65% |
| hsa_circ_0029511 | 4019 | 13 | 0.65% |
| hsa_circ_0077359 | 3710 | 12 | 0.65% |
| hsa_circ_0118879 | 3710 | 12 | 0.65% |
| hsa_circ_0084344 | 3401 | 11 | 0.65% |
| hsa_circ_0111204 | 3401 | 11 | 0.65% |
| hsa_circ_0081508 | 4947 | 16 | 0.65% |
| hsa_circ_0115604 | 3092 | 10 | 0.65% |
| hsa_circ_0029130 | 4020 | 13 | 0.65% |
| hsa_circ_0021489 | 3711 | 12 | 0.65% |
| hsa_circ_0078221 | 3711 | 12 | 0.65% |

| | | | |
|---|---|---|---|
| hsa_circ_0120506 | 4639 | 15 | 0.65% |
| hsa_circ_0076370 | 3402 | 11 | 0.65% |
| hsa_circ_0087987 | 3402 | 11 | 0.65% |
| hsa_circ_0091944 | 3402 | 11 | 0.65% |
| hsa_circ_0098541 | 3402 | 11 | 0.65% |
| hsa_circ_0039656 | 3093 | 10 | 0.65% |
| hsa_circ_0111659 | 3093 | 10 | 0.65% |
| hsa_circ_0040430 | 3712 | 12 | 0.65% |
| hsa_circ_0048639 | 3712 | 12 | 0.65% |
| hsa_circ_0019899 | 3403 | 11 | 0.65% |
| hsa_circ_0094840 | 9281 | 30 | 0.65% |
| hsa_circ_0136302 | 6497 | 21 | 0.65% |
| hsa_circ_0082510 | 4023 | 13 | 0.65% |
| hsa_circ_0128131 | 4023 | 13 | 0.65% |
| hsa_circ_0081248 | 4642 | 15 | 0.65% |
| hsa_circ_0030645 | 3714 | 12 | 0.65% |
| hsa_circ_0036260 | 3714 | 12 | 0.65% |
| hsa_circ_0107354 | 3095 | 10 | 0.65% |
| hsa_circ_0029218 | 9286 | 30 | 0.65% |
| hsa_circ_0057858 | 3405 | 11 | 0.65% |
| hsa_circ_0068597 | 3405 | 11 | 0.65% |
| hsa_circ_0017396 | 4953 | 16 | 0.65% |
| hsa_circ_0051058 | 3715 | 12 | 0.65% |
| hsa_circ_0026108 | 4644 | 15 | 0.65% |
| hsa_circ_0085347 | 4025 | 13 | 0.65% |
| hsa_circ_0046772 | 3406 | 11 | 0.65% |
| hsa_circ_0073453 | 3406 | 11 | 0.65% |
| hsa_circ_0132323 | 6503 | 21 | 0.65% |
| hsa_circ_0004289 | 3716 | 12 | 0.65% |
| hsa_circ_0104131 | 4026 | 13 | 0.65% |
| hsa_circ_0047474 | 17343 | 56 | 0.65% |
| hsa_circ_0126051 | 3097 | 10 | 0.65% |
| hsa_circ_0013746 | 3717 | 12 | 0.65% |
| hsa_circ_0140740 | 3717 | 12 | 0.65% |
| hsa_circ_0018540 | 4027 | 13 | 0.65% |
| hsa_circ_0070490 | 3098 | 10 | 0.65% |
| hsa_circ_0009924 | 3408 | 11 | 0.65% |
| hsa_circ_0090692 | 3408 | 11 | 0.65% |
| hsa_circ_0034647 | 4028 | 13 | 0.65% |
| hsa_circ_0108614 | 4028 | 13 | 0.65% |
| hsa_circ_0028691 | 4338 | 14 | 0.65% |
| hsa_circ_0115812 | 13016 | 42 | 0.65% |
| hsa_circ_0020404 | 3409 | 11 | 0.65% |
| hsa_circ_0057470 | 3409 | 11 | 0.65% |
| hsa_circ_0036481 | 3719 | 12 | 0.65% |
| hsa_circ_0054477 | 3719 | 12 | 0.65% |
| hsa_circ_0134799 | 4650 | 15 | 0.65% |
| hsa_circ_0045079 | 4030 | 13 | 0.65% |
| hsa_circ_0064207 | 3410 | 11 | 0.65% |
| hsa_circ_0077034 | 3410 | 11 | 0.65% |

| | | | |
|---|---|---|---|
| hsa_circ_0084695 | 3410 | 11 | 0.65% |
| hsa_circ_0039807 | 3100 | 10 | 0.65% |
| hsa_circ_0030143 | 4341 | 14 | 0.65% |
| hsa_circ_0133557 | 3721 | 12 | 0.64% |
| hsa_circ_0105840 | 3101 | 10 | 0.64% |
| hsa_circ_0090906 | 4962 | 16 | 0.64% |
| hsa_circ_0129726 | 16127 | 52 | 0.64% |
| hsa_circ_0055208 | 4342 | 14 | 0.64% |
| hsa_circ_0077303 | 4342 | 14 | 0.64% |
| hsa_circ_0047504 | 4343 | 14 | 0.64% |
| hsa_circ_0084885 | 4343 | 14 | 0.64% |
| hsa_circ_0014383 | 3723 | 12 | 0.64% |
| hsa_circ_0051180 | 3413 | 11 | 0.64% |
| hsa_circ_0089937 | 3413 | 11 | 0.64% |
| hsa_circ_0097973 | 4034 | 13 | 0.64% |
| hsa_circ_0053835 | 3724 | 12 | 0.64% |
| hsa_circ_0064062 | 3724 | 12 | 0.64% |
| hsa_circ_0091918 | 3724 | 12 | 0.64% |
| hsa_circ_0081063 | 3414 | 11 | 0.64% |
| hsa_circ_0101667 | 3414 | 11 | 0.64% |
| hsa_circ_0071214 | 3104 | 10 | 0.64% |
| hsa_circ_0069824 | 3415 | 11 | 0.64% |
| hsa_circ_0056906 | 4036 | 13 | 0.64% |
| hsa_circ_0111942 | 4036 | 13 | 0.64% |
| hsa_circ_0114231 | 6520 | 21 | 0.64% |
| hsa_circ_0111821 | 3726 | 12 | 0.64% |
| hsa_circ_0031456 | 3727 | 12 | 0.64% |
| hsa_circ_0109906 | 3106 | 10 | 0.64% |
| hsa_circ_0034175 | 3417 | 11 | 0.64% |
| hsa_circ_0049936 | 3417 | 11 | 0.64% |
| hsa_circ_0082217 | 4349 | 14 | 0.64% |
| hsa_circ_0139774 | 4349 | 14 | 0.64% |
| hsa_circ_0113451 | 17707 | 57 | 0.64% |
| hsa_circ_0091020 | 4039 | 13 | 0.64% |
| hsa_circ_0082667 | 3107 | 10 | 0.64% |
| hsa_circ_0046646 | 6525 | 21 | 0.64% |
| hsa_circ_0034244 | 3418 | 11 | 0.64% |
| hsa_circ_0001306 | 10566 | 34 | 0.64% |
| hsa_circ_0036967 | 4351 | 14 | 0.64% |
| hsa_circ_0032209 | 13676 | 44 | 0.64% |
| hsa_circ_0010317 | 3730 | 12 | 0.64% |
| hsa_circ_0034147 | 3730 | 12 | 0.64% |
| hsa_circ_0058981 | 3730 | 12 | 0.64% |
| hsa_circ_0016004 | 3420 | 11 | 0.64% |
| hsa_circ_0055982 | 3731 | 12 | 0.64% |
| hsa_circ_0122167 | 4354 | 14 | 0.64% |
| hsa_circ_0074446 | 4043 | 13 | 0.64% |
| hsa_circ_0078082 | 3732 | 12 | 0.64% |
| hsa_circ_0041875 | 3421 | 11 | 0.64% |
| hsa_circ_0077630 | 3421 | 11 | 0.64% |

| | | | |
|---|---|---|---|
| hsa_circ_0078009 | 3421 | 11 | 0.64% |
| hsa_circ_0084422 | 3421 | 11 | 0.64% |
| hsa_circ_0122811 | 3421 | 11 | 0.64% |
| hsa_circ_0046925 | 3110 | 10 | 0.64% |
| hsa_circ_0068873 | 3422 | 11 | 0.64% |
| hsa_circ_0134304 | 3422 | 11 | 0.64% |
| hsa_circ_0136263 | 3422 | 11 | 0.64% |
| hsa_circ_0029916 | 3111 | 10 | 0.64% |
| hsa_circ_0023044 | 3423 | 11 | 0.64% |
| hsa_circ_0109819 | 3423 | 11 | 0.64% |
| hsa_circ_0107832 | 7158 | 23 | 0.64% |
| hsa_circ_0028548 | 3735 | 12 | 0.64% |
| hsa_circ_0067298 | 3735 | 12 | 0.64% |
| hsa_circ_0090656 | 3735 | 12 | 0.64% |
| hsa_circ_0131730 | 3735 | 12 | 0.64% |
| hsa_circ_0024086 | 3424 | 11 | 0.64% |
| hsa_circ_0025958 | 3424 | 11 | 0.64% |
| hsa_circ_0041888 | 3424 | 11 | 0.64% |
| hsa_circ_0129606 | 3113 | 10 | 0.64% |
| hsa_circ_0039632 | 4047 | 13 | 0.64% |
| hsa_circ_0045139 | 4047 | 13 | 0.64% |
| hsa_circ_0140165 | 6538 | 21 | 0.64% |
| hsa_circ_0086808 | 3425 | 11 | 0.64% |
| hsa_circ_0110259 | 3425 | 11 | 0.64% |
| hsa_circ_0002196 | 4048 | 13 | 0.64% |
| hsa_circ_0053714 | 4048 | 13 | 0.64% |
| hsa_circ_0013287 | 3737 | 12 | 0.64% |
| hsa_circ_0069949 | 3737 | 12 | 0.64% |
| hsa_circ_0070644 | 3426 | 11 | 0.64% |
| hsa_circ_0072578 | 3426 | 11 | 0.64% |
| hsa_circ_0117834 | 3426 | 11 | 0.64% |
| hsa_circ_0035780 | 4049 | 13 | 0.64% |
| hsa_circ_0117814 | 4049 | 13 | 0.64% |
| hsa_circ_0005503 | 9967 | 32 | 0.64% |
| hsa_circ_0102332 | 12459 | 40 | 0.64% |
| hsa_circ_0118952 | 3738 | 12 | 0.64% |
| hsa_circ_0074105 | 3115 | 10 | 0.64% |
| hsa_circ_0032637 | 4050 | 13 | 0.64% |
| hsa_circ_0087823 | 4050 | 13 | 0.64% |
| hsa_circ_0127197 | 7477 | 24 | 0.64% |
| hsa_circ_0023546 | 3427 | 11 | 0.64% |
| hsa_circ_0075087 | 3427 | 11 | 0.64% |
| hsa_circ_0108520 | 6231 | 20 | 0.64% |
| hsa_circ_0017749 | 3739 | 12 | 0.64% |
| hsa_circ_0069680 | 3739 | 12 | 0.64% |
| hsa_circ_0075063 | 3739 | 12 | 0.64% |
| hsa_circ_0014062 | 3116 | 10 | 0.64% |
| hsa_circ_0012060 | 4051 | 13 | 0.64% |
| hsa_circ_0052726 | 4051 | 13 | 0.64% |
| hsa_circ_0010954 | 3740 | 12 | 0.64% |

| | | | |
|---|---|---|---|
| hsa_circ_0024206 | 3740 | 12 | 0.64% |
| hsa_circ_0061133 | 3740 | 12 | 0.64% |
| hsa_circ_0139816 | 14026 | 45 | 0.64% |
| hsa_circ_0024685 | 3117 | 10 | 0.64% |
| hsa_circ_0027759 | 3117 | 10 | 0.64% |
| hsa_circ_0066224 | 3117 | 10 | 0.64% |
| hsa_circ_0102654 | 3117 | 10 | 0.64% |
| hsa_circ_0106025 | 8416 | 27 | 0.64% |
| hsa_circ_0061590 | 4364 | 14 | 0.64% |
| hsa_circ_0041172 | 3741 | 12 | 0.64% |
| hsa_circ_0066746 | 3741 | 12 | 0.64% |
| hsa_circ_0120777 | 3741 | 12 | 0.64% |
| hsa_circ_0066209 | 5300 | 17 | 0.64% |
| hsa_circ_0102896 | 4053 | 13 | 0.64% |
| hsa_circ_0015469 | 3430 | 11 | 0.64% |
| hsa_circ_0016218 | 3430 | 11 | 0.64% |
| hsa_circ_0049406 | 3430 | 11 | 0.64% |
| hsa_circ_0128145 | 3430 | 11 | 0.64% |
| hsa_circ_0034890 | 3742 | 12 | 0.64% |
| hsa_circ_0018291 | 3743 | 12 | 0.64% |
| hsa_circ_0089426 | 3743 | 12 | 0.64% |
| hsa_circ_0074564 | 4055 | 13 | 0.64% |
| hsa_circ_0010557 | 4368 | 14 | 0.64% |
| hsa_circ_0092565 | 4368 | 14 | 0.64% |
| hsa_circ_0126609 | 4368 | 14 | 0.64% |
| hsa_circ_0025988 | 4056 | 13 | 0.64% |
| hsa_circ_0130411 | 3432 | 11 | 0.64% |
| hsa_circ_0135000 | 3432 | 11 | 0.64% |
| hsa_circ_0029741 | 3120 | 10 | 0.64% |
| hsa_circ_0041861 | 3120 | 10 | 0.64% |
| hsa_circ_0121139 | 4057 | 13 | 0.64% |
| hsa_circ_0013417 | 3745 | 12 | 0.64% |
| hsa_circ_0056725 | 3745 | 12 | 0.64% |
| hsa_circ_0082868 | 3745 | 12 | 0.64% |
| hsa_circ_0042278 | 3433 | 11 | 0.64% |
| hsa_circ_0076723 | 4058 | 13 | 0.64% |
| hsa_circ_0041962 | 3746 | 12 | 0.64% |
| hsa_circ_0101761 | 7180 | 23 | 0.64% |
| hsa_circ_0052599 | 4371 | 14 | 0.64% |
| hsa_circ_0049116 | 3747 | 12 | 0.64% |
| hsa_circ_0049120 | 3747 | 12 | 0.64% |
| hsa_circ_0058347 | 3747 | 12 | 0.64% |
| hsa_circ_0042307 | 3435 | 11 | 0.64% |
| hsa_circ_0103630 | 3435 | 11 | 0.64% |
| hsa_circ_0042001 | 3123 | 10 | 0.64% |
| hsa_circ_0046331 | 3123 | 10 | 0.64% |
| hsa_circ_0082507 | 3123 | 10 | 0.64% |
| hsa_circ_0054325 | 3748 | 12 | 0.64% |
| hsa_circ_0074058 | 3748 | 12 | 0.64% |
| hsa_circ_0084699 | 3436 | 11 | 0.64% |

| | | | |
|---|---|---|---|
| hsa_circ_0069548 | 4061 | 13 | 0.64% |
| hsa_circ_0099640 | 5623 | 18 | 0.64% |
| hsa_circ_0022728 | 3749 | 12 | 0.64% |
| hsa_circ_0032698 | 3749 | 12 | 0.64% |
| hsa_circ_0119811 | 3437 | 11 | 0.64% |
| hsa_circ_0136041 | 3437 | 11 | 0.64% |
| hsa_circ_0055299 | 3750 | 12 | 0.64% |
| hsa_circ_0085373 | 3750 | 12 | 0.64% |
| hsa_circ_0136741 | 3750 | 12 | 0.64% |
| hsa_circ_0063929 | 3125 | 10 | 0.64% |
| hsa_circ_0057565 | 4376 | 14 | 0.64% |
| hsa_circ_0043535 | 3751 | 12 | 0.64% |
| hsa_circ_0019900 | 3439 | 11 | 0.64% |
| hsa_circ_0045031 | 3439 | 11 | 0.64% |
| hsa_circ_0107548 | 3439 | 11 | 0.64% |
| hsa_circ_0073953 | 5315 | 17 | 0.64% |
| hsa_circ_0096572 | 4065 | 13 | 0.64% |
| hsa_circ_0057639 | 3127 | 10 | 0.64% |
| hsa_circ_0073885 | 3753 | 12 | 0.64% |
| hsa_circ_0107267 | 3753 | 12 | 0.64% |
| hsa_circ_0101759 | 7194 | 23 | 0.64% |
| hsa_circ_0011548 | 3128 | 10 | 0.64% |
| hsa_circ_0096391 | 3128 | 10 | 0.64% |
| hsa_circ_0080202 | 3441 | 11 | 0.64% |
| hsa_circ_0035759 | 4067 | 13 | 0.64% |
| hsa_circ_0032613 | 3442 | 11 | 0.64% |
| hsa_circ_0081904 | 3755 | 12 | 0.64% |
| hsa_circ_0046083 | 4068 | 13 | 0.64% |
| hsa_circ_0021487 | 4381 | 14 | 0.64% |
| hsa_circ_0093872 | 8138 | 26 | 0.64% |
| hsa_circ_0013140 | 3756 | 12 | 0.64% |
| hsa_circ_0031513 | 3756 | 12 | 0.64% |
| hsa_circ_0104837 | 3756 | 12 | 0.64% |
| hsa_circ_0069083 | 3443 | 11 | 0.64% |
| hsa_circ_0074938 | 3130 | 10 | 0.64% |
| hsa_circ_0102294 | 5322 | 17 | 0.64% |
| hsa_circ_0044916 | 4070 | 13 | 0.64% |
| hsa_circ_0114694 | 7514 | 24 | 0.64% |
| hsa_circ_0040614 | 3757 | 12 | 0.64% |
| hsa_circ_0024965 | 3444 | 11 | 0.64% |
| hsa_circ_0113995 | 3444 | 11 | 0.64% |
| hsa_circ_0123360 | 3444 | 11 | 0.64% |
| hsa_circ_0103787 | 12838 | 41 | 0.64% |
| hsa_circ_0039974 | 4384 | 14 | 0.64% |
| hsa_circ_0123408 | 4384 | 14 | 0.64% |
| hsa_circ_0018319 | 3758 | 12 | 0.64% |
| hsa_circ_0086082 | 3758 | 12 | 0.64% |
| hsa_circ_0026844 | 3445 | 11 | 0.64% |
| hsa_circ_0062701 | 3445 | 11 | 0.64% |
| hsa_circ_0019161 | 4072 | 13 | 0.64% |

| | | | |
|---|---|---|---|
| hsa_circ_0036890 | 3759 | 12 | 0.64% |
| hsa_circ_0083271 | 3446 | 11 | 0.64% |
| hsa_circ_0066225 | 3133 | 10 | 0.64% |
| hsa_circ_0112186 | 12533 | 40 | 0.64% |
| hsa_circ_0085667 | 3760 | 12 | 0.64% |
| hsa_circ_0084909 | 3447 | 11 | 0.64% |
| hsa_circ_0112579 | 3447 | 11 | 0.64% |
| hsa_circ_0128736 | 3447 | 11 | 0.64% |
| hsa_circ_0109873 | 5954 | 19 | 0.64% |
| hsa_circ_0063270 | 4074 | 13 | 0.64% |
| hsa_circ_0027211 | 3134 | 10 | 0.64% |
| hsa_circ_0030928 | 3761 | 12 | 0.64% |
| hsa_circ_0078109 | 3761 | 12 | 0.64% |
| hsa_circ_0013230 | 3448 | 11 | 0.64% |
| hsa_circ_0029739 | 3448 | 11 | 0.64% |
| hsa_circ_0040717 | 3448 | 11 | 0.64% |
| hsa_circ_0086846 | 3448 | 11 | 0.64% |
| hsa_circ_0100189 | 6897 | 22 | 0.64% |
| hsa_circ_0034282 | 3135 | 10 | 0.64% |
| hsa_circ_0076312 | 3135 | 10 | 0.64% |
| hsa_circ_0014557 | 3136 | 10 | 0.64% |
| hsa_circ_0138975 | 4391 | 14 | 0.64% |
| hsa_circ_0136464 | 6273 | 20 | 0.64% |
| hsa_circ_0033330 | 3764 | 12 | 0.64% |
| hsa_circ_0046159 | 3764 | 12 | 0.64% |
| hsa_circ_0071201 | 3451 | 11 | 0.64% |
| hsa_circ_0112659 | 3451 | 11 | 0.64% |
| hsa_circ_0056203 | 3765 | 12 | 0.64% |
| hsa_circ_0057058 | 3765 | 12 | 0.64% |
| hsa_circ_0087969 | 3765 | 12 | 0.64% |
| hsa_circ_0102867 | 3765 | 12 | 0.64% |
| hsa_circ_0006456 | 6589 | 21 | 0.64% |
| hsa_circ_0030079 | 3452 | 11 | 0.64% |
| hsa_circ_0047142 | 3452 | 11 | 0.64% |
| hsa_circ_0039634 | 4080 | 13 | 0.64% |
| hsa_circ_0124847 | 8475 | 27 | 0.64% |
| hsa_circ_0030368 | 3139 | 10 | 0.64% |
| hsa_circ_0088005 | 3139 | 10 | 0.64% |
| hsa_circ_0136177 | 3139 | 10 | 0.64% |
| hsa_circ_0071617 | 3453 | 11 | 0.64% |
| hsa_circ_0012972 | 3767 | 12 | 0.64% |
| hsa_circ_0030708 | 3767 | 12 | 0.64% |
| hsa_circ_0119351 | 12243 | 39 | 0.64% |
| hsa_circ_0047018 | 4082 | 13 | 0.64% |
| hsa_circ_0083190 | 3768 | 12 | 0.64% |
| hsa_circ_0003370 | 3454 | 11 | 0.64% |
| hsa_circ_0005137 | 3454 | 11 | 0.64% |
| hsa_circ_0086526 | 3454 | 11 | 0.64% |
| hsa_circ_0126199 | 3140 | 10 | 0.64% |
| hsa_circ_0056677 | 3769 | 12 | 0.64% |

| | | | |
|---|---|---|---|
| hsa_circ_0079155 | 3769 | 12 | 0.64% |
| hsa_circ_0120409 | 3769 | 12 | 0.64% |
| hsa_circ_0070562 | 3455 | 11 | 0.64% |
| hsa_circ_0099527 | 3455 | 11 | 0.64% |
| hsa_circ_0018253 | 3141 | 10 | 0.64% |
| hsa_circ_0050671 | 3456 | 11 | 0.64% |
| hsa_circ_0010755 | 3457 | 11 | 0.64% |
| hsa_circ_0034985 | 3457 | 11 | 0.64% |
| hsa_circ_0045689 | 3457 | 11 | 0.64% |
| hsa_circ_0054073 | 3457 | 11 | 0.64% |
| hsa_circ_0076962 | 3457 | 11 | 0.64% |
| hsa_circ_0066066 | 5343 | 17 | 0.64% |
| hsa_circ_0024071 | 3772 | 12 | 0.64% |
| hsa_circ_0004020 | 3458 | 11 | 0.64% |
| hsa_circ_0081372 | 3458 | 11 | 0.64% |
| hsa_circ_0126931 | 12261 | 39 | 0.64% |
| hsa_circ_0056643 | 4087 | 13 | 0.64% |
| hsa_circ_0031454 | 3773 | 12 | 0.64% |
| hsa_circ_0078795 | 3773 | 12 | 0.64% |
| hsa_circ_0026122 | 3459 | 11 | 0.64% |
| hsa_circ_0078705 | 3459 | 11 | 0.64% |
| hsa_circ_0027839 | 3145 | 10 | 0.64% |
| hsa_circ_0044215 | 11952 | 38 | 0.64% |
| hsa_circ_0026329 | 4089 | 13 | 0.64% |
| hsa_circ_0070172 | 3460 | 11 | 0.64% |
| hsa_circ_0104741 | 6291 | 20 | 0.64% |
| hsa_circ_0040476 | 3775 | 12 | 0.64% |
| hsa_circ_0038908 | 11640 | 37 | 0.64% |
| hsa_circ_0084730 | 3461 | 11 | 0.64% |
| hsa_circ_0116031 | 3461 | 11 | 0.64% |
| hsa_circ_0100709 | 14476 | 46 | 0.64% |
| hsa_circ_0019833 | 3462 | 11 | 0.64% |
| hsa_circ_0026514 | 3462 | 11 | 0.64% |
| hsa_circ_0118319 | 3462 | 11 | 0.64% |
| hsa_circ_0026817 | 3777 | 12 | 0.64% |
| hsa_circ_0046333 | 3777 | 12 | 0.64% |
| hsa_circ_0095068 | 3777 | 12 | 0.64% |
| hsa_circ_0066519 | 3148 | 10 | 0.64% |
| hsa_circ_0015841 | 3463 | 11 | 0.64% |
| hsa_circ_0046915 | 3778 | 12 | 0.64% |
| hsa_circ_0133520 | 3778 | 12 | 0.64% |
| hsa_circ_0135924 | 3779 | 12 | 0.64% |
| hsa_circ_0100256 | 11338 | 36 | 0.64% |
| hsa_circ_0049763 | 11654 | 37 | 0.63% |
| hsa_circ_0002845 | 12284 | 39 | 0.63% |
| hsa_circ_0105797 | 4410 | 14 | 0.63% |
| hsa_circ_0064539 | 4095 | 13 | 0.63% |
| hsa_circ_0016838 | 3780 | 12 | 0.63% |
| hsa_circ_0036236 | 3780 | 12 | 0.63% |
| hsa_circ_0014621 | 3465 | 11 | 0.63% |

| | | | |
|---|---|---|---|
| hsa_circ_0062697 | 3465 | 11 | 0.63% |
| hsa_circ_0062992 | 4096 | 13 | 0.63% |
| hsa_circ_0072895 | 3781 | 12 | 0.63% |
| hsa_circ_0032425 | 3466 | 11 | 0.63% |
| hsa_circ_0070894 | 3466 | 11 | 0.63% |
| hsa_circ_0094841 | 9454 | 30 | 0.63% |
| hsa_circ_0060974 | 4412 | 14 | 0.63% |
| hsa_circ_0086437 | 4097 | 13 | 0.63% |
| hsa_circ_0043878 | 3782 | 12 | 0.63% |
| hsa_circ_0029948 | 3467 | 11 | 0.63% |
| hsa_circ_0057777 | 3467 | 11 | 0.63% |
| hsa_circ_0077121 | 3467 | 11 | 0.63% |
| hsa_circ_0109243 | 3467 | 11 | 0.63% |
| hsa_circ_0019514 | 4098 | 13 | 0.63% |
| hsa_circ_0133274 | 7566 | 24 | 0.63% |
| hsa_circ_0040337 | 13556 | 43 | 0.63% |
| hsa_circ_0044609 | 3468 | 11 | 0.63% |
| hsa_circ_0048160 | 3468 | 11 | 0.63% |
| hsa_circ_0062371 | 3468 | 11 | 0.63% |
| hsa_circ_0096562 | 3468 | 11 | 0.63% |
| hsa_circ_0010029 | 4414 | 14 | 0.63% |
| hsa_circ_0013697 | 4099 | 13 | 0.63% |
| hsa_circ_0009660 | 3784 | 12 | 0.63% |
| hsa_circ_0068489 | 3784 | 12 | 0.63% |
| hsa_circ_0057209 | 3154 | 10 | 0.63% |
| hsa_circ_0012244 | 3785 | 12 | 0.63% |
| hsa_circ_0036159 | 3785 | 12 | 0.63% |
| hsa_circ_0061115 | 3785 | 12 | 0.63% |
| hsa_circ_0021032 | 3470 | 11 | 0.63% |
| hsa_circ_0136773 | 3470 | 11 | 0.63% |
| hsa_circ_0080184 | 11042 | 35 | 0.63% |
| hsa_circ_0076840 | 3786 | 12 | 0.63% |
| hsa_circ_0068430 | 4102 | 13 | 0.63% |
| hsa_circ_0136585 | 6313 | 20 | 0.63% |
| hsa_circ_0129123 | 4420 | 14 | 0.63% |
| hsa_circ_0080196 | 3473 | 11 | 0.63% |
| hsa_circ_0018491 | 4736 | 15 | 0.63% |
| hsa_circ_0075700 | 3789 | 12 | 0.63% |
| hsa_circ_0076815 | 3789 | 12 | 0.63% |
| hsa_circ_0063632 | 4105 | 13 | 0.63% |
| hsa_circ_0068556 | 3158 | 10 | 0.63% |
| hsa_circ_0017503 | 3790 | 12 | 0.63% |
| hsa_circ_0080029 | 8844 | 28 | 0.63% |
| hsa_circ_0044780 | 5054 | 16 | 0.63% |
| hsa_circ_0010729 | 3475 | 11 | 0.63% |
| hsa_circ_0010753 | 3475 | 11 | 0.63% |
| hsa_circ_0007616 | 4107 | 13 | 0.63% |
| hsa_circ_0102710 | 4107 | 13 | 0.63% |
| hsa_circ_0108933 | 8215 | 26 | 0.63% |
| hsa_circ_0070970 | 8848 | 28 | 0.63% |

| | | | |
|---|---|---|---|
| hsa_circ_0010597 | 4108 | 13 | 0.63% |
| hsa_circ_0029183 | 3792 | 12 | 0.63% |
| hsa_circ_0058557 | 3792 | 12 | 0.63% |
| hsa_circ_0118405 | 3476 | 11 | 0.63% |
| hsa_circ_0112216 | 3160 | 10 | 0.63% |
| hsa_circ_0140438 | 10431 | 33 | 0.63% |
| hsa_circ_0013052 | 3477 | 11 | 0.63% |
| hsa_circ_0009998 | 3794 | 12 | 0.63% |
| hsa_circ_0063931 | 3478 | 11 | 0.63% |
| hsa_circ_0081509 | 5059 | 16 | 0.63% |
| hsa_circ_0083516 | 4743 | 15 | 0.63% |
| hsa_circ_0099307 | 7906 | 25 | 0.63% |
| hsa_circ_0131852 | 3795 | 12 | 0.63% |
| hsa_circ_0064229 | 3479 | 11 | 0.63% |
| hsa_circ_0080139 | 3479 | 11 | 0.63% |
| hsa_circ_0135441 | 4428 | 14 | 0.63% |
| hsa_circ_0118321 | 16764 | 53 | 0.63% |
| hsa_circ_0101169 | 4112 | 13 | 0.63% |
| hsa_circ_0016176 | 13918 | 44 | 0.63% |
| hsa_circ_0048872 | 4113 | 13 | 0.63% |
| hsa_circ_0055298 | 3797 | 12 | 0.63% |
| hsa_circ_0074352 | 3797 | 12 | 0.63% |
| hsa_circ_0079977 | 3797 | 12 | 0.63% |
| hsa_circ_0096887 | 4430 | 14 | 0.63% |
| hsa_circ_0024974 | 3481 | 11 | 0.63% |
| hsa_circ_0042447 | 3481 | 11 | 0.63% |
| hsa_circ_0086861 | 3481 | 11 | 0.63% |
| hsa_circ_0092263 | 3481 | 11 | 0.63% |
| hsa_circ_0093748 | 3481 | 11 | 0.63% |
| hsa_circ_0029152 | 4114 | 13 | 0.63% |
| hsa_circ_0066210 | 5380 | 17 | 0.63% |
| hsa_circ_0117940 | 3165 | 10 | 0.63% |
| hsa_circ_0140291 | 4115 | 13 | 0.63% |
| hsa_circ_0009222 | 3799 | 12 | 0.63% |
| hsa_circ_0089478 | 3799 | 12 | 0.63% |
| hsa_circ_0073515 | 3800 | 12 | 0.63% |
| hsa_circ_0082869 | 3800 | 12 | 0.63% |
| hsa_circ_0013310 | 9501 | 30 | 0.63% |
| hsa_circ_0000501 | 3484 | 11 | 0.63% |
| hsa_circ_0012539 | 3484 | 11 | 0.63% |
| hsa_circ_0047533 | 3484 | 11 | 0.63% |
| hsa_circ_0086518 | 3484 | 11 | 0.63% |
| hsa_circ_0126575 | 3484 | 11 | 0.63% |
| hsa_circ_0135021 | 3484 | 11 | 0.63% |
| hsa_circ_0034906 | 3801 | 12 | 0.63% |
| hsa_circ_0071984 | 3801 | 12 | 0.63% |
| hsa_circ_0029095 | 4118 | 13 | 0.63% |
| hsa_circ_0075114 | 4118 | 13 | 0.63% |
| hsa_circ_0129067 | 4118 | 13 | 0.63% |
| hsa_circ_0001293 | 3485 | 11 | 0.63% |

| | | | |
|---|---|---|---|
| hsa_circ_0045456 | 3485 | 11 | 0.63% |
| hsa_circ_0053446 | 3485 | 11 | 0.63% |
| hsa_circ_0065196 | 3485 | 11 | 0.63% |
| hsa_circ_0027663 | 3802 | 12 | 0.63% |
| hsa_circ_0033075 | 4436 | 14 | 0.63% |
| hsa_circ_0015846 | 3486 | 11 | 0.63% |
| hsa_circ_0129856 | 3486 | 11 | 0.63% |
| hsa_circ_0029789 | 3803 | 12 | 0.63% |
| hsa_circ_0046913 | 3803 | 12 | 0.63% |
| hsa_circ_0137419 | 3803 | 12 | 0.63% |
| hsa_circ_0070232 | 3487 | 11 | 0.63% |
| hsa_circ_0079983 | 3487 | 11 | 0.63% |
| hsa_circ_0017143 | 3170 | 10 | 0.63% |
| hsa_circ_0139025 | 4439 | 14 | 0.63% |
| hsa_circ_0010062 | 4122 | 13 | 0.63% |
| hsa_circ_0092245 | 3805 | 12 | 0.63% |
| hsa_circ_0119841 | 3805 | 12 | 0.63% |
| hsa_circ_0140668 | 3805 | 12 | 0.63% |
| hsa_circ_0068319 | 3488 | 11 | 0.63% |
| hsa_circ_0112117 | 3488 | 11 | 0.63% |
| hsa_circ_0000492 | 3171 | 10 | 0.63% |
| hsa_circ_0094420 | 3171 | 10 | 0.63% |
| hsa_circ_0085424 | 5391 | 17 | 0.63% |
| hsa_circ_0073278 | 4123 | 13 | 0.63% |
| hsa_circ_0056675 | 3806 | 12 | 0.63% |
| hsa_circ_0070043 | 3172 | 10 | 0.63% |
| hsa_circ_0070297 | 3172 | 10 | 0.63% |
| hsa_circ_0077566 | 8565 | 27 | 0.63% |
| hsa_circ_0021474 | 3807 | 12 | 0.63% |
| hsa_circ_0040799 | 3807 | 12 | 0.63% |
| hsa_circ_0066924 | 3807 | 12 | 0.63% |
| hsa_circ_0084306 | 3807 | 12 | 0.63% |
| hsa_circ_0068966 | 3490 | 11 | 0.63% |
| hsa_circ_0033394 | 3173 | 10 | 0.63% |
| hsa_circ_0046926 | 4125 | 13 | 0.63% |
| hsa_circ_0116480 | 4443 | 14 | 0.63% |
| hsa_circ_0020016 | 3491 | 11 | 0.63% |
| hsa_circ_0065586 | 3491 | 11 | 0.63% |
| hsa_circ_0071907 | 3491 | 11 | 0.63% |
| hsa_circ_0072450 | 3491 | 11 | 0.63% |
| hsa_circ_0081825 | 3491 | 11 | 0.63% |
| hsa_circ_0091385 | 3491 | 11 | 0.63% |
| hsa_circ_0121968 | 19042 | 60 | 0.63% |
| hsa_circ_0039319 | 3809 | 12 | 0.63% |
| hsa_circ_0044864 | 3809 | 12 | 0.63% |
| hsa_circ_0070052 | 4444 | 14 | 0.63% |
| hsa_circ_0086683 | 3492 | 11 | 0.63% |
| hsa_circ_0128977 | 4127 | 13 | 0.63% |
| hsa_circ_0060786 | 3810 | 12 | 0.63% |
| hsa_circ_0073066 | 3810 | 12 | 0.63% |

| | | | |
|---|---|---|---|
| hsa_circ_0085723 | 3493 | 11 | 0.63% |
| hsa_circ_0010621 | 4129 | 13 | 0.63% |
| hsa_circ_0029977 | 4129 | 13 | 0.63% |
| hsa_circ_0053725 | 4129 | 13 | 0.63% |
| hsa_circ_0044453 | 3812 | 12 | 0.63% |
| hsa_circ_0025989 | 4130 | 13 | 0.63% |
| hsa_circ_0031620 | 4130 | 13 | 0.63% |
| hsa_circ_0074305 | 3177 | 10 | 0.63% |
| hsa_circ_0127116 | 3177 | 10 | 0.63% |
| hsa_circ_0038318 | 6990 | 22 | 0.63% |
| hsa_circ_0061357 | 3495 | 11 | 0.63% |
| hsa_circ_0091624 | 3495 | 11 | 0.63% |
| hsa_circ_0046160 | 3813 | 12 | 0.63% |
| hsa_circ_0059636 | 4131 | 13 | 0.63% |
| hsa_circ_0099946 | 3496 | 11 | 0.63% |
| hsa_circ_0063886 | 3814 | 12 | 0.63% |
| hsa_circ_0072580 | 3814 | 12 | 0.63% |
| hsa_circ_0053635 | 4132 | 13 | 0.63% |
| hsa_circ_0103487 | 3497 | 11 | 0.63% |
| hsa_circ_0108440 | 3497 | 11 | 0.63% |
| hsa_circ_0058421 | 3815 | 12 | 0.63% |
| hsa_circ_0128953 | 4451 | 14 | 0.63% |
| hsa_circ_0049766 | 18125 | 57 | 0.63% |
| hsa_circ_0047912 | 3816 | 12 | 0.63% |
| hsa_circ_0022314 | 3498 | 11 | 0.63% |
| hsa_circ_0072231 | 3498 | 11 | 0.63% |
| hsa_circ_0127791 | 5407 | 17 | 0.63% |
| hsa_circ_0045800 | 4135 | 13 | 0.63% |
| hsa_circ_0111818 | 3817 | 12 | 0.63% |
| hsa_circ_0056145 | 3499 | 11 | 0.63% |
| hsa_circ_0073294 | 3499 | 11 | 0.63% |
| hsa_circ_0058012 | 4454 | 14 | 0.63% |
| hsa_circ_0070150 | 4454 | 14 | 0.63% |
| hsa_circ_0037530 | 3500 | 11 | 0.63% |
| hsa_circ_0053753 | 3500 | 11 | 0.63% |
| hsa_circ_0083743 | 3500 | 11 | 0.63% |
| hsa_circ_0122320 | 12410 | 39 | 0.63% |
| hsa_circ_0028444 | 4137 | 13 | 0.63% |
| hsa_circ_0045753 | 4137 | 13 | 0.63% |
| hsa_circ_0090312 | 3819 | 12 | 0.63% |
| hsa_circ_0096642 | 14322 | 45 | 0.63% |
| hsa_circ_0075755 | 3501 | 11 | 0.63% |
| hsa_circ_0123874 | 6684 | 21 | 0.63% |
| hsa_circ_0012893 | 4456 | 14 | 0.63% |
| hsa_circ_0091845 | 4456 | 14 | 0.63% |
| hsa_circ_0047242 | 3183 | 10 | 0.63% |
| hsa_circ_0113533 | 4138 | 13 | 0.63% |
| hsa_circ_0027673 | 3820 | 12 | 0.63% |
| hsa_circ_0089188 | 12416 | 39 | 0.63% |
| hsa_circ_0105172 | 3502 | 11 | 0.63% |

| | | | |
|---|---|---|---|
| hsa_circ_0108441 | 3502 | 11 | 0.63% |
| hsa_circ_0108884 | 3502 | 11 | 0.63% |
| hsa_circ_0056892 | 3184 | 10 | 0.63% |
| hsa_circ_0101584 | 3821 | 12 | 0.63% |
| hsa_circ_0020883 | 3503 | 11 | 0.63% |
| hsa_circ_0122268 | 3503 | 11 | 0.63% |
| hsa_circ_0058815 | 4140 | 13 | 0.63% |
| hsa_circ_0060850 | 4777 | 15 | 0.63% |
| hsa_circ_0001437 | 12105 | 38 | 0.63% |
| hsa_circ_0067559 | 4460 | 14 | 0.63% |
| hsa_circ_0028768 | 3824 | 12 | 0.63% |
| hsa_circ_0063105 | 3824 | 12 | 0.63% |
| hsa_circ_0091842 | 3506 | 11 | 0.63% |
| hsa_circ_0100710 | 14662 | 46 | 0.63% |
| hsa_circ_0031482 | 3825 | 12 | 0.63% |
| hsa_circ_0119313 | 3825 | 12 | 0.63% |
| hsa_circ_0052443 | 3188 | 10 | 0.63% |
| hsa_circ_0133712 | 3188 | 10 | 0.63% |
| hsa_circ_0097786 | 3507 | 11 | 0.63% |
| hsa_circ_0119573 | 9246 | 29 | 0.63% |
| hsa_circ_0082103 | 3508 | 11 | 0.63% |
| hsa_circ_0118108 | 3508 | 11 | 0.63% |
| hsa_circ_0017704 | 3827 | 12 | 0.63% |
| hsa_circ_0088755 | 3827 | 12 | 0.63% |
| hsa_circ_0057225 | 4146 | 13 | 0.63% |
| hsa_circ_0081648 | 15949 | 50 | 0.63% |
| hsa_circ_0098551 | 11484 | 36 | 0.63% |
| hsa_circ_0084061 | 4467 | 14 | 0.63% |
| hsa_circ_0010299 | 3829 | 12 | 0.63% |
| hsa_circ_0047984 | 3829 | 12 | 0.63% |
| hsa_circ_0029623 | 3510 | 11 | 0.63% |
| hsa_circ_0072383 | 3510 | 11 | 0.63% |
| hsa_circ_0032915 | 3191 | 10 | 0.63% |
| hsa_circ_0039660 | 3191 | 10 | 0.63% |
| hsa_circ_0058965 | 3191 | 10 | 0.63% |
| hsa_circ_0005018 | 4469 | 14 | 0.63% |
| hsa_circ_0057507 | 4469 | 14 | 0.63% |
| hsa_circ_0075300 | 4469 | 14 | 0.63% |
| hsa_circ_0043298 | 5746 | 18 | 0.63% |
| hsa_circ_0010682 | 4150 | 13 | 0.63% |
| hsa_circ_0032981 | 3512 | 11 | 0.63% |
| hsa_circ_0045694 | 3512 | 11 | 0.63% |
| hsa_circ_0085182 | 3512 | 11 | 0.63% |
| hsa_circ_0139028 | 13729 | 43 | 0.63% |
| hsa_circ_0009114 | 4470 | 14 | 0.63% |
| hsa_circ_0115101 | 3193 | 10 | 0.63% |
| hsa_circ_0053615 | 4151 | 13 | 0.63% |
| hsa_circ_0071194 | 3513 | 11 | 0.63% |
| hsa_circ_0060683 | 4152 | 13 | 0.63% |
| hsa_circ_0026740 | 3833 | 12 | 0.63% |

| | | | |
|---|---|---|---|
| hsa_circ_0053830 | 3833 | 12 | 0.63% |
| hsa_circ_0064971 | 11500 | 36 | 0.63% |
| hsa_circ_0032636 | 4153 | 13 | 0.63% |
| hsa_circ_0136385 | 10224 | 32 | 0.63% |
| hsa_circ_0106930 | 7029 | 22 | 0.63% |
| hsa_circ_0039808 | 3195 | 10 | 0.63% |
| hsa_circ_0073952 | 5432 | 17 | 0.63% |
| hsa_circ_0069095 | 4154 | 13 | 0.63% |
| hsa_circ_0050359 | 3515 | 11 | 0.63% |
| hsa_circ_0068363 | 3515 | 11 | 0.63% |
| hsa_circ_0123816 | 3515 | 11 | 0.63% |
| hsa_circ_0014731 | 5113 | 16 | 0.63% |
| hsa_circ_0030267 | 4474 | 14 | 0.63% |
| hsa_circ_0011486 | 4155 | 13 | 0.63% |
| hsa_circ_0086915 | 4155 | 13 | 0.63% |
| hsa_circ_0057861 | 3516 | 11 | 0.63% |
| hsa_circ_0124352 | 7991 | 25 | 0.63% |
| hsa_circ_0016605 | 3197 | 10 | 0.63% |
| hsa_circ_0064235 | 3197 | 10 | 0.63% |
| hsa_circ_0081383 | 4476 | 14 | 0.63% |
| hsa_circ_0016260 | 3517 | 11 | 0.63% |
| hsa_circ_0098637 | 3837 | 12 | 0.63% |
| hsa_circ_0026841 | 3518 | 11 | 0.63% |
| hsa_circ_0130103 | 3518 | 11 | 0.63% |
| hsa_circ_0009976 | 3838 | 12 | 0.63% |
| hsa_circ_0034509 | 3838 | 12 | 0.63% |
| hsa_circ_0034240 | 4158 | 13 | 0.63% |
| hsa_circ_0052123 | 4158 | 13 | 0.63% |
| hsa_circ_0139440 | 3199 | 10 | 0.63% |
| hsa_circ_0118259 | 6718 | 21 | 0.63% |
| hsa_circ_0058123 | 3519 | 11 | 0.63% |
| hsa_circ_0073335 | 3519 | 11 | 0.63% |
| hsa_circ_0086880 | 3519 | 11 | 0.63% |
| hsa_circ_0094138 | 3519 | 11 | 0.63% |
| hsa_circ_0078800 | 3839 | 12 | 0.63% |
| hsa_circ_0018930 | 4159 | 13 | 0.63% |
| hsa_circ_0015703 | 4799 | 15 | 0.63% |
| hsa_circ_0056291 | 3840 | 12 | 0.63% |
| hsa_circ_0132964 | 3840 | 12 | 0.63% |
| hsa_circ_0042624 | 3520 | 11 | 0.63% |
| hsa_circ_0052733 | 3200 | 10 | 0.63% |
| hsa_circ_0134517 | 8322 | 26 | 0.62% |
| hsa_circ_0073358 | 4161 | 13 | 0.62% |
| hsa_circ_0024664 | 3841 | 12 | 0.62% |
| hsa_circ_0033215 | 3841 | 12 | 0.62% |
| hsa_circ_0034905 | 3841 | 12 | 0.62% |
| hsa_circ_0127053 | 11844 | 37 | 0.62% |
| hsa_circ_0030488 | 4802 | 15 | 0.62% |
| hsa_circ_0018308 | 6083 | 19 | 0.62% |
| hsa_circ_0003560 | 3522 | 11 | 0.62% |

| | | | |
|---|---|---|---|
| hsa_circ_0006785 | 3522 | 11 | 0.62% |
| hsa_circ_0138018 | 4483 | 14 | 0.62% |
| hsa_circ_0050794 | 3843 | 12 | 0.62% |
| hsa_circ_0072661 | 3523 | 11 | 0.62% |
| hsa_circ_0080007 | 3523 | 11 | 0.62% |
| hsa_circ_0019128 | 3203 | 10 | 0.62% |
| hsa_circ_0067977 | 3203 | 10 | 0.62% |
| hsa_circ_0077349 | 3203 | 10 | 0.62% |
| hsa_circ_0090235 | 3844 | 12 | 0.62% |
| hsa_circ_0070252 | 9290 | 29 | 0.62% |
| hsa_circ_0035070 | 4165 | 13 | 0.62% |
| hsa_circ_0039657 | 3204 | 10 | 0.62% |
| hsa_circ_0039739 | 3204 | 10 | 0.62% |
| hsa_circ_0082108 | 3204 | 10 | 0.62% |
| hsa_circ_0082611 | 3845 | 12 | 0.62% |
| hsa_circ_0069294 | 3525 | 11 | 0.62% |
| hsa_circ_0071299 | 3525 | 11 | 0.62% |
| hsa_circ_0105114 | 3525 | 11 | 0.62% |
| hsa_circ_0066668 | 4807 | 15 | 0.62% |
| hsa_circ_0005999 | 6089 | 19 | 0.62% |
| hsa_circ_0061438 | 5128 | 16 | 0.62% |
| hsa_circ_0015226 | 3205 | 10 | 0.62% |
| hsa_circ_0095674 | 7372 | 23 | 0.62% |
| hsa_circ_0060840 | 4167 | 13 | 0.62% |
| hsa_circ_0023020 | 3847 | 12 | 0.62% |
| hsa_circ_0040688 | 3847 | 12 | 0.62% |
| hsa_circ_0054581 | 3527 | 11 | 0.62% |
| hsa_circ_0135033 | 3527 | 11 | 0.62% |
| hsa_circ_0009810 | 4489 | 14 | 0.62% |
| hsa_circ_0075999 | 4489 | 14 | 0.62% |
| hsa_circ_0013080 | 3848 | 12 | 0.62% |
| hsa_circ_0036834 | 4169 | 13 | 0.62% |
| hsa_circ_0098552 | 11545 | 36 | 0.62% |
| hsa_circ_0014554 | 3207 | 10 | 0.62% |
| hsa_circ_0011127 | 3528 | 11 | 0.62% |
| hsa_circ_0066003 | 3849 | 12 | 0.62% |
| hsa_circ_0077358 | 3849 | 12 | 0.62% |
| hsa_circ_0091496 | 3849 | 12 | 0.62% |
| hsa_circ_0041402 | 3529 | 11 | 0.62% |
| hsa_circ_0010320 | 3850 | 12 | 0.62% |
| hsa_circ_0034648 | 4171 | 13 | 0.62% |
| hsa_circ_0010279 | 3852 | 12 | 0.62% |
| hsa_circ_0058232 | 3852 | 12 | 0.62% |
| hsa_circ_0091962 | 3852 | 12 | 0.62% |
| hsa_circ_0026702 | 3531 | 11 | 0.62% |
| hsa_circ_0045960 | 3531 | 11 | 0.62% |
| hsa_circ_0017327 | 3532 | 11 | 0.62% |
| hsa_circ_0062058 | 3532 | 11 | 0.62% |
| hsa_circ_0015868 | 3211 | 10 | 0.62% |
| hsa_circ_0064935 | 3211 | 10 | 0.62% |

| | | | |
|---|---|---|---|
| hsa_circ_0108435 | 3211 | 10 | 0.62% |
| hsa_circ_0112133 | 12204 | 38 | 0.62% |
| hsa_circ_0073839 | 3533 | 11 | 0.62% |
| hsa_circ_0076015 | 4497 | 14 | 0.62% |
| hsa_circ_0015768 | 3855 | 12 | 0.62% |
| hsa_circ_0025662 | 3534 | 11 | 0.62% |
| hsa_circ_0060074 | 3534 | 11 | 0.62% |
| hsa_circ_0065922 | 3534 | 11 | 0.62% |
| hsa_circ_0125854 | 9960 | 31 | 0.62% |
| hsa_circ_0124844 | 3213 | 10 | 0.62% |
| hsa_circ_0111776 | 6105 | 19 | 0.62% |
| hsa_circ_0025957 | 3535 | 11 | 0.62% |
| hsa_circ_0034732 | 4501 | 14 | 0.62% |
| hsa_circ_0081647 | 16076 | 50 | 0.62% |
| hsa_circ_0133359 | 11575 | 36 | 0.62% |
| hsa_circ_0027888 | 3537 | 11 | 0.62% |
| hsa_circ_0028791 | 3537 | 11 | 0.62% |
| hsa_circ_0085356 | 3537 | 11 | 0.62% |
| hsa_circ_0007278 | 10933 | 34 | 0.62% |
| hsa_circ_0035725 | 3859 | 12 | 0.62% |
| hsa_circ_0054816 | 3859 | 12 | 0.62% |
| hsa_circ_0010349 | 3216 | 10 | 0.62% |
| hsa_circ_0016908 | 3216 | 10 | 0.62% |
| hsa_circ_0035702 | 3538 | 11 | 0.62% |
| hsa_circ_0083160 | 3538 | 11 | 0.62% |
| hsa_circ_0057566 | 4503 | 14 | 0.62% |
| hsa_circ_0014622 | 3217 | 10 | 0.62% |
| hsa_circ_0028447 | 3539 | 11 | 0.62% |
| hsa_circ_0125000 | 4826 | 15 | 0.62% |
| hsa_circ_0105816 | 3861 | 12 | 0.62% |
| hsa_circ_0016307 | 3218 | 10 | 0.62% |
| hsa_circ_0070103 | 3540 | 11 | 0.62% |
| hsa_circ_0091714 | 3540 | 11 | 0.62% |
| hsa_circ_0092084 | 3540 | 11 | 0.62% |
| hsa_circ_0047977 | 3219 | 10 | 0.62% |
| hsa_circ_0104161 | 3219 | 10 | 0.62% |
| hsa_circ_0039809 | 3541 | 11 | 0.62% |
| hsa_circ_0051299 | 3863 | 12 | 0.62% |
| hsa_circ_0085489 | 4185 | 13 | 0.62% |
| hsa_circ_0059550 | 4507 | 14 | 0.62% |
| hsa_circ_0091015 | 4829 | 15 | 0.62% |
| hsa_circ_0090367 | 4508 | 14 | 0.62% |
| hsa_circ_0067307 | 4186 | 13 | 0.62% |
| hsa_circ_0086594 | 4186 | 13 | 0.62% |
| hsa_circ_0035581 | 3864 | 12 | 0.62% |
| hsa_circ_0012080 | 3542 | 11 | 0.62% |
| hsa_circ_0053434 | 3543 | 11 | 0.62% |
| hsa_circ_0025964 | 4510 | 14 | 0.62% |
| hsa_circ_0047301 | 3866 | 12 | 0.62% |
| hsa_circ_0091930 | 3866 | 12 | 0.62% |

| | | | |
|---|---|---|---|
| hsa_circ_0029546 | 3544 | 11 | 0.62% |
| hsa_circ_0033561 | 3544 | 11 | 0.62% |
| hsa_circ_0081510 | 5155 | 16 | 0.62% |
| hsa_circ_0034882 | 3222 | 10 | 0.62% |
| hsa_circ_0106562 | 3222 | 10 | 0.62% |
| hsa_circ_0119043 | 10312 | 32 | 0.62% |
| hsa_circ_0038486 | 7412 | 23 | 0.62% |
| hsa_circ_0130979 | 8379 | 26 | 0.62% |
| hsa_circ_0033076 | 4512 | 14 | 0.62% |
| hsa_circ_0081826 | 3546 | 11 | 0.62% |
| hsa_circ_0011125 | 4191 | 13 | 0.62% |
| hsa_circ_0038733 | 4191 | 13 | 0.62% |
| hsa_circ_0068656 | 4191 | 13 | 0.62% |
| hsa_circ_0026538 | 3869 | 12 | 0.62% |
| hsa_circ_0134193 | 5159 | 16 | 0.62% |
| hsa_circ_0084795 | 4192 | 13 | 0.62% |
| hsa_circ_0024972 | 3225 | 10 | 0.62% |
| hsa_circ_0074023 | 4838 | 15 | 0.62% |
| hsa_circ_0138002 | 10323 | 32 | 0.62% |
| hsa_circ_0085229 | 3226 | 10 | 0.62% |
| hsa_circ_0098504 | 4194 | 13 | 0.62% |
| hsa_circ_0058545 | 3227 | 10 | 0.62% |
| hsa_circ_0073065 | 3227 | 10 | 0.62% |
| hsa_circ_0069498 | 3550 | 11 | 0.62% |
| hsa_circ_0030038 | 3873 | 12 | 0.62% |
| hsa_circ_0067358 | 3873 | 12 | 0.62% |
| hsa_circ_0057713 | 3228 | 10 | 0.62% |
| hsa_circ_0111957 | 3228 | 10 | 0.62% |
| hsa_circ_0015473 | 3551 | 11 | 0.62% |
| hsa_circ_0027213 | 3551 | 11 | 0.62% |
| hsa_circ_0121439 | 3874 | 12 | 0.62% |
| hsa_circ_0028537 | 4197 | 13 | 0.62% |
| hsa_circ_0001731 | 6458 | 20 | 0.62% |
| hsa_circ_0014807 | 3229 | 10 | 0.62% |
| hsa_circ_0044921 | 3552 | 11 | 0.62% |
| hsa_circ_0017255 | 3875 | 12 | 0.62% |
| hsa_circ_0098182 | 3875 | 12 | 0.62% |
| hsa_circ_0017346 | 4198 | 13 | 0.62% |
| hsa_circ_0039633 | 4198 | 13 | 0.62% |
| hsa_circ_0104993 | 4844 | 15 | 0.62% |
| hsa_circ_0057345 | 5490 | 17 | 0.62% |
| hsa_circ_0060975 | 4522 | 14 | 0.62% |
| hsa_circ_0086636 | 4199 | 13 | 0.62% |
| hsa_circ_0053196 | 3876 | 12 | 0.62% |
| hsa_circ_0051178 | 3553 | 11 | 0.62% |
| hsa_circ_0053331 | 3877 | 12 | 0.62% |
| hsa_circ_0041972 | 3554 | 11 | 0.62% |
| hsa_circ_0078572 | 3554 | 11 | 0.62% |
| hsa_circ_0043980 | 6785 | 21 | 0.62% |
| hsa_circ_0066363 | 3231 | 10 | 0.62% |

| | | | |
|---|---|---|---|
| hsa_circ_0074449 | 4201 | 13 | 0.62% |
| hsa_circ_0091012 | 3878 | 12 | 0.62% |
| hsa_circ_0090569 | 5171 | 16 | 0.62% |
| hsa_circ_0110664 | 4525 | 14 | 0.62% |
| hsa_circ_0118961 | 4525 | 14 | 0.62% |
| hsa_circ_0086756 | 4202 | 13 | 0.62% |
| hsa_circ_0044769 | 3879 | 12 | 0.62% |
| hsa_circ_0088911 | 3879 | 12 | 0.62% |
| hsa_circ_0090143 | 3879 | 12 | 0.62% |
| hsa_circ_0133164 | 3556 | 11 | 0.62% |
| hsa_circ_0034907 | 3880 | 12 | 0.62% |
| hsa_circ_0077724 | 3557 | 11 | 0.62% |
| hsa_circ_0068783 | 9378 | 29 | 0.62% |
| hsa_circ_0027884 | 4204 | 13 | 0.62% |
| hsa_circ_0017323 | 3881 | 12 | 0.62% |
| hsa_circ_0060914 | 3881 | 12 | 0.62% |
| hsa_circ_0075299 | 4528 | 14 | 0.62% |
| hsa_circ_0119440 | 3558 | 11 | 0.62% |
| hsa_circ_0122154 | 3558 | 11 | 0.62% |
| hsa_circ_0024615 | 4205 | 13 | 0.62% |
| hsa_circ_0076491 | 3882 | 12 | 0.62% |
| hsa_circ_0113103 | 3882 | 12 | 0.62% |
| hsa_circ_0119569 | 9382 | 29 | 0.62% |
| hsa_circ_0052059 | 3559 | 11 | 0.62% |
| hsa_circ_0096567 | 3559 | 11 | 0.62% |
| hsa_circ_0061233 | 6795 | 21 | 0.62% |
| hsa_circ_0135442 | 4530 | 14 | 0.62% |
| hsa_circ_0082207 | 3883 | 12 | 0.62% |
| hsa_circ_0084853 | 3883 | 12 | 0.62% |
| hsa_circ_0016427 | 3560 | 11 | 0.62% |
| hsa_circ_0056820 | 3560 | 11 | 0.62% |
| hsa_circ_0065486 | 3884 | 12 | 0.62% |
| hsa_circ_0082548 | 3884 | 12 | 0.62% |
| hsa_circ_0113858 | 3884 | 12 | 0.62% |
| hsa_circ_0036638 | 3561 | 11 | 0.62% |
| hsa_circ_0135636 | 4856 | 15 | 0.62% |
| hsa_circ_0108533 | 3885 | 12 | 0.62% |
| hsa_circ_0104657 | 3562 | 11 | 0.62% |
| hsa_circ_0033326 | 3886 | 12 | 0.62% |
| hsa_circ_0103896 | 3239 | 10 | 0.62% |
| hsa_circ_0121773 | 3239 | 10 | 0.62% |
| hsa_circ_0039798 | 13604 | 42 | 0.62% |
| hsa_circ_0063922 | 3563 | 11 | 0.62% |
| hsa_circ_0084910 | 3563 | 11 | 0.62% |
| hsa_circ_0092240 | 3887 | 12 | 0.62% |
| hsa_circ_0121808 | 43085 | 133 | 0.62% |
| hsa_circ_0094728 | 6480 | 20 | 0.62% |
| hsa_circ_0016791 | 3565 | 11 | 0.62% |
| hsa_circ_0026996 | 3565 | 11 | 0.62% |
| hsa_circ_0136387 | 10372 | 32 | 0.62% |

| | | | |
|---|---|---|---|
| hsa_circ_0090050 | 3890 | 12 | 0.62% |
| hsa_circ_0020405 | 3566 | 11 | 0.62% |
| hsa_circ_0080137 | 3566 | 11 | 0.62% |
| hsa_circ_0083511 | 4863 | 15 | 0.62% |
| hsa_circ_0068488 | 3242 | 10 | 0.62% |
| hsa_circ_0093675 | 12320 | 38 | 0.62% |
| hsa_circ_0062351 | 4539 | 14 | 0.62% |
| hsa_circ_0072207 | 4539 | 14 | 0.62% |
| hsa_circ_0080128 | 4215 | 13 | 0.62% |
| hsa_circ_0061354 | 3891 | 12 | 0.62% |
| hsa_circ_0063573 | 3891 | 12 | 0.62% |
| hsa_circ_0044571 | 3567 | 11 | 0.62% |
| hsa_circ_0038917 | 3243 | 10 | 0.62% |
| hsa_circ_0102903 | 3243 | 10 | 0.62% |
| hsa_circ_0062537 | 4216 | 13 | 0.62% |
| hsa_circ_0117621 | 3892 | 12 | 0.62% |
| hsa_circ_0035752 | 4217 | 13 | 0.62% |
| hsa_circ_0030189 | 3569 | 11 | 0.62% |
| hsa_circ_0043821 | 3569 | 11 | 0.62% |
| hsa_circ_0022725 | 3894 | 12 | 0.62% |
| hsa_circ_0081020 | 4219 | 13 | 0.62% |
| hsa_circ_0007143 | 9737 | 30 | 0.62% |
| hsa_circ_0034639 | 3895 | 12 | 0.62% |
| hsa_circ_0007717 | 14282 | 44 | 0.62% |
| hsa_circ_0017330 | 3246 | 10 | 0.62% |
| hsa_circ_0063803 | 3896 | 12 | 0.62% |
| hsa_circ_0069287 | 3896 | 12 | 0.62% |
| hsa_circ_0027757 | 3248 | 10 | 0.62% |
| hsa_circ_0101641 | 5197 | 16 | 0.62% |
| hsa_circ_0000559 | 3573 | 11 | 0.62% |
| hsa_circ_0046332 | 3573 | 11 | 0.62% |
| hsa_circ_0121807 | 43207 | 133 | 0.62% |
| hsa_circ_0099744 | 7148 | 22 | 0.62% |
| hsa_circ_0043705 | 3574 | 11 | 0.62% |
| hsa_circ_0054312 | 3574 | 11 | 0.62% |
| hsa_circ_0069666 | 3574 | 11 | 0.62% |
| hsa_circ_0044451 | 3899 | 12 | 0.62% |
| hsa_circ_0090144 | 3899 | 12 | 0.62% |
| hsa_circ_0134308 | 6499 | 20 | 0.62% |
| hsa_circ_0118160 | 7799 | 24 | 0.62% |
| hsa_circ_0010589 | 4225 | 13 | 0.62% |
| hsa_circ_0054814 | 3900 | 12 | 0.62% |
| hsa_circ_0067395 | 3900 | 12 | 0.62% |
| hsa_circ_0045859 | 3575 | 11 | 0.62% |
| hsa_circ_0052907 | 3575 | 11 | 0.62% |
| hsa_circ_0035514 | 3250 | 10 | 0.62% |
| hsa_circ_0134136 | 4551 | 14 | 0.62% |
| hsa_circ_0017373 | 3901 | 12 | 0.62% |
| hsa_circ_0072664 | 3901 | 12 | 0.62% |
| hsa_circ_0089788 | 3576 | 11 | 0.62% |

| | | | |
|---|---|---|---|
| hsa_circ_0056732 | 4552 | 14 | 0.62% |
| hsa_circ_0030204 | 3902 | 12 | 0.62% |
| hsa_circ_0029518 | 4228 | 13 | 0.61% |
| hsa_circ_0089214 | 3903 | 12 | 0.61% |
| hsa_circ_0131201 | 3903 | 12 | 0.61% |
| hsa_circ_0102902 | 3578 | 11 | 0.61% |
| hsa_circ_0070251 | 9433 | 29 | 0.61% |
| hsa_circ_0115272 | 5530 | 17 | 0.61% |
| hsa_circ_0032469 | 3253 | 10 | 0.61% |
| hsa_circ_0036068 | 4229 | 13 | 0.61% |
| hsa_circ_0061167 | 4229 | 13 | 0.61% |
| hsa_circ_0091945 | 3904 | 12 | 0.61% |
| hsa_circ_0019834 | 3579 | 11 | 0.61% |
| hsa_circ_0029621 | 3579 | 11 | 0.61% |
| hsa_circ_0090895 | 3580 | 11 | 0.61% |
| hsa_circ_0022870 | 4882 | 15 | 0.61% |
| hsa_circ_0012762 | 7486 | 23 | 0.61% |
| hsa_circ_0051458 | 4557 | 14 | 0.61% |
| hsa_circ_0060787 | 3906 | 12 | 0.61% |
| hsa_circ_0130566 | 3906 | 12 | 0.61% |
| hsa_circ_0019432 | 3581 | 11 | 0.61% |
| hsa_circ_0019688 | 3581 | 11 | 0.61% |
| hsa_circ_0032432 | 3581 | 11 | 0.61% |
| hsa_circ_0042625 | 3581 | 11 | 0.61% |
| hsa_circ_0076476 | 3581 | 11 | 0.61% |
| hsa_circ_0057979 | 3907 | 12 | 0.61% |
| hsa_circ_0107316 | 3907 | 12 | 0.61% |
| hsa_circ_0119513 | 8466 | 26 | 0.61% |
| hsa_circ_0016613 | 3582 | 11 | 0.61% |
| hsa_circ_0012562 | 6513 | 20 | 0.61% |
| hsa_circ_0030477 | 3908 | 12 | 0.61% |
| hsa_circ_0066518 | 3908 | 12 | 0.61% |
| hsa_circ_0126356 | 9771 | 30 | 0.61% |
| hsa_circ_0093288 | 3583 | 11 | 0.61% |
| hsa_circ_0112702 | 12380 | 38 | 0.61% |
| hsa_circ_0095370 | 3258 | 10 | 0.61% |
| hsa_circ_0029509 | 3584 | 11 | 0.61% |
| hsa_circ_0051695 | 3584 | 11 | 0.61% |
| hsa_circ_0027194 | 3585 | 11 | 0.61% |
| hsa_circ_0092978 | 3585 | 11 | 0.61% |
| hsa_circ_0128142 | 3585 | 11 | 0.61% |
| hsa_circ_0021252 | 3911 | 12 | 0.61% |
| hsa_circ_0045080 | 3911 | 12 | 0.61% |
| hsa_circ_0010669 | 4238 | 13 | 0.61% |
| hsa_circ_0080366 | 4238 | 13 | 0.61% |
| hsa_circ_0118550 | 4238 | 13 | 0.61% |
| hsa_circ_0090314 | 3586 | 11 | 0.61% |
| hsa_circ_0136055 | 3586 | 11 | 0.61% |
| hsa_circ_0019123 | 3260 | 10 | 0.61% |
| hsa_circ_0116063 | 9456 | 29 | 0.61% |

| | | | |
|---|---|---|---|
| hsa_circ_0034272 | 4239 | 13 | 0.61% |
| hsa_circ_0029182 | 3913 | 12 | 0.61% |
| hsa_circ_0118435 | 3913 | 12 | 0.61% |
| hsa_circ_0116028 | 3587 | 11 | 0.61% |
| hsa_circ_0095371 | 3261 | 10 | 0.61% |
| hsa_circ_0070288 | 3589 | 11 | 0.61% |
| hsa_circ_0071178 | 3589 | 11 | 0.61% |
| hsa_circ_0083685 | 3589 | 11 | 0.61% |
| hsa_circ_0094911 | 3263 | 10 | 0.61% |
| hsa_circ_0099321 | 4242 | 13 | 0.61% |
| hsa_circ_0031953 | 4569 | 14 | 0.61% |
| hsa_circ_0068994 | 3590 | 11 | 0.61% |
| hsa_circ_0093410 | 3590 | 11 | 0.61% |
| hsa_circ_0039740 | 3264 | 10 | 0.61% |
| hsa_circ_0066807 | 4244 | 13 | 0.61% |
| hsa_circ_0032697 | 3918 | 12 | 0.61% |
| hsa_circ_0136868 | 3918 | 12 | 0.61% |
| hsa_circ_0003702 | 3265 | 10 | 0.61% |
| hsa_circ_0139323 | 4245 | 13 | 0.61% |
| hsa_circ_0026839 | 3592 | 11 | 0.61% |
| hsa_circ_0036875 | 3592 | 11 | 0.61% |
| hsa_circ_0126223 | 3592 | 11 | 0.61% |
| hsa_circ_0113532 | 9797 | 30 | 0.61% |
| hsa_circ_0014805 | 3919 | 12 | 0.61% |
| hsa_circ_0034717 | 3919 | 12 | 0.61% |
| hsa_circ_0025990 | 4246 | 13 | 0.61% |
| hsa_circ_0086059 | 3593 | 11 | 0.61% |
| hsa_circ_0125063 | 9146 | 28 | 0.61% |
| hsa_circ_0009510 | 3920 | 12 | 0.61% |
| hsa_circ_0124703 | 3920 | 12 | 0.61% |
| hsa_circ_0012040 | 4247 | 13 | 0.61% |
| hsa_circ_0074113 | 3267 | 10 | 0.61% |
| hsa_circ_0124759 | 7188 | 22 | 0.61% |
| hsa_circ_0022193 | 3594 | 11 | 0.61% |
| hsa_circ_0049732 | 3594 | 11 | 0.61% |
| hsa_circ_0027362 | 3921 | 12 | 0.61% |
| hsa_circ_0066833 | 3921 | 12 | 0.61% |
| hsa_circ_0076270 | 3921 | 12 | 0.61% |
| hsa_circ_0118972 | 3921 | 12 | 0.61% |
| hsa_circ_0135736 | 5556 | 17 | 0.61% |
| hsa_circ_0121016 | 7191 | 22 | 0.61% |
| hsa_circ_0009444 | 3596 | 11 | 0.61% |
| hsa_circ_0052501 | 3596 | 11 | 0.61% |
| hsa_circ_0111052 | 3923 | 12 | 0.61% |
| hsa_circ_0037411 | 4250 | 13 | 0.61% |
| hsa_circ_0045076 | 4250 | 13 | 0.61% |
| hsa_circ_0007021 | 8827 | 27 | 0.61% |
| hsa_circ_0056476 | 4577 | 14 | 0.61% |
| hsa_circ_0121300 | 9481 | 29 | 0.61% |
| hsa_circ_0019977 | 3924 | 12 | 0.61% |

| | | | |
|---|---|---|---|
| hsa_circ_0029300 | 3597 | 11 | 0.61% |
| hsa_circ_0126222 | 3597 | 11 | 0.61% |
| hsa_circ_0049764 | 13081 | 40 | 0.61% |
| hsa_circ_0080668 | 4906 | 15 | 0.61% |
| hsa_circ_0041207 | 3925 | 12 | 0.61% |
| hsa_circ_0096776 | 3925 | 12 | 0.61% |
| hsa_circ_0071616 | 3598 | 11 | 0.61% |
| hsa_circ_0062547 | 3271 | 10 | 0.61% |
| hsa_circ_0053188 | 3926 | 12 | 0.61% |
| hsa_circ_0070741 | 9488 | 29 | 0.61% |
| hsa_circ_0065723 | 3599 | 11 | 0.61% |
| hsa_circ_0090386 | 3599 | 11 | 0.61% |
| hsa_circ_0123946 | 3599 | 11 | 0.61% |
| hsa_circ_0003470 | 12436 | 38 | 0.61% |
| hsa_circ_0112451 | 3600 | 11 | 0.61% |
| hsa_circ_0124350 | 8182 | 25 | 0.61% |
| hsa_circ_0093030 | 11455 | 35 | 0.61% |
| hsa_circ_0025348 | 3273 | 10 | 0.61% |
| hsa_circ_0109382 | 4255 | 13 | 0.61% |
| hsa_circ_0063687 | 3928 | 12 | 0.61% |
| hsa_circ_0021028 | 3601 | 11 | 0.61% |
| hsa_circ_0081640 | 16369 | 50 | 0.61% |
| hsa_circ_0035293 | 4257 | 13 | 0.61% |
| hsa_circ_0081764 | 4257 | 13 | 0.61% |
| hsa_circ_0083007 | 4257 | 13 | 0.61% |
| hsa_circ_0124472 | 8515 | 26 | 0.61% |
| hsa_circ_0064488 | 3930 | 12 | 0.61% |
| hsa_circ_0065743 | 3930 | 12 | 0.61% |
| hsa_circ_0091975 | 3930 | 12 | 0.61% |
| hsa_circ_0043031 | 4258 | 13 | 0.61% |
| hsa_circ_0128517 | 4258 | 13 | 0.61% |
| hsa_circ_0030040 | 3603 | 11 | 0.61% |
| hsa_circ_0058836 | 3603 | 11 | 0.61% |
| hsa_circ_0086897 | 3603 | 11 | 0.61% |
| hsa_circ_0005590 | 3931 | 12 | 0.61% |
| hsa_circ_0104670 | 4914 | 15 | 0.61% |
| hsa_circ_0075754 | 3276 | 10 | 0.61% |
| hsa_circ_0126373 | 3276 | 10 | 0.61% |
| hsa_circ_0056818 | 4259 | 13 | 0.61% |
| hsa_circ_0012531 | 3604 | 11 | 0.61% |
| hsa_circ_0034710 | 3604 | 11 | 0.61% |
| hsa_circ_0043877 | 3932 | 12 | 0.61% |
| hsa_circ_0085910 | 3932 | 12 | 0.61% |
| hsa_circ_0029978 | 4260 | 13 | 0.61% |
| hsa_circ_0016143 | 3277 | 10 | 0.61% |
| hsa_circ_0037529 | 3605 | 11 | 0.61% |
| hsa_circ_0046770 | 3605 | 11 | 0.61% |
| hsa_circ_0102095 | 3605 | 11 | 0.61% |
| hsa_circ_0040800 | 3933 | 12 | 0.61% |
| hsa_circ_0042587 | 3933 | 12 | 0.61% |

| | | | |
|---|---|---|---|
| hsa_circ_0118971 | 3933 | 12 | 0.61% |
| hsa_circ_0000046 | 12456 | 38 | 0.61% |
| hsa_circ_0052689 | 4917 | 15 | 0.61% |
| hsa_circ_0066293 | 3606 | 11 | 0.61% |
| hsa_circ_0090035 | 3606 | 11 | 0.61% |
| hsa_circ_0075148 | 3934 | 12 | 0.61% |
| hsa_circ_0050233 | 8524 | 26 | 0.61% |
| hsa_circ_0020406 | 3607 | 11 | 0.61% |
| hsa_circ_0038252 | 3607 | 11 | 0.61% |
| hsa_circ_0084329 | 3607 | 11 | 0.61% |
| hsa_circ_0015086 | 3935 | 12 | 0.61% |
| hsa_circ_0060050 | 3935 | 12 | 0.61% |
| hsa_circ_0052635 | 4263 | 13 | 0.61% |
| hsa_circ_0030869 | 5248 | 16 | 0.61% |
| hsa_circ_0048645 | 3936 | 12 | 0.61% |
| hsa_circ_0089189 | 3936 | 12 | 0.61% |
| hsa_circ_0092167 | 3608 | 11 | 0.61% |
| hsa_circ_0017222 | 3280 | 10 | 0.61% |
| hsa_circ_0040073 | 4593 | 14 | 0.61% |
| hsa_circ_0031934 | 4265 | 13 | 0.61% |
| hsa_circ_0014620 | 3609 | 11 | 0.61% |
| hsa_circ_0046038 | 3609 | 11 | 0.61% |
| hsa_circ_0046768 | 3609 | 11 | 0.61% |
| hsa_circ_0059079 | 3281 | 10 | 0.61% |
| hsa_circ_0128460 | 8532 | 26 | 0.61% |
| hsa_circ_0048865 | 4266 | 13 | 0.61% |
| hsa_circ_0065104 | 3938 | 12 | 0.61% |
| hsa_circ_0043239 | 3610 | 11 | 0.61% |
| hsa_circ_0045267 | 3610 | 11 | 0.61% |
| hsa_circ_0086847 | 3610 | 11 | 0.61% |
| hsa_circ_0032486 | 3282 | 10 | 0.61% |
| hsa_circ_0139980 | 7222 | 22 | 0.61% |
| hsa_circ_0084594 | 3940 | 12 | 0.61% |
| hsa_circ_0072083 | 4598 | 14 | 0.61% |
| hsa_circ_0094344 | 5255 | 16 | 0.61% |
| hsa_circ_0119616 | 26935 | 82 | 0.61% |
| hsa_circ_0088828 | 15111 | 46 | 0.61% |
| hsa_circ_0026211 | 4271 | 13 | 0.61% |
| hsa_circ_0017269 | 3614 | 11 | 0.61% |
| hsa_circ_0083755 | 3614 | 11 | 0.61% |
| hsa_circ_0027649 | 3943 | 12 | 0.61% |
| hsa_circ_0054542 | 3943 | 12 | 0.61% |
| hsa_circ_0054806 | 3943 | 12 | 0.61% |
| hsa_circ_0081810 | 3943 | 12 | 0.61% |
| hsa_circ_0047403 | 4272 | 13 | 0.61% |
| hsa_circ_0016130 | 5258 | 16 | 0.61% |
| hsa_circ_0074278 | 3615 | 11 | 0.61% |
| hsa_circ_0053805 | 3944 | 12 | 0.61% |
| hsa_circ_0088671 | 3944 | 12 | 0.61% |
| hsa_circ_0133178 | 3944 | 12 | 0.61% |

| | | | |
|---|---|---|---|
| hsa_circ_0031618 | 4273 | 13 | 0.61% |
| hsa_circ_0111955 | 3287 | 10 | 0.61% |
| hsa_circ_0018215 | 3616 | 11 | 0.61% |
| hsa_circ_0090684 | 4274 | 13 | 0.61% |
| hsa_circ_0095718 | 3617 | 11 | 0.61% |
| hsa_circ_0015704 | 4933 | 15 | 0.61% |
| hsa_circ_0031109 | 3947 | 12 | 0.61% |
| hsa_circ_0099212 | 4276 | 13 | 0.61% |
| hsa_circ_0101599 | 4606 | 14 | 0.61% |
| hsa_circ_0115634 | 4277 | 13 | 0.61% |
| hsa_circ_0126828 | 4277 | 13 | 0.61% |
| hsa_circ_0056724 | 3948 | 12 | 0.61% |
| hsa_circ_0015699 | 3619 | 11 | 0.61% |
| hsa_circ_0017268 | 3619 | 11 | 0.61% |
| hsa_circ_0045692 | 3619 | 11 | 0.61% |
| hsa_circ_0053447 | 3619 | 11 | 0.61% |
| hsa_circ_0065587 | 3619 | 11 | 0.61% |
| hsa_circ_0084698 | 3619 | 11 | 0.61% |
| hsa_circ_0073751 | 3290 | 10 | 0.61% |
| hsa_circ_0070147 | 4607 | 14 | 0.61% |
| hsa_circ_0031925 | 4278 | 13 | 0.61% |
| hsa_circ_0035765 | 4278 | 13 | 0.61% |
| hsa_circ_0053665 | 4278 | 13 | 0.61% |
| hsa_circ_0135261 | 53969 | 164 | 0.61% |
| hsa_circ_0123595 | 8227 | 25 | 0.61% |
| hsa_circ_0013376 | 4608 | 14 | 0.61% |
| hsa_circ_0125830 | 6254 | 19 | 0.61% |
| hsa_circ_0089168 | 3950 | 12 | 0.61% |
| hsa_circ_0074827 | 3292 | 10 | 0.61% |
| hsa_circ_0111954 | 3292 | 10 | 0.61% |
| hsa_circ_0125125 | 4280 | 13 | 0.61% |
| hsa_circ_0111377 | 3622 | 11 | 0.61% |
| hsa_circ_0124182 | 9550 | 29 | 0.61% |
| hsa_circ_0068675 | 3952 | 12 | 0.61% |
| hsa_circ_0135432 | 3952 | 12 | 0.61% |
| hsa_circ_0063823 | 4611 | 14 | 0.61% |
| hsa_circ_0019817 | 3623 | 11 | 0.61% |
| hsa_circ_0087657 | 3623 | 11 | 0.61% |
| hsa_circ_0073951 | 5600 | 17 | 0.61% |
| hsa_circ_0063102 | 3953 | 12 | 0.61% |
| hsa_circ_0077059 | 3953 | 12 | 0.61% |
| hsa_circ_0078107 | 3953 | 12 | 0.61% |
| hsa_circ_0131670 | 14165 | 43 | 0.61% |
| hsa_circ_0078108 | 3624 | 11 | 0.61% |
| hsa_circ_0070402 | 4283 | 13 | 0.61% |
| hsa_circ_0104050 | 11205 | 34 | 0.61% |
| hsa_circ_0060851 | 4944 | 15 | 0.61% |
| hsa_circ_0131226 | 13185 | 40 | 0.61% |
| hsa_circ_0032953 | 3626 | 11 | 0.61% |
| hsa_circ_0073716 | 3626 | 11 | 0.61% |

| | | | |
|---|---|---|---|
| hsa_circ_0024891 | 3956 | 12 | 0.61% |
| hsa_circ_0102096 | 4286 | 13 | 0.61% |
| hsa_circ_0136210 | 10222 | 31 | 0.61% |
| hsa_circ_0027247 | 3957 | 12 | 0.61% |
| hsa_circ_0036482 | 3957 | 12 | 0.61% |
| hsa_circ_0022342 | 4287 | 13 | 0.61% |
| hsa_circ_0070044 | 3298 | 10 | 0.61% |
| hsa_circ_0088217 | 3298 | 10 | 0.61% |
| hsa_circ_0027573 | 3628 | 11 | 0.61% |
| hsa_circ_0073511 | 3628 | 11 | 0.61% |
| hsa_circ_0107540 | 3628 | 11 | 0.61% |
| hsa_circ_0028922 | 5607 | 17 | 0.61% |
| hsa_circ_0033345 | 3958 | 12 | 0.61% |
| hsa_circ_0112704 | 3958 | 12 | 0.61% |
| hsa_circ_0105832 | 18144 | 55 | 0.61% |
| hsa_circ_0136918 | 6268 | 19 | 0.61% |
| hsa_circ_0119556 | 9567 | 29 | 0.61% |
| hsa_circ_0036814 | 3299 | 10 | 0.61% |
| hsa_circ_0114737 | 3629 | 11 | 0.61% |
| hsa_circ_0033096 | 4289 | 13 | 0.61% |
| hsa_circ_0038734 | 4289 | 13 | 0.61% |
| hsa_circ_0014688 | 3960 | 12 | 0.61% |
| hsa_circ_0122985 | 12542 | 38 | 0.61% |
| hsa_circ_0077027 | 4621 | 14 | 0.61% |
| hsa_circ_0097104 | 4291 | 13 | 0.61% |
| hsa_circ_0108208 | 3631 | 11 | 0.61% |
| hsa_circ_0130256 | 3631 | 11 | 0.61% |
| hsa_circ_0135637 | 4952 | 15 | 0.61% |
| hsa_circ_0011065 | 4622 | 14 | 0.61% |
| hsa_circ_0082185 | 4622 | 14 | 0.61% |
| hsa_circ_0020225 | 3962 | 12 | 0.61% |
| hsa_circ_0054043 | 3962 | 12 | 0.61% |
| hsa_circ_0034773 | 4623 | 14 | 0.61% |
| hsa_circ_0140610 | 3963 | 12 | 0.61% |
| hsa_circ_0096290 | 4624 | 14 | 0.61% |
| hsa_circ_0124996 | 23453 | 71 | 0.61% |
| hsa_circ_0013706 | 3964 | 12 | 0.61% |
| hsa_circ_0139801 | 3964 | 12 | 0.61% |
| hsa_circ_0043822 | 3634 | 11 | 0.61% |
| hsa_circ_0055817 | 4295 | 13 | 0.61% |
| hsa_circ_0055458 | 3304 | 10 | 0.61% |
| hsa_circ_0009977 | 3965 | 12 | 0.61% |
| hsa_circ_0065063 | 4626 | 14 | 0.61% |
| hsa_circ_0077021 | 3635 | 11 | 0.61% |
| hsa_circ_0129977 | 3635 | 11 | 0.61% |
| hsa_circ_0044865 | 3305 | 10 | 0.61% |
| hsa_circ_0071990 | 3305 | 10 | 0.61% |
| hsa_circ_0115278 | 3636 | 11 | 0.61% |
| hsa_circ_0030142 | 4628 | 14 | 0.61% |
| hsa_circ_0041812 | 3306 | 10 | 0.60% |

| | | | |
|---|---|---|---|
| hsa_circ_0107675 | 3306 | 10 | 0.60% |
| hsa_circ_0063074 | 4298 | 13 | 0.60% |
| hsa_circ_0045627 | 3637 | 11 | 0.60% |
| hsa_circ_0049929 | 3637 | 11 | 0.60% |
| hsa_circ_0012561 | 6613 | 20 | 0.60% |
| hsa_circ_0013415 | 3968 | 12 | 0.60% |
| hsa_circ_0028112 | 3968 | 12 | 0.60% |
| hsa_circ_0070629 | 3307 | 10 | 0.60% |
| hsa_circ_0090682 | 3969 | 12 | 0.60% |
| hsa_circ_0078833 | 10916 | 33 | 0.60% |
| hsa_circ_0030929 | 3308 | 10 | 0.60% |
| hsa_circ_0109071 | 3309 | 10 | 0.60% |
| hsa_circ_0058003 | 3971 | 12 | 0.60% |
| hsa_circ_0107573 | 16878 | 51 | 0.60% |
| hsa_circ_0006445 | 6288 | 19 | 0.60% |
| hsa_circ_0062346 | 4634 | 14 | 0.60% |
| hsa_circ_0101915 | 4634 | 14 | 0.60% |
| hsa_circ_0074464 | 3972 | 12 | 0.60% |
| hsa_circ_0078471 | 3972 | 12 | 0.60% |
| hsa_circ_0036714 | 3641 | 11 | 0.60% |
| hsa_circ_0079213 | 3641 | 11 | 0.60% |
| hsa_circ_0095368 | 3310 | 10 | 0.60% |
| hsa_circ_0043235 | 3642 | 11 | 0.60% |
| hsa_circ_0087447 | 3642 | 11 | 0.60% |
| hsa_circ_0067192 | 3311 | 10 | 0.60% |
| hsa_circ_0089833 | 3311 | 10 | 0.60% |
| hsa_circ_0140709 | 4967 | 15 | 0.60% |
| hsa_circ_0032634 | 4305 | 13 | 0.60% |
| hsa_circ_0135480 | 3974 | 12 | 0.60% |
| hsa_circ_0058544 | 3312 | 10 | 0.60% |
| hsa_circ_0138788 | 4637 | 14 | 0.60% |
| hsa_circ_0065778 | 17886 | 54 | 0.60% |
| hsa_circ_0137293 | 4306 | 13 | 0.60% |
| hsa_circ_0012491 | 3975 | 12 | 0.60% |
| hsa_circ_0036995 | 3644 | 11 | 0.60% |
| hsa_circ_0114427 | 3644 | 11 | 0.60% |
| hsa_circ_0129838 | 6957 | 21 | 0.60% |
| hsa_circ_0035600 | 4638 | 14 | 0.60% |
| hsa_circ_0059300 | 3313 | 10 | 0.60% |
| hsa_circ_0095369 | 3313 | 10 | 0.60% |
| hsa_circ_0046479 | 3645 | 11 | 0.60% |
| hsa_circ_0068344 | 3645 | 11 | 0.60% |
| hsa_circ_0117624 | 3977 | 12 | 0.60% |
| hsa_circ_0027891 | 3646 | 11 | 0.60% |
| hsa_circ_0121507 | 3646 | 11 | 0.60% |
| hsa_circ_0082216 | 4641 | 14 | 0.60% |
| hsa_circ_0082508 | 3315 | 10 | 0.60% |
| hsa_circ_0016585 | 3647 | 11 | 0.60% |
| hsa_circ_0025655 | 3647 | 11 | 0.60% |
| hsa_circ_0106524 | 15252 | 46 | 0.60% |

| | | | |
|---|---|---|---|
| hsa_circ_0034204 | 3316 | 10 | 0.60% |
| hsa_circ_0026700 | 3648 | 11 | 0.60% |
| hsa_circ_0076338 | 3648 | 11 | 0.60% |
| hsa_circ_0091625 | 3648 | 11 | 0.60% |
| hsa_circ_0047041 | 3980 | 12 | 0.60% |
| hsa_circ_0027756 | 3317 | 10 | 0.60% |
| hsa_circ_0080131 | 3649 | 11 | 0.60% |
| hsa_circ_0088567 | 3649 | 11 | 0.60% |
| hsa_circ_0027426 | 3650 | 11 | 0.60% |
| hsa_circ_0028767 | 3650 | 11 | 0.60% |
| hsa_circ_0084370 | 4314 | 13 | 0.60% |
| hsa_circ_0027599 | 4646 | 14 | 0.60% |
| hsa_circ_0017397 | 5310 | 16 | 0.60% |
| hsa_circ_0130936 | 3319 | 10 | 0.60% |
| hsa_circ_0018745 | 3651 | 11 | 0.60% |
| hsa_circ_0087704 | 3651 | 11 | 0.60% |
| hsa_circ_0036235 | 3983 | 12 | 0.60% |
| hsa_circ_0123591 | 8298 | 25 | 0.60% |
| hsa_circ_0040071 | 4647 | 14 | 0.60% |
| hsa_circ_0041423 | 4647 | 14 | 0.60% |
| hsa_circ_0000845 | 11620 | 35 | 0.60% |
| hsa_circ_0073721 | 4980 | 15 | 0.60% |
| hsa_circ_0057567 | 4648 | 14 | 0.60% |
| hsa_circ_0010694 | 3985 | 12 | 0.60% |
| hsa_circ_0040297 | 3985 | 12 | 0.60% |
| hsa_circ_0072163 | 3653 | 11 | 0.60% |
| hsa_circ_0076159 | 3653 | 11 | 0.60% |
| hsa_circ_0086351 | 3321 | 10 | 0.60% |
| hsa_circ_0013069 | 3986 | 12 | 0.60% |
| hsa_circ_0009848 | 3654 | 11 | 0.60% |
| hsa_circ_0011491 | 3654 | 11 | 0.60% |
| hsa_circ_0043706 | 3654 | 11 | 0.60% |
| hsa_circ_0112433 | 9966 | 30 | 0.60% |
| hsa_circ_0110712 | 4319 | 13 | 0.60% |
| hsa_circ_0047517 | 3987 | 12 | 0.60% |
| hsa_circ_0113690 | 3987 | 12 | 0.60% |
| hsa_circ_0136742 | 3987 | 12 | 0.60% |
| hsa_circ_0007598 | 10301 | 31 | 0.60% |
| hsa_circ_0059353 | 4320 | 13 | 0.60% |
| hsa_circ_0054882 | 3988 | 12 | 0.60% |
| hsa_circ_0095404 | 16950 | 51 | 0.60% |
| hsa_circ_0052435 | 4653 | 14 | 0.60% |
| hsa_circ_0054569 | 3656 | 11 | 0.60% |
| hsa_circ_0071195 | 3656 | 11 | 0.60% |
| hsa_circ_0105173 | 3656 | 11 | 0.60% |
| hsa_circ_0047206 | 3324 | 10 | 0.60% |
| hsa_circ_0047243 | 3324 | 10 | 0.60% |
| hsa_circ_0009560 | 3989 | 12 | 0.60% |
| hsa_circ_0010273 | 3989 | 12 | 0.60% |
| hsa_circ_0039020 | 5319 | 16 | 0.60% |

| | | | |
|---|---|---|---|
| hsa_circ_0081205 | 3657 | 11 | 0.60% |
| hsa_circ_0090254 | 4322 | 13 | 0.60% |
| hsa_circ_0065144 | 4987 | 15 | 0.60% |
| hsa_circ_0039625 | 4323 | 13 | 0.60% |
| hsa_circ_0090335 | 4656 | 14 | 0.60% |
| hsa_circ_0065361 | 3991 | 12 | 0.60% |
| hsa_circ_0022192 | 3659 | 11 | 0.60% |
| hsa_circ_0056294 | 3659 | 11 | 0.60% |
| hsa_circ_0068009 | 4990 | 15 | 0.60% |
| hsa_circ_0051059 | 3992 | 12 | 0.60% |
| hsa_circ_0075155 | 3660 | 11 | 0.60% |
| hsa_circ_0004333 | 6655 | 20 | 0.60% |
| hsa_circ_0073777 | 3993 | 12 | 0.60% |
| hsa_circ_0119389 | 9984 | 30 | 0.60% |
| hsa_circ_0105663 | 3994 | 12 | 0.60% |
| hsa_circ_0064094 | 4327 | 13 | 0.60% |
| hsa_circ_0023418 | 7323 | 22 | 0.60% |
| hsa_circ_0010741 | 3662 | 11 | 0.60% |
| hsa_circ_0032295 | 3662 | 11 | 0.60% |
| hsa_circ_0059709 | 3662 | 11 | 0.60% |
| hsa_circ_0065594 | 3662 | 11 | 0.60% |
| hsa_circ_0027634 | 3995 | 12 | 0.60% |
| hsa_circ_0069870 | 3995 | 12 | 0.60% |
| hsa_circ_0025991 | 4328 | 13 | 0.60% |
| hsa_circ_0040504 | 3663 | 11 | 0.60% |
| hsa_circ_0043943 | 3663 | 11 | 0.60% |
| hsa_circ_0047141 | 3663 | 11 | 0.60% |
| hsa_circ_0077118 | 3663 | 11 | 0.60% |
| hsa_circ_0091843 | 3663 | 11 | 0.60% |
| hsa_circ_0078794 | 9658 | 29 | 0.60% |
| hsa_circ_0078812 | 9658 | 29 | 0.60% |
| hsa_circ_0078862 | 9658 | 29 | 0.60% |
| hsa_circ_0078968 | 9658 | 29 | 0.60% |
| hsa_circ_0016027 | 3664 | 11 | 0.60% |
| hsa_circ_0071297 | 3664 | 11 | 0.60% |
| hsa_circ_0041504 | 3331 | 10 | 0.60% |
| hsa_circ_0067978 | 3331 | 10 | 0.60% |
| hsa_circ_0071948 | 3331 | 10 | 0.60% |
| hsa_circ_0057122 | 5996 | 18 | 0.60% |
| hsa_circ_0019513 | 4331 | 13 | 0.60% |
| hsa_circ_0045754 | 4331 | 13 | 0.60% |
| hsa_circ_0034195 | 3665 | 11 | 0.60% |
| hsa_circ_0033102 | 3999 | 12 | 0.60% |
| hsa_circ_0040102 | 3666 | 11 | 0.60% |
| hsa_circ_0075997 | 4666 | 14 | 0.60% |
| hsa_circ_0082039 | 4333 | 13 | 0.60% |
| hsa_circ_0111932 | 4333 | 13 | 0.60% |
| hsa_circ_0122607 | 4333 | 13 | 0.60% |
| hsa_circ_0018825 | 8000 | 24 | 0.60% |
| hsa_circ_0018662 | 4000 | 12 | 0.60% |

| | | | |
|---|---|---|---|
| hsa_circ_0023192 | 4000 | 12 | 0.60% |
| hsa_circ_0038172 | 3667 | 11 | 0.60% |
| hsa_circ_0134167 | 8668 | 26 | 0.60% |
| hsa_circ_0032862 | 4334 | 13 | 0.60% |
| hsa_circ_0136488 | 4001 | 12 | 0.60% |
| hsa_circ_0055980 | 4002 | 12 | 0.60% |
| hsa_circ_0091512 | 4002 | 12 | 0.60% |
| hsa_circ_0031186 | 3335 | 10 | 0.60% |
| hsa_circ_0109899 | 3335 | 10 | 0.60% |
| hsa_circ_0002790 | 3669 | 11 | 0.60% |
| hsa_circ_0067950 | 3669 | 11 | 0.60% |
| hsa_circ_0090713 | 3669 | 11 | 0.60% |
| hsa_circ_0041887 | 4003 | 12 | 0.60% |
| hsa_circ_0019901 | 3670 | 11 | 0.60% |
| hsa_circ_0020863 | 3670 | 11 | 0.60% |
| hsa_circ_0030074 | 3670 | 11 | 0.60% |
| hsa_circ_0024478 | 4004 | 12 | 0.60% |
| hsa_circ_0130473 | 4004 | 12 | 0.60% |
| hsa_circ_0009310 | 15016 | 45 | 0.60% |
| hsa_circ_0061705 | 3671 | 11 | 0.60% |
| hsa_circ_0131127 | 3671 | 11 | 0.60% |
| hsa_circ_0119283 | 4005 | 12 | 0.60% |
| hsa_circ_0120605 | 4005 | 12 | 0.60% |
| hsa_circ_0098111 | 11014 | 33 | 0.60% |
| hsa_circ_0122436 | 4673 | 14 | 0.60% |
| hsa_circ_0001331 | 6342 | 19 | 0.60% |
| hsa_circ_0027823 | 3672 | 11 | 0.60% |
| hsa_circ_0027133 | 4006 | 12 | 0.60% |
| hsa_circ_0034711 | 4006 | 12 | 0.60% |
| hsa_circ_0070671 | 4006 | 12 | 0.60% |
| hsa_circ_0084390 | 4340 | 13 | 0.60% |
| hsa_circ_0116957 | 11352 | 34 | 0.60% |
| hsa_circ_0025671 | 3673 | 11 | 0.60% |
| hsa_circ_0064850 | 4007 | 12 | 0.60% |
| hsa_circ_0084214 | 4007 | 12 | 0.60% |
| hsa_circ_0060852 | 5009 | 15 | 0.60% |
| hsa_circ_0003067 | 10019 | 30 | 0.60% |
| hsa_circ_0015031 | 3674 | 11 | 0.60% |
| hsa_circ_0020046 | 3674 | 11 | 0.60% |
| hsa_circ_0041305 | 3674 | 11 | 0.60% |
| hsa_circ_0084355 | 3674 | 11 | 0.60% |
| hsa_circ_0088963 | 3674 | 11 | 0.60% |
| hsa_circ_0035793 | 3340 | 10 | 0.60% |
| hsa_circ_0076311 | 3340 | 10 | 0.60% |
| hsa_circ_0052118 | 12359 | 37 | 0.60% |
| hsa_circ_0060688 | 4343 | 13 | 0.60% |
| hsa_circ_0086885 | 3675 | 11 | 0.60% |
| hsa_circ_0114491 | 3675 | 11 | 0.60% |
| hsa_circ_0136580 | 6683 | 20 | 0.60% |
| hsa_circ_0081639 | 16708 | 50 | 0.60% |

| | | | |
|---|---|---|---|
| hsa_circ_0054336 | 3676 | 11 | 0.60% |
| hsa_circ_0090893 | 3676 | 11 | 0.60% |
| hsa_circ_0093995 | 10360 | 31 | 0.60% |
| hsa_circ_0054792 | 4011 | 12 | 0.60% |
| hsa_circ_0137126 | 4011 | 12 | 0.60% |
| hsa_circ_0061733 | 3677 | 11 | 0.60% |
| hsa_circ_0004758 | 4346 | 13 | 0.60% |
| hsa_circ_0035224 | 4012 | 12 | 0.60% |
| hsa_circ_0118821 | 4012 | 12 | 0.60% |
| hsa_circ_0073513 | 4347 | 13 | 0.60% |
| hsa_circ_0055457 | 3344 | 10 | 0.60% |
| hsa_circ_0085234 | 4013 | 12 | 0.60% |
| hsa_circ_0115718 | 8695 | 26 | 0.60% |
| hsa_circ_0011828 | 4683 | 14 | 0.60% |
| hsa_circ_0077042 | 4014 | 12 | 0.60% |
| hsa_circ_0075083 | 4349 | 13 | 0.60% |
| hsa_circ_0033105 | 3680 | 11 | 0.60% |
| hsa_circ_0084367 | 3680 | 11 | 0.60% |
| hsa_circ_0045392 | 4684 | 14 | 0.60% |
| hsa_circ_0057509 | 4015 | 12 | 0.60% |
| hsa_circ_0076050 | 4015 | 12 | 0.60% |
| hsa_circ_0042915 | 3681 | 11 | 0.60% |
| hsa_circ_0084338 | 3681 | 11 | 0.60% |
| hsa_circ_0128734 | 3681 | 11 | 0.60% |
| hsa_circ_0027704 | 4685 | 14 | 0.60% |
| hsa_circ_0061313 | 4016 | 12 | 0.60% |
| hsa_circ_0083191 | 4016 | 12 | 0.60% |
| hsa_circ_0057996 | 4351 | 13 | 0.60% |
| hsa_circ_0088845 | 4351 | 13 | 0.60% |
| hsa_circ_0122269 | 3682 | 11 | 0.60% |
| hsa_circ_0029180 | 4017 | 12 | 0.60% |
| hsa_circ_0060788 | 4017 | 12 | 0.60% |
| hsa_circ_0030821 | 3683 | 11 | 0.60% |
| hsa_circ_0100882 | 3683 | 11 | 0.60% |
| hsa_circ_0139579 | 6027 | 18 | 0.60% |
| hsa_circ_0013071 | 4018 | 12 | 0.60% |
| hsa_circ_0015842 | 3684 | 11 | 0.60% |
| hsa_circ_0076475 | 3684 | 11 | 0.60% |
| hsa_circ_0088922 | 4354 | 13 | 0.60% |
| hsa_circ_0052636 | 4355 | 13 | 0.60% |
| hsa_circ_0085357 | 3685 | 11 | 0.60% |
| hsa_circ_0096563 | 3685 | 11 | 0.60% |
| hsa_circ_0116428 | 4021 | 12 | 0.60% |
| hsa_circ_0023994 | 3686 | 11 | 0.60% |
| hsa_circ_0052723 | 3686 | 11 | 0.60% |
| hsa_circ_0093075 | 3351 | 10 | 0.60% |
| hsa_circ_0048866 | 4357 | 13 | 0.60% |
| hsa_circ_0063802 | 4022 | 12 | 0.60% |
| hsa_circ_0041973 | 3687 | 11 | 0.60% |
| hsa_circ_0071179 | 3687 | 11 | 0.60% |

| | | | |
|---|---|---|---|
| hsa_circ_0139011 | 3687 | 11 | 0.60% |
| hsa_circ_0070564 | 5699 | 17 | 0.60% |
| hsa_circ_0136101 | 9722 | 29 | 0.60% |
| hsa_circ_0125645 | 4023 | 12 | 0.60% |
| hsa_circ_0105798 | 13075 | 39 | 0.60% |
| hsa_circ_0117171 | 3688 | 11 | 0.60% |
| hsa_circ_0098745 | 12741 | 38 | 0.60% |
| hsa_circ_0038028 | 10059 | 30 | 0.60% |
| hsa_circ_0011317 | 4359 | 13 | 0.60% |
| hsa_circ_0074631 | 4359 | 13 | 0.60% |
| hsa_circ_0122984 | 12742 | 38 | 0.60% |
| hsa_circ_0017122 | 4024 | 12 | 0.60% |
| hsa_circ_0017366 | 4024 | 12 | 0.60% |
| hsa_circ_0081382 | 4695 | 14 | 0.60% |
| hsa_circ_0035327 | 3689 | 11 | 0.60% |
| hsa_circ_0068878 | 3689 | 11 | 0.60% |
| hsa_circ_0126568 | 5366 | 16 | 0.60% |
| hsa_circ_0015693 | 4360 | 13 | 0.60% |
| hsa_circ_0086903 | 4360 | 13 | 0.60% |
| hsa_circ_0017328 | 3354 | 10 | 0.60% |
| hsa_circ_0036724 | 4025 | 12 | 0.60% |
| hsa_circ_0073316 | 4696 | 14 | 0.60% |
| hsa_circ_0037083 | 3690 | 11 | 0.60% |
| hsa_circ_0111463 | 16438 | 49 | 0.60% |
| hsa_circ_0132012 | 4026 | 12 | 0.60% |
| hsa_circ_0011787 | 3355 | 10 | 0.60% |
| hsa_circ_0136051 | 3355 | 10 | 0.60% |
| hsa_circ_0038533 | 3691 | 11 | 0.60% |
| hsa_circ_0040947 | 3691 | 11 | 0.60% |
| hsa_circ_0056292 | 3691 | 11 | 0.60% |
| hsa_circ_0081217 | 3356 | 10 | 0.60% |
| hsa_circ_0122564 | 7719 | 23 | 0.60% |
| hsa_circ_0113404 | 6713 | 20 | 0.60% |
| hsa_circ_0104815 | 4364 | 13 | 0.60% |
| hsa_circ_0128411 | 24170 | 72 | 0.60% |
| hsa_circ_0002385 | 3693 | 11 | 0.60% |
| hsa_circ_0021574 | 3693 | 11 | 0.60% |
| hsa_circ_0056065 | 3693 | 11 | 0.60% |
| hsa_circ_0063349 | 4701 | 14 | 0.60% |
| hsa_circ_0077018 | 3694 | 11 | 0.60% |
| hsa_circ_0093717 | 3694 | 11 | 0.60% |
| hsa_circ_0056441 | 4030 | 12 | 0.60% |
| hsa_circ_0073544 | 4030 | 12 | 0.60% |
| hsa_circ_0056477 | 4702 | 14 | 0.60% |
| hsa_circ_0116654 | 14111 | 42 | 0.60% |
| hsa_circ_0057172 | 7056 | 21 | 0.60% |
| hsa_circ_0059294 | 4032 | 12 | 0.60% |
| hsa_circ_0062213 | 4032 | 12 | 0.60% |
| hsa_circ_0023419 | 7393 | 22 | 0.60% |
| hsa_circ_0059962 | 4369 | 13 | 0.60% |

| | | | |
|---|---|---|---|
| hsa_circ_0114662 | 10085 | 30 | 0.59% |
| hsa_circ_0118196 | 3698 | 11 | 0.59% |
| hsa_circ_0077354 | 3362 | 10 | 0.59% |
| hsa_circ_0054740 | 4371 | 13 | 0.59% |
| hsa_circ_0086940 | 4035 | 12 | 0.59% |
| hsa_circ_0104067 | 4035 | 12 | 0.59% |
| hsa_circ_0044519 | 3363 | 10 | 0.59% |
| hsa_circ_0023023 | 4036 | 12 | 0.59% |
| hsa_circ_0032087 | 4036 | 12 | 0.59% |
| hsa_circ_0040331 | 3700 | 11 | 0.59% |
| hsa_circ_0075761 | 3700 | 11 | 0.59% |
| hsa_circ_0116894 | 9083 | 27 | 0.59% |
| hsa_circ_0035500 | 3701 | 11 | 0.59% |
| hsa_circ_0056642 | 4374 | 13 | 0.59% |
| hsa_circ_0057854 | 4374 | 13 | 0.59% |
| hsa_circ_0131094 | 4038 | 12 | 0.59% |
| hsa_circ_0136229 | 3365 | 10 | 0.59% |
| hsa_circ_0028780 | 3702 | 11 | 0.59% |
| hsa_circ_0091626 | 3702 | 11 | 0.59% |
| hsa_circ_0062402 | 4039 | 12 | 0.59% |
| hsa_circ_0065070 | 4039 | 12 | 0.59% |
| hsa_circ_0132697 | 10435 | 31 | 0.59% |
| hsa_circ_0054986 | 3703 | 11 | 0.59% |
| hsa_circ_0019714 | 4040 | 12 | 0.59% |
| hsa_circ_0032523 | 10774 | 32 | 0.59% |
| hsa_circ_0084150 | 14141 | 42 | 0.59% |
| hsa_circ_0010571 | 4377 | 13 | 0.59% |
| hsa_circ_0019690 | 3367 | 10 | 0.59% |
| hsa_circ_0023880 | 3704 | 11 | 0.59% |
| hsa_circ_0067571 | 4041 | 12 | 0.59% |
| hsa_circ_0015701 | 5052 | 15 | 0.59% |
| hsa_circ_0114428 | 3705 | 11 | 0.59% |
| hsa_circ_0039128 | 4042 | 12 | 0.59% |
| hsa_circ_0065132 | 4042 | 12 | 0.59% |
| hsa_circ_0046718 | 3369 | 10 | 0.59% |
| hsa_circ_0088983 | 3706 | 11 | 0.59% |
| hsa_circ_0129391 | 4043 | 12 | 0.59% |
| hsa_circ_0077451 | 4380 | 13 | 0.59% |
| hsa_circ_0119563 | 9773 | 29 | 0.59% |
| hsa_circ_0004462 | 6403 | 19 | 0.59% |
| hsa_circ_0042583 | 4044 | 12 | 0.59% |
| hsa_circ_0127815 | 3707 | 11 | 0.59% |
| hsa_circ_0053095 | 4382 | 13 | 0.59% |
| hsa_circ_0127859 | 4382 | 13 | 0.59% |
| hsa_circ_0033519 | 4045 | 12 | 0.59% |
| hsa_circ_0076161 | 4045 | 12 | 0.59% |
| hsa_circ_0107555 | 3708 | 11 | 0.59% |
| hsa_circ_0111672 | 9440 | 28 | 0.59% |
| hsa_circ_0013196 | 4046 | 12 | 0.59% |
| hsa_circ_0043387 | 3709 | 11 | 0.59% |

| | | | |
|---|---|---|---|
| hsa_circ_0007105 | 13827 | 41 | 0.59% |
| hsa_circ_0061625 | 3710 | 11 | 0.59% |
| hsa_circ_0015996 | 7084 | 21 | 0.59% |
| hsa_circ_0009224 | 4048 | 12 | 0.59% |
| hsa_circ_0065105 | 4048 | 12 | 0.59% |
| hsa_circ_0010148 | 7422 | 22 | 0.59% |
| hsa_circ_0140363 | 4386 | 13 | 0.59% |
| hsa_circ_0138859 | 8435 | 25 | 0.59% |
| hsa_circ_0093307 | 4049 | 12 | 0.59% |
| hsa_circ_0123588 | 8436 | 25 | 0.59% |
| hsa_circ_0010598 | 4387 | 13 | 0.59% |
| hsa_circ_0048412 | 4050 | 12 | 0.59% |
| hsa_circ_0116614 | 4050 | 12 | 0.59% |
| hsa_circ_0034214 | 4388 | 13 | 0.59% |
| hsa_circ_0035638 | 3713 | 11 | 0.59% |
| hsa_circ_0039441 | 3713 | 11 | 0.59% |
| hsa_circ_0015657 | 6414 | 19 | 0.59% |
| hsa_circ_0039741 | 3376 | 10 | 0.59% |
| hsa_circ_0128301 | 4389 | 13 | 0.59% |
| hsa_circ_0053791 | 3714 | 11 | 0.59% |
| hsa_circ_0077117 | 3714 | 11 | 0.59% |
| hsa_circ_0125135 | 3714 | 11 | 0.59% |
| hsa_circ_0086902 | 4052 | 12 | 0.59% |
| hsa_circ_0091319 | 4052 | 12 | 0.59% |
| hsa_circ_0056151 | 3377 | 10 | 0.59% |
| hsa_circ_0084292 | 3715 | 11 | 0.59% |
| hsa_circ_0062442 | 4053 | 12 | 0.59% |
| hsa_circ_0022343 | 4391 | 13 | 0.59% |
| hsa_circ_0018216 | 3716 | 11 | 0.59% |
| hsa_circ_0057984 | 4054 | 12 | 0.59% |
| hsa_circ_0004935 | 4392 | 13 | 0.59% |
| hsa_circ_0101760 | 7771 | 23 | 0.59% |
| hsa_circ_0097163 | 8109 | 24 | 0.59% |
| hsa_circ_0017227 | 3717 | 11 | 0.59% |
| hsa_circ_0029291 | 3717 | 11 | 0.59% |
| hsa_circ_0044181 | 12167 | 36 | 0.59% |
| hsa_circ_0100534 | 4056 | 12 | 0.59% |
| hsa_circ_0087250 | 3718 | 11 | 0.59% |
| hsa_circ_0036565 | 3380 | 10 | 0.59% |
| hsa_circ_0072848 | 10479 | 31 | 0.59% |
| hsa_circ_0050027 | 3719 | 11 | 0.59% |
| hsa_circ_0115159 | 8115 | 24 | 0.59% |
| hsa_circ_0086892 | 3720 | 11 | 0.59% |
| hsa_circ_0085885 | 6426 | 19 | 0.59% |
| hsa_circ_0113928 | 4397 | 13 | 0.59% |
| hsa_circ_0016742 | 4059 | 12 | 0.59% |
| hsa_circ_0027161 | 4059 | 12 | 0.59% |
| hsa_circ_0048184 | 4059 | 12 | 0.59% |
| hsa_circ_0082215 | 4059 | 12 | 0.59% |
| hsa_circ_0049373 | 3383 | 10 | 0.59% |

| | | | |
|---|---|---|---|
| hsa_circ_0078934 | 8120 | 24 | 0.59% |
| hsa_circ_0075112 | 4399 | 13 | 0.59% |
| hsa_circ_0032980 | 3384 | 10 | 0.59% |
| hsa_circ_0072822 | 10491 | 31 | 0.59% |
| hsa_circ_0004170 | 6092 | 18 | 0.59% |
| hsa_circ_0021213 | 3723 | 11 | 0.59% |
| hsa_circ_0091902 | 3723 | 11 | 0.59% |
| hsa_circ_0052716 | 3385 | 10 | 0.59% |
| hsa_circ_0046073 | 4401 | 13 | 0.59% |
| hsa_circ_0091414 | 4401 | 13 | 0.59% |
| hsa_circ_0067538 | 3724 | 11 | 0.59% |
| hsa_circ_0071587 | 4063 | 12 | 0.59% |
| hsa_circ_0073638 | 4063 | 12 | 0.59% |
| hsa_circ_0097334 | 9820 | 29 | 0.59% |
| hsa_circ_0015702 | 3725 | 11 | 0.59% |
| hsa_circ_0084330 | 3725 | 11 | 0.59% |
| hsa_circ_0023828 | 4064 | 12 | 0.59% |
| hsa_circ_0028496 | 3726 | 11 | 0.59% |
| hsa_circ_0035591 | 3726 | 11 | 0.59% |
| hsa_circ_0093786 | 3726 | 11 | 0.59% |
| hsa_circ_0100275 | 16259 | 48 | 0.59% |
| hsa_circ_0026836 | 3727 | 11 | 0.59% |
| hsa_circ_0119915 | 3727 | 11 | 0.59% |
| hsa_circ_0128344 | 37948 | 112 | 0.59% |
| hsa_circ_0006902 | 6099 | 18 | 0.59% |
| hsa_circ_0009820 | 4066 | 12 | 0.59% |
| hsa_circ_0013288 | 4066 | 12 | 0.59% |
| hsa_circ_0034494 | 4066 | 12 | 0.59% |
| hsa_circ_0132400 | 4066 | 12 | 0.59% |
| hsa_circ_0074022 | 5083 | 15 | 0.59% |
| hsa_circ_0013816 | 3389 | 10 | 0.59% |
| hsa_circ_0020858 | 3728 | 11 | 0.59% |
| hsa_circ_0064851 | 3728 | 11 | 0.59% |
| hsa_circ_0081772 | 4067 | 12 | 0.59% |
| hsa_circ_0053184 | 4068 | 12 | 0.59% |
| hsa_circ_0021023 | 3729 | 11 | 0.59% |
| hsa_circ_0045787 | 3729 | 11 | 0.59% |
| hsa_circ_0045107 | 21358 | 63 | 0.59% |
| hsa_circ_0063391 | 5086 | 15 | 0.59% |
| hsa_circ_0010869 | 4408 | 13 | 0.59% |
| hsa_circ_0029865 | 4069 | 12 | 0.59% |
| hsa_circ_0073672 | 4069 | 12 | 0.59% |
| hsa_circ_0084849 | 4069 | 12 | 0.59% |
| hsa_circ_0133221 | 4069 | 12 | 0.59% |
| hsa_circ_0010033 | 11190 | 33 | 0.59% |
| hsa_circ_0100152 | 3730 | 11 | 0.59% |
| hsa_circ_0071153 | 3391 | 10 | 0.59% |
| hsa_circ_0044781 | 5426 | 16 | 0.59% |
| hsa_circ_0011677 | 4070 | 12 | 0.59% |
| hsa_circ_0108938 | 3731 | 11 | 0.59% |

| | | | |
|---|---|---|---|
| hsa_circ_0104777 | 9159 | 27 | 0.59% |
| hsa_circ_0059004 | 4410 | 13 | 0.59% |
| hsa_circ_0062534 | 4410 | 13 | 0.59% |
| hsa_circ_0077546 | 10177 | 30 | 0.59% |
| hsa_circ_0061046 | 3732 | 11 | 0.59% |
| hsa_circ_0082511 | 4411 | 13 | 0.59% |
| hsa_circ_0049105 | 4072 | 12 | 0.59% |
| hsa_circ_0070142 | 4751 | 14 | 0.59% |
| hsa_circ_0047701 | 3733 | 11 | 0.59% |
| hsa_circ_0086060 | 3733 | 11 | 0.59% |
| hsa_circ_0131759 | 3733 | 11 | 0.59% |
| hsa_circ_0122175 | 6109 | 18 | 0.59% |
| hsa_circ_0040098 | 4073 | 12 | 0.59% |
| hsa_circ_0090311 | 4073 | 12 | 0.59% |
| hsa_circ_0140611 | 4073 | 12 | 0.59% |
| hsa_circ_0122880 | 9504 | 28 | 0.59% |
| hsa_circ_0032402 | 3734 | 11 | 0.59% |
| hsa_circ_0035639 | 3734 | 11 | 0.59% |
| hsa_circ_0054807 | 4074 | 12 | 0.59% |
| hsa_circ_0083000 | 4416 | 13 | 0.59% |
| hsa_circ_0070486 | 3397 | 10 | 0.59% |
| hsa_circ_0069542 | 4417 | 13 | 0.59% |
| hsa_circ_0085306 | 4417 | 13 | 0.59% |
| hsa_circ_0107927 | 4417 | 13 | 0.59% |
| hsa_circ_0015224 | 3738 | 11 | 0.59% |
| hsa_circ_0024014 | 4078 | 12 | 0.59% |
| hsa_circ_0059354 | 4418 | 13 | 0.59% |
| hsa_circ_0100252 | 5098 | 15 | 0.59% |
| hsa_circ_0050815 | 4079 | 12 | 0.59% |
| hsa_circ_0094295 | 6459 | 19 | 0.59% |
| hsa_circ_0104017 | 13259 | 39 | 0.59% |
| hsa_circ_0113403 | 6800 | 20 | 0.59% |
| hsa_circ_0021572 | 4760 | 14 | 0.59% |
| hsa_circ_0044715 | 4420 | 13 | 0.59% |
| hsa_circ_0037461 | 4080 | 12 | 0.59% |
| hsa_circ_0129957 | 4080 | 12 | 0.59% |
| hsa_circ_0010726 | 3740 | 11 | 0.59% |
| hsa_circ_0030922 | 3740 | 11 | 0.59% |
| hsa_circ_0048735 | 4081 | 12 | 0.59% |
| hsa_circ_0030620 | 3741 | 11 | 0.59% |
| hsa_circ_0027755 | 3401 | 10 | 0.59% |
| hsa_circ_0015085 | 4082 | 12 | 0.59% |
| hsa_circ_0019192 | 4082 | 12 | 0.59% |
| hsa_circ_0061889 | 4082 | 12 | 0.59% |
| hsa_circ_0034338 | 3742 | 11 | 0.59% |
| hsa_circ_0028921 | 5784 | 17 | 0.59% |
| hsa_circ_0030926 | 4083 | 12 | 0.59% |
| hsa_circ_0006197 | 5104 | 15 | 0.59% |
| hsa_circ_0135001 | 3743 | 11 | 0.59% |
| hsa_circ_0045066 | 3403 | 10 | 0.59% |

| | | | |
|---|---|---|---|
| hsa_circ_0073297 | 3403 | 10 | 0.59% |
| hsa_circ_0056739 | 4765 | 14 | 0.59% |
| hsa_circ_0090687 | 3744 | 11 | 0.59% |
| hsa_circ_0117948 | 3404 | 10 | 0.59% |
| hsa_circ_0014034 | 4085 | 12 | 0.59% |
| hsa_circ_0065362 | 4085 | 12 | 0.59% |
| hsa_circ_0002848 | 9532 | 28 | 0.59% |
| hsa_circ_0028792 | 3745 | 11 | 0.59% |
| hsa_circ_0038250 | 3745 | 11 | 0.59% |
| hsa_circ_0094338 | 4767 | 14 | 0.59% |
| hsa_circ_0049439 | 4086 | 12 | 0.59% |
| hsa_circ_0056055 | 4086 | 12 | 0.59% |
| hsa_circ_0109284 | 11578 | 34 | 0.59% |
| hsa_circ_0064227 | 3746 | 11 | 0.59% |
| hsa_circ_0116825 | 4087 | 12 | 0.59% |
| hsa_circ_0042036 | 3747 | 11 | 0.59% |
| hsa_circ_0057121 | 6132 | 18 | 0.59% |
| hsa_circ_0019121 | 3407 | 10 | 0.59% |
| hsa_circ_0054448 | 3407 | 10 | 0.59% |
| hsa_circ_0056322 | 3407 | 10 | 0.59% |
| hsa_circ_0028013 | 3748 | 11 | 0.59% |
| hsa_circ_0139425 | 4089 | 12 | 0.59% |
| hsa_circ_0084371 | 4430 | 13 | 0.59% |
| hsa_circ_0068034 | 4771 | 14 | 0.59% |
| hsa_circ_0012801 | 3749 | 11 | 0.59% |
| hsa_circ_0029088 | 3749 | 11 | 0.59% |
| hsa_circ_0063089 | 3749 | 11 | 0.59% |
| hsa_circ_0072655 | 3749 | 11 | 0.59% |
| hsa_circ_0050795 | 4090 | 12 | 0.59% |
| hsa_circ_0058083 | 4431 | 13 | 0.59% |
| hsa_circ_0028272 | 3750 | 11 | 0.59% |
| hsa_circ_0061397 | 3750 | 11 | 0.59% |
| hsa_circ_0071180 | 3750 | 11 | 0.59% |
| hsa_circ_0010622 | 4432 | 13 | 0.59% |
| hsa_circ_0024130 | 4092 | 12 | 0.59% |
| hsa_circ_0062059 | 3751 | 11 | 0.59% |
| hsa_circ_0119274 | 3751 | 11 | 0.59% |
| hsa_circ_0083662 | 3410 | 10 | 0.59% |
| hsa_circ_0033323 | 4093 | 12 | 0.59% |
| hsa_circ_0046692 | 4093 | 12 | 0.59% |
| hsa_circ_0107541 | 3753 | 11 | 0.59% |
| hsa_circ_0122315 | 6483 | 19 | 0.59% |
| hsa_circ_0076448 | 4436 | 13 | 0.59% |
| hsa_circ_0138197 | 4436 | 13 | 0.59% |
| hsa_circ_0098003 | 5460 | 16 | 0.59% |
| hsa_circ_0040716 | 3754 | 11 | 0.59% |
| hsa_circ_0086848 | 3754 | 11 | 0.59% |
| hsa_circ_0011066 | 4778 | 14 | 0.59% |
| hsa_circ_0019124 | 3413 | 10 | 0.59% |
| hsa_circ_0113818 | 3413 | 10 | 0.59% |

| | | | |
|---|---|---|---|
| hsa_circ_0020843 | 10581 | 31 | 0.59% |
| hsa_circ_0042030 | 3755 | 11 | 0.59% |
| hsa_circ_0063094 | 3755 | 11 | 0.59% |
| hsa_circ_0039643 | 4438 | 13 | 0.59% |
| hsa_circ_0139800 | 4097 | 12 | 0.59% |
| hsa_circ_0114420 | 4098 | 12 | 0.59% |
| hsa_circ_0019330 | 3757 | 11 | 0.59% |
| hsa_circ_0000443 | 7173 | 21 | 0.59% |
| hsa_circ_0020114 | 4782 | 14 | 0.59% |
| hsa_circ_0091903 | 4099 | 12 | 0.59% |
| hsa_circ_0102171 | 4099 | 12 | 0.59% |
| hsa_circ_0017131 | 4100 | 12 | 0.59% |
| hsa_circ_0024803 | 3417 | 10 | 0.59% |
| hsa_circ_0045789 | 3417 | 10 | 0.59% |
| hsa_circ_0108880 | 3417 | 10 | 0.59% |
| hsa_circ_0049733 | 3759 | 11 | 0.59% |
| hsa_circ_0077338 | 4101 | 12 | 0.59% |
| hsa_circ_0091946 | 4101 | 12 | 0.59% |
| hsa_circ_0060474 | 4785 | 14 | 0.59% |
| hsa_circ_0132554 | 8887 | 26 | 0.59% |
| hsa_circ_0138671 | 15725 | 46 | 0.59% |
| hsa_circ_0046161 | 4103 | 12 | 0.58% |
| hsa_circ_0138765 | 17780 | 52 | 0.58% |
| hsa_circ_0064927 | 7523 | 22 | 0.58% |
| hsa_circ_0082006 | 4788 | 14 | 0.58% |
| hsa_circ_0066945 | 4446 | 13 | 0.58% |
| hsa_circ_0090048 | 4104 | 12 | 0.58% |
| hsa_circ_0032902 | 3762 | 11 | 0.58% |
| hsa_circ_0041377 | 3762 | 11 | 0.58% |
| hsa_circ_0071424 | 3762 | 11 | 0.58% |
| hsa_circ_0031207 | 5131 | 15 | 0.58% |
| hsa_circ_0101795 | 13341 | 39 | 0.58% |
| hsa_circ_0041208 | 4105 | 12 | 0.58% |
| hsa_circ_0116037 | 4105 | 12 | 0.58% |
| hsa_circ_0127320 | 3763 | 11 | 0.58% |
| hsa_circ_0135982 | 3763 | 11 | 0.58% |
| hsa_circ_0115411 | 8553 | 25 | 0.58% |
| hsa_circ_0051094 | 4448 | 13 | 0.58% |
| hsa_circ_0074909 | 4106 | 12 | 0.58% |
| hsa_circ_0051175 | 3764 | 11 | 0.58% |
| hsa_circ_0077620 | 5133 | 15 | 0.58% |
| hsa_circ_0039222 | 3422 | 10 | 0.58% |
| hsa_circ_0010641 | 4449 | 13 | 0.58% |
| hsa_circ_0026098 | 4449 | 13 | 0.58% |
| hsa_circ_0039524 | 4449 | 13 | 0.58% |
| hsa_circ_0073034 | 4449 | 13 | 0.58% |
| hsa_circ_0136457 | 4107 | 12 | 0.58% |
| hsa_circ_0093830 | 8899 | 26 | 0.58% |
| hsa_circ_0029086 | 3765 | 11 | 0.58% |
| hsa_circ_0056907 | 4450 | 13 | 0.58% |

| | | | |
|---|---|---|---|
| hsa_circ_0071347 | 4108 | 12 | 0.58% |
| hsa_circ_0009717 | 6163 | 18 | 0.58% |
| hsa_circ_0009990 | 3424 | 10 | 0.58% |
| hsa_circ_0110912 | 5479 | 16 | 0.58% |
| hsa_circ_0120497 | 9246 | 27 | 0.58% |
| hsa_circ_0075701 | 3767 | 11 | 0.58% |
| hsa_circ_0021496 | 4452 | 13 | 0.58% |
| hsa_circ_0103059 | 14042 | 41 | 0.58% |
| hsa_circ_0084956 | 4795 | 14 | 0.58% |
| hsa_circ_0017742 | 4110 | 12 | 0.58% |
| hsa_circ_0066561 | 4110 | 12 | 0.58% |
| hsa_circ_0016609 | 3768 | 11 | 0.58% |
| hsa_circ_0056062 | 3768 | 11 | 0.58% |
| hsa_circ_0058111 | 3768 | 11 | 0.58% |
| hsa_circ_0012919 | 4111 | 12 | 0.58% |
| hsa_circ_0089476 | 4111 | 12 | 0.58% |
| hsa_circ_0098819 | 12676 | 37 | 0.58% |
| hsa_circ_0053666 | 4454 | 13 | 0.58% |
| hsa_circ_0026834 | 3769 | 11 | 0.58% |
| hsa_circ_0090818 | 3770 | 11 | 0.58% |
| hsa_circ_0135820 | 3770 | 11 | 0.58% |
| hsa_circ_0053806 | 4113 | 12 | 0.58% |
| hsa_circ_0086061 | 4113 | 12 | 0.58% |
| hsa_circ_0056635 | 3428 | 10 | 0.58% |
| hsa_circ_0020324 | 16798 | 49 | 0.58% |
| hsa_circ_0036704 | 4114 | 12 | 0.58% |
| hsa_circ_0044752 | 4457 | 13 | 0.58% |
| hsa_circ_0053728 | 4457 | 13 | 0.58% |
| hsa_circ_0047989 | 4115 | 12 | 0.58% |
| hsa_circ_0080191 | 4459 | 13 | 0.58% |
| hsa_circ_0063964 | 4116 | 12 | 0.58% |
| hsa_circ_0076797 | 3773 | 11 | 0.58% |
| hsa_circ_0010662 | 4460 | 13 | 0.58% |
| hsa_circ_0063801 | 4117 | 12 | 0.58% |
| hsa_circ_0087312 | 4117 | 12 | 0.58% |
| hsa_circ_0061027 | 3774 | 11 | 0.58% |
| hsa_circ_0001547 | 5833 | 17 | 0.58% |
| hsa_circ_0066046 | 4118 | 12 | 0.58% |
| hsa_circ_0087433 | 4118 | 12 | 0.58% |
| hsa_circ_0041863 | 3775 | 11 | 0.58% |
| hsa_circ_0072520 | 3775 | 11 | 0.58% |
| hsa_circ_0122155 | 3775 | 11 | 0.58% |
| hsa_circ_0081091 | 3432 | 10 | 0.58% |
| hsa_circ_0115198 | 11669 | 34 | 0.58% |
| hsa_circ_0019160 | 4462 | 13 | 0.58% |
| hsa_circ_0070067 | 4119 | 12 | 0.58% |
| hsa_circ_0042906 | 3776 | 11 | 0.58% |
| hsa_circ_0091168 | 3776 | 11 | 0.58% |
| hsa_circ_0008111 | 4120 | 12 | 0.58% |
| hsa_circ_0071005 | 3777 | 11 | 0.58% |

| | | | |
|---|---|---|---|
| hsa_circ_0053596 | 4464 | 13 | 0.58% |
| hsa_circ_0083968 | 4464 | 13 | 0.58% |
| hsa_circ_0040433 | 4121 | 12 | 0.58% |
| hsa_circ_0089427 | 4121 | 12 | 0.58% |
| hsa_circ_0065588 | 3778 | 11 | 0.58% |
| hsa_circ_0052730 | 4465 | 13 | 0.58% |
| hsa_circ_0117545 | 4465 | 13 | 0.58% |
| hsa_circ_0052711 | 5153 | 15 | 0.58% |
| hsa_circ_0030864 | 5497 | 16 | 0.58% |
| hsa_circ_0064970 | 4810 | 14 | 0.58% |
| hsa_circ_0121027 | 8246 | 24 | 0.58% |
| hsa_circ_0052436 | 4811 | 14 | 0.58% |
| hsa_circ_0115719 | 8935 | 26 | 0.58% |
| hsa_circ_0105085 | 13403 | 39 | 0.58% |
| hsa_circ_0122176 | 6186 | 18 | 0.58% |
| hsa_circ_0136486 | 4124 | 12 | 0.58% |
| hsa_circ_0075082 | 4468 | 13 | 0.58% |
| hsa_circ_0106185 | 5843 | 17 | 0.58% |
| hsa_circ_0053448 | 3781 | 11 | 0.58% |
| hsa_circ_0055599 | 3781 | 11 | 0.58% |
| hsa_circ_0069001 | 3781 | 11 | 0.58% |
| hsa_circ_0105084 | 13406 | 39 | 0.58% |
| hsa_circ_0016936 | 4125 | 12 | 0.58% |
| hsa_circ_0094830 | 11346 | 33 | 0.58% |
| hsa_circ_0085236 | 4126 | 12 | 0.58% |
| hsa_circ_0042840 | 5158 | 15 | 0.58% |
| hsa_circ_0064933 | 3439 | 10 | 0.58% |
| hsa_circ_0077022 | 3783 | 11 | 0.58% |
| hsa_circ_0129965 | 3783 | 11 | 0.58% |
| hsa_circ_0085911 | 4127 | 12 | 0.58% |
| hsa_circ_0115829 | 6880 | 20 | 0.58% |
| hsa_circ_0028783 | 3784 | 11 | 0.58% |
| hsa_circ_0088627 | 5161 | 15 | 0.58% |
| hsa_circ_0027650 | 4129 | 12 | 0.58% |
| hsa_circ_0010750 | 3785 | 11 | 0.58% |
| hsa_circ_0022259 | 3785 | 11 | 0.58% |
| hsa_circ_0039626 | 4474 | 13 | 0.58% |
| hsa_circ_0110336 | 12734 | 37 | 0.58% |
| hsa_circ_0010695 | 4130 | 12 | 0.58% |
| hsa_circ_0042596 | 4130 | 12 | 0.58% |
| hsa_circ_0064377 | 3786 | 11 | 0.58% |
| hsa_circ_0046856 | 3442 | 10 | 0.58% |
| hsa_circ_0099125 | 4475 | 13 | 0.58% |
| hsa_circ_0025100 | 4131 | 12 | 0.58% |
| hsa_circ_0084287 | 4131 | 12 | 0.58% |
| hsa_circ_0122557 | 8951 | 26 | 0.58% |
| hsa_circ_0084368 | 3787 | 11 | 0.58% |
| hsa_circ_0044782 | 5509 | 16 | 0.58% |
| hsa_circ_0060853 | 5165 | 15 | 0.58% |
| hsa_circ_0072890 | 4132 | 12 | 0.58% |

| | | | |
|---|---|---|---|
| hsa_circ_0036233 | 4133 | 12 | 0.58% |
| hsa_circ_0024890 | 4134 | 12 | 0.58% |
| hsa_circ_0027365 | 4134 | 12 | 0.58% |
| hsa_circ_0054458 | 4134 | 12 | 0.58% |
| hsa_circ_0061312 | 4134 | 12 | 0.58% |
| hsa_circ_0010739 | 3790 | 11 | 0.58% |
| hsa_circ_0034314 | 3790 | 11 | 0.58% |
| hsa_circ_0084494 | 4135 | 12 | 0.58% |
| hsa_circ_0022310 | 3791 | 11 | 0.58% |
| hsa_circ_0042367 | 3791 | 11 | 0.58% |
| hsa_circ_0117887 | 13097 | 38 | 0.58% |
| hsa_circ_0029400 | 27919 | 81 | 0.58% |
| hsa_circ_0064095 | 4481 | 13 | 0.58% |
| hsa_circ_0133183 | 3447 | 10 | 0.58% |
| hsa_circ_0001959 | 3792 | 11 | 0.58% |
| hsa_circ_0127328 | 3792 | 11 | 0.58% |
| hsa_circ_0027332 | 4137 | 12 | 0.58% |
| hsa_circ_0085232 | 4137 | 12 | 0.58% |
| hsa_circ_0058589 | 14480 | 42 | 0.58% |
| hsa_circ_0081677 | 13446 | 39 | 0.58% |
| hsa_circ_0010585 | 4482 | 13 | 0.58% |
| hsa_circ_0072816 | 10688 | 31 | 0.58% |
| hsa_circ_0118109 | 7930 | 23 | 0.58% |
| hsa_circ_0093866 | 5172 | 15 | 0.58% |
| hsa_circ_0109907 | 3793 | 11 | 0.58% |
| hsa_circ_0134165 | 6208 | 18 | 0.58% |
| hsa_circ_0119632 | 13106 | 38 | 0.58% |
| hsa_circ_0044610 | 3794 | 11 | 0.58% |
| hsa_circ_0100954 | 19315 | 56 | 0.58% |
| hsa_circ_0076695 | 4484 | 13 | 0.58% |
| hsa_circ_0120236 | 13108 | 38 | 0.58% |
| hsa_circ_0064066 | 4140 | 12 | 0.58% |
| hsa_circ_0041852 | 5866 | 17 | 0.58% |
| hsa_circ_0095826 | 4831 | 14 | 0.58% |
| hsa_circ_0040608 | 4141 | 12 | 0.58% |
| hsa_circ_0131120 | 3796 | 11 | 0.58% |
| hsa_circ_0047068 | 4832 | 14 | 0.58% |
| hsa_circ_0059355 | 4487 | 13 | 0.58% |
| hsa_circ_0067570 | 4142 | 12 | 0.58% |
| hsa_circ_0035328 | 3797 | 11 | 0.58% |
| hsa_circ_0035509 | 3452 | 10 | 0.58% |
| hsa_circ_0018107 | 4833 | 14 | 0.58% |
| hsa_circ_0025757 | 4488 | 13 | 0.58% |
| hsa_circ_0131136 | 17262 | 50 | 0.58% |
| hsa_circ_0000609 | 4143 | 12 | 0.58% |
| hsa_circ_0047195 | 4143 | 12 | 0.58% |
| hsa_circ_0035177 | 3798 | 11 | 0.58% |
| hsa_circ_0127166 | 3798 | 11 | 0.58% |
| hsa_circ_0067633 | 4489 | 13 | 0.58% |
| hsa_circ_0036784 | 13469 | 39 | 0.58% |

| | | | |
|---|---|---|---|
| hsa_circ_0019119 | 3799 | 11 | 0.58% |
| hsa_circ_0021607 | 3799 | 11 | 0.58% |
| hsa_circ_0043242 | 3799 | 11 | 0.58% |
| hsa_circ_0064589 | 3799 | 11 | 0.58% |
| hsa_circ_0032560 | 4490 | 13 | 0.58% |
| hsa_circ_0131121 | 3800 | 11 | 0.58% |
| hsa_circ_0047003 | 4492 | 13 | 0.58% |
| hsa_circ_0073340 | 3801 | 11 | 0.58% |
| hsa_circ_0074948 | 3801 | 11 | 0.58% |
| hsa_circ_0011611 | 13823 | 40 | 0.58% |
| hsa_circ_0051060 | 4147 | 12 | 0.58% |
| hsa_circ_0062443 | 4147 | 12 | 0.58% |
| hsa_circ_0017351 | 6221 | 18 | 0.58% |
| hsa_circ_0041977 | 4493 | 13 | 0.58% |
| hsa_circ_0029206 | 3802 | 11 | 0.58% |
| hsa_circ_0018969 | 4148 | 12 | 0.58% |
| hsa_circ_0042916 | 4494 | 13 | 0.58% |
| hsa_circ_0038739 | 3803 | 11 | 0.58% |
| hsa_circ_0004200 | 7952 | 23 | 0.58% |
| hsa_circ_0028946 | 4149 | 12 | 0.58% |
| hsa_circ_0044448 | 4149 | 12 | 0.58% |
| hsa_circ_0101600 | 4841 | 14 | 0.58% |
| hsa_circ_0023967 | 3804 | 11 | 0.58% |
| hsa_circ_0041552 | 3804 | 11 | 0.58% |
| hsa_circ_0081285 | 3804 | 11 | 0.58% |
| hsa_circ_0086810 | 3804 | 11 | 0.58% |
| hsa_circ_0088964 | 3804 | 11 | 0.58% |
| hsa_circ_0027395 | 5879 | 17 | 0.58% |
| hsa_circ_0026107 | 5188 | 15 | 0.58% |
| hsa_circ_0070327 | 3805 | 11 | 0.58% |
| hsa_circ_0076008 | 4843 | 14 | 0.58% |
| hsa_circ_0095803 | 4843 | 14 | 0.58% |
| hsa_circ_0029399 | 28025 | 81 | 0.58% |
| hsa_circ_0111075 | 4152 | 12 | 0.58% |
| hsa_circ_0050026 | 3806 | 11 | 0.58% |
| hsa_circ_0102609 | 4499 | 13 | 0.58% |
| hsa_circ_0128581 | 4499 | 13 | 0.58% |
| hsa_circ_0086534 | 3807 | 11 | 0.58% |
| hsa_circ_0129771 | 4500 | 13 | 0.58% |
| hsa_circ_0019681 | 3809 | 11 | 0.58% |
| hsa_circ_0119441 | 3809 | 11 | 0.58% |
| hsa_circ_0000442 | 7273 | 21 | 0.58% |
| hsa_circ_0076274 | 5195 | 15 | 0.58% |
| hsa_circ_0024072 | 4156 | 12 | 0.58% |
| hsa_circ_0027578 | 4156 | 12 | 0.58% |
| hsa_circ_0065371 | 3810 | 11 | 0.58% |
| hsa_circ_0079232 | 4503 | 13 | 0.58% |
| hsa_circ_0047244 | 3464 | 10 | 0.58% |
| hsa_circ_0018949 | 4157 | 12 | 0.58% |
| hsa_circ_0053831 | 4157 | 12 | 0.58% |

| | | | |
|---|---|---|---|
| hsa_circ_0069794 | 3811 | 11 | 0.58% |
| hsa_circ_0024224 | 4504 | 13 | 0.58% |
| hsa_circ_0044716 | 4504 | 13 | 0.58% |
| hsa_circ_0081811 | 4158 | 12 | 0.58% |
| hsa_circ_0121967 | 20791 | 60 | 0.58% |
| hsa_circ_0015403 | 5198 | 15 | 0.58% |
| hsa_circ_0065615 | 3812 | 11 | 0.58% |
| hsa_circ_0086559 | 3812 | 11 | 0.58% |
| hsa_circ_0114422 | 4159 | 12 | 0.58% |
| hsa_circ_0032025 | 3466 | 10 | 0.58% |
| hsa_circ_0111530 | 6586 | 19 | 0.58% |
| hsa_circ_0028743 | 3813 | 11 | 0.58% |
| hsa_circ_0100961 | 4854 | 14 | 0.58% |
| hsa_circ_0018233 | 3814 | 11 | 0.58% |
| hsa_circ_0000429 | 9709 | 28 | 0.58% |
| hsa_circ_0105775 | 7976 | 23 | 0.58% |
| hsa_circ_0079117 | 4855 | 14 | 0.58% |
| hsa_circ_0050567 | 3468 | 10 | 0.58% |
| hsa_circ_0064487 | 4162 | 12 | 0.58% |
| hsa_circ_0109054 | 11099 | 32 | 0.58% |
| hsa_circ_0043028 | 4509 | 13 | 0.58% |
| hsa_circ_0048390 | 4856 | 14 | 0.58% |
| hsa_circ_0060789 | 4163 | 12 | 0.58% |
| hsa_circ_0077327 | 4163 | 12 | 0.58% |
| hsa_circ_0096291 | 4857 | 14 | 0.58% |
| hsa_circ_0039973 | 4858 | 14 | 0.58% |
| hsa_circ_0041401 | 4164 | 12 | 0.58% |
| hsa_circ_0102354 | 3817 | 11 | 0.58% |
| hsa_circ_0123256 | 12146 | 35 | 0.58% |
| hsa_circ_0091350 | 5553 | 16 | 0.58% |
| hsa_circ_0012061 | 4512 | 13 | 0.58% |
| hsa_circ_0085310 | 4512 | 13 | 0.58% |
| hsa_circ_0136506 | 4512 | 13 | 0.58% |
| hsa_circ_0115734 | 8678 | 25 | 0.58% |
| hsa_circ_0063097 | 4166 | 12 | 0.58% |
| hsa_circ_0071914 | 4167 | 12 | 0.58% |
| hsa_circ_0043969 | 3820 | 11 | 0.58% |
| hsa_circ_0057855 | 4515 | 13 | 0.58% |
| hsa_circ_0075111 | 4515 | 13 | 0.58% |
| hsa_circ_0082514 | 4515 | 13 | 0.58% |
| hsa_circ_0123180 | 12851 | 37 | 0.58% |
| hsa_circ_0028388 | 4168 | 12 | 0.58% |
| hsa_circ_0031952 | 4863 | 14 | 0.58% |
| hsa_circ_0011807 | 3821 | 11 | 0.58% |
| hsa_circ_0041325 | 3821 | 11 | 0.58% |
| hsa_circ_0113207 | 13895 | 40 | 0.58% |
| hsa_circ_0014770 | 4516 | 13 | 0.58% |
| hsa_circ_0070748 | 4516 | 13 | 0.58% |
| hsa_circ_0125584 | 4864 | 14 | 0.58% |
| hsa_circ_0059009 | 3822 | 11 | 0.58% |

| | | | |
|---|---|---|---|
| hsa_circ_0091912 | 3822 | 11 | 0.58% |
| hsa_circ_0035151 | 4517 | 13 | 0.58% |
| hsa_circ_0037199 | 4517 | 13 | 0.58% |
| hsa_circ_0027662 | 4170 | 12 | 0.58% |
| hsa_circ_0054801 | 4170 | 12 | 0.58% |
| hsa_circ_0089529 | 4170 | 12 | 0.58% |
| hsa_circ_0025651 | 3823 | 11 | 0.58% |
| hsa_circ_0077323 | 3476 | 10 | 0.58% |
| hsa_circ_0097101 | 4519 | 13 | 0.58% |
| hsa_circ_0006799 | 3477 | 10 | 0.58% |
| hsa_circ_0036605 | 3825 | 11 | 0.58% |
| hsa_circ_0073788 | 3825 | 11 | 0.58% |
| hsa_circ_0091886 | 3825 | 11 | 0.58% |
| hsa_circ_0084788 | 5912 | 17 | 0.58% |
| hsa_circ_0090201 | 5217 | 15 | 0.58% |
| hsa_circ_0035341 | 4870 | 14 | 0.57% |
| hsa_circ_0041488 | 4870 | 14 | 0.57% |
| hsa_circ_0042864 | 3479 | 10 | 0.57% |
| hsa_circ_0133208 | 4871 | 14 | 0.57% |
| hsa_circ_0126999 | 10091 | 29 | 0.57% |
| hsa_circ_0134032 | 4524 | 13 | 0.57% |
| hsa_circ_0035111 | 3828 | 11 | 0.57% |
| hsa_circ_0043924 | 3828 | 11 | 0.57% |
| hsa_circ_0068035 | 4873 | 14 | 0.57% |
| hsa_circ_0027753 | 3481 | 10 | 0.57% |
| hsa_circ_0062699 | 3830 | 11 | 0.57% |
| hsa_circ_0071181 | 3830 | 11 | 0.57% |
| hsa_circ_0086916 | 4527 | 13 | 0.57% |
| hsa_circ_0046652 | 9751 | 28 | 0.57% |
| hsa_circ_0130014 | 4179 | 12 | 0.57% |
| hsa_circ_0011784 | 3483 | 10 | 0.57% |
| hsa_circ_0009307 | 15674 | 45 | 0.57% |
| hsa_circ_0113147 | 10102 | 29 | 0.57% |
| hsa_circ_0018076 | 17070 | 49 | 0.57% |
| hsa_circ_0052903 | 3833 | 11 | 0.57% |
| hsa_circ_0068879 | 3833 | 11 | 0.57% |
| hsa_circ_0130926 | 3833 | 11 | 0.57% |
| hsa_circ_0025992 | 4530 | 13 | 0.57% |
| hsa_circ_0092089 | 4530 | 13 | 0.57% |
| hsa_circ_0107528 | 5924 | 17 | 0.57% |
| hsa_circ_0120600 | 4182 | 12 | 0.57% |
| hsa_circ_0137052 | 8713 | 25 | 0.57% |
| hsa_circ_0018063 | 3834 | 11 | 0.57% |
| hsa_circ_0010336 | 4183 | 12 | 0.57% |
| hsa_circ_0060854 | 5230 | 15 | 0.57% |
| hsa_circ_0058542 | 3836 | 11 | 0.57% |
| hsa_circ_0086865 | 3836 | 11 | 0.57% |
| hsa_circ_0016782 | 4185 | 12 | 0.57% |
| hsa_circ_0036833 | 4534 | 13 | 0.57% |
| hsa_circ_0084491 | 4186 | 12 | 0.57% |

| | | | |
|---|---|---|---|
| hsa_circ_0013942 | 4537 | 13 | 0.57% |
| hsa_circ_0136482 | 4537 | 13 | 0.57% |
| hsa_circ_0015627 | 3839 | 11 | 0.57% |
| hsa_circ_0053491 | 3839 | 11 | 0.57% |
| hsa_circ_0042166 | 4887 | 14 | 0.57% |
| hsa_circ_0018321 | 4189 | 12 | 0.57% |
| hsa_circ_0030471 | 3840 | 11 | 0.57% |
| hsa_circ_0135002 | 3840 | 11 | 0.57% |
| hsa_circ_0054285 | 14314 | 41 | 0.57% |
| hsa_circ_0133918 | 35264 | 101 | 0.57% |
| hsa_circ_0130186 | 9777 | 28 | 0.57% |
| hsa_circ_0028392 | 3841 | 11 | 0.57% |
| hsa_circ_0057605 | 3841 | 11 | 0.57% |
| hsa_circ_0003906 | 14317 | 41 | 0.57% |
| hsa_circ_0052690 | 5238 | 15 | 0.57% |
| hsa_circ_0134642 | 9429 | 27 | 0.57% |
| hsa_circ_0045801 | 4540 | 13 | 0.57% |
| hsa_circ_0087820 | 4540 | 13 | 0.57% |
| hsa_circ_0047831 | 5239 | 15 | 0.57% |
| hsa_circ_0129378 | 5240 | 15 | 0.57% |
| hsa_circ_0044192 | 11179 | 32 | 0.57% |
| hsa_circ_0084737 | 3843 | 11 | 0.57% |
| hsa_circ_0105532 | 4193 | 12 | 0.57% |
| hsa_circ_0041487 | 4892 | 14 | 0.57% |
| hsa_circ_0067537 | 3844 | 11 | 0.57% |
| hsa_circ_0107549 | 3844 | 11 | 0.57% |
| hsa_circ_0032861 | 4543 | 13 | 0.57% |
| hsa_circ_0043875 | 4194 | 12 | 0.57% |
| hsa_circ_0084307 | 4194 | 12 | 0.57% |
| hsa_circ_0084314 | 4194 | 12 | 0.57% |
| hsa_circ_0101622 | 4194 | 12 | 0.57% |
| hsa_circ_0050360 | 4195 | 12 | 0.57% |
| hsa_circ_0130097 | 4195 | 12 | 0.57% |
| hsa_circ_0037854 | 4545 | 13 | 0.57% |
| hsa_circ_0086931 | 4545 | 13 | 0.57% |
| hsa_circ_0040077 | 3846 | 11 | 0.57% |
| hsa_circ_0072164 | 3847 | 11 | 0.57% |
| hsa_circ_0067391 | 4197 | 12 | 0.57% |
| hsa_circ_0071928 | 4197 | 12 | 0.57% |
| hsa_circ_0098142 | 4197 | 12 | 0.57% |
| hsa_circ_0004837 | 10493 | 30 | 0.57% |
| hsa_circ_0042626 | 3848 | 11 | 0.57% |
| hsa_circ_0045452 | 3848 | 11 | 0.57% |
| hsa_circ_0098925 | 5947 | 17 | 0.57% |
| hsa_circ_0024672 | 4198 | 12 | 0.57% |
| hsa_circ_0060841 | 4548 | 13 | 0.57% |
| hsa_circ_0052289 | 3849 | 11 | 0.57% |
| hsa_circ_0070563 | 3849 | 11 | 0.57% |
| hsa_circ_0105599 | 3849 | 11 | 0.57% |
| hsa_circ_0080692 | 9448 | 27 | 0.57% |

| | | | |
|---|---|---|---|
| hsa_circ_0136386 | 11198 | 32 | 0.57% |
| hsa_circ_0038317 | 7699 | 22 | 0.57% |
| hsa_circ_0135361 | 9449 | 27 | 0.57% |
| hsa_circ_0037277 | 4900 | 14 | 0.57% |
| hsa_circ_0121177 | 4200 | 12 | 0.57% |
| hsa_circ_0053667 | 4551 | 13 | 0.57% |
| hsa_circ_0069121 | 4201 | 12 | 0.57% |
| hsa_circ_0069686 | 4201 | 12 | 0.57% |
| hsa_circ_0092488 | 3851 | 11 | 0.57% |
| hsa_circ_0028396 | 3852 | 11 | 0.57% |
| hsa_circ_0131066 | 3852 | 11 | 0.57% |
| hsa_circ_0012291 | 4553 | 13 | 0.57% |
| hsa_circ_0098734 | 4553 | 13 | 0.57% |
| hsa_circ_0047515 | 4203 | 12 | 0.57% |
| hsa_circ_0065067 | 4203 | 12 | 0.57% |
| hsa_circ_0037524 | 3853 | 11 | 0.57% |
| hsa_circ_0131449 | 9459 | 27 | 0.57% |
| hsa_circ_0043042 | 3854 | 11 | 0.57% |
| hsa_circ_0076367 | 3854 | 11 | 0.57% |
| hsa_circ_0131714 | 5256 | 15 | 0.57% |
| hsa_circ_0056728 | 3855 | 11 | 0.57% |
| hsa_circ_0099403 | 3855 | 11 | 0.57% |
| hsa_circ_0087435 | 4206 | 12 | 0.57% |
| hsa_circ_0115735 | 8763 | 25 | 0.57% |
| hsa_circ_0063926 | 3856 | 11 | 0.57% |
| hsa_circ_0048413 | 4207 | 12 | 0.57% |
| hsa_circ_0090630 | 3506 | 10 | 0.57% |
| hsa_circ_0053691 | 4558 | 13 | 0.57% |
| hsa_circ_0070842 | 4558 | 13 | 0.57% |
| hsa_circ_0008748 | 14025 | 40 | 0.57% |
| hsa_circ_0032417 | 3857 | 11 | 0.57% |
| hsa_circ_0011067 | 4909 | 14 | 0.57% |
| hsa_circ_0100276 | 16831 | 48 | 0.57% |
| hsa_circ_0122694 | 13326 | 38 | 0.57% |
| hsa_circ_0018490 | 4209 | 12 | 0.57% |
| hsa_circ_0028920 | 5963 | 17 | 0.57% |
| hsa_circ_0019159 | 4560 | 13 | 0.57% |
| hsa_circ_0127793 | 4560 | 13 | 0.57% |
| hsa_circ_0131301 | 9471 | 27 | 0.57% |
| hsa_circ_0059073 | 3508 | 10 | 0.57% |
| hsa_circ_0057008 | 12980 | 37 | 0.57% |
| hsa_circ_0056645 | 3859 | 11 | 0.57% |
| hsa_circ_0130927 | 4210 | 12 | 0.57% |
| hsa_circ_0029459 | 4561 | 13 | 0.57% |
| hsa_circ_0112408 | 16139 | 46 | 0.57% |
| hsa_circ_0078883 | 6316 | 18 | 0.57% |
| hsa_circ_0067644 | 3860 | 11 | 0.57% |
| hsa_circ_0096641 | 15791 | 45 | 0.57% |
| hsa_circ_0101101 | 4211 | 12 | 0.57% |
| hsa_circ_0007957 | 4562 | 13 | 0.57% |

| | | | |
|---|---|---|---|
| hsa_circ_0050339 | 4913 | 14 | 0.57% |
| hsa_circ_0116388 | 11933 | 34 | 0.57% |
| hsa_circ_0023995 | 4212 | 12 | 0.57% |
| hsa_circ_0121524 | 3861 | 11 | 0.57% |
| hsa_circ_0007680 | 3862 | 11 | 0.57% |
| hsa_circ_0057643 | 3511 | 10 | 0.57% |
| hsa_circ_0060211 | 4214 | 12 | 0.57% |
| hsa_circ_0011540 | 3863 | 11 | 0.57% |
| hsa_circ_0054578 | 3863 | 11 | 0.57% |
| hsa_circ_0047017 | 4566 | 13 | 0.57% |
| hsa_circ_0066211 | 5971 | 17 | 0.57% |
| hsa_circ_0049111 | 4215 | 12 | 0.57% |
| hsa_circ_0077115 | 3864 | 11 | 0.57% |
| hsa_circ_0073950 | 5973 | 17 | 0.57% |
| hsa_circ_0052117 | 3865 | 11 | 0.57% |
| hsa_circ_0090714 | 3865 | 11 | 0.57% |
| hsa_circ_0097335 | 3865 | 11 | 0.57% |
| hsa_circ_0128046 | 3865 | 11 | 0.57% |
| hsa_circ_0042912 | 4568 | 13 | 0.57% |
| hsa_circ_0051061 | 4217 | 12 | 0.57% |
| hsa_circ_0052437 | 4920 | 14 | 0.57% |
| hsa_circ_0055452 | 3866 | 11 | 0.57% |
| hsa_circ_0047019 | 4569 | 13 | 0.57% |
| hsa_circ_0113608 | 9841 | 28 | 0.57% |
| hsa_circ_0018108 | 4921 | 14 | 0.57% |
| hsa_circ_0138404 | 3867 | 11 | 0.57% |
| hsa_circ_0044783 | 5626 | 16 | 0.57% |
| hsa_circ_0022297 | 4220 | 12 | 0.57% |
| hsa_circ_0014549 | 3517 | 10 | 0.57% |
| hsa_circ_0109744 | 3869 | 11 | 0.57% |
| hsa_circ_0026582 | 4221 | 12 | 0.57% |
| hsa_circ_0085227 | 4221 | 12 | 0.57% |
| hsa_circ_0137089 | 9147 | 26 | 0.57% |
| hsa_circ_0134002 | 7740 | 22 | 0.57% |
| hsa_circ_0129811 | 3870 | 11 | 0.57% |
| hsa_circ_0109582 | 14073 | 40 | 0.57% |
| hsa_circ_0134155 | 14425 | 41 | 0.57% |
| hsa_circ_0080025 | 4222 | 12 | 0.57% |
| hsa_circ_0039661 | 3519 | 10 | 0.57% |
| hsa_circ_0019081 | 4223 | 12 | 0.57% |
| hsa_circ_0108485 | 4223 | 12 | 0.57% |
| hsa_circ_0120404 | 4223 | 12 | 0.57% |
| hsa_circ_0119812 | 4575 | 13 | 0.57% |
| hsa_circ_0024521 | 4224 | 12 | 0.57% |
| hsa_circ_0030024 | 4224 | 12 | 0.57% |
| hsa_circ_0080132 | 3872 | 11 | 0.57% |
| hsa_circ_0090683 | 3872 | 11 | 0.57% |
| hsa_circ_0130782 | 7041 | 20 | 0.57% |
| hsa_circ_0024414 | 4929 | 14 | 0.57% |
| hsa_circ_0062028 | 4225 | 12 | 0.57% |

| | | | |
|---|---|---|---|
| hsa_circ_0124291 | 4225 | 12 | 0.57% |
| hsa_circ_0054283 | 14436 | 41 | 0.57% |
| hsa_circ_0015401 | 5282 | 15 | 0.57% |
| hsa_circ_0037543 | 4930 | 14 | 0.57% |
| hsa_circ_0094421 | 3522 | 10 | 0.57% |
| hsa_circ_0089475 | 4227 | 12 | 0.57% |
| hsa_circ_0015897 | 3875 | 11 | 0.57% |
| hsa_circ_0038169 | 3875 | 11 | 0.57% |
| hsa_circ_0054065 | 3875 | 11 | 0.57% |
| hsa_circ_0071993 | 3523 | 10 | 0.57% |
| hsa_circ_0043027 | 4580 | 13 | 0.57% |
| hsa_circ_0055635 | 5285 | 15 | 0.57% |
| hsa_circ_0089267 | 5285 | 15 | 0.57% |
| hsa_circ_0042608 | 3876 | 11 | 0.57% |
| hsa_circ_0081203 | 3876 | 11 | 0.57% |
| hsa_circ_0131291 | 12686 | 36 | 0.57% |
| hsa_circ_0054632 | 4229 | 12 | 0.57% |
| hsa_circ_0086849 | 3877 | 11 | 0.57% |
| hsa_circ_0017398 | 5640 | 16 | 0.57% |
| hsa_circ_0133891 | 5640 | 16 | 0.57% |
| hsa_circ_0047898 | 4935 | 14 | 0.57% |
| hsa_circ_0012920 | 4230 | 12 | 0.57% |
| hsa_circ_0064046 | 4230 | 12 | 0.57% |
| hsa_circ_0091936 | 4230 | 12 | 0.57% |
| hsa_circ_0019689 | 3525 | 10 | 0.57% |
| hsa_circ_0036096 | 3878 | 11 | 0.57% |
| hsa_circ_0065189 | 3878 | 11 | 0.57% |
| hsa_circ_0083156 | 3878 | 11 | 0.57% |
| hsa_circ_0136052 | 3878 | 11 | 0.57% |
| hsa_circ_0011212 | 4936 | 14 | 0.57% |
| hsa_circ_0093280 | 4584 | 13 | 0.57% |
| hsa_circ_0108746 | 13401 | 38 | 0.57% |
| hsa_circ_0051174 | 3880 | 11 | 0.57% |
| hsa_circ_0085177 | 4233 | 12 | 0.57% |
| hsa_circ_0111101 | 4233 | 12 | 0.57% |
| hsa_circ_0134433 | 4233 | 12 | 0.57% |
| hsa_circ_0019043 | 5998 | 17 | 0.57% |
| hsa_circ_0010663 | 4588 | 13 | 0.57% |
| hsa_circ_0032370 | 7060 | 20 | 0.57% |
| hsa_circ_0078103 | 4236 | 12 | 0.57% |
| hsa_circ_0099281 | 50134 | 142 | 0.57% |
| hsa_circ_0046920 | 4590 | 13 | 0.57% |
| hsa_circ_0054727 | 4590 | 13 | 0.57% |
| hsa_circ_0104129 | 4590 | 13 | 0.57% |
| hsa_circ_0029507 | 3884 | 11 | 0.57% |
| hsa_circ_0033396 | 3884 | 11 | 0.57% |
| hsa_circ_0011612 | 14126 | 40 | 0.57% |
| hsa_circ_0033207 | 4238 | 12 | 0.57% |
| hsa_circ_0069289 | 3885 | 11 | 0.57% |
| hsa_circ_0031473 | 4239 | 12 | 0.57% |

| | | | |
|---|---|---|---|
| hsa_circ_0077173 | 4594 | 13 | 0.57% |
| hsa_circ_0042305 | 3534 | 10 | 0.57% |
| hsa_circ_0043591 | 4595 | 13 | 0.57% |
| hsa_circ_0111580 | 6010 | 17 | 0.57% |
| hsa_circ_0040435 | 3889 | 11 | 0.57% |
| hsa_circ_0008739 | 4950 | 14 | 0.57% |
| hsa_circ_0036540 | 4243 | 12 | 0.57% |
| hsa_circ_0063961 | 4243 | 12 | 0.57% |
| hsa_circ_0044717 | 4597 | 13 | 0.57% |
| hsa_circ_0028729 | 3890 | 11 | 0.57% |
| hsa_circ_0047138 | 3890 | 11 | 0.57% |
| hsa_circ_0064225 | 3890 | 11 | 0.57% |
| hsa_circ_0071188 | 3890 | 11 | 0.57% |
| hsa_circ_0019284 | 4598 | 13 | 0.57% |
| hsa_circ_0104093 | 10258 | 29 | 0.57% |
| hsa_circ_0046003 | 4599 | 13 | 0.57% |
| hsa_circ_0038436 | 3538 | 10 | 0.57% |
| hsa_circ_0121378 | 8492 | 24 | 0.57% |
| hsa_circ_0027425 | 3539 | 10 | 0.57% |
| hsa_circ_0031457 | 4247 | 12 | 0.57% |
| hsa_circ_0077200 | 4247 | 12 | 0.57% |
| hsa_circ_0086062 | 4247 | 12 | 0.57% |
| hsa_circ_0064540 | 4601 | 13 | 0.57% |
| hsa_circ_0036636 | 4248 | 12 | 0.56% |
| hsa_circ_0123894 | 4248 | 12 | 0.56% |
| hsa_circ_0001125 | 16994 | 48 | 0.56% |
| hsa_circ_0032351 | 4249 | 12 | 0.56% |
| hsa_circ_0085230 | 4249 | 12 | 0.56% |
| hsa_circ_0091947 | 4249 | 12 | 0.56% |
| hsa_circ_0045802 | 4604 | 13 | 0.56% |
| hsa_circ_0014166 | 4250 | 12 | 0.56% |
| hsa_circ_0051062 | 4250 | 12 | 0.56% |
| hsa_circ_0080478 | 3896 | 11 | 0.56% |
| hsa_circ_0131450 | 9563 | 27 | 0.56% |
| hsa_circ_0055456 | 3542 | 10 | 0.56% |
| hsa_circ_0128127 | 4605 | 13 | 0.56% |
| hsa_circ_0016839 | 4251 | 12 | 0.56% |
| hsa_circ_0061045 | 3897 | 11 | 0.56% |
| hsa_circ_0048867 | 4606 | 13 | 0.56% |
| hsa_circ_0043536 | 4252 | 12 | 0.56% |
| hsa_circ_0055088 | 4252 | 12 | 0.56% |
| hsa_circ_0062360 | 4252 | 12 | 0.56% |
| hsa_circ_0068674 | 4252 | 12 | 0.56% |
| hsa_circ_0010272 | 3898 | 11 | 0.56% |
| hsa_circ_0090803 | 4607 | 13 | 0.56% |
| hsa_circ_0076310 | 3544 | 10 | 0.56% |
| hsa_circ_0069871 | 4253 | 12 | 0.56% |
| hsa_circ_0076162 | 4253 | 12 | 0.56% |
| hsa_circ_0050232 | 5672 | 16 | 0.56% |
| hsa_circ_0037063 | 4963 | 14 | 0.56% |

| | | | |
|---|---|---|---|
| hsa_circ_0030566 | 3900 | 11 | 0.56% |
| hsa_circ_0136886 | 3900 | 11 | 0.56% |
| hsa_circ_0018942 | 4255 | 12 | 0.56% |
| hsa_circ_0016584 | 3901 | 11 | 0.56% |
| hsa_circ_0114426 | 3901 | 11 | 0.56% |
| hsa_circ_0120932 | 4256 | 12 | 0.56% |
| hsa_circ_0003910 | 11350 | 32 | 0.56% |
| hsa_circ_0117691 | 4611 | 13 | 0.56% |
| hsa_circ_0059432 | 3902 | 11 | 0.56% |
| hsa_circ_0091927 | 4257 | 12 | 0.56% |
| hsa_circ_0096640 | 15964 | 45 | 0.56% |
| hsa_circ_0022859 | 4612 | 13 | 0.56% |
| hsa_circ_0124471 | 46480 | 131 | 0.56% |
| hsa_circ_0073288 | 4258 | 12 | 0.56% |
| hsa_circ_0012892 | 4970 | 14 | 0.56% |
| hsa_circ_0128715 | 4615 | 13 | 0.56% |
| hsa_circ_0096774 | 4260 | 12 | 0.56% |
| hsa_circ_0115636 | 4260 | 12 | 0.56% |
| hsa_circ_0028781 | 3906 | 11 | 0.56% |
| hsa_circ_0086886 | 3906 | 11 | 0.56% |
| hsa_circ_0006361 | 13495 | 38 | 0.56% |
| hsa_circ_0029997 | 4972 | 14 | 0.56% |
| hsa_circ_0087427 | 7814 | 22 | 0.56% |
| hsa_circ_0021024 | 3907 | 11 | 0.56% |
| hsa_circ_0129198 | 3907 | 11 | 0.56% |
| hsa_circ_0117317 | 3552 | 10 | 0.56% |
| hsa_circ_0081709 | 13854 | 39 | 0.56% |
| hsa_circ_0059961 | 4618 | 13 | 0.56% |
| hsa_circ_0115682 | 8526 | 24 | 0.56% |
| hsa_circ_0023829 | 4263 | 12 | 0.56% |
| hsa_circ_0083002 | 4619 | 13 | 0.56% |
| hsa_circ_0035325 | 4264 | 12 | 0.56% |
| hsa_circ_0067560 | 4264 | 12 | 0.56% |
| hsa_circ_0052637 | 4620 | 13 | 0.56% |
| hsa_circ_0108388 | 5331 | 15 | 0.56% |
| hsa_circ_0029179 | 4265 | 12 | 0.56% |
| hsa_circ_0128469 | 4265 | 12 | 0.56% |
| hsa_circ_0054278 | 14573 | 41 | 0.56% |
| hsa_circ_0128036 | 3910 | 11 | 0.56% |
| hsa_circ_0064060 | 4266 | 12 | 0.56% |
| hsa_circ_0065589 | 3911 | 11 | 0.56% |
| hsa_circ_0057119 | 6401 | 18 | 0.56% |
| hsa_circ_0088923 | 4623 | 13 | 0.56% |
| hsa_circ_0082866 | 4268 | 12 | 0.56% |
| hsa_circ_0095349 | 4268 | 12 | 0.56% |
| hsa_circ_0065768 | 3557 | 10 | 0.56% |
| hsa_circ_0013867 | 8893 | 25 | 0.56% |
| hsa_circ_0016608 | 3913 | 11 | 0.56% |
| hsa_circ_0042448 | 3913 | 11 | 0.56% |
| hsa_circ_0005558 | 9249 | 26 | 0.56% |

| | | | |
|---|---|---|---|
| hsa_circ_0122382 | 7828 | 22 | 0.56% |
| hsa_circ_0050156 | 3914 | 11 | 0.56% |
| hsa_circ_0053506 | 3914 | 11 | 0.56% |
| hsa_circ_0079954 | 3559 | 10 | 0.56% |
| hsa_circ_0010257 | 3915 | 11 | 0.56% |
| hsa_circ_0101811 | 4627 | 13 | 0.56% |
| hsa_circ_0111397 | 5695 | 16 | 0.56% |
| hsa_circ_0139951 | 10680 | 30 | 0.56% |
| hsa_circ_0018927 | 4628 | 13 | 0.56% |
| hsa_circ_0001697 | 4272 | 12 | 0.56% |
| hsa_circ_0047354 | 3916 | 11 | 0.56% |
| hsa_circ_0117633 | 3916 | 11 | 0.56% |
| hsa_circ_0047140 | 3917 | 11 | 0.56% |
| hsa_circ_0049505 | 3561 | 10 | 0.56% |
| hsa_circ_0030923 | 4274 | 12 | 0.56% |
| hsa_circ_0068015 | 4274 | 12 | 0.56% |
| hsa_circ_0084906 | 3918 | 11 | 0.56% |
| hsa_circ_0096525 | 12111 | 34 | 0.56% |
| hsa_circ_0024059 | 4275 | 12 | 0.56% |
| hsa_circ_0086063 | 4275 | 12 | 0.56% |
| hsa_circ_0073484 | 3919 | 11 | 0.56% |
| hsa_circ_0135571 | 3919 | 11 | 0.56% |
| hsa_circ_0139297 | 3920 | 11 | 0.56% |
| hsa_circ_0044753 | 4633 | 13 | 0.56% |
| hsa_circ_0104950 | 8197 | 23 | 0.56% |
| hsa_circ_0063387 | 4277 | 12 | 0.56% |
| hsa_circ_0065366 | 4277 | 12 | 0.56% |
| hsa_circ_0089433 | 4277 | 12 | 0.56% |
| hsa_circ_0117412 | 4277 | 12 | 0.56% |
| hsa_circ_0098281 | 9980 | 28 | 0.56% |
| hsa_circ_0037523 | 3921 | 11 | 0.56% |
| hsa_circ_0038167 | 3921 | 11 | 0.56% |
| hsa_circ_0077071 | 3921 | 11 | 0.56% |
| hsa_circ_0108244 | 13546 | 38 | 0.56% |
| hsa_circ_0057021 | 4278 | 12 | 0.56% |
| hsa_circ_0120353 | 4278 | 12 | 0.56% |
| hsa_circ_0042029 | 3922 | 11 | 0.56% |
| hsa_circ_0024406 | 11767 | 33 | 0.56% |
| hsa_circ_0047198 | 4279 | 12 | 0.56% |
| hsa_circ_0104694 | 7845 | 22 | 0.56% |
| hsa_circ_0075908 | 4636 | 13 | 0.56% |
| hsa_circ_0137141 | 4636 | 13 | 0.56% |
| hsa_circ_0019815 | 3923 | 11 | 0.56% |
| hsa_circ_0046769 | 3923 | 11 | 0.56% |
| hsa_circ_0077725 | 3923 | 11 | 0.56% |
| hsa_circ_0035342 | 4993 | 14 | 0.56% |
| hsa_circ_0049195 | 4280 | 12 | 0.56% |
| hsa_circ_0013344 | 7847 | 22 | 0.56% |
| hsa_circ_0015689 | 4994 | 14 | 0.56% |
| hsa_circ_0016919 | 3924 | 11 | 0.56% |

| | | | |
|---|---|---|---|
| hsa_circ_0126221 | 3924 | 11 | 0.56% |
| hsa_circ_0026691 | 11060 | 31 | 0.56% |
| hsa_circ_0025669 | 3925 | 11 | 0.56% |
| hsa_circ_0080133 | 3925 | 11 | 0.56% |
| hsa_circ_0008738 | 4282 | 12 | 0.56% |
| hsa_circ_0071215 | 3569 | 10 | 0.56% |
| hsa_circ_0065373 | 3926 | 11 | 0.56% |
| hsa_circ_0031175 | 4283 | 12 | 0.56% |
| hsa_circ_0050842 | 4283 | 12 | 0.56% |
| hsa_circ_0018356 | 11779 | 33 | 0.56% |
| hsa_circ_0014771 | 4641 | 13 | 0.56% |
| hsa_circ_0079450 | 4284 | 12 | 0.56% |
| hsa_circ_0007286 | 9284 | 26 | 0.56% |
| hsa_circ_0016836 | 4285 | 12 | 0.56% |
| hsa_circ_0128455 | 4285 | 12 | 0.56% |
| hsa_circ_0105108 | 3928 | 11 | 0.56% |
| hsa_circ_0063075 | 4643 | 13 | 0.56% |
| hsa_circ_0031468 | 4286 | 12 | 0.56% |
| hsa_circ_0068557 | 3572 | 10 | 0.56% |
| hsa_circ_0036783 | 13931 | 39 | 0.56% |
| hsa_circ_0134000 | 7859 | 22 | 0.56% |
| hsa_circ_0039314 | 4644 | 13 | 0.56% |
| hsa_circ_0028769 | 4287 | 12 | 0.56% |
| hsa_circ_0026106 | 3930 | 11 | 0.56% |
| hsa_circ_0092088 | 4645 | 13 | 0.56% |
| hsa_circ_0090764 | 5003 | 14 | 0.56% |
| hsa_circ_0065525 | 3931 | 11 | 0.56% |
| hsa_circ_0093749 | 3931 | 11 | 0.56% |
| hsa_circ_0135560 | 15369 | 43 | 0.56% |
| hsa_circ_0096675 | 9293 | 26 | 0.56% |
| hsa_circ_0047513 | 5004 | 14 | 0.56% |
| hsa_circ_0099528 | 3932 | 11 | 0.56% |
| hsa_circ_0095762 | 8222 | 23 | 0.56% |
| hsa_circ_0020873 | 3933 | 11 | 0.56% |
| hsa_circ_0062444 | 4291 | 12 | 0.56% |
| hsa_circ_0038051 | 3934 | 11 | 0.56% |
| hsa_circ_0076309 | 3934 | 11 | 0.56% |
| hsa_circ_0094901 | 3934 | 11 | 0.56% |
| hsa_circ_0061338 | 15022 | 42 | 0.56% |
| hsa_circ_0137769 | 7511 | 21 | 0.56% |
| hsa_circ_0043995 | 4292 | 12 | 0.56% |
| hsa_circ_0066004 | 4292 | 12 | 0.56% |
| hsa_circ_0042917 | 3577 | 10 | 0.56% |
| hsa_circ_0010090 | 3935 | 11 | 0.56% |
| hsa_circ_0010733 | 3936 | 11 | 0.56% |
| hsa_circ_0019687 | 3936 | 11 | 0.56% |
| hsa_circ_0076160 | 3936 | 11 | 0.56% |
| hsa_circ_0091277 | 3936 | 11 | 0.56% |
| hsa_circ_0011783 | 3579 | 10 | 0.56% |
| hsa_circ_0023536 | 4295 | 12 | 0.56% |

| | | | |
|---|---|---|---|
| hsa_circ_0078410 | 4295 | 12 | 0.56% |
| hsa_circ_0029411 | 5370 | 15 | 0.56% |
| hsa_circ_0067741 | 5370 | 15 | 0.56% |
| hsa_circ_0033116 | 4296 | 12 | 0.56% |
| hsa_circ_0072887 | 3580 | 10 | 0.56% |
| hsa_circ_0030470 | 3939 | 11 | 0.56% |
| hsa_circ_0031816 | 3939 | 11 | 0.56% |
| hsa_circ_0138684 | 5372 | 15 | 0.56% |
| hsa_circ_0100866 | 5014 | 14 | 0.56% |
| hsa_circ_0011449 | 4298 | 12 | 0.56% |
| hsa_circ_0070369 | 4298 | 12 | 0.56% |
| hsa_circ_0048731 | 4657 | 13 | 0.56% |
| hsa_circ_0019126 | 4299 | 12 | 0.56% |
| hsa_circ_0033606 | 4299 | 12 | 0.56% |
| hsa_circ_0071908 | 4299 | 12 | 0.56% |
| hsa_circ_0039109 | 5374 | 15 | 0.56% |
| hsa_circ_0045067 | 3583 | 10 | 0.56% |
| hsa_circ_0076693 | 4658 | 13 | 0.56% |
| hsa_circ_0028774 | 4300 | 12 | 0.56% |
| hsa_circ_0079461 | 5017 | 14 | 0.56% |
| hsa_circ_0090283 | 5017 | 14 | 0.56% |
| hsa_circ_0139521 | 3942 | 11 | 0.56% |
| hsa_circ_0091320 | 4301 | 12 | 0.56% |
| hsa_circ_0009566 | 3943 | 11 | 0.56% |
| hsa_circ_0045448 | 3943 | 11 | 0.56% |
| hsa_circ_0015998 | 3944 | 11 | 0.56% |
| hsa_circ_0042609 | 3944 | 11 | 0.56% |
| hsa_circ_0082517 | 3586 | 10 | 0.56% |
| hsa_circ_0014031 | 3945 | 11 | 0.56% |
| hsa_circ_0041897 | 4304 | 12 | 0.56% |
| hsa_circ_0054633 | 4304 | 12 | 0.56% |
| hsa_circ_0102901 | 3946 | 11 | 0.56% |
| hsa_circ_0017585 | 12197 | 34 | 0.56% |
| hsa_circ_0077547 | 10763 | 30 | 0.56% |
| hsa_circ_0084369 | 3947 | 11 | 0.56% |
| hsa_circ_0106563 | 3947 | 11 | 0.56% |
| hsa_circ_0132001 | 10049 | 28 | 0.56% |
| hsa_circ_0095023 | 9693 | 27 | 0.56% |
| hsa_circ_0121433 | 5026 | 14 | 0.56% |
| hsa_circ_0095433 | 4308 | 12 | 0.56% |
| hsa_circ_0039890 | 3949 | 11 | 0.56% |
| hsa_circ_0050296 | 3949 | 11 | 0.56% |
| hsa_circ_0081972 | 3949 | 11 | 0.56% |
| hsa_circ_0104904 | 10412 | 29 | 0.56% |
| hsa_circ_0136375 | 12567 | 35 | 0.56% |
| hsa_circ_0011068 | 5027 | 14 | 0.56% |
| hsa_circ_0044438 | 5027 | 14 | 0.56% |
| hsa_circ_0019511 | 4668 | 13 | 0.56% |
| hsa_circ_0123757 | 4309 | 12 | 0.56% |
| hsa_circ_0108839 | 12928 | 36 | 0.56% |

| | | | |
|---|---|---|---|
| hsa_circ_0049152 | 4310 | 12 | 0.56% |
| hsa_circ_0017223 | 3952 | 11 | 0.56% |
| hsa_circ_0090255 | 4672 | 13 | 0.56% |
| hsa_circ_0093880 | 7907 | 22 | 0.56% |
| hsa_circ_0077116 | 3954 | 11 | 0.56% |
| hsa_circ_0091737 | 6112 | 17 | 0.56% |
| hsa_circ_0062403 | 4315 | 12 | 0.56% |
| hsa_circ_0084848 | 4315 | 12 | 0.56% |
| hsa_circ_0128342 | 7912 | 22 | 0.56% |
| hsa_circ_0044784 | 5755 | 16 | 0.56% |
| hsa_circ_0046922 | 4676 | 13 | 0.56% |
| hsa_circ_0076009 | 5036 | 14 | 0.56% |
| hsa_circ_0018660 | 4317 | 12 | 0.56% |
| hsa_circ_0020554 | 4317 | 12 | 0.56% |
| hsa_circ_0034484 | 4317 | 12 | 0.56% |
| hsa_circ_0078691 | 4677 | 13 | 0.56% |
| hsa_circ_0091844 | 3958 | 11 | 0.56% |
| hsa_circ_0116925 | 3958 | 11 | 0.56% |
| hsa_circ_0038898 | 9716 | 27 | 0.56% |
| hsa_circ_0072119 | 4319 | 12 | 0.56% |
| hsa_circ_0068036 | 5039 | 14 | 0.56% |
| hsa_circ_0062585 | 5399 | 15 | 0.56% |
| hsa_circ_0052727 | 4680 | 13 | 0.56% |
| hsa_circ_0077444 | 4680 | 13 | 0.56% |
| hsa_circ_0089215 | 4320 | 12 | 0.56% |
| hsa_circ_0030087 | 3960 | 11 | 0.56% |
| hsa_circ_0110924 | 3600 | 10 | 0.56% |
| hsa_circ_0026923 | 5761 | 16 | 0.56% |
| hsa_circ_0115255 | 5041 | 14 | 0.56% |
| hsa_circ_0015479 | 4321 | 12 | 0.56% |
| hsa_circ_0051170 | 3961 | 11 | 0.56% |
| hsa_circ_0091803 | 5042 | 14 | 0.56% |
| hsa_circ_0072896 | 4322 | 12 | 0.56% |
| hsa_circ_0072177 | 3962 | 11 | 0.56% |
| hsa_circ_0121295 | 16574 | 46 | 0.56% |
| hsa_circ_0118823 | 11891 | 33 | 0.56% |
| hsa_circ_0115464 | 13693 | 38 | 0.56% |
| hsa_circ_0012626 | 4685 | 13 | 0.55% |
| hsa_circ_0012889 | 5046 | 14 | 0.55% |
| hsa_circ_0068880 | 3965 | 11 | 0.55% |
| hsa_circ_0046349 | 4686 | 13 | 0.55% |
| hsa_circ_0117280 | 10093 | 28 | 0.55% |
| hsa_circ_0130859 | 10094 | 28 | 0.55% |
| hsa_circ_0127391 | 10816 | 30 | 0.55% |
| hsa_circ_0085104 | 3966 | 11 | 0.55% |
| hsa_circ_0035343 | 5048 | 14 | 0.55% |
| hsa_circ_0033319 | 4327 | 12 | 0.55% |
| hsa_circ_0066062 | 4327 | 12 | 0.55% |
| hsa_circ_0066011 | 4688 | 13 | 0.55% |
| hsa_circ_0036097 | 3967 | 11 | 0.55% |

| | | | |
|---|---|---|---|
| hsa_circ_0070324 | 3967 | 11 | 0.55% |
| hsa_circ_0086850 | 3967 | 11 | 0.55% |
| hsa_circ_0117295 | 3967 | 11 | 0.55% |
| hsa_circ_0073949 | 6131 | 17 | 0.55% |
| hsa_circ_0054473 | 4328 | 12 | 0.55% |
| hsa_circ_0083192 | 4328 | 12 | 0.55% |
| hsa_circ_0134198 | 7214 | 20 | 0.55% |
| hsa_circ_0021570 | 3968 | 11 | 0.55% |
| hsa_circ_0094644 | 4329 | 12 | 0.55% |
| hsa_circ_0090883 | 3970 | 11 | 0.55% |
| hsa_circ_0049675 | 4331 | 12 | 0.55% |
| hsa_circ_0053821 | 4331 | 12 | 0.55% |
| hsa_circ_0010580 | 4692 | 13 | 0.55% |
| hsa_circ_0016774 | 4692 | 13 | 0.55% |
| hsa_circ_0035766 | 4692 | 13 | 0.55% |
| hsa_circ_0112380 | 11912 | 33 | 0.55% |
| hsa_circ_0010948 | 4332 | 12 | 0.55% |
| hsa_circ_0113323 | 3610 | 10 | 0.55% |
| hsa_circ_0136110 | 5416 | 15 | 0.55% |
| hsa_circ_0017367 | 4333 | 12 | 0.55% |
| hsa_circ_0021029 | 3972 | 11 | 0.55% |
| hsa_circ_0077460 | 4695 | 13 | 0.55% |
| hsa_circ_0029177 | 4334 | 12 | 0.55% |
| hsa_circ_0086464 | 3973 | 11 | 0.55% |
| hsa_circ_0076016 | 5057 | 14 | 0.55% |
| hsa_circ_0116752 | 15172 | 42 | 0.55% |
| hsa_circ_0016529 | 4335 | 12 | 0.55% |
| hsa_circ_0056566 | 3974 | 11 | 0.55% |
| hsa_circ_0020057 | 5420 | 15 | 0.55% |
| hsa_circ_0040477 | 4336 | 12 | 0.55% |
| hsa_circ_0091278 | 3975 | 11 | 0.55% |
| hsa_circ_0137566 | 4698 | 13 | 0.55% |
| hsa_circ_0016607 | 3976 | 11 | 0.55% |
| hsa_circ_0024123 | 3976 | 11 | 0.55% |
| hsa_circ_0119819 | 3976 | 11 | 0.55% |
| hsa_circ_0120977 | 6868 | 19 | 0.55% |
| hsa_circ_0117311 | 9039 | 25 | 0.55% |
| hsa_circ_0072208 | 5062 | 14 | 0.55% |
| hsa_circ_0087313 | 4339 | 12 | 0.55% |
| hsa_circ_0016008 | 3616 | 10 | 0.55% |
| hsa_circ_0085222 | 3616 | 10 | 0.55% |
| hsa_circ_0014689 | 4340 | 12 | 0.55% |
| hsa_circ_0045803 | 4702 | 13 | 0.55% |
| hsa_circ_0119325 | 4341 | 12 | 0.55% |
| hsa_circ_0132726 | 4703 | 13 | 0.55% |
| hsa_circ_0076018 | 4342 | 12 | 0.55% |
| hsa_circ_0084621 | 4704 | 13 | 0.55% |
| hsa_circ_0066298 | 3981 | 11 | 0.55% |
| hsa_circ_0087202 | 3981 | 11 | 0.55% |
| hsa_circ_0128855 | 14839 | 41 | 0.55% |

| | | | |
|---|---|---|---|
| hsa_circ_0123984 | 12307 | 34 | 0.55% |
| hsa_circ_0079611 | 13393 | 37 | 0.55% |
| hsa_circ_0054969 | 5792 | 16 | 0.55% |
| hsa_circ_0048414 | 4344 | 12 | 0.55% |
| hsa_circ_0073886 | 4344 | 12 | 0.55% |
| hsa_circ_0068967 | 3982 | 11 | 0.55% |
| hsa_circ_0080479 | 3620 | 10 | 0.55% |
| hsa_circ_0035193 | 5069 | 14 | 0.55% |
| hsa_circ_0070140 | 5069 | 14 | 0.55% |
| hsa_circ_0053586 | 4707 | 13 | 0.55% |
| hsa_circ_0057859 | 3983 | 11 | 0.55% |
| hsa_circ_0030857 | 5794 | 16 | 0.55% |
| hsa_circ_0029889 | 4346 | 12 | 0.55% |
| hsa_circ_0089474 | 4346 | 12 | 0.55% |
| hsa_circ_0021608 | 3984 | 11 | 0.55% |
| hsa_circ_0015035 | 6520 | 18 | 0.55% |
| hsa_circ_0073445 | 4347 | 12 | 0.55% |
| hsa_circ_0078222 | 4347 | 12 | 0.55% |
| hsa_circ_0116778 | 10144 | 28 | 0.55% |
| hsa_circ_0121592 | 4710 | 13 | 0.55% |
| hsa_circ_0059493 | 4348 | 12 | 0.55% |
| hsa_circ_0080884 | 4348 | 12 | 0.55% |
| hsa_circ_0019704 | 3986 | 11 | 0.55% |
| hsa_circ_0034354 | 3986 | 11 | 0.55% |
| hsa_circ_0055912 | 3986 | 11 | 0.55% |
| hsa_circ_0078003 | 3986 | 11 | 0.55% |
| hsa_circ_0024223 | 4711 | 13 | 0.55% |
| hsa_circ_0036227 | 4349 | 12 | 0.55% |
| hsa_circ_0117622 | 4349 | 12 | 0.55% |
| hsa_circ_0021629 | 5437 | 15 | 0.55% |
| hsa_circ_0060842 | 4713 | 13 | 0.55% |
| hsa_circ_0122628 | 11964 | 33 | 0.55% |
| hsa_circ_0036606 | 3988 | 11 | 0.55% |
| hsa_circ_0070122 | 3988 | 11 | 0.55% |
| hsa_circ_0073313 | 3988 | 11 | 0.55% |
| hsa_circ_0082881 | 4714 | 13 | 0.55% |
| hsa_circ_0116595 | 8703 | 24 | 0.55% |
| hsa_circ_0017861 | 4352 | 12 | 0.55% |
| hsa_circ_0023996 | 4352 | 12 | 0.55% |
| hsa_circ_0008587 | 3627 | 10 | 0.55% |
| hsa_circ_0109963 | 15959 | 44 | 0.55% |
| hsa_circ_0032415 | 3990 | 11 | 0.55% |
| hsa_circ_0041804 | 3990 | 11 | 0.55% |
| hsa_circ_0043957 | 3990 | 11 | 0.55% |
| hsa_circ_0121193 | 4353 | 12 | 0.55% |
| hsa_circ_0035781 | 4716 | 13 | 0.55% |
| hsa_circ_0037025 | 3628 | 10 | 0.55% |
| hsa_circ_0026692 | 11247 | 31 | 0.55% |
| hsa_circ_0064846 | 4354 | 12 | 0.55% |
| hsa_circ_0021485 | 5080 | 14 | 0.55% |

| | | | |
|---|---|---|---|
| hsa_circ_0015843 | 3992 | 11 | 0.55% |
| hsa_circ_0095719 | 3992 | 11 | 0.55% |
| hsa_circ_0101003 | 4719 | 13 | 0.55% |
| hsa_circ_0028770 | 4356 | 12 | 0.55% |
| hsa_circ_0125395 | 4356 | 12 | 0.55% |
| hsa_circ_0010456 | 4357 | 12 | 0.55% |
| hsa_circ_0139520 | 5447 | 15 | 0.55% |
| hsa_circ_0042990 | 15252 | 42 | 0.55% |
| hsa_circ_0071877 | 4358 | 12 | 0.55% |
| hsa_circ_0088831 | 4358 | 12 | 0.55% |
| hsa_circ_0124500 | 3995 | 11 | 0.55% |
| hsa_circ_0016935 | 4359 | 12 | 0.55% |
| hsa_circ_0084587 | 4359 | 12 | 0.55% |
| hsa_circ_0134774 | 4359 | 12 | 0.55% |
| hsa_circ_0042943 | 6902 | 19 | 0.55% |
| hsa_circ_0028404 | 3996 | 11 | 0.55% |
| hsa_circ_0071913 | 3996 | 11 | 0.55% |
| hsa_circ_0062410 | 6540 | 18 | 0.55% |
| hsa_circ_0140653 | 4360 | 12 | 0.55% |
| hsa_circ_0024413 | 5088 | 14 | 0.55% |
| hsa_circ_0062339 | 5089 | 14 | 0.55% |
| hsa_circ_0086898 | 4362 | 12 | 0.55% |
| hsa_circ_0129427 | 3999 | 11 | 0.55% |
| hsa_circ_0120687 | 10180 | 28 | 0.55% |
| hsa_circ_0110625 | 5090 | 14 | 0.55% |
| hsa_circ_0060806 | 4727 | 13 | 0.55% |
| hsa_circ_0108049 | 4000 | 11 | 0.55% |
| hsa_circ_0075909 | 4728 | 13 | 0.55% |
| hsa_circ_0090696 | 4728 | 13 | 0.55% |
| hsa_circ_0066497 | 9821 | 27 | 0.55% |
| hsa_circ_0020432 | 4002 | 11 | 0.55% |
| hsa_circ_0082021 | 4366 | 12 | 0.55% |
| hsa_circ_0089222 | 4366 | 12 | 0.55% |
| hsa_circ_0041286 | 4003 | 11 | 0.55% |
| hsa_circ_0091126 | 4003 | 11 | 0.55% |
| hsa_circ_0044862 | 4367 | 12 | 0.55% |
| hsa_circ_0082845 | 4367 | 12 | 0.55% |
| hsa_circ_0093238 | 5460 | 15 | 0.55% |
| hsa_circ_0109352 | 4732 | 13 | 0.55% |
| hsa_circ_0122630 | 4732 | 13 | 0.55% |
| hsa_circ_0054050 | 4368 | 12 | 0.55% |
| hsa_circ_0105784 | 4004 | 11 | 0.55% |
| hsa_circ_0000964 | 11650 | 32 | 0.55% |
| hsa_circ_0082005 | 5097 | 14 | 0.55% |
| hsa_circ_0033336 | 4369 | 12 | 0.55% |
| hsa_circ_0111651 | 14928 | 41 | 0.55% |
| hsa_circ_0096526 | 12381 | 34 | 0.55% |
| hsa_circ_0027567 | 4006 | 11 | 0.55% |
| hsa_circ_0084597 | 4371 | 12 | 0.55% |
| hsa_circ_0127392 | 10929 | 30 | 0.55% |

| | | | |
|---|---|---|---|
| hsa_circ_0072369 | 4736 | 13 | 0.55% |
| hsa_circ_0044785 | 5829 | 16 | 0.55% |
| hsa_circ_0060855 | 5465 | 15 | 0.55% |
| hsa_circ_0074307 | 4008 | 11 | 0.55% |
| hsa_circ_0109654 | 15304 | 42 | 0.55% |
| hsa_circ_0053379 | 4373 | 12 | 0.55% |
| hsa_circ_0090854 | 4373 | 12 | 0.55% |
| hsa_circ_0135552 | 6924 | 19 | 0.55% |
| hsa_circ_0113270 | 12026 | 33 | 0.55% |
| hsa_circ_0019285 | 4738 | 13 | 0.55% |
| hsa_circ_0052253 | 5103 | 14 | 0.55% |
| hsa_circ_0073485 | 4374 | 12 | 0.55% |
| hsa_circ_0012694 | 4739 | 13 | 0.55% |
| hsa_circ_0105785 | 4010 | 11 | 0.55% |
| hsa_circ_0063095 | 4375 | 12 | 0.55% |
| hsa_circ_0083375 | 5469 | 15 | 0.55% |
| hsa_circ_0035501 | 4011 | 11 | 0.55% |
| hsa_circ_0036483 | 4376 | 12 | 0.55% |
| hsa_circ_0054509 | 4376 | 12 | 0.55% |
| hsa_circ_0062579 | 4012 | 11 | 0.55% |
| hsa_circ_0015208 | 4377 | 12 | 0.55% |
| hsa_circ_0056478 | 5107 | 14 | 0.55% |
| hsa_circ_0073480 | 3648 | 10 | 0.55% |
| hsa_circ_0020407 | 4013 | 11 | 0.55% |
| hsa_circ_0000090 | 10580 | 29 | 0.55% |
| hsa_circ_0010220 | 4378 | 12 | 0.55% |
| hsa_circ_0133990 | 8028 | 22 | 0.55% |
| hsa_circ_0000943 | 4014 | 11 | 0.55% |
| hsa_circ_0012910 | 5839 | 16 | 0.55% |
| hsa_circ_0092087 | 4745 | 13 | 0.55% |
| hsa_circ_0051646 | 4746 | 13 | 0.55% |
| hsa_circ_0053668 | 4746 | 13 | 0.55% |
| hsa_circ_0073656 | 4381 | 12 | 0.55% |
| hsa_circ_0038165 | 4016 | 11 | 0.55% |
| hsa_circ_0111305 | 7667 | 21 | 0.55% |
| hsa_circ_0097208 | 4747 | 13 | 0.55% |
| hsa_circ_0108840 | 13146 | 36 | 0.55% |
| hsa_circ_0027838 | 4382 | 12 | 0.55% |
| hsa_circ_0077111 | 4017 | 11 | 0.55% |
| hsa_circ_0062752 | 4018 | 11 | 0.55% |
| hsa_circ_0013705 | 4384 | 12 | 0.55% |
| hsa_circ_0016070 | 4384 | 12 | 0.55% |
| hsa_circ_0017123 | 4384 | 12 | 0.55% |
| hsa_circ_0050832 | 3654 | 10 | 0.55% |
| hsa_circ_0010846 | 4385 | 12 | 0.55% |
| hsa_circ_0041209 | 4385 | 12 | 0.55% |
| hsa_circ_0115209 | 5116 | 14 | 0.55% |
| hsa_circ_0068920 | 4020 | 11 | 0.55% |
| hsa_circ_0046715 | 4751 | 13 | 0.55% |
| hsa_circ_0129835 | 9868 | 27 | 0.55% |

| | | | |
|---|---|---|---|
| hsa_circ_0088826 | 4386 | 12 | 0.55% |
| hsa_circ_0030489 | 5483 | 15 | 0.55% |
| hsa_circ_0025087 | 4752 | 13 | 0.55% |
| hsa_circ_0039315 | 4752 | 13 | 0.55% |
| hsa_circ_0068485 | 4021 | 11 | 0.55% |
| hsa_circ_0023716 | 6581 | 18 | 0.55% |
| hsa_circ_0106401 | 11702 | 32 | 0.55% |
| hsa_circ_0108254 | 4755 | 13 | 0.55% |
| hsa_circ_0070726 | 4024 | 11 | 0.55% |
| hsa_circ_0091981 | 4391 | 12 | 0.55% |
| hsa_circ_0123858 | 4758 | 13 | 0.55% |
| hsa_circ_0077199 | 4392 | 12 | 0.55% |
| hsa_circ_0128660 | 15373 | 42 | 0.55% |
| hsa_circ_0066943 | 4759 | 13 | 0.55% |
| hsa_circ_0028361 | 4393 | 12 | 0.55% |
| hsa_circ_0050113 | 4027 | 11 | 0.55% |
| hsa_circ_0013864 | 9153 | 25 | 0.55% |
| hsa_circ_0064096 | 4760 | 13 | 0.55% |
| hsa_circ_0069540 | 4760 | 13 | 0.55% |
| hsa_circ_0104128 | 4760 | 13 | 0.55% |
| hsa_circ_0041682 | 4394 | 12 | 0.55% |
| hsa_circ_0037522 | 4028 | 11 | 0.55% |
| hsa_circ_0042627 | 4028 | 11 | 0.55% |
| hsa_circ_0051047 | 5493 | 15 | 0.55% |
| hsa_circ_0028685 | 5127 | 14 | 0.55% |
| hsa_circ_0058097 | 4761 | 13 | 0.55% |
| hsa_circ_0082499 | 4761 | 13 | 0.55% |
| hsa_circ_0112200 | 4395 | 12 | 0.55% |
| hsa_circ_0041708 | 5128 | 14 | 0.55% |
| hsa_circ_0025103 | 4396 | 12 | 0.55% |
| hsa_circ_0093860 | 4396 | 12 | 0.55% |
| hsa_circ_0076689 | 4763 | 13 | 0.55% |
| hsa_circ_0010437 | 4031 | 11 | 0.55% |
| hsa_circ_0060963 | 4031 | 11 | 0.55% |
| hsa_circ_0048723 | 4764 | 13 | 0.55% |
| hsa_circ_0062334 | 5497 | 15 | 0.55% |
| hsa_circ_0024240 | 48012 | 131 | 0.55% |
| hsa_circ_0111847 | 4765 | 13 | 0.55% |
| hsa_circ_0047247 | 4399 | 12 | 0.55% |
| hsa_circ_0073010 | 4399 | 12 | 0.55% |
| hsa_circ_0030140 | 4766 | 13 | 0.55% |
| hsa_circ_0065407 | 4766 | 13 | 0.55% |
| hsa_circ_0070780 | 4766 | 13 | 0.55% |
| hsa_circ_0030460 | 4033 | 11 | 0.55% |
| hsa_circ_0047152 | 4033 | 11 | 0.55% |
| hsa_circ_0063720 | 5134 | 14 | 0.55% |
| hsa_circ_0024931 | 4768 | 13 | 0.55% |
| hsa_circ_0064217 | 4035 | 11 | 0.55% |
| hsa_circ_0070310 | 4035 | 11 | 0.55% |
| hsa_circ_0127157 | 4035 | 11 | 0.55% |

| | | | |
|---|---|---|---|
| hsa_circ_0132937 | 11007 | 30 | 0.55% |
| hsa_circ_0019814 | 4036 | 11 | 0.55% |
| hsa_circ_0090855 | 4403 | 12 | 0.55% |
| hsa_circ_0110324 | 5137 | 14 | 0.55% |
| hsa_circ_0107672 | 24221 | 66 | 0.54% |
| hsa_circ_0063086 | 4037 | 11 | 0.54% |
| hsa_circ_0081218 | 3670 | 10 | 0.54% |
| hsa_circ_0016606 | 4038 | 11 | 0.54% |
| hsa_circ_0101782 | 4773 | 13 | 0.54% |
| hsa_circ_0123313 | 4773 | 13 | 0.54% |
| hsa_circ_0052679 | 4406 | 12 | 0.54% |
| hsa_circ_0019320 | 4039 | 11 | 0.54% |
| hsa_circ_0030856 | 5875 | 16 | 0.54% |
| hsa_circ_0076493 | 4408 | 12 | 0.54% |
| hsa_circ_0068598 | 4041 | 11 | 0.54% |
| hsa_circ_0048868 | 4776 | 13 | 0.54% |
| hsa_circ_0006727 | 16165 | 44 | 0.54% |
| hsa_circ_0103866 | 12859 | 35 | 0.54% |
| hsa_circ_0123572 | 9186 | 25 | 0.54% |
| hsa_circ_0029506 | 4777 | 13 | 0.54% |
| hsa_circ_0047514 | 5145 | 14 | 0.54% |
| hsa_circ_0058196 | 4778 | 13 | 0.54% |
| hsa_circ_0087810 | 4043 | 11 | 0.54% |
| hsa_circ_0076000 | 5146 | 14 | 0.54% |
| hsa_circ_0018320 | 4411 | 12 | 0.54% |
| hsa_circ_0098326 | 4411 | 12 | 0.54% |
| hsa_circ_0108961 | 4411 | 12 | 0.54% |
| hsa_circ_0054009 | 4044 | 11 | 0.54% |
| hsa_circ_0136046 | 4044 | 11 | 0.54% |
| hsa_circ_0036782 | 14338 | 39 | 0.54% |
| hsa_circ_0016376 | 4412 | 12 | 0.54% |
| hsa_circ_0038677 | 4780 | 13 | 0.54% |
| hsa_circ_0019451 | 4045 | 11 | 0.54% |
| hsa_circ_0128829 | 4045 | 11 | 0.54% |
| hsa_circ_0104685 | 5516 | 15 | 0.54% |
| hsa_circ_0108841 | 13239 | 36 | 0.54% |
| hsa_circ_0089440 | 4413 | 12 | 0.54% |
| hsa_circ_0005408 | 8091 | 22 | 0.54% |
| hsa_circ_0000444 | 9930 | 27 | 0.54% |
| hsa_circ_0128676 | 5149 | 14 | 0.54% |
| hsa_circ_0004555 | 5517 | 15 | 0.54% |
| hsa_circ_0071982 | 3678 | 10 | 0.54% |
| hsa_circ_0117712 | 3678 | 10 | 0.54% |
| hsa_circ_0071166 | 4414 | 12 | 0.54% |
| hsa_circ_0087314 | 4414 | 12 | 0.54% |
| hsa_circ_0009978 | 4415 | 12 | 0.54% |
| hsa_circ_0081708 | 22443 | 61 | 0.54% |
| hsa_circ_0122653 | 11408 | 31 | 0.54% |
| hsa_circ_0074784 | 4416 | 12 | 0.54% |
| hsa_circ_0085092 | 3680 | 10 | 0.54% |

| | | | |
|---|---|---|---|
| hsa_circ_0116846 | 15825 | 43 | 0.54% |
| hsa_circ_0086917 | 4785 | 13 | 0.54% |
| hsa_circ_0047161 | 4049 | 11 | 0.54% |
| hsa_circ_0085209 | 4786 | 13 | 0.54% |
| hsa_circ_0090744 | 4786 | 13 | 0.54% |
| hsa_circ_0072178 | 4418 | 12 | 0.54% |
| hsa_circ_0076051 | 4418 | 12 | 0.54% |
| hsa_circ_0035676 | 4787 | 13 | 0.54% |
| hsa_circ_0046345 | 4787 | 13 | 0.54% |
| hsa_circ_0086893 | 4787 | 13 | 0.54% |
| hsa_circ_0047986 | 4419 | 12 | 0.54% |
| hsa_circ_0071606 | 4419 | 12 | 0.54% |
| hsa_circ_0075702 | 4419 | 12 | 0.54% |
| hsa_circ_0091976 | 4419 | 12 | 0.54% |
| hsa_circ_0084903 | 4051 | 11 | 0.54% |
| hsa_circ_0084957 | 5156 | 14 | 0.54% |
| hsa_circ_0101232 | 4788 | 13 | 0.54% |
| hsa_circ_0004650 | 4052 | 11 | 0.54% |
| hsa_circ_0046281 | 4789 | 13 | 0.54% |
| hsa_circ_0091101 | 4789 | 13 | 0.54% |
| hsa_circ_0084619 | 4790 | 13 | 0.54% |
| hsa_circ_0010547 | 5159 | 14 | 0.54% |
| hsa_circ_0085863 | 4791 | 13 | 0.54% |
| hsa_circ_0062205 | 4423 | 12 | 0.54% |
| hsa_circ_0069549 | 5898 | 16 | 0.54% |
| hsa_circ_0103217 | 4055 | 11 | 0.54% |
| hsa_circ_0128467 | 14747 | 40 | 0.54% |
| hsa_circ_0035344 | 5162 | 14 | 0.54% |
| hsa_circ_0044756 | 4056 | 11 | 0.54% |
| hsa_circ_0052691 | 5531 | 15 | 0.54% |
| hsa_circ_0070862 | 4425 | 12 | 0.54% |
| hsa_circ_0092250 | 4794 | 13 | 0.54% |
| hsa_circ_0024098 | 4426 | 12 | 0.54% |
| hsa_circ_0000726 | 7378 | 20 | 0.54% |
| hsa_circ_0117693 | 3689 | 10 | 0.54% |
| hsa_circ_0036957 | 5167 | 14 | 0.54% |
| hsa_circ_0067374 | 4060 | 11 | 0.54% |
| hsa_circ_0033093 | 4799 | 13 | 0.54% |
| hsa_circ_0086851 | 4061 | 11 | 0.54% |
| hsa_circ_0056901 | 10707 | 29 | 0.54% |
| hsa_circ_0090761 | 5169 | 14 | 0.54% |
| hsa_circ_0097096 | 4800 | 13 | 0.54% |
| hsa_circ_0068525 | 5908 | 16 | 0.54% |
| hsa_circ_0026997 | 4062 | 11 | 0.54% |
| hsa_circ_0035304 | 4062 | 11 | 0.54% |
| hsa_circ_0059152 | 4062 | 11 | 0.54% |
| hsa_circ_0134623 | 3693 | 10 | 0.54% |
| hsa_circ_0127288 | 4432 | 12 | 0.54% |
| hsa_circ_0005459 | 3694 | 10 | 0.54% |
| hsa_circ_0086064 | 4433 | 12 | 0.54% |

| | | | |
|---|---|---|---|
| hsa_circ_0104688 | 4433 | 12 | 0.54% |
| hsa_circ_0120765 | 4433 | 12 | 0.54% |
| hsa_circ_0084690 | 4064 | 11 | 0.54% |
| hsa_circ_0066138 | 4065 | 11 | 0.54% |
| hsa_circ_0055642 | 5544 | 15 | 0.54% |
| hsa_circ_0028684 | 5175 | 14 | 0.54% |
| hsa_circ_0073944 | 6284 | 17 | 0.54% |
| hsa_circ_0041367 | 4067 | 11 | 0.54% |
| hsa_circ_0075162 | 4067 | 11 | 0.54% |
| hsa_circ_0093750 | 4067 | 11 | 0.54% |
| hsa_circ_0096727 | 10724 | 29 | 0.54% |
| hsa_circ_0025952 | 4068 | 11 | 0.54% |
| hsa_circ_0084495 | 4438 | 12 | 0.54% |
| hsa_circ_0011780 | 3699 | 10 | 0.54% |
| hsa_circ_0131627 | 4069 | 11 | 0.54% |
| hsa_circ_0027010 | 4439 | 12 | 0.54% |
| hsa_circ_0077195 | 4439 | 12 | 0.54% |
| hsa_circ_0093342 | 11099 | 30 | 0.54% |
| hsa_circ_0046748 | 4440 | 12 | 0.54% |
| hsa_circ_0070066 | 4440 | 12 | 0.54% |
| hsa_circ_0011069 | 5181 | 14 | 0.54% |
| hsa_circ_0112855 | 4072 | 11 | 0.54% |
| hsa_circ_0020058 | 5554 | 15 | 0.54% |
| hsa_circ_0049445 | 4073 | 11 | 0.54% |
| hsa_circ_0054874 | 4444 | 12 | 0.54% |
| hsa_circ_0054508 | 4445 | 12 | 0.54% |
| hsa_circ_0089428 | 4445 | 12 | 0.54% |
| hsa_circ_0124965 | 4445 | 12 | 0.54% |
| hsa_circ_0016011 | 3705 | 10 | 0.54% |
| hsa_circ_0036566 | 3705 | 10 | 0.54% |
| hsa_circ_0067128 | 11486 | 31 | 0.54% |
| hsa_circ_0071897 | 4076 | 11 | 0.54% |
| hsa_circ_0073330 | 4076 | 11 | 0.54% |
| hsa_circ_0056479 | 5188 | 14 | 0.54% |
| hsa_circ_0013418 | 4447 | 12 | 0.54% |
| hsa_circ_0128753 | 7412 | 20 | 0.54% |
| hsa_circ_0030855 | 5930 | 16 | 0.54% |
| hsa_circ_0038080 | 4077 | 11 | 0.54% |
| hsa_circ_0029966 | 5189 | 14 | 0.54% |
| hsa_circ_0019222 | 4448 | 12 | 0.54% |
| hsa_circ_0063103 | 4449 | 12 | 0.54% |
| hsa_circ_0091702 | 4449 | 12 | 0.54% |
| hsa_circ_0052307 | 22248 | 60 | 0.54% |
| hsa_circ_0095434 | 4450 | 12 | 0.54% |
| hsa_circ_0036735 | 3709 | 10 | 0.54% |
| hsa_circ_0007523 | 7789 | 21 | 0.54% |
| hsa_circ_0031176 | 4451 | 12 | 0.54% |
| hsa_circ_0077477 | 4451 | 12 | 0.54% |
| hsa_circ_0060856 | 5564 | 15 | 0.54% |
| hsa_circ_0036863 | 4452 | 12 | 0.54% |

| | | | |
|---|---|---|---|
| hsa_circ_0123387 | 4452 | 12 | 0.54% |
| hsa_circ_0088607 | 5937 | 16 | 0.54% |
| hsa_circ_0028730 | 4453 | 12 | 0.54% |
| hsa_circ_0027427 | 4083 | 11 | 0.54% |
| hsa_circ_0030472 | 4083 | 11 | 0.54% |
| hsa_circ_0095813 | 9651 | 26 | 0.54% |
| hsa_circ_0103215 | 12250 | 33 | 0.54% |
| hsa_circ_0111070 | 16335 | 44 | 0.54% |
| hsa_circ_0064844 | 4455 | 12 | 0.54% |
| hsa_circ_0125341 | 43815 | 118 | 0.54% |
| hsa_circ_0085483 | 4828 | 13 | 0.54% |
| hsa_circ_0028428 | 5571 | 15 | 0.54% |
| hsa_circ_0055606 | 9657 | 26 | 0.54% |
| hsa_circ_0003811 | 10030 | 27 | 0.54% |
| hsa_circ_0060790 | 4458 | 12 | 0.54% |
| hsa_circ_0049366 | 4087 | 11 | 0.54% |
| hsa_circ_0047139 | 4088 | 11 | 0.54% |
| hsa_circ_0052998 | 4088 | 11 | 0.54% |
| hsa_circ_0087988 | 4088 | 11 | 0.54% |
| hsa_circ_0036484 | 4460 | 12 | 0.54% |
| hsa_circ_0035675 | 4089 | 11 | 0.54% |
| hsa_circ_0061719 | 4089 | 11 | 0.54% |
| hsa_circ_0130619 | 33084 | 89 | 0.54% |
| hsa_circ_0091736 | 6320 | 17 | 0.54% |
| hsa_circ_0110839 | 22309 | 60 | 0.54% |
| hsa_circ_0074465 | 4462 | 12 | 0.54% |
| hsa_circ_0101591 | 4462 | 12 | 0.54% |
| hsa_circ_0019904 | 4834 | 13 | 0.54% |
| hsa_circ_0131054 | 20452 | 55 | 0.54% |
| hsa_circ_0027748 | 4091 | 11 | 0.54% |
| hsa_circ_0043662 | 16736 | 45 | 0.54% |
| hsa_circ_0057982 | 4463 | 12 | 0.54% |
| hsa_circ_0002246 | 4835 | 13 | 0.54% |
| hsa_circ_0114723 | 18227 | 49 | 0.54% |
| hsa_circ_0086871 | 4836 | 13 | 0.54% |
| hsa_circ_0016005 | 4092 | 11 | 0.54% |
| hsa_circ_0064928 | 3720 | 10 | 0.54% |
| hsa_circ_0057860 | 4093 | 11 | 0.54% |
| hsa_circ_0070340 | 4093 | 11 | 0.54% |
| hsa_circ_0123607 | 14513 | 39 | 0.54% |
| hsa_circ_0070137 | 5210 | 14 | 0.54% |
| hsa_circ_0094882 | 4466 | 12 | 0.54% |
| hsa_circ_0071147 | 3722 | 10 | 0.54% |
| hsa_circ_0129950 | 4839 | 13 | 0.54% |
| hsa_circ_0102065 | 4467 | 12 | 0.54% |
| hsa_circ_0090992 | 5584 | 15 | 0.54% |
| hsa_circ_0028393 | 4095 | 11 | 0.54% |
| hsa_circ_0021578 | 5212 | 14 | 0.54% |
| hsa_circ_0124564 | 8563 | 23 | 0.54% |
| hsa_circ_0033313 | 4468 | 12 | 0.54% |

| | | | |
|---|---|---|---|
| hsa_circ_0116390 | 16757 | 45 | 0.54% |
| hsa_circ_0138428 | 4469 | 12 | 0.54% |
| hsa_circ_0005905 | 4470 | 12 | 0.54% |
| hsa_circ_0053807 | 4470 | 12 | 0.54% |
| hsa_circ_0073468 | 4470 | 12 | 0.54% |
| hsa_circ_0041881 | 4471 | 12 | 0.54% |
| hsa_circ_0050766 | 7452 | 20 | 0.54% |
| hsa_circ_0037315 | 3726 | 10 | 0.54% |
| hsa_circ_0029174 | 4844 | 13 | 0.54% |
| hsa_circ_0021064 | 4099 | 11 | 0.54% |
| hsa_circ_0101717 | 19750 | 53 | 0.54% |
| hsa_circ_0065064 | 5217 | 14 | 0.54% |
| hsa_circ_0097330 | 10807 | 29 | 0.54% |
| hsa_circ_0029504 | 4845 | 13 | 0.54% |
| hsa_circ_0075703 | 4473 | 12 | 0.54% |
| hsa_circ_0138392 | 5219 | 14 | 0.54% |
| hsa_circ_0109270 | 14166 | 38 | 0.54% |
| hsa_circ_0049112 | 4474 | 12 | 0.54% |
| hsa_circ_0073289 | 4474 | 12 | 0.54% |
| hsa_circ_0005213 | 16407 | 44 | 0.54% |
| hsa_circ_0039071 | 4848 | 13 | 0.54% |
| hsa_circ_0078422 | 7087 | 19 | 0.54% |
| hsa_circ_0001209 | 4849 | 13 | 0.54% |
| hsa_circ_0104500 | 4849 | 13 | 0.54% |
| hsa_circ_0012772 | 4103 | 11 | 0.54% |
| hsa_circ_0112546 | 4103 | 11 | 0.54% |
| hsa_circ_0111638 | 3730 | 10 | 0.54% |
| hsa_circ_0061250 | 9699 | 26 | 0.54% |
| hsa_circ_0127338 | 13431 | 36 | 0.54% |
| hsa_circ_0065590 | 4104 | 11 | 0.54% |
| hsa_circ_0092079 | 4104 | 11 | 0.54% |
| hsa_circ_0027750 | 3731 | 10 | 0.54% |
| hsa_circ_0035782 | 4851 | 13 | 0.54% |
| hsa_circ_0089136 | 4478 | 12 | 0.54% |
| hsa_circ_0103825 | 4105 | 11 | 0.54% |
| hsa_circ_0088617 | 5972 | 16 | 0.54% |
| hsa_circ_0071915 | 4479 | 12 | 0.54% |
| hsa_circ_0085190 | 5226 | 14 | 0.54% |
| hsa_circ_0052638 | 4853 | 13 | 0.54% |
| hsa_circ_0013834 | 4480 | 12 | 0.54% |
| hsa_circ_0094641 | 4480 | 12 | 0.54% |
| hsa_circ_0046282 | 4854 | 13 | 0.54% |
| hsa_circ_0065414 | 4481 | 12 | 0.54% |
| hsa_circ_0078105 | 4481 | 12 | 0.54% |
| hsa_circ_0119049 | 4855 | 13 | 0.54% |
| hsa_circ_0028919 | 6349 | 17 | 0.54% |
| hsa_circ_0107099 | 11952 | 32 | 0.54% |
| hsa_circ_0015083 | 4482 | 12 | 0.54% |
| hsa_circ_0134713 | 7471 | 20 | 0.54% |
| hsa_circ_0131162 | 6724 | 18 | 0.54% |

| | | | |
|---|---|---|---|
| hsa_circ_0006923 | 4110 | 11 | 0.54% |
| hsa_circ_0098126 | 11958 | 32 | 0.54% |
| hsa_circ_0025495 | 4858 | 13 | 0.54% |
| hsa_circ_0061140 | 3737 | 10 | 0.54% |
| hsa_circ_0052997 | 4111 | 11 | 0.54% |
| hsa_circ_0045908 | 4859 | 13 | 0.54% |
| hsa_circ_0041854 | 3738 | 10 | 0.54% |
| hsa_circ_0117937 | 3738 | 10 | 0.54% |
| hsa_circ_0019813 | 4112 | 11 | 0.54% |
| hsa_circ_0020015 | 4112 | 11 | 0.54% |
| hsa_circ_0087201 | 4112 | 11 | 0.54% |
| hsa_circ_0110788 | 4486 | 12 | 0.53% |
| hsa_circ_0107098 | 11963 | 32 | 0.53% |
| hsa_circ_0023223 | 4860 | 13 | 0.53% |
| hsa_circ_0071182 | 4860 | 13 | 0.53% |
| hsa_circ_0055605 | 9721 | 26 | 0.53% |
| hsa_circ_0084279 | 4113 | 11 | 0.53% |
| hsa_circ_0091887 | 4113 | 11 | 0.53% |
| hsa_circ_0012042 | 4861 | 13 | 0.53% |
| hsa_circ_0015708 | 5610 | 15 | 0.53% |
| hsa_circ_0090669 | 4489 | 12 | 0.53% |
| hsa_circ_0012980 | 4115 | 11 | 0.53% |
| hsa_circ_0112145 | 4115 | 11 | 0.53% |
| hsa_circ_0130542 | 4865 | 13 | 0.53% |
| hsa_circ_0034479 | 4491 | 12 | 0.53% |
| hsa_circ_0054809 | 4491 | 12 | 0.53% |
| hsa_circ_0069458 | 4117 | 11 | 0.53% |
| hsa_circ_0002734 | 43416 | 116 | 0.53% |
| hsa_circ_0070472 | 5240 | 14 | 0.53% |
| hsa_circ_0000476 | 12352 | 33 | 0.53% |
| hsa_circ_0111525 | 7112 | 19 | 0.53% |
| hsa_circ_0136688 | 72620 | 194 | 0.53% |
| hsa_circ_0030641 | 4492 | 12 | 0.53% |
| hsa_circ_0075986 | 5241 | 14 | 0.53% |
| hsa_circ_0068037 | 5242 | 14 | 0.53% |
| hsa_circ_0117129 | 10859 | 29 | 0.53% |
| hsa_circ_0072368 | 4868 | 13 | 0.53% |
| hsa_circ_0084710 | 4494 | 12 | 0.53% |
| hsa_circ_0086065 | 4494 | 12 | 0.53% |
| hsa_circ_0103216 | 4120 | 11 | 0.53% |
| hsa_circ_0138629 | 7117 | 19 | 0.53% |
| hsa_circ_0023187 | 4495 | 12 | 0.53% |
| hsa_circ_0032997 | 4495 | 12 | 0.53% |
| hsa_circ_0072174 | 4496 | 12 | 0.53% |
| hsa_circ_0084729 | 4871 | 13 | 0.53% |
| hsa_circ_0070050 | 4122 | 11 | 0.53% |
| hsa_circ_0014736 | 5621 | 15 | 0.53% |
| hsa_circ_0125968 | 4872 | 13 | 0.53% |
| hsa_circ_0033435 | 4123 | 11 | 0.53% |
| hsa_circ_0036874 | 4123 | 11 | 0.53% |

| | | | |
|---|---|---|---|
| hsa_circ_0042942 | 7122 | 19 | 0.53% |
| hsa_circ_0071203 | 4873 | 13 | 0.53% |
| hsa_circ_0028784 | 4124 | 11 | 0.53% |
| hsa_circ_0039742 | 4124 | 11 | 0.53% |
| hsa_circ_0065174 | 4499 | 12 | 0.53% |
| hsa_circ_0029099 | 4874 | 13 | 0.53% |
| hsa_circ_0054725 | 4875 | 13 | 0.53% |
| hsa_circ_0085568 | 4501 | 12 | 0.53% |
| hsa_circ_0028403 | 4126 | 11 | 0.53% |
| hsa_circ_0030622 | 4126 | 11 | 0.53% |
| hsa_circ_0055963 | 4126 | 11 | 0.53% |
| hsa_circ_0040091 | 4502 | 12 | 0.53% |
| hsa_circ_0017167 | 4127 | 11 | 0.53% |
| hsa_circ_0052700 | 5628 | 15 | 0.53% |
| hsa_circ_0033303 | 4503 | 12 | 0.53% |
| hsa_circ_0073668 | 4504 | 12 | 0.53% |
| hsa_circ_0081970 | 4504 | 12 | 0.53% |
| hsa_circ_0089206 | 4504 | 12 | 0.53% |
| hsa_circ_0029503 | 4505 | 12 | 0.53% |
| hsa_circ_0062401 | 4505 | 12 | 0.53% |
| hsa_circ_0092243 | 4505 | 12 | 0.53% |
| hsa_circ_0026100 | 4130 | 11 | 0.53% |
| hsa_circ_0075851 | 4881 | 13 | 0.53% |
| hsa_circ_0127690 | 13895 | 37 | 0.53% |
| hsa_circ_0028442 | 4883 | 13 | 0.53% |
| hsa_circ_0082100 | 4132 | 11 | 0.53% |
| hsa_circ_0002977 | 12772 | 34 | 0.53% |
| hsa_circ_0019601 | 4508 | 12 | 0.53% |
| hsa_circ_0034771 | 4884 | 13 | 0.53% |
| hsa_circ_0015594 | 4509 | 12 | 0.53% |
| hsa_circ_0028362 | 4509 | 12 | 0.53% |
| hsa_circ_0041951 | 4885 | 13 | 0.53% |
| hsa_circ_0074268 | 4885 | 13 | 0.53% |
| hsa_circ_0020854 | 4134 | 11 | 0.53% |
| hsa_circ_0054507 | 4510 | 12 | 0.53% |
| hsa_circ_0138597 | 4886 | 13 | 0.53% |
| hsa_circ_0091804 | 5263 | 14 | 0.53% |
| hsa_circ_0071078 | 4889 | 13 | 0.53% |
| hsa_circ_0030490 | 5642 | 15 | 0.53% |
| hsa_circ_0042662 | 4890 | 13 | 0.53% |
| hsa_circ_0126681 | 21065 | 56 | 0.53% |
| hsa_circ_0050841 | 4514 | 12 | 0.53% |
| hsa_circ_0075152 | 4514 | 12 | 0.53% |
| hsa_circ_0083193 | 4514 | 12 | 0.53% |
| hsa_circ_0034337 | 4138 | 11 | 0.53% |
| hsa_circ_0051167 | 4138 | 11 | 0.53% |
| hsa_circ_0081517 | 4138 | 11 | 0.53% |
| hsa_circ_0111269 | 5267 | 14 | 0.53% |
| hsa_circ_0025088 | 4891 | 13 | 0.53% |
| hsa_circ_0056140 | 4515 | 12 | 0.53% |

| | | | |
|---|---|---|---|
| hsa_circ_0129480 | 4515 | 12 | 0.53% |
| hsa_circ_0022613 | 4516 | 12 | 0.53% |
| hsa_circ_0068599 | 4140 | 11 | 0.53% |
| hsa_circ_0081315 | 4140 | 11 | 0.53% |
| hsa_circ_0128973 | 4893 | 13 | 0.53% |
| hsa_circ_0056051 | 4517 | 12 | 0.53% |
| hsa_circ_0098143 | 4517 | 12 | 0.53% |
| hsa_circ_0126853 | 4517 | 12 | 0.53% |
| hsa_circ_0122784 | 12422 | 33 | 0.53% |
| hsa_circ_0057048 | 5271 | 14 | 0.53% |
| hsa_circ_0066610 | 4518 | 12 | 0.53% |
| hsa_circ_0069935 | 4518 | 12 | 0.53% |
| hsa_circ_0015881 | 4142 | 11 | 0.53% |
| hsa_circ_0052724 | 4142 | 11 | 0.53% |
| hsa_circ_0069975 | 4142 | 11 | 0.53% |
| hsa_circ_0011832 | 4519 | 12 | 0.53% |
| hsa_circ_0113027 | 4519 | 12 | 0.53% |
| hsa_circ_0048185 | 4520 | 12 | 0.53% |
| hsa_circ_0119000 | 7534 | 20 | 0.53% |
| hsa_circ_0093861 | 4521 | 12 | 0.53% |
| hsa_circ_0122877 | 47095 | 125 | 0.53% |
| hsa_circ_0058197 | 4898 | 13 | 0.53% |
| hsa_circ_0070325 | 4145 | 11 | 0.53% |
| hsa_circ_0077355 | 4146 | 11 | 0.53% |
| hsa_circ_0115110 | 4146 | 11 | 0.53% |
| hsa_circ_0056132 | 10932 | 29 | 0.53% |
| hsa_circ_0034222 | 4901 | 13 | 0.53% |
| hsa_circ_0124905 | 4148 | 11 | 0.53% |
| hsa_circ_0009950 | 4526 | 12 | 0.53% |
| hsa_circ_0057585 | 21501 | 57 | 0.53% |
| hsa_circ_0084588 | 4527 | 12 | 0.53% |
| hsa_circ_0091144 | 4527 | 12 | 0.53% |
| hsa_circ_0025954 | 4150 | 11 | 0.53% |
| hsa_circ_0125912 | 3773 | 10 | 0.53% |
| hsa_circ_0020059 | 5660 | 15 | 0.53% |
| hsa_circ_0062928 | 4528 | 12 | 0.53% |
| hsa_circ_0043537 | 4529 | 12 | 0.53% |
| hsa_circ_0124298 | 4152 | 11 | 0.53% |
| hsa_circ_0024124 | 4530 | 12 | 0.53% |
| hsa_circ_0047248 | 4530 | 12 | 0.53% |
| hsa_circ_0071791 | 4530 | 12 | 0.53% |
| hsa_circ_0084215 | 4530 | 12 | 0.53% |
| hsa_circ_0119958 | 4530 | 12 | 0.53% |
| hsa_circ_0121590 | 10193 | 27 | 0.53% |
| hsa_circ_0042202 | 4154 | 11 | 0.53% |
| hsa_circ_0024412 | 5287 | 14 | 0.53% |
| hsa_circ_0002953 | 9819 | 26 | 0.53% |
| hsa_circ_0058222 | 4532 | 12 | 0.53% |
| hsa_circ_0020408 | 4155 | 11 | 0.53% |
| hsa_circ_0077156 | 4155 | 11 | 0.53% |

| | | | |
|---|---|---|---|
| hsa_circ_0087984 | 4155 | 11 | 0.53% |
| hsa_circ_0041458 | 4533 | 12 | 0.53% |
| hsa_circ_0082512 | 4911 | 13 | 0.53% |
| hsa_circ_0082004 | 5289 | 14 | 0.53% |
| hsa_circ_0003145 | 3778 | 10 | 0.53% |
| hsa_circ_0088612 | 4534 | 12 | 0.53% |
| hsa_circ_0045453 | 4157 | 11 | 0.53% |
| hsa_circ_0002699 | 4535 | 12 | 0.53% |
| hsa_circ_0070682 | 4535 | 12 | 0.53% |
| hsa_circ_0069914 | 5291 | 14 | 0.53% |
| hsa_circ_0113765 | 5669 | 15 | 0.53% |
| hsa_circ_0128052 | 13608 | 36 | 0.53% |
| hsa_circ_0015220 | 4158 | 11 | 0.53% |
| hsa_circ_0074106 | 3780 | 10 | 0.53% |
| hsa_circ_0112807 | 8317 | 22 | 0.53% |
| hsa_circ_0053449 | 4159 | 11 | 0.53% |
| hsa_circ_0071202 | 4159 | 11 | 0.53% |
| hsa_circ_0103250 | 18528 | 49 | 0.53% |
| hsa_circ_0034229 | 4916 | 13 | 0.53% |
| hsa_circ_0085864 | 4916 | 13 | 0.53% |
| hsa_circ_0002851 | 4538 | 12 | 0.53% |
| hsa_circ_0053565 | 4160 | 11 | 0.53% |
| hsa_circ_0136211 | 11724 | 31 | 0.53% |
| hsa_circ_0032979 | 4917 | 13 | 0.53% |
| hsa_circ_0047978 | 4539 | 12 | 0.53% |
| hsa_circ_0009565 | 4161 | 11 | 0.53% |
| hsa_circ_0081680 | 13998 | 37 | 0.53% |
| hsa_circ_0049106 | 4540 | 12 | 0.53% |
| hsa_circ_0052115 | 4540 | 12 | 0.53% |
| hsa_circ_0063721 | 5297 | 14 | 0.53% |
| hsa_circ_0005147 | 4162 | 11 | 0.53% |
| hsa_circ_0049031 | 4920 | 13 | 0.53% |
| hsa_circ_0088832 | 4542 | 12 | 0.53% |
| hsa_circ_0040801 | 4543 | 12 | 0.53% |
| hsa_circ_0073481 | 3787 | 10 | 0.53% |
| hsa_circ_0116206 | 4166 | 11 | 0.53% |
| hsa_circ_0096738 | 4545 | 12 | 0.53% |
| hsa_circ_0034983 | 3788 | 10 | 0.53% |
| hsa_circ_0135881 | 4167 | 11 | 0.53% |
| hsa_circ_0069393 | 4546 | 12 | 0.53% |
| hsa_circ_0124922 | 4926 | 13 | 0.53% |
| hsa_circ_0094642 | 4927 | 13 | 0.53% |
| hsa_circ_0089161 | 4548 | 12 | 0.53% |
| hsa_circ_0044622 | 4169 | 11 | 0.53% |
| hsa_circ_0102435 | 17816 | 47 | 0.53% |
| hsa_circ_0009807 | 5307 | 14 | 0.53% |
| hsa_circ_0016792 | 4170 | 11 | 0.53% |
| hsa_circ_0030078 | 4170 | 11 | 0.53% |
| hsa_circ_0030709 | 4170 | 11 | 0.53% |
| hsa_circ_0072156 | 4170 | 11 | 0.53% |

| | | | |
|---|---|---|---|
| hsa_circ_0076308 | 4170 | 11 | 0.53% |
| hsa_circ_0091904 | 4551 | 12 | 0.53% |
| hsa_circ_0036826 | 4931 | 13 | 0.53% |
| hsa_circ_0038615 | 4931 | 13 | 0.53% |
| hsa_circ_0053572 | 4931 | 13 | 0.53% |
| hsa_circ_0016278 | 4552 | 12 | 0.53% |
| hsa_circ_0122499 | 6830 | 18 | 0.53% |
| hsa_circ_0090665 | 4933 | 13 | 0.53% |
| hsa_circ_0085055 | 5692 | 15 | 0.53% |
| hsa_circ_0063954 | 4554 | 12 | 0.53% |
| hsa_circ_0037148 | 4934 | 13 | 0.53% |
| hsa_circ_0122781 | 12525 | 33 | 0.53% |
| hsa_circ_0027212 | 4175 | 11 | 0.53% |
| hsa_circ_0047196 | 4555 | 12 | 0.53% |
| hsa_circ_0087428 | 8351 | 22 | 0.53% |
| hsa_circ_0024010 | 5694 | 15 | 0.53% |
| hsa_circ_0026924 | 6074 | 16 | 0.53% |
| hsa_circ_0041399 | 4557 | 12 | 0.53% |
| hsa_circ_0091846 | 5317 | 14 | 0.53% |
| hsa_circ_0100960 | 4558 | 12 | 0.53% |
| hsa_circ_0039616 | 4938 | 13 | 0.53% |
| hsa_circ_0036872 | 4180 | 11 | 0.53% |
| hsa_circ_0008650 | 7221 | 19 | 0.53% |
| hsa_circ_0080517 | 4941 | 13 | 0.53% |
| hsa_circ_0136863 | 4941 | 13 | 0.53% |
| hsa_circ_0021542 | 12545 | 33 | 0.53% |
| hsa_circ_0011360 | 4562 | 12 | 0.53% |
| hsa_circ_0025471 | 9886 | 26 | 0.53% |
| hsa_circ_0037520 | 4183 | 11 | 0.53% |
| hsa_circ_0087562 | 4183 | 11 | 0.53% |
| hsa_circ_0014590 | 4564 | 12 | 0.53% |
| hsa_circ_0125970 | 4564 | 12 | 0.53% |
| hsa_circ_0011770 | 4184 | 11 | 0.53% |
| hsa_circ_0064097 | 4945 | 13 | 0.53% |
| hsa_circ_0125152 | 7608 | 20 | 0.53% |
| hsa_circ_0127484 | 4946 | 13 | 0.53% |
| hsa_circ_0016425 | 4947 | 13 | 0.53% |
| hsa_circ_0053669 | 4947 | 13 | 0.53% |
| hsa_circ_0125975 | 4947 | 13 | 0.53% |
| hsa_circ_0062695 | 4186 | 11 | 0.53% |
| hsa_circ_0074630 | 4567 | 12 | 0.53% |
| hsa_circ_0011779 | 3806 | 10 | 0.53% |
| hsa_circ_0027204 | 3806 | 10 | 0.53% |
| hsa_circ_0089772 | 15226 | 40 | 0.53% |
| hsa_circ_0028724 | 4568 | 12 | 0.53% |
| hsa_circ_0030139 | 4949 | 13 | 0.53% |
| hsa_circ_0047584 | 4188 | 11 | 0.53% |
| hsa_circ_0074308 | 4188 | 11 | 0.53% |
| hsa_circ_0053326 | 4569 | 12 | 0.53% |
| hsa_circ_0063098 | 4569 | 12 | 0.53% |

| | | | |
|---|---|---|---|
| hsa_circ_0078789 | 4950 | 13 | 0.53% |
| hsa_circ_0038435 | 3808 | 10 | 0.53% |
| hsa_circ_0125151 | 7618 | 20 | 0.53% |
| hsa_circ_0068881 | 4571 | 12 | 0.53% |
| hsa_circ_0016476 | 7238 | 19 | 0.53% |
| hsa_circ_0122954 | 4953 | 13 | 0.52% |
| hsa_circ_0100707 | 4572 | 12 | 0.52% |
| hsa_circ_0068341 | 4191 | 11 | 0.52% |
| hsa_circ_0081704 | 23247 | 61 | 0.52% |
| hsa_circ_0123380 | 8004 | 21 | 0.52% |
| hsa_circ_0083292 | 5336 | 14 | 0.52% |
| hsa_circ_0029512 | 4955 | 13 | 0.52% |
| hsa_circ_0032354 | 4956 | 13 | 0.52% |
| hsa_circ_0114240 | 16775 | 44 | 0.52% |
| hsa_circ_0039663 | 3813 | 10 | 0.52% |
| hsa_circ_0092654 | 4957 | 13 | 0.52% |
| hsa_circ_0042203 | 4576 | 12 | 0.52% |
| hsa_circ_0137181 | 6102 | 16 | 0.52% |
| hsa_circ_0117505 | 4958 | 13 | 0.52% |
| hsa_circ_0010318 | 4577 | 12 | 0.52% |
| hsa_circ_0061767 | 5340 | 14 | 0.52% |
| hsa_circ_0128958 | 5340 | 14 | 0.52% |
| hsa_circ_0053554 | 4196 | 11 | 0.52% |
| hsa_circ_0045069 | 4959 | 13 | 0.52% |
| hsa_circ_0138769 | 9537 | 25 | 0.52% |
| hsa_circ_0138772 | 9537 | 25 | 0.52% |
| hsa_circ_0128738 | 4197 | 11 | 0.52% |
| hsa_circ_0091408 | 16791 | 44 | 0.52% |
| hsa_circ_0039743 | 4198 | 11 | 0.52% |
| hsa_circ_0039636 | 4962 | 13 | 0.52% |
| hsa_circ_0049426 | 4581 | 12 | 0.52% |
| hsa_circ_0067629 | 4963 | 13 | 0.52% |
| hsa_circ_0024674 | 4200 | 11 | 0.52% |
| hsa_circ_0046852 | 4964 | 13 | 0.52% |
| hsa_circ_0068601 | 4964 | 13 | 0.52% |
| hsa_circ_0082500 | 4965 | 13 | 0.52% |
| hsa_circ_0074712 | 5729 | 15 | 0.52% |
| hsa_circ_0079338 | 4202 | 11 | 0.52% |
| hsa_circ_0085414 | 4202 | 11 | 0.52% |
| hsa_circ_0119275 | 4202 | 11 | 0.52% |
| hsa_circ_0086866 | 4203 | 11 | 0.52% |
| hsa_circ_0037462 | 4968 | 13 | 0.52% |
| hsa_circ_0117909 | 4968 | 13 | 0.52% |
| hsa_circ_0130745 | 8026 | 21 | 0.52% |
| hsa_circ_0030495 | 5733 | 15 | 0.52% |
| hsa_circ_0070135 | 5351 | 14 | 0.52% |
| hsa_circ_0053587 | 4969 | 13 | 0.52% |
| hsa_circ_0011495 | 4587 | 12 | 0.52% |
| hsa_circ_0089207 | 4587 | 12 | 0.52% |
| hsa_circ_0042941 | 7263 | 19 | 0.52% |

| | | | |
|---|---|---|---|
| hsa_circ_0012094 | 4205 | 11 | 0.52% |
| hsa_circ_0015853 | 4205 | 11 | 0.52% |
| hsa_circ_0137462 | 4205 | 11 | 0.52% |
| hsa_circ_0060857 | 5735 | 15 | 0.52% |
| hsa_circ_0108958 | 4588 | 12 | 0.52% |
| hsa_circ_0013861 | 9561 | 25 | 0.52% |
| hsa_circ_0039039 | 4972 | 13 | 0.52% |
| hsa_circ_0041485 | 5355 | 14 | 0.52% |
| hsa_circ_0024732 | 4590 | 12 | 0.52% |
| hsa_circ_0066324 | 4590 | 12 | 0.52% |
| hsa_circ_0046763 | 3825 | 10 | 0.52% |
| hsa_circ_0086888 | 4973 | 13 | 0.52% |
| hsa_circ_0041553 | 4208 | 11 | 0.52% |
| hsa_circ_0086387 | 4208 | 11 | 0.52% |
| hsa_circ_0042908 | 4974 | 13 | 0.52% |
| hsa_circ_0077452 | 4974 | 13 | 0.52% |
| hsa_circ_0113242 | 4209 | 11 | 0.52% |
| hsa_circ_0122265 | 4209 | 11 | 0.52% |
| hsa_circ_0015119 | 6505 | 17 | 0.52% |
| hsa_circ_0073407 | 4210 | 11 | 0.52% |
| hsa_circ_0048389 | 5741 | 15 | 0.52% |
| hsa_circ_0060336 | 4211 | 11 | 0.52% |
| hsa_circ_0125032 | 4595 | 12 | 0.52% |
| hsa_circ_0091672 | 4979 | 13 | 0.52% |
| hsa_circ_0004933 | 4596 | 12 | 0.52% |
| hsa_circ_0063799 | 4596 | 12 | 0.52% |
| hsa_circ_0076774 | 4596 | 12 | 0.52% |
| hsa_circ_0032292 | 4213 | 11 | 0.52% |
| hsa_circ_0045785 | 4214 | 11 | 0.52% |
| hsa_circ_0084779 | 4214 | 11 | 0.52% |
| hsa_circ_0082509 | 3831 | 10 | 0.52% |
| hsa_circ_0120978 | 9195 | 24 | 0.52% |
| hsa_circ_0075988 | 5364 | 14 | 0.52% |
| hsa_circ_0053715 | 4981 | 13 | 0.52% |
| hsa_circ_0082194 | 4598 | 12 | 0.52% |
| hsa_circ_0087996 | 4215 | 11 | 0.52% |
| hsa_circ_0111270 | 5365 | 14 | 0.52% |
| hsa_circ_0075932 | 4982 | 13 | 0.52% |
| hsa_circ_0135391 | 45994 | 120 | 0.52% |
| hsa_circ_0033320 | 4600 | 12 | 0.52% |
| hsa_circ_0107100 | 12267 | 32 | 0.52% |
| hsa_circ_0010565 | 5367 | 14 | 0.52% |
| hsa_circ_0025953 | 4217 | 11 | 0.52% |
| hsa_circ_0139495 | 6134 | 16 | 0.52% |
| hsa_circ_0024225 | 4984 | 13 | 0.52% |
| hsa_circ_0027749 | 3834 | 10 | 0.52% |
| hsa_circ_0072341 | 4601 | 12 | 0.52% |
| hsa_circ_0125944 | 4601 | 12 | 0.52% |
| hsa_circ_0059994 | 4218 | 11 | 0.52% |
| hsa_circ_0066935 | 4986 | 13 | 0.52% |

| | | | |
|---|---|---|---|
| hsa_circ_0130254 | 4986 | 13 | 0.52% |
| hsa_circ_0075159 | 4219 | 11 | 0.52% |
| hsa_circ_0033305 | 4603 | 12 | 0.52% |
| hsa_circ_0065116 | 4603 | 12 | 0.52% |
| hsa_circ_0071183 | 4987 | 13 | 0.52% |
| hsa_circ_0138133 | 26472 | 69 | 0.52% |
| hsa_circ_0053792 | 4604 | 12 | 0.52% |
| hsa_circ_0088131 | 4221 | 11 | 0.52% |
| hsa_circ_0021020 | 4222 | 11 | 0.52% |
| hsa_circ_0025667 | 4222 | 11 | 0.52% |
| hsa_circ_0045441 | 4222 | 11 | 0.52% |
| hsa_circ_0035233 | 4606 | 12 | 0.52% |
| hsa_circ_0084497 | 4606 | 12 | 0.52% |
| hsa_circ_0091584 | 4606 | 12 | 0.52% |
| hsa_circ_0136047 | 4223 | 11 | 0.52% |
| hsa_circ_0035663 | 4224 | 11 | 0.52% |
| hsa_circ_0073299 | 4224 | 11 | 0.52% |
| hsa_circ_0139389 | 4224 | 11 | 0.52% |
| hsa_circ_0028006 | 4609 | 12 | 0.52% |
| hsa_circ_0024968 | 4225 | 11 | 0.52% |
| hsa_circ_0035301 | 4225 | 11 | 0.52% |
| hsa_circ_0104806 | 12676 | 33 | 0.52% |
| hsa_circ_0054506 | 4611 | 12 | 0.52% |
| hsa_circ_0071980 | 4227 | 11 | 0.52% |
| hsa_circ_0110666 | 4227 | 11 | 0.52% |
| hsa_circ_0020756 | 4612 | 12 | 0.52% |
| hsa_circ_0066307 | 4612 | 12 | 0.52% |
| hsa_circ_0028426 | 5766 | 15 | 0.52% |
| hsa_circ_0028776 | 4229 | 11 | 0.52% |
| hsa_circ_0040594 | 4229 | 11 | 0.52% |
| hsa_circ_0047048 | 4614 | 12 | 0.52% |
| hsa_circ_0073443 | 4614 | 12 | 0.52% |
| hsa_circ_0055962 | 4231 | 11 | 0.52% |
| hsa_circ_0104367 | 11924 | 31 | 0.52% |
| hsa_circ_0039099 | 4616 | 12 | 0.52% |
| hsa_circ_0069978 | 5771 | 15 | 0.52% |
| hsa_circ_0056128 | 4617 | 12 | 0.52% |
| hsa_circ_0129869 | 8467 | 22 | 0.52% |
| hsa_circ_0020409 | 4234 | 11 | 0.52% |
| hsa_circ_0009961 | 4619 | 12 | 0.52% |
| hsa_circ_0118315 | 5389 | 14 | 0.52% |
| hsa_circ_0137565 | 13474 | 35 | 0.52% |
| hsa_circ_0081676 | 15015 | 39 | 0.52% |
| hsa_circ_0074242 | 5391 | 14 | 0.52% |
| hsa_circ_0130849 | 4621 | 12 | 0.52% |
| hsa_circ_0022010 | 13093 | 34 | 0.52% |
| hsa_circ_0010267 | 5777 | 15 | 0.52% |
| hsa_circ_0025663 | 4237 | 11 | 0.52% |
| hsa_circ_0042613 | 4237 | 11 | 0.52% |
| hsa_circ_0051297 | 4623 | 12 | 0.52% |

| | | | |
|---|---|---|---|
| hsa_circ_0095551 | 5779 | 15 | 0.52% |
| hsa_circ_0030088 | 4239 | 11 | 0.52% |
| hsa_circ_0051164 | 4239 | 11 | 0.52% |
| hsa_circ_0045074 | 5010 | 13 | 0.52% |
| hsa_circ_0027747 | 4240 | 11 | 0.52% |
| hsa_circ_0081907 | 5011 | 13 | 0.52% |
| hsa_circ_0106680 | 8095 | 21 | 0.52% |
| hsa_circ_0009817 | 4626 | 12 | 0.52% |
| hsa_circ_0012923 | 4626 | 12 | 0.52% |
| hsa_circ_0026585 | 4627 | 12 | 0.52% |
| hsa_circ_0090598 | 4627 | 12 | 0.52% |
| hsa_circ_0024411 | 5399 | 14 | 0.52% |
| hsa_circ_0065172 | 4628 | 12 | 0.52% |
| hsa_circ_0082795 | 4628 | 12 | 0.52% |
| hsa_circ_0140161 | 5014 | 13 | 0.52% |
| hsa_circ_0038233 | 5400 | 14 | 0.52% |
| hsa_circ_0086586 | 4629 | 12 | 0.52% |
| hsa_circ_0072367 | 5015 | 13 | 0.52% |
| hsa_circ_0118479 | 4630 | 12 | 0.52% |
| hsa_circ_0089796 | 23925 | 62 | 0.52% |
| hsa_circ_0029502 | 4631 | 12 | 0.52% |
| hsa_circ_0043538 | 4631 | 12 | 0.52% |
| hsa_circ_0001800 | 9649 | 25 | 0.52% |
| hsa_circ_0032857 | 5018 | 13 | 0.52% |
| hsa_circ_0053484 | 4246 | 11 | 0.52% |
| hsa_circ_0104807 | 12739 | 33 | 0.52% |
| hsa_circ_0058105 | 5019 | 13 | 0.52% |
| hsa_circ_0041397 | 4633 | 12 | 0.52% |
| hsa_circ_0017316 | 4247 | 11 | 0.52% |
| hsa_circ_0087983 | 4247 | 11 | 0.52% |
| hsa_circ_0140242 | 4247 | 11 | 0.52% |
| hsa_circ_0016475 | 7336 | 19 | 0.52% |
| hsa_circ_0009806 | 5406 | 14 | 0.52% |
| hsa_circ_0056480 | 5406 | 14 | 0.52% |
| hsa_circ_0036643 | 4635 | 12 | 0.52% |
| hsa_circ_0048415 | 4635 | 12 | 0.52% |
| hsa_circ_0096773 | 4635 | 12 | 0.52% |
| hsa_circ_0073483 | 4249 | 11 | 0.52% |
| hsa_circ_0076472 | 4249 | 11 | 0.52% |
| hsa_circ_0110309 | 4249 | 11 | 0.52% |
| hsa_circ_0010677 | 5022 | 13 | 0.52% |
| hsa_circ_0114484 | 12362 | 32 | 0.52% |
| hsa_circ_0090758 | 5023 | 13 | 0.52% |
| hsa_circ_0038270 | 4251 | 11 | 0.52% |
| hsa_circ_0035760 | 5024 | 13 | 0.52% |
| hsa_circ_0021581 | 4638 | 12 | 0.52% |
| hsa_circ_0110917 | 11209 | 29 | 0.52% |
| hsa_circ_0058088 | 5025 | 13 | 0.52% |
| hsa_circ_0038266 | 4252 | 11 | 0.52% |
| hsa_circ_0092086 | 5026 | 13 | 0.52% |

| | | | |
|---|---|---|---|
| hsa_circ_0100140 | 6961 | 18 | 0.52% |
| hsa_circ_0052290 | 3868 | 10 | 0.52% |
| hsa_circ_0008632 | 12379 | 32 | 0.52% |
| hsa_circ_0025666 | 5029 | 13 | 0.52% |
| hsa_circ_0013640 | 14701 | 38 | 0.52% |
| hsa_circ_0070230 | 4256 | 11 | 0.52% |
| hsa_circ_0094907 | 4256 | 11 | 0.52% |
| hsa_circ_0025091 | 5031 | 13 | 0.52% |
| hsa_circ_0064054 | 4644 | 12 | 0.52% |
| hsa_circ_0065173 | 4644 | 12 | 0.52% |
| hsa_circ_0077201 | 4644 | 12 | 0.52% |
| hsa_circ_0079115 | 5419 | 14 | 0.52% |
| hsa_circ_0116815 | 4645 | 12 | 0.52% |
| hsa_circ_0033287 | 4646 | 12 | 0.52% |
| hsa_circ_0042154 | 4646 | 12 | 0.52% |
| hsa_circ_0083194 | 4646 | 12 | 0.52% |
| hsa_circ_0054293 | 4259 | 11 | 0.52% |
| hsa_circ_0124733 | 8519 | 22 | 0.52% |
| hsa_circ_0031177 | 4647 | 12 | 0.52% |
| hsa_circ_0062335 | 5809 | 15 | 0.52% |
| hsa_circ_0020282 | 3873 | 10 | 0.52% |
| hsa_circ_0062399 | 4261 | 11 | 0.52% |
| hsa_circ_0088210 | 4262 | 11 | 0.52% |
| hsa_circ_0111271 | 5425 | 14 | 0.52% |
| hsa_circ_0089530 | 4650 | 12 | 0.52% |
| hsa_circ_0071400 | 5038 | 13 | 0.52% |
| hsa_circ_0012557 | 7751 | 20 | 0.52% |
| hsa_circ_0116005 | 5426 | 14 | 0.52% |
| hsa_circ_0049187 | 4651 | 12 | 0.52% |
| hsa_circ_0053692 | 5039 | 13 | 0.52% |
| hsa_circ_0071077 | 5039 | 13 | 0.52% |
| hsa_circ_0033445 | 4652 | 12 | 0.52% |
| hsa_circ_0080195 | 4652 | 12 | 0.52% |
| hsa_circ_0035667 | 5040 | 13 | 0.52% |
| hsa_circ_0007645 | 4265 | 11 | 0.52% |
| hsa_circ_0044847 | 4653 | 12 | 0.52% |
| hsa_circ_0071144 | 3878 | 10 | 0.52% |
| hsa_circ_0032996 | 4655 | 12 | 0.52% |
| hsa_circ_0090730 | 5043 | 13 | 0.52% |
| hsa_circ_0109358 | 5043 | 13 | 0.52% |
| hsa_circ_0060858 | 5819 | 15 | 0.52% |
| hsa_circ_0083293 | 5432 | 14 | 0.52% |
| hsa_circ_0037408 | 5045 | 13 | 0.52% |
| hsa_circ_0032344 | 4657 | 12 | 0.52% |
| hsa_circ_0036485 | 4657 | 12 | 0.52% |
| hsa_circ_0041210 | 4657 | 12 | 0.52% |
| hsa_circ_0094863 | 4269 | 11 | 0.52% |
| hsa_circ_0020844 | 12032 | 31 | 0.52% |
| hsa_circ_0046339 | 5046 | 13 | 0.52% |
| hsa_circ_0101261 | 7375 | 19 | 0.52% |

| | | | |
|---|---|---|---|
| hsa_circ_0047907 | 5047 | 13 | 0.52% |
| hsa_circ_0027141 | 4659 | 12 | 0.52% |
| hsa_circ_0033223 | 4659 | 12 | 0.52% |
| hsa_circ_0074694 | 4659 | 12 | 0.52% |
| hsa_circ_0017218 | 4271 | 11 | 0.52% |
| hsa_circ_0115287 | 4271 | 11 | 0.52% |
| hsa_circ_0056900 | 11264 | 29 | 0.51% |
| hsa_circ_0067565 | 4661 | 12 | 0.51% |
| hsa_circ_0126496 | 4273 | 11 | 0.51% |
| hsa_circ_0078982 | 6216 | 16 | 0.51% |
| hsa_circ_0051116 | 3885 | 10 | 0.51% |
| hsa_circ_0130527 | 12822 | 33 | 0.51% |
| hsa_circ_0103864 | 23315 | 60 | 0.51% |
| hsa_circ_0074628 | 5052 | 13 | 0.51% |
| hsa_circ_0073636 | 5441 | 14 | 0.51% |
| hsa_circ_0010457 | 4664 | 12 | 0.51% |
| hsa_circ_0023533 | 4664 | 12 | 0.51% |
| hsa_circ_0131653 | 4276 | 11 | 0.51% |
| hsa_circ_0078077 | 4665 | 12 | 0.51% |
| hsa_circ_0086887 | 4665 | 12 | 0.51% |
| hsa_circ_0125577 | 40431 | 104 | 0.51% |
| hsa_circ_0015470 | 5054 | 13 | 0.51% |
| hsa_circ_0026833 | 4277 | 11 | 0.51% |
| hsa_circ_0043956 | 4277 | 11 | 0.51% |
| hsa_circ_0014588 | 4666 | 12 | 0.51% |
| hsa_circ_0015750 | 4666 | 12 | 0.51% |
| hsa_circ_0135833 | 4666 | 12 | 0.51% |
| hsa_circ_0072347 | 5055 | 13 | 0.51% |
| hsa_circ_0100752 | 5055 | 13 | 0.51% |
| hsa_circ_0075995 | 5446 | 14 | 0.51% |
| hsa_circ_0034284 | 3890 | 10 | 0.51% |
| hsa_circ_0140419 | 12450 | 32 | 0.51% |
| hsa_circ_0084216 | 4669 | 12 | 0.51% |
| hsa_circ_0138131 | 26849 | 69 | 0.51% |
| hsa_circ_0122368 | 6226 | 16 | 0.51% |
| hsa_circ_0055089 | 4670 | 12 | 0.51% |
| hsa_circ_0078699 | 4670 | 12 | 0.51% |
| hsa_circ_0013144 | 11286 | 29 | 0.51% |
| hsa_circ_0035147 | 5060 | 13 | 0.51% |
| hsa_circ_0027129 | 4671 | 12 | 0.51% |
| hsa_circ_0042663 | 4671 | 12 | 0.51% |
| hsa_circ_0128405 | 4671 | 12 | 0.51% |
| hsa_circ_0049425 | 4282 | 11 | 0.51% |
| hsa_circ_0029500 | 5061 | 13 | 0.51% |
| hsa_circ_0060481 | 5452 | 14 | 0.51% |
| hsa_circ_0076002 | 5452 | 14 | 0.51% |
| hsa_circ_0101455 | 4284 | 11 | 0.51% |
| hsa_circ_0007433 | 5063 | 13 | 0.51% |
| hsa_circ_0010287 | 4674 | 12 | 0.51% |
| hsa_circ_0092450 | 11687 | 30 | 0.51% |

| | | | |
|---|---|---|---|
| hsa_circ_0024733 | 4676 | 12 | 0.51% |
| hsa_circ_0019224 | 4287 | 11 | 0.51% |
| hsa_circ_0024094 | 4677 | 12 | 0.51% |
| hsa_circ_0112715 | 5457 | 14 | 0.51% |
| hsa_circ_0003755 | 4678 | 12 | 0.51% |
| hsa_circ_0063052 | 10139 | 26 | 0.51% |
| hsa_circ_0040423 | 5460 | 14 | 0.51% |
| hsa_circ_0075985 | 4680 | 12 | 0.51% |
| hsa_circ_0034157 | 4290 | 11 | 0.51% |
| hsa_circ_0021571 | 4291 | 11 | 0.51% |
| hsa_circ_0004347 | 13264 | 34 | 0.51% |
| hsa_circ_0100701 | 5072 | 13 | 0.51% |
| hsa_circ_0039615 | 5073 | 13 | 0.51% |
| hsa_circ_0094079 | 5073 | 13 | 0.51% |
| hsa_circ_0062445 | 4683 | 12 | 0.51% |
| hsa_circ_0095276 | 4684 | 12 | 0.51% |
| hsa_circ_0045755 | 5075 | 13 | 0.51% |
| hsa_circ_0128472 | 16008 | 41 | 0.51% |
| hsa_circ_0007029 | 30064 | 77 | 0.51% |
| hsa_circ_0137176 | 4686 | 12 | 0.51% |
| hsa_circ_0061998 | 3905 | 10 | 0.51% |
| hsa_circ_0061595 | 5077 | 13 | 0.51% |
| hsa_circ_0062698 | 4296 | 11 | 0.51% |
| hsa_circ_0071169 | 4296 | 11 | 0.51% |
| hsa_circ_0137341 | 7811 | 20 | 0.51% |
| hsa_circ_0087132 | 9764 | 25 | 0.51% |
| hsa_circ_0094595 | 9764 | 25 | 0.51% |
| hsa_circ_0087315 | 4687 | 12 | 0.51% |
| hsa_circ_0028567 | 3906 | 10 | 0.51% |
| hsa_circ_0035278 | 5078 | 13 | 0.51% |
| hsa_circ_0053815 | 4688 | 12 | 0.51% |
| hsa_circ_0091905 | 4688 | 12 | 0.51% |
| hsa_circ_0055509 | 4689 | 12 | 0.51% |
| hsa_circ_0138773 | 9769 | 25 | 0.51% |
| hsa_circ_0013765 | 5080 | 13 | 0.51% |
| hsa_circ_0055961 | 4299 | 11 | 0.51% |
| hsa_circ_0085865 | 5081 | 13 | 0.51% |
| hsa_circ_0112693 | 7036 | 18 | 0.51% |
| hsa_circ_0109314 | 3909 | 10 | 0.51% |
| hsa_circ_0019835 | 4300 | 11 | 0.51% |
| hsa_circ_0019902 | 4300 | 11 | 0.51% |
| hsa_circ_0121856 | 4300 | 11 | 0.51% |
| hsa_circ_0053647 | 5082 | 13 | 0.51% |
| hsa_circ_0013961 | 5084 | 13 | 0.51% |
| hsa_circ_0052639 | 5084 | 13 | 0.51% |
| hsa_circ_0024058 | 4693 | 12 | 0.51% |
| hsa_circ_0026319 | 4302 | 11 | 0.51% |
| hsa_circ_0026104 | 5867 | 15 | 0.51% |
| hsa_circ_0091432 | 5085 | 13 | 0.51% |
| hsa_circ_0053824 | 4694 | 12 | 0.51% |

| | | | |
|---|---|---|---|
| hsa_circ_0068985 | 4303 | 11 | 0.51% |
| hsa_circ_0078585 | 4303 | 11 | 0.51% |
| hsa_circ_0059777 | 4304 | 11 | 0.51% |
| hsa_circ_0044718 | 5087 | 13 | 0.51% |
| hsa_circ_0020473 | 4305 | 11 | 0.51% |
| hsa_circ_0047245 | 3914 | 10 | 0.51% |
| hsa_circ_0125971 | 4697 | 12 | 0.51% |
| hsa_circ_0103568 | 8220 | 21 | 0.51% |
| hsa_circ_0086698 | 14876 | 38 | 0.51% |
| hsa_circ_0032524 | 21925 | 56 | 0.51% |
| hsa_circ_0117536 | 15270 | 39 | 0.51% |
| hsa_circ_0028771 | 4699 | 12 | 0.51% |
| hsa_circ_0029891 | 6657 | 17 | 0.51% |
| hsa_circ_0010515 | 4700 | 12 | 0.51% |
| hsa_circ_0031462 | 4700 | 12 | 0.51% |
| hsa_circ_0056050 | 4700 | 12 | 0.51% |
| hsa_circ_0042160 | 3917 | 10 | 0.51% |
| hsa_circ_0074108 | 4701 | 12 | 0.51% |
| hsa_circ_0009815 | 5485 | 14 | 0.51% |
| hsa_circ_0105296 | 8228 | 21 | 0.51% |
| hsa_circ_0109316 | 4310 | 11 | 0.51% |
| hsa_circ_0070900 | 5094 | 13 | 0.51% |
| hsa_circ_0062990 | 5486 | 14 | 0.51% |
| hsa_circ_0071009 | 5486 | 14 | 0.51% |
| hsa_circ_0101638 | 12148 | 31 | 0.51% |
| hsa_circ_0068153 | 10189 | 26 | 0.51% |
| hsa_circ_0100965 | 10189 | 26 | 0.51% |
| hsa_circ_0137336 | 7838 | 20 | 0.51% |
| hsa_circ_0011777 | 3919 | 10 | 0.51% |
| hsa_circ_0038164 | 4311 | 11 | 0.51% |
| hsa_circ_0017579 | 13327 | 34 | 0.51% |
| hsa_circ_0132755 | 5880 | 15 | 0.51% |
| hsa_circ_0135134 | 5880 | 15 | 0.51% |
| hsa_circ_0011070 | 5489 | 14 | 0.51% |
| hsa_circ_0060791 | 4706 | 12 | 0.51% |
| hsa_circ_0053525 | 4314 | 11 | 0.51% |
| hsa_circ_0086904 | 5099 | 13 | 0.51% |
| hsa_circ_0084589 | 4707 | 12 | 0.51% |
| hsa_circ_0125218 | 4707 | 12 | 0.51% |
| hsa_circ_0081273 | 4316 | 11 | 0.51% |
| hsa_circ_0128470 | 16088 | 41 | 0.51% |
| hsa_circ_0112691 | 8633 | 22 | 0.51% |
| hsa_circ_0054810 | 4709 | 12 | 0.51% |
| hsa_circ_0081377 | 5494 | 14 | 0.51% |
| hsa_circ_0128963 | 5494 | 14 | 0.51% |
| hsa_circ_0048499 | 4317 | 11 | 0.51% |
| hsa_circ_0020410 | 4319 | 11 | 0.51% |
| hsa_circ_0100515 | 5105 | 13 | 0.51% |
| hsa_circ_0063957 | 4713 | 12 | 0.51% |
| hsa_circ_0099605 | 4713 | 12 | 0.51% |

| | | | |
|---|---|---|---|
| hsa_circ_0039391 | 3928 | 10 | 0.51% |
| hsa_circ_0132641 | 14142 | 36 | 0.51% |
| hsa_circ_0057998 | 5107 | 13 | 0.51% |
| hsa_circ_0020060 | 5893 | 15 | 0.51% |
| hsa_circ_0042939 | 7467 | 19 | 0.51% |
| hsa_circ_0024410 | 5502 | 14 | 0.51% |
| hsa_circ_0071184 | 5109 | 13 | 0.51% |
| hsa_circ_0049113 | 4716 | 12 | 0.51% |
| hsa_circ_0104157 | 4716 | 12 | 0.51% |
| hsa_circ_0089938 | 4323 | 11 | 0.51% |
| hsa_circ_0011820 | 3930 | 10 | 0.51% |
| hsa_circ_0100253 | 5896 | 15 | 0.51% |
| hsa_circ_0019905 | 5110 | 13 | 0.51% |
| hsa_circ_0092418 | 8648 | 22 | 0.51% |
| hsa_circ_0122636 | 12974 | 33 | 0.51% |
| hsa_circ_0066933 | 5111 | 13 | 0.51% |
| hsa_circ_0084357 | 5111 | 13 | 0.51% |
| hsa_circ_0133914 | 5111 | 13 | 0.51% |
| hsa_circ_0084360 | 4325 | 11 | 0.51% |
| hsa_circ_0087806 | 4325 | 11 | 0.51% |
| hsa_circ_0054042 | 5112 | 13 | 0.51% |
| hsa_circ_0047512 | 4719 | 12 | 0.51% |
| hsa_circ_0010350 | 3933 | 10 | 0.51% |
| hsa_circ_0112701 | 14947 | 38 | 0.51% |
| hsa_circ_0122741 | 7474 | 19 | 0.51% |
| hsa_circ_0091276 | 6294 | 16 | 0.51% |
| hsa_circ_0091100 | 5114 | 13 | 0.51% |
| hsa_circ_0038433 | 3934 | 10 | 0.51% |
| hsa_circ_0087010 | 4328 | 11 | 0.51% |
| hsa_circ_0064040 | 4722 | 12 | 0.51% |
| hsa_circ_0113437 | 4722 | 12 | 0.51% |
| hsa_circ_0137468 | 5116 | 13 | 0.51% |
| hsa_circ_0064212 | 4329 | 11 | 0.51% |
| hsa_circ_0085223 | 4329 | 11 | 0.51% |
| hsa_circ_0118772 | 4329 | 11 | 0.51% |
| hsa_circ_0103310 | 20466 | 52 | 0.51% |
| hsa_circ_0124653 | 12991 | 33 | 0.51% |
| hsa_circ_0005369 | 4331 | 11 | 0.51% |
| hsa_circ_0117418 | 7481 | 19 | 0.51% |
| hsa_circ_0111023 | 4725 | 12 | 0.51% |
| hsa_circ_0079331 | 5120 | 13 | 0.51% |
| hsa_circ_0047979 | 4727 | 12 | 0.51% |
| hsa_circ_0066320 | 4727 | 12 | 0.51% |
| hsa_circ_0001996 | 15759 | 40 | 0.51% |
| hsa_circ_0115206 | 15759 | 40 | 0.51% |
| hsa_circ_0041495 | 5516 | 14 | 0.51% |
| hsa_circ_0119725 | 9063 | 23 | 0.51% |
| hsa_circ_0075784 | 5517 | 14 | 0.51% |
| hsa_circ_0060859 | 5912 | 15 | 0.51% |
| hsa_circ_0061088 | 4730 | 12 | 0.51% |

| | | | |
|---|---|---|---|
| hsa_circ_0039744 | 4336 | 11 | 0.51% |
| hsa_circ_0071076 | 5125 | 13 | 0.51% |
| hsa_circ_0053093 | 5126 | 13 | 0.51% |
| hsa_circ_0031752 | 5521 | 14 | 0.51% |
| hsa_circ_0022260 | 4338 | 11 | 0.51% |
| hsa_circ_0058783 | 4338 | 11 | 0.51% |
| hsa_circ_0001924 | 24451 | 62 | 0.51% |
| hsa_circ_0033254 | 4733 | 12 | 0.51% |
| hsa_circ_0046379 | 4339 | 11 | 0.51% |
| hsa_circ_0133541 | 4339 | 11 | 0.51% |
| hsa_circ_0061672 | 4734 | 12 | 0.51% |
| hsa_circ_0074629 | 4734 | 12 | 0.51% |
| hsa_circ_0140741 | 4734 | 12 | 0.51% |
| hsa_circ_0084515 | 9863 | 25 | 0.51% |
| hsa_circ_0080684 | 5129 | 13 | 0.51% |
| hsa_circ_0122433 | 4340 | 11 | 0.51% |
| hsa_circ_0069547 | 5130 | 13 | 0.51% |
| hsa_circ_0091701 | 4736 | 12 | 0.51% |
| hsa_circ_0010558 | 5526 | 14 | 0.51% |
| hsa_circ_0009564 | 4342 | 11 | 0.51% |
| hsa_circ_0078385 | 4342 | 11 | 0.51% |
| hsa_circ_0112502 | 38303 | 97 | 0.51% |
| hsa_circ_0074495 | 7108 | 18 | 0.51% |
| hsa_circ_0052904 | 4344 | 11 | 0.51% |
| hsa_circ_0042485 | 4739 | 12 | 0.51% |
| hsa_circ_0058914 | 4739 | 12 | 0.51% |
| hsa_circ_0132365 | 5134 | 13 | 0.51% |
| hsa_circ_0006759 | 78995 | 200 | 0.51% |
| hsa_circ_0091772 | 5530 | 14 | 0.51% |
| hsa_circ_0042592 | 4740 | 12 | 0.51% |
| hsa_circ_0080980 | 4740 | 12 | 0.51% |
| hsa_circ_0102290 | 9877 | 25 | 0.51% |
| hsa_circ_0123601 | 4346 | 11 | 0.51% |
| hsa_circ_0117944 | 3951 | 10 | 0.51% |
| hsa_circ_0073012 | 47022 | 119 | 0.51% |
| hsa_circ_0072348 | 5137 | 13 | 0.51% |
| hsa_circ_0011772 | 4347 | 11 | 0.51% |
| hsa_circ_0090754 | 5533 | 14 | 0.51% |
| hsa_circ_0073966 | 5534 | 14 | 0.51% |
| hsa_circ_0118071 | 5534 | 14 | 0.51% |
| hsa_circ_0118723 | 15813 | 40 | 0.51% |
| hsa_circ_0074709 | 5930 | 15 | 0.51% |
| hsa_circ_0008444 | 10675 | 27 | 0.51% |
| hsa_circ_0075699 | 4745 | 12 | 0.51% |
| hsa_circ_0114855 | 11072 | 28 | 0.51% |
| hsa_circ_0054565 | 4350 | 11 | 0.51% |
| hsa_circ_0071133 | 4350 | 11 | 0.51% |
| hsa_circ_0086529 | 4350 | 11 | 0.51% |
| hsa_circ_0071161 | 4746 | 12 | 0.51% |
| hsa_circ_0123102 | 4746 | 12 | 0.51% |

| | | | |
|---|---|---|---|
| hsa_circ_0043958 | 4351 | 11 | 0.51% |
| hsa_circ_0124294 | 4352 | 11 | 0.51% |
| hsa_circ_0056138 | 4748 | 12 | 0.51% |
| hsa_circ_0094949 | 13058 | 33 | 0.51% |
| hsa_circ_0047207 | 3957 | 10 | 0.51% |
| hsa_circ_0014746 | 5540 | 14 | 0.51% |
| hsa_circ_0017363 | 4353 | 11 | 0.51% |
| hsa_circ_0047925 | 5146 | 13 | 0.51% |
| hsa_circ_0077621 | 5938 | 15 | 0.51% |
| hsa_circ_0132569 | 19003 | 48 | 0.51% |
| hsa_circ_0111641 | 4752 | 12 | 0.51% |
| hsa_circ_0123914 | 4752 | 12 | 0.51% |
| hsa_circ_0053794 | 5149 | 13 | 0.50% |
| hsa_circ_0076724 | 5149 | 13 | 0.50% |
| hsa_circ_0029063 | 4753 | 12 | 0.50% |
| hsa_circ_0081968 | 4753 | 12 | 0.50% |
| hsa_circ_0049314 | 5151 | 13 | 0.50% |
| hsa_circ_0059413 | 5151 | 13 | 0.50% |
| hsa_circ_0054964 | 5548 | 14 | 0.50% |
| hsa_circ_0073482 | 3963 | 10 | 0.50% |
| hsa_circ_0137520 | 4756 | 12 | 0.50% |
| hsa_circ_0105176 | 15061 | 38 | 0.50% |
| hsa_circ_0042156 | 4360 | 11 | 0.50% |
| hsa_circ_0045073 | 5153 | 13 | 0.50% |
| hsa_circ_0077445 | 5153 | 13 | 0.50% |
| hsa_circ_0053189 | 4757 | 12 | 0.50% |
| hsa_circ_0028777 | 4361 | 11 | 0.50% |
| hsa_circ_0088211 | 4361 | 11 | 0.50% |
| hsa_circ_0016843 | 10309 | 26 | 0.50% |
| hsa_circ_0072886 | 4758 | 12 | 0.50% |
| hsa_circ_0110611 | 5155 | 13 | 0.50% |
| hsa_circ_0054300 | 4362 | 11 | 0.50% |
| hsa_circ_0053822 | 4760 | 12 | 0.50% |
| hsa_circ_0129834 | 4760 | 12 | 0.50% |
| hsa_circ_0065399 | 5157 | 13 | 0.50% |
| hsa_circ_0078813 | 11507 | 29 | 0.50% |
| hsa_circ_0091583 | 4762 | 12 | 0.50% |
| hsa_circ_0074260 | 5556 | 14 | 0.50% |
| hsa_circ_0070116 | 4366 | 11 | 0.50% |
| hsa_circ_0081289 | 4366 | 11 | 0.50% |
| hsa_circ_0070683 | 4763 | 12 | 0.50% |
| hsa_circ_0112694 | 7145 | 18 | 0.50% |
| hsa_circ_0104047 | 11512 | 29 | 0.50% |
| hsa_circ_0132638 | 14291 | 36 | 0.50% |
| hsa_circ_0062461 | 79001 | 199 | 0.50% |
| hsa_circ_0079395 | 4367 | 11 | 0.50% |
| hsa_circ_0058713 | 15090 | 38 | 0.50% |
| hsa_circ_0032343 | 4766 | 12 | 0.50% |
| hsa_circ_0002370 | 79436 | 200 | 0.50% |
| hsa_circ_0045439 | 4370 | 11 | 0.50% |

| | | | |
|---|---|---|---|
| hsa_circ_0075558 | 3973 | 10 | 0.50% |
| hsa_circ_0019565 | 5960 | 15 | 0.50% |
| hsa_circ_0065170 | 4768 | 12 | 0.50% |
| hsa_circ_0040424 | 5563 | 14 | 0.50% |
| hsa_circ_0112692 | 8742 | 22 | 0.50% |
| hsa_circ_0046766 | 3974 | 10 | 0.50% |
| hsa_circ_0057057 | 4769 | 12 | 0.50% |
| hsa_circ_0051308 | 4770 | 12 | 0.50% |
| hsa_circ_0054582 | 4770 | 12 | 0.50% |
| hsa_circ_0035783 | 5168 | 13 | 0.50% |
| hsa_circ_0055643 | 5965 | 15 | 0.50% |
| hsa_circ_0018292 | 4772 | 12 | 0.50% |
| hsa_circ_0029875 | 5170 | 13 | 0.50% |
| hsa_circ_0081901 | 5170 | 13 | 0.50% |
| hsa_circ_0136050 | 5170 | 13 | 0.50% |
| hsa_circ_0068981 | 3977 | 10 | 0.50% |
| hsa_circ_0025955 | 4375 | 11 | 0.50% |
| hsa_circ_0121204 | 4375 | 11 | 0.50% |
| hsa_circ_0058826 | 5171 | 13 | 0.50% |
| hsa_circ_0106517 | 5171 | 13 | 0.50% |
| hsa_circ_0098920 | 13923 | 35 | 0.50% |
| hsa_circ_0002108 | 11539 | 29 | 0.50% |
| hsa_circ_0055525 | 4377 | 11 | 0.50% |
| hsa_circ_0056052 | 4775 | 12 | 0.50% |
| hsa_circ_0108955 | 4775 | 12 | 0.50% |
| hsa_circ_0090227 | 5173 | 13 | 0.50% |
| hsa_circ_0029956 | 5572 | 14 | 0.50% |
| hsa_circ_0076001 | 5572 | 14 | 0.50% |
| hsa_circ_0123743 | 4776 | 12 | 0.50% |
| hsa_circ_0041965 | 4378 | 11 | 0.50% |
| hsa_circ_0040152 | 5175 | 13 | 0.50% |
| hsa_circ_0047193 | 4777 | 12 | 0.50% |
| hsa_circ_0029403 | 14731 | 37 | 0.50% |
| hsa_circ_0102338 | 4778 | 12 | 0.50% |
| hsa_circ_0140318 | 4380 | 11 | 0.50% |
| hsa_circ_0037084 | 4779 | 12 | 0.50% |
| hsa_circ_0073341 | 4779 | 12 | 0.50% |
| hsa_circ_0093066 | 4381 | 11 | 0.50% |
| hsa_circ_0118142 | 7169 | 18 | 0.50% |
| hsa_circ_0107605 | 4382 | 11 | 0.50% |
| hsa_circ_0097356 | 4781 | 12 | 0.50% |
| hsa_circ_0024409 | 5578 | 14 | 0.50% |
| hsa_circ_0079707 | 5578 | 14 | 0.50% |
| hsa_circ_0065175 | 4383 | 11 | 0.50% |
| hsa_circ_0095390 | 5977 | 15 | 0.50% |
| hsa_circ_0136102 | 11556 | 29 | 0.50% |
| hsa_circ_0056054 | 4782 | 12 | 0.50% |
| hsa_circ_0032856 | 5181 | 13 | 0.50% |
| hsa_circ_0055960 | 4384 | 11 | 0.50% |
| hsa_circ_0103034 | 7573 | 19 | 0.50% |

| | | | |
|---|---|---|---|
| hsa_circ_0076201 | 6378 | 16 | 0.50% |
| hsa_circ_0016625 | 4385 | 11 | 0.50% |
| hsa_circ_0093751 | 4784 | 12 | 0.50% |
| hsa_circ_0033321 | 4785 | 12 | 0.50% |
| hsa_circ_0071909 | 4785 | 12 | 0.50% |
| hsa_circ_0110141 | 21136 | 53 | 0.50% |
| hsa_circ_0063238 | 4387 | 11 | 0.50% |
| hsa_circ_0028385 | 4786 | 12 | 0.50% |
| hsa_circ_0081173 | 5185 | 13 | 0.50% |
| hsa_circ_0034208 | 5186 | 13 | 0.50% |
| hsa_circ_0084361 | 5186 | 13 | 0.50% |
| hsa_circ_0135964 | 10771 | 27 | 0.50% |
| hsa_circ_0071981 | 4389 | 11 | 0.50% |
| hsa_circ_0133269 | 13567 | 34 | 0.50% |
| hsa_circ_0016656 | 12370 | 31 | 0.50% |
| hsa_circ_0062060 | 4390 | 11 | 0.50% |
| hsa_circ_0015463 | 5189 | 13 | 0.50% |
| hsa_circ_0121434 | 5589 | 14 | 0.50% |
| hsa_circ_0009173 | 4393 | 11 | 0.50% |
| hsa_circ_0130439 | 10784 | 27 | 0.50% |
| hsa_circ_0035664 | 5193 | 13 | 0.50% |
| hsa_circ_0037464 | 5193 | 13 | 0.50% |
| hsa_circ_0007169 | 5992 | 15 | 0.50% |
| hsa_circ_0018228 | 5593 | 14 | 0.50% |
| hsa_circ_0102062 | 4794 | 12 | 0.50% |
| hsa_circ_0079341 | 3995 | 10 | 0.50% |
| hsa_circ_0067001 | 4795 | 12 | 0.50% |
| hsa_circ_0130328 | 19980 | 50 | 0.50% |
| hsa_circ_0010642 | 5195 | 13 | 0.50% |
| hsa_circ_0070335 | 4396 | 11 | 0.50% |
| hsa_circ_0073269 | 5196 | 13 | 0.50% |
| hsa_circ_0077346 | 5196 | 13 | 0.50% |
| hsa_circ_0076010 | 5596 | 14 | 0.50% |
| hsa_circ_0091913 | 4797 | 12 | 0.50% |
| hsa_circ_0088965 | 4398 | 11 | 0.50% |
| hsa_circ_0014846 | 4798 | 12 | 0.50% |
| hsa_circ_0137608 | 9997 | 25 | 0.50% |
| hsa_circ_0102252 | 13996 | 35 | 0.50% |
| hsa_circ_0027447 | 4399 | 11 | 0.50% |
| hsa_circ_0072165 | 4399 | 11 | 0.50% |
| hsa_circ_0081264 | 5999 | 15 | 0.50% |
| hsa_circ_0073200 | 5200 | 13 | 0.50% |
| hsa_circ_0020850 | 4401 | 11 | 0.50% |
| hsa_circ_0112827 | 13204 | 33 | 0.50% |
| hsa_circ_0088938 | 5202 | 13 | 0.50% |
| hsa_circ_0049813 | 5203 | 13 | 0.50% |
| hsa_circ_0122260 | 5203 | 13 | 0.50% |
| hsa_circ_0123084 | 12007 | 30 | 0.50% |
| hsa_circ_0085724 | 4403 | 11 | 0.50% |
| hsa_circ_0091645 | 4804 | 12 | 0.50% |

| | | | |
|---|---|---|---|
| hsa_circ_0091847 | 5605 | 14 | 0.50% |
| hsa_circ_0005055 | 4404 | 11 | 0.50% |
| hsa_circ_0104468 | 9609 | 24 | 0.50% |
| hsa_circ_0058089 | 5205 | 13 | 0.50% |
| hsa_circ_0034111 | 5606 | 14 | 0.50% |
| hsa_circ_0075482 | 4806 | 12 | 0.50% |
| hsa_circ_0094883 | 4806 | 12 | 0.50% |
| hsa_circ_0097111 | 16422 | 41 | 0.50% |
| hsa_circ_0020411 | 4406 | 11 | 0.50% |
| hsa_circ_0003031 | 14822 | 37 | 0.50% |
| hsa_circ_0034602 | 5208 | 13 | 0.50% |
| hsa_circ_0046851 | 5208 | 13 | 0.50% |
| hsa_circ_0087186 | 5208 | 13 | 0.50% |
| hsa_circ_0039442 | 4407 | 11 | 0.50% |
| hsa_circ_0091431 | 5209 | 13 | 0.50% |
| hsa_circ_0038432 | 4007 | 10 | 0.50% |
| hsa_circ_0018831 | 16029 | 40 | 0.50% |
| hsa_circ_0011771 | 4409 | 11 | 0.50% |
| hsa_circ_0061583 | 4810 | 12 | 0.50% |
| hsa_circ_0058094 | 5211 | 13 | 0.50% |
| hsa_circ_0130329 | 20043 | 50 | 0.50% |
| hsa_circ_0100577 | 13631 | 34 | 0.50% |
| hsa_circ_0090759 | 5212 | 13 | 0.50% |
| hsa_circ_0099604 | 14434 | 36 | 0.50% |
| hsa_circ_0091906 | 4812 | 12 | 0.50% |
| hsa_circ_0015624 | 4411 | 11 | 0.50% |
| hsa_circ_0083152 | 4411 | 11 | 0.50% |
| hsa_circ_0123448 | 5615 | 14 | 0.50% |
| hsa_circ_0124604 | 9627 | 24 | 0.50% |
| hsa_circ_0028782 | 4413 | 11 | 0.50% |
| hsa_circ_0120349 | 12839 | 32 | 0.50% |
| hsa_circ_0030137 | 5216 | 13 | 0.50% |
| hsa_circ_0037463 | 5216 | 13 | 0.50% |
| hsa_circ_0046918 | 5216 | 13 | 0.50% |
| hsa_circ_0053640 | 5216 | 13 | 0.50% |
| hsa_circ_0016778 | 4815 | 12 | 0.50% |
| hsa_circ_0139433 | 4815 | 12 | 0.50% |
| hsa_circ_0078078 | 4816 | 12 | 0.50% |
| hsa_circ_0063742 | 4415 | 11 | 0.50% |
| hsa_circ_0101257 | 7626 | 19 | 0.50% |
| hsa_circ_0118727 | 16059 | 40 | 0.50% |
| hsa_circ_0098932 | 4818 | 12 | 0.50% |
| hsa_circ_0135427 | 4818 | 12 | 0.50% |
| hsa_circ_0125351 | 5220 | 13 | 0.50% |
| hsa_circ_0117503 | 4417 | 11 | 0.50% |
| hsa_circ_0054757 | 4819 | 12 | 0.50% |
| hsa_circ_0053610 | 5221 | 13 | 0.50% |
| hsa_circ_0032328 | 4820 | 12 | 0.50% |
| hsa_circ_0115744 | 44587 | 111 | 0.50% |
| hsa_circ_0059949 | 5222 | 13 | 0.50% |

| | | | |
|---|---|---|---|
| hsa_circ_0072366 | 5222 | 13 | 0.50% |
| hsa_circ_0033283 | 4419 | 11 | 0.50% |
| hsa_circ_0060792 | 4821 | 12 | 0.50% |
| hsa_circ_0103253 | 5223 | 13 | 0.50% |
| hsa_circ_0054924 | 80359 | 200 | 0.50% |
| hsa_circ_0024969 | 4420 | 11 | 0.50% |
| hsa_circ_0024407 | 5628 | 14 | 0.50% |
| hsa_circ_0038234 | 5628 | 14 | 0.50% |
| hsa_circ_0048029 | 4824 | 12 | 0.50% |
| hsa_circ_0020855 | 4423 | 11 | 0.50% |
| hsa_circ_0125860 | 4826 | 12 | 0.50% |
| hsa_circ_0128186 | 17295 | 43 | 0.50% |
| hsa_circ_0052640 | 5229 | 13 | 0.50% |
| hsa_circ_0056136 | 4827 | 12 | 0.50% |
| hsa_circ_0124903 | 4425 | 11 | 0.50% |
| hsa_circ_0073781 | 4023 | 10 | 0.50% |
| hsa_circ_0063722 | 5633 | 14 | 0.50% |
| hsa_circ_0042905 | 4426 | 11 | 0.50% |
| hsa_circ_0042910 | 4426 | 11 | 0.50% |
| hsa_circ_0069677 | 4829 | 12 | 0.50% |
| hsa_circ_0085224 | 4829 | 12 | 0.50% |
| hsa_circ_0015857 | 12475 | 31 | 0.50% |
| hsa_circ_0120877 | 15696 | 39 | 0.50% |
| hsa_circ_0044222 | 5232 | 13 | 0.50% |
| hsa_circ_0118383 | 14087 | 35 | 0.50% |
| hsa_circ_0053450 | 4428 | 11 | 0.50% |
| hsa_circ_0088191 | 4428 | 11 | 0.50% |
| hsa_circ_0130999 | 14092 | 35 | 0.50% |
| hsa_circ_0096635 | 4832 | 12 | 0.50% |
| hsa_circ_0128620 | 11678 | 29 | 0.50% |
| hsa_circ_0135408 | 11276 | 28 | 0.50% |
| hsa_circ_0104251 | 22956 | 57 | 0.50% |
| hsa_circ_0021353 | 4833 | 12 | 0.50% |
| hsa_circ_0101240 | 5639 | 14 | 0.50% |
| hsa_circ_0028569 | 4028 | 10 | 0.50% |
| hsa_circ_0036851 | 4834 | 12 | 0.50% |
| hsa_circ_0046039 | 4834 | 12 | 0.50% |
| hsa_circ_0063076 | 5237 | 13 | 0.50% |
| hsa_circ_0056277 | 5238 | 13 | 0.50% |
| hsa_circ_0009032 | 5641 | 14 | 0.50% |
| hsa_circ_0019906 | 5239 | 13 | 0.50% |
| hsa_circ_0069007 | 4836 | 12 | 0.50% |
| hsa_circ_0089134 | 4836 | 12 | 0.50% |
| hsa_circ_0018830 | 16122 | 40 | 0.50% |
| hsa_circ_0126134 | 6046 | 15 | 0.50% |
| hsa_circ_0128800 | 6046 | 15 | 0.50% |
| hsa_circ_0044757 | 5643 | 14 | 0.50% |
| hsa_circ_0090735 | 5240 | 13 | 0.50% |
| hsa_circ_0091931 | 4837 | 12 | 0.50% |
| hsa_circ_0101205 | 12096 | 30 | 0.50% |

| | | | |
|---|---|---|---|
| hsa_circ_0036825 | 5242 | 13 | 0.50% |
| hsa_circ_0010727 | 4436 | 11 | 0.50% |
| hsa_circ_0105297 | 8469 | 21 | 0.50% |
| hsa_circ_0118724 | 16132 | 40 | 0.50% |
| hsa_circ_0127699 | 11293 | 28 | 0.50% |
| hsa_circ_0042599 | 4437 | 11 | 0.50% |
| hsa_circ_0052709 | 6053 | 15 | 0.50% |
| hsa_circ_0107013 | 5246 | 13 | 0.50% |
| hsa_circ_0050958 | 15336 | 38 | 0.50% |
| hsa_circ_0077110 | 4441 | 11 | 0.50% |
| hsa_circ_0111287 | 4845 | 12 | 0.50% |
| hsa_circ_0062694 | 4442 | 11 | 0.50% |
| hsa_circ_0138316 | 12116 | 30 | 0.50% |
| hsa_circ_0133659 | 6462 | 16 | 0.50% |
| hsa_circ_0022009 | 13733 | 34 | 0.50% |
| hsa_circ_0109055 | 12928 | 32 | 0.50% |
| hsa_circ_0016926 | 4848 | 12 | 0.50% |
| hsa_circ_0019444 | 4444 | 11 | 0.50% |
| hsa_circ_0043373 | 4444 | 11 | 0.50% |
| hsa_circ_0100971 | 8889 | 22 | 0.49% |
| hsa_circ_0039495 | 5253 | 13 | 0.49% |
| hsa_circ_0066139 | 4445 | 11 | 0.49% |
| hsa_circ_0100972 | 8892 | 22 | 0.49% |
| hsa_circ_0015999 | 4446 | 11 | 0.49% |
| hsa_circ_0098921 | 14147 | 35 | 0.49% |
| hsa_circ_0028571 | 4042 | 10 | 0.49% |
| hsa_circ_0092082 | 5256 | 13 | 0.49% |
| hsa_circ_0133594 | 12131 | 30 | 0.49% |
| hsa_circ_0014843 | 4853 | 12 | 0.49% |
| hsa_circ_0027698 | 5663 | 14 | 0.49% |
| hsa_circ_0034650 | 4450 | 11 | 0.49% |
| hsa_circ_0050114 | 4450 | 11 | 0.49% |
| hsa_circ_0093929 | 4855 | 12 | 0.49% |
| hsa_circ_0092434 | 14971 | 37 | 0.49% |
| hsa_circ_0078370 | 4451 | 11 | 0.49% |
| hsa_circ_0040478 | 4856 | 12 | 0.49% |
| hsa_circ_0096737 | 4856 | 12 | 0.49% |
| hsa_circ_0019183 | 4857 | 12 | 0.49% |
| hsa_circ_0020757 | 4857 | 12 | 0.49% |
| hsa_circ_0113567 | 11738 | 29 | 0.49% |
| hsa_circ_0092225 | 4858 | 12 | 0.49% |
| hsa_circ_0059357 | 5263 | 13 | 0.49% |
| hsa_circ_0128870 | 17005 | 42 | 0.49% |
| hsa_circ_0073231 | 4454 | 11 | 0.49% |
| hsa_circ_0047980 | 4859 | 12 | 0.49% |
| hsa_circ_0066611 | 4859 | 12 | 0.49% |
| hsa_circ_0103531 | 14982 | 37 | 0.49% |
| hsa_circ_0077339 | 5264 | 13 | 0.49% |
| hsa_circ_0008254 | 11743 | 29 | 0.49% |
| hsa_circ_0060860 | 6074 | 15 | 0.49% |

| | | | |
|---|---|---|---|
| hsa_circ_0105020 | 15793 | 39 | 0.49% |
| hsa_circ_0126491 | 12554 | 31 | 0.49% |
| hsa_circ_0020763 | 5265 | 13 | 0.49% |
| hsa_circ_0027635 | 4860 | 12 | 0.49% |
| hsa_circ_0132417 | 4860 | 12 | 0.49% |
| hsa_circ_0084731 | 4455 | 11 | 0.49% |
| hsa_circ_0072883 | 4050 | 10 | 0.49% |
| hsa_circ_0121823 | 18631 | 46 | 0.49% |
| hsa_circ_0120350 | 12961 | 32 | 0.49% |
| hsa_circ_0011071 | 5671 | 14 | 0.49% |
| hsa_circ_0007913 | 4861 | 12 | 0.49% |
| hsa_circ_0024131 | 4861 | 12 | 0.49% |
| hsa_circ_0043044 | 4456 | 11 | 0.49% |
| hsa_circ_0059153 | 4456 | 11 | 0.49% |
| hsa_circ_0139815 | 18231 | 45 | 0.49% |
| hsa_circ_0067105 | 8508 | 21 | 0.49% |
| hsa_circ_0045208 | 5267 | 13 | 0.49% |
| hsa_circ_0073943 | 6888 | 17 | 0.49% |
| hsa_circ_0098428 | 14587 | 36 | 0.49% |
| hsa_circ_0112619 | 7701 | 19 | 0.49% |
| hsa_circ_0059358 | 12565 | 31 | 0.49% |
| hsa_circ_0071603 | 4864 | 12 | 0.49% |
| hsa_circ_0084845 | 4864 | 12 | 0.49% |
| hsa_circ_0090922 | 8107 | 20 | 0.49% |
| hsa_circ_0106288 | 5675 | 14 | 0.49% |
| hsa_circ_0083203 | 21079 | 52 | 0.49% |
| hsa_circ_0005620 | 4865 | 12 | 0.49% |
| hsa_circ_0082890 | 14190 | 35 | 0.49% |
| hsa_circ_0112372 | 5676 | 14 | 0.49% |
| hsa_circ_0100415 | 4460 | 11 | 0.49% |
| hsa_circ_0013817 | 5271 | 13 | 0.49% |
| hsa_circ_0020555 | 4866 | 12 | 0.49% |
| hsa_circ_0065330 | 5272 | 13 | 0.49% |
| hsa_circ_0123662 | 4461 | 11 | 0.49% |
| hsa_circ_0019903 | 4462 | 11 | 0.49% |
| hsa_circ_0051161 | 4462 | 11 | 0.49% |
| hsa_circ_0008737 | 13387 | 33 | 0.49% |
| hsa_circ_0022281 | 4868 | 12 | 0.49% |
| hsa_circ_0070063 | 4868 | 12 | 0.49% |
| hsa_circ_0070901 | 5274 | 13 | 0.49% |
| hsa_circ_0077446 | 5274 | 13 | 0.49% |
| hsa_circ_0130800 | 5680 | 14 | 0.49% |
| hsa_circ_0028757 | 4463 | 11 | 0.49% |
| hsa_circ_0088130 | 4463 | 11 | 0.49% |
| hsa_circ_0043115 | 15418 | 38 | 0.49% |
| hsa_circ_0034234 | 5275 | 13 | 0.49% |
| hsa_circ_0086809 | 5275 | 13 | 0.49% |
| hsa_circ_0035363 | 4464 | 11 | 0.49% |
| hsa_circ_0138127 | 28002 | 69 | 0.49% |
| hsa_circ_0002224 | 13394 | 33 | 0.49% |

| | | | |
|---|---|---|---|
| hsa_circ_0059433 | 4465 | 11 | 0.49% |
| hsa_circ_0066002 | 4871 | 12 | 0.49% |
| hsa_circ_0091953 | 4871 | 12 | 0.49% |
| hsa_circ_0123901 | 4871 | 12 | 0.49% |
| hsa_circ_0029876 | 5279 | 13 | 0.49% |
| hsa_circ_0015536 | 4873 | 12 | 0.49% |
| hsa_circ_0105834 | 4467 | 11 | 0.49% |
| hsa_circ_0023544 | 4468 | 11 | 0.49% |
| hsa_circ_0138198 | 5281 | 13 | 0.49% |
| hsa_circ_0035629 | 4875 | 12 | 0.49% |
| hsa_circ_0038738 | 4875 | 12 | 0.49% |
| hsa_circ_0054798 | 4875 | 12 | 0.49% |
| hsa_circ_0064049 | 4875 | 12 | 0.49% |
| hsa_circ_0084356 | 4469 | 11 | 0.49% |
| hsa_circ_0089848 | 4876 | 12 | 0.49% |
| hsa_circ_0046015 | 4877 | 12 | 0.49% |
| hsa_circ_0055993 | 5690 | 14 | 0.49% |
| hsa_circ_0037458 | 4878 | 12 | 0.49% |
| hsa_circ_0067485 | 17887 | 44 | 0.49% |
| hsa_circ_0114936 | 6098 | 15 | 0.49% |
| hsa_circ_0046712 | 5285 | 13 | 0.49% |
| hsa_circ_0076686 | 5285 | 13 | 0.49% |
| hsa_circ_0138387 | 5692 | 14 | 0.49% |
| hsa_circ_0065591 | 4473 | 11 | 0.49% |
| hsa_circ_0120045 | 6100 | 15 | 0.49% |
| hsa_circ_0110690 | 4880 | 12 | 0.49% |
| hsa_circ_0130801 | 8948 | 22 | 0.49% |
| hsa_circ_0033436 | 4474 | 11 | 0.49% |
| hsa_circ_0137983 | 6101 | 15 | 0.49% |
| hsa_circ_0058536 | 4475 | 11 | 0.49% |
| hsa_circ_0090670 | 5289 | 13 | 0.49% |
| hsa_circ_0091848 | 5696 | 14 | 0.49% |
| hsa_circ_0098119 | 14648 | 36 | 0.49% |
| hsa_circ_0021211 | 4476 | 11 | 0.49% |
| hsa_circ_0067736 | 4476 | 11 | 0.49% |
| hsa_circ_0078981 | 4883 | 12 | 0.49% |
| hsa_circ_0077525 | 5291 | 13 | 0.49% |
| hsa_circ_0082798 | 5291 | 13 | 0.49% |
| hsa_circ_0043990 | 4884 | 12 | 0.49% |
| hsa_circ_0105491 | 16283 | 40 | 0.49% |
| hsa_circ_0074693 | 4885 | 12 | 0.49% |
| hsa_circ_0081643 | 20360 | 50 | 0.49% |
| hsa_circ_0102692 | 5701 | 14 | 0.49% |
| hsa_circ_0060820 | 4887 | 12 | 0.49% |
| hsa_circ_0026103 | 6109 | 15 | 0.49% |
| hsa_circ_0098271 | 6110 | 15 | 0.49% |
| hsa_circ_0035784 | 5296 | 13 | 0.49% |
| hsa_circ_0021580 | 4889 | 12 | 0.49% |
| hsa_circ_0138122 | 28112 | 69 | 0.49% |
| hsa_circ_0038735 | 5297 | 13 | 0.49% |

| | | | |
|---|---|---|---|
| hsa_circ_0019077 | 5298 | 13 | 0.49% |
| hsa_circ_0042606 | 4483 | 11 | 0.49% |
| hsa_circ_0007988 | 5706 | 14 | 0.49% |
| hsa_circ_0013131 | 5706 | 14 | 0.49% |
| hsa_circ_0056142 | 4484 | 11 | 0.49% |
| hsa_circ_0111503 | 4484 | 11 | 0.49% |
| hsa_circ_0130210 | 4484 | 11 | 0.49% |
| hsa_circ_0041876 | 4892 | 12 | 0.49% |
| hsa_circ_0110143 | 6932 | 17 | 0.49% |
| hsa_circ_0121644 | 10602 | 26 | 0.49% |
| hsa_circ_0128020 | 13458 | 33 | 0.49% |
| hsa_circ_0070321 | 4486 | 11 | 0.49% |
| hsa_circ_0127869 | 12645 | 31 | 0.49% |
| hsa_circ_0042588 | 4895 | 12 | 0.49% |
| hsa_circ_0076725 | 5303 | 13 | 0.49% |
| hsa_circ_0009804 | 5712 | 14 | 0.49% |
| hsa_circ_0066420 | 4896 | 12 | 0.49% |
| hsa_circ_0077054 | 4896 | 12 | 0.49% |
| hsa_circ_0077197 | 4896 | 12 | 0.49% |
| hsa_circ_0010670 | 5305 | 13 | 0.49% |
| hsa_circ_0063087 | 4897 | 12 | 0.49% |
| hsa_circ_0097684 | 31015 | 76 | 0.49% |
| hsa_circ_0060988 | 4081 | 10 | 0.49% |
| hsa_circ_0122563 | 7346 | 18 | 0.49% |
| hsa_circ_0111777 | 10611 | 26 | 0.49% |
| hsa_circ_0052258 | 9387 | 23 | 0.49% |
| hsa_circ_0096564 | 4490 | 11 | 0.49% |
| hsa_circ_0022862 | 6125 | 15 | 0.49% |
| hsa_circ_0029203 | 6942 | 17 | 0.49% |
| hsa_circ_0077454 | 5309 | 13 | 0.49% |
| hsa_circ_0102465 | 8577 | 21 | 0.49% |
| hsa_circ_0010555 | 5719 | 14 | 0.49% |
| hsa_circ_0114129 | 4494 | 11 | 0.49% |
| hsa_circ_0045072 | 5312 | 13 | 0.49% |
| hsa_circ_0081996 | 5312 | 13 | 0.49% |
| hsa_circ_0062925 | 20840 | 51 | 0.49% |
| hsa_circ_0114424 | 5721 | 14 | 0.49% |
| hsa_circ_0126721 | 6130 | 15 | 0.49% |
| hsa_circ_0038208 | 15122 | 37 | 0.49% |
| hsa_circ_0042449 | 4496 | 11 | 0.49% |
| hsa_circ_0070336 | 4496 | 11 | 0.49% |
| hsa_circ_0118072 | 5723 | 14 | 0.49% |
| hsa_circ_0016543 | 4089 | 10 | 0.49% |
| hsa_circ_0046765 | 4089 | 10 | 0.49% |
| hsa_circ_0100040 | 8996 | 22 | 0.49% |
| hsa_circ_0064271 | 4907 | 12 | 0.49% |
| hsa_circ_0005407 | 6952 | 17 | 0.49% |
| hsa_circ_0067106 | 8589 | 21 | 0.49% |
| hsa_circ_0034461 | 4908 | 12 | 0.49% |
| hsa_circ_0047064 | 5727 | 14 | 0.49% |

| | | | |
|---|---|---|---|
| hsa_circ_0072166 | 4500 | 11 | 0.49% |
| hsa_circ_0066294 | 4910 | 12 | 0.49% |
| hsa_circ_0121572 | 8593 | 21 | 0.49% |
| hsa_circ_0075556 | 4092 | 10 | 0.49% |
| hsa_circ_0068038 | 5729 | 14 | 0.49% |
| hsa_circ_0018816 | 5321 | 13 | 0.49% |
| hsa_circ_0087448 | 4503 | 11 | 0.49% |
| hsa_circ_0107160 | 4503 | 11 | 0.49% |
| hsa_circ_0030138 | 5322 | 13 | 0.49% |
| hsa_circ_0015709 | 6141 | 15 | 0.49% |
| hsa_circ_0108949 | 4913 | 12 | 0.49% |
| hsa_circ_0112196 | 4913 | 12 | 0.49% |
| hsa_circ_0061302 | 15149 | 37 | 0.49% |
| hsa_circ_0013969 | 10236 | 25 | 0.49% |
| hsa_circ_0007033 | 9827 | 24 | 0.49% |
| hsa_circ_0105021 | 15971 | 39 | 0.49% |
| hsa_circ_0078974 | 11876 | 29 | 0.49% |
| hsa_circ_0021176 | 5324 | 13 | 0.49% |
| hsa_circ_0042600 | 4505 | 11 | 0.49% |
| hsa_circ_0112098 | 5734 | 14 | 0.49% |
| hsa_circ_0103822 | 4915 | 12 | 0.49% |
| hsa_circ_0062400 | 5325 | 13 | 0.49% |
| hsa_circ_0134157 | 24987 | 61 | 0.49% |
| hsa_circ_0053197 | 5326 | 13 | 0.49% |
| hsa_circ_0079107 | 5736 | 14 | 0.49% |
| hsa_circ_0068600 | 4507 | 11 | 0.49% |
| hsa_circ_0091372 | 4507 | 11 | 0.49% |
| hsa_circ_0098922 | 14342 | 35 | 0.49% |
| hsa_circ_0025475 | 4098 | 10 | 0.49% |
| hsa_circ_0047246 | 4098 | 10 | 0.49% |
| hsa_circ_0052543 | 6967 | 17 | 0.49% |
| hsa_circ_0052438 | 5739 | 14 | 0.49% |
| hsa_circ_0133660 | 6559 | 16 | 0.49% |
| hsa_circ_0032623 | 4920 | 12 | 0.49% |
| hsa_circ_0041294 | 4510 | 11 | 0.49% |
| hsa_circ_0100443 | 21734 | 53 | 0.49% |
| hsa_circ_0086584 | 4921 | 12 | 0.49% |
| hsa_circ_0013367 | 4511 | 11 | 0.49% |
| hsa_circ_0038913 | 10664 | 26 | 0.49% |
| hsa_circ_0045442 | 4512 | 11 | 0.49% |
| hsa_circ_0053280 | 4512 | 11 | 0.49% |
| hsa_circ_0001241 | 55789 | 136 | 0.49% |
| hsa_circ_0036824 | 5333 | 13 | 0.49% |
| hsa_circ_0076820 | 4923 | 12 | 0.49% |
| hsa_circ_0105117 | 4513 | 11 | 0.49% |
| hsa_circ_0094247 | 10259 | 25 | 0.49% |
| hsa_circ_0047889 | 4925 | 12 | 0.49% |
| hsa_circ_0063956 | 4925 | 12 | 0.49% |
| hsa_circ_0135407 | 43095 | 105 | 0.49% |
| hsa_circ_0052976 | 7388 | 18 | 0.49% |

| | | | |
|---|---|---|---|
| hsa_circ_0022605 | 4926 | 12 | 0.49% |
| hsa_circ_0023534 | 4926 | 12 | 0.49% |
| hsa_circ_0038406 | 22578 | 55 | 0.49% |
| hsa_circ_0033309 | 4927 | 12 | 0.49% |
| hsa_circ_0067630 | 5338 | 13 | 0.49% |
| hsa_circ_0034981 | 4517 | 11 | 0.49% |
| hsa_circ_0024183 | 4928 | 12 | 0.49% |
| hsa_circ_0035927 | 5339 | 13 | 0.49% |
| hsa_circ_0065392 | 5339 | 13 | 0.49% |
| hsa_circ_0064059 | 4929 | 12 | 0.49% |
| hsa_circ_0016716 | 6162 | 15 | 0.49% |
| hsa_circ_0057603 | 4519 | 11 | 0.49% |
| hsa_circ_0084331 | 4519 | 11 | 0.49% |
| hsa_circ_0086852 | 4519 | 11 | 0.49% |
| hsa_circ_0054866 | 4930 | 12 | 0.49% |
| hsa_circ_0019286 | 5341 | 13 | 0.49% |
| hsa_circ_0094071 | 4931 | 12 | 0.49% |
| hsa_circ_0136376 | 14385 | 35 | 0.49% |
| hsa_circ_0037652 | 4932 | 12 | 0.49% |
| hsa_circ_0019907 | 5344 | 13 | 0.49% |
| hsa_circ_0078790 | 5344 | 13 | 0.49% |
| hsa_circ_0058284 | 4522 | 11 | 0.49% |
| hsa_circ_0111303 | 4522 | 11 | 0.49% |
| hsa_circ_0103231 | 7400 | 18 | 0.49% |
| hsa_circ_0109359 | 5346 | 13 | 0.49% |
| hsa_circ_0091892 | 4935 | 12 | 0.49% |
| hsa_circ_0138573 | 4935 | 12 | 0.49% |
| hsa_circ_0018826 | 16454 | 40 | 0.49% |
| hsa_circ_0038261 | 4525 | 11 | 0.49% |
| hsa_circ_0124734 | 4937 | 12 | 0.49% |
| hsa_circ_0027140 | 4938 | 12 | 0.49% |
| hsa_circ_0053793 | 4938 | 12 | 0.49% |
| hsa_circ_0068498 | 5762 | 14 | 0.49% |
| hsa_circ_0019874 | 4939 | 12 | 0.49% |
| hsa_circ_0042178 | 4939 | 12 | 0.49% |
| hsa_circ_0028778 | 4528 | 11 | 0.49% |
| hsa_circ_0091933 | 4940 | 12 | 0.49% |
| hsa_circ_0047004 | 5352 | 13 | 0.49% |
| hsa_circ_0020061 | 6176 | 15 | 0.49% |
| hsa_circ_0066468 | 4530 | 11 | 0.49% |
| hsa_circ_0029979 | 5354 | 13 | 0.49% |
| hsa_circ_0126334 | 5354 | 13 | 0.49% |
| hsa_circ_0006257 | 9473 | 23 | 0.49% |
| hsa_circ_0045804 | 5355 | 13 | 0.49% |
| hsa_circ_0009642 | 7827 | 19 | 0.49% |
| hsa_circ_0092268 | 4532 | 11 | 0.49% |
| hsa_circ_0109885 | 13599 | 33 | 0.49% |
| hsa_circ_0027889 | 4533 | 11 | 0.49% |
| hsa_circ_0027289 | 7006 | 17 | 0.49% |
| hsa_circ_0135281 | 9479 | 23 | 0.49% |

| | | | |
|---|---|---|---|
| hsa_circ_0001600 | 4534 | 11 | 0.49% |
| hsa_circ_0094095 | 26380 | 64 | 0.49% |
| hsa_circ_0002813 | 13191 | 32 | 0.49% |
| hsa_circ_0135851 | 4947 | 12 | 0.49% |
| hsa_circ_0108988 | 5361 | 13 | 0.48% |
| hsa_circ_0085225 | 4949 | 12 | 0.48% |
| hsa_circ_0042150 | 5774 | 14 | 0.48% |
| hsa_circ_0128702 | 5362 | 13 | 0.48% |
| hsa_circ_0010265 | 6187 | 15 | 0.48% |
| hsa_circ_0034980 | 4125 | 10 | 0.48% |
| hsa_circ_0036789 | 4125 | 10 | 0.48% |
| hsa_circ_0053636 | 5364 | 13 | 0.48% |
| hsa_circ_0104578 | 7015 | 17 | 0.48% |
| hsa_circ_0121656 | 7428 | 18 | 0.48% |
| hsa_circ_0040479 | 4952 | 12 | 0.48% |
| hsa_circ_0074720 | 4540 | 11 | 0.48% |
| hsa_circ_0029499 | 4953 | 12 | 0.48% |
| hsa_circ_0073840 | 4953 | 12 | 0.48% |
| hsa_circ_0035736 | 5366 | 13 | 0.48% |
| hsa_circ_0051279 | 5366 | 13 | 0.48% |
| hsa_circ_0047250 | 4129 | 10 | 0.48% |
| hsa_circ_0114425 | 5782 | 14 | 0.48% |
| hsa_circ_0108964 | 20653 | 50 | 0.48% |
| hsa_circ_0107279 | 16110 | 39 | 0.48% |
| hsa_circ_0121151 | 5370 | 13 | 0.48% |
| hsa_circ_0028748 | 4544 | 11 | 0.48% |
| hsa_circ_0043217 | 4544 | 11 | 0.48% |
| hsa_circ_0058247 | 5784 | 14 | 0.48% |
| hsa_circ_0049109 | 4958 | 12 | 0.48% |
| hsa_circ_0061139 | 4132 | 10 | 0.48% |
| hsa_circ_0100514 | 5372 | 13 | 0.48% |
| hsa_circ_0081590 | 4960 | 12 | 0.48% |
| hsa_circ_0084293 | 4960 | 12 | 0.48% |
| hsa_circ_0085210 | 4960 | 12 | 0.48% |
| hsa_circ_0053808 | 4961 | 12 | 0.48% |
| hsa_circ_0088101 | 16954 | 41 | 0.48% |
| hsa_circ_0099722 | 7030 | 17 | 0.48% |
| hsa_circ_0033095 | 5376 | 13 | 0.48% |
| hsa_circ_0041734 | 5376 | 13 | 0.48% |
| hsa_circ_0019678 | 4549 | 11 | 0.48% |
| hsa_circ_0124900 | 4549 | 11 | 0.48% |
| hsa_circ_0094283 | 4963 | 12 | 0.48% |
| hsa_circ_0104135 | 4963 | 12 | 0.48% |
| hsa_circ_0125652 | 5377 | 13 | 0.48% |
| hsa_circ_0018232 | 18200 | 44 | 0.48% |
| hsa_circ_0124605 | 9930 | 24 | 0.48% |
| hsa_circ_0024216 | 4965 | 12 | 0.48% |
| hsa_circ_0086175 | 4965 | 12 | 0.48% |
| hsa_circ_0073455 | 5379 | 13 | 0.48% |
| hsa_circ_0076513 | 4552 | 11 | 0.48% |

| | | | |
|---|---|---|---|
| hsa_circ_0055254 | 4966 | 12 | 0.48% |
| hsa_circ_0093858 | 4966 | 12 | 0.48% |
| hsa_circ_0061285 | 7450 | 18 | 0.48% |
| hsa_circ_0056162 | 4967 | 12 | 0.48% |
| hsa_circ_0138196 | 5381 | 13 | 0.48% |
| hsa_circ_0010552 | 5795 | 14 | 0.48% |
| hsa_circ_0010572 | 5383 | 13 | 0.48% |
| hsa_circ_0053754 | 4555 | 11 | 0.48% |
| hsa_circ_0020062 | 6212 | 15 | 0.48% |
| hsa_circ_0074838 | 6213 | 15 | 0.48% |
| hsa_circ_0060793 | 4971 | 12 | 0.48% |
| hsa_circ_0128026 | 5386 | 13 | 0.48% |
| hsa_circ_0054800 | 4972 | 12 | 0.48% |
| hsa_circ_0100992 | 4972 | 12 | 0.48% |
| hsa_circ_0013086 | 5387 | 13 | 0.48% |
| hsa_circ_0131482 | 5387 | 13 | 0.48% |
| hsa_circ_0111288 | 4973 | 12 | 0.48% |
| hsa_circ_0005102 | 10361 | 25 | 0.48% |
| hsa_circ_0028779 | 4559 | 11 | 0.48% |
| hsa_circ_0071885 | 4559 | 11 | 0.48% |
| hsa_circ_0029501 | 5388 | 13 | 0.48% |
| hsa_circ_0073267 | 4974 | 12 | 0.48% |
| hsa_circ_0138574 | 4974 | 12 | 0.48% |
| hsa_circ_0132543 | 7876 | 19 | 0.48% |
| hsa_circ_0019908 | 5389 | 13 | 0.48% |
| hsa_circ_0067640 | 4560 | 11 | 0.48% |
| hsa_circ_0138799 | 4560 | 11 | 0.48% |
| hsa_circ_0028725 | 4975 | 12 | 0.48% |
| hsa_circ_0049500 | 20734 | 50 | 0.48% |
| hsa_circ_0140123 | 13273 | 32 | 0.48% |
| hsa_circ_0041668 | 4564 | 11 | 0.48% |
| hsa_circ_0054054 | 4979 | 12 | 0.48% |
| hsa_circ_0119562 | 4979 | 12 | 0.48% |
| hsa_circ_0027455 | 4980 | 12 | 0.48% |
| hsa_circ_0059454 | 4566 | 11 | 0.48% |
| hsa_circ_0050838 | 5812 | 14 | 0.48% |
| hsa_circ_0090141 | 4982 | 12 | 0.48% |
| hsa_circ_0030491 | 6228 | 15 | 0.48% |
| hsa_circ_0063723 | 5813 | 14 | 0.48% |
| hsa_circ_0084372 | 5398 | 13 | 0.48% |
| hsa_circ_0064058 | 4983 | 12 | 0.48% |
| hsa_circ_0061300 | 7475 | 18 | 0.48% |
| hsa_circ_0138630 | 7891 | 19 | 0.48% |
| hsa_circ_0009983 | 4984 | 12 | 0.48% |
| hsa_circ_0034283 | 4984 | 12 | 0.48% |
| hsa_circ_0034446 | 4984 | 12 | 0.48% |
| hsa_circ_0112856 | 4569 | 11 | 0.48% |
| hsa_circ_0124339 | 4569 | 11 | 0.48% |
| hsa_circ_0010633 | 5400 | 13 | 0.48% |
| hsa_circ_0053597 | 5400 | 13 | 0.48% |

| | | | |
|---|---|---|---|
| hsa_circ_0015592 | 4985 | 12 | 0.48% |
| hsa_circ_0071171 | 5401 | 13 | 0.48% |
| hsa_circ_0010599 | 5403 | 13 | 0.48% |
| hsa_circ_0135003 | 5404 | 13 | 0.48% |
| hsa_circ_0136923 | 41585 | 100 | 0.48% |
| hsa_circ_0049501 | 20795 | 50 | 0.48% |
| hsa_circ_0076273 | 4575 | 11 | 0.48% |
| hsa_circ_0101506 | 5407 | 13 | 0.48% |
| hsa_circ_0107935 | 5408 | 13 | 0.48% |
| hsa_circ_0055126 | 4992 | 12 | 0.48% |
| hsa_circ_0084075 | 4576 | 11 | 0.48% |
| hsa_circ_0009777 | 5825 | 14 | 0.48% |
| hsa_circ_0065400 | 5409 | 13 | 0.48% |
| hsa_circ_0010643 | 5410 | 13 | 0.48% |
| hsa_circ_0101589 | 4994 | 12 | 0.48% |
| hsa_circ_0028714 | 4578 | 11 | 0.48% |
| hsa_circ_0089468 | 4578 | 11 | 0.48% |
| hsa_circ_0089813 | 17901 | 43 | 0.48% |
| hsa_circ_0036156 | 4996 | 12 | 0.48% |
| hsa_circ_0074625 | 5413 | 13 | 0.48% |
| hsa_circ_0096547 | 5413 | 13 | 0.48% |
| hsa_circ_0116383 | 30821 | 74 | 0.48% |
| hsa_circ_0013210 | 4998 | 12 | 0.48% |
| hsa_circ_0081900 | 5415 | 13 | 0.48% |
| hsa_circ_0042907 | 4582 | 11 | 0.48% |
| hsa_circ_0126238 | 5832 | 14 | 0.48% |
| hsa_circ_0020412 | 4583 | 11 | 0.48% |
| hsa_circ_0045784 | 4583 | 11 | 0.48% |
| hsa_circ_0065511 | 4167 | 10 | 0.48% |
| hsa_circ_0004886 | 8751 | 21 | 0.48% |
| hsa_circ_0045444 | 4584 | 11 | 0.48% |
| hsa_circ_0060351 | 6251 | 15 | 0.48% |
| hsa_circ_0013840 | 4585 | 11 | 0.48% |
| hsa_circ_0094136 | 11255 | 27 | 0.48% |
| hsa_circ_0096771 | 16258 | 39 | 0.48% |
| hsa_circ_0064210 | 4586 | 11 | 0.48% |
| hsa_circ_0099006 | 15012 | 36 | 0.48% |
| hsa_circ_0073232 | 4587 | 11 | 0.48% |
| hsa_circ_0078356 | 4587 | 11 | 0.48% |
| hsa_circ_0119633 | 17515 | 42 | 0.48% |
| hsa_circ_0130392 | 38378 | 92 | 0.48% |
| hsa_circ_0007514 | 5423 | 13 | 0.48% |
| hsa_circ_0015227 | 4172 | 10 | 0.48% |
| hsa_circ_0114582 | 5424 | 13 | 0.48% |
| hsa_circ_0086288 | 4590 | 11 | 0.48% |
| hsa_circ_0135026 | 4590 | 11 | 0.48% |
| hsa_circ_0067671 | 4591 | 11 | 0.48% |
| hsa_circ_0102871 | 16697 | 40 | 0.48% |
| hsa_circ_0016169 | 12106 | 29 | 0.48% |
| hsa_circ_0132840 | 5427 | 13 | 0.48% |

| | | | |
|---|---|---|---|
| hsa_circ_0105799 | 17118 | 41 | 0.48% |
| hsa_circ_0039645 | 5428 | 13 | 0.48% |
| hsa_circ_0133881 | 6681 | 16 | 0.48% |
| hsa_circ_0023539 | 5012 | 12 | 0.48% |
| hsa_circ_0138120 | 28820 | 69 | 0.48% |
| hsa_circ_0052316 | 5848 | 14 | 0.48% |
| hsa_circ_0035588 | 4595 | 11 | 0.48% |
| hsa_circ_0033534 | 5431 | 13 | 0.48% |
| hsa_circ_0063077 | 5431 | 13 | 0.48% |
| hsa_circ_0101233 | 6686 | 16 | 0.48% |
| hsa_circ_0123484 | 7104 | 17 | 0.48% |
| hsa_circ_0011773 | 4179 | 10 | 0.48% |
| hsa_circ_0053797 | 5015 | 12 | 0.48% |
| hsa_circ_0042450 | 4598 | 11 | 0.48% |
| hsa_circ_0090251 | 5435 | 13 | 0.48% |
| hsa_circ_0092247 | 5435 | 13 | 0.48% |
| hsa_circ_0031514 | 5017 | 12 | 0.48% |
| hsa_circ_0081286 | 4599 | 11 | 0.48% |
| hsa_circ_0140566 | 4599 | 11 | 0.48% |
| hsa_circ_0117385 | 15889 | 38 | 0.48% |
| hsa_circ_0026583 | 5018 | 12 | 0.48% |
| hsa_circ_0134137 | 5855 | 14 | 0.48% |
| hsa_circ_0079533 | 6692 | 16 | 0.48% |
| hsa_circ_0009443 | 4601 | 11 | 0.48% |
| hsa_circ_0047797 | 5438 | 13 | 0.48% |
| hsa_circ_0045738 | 15896 | 38 | 0.48% |
| hsa_circ_0027333 | 5020 | 12 | 0.48% |
| hsa_circ_0091849 | 5857 | 14 | 0.48% |
| hsa_circ_0058722 | 21338 | 51 | 0.48% |
| hsa_circ_0025727 | 5022 | 12 | 0.48% |
| hsa_circ_0040390 | 17996 | 43 | 0.48% |
| hsa_circ_0108122 | 5441 | 13 | 0.48% |
| hsa_circ_0042658 | 5442 | 13 | 0.48% |
| hsa_circ_0057044 | 5862 | 14 | 0.48% |
| hsa_circ_0139786 | 5025 | 12 | 0.48% |
| hsa_circ_0134957 | 7119 | 17 | 0.48% |
| hsa_circ_0136838 | 15496 | 37 | 0.48% |
| hsa_circ_0050045 | 5026 | 12 | 0.48% |
| hsa_circ_0025084 | 5445 | 13 | 0.48% |
| hsa_circ_0122706 | 9216 | 22 | 0.48% |
| hsa_circ_0029877 | 5446 | 13 | 0.48% |
| hsa_circ_0104138 | 5446 | 13 | 0.48% |
| hsa_circ_0060861 | 6287 | 15 | 0.48% |
| hsa_circ_0071159 | 5030 | 12 | 0.48% |
| hsa_circ_0010671 | 5450 | 13 | 0.48% |
| hsa_circ_0041457 | 5450 | 13 | 0.48% |
| hsa_circ_0123915 | 5031 | 12 | 0.48% |
| hsa_circ_0062583 | 6289 | 15 | 0.48% |
| hsa_circ_0086519 | 4612 | 11 | 0.48% |
| hsa_circ_0127878 | 13417 | 32 | 0.48% |

| | | | |
|---|---|---|---|
| hsa_circ_0091805 | 5870 | 14 | 0.48% |
| hsa_circ_0101639 | 5870 | 14 | 0.48% |
| hsa_circ_0102237 | 15097 | 36 | 0.48% |
| hsa_circ_0019909 | 5452 | 13 | 0.48% |
| hsa_circ_0031850 | 15518 | 37 | 0.48% |
| hsa_circ_0064098 | 5454 | 13 | 0.48% |
| hsa_circ_0010548 | 5874 | 14 | 0.48% |
| hsa_circ_0053451 | 4616 | 11 | 0.48% |
| hsa_circ_0016274 | 5875 | 14 | 0.48% |
| hsa_circ_0085198 | 5875 | 14 | 0.48% |
| hsa_circ_0129517 | 5036 | 12 | 0.48% |
| hsa_circ_0046850 | 5456 | 13 | 0.48% |
| hsa_circ_0063365 | 5037 | 12 | 0.48% |
| hsa_circ_0009181 | 11334 | 27 | 0.48% |
| hsa_circ_0020136 | 4618 | 11 | 0.48% |
| hsa_circ_0103692 | 5458 | 13 | 0.48% |
| hsa_circ_0059609 | 4619 | 11 | 0.48% |
| hsa_circ_0091876 | 4619 | 11 | 0.48% |
| hsa_circ_0033048 | 5039 | 12 | 0.48% |
| hsa_circ_0046847 | 5459 | 13 | 0.48% |
| hsa_circ_0017860 | 6300 | 15 | 0.48% |
| hsa_circ_0018465 | 5040 | 12 | 0.48% |
| hsa_circ_0102488 | 18483 | 44 | 0.48% |
| hsa_circ_0010683 | 5461 | 13 | 0.48% |
| hsa_circ_0025642 | 5041 | 12 | 0.48% |
| hsa_circ_0030492 | 6302 | 15 | 0.48% |
| hsa_circ_0053631 | 5462 | 13 | 0.48% |
| hsa_circ_0078299 | 5042 | 12 | 0.48% |
| hsa_circ_0111583 | 5042 | 12 | 0.48% |
| hsa_circ_0013809 | 10928 | 26 | 0.48% |
| hsa_circ_0130721 | 7986 | 19 | 0.48% |
| hsa_circ_0015896 | 5885 | 14 | 0.48% |
| hsa_circ_0020758 | 5045 | 12 | 0.48% |
| hsa_circ_0096708 | 10511 | 25 | 0.48% |
| hsa_circ_0027134 | 5046 | 12 | 0.48% |
| hsa_circ_0044733 | 5046 | 12 | 0.48% |
| hsa_circ_0072891 | 5046 | 12 | 0.48% |
| hsa_circ_0121921 | 5047 | 12 | 0.48% |
| hsa_circ_0018109 | 5889 | 14 | 0.48% |
| hsa_circ_0035141 | 5469 | 13 | 0.48% |
| hsa_circ_0100984 | 7574 | 18 | 0.48% |
| hsa_circ_0041274 | 4629 | 11 | 0.48% |
| hsa_circ_0084900 | 4629 | 11 | 0.48% |
| hsa_circ_0120308 | 12632 | 30 | 0.47% |
| hsa_circ_0115275 | 5474 | 13 | 0.47% |
| hsa_circ_0082971 | 4632 | 11 | 0.47% |
| hsa_circ_0128417 | 16424 | 39 | 0.47% |
| hsa_circ_0108510 | 13056 | 31 | 0.47% |
| hsa_circ_0058782 | 5055 | 12 | 0.47% |
| hsa_circ_0065166 | 5055 | 12 | 0.47% |

| | | | |
|---|---|---|---|
| hsa_circ_0070062 | 5055 | 12 | 0.47% |
| hsa_circ_0140565 | 4634 | 11 | 0.47% |
| hsa_circ_0093857 | 5056 | 12 | 0.47% |
| hsa_circ_0119092 | 19803 | 47 | 0.47% |
| hsa_circ_0005798 | 8428 | 20 | 0.47% |
| hsa_circ_0078104 | 5057 | 12 | 0.47% |
| hsa_circ_0061598 | 5479 | 13 | 0.47% |
| hsa_circ_0028744 | 4637 | 11 | 0.47% |
| hsa_circ_0085201 | 5902 | 14 | 0.47% |
| hsa_circ_0054110 | 5059 | 12 | 0.47% |
| hsa_circ_0065402 | 5481 | 13 | 0.47% |
| hsa_circ_0095671 | 5481 | 13 | 0.47% |
| hsa_circ_0128521 | 5481 | 13 | 0.47% |
| hsa_circ_0098923 | 14758 | 35 | 0.47% |
| hsa_circ_0098120 | 12652 | 30 | 0.47% |
| hsa_circ_0107156 | 13498 | 32 | 0.47% |
| hsa_circ_0045685 | 4640 | 11 | 0.47% |
| hsa_circ_0053670 | 5484 | 13 | 0.47% |
| hsa_circ_0074632 | 5484 | 13 | 0.47% |
| hsa_circ_0124576 | 10547 | 25 | 0.47% |
| hsa_circ_0012541 | 5063 | 12 | 0.47% |
| hsa_circ_0017173 | 5063 | 12 | 0.47% |
| hsa_circ_0028749 | 5063 | 12 | 0.47% |
| hsa_circ_0014194 | 5907 | 14 | 0.47% |
| hsa_circ_0096407 | 5907 | 14 | 0.47% |
| hsa_circ_0127333 | 8863 | 21 | 0.47% |
| hsa_circ_0052439 | 5909 | 14 | 0.47% |
| hsa_circ_0065891 | 5909 | 14 | 0.47% |
| hsa_circ_0021716 | 5065 | 12 | 0.47% |
| hsa_circ_0027065 | 14351 | 34 | 0.47% |
| hsa_circ_0028568 | 4221 | 10 | 0.47% |
| hsa_circ_0052701 | 6332 | 15 | 0.47% |
| hsa_circ_0009543 | 4644 | 11 | 0.47% |
| hsa_circ_0027187 | 4644 | 11 | 0.47% |
| hsa_circ_0111504 | 4644 | 11 | 0.47% |
| hsa_circ_0021992 | 6333 | 15 | 0.47% |
| hsa_circ_0128974 | 5489 | 13 | 0.47% |
| hsa_circ_0002090 | 5490 | 13 | 0.47% |
| hsa_circ_0052641 | 5490 | 13 | 0.47% |
| hsa_circ_0111975 | 5068 | 12 | 0.47% |
| hsa_circ_0027600 | 5913 | 14 | 0.47% |
| hsa_circ_0077155 | 4646 | 11 | 0.47% |
| hsa_circ_0039653 | 4224 | 10 | 0.47% |
| hsa_circ_0019430 | 4647 | 11 | 0.47% |
| hsa_circ_0055453 | 4647 | 11 | 0.47% |
| hsa_circ_0065167 | 5071 | 12 | 0.47% |
| hsa_circ_0032989 | 5072 | 12 | 0.47% |
| hsa_circ_0137781 | 5072 | 12 | 0.47% |
| hsa_circ_0050973 | 5495 | 13 | 0.47% |
| hsa_circ_0140564 | 4650 | 11 | 0.47% |

| | | | |
|---|---|---|---|
| hsa_circ_0053641 | 5496 | 13 | 0.47% |
| hsa_circ_0053730 | 5496 | 13 | 0.47% |
| hsa_circ_0002753 | 10571 | 25 | 0.47% |
| hsa_circ_0036823 | 5497 | 13 | 0.47% |
| hsa_circ_0059563 | 4652 | 11 | 0.47% |
| hsa_circ_0045070 | 5498 | 13 | 0.47% |
| hsa_circ_0025664 | 5499 | 13 | 0.47% |
| hsa_circ_0078578 | 4653 | 11 | 0.47% |
| hsa_circ_0003447 | 4654 | 11 | 0.47% |
| hsa_circ_0119700 | 5924 | 14 | 0.47% |
| hsa_circ_0086867 | 5501 | 13 | 0.47% |
| hsa_circ_0064055 | 5079 | 12 | 0.47% |
| hsa_circ_0090926 | 5079 | 12 | 0.47% |
| hsa_circ_0132968 | 4656 | 11 | 0.47% |
| hsa_circ_0105386 | 6350 | 15 | 0.47% |
| hsa_circ_0027651 | 5080 | 12 | 0.47% |
| hsa_circ_0028007 | 5080 | 12 | 0.47% |
| hsa_circ_0036639 | 5080 | 12 | 0.47% |
| hsa_circ_0008149 | 4657 | 11 | 0.47% |
| hsa_circ_0000576 | 18630 | 44 | 0.47% |
| hsa_circ_0119372 | 5928 | 14 | 0.47% |
| hsa_circ_0098617 | 4658 | 11 | 0.47% |
| hsa_circ_0101691 | 5082 | 12 | 0.47% |
| hsa_circ_0014591 | 5083 | 12 | 0.47% |
| hsa_circ_0033306 | 5084 | 12 | 0.47% |
| hsa_circ_0140143 | 14405 | 34 | 0.47% |
| hsa_circ_0076841 | 5508 | 13 | 0.47% |
| hsa_circ_0027697 | 5933 | 14 | 0.47% |
| hsa_circ_0063724 | 5933 | 14 | 0.47% |
| hsa_circ_0028232 | 5086 | 12 | 0.47% |
| hsa_circ_0067606 | 5510 | 13 | 0.47% |
| hsa_circ_0108509 | 13141 | 31 | 0.47% |
| hsa_circ_0064590 | 4663 | 11 | 0.47% |
| hsa_circ_0071900 | 4663 | 11 | 0.47% |
| hsa_circ_0013111 | 5087 | 12 | 0.47% |
| hsa_circ_0061880 | 5087 | 12 | 0.47% |
| hsa_circ_0098620 | 5087 | 12 | 0.47% |
| hsa_circ_0001242 | 57654 | 136 | 0.47% |
| hsa_circ_0098194 | 6783 | 16 | 0.47% |
| hsa_circ_0128187 | 46636 | 110 | 0.47% |
| hsa_circ_0035826 | 6360 | 15 | 0.47% |
| hsa_circ_0061083 | 5512 | 13 | 0.47% |
| hsa_circ_0122911 | 5512 | 13 | 0.47% |
| hsa_circ_0062942 | 4240 | 10 | 0.47% |
| hsa_circ_0126091 | 7209 | 17 | 0.47% |
| hsa_circ_0074257 | 5937 | 14 | 0.47% |
| hsa_circ_0072167 | 5089 | 12 | 0.47% |
| hsa_circ_0138992 | 5090 | 12 | 0.47% |
| hsa_circ_0020413 | 4667 | 11 | 0.47% |
| hsa_circ_0099124 | 19941 | 47 | 0.47% |

| | | | |
|---|---|---|---|
| hsa_circ_0101510 | 49644 | 117 | 0.47% |
| hsa_circ_0015817 | 5092 | 12 | 0.47% |
| hsa_circ_0114711 | 10611 | 25 | 0.47% |
| hsa_circ_0004790 | 5518 | 13 | 0.47% |
| hsa_circ_0098295 | 13159 | 31 | 0.47% |
| hsa_circ_0081232 | 6368 | 15 | 0.47% |
| hsa_circ_0042657 | 5519 | 13 | 0.47% |
| hsa_circ_0086499 | 4670 | 11 | 0.47% |
| hsa_circ_0000946 | 19107 | 45 | 0.47% |
| hsa_circ_0025738 | 5520 | 13 | 0.47% |
| hsa_circ_0116958 | 4671 | 11 | 0.47% |
| hsa_circ_0125745 | 7222 | 17 | 0.47% |
| hsa_circ_0053185 | 5098 | 12 | 0.47% |
| hsa_circ_0062446 | 5098 | 12 | 0.47% |
| hsa_circ_0085213 | 5098 | 12 | 0.47% |
| hsa_circ_0020669 | 5523 | 13 | 0.47% |
| hsa_circ_0016657 | 13172 | 31 | 0.47% |
| hsa_circ_0044838 | 5099 | 12 | 0.47% |
| hsa_circ_0028758 | 5100 | 12 | 0.47% |
| hsa_circ_0128323 | 4675 | 11 | 0.47% |
| hsa_circ_0081255 | 6376 | 15 | 0.47% |
| hsa_circ_0019076 | 5526 | 13 | 0.47% |
| hsa_circ_0052977 | 7653 | 18 | 0.47% |
| hsa_circ_0133090 | 43369 | 102 | 0.47% |
| hsa_circ_0043219 | 5528 | 13 | 0.47% |
| hsa_circ_0027811 | 14459 | 34 | 0.47% |
| hsa_circ_0051586 | 14034 | 33 | 0.47% |
| hsa_circ_0024673 | 4678 | 11 | 0.47% |
| hsa_circ_0078140 | 4253 | 10 | 0.47% |
| hsa_circ_0072502 | 6805 | 16 | 0.47% |
| hsa_circ_0026624 | 7657 | 18 | 0.47% |
| hsa_circ_0081694 | 25950 | 61 | 0.47% |
| hsa_circ_0135842 | 14039 | 33 | 0.47% |
| hsa_circ_0082495 | 5531 | 13 | 0.47% |
| hsa_circ_0102635 | 17020 | 40 | 0.47% |
| hsa_circ_0047966 | 5106 | 12 | 0.47% |
| hsa_circ_0054579 | 5106 | 12 | 0.47% |
| hsa_circ_0053588 | 5532 | 13 | 0.47% |
| hsa_circ_0070124 | 5958 | 14 | 0.47% |
| hsa_circ_0099829 | 28515 | 67 | 0.47% |
| hsa_circ_0138499 | 21283 | 50 | 0.47% |
| hsa_circ_0024184 | 5108 | 12 | 0.47% |
| hsa_circ_0085226 | 5108 | 12 | 0.47% |
| hsa_circ_0090258 | 5534 | 13 | 0.47% |
| hsa_circ_0011492 | 5109 | 12 | 0.47% |
| hsa_circ_0020316 | 8941 | 21 | 0.47% |
| hsa_circ_0110144 | 7238 | 17 | 0.47% |
| hsa_circ_0128600 | 15335 | 36 | 0.47% |
| hsa_circ_0026099 | 5538 | 13 | 0.47% |
| hsa_circ_0123264 | 10651 | 25 | 0.47% |

| | | | |
|---|---|---|---|
| hsa_circ_0053755 | 4687 | 11 | 0.47% |
| hsa_circ_0068982 | 4687 | 11 | 0.47% |
| hsa_circ_0009641 | 8097 | 19 | 0.47% |
| hsa_circ_0031178 | 5114 | 12 | 0.47% |
| hsa_circ_0025726 | 5115 | 12 | 0.47% |
| hsa_circ_0049922 | 4689 | 11 | 0.47% |
| hsa_circ_0123121 | 4689 | 11 | 0.47% |
| hsa_circ_0118351 | 5542 | 13 | 0.47% |
| hsa_circ_0135715 | 12363 | 29 | 0.47% |
| hsa_circ_0043539 | 5116 | 12 | 0.47% |
| hsa_circ_0104952 | 14922 | 35 | 0.47% |
| hsa_circ_0040365 | 5969 | 14 | 0.47% |
| hsa_circ_0020851 | 4690 | 11 | 0.47% |
| hsa_circ_0085866 | 5543 | 13 | 0.47% |
| hsa_circ_0090243 | 5543 | 13 | 0.47% |
| hsa_circ_0125450 | 20471 | 48 | 0.47% |
| hsa_circ_0001509 | 5118 | 12 | 0.47% |
| hsa_circ_0046334 | 5118 | 12 | 0.47% |
| hsa_circ_0066325 | 5118 | 12 | 0.47% |
| hsa_circ_0055636 | 6399 | 15 | 0.47% |
| hsa_circ_0063078 | 5546 | 13 | 0.47% |
| hsa_circ_0105970 | 15785 | 37 | 0.47% |
| hsa_circ_0060862 | 6400 | 15 | 0.47% |
| hsa_circ_0060794 | 5120 | 12 | 0.47% |
| hsa_circ_0114021 | 5120 | 12 | 0.47% |
| hsa_circ_0013935 | 15789 | 37 | 0.47% |
| hsa_circ_0137352 | 8108 | 19 | 0.47% |
| hsa_circ_0137527 | 8535 | 20 | 0.47% |
| hsa_circ_0078106 | 5121 | 12 | 0.47% |
| hsa_circ_0084272 | 4695 | 11 | 0.47% |
| hsa_circ_0101587 | 5122 | 12 | 0.47% |
| hsa_circ_0001704 | 4696 | 11 | 0.47% |
| hsa_circ_0017085 | 4696 | 11 | 0.47% |
| hsa_circ_0055101 | 5123 | 12 | 0.47% |
| hsa_circ_0070860 | 5123 | 12 | 0.47% |
| hsa_circ_0073841 | 5123 | 12 | 0.47% |
| hsa_circ_0104905 | 12381 | 29 | 0.47% |
| hsa_circ_0102481 | 5551 | 13 | 0.47% |
| hsa_circ_0036637 | 5124 | 12 | 0.47% |
| hsa_circ_0040425 | 5979 | 14 | 0.47% |
| hsa_circ_0024805 | 5552 | 13 | 0.47% |
| hsa_circ_0070117 | 5980 | 14 | 0.47% |
| hsa_circ_0103004 | 5553 | 13 | 0.47% |
| hsa_circ_0126335 | 5554 | 13 | 0.47% |
| hsa_circ_0032262 | 5127 | 12 | 0.47% |
| hsa_circ_0043218 | 4700 | 11 | 0.47% |
| hsa_circ_0113859 | 5555 | 13 | 0.47% |
| hsa_circ_0058662 | 16667 | 39 | 0.47% |
| hsa_circ_0126560 | 17096 | 40 | 0.47% |
| hsa_circ_0014772 | 5557 | 13 | 0.47% |

| | | | |
|---|---|---|---|
| hsa_circ_0084348 | 5557 | 13 | 0.47% |
| hsa_circ_0003306 | 5130 | 12 | 0.47% |
| hsa_circ_0077032 | 5130 | 12 | 0.47% |
| hsa_circ_0113856 | 5559 | 13 | 0.47% |
| hsa_circ_0136113 | 10264 | 24 | 0.47% |
| hsa_circ_0045028 | 5132 | 12 | 0.47% |
| hsa_circ_0038029 | 14542 | 34 | 0.47% |
| hsa_circ_0012807 | 4705 | 11 | 0.47% |
| hsa_circ_0099164 | 7272 | 17 | 0.47% |
| hsa_circ_0027428 | 4706 | 11 | 0.47% |
| hsa_circ_0024203 | 5134 | 12 | 0.47% |
| hsa_circ_0126816 | 9414 | 22 | 0.47% |
| hsa_circ_0127424 | 5563 | 13 | 0.47% |
| hsa_circ_0104816 | 5564 | 13 | 0.47% |
| hsa_circ_0009015 | 15409 | 36 | 0.47% |
| hsa_circ_0057999 | 5565 | 13 | 0.47% |
| hsa_circ_0050023 | 5566 | 13 | 0.47% |
| hsa_circ_0135244 | 5139 | 12 | 0.47% |
| hsa_circ_0091919 | 5140 | 12 | 0.47% |
| hsa_circ_0083288 | 4712 | 11 | 0.47% |
| hsa_circ_0039619 | 5569 | 13 | 0.47% |
| hsa_circ_0079112 | 5998 | 14 | 0.47% |
| hsa_circ_0101636 | 54846 | 128 | 0.47% |
| hsa_circ_0019822 | 4714 | 11 | 0.47% |
| hsa_circ_0054794 | 5143 | 12 | 0.47% |
| hsa_circ_0088842 | 5572 | 13 | 0.47% |
| hsa_circ_0028557 | 4715 | 11 | 0.47% |
| hsa_circ_0046937 | 20148 | 47 | 0.47% |
| hsa_circ_0137175 | 5573 | 13 | 0.47% |
| hsa_circ_0074309 | 4716 | 11 | 0.47% |
| hsa_circ_0011830 | 6003 | 14 | 0.47% |
| hsa_circ_0025831 | 4717 | 11 | 0.47% |
| hsa_circ_0078573 | 4718 | 11 | 0.47% |
| hsa_circ_0053598 | 5576 | 13 | 0.47% |
| hsa_circ_0098505 | 7721 | 18 | 0.47% |
| hsa_circ_0065065 | 6007 | 14 | 0.47% |
| hsa_circ_0045063 | 5578 | 13 | 0.47% |
| hsa_circ_0028374 | 5149 | 12 | 0.47% |
| hsa_circ_0116208 | 5149 | 12 | 0.47% |
| hsa_circ_0131742 | 13306 | 31 | 0.47% |
| hsa_circ_0091907 | 5151 | 12 | 0.47% |
| hsa_circ_0087316 | 5152 | 12 | 0.47% |
| hsa_circ_0128322 | 5152 | 12 | 0.47% |
| hsa_circ_0091916 | 5154 | 12 | 0.47% |
| hsa_circ_0010261 | 6443 | 15 | 0.47% |
| hsa_circ_0139742 | 5584 | 13 | 0.47% |
| hsa_circ_0087801 | 5155 | 12 | 0.47% |
| hsa_circ_0109315 | 4296 | 10 | 0.47% |
| hsa_circ_0132541 | 8164 | 19 | 0.47% |
| hsa_circ_0009979 | 5157 | 12 | 0.47% |

| | | | |
|---|---|---|---|
| hsa_circ_0033324 | 5158 | 12 | 0.47% |
| hsa_circ_0089208 | 5588 | 13 | 0.47% |
| hsa_circ_0019186 | 5160 | 12 | 0.47% |
| hsa_circ_0130846 | 5160 | 12 | 0.47% |
| hsa_circ_0045684 | 4730 | 11 | 0.47% |
| hsa_circ_0032728 | 5162 | 12 | 0.46% |
| hsa_circ_0108528 | 4732 | 11 | 0.46% |
| hsa_circ_0053671 | 5593 | 13 | 0.46% |
| hsa_circ_0139741 | 5593 | 13 | 0.46% |
| hsa_circ_0045735 | 16349 | 38 | 0.46% |
| hsa_circ_0075153 | 5163 | 12 | 0.46% |
| hsa_circ_0027613 | 5594 | 13 | 0.46% |
| hsa_circ_0139877 | 5594 | 13 | 0.46% |
| hsa_circ_0128877 | 11619 | 27 | 0.46% |
| hsa_circ_0025836 | 5164 | 12 | 0.46% |
| hsa_circ_0054912 | 5164 | 12 | 0.46% |
| hsa_circ_0074197 | 5164 | 12 | 0.46% |
| hsa_circ_0118970 | 5164 | 12 | 0.46% |
| hsa_circ_0139962 | 9898 | 23 | 0.46% |
| hsa_circ_0122644 | 6456 | 15 | 0.46% |
| hsa_circ_0058498 | 5165 | 12 | 0.46% |
| hsa_circ_0137257 | 21521 | 50 | 0.46% |
| hsa_circ_0045207 | 5596 | 13 | 0.46% |
| hsa_circ_0124702 | 6457 | 15 | 0.46% |
| hsa_circ_0085867 | 5597 | 13 | 0.46% |
| hsa_circ_0126786 | 5597 | 13 | 0.46% |
| hsa_circ_0125433 | 14209 | 33 | 0.46% |
| hsa_circ_0010311 | 5167 | 12 | 0.46% |
| hsa_circ_0051159 | 4737 | 11 | 0.46% |
| hsa_circ_0099618 | 4737 | 11 | 0.46% |
| hsa_circ_0010544 | 6029 | 14 | 0.46% |
| hsa_circ_0096966 | 5168 | 12 | 0.46% |
| hsa_circ_0111289 | 5168 | 12 | 0.46% |
| hsa_circ_0140724 | 37904 | 88 | 0.46% |
| hsa_circ_0019675 | 4738 | 11 | 0.46% |
| hsa_circ_0130935 | 5600 | 13 | 0.46% |
| hsa_circ_0030188 | 6462 | 15 | 0.46% |
| hsa_circ_0128511 | 6032 | 14 | 0.46% |
| hsa_circ_0020856 | 4740 | 11 | 0.46% |
| hsa_circ_0094645 | 5171 | 12 | 0.46% |
| hsa_circ_0033228 | 5172 | 12 | 0.46% |
| hsa_circ_0052097 | 4741 | 11 | 0.46% |
| hsa_circ_0016275 | 6035 | 14 | 0.46% |
| hsa_circ_0085195 | 6035 | 14 | 0.46% |
| hsa_circ_0108085 | 4742 | 11 | 0.46% |
| hsa_circ_0062693 | 5605 | 13 | 0.46% |
| hsa_circ_0015472 | 5174 | 12 | 0.46% |
| hsa_circ_0016730 | 6469 | 15 | 0.46% |
| hsa_circ_0043374 | 4744 | 11 | 0.46% |
| hsa_circ_0060712 | 6038 | 14 | 0.46% |

| | | | |
|---|---|---|---|
| hsa_circ_0054587 | 4745 | 11 | 0.46% |
| hsa_circ_0084901 | 4745 | 11 | 0.46% |
| hsa_circ_0076920 | 10785 | 25 | 0.46% |
| hsa_circ_0023537 | 5177 | 12 | 0.46% |
| hsa_circ_0075949 | 5609 | 13 | 0.46% |
| hsa_circ_0046917 | 5610 | 13 | 0.46% |
| hsa_circ_0073126 | 5610 | 13 | 0.46% |
| hsa_circ_0015995 | 8632 | 20 | 0.46% |
| hsa_circ_0132695 | 13382 | 31 | 0.46% |
| hsa_circ_0083912 | 6044 | 14 | 0.46% |
| hsa_circ_0053778 | 4749 | 11 | 0.46% |
| hsa_circ_0003656 | 12525 | 29 | 0.46% |
| hsa_circ_0021019 | 4751 | 11 | 0.46% |
| hsa_circ_0137097 | 5184 | 12 | 0.46% |
| hsa_circ_0070902 | 5617 | 13 | 0.46% |
| hsa_circ_0014589 | 5185 | 12 | 0.46% |
| hsa_circ_0073657 | 5185 | 12 | 0.46% |
| hsa_circ_0087802 | 4753 | 11 | 0.46% |
| hsa_circ_0027182 | 4755 | 11 | 0.46% |
| hsa_circ_0107924 | 5620 | 13 | 0.46% |
| hsa_circ_0138230 | 5620 | 13 | 0.46% |
| hsa_circ_0103689 | 20319 | 47 | 0.46% |
| hsa_circ_0009640 | 8215 | 19 | 0.46% |
| hsa_circ_0035727 | 5621 | 13 | 0.46% |
| hsa_circ_0036819 | 5622 | 13 | 0.46% |
| hsa_circ_0086853 | 5622 | 13 | 0.46% |
| hsa_circ_0089811 | 18596 | 43 | 0.46% |
| hsa_circ_0066315 | 5190 | 12 | 0.46% |
| hsa_circ_0138686 | 5190 | 12 | 0.46% |
| hsa_circ_0132751 | 5623 | 13 | 0.46% |
| hsa_circ_0013249 | 6056 | 14 | 0.46% |
| hsa_circ_0013704 | 5191 | 12 | 0.46% |
| hsa_circ_0044186 | 16006 | 37 | 0.46% |
| hsa_circ_0057778 | 4759 | 11 | 0.46% |
| hsa_circ_0009661 | 5192 | 12 | 0.46% |
| hsa_circ_0034599 | 5625 | 13 | 0.46% |
| hsa_circ_0117318 | 4760 | 11 | 0.46% |
| hsa_circ_0071945 | 4328 | 10 | 0.46% |
| hsa_circ_0114660 | 11253 | 26 | 0.46% |
| hsa_circ_0108963 | 21646 | 50 | 0.46% |
| hsa_circ_0052642 | 5628 | 13 | 0.46% |
| hsa_circ_0108812 | 5628 | 13 | 0.46% |
| hsa_circ_0114633 | 7794 | 18 | 0.46% |
| hsa_circ_0025656 | 4763 | 11 | 0.46% |
| hsa_circ_0113388 | 7796 | 18 | 0.46% |
| hsa_circ_0102289 | 10828 | 25 | 0.46% |
| hsa_circ_0073404 | 4765 | 11 | 0.46% |
| hsa_circ_0086440 | 5632 | 13 | 0.46% |
| hsa_circ_0007313 | 20796 | 48 | 0.46% |
| hsa_circ_0078592 | 4766 | 11 | 0.46% |

| | | | |
|---|---|---|---|
| hsa_circ_0085631 | 5200 | 12 | 0.46% |
| hsa_circ_0052702 | 6502 | 15 | 0.46% |
| hsa_circ_0116937 | 19940 | 46 | 0.46% |
| hsa_circ_0132717 | 12138 | 28 | 0.46% |
| hsa_circ_0102355 | 18209 | 42 | 0.46% |
| hsa_circ_0033224 | 5203 | 12 | 0.46% |
| hsa_circ_0054741 | 5637 | 13 | 0.46% |
| hsa_circ_0020002 | 4770 | 11 | 0.46% |
| hsa_circ_0015700 | 6505 | 15 | 0.46% |
| hsa_circ_0124534 | 16046 | 37 | 0.46% |
| hsa_circ_0018442 | 4337 | 10 | 0.46% |
| hsa_circ_0042578 | 5205 | 12 | 0.46% |
| hsa_circ_0034601 | 5639 | 13 | 0.46% |
| hsa_circ_0101511 | 50761 | 117 | 0.46% |
| hsa_circ_0140136 | 12148 | 28 | 0.46% |
| hsa_circ_0112088 | 7810 | 18 | 0.46% |
| hsa_circ_0033259 | 5207 | 12 | 0.46% |
| hsa_circ_0054051 | 5641 | 13 | 0.46% |
| hsa_circ_0025095 | 5642 | 13 | 0.46% |
| hsa_circ_0027636 | 5208 | 12 | 0.46% |
| hsa_circ_0073268 | 5208 | 12 | 0.46% |
| hsa_circ_0073337 | 4774 | 11 | 0.46% |
| hsa_circ_0035677 | 5643 | 13 | 0.46% |
| hsa_circ_0064541 | 5643 | 13 | 0.46% |
| hsa_circ_0062936 | 5209 | 12 | 0.46% |
| hsa_circ_0013841 | 4775 | 11 | 0.46% |
| hsa_circ_0124581 | 4775 | 11 | 0.46% |
| hsa_circ_0134139 | 14761 | 34 | 0.46% |
| hsa_circ_0033533 | 5644 | 13 | 0.46% |
| hsa_circ_0113350 | 21709 | 50 | 0.46% |
| hsa_circ_0088127 | 4778 | 11 | 0.46% |
| hsa_circ_0115301 | 5648 | 13 | 0.46% |
| hsa_circ_0010306 | 5214 | 12 | 0.46% |
| hsa_circ_0053961 | 5214 | 12 | 0.46% |
| hsa_circ_0091093 | 6084 | 14 | 0.46% |
| hsa_circ_0047046 | 5215 | 12 | 0.46% |
| hsa_circ_0063725 | 6085 | 14 | 0.46% |
| hsa_circ_0038271 | 4782 | 11 | 0.46% |
| hsa_circ_0133151 | 5217 | 12 | 0.46% |
| hsa_circ_0029298 | 4783 | 11 | 0.46% |
| hsa_circ_0026480 | 22178 | 51 | 0.46% |
| hsa_circ_0126918 | 27402 | 63 | 0.46% |
| hsa_circ_0035737 | 5655 | 13 | 0.46% |
| hsa_circ_0007666 | 6091 | 14 | 0.46% |
| hsa_circ_0130718 | 8267 | 19 | 0.46% |
| hsa_circ_0074494 | 7832 | 18 | 0.46% |
| hsa_circ_0029456 | 5657 | 13 | 0.46% |
| hsa_circ_0053779 | 5659 | 13 | 0.46% |
| hsa_circ_0056977 | 6965 | 16 | 0.46% |
| hsa_circ_0020063 | 6530 | 15 | 0.46% |

| | | | |
|---|---|---|---|
| hsa_circ_0029491 | 5225 | 12 | 0.46% |
| hsa_circ_0102054 | 4790 | 11 | 0.46% |
| hsa_circ_0099568 | 5661 | 13 | 0.46% |
| hsa_circ_0012914 | 6532 | 15 | 0.46% |
| hsa_circ_0056053 | 5226 | 12 | 0.46% |
| hsa_circ_0108526 | 4791 | 11 | 0.46% |
| hsa_circ_0096628 | 6534 | 15 | 0.46% |
| hsa_circ_0053809 | 5228 | 12 | 0.46% |
| hsa_circ_0056720 | 5228 | 12 | 0.46% |
| hsa_circ_0031356 | 16122 | 37 | 0.46% |
| hsa_circ_0010715 | 5229 | 12 | 0.46% |
| hsa_circ_0115972 | 5229 | 12 | 0.46% |
| hsa_circ_0015381 | 5665 | 13 | 0.46% |
| hsa_circ_0135888 | 20917 | 48 | 0.46% |
| hsa_circ_0104821 | 18303 | 42 | 0.46% |
| hsa_circ_0002818 | 16560 | 38 | 0.46% |
| hsa_circ_0027339 | 16126 | 37 | 0.46% |
| hsa_circ_0045071 | 5667 | 13 | 0.46% |
| hsa_circ_0069720 | 5667 | 13 | 0.46% |
| hsa_circ_0024438 | 5232 | 12 | 0.46% |
| hsa_circ_0042177 | 5232 | 12 | 0.46% |
| hsa_circ_0030468 | 4796 | 11 | 0.46% |
| hsa_circ_0119794 | 20493 | 47 | 0.46% |
| hsa_circ_0113329 | 5669 | 13 | 0.46% |
| hsa_circ_0027153 | 5233 | 12 | 0.46% |
| hsa_circ_0053979 | 4797 | 11 | 0.46% |
| hsa_circ_0120160 | 14394 | 33 | 0.46% |
| hsa_circ_0031469 | 6107 | 14 | 0.46% |
| hsa_circ_0043951 | 4799 | 11 | 0.46% |
| hsa_circ_0138786 | 11344 | 26 | 0.46% |
| hsa_circ_0046845 | 5672 | 13 | 0.46% |
| hsa_circ_0096811 | 5672 | 13 | 0.46% |
| hsa_circ_0082877 | 6546 | 15 | 0.46% |
| hsa_circ_0041289 | 4801 | 11 | 0.46% |
| hsa_circ_0077343 | 5674 | 13 | 0.46% |
| hsa_circ_0081236 | 6547 | 15 | 0.46% |
| hsa_circ_0029405 | 12222 | 28 | 0.46% |
| hsa_circ_0107542 | 4802 | 11 | 0.46% |
| hsa_circ_0066036 | 5239 | 12 | 0.46% |
| hsa_circ_0057821 | 4803 | 11 | 0.46% |
| hsa_circ_0084334 | 5677 | 13 | 0.46% |
| hsa_circ_0020961 | 5241 | 12 | 0.46% |
| hsa_circ_0020414 | 4805 | 11 | 0.46% |
| hsa_circ_0129915 | 5679 | 13 | 0.46% |
| hsa_circ_0008127 | 6116 | 14 | 0.46% |
| hsa_circ_0051649 | 6116 | 14 | 0.46% |
| hsa_circ_0120161 | 14417 | 33 | 0.46% |
| hsa_circ_0072951 | 5680 | 13 | 0.46% |
| hsa_circ_0132757 | 8740 | 20 | 0.46% |
| hsa_circ_0003610 | 6118 | 14 | 0.46% |

| | | | |
|---|---|---|---|
| hsa_circ_0082797 | 5681 | 13 | 0.46% |
| hsa_circ_0060169 | 6119 | 14 | 0.46% |
| hsa_circ_0034148 | 5245 | 12 | 0.46% |
| hsa_circ_0121862 | 13550 | 31 | 0.46% |
| hsa_circ_0121863 | 13553 | 31 | 0.46% |
| hsa_circ_0086864 | 5684 | 13 | 0.46% |
| hsa_circ_0035712 | 5247 | 12 | 0.46% |
| hsa_circ_0042593 | 5247 | 12 | 0.46% |
| hsa_circ_0010600 | 5685 | 13 | 0.46% |
| hsa_circ_0028359 | 5248 | 12 | 0.46% |
| hsa_circ_0030834 | 6998 | 16 | 0.46% |
| hsa_circ_0033437 | 5249 | 12 | 0.46% |
| hsa_circ_0058080 | 5687 | 13 | 0.46% |
| hsa_circ_0084362 | 5687 | 13 | 0.46% |
| hsa_circ_0091850 | 6125 | 14 | 0.46% |
| hsa_circ_0049110 | 5250 | 12 | 0.46% |
| hsa_circ_0034740 | 6127 | 14 | 0.46% |
| hsa_circ_0037739 | 6127 | 14 | 0.46% |
| hsa_circ_0123072 | 5252 | 12 | 0.46% |
| hsa_circ_0088839 | 5690 | 13 | 0.46% |
| hsa_circ_0114875 | 17946 | 41 | 0.46% |
| hsa_circ_0123902 | 5253 | 12 | 0.46% |
| hsa_circ_0008561 | 5691 | 13 | 0.46% |
| hsa_circ_0086368 | 6567 | 15 | 0.46% |
| hsa_circ_0044182 | 16200 | 37 | 0.46% |
| hsa_circ_0010327 | 5255 | 12 | 0.46% |
| hsa_circ_0078099 | 5256 | 12 | 0.46% |
| hsa_circ_0046764 | 4818 | 11 | 0.46% |
| hsa_circ_0016952 | 22778 | 52 | 0.46% |
| hsa_circ_0098506 | 7885 | 18 | 0.46% |
| hsa_circ_0135390 | 52569 | 120 | 0.46% |
| hsa_circ_0004634 | 15773 | 36 | 0.46% |
| hsa_circ_0042893 | 5697 | 13 | 0.46% |
| hsa_circ_0095600 | 9644 | 22 | 0.46% |
| hsa_circ_0089429 | 5261 | 12 | 0.46% |
| hsa_circ_0073237 | 5262 | 12 | 0.46% |
| hsa_circ_0067988 | 6578 | 15 | 0.46% |
| hsa_circ_0138229 | 5701 | 13 | 0.46% |
| hsa_circ_0027338 | 16228 | 37 | 0.46% |
| hsa_circ_0028752 | 5265 | 12 | 0.46% |
| hsa_circ_0034395 | 5267 | 12 | 0.46% |
| hsa_circ_0079090 | 21510 | 49 | 0.46% |
| hsa_circ_0019097 | 6146 | 14 | 0.46% |
| hsa_circ_0061584 | 5269 | 12 | 0.46% |
| hsa_circ_0113238 | 12295 | 28 | 0.46% |
| hsa_circ_0067844 | 7466 | 17 | 0.46% |
| hsa_circ_0109317 | 4392 | 10 | 0.46% |
| hsa_circ_0027168 | 5271 | 12 | 0.46% |
| hsa_circ_0036802 | 4393 | 10 | 0.46% |
| hsa_circ_0050791 | 5711 | 13 | 0.46% |

| | | | |
|---|---|---|---|
| hsa_circ_0129171 | 15816 | 36 | 0.46% |
| hsa_circ_0083336 | 6590 | 15 | 0.46% |
| hsa_circ_0028148 | 4833 | 11 | 0.46% |
| hsa_circ_0025080 | 5713 | 13 | 0.46% |
| hsa_circ_0091893 | 5274 | 12 | 0.46% |
| hsa_circ_0045513 | 4395 | 10 | 0.46% |
| hsa_circ_0070287 | 4835 | 11 | 0.46% |
| hsa_circ_0112312 | 8792 | 20 | 0.45% |
| hsa_circ_0045205 | 5715 | 13 | 0.45% |
| hsa_circ_0128584 | 11870 | 27 | 0.45% |
| hsa_circ_0001431 | 4836 | 11 | 0.45% |
| hsa_circ_0037112 | 4836 | 11 | 0.45% |
| hsa_circ_0045446 | 4836 | 11 | 0.45% |
| hsa_circ_0072814 | 16707 | 38 | 0.45% |
| hsa_circ_0123936 | 11436 | 26 | 0.45% |
| hsa_circ_0016629 | 5718 | 13 | 0.45% |
| hsa_circ_0115530 | 5718 | 13 | 0.45% |
| hsa_circ_0009800 | 6158 | 14 | 0.45% |
| hsa_circ_0001822 | 7919 | 18 | 0.45% |
| hsa_circ_0044404 | 7040 | 16 | 0.45% |
| hsa_circ_0071604 | 5280 | 12 | 0.45% |
| hsa_circ_0139044 | 5280 | 12 | 0.45% |
| hsa_circ_0068974 | 4840 | 11 | 0.45% |
| hsa_circ_0116839 | 5721 | 13 | 0.45% |
| hsa_circ_0002313 | 24648 | 56 | 0.45% |
| hsa_circ_0116462 | 5282 | 12 | 0.45% |
| hsa_circ_0053770 | 4842 | 11 | 0.45% |
| hsa_circ_0075305 | 5723 | 13 | 0.45% |
| hsa_circ_0122489 | 5283 | 12 | 0.45% |
| hsa_circ_0001517 | 6604 | 15 | 0.45% |
| hsa_circ_0118141 | 11450 | 26 | 0.45% |
| hsa_circ_0058374 | 5725 | 13 | 0.45% |
| hsa_circ_0081261 | 6606 | 15 | 0.45% |
| hsa_circ_0081675 | 17177 | 39 | 0.45% |
| hsa_circ_0118098 | 4845 | 11 | 0.45% |
| hsa_circ_0012627 | 5726 | 13 | 0.45% |
| hsa_circ_0084349 | 5726 | 13 | 0.45% |
| hsa_circ_0078901 | 17619 | 40 | 0.45% |
| hsa_circ_0099593 | 11016 | 25 | 0.45% |
| hsa_circ_0052643 | 5729 | 13 | 0.45% |
| hsa_circ_0126459 | 5729 | 13 | 0.45% |
| hsa_circ_0121897 | 4408 | 10 | 0.45% |
| hsa_circ_0136042 | 4849 | 11 | 0.45% |
| hsa_circ_0048387 | 6613 | 15 | 0.45% |
| hsa_circ_0123122 | 4850 | 11 | 0.45% |
| hsa_circ_0134989 | 4850 | 11 | 0.45% |
| hsa_circ_0085300 | 25132 | 57 | 0.45% |
| hsa_circ_0011833 | 5291 | 12 | 0.45% |
| hsa_circ_0069933 | 6173 | 14 | 0.45% |
| hsa_circ_0026925 | 7055 | 16 | 0.45% |

| | | | |
|---|---|---|---|
| hsa_circ_0047054 | 5293 | 12 | 0.45% |
| hsa_circ_0041292 | 4852 | 11 | 0.45% |
| hsa_circ_0053756 | 4852 | 11 | 0.45% |
| hsa_circ_0101239 | 6176 | 14 | 0.45% |
| hsa_circ_0071006 | 4854 | 11 | 0.45% |
| hsa_circ_0033032 | 6180 | 14 | 0.45% |
| hsa_circ_0027205 | 4856 | 11 | 0.45% |
| hsa_circ_0129669 | 19867 | 45 | 0.45% |
| hsa_circ_0071610 | 5298 | 12 | 0.45% |
| hsa_circ_0020320 | 5740 | 13 | 0.45% |
| hsa_circ_0084217 | 5300 | 12 | 0.45% |
| hsa_circ_0134622 | 22088 | 50 | 0.45% |
| hsa_circ_0109707 | 14137 | 32 | 0.45% |
| hsa_circ_0084339 | 4860 | 11 | 0.45% |
| hsa_circ_0044719 | 5744 | 13 | 0.45% |
| hsa_circ_0065592 | 4861 | 11 | 0.45% |
| hsa_circ_0071062 | 5303 | 12 | 0.45% |
| hsa_circ_0088326 | 5745 | 13 | 0.45% |
| hsa_circ_0131157 | 11934 | 27 | 0.45% |
| hsa_circ_0091353 | 9282 | 21 | 0.45% |
| hsa_circ_0054504 | 5304 | 12 | 0.45% |
| hsa_circ_0075483 | 5304 | 12 | 0.45% |
| hsa_circ_0016953 | 22985 | 52 | 0.45% |
| hsa_circ_0056905 | 5747 | 13 | 0.45% |
| hsa_circ_0071436 | 5305 | 12 | 0.45% |
| hsa_circ_0138485 | 21665 | 49 | 0.45% |
| hsa_circ_0130326 | 5748 | 13 | 0.45% |
| hsa_circ_0118904 | 13709 | 31 | 0.45% |
| hsa_circ_0031067 | 5749 | 13 | 0.45% |
| hsa_circ_0049820 | 4865 | 11 | 0.45% |
| hsa_circ_0089151 | 4865 | 11 | 0.45% |
| hsa_circ_0117136 | 5752 | 13 | 0.45% |
| hsa_circ_0085211 | 5310 | 12 | 0.45% |
| hsa_circ_0070742 | 12833 | 29 | 0.45% |
| hsa_circ_0053642 | 5753 | 13 | 0.45% |
| hsa_circ_0036813 | 6196 | 14 | 0.45% |
| hsa_circ_0016922 | 5311 | 12 | 0.45% |
| hsa_circ_0052440 | 6197 | 14 | 0.45% |
| hsa_circ_0136230 | 6640 | 15 | 0.45% |
| hsa_circ_0041667 | 5312 | 12 | 0.45% |
| hsa_circ_0061138 | 4427 | 10 | 0.45% |
| hsa_circ_0053526 | 4870 | 11 | 0.45% |
| hsa_circ_0053541 | 4870 | 11 | 0.45% |
| hsa_circ_0027177 | 5313 | 12 | 0.45% |
| hsa_circ_0056729 | 5313 | 12 | 0.45% |
| hsa_circ_0091928 | 5313 | 12 | 0.45% |
| hsa_circ_0113330 | 5758 | 13 | 0.45% |
| hsa_circ_0096747 | 13289 | 30 | 0.45% |
| hsa_circ_0129907 | 5316 | 12 | 0.45% |
| hsa_circ_0105295 | 35893 | 81 | 0.45% |

| | | | |
|---|---|---|---|
| hsa_circ_0015762 | 6647 | 15 | 0.45% |
| hsa_circ_0027644 | 5318 | 12 | 0.45% |
| hsa_circ_0054795 | 5318 | 12 | 0.45% |
| hsa_circ_0130211 | 4876 | 11 | 0.45% |
| hsa_circ_0071162 | 5320 | 12 | 0.45% |
| hsa_circ_0024854 | 6208 | 14 | 0.45% |
| hsa_circ_0058112 | 4878 | 11 | 0.45% |
| hsa_circ_0019433 | 4879 | 11 | 0.45% |
| hsa_circ_0112086 | 7984 | 18 | 0.45% |
| hsa_circ_0033304 | 5323 | 12 | 0.45% |
| hsa_circ_0011111 | 5767 | 13 | 0.45% |
| hsa_circ_0029878 | 5767 | 13 | 0.45% |
| hsa_circ_0070333 | 4880 | 11 | 0.45% |
| hsa_circ_0101585 | 5324 | 12 | 0.45% |
| hsa_circ_0061392 | 5769 | 13 | 0.45% |
| hsa_circ_0094643 | 5769 | 13 | 0.45% |
| hsa_circ_0014195 | 6213 | 14 | 0.45% |
| hsa_circ_0032508 | 5326 | 12 | 0.45% |
| hsa_circ_0043540 | 5326 | 12 | 0.45% |
| hsa_circ_0065164 | 5326 | 12 | 0.45% |
| hsa_circ_0125488 | 5770 | 13 | 0.45% |
| hsa_circ_0094273 | 7546 | 17 | 0.45% |
| hsa_circ_0123822 | 13761 | 31 | 0.45% |
| hsa_circ_0063728 | 6215 | 14 | 0.45% |
| hsa_circ_0120476 | 7104 | 16 | 0.45% |
| hsa_circ_0029492 | 5328 | 12 | 0.45% |
| hsa_circ_0074621 | 5773 | 13 | 0.45% |
| hsa_circ_0081318 | 5329 | 12 | 0.45% |
| hsa_circ_0035930 | 4885 | 11 | 0.45% |
| hsa_circ_0140458 | 5774 | 13 | 0.45% |
| hsa_circ_0036012 | 5330 | 12 | 0.45% |
| hsa_circ_0120475 | 7107 | 16 | 0.45% |
| hsa_circ_0038605 | 9774 | 22 | 0.45% |
| hsa_circ_0091358 | 5332 | 12 | 0.45% |
| hsa_circ_0138498 | 5333 | 12 | 0.45% |
| hsa_circ_0034201 | 5778 | 13 | 0.45% |
| hsa_circ_0126006 | 7556 | 17 | 0.45% |
| hsa_circ_0042589 | 5334 | 12 | 0.45% |
| hsa_circ_0090722 | 5779 | 13 | 0.45% |
| hsa_circ_0074057 | 5335 | 12 | 0.45% |
| hsa_circ_0111290 | 5335 | 12 | 0.45% |
| hsa_circ_0042610 | 4891 | 11 | 0.45% |
| hsa_circ_0073338 | 4891 | 11 | 0.45% |
| hsa_circ_0013248 | 6225 | 14 | 0.45% |
| hsa_circ_0071256 | 4892 | 11 | 0.45% |
| hsa_circ_0130243 | 5337 | 12 | 0.45% |
| hsa_circ_0036818 | 5783 | 13 | 0.45% |
| hsa_circ_0137071 | 15572 | 35 | 0.45% |
| hsa_circ_0025085 | 5784 | 13 | 0.45% |
| hsa_circ_0124860 | 8009 | 18 | 0.45% |

| | | | |
|---|---|---|---|
| hsa_circ_0078685 | 5785 | 13 | 0.45% |
| hsa_circ_0068039 | 6231 | 14 | 0.45% |
| hsa_circ_0038426 | 4896 | 11 | 0.45% |
| hsa_circ_0072888 | 5343 | 12 | 0.45% |
| hsa_circ_0075064 | 5343 | 12 | 0.45% |
| hsa_circ_0000955 | 19592 | 44 | 0.45% |
| hsa_circ_0110145 | 7571 | 17 | 0.45% |
| hsa_circ_0101881 | 6236 | 14 | 0.45% |
| hsa_circ_0114140 | 5791 | 13 | 0.45% |
| hsa_circ_0130581 | 6237 | 14 | 0.45% |
| hsa_circ_0086176 | 5347 | 12 | 0.45% |
| hsa_circ_0055239 | 6239 | 14 | 0.45% |
| hsa_circ_0070120 | 6240 | 14 | 0.45% |
| hsa_circ_0075156 | 8023 | 18 | 0.45% |
| hsa_circ_0103277 | 18721 | 42 | 0.45% |
| hsa_circ_0114655 | 5349 | 12 | 0.45% |
| hsa_circ_0120970 | 5349 | 12 | 0.45% |
| hsa_circ_0109536 | 15156 | 34 | 0.45% |
| hsa_circ_0084294 | 5350 | 12 | 0.45% |
| hsa_circ_0112192 | 5350 | 12 | 0.45% |
| hsa_circ_0020326 | 21848 | 49 | 0.45% |
| hsa_circ_0001243 | 60646 | 136 | 0.45% |
| hsa_circ_0075946 | 7581 | 17 | 0.45% |
| hsa_circ_0098253 | 20069 | 45 | 0.45% |
| hsa_circ_0053573 | 5798 | 13 | 0.45% |
| hsa_circ_0138988 | 5352 | 12 | 0.45% |
| hsa_circ_0116949 | 4906 | 11 | 0.45% |
| hsa_circ_0081249 | 6691 | 15 | 0.45% |
| hsa_circ_0019096 | 6245 | 14 | 0.45% |
| hsa_circ_0066309 | 5353 | 12 | 0.45% |
| hsa_circ_0066295 | 5355 | 12 | 0.45% |
| hsa_circ_0073842 | 5356 | 12 | 0.45% |
| hsa_circ_0082996 | 4910 | 11 | 0.45% |
| hsa_circ_0069979 | 6696 | 15 | 0.45% |
| hsa_circ_0038237 | 4912 | 11 | 0.45% |
| hsa_circ_0069678 | 5359 | 12 | 0.45% |
| hsa_circ_0024665 | 5361 | 12 | 0.45% |
| hsa_circ_0056722 | 5361 | 12 | 0.45% |
| hsa_circ_0047981 | 5364 | 12 | 0.45% |
| hsa_circ_0123803 | 12965 | 29 | 0.45% |
| hsa_circ_0028363 | 5366 | 12 | 0.45% |
| hsa_circ_0108950 | 5814 | 13 | 0.45% |
| hsa_circ_0067760 | 5367 | 12 | 0.45% |
| hsa_circ_0099743 | 16553 | 37 | 0.45% |
| hsa_circ_0034235 | 5816 | 13 | 0.45% |
| hsa_circ_0055816 | 11185 | 25 | 0.45% |
| hsa_circ_0127739 | 5369 | 12 | 0.45% |
| hsa_circ_0119194 | 10739 | 24 | 0.45% |
| hsa_circ_0129099 | 5817 | 13 | 0.45% |
| hsa_circ_0016276 | 6265 | 14 | 0.45% |

| | | | |
|---|---|---|---|
| hsa_circ_0041864 | 5371 | 12 | 0.45% |
| hsa_circ_0046844 | 5820 | 13 | 0.45% |
| hsa_circ_0039090 | 5821 | 13 | 0.45% |
| hsa_circ_0096923 | 5821 | 13 | 0.45% |
| hsa_circ_0003257 | 16568 | 37 | 0.45% |
| hsa_circ_0004383 | 5822 | 13 | 0.45% |
| hsa_circ_0084332 | 4927 | 11 | 0.45% |
| hsa_circ_0054611 | 5375 | 12 | 0.45% |
| hsa_circ_0075990 | 6271 | 14 | 0.45% |
| hsa_circ_0103958 | 5824 | 13 | 0.45% |
| hsa_circ_0077385 | 5376 | 12 | 0.45% |
| hsa_circ_0110543 | 7169 | 16 | 0.45% |
| hsa_circ_0016626 | 5825 | 13 | 0.45% |
| hsa_circ_0091869 | 4929 | 11 | 0.45% |
| hsa_circ_0031066 | 5826 | 13 | 0.45% |
| hsa_circ_0058090 | 5826 | 13 | 0.45% |
| hsa_circ_0136876 | 12997 | 29 | 0.45% |
| hsa_circ_0047959 | 4930 | 11 | 0.45% |
| hsa_circ_0124266 | 7172 | 16 | 0.45% |
| hsa_circ_0064722 | 5379 | 12 | 0.45% |
| hsa_circ_0081877 | 5379 | 12 | 0.45% |
| hsa_circ_0112413 | 23314 | 52 | 0.45% |
| hsa_circ_0050454 | 4932 | 11 | 0.45% |
| hsa_circ_0089467 | 4932 | 11 | 0.45% |
| hsa_circ_0058098 | 5829 | 13 | 0.45% |
| hsa_circ_0108593 | 16593 | 37 | 0.45% |
| hsa_circ_0072209 | 6279 | 14 | 0.45% |
| hsa_circ_0104300 | 5831 | 13 | 0.45% |
| hsa_circ_0074493 | 8075 | 18 | 0.45% |
| hsa_circ_0084340 | 5832 | 13 | 0.45% |
| hsa_circ_0036812 | 6281 | 14 | 0.45% |
| hsa_circ_0057835 | 5833 | 13 | 0.45% |
| hsa_circ_0138917 | 5833 | 13 | 0.45% |
| hsa_circ_0027201 | 4487 | 10 | 0.45% |
| hsa_circ_0049914 | 4936 | 11 | 0.45% |
| hsa_circ_0033532 | 5834 | 13 | 0.45% |
| hsa_circ_0024093 | 5386 | 12 | 0.45% |
| hsa_circ_0139937 | 16162 | 36 | 0.45% |
| hsa_circ_0038447 | 24693 | 55 | 0.45% |
| hsa_circ_0128585 | 12124 | 27 | 0.45% |
| hsa_circ_0076368 | 6736 | 15 | 0.45% |
| hsa_circ_0063726 | 6287 | 14 | 0.45% |
| hsa_circ_0102640 | 22461 | 50 | 0.45% |
| hsa_circ_0016954 | 23360 | 52 | 0.45% |
| hsa_circ_0020319 | 5391 | 12 | 0.45% |
| hsa_circ_0059682 | 10783 | 24 | 0.45% |
| hsa_circ_0010259 | 6740 | 15 | 0.45% |
| hsa_circ_0091908 | 5392 | 12 | 0.45% |
| hsa_circ_0123099 | 5392 | 12 | 0.45% |
| hsa_circ_0122627 | 5842 | 13 | 0.45% |

| | | | |
|---|---|---|---|
| hsa_circ_0042601 | 4944 | 11 | 0.44% |
| hsa_circ_0066743 | 5843 | 13 | 0.44% |
| hsa_circ_0033094 | 5844 | 13 | 0.44% |
| hsa_circ_0071396 | 5395 | 12 | 0.44% |
| hsa_circ_0110146 | 7643 | 17 | 0.44% |
| hsa_circ_0033231 | 5396 | 12 | 0.44% |
| hsa_circ_0074056 | 5397 | 12 | 0.44% |
| hsa_circ_0023535 | 5398 | 12 | 0.44% |
| hsa_circ_0017079 | 5848 | 13 | 0.44% |
| hsa_circ_0112321 | 13048 | 29 | 0.44% |
| hsa_circ_0082997 | 5850 | 13 | 0.44% |
| hsa_circ_0049188 | 5400 | 12 | 0.44% |
| hsa_circ_0061680 | 5400 | 12 | 0.44% |
| hsa_circ_0020415 | 4951 | 11 | 0.44% |
| hsa_circ_0111978 | 4951 | 11 | 0.44% |
| hsa_circ_0001575 | 20708 | 46 | 0.44% |
| hsa_circ_0040867 | 6753 | 15 | 0.44% |
| hsa_circ_0077192 | 5403 | 12 | 0.44% |
| hsa_circ_0079214 | 5403 | 12 | 0.44% |
| hsa_circ_0061186 | 4953 | 11 | 0.44% |
| hsa_circ_0015877 | 17565 | 39 | 0.44% |
| hsa_circ_0046919 | 5855 | 13 | 0.44% |
| hsa_circ_0033339 | 5405 | 12 | 0.44% |
| hsa_circ_0035728 | 5857 | 13 | 0.44% |
| hsa_circ_0134550 | 5857 | 13 | 0.44% |
| hsa_circ_0110881 | 5408 | 12 | 0.44% |
| hsa_circ_0064048 | 5409 | 12 | 0.44% |
| hsa_circ_0139194 | 13974 | 31 | 0.44% |
| hsa_circ_0127718 | 16682 | 37 | 0.44% |
| hsa_circ_0111629 | 5411 | 12 | 0.44% |
| hsa_circ_0091143 | 5412 | 12 | 0.44% |
| hsa_circ_0102052 | 4961 | 11 | 0.44% |
| hsa_circ_0133926 | 4961 | 11 | 0.44% |
| hsa_circ_0095329 | 10826 | 24 | 0.44% |
| hsa_circ_0021479 | 5413 | 12 | 0.44% |
| hsa_circ_0054572 | 4962 | 11 | 0.44% |
| hsa_circ_0049674 | 5414 | 12 | 0.44% |
| hsa_circ_0134082 | 5414 | 12 | 0.44% |
| hsa_circ_0012532 | 4963 | 11 | 0.44% |
| hsa_circ_0124585 | 11732 | 26 | 0.44% |
| hsa_circ_0064591 | 4964 | 11 | 0.44% |
| hsa_circ_0113331 | 18955 | 42 | 0.44% |
| hsa_circ_0109528 | 17603 | 39 | 0.44% |
| hsa_circ_0074310 | 4965 | 11 | 0.44% |
| hsa_circ_0042911 | 5868 | 13 | 0.44% |
| hsa_circ_0070182 | 5868 | 13 | 0.44% |
| hsa_circ_0070999 | 5868 | 13 | 0.44% |
| hsa_circ_0008552 | 10382 | 23 | 0.44% |
| hsa_circ_0070848 | 6320 | 14 | 0.44% |
| hsa_circ_0130580 | 6320 | 14 | 0.44% |

| | | | |
|---|---|---|---|
| hsa_circ_0114143 | 5869 | 13 | 0.44% |
| hsa_circ_0015591 | 5420 | 12 | 0.44% |
| hsa_circ_0134554 | 5420 | 12 | 0.44% |
| hsa_circ_0057841 | 5872 | 13 | 0.44% |
| hsa_circ_0098269 | 6777 | 15 | 0.44% |
| hsa_circ_0109318 | 4519 | 10 | 0.44% |
| hsa_circ_0139748 | 42933 | 95 | 0.44% |
| hsa_circ_0136197 | 14914 | 33 | 0.44% |
| hsa_circ_0096947 | 20340 | 45 | 0.44% |
| hsa_circ_0048730 | 5876 | 13 | 0.44% |
| hsa_circ_0101219 | 5424 | 12 | 0.44% |
| hsa_circ_0060863 | 6781 | 15 | 0.44% |
| hsa_circ_0137317 | 5425 | 12 | 0.44% |
| hsa_circ_0071007 | 4973 | 11 | 0.44% |
| hsa_circ_0130640 | 8590 | 19 | 0.44% |
| hsa_circ_0086889 | 5878 | 13 | 0.44% |
| hsa_circ_0108676 | 19443 | 43 | 0.44% |
| hsa_circ_0061585 | 5426 | 12 | 0.44% |
| hsa_circ_0007274 | 4974 | 11 | 0.44% |
| hsa_circ_0009684 | 5879 | 13 | 0.44% |
| hsa_circ_0045756 | 5879 | 13 | 0.44% |
| hsa_circ_0028750 | 5427 | 12 | 0.44% |
| hsa_circ_0090866 | 4975 | 11 | 0.44% |
| hsa_circ_0114325 | 15832 | 35 | 0.44% |
| hsa_circ_0055235 | 6333 | 14 | 0.44% |
| hsa_circ_0068040 | 6334 | 14 | 0.44% |
| hsa_circ_0054787 | 5430 | 12 | 0.44% |
| hsa_circ_0037460 | 5883 | 13 | 0.44% |
| hsa_circ_0042730 | 5883 | 13 | 0.44% |
| hsa_circ_0093239 | 19912 | 44 | 0.44% |
| hsa_circ_0130790 | 11768 | 26 | 0.44% |
| hsa_circ_0056814 | 5887 | 13 | 0.44% |
| hsa_circ_0105395 | 23555 | 52 | 0.44% |
| hsa_circ_0123189 | 13137 | 29 | 0.44% |
| hsa_circ_0070119 | 6342 | 14 | 0.44% |
| hsa_circ_0101220 | 5436 | 12 | 0.44% |
| hsa_circ_0085200 | 6343 | 14 | 0.44% |
| hsa_circ_0015474 | 5890 | 13 | 0.44% |
| hsa_circ_0034322 | 5890 | 13 | 0.44% |
| hsa_circ_0053611 | 5890 | 13 | 0.44% |
| hsa_circ_0078921 | 5438 | 12 | 0.44% |
| hsa_circ_0082203 | 5438 | 12 | 0.44% |
| hsa_circ_0093993 | 11785 | 26 | 0.44% |
| hsa_circ_0119346 | 18132 | 40 | 0.44% |
| hsa_circ_0111400 | 53040 | 117 | 0.44% |
| hsa_circ_0052333 | 21773 | 48 | 0.44% |
| hsa_circ_0069943 | 5444 | 12 | 0.44% |
| hsa_circ_0077330 | 5447 | 12 | 0.44% |
| hsa_circ_0132418 | 5448 | 12 | 0.44% |
| hsa_circ_0010089 | 5450 | 12 | 0.44% |

| | | | |
|---|---|---|---|
| hsa_circ_0017527 | 24980 | 55 | 0.44% |
| hsa_circ_0019571 | 5452 | 12 | 0.44% |
| hsa_circ_0047214 | 5452 | 12 | 0.44% |
| hsa_circ_0089445 | 7270 | 16 | 0.44% |
| hsa_circ_0029879 | 5907 | 13 | 0.44% |
| hsa_circ_0138768 | 11361 | 25 | 0.44% |
| hsa_circ_0026475 | 23181 | 51 | 0.44% |
| hsa_circ_0099442 | 5909 | 13 | 0.44% |
| hsa_circ_0139195 | 14091 | 31 | 0.44% |
| hsa_circ_0042061 | 5910 | 13 | 0.44% |
| hsa_circ_0056509 | 5001 | 11 | 0.44% |
| hsa_circ_0106906 | 5001 | 11 | 0.44% |
| hsa_circ_0103377 | 5911 | 13 | 0.44% |
| hsa_circ_0066944 | 5912 | 13 | 0.44% |
| hsa_circ_0052665 | 24104 | 53 | 0.44% |
| hsa_circ_0060576 | 5003 | 11 | 0.44% |
| hsa_circ_0059525 | 6823 | 15 | 0.44% |
| hsa_circ_0053542 | 5004 | 11 | 0.44% |
| hsa_circ_0078355 | 5004 | 11 | 0.44% |
| hsa_circ_0072181 | 5459 | 12 | 0.44% |
| hsa_circ_0118862 | 5460 | 12 | 0.44% |
| hsa_circ_0130189 | 12286 | 27 | 0.44% |
| hsa_circ_0030317 | 6826 | 15 | 0.44% |
| hsa_circ_0071935 | 5006 | 11 | 0.44% |
| hsa_circ_0086854 | 5917 | 13 | 0.44% |
| hsa_circ_0068882 | 5462 | 12 | 0.44% |
| hsa_circ_0025973 | 6373 | 14 | 0.44% |
| hsa_circ_0132364 | 10017 | 22 | 0.44% |
| hsa_circ_0020130 | 12749 | 28 | 0.44% |
| hsa_circ_0025665 | 5920 | 13 | 0.44% |
| hsa_circ_0125868 | 7742 | 17 | 0.44% |
| hsa_circ_0044714 | 6376 | 14 | 0.44% |
| hsa_circ_0028179 | 5010 | 11 | 0.44% |
| hsa_circ_0073125 | 5921 | 13 | 0.44% |
| hsa_circ_0137780 | 20496 | 45 | 0.44% |
| hsa_circ_0110544 | 7288 | 16 | 0.44% |
| hsa_circ_0032458 | 6378 | 14 | 0.44% |
| hsa_circ_0114575 | 5467 | 12 | 0.44% |
| hsa_circ_0071075 | 5923 | 13 | 0.44% |
| hsa_circ_0070231 | 13213 | 29 | 0.44% |
| hsa_circ_0034810 | 5012 | 11 | 0.44% |
| hsa_circ_0131370 | 14126 | 31 | 0.44% |
| hsa_circ_0037931 | 5924 | 13 | 0.44% |
| hsa_circ_0033331 | 5469 | 12 | 0.44% |
| hsa_circ_0098125 | 46490 | 102 | 0.44% |
| hsa_circ_0112286 | 6381 | 14 | 0.44% |
| hsa_circ_0107261 | 44212 | 97 | 0.44% |
| hsa_circ_0100703 | 17776 | 39 | 0.44% |
| hsa_circ_0086872 | 5926 | 13 | 0.44% |
| hsa_circ_0058084 | 5928 | 13 | 0.44% |

| | | | |
|---|---|---|---|
| hsa_circ_0005461 | 4560 | 10 | 0.44% |
| hsa_circ_0010644 | 5929 | 13 | 0.44% |
| hsa_circ_0103707 | 5473 | 12 | 0.44% |
| hsa_circ_0062696 | 5017 | 11 | 0.44% |
| hsa_circ_0074107 | 4561 | 10 | 0.44% |
| hsa_circ_0084590 | 5474 | 12 | 0.44% |
| hsa_circ_0019181 | 6843 | 15 | 0.44% |
| hsa_circ_0049818 | 5475 | 12 | 0.44% |
| hsa_circ_0078999 | 6844 | 15 | 0.44% |
| hsa_circ_0035789 | 4563 | 10 | 0.44% |
| hsa_circ_0078144 | 4563 | 10 | 0.44% |
| hsa_circ_0132667 | 5476 | 12 | 0.44% |
| hsa_circ_0029493 | 5477 | 12 | 0.44% |
| hsa_circ_0126930 | 13695 | 30 | 0.44% |
| hsa_circ_0093580 | 10956 | 24 | 0.44% |
| hsa_circ_0040769 | 5023 | 11 | 0.44% |
| hsa_circ_0019321 | 5480 | 12 | 0.44% |
| hsa_circ_0027637 | 5481 | 12 | 0.44% |
| hsa_circ_0107543 | 5025 | 11 | 0.44% |
| hsa_circ_0127570 | 5025 | 11 | 0.44% |
| hsa_circ_0129964 | 5025 | 11 | 0.44% |
| hsa_circ_0135312 | 16905 | 37 | 0.44% |
| hsa_circ_0049419 | 5026 | 11 | 0.44% |
| hsa_circ_0053757 | 5026 | 11 | 0.44% |
| hsa_circ_0033299 | 5483 | 12 | 0.44% |
| hsa_circ_0052933 | 5940 | 13 | 0.44% |
| hsa_circ_0062088 | 5940 | 13 | 0.44% |
| hsa_circ_0042239 | 8225 | 18 | 0.44% |
| hsa_circ_0097473 | 9597 | 21 | 0.44% |
| hsa_circ_0027652 | 5485 | 12 | 0.44% |
| hsa_circ_0025827 | 5486 | 12 | 0.44% |
| hsa_circ_0008919 | 17831 | 39 | 0.44% |
| hsa_circ_0009994 | 5487 | 12 | 0.44% |
| hsa_circ_0061644 | 5487 | 12 | 0.44% |
| hsa_circ_0073670 | 5487 | 12 | 0.44% |
| hsa_circ_0120947 | 6402 | 14 | 0.44% |
| hsa_circ_0002825 | 15549 | 34 | 0.44% |
| hsa_circ_0045029 | 5488 | 12 | 0.44% |
| hsa_circ_0082817 | 5488 | 12 | 0.44% |
| hsa_circ_0053452 | 5031 | 11 | 0.44% |
| hsa_circ_0053738 | 5031 | 11 | 0.44% |
| hsa_circ_0094205 | 5946 | 13 | 0.44% |
| hsa_circ_0086388 | 5032 | 11 | 0.44% |
| hsa_circ_0036815 | 5947 | 13 | 0.44% |
| hsa_circ_0073238 | 5490 | 12 | 0.44% |
| hsa_circ_0082193 | 5490 | 12 | 0.44% |
| hsa_circ_0001587 | 25626 | 56 | 0.44% |
| hsa_circ_0012046 | 5949 | 13 | 0.44% |
| hsa_circ_0134761 | 16017 | 35 | 0.44% |
| hsa_circ_0084333 | 5034 | 11 | 0.44% |

| | | | |
|---|---|---|---|
| hsa_circ_0026634 | 19221 | 42 | 0.44% |
| hsa_circ_0042900 | 5035 | 11 | 0.44% |
| hsa_circ_0053192 | 5951 | 13 | 0.44% |
| hsa_circ_0103617 | 25179 | 55 | 0.44% |
| hsa_circ_0087978 | 5036 | 11 | 0.44% |
| hsa_circ_0068041 | 6410 | 14 | 0.44% |
| hsa_circ_0055368 | 7327 | 16 | 0.44% |
| hsa_circ_0049511 | 24274 | 53 | 0.44% |
| hsa_circ_0067979 | 6870 | 15 | 0.44% |
| hsa_circ_0052441 | 6413 | 14 | 0.44% |
| hsa_circ_0135027 | 5039 | 11 | 0.44% |
| hsa_circ_0029087 | 5956 | 13 | 0.44% |
| hsa_circ_0068188 | 5499 | 12 | 0.44% |
| hsa_circ_0014882 | 5044 | 11 | 0.44% |
| hsa_circ_0042604 | 5044 | 11 | 0.44% |
| hsa_circ_0134043 | 8254 | 18 | 0.44% |
| hsa_circ_0071886 | 5046 | 11 | 0.44% |
| hsa_circ_0048941 | 7340 | 16 | 0.44% |
| hsa_circ_0139512 | 22025 | 48 | 0.44% |
| hsa_circ_0010256 | 6883 | 15 | 0.44% |
| hsa_circ_0068975 | 5048 | 11 | 0.44% |
| hsa_circ_0085532 | 5048 | 11 | 0.44% |
| hsa_circ_0100028 | 11015 | 24 | 0.44% |
| hsa_circ_0070118 | 6426 | 14 | 0.44% |
| hsa_circ_0131251 | 15149 | 33 | 0.44% |
| hsa_circ_0052935 | 7345 | 16 | 0.44% |
| hsa_circ_0122610 | 6427 | 14 | 0.44% |
| hsa_circ_0028735 | 5051 | 11 | 0.44% |
| hsa_circ_0009796 | 6429 | 14 | 0.44% |
| hsa_circ_0091909 | 5511 | 12 | 0.44% |
| hsa_circ_0136797 | 5511 | 12 | 0.44% |
| hsa_circ_0136423 | 14698 | 32 | 0.44% |
| hsa_circ_0015760 | 6890 | 15 | 0.44% |
| hsa_circ_0056534 | 5973 | 13 | 0.44% |
| hsa_circ_0120841 | 5973 | 13 | 0.44% |
| hsa_circ_0094392 | 12406 | 27 | 0.44% |
| hsa_circ_0078112 | 6433 | 14 | 0.44% |
| hsa_circ_0069499 | 5055 | 11 | 0.44% |
| hsa_circ_0075557 | 5055 | 11 | 0.44% |
| hsa_circ_0077722 | 5055 | 11 | 0.44% |
| hsa_circ_0090654 | 5515 | 12 | 0.44% |
| hsa_circ_0129169 | 16087 | 35 | 0.44% |
| hsa_circ_0125241 | 5056 | 11 | 0.44% |
| hsa_circ_0054734 | 5977 | 13 | 0.44% |
| hsa_circ_0114142 | 5977 | 13 | 0.44% |
| hsa_circ_0135216 | 12874 | 28 | 0.43% |
| hsa_circ_0044403 | 7358 | 16 | 0.43% |
| hsa_circ_0092080 | 5979 | 13 | 0.43% |
| hsa_circ_0028745 | 5520 | 12 | 0.43% |
| hsa_circ_0127255 | 5520 | 12 | 0.43% |

| | | | |
|---|---|---|---|
| hsa_circ_0140234 | 5981 | 13 | 0.43% |
| hsa_circ_0108677 | 19785 | 43 | 0.43% |
| hsa_circ_0018357 | 15185 | 33 | 0.43% |
| hsa_circ_0132730 | 5522 | 12 | 0.43% |
| hsa_circ_0129906 | 6903 | 15 | 0.43% |
| hsa_circ_0084351 | 5983 | 13 | 0.43% |
| hsa_circ_0032888 | 6904 | 15 | 0.43% |
| hsa_circ_0106053 | 9206 | 20 | 0.43% |
| hsa_circ_0084324 | 5984 | 13 | 0.43% |
| hsa_circ_0093643 | 10128 | 22 | 0.43% |
| hsa_circ_0076721 | 5985 | 13 | 0.43% |
| hsa_circ_0101817 | 11051 | 24 | 0.43% |
| hsa_circ_0104761 | 5986 | 13 | 0.43% |
| hsa_circ_0054785 | 5526 | 12 | 0.43% |
| hsa_circ_0116461 | 5526 | 12 | 0.43% |
| hsa_circ_0045434 | 5068 | 11 | 0.43% |
| hsa_circ_0066296 | 5529 | 12 | 0.43% |
| hsa_circ_0042161 | 5069 | 11 | 0.43% |
| hsa_circ_0083277 | 5069 | 11 | 0.43% |
| hsa_circ_0103108 | 7834 | 17 | 0.43% |
| hsa_circ_0009952 | 5992 | 13 | 0.43% |
| hsa_circ_0009948 | 5993 | 13 | 0.43% |
| hsa_circ_0061185 | 5071 | 11 | 0.43% |
| hsa_circ_0002856 | 18442 | 40 | 0.43% |
| hsa_circ_0038912 | 6455 | 14 | 0.43% |
| hsa_circ_0077331 | 5994 | 13 | 0.43% |
| hsa_circ_0033314 | 5533 | 12 | 0.43% |
| hsa_circ_0114782 | 15677 | 34 | 0.43% |
| hsa_circ_0047005 | 5997 | 13 | 0.43% |
| hsa_circ_0116936 | 5997 | 13 | 0.43% |
| hsa_circ_0098113 | 6920 | 15 | 0.43% |
| hsa_circ_0024666 | 5536 | 12 | 0.43% |
| hsa_circ_0078073 | 5536 | 12 | 0.43% |
| hsa_circ_0036808 | 6460 | 14 | 0.43% |
| hsa_circ_0026926 | 7383 | 16 | 0.43% |
| hsa_circ_0035143 | 5999 | 13 | 0.43% |
| hsa_circ_0123190 | 20309 | 44 | 0.43% |
| hsa_circ_0127548 | 5539 | 12 | 0.43% |
| hsa_circ_0084341 | 6001 | 13 | 0.43% |
| hsa_circ_0063393 | 5540 | 12 | 0.43% |
| hsa_circ_0000136 | 11542 | 25 | 0.43% |
| hsa_circ_0105482 | 5541 | 12 | 0.43% |
| hsa_circ_0139858 | 15239 | 33 | 0.43% |
| hsa_circ_0021016 | 5080 | 11 | 0.43% |
| hsa_circ_0029686 | 5542 | 12 | 0.43% |
| hsa_circ_0032460 | 16628 | 36 | 0.43% |
| hsa_circ_0008447 | 6467 | 14 | 0.43% |
| hsa_circ_0139856 | 15244 | 33 | 0.43% |
| hsa_circ_0138543 | 15246 | 33 | 0.43% |
| hsa_circ_0110147 | 7854 | 17 | 0.43% |

| | | | |
|---|---|---|---|
| hsa_circ_0086890 | 6006 | 13 | 0.43% |
| hsa_circ_0111717 | 6006 | 13 | 0.43% |
| hsa_circ_0015221 | 5082 | 11 | 0.43% |
| hsa_circ_0045783 | 5083 | 11 | 0.43% |
| hsa_circ_0090723 | 6008 | 13 | 0.43% |
| hsa_circ_0031403 | 7857 | 17 | 0.43% |
| hsa_circ_0071008 | 5084 | 11 | 0.43% |
| hsa_circ_0073331 | 5084 | 11 | 0.43% |
| hsa_circ_0090184 | 5085 | 11 | 0.43% |
| hsa_circ_0102110 | 7859 | 17 | 0.43% |
| hsa_circ_0137424 | 6011 | 13 | 0.43% |
| hsa_circ_0123652 | 6012 | 13 | 0.43% |
| hsa_circ_0066306 | 5550 | 12 | 0.43% |
| hsa_circ_0090927 | 5551 | 12 | 0.43% |
| hsa_circ_0083652 | 12953 | 28 | 0.43% |
| hsa_circ_0086811 | 5552 | 12 | 0.43% |
| hsa_circ_0034226 | 6015 | 13 | 0.43% |
| hsa_circ_0096446 | 10642 | 23 | 0.43% |
| hsa_circ_0049920 | 5090 | 11 | 0.43% |
| hsa_circ_0033249 | 5553 | 12 | 0.43% |
| hsa_circ_0104990 | 9256 | 20 | 0.43% |
| hsa_circ_0027645 | 5554 | 12 | 0.43% |
| hsa_circ_0080166 | 6480 | 14 | 0.43% |
| hsa_circ_0024011 | 6943 | 15 | 0.43% |
| hsa_circ_0070055 | 5555 | 12 | 0.43% |
| hsa_circ_0132003 | 7407 | 16 | 0.43% |
| hsa_circ_0133722 | 7408 | 16 | 0.43% |
| hsa_circ_0018536 | 5557 | 12 | 0.43% |
| hsa_circ_0044183 | 17135 | 37 | 0.43% |
| hsa_circ_0073843 | 5558 | 12 | 0.43% |
| hsa_circ_0076195 | 10190 | 22 | 0.43% |
| hsa_circ_0052442 | 6485 | 14 | 0.43% |
| hsa_circ_0071662 | 6022 | 13 | 0.43% |
| hsa_circ_0081865 | 13897 | 30 | 0.43% |
| hsa_circ_0005187 | 7875 | 17 | 0.43% |
| hsa_circ_0128586 | 12508 | 27 | 0.43% |
| hsa_circ_0138881 | 26871 | 58 | 0.43% |
| hsa_circ_0069071 | 7414 | 16 | 0.43% |
| hsa_circ_0061596 | 6024 | 13 | 0.43% |
| hsa_circ_0072679 | 6024 | 13 | 0.43% |
| hsa_circ_0118667 | 21781 | 47 | 0.43% |
| hsa_circ_0096812 | 6025 | 13 | 0.43% |
| hsa_circ_0093685 | 5563 | 12 | 0.43% |
| hsa_circ_0025094 | 6027 | 13 | 0.43% |
| hsa_circ_0002258 | 16695 | 36 | 0.43% |
| hsa_circ_0010328 | 5565 | 12 | 0.43% |
| hsa_circ_0027178 | 5565 | 12 | 0.43% |
| hsa_circ_0036642 | 5566 | 12 | 0.43% |
| hsa_circ_0137137 | 6030 | 13 | 0.43% |
| hsa_circ_0086873 | 6031 | 13 | 0.43% |

| | | | |
|---|---|---|---|
| hsa_circ_0114318 | 21806 | 47 | 0.43% |
| hsa_circ_0094100 | 24132 | 52 | 0.43% |
| hsa_circ_0110942 | 17635 | 38 | 0.43% |
| hsa_circ_0034200 | 5105 | 11 | 0.43% |
| hsa_circ_0105062 | 18564 | 40 | 0.43% |
| hsa_circ_0058081 | 6035 | 13 | 0.43% |
| hsa_circ_0078005 | 5572 | 12 | 0.43% |
| hsa_circ_0043541 | 5573 | 12 | 0.43% |
| hsa_circ_0028561 | 5109 | 11 | 0.43% |
| hsa_circ_0090622 | 6038 | 13 | 0.43% |
| hsa_circ_0072720 | 6503 | 14 | 0.43% |
| hsa_circ_0046010 | 6039 | 13 | 0.43% |
| hsa_circ_0075991 | 6504 | 14 | 0.43% |
| hsa_circ_0121432 | 9292 | 20 | 0.43% |
| hsa_circ_0097477 | 9758 | 21 | 0.43% |
| hsa_circ_0025081 | 6041 | 13 | 0.43% |
| hsa_circ_0140311 | 8366 | 18 | 0.43% |
| hsa_circ_0131048 | 5578 | 12 | 0.43% |
| hsa_circ_0097474 | 9762 | 21 | 0.43% |
| hsa_circ_0035785 | 6044 | 13 | 0.43% |
| hsa_circ_0049814 | 6044 | 13 | 0.43% |
| hsa_circ_0103145 | 7439 | 16 | 0.43% |
| hsa_circ_0038027 | 6046 | 13 | 0.43% |
| hsa_circ_0123091 | 49768 | 107 | 0.43% |
| hsa_circ_0009791 | 6512 | 14 | 0.43% |
| hsa_circ_0063079 | 6047 | 13 | 0.43% |
| hsa_circ_0123731 | 5582 | 12 | 0.43% |
| hsa_circ_0138398 | 5117 | 11 | 0.43% |
| hsa_circ_0021993 | 6979 | 15 | 0.43% |
| hsa_circ_0007780 | 5584 | 12 | 0.43% |
| hsa_circ_0108823 | 6515 | 14 | 0.43% |
| hsa_circ_0057849 | 6050 | 13 | 0.43% |
| hsa_circ_0088466 | 6516 | 14 | 0.43% |
| hsa_circ_0030126 | 15365 | 33 | 0.43% |
| hsa_circ_0049502 | 23283 | 50 | 0.43% |
| hsa_circ_0127483 | 6985 | 15 | 0.43% |
| hsa_circ_0035992 | 20024 | 43 | 0.43% |
| hsa_circ_0081765 | 6055 | 13 | 0.43% |
| hsa_circ_0116986 | 8385 | 18 | 0.43% |
| hsa_circ_0128833 | 27020 | 58 | 0.43% |
| hsa_circ_0098114 | 6988 | 15 | 0.43% |
| hsa_circ_0114874 | 20500 | 44 | 0.43% |
| hsa_circ_0062244 | 18171 | 39 | 0.43% |
| hsa_circ_0130627 | 5592 | 12 | 0.43% |
| hsa_circ_0062631 | 5126 | 11 | 0.43% |
| hsa_circ_0138800 | 5126 | 11 | 0.43% |
| hsa_circ_0096943 | 20973 | 45 | 0.43% |
| hsa_circ_0027429 | 5128 | 11 | 0.43% |
| hsa_circ_0112201 | 18183 | 39 | 0.43% |
| hsa_circ_0011342 | 18652 | 40 | 0.43% |

| | | | |
|---|---|---|---|
| hsa_circ_0082501 | 6063 | 13 | 0.43% |
| hsa_circ_0115822 | 10261 | 22 | 0.43% |
| hsa_circ_0112284 | 6530 | 14 | 0.43% |
| hsa_circ_0001812 | 9329 | 20 | 0.43% |
| hsa_circ_0037454 | 6064 | 13 | 0.43% |
| hsa_circ_0083345 | 6064 | 13 | 0.43% |
| hsa_circ_0094203 | 6064 | 13 | 0.43% |
| hsa_circ_0027383 | 7930 | 17 | 0.43% |
| hsa_circ_0053546 | 6065 | 13 | 0.43% |
| hsa_circ_0026927 | 7466 | 16 | 0.43% |
| hsa_circ_0104982 | 7935 | 17 | 0.43% |
| hsa_circ_0000540 | 22406 | 48 | 0.43% |
| hsa_circ_0138332 | 12604 | 27 | 0.43% |
| hsa_circ_0053492 | 5135 | 11 | 0.43% |
| hsa_circ_0053800 | 5602 | 12 | 0.43% |
| hsa_circ_0035738 | 6069 | 13 | 0.43% |
| hsa_circ_0059119 | 5136 | 11 | 0.43% |
| hsa_circ_0013703 | 5605 | 12 | 0.43% |
| hsa_circ_0134324 | 5605 | 12 | 0.43% |
| hsa_circ_0121861 | 19620 | 42 | 0.43% |
| hsa_circ_0047211 | 5606 | 12 | 0.43% |
| hsa_circ_0056636 | 5139 | 11 | 0.43% |
| hsa_circ_0020064 | 7008 | 15 | 0.43% |
| hsa_circ_0002802 | 18690 | 40 | 0.43% |
| hsa_circ_0010254 | 7009 | 15 | 0.43% |
| hsa_circ_0139668 | 15420 | 33 | 0.43% |
| hsa_circ_0016568 | 5141 | 11 | 0.43% |
| hsa_circ_0122623 | 6076 | 13 | 0.43% |
| hsa_circ_0051293 | 5609 | 12 | 0.43% |
| hsa_circ_0089209 | 6077 | 13 | 0.43% |
| hsa_circ_0086804 | 5610 | 12 | 0.43% |
| hsa_circ_0129553 | 12156 | 26 | 0.43% |
| hsa_circ_0028399 | 5143 | 11 | 0.43% |
| hsa_circ_0047967 | 5611 | 12 | 0.43% |
| hsa_circ_0053952 | 5611 | 12 | 0.43% |
| hsa_circ_0089275 | 6547 | 14 | 0.43% |
| hsa_circ_0110545 | 13096 | 28 | 0.43% |
| hsa_circ_0016289 | 5614 | 12 | 0.43% |
| hsa_circ_0031669 | 6082 | 13 | 0.43% |
| hsa_circ_0095450 | 6082 | 13 | 0.43% |
| hsa_circ_0068043 | 6550 | 14 | 0.43% |
| hsa_circ_0112399 | 14506 | 31 | 0.43% |
| hsa_circ_0015363 | 14975 | 32 | 0.43% |
| hsa_circ_0093084 | 6084 | 13 | 0.43% |
| hsa_circ_0011839 | 5616 | 12 | 0.43% |
| hsa_circ_0069124 | 6085 | 13 | 0.43% |
| hsa_circ_0136533 | 5618 | 12 | 0.43% |
| hsa_circ_0138602 | 7960 | 17 | 0.43% |
| hsa_circ_0078049 | 5619 | 12 | 0.43% |
| hsa_circ_0104718 | 5619 | 12 | 0.43% |

| | | | |
|---|---|---|---|
| hsa_circ_0093688 | 61814 | 132 | 0.43% |
| hsa_circ_0115422 | 14986 | 32 | 0.43% |
| hsa_circ_0119279 | 16861 | 36 | 0.43% |
| hsa_circ_0044401 | 7494 | 16 | 0.43% |
| hsa_circ_0053574 | 6090 | 13 | 0.43% |
| hsa_circ_0076810 | 5623 | 12 | 0.43% |
| hsa_circ_0061905 | 6561 | 14 | 0.43% |
| hsa_circ_0119752 | 13591 | 29 | 0.43% |
| hsa_circ_0061432 | 6094 | 13 | 0.43% |
| hsa_circ_0088618 | 6563 | 14 | 0.43% |
| hsa_circ_0113126 | 5626 | 12 | 0.43% |
| hsa_circ_0031183 | 6564 | 14 | 0.43% |
| hsa_circ_0057030 | 6564 | 14 | 0.43% |
| hsa_circ_0060864 | 7033 | 15 | 0.43% |
| hsa_circ_0133239 | 5629 | 12 | 0.43% |
| hsa_circ_0097475 | 9851 | 21 | 0.43% |
| hsa_circ_0032525 | 26272 | 56 | 0.43% |
| hsa_circ_0028147 | 4692 | 10 | 0.43% |
| hsa_circ_0094743 | 6100 | 13 | 0.43% |
| hsa_circ_0091851 | 6570 | 14 | 0.43% |
| hsa_circ_0114803 | 9386 | 20 | 0.43% |
| hsa_circ_0120136 | 9387 | 20 | 0.43% |
| hsa_circ_0007261 | 23937 | 51 | 0.43% |
| hsa_circ_0013936 | 17367 | 37 | 0.43% |
| hsa_circ_0136818 | 6572 | 14 | 0.43% |
| hsa_circ_0053771 | 6103 | 13 | 0.43% |
| hsa_circ_0087253 | 5634 | 12 | 0.43% |
| hsa_circ_0021403 | 6104 | 13 | 0.43% |
| hsa_circ_0077453 | 6104 | 13 | 0.43% |
| hsa_circ_0036805 | 6574 | 14 | 0.43% |
| hsa_circ_0045682 | 5166 | 11 | 0.43% |
| hsa_circ_0027690 | 6575 | 14 | 0.43% |
| hsa_circ_0138745 | 21134 | 45 | 0.43% |
| hsa_circ_0135215 | 13153 | 28 | 0.43% |
| hsa_circ_0119337 | 6110 | 13 | 0.43% |
| hsa_circ_0015590 | 5642 | 12 | 0.43% |
| hsa_circ_0027382 | 7993 | 17 | 0.43% |
| hsa_circ_0138193 | 6113 | 13 | 0.43% |
| hsa_circ_0125867 | 7994 | 17 | 0.43% |
| hsa_circ_0100699 | 17870 | 38 | 0.43% |
| hsa_circ_0056727 | 8936 | 19 | 0.43% |
| hsa_circ_0009788 | 6585 | 14 | 0.43% |
| hsa_circ_0038279 | 5174 | 11 | 0.43% |
| hsa_circ_0088609 | 5174 | 11 | 0.43% |
| hsa_circ_0140769 | 6115 | 13 | 0.43% |
| hsa_circ_0093765 | 14115 | 30 | 0.43% |
| hsa_circ_0036151 | 5646 | 12 | 0.43% |
| hsa_circ_0006439 | 12234 | 26 | 0.43% |
| hsa_circ_0013837 | 5176 | 11 | 0.43% |
| hsa_circ_0053514 | 5176 | 11 | 0.43% |

| | | | |
|---|---|---|---|
| hsa_circ_0123126 | 7059 | 15 | 0.42% |
| hsa_circ_0096802 | 24002 | 51 | 0.42% |
| hsa_circ_0031515 | 5649 | 12 | 0.42% |
| hsa_circ_0071607 | 5649 | 12 | 0.42% |
| hsa_circ_0096332 | 16949 | 36 | 0.42% |
| hsa_circ_0071134 | 5651 | 12 | 0.42% |
| hsa_circ_0013605 | 6593 | 14 | 0.42% |
| hsa_circ_0126936 | 7065 | 15 | 0.42% |
| hsa_circ_0058248 | 6594 | 14 | 0.42% |
| hsa_circ_0053531 | 6124 | 13 | 0.42% |
| hsa_circ_0048724 | 6125 | 13 | 0.42% |
| hsa_circ_0074055 | 5654 | 12 | 0.42% |
| hsa_circ_0018755 | 23096 | 49 | 0.42% |
| hsa_circ_0075974 | 6599 | 14 | 0.42% |
| hsa_circ_0060396 | 20741 | 44 | 0.42% |
| hsa_circ_0033226 | 5657 | 12 | 0.42% |
| hsa_circ_0027185 | 6129 | 13 | 0.42% |
| hsa_circ_0127033 | 7072 | 15 | 0.42% |
| hsa_circ_0002641 | 14145 | 30 | 0.42% |
| hsa_circ_0029880 | 6130 | 13 | 0.42% |
| hsa_circ_0035071 | 6130 | 13 | 0.42% |
| hsa_circ_0081297 | 5187 | 11 | 0.42% |
| hsa_circ_0104708 | 5187 | 11 | 0.42% |
| hsa_circ_0042176 | 5659 | 12 | 0.42% |
| hsa_circ_0056533 | 6131 | 13 | 0.42% |
| hsa_circ_0136754 | 5660 | 12 | 0.42% |
| hsa_circ_0129248 | 8963 | 19 | 0.42% |
| hsa_circ_0081969 | 5661 | 12 | 0.42% |
| hsa_circ_0099467 | 5661 | 12 | 0.42% |
| hsa_circ_0001213 | 15097 | 32 | 0.42% |
| hsa_circ_0070047 | 5662 | 12 | 0.42% |
| hsa_circ_0123087 | 6606 | 14 | 0.42% |
| hsa_circ_0019095 | 6607 | 14 | 0.42% |
| hsa_circ_0084560 | 6136 | 13 | 0.42% |
| hsa_circ_0028738 | 5664 | 12 | 0.42% |
| hsa_circ_0105294 | 25961 | 55 | 0.42% |
| hsa_circ_0028562 | 5193 | 11 | 0.42% |
| hsa_circ_0135031 | 5193 | 11 | 0.42% |
| hsa_circ_0137350 | 8973 | 19 | 0.42% |
| hsa_circ_0035786 | 6140 | 13 | 0.42% |
| hsa_circ_0132716 | 6140 | 13 | 0.42% |
| hsa_circ_0034225 | 6141 | 13 | 0.42% |
| hsa_circ_0125667 | 6615 | 14 | 0.42% |
| hsa_circ_0025885 | 6144 | 13 | 0.42% |
| hsa_circ_0077312 | 6144 | 13 | 0.42% |
| hsa_circ_0093310 | 15124 | 32 | 0.42% |
| hsa_circ_0137645 | 7562 | 16 | 0.42% |
| hsa_circ_0110632 | 23165 | 49 | 0.42% |
| hsa_circ_0098810 | 6147 | 13 | 0.42% |
| hsa_circ_0121587 | 12768 | 27 | 0.42% |

| | | | |
|---|---|---|---|
| hsa_circ_0090623 | 6149 | 13 | 0.42% |
| hsa_circ_0024889 | 5676 | 12 | 0.42% |
| hsa_circ_0078998 | 6623 | 14 | 0.42% |
| hsa_circ_0012684 | 6150 | 13 | 0.42% |
| hsa_circ_0112282 | 6624 | 14 | 0.42% |
| hsa_circ_0115326 | 15141 | 32 | 0.42% |
| hsa_circ_0031179 | 5679 | 12 | 0.42% |
| hsa_circ_0084073 | 5206 | 11 | 0.42% |
| hsa_circ_0131209 | 21298 | 45 | 0.42% |
| hsa_circ_0016777 | 6153 | 13 | 0.42% |
| hsa_circ_0019141 | 5680 | 12 | 0.42% |
| hsa_circ_0080167 | 6629 | 14 | 0.42% |
| hsa_circ_0091138 | 6156 | 13 | 0.42% |
| hsa_circ_0114416 | 5683 | 12 | 0.42% |
| hsa_circ_0132069 | 5683 | 12 | 0.42% |
| hsa_circ_0025972 | 7104 | 15 | 0.42% |
| hsa_circ_0114777 | 6157 | 13 | 0.42% |
| hsa_circ_0000209 | 14683 | 31 | 0.42% |
| hsa_circ_0051075 | 15157 | 32 | 0.42% |
| hsa_circ_0128146 | 5211 | 11 | 0.42% |
| hsa_circ_0027458 | 6159 | 13 | 0.42% |
| hsa_circ_0058820 | 6159 | 13 | 0.42% |
| hsa_circ_0119526 | 6159 | 13 | 0.42% |
| hsa_circ_0035753 | 6160 | 13 | 0.42% |
| hsa_circ_0018817 | 6163 | 13 | 0.42% |
| hsa_circ_0077805 | 5689 | 12 | 0.42% |
| hsa_circ_0090248 | 6165 | 13 | 0.42% |
| hsa_circ_0016001 | 5691 | 12 | 0.42% |
| hsa_circ_0084218 | 5691 | 12 | 0.42% |
| hsa_circ_0018729 | 6640 | 14 | 0.42% |
| hsa_circ_0063080 | 6166 | 13 | 0.42% |
| hsa_circ_0139740 | 6166 | 13 | 0.42% |
| hsa_circ_0089512 | 5692 | 12 | 0.42% |
| hsa_circ_0036803 | 6642 | 14 | 0.42% |
| hsa_circ_0042349 | 19452 | 41 | 0.42% |
| hsa_circ_0042157 | 5219 | 11 | 0.42% |
| hsa_circ_0131305 | 5219 | 11 | 0.42% |
| hsa_circ_0131210 | 21354 | 45 | 0.42% |
| hsa_circ_0045736 | 18035 | 38 | 0.42% |
| hsa_circ_0128041 | 6170 | 13 | 0.42% |
| hsa_circ_0084899 | 5221 | 11 | 0.42% |
| hsa_circ_0031502 | 5697 | 12 | 0.42% |
| hsa_circ_0053758 | 5223 | 11 | 0.42% |
| hsa_circ_0063392 | 5699 | 12 | 0.42% |
| hsa_circ_0000228 | 14249 | 30 | 0.42% |
| hsa_circ_0081239 | 7125 | 15 | 0.42% |
| hsa_circ_0090704 | 6175 | 13 | 0.42% |
| hsa_circ_0039630 | 6177 | 13 | 0.42% |
| hsa_circ_0091142 | 5702 | 12 | 0.42% |
| hsa_circ_0033220 | 5703 | 12 | 0.42% |

| | | | |
|---|---|---|---|
| hsa_circ_0040861 | 7129 | 15 | 0.42% |
| hsa_circ_0010343 | 4753 | 10 | 0.42% |
| hsa_circ_0137661 | 6179 | 13 | 0.42% |
| hsa_circ_0112384 | 7130 | 15 | 0.42% |
| hsa_circ_0094884 | 5704 | 12 | 0.42% |
| hsa_circ_0063298 | 20444 | 43 | 0.42% |
| hsa_circ_0060915 | 5706 | 12 | 0.42% |
| hsa_circ_0117961 | 6658 | 14 | 0.42% |
| hsa_circ_0056159 | 20452 | 43 | 0.42% |
| hsa_circ_0099807 | 9037 | 19 | 0.42% |
| hsa_circ_0064592 | 5232 | 11 | 0.42% |
| hsa_circ_0105272 | 8562 | 18 | 0.42% |
| hsa_circ_0086855 | 6184 | 13 | 0.42% |
| hsa_circ_0125807 | 15699 | 33 | 0.42% |
| hsa_circ_0130391 | 14748 | 31 | 0.42% |
| hsa_circ_0027646 | 5709 | 12 | 0.42% |
| hsa_circ_0055990 | 6661 | 14 | 0.42% |
| hsa_circ_0113334 | 20459 | 43 | 0.42% |
| hsa_circ_0119213 | 6662 | 14 | 0.42% |
| hsa_circ_0104269 | 24746 | 52 | 0.42% |
| hsa_circ_0052692 | 7139 | 15 | 0.42% |
| hsa_circ_0042590 | 5712 | 12 | 0.42% |
| hsa_circ_0106404 | 14281 | 30 | 0.42% |
| hsa_circ_0082656 | 12377 | 26 | 0.42% |
| hsa_circ_0044398 | 7617 | 16 | 0.42% |
| hsa_circ_0002379 | 6189 | 13 | 0.42% |
| hsa_circ_0024118 | 6189 | 13 | 0.42% |
| hsa_circ_0019184 | 5714 | 12 | 0.42% |
| hsa_circ_0035124 | 5714 | 12 | 0.42% |
| hsa_circ_0036640 | 5714 | 12 | 0.42% |
| hsa_circ_0106432 | 7620 | 16 | 0.42% |
| hsa_circ_0056510 | 5239 | 11 | 0.42% |
| hsa_circ_0090294 | 6668 | 14 | 0.42% |
| hsa_circ_0071939 | 4763 | 10 | 0.42% |
| hsa_circ_0128587 | 12862 | 27 | 0.42% |
| hsa_circ_0070739 | 15723 | 33 | 0.42% |
| hsa_circ_0042898 | 5242 | 11 | 0.42% |
| hsa_circ_0049026 | 5719 | 12 | 0.42% |
| hsa_circ_0004632 | 10009 | 21 | 0.42% |
| hsa_circ_0111260 | 6198 | 13 | 0.42% |
| hsa_circ_0075790 | 8583 | 18 | 0.42% |
| hsa_circ_0053453 | 5246 | 11 | 0.42% |
| hsa_circ_0028746 | 5724 | 12 | 0.42% |
| hsa_circ_0074311 | 5247 | 11 | 0.42% |
| hsa_circ_0132744 | 46271 | 97 | 0.42% |
| hsa_circ_0035739 | 6202 | 13 | 0.42% |
| hsa_circ_0139554 | 20515 | 43 | 0.42% |
| hsa_circ_0015873 | 18609 | 39 | 0.42% |
| hsa_circ_0009953 | 6203 | 13 | 0.42% |
| hsa_circ_0037930 | 6203 | 13 | 0.42% |

| | | | |
|---|---|---|---|
| hsa_circ_0028720 | 5726 | 12 | 0.42% |
| hsa_circ_0004995 | 5249 | 11 | 0.42% |
| hsa_circ_0058543 | 5727 | 12 | 0.42% |
| hsa_circ_0093689 | 63000 | 132 | 0.42% |
| hsa_circ_0059995 | 5250 | 11 | 0.42% |
| hsa_circ_0128461 | 14319 | 30 | 0.42% |
| hsa_circ_0024220 | 6205 | 13 | 0.42% |
| hsa_circ_0109961 | 20047 | 42 | 0.42% |
| hsa_circ_0056532 | 6206 | 13 | 0.42% |
| hsa_circ_0106642 | 5731 | 12 | 0.42% |
| hsa_circ_0060865 | 7165 | 15 | 0.42% |
| hsa_circ_0073239 | 5732 | 12 | 0.42% |
| hsa_circ_0084256 | 5732 | 12 | 0.42% |
| hsa_circ_0072652 | 6688 | 14 | 0.42% |
| hsa_circ_0007495 | 18633 | 39 | 0.42% |
| hsa_circ_0010657 | 6211 | 13 | 0.42% |
| hsa_circ_0126982 | 12900 | 27 | 0.42% |
| hsa_circ_0101814 | 67847 | 142 | 0.42% |
| hsa_circ_0065403 | 6213 | 13 | 0.42% |
| hsa_circ_0112202 | 18642 | 39 | 0.42% |
| hsa_circ_0053776 | 5736 | 12 | 0.42% |
| hsa_circ_0028565 | 4780 | 10 | 0.42% |
| hsa_circ_0133985 | 9561 | 20 | 0.42% |
| hsa_circ_0091852 | 6694 | 14 | 0.42% |
| hsa_circ_0120867 | 6694 | 14 | 0.42% |
| hsa_circ_0036147 | 5739 | 12 | 0.42% |
| hsa_circ_0059790 | 5739 | 12 | 0.42% |
| hsa_circ_0078479 | 8131 | 17 | 0.42% |
| hsa_circ_0110823 | 6218 | 13 | 0.42% |
| hsa_circ_0116384 | 35395 | 74 | 0.42% |
| hsa_circ_0002283 | 17699 | 37 | 0.42% |
| hsa_circ_0119743 | 11003 | 23 | 0.42% |
| hsa_circ_0072175 | 5741 | 12 | 0.42% |
| hsa_circ_0058889 | 6698 | 14 | 0.42% |
| hsa_circ_0045681 | 5263 | 11 | 0.42% |
| hsa_circ_0086401 | 6223 | 13 | 0.42% |
| hsa_circ_0011893 | 5745 | 12 | 0.42% |
| hsa_circ_0016095 | 6224 | 13 | 0.42% |
| hsa_circ_0043222 | 6225 | 13 | 0.42% |
| hsa_circ_0093311 | 16760 | 35 | 0.42% |
| hsa_circ_0092266 | 5268 | 11 | 0.42% |
| hsa_circ_0054731 | 6226 | 13 | 0.42% |
| hsa_circ_0132258 | 11495 | 24 | 0.42% |
| hsa_circ_0114400 | 16764 | 35 | 0.42% |
| hsa_circ_0024276 | 5270 | 11 | 0.42% |
| hsa_circ_0111770 | 11501 | 24 | 0.42% |
| hsa_circ_0112444 | 7189 | 15 | 0.42% |
| hsa_circ_0029296 | 5273 | 11 | 0.42% |
| hsa_circ_0139860 | 15822 | 33 | 0.42% |
| hsa_circ_0020065 | 7192 | 15 | 0.42% |

| | | | |
|---|---|---|---|
| hsa_circ_0080298 | 5754 | 12 | 0.42% |
| hsa_circ_0098467 | 6234 | 13 | 0.42% |
| hsa_circ_0015588 | 5755 | 12 | 0.42% |
| hsa_circ_0058892 | 5755 | 12 | 0.42% |
| hsa_circ_0067668 | 5755 | 12 | 0.42% |
| hsa_circ_0131487 | 27338 | 57 | 0.42% |
| hsa_circ_0098468 | 6235 | 13 | 0.42% |
| hsa_circ_0134745 | 6235 | 13 | 0.42% |
| hsa_circ_0104983 | 8154 | 17 | 0.42% |
| hsa_circ_0139590 | 6716 | 14 | 0.42% |
| hsa_circ_0133038 | 5277 | 11 | 0.42% |
| hsa_circ_0065240 | 5757 | 12 | 0.42% |
| hsa_circ_0053716 | 6237 | 13 | 0.42% |
| hsa_circ_0117101 | 11035 | 23 | 0.42% |
| hsa_circ_0129860 | 5758 | 12 | 0.42% |
| hsa_circ_0010601 | 6238 | 13 | 0.42% |
| hsa_circ_0061904 | 6719 | 14 | 0.42% |
| hsa_circ_0070112 | 6720 | 14 | 0.42% |
| hsa_circ_0027203 | 4800 | 10 | 0.42% |
| hsa_circ_0054735 | 6246 | 13 | 0.42% |
| hsa_circ_0002644 | 17298 | 36 | 0.42% |
| hsa_circ_0054782 | 5766 | 12 | 0.42% |
| hsa_circ_0073844 | 5766 | 12 | 0.42% |
| hsa_circ_0094291 | 13936 | 29 | 0.42% |
| hsa_circ_0012359 | 5767 | 12 | 0.42% |
| hsa_circ_0127854 | 11054 | 23 | 0.42% |
| hsa_circ_0068564 | 21148 | 44 | 0.42% |
| hsa_circ_0132254 | 8171 | 17 | 0.42% |
| hsa_circ_0027964 | 5768 | 12 | 0.42% |
| hsa_circ_0053772 | 6249 | 13 | 0.42% |
| hsa_circ_0138603 | 8172 | 17 | 0.42% |
| hsa_circ_0053507 | 5288 | 11 | 0.42% |
| hsa_circ_0070370 | 5769 | 12 | 0.42% |
| hsa_circ_0027207 | 6250 | 13 | 0.42% |
| hsa_circ_0072033 | 7212 | 15 | 0.42% |
| hsa_circ_0134564 | 25002 | 52 | 0.42% |
| hsa_circ_0001720 | 8655 | 18 | 0.42% |
| hsa_circ_0112355 | 5770 | 12 | 0.42% |
| hsa_circ_0093754 | 17311 | 36 | 0.42% |
| hsa_circ_0129715 | 9139 | 19 | 0.42% |
| hsa_circ_0028564 | 5292 | 11 | 0.42% |
| hsa_circ_0073785 | 5292 | 11 | 0.42% |
| hsa_circ_0006140 | 15877 | 33 | 0.42% |
| hsa_circ_0115810 | 6736 | 14 | 0.42% |
| hsa_circ_0136751 | 5774 | 12 | 0.42% |
| hsa_circ_0094376 | 7700 | 16 | 0.42% |
| hsa_circ_0103319 | 14440 | 30 | 0.42% |
| hsa_circ_0020759 | 5776 | 12 | 0.42% |
| hsa_circ_0109060 | 25512 | 53 | 0.42% |
| hsa_circ_0107925 | 6258 | 13 | 0.42% |

| | | | |
|---|---|---|---|
| hsa_circ_0026668 | 5296 | 11 | 0.42% |
| hsa_circ_0093985 | 22147 | 46 | 0.42% |
| hsa_circ_0028376 | 6261 | 13 | 0.42% |
| hsa_circ_0045757 | 6261 | 13 | 0.42% |
| hsa_circ_0063081 | 6262 | 13 | 0.42% |
| hsa_circ_0111950 | 8189 | 17 | 0.42% |
| hsa_circ_0131211 | 21678 | 45 | 0.42% |
| hsa_circ_0054777 | 5781 | 12 | 0.42% |
| hsa_circ_0084373 | 6263 | 13 | 0.42% |
| hsa_circ_0096937 | 21682 | 45 | 0.42% |
| hsa_circ_0131972 | 6264 | 13 | 0.42% |
| hsa_circ_0089763 | 5783 | 12 | 0.42% |
| hsa_circ_0109704 | 15424 | 32 | 0.41% |
| hsa_circ_0077336 | 6266 | 13 | 0.41% |
| hsa_circ_0130112 | 6266 | 13 | 0.41% |
| hsa_circ_0041282 | 5784 | 12 | 0.41% |
| hsa_circ_0046840 | 6267 | 13 | 0.41% |
| hsa_circ_0047796 | 6267 | 13 | 0.41% |
| hsa_circ_0013827 | 5786 | 12 | 0.41% |
| hsa_circ_0020416 | 5304 | 11 | 0.41% |
| hsa_circ_0018305 | 21226 | 44 | 0.41% |
| hsa_circ_0130422 | 13508 | 28 | 0.41% |
| hsa_circ_0047215 | 5307 | 11 | 0.41% |
| hsa_circ_0087282 | 11581 | 24 | 0.41% |
| hsa_circ_0024706 | 5309 | 11 | 0.41% |
| hsa_circ_0084493 | 6275 | 13 | 0.41% |
| hsa_circ_0074548 | 6277 | 13 | 0.41% |
| hsa_circ_0080969 | 7727 | 16 | 0.41% |
| hsa_circ_0010274 | 5796 | 12 | 0.41% |
| hsa_circ_0066302 | 5796 | 12 | 0.41% |
| hsa_circ_0078822 | 20770 | 43 | 0.41% |
| hsa_circ_0104616 | 26567 | 55 | 0.41% |
| hsa_circ_0090671 | 6280 | 13 | 0.41% |
| hsa_circ_0005554 | 8214 | 17 | 0.41% |
| hsa_circ_0047729 | 5799 | 12 | 0.41% |
| hsa_circ_0077344 | 6283 | 13 | 0.41% |
| hsa_circ_0060866 | 7253 | 15 | 0.41% |
| hsa_circ_0112203 | 18858 | 39 | 0.41% |
| hsa_circ_0005476 | 5319 | 11 | 0.41% |
| hsa_circ_0111042 | 5319 | 11 | 0.41% |
| hsa_circ_0053761 | 6287 | 13 | 0.41% |
| hsa_circ_0089081 | 6287 | 13 | 0.41% |
| hsa_circ_0125250 | 6287 | 13 | 0.41% |
| hsa_circ_0021006 | 7255 | 15 | 0.41% |
| hsa_circ_0068421 | 5804 | 12 | 0.41% |
| hsa_circ_0066013 | 6288 | 13 | 0.41% |
| hsa_circ_0053330 | 30480 | 63 | 0.41% |
| hsa_circ_0045435 | 5322 | 11 | 0.41% |
| hsa_circ_0025059 | 5806 | 12 | 0.41% |
| hsa_circ_0095591 | 54679 | 113 | 0.41% |

| | | | |
|---|---|---|---|
| hsa_circ_0062937 | 5807 | 12 | 0.41% |
| hsa_circ_0018298 | 21296 | 44 | 0.41% |
| hsa_circ_0065593 | 5808 | 12 | 0.41% |
| hsa_circ_0077317 | 5808 | 12 | 0.41% |
| hsa_circ_0028753 | 5809 | 12 | 0.41% |
| hsa_circ_0050799 | 5809 | 12 | 0.41% |
| hsa_circ_0011788 | 6294 | 13 | 0.41% |
| hsa_circ_0102023 | 18398 | 38 | 0.41% |
| hsa_circ_0102165 | 4842 | 10 | 0.41% |
| hsa_circ_0041275 | 5327 | 11 | 0.41% |
| hsa_circ_0047411 | 7265 | 15 | 0.41% |
| hsa_circ_0126611 | 18405 | 38 | 0.41% |
| hsa_circ_0138604 | 8235 | 17 | 0.41% |
| hsa_circ_0088260 | 5813 | 12 | 0.41% |
| hsa_circ_0084561 | 6298 | 13 | 0.41% |
| hsa_circ_0139802 | 15503 | 32 | 0.41% |
| hsa_circ_0100759 | 8238 | 17 | 0.41% |
| hsa_circ_0060617 | 10177 | 21 | 0.41% |
| hsa_circ_0073786 | 5817 | 12 | 0.41% |
| hsa_circ_0086012 | 16970 | 35 | 0.41% |
| hsa_circ_0116669 | 21336 | 44 | 0.41% |
| hsa_circ_0015850 | 5819 | 12 | 0.41% |
| hsa_circ_0033255 | 5819 | 12 | 0.41% |
| hsa_circ_0132560 | 6304 | 13 | 0.41% |
| hsa_circ_0101653 | 5335 | 11 | 0.41% |
| hsa_circ_0067625 | 6306 | 13 | 0.41% |
| hsa_circ_0086995 | 14068 | 29 | 0.41% |
| hsa_circ_0037557 | 6307 | 13 | 0.41% |
| hsa_circ_0109266 | 18928 | 39 | 0.41% |
| hsa_circ_0087251 | 5824 | 12 | 0.41% |
| hsa_circ_0127535 | 7766 | 16 | 0.41% |
| hsa_circ_0036797 | 6797 | 14 | 0.41% |
| hsa_circ_0047958 | 4855 | 10 | 0.41% |
| hsa_circ_0105061 | 24764 | 51 | 0.41% |
| hsa_circ_0011892 | 5827 | 12 | 0.41% |
| hsa_circ_0131456 | 5827 | 12 | 0.41% |
| hsa_circ_0034092 | 6314 | 13 | 0.41% |
| hsa_circ_0036212 | 19430 | 40 | 0.41% |
| hsa_circ_0074419 | 6315 | 13 | 0.41% |
| hsa_circ_0138492 | 5344 | 11 | 0.41% |
| hsa_circ_0122629 | 6316 | 13 | 0.41% |
| hsa_circ_0101650 | 9717 | 20 | 0.41% |
| hsa_circ_0042146 | 6803 | 14 | 0.41% |
| hsa_circ_0061928 | 6318 | 13 | 0.41% |
| hsa_circ_0006541 | 13123 | 27 | 0.41% |
| hsa_circ_0061903 | 6805 | 14 | 0.41% |
| hsa_circ_0095449 | 6319 | 13 | 0.41% |
| hsa_circ_0114016 | 17015 | 35 | 0.41% |
| hsa_circ_0094292 | 14099 | 29 | 0.41% |
| hsa_circ_0122990 | 21392 | 44 | 0.41% |

| | | | |
|---|---|---|---|
| hsa_circ_0016000 | 5349 | 11 | 0.41% |
| hsa_circ_0104346 | 5349 | 11 | 0.41% |
| hsa_circ_0082208 | 6808 | 14 | 0.41% |
| hsa_circ_0128424 | 18482 | 38 | 0.41% |
| hsa_circ_0056531 | 6323 | 13 | 0.41% |
| hsa_circ_0117315 | 9728 | 20 | 0.41% |
| hsa_circ_0140432 | 9729 | 20 | 0.41% |
| hsa_circ_0023417 | 5351 | 11 | 0.41% |
| hsa_circ_0075041 | 7297 | 15 | 0.41% |
| hsa_circ_0013519 | 5838 | 12 | 0.41% |
| hsa_circ_0033264 | 5838 | 12 | 0.41% |
| hsa_circ_0032111 | 5352 | 11 | 0.41% |
| hsa_circ_0001476 | 9731 | 20 | 0.41% |
| hsa_circ_0036148 | 5839 | 12 | 0.41% |
| hsa_circ_0069938 | 5841 | 12 | 0.41% |
| hsa_circ_0001592 | 6330 | 13 | 0.41% |
| hsa_circ_0126734 | 7305 | 15 | 0.41% |
| hsa_circ_0094775 | 6818 | 14 | 0.41% |
| hsa_circ_0014192 | 6333 | 13 | 0.41% |
| hsa_circ_0046647 | 22899 | 47 | 0.41% |
| hsa_circ_0039075 | 6336 | 13 | 0.41% |
| hsa_circ_0122316 | 5849 | 12 | 0.41% |
| hsa_circ_0111384 | 18523 | 38 | 0.41% |
| hsa_circ_0073845 | 5850 | 12 | 0.41% |
| hsa_circ_0124525 | 6339 | 13 | 0.41% |
| hsa_circ_0087133 | 12191 | 25 | 0.41% |
| hsa_circ_0014844 | 5852 | 12 | 0.41% |
| hsa_circ_0138787 | 7317 | 15 | 0.41% |
| hsa_circ_0131873 | 12683 | 26 | 0.41% |
| hsa_circ_0062460 | 97076 | 199 | 0.41% |
| hsa_circ_0062459 | 97097 | 199 | 0.41% |
| hsa_circ_0111779 | 24888 | 51 | 0.41% |
| hsa_circ_0091853 | 6832 | 14 | 0.41% |
| hsa_circ_0077649 | 5856 | 12 | 0.41% |
| hsa_circ_0082722 | 5368 | 11 | 0.41% |
| hsa_circ_0058888 | 6833 | 14 | 0.41% |
| hsa_circ_0040455 | 6345 | 13 | 0.41% |
| hsa_circ_0132842 | 17087 | 35 | 0.41% |
| hsa_circ_0090749 | 6835 | 14 | 0.41% |
| hsa_circ_0043412 | 6836 | 14 | 0.41% |
| hsa_circ_0053599 | 6349 | 13 | 0.41% |
| hsa_circ_0020066 | 7327 | 15 | 0.41% |
| hsa_circ_0027130 | 5862 | 12 | 0.41% |
| hsa_circ_0054773 | 5862 | 12 | 0.41% |
| hsa_circ_0139728 | 6351 | 13 | 0.41% |
| hsa_circ_0004642 | 5863 | 12 | 0.41% |
| hsa_circ_0025819 | 5863 | 12 | 0.41% |
| hsa_circ_0053543 | 5375 | 11 | 0.41% |
| hsa_circ_0058249 | 6841 | 14 | 0.41% |
| hsa_circ_0140440 | 14173 | 29 | 0.41% |

| | | | |
|---|---|---|---|
| hsa_circ_0037949 | 5865 | 12 | 0.41% |
| hsa_circ_0027813 | 17596 | 36 | 0.41% |
| hsa_circ_0059356 | 7821 | 16 | 0.41% |
| hsa_circ_0035740 | 6355 | 13 | 0.41% |
| hsa_circ_0044712 | 6844 | 14 | 0.41% |
| hsa_circ_0028550 | 5378 | 11 | 0.41% |
| hsa_circ_0134746 | 6357 | 13 | 0.41% |
| hsa_circ_0053825 | 5868 | 12 | 0.41% |
| hsa_circ_0008441 | 5380 | 11 | 0.41% |
| hsa_circ_0113344 | 8805 | 18 | 0.41% |
| hsa_circ_0006300 | 14677 | 30 | 0.41% |
| hsa_circ_0111943 | 8318 | 17 | 0.41% |
| hsa_circ_0016634 | 6361 | 13 | 0.41% |
| hsa_circ_0061901 | 6852 | 14 | 0.41% |
| hsa_circ_0028434 | 6363 | 13 | 0.41% |
| hsa_circ_0061297 | 8812 | 18 | 0.41% |
| hsa_circ_0044178 | 17625 | 36 | 0.41% |
| hsa_circ_0056828 | 5875 | 12 | 0.41% |
| hsa_circ_0123183 | 19094 | 39 | 0.41% |
| hsa_circ_0096472 | 12731 | 26 | 0.41% |
| hsa_circ_0077047 | 5876 | 12 | 0.41% |
| hsa_circ_0112742 | 29870 | 61 | 0.41% |
| hsa_circ_0073240 | 5877 | 12 | 0.41% |
| hsa_circ_0053717 | 6367 | 13 | 0.41% |
| hsa_circ_0034741 | 6857 | 14 | 0.41% |
| hsa_circ_0101116 | 5878 | 12 | 0.41% |
| hsa_circ_0132412 | 5878 | 12 | 0.41% |
| hsa_circ_0120951 | 6858 | 14 | 0.41% |
| hsa_circ_0076289 | 5879 | 12 | 0.41% |
| hsa_circ_0135557 | 8330 | 17 | 0.41% |
| hsa_circ_0055207 | 6370 | 13 | 0.41% |
| hsa_circ_0089125 | 5881 | 12 | 0.41% |
| hsa_circ_0057754 | 5391 | 11 | 0.41% |
| hsa_circ_0000353 | 14703 | 30 | 0.41% |
| hsa_circ_0047011 | 6373 | 13 | 0.41% |
| hsa_circ_0124921 | 5884 | 12 | 0.41% |
| hsa_circ_0138860 | 12749 | 26 | 0.41% |
| hsa_circ_0012882 | 6375 | 13 | 0.41% |
| hsa_circ_0084317 | 6375 | 13 | 0.41% |
| hsa_circ_0100076 | 8337 | 17 | 0.41% |
| hsa_circ_0006669 | 12261 | 25 | 0.41% |
| hsa_circ_0117789 | 19128 | 39 | 0.41% |
| hsa_circ_0135558 | 8339 | 17 | 0.41% |
| hsa_circ_0023591 | 6868 | 14 | 0.41% |
| hsa_circ_0131212 | 22082 | 45 | 0.41% |
| hsa_circ_0042350 | 17670 | 36 | 0.41% |
| hsa_circ_0139383 | 6383 | 13 | 0.41% |
| hsa_circ_0020962 | 5892 | 12 | 0.41% |
| hsa_circ_0138399 | 5401 | 11 | 0.41% |
| hsa_circ_0026043 | 6384 | 13 | 0.41% |

| | | | |
|---|---|---|---|
| hsa_circ_0048709 | 8840 | 18 | 0.41% |
| hsa_circ_0119226 | 21118 | 43 | 0.41% |
| hsa_circ_0063867 | 18173 | 37 | 0.41% |
| hsa_circ_0069500 | 5403 | 11 | 0.41% |
| hsa_circ_0114141 | 6387 | 13 | 0.41% |
| hsa_circ_0101202 | 14740 | 30 | 0.41% |
| hsa_circ_0098264 | 7370 | 15 | 0.41% |
| hsa_circ_0025837 | 5896 | 12 | 0.41% |
| hsa_circ_0028552 | 5405 | 11 | 0.41% |
| hsa_circ_0068979 | 5405 | 11 | 0.41% |
| hsa_circ_0097432 | 8354 | 17 | 0.41% |
| hsa_circ_0016179 | 5897 | 12 | 0.41% |
| hsa_circ_0024462 | 5407 | 11 | 0.41% |
| hsa_circ_0023414 | 6882 | 14 | 0.41% |
| hsa_circ_0070159 | 6882 | 14 | 0.41% |
| hsa_circ_0127663 | 5408 | 11 | 0.41% |
| hsa_circ_0136237 | 12783 | 26 | 0.41% |
| hsa_circ_0073782 | 4917 | 10 | 0.41% |
| hsa_circ_0071921 | 5901 | 12 | 0.41% |
| hsa_circ_0086856 | 6393 | 13 | 0.41% |
| hsa_circ_0027614 | 6395 | 13 | 0.41% |
| hsa_circ_0048725 | 6395 | 13 | 0.41% |
| hsa_circ_0040863 | 6889 | 14 | 0.41% |
| hsa_circ_0116945 | 5905 | 12 | 0.41% |
| hsa_circ_0120235 | 10336 | 21 | 0.41% |
| hsa_circ_0045449 | 6399 | 13 | 0.41% |
| hsa_circ_0053760 | 5907 | 12 | 0.41% |
| hsa_circ_0036055 | 28064 | 57 | 0.41% |
| hsa_circ_0052644 | 6401 | 13 | 0.41% |
| hsa_circ_0088259 | 5912 | 12 | 0.41% |
| hsa_circ_0083346 | 6405 | 13 | 0.41% |
| hsa_circ_0003793 | 14781 | 30 | 0.41% |
| hsa_circ_0030481 | 7392 | 15 | 0.41% |
| hsa_circ_0100708 | 5915 | 12 | 0.41% |
| hsa_circ_0053693 | 6408 | 13 | 0.41% |
| hsa_circ_0120783 | 17255 | 35 | 0.41% |
| hsa_circ_0067220 | 32540 | 66 | 0.41% |
| hsa_circ_0018657 | 6903 | 14 | 0.41% |
| hsa_circ_0086881 | 6410 | 13 | 0.41% |
| hsa_circ_0082561 | 5917 | 12 | 0.41% |
| hsa_circ_0028364 | 5919 | 12 | 0.41% |
| hsa_circ_0045680 | 5919 | 12 | 0.41% |
| hsa_circ_0058893 | 5919 | 12 | 0.41% |
| hsa_circ_0132665 | 5920 | 12 | 0.41% |
| hsa_circ_0013835 | 5428 | 11 | 0.41% |
| hsa_circ_0134747 | 6415 | 13 | 0.41% |
| hsa_circ_0091139 | 5922 | 12 | 0.41% |
| hsa_circ_0063927 | 5430 | 11 | 0.41% |
| hsa_circ_0128956 | 6911 | 14 | 0.41% |
| hsa_circ_0026671 | 6418 | 13 | 0.41% |

| | | | |
|---|---|---|---|
| hsa_circ_0042235 | 8888 | 18 | 0.41% |
| hsa_circ_0103922 | 25188 | 51 | 0.40% |
| hsa_circ_0101196 | 14818 | 30 | 0.40% |
| hsa_circ_0011671 | 37047 | 75 | 0.40% |
| hsa_circ_0081237 | 7410 | 15 | 0.40% |
| hsa_circ_0108394 | 6916 | 14 | 0.40% |
| hsa_circ_0053530 | 5434 | 11 | 0.40% |
| hsa_circ_0069872 | 5930 | 12 | 0.40% |
| hsa_circ_0114490 | 18783 | 38 | 0.40% |
| hsa_circ_0101197 | 14830 | 30 | 0.40% |
| hsa_circ_0137235 | 8898 | 18 | 0.40% |
| hsa_circ_0061271 | 7415 | 15 | 0.40% |
| hsa_circ_0070104 | 6921 | 14 | 0.40% |
| hsa_circ_0000135 | 19777 | 40 | 0.40% |
| hsa_circ_0028736 | 5934 | 12 | 0.40% |
| hsa_circ_0073784 | 5934 | 12 | 0.40% |
| hsa_circ_0121317 | 25715 | 52 | 0.40% |
| hsa_circ_0140303 | 25716 | 52 | 0.40% |
| hsa_circ_0066666 | 8902 | 18 | 0.40% |
| hsa_circ_0009561 | 5937 | 12 | 0.40% |
| hsa_circ_0120948 | 6927 | 14 | 0.40% |
| hsa_circ_0008544 | 12866 | 26 | 0.40% |
| hsa_circ_0031180 | 5939 | 12 | 0.40% |
| hsa_circ_0085214 | 5939 | 12 | 0.40% |
| hsa_circ_0111931 | 8414 | 17 | 0.40% |
| hsa_circ_0034202 | 6435 | 13 | 0.40% |
| hsa_circ_0045445 | 6435 | 13 | 0.40% |
| hsa_circ_0032505 | 5941 | 12 | 0.40% |
| hsa_circ_0070312 | 5446 | 11 | 0.40% |
| hsa_circ_0047006 | 6437 | 13 | 0.40% |
| hsa_circ_0131961 | 6437 | 13 | 0.40% |
| hsa_circ_0073846 | 5942 | 12 | 0.40% |
| hsa_circ_0056678 | 5943 | 12 | 0.40% |
| hsa_circ_0076816 | 5943 | 12 | 0.40% |
| hsa_circ_0058004 | 6439 | 13 | 0.40% |
| hsa_circ_0016566 | 5449 | 11 | 0.40% |
| hsa_circ_0090039 | 5946 | 12 | 0.40% |
| hsa_circ_0118234 | 8424 | 17 | 0.40% |
| hsa_circ_0088840 | 6442 | 13 | 0.40% |
| hsa_circ_0021017 | 5451 | 11 | 0.40% |
| hsa_circ_0114144 | 13876 | 28 | 0.40% |
| hsa_circ_0064836 | 5947 | 12 | 0.40% |
| hsa_circ_0054770 | 5948 | 12 | 0.40% |
| hsa_circ_0103139 | 9419 | 19 | 0.40% |
| hsa_circ_0104380 | 8428 | 17 | 0.40% |
| hsa_circ_0031503 | 5950 | 12 | 0.40% |
| hsa_circ_0053784 | 6446 | 13 | 0.40% |
| hsa_circ_0097476 | 10413 | 21 | 0.40% |
| hsa_circ_0132556 | 30250 | 61 | 0.40% |
| hsa_circ_0122820 | 10911 | 22 | 0.40% |

| | | | |
|---|---|---|---|
| hsa_circ_0020417 | 5456 | 11 | 0.40% |
| hsa_circ_0068940 | 9425 | 19 | 0.40% |
| hsa_circ_0129059 | 9922 | 20 | 0.40% |
| hsa_circ_0120037 | 15381 | 31 | 0.40% |
| hsa_circ_0018167 | 5458 | 11 | 0.40% |
| hsa_circ_0013818 | 10420 | 21 | 0.40% |
| hsa_circ_0068262 | 13895 | 28 | 0.40% |
| hsa_circ_0070851 | 6948 | 14 | 0.40% |
| hsa_circ_0136668 | 5958 | 12 | 0.40% |
| hsa_circ_0138576 | 5958 | 12 | 0.40% |
| hsa_circ_0128412 | 35750 | 72 | 0.40% |
| hsa_circ_0041276 | 5462 | 11 | 0.40% |
| hsa_circ_0111043 | 5462 | 11 | 0.40% |
| hsa_circ_0074255 | 6952 | 14 | 0.40% |
| hsa_circ_0135841 | 16387 | 33 | 0.40% |
| hsa_circ_0113285 | 5959 | 12 | 0.40% |
| hsa_circ_0036795 | 6954 | 14 | 0.40% |
| hsa_circ_0033300 | 5964 | 12 | 0.40% |
| hsa_circ_0071608 | 5964 | 12 | 0.40% |
| hsa_circ_0070843 | 6959 | 14 | 0.40% |
| hsa_circ_0102111 | 8451 | 17 | 0.40% |
| hsa_circ_0033033 | 6960 | 14 | 0.40% |
| hsa_circ_0002722 | 17403 | 35 | 0.40% |
| hsa_circ_0059785 | 6464 | 13 | 0.40% |
| hsa_circ_0066316 | 5967 | 12 | 0.40% |
| hsa_circ_0128782 | 6962 | 14 | 0.40% |
| hsa_circ_0044196 | 20390 | 41 | 0.40% |
| hsa_circ_0121335 | 8455 | 17 | 0.40% |
| hsa_circ_0025801 | 6964 | 14 | 0.40% |
| hsa_circ_0098137 | 5472 | 11 | 0.40% |
| hsa_circ_0112809 | 13432 | 27 | 0.40% |
| hsa_circ_0028231 | 6468 | 13 | 0.40% |
| hsa_circ_0029987 | 6468 | 13 | 0.40% |
| hsa_circ_0135172 | 12937 | 26 | 0.40% |
| hsa_circ_0131273 | 16423 | 33 | 0.40% |
| hsa_circ_0124559 | 10451 | 21 | 0.40% |
| hsa_circ_0015849 | 5972 | 12 | 0.40% |
| hsa_circ_0056530 | 6470 | 13 | 0.40% |
| hsa_circ_0068559 | 25887 | 52 | 0.40% |
| hsa_circ_0013702 | 5974 | 12 | 0.40% |
| hsa_circ_0127256 | 5974 | 12 | 0.40% |
| hsa_circ_0125801 | 15438 | 31 | 0.40% |
| hsa_circ_0062015 | 6972 | 14 | 0.40% |
| hsa_circ_0124746 | 10459 | 21 | 0.40% |
| hsa_circ_0034600 | 6475 | 13 | 0.40% |
| hsa_circ_0039620 | 6475 | 13 | 0.40% |
| hsa_circ_0016288 | 5977 | 12 | 0.40% |
| hsa_circ_0100073 | 8468 | 17 | 0.40% |
| hsa_circ_0027647 | 5978 | 12 | 0.40% |
| hsa_circ_0087340 | 8469 | 17 | 0.40% |

| | | | |
|---|---|---|---|
| hsa_circ_0126790 | 5480 | 11 | 0.40% |
| hsa_circ_0134431 | 5480 | 11 | 0.40% |
| hsa_circ_0085196 | 6975 | 14 | 0.40% |
| hsa_circ_0114704 | 13453 | 27 | 0.40% |
| hsa_circ_0012843 | 13952 | 28 | 0.40% |
| hsa_circ_0037459 | 6478 | 13 | 0.40% |
| hsa_circ_0123188 | 21927 | 44 | 0.40% |
| hsa_circ_0053493 | 5482 | 11 | 0.40% |
| hsa_circ_0075638 | 5981 | 12 | 0.40% |
| hsa_circ_0025658 | 6480 | 13 | 0.40% |
| hsa_circ_0098124 | 30408 | 61 | 0.40% |
| hsa_circ_0018322 | 5982 | 12 | 0.40% |
| hsa_circ_0101512 | 58329 | 117 | 0.40% |
| hsa_circ_0011623 | 6481 | 13 | 0.40% |
| hsa_circ_0091646 | 5983 | 12 | 0.40% |
| hsa_circ_0112121 | 6482 | 13 | 0.40% |
| hsa_circ_0016002 | 5985 | 12 | 0.40% |
| hsa_circ_0004451 | 19952 | 40 | 0.40% |
| hsa_circ_0113739 | 7482 | 15 | 0.40% |
| hsa_circ_0066287 | 5986 | 12 | 0.40% |
| hsa_circ_0124378 | 5986 | 12 | 0.40% |
| hsa_circ_0027169 | 6485 | 13 | 0.40% |
| hsa_circ_0130390 | 15465 | 31 | 0.40% |
| hsa_circ_0046045 | 5987 | 12 | 0.40% |
| hsa_circ_0042791 | 6985 | 14 | 0.40% |
| hsa_circ_0044411 | 9980 | 20 | 0.40% |
| hsa_circ_0021994 | 7485 | 15 | 0.40% |
| hsa_circ_0084315 | 6488 | 13 | 0.40% |
| hsa_circ_0128489 | 17967 | 36 | 0.40% |
| hsa_circ_0065162 | 5989 | 12 | 0.40% |
| hsa_circ_0100833 | 15973 | 32 | 0.40% |
| hsa_circ_0032403 | 5491 | 11 | 0.40% |
| hsa_circ_0030053 | 15977 | 32 | 0.40% |
| hsa_circ_0073663 | 5992 | 12 | 0.40% |
| hsa_circ_0103917 | 16978 | 34 | 0.40% |
| hsa_circ_0036729 | 5493 | 11 | 0.40% |
| hsa_circ_0092516 | 6493 | 13 | 0.40% |
| hsa_circ_0134221 | 12988 | 26 | 0.40% |
| hsa_circ_0010645 | 6494 | 13 | 0.40% |
| hsa_circ_0035659 | 6494 | 13 | 0.40% |
| hsa_circ_0129189 | 12489 | 25 | 0.40% |
| hsa_circ_0010305 | 5995 | 12 | 0.40% |
| hsa_circ_0089789 | 5995 | 12 | 0.40% |
| hsa_circ_0061272 | 7495 | 15 | 0.40% |
| hsa_circ_0074054 | 5996 | 12 | 0.40% |
| hsa_circ_0127389 | 5996 | 12 | 0.40% |
| hsa_circ_0130729 | 5498 | 11 | 0.40% |
| hsa_circ_0140390 | 18994 | 38 | 0.40% |
| hsa_circ_0089776 | 7998 | 16 | 0.40% |
| hsa_circ_0078138 | 4999 | 10 | 0.40% |

| | | | |
|---|---|---|---|
| hsa_circ_0057324 | 5999 | 12 | 0.40% |
| hsa_circ_0033776 | 6499 | 13 | 0.40% |
| hsa_circ_0086857 | 6502 | 13 | 0.40% |
| hsa_circ_0029494 | 6002 | 12 | 0.40% |
| hsa_circ_0059996 | 5502 | 11 | 0.40% |
| hsa_circ_0117843 | 8004 | 16 | 0.40% |
| hsa_circ_0094766 | 6003 | 12 | 0.40% |
| hsa_circ_0059997 | 23015 | 46 | 0.40% |
| hsa_circ_0059791 | 6005 | 12 | 0.40% |
| hsa_circ_0099823 | 6005 | 12 | 0.40% |
| hsa_circ_0113522 | 13011 | 26 | 0.40% |
| hsa_circ_0122611 | 7006 | 14 | 0.40% |
| hsa_circ_0074312 | 5505 | 11 | 0.40% |
| hsa_circ_0135717 | 14014 | 28 | 0.40% |
| hsa_circ_0078476 | 8511 | 17 | 0.40% |
| hsa_circ_0063056 | 6008 | 12 | 0.40% |
| hsa_circ_0112796 | 12520 | 25 | 0.40% |
| hsa_circ_0060867 | 7512 | 15 | 0.40% |
| hsa_circ_0042575 | 6010 | 12 | 0.40% |
| hsa_circ_0130628 | 6010 | 12 | 0.40% |
| hsa_circ_0098924 | 7014 | 14 | 0.40% |
| hsa_circ_0016287 | 6514 | 13 | 0.40% |
| hsa_circ_0061924 | 6515 | 13 | 0.40% |
| hsa_circ_0114700 | 13533 | 27 | 0.40% |
| hsa_circ_0114634 | 9022 | 18 | 0.40% |
| hsa_circ_0032284 | 6517 | 13 | 0.40% |
| hsa_circ_0073847 | 6017 | 12 | 0.40% |
| hsa_circ_0028563 | 5516 | 11 | 0.40% |
| hsa_circ_0045436 | 5516 | 11 | 0.40% |
| hsa_circ_0064542 | 6519 | 13 | 0.40% |
| hsa_circ_0128122 | 7525 | 15 | 0.40% |
| hsa_circ_0009782 | 7024 | 14 | 0.40% |
| hsa_circ_0055233 | 7025 | 14 | 0.40% |
| hsa_circ_0058282 | 5520 | 11 | 0.40% |
| hsa_circ_0025057 | 6022 | 12 | 0.40% |
| hsa_circ_0058478 | 6022 | 12 | 0.40% |
| hsa_circ_0082646 | 13049 | 26 | 0.40% |
| hsa_circ_0031181 | 6023 | 12 | 0.40% |
| hsa_circ_0100072 | 8534 | 17 | 0.40% |
| hsa_circ_0059789 | 5522 | 11 | 0.40% |
| hsa_circ_0001591 | 14057 | 28 | 0.40% |
| hsa_circ_0139417 | 6527 | 13 | 0.40% |
| hsa_circ_0053759 | 5524 | 11 | 0.40% |
| hsa_circ_0086891 | 6530 | 13 | 0.40% |
| hsa_circ_0027653 | 6028 | 12 | 0.40% |
| hsa_circ_0091854 | 7033 | 14 | 0.40% |
| hsa_circ_0107952 | 7033 | 14 | 0.40% |
| hsa_circ_0087123 | 19594 | 39 | 0.40% |
| hsa_circ_0016630 | 6532 | 13 | 0.40% |
| hsa_circ_0132631 | 10051 | 20 | 0.40% |

| | | | |
|---|---|---|---|
| hsa_circ_0059992 | 23121 | 46 | 0.40% |
| hsa_circ_0137534 | 7540 | 15 | 0.40% |
| hsa_circ_0059614 | 6535 | 13 | 0.40% |
| hsa_circ_0068500 | 18098 | 36 | 0.40% |
| hsa_circ_0027162 | 6033 | 12 | 0.40% |
| hsa_circ_0103378 | 8047 | 16 | 0.40% |
| hsa_circ_0131451 | 16598 | 33 | 0.40% |
| hsa_circ_0053762 | 6544 | 13 | 0.40% |
| hsa_circ_0088837 | 6544 | 13 | 0.40% |
| hsa_circ_0113203 | 20137 | 40 | 0.40% |
| hsa_circ_0025863 | 9566 | 19 | 0.40% |
| hsa_circ_0069873 | 6042 | 12 | 0.40% |
| hsa_circ_0017858 | 7553 | 15 | 0.40% |
| hsa_circ_0102086 | 7555 | 15 | 0.40% |
| hsa_circ_0001961 | 9067 | 18 | 0.40% |
| hsa_circ_0061273 | 7556 | 15 | 0.40% |
| hsa_circ_0038251 | 6046 | 12 | 0.40% |
| hsa_circ_0132045 | 7559 | 15 | 0.40% |
| hsa_circ_0139119 | 7559 | 15 | 0.40% |
| hsa_circ_0083253 | 8063 | 16 | 0.40% |
| hsa_circ_0094384 | 14114 | 28 | 0.40% |
| hsa_circ_0138188 | 6553 | 13 | 0.40% |
| hsa_circ_0047045 | 6049 | 12 | 0.40% |
| hsa_circ_0098728 | 18652 | 37 | 0.40% |
| hsa_circ_0044208 | 20670 | 41 | 0.40% |
| hsa_circ_0017572 | 6554 | 13 | 0.40% |
| hsa_circ_0072168 | 6050 | 12 | 0.40% |
| hsa_circ_0123177 | 19666 | 39 | 0.40% |
| hsa_circ_0073783 | 5547 | 11 | 0.40% |
| hsa_circ_0099394 | 31265 | 62 | 0.40% |
| hsa_circ_0090745 | 7060 | 14 | 0.40% |
| hsa_circ_0053527 | 6556 | 13 | 0.40% |
| hsa_circ_0077382 | 6556 | 13 | 0.40% |
| hsa_circ_0049937 | 6052 | 12 | 0.40% |
| hsa_circ_0125153 | 6052 | 12 | 0.40% |
| hsa_circ_0043948 | 5549 | 11 | 0.40% |
| hsa_circ_0053798 | 6054 | 12 | 0.40% |
| hsa_circ_0103918 | 23713 | 47 | 0.40% |
| hsa_circ_0137987 | 14635 | 29 | 0.40% |
| hsa_circ_0048698 | 9084 | 18 | 0.40% |
| hsa_circ_0107926 | 6561 | 13 | 0.40% |
| hsa_circ_0134762 | 7066 | 14 | 0.40% |
| hsa_circ_0077202 | 6562 | 13 | 0.40% |
| hsa_circ_0015216 | 5553 | 11 | 0.40% |
| hsa_circ_0118242 | 6058 | 12 | 0.40% |
| hsa_circ_0133728 | 6563 | 13 | 0.40% |
| hsa_circ_0015848 | 6060 | 12 | 0.40% |
| hsa_circ_0077196 | 6060 | 12 | 0.40% |
| hsa_circ_0095207 | 13638 | 27 | 0.40% |
| hsa_circ_0123804 | 21721 | 43 | 0.40% |

| | | | |
|---|---|---|---|
| hsa_circ_0112188 | 6062 | 12 | 0.40% |
| hsa_circ_0114697 | 13640 | 27 | 0.40% |
| hsa_circ_0063082 | 6569 | 13 | 0.40% |
| hsa_circ_0060868 | 7580 | 15 | 0.40% |
| hsa_circ_0034733 | 7075 | 14 | 0.40% |
| hsa_circ_0042524 | 5559 | 11 | 0.40% |
| hsa_circ_0029495 | 6570 | 13 | 0.40% |
| hsa_circ_0132567 | 24259 | 48 | 0.40% |
| hsa_circ_0078094 | 6065 | 12 | 0.40% |
| hsa_circ_0092748 | 19718 | 39 | 0.40% |
| hsa_circ_0099909 | 7586 | 15 | 0.40% |
| hsa_circ_0098266 | 6070 | 12 | 0.40% |
| hsa_circ_0085791 | 6576 | 13 | 0.40% |
| hsa_circ_0035741 | 6577 | 13 | 0.40% |
| hsa_circ_0131192 | 19734 | 39 | 0.40% |
| hsa_circ_0053773 | 6578 | 13 | 0.40% |
| hsa_circ_0073891 | 6073 | 12 | 0.40% |
| hsa_circ_0083368 | 17715 | 35 | 0.40% |
| hsa_circ_0087849 | 7086 | 14 | 0.40% |
| hsa_circ_0096634 | 6580 | 13 | 0.40% |
| hsa_circ_0127235 | 6075 | 12 | 0.40% |
| hsa_circ_0085812 | 9114 | 18 | 0.39% |
| hsa_circ_0066439 | 6076 | 12 | 0.39% |
| hsa_circ_0119823 | 6076 | 12 | 0.39% |
| hsa_circ_0003654 | 14684 | 29 | 0.39% |
| hsa_circ_0100757 | 8608 | 17 | 0.39% |
| hsa_circ_0018656 | 7089 | 14 | 0.39% |
| hsa_circ_0010602 | 6583 | 13 | 0.39% |
| hsa_circ_0026672 | 6584 | 13 | 0.39% |
| hsa_circ_0071782 | 6078 | 12 | 0.39% |
| hsa_circ_0136130 | 6585 | 13 | 0.39% |
| hsa_circ_0053733 | 5573 | 11 | 0.39% |
| hsa_circ_0136865 | 66880 | 132 | 0.39% |
| hsa_circ_0034107 | 17231 | 34 | 0.39% |
| hsa_circ_0016286 | 6589 | 13 | 0.39% |
| hsa_circ_0102427 | 19769 | 39 | 0.39% |
| hsa_circ_0024458 | 5576 | 11 | 0.39% |
| hsa_circ_0128419 | 6088 | 12 | 0.39% |
| hsa_circ_0115823 | 11162 | 22 | 0.39% |
| hsa_circ_0068670 | 6089 | 12 | 0.39% |
| hsa_circ_0108511 | 15736 | 31 | 0.39% |
| hsa_circ_0138186 | 7107 | 14 | 0.39% |
| hsa_circ_0085899 | 6600 | 13 | 0.39% |
| hsa_circ_0125121 | 8124 | 16 | 0.39% |
| hsa_circ_0100070 | 8632 | 17 | 0.39% |
| hsa_circ_0015637 | 61440 | 121 | 0.39% |
| hsa_circ_0107321 | 17772 | 35 | 0.39% |
| hsa_circ_0098397 | 5080 | 10 | 0.39% |
| hsa_circ_0053600 | 6606 | 13 | 0.39% |
| hsa_circ_0100062 | 20837 | 41 | 0.39% |

| | | | |
|---|---|---|---|
| hsa_circ_0010581 | 6607 | 13 | 0.39% |
| hsa_circ_0127530 | 8132 | 16 | 0.39% |
| hsa_circ_0053544 | 5591 | 11 | 0.39% |
| hsa_circ_0009562 | 6100 | 12 | 0.39% |
| hsa_circ_0077325 | 6100 | 12 | 0.39% |
| hsa_circ_0018685 | 6609 | 13 | 0.39% |
| hsa_circ_0115737 | 25930 | 51 | 0.39% |
| hsa_circ_0109964 | 22377 | 44 | 0.39% |
| hsa_circ_0108393 | 27973 | 55 | 0.39% |
| hsa_circ_0137980 | 9664 | 19 | 0.39% |
| hsa_circ_0077651 | 6614 | 13 | 0.39% |
| hsa_circ_0044710 | 7124 | 14 | 0.39% |
| hsa_circ_0081172 | 6617 | 13 | 0.39% |
| hsa_circ_0067884 | 12224 | 24 | 0.39% |
| hsa_circ_0134222 | 13243 | 26 | 0.39% |
| hsa_circ_0094374 | 8151 | 16 | 0.39% |
| hsa_circ_0136111 | 21908 | 43 | 0.39% |
| hsa_circ_0111299 | 6114 | 12 | 0.39% |
| hsa_circ_0053616 | 6624 | 13 | 0.39% |
| hsa_circ_0129913 | 9682 | 19 | 0.39% |
| hsa_circ_0070859 | 6115 | 12 | 0.39% |
| hsa_circ_0035767 | 6625 | 13 | 0.39% |
| hsa_circ_0114692 | 13760 | 27 | 0.39% |
| hsa_circ_0054763 | 6117 | 12 | 0.39% |
| hsa_circ_0039622 | 6627 | 13 | 0.39% |
| hsa_circ_0105524 | 6627 | 13 | 0.39% |
| hsa_circ_0101779 | 6118 | 12 | 0.39% |
| hsa_circ_0010559 | 7138 | 14 | 0.39% |
| hsa_circ_0033270 | 6119 | 12 | 0.39% |
| hsa_circ_0046822 | 13258 | 26 | 0.39% |
| hsa_circ_0078312 | 7649 | 15 | 0.39% |
| hsa_circ_0134755 | 7140 | 14 | 0.39% |
| hsa_circ_0133727 | 6632 | 13 | 0.39% |
| hsa_circ_0093564 | 42343 | 83 | 0.39% |
| hsa_circ_0058835 | 6123 | 12 | 0.39% |
| hsa_circ_0112517 | 15818 | 31 | 0.39% |
| hsa_circ_0033294 | 6124 | 12 | 0.39% |
| hsa_circ_0082785 | 6124 | 12 | 0.39% |
| hsa_circ_0024199 | 6125 | 12 | 0.39% |
| hsa_circ_0108711 | 6636 | 13 | 0.39% |
| hsa_circ_0127314 | 13273 | 26 | 0.39% |
| hsa_circ_0100071 | 8679 | 17 | 0.39% |
| hsa_circ_0128251 | 6637 | 13 | 0.39% |
| hsa_circ_0009180 | 8170 | 16 | 0.39% |
| hsa_circ_0055495 | 6130 | 12 | 0.39% |
| hsa_circ_0071916 | 6132 | 12 | 0.39% |
| hsa_circ_0075987 | 7158 | 14 | 0.39% |
| hsa_circ_0135563 | 44482 | 87 | 0.39% |
| hsa_circ_0091099 | 6648 | 13 | 0.39% |
| hsa_circ_0069874 | 6137 | 12 | 0.39% |

| | | | |
|---|---|---|---|
| hsa_circ_0124957 | 7164 | 14 | 0.39% |
| hsa_circ_0134054 | 9211 | 18 | 0.39% |
| hsa_circ_0097319 | 6654 | 13 | 0.39% |
| hsa_circ_0124056 | 6654 | 13 | 0.39% |
| hsa_circ_0125537 | 22523 | 44 | 0.39% |
| hsa_circ_0134029 | 21500 | 42 | 0.39% |
| hsa_circ_0108751 | 15359 | 30 | 0.39% |
| hsa_circ_0023579 | 7168 | 14 | 0.39% |
| hsa_circ_0047216 | 5632 | 11 | 0.39% |
| hsa_circ_0114689 | 13826 | 27 | 0.39% |
| hsa_circ_0117861 | 6658 | 13 | 0.39% |
| hsa_circ_0059990 | 5634 | 11 | 0.39% |
| hsa_circ_0129981 | 5635 | 11 | 0.39% |
| hsa_circ_0052003 | 6148 | 12 | 0.39% |
| hsa_circ_0130166 | 7173 | 14 | 0.39% |
| hsa_circ_0123928 | 23571 | 46 | 0.39% |
| hsa_circ_0030293 | 6662 | 13 | 0.39% |
| hsa_circ_0063083 | 6662 | 13 | 0.39% |
| hsa_circ_0027202 | 7687 | 15 | 0.39% |
| hsa_circ_0097970 | 19987 | 39 | 0.39% |
| hsa_circ_0123929 | 23576 | 46 | 0.39% |
| hsa_circ_0114684 | 13839 | 27 | 0.39% |
| hsa_circ_0075977 | 7176 | 14 | 0.39% |
| hsa_circ_0135909 | 6664 | 13 | 0.39% |
| hsa_circ_0084358 | 6665 | 13 | 0.39% |
| hsa_circ_0079763 | 457371 | 892 | 0.39% |
| hsa_circ_0032020 | 6666 | 13 | 0.39% |
| hsa_circ_0047007 | 6667 | 13 | 0.39% |
| hsa_circ_0028733 | 5642 | 11 | 0.39% |
| hsa_circ_0108464 | 6668 | 13 | 0.39% |
| hsa_circ_0058894 | 7694 | 15 | 0.39% |
| hsa_circ_0116245 | 31804 | 62 | 0.39% |
| hsa_circ_0089547 | 9234 | 18 | 0.39% |
| hsa_circ_0127493 | 6670 | 13 | 0.39% |
| hsa_circ_0045450 | 6671 | 13 | 0.39% |
| hsa_circ_0005200 | 6158 | 12 | 0.39% |
| hsa_circ_0130872 | 6158 | 12 | 0.39% |
| hsa_circ_0029933 | 7700 | 15 | 0.39% |
| hsa_circ_0013701 | 6160 | 12 | 0.39% |
| hsa_circ_0028731 | 6160 | 12 | 0.39% |
| hsa_circ_0137297 | 9241 | 18 | 0.39% |
| hsa_circ_0027829 | 7191 | 14 | 0.39% |
| hsa_circ_0124892 | 6164 | 12 | 0.39% |
| hsa_circ_0053561 | 6678 | 13 | 0.39% |
| hsa_circ_0108386 | 6679 | 13 | 0.39% |
| hsa_circ_0025795 | 7193 | 14 | 0.39% |
| hsa_circ_0019875 | 6166 | 12 | 0.39% |
| hsa_circ_0039951 | 6166 | 12 | 0.39% |
| hsa_circ_0078475 | 8736 | 17 | 0.39% |
| hsa_circ_0025862 | 9764 | 19 | 0.39% |

| | | | |
|---|---|---|---|
| hsa_circ_0039621 | 6682 | 13 | 0.39% |
| hsa_circ_0053694 | 6682 | 13 | 0.39% |
| hsa_circ_0006920 | 6168 | 12 | 0.39% |
| hsa_circ_0066297 | 6168 | 12 | 0.39% |
| hsa_circ_0123120 | 5654 | 11 | 0.39% |
| hsa_circ_0043411 | 7197 | 14 | 0.39% |
| hsa_circ_0092747 | 20049 | 39 | 0.39% |
| hsa_circ_0012336 | 7199 | 14 | 0.39% |
| hsa_circ_0010312 | 6172 | 12 | 0.39% |
| hsa_circ_0085338 | 6687 | 13 | 0.39% |
| hsa_circ_0028559 | 5659 | 11 | 0.39% |
| hsa_circ_0010417 | 6174 | 12 | 0.39% |
| hsa_circ_0070298 | 5661 | 11 | 0.39% |
| hsa_circ_0014863 | 6691 | 13 | 0.39% |
| hsa_circ_0090741 | 6693 | 13 | 0.39% |
| hsa_circ_0009077 | 27293 | 53 | 0.39% |
| hsa_circ_0013419 | 6182 | 12 | 0.39% |
| hsa_circ_0077345 | 6699 | 13 | 0.39% |
| hsa_circ_0006119 | 23706 | 46 | 0.39% |
| hsa_circ_0027191 | 5669 | 11 | 0.39% |
| hsa_circ_0139474 | 59784 | 116 | 0.39% |
| hsa_circ_0037929 | 6700 | 13 | 0.39% |
| hsa_circ_0003415 | 6701 | 13 | 0.39% |
| hsa_circ_0005799 | 6701 | 13 | 0.39% |
| hsa_circ_0090724 | 6701 | 13 | 0.39% |
| hsa_circ_0135440 | 7733 | 15 | 0.39% |
| hsa_circ_0069023 | 6702 | 13 | 0.39% |
| hsa_circ_0135162 | 6187 | 12 | 0.39% |
| hsa_circ_0129693 | 6703 | 13 | 0.39% |
| hsa_circ_0129251 | 6704 | 13 | 0.39% |
| hsa_circ_0070863 | 6189 | 12 | 0.39% |
| hsa_circ_0107068 | 21664 | 42 | 0.39% |
| hsa_circ_0095205 | 13928 | 27 | 0.39% |
| hsa_circ_0008374 | 18571 | 36 | 0.39% |
| hsa_circ_0073500 | 11865 | 23 | 0.39% |
| hsa_circ_0096751 | 5676 | 11 | 0.39% |
| hsa_circ_0127201 | 12902 | 25 | 0.39% |
| hsa_circ_0084295 | 6193 | 12 | 0.39% |
| hsa_circ_0033961 | 51611 | 100 | 0.39% |
| hsa_circ_0107611 | 16516 | 32 | 0.39% |
| hsa_circ_0069004 | 6194 | 12 | 0.39% |
| hsa_circ_0099910 | 7745 | 15 | 0.39% |
| hsa_circ_0092759 | 6713 | 13 | 0.39% |
| hsa_circ_0079523 | 11361 | 22 | 0.39% |
| hsa_circ_0031182 | 6197 | 12 | 0.39% |
| hsa_circ_0139318 | 6197 | 12 | 0.39% |
| hsa_circ_0123972 | 20142 | 39 | 0.39% |
| hsa_circ_0043311 | 21178 | 41 | 0.39% |
| hsa_circ_0010573 | 6716 | 13 | 0.39% |
| hsa_circ_0037111 | 6716 | 13 | 0.39% |

| | | | |
|---|---|---|---|
| hsa_circ_0118585 | 13950 | 27 | 0.39% |
| hsa_circ_0096599 | 13954 | 27 | 0.39% |
| hsa_circ_0033438 | 6202 | 12 | 0.39% |
| hsa_circ_0124706 | 52207 | 101 | 0.39% |
| hsa_circ_0037432 | 6720 | 13 | 0.39% |
| hsa_circ_0067061 | 7238 | 14 | 0.39% |
| hsa_circ_0078000 | 6205 | 12 | 0.39% |
| hsa_circ_0053179 | 6724 | 13 | 0.39% |
| hsa_circ_0081334 | 23276 | 45 | 0.39% |
| hsa_circ_0055763 | 5690 | 11 | 0.39% |
| hsa_circ_0027638 | 6209 | 12 | 0.39% |
| hsa_circ_0075130 | 24319 | 47 | 0.39% |
| hsa_circ_0015611 | 10867 | 21 | 0.39% |
| hsa_circ_0050775 | 6210 | 12 | 0.39% |
| hsa_circ_0126485 | 37786 | 73 | 0.39% |
| hsa_circ_0027206 | 6730 | 13 | 0.39% |
| hsa_circ_0032404 | 5697 | 11 | 0.39% |
| hsa_circ_0080192 | 6734 | 13 | 0.39% |
| hsa_circ_0031107 | 6216 | 12 | 0.39% |
| hsa_circ_0082202 | 7253 | 14 | 0.39% |
| hsa_circ_0072884 | 6217 | 12 | 0.39% |
| hsa_circ_0053739 | 5699 | 11 | 0.39% |
| hsa_circ_0057843 | 6736 | 13 | 0.39% |
| hsa_circ_0088325 | 6737 | 13 | 0.39% |
| hsa_circ_0077352 | 6219 | 12 | 0.39% |
| hsa_circ_0070076 | 7257 | 14 | 0.39% |
| hsa_circ_0053672 | 6740 | 13 | 0.39% |
| hsa_circ_0033285 | 6222 | 12 | 0.39% |
| hsa_circ_0029980 | 6741 | 13 | 0.39% |
| hsa_circ_0029919 | 5704 | 11 | 0.39% |
| hsa_circ_0121946 | 39421 | 76 | 0.39% |
| hsa_circ_0033291 | 6225 | 12 | 0.39% |
| hsa_circ_0113423 | 6744 | 13 | 0.39% |
| hsa_circ_0092076 | 6746 | 13 | 0.39% |
| hsa_circ_0005688 | 8304 | 16 | 0.39% |
| hsa_circ_0124391 | 8304 | 16 | 0.39% |
| hsa_circ_0000502 | 7785 | 15 | 0.39% |
| hsa_circ_0033275 | 6228 | 12 | 0.39% |
| hsa_circ_0071138 | 6228 | 12 | 0.39% |
| hsa_circ_0040456 | 6750 | 13 | 0.39% |
| hsa_circ_0125267 | 11944 | 23 | 0.39% |
| hsa_circ_0016284 | 6753 | 13 | 0.39% |
| hsa_circ_0062512 | 6753 | 13 | 0.39% |
| hsa_circ_0108521 | 6753 | 13 | 0.39% |
| hsa_circ_0126351 | 23376 | 45 | 0.39% |
| hsa_circ_0117108 | 7273 | 14 | 0.38% |
| hsa_circ_0081331 | 23381 | 45 | 0.38% |
| hsa_circ_0063496 | 6235 | 12 | 0.38% |
| hsa_circ_0127695 | 20787 | 40 | 0.38% |
| hsa_circ_0035694 | 6756 | 13 | 0.38% |

| | | | |
|---|---|---|---|
| hsa_circ_0063084 | 6756 | 13 | 0.38% |
| hsa_circ_0107808 | 6758 | 13 | 0.38% |
| hsa_circ_0122782 | 18715 | 36 | 0.38% |
| hsa_circ_0104820 | 21835 | 42 | 0.38% |
| hsa_circ_0093043 | 21317 | 41 | 0.38% |
| hsa_circ_0114606 | 16638 | 32 | 0.38% |
| hsa_circ_0016440 | 9883 | 19 | 0.38% |
| hsa_circ_0073669 | 6242 | 12 | 0.38% |
| hsa_circ_0071745 | 5722 | 11 | 0.38% |
| hsa_circ_0101103 | 27050 | 52 | 0.38% |
| hsa_circ_0139475 | 60348 | 116 | 0.38% |
| hsa_circ_0078100 | 6243 | 12 | 0.38% |
| hsa_circ_0056711 | 6244 | 12 | 0.38% |
| hsa_circ_0070409 | 6244 | 12 | 0.38% |
| hsa_circ_0007339 | 6765 | 13 | 0.38% |
| hsa_circ_0126201 | 6765 | 13 | 0.38% |
| hsa_circ_0059788 | 5725 | 11 | 0.38% |
| hsa_circ_0137902 | 13014 | 25 | 0.38% |
| hsa_circ_0137901 | 13016 | 25 | 0.38% |
| hsa_circ_0011950 | 5728 | 11 | 0.38% |
| hsa_circ_0082645 | 13539 | 26 | 0.38% |
| hsa_circ_0019389 | 23433 | 45 | 0.38% |
| hsa_circ_0012880 | 6770 | 13 | 0.38% |
| hsa_circ_0027200 | 5208 | 10 | 0.38% |
| hsa_circ_0084342 | 6771 | 13 | 0.38% |
| hsa_circ_0101210 | 6771 | 13 | 0.38% |
| hsa_circ_0083278 | 5730 | 11 | 0.38% |
| hsa_circ_0127529 | 8335 | 16 | 0.38% |
| hsa_circ_0070870 | 6774 | 13 | 0.38% |
| hsa_circ_0034976 | 5732 | 11 | 0.38% |
| hsa_circ_0098790 | 5732 | 11 | 0.38% |
| hsa_circ_0063923 | 6255 | 12 | 0.38% |
| hsa_circ_0011485 | 7299 | 14 | 0.38% |
| hsa_circ_0046767 | 5735 | 11 | 0.38% |
| hsa_circ_0035754 | 6778 | 13 | 0.38% |
| hsa_circ_0066299 | 6257 | 12 | 0.38% |
| hsa_circ_0137190 | 71959 | 138 | 0.38% |
| hsa_circ_0073332 | 5736 | 11 | 0.38% |
| hsa_circ_0094739 | 6779 | 13 | 0.38% |
| hsa_circ_0028365 | 6258 | 12 | 0.38% |
| hsa_circ_0081815 | 6258 | 12 | 0.38% |
| hsa_circ_0116504 | 23994 | 46 | 0.38% |
| hsa_circ_0121964 | 44860 | 86 | 0.38% |
| hsa_circ_0132099 | 9914 | 19 | 0.38% |
| hsa_circ_0085636 | 6263 | 12 | 0.38% |
| hsa_circ_0101242 | 37582 | 72 | 0.38% |
| hsa_circ_0023590 | 7308 | 14 | 0.38% |
| hsa_circ_0047718 | 6786 | 13 | 0.38% |
| hsa_circ_0123545 | 24014 | 46 | 0.38% |
| hsa_circ_0046914 | 6787 | 13 | 0.38% |

| | | | |
|---|---|---|---|
| hsa_circ_0104046 | 15144 | 29 | 0.38% |
| hsa_circ_0072176 | 6268 | 12 | 0.38% |
| hsa_circ_0009949 | 6791 | 13 | 0.38% |
| hsa_circ_0084288 | 6269 | 12 | 0.38% |
| hsa_circ_0085193 | 7315 | 14 | 0.38% |
| hsa_circ_0113276 | 6270 | 12 | 0.38% |
| hsa_circ_0090725 | 6793 | 13 | 0.38% |
| hsa_circ_0002911 | 24562 | 47 | 0.38% |
| hsa_circ_0010288 | 6273 | 12 | 0.38% |
| hsa_circ_0033216 | 6274 | 12 | 0.38% |
| hsa_circ_0017562 | 6797 | 13 | 0.38% |
| hsa_circ_0086858 | 6797 | 13 | 0.38% |
| hsa_circ_0031504 | 7320 | 14 | 0.38% |
| hsa_circ_0063727 | 7320 | 14 | 0.38% |
| hsa_circ_0112278 | 7321 | 14 | 0.38% |
| hsa_circ_0034321 | 6800 | 13 | 0.38% |
| hsa_circ_0139476 | 60687 | 116 | 0.38% |
| hsa_circ_0015217 | 5755 | 11 | 0.38% |
| hsa_circ_0038267 | 5755 | 11 | 0.38% |
| hsa_circ_0004778 | 6803 | 13 | 0.38% |
| hsa_circ_0057755 | 6280 | 12 | 0.38% |
| hsa_circ_0073029 | 28789 | 55 | 0.38% |
| hsa_circ_0026928 | 8376 | 16 | 0.38% |
| hsa_circ_0123704 | 7329 | 14 | 0.38% |
| hsa_circ_0092702 | 15714 | 30 | 0.38% |
| hsa_circ_0042878 | 6810 | 13 | 0.38% |
| hsa_circ_0081329 | 22002 | 42 | 0.38% |
| hsa_circ_0111995 | 7337 | 14 | 0.38% |
| hsa_circ_0131290 | 19391 | 37 | 0.38% |
| hsa_circ_0053695 | 6814 | 13 | 0.38% |
| hsa_circ_0056634 | 5766 | 11 | 0.38% |
| hsa_circ_0138819 | 24637 | 47 | 0.38% |
| hsa_circ_0015222 | 5768 | 11 | 0.38% |
| hsa_circ_0132207 | 6817 | 13 | 0.38% |
| hsa_circ_0130694 | 23075 | 44 | 0.38% |
| hsa_circ_0112700 | 19929 | 38 | 0.38% |
| hsa_circ_0039949 | 6295 | 12 | 0.38% |
| hsa_circ_0035768 | 6820 | 13 | 0.38% |
| hsa_circ_0132205 | 6820 | 13 | 0.38% |
| hsa_circ_0067607 | 6821 | 13 | 0.38% |
| hsa_circ_0082643 | 13643 | 26 | 0.38% |
| hsa_circ_0072825 | 18366 | 35 | 0.38% |
| hsa_circ_0012044 | 6823 | 13 | 0.38% |
| hsa_circ_0132675 | 6299 | 12 | 0.38% |
| hsa_circ_0104146 | 6824 | 13 | 0.38% |
| hsa_circ_0016160 | 22572 | 43 | 0.38% |
| hsa_circ_0106436 | 25200 | 48 | 0.38% |
| hsa_circ_0122626 | 7350 | 14 | 0.38% |
| hsa_circ_0077332 | 6825 | 13 | 0.38% |
| hsa_circ_0090652 | 6825 | 13 | 0.38% |

| | | | |
|---|---|---|---|
| hsa_circ_0024993 | 9453 | 18 | 0.38% |
| hsa_circ_0065130 | 6302 | 12 | 0.38% |
| hsa_circ_0027615 | 6828 | 13 | 0.38% |
| hsa_circ_0053575 | 6828 | 13 | 0.38% |
| hsa_circ_0007531 | 11030 | 21 | 0.38% |
| hsa_circ_0076372 | 23113 | 44 | 0.38% |
| hsa_circ_0055291 | 18912 | 36 | 0.38% |
| hsa_circ_0009059 | 6304 | 12 | 0.38% |
| hsa_circ_0078575 | 5779 | 11 | 0.38% |
| hsa_circ_0077194 | 6306 | 12 | 0.38% |
| hsa_circ_0029280 | 5255 | 10 | 0.38% |
| hsa_circ_0068668 | 6307 | 12 | 0.38% |
| hsa_circ_0067194 | 16820 | 32 | 0.38% |
| hsa_circ_0009563 | 6308 | 12 | 0.38% |
| hsa_circ_0089361 | 6309 | 12 | 0.38% |
| hsa_circ_0105641 | 6835 | 13 | 0.38% |
| hsa_circ_0114562 | 13145 | 25 | 0.38% |
| hsa_circ_0010313 | 6310 | 12 | 0.38% |
| hsa_circ_0073286 | 6310 | 12 | 0.38% |
| hsa_circ_0025893 | 6837 | 13 | 0.38% |
| hsa_circ_0101912 | 12624 | 24 | 0.38% |
| hsa_circ_0092943 | 5786 | 11 | 0.38% |
| hsa_circ_0117494 | 25778 | 49 | 0.38% |
| hsa_circ_0130629 | 6313 | 12 | 0.38% |
| hsa_circ_0101705 | 26313 | 50 | 0.38% |
| hsa_circ_0029297 | 5790 | 11 | 0.38% |
| hsa_circ_0031184 | 7370 | 14 | 0.38% |
| hsa_circ_0015531 | 7371 | 14 | 0.38% |
| hsa_circ_0082717 | 6320 | 12 | 0.38% |
| hsa_circ_0048253 | 13695 | 26 | 0.38% |
| hsa_circ_0138776 | 10008 | 19 | 0.38% |
| hsa_circ_0010646 | 6848 | 13 | 0.38% |
| hsa_circ_0125438 | 6848 | 13 | 0.38% |
| hsa_circ_0109629 | 21071 | 40 | 0.38% |
| hsa_circ_0084219 | 6322 | 12 | 0.38% |
| hsa_circ_0119859 | 7376 | 14 | 0.38% |
| hsa_circ_0130832 | 15807 | 30 | 0.38% |
| hsa_circ_0129925 | 29510 | 56 | 0.38% |
| hsa_circ_0010541 | 7378 | 14 | 0.38% |
| hsa_circ_0006820 | 6325 | 12 | 0.38% |
| hsa_circ_0125360 | 24249 | 46 | 0.38% |
| hsa_circ_0042204 | 6854 | 13 | 0.38% |
| hsa_circ_0137463 | 58528 | 111 | 0.38% |
| hsa_circ_0096625 | 6855 | 13 | 0.38% |
| hsa_circ_0130690 | 23207 | 44 | 0.38% |
| hsa_circ_0053454 | 5802 | 11 | 0.38% |
| hsa_circ_0090736 | 7385 | 14 | 0.38% |
| hsa_circ_0030282 | 22156 | 42 | 0.38% |
| hsa_circ_0120913 | 5803 | 11 | 0.38% |
| hsa_circ_0116359 | 17938 | 34 | 0.38% |

| | | | |
|---|---|---|---|
| hsa_circ_0017168 | 7914 | 15 | 0.38% |
| hsa_circ_0134317 | 20577 | 39 | 0.38% |
| hsa_circ_0114337 | 18469 | 35 | 0.38% |
| hsa_circ_0139115 | 6333 | 12 | 0.38% |
| hsa_circ_0053176 | 6864 | 13 | 0.38% |
| hsa_circ_0091855 | 7393 | 14 | 0.38% |
| hsa_circ_0070849 | 7394 | 14 | 0.38% |
| hsa_circ_0123820 | 14790 | 28 | 0.38% |
| hsa_circ_0091806 | 7395 | 14 | 0.38% |
| hsa_circ_0097964 | 23779 | 45 | 0.38% |
| hsa_circ_0039087 | 6870 | 13 | 0.38% |
| hsa_circ_0053673 | 6870 | 13 | 0.38% |
| hsa_circ_0021630 | 20083 | 38 | 0.38% |
| hsa_circ_0055069 | 17969 | 34 | 0.38% |
| hsa_circ_0118519 | 8985 | 17 | 0.38% |
| hsa_circ_0123124 | 15328 | 29 | 0.38% |
| hsa_circ_0012483 | 12159 | 23 | 0.38% |
| hsa_circ_0124357 | 6874 | 13 | 0.38% |
| hsa_circ_0028558 | 5818 | 11 | 0.38% |
| hsa_circ_0058833 | 6348 | 12 | 0.38% |
| hsa_circ_0015692 | 6879 | 13 | 0.38% |
| hsa_circ_0116391 | 26988 | 51 | 0.38% |
| hsa_circ_0139950 | 17464 | 33 | 0.38% |
| hsa_circ_0071968 | 6880 | 13 | 0.38% |
| hsa_circ_0084343 | 6880 | 13 | 0.38% |
| hsa_circ_0012879 | 6881 | 13 | 0.38% |
| hsa_circ_0005390 | 25938 | 49 | 0.38% |
| hsa_circ_0110140 | 28066 | 53 | 0.38% |
| hsa_circ_0050779 | 6355 | 12 | 0.38% |
| hsa_circ_0125340 | 6355 | 12 | 0.38% |
| hsa_circ_0126933 | 20655 | 39 | 0.38% |
| hsa_circ_0024450 | 6356 | 12 | 0.38% |
| hsa_circ_0107525 | 7416 | 14 | 0.38% |
| hsa_circ_0115537 | 28076 | 53 | 0.38% |
| hsa_circ_0103585 | 5828 | 11 | 0.38% |
| hsa_circ_0005144 | 29671 | 56 | 0.38% |
| hsa_circ_0056529 | 6888 | 13 | 0.38% |
| hsa_circ_0024013 | 6360 | 12 | 0.38% |
| hsa_circ_0028553 | 5830 | 11 | 0.38% |
| hsa_circ_0069432 | 5830 | 11 | 0.38% |
| hsa_circ_0132749 | 9014 | 17 | 0.38% |
| hsa_circ_0134124 | 20681 | 39 | 0.38% |
| hsa_circ_0119893 | 7424 | 14 | 0.38% |
| hsa_circ_0028732 | 6364 | 12 | 0.38% |
| hsa_circ_0010258 | 6365 | 12 | 0.38% |
| hsa_circ_0072182 | 6365 | 12 | 0.38% |
| hsa_circ_0034108 | 18037 | 34 | 0.38% |
| hsa_circ_0108288 | 7429 | 14 | 0.38% |
| hsa_circ_0124713 | 10083 | 19 | 0.38% |
| hsa_circ_0082998 | 6899 | 13 | 0.38% |

| | | | |
|---|---|---|---|
| hsa_circ_0097830 | 9555 | 18 | 0.38% |
| hsa_circ_0118634 | 15929 | 30 | 0.38% |
| hsa_circ_0028721 | 6372 | 12 | 0.38% |
| hsa_circ_0074051 | 6904 | 13 | 0.38% |
| hsa_circ_0132100 | 10092 | 19 | 0.38% |
| hsa_circ_0076069 | 7968 | 15 | 0.38% |
| hsa_circ_0127842 | 14875 | 28 | 0.38% |
| hsa_circ_0114338 | 18595 | 35 | 0.38% |
| hsa_circ_0071324 | 6907 | 13 | 0.38% |
| hsa_circ_0038448 | 29226 | 55 | 0.38% |
| hsa_circ_0089846 | 6377 | 12 | 0.38% |
| hsa_circ_0121691 | 14880 | 28 | 0.38% |
| hsa_circ_0028354 | 6378 | 12 | 0.38% |
| hsa_circ_0078473 | 9038 | 17 | 0.38% |
| hsa_circ_0073848 | 6380 | 12 | 0.38% |
| hsa_circ_0112450 | 6915 | 13 | 0.38% |
| hsa_circ_0093691 | 19685 | 37 | 0.38% |
| hsa_circ_0035397 | 6917 | 13 | 0.38% |
| hsa_circ_0073333 | 5853 | 11 | 0.38% |
| hsa_circ_0074256 | 7450 | 14 | 0.38% |
| hsa_circ_0020035 | 6387 | 12 | 0.38% |
| hsa_circ_0104692 | 24484 | 46 | 0.38% |
| hsa_circ_0131445 | 28743 | 54 | 0.38% |
| hsa_circ_0124842 | 8517 | 16 | 0.38% |
| hsa_circ_0073336 | 6388 | 12 | 0.38% |
| hsa_circ_0131269 | 7453 | 14 | 0.38% |
| hsa_circ_0126272 | 21831 | 41 | 0.38% |
| hsa_circ_0073664 | 6392 | 12 | 0.38% |
| hsa_circ_0136796 | 35693 | 67 | 0.38% |
| hsa_circ_0027154 | 6393 | 12 | 0.38% |
| hsa_circ_0130493 | 6393 | 12 | 0.38% |
| hsa_circ_0111994 | 7459 | 14 | 0.38% |
| hsa_circ_0139747 | 6927 | 13 | 0.38% |
| hsa_circ_0070077 | 7460 | 14 | 0.38% |
| hsa_circ_0125189 | 10125 | 19 | 0.38% |
| hsa_circ_0045907 | 6929 | 13 | 0.38% |
| hsa_circ_0041524 | 7463 | 14 | 0.38% |
| hsa_circ_0103982 | 18660 | 35 | 0.38% |
| hsa_circ_0084267 | 6399 | 12 | 0.38% |
| hsa_circ_0125326 | 7466 | 14 | 0.38% |
| hsa_circ_0058890 | 5867 | 11 | 0.37% |
| hsa_circ_0139812 | 24541 | 46 | 0.37% |
| hsa_circ_0070091 | 7469 | 14 | 0.37% |
| hsa_circ_0109102 | 6402 | 12 | 0.37% |
| hsa_circ_0130313 | 18677 | 35 | 0.37% |
| hsa_circ_0139030 | 14943 | 28 | 0.37% |
| hsa_circ_0011901 | 6405 | 12 | 0.37% |
| hsa_circ_0070306 | 5872 | 11 | 0.37% |
| hsa_circ_0027639 | 6406 | 12 | 0.37% |
| hsa_circ_0101504 | 8542 | 16 | 0.37% |

| | | | |
|---|---|---|---|
| hsa_circ_0022181 | 5873 | 11 | 0.37% |
| hsa_circ_0082639 | 13883 | 26 | 0.37% |
| hsa_circ_0099819 | 6408 | 12 | 0.37% |
| hsa_circ_0004515 | 26167 | 49 | 0.37% |
| hsa_circ_0093692 | 19759 | 37 | 0.37% |
| hsa_circ_0012484 | 12284 | 23 | 0.37% |
| hsa_circ_0103983 | 18694 | 35 | 0.37% |
| hsa_circ_0076842 | 6944 | 13 | 0.37% |
| hsa_circ_0010314 | 6410 | 12 | 0.37% |
| hsa_circ_0086996 | 15491 | 29 | 0.37% |
| hsa_circ_0057110 | 29381 | 55 | 0.37% |
| hsa_circ_0066886 | 6411 | 12 | 0.37% |
| hsa_circ_0075506 | 7482 | 14 | 0.37% |
| hsa_circ_0014431 | 6414 | 12 | 0.37% |
| hsa_circ_0131288 | 10156 | 19 | 0.37% |
| hsa_circ_0124991 | 6951 | 13 | 0.37% |
| hsa_circ_0130558 | 6951 | 13 | 0.37% |
| hsa_circ_0034596 | 6952 | 13 | 0.37% |
| hsa_circ_0044732 | 6419 | 12 | 0.37% |
| hsa_circ_0134226 | 27281 | 51 | 0.37% |
| hsa_circ_0065990 | 7489 | 14 | 0.37% |
| hsa_circ_0015586 | 6422 | 12 | 0.37% |
| hsa_circ_0025318 | 20872 | 39 | 0.37% |
| hsa_circ_0043946 | 6958 | 13 | 0.37% |
| hsa_circ_0049815 | 6959 | 13 | 0.37% |
| hsa_circ_0071148 | 6959 | 13 | 0.37% |
| hsa_circ_0124786 | 8031 | 15 | 0.37% |
| hsa_circ_0071890 | 6425 | 12 | 0.37% |
| hsa_circ_0034203 | 6962 | 13 | 0.37% |
| hsa_circ_0021033 | 5891 | 11 | 0.37% |
| hsa_circ_0100632 | 53555 | 100 | 0.37% |
| hsa_circ_0126273 | 21960 | 41 | 0.37% |
| hsa_circ_0093640 | 14463 | 27 | 0.37% |
| hsa_circ_0116813 | 16609 | 31 | 0.37% |
| hsa_circ_0095556 | 7501 | 14 | 0.37% |
| hsa_circ_0089791 | 49839 | 93 | 0.37% |
| hsa_circ_0077324 | 6431 | 12 | 0.37% |
| hsa_circ_0056014 | 18225 | 34 | 0.37% |
| hsa_circ_0135460 | 15548 | 29 | 0.37% |
| hsa_circ_0029881 | 6970 | 13 | 0.37% |
| hsa_circ_0099186 | 6970 | 13 | 0.37% |
| hsa_circ_0131637 | 28953 | 54 | 0.37% |
| hsa_circ_0047209 | 6434 | 12 | 0.37% |
| hsa_circ_0004314 | 7507 | 14 | 0.37% |
| hsa_circ_0119093 | 12335 | 23 | 0.37% |
| hsa_circ_0015847 | 6436 | 12 | 0.37% |
| hsa_circ_0127204 | 31645 | 59 | 0.37% |
| hsa_circ_0000750 | 32184 | 60 | 0.37% |
| hsa_circ_0069501 | 5901 | 11 | 0.37% |
| hsa_circ_0056805 | 6974 | 13 | 0.37% |

| | | | |
|---|---|---|---|
| hsa_circ_0016277 | 7511 | 14 | 0.37% |
| hsa_circ_0057315 | 6440 | 12 | 0.37% |
| hsa_circ_0077321 | 6440 | 12 | 0.37% |
| hsa_circ_0111905 | 6444 | 12 | 0.37% |
| hsa_circ_0121821 | 6982 | 13 | 0.37% |
| hsa_circ_0081768 | 6985 | 13 | 0.37% |
| hsa_circ_0096077 | 18807 | 35 | 0.37% |
| hsa_circ_0122878 | 67170 | 125 | 0.37% |
| hsa_circ_0007815 | 11287 | 21 | 0.37% |
| hsa_circ_0080032 | 14513 | 27 | 0.37% |
| hsa_circ_0026579 | 6451 | 12 | 0.37% |
| hsa_circ_0087086 | 6989 | 13 | 0.37% |
| hsa_circ_0028554 | 5915 | 11 | 0.37% |
| hsa_circ_0045437 | 5915 | 11 | 0.37% |
| hsa_circ_0033243 | 6453 | 12 | 0.37% |
| hsa_circ_0077652 | 5916 | 11 | 0.37% |
| hsa_circ_0103146 | 16135 | 30 | 0.37% |
| hsa_circ_0018655 | 7531 | 14 | 0.37% |
| hsa_circ_0057107 | 29591 | 55 | 0.37% |
| hsa_circ_0031059 | 6995 | 13 | 0.37% |
| hsa_circ_0096933 | 24214 | 45 | 0.37% |
| hsa_circ_0126526 | 11840 | 22 | 0.37% |
| hsa_circ_0054571 | 5922 | 11 | 0.37% |
| hsa_circ_0097651 | 47379 | 88 | 0.37% |
| hsa_circ_0035398 | 7000 | 13 | 0.37% |
| hsa_circ_0009013 | 21001 | 39 | 0.37% |
| hsa_circ_0053674 | 7002 | 13 | 0.37% |
| hsa_circ_0053763 | 7002 | 13 | 0.37% |
| hsa_circ_0035742 | 7004 | 13 | 0.37% |
| hsa_circ_0127849 | 15087 | 28 | 0.37% |
| hsa_circ_0053648 | 7005 | 13 | 0.37% |
| hsa_circ_0118921 | 16167 | 30 | 0.37% |
| hsa_circ_0053545 | 5931 | 11 | 0.37% |
| hsa_circ_0083449 | 5931 | 11 | 0.37% |
| hsa_circ_0078101 | 6471 | 12 | 0.37% |
| hsa_circ_0026102 | 5933 | 11 | 0.37% |
| hsa_circ_0131871 | 17265 | 32 | 0.37% |
| hsa_circ_0134091 | 12949 | 24 | 0.37% |
| hsa_circ_0082786 | 6476 | 12 | 0.37% |
| hsa_circ_0078677 | 7016 | 13 | 0.37% |
| hsa_circ_0082562 | 6477 | 12 | 0.37% |
| hsa_circ_0084298 | 6477 | 12 | 0.37% |
| hsa_circ_0016610 | 7017 | 13 | 0.37% |
| hsa_circ_0124753 | 14035 | 26 | 0.37% |
| hsa_circ_0028726 | 6479 | 12 | 0.37% |
| hsa_circ_0078675 | 7019 | 13 | 0.37% |
| hsa_circ_0132976 | 7020 | 13 | 0.37% |
| hsa_circ_0111929 | 6481 | 12 | 0.37% |
| hsa_circ_0093463 | 11882 | 22 | 0.37% |
| hsa_circ_0073671 | 6483 | 12 | 0.37% |

| | | | |
|---|---|---|---|
| hsa_circ_0103147 | 16209 | 30 | 0.37% |
| hsa_circ_0095243 | 9190 | 17 | 0.37% |
| hsa_circ_0024308 | 7569 | 14 | 0.37% |
| hsa_circ_0109940 | 23252 | 43 | 0.37% |
| hsa_circ_0057102 | 29746 | 55 | 0.37% |
| hsa_circ_0071163 | 7032 | 13 | 0.37% |
| hsa_circ_0013822 | 19475 | 36 | 0.37% |
| hsa_circ_0026929 | 8656 | 16 | 0.37% |
| hsa_circ_0053718 | 7035 | 13 | 0.37% |
| hsa_circ_0091910 | 6495 | 12 | 0.37% |
| hsa_circ_0112274 | 7578 | 14 | 0.37% |
| hsa_circ_0123116 | 6496 | 12 | 0.37% |
| hsa_circ_0058534 | 7038 | 13 | 0.37% |
| hsa_circ_0061386 | 41146 | 76 | 0.37% |
| hsa_circ_0096885 | 17325 | 32 | 0.37% |
| hsa_circ_0139285 | 7041 | 13 | 0.37% |
| hsa_circ_0089784 | 30334 | 56 | 0.37% |
| hsa_circ_0010658 | 7042 | 13 | 0.37% |
| hsa_circ_0122622 | 7584 | 14 | 0.37% |
| hsa_circ_0061675 | 6502 | 12 | 0.37% |
| hsa_circ_0139552 | 8130 | 15 | 0.37% |
| hsa_circ_0065363 | 7046 | 13 | 0.37% |
| hsa_circ_0107590 | 7047 | 13 | 0.37% |
| hsa_circ_0081293 | 5964 | 11 | 0.37% |
| hsa_circ_0139400 | 16808 | 31 | 0.37% |
| hsa_circ_0010703 | 6507 | 12 | 0.37% |
| hsa_circ_0136276 | 20606 | 38 | 0.37% |
| hsa_circ_0078472 | 9219 | 17 | 0.37% |
| hsa_circ_0033288 | 6508 | 12 | 0.37% |
| hsa_circ_0034143 | 6508 | 12 | 0.37% |
| hsa_circ_0103172 | 18440 | 34 | 0.37% |
| hsa_circ_0107544 | 7051 | 13 | 0.37% |
| hsa_circ_0025820 | 6509 | 12 | 0.37% |
| hsa_circ_0101654 | 10306 | 19 | 0.37% |
| hsa_circ_0073124 | 7053 | 13 | 0.37% |
| hsa_circ_0096616 | 7053 | 13 | 0.37% |
| hsa_circ_0086171 | 15193 | 28 | 0.37% |
| hsa_circ_0026669 | 5970 | 11 | 0.37% |
| hsa_circ_0076811 | 6514 | 12 | 0.37% |
| hsa_circ_0114594 | 13034 | 24 | 0.37% |
| hsa_circ_0111989 | 7607 | 14 | 0.37% |
| hsa_circ_0008721 | 11411 | 21 | 0.37% |
| hsa_circ_0016180 | 6521 | 12 | 0.37% |
| hsa_circ_0025055 | 6522 | 12 | 0.37% |
| hsa_circ_0029882 | 7070 | 13 | 0.37% |
| hsa_circ_0035732 | 7070 | 13 | 0.37% |
| hsa_circ_0057756 | 6527 | 12 | 0.37% |
| hsa_circ_0127870 | 19583 | 36 | 0.37% |
| hsa_circ_0125435 | 6528 | 12 | 0.37% |
| hsa_circ_0058895 | 8162 | 15 | 0.37% |

| | | | |
|---|---|---|---|
| hsa_circ_0129960 | 8707 | 16 | 0.37% |
| hsa_circ_0113699 | 10884 | 20 | 0.37% |
| hsa_circ_0073302 | 7619 | 14 | 0.37% |
| hsa_circ_0087845 | 7075 | 13 | 0.37% |
| hsa_circ_0126274 | 20142 | 37 | 0.37% |
| hsa_circ_0017563 | 7077 | 13 | 0.37% |
| hsa_circ_0096119 | 17965 | 33 | 0.37% |
| hsa_circ_0119099 | 10892 | 20 | 0.37% |
| hsa_circ_0022179 | 5992 | 11 | 0.37% |
| hsa_circ_0116497 | 5992 | 11 | 0.37% |
| hsa_circ_0042896 | 7082 | 13 | 0.37% |
| hsa_circ_0102433 | 26698 | 49 | 0.37% |
| hsa_circ_0086172 | 15256 | 28 | 0.37% |
| hsa_circ_0129201 | 15802 | 29 | 0.37% |
| hsa_circ_0056723 | 7085 | 13 | 0.37% |
| hsa_circ_0019125 | 6540 | 12 | 0.37% |
| hsa_circ_0136840 | 14171 | 26 | 0.37% |
| hsa_circ_0132622 | 25617 | 47 | 0.37% |
| hsa_circ_0125238 | 19622 | 36 | 0.37% |
| hsa_circ_0037014 | 19081 | 35 | 0.37% |
| hsa_circ_0041287 | 5998 | 11 | 0.37% |
| hsa_circ_0001655 | 20725 | 38 | 0.37% |
| hsa_circ_0095240 | 9272 | 17 | 0.37% |
| hsa_circ_0039617 | 7091 | 13 | 0.37% |
| hsa_circ_0094208 | 18006 | 33 | 0.37% |
| hsa_circ_0091856 | 7639 | 14 | 0.37% |
| hsa_circ_0025644 | 6548 | 12 | 0.37% |
| hsa_circ_0124889 | 6548 | 12 | 0.37% |
| hsa_circ_0068169 | 7094 | 13 | 0.37% |
| hsa_circ_0102503 | 6549 | 12 | 0.37% |
| hsa_circ_0074048 | 7095 | 13 | 0.37% |
| hsa_circ_0101230 | 8733 | 16 | 0.37% |
| hsa_circ_0035787 | 7096 | 13 | 0.37% |
| hsa_circ_0053740 | 6552 | 12 | 0.37% |
| hsa_circ_0113279 | 6552 | 12 | 0.37% |
| hsa_circ_0104406 | 7100 | 13 | 0.37% |
| hsa_circ_0084296 | 6557 | 12 | 0.37% |
| hsa_circ_0023580 | 7650 | 14 | 0.37% |
| hsa_circ_0108848 | 51927 | 95 | 0.37% |
| hsa_circ_0039316 | 7106 | 13 | 0.37% |
| hsa_circ_0070300 | 6013 | 11 | 0.37% |
| hsa_circ_0038736 | 7107 | 13 | 0.37% |
| hsa_circ_0054774 | 6561 | 12 | 0.37% |
| hsa_circ_0053555 | 7108 | 13 | 0.37% |
| hsa_circ_0092797 | 7108 | 13 | 0.37% |
| hsa_circ_0067083 | 6015 | 11 | 0.37% |
| hsa_circ_0088104 | 7110 | 13 | 0.37% |
| hsa_circ_0095799 | 16958 | 31 | 0.37% |
| hsa_circ_0025082 | 7112 | 13 | 0.37% |
| hsa_circ_0108365 | 12038 | 22 | 0.37% |

| | | | |
|---|---|---|---|
| hsa_circ_0057094 | 30099 | 55 | 0.37% |
| hsa_circ_0110446 | 32291 | 59 | 0.37% |
| hsa_circ_0010560 | 7664 | 14 | 0.37% |
| hsa_circ_0053455 | 6022 | 11 | 0.37% |
| hsa_circ_0033311 | 6570 | 12 | 0.37% |
| hsa_circ_0082191 | 6570 | 12 | 0.37% |
| hsa_circ_0104939 | 12049 | 22 | 0.37% |
| hsa_circ_0024724 | 17526 | 32 | 0.37% |
| hsa_circ_0126748 | 31226 | 57 | 0.37% |
| hsa_circ_0032378 | 23010 | 42 | 0.37% |
| hsa_circ_0119553 | 18080 | 33 | 0.37% |
| hsa_circ_0010691 | 7671 | 14 | 0.37% |
| hsa_circ_0078313 | 8220 | 15 | 0.36% |
| hsa_circ_0092241 | 7125 | 13 | 0.36% |
| hsa_circ_0072169 | 6577 | 12 | 0.36% |
| hsa_circ_0080208 | 6577 | 12 | 0.36% |
| hsa_circ_0108827 | 11510 | 21 | 0.36% |
| hsa_circ_0133873 | 14799 | 27 | 0.36% |
| hsa_circ_0034106 | 18641 | 34 | 0.36% |
| hsa_circ_0095996 | 17002 | 31 | 0.36% |
| hsa_circ_0086859 | 7132 | 13 | 0.36% |
| hsa_circ_0134838 | 15363 | 28 | 0.36% |
| hsa_circ_0020760 | 7133 | 13 | 0.36% |
| hsa_circ_0001759 | 13171 | 24 | 0.36% |
| hsa_circ_0004836 | 6586 | 12 | 0.36% |
| hsa_circ_0136401 | 7135 | 13 | 0.36% |
| hsa_circ_0018087 | 74114 | 135 | 0.36% |
| hsa_circ_0114706 | 7686 | 14 | 0.36% |
| hsa_circ_0019282 | 7688 | 14 | 0.36% |
| hsa_circ_0090716 | 7139 | 13 | 0.36% |
| hsa_circ_0129252 | 7139 | 13 | 0.36% |
| hsa_circ_0114239 | 24177 | 44 | 0.36% |
| hsa_circ_0084309 | 7145 | 13 | 0.36% |
| hsa_circ_0034149 | 6596 | 12 | 0.36% |
| hsa_circ_0061536 | 6596 | 12 | 0.36% |
| hsa_circ_0127531 | 13742 | 25 | 0.36% |
| hsa_circ_0057092 | 30241 | 55 | 0.36% |
| hsa_circ_0038909 | 21444 | 39 | 0.36% |
| hsa_circ_0091073 | 7698 | 14 | 0.36% |
| hsa_circ_0034220 | 7149 | 13 | 0.36% |
| hsa_circ_0114533 | 10456 | 19 | 0.36% |
| hsa_circ_0091807 | 7705 | 14 | 0.36% |
| hsa_circ_0133135 | 18164 | 33 | 0.36% |
| hsa_circ_0068166 | 7157 | 13 | 0.36% |
| hsa_circ_0110166 | 15417 | 28 | 0.36% |
| hsa_circ_0018142 | 9911 | 18 | 0.36% |
| hsa_circ_0101505 | 8810 | 16 | 0.36% |
| hsa_circ_0053576 | 7159 | 13 | 0.36% |
| hsa_circ_0114592 | 13219 | 24 | 0.36% |
| hsa_circ_0104729 | 9364 | 17 | 0.36% |

| | | | |
|---|---|---|---|
| hsa_circ_0093044 | 22035 | 40 | 0.36% |
| hsa_circ_0044855 | 6611 | 12 | 0.36% |
| hsa_circ_0119881 | 8816 | 16 | 0.36% |
| hsa_circ_0036421 | 7164 | 13 | 0.36% |
| hsa_circ_0091870 | 6613 | 12 | 0.36% |
| hsa_circ_0018537 | 7165 | 13 | 0.36% |
| hsa_circ_0073849 | 6614 | 12 | 0.36% |
| hsa_circ_0133611 | 19292 | 35 | 0.36% |
| hsa_circ_0035139 | 7166 | 13 | 0.36% |
| hsa_circ_0132722 | 6064 | 11 | 0.36% |
| hsa_circ_0053675 | 7167 | 13 | 0.36% |
| hsa_circ_0082201 | 7719 | 14 | 0.36% |
| hsa_circ_0018994 | 6065 | 11 | 0.36% |
| hsa_circ_0003824 | 7168 | 13 | 0.36% |
| hsa_circ_0136651 | 8824 | 16 | 0.36% |
| hsa_circ_0076837 | 7171 | 13 | 0.36% |
| hsa_circ_0084322 | 7173 | 13 | 0.36% |
| hsa_circ_0063468 | 30354 | 55 | 0.36% |
| hsa_circ_0110862 | 13799 | 25 | 0.36% |
| hsa_circ_0035743 | 7176 | 13 | 0.36% |
| hsa_circ_0071149 | 7177 | 13 | 0.36% |
| hsa_circ_0124840 | 8837 | 16 | 0.36% |
| hsa_circ_0091911 | 6628 | 12 | 0.36% |
| hsa_circ_0009980 | 6629 | 12 | 0.36% |
| hsa_circ_0071063 | 6629 | 12 | 0.36% |
| hsa_circ_0102666 | 29279 | 53 | 0.36% |
| hsa_circ_0102476 | 21551 | 39 | 0.36% |
| hsa_circ_0060075 | 7185 | 13 | 0.36% |
| hsa_circ_0065411 | 6634 | 12 | 0.36% |
| hsa_circ_0120285 | 7188 | 13 | 0.36% |
| hsa_circ_0133630 | 12720 | 23 | 0.36% |
| hsa_circ_0053764 | 7191 | 13 | 0.36% |
| hsa_circ_0137707 | 21576 | 39 | 0.36% |
| hsa_circ_0010623 | 7192 | 13 | 0.36% |
| hsa_circ_0115989 | 6639 | 12 | 0.36% |
| hsa_circ_0062099 | 8299 | 15 | 0.36% |
| hsa_circ_0056527 | 7193 | 13 | 0.36% |
| hsa_circ_0061537 | 6641 | 12 | 0.36% |
| hsa_circ_0035399 | 7195 | 13 | 0.36% |
| hsa_circ_0124101 | 12730 | 23 | 0.36% |
| hsa_circ_0007081 | 18266 | 33 | 0.36% |
| hsa_circ_0137178 | 22701 | 41 | 0.36% |
| hsa_circ_0096892 | 11076 | 20 | 0.36% |
| hsa_circ_0094486 | 20501 | 37 | 0.36% |
| hsa_circ_0082195 | 6650 | 12 | 0.36% |
| hsa_circ_0037542 | 21060 | 38 | 0.36% |
| hsa_circ_0082061 | 6651 | 12 | 0.36% |
| hsa_circ_0090737 | 7760 | 14 | 0.36% |
| hsa_circ_0118520 | 9423 | 17 | 0.36% |
| hsa_circ_0048726 | 7207 | 13 | 0.36% |

| | | | |
|---|---|---|---|
| hsa_circ_0099391 | 10534 | 19 | 0.36% |
| hsa_circ_0108294 | 7762 | 14 | 0.36% |
| hsa_circ_0127524 | 8874 | 16 | 0.36% |
| hsa_circ_0098906 | 8875 | 16 | 0.36% |
| hsa_circ_0113419 | 6657 | 12 | 0.36% |
| hsa_circ_0008436 | 21084 | 38 | 0.36% |
| hsa_circ_0128802 | 50499 | 91 | 0.36% |
| hsa_circ_0005266 | 9990 | 18 | 0.36% |
| hsa_circ_0019527 | 6660 | 12 | 0.36% |
| hsa_circ_0134238 | 39962 | 72 | 0.36% |
| hsa_circ_0021861 | 24981 | 45 | 0.36% |
| hsa_circ_0060167 | 29978 | 54 | 0.36% |
| hsa_circ_0053459 | 7217 | 13 | 0.36% |
| hsa_circ_0138862 | 22208 | 40 | 0.36% |
| hsa_circ_0035788 | 7218 | 13 | 0.36% |
| hsa_circ_0032110 | 7774 | 14 | 0.36% |
| hsa_circ_0131160 | 15551 | 28 | 0.36% |
| hsa_circ_0102514 | 11108 | 20 | 0.36% |
| hsa_circ_0102998 | 28887 | 52 | 0.36% |
| hsa_circ_0130385 | 20555 | 37 | 0.36% |
| hsa_circ_0024451 | 6667 | 12 | 0.36% |
| hsa_circ_0073123 | 7223 | 13 | 0.36% |
| hsa_circ_0099901 | 26117 | 47 | 0.36% |
| hsa_circ_0035135 | 7224 | 13 | 0.36% |
| hsa_circ_0078582 | 6113 | 11 | 0.36% |
| hsa_circ_0103099 | 11671 | 21 | 0.36% |
| hsa_circ_0080464 | 24460 | 44 | 0.36% |
| hsa_circ_0055272 | 7783 | 14 | 0.36% |
| hsa_circ_0004739 | 16678 | 30 | 0.36% |
| hsa_circ_0123110 | 8339 | 15 | 0.36% |
| hsa_circ_0054758 | 6673 | 12 | 0.36% |
| hsa_circ_0027517 | 6675 | 12 | 0.36% |
| hsa_circ_0092760 | 7232 | 13 | 0.36% |
| hsa_circ_0129384 | 13911 | 25 | 0.36% |
| hsa_circ_0008147 | 11130 | 20 | 0.36% |
| hsa_circ_0090279 | 7791 | 14 | 0.36% |
| hsa_circ_0010603 | 7235 | 13 | 0.36% |
| hsa_circ_0115645 | 7792 | 14 | 0.36% |
| hsa_circ_0023604 | 6680 | 12 | 0.36% |
| hsa_circ_0029542 | 25607 | 46 | 0.36% |
| hsa_circ_0137975 | 7238 | 13 | 0.36% |
| hsa_circ_0069785 | 6125 | 11 | 0.36% |
| hsa_circ_0011617 | 7240 | 13 | 0.36% |
| hsa_circ_0010255 | 8355 | 15 | 0.36% |
| hsa_circ_0034227 | 7243 | 13 | 0.36% |
| hsa_circ_0135544 | 18392 | 33 | 0.36% |
| hsa_circ_0038240 | 6688 | 12 | 0.36% |
| hsa_circ_0096441 | 23966 | 43 | 0.36% |
| hsa_circ_0123479 | 7246 | 13 | 0.36% |
| hsa_circ_0114675 | 9476 | 17 | 0.36% |

| | | | |
|---|---|---|---|
| hsa_circ_0109058 | 29544 | 53 | 0.36% |
| hsa_circ_0084220 | 6691 | 12 | 0.36% |
| hsa_circ_0134239 | 6691 | 12 | 0.36% |
| hsa_circ_0002384 | 10596 | 19 | 0.36% |
| hsa_circ_0085900 | 7251 | 13 | 0.36% |
| hsa_circ_0028370 | 6698 | 12 | 0.36% |
| hsa_circ_0084284 | 6698 | 12 | 0.36% |
| hsa_circ_0071139 | 7258 | 13 | 0.36% |
| hsa_circ_0065408 | 7259 | 13 | 0.36% |
| hsa_circ_0012485 | 12844 | 23 | 0.36% |
| hsa_circ_0042792 | 7820 | 14 | 0.36% |
| hsa_circ_0033286 | 6703 | 12 | 0.36% |
| hsa_circ_0084299 | 6704 | 12 | 0.36% |
| hsa_circ_0070322 | 6146 | 11 | 0.36% |
| hsa_circ_0117361 | 17321 | 31 | 0.36% |
| hsa_circ_0062089 | 7266 | 13 | 0.36% |
| hsa_circ_0138306 | 7826 | 14 | 0.36% |
| hsa_circ_0024459 | 6150 | 11 | 0.36% |
| hsa_circ_0071917 | 6710 | 12 | 0.36% |
| hsa_circ_0118861 | 6712 | 12 | 0.36% |
| hsa_circ_0028555 | 6153 | 11 | 0.36% |
| hsa_circ_0027928 | 6154 | 11 | 0.36% |
| hsa_circ_0135966 | 15665 | 28 | 0.36% |
| hsa_circ_0064050 | 6715 | 12 | 0.36% |
| hsa_circ_0085183 | 7835 | 14 | 0.36% |
| hsa_circ_0114589 | 13435 | 24 | 0.36% |
| hsa_circ_0034156 | 6158 | 11 | 0.36% |
| hsa_circ_0136777 | 22956 | 41 | 0.36% |
| hsa_circ_0099507 | 12318 | 22 | 0.36% |
| hsa_circ_0024277 | 6159 | 11 | 0.36% |
| hsa_circ_0003170 | 14559 | 26 | 0.36% |
| hsa_circ_0070078 | 7840 | 14 | 0.36% |
| hsa_circ_0060987 | 7282 | 13 | 0.36% |
| hsa_circ_0061535 | 6722 | 12 | 0.36% |
| hsa_circ_0046912 | 7283 | 13 | 0.36% |
| hsa_circ_0021862 | 25221 | 45 | 0.36% |
| hsa_circ_0108898 | 11213 | 20 | 0.36% |
| hsa_circ_0079547 | 7290 | 13 | 0.36% |
| hsa_circ_0094487 | 20752 | 37 | 0.36% |
| hsa_circ_0034144 | 8974 | 16 | 0.36% |
| hsa_circ_0069449 | 6731 | 12 | 0.36% |
| hsa_circ_0122097 | 12902 | 23 | 0.36% |
| hsa_circ_0063085 | 7293 | 13 | 0.36% |
| hsa_circ_0094118 | 22443 | 40 | 0.36% |
| hsa_circ_0083800 | 6172 | 11 | 0.36% |
| hsa_circ_0006062 | 10102 | 18 | 0.36% |
| hsa_circ_0011814 | 7858 | 14 | 0.36% |
| hsa_circ_0084310 | 7298 | 13 | 0.36% |
| hsa_circ_0084312 | 7298 | 13 | 0.36% |
| hsa_circ_0099269 | 7298 | 13 | 0.36% |

| | | | |
|---|---|---|---|
| hsa_circ_0098285 | 15164 | 27 | 0.36% |
| hsa_circ_0029204 | 9549 | 17 | 0.36% |
| hsa_circ_0103368 | 7303 | 13 | 0.36% |
| hsa_circ_0077441 | 7304 | 13 | 0.36% |
| hsa_circ_0027174 | 7311 | 13 | 0.36% |
| hsa_circ_0118318 | 11249 | 20 | 0.36% |
| hsa_circ_0101264 | 22499 | 40 | 0.36% |
| hsa_circ_0093874 | 15187 | 27 | 0.36% |
| hsa_circ_0033279 | 6751 | 12 | 0.36% |
| hsa_circ_0071596 | 6751 | 12 | 0.36% |
| hsa_circ_0104051 | 19129 | 34 | 0.36% |
| hsa_circ_0033042 | 6755 | 12 | 0.36% |
| hsa_circ_0134389 | 6755 | 12 | 0.36% |
| hsa_circ_0018141 | 10133 | 18 | 0.36% |
| hsa_circ_0118373 | 13513 | 24 | 0.36% |
| hsa_circ_0053566 | 7320 | 13 | 0.36% |
| hsa_circ_0001611 | 38290 | 68 | 0.36% |
| hsa_circ_0137705 | 21962 | 39 | 0.36% |
| hsa_circ_0127983 | 18585 | 33 | 0.36% |
| hsa_circ_0020761 | 7322 | 13 | 0.36% |
| hsa_circ_0110665 | 28734 | 51 | 0.35% |
| hsa_circ_0053765 | 7326 | 13 | 0.35% |
| hsa_circ_0012105 | 20295 | 36 | 0.35% |
| hsa_circ_0019317 | 10150 | 18 | 0.35% |
| hsa_circ_0010654 | 7331 | 13 | 0.35% |
| hsa_circ_0108290 | 7895 | 14 | 0.35% |
| hsa_circ_0031676 | 11843 | 21 | 0.35% |
| hsa_circ_0034134 | 7333 | 13 | 0.35% |
| hsa_circ_0065103 | 13539 | 24 | 0.35% |
| hsa_circ_0053494 | 6207 | 11 | 0.35% |
| hsa_circ_0098238 | 20879 | 37 | 0.35% |
| hsa_circ_0122638 | 14108 | 25 | 0.35% |
| hsa_circ_0134487 | 8467 | 15 | 0.35% |
| hsa_circ_0132017 | 24275 | 43 | 0.35% |
| hsa_circ_0016308 | 6775 | 12 | 0.35% |
| hsa_circ_0071743 | 6211 | 11 | 0.35% |
| hsa_circ_0010582 | 7341 | 13 | 0.35% |
| hsa_circ_0134863 | 39530 | 70 | 0.35% |
| hsa_circ_0029883 | 7343 | 13 | 0.35% |
| hsa_circ_0127516 | 7345 | 13 | 0.35% |
| hsa_circ_0073328 | 6216 | 11 | 0.35% |
| hsa_circ_0122098 | 12998 | 23 | 0.35% |
| hsa_circ_0026096 | 7347 | 13 | 0.35% |
| hsa_circ_0053696 | 7350 | 13 | 0.35% |
| hsa_circ_0033301 | 6785 | 12 | 0.35% |
| hsa_circ_0110859 | 9047 | 16 | 0.35% |
| hsa_circ_0010315 | 6786 | 12 | 0.35% |
| hsa_circ_0025963 | 6787 | 12 | 0.35% |
| hsa_circ_0117103 | 9615 | 17 | 0.35% |
| hsa_circ_0123355 | 11315 | 20 | 0.35% |

| | | | |
|---|---|---|---|
| hsa_circ_0021863 | 25464 | 45 | 0.35% |
| hsa_circ_0132038 | 84324 | 149 | 0.35% |
| hsa_circ_0132832 | 18676 | 33 | 0.35% |
| hsa_circ_0023581 | 7926 | 14 | 0.35% |
| hsa_circ_0003698 | 14157 | 25 | 0.35% |
| hsa_circ_0053795 | 7362 | 13 | 0.35% |
| hsa_circ_0122542 | 16423 | 29 | 0.35% |
| hsa_circ_0008679 | 10196 | 18 | 0.35% |
| hsa_circ_0027155 | 7364 | 13 | 0.35% |
| hsa_circ_0034196 | 7364 | 13 | 0.35% |
| hsa_circ_0113737 | 8497 | 15 | 0.35% |
| hsa_circ_0017564 | 7365 | 13 | 0.35% |
| hsa_circ_0053508 | 7367 | 13 | 0.35% |
| hsa_circ_0011487 | 6801 | 12 | 0.35% |
| hsa_circ_0110483 | 31174 | 55 | 0.35% |
| hsa_circ_0086862 | 7370 | 13 | 0.35% |
| hsa_circ_0102345 | 7937 | 14 | 0.35% |
| hsa_circ_0094488 | 20977 | 37 | 0.35% |
| hsa_circ_0135575 | 38559 | 68 | 0.35% |
| hsa_circ_0113695 | 7372 | 13 | 0.35% |
| hsa_circ_0057836 | 7374 | 13 | 0.35% |
| hsa_circ_0132018 | 24392 | 43 | 0.35% |
| hsa_circ_0105968 | 32335 | 57 | 0.35% |
| hsa_circ_0050839 | 7942 | 14 | 0.35% |
| hsa_circ_0016611 | 7377 | 13 | 0.35% |
| hsa_circ_0009969 | 6810 | 12 | 0.35% |
| hsa_circ_0126145 | 32360 | 57 | 0.35% |
| hsa_circ_0120730 | 7381 | 13 | 0.35% |
| hsa_circ_0123195 | 7381 | 13 | 0.35% |
| hsa_circ_0117195 | 20443 | 36 | 0.35% |
| hsa_circ_0140436 | 13063 | 23 | 0.35% |
| hsa_circ_0120058 | 18176 | 32 | 0.35% |
| hsa_circ_0054025 | 11928 | 21 | 0.35% |
| hsa_circ_0019313 | 10224 | 18 | 0.35% |
| hsa_circ_0129083 | 28974 | 51 | 0.35% |
| hsa_circ_0002172 | 9090 | 16 | 0.35% |
| hsa_circ_0137982 | 8522 | 15 | 0.35% |
| hsa_circ_0082199 | 7960 | 14 | 0.35% |
| hsa_circ_0005715 | 18196 | 32 | 0.35% |
| hsa_circ_0101237 | 40950 | 72 | 0.35% |
| hsa_circ_0043220 | 7395 | 13 | 0.35% |
| hsa_circ_0018409 | 7964 | 14 | 0.35% |
| hsa_circ_0013120 | 6827 | 12 | 0.35% |
| hsa_circ_0035749 | 7396 | 13 | 0.35% |
| hsa_circ_0038427 | 20483 | 36 | 0.35% |
| hsa_circ_0070294 | 6259 | 11 | 0.35% |
| hsa_circ_0122538 | 16502 | 29 | 0.35% |
| hsa_circ_0077309 | 7398 | 13 | 0.35% |
| hsa_circ_0134332 | 6829 | 12 | 0.35% |
| hsa_circ_0053435 | 6261 | 11 | 0.35% |

| | | | |
|---|---|---|---|
| hsa_circ_0133365 | 19922 | 35 | 0.35% |
| hsa_circ_0081230 | 8538 | 15 | 0.35% |
| hsa_circ_0037947 | 7400 | 13 | 0.35% |
| hsa_circ_0053741 | 7400 | 13 | 0.35% |
| hsa_circ_0078090 | 6831 | 12 | 0.35% |
| hsa_circ_0124902 | 10248 | 18 | 0.35% |
| hsa_circ_0071918 | 6832 | 12 | 0.35% |
| hsa_circ_0028717 | 6834 | 12 | 0.35% |
| hsa_circ_0100949 | 14808 | 26 | 0.35% |
| hsa_circ_0026670 | 6266 | 11 | 0.35% |
| hsa_circ_0090685 | 7407 | 13 | 0.35% |
| hsa_circ_0007596 | 20515 | 36 | 0.35% |
| hsa_circ_0058828 | 6839 | 12 | 0.35% |
| hsa_circ_0094448 | 22228 | 39 | 0.35% |
| hsa_circ_0136434 | 7412 | 13 | 0.35% |
| hsa_circ_0024452 | 6842 | 12 | 0.35% |
| hsa_circ_0140166 | 23377 | 41 | 0.35% |
| hsa_circ_0087135 | 17107 | 30 | 0.35% |
| hsa_circ_0010300 | 6843 | 12 | 0.35% |
| hsa_circ_0020212 | 6843 | 12 | 0.35% |
| hsa_circ_0084221 | 6843 | 12 | 0.35% |
| hsa_circ_0069212 | 6844 | 12 | 0.35% |
| hsa_circ_0093561 | 41069 | 72 | 0.35% |
| hsa_circ_0132206 | 7416 | 13 | 0.35% |
| hsa_circ_0005433 | 18826 | 33 | 0.35% |
| hsa_circ_0115586 | 30814 | 54 | 0.35% |
| hsa_circ_0078085 | 6848 | 12 | 0.35% |
| hsa_circ_0129642 | 19974 | 35 | 0.35% |
| hsa_circ_0003052 | 9703 | 17 | 0.35% |
| hsa_circ_0008972 | 18265 | 32 | 0.35% |
| hsa_circ_0070079 | 7995 | 14 | 0.35% |
| hsa_circ_0070846 | 7995 | 14 | 0.35% |
| hsa_circ_0035587 | 6853 | 12 | 0.35% |
| hsa_circ_0139029 | 30271 | 53 | 0.35% |
| hsa_circ_0068893 | 7425 | 13 | 0.35% |
| hsa_circ_0106127 | 7425 | 13 | 0.35% |
| hsa_circ_0025078 | 7427 | 13 | 0.35% |
| hsa_circ_0056525 | 7427 | 13 | 0.35% |
| hsa_circ_0009085 | 7999 | 14 | 0.35% |
| hsa_circ_0033077 | 7999 | 14 | 0.35% |
| hsa_circ_0012221 | 24000 | 42 | 0.35% |
| hsa_circ_0122565 | 26287 | 46 | 0.35% |
| hsa_circ_0122535 | 16573 | 29 | 0.35% |
| hsa_circ_0005447 | 19432 | 34 | 0.35% |
| hsa_circ_0102747 | 7434 | 13 | 0.35% |
| hsa_circ_0033295 | 6867 | 12 | 0.35% |
| hsa_circ_0136092 | 14307 | 25 | 0.35% |
| hsa_circ_0035610 | 7440 | 13 | 0.35% |
| hsa_circ_0126636 | 26327 | 46 | 0.35% |
| hsa_circ_0101236 | 16599 | 29 | 0.35% |

| | | | |
|---|---|---|---|
| hsa_circ_0007740 | 8015 | 14 | 0.35% |
| hsa_circ_0123352 | 11452 | 20 | 0.35% |
| hsa_circ_0082975 | 7444 | 13 | 0.35% |
| hsa_circ_0106090 | 13171 | 23 | 0.35% |
| hsa_circ_0045390 | 6872 | 12 | 0.35% |
| hsa_circ_0035590 | 6875 | 12 | 0.35% |
| hsa_circ_0017171 | 8594 | 15 | 0.35% |
| hsa_circ_0103369 | 7449 | 13 | 0.35% |
| hsa_circ_0065156 | 6876 | 12 | 0.35% |
| hsa_circ_0010604 | 7450 | 13 | 0.35% |
| hsa_circ_0080701 | 6305 | 11 | 0.35% |
| hsa_circ_0027812 | 20642 | 36 | 0.35% |
| hsa_circ_0035708 | 7455 | 13 | 0.35% |
| hsa_circ_0130978 | 16061 | 28 | 0.35% |
| hsa_circ_0058250 | 8032 | 14 | 0.35% |
| hsa_circ_0008959 | 9180 | 16 | 0.35% |
| hsa_circ_0035755 | 7459 | 13 | 0.35% |
| hsa_circ_0020969 | 12624 | 22 | 0.35% |
| hsa_circ_0127422 | 14346 | 25 | 0.35% |
| hsa_circ_0090624 | 7460 | 13 | 0.35% |
| hsa_circ_0127532 | 14347 | 25 | 0.35% |
| hsa_circ_0029294 | 6313 | 11 | 0.35% |
| hsa_circ_0085074 | 8609 | 15 | 0.35% |
| hsa_circ_0093026 | 23535 | 41 | 0.35% |
| hsa_circ_0035678 | 7463 | 13 | 0.35% |
| hsa_circ_0027188 | 6315 | 11 | 0.35% |
| hsa_circ_0028551 | 6315 | 11 | 0.35% |
| hsa_circ_0004674 | 13779 | 24 | 0.35% |
| hsa_circ_0071618 | 6316 | 11 | 0.35% |
| hsa_circ_0003727 | 7465 | 13 | 0.35% |
| hsa_circ_0034140 | 7465 | 13 | 0.35% |
| hsa_circ_0046036 | 6891 | 12 | 0.35% |
| hsa_circ_0000233 | 10337 | 18 | 0.35% |
| hsa_circ_0029496 | 7466 | 13 | 0.35% |
| hsa_circ_0126406 | 16655 | 29 | 0.35% |
| hsa_circ_0065157 | 6892 | 12 | 0.35% |
| hsa_circ_0016167 | 24712 | 43 | 0.35% |
| hsa_circ_0004353 | 9770 | 17 | 0.35% |
| hsa_circ_0113691 | 14943 | 26 | 0.35% |
| hsa_circ_0000003 | 29313 | 51 | 0.35% |
| hsa_circ_0027179 | 7473 | 13 | 0.35% |
| hsa_circ_0090705 | 7473 | 13 | 0.35% |
| hsa_circ_0053649 | 7476 | 13 | 0.35% |
| hsa_circ_0015223 | 6326 | 11 | 0.35% |
| hsa_circ_0140002 | 6326 | 11 | 0.35% |
| hsa_circ_0094372 | 8052 | 14 | 0.35% |
| hsa_circ_0053742 | 7478 | 13 | 0.35% |
| hsa_circ_0024704 | 6328 | 11 | 0.35% |
| hsa_circ_0102279 | 26463 | 46 | 0.35% |
| hsa_circ_0061821 | 7479 | 13 | 0.35% |

| | | | |
|---|---|---|---|
| hsa_circ_0073385 | 6904 | 12 | 0.35% |
| hsa_circ_0056730 | 13809 | 24 | 0.35% |
| hsa_circ_0111437 | 8632 | 15 | 0.35% |
| hsa_circ_0112843 | 20717 | 36 | 0.35% |
| hsa_circ_0089861 | 6906 | 12 | 0.35% |
| hsa_circ_0057837 | 7484 | 13 | 0.35% |
| hsa_circ_0011836 | 6909 | 12 | 0.35% |
| hsa_circ_0078840 | 12675 | 22 | 0.35% |
| hsa_circ_0060073 | 7491 | 13 | 0.35% |
| hsa_circ_0072663 | 7491 | 13 | 0.35% |
| hsa_circ_0077819 | 7491 | 13 | 0.35% |
| hsa_circ_0015212 | 6916 | 12 | 0.35% |
| hsa_circ_0129905 | 59950 | 104 | 0.35% |
| hsa_circ_0007119 | 35166 | 61 | 0.35% |
| hsa_circ_0096330 | 89951 | 156 | 0.35% |
| hsa_circ_0076828 | 7497 | 13 | 0.35% |
| hsa_circ_0114695 | 20761 | 36 | 0.35% |
| hsa_circ_0114742 | 17302 | 30 | 0.35% |
| hsa_circ_0035429 | 12112 | 21 | 0.35% |
| hsa_circ_0092851 | 12112 | 21 | 0.35% |
| hsa_circ_0069125 | 7499 | 13 | 0.35% |
| hsa_circ_0007533 | 21344 | 37 | 0.35% |
| hsa_circ_0030651 | 9809 | 17 | 0.35% |
| hsa_circ_0046033 | 9234 | 16 | 0.35% |
| hsa_circ_0125370 | 28868 | 50 | 0.35% |
| hsa_circ_0103082 | 27136 | 47 | 0.35% |
| hsa_circ_0123349 | 11550 | 20 | 0.35% |
| hsa_circ_0120378 | 15593 | 27 | 0.35% |
| hsa_circ_0030474 | 8666 | 15 | 0.35% |
| hsa_circ_0070289 | 6934 | 12 | 0.35% |
| hsa_circ_0073028 | 24847 | 43 | 0.35% |
| hsa_circ_0137782 | 16758 | 29 | 0.35% |
| hsa_circ_0077314 | 7513 | 13 | 0.35% |
| hsa_circ_0113280 | 6936 | 12 | 0.35% |
| hsa_circ_0104663 | 11562 | 20 | 0.35% |
| hsa_circ_0037450 | 6938 | 12 | 0.35% |
| hsa_circ_0100274 | 23127 | 40 | 0.35% |
| hsa_circ_0053623 | 7517 | 13 | 0.35% |
| hsa_circ_0126819 | 20818 | 36 | 0.35% |
| hsa_circ_0077322 | 6940 | 12 | 0.35% |
| hsa_circ_0005044 | 7521 | 13 | 0.35% |
| hsa_circ_0013823 | 20834 | 36 | 0.35% |
| hsa_circ_0127811 | 21995 | 38 | 0.35% |
| hsa_circ_0120952 | 8104 | 14 | 0.35% |
| hsa_circ_0065401 | 7532 | 13 | 0.35% |
| hsa_circ_0053518 | 6374 | 11 | 0.35% |
| hsa_circ_0127844 | 8694 | 15 | 0.35% |
| hsa_circ_0009962 | 6957 | 12 | 0.34% |
| hsa_circ_0106991 | 27250 | 47 | 0.34% |
| hsa_circ_0066904 | 6958 | 12 | 0.34% |

| | | | |
|---|---|---|---|
| hsa_circ_0053676 | 7538 | 13 | 0.34% |
| hsa_circ_0078089 | 6961 | 12 | 0.34% |
| hsa_circ_0128476 | 14506 | 25 | 0.34% |
| hsa_circ_0093722 | 11605 | 20 | 0.34% |
| hsa_circ_0108732 | 15676 | 27 | 0.34% |
| hsa_circ_0094560 | 23226 | 40 | 0.34% |
| hsa_circ_0008825 | 29034 | 50 | 0.34% |
| hsa_circ_0135584 | 9291 | 16 | 0.34% |
| hsa_circ_0034302 | 7550 | 13 | 0.34% |
| hsa_circ_0104480 | 21489 | 37 | 0.34% |
| hsa_circ_0032452 | 8135 | 14 | 0.34% |
| hsa_circ_0030478 | 8720 | 15 | 0.34% |
| hsa_circ_0053456 | 6395 | 11 | 0.34% |
| hsa_circ_0005858 | 9302 | 16 | 0.34% |
| hsa_circ_0099157 | 11050 | 19 | 0.34% |
| hsa_circ_0099009 | 20356 | 35 | 0.34% |
| hsa_circ_0046335 | 7563 | 13 | 0.34% |
| hsa_circ_0084262 | 6983 | 12 | 0.34% |
| hsa_circ_0014742 | 7565 | 13 | 0.34% |
| hsa_circ_0076869 | 29105 | 50 | 0.34% |
| hsa_circ_0137040 | 6986 | 12 | 0.34% |
| hsa_circ_0004561 | 8151 | 14 | 0.34% |
| hsa_circ_0086860 | 7569 | 13 | 0.34% |
| hsa_circ_0084222 | 6987 | 12 | 0.34% |
| hsa_circ_0095784 | 7570 | 13 | 0.34% |
| hsa_circ_0004419 | 6988 | 12 | 0.34% |
| hsa_circ_0123346 | 7571 | 13 | 0.34% |
| hsa_circ_0132754 | 11649 | 20 | 0.34% |
| hsa_circ_0000354 | 7575 | 13 | 0.34% |
| hsa_circ_0121848 | 6993 | 12 | 0.34% |
| hsa_circ_0029408 | 19231 | 33 | 0.34% |
| hsa_circ_0118860 | 6994 | 12 | 0.34% |
| hsa_circ_0100950 | 15161 | 26 | 0.34% |
| hsa_circ_0135583 | 9330 | 16 | 0.34% |
| hsa_circ_0123083 | 21579 | 37 | 0.34% |
| hsa_circ_0129253 | 7584 | 13 | 0.34% |
| hsa_circ_0027195 | 8756 | 15 | 0.34% |
| hsa_circ_0024200 | 7005 | 12 | 0.34% |
| hsa_circ_0054745 | 7589 | 13 | 0.34% |
| hsa_circ_0130877 | 14013 | 24 | 0.34% |
| hsa_circ_0134078 | 21021 | 36 | 0.34% |
| hsa_circ_0129324 | 21606 | 37 | 0.34% |
| hsa_circ_0010647 | 7592 | 13 | 0.34% |
| hsa_circ_0114712 | 14602 | 25 | 0.34% |
| hsa_circ_0103731 | 16941 | 29 | 0.34% |
| hsa_circ_0115525 | 9931 | 17 | 0.34% |
| hsa_circ_0108287 | 8179 | 14 | 0.34% |
| hsa_circ_0126175 | 15192 | 26 | 0.34% |
| hsa_circ_0045389 | 7013 | 12 | 0.34% |
| hsa_circ_0088170 | 7598 | 13 | 0.34% |

| | | | |
|---|---|---|---|
| hsa_circ_0092947 | 8770 | 15 | 0.34% |
| hsa_circ_0065158 | 7016 | 12 | 0.34% |
| hsa_circ_0070895 | 7017 | 12 | 0.34% |
| hsa_circ_0107496 | 7017 | 12 | 0.34% |
| hsa_circ_0129520 | 8772 | 15 | 0.34% |
| hsa_circ_0000627 | 21638 | 37 | 0.34% |
| hsa_circ_0020318 | 7605 | 13 | 0.34% |
| hsa_circ_0113277 | 7020 | 12 | 0.34% |
| hsa_circ_0003365 | 10531 | 18 | 0.34% |
| hsa_circ_0132570 | 28085 | 48 | 0.34% |
| hsa_circ_0137572 | 10532 | 18 | 0.34% |
| hsa_circ_0118863 | 7022 | 12 | 0.34% |
| hsa_circ_0132214 | 8193 | 14 | 0.34% |
| hsa_circ_0025894 | 12880 | 22 | 0.34% |
| hsa_circ_0024326 | 34543 | 59 | 0.34% |
| hsa_circ_0055236 | 8198 | 14 | 0.34% |
| hsa_circ_0084275 | 7029 | 12 | 0.34% |
| hsa_circ_0137649 | 6445 | 11 | 0.34% |
| hsa_circ_0024192 | 7617 | 13 | 0.34% |
| hsa_circ_0032107 | 8204 | 14 | 0.34% |
| hsa_circ_0017169 | 8791 | 15 | 0.34% |
| hsa_circ_0092848 | 12311 | 21 | 0.34% |
| hsa_circ_0066221 | 7035 | 12 | 0.34% |
| hsa_circ_0138904 | 27556 | 47 | 0.34% |
| hsa_circ_0061801 | 6450 | 11 | 0.34% |
| hsa_circ_0093243 | 28149 | 48 | 0.34% |
| hsa_circ_0119040 | 9387 | 16 | 0.34% |
| hsa_circ_0020762 | 7627 | 13 | 0.34% |
| hsa_circ_0125225 | 7629 | 13 | 0.34% |
| hsa_circ_0020208 | 7043 | 12 | 0.34% |
| hsa_circ_0124958 | 8218 | 14 | 0.34% |
| hsa_circ_0070080 | 8222 | 14 | 0.34% |
| hsa_circ_0105161 | 19383 | 33 | 0.34% |
| hsa_circ_0129245 | 15859 | 27 | 0.34% |
| hsa_circ_0113499 | 9986 | 17 | 0.34% |
| hsa_circ_0115626 | 19392 | 33 | 0.34% |
| hsa_circ_0138734 | 24688 | 42 | 0.34% |
| hsa_circ_0127777 | 7642 | 13 | 0.34% |
| hsa_circ_0126590 | 61730 | 105 | 0.34% |
| hsa_circ_0132189 | 7057 | 12 | 0.34% |
| hsa_circ_0070292 | 6472 | 11 | 0.34% |
| hsa_circ_0060795 | 7061 | 12 | 0.34% |
| hsa_circ_0013142 | 12358 | 21 | 0.34% |
| hsa_circ_0033260 | 7062 | 12 | 0.34% |
| hsa_circ_0129416 | 10005 | 17 | 0.34% |
| hsa_circ_0096469 | 11184 | 19 | 0.34% |
| hsa_circ_0119570 | 30021 | 51 | 0.34% |
| hsa_circ_0033244 | 7064 | 12 | 0.34% |
| hsa_circ_0033276 | 7065 | 12 | 0.34% |
| hsa_circ_0114708 | 14719 | 25 | 0.34% |

| | | | |
|---|---|---|---|
| hsa_circ_0129100 | 28261 | 48 | 0.34% |
| hsa_circ_0055070 | 20022 | 34 | 0.34% |
| hsa_circ_0123717 | 17078 | 29 | 0.34% |
| hsa_circ_0087146 | 24751 | 42 | 0.34% |
| hsa_circ_0122174 | 10608 | 18 | 0.34% |
| hsa_circ_0087252 | 7072 | 12 | 0.34% |
| hsa_circ_0069545 | 7664 | 13 | 0.34% |
| hsa_circ_0055289 | 21229 | 36 | 0.34% |
| hsa_circ_0089779 | 14743 | 25 | 0.34% |
| hsa_circ_0027142 | 7077 | 12 | 0.34% |
| hsa_circ_0054742 | 7668 | 13 | 0.34% |
| hsa_circ_0124904 | 8260 | 14 | 0.34% |
| hsa_circ_0039097 | 8852 | 15 | 0.34% |
| hsa_circ_0016164 | 25381 | 43 | 0.34% |
| hsa_circ_0035687 | 7674 | 13 | 0.34% |
| hsa_circ_0090129 | 7674 | 13 | 0.34% |
| hsa_circ_0071598 | 7085 | 12 | 0.34% |
| hsa_circ_0073311 | 8266 | 14 | 0.34% |
| hsa_circ_0120949 | 8267 | 14 | 0.34% |
| hsa_circ_0138663 | 25399 | 43 | 0.34% |
| hsa_circ_0010648 | 7679 | 13 | 0.34% |
| hsa_circ_0069502 | 6498 | 11 | 0.34% |
| hsa_circ_0052335 | 7089 | 12 | 0.34% |
| hsa_circ_0132188 | 7089 | 12 | 0.34% |
| hsa_circ_0065404 | 7681 | 13 | 0.34% |
| hsa_circ_0126460 | 18324 | 31 | 0.34% |
| hsa_circ_0093244 | 28375 | 48 | 0.34% |
| hsa_circ_0070316 | 6503 | 11 | 0.34% |
| hsa_circ_0052894 | 7687 | 13 | 0.34% |
| hsa_circ_0138664 | 25430 | 43 | 0.34% |
| hsa_circ_0077350 | 7097 | 12 | 0.34% |
| hsa_circ_0113676 | 19519 | 33 | 0.34% |
| hsa_circ_0095786 | 7690 | 13 | 0.34% |
| hsa_circ_0054537 | 7099 | 12 | 0.34% |
| hsa_circ_0109860 | 25439 | 43 | 0.34% |
| hsa_circ_0127357 | 19526 | 33 | 0.34% |
| hsa_circ_0085334 | 7693 | 13 | 0.34% |
| hsa_circ_0105984 | 37288 | 63 | 0.34% |
| hsa_circ_0053495 | 7695 | 13 | 0.34% |
| hsa_circ_0110860 | 9471 | 16 | 0.34% |
| hsa_circ_0124092 | 12431 | 21 | 0.34% |
| hsa_circ_0134593 | 26644 | 45 | 0.34% |
| hsa_circ_0106405 | 6517 | 11 | 0.34% |
| hsa_circ_0000938 | 34957 | 59 | 0.34% |
| hsa_circ_0096967 | 20146 | 34 | 0.34% |
| hsa_circ_0002203 | 19558 | 33 | 0.34% |
| hsa_circ_0124916 | 17783 | 30 | 0.34% |
| hsa_circ_0000727 | 13043 | 22 | 0.34% |
| hsa_circ_0131477 | 16603 | 28 | 0.34% |
| hsa_circ_0100418 | 16012 | 27 | 0.34% |

| | | | |
|---|---|---|---|
| hsa_circ_0094454 | 23131 | 39 | 0.34% |
| hsa_circ_0090717 | 7713 | 13 | 0.34% |
| hsa_circ_0053457 | 6527 | 11 | 0.34% |
| hsa_circ_0045388 | 7121 | 12 | 0.34% |
| hsa_circ_0091128 | 7718 | 13 | 0.34% |
| hsa_circ_0122532 | 17218 | 29 | 0.34% |
| hsa_circ_0094116 | 23749 | 40 | 0.34% |
| hsa_circ_0133307 | 24344 | 41 | 0.34% |
| hsa_circ_0053719 | 7719 | 13 | 0.34% |
| hsa_circ_0132594 | 7720 | 13 | 0.34% |
| hsa_circ_0112185 | 7128 | 12 | 0.34% |
| hsa_circ_0123329 | 19603 | 33 | 0.34% |
| hsa_circ_0018316 | 7130 | 12 | 0.34% |
| hsa_circ_0109841 | 21990 | 37 | 0.34% |
| hsa_circ_0122616 | 7729 | 13 | 0.34% |
| hsa_circ_0035714 | 7732 | 13 | 0.34% |
| hsa_circ_0095855 | 8923 | 15 | 0.34% |
| hsa_circ_0055071 | 20226 | 34 | 0.34% |
| hsa_circ_0106878 | 45818 | 77 | 0.34% |
| hsa_circ_0125699 | 13091 | 22 | 0.34% |
| hsa_circ_0034109 | 14282 | 24 | 0.34% |
| hsa_circ_0038411 | 32731 | 55 | 0.34% |
| hsa_circ_0029292 | 6547 | 11 | 0.34% |
| hsa_circ_0114060 | 7741 | 13 | 0.34% |
| hsa_circ_0066288 | 7146 | 12 | 0.34% |
| hsa_circ_0126939 | 18462 | 31 | 0.34% |
| hsa_circ_0013824 | 21448 | 36 | 0.34% |
| hsa_circ_0128801 | 17880 | 30 | 0.34% |
| hsa_circ_0011806 | 8347 | 14 | 0.34% |
| hsa_circ_0035679 | 7751 | 13 | 0.34% |
| hsa_circ_0017186 | 46511 | 78 | 0.34% |
| hsa_circ_0001351 | 7752 | 13 | 0.34% |
| hsa_circ_0092553 | 7752 | 13 | 0.34% |
| hsa_circ_0105611 | 16697 | 28 | 0.34% |
| hsa_circ_0129943 | 7753 | 13 | 0.34% |
| hsa_circ_0132743 | 57852 | 97 | 0.34% |
| hsa_circ_0017571 | 7754 | 13 | 0.34% |
| hsa_circ_0021030 | 7158 | 12 | 0.34% |
| hsa_circ_0015312 | 10141 | 17 | 0.34% |
| hsa_circ_0140725 | 24460 | 41 | 0.34% |
| hsa_circ_0082081 | 10742 | 18 | 0.34% |
| hsa_circ_0096701 | 85346 | 143 | 0.34% |
| hsa_circ_0102439 | 9551 | 16 | 0.34% |
| hsa_circ_0013825 | 21500 | 36 | 0.33% |
| hsa_circ_0108247 | 22697 | 38 | 0.33% |
| hsa_circ_0034044 | 17323 | 29 | 0.33% |
| hsa_circ_0118442 | 7766 | 13 | 0.33% |
| hsa_circ_0090719 | 7767 | 13 | 0.33% |
| hsa_circ_0060796 | 7170 | 12 | 0.33% |
| hsa_circ_0113281 | 7170 | 12 | 0.33% |

| | | | |
|---|---|---|---|
| hsa_circ_0035697 | 7769 | 13 | 0.33% |
| hsa_circ_0028355 | 7172 | 12 | 0.33% |
| hsa_circ_0016156 | 25700 | 43 | 0.33% |
| hsa_circ_0122429 | 31081 | 52 | 0.33% |
| hsa_circ_0104481 | 22116 | 37 | 0.33% |
| hsa_circ_0035689 | 7771 | 13 | 0.33% |
| hsa_circ_0094472 | 22720 | 38 | 0.33% |
| hsa_circ_0062818 | 8371 | 14 | 0.33% |
| hsa_circ_0034112 | 8373 | 14 | 0.33% |
| hsa_circ_0103997 | 10767 | 18 | 0.33% |
| hsa_circ_0004185 | 23927 | 40 | 0.33% |
| hsa_circ_0126857 | 11964 | 20 | 0.33% |
| hsa_circ_0091866 | 7179 | 12 | 0.33% |
| hsa_circ_0133769 | 7778 | 13 | 0.33% |
| hsa_circ_0091857 | 8377 | 14 | 0.33% |
| hsa_circ_0037449 | 7182 | 12 | 0.33% |
| hsa_circ_0126858 | 11972 | 20 | 0.33% |
| hsa_circ_0117964 | 18561 | 31 | 0.33% |
| hsa_circ_0053726 | 7784 | 13 | 0.33% |
| hsa_circ_0124901 | 8384 | 14 | 0.33% |
| hsa_circ_0138644 | 25756 | 43 | 0.33% |
| hsa_circ_0033296 | 7189 | 12 | 0.33% |
| hsa_circ_0041277 | 7192 | 12 | 0.33% |
| hsa_circ_0075066 | 17985 | 30 | 0.33% |
| hsa_circ_0118958 | 8994 | 15 | 0.33% |
| hsa_circ_0010605 | 7795 | 13 | 0.33% |
| hsa_circ_0085079 | 8996 | 15 | 0.33% |
| hsa_circ_0002741 | 7198 | 12 | 0.33% |
| hsa_circ_0138861 | 15596 | 26 | 0.33% |
| hsa_circ_0025086 | 7798 | 13 | 0.33% |
| hsa_circ_0045387 | 7202 | 12 | 0.33% |
| hsa_circ_0059386 | 8403 | 14 | 0.33% |
| hsa_circ_0134140 | 20408 | 34 | 0.33% |
| hsa_circ_0037444 | 7204 | 12 | 0.33% |
| hsa_circ_0069060 | 11410 | 19 | 0.33% |
| hsa_circ_0050831 | 7807 | 13 | 0.33% |
| hsa_circ_0101509 | 39038 | 65 | 0.33% |
| hsa_circ_0042221 | 10811 | 18 | 0.33% |
| hsa_circ_0053624 | 7814 | 13 | 0.33% |
| hsa_circ_0071881 | 8417 | 14 | 0.33% |
| hsa_circ_0000766 | 46301 | 77 | 0.33% |
| hsa_circ_0073183 | 7818 | 13 | 0.33% |
| hsa_circ_0101644 | 31875 | 53 | 0.33% |
| hsa_circ_0127242 | 22859 | 38 | 0.33% |
| hsa_circ_0033281 | 7219 | 12 | 0.33% |
| hsa_circ_0132190 | 7219 | 12 | 0.33% |
| hsa_circ_0127243 | 22864 | 38 | 0.33% |
| hsa_circ_0053496 | 7823 | 13 | 0.33% |
| hsa_circ_0129337 | 7823 | 13 | 0.33% |
| hsa_circ_0091808 | 8426 | 14 | 0.33% |

| | | | |
|---|---|---|---|
| hsa_circ_0123764 | 27697 | 46 | 0.33% |
| hsa_circ_0115777 | 10838 | 18 | 0.33% |
| hsa_circ_0021034 | 9033 | 15 | 0.33% |
| hsa_circ_0120494 | 8432 | 14 | 0.33% |
| hsa_circ_0016073 | 7830 | 13 | 0.33% |
| hsa_circ_0006263 | 22891 | 38 | 0.33% |
| hsa_circ_0011902 | 7229 | 12 | 0.33% |
| hsa_circ_0094471 | 22893 | 38 | 0.33% |
| hsa_circ_0033280 | 7232 | 12 | 0.33% |
| hsa_circ_0111537 | 11451 | 19 | 0.33% |
| hsa_circ_0075818 | 25316 | 42 | 0.33% |
| hsa_circ_0037440 | 7836 | 13 | 0.33% |
| hsa_circ_0055237 | 8440 | 14 | 0.33% |
| hsa_circ_0132019 | 74771 | 124 | 0.33% |
| hsa_circ_0029497 | 7840 | 13 | 0.33% |
| hsa_circ_0114165 | 12065 | 20 | 0.33% |
| hsa_circ_0092946 | 9050 | 15 | 0.33% |
| hsa_circ_0033034 | 8448 | 14 | 0.33% |
| hsa_circ_0046719 | 7845 | 13 | 0.33% |
| hsa_circ_0132020 | 74830 | 124 | 0.33% |
| hsa_circ_0114009 | 21125 | 35 | 0.33% |
| hsa_circ_0000143 | 73036 | 121 | 0.33% |
| hsa_circ_0010245 | 6640 | 11 | 0.33% |
| hsa_circ_0053589 | 7850 | 13 | 0.33% |
| hsa_circ_0012655 | 9058 | 15 | 0.33% |
| hsa_circ_0001597 | 62204 | 103 | 0.33% |
| hsa_circ_0109051 | 27177 | 45 | 0.33% |
| hsa_circ_0089778 | 15099 | 25 | 0.33% |
| hsa_circ_0044184 | 32010 | 53 | 0.33% |
| hsa_circ_0102430 | 24164 | 40 | 0.33% |
| hsa_circ_0010294 | 7253 | 12 | 0.33% |
| hsa_circ_0008551 | 10276 | 17 | 0.33% |
| hsa_circ_0010289 | 7254 | 12 | 0.33% |
| hsa_circ_0105529 | 16322 | 27 | 0.33% |
| hsa_circ_0108733 | 16322 | 27 | 0.33% |
| hsa_circ_0108424 | 16930 | 28 | 0.33% |
| hsa_circ_0090873 | 7256 | 12 | 0.33% |
| hsa_circ_0112114 | 25398 | 42 | 0.33% |
| hsa_circ_0101262 | 11493 | 19 | 0.33% |
| hsa_circ_0031409 | 13308 | 22 | 0.33% |
| hsa_circ_0017918 | 12707 | 21 | 0.33% |
| hsa_circ_0125416 | 8472 | 14 | 0.33% |
| hsa_circ_0132698 | 45389 | 75 | 0.33% |
| hsa_circ_0094348 | 7868 | 13 | 0.33% |
| hsa_circ_0076267 | 9079 | 15 | 0.33% |
| hsa_circ_0112112 | 25426 | 42 | 0.33% |
| hsa_circ_0117362 | 21190 | 35 | 0.33% |
| hsa_circ_0085071 | 9082 | 15 | 0.33% |
| hsa_circ_0125339 | 71449 | 118 | 0.33% |
| hsa_circ_0138005 | 18165 | 30 | 0.33% |

| | | | |
|---|---|---|---|
| hsa_circ_0035769 | 7872 | 13 | 0.33% |
| hsa_circ_0067139 | 7872 | 13 | 0.33% |
| hsa_circ_0076833 | 7872 | 13 | 0.33% |
| hsa_circ_0085102 | 6661 | 11 | 0.33% |
| hsa_circ_0111778 | 30889 | 51 | 0.33% |
| hsa_circ_0037455 | 7874 | 13 | 0.33% |
| hsa_circ_0113359 | 7874 | 13 | 0.33% |
| hsa_circ_0127838 | 8480 | 14 | 0.33% |
| hsa_circ_0128858 | 30904 | 51 | 0.33% |
| hsa_circ_0093554 | 7878 | 13 | 0.33% |
| hsa_circ_0125512 | 20605 | 34 | 0.33% |
| hsa_circ_0105527 | 16363 | 27 | 0.33% |
| hsa_circ_0016775 | 7881 | 13 | 0.33% |
| hsa_circ_0064995 | 10913 | 18 | 0.33% |
| hsa_circ_0051048 | 9096 | 15 | 0.33% |
| hsa_circ_0085066 | 9099 | 15 | 0.33% |
| hsa_circ_0123323 | 41858 | 69 | 0.33% |
| hsa_circ_0081287 | 7281 | 12 | 0.33% |
| hsa_circ_0053720 | 7888 | 13 | 0.33% |
| hsa_circ_0026093 | 7889 | 13 | 0.33% |
| hsa_circ_0121675 | 7889 | 13 | 0.33% |
| hsa_circ_0070105 | 8496 | 14 | 0.33% |
| hsa_circ_0055271 | 8499 | 14 | 0.33% |
| hsa_circ_0035660 | 8500 | 14 | 0.33% |
| hsa_circ_0008468 | 23682 | 39 | 0.33% |
| hsa_circ_0109052 | 26111 | 43 | 0.33% |
| hsa_circ_0123324 | 41913 | 69 | 0.33% |
| hsa_circ_0121611 | 7290 | 12 | 0.33% |
| hsa_circ_0100973 | 13368 | 22 | 0.33% |
| hsa_circ_0125735 | 23092 | 38 | 0.33% |
| hsa_circ_0114866 | 9724 | 16 | 0.33% |
| hsa_circ_0120950 | 8509 | 14 | 0.33% |
| hsa_circ_0118583 | 16412 | 27 | 0.33% |
| hsa_circ_0091858 | 8510 | 14 | 0.33% |
| hsa_circ_0027175 | 7904 | 13 | 0.33% |
| hsa_circ_0054743 | 7905 | 13 | 0.33% |
| hsa_circ_0134599 | 33445 | 55 | 0.33% |
| hsa_circ_0088827 | 7907 | 13 | 0.33% |
| hsa_circ_0078102 | 8517 | 14 | 0.33% |
| hsa_circ_0027151 | 7909 | 13 | 0.33% |
| hsa_circ_0111541 | 18862 | 31 | 0.33% |
| hsa_circ_0057201 | 7302 | 12 | 0.33% |
| hsa_circ_0033071 | 21909 | 36 | 0.33% |
| hsa_circ_0084273 | 7303 | 12 | 0.33% |
| hsa_circ_0140473 | 37736 | 62 | 0.33% |
| hsa_circ_0127685 | 9132 | 15 | 0.33% |
| hsa_circ_0027881 | 8528 | 14 | 0.33% |
| hsa_circ_0033256 | 7310 | 12 | 0.33% |
| hsa_circ_0070326 | 6705 | 11 | 0.33% |
| hsa_circ_0078576 | 6706 | 11 | 0.33% |

| | | | |
|---|---|---|---|
| hsa_circ_0114663 | 7317 | 12 | 0.33% |
| hsa_circ_0077340 | 7938 | 13 | 0.33% |
| hsa_circ_0129060 | 21985 | 36 | 0.33% |
| hsa_circ_0003960 | 8550 | 14 | 0.33% |
| hsa_circ_0053577 | 7940 | 13 | 0.33% |
| hsa_circ_0109551 | 26876 | 44 | 0.33% |
| hsa_circ_0013749 | 7942 | 13 | 0.33% |
| hsa_circ_0073606 | 7942 | 13 | 0.33% |
| hsa_circ_0121205 | 8555 | 14 | 0.33% |
| hsa_circ_0091667 | 8556 | 14 | 0.33% |
| hsa_circ_0120274 | 15279 | 25 | 0.33% |
| hsa_circ_0128184 | 37896 | 62 | 0.33% |
| hsa_circ_0140171 | 9781 | 16 | 0.33% |
| hsa_circ_0129225 | 7949 | 13 | 0.33% |
| hsa_circ_0108291 | 16514 | 27 | 0.33% |
| hsa_circ_0045386 | 7341 | 12 | 0.33% |
| hsa_circ_0072170 | 7342 | 12 | 0.33% |
| hsa_circ_0115811 | 7954 | 13 | 0.33% |
| hsa_circ_0006770 | 17749 | 29 | 0.33% |
| hsa_circ_0024201 | 7957 | 13 | 0.33% |
| hsa_circ_0053485 | 7961 | 13 | 0.33% |
| hsa_circ_0105269 | 17147 | 28 | 0.33% |
| hsa_circ_0113750 | 8575 | 14 | 0.33% |
| hsa_circ_0127871 | 22053 | 36 | 0.33% |
| hsa_circ_0015836 | 7351 | 12 | 0.33% |
| hsa_circ_0035136 | 7965 | 13 | 0.33% |
| hsa_circ_0056628 | 6740 | 11 | 0.33% |
| hsa_circ_0064045 | 7353 | 12 | 0.33% |
| hsa_circ_0101518 | 71696 | 117 | 0.33% |
| hsa_circ_0028358 | 7354 | 12 | 0.33% |
| hsa_circ_0117199 | 28805 | 47 | 0.33% |
| hsa_circ_0053458 | 6743 | 11 | 0.33% |
| hsa_circ_0123109 | 9196 | 15 | 0.33% |
| hsa_circ_0084300 | 7970 | 13 | 0.33% |
| hsa_circ_0094109 | 21461 | 35 | 0.33% |
| hsa_circ_0073324 | 6745 | 11 | 0.33% |
| hsa_circ_0070081 | 8585 | 14 | 0.33% |
| hsa_circ_0060071 | 7974 | 13 | 0.33% |
| hsa_circ_0016622 | 7975 | 13 | 0.33% |
| hsa_circ_0046037 | 7362 | 12 | 0.33% |
| hsa_circ_0095383 | 27622 | 45 | 0.33% |
| hsa_circ_0015213 | 7366 | 12 | 0.33% |
| hsa_circ_0081633 | 9822 | 16 | 0.33% |
| hsa_circ_0077333 | 7982 | 13 | 0.33% |
| hsa_circ_0101514 | 71871 | 117 | 0.33% |
| hsa_circ_0127567 | 22118 | 36 | 0.33% |
| hsa_circ_0101515 | 71885 | 117 | 0.33% |
| hsa_circ_0017124 | 7376 | 12 | 0.33% |
| hsa_circ_0140648 | 22745 | 37 | 0.33% |
| hsa_circ_0053601 | 7992 | 13 | 0.33% |

| | | | |
|---|---|---|---|
| hsa_circ_0028718 | 7378 | 12 | 0.33% |
| hsa_circ_0112183 | 7379 | 12 | 0.33% |
| hsa_circ_0024306 | 8610 | 14 | 0.33% |
| hsa_circ_0139320 | 10462 | 17 | 0.32% |
| hsa_circ_0028022 | 14770 | 24 | 0.32% |
| hsa_circ_0128798 | 18463 | 30 | 0.32% |
| hsa_circ_0070491 | 6770 | 11 | 0.32% |
| hsa_circ_0035744 | 8002 | 13 | 0.32% |
| hsa_circ_0090715 | 8002 | 13 | 0.32% |
| hsa_circ_0104018 | 27701 | 45 | 0.32% |
| hsa_circ_0068067 | 8003 | 13 | 0.32% |
| hsa_circ_0054736 | 8008 | 13 | 0.32% |
| hsa_circ_0102691 | 8625 | 14 | 0.32% |
| hsa_circ_0073665 | 7394 | 12 | 0.32% |
| hsa_circ_0122949 | 29578 | 48 | 0.32% |
| hsa_circ_0113282 | 7395 | 12 | 0.32% |
| hsa_circ_0035680 | 8014 | 13 | 0.32% |
| hsa_circ_0053460 | 8015 | 13 | 0.32% |
| hsa_circ_0090697 | 8016 | 13 | 0.32% |
| hsa_circ_0116380 | 45630 | 74 | 0.32% |
| hsa_circ_0126740 | 8020 | 13 | 0.32% |
| hsa_circ_0027170 | 8021 | 13 | 0.32% |
| hsa_circ_0137142 | 13580 | 22 | 0.32% |
| hsa_circ_0136039 | 25312 | 41 | 0.32% |
| hsa_circ_0035717 | 8026 | 13 | 0.32% |
| hsa_circ_0108370 | 17908 | 29 | 0.32% |
| hsa_circ_0109446 | 16673 | 27 | 0.32% |
| hsa_circ_0130331 | 11734 | 19 | 0.32% |
| hsa_circ_0070143 | 8647 | 14 | 0.32% |
| hsa_circ_0037439 | 8031 | 13 | 0.32% |
| hsa_circ_0099827 | 12976 | 21 | 0.32% |
| hsa_circ_0051309 | 8033 | 13 | 0.32% |
| hsa_circ_0099697 | 8651 | 14 | 0.32% |
| hsa_circ_0105528 | 16695 | 27 | 0.32% |
| hsa_circ_0092155 | 7421 | 12 | 0.32% |
| hsa_circ_0006378 | 32160 | 52 | 0.32% |
| hsa_circ_0118858 | 7423 | 12 | 0.32% |
| hsa_circ_0124920 | 8043 | 13 | 0.32% |
| hsa_circ_0100704 | 21658 | 35 | 0.32% |
| hsa_circ_0106523 | 27228 | 44 | 0.32% |
| hsa_circ_0009957 | 7426 | 12 | 0.32% |
| hsa_circ_0093551 | 8045 | 13 | 0.32% |
| hsa_circ_0055800 | 15475 | 25 | 0.32% |
| hsa_circ_0037570 | 7428 | 12 | 0.32% |
| hsa_circ_0038262 | 6812 | 11 | 0.32% |
| hsa_circ_0117889 | 8051 | 13 | 0.32% |
| hsa_circ_0008559 | 19819 | 32 | 0.32% |
| hsa_circ_0055601 | 8053 | 13 | 0.32% |
| hsa_circ_0125417 | 8673 | 14 | 0.32% |
| hsa_circ_0103083 | 27261 | 44 | 0.32% |

| | | | |
|---|---|---|---|
| hsa_circ_0058694 | 12398 | 20 | 0.32% |
| hsa_circ_0034301 | 8060 | 13 | 0.32% |
| hsa_circ_0084252 | 7440 | 12 | 0.32% |
| hsa_circ_0046336 | 8061 | 13 | 0.32% |
| hsa_circ_0128778 | 18603 | 30 | 0.32% |
| hsa_circ_0136808 | 20464 | 33 | 0.32% |
| hsa_circ_0132247 | 8684 | 14 | 0.32% |
| hsa_circ_0055985 | 8688 | 14 | 0.32% |
| hsa_circ_0009148 | 14897 | 24 | 0.32% |
| hsa_circ_0137342 | 12417 | 20 | 0.32% |
| hsa_circ_0127240 | 23600 | 38 | 0.32% |
| hsa_circ_0100990 | 8696 | 14 | 0.32% |
| hsa_circ_0027180 | 8076 | 13 | 0.32% |
| hsa_circ_0130592 | 15531 | 25 | 0.32% |
| hsa_circ_0123358 | 30445 | 49 | 0.32% |
| hsa_circ_0130327 | 11810 | 19 | 0.32% |
| hsa_circ_0104581 | 17409 | 28 | 0.32% |
| hsa_circ_0046240 | 8084 | 13 | 0.32% |
| hsa_circ_0129267 | 10572 | 17 | 0.32% |
| hsa_circ_0120000 | 8085 | 13 | 0.32% |
| hsa_circ_0008629 | 8086 | 13 | 0.32% |
| hsa_circ_0132750 | 18040 | 29 | 0.32% |
| hsa_circ_0104617 | 23640 | 38 | 0.32% |
| hsa_circ_0117126 | 10578 | 17 | 0.32% |
| hsa_circ_0061659 | 7467 | 12 | 0.32% |
| hsa_circ_0059189 | 16179 | 26 | 0.32% |
| hsa_circ_0018940 | 20538 | 33 | 0.32% |
| hsa_circ_0107165 | 10581 | 17 | 0.32% |
| hsa_circ_0046927 | 8093 | 13 | 0.32% |
| hsa_circ_0125797 | 19300 | 31 | 0.32% |
| hsa_circ_0068991 | 7472 | 12 | 0.32% |
| hsa_circ_0075975 | 8718 | 14 | 0.32% |
| hsa_circ_0027196 | 7473 | 12 | 0.32% |
| hsa_circ_0016456 | 8097 | 13 | 0.32% |
| hsa_circ_0053721 | 8099 | 13 | 0.32% |
| hsa_circ_0117405 | 6854 | 11 | 0.32% |
| hsa_circ_0053567 | 8101 | 13 | 0.32% |
| hsa_circ_0126523 | 19320 | 31 | 0.32% |
| hsa_circ_0061433 | 8102 | 13 | 0.32% |
| hsa_circ_0110861 | 9981 | 16 | 0.32% |
| hsa_circ_0040925 | 7487 | 12 | 0.32% |
| hsa_circ_0072171 | 7489 | 12 | 0.32% |
| hsa_circ_0104579 | 17475 | 28 | 0.32% |
| hsa_circ_0099478 | 19973 | 32 | 0.32% |
| hsa_circ_0125796 | 19357 | 31 | 0.32% |
| hsa_circ_0129202 | 14363 | 23 | 0.32% |
| hsa_circ_0097320 | 22484 | 36 | 0.32% |
| hsa_circ_0104681 | 69961 | 112 | 0.32% |
| hsa_circ_0045384 | 7496 | 12 | 0.32% |
| hsa_circ_0061080 | 8122 | 13 | 0.32% |

| | | | |
|---|---|---|---|
| hsa_circ_0050112 | 6873 | 11 | 0.32% |
| hsa_circ_0113367 | 17495 | 28 | 0.32% |
| hsa_circ_0113919 | 23744 | 38 | 0.32% |
| hsa_circ_0053590 | 8124 | 13 | 0.32% |
| hsa_circ_0053602 | 8124 | 13 | 0.32% |
| hsa_circ_0000547 | 33763 | 54 | 0.32% |
| hsa_circ_0020793 | 10006 | 16 | 0.32% |
| hsa_circ_0055072 | 21265 | 34 | 0.32% |
| hsa_circ_0037568 | 7506 | 12 | 0.32% |
| hsa_circ_0128227 | 16280 | 26 | 0.32% |
| hsa_circ_0037445 | 7514 | 12 | 0.32% |
| hsa_circ_0101494 | 18786 | 30 | 0.32% |
| hsa_circ_0030465 | 9394 | 15 | 0.32% |
| hsa_circ_0093168 | 9394 | 15 | 0.32% |
| hsa_circ_0042793 | 8768 | 14 | 0.32% |
| hsa_circ_0129894 | 8143 | 13 | 0.32% |
| hsa_circ_0017691 | 33829 | 54 | 0.32% |
| hsa_circ_0101808 | 77692 | 124 | 0.32% |
| hsa_circ_0070082 | 8772 | 14 | 0.32% |
| hsa_circ_0016776 | 8151 | 13 | 0.32% |
| hsa_circ_0073779 | 7524 | 12 | 0.32% |
| hsa_circ_0102094 | 11914 | 19 | 0.32% |
| hsa_circ_0010606 | 8152 | 13 | 0.32% |
| hsa_circ_0011801 | 8781 | 14 | 0.32% |
| hsa_circ_0114701 | 23835 | 38 | 0.32% |
| hsa_circ_0107955 | 13174 | 21 | 0.32% |
| hsa_circ_0118395 | 11292 | 18 | 0.32% |
| hsa_circ_0035665 | 8157 | 13 | 0.32% |
| hsa_circ_0028170 | 7532 | 12 | 0.32% |
| hsa_circ_0037914 | 8164 | 13 | 0.32% |
| hsa_circ_0107956 | 8164 | 13 | 0.32% |
| hsa_circ_0101231 | 45222 | 72 | 0.32% |
| hsa_circ_0070083 | 8794 | 14 | 0.32% |
| hsa_circ_0000578 | 27646 | 44 | 0.32% |
| hsa_circ_0037438 | 8171 | 13 | 0.32% |
| hsa_circ_0070492 | 6914 | 11 | 0.32% |
| hsa_circ_0062822 | 7543 | 12 | 0.32% |
| hsa_circ_0094459 | 23887 | 38 | 0.32% |
| hsa_circ_0025374 | 8172 | 13 | 0.32% |
| hsa_circ_0055270 | 8802 | 14 | 0.32% |
| hsa_circ_0033277 | 7546 | 12 | 0.32% |
| hsa_circ_0095063 | 23268 | 37 | 0.32% |
| hsa_circ_0035968 | 33960 | 54 | 0.32% |
| hsa_circ_0037013 | 22012 | 35 | 0.32% |
| hsa_circ_0034281 | 25157 | 40 | 0.32% |
| hsa_circ_0113360 | 8180 | 13 | 0.32% |
| hsa_circ_0009033 | 20768 | 33 | 0.32% |
| hsa_circ_0138993 | 25803 | 41 | 0.32% |
| hsa_circ_0139500 | 21403 | 34 | 0.32% |
| hsa_circ_0061239 | 22663 | 36 | 0.32% |

| | | | |
|---|---|---|---|
| hsa_circ_0088126 | 51629 | 82 | 0.32% |
| hsa_circ_0139942 | 10074 | 16 | 0.32% |
| hsa_circ_0046035 | 7556 | 12 | 0.32% |
| hsa_circ_0073317 | 8818 | 14 | 0.32% |
| hsa_circ_0095764 | 8821 | 14 | 0.32% |
| hsa_circ_0060980 | 8191 | 13 | 0.32% |
| hsa_circ_0054570 | 6931 | 11 | 0.32% |
| hsa_circ_0005534 | 30249 | 48 | 0.32% |
| hsa_circ_0117595 | 6933 | 11 | 0.32% |
| hsa_circ_0127336 | 26475 | 42 | 0.32% |
| hsa_circ_0035745 | 8197 | 13 | 0.32% |
| hsa_circ_0065041 | 11981 | 19 | 0.32% |
| hsa_circ_0068987 | 7570 | 12 | 0.32% |
| hsa_circ_0052305 | 37237 | 59 | 0.32% |
| hsa_circ_0121332 | 37241 | 59 | 0.32% |
| hsa_circ_0132681 | 8838 | 14 | 0.32% |
| hsa_circ_0115630 | 11364 | 18 | 0.32% |
| hsa_circ_0117782 | 18941 | 30 | 0.32% |
| hsa_circ_0035681 | 8208 | 13 | 0.32% |
| hsa_circ_0117588 | 15794 | 25 | 0.32% |
| hsa_circ_0021035 | 9477 | 15 | 0.32% |
| hsa_circ_0127845 | 10111 | 16 | 0.32% |
| hsa_circ_0016177 | 27817 | 44 | 0.32% |
| hsa_circ_0093697 | 37301 | 59 | 0.32% |
| hsa_circ_0048578 | 14546 | 23 | 0.32% |
| hsa_circ_0129857 | 8223 | 13 | 0.32% |
| hsa_circ_0038412 | 34794 | 55 | 0.32% |
| hsa_circ_0078079 | 7593 | 12 | 0.32% |
| hsa_circ_0024985 | 43029 | 68 | 0.32% |
| hsa_circ_0069672 | 7601 | 12 | 0.32% |
| hsa_circ_0137464 | 15840 | 25 | 0.32% |
| hsa_circ_0077337 | 8238 | 13 | 0.32% |
| hsa_circ_0097851 | 6973 | 11 | 0.32% |
| hsa_circ_0046031 | 10143 | 16 | 0.32% |
| hsa_circ_0134028 | 26626 | 42 | 0.32% |
| hsa_circ_0054737 | 8245 | 13 | 0.32% |
| hsa_circ_0139533 | 14588 | 23 | 0.32% |
| hsa_circ_0114609 | 16494 | 26 | 0.32% |
| hsa_circ_0082978 | 8248 | 13 | 0.32% |
| hsa_circ_0004600 | 13331 | 21 | 0.32% |
| hsa_circ_0035698 | 8254 | 13 | 0.31% |
| hsa_circ_0125792 | 24131 | 38 | 0.31% |
| hsa_circ_0000579 | 27943 | 44 | 0.31% |
| hsa_circ_0111810 | 8257 | 13 | 0.31% |
| hsa_circ_0132564 | 7623 | 12 | 0.31% |
| hsa_circ_0130147 | 9529 | 15 | 0.31% |
| hsa_circ_0114685 | 24141 | 38 | 0.31% |
| hsa_circ_0037574 | 6989 | 11 | 0.31% |
| hsa_circ_0139562 | 8896 | 14 | 0.31% |
| hsa_circ_0093698 | 37499 | 59 | 0.31% |

| | | | |
|---|---|---|---|
| hsa_circ_0105489 | 25426 | 40 | 0.31% |
| hsa_circ_0117585 | 15893 | 25 | 0.31% |
| hsa_circ_0021031 | 7630 | 12 | 0.31% |
| hsa_circ_0024881 | 9538 | 15 | 0.31% |
| hsa_circ_0132039 | 94747 | 149 | 0.31% |
| hsa_circ_0136736 | 17170 | 27 | 0.31% |
| hsa_circ_0015218 | 6999 | 11 | 0.31% |
| hsa_circ_0077315 | 8272 | 13 | 0.31% |
| hsa_circ_0011965 | 15909 | 25 | 0.31% |
| hsa_circ_0070177 | 8910 | 14 | 0.31% |
| hsa_circ_0127239 | 24185 | 38 | 0.31% |
| hsa_circ_0114316 | 20367 | 32 | 0.31% |
| hsa_circ_0130683 | 23555 | 37 | 0.31% |
| hsa_circ_0129113 | 9550 | 15 | 0.31% |
| hsa_circ_0124292 | 7641 | 12 | 0.31% |
| hsa_circ_0010664 | 8915 | 14 | 0.31% |
| hsa_circ_0108801 | 21016 | 33 | 0.31% |
| hsa_circ_0115632 | 17832 | 28 | 0.31% |
| hsa_circ_0101198 | 19108 | 30 | 0.31% |
| hsa_circ_0108765 | 24206 | 38 | 0.31% |
| hsa_circ_0065042 | 12106 | 19 | 0.31% |
| hsa_circ_0083298 | 8921 | 14 | 0.31% |
| hsa_circ_0037437 | 8284 | 13 | 0.31% |
| hsa_circ_0028366 | 7647 | 12 | 0.31% |
| hsa_circ_0119282 | 14657 | 23 | 0.31% |
| hsa_circ_0010553 | 8922 | 14 | 0.31% |
| hsa_circ_0127780 | 13384 | 21 | 0.31% |
| hsa_circ_0117960 | 27409 | 43 | 0.31% |
| hsa_circ_0007600 | 8929 | 14 | 0.31% |
| hsa_circ_0033265 | 7656 | 12 | 0.31% |
| hsa_circ_0118855 | 7660 | 12 | 0.31% |
| hsa_circ_0096673 | 20432 | 32 | 0.31% |
| hsa_circ_0128318 | 8939 | 14 | 0.31% |
| hsa_circ_0137410 | 18520 | 29 | 0.31% |
| hsa_circ_0031073 | 16605 | 26 | 0.31% |
| hsa_circ_0134158 | 38959 | 61 | 0.31% |
| hsa_circ_0084263 | 7667 | 12 | 0.31% |
| hsa_circ_0105454 | 8308 | 13 | 0.31% |
| hsa_circ_0106879 | 49212 | 77 | 0.31% |
| hsa_circ_0045381 | 7670 | 12 | 0.31% |
| hsa_circ_0101195 | 19180 | 30 | 0.31% |
| hsa_circ_0015204 | 7673 | 12 | 0.31% |
| hsa_circ_0085093 | 7674 | 12 | 0.31% |
| hsa_circ_0129959 | 7675 | 12 | 0.31% |
| hsa_circ_0005934 | 15991 | 25 | 0.31% |
| hsa_circ_0042794 | 8955 | 14 | 0.31% |
| hsa_circ_0021345 | 8957 | 14 | 0.31% |
| hsa_circ_0016621 | 8320 | 13 | 0.31% |
| hsa_circ_0132044 | 9602 | 15 | 0.31% |
| hsa_circ_0060070 | 8324 | 13 | 0.31% |

| | | | |
|---|---|---|---|
| hsa_circ_0010845 | 8328 | 13 | 0.31% |
| hsa_circ_0140727 | 19222 | 30 | 0.31% |
| hsa_circ_0100957 | 26913 | 42 | 0.31% |
| hsa_circ_0087885 | 7049 | 11 | 0.31% |
| hsa_circ_0102410 | 28840 | 45 | 0.31% |
| hsa_circ_0054728 | 8337 | 13 | 0.31% |
| hsa_circ_0027146 | 7697 | 12 | 0.31% |
| hsa_circ_0116000 | 8340 | 13 | 0.31% |
| hsa_circ_0127709 | 8342 | 13 | 0.31% |
| hsa_circ_0010114 | 8984 | 14 | 0.31% |
| hsa_circ_0034811 | 7701 | 12 | 0.31% |
| hsa_circ_0013819 | 23104 | 36 | 0.31% |
| hsa_circ_0077326 | 8345 | 13 | 0.31% |
| hsa_circ_0035583 | 7704 | 12 | 0.31% |
| hsa_circ_0137408 | 18620 | 29 | 0.31% |
| hsa_circ_0046337 | 8348 | 13 | 0.31% |
| hsa_circ_0073877 | 8348 | 13 | 0.31% |
| hsa_circ_0013662 | 8349 | 13 | 0.31% |
| hsa_circ_0091121 | 8349 | 13 | 0.31% |
| hsa_circ_0138968 | 17985 | 28 | 0.31% |
| hsa_circ_0088123 | 52693 | 82 | 0.31% |
| hsa_circ_0038238 | 7713 | 12 | 0.31% |
| hsa_circ_0119697 | 23782 | 37 | 0.31% |
| hsa_circ_0008481 | 21214 | 33 | 0.31% |
| hsa_circ_0010549 | 9001 | 14 | 0.31% |
| hsa_circ_0068968 | 7716 | 12 | 0.31% |
| hsa_circ_0121850 | 28945 | 45 | 0.31% |
| hsa_circ_0136419 | 20584 | 32 | 0.31% |
| hsa_circ_0014873 | 24444 | 38 | 0.31% |
| hsa_circ_0128218 | 10293 | 16 | 0.31% |
| hsa_circ_0126462 | 30239 | 47 | 0.31% |
| hsa_circ_0126463 | 30242 | 47 | 0.31% |
| hsa_circ_0077647 | 8365 | 13 | 0.31% |
| hsa_circ_0132368 | 19305 | 30 | 0.31% |
| hsa_circ_0137179 | 12870 | 20 | 0.31% |
| hsa_circ_0053509 | 8368 | 13 | 0.31% |
| hsa_circ_0109831 | 12232 | 19 | 0.31% |
| hsa_circ_0077266 | 9657 | 15 | 0.31% |
| hsa_circ_0033271 | 7730 | 12 | 0.31% |
| hsa_circ_0034329 | 21262 | 33 | 0.31% |
| hsa_circ_0016756 | 8376 | 13 | 0.31% |
| hsa_circ_0135579 | 9665 | 15 | 0.31% |
| hsa_circ_0029409 | 21267 | 33 | 0.31% |
| hsa_circ_0077900 | 25780 | 40 | 0.31% |
| hsa_circ_0035578 | 7734 | 12 | 0.31% |
| hsa_circ_0042935 | 8383 | 13 | 0.31% |
| hsa_circ_0041666 | 7739 | 12 | 0.31% |
| hsa_circ_0096928 | 24516 | 38 | 0.31% |
| hsa_circ_0035721 | 8388 | 13 | 0.31% |
| hsa_circ_0084132 | 29683 | 46 | 0.31% |

| | | | |
|---|---|---|---|
| hsa_circ_0029911 | 7100 | 11 | 0.31% |
| hsa_circ_0053677 | 8391 | 13 | 0.31% |
| hsa_circ_0108899 | 33565 | 52 | 0.31% |
| hsa_circ_0099468 | 16140 | 25 | 0.31% |
| hsa_circ_0136032 | 7748 | 12 | 0.31% |
| hsa_circ_0068068 | 8394 | 13 | 0.31% |
| hsa_circ_0082972 | 8396 | 13 | 0.31% |
| hsa_circ_0095672 | 20677 | 32 | 0.31% |
| hsa_circ_0137314 | 14218 | 22 | 0.31% |
| hsa_circ_0021025 | 7758 | 12 | 0.31% |
| hsa_circ_0062093 | 9700 | 15 | 0.31% |
| hsa_circ_0136529 | 8409 | 13 | 0.31% |
| hsa_circ_0072172 | 7764 | 12 | 0.31% |
| hsa_circ_0006329 | 16176 | 25 | 0.31% |
| hsa_circ_0125579 | 70539 | 109 | 0.31% |
| hsa_circ_0082994 | 8413 | 13 | 0.31% |
| hsa_circ_0053697 | 8414 | 13 | 0.31% |
| hsa_circ_0136653 | 17479 | 27 | 0.31% |
| hsa_circ_0096265 | 8416 | 13 | 0.31% |
| hsa_circ_0129841 | 19425 | 30 | 0.31% |
| hsa_circ_0105031 | 14895 | 23 | 0.31% |
| hsa_circ_0090693 | 8419 | 13 | 0.31% |
| hsa_circ_0010607 | 8420 | 13 | 0.31% |
| hsa_circ_0112285 | 9720 | 15 | 0.31% |
| hsa_circ_0118725 | 40831 | 63 | 0.31% |
| hsa_circ_0037436 | 8426 | 13 | 0.31% |
| hsa_circ_0037564 | 7779 | 12 | 0.31% |
| hsa_circ_0008669 | 11669 | 18 | 0.31% |
| hsa_circ_0126268 | 26582 | 41 | 0.31% |
| hsa_circ_0140789 | 18155 | 28 | 0.31% |
| hsa_circ_0096331 | 63546 | 98 | 0.31% |
| hsa_circ_0034217 | 8432 | 13 | 0.31% |
| hsa_circ_0037453 | 8432 | 13 | 0.31% |
| hsa_circ_0047412 | 26604 | 41 | 0.31% |
| hsa_circ_0133226 | 18169 | 28 | 0.31% |
| hsa_circ_0125978 | 12979 | 20 | 0.31% |
| hsa_circ_0055230 | 9086 | 14 | 0.31% |
| hsa_circ_0124745 | 13630 | 21 | 0.31% |
| hsa_circ_0132613 | 17525 | 27 | 0.31% |
| hsa_circ_0105032 | 23380 | 36 | 0.31% |
| hsa_circ_0119138 | 21432 | 33 | 0.31% |
| hsa_circ_0028371 | 7794 | 12 | 0.31% |
| hsa_circ_0139514 | 11692 | 18 | 0.31% |
| hsa_circ_0124804 | 51980 | 80 | 0.31% |
| hsa_circ_0008720 | 26641 | 41 | 0.31% |
| hsa_circ_0027148 | 8448 | 13 | 0.31% |
| hsa_circ_0015199 | 7799 | 12 | 0.31% |
| hsa_circ_0107498 | 34450 | 53 | 0.31% |
| hsa_circ_0130461 | 18852 | 29 | 0.31% |
| hsa_circ_0033282 | 7801 | 12 | 0.31% |

| | | | |
|---|---|---|---|
| hsa_circ_0091867 | 7801 | 12 | 0.31% |
| hsa_circ_0128830 | 22106 | 34 | 0.31% |
| hsa_circ_0126269 | 26665 | 41 | 0.31% |
| hsa_circ_0010290 | 7806 | 12 | 0.31% |
| hsa_circ_0000833 | 14962 | 23 | 0.31% |
| hsa_circ_0046338 | 8458 | 13 | 0.31% |
| hsa_circ_0119385 | 17569 | 27 | 0.31% |
| hsa_circ_0127032 | 13666 | 21 | 0.31% |
| hsa_circ_0135401 | 18876 | 29 | 0.31% |
| hsa_circ_0080940 | 7812 | 12 | 0.31% |
| hsa_circ_0046027 | 10419 | 16 | 0.31% |
| hsa_circ_0137347 | 28011 | 43 | 0.31% |
| hsa_circ_0028023 | 9125 | 14 | 0.31% |
| hsa_circ_0138639 | 9128 | 14 | 0.31% |
| hsa_circ_0100988 | 7824 | 12 | 0.31% |
| hsa_circ_0126461 | 20216 | 31 | 0.31% |
| hsa_circ_0054726 | 8478 | 13 | 0.31% |
| hsa_circ_0069503 | 7176 | 11 | 0.31% |
| hsa_circ_0018304 | 24790 | 38 | 0.31% |
| hsa_circ_0126798 | 9134 | 14 | 0.31% |
| hsa_circ_0114613 | 16966 | 26 | 0.31% |
| hsa_circ_0011785 | 8484 | 13 | 0.31% |
| hsa_circ_0082192 | 9138 | 14 | 0.31% |
| hsa_circ_0126270 | 26765 | 41 | 0.31% |
| hsa_circ_0070852 | 9140 | 14 | 0.31% |
| hsa_circ_0087238 | 8488 | 13 | 0.31% |
| hsa_circ_0101241 | 18936 | 29 | 0.31% |
| hsa_circ_0036129 | 9145 | 14 | 0.31% |
| hsa_circ_0132215 | 9145 | 14 | 0.31% |
| hsa_circ_0095667 | 20903 | 32 | 0.31% |
| hsa_circ_0061236 | 23519 | 36 | 0.31% |
| hsa_circ_0128477 | 16333 | 25 | 0.31% |
| hsa_circ_0010303 | 7840 | 12 | 0.31% |
| hsa_circ_0110501 | 17643 | 27 | 0.31% |
| hsa_circ_0129716 | 12416 | 19 | 0.31% |
| hsa_circ_0000230 | 88220 | 135 | 0.31% |
| hsa_circ_0140162 | 28754 | 44 | 0.31% |
| hsa_circ_0003804 | 10456 | 16 | 0.31% |
| hsa_circ_0101895 | 10456 | 16 | 0.31% |
| hsa_circ_0104715 | 28755 | 44 | 0.31% |
| hsa_circ_0131894 | 26799 | 41 | 0.31% |
| hsa_circ_0053766 | 8500 | 13 | 0.31% |
| hsa_circ_0047603 | 9810 | 15 | 0.31% |
| hsa_circ_0084257 | 7850 | 12 | 0.31% |
| hsa_circ_0082990 | 8505 | 13 | 0.31% |
| hsa_circ_0112283 | 9814 | 15 | 0.31% |
| hsa_circ_0103275 | 30754 | 47 | 0.31% |
| hsa_circ_0008593 | 11780 | 18 | 0.31% |
| hsa_circ_0140045 | 32724 | 50 | 0.31% |
| hsa_circ_0090688 | 8509 | 13 | 0.31% |

| | | | |
|---|---|---|---|
| hsa_circ_0053578 | 8511 | 13 | 0.31% |
| hsa_circ_0042795 | 9166 | 14 | 0.31% |
| hsa_circ_0098512 | 9824 | 15 | 0.31% |
| hsa_circ_0004126 | 88445 | 135 | 0.31% |
| hsa_circ_0082879 | 9829 | 15 | 0.31% |
| hsa_circ_0139743 | 11796 | 18 | 0.31% |
| hsa_circ_0046030 | 10486 | 16 | 0.31% |
| hsa_circ_0033274 | 7867 | 12 | 0.31% |
| hsa_circ_0127781 | 13768 | 21 | 0.31% |
| hsa_circ_0010624 | 8525 | 13 | 0.30% |
| hsa_circ_0053650 | 8526 | 13 | 0.30% |
| hsa_circ_0126271 | 26894 | 41 | 0.30% |
| hsa_circ_0127719 | 15088 | 23 | 0.30% |
| hsa_circ_0109861 | 22314 | 34 | 0.30% |
| hsa_circ_0094371 | 22976 | 35 | 0.30% |
| hsa_circ_0084223 | 7881 | 12 | 0.30% |
| hsa_circ_0114611 | 36136 | 55 | 0.30% |
| hsa_circ_0037940 | 8542 | 13 | 0.30% |
| hsa_circ_0101206 | 8545 | 13 | 0.30% |
| hsa_circ_0110500 | 17748 | 27 | 0.30% |
| hsa_circ_0115701 | 46678 | 71 | 0.30% |
| hsa_circ_0015214 | 7232 | 11 | 0.30% |
| hsa_circ_0034210 | 8547 | 13 | 0.30% |
| hsa_circ_0081201 | 8549 | 13 | 0.30% |
| hsa_circ_0122767 | 10522 | 16 | 0.30% |
| hsa_circ_0073318 | 9208 | 14 | 0.30% |
| hsa_circ_0037434 | 8551 | 13 | 0.30% |
| hsa_circ_0068988 | 7894 | 12 | 0.30% |
| hsa_circ_0090691 | 8552 | 13 | 0.30% |
| hsa_circ_0033261 | 7896 | 12 | 0.30% |
| hsa_circ_0114063 | 18425 | 28 | 0.30% |
| hsa_circ_0077028 | 9216 | 14 | 0.30% |
| hsa_circ_0126073 | 29629 | 45 | 0.30% |
| hsa_circ_0003760 | 8560 | 13 | 0.30% |
| hsa_circ_0053532 | 8562 | 13 | 0.30% |
| hsa_circ_0104980 | 11198 | 17 | 0.30% |
| hsa_circ_0019035 | 16474 | 25 | 0.30% |
| hsa_circ_0127327 | 9888 | 15 | 0.30% |
| hsa_circ_0056627 | 7911 | 12 | 0.30% |
| hsa_circ_0070318 | 7252 | 11 | 0.30% |
| hsa_circ_0108821 | 9890 | 15 | 0.30% |
| hsa_circ_0117788 | 35607 | 54 | 0.30% |
| hsa_circ_0015774 | 7913 | 12 | 0.30% |
| hsa_circ_0116952 | 22424 | 34 | 0.30% |
| hsa_circ_0034223 | 8577 | 13 | 0.30% |
| hsa_circ_0108849 | 62730 | 95 | 0.30% |
| hsa_circ_0071609 | 9247 | 14 | 0.30% |
| hsa_circ_0108422 | 7928 | 12 | 0.30% |
| hsa_circ_0061434 | 10573 | 16 | 0.30% |
| hsa_circ_0106509 | 35030 | 53 | 0.30% |

| | | | |
|---|---|---|---|
| hsa_circ_0138843 | 23134 | 35 | 0.30% |
| hsa_circ_0047093 | 37028 | 56 | 0.30% |
| hsa_circ_0131741 | 30429 | 46 | 0.30% |
| hsa_circ_0053562 | 8601 | 13 | 0.30% |
| hsa_circ_0034138 | 7278 | 11 | 0.30% |
| hsa_circ_0024121 | 8602 | 13 | 0.30% |
| hsa_circ_0053678 | 8602 | 13 | 0.30% |
| hsa_circ_0008578 | 8605 | 13 | 0.30% |
| hsa_circ_0093547 | 8606 | 13 | 0.30% |
| hsa_circ_0029917 | 7945 | 12 | 0.30% |
| hsa_circ_0121549 | 8609 | 13 | 0.30% |
| hsa_circ_0077644 | 7947 | 12 | 0.30% |
| hsa_circ_0131453 | 28477 | 43 | 0.30% |
| hsa_circ_0121534 | 13913 | 21 | 0.30% |
| hsa_circ_0002312 | 11272 | 17 | 0.30% |
| hsa_circ_0137584 | 10610 | 16 | 0.30% |
| hsa_circ_0134185 | 49078 | 74 | 0.30% |
| hsa_circ_0097292 | 9951 | 15 | 0.30% |
| hsa_circ_0000062 | 8625 | 13 | 0.30% |
| hsa_circ_0093160 | 8627 | 13 | 0.30% |
| hsa_circ_0000004 | 33847 | 51 | 0.30% |
| hsa_circ_0034998 | 9958 | 15 | 0.30% |
| hsa_circ_0115288 | 8632 | 13 | 0.30% |
| hsa_circ_0106037 | 17931 | 27 | 0.30% |
| hsa_circ_0000169 | 11954 | 18 | 0.30% |
| hsa_circ_0053724 | 8636 | 13 | 0.30% |
| hsa_circ_0082979 | 8637 | 13 | 0.30% |
| hsa_circ_0095261 | 46511 | 70 | 0.30% |
| hsa_circ_0138826 | 24589 | 37 | 0.30% |
| hsa_circ_0114872 | 34564 | 52 | 0.30% |
| hsa_circ_0000767 | 51184 | 77 | 0.30% |
| hsa_circ_0082878 | 9972 | 15 | 0.30% |
| hsa_circ_0035054 | 31913 | 48 | 0.30% |
| hsa_circ_0006355 | 13300 | 20 | 0.30% |
| hsa_circ_0099061 | 15296 | 23 | 0.30% |
| hsa_circ_0010649 | 8646 | 13 | 0.30% |
| hsa_circ_0033272 | 7981 | 12 | 0.30% |
| hsa_circ_0134182 | 49219 | 74 | 0.30% |
| hsa_circ_0011484 | 9313 | 14 | 0.30% |
| hsa_circ_0027246 | 35257 | 53 | 0.30% |
| hsa_circ_0115695 | 8649 | 13 | 0.30% |
| hsa_circ_0003301 | 20646 | 31 | 0.30% |
| hsa_circ_0139471 | 9324 | 14 | 0.30% |
| hsa_circ_0073322 | 7326 | 11 | 0.30% |
| hsa_circ_0053603 | 8660 | 13 | 0.30% |
| hsa_circ_0118999 | 15992 | 24 | 0.30% |
| hsa_circ_0126074 | 29989 | 45 | 0.30% |
| hsa_circ_0081030 | 10663 | 16 | 0.30% |
| hsa_circ_0079961 | 23327 | 35 | 0.30% |
| hsa_circ_0081252 | 9999 | 15 | 0.30% |

| | | | |
|---|---|---|---|
| hsa_circ_0134180 | 49339 | 74 | 0.30% |
| hsa_circ_0027135 | 8001 | 12 | 0.30% |
| hsa_circ_0128789 | 19336 | 29 | 0.30% |
| hsa_circ_0120784 | 16004 | 24 | 0.30% |
| hsa_circ_0038907 | 26007 | 39 | 0.30% |
| hsa_circ_0095604 | 23345 | 35 | 0.30% |
| hsa_circ_0112410 | 17342 | 26 | 0.30% |
| hsa_circ_0084285 | 8673 | 13 | 0.30% |
| hsa_circ_0037565 | 8007 | 12 | 0.30% |
| hsa_circ_0138967 | 20022 | 30 | 0.30% |
| hsa_circ_0002525 | 18691 | 28 | 0.30% |
| hsa_circ_0125239 | 8011 | 12 | 0.30% |
| hsa_circ_0132742 | 64776 | 97 | 0.30% |
| hsa_circ_0033257 | 8016 | 12 | 0.30% |
| hsa_circ_0116922 | 8016 | 12 | 0.30% |
| hsa_circ_0134176 | 49442 | 74 | 0.30% |
| hsa_circ_0039198 | 8690 | 13 | 0.30% |
| hsa_circ_0103045 | 9359 | 14 | 0.30% |
| hsa_circ_0001218 | 34765 | 52 | 0.30% |
| hsa_circ_0101815 | 11370 | 17 | 0.30% |
| hsa_circ_0103079 | 21403 | 32 | 0.30% |
| hsa_circ_0056160 | 8696 | 13 | 0.30% |
| hsa_circ_0073283 | 8029 | 12 | 0.30% |
| hsa_circ_0118722 | 42170 | 63 | 0.30% |
| hsa_circ_0119750 | 12049 | 18 | 0.30% |
| hsa_circ_0016773 | 8703 | 13 | 0.30% |
| hsa_circ_0133229 | 24772 | 37 | 0.30% |
| hsa_circ_0053625 | 8704 | 13 | 0.30% |
| hsa_circ_0000768 | 51558 | 77 | 0.30% |
| hsa_circ_0089253 | 30801 | 46 | 0.30% |
| hsa_circ_0053461 | 8706 | 13 | 0.30% |
| hsa_circ_0016631 | 8707 | 13 | 0.30% |
| hsa_circ_0029692 | 14739 | 22 | 0.30% |
| hsa_circ_0045377 | 8040 | 12 | 0.30% |
| hsa_circ_0112228 | 15411 | 23 | 0.30% |
| hsa_circ_0010268 | 10051 | 15 | 0.30% |
| hsa_circ_0073320 | 7372 | 11 | 0.30% |
| hsa_circ_0039098 | 10053 | 15 | 0.30% |
| hsa_circ_0047094 | 37535 | 56 | 0.30% |
| hsa_circ_0109855 | 28826 | 43 | 0.30% |
| hsa_circ_0109867 | 22795 | 34 | 0.30% |
| hsa_circ_0024325 | 39560 | 59 | 0.30% |
| hsa_circ_0000769 | 51641 | 77 | 0.30% |
| hsa_circ_0055941 | 9390 | 14 | 0.30% |
| hsa_circ_0136422 | 21464 | 32 | 0.30% |
| hsa_circ_0117728 | 18784 | 28 | 0.30% |
| hsa_circ_0076514 | 7380 | 11 | 0.30% |
| hsa_circ_0138519 | 18788 | 28 | 0.30% |
| hsa_circ_0121622 | 9396 | 14 | 0.30% |
| hsa_circ_0101516 | 78560 | 117 | 0.30% |

| | | | |
|---|---|---|---|
| hsa_circ_0095260 | 47010 | 70 | 0.30% |
| hsa_circ_0125538 | 10074 | 15 | 0.30% |
| hsa_circ_0128409 | 48372 | 72 | 0.30% |
| hsa_circ_0114233 | 30919 | 46 | 0.30% |
| hsa_circ_0104808 | 10084 | 15 | 0.30% |
| hsa_circ_0131930 | 8069 | 12 | 0.30% |
| hsa_circ_0127866 | 29590 | 44 | 0.30% |
| hsa_circ_0126859 | 17485 | 26 | 0.30% |
| hsa_circ_0116953 | 22866 | 34 | 0.30% |
| hsa_circ_0077310 | 8744 | 13 | 0.30% |
| hsa_circ_0134250 | 78042 | 116 | 0.30% |
| hsa_circ_0053679 | 8748 | 13 | 0.30% |
| hsa_circ_0053515 | 8749 | 13 | 0.30% |
| hsa_circ_0133615 | 14134 | 21 | 0.30% |
| hsa_circ_0081250 | 10096 | 15 | 0.30% |
| hsa_circ_0113744 | 32312 | 48 | 0.30% |
| hsa_circ_0011514 | 25581 | 38 | 0.30% |
| hsa_circ_0011826 | 9426 | 14 | 0.30% |
| hsa_circ_0132183 | 34338 | 51 | 0.30% |
| hsa_circ_0113753 | 28959 | 43 | 0.30% |
| hsa_circ_0009087 | 22903 | 34 | 0.30% |
| hsa_circ_0120269 | 8084 | 12 | 0.30% |
| hsa_circ_0084224 | 8089 | 12 | 0.30% |
| hsa_circ_0130001 | 22925 | 34 | 0.30% |
| hsa_circ_0131969 | 8767 | 13 | 0.30% |
| hsa_circ_0056629 | 8768 | 13 | 0.30% |
| hsa_circ_0076870 | 33729 | 50 | 0.30% |
| hsa_circ_0138516 | 18892 | 28 | 0.30% |
| hsa_circ_0125740 | 15520 | 23 | 0.30% |
| hsa_circ_0047475 | 76257 | 113 | 0.30% |
| hsa_circ_0101235 | 9451 | 14 | 0.30% |
| hsa_circ_0106637 | 23630 | 35 | 0.30% |
| hsa_circ_0117623 | 12828 | 19 | 0.30% |
| hsa_circ_0073323 | 7427 | 11 | 0.30% |
| hsa_circ_0114873 | 35118 | 52 | 0.30% |
| hsa_circ_0129803 | 17559 | 26 | 0.30% |
| hsa_circ_0054721 | 8782 | 13 | 0.30% |
| hsa_circ_0035682 | 8783 | 13 | 0.30% |
| hsa_circ_0109852 | 29053 | 43 | 0.30% |
| hsa_circ_0047476 | 76366 | 113 | 0.30% |
| hsa_circ_0010295 | 8110 | 12 | 0.30% |
| hsa_circ_0052897 | 8789 | 13 | 0.30% |
| hsa_circ_0073285 | 8113 | 12 | 0.30% |
| hsa_circ_0095201 | 10142 | 15 | 0.30% |
| hsa_circ_0033078 | 9466 | 14 | 0.30% |
| hsa_circ_0074699 | 34501 | 51 | 0.30% |
| hsa_circ_0058199 | 9471 | 14 | 0.30% |
| hsa_circ_0056016 | 18943 | 28 | 0.30% |
| hsa_circ_0116954 | 23003 | 34 | 0.30% |
| hsa_circ_0119192 | 16239 | 24 | 0.30% |

| | | | |
|---|---|---|---|
| hsa_circ_0084274 | 8120 | 12 | 0.30% |
| hsa_circ_0126799 | 9474 | 14 | 0.30% |
| hsa_circ_0082690 | 8123 | 12 | 0.30% |
| hsa_circ_0027540 | 8124 | 12 | 0.30% |
| hsa_circ_0134865 | 15573 | 23 | 0.30% |
| hsa_circ_0071605 | 9481 | 14 | 0.30% |
| hsa_circ_0103064 | 10159 | 15 | 0.30% |
| hsa_circ_0105444 | 15581 | 23 | 0.30% |
| hsa_circ_0053637 | 8808 | 13 | 0.30% |
| hsa_circ_0000934 | 27103 | 40 | 0.30% |
| hsa_circ_0109858 | 23039 | 34 | 0.30% |
| hsa_circ_0000006 | 43368 | 64 | 0.30% |
| hsa_circ_0026092 | 8811 | 13 | 0.30% |
| hsa_circ_0109859 | 23051 | 34 | 0.29% |
| hsa_circ_0010262 | 10170 | 15 | 0.29% |
| hsa_circ_0053722 | 8814 | 13 | 0.29% |
| hsa_circ_0053011 | 7458 | 11 | 0.29% |
| hsa_circ_0122774 | 24409 | 36 | 0.29% |
| hsa_circ_0033266 | 8137 | 12 | 0.29% |
| hsa_circ_0070313 | 7461 | 11 | 0.29% |
| hsa_circ_0113563 | 15603 | 23 | 0.29% |
| hsa_circ_0094379 | 29850 | 44 | 0.29% |
| hsa_circ_0008528 | 27149 | 40 | 0.29% |
| hsa_circ_0084591 | 8145 | 12 | 0.29% |
| hsa_circ_0034997 | 10182 | 15 | 0.29% |
| hsa_circ_0010542 | 9508 | 14 | 0.29% |
| hsa_circ_0055935 | 9510 | 14 | 0.29% |
| hsa_circ_0110607 | 8832 | 13 | 0.29% |
| hsa_circ_0113275 | 8154 | 12 | 0.29% |
| hsa_circ_0140728 | 24468 | 36 | 0.29% |
| hsa_circ_0026097 | 8837 | 13 | 0.29% |
| hsa_circ_0082016 | 8158 | 12 | 0.29% |
| hsa_circ_0117977 | 10199 | 15 | 0.29% |
| hsa_circ_0007124 | 12241 | 18 | 0.29% |
| hsa_circ_0091180 | 76188 | 112 | 0.29% |
| hsa_circ_0091565 | 21090 | 31 | 0.29% |
| hsa_circ_0018435 | 9525 | 14 | 0.29% |
| hsa_circ_0134965 | 10886 | 16 | 0.29% |
| hsa_circ_0030912 | 15649 | 23 | 0.29% |
| hsa_circ_0042771 | 9526 | 14 | 0.29% |
| hsa_circ_0134986 | 9528 | 14 | 0.29% |
| hsa_circ_0111044 | 34034 | 50 | 0.29% |
| hsa_circ_0034039 | 19745 | 29 | 0.29% |
| hsa_circ_0046028 | 10896 | 16 | 0.29% |
| hsa_circ_0121641 | 14986 | 22 | 0.29% |
| hsa_circ_0076825 | 8856 | 13 | 0.29% |
| hsa_circ_0095666 | 9538 | 14 | 0.29% |
| hsa_circ_0053510 | 8861 | 13 | 0.29% |
| hsa_circ_0025975 | 27952 | 41 | 0.29% |
| hsa_circ_0099615 | 34101 | 50 | 0.29% |

| | | | |
|---|---|---|---|
| hsa_circ_0061248 | 25237 | 37 | 0.29% |
| hsa_circ_0101833 | 17054 | 25 | 0.29% |
| hsa_circ_0006158 | 21151 | 31 | 0.29% |
| hsa_circ_0075637 | 8874 | 13 | 0.29% |
| hsa_circ_0034215 | 8876 | 13 | 0.29% |
| hsa_circ_0010583 | 8877 | 13 | 0.29% |
| hsa_circ_0096977 | 16392 | 24 | 0.29% |
| hsa_circ_0104504 | 24591 | 36 | 0.29% |
| hsa_circ_0045376 | 8197 | 12 | 0.29% |
| hsa_circ_0098050 | 22544 | 33 | 0.29% |
| hsa_circ_0090686 | 8883 | 13 | 0.29% |
| hsa_circ_0005893 | 8201 | 12 | 0.29% |
| hsa_circ_0035699 | 8886 | 13 | 0.29% |
| hsa_circ_0080702 | 7523 | 11 | 0.29% |
| hsa_circ_0131966 | 8891 | 13 | 0.29% |
| hsa_circ_0131631 | 9576 | 14 | 0.29% |
| hsa_circ_0066963 | 8210 | 12 | 0.29% |
| hsa_circ_0132552 | 37631 | 55 | 0.29% |
| hsa_circ_0037571 | 8211 | 12 | 0.29% |
| hsa_circ_0132164 | 26002 | 38 | 0.29% |
| hsa_circ_0082989 | 8901 | 13 | 0.29% |
| hsa_circ_0034996 | 10275 | 15 | 0.29% |
| hsa_circ_0140649 | 20556 | 30 | 0.29% |
| hsa_circ_0139859 | 22615 | 33 | 0.29% |
| hsa_circ_0020490 | 8225 | 12 | 0.29% |
| hsa_circ_0021036 | 10283 | 15 | 0.29% |
| hsa_circ_0014674 | 32907 | 48 | 0.29% |
| hsa_circ_0009176 | 13714 | 20 | 0.29% |
| hsa_circ_0125750 | 19886 | 29 | 0.29% |
| hsa_circ_0033079 | 9601 | 14 | 0.29% |
| hsa_circ_0111944 | 29489 | 43 | 0.29% |
| hsa_circ_0138337 | 15091 | 22 | 0.29% |
| hsa_circ_0027138 | 8232 | 12 | 0.29% |
| hsa_circ_0132203 | 24697 | 36 | 0.29% |
| hsa_circ_0030467 | 10291 | 15 | 0.29% |
| hsa_circ_0054568 | 7547 | 11 | 0.29% |
| hsa_circ_0135865 | 25386 | 37 | 0.29% |
| hsa_circ_0134141 | 36365 | 53 | 0.29% |
| hsa_circ_0015168 | 12364 | 18 | 0.29% |
| hsa_circ_0103176 | 8243 | 12 | 0.29% |
| hsa_circ_0140249 | 8932 | 13 | 0.29% |
| hsa_circ_0027189 | 8245 | 12 | 0.29% |
| hsa_circ_0124401 | 8246 | 12 | 0.29% |
| hsa_circ_0115591 | 23366 | 34 | 0.29% |
| hsa_circ_0140431 | 15122 | 22 | 0.29% |
| hsa_circ_0047752 | 8936 | 13 | 0.29% |
| hsa_circ_0073279 | 8251 | 12 | 0.29% |
| hsa_circ_0055614 | 17881 | 26 | 0.29% |
| hsa_circ_0096674 | 8942 | 13 | 0.29% |
| hsa_circ_0040923 | 8267 | 12 | 0.29% |

| | | | |
|---|---|---|---|
| hsa_circ_0074700 | 35139 | 51 | 0.29% |
| hsa_circ_0089268 | 10335 | 15 | 0.29% |
| hsa_circ_0054722 | 8957 | 13 | 0.29% |
| hsa_circ_0133912 | 11714 | 17 | 0.29% |
| hsa_circ_0126820 | 68218 | 99 | 0.29% |
| hsa_circ_0129800 | 17916 | 26 | 0.29% |
| hsa_circ_0106616 | 28258 | 41 | 0.29% |
| hsa_circ_0101809 | 13096 | 19 | 0.29% |
| hsa_circ_0104903 | 22754 | 33 | 0.29% |
| hsa_circ_0107949 | 15861 | 23 | 0.29% |
| hsa_circ_0035713 | 8965 | 13 | 0.29% |
| hsa_circ_0011823 | 9657 | 14 | 0.29% |
| hsa_circ_0118093 | 35194 | 51 | 0.29% |
| hsa_circ_0002184 | 19326 | 28 | 0.29% |
| hsa_circ_0007582 | 24864 | 36 | 0.29% |
| hsa_circ_0001826 | 12437 | 18 | 0.29% |
| hsa_circ_0080703 | 7604 | 11 | 0.29% |
| hsa_circ_0073235 | 8987 | 13 | 0.29% |
| hsa_circ_0089761 | 8302 | 12 | 0.29% |
| hsa_circ_0098193 | 37363 | 54 | 0.29% |
| hsa_circ_0115404 | 9687 | 14 | 0.29% |
| hsa_circ_0139957 | 24225 | 35 | 0.29% |
| hsa_circ_0053612 | 9000 | 13 | 0.29% |
| hsa_circ_0093558 | 49850 | 72 | 0.29% |
| hsa_circ_0100074 | 16617 | 24 | 0.29% |
| hsa_circ_0090678 | 9002 | 13 | 0.29% |
| hsa_circ_0028549 | 8314 | 12 | 0.29% |
| hsa_circ_0083104 | 9009 | 13 | 0.29% |
| hsa_circ_0134107 | 23573 | 34 | 0.29% |
| hsa_circ_0045374 | 8326 | 12 | 0.29% |
| hsa_circ_0106218 | 34692 | 50 | 0.29% |
| hsa_circ_0096748 | 20817 | 30 | 0.29% |
| hsa_circ_0099151 | 27065 | 39 | 0.29% |
| hsa_circ_0115463 | 37478 | 54 | 0.29% |
| hsa_circ_0017433 | 9719 | 14 | 0.29% |
| hsa_circ_0021311 | 8331 | 12 | 0.29% |
| hsa_circ_0034213 | 9030 | 13 | 0.29% |
| hsa_circ_0075476 | 24312 | 35 | 0.29% |
| hsa_circ_0070307 | 7641 | 11 | 0.29% |
| hsa_circ_0074696 | 35428 | 51 | 0.29% |
| hsa_circ_0031702 | 29179 | 42 | 0.29% |
| hsa_circ_0106089 | 12506 | 18 | 0.29% |
| hsa_circ_0038818 | 29185 | 42 | 0.29% |
| hsa_circ_0139784 | 25715 | 37 | 0.29% |
| hsa_circ_0096394 | 20854 | 30 | 0.29% |
| hsa_circ_0124402 | 8342 | 12 | 0.29% |
| hsa_circ_0087436 | 8348 | 12 | 0.29% |
| hsa_circ_0082665 | 11828 | 17 | 0.29% |
| hsa_circ_0110605 | 9046 | 13 | 0.29% |
| hsa_circ_0121610 | 7658 | 11 | 0.29% |

| | | | |
|---|---|---|---|
| hsa_circ_0053698 | 9051 | 13 | 0.29% |
| hsa_circ_0055618 | 26457 | 38 | 0.29% |
| hsa_circ_0035683 | 9052 | 13 | 0.29% |
| hsa_circ_0025043 | 34828 | 50 | 0.29% |
| hsa_circ_0105704 | 9754 | 14 | 0.29% |
| hsa_circ_0054567 | 7665 | 11 | 0.29% |
| hsa_circ_0126366 | 21602 | 31 | 0.29% |
| hsa_circ_0128510 | 17427 | 25 | 0.29% |
| hsa_circ_0130590 | 48102 | 69 | 0.29% |
| hsa_circ_0094657 | 23008 | 33 | 0.29% |
| hsa_circ_0128509 | 17431 | 25 | 0.29% |
| hsa_circ_0071922 | 8367 | 12 | 0.29% |
| hsa_circ_0104908 | 17435 | 25 | 0.29% |
| hsa_circ_0089777 | 17437 | 25 | 0.29% |
| hsa_circ_0070301 | 7673 | 11 | 0.29% |
| hsa_circ_0137151 | 40458 | 58 | 0.29% |
| hsa_circ_0010650 | 9072 | 13 | 0.29% |
| hsa_circ_0066962 | 8376 | 12 | 0.29% |
| hsa_circ_0110608 | 9075 | 13 | 0.29% |
| hsa_circ_0033262 | 8377 | 12 | 0.29% |
| hsa_circ_0087155 | 9077 | 13 | 0.29% |
| hsa_circ_0033210 | 8381 | 12 | 0.29% |
| hsa_circ_0027192 | 8384 | 12 | 0.29% |
| hsa_circ_0111396 | 29346 | 42 | 0.29% |
| hsa_circ_0084225 | 8385 | 12 | 0.29% |
| hsa_circ_0126367 | 21672 | 31 | 0.29% |
| hsa_circ_0034154 | 9089 | 13 | 0.29% |
| hsa_circ_0068892 | 8390 | 12 | 0.29% |
| hsa_circ_0117737 | 44052 | 63 | 0.29% |
| hsa_circ_0077642 | 8391 | 12 | 0.29% |
| hsa_circ_0097086 | 27280 | 39 | 0.29% |
| hsa_circ_0078945 | 8395 | 12 | 0.29% |
| hsa_circ_0078315 | 9795 | 14 | 0.29% |
| hsa_circ_0102489 | 27287 | 39 | 0.29% |
| hsa_circ_0096978 | 16793 | 24 | 0.29% |
| hsa_circ_0030920 | 21696 | 31 | 0.29% |
| hsa_circ_0131366 | 23802 | 34 | 0.29% |
| hsa_circ_0131970 | 9104 | 13 | 0.29% |
| hsa_circ_0132571 | 33620 | 48 | 0.29% |
| hsa_circ_0095067 | 23114 | 33 | 0.29% |
| hsa_circ_0008875 | 17516 | 25 | 0.29% |
| hsa_circ_0121965 | 9109 | 13 | 0.29% |
| hsa_circ_0112279 | 10511 | 15 | 0.29% |
| hsa_circ_0140651 | 9111 | 13 | 0.29% |
| hsa_circ_0102378 | 8416 | 12 | 0.29% |
| hsa_circ_0136428 | 20340 | 29 | 0.29% |
| hsa_circ_0092424 | 9120 | 13 | 0.29% |
| hsa_circ_0077628 | 28064 | 40 | 0.29% |
| hsa_circ_0010291 | 8420 | 12 | 0.29% |
| hsa_circ_0101247 | 13332 | 19 | 0.29% |

| | | | |
|---|---|---|---|
| hsa_circ_0082992 | 9124 | 13 | 0.28% |
| hsa_circ_0110606 | 9126 | 13 | 0.28% |
| hsa_circ_0110546 | 21776 | 31 | 0.28% |
| hsa_circ_0078579 | 8430 | 12 | 0.28% |
| hsa_circ_0101226 | 50581 | 72 | 0.28% |
| hsa_circ_0110547 | 21779 | 31 | 0.28% |
| hsa_circ_0107584 | 11242 | 16 | 0.28% |
| hsa_circ_0090679 | 9136 | 13 | 0.28% |
| hsa_circ_0116733 | 9139 | 13 | 0.28% |
| hsa_circ_0045373 | 8438 | 12 | 0.28% |
| hsa_circ_0132216 | 9845 | 14 | 0.28% |
| hsa_circ_0039095 | 10550 | 15 | 0.28% |
| hsa_circ_0110828 | 10553 | 15 | 0.28% |
| hsa_circ_0131200 | 14776 | 21 | 0.28% |
| hsa_circ_0055619 | 26739 | 38 | 0.28% |
| hsa_circ_0119739 | 19703 | 28 | 0.28% |
| hsa_circ_0070125 | 9854 | 14 | 0.28% |
| hsa_circ_0079367 | 37305 | 53 | 0.28% |
| hsa_circ_0004520 | 13377 | 19 | 0.28% |
| hsa_circ_0065481 | 9858 | 14 | 0.28% |
| hsa_circ_0005259 | 30988 | 44 | 0.28% |
| hsa_circ_0118582 | 19017 | 27 | 0.28% |
| hsa_circ_0084245 | 8453 | 12 | 0.28% |
| hsa_circ_0138368 | 16204 | 23 | 0.28% |
| hsa_circ_0010608 | 9160 | 13 | 0.28% |
| hsa_circ_0074697 | 35937 | 51 | 0.28% |
| hsa_circ_0072907 | 11979 | 17 | 0.28% |
| hsa_circ_0002262 | 21854 | 31 | 0.28% |
| hsa_circ_0111259 | 26790 | 38 | 0.28% |
| hsa_circ_0013846 | 9165 | 13 | 0.28% |
| hsa_circ_0071602 | 9871 | 14 | 0.28% |
| hsa_circ_0002128 | 21865 | 31 | 0.28% |
| hsa_circ_0038813 | 29624 | 42 | 0.28% |
| hsa_circ_0053462 | 9174 | 13 | 0.28% |
| hsa_circ_0096813 | 9174 | 13 | 0.28% |
| hsa_circ_0099203 | 9174 | 13 | 0.28% |
| hsa_circ_0070477 | 9179 | 13 | 0.28% |
| hsa_circ_0101781 | 38840 | 55 | 0.28% |
| hsa_circ_0046025 | 11300 | 16 | 0.28% |
| hsa_circ_0135465 | 24023 | 34 | 0.28% |
| hsa_circ_0104981 | 12013 | 17 | 0.28% |
| hsa_circ_0002129 | 19083 | 27 | 0.28% |
| hsa_circ_0033080 | 9895 | 14 | 0.28% |
| hsa_circ_0100991 | 12024 | 17 | 0.28% |
| hsa_circ_0125536 | 12733 | 18 | 0.28% |
| hsa_circ_0002769 | 9906 | 14 | 0.28% |
| hsa_circ_0115426 | 24060 | 34 | 0.28% |
| hsa_circ_0053617 | 9200 | 13 | 0.28% |
| hsa_circ_0073314 | 9911 | 14 | 0.28% |
| hsa_circ_0117200 | 33276 | 47 | 0.28% |

| | | | |
|---|---|---|---|
| hsa_circ_0113972 | 29029 | 41 | 0.28% |
| hsa_circ_0033258 | 8497 | 12 | 0.28% |
| hsa_circ_0133032 | 30454 | 43 | 0.28% |
| hsa_circ_0121299 | 20542 | 29 | 0.28% |
| hsa_circ_0124739 | 12753 | 18 | 0.28% |
| hsa_circ_0099274 | 20550 | 29 | 0.28% |
| hsa_circ_0035716 | 9214 | 13 | 0.28% |
| hsa_circ_0010625 | 9216 | 13 | 0.28% |
| hsa_circ_0130876 | 17015 | 24 | 0.28% |
| hsa_circ_0127904 | 19853 | 28 | 0.28% |
| hsa_circ_0028367 | 9218 | 13 | 0.28% |
| hsa_circ_0120936 | 23408 | 33 | 0.28% |
| hsa_circ_0130317 | 9932 | 14 | 0.28% |
| hsa_circ_0035691 | 9224 | 13 | 0.28% |
| hsa_circ_0134871 | 11356 | 16 | 0.28% |
| hsa_circ_0045368 | 8521 | 12 | 0.28% |
| hsa_circ_0120364 | 20593 | 29 | 0.28% |
| hsa_circ_0095362 | 41905 | 59 | 0.28% |
| hsa_circ_0132756 | 14207 | 20 | 0.28% |
| hsa_circ_0056769 | 76010 | 107 | 0.28% |
| hsa_circ_0132052 | 9946 | 14 | 0.28% |
| hsa_circ_0120935 | 23445 | 33 | 0.28% |
| hsa_circ_0137851 | 22030 | 31 | 0.28% |
| hsa_circ_0035584 | 8528 | 12 | 0.28% |
| hsa_circ_0001586 | 39800 | 56 | 0.28% |
| hsa_circ_0140227 | 25594 | 36 | 0.28% |
| hsa_circ_0129890 | 29151 | 41 | 0.28% |
| hsa_circ_0090625 | 9245 | 13 | 0.28% |
| hsa_circ_0094594 | 12090 | 17 | 0.28% |
| hsa_circ_0025763 | 27739 | 39 | 0.28% |
| hsa_circ_0140621 | 51224 | 72 | 0.28% |
| hsa_circ_0053626 | 9249 | 13 | 0.28% |
| hsa_circ_0058822 | 9250 | 13 | 0.28% |
| hsa_circ_0005822 | 19216 | 27 | 0.28% |
| hsa_circ_0118567 | 16374 | 23 | 0.28% |
| hsa_circ_0134416 | 9257 | 13 | 0.28% |
| hsa_circ_0024120 | 9972 | 14 | 0.28% |
| hsa_circ_0045365 | 8553 | 12 | 0.28% |
| hsa_circ_0095812 | 18554 | 26 | 0.28% |
| hsa_circ_0125494 | 39251 | 55 | 0.28% |
| hsa_circ_0114414 | 22837 | 32 | 0.28% |
| hsa_circ_0056708 | 9286 | 13 | 0.28% |
| hsa_circ_0130006 | 9286 | 13 | 0.28% |
| hsa_circ_0077821 | 20003 | 28 | 0.28% |
| hsa_circ_0062504 | 25006 | 35 | 0.28% |
| hsa_circ_0083869 | 16435 | 23 | 0.28% |
| hsa_circ_0093306 | 16442 | 23 | 0.28% |
| hsa_circ_0107916 | 9295 | 13 | 0.28% |
| hsa_circ_0052895 | 9299 | 13 | 0.28% |
| hsa_circ_0134864 | 50080 | 70 | 0.28% |

| | | | |
|---|---|---|---|
| hsa_circ_0134872 | 11447 | 16 | 0.28% |
| hsa_circ_0006671 | 25759 | 36 | 0.28% |
| hsa_circ_0113975 | 11450 | 16 | 0.28% |
| hsa_circ_0004915 | 23623 | 33 | 0.28% |
| hsa_circ_0004147 | 20763 | 29 | 0.28% |
| hsa_circ_0077625 | 28639 | 40 | 0.28% |
| hsa_circ_0119449 | 17901 | 25 | 0.28% |
| hsa_circ_0019046 | 10743 | 15 | 0.28% |
| hsa_circ_0053591 | 9311 | 13 | 0.28% |
| hsa_circ_0008763 | 29368 | 41 | 0.28% |
| hsa_circ_0116507 | 18624 | 26 | 0.28% |
| hsa_circ_0038802 | 30092 | 42 | 0.28% |
| hsa_circ_0132053 | 10032 | 14 | 0.28% |
| hsa_circ_0101203 | 21498 | 30 | 0.28% |
| hsa_circ_0100545 | 22218 | 31 | 0.28% |
| hsa_circ_0016632 | 9318 | 13 | 0.28% |
| hsa_circ_0135880 | 12902 | 18 | 0.28% |
| hsa_circ_0120257 | 23655 | 33 | 0.28% |
| hsa_circ_0026133 | 38709 | 54 | 0.28% |
| hsa_circ_0078314 | 9319 | 13 | 0.28% |
| hsa_circ_0093562 | 51616 | 72 | 0.28% |
| hsa_circ_0134427 | 8603 | 12 | 0.28% |
| hsa_circ_0046021 | 11471 | 16 | 0.28% |
| hsa_circ_0130408 | 20076 | 28 | 0.28% |
| hsa_circ_0048716 | 50191 | 70 | 0.28% |
| hsa_circ_0078322 | 12192 | 17 | 0.28% |
| hsa_circ_0035718 | 9325 | 13 | 0.28% |
| hsa_circ_0033250 | 8608 | 12 | 0.28% |
| hsa_circ_0126988 | 15785 | 22 | 0.28% |
| hsa_circ_0002062 | 31572 | 44 | 0.28% |
| hsa_circ_0033232 | 8618 | 12 | 0.28% |
| hsa_circ_0125362 | 34477 | 48 | 0.28% |
| hsa_circ_0101810 | 9339 | 13 | 0.28% |
| hsa_circ_0046023 | 11498 | 16 | 0.28% |
| hsa_circ_0015845 | 10063 | 14 | 0.28% |
| hsa_circ_0053643 | 9346 | 13 | 0.28% |
| hsa_circ_0074698 | 36669 | 51 | 0.28% |
| hsa_circ_0104683 | 10068 | 14 | 0.28% |
| hsa_circ_0130755 | 23733 | 33 | 0.28% |
| hsa_circ_0037451 | 9353 | 13 | 0.28% |
| hsa_circ_0124473 | 32382 | 45 | 0.28% |
| hsa_circ_0101199 | 21588 | 30 | 0.28% |
| hsa_circ_0134084 | 25187 | 35 | 0.28% |
| hsa_circ_0011792 | 10075 | 14 | 0.28% |
| hsa_circ_0010250 | 10797 | 15 | 0.28% |
| hsa_circ_0096597 | 51830 | 72 | 0.28% |
| hsa_circ_0048717 | 34556 | 48 | 0.28% |
| hsa_circ_0010022 | 36723 | 51 | 0.28% |
| hsa_circ_0093986 | 35295 | 49 | 0.28% |
| hsa_circ_0084258 | 8644 | 12 | 0.28% |

| | | | |
|---|---|---|---|
| hsa_circ_0096500 | 22333 | 31 | 0.28% |
| hsa_circ_0102633 | 10812 | 15 | 0.28% |
| hsa_circ_0035746 | 9371 | 13 | 0.28% |
| hsa_circ_0053651 | 9374 | 13 | 0.28% |
| hsa_circ_0053463 | 9375 | 13 | 0.28% |
| hsa_circ_0124546 | 23802 | 33 | 0.28% |
| hsa_circ_0103690 | 28130 | 39 | 0.28% |
| hsa_circ_0009958 | 8656 | 12 | 0.28% |
| hsa_circ_0114415 | 23086 | 32 | 0.28% |
| hsa_circ_0121296 | 20922 | 29 | 0.28% |
| hsa_circ_0087156 | 9382 | 13 | 0.28% |
| hsa_circ_0124505 | 21663 | 30 | 0.28% |
| hsa_circ_0029649 | 9393 | 13 | 0.28% |
| hsa_circ_0132184 | 33237 | 46 | 0.28% |
| hsa_circ_0102874 | 25290 | 35 | 0.28% |
| hsa_circ_0088223 | 8671 | 12 | 0.28% |
| hsa_circ_0133610 | 27465 | 38 | 0.28% |
| hsa_circ_0029931 | 10843 | 15 | 0.28% |
| hsa_circ_0084268 | 9398 | 13 | 0.28% |
| hsa_circ_0076862 | 11567 | 16 | 0.28% |
| hsa_circ_0084226 | 8676 | 12 | 0.28% |
| hsa_circ_0015684 | 10128 | 14 | 0.28% |
| hsa_circ_0104937 | 15916 | 22 | 0.28% |
| hsa_circ_0005968 | 20266 | 28 | 0.28% |
| hsa_circ_0134873 | 11584 | 16 | 0.28% |
| hsa_circ_0038268 | 7965 | 11 | 0.28% |
| hsa_circ_0005431 | 22450 | 31 | 0.28% |
| hsa_circ_0008663 | 22461 | 31 | 0.28% |
| hsa_circ_0056839 | 12320 | 17 | 0.28% |
| hsa_circ_0075967 | 18850 | 26 | 0.28% |
| hsa_circ_0132734 | 11600 | 16 | 0.28% |
| hsa_circ_0101216 | 52212 | 72 | 0.28% |
| hsa_circ_0117087 | 32634 | 45 | 0.28% |
| hsa_circ_0082980 | 9428 | 13 | 0.28% |
| hsa_circ_0140243 | 7978 | 11 | 0.28% |
| hsa_circ_0027654 | 8705 | 12 | 0.28% |
| hsa_circ_0077316 | 9433 | 13 | 0.28% |
| hsa_circ_0081031 | 11612 | 16 | 0.28% |
| hsa_circ_0033267 | 8709 | 12 | 0.28% |
| hsa_circ_0130316 | 10161 | 14 | 0.28% |
| hsa_circ_0124594 | 33398 | 46 | 0.28% |
| hsa_circ_0081763 | 9439 | 13 | 0.28% |
| hsa_circ_0130879 | 33405 | 46 | 0.28% |
| hsa_circ_0113990 | 21068 | 29 | 0.28% |
| hsa_circ_0116951 | 25436 | 35 | 0.28% |
| hsa_circ_0133680 | 37074 | 51 | 0.28% |
| hsa_circ_0006488 | 12358 | 17 | 0.28% |
| hsa_circ_0124697 | 8724 | 12 | 0.28% |
| hsa_circ_0046761 | 10906 | 15 | 0.28% |
| hsa_circ_0053699 | 9453 | 13 | 0.28% |

| | | | |
|---|---|---|---|
| hsa_circ_0087948 | 37813 | 52 | 0.28% |
| hsa_circ_0132014 | 20362 | 28 | 0.28% |
| hsa_circ_0090662 | 9454 | 13 | 0.28% |
| hsa_circ_0028324 | 22545 | 31 | 0.28% |
| hsa_circ_0134848 | 21821 | 30 | 0.27% |
| hsa_circ_0134839 | 20370 | 28 | 0.27% |
| hsa_circ_0095200 | 10914 | 15 | 0.27% |
| hsa_circ_0030461 | 10915 | 15 | 0.27% |
| hsa_circ_0133212 | 9460 | 13 | 0.27% |
| hsa_circ_0121478 | 16010 | 22 | 0.27% |
| hsa_circ_0076515 | 8006 | 11 | 0.27% |
| hsa_circ_0105925 | 22563 | 31 | 0.27% |
| hsa_circ_0076826 | 9462 | 13 | 0.27% |
| hsa_circ_0093563 | 52410 | 72 | 0.27% |
| hsa_circ_0021295 | 8009 | 11 | 0.27% |
| hsa_circ_0007515 | 8740 | 12 | 0.27% |
| hsa_circ_0027814 | 29862 | 41 | 0.27% |
| hsa_circ_0004710 | 16024 | 22 | 0.27% |
| hsa_circ_0113283 | 8742 | 12 | 0.27% |
| hsa_circ_0033289 | 8743 | 12 | 0.27% |
| hsa_circ_0137161 | 8744 | 12 | 0.27% |
| hsa_circ_0010609 | 9474 | 13 | 0.27% |
| hsa_circ_0057224 | 10935 | 15 | 0.27% |
| hsa_circ_0055238 | 10206 | 14 | 0.27% |
| hsa_circ_0073315 | 10207 | 14 | 0.27% |
| hsa_circ_0116500 | 39386 | 54 | 0.27% |
| hsa_circ_0098902 | 24822 | 34 | 0.27% |
| hsa_circ_0011781 | 10221 | 14 | 0.27% |
| hsa_circ_0010017 | 37238 | 51 | 0.27% |
| hsa_circ_0119863 | 14604 | 20 | 0.27% |
| hsa_circ_0103174 | 8763 | 12 | 0.27% |
| hsa_circ_0124880 | 18989 | 26 | 0.27% |
| hsa_circ_0094385 | 26293 | 36 | 0.27% |
| hsa_circ_0070435 | 18260 | 25 | 0.27% |
| hsa_circ_0107574 | 34336 | 47 | 0.27% |
| hsa_circ_0045364 | 8770 | 12 | 0.27% |
| hsa_circ_0104818 | 9502 | 13 | 0.27% |
| hsa_circ_0125880 | 27058 | 37 | 0.27% |
| hsa_circ_0085050 | 10971 | 15 | 0.27% |
| hsa_circ_0111352 | 19748 | 27 | 0.27% |
| hsa_circ_0084264 | 8778 | 12 | 0.27% |
| hsa_circ_0053604 | 9513 | 13 | 0.27% |
| hsa_circ_0033247 | 8785 | 12 | 0.27% |
| hsa_circ_0100849 | 30020 | 41 | 0.27% |
| hsa_circ_0071592 | 8788 | 12 | 0.27% |
| hsa_circ_0112400 | 35153 | 48 | 0.27% |
| hsa_circ_0068969 | 8791 | 12 | 0.27% |
| hsa_circ_0033092 | 9527 | 13 | 0.27% |
| hsa_circ_0124700 | 41776 | 57 | 0.27% |
| hsa_circ_0069223 | 8795 | 12 | 0.27% |

| | | | |
|---|---|---|---|
| hsa_circ_0097205 | 21258 | 29 | 0.27% |
| hsa_circ_0037573 | 8064 | 11 | 0.27% |
| hsa_circ_0056626 | 9532 | 13 | 0.27% |
| hsa_circ_0111531 | 17598 | 24 | 0.27% |
| hsa_circ_0115922 | 41075 | 56 | 0.27% |
| hsa_circ_0134861 | 51366 | 70 | 0.27% |
| hsa_circ_0044188 | 27153 | 37 | 0.27% |
| hsa_circ_0094087 | 30836 | 42 | 0.27% |
| hsa_circ_0096636 | 10281 | 14 | 0.27% |
| hsa_circ_0111441 | 10281 | 14 | 0.27% |
| hsa_circ_0089269 | 11016 | 15 | 0.27% |
| hsa_circ_0126860 | 31583 | 43 | 0.27% |
| hsa_circ_0139324 | 30118 | 41 | 0.27% |
| hsa_circ_0129705 | 22042 | 30 | 0.27% |
| hsa_circ_0128751 | 41149 | 56 | 0.27% |
| hsa_circ_0038786 | 30872 | 42 | 0.27% |
| hsa_circ_0005618 | 27937 | 38 | 0.27% |
| hsa_circ_0130368 | 24998 | 34 | 0.27% |
| hsa_circ_0094263 | 15441 | 21 | 0.27% |
| hsa_circ_0132217 | 10297 | 14 | 0.27% |
| hsa_circ_0053568 | 9562 | 13 | 0.27% |
| hsa_circ_0115592 | 16183 | 22 | 0.27% |
| hsa_circ_0104733 | 21339 | 29 | 0.27% |
| hsa_circ_0003591 | 9567 | 13 | 0.27% |
| hsa_circ_0097971 | 29437 | 40 | 0.27% |
| hsa_circ_0118575 | 16929 | 23 | 0.27% |
| hsa_circ_0129336 | 16934 | 23 | 0.27% |
| hsa_circ_0062692 | 11795 | 16 | 0.27% |
| hsa_circ_0005507 | 18431 | 25 | 0.27% |
| hsa_circ_0021307 | 8850 | 12 | 0.27% |
| hsa_circ_0128416 | 32461 | 44 | 0.27% |
| hsa_circ_0000983 | 20657 | 28 | 0.27% |
| hsa_circ_0133587 | 28035 | 38 | 0.27% |
| hsa_circ_0140495 | 15494 | 21 | 0.27% |
| hsa_circ_0028368 | 9592 | 13 | 0.27% |
| hsa_circ_0053497 | 9592 | 13 | 0.27% |
| hsa_circ_0084301 | 9592 | 13 | 0.27% |
| hsa_circ_0020486 | 10331 | 14 | 0.27% |
| hsa_circ_0128352 | 14024 | 19 | 0.27% |
| hsa_circ_0087157 | 9596 | 13 | 0.27% |
| hsa_circ_0139519 | 64977 | 88 | 0.27% |
| hsa_circ_0139326 | 30274 | 41 | 0.27% |
| hsa_circ_0019978 | 8863 | 12 | 0.27% |
| hsa_circ_0084794 | 9607 | 13 | 0.27% |
| hsa_circ_0131514 | 24388 | 33 | 0.27% |
| hsa_circ_0113743 | 35477 | 48 | 0.27% |
| hsa_circ_0107499 | 54694 | 74 | 0.27% |
| hsa_circ_0114326 | 25872 | 35 | 0.27% |
| hsa_circ_0097962 | 29569 | 40 | 0.27% |
| hsa_circ_0007873 | 22178 | 30 | 0.27% |

| | | | |
|---|---|---|---|
| hsa_circ_0070293 | 8132 | 11 | 0.27% |
| hsa_circ_0123319 | 51037 | 69 | 0.27% |
| hsa_circ_0084227 | 8880 | 12 | 0.27% |
| hsa_circ_0091567 | 10363 | 14 | 0.27% |
| hsa_circ_0010610 | 9623 | 13 | 0.27% |
| hsa_circ_0094191 | 11105 | 15 | 0.27% |
| hsa_circ_0017526 | 40727 | 55 | 0.27% |
| hsa_circ_0118568 | 17032 | 23 | 0.27% |
| hsa_circ_0071599 | 10368 | 14 | 0.27% |
| hsa_circ_0066593 | 9629 | 13 | 0.27% |
| hsa_circ_0101227 | 53356 | 72 | 0.27% |
| hsa_circ_0111596 | 31146 | 42 | 0.27% |
| hsa_circ_0053680 | 9641 | 13 | 0.27% |
| hsa_circ_0087951 | 9641 | 13 | 0.27% |
| hsa_circ_0010626 | 9642 | 13 | 0.27% |
| hsa_circ_0118980 | 15577 | 21 | 0.27% |
| hsa_circ_0140496 | 15579 | 21 | 0.27% |
| hsa_circ_0136029 | 13356 | 18 | 0.27% |
| hsa_circ_0139332 | 31166 | 42 | 0.27% |
| hsa_circ_0002175 | 32653 | 44 | 0.27% |
| hsa_circ_0110514 | 9648 | 13 | 0.27% |
| hsa_circ_0101513 | 86852 | 117 | 0.27% |
| hsa_circ_0047408 | 11138 | 15 | 0.27% |
| hsa_circ_0009093 | 26733 | 36 | 0.27% |
| hsa_circ_0140427 | 34902 | 47 | 0.27% |
| hsa_circ_0010270 | 11141 | 15 | 0.27% |
| hsa_circ_0076276 | 9657 | 13 | 0.27% |
| hsa_circ_0078070 | 8916 | 12 | 0.27% |
| hsa_circ_0116876 | 24525 | 33 | 0.27% |
| hsa_circ_0125490 | 23044 | 31 | 0.27% |
| hsa_circ_0011535 | 33454 | 45 | 0.27% |
| hsa_circ_0033236 | 8922 | 12 | 0.27% |
| hsa_circ_0112675 | 8922 | 12 | 0.27% |
| hsa_circ_0033245 | 8926 | 12 | 0.27% |
| hsa_circ_0098121 | 24553 | 33 | 0.27% |
| hsa_circ_0090706 | 9673 | 13 | 0.27% |
| hsa_circ_0111532 | 17862 | 24 | 0.27% |
| hsa_circ_0096886 | 12653 | 17 | 0.27% |
| hsa_circ_0104052 | 25337 | 34 | 0.27% |
| hsa_circ_0047472 | 21615 | 29 | 0.27% |
| hsa_circ_0111442 | 10445 | 14 | 0.27% |
| hsa_circ_0099428 | 18656 | 25 | 0.27% |
| hsa_circ_0071588 | 8955 | 12 | 0.27% |
| hsa_circ_0136112 | 32097 | 43 | 0.27% |
| hsa_circ_0132747 | 29858 | 40 | 0.27% |
| hsa_circ_0035705 | 9705 | 13 | 0.27% |
| hsa_circ_0011786 | 10452 | 14 | 0.27% |
| hsa_circ_0138130 | 9708 | 13 | 0.27% |
| hsa_circ_0035684 | 9710 | 13 | 0.27% |
| hsa_circ_0140225 | 26897 | 36 | 0.27% |

| | | | |
|---|---|---|---|
| hsa_circ_0010269 | 11217 | 15 | 0.27% |
| hsa_circ_0125829 | 17202 | 23 | 0.27% |
| hsa_circ_0084793 | 9725 | 13 | 0.27% |
| hsa_circ_0090626 | 9725 | 13 | 0.27% |
| hsa_circ_0099434 | 26936 | 36 | 0.27% |
| hsa_circ_0053579 | 9735 | 13 | 0.27% |
| hsa_circ_0121206 | 10489 | 14 | 0.27% |
| hsa_circ_0073234 | 9741 | 13 | 0.27% |
| hsa_circ_0102946 | 29976 | 40 | 0.27% |
| hsa_circ_0090707 | 10492 | 14 | 0.27% |
| hsa_circ_0060311 | 8244 | 11 | 0.27% |
| hsa_circ_0035116 | 9744 | 13 | 0.27% |
| hsa_circ_0137547 | 32984 | 44 | 0.27% |
| hsa_circ_0137548 | 32988 | 44 | 0.27% |
| hsa_circ_0035704 | 9748 | 13 | 0.27% |
| hsa_circ_0109310 | 23999 | 32 | 0.27% |
| hsa_circ_0073778 | 9753 | 13 | 0.27% |
| hsa_circ_0078326 | 9003 | 12 | 0.27% |
| hsa_circ_0109311 | 24013 | 32 | 0.27% |
| hsa_circ_0054564 | 8255 | 11 | 0.27% |
| hsa_circ_0125368 | 37549 | 50 | 0.27% |
| hsa_circ_0140244 | 8262 | 11 | 0.27% |
| hsa_circ_0089251 | 18030 | 24 | 0.27% |
| hsa_circ_0104834 | 20284 | 27 | 0.27% |
| hsa_circ_0081174 | 10519 | 14 | 0.27% |
| hsa_circ_0100859 | 21042 | 28 | 0.27% |
| hsa_circ_0125345 | 27058 | 36 | 0.27% |
| hsa_circ_0126369 | 27058 | 36 | 0.27% |
| hsa_circ_0112888 | 34577 | 46 | 0.27% |
| hsa_circ_0087154 | 9773 | 13 | 0.27% |
| hsa_circ_0007972 | 18795 | 25 | 0.27% |
| hsa_circ_0052882 | 9777 | 13 | 0.27% |
| hsa_circ_0104053 | 25580 | 34 | 0.27% |
| hsa_circ_0059686 | 45915 | 61 | 0.27% |
| hsa_circ_0083095 | 9786 | 13 | 0.27% |
| hsa_circ_0118092 | 38393 | 51 | 0.27% |
| hsa_circ_0135387 | 27854 | 37 | 0.27% |
| hsa_circ_0097206 | 21835 | 29 | 0.27% |
| hsa_circ_0076835 | 9790 | 13 | 0.27% |
| hsa_circ_0084228 | 9039 | 12 | 0.27% |
| hsa_circ_0101485 | 9797 | 13 | 0.27% |
| hsa_circ_0033273 | 9046 | 12 | 0.27% |
| hsa_circ_0110400 | 29400 | 39 | 0.27% |
| hsa_circ_0131509 | 9803 | 13 | 0.27% |
| hsa_circ_0097354 | 39214 | 52 | 0.27% |
| hsa_circ_0018436 | 10564 | 14 | 0.27% |
| hsa_circ_0095340 | 45275 | 60 | 0.27% |
| hsa_circ_0063388 | 50571 | 67 | 0.26% |
| hsa_circ_0091568 | 10568 | 14 | 0.26% |
| hsa_circ_0010292 | 9060 | 12 | 0.26% |

| | | | |
|---|---|---|---|
| hsa_circ_0134842 | 61161 | 81 | 0.26% |
| hsa_circ_0098749 | 41531 | 55 | 0.26% |
| hsa_circ_0094181 | 30214 | 40 | 0.26% |
| hsa_circ_0132037 | 9823 | 13 | 0.26% |
| hsa_circ_0075686 | 9825 | 13 | 0.26% |
| hsa_circ_0122870 | 21920 | 29 | 0.26% |
| hsa_circ_0007851 | 17385 | 23 | 0.26% |
| hsa_circ_0084253 | 9071 | 12 | 0.26% |
| hsa_circ_0033239 | 9073 | 12 | 0.26% |
| hsa_circ_0056015 | 21171 | 28 | 0.26% |
| hsa_circ_0123123 | 11343 | 15 | 0.26% |
| hsa_circ_0053436 | 9834 | 13 | 0.26% |
| hsa_circ_0083091 | 9835 | 13 | 0.26% |
| hsa_circ_0136381 | 9837 | 13 | 0.26% |
| hsa_circ_0013934 | 28000 | 37 | 0.26% |
| hsa_circ_0134249 | 87789 | 116 | 0.26% |
| hsa_circ_0118559 | 17414 | 23 | 0.26% |
| hsa_circ_0135384 | 19691 | 26 | 0.26% |
| hsa_circ_0086009 | 11361 | 15 | 0.26% |
| hsa_circ_0091275 | 43935 | 58 | 0.26% |
| hsa_circ_0094559 | 30313 | 40 | 0.26% |
| hsa_circ_0010574 | 9853 | 13 | 0.26% |
| hsa_circ_0119512 | 38672 | 51 | 0.26% |
| hsa_circ_0037448 | 9861 | 13 | 0.26% |
| hsa_circ_0028369 | 9865 | 13 | 0.26% |
| hsa_circ_0093716 | 11383 | 15 | 0.26% |
| hsa_circ_0064024 | 293746 | 387 | 0.26% |
| hsa_circ_0085333 | 18219 | 24 | 0.26% |
| hsa_circ_0053605 | 9870 | 13 | 0.26% |
| hsa_circ_0140124 | 28855 | 38 | 0.26% |
| hsa_circ_0053547 | 9872 | 13 | 0.26% |
| hsa_circ_0064025 | 293975 | 387 | 0.26% |
| hsa_circ_0138048 | 15197 | 20 | 0.26% |
| hsa_circ_0061581 | 11400 | 15 | 0.26% |
| hsa_circ_0016627 | 9888 | 13 | 0.26% |
| hsa_circ_0130685 | 28145 | 37 | 0.26% |
| hsa_circ_0044176 | 25109 | 33 | 0.26% |
| hsa_circ_0004422 | 28165 | 37 | 0.26% |
| hsa_circ_0035726 | 9896 | 13 | 0.26% |
| hsa_circ_0098186 | 22076 | 29 | 0.26% |
| hsa_circ_0134108 | 12946 | 17 | 0.26% |
| hsa_circ_0064026 | 294737 | 387 | 0.26% |
| hsa_circ_0127423 | 16761 | 22 | 0.26% |
| hsa_circ_0123343 | 9144 | 12 | 0.26% |
| hsa_circ_0121533 | 27433 | 36 | 0.26% |
| hsa_circ_0007400 | 25910 | 34 | 0.26% |
| hsa_circ_0052415 | 9146 | 12 | 0.26% |
| hsa_circ_0053464 | 9912 | 13 | 0.26% |
| hsa_circ_0050837 | 10676 | 14 | 0.26% |
| hsa_circ_0056622 | 9155 | 12 | 0.26% |

| | | | |
|---|---|---|---|
| hsa_circ_0121854 | 9922 | 13 | 0.26% |
| hsa_circ_0099465 | 22138 | 29 | 0.26% |
| hsa_circ_0121855 | 9925 | 13 | 0.26% |
| hsa_circ_0134789 | 11454 | 15 | 0.26% |
| hsa_circ_0122036 | 16036 | 21 | 0.26% |
| hsa_circ_0109951 | 29791 | 39 | 0.26% |
| hsa_circ_0125661 | 32120 | 42 | 0.26% |
| hsa_circ_0008065 | 14531 | 19 | 0.26% |
| hsa_circ_0090140 | 11473 | 15 | 0.26% |
| hsa_circ_0034207 | 9946 | 13 | 0.26% |
| hsa_circ_0035685 | 9946 | 13 | 0.26% |
| hsa_circ_0027147 | 9947 | 13 | 0.26% |
| hsa_circ_0006938 | 32914 | 43 | 0.26% |
| hsa_circ_0006038 | 35976 | 47 | 0.26% |
| hsa_circ_0134980 | 13013 | 17 | 0.26% |
| hsa_circ_0007114 | 10717 | 14 | 0.26% |
| hsa_circ_0066014 | 9952 | 13 | 0.26% |
| hsa_circ_0129859 | 12251 | 16 | 0.26% |
| hsa_circ_0078357 | 32162 | 42 | 0.26% |
| hsa_circ_0007253 | 31400 | 41 | 0.26% |
| hsa_circ_0008048 | 27576 | 36 | 0.26% |
| hsa_circ_0111526 | 18388 | 24 | 0.26% |
| hsa_circ_0138307 | 12259 | 16 | 0.26% |
| hsa_circ_0115679 | 16859 | 22 | 0.26% |
| hsa_circ_0113424 | 16861 | 22 | 0.26% |
| hsa_circ_0127007 | 25302 | 33 | 0.26% |
| hsa_circ_0095669 | 31440 | 41 | 0.26% |
| hsa_circ_0139746 | 10736 | 14 | 0.26% |
| hsa_circ_0046017 | 12274 | 16 | 0.26% |
| hsa_circ_0010263 | 11507 | 15 | 0.26% |
| hsa_circ_0122005 | 19182 | 25 | 0.26% |
| hsa_circ_0008971 | 30712 | 40 | 0.26% |
| hsa_circ_0027871 | 17663 | 23 | 0.26% |
| hsa_circ_0101502 | 33803 | 44 | 0.26% |
| hsa_circ_0084247 | 9989 | 13 | 0.26% |
| hsa_circ_0120154 | 40735 | 53 | 0.26% |
| hsa_circ_0059793 | 37703 | 49 | 0.26% |
| hsa_circ_0120934 | 25412 | 33 | 0.26% |
| hsa_circ_0090698 | 10782 | 14 | 0.26% |
| hsa_circ_0016763 | 10012 | 13 | 0.26% |
| hsa_circ_0035700 | 10013 | 13 | 0.26% |
| hsa_circ_0117593 | 11557 | 15 | 0.26% |
| hsa_circ_0045193 | 10787 | 14 | 0.26% |
| hsa_circ_0010575 | 10017 | 13 | 0.26% |
| hsa_circ_0117594 | 11561 | 15 | 0.26% |
| hsa_circ_0053465 | 10021 | 13 | 0.26% |
| hsa_circ_0002797 | 10795 | 14 | 0.26% |
| hsa_circ_0034742 | 10797 | 14 | 0.26% |
| hsa_circ_0034113 | 10803 | 14 | 0.26% |
| hsa_circ_0070898 | 10803 | 14 | 0.26% |

| | | | |
|---|---|---|---|
| hsa_circ_0109306 | 36268 | 47 | 0.26% |
| hsa_circ_0109369 | 20069 | 26 | 0.26% |
| hsa_circ_0023997 | 9264 | 12 | 0.26% |
| hsa_circ_0112162 | 19305 | 25 | 0.26% |
| hsa_circ_0090672 | 10039 | 13 | 0.26% |
| hsa_circ_0034328 | 25491 | 33 | 0.26% |
| hsa_circ_0033278 | 9270 | 12 | 0.26% |
| hsa_circ_0035688 | 10043 | 13 | 0.26% |
| hsa_circ_0095670 | 31689 | 41 | 0.26% |
| hsa_circ_0099425 | 19323 | 25 | 0.26% |
| hsa_circ_0053618 | 10048 | 13 | 0.26% |
| hsa_circ_0128750 | 34010 | 44 | 0.26% |
| hsa_circ_0082986 | 10049 | 13 | 0.26% |
| hsa_circ_0074394 | 20111 | 26 | 0.26% |
| hsa_circ_0134236 | 20114 | 26 | 0.26% |
| hsa_circ_0007872 | 10060 | 13 | 0.26% |
| hsa_circ_0112163 | 19348 | 25 | 0.26% |
| hsa_circ_0127359 | 23220 | 30 | 0.26% |
| hsa_circ_0064817 | 9290 | 12 | 0.26% |
| hsa_circ_0102534 | 21681 | 28 | 0.26% |
| hsa_circ_0115690 | 23236 | 30 | 0.26% |
| hsa_circ_0105608 | 10070 | 13 | 0.26% |
| hsa_circ_0100997 | 10848 | 14 | 0.26% |
| hsa_circ_0106392 | 29447 | 38 | 0.26% |
| hsa_circ_0016602 | 10851 | 14 | 0.26% |
| hsa_circ_0111016 | 22482 | 29 | 0.26% |
| hsa_circ_0099828 | 16281 | 21 | 0.26% |
| hsa_circ_0106221 | 10856 | 14 | 0.26% |
| hsa_circ_0016772 | 10083 | 13 | 0.26% |
| hsa_circ_0101822 | 11635 | 15 | 0.26% |
| hsa_circ_0122152 | 37233 | 48 | 0.26% |
| hsa_circ_0062929 | 10088 | 13 | 0.26% |
| hsa_circ_0123389 | 40354 | 52 | 0.26% |
| hsa_circ_0094110 | 27163 | 35 | 0.26% |
| hsa_circ_0010285 | 9314 | 12 | 0.26% |
| hsa_circ_0105920 | 28719 | 37 | 0.26% |
| hsa_circ_0078358 | 32609 | 42 | 0.26% |
| hsa_circ_0079910 | 20963 | 27 | 0.26% |
| hsa_circ_0124643 | 20965 | 27 | 0.26% |
| hsa_circ_0070853 | 9318 | 12 | 0.26% |
| hsa_circ_0111163 | 38828 | 50 | 0.26% |
| hsa_circ_0004284 | 16314 | 21 | 0.26% |
| hsa_circ_0070162 | 10876 | 14 | 0.26% |
| hsa_circ_0078342 | 112651 | 145 | 0.26% |
| hsa_circ_0139677 | 10877 | 14 | 0.26% |
| hsa_circ_0074393 | 20203 | 26 | 0.26% |
| hsa_circ_0122330 | 16322 | 21 | 0.26% |
| hsa_circ_0007487 | 10884 | 14 | 0.26% |
| hsa_circ_0093723 | 14772 | 19 | 0.26% |
| hsa_circ_0101200 | 30335 | 39 | 0.26% |

| | | | |
|---|---|---|---|
| hsa_circ_0104628 | 9334 | 12 | 0.26% |
| hsa_circ_0063469 | 42792 | 55 | 0.26% |
| hsa_circ_0035666 | 10116 | 13 | 0.26% |
| hsa_circ_0123449 | 24124 | 31 | 0.26% |
| hsa_circ_0084265 | 10117 | 13 | 0.26% |
| hsa_circ_0078341 | 112856 | 145 | 0.26% |
| hsa_circ_0026090 | 10120 | 13 | 0.26% |
| hsa_circ_0121851 | 35031 | 45 | 0.26% |
| hsa_circ_0138774 | 19469 | 25 | 0.26% |
| hsa_circ_0105731 | 24144 | 31 | 0.26% |
| hsa_circ_0111410 | 32721 | 42 | 0.26% |
| hsa_circ_0131123 | 34289 | 44 | 0.26% |
| hsa_circ_0115689 | 10132 | 13 | 0.26% |
| hsa_circ_0120041 | 17152 | 22 | 0.26% |
| hsa_circ_0010611 | 10915 | 14 | 0.26% |
| hsa_circ_0020118 | 92807 | 119 | 0.26% |
| hsa_circ_0107201 | 60832 | 78 | 0.26% |
| hsa_circ_0080030 | 21840 | 28 | 0.26% |
| hsa_circ_0096434 | 14820 | 19 | 0.26% |
| hsa_circ_0099351 | 11700 | 15 | 0.26% |
| hsa_circ_0090699 | 10920 | 14 | 0.26% |
| hsa_circ_0114583 | 10140 | 13 | 0.26% |
| hsa_circ_0076889 | 10143 | 13 | 0.26% |
| hsa_circ_0078359 | 32786 | 42 | 0.26% |
| hsa_circ_0115572 | 14839 | 19 | 0.26% |
| hsa_circ_0055617 | 20318 | 26 | 0.26% |
| hsa_circ_0078338 | 113315 | 145 | 0.26% |
| hsa_circ_0124256 | 73471 | 94 | 0.26% |
| hsa_circ_0062930 | 10161 | 13 | 0.26% |
| hsa_circ_0139548 | 21107 | 27 | 0.26% |
| hsa_circ_0084229 | 9381 | 12 | 0.26% |
| hsa_circ_0078031 | 290905 | 372 | 0.26% |
| hsa_circ_0127643 | 25810 | 33 | 0.26% |
| hsa_circ_0009965 | 11737 | 15 | 0.26% |
| hsa_circ_0073306 | 10964 | 14 | 0.26% |
| hsa_circ_0001859 | 34461 | 44 | 0.26% |
| hsa_circ_0069222 | 10182 | 13 | 0.26% |
| hsa_circ_0070856 | 9399 | 12 | 0.26% |
| hsa_circ_0114404 | 22716 | 29 | 0.26% |
| hsa_circ_0010578 | 10968 | 14 | 0.26% |
| hsa_circ_0089797 | 48581 | 62 | 0.26% |
| hsa_circ_0033237 | 9403 | 12 | 0.26% |
| hsa_circ_0104812 | 15674 | 20 | 0.26% |
| hsa_circ_0081371 | 109736 | 140 | 0.26% |
| hsa_circ_0070299 | 8627 | 11 | 0.26% |
| hsa_circ_0082666 | 13335 | 17 | 0.25% |
| hsa_circ_0078065 | 9415 | 12 | 0.25% |
| hsa_circ_0139549 | 21186 | 27 | 0.25% |
| hsa_circ_0033217 | 9418 | 12 | 0.25% |
| hsa_circ_0121906 | 21192 | 27 | 0.25% |

| | | | |
|---|---|---|---|
| hsa_circ_0078329 | 10206 | 13 | 0.25% |
| hsa_circ_0140245 | 8636 | 11 | 0.25% |
| hsa_circ_0133358 | 28264 | 36 | 0.25% |
| hsa_circ_0134266 | 27486 | 35 | 0.25% |
| hsa_circ_0139557 | 22777 | 29 | 0.25% |
| hsa_circ_0017174 | 10212 | 13 | 0.25% |
| hsa_circ_0016779 | 11000 | 14 | 0.25% |
| hsa_circ_0134565 | 11011 | 14 | 0.25% |
| hsa_circ_0127855 | 18095 | 23 | 0.25% |
| hsa_circ_0128880 | 33831 | 43 | 0.25% |
| hsa_circ_0116923 | 11016 | 14 | 0.25% |
| hsa_circ_0023531 | 22821 | 29 | 0.25% |
| hsa_circ_0010612 | 11018 | 14 | 0.25% |
| hsa_circ_0055234 | 11019 | 14 | 0.25% |
| hsa_circ_0133600 | 10232 | 13 | 0.25% |
| hsa_circ_0084685 | 27551 | 35 | 0.25% |
| hsa_circ_0106016 | 29127 | 37 | 0.25% |
| hsa_circ_0111059 | 29137 | 37 | 0.25% |
| hsa_circ_0103728 | 11025 | 14 | 0.25% |
| hsa_circ_0039088 | 11814 | 15 | 0.25% |
| hsa_circ_0113985 | 10241 | 13 | 0.25% |
| hsa_circ_0123073 | 9454 | 12 | 0.25% |
| hsa_circ_0046016 | 9455 | 12 | 0.25% |
| hsa_circ_0119417 | 22071 | 28 | 0.25% |
| hsa_circ_0098609 | 37843 | 48 | 0.25% |
| hsa_circ_0126973 | 10250 | 13 | 0.25% |
| hsa_circ_0113756 | 18928 | 24 | 0.25% |
| hsa_circ_0126360 | 34704 | 44 | 0.25% |
| hsa_circ_0075642 | 39449 | 50 | 0.25% |
| hsa_circ_0134417 | 10258 | 13 | 0.25% |
| hsa_circ_0095248 | 29202 | 37 | 0.25% |
| hsa_circ_0103956 | 32385 | 41 | 0.25% |
| hsa_circ_0120245 | 20540 | 26 | 0.25% |
| hsa_circ_0114702 | 30039 | 38 | 0.25% |
| hsa_circ_0139325 | 32417 | 41 | 0.25% |
| hsa_circ_0075687 | 10279 | 13 | 0.25% |
| hsa_circ_0127452 | 18984 | 24 | 0.25% |
| hsa_circ_0037446 | 10284 | 13 | 0.25% |
| hsa_circ_0057897 | 12658 | 16 | 0.25% |
| hsa_circ_0053627 | 10288 | 13 | 0.25% |
| hsa_circ_0023998 | 9498 | 12 | 0.25% |
| hsa_circ_0114131 | 18209 | 23 | 0.25% |
| hsa_circ_0130004 | 19002 | 24 | 0.25% |
| hsa_circ_0117787 | 42756 | 54 | 0.25% |
| hsa_circ_0093673 | 14252 | 18 | 0.25% |
| hsa_circ_0102752 | 13465 | 17 | 0.25% |
| hsa_circ_0082725 | 39624 | 50 | 0.25% |
| hsa_circ_0016037 | 31702 | 40 | 0.25% |
| hsa_circ_0114341 | 26160 | 33 | 0.25% |
| hsa_circ_0119371 | 25372 | 32 | 0.25% |

| | | | |
|---|---|---|---|
| hsa_circ_0131498 | 23794 | 30 | 0.25% |
| hsa_circ_0137885 | 38876 | 49 | 0.25% |
| hsa_circ_0033229 | 9521 | 12 | 0.25% |
| hsa_circ_0028372 | 10322 | 13 | 0.25% |
| hsa_circ_0073321 | 9529 | 12 | 0.25% |
| hsa_circ_0140246 | 8735 | 11 | 0.25% |
| hsa_circ_0133725 | 19859 | 25 | 0.25% |
| hsa_circ_0127573 | 27810 | 35 | 0.25% |
| hsa_circ_0011808 | 11126 | 14 | 0.25% |
| hsa_circ_0126000 | 99373 | 125 | 0.25% |
| hsa_circ_0105097 | 31833 | 40 | 0.25% |
| hsa_circ_0129889 | 32638 | 41 | 0.25% |
| hsa_circ_0123613 | 9553 | 12 | 0.25% |
| hsa_circ_0094786 | 27069 | 34 | 0.25% |
| hsa_circ_0102646 | 47775 | 60 | 0.25% |
| hsa_circ_0016208 | 11149 | 14 | 0.25% |
| hsa_circ_0073312 | 11150 | 14 | 0.25% |
| hsa_circ_0131172 | 30273 | 38 | 0.25% |
| hsa_circ_0090666 | 10358 | 13 | 0.25% |
| hsa_circ_0104691 | 10359 | 13 | 0.25% |
| hsa_circ_0119280 | 40641 | 51 | 0.25% |
| hsa_circ_0138957 | 28688 | 36 | 0.25% |
| hsa_circ_0053606 | 10361 | 13 | 0.25% |
| hsa_circ_0130375 | 41445 | 52 | 0.25% |
| hsa_circ_0074633 | 10367 | 13 | 0.25% |
| hsa_circ_0108846 | 11962 | 15 | 0.25% |
| hsa_circ_0094139 | 35093 | 44 | 0.25% |
| hsa_circ_0108138 | 30315 | 38 | 0.25% |
| hsa_circ_0139511 | 11967 | 15 | 0.25% |
| hsa_circ_0057898 | 12767 | 16 | 0.25% |
| hsa_circ_0034197 | 10374 | 13 | 0.25% |
| hsa_circ_0125668 | 23942 | 30 | 0.25% |
| hsa_circ_0025793 | 19954 | 25 | 0.25% |
| hsa_circ_0117127 | 30353 | 38 | 0.25% |
| hsa_circ_0114076 | 38341 | 48 | 0.25% |
| hsa_circ_0108303 | 28759 | 36 | 0.25% |
| hsa_circ_0035706 | 10386 | 13 | 0.25% |
| hsa_circ_0117973 | 15180 | 19 | 0.25% |
| hsa_circ_0094523 | 30366 | 38 | 0.25% |
| hsa_circ_0115510 | 14384 | 18 | 0.25% |
| hsa_circ_0110497 | 27170 | 34 | 0.25% |
| hsa_circ_0021354 | 10400 | 13 | 0.25% |
| hsa_circ_0134422 | 23204 | 29 | 0.25% |
| hsa_circ_0094234 | 28014 | 35 | 0.25% |
| hsa_circ_0084230 | 9605 | 12 | 0.25% |
| hsa_circ_0132350 | 37637 | 47 | 0.25% |
| hsa_circ_0131710 | 14419 | 18 | 0.25% |
| hsa_circ_0105615 | 10417 | 13 | 0.25% |
| hsa_circ_0137435 | 20843 | 26 | 0.25% |
| hsa_circ_0108855 | 49710 | 62 | 0.25% |

| | | | |
|---|---|---|---|
| hsa_circ_0095662 | 28079 | 35 | 0.25% |
| hsa_circ_0018916 | 12837 | 16 | 0.25% |
| hsa_circ_0122215 | 34504 | 43 | 0.25% |
| hsa_circ_0103729 | 11236 | 14 | 0.25% |
| hsa_circ_0010550 | 11237 | 14 | 0.25% |
| hsa_circ_0119193 | 29711 | 37 | 0.25% |
| hsa_circ_0134276 | 32137 | 40 | 0.25% |
| hsa_circ_0005675 | 32951 | 41 | 0.25% |
| hsa_circ_0104095 | 29748 | 37 | 0.25% |
| hsa_circ_0061246 | 36186 | 45 | 0.25% |
| hsa_circ_0140429 | 28963 | 36 | 0.25% |
| hsa_circ_0010576 | 10461 | 13 | 0.25% |
| hsa_circ_0140221 | 41044 | 51 | 0.25% |
| hsa_circ_0116781 | 10466 | 13 | 0.25% |
| hsa_circ_0061245 | 36244 | 45 | 0.25% |
| hsa_circ_0033756 | 14500 | 18 | 0.25% |
| hsa_circ_0111454 | 26584 | 33 | 0.25% |
| hsa_circ_0134978 | 10473 | 13 | 0.25% |
| hsa_circ_0095430 | 26587 | 33 | 0.25% |
| hsa_circ_0122882 | 20144 | 25 | 0.25% |
| hsa_circ_0039246 | 20955 | 26 | 0.25% |
| hsa_circ_0130684 | 29832 | 37 | 0.25% |
| hsa_circ_0077334 | 10485 | 13 | 0.25% |
| hsa_circ_0024117 | 11294 | 14 | 0.25% |
| hsa_circ_0026087 | 10495 | 13 | 0.25% |
| hsa_circ_0139177 | 10496 | 13 | 0.25% |
| hsa_circ_0039327 | 21800 | 27 | 0.25% |
| hsa_circ_0044213 | 39573 | 49 | 0.25% |
| hsa_circ_0055603 | 11307 | 14 | 0.25% |
| hsa_circ_0104094 | 13739 | 17 | 0.25% |
| hsa_circ_0125122 | 17784 | 22 | 0.25% |
| hsa_circ_0136198 | 26687 | 33 | 0.25% |
| hsa_circ_0123390 | 42060 | 52 | 0.25% |
| hsa_circ_0029404 | 33166 | 41 | 0.25% |
| hsa_circ_0111996 | 29933 | 37 | 0.25% |
| hsa_circ_0000229 | 47732 | 59 | 0.25% |
| hsa_circ_0021355 | 10518 | 13 | 0.25% |
| hsa_circ_0055987 | 11333 | 14 | 0.25% |
| hsa_circ_0090655 | 10526 | 13 | 0.25% |
| hsa_circ_0114674 | 23493 | 29 | 0.25% |
| hsa_circ_0015096 | 28357 | 35 | 0.25% |
| hsa_circ_0010251 | 12155 | 15 | 0.25% |
| hsa_circ_0034205 | 10538 | 13 | 0.25% |
| hsa_circ_0008233 | 17837 | 22 | 0.25% |
| hsa_circ_0125120 | 17839 | 22 | 0.25% |
| hsa_circ_0117587 | 20275 | 25 | 0.25% |
| hsa_circ_0097853 | 10550 | 13 | 0.25% |
| hsa_circ_0101501 | 34086 | 42 | 0.25% |
| hsa_circ_0076821 | 10551 | 13 | 0.25% |
| hsa_circ_0125861 | 11365 | 14 | 0.25% |

| | | | |
|---|---|---|---|
| hsa_circ_0010570 | 10560 | 13 | 0.25% |
| hsa_circ_0000585 | 10562 | 13 | 0.25% |
| hsa_circ_0002527 | 12187 | 15 | 0.25% |
| hsa_circ_0134979 | 10568 | 13 | 0.25% |
| hsa_circ_0094758 | 25216 | 31 | 0.25% |
| hsa_circ_0010551 | 11389 | 14 | 0.25% |
| hsa_circ_0010282 | 9763 | 12 | 0.25% |
| hsa_circ_0004417 | 30103 | 37 | 0.25% |
| hsa_circ_0096889 | 16272 | 20 | 0.25% |
| hsa_circ_0057199 | 13018 | 16 | 0.25% |
| hsa_circ_0095867 | 26854 | 33 | 0.25% |
| hsa_circ_0094293 | 45571 | 56 | 0.25% |
| hsa_circ_0124704 | 12209 | 15 | 0.25% |
| hsa_circ_0053580 | 10583 | 13 | 0.25% |
| hsa_circ_0053613 | 10584 | 13 | 0.25% |
| hsa_circ_0082882 | 11403 | 14 | 0.25% |
| hsa_circ_0111544 | 34214 | 42 | 0.25% |
| hsa_circ_0133321 | 39109 | 48 | 0.25% |
| hsa_circ_0084248 | 10594 | 13 | 0.25% |
| hsa_circ_0118576 | 33413 | 41 | 0.25% |
| hsa_circ_0118870 | 11411 | 14 | 0.25% |
| hsa_circ_0125690 | 32610 | 40 | 0.25% |
| hsa_circ_0090139 | 12230 | 15 | 0.25% |
| hsa_circ_0023415 | 31801 | 39 | 0.25% |
| hsa_circ_0055231 | 11419 | 14 | 0.25% |
| hsa_circ_0108639 | 11425 | 14 | 0.25% |
| hsa_circ_0111990 | 30203 | 37 | 0.25% |
| hsa_circ_0107954 | 29392 | 36 | 0.24% |
| hsa_circ_0131344 | 17148 | 21 | 0.24% |
| hsa_circ_0127895 | 34307 | 42 | 0.24% |
| hsa_circ_0098511 | 13076 | 16 | 0.24% |
| hsa_circ_0118569 | 33516 | 41 | 0.24% |
| hsa_circ_0140134 | 23709 | 29 | 0.24% |
| hsa_circ_0123461 | 9812 | 12 | 0.24% |
| hsa_circ_0125862 | 11459 | 14 | 0.24% |
| hsa_circ_0010271 | 9825 | 12 | 0.24% |
| hsa_circ_0053511 | 10654 | 13 | 0.24% |
| hsa_circ_0101780 | 17215 | 21 | 0.24% |
| hsa_circ_0098992 | 18035 | 22 | 0.24% |
| hsa_circ_0052883 | 10661 | 13 | 0.24% |
| hsa_circ_0053581 | 10661 | 13 | 0.24% |
| hsa_circ_0124254 | 77109 | 94 | 0.24% |
| hsa_circ_0035696 | 10668 | 13 | 0.24% |
| hsa_circ_0053516 | 10672 | 13 | 0.24% |
| hsa_circ_0084231 | 9853 | 12 | 0.24% |
| hsa_circ_0080465 | 36139 | 44 | 0.24% |
| hsa_circ_0071595 | 11500 | 14 | 0.24% |
| hsa_circ_0124255 | 77244 | 94 | 0.24% |
| hsa_circ_0123444 | 31239 | 38 | 0.24% |
| hsa_circ_0045185 | 11513 | 14 | 0.24% |

| | | | |
|---|---|---|---|
| hsa_circ_0013811 | 22226 | 27 | 0.24% |
| hsa_circ_0122159 | 22228 | 27 | 0.24% |
| hsa_circ_0131485 | 27173 | 33 | 0.24% |
| hsa_circ_0009959 | 9884 | 12 | 0.24% |
| hsa_circ_0033251 | 9885 | 12 | 0.24% |
| hsa_circ_0058696 | 19771 | 24 | 0.24% |
| hsa_circ_0067710 | 46135 | 56 | 0.24% |
| hsa_circ_0049510 | 43665 | 53 | 0.24% |
| hsa_circ_0038239 | 9887 | 12 | 0.24% |
| hsa_circ_0023999 | 9888 | 12 | 0.24% |
| hsa_circ_0010577 | 10713 | 13 | 0.24% |
| hsa_circ_0120767 | 11540 | 14 | 0.24% |
| hsa_circ_0101228 | 10717 | 13 | 0.24% |
| hsa_circ_0116253 | 36286 | 44 | 0.24% |
| hsa_circ_0015946 | 9898 | 12 | 0.24% |
| hsa_circ_0077318 | 10729 | 13 | 0.24% |
| hsa_circ_0020327 | 45397 | 55 | 0.24% |
| hsa_circ_0093301 | 48700 | 59 | 0.24% |
| hsa_circ_0049097 | 10731 | 13 | 0.24% |
| hsa_circ_0055803 | 37984 | 46 | 0.24% |
| hsa_circ_0124719 | 11567 | 14 | 0.24% |
| hsa_circ_0034135 | 10743 | 13 | 0.24% |
| hsa_circ_0054944 | 14049 | 17 | 0.24% |
| hsa_circ_0134430 | 21494 | 26 | 0.24% |
| hsa_circ_0098793 | 11579 | 14 | 0.24% |
| hsa_circ_0098791 | 11583 | 14 | 0.24% |
| hsa_circ_0111999 | 17375 | 21 | 0.24% |
| hsa_circ_0119258 | 10758 | 13 | 0.24% |
| hsa_circ_0053592 | 10760 | 13 | 0.24% |
| hsa_circ_0053607 | 10763 | 13 | 0.24% |
| hsa_circ_0128876 | 44728 | 54 | 0.24% |
| hsa_circ_0115094 | 32304 | 39 | 0.24% |
| hsa_circ_0108935 | 21538 | 26 | 0.24% |
| hsa_circ_0015682 | 11599 | 14 | 0.24% |
| hsa_circ_0120715 | 25692 | 31 | 0.24% |
| hsa_circ_0065482 | 11603 | 14 | 0.24% |
| hsa_circ_0006963 | 25693 | 31 | 0.24% |
| hsa_circ_0139736 | 40617 | 49 | 0.24% |
| hsa_circ_0132745 | 80457 | 97 | 0.24% |
| hsa_circ_0011810 | 11613 | 14 | 0.24% |
| hsa_circ_0090137 | 12444 | 15 | 0.24% |
| hsa_circ_0131479 | 27377 | 33 | 0.24% |
| hsa_circ_0127500 | 26560 | 32 | 0.24% |
| hsa_circ_0094262 | 10793 | 13 | 0.24% |
| hsa_circ_0134757 | 30719 | 37 | 0.24% |
| hsa_circ_0119191 | 23250 | 28 | 0.24% |
| hsa_circ_0093089 | 48166 | 58 | 0.24% |
| hsa_circ_0111302 | 40713 | 49 | 0.24% |
| hsa_circ_0026476 | 46554 | 56 | 0.24% |
| hsa_circ_0081611 | 13303 | 16 | 0.24% |

| | | | |
|---|---|---|---|
| hsa_circ_0093582 | 18295 | 22 | 0.24% |
| hsa_circ_0053681 | 10815 | 13 | 0.24% |
| hsa_circ_0013865 | 23294 | 28 | 0.24% |
| hsa_circ_0115830 | 26624 | 32 | 0.24% |
| hsa_circ_0129339 | 28301 | 34 | 0.24% |
| hsa_circ_0082987 | 10821 | 13 | 0.24% |
| hsa_circ_0103283 | 25811 | 31 | 0.24% |
| hsa_circ_0133140 | 87433 | 105 | 0.24% |
| hsa_circ_0115954 | 33314 | 40 | 0.24% |
| hsa_circ_0101519 | 39145 | 47 | 0.24% |
| hsa_circ_0111249 | 40816 | 49 | 0.24% |
| hsa_circ_0133677 | 42484 | 51 | 0.24% |
| hsa_circ_0019374 | 46658 | 56 | 0.24% |
| hsa_circ_0033233 | 9999 | 12 | 0.24% |
| hsa_circ_0073599 | 10000 | 12 | 0.24% |
| hsa_circ_0005940 | 44169 | 53 | 0.24% |
| hsa_circ_0139169 | 10834 | 13 | 0.24% |
| hsa_circ_0108325 | 21669 | 26 | 0.24% |
| hsa_circ_0016172 | 11683 | 14 | 0.24% |
| hsa_circ_0000694 | 32548 | 39 | 0.24% |
| hsa_circ_0053582 | 10850 | 13 | 0.24% |
| hsa_circ_0127522 | 15858 | 19 | 0.24% |
| hsa_circ_0120477 | 11685 | 14 | 0.24% |
| hsa_circ_0082966 | 10851 | 13 | 0.24% |
| hsa_circ_0099432 | 27546 | 33 | 0.24% |
| hsa_circ_0105792 | 50085 | 60 | 0.24% |
| hsa_circ_0078330 | 10852 | 13 | 0.24% |
| hsa_circ_0139498 | 21706 | 26 | 0.24% |
| hsa_circ_0034729 | 40091 | 48 | 0.24% |
| hsa_circ_0117111 | 53464 | 64 | 0.24% |
| hsa_circ_0111235 | 40962 | 49 | 0.24% |
| hsa_circ_0132863 | 71068 | 85 | 0.24% |
| hsa_circ_0052884 | 10872 | 13 | 0.24% |
| hsa_circ_0010280 | 10036 | 12 | 0.24% |
| hsa_circ_0033755 | 12554 | 15 | 0.24% |
| hsa_circ_0032355 | 41861 | 50 | 0.24% |
| hsa_circ_0101651 | 31820 | 38 | 0.24% |
| hsa_circ_0053519 | 10886 | 13 | 0.24% |
| hsa_circ_0099966 | 53606 | 64 | 0.24% |
| hsa_circ_0108922 | 24292 | 29 | 0.24% |
| hsa_circ_0102179 | 33518 | 40 | 0.24% |
| hsa_circ_0118563 | 34358 | 41 | 0.24% |
| hsa_circ_0119861 | 11732 | 14 | 0.24% |
| hsa_circ_0133771 | 49452 | 59 | 0.24% |
| hsa_circ_0101832 | 20956 | 25 | 0.24% |
| hsa_circ_0010275 | 10059 | 12 | 0.24% |
| hsa_circ_0134330 | 31859 | 38 | 0.24% |
| hsa_circ_0123705 | 20961 | 25 | 0.24% |
| hsa_circ_0000693 | 32712 | 39 | 0.24% |
| hsa_circ_0093559 | 60397 | 72 | 0.24% |

| | | | |
|---|---|---|---|
| hsa_circ_0099964 | 53702 | 64 | 0.24% |
| hsa_circ_0111981 | 20142 | 24 | 0.24% |
| hsa_circ_0117969 | 15948 | 19 | 0.24% |
| hsa_circ_0108300 | 22671 | 27 | 0.24% |
| hsa_circ_0075695 | 15118 | 18 | 0.24% |
| hsa_circ_0114417 | 31918 | 38 | 0.24% |
| hsa_circ_0126508 | 22685 | 27 | 0.24% |
| hsa_circ_0113057 | 51253 | 61 | 0.24% |
| hsa_circ_0077311 | 10923 | 13 | 0.24% |
| hsa_circ_0131486 | 47901 | 57 | 0.24% |
| hsa_circ_0108299 | 22691 | 27 | 0.24% |
| hsa_circ_0099451 | 24372 | 29 | 0.24% |
| hsa_circ_0101652 | 31941 | 38 | 0.24% |
| hsa_circ_0111187 | 41189 | 49 | 0.24% |
| hsa_circ_0054435 | 20178 | 24 | 0.24% |
| hsa_circ_0094776 | 11771 | 14 | 0.24% |
| hsa_circ_0139496 | 21870 | 26 | 0.24% |
| hsa_circ_0117517 | 11783 | 14 | 0.24% |
| hsa_circ_0120508 | 41255 | 49 | 0.24% |
| hsa_circ_0001781 | 17682 | 21 | 0.24% |
| hsa_circ_0030298 | 27791 | 33 | 0.24% |
| hsa_circ_0050836 | 11791 | 14 | 0.24% |
| hsa_circ_0074634 | 10949 | 13 | 0.24% |
| hsa_circ_0102718 | 10949 | 13 | 0.24% |
| hsa_circ_0129839 | 60649 | 72 | 0.24% |
| hsa_circ_0062967 | 12639 | 15 | 0.24% |
| hsa_circ_0102754 | 28659 | 34 | 0.24% |
| hsa_circ_0122758 | 30345 | 36 | 0.24% |
| hsa_circ_0125615 | 52276 | 62 | 0.24% |
| hsa_circ_0082983 | 10963 | 13 | 0.24% |
| hsa_circ_0098187 | 24456 | 29 | 0.24% |
| hsa_circ_0099963 | 53984 | 64 | 0.24% |
| hsa_circ_0098184 | 28679 | 34 | 0.24% |
| hsa_circ_0139494 | 21933 | 26 | 0.24% |
| hsa_circ_0102743 | 10969 | 13 | 0.24% |
| hsa_circ_0061997 | 10128 | 12 | 0.24% |
| hsa_circ_0100863 | 17727 | 21 | 0.24% |
| hsa_circ_0122589 | 32922 | 39 | 0.24% |
| hsa_circ_0073307 | 11820 | 14 | 0.24% |
| hsa_circ_0132557 | 51516 | 61 | 0.24% |
| hsa_circ_0128856 | 43072 | 51 | 0.24% |
| hsa_circ_0057899 | 13513 | 16 | 0.24% |
| hsa_circ_0127398 | 43922 | 52 | 0.24% |
| hsa_circ_0130421 | 24506 | 29 | 0.24% |
| hsa_circ_0133137 | 38889 | 46 | 0.24% |
| hsa_circ_0033725 | 15221 | 18 | 0.24% |
| hsa_circ_0130933 | 22835 | 27 | 0.24% |
| hsa_circ_0122785 | 32984 | 39 | 0.24% |
| hsa_circ_0130934 | 22840 | 27 | 0.24% |
| hsa_circ_0137895 | 30455 | 36 | 0.24% |

| | | | |
|---|---|---|---|
| hsa_circ_0021111 | 32150 | 38 | 0.24% |
| hsa_circ_0024668 | 10154 | 12 | 0.24% |
| hsa_circ_0134156 | 34714 | 41 | 0.24% |
| hsa_circ_0138320 | 31328 | 37 | 0.24% |
| hsa_circ_0001515 | 14395 | 17 | 0.24% |
| hsa_circ_0020328 | 46578 | 55 | 0.24% |
| hsa_circ_0006100 | 32207 | 38 | 0.24% |
| hsa_circ_0082963 | 11020 | 13 | 0.24% |
| hsa_circ_0088212 | 10174 | 12 | 0.24% |
| hsa_circ_0139488 | 30525 | 36 | 0.24% |
| hsa_circ_0128373 | 24601 | 29 | 0.24% |
| hsa_circ_0073303 | 11878 | 14 | 0.24% |
| hsa_circ_0124562 | 26305 | 31 | 0.24% |
| hsa_circ_0034299 | 42439 | 50 | 0.24% |
| hsa_circ_0105023 | 40752 | 48 | 0.24% |
| hsa_circ_0011778 | 11887 | 14 | 0.24% |
| hsa_circ_0035686 | 11042 | 13 | 0.24% |
| hsa_circ_0030462 | 12741 | 15 | 0.24% |
| hsa_circ_0034153 | 11043 | 13 | 0.24% |
| hsa_circ_0130596 | 31431 | 37 | 0.24% |
| hsa_circ_0093560 | 61191 | 72 | 0.24% |
| hsa_circ_0101211 | 11049 | 13 | 0.24% |
| hsa_circ_0096913 | 28900 | 34 | 0.24% |
| hsa_circ_0132732 | 13605 | 16 | 0.24% |
| hsa_circ_0029406 | 30657 | 36 | 0.23% |
| hsa_circ_0136181 | 37479 | 44 | 0.23% |
| hsa_circ_0090134 | 12779 | 15 | 0.23% |
| hsa_circ_0096914 | 28976 | 34 | 0.23% |
| hsa_circ_0131624 | 20456 | 24 | 0.23% |
| hsa_circ_0066015 | 11087 | 13 | 0.23% |
| hsa_circ_0046018 | 13651 | 16 | 0.23% |
| hsa_circ_0034198 | 11092 | 13 | 0.23% |
| hsa_circ_0034099 | 35841 | 42 | 0.23% |
| hsa_circ_0125441 | 12803 | 15 | 0.23% |
| hsa_circ_0139485 | 22200 | 26 | 0.23% |
| hsa_circ_0077320 | 11103 | 13 | 0.23% |
| hsa_circ_0092465 | 15375 | 18 | 0.23% |
| hsa_circ_0125967 | 26483 | 31 | 0.23% |
| hsa_circ_0076829 | 11106 | 13 | 0.23% |
| hsa_circ_0005123 | 40163 | 47 | 0.23% |
| hsa_circ_0118552 | 11964 | 14 | 0.23% |
| hsa_circ_0133224 | 18802 | 22 | 0.23% |
| hsa_circ_0124805 | 15387 | 18 | 0.23% |
| hsa_circ_0112557 | 43606 | 51 | 0.23% |
| hsa_circ_0135400 | 11974 | 14 | 0.23% |
| hsa_circ_0136420 | 27371 | 32 | 0.23% |
| hsa_circ_0053556 | 11120 | 13 | 0.23% |
| hsa_circ_0102716 | 11121 | 13 | 0.23% |
| hsa_circ_0083366 | 17110 | 20 | 0.23% |
| hsa_circ_0128874 | 12834 | 15 | 0.23% |

| | | | |
|---|---|---|---|
| hsa_circ_0140226 | 12848 | 15 | 0.23% |
| hsa_circ_0035132 | 11136 | 13 | 0.23% |
| hsa_circ_0066823 | 13707 | 16 | 0.23% |
| hsa_circ_0070844 | 11994 | 14 | 0.23% |
| hsa_circ_0121970 | 65131 | 76 | 0.23% |
| hsa_circ_0094774 | 12000 | 14 | 0.23% |
| hsa_circ_0064611 | 30859 | 36 | 0.23% |
| hsa_circ_0082959 | 11144 | 13 | 0.23% |
| hsa_circ_0098191 | 27442 | 32 | 0.23% |
| hsa_circ_0047477 | 96979 | 113 | 0.23% |
| hsa_circ_0113977 | 11161 | 13 | 0.23% |
| hsa_circ_0128601 | 35202 | 41 | 0.23% |
| hsa_circ_0123765 | 39498 | 46 | 0.23% |
| hsa_circ_0102155 | 44653 | 52 | 0.23% |
| hsa_circ_0136226 | 22330 | 26 | 0.23% |
| hsa_circ_0121032 | 26625 | 31 | 0.23% |
| hsa_circ_0053466 | 11168 | 13 | 0.23% |
| hsa_circ_0094947 | 27504 | 32 | 0.23% |
| hsa_circ_0017917 | 40399 | 47 | 0.23% |
| hsa_circ_0052885 | 11177 | 13 | 0.23% |
| hsa_circ_0098599 | 51596 | 60 | 0.23% |
| hsa_circ_0127022 | 41286 | 48 | 0.23% |
| hsa_circ_0033777 | 15498 | 18 | 0.23% |
| hsa_circ_0126458 | 33581 | 39 | 0.23% |
| hsa_circ_0103144 | 35309 | 41 | 0.23% |
| hsa_circ_0113496 | 29281 | 34 | 0.23% |
| hsa_circ_0131937 | 36189 | 42 | 0.23% |
| hsa_circ_0134846 | 18096 | 21 | 0.23% |
| hsa_circ_0085897 | 12931 | 15 | 0.23% |
| hsa_circ_0131603 | 32762 | 38 | 0.23% |
| hsa_circ_0121843 | 40526 | 47 | 0.23% |
| hsa_circ_0120064 | 25875 | 30 | 0.23% |
| hsa_circ_0124629 | 43142 | 50 | 0.23% |
| hsa_circ_0042748 | 12081 | 14 | 0.23% |
| hsa_circ_0001991 | 38843 | 45 | 0.23% |
| hsa_circ_0121844 | 40579 | 47 | 0.23% |
| hsa_circ_0082880 | 12955 | 15 | 0.23% |
| hsa_circ_0121512 | 21598 | 25 | 0.23% |
| hsa_circ_0100861 | 18147 | 21 | 0.23% |
| hsa_circ_0090673 | 11234 | 13 | 0.23% |
| hsa_circ_0134859 | 60508 | 70 | 0.23% |
| hsa_circ_0108297 | 23352 | 27 | 0.23% |
| hsa_circ_0137585 | 13842 | 16 | 0.23% |
| hsa_circ_0055600 | 11250 | 13 | 0.23% |
| hsa_circ_0123085 | 32889 | 38 | 0.23% |
| hsa_circ_0130409 | 24237 | 28 | 0.23% |
| hsa_circ_0137465 | 21643 | 25 | 0.23% |
| hsa_circ_0116544 | 42426 | 49 | 0.23% |
| hsa_circ_0112553 | 44158 | 51 | 0.23% |
| hsa_circ_0139505 | 22513 | 26 | 0.23% |

| | | | |
|---|---|---|---|
| hsa_circ_0105929 | 18193 | 21 | 0.23% |
| hsa_circ_0035692 | 11263 | 13 | 0.23% |
| hsa_circ_0073308 | 12134 | 14 | 0.23% |
| hsa_circ_0008281 | 52877 | 61 | 0.23% |
| hsa_circ_0052890 | 11269 | 13 | 0.23% |
| hsa_circ_0002939 | 32955 | 38 | 0.23% |
| hsa_circ_0094693 | 23419 | 27 | 0.23% |
| hsa_circ_0094941 | 33850 | 39 | 0.23% |
| hsa_circ_0096915 | 29511 | 34 | 0.23% |
| hsa_circ_0064994 | 11287 | 13 | 0.23% |
| hsa_circ_0097298 | 23445 | 27 | 0.23% |
| hsa_circ_0082976 | 11290 | 13 | 0.23% |
| hsa_circ_0102621 | 35612 | 41 | 0.23% |
| hsa_circ_0078360 | 36484 | 42 | 0.23% |
| hsa_circ_0113274 | 10424 | 12 | 0.23% |
| hsa_circ_0108828 | 35631 | 41 | 0.23% |
| hsa_circ_0117095 | 13042 | 15 | 0.23% |
| hsa_circ_0122322 | 53921 | 62 | 0.23% |
| hsa_circ_0085059 | 13046 | 15 | 0.23% |
| hsa_circ_0118540 | 29584 | 34 | 0.23% |
| hsa_circ_0056227 | 12185 | 14 | 0.23% |
| hsa_circ_0116501 | 47009 | 54 | 0.23% |
| hsa_circ_0097847 | 41805 | 48 | 0.23% |
| hsa_circ_0005076 | 22647 | 26 | 0.23% |
| hsa_circ_0127517 | 16558 | 19 | 0.23% |
| hsa_circ_0111538 | 26152 | 30 | 0.23% |
| hsa_circ_0053548 | 11333 | 13 | 0.23% |
| hsa_circ_0034797 | 27898 | 32 | 0.23% |
| hsa_circ_0108305 | 27032 | 31 | 0.23% |
| hsa_circ_0020487 | 12208 | 14 | 0.23% |
| hsa_circ_0111418 | 41873 | 48 | 0.23% |
| hsa_circ_0076407 | 41006 | 47 | 0.23% |
| hsa_circ_0047395 | 15708 | 18 | 0.23% |
| hsa_circ_0018835 | 31417 | 36 | 0.23% |
| hsa_circ_0138591 | 26183 | 30 | 0.23% |
| hsa_circ_0139395 | 14839 | 17 | 0.23% |
| hsa_circ_0094955 | 13968 | 16 | 0.23% |
| hsa_circ_0115923 | 48903 | 56 | 0.23% |
| hsa_circ_0033234 | 10480 | 12 | 0.23% |
| hsa_circ_0051491 | 55898 | 64 | 0.23% |
| hsa_circ_0130114 | 54158 | 62 | 0.23% |
| hsa_circ_0137994 | 11357 | 13 | 0.23% |
| hsa_circ_0110855 | 12234 | 14 | 0.23% |
| hsa_circ_0114560 | 90889 | 104 | 0.23% |
| hsa_circ_0135278 | 23597 | 27 | 0.23% |
| hsa_circ_0103798 | 45464 | 52 | 0.23% |
| hsa_circ_0082981 | 11370 | 13 | 0.23% |
| hsa_circ_0000294 | 24491 | 28 | 0.23% |
| hsa_circ_0138590 | 34123 | 39 | 0.23% |
| hsa_circ_0021356 | 11376 | 13 | 0.23% |

| | | | |
|---|---|---|---|
| hsa_circ_0068781 | 29759 | 34 | 0.23% |
| hsa_circ_0021793 | 31526 | 36 | 0.23% |
| hsa_circ_0139399 | 20150 | 23 | 0.23% |
| hsa_circ_0004199 | 25408 | 29 | 0.23% |
| hsa_circ_0070854 | 10515 | 12 | 0.23% |
| hsa_circ_0084232 | 10517 | 12 | 0.23% |
| hsa_circ_0052891 | 11394 | 13 | 0.23% |
| hsa_circ_0093690 | 32431 | 37 | 0.23% |
| hsa_circ_0122757 | 31561 | 36 | 0.23% |
| hsa_circ_0028356 | 11399 | 13 | 0.23% |
| hsa_circ_0122754 | 22818 | 26 | 0.23% |
| hsa_circ_0118580 | 35985 | 41 | 0.23% |
| hsa_circ_0118388 | 21070 | 24 | 0.23% |
| hsa_circ_0100801 | 19316 | 22 | 0.23% |
| hsa_circ_0046009 | 14052 | 16 | 0.23% |
| hsa_circ_0129220 | 40419 | 46 | 0.23% |
| hsa_circ_0099832 | 12308 | 14 | 0.23% |
| hsa_circ_0124289 | 21101 | 24 | 0.23% |
| hsa_circ_0053467 | 11430 | 13 | 0.23% |
| hsa_circ_0101715 | 48362 | 55 | 0.23% |
| hsa_circ_0134857 | 21104 | 24 | 0.23% |
| hsa_circ_0131510 | 11438 | 13 | 0.23% |
| hsa_circ_0035190 | 27280 | 31 | 0.23% |
| hsa_circ_0032387 | 12320 | 14 | 0.23% |
| hsa_circ_0021357 | 11445 | 13 | 0.23% |
| hsa_circ_0138746 | 39624 | 45 | 0.23% |
| hsa_circ_0056625 | 11448 | 13 | 0.23% |
| hsa_circ_0091210 | 118901 | 135 | 0.23% |
| hsa_circ_0002571 | 14092 | 16 | 0.23% |
| hsa_circ_0137995 | 11452 | 13 | 0.23% |
| hsa_circ_0078684 | 13217 | 15 | 0.23% |
| hsa_circ_0137034 | 58155 | 66 | 0.23% |
| hsa_circ_0013638 | 40553 | 46 | 0.23% |
| hsa_circ_0114168 | 34385 | 39 | 0.23% |
| hsa_circ_0001760 | 21163 | 24 | 0.23% |
| hsa_circ_0122539 | 25572 | 29 | 0.23% |
| hsa_circ_0116814 | 44976 | 51 | 0.23% |
| hsa_circ_0039926 | 34421 | 39 | 0.23% |
| hsa_circ_0104048 | 30026 | 34 | 0.23% |
| hsa_circ_0057900 | 14136 | 16 | 0.23% |
| hsa_circ_0008437 | 19451 | 22 | 0.23% |
| hsa_circ_0021358 | 11499 | 13 | 0.23% |
| hsa_circ_0071593 | 12386 | 14 | 0.23% |
| hsa_circ_0020488 | 12389 | 14 | 0.23% |
| hsa_circ_0061238 | 39831 | 45 | 0.23% |
| hsa_circ_0140430 | 52257 | 59 | 0.23% |
| hsa_circ_0122775 | 37219 | 42 | 0.23% |
| hsa_circ_0101527 | 32801 | 37 | 0.23% |
| hsa_circ_0094313 | 29255 | 33 | 0.23% |
| hsa_circ_0080234 | 31917 | 36 | 0.23% |

| | | | |
|---|---|---|---|
| hsa_circ_0112002 | 15076 | 17 | 0.23% |
| hsa_circ_0100798 | 19513 | 22 | 0.23% |
| hsa_circ_0134142 | 47012 | 53 | 0.23% |
| hsa_circ_0133735 | 26625 | 30 | 0.23% |
| hsa_circ_0002495 | 17764 | 20 | 0.23% |
| hsa_circ_0013047 | 39089 | 44 | 0.23% |
| hsa_circ_0119876 | 18661 | 21 | 0.23% |
| hsa_circ_0102742 | 19551 | 22 | 0.23% |
| hsa_circ_0120727 | 31998 | 36 | 0.23% |
| hsa_circ_0112068 | 38223 | 43 | 0.22% |
| hsa_circ_0093373 | 13335 | 15 | 0.22% |
| hsa_circ_0112060 | 28452 | 32 | 0.22% |
| hsa_circ_0000962 | 37347 | 42 | 0.22% |
| hsa_circ_0121178 | 16017 | 18 | 0.22% |
| hsa_circ_0056623 | 10680 | 12 | 0.22% |
| hsa_circ_0082982 | 11571 | 13 | 0.22% |
| hsa_circ_0112614 | 23143 | 26 | 0.22% |
| hsa_circ_0140772 | 20474 | 23 | 0.22% |
| hsa_circ_0112061 | 28506 | 32 | 0.22% |
| hsa_circ_0034139 | 21382 | 24 | 0.22% |
| hsa_circ_0021359 | 11586 | 13 | 0.22% |
| hsa_circ_0093303 | 11589 | 13 | 0.22% |
| hsa_circ_0053468 | 11595 | 13 | 0.22% |
| hsa_circ_0034194 | 11597 | 13 | 0.22% |
| hsa_circ_0016612 | 11599 | 13 | 0.22% |
| hsa_circ_0127684 | 13384 | 15 | 0.22% |
| hsa_circ_0033778 | 16069 | 18 | 0.22% |
| hsa_circ_0134843 | 72320 | 81 | 0.22% |
| hsa_circ_0131358 | 50903 | 57 | 0.22% |
| hsa_circ_0097147 | 24114 | 27 | 0.22% |
| hsa_circ_0098584 | 16083 | 18 | 0.22% |
| hsa_circ_0056230 | 12512 | 14 | 0.22% |
| hsa_circ_0108456 | 25030 | 28 | 0.22% |
| hsa_circ_0127970 | 39350 | 44 | 0.22% |
| hsa_circ_0121171 | 16113 | 18 | 0.22% |
| hsa_circ_0082967 | 11642 | 13 | 0.22% |
| hsa_circ_0107958 | 34931 | 39 | 0.22% |
| hsa_circ_0077319 | 11650 | 13 | 0.22% |
| hsa_circ_0113717 | 11650 | 13 | 0.22% |
| hsa_circ_0002133 | 35847 | 40 | 0.22% |
| hsa_circ_0056624 | 11653 | 13 | 0.22% |
| hsa_circ_0139418 | 11654 | 13 | 0.22% |
| hsa_circ_0042936 | 37653 | 42 | 0.22% |
| hsa_circ_0104366 | 27804 | 31 | 0.22% |
| hsa_circ_0026094 | 12559 | 14 | 0.22% |
| hsa_circ_0110274 | 39476 | 44 | 0.22% |
| hsa_circ_0089843 | 36795 | 41 | 0.22% |
| hsa_circ_0110999 | 19749 | 22 | 0.22% |
| hsa_circ_0117699 | 20654 | 23 | 0.22% |
| hsa_circ_0027149 | 11674 | 13 | 0.22% |

| | | | |
|---|---|---|---|
| hsa_circ_0137582 | 29652 | 33 | 0.22% |
| hsa_circ_0110296 | 13485 | 15 | 0.22% |
| hsa_circ_0138575 | 22487 | 25 | 0.22% |
| hsa_circ_0123117 | 33284 | 37 | 0.22% |
| hsa_circ_0134594 | 47708 | 53 | 0.22% |
| hsa_circ_0076831 | 11704 | 13 | 0.22% |
| hsa_circ_0050564 | 81033 | 90 | 0.22% |
| hsa_circ_0064627 | 27926 | 31 | 0.22% |
| hsa_circ_0049508 | 47753 | 53 | 0.22% |
| hsa_circ_0137087 | 27044 | 30 | 0.22% |
| hsa_circ_0126898 | 18935 | 21 | 0.22% |
| hsa_circ_0006294 | 37884 | 42 | 0.22% |
| hsa_circ_0013799 | 12637 | 14 | 0.22% |
| hsa_circ_0124499 | 16252 | 18 | 0.22% |
| hsa_circ_0098127 | 28902 | 32 | 0.22% |
| hsa_circ_0119067 | 40661 | 45 | 0.22% |
| hsa_circ_0002930 | 16266 | 18 | 0.22% |
| hsa_circ_0082977 | 11750 | 13 | 0.22% |
| hsa_circ_0137766 | 37072 | 41 | 0.22% |
| hsa_circ_0053437 | 11757 | 13 | 0.22% |
| hsa_circ_0101222 | 26230 | 29 | 0.22% |
| hsa_circ_0137767 | 37109 | 41 | 0.22% |
| hsa_circ_0133322 | 48887 | 54 | 0.22% |
| hsa_circ_0052896 | 11782 | 13 | 0.22% |
| hsa_circ_0135986 | 22659 | 25 | 0.22% |
| hsa_circ_0010246 | 13601 | 15 | 0.22% |
| hsa_circ_0009960 | 11797 | 13 | 0.22% |
| hsa_circ_0127546 | 33585 | 37 | 0.22% |
| hsa_circ_0105037 | 34506 | 38 | 0.22% |
| hsa_circ_0133770 | 11806 | 13 | 0.22% |
| hsa_circ_0129888 | 37235 | 41 | 0.22% |
| hsa_circ_0135382 | 50891 | 56 | 0.22% |
| hsa_circ_0035690 | 11815 | 13 | 0.22% |
| hsa_circ_0135716 | 16360 | 18 | 0.22% |
| hsa_circ_0082974 | 11817 | 13 | 0.22% |
| hsa_circ_0128099 | 47286 | 52 | 0.22% |
| hsa_circ_0119749 | 15460 | 17 | 0.22% |
| hsa_circ_0084235 | 11828 | 13 | 0.22% |
| hsa_circ_0094632 | 36405 | 40 | 0.22% |
| hsa_circ_0129122 | 22756 | 25 | 0.22% |
| hsa_circ_0122768 | 24580 | 27 | 0.22% |
| hsa_circ_0084233 | 10925 | 12 | 0.22% |
| hsa_circ_0073309 | 12747 | 14 | 0.22% |
| hsa_circ_0006330 | 38248 | 42 | 0.22% |
| hsa_circ_0102334 | 26411 | 29 | 0.22% |
| hsa_circ_0016900 | 46469 | 51 | 0.22% |
| hsa_circ_0026095 | 13671 | 15 | 0.22% |
| hsa_circ_0127006 | 36460 | 40 | 0.22% |
| hsa_circ_0033235 | 10938 | 12 | 0.22% |
| hsa_circ_0098912 | 13676 | 15 | 0.22% |

| | | | |
|---|---|---|---|
| hsa_circ_0098331 | 13685 | 15 | 0.22% |
| hsa_circ_0096700 | 37418 | 41 | 0.22% |
| hsa_circ_0132032 | 16428 | 18 | 0.22% |
| hsa_circ_0132568 | 10953 | 12 | 0.22% |
| hsa_circ_0132572 | 10953 | 12 | 0.22% |
| hsa_circ_0134858 | 21907 | 24 | 0.22% |
| hsa_circ_0140224 | 13692 | 15 | 0.22% |
| hsa_circ_0105974 | 52058 | 57 | 0.22% |
| hsa_circ_0056229 | 34706 | 38 | 0.22% |
| hsa_circ_0007971 | 38370 | 42 | 0.22% |
| hsa_circ_0118553 | 12791 | 14 | 0.22% |
| hsa_circ_0053549 | 11878 | 13 | 0.22% |
| hsa_circ_0122769 | 24671 | 27 | 0.22% |
| hsa_circ_0004875 | 11879 | 13 | 0.22% |
| hsa_circ_0101044 | 17362 | 19 | 0.22% |
| hsa_circ_0097297 | 24675 | 27 | 0.22% |
| hsa_circ_0087746 | 12795 | 14 | 0.22% |
| hsa_circ_0082960 | 11882 | 13 | 0.22% |
| hsa_circ_0107508 | 33820 | 37 | 0.22% |
| hsa_circ_0033227 | 10980 | 12 | 0.22% |
| hsa_circ_0132565 | 26546 | 29 | 0.22% |
| hsa_circ_0124252 | 50364 | 55 | 0.22% |
| hsa_circ_0140370 | 29307 | 32 | 0.22% |
| hsa_circ_0125575 | 30223 | 33 | 0.22% |
| hsa_circ_0125695 | 35721 | 39 | 0.22% |
| hsa_circ_0137549 | 40311 | 44 | 0.22% |
| hsa_circ_0140437 | 39396 | 43 | 0.22% |
| hsa_circ_0133624 | 17411 | 19 | 0.22% |
| hsa_circ_0138592 | 26581 | 29 | 0.22% |
| hsa_circ_0122156 | 19250 | 21 | 0.22% |
| hsa_circ_0114374 | 12842 | 14 | 0.22% |
| hsa_circ_0123108 | 35776 | 39 | 0.22% |
| hsa_circ_0005025 | 22935 | 25 | 0.22% |
| hsa_circ_0140773 | 21108 | 23 | 0.22% |
| hsa_circ_0128140 | 26616 | 29 | 0.22% |
| hsa_circ_0108192 | 19275 | 21 | 0.22% |
| hsa_circ_0015337 | 31212 | 34 | 0.22% |
| hsa_circ_0136710 | 22036 | 24 | 0.22% |
| hsa_circ_0090657 | 12855 | 14 | 0.22% |
| hsa_circ_0129421 | 43158 | 47 | 0.22% |
| hsa_circ_0107159 | 20206 | 22 | 0.22% |
| hsa_circ_0010276 | 11026 | 12 | 0.22% |
| hsa_circ_0069983 | 17467 | 19 | 0.22% |
| hsa_circ_0089775 | 22987 | 25 | 0.22% |
| hsa_circ_0084236 | 11954 | 13 | 0.22% |
| hsa_circ_0094395 | 11958 | 13 | 0.22% |
| hsa_circ_0125693 | 35883 | 39 | 0.22% |
| hsa_circ_0116788 | 13802 | 15 | 0.22% |
| hsa_circ_0119418 | 25764 | 28 | 0.22% |
| hsa_circ_0093181 | 68099 | 74 | 0.22% |

| | | | |
|---|---|---|---|
| hsa_circ_0140690 | 32211 | 35 | 0.22% |
| hsa_circ_0092676 | 71788 | 78 | 0.22% |
| hsa_circ_0052892 | 11966 | 13 | 0.22% |
| hsa_circ_0053469 | 11966 | 13 | 0.22% |
| hsa_circ_0133231 | 34991 | 38 | 0.22% |
| hsa_circ_0070737 | 30390 | 33 | 0.22% |
| hsa_circ_0122435 | 22115 | 24 | 0.22% |
| hsa_circ_0140652 | 56216 | 61 | 0.22% |
| hsa_circ_0108250 | 13824 | 15 | 0.22% |
| hsa_circ_0123445 | 35060 | 38 | 0.22% |
| hsa_circ_0132023 | 16608 | 18 | 0.22% |
| hsa_circ_0126937 | 20305 | 22 | 0.22% |
| hsa_circ_0121176 | 19383 | 21 | 0.22% |
| hsa_circ_0131122 | 31389 | 34 | 0.22% |
| hsa_circ_0052893 | 12005 | 13 | 0.22% |
| hsa_circ_0131199 | 23093 | 25 | 0.22% |
| hsa_circ_0017170 | 13857 | 15 | 0.22% |
| hsa_circ_0003466 | 36032 | 39 | 0.22% |
| hsa_circ_0062519 | 37886 | 41 | 0.22% |
| hsa_circ_0125611 | 36067 | 39 | 0.22% |
| hsa_circ_0123535 | 28683 | 31 | 0.22% |
| hsa_circ_0115090 | 35162 | 38 | 0.22% |
| hsa_circ_0100428 | 22208 | 24 | 0.22% |
| hsa_circ_0134853 | 64783 | 70 | 0.22% |
| hsa_circ_0026085 | 12032 | 13 | 0.22% |
| hsa_circ_0057539 | 14810 | 16 | 0.22% |
| hsa_circ_0095249 | 34252 | 37 | 0.22% |
| hsa_circ_0028357 | 12036 | 13 | 0.22% |
| hsa_circ_0002545 | 12038 | 13 | 0.22% |
| hsa_circ_0046004 | 12040 | 13 | 0.22% |
| hsa_circ_0044175 | 39827 | 43 | 0.22% |
| hsa_circ_0133346 | 37986 | 41 | 0.22% |
| hsa_circ_0131469 | 40792 | 44 | 0.22% |
| hsa_circ_0128147 | 43584 | 47 | 0.22% |
| hsa_circ_0134256 | 31536 | 34 | 0.22% |
| hsa_circ_0118357 | 21338 | 23 | 0.22% |
| hsa_circ_0122765 | 23194 | 25 | 0.22% |
| hsa_circ_0103686 | 15781 | 17 | 0.22% |
| hsa_circ_0006453 | 12068 | 13 | 0.22% |
| hsa_circ_0103683 | 15784 | 17 | 0.22% |
| hsa_circ_0076077 | 39005 | 42 | 0.22% |
| hsa_circ_0075817 | 40876 | 44 | 0.22% |
| hsa_circ_0128860 | 56709 | 61 | 0.22% |
| hsa_circ_0099363 | 23242 | 25 | 0.22% |
| hsa_circ_0139419 | 38140 | 41 | 0.21% |
| hsa_circ_0033230 | 11166 | 12 | 0.21% |
| hsa_circ_0033225 | 11167 | 12 | 0.21% |
| hsa_circ_0134434 | 69804 | 75 | 0.21% |
| hsa_circ_0132375 | 22339 | 24 | 0.21% |
| hsa_circ_0132566 | 11172 | 12 | 0.21% |

| | | | |
|---|---|---|---|
| hsa_circ_0110940 | 48427 | 52 | 0.21% |
| hsa_circ_0099596 | 71710 | 77 | 0.21% |
| hsa_circ_0140228 | 32608 | 35 | 0.21% |
| hsa_circ_0125787 | 34494 | 37 | 0.21% |
| hsa_circ_0132343 | 68081 | 73 | 0.21% |
| hsa_circ_0010252 | 13990 | 15 | 0.21% |
| hsa_circ_0132723 | 16794 | 18 | 0.21% |
| hsa_circ_0097355 | 48553 | 52 | 0.21% |
| hsa_circ_0109313 | 17744 | 19 | 0.21% |
| hsa_circ_0093085 | 54175 | 58 | 0.21% |
| hsa_circ_0000397 | 12144 | 13 | 0.21% |
| hsa_circ_0118577 | 38327 | 41 | 0.21% |
| hsa_circ_0110941 | 48632 | 52 | 0.21% |
| hsa_circ_0074240 | 40230 | 43 | 0.21% |
| hsa_circ_0033779 | 16841 | 18 | 0.21% |
| hsa_circ_0017172 | 14035 | 15 | 0.21% |
| hsa_circ_0052886 | 12164 | 13 | 0.21% |
| hsa_circ_0140688 | 32756 | 35 | 0.21% |
| hsa_circ_0121845 | 46813 | 50 | 0.21% |
| hsa_circ_0137432 | 19673 | 21 | 0.21% |
| hsa_circ_0118570 | 38430 | 41 | 0.21% |
| hsa_circ_0139416 | 12188 | 13 | 0.21% |
| hsa_circ_0080991 | 12190 | 13 | 0.21% |
| hsa_circ_0006880 | 35637 | 38 | 0.21% |
| hsa_circ_0099594 | 45018 | 48 | 0.21% |
| hsa_circ_0033571 | 30953 | 33 | 0.21% |
| hsa_circ_0003265 | 12194 | 13 | 0.21% |
| hsa_circ_0128302 | 30955 | 33 | 0.21% |
| hsa_circ_0026091 | 14075 | 15 | 0.21% |
| hsa_circ_0108454 | 19712 | 21 | 0.21% |
| hsa_circ_0098687 | 55384 | 59 | 0.21% |
| hsa_circ_0117425 | 27230 | 29 | 0.21% |
| hsa_circ_0124630 | 12208 | 13 | 0.21% |
| hsa_circ_0117533 | 24425 | 26 | 0.21% |
| hsa_circ_0050830 | 14094 | 15 | 0.21% |
| hsa_circ_0139515 | 31026 | 33 | 0.21% |
| hsa_circ_0062520 | 38553 | 41 | 0.21% |
| hsa_circ_0108249 | 32936 | 35 | 0.21% |
| hsa_circ_0000400 | 55536 | 59 | 0.21% |
| hsa_circ_0134854 | 22594 | 24 | 0.21% |
| hsa_circ_0115627 | 31071 | 33 | 0.21% |
| hsa_circ_0124650 | 40487 | 43 | 0.21% |
| hsa_circ_0126017 | 14125 | 15 | 0.21% |
| hsa_circ_0099831 | 16012 | 17 | 0.21% |
| hsa_circ_0133900 | 14131 | 15 | 0.21% |
| hsa_circ_0033240 | 11308 | 12 | 0.21% |
| hsa_circ_0019100 | 22623 | 24 | 0.21% |
| hsa_circ_0084234 | 11312 | 12 | 0.21% |
| hsa_circ_0112810 | 28284 | 30 | 0.21% |
| hsa_circ_0053470 | 12267 | 13 | 0.21% |

| | | | |
|---|---|---|---|
| hsa_circ_0013090 | 31143 | 33 | 0.21% |
| hsa_circ_0134197 | 21706 | 23 | 0.21% |
| hsa_circ_0005119 | 35869 | 38 | 0.21% |
| hsa_circ_0118310 | 12272 | 13 | 0.21% |
| hsa_circ_0104535 | 31165 | 33 | 0.21% |
| hsa_circ_0112808 | 28333 | 30 | 0.21% |
| hsa_circ_0117532 | 24556 | 26 | 0.21% |
| hsa_circ_0103687 | 16059 | 17 | 0.21% |
| hsa_circ_0098138 | 26455 | 28 | 0.21% |
| hsa_circ_0103684 | 16062 | 17 | 0.21% |
| hsa_circ_0086001 | 14175 | 15 | 0.21% |
| hsa_circ_0102542 | 11364 | 12 | 0.21% |
| hsa_circ_0136707 | 16101 | 17 | 0.21% |
| hsa_circ_0103730 | 27476 | 29 | 0.21% |
| hsa_circ_0102357 | 40747 | 43 | 0.21% |
| hsa_circ_0053520 | 12321 | 13 | 0.21% |
| hsa_circ_0127299 | 12321 | 13 | 0.21% |
| hsa_circ_0124257 | 89124 | 94 | 0.21% |
| hsa_circ_0097253 | 51207 | 54 | 0.21% |
| hsa_circ_0024000 | 11382 | 12 | 0.21% |
| hsa_circ_0089192 | 36047 | 38 | 0.21% |
| hsa_circ_0084237 | 12338 | 13 | 0.21% |
| hsa_circ_0007908 | 23729 | 25 | 0.21% |
| hsa_circ_0097254 | 51257 | 54 | 0.21% |
| hsa_circ_0021364 | 13289 | 14 | 0.21% |
| hsa_circ_0103142 | 37044 | 39 | 0.21% |
| hsa_circ_0018306 | 42752 | 45 | 0.21% |
| hsa_circ_0122157 | 19952 | 21 | 0.21% |
| hsa_circ_0040922 | 20908 | 22 | 0.21% |
| hsa_circ_0126855 | 44685 | 47 | 0.21% |
| hsa_circ_0103046 | 39947 | 42 | 0.21% |
| hsa_circ_0138585 | 92268 | 97 | 0.21% |
| hsa_circ_0080214 | 23785 | 25 | 0.21% |
| hsa_circ_0093633 | 29507 | 31 | 0.21% |
| hsa_circ_0033221 | 11423 | 12 | 0.21% |
| hsa_circ_0136703 | 16184 | 17 | 0.21% |
| hsa_circ_0129320 | 25709 | 27 | 0.21% |
| hsa_circ_0128693 | 43830 | 46 | 0.21% |
| hsa_circ_0089193 | 36225 | 38 | 0.21% |
| hsa_circ_0000218 | 70570 | 74 | 0.21% |
| hsa_circ_0137173 | 16216 | 17 | 0.21% |
| hsa_circ_0137262 | 74417 | 78 | 0.21% |
| hsa_circ_0097244 | 51526 | 54 | 0.21% |
| hsa_circ_0127499 | 30539 | 32 | 0.21% |
| hsa_circ_0000005 | 49639 | 52 | 0.21% |
| hsa_circ_0115894 | 16231 | 17 | 0.21% |
| hsa_circ_0053533 | 12412 | 13 | 0.21% |
| hsa_circ_0071435 | 27705 | 29 | 0.21% |
| hsa_circ_0000386 | 31530 | 33 | 0.21% |
| hsa_circ_0099278 | 18154 | 19 | 0.21% |

| | | | |
|---|---|---|---|
| hsa_circ_0114317 | 30587 | 32 | 0.21% |
| hsa_circ_0116950 | 24852 | 26 | 0.21% |
| hsa_circ_0027139 | 12426 | 13 | 0.21% |
| hsa_circ_0138338 | 49718 | 52 | 0.21% |
| hsa_circ_0000526 | 32519 | 34 | 0.21% |
| hsa_circ_0033081 | 13391 | 14 | 0.21% |
| hsa_circ_0128407 | 53587 | 56 | 0.21% |
| hsa_circ_0131242 | 12440 | 13 | 0.21% |
| hsa_circ_0139813 | 53590 | 56 | 0.21% |
| hsa_circ_0137768 | 39252 | 41 | 0.21% |
| hsa_circ_0110297 | 14361 | 15 | 0.21% |
| hsa_circ_0098283 | 51718 | 54 | 0.21% |
| hsa_circ_0129266 | 49809 | 52 | 0.21% |
| hsa_circ_0118304 | 17246 | 18 | 0.21% |
| hsa_circ_0138502 | 13417 | 14 | 0.21% |
| hsa_circ_0086248 | 12459 | 13 | 0.21% |
| hsa_circ_0096587 | 42176 | 44 | 0.21% |
| hsa_circ_0120137 | 44096 | 46 | 0.21% |
| hsa_circ_0125142 | 12462 | 13 | 0.21% |
| hsa_circ_0015385 | 17259 | 18 | 0.21% |
| hsa_circ_0132245 | 15342 | 16 | 0.21% |
| hsa_circ_0099602 | 34522 | 36 | 0.21% |
| hsa_circ_0095654 | 12467 | 13 | 0.21% |
| hsa_circ_0121061 | 43194 | 45 | 0.21% |
| hsa_circ_0085898 | 14404 | 15 | 0.21% |
| hsa_circ_0053471 | 12485 | 13 | 0.21% |
| hsa_circ_0084238 | 12485 | 13 | 0.21% |
| hsa_circ_0100633 | 43226 | 45 | 0.21% |
| hsa_circ_0101215 | 27861 | 29 | 0.21% |
| hsa_circ_0092670 | 13454 | 14 | 0.21% |
| hsa_circ_0136439 | 12496 | 13 | 0.21% |
| hsa_circ_0087834 | 50009 | 52 | 0.21% |
| hsa_circ_0112840 | 33662 | 35 | 0.21% |
| hsa_circ_0009741 | 42320 | 44 | 0.21% |
| hsa_circ_0021365 | 13467 | 14 | 0.21% |
| hsa_circ_0099273 | 18278 | 19 | 0.21% |
| hsa_circ_0032162 | 13473 | 14 | 0.21% |
| hsa_circ_0103743 | 20213 | 21 | 0.21% |
| hsa_circ_0137319 | 14443 | 15 | 0.21% |
| hsa_circ_0108544 | 48153 | 50 | 0.21% |
| hsa_circ_0108545 | 48156 | 50 | 0.21% |
| hsa_circ_0126788 | 32761 | 34 | 0.21% |
| hsa_circ_0002287 | 36617 | 38 | 0.21% |
| hsa_circ_0011793 | 13496 | 14 | 0.21% |
| hsa_circ_0101213 | 15427 | 16 | 0.21% |
| hsa_circ_0033780 | 17359 | 18 | 0.21% |
| hsa_circ_0095979 | 46293 | 48 | 0.21% |
| hsa_circ_0134190 | 32798 | 34 | 0.21% |
| hsa_circ_0123478 | 39561 | 41 | 0.21% |
| hsa_circ_0115711 | 35733 | 37 | 0.21% |

| | | | |
|---|---|---|---|
| hsa_circ_0033218 | 11597 | 12 | 0.21% |
| hsa_circ_0108626 | 28033 | 29 | 0.21% |
| hsa_circ_0010253 | 14500 | 15 | 0.21% |
| hsa_circ_0112795 | 39638 | 41 | 0.21% |
| hsa_circ_0122759 | 19337 | 20 | 0.21% |
| hsa_circ_0136424 | 34810 | 36 | 0.21% |
| hsa_circ_0034832 | 12578 | 13 | 0.21% |
| hsa_circ_0026088 | 14515 | 15 | 0.21% |
| hsa_circ_0087832 | 50332 | 52 | 0.21% |
| hsa_circ_0033208 | 11624 | 12 | 0.21% |
| hsa_circ_0108369 | 31008 | 32 | 0.21% |
| hsa_circ_0139040 | 28104 | 29 | 0.21% |
| hsa_circ_0006742 | 31997 | 33 | 0.21% |
| hsa_circ_0132856 | 41697 | 43 | 0.21% |
| hsa_circ_0100429 | 28143 | 29 | 0.21% |
| hsa_circ_0114238 | 25234 | 26 | 0.21% |
| hsa_circ_0092671 | 13592 | 14 | 0.21% |
| hsa_circ_0094526 | 76778 | 79 | 0.21% |
| hsa_circ_0113925 | 26242 | 27 | 0.21% |
| hsa_circ_0133230 | 36954 | 38 | 0.21% |
| hsa_circ_0084239 | 12643 | 13 | 0.21% |
| hsa_circ_0116387 | 35996 | 37 | 0.21% |
| hsa_circ_0024001 | 11675 | 12 | 0.21% |
| hsa_circ_0120842 | 43790 | 45 | 0.21% |
| hsa_circ_0137127 | 14597 | 15 | 0.21% |
| hsa_circ_0080880 | 43818 | 45 | 0.21% |
| hsa_circ_0033211 | 11685 | 12 | 0.21% |
| hsa_circ_0130928 | 22397 | 23 | 0.21% |
| hsa_circ_0080036 | 55563 | 57 | 0.21% |
| hsa_circ_0084240 | 12674 | 13 | 0.21% |
| hsa_circ_0139265 | 40951 | 42 | 0.21% |
| hsa_circ_0111228 | 37059 | 38 | 0.21% |
| hsa_circ_0131467 | 42934 | 44 | 0.20% |
| hsa_circ_0125669 | 41960 | 43 | 0.20% |
| hsa_circ_0103077 | 12689 | 13 | 0.20% |
| hsa_circ_0101499 | 39054 | 40 | 0.20% |
| hsa_circ_0117097 | 34173 | 35 | 0.20% |
| hsa_circ_0020290 | 28326 | 29 | 0.20% |
| hsa_circ_0131468 | 43018 | 44 | 0.20% |
| hsa_circ_0013763 | 33259 | 34 | 0.20% |
| hsa_circ_0100924 | 22500 | 23 | 0.20% |
| hsa_circ_0082964 | 12722 | 13 | 0.20% |
| hsa_circ_0125491 | 33288 | 34 | 0.20% |
| hsa_circ_0112799 | 40157 | 41 | 0.20% |
| hsa_circ_0104411 | 63672 | 65 | 0.20% |
| hsa_circ_0128508 | 48993 | 50 | 0.20% |
| hsa_circ_0007138 | 17643 | 18 | 0.20% |
| hsa_circ_0034836 | 12743 | 13 | 0.20% |
| hsa_circ_0129117 | 34319 | 35 | 0.20% |
| hsa_circ_0124928 | 29417 | 30 | 0.20% |

| | | | |
|---|---|---|---|
| hsa_circ_0128507 | 49030 | 50 | 0.20% |
| hsa_circ_0065479 | 13733 | 14 | 0.20% |
| hsa_circ_0130724 | 15697 | 16 | 0.20% |
| hsa_circ_0021360 | 12756 | 13 | 0.20% |
| hsa_circ_0101706 | 53969 | 55 | 0.20% |
| hsa_circ_0137277 | 12759 | 13 | 0.20% |
| hsa_circ_0120504 | 28470 | 29 | 0.20% |
| hsa_circ_0132673 | 43199 | 44 | 0.20% |
| hsa_circ_0113284 | 11787 | 12 | 0.20% |
| hsa_circ_0125911 | 28488 | 29 | 0.20% |
| hsa_circ_0006198 | 49119 | 50 | 0.20% |
| hsa_circ_0111309 | 19651 | 20 | 0.20% |
| hsa_circ_0064498 | 35373 | 36 | 0.20% |
| hsa_circ_0122604 | 20637 | 21 | 0.20% |
| hsa_circ_0001056 | 32434 | 33 | 0.20% |
| hsa_circ_0034150 | 13772 | 14 | 0.20% |
| hsa_circ_0103189 | 13777 | 14 | 0.20% |
| hsa_circ_0124929 | 29544 | 30 | 0.20% |
| hsa_circ_0094656 | 49252 | 50 | 0.20% |
| hsa_circ_0100925 | 22656 | 23 | 0.20% |
| hsa_circ_0123105 | 42373 | 43 | 0.20% |
| hsa_circ_0140238 | 35481 | 36 | 0.20% |
| hsa_circ_0124642 | 26611 | 27 | 0.20% |
| hsa_circ_0105184 | 37468 | 38 | 0.20% |
| hsa_circ_0135573 | 67051 | 68 | 0.20% |
| hsa_circ_0115710 | 22682 | 23 | 0.20% |
| hsa_circ_0138607 | 39447 | 40 | 0.20% |
| hsa_circ_0136817 | 30575 | 31 | 0.20% |
| hsa_circ_0123357 | 42415 | 43 | 0.20% |
| hsa_circ_0098550 | 38480 | 39 | 0.20% |
| hsa_circ_0101901 | 37503 | 38 | 0.20% |
| hsa_circ_0033212 | 11846 | 12 | 0.20% |
| hsa_circ_0103702 | 30613 | 31 | 0.20% |
| hsa_circ_0114510 | 40523 | 41 | 0.20% |
| hsa_circ_0098189 | 36574 | 37 | 0.20% |
| hsa_circ_0000695 | 42512 | 43 | 0.20% |
| hsa_circ_0021361 | 12857 | 13 | 0.20% |
| hsa_circ_0033781 | 17818 | 18 | 0.20% |
| hsa_circ_0052887 | 12869 | 13 | 0.20% |
| hsa_circ_0108460 | 16835 | 17 | 0.20% |
| hsa_circ_0009169 | 14862 | 15 | 0.20% |
| hsa_circ_0111979 | 40623 | 41 | 0.20% |
| hsa_circ_0130997 | 28737 | 29 | 0.20% |
| hsa_circ_0000577 | 43670 | 44 | 0.20% |
| hsa_circ_0126365 | 56575 | 57 | 0.20% |
| hsa_circ_0131135 | 80464 | 81 | 0.20% |
| hsa_circ_0053550 | 12917 | 13 | 0.20% |
| hsa_circ_0135322 | 54649 | 55 | 0.20% |
| hsa_circ_0120557 | 38761 | 39 | 0.20% |
| hsa_circ_0108740 | 27829 | 28 | 0.20% |

| | | | |
|---|---|---|---|
| hsa_circ_0117493 | 33794 | 34 | 0.20% |
| hsa_circ_0137484 | 68611 | 69 | 0.20% |
| hsa_circ_0127701 | 14917 | 15 | 0.20% |
| hsa_circ_0021362 | 12930 | 13 | 0.20% |
| hsa_circ_0024441 | 12937 | 13 | 0.20% |
| hsa_circ_0084241 | 12943 | 13 | 0.20% |
| hsa_circ_0103158 | 39827 | 40 | 0.20% |
| hsa_circ_0030237 | 29878 | 30 | 0.20% |
| hsa_circ_0053498 | 12967 | 13 | 0.20% |
| hsa_circ_0122617 | 13969 | 14 | 0.20% |
| hsa_circ_0053534 | 12976 | 13 | 0.20% |
| hsa_circ_0103367 | 23962 | 24 | 0.20% |
| hsa_circ_0137263 | 77902 | 78 | 0.20% |
| hsa_circ_0035349 | 40975 | 41 | 0.20% |
| hsa_circ_0100970 | 23987 | 24 | 0.20% |
| hsa_circ_0127476 | 38979 | 39 | 0.20% |
| hsa_circ_0130586 | 17993 | 18 | 0.20% |
| hsa_circ_0104825 | 33041 | 33 | 0.20% |
| hsa_circ_0030463 | 15025 | 15 | 0.20% |
| hsa_circ_0130929 | 39073 | 39 | 0.20% |
| hsa_circ_0101229 | 29071 | 29 | 0.20% |
| hsa_circ_0021363 | 13035 | 13 | 0.20% |
| hsa_circ_0122786 | 49154 | 49 | 0.20% |
| hsa_circ_0110597 | 24079 | 24 | 0.20% |
| hsa_circ_0104938 | 39129 | 39 | 0.20% |
| hsa_circ_0002503 | 24089 | 24 | 0.20% |
| hsa_circ_0127390 | 45168 | 45 | 0.20% |
| hsa_circ_0125482 | 49183 | 49 | 0.20% |
| hsa_circ_0073295 | 14054 | 14 | 0.20% |
| hsa_circ_0107384 | 52206 | 52 | 0.20% |
| hsa_circ_0099286 | 28114 | 28 | 0.20% |
| hsa_circ_0058602 | 16067 | 16 | 0.20% |
| hsa_circ_0104738 | 38160 | 38 | 0.20% |
| hsa_circ_0117502 | 12057 | 12 | 0.20% |
| hsa_circ_0034300 | 50251 | 50 | 0.20% |
| hsa_circ_0103188 | 14074 | 14 | 0.20% |
| hsa_circ_0107383 | 52278 | 52 | 0.20% |
| hsa_circ_0130725 | 17091 | 17 | 0.20% |
| hsa_circ_0033726 | 18098 | 18 | 0.20% |
| hsa_circ_0087964 | 38210 | 38 | 0.20% |
| hsa_circ_0140472 | 62352 | 62 | 0.20% |
| hsa_circ_0103047 | 32185 | 32 | 0.20% |
| hsa_circ_0131515 | 16097 | 16 | 0.20% |
| hsa_circ_0094284 | 27167 | 27 | 0.20% |
| hsa_circ_0070126 | 14113 | 14 | 0.20% |
| hsa_circ_0115268 | 43381 | 43 | 0.20% |
| hsa_circ_0137095 | 16148 | 16 | 0.20% |
| hsa_circ_0087961 | 38358 | 38 | 0.20% |
| hsa_circ_0103379 | 26250 | 26 | 0.20% |
| hsa_circ_0095055 | 14138 | 14 | 0.20% |

| | | | |
|---|---|---|---|
| hsa_circ_0136655 | 43427 | 43 | 0.20% |
| hsa_circ_0139408 | 13130 | 13 | 0.20% |
| hsa_circ_0057220 | 13134 | 13 | 0.20% |
| hsa_circ_0110957 | 16175 | 16 | 0.20% |
| hsa_circ_0133237 | 15166 | 15 | 0.20% |
| hsa_circ_0125132 | 50564 | 50 | 0.20% |
| hsa_circ_0001210 | 25285 | 25 | 0.20% |
| hsa_circ_0093207 | 50587 | 50 | 0.20% |
| hsa_circ_0123600 | 14165 | 14 | 0.20% |
| hsa_circ_0130593 | 15187 | 15 | 0.20% |
| hsa_circ_0020321 | 15190 | 15 | 0.20% |
| hsa_circ_0001217 | 52676 | 52 | 0.20% |
| hsa_circ_0034833 | 13175 | 13 | 0.20% |
| hsa_circ_0121849 | 25338 | 25 | 0.20% |
| hsa_circ_0116001 | 15210 | 15 | 0.20% |
| hsa_circ_0082970 | 13182 | 13 | 0.20% |
| hsa_circ_0099666 | 49691 | 49 | 0.20% |
| hsa_circ_0128862 | 29412 | 29 | 0.20% |
| hsa_circ_0004521 | 64938 | 64 | 0.20% |
| hsa_circ_0021776 | 22330 | 22 | 0.20% |
| hsa_circ_0001761 | 27413 | 27 | 0.20% |
| hsa_circ_0109367 | 22361 | 22 | 0.20% |
| hsa_circ_0117501 | 12199 | 12 | 0.20% |
| hsa_circ_0108298 | 27454 | 27 | 0.20% |
| hsa_circ_0100266 | 37624 | 37 | 0.20% |
| hsa_circ_0084242 | 13220 | 13 | 0.20% |
| hsa_circ_0140368 | 13224 | 13 | 0.20% |
| hsa_circ_0103048 | 32554 | 32 | 0.20% |
| hsa_circ_0099475 | 48842 | 48 | 0.20% |
| hsa_circ_0053512 | 13231 | 13 | 0.20% |
| hsa_circ_0098323 | 71246 | 70 | 0.20% |
| hsa_circ_0095354 | 27487 | 27 | 0.20% |
| hsa_circ_0121400 | 29542 | 29 | 0.20% |
| hsa_circ_0005723 | 37705 | 37 | 0.20% |
| hsa_circ_0104730 | 38739 | 38 | 0.20% |
| hsa_circ_0138918 | 42818 | 42 | 0.20% |
| hsa_circ_0015339 | 37725 | 37 | 0.20% |
| hsa_circ_0099275 | 19389 | 19 | 0.20% |
| hsa_circ_0053438 | 13278 | 13 | 0.20% |
| hsa_circ_0124803 | 81723 | 80 | 0.20% |
| hsa_circ_0001711 | 50064 | 49 | 0.20% |
| hsa_circ_0101209 | 29635 | 29 | 0.20% |
| hsa_circ_0053472 | 13287 | 13 | 0.20% |
| hsa_circ_0102828 | 16355 | 16 | 0.20% |
| hsa_circ_0133096 | 35800 | 35 | 0.20% |
| hsa_circ_0111556 | 28647 | 28 | 0.20% |
| hsa_circ_0119978 | 42989 | 42 | 0.20% |
| hsa_circ_0101217 | 73700 | 72 | 0.20% |
| hsa_circ_0136194 | 16381 | 16 | 0.20% |
| hsa_circ_0103699 | 26622 | 26 | 0.20% |

| | | | |
|---|---|---|---|
| hsa_circ_0103137 | 15363 | 15 | 0.20% |
| hsa_circ_0082968 | 13325 | 13 | 0.20% |
| hsa_circ_0104693 | 47174 | 46 | 0.20% |
| hsa_circ_0099158 | 47179 | 46 | 0.20% |
| hsa_circ_0134918 | 22580 | 22 | 0.19% |
| hsa_circ_0100550 | 36950 | 36 | 0.19% |
| hsa_circ_0100248 | 51348 | 50 | 0.19% |
| hsa_circ_0094393 | 14378 | 14 | 0.19% |
| hsa_circ_0130419 | 33907 | 33 | 0.19% |
| hsa_circ_0100927 | 23634 | 23 | 0.19% |
| hsa_circ_0134763 | 18502 | 18 | 0.19% |
| hsa_circ_0126854 | 48315 | 47 | 0.19% |
| hsa_circ_0111930 | 13369 | 13 | 0.19% |
| hsa_circ_0004928 | 39106 | 38 | 0.19% |
| hsa_circ_0093304 | 60763 | 59 | 0.19% |
| hsa_circ_0129893 | 16479 | 16 | 0.19% |
| hsa_circ_0136455 | 36057 | 35 | 0.19% |
| hsa_circ_0077432 | 53577 | 52 | 0.19% |
| hsa_circ_0112452 | 50487 | 49 | 0.19% |
| hsa_circ_0117107 | 23699 | 23 | 0.19% |
| hsa_circ_0005156 | 38151 | 37 | 0.19% |
| hsa_circ_0002031 | 38175 | 37 | 0.19% |
| hsa_circ_0115766 | 44380 | 43 | 0.19% |
| hsa_circ_0115761 | 32000 | 31 | 0.19% |
| hsa_circ_0098458 | 14463 | 14 | 0.19% |
| hsa_circ_0020322 | 15497 | 15 | 0.19% |
| hsa_circ_0098459 | 14464 | 14 | 0.19% |
| hsa_circ_0061711 | 14466 | 14 | 0.19% |
| hsa_circ_0098052 | 45519 | 44 | 0.19% |
| hsa_circ_0103693 | 66236 | 64 | 0.19% |
| hsa_circ_0132753 | 17595 | 17 | 0.19% |
| hsa_circ_0034834 | 13458 | 13 | 0.19% |
| hsa_circ_0003383 | 24850 | 24 | 0.19% |
| hsa_circ_0126170 | 57997 | 56 | 0.19% |
| hsa_circ_0132752 | 17626 | 17 | 0.19% |
| hsa_circ_0133874 | 33184 | 32 | 0.19% |
| hsa_circ_0129419 | 34229 | 33 | 0.19% |
| hsa_circ_0130565 | 14528 | 14 | 0.19% |
| hsa_circ_0111951 | 55008 | 53 | 0.19% |
| hsa_circ_0053473 | 13505 | 13 | 0.19% |
| hsa_circ_0093299 | 61294 | 59 | 0.19% |
| hsa_circ_0099578 | 43639 | 42 | 0.19% |
| hsa_circ_0084243 | 13508 | 13 | 0.19% |
| hsa_circ_0115709 | 15596 | 15 | 0.19% |
| hsa_circ_0138445 | 37436 | 36 | 0.19% |
| hsa_circ_0136391 | 28082 | 27 | 0.19% |
| hsa_circ_0082961 | 13522 | 13 | 0.19% |
| hsa_circ_0100257 | 47849 | 46 | 0.19% |
| hsa_circ_0121841 | 48897 | 47 | 0.19% |
| hsa_circ_0114413 | 39545 | 38 | 0.19% |

| | | | |
|---|---|---|---|
| hsa_circ_0101212 | 13529 | 13 | 0.19% |
| hsa_circ_0138364 | 44774 | 43 | 0.19% |
| hsa_circ_0131452 | 39576 | 38 | 0.19% |
| hsa_circ_0061712 | 14591 | 14 | 0.19% |
| hsa_circ_0099694 | 13550 | 13 | 0.19% |
| hsa_circ_0029715 | 14593 | 14 | 0.19% |
| hsa_circ_0139407 | 13555 | 13 | 0.19% |
| hsa_circ_0101201 | 40716 | 39 | 0.19% |
| hsa_circ_0001763 | 28199 | 27 | 0.19% |
| hsa_circ_0006783 | 53300 | 51 | 0.19% |
| hsa_circ_0107387 | 42870 | 41 | 0.19% |
| hsa_circ_0127302 | 41825 | 40 | 0.19% |
| hsa_circ_0111923 | 13606 | 13 | 0.19% |
| hsa_circ_0109304 | 40821 | 39 | 0.19% |
| hsa_circ_0033209 | 12563 | 12 | 0.19% |
| hsa_circ_0052888 | 13610 | 13 | 0.19% |
| hsa_circ_0084791 | 13617 | 13 | 0.19% |
| hsa_circ_0108803 | 34567 | 33 | 0.19% |
| hsa_circ_0088336 | 55519 | 53 | 0.19% |
| hsa_circ_0129752 | 39818 | 38 | 0.19% |
| hsa_circ_0137174 | 18869 | 18 | 0.19% |
| hsa_circ_0136006 | 14679 | 14 | 0.19% |
| hsa_circ_0139894 | 15738 | 15 | 0.19% |
| hsa_circ_0129411 | 17837 | 17 | 0.19% |
| hsa_circ_0002858 | 27282 | 26 | 0.19% |
| hsa_circ_0033782 | 18892 | 18 | 0.19% |
| hsa_circ_0117592 | 75601 | 72 | 0.19% |
| hsa_circ_0107388 | 43057 | 41 | 0.19% |
| hsa_circ_0101223 | 30461 | 29 | 0.19% |
| hsa_circ_0053474 | 13667 | 13 | 0.19% |
| hsa_circ_0003186 | 27346 | 26 | 0.19% |
| hsa_circ_0113748 | 24195 | 23 | 0.19% |
| hsa_circ_0003358 | 34728 | 33 | 0.19% |
| hsa_circ_0099695 | 13687 | 13 | 0.19% |
| hsa_circ_0101224 | 30548 | 29 | 0.19% |
| hsa_circ_0138919 | 44254 | 42 | 0.19% |
| hsa_circ_0121552 | 27405 | 26 | 0.19% |
| hsa_circ_0137278 | 28465 | 27 | 0.19% |
| hsa_circ_0121806 | 25318 | 24 | 0.19% |
| hsa_circ_0009001 | 37979 | 36 | 0.19% |
| hsa_circ_0056231 | 14775 | 14 | 0.19% |
| hsa_circ_0080989 | 25335 | 24 | 0.19% |
| hsa_circ_0075525 | 44365 | 42 | 0.19% |
| hsa_circ_0122881 | 29579 | 28 | 0.19% |
| hsa_circ_0093687 | 43317 | 41 | 0.19% |
| hsa_circ_0096338 | 26415 | 25 | 0.19% |
| hsa_circ_0107389 | 43321 | 41 | 0.19% |
| hsa_circ_0125687 | 32789 | 31 | 0.19% |
| hsa_circ_0052889 | 13756 | 13 | 0.19% |
| hsa_circ_0099638 | 13759 | 13 | 0.19% |

| | | | |
|---|---|---|---|
| hsa_circ_0134849 | 37070 | 35 | 0.19% |
| hsa_circ_0096337 | 26504 | 25 | 0.19% |
| hsa_circ_0121498 | 27568 | 26 | 0.19% |
| hsa_circ_0139393 | 41361 | 39 | 0.19% |
| hsa_circ_0113718 | 32883 | 31 | 0.19% |
| hsa_circ_0004141 | 58368 | 55 | 0.19% |
| hsa_circ_0093475 | 27599 | 26 | 0.19% |
| hsa_circ_0076812 | 13804 | 13 | 0.19% |
| hsa_circ_0113754 | 47785 | 45 | 0.19% |
| hsa_circ_0133761 | 29739 | 28 | 0.19% |
| hsa_circ_0123672 | 13809 | 13 | 0.19% |
| hsa_circ_0093273 | 40382 | 38 | 0.19% |
| hsa_circ_0094948 | 36134 | 34 | 0.19% |
| hsa_circ_0100999 | 25510 | 24 | 0.19% |
| hsa_circ_0137198 | 26617 | 25 | 0.19% |
| hsa_circ_0108497 | 37301 | 35 | 0.19% |
| hsa_circ_0132740 | 31995 | 30 | 0.19% |
| hsa_circ_0111170 | 14934 | 14 | 0.19% |
| hsa_circ_0026086 | 16003 | 15 | 0.19% |
| hsa_circ_0130858 | 14937 | 14 | 0.19% |
| hsa_circ_0125338 | 26689 | 25 | 0.19% |
| hsa_circ_0004415 | 33098 | 31 | 0.19% |
| hsa_circ_0123821 | 37394 | 35 | 0.19% |
| hsa_circ_0058033 | 24574 | 23 | 0.19% |
| hsa_circ_0129228 | 33155 | 31 | 0.19% |
| hsa_circ_0096586 | 47075 | 44 | 0.19% |
| hsa_circ_0136096 | 47076 | 44 | 0.19% |
| hsa_circ_0129948 | 53520 | 50 | 0.19% |
| hsa_circ_0078837 | 52452 | 49 | 0.19% |
| hsa_circ_0093087 | 43892 | 41 | 0.19% |
| hsa_circ_0003519 | 32120 | 30 | 0.19% |
| hsa_circ_0000227 | 32123 | 30 | 0.19% |
| hsa_circ_0103393 | 32149 | 30 | 0.19% |
| hsa_circ_0001762 | 28940 | 27 | 0.19% |
| hsa_circ_0002189 | 47166 | 44 | 0.19% |
| hsa_circ_0137117 | 86878 | 81 | 0.19% |
| hsa_circ_0104888 | 13944 | 13 | 0.19% |
| hsa_circ_0120299 | 13947 | 13 | 0.19% |
| hsa_circ_0098911 | 15023 | 14 | 0.19% |
| hsa_circ_0124625 | 53683 | 50 | 0.19% |
| hsa_circ_0080879 | 62276 | 58 | 0.19% |
| hsa_circ_0034326 | 34386 | 32 | 0.19% |
| hsa_circ_0016173 | 16128 | 15 | 0.19% |
| hsa_circ_0128792 | 27956 | 26 | 0.19% |
| hsa_circ_0121504 | 50546 | 47 | 0.19% |
| hsa_circ_0005124 | 48414 | 45 | 0.19% |
| hsa_circ_0080878 | 62414 | 58 | 0.19% |
| hsa_circ_0093088 | 44125 | 41 | 0.19% |
| hsa_circ_0134890 | 40917 | 38 | 0.19% |
| hsa_circ_0004938 | 30172 | 28 | 0.19% |

| | | | |
|---|---|---|---|
| hsa_circ_0109366 | 16169 | 15 | 0.19% |
| hsa_circ_0140474 | 16170 | 15 | 0.19% |
| hsa_circ_0008180 | 38818 | 36 | 0.19% |
| hsa_circ_0140766 | 16177 | 15 | 0.19% |
| hsa_circ_0111548 | 60429 | 56 | 0.19% |
| hsa_circ_0070094 | 15108 | 14 | 0.19% |
| hsa_circ_0100408 | 31300 | 29 | 0.19% |
| hsa_circ_0124502 | 31302 | 29 | 0.19% |
| hsa_circ_0024002 | 12954 | 12 | 0.19% |
| hsa_circ_0132608 | 41032 | 38 | 0.19% |
| hsa_circ_0012283 | 42139 | 39 | 0.19% |
| hsa_circ_0119994 | 46475 | 43 | 0.19% |
| hsa_circ_0134850 | 75671 | 70 | 0.19% |
| hsa_circ_0080877 | 62719 | 58 | 0.18% |
| hsa_circ_0109312 | 20556 | 19 | 0.18% |
| hsa_circ_0129226 | 33549 | 31 | 0.18% |
| hsa_circ_0012284 | 42215 | 39 | 0.18% |
| hsa_circ_0048968 | 59552 | 55 | 0.18% |
| hsa_circ_0138599 | 43324 | 40 | 0.18% |
| hsa_circ_0130655 | 30328 | 28 | 0.18% |
| hsa_circ_0122776 | 19512 | 18 | 0.18% |
| hsa_circ_0137256 | 24933 | 23 | 0.18% |
| hsa_circ_0099833 | 47704 | 44 | 0.18% |
| hsa_circ_0137149 | 47713 | 44 | 0.18% |
| hsa_circ_0078323 | 18444 | 17 | 0.18% |
| hsa_circ_0059419 | 18445 | 17 | 0.18% |
| hsa_circ_0111533 | 32563 | 30 | 0.18% |
| hsa_circ_0135711 | 17367 | 16 | 0.18% |
| hsa_circ_0138596 | 27138 | 25 | 0.18% |
| hsa_circ_0136453 | 73903 | 68 | 0.18% |
| hsa_circ_0094085 | 45652 | 42 | 0.18% |
| hsa_circ_0139506 | 41321 | 38 | 0.18% |
| hsa_circ_0099767 | 44589 | 41 | 0.18% |
| hsa_circ_0002865 | 51137 | 47 | 0.18% |
| hsa_circ_0065483 | 15236 | 14 | 0.18% |
| hsa_circ_0126809 | 46797 | 43 | 0.18% |
| hsa_circ_0002765 | 32652 | 30 | 0.18% |
| hsa_circ_0007045 | 32655 | 30 | 0.18% |
| hsa_circ_0024003 | 13068 | 12 | 0.18% |
| hsa_circ_0111579 | 46839 | 43 | 0.18% |
| hsa_circ_0105922 | 17441 | 16 | 0.18% |
| hsa_circ_0006416 | 19628 | 18 | 0.18% |
| hsa_circ_0105687 | 15267 | 14 | 0.18% |
| hsa_circ_0042443 | 24000 | 22 | 0.18% |
| hsa_circ_0134852 | 26182 | 24 | 0.18% |
| hsa_circ_0130584 | 19640 | 18 | 0.18% |
| hsa_circ_0001191 | 36010 | 33 | 0.18% |
| hsa_circ_0000903 | 24041 | 22 | 0.18% |
| hsa_circ_0130598 | 24049 | 22 | 0.18% |
| hsa_circ_0137884 | 17495 | 16 | 0.18% |

| | | | |
|---|---|---|---|
| hsa_circ_0056228 | 15309 | 14 | 0.18% |
| hsa_circ_0069216 | 38286 | 35 | 0.18% |
| hsa_circ_0136002 | 35026 | 32 | 0.18% |
| hsa_circ_0123333 | 42702 | 39 | 0.18% |
| hsa_circ_0118919 | 38334 | 35 | 0.18% |
| hsa_circ_0094655 | 54766 | 50 | 0.18% |
| hsa_circ_0099975 | 87668 | 80 | 0.18% |
| hsa_circ_0013810 | 32886 | 30 | 0.18% |
| hsa_circ_0134191 | 37273 | 34 | 0.18% |
| hsa_circ_0122844 | 72357 | 66 | 0.18% |
| hsa_circ_0093305 | 64727 | 59 | 0.18% |
| hsa_circ_0033783 | 19749 | 18 | 0.18% |
| hsa_circ_0095428 | 40602 | 37 | 0.18% |
| hsa_circ_0100555 | 15367 | 14 | 0.18% |
| hsa_circ_0122762 | 50502 | 46 | 0.18% |
| hsa_circ_0080203 | 31852 | 29 | 0.18% |
| hsa_circ_0006005 | 38456 | 35 | 0.18% |
| hsa_circ_0117312 | 19794 | 18 | 0.18% |
| hsa_circ_0105927 | 24197 | 22 | 0.18% |
| hsa_circ_0036955 | 47297 | 43 | 0.18% |
| hsa_circ_0118920 | 47303 | 43 | 0.18% |
| hsa_circ_0095457 | 27511 | 25 | 0.18% |
| hsa_circ_0132603 | 33029 | 30 | 0.18% |
| hsa_circ_0107146 | 35234 | 32 | 0.18% |
| hsa_circ_0118867 | 48451 | 44 | 0.18% |
| hsa_circ_0003523 | 18726 | 17 | 0.18% |
| hsa_circ_0054279 | 48484 | 44 | 0.18% |
| hsa_circ_0138626 | 41877 | 38 | 0.18% |
| hsa_circ_0118866 | 48490 | 44 | 0.18% |
| hsa_circ_0012285 | 43032 | 39 | 0.18% |
| hsa_circ_0011166 | 52987 | 48 | 0.18% |
| hsa_circ_0096871 | 37536 | 34 | 0.18% |
| hsa_circ_0082957 | 15459 | 14 | 0.18% |
| hsa_circ_0100268 | 40860 | 37 | 0.18% |
| hsa_circ_0130784 | 23201 | 21 | 0.18% |
| hsa_circ_0029834 | 19888 | 18 | 0.18% |
| hsa_circ_0074984 | 191233 | 173 | 0.18% |
| hsa_circ_0004188 | 19899 | 18 | 0.18% |
| hsa_circ_0115985 | 45334 | 41 | 0.18% |
| hsa_circ_0054275 | 48673 | 44 | 0.18% |
| hsa_circ_0130723 | 21020 | 19 | 0.18% |
| hsa_circ_0001468 | 28771 | 26 | 0.18% |
| hsa_circ_0120940 | 42056 | 38 | 0.18% |
| hsa_circ_0124583 | 90765 | 82 | 0.18% |
| hsa_circ_0004182 | 27680 | 25 | 0.18% |
| hsa_circ_0117630 | 28791 | 26 | 0.18% |
| hsa_circ_0092576 | 45419 | 41 | 0.18% |
| hsa_circ_0125401 | 28808 | 26 | 0.18% |
| hsa_circ_0054271 | 48773 | 44 | 0.18% |
| hsa_circ_0121446 | 35484 | 32 | 0.18% |

| | | | |
|---|---|---|---|
| hsa_circ_0103073 | 21072 | 19 | 0.18% |
| hsa_circ_0101221 | 17751 | 16 | 0.18% |
| hsa_circ_0120190 | 19973 | 18 | 0.18% |
| hsa_circ_0033757 | 19974 | 18 | 0.18% |
| hsa_circ_0138371 | 25526 | 23 | 0.18% |
| hsa_circ_0111273 | 19978 | 18 | 0.18% |
| hsa_circ_0136374 | 15539 | 14 | 0.18% |
| hsa_circ_0029417 | 15546 | 14 | 0.18% |
| hsa_circ_0119047 | 38874 | 35 | 0.18% |
| hsa_circ_0054269 | 48876 | 44 | 0.18% |
| hsa_circ_0115325 | 53350 | 48 | 0.18% |
| hsa_circ_0083181 | 52259 | 47 | 0.18% |
| hsa_circ_0110854 | 27801 | 25 | 0.18% |
| hsa_circ_0139956 | 55611 | 50 | 0.18% |
| hsa_circ_0134192 | 37835 | 34 | 0.18% |
| hsa_circ_0114563 | 35615 | 32 | 0.18% |
| hsa_circ_0126938 | 24491 | 22 | 0.18% |
| hsa_circ_0134666 | 45646 | 41 | 0.18% |
| hsa_circ_0000892 | 72393 | 65 | 0.18% |
| hsa_circ_0138594 | 36762 | 33 | 0.18% |
| hsa_circ_0100958 | 66887 | 60 | 0.18% |
| hsa_circ_0130495 | 37904 | 34 | 0.18% |
| hsa_circ_0093875 | 30129 | 27 | 0.18% |
| hsa_circ_0016727 | 32366 | 29 | 0.18% |
| hsa_circ_0102509 | 41308 | 37 | 0.18% |
| hsa_circ_0116506 | 29028 | 26 | 0.18% |
| hsa_circ_0103072 | 21213 | 19 | 0.18% |
| hsa_circ_0109703 | 86014 | 77 | 0.18% |
| hsa_circ_0096736 | 45801 | 41 | 0.18% |
| hsa_circ_0093825 | 42458 | 38 | 0.18% |
| hsa_circ_0001658 | 48071 | 43 | 0.18% |
| hsa_circ_0136456 | 44736 | 40 | 0.18% |
| hsa_circ_0002132 | 49225 | 44 | 0.18% |
| hsa_circ_0121971 | 30207 | 27 | 0.18% |
| hsa_circ_0127843 | 35807 | 32 | 0.18% |
| hsa_circ_0130138 | 30217 | 27 | 0.18% |
| hsa_circ_0111540 | 34714 | 31 | 0.18% |
| hsa_circ_0004019 | 48161 | 43 | 0.18% |
| hsa_circ_0124390 | 59402 | 53 | 0.18% |
| hsa_circ_0103364 | 82941 | 74 | 0.18% |
| hsa_circ_0134847 | 78481 | 70 | 0.18% |
| hsa_circ_0119879 | 28044 | 25 | 0.18% |
| hsa_circ_0001108 | 41507 | 37 | 0.18% |
| hsa_circ_0033784 | 20195 | 18 | 0.18% |
| hsa_circ_0119551 | 43782 | 39 | 0.18% |
| hsa_circ_0111539 | 34816 | 31 | 0.18% |
| hsa_circ_0125872 | 19101 | 17 | 0.18% |
| hsa_circ_0138369 | 25851 | 23 | 0.18% |
| hsa_circ_0004701 | 48341 | 43 | 0.18% |
| hsa_circ_0139814 | 50591 | 45 | 0.18% |

| | | | |
|---|---|---|---|
| hsa_circ_0101225 | 32605 | 29 | 0.18% |
| hsa_circ_0124579 | 61877 | 55 | 0.18% |
| hsa_circ_0123115 | 18016 | 16 | 0.18% |
| hsa_circ_0132731 | 46167 | 41 | 0.18% |
| hsa_circ_0001107 | 41676 | 37 | 0.18% |
| hsa_circ_0100269 | 41681 | 37 | 0.18% |
| hsa_circ_0128580 | 60854 | 54 | 0.18% |
| hsa_circ_0094630 | 45080 | 40 | 0.18% |
| hsa_circ_0099619 | 37194 | 33 | 0.18% |
| hsa_circ_0093773 | 32687 | 29 | 0.18% |
| hsa_circ_0079083 | 23682 | 21 | 0.18% |
| hsa_circ_0072846 | 41730 | 37 | 0.18% |
| hsa_circ_0121969 | 85748 | 76 | 0.18% |
| hsa_circ_0092596 | 46267 | 41 | 0.18% |
| hsa_circ_0133094 | 39501 | 35 | 0.18% |
| hsa_circ_0099976 | 90371 | 80 | 0.18% |
| hsa_circ_0130682 | 24867 | 22 | 0.18% |
| hsa_circ_0131274 | 53164 | 47 | 0.18% |
| hsa_circ_0137783 | 14713 | 13 | 0.18% |
| hsa_circ_0125869 | 37369 | 33 | 0.18% |
| hsa_circ_0002335 | 56621 | 50 | 0.18% |
| hsa_circ_0137183 | 49842 | 44 | 0.18% |
| hsa_circ_0112800 | 46453 | 41 | 0.18% |
| hsa_circ_0046642 | 53268 | 47 | 0.18% |
| hsa_circ_0112161 | 26076 | 23 | 0.18% |
| hsa_circ_0130644 | 62424 | 55 | 0.18% |
| hsa_circ_0131625 | 69263 | 61 | 0.18% |
| hsa_circ_0122544 | 52244 | 46 | 0.18% |
| hsa_circ_0074987 | 61368 | 54 | 0.18% |
| hsa_circ_0106383 | 69369 | 61 | 0.18% |
| hsa_circ_0139737 | 89844 | 79 | 0.18% |
| hsa_circ_0130642 | 62599 | 55 | 0.18% |
| hsa_circ_0093743 | 19355 | 17 | 0.18% |
| hsa_circ_0120138 | 52381 | 46 | 0.18% |
| hsa_circ_0093357 | 44413 | 39 | 0.18% |
| hsa_circ_0099284 | 35307 | 31 | 0.18% |
| hsa_circ_0059420 | 19366 | 17 | 0.18% |
| hsa_circ_0099977 | 91141 | 80 | 0.18% |
| hsa_circ_0084559 | 55845 | 49 | 0.18% |
| hsa_circ_0058208 | 59266 | 52 | 0.18% |
| hsa_circ_0135835 | 58127 | 51 | 0.18% |
| hsa_circ_0130450 | 37619 | 33 | 0.18% |
| hsa_circ_0095761 | 39914 | 35 | 0.18% |
| hsa_circ_0099267 | 17118 | 15 | 0.18% |
| hsa_circ_0008395 | 37664 | 33 | 0.18% |
| hsa_circ_0115986 | 46816 | 41 | 0.18% |
| hsa_circ_0004336 | 28547 | 25 | 0.18% |
| hsa_circ_0099978 | 91370 | 80 | 0.18% |
| hsa_circ_0103150 | 47971 | 42 | 0.18% |
| hsa_circ_0139785 | 43413 | 38 | 0.18% |

| | | | |
|---|---|---|---|
| hsa_circ_0100974 | 25134 | 22 | 0.18% |
| hsa_circ_0129120 | 17138 | 15 | 0.18% |
| hsa_circ_0021754 | 43438 | 38 | 0.17% |
| hsa_circ_0128420 | 35457 | 31 | 0.17% |
| hsa_circ_0034021 | 61802 | 54 | 0.17% |
| hsa_circ_0131463 | 74396 | 65 | 0.17% |
| hsa_circ_0111964 | 85891 | 75 | 0.17% |
| hsa_circ_0132080 | 34377 | 30 | 0.17% |
| hsa_circ_0079078 | 24067 | 21 | 0.17% |
| hsa_circ_0125156 | 30956 | 27 | 0.17% |
| hsa_circ_0114373 | 32121 | 28 | 0.17% |
| hsa_circ_0119082 | 22950 | 20 | 0.17% |
| hsa_circ_0118301 | 18360 | 16 | 0.17% |
| hsa_circ_0053499 | 14923 | 13 | 0.17% |
| hsa_circ_0131669 | 56250 | 49 | 0.17% |
| hsa_circ_0109349 | 14929 | 13 | 0.17% |
| hsa_circ_0136095 | 50565 | 44 | 0.17% |
| hsa_circ_0095641 | 43685 | 38 | 0.17% |
| hsa_circ_0027245 | 86232 | 75 | 0.17% |
| hsa_circ_0114078 | 33355 | 29 | 0.17% |
| hsa_circ_0001676 | 14956 | 13 | 0.17% |
| hsa_circ_0003841 | 42579 | 37 | 0.17% |
| hsa_circ_0025892 | 40280 | 35 | 0.17% |
| hsa_circ_0108496 | 50641 | 44 | 0.17% |
| hsa_circ_0072804 | 42592 | 37 | 0.17% |
| hsa_circ_0096037 | 17268 | 15 | 0.17% |
| hsa_circ_0139734 | 90966 | 79 | 0.17% |
| hsa_circ_0107478 | 48384 | 42 | 0.17% |
| hsa_circ_0108420 | 78355 | 68 | 0.17% |
| hsa_circ_0133139 | 33474 | 29 | 0.17% |
| hsa_circ_0006261 | 19630 | 17 | 0.17% |
| hsa_circ_0137485 | 79686 | 69 | 0.17% |
| hsa_circ_0102253 | 40431 | 35 | 0.17% |
| hsa_circ_0122618 | 16181 | 14 | 0.17% |
| hsa_circ_0027242 | 86692 | 75 | 0.17% |
| hsa_circ_0137260 | 90177 | 78 | 0.17% |
| hsa_circ_0095246 | 34699 | 30 | 0.17% |
| hsa_circ_0136114 | 27806 | 24 | 0.17% |
| hsa_circ_0102629 | 20870 | 18 | 0.17% |
| hsa_circ_0026148 | 74223 | 64 | 0.17% |
| hsa_circ_0095361 | 19717 | 17 | 0.17% |
| hsa_circ_0104942 | 32477 | 28 | 0.17% |
| hsa_circ_0110678 | 70771 | 61 | 0.17% |
| hsa_circ_0098287 | 20909 | 18 | 0.17% |
| hsa_circ_0125157 | 31364 | 27 | 0.17% |
| hsa_circ_0101903 | 40661 | 35 | 0.17% |
| hsa_circ_0093271 | 33691 | 29 | 0.17% |
| hsa_circ_0112159 | 61599 | 53 | 0.17% |
| hsa_circ_0044179 | 61622 | 53 | 0.17% |
| hsa_circ_0006630 | 19778 | 17 | 0.17% |

| | | | |
|---|---|---|---|
| hsa_circ_0118541 | 39561 | 34 | 0.17% |
| hsa_circ_0131513 | 38407 | 33 | 0.17% |
| hsa_circ_0139431 | 70996 | 61 | 0.17% |
| hsa_circ_0101646 | 61756 | 53 | 0.17% |
| hsa_circ_0115197 | 88582 | 76 | 0.17% |
| hsa_circ_0132114 | 18649 | 16 | 0.17% |
| hsa_circ_0122887 | 55970 | 48 | 0.17% |
| hsa_circ_0104940 | 32659 | 28 | 0.17% |
| hsa_circ_0134076 | 36164 | 31 | 0.17% |
| hsa_circ_0134432 | 29168 | 25 | 0.17% |
| hsa_circ_0002711 | 23346 | 20 | 0.17% |
| hsa_circ_0094062 | 59542 | 51 | 0.17% |
| hsa_circ_0094063 | 59545 | 51 | 0.17% |
| hsa_circ_0020027 | 16349 | 14 | 0.17% |
| hsa_circ_0113884 | 24544 | 21 | 0.17% |
| hsa_circ_0104912 | 25714 | 22 | 0.17% |
| hsa_circ_0138522 | 44418 | 38 | 0.17% |
| hsa_circ_0094386 | 42081 | 36 | 0.17% |
| hsa_circ_0132110 | 18718 | 16 | 0.17% |
| hsa_circ_0104349 | 47976 | 41 | 0.17% |
| hsa_circ_0133768 | 18723 | 16 | 0.17% |
| hsa_circ_0114307 | 31602 | 27 | 0.17% |
| hsa_circ_0105985 | 73760 | 63 | 0.17% |
| hsa_circ_0137194 | 37508 | 32 | 0.17% |
| hsa_circ_0102875 | 41025 | 35 | 0.17% |
| hsa_circ_0076843 | 16411 | 14 | 0.17% |
| hsa_circ_0111543 | 49234 | 42 | 0.17% |
| hsa_circ_0003968 | 45723 | 39 | 0.17% |
| hsa_circ_0122602 | 15252 | 13 | 0.17% |
| hsa_circ_0121180 | 29343 | 25 | 0.17% |
| hsa_circ_0130423 | 32903 | 28 | 0.17% |
| hsa_circ_0033727 | 21168 | 18 | 0.17% |
| hsa_circ_0032834 | 29401 | 25 | 0.17% |
| hsa_circ_0128197 | 29404 | 25 | 0.17% |
| hsa_circ_0107959 | 45876 | 39 | 0.17% |
| hsa_circ_0034119 | 15294 | 13 | 0.17% |
| hsa_circ_0101207 | 15303 | 13 | 0.17% |
| hsa_circ_0121173 | 29439 | 25 | 0.17% |
| hsa_circ_0101683 | 25912 | 22 | 0.17% |
| hsa_circ_0109477 | 55358 | 47 | 0.17% |
| hsa_circ_0109476 | 55364 | 47 | 0.17% |
| hsa_circ_0100851 | 48299 | 41 | 0.17% |
| hsa_circ_0135749 | 48319 | 41 | 0.17% |
| hsa_circ_0139305 | 42430 | 36 | 0.17% |
| hsa_circ_0105683 | 28308 | 24 | 0.17% |
| hsa_circ_0076844 | 16519 | 14 | 0.17% |
| hsa_circ_0102240 | 16522 | 14 | 0.17% |
| hsa_circ_0042113 | 40167 | 34 | 0.17% |
| hsa_circ_0114495 | 38986 | 33 | 0.17% |
| hsa_circ_0033566 | 17731 | 15 | 0.17% |

| | | | |
|---|---|---|---|
| hsa_circ_0113692 | 34295 | 29 | 0.17% |
| hsa_circ_0106545 | 52074 | 44 | 0.17% |
| hsa_circ_0125995 | 15391 | 13 | 0.17% |
| hsa_circ_0070448 | 48572 | 41 | 0.17% |
| hsa_circ_0134860 | 15401 | 13 | 0.17% |
| hsa_circ_0114846 | 41480 | 35 | 0.17% |
| hsa_circ_0138605 | 47407 | 40 | 0.17% |
| hsa_circ_0108459 | 32010 | 27 | 0.17% |
| hsa_circ_0111752 | 46249 | 39 | 0.17% |
| hsa_circ_0076845 | 16603 | 14 | 0.17% |
| hsa_circ_0100116 | 53372 | 45 | 0.17% |
| hsa_circ_0100753 | 72357 | 61 | 0.17% |
| hsa_circ_0092577 | 28470 | 24 | 0.17% |
| hsa_circ_0139874 | 30859 | 26 | 0.17% |
| hsa_circ_0107953 | 46298 | 39 | 0.17% |
| hsa_circ_0132951 | 41566 | 35 | 0.17% |
| hsa_circ_0130141 | 43946 | 37 | 0.17% |
| hsa_circ_0132558 | 60587 | 51 | 0.17% |
| hsa_circ_0126015 | 24949 | 21 | 0.17% |
| hsa_circ_0036948 | 62990 | 53 | 0.17% |
| hsa_circ_0137917 | 48749 | 41 | 0.17% |
| hsa_circ_0129892 | 29726 | 25 | 0.17% |
| hsa_circ_0139671 | 73759 | 62 | 0.17% |
| hsa_circ_0002033 | 29743 | 25 | 0.17% |
| hsa_circ_0123033 | 42839 | 36 | 0.17% |
| hsa_circ_0102709 | 65465 | 55 | 0.17% |
| hsa_circ_0101752 | 58327 | 49 | 0.17% |
| hsa_circ_0135985 | 19046 | 16 | 0.17% |
| hsa_circ_0132724 | 21436 | 18 | 0.17% |
| hsa_circ_0088433 | 27391 | 23 | 0.17% |
| hsa_circ_0125871 | 32184 | 27 | 0.17% |
| hsa_circ_0033567 | 17882 | 15 | 0.17% |
| hsa_circ_0099348 | 17891 | 15 | 0.17% |
| hsa_circ_0129229 | 47711 | 40 | 0.17% |
| hsa_circ_0094278 | 37002 | 31 | 0.17% |
| hsa_circ_0121511 | 44164 | 37 | 0.17% |
| hsa_circ_0073233 | 52540 | 44 | 0.17% |
| hsa_circ_0019246 | 40607 | 34 | 0.17% |
| hsa_circ_0103370 | 17928 | 15 | 0.17% |
| hsa_circ_0137563 | 58578 | 49 | 0.17% |
| hsa_circ_0124486 | 37080 | 31 | 0.17% |
| hsa_circ_0055602 | 16756 | 14 | 0.17% |
| hsa_circ_0124560 | 44285 | 37 | 0.17% |
| hsa_circ_0140099 | 26338 | 22 | 0.17% |
| hsa_circ_0127847 | 31136 | 26 | 0.17% |
| hsa_circ_0080033 | 68272 | 57 | 0.17% |
| hsa_circ_0033568 | 17984 | 15 | 0.17% |
| hsa_circ_0067972 | 53959 | 45 | 0.17% |
| hsa_circ_0126016 | 25198 | 21 | 0.17% |
| hsa_circ_0127749 | 25204 | 21 | 0.17% |

| | | | |
|---|---|---|---|
| hsa_circ_0101824 | 34852 | 29 | 0.17% |
| hsa_circ_0132741 | 26443 | 22 | 0.17% |
| hsa_circ_0140237 | 16828 | 14 | 0.17% |
| hsa_circ_0067966 | 54103 | 45 | 0.17% |
| hsa_circ_0045945 | 51705 | 43 | 0.17% |
| hsa_circ_0127713 | 63751 | 53 | 0.17% |
| hsa_circ_0101102 | 58943 | 49 | 0.17% |
| hsa_circ_0096627 | 19257 | 16 | 0.17% |
| hsa_circ_0024151 | 127592 | 106 | 0.17% |
| hsa_circ_0135152 | 36126 | 30 | 0.17% |
| hsa_circ_0122639 | 42157 | 35 | 0.17% |
| hsa_circ_0105924 | 42170 | 35 | 0.17% |
| hsa_circ_0023301 | 93983 | 78 | 0.17% |
| hsa_circ_0033569 | 18083 | 15 | 0.17% |
| hsa_circ_0131194 | 47035 | 39 | 0.17% |
| hsa_circ_0121910 | 16896 | 14 | 0.17% |
| hsa_circ_0100975 | 26561 | 22 | 0.17% |
| hsa_circ_0135403 | 44673 | 37 | 0.17% |
| hsa_circ_0122640 | 42263 | 35 | 0.17% |
| hsa_circ_0013805 | 44688 | 37 | 0.17% |
| hsa_circ_0129097 | 35039 | 29 | 0.17% |
| hsa_circ_0118341 | 49549 | 41 | 0.17% |
| hsa_circ_0033570 | 18142 | 15 | 0.17% |
| hsa_circ_0101214 | 19382 | 16 | 0.17% |
| hsa_circ_0076846 | 16960 | 14 | 0.17% |
| hsa_circ_0022426 | 18173 | 15 | 0.17% |
| hsa_circ_0078407 | 52122 | 43 | 0.16% |
| hsa_circ_0114673 | 35178 | 29 | 0.16% |
| hsa_circ_0139489 | 63137 | 52 | 0.16% |
| hsa_circ_0132672 | 42498 | 35 | 0.16% |
| hsa_circ_0095080 | 64381 | 53 | 0.16% |
| hsa_circ_0133068 | 38877 | 32 | 0.16% |
| hsa_circ_0033493 | 72902 | 60 | 0.16% |
| hsa_circ_0115631 | 36467 | 30 | 0.16% |
| hsa_circ_0088434 | 27959 | 23 | 0.16% |
| hsa_circ_0125411 | 79035 | 65 | 0.16% |
| hsa_circ_0108272 | 18239 | 15 | 0.16% |
| hsa_circ_0134120 | 41356 | 34 | 0.16% |
| hsa_circ_0034117 | 17046 | 14 | 0.16% |
| hsa_circ_0108266 | 19492 | 16 | 0.16% |
| hsa_circ_0094264 | 29255 | 24 | 0.16% |
| hsa_circ_0000552 | 68264 | 56 | 0.16% |
| hsa_circ_0139999 | 35354 | 29 | 0.16% |
| hsa_circ_0134418 | 25602 | 21 | 0.16% |
| hsa_circ_0118317 | 57372 | 47 | 0.16% |
| hsa_circ_0130591 | 50050 | 41 | 0.16% |
| hsa_circ_0033572 | 18317 | 15 | 0.16% |
| hsa_circ_0076847 | 17099 | 14 | 0.16% |
| hsa_circ_0139508 | 46439 | 38 | 0.16% |
| hsa_circ_0001446 | 35445 | 29 | 0.16% |

| | | | |
|---|---|---|---|
| hsa_circ_0139555 | 52582 | 43 | 0.16% |
| hsa_circ_0134253 | 63591 | 52 | 0.16% |
| hsa_circ_0123612 | 17134 | 14 | 0.16% |
| hsa_circ_0132748 | 34271 | 28 | 0.16% |
| hsa_circ_0022427 | 18361 | 15 | 0.16% |
| hsa_circ_0130889 | 29384 | 24 | 0.16% |
| hsa_circ_0113694 | 40404 | 33 | 0.16% |
| hsa_circ_0006205 | 40452 | 33 | 0.16% |
| hsa_circ_0118336 | 44159 | 36 | 0.16% |
| hsa_circ_0034151 | 17184 | 14 | 0.16% |
| hsa_circ_0055804 | 56484 | 46 | 0.16% |
| hsa_circ_0125736 | 55270 | 45 | 0.16% |
| hsa_circ_0133225 | 36861 | 30 | 0.16% |
| hsa_circ_0096706 | 24576 | 20 | 0.16% |
| hsa_circ_0125145 | 17222 | 14 | 0.16% |
| hsa_circ_0107960 | 48001 | 39 | 0.16% |
| hsa_circ_0046641 | 57850 | 47 | 0.16% |
| hsa_circ_0139255 | 46776 | 38 | 0.16% |
| hsa_circ_0129871 | 32010 | 26 | 0.16% |
| hsa_circ_0114164 | 35711 | 29 | 0.16% |
| hsa_circ_0125702 | 18476 | 15 | 0.16% |
| hsa_circ_0073281 | 17247 | 14 | 0.16% |
| hsa_circ_0133612 | 20951 | 17 | 0.16% |
| hsa_circ_0137233 | 46832 | 38 | 0.16% |
| hsa_circ_0065484 | 18493 | 15 | 0.16% |
| hsa_circ_0003462 | 49323 | 40 | 0.16% |
| hsa_circ_0130980 | 56738 | 46 | 0.16% |
| hsa_circ_0108428 | 71548 | 58 | 0.16% |
| hsa_circ_0076848 | 17278 | 14 | 0.16% |
| hsa_circ_0022428 | 18515 | 15 | 0.16% |
| hsa_circ_0123391 | 85172 | 69 | 0.16% |
| hsa_circ_0120505 | 82749 | 67 | 0.16% |
| hsa_circ_0127481 | 46936 | 38 | 0.16% |
| hsa_circ_0025824 | 96408 | 78 | 0.16% |
| hsa_circ_0127068 | 49447 | 40 | 0.16% |
| hsa_circ_0093730 | 17310 | 14 | 0.16% |
| hsa_circ_0133671 | 61823 | 50 | 0.16% |
| hsa_circ_0047050 | 39600 | 32 | 0.16% |
| hsa_circ_0138309 | 23522 | 19 | 0.16% |
| hsa_circ_0129942 | 21051 | 17 | 0.16% |
| hsa_circ_0093076 | 24769 | 20 | 0.16% |
| hsa_circ_0137647 | 59474 | 48 | 0.16% |
| hsa_circ_0130113 | 47093 | 38 | 0.16% |
| hsa_circ_0108273 | 18599 | 15 | 0.16% |
| hsa_circ_0133829 | 44646 | 36 | 0.16% |
| hsa_circ_0020025 | 32250 | 26 | 0.16% |
| hsa_circ_0047774 | 64512 | 52 | 0.16% |
| hsa_circ_0022429 | 18614 | 15 | 0.16% |
| hsa_circ_0134601 | 24824 | 20 | 0.16% |
| hsa_circ_0119255 | 44687 | 36 | 0.16% |

| | | | |
|---|---|---|---|
| hsa_circ_0128048 | 50917 | 41 | 0.16% |
| hsa_circ_0108702 | 49711 | 40 | 0.16% |
| hsa_circ_0103044 | 19906 | 16 | 0.16% |
| hsa_circ_0102711 | 46044 | 37 | 0.16% |
| hsa_circ_0129272 | 52267 | 42 | 0.16% |
| hsa_circ_0016170 | 56004 | 45 | 0.16% |
| hsa_circ_0098060 | 52296 | 42 | 0.16% |
| hsa_circ_0102746 | 34865 | 28 | 0.16% |
| hsa_circ_0099352 | 21170 | 17 | 0.16% |
| hsa_circ_0076849 | 17436 | 14 | 0.16% |
| hsa_circ_0099982 | 48603 | 39 | 0.16% |
| hsa_circ_0097087 | 52349 | 42 | 0.16% |
| hsa_circ_0123137 | 32413 | 26 | 0.16% |
| hsa_circ_0124781 | 38686 | 31 | 0.16% |
| hsa_circ_0023664 | 58654 | 47 | 0.16% |
| hsa_circ_0132671 | 26226 | 21 | 0.16% |
| hsa_circ_0115532 | 22489 | 18 | 0.16% |
| hsa_circ_0094088 | 79995 | 64 | 0.16% |
| hsa_circ_0108274 | 18763 | 15 | 0.16% |
| hsa_circ_0099353 | 21269 | 17 | 0.16% |
| hsa_circ_0128045 | 51309 | 41 | 0.16% |
| hsa_circ_0140235 | 17527 | 14 | 0.16% |
| hsa_circ_0125146 | 17537 | 14 | 0.16% |
| hsa_circ_0117690 | 26306 | 21 | 0.16% |
| hsa_circ_0044189 | 77673 | 62 | 0.16% |
| hsa_circ_0122380 | 52642 | 42 | 0.16% |
| hsa_circ_0092473 | 52756 | 42 | 0.16% |
| hsa_circ_0108357 | 57788 | 46 | 0.16% |
| hsa_circ_0012863 | 74122 | 59 | 0.16% |
| hsa_circ_0130594 | 46495 | 37 | 0.16% |
| hsa_circ_0008885 | 22623 | 18 | 0.16% |
| hsa_circ_0094770 | 32692 | 26 | 0.16% |
| hsa_circ_0076850 | 17620 | 14 | 0.16% |
| hsa_circ_0127028 | 41538 | 33 | 0.16% |
| hsa_circ_0095971 | 36512 | 29 | 0.16% |
| hsa_circ_0107951 | 49115 | 39 | 0.16% |
| hsa_circ_0109954 | 68022 | 54 | 0.16% |
| hsa_circ_0121541 | 37791 | 30 | 0.16% |
| hsa_circ_0122023 | 17638 | 14 | 0.16% |
| hsa_circ_0095057 | 22696 | 18 | 0.16% |
| hsa_circ_0096336 | 31554 | 25 | 0.16% |
| hsa_circ_0099479 | 51758 | 41 | 0.16% |
| hsa_circ_0134841 | 54405 | 43 | 0.16% |
| hsa_circ_0116141 | 22778 | 18 | 0.16% |
| hsa_circ_0132043 | 18983 | 15 | 0.16% |
| hsa_circ_0137429 | 43046 | 34 | 0.16% |
| hsa_circ_0115712 | 67136 | 53 | 0.16% |
| hsa_circ_0118868 | 29141 | 23 | 0.16% |
| hsa_circ_0098990 | 46886 | 37 | 0.16% |
| hsa_circ_0114623 | 20278 | 16 | 0.16% |

| | | | |
|---|---|---|---|
| hsa_circ_0137886 | 62106 | 49 | 0.16% |
| hsa_circ_0103022 | 34227 | 27 | 0.16% |
| hsa_circ_0114409 | 15215 | 12 | 0.16% |
| hsa_circ_0099840 | 29168 | 23 | 0.16% |
| hsa_circ_0073287 | 17760 | 14 | 0.16% |
| hsa_circ_0100780 | 27912 | 22 | 0.16% |
| hsa_circ_0104414 | 48220 | 38 | 0.16% |
| hsa_circ_0139398 | 54571 | 43 | 0.16% |
| hsa_circ_0107477 | 53334 | 42 | 0.16% |
| hsa_circ_0130556 | 33027 | 26 | 0.16% |
| hsa_circ_0130555 | 33045 | 26 | 0.16% |
| hsa_circ_0119549 | 58475 | 46 | 0.16% |
| hsa_circ_0137168 | 33052 | 26 | 0.16% |
| hsa_circ_0114843 | 55953 | 44 | 0.16% |
| hsa_circ_0092555 | 20353 | 16 | 0.16% |
| hsa_circ_0117959 | 59833 | 47 | 0.16% |
| hsa_circ_0139547 | 47112 | 37 | 0.16% |
| hsa_circ_0099354 | 21651 | 17 | 0.16% |
| hsa_circ_0100267 | 47153 | 37 | 0.16% |
| hsa_circ_0115489 | 88041 | 69 | 0.16% |
| hsa_circ_0112697 | 44664 | 35 | 0.16% |
| hsa_circ_0096612 | 68958 | 54 | 0.16% |
| hsa_circ_0073273 | 24264 | 19 | 0.16% |
| hsa_circ_0139670 | 61326 | 48 | 0.16% |
| hsa_circ_0124637 | 42162 | 33 | 0.16% |
| hsa_circ_0129118 | 60077 | 47 | 0.16% |
| hsa_circ_0133256 | 60089 | 47 | 0.16% |
| hsa_circ_0076851 | 17902 | 14 | 0.16% |
| hsa_circ_0124651 | 55007 | 43 | 0.16% |
| hsa_circ_0099344 | 89634 | 70 | 0.16% |
| hsa_circ_0033758 | 23054 | 18 | 0.16% |
| hsa_circ_0004173 | 32035 | 25 | 0.16% |
| hsa_circ_0139161 | 47427 | 37 | 0.16% |
| hsa_circ_0108699 | 69258 | 54 | 0.16% |
| hsa_circ_0114103 | 41070 | 32 | 0.16% |
| hsa_circ_0114102 | 41085 | 32 | 0.16% |
| hsa_circ_0102084 | 65528 | 51 | 0.16% |
| hsa_circ_0129241 | 28268 | 22 | 0.16% |
| hsa_circ_0092486 | 80957 | 63 | 0.16% |
| hsa_circ_0130716 | 77118 | 60 | 0.16% |
| hsa_circ_0098128 | 61739 | 48 | 0.16% |
| hsa_circ_0127195 | 45033 | 35 | 0.16% |
| hsa_circ_0136717 | 36040 | 28 | 0.16% |
| hsa_circ_0069067 | 20603 | 16 | 0.16% |
| hsa_circ_0093695 | 76016 | 59 | 0.16% |
| hsa_circ_0123610 | 19336 | 15 | 0.16% |
| hsa_circ_0076852 | 18055 | 14 | 0.16% |
| hsa_circ_0077820 | 42563 | 33 | 0.16% |
| hsa_circ_0003495 | 63222 | 49 | 0.16% |
| hsa_circ_0093672 | 55544 | 43 | 0.15% |

| | | | |
|---|---|---|---|
| hsa_circ_0002144 | 76214 | 59 | 0.15% |
| hsa_circ_0103745 | 46509 | 36 | 0.15% |
| hsa_circ_0076871 | 64649 | 50 | 0.15% |
| hsa_circ_0129414 | 23276 | 18 | 0.15% |
| hsa_circ_0114128 | 20708 | 16 | 0.15% |
| hsa_circ_0133136 | 82850 | 64 | 0.15% |
| hsa_circ_0026083 | 19419 | 15 | 0.15% |
| hsa_circ_0130654 | 38865 | 30 | 0.15% |
| hsa_circ_0099051 | 20736 | 16 | 0.15% |
| hsa_circ_0129230 | 51855 | 40 | 0.15% |
| hsa_circ_0125347 | 37603 | 29 | 0.15% |
| hsa_circ_0133953 | 23365 | 18 | 0.15% |
| hsa_circ_0109364 | 36350 | 28 | 0.15% |
| hsa_circ_0137318 | 25965 | 20 | 0.15% |
| hsa_circ_0076853 | 18181 | 14 | 0.15% |
| hsa_circ_0055550 | 592527 | 456 | 0.15% |
| hsa_circ_0136718 | 36442 | 28 | 0.15% |
| hsa_circ_0125523 | 39085 | 30 | 0.15% |
| hsa_circ_0126171 | 72972 | 56 | 0.15% |
| hsa_circ_0032160 | 18251 | 14 | 0.15% |
| hsa_circ_0097293 | 19565 | 15 | 0.15% |
| hsa_circ_0133482 | 73058 | 56 | 0.15% |
| hsa_circ_0130369 | 44360 | 34 | 0.15% |
| hsa_circ_0096335 | 32626 | 25 | 0.15% |
| hsa_circ_0033785 | 23493 | 18 | 0.15% |
| hsa_circ_0076854 | 18277 | 14 | 0.15% |
| hsa_circ_0139024 | 63985 | 49 | 0.15% |
| hsa_circ_0097433 | 26126 | 20 | 0.15% |
| hsa_circ_0125524 | 39210 | 30 | 0.15% |
| hsa_circ_0137431 | 36596 | 28 | 0.15% |
| hsa_circ_0139718 | 60151 | 46 | 0.15% |
| hsa_circ_0096909 | 54932 | 42 | 0.15% |
| hsa_circ_0119862 | 61527 | 47 | 0.15% |
| hsa_circ_0101678 | 40587 | 31 | 0.15% |
| hsa_circ_0114614 | 20968 | 16 | 0.15% |
| hsa_circ_0136647 | 36709 | 28 | 0.15% |
| hsa_circ_0114637 | 20984 | 16 | 0.15% |
| hsa_circ_0052591 | 43280 | 33 | 0.15% |
| hsa_circ_0138443 | 52512 | 40 | 0.15% |
| hsa_circ_0095818 | 36764 | 28 | 0.15% |
| hsa_circ_0076855 | 18382 | 14 | 0.15% |
| hsa_circ_0139437 | 49905 | 38 | 0.15% |
| hsa_circ_0112398 | 19727 | 15 | 0.15% |
| hsa_circ_0140698 | 43412 | 33 | 0.15% |
| hsa_circ_0133222 | 64469 | 49 | 0.15% |
| hsa_circ_0025766 | 92170 | 70 | 0.15% |
| hsa_circ_0060397 | 55350 | 42 | 0.15% |
| hsa_circ_0121179 | 23726 | 18 | 0.15% |
| hsa_circ_0134780 | 55364 | 42 | 0.15% |
| hsa_circ_0108438 | 46152 | 35 | 0.15% |

| | | | |
|---|---|---|---|
| hsa_circ_0108437 | 46154 | 35 | 0.15% |
| hsa_circ_0130361 | 44891 | 34 | 0.15% |
| hsa_circ_0140765 | 29081 | 22 | 0.15% |
| hsa_circ_0101208 | 21156 | 16 | 0.15% |
| hsa_circ_0121172 | 23822 | 18 | 0.15% |
| hsa_circ_0033786 | 23824 | 18 | 0.15% |
| hsa_circ_0131193 | 51624 | 39 | 0.15% |
| hsa_circ_0004153 | 48995 | 37 | 0.15% |
| hsa_circ_0137564 | 66215 | 50 | 0.15% |
| hsa_circ_0076856 | 18544 | 14 | 0.15% |
| hsa_circ_0034137 | 31805 | 24 | 0.15% |
| hsa_circ_0098510 | 38480 | 29 | 0.15% |
| hsa_circ_0113376 | 21236 | 16 | 0.15% |
| hsa_circ_0094946 | 18595 | 14 | 0.15% |
| hsa_circ_0080979 | 106311 | 80 | 0.15% |
| hsa_circ_0006210 | 54499 | 41 | 0.15% |
| hsa_circ_0131089 | 49184 | 37 | 0.15% |
| hsa_circ_0135324 | 41214 | 31 | 0.15% |
| hsa_circ_0131359 | 38573 | 29 | 0.15% |
| hsa_circ_0117331 | 49217 | 37 | 0.15% |
| hsa_circ_0132284 | 50574 | 38 | 0.15% |
| hsa_circ_0134840 | 45300 | 34 | 0.15% |
| hsa_circ_0136435 | 21319 | 16 | 0.15% |
| hsa_circ_0098311 | 58632 | 44 | 0.15% |
| hsa_circ_0096609 | 54665 | 41 | 0.15% |
| hsa_circ_0093865 | 74772 | 56 | 0.15% |
| hsa_circ_0132363 | 29390 | 22 | 0.15% |
| hsa_circ_0004583 | 45422 | 34 | 0.15% |
| hsa_circ_0112396 | 20050 | 15 | 0.15% |
| hsa_circ_0116068 | 53468 | 40 | 0.15% |
| hsa_circ_0139722 | 26748 | 20 | 0.15% |
| hsa_circ_0100976 | 46851 | 35 | 0.15% |
| hsa_circ_0076857 | 18751 | 14 | 0.15% |
| hsa_circ_0134771 | 56271 | 42 | 0.15% |
| hsa_circ_0113259 | 58965 | 44 | 0.15% |
| hsa_circ_0115639 | 26805 | 20 | 0.15% |
| hsa_circ_0001585 | 91153 | 68 | 0.15% |
| hsa_circ_0055147 | 60325 | 45 | 0.15% |
| hsa_circ_0120787 | 36219 | 27 | 0.15% |
| hsa_circ_0129413 | 28176 | 21 | 0.15% |
| hsa_circ_0080235 | 51012 | 38 | 0.15% |
| hsa_circ_0095635 | 59191 | 44 | 0.15% |
| hsa_circ_0063302 | 82111 | 61 | 0.15% |
| hsa_circ_0033787 | 24268 | 18 | 0.15% |
| hsa_circ_0063299 | 82260 | 61 | 0.15% |
| hsa_circ_0129947 | 67430 | 50 | 0.15% |
| hsa_circ_0121782 | 39121 | 29 | 0.15% |
| hsa_circ_0125481 | 59359 | 44 | 0.15% |
| hsa_circ_0026082 | 18909 | 14 | 0.15% |
| hsa_circ_0101818 | 48648 | 36 | 0.15% |

| | | | |
|---|---|---|---|
| hsa_circ_0097089 | 94604 | 70 | 0.15% |
| hsa_circ_0137430 | 37843 | 28 | 0.15% |
| hsa_circ_0126464 | 75691 | 56 | 0.15% |
| hsa_circ_0131503 | 36522 | 27 | 0.15% |
| hsa_circ_0135121 | 39234 | 29 | 0.15% |
| hsa_circ_0096883 | 46027 | 34 | 0.15% |
| hsa_circ_0108543 | 46031 | 34 | 0.15% |
| hsa_circ_0102538 | 62324 | 46 | 0.15% |
| hsa_circ_0131605 | 33877 | 25 | 0.15% |
| hsa_circ_0100260 | 89482 | 66 | 0.15% |
| hsa_circ_0083120 | 20338 | 15 | 0.15% |
| hsa_circ_0106679 | 51538 | 38 | 0.15% |
| hsa_circ_0017176 | 40714 | 30 | 0.15% |
| hsa_circ_0119074 | 25789 | 19 | 0.15% |
| hsa_circ_0117147 | 52951 | 39 | 0.15% |
| hsa_circ_0032844 | 42096 | 31 | 0.15% |
| hsa_circ_0122314 | 32592 | 24 | 0.15% |
| hsa_circ_0115638 | 35346 | 26 | 0.15% |
| hsa_circ_0132124 | 24480 | 18 | 0.15% |
| hsa_circ_0129891 | 21771 | 16 | 0.15% |
| hsa_circ_0009066 | 65335 | 48 | 0.15% |
| hsa_circ_0126897 | 28611 | 21 | 0.15% |
| hsa_circ_0103685 | 64048 | 47 | 0.15% |
| hsa_circ_0116505 | 59971 | 44 | 0.15% |
| hsa_circ_0097849 | 24558 | 18 | 0.15% |
| hsa_circ_0105180 | 60055 | 44 | 0.15% |
| hsa_circ_0095358 | 20485 | 15 | 0.15% |
| hsa_circ_0140686 | 40982 | 30 | 0.15% |
| hsa_circ_0126013 | 56067 | 41 | 0.15% |
| hsa_circ_0139490 | 71110 | 52 | 0.15% |
| hsa_circ_0103688 | 50598 | 37 | 0.15% |
| hsa_circ_0126465 | 76702 | 56 | 0.15% |
| hsa_circ_0120039 | 35623 | 26 | 0.15% |
| hsa_circ_0135175 | 53484 | 39 | 0.15% |
| hsa_circ_0131289 | 28813 | 21 | 0.15% |
| hsa_circ_0076858 | 19214 | 14 | 0.15% |
| hsa_circ_0100258 | 85215 | 62 | 0.15% |
| hsa_circ_0106212 | 52282 | 38 | 0.15% |
| hsa_circ_0125689 | 56427 | 41 | 0.15% |
| hsa_circ_0117752 | 34477 | 25 | 0.15% |
| hsa_circ_0093877 | 22072 | 16 | 0.14% |
| hsa_circ_0118596 | 23474 | 17 | 0.14% |
| hsa_circ_0119136 | 59407 | 43 | 0.14% |
| hsa_circ_0115687 | 59442 | 43 | 0.14% |
| hsa_circ_0131464 | 89888 | 65 | 0.14% |
| hsa_circ_0006975 | 44263 | 32 | 0.14% |
| hsa_circ_0004328 | 22133 | 16 | 0.14% |
| hsa_circ_0076859 | 19369 | 14 | 0.14% |
| hsa_circ_0127385 | 70569 | 51 | 0.14% |
| hsa_circ_0102440 | 62358 | 45 | 0.14% |

| | | | |
|---|---|---|---|
| hsa_circ_0138845 | 62376 | 45 | 0.14% |
| hsa_circ_0100138 | 59625 | 43 | 0.14% |
| hsa_circ_0110890 | 88755 | 64 | 0.14% |
| hsa_circ_0058908 | 62442 | 45 | 0.14% |
| hsa_circ_0139507 | 52762 | 38 | 0.14% |
| hsa_circ_0033738 | 216637 | 156 | 0.14% |
| hsa_circ_0137616 | 47217 | 34 | 0.14% |
| hsa_circ_0100151 | 48639 | 35 | 0.14% |
| hsa_circ_0103135 | 23632 | 17 | 0.14% |
| hsa_circ_0129240 | 30588 | 22 | 0.14% |
| hsa_circ_0128806 | 62569 | 45 | 0.14% |
| hsa_circ_0135433 | 79318 | 57 | 0.14% |
| hsa_circ_0097874 | 50143 | 36 | 0.14% |
| hsa_circ_0129984 | 19508 | 14 | 0.14% |
| hsa_circ_0104949 | 50196 | 36 | 0.14% |
| hsa_circ_0128425 | 59980 | 43 | 0.14% |
| hsa_circ_0034114 | 19539 | 14 | 0.14% |
| hsa_circ_0097090 | 97701 | 70 | 0.14% |
| hsa_circ_0101648 | 69940 | 50 | 0.14% |
| hsa_circ_0133585 | 53179 | 38 | 0.14% |
| hsa_circ_0033788 | 25207 | 18 | 0.14% |
| hsa_circ_0125492 | 58933 | 42 | 0.14% |
| hsa_circ_0018084 | 66132 | 47 | 0.14% |
| hsa_circ_0129124 | 63328 | 45 | 0.14% |
| hsa_circ_0094596 | 47858 | 34 | 0.14% |
| hsa_circ_0033764 | 219601 | 156 | 0.14% |
| hsa_circ_0121783 | 40838 | 29 | 0.14% |
| hsa_circ_0124561 | 56342 | 40 | 0.14% |
| hsa_circ_0130880 | 64803 | 46 | 0.14% |
| hsa_circ_0120503 | 40871 | 29 | 0.14% |
| hsa_circ_0005114 | 80337 | 57 | 0.14% |
| hsa_circ_0076860 | 19738 | 14 | 0.14% |
| hsa_circ_0133626 | 46527 | 33 | 0.14% |
| hsa_circ_0099443 | 53623 | 38 | 0.14% |
| hsa_circ_0134602 | 35286 | 25 | 0.14% |
| hsa_circ_0136603 | 69175 | 49 | 0.14% |
| hsa_circ_0003918 | 26823 | 19 | 0.14% |
| hsa_circ_0139731 | 98860 | 70 | 0.14% |
| hsa_circ_0127480 | 53679 | 38 | 0.14% |
| hsa_circ_0107957 | 26868 | 19 | 0.14% |
| hsa_circ_0103865 | 93363 | 66 | 0.14% |
| hsa_circ_0132737 | 66535 | 47 | 0.14% |
| hsa_circ_0120383 | 70795 | 50 | 0.14% |
| hsa_circ_0102249 | 46769 | 33 | 0.14% |
| hsa_circ_0033905 | 221134 | 156 | 0.14% |
| hsa_circ_0113755 | 75157 | 53 | 0.14% |
| hsa_circ_0003955 | 90766 | 64 | 0.14% |
| hsa_circ_0002816 | 96450 | 68 | 0.14% |
| hsa_circ_0132736 | 66691 | 47 | 0.14% |
| hsa_circ_0085302 | 55378 | 39 | 0.14% |

| | | | |
|---|---|---|---|
| hsa_circ_0104943 | 83831 | 59 | 0.14% |
| hsa_circ_0033789 | 25584 | 18 | 0.14% |
| hsa_circ_0105600 | 45498 | 32 | 0.14% |
| hsa_circ_0078641 | 39817 | 28 | 0.14% |
| hsa_circ_0133347 | 58325 | 41 | 0.14% |
| hsa_circ_0140476 | 22763 | 16 | 0.14% |
| hsa_circ_0020970 | 236325 | 166 | 0.14% |
| hsa_circ_0129412 | 29922 | 21 | 0.14% |
| hsa_circ_0101416 | 61277 | 43 | 0.14% |
| hsa_circ_0137298 | 91256 | 64 | 0.14% |
| hsa_circ_0100781 | 21393 | 15 | 0.14% |
| hsa_circ_0137345 | 98445 | 69 | 0.14% |
| hsa_circ_0140475 | 22828 | 16 | 0.14% |
| hsa_circ_0033893 | 25686 | 18 | 0.14% |
| hsa_circ_0033739 | 222884 | 156 | 0.14% |
| hsa_circ_0069996 | 162888 | 114 | 0.14% |
| hsa_circ_0108267 | 22877 | 16 | 0.14% |
| hsa_circ_0100259 | 88661 | 62 | 0.14% |
| hsa_circ_0115629 | 65793 | 46 | 0.14% |
| hsa_circ_0093876 | 54459 | 38 | 0.14% |
| hsa_circ_0123719 | 57328 | 40 | 0.14% |
| hsa_circ_0008273 | 55932 | 39 | 0.14% |
| hsa_circ_0118471 | 60244 | 42 | 0.14% |
| hsa_circ_0125749 | 66006 | 46 | 0.14% |
| hsa_circ_0002842 | 22960 | 16 | 0.14% |
| hsa_circ_0100261 | 94744 | 66 | 0.14% |
| hsa_circ_0055801 | 66057 | 46 | 0.14% |
| hsa_circ_0116097 | 33031 | 23 | 0.14% |
| hsa_circ_0122430 | 74696 | 52 | 0.14% |
| hsa_circ_0114615 | 54590 | 38 | 0.14% |
| hsa_circ_0060046 | 71863 | 50 | 0.14% |
| hsa_circ_0007170 | 56058 | 39 | 0.14% |
| hsa_circ_0104615 | 79072 | 55 | 0.14% |
| hsa_circ_0003156 | 46011 | 32 | 0.14% |
| hsa_circ_0068511 | 57547 | 40 | 0.14% |
| hsa_circ_0006219 | 24474 | 17 | 0.14% |
| hsa_circ_0033765 | 224592 | 156 | 0.14% |
| hsa_circ_0060045 | 71988 | 50 | 0.14% |
| hsa_circ_0134790 | 63388 | 44 | 0.14% |
| hsa_circ_0121530 | 77827 | 54 | 0.14% |
| hsa_circ_0107381 | 46194 | 32 | 0.14% |
| hsa_circ_0101821 | 23099 | 16 | 0.14% |
| hsa_circ_0111780 | 73776 | 51 | 0.14% |
| hsa_circ_0113701 | 23171 | 16 | 0.14% |
| hsa_circ_0133232 | 76804 | 53 | 0.14% |
| hsa_circ_0126744 | 23191 | 16 | 0.14% |
| hsa_circ_0095054 | 23205 | 16 | 0.14% |
| hsa_circ_0114071 | 21761 | 15 | 0.14% |
| hsa_circ_0120038 | 20321 | 14 | 0.14% |
| hsa_circ_0034133 | 34864 | 24 | 0.14% |

| | | | |
|---|---|---|---|
| hsa_circ_0105614 | 21790 | 15 | 0.14% |
| hsa_circ_0018083 | 68354 | 47 | 0.14% |
| hsa_circ_0117629 | 37823 | 26 | 0.14% |
| hsa_circ_0097969 | 62591 | 43 | 0.14% |
| hsa_circ_0116728 | 20394 | 14 | 0.14% |
| hsa_circ_0111230 | 21863 | 15 | 0.14% |
| hsa_circ_0121398 | 52500 | 36 | 0.14% |
| hsa_circ_0097088 | 61255 | 42 | 0.14% |
| hsa_circ_0033954 | 59809 | 41 | 0.14% |
| hsa_circ_0122022 | 55460 | 38 | 0.14% |
| hsa_circ_0130410 | 49632 | 34 | 0.14% |
| hsa_circ_0131167 | 52579 | 36 | 0.14% |
| hsa_circ_0033790 | 26290 | 18 | 0.14% |
| hsa_circ_0034132 | 18995 | 13 | 0.14% |
| hsa_circ_0125337 | 36543 | 25 | 0.14% |
| hsa_circ_0018082 | 68772 | 47 | 0.14% |
| hsa_circ_0117958 | 68785 | 47 | 0.14% |
| hsa_circ_0095453 | 35139 | 24 | 0.14% |
| hsa_circ_0114617 | 55659 | 38 | 0.14% |
| hsa_circ_0092580 | 55661 | 38 | 0.14% |
| hsa_circ_0098312 | 64464 | 44 | 0.14% |
| hsa_circ_0017528 | 30778 | 21 | 0.14% |
| hsa_circ_0103070 | 36728 | 25 | 0.14% |
| hsa_circ_0114235 | 66135 | 45 | 0.14% |
| hsa_circ_0119550 | 67636 | 46 | 0.14% |
| hsa_circ_0052300 | 72059 | 49 | 0.14% |
| hsa_circ_0061306 | 70602 | 48 | 0.14% |
| hsa_circ_0098914 | 33853 | 23 | 0.14% |
| hsa_circ_0136848 | 63360 | 43 | 0.14% |
| hsa_circ_0122755 | 91480 | 62 | 0.14% |
| hsa_circ_0132033 | 26578 | 18 | 0.14% |
| hsa_circ_0138427 | 28056 | 19 | 0.14% |
| hsa_circ_0100313 | 35456 | 24 | 0.14% |
| hsa_circ_0117725 | 47307 | 32 | 0.14% |
| hsa_circ_0135747 | 68143 | 46 | 0.14% |
| hsa_circ_0069995 | 168898 | 114 | 0.13% |
| hsa_circ_0101500 | 45960 | 31 | 0.13% |
| hsa_circ_0006447 | 68245 | 46 | 0.13% |
| hsa_circ_0120286 | 90511 | 61 | 0.13% |
| hsa_circ_0096739 | 71251 | 48 | 0.13% |
| hsa_circ_0138883 | 44587 | 30 | 0.13% |
| hsa_circ_0132024 | 26758 | 18 | 0.13% |
| hsa_circ_0068063 | 98116 | 66 | 0.13% |
| hsa_circ_0102827 | 62439 | 42 | 0.13% |
| hsa_circ_0004176 | 26767 | 18 | 0.13% |
| hsa_circ_0020985 | 87810 | 59 | 0.13% |
| hsa_circ_0130719 | 37218 | 25 | 0.13% |
| hsa_circ_0125534 | 58079 | 39 | 0.13% |
| hsa_circ_0121515 | 46207 | 31 | 0.13% |
| hsa_circ_0111642 | 31309 | 21 | 0.13% |

| | | | |
|---|---|---|---|
| hsa_circ_0137273 | 38808 | 26 | 0.13% |
| hsa_circ_0137274 | 38813 | 26 | 0.13% |
| hsa_circ_0140723 | 76158 | 51 | 0.13% |
| hsa_circ_0069993 | 170296 | 114 | 0.13% |
| hsa_circ_0134077 | 58273 | 39 | 0.13% |
| hsa_circ_0017529 | 31404 | 21 | 0.13% |
| hsa_circ_0140133 | 43388 | 29 | 0.13% |
| hsa_circ_0120284 | 91278 | 61 | 0.13% |
| hsa_circ_0136825 | 23946 | 16 | 0.13% |
| hsa_circ_0004135 | 62869 | 42 | 0.13% |
| hsa_circ_0130597 | 55393 | 37 | 0.13% |
| hsa_circ_0095673 | 23959 | 16 | 0.13% |
| hsa_circ_0067092 | 67417 | 45 | 0.13% |
| hsa_circ_0133223 | 73432 | 49 | 0.13% |
| hsa_circ_0124862 | 39012 | 26 | 0.13% |
| hsa_circ_0034122 | 30028 | 20 | 0.13% |
| hsa_circ_0102826 | 63107 | 42 | 0.13% |
| hsa_circ_0005890 | 94738 | 63 | 0.13% |
| hsa_circ_0096905 | 55645 | 37 | 0.13% |
| hsa_circ_0124923 | 66183 | 44 | 0.13% |
| hsa_circ_0101415 | 64752 | 43 | 0.13% |
| hsa_circ_0125335 | 75360 | 50 | 0.13% |
| hsa_circ_0100963 | 48238 | 32 | 0.13% |
| hsa_circ_0122872 | 22614 | 15 | 0.13% |
| hsa_circ_0098792 | 43725 | 29 | 0.13% |
| hsa_circ_0136427 | 51293 | 34 | 0.13% |
| hsa_circ_0138789 | 70951 | 47 | 0.13% |
| hsa_circ_0104044 | 60423 | 40 | 0.13% |
| hsa_circ_0130406 | 60424 | 40 | 0.13% |
| hsa_circ_0138308 | 33244 | 22 | 0.13% |
| hsa_circ_0130424 | 66555 | 44 | 0.13% |
| hsa_circ_0132050 | 40847 | 27 | 0.13% |
| hsa_circ_0033791 | 27246 | 18 | 0.13% |
| hsa_circ_0018467 | 78719 | 52 | 0.13% |
| hsa_circ_0004362 | 63617 | 42 | 0.13% |
| hsa_circ_0103141 | 33355 | 22 | 0.13% |
| hsa_circ_0007502 | 89460 | 59 | 0.13% |
| hsa_circ_0006562 | 89463 | 59 | 0.13% |
| hsa_circ_0130833 | 72862 | 48 | 0.13% |
| hsa_circ_0105698 | 63758 | 42 | 0.13% |
| hsa_circ_0122431 | 78951 | 52 | 0.13% |
| hsa_circ_0140239 | 60765 | 40 | 0.13% |
| hsa_circ_0013800 | 621610 | 409 | 0.13% |
| hsa_circ_0095849 | 53214 | 35 | 0.13% |
| hsa_circ_0067089 | 68514 | 45 | 0.13% |
| hsa_circ_0108378 | 51779 | 34 | 0.13% |
| hsa_circ_0098190 | 63970 | 42 | 0.13% |
| hsa_circ_0034124 | 36559 | 24 | 0.13% |
| hsa_circ_0134268 | 44190 | 29 | 0.13% |
| hsa_circ_0130881 | 70124 | 46 | 0.13% |

| | | | |
|---|---|---|---|
| hsa_circ_0077744 | 35083 | 23 | 0.13% |
| hsa_circ_0032941 | 59490 | 39 | 0.13% |
| hsa_circ_0122756 | 94600 | 62 | 0.13% |
| hsa_circ_0131438 | 53486 | 35 | 0.13% |
| hsa_circ_0134977 | 64229 | 42 | 0.13% |
| hsa_circ_0119880 | 85682 | 56 | 0.13% |
| hsa_circ_0095334 | 27556 | 18 | 0.13% |
| hsa_circ_0077745 | 35222 | 23 | 0.13% |
| hsa_circ_0114717 | 56671 | 37 | 0.13% |
| hsa_circ_0139882 | 49024 | 32 | 0.13% |
| hsa_circ_0020259 | 65884 | 43 | 0.13% |
| hsa_circ_0097301 | 55182 | 36 | 0.13% |
| hsa_circ_0097302 | 55182 | 36 | 0.13% |
| hsa_circ_0102872 | 53710 | 35 | 0.13% |
| hsa_circ_0095853 | 42996 | 28 | 0.13% |
| hsa_circ_0033720 | 27648 | 18 | 0.13% |
| hsa_circ_0011507 | 76814 | 50 | 0.13% |
| hsa_circ_0122605 | 27670 | 18 | 0.13% |
| hsa_circ_0111534 | 47656 | 31 | 0.13% |
| hsa_circ_0105612 | 39977 | 26 | 0.13% |
| hsa_circ_0093775 | 38459 | 25 | 0.13% |
| hsa_circ_0130602 | 70804 | 46 | 0.13% |
| hsa_circ_0139672 | 95445 | 62 | 0.13% |
| hsa_circ_0112740 | 26185 | 17 | 0.13% |
| hsa_circ_0114616 | 58543 | 38 | 0.13% |
| hsa_circ_0115713 | 81801 | 53 | 0.13% |
| hsa_circ_0117711 | 71067 | 46 | 0.13% |
| hsa_circ_0117701 | 35549 | 23 | 0.13% |
| hsa_circ_0112690 | 54115 | 35 | 0.13% |
| hsa_circ_0124590 | 40213 | 26 | 0.13% |
| hsa_circ_0124740 | 41760 | 27 | 0.13% |
| hsa_circ_0136815 | 40222 | 26 | 0.13% |
| hsa_circ_0139162 | 57254 | 37 | 0.13% |
| hsa_circ_0096911 | 63478 | 41 | 0.13% |
| hsa_circ_0119137 | 66594 | 43 | 0.13% |
| hsa_circ_0137528 | 32544 | 21 | 0.13% |
| hsa_circ_0121405 | 65097 | 42 | 0.13% |
| hsa_circ_0131511 | 69755 | 45 | 0.13% |
| hsa_circ_0104045 | 62010 | 40 | 0.13% |
| hsa_circ_0099550 | 82174 | 53 | 0.13% |
| hsa_circ_0112689 | 76058 | 49 | 0.13% |
| hsa_circ_0013940 | 40384 | 26 | 0.13% |
| hsa_circ_0129235 | 62134 | 40 | 0.13% |
| hsa_circ_0111527 | 48182 | 31 | 0.13% |
| hsa_circ_0034115 | 21768 | 14 | 0.13% |
| hsa_circ_0096912 | 63775 | 41 | 0.13% |
| hsa_circ_0096744 | 31115 | 20 | 0.13% |
| hsa_circ_0094534 | 84023 | 54 | 0.13% |
| hsa_circ_0121181 | 38925 | 25 | 0.13% |
| hsa_circ_0099627 | 52952 | 34 | 0.13% |

| | | | |
|---|---|---|---|
| hsa_circ_0115526 | 82607 | 53 | 0.13% |
| hsa_circ_0131495 | 48348 | 31 | 0.13% |
| hsa_circ_0103138 | 35890 | 23 | 0.13% |
| hsa_circ_0130603 | 71837 | 46 | 0.13% |
| hsa_circ_0093272 | 59344 | 38 | 0.13% |
| hsa_circ_0130604 | 71852 | 46 | 0.13% |
| hsa_circ_0139883 | 50045 | 32 | 0.13% |
| hsa_circ_0139880 | 50057 | 32 | 0.13% |
| hsa_circ_0119046 | 65751 | 42 | 0.13% |
| hsa_circ_0108524 | 67336 | 43 | 0.13% |
| hsa_circ_0130455 | 48559 | 31 | 0.13% |
| hsa_circ_0127360 | 53313 | 34 | 0.13% |
| hsa_circ_0139881 | 50178 | 32 | 0.13% |
| hsa_circ_0101000 | 64368 | 41 | 0.13% |
| hsa_circ_0093686 | 26701 | 17 | 0.13% |
| hsa_circ_0034123 | 31417 | 20 | 0.13% |
| hsa_circ_0107950 | 40961 | 26 | 0.13% |
| hsa_circ_0095850 | 55199 | 35 | 0.13% |
| hsa_circ_0113749 | 71007 | 45 | 0.13% |
| hsa_circ_0138500 | 74253 | 47 | 0.13% |
| hsa_circ_0068042 | 104347 | 66 | 0.13% |
| hsa_circ_0006252 | 52176 | 33 | 0.13% |
| hsa_circ_0118226 | 45961 | 29 | 0.13% |
| hsa_circ_0132715 | 99880 | 63 | 0.13% |
| hsa_circ_0044190 | 98347 | 62 | 0.13% |
| hsa_circ_0119016 | 50771 | 32 | 0.13% |
| hsa_circ_0139739 | 49287 | 31 | 0.13% |
| hsa_circ_0140767 | 34983 | 22 | 0.13% |
| hsa_circ_0132735 | 74743 | 47 | 0.13% |
| hsa_circ_0122426 | 82766 | 52 | 0.13% |
| hsa_circ_0062498 | 70088 | 44 | 0.13% |
| hsa_circ_0099692 | 47836 | 30 | 0.13% |
| hsa_circ_0125436 | 47841 | 30 | 0.13% |
| hsa_circ_0118340 | 65402 | 41 | 0.13% |
| hsa_circ_0128047 | 65483 | 41 | 0.13% |
| hsa_circ_0110100 | 75087 | 47 | 0.13% |
| hsa_circ_0098908 | 27169 | 17 | 0.13% |
| hsa_circ_0099206 | 36760 | 23 | 0.13% |
| hsa_circ_0099693 | 47973 | 30 | 0.13% |
| hsa_circ_0098909 | 27186 | 17 | 0.13% |
| hsa_circ_0113688 | 71975 | 45 | 0.13% |
| hsa_circ_0116502 | 86372 | 54 | 0.13% |
| hsa_circ_0114070 | 35218 | 22 | 0.12% |
| hsa_circ_0006071 | 75251 | 47 | 0.12% |
| hsa_circ_0139961 | 88262 | 55 | 0.12% |
| hsa_circ_0031701 | 229813 | 143 | 0.12% |
| hsa_circ_0077989 | 69174 | 43 | 0.12% |
| hsa_circ_0093746 | 28991 | 18 | 0.12% |
| hsa_circ_0129236 | 64439 | 40 | 0.12% |
| hsa_circ_0130005 | 48414 | 30 | 0.12% |

| | | | |
|---|---|---|---|
| hsa_circ_0115688 | 69453 | 43 | 0.12% |
| hsa_circ_0130420 | 53302 | 33 | 0.12% |
| hsa_circ_0138848 | 30697 | 19 | 0.12% |
| hsa_circ_0130722 | 35553 | 22 | 0.12% |
| hsa_circ_0055856 | 84093 | 52 | 0.12% |
| hsa_circ_0123107 | 63090 | 39 | 0.12% |
| hsa_circ_0127319 | 29161 | 18 | 0.12% |
| hsa_circ_0108429 | 56702 | 35 | 0.12% |
| hsa_circ_0034125 | 38883 | 24 | 0.12% |
| hsa_circ_0122740 | 72909 | 45 | 0.12% |
| hsa_circ_0136716 | 42192 | 26 | 0.12% |
| hsa_circ_0138600 | 64946 | 40 | 0.12% |
| hsa_circ_0104941 | 95797 | 59 | 0.12% |
| hsa_circ_0137195 | 66628 | 41 | 0.12% |
| hsa_circ_0138601 | 65009 | 40 | 0.12% |
| hsa_circ_0125574 | 43897 | 27 | 0.12% |
| hsa_circ_0076612 | 78213 | 48 | 0.12% |
| hsa_circ_0066708 | 206955 | 127 | 0.12% |
| hsa_circ_0136007 | 66843 | 41 | 0.12% |
| hsa_circ_0136646 | 61999 | 38 | 0.12% |
| hsa_circ_0017916 | 76707 | 47 | 0.12% |
| hsa_circ_0055857 | 84870 | 52 | 0.12% |
| hsa_circ_0132733 | 76748 | 47 | 0.12% |
| hsa_circ_0115628 | 53890 | 33 | 0.12% |
| hsa_circ_0085047 | 117789 | 72 | 0.12% |
| hsa_circ_0111993 | 60533 | 37 | 0.12% |
| hsa_circ_0136382 | 81818 | 50 | 0.12% |
| hsa_circ_0135040 | 65483 | 40 | 0.12% |
| hsa_circ_0122163 | 37697 | 23 | 0.12% |
| hsa_circ_0122651 | 81995 | 50 | 0.12% |
| hsa_circ_0045369 | 86948 | 53 | 0.12% |
| hsa_circ_0126011 | 21339 | 13 | 0.12% |
| hsa_circ_0112688 | 80448 | 49 | 0.12% |
| hsa_circ_0077662 | 55858 | 34 | 0.12% |
| hsa_circ_0136425 | 59159 | 36 | 0.12% |
| hsa_circ_0140135 | 55892 | 34 | 0.12% |
| hsa_circ_0008029 | 60853 | 37 | 0.12% |
| hsa_circ_0111300 | 70766 | 43 | 0.12% |
| hsa_circ_0002270 | 41146 | 25 | 0.12% |
| hsa_circ_0134228 | 92280 | 56 | 0.12% |
| hsa_circ_0103887 | 88987 | 54 | 0.12% |
| hsa_circ_0077660 | 56094 | 34 | 0.12% |
| hsa_circ_0108446 | 57782 | 35 | 0.12% |
| hsa_circ_0000391 | 57799 | 35 | 0.12% |
| hsa_circ_0033728 | 29726 | 18 | 0.12% |
| hsa_circ_0077658 | 56191 | 34 | 0.12% |
| hsa_circ_0115714 | 87603 | 53 | 0.12% |
| hsa_circ_0138370 | 46315 | 28 | 0.12% |
| hsa_circ_0108447 | 57929 | 35 | 0.12% |
| hsa_circ_0100962 | 61286 | 37 | 0.12% |

| | | | |
|---|---|---|---|
| hsa_circ_0137129 | 64601 | 39 | 0.12% |
| hsa_circ_0100650 | 29821 | 18 | 0.12% |
| hsa_circ_0134987 | 38160 | 23 | 0.12% |
| hsa_circ_0100191 | 51509 | 31 | 0.12% |
| hsa_circ_0045370 | 88072 | 53 | 0.12% |
| hsa_circ_0138587 | 68135 | 41 | 0.12% |
| hsa_circ_0137912 | 69908 | 42 | 0.12% |
| hsa_circ_0077657 | 56625 | 34 | 0.12% |
| hsa_circ_0133093 | 71633 | 43 | 0.12% |
| hsa_circ_0111535 | 51684 | 31 | 0.12% |
| hsa_circ_0011488 | 83407 | 50 | 0.12% |
| hsa_circ_0033839 | 260238 | 156 | 0.12% |
| hsa_circ_0111466 | 45043 | 27 | 0.12% |
| hsa_circ_0101671 | 65117 | 39 | 0.12% |
| hsa_circ_0033822 | 78475 | 47 | 0.12% |
| hsa_circ_0107608 | 81872 | 49 | 0.12% |
| hsa_circ_0101672 | 65168 | 39 | 0.12% |
| hsa_circ_0125684 | 66881 | 40 | 0.12% |
| hsa_circ_0030440 | 77000 | 46 | 0.12% |
| hsa_circ_0034126 | 40177 | 24 | 0.12% |
| hsa_circ_0132692 | 67054 | 40 | 0.12% |
| hsa_circ_0132118 | 75464 | 45 | 0.12% |
| hsa_circ_0101193 | 65505 | 39 | 0.12% |
| hsa_circ_0111997 | 62174 | 37 | 0.12% |
| hsa_circ_0033894 | 30262 | 18 | 0.12% |
| hsa_circ_0076868 | 84158 | 50 | 0.12% |
| hsa_circ_0130094 | 45463 | 27 | 0.12% |
| hsa_circ_0132720 | 77500 | 46 | 0.12% |
| hsa_circ_0083793 | 97732 | 58 | 0.12% |
| hsa_circ_0033896 | 79438 | 47 | 0.12% |
| hsa_circ_0055538 | 368628 | 218 | 0.12% |
| hsa_circ_0001548 | 42282 | 25 | 0.12% |
| hsa_circ_0134591 | 76113 | 45 | 0.12% |
| hsa_circ_0036002 | 86265 | 51 | 0.12% |
| hsa_circ_0126672 | 76149 | 45 | 0.12% |
| hsa_circ_0034127 | 40627 | 24 | 0.12% |
| hsa_circ_0132165 | 76222 | 45 | 0.12% |
| hsa_circ_0137321 | 44041 | 26 | 0.12% |
| hsa_circ_0132718 | 77976 | 46 | 0.12% |
| hsa_circ_0013937 | 62770 | 37 | 0.12% |
| hsa_circ_0124503 | 81529 | 48 | 0.12% |
| hsa_circ_0117891 | 54364 | 32 | 0.12% |
| hsa_circ_0139048 | 57792 | 34 | 0.12% |
| hsa_circ_0033766 | 265174 | 156 | 0.12% |
| hsa_circ_0132691 | 47626 | 28 | 0.12% |
| hsa_circ_0130975 | 95383 | 56 | 0.12% |
| hsa_circ_0125576 | 68187 | 40 | 0.12% |
| hsa_circ_0123035 | 61396 | 36 | 0.12% |
| hsa_circ_0123330 | 76888 | 45 | 0.12% |
| hsa_circ_0125798 | 78606 | 46 | 0.12% |

| | | | |
|---|---|---|---|
| hsa_circ_0131502 | 68371 | 40 | 0.12% |
| hsa_circ_0092834 | 25645 | 15 | 0.12% |
| hsa_circ_0001908 | 23944 | 14 | 0.12% |
| hsa_circ_0099525 | 61638 | 36 | 0.12% |
| hsa_circ_0129238 | 58237 | 34 | 0.12% |
| hsa_circ_0104054 | 78814 | 46 | 0.12% |
| hsa_circ_0003138 | 42852 | 25 | 0.12% |
| hsa_circ_0003993 | 42866 | 25 | 0.12% |
| hsa_circ_0131158 | 84072 | 49 | 0.12% |
| hsa_circ_0130589 | 94418 | 55 | 0.12% |
| hsa_circ_0103136 | 56681 | 33 | 0.12% |
| hsa_circ_0108271 | 56707 | 33 | 0.12% |
| hsa_circ_0099526 | 61863 | 36 | 0.12% |
| hsa_circ_0089804 | 75778 | 44 | 0.12% |
| hsa_circ_0130416 | 56841 | 33 | 0.12% |
| hsa_circ_0033767 | 269157 | 156 | 0.12% |
| hsa_circ_0125437 | 48336 | 28 | 0.12% |
| hsa_circ_0034118 | 34564 | 20 | 0.12% |
| hsa_circ_0134751 | 24270 | 14 | 0.12% |
| hsa_circ_0033840 | 271017 | 156 | 0.12% |
| hsa_circ_0113697 | 86867 | 50 | 0.12% |
| hsa_circ_0000249 | 71234 | 41 | 0.12% |
| hsa_circ_0089801 | 133803 | 77 | 0.12% |
| hsa_circ_0135381 | 99061 | 57 | 0.12% |
| hsa_circ_0127757 | 74767 | 43 | 0.12% |
| hsa_circ_0135293 | 69567 | 40 | 0.11% |
| hsa_circ_0101819 | 38280 | 22 | 0.11% |
| hsa_circ_0130418 | 57525 | 33 | 0.11% |
| hsa_circ_0130417 | 57529 | 33 | 0.11% |
| hsa_circ_0096606 | 71521 | 41 | 0.11% |
| hsa_circ_0109905 | 71536 | 41 | 0.11% |
| hsa_circ_0132025 | 31409 | 18 | 0.11% |
| hsa_circ_0132026 | 31421 | 18 | 0.11% |
| hsa_circ_0122024 | 61104 | 35 | 0.11% |
| hsa_circ_0062183 | 96138 | 55 | 0.11% |
| hsa_circ_0140240 | 76938 | 44 | 0.11% |
| hsa_circ_0034128 | 41992 | 24 | 0.11% |
| hsa_circ_0132344 | 61327 | 35 | 0.11% |
| hsa_circ_0113740 | 52593 | 30 | 0.11% |
| hsa_circ_0114791 | 47348 | 27 | 0.11% |
| hsa_circ_0134555 | 73719 | 42 | 0.11% |
| hsa_circ_0101674 | 54487 | 31 | 0.11% |
| hsa_circ_0137779 | 79112 | 45 | 0.11% |
| hsa_circ_0102753 | 36929 | 21 | 0.11% |
| hsa_circ_0138994 | 82670 | 47 | 0.11% |
| hsa_circ_0101675 | 54538 | 31 | 0.11% |
| hsa_circ_0130413 | 44021 | 25 | 0.11% |
| hsa_circ_0013971 | 17615 | 10 | 0.11% |
| hsa_circ_0130720 | 44038 | 25 | 0.11% |
| hsa_circ_0097337 | 86348 | 49 | 0.11% |

| | | | |
|---|---|---|---|
| hsa_circ_0034129 | 42304 | 24 | 0.11% |
| hsa_circ_0101647 | 93432 | 53 | 0.11% |
| hsa_circ_0072852 | 61717 | 35 | 0.11% |
| hsa_circ_0130414 | 44086 | 25 | 0.11% |
| hsa_circ_0112687 | 86429 | 49 | 0.11% |
| hsa_circ_0017821 | 114663 | 65 | 0.11% |
| hsa_circ_0135868 | 58220 | 33 | 0.11% |
| hsa_circ_0113741 | 84691 | 48 | 0.11% |
| hsa_circ_0076872 | 173025 | 98 | 0.11% |
| hsa_circ_0119286 | 63574 | 36 | 0.11% |
| hsa_circ_0101649 | 88519 | 50 | 0.11% |
| hsa_circ_0126706 | 85092 | 48 | 0.11% |
| hsa_circ_0127762 | 53186 | 30 | 0.11% |
| hsa_circ_0033792 | 31935 | 18 | 0.11% |
| hsa_circ_0129247 | 46133 | 26 | 0.11% |
| hsa_circ_0107141 | 86950 | 49 | 0.11% |
| hsa_circ_0129232 | 78199 | 44 | 0.11% |
| hsa_circ_0129233 | 78199 | 44 | 0.11% |
| hsa_circ_0105602 | 74691 | 42 | 0.11% |
| hsa_circ_0111652 | 48048 | 27 | 0.11% |
| hsa_circ_0127763 | 53400 | 30 | 0.11% |
| hsa_circ_0130861 | 80276 | 45 | 0.11% |
| hsa_circ_0106639 | 53539 | 30 | 0.11% |
| hsa_circ_0033883 | 32136 | 18 | 0.11% |
| hsa_circ_0008136 | 48233 | 27 | 0.11% |
| hsa_circ_0102712 | 66111 | 37 | 0.11% |
| hsa_circ_0001105 | 76833 | 43 | 0.11% |
| hsa_circ_0111639 | 75048 | 42 | 0.11% |
| hsa_circ_0033841 | 279731 | 156 | 0.11% |
| hsa_circ_0119344 | 68158 | 38 | 0.11% |
| hsa_circ_0033842 | 279896 | 156 | 0.11% |
| hsa_circ_0033740 | 279948 | 156 | 0.11% |
| hsa_circ_0000146 | 73585 | 41 | 0.11% |
| hsa_circ_0013938 | 46683 | 26 | 0.11% |
| hsa_circ_0132687 | 26947 | 15 | 0.11% |
| hsa_circ_0125870 | 64718 | 36 | 0.11% |
| hsa_circ_0033689 | 280496 | 156 | 0.11% |
| hsa_circ_0072812 | 62977 | 35 | 0.11% |
| hsa_circ_0133095 | 71999 | 40 | 0.11% |
| hsa_circ_0088596 | 81016 | 45 | 0.11% |
| hsa_circ_0011416 | 109865 | 61 | 0.11% |
| hsa_circ_0034130 | 43255 | 24 | 0.11% |
| hsa_circ_0133215 | 59504 | 33 | 0.11% |
| hsa_circ_0011417 | 110123 | 61 | 0.11% |
| hsa_circ_0122313 | 59607 | 33 | 0.11% |
| hsa_circ_0000376 | 48782 | 27 | 0.11% |
| hsa_circ_0095805 | 70481 | 39 | 0.11% |
| hsa_circ_0114672 | 56038 | 31 | 0.11% |
| hsa_circ_0038894 | 112088 | 62 | 0.11% |
| hsa_circ_0129246 | 47134 | 26 | 0.11% |

| | | | |
|---|---|---|---|
| hsa_circ_0125535 | 70764 | 39 | 0.11% |
| hsa_circ_0111047 | 25403 | 14 | 0.11% |
| hsa_circ_0034131 | 43613 | 24 | 0.11% |
| hsa_circ_0111093 | 92708 | 51 | 0.11% |
| hsa_circ_0127764 | 54566 | 30 | 0.11% |
| hsa_circ_0140771 | 56408 | 31 | 0.11% |
| hsa_circ_0139148 | 78310 | 43 | 0.11% |
| hsa_circ_0080978 | 146122 | 80 | 0.11% |
| hsa_circ_0137604 | 65881 | 36 | 0.11% |
| hsa_circ_0140729 | 80835 | 44 | 0.11% |
| hsa_circ_0034184 | 167259 | 91 | 0.11% |
| hsa_circ_0071637 | 99331 | 54 | 0.11% |
| hsa_circ_0034275 | 82839 | 45 | 0.11% |
| hsa_circ_0135852 | 33159 | 18 | 0.11% |
| hsa_circ_0100977 | 68234 | 37 | 0.11% |
| hsa_circ_0049136 | 27663 | 15 | 0.11% |
| hsa_circ_0130415 | 46143 | 25 | 0.11% |
| hsa_circ_0095780 | 95992 | 52 | 0.11% |
| hsa_circ_0104810 | 66484 | 36 | 0.11% |
| hsa_circ_0104811 | 66488 | 36 | 0.11% |
| hsa_circ_0137617 | 83115 | 45 | 0.11% |
| hsa_circ_0094480 | 62817 | 34 | 0.11% |
| hsa_circ_0033843 | 288250 | 156 | 0.11% |
| hsa_circ_0101645 | 97936 | 53 | 0.11% |
| hsa_circ_0129912 | 62872 | 34 | 0.11% |
| hsa_circ_0132693 | 74048 | 40 | 0.11% |
| hsa_circ_0101192 | 72217 | 39 | 0.11% |
| hsa_circ_0129911 | 62973 | 34 | 0.11% |
| hsa_circ_0137789 | 51950 | 28 | 0.11% |
| hsa_circ_0075549 | 68732 | 37 | 0.11% |
| hsa_circ_0076867 | 92917 | 50 | 0.11% |
| hsa_circ_0108296 | 83781 | 45 | 0.11% |
| hsa_circ_0096872 | 83793 | 45 | 0.11% |
| hsa_circ_0108295 | 83807 | 45 | 0.11% |
| hsa_circ_0131604 | 70795 | 38 | 0.11% |
| hsa_circ_0135799 | 81979 | 44 | 0.11% |
| hsa_circ_0001106 | 80181 | 43 | 0.11% |
| hsa_circ_0115527 | 72731 | 39 | 0.11% |
| hsa_circ_0109365 | 52218 | 28 | 0.11% |
| hsa_circ_0102713 | 69047 | 37 | 0.11% |
| hsa_circ_0122865 | 50408 | 27 | 0.11% |
| hsa_circ_0101816 | 93373 | 50 | 0.11% |
| hsa_circ_0044191 | 130815 | 70 | 0.11% |
| hsa_circ_0034120 | 24308 | 13 | 0.11% |
| hsa_circ_0137887 | 91726 | 49 | 0.11% |
| hsa_circ_0098516 | 44965 | 24 | 0.11% |
| hsa_circ_0122158 | 65578 | 35 | 0.11% |
| hsa_circ_0033844 | 292464 | 156 | 0.11% |
| hsa_circ_0125483 | 48776 | 26 | 0.11% |
| hsa_circ_0055598 | 30026 | 16 | 0.11% |

| | | | |
|---|---|---|---|
| hsa_circ_0033890 | 88225 | 47 | 0.11% |
| hsa_circ_0098513 | 45088 | 24 | 0.11% |
| hsa_circ_0098514 | 45092 | 24 | 0.11% |
| hsa_circ_0094765 | 41389 | 22 | 0.11% |
| hsa_circ_0109124 | 94090 | 50 | 0.11% |
| hsa_circ_0093098 | 43288 | 23 | 0.11% |
| hsa_circ_0001764 | 26359 | 14 | 0.11% |
| hsa_circ_0094773 | 37702 | 20 | 0.11% |
| hsa_circ_0094764 | 41487 | 22 | 0.11% |
| hsa_circ_0006934 | 73592 | 39 | 0.11% |
| hsa_circ_0133934 | 62303 | 33 | 0.11% |
| hsa_circ_0033741 | 294699 | 156 | 0.11% |
| hsa_circ_0033845 | 294817 | 156 | 0.11% |
| hsa_circ_0125138 | 35946 | 19 | 0.11% |
| hsa_circ_0038895 | 117311 | 62 | 0.11% |
| hsa_circ_0132209 | 70071 | 37 | 0.11% |
| hsa_circ_0124638 | 70075 | 37 | 0.11% |
| hsa_circ_0120040 | 75791 | 40 | 0.11% |
| hsa_circ_0122312 | 45475 | 24 | 0.11% |
| hsa_circ_0092693 | 32224 | 17 | 0.11% |
| hsa_circ_0100547 | 72033 | 38 | 0.11% |
| hsa_circ_0033793 | 34121 | 18 | 0.11% |
| hsa_circ_0004993 | 91073 | 48 | 0.11% |
| hsa_circ_0135768 | 56957 | 30 | 0.11% |
| hsa_circ_0033823 | 89349 | 47 | 0.11% |
| hsa_circ_0002295 | 79940 | 42 | 0.11% |
| hsa_circ_0130595 | 70457 | 37 | 0.11% |
| hsa_circ_0111171 | 72523 | 38 | 0.10% |
| hsa_circ_0136430 | 57409 | 30 | 0.10% |
| hsa_circ_0127350 | 30627 | 16 | 0.10% |
| hsa_circ_0108448 | 93870 | 49 | 0.10% |
| hsa_circ_0033846 | 298897 | 156 | 0.10% |
| hsa_circ_0137529 | 40245 | 21 | 0.10% |
| hsa_circ_0132721 | 88230 | 46 | 0.10% |
| hsa_circ_0104055 | 88431 | 46 | 0.10% |
| hsa_circ_0123356 | 55773 | 29 | 0.10% |
| hsa_circ_0126934 | 42336 | 22 | 0.10% |
| hsa_circ_0131506 | 78920 | 41 | 0.10% |
| hsa_circ_0123353 | 55910 | 29 | 0.10% |
| hsa_circ_0132719 | 88706 | 46 | 0.10% |
| hsa_circ_0139269 | 98356 | 51 | 0.10% |
| hsa_circ_0123350 | 56008 | 29 | 0.10% |
| hsa_circ_0062505 | 67603 | 35 | 0.10% |
| hsa_circ_0109903 | 77409 | 40 | 0.10% |
| hsa_circ_0128512 | 91008 | 47 | 0.10% |
| hsa_circ_0126610 | 91113 | 47 | 0.10% |
| hsa_circ_0130587 | 62046 | 32 | 0.10% |
| hsa_circ_0033794 | 34978 | 18 | 0.10% |
| hsa_circ_0101750 | 50551 | 26 | 0.10% |
| hsa_circ_0037856 | 97237 | 50 | 0.10% |

| | | | |
|---|---|---|---|
| hsa_circ_0132054 | 83651 | 43 | 0.10% |
| hsa_circ_0129231 | 85666 | 44 | 0.10% |
| hsa_circ_0117589 | 62440 | 32 | 0.10% |
| hsa_circ_0131500 | 56671 | 29 | 0.10% |
| hsa_circ_0117586 | 62539 | 32 | 0.10% |
| hsa_circ_0033795 | 35193 | 18 | 0.10% |
| hsa_circ_0129227 | 86060 | 44 | 0.10% |
| hsa_circ_0033796 | 35213 | 18 | 0.10% |
| hsa_circ_0121554 | 46951 | 24 | 0.10% |
| hsa_circ_0125349 | 52832 | 27 | 0.10% |
| hsa_circ_0132055 | 84144 | 43 | 0.10% |
| hsa_circ_0140248 | 76355 | 39 | 0.10% |
| hsa_circ_0121550 | 80377 | 41 | 0.10% |
| hsa_circ_0128120 | 62760 | 32 | 0.10% |
| hsa_circ_0138536 | 94156 | 48 | 0.10% |
| hsa_circ_0124276 | 64817 | 33 | 0.10% |
| hsa_circ_0038892 | 108457 | 55 | 0.10% |
| hsa_circ_0124275 | 65141 | 33 | 0.10% |
| hsa_circ_0097577 | 96727 | 49 | 0.10% |
| hsa_circ_0013091 | 146152 | 74 | 0.10% |
| hsa_circ_0025468 | 225177 | 114 | 0.10% |
| hsa_circ_0127203 | 96814 | 49 | 0.10% |
| hsa_circ_0101670 | 98864 | 50 | 0.10% |
| hsa_circ_0068512 | 108948 | 55 | 0.10% |
| hsa_circ_0130932 | 75548 | 38 | 0.10% |
| hsa_circ_0130585 | 63693 | 32 | 0.10% |
| hsa_circ_0136421 | 71832 | 36 | 0.10% |
| hsa_circ_0102250 | 81898 | 41 | 0.10% |
| hsa_circ_0092597 | 73937 | 37 | 0.10% |
| hsa_circ_0118225 | 38052 | 19 | 0.10% |
| hsa_circ_0115528 | 78128 | 39 | 0.10% |
| hsa_circ_0002202 | 98209 | 49 | 0.10% |
| hsa_circ_0035406 | 116285 | 58 | 0.10% |
| hsa_circ_0038893 | 110279 | 55 | 0.10% |
| hsa_circ_0113369 | 88305 | 44 | 0.10% |
| hsa_circ_0092598 | 94392 | 47 | 0.10% |
| hsa_circ_0003737 | 68287 | 34 | 0.10% |
| hsa_circ_0033824 | 94477 | 47 | 0.10% |
| hsa_circ_0130588 | 64470 | 32 | 0.10% |
| hsa_circ_0124811 | 66499 | 33 | 0.10% |
| hsa_circ_0033847 | 314555 | 156 | 0.10% |
| hsa_circ_0096890 | 82675 | 41 | 0.10% |
| hsa_circ_0124809 | 66655 | 33 | 0.10% |
| hsa_circ_0125799 | 93281 | 46 | 0.10% |
| hsa_circ_0018067 | 103543 | 51 | 0.10% |
| hsa_circ_0093095 | 58886 | 29 | 0.10% |
| hsa_circ_0074783 | 87434 | 43 | 0.10% |
| hsa_circ_0114597 | 73235 | 36 | 0.10% |
| hsa_circ_0102444 | 67165 | 33 | 0.10% |
| hsa_circ_0038896 | 126424 | 62 | 0.10% |

| | | | |
|---|---|---|---|
| hsa_circ_0119744 | 32640 | 16 | 0.10% |
| hsa_circ_0093878 | 53075 | 26 | 0.10% |
| hsa_circ_0033797 | 36777 | 18 | 0.10% |
| hsa_circ_0121682 | 85855 | 42 | 0.10% |
| hsa_circ_0095815 | 71651 | 35 | 0.10% |
| hsa_circ_0124633 | 73715 | 36 | 0.10% |
| hsa_circ_0033825 | 96389 | 47 | 0.10% |
| hsa_circ_0099477 | 47207 | 23 | 0.10% |
| hsa_circ_0033826 | 96491 | 47 | 0.10% |
| hsa_circ_0139732 | 82145 | 40 | 0.10% |
| hsa_circ_0140164 | 78094 | 38 | 0.10% |
| hsa_circ_0124712 | 36994 | 18 | 0.10% |
| hsa_circ_0133920 | 30926 | 15 | 0.10% |
| hsa_circ_0013939 | 53648 | 26 | 0.10% |
| hsa_circ_0001488 | 86861 | 42 | 0.10% |
| hsa_circ_0100556 | 88947 | 43 | 0.10% |
| hsa_circ_0100969 | 91108 | 44 | 0.10% |
| hsa_circ_0122862 | 45556 | 22 | 0.10% |
| hsa_circ_0124455 | 31077 | 15 | 0.10% |
| hsa_circ_0074028 | 140893 | 68 | 0.10% |
| hsa_circ_0095668 | 85006 | 41 | 0.10% |
| hsa_circ_0033848 | 324114 | 156 | 0.10% |
| hsa_circ_0101778 | 33244 | 16 | 0.10% |
| hsa_circ_0109904 | 85237 | 41 | 0.10% |
| hsa_circ_0120969 | 62377 | 30 | 0.10% |
| hsa_circ_0033798 | 37430 | 18 | 0.10% |
| hsa_circ_0033849 | 325703 | 156 | 0.10% |
| hsa_circ_0033799 | 37596 | 18 | 0.10% |
| hsa_circ_0098515 | 52248 | 25 | 0.10% |
| hsa_circ_0034116 | 50259 | 24 | 0.10% |
| hsa_circ_0089802 | 92230 | 44 | 0.10% |
| hsa_circ_0094475 | 73568 | 35 | 0.10% |
| hsa_circ_0094476 | 73623 | 35 | 0.10% |
| hsa_circ_0135323 | 75843 | 36 | 0.09% |
| hsa_circ_0007377 | 92780 | 44 | 0.09% |
| hsa_circ_0133338 | 80209 | 38 | 0.09% |
| hsa_circ_0064610 | 86618 | 41 | 0.09% |
| hsa_circ_0137434 | 63471 | 30 | 0.09% |
| hsa_circ_0001031 | 63518 | 30 | 0.09% |
| hsa_circ_0036567 | 84887 | 40 | 0.09% |
| hsa_circ_0114676 | 87022 | 41 | 0.09% |
| hsa_circ_0033742 | 331351 | 156 | 0.09% |
| hsa_circ_0096910 | 89447 | 42 | 0.09% |
| hsa_circ_0123710 | 44733 | 21 | 0.09% |
| hsa_circ_0080432 | 110830 | 52 | 0.09% |
| hsa_circ_0137433 | 68245 | 32 | 0.09% |
| hsa_circ_0001909 | 29867 | 14 | 0.09% |
| hsa_circ_0033800 | 38474 | 18 | 0.09% |
| hsa_circ_0139673 | 34233 | 16 | 0.09% |
| hsa_circ_0033801 | 38560 | 18 | 0.09% |

| | | | |
|---|---|---|---|
| hsa_circ_0100557 | 94359 | 44 | 0.09% |
| hsa_circ_0033802 | 38658 | 18 | 0.09% |
| hsa_circ_0101777 | 34376 | 16 | 0.09% |
| hsa_circ_0033803 | 38765 | 18 | 0.09% |
| hsa_circ_0102181 | 71223 | 33 | 0.09% |
| hsa_circ_0132034 | 51939 | 24 | 0.09% |
| hsa_circ_0132030 | 52022 | 24 | 0.09% |
| hsa_circ_0132029 | 52056 | 24 | 0.09% |
| hsa_circ_0018757 | 119318 | 55 | 0.09% |
| hsa_circ_0132027 | 52119 | 24 | 0.09% |
| hsa_circ_0133586 | 87149 | 40 | 0.09% |
| hsa_circ_0033743 | 340008 | 156 | 0.09% |
| hsa_circ_0108268 | 34876 | 16 | 0.09% |
| hsa_circ_0086243 | 93781 | 43 | 0.09% |
| hsa_circ_0033850 | 340451 | 156 | 0.09% |
| hsa_circ_0018758 | 120076 | 55 | 0.09% |
| hsa_circ_0033827 | 102654 | 47 | 0.09% |
| hsa_circ_0122037 | 80831 | 37 | 0.09% |
| hsa_circ_0128414 | 83016 | 38 | 0.09% |
| hsa_circ_0034121 | 28417 | 13 | 0.09% |
| hsa_circ_0124584 | 65722 | 30 | 0.09% |
| hsa_circ_0033768 | 341806 | 156 | 0.09% |
| hsa_circ_0086244 | 94314 | 43 | 0.09% |
| hsa_circ_0133609 | 57037 | 26 | 0.09% |
| hsa_circ_0132035 | 63720 | 29 | 0.09% |
| hsa_circ_0033734 | 103514 | 47 | 0.09% |
| hsa_circ_0122038 | 92716 | 42 | 0.09% |
| hsa_circ_0118195 | 30912 | 14 | 0.09% |
| hsa_circ_0100558 | 97317 | 44 | 0.09% |
| hsa_circ_0100554 | 97328 | 44 | 0.09% |
| hsa_circ_0100779 | 64158 | 29 | 0.09% |
| hsa_circ_0101894 | 91001 | 41 | 0.09% |
| hsa_circ_0094375 | 99929 | 45 | 0.09% |
| hsa_circ_0096735 | 91077 | 41 | 0.09% |
| hsa_circ_0033994 | 82203 | 37 | 0.09% |
| hsa_circ_0047400 | 73330 | 33 | 0.09% |
| hsa_circ_0121012 | 86736 | 39 | 0.09% |
| hsa_circ_0129638 | 71345 | 32 | 0.09% |
| hsa_circ_0025622 | 84922 | 38 | 0.09% |
| hsa_circ_0033761 | 105368 | 47 | 0.09% |
| hsa_circ_0100546 | 69565 | 31 | 0.09% |
| hsa_circ_0044187 | 157268 | 70 | 0.09% |
| hsa_circ_0008221 | 74161 | 33 | 0.09% |
| hsa_circ_0006957 | 74332 | 33 | 0.09% |
| hsa_circ_0125343 | 90245 | 40 | 0.09% |
| hsa_circ_0131728 | 85800 | 38 | 0.09% |
| hsa_circ_0130087 | 92619 | 41 | 0.09% |
| hsa_circ_0140768 | 79124 | 35 | 0.09% |
| hsa_circ_0121509 | 83650 | 37 | 0.09% |
| hsa_circ_0025621 | 86462 | 38 | 0.09% |

| | | | |
|---|---|---|---|
| hsa_circ_0121539 | 65991 | 29 | 0.09% |
| hsa_circ_0017915 | 109416 | 48 | 0.09% |
| hsa_circ_0033995 | 242002 | 106 | 0.09% |
| hsa_circ_0130599 | 84839 | 37 | 0.09% |
| hsa_circ_0126012 | 48170 | 21 | 0.09% |
| hsa_circ_0033804 | 41302 | 18 | 0.09% |
| hsa_circ_0133339 | 87205 | 38 | 0.09% |
| hsa_circ_0013806 | 112501 | 49 | 0.09% |
| hsa_circ_0096732 | 94221 | 41 | 0.09% |
| hsa_circ_0095053 | 39101 | 17 | 0.09% |
| hsa_circ_0033769 | 359874 | 156 | 0.09% |
| hsa_circ_0121041 | 85365 | 37 | 0.09% |
| hsa_circ_0129237 | 92619 | 40 | 0.09% |
| hsa_circ_0137322 | 60233 | 26 | 0.09% |
| hsa_circ_0111622 | 81319 | 35 | 0.09% |
| hsa_circ_0136107 | 95868 | 41 | 0.09% |
| hsa_circ_0121042 | 86518 | 37 | 0.09% |
| hsa_circ_0009010 | 77168 | 33 | 0.09% |
| hsa_circ_0097336 | 77304 | 33 | 0.09% |
| hsa_circ_0100967 | 82275 | 35 | 0.09% |
| hsa_circ_0078719 | 115442 | 49 | 0.08% |
| hsa_circ_0089798 | 182282 | 77 | 0.08% |
| hsa_circ_0033828 | 111657 | 47 | 0.08% |
| hsa_circ_0054436 | 142607 | 60 | 0.08% |
| hsa_circ_0087105 | 107131 | 45 | 0.08% |
| hsa_circ_0100968 | 83478 | 35 | 0.08% |
| hsa_circ_0117104 | 57252 | 24 | 0.08% |
| hsa_circ_0111162 | 50122 | 21 | 0.08% |
| hsa_circ_0111157 | 50174 | 21 | 0.08% |
| hsa_circ_0033829 | 112669 | 47 | 0.08% |
| hsa_circ_0033830 | 112869 | 47 | 0.08% |
| hsa_circ_0136429 | 81695 | 34 | 0.08% |
| hsa_circ_0107092 | 94019 | 39 | 0.08% |
| hsa_circ_0017914 | 116384 | 48 | 0.08% |
| hsa_circ_0094233 | 99637 | 41 | 0.08% |
| hsa_circ_0132729 | 73312 | 30 | 0.08% |
| hsa_circ_0033750 | 382143 | 156 | 0.08% |
| hsa_circ_0131494 | 93211 | 38 | 0.08% |
| hsa_circ_0131501 | 71730 | 29 | 0.08% |
| hsa_circ_0049100 | 42149 | 17 | 0.08% |
| hsa_circ_0114670 | 77041 | 31 | 0.08% |
| hsa_circ_0140588 | 92322 | 37 | 0.08% |
| hsa_circ_0033759 | 45017 | 18 | 0.08% |
| hsa_circ_0114669 | 77597 | 31 | 0.08% |
| hsa_circ_0073756 | 67596 | 27 | 0.08% |
| hsa_circ_0137272 | 65591 | 26 | 0.08% |
| hsa_circ_0111159 | 53184 | 21 | 0.08% |
| hsa_circ_0121404 | 78820 | 31 | 0.08% |
| hsa_circ_0138365 | 81627 | 32 | 0.08% |
| hsa_circ_0091531 | 112893 | 44 | 0.08% |

| | | | |
|---|---|---|---|
| hsa_circ_0017177 | 148896 | 58 | 0.08% |
| hsa_circ_0033729 | 46353 | 18 | 0.08% |
| hsa_circ_0086199 | 170940 | 66 | 0.08% |
| hsa_circ_0083359 | 111474 | 43 | 0.08% |
| hsa_circ_0063775 | 41671 | 16 | 0.08% |
| hsa_circ_0033955 | 261429 | 100 | 0.08% |
| hsa_circ_0108439 | 44707 | 17 | 0.08% |
| hsa_circ_0132727 | 78904 | 30 | 0.08% |
| hsa_circ_0100966 | 92378 | 35 | 0.08% |
| hsa_circ_0033805 | 47766 | 18 | 0.08% |
| hsa_circ_0062491 | 116861 | 44 | 0.08% |
| hsa_circ_0095993 | 79875 | 30 | 0.08% |
| hsa_circ_0032124 | 136105 | 51 | 0.07% |
| hsa_circ_0096826 | 37554 | 14 | 0.07% |
| hsa_circ_0094455 | 93936 | 35 | 0.07% |
| hsa_circ_0108387 | 91296 | 34 | 0.07% |
| hsa_circ_0033730 | 48449 | 18 | 0.07% |
| hsa_circ_0033878 | 48762 | 18 | 0.07% |
| hsa_circ_0035409 | 157957 | 58 | 0.07% |
| hsa_circ_0080357 | 158037 | 58 | 0.07% |
| hsa_circ_0095350 | 70921 | 26 | 0.07% |
| hsa_circ_0031772 | 218630 | 80 | 0.07% |
| hsa_circ_0033806 | 49249 | 18 | 0.07% |
| hsa_circ_0035407 | 158971 | 58 | 0.07% |
| hsa_circ_0108269 | 43929 | 16 | 0.07% |
| hsa_circ_0062496 | 122441 | 44 | 0.07% |
| hsa_circ_0025580 | 161704 | 58 | 0.07% |
| hsa_circ_0047501 | 122940 | 44 | 0.07% |
| hsa_circ_0033807 | 50454 | 18 | 0.07% |
| hsa_circ_0108270 | 45005 | 16 | 0.07% |
| hsa_circ_0033831 | 146336 | 52 | 0.07% |
| hsa_circ_0125137 | 53745 | 19 | 0.07% |
| hsa_circ_0033808 | 51042 | 18 | 0.07% |
| hsa_circ_0029734 | 184927 | 65 | 0.07% |
| hsa_circ_0033832 | 148936 | 52 | 0.07% |
| hsa_circ_0103187 | 40282 | 14 | 0.07% |
| hsa_circ_0078382 | 233176 | 81 | 0.07% |
| hsa_circ_0080355 | 167218 | 58 | 0.07% |
| hsa_circ_0033760 | 52018 | 18 | 0.07% |
| hsa_circ_0033809 | 52230 | 18 | 0.07% |
| hsa_circ_0118198 | 62058 | 21 | 0.07% |
| hsa_circ_0033960 | 295824 | 100 | 0.07% |
| hsa_circ_0033735 | 154589 | 52 | 0.07% |
| hsa_circ_0033810 | 53796 | 18 | 0.07% |
| hsa_circ_0033833 | 155545 | 52 | 0.07% |
| hsa_circ_0033736 | 155775 | 52 | 0.07% |
| hsa_circ_0033762 | 157610 | 52 | 0.07% |
| hsa_circ_0089799 | 234235 | 77 | 0.07% |
| hsa_circ_0074856 | 122913 | 40 | 0.07% |
| hsa_circ_0033811 | 55501 | 18 | 0.06% |

| | | | |
|---|---|---|---|
| hsa_circ_0069858 | 296417 | 96 | 0.06% |
| hsa_circ_0085301 | 176546 | 57 | 0.06% |
| hsa_circ_0033744 | 488837 | 156 | 0.06% |
| hsa_circ_0111160 | 78531 | 25 | 0.06% |
| hsa_circ_0132725 | 95637 | 30 | 0.06% |
| hsa_circ_0033834 | 167788 | 52 | 0.06% |
| hsa_circ_0077693 | 187265 | 57 | 0.06% |
| hsa_circ_0033737 | 171337 | 52 | 0.06% |
| hsa_circ_0033745 | 517852 | 156 | 0.06% |
| hsa_circ_0066635 | 207725 | 62 | 0.06% |
| hsa_circ_0033835 | 174551 | 52 | 0.06% |
| hsa_circ_0033812 | 60450 | 18 | 0.06% |
| hsa_circ_0033996 | 356025 | 106 | 0.06% |
| hsa_circ_0033731 | 60552 | 18 | 0.06% |
| hsa_circ_0034327 | 215486 | 64 | 0.06% |
| hsa_circ_0013941 | 47357 | 14 | 0.06% |
| hsa_circ_0033836 | 181587 | 52 | 0.06% |
| hsa_circ_0033837 | 182863 | 52 | 0.06% |
| hsa_circ_0033763 | 183023 | 52 | 0.06% |
| hsa_circ_0033813 | 63396 | 18 | 0.06% |
| hsa_circ_0034000 | 375533 | 106 | 0.06% |
| hsa_circ_0033732 | 63811 | 18 | 0.06% |
| hsa_circ_0086197 | 236715 | 66 | 0.06% |
| hsa_circ_0086198 | 237262 | 66 | 0.06% |
| hsa_circ_0033814 | 64968 | 18 | 0.06% |
| hsa_circ_0033838 | 188308 | 52 | 0.06% |
| hsa_circ_0033980 | 385212 | 106 | 0.06% |
| hsa_circ_0033815 | 66496 | 18 | 0.05% |
| hsa_circ_0033816 | 67400 | 18 | 0.05% |
| hsa_circ_0033817 | 68076 | 18 | 0.05% |
| hsa_circ_0033895 | 68248 | 18 | 0.05% |
| hsa_circ_0033733 | 68353 | 18 | 0.05% |
| hsa_circ_0033818 | 68780 | 18 | 0.05% |
| hsa_circ_0033721 | 199357 | 52 | 0.05% |
| hsa_circ_0033819 | 70094 | 18 | 0.05% |
| hsa_circ_0033746 | 609531 | 156 | 0.05% |
| hsa_circ_0033820 | 70407 | 18 | 0.05% |
| hsa_circ_0057213 | 59880 | 15 | 0.05% |
| hsa_circ_0043601 | 240662 | 59 | 0.05% |
| hsa_circ_0043602 | 242289 | 59 | 0.05% |
| hsa_circ_0043603 | 242759 | 59 | 0.05% |
| hsa_circ_0043605 | 244563 | 59 | 0.05% |
| hsa_circ_0043604 | 244944 | 59 | 0.05% |
| hsa_circ_0033997 | 442732 | 106 | 0.05% |
| hsa_circ_0074029 | 284486 | 68 | 0.05% |
| hsa_circ_0033722 | 658353 | 156 | 0.05% |
| hsa_circ_0033988 | 453671 | 106 | 0.05% |
| hsa_circ_0033821 | 77170 | 18 | 0.05% |
| hsa_circ_0062480 | 242132 | 55 | 0.05% |
| hsa_circ_0033747 | 711581 | 156 | 0.04% |

| | | | |
|---|---|---|---|
| hsa_circ_0062481 | 235696 | 50 | 0.04% |
| hsa_circ_0069749 | 826926 | 174 | 0.04% |
| hsa_circ_0033929 | 755115 | 156 | 0.04% |
| hsa_circ_0033925 | 757704 | 156 | 0.04% |
| hsa_circ_0033913 | 766508 | 156 | 0.04% |
| hsa_circ_0013829 | 541092 | 109 | 0.04% |
| hsa_circ_0013828 | 314809 | 62 | 0.04% |
| hsa_circ_0033967 | 544476 | 106 | 0.04% |
| hsa_circ_0033748 | 828200 | 156 | 0.04% |
| hsa_circ_0062465 | 611623 | 94 | 0.03% |
| hsa_circ_0062466 | 676065 | 94 | 0.03% |
| hsa_circ_0038910 | 647007 | 87 | 0.03% |
| hsa_circ_0062469 | 504699 | 66 | 0.03% |
| hsa_circ_0062467 | 507204 | 66 | 0.03% |
| hsa_circ_0062472 | 529314 | 66 | 0.02% |
| hsa_circ_0061589 | 905233 | 96 | 0.02% |
